## Supplementary Table 5 for "Distinct roles for MNK1 and MNK2 in social and cognitive behavior through kinase-specific regulation of the synaptic proteome and phosphoproteome"

**TMT all phosphosites cortex**

| phosphosite CX | normalized protein expression CX | | | | | | | | | | | | | | | | LogFC over WT CX | | | p-value / WT CX | | | adjusted p-value / WT CX | | |
|---|---|---|---|---|---|---|---|---|---|---|---|---|---|---|---|---|---|---|---|---|---|---|---|---|---|
| id | CX_WT | CX_WT | CX_WT | CX_WT | CX_MNK1 | CX_MNK1 | CX_MNK1 | CX_MNK1 | CX_MNK2 | CX_MNK2 | CX_MNK2 | CX_MNK2 | CX_MNKD | CX_MNKD | CX_MNKD | CX_MNKD | logFC.CX_ | logFC.CX_ | logFC.CX_ | P.Value.CX | P.Value.CX | P.Value.CX | adj.P.Val.C | adj.P.Val.C | adj.P.Val.CX_MNKDKO.over.CX_WT |
| 0610010F05Rik_M0QWF3_S116s_1 | 1.25 | -0.61 | -0.47 | 1.76 | 1.12 | -1.02 | 1.12 | 1.45 | 0.48 | -0.02 | 1.29 | 0.35 | -1.26 | -1.61 | -0.64 | -0.9 | 0.185 | 0.0425 | -1.585 | 0.020393 | 0.79955 | 0.943631 | 1 | 0.999631 | 0.393617 |
| 0610011F06Rik_G5E8X1_S149s_1 | -0.57 | -0.3 | -0.24 | -0.91 | -0.51 | -0.44 | -0.66 | -0.22 | -0.32 | -0.42 | -0.29 | -0.17 | 0.67 | 0.19 | 0.02 | 0.19 | 0.0475 | 0.205 | 0.7725 | 0.02807 | 0.860484 | 0.455674 | 1 | 0.999631 | 0.423977 |
| 1110002E22Rik_E0CYV9_S1338s_1 | 0.34 | 0.08 | -0.72 | -0.32 | 0.15 | -0.25 | 0.14 | -0.14 | -0.06 | 0.25 | -0.29 | -0.28 | -1.29 | -1.66 | -1.45 | -0.61 | 0.13 | 0.06 | -1.0975 | 0.012958 | 0.67527 | 0.852349 | 1 | 0.999631 | 0.385092 |
| 1110037F02Rik_E9PZY8_S1628s_1 | 0.24 | 0.09 | -0.04 | -0.44 | 0.04 | 0.05 | -0.35 | -0.81 | 0.52 | 0.82 | -0.17 | -0.19 | 1 | 0.96 | -0.48 | -0.21 | -0.23 | 0.2825 | 0.355 | 0.411353 | 0.461068 | 0.412655 | 1 | 0.999631 | 0.767937 |
| 1700021K19Rik_D3Z7B1_S137s_1 | 0.32 | 0.43 | 0.68 | 0.5 | 0.1 | 0.51 | 0.47 | 1.27 | 0.41 | 1.21 | 0.84 | 0.91 | 0.67 | 0.89 | 1.52 | 0.72 | 0.105 | 0.36 | 0.4675 | 0.150775 | 0.736425 | 0.217359 | 1 | 0.999631 | 0.623336 |
| 1700112E06Rik_E0CYG4_T63t_1 | -0.22 | -0.87 | -0.28 | -1.34 | -0.21 | -0.85 | -0.53 | -0.11 | -1.34 | -0.4 | -0.23 | -0.49 | 0.39 | -0.43 | -0.63 | -1 | 0.2525 | 0.0625 | 0.26 | 0.533066 | 0.471173 | 0.870449 | 1 | 0.999631 | 0.827798 |
| 2010300C02Rik_E9Q3M9_S1094s_1 | 1.71 | 1.3 | 0.84 | 1.59 | 1.88 | 1.26 | 2.08 | 1.19 | 2.06 | 0.85 | 2.29 | 1.42 | -0.13 | 1.05 | -0.36 | 2.47 | 0.2425 | 0.295 | -0.6025 | 0.33879 | 0.473579 | 0.46057 | 1 | 0.999631 | 0.731958 |
| 2010300C02Rik_E9Q3M9_S1123s_1 | 0.99 | 2.14 | 0.4 | 1.35 | 2.11 | 1.79 | 0.93 | 0.65 | 2.46 | 1.16 | 0.71 | 0.97 | -0.23 | 1.56 | 0.15 | 1.33 | 0.15 | 0.105 | -0.5175 | 0.344494 | 0.753449 | 0.83514 | 1 | 0.999631 | 0.734049 |
| 2010300C02Rik_E9Q3M9_S113s_1 | 1.82 | 0.21 | -1.52 | 1.8 | 2.3 | 1.09 | 1.58 | 1.76 | 1.34 | -0.11 | 0.93 | -0.63 | -4.29 | 1.37 | 0.45 | 0.17 | 1.105 | -0.195 | -1.1525 | 0.371189 | 0.155186 | 0.808331 | 1 | 0.999631 | 0.746338 |
| 2010300C02Rik_E9Q3M9_S162s_1 | -0.44 | -0.3 | -0.72 | -0.94 | -0.97 | -0.2 | -0.73 | -0.83 | -0.85 | -0.25 | -1.32 | -1.07 | -0.31 | -0.08 | -0.74 | -0.56 | -0.0825 | -0.2725 | 0.1775 | 0.562729 | 0.777339 | 0.411045 | 1 | 0.999631 | 0.841038 |
| 2010300C02Rik_E9Q3M9_S189s_1 | 0.39 | -0.97 | -0.8 | -0.81 | -0.93 | -0.72 | -0.78 | -1.08 | -0.96 | 0.06 | -1.52 | -1.11 | -1.44 | -0.48 | -0.9 | -0.98 | -0.33 | -0.335 | -0.4025 | 0.32378 | 0.360785 | 0.464307 | 1 | 0.999631 | 0.724485 |
| 2010300C02Rik_E9Q3M9_S325s_1 | 0.28 | 0.32 | -1.73 | -0.87 | 0.49 | 0.28 | -0.41 | -0.32 | 0.53 | 0.73 | -1.07 | -1.3 | 0.83 | -0.11 | -0.66 | -0.9 | 0.51 | 0.2225 | 0.29 | 0.616648 | 0.326753 | 0.732801 | 1 | 0.999631 | 0.861203 |
| 2010300C02Rik_E9Q3M9_S465s_1 | -0.86 | -1.61 | -4.73 | -2.88 | -1.47 | -1.82 | -4.43 | -4.65 | -1.3 | -0.18 | -1.91 | -3.27 | -1.22 | -1.69 | -5.67 | -4.46 | -0.5725 | 0.855 | -0.74 | 0.52777 | 0.579301 | 0.365307 | 1 | 0.999631 | 0.824665 |
| 2010300C02Rik_E9Q3M9_S47s_1 | 0.42 | 0.48 | -0.68 | 0.71 | 0.4 | 0.48 | 0.87 | 0.87 | 0.55 | 0.39 | 0.57 | 0.15 | -0.93 | 0.77 | 0.87 | 0.62 | 0.4225 | 0.1825 | 0.1 | 0.842242 | 0.256576 | 0.613613 | 1 | 0.999631 | 0.947766 |
| 2010300C02Rik_E9Q3M9_S507s_1 | -1.39 | 1.53 | 1.41 | -0.46 | 1.49 | 0.52 | 0.48 | 1.21 | 1.48 | -1.18 | 1.9 | 2.34 | 2.37 | 0.36 | 1.44 | 0.6 | 0.6525 | 0.8625 | 0.92 | 0.236198 | 0.346176 | 0.363186 | 1 | 0.999631 | 0.683301 |
| 2010300C02Rik_E9Q3M9_S570s_1 | -1.25 | 0.27 | 1.33 | -0.24 | 0.82 | -0.66 | 0.41 | 0.59 | 0.5 | -1.42 | 1.62 | 1.59 | 1.25 | -0.08 | 1.59 | 0.42 | 0.2625 | 0.545 | 0.7675 | 0.220141 | 0.648403 | 0.493103 | 1 | 0.999631 | 0.670699 |
| 2010300C02Rik_E9Q3M9_S572s_1 | -0.8 | 0.09 | 0.75 | 0.09 | 0.41 | -0.21 | 0.85 | 0.11 | 0.26 | -1.29 | 1.11 | 1.57 | 0.86 | -0.23 | 1.57 | 0.52 | 0.2575 | 0.38 | 0.6475 | 0.196027 | 0.52018 | 0.553884 | 1 | 0.999631 | 0.653373 |
| 2010300C02Rik_E9Q3M9_S612s_1 | -2.15 | 1.1 | 3.18 | -0.39 | 1.73 | -0.04 | 0.32 | 2.11 | 1.25 | -2.05 | 1.18 | 3.36 | 2.04 | 0.07 | 2.74 | 2.06 | 0.595 | 0.5 | 1.2925 | 0.248653 | 0.580695 | 0.71283 | 1 | 0.999631 | 0.685824 |
| 2010300C02Rik_E9Q3M9_S712s_1 | 0.26 | 0.3 | 0.17 | 0.47 | 0.28 | 0.52 | -0.23 | 0.61 | 0.11 | 0.05 | -0.03 | 0.89 | -0.07 | 0.98 | 0.24 | 1.05 | -0.005 | -0.045 | 0.25 | 0.474263 | 0.985738 | 0.880533 | 1 | 0.999631 | 0.799827 |
| 2010300C02Rik_E9Q3M9_S730s_1 | 0.75 | 1.55 | -0.04 | -0.47 | -0.04 | 1.36 | -1.3 | -0.16 | 0.96 | 1.66 | -1.12 | 0.09 | -1.01 | 2.62 | 0.66 | -0.71 | -0.4825 | -0.05 | -0.0575 | 0.943876 | 0.451684 | 0.940374 | 1 | 0.999631 | 0.981629 |
| 2010300C02Rik_E9Q3M9_S898s_1 | 2 | 0.8 | -0.57 | 1.82 | 2.62 | 1.82 | 1.49 | 1.83 | 1.8 | 0.55 | 1.09 | 0.28 | -1.46 | 1.57 | 0.96 | 1.13 | 0.9275 | -0.0825 | -0.4625 | 0.560425 | 0.133512 | 0.893333 | 1 | 0.999631 | 0.839898 |
| 2010300C02Rik_E9Q3M9_S918s_1 | 1.78 | -0.73 | -1.97 | 1 | 1.38 | 0.3 | 1.61 | 0.71 | 0.66 | -0.01 | 0.74 | -1.32 | -1.85 | -0.34 | -0.38 | -1.33 | 0.98 | -0.0025 | -0.995 | 0.234353 | 0.230978 | 0.997645 | 1 | 1 | 0.683223 |
| 2010300C02Rik_E9Q3M9_S92s_1 | 0.38 | 0.2 | -0.68 | 0.12 | 0.45 | -0.43 | -0.39 | 0.26 | 0.02 | 0.13 | -0.16 | -0.25 | -1.23 | 0.02 | -0.11 | 0.34 | -0.0325 | -0.07 | -0.25 | 0.560042 | 0.926507 | 0.823765 | 1 | 0.999631 | 0.839805 |
| 2010300C02Rik_E9Q3M9_S95s_1 | 0.27 | -0.61 | -1.39 | 0.26 | 0.81 | -1.28 | -0.5 | -0.23 | -0.16 | -1.01 | -0.51 | 0.19 | -2.24 | -0.15 | -0.55 | -0.35 | 0.0675 | -0.005 | -0.455 | 0.435396 | 0.900649 | 0.991333 | 1 | 0.999631 | 0.780052 |
| 2010300C02Rik_E9Q3M9_S961s_1 | -0.01 | 0.17 | 0.05 | 0.14 | -0.24 | 0.2 | -0.19 | 0.35 | -0.01 | 0.32 | 0.24 | 0.54 | 0.61 | 0.28 | 0.48 | 0.65 | -0.0575 | 0.185 | 0.4175 | 0.138897 | 0.824015 | 0.475156 | 1 | 0.999631 | 0.611718 |
| 2010300C02Rik_E9Q3M9_T660t_1 | -0.69 | 0.47 | -0.11 | -0.54 | 0.04 | 0.01 | -0.56 | -0.13 | -0.13 | -0.26 | -0.25 | 0.03 | 0.46 | 0.35 | 0 | 0.05 | 0.0575 | 0.065 | 0.4325 | 0.228117 | 0.863167 | 0.841413 | 1 | 0.999631 | 0.679038 |
| 2010300C02Rik_E9Q3M9_T959t_1 | -0.22 | -0.25 | 0.15 | -0.21 | -0.2 | -0.82 | -0.64 | 0.14 | -0.65 | -0.08 | -0.25 | 0.54 | -0.32 | -0.21 | 0.2 | 0.21 | -0.2475 | 0.0225 | 0.1025 | 0.723321 | 0.417472 | 0.94435 | 1 | 0.999631 | 0.905674 |
| 2010300C02Rik_F7B0R9_S599s_1 | 1.29 | 0.6 | 0.77 | 1.65 | 1.36 | 0.79 | 1.9 | 1.61 | 1.4 | 0.11 | 1.84 | 1.04 | -0.66 | 0.87 | -0.07 | 1.92 | 0.3375 | 0.02 | -0.5625 | 0.333282 | 0.36032 | 0.963394 | 1 | 0.999631 | 0.728079 |
| 2310035C23Rik_G3X9J4_S141s_1 | -0.8 | 0.34 | 0.15 | -0.64 | -0.75 | -0.36 | -0.01 | 0.68 | -0.48 | 0.03 | 0.1 | 0.35 | 0.16 | -0.05 | -0.13 | 0.1 | 0.1275 | 0.2375 | 0.2575 | 0.46684 | 0.758322 | 0.51831 | 1 | 0.999631 | 0.795267 |
| 2310035C23Rik_G3X9J4_S180s_1 | 0 | 0.16 | -0.77 | -0.55 | 0.13 | 0.24 | -0.68 | -1.18 | 0.1 | -0.22 | -0.4 | -0.84 | -0.56 | -0.1 | -0.84 | 0.22 | -0.0825 | -0.05 | -0.03 | 0.934919 | 0.838947 | 0.883224 | 1 | 0.999631 | 0.979037 |
| 2310035C23Rik_G3X9J4_S182s_1 | 0.31 | 0.47 | -0.19 | 0.43 | 0.52 | 0.2 | 0.02 | 0.37 | 0.43 | 0.21 | 0.64 | 0.67 | -0.29 | 0.3 | 0.27 | 0.79 | 0.0225 | 0.2325 | 0.0125 | 0.970336 | 0.934545 | 0.417243 | 1 | 0.999631 | 0.989586 |
| 2310035C23Rik_G3X9J4_S193s_1 | 1.16 | -0.06 | 1.36 | 1.3 | -0.57 | 0.35 | 2.98 | -0.28 | 0.38 | 1.43 | 1.25 | 0.73 | -1 | -1.28 | 0.16 | -0.58 | -0.32 | 0.0075 | -1.615 | 0.005095 | 0.678257 | 0.985658 | 1 | 0.999631 | 0.351308 |
| 2310035C23Rik_G3X9J4_S243s_1 | -0.23 | 0.58 | 0 | -0.14 | 0.15 | 0.34 | -0.55 | 0.26 | 0.29 | 0.18 | -0.05 | 0.14 | 0.79 | 0.63 | -0.06 | 0.09 | -0.0025 | 0.0875 | 0.31 | 0.372106 | 0.993776 | 0.759978 | 1 | 0.999631 | 0.746839 |
| 2310035C23Rik_G3X9J4_S244s_1 | -0.23 | 0.58 | 0.25 | 0.15 | 0.15 | 0.34 | -0.36 | 0.49 | 0.29 | 0.18 | 0.36 | 0.54 | 0.79 | 0.63 | 0.28 | 0.57 | -0.0325 | 0.155 | 0.38 | 0.226437 | 0.915606 | 0.583017 | 1 | 0.999631 | 0.677523 |
| 2310035C23Rik_G3X9J4_S422s_1 | -0.64 | 1.39 | 2.22 | 1.96 | 1.8 | 1.88 | 2.7 | 2.16 | 2.22 | -0.15 | 2.29 | 2.5 | -0.19 | 0.8 | 1.61 | 0.84 | 0.9025 | 0.4825 | -0.4675 | 0.488327 | 0.163108 | 0.545599 | 1 | 0.999631 | 0.80665 |
| 2310035C23Rik_G3X9J4_S56s_1 | -0.68 | 0.01 | -0.46 | -1.03 | -0.24 | -0.54 | -0.48 | -0.7 | -0.22 | -0.81 | -0.46 | -0.22 | 0.04 | 0.02 | -1.11 | -0.84 | 0.05 | 0.1125 | 0.0675 | 0.864112 | 0.8678 | 0.725019 | 1 | 0.999631 | 0.955131 |
| 2310035C23Rik_G3X9J4_T143t_1 | -0.08 | -0.05 | 0.24 | -0.21 | -0.09 | -0.04 | -0.57 | -0.52 | -0.2 | -0.35 | -0.28 | -0.14 | 0.56 | -0.04 | 0.04 | -0.02 | -0.28 | -0.2175 | 0.16 | 0.584648 | 0.306579 | 0.395128 | 1 | 0.999631 | 0.849205 |
| 4833439L19Rik_D3Z1F7_S164s_1 | 0.15 | 0.73 | 1.67 | 0.65 | 0.15 | 0.48 | 0.69 | 1.65 | 0.15 | 0.92 | 1.73 | 1.49 | 0.65 | 1.27 | 2.17 | 1.52 | -0.0575 | 0.2725 | 0.6025 | 0.198039 | 0.895997 | 0.557584 | 1 | 0.999631 | 0.65543 |
| 4931406P16Rik_Q8C5X1_S662s_1 | 1.32 | 1.45 | 0.97 | 0.71 | 1.04 | 1.34 | 0.84 | 0.88 | 1.36 | 1.53 | 1.48 | 0.74 | 1.21 | 1.52 | 0.28 | 2.3 | -0.0875 | 0.165 | 0.215 | 0.634233 | 0.756882 | 0.600912 | 1 | 0.999631 | 0.868569 |
| 5031439G07Rik_B1APX2_S30s_1 | 1.99 | -0.49 | -1.41 | 1.51 | 1.36 | 0.48 | 1.35 | 0.44 | 1.06 | 0.03 | 0.3 | -0.71 | -1.7 | -0.4 | -0.93 | -0.71 | 0.5075 | -0.23 | -1.335 | 0.099504 | 0.501172 | 0.76845 | 1 | 0.999631 | 0.562298 |
| 5031439G07Rik_B1APX2_S311s_2 | -1.89 | -1.4 | -0.58 | -1.81 | -0.81 | -0.93 | -1.76 | -1.88 | -0.57 | -1.73 | -0.94 | -1.66 | -0.62 | -0.35 | -1.11 | -2.21 | 0.075 | 0.195 | 0.3475 | 0.483992 | 0.854329 | 0.641511 | 1 | 0.999631 | 0.803422 |
| 5031439G07Rik_B1APX2_S322s_1 | -0.48 | 0.11 | -1.9 | -1.76 | -0.7 | -0.2 | -0.88 | -1.46 | -0.29 | -0.92 | -4.12 | -0.88 | -0.97 | -0.97 | 0.17 | -0.43 | 0.1975 | -0.545 | 0.4575 | 0.39542 | 0.703138 | 0.534775 | 1 | 0.999631 | 0.760219 |
| 5031439G07Rik_B1APX2_S322s_2 | -1.64 | -0.99 | -1.15 | -1.95 | -0.43 | -0.58 | -1.69 | -1.54 | -0.66 | -1.58 | -1 | -1.39 | -0.77 | -0.56 | -1.35 | -2.27 | 0.3725 | 0.275 | 0.195 | 0.661703 | 0.359651 | 0.432314 | 1 | 0.999631 | 0.88095 |
| 5031439G07Rik_B1APX2_S347s_1 | 0 | 0.98 | 1.39 | 0.79 | 0.65 | 0.95 | 0.53 | 0.78 | 1.16 | 0.99 | -0.39 | 0.73 | 1.1 | 1.59 | 1.05 | 1.51 | -0.0625 | -0.1675 | 0.5225 | 0.176565 | 0.85425 | 0.708785 | 1 | 0.999631 | 0.642116 |
| 5031439G07Rik_B1APX2_S405s_1 | -1.17 | 0.47 | -0.18 | 0.09 | 0.38 | 0.29 | -0.23 | 0.28 | 0.64 | -0.51 | 0.6 | 0.54 | 0.74 | 0.94 | 0.36 | 0.56 | 0.3775 | 0.515 | 0.8475 | 0.055007 | 0.34161 | 0.260332 | 1 | 0.999631 | 0.492066 |
| 5031439G07Rik_B1APX2_T336t_1 | -1.11 | -0.01 | 0.14 | -0.73 | -0.35 | -0.38 | -0.75 | -0.18 | -0.18 | -0.21 | 0 | 0.38 | 0.38 | -0.21 | 0.33 | 0.44 | 0.0125 | 0.425 | 0.6625 | 0.099919 | 0.971435 | 0.2545 | 1 | 0.999631 | 0.562917 |
| 5730455P16Rik_J3KMQ6_S19s_1 | 0.53 | 0.7 | 0.9 | 1.06 | 0.62 | 0.51 | 0.41 | 0.94 | 0.71 | 1.48 | 0.67 | 0.85 | 0.19 | 0.25 | 0.5 | 0.17 | -0.1775 | 0.13 | -0.52 | 0.085786 | 0.50567 | 0.664679 | 1 | 0.999631 | 0.551251 |
| 6330403A02Rik_J3QP43_S27s_1 | -0.56 | -0.62 | -0.59 | -0.53 | -0.47 | -0.07 | -0.75 | -0.57 | -0.59 | -0.1 | -0.51 | -0.17 | 0.1 | -0.59 | -0.29 | -0.05 | 0.11 | 0.2325 | 0.3675 | 0.213645 | 0.668955 | 0.374863 | 1 | 0.999631 | 0.667153 |
| 6330403A02Rik_J3QP43_S28s_1 | -0.33 | -0.93 | -1.01 | -0.94 | -0.67 | -0.83 | -1.26 | -1.12 | -0.63 | -0.45 | -0.99 | -1.02 | -0.45 | -0.76 | -0.89 | -0.9 | -0.1675 | 0.03 | 0.0525 | 0.860302 | 0.559591 | 0.918552 | 1 | 0.999631 | 0.953592 |
| 6330403A02Rik_J3QP43_S29s_1 | -0.78 | -1.34 | 0.99 | -0.87 | -1.13 | -0.89 | -1.34 | 0.28 | -1.03 | -0.37 | -0.84 | -0.1 | 0.34 | -1.53 | -0.23 | -0.2 | -0.27 | -0.085 | 0.095 | 0.872491 | 0.638161 | 0.870202 | 1 | 0.999631 | 0.959468 |
| 6430548M08Rik_Q3TA40_S208s_1 | 0.67 | -1.4 | -1.08 | 0.41 | 0.65 | 0.24 | -0.08 | -0.54 | -2.03 | -1.63 | -0.42 | -1.01 | -1.12 | -0.62 | -0.74 | -0.14 | 0.4175 | -0.9225 | -0.305 | 0.565424 | 0.441724 | 0.134488 | 1 | 0.999631 | 0.84139 |
| 6430548M08Rik_Q3TA40_S275s_1 | -1.43 | -1.08 | -0.84 | -1.21 | -1.17 | -1.32 | -1.25 | -1.16 | -1.21 | -1.05 | -1.09 | -0.96 | -0.46 | -0.92 | -1.09 | -0.86 | -0.085 | 0.0625 | 0.3075 | 0.310388 | 0.73451 | 0.810163 | 1 | 0.999631 | 0.71924 |
| 6430548M08Rik_Q3TA40_S72s_1 | -0.62 | 0.99 | 0.35 | 0.7 | 0.37 | 0.46 | -0.5 | 1.33 | 0.71 | 0.87 | 0.42 | 0.02 | 0.88 | 0.58 | 0.75 | 0.81 | 0.06 | 0.15 | 0.4 | 0.315539 | 0.901369 | 0.712875 | 1 | 0.999631 | 0.72069 |
| 8030462N17Rik_Q0VAW6_S145s_1 | 0.29 | 0.35 | 1.06 | 1.51 | 0.27 | 0.04 | 1.18 | 0.91 | 0 | 0.51 | 0.88 | 0.65 | 0.01 | -1.22 | 1.07 | -0.19 | -0.2025 | -0.2925 | -0.885 | 0.114612 | 0.615596 | 0.440943 | 1 | 0.999631 | 0.579001 |
| 8030462N17Rik_Q0VAW6_S66s_1 | 0.35 | 1.58 | 0.59 | 1.64 | 0.81 | 0.93 | 0.62 | 0.38 | 1.11 | 0.84 | 0.55 | 0.97 | 1 | 1.33 | 1.07 | 0.52 | -0.355 | -0.1725 | -0.06 | 0.880299 | 0.351125 | 0.649214 | 1 | 0.999631 | 0.963151 |
| A0A0G2JDV2_S170s_1 | 0.45 | 0.89 | 0.74 | 0.46 | 1.14 | 1.24 | -0.22 | 0.24 | 1.21 | 1.68 | -1.81 | -0.51 | 1.08 | 1.33 | 0.54 | -0.32 | -0.035 | -0.4925 | 0.0225 | 0.955477 | 0.92704 | 0.498868 | 1 | 0.999631 | 0.984649 |
| A0A0G2JFH6_S27s_1 | 0.36 | -0.78 | 0.01 | -0.8 | -1.66 | -0.1 | 0.97 | -0.28 | -0.29 | 1.18 | -0.08 | 1.15 | 0.36 | 0.28 | -0.22 | -0.49 | 0.035 | 0.7925 | 0.285 | 0.46729 | 0.950174 | 0.117492 | 1 | 0.999631 | 0.795438 |
| A0A0G2JFH6_S28s_1 | 1.86 | -0.75 | 0.95 | -0.1 | 1.2 | -0.04 | -1.06 | 1.3 | 0.13 | 1.63 | -0.03 | 0.62 | -1.1 | 0.24 | -0.23 | -2.25 | -0.14 | 0.0975 | -1.325 | 0.08388 | 0.843595 | 0.87532 | 1 | 0.999631 | 0.549732 |
| A0A0G2JGJ4_S77s_1 | 1.11 | -0.27 | 2.22 | 1.14 | -0.93 | -0.59 | 1.18 | 0.23 | -0.38 | 0.21 | 0.3 | 0.03 | -0.4 | -0.6 | 0.74 | 0.52 | -1.0775 | -1.01 | -0.985 | 0.104539 | 0.108834 | 0.069049 | 1 | 0.999631 | 0.568213 |
| A0A0J9YUM2_S765s_1 | 0.84 | -0.27 | -0.17 | 0.41 | 1.54 | 0.17 | 1.05 | 0.27 | 0.47 | -1.63 | 0.65 | 0.16 | -1.96 | -0.14 | -0.85 | -0.02 | 0.555 | -0.29 | -0.945 | 0.080475 | 0.202198 | 0.596329 | 1 | 0.999631 | 0.545579 |
| A0A0J9YUM2_S766s_1 | -0.13 | -0.44 | 0.3 | 0.83 | 0.99 | -0.09 | 1.4 | 0.62 | 0.06 | -1.9 | 1.16 | 0.66 | -1.03 | -0.61 | -0.29 | 0.67 | 0.59 | -0.145 | -0.455 | 0.327288 | 0.178953 | 0.824011 | 1 | 0.999631 | 0.724678 |
| A0A0J9YVH8_S1262s_1 | -0.51 | 0.13 | 0.32 | -0.79 | 1.23 | -0.31 | -0.37 | 0.15 | -1.44 | -2.85 | 0.76 | -0.67 | -1.07 | -1.63 | -0.02 | -0.4 | 0.3875 | -0.8375 | -0.5675 | 0.220502 | 0.389529 | 0.255125 | 1 | 0.999631 | 0.670752 |
| A0A0N4SUT6_T239t_1 | -0.12 | 0.49 | 0.04 | -0.33 | 0.76 | 0.24 | 0.08 | -0.07 | -0.31 | -0.86 | -0.29 | 0.03 | 0.91 | 0.53 | 1.58 | 1.54 | 0.2325 | -0.3775 | 1.12 | 0.010187 | 0.456778 | 0.250345 | 1 | 0.999631 | 0.383133 |
| A0A0N4SW73_S1188s_1 | 1.03 | -0.06 | 1.1 | 0.87 | 0.23 | 2.08 | 0.99 | 0.61 | 0.06 | 1.69 | -0.42 | -0.11 | -0.49 | 2.4 | 0.81 | 0.98 | 0.2425 | -0.43 | 0.19 | 0.749337 | 0.6017 | 0.4102 | 1 | 0.999631 | 0.912137 |
| A0A0N4SW93_S866s_1 | 1.16 | 0.08 | -0.42 | 0.57 | 1.38 | 0.04 | 0.34 | 0.16 | 0.28 | -0.58 | -0.03 | -0.32 | -0.98 | 0.08 | -0.75 | -0.09 | 0.1325 | -0.51 | -0.7825 | 0.093843 | 0.765208 | 0.220314 | 1 | 0.999631 | 0.558128 |
| A0A0R4J006_S143s_1 | -0.5 | 0.36 | -0.15 | -0.08 | 0.1 | -0.83 | -0.52 | -0.54 | -0.3 | 0.44 | -0.07 | 0.32 | -0.4 | 0.41 | 0.32 | -0.04 | -0.355 | 0.19 | 0.165 | 0.61906 | 0.275986 | 0.546485 | 1 | 0.999631 | 0.861888 |
| A0A0R4J0J1_S543s_1 | -0.32 | -1.2 | -1.31 | -0.44 | -0.64 | -0.83 | -0.37 | -1.01 | -0.62 | -0.71 | -1.19 | -1.65 | -0.81 | -0.32 | -1.49 | -0.86 | 0.105 | -0.225 | -0.0525 | 0.891183 | 0.7511 | 0.551233 | 1 | 0.999631 | 0.96415 |
| A0A0R4J0K0_T176t_1 | 0.13 | -0.19 | -0.43 | 0.13 | 0.01 | -0.09 | -0.09 | 0.28 | 0.13 | -0.6 | -0.22 | -0.25 | -0.71 | -0.3 | 0.05 | 0.59 | 0.1175 | -0.145 | -0.0025 | 0.994429 | 0.656565 | 0.618128 | 1 | 0.999631 | 0.997705 |
| A0A0R4J0M1_S162s_1 | -1.2 | -0.42 | 0.67 | -1.43 | -0.24 | -0.47 | -0.83 | 0.28 | -0.6 | -0.63 | -0.38 | -0.92 | -0.04 | -0.19 | -0.77 | 0.37 | 0.28 | -0.0375 | 0.4375 | 0.395656 | 0.575395 | 0.936384 | 1 | 0.999631 | 0.760249 |
| A0A0R4J1E3_S613s_1 | -0.69 | -0.35 | -0.62 | -0.53 | -0.41 | -0.62 | -0.82 | -0.71 | -0.27 | -0.5 | -0.73 | -0.3 | 0.13 | 0.01 | -0.55 | -0.21 | -0.0925 | 0.0975 | 0.3925 | 0.194317 | 0.706516 | 0.706815 | 1 | 0.999631 | 0.65176 |
| A0A0R4J216_S30s_1 | 0.25 | 0.22 | -0.07 | 0.38 | -0.23 | 0.18 | -0.25 | 0.01 | -0.49 | 1.49 | 0 | -1.04 | -0.16 | -0.04 | -0.11 | -0.64 | -0.2675 | -0.205 | -0.4325 | 0.156375 | 0.306673 | 0.693463 | 1 | 0.999631 | 0.626452 |
| A0A0R4J216_T23t_1 | 1.17 | -0.22 | -1.39 | -0.72 | -0.33 | -0.79 | -0.46 | -2.3 | -1.42 | 0.28 | 0.22 | -2.17 | -0.35 | -1.39 | -1.34 | -1.19 | -0.68 | -0.4825 | -0.7775 | 0.180805 | 0.298073 | 0.510063 | 1 | 0.999631 | 0.64376 |
| A0A0U1RPY2_S57s_1 | 1.7 | 0.71 | 0.79 | 0.44 | -0.95 | 1.08 | 2.24 | -1.05 | -0.28 | 1.57 | -1.73 | -1.04 | -0.36 | -0.62 | -0.66 | -0.96 | -0.58 | -1.28 | -1.56 | 0.001293 | 0.444355 | 0.085056 | 1 | 0.999631 | 0.351308 |
| A0A140LHX7_S67s_1 | -0.29 | 0.45 | -0.27 | -0.6 | -0.29 | 0.27 | -1.02 | -0.59 | -0.01 | 0.28 | -0.61 | -0.13 | 0.87 | 0.41 | -0.58 | 0.22 | -0.23 | 0.06 | 0.4075 | 0.321151 | 0.537613 | 0.858704 | 1 | 0.999631 | 0.72408 |
| A0A140LHX7_S68s_1 | -0.61 | -1.18 | 0.47 | -0.32 | -1.6 | -1.75 | -0.55 | -0.18 | -1.58 | -1.04 | 0.16 | 0.34 | -0.2 | -1.01 | 0 | 0.8 | -0.61 | -0.12 | 0.3075 | 0.533173 | 0.230323 | 0.823697 | 1 | 0.999631 | 0.827798 |
| A0A140LI67_S906s_1 | 1.5 | 0.28 | 0.44 | -0.45 | 0 | -0.36 | -0.13 | -0.24 | 0.35 | 1.4 | -0.4 | -0.18 | -0.88 | -1.05 | 1.05 | -1.26 | -0.625 | -0.15 | -0.9775 | 0.131544 | 0.150697 | 0.777792 | 1 | 0.999631 | 0.603456 |
| A0A140LIW3_S1289s_1 | 0.87 | -0.63 | -0.29 | 0.05 | -0.82 | -0.19 | 0.48 | 0.46 | -1.77 | 0.07 | -0.11 | -0.1 | -1.02 | -0.17 | 0.01 | 0.13 | -0.0175 | -0.4775 | -0.2625 | 0.540301 | 0.967943 | 0.35957 | 1 | 0.999631 | 0.831295 |
| A0A140LIW3_S1290s_1 | 0.87 | -0.63 | -0.25 | 0.33 | -0.82 | -0.19 | 0.82 | 0.57 | -1.77 | 0.07 | 0.15 | -0.23 | -1.02 | -0.17 | -0.47 | 0 | 0.015 | -0.525 | -0.495 | 0.249519 | 0.974691 | 0.327895 | 1 | 0.999631 | 0.686359 |
| A0A140LIW3_S1350s_1 | -0.41 | -0.39 | 0.1 | -0.67 | 0.31 | 0.16 | -0.26 | -1.29 | 0.06 | -0.19 | -2 | -3.07 | 0.52 | -0.2 | -1 | -0.24 | 0.0725 | -0.9575 | 0.1125 | 0.769682 | 0.855984 | 0.182921 | 1 | 0.999631 | 0.918554 |
| A0A140LIW3_S1450s_1 | -0.71 | -0.56 | 0.14 | -0.31 | -0.62 | -0.55 | -0.66 | 0.32 | -0.84 | -0.27 | -0.32 | 0.25 | 0.9 | 0.07 | -0.11 | -0.18 | -0.0175 | 0.065 | 0.53 | 0.163575 | 0.958444 | 0.847268 | 1 | 0.999631 | 0.631937 |
| A0A140LJD2_S105s_1 | -0.43 | -0.12 | -0.17 | -0.5 | -0.06 | 0.11 | -0.74 | -0.24 | 0.12 | 0.17 | -0.29 | -0.02 | -0.03 | 0.64 | -0.34 | 0.34 | 0.0725 | 0.3 | 0.4575 | 0.172639 | 0.798587 | 0.268874 | 1 | 0.999631 | 0.642116 |
| A0A1B0GQW2_S50s_1 | 0.72 | 0.17 | 0.34 | 0 | -0.06 | -0.04 | 0.19 | 0.35 | 0.02 | 1.03 | 0.58 | 0.78 | 0.12 | 0.37 | 0.54 | 0.58 | -0.1975 | 0.295 | 0.095 | 0.749252 | 0.474403 | 0.370725 | 1 | 0.999631 | 0.912137 |
| A0A1B0GR85_S1132s_1 | -0.61 | 0.24 | -0.4 | 0.03 | 0.38 | 0.58 | -0.25 | -0.48 | 0.16 | -0.33 | 0.61 | -0.64 | 0.05 | 0.58 | -0.31 | 0.65 | 0.2425 | 0.135 | 0.4275 | 0.244466 | 0.49186 | 0.713291 | 1 | 0.999631 | 0.685751 |
| A0A1B0GR85_S1194s_1 | 0.08 | -1.02 | -1.27 | -1.1 | -0.37 | 0.01 | -0.47 | -2.68 | -1.68 | -1.67 | -1.48 | -0.57 | -0.22 | -0.67 | -0.66 | -1.63 | -0.05 | -0.5225 | 0.0325 | 0.94025 | 0.935092 | 0.224079 | 1 | 0.999631 | 0.981146 |
| A0A1B0GR85_S1194s_2 | -0.22 | -0.94 | 0.37 | 1.09 | -0.85 | -1.52 | 1.14 | 0.47 | -0.81 | -0.06 | 0.03 | -0.27 | -2.89 | -2.18 | -0.58 | -0.07 | -0.265 | -0.3525 | -1.505 | 0.053793 | 0.690314 | 0.449379 | 1 | 0.999631 | 0.4909 |
| A0A1B0GR85_S1202s_2 | -0.22 | -0.94 | 0.37 | 1.09 | -0.85 | -1.52 | 1.14 | 0.47 | -0.81 | -0.06 | 0.03 | -0.27 | -2.89 | -2.18 | -0.58 | -0.07 | -0.265 | -0.3525 | -1.505 | 0.053793 | 0.690314 | 0.449379 | 1 | 0.999631 | 0.4909 |
| A0A1B0GR85_S1331s_1 | -0.2 | 0.11 | 0.31 | -0.32 | -0.02 | 0.28 | -0.54 | 0.09 | -0.53 | 0.26 | -0.07 | 0.05 | 0.39 | 0.16 | -0.52 | 0.17 | -0.0225 | -0.0475 | 0.075 | 0.81695 | 0.939066 | 0.873474 | 1 | 0.999631 | 0.937777 |
| A0A1B0GR85_S1454s_1 | -0.09 | 0.52 | 0.37 | 0.23 | -0.07 | 0.77 | -0.02 | 0.31 | 0.1 | 1.11 | 0.12 | 0.31 | 0.04 | 1.13 | -0.09 | 0.56 | -0.01 | 0.1525 | 0.1525 | 0.671567 | 0.973203 | 0.642008 | 1 | 0.999631 | 0.884779 |
| A0A1B0GS09_S77s_1 | 0.46 | -0.13 | -0.31 | -0.27 | 0.29 | 0.52 | -0.48 | -0.09 | 0 | 0.23 | -1.22 | 0.39 | 0.54 | 1.25 | -0.32 | -0.39 | 0.1225 | -0.0875 | 0.3325 | 0.450834 | 0.708434 | 0.832342 | 1 | 0.999631 | 0.787797 |
| A0A1D5RLB2_S38s_1 | 0.34 | -0.37 | -0.17 | 0.26 | -0.63 | -0.17 | -0.03 | -0.42 | -0.29 | 0.59 | -0.41 | 0.19 | 0.06 | -0.75 | 0.01 | -0.24 | -0.3275 | 0.005 | -0.245 | 0.461164 | 0.273127 | 0.988083 | 1 | 0.999631 | 0.792226 |
| A0A1D5RLL0_S106s_1 | -0.11 | 0.42 | 0.4 | -0.23 | -0.07 | 0.2 | 0.13 | 0.48 | 0.38 | 0.36 | 0.73 | 0.31 | 1.02 | 0.24 | -0.21 | -0.06 | 0.065 | 0.325 | 0.1275 | 0.728938 | 0.818226 | 0.271414 | 1 | 0.999631 | 0.905923 |
| A0A1W2P772_S197s_1 | -0.41 | -0.79 | -0.73 | -0.66 | -0.62 | -0.43 | -1.45 | -0.63 | -1.69 | -0.58 | -0.29 | -0.51 | -0.39 | 0.08 | -0.28 | -0.82 | -0.135 | -0.12 | 0.295 | 0.343818 | 0.657297 | 0.738504 | 1 | 0.999631 | 0.733852 |
| A0A1W2P772_S207s_1 | 0.7 | 0.09 | 0.3 | 0.77 | 0.24 | 0.15 | 0.49 | 0.52 | -0.23 | -0.01 | 0.39 | -0.77 | -1.35 | 0.07 | -1.08 | 0.32 | -0.115 | -0.62 | -0.975 | 0.050085 | 0.676031 | 0.091486 | 1 | 0.999631 | 0.488436 |
| A0A1W2P772_S210s_1 | -0.49 | -3.81 | 0.17 | 0.91 | -0.91 | -3.55 | 0.52 | 0.95 | -2.67 | -3.17 | -0.45 | 1.2 | -2.97 | -3.34 | 0.54 | 0.37 | 0.0575 | -0.4675 | -0.545 | 0.663414 | 0.962976 | 0.707705 | 1 | 0.999631 | 0.881968 |
| A0A1W2P832_S760s_1 | 0.39 | 0.03 | 0.09 | 0.49 | 0.37 | 0.27 | 0.24 | 0.47 | 0 | 0.15 | -0.18 | 0.78 | -0.51 | 0.98 | 0.65 | 0.4 | 0.0875 | -0.0625 | 0.13 | 0.73087 | 0.725155 | 0.838758 | 1 | 0.999631 | 0.906897 |
| A0A1W2P832_S776s_1 | 1.06 | 2.34 | 1.38 | 1.34 | 1.73 | 1.71 | 0.47 | 1.46 | 1.95 | 1.76 | 1.09 | 1.48 | 2.06 | 1.84 | 1.88 | 1.96 | -0.1875 | 0.04 | 0.405 | 0.256881 | 0.6479 | 0.912899 | 1 | 0.999631 | 0.691567 |
| A0A1Y7VKX4_S609s_1 | 0.54 | 0.25 | 1.69 | 0.79 | -0.1 | 0.45 | 0.81 | 1.02 | 0.47 | 0.54 | 1.14 | 1.01 | -0.01 | -0.13 | 1.09 | 1.09 | -0.2725 | -0.0275 | -0.3075 | 0.503738 | 0.50117 | 0.941807 | 1 | 0.999631 | 0.814226 |
| A0A1Y7VLQ2_S103s_2 | -0.14 | 1.36 | -0.1 | -0.04 | -0.73 | 0.74 | -1.28 | -0.9 | -0.57 | -0.67 | 1.37 | -0.77 | -0.64 | 1.81 | 0.46 | -0.02 | -0.8125 | -0.43 | 0.1325 | 0.820167 | 0.148986 | 0.462235 | 1 | 0.999631 | 0.938303 |
| A0A1Y7VLQ2_S95s_2 | -0.14 | 1.36 | -0.1 | -0.04 | -0.73 | 0.74 | -1.28 | -0.9 | -0.57 | -0.67 | 1.37 | -0.77 | -0.64 | 1.81 | 0.46 | -0.02 | -0.8125 | -0.43 | 0.1325 | 0.820167 | 0.148986 | 0.462235 | 1 | 0.999631 | 0.938303 |
| A0A1Y7VNP1_S418s_2 | 0.09 | 0.27 | 0.22 | 0.31 | 0.21 | -0.12 | 0.6 | -0.38 | 0 | -0.16 | 0.27 | -0.48 | -0.24 | 0.58 | 0.45 | 0.88 | -0.145 | -0.315 | 0.195 | 0.549203 | 0.618609 | 0.263204 | 1 | 0.999631 | 0.83507 |
| A0A1Y7VNP1_S421s_2 | -0.05 | 0.2 | 0.22 | 0.31 | 0.31 | -0.81 | 0.6 | -0.38 | 0.06 | -0.44 | 0.27 | -0.48 | -1.04 | 0.58 | 0.45 | 0.88 | -0.24 | -0.3175 | 0.0475 | 0.913871 | 0.505392 | 0.286702 | 1 | 0.999631 | 0.972551 |
| A0A1Y7VNZ6_S556s_1 | 1.03 | 0.36 | 0.62 | 0.34 | 1.54 | 0.5 | 0.58 | 0.8 | 0.52 | 1.19 | 1.05 | 0.39 | -0.78 | 0.56 | 0.11 | 0.88 | 0.2675 | 0.2 | -0.395 | 0.349499 | 0.423793 | 0.531168 | 1 | 0.999631 | 0.736887 |
| A0A286YCB8_S126s_1 | 0.16 | 0.69 | 0.64 | -0.08 | 0.98 | 0.1 | -0.17 | -0.23 | 1.03 | 0.36 | -0.02 | 0.94 | 0.27 | 1.39 | -1.99 | 0.49 | -0.1825 | 0.225 | -0.3125 | 0.639687 | 0.614509 | 0.525003 | 1 | 0.999631 | 0.870995 |
| A0A286YCB8_S127s_1 | 1.34 | 3.73 | 0.79 | 0.67 | 2.64 | 3.21 | -0.15 | -0.03 | -0.06 | 2.54 | 0.42 | 0.82 | 4.12 | 3.36 | 0.61 | 1.13 | -0.215 | -0.7025 | 0.6725 | 0.485638 | 0.825576 | 0.389649 | 1 | 0.999631 | 0.804287 |
| A0A286YCB8_S131s_1 | -0.97 | 0 | 0.36 | -0.26 | -0.21 | -0.02 | -0.7 | 0.13 | -0.94 | -0.99 | 0.52 | -0.12 | 0.99 | 0.39 | 0.1 | -0.14 | 0.0175 | -0.165 | 0.5525 | 0.185849 | 0.961054 | 0.714149 | 1 | 0.999631 | 0.646224 |
| A0A286YCX6_S971s_1 | -0.1 | -1.26 | -1.42 | -1.29 | -1.03 | -1.75 | -1.16 | -0.93 | -1.17 | -2.15 | -1.63 | -1.23 | 0.22 | -1.21 | -0.55 | -1.25 | -0.2 | -0.5275 | 0.32 | 0.490202 | 0.596059 | 0.203644 | 1 | 0.999631 | 0.807727 |
| A0A286YCX6_T968t_1 | -1 | -0.82 | -0.05 | -0.48 | -0.43 | -0.39 | -1.3 | 0.28 | -0.9 | 0.11 | -1.07 | -1.62 | -0.41 | 0.07 | 0.78 | 0.91 | 0.1275 | -0.2825 | 0.925 | 0.040002 | 0.746106 | 0.506716 | 1 | 0.999631 | 0.472195 |
| A0A286YD83_S49s_1 | -1.15 | -1.34 | -2.79 | -2.16 | -1.58 | -1.76 | -3.16 | -3.15 | -1.55 | -0.47 | -1.87 | -2.14 | -0.56 | -1.72 | -1.86 | -1.98 | -0.5525 | 0.3525 | 0.33 | 0.50155 | 0.311676 | 0.486849 | 1 | 0.999631 | 0.814092 |
| A0A286YDE4_S719s_1 | 1.89 | -0.28 | -0.28 | 0.94 | 0.54 | 0.37 | 1.73 | 0.2 | 0.6 | 0.93 | 0.2 | -0.28 | -1.18 | -0.74 | 0.12 | -0.4 | 0.1425 | -0.205 | -1.1175 | 0.066145 | 0.804067 | 0.707914 | 1 | 0.999631 | 0.516933 |
| A0A286YDL2_S718s_1 | 0.2 | 1.16 | 0.67 | 0.02 | 0.4 | 0.27 | -0.59 | -0.46 | 0.21 | 0.77 | 0.79 | -0.12 | 0.21 | 0.99 | 0.03 | 0.24 | -0.6075 | -0.1 | -0.145 | 0.698634 | 0.129146 | 0.785985 | 1 | 0.999631 | 0.895842 |
| A0A286YDL2_S741s_1 | 1.01 | 0.07 | -0.63 | 0.7 | 0.69 | -0.47 | 0.81 | 0.1 | -0.12 | -0.46 | 1.04 | -0.47 | -0.81 | -0.02 | -1 | 0.28 | -0.005 | -0.29 | -0.675 | 0.170543 | 0.991155 | 0.554907 | 1 | 0.999631 | 0.64073 |
| A0A286YDL2_S741s_2 | 1.36 | 0.74 | -0.29 | 1.69 | 1.41 | 0.47 | 1.07 | 1.09 | 0.68 | -0.06 | 1.12 | 0.2 | -1.49 | 1.31 | 0.66 | 0.9 | 0.135 | -0.39 | -0.53 | 0.442541 | 0.768826 | 0.430014 | 1 | 0.999631 | 0.782367 |
| A0A286YDL2_S745s_1 | 0.14 | 0.61 | 0.14 | 0.54 | -0.27 | 0.4 | -0.21 | 0.38 | 0.26 | 0.69 | 0.69 | 0.47 | -0.28 | 0.8 | 1.22 | 0.89 | -0.2825 | 0.17 | 0.3 | 0.44103 | 0.345114 | 0.535266 | 1 | 0.999631 | 0.781983 |
| A0A286YDL2_S745s_2 | 1.36 | 0.74 | -1.14 | 0.26 | 1.41 | 0.47 | -0.03 | -0.15 | 0.68 | -0.06 | 0.12 | -1.19 | -1.49 | 1.31 | -0.38 | -0.36 | 0.12 | -0.4175 | -0.535 | 0.449298 | 0.836321 | 0.493241 | 1 | 0.999631 | 0.786904 |
| A0A286YDL2_S750s_1 | -0.85 | -3.01 | -1.83 | -2.53 | -2.23 | -2.83 | -1.55 | -2.19 | -1.61 | -1.04 | -1.25 | -1.91 | -1.74 | -2.11 | -2.7 | -4.08 | -0.145 | 0.6025 | -0.6025 | 0.349073 | 0.773342 | 0.233258 | 1 | 0.999631 | 0.736748 |
| A0A2C9F2C1_S114s_1 | -0.71 | -1.08 | -0.49 | 0.39 | 0.39 | -0.54 | 0.43 | -0.56 | -0.5 | -0.74 | -0.01 | 0.96 | -0.82 | 0.25 | -1.11 | 0.12 | 0.4025 | 0.4 | 0.0825 | 0.856542 | 0.344598 | 0.405655 | 1 | 0.999631 | 0.951817 |
| A0A2I3BPI6_T9t_1 | -0.85 | -1.15 | 0.42 | 0.15 | -0.74 | -0.82 | 0.51 | 0.28 | 0.41 | -1.19 | 0.29 | 0.36 | 0.21 | -0.72 | 0.22 | -0.15 | 0.165 | 0.325 | 0.2475 | 0.578519 | 0.733718 | 0.529315 | 1 | 0.999631 | 0.84622 |
| A0A2I3BQ39_S187s_1 | 0.63 | -1.01 | 0.19 | 1.23 | 1.34 | 0.33 | 0.9 | 1.11 | 0.57 | 0.21 | -0.15 | 0.5 | -1.05 | -0.76 | -0.07 | 0.01 | 0.66 | 0.0225 | -0.7275 | 0.177962 | 0.19958 | 0.962664 | 1 | 0.999631 | 0.642395 |
| A0A2I3BQP6_S426s_1 | 0.18 | 0.22 | -0.07 | 0.49 | 0.41 | -0.2 | 0.09 | 0.2 | -0.05 | -0.19 | 0.05 | 0.02 | -0.15 | -0.03 | -0.42 | 0.09 | -0.08 | -0.2475 | -0.3325 | 0.260614 | 0.764903 | 0.34818 | 1 | 0.999631 | 0.692669 |
| A0A2I3BRN2_S374s_1 | 2.27 | -1.58 | -0.1 | 0.1 | -1.04 | -0.14 | 1.83 | -0.71 | -1.06 | 1.43 | -0.84 | -1.18 | -0.62 | -1.05 | -0.14 | -1.74 | -0.1875 | -0.585 | -1.06 | 0.18281 | 0.832181 | 0.509259 | 1 | 0.999631 | 0.643861 |
| A0A2R8VHB6_S125s_1 | 0.41 | 0.39 | -0.48 | -0.51 | 0.69 | 0.07 | -0.67 | -0.68 | 0.59 | 0.17 | -0.46 | 0.3 | 0.33 | 0.5 | 0.19 | -0.18 | -0.1 | 0.1975 | 0.2575 | 0.470527 | 0.810329 | 0.594882 | 1 | 0.999631 | 0.797463 |
| A0A2R8VHB6_S130s_1 | -0.13 | 0.78 | 0.6 | 0.95 | 1.62 | 1.15 | 0.49 | 1.23 | -0.16 | 0.86 | 0.99 | 1.2 | 1.26 | 0.22 | 1.24 | 1.04 | 0.5725 | 0.1725 | 0.39 | 0.316285 | 0.134389 | 0.666083 | 1 | 0.999631 | 0.720734 |
| A0A2R8VHB6_S148s_1 | 1.58 | -0.63 | 0.4 | -0.41 | -1.64 | 0.26 | 2.66 | -1.84 | -1.14 | 1.96 | -0.99 | -0.9 | -0.62 | -1.51 | -1.63 | -1.41 | -0.375 | -0.5025 | -1.5275 | 0.013988 | 0.707886 | 0.528108 | 1 | 0.999631 | 0.385092 |
| A0A2R8VHB6_S148s_2 | -1.9 | 0.47 | -1.1 | -1.69 | -0.76 | 0.12 | -0.93 | -1.46 | -0.53 | -0.16 | -1.52 | -0.92 | 1.44 | -0.11 | -1.89 | -1.82 | 0.2975 | 0.2725 | 0.46 | 0.582827 | 0.605651 | 0.629926 | 1 | 0.999631 | 0.848004 |
| A0A2R8W6P0_S335s_1 | -0.2 | -0.44 | -0.63 | -0.33 | -0.27 | -0.6 | -0.67 | -0.39 | -0.7 | -0.52 | -0.68 | -0.71 | 0.36 | -0.5 | -0.64 | -0.27 | -0.0825 | -0.2525 | 0.1375 | 0.669957 | 0.742284 | 0.32476 | 1 | 0.999631 | 0.884054 |
| A0A3B2WBC6_S10s_1 | -0.55 | 0.22 | -0.4 | -0.08 | 0.18 | -0.56 | -0.76 | -0.69 | -0.3 | 0.26 | -0.38 | -0.26 | 0.95 | 0.67 | -0.14 | -0.03 | -0.255 | 0.0325 | 0.565 | 0.143291 | 0.436811 | 0.91388 | 1 | 0.999631 | 0.614578 |
| A0A3B2WCD2_S1010s_1 | 0.81 | 0.03 | -0.28 | 0.7 | 0.24 | 0.51 | 1.46 | -0.35 | -0.99 | 1.41 | 0.02 | -0.88 | -0.77 | -0.75 | -0.79 | -1.31 | 0.15 | -0.425 | -1.22 | 0.005678 | 0.73696 | 0.459196 | 1 | 0.999631 | 0.351545 |
| A0A3B2WCD2_S1019s_1 | -0.52 | 0.64 | -0.14 | -0.29 | 0.09 | 0.46 | 0.42 | 0.07 | 0.83 | 0.25 | 0.79 | 0.67 | 1.13 | 0.53 | 0.59 | 0.35 | 0.3375 | 0.7125 | 0.7275 | 0.063606 | 0.306171 | 0.055541 | 1 | 0.999631 | 0.513165 |
| A0A3B2WCD2_S1217s_1 | 1.47 | -0.74 | 0.14 | -0.18 | -1.99 | 0.2 | 1.24 | -1.72 | -1.66 | 1.58 | -0.21 | -1.52 | -0.89 | -1.02 | -1.07 | -1.77 | -0.74 | -0.625 | -1.36 | 0.018559 | 0.361234 | 0.430865 | 1 | 0.999631 | 0.391024 |
| A0A3B2WCD2_S1301s_1 | 0.8 | 0.87 | 1.72 | 1.59 | 0.07 | 0.6 | 1.5 | 1.3 | 0.85 | 0.61 | 0.36 | 2.32 | 1.31 | 0.8 | 2.03 | 0.99 | -0.3775 | -0.21 | 0.0375 | 0.923458 | 0.364916 | 0.663393 | 1 | 0.999631 | 0.975032 |
| A0A3B2WCD2_S1379s_1 | 0.09 | 0.01 | -1.19 | 0.77 | 0.7 | -0.82 | 0.77 | -0.15 | -0.22 | -1.31 | 0.71 | -0.56 | 0.02 | -0.32 | -0.93 | 0.25 | 0.205 | -0.265 | -0.165 | 0.726849 | 0.692696 | 0.627604 | 1 | 0.999631 | 0.905774 |
| A0A3B2WCD2_S877s_1 | -1.07 | 1.07 | 0.65 | -0.35 | 1.07 | 0.35 | 0.54 | 0.42 | 0.91 | -0.35 | 1.38 | 1.63 | 1.83 | 0.22 | 1.53 | 0.42 | 0.52 | 0.8175 | 0.925 | 0.132721 | 0.29673 | 0.19217 | 1 | 0.999631 | 0.604583 |
| A0A3B2WCD2_S898s_1 | -0.62 | -0.52 | -0.01 | -0.28 | -0.25 | 0.02 | -0.59 | -0.99 | -1.02 | -0.53 | -0.17 | 0.22 | 0.09 | -0.23 | -1.1 | 0.49 | -0.095 | -0.0175 | 0.17 | 0.666528 | 0.760749 | 0.9591 | 1 | 0.999631 | 0.88264 |
| A0A3B2WCD2_S951s_1 | -1.47 | 0.46 | -0.05 | -0.01 | 0.23 | 0.13 | 0.19 | -0.03 | 0.47 | -0.33 | 0.91 | 0.53 | 1 | 0.67 | 0.93 | 0.7 | 0.3975 | 0.6625 | 1.0925 | 0.028116 | 0.353096 | 0.184217 | 1 | 0.999631 | 0.423977 |
| A0A3Q4EBK4_S301s_1 | -0.54 | 0.23 | 1.42 | -0.17 | 0.31 | -0.13 | 0.01 | 0.21 | -0.64 | -1.41 | 0.18 | -0.35 | -1.01 | 0.26 | -0.55 | 1.3 | -0.135 | -0.79 | -0.235 | 0.697371 | 0.753361 | 0.144719 | 1 | 0.999631 | 0.895731 |
| A0A494B917_S42s_1 | -0.2 | -0.46 | -0.12 | -0.75 | 0.97 | -1.1 | -0.27 | 0.84 | 0.09 | -0.56 | 0.87 | 1.01 | -1.32 | 0.13 | 0.33 | 0.42 | 0.4925 | 0.735 | 0.2725 | 0.536924 | 0.321078 | 0.09325 | 1 | 0.999631 | 0.830094 |
| A0A494BAF6_S191s_1 | 0.87 | 1.5 | 1.3 | 0.77 | 0.02 | 0.49 | 0.53 | 0.72 | 0.82 | 0.51 | 1.04 | 1.3 | 1.3 | -0.06 | 0.17 | 0.14 | -0.67 | -0.1925 | -0.7225 | 0.085468 | 0.044182 | 0.537878 | 1 | 0.999631 | 0.551124 |
| A0A498WGM0_S167s_1 | 0.49 | -0.32 | -0.38 | -0.56 | -0.24 | 0.34 | 0.02 | 0.04 | -1.68 | -0.16 | -0.76 | -0.9 | -0.51 | -0.17 | -0.32 | -0.73 | 0.2325 | -0.6825 | -0.24 | 0.477636 | 0.465408 | 0.114194 | 1 | 0.999631 | 0.800237 |
| A0A571BDA2_S895s_1 | 0.35 | 0.37 | -0.52 | -0.51 | -0.1 | 0.48 | -0.8 | -0.17 | -0.25 | -0.27 | -0.1 | -0.48 | 1.26 | 0.35 | 0.21 | 0.45 | -0.07 | -0.1975 | 0.645 | 0.114761 | 0.853172 | 0.543991 | 1 | 0.999631 | 0.579186 |
| A0A571BDA2_S971s_1 | -1.42 | -1.18 | -0.14 | -0.88 | -1.39 | -1.14 | -0.63 | -0.95 | -1.55 | -1.45 | -1.21 | -1.02 | 0.16 | -0.83 | -0.91 | 0.02 | -0.1225 | -0.4025 | 0.515 | 0.229408 | 0.727065 | 0.260248 | 1 | 0.999631 | 0.679851 |
| A0A571BEK8_S23s_1 | 2.24 | -0.95 | 1.01 | 2.3 | 1.42 | 0.97 | 2.27 | 1.42 | -0.95 | 0.34 | 1.73 | -0.98 | -2.52 | -0.59 | -0.53 | 0.19 | 0.37 | -1.115 | -2.0125 | 0.033624 | 0.606246 | 0.218925 | 1 | 0.999631 | 0.443263 |
| A0A571BGD8_S21s_1 | -0.24 | -0.44 | -0.33 | -0.86 | 1.9 | 0.15 | -0.25 | -0.73 | 0.09 | 0.92 | 0.62 | 1.23 | -0.37 | -0.16 | -0.42 | -0.04 | 0.735 | 1.1825 | 0.22 | 0.450924 | 0.196479 | 0.004988 | 1 | 0.999631 | 0.787797 |
| A0A5F8MPK1_S6s_1 | 1.52 | -0.21 | -0.71 | 0.2 | 0.43 | 0.97 | 0.55 | 0.79 | -0.33 | 0.64 | 0.53 | -0.22 | -0.44 | 0.61 | 0.31 | 0.35 | 0.485 | -0.045 | 0.0075 | 0.988126 | 0.315689 | 0.929519 | 1 | 0.999631 | 0.995409 |
| A0A5F8MQ17_S313s_1 | -0.12 | 0.7 | 0.54 | 0.29 | 0.38 | 0.21 | -0.38 | -0.38 | 0.34 | 0.35 | -0.07 | 0.62 | 0.89 | 0.8 | 0.49 | 0.85 | -0.395 | -0.0425 | 0.405 | 0.2013 | 0.231383 | 0.888232 | 1 | 0.999631 | 0.658952 |
| A0A5F8MQ82_S377s_1 | -0.09 | -0.22 | -0.67 | 0.46 | -0.15 | -0.5 | -0.66 | 0.31 | 0.19 | -0.06 | -0.54 | -0.38 | -1.03 | -0.02 | 0.37 | 0.22 | -0.12 | -0.0675 | 0.015 | 0.970821 | 0.730478 | 0.839814 | 1 | 0.999631 | 0.989618 |
| A0A5H1ZRL1_S6s_1 | 0.18 | 1.78 | 0.91 | -2.3 | 1.99 | 2.11 | -0.26 | 0.68 | 1.45 | 0.56 | 0.73 | 1.03 | 2.47 | 2.4 | -0.15 | -0.36 | 0.9875 | 0.8 | 0.9475 | 0.355712 | 0.28955 | 0.32445 | 1 | 0.999631 | 0.74034 |
| A0A6I8MWZ7_S543s_1 | 0.54 | -0.11 | 0.09 | 0.01 | 0.27 | 0.25 | 0.1 | 0.86 | -0.18 | 0.36 | 0.76 | 0.21 | 0.53 | 0.21 | 0.84 | 0.96 | 0.2375 | 0.155 | 0.5025 | 0.130787 | 0.422212 | 0.618176 | 1 | 0.999631 | 0.602177 |
| A0A7N9VR94_S286s_1 | 0.94 | 0.98 | -0.31 | -1.03 | -0.01 | 0.35 | -0.11 | -0.44 | 1.08 | -0.24 | 0.76 | 0.41 | 1.97 | -0.19 | 0.2 | -1.12 | -0.1975 | 0.3575 | 0.07 | 0.922355 | 0.687622 | 0.505858 | 1 | 0.999631 | 0.975032 |
| A0A7N9VR94_S3383s_1 | 0.55 | 0.97 | -0.88 | 0.19 | 0.71 | -0.28 | 0.37 | 0.28 | 0.28 | 1.3 | 0.35 | -0.5 | 0.17 | 0.51 | 0.26 | -0.16 | 0.0625 | 0.15 | -0.0125 | 0.976712 | 0.885901 | 0.769169 | 1 | 0.999631 | 0.992016 |
| A0A7N9VR94_S55s_1 | -0.92 | -0.73 | -0.61 | -0.56 | -0.5 | 0.62 | -0.71 | -0.27 | 0.31 | 0.64 | -0.51 | -0.53 | 0.39 | 0.74 | 0.44 | 0.22 | 0.49 | 0.6825 | 1.1525 | 0.001885 | 0.17078 | 0.073296 | 1 | 0.999631 | 0.351308 |
| A230065H16Rik_G3UW55_S11s_1 | -0.26 | 0.3 | 0.35 | -0.05 | 0 | 0.5 | -0.39 | -0.09 | 0.06 | 1.07 | 0.79 | -0.03 | 0.53 | -0.53 | -0.09 | 0.22 | -0.08 | 0.3875 | -0.0525 | 0.87612 | 0.789319 | 0.281152 | 1 | 0.999631 | 0.961046 |
| A2AG58-2_A2AG58_S205s_1 | -0.31 | 0.05 | -0.83 | -0.63 | 0.06 | -0.28 | -0.78 | -1.73 | -0.17 | 0.43 | -0.34 | -0.46 | -0.22 | -0.26 | 0.31 | -0.09 | -0.2525 | 0.295 | 0.365 | 0.267064 | 0.558179 | 0.380011 | 1 | 0.999631 | 0.695712 |
| A830010M20Rik_Q3TEG0_T427t_1 | 0.01 | -0.23 | -0.06 | 0.17 | -0.73 | -0.23 | -0.26 | -0.11 | -0.57 | 0.39 | 0.25 | 0.27 | -0.19 | -0.58 | -0.19 | 0.65 | -0.305 | 0.1125 | -0.05 | 0.882596 | 0.261418 | 0.714226 | 1 | 0.999631 | 0.963505 |
| Aagab_Q8R2R3_S211s_1 | -0.26 | -0.77 | -0.4 | 0.19 | -1.86 | -0.88 | -0.13 | -1.13 | -2.4 | 0.38 | -0.67 | 0.5 | -1.46 | -1.08 | -0.09 | 0.1 | -0.69 | -0.2375 | -0.3225 | 0.463446 | 0.116222 | 0.707963 | 1 | 0.999631 | 0.793615 |
| Aak1_Q3UHJ0_S18s_1 | -0.29 | 0.4 | -0.52 | -1.03 | 0.13 | -0.59 | -0.73 | -0.88 | -0.44 | -0.5 | -0.95 | -0.32 | 0.43 | 0 | -0.73 | -1.15 | -0.1575 | -0.1925 | -0.0025 | 0.995643 | 0.682502 | 0.595782 | 1 | 0.999631 | 0.997705 |
| Aak1_Q3UHJ0_S21s_1 | 0.11 | 0.39 | -0.47 | -0.88 | -0.2 | 0.33 | -0.68 | -0.29 | 0.06 | -0.67 | -0.31 | -0.5 | 1.15 | 0.28 | -0.98 | -0.83 | 0.0025 | -0.1425 | 0.1175 | 0.827669 | 0.994622 | 0.692772 | 1 | 0.999631 | 0.941202 |
| Aak1_Q3UHJ0_S621s_2 | 0.13 | -0.15 | -0.31 | -0.09 | 0.48 | 0.06 | -0.22 | -0.05 | -0.36 | 0.27 | -0.56 | -0.52 | -0.16 | -0.12 | -0.43 | 0.18 | 0.1725 | -0.1875 | -0.0275 | 0.922834 | 0.525287 | 0.528679 | 1 | 0.999631 | 0.975032 |
| Aak1_Q3UHJ0_S622s_1 | -0.16 | 0.11 | -0.36 | -0.07 | -0.89 | -1.82 | -0.88 | -0.61 | -0.24 | 0.2 | -0.56 | -0.34 | 0.78 | -0.11 | -0.35 | -0.67 | -0.93 | -0.115 | 0.0325 | 0.92985 | 0.016427 | 0.684621 | 1 | 0.999631 | 0.97754 |
| Aak1_Q3UHJ0_S622s_2 | 0.12 | 0.16 | -0.04 | -0.15 | 0.19 | 0.07 | -0.21 | 0.07 | -0.05 | -0.08 | -0.42 | -0.07 | -0.17 | 0.39 | 0.05 | 0.2 | 0.0075 | -0.1775 | 0.095 | 0.73357 | 0.975462 | 0.489149 | 1 | 0.999631 | 0.907462 |
| Aak1_Q3UHJ0_S635s_1 | 1.3 | -0.87 | -1.26 | 0.9 | 1.66 | 0.69 | 0.87 | 0.88 | -0.24 | -0.39 | -0.39 | -0.53 | -1.92 | -0.14 | -0.47 | -0.74 | 1.0075 | -0.405 | -0.835 | 0.228017 | 0.120851 | 0.493617 | 1 | 0.999631 | 0.679038 |
| Aak1_Q3UHJ0_S650s_1 | 1.69 | 0.02 | -0.43 | 1.83 | 2.4 | 1.43 | 1.86 | 2.07 | 1.17 | -0.01 | 1.59 | 0.66 | -1.42 | 1.26 | 0.93 | 0.43 | 1.1625 | 0.075 | -0.4775 | 0.51839 | 0.060642 | 0.901997 | 1 | 0.999631 | 0.819449 |
| Aak1_Q3UHJ0_S668s_1 | 2.19 | 0.7 | 0.78 | 1.27 | 0.97 | 0.99 | 1.42 | 1.18 | 0.72 | 0.89 | 0.2 | -0.79 | -1.56 | 0.24 | 0.42 | -0.58 | -0.095 | -0.98 | -1.605 | 0.012321 | 0.799841 | 0.068412 | 1 | 0.999631 | 0.385092 |
| Aak1_Q3UHJ0_S668s_2 | 3.14 | 0.52 | 0.64 | 2.24 | 1.83 | 1.15 | 2.57 | 1.28 | 1.32 | 1.87 | 0.27 | -0.14 | -1.85 | 0.35 | 0.69 | -0.42 | 0.0725 | -0.805 | -1.9425 | 0.024914 | 0.909613 | 0.267569 | 1 | 0.999631 | 0.418003 |
| Aak1_Q3UHJ0_S676s_1 | 0.98 | 0.1 | 0.56 | 0.55 | 0.07 | 0.53 | 0.98 | -0.04 | 0.62 | 0.67 | -0.51 | -0.71 | -0.78 | -0.66 | -0.15 | -0.69 | -0.1625 | -0.53 | -1.1175 | 0.005124 | 0.62834 | 0.219236 | 1 | 0.999631 | 0.351308 |
| Aak1_Q3UHJ0_S676s_2 | 3.33 | -0.71 | -0.06 | 1.42 | 1.56 | 0.53 | 2.1 | 0.57 | 0.56 | 0.72 | -0.3 | -0.62 | -2.29 | -0.63 | -0.22 | -0.63 | 0.195 | -0.905 | -1.9375 | 0.047176 | 0.818338 | 0.293287 | 1 | 0.999631 | 0.480102 |
| Aak1_Q3UHJ0_S680s_1 | -0.2 | -0.46 | -1.2 | -0.86 | 0.39 | -0.12 | -0.28 | -0.64 | -0.42 | -1.42 | -0.58 | -0.86 | -1 | -0.18 | -1.32 | -0.6 | 0.5175 | -0.14 | -0.095 | 0.798781 | 0.152817 | 0.690362 | 1 | 0.999631 | 0.93091 |

| | | | | | | | | | | | | | | | | | | | | | | | | |
|---|---|---|---|---|---|---|---|---|---|---|---|---|---|---|---|---|---|---|---|---|---|---|---|---|
| Aak1_Q3UHJ0_S729s_1 | -0.05 | -0.06 | -0.2 | 0.2 | 0.52 | -0.02 | 0 | 0.29 | -0.1 | -0.85 | 0.01 | -0.16 | -0.38 | 0.11 | -0.07 | 0.17 | 0.225 | -0.2475 | -0.015 | 0.957552 | 0.395225 | 0.408718 | 1 | 0.999631 | 0.98529 |
| Aak1_Q3UHJ0_S795s_1 | 1.71 | 0.47 | -0.88 | 1.97 | 2.22 | 1.29 | 1.82 | 1.65 | 0.89 | -0.41 | 0.79 | 0.1 | -1.77 | 1.4 | 0.21 | 1.16 | 0.9275 | -0.475 | -0.5675 | 0.505995 | 0.154156 | 0.470756 | 1 | 0.999631 | 0.814836 |
| Aak1_Q3UHJ0_S848s_1 | 0.72 | -0.48 | -1.17 | 0.92 | 1.16 | -0.96 | 0.35 | -0.54 | -0.12 | -1.58 | -0.26 | -1.23 | 0.1 | -0.44 | -1.15 | -0.93 | 0.005 | -0.795 | -0.6025 | 0.275102 | 0.993528 | 0.181523 | 1 | 0.999631 | 0.700187 |
| Aak1_Q3UHJ0_S936s_1 | -1.23 | -0.6 | -0.43 | -0.53 | -0.31 | -1.01 | -0.38 | -0.6 | -0.33 | -1.53 | -0.02 | 0 | 0.14 | -1.04 | -0.73 | -0.52 | 0.1225 | 0.2275 | 0.16 | 0.657332 | 0.685816 | 0.583153 | 1 | 0.999631 | 0.879224 |
| Aak1_Q3UHJ0_T604t_1 | -0.07 | 0.19 | 0.03 | -0.18 | -0.05 | 0.11 | -0.58 | 0.17 | 0.22 | 0.13 | 0.08 | -0.09 | 0.57 | 0.51 | 0.15 | 0.25 | -0.08 | 0.0925 | 0.3775 | 0.190555 | 0.771272 | 0.711916 | 1 | 0.999631 | 0.648674 |
| Aak1_Q3UHJ0_T618t_1 | -1.23 | -0.69 | 0.18 | -1.46 | -1.91 | -1.38 | -1.71 | -1.55 | -1.43 | -0.58 | -0.98 | -0.59 | 0.86 | -0.61 | -0.89 | -0.57 | -0.8375 | -0.095 | 0.4975 | 0.341613 | 0.054311 | 0.82105 | 1 | 0.999631 | 0.73317 |
| Aak1_Q3UHJ0_T618t_2 | 0.12 | 0.09 | -0.12 | -0.13 | 0.26 | 0.07 | -0.22 | 0.03 | -0.12 | 0 | -0.46 | -0.2 | -0.17 | 0.28 | -0.08 | 0.19 | 0.045 | -0.185 | 0.065 | 0.812902 | 0.85523 | 0.47365 | 1 | 0.999631 | 0.937041 |
| Aak1_Q3UHJ0_T672t_2 | 3.47 | -2.08 | -0.28 | 0.51 | 1.45 | 0.15 | 1.77 | -0.22 | -0.43 | -0.82 | -0.33 | -1.36 | -2.02 | -1.48 | -1.12 | -0.53 | 0.3825 | -1.14 | -1.6925 | 0.122071 | 0.722692 | 0.278741 | 1 | 0.999631 | 0.590723 |
| Aak1_Q3UHJ0_T679t_1 | -0.2 | -0.46 | -1.2 | -0.86 | 0.39 | -0.12 | -0.28 | -0.64 | -0.42 | -1.42 | -0.58 | -0.86 | -1 | -0.18 | -1.32 | -0.6 | 0.5175 | -0.14 | -0.095 | 0.798781 | 0.152817 | 0.690362 | 1 | 0.999631 | 0.93091 |
| Aatk_B1AZF9_S662s_1 | -1.96 | 4.09 | 3.49 | 0.47 | 1.88 | 3.44 | 1.12 | 1.89 | 3.39 | 0.4 | 2.69 | 3.86 | 4.34 | 3.29 | 2.69 | 2.34 | 0.56 | 1.0625 | 1.6425 | 0.206608 | 0.659941 | 0.443588 | 1 | 0.999631 | 0.6605 |
| Aatk_B1AZF9_S670s_1 | 0 | -0.18 | -0.54 | 0.61 | 0.32 | -0.42 | 0.32 | -0.93 | -0.27 | 0.31 | -1.1 | -1.6 | 0.11 | 0.78 | 0.08 | 0.79 | -0.15 | -0.6375 | 0.4675 | 0.214886 | 0.70515 | 0.197556 | 1 | 0.999631 | 0.667764 |
| Aatk_B1AZF9_S987s_1 | 0.25 | -0.5 | -0.38 | -0.29 | -0.29 | -0.47 | -0.13 | 0.04 | -0.41 | -0.11 | -0.31 | -0.71 | -0.29 | 0.36 | -0.47 | -0.44 | 0.0175 | -0.155 | 0.02 | 0.951565 | 0.950105 | 0.597925 | 1 | 0.999631 | 0.984079 |
| Abca2_P41234_S2381s_1 | -2.29 | -0.71 | 0.55 | 0.09 | -0.95 | -0.82 | 0.02 | 0.7 | 0.43 | -0.97 | -0.07 | 1.07 | 0.82 | -0.12 | 0.31 | 1.14 | 0.3275 | 0.705 | 1.1275 | 0.092634 | 0.618916 | 0.312089 | 1 | 0.999631 | 0.557279 |
| Abcb1a_P21447_S667s_1 | 0.88 | 1.27 | 1.2 | 1.07 | 1.24 | 1.53 | 0.86 | 0.4 | 2.14 | 2.39 | 0.63 | 1.5 | 0.79 | 0.68 | 1.07 | 0.71 | -0.0975 | 0.56 | -0.2925 | 0.299252 | 0.755517 | 0.193152 | 1 | 0.999631 | 0.712604 |
| Abcc4_E9Q236_T561t_1 | -0.34 | 0.35 | -0.45 | -0.36 | -0.31 | -0.45 | -0.75 | -1.19 | -0.32 | 0.23 | -1.18 | -0.27 | 0.73 | -0.4 | -0.4 | -0.32 | -0.475 | -0.185 | 0.1025 | 0.783918 | 0.158284 | 0.621524 | 1 | 0.999631 | 0.922628 |
| Abcd2_Q61285_S58s_1 | -0.26 | -1.56 | -0.54 | -0.62 | -0.94 | -0.54 | -0.53 | -1.06 | -0.28 | 0.2 | -0.57 | -0.49 | -0.35 | -1.49 | -0.73 | -1.13 | -0.0225 | 0.46 | -0.18 | 0.653869 | 0.948042 | 0.22412 | 1 | 0.999631 | 0.878409 |
| Abcd3_P55096_S424s_1 | -1.09 | -1.07 | -0.88 | -1.61 | -0.75 | -1.41 | -1.31 | -1.64 | 0.02 | -0.36 | -0.81 | -0.57 | -0.19 | -0.43 | -0.23 | -0.77 | -0.115 | 0.7325 | 0.7575 | 0.030097 | 0.706716 | 0.035854 | 1 | 0.999631 | 0.436111 |
| Abcf1_Q6P542_S194s_1 | -1.44 | 0.43 | 0.31 | 0.57 | 0.34 | 0.2 | -0.3 | 0.29 | 0.35 | 0.26 | 0.46 | -0.58 | 0.47 | 0.16 | 0.41 | -1.29 | 0.165 | 0.155 | -0.03 | 0.958542 | 0.726834 | 0.757537 | 1 | 0.999631 | 0.98564 |
| Abhd17b_Q7M759_S282s_1 | -0.71 | -0.28 | 0.44 | 0 | -0.26 | -0.83 | -0.76 | 0.04 | -0.52 | -0.22 | -0.48 | 0.82 | 0.66 | -0.14 | -0.11 | 1.08 | -0.315 | 0.0375 | 0.51 | 0.227625 | 0.376562 | 0.926318 | 1 | 0.999631 | 0.678949 |
| Abi1_B7ZCU4_S183s_1 | -0.16 | -0.22 | -0.02 | -0.54 | -0.58 | -0.55 | -0.74 | -0.42 | -0.27 | 0.1 | -0.33 | -0.16 | 0.47 | -0.03 | -0.15 | -0.06 | -0.3375 | 0.07 | 0.2925 | 0.330867 | 0.197475 | 0.791337 | 1 | 0.999631 | 0.727302 |
| Abi1_B7ZCU4_S222s_1 | 2.98 | 0.19 | -0.76 | 1.37 | 2.53 | 2.67 | 2.27 | 1.69 | 1.9 | 0.47 | 1.18 | 0.1 | -1.2 | 0.25 | 1.15 | -0.56 | 1.345 | -0.0325 | -1.035 | 0.229835 | 0.092114 | 0.967161 | 1 | 0.999631 | 0.680524 |
| Abi1_B7ZCU4_S225s_1 | 2.1 | -1.51 | -0.24 | 1.61 | 0.33 | -0.33 | 1.25 | 0.52 | -0.51 | 0.32 | 0.14 | -0.56 | -1.43 | -1.3 | -0.33 | -0.92 | -0.0475 | -0.6425 | -1.485 | 0.075667 | 0.951663 | 0.408621 | 1 | 0.999631 | 0.541044 |
| Abi1_B7ZCU4_S231s_1 | 3.1 | 0.8 | 0.39 | 1.52 | 2.52 | 2.77 | 2.23 | 1.72 | 1.94 | 1.3 | 1.17 | 0.67 | -1.17 | 1.22 | 1.25 | 0.88 | 0.8575 | -0.1825 | -0.9075 | 0.233357 | 0.162197 | 0.759276 | 1 | 0.999631 | 0.683077 |
| Abi2_A0A087WRS3_S191s_1 | 0.11 | 1.48 | -3.03 | -1.91 | -1.03 | -1.89 | -0.32 | -3.2 | -0.98 | 0.6 | -3.69 | -3.6 | -1.05 | -1.31 | -3.94 | -4.95 | -0.7725 | -1.08 | -1.975 | 0.119269 | 0.45479 | 0.394943 | 1 | 0.999631 | 0.586867 |
| Abi2_A0A087WRS3_S195s_1 | 1.02 | -2.21 | -2.33 | -1.05 | -3.49 | -1.64 | 0.07 | -2.22 | -2.83 | 1.73 | -2.47 | -2.89 | -2.46 | -2.08 | -2.15 | -1.77 | -0.6775 | -0.4725 | -0.9725 | 0.185871 | 0.470736 | 0.686728 | 1 | 0.999631 | 0.646224 |
| Abi2_Q6AXH6_S183s_1 | -1.01 | -0.84 | -0.47 | -0.78 | -1.04 | -1.13 | -1.05 | -0.81 | -1.05 | -0.42 | -0.91 | -0.36 | -0.18 | -0.9 | -0.53 | -0.34 | -0.2325 | 0.09 | 0.2875 | 0.34779 | 0.365563 | 0.757823 | 1 | 0.999631 | 0.736498 |
| Abi2_Q6AXH6_S216s_1 | 1.58 | -1.41 | -1.5 | 0.82 | 1.53 | 0.2 | 1.33 | 1.31 | -0.37 | -0.66 | -0.17 | -1.69 | -2.79 | -0.39 | -1.11 | -0.75 | 1.22 | -0.595 | -1.1325 | 0.189115 | 0.122816 | 0.436749 | 1 | 0.999631 | 0.648556 |
| Abi2_Q6AXH6_S227s_1 | 0.85 | 0.65 | 0.64 | 0.44 | 0.5 | 0.51 | 0.51 | 1.03 | 0.86 | 0.84 | 0.53 | 0.2 | 0.12 | 0.73 | 0.35 | 0.74 | -0.0075 | -0.0375 | -0.16 | 0.585656 | 0.976991 | 0.892353 | 1 | 0.999631 | 0.849524 |
| Abl1_P00520_S709s_1 | 0.68 | 1.29 | 0.09 | 1.4 | 0.49 | 0.87 | 1.15 | 1.7 | 1.24 | 1.55 | 0.64 | 0.67 | -0.45 | 0.06 | 1.46 | 0.1 | 0.1875 | 0.16 | -0.5725 | 0.259838 | 0.641108 | 0.684481 | 1 | 0.999631 | 0.692624 |
| Abl1_P00520_S713s_1 | 2.88 | -0.67 | 0.88 | 1.46 | 0.4 | 0.87 | 2.04 | 1.22 | 0.43 | 1.5 | 1.18 | 0.04 | -1.9 | 0.06 | 0.47 | -0.21 | -0.005 | -0.35 | -1.5325 | 0.075152 | 0.994427 | 0.627397 | 1 | 0.999631 | 0.541044 |
| Abl1_P00520_S903s_1 | -0.59 | -0.3 | -2.25 | -3.08 | -0.8 | -2.01 | -1.95 | -2.57 | -0.17 | -0.07 | -2.29 | -2.13 | -0.67 | -0.06 | -2.69 | -2.56 | -0.2775 | 0.39 | 0.06 | 0.941478 | 0.683034 | 0.623691 | 1 | 0.999631 | 0.981286 |
| Abl1_P00520_Y179y_1 | 0.29 | 0.78 | 0.42 | -0.51 | -0.45 | 0.26 | -0.21 | -0.09 | 0.26 | 0.8 | -0.71 | -0.26 | 0.3 | -0.02 | -0.55 | -0.04 | -0.3675 | -0.2225 | -0.3225 | 0.390455 | 0.298718 | 0.603848 | 1 | 0.999631 | 0.758901 |
| Abl2_F8VQH0_S632s_1 | 0.08 | -0.2 | -0.28 | 0.3 | -0.06 | 0.66 | 0.12 | 0.89 | -0.12 | -0.59 | 0.06 | 0.16 | -1.12 | 0.03 | -0.45 | 0.21 | 0.4275 | -0.0975 | -0.3075 | 0.413198 | 0.19304 | 0.741154 | 1 | 0.999631 | 0.768474 |
| Abl2_F8VQH0_S785s_1 | -0.94 | -0.22 | -0.13 | 0.59 | -0.62 | -0.48 | 0.93 | 0.46 | 0.34 | 0.1 | 0.64 | -0.57 | -0.5 | -0.73 | 0.21 | 0.9 | 0.2475 | 0.3025 | 0.145 | 0.759457 | 0.597544 | 0.469259 | 1 | 0.999631 | 0.914614 |
| Abl2_F8VQH0_S819s_1 | 0.82 | -0.38 | 0.04 | 1.45 | -0.25 | -0.47 | 1.83 | 0.57 | -0.41 | 0.57 | 1.54 | 0.97 | -0.93 | -0.25 | -0.44 | 1.55 | -0.0625 | 0.185 | -0.5 | 0.423925 | 0.916641 | 0.731803 | 1 | 0.999631 | 0.773835 |
| Abl2_F8VQH0_S822s_1 | 0.25 | 0.48 | -0.67 | 0.31 | 0.56 | 0.01 | 0.32 | 0.02 | 0.03 | 0.69 | 0.49 | 0.65 | -0.14 | 0.42 | -0.43 | 0.2 | 0.135 | 0.3725 | -0.08 | 0.826793 | 0.684706 | 0.292852 | 1 | 0.999631 | 0.940834 |
| Abl2_F8VQH0_S936s_1 | 0.8 | -0.27 | -2.57 | 0.76 | 1.33 | 0.63 | 1.16 | 0.12 | 0.76 | -0.19 | -0.28 | -1.39 | -0.96 | 0.55 | -0.91 | 0.27 | 1.13 | 0.045 | 0.0575 | 0.940576 | 0.148329 | 0.954769 | 1 | 0.999631 | 0.981146 |
| Abl2_F8VQH0_Y439y_1 | -1.04 | -1.18 | -1.73 | -0.97 | -0.77 | -0.55 | -0.83 | -1.1 | -1.01 | -1.57 | -1.27 | -0.79 | -0.56 | -1.05 | -1.02 | -2.96 | 0.4175 | 0.07 | -0.1675 | 0.751158 | 0.163987 | 0.820959 | 1 | 0.999631 | 0.912766 |
| Ablim1_E9Q9C4_S304s_1 | 0.95 | 0.22 | -0.54 | 1.73 | 0.59 | 0.51 | 0.89 | 1.75 | 0.19 | 0.8 | 0.81 | 0.11 | -1.32 | 0.06 | -0.02 | 1.57 | 0.345 | -0.1125 | -0.5175 | 0.454453 | 0.513993 | 0.820864 | 1 | 0.999631 | 0.789715 |
| Ablim1_E9Q9C4_S319s_1 | -0.6 | -1.97 | -0.77 | -0.29 | -0.52 | -0.77 | -0.19 | -0.35 | -0.99 | -1.43 | -1.56 | -1.3 | -1.5 | -0.88 | -0.34 | -0.43 | 0.45 | -0.4125 | 0.12 | 0.790696 | 0.270765 | 0.316155 | 1 | 0.999631 | 0.926419 |
| Ablim1_E9Q9C4_S319s_2 | -0.38 | -2.2 | -2.16 | -2.82 | -0.86 | -2.08 | -1.69 | -2.22 | -1.78 | -0.64 | -2.85 | -2.72 | -0.58 | -1.82 | -4.28 | -3.81 | 0.1775 | -0.1075 | -0.7325 | 0.410683 | 0.750256 | 0.870259 | 1 | 0.999631 | 0.767937 |
| Ablim1_E9Q9C4_S378s_1 | -0.32 | -0.33 | 0.42 | 0.11 | 0.16 | -0.07 | 0.8 | -0.35 | 0.4 | -0.42 | 0.21 | 0.11 | -0.61 | -0.82 | -0.36 | -0.85 | 0.165 | 0.105 | -0.63 | 0.064474 | 0.629562 | 0.740004 | 1 | 0.999631 | 0.514485 |
| Ablim1_E9Q9C4_S383s_1 | -0.41 | -0.24 | -0.82 | -1.14 | -0.61 | -0.59 | -1.09 | -0.73 | -0.68 | -0.29 | -0.97 | -0.51 | 1.02 | -0.55 | -0.95 | 0.02 | -0.1025 | 0.04 | 0.5375 | 0.264586 | 0.731761 | 0.898398 | 1 | 0.999631 | 0.6935 |
| Ablim1_E9Q9C4_S387s_1 | -1.02 | -0.56 | 0.5 | 0.44 | 0.61 | -2.68 | -0.64 | 0.22 | -1.4 | -2.17 | -1.44 | -0.85 | 1.05 | 0.1 | -0.35 | -0.45 | -0.4625 | -1.305 | 0.2475 | 0.615802 | 0.528764 | 0.015869 | 1 | 0.999631 | 0.861203 |
| Ablim1_E9Q9C4_S404s_1 | 0.66 | 0.36 | -0.37 | 0.89 | 1.18 | 0.64 | 0.38 | 0.63 | 0.65 | 0.67 | 0.06 | 0.62 | 0.5 | 0.73 | -0.03 | 0.02 | 0.3225 | 0.115 | -0.08 | 0.829916 | 0.368543 | 0.743124 | 1 | 0.999631 | 0.942187 |
| Ablim1_E9Q9C4_S407s_1 | -0.98 | 0.51 | -0.5 | -0.24 | 0.04 | -0.24 | -0.61 | -0.98 | 0.5 | 0.02 | -0.62 | -0.24 | -0.16 | -0.22 | -0.45 | -0.38 | -0.145 | 0.2175 | -4.9E-17 | 1 | 0.711351 | 0.590941 | 1 | 0.999631 | 1 |
| Ablim1_E9Q9C4_S410s_1 | -0.78 | -0.87 | -0.17 | 0.6 | -1.03 | -0.4 | -0.64 | 0.14 | -0.2 | -0.62 | 0.17 | -0.18 | 0.09 | 0.25 | -1.15 | -0.47 | -0.1775 | 0.0975 | -0.015 | 0.973914 | 0.670897 | 0.803771 | 1 | 0.999631 | 0.990749 |
| Ablim1_E9Q9C4_S425s_1 | 0.82 | -0.68 | -1.08 | -0.12 | -0.24 | -0.05 | 0.46 | -0.23 | -1.09 | 0.01 | -0.59 | -1.07 | -1.35 | -0.65 | -0.64 | -0.36 | 0.25 | -0.42 | -0.485 | 0.303733 | 0.567661 | 0.381789 | 1 | 0.999631 | 0.715879 |
| Ablim1_E9Q9C4_S442s_1 | 0.58 | 1.32 | 0.37 | 0.83 | 1.11 | 2.17 | 0.74 | 1.08 | 1.29 | 1.07 | 0.99 | 0.25 | 0.34 | 1.47 | 0.62 | 1.25 | 0.5 | 0.125 | 0.145 | 0.699388 | 0.212092 | 0.718828 | 1 | 0.999631 | 0.896612 |
| Ablim1_E9Q9C4_S491s_1 | -0.74 | -0.11 | -0.04 | 0.1 | -0.13 | 0.1 | -0.19 | 0.28 | -0.73 | -0.05 | -0.61 | 0 | -0.53 | -0.35 | -0.16 | -0.23 | 0.2125 | -0.15 | -0.12 | 0.693999 | 0.467616 | 0.643484 | 1 | 0.999631 | 0.895131 |
| Ablim1_E9Q9C4_S492s_1 | -1.08 | -0.22 | -0.84 | -0.61 | 0 | -0.43 | -0.67 | -0.54 | -0.04 | -0.67 | -1.15 | -0.41 | 0.19 | -0.2 | -1.01 | -1.28 | 0.2775 | 0.12 | 0.1125 | 0.783223 | 0.363113 | 0.725782 | 1 | 0.999631 | 0.922628 |
| Ablim1_E9Q9C4_S544s_1 | -0.36 | -0.38 | -0.37 | -0.39 | -0.19 | 1.35 | 0.93 | -0.13 | 0.19 | -0.7 | 0.19 | 0.07 | -1.12 | 0.2 | 0.2 | -0.19 | 0.865 | 0.3125 | 0.1475 | 0.683785 | 0.049404 | 0.304292 | 1 | 0.999631 | 0.889602 |
| Ablim1_E9Q9C4_S559s_1 | 0.46 | 0.92 | 0.22 | 0.18 | 0.67 | 1.54 | 0.04 | 0.24 | 1.13 | 0.84 | 0.12 | 0.42 | -0.06 | 1.41 | 0.11 | 0.49 | 0.1775 | 0.1825 | 0.0425 | 0.914528 | 0.646402 | 0.585128 | 1 | 0.999631 | 0.972695 |
| Ablim1_E9Q9C4_S573s_1 | -1.02 | 1.03 | 0.36 | -0.3 | 1.03 | 1.14 | 0.76 | 1.76 | 1.6 | 0.08 | 1.43 | 1.97 | 1.61 | 1.18 | 0.83 | 0.81 | 1.155 | 1.2525 | 1.09 | 0.039759 | 0.031133 | 0.045939 | 1 | 0.999631 | 0.471305 |
| Ablim1_E9Q9C4_S610s_1 | 0.77 | 0.46 | 0.76 | 1.68 | 0.85 | 1 | 1 | 1.68 | 0.48 | 0.54 | 1.18 | 0.92 | -0.48 | 1.42 | 1.05 | 1.43 | 0.215 | -0.1375 | -0.0625 | 0.900865 | 0.544633 | 0.695815 | 1 | 0.999631 | 0.968153 |
| Ablim1_E9Q9C4_T303t_1 | 0.1 | 0.58 | 0.2 | 0.39 | 0.15 | 0.73 | -0.05 | 0.12 | 0.02 | 0.56 | 0.43 | 0.68 | 0.35 | 0.45 | 0.04 | -0.01 | -0.08 | 0.105 | -0.11 | 0.699847 | 0.776095 | 0.706654 | 1 | 0.999631 | 0.896612 |
| Ablim2_E9Q0W6_S313s_1 | -2.45 | -0.54 | -0.63 | -2.03 | -1.21 | -1.23 | -1.61 | -1.75 | -1.2 | -1.97 | -1.38 | -1.01 | 0.57 | -1.19 | -1.07 | -1.74 | -0.0375 | 0.0225 | 0.555 | 0.386527 | 0.937471 | 0.964137 | 1 | 0.999631 | 0.756141 |
| Ablim2_E9Q0W6_S366s_1 | 1.66 | 0.28 | 2.17 | 1.4 | -0.04 | 0.99 | 1.89 | 0.26 | 0.57 | 2.48 | 0.71 | 0.57 | -1.02 | -0.66 | 0.81 | -0.73 | -0.6025 | -0.295 | -1.7775 | 0.007476 | 0.280763 | 0.604691 | 1 | 0.999631 | 0.358173 |
| Ablim2_E9Q0W6_S402s_1 | 1.69 | -0.87 | -0.06 | 0.45 | -0.96 | -0.34 | 0.72 | -0.84 | -1.27 | 1.15 | -0.88 | -1.47 | -1.52 | -1.58 | -2.19 | -2.13 | -0.6575 | -0.92 | -2.1575 | 0.002231 | 0.285513 | 0.218585 | 1 | 0.999631 | 0.351308 |
| Ablim2_E9Q0W6_S457s_1 | -0.63 | -2.46 | 0.24 | 0 | -2.33 | -1.68 | -0.86 | -0.59 | -2.32 | -0.47 | -1.17 | -1.14 | -0.77 | -1.7 | -0.49 | -0.42 | -0.6525 | -0.5625 | -0.1325 | 0.829465 | 0.328562 | 0.395599 | 1 | 0.999631 | 0.942058 |
| Ablim2_E9Q0W6_S486s_1 | -1.02 | 0.32 | -0.97 | -0.95 | 1.39 | -0.07 | -0.6 | -0.79 | 0.46 | -1.29 | -0.74 | -0.1 | 0.08 | 0.18 | -0.3 | -1.23 | 0.6375 | 0.2375 | 0.3375 | 0.463849 | 0.258141 | 0.623138 | 1 | 0.999631 | 0.793747 |
| Ablim2_E9Q0W6_S487s_1 | -1.02 | 0.32 | 0.27 | 0.57 | 1.39 | -0.07 | 0.52 | 0.63 | 0.46 | -1.29 | -0.13 | 0.6 | 0.08 | 0.18 | 0.26 | 1.03 | 0.5825 | -0.125 | 0.3525 | 0.418344 | 0.216208 | 0.811669 | 1 | 0.999631 | 0.771707 |
| Ablim2_E9Q0W6_S511s_1 | -0.51 | 0.36 | -0.23 | -0.65 | 0.15 | 0.19 | -0.76 | -0.45 | -0.04 | -0.07 | -0.73 | -0.22 | 0.53 | 0.38 | -0.57 | -0.21 | 0.04 | -0.0075 | 0.29 | 0.449868 | 0.90929 | 0.981762 | 1 | 0.999631 | 0.78723 |
| Ablim2_E9Q0W6_T397t_1 | -0.25 | -2.07 | -1.58 | -1.32 | -2.65 | -2.3 | -0.8 | -1.12 | -2.67 | -1.09 | -1.35 | -1.34 | -2.89 | -2.05 | -0.87 | -1.49 | -0.4125 | -0.3075 | -0.52 | 0.346407 | 0.454486 | 0.541384 | 1 | 0.999631 | 0.735175 |
| Ablim3_Q69ZX8_S282s_1 | -1.75 | -0.71 | -0.96 | -1.76 | -0.9 | -1.13 | 0.37 | -1.42 | -0.2 | -2.18 | -0.01 | 0.57 | 1.26 | -1.33 | -1.06 | -0.99 | 0.525 | 0.84 | 0.765 | 0.221867 | 0.269518 | 0.182283 | 1 | 0.999631 | 0.672466 |
| Ablim3_Q69ZX8_S337s_1 | -0.24 | -1.17 | -0.51 | -0.44 | -0.79 | -0.7 | -0.6 | -0.09 | -0.73 | -0.19 | -0.1 | -0.19 | -0.54 | -0.44 | 0.05 | -0.19 | 0.045 | 0.2875 | 0.31 | 0.347705 | 0.88466 | 0.369074 | 1 | 0.999631 | 0.736498 |
| Ablim3_Q69ZX8_S372s_1 | 0.86 | 0.61 | 0.25 | 0.25 | 0.47 | 0.67 | 0.4 | 0.28 | 0.33 | 0.22 | 0.09 | 1 | 1.38 | 0.61 | 1.12 | 0.62 | -0.0375 | -0.0825 | 0.44 | 0.193437 | 0.888079 | 0.793651 | 1 | 0.999631 | 0.650087 |
| Ablim3_Q69ZX8_S373s_1 | 0.73 | 0.13 | -0.96 | -0.35 | 0.05 | 0.06 | -0.59 | 0.18 | 0.22 | -0.91 | -0.84 | -1.19 | -0.6 | 0 | -0.17 | -0.92 | 0.0375 | -0.5675 | -0.31 | 0.472294 | 0.92564 | 0.236761 | 1 | 0.999631 | 0.79887 |
| Ablim3_Q69ZX8_S408s_1 | -0.15 | -1.54 | -1.13 | -0.11 | -0.36 | 0.08 | -0.97 | -0.58 | 0.05 | 0.06 | -1.35 | -1.23 | -0.82 | -0.28 | -0.03 | 0.01 | 0.275 | 0.115 | 0.4525 | 0.295166 | 0.514182 | 0.818606 | 1 | 0.999631 | 0.708769 |
| Ablim3_Q69ZX8_S502s_1 | -0.57 | -0.77 | -0.76 | -0.51 | -0.04 | -0.5 | -0.86 | -0.48 | -0.42 | 0.34 | -0.37 | 0.02 | -0.35 | -0.47 | -0.62 | -0.46 | 0.1825 | 0.545 | 0.1775 | 0.505924 | 0.506154 | 0.079102 | 1 | 0.999631 | 0.814836 |
| Ablim3_Q69ZX8_T542t_1 | -2.2 | -0.27 | -1.06 | -0.83 | 0.91 | 0.26 | -0.83 | -2.02 | -0.18 | -0.61 | -1.01 | -0.67 | 1.27 | 0.94 | -0.74 | -0.58 | 0.67 | 0.4725 | 1.3125 | 0.04958 | 0.333875 | 0.294111 | 1 | 0.999631 | 0.487521 |
| Abr_Q5SSL4_S53s_1 | 0.22 | -0.25 | -0.55 | -0.8 | 0.33 | 0.42 | -0.95 | -0.73 | -0.29 | -0.53 | -0.67 | -0.43 | 0.62 | 0.15 | -0.64 | -0.28 | 0.1125 | -0.135 | 0.3075 | 0.430518 | 0.786369 | 0.659766 | 1 | 0.999631 | 0.778235 |
| Abr_Q5SSL4_S72s_1 | -1.04 | -0.43 | -0.22 | -1.47 | 0.03 | -0.69 | 0.28 | -1.36 | -0.32 | -1.76 | 0.12 | -0.81 | -0.77 | -0.94 | -0.9 | -1.86 | 0.355 | 0.0975 | -0.3275 | 0.423872 | 0.439361 | 0.837678 | 1 | 0.999631 | 0.773835 |
| Abr_Q5SSL4_T294t_1 | -1.28 | -4.43 | -0.78 | -0.73 | -3.58 | -3.67 | 0.05 | -2 | -4.04 | -2.5 | -0.83 | -0.71 | -3.76 | -3.35 | -1.04 | -1.4 | -0.495 | -0.215 | -0.5825 | 0.546874 | 0.643737 | 0.833273 | 1 | 0.999631 | 0.834213 |
| Acap2_D4AFX6_S386s_1 | 0.24 | 0.12 | 0.11 | -0.6 | -1.08 | 0.55 | 0.29 | -0.21 | 0.71 | 1.41 | -0.1 | -0.11 | 1.11 | 1.12 | 0.41 | 0.75 | -0.08 | 0.51 | 0.88 | 0.021931 | 0.844502 | 0.240002 | 1 | 0.999631 | 0.405544 |
| Acap2_D4AFX6_S388s_2 | 0.06 | 0.6 | 0.29 | 0.57 | 0.2 | 0.34 | -0.05 | -0.59 | 0.24 | 0.91 | 0.78 | 0.53 | -0.53 | 0.32 | 0.4 | -0.64 | -0.405 | 0.235 | -0.4925 | 0.186915 | 0.201191 | 0.419307 | 1 | 0.999631 | 0.646474 |
| Acap2_D4AFX6_S391s_1 | -0.37 | -0.25 | 0.19 | -0.06 | -0.98 | 0.1 | -0.5 | -0.7 | -0.09 | -0.64 | -0.06 | -0.2 | -0.18 | -0.76 | -0.06 | -0.41 | -0.3975 | -0.125 | -0.23 | 0.451813 | 0.223015 | 0.656195 | 1 | 0.999631 | 0.788148 |
| Acap2_D4AFX6_S391s_2 | -0.34 | -0.24 | 0.29 | 0.57 | 0.55 | 0.06 | -0.05 | -0.59 | -0.47 | 0.44 | 0.78 | 0.53 | -2.37 | 0.06 | 0.4 | -0.64 | -0.0775 | 0.25 | -0.7075 | 0.252745 | 0.824234 | 0.508721 | 1 | 0.999631 | 0.688432 |
| Acap2_D4AFX6_S528s_1 | -0.84 | 0.48 | -1.68 | -2.18 | 0.32 | 0.07 | -1.74 | -2.43 | 0.06 | -0.4 | -1.45 | -1.49 | 0.88 | 0.42 | -1.57 | -1.54 | 0.11 | 0.235 | 0.6025 | 0.4357 | 0.887772 | 0.710447 | 1 | 0.999631 | 0.780324 |
| Acap2_D4AFX6_S559s_1 | -0.16 | -0.51 | 0.16 | 2.03 | 2.88 | -0.05 | 2.56 | 1.46 | 1.33 | -2.19 | 2.48 | 2.19 | -1.76 | -0.49 | -0.59 | 0.37 | 1.3325 | 0.5725 | -0.9975 | 0.144532 | 0.106068 | 0.587443 | 1 | 0.999631 | 0.616037 |
| Acap2_D4AFX6_S562s_1 | 0.83 | 0.49 | -0.48 | -0.34 | -0.29 | 0.16 | 0.24 | -0.83 | -0.4 | 0.6 | -0.77 | -0.65 | -1.71 | -0.97 | -2.7 | -0.75 | -0.305 | -0.43 | -1.6575 | 0.00872 | 0.457508 | 0.340948 | 1 | 0.999631 | 0.365244 |
| Acap2_D4AFX6_S568s_1 | 0.15 | -0.13 | 0.28 | 0.78 | 1.55 | 0.89 | 1.95 | 0.65 | 0.99 | -0.59 | 0.68 | 0.99 | -1.3 | -1.05 | -0.45 | -0.92 | 0.99 | 0.2475 | -1.2 | 0.004522 | 0.022594 | 0.562517 | 1 | 0.999631 | 0.351308 |
| Acap2_D4AFX6_S580s_1 | -0.57 | 0.41 | 1.43 | 1.01 | -0.61 | 0.17 | 0.03 | 1.38 | 0.41 | -0.68 | 0.99 | -1.05 | -0.02 | 0.87 | 1.93 | -1.51 | -0.3275 | -0.6525 | -0.2525 | 0.735231 | 0.555891 | 0.282228 | 1 | 0.999631 | 0.90759 |
| Acap3_Q6NXL5_S383s_1 | 0.38 | -0.02 | 0.2 | 0.33 | -0.02 | 0.32 | 0.16 | 0.53 | 0.6 | 1 | 0.84 | 0.45 | 1 | 0.63 | 0.57 | 0.82 | 0.025 | 0.5 | 0.5325 | 0.077718 | 0.922444 | 0.085519 | 1 | 0.999631 | 0.543121 |
| Acap3_Q6NXL5_S383s_2 | -0.6 | -0.23 | -0.3 | -0.52 | -0.63 | 0.73 | 0.06 | 0.12 | 0.74 | 0.75 | 0.37 | 0.28 | 0.09 | 0.48 | 0.22 | 0.34 | 0.4825 | 0.9475 | 0.695 | 0.027021 | 0.16854 | 0.005152 | 1 | 0.999631 | 0.422893 |
| Acap3_Q6NXL5_S388s_1 | -0.48 | 1.01 | 1.16 | 0.18 | 0.02 | 0.71 | -0.1 | -0.51 | 0.52 | 0.87 | 0.28 | 0.05 | 0.75 | 0.76 | 1.07 | 1.12 | -0.4375 | -0.0375 | 0.4575 | 0.282463 | 0.337908 | 0.929282 | 1 | 0.999631 | 0.70392 |
| Acin1_F6RJ39_S414s_1 | 0.9 | 0.67 | 0.98 | 0.2 | 0.84 | 0.43 | 0.33 | -0.69 | 1.12 | 1.19 | 0.97 | 2.05 | 1.82 | 0.48 | 2.11 | 1.38 | -0.46 | 0.645 | 0.76 | 0.090203 | 0.246352 | 0.085019 | 1 | 0.999631 | 0.557085 |
| Acin1_F6RJ39_S425s_1 | -0.27 | 0.44 | -0.12 | 0.07 | 0.51 | 0.05 | -0.42 | 0.15 | 0.27 | 0.42 | -0.29 | -0.16 | 0.98 | 0.58 | -0.16 | 0.12 | 0.0425 | 0.03 | 0.35 | 0.331208 | 0.889038 | 0.92097 | 1 | 0.999631 | 0.727302 |
| Acin1_F6RJ39_S437s_1 | 2.54 | -2.36 | -0.19 | -0.35 | -2.7 | -0.62 | 2.87 | -2.41 | -5.6 | 2.41 | -1.61 | -1.93 | -3 | -3.45 | -1.98 | -2.91 | -0.625 | -1.5925 | -2.745 | 0.01201 | 0.653023 | 0.341089 | 1 | 0.999631 | 0.385092 |
| Acin1_F6RJ39_S541s_1 | -0.52 | 0.41 | 0.21 | -0.79 | 0.39 | -0.7 | -0.93 | -0.01 | -0.26 | 0.44 | -0.54 | -0.24 | 1.01 | -0.02 | -1.98 | -0.67 | -0.14 | 0.0225 | -0.2425 | 0.697941 | 0.737153 | 0.95248 | 1 | 0.999631 | 0.895731 |
| Acin1_F6RJ39_S601s_1 | 0.65 | 1.02 | 1.08 | 0.98 | 0 | 0.52 | 0.38 | 0.24 | -0.6 | 1.38 | 1.38 | 0.2 | 0.33 | 0.76 | 1.18 | 1.83 | -0.6475 | -0.3425 | 0.0925 | 0.804833 | 0.028747 | 0.477488 | 1 | 0.999631 | 0.934231 |
| Acin1_F6RJ39_S603s_1 | 0.5 | 0.48 | 0.6 | 0.42 | 0.2 | 0.34 | 0.11 | -0.16 | 0.57 | 0.94 | 0.53 | 0.26 | 1.15 | 0.43 | 0.34 | -0.02 | -0.3775 | 0.075 | -0.025 | 0.938787 | 0.145863 | 0.776672 | 1 | 0.999631 | 0.980279 |
| Acin1_F6RJ39_S675s_1 | 1.59 | 0.34 | 1.29 | 0.6 | 0.68 | 0.69 | 0.91 | -0.01 | 1.29 | 2.11 | 0.19 | 0.58 | 0.71 | 0.81 | 0.96 | 0.09 | -0.3875 | 0.0875 | -0.3125 | 0.422613 | 0.310533 | 0.858204 | 1 | 0.999631 | 0.773293 |
| Acin1_F6RJ39_S784s_1 | 1.37 | 0.15 | 1.23 | 0.51 | 0.2 | 0.67 | 0.75 | 0.61 | 1.58 | 1.76 | 0.99 | 0.82 | 0.42 | 0.65 | 0.9 | -0.21 | -0.2575 | 0.4725 | -0.375 | 0.359491 | 0.464312 | 0.240752 | 1 | 0.999631 | 0.741182 |
| Acin1_F6RJ39_S832s_1 | 1.68 | 1.07 | 2.77 | 1.51 | 0.28 | 2.14 | 2.18 | 0.88 | 1.74 | 3.12 | 1.58 | 1.44 | 0.42 | 0.24 | 1.67 | 1.31 | -0.3875 | 0.2125 | -0.8475 | 0.104167 | 0.48401 | 0.673581 | 1 | 0.999631 | 0.568213 |
| Acin1_F6RJ39_S923s_1 | 0.24 | 0.03 | 0.08 | 0.34 | 0.19 | -0.07 | 0.78 | -0.05 | -0.08 | 0.01 | 0.84 | 0.11 | -0.81 | 1.02 | -0.47 | 1.25 | 0.04 | 0.0475 | 0.075 | 0.881048 | 0.888763 | 0.873845 | 1 | 0.999631 | 0.963364 |
| Acin1_F6RJ39_S937s_1 | 0.02 | -0.05 | 0.25 | -0.17 | -0.2 | -0.27 | -0.52 | -0.15 | -0.27 | 0.37 | -0.13 | 0.01 | 0.24 | 0.23 | 0.17 | 0.51 | -0.2975 | -0.0175 | 0.275 | 0.323914 | 0.246517 | 0.948572 | 1 | 0.999631 | 0.724485 |
| Acin1_Q9JIX8_S858s_1 | 1.84 | 1.28 | 2.79 | 1.46 | 0.38 | 2.36 | 2.15 | 0.87 | 1.94 | 3.36 | 1.49 | 1.45 | 0.47 | 0.32 | 1.59 | 1.32 | -0.4025 | 0.2175 | -0.9175 | 0.067386 | 0.464197 | 0.679635 | 1 | 0.999631 | 0.521449 |
| Acly_Q3TS02_S455s_1 | 1.12 | 0.43 | 0.48 | 1.15 | 0.4 | 1.03 | 1.14 | 0.55 | 1.21 | 1.22 | 0.81 | 0.74 | -0.51 | 0.4 | 0.64 | 0.03 | -0.015 | 0.2 | -0.655 | 0.096765 | 0.962218 | 0.517741 | 1 | 0.999631 | 0.55886 |
| Acly_Q3TS02_S481s_1 | 0.4 | 0.63 | 0.67 | 0.56 | 0.49 | 0.39 | 0.05 | 0.75 | 0.52 | 0.67 | 0.45 | 1.09 | 0.58 | 1.2 | 0.87 | 1.05 | -0.145 | 0.1175 | 0.36 | 0.219187 | 0.581163 | 0.662722 | 1 | 0.999631 | 0.669859 |
| Aco2_Q99KI0_S559s_1 | -0.12 | -0.55 | -0.16 | -1.22 | -0.58 | -0.58 | -0.12 | -0.4 | -0.14 | 0.36 | -0.24 | -0.13 | -0.25 | -0.27 | -0.54 | -0.33 | 0.0925 | 0.475 | 0.165 | 0.623271 | 0.774981 | 0.179928 | 1 | 0.999631 | 0.862913 |
| Acot1_O55137_S416s_1 | -0.28 | 0.82 | 1.54 | 0 | 0.09 | -0.38 | 0.7 | 0.93 | 1.03 | 1.08 | 0.77 | 1.94 | 1.25 | 1.75 | 1.85 | 1.37 | -0.185 | 0.685 | 1.035 | 0.03804 | 0.701592 | 0.166757 | 1 | 0.999631 | 0.459475 |
| Acot1_O55137_S56s_1 | -0.14 | 0.84 | 0.35 | -0.22 | 0.24 | 0.06 | 0.15 | 0.75 | 0.55 | 0.49 | -0.05 | 0.65 | 1.26 | 0.31 | 0.57 | 0.08 | 0.0925 | 0.2025 | 0.3475 | 0.378722 | 0.780136 | 0.552989 | 1 | 0.999631 | 0.751637 |
| Acot11_F6SS52_S14s_1 | 0.19 | 1.03 | 0.87 | 0.82 | 0.82 | 1.43 | 0.51 | 1.78 | 0.99 | 0.92 | 1.22 | 1.15 | 0.39 | 1.6 | 1.27 | 1.8 | 0.4075 | 0.3425 | 0.5375 | 0.190465 | 0.278353 | 0.250096 | 1 | 0.999631 | 0.648674 |
| Acot11_F6SS52_S20s_1 | -0.81 | -1.09 | 0.52 | -0.01 | -0.18 | -1.07 | -1.13 | -0.03 | -1.02 | 0.04 | -0.21 | -0.87 | -0.24 | -1.02 | 0.14 | 0.38 | -0.255 | -0.1675 | 0.1625 | 0.729972 | 0.575817 | 0.706027 | 1 | 0.999631 | 0.906325 |
| Acot7_Q91V12_S285s_1 | -0.29 | 1.18 | -0.35 | -0.81 | 0.08 | 1.24 | -0.14 | 0.53 | 0.78 | 0.24 | 0.44 | 0.88 | 0.24 | 0.47 | 1.2 | 0.46 | 0.495 | 0.6525 | 0.66 | 0.181715 | 0.331409 | 0.168525 | 1 | 0.999631 | 0.643861 |
| Acot7_Q91V12_S2s_1 | -0.93 | -0.15 | -0.16 | -0.21 | -0.36 | -1.22 | -0.35 | 0.29 | -0.56 | -1.85 | -1.03 | -0.15 | -1.14 | -0.53 | 0.46 | -0.64 | -0.0475 | -0.535 | -0.1 | 0.805542 | 0.899917 | 0.218535 | 1 | 0.999631 | 0.934743 |
| Acsbg1_Q99PU5_S55s_1 | 0 | 0.57 | 0.07 | -0.05 | 0.25 | -0.3 | -0.53 | 0.61 | 0.39 | 0.69 | 0.3 | 0.77 | 0.85 | 0.29 | 0.4 | 0.94 | -0.14 | 0.39 | 0.4725 | 0.150094 | 0.676765 | 0.185353 | 1 | 0.999631 | 0.622782 |
| Acsbg1_Q99PU5_S70s_1 | -0.61 | 0.35 | -0.24 | -0.2 | -0.11 | 0.86 | -0.48 | 0.12 | -0.15 | -0.48 | -0.2 | -0.51 | 0.47 | 1.11 | 0.23 | 0.07 | 0.2725 | -0.16 | 0.645 | 0.093759 | 0.462775 | 0.593333 | 1 | 0.999631 | 0.558128 |
| Acsl6_Q5F2C5_S364s_1 | -1.41 | -0.1 | -0.37 | -0.79 | -0.52 | -0.45 | -1.57 | -1.66 | -0.23 | -1.2 | -0.98 | -0.38 | 0.32 | 0.24 | 0.08 | -0.09 | -0.3825 | -0.03 | 0.805 | 0.043055 | 0.380646 | 0.938171 | 1 | 0.999631 | 0.475443 |
| Acss2_Q9QXG4_S265s_1 | 0.24 | 0.74 | 0.24 | -0.23 | 0.12 | 0.5 | -0.32 | 0.02 | 0.42 | 0.23 | 0.14 | 0.25 | 1.15 | 1 | -0.35 | 0.31 | -0.1675 | 0.0125 | 0.28 | 0.504068 | 0.595355 | 0.965743 | 1 | 0.999631 | 0.814292 |
| Acss2_Q9QXG4_S267s_1 | -0.21 | -0.44 | 0.8 | -0.47 | -0.53 | -0.45 | -0.62 | -0.41 | -0.24 | -0.67 | 0.42 | 0.38 | -0.26 | -0.08 | 0 | 1.02 | -0.4225 | 0.0525 | 0.25 | 0.560633 | 0.230967 | 0.897286 | 1 | 0.999631 | 0.839977 |
| Acss2_Q9QXG4_S30s_1 | -0.09 | 0.77 | 0.76 | -0.03 | 0.14 | 0.57 | -0.15 | 0.52 | 0.75 | 0.6 | 1.01 | 0.86 | 1.27 | 1.25 | 0.78 | 1.19 | -0.0825 | 0.4525 | 0.77 | 0.041229 | 0.804219 | 0.174789 | 1 | 0.999631 | 0.474979 |
| Actb_P60710_S233s_1 | 0.1 | 0.16 | 0.72 | -0.39 | 0.13 | 0.52 | -0.57 | 0.12 | -0.1 | -0.08 | -0.75 | 0.03 | 0.78 | 0.67 | 0.36 | 0.1 | -0.0975 | -0.3725 | 0.33 | 0.343319 | 0.780938 | 0.28351 | 1 | 0.999631 | 0.73348 |
| Actb_P60710_S239s_1 | -0.85 | -0.2 | -0.16 | -0.46 | -0.58 | 0.19 | -0.8 | -1.32 | -0.08 | -0.2 | -2.06 | -2.08 | 0.77 | 0.18 | -0.02 | -0.29 | -0.21 | -0.6875 | 0.5775 | 0.113776 | 0.573475 | 0.22229 | 1 | 0.999631 | 0.579001 |
| Actb_P60710_S52s_1 | -0.59 | 3.13 | 1.72 | -0.76 | 2.67 | 3.3 | 1.96 | -0.27 | 1.36 | 0.55 | 0.29 | 1.09 | 1.96 | 0.41 | 0.72 | -2.18 | 1.04 | -0.0525 | -0.6475 | 0.555786 | 0.333575 | 0.950638 | 1 | 0.999631 | 0.838305 |
| Actb_P60710_T203t_1 | -0.35 | 0.15 | 0.04 | -0.26 | -0.44 | -0.25 | -0.67 | -0.4 | -0.31 | -0.07 | 0.04 | -0.02 | -0.11 | -0.1 | -0.62 | -0.73 | -0.335 | 0.015 | -0.285 | 0.360366 | 0.212451 | 0.954457 | 1 | 0.999631 | 0.741182 |
| Actb_P60710_Y198y_1 | 1.43 | -1.59 | -0.97 | -1.2 | -1.78 | -2.39 | 1.39 | -1.12 | -1.69 | -0.69 | -1.09 | -0.97 | -0.75 | -1.78 | -0.9 | -1.39 | -0.3925 | -0.5275 | -0.6225 | 0.348687 | 0.673746 | 0.420887 | 1 | 0.999631 | 0.736697 |
| Actr3_Q99JY9_S418s_1 | -0.24 | 1.46 | 1.48 | -0.42 | 2.87 | 0.45 | 1.84 | -0.65 | 4.27 | 0.11 | 3.26 | 4.1 | 5.01 | 0.67 | 0.33 | -0.03 | 0.5575 | 2.365 | 0.925 | 0.407724 | 0.498002 | 0.030774 | 1 | 0.999631 | 0.766996 |
| Acvr2a_P27038_S184s_1 | 0.28 | 1.12 | 0.81 | 0.72 | 0.2 | 1.21 | -0.01 | 0.16 | 0.16 | 0.48 | 0.71 | 0.28 | 1.39 | 1.4 | 0.23 | 0.85 | -0.3425 | -0.325 | 0.235 | 0.529615 | 0.345267 | 0.28398 | 1 | 0.999631 | 0.826087 |
| Acy3_Q91XE4_S7s_1 | 2.16 | -1.05 | -1.53 | -0.74 | -1.07 | -1.78 | 0.98 | -1.52 | 0.49 | 0.37 | -1.38 | 0.79 | -1.89 | -0.7 | -2.27 | -3.44 | -0.5575 | 0.3575 | -1.785 | 0.063139 | 0.540483 | 0.673281 | 1 | 0.999631 | 0.512978 |
| Adam22_Q9R1V6_S772s_1 | -0.53 | 0.13 | -0.5 | -1.27 | -0.67 | -0.36 | -0.55 | -0.61 | 0.06 | -0.8 | 0.01 | 0.22 | 0.52 | -0.77 | -0.44 | -1.38 | -0.005 | 0.415 | 0.025 | 0.958138 | 0.988021 | 0.296776 | 1 | 0.999631 | 0.985497 |
| Adam22_Q9R1V6_S774s_1 | -0.28 | 0.4 | 0.28 | 0.17 | 1.19 | 0.47 | 0.84 | 0.61 | 1.14 | -0.17 | 1.35 | 1.36 | 1.29 | 0.22 | 1.06 | -0.35 | 0.635 | 0.7775 | 0.4125 | 0.338683 | 0.049738 | 0.079916 | 1 | 0.999631 | 0.731883 |
| Adam22_Q9R1V6_S796s_1 | 0.38 | 0 | -0.78 | -0.95 | -0.56 | 0.2 | -0.85 | -0.75 | 0.16 | 0.26 | -0.53 | -0.57 | 0.55 | 0.23 | -0.78 | -0.85 | -0.1525 | 0.1675 | 0.125 | 0.788967 | 0.703876 | 0.675554 | 1 | 0.999631 | 0.925513 |
| Adam22_Q9R1V6_S813s_1 | -0.56 | -0.63 | 0.22 | -0.37 | -0.66 | -0.35 | -0.43 | -0.55 | 0.44 | 0.29 | -0.3 | 0.18 | 0.21 | -0.86 | -0.24 | 0.24 | -0.1625 | 0.4875 | 0.1725 | 0.639805 | 0.569068 | 0.144962 | 1 | 0.999631 | 0.870995 |
| Adam22_Q9R1V6_S819s_1 | -0.81 | -0.64 | -0.94 | -0.99 | -0.41 | -0.46 | -1.04 | -0.65 | -0.49 | -0.94 | -1.27 | -0.53 | -0.21 | -0.09 | -0.98 | -0.66 | 0.205 | 0.0375 | 0.36 | 0.264655 | 0.44388 | 0.896494 | 1 | 0.999631 | 0.6935 |
| Adam22_Q9R1V6_S867s_1 | -0.87 | -1.9 | -0.09 | 0.17 | -2.2 | -1.56 | 0.04 | -0.14 | -2.04 | -0.98 | 0.93 | 0.33 | 0.3 | -1.29 | 0.36 | -0.18 | -0.2925 | 0.2325 | 0.47 | 0.408255 | 0.6499 | 0.74749 | 1 | 0.999631 | 0.766996 |
| Adam22_Q9R1V6_S867s_2 | -0.77 | -0.97 | -0.53 | -2.11 | -2.42 | -1.64 | -1.43 | -1.37 | -1.97 | -1.07 | -0.01 | -1.72 | 1.79 | -2.32 | -1.47 | -1.3 | -0.62 | -0.0975 | 0.27 | 0.747653 | 0.163809 | 0.852635 | 1 | 0.999631 | 0.911931 |
| Adam22_Q9R1V6_S871s_2 | -0.88 | -1.48 | -0.53 | -2.11 | -2.72 | -1.71 | -1.43 | -1.37 | -2.23 | -1.01 | -0.01 | -1.72 | 1.3 | -2.17 | -1.47 | -1.3 | -0.5575 | 0.0075 | 0.34 | 0.645775 | 0.235487 | 0.989107 | 1 | 0.999631 | 0.873384 |
| Adam22_Q9R1V6_S874s_1 | 0.38 | -0.36 | -0.12 | -0.15 | -0.41 | -0.64 | -0.14 | 0.08 | -0.59 | -1.13 | -0.25 | 0.29 | -1.02 | -1.32 | 0.08 | -0.11 | -0.215 | -0.3575 | -0.53 | 0.204998 | 0.467796 | 0.341477 | 1 | 0.999631 | 0.6605 |
| Adcy2_Q3V1Q3_S1089s_1 | 0.17 | 0.7 | -0.24 | -0.01 | 0.02 | 0.36 | -0.32 | 0.2 | -0.01 | -0.12 | -0.29 | -0.35 | 0.29 | -0.13 | -0.83 | 0.14 | -0.09 | -0.3475 | -0.2875 | 0.438362 | 0.769008 | 0.256761 | 1 | 0.999631 | 0.781942 |
| Adcy2_Q3V1Q3_S476s_1 | -1.28 | -0.25 | -0.64 | -1.03 | -0.64 | -0.39 | -1.04 | -0.78 | -0.79 | -0.39 | -0.58 | -0.6 | 0.2 | -0.54 | -0.56 | -0.64 | 0.0875 | 0.21 | 0.415 | 0.257278 | 0.781454 | 0.501591 | 1 | 0.999631 | 0.691981 |
| Adcy5_P84309_S137s_1 | 0.16 | -0.28 | -0.5 | 0.02 | -0.06 | -0.88 | -1.08 | -0.95 | -0.36 | -0.26 | -1.31 | 0 | 0.28 | -0.2 | -0.53 | 0.08 | -0.5925 | -0.3325 | 0.0575 | 0.856 | 0.090221 | 0.363013 | 1 | 0.999631 | 0.951817 |
| Adcy5_P84309_S156s_1 | -0.08 | 0.37 | -0.82 | 0.73 | 0.25 | -0.17 | -0.24 | 0.68 | 0.07 | -0.27 | -0.12 | 0.62 | -0.27 | -0.01 | 1.09 | 0.32 | 0.08 | 0.025 | 0.2325 | 0.604739 | 0.841612 | 0.950031 | 1 | 0.999631 | 0.85817 |
| Adcy5_P84309_S49s_1 | 1.2 | -0.65 | -1.6 | 0.06 | 0.87 | -0.43 | 0.26 | -0.93 | 0.38 | 0.9 | -0.69 | -0.33 | -1.03 | -0.99 | -0.06 | -1.09 | 0.19 | 0.3125 | -0.545 | 0.362891 | 0.76535 | 0.618683 | 1 | 0.999631 | 0.74343 |
| Adcy5_P84309_S8s_1 | -0.71 | -0.86 | -0.81 | -0.6 | -0.85 | -1.35 | -0.96 | -0.34 | -0.75 | -0.88 | -0.59 | -0.48 | 0.1 | -0.54 | -0.31 | -0.42 | -0.13 | 0.07 | 0.4525 | 0.132845 | 0.654147 | 0.780291 | 1 | 0.999631 | 0.604808 |
| Adcy5_P84309_T62t_1 | 2.76 | -0.92 | -0.46 | 1.1 | 1.53 | -0.06 | 1.46 | 0.09 | -0.1 | 1.02 | -0.46 | -1.26 | -2.81 | -1.82 | 0.01 | -1.28 | 0.135 | -0.82 | -2.095 | 0.036312 | 0.868793 | 0.340429 | 1 | 0.999631 | 0.453963 |
| Adcy5_P84309_T665t_1 | -0.71 | -0.29 | -0.16 | 1.15 | 0.93 | 0 | 0.93 | 2.01 | 0.44 | -0.54 | 0.72 | 0.8 | -0.54 | 0.23 | 1.09 | 1.16 | 0.97 | 0.3575 | 0.4875 | 0.371003 | 0.092785 | 0.467169 | 1 | 0.999631 | 0.746338 |
| Adcy6_F8VQ52_S53s_1 | 1.04 | -0.77 | -0.99 | -0.32 | 0.08 | -0.6 | -0.28 | -0.63 | -0.12 | 0.52 | -0.45 | -0.91 | -0.49 | -0.3 | -1.16 | -0.4 | -0.0975 | 0.02 | -0.3275 | 0.500839 | 0.833524 | 0.968922 | 1 | 0.999631 | 0.814043 |
| Adcy6_F8VQ52_S576s_1 | -0.22 | -0.85 | -0.49 | -0.14 | 0.02 | -0.16 | -0.19 | -0.33 | -0.2 | -0.05 | -0.19 | -0.37 | -0.4 | 0.2 | -0.4 | -0.1 | 0.26 | 0.2225 | 0.25 | 0.427202 | 0.345917 | 0.427653 | 1 | 0.999631 | 0.775206 |
| Adcy8_P97490_S67s_1 | -0.63 | -0.57 | -1.96 | -1.82 | -1.04 | -1.09 | -0.95 | -1.74 | -1.4 | -1.67 | -1.92 | -2.19 | -0.55 | -0.64 | -2.16 | -1.69 | 0.04 | -0.55 | -0.015 | 0.976792 | 0.9229 | 0.206261 | 1 | 0.999631 | 0.992016 |
| Adcy9_P51830_S1273s_1 | -0.09 | 0.34 | 0.31 | 0.23 | 0.15 | -0.14 | -0.19 | 0.33 | -0.27 | -0.31 | 0.03 | -0.1 | -0.45 | 0.03 | 0.11 | 0.66 | -0.16 | -0.36 | -0.11 | 0.73617 | 0.544119 | 0.18355 | 1 | 0.999631 | 0.90759 |
| Adcy9_P51830_S354s_1 | 1.24 | 0.12 | -4.32 | -3.13 | -0.58 | 0.49 | -2.45 | -4.2 | -0.73 | 1.34 | -3.68 | -4.49 | 0.04 | -0.88 | -5.91 | -6.92 | -0.1625 | -0.3675 | -1.895 | 0.314027 | 0.908877 | 0.818128 | 1 | 0.999631 | 0.719959 |
| Adcy9_P51830_S39s_1 | 2.78 | -0.87 | 0.19 | 1.5 | 0.75 | 0.86 | 2.11 | 1 | 0.18 | 0.18 | -0.11 | -1.42 | -2.5 | -0.89 | 0.25 | -0.92 | 0.28 | -1.1925 | -1.915 | 0.043647 | 0.709271 | 0.147469 | 1 | 0.999631 | 0.475443 |
| Adcy9_P51830_S49s_1 | -0.15 | 0.26 | -0.3 | 0.09 | -0.25 | -0.41 | 0.33 | 0.37 | -0.09 | -0.67 | -0.04 | -0.03 | -0.63 | -0.1 | 0.3 | 0.08 | 0.035 | -0.1825 | -0.0625 | 0.844622 | 0.906724 | 0.529768 | 1 | 0.999631 | 0.94856 |
| Adcy9_P51830_S610s_1 | -1.21 | -1.32 | -1.36 | -1.61 | -1.51 | -1.93 | -1.84 | -1.77 | -1.91 | -1.43 | -1.58 | -1.45 | -0.53 | -2.11 | -2 | -1.47 | -0.3875 | -0.2175 | -0.1525 | 0.701562 | 0.142261 | 0.415103 | 1 | 0.999631 | 0.896835 |
| Adcy9_P51830_S613s_1 | -0.41 | -0.74 | -0.35 | -0.3 | -0.61 | -0.98 | -0.84 | -0.15 | -1.31 | -1.22 | -0.46 | -0.69 | -0.87 | -1.16 | -0.47 | -0.59 | -0.195 | -0.47 | -0.3225 | 0.289671 | 0.498002 | 0.143921 | 1 | 0.999631 | 0.706164 |
| Adcy9_P51830_S678s_1 | -1.6 | -1.18 | -1.48 | -2.27 | -0.02 | -1.23 | -0.73 | -2.42 | 0.31 | -1.75 | -0.74 | -0.71 | 1.86 | -2.33 | -2.38 | -2.56 | 0.5325 | 0.91 | 0.28 | 0.766496 | 0.316174 | 0.074794 | 1 | 0.999631 | 0.916961 |
| Adcy9_P51830_T1275t_1 | -0.47 | -0.55 | 0.89 | 0.34 | -0.87 | -0.81 | -0.12 | 1.28 | -1.26 | -1.32 | 0.62 | 0.71 | -0.58 | -0.53 | -0.19 | 0.96 | -0.1825 | -0.365 | -0.1375 | 0.775824 | 0.743143 | 0.548212 | 1 | 0.999631 | 0.920998 |
| Adcyap1r1_E9Q3E8_S438s_1 | -1.42 | -0.23 | 0.45 | -0.41 | -0.57 | 0.02 | -0.74 | 0.03 | -0.07 | -1.22 | -1 | 0.17 | -2.09 | -0.03 | -0.14 | -1.36 | 0.0875 | -0.1275 | -0.5025 | 0.394273 | 0.837917 | 0.795528 | 1 | 0.999631 | 0.759303 |
| Add1_F8WGR0_S12s_1 | -0.6 | -0.87 | -0.3 | -0.75 | -1.16 | -1.11 | -0.82 | -0.93 | -1.28 | -0.5 | -0.74 | -0.6 | -0.48 | -1.14 | -0.43 | -0.4 | -0.375 | -0.15 | 0.0175 | 0.954898 | 0.166538 | 0.612853 | 1 | 0.999631 | 0.984441 |
| Add1_F8WGR0_S353s_2 | 0.02 | -0.92 | -0.08 | -1 | -1.37 | -0.52 | -0.52 | -0.44 | -0.68 | 0.49 | -1.49 | -0.93 | -0.28 | -1.1 | -0.68 | -1.56 | -0.2175 | -0.1575 | -0.41 | 0.325509 | 0.557768 | 0.742011 | 1 | 0.999631 | 0.724678 |
| Add1_F8WGR0_S355s_1 | -0.17 | -0.91 | -0.71 | -1 | -1.05 | -1.12 | -0.92 | -0.69 | -1.31 | -0.33 | -0.66 | -0.83 | -0.3 | -0.42 | -0.74 | -1.09 | -0.2475 | -0.085 | 0.06 | 0.855906 | 0.394685 | 0.796415 | 1 | 0.999631 | 0.951817 |
| Add1_F8WGR0_S355s_2 | -0.32 | -0.89 | -0.06 | -0.68 | -1.08 | -0.57 | -0.48 | -0.07 | -0.65 | 0 | -1.13 | -0.77 | -0.35 | -1.16 | -0.43 | -1.15 | -0.0625 | -0.15 | -0.285 | 0.420345 | 0.84684 | 0.663398 | 1 | 0.999631 | 0.77267 |
| Add1_F8WGR0_S366s_1 | -0.25 | -0.86 | 0.09 | -0.4 | -0.73 | -0.65 | -0.69 | -0.36 | -0.47 | -0.57 | -0.34 | -0.3 | -0.47 | -0.75 | -0.24 | -0.04 | -0.2525 | -0.065 | -0.02 | 0.95112 | 0.390191 | 0.823532 | 1 | 0.999631 | 0.984079 |
| Add1_F8WGR0_S457s_2 | 0.72 | -0.96 | 0.45 | -0.96 | -1.65 | -0.19 | 0.52 | -1.26 | -0.63 | 0.85 | -1.79 | -1.5 | -0.04 | -1.38 | -1.81 | -1.45 | -0.4575 | -0.58 | -0.9825 | 0.099543 | 0.455731 | 0.393904 | 1 | 0.999631 | 0.562298 |
| Add1_F8WGR0_S464s_1 | 1.42 | -1.2 | 0.44 | -0.94 | -1.49 | 0.64 | 0.37 | -1.3 | -0.58 | 1.03 | -0.19 | -1.92 | -0.54 | -1.6 | -1.05 | -1.57 | -0.375 | -0.345 | -1.12 | 0.087646 | 0.608555 | 0.652112 | 1 | 0.999631 | 0.553787 |
| Add1_F8WGR0_S465s_1 | 1.34 | -1.73 | -0.15 | 0.13 | -1.62 | 0.74 | 1.16 | -1.38 | -0.56 | 0.96 | -1.08 | -0.51 | -0.28 | -2.03 | -0.04 | -1.16 | -0.1725 | -0.195 | -0.775 | 0.277487 | 0.835394 | 0.776612 | 1 | 0.999631 | 0.700874 |
| Add1_F8WGR0_S465s_2 | 0.72 | -0.96 | 0.45 | -0.96 | -1.65 | -0.19 | 0.52 | -1.26 | -0.63 | 0.85 | -1.79 | -1.5 | -0.04 | -1.38 | -1.81 | -1.45 | -0.4575 | -0.58 | -0.9825 | 0.099543 | 0.455731 | 0.393904 | 1 | 0.999631 | 0.562298 |
| Add1_F8WGR0_S617s_1 | 0.36 | -0.4 | -0.56 | -0.8 | -0.64 | -0.15 | -0.58 | -0.96 | -0.21 | 0.77 | -0.83 | -0.91 | -0.03 | -0.62 | -0.91 | -1.23 | -0.2325 | 0.055 | -0.3475 | 0.381007 | 0.496289 | 0.903143 | 1 | 0.999631 | 0.752619 |
| Add1_F8WGR0_S631s_1 | -1.33 | -0.33 | -0.69 | -2.54 | 0.1 | -0.95 | -2.06 | -2.37 | -0.63 | -1.24 | -2.07 | -1.43 | 1.89 | -0.4 | -2.63 | -1.11 | -0.0975 | -0.12 | 0.66 | 0.474857 | 0.882895 | 0.8214 | 1 | 0.999631 | 0.799827 |
| Add1_F8WGR0_T358t_1 | -0.27 | -0.39 | -0.06 | -0.16 | -0.96 | -1 | -0.51 | -0.28 | -0.58 | -0.61 | -0.14 | -0.51 | -0.13 | -0.71 | -0.53 | -0.98 | -0.4675 | -0.24 | -0.3675 | 0.237166 | 0.114363 | 0.364364 | 1 | 0.999631 | 0.683301 |
| Add1_F8WGR0_T358t_2 | -0.35 | -0.96 | -0.06 | -0.68 | -1.08 | -0.57 | -0.48 | -0.07 | -0.68 | -0.02 | -1.13 | -0.77 | -0.35 | -1.14 | -0.43 | -1.15 | -0.0375 | -0.1375 | -0.255 | 0.472767 | 0.908912 | 0.691492 | 1 | 0.999631 | 0.799014 |
| Add1_F8WGR0_T641t_1 | -1.32 | -1.44 | -0.28 | -1.23 | -1.81 | -1.9 | -1.55 | -1.02 | -1.66 | -1.14 | -0.98 | -0.59 | -0.39 | -1.17 | -0.83 | -0.53 | -0.5025 | -0.025 | 0.3375 | 0.364781 | 0.181036 | 0.946448 | 1 | 0.999631 | 0.745299 |
| Add1_F8WGR0_T641t_2 | 0.13 | 0.68 | 0.77 | 0.01 | 0.01 | 0.61 | -0.04 | -0.32 | 0.08 | 0.83 | -0.46 | -0.04 | 0 | 0.12 | 0.11 | 0.7 | -0.3325 | -0.295 | -0.165 | 0.614983 | 0.314179 | 0.423407 | 1 | 0.999631 | 0.861203 |
| Add1_F8WGR0_T645t_1 | -1.27 | -0.24 | 0.53 | -0.85 | -1.39 | 0.03 | -0.99 | -0.48 | 0.13 | -0.42 | 0.16 | -0.01 | 0.27 | 0.31 | 0.04 | 0.63 | -0.25 | 0.4225 | 0.77 | 0.092311 | 0.600289 | 0.32515 | 1 | 0.999631 | 0.557085 |
| Add1_F8WGR0_T645t_2 | 0.13 | 0.68 | 0.55 | -0.1 | 0.01 | 0.61 | -0.14 | -0.17 | 0.08 | 0.83 | -0.51 | 0 | 0 | 0.12 | 0 | 0.82 | -0.2375 | -0.215 | -0.08 | 0.812079 | 0.452334 | 0.556541 | 1 | 0.999631 | 0.936949 |
| Add1_F8WHZ9_S631s_1 | -1.33 | -0.33 | -0.69 | -2.54 | 0.1 | -0.95 | -2.06 | -2.37 | -0.63 | -1.24 | -2.07 | -1.43 | 1.89 | -0.4 | -2.63 | -1.11 | -0.0975 | -0.12 | 0.66 | 0.474857 | 0.882895 | 0.8214 | 1 | 0.999631 | 0.799827 |
| Add1_F8WHZ9_T641t_1 | -1.26 | -1.32 | 0.19 | -0.93 | -1.67 | -1.66 | -1.29 | -0.71 | -1.41 | -0.88 | -0.6 | -0.08 | 0.04 | -0.93 | -0.41 | -0.03 | -0.5025 | 0.0875 | 0.4975 | 0.259255 | 0.243955 | 0.842988 | 1 | 0.999631 | 0.692624 |

| | | | | | | | | | | | | | | | | | | | | | | | | | | |
|---|---|---|---|---|---|---|---|---|---|---|---|---|---|---|---|---|---|---|---|---|---|---|---|---|---|---|
| Add1_F8WHZ9_T641t_2 | 0.13 | 0.68 | 0.77 | 0.01 | 0.01 | 0.61 | -0.04 | -0.32 | 0.08 | 0.83 | -0.46 | -0.04 | 0 | 0.12 | 0.11 | 0.7 | -0.3325 | -0.295 | -0.165 | 0.614983 | 0.314179 | 0.423407 | | 1 | 0.999631 | 0.861203 |
| Add1_F8WHZ9_T645t_1 | -1.27 | -0.24 | 0.53 | -0.85 | -1.39 | 0.03 | -0.99 | -0.48 | 0.13 | -0.42 | 0.16 | -0.01 | 0.27 | 0.31 | 0.04 | 0.63 | -0.25 | 0.4225 | 0.77 | 0.092311 | 0.600289 | 0.32515 | | 1 | 0.999631 | 0.557085 |
| Add1_F8WHZ9_T645t_2 | 0.13 | 0.68 | 0.55 | -0.1 | 0.01 | 0.61 | -0.14 | -0.17 | 0.08 | 0.83 | -0.51 | 0 | 0 | 0.12 | 0 | 0.82 | -0.2375 | -0.215 | -0.08 | 0.812079 | 0.452334 | 0.556541 | | 1 | 0.999631 | 0.936949 |
| Add1_Q9QYC0_S600s_1 | -1.33 | -0.33 | -0.69 | -2.54 | 0.1 | -0.95 | -2.06 | -2.37 | -0.63 | -1.24 | -2.07 | -1.43 | 1.89 | -0.4 | -2.63 | -1.11 | -0.0975 | -0.12 | 0.66 | 0.474857 | 0.882895 | 0.8214 | | 1 | 0.999631 | 0.799827 |
| Add1_Q9QYC0_S628s_1 | 0.08 | 1.59 | 2.05 | 0.98 | 0.48 | 1.83 | 0.34 | 1.05 | 1.19 | 1.65 | 0.92 | 1.47 | 1.56 | 1.6 | 1.45 | 2.05 | -0.25 | 0.1325 | 0.49 | 0.287286 | 0.624639 | 0.767368 | | 1 | 0.999631 | 0.705972 |
| Add1_Q9QYC0_T610t_1 | -1.64 | -1.97 | -2.92 | -3.58 | -2.07 | -2.24 | -3.02 | -3.6 | -2.06 | -1.6 | -2.67 | -3.24 | -0.73 | -1.72 | -3.63 | -3.3 | -0.205 | 0.135 | 0.1825 | 0.799486 | 0.697946 | 0.799837 | | 1 | 0.999631 | 0.931051 |
| Add1_Q9QYC0_T610t_2 | 0.13 | 0.68 | 0.12 | -0.3 | 0.01 | 0.61 | -0.58 | -0.39 | 0.08 | 0.83 | -0.76 | -0.17 | 0 | 0.12 | -0.26 | 0.07 | -0.245 | -0.1625 | -0.175 | 0.576566 | 0.49738 | 0.68479 | | 1 | 0.999631 | 0.846204 |
| Add1_Q9QYC0_T614t_1 | -1.27 | -0.24 | 0.53 | -0.85 | -1.39 | 0.03 | -0.99 | -0.48 | 0.13 | -0.42 | 0.16 | -0.01 | 0.27 | 0.31 | 0.04 | 0.63 | -0.25 | 0.4225 | 0.77 | 0.092311 | 0.600289 | 0.32515 | | 1 | 0.999631 | 0.557085 |
| Add1_Q9QYC0_T614t_2 | 0.13 | 0.68 | -0.14 | -0.41 | 0.01 | 0.61 | -0.78 | -0.37 | 0.08 | 0.83 | -0.86 | -0.2 | 0 | 0.12 | -0.39 | -0.07 | -0.1975 | -0.1025 | -0.15 | 0.651497 | 0.611683 | 0.808501 | | 1 | 0.999631 | 0.876488 |
| Add2_Q9QYB8_S11s_1 | -0.75 | -0.46 | -0.08 | -0.66 | -0.55 | -1.02 | -0.91 | -0.68 | -0.63 | -0.88 | -0.4 | -0.6 | 0.02 | -0.62 | -0.56 | -0.56 | -0.3025 | -0.14 | 0.0575 | 0.851921 | 0.282672 | 0.616805 | | 1 | 0.999631 | 0.950099 |
| Add2_Q9QYB8_S528s_1 | 1.17 | -1.05 | -1.22 | 0.04 | -0.39 | -0.78 | -0.56 | -0.63 | -0.86 | -0.65 | -1.26 | -1.08 | -0.25 | -0.01 | -0.62 | -0.68 | -0.325 | -0.6975 | -0.125 | 0.816688 | 0.535994 | 0.210018 | | 1 | 0.999631 | 0.937777 |
| Add2_Q9QYB8_S530s_1 | -0.19 | -0.59 | -1.4 | 0.13 | 0.14 | -0.32 | -0.2 | -0.34 | -0.31 | -0.13 | -0.97 | -1.09 | -0.46 | -0.52 | 0 | 0.01 | 0.3325 | -0.1125 | 0.27 | 0.494342 | 0.376755 | 0.785825 | | 1 | 0.999631 | 0.81007 |
| Add2_Q9QYB8_S532s_1 | 0.9 | 0.07 | -0.02 | 0.09 | 0.64 | -0.14 | 0.4 | -0.64 | 0.49 | 0.57 | -0.16 | 0.88 | -0.67 | -0.03 | 0.02 | 0.75 | -0.195 | 0.185 | -0.2425 | 0.538851 | 0.604647 | 0.596827 | | 1 | 0.999631 | 0.830899 |
| Add2_Q9QYB8_S539s_1 | 0.71 | 0.41 | 0.82 | 0.32 | -0.14 | 0.79 | -0.08 | 1.1 | 0.62 | 0.81 | -0.73 | 1.1 | 0.63 | 0.76 | 0.99 | 1.31 | -0.1475 | -0.115 | 0.3575 | 0.248465 | 0.681588 | 0.786604 | | 1 | 0.999631 | 0.685751 |
| Add2_Q9QYB8_S594s_1 | -0.88 | 1.18 | -0.22 | 0.16 | 0.23 | 0.98 | 0 | 0.32 | 1.35 | 0.66 | 0.59 | 0.92 | 2.29 | 1.13 | 0.17 | 1 | 0.3225 | 0.82 | 1.0875 | 0.078775 | 0.488822 | 0.095766 | | 1 | 0.999631 | 0.545479 |
| Add2_Q9QYB8_S598s_1 | 0.8 | 1.77 | 1.03 | 1.11 | 1.49 | 2.12 | 0.04 | 0.58 | 1.24 | 1.91 | -0.06 | 0.08 | 1.79 | 1.99 | -0.01 | -0.45 | -0.12 | -0.385 | -0.3475 | 0.563631 | 0.803237 | 0.441665 | | 1 | 0.999631 | 0.841252 |
| Add2_Q9QYB8_S598s_2 | 0.75 | 1.18 | 0.28 | -0.32 | 1.07 | 1.08 | -0.5 | 0.3 | 0.88 | 0.71 | -0.27 | 0.02 | 0.32 | 1.42 | -0.18 | 0.07 | 0.015 | -0.1375 | -0.065 | 0.889565 | 0.974605 | 0.746477 | | 1 | 0.999631 | 0.963505 |
| Add2_Q9QYB8_S598s_3 | 0.42 | 1.37 | 1.27 | 1.68 | 1.22 | 1.3 | 0.99 | 1.63 | 1.04 | 1.01 | 1.17 | 0.88 | 0.1 | 0.96 | 1.15 | 1.62 | 0.1 | -0.16 | -0.2275 | 0.596475 | 0.767161 | 0.628781 | | 1 | 0.999631 | 0.854508 |
| Add2_Q9QYB8_S602s_1 | -0.83 | -1.07 | 0.71 | -1.12 | -1.92 | -2.09 | -0.84 | -0.57 | -2.13 | -0.6 | -0.17 | -0.11 | 0.54 | -0.95 | -1.12 | -0.46 | -0.7775 | -0.175 | 0.08 | 0.881341 | 0.167586 | 0.765759 | | 1 | 0.999631 | 0.963364 |
| Add2_Q9QYB8_S602s_2 | -1.14 | -0.59 | -0.44 | -1.62 | -1.78 | -1 | -1.47 | -1.46 | -1.26 | -0.4 | -1.32 | -0.85 | 0.48 | -1.08 | -1.08 | -0.89 | -0.48 | -0.01 | 0.305 | 0.510843 | 0.18674 | 0.97848 | | 1 | 0.999631 | 0.817687 |
| Add2_Q9QYB8_S602s_3 | 0.42 | 1.37 | 1.27 | 1.68 | 1.22 | 1.3 | 0.99 | 1.63 | 1.04 | 1.01 | 1.17 | 0.88 | 0.1 | 0.96 | 1.15 | 1.62 | 0.1 | -0.16 | -0.2275 | 0.596475 | 0.767161 | 0.628781 | | 1 | 0.999631 | 0.854508 |
| Add2_Q9QYB8_S606s_2 | -1.48 | -0.84 | -0.12 | -1.07 | -2.26 | -1.28 | -1.05 | -0.61 | -1.47 | -0.45 | -0.89 | -0.58 | 0.42 | -1.31 | -0.75 | -0.48 | -0.4225 | 0.03 | 0.3475 | 0.452843 | 0.34771 | 0.937753 | | 1 | 0.999631 | 0.788525 |
| Add2_Q9QYB8_S606s_3 | 0.42 | 1.37 | 1.27 | 1.68 | 1.22 | 1.3 | 0.99 | 1.63 | 1.04 | 1.01 | 1.17 | 0.88 | 0.1 | 0.96 | 1.15 | 1.62 | 0.1 | -0.16 | -0.2275 | 0.596475 | 0.767161 | 0.628781 | | 1 | 0.999631 | 0.854508 |
| Add2_Q9QYB8_S60s_1 | 0.08 | 0.64 | 0.38 | -0.26 | -0.07 | 0.91 | -0.36 | 0.2 | 0.03 | 0.33 | -0.01 | 0.08 | 0.49 | 0.78 | 0.58 | 0.72 | -0.04 | -0.1025 | 0.4325 | 0.177456 | 0.911121 | 0.726521 | | 1 | 0.999631 | 0.642116 |
| Add2_Q9QYB8_S614s_2 | -4.03 | -6.59 | -0.08 | 0.22 | -6.49 | -3.53 | 0.43 | -1.35 | -7.22 | -6.29 | -1.07 | -0.93 | -7.96 | -3.41 | -1.08 | -1.39 | -0.115 | -1.2575 | -0.84 | 0.660613 | 0.950956 | 0.529876 | | 1 | 0.999631 | 0.880315 |
| Add2_Q9QYB8_S618s_1 | -0.41 | -0.13 | 1.12 | -1.67 | -3.16 | -0.94 | -1.47 | -3.29 | -1.94 | 0.64 | -1.6 | -1.05 | 1.87 | -0.67 | -0.99 | -1.38 | -1.9425 | -0.715 | -0.02 | 0.980308 | 0.022365 | 0.33261 | | 1 | 0.999631 | 0.994107 |
| Add2_Q9QYB8_S618s_2 | -4.03 | -6.59 | -0.08 | 0.22 | -6.49 | -3.53 | 0.43 | -1.35 | -7.22 | -6.29 | -1.07 | -0.93 | -7.96 | -3.41 | -1.08 | -1.39 | -0.115 | -1.2575 | -0.84 | 0.660613 | 0.950956 | 0.529876 | | 1 | 0.999631 | 0.880315 |
| Add2_Q9QYB8_S623s_1 | -1.61 | -4.54 | -3.78 | -4.11 | -3.37 | -3.75 | -2.92 | -3.21 | -3.05 | -2.3 | -2.79 | -4.12 | -1.95 | -3.9 | -4.72 | -4.22 | 0.1975 | 0.445 | -0.1875 | 0.809835 | 0.745936 | 0.51577 | | 1 | 0.999631 | 0.936643 |
| Add2_Q9QYB8_S62s_1 | -0.29 | -0.4 | 0.38 | -0.15 | -0.62 | -0.33 | -0.7 | -0.6 | -1.35 | -0.13 | -0.69 | -0.44 | -0.6 | 0.11 | 0.44 | 0.27 | -0.4475 | -0.5375 | 0.17 | 0.625676 | 0.129327 | 0.148682 | | 1 | 0.999631 | 0.864021 |
| Add2_Q9QYB8_S692s_1 | -0.72 | 0.21 | 0.36 | -0.16 | -0.34 | -0.19 | -0.55 | -0.2 | -0.05 | 0.18 | -0.13 | 0.35 | 1.25 | 0.34 | 0.17 | 0.37 | -0.2425 | 0.165 | 0.61 | 0.131118 | 0.441416 | 0.610891 | | 1 | 0.999631 | 0.602916 |
| Add2_Q9QYB8_S692s_2 | -3.1 | -3.4 | 0.85 | 0.04 | -3.67 | -3.36 | 0.23 | 0.2 | -4.86 | -2.22 | 0.43 | 0.53 | -4.57 | -3.29 | 0.17 | 1.08 | -0.2475 | -0.1275 | -0.25 | 0.863583 | 0.84899 | 0.928705 | | 1 | 0.999631 | 0.955131 |
| Add2_Q9QYB8_S696s_1 | -0.19 | 0.88 | 1.09 | 0.16 | 0.25 | 1.1 | -0.4 | 0.62 | 0.44 | 0.64 | 0.11 | 0.68 | 1.92 | 0.93 | 0.67 | 0.94 | -0.0925 | -0.0175 | 0.63 | 0.156924 | 0.829228 | 0.961246 | | 1 | 0.999631 | 0.626839 |
| Add2_Q9QYB8_S700s_1 | -2.89 | -5.49 | -3.01 | -2.42 | -4.08 | -5.08 | -2.53 | -2.43 | -4.81 | -3.79 | -2.55 | -2.6 | -3.4 | -4.76 | -2.83 | -2.66 | -0.0775 | 0.015 | 0.04 | 0.956771 | 0.924712 | 0.984489 | | 1 | 0.999631 | 0.984803 |
| Add2_Q9QYB8_T561t_1 | -0.74 | -0.11 | -0.66 | -0.86 | -0.71 | -0.62 | -0.72 | -0.57 | -0.54 | -1 | -0.32 | 0.13 | 0.18 | -0.24 | -0.62 | -0.27 | -0.0625 | 0.16 | 0.355 | 0.280069 | 0.814719 | 0.635777 | | 1 | 0.999631 | 0.701922 |
| Add3_Q9QYB5_S42s_1 | -0.39 | 0.23 | -0.1 | -0.04 | -0.28 | 0.34 | -0.16 | 0.08 | 0.44 | 0.5 | -0.02 | -0.04 | 0.47 | 0.81 | -0.05 | 0.25 | 0.07 | 0.295 | 0.445 | 0.176122 | 0.798844 | 0.313877 | | 1 | 0.999631 | 0.642116 |
| Add3_Q9QYB5_S461s_1 | 0.67 | -1.18 | 0.4 | -0.06 | -1.96 | -0.33 | 1.7 | -1.54 | -1.52 | 0.89 | -1.18 | -1.37 | -1.61 | -1.77 | -1.34 | -1.89 | -0.49 | -0.7525 | -1.61 | 0.00379 | 0.544191 | 0.250434 | | 1 | 0.999631 | 0.351308 |
| Add3_Q9QYB5_S641s_1 | -0.88 | -1.04 | -0.81 | -1.09 | -0.95 | -0.9 | -0.9 | -0.75 | -0.61 | -0.41 | -0.59 | -0.77 | -0.49 | -0.65 | -0.49 | -0.27 | 0.08 | 0.36 | 0.48 | 0.097 | 0.734275 | 0.170467 | | 1 | 0.999631 | 0.55886 |
| Add3_Q9QYB5_S645s_1 | 0.06 | 0.8 | 0.62 | 0.19 | 0.25 | 0.34 | -0.11 | 0.24 | 0.17 | 0.86 | 0.34 | 0.54 | 1.08 | 0.52 | 0.11 | 0.65 | -0.2375 | 0.06 | 0.1725 | 0.608923 | 0.407241 | 0.84303 | | 1 | 0.999631 | 0.858851 |
| Add3_Q9QYB5_S649s_1 | -1.85 | -3.52 | -3.05 | -1.96 | -2.77 | -4.36 | -1.77 | -2.29 | -3.93 | -3.27 | -1.8 | -2.68 | -2.52 | -3.66 | -2.18 | -2.58 | -0.2025 | -0.325 | -0.14 | 0.779316 | 0.745741 | 0.567195 | | 1 | 0.999631 | 0.921416 |
| Add3_Q9QYB5_S64s_1 | 0.33 | 1.36 | 1.37 | 0.72 | 1.03 | 0.94 | 0.35 | 1.1 | 0.95 | 1.41 | 0.62 | 1.09 | 0.33 | 0.86 | 1.58 | 1.69 | -0.09 | 0.0725 | 0.17 | 0.68798 | 0.79245 | 0.833788 | | 1 | 0.999631 | 0.891733 |
| Add3_Q9QYB5_T12t_1 | -0.88 | -0.55 | -0.55 | -0.96 | -0.92 | -0.92 | -1.13 | -0.84 | -1.19 | -0.48 | -1.3 | -0.95 | 0.11 | -0.81 | -0.81 | -0.35 | -0.2175 | -0.245 | 0.27 | 0.413628 | 0.391112 | 0.413095 | | 1 | 0.999631 | 0.768649 |
| Adgra1_Q8C4G9_S537s_1 | 0.11 | -0.9 | -0.35 | 1.78 | 0.44 | -2.02 | -0.21 | 0.46 | 0.32 | -0.56 | -0.04 | -0.08 | -0.82 | -0.14 | -1.29 | 0.43 | -0.4925 | -0.25 | -0.615 | 0.336856 | 0.501642 | 0.656433 | | 1 | 0.999631 | 0.730499 |
| Adgra1_Q8C4G9_T239t_1 | -1.44 | 2.22 | 0.6 | -0.98 | 2.03 | 1.15 | 0.7 | 0.46 | 3.86 | 0.21 | 0.96 | 1.53 | 3.19 | 1.26 | 1.05 | -0.21 | 0.985 | 1.54 | 1.2225 | 0.210561 | 0.229755 | 0.143603 | | 1 | 0.999631 | 0.664726 |
| Adgrb1_E9Q4H5_S336s_1 | -0.01 | -0.05 | -0.24 | -0.23 | 0.33 | 0.73 | -0.6 | -0.23 | -0.29 | -0.13 | 0.57 | -0.63 | -0.12 | 0.06 | -0.34 | 0.84 | 0.19 | 0.0125 | 0.2425 | 0.473768 | 0.576403 | 0.968767 | | 1 | 0.999631 | 0.799779 |
| Adgrb1_E9Q4H5_S349s_1 | -0.57 | 0.87 | 0.33 | -1.34 | 1.29 | 0.67 | 1.57 | -0.15 | 1.52 | -0.59 | 1.44 | 1.24 | 2.32 | 0.01 | -0.43 | -1.1 | 1.0225 | 1.08 | 0.3775 | 0.629105 | 0.095571 | 0.112409 | | 1 | 0.999631 | 0.866534 |
| Adgrb1_E9Q4H5_S364s_1 | 0.4 | 0.82 | 1 | 1.13 | 1.09 | 1.25 | 0.33 | 1.08 | 0.8 | 0.13 | 0.46 | 1.47 | 0.3 | 0.86 | 1.48 | 1.78 | 0.1 | -0.1225 | 0.2675 | 0.502123 | 0.750058 | 0.73467 | | 1 | 0.999631 | 0.814092 |
| Adgrb1_E9Q4H5_S527s_1 | -1.28 | 0.51 | -0.37 | -1.4 | 0.46 | -0.26 | -0.38 | -1.16 | 0.1 | -0.98 | -0.45 | 0.22 | -0.48 | -0.88 | -0.95 | -1.15 | 0.3 | 0.3575 | -0.23 | 0.6201 | 0.564845 | 0.478683 | | 1 | 0.999631 | 0.861888 |
| Adgrb1_E9Q4H5_T308t_1 | 0.67 | 0.81 | -0.42 | 1.59 | 1.01 | 1.02 | 1.14 | 1.67 | -0.65 | -1.63 | 1.2 | -0.6 | -0.31 | 0.34 | -0.21 | 0.92 | 0.5475 | -1.0825 | -0.4775 | 0.337382 | 0.225334 | 0.11977 | | 1 | 0.999631 | 0.730499 |
| Adgrb2_B1ASC0_S1131s_1 | 1.6 | -0.84 | 0.62 | 0.78 | -1.3 | 0.74 | 1.14 | 0.34 | 0.19 | 0.06 | 0 | -0.44 | 0.33 | -1.06 | 0.66 | -0.62 | -0.31 | -0.5875 | -0.7125 | 0.246037 | 0.639268 | 0.256487 | | 1 | 0.999631 | 0.685751 |
| Adgrb2_B1ASC0_S1415s_1 | -0.86 | -1.54 | -1.8 | -1.33 | -1.63 | -1.06 | -1.63 | -1.38 | -1.69 | -0.74 | -1.28 | -1.7 | -0.83 | -1.13 | -1.35 | -1.4 | -0.0425 | 0.03 | 0.205 | 0.526697 | 0.888045 | 0.930501 | | 1 | 0.999631 | 0.824261 |
| Adgrl1_H7BX15_S1128s_1 | 0.52 | -0.97 | -0.86 | -0.35 | -0.88 | -0.43 | -0.22 | -0.72 | -0.67 | 0.18 | -0.51 | -0.72 | -0.66 | -0.75 | -0.89 | -0.64 | -0.1475 | -0.015 | -0.32 | 0.408933 | 0.700761 | 0.970604 | | 1 | 0.999631 | 0.766996 |
| Adgrl1_H7BX15_S1146s_1 | -0.33 | 0.97 | 0.33 | 1.64 | 1.87 | 1.76 | 2.41 | 1.55 | 2.47 | -0.65 | 2.26 | 2.08 | -1.19 | -0.04 | 1.51 | -0.68 | 1.245 | 0.8875 | -0.7525 | 0.262568 | 0.018398 | 0.253485 | | 1 | 0.999631 | 0.693034 |
| Adgrl1_H7BX15_S1171s_1 | -0.76 | 0.09 | -0.66 | -0.55 | 0.17 | -0.04 | -0.73 | -0.34 | 0.06 | -0.16 | -0.4 | -0.6 | 0.59 | 0.32 | -1.28 | -0.91 | 0.235 | 0.195 | 0.15 | 0.75563 | 0.470382 | 0.531335 | | 1 | 0.999631 | 0.913663 |
| Adgrl1_H7BX15_S1213s_1 | -0.95 | -0.41 | -2.58 | -2.15 | -0.67 | -0.66 | -2.23 | -1.99 | -0.16 | -1.21 | -2.28 | -2.21 | -0.32 | -0.49 | -1.26 | -3.28 | 0.135 | 0.0575 | 0.185 | 0.803955 | 0.820667 | 0.928442 | | 1 | 0.999631 | 0.93396 |
| Adgrl1_H7BX15_S1219s_1 | -1.08 | -1.58 | -1.06 | -0.53 | -1.36 | -2.82 | -1.32 | -1.28 | -1.62 | -2.11 | -0.81 | -1.65 | -0.54 | -2.78 | -0.92 | -1.25 | -0.6325 | -0.485 | -0.31 | 0.546866 | 0.160579 | 0.211981 | | 1 | 0.999631 | 0.834213 |
| Adgrl1_H7BX15_S1222s_1 | -0.02 | -1.72 | -0.86 | -0.29 | -0.52 | -1.34 | -0.41 | -0.47 | -1.27 | -1.61 | -0.24 | 0.08 | -0.63 | -1.06 | 0.02 | -0.6 | 0.0375 | -0.0375 | 0.155 | 0.72536 | 0.930115 | 0.942292 | | 1 | 0.999631 | 0.90568 |
| Adgrl1_H7BX15_S1374s_1 | -0.27 | -1.14 | 0.4 | -0.37 | -0.53 | -0.15 | -0.5 | -0.8 | -0.29 | -0.07 | -1.19 | -0.14 | 0.35 | 0.24 | -0.11 | -0.1 | -0.15 | -0.0775 | 0.44 | 0.258236 | 0.68146 | 0.851693 | | 1 | 0.999631 | 0.692624 |
| Adgrl1_H7BX15_S1384s_1 | 0.23 | 1.1 | -1.75 | 0.54 | 2.46 | 0.28 | 0.6 | 0.42 | 1.76 | -1.04 | 1.22 | 0.22 | -0.56 | 1.17 | -1.29 | 0.07 | 0.91 | 0.51 | -0.1825 | 0.798937 | 0.219495 | 0.512554 | | 1 | 0.999631 | 0.930937 |
| Adgrl1_H7BX15_S1444s_1 | -0.49 | 0 | -0.22 | -0.17 | -0.35 | -0.67 | -0.3 | -0.41 | -0.84 | -1.06 | -0.66 | -0.76 | 0.09 | -0.34 | 0.14 | -0.22 | -0.2125 | -0.61 | 0.1375 | 0.630487 | 0.405587 | 0.038451 | | 1 | 0.999631 | 0.867509 |
| Adgrl1_H7BX15_T1172t_1 | -0.91 | -1.82 | -0.22 | 0.19 | -1.14 | -1.85 | -0.52 | -0.05 | -1.9 | -2.92 | -0.51 | -0.21 | -0.19 | -1.14 | 0.32 | -0.41 | -0.2 | -0.695 | 0.335 | 0.514693 | 0.712562 | 0.324373 | | 1 | 0.999631 | 0.819375 |
| Adgrl2_Q0P6A0_S145s_1 | -0.59 | 0.16 | -0.09 | -0.03 | 0.23 | -0.24 | -0.58 | -0.07 | 0.01 | -0.39 | 0.09 | 0.55 | 0.68 | 0.39 | -0.04 | 0.42 | -0.0275 | 0.2025 | 0.5 | 0.131792 | 0.926357 | 0.524105 | | 1 | 0.999631 | 0.603456 |
| Adgrl2_Q0P6A0_S175s_1 | -1.38 | -0.24 | -0.12 | -0.42 | 0.42 | -0.25 | -0.59 | 0.09 | -0.02 | -0.34 | 0.12 | 0.07 | 0.07 | 0.56 | -0.45 | 0.15 | 0.4575 | 0.4975 | 0.6225 | 0.130719 | 0.241395 | 0.173178 | | 1 | 0.999631 | 0.602177 |
| Adgrl2_Q0P6A0_S180s_1 | -0.83 | 0.33 | -0.72 | -0.3 | -0.14 | 0.04 | -0.31 | -0.59 | -0.45 | -0.01 | -0.04 | 0.19 | 0.18 | 0.17 | 0.34 | 0.43 | 0.13 | 0.3025 | 0.66 | 0.07414 | 0.698 | 0.384187 | | 1 | 0.999631 | 0.539897 |
| Adgrl3_F6WMY5_S558s_1 | 1.75 | -2.55 | -1.91 | -0.27 | -0.68 | -0.97 | 0.56 | -1.88 | -1.53 | -0.71 | -1.61 | -2.09 | -1.64 | -2.05 | -1.59 | -3.87 | 0.0025 | -0.74 | -1.5425 | 0.126492 | 0.997867 | 0.405769 | | 1 | 0.999631 | 0.596745 |
| Adgrl3_F6WMY5_S576s_1 | -0.03 | -0.27 | 0.6 | 0.8 | 0.58 | 0.54 | 2.84 | 0.33 | 0.66 | -1.54 | 0.82 | 0.94 | -1.74 | -1.99 | 0.11 | -1.24 | 0.7975 | -0.055 | -1.49 | 0.014033 | 0.193551 | 0.925289 | | 1 | 0.999631 | 0.385092 |
| Adgrl3_F6WMY5_S590s_1 | -1.06 | -1.09 | -0.88 | -0.47 | -0.74 | -1.47 | -0.67 | -0.96 | -1.17 | -1.27 | -1.51 | -0.63 | -1.41 | -0.95 | 0.35 | -0.37 | -0.085 | -0.27 | 0.28 | 0.509451 | 0.77465 | 0.386953 | | 1 | 0.999631 | 0.816932 |
| Adgrl3_F6WMY5_S776s_1 | -0.29 | 1.43 | -0.33 | -0.3 | 0.75 | 1.04 | -0.33 | -0.82 | 1.2 | 0.05 | 0.44 | 0.03 | 0.1 | 1.7 | -0.98 | -1.28 | 0.0325 | 0.3025 | -0.2425 | 0.733038 | 0.954048 | 0.540807 | | 1 | 0.999631 | 0.907434 |
| Adgrl3_F6WMY5_S821s_1 | 0.68 | 1.32 | 0.72 | 1.34 | 1.19 | 1.85 | 0.5 | 1.51 | 1.49 | 1.46 | 1.13 | 0.72 | -0.03 | 1.75 | 1.31 | 0.96 | 0.2475 | 0.185 | -0.0175 | 0.967439 | 0.500299 | 0.563404 | | 1 | 0.999631 | 0.988504 |
| Adgrl3_F6WMY5_S858s_1 | -0.32 | 1.72 | 2.85 | 1.34 | 0.42 | 2.33 | 3.5 | 1.48 | 1.25 | -0.07 | 1.99 | 2.41 | 2.13 | 0.48 | 2.81 | 0.5 | 0.535 | -0.0025 | 0.0825 | 0.91477 | 0.512421 | 0.997347 | | 1 | 1 | 0.972775 |
| Adh6a_E9Q4Y1_T3t_3 | -1.92 | -0.59 | -0.61 | -1.55 | -0.11 | -1.11 | -0.75 | -1.53 | -0.57 | -1.8 | -0.72 | -0.04 | 0.58 | -1.53 | -0.55 | -0.63 | 0.2925 | 0.385 | 0.635 | 0.23856 | 0.510481 | 0.430038 | | 1 | 0.999631 | 0.683403 |
| Adh6a_E9Q4Y1_T7t_3 | -1.92 | -0.59 | -0.61 | -1.55 | -0.11 | -1.11 | -0.75 | -1.53 | -0.57 | -1.8 | -0.72 | -0.04 | 0.58 | -1.53 | -0.55 | -0.63 | 0.2925 | 0.385 | 0.635 | 0.23856 | 0.510481 | 0.430038 | | 1 | 0.999631 | 0.683403 |
| Adh6a_E9Q4Y1_T9t_3 | -1.92 | -0.59 | -0.61 | -1.55 | -0.11 | -1.11 | -0.75 | -1.53 | -0.57 | -1.8 | -0.72 | -0.04 | 0.58 | -1.53 | -0.55 | -0.63 | 0.2925 | 0.385 | 0.635 | 0.23856 | 0.510481 | 0.430038 | | 1 | 0.999631 | 0.683403 |
| Adra1d_A2ANQ2_S444s_1 | 1.71 | 0.34 | 0.19 | 0.73 | 0.06 | 1.16 | 1.17 | 0.63 | 0.35 | 1.07 | -0.32 | 0.09 | -1.28 | -0.77 | -0.33 | -0.4 | 0.0125 | -0.445 | -1.4375 | 0.00611 | 0.976544 | 0.328795 | | 1 | 0.999631 | 0.351545 |
| Adra1d_P97714_S281s_1 | 1.05 | 0.14 | -0.14 | 1.28 | 0.82 | 0 | 0.99 | 0.72 | 0.53 | -0.08 | 0.86 | 0.55 | -1.06 | 0.32 | -0.16 | 0.47 | 0.05 | -0.1175 | -0.69 | 0.168693 | 0.902491 | 0.772684 | | 1 | 0.999631 | 0.637839 |
| Adra2a_Q01338_S255s_1 | 0.42 | -0.48 | -0.24 | -0.29 | -0.55 | -0.4 | -0.16 | 0.68 | -0.25 | 0.05 | -0.25 | -0.85 | -0.46 | -0.48 | 0.03 | -0.86 | 0.04 | -0.1775 | -0.295 | 0.389767 | 0.911658 | 0.589815 | | 1 | 0.999631 | 0.758766 |
| Adra2a_Q01338_S339s_1 | 0.06 | -0.17 | -0.07 | 0.48 | 0.36 | -0.43 | 0.26 | 0.47 | -0.29 | -0.53 | 0.63 | 0.07 | 0.16 | -0.19 | 0.18 | 1.01 | 0.09 | -0.105 | 0.215 | 0.542411 | 0.769242 | 0.755996 | | 1 | 0.999631 | 0.832881 |
| Adra2a_Q01338_S346s_1 | -0.64 | -0.91 | -1.2 | -1.33 | -1.29 | -1.05 | -1.19 | -0.97 | -0.62 | 0.16 | -1.94 | -1.07 | 0.09 | -0.56 | -0.43 | -1.37 | -0.105 | 0.1525 | 0.4525 | 0.247699 | 0.694601 | 0.737758 | | 1 | 0.999631 | 0.685751 |
| Adra2a_Q01338_S360s_1 | 0.47 | -2.21 | -0.76 | -0.46 | -1.44 | -2.17 | 0.17 | -0.18 | -2.02 | -0.29 | -1.13 | -1.31 | -1.32 | -2.09 | -0.8 | -1.56 | -0.165 | -0.4475 | -0.7025 | 0.235187 | 0.811103 | 0.462978 | | 1 | 0.999631 | 0.683301 |
| Adra2a_Q01338_S362s_1 | 0.46 | -1.77 | -0.65 | -1.59 | -0.58 | -2.35 | 0.27 | 0.54 | -1.57 | 0.36 | 0.57 | 0.6 | -1.17 | -1.6 | 0.17 | 0.37 | 0.3575 | 0.8775 | 0.33 | 0.606612 | 0.62638 | 0.202519 | | 1 | 0.999631 | 0.858318 |
| Adra2c_Q01337_S257s_1 | -0.77 | -1.35 | -0.51 | -0.95 | -1.42 | -1.29 | -1.13 | -0.48 | -1.38 | -0.91 | -0.77 | -0.77 | -0.59 | -1.36 | -0.9 | -0.55 | -0.185 | -0.0625 | 0.045 | 0.891698 | 0.567445 | 0.83631 | | 1 | 0.999631 | 0.964276 |
| Adra2c_Q01337_S287s_1 | 0 | -0.49 | -3.95 | -2.92 | 0.08 | -0.87 | -1.76 | -2.54 | 0.16 | 0.33 | -1.15 | -4.27 | -0.81 | -0.18 | -2.74 | -3.18 | 0.5675 | 0.6075 | 0.1125 | 0.912296 | 0.555278 | 0.62094 | | 1 | 0.999631 | 0.972165 |
| Adrb1_P34971_S274s_1 | -0.97 | 0.69 | -0.98 | -2.08 | -0.23 | 0.49 | -2.01 | -1.34 | -0.13 | -0.22 | -1.83 | -1.01 | 0.98 | 0.19 | -1.47 | -1.14 | 0.0625 | 0.0375 | 0.475 | 0.511308 | 0.929411 | 0.952398 | | 1 | 0.999631 | 0.817687 |
| Adrbk1_Q7TS64_S628s_1 | 0.43 | 0.26 | 0.58 | 0.1 | 0.4 | 0.39 | 0.11 | 0.46 | 0.43 | 0.3 | 0.27 | 0.44 | -0.49 | 0.89 | 0.63 | 0.65 | -0.0025 | 0.0175 | 0.0775 | 0.833883 | 0.992029 | 0.944798 | | 1 | 0.999631 | 0.943862 |
| AF529169_Q8K3V7_S118s_1 | 0.7 | -0.1 | -0.06 | 0.55 | 0.33 | -0.11 | 0.72 | 0.19 | 0.37 | 0.8 | 0.11 | 1.18 | -0.17 | 0.25 | -0.19 | -0.15 | 0.01 | 0.3425 | -0.3375 | 0.301711 | 0.974892 | 0.342121 | | 1 | 0.999631 | 0.714303 |
| Afap1_Q80YS6_S549s_1 | 1.15 | 0.05 | 0.73 | 0.81 | 0 | 0.62 | 0.9 | 0.15 | 0.37 | 1.35 | 0.31 | 0.72 | -0.63 | -0.88 | 0.98 | -0.31 | -0.2675 | 0.0025 | -0.895 | 0.078519 | 0.443508 | 0.994502 | | 1 | 0.999631 | 0.544927 |
| Afap1_Q80YS6_S669s_1 | -0.02 | -0.07 | 0.11 | 0.24 | 0.19 | 0.32 | -0.1 | -0.09 | 0.5 | 0.24 | 0.07 | 0.05 | 0.13 | 0.12 | -0.32 | -0.04 | 0.015 | 0.15 | -0.0925 | 0.736589 | 0.951893 | 0.562856 | | 1 | 0.999631 | 0.90765 |
| Afap1l1_Q8BZI0_S329s_1 | 1.36 | 0.83 | 1.1 | 1.56 | 1.56 | 1.55 | 1.2 | 1.38 | 1.26 | -0.03 | 1.15 | 0.64 | -0.13 | 1.75 | 1.19 | 0.79 | 0.21 | -0.4575 | -0.3125 | 0.476574 | 0.445647 | 0.228927 | | 1 | 0.999631 | 0.799947 |
| Afap1l1_Q8BZI0_S747s_1 | 1.2 | 1.22 | 0.49 | 0.78 | 1.37 | 1.26 | 0.13 | 0.06 | 1.29 | 1.2 | 0.37 | 0.65 | 1.17 | 1.54 | 0.5 | 0.2 | -0.2175 | -0.045 | -0.07 | 0.856045 | 0.588792 | 0.892381 | | 1 | 0.999631 | 0.951817 |
| Afap1l2_Q5DTU0_S484s_1 | 0.01 | 1.03 | 0.25 | 0.11 | 0.65 | 0.79 | -0.38 | -0.27 | 0.7 | 0.76 | 0.59 | 0.98 | 1.23 | 0.76 | 1.04 | 0.74 | -0.1525 | 0.4075 | 0.5925 | 0.098615 | 0.6973 | 0.211625 | | 1 | 0.999631 | 0.562298 |
| Aff4_Q9ESC8_S1059s_1 | -0.24 | 0.35 | 0.34 | -0.22 | -0.48 | 0.07 | -0.54 | -0.81 | -0.46 | -0.18 | -0.08 | 0.28 | 0.53 | 0.48 | 0.21 | -0.64 | -0.4975 | -0.1675 | 0.0875 | 0.810421 | 0.128655 | 0.580968 | | 1 | 0.999631 | 0.936878 |
| Aff4_Q9ESC8_S544s_1 | 2.68 | -0.05 | 1.5 | 1.36 | 0 | -0.03 | 2.29 | 0.08 | 0.6 | 1.97 | 0.43 | -0.21 | 0.52 | -1.29 | -0.1 | -0.69 | -0.7875 | -0.675 | -1.7625 | 0.016054 | 0.279916 | 0.313193 | | 1 | 0.999631 | 0.388832 |
| Aff4_Q9ESC8_S614s_1 | 2.57 | 1.58 | 1.94 | 1.16 | 0.78 | 1.62 | 1.44 | 0.63 | 1.11 | 1.78 | 0.15 | 1.26 | 0.26 | 0.93 | 0.26 | -0.06 | -0.695 | -0.7375 | -1.465 | 0.003719 | 0.101524 | 0.12269 | | 1 | 0.999631 | 0.351308 |
| Aff4_Q9ESC8_S615s_1 | 2.57 | 1.58 | 1.94 | 1.16 | 0.78 | 1.62 | 1.44 | 0.63 | 1.11 | 1.78 | 0.15 | 1.26 | 0.26 | 0.93 | 0.26 | -0.06 | -0.695 | -0.7375 | -1.465 | 0.003719 | 0.101524 | 0.12269 | | 1 | 0.999631 | 0.351308 |
| Aftph_H3BKM5_S240s_1 | -0.52 | -0.7 | -0.54 | -0.1 | -0.69 | -0.5 | -0.22 | -0.57 | -0.87 | -1.23 | -0.12 | 0.34 | 0.07 | -0.05 | 0.02 | -0.16 | -0.03 | -0.005 | 0.435 | 0.140993 | 0.909219 | 0.98985 | | 1 | 0.999631 | 0.61245 |
| Aftph_H3BKM5_S525s_1 | -0.35 | 4.68 | 3.52 | 0.21 | 5.67 | 3.16 | 3.69 | 1.65 | 7.62 | 0.12 | 4.92 | 6.74 | 8.15 | 2.43 | 2.81 | 0.81 | 1.5275 | 2.835 | 1.535 | 0.372823 | 0.245542 | 0.131942 | | 1 | 0.999631 | 0.747035 |
| Agap1_E9Q3K3_S124s_1 | -0.72 | -0.45 | 0.13 | -1.23 | -0.76 | -0.61 | -1.2 | -0.1 | -1.83 | -0.32 | -1.56 | 0.3 | -0.23 | -0.77 | -0.63 | -0.3 | -0.1 | -0.285 | 0.085 | 0.813864 | 0.79135 | 0.599835 | | 1 | 0.999631 | 0.937041 |
| Agap1_E9Q3K3_S152s_1 | 2.69 | -0.87 | -0.09 | 1.41 | 1.1 | 0.52 | 2.12 | 0.86 | 0.41 | -0.44 | 0.38 | -1.03 | -1.45 | -0.89 | -1.33 | -0.54 | 0.365 | -0.955 | -1.8375 | 0.027444 | 0.63197 | 0.229647 | | 1 | 0.999631 | 0.423809 |
| Agap1_E9Q3K3_S292s_1 | 0.3 | -1.84 | -1.54 | -1.81 | -1.3 | -1.07 | -1.73 | -2.17 | -0.7 | -0.96 | -2.77 | -1.77 | -1.88 | -2.34 | -1.83 | -2.58 | -0.345 | -0.3275 | -0.935 | 0.093888 | 0.515955 | 0.603516 | | 1 | 0.999631 | 0.558128 |
| Agap1_E9Q3K3_S295s_1 | 0.79 | -1.56 | -1.81 | -0.34 | -0.72 | -1.04 | -0.16 | -0.35 | -0.91 | 0.65 | -0.87 | -2.12 | -1.28 | -1.21 | -1.8 | -1.1 | 0.1625 | -0.0825 | -0.6175 | 0.294233 | 0.777525 | 0.910128 | | 1 | 0.999631 | 0.708293 |
| Agap1_E9Q3K3_S295s_2 | -0.19 | -4.5 | -0.43 | -0.16 | 0.04 | -2.9 | -0.05 | 0.4 | -1.36 | -0.24 | -0.3 | -0.39 | -1.77 | -2.25 | -0.42 | -0.44 | 0.6925 | 0.7475 | 0.1 | 0.919652 | 0.540072 | 0.437973 | | 1 | 0.999631 | 0.974724 |
| Agap1_E9Q3K3_S57s_1 | 3.11 | -0.49 | -0.59 | 1.52 | 1.34 | 0.59 | 2.15 | 0.72 | 0.65 | 0.89 | 0.34 | -0.59 | -1.98 | -0.1 | 0.07 | -1.1 | 0.3125 | -0.565 | -1.665 | 0.079603 | 0.707852 | 0.499188 | | 1 | 0.999631 | 0.545579 |
| Agap1_E9Q3K3_S591s_1 | 2.41 | 0.46 | 0.96 | 1.52 | 0.56 | 1.33 | 2.13 | 0.77 | 1.16 | 1.66 | 1.12 | 0.63 | -1.02 | 0.04 | 0.93 | 0.5 | -0.14 | -0.195 | -1.225 | 0.047123 | 0.783619 | 0.670674 | | 1 | 0.999631 | 0.480102 |
| Agap1_E9Q3K3_S674s_1 | -1.26 | -0.59 | -0.29 | -0.21 | -0.36 | -0.26 | 0.23 | 1.06 | -0.56 | -0.93 | 0.6 | 0.45 | -0.03 | 0 | 1.35 | 0.46 | 0.755 | 0.4775 | 1.0325 | 0.030323 | 0.08695 | 0.293126 | | 1 | 0.999631 | 0.43753 |
| Agap1_E9Q3K3_S720s_1 | 0.85 | 0 | -0.18 | 0.63 | 0.62 | 0.85 | 0.34 | 1.13 | 0.19 | 0.43 | -0.11 | 0.02 | -0.58 | 0.73 | 0.39 | 0.22 | 0.41 | -0.1925 | -0.135 | 0.734574 | 0.240534 | 0.559723 | | 1 | 0.999631 | 0.907462 |
| Agap1_E9Q3K3_S92s_1 | 0.96 | 0.08 | -0.39 | 1.24 | 1.28 | 0.81 | 0.61 | 0.99 | 1.71 | 0.84 | 0.72 | 0.72 | -0.53 | 1.02 | 0.97 | 0.92 | 0.45 | 0.525 | 0.1225 | 0.809643 | 0.283847 | 0.253085 | | 1 | 0.999631 | 0.936634 |
| Agap1_E9Q3K3_T1035t_1 | 0.61 | -1.34 | -1.13 | 1.23 | 1.96 | -1.1 | 1.17 | 0.85 | 0.4 | -0.79 | 0.3 | -1.04 | -2.77 | -0.18 | -0.26 | 0.44 | 0.8775 | -0.125 | -0.535 | 0.522614 | 0.285096 | 0.849834 | | 1 | 0.999631 | 0.821224 |
| Agap1_E9Q3K3_T224t_1 | 0.33 | 0.53 | 0.48 | 1.15 | 1.02 | 0.83 | 0.56 | 1.21 | 1.41 | 0.75 | 1.59 | 1.15 | 0.48 | 1.37 | 1.02 | 1.18 | 0.2825 | 0.6025 | 0.39 | 0.259314 | 0.350206 | 0.082736 | | 1 | 0.999631 | 0.692624 |
| Agap2_Q3UHD9_S1148s_1 | -0.53 | 0.71 | 0.94 | -0.46 | 0.33 | 0.34 | -0.12 | -0.1 | 0.41 | -0.31 | -0.53 | -1.24 | 0.2 | 1.27 | 0.86 | 0.84 | -0.0525 | -0.5825 | 0.6275 | 0.179617 | 0.897216 | 0.251796 | | 1 | 0.999631 | 0.64376 |
| Agap2_Q3UHD9_S1171s_1 | 0.92 | 0.1 | 0.93 | 1.6 | 0.13 | 0.37 | 1.45 | 1 | 0.32 | 1.02 | 1.33 | 0.54 | -1.23 | -0.14 | 1.67 | 1.28 | -0.15 | -0.085 | -0.4925 | 0.460312 | 0.723987 | 0.830402 | | 1 | 0.999631 | 0.791962 |
| Agap2_Q3UHD9_S1171s_2 | 1.67 | 0.83 | -1.12 | 1.99 | 2.4 | 2.04 | 2 | 1.5 | 1.72 | 1.34 | 1.39 | -0.12 | -2.42 | 0.52 | 0.19 | 0.77 | 1.1425 | 0.24 | -1.0775 | 0.237591 | 0.101851 | 0.738978 | | 1 | 0.999631 | 0.683301 |
| Agap2_Q3UHD9_S1175s_2 | 1.69 | 0.89 | -1.12 | 1.99 | 2.55 | 2.16 | 2 | 1.5 | 1.66 | 1.35 | 1.39 | -0.12 | -2.93 | 0.51 | 0.19 | 0.77 | 1.19 | 0.2075 | -1.2275 | 0.217344 | 0.093931 | 0.772673 | | 1 | 0.999631 | 0.669707 |
| Agap2_Q3UHD9_S149s_1 | -1 | 0.55 | 0.82 | -0.86 | -0.82 | -1.07 | -0.22 | -0.91 | -0.57 | -0.14 | 0.31 | -0.17 | 1.22 | -0.54 | -1.16 | 0.63 | -0.6325 | -0.02 | 0.16 | 0.804092 | 0.207455 | 0.966914 | | 1 | 0.999631 | 0.934014 |
| Agap2_Q3UHD9_S168s_1 | 2.39 | -2.69 | -0.68 | 0.26 | 0.95 | -1.47 | 1.5 | -0.47 | -1.27 | -1.88 | -1.5 | -2.78 | -2.78 | -2.35 | -0.6 | -1.63 | 0.3075 | -1.6775 | -1.66 | 0.117846 | 0.774829 | 0.104836 | | 1 | 0.999631 | 0.585087 |
| Agap2_Q3UHD9_S211s_1 | 0.11 | 0.71 | 0.24 | -0.07 | 0.3 | 1.33 | -0.14 | 0.44 | 0.65 | 0.71 | 0.45 | 0.17 | 1.11 | 1.37 | 0.25 | 0.89 | 0.235 | 0.2475 | 0.6575 | 0.083841 | 0.528985 | 0.404551 | | 1 | 0.999631 | 0.549732 |
| Agap2_Q3UHD9_S26s_1 | -0.33 | -0.34 | -0.51 | -1.42 | 0.25 | -0.08 | -0.47 | -1.21 | -0.25 | 0.26 | -1.04 | -1.2 | -0.22 | 0.52 | 0.16 | -0.89 | 0.2725 | 0.0925 | 0.5425 | 0.210164 | 0.509588 | 0.82957 | | 1 | 0.999631 | 0.66441 |
| Agap2_Q3UHD9_S33s_1 | 0.5 | -0.3 | -0.61 | 0.19 | 0.83 | -0.08 | 0.66 | 0.34 | -0.16 | -0.35 | -0.05 | -0.43 | -0.35 | -0.16 | -0.44 | -0.42 | 0.4925 | -0.1925 | -0.2875 | 0.392722 | 0.17964 | 0.552652 | | 1 | 0.999631 | 0.758901 |
| Agap2_Q3UHD9_S349s_1 | -0.41 | 1.58 | 1.53 | -0.47 | 0.07 | -0.01 | -0.29 | -0.79 | 0.32 | 0.51 | 0.76 | 1.6 | 1.6 | -0.12 | -0.23 | 1.83 | -0.8125 | 0.24 | 0.2125 | 0.763275 | 0.167233 | 0.683722 | | 1 | 0.999631 | 0.915608 |
| Agap2_Q3UHD9_S368s_1 | -0.31 | 0.06 | -1.97 | 0.52 | 0.18 | 0.01 | 0.17 | 0.86 | -0.27 | -0.84 | 0.18 | 0.36 | -1.89 | -0.37 | -1.18 | -0.38 | 0.73 | 0.2825 | -0.53 | 0.384035 | 0.190609 | 0.615278 | | 1 | 0.999631 | 0.754721 |
| Agap2_Q3UHD9_S424s_1 | -1.14 | -0.72 | -0.91 | -0.8 | -0.36 | -0.39 | -1.05 | -0.24 | -0.17 | -1.13 | -0.38 | 0.09 | -0.21 | -0.27 | 0.05 | -0.05 | 0.3825 | 0.495 | 0.7725 | 0.016262 | 0.197223 | 0.156939 | | 1 | 0.999631 | 0.388832 |
| Agap2_Q3UHD9_S632s_1 | 0.29 | 1.16 | -0.2 | -0.23 | 1.38 | 0.14 | 0.51 | 0.28 | -1.46 | -0.81 | 0.25 | 0.93 | 0.51 | 1.4 | 0.87 | 1.35 | 0.3225 | -0.5275 | 0.7775 | 0.080164 | 0.451998 | 0.368772 | | 1 | 0.999631 | 0.545579 |
| Agap2_Q3UHD9_S642s_1 | 2.12 | 0.88 | 0.85 | 2.31 | 2.71 | 1.02 | 2.61 | 2.51 | 1.45 | 0.23 | 2.03 | 1.54 | -0.1 | 1.11 | 0.41 | 1.72 | 0.6725 | -0.2275 | -0.755 | 0.173302 | 0.215635 | 0.660364 | | 1 | 0.999631 | 0.642116 |
| Agap2_Q3UHD9_S650s_1 | 2.37 | -0.35 | 0.09 | -0.62 | -1.27 | 0.46 | 3.25 | -3.9 | -3.04 | 1.54 | -2.49 | -0.25 | -0.08 | -2.33 | -0.73 | -0.8 | -0.7375 | -1.4325 | -1.3575 | 0.088278 | 0.60106 | 0.205745 | | 1 | 0.999631 | 0.553858 |
| Agap2_Q3UHD9_S921s_1 | -1 | -0.3 | -0.85 | -0.79 | -0.02 | -0.4 | -1.02 | -0.59 | -0.6 | -1.15 | -0.71 | -0.7 | -0.29 | -0.36 | -0.56 | -0.46 | 0.2275 | -0.055 | 0.3175 | 0.283911 | 0.471376 | 0.847355 | | 1 | 0.999631 | 0.704942 |
| Agap2_Q3UHD9_S99s_1 | 0.86 | -1.53 | -3.62 | -1.03 | 0.29 | -1.36 | -0.34 | 0.83 | 0.78 | -0.93 | -0.67 | -0.58 | -0.53 | 0.12 | -0.35 | -2.11 | 1.185 | 0.98 | 0.6125 | 0.499668 | 0.206095 | 0.275865 | | 1 | 0.999631 | 0.813279 |
| Agap2_Q3UHD9_T350t_1 | -0.26 | -1.53 | -0.44 | -0.53 | -0.29 | -0.6 | -0.94 | -0.5 | -0.28 | 0.21 | -0.56 | 0.09 | -0.77 | -0.64 | -0.51 | 0.11 | 0.1075 | 0.555 | 0.2375 | 0.537359 | 0.75721 | 0.152355 | | 1 | 0.999631 | 0.830391 |
| Agap2_Q3UHD9_T629t_1 | 0.45 | -1.14 | -1.98 | -0.76 | 0.36 | -1.4 | -1.11 | -0.68 | -1.08 | -0.75 | -1.4 | -0.83 | 0.2 | -1.22 | -1.38 | -2.01 | 0.15 | -0.1575 | -0.245 | 0.69461 | 0.795283 | 0.752725 | | 1 | 0.999631 | 0.895302 |
| Agap2_Q3UHD9_T803t_1 | 0.08 | 1.2 | 0.54 | -0.12 | 0.8 | 0.26 | -0.33 | -0.23 | 1.06 | -0.12 | -0.07 | 0.29 | 2.05 | 0.83 | -0.3 | 0.9 | -0.3 | -0.135 | 0.445 | 0.408797 | 0.456581 | 0.741619 | | 1 | 0.999631 | 0.766996 |
| Agap3_Q8VHH5_S395s_1 | 1.93 | -0.1 | -0.89 | 2.37 | 1.24 | 0.37 | 1.91 | 1.19 | 0.5 | 0.26 | 2.03 | -0.03 | -2.17 | -0.59 | -0.66 | -0.41 | 0.35 | -0.1375 | -1.785 | 0.041113 | 0.639878 | 0.863345 | | 1 | 0.999631 | 0.474979 |
| Agap3_Q8VHH5_S478s_1 | 1.94 | -0.58 | -1.33 | 0.98 | 2.59 | 0.54 | 1.45 | 1.53 | 0.81 | -0.12 | 0.76 | -1.31 | -2.98 | 0.29 | -0.65 | -1.23 | 1.275 | -0.2175 | -1.395 | 0.133859 | 0.112971 | 0.781689 | | 1 | 0.999631 | 0.60492 |
| Agap3_Q8VHH5_S573s_1 | 0.95 | 0.79 | 0.13 | 0.64 | 0.61 | 1.13 | 0.32 | 0.59 | 1.35 | 1.19 | -0.01 | 0.77 | 0.14 | 0.85 | 0.83 | 0.45 | 0.035 | 0.1975 | -0.06 | 0.853334 | 0.908487 | 0.602171 | | 1 | 0.999631 | 0.950701 |
| Agfg1_Q8K2K6_T177t_2 | 2.3 | 0.85 | -0.39 | 2.67 | 2.45 | 2.05 | 2.77 | 2.26 | 1.96 | 0.65 | 2.22 | 0.9 | -2.67 | 1.38 | 0.66 | 0.76 | 1.025 | 0.075 | -1.325 | 0.199843 | 0.135176 | 0.916112 | | 1 | 0.999631 | 0.657804 |
| Agps_Q8C0I1_S52s_1 | -1.11 | -1.98 | -0.91 | -0.34 | -1.82 | -2.21 | -0.58 | -0.26 | -2.17 | -1.72 | -0.29 | 0.11 | -0.54 | -1.84 | -0.53 | -1.13 | -0.1325 | 0.0675 | 0.075 | 0.869427 | 0.805061 | 0.908918 | | 1 | 0.999631 | 0.957915 |
| Agps_Q8C0I1_S57s_1 | -0.74 | -0.64 | -0.05 | -0.49 | -0.45 | -0.55 | -0.6 | -0.26 | -1.04 | -1.42 | -0.33 | -0.05 | -0.41 | -0.61 | -0.25 | -0.38 | 0.015 | -0.23 | 0.0675 | 0.815929 | 0.955074 | 0.543621 | | 1 | 0.999631 | 0.93761 |
| Ahctf1_Q8CJF7_S1218s_1 | -0.48 | -0.68 | -0.6 | -1.09 | -0.46 | -1.38 | -0.62 | -0.76 | -0.64 | -0.36 | 0.34 | -0.1 | 0.64 | -0.34 | 0.13 | 0.7 | -0.0925 | 0.5225 | 0.995 | 0.01442 | 0.759939 | 0.119066 | | 1 | 0.999631 | 0.385092 |
| Ahctf1_Q8CJF7_S1381s_1 | 1.03 | 1.57 | -1.88 | -1.03 | 0.7 | 2.08 | -1.39 | -2.92 | 1.64 | 1.98 | -1.12 | -2.13 | 1.93 | 1.65 | -2.89 | -1.58 | -0.305 | 0.17 | -0.145 | 0.905729 | 0.795682 | 0.879442 | | 1 | 0.999631 | 0.969317 |
| Ahctf1_Q8CJF7_S1507s_1 | -0.63 | -0.23 | -0.97 | -0.75 | -0.85 | -1.11 | -1.06 | -0.64 | -1.1 | -0.86 | -1.57 | -2.73 | -0.11 | -0.96 | -1.23 | -0.67 | -0.27 | -0.92 | -0.0975 | 0.778426 | 0.338983 | 0.05939 | | 1 | 0.999631 | 0.921416 |
| Ahcyl2_F8WGT1_S109s_1 | -2.05 | -0.21 | -0.04 | -0.52 | -0.37 | -0.13 | -0.4 | 0.06 | -0.3 | -0.9 | -0.32 | 0.32 | 1.05 | 0.41 | 0.25 | 1.73 | 0.495 | 0.405 | 1.565 | 0.014233 | 0.291266 | 0.423013 | | 1 | 0.999631 | 0.385092 |
| Ahdc1_Q6PAL7_S1060s_1 | -0.18 | -0.51 | -0.33 | -0.03 | -0.09 | -0.51 | -1.17 | -0.86 | -1.1 | 0.2 | -0.3 | -0.19 | 0.73 | -0.15 | -0.36 | -0.31 | -0.395 | -0.085 | 0.24 | 0.485838 | 0.221404 | 0.801755 | | 1 | 0.999631 | 0.804488 |
| Ahdc1_Q6PAL7_S1180s_1 | -0.41 | 0.18 | 0.06 | 0.03 | -0.1 | 0.03 | -0.44 | 0.27 | 0.13 | -0.02 | -0.16 | 0.91 | -0.08 | 0.91 | 0.27 | -0.36 | -0.025 | 0.25 | 0.22 | 0.540225 | 0.928535 | 0.450976 | | 1 | 0.999631 | 0.831295 |
| Ahdc1_Q6PAL7_S1467s_1 | 1.23 | -0.03 | 0.58 | 0.58 | -0.31 | -0.2 | 0.5 | -0.09 | -0.18 | 0.97 | 0.06 | -0.39 | -0.4 | -0.55 | -0.07 | -0.93 | -0.615 | -0.475 | -1.0775 | 0.012916 | 0.100292 | 0.258084 | | 1 | 0.999631 | 0.385092 |
| Ahdc1_Q6PAL7_S892s_1 | -0.29 | -0.1 | -0.22 | -0.59 | -0.08 | -1.13 | -0.4 | -0.3 | -1.6 | -0.52 | 0.24 | 0.65 | 0.59 | -0.83 | 0.13 | 0.2 | -0.1775 | -0.0075 | 0.3225 | 0.386934 | 0.567068 | 0.987621 | | 1 | 0.999631 | 0.756701 |
| Ahdc1_Q6PAL7_S99s_1 | 0.44 | -0.71 | 0.38 | 1.42 | -0.6 | -0.7 | 0.78 | 0.72 | -0.74 | -0.32 | 0.09 | 1.39 | -0.24 | -1.45 | 0.48 | 1.08 | -0.3325 | -0.2775 | -0.415 | 0.514109 | 0.546589 | 0.634712 | | 1 | 0.999631 | 0.81893 |
| Ahi1_E9QP54_S26s_1 | 2.58 | -0.07 | 2.53 | 2 | 0.46 | 1.39 | 2.45 | 1.68 | 1.6 | 2.37 | -0.73 | 1.08 | 0.97 | 1.17 | 2.16 | 0.45 | -0.265 | -0.68 | -0.5725 | 0.385369 | 0.69141 | 0.402659 | | 1 | 0.999631 | 0.756021 |
| Ahi1_E9QP54_S974s_1 | 0.26 | -1.47 | 1.04 | -0.36 | -0.51 | -0.51 | 0.03 | 0.8 | -0.06 | 0.05 | -0.83 | 1.08 | 0.46 | 0.45 | 1.64 | 0.5 | 0.085 | 0.1925 | 0.895 | 0.134068 | 0.879564 | 0.749057 | | 1 | 0.999631 | 0.60492 |
| Ahi1_E9QP54_T122t_1 | 0.02 | -2.8 | 0.37 | -0.88 | -1.9 | -1.89 | -0.13 | 0.38 | -1.36 | -0.53 | -1.66 | 0.06 | -0.69 | -0.55 | 1.07 | -0.45 | -0.0625 | -0.05 | 0.6675 | 0.367124 | 0.938196 | 0.944561 | | 1 | 0.999631 | 0.745796 |
| Ahi1_E9QP54_T5t_1 | 0.97 | -0.15 | -1.85 | 1.22 | 0.19 | 0.29 | -1.08 | -0.75 | 0.66 | 0.96 | -3.21 | -1.6 | 0.71 | 1.71 | -0.49 | -0.8 | -0.385 | -0.845 | 0.235 | 0.766716 | 0.579526 | 0.425453 | | 1 | 0.999631 | 0.916961 |
| Ahnak_E9Q616_S136s_1 | 0.68 | 1.08 | -0.06 | -0.56 | 0.97 | 0.47 | -0.56 | 0.39 | 0.26 | -0.47 | 0.08 | 0.02 | -0.78 | 0.33 | 1.1 | 0.09 | 0.0325 | -0.3125 | -0.1 | 0.844065 | 0.944295 | 0.450852 | | 1 | 0.999631 | 0.948381 |
| Ahnak_E9Q616_S5504s_1 | -0.68 | 0.02 | -0.14 | -0.04 | -0.56 | -0.1 | -0.23 | -0.55 | 0 | 0.58 | -0.92 | 0.66 | -0.79 | -0.43 | 0.45 | -0.05 | -0.15 | 0.29 | 0.005 | 0.988908 | 0.594126 | 0.482154 | | 1 | 0.999631 | 0.995427 |
| Ahsa2_F6S1R2_S175s_1 | 1.48 | -0.46 | -0.85 | 0.89 | 2.17 | 0.82 | 1.31 | 0.77 | 0.26 | -1.14 | -0.56 | -1.02 | -1.06 | 0.82 | -0.76 | -0.24 | 1.0025 | -0.88 | -0.575 | 0.366322 | 0.109173 | 0.154142 | | 1 | 0.999631 | 0.745796 |
| Ahsg_P29699_S138s_1 | 1.74 | 2.39 | 1.52 | 2.29 | 2.04 | 3.08 | 1.73 | 1.71 | 1.98 | 1.35 | 1.94 | 2.23 | 2.45 | 3.31 | 2.69 | 2.75 | 0.155 | -0.11 | 0.815 | 0.035815 | 0.694078 | 0.74111 | | 1 | 0.999631 | 0.453811 |
| Aida_Q8C4Q6_S161s_1 | -0.85 | -0.86 | -0.43 | -0.83 | -1 | -1.2 | -0.14 | -0.51 | -0.96 | -1.23 | -0.71 | -1.19 | -0.32 | -1.06 | -0.83 | -0.54 | 0.03 | -0.28 | 0.055 | 0.85432 | 0.923792 | 0.31303 | | 1 | 0.999631 | 0.951263 |
| Aimp1_Q3UZG4_S147s_1 | 0.68 | 1.48 | 1.04 | 0.7 | 0.93 | 1.49 | 0.45 | 0.37 | 1.43 | 2.06 | 0.81 | 1.4 | 0.04 | 2.04 | 1.2 | 1.1 | -0.165 | 0.45 | 0.12 | 0.789858 | 0.638198 | 0.222712 | | 1 | 0.999631 | 0.925678 |
| Aimp2_Q8R010_S36s_1 | -0.25 | 0.18 | -1.41 | -0.73 | 0.27 | -0.37 | -0.85 | -1 | 0.11 | -0.17 | -1.11 | -0.39 | 0.36 | 0.3 | -0.49 | 0.02 | 0.065 | 0.1625 | 0.6 | 0.166102 | 0.88117 | 0.705758 | | 1 | 0.999631 | 0.63587 |
| Akap1_O08715_S101s_1 | 0.04 | -0.03 | -0.38 | 0.39 | 0.86 | -0.12 | 0.65 | 0.23 | 0.45 | -0.2 | 0.7 | 0 | 0.73 | 0.64 | -0.73 | 0.21 | 0.4 | 0.2325 | 0.2075 | 0.602761 | 0.22833 | 0.472745 | | 1 | 0.999631 | 0.857278 |
| Akap1_O08715_S428s_1 | 1.38 | 0.5 | -1.89 | -0.29 | 0.37 | 0.4 | -0.22 | -1.08 | 0.98 | 1.39 | -1.11 | -1.18 | 2.25 | 0.3 | -1.66 | -0.62 | -0.0575 | 0.095 | 0.1425 | 0.878235 | 0.933354 | 0.910709 | | 1 | 0.999631 | 0.962347 |
| Akap1_O08715_S55s_1 | 0.26 | -0.15 | -0.04 | 0.37 | 0.44 | -0.11 | 0.31 | 0.54 | 0.27 | -0.34 | 0.12 | 0.28 | 0.19 | 0.53 | 0.18 | 0.44 | 0.185 | -0.0275 | 0.225 | 0.433859 | 0.506156 | 0.922667 | | 1 | 0.999631 | 0.779621 |
| Akap1_O08715_T487t_1 | -0.42 | -1.03 | -0.24 | -0.59 | -0.3 | -1.05 | -0.59 | -0.37 | -0.69 | -0.55 | -0.51 | -0.44 | 0.32 | -0.4 | -0.69 | 0.22 | -0.0075 | 0.0225 | 0.4325 | 0.237293 | 0.980123 | 0.93533 | | 1 | 0.999631 | 0.683301 |
| Akap10_O88845_S187s_1 | 0.07 | 0.08 | -0.24 | 0.48 | 0.82 | 0.8 | 0.61 | 0.45 | 0.69 | -0.24 | 0.76 | -0.58 | -0.76 | 0.47 | 0.35 | -0.07 | 0.5725 | 0.06 | -0.1 | 0.782967 | 0.054592 | 0.876326 | | 1 | 0.999631 | 0.92257 |
| Akap10_O88845_S281s_1 | 1.18 | 1.62 | -0.33 | 0.87 | 1.96 | 1.76 | 1.8 | 1.23 | 1.83 | 1.04 | 0.83 | 0.09 | -0.01 | 1.82 | 0.74 | -0.76 | 0.8525 | 0.1125 | -0.3875 | 0.539015 | 0.07505 | 0.827859 | | 1 | 0.999631 | 0.830899 |
| Akap11_E9Q777_S910s_1 | 0.54 | -0.11 | -1.56 | -0.97 | 0.06 | -1.01 | -1.12 | -0.73 | 0.42 | 0.37 | -0.82 | -0.85 | 1.03 | 0.48 | -0.15 | -1.31 | -0.175 | 0.305 | 0.5375 | 0.394469 | 0.727404 | 0.577182 | | 1 | 0.999631 | 0.759319 |
| Akap11_E9Q777_S912s_1 | 0.54 | -0.11 | -1.56 | -0.97 | 0.06 | -1.01 | -1.12 | -0.73 | 0.42 | 0.37 | -0.82 | -0.85 | 1.03 | 0.48 | -0.15 | -1.31 | -0.175 | 0.305 | 0.5375 | 0.394469 | 0.727404 | 0.577182 | | 1 | 0.999631 | 0.759319 |
| Akap12_Q9WTQ5_S1292s_1 | -0.46 | -0.62 | 0.87 | 0 | -1.14 | -0.95 | 0.28 | 0.43 | -0.28 | -0.41 | -0.82 | 0.73 | 0.62 | -0.55 | 1.16 | -0.28 | -0.2925 | -0.1425 | 0.29 | 0.563207 | 0.558238 | 0.755892 | | 1 | 0.999631 | 0.841038 |
| Akap12_Q9WTQ5_S1645s_1 | 0.24 | -0.74 | 0.38 | -0.02 | -0.89 | -0.41 | 0.69 | -0.28 | -0.77 | 0.15 | -0.9 | -0.42 | 0.15 | -0.7 | 0.25 | 0.11 | -0.1875 | -0.45 | -0.0125 | 0.973244 | 0.651593 | 0.242036 | | 1 | 0.999631 | 0.990577 |
| Akap12_Q9WTQ5_S234s_1 | -0.48 | 0.71 | 0.5 | 0.88 | 0.55 | 0 | 0.34 | 0.92 | 0.51 | -0.03 | 1 | 1.69 | 0.73 | 0.51 | 1.25 | 1.8 | 0.05 | 0.39 | 0.67 | 0.141 | 0.894099 | 0.407825 | | 1 | 0.999631 | 0.61245 |
| Akap12_Q9WTQ5_S270s_1 | -1.76 | -2.89 | -0.07 | -0.08 | -2.41 | -2.52 | -0.39 | -0.04 | -2.61 | -2.01 | 0.1 | 0.76 | -0.22 | -2.01 | 1.33 | 0.93 | -0.14 | 0.26 | 1.2075 | 0.190335 | 0.865601 | 0.778711 | | 1 | 0.999631 | 0.648674 |
| Akap12_Q9WTQ5_S350s_1 | -0.11 | 0.33 | 0.43 | 0.12 | 0.28 | 0.15 | 0.13 | 0.07 | 0.42 | 0.4 | 0.48 | 0.69 | 0.98 | 0.64 | 0.78 | 0.51 | -0.035 | 0.305 | 0.535 | 0.084238 | 0.888437 | 0.259844 | | 1 | 0.999631 | 0.549732 |
| Akap12_Q9WTQ5_S467s_1 | -0.97 | -1.04 | -1.92 | -1.67 | -0.76 | -1.03 | -1.16 | -1.59 | -1.13 | -0.83 | -1.68 | -1.04 | -0.38 | -0.91 | -1.36 | -0.98 | 0.265 | 0.23 | 0.4925 | 0.191247 | 0.43193 | 0.50509 | | 1 | 0.999631 | 0.648674 |
| Akap12_Q9WTQ5_S507s_1 | 0.05 | 0.92 | -0.34 | 0.53 | 1.98 | 1.27 | 0.75 | 0.5 | 1.68 | -0.55 | 0.99 | 0.61 | -1.13 | 2.38 | 0.93 | 0.6 | 0.835 | 0.3925 | 0.405 | 0.558484 | 0.072217 | 0.449596 | | 1 | 0.999631 | 0.838464 |
| Akap12_Q9WTQ5_S543s_1 | -0.39 | 0.68 | 1.82 | 1.06 | 0.81 | 0.91 | 0.65 | 1.24 | 0.72 | 1.73 | 0.72 | 2.16 | 0.05 | 1.25 | 2.39 | 2.3 | 0.11 | 0.54 | 0.705 | 0.287853 | 0.810266 | 0.334411 | | 1 | 0.999631 | 0.705972 |

| | | | | | | | | | | | | | | | | | | | | | | | | | |
|---|---|---|---|---|---|---|---|---|---|---|---|---|---|---|---|---|---|---|---|---|---|---|---|---|---|
| Akap12_Q9WTQ5_S584s_1 | -0.76 | 0.01 | -0.2 | -1.24 | -0.29 | -0.33 | -0.69 | -0.33 | -0.19 | -0.15 | -0.65 | 0.07 | -0.22 | -0.21 | 0.13 | -0.28 | 0.1375 | 0.3175 | 0.4025 | 0.269565 | 0.683036 | 0.38245 | 1 | 0.999631 | 0.697839 |
| Akap12_Q9WTQ5_S598s_1 | 1.07 | 1.95 | 1.29 | 1.75 | 2.52 | 1.41 | 1.35 | 2.31 | 2.75 | 1.21 | 2.47 | 3.35 | 0.81 | 3.08 | 3.48 | 2.88 | 0.3825 | 0.93 | 1.0475 | 0.095021 | 0.324989 | 0.075105 | 1 | 0.999631 | 0.558128 |
| Akap12_Q9WTQ5_S613s_1 | -0.36 | 0.63 | -0.13 | -1.07 | 0.55 | 0.34 | -0.98 | -0.38 | 1.36 | 1.32 | -1.07 | -0.89 | 1.54 | 1.17 | 0.44 | -1.08 | 0.115 | 0.4125 | 0.75 | 0.239949 | 0.807416 | 0.546052 | 1 | 0.999631 | 0.683719 |
| Akap12_Q9WTQ5_S682s_1 | 2.62 | 1.05 | 0.97 | 1.05 | 0.36 | 1.59 | 2.27 | 0.67 | 0.41 | 2.86 | 0.01 | -0.39 | 0.82 | 0.33 | 0.4 | -0.91 | -0.2 | -0.7 | -1.2625 | 0.032676 | 0.71393 | 0.354061 | 1 | 0.999631 | 0.44294 |
| Akap12_Q9WTQ5_S684s_1 | 1.87 | -0.71 | 1.06 | 0.42 | 0.07 | 0.32 | 1.64 | -0.44 | -0.01 | 2.29 | -0.75 | -0.64 | 0.17 | -0.37 | 0.15 | -0.09 | -0.2625 | -0.4375 | -0.695 | 0.206842 | 0.678042 | 0.580552 | 1 | 0.999631 | 0.660763 |
| Akap12_Q9WTQ5_S786s_1 | 1.09 | 1.26 | 0.58 | 1.88 | 2.42 | 2.12 | 1.43 | 1.76 | 1.99 | 1 | 1.46 | 2.35 | -0.08 | 2.02 | 1.99 | 1.98 | 0.73 | 0.4975 | 0.275 | 0.616063 | 0.069308 | 0.24138 | 1 | 0.999631 | 0.861203 |
| Akap12_Q9WTQ5_S885s_1 | 2.38 | -0.65 | 0.52 | -0.28 | -0.43 | -0.37 | 0.61 | -0.37 | -0.03 | 0.78 | 0.03 | -0.02 | 1.96 | -1.41 | 0.16 | -0.61 | -0.6325 | -0.3025 | -0.4675 | 0.587106 | 0.338217 | 0.635485 | 1 | 0.999631 | 0.849999 |
| Akap5_H3BIV5_S149s_1 | -0.02 | -0.34 | -0.89 | -0.39 | -1.02 | 1.47 | 0.02 | 0.26 | -2.01 | -1.01 | -0.4 | -4.4 | -2.19 | 0.34 | -2.35 | -1.83 | 0.5925 | -1.545 | -1.0975 | 0.089096 | 0.259202 | 0.074784 | 1 | 0.999631 | 0.554748 |
| Akap5_H3BIV5_S178s_1 | -1 | -0.68 | -1.14 | -0.26 | -0.42 | -0.8 | -0.68 | -0.38 | -0.22 | -0.98 | -1.14 | -1.04 | -0.34 | -0.35 | -0.43 | -0.62 | 0.2 | -0.075 | 0.335 | 0.287425 | 0.497218 | 0.823374 | 1 | 0.999631 | 0.705972 |
| Akap5_H3BIV5_S249s_1 | 1.07 | -1.49 | -1 | -0.02 | 0.95 | -0.68 | 0.27 | -0.41 | -0.8 | -1.21 | 0.04 | -0.52 | -0.79 | -1.03 | -1.06 | -0.54 | 0.3925 | -0.2625 | -0.495 | 0.370781 | 0.522838 | 0.648745 | 1 | 0.999631 | 0.746338 |
| Akap5_H3BIV5_S258s_1 | -0.13 | -0.15 | -0.24 | -0.2 | -0.27 | -0.31 | -0.79 | -0.03 | -0.31 | -0.08 | -0.56 | -0.11 | -0.43 | 0.28 | -0.05 | -0.16 | -0.17 | -0.085 | 0.09 | 0.750144 | 0.522083 | 0.73765 | 1 | 0.999631 | 0.91245 |
| Akap5_H3BIV5_S269s_1 | 1.26 | -2.13 | -2.25 | 0.05 | 0.05 | -0.78 | 0.4 | -1.39 | -0.43 | -1.3 | -0.01 | -1.51 | -1.13 | -1.04 | -1.74 | -2.49 | 0.3375 | -0.045 | -0.8325 | 0.313131 | 0.684018 | 0.955865 | 1 | 0.999631 | 0.719861 |
| Akap5_H3BIV5_S277s_1 | -0.76 | -1.3 | -1.81 | -1.46 | -1.33 | -1.53 | -1.38 | -2.14 | -1.32 | -1.11 | -2.15 | -2.07 | -0.85 | -0.79 | -2.47 | -1.94 | -0.2625 | -0.33 | -0.18 | 0.697833 | 0.433036 | 0.381848 | 1 | 0.999631 | 0.895731 |
| Akap5_H3BIV5_S29s_1 | 0.68 | -0.68 | -1.04 | -1.13 | -0.67 | -0.52 | -0.78 | -1.57 | -0.09 | 0.8 | -1.45 | -1.67 | -0.2 | -0.54 | -1.08 | -1.61 | -0.3425 | -0.06 | -0.315 | 0.532026 | 0.463583 | 0.926018 | 1 | 0.999631 | 0.827588 |
| Akap5_H3BIV5_S75s_1 | 2.33 | -0.82 | 2.25 | 0.12 | 1.49 | -0.66 | 4.85 | -2.52 | 1.43 | 0.55 | 3.01 | 4.1 | 1.47 | -1.78 | 1.63 | -0.06 | -0.18 | 1.3025 | -0.655 | 0.499713 | 0.90429 | 0.198887 | 1 | 0.999631 | 0.813279 |
| Akap5_H3BIV5_T227t_1 | -1.08 | -0.72 | -0.85 | -0.51 | -0.74 | -1.01 | -1.08 | -0.63 | -0.8 | -0.79 | -1.13 | -0.29 | -0.38 | -0.28 | -0.5 | -0.47 | -0.075 | 0.0375 | 0.3825 | 0.185867 | 0.775359 | 0.898195 | 1 | 0.999631 | 0.646224 |
| Akap6_E9Q9K8_S1070s_1 | -1.24 | -0.12 | -0.26 | -0.61 | 0.1 | -0.31 | -0.54 | 0.16 | -0.07 | -0.51 | 0.1 | 0.09 | 0.29 | -0.16 | -0.16 | 0.32 | 0.41 | 0.46 | 0.63 | 0.091867 | 0.242421 | 0.19223 | 1 | 0.999631 | 0.557085 |
| Akap6_E9Q9K8_S423s_1 | -0.5 | -1.85 | -0.7 | 1.31 | -1.2 | -1.63 | 1.02 | 1.53 | -1.13 | -1.2 | 1.35 | 1.29 | -2.73 | -1.12 | 0.72 | 1.29 | 0.365 | 0.5125 | -0.025 | 0.979282 | 0.681906 | 0.549194 | 1 | 0.999631 | 0.993455 |
| Akap6_E9Q9K8_S480s_1 | 1.37 | 1.04 | -0.31 | 1.74 | 1.8 | 1.37 | 1.24 | 1.47 | 1.75 | 0.94 | 1.76 | 0.99 | -1.66 | 1.41 | 1.26 | 1.18 | 0.51 | 0.4 | -0.4125 | 0.588293 | 0.271077 | 0.414582 | 1 | 0.999631 | 0.850634 |
| Akap6_E9Q9K8_S533s_1 | 0.61 | -0.27 | -0.93 | -0.01 | 0.5 | -0.27 | 0.41 | 0.46 | -0.49 | -0.07 | -0.46 | -1.53 | -0.57 | 0.47 | 0.26 | -0.44 | 0.425 | -0.4875 | 0.08 | 0.849524 | 0.280006 | 0.284852 | 1 | 0.999631 | 0.949525 |
| Akap6_E9Q9K8_T2234t_1 | 0.37 | 0.12 | 0.12 | 0.41 | -0.45 | 0.37 | -0.27 | 0.73 | -0.5 | 0.32 | 0.13 | -0.18 | -0.36 | 0.31 | 0.91 | 0.35 | -0.16 | -0.3125 | 0.0475 | 0.888268 | 0.628257 | 0.290268 | 1 | 0.999631 | 0.963505 |
| Akap9_E9QQ10_T361t_1 | -0.64 | -0.78 | -0.73 | -1.46 | -0.53 | -0.38 | -0.34 | 0.02 | -0.67 | -0.53 | -0.85 | 0.03 | 0.16 | -0.42 | -0.16 | 0.36 | 0.595 | 0.3975 | 0.8875 | 0.021013 | 0.064798 | 0.238294 | 1 | 0.999631 | 0.400544 |
| Akt1s1_E9QKI5_S118s_1 | -1.33 | -0.01 | -0.3 | -1.92 | -2.43 | -2.24 | -1.52 | -1.48 | -2.35 | -2.47 | -1.58 | -1.29 | 0.37 | -1.48 | -0.31 | -1.11 | -1.0275 | -1.0325 | 0.2575 | 0.648606 | 0.055326 | 0.063601 | 1 | 0.999631 | 0.874615 |
| Akt1s1_E9QKI5_S143s_1 | -0.96 | -0.2 | -0.53 | -0.54 | -1.34 | -0.16 | -0.46 | -0.56 | -0.41 | -0.35 | -1.23 | -0.39 | 0.17 | -0.36 | -0.15 | -0.79 | -0.0725 | -0.0375 | 0.275 | 0.411873 | 0.828425 | 0.907109 | 1 | 0.999631 | 0.767937 |
| Akt1s1_E9QKI5_S229s_1 | -1.22 | -0.84 | -0.61 | -1.57 | -1.76 | -1.2 | -1.14 | -1.71 | -1.54 | -0.9 | 0.2 | 0.45 | 1.16 | -0.91 | -1.16 | -1.12 | -0.3925 | 0.6125 | 0.5525 | 0.331587 | 0.234833 | 0.23012 | 1 | 0.999631 | 0.727302 |
| Akt1s1_E9QKI5_S230s_1 | -1.22 | -0.84 | -0.61 | -1.57 | -1.76 | -1.2 | -1.14 | -1.71 | -1.54 | -0.9 | 0.2 | 0.45 | 1.16 | -0.91 | -1.16 | -1.12 | -0.3925 | 0.6125 | 0.5525 | 0.331587 | 0.234833 | 0.23012 | 1 | 0.999631 | 0.727302 |
| Akt1s1_E9QKI5_S239s_1 | -0.52 | 0.77 | 1.05 | 0.15 | 0.83 | -0.02 | 0.55 | -0.58 | 1 | 0.2 | 0.48 | 1.83 | 1.16 | 0.16 | 0.25 | -0.66 | -0.1675 | 0.515 | -0.135 | 0.783823 | 0.71198 | 0.298279 | 1 | 0.999631 | 0.922628 |
| Akt3_Q9WUA6_S34s_1 | 0.08 | 0.46 | 1.02 | 0.4 | 0.07 | 0.73 | -0.08 | 0.18 | -0.1 | 0.36 | 0.56 | -0.53 | 0.83 | 0.61 | 0.96 | 1.05 | -0.265 | -0.4175 | 0.3725 | 0.247848 | 0.408945 | 0.251838 | 1 | 0.999631 | 0.685751 |
| Akt3_Q9WUA6_T447t_1 | -0.66 | 1.23 | 0.61 | -0.9 | 1.54 | -0.32 | 1.24 | -0.49 | 2.42 | -1.14 | 1.42 | 2.63 | 2.71 | -0.42 | -0.41 | -0.44 | 0.4225 | 1.2625 | 0.29 | 0.722005 | 0.521258 | 0.171614 | 1 | 0.999631 | 0.905674 |
| Alad_P10518_S215s_1 | -0.83 | -0.84 | 0.79 | 0.26 | -0.48 | -0.64 | 0.19 | 0.45 | 0 | -0.09 | 0.47 | 1.05 | 0.23 | -0.28 | 0.79 | 0.82 | 0.035 | 0.5125 | 0.545 | 0.265454 | 0.93974 | 0.289138 | 1 | 0.999631 | 0.694352 |
| Aldh1a1_P24549_S2s_1 | -0.44 | 0.01 | -0.49 | -0.71 | 0.14 | -0.78 | -1.05 | -1.11 | -0.72 | -0.82 | -0.57 | -1.23 | -0.18 | -0.65 | -0.16 | -0.06 | -0.2925 | -0.4275 | 0.145 | 0.633959 | 0.416441 | 0.16525 | 1 | 0.999631 | 0.868569 |
| Aldh1a1_P24549_S3s_1 | -0.44 | 0.01 | -0.49 | -0.71 | 0.14 | -0.78 | -1.05 | -1.11 | -0.72 | -0.82 | -0.57 | -1.23 | -0.18 | -0.65 | -0.16 | -0.06 | -0.2925 | -0.4275 | 0.145 | 0.633959 | 0.416441 | 0.16525 | 1 | 0.999631 | 0.868569 |
| Aldh1l1_Q8R0Y6_T544t_1 | -0.16 | 0.78 | -0.15 | -1.61 | -1.39 | 1.01 | -1.14 | -1.29 | 0.03 | 0.28 | -1.31 | -1.38 | 0.18 | 1.89 | -0.9 | -0.92 | -0.4175 | -0.31 | 0.3475 | 0.634767 | 0.53876 | 0.607878 | 1 | 0.999631 | 0.868886 |
| Aldoa_P05064_S39s_1 | -0.54 | -1.03 | 0.77 | 0.28 | -0.69 | -0.86 | -0.2 | 0.56 | -1.05 | -0.86 | 0.68 | 0.55 | -0.59 | -0.44 | 0.48 | 1.03 | -0.1675 | -0.04 | 0.25 | 0.636108 | 0.730782 | 0.943181 | 1 | 0.999631 | 0.869397 |
| Aldoa_P05064_S46s_1 | -0.07 | 0.78 | 0.7 | 0.29 | 0.28 | 0.5 | 0.21 | 0.19 | 0.16 | 0.11 | 0.45 | 0.46 | 0.15 | 0.58 | 0.21 | 0.64 | -0.13 | -0.13 | -0.03 | 0.924887 | 0.650306 | 0.662964 | 1 | 0.999631 | 0.975422 |
| Aldoc_P05063_S36s_1 | -0.22 | 0.93 | 0.95 | 0.49 | 0.79 | 0.35 | -0.13 | 0.04 | 0.39 | -0.06 | 0.12 | 0.63 | 0.24 | -0.02 | 0.47 | 0.16 | -0.275 | -0.2675 | -0.325 | 0.363407 | 0.455142 | 0.453443 | 1 | 0.999631 | 0.743747 |
| Aldoc_P05063_S39s_1 | 0.6 | 0.79 | 0.18 | -0.6 | 0.19 | 0.07 | -0.89 | -0.43 | -0.56 | 0.06 | -0.22 | 0.17 | 0.02 | 0.2 | -0.15 | 0.5 | -0.5075 | -0.38 | -0.1 | 0.790903 | 0.223846 | 0.323107 | 1 | 0.999631 | 0.926419 |
| Aldoc_P05063_S45s_1 | 0.22 | 0.74 | 0.67 | -0.36 | 0.09 | 0.55 | -0.03 | -0.71 | 1.03 | 0.46 | 0.57 | 0.67 | 0.95 | 0.52 | 0.42 | 0.67 | -0.3425 | 0.365 | 0.3225 | 0.356892 | 0.37523 | 0.28776 | 1 | 0.999631 | 0.740719 |
| Alkbh3_Q8K1E6_S28s_1 | 1.51 | 0.46 | 0.57 | 0.99 | 2.28 | 1.88 | 1.92 | 0.51 | 2.13 | -0.39 | 0.44 | -0.69 | -3.16 | -0.47 | 0.16 | -1.92 | 0.765 | -0.51 | -2.23 | 0.008827 | 0.109483 | 0.414526 | 1 | 0.999631 | 0.365244 |
| Alkbh5_Q3TSG4_S362s_1 | 0.79 | 2.37 | 1.63 | 1.02 | 0.66 | 2.32 | 1.07 | 1.34 | 2.01 | 2.01 | 2.02 | 0.98 | 1.65 | 2.52 | 1.37 | 2.98 | -0.105 | 0.3025 | 0.6775 | 0.191804 | 0.825064 | 0.492404 | 1 | 0.999631 | 0.648674 |
| Alkbh5_Q3TSG4_S385s_1 | 0.46 | 1.68 | 0.39 | 0.43 | 0.65 | 1.45 | 0.33 | 0.47 | 0.97 | 1.34 | 0.71 | 0.71 | 2.12 | 1.66 | 0.94 | 1.14 | -0.015 | 0.1925 | 0.725 | 0.110805 | 0.970351 | 0.608336 | 1 | 0.999631 | 0.576275 |
| Alkbh5_Q3TSG4_S65s_1 | 2.5 | 2.54 | 3.12 | 2.44 | 1.2 | 3 | 2.22 | 1.07 | 2.3 | 3.64 | 1.66 | 1.34 | 1.4 | 1.65 | 1.73 | 0.79 | -0.7775 | -0.415 | -1.2575 | 0.003559 | 0.117154 | 0.419145 | 1 | 0.999631 | 0.351308 |
| Als2_Q920R0_S1016s_1 | -1.1 | -0.16 | -0.49 | -0.27 | 0.15 | -0.48 | -0.82 | -0.45 | -0.27 | -0.33 | -0.69 | -0.58 | 0.54 | 0.06 | -0.48 | -0.69 | 0.105 | 0.0375 | 0.3625 | 0.35432 | 0.751894 | 0.901991 | 1 | 0.999631 | 0.739951 |
| Als2_Q920R0_S1329s_1 | -1.76 | 0.07 | 1.04 | -0.22 | 0.96 | 0.14 | 1.31 | -0.67 | 2.39 | -1.46 | 1.45 | 3.29 | 2.92 | -0.81 | 0.78 | 0.16 | 0.6525 | 1.635 | 0.98 | 0.264967 | 0.329544 | 0.133368 | 1 | 0.999631 | 0.693784 |
| Als2_Q920R0_S1458s_1 | -0.91 | -0.57 | -1.05 | -1.2 | -0.98 | -0.15 | -0.86 | -1.06 | -0.61 | -0.63 | -2.07 | -0.78 | -2.15 | 0.21 | -0.19 | -0.34 | 0.17 | -0.09 | 0.315 | 0.543346 | 0.58166 | 0.818612 | 1 | 0.999631 | 0.832881 |
| Als2_Q920R0_S460s_1 | -0.11 | 1.03 | 0.97 | 0.88 | 0.83 | 1.06 | 0.86 | 0.82 | 0.85 | 0.88 | 1.03 | 0.79 | 0.38 | 1.29 | 0.75 | 1.7 | 0.2 | 0.195 | 0.3375 | 0.422369 | 0.538457 | 0.555658 | 1 | 0.999631 | 0.773293 |
| Als2_Q920R0_S477s_1 | -1.29 | -0.23 | 0.97 | 0.32 | -0.54 | 0.06 | -1.17 | -0.87 | 0.71 | -1.52 | -2.61 | 0.64 | 0.25 | -0.82 | 0.76 | 0.9 | -0.5725 | -0.6375 | 0.33 | 0.565462 | 0.277387 | 0.45055 | 1 | 0.999631 | 0.84139 |
| Als2_Q920R0_S486s_1 | -1.55 | -0.78 | -0.88 | -1.03 | -1.62 | -0.93 | -1.3 | -0.81 | -1.8 | -0.77 | -0.59 | -0.28 | -0.26 | -0.75 | -1.04 | 0.01 | -0.105 | 0.2 | 0.55 | 0.138983 | 0.733864 | 0.609409 | 1 | 0.999631 | 0.611718 |
| Als2_Q920R0_T504t_1 | -0.75 | -0.87 | -0.4 | -0.63 | -1.01 | -0.86 | -0.41 | -0.33 | -0.72 | -0.89 | -0.28 | 0.34 | 0.05 | -0.33 | 0.04 | -0.1 | 0.01 | 0.275 | 0.5775 | 0.059439 | 0.971283 | 0.424894 | 1 | 0.999631 | 0.50521 |
| Amer2_Q8CCJ4_S223s_1 | -0.09 | 2.23 | 1.78 | 1.05 | 0.59 | 2.1 | 0.38 | 1.52 | 1.72 | 1.82 | 0.95 | 2.24 | -0.33 | 2.72 | 1.69 | 2.23 | -0.095 | 0.44 | 0.335 | 0.651824 | 0.871342 | 0.419032 | 1 | 0.999631 | 0.876813 |
| Amer2_Q8CCJ4_S227s_1 | 0.05 | 0.8 | 0.36 | 0.35 | 0.69 | 0.81 | -0.18 | -0.1 | 0.5 | 0.26 | 0.67 | 0.47 | 1.17 | 0.92 | 0.02 | 0.73 | -0.085 | 0.085 | 0.32 | 0.369902 | 0.801476 | 0.759294 | 1 | 0.999631 | 0.746338 |
| Amer2_Q8CCJ4_S244s_1 | 2.12 | 0.08 | 1.31 | 0.96 | -0.27 | -0.13 | 1.16 | 0.28 | -0.2 | 1.48 | -0.32 | -0.27 | -0.55 | -0.91 | -0.02 | -0.16 | -0.8575 | -0.945 | -1.5275 | 0.00752 | 0.110305 | 0.116882 | 1 | 0.999631 | 0.358173 |
| Amer2_Q8CCJ4_S269s_1 | 1.32 | 2.13 | 0.92 | -0.45 | 2.1 | 1.47 | 1.69 | 0.42 | 1.85 | -0.74 | 2.18 | 1.3 | 1.93 | 0.48 | 0.17 | -1.48 | 0.44 | 0.1675 | -0.705 | 0.372031 | 0.46007 | 0.823199 | 1 | 0.999631 | 0.746837 |
| Amer2_Q8CCJ4_S280s_1 | -0.68 | -0.86 | -0.61 | -0.9 | -0.76 | -1.09 | -0.97 | -0.82 | -1.14 | -0.71 | -0.64 | -0.48 | -0.06 | -0.68 | -0.71 | -0.45 | -0.1475 | 0.02 | 0.2875 | 0.330942 | 0.544836 | 0.94069 | 1 | 0.999631 | 0.727302 |
| Amer2_Q8CCJ4_S71s_1 | -0.14 | -0.29 | -0.85 | -0.06 | 0.52 | 0.34 | -0.26 | -0.22 | 0.12 | -0.42 | -0.33 | 0.15 | -0.27 | 0.39 | -0.11 | 0.04 | 0.43 | 0.215 | 0.3475 | 0.285481 | 0.195049 | 0.485693 | 1 | 0.999631 | 0.705851 |
| Amer2_Q8CCJ4_S98s_1 | -0.69 | -0.08 | -1.15 | -0.02 | -0.19 | -0.7 | -0.25 | -0.36 | -0.85 | -0.77 | -0.85 | -0.49 | -1.33 | 0 | -0.83 | -0.47 | 0.11 | -0.255 | -0.1725 | 0.674006 | 0.74082 | 0.450034 | 1 | 0.999631 | 0.885515 |
| Amer2_Q8CCJ4_T100t_1 | -0.69 | -0.08 | -1.15 | -0.02 | -0.19 | -0.7 | -0.25 | -0.36 | -0.85 | -0.77 | -0.85 | -0.49 | -1.33 | 0 | -0.83 | -0.47 | 0.11 | -0.255 | -0.1725 | 0.674006 | 0.74082 | 0.450034 | 1 | 0.999631 | 0.885515 |
| Amer3_Q6NS69_S17s_1 | 1.13 | 0.17 | -0.35 | 1.22 | 1.43 | 0.62 | 1.39 | 0.43 | 0.5 | -0.63 | 0.48 | -0.35 | -2.15 | 0.38 | -0.73 | 0.07 | 0.425 | -0.5425 | -1.15 | 0.086704 | 0.352295 | 0.259351 | 1 | 0.999631 | 0.553514 |
| Amot_Q8VHG2_S1083s_1 | 1.06 | 0 | -0.43 | -0.4 | -0.75 | 0.38 | -0.02 | 0.01 | -0.19 | -0.8 | 0.14 | 0.19 | -0.05 | 0.15 | 0.48 | 0.79 | -0.1525 | -0.2225 | 0.285 | 0.494356 | 0.715855 | 0.598646 | 1 | 0.999631 | 0.81007 |
| Amot_Q8VHG2_S149s_1 | 1.2 | 0.18 | -0.28 | 0.41 | 0.55 | 0.61 | 1.5 | 0.44 | 0.4 | 1.27 | -0.14 | -0.13 | -1.33 | -0.5 | 0.16 | -1.15 | 0.3975 | -0.0275 | -1.0825 | 0.036348 | 0.331859 | 0.950821 | 1 | 0.999631 | 0.453963 |
| Amot_Q8VHG2_S693s_1 | -0.87 | -0.18 | -0.88 | -1.08 | -0.18 | -0.34 | -1 | -1.14 | -0.44 | -0.57 | -0.78 | 0.1 | 0.34 | 0.43 | -1.04 | 0.29 | 0.0875 | 0.33 | 0.7575 | 0.093153 | 0.798514 | 0.325583 | 1 | 0.999631 | 0.558128 |
| Amotl1_Q9D4H4_S645s_1 | 1.4 | -0.67 | -0.05 | -0.02 | -0.48 | 0.23 | 0.9 | -2.12 | -0.74 | 1.2 | -1.35 | 0.05 | -0.15 | -0.81 | 0.15 | -0.91 | -0.5325 | -0.375 | -0.595 | 0.242097 | 0.449532 | 0.558351 | 1 | 0.999631 | 0.684914 |
| Amotl2_B8JK81_S50s_1 | 0.69 | -0.24 | 0.7 | 0.56 | -0.15 | -0.05 | 0.16 | 0.02 | -0.03 | 1.09 | -0.11 | 0.37 | 0.31 | -0.04 | 0.66 | 0.37 | -0.4325 | -0.0975 | -0.1025 | 0.760503 | 0.169744 | 0.796611 | 1 | 0.999631 | 0.914614 |
| Amotl2_B8JK81_S51s_1 | 0.69 | -0.24 | 0.7 | 0.56 | -0.15 | -0.05 | 0.16 | 0.02 | -0.03 | 1.09 | -0.11 | 0.37 | 0.31 | -0.04 | 0.66 | 0.37 | -0.4325 | -0.0975 | -0.1025 | 0.760503 | 0.169744 | 0.796611 | 1 | 0.999631 | 0.914614 |
| Ampd2_A2AE27_S3s_1 | -0.22 | -1.23 | -1.16 | -1.54 | -1.23 | -0.28 | -0.65 | -3.78 | -1.72 | 1.02 | -1.6 | -1.59 | -1.37 | -0.11 | -0.62 | 0.35 | -0.4475 | 0.065 | 0.6 | 0.211929 | 0.548295 | 0.920339 | 1 | 0.999631 | 0.665857 |
| Ampd2_Q9DBT5_S12s_1 | 0.01 | -0.38 | 0.3 | 0.8 | 0.69 | 0.1 | 0.48 | 1.33 | 0.15 | -0.55 | 0.64 | 1.01 | -0.51 | 0.78 | 1.26 | 0.1 | 0.4675 | 0.13 | 0.225 | 0.62405 | 0.230466 | 0.758253 | 1 | 0.999631 | 0.863524 |
| Ampd2_Q9DBT5_S19s_1 | 1.29 | 0.21 | -0.24 | 1.43 | 1.94 | 0.91 | 1.34 | 1.2 | 0.86 | -0.05 | 0.73 | 0.45 | -1.33 | 0.86 | 0.69 | 0.72 | 0.675 | -0.175 | -0.4375 | 0.474813 | 0.154968 | 0.696834 | 1 | 0.999631 | 0.799827 |
| Ampd2_Q9DBT5_S53s_1 | 2.25 | 1.91 | 0.2 | 2.81 | 2.92 | 2.18 | 2.05 | 2.29 | 2.75 | 1.57 | 1.95 | 1.54 | -1.75 | 1.97 | 1.58 | 1.88 | 0.5675 | 0.16 | -0.8725 | 0.348881 | 0.31285 | 0.781873 | 1 | 0.999631 | 0.736697 |
| Ampd2_Q9DBT5_S87s_1 | 1.18 | 1.07 | 1.37 | 1.22 | 0.82 | 0.91 | 0.57 | 0.51 | 0.95 | 1.41 | 0.92 | 0.99 | 0.05 | 0.63 | 0.46 | 0.86 | -0.5075 | -0.1425 | -0.71 | 0.034274 | 0.061656 | 0.584921 | 1 | 0.999631 | 0.446273 |
| Ampd3_O08739_S85s_1 | 0 | 1.76 | 0.95 | 0.85 | 0.69 | 0.95 | 0.25 | 0.61 | 1.1 | 0.51 | 0.63 | 1.49 | 0.38 | 1.81 | 0.37 | 0.87 | -0.265 | 0.0425 | -0.0325 | 0.945963 | 0.506834 | 0.920414 | 1 | 0.999631 | 0.982217 |
| Amph_Q7TQF7_S244s_1 | 0.5 | 1.2 | -0.26 | 0.95 | 0.85 | 0.69 | 0.57 | 1.67 | 0.52 | -0.22 | 1.27 | 0.12 | -0.67 | 0.39 | 0.15 | 0.87 | 0.3475 | -0.175 | -0.4125 | 0.371972 | 0.402159 | 0.695781 | 1 | 0.999631 | 0.746837 |
| Amph_Q7TQF7_S250s_1 | -0.66 | -0.59 | -0.47 | -0.35 | -0.2 | -0.66 | -0.82 | -0.52 | -0.43 | -1.01 | -0.53 | 0.02 | -0.1 | -0.91 | -0.92 | -0.06 | -0.0325 | 0.03 | 0.02 | 0.951219 | 0.899495 | 0.920074 | 1 | 0.999631 | 0.984079 |
| Amph_Q7TQF7_S252s_1 | -0.75 | -0.16 | -0.44 | -0.46 | -0.34 | -0.32 | -0.82 | -0.19 | -0.85 | -0.75 | -0.36 | -0.11 | -0.33 | -0.3 | -0.36 | 0.26 | 0.035 | -0.065 | 0.27 | 0.376052 | 0.897821 | 0.824581 | 1 | 0.999631 | 0.748307 |
| Amph_Q7TQF7_S262s_1 | -0.88 | 0.23 | -0.43 | -3.43 | -1.46 | -0.95 | -3.27 | -2.86 | -0.89 | -0.47 | -2.59 | -1.19 | 1.39 | -0.43 | -1.51 | -0.96 | -1.0075 | -0.1575 | 0.75 | 0.402595 | 0.251404 | 0.845727 | 1 | 0.999631 | 0.763968 |
| Amph_Q7TQF7_S262s_2 | -1.4 | -0.18 | -0.46 | -0.62 | -0.9 | -0.52 | -0.86 | -0.44 | -0.56 | -0.86 | -0.73 | 0.19 | -0.25 | -0.27 | -0.2 | -0.3 | -0.015 | 0.175 | 0.41 | 0.236527 | 0.963559 | 0.64321 | 1 | 0.999631 | 0.683301 |
| Amph_Q7TQF7_S272s_1 | -0.37 | -0.32 | -0.78 | -0.23 | -0.24 | -0.55 | -0.36 | -1.2 | -0.42 | -0.86 | -1.5 | -0.52 | -0.48 | -0.55 | -1.05 | -0.29 | -0.1625 | -0.4 | -0.1675 | 0.584345 | 0.596979 | 0.239673 | 1 | 0.999631 | 0.849121 |
| Amph_Q7TQF7_S272s_2 | -1.45 | -0.33 | -0.56 | -0.67 | -0.91 | -0.65 | -0.88 | -0.5 | -0.59 | -0.78 | -0.65 | 0.15 | -0.28 | -0.29 | -0.26 | -0.3 | 0.0175 | 0.285 | 0.47 | 0.168047 | 0.955535 | 0.431687 | 1 | 0.999631 | 0.637399 |
| Amph_Q7TQF7_S500s_1 | -0.38 | -0.72 | -0.47 | -0.78 | -0.94 | -0.69 | -0.88 | -0.88 | -0.88 | -0.58 | -0.73 | -0.63 | 0.06 | -0.6 | -0.58 | -0.19 | -0.26 | -0.1175 | 0.26 | 0.390567 | 0.300877 | 0.645667 | 1 | 0.999631 | 0.758901 |
| Amph_Q7TQF7_T260t_1 | 0.44 | 1.12 | -1.58 | -2.18 | 1.02 | 0.06 | -2.67 | -1.94 | 1.34 | 1.67 | -1.08 | -1.25 | 2.15 | 2.09 | -1.56 | -1.33 | -0.3325 | 0.72 | 0.8875 | 0.428923 | 0.740719 | 0.458962 | 1 | 0.999631 | 0.776616 |
| Anapc5_Q3TWF7_T217t_1 | 0.2 | -0.06 | -1.31 | -0.15 | 0.01 | -0.47 | -0.57 | 0.57 | 0.39 | -0.05 | -0.65 | 0.02 | 0.98 | -0.28 | 0.4 | 0.1 | 0.215 | 0.2575 | 0.63 | 0.168797 | 0.611379 | 0.531543 | 1 | 0.999631 | 0.637839 |
| Ank1_Q02357_S1804s_1 | 1.56 | -0.22 | 0.38 | 0.47 | -0.63 | -0.17 | 0.72 | -0.41 | -0.84 | 0.59 | -1.08 | -1.18 | -0.74 | -1.18 | -0.4 | -1.1 | -0.67 | -1.175 | -1.4025 | 0.007426 | 0.164972 | 0.045166 | 1 | 0.999631 | 0.358173 |
| Ank1_Q02357_S897s_1 | 0.27 | 0.13 | -0.59 | -0.17 | -0.03 | 0.26 | -0.27 | -0.09 | -0.22 | -0.26 | -0.72 | -1.17 | -1.71 | 0.08 | -0.65 | -0.45 | 0.0575 | -0.5025 | -0.5925 | 0.188791 | 0.843287 | 0.16172 | 1 | 0.999631 | 0.648316 |
| Ank1_Q02357_T1421t_1 | 1.39 | 1.48 | -1.53 | 0.61 | 0.85 | -0.02 | -0.01 | 0.33 | 0.46 | -0.64 | -0.84 | -0.45 | 0.04 | -0.11 | -1.08 | -0.04 | -0.2 | -0.855 | -0.785 | 0.256862 | 0.759168 | 0.225895 | 1 | 0.999631 | 0.691567 |
| Ank1_Q02357_T907t_1 | -1 | -0.89 | -1.73 | -0.97 | -0.5 | -1.41 | -0.58 | -0.97 | -1.05 | -0.57 | -1.07 | -0.99 | -0.31 | 0.01 | -1.53 | -1.17 | 0.2825 | 0.2275 | 0.3975 | 0.357441 | 0.39895 | 0.458701 | 1 | 0.999631 | 0.740986 |
| Ank2_Q8C8R3_S1454s_1 | -0.68 | 0.18 | -0.04 | -0.17 | -0.34 | -0.61 | -0.53 | 0.36 | -0.45 | -0.2 | -0.32 | 0.37 | 0.05 | -0.38 | -0.19 | 0.46 | -0.1025 | 0.0275 | 0.1625 | 0.624985 | 0.755559 | 0.931032 | 1 | 0.999631 | 0.863768 |
| Ank2_Q8C8R3_S1456s_1 | -0.75 | 0.15 | 0.04 | -0.15 | -0.25 | -0.1 | -0.28 | 0.14 | -0.43 | 0 | -0.11 | 0.39 | 0.47 | 0.02 | -0.11 | 0.54 | 0.055 | 0.14 | 0.4075 | 0.236038 | 0.850702 | 0.664357 | 1 | 0.999631 | 0.683301 |
| Ank2_Q8C8R3_S1727s_1 | 0.15 | -1.42 | -0.07 | -0.55 | -2.76 | -1.24 | -0.31 | -1.8 | -1.23 | 0.51 | -0.54 | -1.27 | -1.53 | -2.56 | -1.05 | -2.08 | -1.055 | -0.16 | -1.3325 | 0.016473 | 0.087116 | 0.754853 | 1 | 0.999631 | 0.389192 |
| Ank2_Q8C8R3_S1727s_2 | -2.01 | -2 | 0.21 | -2.03 | -2.61 | -1.52 | -2.25 | -2.81 | -1.35 | -0.1 | -1.4 | -1.26 | 0.36 | -2.21 | -0.66 | -2.86 | -0.84 | 0.43 | 0.115 | 0.885755 | 0.162984 | 0.467329 | 1 | 0.999631 | 0.963505 |
| Ank2_Q8C8R3_S1728s_1 | 0.27 | -1.56 | -1.53 | -0.66 | -1.38 | -0.72 | -1.2 | -1.16 | -1.22 | 0.88 | -2.05 | -2.47 | -0.74 | -2.11 | -3.23 | -2.5 | -0.245 | -0.345 | -1.275 | 0.061206 | 0.582847 | 0.651652 | 1 | 0.999631 | 0.509355 |
| Ank2_Q8C8R3_S1728s_2 | -2.01 | -2 | 0.21 | -2.03 | -2.61 | -1.52 | -2.25 | -2.81 | -1.35 | -0.1 | -1.4 | -1.26 | 0.36 | -2.21 | -0.66 | -2.86 | -0.84 | 0.43 | 0.115 | 0.885755 | 0.162984 | 0.467329 | 1 | 0.999631 | 0.963505 |
| Ank2_Q8C8R3_S1731s_1 | 1.91 | 0.14 | 0.7 | 1.09 | 0.18 | 1.06 | 1.61 | 0.22 | 0.76 | 1.87 | 0.29 | -0.28 | -1.78 | -0.74 | 0.2 | -0.92 | -0.1925 | -0.3 | -1.77 | 0.006456 | 0.690227 | 0.585664 | 1 | 0.999631 | 0.351545 |
| Ank2_Q8C8R3_S1750s_1 | 1.1 | -0.87 | -1.89 | 0.79 | 1.41 | -0.42 | 0.67 | 0.62 | 0.39 | -0.87 | -0.07 | -0.8 | -1.91 | -0.65 | -0.88 | -0.77 | 0.7875 | -0.12 | -0.835 | 0.237175 | 0.280977 | 0.861005 | 1 | 0.999631 | 0.683301 |
| Ank2_Q8C8R3_S1805s_1 | -1.09 | 1.49 | 2.59 | -0.22 | 2.36 | 1.04 | 2.81 | 1.22 | 3.18 | -0.31 | 3.18 | 4.31 | 2.31 | -0.71 | 1.62 | 0.01 | 1.165 | 1.8975 | 0.115 | 0.901708 | 0.175175 | 0.114381 | 1 | 0.999631 | 0.968183 |
| Ank2_Q8C8R3_S1821s_1 | 0.22 | 2.53 | 1.63 | 0.66 | 6.15 | 2.83 | 5.19 | 3.32 | 6.7 | -0.46 | 6.03 | 6.74 | 3.68 | 0.77 | 3.35 | 0.3 | 3.1125 | 3.4925 | 0.765 | 0.390171 | 0.003449 | 0.045098 | 1 | 0.999631 | 0.758901 |
| Ank2_Q8C8R3_S1835s_1 | -0.16 | 3.34 | 2.11 | -0.1 | 4.87 | 1.7 | 3.25 | 0.93 | 6.23 | 0.01 | 3.33 | 4.5 | 5.13 | 1.44 | 1.01 | 1.45 | 1.39 | 2.22 | 0.96 | 0.391567 | 0.207045 | 0.122068 | 1 | 0.999631 | 0.758901 |
| Ank2_Q8C8R3_S1844s_1 | -0.4 | -0.56 | -1.11 | -0.76 | -0.09 | -0.21 | -0.58 | -0.83 | -0.43 | -0.36 | -0.44 | -0.54 | 0.03 | -0.1 | -0.5 | -0.44 | 0.28 | 0.265 | 0.455 | 0.155417 | 0.354887 | 0.337171 | 1 | 0.999631 | 0.626444 |
| Ank2_Q8C8R3_S1847s_1 | -0.09 | -0.56 | -0.52 | 0.07 | -0.06 | -0.47 | -0.09 | -0.31 | -0.86 | -0.25 | 0.14 | -0.18 | -0.35 | -0.32 | 0.18 | 0.37 | 0.0425 | -0.0125 | 0.245 | 0.452374 | 0.876278 | 0.968854 | 1 | 0.999631 | 0.788364 |
| Ank2_Q8C8R3_S1856s_1 | 1.31 | 0.2 | -0.98 | -0.24 | 0.6 | 0.23 | -0.11 | -0.02 | 1.22 | 1.53 | -0.2 | 0.18 | 0.82 | 0.32 | -0.25 | -0.38 | 0.1025 | 0.61 | 0.055 | 0.916062 | 0.830472 | 0.30677 | 1 | 0.999631 | 0.973495 |
| Ank2_Q8C8R3_S1870s_1 | 0.85 | 3.23 | 4.78 | 2.54 | 5.06 | 2.98 | 6.65 | -1.17 | 5.87 | 0.04 | 5.39 | 6.89 | 6.79 | -0.4 | 5.62 | -0.11 | 0.53 | 1.6975 | 0.125 | 0.94184 | 0.739547 | 0.267414 | 1 | 0.999631 | 0.981463 |
| Ank2_Q8C8R3_S1879s_1 | -1.79 | -0.04 | -0.68 | -1.11 | 0.18 | -0.25 | -0.18 | -0.03 | 1.06 | -1.67 | 0.83 | 1.26 | 1.7 | -0.26 | 0.51 | -0.43 | 0.835 | 1.275 | 1.285 | 0.042796 | 0.054373 | 0.091076 | 1 | 0.999631 | 0.475443 |
| Ank2_Q8C8R3_S1893s_1 | 1.98 | 2.36 | 1.02 | -0.96 | 3.81 | 2.32 | 4.48 | -1.63 | 5.54 | 1.75 | 0.95 | 2.22 | 4.12 | -0.55 | -1.32 | -2.5 | 1.145 | 1.515 | -1.1625 | 0.405499 | 0.396629 | 0.181928 | 1 | 0.999631 | 0.765965 |
| Ank2_Q8C8R3_S1953s_1 | -0.51 | -0.41 | 1.09 | -0.19 | 1.21 | -0.36 | 1.29 | -0.93 | 1.96 | -0.46 | 1.05 | 2.6 | 1.13 | -0.37 | 1.23 | -0.15 | 0.3075 | 1.2925 | 0.465 | 0.387079 | 0.613953 | 0.081986 | 1 | 0.999631 | 0.756701 |
| Ank2_Q8C8R3_S1964s_1 | 0.56 | -0.36 | -0.91 | 0.19 | 0.42 | -0.27 | 0.3 | 0.17 | 0.03 | -0.25 | 0.1 | 0.1 | -1.2 | -0.16 | 0.06 | -0.01 | 0.285 | 0.125 | -0.1975 | 0.654781 | 0.451001 | 0.731494 | 1 | 0.999631 | 0.878941 |
| Ank2_Q8C8R3_S2118s_1 | -1.88 | -0.57 | 0.62 | -0.49 | -1.27 | -1.48 | -0.26 | -0.55 | -1.1 | -1.76 | 0.44 | 0.99 | -1 | -1.29 | 0.35 | 0.08 | -0.31 | 0.2225 | 0.115 | 0.846398 | 0.569915 | 0.760439 | 1 | 0.999631 | 0.948838 |
| Ank2_Q8C8R3_S2163s_1 | 2.15 | -0.38 | -1.44 | 1.14 | 1.57 | -0.54 | 1.62 | 0.94 | 0.12 | -1.43 | -0.2 | -0.91 | -1.03 | -0.21 | -0.87 | -0.67 | 0.53 | -0.9725 | -1.0625 | 0.162551 | 0.522907 | 0.223769 | 1 | 0.999631 | 0.631442 |
| Ank2_Q8C8R3_S2204s_1 | 1.16 | 0.96 | -0.21 | 0.35 | 0.54 | -0.01 | -0.22 | -0.13 | 0.23 | 0.06 | -0.16 | -0.78 | -0.45 | 0.63 | 0.24 | 0.17 | -0.52 | -0.7275 | -0.4175 | 0.315179 | 0.183702 | 0.090769 | 1 | 0.999631 | 0.72069 |
| Ank2_Q8C8R3_S2266s_1 | -0.03 | 0.1 | -0.41 | -0.71 | -0.49 | -0.03 | -0.73 | -0.63 | -0.31 | 0.22 | -0.66 | -0.29 | 0.6 | -0.04 | -0.21 | -0.47 | -0.2075 | 0.0025 | 0.2325 | 0.511482 | 0.497417 | 0.993739 | 1 | 0.999631 | 0.817687 |
| Ank2_Q8C8R3_S2392s_1 | -0.68 | -0.72 | -0.9 | -0.95 | -0.66 | -0.87 | -1.14 | -0.74 | -0.52 | -0.29 | -0.69 | -0.45 | 0.18 | -0.28 | -0.49 | -0.39 | -0.04 | 0.325 | 0.5675 | 0.071353 | 0.872046 | 0.215105 | 1 | 0.999631 | 0.531405 |
| Ank2_Q8C8R3_S2450s_1 | -0.65 | -0.21 | -2.19 | -3.78 | -0.17 | -0.33 | -2.3 | -3.37 | 0.04 | 0.91 | -1.64 | -3.84 | 1.38 | 0.27 | -4.46 | -3.24 | 0.165 | 0.575 | 0.195 | 0.885589 | 0.86469 | 0.614922 | 1 | 0.999631 | 0.963505 |
| Ank2_Q8C8R3_S2455s_2 | -1.09 | 0.22 | -0.02 | -1.18 | -1.42 | -0.73 | -0.86 | 0.17 | -0.2 | -0.61 | -0.52 | 0.74 | 0.95 | -0.57 | -0.9 | 0 | -0.1925 | 0.37 | 0.3875 | 0.457856 | 0.68099 | 0.4305 | 1 | 0.999631 | 0.790722 |
| Ank2_Q8C8R3_S2458s_2 | -1.09 | 0.22 | -0.02 | -1.18 | -1.42 | -0.73 | -0.86 | 0.17 | -0.2 | -0.61 | -0.52 | 0.74 | 0.95 | -0.57 | -0.9 | 0 | -0.1925 | 0.37 | 0.3875 | 0.457856 | 0.68099 | 0.4305 | 1 | 0.999631 | 0.790722 |
| Ank2_Q8C8R3_S2524s_1 | -1.53 | -1.36 | -2.05 | -2 | -1.16 | -1.43 | -1.64 | -1.86 | -1.34 | -1.24 | -1.6 | -1.64 | -0.3 | -0.82 | -1.67 | -1.62 | 0.2125 | 0.28 | 0.6325 | 0.135524 | 0.477001 | 0.341149 | 1 | 0.999631 | 0.607806 |
| Ank2_Q8C8R3_S2739s_1 | 0.7 | 0.95 | 0.09 | 0.39 | 1.08 | -0.21 | 0.41 | -1.45 | 1.63 | -0.1 | 1.12 | 1.75 | 0.37 | 0.84 | -0.46 | -0.48 | -0.575 | 0.5675 | -0.465 | 0.260818 | 0.290744 | 0.228893 | 1 | 0.999631 | 0.69271 |
| Ank2_Q8C8R3_S2746s_1 | 0.6 | 0.35 | 1.19 | 0.65 | 0.87 | 0.57 | 0.58 | 1.17 | 1.6 | 1.07 | 1.93 | 2.3 | 1.17 | 0.17 | 1.97 | 0.99 | 0.1 | 1.0275 | 0.3775 | 0.380813 | 0.73489 | 0.014865 | 1 | 0.999631 | 0.752619 |
| Ank2_Q8C8R3_S2769s_1 | 0.08 | -2.3 | -0.79 | -1.07 | -1.98 | -2.73 | -1.15 | -1.17 | -2.15 | -0.85 | -1.89 | -1.05 | -1.58 | -1.5 | 0.35 | -2.11 | -0.7375 | -0.465 | -0.19 | 0.771456 | 0.212651 | 0.401358 | 1 | 0.999631 | 0.919413 |
| Ank2_Q8C8R3_S2783s_1 | -1.71 | -2.47 | -2.06 | -2.18 | -2.45 | -2.16 | -1.69 | -2.04 | -2.29 | -0.98 | -2.76 | -1.63 | -1.01 | -1.95 | -2.45 | -1.37 | 0.02 | 0.19 | 0.41 | 0.302932 | 0.945388 | 0.653925 | 1 | 0.999631 | 0.714978 |
| Ank2_Q8C8R3_S27s_1 | -0.64 | 0.4 | 0.6 | -0.23 | 0.35 | 0.51 | -0.63 | 0.27 | 0.33 | 0.39 | -0.28 | 0.46 | 0.97 | 0.43 | 0.37 | 0.53 | 0.0925 | 0.1925 | 0.5425 | 0.158094 | 0.813494 | 0.599168 | 1 | 0.999631 | 0.628615 |
| Ank2_Q8C8R3_S2852s_1 | -0.13 | -0.3 | -1.38 | -0.41 | -0.2 | -0.64 | -0.6 | -0.28 | -0.27 | -0.29 | -0.29 | -0.41 | -0.15 | -0.35 | -1.2 | -0.77 | 0.125 | 0.24 | -0.0625 | 0.87373 | 0.712621 | 0.477165 | 1 | 0.999631 | 0.959802 |
| Ank2_Q8C8R3_S2852s_2 | -0.9 | -0.26 | 0.8 | 1.01 | 0.19 | -0.52 | 0.53 | 1.85 | -0.29 | -0.86 | 1.49 | 2.41 | 0.13 | -0.19 | 2.03 | 2.03 | 0.35 | 0.525 | 0.8375 | 0.229 | 0.566158 | 0.505244 | 1 | 0.999631 | 0.679627 |
| Ank2_Q8C8R3_S2855s_1 | 0.01 | 1.07 | 0.76 | 0.09 | 0.36 | 0.39 | -0.38 | 0.5 | 0.39 | 0.89 | 0.53 | 1.08 | 2.44 | 0.99 | 1.06 | 1.04 | -0.265 | 0.24 | 0.9 | 0.064053 | 0.461126 | 0.492553 | 1 | 0.999631 | 0.513532 |
| Ank2_Q8C8R3_S29s_1 | -0.55 | 0.14 | 0.02 | -0.51 | -0.4 | 0.13 | -0.57 | -0.34 | 0.05 | 0.31 | -0.35 | 0.06 | 0.73 | 0.2 | -0.19 | 0.12 | -0.07 | 0.2425 | 0.44 | 0.205838 | 0.814206 | 0.427436 | 1 | 0.999631 | 0.6605 |
| Ank2_Q8C8R3_S3060s_1 | -0.59 | -0.43 | -1.41 | -0.73 | 0.97 | 0.21 | -1.01 | -0.99 | 0.88 | -1.32 | -0.97 | -0.29 | -0.26 | -0.38 | -0.82 | -0.29 | 0.585 | 0.365 | 0.3525 | 0.296504 | 0.257169 | 0.473873 | 1 | 0.999631 | 0.709528 |
| Ank2_Q8C8R3_S30s_1 | -2.04 | -0.19 | 0.11 | -0.56 | -0.41 | -0.56 | -0.58 | 0.15 | -0.26 | -1 | 0.1 | 0.04 | 0.31 | -0.5 | 0.41 | -0.26 | 0.32 | 0.39 | 0.66 | 0.208555 | 0.509892 | 0.450938 | 1 | 0.999631 | 0.66217 |
| Ank2_Q8C8R3_S3218s_1 | -0.2 | -0.87 | -0.69 | 0.06 | 0.15 | -0.62 | -0.43 | 0.23 | -0.19 | -1.25 | -1.38 | 0.34 | -0.32 | -0.53 | -0.64 | -0.26 | 0.2575 | -0.195 | -0.0125 | 0.968619 | 0.452537 | 0.671009 | 1 | 0.999631 | 0.988771 |
| Ank2_Q8C8R3_S3254s_1 | -0.42 | -1.07 | -0.25 | -0.64 | -0.76 | -0.73 | -1.67 | -0.35 | -0.4 | -0.71 | -0.32 | -2.21 | 0.69 | -0.45 | -0.62 | -0.33 | -0.2825 | -0.315 | 0.4175 | 0.287427 | 0.436345 | 0.500857 | 1 | 0.999631 | 0.705972 |
| Ank2_Q8C8R3_S3264s_1 | 0.83 | 0.56 | -0.39 | 1.17 | 1.22 | 0.99 | 0.53 | 1.17 | 1.16 | 0.56 | 0.41 | 0.48 | -0.06 | 1.13 | 0.43 | 1.03 | 0.435 | 0.11 | 0.09 | 0.837653 | 0.271579 | 0.779551 | 1 | 0.999631 | 0.946036 |
| Ank2_Q8C8R3_S3343s_1 | -0.3 | -0.08 | -0.79 | -0.56 | 0.31 | -0.57 | -0.14 | -0.03 | -0.86 | -0.37 | 0.17 | -0.09 | -0.77 | -0.5 | -1.2 | -0.07 | 0.325 | 0.145 | -0.2025 | 0.560096 | 0.294728 | 0.657016 | 1 | 0.999631 | 0.839805 |
| Ank2_Q8C8R3_S3371s_1 | -0.41 | -0.77 | -0.07 | -0.64 | -1.28 | -0.69 | -0.46 | -1.12 | -0.66 | -0.14 | -0.77 | -1.26 | -0.5 | -1.6 | -0.28 | -0.68 | -0.415 | -0.235 | -0.2925 | 0.437421 | 0.193001 | 0.481212 | 1 | 0.999631 | 0.781057 |
| Ank2_Q8C8R3_S3390s_1 | -0.2 | 0.6 | 0.26 | -0.26 | -0.36 | 0.18 | -0.05 | -0.12 | 0.77 | 0.82 | 0.8 | 0.75 | 1.31 | 0.24 | 0.73 | 0.67 | -0.1875 | 0.685 | 0.6375 | 0.09569 | 0.532201 | 0.037679 | 1 | 0.999631 | 0.558128 |
| Ank2_Q8C8R3_S3407s_1 | 0.15 | -0.3 | -0.71 | 0 | 0.14 | -0.14 | -0.4 | -0.38 | 0 | -0.09 | -0.4 | -0.07 | -0.14 | -0.27 | -0.64 | -0.14 | 0.02 | 0.075 | -0.0825 | 0.792695 | 0.945925 | 0.798 | 1 | 0.999631 | 0.927581 |
| Ank2_Q8C8R3_S3439s_1 | 0.71 | -2.27 | -0.83 | -0.83 | -1.53 | -1.78 | 0.45 | -1.69 | -1.69 | -0.83 | -1.74 | -1.55 | -2.43 | -3.06 | -1.67 | -3.24 | -0.3325 | -0.6475 | -1.795 | 0.016893 | 0.642918 | 0.287019 | 1 | 0.999631 | 0.391024 |
| Ank2_Q8C8R3_S34s_1 | -0.91 | 0.15 | 0.19 | -0.2 | -0.06 | -0.09 | -0.58 | -0.04 | 0.12 | -0.19 | -0.15 | 0.36 | 0.66 | -0.22 | 0.2 | 0.08 | 9.81E-18 | 0.2275 | 0.3725 | 0.316661 | 1 | 0.501594 | 1 | 0.999631 | 0.720734 |
| Ank2_Q8C8R3_S34s_2 | -1.24 | 1.46 | 0.46 | 0.75 | 0.84 | 1.01 | -0.16 | 0.9 | 1.28 | 0.3 | 0.58 | 1.05 | 1.7 | 1.18 | 0.79 | 0.99 | 0.29 | 0.445 | 0.8075 | 0.17143 | 0.61705 | 0.439033 | 1 | 0.999631 | 0.641657 |
| Ank2_Q8C8R3_S3714s_1 | -1.5 | -0.8 | -0.33 | -0.99 | -1.11 | -1.48 | -0.76 | -0.68 | -1.25 | -1.54 | -0.53 | -0.29 | -0.31 | -0.85 | -0.28 | -0.12 | -0.1025 | 0.0025 | 0.515 | 0.161449 | 0.762879 | 0.994954 | 1 | 0.999631 | 0.631442 |
| Ank2_Q8C8R3_S3720s_1 | -0.56 | -0.84 | -0.87 | -1.08 | -0.4 | -0.6 | -0.84 | -0.49 | -0.7 | -0.88 | -0.78 | -0.64 | -0.35 | -0.14 | -0.77 | -0.61 | 0.255 | 0.0875 | 0.37 | 0.22417 | 0.330677 | 0.731966 | 1 | 0.999631 | 0.67471 |
| Ank2_Q8C8R3_S3792s_1 | -0.53 | 0.37 | 0.83 | -0.29 | -0.51 | 0.28 | 0.03 | 0.19 | -0.04 | 0.31 | 0.72 | 0.51 | 1.06 | 0.41 | 1.08 | 1.3 | -0.0975 | 0.28 | 0.8675 | 0.048438 | 0.794388 | 0.461669 | 1 | 0.999631 | 0.48343 |
| Ank2_Q8C8R3_S3808s_1 | 0.23 | -3.04 | -0.58 | 0.06 | -3.31 | -2.83 | 2.94 | -3.05 | -3.77 | -0.86 | -2.93 | -1.48 | -2.43 | -3 | -0.91 | -2.38 | -0.73 | -1.4275 | -1.3475 | 0.104052 | 0.611772 | 0.128149 | 1 | 0.999631 | 0.568213 |
| Ank2_Q8C8R3_S3814s_1 | -0.69 | -0.73 | -0.78 | -0.66 | -0.64 | -0.79 | -0.98 | -0.61 | -0.89 | -0.88 | -0.73 | -0.69 | -0.13 | -0.63 | -0.76 | -0.44 | -0.04 | -0.0825 | 0.225 | 0.427235 | 0.86625 | 0.731224 | 1 | 0.999631 | 0.775206 |
| Ank2_Q8C8R3_S3878s_1 | 2.33 | 1.12 | 0.18 | 1.87 | 0.56 | 1.06 | 1.1 | 0.4 | 0.64 | 1.27 | 0.24 | 0.41 | -0.08 | 0.08 | -0.04 | -0.73 | -0.595 | -0.735 | -1.5675 | 0.008744 | 0.230877 | 0.161327 | 1 | 0.999631 | 0.365244 |
| Ank2_Q8C8R3_S3879s_1 | 2.77 | 1.85 | 1.23 | 2.5 | 0.92 | 1.84 | 2.05 | 1.13 | 1.31 | 1.99 | 1.11 | 1.2 | 0.26 | 0.48 | 1.12 | 0.72 | -0.6025 | -0.685 | -1.4425 | 0.005257 | 0.184017 | 0.117746 | 1 | 0.999631 | 0.351308 |
| Ank2_Q8C8R3_S3921s_1 | 1.76 | -3.35 | -1.8 | -0.96 | -1.9 | -2.01 | 2.02 | -3.14 | -2.3 | -0.17 | -2.92 | -2.37 | -2.45 | -3.18 | -2.92 | -2.45 | -0.17 | -0.8525 | -1.6625 | 0.099155 | 0.897685 | 0.428833 | 1 | 0.999631 | 0.562298 |
| Ank2_Q8C8R3_S3s_2 | -0.19 | 0.5 | -0.73 | -3.02 | -0.39 | -0.28 | -0.01 | 0.68 | -0.32 | 0.35 | -0.82 | -0.05 | 0.16 | -0.89 | -2.02 | -0.08 | 0.86 | 0.65 | 0.1525 | 0.847315 | 0.241975 | 0.371992 | 1 | 0.999631 | 0.948838 |
| Ank2_Q8C8R3_S44s_1 | -0.64 | 0.11 | -0.13 | -0.7 | -0.45 | -0.04 | -0.85 | -0.7 | -0.02 | 0.09 | -0.3 | -0.01 | 0.61 | 0.14 | -0.34 | -0.34 | -0.17 | 0.28 | 0.3575 | 0.326305 | 0.593183 | 0.354315 | 1 | 0.999631 | 0.724678 |
| Ank2_Q8C8R3_S49s_1 | -0.91 | 0.15 | 0.19 | -0.2 | -0.06 | -0.09 | -0.58 | -0.04 | 0.12 | -0.19 | -0.15 | 0.36 | 0.66 | -0.22 | 0.2 | 0.08 | 9.81E-18 | 0.2275 | 0.3725 | 0.316661 | 1 | 0.501594 | 1 | 0.999631 | 0.720734 |
| Ank2_Q8C8R3_S49s_2 | -1.24 | 1.46 | 0.46 | 0.75 | 0.84 | 1.01 | -0.16 | 0.9 | 1.28 | 0.3 | 0.58 | 1.05 | 1.7 | 1.18 | 0.79 | 0.99 | 0.29 | 0.445 | 0.8075 | 0.17143 | 0.61705 | 0.439033 | 1 | 0.999631 | 0.641657 |
| Ank2_Q8C8R3_S588s_1 | -0.68 | 0.18 | -0.14 | -0.25 | -0.34 | -0.61 | -0.72 | 0.03 | -0.45 | -0.2 | -0.51 | 0.05 | 0.05 | -0.38 | -0.32 | 0.17 | -0.1875 | -0.055 | 0.1025 | 0.744291 | 0.54116 | 0.85341 | 1 | 0.999631 | 0.910233 |
| Ank2_Q8C8R3_S590s_1 | -0.82 | 0.02 | -0.14 | -0.41 | -0.4 | -0.24 | -0.4 | -0.16 | -0.58 | -0.09 | -0.35 | 0.09 | 0.26 | -0.15 | -0.35 | 0.25 | 0.0375 | 0.105 | 0.34 | 0.303114 | 0.892413 | 0.732217 | 1 | 0.999631 | 0.714978 |
| Ank2_Q8C8R3_S6s_2 | -0.19 | 0.5 | -0.73 | -3.02 | -0.39 | -0.28 | -0.01 | 0.68 | -0.32 | 0.35 | -0.82 | -0.05 | 0.16 | -0.89 | -2.02 | -0.08 | 0.86 | 0.65 | 0.1525 | 0.847315 | 0.241975 | 0.371992 | 1 | 0.999631 | 0.948838 |
| Ank2_Q8C8R3_S7s_1 | 2.16 | 2.15 | 0.49 | 2.07 | 2.31 | 3.06 | 1.75 | 2.55 | 2.36 | 1.44 | 2.36 | 1.39 | -0.98 | 2.56 | 1.98 | 1.32 | 0.7 | 0.17 | -0.4975 | 0.521751 | 0.15884 | 0.72079 | 1 | 0.999631 | 0.820623 |
| Ank2_Q8C8R3_S892s_1 | 1.69 | 0.32 | -0.93 | 2.22 | 1.95 | 1.63 | 1.89 | 0.87 | 1.4 | -0.14 | 0.98 | -0.77 | -1.96 | 0.87 | -0.16 | -0.24 | 0.76 | -0.4575 | -1.1975 | 0.15797 | 0.271801 | 0.553219 | 1 | 0.999631 | 0.628615 |
| Ank2_Q8C8R3_S939s_1 | 3.16 | -1.87 | -2.76 | 2.1 | 1.26 | -0.04 | 2.59 | -0.31 | 0.22 | -0.69 | 0.47 | -1.56 | -3.77 | -1.86 | -1.46 | -2.55 | 0.7175 | -0.5475 | -2.5675 | 0.07183 | 0.600453 | 0.675707 | 1 | 0.999631 | 0.532005 |
| Ank2_Q8C8R3_T2123t_1 | -0.19 | -0.28 | -1.7 | 0.84 | 0.35 | -0.67 | -0.18 | -0.48 | 0.16 | -1.07 | -1.17 | -0.81 | -0.26 | 0.43 | -1.22 | -0.5 | 0.0875 | -0.39 | -0.055 | 0.923564 | 0.867759 | 0.490147 | 1 | 0.999631 | 0.975032 |
| Ank2_Q8C8R3_T2260t_1 | -0.21 | -0.5 | -0.28 | -0.59 | -0.92 | -0.64 | -0.75 | -0.48 | -0.85 | -0.3 | -0.51 | -0.01 | -0.07 | -0.15 | -0.2 | -0.06 | -0.3025 | -0.0225 | 0.275 | 0.314499 | 0.244316 | 0.937533 | 1 | 0.999631 | 0.720156 |
| Ank2_Q8C8R3_T2361t_1 | -1.57 | -0.83 | -1.27 | -0.7 | -0.49 | -1.34 | -0.45 | -0.69 | -0.83 | -1.62 | -0.51 | -0.57 | -1.03 | -1.02 | -1.16 | -0.86 | 0.35 | 0.21 | 0.075 | 0.807194 | 0.302284 | 0.561751 | 1 | 0.999631 | 0.935599 |
| Ank2_Q8C8R3_T2570t_1 | -0.79 | -0.12 | 0.09 | -0.01 | -0.26 | -0.18 | -0.59 | -0.17 | -0.06 | 0.15 | -0.36 | 0.21 | 0.88 | 0.1 | 0.03 | 0.33 | -0.0925 | 0.1925 | 0.5425 | 0.135254 | 0.751856 | 0.534839 | 1 | 0.999631 | 0.607096 |
| Ank2_Q8C8R3_T2805t_1 | 1.55 | 0.45 | 0.73 | 2.21 | 0.84 | 1.41 | 1.56 | 0.24 | 1.17 | 2.29 | 0.93 | 1.05 | -1.68 | -0.04 | 0.48 | 1.14 | -0.2225 | 0.125 | -1.26 | 0.076087 | 0.641595 | 0.797325 | 1 | 0.999631 | 0.542138 |
| Ank2_Q8C8R3_T3063t_1 | 0.43 | 0.66 | 0.21 | 0.78 | -0.49 | 0.07 | 1.59 | 0.81 | 1.51 | -0.44 | 0.61 | 0.67 | -3.22 | -1.63 | 0.74 | -0.56 | -0.025 | 0.0675 | -1.6875 | 0.044093 | 0.955896 | 0.872783 | 1 | 0.999631 | 0.475443 |
| Ank2_Q8C8R3_T3078t_1 | -0.41 | -0.14 | -0.61 | -0.41 | -0.05 | -0.57 | -0.79 | -0.46 | -0.08 | -0.19 | -0.38 | -0.01 | -0.23 | -0.04 | -0.55 | -0.06 | -0.075 | 0.2275 | 0.1725 | 0.546326 | 0.783294 | 0.386262 | 1 | 0.999631 | 0.834213 |
| Ank2_Q8C8R3_T3746t_1 | -1.45 | -0.27 | -0.62 | -0.8 | 0.24 | -0.7 | 0.01 | -0.45 | 0.4 | -1.49 | -0.3 | -0.02 | 0.28 | -0.62 | -0.33 | -0.55 | 0.56 | 0.4325 | 0.48 | 0.209827 | 0.137666 | 0.360315 | 1 | 0.999631 | 0.66441 |
| Ank2_Q8C8R3_T3767t_1 | 0.19 | 0.94 | 0.4 | 0.4 | 0.74 | 0.3 | -0.22 | 0.43 | 0.83 | 0.74 | 0.21 | 1.08 | 1.75 | 1.01 | 0.65 | 1 | -0.17 | 0.2325 | 0.62 | 0.095464 | 0.587396 | 0.460677 | 1 | 0.999631 | 0.558128 |
| Ank2_Q8C8R3_T3773t_1 | -1 | -0.77 | 0.42 | -0.49 | -1.2 | -1.34 | -0.69 | -0.4 | -1.08 | -0.73 | -0.11 | 0.11 | 0.51 | -0.76 | 0.11 | 0.27 | -0.4475 | 0.0075 | 0.4925 | 0.264639 | 0.267487 | 0.985705 | 1 | 0.999631 | 0.6935 |
| Ank2_Q8C8R3_T3784t_1 | -0.03 | 0.03 | -0.51 | -0.83 | -0.38 | 0.35 | -0.89 | -0.9 | 0.04 | 0.45 | -0.94 | -0.52 | 0.39 | 0.4 | -0.37 | -0.11 | -0.12 | 0.0925 | 0.4125 | 0.246168 | 0.749268 | 0.81111 | 1 | 0.999631 | 0.685751 |
| Ank2_Q8C8R3_T3813t_1 | -0.91 | -0.74 | -1.16 | -0.85 | -1.01 | -1.26 | -1.03 | -0.88 | -1.33 | -1.25 | -1.06 | -0.68 | -0.59 | -0.95 | -0.98 | -0.58 | -0.13 | -0.165 | 0.14 | 0.618445 | 0.600581 | 0.551266 | 1 | 0.999631 | 0.861888 |
| Ank2_Q8C8R3_T5t_1 | -0.13 | -1.54 | -0.42 | -2.36 | -0.48 | -0.88 | -1.01 | -1.74 | -0.79 | 0.55 | -0.79 | -0.61 | -1.7 | -0.26 | -0.48 | -0.14 | 0.085 | 0.7025 | 0.4675 | 0.425778 | 0.873492 | 0.229627 | 1 | 0.999631 | 0.774624 |
| Ank2_Q8C8R3_T5t_2 | -0.19 | 0.5 | -0.73 | -3.02 | -0.39 | -0.28 | -0.01 | 0.68 | -0.32 | 0.35 | -0.82 | -0.05 | 0.16 | -0.89 | -2.02 | -0.08 | 0.86 | 0.65 | 0.1525 | 0.847315 | 0.241975 | 0.371992 | 1 | 0.999631 | 0.948838 |
| Ank3_A0A087WNU5_S1186s_1 | -1.59 | -3.07 | -0.5 | -0.39 | -2.09 | -3.1 | -0.14 | -1.44 | -2.57 | -2.2 | -1.79 | -1.37 | -2.06 | -3.46 | -0.99 | -0.63 | -0.305 | -0.595 | -0.3975 | 0.611702 | 0.693055 | 0.343439 | 1 | 0.999631 | 0.859812 |
| Ank3_A0A087WNU5_S1316s_1 | -1.6 | -0.15 | 0.01 | -0.81 | -0.49 | -0.69 | -0.81 | -0.32 | -0.39 | -1.08 | -0.35 | -0.14 | 0.4 | -0.59 | -0.42 | -0.98 | 0.06 | 0.1475 | 0.24 | 0.605505 | 0.877931 | 0.727479 | 1 | 0.999631 | 0.85817 |
| Ank3_A0A087WNU5_S1338s_1 | -0.05 | 0.84 | 0.31 | -0.34 | -0.39 | -0.43 | 0.38 | -1.22 | -0.19 | -0.09 | -1.02 | -0.27 | 0.42 | -0.78 | 0.21 | -1.28 | -0.605 | -0.5825 | -0.5475 | 0.258945 | 0.165612 | 0.132686 | 1 | 0.999631 | 0.692624 |
| Ank3_A0A087WNU5_S1378s_1 | -1.93 | -4.5 | -5.11 | -5.55 | -3.82 | -4.35 | -3.75 | -8.02 | -4.27 | -2.64 | -4.98 | -7.17 | -3.85 | -4.38 | -6.98 | -7.24 | -0.7125 | -0.4925 | -1.34 | 0.20755 | 0.527208 | 0.646218 | 1 | 0.999631 | 0.6614 |

| | | | | | | | | | | | | | | | | | | | | | | | | | |
|---|---|---|---|---|---|---|---|---|---|---|---|---|---|---|---|---|---|---|---|---|---|---|---|---|---|
| Ank3_A0A087WNU5_S1380s_1 | 1.04 | -3.12 | -1.96 | -1.36 | -2.65 | -1.59 | 1.82 | -4.05 | -2.8 | 0.88 | -2.04 | -2.61 | -1.53 | -2.74 | -2.87 | -3.65 | -0.2675 | -0.2925 | -1.3475 | 0.136067 | 0.837064 | 0.781848 | 1 | 0.999631 | 0.608909 |
| Ank3_A0A087WNU5_S1390s_1 | -0.43 | 0.49 | 0.89 | 0.56 | 0 | -0.13 | -0.09 | 0.4 | 0.44 | -0.17 | -0.4 | 0.23 | 0.75 | -0.34 | 0.83 | 0.94 | -0.3325 | -0.3525 | 0.1675 | 0.692911 | 0.342944 | 0.35112 | 1 | 0.999631 | 0.894897 |
| Ank3_A0A087WNU5_S2266s_1 | 1.17 | 2.33 | 2.14 | 0.77 | 1.48 | 2.65 | 1.06 | 1.13 | 1.7 | 2.32 | 1.74 | 1.2 | 1.35 | 1.62 | 1.61 | 2.65 | -0.0225 | 0.1375 | 0.205 | 0.661204 | 0.9637 | 0.75344 | 1 | 0.999631 | 0.880543 |
| Ank3_A0A087WNU5_S2266s_2 | 1.48 | 1.45 | 0.41 | 1.29 | 1.44 | 1.01 | 1.64 | 0.88 | 1.05 | 1.29 | 1.72 | -0.12 | -2.46 | 0.18 | -0.05 | 0.99 | 0.085 | -0.1725 | -1.4925 | 0.054085 | 0.804348 | 0.706365 | 1 | 0.999631 | 0.491994 |
| Ank3_A0A087WNU5_S2704s_1 | -0.14 | 2.11 | 0.32 | 1.06 | 1.86 | 0.25 | 0.61 | 0.85 | 1.89 | -0.21 | 0.78 | 1 | 2.3 | 0.52 | 0.43 | 0.76 | 0.055 | 0.0275 | 0.165 | 0.783474 | 0.920386 | 0.963166 | 1 | 0.999631 | 0.922628 |
| Ank3_A0A087WNU5_T2270t_1 | -0.03 | 0.21 | -0.41 | -0.29 | 0.12 | -0.34 | 0.16 | -0.5 | -0.18 | 0.13 | 0.17 | 0.27 | 0.32 | -0.25 | -0.63 | 0.02 | -0.01 | 0.2275 | -0.005 | 0.987682 | 0.972326 | 0.414795 | 1 | 0.999631 | 0.995409 |
| Ank3_A0A087WNU5_T2270t_2 | 1.48 | 1.45 | 0.41 | 1.29 | 1.44 | 1.01 | 1.64 | 0.88 | 1.05 | 1.29 | 1.72 | -0.12 | -2.46 | 0.18 | -0.05 | 0.99 | 0.085 | -0.1725 | -1.4925 | 0.054085 | 0.804348 | 0.706365 | 1 | 0.999631 | 0.491994 |
| Ank3_G5E8K5_S1648s_1 | 0.31 | -1.2 | -1.09 | 0.29 | -0.35 | -1.62 | 0.15 | -0.09 | -1.02 | -1.64 | -1.04 | -1.43 | -1.82 | -0.38 | -0.88 | -0.73 | -0.055 | -0.86 | -0.53 | 0.301008 | 0.917453 | 0.075047 | 1 | 0.999631 | 0.714243 |
| Ank3_G5E8K5_S1656s_1 | 2.44 | 2.06 | -0.64 | 2.25 | 2.59 | 2.83 | 1.96 | 1.04 | 2.71 | 2.68 | 0.94 | 0.39 | 1.41 | 1.94 | 0.85 | 0.06 | 0.5775 | 0.1525 | -0.4625 | 0.531559 | 0.43676 | 0.853035 | 1 | 0.999631 | 0.827175 |
| Ank3_G5E8K5_S27s_1 | 0.13 | 1.05 | 1.01 | -0.04 | 0.35 | 1.67 | -0.22 | 0.26 | 0.9 | 0.79 | 0.1 | 0.85 | 0.43 | 1.35 | 0.72 | 0.97 | -0.0225 | 0.1225 | 0.33 | 0.39662 | 0.96196 | 0.741304 | 1 | 0.999631 | 0.761029 |
| Ank3_G5E8K5_S507s_1 | 0.07 | 0.65 | -0.35 | -0.32 | 0.05 | 0.89 | 0 | 1.23 | 0.19 | 0.07 | -0.69 | 0.36 | 1.05 | 1.05 | 0.56 | 0 | 0.53 | -0.03 | 0.6525 | 0.109139 | 0.198205 | 0.933778 | 1 | 0.999631 | 0.572424 |
| Ank3_G5E8K5_S606s_1 | -0.93 | -1.01 | -0.37 | -1.5 | -1.47 | -1.11 | -1.77 | -2.15 | -0.7 | -1.41 | -1.85 | -1.08 | 0.19 | -1.23 | -1.9 | -1.07 | -0.6725 | -0.3075 | -0.05 | 0.916962 | 0.078758 | 0.409973 | 1 | 0.999631 | 0.973513 |
| Ank3_G5E8K5_S919s_1 | -2.05 | 0.61 | 0.03 | -1.95 | 0.02 | -0.04 | -0.92 | -2.58 | 0.53 | -0.95 | -0.91 | -0.16 | 0.38 | -0.17 | -0.78 | -0.64 | -0.04 | 0.4675 | 0.5375 | 0.419148 | 0.959885 | 0.499558 | 1 | 0.999631 | 0.772152 |
| Ank3_G5E8K5_S921s_1 | -0.76 | 0.93 | -0.21 | 0.88 | 2.18 | 0.33 | 2.19 | 0.5 | 1.91 | -2.57 | 1.08 | 0.97 | -0.24 | -0.04 | -0.05 | 0.68 | 1.09 | 0.1375 | -0.1225 | 0.789604 | 0.092891 | 0.883081 | 1 | 0.999631 | 0.925593 |
| Ank3_G5E8K5_T1201t_1 | 0.32 | 0.02 | 0.65 | -0.37 | -0.87 | -0.36 | -0.38 | -0.13 | -0.71 | -0.13 | -0.14 | -0.06 | 0.13 | -0.07 | 0.17 | 0.33 | -0.59 | -0.415 | -0.015 | 0.962331 | 0.087581 | 0.217738 | 1 | 0.999631 | 0.986642 |
| Ank3_S4R162_S33s_1 | 0.57 | 1.94 | 1.14 | 0.12 | 1.11 | 2.71 | 1.21 | 0.44 | 2.19 | 1.1 | 0.75 | 1.01 | 1.53 | 0.69 | 0.58 | -0.84 | 0.425 | 0.32 | -0.4525 | 0.441046 | 0.458979 | 0.513204 | 1 | 0.999631 | 0.781983 |
| Ank3_S4R162_S73s_1 | 0.39 | -0.64 | -1 | -0.95 | -1.21 | -0.12 | -0.37 | -2.34 | -1.65 | 0.12 | -1.87 | -2.24 | -0.67 | -1.58 | -1.39 | -1.75 | -0.46 | -0.86 | -0.7975 | 0.079385 | 0.409719 | 0.153081 | 1 | 0.999631 | 0.545579 |
| Ank3_S4R162_S87s_1 | 0.73 | 0.23 | -0.51 | 0.71 | 1.65 | -0.17 | 0.5 | 0.85 | 0.62 | -0.05 | 0.66 | 0.47 | -0.46 | 0.26 | 0.09 | 0.59 | 0.4175 | 0.135 | -0.17 | 0.666463 | 0.373172 | 0.711782 | 1 | 0.999631 | 0.88264 |
| Ank3_S4R162_T32t_1 | -0.59 | -2.45 | -1.54 | -2.24 | -0.4 | -1.93 | -0.64 | -1.38 | -0.25 | -1.05 | -1.27 | -0.29 | -0.18 | -2.81 | -2.37 | -4.22 | 0.6175 | 0.99 | -0.69 | 0.405765 | 0.243827 | 0.060797 | 1 | 0.999631 | 0.765969 |
| Ank3_S4R1S2_S1411s_1 | 0.73 | 0.23 | -0.51 | 0.71 | 1.65 | -0.17 | 0.5 | 0.85 | 0.62 | -0.05 | 0.66 | 0.47 | -0.46 | 0.26 | 0.09 | 0.59 | 0.4175 | 0.135 | -0.17 | 0.666463 | 0.373172 | 0.711782 | 1 | 0.999631 | 0.88264 |
| Ank3_S4R1S2_S1421s_1 | -2.06 | -1.44 | -1.27 | -0.29 | -1.27 | -1.89 | -1.03 | -0.43 | -2.1 | -2.06 | -0.67 | -0.6 | -1.6 | -1.94 | -1 | -1.1 | 0.11 | -0.0925 | -0.145 | 0.738893 | 0.809836 | 0.859172 | 1 | 0.999631 | 0.908082 |
| Ank3_S4R2K9_S1645s_1 | -0.69 | -1.2 | -1.11 | -0.91 | -1.01 | -1.23 | -1.01 | -0.81 | -1.06 | -1.17 | -0.9 | -0.44 | 0.14 | -0.67 | -0.54 | -0.26 | -0.0375 | 0.085 | 0.645 | 0.058729 | 0.883046 | 0.767086 | 1 | 0.999631 | 0.501503 |
| Ank3_S4R2K9_S1679s_1 | 0.68 | -0.85 | -0.09 | -0.61 | 0.09 | 0.11 | -0.04 | -0.51 | 0.16 | 0.49 | -0.75 | -1.17 | -1.5 | -0.11 | -0.15 | -0.39 | 0.13 | -0.1 | -0.32 | 0.495281 | 0.733693 | 0.838422 | 1 | 0.999631 | 0.810677 |
| Ank3_S4R2K9_S1853s_1 | 0.99 | -1.42 | -2.57 | 0.11 | 1.54 | 0.5 | 0.93 | 0.12 | 0.19 | -1.67 | -0.32 | -1.33 | -1.94 | -0.3 | -1.4 | -1.22 | 1.495 | -0.06 | -0.4925 | 0.519578 | 0.069019 | 0.939623 | 1 | 0.999631 | 0.819456 |
| Ank3_S4R2K9_S2113s_1 | 1.04 | 1.11 | 0.18 | 0.79 | 1.76 | 1.67 | 1.04 | 1.49 | 1.27 | 0.84 | 0.85 | 1.22 | 0.52 | 2.08 | 0.39 | 1.54 | 0.71 | 0.265 | 0.3525 | 0.445952 | 0.045817 | 0.400834 | 1 | 0.999631 | 0.784402 |
| Ank3_S4R2K9_S2215s_1 | 1.77 | -1.02 | -0.54 | -0.04 | -1.85 | -0.7 | 2.43 | -2.25 | -2.11 | 0.82 | -1.3 | -1.82 | -2.22 | -2.85 | -1.99 | -2.57 | -0.635 | -1.145 | -2.45 | 0.001845 | 0.548775 | 0.170469 | 1 | 0.999631 | 0.351308 |
| Ank3_S4R2K9_S2246s_1 | -0.17 | -0.9 | -0.06 | -0.32 | -0.92 | -0.63 | -0.21 | -0.84 | -0.74 | 0.06 | -0.49 | 0.01 | 0.5 | -0.6 | -0.32 | 0.05 | -0.2875 | 0.0725 | 0.27 | 0.454562 | 0.356747 | 0.823826 | 1 | 0.999631 | 0.789731 |
| Ank3_S4R2K9_S2330s_1 | 2.91 | 0.29 | 0 | 1.33 | 0.17 | 1.01 | 1.82 | 0.61 | 0.14 | 1.44 | -0.32 | -0.59 | -1.4 | -0.55 | 0.1 | -0.84 | -0.23 | -0.965 | -1.805 | 0.018766 | 0.729587 | 0.19458 | 1 | 0.999631 | 0.391024 |
| Ankhd1_F7DE17_S168s_1 | 0.66 | 1.17 | 1.78 | 0.92 | 1.01 | 1.38 | 1.1 | 1.97 | 1.16 | 0.44 | 2.62 | 2.29 | 0.62 | 1.77 | 3.04 | 2.11 | 0.2325 | 0.495 | 0.7525 | 0.172277 | 0.513084 | 0.35589 | 1 | 0.999631 | 0.642116 |
| Ankhd1_F7DE17_T29t_1 | 1.23 | 0.75 | 0.2 | 0.47 | 0.58 | 1.73 | 1.15 | 0.46 | 0.95 | 1.12 | 1.43 | 1.15 | -0.2 | 1.38 | 1.01 | 0.21 | 0.3175 | 0.5 | -0.0625 | 0.884686 | 0.411282 | 0.13049 | 1 | 0.999631 | 0.963505 |
| Ankib1_Q6ZPS6_S738s_1 | -0.14 | -0.25 | -0.92 | -1.13 | -0.77 | -0.59 | -1.18 | -0.99 | -0.54 | 0.08 | -0.63 | -0.84 | 0.38 | -0.31 | -0.94 | -0.39 | -0.2725 | 0.1275 | 0.295 | 0.459479 | 0.407063 | 0.71816 | 1 | 0.999631 | 0.79152 |
| Ankrd11_E9Q4F7_S1676s_1 | 0.64 | 1.27 | 0.93 | 0.52 | 0.48 | 0.97 | 0.21 | 0.3 | 0.61 | 1.33 | 0.67 | 0.63 | 0.65 | 1.08 | 0.67 | 1.56 | -0.35 | -0.03 | 0.15 | 0.659035 | 0.261321 | 0.922688 | 1 | 0.999631 | 0.880034 |
| Ankrd12_G5E893_T1349t_1 | 1.37 | -0.02 | 1.63 | 2.19 | 0.31 | -0.18 | 0.94 | 1.94 | 0.27 | 2.48 | 1.05 | 3.11 | 0.04 | 0.76 | 2.22 | 2.2 | -0.54 | 0.435 | 0.0125 | 0.984525 | 0.373471 | 0.545241 | 1 | 0.999631 | 0.995133 |
| Ankrd13c_Q3UX43_S411s_1 | -0.61 | 0.14 | -0.2 | -0.58 | -0.41 | 0.15 | -0.88 | -0.48 | -0.04 | -0.54 | -0.46 | -0.06 | 0.08 | 0.13 | -0.7 | -0.07 | -0.0925 | 0.0375 | 0.1725 | 0.606363 | 0.774672 | 0.899938 | 1 | 0.999631 | 0.858185 |
| Ankrd13d_Q6PD24_T556t_1 | -0.89 | -0.85 | 0.23 | -0.53 | -0.92 | -0.69 | -0.73 | -0.77 | -0.76 | -0.33 | -0.09 | -0.06 | -0.72 | -0.57 | 0.09 | 0.11 | -0.2675 | 0.2 | 0.2375 | 0.532682 | 0.405522 | 0.56756 | 1 | 0.999631 | 0.8277 |
| Ankrd17_E9QKG6_S1804s_1 | -0.03 | -0.04 | 0.69 | 0.1 | 0.35 | 0.27 | 0.1 | 0.74 | 0.51 | 0.54 | -0.4 | 0.66 | 1.06 | 1.04 | -0.04 | 0.7 | 0.185 | 0.1475 | 0.51 | 0.177795 | 0.528913 | 0.668671 | 1 | 0.999631 | 0.642116 |
| Ankrd17_E9QKG6_S2150s_1 | 1.05 | 0.37 | -1.97 | -1.29 | 0.68 | 0.1 | 0.1 | -1.26 | 0.33 | -0.25 | -1.35 | -0.36 | -0.69 | -0.33 | -2.51 | -0.22 | 0.365 | 0.0525 | -0.4775 | 0.540526 | 0.613983 | 0.940098 | 1 | 0.999631 | 0.831391 |
| Ankrd27_Q3UMR0_S634s_1 | -0.39 | -1.75 | -0.38 | -0.64 | -1.31 | -1.52 | -0.62 | -0.86 | -1.14 | -0.76 | -1.08 | -0.29 | -0.86 | -0.33 | -0.47 | -0.52 | -0.2875 | -0.0275 | 0.245 | 0.524059 | 0.470279 | 0.944365 | 1 | 0.999631 | 0.822727 |
| Ankrd34a_B2RW11_S149s_1 | -0.91 | -1.41 | -0.25 | -0.5 | -1.18 | -1.65 | -0.71 | -0.57 | -1.22 | -1.93 | -0.08 | -0.29 | -0.35 | -0.74 | -0.07 | 0.09 | -0.26 | -0.1125 | 0.5 | 0.187198 | 0.488873 | 0.814315 | 1 | 0.999631 | 0.646474 |
| Ankrd34a_B2RW11_S183s_1 | -0.92 | -0.77 | -0.36 | -0.48 | -0.53 | -0.84 | -1.15 | -0.13 | -0.94 | -0.86 | -0.44 | 0.01 | 0.41 | -0.45 | 0.14 | -0.14 | -0.03 | 0.075 | 0.6225 | 0.071089 | 0.92264 | 0.81269 | 1 | 0.999631 | 0.531307 |
| Ankrd34a_B2RW11_S225s_1 | -0.68 | -0.65 | -0.9 | 0.57 | 0.36 | 0.02 | 0.16 | 0.77 | -0.02 | -0.61 | -0.12 | -0.22 | -1.77 | -0.8 | -0.02 | -0.02 | 0.7425 | 0.1725 | -0.2375 | 0.641406 | 0.076857 | 0.651172 | 1 | 0.999631 | 0.871335 |
| Ankrd34a_B2RW11_S246s_1 | 0.02 | 0.47 | 0.3 | -0.01 | 0.2 | 0.34 | 0.01 | -0.06 | 0.6 | -0.01 | 0.17 | 0.01 | 0.48 | 1 | -0.05 | 0.24 | -0.0725 | -0.0025 | 0.2225 | 0.501371 | 0.777993 | 0.992868 | 1 | 0.999631 | 0.814092 |
| Ankrd34a_B2RW11_S391s_1 | -0.07 | 0.04 | -0.74 | 0.25 | 1.05 | 0.35 | -0.1 | -0.03 | 0.76 | -0.93 | 0.07 | -0.49 | -0.97 | 0.55 | -0.23 | 0.13 | 0.4475 | -0.0175 | 1.57E-16 | 1 | 0.234994 | 0.967102 | 1 | 0.999631 | 1 |
| Ankrd34a_B2RW11_S408s_1 | -0.58 | -1.73 | -0.4 | -0.15 | -1.24 | -1.05 | 0.26 | 0.13 | -1.59 | -0.92 | 0.73 | 0.04 | -2.95 | -1.04 | -0.29 | 0.82 | 0.24 | 0.28 | -0.15 | 0.843564 | 0.628145 | 0.625687 | 1 | 0.999631 | 0.94821 |
| Ankrd34a_B2RW11_S460s_1 | -0.02 | 1.09 | -0.21 | 0.53 | 0.8 | 0.77 | 0.35 | 0.49 | 0.91 | -0.07 | 0.76 | 0.68 | 0.51 | 1.27 | 0.91 | 1 | 0.255 | 0.2225 | 0.575 | 0.146525 | 0.466247 | 0.566829 | 1 | 0.999631 | 0.618378 |
| Ankrd34a_B2RW11_S472s_1 | -0.14 | 3.68 | 2.88 | -0.56 | 5.31 | 1.98 | 4.69 | 1.4 | 7.23 | -0.44 | 4.31 | 5.74 | 6.38 | 0.94 | 2.32 | -0.1 | 1.88 | 2.745 | 0.92 | 0.540899 | 0.149564 | 0.125496 | 1 | 0.999631 | 0.831715 |
| Ankrd34a_B2RW11_T315t_1 | -0.35 | 0.82 | 1.45 | 0.37 | 0.34 | 0.42 | 0.5 | 1.04 | 0.51 | 0.24 | 1.22 | 1.05 | 0.65 | 0.77 | 1.3 | 1.46 | 0.0025 | 0.1825 | 0.4725 | 0.290965 | 0.995137 | 0.678511 | 1 | 0.999631 | 0.706164 |
| Ankrd34b_Q3UUF8_S293s_1 | -0.29 | 3.01 | 0.32 | -0.97 | 4.13 | 0.98 | 2.14 | -0.13 | 6.32 | 0.23 | 2 | 2.57 | 4.38 | 1.4 | 0.88 | -0.25 | 1.2625 | 2.2625 | 1.085 | 0.343949 | 0.259556 | 0.111971 | 1 | 0.999631 | 0.733852 |
| Ankrd34c_Q8BLB8_S231s_1 | -0.17 | -0.05 | 0.62 | -0.47 | 0.43 | -0.55 | 0.27 | -0.73 | 0.19 | -0.32 | 0.53 | 0.09 | -0.03 | -0.77 | 0.53 | 0.56 | -0.1275 | 0.14 | 0.09 | 0.825338 | 0.739461 | 0.67886 | 1 | 0.999631 | 0.940329 |
| Ankrd34c_Q8BLB8_T347t_1 | 1 | 1.68 | -0.53 | 0.52 | 0.91 | 1.5 | 1.33 | 0.87 | 1.22 | 0.55 | 1.01 | 0 | -0.4 | 1.65 | -0.79 | -0.21 | 0.485 | 0.0275 | -0.605 | 0.357527 | 0.313757 | 0.95679 | 1 | 0.999631 | 0.740986 |
| Ankrd50_F7BE84_S1101s_1 | -1.48 | -0.5 | -0.41 | 0.21 | -1.08 | -1.22 | -1.17 | 1.45 | -0.77 | -0.71 | 1.1 | -0.13 | -0.61 | -0.61 | -0.27 | -0.55 | 0.04 | 0.4175 | 0.035 | 0.928314 | 0.951592 | 0.432079 | 1 | 0.999631 | 0.976721 |
| Ankrd50_F7BE84_S1279s_1 | 1.67 | -0.7 | 0.05 | -0.44 | -1.44 | 0.17 | 2.13 | -2.12 | -2.46 | 1.35 | -1.47 | -1.61 | -1.13 | -1.96 | -1.48 | -1.86 | -0.46 | -1.1925 | -1.7525 | 0.007586 | 0.625452 | 0.186788 | 1 | 0.999631 | 0.358173 |
| Ankrd63_A2ARS0_S193s_1 | -1.2 | -0.15 | -0.15 | -2 | -3.33 | -2.6 | -3.39 | -1.75 | -1.13 | 0.21 | -2.02 | -0.56 | 1.09 | -0.48 | 0.07 | -0.33 | -1.8925 | 3.14E-16 | 0.9625 | 0.096893 | 0.005866 | 1 | 1 | 1 | 0.55886 |
| Ankrd63_A2ARS0_S193s_2 | 0.57 | -0.65 | -1.45 | 0.78 | 0.52 | -1.23 | -0.23 | 0.82 | -0.2 | -0.4 | -0.78 | -0.52 | -2.17 | -0.48 | -0.02 | -0.45 | 0.1575 | -0.2875 | -0.5925 | 0.363898 | 0.801326 | 0.575581 | 1 | 0.999631 | 0.743941 |
| Ankrd63_A2ARS0_S203s_1 | 0.85 | -1.2 | -1.87 | -0.18 | -0.15 | -1.23 | -0.32 | 0 | -0.87 | -0.52 | -1.44 | -0.96 | -2.35 | -1.37 | -0.91 | -1.64 | 0.175 | -0.3475 | -0.9675 | 0.134463 | 0.768591 | 0.548652 | 1 | 0.999631 | 0.605974 |
| Ankrd63_A2ARS0_S203s_2 | 0.56 | -0.71 | -1.47 | 0.81 | 0.5 | -1.23 | -0.15 | 0.85 | -0.25 | -0.4 | -0.72 | -0.54 | -2.13 | -0.34 | -0.01 | -0.59 | 0.195 | -0.275 | -0.565 | 0.388433 | 0.758315 | 0.595966 | 1 | 0.999631 | 0.757772 |
| Ankrd63_A2ARS0_S268s_1 | -0.56 | 0.12 | -0.43 | 0.66 | 0.15 | -0.52 | -0.34 | 0.08 | -0.13 | -0.96 | -0.09 | -0.46 | -1.25 | -0.3 | -0.36 | -1.05 | -0.105 | -0.3575 | -0.6875 | 0.106416 | 0.765581 | 0.347548 | 1 | 0.999631 | 0.568797 |
| Ankrd63_A2ARS0_S293s_1 | 1.77 | -2.66 | -4.02 | 0.01 | 1.01 | -2.95 | 0.24 | 0.23 | -0.59 | -3.44 | -1.39 | -2.02 | -3 | -1.7 | -2.07 | -3.68 | 0.8575 | -0.635 | -1.3875 | 0.251889 | 0.525708 | 0.607418 | 1 | 0.999631 | 0.687703 |
| Ankrd63_A2ARS0_S341s_1 | 1.24 | 0.47 | -1.31 | 2.27 | 2.89 | -0.29 | 0.63 | 2.11 | 1.5 | -0.08 | 0.42 | 1.23 | 0.13 | 0.66 | 1.27 | 0.72 | 0.6675 | 0.1 | 0.0275 | 0.968667 | 0.465238 | 0.892678 | 1 | 0.999631 | 0.988771 |
| Ankrd63_A2ARS0_T342t_1 | 1.24 | 0.47 | -1.31 | 2.27 | 2.89 | -0.29 | 0.63 | 2.11 | 1.5 | -0.08 | 0.42 | 1.23 | 0.13 | 0.66 | 1.27 | 0.72 | 0.6675 | 0.1 | 0.0275 | 0.968667 | 0.465238 | 0.892678 | 1 | 0.999631 | 0.988771 |
| Anks1_Q3UHP6_S663s_1 | 0.25 | 0.92 | 0.37 | -0.84 | 0.77 | 1.13 | -0.58 | 0.41 | 1.05 | 0.84 | -0.92 | -1.19 | 1.57 | 1.04 | -0.15 | -0.72 | 0.2575 | -0.23 | 0.26 | 0.660036 | 0.6022 | 0.712895 | 1 | 0.999631 | 0.880315 |
| Anks1_Q3UHP6_S903s_1 | -0.81 | -1.24 | 0.6 | -1.47 | 0.92 | -1.32 | 1.85 | 0.1 | 0.92 | -0.27 | 2.03 | 2.77 | 1.66 | -2.49 | 0.5 | 0.54 | 1.1175 | 2.0925 | 0.7825 | 0.376985 | 0.144721 | 0.015046 | 1 | 0.999631 | 0.749236 |
| Anks1b_S4R286_S1222s_1 | 1.61 | -0.72 | 0.19 | 0.88 | 1.78 | 0.05 | 1.13 | 0.6 | 0.01 | -0.91 | 0.91 | -1.03 | -1.96 | -0.29 | -1.26 | 0.13 | 0.4 | -0.745 | -1.335 | 0.053349 | 0.484931 | 0.240672 | 1 | 0.999631 | 0.490856 |
| Anks1b_S4R286_S364s_1 | -0.17 | 0.34 | 1.73 | 1.69 | 0.09 | 0.47 | 1.07 | 0.93 | 0.52 | 1.14 | 2.06 | 2.57 | 0.31 | 0.92 | 2.72 | 2.76 | -0.2575 | 0.675 | 0.78 | 0.28048 | 0.607189 | 0.280184 | 1 | 0.999631 | 0.701922 |
| Anks1b_S4R286_S510s_1 | -1.23 | -0.04 | -0.2 | -0.17 | 0.03 | 0.03 | -0.88 | -0.48 | -0.11 | -0.63 | -0.2 | 0.17 | 0.35 | -0.18 | -0.06 | -0.25 | 0.085 | 0.2175 | 0.375 | 0.308409 | 0.818884 | 0.545922 | 1 | 0.999631 | 0.718521 |
| Anks1b_S4R286_S589s_1 | 0.32 | 5.62 | 2.3 | 0.69 | 4.8 | 4.38 | 1.84 | 1.5 | 5.05 | 1.83 | 3.05 | 4.07 | 5.55 | 3.17 | 2.32 | 1.35 | 0.8975 | 1.2675 | 0.865 | 0.501166 | 0.481681 | 0.302075 | 1 | 0.999631 | 0.814092 |
| Anks1b_S4R286_T968t_1 | -1.99 | -2.53 | -0.96 | -0.98 | -2.85 | -2.27 | -1.69 | -1.33 | -2.72 | -1.66 | -1.57 | -1.14 | -1.6 | -2.21 | -1.65 | -0.43 | -0.42 | -0.1575 | 0.1425 | 0.781105 | 0.395086 | 0.748279 | 1 | 0.999631 | 0.921543 |
| Anln_Q8K298_S637s_1 | -0.3 | 0.39 | 1.87 | 0.56 | 0.46 | 0.72 | 1.08 | 0.57 | 1.1 | 0.21 | 1.35 | 2.07 | 0.29 | 0.44 | 1.6 | 0.56 | 0.0775 | 0.5525 | 0.0925 | 0.857285 | 0.864809 | 0.328505 | 1 | 0.999631 | 0.952021 |
| Anln_Q8K298_S656s_1 | 0.5 | 1.05 | 1.71 | 0.49 | 0.21 | 0.77 | -0.25 | -0.32 | 0.83 | 1.9 | 0.46 | 0.64 | 2.04 | 0.47 | 0.91 | 0.1 | -0.835 | 0.02 | -0.0575 | 0.906809 | 0.055453 | 0.963034 | 1 | 0.999631 | 0.969801 |
| Ano8_Q6PB70_S637s_1 | -0.23 | 0.11 | -0.15 | -0.55 | -0.03 | -0.23 | -0.73 | -0.41 | -0.33 | -0.02 | -0.2 | -0.33 | 0.57 | -0.25 | -0.19 | 0.1 | -0.145 | -0.015 | 0.2625 | 0.416471 | 0.608703 | 0.955309 | 1 | 0.999631 | 0.770712 |
| Antxr1_Q9CZ52_S561s_1 | -0.34 | -0.52 | -0.59 | -0.75 | -0.32 | -0.74 | -0.36 | -0.36 | -0.86 | -0.43 | -1.04 | -0.95 | 0.16 | -0.64 | -0.72 | -0.78 | 0.105 | -0.27 | 0.055 | 0.863417 | 0.676383 | 0.330081 | 1 | 0.999631 | 0.955107 |
| Anxa7_Q07076_S311s_1 | 0.49 | -0.41 | 0.68 | -0.08 | -0.49 | -0.27 | -0.01 | -0.28 | -0.11 | 0.74 | 0.4 | 0.4 | -0.23 | -0.61 | 0.07 | -0.4 | -0.4325 | 0.1875 | -0.4625 | 0.204353 | 0.196251 | 0.592306 | 1 | 0.999631 | 0.659876 |
| Ap1g1_Q8CBB7_S235s_1 | -0.5 | -0.52 | -0.3 | -0.45 | -0.63 | -0.62 | -0.64 | -0.6 | -0.84 | -0.48 | -0.67 | -0.02 | -0.21 | -0.24 | -0.35 | -0.08 | -0.18 | -0.06 | 0.2225 | 0.403394 | 0.440538 | 0.830656 | 1 | 0.999631 | 0.764348 |
| Ap2a1_P17426_S517s_1 | -0.74 | -2.04 | -0.8 | -2.11 | -1.56 | -1.86 | -0.94 | -0.67 | -2.21 | -2.11 | -1.55 | -0.21 | -1.27 | -2.54 | -0.67 | -1.11 | 0.165 | -0.0975 | 0.025 | 0.961664 | 0.714855 | 0.859582 | 1 | 0.999631 | 0.986594 |
| Ap2a1_P17426_S518s_1 | -0.74 | -2.04 | -0.8 | -2.11 | -1.56 | -1.86 | -0.94 | -0.67 | -2.21 | -2.11 | -1.55 | -0.21 | -1.27 | -2.54 | -0.67 | -1.11 | 0.165 | -0.0975 | 0.025 | 0.961664 | 0.714855 | 0.859582 | 1 | 0.999631 | 0.986594 |
| Ap2a1_P17426_S611s_1 | 0.8 | 0.76 | 0.03 | 0.02 | 0.58 | 0.58 | 0.19 | 0.89 | 0.03 | 0.56 | 0.86 | -0.18 | -0.84 | 0.87 | 0.19 | 0.91 | 0.1575 | -0.085 | -0.12 | 0.793791 | 0.617715 | 0.812362 | 1 | 0.999631 | 0.928267 |
| Ap2a1_P17426_S626s_1 | 0.14 | 0.3 | 0.64 | -0.13 | -0.91 | 0.06 | -0.14 | -0.21 | -0.78 | -0.17 | -0.11 | -0.3 | -0.19 | 0.15 | 0 | 0.35 | -0.5375 | -0.5775 | -0.16 | 0.598166 | 0.11386 | 0.077645 | 1 | 0.999631 | 0.855276 |
| Ap2a1_P17426_S631s_1 | 0.96 | -0.15 | 0.3 | 0.5 | 0.64 | 0.91 | 0.51 | 1.07 | 0.04 | 0 | 0.55 | 0.09 | 0.26 | 1.09 | 0.43 | 0.38 | 0.38 | -0.2325 | 0.1375 | 0.697153 | 0.245205 | 0.476013 | 1 | 0.999631 | 0.895731 |
| Ap2a1_P17426_S655s_1 | -0.72 | -0.09 | 2.27 | 0.97 | 0 | -0.41 | 0.04 | 0.22 | -0.68 | -1.35 | 0.66 | 1.95 | 0.67 | 0.48 | 1.53 | 0.51 | -0.645 | -0.4625 | 0.19 | 0.763879 | 0.299608 | 0.592282 | 1 | 0.999631 | 0.915728 |
| Ap2m1_P84091_S45s_1 | -0.33 | -0.24 | -0.14 | -0.68 | -0.97 | -0.77 | -0.94 | -1.1 | -1.14 | -0.64 | -0.87 | -1.17 | -0.66 | -0.85 | -1.46 | -1.21 | -0.5975 | -0.6075 | -0.6975 | 0.044513 | 0.037374 | 0.048811 | 1 | 0.999631 | 0.475443 |
| Ap2m1_P84091_T156t_1 | -0.17 | 0.24 | -0.33 | 0.32 | 0.23 | -0.01 | 0.08 | 1.26 | 0.18 | -0.32 | 0.63 | 0.36 | 0.7 | 0.55 | 0.42 | 0.33 | 0.375 | 0.1975 | 0.485 | 0.121387 | 0.309299 | 0.537038 | 1 | 0.999631 | 0.589748 |
| Ap3b2_Q9JME5_S272s_1 | -0.48 | 0.65 | 0.71 | 0.12 | 0.37 | 0.35 | -0.09 | 0.88 | 0.34 | 0.21 | 0.37 | 0.75 | 1.31 | 0.56 | 0.78 | 0.51 | 0.1275 | 0.1675 | 0.54 | 0.169598 | 0.726054 | 0.627987 | 1 | 0.999631 | 0.639508 |
| Ap3b2_Q9JME5_S8s_1 | 2.1 | 1.14 | 0.19 | 1.85 | 2.02 | 0.94 | 1.63 | 1.44 | 1.84 | 1.81 | 0.78 | 0.77 | 0.49 | 1.13 | 1.57 | 0.71 | 0.1875 | -0.02 | -0.345 | 0.47567 | 0.686512 | 0.967944 | 1 | 0.999631 | 0.799882 |
| Ap3d1_O54774_S760s_1 | 0.31 | 2.72 | 1.75 | -0.78 | 3.34 | 1.49 | 1.41 | 0.78 | 3.95 | 0.19 | 2.96 | 3.75 | 5.03 | 0.79 | 1.67 | 0.56 | 0.755 | 1.7125 | 1.0125 | 0.366783 | 0.371857 | 0.112783 | 1 | 0.999631 | 0.745796 |
| Ap3d1_O54774_S825s_1 | 0.66 | -0.53 | -0.85 | -0.62 | -0.46 | -0.58 | -0.02 | -0.83 | 0.08 | -0.09 | -1.07 | -1.28 | 0.46 | -1.16 | -0.92 | -1.51 | -0.1375 | -0.255 | -0.4475 | 0.39758 | 0.723843 | 0.587355 | 1 | 0.999631 | 0.761181 |
| Ap4e1_Q80V94_S675s_1 | -0.66 | -0.51 | -0.42 | -0.38 | -0.55 | -0.76 | -0.42 | -0.54 | -0.74 | 0.08 | -0.15 | 0.21 | 0.22 | -0.25 | -0.39 | 0.61 | -0.075 | 0.3425 | 0.54 | 0.115587 | 0.754094 | 0.266647 | 1 | 0.999631 | 0.580776 |
| Apba1_B2RUJ5_S246s_1 | 1.15 | 1.13 | 1.71 | 1.13 | 0.07 | 1.56 | 2.6 | 0.15 | -0.03 | 1.17 | 0.87 | 1.03 | 0.49 | 0.12 | 1.74 | 0.53 | -0.185 | -0.52 | -0.56 | 0.185706 | 0.745562 | 0.157282 | 1 | 0.999631 | 0.646224 |
| Apba1_B2RUJ5_S289s_1 | -1.06 | -1.13 | -0.2 | -1.41 | -1.46 | -1.46 | -0.41 | -2 | -1.12 | -0.52 | -0.79 | -0.4 | 0.25 | -1.77 | -0.74 | -1.46 | -0.3825 | 0.2425 | 0.02 | 0.967922 | 0.372195 | 0.490467 | 1 | 0.999631 | 0.988771 |
| Apba1_B2RUJ5_S317s_1 | -0.65 | -1.48 | -1.16 | -1.69 | -2.04 | -1.86 | -1.08 | -1.67 | -1.89 | -0.74 | -1.62 | -1.61 | -0.39 | -1.3 | -1.85 | -1.42 | -0.4175 | -0.22 | 0.005 | 0.990107 | 0.24007 | 0.552331 | 1 | 0.999631 | 0.995892 |
| Apba1_B2RUJ5_S317s_2 | -0.68 | -0.61 | 0.17 | 0.3 | -0.09 | -0.78 | 0.04 | 0.42 | -0.25 | -0.63 | 0.94 | 1.61 | 0.54 | 0.04 | 1.22 | 0.96 | 0.1025 | 0.6225 | 0.895 | 0.042586 | 0.784472 | 0.265045 | 1 | 0.999631 | 0.475443 |
| Apba1_B2RUJ5_S372s_1 | -0.19 | -0.46 | -0.32 | 1.45 | 0.51 | 0.56 | 0.53 | 1.55 | 0.83 | 0.43 | 1.71 | 1.47 | -2.11 | 1.42 | 1.38 | 1.21 | 0.6675 | 0.99 | 0.355 | 0.6747 | 0.190747 | 0.074588 | 1 | 0.999631 | 0.885515 |
| Apba1_B2RUJ5_S391s_1 | 0.74 | -0.47 | -0.02 | -0.29 | -0.34 | -0.14 | -0.04 | -0.82 | -0.14 | 1.01 | -0.49 | -0.57 | -0.46 | -0.84 | -0.74 | -0.9 | -0.325 | -0.0375 | -0.725 | 0.057894 | 0.361807 | 0.932782 | 1 | 0.999631 | 0.49853 |
| Apba1_B2RUJ5_S573s_1 | 0.61 | 0.02 | -0.15 | 0 | -0.05 | 0.32 | 0.31 | -0.36 | 0.48 | 0.95 | -0.35 | -0.66 | -0.34 | -0.42 | -0.5 | -0.88 | -0.065 | -0.015 | -0.655 | 0.053747 | 0.827582 | 0.971009 | 1 | 0.999631 | 0.4909 |
| Apba1_B2RUJ5_T309t_1 | 0 | -0.28 | -0.57 | -0.39 | -0.62 | -0.57 | -0.4 | -0.72 | -0.2 | 0.3 | -0.6 | -0.02 | 0.03 | -0.03 | -1.18 | -0.03 | -0.2675 | 0.18 | 0.0075 | 0.983528 | 0.304556 | 0.551408 | 1 | 0.999631 | 0.995133 |
| Apba1_B2RUJ5_T309t_2 | -0.68 | -0.61 | 0.17 | 0.3 | -0.09 | -0.78 | 0.04 | 0.42 | -0.25 | -0.63 | 0.94 | 1.61 | 0.54 | 0.04 | 1.22 | 0.96 | 0.1025 | 0.6225 | 0.895 | 0.042586 | 0.784472 | 0.265045 | 1 | 0.999631 | 0.475443 |
| Apba1_B2RUJ5_T333t_1 | 0.22 | -0.45 | -1.3 | 0.13 | -0.21 | -0.17 | 0.12 | 0.14 | -0.33 | -0.05 | 0.03 | 0 | -0.97 | -0.42 | -0.37 | -0.25 | 0.32 | 0.2625 | -0.1525 | 0.707821 | 0.404878 | 0.496279 | 1 | 0.999631 | 0.898881 |
| Apba1_B2RUJ5_T375t_1 | -0.19 | 2.38 | 1.12 | 0.3 | 0.34 | 1.77 | 0.04 | 0.67 | 1.7 | 1.76 | 0.75 | 1.31 | 1.44 | 2.28 | 0.8 | 1.71 | -0.1975 | 0.4775 | 0.655 | 0.2796 | 0.746603 | 0.403875 | 1 | 0.999631 | 0.701922 |
| Apba2_P98084_S479s_1 | 0 | 0.57 | -0.26 | -0.62 | 0.11 | 0.46 | -0.46 | 0.86 | -0.08 | 0.55 | 0.22 | -0.2 | -0.26 | 0.81 | -0.27 | -0.21 | 0.32 | 0.2 | 0.095 | 0.809032 | 0.41579 | 0.564623 | 1 | 0.999631 | 0.93635 |
| Apba2_P98084_S481s_1 | 0.1 | -0.59 | -1.97 | -0.56 | 0.09 | -0.25 | -0.73 | -0.5 | -0.24 | -0.49 | -0.7 | -0.91 | -1.57 | -0.24 | -0.68 | -0.67 | 0.4075 | 0.17 | -0.035 | 0.943746 | 0.378292 | 0.706244 | 1 | 0.999631 | 0.981629 |
| Apba2_P98084_T307t_1 | 2.59 | -0.41 | -0.06 | 1.74 | 0.03 | 0.79 | 2.69 | 0.48 | -0.54 | 0.15 | 0.4 | -0.56 | -2.86 | -1.02 | 0.11 | -0.61 | 0.0325 | -1.1025 | -2.06 | 0.030551 | 0.967889 | 0.128418 | 1 | 0.999631 | 0.438336 |
| Apbb1ip_Q8R5A3_S507s_1 | 0.47 | 0.89 | 0.1 | -0.65 | -0.04 | 0.54 | 0.73 | 0.15 | 0.56 | 1.3 | -0.18 | 1.84 | 1.07 | 0.12 | 0.48 | -0.75 | 0.1425 | 0.6775 | 0.0275 | 0.954792 | 0.710851 | 0.209394 | 1 | 0.999631 | 0.984441 |
| Apbb2_F6V8Y6_S413s_1 | -0.59 | 0.95 | 0.84 | -0.73 | 0.04 | 0.68 | -0.45 | -0.1 | 0.45 | 0.12 | -0.09 | 0.69 | 2.66 | 0.42 | 0.84 | -0.6 | -0.075 | 0.175 | 0.7125 | 0.335202 | 0.876177 | 0.708805 | 1 | 0.999631 | 0.730219 |
| Apbb3_Q8R1C9_S111s_1 | 0.79 | 0.27 | 0.61 | 0.81 | -0.04 | 0.66 | 1.32 | 0.35 | 0.61 | 1.18 | 1.17 | 1.02 | 0.17 | 0.12 | 0.46 | 0.32 | -0.0475 | 0.375 | -0.3525 | 0.228614 | 0.890574 | 0.201306 | 1 | 0.999631 | 0.679038 |
| Apc_B2RUG9_S1030s_1 | 0.57 | 0.6 | 0.42 | 0.69 | 0.76 | 0.62 | 0.81 | 0.73 | 1.5 | 0.33 | 1.18 | 0.97 | -0.16 | 0.01 | 0.91 | 0.59 | 0.16 | 0.425 | -0.2325 | 0.486333 | 0.497522 | 0.19952 | 1 | 0.999631 | 0.804724 |
| Apc_B2RUG9_S1040s_1 | 0.18 | -3.5 | 2.33 | 0.76 | -2.83 | -2.38 | 2.91 | 0.14 | -2.01 | -0.16 | 0.21 | 0.13 | -2.02 | -3.64 | 0.12 | -0.31 | -0.4825 | -0.4 | -1.405 | 0.283861 | 0.753413 | 0.728551 | 1 | 0.999631 | 0.704942 |
| Apc_B2RUG9_S1370s_1 | -1.09 | -1.48 | -1.22 | -1.74 | -1.41 | -1.8 | -1.36 | -1.7 | -1.73 | -1.42 | -1.16 | 0.1 | -0.91 | -1.52 | -1.3 | -1.33 | -0.185 | 0.33 | 0.1175 | 0.695761 | 0.50021 | 0.446943 | 1 | 0.999631 | 0.895683 |
| Apc_B2RUG9_S1435s_1 | -0.57 | -0.3 | 0.15 | -0.1 | -0.34 | -0.84 | -0.86 | -0.06 | -0.76 | -0.24 | -0.71 | 0.38 | -0.02 | -0.03 | 0.6 | -0.08 | -0.32 | -0.1275 | 0.3225 | 0.315947 | 0.310381 | 0.713957 | 1 | 0.999631 | 0.720734 |
| Apc_B2RUG9_S1772s_1 | 0.89 | -0.73 | -1.29 | -1.24 | -0.82 | -0.77 | -0.75 | -0.68 | -0.33 | -0.25 | -0.52 | 0.76 | 0.59 | -0.33 | -0.21 | -0.95 | -0.1625 | 0.5075 | 0.3675 | 0.513895 | 0.737251 | 0.362012 | 1 | 0.999631 | 0.81893 |
| Apc_B2RUG9_S2260s_1 | 0.54 | -0.23 | -0.28 | 0.46 | 0.05 | 0.12 | 0.14 | 0.23 | 0.29 | 0.6 | 0.27 | 0.01 | -0.76 | -0.22 | 0.03 | 0.15 | 0.0125 | 0.17 | -0.3225 | 0.371101 | 0.965816 | 0.591142 | 1 | 0.999631 | 0.746338 |
| Apc_B2RUG9_S2283s_1 | 1.16 | 0.49 | 1.11 | 0.5 | 0.44 | 1.22 | 0.37 | 0.83 | 0.49 | 1.23 | 0.65 | 0.92 | -0.06 | 0.65 | 1.18 | 1.19 | -0.1 | 0.0075 | -0.075 | 0.84471 | 0.754199 | 0.980807 | 1 | 0.999631 | 0.94856 |
| Apc_B2RUG9_S2290s_1 | 2.23 | 0.51 | 2.53 | 2.45 | 0.07 | 0.88 | 2.47 | 1.84 | 0.08 | 0.93 | 1.94 | 1.63 | -0.27 | 0.53 | 1.23 | 1.25 | -0.615 | -0.785 | -1.245 | 0.045696 | 0.346754 | 0.196427 | 1 | 0.999631 | 0.477252 |
| Apc_B2RUG9_S2449s_1 | 0.5 | 0.43 | -1.49 | -0.59 | 0.49 | 0.46 | -0.4 | -0.59 | 0.82 | 0.85 | 0 | -0.59 | -0.51 | 1.01 | -0.65 | -0.27 | 0.2775 | 0.5575 | 0.1825 | 0.74538 | 0.59103 | 0.319474 | 1 | 0.999631 | 0.910585 |
| Apc_B2RUG9_S2674s_1 | -0.04 | -0.25 | -0.02 | 0.09 | 0.13 | -0.55 | -0.48 | 0.52 | -0.15 | -0.26 | 0.02 | -0.35 | -0.15 | -0.22 | 1.32 | -0.08 | -0.04 | -0.13 | 0.2725 | 0.500601 | 0.899259 | 0.607044 | 1 | 0.999631 | 0.813915 |
| Apc_B2RUG9_S2710s_1 | -0.27 | 0.03 | 0.06 | -0.27 | 0.08 | -0.7 | -0.3 | 0.28 | 0.55 | 0 | 0.16 | 0.35 | 1.28 | -0.82 | -0.93 | 0.13 | -0.0475 | 0.3775 | 0.0275 | 0.956055 | 0.873711 | 0.177972 | 1 | 0.999631 | 0.984741 |
| Apc_B2RUG9_S2771s_1 | -1.16 | 2.52 | 2.82 | 0.25 | 4.89 | 1.51 | 3.76 | 0.46 | 6.93 | -1.15 | 3.25 | 4.92 | 6.35 | -0.28 | 0.22 | 0.32 | 1.5475 | 2.38 | 0.545 | 0.724019 | 0.212046 | 0.175455 | 1 | 0.999631 | 0.905674 |
| Apc_B2RUG9_S2829s_1 | 1.81 | 0.09 | 0.88 | 0.48 | -1.51 | 1.09 | 1.76 | -0.87 | -0.17 | 0.81 | -1.52 | 0.64 | 0.03 | -0.83 | -0.03 | -0.34 | -0.6975 | -0.875 | -1.1075 | 0.02519 | 0.36704 | 0.161925 | 1 | 0.999631 | 0.421067 |
| Apc_B2RUG9_S742s_1 | -1.32 | -0.71 | -0.65 | -1.01 | -1.16 | -0.38 | -0.37 | -1.2 | -0.66 | -0.39 | -0.51 | -0.17 | -0.75 | -0.16 | -1.74 | -0.25 | 0.145 | 0.49 | 0.1975 | 0.633246 | 0.656228 | 0.106413 | 1 | 0.999631 | 0.868569 |
| Apc_B2RUG9_S838s_1 | 2.69 | -0.91 | -0.63 | 1.27 | 0.31 | -0.31 | 2.55 | -0.56 | -0.4 | 0.51 | 0.06 | -1.1 | -2.46 | -1.47 | -1.2 | -1.52 | -0.1075 | -0.8375 | -2.2675 | 0.014619 | 0.910067 | 0.310578 | 1 | 0.999631 | 0.386443 |
| Apc_B2RUG9_S983s_1 | 0.36 | -0.78 | -0.27 | 1.6 | 0.42 | 0.3 | 1.14 | 1.15 | 0.48 | 0.34 | 0.16 | 1.08 | -0.89 | 0.2 | 0.3 | 0.18 | 0.525 | 0.2875 | -0.28 | 0.61008 | 0.327103 | 0.581692 | 1 | 0.999631 | 0.859048 |
| Apc_B2RUG9_T1437t_1 | 0.89 | 1.21 | 1.04 | -1.06 | 1.14 | 1.56 | -0.03 | 0.07 | 1.46 | 1.17 | 1.43 | 0.55 | 1.33 | 1.94 | 0.16 | 0.27 | 0.165 | 0.6325 | 0.405 | 0.516962 | 0.782715 | 0.253076 | 1 | 0.999631 | 0.819449 |
| Apc2_D3YTR0_S1230s_1 | -0.56 | 0.14 | -1.06 | -0.79 | -0.35 | -0.47 | -1 | 0.05 | -0.53 | -1.02 | 0.16 | -0.31 | 0.07 | -0.07 | 0.08 | -0.67 | 0.125 | 0.1425 | 0.42 | 0.261507 | 0.72901 | 0.707258 | 1 | 0.999631 | 0.69271 |
| Apc2_D3YTR0_S1834s_1 | 1.71 | -0.8 | 0.14 | -0.21 | -0.98 | -0.21 | 0.56 | -0.79 | -1.16 | 0.78 | -0.67 | -1.18 | -0.88 | -1.59 | -0.95 | -1.4 | -0.565 | -0.7675 | -1.415 | 0.02201 | 0.342266 | 0.246974 | 1 | 0.999631 | 0.405544 |
| Apc2_D3YTR0_S1880s_1 | -0.17 | -3.4 | -0.58 | -0.74 | -1.02 | -0.6 | -2.09 | -2.04 | -2.55 | -0.19 | -1.13 | -1.85 | -1.25 | -0.06 | -1.31 | -0.64 | -0.215 | -0.2075 | 0.4075 | 0.564912 | 0.767662 | 0.791639 | 1 | 0.999631 | 0.84139 |
| Apc2_D3YTR0_S1890s_1 | -0.77 | -0.48 | -0.89 | -0.34 | 0.14 | -0.41 | -0.73 | -0.45 | -0.18 | -0.07 | -0.38 | -0.59 | 0.23 | 0.27 | -0.24 | 0.32 | 0.2575 | 0.315 | 0.765 | 0.024826 | 0.387533 | 0.268227 | 1 | 0.999631 | 0.418003 |
| Apc2_D3YTR0_S2010s_1 | 1.13 | 0.96 | 1.15 | 1.46 | 1.37 | 0.98 | 1.16 | 1.28 | 1.8 | 0.29 | 0.71 | 0.7 | 0.92 | 1.3 | -0.02 | 1.02 | 0.0225 | -0.3 | -0.37 | 0.313346 | 0.928771 | 0.423598 | 1 | 0.999631 | 0.719861 |
| Apc2_D3YTR0_S2128s_1 | 1.24 | -1.05 | -0.49 | -0.48 | -0.91 | -0.85 | 1.05 | -1.05 | -1.26 | 0.59 | -1.44 | -0.99 | -0.49 | -1.35 | -1.82 | -0.58 | -0.245 | -0.58 | -0.865 | 0.138756 | 0.698706 | 0.358213 | 1 | 0.999631 | 0.611577 |
| Apc2_D3YTR0_S2172s_1 | 1 | 1.34 | -0.41 | -0.94 | 1.09 | 0.9 | -0.89 | -0.4 | 1.17 | 2.21 | -0.35 | 0.13 | 0.3 | 1.39 | -0.02 | 0.21 | -0.0725 | 0.5425 | 0.2225 | 0.702126 | 0.911568 | 0.447802 | 1 | 0.999631 | 0.896857 |
| Apc2_D3YTR0_S2205s_1 | -0.67 | -0.63 | -0.32 | -1.6 | -0.3 | -0.85 | -0.54 | -0.5 | -0.1 | -1.15 | -0.15 | -0.36 | 0.5 | -1.15 | -0.64 | -1.09 | 0.2575 | 0.365 | 0.21 | 0.652658 | 0.450814 | 0.35689 | 1 | 0.999631 | 0.877704 |
| Apc2_D3YTR0_T128t_1 | -0.25 | -0.9 | -0.35 | -0.03 | -1.22 | -1.45 | -0.26 | 0.09 | -1.15 | -0.75 | -0.84 | -0.5 | -0.97 | -0.85 | -0.51 | -0.54 | -0.3275 | -0.4275 | -0.335 | 0.295254 | 0.435616 | 0.179705 | 1 | 0.999631 | 0.708769 |
| Api5_O35841_S464s_1 | 2.44 | 1.5 | 3.45 | 1.46 | 1.25 | 1.43 | 2.13 | 0.65 | 1.29 | 2.02 | 1.29 | 0.96 | 0.09 | 0.78 | 1.39 | 0.04 | -0.8475 | -0.8225 | -1.6375 | 0.011245 | 0.128025 | 0.121811 | 1 | 0.999631 | 0.385092 |
| Aplf_Q9D842_S23s_1 | 0.51 | 1.44 | 1.24 | 1.27 | 0.97 | 1.25 | 0.79 | 1.37 | 1.38 | 1.21 | 0.98 | 1.81 | 1.51 | 1.53 | 1.86 | 1.38 | -0.02 | 0.23 | 0.455 | 0.172807 | 0.947578 | 0.486237 | 1 | 0.999631 | 0.642116 |
| Aplp2_Q60709_T724t_1 | -0.26 | 0.12 | -0.58 | 0.16 | -0.48 | -0.69 | -0.73 | 0.19 | -0.4 | -0.7 | -0.93 | -0.5 | -0.27 | -0.2 | -0.45 | 0.15 | -0.2875 | -0.4925 | -0.0525 | 0.865332 | 0.382099 | 0.117634 | 1 | 0.999631 | 0.955768 |
| Apoe_P08226_T140t_1 | -0.5 | 1.63 | 1.07 | 0.62 | 1.26 | 1.02 | 0.25 | 0.39 | 0.83 | 0.26 | 1.48 | 1.3 | 2.09 | 0.39 | 2.56 | 0.9 | 0.025 | 0.2625 | 0.78 | 0.223361 | 0.958841 | 0.602424 | 1 | 0.999631 | 0.67471 |
| App_Q3TWF3_S441s_1 | -0.17 | 1.16 | 0.3 | -0.01 | 0.2 | 0.25 | -0.22 | 0.74 | 0.5 | 0.71 | 0.3 | 0.85 | 1.23 | 1.47 | 1.75 | 1.1 | -0.0775 | 0.27 | 1.0675 | 0.015413 | 0.835371 | 0.454818 | 1 | 0.999631 | 0.388832 |
| App_Q3TWF3_T725t_1 | 0.11 | -0.22 | -0.71 | -1.48 | -0.49 | -0.03 | -0.86 | -0.63 | -0.72 | 0.36 | -0.96 | -0.64 | 0.09 | 0.47 | -1.12 | -0.93 | 0.0725 | 0.085 | 0.2025 | 0.684984 | 0.854078 | 0.848579 | 1 | 0.999631 | 0.890166 |
| Appl1_Q8K3H0_S401s_1 | 0.2 | 0.08 | -0.77 | 0.52 | 0.56 | 0.33 | 0.8 | 0.51 | 0.46 | -1.01 | 0.68 | 0.22 | -2.06 | 0.61 | 0.35 | 0.3 | 0.5425 | 0.08 | -0.2075 | 0.737481 | 0.126609 | 0.860795 | 1 | 0.999631 | 0.907787 |
| Aqp4_P55088_S254s_1 | -0.98 | -1.09 | -0.29 | -0.89 | -1.53 | -0.45 | -0.69 | -1.33 | -1.36 | -0.06 | -0.39 | -1.66 | -0.01 | -0.49 | -0.86 | -0.43 | -0.1875 | -0.055 | 0.365 | 0.279204 | 0.589439 | 0.896666 | 1 | 0.999631 | 0.70145 |
| Aqp4_P55088_S263s_1 | 0.22 | 1.24 | 1.49 | 0.22 | 0.25 | 1.59 | 0.15 | -0.22 | 0.75 | 1.25 | 0.03 | 0.6 | 1.09 | 1.36 | 0.65 | 1.6 | -0.35 | -0.135 | 0.3825 | 0.361936 | 0.479717 | 0.749058 | 1 | 0.999631 | 0.742006 |
| Araf_P04627_S255s_1 | 0.92 | 0.57 | 0.72 | 0.18 | 0.3 | 0.24 | -0.07 | 0.33 | 0.35 | 0.92 | 1.09 | 0.41 | -0.45 | 0.15 | 0.42 | 0.81 | -0.3975 | 0.095 | -0.365 | 0.321609 | 0.166302 | 0.759391 | 1 | 0.999631 | 0.724092 |
| Araf_P04627_S580s_1 | 0.44 | -0.13 | 0.44 | -0.33 | -0.14 | -0.15 | -0.03 | -0.25 | -0.14 | 0.63 | -0.84 | 0.24 | 0.2 | 0.32 | -0.52 | 0.05 | -0.2475 | -0.1325 | -0.0925 | 0.784695 | 0.389904 | 0.733567 | 1 | 0.999631 | 0.922904 |
| Arcn1_Q5XJY5_S493s_1 | 0.18 | 0.25 | 0.04 | -0.37 | -0.24 | 0.39 | -0.25 | -0.13 | 0.44 | 0.64 | 0.33 | 0.1 | 0.35 | 0.83 | 0.63 | 0.41 | -0.0825 | 0.3525 | 0.53 | 0.094359 | 0.771334 | 0.224356 | 1 | 0.999631 | 0.558128 |
| Arel1_E9PZX5_S337s_1 | -1.52 | -0.86 | -0.12 | -0.19 | -0.89 | -0.94 | -0.66 | -0.51 | -0.84 | -0.6 | -0.9 | -0.6 | 0.06 | -0.2 | 0.58 | 0.27 | -0.0775 | -0.0625 | 0.85 | 0.053 | 0.831687 | 0.864696 | 1 | 0.999631 | 0.489777 |
| Arf6_P62331_S38s_1 | -0.8 | -0.13 | 0.03 | -0.35 | -0.64 | 0.75 | -0.57 | 0.85 | -0.51 | -0.62 | 0.52 | -0.25 | -1.84 | 1.29 | -1.71 | -0.65 | 0.41 | 0.0975 | -0.415 | 0.537317 | 0.356924 | 0.782018 | 1 | 0.999631 | 0.830391 |
| Arfgap1_V9GWV1_S150s_1 | 1.94 | -0.74 | 1.05 | 1.23 | -0.51 | 0.17 | 1.7 | 1.29 | -0.74 | 0.46 | -0.75 | 0.24 | -0.27 | -0.56 | 0.92 | 0.2 | -0.2075 | -1.0675 | -0.7975 | 0.202319 | 0.759866 | 0.098356 | 1 | 0.999631 | 0.659062 |
| Arfgap1_V9GWV1_S159s_1 | 0.45 | -0.51 | -0.5 | 0.3 | -0.24 | -1.11 | 1.13 | -0.47 | -1.27 | -1.54 | -0.82 | -0.85 | -0.93 | -1.25 | -0.14 | -0.88 | -0.1075 | -1.055 | -0.735 | 0.078258 | 0.830614 | 0.01237 | 1 | 0.999631 | 0.543873 |
| Arfgap1_V9GWV1_S256s_1 | -1.1 | -0.42 | -0.73 | -0.89 | 0.2 | 0.01 | -0.35 | 0.35 | -0.81 | -2.33 | 0.27 | -0.22 | -1.41 | -1.77 | -0.14 | -0.79 | 0.8375 | 0.0125 | -0.2425 | 0.554459 | 0.014155 | 0.98156 | 1 | 0.999631 | 0.838079 |
| Arfgap1_V9GWV1_S330s_1 | -0.34 | -1.1 | -1.57 | -0.27 | -0.76 | -0.63 | -0.6 | -0.7 | -0.45 | -0.66 | -0.99 | -0.71 | -1.03 | -1 | -1.31 | -0.94 | 0.1475 | 0.1175 | -0.25 | 0.501736 | 0.671454 | 0.746885 | 1 | 0.999631 | 0.814092 |
| Arfgap1_V9GWV1_S330s_2 | -0.59 | 0.33 | 0.29 | 0.07 | 0.37 | -0.21 | 0.01 | 0.68 | 0.06 | -0.87 | -0.75 | -0.5 | -0.26 | 0.52 | -0.14 | 0.78 | 0.1875 | -0.54 | 0.2 | 0.593245 | 0.573767 | 0.13768 | 1 | 0.999631 | 0.853305 |
| Arfgap1_V9GWV1_S335s_2 | -0.59 | 0.33 | 0.56 | 0.29 | 0.37 | -0.21 | 0.19 | 0.35 | 0.06 | -0.87 | -1.13 | -0.31 | -0.26 | 0.52 | -0.3 | 1.43 | 0.0275 | -0.71 | 0.2 | 0.671533 | 0.933279 | 0.091865 | 1 | 0.999631 | 0.884779 |
| Arfgap1_V9GWV1_S347s_1 | -1.25 | -1.37 | -0.69 | -0.57 | -0.42 | -1.86 | -0.05 | -0.43 | -0.74 | -2.06 | -0.42 | -0.94 | -0.27 | -0.76 | -0.87 | -0.88 | 0.28 | -0.07 | 0.275 | 0.405896 | 0.525446 | 0.865824 | 1 | 0.999631 | 0.765969 |
| Arfgap1_V9GWV1_T135t_1 | -1.25 | -0.52 | -0.85 | -1.12 | -0.59 | -0.51 | -0.94 | -0.53 | -0.44 | -1.04 | -0.8 | -0.69 | 0.48 | 0.12 | -0.54 | -0.5 | 0.2925 | 0.1925 | 0.825 | 0.038214 | 0.303483 | 0.511742 | 1 | 0.999631 | 0.459475 |
| Arfgap3_Q9D8S3_S403s_1 | 2.57 | -2.42 | -0.88 | 0.56 | -1.12 | -0.46 | 1.51 | -0.8 | -1.39 | 0.23 | -0.48 | -1.57 | -3.05 | -2.24 | -1.25 | -2.79 | -0.175 | -0.76 | -2.29 | 0.037697 | 0.866805 | 0.4484 | 1 | 0.999631 | 0.458331 |
| Arfgef1_G3X9K3_S1076s_1 | 1.85 | 0.76 | -3.54 | -1.01 | 0.39 | 1.5 | -1.59 | -1.35 | 1.21 | 1.7 | -1.12 | -2.59 | 0.19 | 0.69 | -2.95 | -4.72 | 0.2225 | 0.285 | -1.2125 | 0.416614 | 0.850972 | 0.82925 | 1 | 0.999631 | 0.770838 |
| Arfgef1_G3X9K3_S1550s_1 | -2.21 | -2.24 | -0.02 | -2 | -1.77 | -2.59 | -0.22 | -2.28 | -1.45 | -2.96 | -1.12 | -0.08 | -0.65 | -3.25 | -0.55 | -1.69 | -0.0975 | 0.215 | 0.0825 | 0.909604 | 0.883883 | 0.762919 | 1 | 0.999631 | 0.970597 |
| Arfgef1_G3X9K3_S1563s_1 | -0.88 | 1.14 | -0.06 | -1.14 | 2.56 | 0.41 | 2.11 | -0.49 | 2.86 | -1.45 | 1.59 | 2.47 | -0.42 | -0.81 | -0.38 | -0.48 | 1.3825 | 1.6025 | -0.2875 | 0.568452 | 0.098096 | 0.118155 | 1 | 0.999631 | 0.843185 |
| Arfgef1_G3X9K3_S1566s_1 | 0.18 | 1.41 | -2.02 | 0.01 | 0.6 | 0.85 | 0.26 | -1.01 | 0.87 | -0.57 | 0.07 | -0.9 | -0.83 | -0.25 | -0.43 | -0.5 | 0.28 | -0.0275 | -0.3975 | 0.542973 | 0.700058 | 0.969477 | 1 | 0.999631 | 0.832881 |
| Arfgef1_G3X9K3_S234s_1 | -0.63 | -1.64 | -0.34 | -0.97 | -0.91 | -0.79 | 0.09 | 0.26 | -0.14 | -1.65 | 0.08 | -1.04 | -1.5 | -1.15 | 0.14 | -0.78 | 0.5575 | 0.2075 | 0.0725 | 0.871826 | 0.194682 | 0.662009 | 1 | 0.999631 | 0.959119 |
| Arfgef1_G3X9K3_S52s_1 | 1.6 | 0.14 | 1.64 | 1 | -0.32 | 0.68 | 1.28 | 0.47 | 0.35 | 1.32 | 0.73 | 0.98 | -0.31 | -0.46 | 0.53 | 0.55 | -0.5675 | -0.25 | -1.0175 | 0.041821 | 0.237542 | 0.548263 | 1 | 0.999631 | 0.475443 |
| Arfgef1_G3X9K3_S661s_1 | -1.22 | -0.65 | -0.64 | -0.77 | -0.37 | -0.57 | -1.05 | -0.5 | -0.54 | -0.99 | -0.51 | 0.25 | -0.67 | -0.31 | -1.21 | -0.79 | 0.1975 | 0.3725 | 0.075 | 0.813061 | 0.488582 | 0.285901 | 1 | 0.999631 | 0.937041 |
| Arfgef2_A2A5R2_S621s_1 | -1.16 | -0.34 | -1.19 | -1.06 | -1.26 | -1.16 | -1.02 | -1.52 | -1.3 | -0.96 | -1.62 | -0.37 | 0.59 | -0.88 | -0.54 | -0.41 | -0.3025 | -0.125 | 0.6275 | 0.139362 | 0.318141 | 0.732586 | 1 | 0.999631 | 0.611718 |
| Arfgef2_F6YCJ0_S129s_1 | -0.82 | -0.3 | -2.38 | -3.64 | -0.28 | 0.05 | -2.26 | -2.91 | 0.21 | -1.67 | -1.7 | -2.68 | 0.26 | -0.57 | -1.41 | -1.98 | 0.435 | 0.325 | 0.86 | 0.290752 | 0.634701 | 0.702421 | 1 | 0.999631 | 0.706164 |
| Arfgef2_F6YCJ0_S138s_1 | -0.47 | -1.21 | -0.58 | -0.62 | -1.19 | -0.84 | -1.17 | -1.09 | -0.71 | -0.47 | -1.98 | -0.98 | -0.25 | -0.22 | -0.74 | -0.97 | -0.3525 | -0.315 | 0.175 | 0.594323 | 0.216903 | 0.426047 | 1 | 0.999631 | 0.85392 |
| Arfgef2_F6YCJ0_S187s_1 | 2.14 | 0.27 | 1.51 | 1.08 | -0.79 | 1.33 | 2.44 | -0.31 | -0.3 | 2.19 | -0.07 | -0.05 | -0.06 | -0.63 | -0.06 | -0.23 | -0.5825 | -0.8075 | -1.495 | 0.005336 | 0.439105 | 0.223653 | 1 | 0.999631 | 0.351308 |
| Arfgef2_F6YCJ0_T155t_1 | 0.3 | 1.1 | 2.02 | 1.43 | 0.96 | 0.83 | 1.34 | 1 | 1.72 | 0.58 | 0.73 | 0.99 | 0.85 | -0.36 | 0.85 | -0.58 | -0.18 | -0.2075 | -1.0225 | 0.064925 | 0.64248 | 0.635916 | 1 | 0.999631 | 0.515898 |
| Arfgef3_Q3UGY8_S1736s_1 | 2.03 | 1.29 | 2.53 | 1.77 | 0 | 1.32 | 2.11 | 1.05 | 0.47 | 1.44 | 0.66 | 0.97 | 0.82 | 0.13 | 0.85 | 0.38 | -0.785 | -1.02 | -1.36 | 0.003516 | 0.126683 | 0.018086 | 1 | 0.999631 | 0.351308 |
| Arfgef3_Q3UGY8_S1957s_1 | 1.37 | 1.2 | 0.08 | 0.66 | -0.25 | 1.08 | 0.99 | 0.08 | -0.14 | 0.88 | -0.71 | -0.27 | -1.02 | -0.5 | 0.87 | -1.82 | -0.3525 | -0.8875 | -1.445 | 0.03066 | 0.422772 | 0.067799 | 1 | 0.999631 | 0.438906 |
| Arfgef3_Q3UGY8_S2054s_1 | 0.2 | 0.57 | 0.59 | 0.91 | 0.11 | 0.42 | 0.3 | 0.85 | 0.34 | 0.34 | 0.95 | 0.5 | -0.47 | 0.63 | 0.64 | 1.1 | -0.1475 | -0.035 | -0.0925 | 0.813796 | 0.61045 | 0.904011 | 1 | 0.999631 | 0.937041 |
| Arfgef3_Q3UGY8_S2088s_1 | -0.21 | 0.2 | -0.37 | 0.2 | 0.07 | -0.12 | 0.13 | 0.73 | 0.33 | 0.65 | 0.4 | 0.5 | 0.14 | 0.32 | 1.14 | 0.3 | 0.2475 | 0.515 | 0.52 | 0.145172 | 0.415538 | 0.081476 | 1 | 0.999631 | 0.616182 |
| Arfgef3_Q3UGY8_S471s_1 | -0.26 | 0.73 | 1.12 | 0.11 | 0.82 | 0.38 | 0.42 | 0.86 | 0.81 | -0.67 | 1.38 | 1.54 | 0.86 | 1.02 | 1.31 | 1.2 | 0.195 | 0.34 | 0.6725 | 0.092048 | 0.589412 | 0.538876 | 1 | 0.999631 | 0.557085 |

| | | | | | | | | | | | | | | | | | | | | | | | | | |
|---|---|---|---|---|---|---|---|---|---|---|---|---|---|---|---|---|---|---|---|---|---|---|---|---|---|
| Arglu1_Q3UL36_S75s_1 | 1.68 | -1.28 | 0.41 | 0.48 | -1.5 | -1.15 | 1.38 | -0.38 | -2.55 | 0.83 | -0.38 | -0.45 | -2 | -2.06 | -0.16 | -0.34 | -0.735 | -0.96 | -1.4625 | 0.061184 | 0.35416 | 0.255805 | 1 | 0.999631 | 0.509355 |
| Arhgap1_Q5FWK3_S223s_1 | -2.06 | -6.81 | 1.21 | 1.53 | -3.43 | -4.29 | 1.76 | 0.76 | -7.52 | -3.13 | 0.52 | 0.76 | -4.39 | -4.27 | 2.03 | 0.04 | 0.2325 | -0.81 | -0.115 | 0.956181 | 0.910403 | 0.727128 | 1 | 0.999631 | 0.984772 |
| Arhgap1_Q5FWK3_S27s_1 | 0.61 | 0.49 | -0.45 | 0.71 | 0.63 | 0.05 | 0 | -0.04 | 0.31 | 0.05 | 0.08 | 0.45 | -0.81 | 0.24 | -0.15 | 0.36 | -0.18 | -0.1175 | -0.43 | 0.296452 | 0.602655 | 0.72535 | 1 | 0.999631 | 0.709528 |
| Arhgap1_Q5FWK3_S51s_1 | -0.18 | -0.02 | -0.39 | -0.66 | -0.37 | -0.6 | -0.83 | -0.11 | -0.24 | 0.51 | 0.28 | 0.3 | 0.69 | 0.38 | 0.19 | 0.09 | -0.165 | 0.525 | 0.65 | 0.05106 | 0.565092 | 0.098679 | 1 | 0.999631 | 0.489558 |
| Arhgap10_Q6Y5D8_S488s_1 | -0.14 | 0.19 | -0.05 | -0.65 | -0.43 | 0.13 | -0.48 | -0.43 | -0.56 | -0.06 | -0.32 | 0.02 | 0.03 | -0.27 | -0.59 | -0.19 | -0.14 | -0.0675 | -0.0925 | 0.766644 | 0.637385 | 0.821128 | 1 | 0.999631 | 0.916961 |
| Arhgap12_B2RUJ8_S199s_1 | 0.08 | 0.78 | 0.83 | 0.48 | 0.44 | 0.78 | 0.2 | -0.31 | 1.16 | 0.93 | -0.22 | -0.19 | 1.76 | -0.42 | 1.45 | 0.46 | -0.265 | -0.1225 | 0.27 | 0.593184 | 0.428363 | 0.7657 | 1 | 0.999631 | 0.853305 |
| Arhgap12_B2RUJ8_S213s_1 | -0.25 | 0.35 | 0.65 | 0.56 | -0.38 | -0.22 | 0.13 | 0.85 | 0.41 | 0.22 | -0.47 | 0.34 | 0.44 | -0.21 | 1.15 | 0.22 | -0.2325 | -0.2025 | 0.0725 | 0.849972 | 0.525749 | 0.549334 | 1 | 0.999631 | 0.949525 |
| Arhgap17_E9QAJ9_S669s_1 | -0.75 | 0.59 | 1.17 | 0.58 | 0.24 | -0.48 | 0.16 | -0.14 | 1.22 | 0.89 | 0.48 | 0.71 | 0.61 | 0.45 | 0.07 | 0.43 | -0.4525 | 0.4275 | -0.0075 | 0.986086 | 0.30704 | 0.336601 | 1 | 0.999631 | 0.995133 |
| Arhgap17_E9QAJ9_S671s_1 | -0.1 | 0.4 | 1.17 | 0.58 | 0.06 | -0.34 | 0.19 | -0.07 | 0.89 | 1.08 | 0.62 | 0.76 | 0.66 | 0.77 | 0.42 | 0.43 | -0.5525 | 0.325 | 0.0575 | 0.865991 | 0.116872 | 0.338933 | 1 | 0.999631 | 0.955982 |
| Arhgap20_Q6IFT4_S46s_1 | 0.8 | 1.31 | 0.55 | 1.33 | 1.35 | 1.46 | 0.98 | 1.6 | 1.72 | 0.89 | 1.81 | 1.35 | 1.03 | 1.88 | 0.89 | 1.7 | 0.35 | 0.445 | 0.3775 | 0.308893 | 0.258249 | 0.20363 | 1 | 0.999631 | 0.718845 |
| Arhgap21_B7ZCJ0_S1090s_2 | 0.81 | -0.64 | -0.31 | 1.06 | 1.44 | -0.09 | 1.35 | 1 | 0.99 | -0.27 | 0.4 | 0.51 | -1.96 | -1.84 | -0.12 | -0.34 | 0.695 | 0.1775 | -1.295 | 0.048137 | 0.192836 | 0.708193 | 1 | 0.999631 | 0.48343 |
| Arhgap21_B7ZCJ0_S1093s_2 | 0.81 | -0.64 | -0.31 | 1.06 | 1.44 | -0.09 | 1.35 | 1 | 0.99 | -0.27 | 0.4 | 0.51 | -1.96 | -1.84 | -0.12 | -0.34 | 0.695 | 0.1775 | -1.295 | 0.048137 | 0.192836 | 0.708193 | 1 | 0.999631 | 0.48343 |
| Arhgap21_B7ZCJ0_S1502s_1 | 0.74 | 1.19 | 0.33 | -0.04 | 0.6 | 1.4 | 0.44 | -0.06 | 1.15 | 1.21 | 0.67 | 0.52 | 0.5 | 0.92 | 0.68 | 0.63 | 0.04 | 0.3325 | 0.1275 | 0.71043 | 0.921212 | 0.35783 | 1 | 0.999631 | 0.899851 |
| Arhgap21_B7ZCJ0_S1729s_1 | 0.81 | 1.37 | 0.42 | 0.78 | 0.92 | 0.32 | -0.2 | 0.92 | 0.59 | 0.71 | 0.62 | 0.58 | -0.05 | 0.72 | 0.3 | 1.17 | -0.355 | -0.22 | -0.31 | 0.411834 | 0.33448 | 0.450922 | 1 | 0.999631 | 0.767937 |
| Arhgap21_B7ZCJ0_S42s_1 | -0.7 | -0.12 | -0.14 | -1.08 | -0.06 | -0.19 | -0.81 | -0.8 | -0.45 | -0.73 | -0.49 | 0.04 | 0.56 | 0.12 | -0.6 | -0.41 | 0.045 | 0.1025 | 0.4275 | 0.281562 | 0.894817 | 0.758869 | 1 | 0.999631 | 0.702855 |
| Arhgap21_B7ZCJ0_S436s_1 | -1.5 | -1.43 | -2.99 | -1.86 | -2.28 | -2.61 | -1.65 | -2.53 | -2.18 | -1.59 | -2.3 | -3.45 | -1.03 | -1.91 | -2.6 | -3.19 | -0.3225 | -0.435 | -0.2375 | 0.66609 | 0.447585 | 0.396283 | 1 | 0.999631 | 0.88264 |
| Arhgap21_B7ZCJ0_S470s_1 | -0.14 | 1 | 0.04 | -0.38 | 0.27 | -0.82 | -0.65 | 0.27 | 0.75 | -1.23 | -0.06 | 0.01 | 0.55 | 0.47 | 1.17 | 0.52 | -0.3625 | -0.2625 | 0.5475 | 0.172049 | 0.394733 | 0.592241 | 1 | 0.999631 | 0.641657 |
| Arhgap21_B7ZCJ0_S478s_1 | -1.33 | -0.72 | -0.91 | -1.01 | -1.14 | -1.4 | -1.25 | -1.36 | -0.94 | -0.4 | -1.79 | -1.68 | -0.22 | -0.22 | -1.86 | -1.39 | -0.295 | -0.21 | 0.07 | 0.87346 | 0.262818 | 0.579331 | 1 | 0.999631 | 0.959788 |
| Arhgap21_B7ZCJ0_S687s_1 | -1.09 | -1.05 | -0.64 | -1.02 | -1.19 | -1.43 | -0.84 | -0.81 | -1.37 | -1.16 | -0.55 | -0.31 | -1.28 | -0.78 | -0.9 | -0.86 | -0.1175 | 0.1025 | -0.005 | 0.985907 | 0.666871 | 0.753578 | 1 | 0.999631 | 0.995133 |
| Arhgap21_B7ZCJ0_T62t_1 | -1.72 | 0.65 | -0.53 | -0.52 | -0.37 | -0.53 | -0.43 | -1.01 | 0.47 | -1.29 | 1.12 | 0.02 | 0.5 | 0.46 | 0.5 | -0.38 | -0.055 | 0.61 | 0.8 | 0.137279 | 0.908529 | 0.348106 | 1 | 0.999631 | 0.610369 |
| Arhgap21_B7ZCJ0_T686t_1 | -1.09 | -1.05 | -0.64 | -1.02 | -1.19 | -1.43 | -0.84 | -0.81 | -1.37 | -1.16 | -0.55 | -0.31 | -1.28 | -0.78 | -0.9 | -0.86 | -0.1175 | 0.1025 | -0.005 | 0.985907 | 0.666871 | 0.753578 | 1 | 0.999631 | 0.995133 |
| Arhgap22_Q8BL80_S22s_1 | 0.52 | 1.19 | 0.78 | 0.82 | 0.22 | 0.69 | 0.66 | 0.34 | 0.63 | 1.15 | 0.73 | 0.64 | -0.12 | 0.58 | 0 | 0.76 | -0.35 | -0.04 | -0.5225 | 0.136537 | 0.216951 | 0.886736 | 1 | 0.999631 | 0.609945 |
| Arhgap22_Q8BL80_S387s_1 | 0.95 | 1.07 | -0.88 | -1.11 | 0.13 | 0.9 | -0.27 | 0.25 | 0.74 | 1.73 | 0.59 | 1 | 0.62 | 0.92 | -1.5 | 0.16 | 0.245 | 1.0075 | 0.0425 | 0.951745 | 0.670563 | 0.107932 | 1 | 0.999631 | 0.984166 |
| Arhgap23_F8VQ11_S1182s_1 | 0.24 | 0.53 | 0.1 | 0.24 | 0.59 | 0.04 | -0.09 | 0.3 | 0.17 | 0.69 | 0.07 | 0.18 | 0.86 | 0.61 | -0.05 | 0.19 | -0.0675 | 0 | 0.125 | 0.691289 | 0.802061 | 1 | 1 | 1 | 0.893997 |
| Arhgap23_F8VQ11_S145s_1 | 0.14 | 0.3 | 0.43 | 0.2 | 0.33 | 0.11 | 0.03 | 0.3 | 0.73 | 0.49 | 0.13 | 0.35 | 0.56 | 0.75 | 0.3 | 0.21 | -0.075 | 0.1575 | 0.1875 | 0.505623 | 0.754788 | 0.552328 | 1 | 0.999631 | 0.814836 |
| Arhgap23_F8VQ11_S155s_1 | -0.79 | 0.05 | 0.2 | -0.61 | -0.31 | -0.59 | -0.66 | -0.51 | -0.5 | -0.37 | -0.35 | -0.05 | 0.16 | -0.02 | -0.01 | 0.48 | -0.23 | -0.03 | 0.44 | 0.211389 | 0.464369 | 0.925331 | 1 | 0.999631 | 0.665474 |
| Arhgap23_F8VQ11_S166s_1 | -1.13 | -0.48 | -0.34 | -0.55 | -0.81 | -0.84 | -0.94 | -0.22 | -0.77 | -0.9 | -0.32 | -0.56 | -0.38 | -0.23 | -0.36 | -0.21 | -0.0775 | -0.0125 | 0.33 | 0.276808 | 0.796929 | 0.966253 | 1 | 0.999631 | 0.700187 |
| Arhgap23_F8VQ11_S215s_1 | 1.09 | 0.15 | 0.6 | 1.26 | 0.93 | 0.31 | 0.79 | 1.24 | 0.44 | -0.03 | 1.15 | 0.99 | -0.75 | 0.48 | 0.87 | 0.6 | 0.0425 | -0.1375 | -0.475 | 0.296214 | 0.902642 | 0.723043 | 1 | 0.999631 | 0.709528 |
| Arhgap23_F8VQ11_S344s_1 | -0.72 | -0.42 | -0.3 | -0.04 | 0.39 | -0.26 | 0.28 | 0.74 | -0.19 | -0.67 | -1.38 | 0.31 | -0.17 | -0.17 | 0.9 | 0.4 | 0.6575 | -0.1125 | 0.61 | 0.106068 | 0.0551 | 0.778424 | 1 | 0.999631 | 0.568797 |
| Arhgap23_F8VQ11_S388s_1 | 0.16 | 1.85 | 1.89 | 0.95 | 3.02 | 1.27 | 3.03 | 1.8 | 4.66 | 0.18 | 2.93 | 4.08 | 3.39 | 0.76 | 2.11 | 0.61 | 1.0675 | 1.75 | 0.505 | 0.467827 | 0.078986 | 0.08585 | 1 | 0.999631 | 0.795565 |
| Arhgap23_F8VQ11_S400s_1 | 0.79 | 0.54 | 0.48 | 0.36 | 0.71 | 0.8 | 0.61 | 0.58 | 0.92 | 0.43 | 0.62 | 0.6 | -0.27 | 0.96 | 0.69 | 0.74 | 0.1325 | 0.1 | -0.0125 | 0.971254 | 0.586373 | 0.702429 | 1 | 0.999631 | 0.989721 |
| Arhgap23_F8VQ11_S413s_1 | -0.36 | -0.3 | 0.66 | 0.21 | 0.79 | -0.13 | 0.97 | 0.8 | 0.1 | -0.38 | 1.17 | 1.93 | 0.33 | 0.31 | 1.37 | 0.53 | 0.555 | 0.6525 | 0.5825 | 0.149749 | 0.15297 | 0.240971 | 1 | 0.999631 | 0.621978 |
| Arhgap23_F8VQ11_T127t_1 | -1.08 | 0.26 | -0.82 | -0.5 | -0.15 | -0.2 | 0.06 | -0.24 | 0.08 | -1.29 | 0.17 | 0.41 | 0.41 | 0.02 | -0.06 | 0.27 | 0.4025 | 0.3775 | 0.695 | 0.077248 | 0.248283 | 0.426347 | 1 | 0.999631 | 0.542683 |
| Arhgap23_F8VQ11_T897t_1 | 0.75 | 0.71 | 2.27 | 1.03 | -0.24 | 1.22 | 1.2 | 0.24 | 0.26 | 1.24 | 0.35 | 0.6 | 0.56 | 0.24 | 0.59 | 1.16 | -0.585 | -0.5775 | -0.5525 | 0.212951 | 0.24764 | 0.198766 | 1 | 0.999631 | 0.666787 |
| Arhgap24_D3Z5T4_T241t_1 | -0.38 | -0.44 | -1.69 | -2.17 | -2.1 | -2.48 | -1.08 | -2.35 | -3.17 | -2.08 | -1.71 | -1.98 | -2.01 | -2.61 | -2.57 | -2.58 | -0.8325 | -1.065 | -1.2725 | 0.019662 | 0.129073 | 0.063952 | 1 | 0.999631 | 0.391668 |
| Arhgap25_Q8BYW1_S274s_1 | -1.08 | -0.92 | -0.41 | -1.21 | -0.97 | -0.7 | -0.93 | -1.13 | -0.92 | -1.52 | -0.26 | -0.82 | -0.55 | -0.65 | -0.66 | -0.61 | -0.0275 | 0.025 | 0.2875 | 0.33827 | 0.921423 | 0.943136 | 1 | 0.999631 | 0.731571 |
| Arhgap26_F7D661_S18s_1 | -1.38 | 0.49 | -0.16 | -0.02 | 1.21 | 0.22 | 1.04 | -1.14 | 0.63 | -1.25 | 0.9 | 1.51 | -0.36 | 0.12 | -1.31 | 0.35 | 0.6 | 0.715 | -0.0325 | 0.949463 | 0.331866 | 0.282589 | 1 | 0.999631 | 0.983795 |
| Arhgap31_A6X8Z5_S1163s_1 | 0.53 | 1.39 | -0.26 | 0.11 | 1.22 | 0.7 | -0.41 | -0.17 | 1.13 | 0.92 | -0.48 | 0.04 | 1.15 | 1.73 | 0.07 | -0.03 | -0.1075 | -0.04 | 0.2875 | 0.585872 | 0.826238 | 0.935297 | 1 | 0.999631 | 0.849524 |
| Arhgap31_A6X8Z5_S1242s_1 | -0.52 | 0.58 | 1.26 | -0.24 | 1.77 | 0.11 | 0.68 | 1.66 | 0.98 | -0.67 | 0.88 | 1.79 | 2.01 | 0.17 | 2.04 | -0.38 | 0.785 | 0.475 | 0.69 | 0.313231 | 0.158936 | 0.433468 | 1 | 0.999631 | 0.719861 |
| Arhgap31_A6X8Z5_S765s_1 | -0.82 | -0.23 | -0.26 | -1.08 | 0.35 | -0.2 | -0.76 | -1.14 | 0 | -0.44 | -0.78 | -0.51 | 0.26 | 0.42 | -0.83 | 0.29 | 0.16 | 0.165 | 0.6325 | 0.127915 | 0.686237 | 0.611657 | 1 | 0.999631 | 0.598425 |
| Arhgap31_A6X8Z5_S850s_1 | -0.2 | -0.69 | 0.36 | -0.36 | -0.48 | 0.2 | -0.12 | -0.67 | -0.51 | 0.21 | 0.3 | 0.22 | -0.39 | 0.33 | 0.13 | 0.62 | -0.045 | 0.2775 | 0.395 | 0.284128 | 0.892212 | 0.418934 | 1 | 0.999631 | 0.705132 |
| Arhgap32_Q811P8_S1405s_1 | 0.26 | 0.59 | -0.2 | 0.48 | 0.6 | 0.29 | -0.07 | 0.33 | 0.5 | -0.03 | 0.09 | 0.62 | -0.09 | 0.47 | 0.33 | 0.34 | 0.005 | 0.0125 | -0.02 | 0.948292 | 0.986337 | 0.967383 | 1 | 0.999631 | 0.983277 |
| Arhgap32_Q811P8_S1588s_1 | 1.06 | 1.04 | 0.4 | 1.19 | 0.9 | 0.17 | 0.88 | 0.67 | -0.13 | -1.63 | 0.92 | -0.24 | -0.47 | 0.56 | 0.35 | 0.75 | -0.2675 | -1.1925 | -0.625 | 0.112527 | 0.389833 | 0.042278 | 1 | 0.999631 | 0.577306 |
| Arhgap32_Q811P8_S1823s_1 | 1.57 | -0.97 | -0.17 | 0.66 | -0.68 | -0.21 | 1.14 | -0.34 | -0.41 | 0.71 | 0.03 | -0.29 | -1.82 | -1.15 | -0.08 | -1.05 | -0.295 | -0.2625 | -1.2975 | 0.051116 | 0.630486 | 0.638378 | 1 | 0.999631 | 0.489558 |
| Arhgap32_Q811P8_S1876s_1 | 0.62 | 1.8 | 1.67 | -1.03 | 2.52 | 1.33 | 3.73 | 0.38 | 3.68 | 0.76 | 3 | 3.68 | 2.91 | -0.22 | 0.21 | -0.52 | 1.225 | 2.015 | -0.17 | 0.8473 | 0.171918 | 0.034422 | 1 | 0.999631 | 0.948838 |
| Arhgap32_Q811P8_S47s_1 | -0.9 | -0.36 | -0.29 | -0.01 | 0.56 | -0.55 | -0.29 | 0.3 | 0.79 | 0.1 | -0.26 | 0.42 | -0.74 | -0.41 | 0.29 | 0.5 | 0.395 | 0.6525 | 0.3 | 0.438602 | 0.272102 | 0.078018 | 1 | 0.999631 | 0.781983 |
| Arhgap32_Q811P8_S613s_1 | 0.74 | -0.99 | -1.35 | 0.49 | 1.46 | -0.57 | 1.12 | 0.07 | -0.03 | -1.28 | 0.22 | -0.64 | -0.91 | -0.38 | -1.12 | -0.68 | 0.7975 | -0.155 | -0.495 | 0.350314 | 0.224764 | 0.786368 | 1 | 0.999631 | 0.737237 |
| Arhgap32_Q811P8_S655s_1 | 0.82 | -0.59 | -2.18 | 0.77 | 1.2 | 0.04 | 0.33 | 0.18 | 0.91 | -0.46 | -0.15 | -0.62 | -1.46 | 0.15 | -1.83 | -0.78 | 0.7325 | 0.215 | -0.685 | 0.360494 | 0.290334 | 0.759458 | 1 | 0.999631 | 0.741182 |
| Arhgap32_Q811P8_S682s_1 | 0.38 | 1.36 | 1.98 | 1.75 | 1.02 | 1.35 | 1.68 | 1.87 | 1.64 | 0.14 | 1.94 | 1.95 | 0.71 | 0.36 | 2.22 | 2.11 | 0.1125 | 0.05 | -0.0175 | 0.974661 | 0.780297 | 0.923792 | 1 | 0.999631 | 0.990998 |
| Arhgap32_Q811P8_S856s_1 | -2.27 | -1.02 | 0.73 | -0.06 | -1.13 | -1.69 | -0.42 | -0.19 | -1.72 | -2.32 | 0.73 | 0.72 | 0.28 | -0.9 | 0.07 | 0.54 | -0.2025 | 0.0075 | 0.6525 | 0.325123 | 0.756782 | 0.993313 | 1 | 0.999631 | 0.72463 |
| Arhgap32_Q811P8_S892s_1 | 1.71 | -0.54 | -2.25 | 1.09 | 2.62 | 0.32 | 1.3 | 0.9 | 0.53 | -1.84 | 0.02 | -1.64 | -2.46 | -0.16 | -0.92 | -0.43 | 1.2825 | -0.735 | -0.995 | 0.276851 | 0.167244 | 0.435232 | 1 | 0.999631 | 0.700187 |
| Arhgap32_Q811P8_S935s_1 | -0.36 | 0.33 | -0.94 | -0.51 | 0.07 | 0 | -0.57 | -0.34 | -0.2 | -1.06 | -1.15 | -0.04 | -0.52 | 0.33 | 0.04 | -0.5 | 0.16 | -0.2425 | 0.2075 | 0.583652 | 0.638792 | 0.549526 | 1 | 0.999631 | 0.848843 |
| Arhgap32_Q811P8_S952s_1 | -0.95 | -0.46 | -1.06 | -0.69 | -0.47 | -1.13 | -0.94 | -1.07 | -0.44 | -1.25 | -0.46 | -0.65 | -0.32 | -0.64 | -0.77 | -0.85 | -0.1125 | 0.09 | 0.145 | 0.623049 | 0.69025 | 0.76797 | 1 | 0.999631 | 0.862913 |
| Arhgap33_Q80YF9_S1144s_1 | 0.85 | 0.06 | -0.63 | 0.11 | 0.75 | 0.11 | 0.73 | -0.24 | 0.51 | 1.27 | 2.08 | 0.21 | 1.48 | 0.62 | -0.01 | 0.7 | 0.24 | 0.92 | 0.6 | 0.18921 | 0.546956 | 0.086463 | 1 | 0.999631 | 0.648664 |
| Arhgap33_Q80YF9_S594s_1 | 0.12 | -0.91 | -0.77 | -0.45 | -0.71 | -0.6 | -0.38 | -0.36 | -0.43 | -0.05 | -0.71 | -0.85 | -0.71 | -0.68 | -0.65 | -0.31 | -0.01 | -0.0075 | -0.085 | 0.794079 | 0.973737 | 0.982211 | 1 | 0.999631 | 0.928498 |
| Arhgap33_Q80YF9_S660s_1 | -0.97 | -0.85 | -0.01 | -0.03 | -0.27 | -1.09 | -0.51 | 0.08 | -0.43 | -0.16 | 0.34 | 0.7 | 0.27 | -0.61 | 0.26 | 0.77 | 0.0175 | 0.5775 | 0.6375 | 0.140128 | 0.962588 | 0.154225 | 1 | 0.999631 | 0.611718 |
| Arhgap33_Q80YF9_S887s_1 | -0.22 | 0.08 | -0.63 | -0.32 | -0.17 | 0.3 | -0.88 | -0.68 | -0.1 | -0.27 | -0.5 | -0.53 | -0.39 | 0.17 | -0.2 | -0.05 | -0.085 | -0.0775 | 0.155 | 0.604787 | 0.802105 | 0.780649 | 1 | 0.999631 | 0.85817 |
| Arhgap33_Q80YF9_S949s_1 | 0.94 | -0.87 | -1.17 | 0.37 | 0.58 | -0.18 | 0.32 | -0.01 | -0.29 | 0.11 | -0.55 | -1.1 | -1.3 | -0.38 | -0.64 | -0.82 | 0.36 | -0.275 | -0.6025 | 0.255232 | 0.474778 | 0.603503 | 1 | 0.999631 | 0.690381 |
| Arhgap33_Q80YF9_S949s_2 | 0.54 | 0.63 | -1.32 | -0.68 | 1.75 | 1.03 | -0.93 | -1.01 | 1.12 | 1.03 | -1.12 | -1.77 | 1.22 | 0.73 | -0.85 | -1 | 0.4175 | 0.0225 | 0.2325 | 0.724215 | 0.576901 | 0.976804 | 1 | 0.999631 | 0.905674 |
| Arhgap33_Q80YF9_S998s_1 | -0.5 | -0.34 | -1.84 | -0.65 | 0.14 | -1.41 | 0.49 | -0.68 | 0.14 | -2.18 | 1.01 | -0.13 | -1.36 | -1.8 | -2.13 | -1.99 | 0.4675 | 0.5425 | -0.9875 | 0.031984 | 0.369705 | 0.430776 | 1 | 0.999631 | 0.442395 |
| Arhgap33_Q80YF9_T943t_1 | -1.94 | 0.13 | -0.66 | -1.66 | -1.23 | -1.14 | -1.42 | -0.64 | -0.7 | -0.01 | -1.19 | -0.46 | 0.54 | -0.56 | -0.78 | -0.77 | -0.075 | 0.4425 | 0.64 | 0.248559 | 0.875183 | 0.391412 | 1 | 0.999631 | 0.685751 |
| Arhgap33_Q80YF9_T943t_2 | 0.54 | 0.63 | -1.32 | -0.68 | 1.75 | 1.03 | -0.93 | -1.01 | 1.12 | 1.03 | -1.12 | -1.77 | 1.22 | 0.73 | -0.85 | -1 | 0.4175 | 0.0225 | 0.2325 | 0.724215 | 0.576901 | 0.976804 | 1 | 0.999631 | 0.905674 |
| Arhgap35_Q91YM2_S1134s_1 | -2.15 | 0.15 | -0.18 | -1.72 | -0.28 | 0.38 | -0.37 | -1.62 | -0.26 | -1.41 | -0.88 | -0.18 | -0.2 | -0.91 | -1.27 | -0.94 | 0.5025 | 0.2925 | 0.145 | 0.796868 | 0.432992 | 0.617174 | 1 | 0.999631 | 0.929952 |
| Arhgap35_Q91YM2_S1134s_2 | -1.27 | -1.39 | -0.78 | 0.3 | 0.23 | -0.64 | 0.79 | -0.65 | 0.29 | -0.07 | 1.04 | 1.08 | -2.8 | -1.07 | 0.73 | 0.36 | 0.7175 | 1.37 | 0.09 | 0.907828 | 0.167144 | 0.014073 | 1 | 0.999631 | 0.96989 |
| Arhgap35_Q91YM2_S1142s_1 | -0.61 | -0.96 | -0.81 | -1.35 | -0.83 | -1.12 | -0.56 | -0.38 | -0.89 | -1.67 | -0.75 | -0.15 | -0.58 | -1.1 | -0.19 | -2.45 | 0.21 | 0.0675 | -0.1475 | 0.766809 | 0.48028 | 0.857048 | 1 | 0.999631 | 0.916961 |
| Arhgap35_Q91YM2_S1142s_2 | -1.27 | -1.39 | -0.78 | 0.3 | 0.23 | -0.64 | 0.79 | -0.65 | 0.29 | -0.07 | 1.04 | 1.08 | -2.8 | -1.07 | 0.73 | 0.36 | 0.7175 | 1.37 | 0.09 | 0.907828 | 0.167144 | 0.014073 | 1 | 0.999631 | 0.96989 |
| Arhgap35_Q91YM2_S975s_1 | -1.07 | -0.29 | -0.46 | -0.26 | -0.4 | -0.18 | -0.79 | -0.21 | -0.48 | -0.48 | -0.19 | -0.07 | 0.89 | -0.13 | -0.36 | -0.68 | 0.125 | 0.215 | 0.45 | 0.285131 | 0.677142 | 0.47339 | 1 | 0.999631 | 0.705851 |
| Arhgap39_H7BX46_S117s_1 | 1.56 | -1.52 | 0.92 | -0.56 | -2.57 | 0.31 | 2.28 | -2.13 | -1.76 | 2.02 | -0.63 | -1.96 | -0.87 | -2.37 | -0.7 | -2.28 | -0.6275 | -0.6825 | -1.655 | 0.044092 | 0.583691 | 0.498116 | 1 | 0.999631 | 0.475443 |
| Arhgap39_H7BX46_S125s_1 | 1.74 | -3.18 | -0.88 | -1.66 | -3.19 | -1.66 | 2.98 | -4.3 | -8.51 | 1.33 | -1.64 | -2.77 | -2.04 | -3.78 | -3.12 | -3.24 | -0.5475 | -1.9025 | -2.05 | 0.050367 | 0.734685 | 0.341258 | 1 | 0.999631 | 0.488571 |
| Arhgap39_H7BX46_S171s_1 | -0.67 | -0.22 | 0.01 | -0.58 | -0.82 | -0.09 | -0.81 | -0.57 | -0.55 | -0.5 | -0.25 | 0.02 | 0.22 | -0.04 | -0.19 | -0.05 | -0.2075 | 0.045 | 0.35 | 0.25099 | 0.491553 | 0.876996 | 1 | 0.999631 | 0.686822 |
| Arhgap39_H7BX46_S297s_1 | 2.51 | 0.63 | 1.61 | 1.58 | 0.11 | 1.84 | 1.5 | 0.4 | 1.43 | 1.71 | 0.16 | -0.12 | 0.47 | 0.06 | 0.05 | -0.3 | -0.62 | -0.7875 | -1.5125 | 0.004988 | 0.255978 | 0.175546 | 1 | 0.999631 | 0.351308 |
| Arhgap39_H7BX46_S364s_1 | 0.48 | 0.29 | -0.43 | 0.38 | 1.65 | 0.88 | 0.31 | 0.4 | 0.6 | -0.67 | 0.37 | -0.9 | 0.3 | -0.18 | -1.54 | 0.06 | 0.63 | -0.33 | -0.52 | 0.270987 | 0.123941 | 0.446712 | 1 | 0.999631 | 0.698674 |
| Arhgap39_H7BX46_S382s_1 | 0.2 | 0 | -0.36 | 0.2 | 0.36 | -0.36 | 0.31 | -0.57 | -0.57 | -0.55 | -0.02 | 0.03 | -0.53 | -0.35 | -1.18 | 0.16 | -0.075 | -0.2875 | -0.485 | 0.19588 | 0.814049 | 0.339033 | 1 | 0.999631 | 0.653145 |
| Arhgap39_H7BX46_S599s_1 | -0.54 | 0.02 | 0.12 | 0.68 | 0.24 | -0.27 | 0.28 | -0.18 | -0.86 | -0.34 | 0.39 | 0.28 | -0.01 | -0.06 | 0.26 | 0.85 | -0.0525 | -0.2025 | 0.19 | 0.609092 | 0.873234 | 0.612487 | 1 | 0.999631 | 0.858851 |
| Arhgap39_H7BX46_S639s_1 | 0.03 | -0.86 | -1.33 | -0.28 | -0.68 | -1.01 | -0.32 | -0.42 | -1.65 | -0.6 | -0.11 | 0.69 | -1.49 | -0.42 | 0.69 | -0.31 | 0.0025 | 0.1925 | 0.2275 | 0.658184 | 0.994488 | 0.720367 | 1 | 0.999631 | 0.879726 |
| Arhgap39_H7BX46_S685s_1 | 0.01 | -0.82 | -0.3 | 0.31 | -0.11 | -0.36 | -0.2 | 0.45 | 0.07 | -0.93 | 0.5 | -0.76 | -1.49 | 0.16 | 0.18 | 0.44 | 0.145 | -0.08 | 0.0225 | 0.963102 | 0.667631 | 0.84896 | 1 | 0.999631 | 0.987129 |
| Arhgap39_H7BX46_T532t_1 | 2.48 | 0.78 | -0.33 | 1.23 | 2.65 | 1.57 | 1.58 | 0.95 | 1.3 | 0.13 | 1.34 | -0.83 | -1.26 | 1.37 | -0.3 | 0.76 | 0.6475 | -0.555 | -0.8975 | 0.234092 | 0.30482 | 0.432602 | 1 | 0.999631 | 0.683077 |
| Arhgap44_Q5SSM3_S598s_1 | 0.27 | -2.5 | 1.01 | 1.05 | -1.27 | -1.05 | 1.3 | 0.79 | -2.05 | -1.1 | 0.77 | 0.49 | -2.96 | -2.22 | 1.31 | -0.26 | -0.015 | -0.43 | -0.99 | 0.373299 | 0.986915 | 0.645497 | 1 | 0.999631 | 0.747035 |
| Arhgap44_Q5SSM3_S606s_1 | -1.87 | -0.79 | -0.27 | -0.98 | -1.25 | -0.32 | -1.52 | -1.19 | -1.49 | -0.39 | -0.94 | -0.77 | -1.04 | -0.41 | -0.36 | -1.21 | -0.0925 | 0.08 | 0.2225 | 0.593614 | 0.825342 | 0.845805 | 1 | 0.999631 | 0.853502 |
| Arhgap44_Q5SSM3_S636s_1 | 0.58 | -0.74 | -0.73 | 0.24 | 0.86 | 0.01 | 0.3 | 0.64 | -0.13 | -0.99 | 0.09 | -0.35 | -0.85 | -0.14 | -0.01 | -0.13 | 0.615 | -0.1825 | -0.12 | 0.769035 | 0.141864 | 0.662083 | 1 | 0.999631 | 0.918318 |
| Arhgap44_Q5SSM3_S805s_1 | 0.53 | 1.83 | 1.61 | 1.31 | 1.36 | 1.71 | 1.08 | 1.34 | 1.49 | 1.12 | 1.58 | 1.39 | 1.29 | 1.45 | 1.2 | 1.62 | 0.0525 | 0.075 | 0.07 | 0.843038 | 0.878874 | 0.827486 | 1 | 0.999631 | 0.948139 |
| Arhgap5_E9PYT0_S1219s_1 | -0.76 | -0.8 | -1.07 | -0.67 | -0.81 | -1.42 | -0.73 | -0.05 | -0.77 | -1.44 | -0.85 | -0.84 | -0.74 | -0.87 | -0.2 | -0.12 | 0.0725 | -0.15 | 0.3425 | 0.278524 | 0.827638 | 0.592159 | 1 | 0.999631 | 0.70098 |
| Arhgap5_E9PYT0_S968s_1 | -0.29 | 0.47 | 0.35 | 0.51 | 0.27 | -0.27 | 0.33 | 0.9 | 0.01 | -1.27 | 0.73 | -0.71 | 0.39 | 0.44 | 0.59 | 0.6 | 0.0475 | -0.57 | 0.245 | 0.422029 | 0.888539 | 0.234318 | 1 | 0.999631 | 0.773027 |
| Arhgap5_E9PYT0_T1218t_1 | -0.76 | -0.8 | 0 | -0.27 | -0.81 | -1.42 | -0.25 | -0.14 | -0.77 | -1.44 | -0.66 | -0.11 | -0.74 | -0.87 | -0.1 | 0.36 | -0.1975 | -0.2875 | 0.12 | 0.753768 | 0.596845 | 0.438831 | 1 | 0.999631 | 0.913377 |
| Arhgdia_Q99PT1_S34s_1 | 0.61 | 1.75 | -0.51 | 0.66 | 0.85 | 1.02 | 0 | 0.98 | 1.04 | 0.54 | 0.47 | 0.45 | -0.96 | 1.2 | -0.02 | 0.58 | 0.085 | -0.0025 | -0.4275 | 0.478254 | 0.862322 | 0.995724 | 1 | 0.999631 | 0.800612 |
| Arhgdia_Q99PT1_Y133y_1 | -0.14 | -1.75 | -0.7 | -0.4 | -0.98 | -1.5 | -0.55 | -0.72 | -1.08 | -0.22 | -1.37 | -1.87 | -1.63 | -3.51 | -1.23 | -2.41 | -0.19 | -0.3875 | -1.4475 | 0.026916 | 0.644556 | 0.423721 | 1 | 0.999631 | 0.422772 |
| Arhgef1_E9PUF7_S961s_1 | -1.33 | 0.38 | 0.21 | -0.5 | 0.34 | -0.6 | 0.15 | 0.38 | 0.03 | -0.1 | 0.24 | 0.63 | -0.39 | -0.31 | -1.45 | 0.25 | 0.3775 | 0.51 | -0.165 | 0.743638 | 0.399878 | 0.246275 | 1 | 0.999631 | 0.90976 |
| Arhgef1_E9PUF7_T429t_1 | 1.16 | 0.19 | -0.04 | 0.98 | 0.82 | -0.1 | 0.83 | 0.55 | 0.61 | 0.61 | 0.64 | 0.53 | -0.84 | -0.01 | 0.32 | 0.36 | -0.0475 | 0.025 | -0.615 | 0.164077 | 0.900326 | 0.941327 | 1 | 0.999631 | 0.632208 |
| Arhgef10_Q8C033_S7s_1 | 0.32 | 0.17 | -0.43 | -0.42 | -0.91 | -1.01 | -0.71 | -0.85 | -0.91 | 0.37 | -1.2 | -0.37 | 0.43 | -0.25 | -0.87 | -0.04 | -0.78 | -0.4375 | -0.0925 | 0.804683 | 0.019457 | 0.296486 | 1 | 0.999631 | 0.934231 |
| Arhgef10_Q8C033_S852s_1 | -0.43 | -0.62 | -1.29 | -1.27 | 0.08 | -0.17 | -2.11 | -1.62 | 0 | -0.22 | -1.57 | -1.93 | 0.61 | -0.58 | 0.32 | -0.89 | -0.0525 | -0.0275 | 0.7675 | 0.097948 | 0.921931 | 0.956272 | 1 | 0.999631 | 0.560722 |
| Arhgef11_Q68FM7_S1125s_1 | -0.49 | -0.22 | 1.02 | 0.04 | 0.28 | -0.8 | 0.86 | -0.45 | 1.25 | -1.02 | 0.95 | 2.34 | 1.54 | -0.88 | 0.69 | -0.32 | -0.115 | 0.7925 | 0.17 | 0.769248 | 0.807851 | 0.26758 | 1 | 0.999631 | 0.918358 |
| Arhgef11_Q68FM7_S1353s_1 | 1.82 | 0.74 | -0.44 | -0.17 | 2.32 | 1.25 | 0.62 | 0.6 | 1.62 | 1.02 | 0.62 | 0.57 | 0.6 | 0.76 | -1.55 | -0.34 | 0.71 | 0.47 | -0.62 | 0.358357 | 0.246683 | 0.384268 | 1 | 0.999631 | 0.741182 |
| Arhgef11_Q68FM7_S1488s_1 | -0.78 | -0.24 | -0.63 | -0.65 | -0.25 | -0.3 | -0.5 | -0.74 | -0.27 | -0.25 | -0.52 | -0.11 | -0.12 | 0.02 | -0.27 | -0.19 | 0.1275 | 0.2875 | 0.435 | 0.138593 | 0.629491 | 0.291122 | 1 | 0.999631 | 0.611124 |
| Arhgef11_Q68FM7_S253s_1 | -1.27 | -0.85 | -0.15 | 0.62 | 0.66 | -1.78 | 0.88 | 0.17 | -0.05 | -2.98 | 0.93 | 0.81 | -0.77 | -2.03 | 0.2 | 0.09 | 0.395 | 0.09 | -0.215 | 0.721812 | 0.550164 | 0.917847 | 1 | 0.999631 | 0.905674 |
| Arhgef11_Q68FM7_S295s_1 | -0.51 | -0.43 | -0.27 | 0.21 | -0.18 | -0.7 | -0.71 | 0.05 | -0.68 | -0.55 | -0.21 | 0.07 | 0.49 | 0.11 | -0.43 | 0.31 | -0.135 | -0.0925 | 0.37 | 0.277768 | 0.661717 | 0.762401 | 1 | 0.999631 | 0.700874 |
| Arhgef11_Q68FM7_S311s_1 | 0.61 | -0.77 | -1.01 | 0.35 | 0.98 | -0.22 | 0.11 | 0.22 | 0.09 | -0.22 | 0.24 | 0.1 | -0.64 | 0.35 | -0.17 | 0.29 | 0.4775 | 0.2575 | 0.1625 | 0.723752 | 0.310208 | 0.541815 | 1 | 0.999631 | 0.905674 |
| Arhgef11_Q68FM7_S585s_1 | 1.06 | 0.22 | -1.16 | 2.1 | 3.52 | 1.34 | 1.72 | 1.45 | 2.35 | -0.88 | 1.07 | 0.9 | -0.37 | 1.32 | 1.2 | 0.23 | 1.4525 | 0.305 | 0.04 | 0.954843 | 0.078257 | 0.716232 | 1 | 0.999631 | 0.984441 |
| Arhgef11_Q68FM7_S663s_1 | -1.33 | -0.09 | -0.65 | 0.64 | 0.26 | 0.42 | -0.12 | 0.21 | 0.88 | -0.1 | -0.14 | 0.79 | -0.33 | -0.13 | 0.67 | 0.22 | 0.55 | 0.715 | 0.465 | 0.329569 | 0.217038 | 0.160341 | 1 | 0.999631 | 0.726614 |
| Arhgef11_Q68FM7_S665s_1 | -0.27 | 0.07 | -0.16 | 0.3 | 0.04 | -0.11 | -0.17 | 0.45 | -0.23 | -0.22 | 0.05 | 0.25 | -0.92 | 0.71 | 0.3 | 0.42 | 0.0675 | -0.0225 | 0.1425 | 0.724696 | 0.806336 | 0.93475 | 1 | 0.999631 | 0.905674 |
| Arhgef12_F8VQN6_S1177s_1 | -2.16 | -1.35 | -4.9 | -3.08 | -1.28 | -1.69 | -3.62 | -4.05 | -2.6 | -1.17 | -3.41 | -5.14 | -2.67 | -0.94 | -3.23 | -1.81 | 0.2125 | -0.2075 | 0.71 | 0.382286 | 0.811478 | 0.831308 | 1 | 0.999631 | 0.753668 |
| Arhgef12_F8VQN6_S1328s_1 | -0.28 | 0.49 | 0.05 | 0.26 | 0.12 | 0.31 | 0.52 | -0.32 | 0.26 | 0 | 0.42 | 0.72 | 0.43 | 0 | 0.81 | 0.55 | 0.0275 | 0.22 | 0.3175 | 0.332386 | 0.927645 | 0.468739 | 1 | 0.999631 | 0.728079 |
| Arhgef12_F8VQN6_S190s_1 | -0.29 | -0.8 | 0.13 | -0.53 | -0.67 | -0.43 | -0.08 | -0.27 | -0.05 | 0.12 | 0.29 | 0.38 | 0.44 | -0.09 | -0.47 | 0.04 | 0.01 | 0.5575 | 0.3525 | 0.311444 | 0.97322 | 0.085705 | 1 | 0.999631 | 0.71924 |
| Arhgef12_F8VQN6_S342s_1 | -0.76 | 0 | -1.72 | -1.79 | -0.85 | -1.08 | -0.83 | -1.67 | -0.74 | -0.29 | -0.97 | -1.07 | -0.46 | -1.93 | -0.82 | -0.67 | -0.04 | 0.3 | 0.0975 | 0.848834 | 0.929519 | 0.511018 | 1 | 0.999631 | 0.949525 |
| Arhgef12_F8VQN6_T737t_1 | 2.67 | 1.02 | 0.65 | 0.96 | 0.54 | 2.1 | 2.11 | 0.13 | 0.72 | 1.01 | 0.77 | 0.33 | -1.12 | -0.16 | 0.46 | -0.17 | -0.105 | -0.6175 | -1.5725 | 0.013002 | 0.865494 | 0.203218 | 1 | 0.999631 | 0.385092 |
| Arhgef17_Q80U35_S142s_1 | -1.01 | 0.17 | 0.64 | -0.42 | 0.01 | 0.39 | -0.4 | -0.6 | -0.21 | 0.47 | -0.08 | -1.49 | -0.14 | -0.51 | -0.93 | 0 | 0.005 | -0.1725 | -0.24 | 0.575408 | 0.990433 | 0.738487 | 1 | 0.999631 | 0.846085 |
| Arhgef17_Q80U35_S152s_1 | -0.52 | -1.34 | 0.95 | -1.89 | -1.23 | -0.82 | -0.7 | -1.11 | -1.49 | -0.13 | -0.02 | -0.11 | 0.66 | -0.36 | 0.55 | 0.66 | -0.265 | 0.2625 | 1.0775 | 0.099507 | 0.645798 | 0.683198 | 1 | 0.999631 | 0.562298 |
| Arhgef17_Q80U35_S2005s_1 | -0.05 | 0.24 | 0.11 | -0.32 | -1.08 | 0.1 | -0.82 | 0.58 | -0.09 | 0.05 | -2.01 | 0.09 | 0.18 | 0.41 | 0.68 | -0.2 | -0.3 | -0.485 | 0.2725 | 0.393026 | 0.468428 | 0.340956 | 1 | 0.999631 | 0.758901 |
| Arhgef18_Q6P9R4_S577s_1 | -0.16 | 0.48 | 0.41 | -0.31 | -0.06 | 0.33 | -0.41 | 0.23 | 0.35 | 0.45 | 0.15 | 0.71 | 0.47 | 0.73 | 0.54 | 0.56 | -0.0825 | 0.31 | 0.47 | 0.147134 | 0.792125 | 0.32081 | 1 | 0.999631 | 0.61916 |
| Arhgef2_H3BJU7_S617s_1 | 0.17 | 0.58 | -0.17 | 0.29 | 0.44 | 0.09 | 0.38 | 0.23 | 0.38 | 0.17 | 0.58 | 0.42 | 0.68 | 0.3 | -0.52 | 0.62 | 0.0675 | 0.17 | 0.0525 | 0.885268 | 0.801572 | 0.543731 | 1 | 0.999631 | 0.963505 |
| Arhgef2_H3BJU7_S856s_1 | 0.68 | 0.87 | 0.25 | 1.06 | 1.37 | 0.22 | 0.92 | 0.72 | 0.59 | -0.24 | 0.93 | 1.27 | -0.82 | 0.54 | 0.53 | -0.37 | 0.0925 | -0.0775 | -0.745 | 0.088736 | 0.781937 | 0.841021 | 1 | 0.999631 | 0.553858 |
| Arhgef2_H3BJU7_S902s_1 | 0.28 | 0.66 | -0.26 | -0.13 | 0.16 | 0.33 | -0.03 | -0.47 | 0.71 | 0.8 | 0.15 | -0.19 | 1.05 | 0.16 | -0.7 | -0.94 | -0.14 | 0.23 | -0.245 | 0.614144 | 0.662459 | 0.518385 | 1 | 0.999631 | 0.860996 |
| Arhgef2_H3BJU7_S911s_1 | -1.57 | 0.97 | 0.44 | -1.82 | -1.6 | 1.1 | -0.72 | -1.04 | 0.74 | 0.7 | -0.57 | 0.44 | 3.2 | 1.41 | 0.66 | 0.46 | -0.07 | 0.8225 | 1.9275 | 0.037561 | 0.930071 | 0.24773 | 1 | 0.999631 | 0.458256 |
| Arhgef2_H3BJU7_S926s_1 | -0.33 | 1.52 | 1.79 | -0.28 | -0.96 | 1.27 | -0.65 | 0.35 | 0.55 | 1.74 | 0.48 | 1.02 | 1.79 | 1.49 | 1.02 | 1.63 | -0.6725 | 0.2725 | 0.8075 | 0.165049 | 0.332944 | 0.642737 | 1 | 0.999631 | 0.633483 |
| Arhgef2_H3BJU7_S926s_2 | 1.69 | 1.23 | -0.82 | 1.59 | 2.37 | 1.31 | 1.63 | 1.34 | 1.44 | 0.91 | 1.85 | -0.25 | -1.88 | 1.16 | 0.03 | 1.01 | 0.74 | 0.065 | -0.8425 | 0.304858 | 0.221828 | 0.922328 | 1 | 0.999631 | 0.716239 |
| Arhgef2_H3BJU7_S930s_2 | 1.69 | 1.23 | -0.82 | 1.59 | 2.37 | 1.31 | 1.63 | 1.34 | 1.44 | 0.91 | 1.85 | -0.25 | -1.88 | 1.16 | 0.03 | 1.01 | 0.74 | 0.065 | -0.8425 | 0.304858 | 0.221828 | 0.922328 | 1 | 0.999631 | 0.716239 |
| Arhgef2_H3BJU7_T766t_1 | -0.3 | -0.64 | 0.02 | 0.17 | 0.02 | -0.69 | 0.03 | 0.36 | -0.45 | -0.38 | 0.4 | 0.14 | -0.36 | 0.12 | -0.03 | 0.99 | 0.1175 | 0.115 | 0.3675 | 0.342197 | 0.720831 | 0.72595 | 1 | 0.999631 | 0.73347 |
| Arhgef25_G5E825_T603t_1 | -0.74 | -0.69 | -1.92 | -0.76 | 0.19 | -0.45 | -0.99 | -1.09 | 0.07 | -0.83 | -1.05 | -0.71 | 0.05 | -0.04 | -2.21 | 0.72 | 0.4425 | 0.3975 | 0.6575 | 0.312852 | 0.302066 | 0.330978 | 1 | 0.999631 | 0.719861 |
| Arhgef26_D3YYY8_S123s_1 | -0.93 | 0.96 | -0.34 | 0.9 | -0.19 | -0.03 | 0.08 | 0.78 | 0.48 | 0.49 | 1.11 | 1.24 | 0.29 | 0.24 | 1.55 | 0.65 | 0.0125 | 0.6825 | 0.535 | 0.321597 | 0.979452 | 0.182834 | 1 | 0.999631 | 0.724092 |
| Arhgef26_D3YYY8_S199s_1 | -1.7 | -0.51 | 0.21 | 0.16 | -0.56 | -0.66 | -0.87 | -0.95 | -1.1 | -0.54 | -0.93 | -0.94 | -0.65 | 0.06 | -0.94 | -0.53 | -0.3 | -0.4175 | -0.055 | 0.908431 | 0.50187 | 0.365824 | 1 | 0.999631 | 0.969941 |
| Arhgef26_D3YYY8_S202s_1 | -1.7 | -0.51 | 0.21 | 0.16 | -0.56 | -0.66 | -0.87 | -0.95 | -1.1 | -0.54 | -0.93 | -0.94 | -0.65 | 0.06 | -0.94 | -0.53 | -0.3 | -0.4175 | -0.055 | 0.908431 | 0.50187 | 0.365824 | 1 | 0.999631 | 0.969941 |
| Arhgef26_D3YYY8_S221s_1 | -0.27 | -0.35 | -0.77 | -0.9 | 0.28 | -0.44 | 0.18 | 0.79 | -0.53 | -1.15 | 0.03 | -0.56 | 0.11 | -0.39 | -0.36 | -0.03 | 0.775 | 0.02 | 0.405 | 0.199676 | 0.040677 | 0.952378 | 1 | 0.999631 | 0.657804 |
| Arhgef28_G5E8P2_S1198s_1 | 0.8 | -0.64 | -0.07 | -0.04 | -0.82 | -0.55 | 1.53 | -0.51 | -0.91 | 1.55 | -0.09 | -0.75 | -0.32 | -1.93 | -0.22 | -1.53 | -0.1 | -0.0625 | -1.0125 | 0.06613 | 0.859571 | 0.914456 | 1 | 0.999631 | 0.516933 |
| Arhgef4_E9PYK8_S1277s_1 | 0.23 | 0.3 | -0.02 | -0.21 | 0.09 | -0.46 | -0.05 | -0.3 | 0.14 | 0.17 | -0.32 | -0.05 | -0.13 | 0.55 | 0.51 | 0.35 | -0.255 | -0.09 | 0.245 | 0.422925 | 0.350616 | 0.74057 | 1 | 0.999631 | 0.773293 |
| Arhgef40_S4R189_S1089s_1 | 1.75 | -0.22 | 1.09 | 0.58 | 0.34 | 1.3 | 0.98 | -0.35 | 0.8 | 1.34 | 0.57 | -0.02 | -0.16 | -0.34 | 0.51 | 0.1 | -0.2325 | -0.1275 | -0.7725 | 0.112153 | 0.653316 | 0.791456 | 1 | 0.999631 | 0.577068 |
| Arhgef40_S4R189_S1494s_1 | -1.31 | -0.57 | -0.3 | -0.33 | -1.37 | -0.91 | -0.76 | -0.63 | -1.16 | -0.78 | -0.95 | -0.09 | -0.85 | -0.86 | 0.37 | -1.5 | -0.29 | -0.1175 | -0.0825 | 0.855653 | 0.386538 | 0.746202 | 1 | 0.999631 | 0.951817 |
| Arhgef6_F6WMJ3_S248s_1 | -0.25 | -0.06 | 0.03 | -0.53 | -0.65 | -0.66 | -0.66 | -0.18 | -0.54 | -1.01 | -0.34 | -0.29 | -0.05 | -0.21 | 0.16 | -0.14 | -0.335 | -0.3425 | 0.1425 | 0.614969 | 0.2292 | 0.256256 | 1 | 0.999631 | 0.861203 |
| Arhgef6_F6WMJ3_S511s_1 | -0.9 | 0.12 | -0.04 | -0.19 | -0.39 | 0.2 | -0.79 | -0.13 | -0.01 | -0.47 | -0.33 | 0.2 | 0.44 | 0.05 | 0.28 | 0.25 | -0.025 | 0.1 | 0.5075 | 0.137357 | 0.941591 | 0.759703 | 1 | 0.999631 | 0.610409 |
| Arhgef6_F6WMJ3_S663s_1 | -0.18 | 0.11 | -0.12 | 0.06 | -0.12 | 0.2 | -0.81 | 0.24 | 0.24 | 0.22 | -0.8 | -0.33 | 0.73 | 0.43 | -0.54 | -0.39 | -0.09 | -0.135 | 0.09 | 0.804918 | 0.771078 | 0.673324 | 1 | 0.999631 | 0.934231 |
| Arhgef6_F6WMJ3_S703s_1 | -0.54 | 0.73 | 0.67 | 0.14 | 0.07 | -0.33 | -0.31 | 0.1 | -0.1 | -0.49 | -0.29 | 0.01 | 0.55 | 0.73 | 0.16 | 0.55 | -0.3675 | -0.4675 | 0.2475 | 0.50257 | 0.304964 | 0.205144 | 1 | 0.999631 | 0.814092 |
| Arhgef7_Q9ES28_S103s_1 | -0.78 | 0.33 | -0.45 | -0.04 | 0.36 | 0.02 | -0.14 | 0.02 | 0.31 | -0.61 | 0.15 | 0.29 | -0.03 | 0.33 | -0.27 | 0.34 | 0.3 | 0.27 | 0.3275 | 0.353505 | 0.352645 | 0.458907 | 1 | 0.999631 | 0.73949 |
| Arhgef7_Q9ES28_S176s_1 | 0.38 | -0.54 | 0.71 | 0.37 | 0.01 | 0.38 | 0.1 | 0.02 | 0.1 | 0.67 | -0.79 | -0.06 | 0.14 | -0.02 | 0.37 | -0.42 | -0.1025 | -0.25 | -0.2125 | 0.562042 | 0.753732 | 0.546579 | 1 | 0.999631 | 0.84072 |
| Arhgef7_Q9ES28_S445s_1 | -0.38 | -0.38 | -0.09 | -0.58 | -0.68 | -0.54 | -0.71 | -0.75 | -0.83 | -0.35 | -0.41 | -0.24 | -0.09 | -0.33 | -0.45 | -0.07 | -0.3125 | -0.1 | 0.1225 | 0.660853 | 0.219614 | 0.713647 | 1 | 0.999631 | 0.880315 |
| Arhgef7_Q9ES28_S621s_1 | -0.42 | -0.22 | -0.66 | -0.23 | -0.44 | -0.48 | -0.71 | -0.35 | -0.4 | -0.2 | -0.02 | -0.06 | -0.53 | -0.49 | -0.3 | 0.1 | -0.1125 | 0.2125 | 0.0775 | 0.791546 | 0.654196 | 0.422296 | 1 | 0.999631 | 0.926807 |
| Arhgef7_Q9ES28_S665s_1 | 0.15 | -0.5 | -0.79 | 0.51 | 0.36 | -0.84 | 0.37 | 0.12 | -0.53 | -0.37 | 0.37 | -0.08 | -0.69 | -0.1 | -0.68 | 0.03 | 0.16 | 0.005 | -0.2025 | 0.601629 | 0.698676 | 0.98947 | 1 | 0.999631 | 0.856765 |
| Arhgef7_Q9ES28_S719s_1 | 0.99 | 0.29 | 0.51 | 0.46 | -0.53 | 0.4 | 0.87 | -0.49 | 0.77 | 1.05 | 0.34 | 0.06 | 0.12 | -0.38 | -0.51 | -0.91 | -0.5 | -0.0075 | -0.9825 | 0.013376 | 0.215222 | 0.981491 | 1 | 0.999631 | 0.385092 |
| Arhgef7_Q9ES28_S80s_1 | 0.04 | -1.32 | 0.04 | 0.13 | -1.82 | -1.4 | 0.71 | -0.68 | -1.4 | -0.96 | -0.48 | -0.9 | -1.82 | -1.9 | 0.13 | -0.29 | -0.52 | -0.6575 | -0.6925 | 0.247108 | 0.390306 | 0.129545 | 1 | 0.999631 | 0.685751 |
| Arhgef7_Q9ES28_S83s_1 | 0.1 | 0.61 | -0.52 | -1.05 | 0.02 | 1.03 | -0.94 | 0.28 | 0.5 | 0.46 | -0.81 | -1.01 | 0.02 | 0.06 | -3.62 | 0.12 | 0.3125 | -7.9E-17 | -0.64 | 0.461755 | 0.542599 | 1 | 1 | 1 | 0.792786 |
| Arid1a_E9QAQ7_S1760s_1 | -0.35 | 0.91 | 1.11 | 0.42 | 0.46 | 0.62 | 0.16 | 0.5 | 0.63 | 1.03 | 0.94 | 1.08 | 0.12 | 0.95 | 1.11 | 0.68 | -0.0875 | 0.3975 | 0.1925 | 0.640262 | 0.809156 | 0.295769 | 1 | 0.999631 | 0.870995 |
| Arid1a_E9QAQ7_S365s_1 | 3.38 | 0 | -0.19 | -0.62 | -1.94 | -0.64 | 0.87 | -0.86 | -1.07 | 2.72 | -0.1 | 0.49 | -1.8 | -0.19 | -1.09 | -0.95 | -1.285 | -0.1325 | -1.65 | 0.075605 | 0.192716 | 0.899823 | 1 | 0.999631 | 0.541044 |
| Arid1a_E9QAQ7_S770s_1 | 0.04 | 1.2 | 0.49 | -0.42 | 0.44 | 0.53 | 0.16 | -0.39 | 0.89 | 0.91 | 0.02 | 0.76 | 1.19 | 0.99 | 0.29 | 0.39 | -0.1425 | 0.3175 | 0.3875 | 0.369243 | 0.725491 | 0.44771 | 1 | 0.999631 | 0.746224 |
| Arid1b_E9Q6R4_S1549s_1 | 0.78 | 1.45 | 1 | 0.93 | 0.93 | 0.79 | 0.62 | 1.23 | 1.23 | 1.29 | 1.09 | 1.23 | 1.46 | 1.51 | 1.08 | 1.35 | -0.1475 | 0.17 | 0.31 | 0.301286 | 0.597674 | 0.527455 | 1 | 0.999631 | 0.714243 |
| Arid1b_E9Q6R4_S1755s_1 | 0.65 | 1.61 | 1.47 | 0.9 | 1.52 | 1.22 | 0.42 | 1.89 | 1.42 | 1.15 | 0.98 | 2.1 | 1.2 | 1.61 | 0.77 | 1.48 | 0.105 | 0.255 | 0.1075 | 0.759819 | 0.790099 | 0.491431 | 1 | 0.999631 | 0.914614 |
| Arid1b_E9Q6R4_S1756s_1 | 0.65 | 1.61 | 1.47 | 0.9 | 1.52 | 1.22 | 0.42 | 1.89 | 1.42 | 1.15 | 0.98 | 2.1 | 1.2 | 1.61 | 0.77 | 1.48 | 0.105 | 0.255 | 0.1075 | 0.759819 | 0.790099 | 0.491431 | 1 | 0.999631 | 0.914614 |
| Arid4a_F8VPQ2_S867s_1 | -0.03 | 1.48 | -0.04 | 0.17 | 0.03 | 0.19 | -0.13 | -0.03 | 0.2 | 0.6 | 0.6 | -0.5 | 0.88 | 0.79 | -0.16 | 0.29 | -0.38 | -0.17 | 0.055 | 0.900623 | 0.333821 | 0.70152 | 1 | 0.999631 | 0.967994 |
| Arid5b_Q8BM75_S789s_1 | -0.19 | -0.39 | 0.07 | -0.39 | -0.51 | -0.66 | -0.45 | -0.02 | 0.08 | -0.65 | 0.17 | 0.49 | -0.15 | -0.28 | -0.24 | 0.08 | -0.185 | 0.2475 | 0.0775 | 0.78027 | 0.496908 | 0.449613 | 1 | 0.999631 | 0.921416 |
| Arl13b_Q640N2_S328s_1 | -0.31 | -0.2 | 0.28 | -0.37 | -0.58 | -0.93 | -0.52 | -0.73 | -0.92 | -0.09 | 0 | 0.6 | -0.45 | -0.31 | -0.32 | 0.11 | -0.54 | 0.0475 | -0.0925 | 0.757342 | 0.067798 | 0.898386 | 1 | 0.999631 | 0.914449 |
| Arl2_Q9D0J4_S45s_1 | -0.46 | 0.57 | 0.54 | 0.19 | 0.4 | 0.27 | -0.56 | 0.19 | -0.4 | 0.26 | 0.12 | 0.36 | 0.96 | 0.87 | 0.55 | 0.66 | -0.135 | -0.125 | 0.55 | 0.120105 | 0.701583 | 0.713274 | 1 | 0.999631 | 0.587703 |
| Arl6ip6_Q8BH07_T19t_1 | 1.35 | 0.84 | 1.72 | 1.64 | 0.63 | 1.2 | 0.85 | 1.27 | -0.06 | 2.06 | 1.35 | 0.51 | -0.24 | 1.02 | 1.19 | 1.49 | -0.4 | -0.4225 | -0.5225 | 0.247662 | 0.212624 | 0.393238 | 1 | 0.999631 | 0.685751 |
| Armcx4_E9PWM3_S34s_1 | 1.1 | -2.97 | 0.63 | -1.03 | -1.91 | -0.91 | 0.78 | -0.46 | -1.86 | -0.34 | -2.42 | -1.72 | -1.52 | -1.95 | -4.67 | -3.1 | -0.0575 | -1.0175 | -2.2425 | 0.044419 | 0.950695 | 0.269311 | 1 | 0.999631 | 0.475443 |
| Arpc1a_Q9R0Q6_S236s_1 | -0.01 | 0.1 | 1.1 | 0.64 | -0.23 | 0.55 | 0.04 | 0.52 | -0.01 | 0.46 | 0.75 | 0.54 | 0.31 | 0.69 | 0.22 | 0.48 | -0.2375 | -0.0225 | -0.0325 | 0.924082 | 0.501802 | 0.947988 | 1 | 0.999631 | 0.975324 |
| Arpc5_Q9CPW4_S77s_1 | -0.41 | 0.88 | -0.19 | 0.19 | 0.04 | 0.17 | -0.43 | -0.65 | 0.54 | 0.47 | 0.12 | 0.01 | 0.87 | 0.94 | -0.09 | 0.05 | -0.335 | 0.1675 | 0.325 | 0.43471 | 0.368353 | 0.63436 | 1 | 0.999631 | 0.779717 |
| Arpc5l_Q9D898_S64s_1 | 1.35 | 1.69 | 0.96 | 0.64 | 0.19 | 1.52 | 1.19 | -0.16 | 0.95 | 1.32 | 0.26 | 0.17 | 1.34 | 0.99 | 0.01 | 0.1 | -0.475 | -0.485 | -0.55 | 0.206057 | 0.301232 | 0.222271 | 1 | 0.999631 | 0.6605 |
| Arpp19_P56212_S2s_1 | -0.03 | 0.81 | 0.83 | 0.56 | 1 | 0.48 | 0.65 | 0.53 | 0.34 | -0.53 | 1.11 | 0.6 | 0.35 | 0.17 | 1.02 | 0.42 | 0.1225 | -0.1625 | -0.0525 | 0.876795 | 0.681642 | 0.690478 | 1 | 0.999631 | 0.961373 |
| Arpp19_P56212_S7s_1 | -2.52 | -4.09 | -3.8 | -4.35 | -4.79 | -4 | -3.97 | -4.48 | -5.45 | -2.99 | -4.73 | -6.09 | -3.17 | -4.58 | -5.09 | -3.48 | -0.62 | -1.125 | -0.39 | 0.493389 | 0.180596 | 0.131016 | 1 | 0.999631 | 0.80955 |
| Arpp19_P56212_T22t_1 | -2.41 | -1.24 | -2.29 | -3.03 | -1.93 | -1.84 | -1.76 | -2.8 | -1.91 | -2.26 | -1.44 | -1.21 | -0.95 | -0.62 | -3 | -2.31 | 0.16 | 0.5375 | 0.5225 | 0.401829 | 0.712937 | 0.236805 | 1 | 0.999631 | 0.763221 |
| Arpp19_P56212_T6t_1 | -2.41 | -1.24 | -2.29 | -3.03 | -1.93 | -1.84 | -1.76 | -2.8 | -1.91 | -2.26 | -1.44 | -1.21 | -0.95 | -0.62 | -3 | -2.31 | 0.16 | 0.5375 | 0.5225 | 0.401829 | 0.712937 | 0.236805 | 1 | 0.999631 | 0.763221 |
| Arpp21_E0CYU3_S32s_1 | 1.04 | 1.43 | 0.92 | 0.62 | 1.09 | 1.52 | 0.94 | 0.33 | 1.76 | 1.93 | 1.01 | 1.16 | 0.87 | 0.97 | 1.58 | 0.54 | -0.0325 | 0.4625 | -0.0125 | 0.970627 | 0.922822 | 0.184514 | 1 | 0.999631 | 0.989618 |
| Arpp21_Q9DCB4_S346s_1 | 1.49 | 0.51 | 1.06 | 1.32 | 0.71 | 1.19 | 1.51 | 1.33 | 1.05 | 1.65 | 1.33 | 1.08 | -0.26 | 0.29 | 1.01 | 0.18 | 0.09 | 0.1825 | -0.79 | 0.058723 | 0.77972 | 0.568312 | 1 | 0.999631 | 0.501503 |
| Arpp21_Q9DCB4_S362s_1 | 1.75 | 0.55 | 1.03 | 0.34 | 1.52 | 1.75 | 0.7 | 0.26 | 1.92 | 1.98 | -0.13 | 0.39 | 0.18 | 0.74 | 0.06 | -0.71 | 0.14 | 0.1225 | -0.85 | 0.075706 | 0.757118 | 0.830141 | 1 | 0.999631 | 0.541044 |
| Arpp21_Q9DCB4_S381s_1 | 0.6 | 1.06 | 1.08 | 0.84 | 0.8 | 0.72 | 0.55 | 1.19 | 0.94 | 1.24 | 1.2 | 1.78 | 0.89 | 1.3 | 1.36 | 1.71 | -0.08 | 0.395 | 0.42 | 0.187489 | 0.766745 | 0.197954 | 1 | 0.999631 | 0.646474 |
| Arpp21_Q9DCB4_S408s_1 | 1.16 | 1.38 | 1.13 | 1.02 | 1.49 | 0.67 | 0.97 | 1.06 | 2.53 | 1.06 | 0.9 | 1.29 | 0 | 0.3 | 1.24 | 0.12 | -0.125 | 0.2725 | -0.7575 | 0.05173 | 0.64947 | 0.494088 | 1 | 0.999631 | 0.489777 |
| Arpp21_Q9DCB4_T87t_1 | 1.76 | 0.54 | 1.31 | 1.96 | 1.03 | 0.91 | 1.96 | 1.72 | 1.17 | 1.68 | 1.59 | 1.71 | -0.79 | 0.8 | 1.7 | 1.08 | 0.0125 | 0.145 | -0.695 | 0.239865 | 0.975513 | 0.694428 | 1 | 0.999631 | 0.683719 |
| Arrb1_J3QNU6_S413s_1 | -1.35 | 1.19 | 3.31 | 0.49 | 1.12 | -0.2 | 2.68 | 1.53 | 2.19 | -1.41 | 3.65 | 4.36 | 1.69 | -1.42 | 2.69 | 1.22 | 0.3725 | 1.2875 | 0.135 | 0.902846 | 0.703212 | 0.358102 | 1 | 0.999631 | 0.968183 |
| Arvcf_P98203_S273s_1 | -1.01 | -1.1 | -0.35 | -1.09 | -1.34 | -0.86 | -1.79 | -1.34 | -1.4 | -0.4 | -0.93 | -0.96 | 0.15 | -0.75 | -0.74 | -0.36 | -0.445 | -0.035 | 0.4625 | 0.196637 | 0.178552 | 0.914744 | 1 | 0.999631 | 0.653373 |
| Arvcf_P98203_S281s_1 | 1.39 | -0.98 | -1.35 | -0.88 | -1.5 | -0.05 | -0.38 | -1.49 | -0.94 | 1.35 | -1.16 | -1.69 | -0.39 | -1.1 | -1.23 | -1.95 | -0.4 | -0.155 | -0.7125 | 0.275201 | 0.542149 | 0.847166 | 1 | 0.999631 | 0.700187 |
| Arvcf_P98203_S283s_1 | 1.3 | -1.11 | -0.81 | -0.4 | -1.09 | -0.35 | -0.01 | -1.31 | -1.13 | 0.92 | -1.16 | -1.35 | -0.8 | -1.89 | -1.31 | -1.64 | -0.435 | -0.425 | -1.155 | 0.057319 | 0.448716 | 0.535455 | 1 | 0.999631 | 0.49679 |
| Arvcf_P98203_S873s_1 | 1.52 | 0.32 | -0.39 | 0.32 | 0.25 | 0.7 | 0.58 | 0.11 | 0.33 | -0.2 | 0.36 | -1.09 | -1.08 | 0.64 | -0.39 | 0.06 | -0.0325 | -0.5925 | -0.635 | 0.232413 | 0.937795 | 0.249846 | 1 | 0.999631 | 0.682449 |
| Arvcf_P98203_T573t_1 | -1.65 | 1.15 | 1.27 | -0.6 | 0.32 | 1.42 | 0.38 | 0.97 | 1.75 | -2.04 | 0.49 | 2.34 | 1.96 | 0.02 | 0.94 | 0.87 | 0.73 | 0.5925 | 0.905 | 0.225725 | 0.291669 | 0.570608 | 1 | 0.999631 | 0.676382 |
| Asap1_E9QMJ1_S1010s_1 | 0.53 | -0.04 | -0.45 | 1.24 | 3.56 | 0.28 | 5.24 | 1.24 | 3.54 | 0.62 | 3.33 | 2.68 | 0.46 | -0.26 | 0.52 | -0.09 | 2.26 | 2.2225 | -0.1625 | 0.703702 | 0.047567 | 0.008322 | 1 | 0.999631 | 0.897149 |

| | | | | | | | | | | | | | | | | | | | | | | | | | |
|---|---|---|---|---|---|---|---|---|---|---|---|---|---|---|---|---|---|---|---|---|---|---|---|---|---|
| Asap1_E9QMJ1_S1024s_1 | -0.08 | 0.67 | -0.35 | -0.01 | 0.26 | 0.41 | -0.4 | -0.17 | 0.23 | -0.13 | -0.24 | 0.22 | 0.32 | -0.08 | -0.49 | 0.35 | -0.0325 | -0.0375 | -0.0325 | 0.926064 | 0.921233 | 0.904636 | 1 | 0.999631 | 0.97588 |
| Asap1_E9QMJ1_S1111s_1 | -1.26 | -3.01 | -2.21 | -1.25 | -2.97 | -2.43 | -1.87 | -3.76 | -2.48 | -2.51 | -1.43 | -2.1 | -2.02 | -1.77 | -3.8 | -1.04 | -0.825 | -0.1975 | -0.225 | 0.729434 | 0.149409 | 0.678598 | 1 | 0.999631 | 0.905923 |
| Asap1_E9QMJ1_S505s_1 | -0.66 | -0.39 | -0.36 | -0.49 | -0.5 | -0.39 | -0.83 | -0.65 | -0.67 | -0.64 | -0.62 | -0.22 | 0.2 | -0.3 | -0.28 | -0.1 | -0.1175 | -0.0625 | 0.355 | 0.218959 | 0.634674 | 0.808193 | 1 | 0.999631 | 0.669859 |
| Asap1_E9QMJ1_S819s_1 | -1.35 | 0.38 | -0.23 | -0.41 | -0.99 | -0.29 | -1.13 | -0.65 | -0.25 | -0.3 | -0.64 | -0.28 | -0.06 | -0.36 | -0.54 | 0.2 | -0.3625 | 0.035 | 0.2125 | 0.607996 | 0.382179 | 0.928058 | 1 | 0.999631 | 0.858527 |
| Asap1_E9QMJ1_S823s_1 | 2.01 | -1.37 | -1.53 | 1.41 | 2.76 | 0.24 | 1.75 | 0.77 | 0.67 | -0.95 | -0.07 | -1.04 | -1.32 | -0.5 | -0.73 | -1.03 | 1.25 | -0.4775 | -1.025 | 0.227756 | 0.199572 | 0.588489 | 1 | 0.999631 | 0.678949 |
| Asap2_D3YX85_S712s_1 | 2.34 | -0.91 | -0.46 | 1.5 | 1.36 | 0.84 | 2 | 0.89 | 0.29 | 1.08 | 1 | -0.49 | -3.38 | -0.24 | 0.27 | -0.22 | 0.655 | -0.1475 | -1.51 | 0.145889 | 0.376487 | 0.845672 | 1 | 0.999631 | 0.616968 |
| Asb3_D6RDR7_S35s_1 | 0.92 | 0.76 | 1.25 | 1.69 | 0.69 | 0.76 | 0.76 | 1.63 | 0.85 | 1.3 | 1.31 | 1.22 | -0.33 | 0.95 | 1.52 | 1.22 | -0.195 | 0.015 | -0.315 | 0.492509 | 0.568896 | 0.96069 | 1 | 0.999631 | 0.80955 |
| Ash1l_Q99MY8_S1678s_1 | 1.43 | -0.84 | 0.64 | -0.25 | -2.18 | -1.4 | 0.83 | -1.44 | -1.61 | 0.45 | -0.94 | -0.54 | -1.22 | -1.79 | -1.22 | -2.08 | -1.2925 | -0.905 | -1.8225 | 0.005143 | 0.09882 | 0.155157 | 1 | 0.999631 | 0.351308 |
| Asic1_Q6NXK8_S497s_1 | 1.51 | 2.9 | -1.62 | -0.35 | 4.72 | 1.29 | 3.05 | 0.63 | 5.13 | 0.97 | 2.53 | 3.07 | 3.59 | 1.45 | 1.76 | -0.38 | 1.8125 | 2.315 | 0.995 | 0.373508 | 0.14218 | 0.064257 | 1 | 0.999631 | 0.747035 |
| Aspscr1_Q8VBT9_S116s_1 | -1.44 | -1.51 | -1.54 | -1.44 | -1.11 | -1.36 | -1.18 | -1.74 | -1.75 | -1.22 | -1.44 | -1.31 | -1 | -1.34 | -1.06 | -1.1 | 0.135 | 0.0525 | 0.3575 | 0.193274 | 0.599679 | 0.836787 | 1 | 0.999631 | 0.649965 |
| Aspscr1_Q8VBT9_S202s_1 | 1.05 | -0.05 | -0.47 | 0.3 | 0.33 | 1.85 | 0 | 0.35 | 0.32 | 2.3 | 0.01 | -0.06 | -1.82 | 1.66 | 0.35 | 0.82 | 0.425 | 0.435 | 0.045 | 0.949776 | 0.39816 | 0.466901 | 1 | 0.999631 | 0.98392 |
| Astn1_Q61137_S192s_1 | -1.28 | -1.12 | -1.75 | -2.64 | -1.01 | -1.16 | -1.39 | -2.61 | -0.67 | -1.06 | -2.01 | -1.46 | 0.16 | -0.49 | -1 | -0.7 | 0.155 | 0.3975 | 1.19 | 0.018505 | 0.744511 | 0.377415 | 1 | 0.999631 | 0.391024 |
| Astn1_Q61137_S274s_1 | -0.05 | 0.29 | -0.64 | 0.2 | 0.06 | -0.02 | 0.03 | -0.03 | 0.05 | -0.47 | -0.25 | -0.25 | -0.7 | 0.01 | -0.51 | -0.03 | 0.06 | -0.18 | -0.2575 | 0.454794 | 0.833868 | 0.559916 | 1 | 0.999631 | 0.789905 |
| Astn1_Q61137_S893s_1 | 1.3 | -0.51 | 0.24 | 0.3 | -1.29 | -0.62 | 0.44 | -0.92 | 0.34 | 1.42 | -0.64 | -1.25 | -0.2 | -2.59 | -0.86 | -2.02 | -0.93 | -0.365 | -1.75 | 0.014427 | 0.084683 | 0.563518 | 1 | 0.999631 | 0.385092 |
| Astn1_Q61137_S898s_1 | 1.27 | -0.89 | -0.08 | -0.23 | -1.08 | 0.26 | 0.57 | -0.71 | 0.14 | 1.5 | -0.57 | -0.83 | -0.35 | -2.12 | -1.15 | -1.67 | -0.2575 | 0.0425 | -1.34 | 0.032911 | 0.640971 | 0.945766 | 1 | 0.999631 | 0.44294 |
| Atad1_Q9D5T0_S322s_1 | -1.04 | -0.48 | -1.16 | -0.63 | -0.94 | -0.43 | -0.61 | -0.95 | -0.82 | -0.36 | 0.06 | -0.51 | -0.58 | 0.08 | -0.17 | -1.63 | 0.095 | 0.42 | 0.2525 | 0.553714 | 0.738578 | 0.197956 | 1 | 0.999631 | 0.837943 |
| Atat1_Q8K341_S249s_1 | 1.78 | -0.63 | 0.2 | 1.36 | 0.44 | 0.61 | 1.62 | 0.86 | 0.66 | 1.09 | 1.23 | 0.35 | -0.91 | -0.38 | 0.92 | 0.05 | 0.205 | 0.155 | -0.7575 | 0.232775 | 0.713413 | 0.775807 | 1 | 0.999631 | 0.682707 |
| Atat1_Q8K341_S253s_1 | 1.56 | 1.56 | 0.93 | 0.98 | 1.01 | 2.06 | 1.6 | 0.51 | 2.29 | 2.36 | 0.93 | 0.77 | 0.64 | 0.59 | 0.84 | 0.15 | 0.0375 | 0.33 | -0.7025 | 0.046788 | 0.923336 | 0.471768 | 1 | 0.999631 | 0.480102 |
| Atat1_Q8K341_T238t_1 | 0.42 | 0.54 | 0.33 | -0.04 | 0.25 | 0.31 | 0.25 | 0.46 | 0.43 | 0.52 | 0.22 | 0.1 | 0.03 | 0.65 | 0.43 | 0.31 | 0.005 | 0.005 | 0.0425 | 0.885021 | 0.984162 | 0.985166 | 1 | 0.999631 | 0.963505 |
| Atcay_Q8BHE3_S370s_1 | 0.88 | 0.1 | -0.77 | -0.89 | 0.14 | 0.11 | -0.09 | -2.24 | 0.54 | 1.07 | -1.61 | -1.45 | -0.2 | -0.43 | -1.84 | -2.39 | -0.35 | -0.1925 | -1.045 | 0.112429 | 0.584411 | 0.786573 | 1 | 0.999631 | 0.577306 |
| Atcay_Q8BHE3_S46s_1 | -0.91 | -1.02 | -1.38 | -1.35 | -1.3 | -1.02 | -1.43 | -1.44 | -0.97 | -0.84 | -1.77 | -1.22 | -0.01 | -0.62 | -1.46 | -1.04 | -0.1325 | -0.035 | 0.3825 | 0.317523 | 0.611757 | 0.909215 | 1 | 0.999631 | 0.721033 |
| Atf2_P16951_S149s_1 | -0.77 | 0.63 | -0.96 | -0.08 | 1.56 | -0.29 | -0.41 | -0.69 | -0.19 | -0.34 | -0.85 | -0.18 | 0.97 | 0.41 | -0.11 | 0.35 | 0.3375 | -0.095 | 0.7 | 0.128655 | 0.558454 | 0.814553 | 1 | 0.999631 | 0.599259 |
| Atf2_P16951_S150s_1 | -0.77 | 0.63 | -0.96 | -0.08 | 1.56 | -0.29 | -0.41 | -0.69 | -0.19 | -0.34 | -0.85 | -0.18 | 0.97 | 0.41 | -0.11 | 0.35 | 0.3375 | -0.095 | 0.7 | 0.128655 | 0.558454 | 0.814553 | 1 | 0.999631 | 0.599259 |
| Atf2_P16951_S212s_2 | 0.05 | 1.08 | 0.48 | 0.24 | -0.46 | 0.09 | 0.32 | 1.38 | -0.12 | 0.09 | 0.59 | 1.08 | -0.23 | 0.33 | 0.33 | 0.76 | -0.13 | -0.0525 | -0.165 | 0.644139 | 0.765716 | 0.888507 | 1 | 0.999631 | 0.872436 |
| Atf2_P16951_S94s_1 | 0.24 | 1.13 | 1.05 | 0.6 | 0.75 | 0.74 | -0.1 | 0.7 | 1.15 | 0.92 | 0.7 | 1.16 | 0.88 | 1.53 | 0.91 | 1.52 | -0.2325 | 0.2275 | 0.455 | 0.201811 | 0.490746 | 0.463414 | 1 | 0.999631 | 0.658952 |
| Atf2_P16951_T53t_1 | -0.62 | 0.49 | 1.05 | -0.1 | 0.03 | 0.93 | 0.64 | 1.06 | 0.3 | -1.99 | 2.16 | 2.04 | 1.01 | -0.43 | 0.92 | 2.64 | 0.46 | 0.4225 | 0.83 | 0.221022 | 0.291734 | 0.640682 | 1 | 0.999631 | 0.671424 |
| Atg16l1_Q8C0J2_S303s_1 | 1.11 | -0.26 | 0.32 | 0.6 | 0.5 | -0.32 | 0.8 | 0.56 | -0.03 | 0.07 | 0.75 | 1 | 0.01 | -0.13 | 0.18 | 0.82 | -0.0575 | 0.005 | -0.2225 | 0.568162 | 0.88158 | 0.989912 | 1 | 0.999631 | 0.843185 |
| Atg2a_Q6P4T0_S1243s_1 | -0.99 | 0.67 | 1.2 | 1.1 | -0.12 | 0.3 | -0.26 | -0.81 | 0.03 | -1.06 | 0.66 | 1.14 | 1.55 | 1.6 | -0.04 | 0.15 | -0.7175 | -0.3025 | 0.32 | 0.602756 | 0.187782 | 0.632495 | 1 | 0.999631 | 0.857278 |
| Atg2b_Q80XK6_S1571s_1 | -0.61 | 0.38 | 1.54 | 1.33 | 0.21 | 0.96 | 0.5 | 0.85 | 0.05 | 0.14 | 0.69 | 1.19 | -1.09 | 0.31 | 1.1 | 1.54 | -0.03 | -0.1425 | -0.195 | 0.773511 | 0.95126 | 0.786047 | 1 | 0.999631 | 0.92076 |
| Atg2b_Q80XK6_S1581s_1 | -0.13 | 0.28 | 1.28 | 0.25 | -0.54 | -1 | -0.29 | 0.11 | -0.24 | 0.03 | -0.34 | -1.55 | -1.37 | 0.17 | 1.22 | -1.49 | -0.85 | -0.945 | -0.7875 | 0.239522 | 0.051176 | 0.060127 | 1 | 0.999631 | 0.683719 |
| Atg2b_Q80XK6_S1741s_1 | 0.53 | 0.75 | 0.09 | 0.34 | 0.85 | 0.47 | 0.64 | 0.96 | 0.8 | 0.22 | 0.26 | 0.76 | 0.33 | 0.81 | 0.81 | 0.56 | 0.3025 | 0.0825 | 0.2 | 0.502393 | 0.278522 | 0.778605 | 1 | 0.999631 | 0.814092 |
| Atg2b_Q80XK6_S239s_1 | 0.28 | 0.9 | 0.04 | -0.22 | 0.99 | 0.64 | -0.11 | 0.4 | 1.14 | 0.27 | 1.16 | 0.74 | 1.08 | 1.14 | 0.47 | 1.08 | 0.23 | 0.5775 | 0.6925 | 0.069097 | 0.524733 | 0.127842 | 1 | 0.999631 | 0.524251 |
| Atg2b_Q80XK6_S240s_1 | 0.28 | 0.9 | 0.04 | -0.22 | 0.99 | 0.64 | -0.11 | 0.4 | 1.14 | 0.27 | 1.16 | 0.74 | 1.08 | 1.14 | 0.47 | 1.08 | 0.23 | 0.5775 | 0.6925 | 0.069097 | 0.524733 | 0.127842 | 1 | 0.999631 | 0.524251 |
| Atg2b_Q80XK6_S255s_1 | 0.62 | 0.57 | -0.62 | 0.44 | 0.49 | 0.1 | 0.6 | 0.56 | 0.93 | 0.12 | 0.37 | -0.25 | -1.63 | 0.22 | 0.31 | 0.29 | 0.185 | 0.04 | -0.455 | 0.396895 | 0.595903 | 0.919792 | 1 | 0.999631 | 0.761093 |
| Atg2b_Q80XK6_S379s_1 | -0.01 | -0.04 | -0.22 | 2.04 | 1.44 | 0.94 | 1.49 | 1.46 | 0.74 | -0.24 | 1.53 | 0.44 | -1.89 | 0.82 | 0.76 | 0.56 | 0.89 | 0.175 | -0.38 | 0.610985 | 0.10763 | 0.767891 | 1 | 0.999631 | 0.859461 |
| Atg2b_Q80XK6_S885s_1 | -0.54 | 2.03 | 1.93 | 1.79 | 0.83 | 0.95 | 1.33 | 2.47 | 0.88 | 0.83 | 1.39 | 2.59 | 1.46 | 1.8 | 2.4 | 2.31 | 0.0925 | 0.12 | 0.69 | 0.264416 | 0.885798 | 0.856431 | 1 | 0.999631 | 0.6935 |
| Atg9a_Q68FE2_S735s_1 | 0.06 | -0.79 | -0.28 | -0.47 | -1.74 | -0.41 | -0.03 | -0.32 | -1.08 | 0.05 | -0.39 | -0.49 | -1.58 | -1.9 | -0.66 | -1.01 | -0.255 | -0.1075 | -0.9175 | 0.031053 | 0.54476 | 0.751625 | 1 | 0.999631 | 0.44065 |
| Atg9a_Q68FE2_S735s_2 | -0.15 | -0.45 | 0.5 | -0.52 | -1.2 | -0.3 | 0.05 | 0.13 | -0.88 | 0.11 | 0.07 | -0.01 | -0.52 | -0.62 | -1.17 | -1.1 | -0.175 | -0.0225 | -0.6975 | 0.066766 | 0.656365 | 0.950498 | 1 | 0.999631 | 0.519446 |
| Atg9a_Q68FE2_S738s_2 | -0.49 | -0.37 | 0.59 | -0.69 | -1.3 | -0.22 | 0.01 | 0.11 | -0.67 | 0.29 | -0.16 | -0.23 | -0.56 | -0.54 | -0.94 | -1.28 | -0.11 | 0.0475 | -0.59 | 0.138447 | 0.795988 | 0.89839 | 1 | 0.999631 | 0.611124 |
| Atg9a_Q68FE2_S741s_2 | 0.5 | -0.65 | 0.25 | -0.13 | -0.96 | -0.49 | 0.16 | 0.18 | -1.49 | -0.4 | 0.6 | 0.51 | -0.42 | -0.83 | -1.85 | -0.68 | -0.27 | -0.1875 | -0.9375 | 0.044915 | 0.48758 | 0.716799 | 1 | 0.999631 | 0.475465 |
| Atg9a_Q68FE2_S761s_1 | 0.59 | 0.06 | 0.18 | 0.9 | -0.07 | -0.05 | 0.67 | 0.76 | 0.26 | -0.06 | 0.38 | 1.05 | -0.48 | 0.43 | 1.04 | 0.91 | -0.105 | -0.025 | 0.0425 | 0.917823 | 0.754034 | 0.942197 | 1 | 0.999631 | 0.97388 |
| Atl1_Q8BH66_S10s_1 | -1.3 | 0.18 | -0.66 | 0.67 | 1.21 | 0.33 | 0.53 | 0.68 | 1.05 | -1.08 | 0.62 | 0.36 | 0.01 | -0.19 | 0.52 | 0.96 | 0.965 | 0.515 | 0.6025 | 0.23727 | 0.058106 | 0.386746 | 1 | 0.999631 | 0.683301 |
| Atl1_Q8BH66_S15s_1 | 0.22 | 4.09 | 0.43 | 0.05 | 3.02 | 4.16 | 0.74 | 0.67 | 3.98 | 0.92 | 1.15 | 1.17 | 3.4 | 3.93 | 1.04 | 0.39 | 0.95 | 0.6075 | 0.9925 | 0.379023 | 0.401548 | 0.563544 | 1 | 0.999631 | 0.751637 |
| Atl1_Q8BH66_S18s_1 | -2.6 | 0.78 | 0.77 | -1.07 | 2.54 | 0.57 | 0.41 | -0.74 | 2.34 | -1.38 | 0.54 | 1.48 | 1.33 | 0.42 | 0.63 | 0.08 | 1.225 | 1.275 | 1.145 | 0.151198 | 0.202954 | 0.216047 | 1 | 0.999631 | 0.624065 |
| Atn1_O35126_S646s_1 | 0.35 | 1.27 | 1.78 | 0.92 | 1.17 | 0.59 | 0.34 | 1.19 | 1.46 | 1.44 | 1.18 | 1.69 | 1.49 | 1.79 | 1.69 | 1.9 | -0.2575 | 0.3625 | 0.6375 | 0.102139 | 0.50446 | 0.320127 | 1 | 0.999631 | 0.564448 |
| Atn1_O35126_T654t_1 | 0.9 | 2.69 | 2.31 | 1.57 | 1.31 | 2.31 | 1.35 | 2.17 | 2.2 | 1.63 | 2.28 | 2.35 | 2.32 | 2.5 | 2.08 | 2.23 | -0.0825 | 0.2475 | 0.415 | 0.335789 | 0.857218 | 0.567959 | 1 | 0.999631 | 0.730466 |
| Atp11c_F6Q8D3_S1105s_1 | 0.68 | 0.24 | 1.28 | 0.53 | -0.69 | -0.45 | 0.67 | 0.09 | -0.11 | 0.43 | -1.02 | 0.52 | -0.01 | -1.41 | 0.43 | 0.21 | -0.7775 | -0.7275 | -0.8775 | 0.08196 | 0.067103 | 0.107962 | 1 | 0.999631 | 0.54792 |
| Atp13a1_Q9EPE9_S22s_1 | 0.21 | 0.26 | 1.65 | 0.67 | -0.29 | 0.39 | 2.42 | 0.93 | 0.05 | -0.56 | 1.17 | 0.81 | 0.23 | -0.72 | 1.12 | 0.04 | 0.165 | -0.33 | -0.53 | 0.291533 | 0.784035 | 0.504375 | 1 | 0.999631 | 0.706164 |
| Atp13a1_Q9EPE9_S896s_1 | -0.98 | -2.46 | 1.56 | -0.05 | -1.26 | -3.94 | 0.97 | 1.55 | -0.6 | -0.68 | 2.14 | 2.47 | -0.03 | -1.38 | 2.19 | 1.52 | -0.1875 | 1.315 | 1.0575 | 0.301569 | 0.883618 | 0.222771 | 1 | 0.999631 | 0.714243 |
| Atp1a1_Q8VDN2_S16s_1 | 0.47 | 0.6 | 1.49 | 0.44 | 0.26 | 1.8 | 1.01 | 1.4 | 0.58 | 0.48 | 1.31 | 1.59 | 0.97 | 0.73 | 1.46 | 1.27 | 0.3675 | 0.24 | 0.3575 | 0.323079 | 0.381898 | 0.539779 | 1 | 0.999631 | 0.724485 |
| Atp1a1_Q8VDN2_S207s_1 | -0.64 | -1.46 | -1.16 | -1.68 | -2.44 | -1.29 | -1.33 | -1.15 | -1.84 | -1.01 | -1.96 | -1.52 | -1.85 | -1.89 | -1.75 | -1.04 | -0.3175 | -0.3475 | -0.3975 | 0.277892 | 0.417891 | 0.332349 | 1 | 0.999631 | 0.700874 |
| Atp1a1_Q8VDN2_S228s_1 | -1.08 | -0.25 | -0.74 | -0.95 | -0.48 | -0.39 | -1.36 | -1.09 | -0.78 | -0.52 | -1.01 | -1.07 | -0.34 | -0.23 | -1.1 | -0.5 | -0.075 | -0.09 | 0.2125 | 0.530498 | 0.823242 | 0.763559 | 1 | 0.999631 | 0.826702 |
| Atp1a1_Q8VDN2_S40s_1 | 1.4 | -0.38 | 0.95 | 0.75 | -0.83 | 0.17 | 0.81 | -0.79 | -0.58 | 0.48 | 0.39 | 0.46 | -1.03 | -1.22 | 0 | -0.64 | -0.84 | -0.4925 | -1.4025 | 0.011513 | 0.125291 | 0.288135 | 1 | 0.999631 | 0.385092 |
| Atp1a1_Q8VDN2_T485t_1 | -0.46 | -0.46 | -0.1 | -1.3 | -0.29 | -0.55 | -0.79 | -0.67 | -0.42 | -0.76 | -0.7 | 0.1 | -0.21 | -0.26 | -0.96 | -0.15 | 0.005 | 0.135 | 0.185 | 0.613798 | 0.987618 | 0.705922 | 1 | 0.999631 | 0.860866 |
| Atp1a1_Q8VDN2_Y684y_1 | -0.96 | 0.2 | 0.8 | 0.08 | 0.61 | -0.66 | 1.04 | -1.2 | 1.72 | -1.56 | -0.13 | 1.01 | 1.99 | -1.3 | 0.13 | -0.88 | -0.0825 | 0.23 | -0.045 | 0.950309 | 0.887175 | 0.74799 | 1 | 0.999631 | 0.984079 |
| Atp1a2_Q6PIE5_S10s_1 | -0.25 | -1.23 | -1 | -1.01 | -1.59 | -0.52 | -0.92 | -1.22 | -0.95 | -0.43 | -0.53 | -0.16 | 0.03 | 0.07 | -0.57 | -0.4 | -0.19 | 0.355 | 0.655 | 0.074486 | 0.584179 | 0.290456 | 1 | 0.999631 | 0.540038 |
| Atp1a2_Q6PIE5_S439s_1 | -1.13 | -1.53 | -1.07 | -1.34 | -1.72 | -1.14 | -1.27 | -1.75 | -1.21 | -0.95 | -0.38 | -0.3 | -0.11 | -0.85 | -0.36 | -0.81 | -0.2025 | 0.5575 | 0.735 | 0.034741 | 0.467275 | 0.097993 | 1 | 0.999631 | 0.449301 |
| Atp1a2_Q6PIE5_S464s_1 | 0.25 | 1.83 | 1.68 | 0.98 | 0.81 | 1.53 | 0.11 | 0.98 | 1.11 | 1.13 | 0.3 | -0.81 | 1.38 | 1.9 | 1.5 | 1.33 | -0.3275 | -0.7525 | 0.3425 | 0.407421 | 0.473143 | 0.187071 | 1 | 0.999631 | 0.766996 |
| Atp1a2_Q6PIE5_S496s_1 | -0.56 | -0.08 | -0.32 | -0.78 | -0.4 | -0.13 | -1.3 | -0.57 | -0.47 | -0.27 | -0.67 | -0.04 | 0.36 | 0.08 | -0.19 | -0.04 | -0.165 | 0.0725 | 0.4875 | 0.12628 | 0.62164 | 0.80235 | 1 | 0.999631 | 0.596745 |
| Atp1a2_Q6PIE5_S58s_1 | -1.03 | -0.24 | 0.67 | 0.08 | -0.76 | 0.49 | -0.11 | -0.99 | -0.84 | 0.13 | 0.32 | 0.97 | -0.55 | -0.56 | 2 | 0.33 | -0.2125 | 0.275 | 0.435 | 0.495544 | 0.649752 | 0.578766 | 1 | 0.999631 | 0.810848 |
| Atp1a3_Q6PIC6_S206s_1 | -0.69 | -0.4 | -0.83 | -1.68 | -1.08 | 0.06 | -1.22 | -1.25 | -0.37 | 0.22 | -1.07 | -1.42 | 0.59 | -0.25 | -1.65 | -1.32 | 0.0275 | 0.24 | 0.2425 | 0.656511 | 0.947086 | 0.595734 | 1 | 0.999631 | 0.879056 |
| Atp1a3_Q6PIC6_S207s_1 | -0.63 | -0.76 | -0.16 | -0.96 | -1.14 | -0.41 | -1.02 | -1.1 | -0.37 | 0.14 | -0.48 | -0.75 | -0.36 | -0.42 | -0.5 | -0.01 | -0.29 | 0.2625 | 0.305 | 0.327222 | 0.349211 | 0.413882 | 1 | 0.999631 | 0.724678 |
| Atp1a3_Q6PIC6_S212s_1 | 0.34 | 0.04 | -0.91 | -0.17 | -0.91 | -0.35 | -0.92 | -1.24 | 0.1 | 0.41 | -0.43 | -0.06 | 0.66 | -0.52 | -0.31 | 0.16 | -0.68 | 0.18 | 0.1725 | 0.666848 | 0.077948 | 0.614288 | 1 | 0.999631 | 0.88264 |
| Atp1a3_Q6PIC6_S218s_1 | -1.15 | -1.05 | -0.63 | -0.65 | -1.09 | -0.9 | -0.84 | -0.74 | -1.07 | -0.71 | -0.91 | 0.02 | -0.04 | -0.32 | -0.15 | -0.61 | -0.0225 | 0.2025 | 0.59 | 0.068957 | 0.930836 | 0.542015 | 1 | 0.999631 | 0.52409 |
| Atp1a3_Q6PIC6_S37s_1 | 0.85 | 0.96 | 0.94 | 0.69 | 0.32 | 1.19 | 0.47 | 0.56 | 0.6 | 0.88 | 0.44 | 0.69 | 1.12 | 1.07 | 0.92 | 1.01 | -0.225 | -0.2075 | 0.17 | 0.520027 | 0.431533 | 0.419071 | 1 | 0.999631 | 0.819528 |
| Atp1a3_Q6PIC6_S442s_1 | -1.08 | 0.6 | 0.46 | 0.11 | 0.42 | -0.19 | -0.04 | 0.87 | -0.1 | -0.42 | 0.32 | 0.23 | 0.14 | -0.04 | 1.38 | 1.41 | 0.2425 | -0.015 | 0.7 | 0.19628 | 0.583676 | 0.97152 | 1 | 0.999631 | 0.653373 |
| Atp1a3_Q6PIC6_S474s_1 | -0.49 | 0.2 | 1.31 | 0.15 | -0.12 | -0.38 | -0.4 | 0.12 | -0.05 | -0.57 | -1.45 | 0.9 | 0.43 | -0.07 | -0.23 | 0.83 | -0.4875 | -0.585 | -0.0525 | 0.906223 | 0.239507 | 0.31701 | 1 | 0.999631 | 0.969391 |
| Atp1a3_Q6PIC6_S56s_1 | -0.37 | 0.94 | 1.63 | 0.45 | 0.09 | 1.37 | 0.53 | 0.71 | 0.73 | -0.08 | 0.46 | 1.44 | 1.36 | 1.57 | 1.92 | 1.39 | 0.0125 | -0.025 | 0.8975 | 0.065638 | 0.978915 | 0.960039 | 1 | 0.999631 | 0.516933 |
| Atp1a3_Q6PIC6_S643s_1 | 0.02 | 1.41 | 0.22 | 0.09 | 0.49 | 0.67 | -0.72 | -0.45 | 0.49 | 0.4 | -0.01 | 0.09 | 0.88 | 1.17 | -0.2 | -0.36 | -0.4375 | -0.1925 | -0.0625 | 0.897814 | 0.349367 | 0.609373 | 1 | 0.999631 | 0.966758 |
| Atp1a3_Q6PIC6_S712s_1 | -0.21 | -0.31 | -0.53 | -0.59 | -0.51 | -0.49 | -0.68 | -0.72 | -0.56 | -1.44 | -0.21 | -0.2 | 0.08 | -0.1 | -0.37 | -0.31 | -0.19 | -0.1925 | 0.235 | 0.40623 | 0.441278 | 0.581402 | 1 | 0.999631 | 0.766128 |
| Atp1a3_Q6PIC6_T368t_1 | 1.89 | 1.58 | 2.3 | 1.54 | -0.49 | 1.91 | 1.05 | 1.13 | -0.21 | 1.57 | 1.03 | -2.03 | -0.87 | 0.87 | 0.5 | -0.05 | -0.9275 | -1.7375 | -1.715 | 0.002379 | 0.088018 | 0.035825 | 1 | 0.999631 | 0.351308 |
| Atp1a3_Q6PIC6_T370t_1 | 0.89 | 0.01 | 1.19 | 1.42 | -0.43 | 1.43 | 1.47 | 1.4 | -0.04 | 0.17 | 1.11 | -0.71 | 0.41 | 0.57 | 1.12 | 0.18 | 0.09 | -0.745 | -0.3075 | 0.44356 | 0.862416 | 0.137386 | 1 | 0.999631 | 0.782783 |
| Atp1a3_Q6PIC6_T71t_1 | -2.82 | -1.23 | 0.76 | -1.09 | -1.22 | -0.22 | -0.32 | -0.56 | -0.54 | -1.64 | 0.18 | 1.14 | 0.96 | 0.05 | 0.57 | 1.25 | 0.515 | 0.88 | 1.8025 | 0.0244 | 0.455775 | 0.298922 | 1 | 0.999631 | 0.416717 |
| Atp2a2_O55143_S553s_1 | -0.61 | -0.45 | -0.21 | -0.9 | -0.56 | -0.43 | -1.5 | 0.69 | -0.65 | -0.44 | 1.19 | 0.88 | 0.49 | -0.16 | -0.51 | 0.09 | 0.0925 | 0.7875 | 0.52 | 0.138098 | 0.838625 | 0.117852 | 1 | 0.999631 | 0.611124 |
| Atp2a2_O55143_S661s_1 | -1.79 | -0.6 | -0.86 | -2.7 | -0.23 | -0.84 | -0.37 | -1.15 | 0.23 | -1.27 | -0.1 | 0.9 | 0.66 | -0.13 | -1.72 | -0.8 | 0.84 | 1.4275 | 0.99 | 0.13852 | 0.1129 | 0.036951 | 1 | 0.999631 | 0.611124 |
| Atp2a2_O55143_S663s_1 | -3.2 | -0.34 | -0.8 | -1.82 | 0.34 | -1.61 | 0.5 | -1.45 | 0.62 | -2.69 | 0.39 | 0.85 | 0.85 | -1.84 | -1.68 | -1.03 | 0.985 | 1.3325 | 0.615 | 0.434406 | 0.207255 | 0.165762 | 1 | 0.999631 | 0.779717 |
| Atp2b1_G5E829_S1139s_1 | 0.59 | -0.69 | -1.16 | 0.02 | -0.05 | -0.89 | 0.29 | -0.14 | -0.83 | -1.03 | -1.24 | -1.33 | -0.71 | -0.73 | -1.14 | -0.51 | 0.1125 | -0.7975 | -0.4625 | 0.286751 | 0.800622 | 0.076064 | 1 | 0.999631 | 0.705972 |
| Atp2b1_G5E829_S1140s_1 | 0.59 | -0.69 | -1.16 | 0.02 | -0.05 | -0.89 | 0.29 | -0.14 | -0.83 | -1.03 | -1.24 | -1.33 | -0.71 | -0.73 | -1.14 | -0.51 | 0.1125 | -0.7975 | -0.4625 | 0.286751 | 0.800622 | 0.076064 | 1 | 0.999631 | 0.705972 |
| Atp2b1_G5E829_S1177s_1 | -0.26 | -1.4 | -0.57 | -0.93 | -1.8 | -1.43 | -0.89 | -1.28 | -1.32 | 0.19 | -1.84 | -1.34 | -0.92 | -1.97 | -1.2 | -1.54 | -0.56 | -0.2875 | -0.6175 | 0.12253 | 0.126585 | 0.554716 | 1 | 0.999631 | 0.591264 |
| Atp2b1_G5E829_S1178s_1 | -1.04 | -1.21 | 0.06 | -0.93 | -2.49 | -1.28 | -1.28 | -1.24 | -1.59 | 0.16 | -1.49 | -1.02 | -0.46 | -1.95 | -0.89 | -1.04 | -0.7925 | -0.205 | -0.305 | 0.48553 | 0.081516 | 0.667234 | 1 | 0.999631 | 0.804287 |
| Atp2b1_G5E829_S1178s_2 | 2.4 | -0.27 | -0.19 | 1.09 | 1.55 | 0.28 | 1.32 | 0.28 | 0.14 | -1.1 | -0.54 | -0.74 | -0.86 | -0.87 | -0.17 | -0.81 | 0.1 | -1.3175 | -1.435 | 0.034419 | 0.87566 | 0.055848 | 1 | 0.999631 | 0.446273 |
| Atp2b1_G5E829_S1182s_2 | 2.4 | -0.27 | -0.19 | 1.09 | 1.55 | 0.28 | 1.32 | 0.28 | 0.14 | -1.1 | -0.54 | -0.74 | -0.86 | -0.87 | -0.17 | -0.81 | 0.1 | -1.3175 | -1.435 | 0.034419 | 0.87566 | 0.055848 | 1 | 0.999631 | 0.446273 |
| Atp2b1_G5E829_S17s_1 | -1.35 | -1.62 | -1.21 | -1.97 | -1.56 | -2.15 | -1.31 | -1.92 | -1.77 | -1.22 | -1.69 | -1.62 | -1.16 | -2.11 | -2.8 | -1.83 | -0.1975 | -0.0375 | -0.4375 | 0.29104 | 0.525383 | 0.89766 | 1 | 0.999631 | 0.706164 |
| Atp2b1_G5E829_S338s_1 | -1.12 | -2.15 | -0.74 | -1.16 | -0.92 | -2.37 | -0.22 | -0.98 | -1.33 | -3.1 | 0.17 | -0.61 | -0.74 | -1.56 | -0.75 | -1.07 | 0.17 | 0.075 | 0.2625 | 0.504222 | 0.737563 | 0.912077 | 1 | 0.999631 | 0.814292 |
| Atp2b2_Q3UHH0_S1100s_1 | -0.47 | -0.93 | -1.28 | -0.91 | -1.24 | -2.18 | -1.16 | -2.82 | -2.19 | -1.89 | -1.29 | -2.52 | -0.89 | -1.13 | -2.76 | -1.62 | -0.9525 | -1.075 | -0.7025 | 0.139809 | 0.046302 | 0.011277 | 1 | 0.999631 | 0.611718 |
| Atp2b2_Q3UHH0_S1101s_1 | -0.47 | -0.93 | -0.9 | -1.25 | -1.24 | -2.18 | -1.29 | -1.61 | -2.19 | -1.89 | -0.85 | -1.69 | -0.89 | -1.13 | -1.92 | -1.44 | -0.6925 | -0.7675 | -0.4575 | 0.198184 | 0.051607 | 0.056419 | 1 | 0.999631 | 0.655701 |
| Atp2b2_Q3UHH0_S1111s_1 | -0.69 | 0.68 | 0.08 | 0.11 | 0.21 | 0.26 | -0.61 | 0.11 | 0.21 | -0.23 | -0.48 | -0.03 | 0.87 | 0.49 | -0.25 | 0.19 | -0.0525 | -0.1775 | 0.28 | 0.483958 | 0.886217 | 0.618247 | 1 | 0.999631 | 0.803422 |
| Atp2b2_Q3UHH0_S1114s_1 | -0.86 | 1.78 | 1.4 | 0.36 | 1.19 | 1.08 | 0.42 | 0.96 | 0.88 | -0.42 | 1.18 | 1.24 | 1.72 | 1.12 | 0.6 | 1.05 | 0.2425 | 0.05 | 0.4525 | 0.444389 | 0.66788 | 0.937503 | 1 | 0.999631 | 0.78327 |
| Atp2b2_Q3UHH0_S1151s_1 | -0.07 | -0.78 | -1.18 | -0.93 | -0.23 | 0.31 | -1.1 | -1.95 | 0.06 | 0.41 | -1.03 | -1.2 | 1 | 0.51 | -1.75 | -2.06 | -0.0025 | 0.3 | 0.165 | 0.818316 | 0.996107 | 0.513102 | 1 | 0.999631 | 0.938303 |
| Atp2b2_Q3UHH0_S1152s_1 | -0.16 | -0.37 | -0.66 | -0.99 | -0.77 | -0.13 | -1.11 | -1.47 | -0.34 | -0.5 | -0.97 | -1.04 | 0.03 | -0.72 | -1.44 | -0.84 | -0.325 | -0.1675 | -0.1975 | 0.610161 | 0.377875 | 0.596333 | 1 | 0.999631 | 0.859048 |
| Atp2b2_Q3UHH0_S1156s_1 | -0.51 | -0.62 | 0.36 | -2.46 | -0.73 | -1.09 | -0.18 | -0.52 | -0.94 | -2.78 | -0.05 | -0.23 | -1.1 | -2.48 | -1.48 | -2.49 | 0.1775 | -0.1925 | -1.08 | 0.107921 | 0.755099 | 0.800003 | 1 | 0.999631 | 0.569416 |
| Atp2b2_Q3UHH0_S1162s_1 | -0.12 | -1.13 | -1.34 | -0.51 | 1.41 | 0.46 | 0.14 | 0.46 | 0.07 | -1.18 | -0.36 | 0.5 | -0.15 | -0.24 | -1.29 | -0.16 | 1.3925 | 0.5325 | 0.315 | 0.451503 | 0.005661 | 0.251744 | 1 | 0.999631 | 0.788148 |
| Atp2b2_Q3UHH0_S1175s_1 | -0.51 | -0.56 | -0.69 | -0.81 | -0.41 | -0.32 | -1.01 | -0.74 | -0.4 | -1.5 | -0.87 | -0.23 | 0.88 | -0.43 | -1.14 | -0.53 | 0.0225 | -0.1075 | 0.3375 | 0.445833 | 0.932918 | 0.751383 | 1 | 0.999631 | 0.784402 |
| Atp2b2_Q3UHH0_S18s_1 | -0.68 | 1.21 | 0.95 | 0.64 | 1.8 | 1.08 | 1.04 | 1.44 | 0.66 | -2.03 | 0.8 | 1.4 | 1.11 | 0.78 | 1.36 | 0.63 | 0.81 | -0.3225 | 0.44 | 0.33996 | 0.092338 | 0.675573 | 1 | 0.999631 | 0.73263 |
| Atp2b2_Q3UHH0_S19s_1 | 0.6 | 0.34 | 0.23 | -0.04 | 1.86 | 1.13 | 1.15 | 0.88 | 0.6 | -0.58 | 0.09 | 0.7 | 0.76 | 0.76 | 0.54 | -0.21 | 0.9725 | -0.08 | 0.18 | 0.593704 | 0.009644 | 0.822673 | 1 | 0.999631 | 0.853512 |
| Atp2b2_Q3UHH0_T1113t_1 | -0.87 | 1.94 | 1.29 | 0.36 | 1 | 1 | 0.31 | 0.59 | 0.97 | -0.14 | 0.8 | 1.35 | 1.95 | 1.27 | 0.62 | 1.13 | 0.045 | 0.065 | 0.5625 | 0.365463 | 0.937659 | 0.916751 | 1 | 0.999631 | 0.74565 |
| Atp2b2_S4R1C4_S1107s_1 | -1.73 | -1.22 | -1.35 | -1.88 | -1.69 | -0.82 | -1.79 | -1.4 | -1.39 | -1.73 | -1.32 | -1.85 | 1.01 | -1.1 | -1.4 | -1.7 | 0.12 | -0.0275 | 0.7475 | 0.218248 | 0.706382 | 0.924148 | 1 | 0.999631 | 0.669859 |
| Atp2b2_S4R1C4_S1116s_2 | -1.42 | -0.8 | 0.7 | 0.64 | -0.91 | -0.84 | -1.06 | -0.49 | -0.76 | -0.92 | 0.21 | 0.21 | 0.35 | 0.04 | 1.29 | -0.23 | -0.605 | -0.095 | 0.5825 | 0.323669 | 0.250666 | 0.866045 | 1 | 0.999631 | 0.724485 |
| Atp2b2_S4R1C4_S1117s_2 | -1.12 | -0.89 | -0.09 | -0.19 | -0.85 | -0.39 | -0.29 | -0.54 | -0.6 | -0.8 | -0.53 | -1.04 | 0.74 | -0.17 | 0.06 | -1.15 | 0.055 | -0.17 | 0.4425 | 0.348622 | 0.866176 | 0.609941 | 1 | 0.999631 | 0.736697 |
| Atp2b2_S4R1C4_S1130s_1 | -1.85 | -0.88 | -0.39 | -0.75 | -1.34 | -0.6 | -0.85 | -2.03 | -1.53 | -0.67 | 0.2 | -0.48 | 1.35 | -0.36 | -0.72 | -0.09 | -0.2375 | 0.3475 | 1.0125 | 0.075579 | 0.586828 | 0.457443 | 1 | 0.999631 | 0.541044 |
| Atp2b2_S4R1C4_S1130s_2 | -0.7 | -1.06 | -0.09 | -0.19 | -0.77 | 0.28 | -0.29 | -0.54 | -0.34 | -0.6 | -0.53 | -1.04 | 1.32 | -0.56 | 0.06 | -1.15 | 0.18 | -0.1175 | 0.4275 | 0.434762 | 0.608981 | 0.719317 | 1 | 0.999631 | 0.779717 |
| Atp2b2_S4R1C4_S1135s_1 | -0.14 | 0.05 | -0.17 | 0.84 | 0.26 | 0.6 | 0.68 | -0.16 | -0.15 | 0.17 | 1.6 | 2.02 | 0.75 | 1.01 | -2.21 | 0.48 | 0.2 | 0.765 | -0.1375 | 0.843308 | 0.559574 | 0.179269 | 1 | 0.999631 | 0.94821 |
| Atp2b2_S4R1C4_S1142s_1 | -1.72 | -0.15 | -1.35 | -1.61 | -1.07 | -0.72 | -1.49 | -0.76 | -0.82 | -1.24 | -1.76 | -1.04 | -0.02 | -0.44 | -0.61 | -0.68 | 0.1975 | -0.0075 | 0.77 | 0.083931 | 0.62759 | 0.985657 | 1 | 0.999631 | 0.549732 |
| Atp2b3_A2ALL9_S1124s_1 | -0.63 | -0.23 | 0.02 | -0.58 | -0.88 | -0.47 | -0.31 | -0.51 | -0.31 | -0.5 | -0.8 | -1.17 | 0.63 | 0.12 | -0.16 | -0.05 | -0.1875 | -0.34 | 0.49 | 0.146366 | 0.505503 | 0.28812 | 1 | 0.999631 | 0.618217 |
| Atp2b3_A2ALL9_S1126s_1 | -0.56 | -0.87 | 0.58 | 0.1 | -1.04 | -0.56 | -0.56 | 0.38 | -1.04 | -0.67 | 0.6 | 0.56 | -0.23 | -0.51 | 0.84 | 0.73 | -0.2575 | 0.05 | 0.395 | 0.40298 | 0.555053 | 0.920949 | 1 | 0.999631 | 0.764348 |
| Atp2b3_Q8C048_S315s_1 | -0.85 | -3.07 | 0.57 | -0.82 | -1.02 | -1.14 | 1.33 | -4.45 | -0.61 | -2.16 | -2.27 | -0.95 | -1.57 | -3.36 | -2.07 | -4.4 | -0.2775 | -0.455 | -1.8075 | 0.056936 | 0.817256 | 0.553356 | 1 | 0.999631 | 0.496556 |
| Atp2b3_Q8C048_S316s_1 | 0.98 | -2 | -1.14 | -0.16 | -0.36 | -1.14 | 1.76 | -1.25 | -0.22 | -1.85 | 0.49 | -0.58 | -1.87 | -3.1 | -2.96 | -3.42 | 0.3325 | 0.04 | -2.2575 | 0.005746 | 0.688254 | 0.955459 | 1 | 0.999631 | 0.351545 |
| Atp2b3_Q8C048_S336s_1 | -0.74 | -0.02 | -0.44 | 0.32 | -0.28 | 0.49 | 0.45 | -0.48 | 0.3 | 0.65 | 0 | 0.46 | 0.31 | -1.08 | -0.02 | -2.55 | 0.265 | 0.5725 | -0.615 | 0.333589 | 0.471581 | 0.103729 | 1 | 0.999631 | 0.728079 |
| Atp2b3_Q8C048_S578s_1 | -0.19 | 1.51 | 0.73 | 0.37 | 1.74 | 1 | 0.01 | 0.33 | 0.69 | 0.01 | 0.54 | 1.12 | 1.45 | 1.5 | 0.38 | 1.02 | 0.165 | -0.015 | 0.4825 | 0.290625 | 0.738185 | 0.97181 | 1 | 0.999631 | 0.706164 |
| Atp2b3_Q8C048_S8s_1 | -2.41 | -3.73 | -0.63 | -1.05 | -2.15 | -3.37 | -0.98 | -0.68 | -3.07 | -2.65 | -1.02 | -1.15 | -2.14 | -2.99 | -1.03 | -0.63 | 0.16 | -0.0175 | 0.2575 | 0.73983 | 0.844116 | 0.981862 | 1 | 0.999631 | 0.908631 |
| Atp2b3_Q8C048_T1079t_1 | -2.11 | -0.71 | -1.43 | -1.15 | -0.8 | -1.19 | -1.02 | -0.99 | -0.82 | -1.25 | -0.78 | -0.81 | -0.04 | -0.61 | -0.6 | -1.45 | 0.35 | 0.435 | 0.675 | 0.135231 | 0.316315 | 0.235258 | 1 | 0.999631 | 0.607096 |
| Atp2b4_E9Q828_S1065s_1 | 0.16 | -1.46 | -0.94 | -0.94 | -1.17 | -0.67 | -0.65 | -1.21 | -0.42 | 0.41 | -1.43 | -1.59 | -0.69 | -2 | -1.49 | -2.19 | -0.13 | 0.0375 | -0.7975 | 0.113284 | 0.736255 | 0.944464 | 1 | 0.999631 | 0.577428 |
| Atp2b4_E9Q828_S329s_1 | -1.33 | -0.52 | -0.25 | -1.22 | -0.18 | -0.94 | -0.13 | -0.64 | -0.32 | -1.41 | -0.25 | 0.43 | 0.56 | -1.08 | -0.66 | -0.66 | 0.3575 | 0.4425 | 0.37 | 0.412536 | 0.325787 | 0.341059 | 1 | 0.999631 | 0.768114 |
| Atp2c1_Q3UZR5_S944s_1 | -0.53 | -1.56 | -1.49 | -0.02 | -0.94 | -1.14 | -0.66 | 0.92 | -1.5 | 0.23 | -1.92 | -0.43 | 0.47 | 0.02 | -0.99 | -0.35 | 0.445 | -0.005 | 0.6875 | 0.169558 | 0.4275 | 0.992985 | 1 | 0.999631 | 0.639508 |
| Atp6v1a_P50516_S384s_1 | -0.76 | -0.19 | -0.73 | -1.07 | -1.11 | -0.67 | -1.08 | -0.68 | -0.25 | -0.51 | -1.09 | -0.69 | 0.09 | 0 | -0.77 | -0.16 | -0.1975 | 0.0525 | 0.4775 | 0.176135 | 0.501927 | 0.868228 | 1 | 0.999631 | 0.642116 |
| Atp6v1a_P50516_S568s_1 | -0.95 | -0.27 | -0.58 | -0.76 | -0.55 | -0.79 | -0.9 | -0.33 | -0.9 | -0.25 | -0.92 | -0.8 | 0 | 0.04 | -0.6 | -0.49 | -0.0025 | -0.0775 | 0.3775 | 0.243228 | 0.992794 | 0.793104 | 1 | 0.999631 | 0.685637 |
| Atp6v1b2_P62814_S498s_1 | -0.65 | -0.47 | 0.09 | -0.42 | -0.25 | -0.4 | -0.82 | -0.79 | -1.05 | -0.14 | -0.77 | -1.26 | -0.13 | -0.21 | -0.68 | -0.16 | -0.2025 | -0.4425 | 0.0675 | 0.824828 | 0.486541 | 0.210184 | 1 | 0.999631 | 0.940061 |
| Atp6v1e1_P50518_T195t_1 | 0.7 | -0.01 | 0.27 | -0.06 | -0.19 | 0.3 | 0.42 | 0.27 | 0.11 | 0.89 | 0.79 | 1.39 | 0.31 | 0.62 | 1.13 | 0.61 | -0.025 | 0.57 | 0.4425 | 0.191997 | 0.931438 | 0.130782 | 1 | 0.999631 | 0.648674 |
| Atp6v1g2_Q9WTT4_S65s_1 | -0.52 | 0.18 | -1.57 | -1.38 | -0.44 | -0.23 | -0.61 | -1.1 | -0.71 | -1.11 | -1.25 | -1.95 | -0.18 | 0.27 | -0.24 | -0.95 | 0.2275 | -0.4325 | 0.5475 | 0.2588 | 0.603551 | 0.363993 | 1 | 0.999631 | 0.692624 |
| Atp6v1h_Q8BVE3_S483s_1 | 0.14 | -2.31 | -3.8 | -3.66 | -2.2 | -2.84 | -3.37 | -3.88 | -2.58 | -1.22 | -2.17 | -3.28 | -3.36 | -2.57 | -3.71 | -5.12 | -0.665 | 0.095 | -1.2825 | 0.180417 | 0.442928 | 0.913893 | 1 | 0.999631 | 0.64376 |
| Atp6v1h_Q8BVE3_S58s_1 | -0.17 | -1.36 | -1.6 | -1.01 | -0.67 | -1.19 | -1.35 | -1.02 | -0.4 | -0.59 | -1.33 | -0.89 | 0.8 | -0.41 | 0.21 | -1.71 | -0.0225 | 0.2325 | 0.7575 | 0.20607 | 0.950905 | 0.554632 | 1 | 0.999631 | 0.6605 |
| Atp6v1h_Q8BVE3_S59s_1 | -0.17 | -1.36 | -1.47 | -1.02 | -0.67 | -1.19 | -1.42 | -0.91 | -0.4 | -0.59 | -1.16 | -0.9 | 0.8 | -0.41 | 0.38 | -1.78 | -0.0425 | 0.2425 | 0.7525 | 0.218239 | 0.90605 | 0.514401 | 1 | 0.999631 | 0.669859 |
| Atp7a_Q64430_S356s_1 | -0.5 | -0.47 | -0.19 | -0.4 | -0.63 | -1.53 | -0.68 | -0.02 | -1.67 | -0.41 | 0.15 | -1.28 | 1.4 | -0.46 | -2.23 | -0.08 | -0.325 | -0.4125 | 0.0475 | 0.943369 | 0.36196 | 0.339923 | 1 | 0.999631 | 0.981629 |
| Atp8a1_P70704_S1109s_1 | -0.47 | -0.87 | 1.82 | -1.72 | -1.05 | -0.75 | -0.26 | 1.58 | -1.18 | -1.42 | 1.3 | 1.45 | -0.78 | -0.68 | 2.44 | 2 | 0.19 | 0.3475 | 1.055 | 0.295471 | 0.819992 | 0.711963 | 1 | 0.999631 | 0.708945 |
| Atp8a1_P70704_S1111s_1 | -0.7 | -0.73 | -0.51 | -1.02 | -0.27 | -1.1 | -0.94 | -0.75 | -0.9 | -0.99 | -1.28 | -0.59 | -1.27 | -0.81 | -0.65 | -0.67 | -0.025 | -0.2 | -0.11 | 0.708685 | 0.930027 | 0.476887 | 1 | 0.999631 | 0.898927 |
| Atp8a1_P70704_S29s_1 | -0.65 | -0.62 | -4.09 | -2.5 | -0.75 | -0.76 | -1.61 | -2.27 | 0.31 | 0.54 | 0.3 | -2.27 | 0.82 | -0.35 | -2.18 | -3.1 | 0.6175 | 1.685 | 0.7625 | 0.468094 | 0.445175 | 0.093779 | 1 | 0.999631 | 0.795565 |
| Atp8a1_P70704_T28t_1 | 0.31 | 1.05 | 0.22 | -0.21 | 1.05 | 0.99 | 0.35 | -0.06 | 2.06 | 2.59 | 0.6 | 1.24 | 2.73 | 1.1 | 0.73 | 1.17 | 0.24 | 1.28 | 1.09 | 0.048873 | 0.53683 | 0.024532 | 1 | 0.999631 | 0.48544 |
| Atp8a2_P98200_S421s_1 | 0.11 | 0.07 | -0.3 | -0.35 | -0.39 | 0.37 | -0.25 | -0.4 | 0.27 | 0.94 | -0.11 | -0.23 | -0.01 | -0.1 | -0.23 | -0.93 | -0.05 | 0.335 | -0.2 | 0.53898 | 0.863191 | 0.333038 | 1 | 0.999631 | 0.830899 |
| Atp8a2_P98200_S422s_1 | -0.3 | -1.14 | -0.3 | -0.35 | -1.21 | -0.52 | -0.25 | -0.4 | -0.4 | 0.24 | -0.11 | -0.23 | -0.06 | -0.71 | -0.23 | -0.93 | -0.0725 | 0.3975 | 0.04 | 0.908467 | 0.828482 | 0.22257 | 1 | 0.999631 | 0.969941 |
| Atp8a2_P98200_S435s_1 | 0.99 | 0.12 | 1.09 | 0.15 | -0.31 | 0.05 | 0.59 | -0.03 | 0.23 | 0.26 | 0.2 | 0.57 | 0.54 | 0.25 | 0.09 | -0.33 | -0.5125 | -0.2725 | -0.45 | 0.2358 | 0.167209 | 0.415943 | 1 | 0.999631 | 0.683301 |
| Atp8a2_P98200_S6s_1 | -0.45 | -0.98 | -1.64 | -1.53 | -0.38 | -0.87 | -1.2 | -1.6 | -0.62 | -1.02 | -1.35 | -0.43 | -0.38 | -0.64 | -0.4 | -0.44 | 0.1375 | 0.295 | 0.685 | 0.069296 | 0.723093 | 0.433523 | 1 | 0.999631 | 0.525113 |
| Atp8a2_P98200_T1108t_1 | -0.43 | -0.99 | -1.11 | -0.31 | 0.19 | -0.57 | -0.41 | -1.05 | -0.15 | -0.47 | -0.92 | -0.64 | 0.38 | -0.51 | -0.77 | -0.25 | 0.25 | 0.165 | 0.4225 | 0.262235 | 0.483747 | 0.60595 | 1 | 0.999631 | 0.69292 |
| Atrx_Q61687_S1290s_1 | -0.38 | -0.57 | 0.58 | -0.28 | -0.42 | -0.43 | -0.33 | -0.18 | -0.27 | 0.76 | -0.14 | 0.04 | 0.4 | -0.06 | 0.52 | 0.37 | -0.1775 | 0.26 | 0.47 | 0.194105 | 0.574658 | 0.48841 | 1 | 0.999631 | 0.651261 |
| Atrx_Q61687_S315s_1 | -3.27 | -1.73 | -2.48 | -2.34 | -0.35 | -3.11 | -0.75 | -1.86 | 0.72 | -3.39 | -1.08 | -0.52 | 1.54 | -1.99 | -1.31 | -2.96 | 0.9375 | 1.3875 | 1.275 | 0.169214 | 0.156815 | 0.109663 | 1 | 0.999631 | 0.638979 |
| Atrx_Q61687_S590s_1 | -1.14 | -0.96 | -1.1 | -1.82 | -1.12 | -1.16 | -1.33 | -1.64 | -1.09 | -1.41 | -1.36 | -0.53 | -0.49 | -0.85 | -1.26 | -0.4 | -0.0575 | 0.1575 | 0.505 | 0.159671 | 0.844938 | 0.635689 | 1 | 0.999631 | 0.629764 |
| Atrx_Q61687_S665s_1 | 0.82 | 0.66 | 0.06 | -0.96 | -0.01 | -0.49 | -0.67 | -0.98 | 0.11 | 0.71 | -1.43 | -0.91 | 0.73 | 0.75 | -0.89 | -0.85 | -0.6825 | -0.525 | -0.21 | 0.712311 | 0.144283 | 0.374186 | 1 | 0.999631 | 0.9008 |
| Atrx_Q61687_S801s_1 | 0.28 | 1.35 | 0.19 | -0.46 | 0.62 | 1.22 | -0.76 | -0.97 | 0.57 | 0.74 | -0.04 | 0.95 | 1.55 | 1.6 | 0.63 | 0.21 | -0.3125 | 0.215 | 0.6575 | 0.200561 | 0.598503 | 0.619526 | 1 | 0.999631 | 0.658263 |
| Atrx_Q61687_S912s_1 | 0.67 | -0.92 | 1.2 | 0.88 | -1.35 | -1.22 | 1.06 | -0.26 | -0.38 | 0.43 | 0.03 | -0.2 | -0.89 | -1.87 | 0.3 | -0.72 | -0.9 | -0.4875 | -1.2525 | 0.056888 | 0.188729 | 0.325458 | 1 | 0.999631 | 0.496556 |
| Atxn1_J3QPR1_S213s_1 | -0.12 | 0.4 | 0.37 | 0.81 | -0.1 | 0.41 | -0.1 | 0.74 | -0.08 | 0.15 | 0.45 | 0.16 | 0.71 | 1.14 | 0.06 | 0.7 | -0.1275 | -0.195 | 0.2875 | 0.419589 | 0.695965 | 0.517081 | 1 | 0.999631 | 0.772152 |
| Atxn1_J3QPR1_S759s_1 | 0.95 | 2.46 | 2.08 | 1.42 | 1.64 | 1.59 | 0.9 | 1.84 | 1.9 | 1.77 | 2.09 | 2.04 | 1.7 | 2.43 | 1.87 | 2.13 | -0.235 | 0.2225 | 0.305 | 0.450272 | 0.559229 | 0.552638 | 1 | 0.999631 | 0.787533 |
| Atxn1_J3QPR1_S81s_1 | -0.72 | 0.59 | 0.45 | 0.44 | 0.2 | 0.51 | -0.05 | 0.23 | 0.42 | 0.31 | -0.03 | 0.31 | 0.46 | 0.69 | 0.52 | 1.09 | 0.0325 | 0.0625 | 0.5 | 0.20332 | 0.926685 | 0.860663 | 1 | 0.999631 | 0.659781 |
| Atxn1l_P0C7T6_T328t_1 | 0.38 | 0.65 | 0.91 | 0.65 | -0.03 | 0.11 | 0.71 | 0.84 | 0.79 | 0.76 | 0.96 | 0.84 | 0.53 | 0.79 | 0.98 | 1.43 | -0.24 | 0.19 | 0.285 | 0.370102 | 0.435148 | 0.461442 | 1 | 0.999631 | 0.746338 |
| Atxn2_F6U2C2_S190s_1 | 1.82 | 0.01 | -0.29 | 0.86 | -0.24 | 0.13 | 1.71 | 0.76 | -1.99 | -0.02 | 0.62 | -0.4 | -1.69 | -1.52 | -0.44 | 0 | -0.01 | -1.0475 | -1.5125 | 0.023969 | 0.98622 | 0.134689 | 1 | 0.999631 | 0.415644 |
| Atxn2_F6U2C2_S276s_1 | 0.76 | 1.19 | 0.49 | 2.08 | 1.42 | 1.49 | 2.05 | 1 | 1.87 | 1.2 | 1.32 | 0.75 | -2.15 | 1.13 | 0.78 | 0.08 | 0.36 | 0.155 | -1.17 | 0.127797 | 0.38924 | 0.713249 | 1 | 0.999631 | 0.598337 |
| Atxn2_F6U2C2_S356s_1 | -1.45 | -0.94 | 0.8 | 1.83 | -0.79 | -0.38 | 0.63 | 1.34 | -0.99 | -1.52 | 1.53 | 0.94 | -0.46 | -1.06 | 1.34 | 1.33 | 0.14 | -0.07 | 0.2275 | 0.788993 | 0.859075 | 0.939292 | 1 | 0.999631 | 0.925513 |
| Atxn2_F6U2C2_S388s_1 | -0.1 | 0.29 | -0.35 | 0.32 | 0.96 | 0.5 | -0.06 | 0.46 | 0.38 | -1.53 | 0.49 | 0.27 | 0.77 | 0.09 | 0.81 | 0.41 | 0.425 | -0.1375 | 0.48 | 0.154611 | 0.198358 | 0.776839 | 1 | 0.999631 | 0.625677 |
| Atxn2_F6U2C2_S403s_1 | 1.27 | -0.12 | 0.15 | 1.14 | 1.11 | 0.56 | 1.44 | 0.77 | 0.45 | -0.38 | 1.8 | 1.26 | -0.95 | 0.29 | -0.23 | 0.87 | 0.36 | 0.1725 | -0.615 | 0.236084 | 0.380966 | 0.753315 | 1 | 0.999631 | 0.683301 |
| Atxn2_F6U2C2_S444s_1 | 1.42 | 0.84 | 0.33 | 0.53 | 0.05 | 0.69 | 0.62 | 0.34 | 0.8 | 1.5 | 0.9 | 0.96 | 0.03 | 1.04 | 1.13 | 0.62 | -0.355 | 0.26 | -0.075 | 0.844087 | 0.288792 | 0.443949 | 1 | 0.999631 | 0.948381 |
| Atxn2_F6U2C2_S519s_1 | -0.5 | 0.18 | -0.04 | -0.23 | 0.36 | 0.47 | 0.56 | 0.35 | 0.51 | -0.61 | 0.97 | 0.8 | -0.2 | 0.77 | 1.16 | 0.78 | 0.5825 | 0.565 | 0.775 | 0.057636 | 0.045705 | 0.177229 | 1 | 0.999631 | 0.497148 |
| Atxn2_F6U2C2_S551s_1 | 0.41 | 1.71 | 1.27 | 0.6 | -0.1 | 1.32 | 0.9 | 0.97 | 0.74 | 1.52 | 0.89 | 0.91 | 1.57 | 1.53 | 1.54 | 0.88 | -0.225 | 0.0175 | 0.3825 | 0.32704 | 0.597354 | 0.962526 | 1 | 0.999631 | 0.724678 |
| Atxn2_F6U2C2_T240t_1 | 2.15 | 1.44 | 0.54 | 1.54 | 1.83 | 3.35 | 1.5 | 2.1 | 1.4 | 1.36 | 1.85 | -0.42 | -1.22 | 2.01 | 0.71 | 1.84 | 0.7775 | -0.37 | -0.5825 | 0.42576 | 0.138361 | 0.510777 | 1 | 0.999631 | 0.774624 |
| Atxn2_F6U2C2_T326t_1 | -0.66 | 0.74 | -0.61 | 0.21 | 0.09 | 0.3 | -0.25 | 0.91 | 0.75 | -0.08 | 0.03 | 0.73 | 0.34 | 0.76 | 0.36 | 0.61 | 0.3425 | 0.4375 | 0.5975 | 0.144673 | 0.417621 | 0.302564 | 1 | 0.999631 | 0.616037 |
| Atxn2_F6U2C2_T401t_1 | 1.27 | -0.12 | 0.15 | 1.14 | 1.11 | 0.56 | 1.44 | 0.77 | 0.45 | -0.38 | 1.8 | 1.26 | -0.95 | 0.29 | -0.23 | 0.87 | 0.36 | 0.1725 | -0.615 | 0.236084 | 0.380966 | 0.753315 | 1 | 0.999631 | 0.683301 |
| Atxn2_F6U2C2_T431t_1 | 0.63 | 1.02 | 0.11 | 0.14 | 0.78 | 0.92 | 0.25 | 0.83 | 0.47 | 1 | 0.74 | 1.13 | -0.06 | 1.26 | -0.64 | 1.17 | 0.22 | 0.36 | -0.0425 | 0.93155 | 0.491468 | 0.277846 | 1 | 0.999631 | 0.978006 |
| Atxn2_F6U2C2_T521t_1 | -0.84 | 0.02 | 0.45 | 0.29 | -1.1 | -0.51 | 0.16 | 0.21 | 0.49 | 0.16 | 0.19 | 1.57 | 0.71 | 0.4 | 1.24 | 1.06 | -0.29 | 0.6225 | 0.8725 | 0.041312 | 0.494446 | 0.175281 | 1 | 0.999631 | 0.474979 |
| Atxn2l_Q7TQH0_S109s_1 | -0.06 | 1.02 | -0.02 | 0.08 | 0.57 | -0.01 | -0.15 | -0.36 | 0.08 | -0.21 | 0.09 | -0.34 | -0.04 | 0.5 | 0.59 | 0.87 | -0.2425 | -0.35 | 0.225 | 0.542097 | 0.497115 | 0.303734 | 1 | 0.999631 | 0.83268 |
| Atxn2l_Q7TQH0_S304s_1 | 1.79 | 0.58 | 0.93 | 0.9 | 1.23 | 1.52 | 0.79 | 1.28 | 1.44 | 2.46 | 1.14 | 1.37 | -0.36 | 2.09 | 1.68 | 0.89 | 0.155 | 0.5525 | 0.025 | 0.964058 | 0.6475 | 0.191201 | 1 | 0.999631 | 0.987331 |
| Atxn2l_Q7TQH0_S337s_1 | 0.87 | 1.95 | 1.47 | 0.32 | 1.51 | 1.61 | 2.07 | -0.1 | 1.66 | 1.16 | 0.77 | 1.07 | 0.52 | 1.64 | 0.83 | 0.6 | 0.12 | 0.0125 | -0.255 | 0.566092 | 0.825517 | 0.975566 | 1 | 0.999631 | 0.841733 |
| Atxn2l_Q7TQH0_S388s_1 | 1.11 | 0.54 | 1.22 | 1 | 0.36 | 1.9 | 1.16 | 0.68 | 0.62 | 0.89 | 0.99 | 1.17 | -2.05 | 2.13 | 1.47 | 1.39 | 0.0575 | -0.05 | -0.2325 | 0.779427 | 0.879729 | 0.859579 | 1 | 0.999631 | 0.921416 |
| Atxn2l_Q7TQH0_S591s_1 | -0.97 | 0.24 | 0.61 | 0.28 | 0.12 | -0.09 | 0 | 0.32 | 0.24 | -0.41 | 0.5 | 0.69 | 0.62 | -0.01 | 0.56 | 0.63 | 0.0475 | 0.215 | 0.41 | 0.321238 | 0.898957 | 0.614095 | 1 | 0.999631 | 0.72408 |
| Atxn2l_Q7TQH0_S631s_1 | 0.09 | 1.57 | 1.39 | 0.74 | 0.46 | 0.73 | 0.31 | 0.82 | 1.03 | 0.8 | 1.28 | 1.6 | 1.41 | 1.07 | 0.83 | 1.53 | -0.3675 | 0.23 | 0.2625 | 0.51441 | 0.337748 | 0.56234 | 1 | 0.999631 | 0.819052 |
| Atxn2l_Q7TQH0_S681s_1 | 0.98 | 0.56 | 0.73 | 0.3 | 0.98 | 0.26 | 0.72 | 0.23 | 1.43 | 1.06 | 0.59 | 0.82 | 0.12 | -0.25 | 0.64 | 0.85 | -0.095 | 0.3325 | -0.3025 | 0.392792 | 0.749826 | 0.287961 | 1 | 0.999631 | 0.758901 |
| Atxn7l2_B7ZCC4_S514s_1 | 0.34 | 1.23 | 0.57 | 0.46 | 0.37 | 0.75 | -0.37 | -0.8 | 0.85 | 0.83 | 0.23 | 0.83 | 0.84 | 1.12 | 0.6 | 0.7 | -0.6625 | 0.035 | 0.165 | 0.604146 | 0.12659 | 0.911286 | 1 | 0.999631 | 0.85817 |
| Aurkaip1_Q9DCJ7_S51s_1 | 0.75 | 2.44 | 1.52 | 1.42 | 1.42 | 1.54 | 0.98 | 1.85 | 2.12 | 1.88 | 2.2 | 1.9 | 0.84 | 2.42 | 1.35 | 1.95 | -0.085 | 0.4925 | 0.1075 | 0.821749 | 0.830548 | 0.213971 | 1 | 0.999631 | 0.938464 |
| Avl9_Q80U56_S368s_1 | -0.24 | -0.03 | -0.58 | -0.32 | -0.08 | -0.09 | -0.91 | -0.11 | -0.24 | -0.53 | -0.01 | 0.18 | -0.06 | -0.34 | -0.26 | -0.26 | -0.005 | 0.1425 | 0.0625 | 0.820397 | 0.986614 | 0.617614 | 1 | 0.999631 | 0.938379 |
| Axin1_F6XWC2_S445s_1 | 2.02 | -0.19 | 1.37 | 0.71 | -1.06 | 0.38 | 2.26 | 0.3 | -1.02 | 0.98 | 0.3 | -0.36 | -0.54 | -0.9 | 0.29 | 0.01 | -0.5075 | -1.0025 | -1.2625 | 0.031997 | 0.492771 | 0.111375 | 1 | 0.999631 | 0.442395 |
| Axin2_O88566_S454s_1 | 1.74 | 0.33 | 0.35 | 1.22 | 1.37 | 1.49 | 1.16 | 0.93 | 1.14 | 0.89 | 0.18 | 0.14 | -0.31 | 0.05 | -0.66 | 0.61 | 0.3275 | -0.3225 | -0.9875 | 0.046345 | 0.398589 | 0.458065 | 1 | 0.999631 | 0.479949 |
| B630019K06Rik_Q7TNS5_S41s_1 | -0.47 | -2.73 | -0.44 | -0.81 | -2.14 | -1.82 | -0.77 | -0.94 | -2.39 | -1.01 | -0.84 | -1.22 | -1.61 | -2.32 | -0.47 | -1.11 | -0.305 | -0.2525 | -0.265 | 0.665292 | 0.60199 | 0.671151 | 1 | 0.999631 | 0.882566 |
| B630019K06Rik_Q7TNS5_S42s_1 | 0.16 | -0.77 | -0.16 | -1.03 | -0.78 | -0.64 | -0.41 | -0.96 | -0.54 | 0.17 | -1.24 | -0.7 | 0.31 | -1.12 | 0.31 | -1.74 | -0.2475 | -0.1275 | -0.11 | 0.840351 | 0.466437 | 0.754948 | 1 | 0.999631 | 0.946678 |

| | | | | | | | | | | | | | | | | | | | | | | | | | |
|---|---|---|---|---|---|---|---|---|---|---|---|---|---|---|---|---|---|---|---|---|---|---|---|---|---|
| B630019K06Rik_Q7TNS5_S97s_1 | -0.68 | -0.82 | -0.26 | 0.32 | 0.2 | 0.23 | 0.02 | 1.34 | 0.45 | -1.27 | 1.18 | 0.64 | 0.56 | 0.53 | 0.97 | 0.04 | 0.8075 | 0.61 | 0.885 | 0.033404 | 0.067136 | 0.279957 | 1 | 0.999631 | 0.44294 |
| Babam1_Q3UI43_S48s_1 | 1.2 | 0.25 | -0.48 | 0.88 | 1.12 | 1.12 | 0.65 | 0.71 | 0.79 | 0.83 | 0.25 | 0.17 | 0.85 | 0.76 | 0.52 | 1.08 | 0.4375 | 0.0475 | 0.34 | 0.413415 | 0.286697 | 0.908896 | 1 | 0.999631 | 0.768474 |
| Bad_Q3U9H3_S128s_1 | 0.21 | -0.07 | 0.67 | 0.3 | -0.3 | 0.48 | 0.87 | 0.12 | 0.69 | 1.32 | 0.27 | -0.48 | -0.08 | -0.44 | 0.48 | -0.93 | 0.015 | 0.1725 | -0.52 | 0.185733 | 0.964 | 0.677887 | 1 | 0.999631 | 0.646224 |
| Bad_Q3U9H3_S94s_1 | -0.18 | 0.42 | 0.53 | 0.03 | 0.43 | 0.22 | -0.08 | 0.7 | 0.47 | 0.15 | 0.44 | -0.1 | 0.94 | 1.03 | 0.05 | 0.37 | 0.1175 | 0.04 | 0.3975 | 0.267573 | 0.694785 | 0.892038 | 1 | 0.999631 | 0.695976 |
| Bag2_Q91YN9_S20s_1 | 0.84 | 1.33 | -0.27 | 0.77 | 0.09 | 1.29 | 2.8 | -0.56 | 0.75 | 1.9 | 0.11 | -0.22 | 0.05 | 0.37 | 0.31 | -0.36 | 0.2375 | -0.0325 | -0.575 | 0.172043 | 0.740802 | 0.95166 | 1 | 0.999631 | 0.641657 |
| Bag3_Q9JLV1_S179s_1 | -1.16 | -0.12 | 0.69 | -0.06 | 0.17 | -0.24 | -0.12 | 0.17 | 0.2 | -0.17 | -0.91 | 0.88 | -1.78 | 1.12 | 0.82 | 0.79 | 0.1575 | 0.1625 | 0.4 | 0.565211 | 0.693986 | 0.747739 | 1 | 0.999631 | 0.84139 |
| Bag3_Q9JLV1_S297s_1 | 1 | -0.24 | 0.37 | 0.23 | 0.52 | 0.24 | 0.03 | 0.31 | 0.43 | 1.25 | -0.3 | -0.11 | -1.18 | 0.93 | 0.45 | 0.05 | -0.065 | -0.0225 | -0.2775 | 0.580045 | 0.839977 | 0.958268 | 1 | 0.999631 | 0.846727 |
| Bag3_Q9JLV1_S390s_1 | -0.47 | -0.16 | -0.33 | -0.42 | -0.09 | 0.03 | -0.49 | -0.22 | -0.15 | -0.45 | -0.02 | 0.29 | 0.85 | 0.31 | -0.27 | -0.68 | 0.1525 | 0.2625 | 0.3975 | 0.307571 | 0.546784 | 0.348843 | 1 | 0.999631 | 0.717856 |
| Bag4_Q8CI61_S175s_1 | 1.89 | 0.48 | 0.18 | 1.47 | 0.92 | -0.28 | 0.72 | 0.9 | 0.39 | 0.7 | 0.53 | 0.27 | -0.06 | 0.48 | -0.45 | 0.47 | -0.44 | -0.5325 | -0.895 | 0.074341 | 0.361369 | 0.22264 | 1 | 0.999631 | 0.539897 |
| Bag4_Q8CI61_S175s_2 | 1.52 | 1.77 | 1.48 | 2.15 | 1.7 | 0.64 | 1.45 | 2.51 | 1.54 | 0.92 | 1.88 | 1.93 | -0.86 | 1.46 | 2.51 | 1.07 | -0.155 | -0.1625 | -0.685 | 0.303961 | 0.708542 | 0.62528 | 1 | 0.999631 | 0.715879 |
| Bag4_Q8CI61_S179s_2 | 1.52 | 1.77 | 1.48 | 2.15 | 1.7 | 0.64 | 1.45 | 2.51 | 1.54 | 0.92 | 1.88 | 1.93 | -0.86 | 1.46 | 2.51 | 1.07 | -0.155 | -0.1625 | -0.685 | 0.303961 | 0.708542 | 0.62528 | 1 | 0.999631 | 0.715879 |
| Bag6_Q3UF95_S1085s_1 | -0.53 | -1.58 | -0.2 | -0.47 | -0.76 | -0.96 | -0.92 | -0.73 | -0.76 | -0.57 | -1.43 | -0.31 | -0.16 | -0.02 | -0.13 | -0.37 | -0.1475 | -0.0725 | 0.525 | 0.168136 | 0.668156 | 0.855926 | 1 | 0.999631 | 0.637399 |
| Bag6_Q3UF95_S968s_2 | -2.67 | -5.99 | -4.48 | -4.74 | -4.88 | -4.91 | -5.19 | -4.03 | -5.94 | -3.8 | -3.97 | -2.38 | -3.76 | -5.17 | -4.46 | -3.13 | -0.2825 | 0.4475 | 0.34 | 0.638072 | 0.665655 | 0.610646 | 1 | 0.999631 | 0.87033 |
| Bag6_Q3UF95_S977s_1 | 0.22 | 0.87 | 0.55 | 0.3 | 0.55 | 0.33 | 0.18 | 0.38 | 0.32 | 0.31 | 0.8 | 0.92 | 0.72 | 0.5 | 0.52 | 0.78 | -0.125 | 0.1025 | 0.145 | 0.61562 | 0.638181 | 0.729959 | 1 | 0.999631 | 0.861203 |
| Bag6_Q3UF95_S977s_2 | -2.67 | -5.99 | -4.48 | -4.74 | -4.88 | -4.91 | -5.19 | -4.03 | -5.94 | -3.8 | -3.97 | -2.38 | -3.76 | -5.17 | -4.46 | -3.13 | -0.2825 | 0.4475 | 0.34 | 0.638072 | 0.665655 | 0.610646 | 1 | 0.999631 | 0.87033 |
| Bag6_Q3UF95_T967t_1 | 1.59 | -1.32 | 1.05 | 0.68 | -1.71 | -1.43 | 1.6 | -0.22 | -1.17 | 0.79 | 0.15 | -0.12 | -1.4 | -0.95 | -0.12 | -0.45 | -0.94 | -0.5875 | -1.23 | 0.073593 | 0.287276 | 0.39382 | 1 | 0.999631 | 0.539296 |
| Bai3_Q80ZF8_S1220s_1 | -0.49 | 0.38 | 1.68 | 1.61 | 0.16 | -0.15 | 1.42 | 1.66 | -0.13 | -0.18 | 1.7 | 2.19 | 0.63 | 0.07 | 2.12 | 2.29 | -0.0225 | 0.1 | 0.4825 | 0.480693 | 0.971014 | 0.888883 | 1 | 0.999631 | 0.801848 |
| Bai3_Q80ZF8_S1411s_1 | -1.35 | -1.02 | 0.1 | -1 | -1.32 | -1.26 | -1.13 | -0.19 | -1.69 | -1.37 | -0.06 | 0.19 | -0.49 | -1.04 | -0.01 | -0.16 | -0.1575 | 0.085 | 0.3925 | 0.348905 | 0.702938 | 0.87212 | 1 | 0.999631 | 0.736697 |
| Bai3_Q80ZF8_S751s_1 | -1.18 | -0.2 | -0.17 | -0.2 | -0.85 | -0.59 | -0.73 | -0.62 | -0.26 | -0.98 | 0.1 | -0.12 | 1.06 | 0.04 | -1.84 | -0.08 | -0.26 | 0.1225 | 0.2325 | 0.696687 | 0.410228 | 0.740327 | 1 | 0.999631 | 0.895731 |
| Bai3_Q80ZF8_T1215t_1 | 1.41 | 1.24 | -0.14 | 1.57 | 1.37 | 1.49 | 1.98 | 1.3 | 0.03 | -0.31 | 1.88 | -1.1 | -1.86 | 0.64 | -0.83 | 0.11 | 0.515 | -0.895 | -1.505 | 0.03185 | 0.236591 | 0.200038 | 1 | 0.999631 | 0.442395 |
| Baiap2_Q8BKX1_S324s_1 | -0.16 | -3.64 | -0.97 | -0.93 | -1.25 | -1.49 | -1.02 | -0.76 | -1 | -0.48 | -1.58 | -0.65 | -1.52 | -1.28 | -0.76 | -0.85 | 0.295 | 0.4975 | 0.3225 | 0.644091 | 0.669895 | 0.489411 | 1 | 0.999631 | 0.872436 |
| Baiap2_Q8BKX1_S326s_1 | -0.41 | -0.67 | -0.44 | -1.11 | -0.8 | -0.64 | -0.96 | -0.79 | -1.08 | -0.14 | -0.29 | -0.25 | 0.3 | -0.5 | -0.76 | -0.51 | -0.14 | 0.2175 | 0.29 | 0.407489 | 0.604562 | 0.508661 | 1 | 0.999631 | 0.766996 |
| Baiap2_Q8BKX1_S367s_1 | 1.96 | 1.29 | -0.71 | 2.33 | 2.45 | 2.03 | 2.1 | 2.49 | 1.75 | 0.37 | 2.03 | -0.39 | -1.61 | 1.94 | 0.16 | 0.54 | 1.05 | -0.2775 | -0.96 | 0.280906 | 0.114467 | 0.722945 | 1 | 0.999631 | 0.701922 |
| Baiap2_Q8BKX1_S396s_1 | -0.32 | 0.66 | -0.33 | -0.06 | -0.33 | 0.18 | -0.02 | 0.54 | -0.22 | 0.09 | -0.65 | 0.73 | -0.02 | 0.53 | -0.56 | 0.58 | 0.105 | 1.96E-17 | 0.145 | 0.708681 | 0.753924 | 1 | 1 | 1 | 0.898927 |
| Baiap2_Q8BKX1_S515s_1 | -0.21 | 0.75 | 0.68 | 0.5 | 2.32 | 0.86 | 0.73 | 1.09 | 1.2 | -1.12 | 1.75 | 1.56 | 0.8 | 0.72 | 1.28 | 0.23 | 0.82 | 0.4175 | 0.3275 | 0.371199 | 0.075809 | 0.51436 | 1 | 0.999631 | 0.746338 |
| Baiap2_Q8BKX1_T341t_1 | 0.12 | 1.81 | -0.82 | 0.84 | 2 | 2.17 | 1.03 | 1.34 | 1.89 | -0.28 | 1.04 | -0.2 | -1.15 | 1.16 | 0.17 | 0.33 | 1.1475 | 0.125 | -0.36 | 0.586574 | 0.065961 | 0.854542 | 1 | 0.999631 | 0.849724 |
| Baiap2_Q8BKX1_T521t_1 | -0.54 | -0.13 | -0.55 | -0.98 | -0.21 | -0.38 | -0.71 | -0.54 | -0.42 | -0.45 | -0.75 | -0.58 | 0.24 | -0.2 | -0.49 | -0.55 | 0.09 | 1.96E-17 | 0.3 | 0.369319 | 0.750658 | 1 | 1 | 1 | 0.746224 |
| Baiap2l2_Q80Y61_S459s_1 | 1.74 | 1.1 | -1.99 | -1.12 | 2.59 | 1.51 | -1.12 | -1.69 | 1.75 | 1.82 | -2.54 | -1.08 | 3.22 | 1.7 | -1.59 | -1.06 | 0.39 | 0.055 | 0.635 | 0.60564 | 0.737282 | 0.963296 | 1 | 0.999631 | 0.85817 |
| Baiap3_S4R1K2_S215s_1 | 0.99 | 1.24 | 3 | 1.36 | 0.4 | 1.19 | 1.97 | 1.31 | 1.99 | 1.64 | 2.45 | 2.72 | 1.3 | 0.95 | 2.41 | 2.68 | -0.43 | 0.5525 | 0.1875 | 0.743559 | 0.416416 | 0.277704 | 1 | 0.999631 | 0.90976 |
| Basp1_Q91XV3_S128s_1 | 0.01 | -0.14 | 0.47 | 0.12 | 0.45 | -0.03 | -0.43 | 0.42 | -1.08 | 0.2 | 0.4 | 0.75 | -0.69 | 0.76 | 0.43 | 0.64 | -0.0125 | -0.0475 | 0.17 | 0.662663 | 0.967272 | 0.910358 | 1 | 0.999631 | 0.881311 |
| Basp1_Q91XV3_S128s_2 | -0.71 | -0.65 | -0.34 | -0.39 | -0.83 | -0.4 | -0.97 | -0.84 | -0.56 | -0.33 | -1.07 | -0.66 | -0.27 | -0.43 | -0.49 | 0 | -0.2375 | -0.1325 | 0.225 | 0.429401 | 0.371642 | 0.636479 | 1 | 0.999631 | 0.776658 |
| Basp1_Q91XV3_S131s_1 | -0.5 | -0.72 | -0.53 | -0.21 | -1.13 | -1.11 | -0.32 | -0.7 | -1.09 | -0.67 | 0.04 | -0.65 | -0.13 | -0.74 | -0.51 | 0.53 | -0.325 | -0.1025 | 0.2775 | 0.438336 | 0.279193 | 0.747603 | 1 | 0.999631 | 0.781942 |
| Basp1_Q91XV3_S131s_2 | -0.71 | -0.65 | -0.15 | -0.05 | -0.83 | -0.4 | -0.48 | -0.2 | -0.56 | -0.33 | -0.66 | 0.01 | -0.27 | -0.43 | -0.17 | 0.27 | -0.0875 | 0.005 | 0.24 | 0.452578 | 0.761562 | 0.986726 | 1 | 0.999631 | 0.788364 |
| Basp1_Q91XV3_S167s_1 | -0.59 | -0.84 | -0.7 | -0.23 | -0.81 | -1.08 | -0.59 | -0.21 | -1.31 | -1.01 | -0.62 | -0.42 | -0.34 | -0.81 | -0.78 | 0.15 | -0.0825 | -0.25 | 0.145 | 0.664038 | 0.779145 | 0.424411 | 1 | 0.999631 | 0.881993 |
| Basp1_Q91XV3_S169s_2 | -0.83 | -1.08 | -0.54 | -0.49 | -0.73 | -1.08 | -0.25 | -1.16 | -1.22 | -0.92 | -0.23 | -0.3 | 0.42 | -0.43 | -0.54 | 0.09 | -0.07 | 0.0675 | 0.62 | 0.087467 | 0.819448 | 0.837824 | 1 | 0.999631 | 0.553616 |
| Basp1_Q91XV3_S191s_1 | -0.67 | -1.02 | 0.95 | -0.21 | -1.59 | -1.71 | -0.29 | -1.81 | -1.11 | -1.6 | -0.13 | -0.46 | -4.02 | -1.32 | -0.17 | 0.29 | -1.1125 | -0.5875 | -1.0675 | 0.257758 | 0.054457 | 0.266325 | 1 | 0.999631 | 0.692624 |
| Basp1_Q91XV3_S218s_1 | -1.12 | -1.79 | -1.84 | -1.41 | -1.32 | -2.05 | -1.74 | -1.77 | -1.87 | -1.85 | -1.46 | -1.96 | -0.53 | -1.49 | -1.44 | -1.54 | -0.18 | -0.245 | 0.29 | 0.417064 | 0.545018 | 0.406321 | 1 | 0.999631 | 0.771432 |
| Basp1_Q91XV3_S40s_1 | -0.33 | 0.76 | 1.47 | -0.28 | -0.18 | 0.4 | -0.06 | 1.15 | -1.11 | -0.13 | -0.21 | -0.73 | -0.1 | -0.24 | 0.89 | 0.48 | -0.0775 | -0.95 | -0.1475 | 0.763648 | 0.876 | 0.069325 | 1 | 0.999631 | 0.915694 |
| Basp1_Q91XV3_S40s_2 | -1.37 | 1.23 | -1.8 | -2.1 | 1.14 | 0.86 | -1.6 | -1.86 | 1.17 | -0.51 | -1.23 | -1.61 | 1.49 | 0.43 | -1.2 | -1.24 | 0.645 | 0.465 | 0.88 | 0.326696 | 0.501433 | 0.589176 | 1 | 0.999631 | 0.724678 |
| Basp1_Q91XV3_S92s_1 | -0.4 | 0.92 | 0.55 | 0.05 | 0.28 | 0.65 | -0.36 | 0.7 | 0.49 | 0.24 | 0.17 | 0.61 | 1.17 | 0.8 | 0.41 | 0.99 | 0.0375 | 0.0975 | 0.5625 | 0.153649 | 0.922924 | 0.779198 | 1 | 0.999631 | 0.625198 |
| Basp1_Q91XV3_T132t_1 | -0.49 | -0.28 | -0.69 | 0.04 | -0.05 | -0.83 | -1.26 | -0.23 | -0.61 | -1.17 | -0.1 | -0.86 | 0.27 | -1.17 | 0.57 | 0.14 | -0.2375 | -0.33 | 0.3075 | 0.476097 | 0.501157 | 0.3278 | 1 | 0.999631 | 0.799882 |
| Basp1_Q91XV3_T132t_2 | -0.82 | 0.58 | -0.29 | -0.52 | 1.03 | 0.3 | -1.08 | -1.04 | 0.71 | -0.14 | -1.1 | -0.77 | 1 | 0.48 | -0.51 | -0.08 | 0.065 | -0.0625 | 0.485 | 0.29206 | 0.906055 | 0.896246 | 1 | 0.999631 | 0.706164 |
| Basp1_Q91XV3_T183t_1 | 0.21 | 0.85 | 2.37 | 1.66 | 0.22 | 0.9 | 1.54 | 1.79 | 0.5 | 0.44 | 1.82 | 1.61 | 0.31 | 0.44 | 1.75 | 1.62 | -0.16 | -0.18 | -0.2425 | 0.666563 | 0.767843 | 0.744214 | 1 | 0.999631 | 0.88264 |
| Basp1_Q91XV3_T31t_1 | -0.33 | -3.27 | 0.11 | -0.2 | -2.24 | -2.35 | 0.2 | -1.03 | -2.67 | -1.7 | -1.19 | -0.85 | -1.68 | -3.59 | -0.71 | -1.98 | -0.4325 | -0.68 | -1.0675 | 0.23106 | 0.616887 | 0.390888 | 1 | 0.999631 | 0.681463 |
| Basp1_Q91XV3_T31t_2 | 1.07 | -0.85 | -0.49 | -1.05 | -0.97 | -0.16 | 0.04 | -1.64 | -0.86 | 0.67 | -1.57 | -1.76 | -0.68 | -1.27 | -1.87 | -2.12 | -0.3525 | -0.55 | -1.155 | 0.054922 | 0.535846 | 0.411444 | 1 | 0.999631 | 0.492066 |
| Basp1_Q91XV3_T36t_1 | 0.5 | -1.6 | -0.53 | -0.96 | -1.78 | -0.73 | -0.29 | -1.41 | -1.25 | 0.08 | -1.12 | -1.35 | -0.98 | -2.12 | -1.27 | -1.75 | -0.405 | -0.2625 | -0.8825 | 0.09621 | 0.439387 | 0.615506 | 1 | 0.999631 | 0.55886 |
| Basp1_Q91XV3_T36t_2 | 0.5 | 0.05 | -0.9 | -1.39 | -0.04 | 0.25 | -0.41 | -1.71 | 0.02 | 0.3 | -1.45 | -1.71 | 0.27 | -0.56 | -1.61 | -1.78 | -0.0425 | -0.275 | -0.485 | 0.419514 | 0.939356 | 0.651864 | 1 | 0.999631 | 0.772152 |
| Baz1b_Q9Z277_S1464s_1 | -1.37 | -1.64 | -0.57 | -1.95 | -1.56 | -1.72 | -1.02 | -1.76 | -1.07 | -2.04 | -0.78 | -2.25 | 0.7 | -1.5 | -2.06 | -1.84 | -0.1325 | -0.1525 | 0.2075 | 0.743253 | 0.716111 | 0.738039 | 1 | 0.999631 | 0.90972 |
| Baz1b_Q9Z277_S1464s_2 | -1.08 | 0.4 | -0.96 | -0.8 | -0.19 | -0.85 | -0.64 | -1.42 | -0.56 | -0.59 | -0.69 | -1.05 | -0.55 | -1.08 | -1.5 | -0.9 | -0.165 | -0.1125 | -0.3975 | 0.346459 | 0.696511 | 0.768274 | 1 | 0.999631 | 0.735175 |
| Baz1b_Q9Z277_S1468s_2 | -1.08 | 0.4 | -0.96 | -0.8 | -0.19 | -0.85 | -0.64 | -1.42 | -0.56 | -0.59 | -0.69 | -1.05 | -0.55 | -1.08 | -1.5 | -0.9 | -0.165 | -0.1125 | -0.3975 | 0.346459 | 0.696511 | 0.768274 | 1 | 0.999631 | 0.735175 |
| BC023829_Q3TYW3_S305s_1 | -0.44 | -1.23 | -0.81 | -0.76 | -1.03 | -0.42 | -1.05 | -0.87 | -0.85 | 0.08 | -0.64 | -0.58 | 0.47 | -0.09 | -0.56 | -0.44 | -0.0325 | 0.3125 | 0.655 | 0.08085 | 0.910768 | 0.33939 | 1 | 0.999631 | 0.545579 |
| Bcam_Q9R069_S615s_1 | 1.19 | 2.09 | -0.9 | 0.7 | 1.31 | 1.04 | 0.17 | -0.54 | 0.84 | 2.2 | 0.78 | 0.18 | 2.09 | 0.69 | 0.18 | 0.12 | -0.275 | 0.23 | -5.1E-16 | 1 | 0.683437 | 0.735012 | 1 | 0.999631 | 1 |
| Bcan_Q61361_S29s_1 | 2 | 0.8 | 0.9 | 1.45 | -0.47 | 1.64 | 3.84 | -1.49 | -0.91 | 2.91 | 0.23 | 2.1 | 0.88 | 0.2 | 1.95 | 2.15 | -0.4075 | -0.205 | 0.0075 | 0.988261 | 0.696869 | 0.798174 | 1 | 0.999631 | 0.995409 |
| Bcan_Q61361_S587s_1 | -0.15 | 0.22 | 0.7 | 0.6 | 0.02 | 0.39 | 0.96 | 0 | -0.19 | 0.19 | 1.1 | 1.98 | 0.28 | 0.39 | 1.03 | 1.19 | 3.93E-17 | 0.4275 | 0.38 | 0.296686 | 1 | 0.399085 | 1 | 0.999631 | 0.7098 |
| Bcar1_Q61140_S572s_1 | -0.85 | 0.35 | -0.54 | -0.75 | -0.47 | -0.89 | -1.3 | -0.72 | -1.94 | -0.13 | -0.69 | -0.64 | -0.83 | -0.49 | 0.04 | 0.4 | -0.3975 | -0.4025 | 0.2275 | 0.579217 | 0.275413 | 0.390954 | 1 | 0.999631 | 0.846484 |
| Bcar1_Q61140_S643s_1 | -0.34 | -2.19 | -0.4 | -0.37 | -0.61 | -0.79 | -0.96 | -0.77 | -0.72 | -0.44 | -1.56 | -0.46 | -0.58 | 0.1 | -0.15 | -0.56 | 0.0425 | 0.03 | 0.5275 | 0.280381 | 0.924091 | 0.952012 | 1 | 0.999631 | 0.701922 |
| Bcar3_Q9QZK2_S164s_1 | -0.55 | -0.45 | -0.68 | -0.24 | -0.45 | -0.95 | -0.43 | -0.53 | -0.27 | -0.24 | -0.91 | -0.22 | -0.35 | -0.37 | -1.6 | 0.2 | -0.11 | 0.07 | -0.05 | 0.902928 | 0.672834 | 0.80537 | 1 | 0.999631 | 0.968183 |
| Bcar3_Q9QZK2_S245s_1 | -0.62 | 0.5 | 0.21 | -0.43 | -0.03 | -0.42 | -0.2 | 0.41 | 0.44 | -0.08 | 0.8 | -0.22 | 1.08 | -0.15 | 0.57 | 1.14 | 0.025 | 0.32 | 0.745 | 0.096672 | 0.942626 | 0.405506 | 1 | 0.999631 | 0.55886 |
| Bcar3_Q9QZK2_S250s_1 | 0.57 | 1.1 | -1.05 | 0.14 | 0.47 | -0.47 | -0.03 | -0.46 | 0.64 | -0.89 | 0.72 | -0.1 | -0.36 | -0.33 | 0.61 | 0.87 | -0.3125 | -0.0975 | 0.0075 | 0.988594 | 0.521091 | 0.859042 | 1 | 0.999631 | 0.995409 |
| Bcas1_Q80YN3_S316s_1 | 1.37 | -0.04 | 0.81 | 1 | -0.41 | 0.73 | 0.7 | -0.76 | -1.91 | 0.75 | -0.71 | -0.83 | 0.17 | -0.28 | 0.15 | -1.23 | -0.72 | -1.46 | -1.0825 | 0.032913 | 0.144659 | 0.027535 | 1 | 0.999631 | 0.44294 |
| Bcas1_Q80YN3_S388s_1 | -0.1 | -0.48 | -0.18 | -0.39 | 0.14 | -0.72 | -1.02 | -1.59 | -0.59 | -0.35 | -0.67 | -0.6 | 0.93 | 0.58 | -1.31 | -0.76 | -0.51 | -0.265 | 0.1475 | 0.774194 | 0.204037 | 0.308881 | 1 | 0.999631 | 0.920998 |
| Bcas1_Q80YN3_S390s_1 | -0.1 | -0.48 | -0.18 | -0.39 | 0.14 | -0.72 | -1.02 | -1.59 | -0.59 | -0.35 | -0.67 | -0.6 | 0.93 | 0.58 | -1.31 | -0.76 | -0.51 | -0.265 | 0.1475 | 0.774194 | 0.204037 | 0.308881 | 1 | 0.999631 | 0.920998 |
| Bcas1_Q80YN3_S414s_1 | -0.73 | -0.69 | 0.46 | 0.13 | -0.98 | -0.81 | -1.01 | -1.32 | -0.93 | 0.09 | -0.22 | 0.4 | 0.31 | -1.42 | 0.42 | 0.31 | -0.8225 | 0.0425 | 0.1125 | 0.824819 | 0.037223 | 0.918748 | 1 | 0.999631 | 0.940061 |
| Bcas1_Q80YN3_S425s_1 | -1.05 | -1.28 | -1.58 | -1.65 | -0.72 | -1.22 | -1.32 | -1.01 | -1.23 | -1.3 | -1.08 | -0.39 | 0.27 | -1.22 | -1.92 | -0.96 | 0.3225 | 0.39 | 0.4325 | 0.366335 | 0.260896 | 0.233598 | 1 | 0.999631 | 0.745796 |
| Bcas1_Q80YN3_S443s_1 | -2.87 | 1.94 | 0.36 | 0.16 | 1.52 | 0.38 | -1.51 | 0.68 | 0.47 | -0.67 | -0.17 | 0.97 | 0.57 | 1.97 | 1.31 | 0.62 | 0.37 | 0.2525 | 1.22 | 0.200679 | 0.718435 | 0.785316 | 1 | 0.999631 | 0.658263 |
| Bcas1_Q80YN3_S497s_1 | -3.51 | -1.77 | -0.33 | -2.07 | -1.29 | -1.85 | -0.5 | -2.24 | -1.07 | -2.92 | -0.83 | -0.7 | 0.28 | -3.68 | -1.23 | -2.31 | 0.45 | 0.54 | 0.185 | 0.839617 | 0.506916 | 0.470396 | 1 | 0.999631 | 0.946365 |
| Bcas1_Q80YN3_S525s_1 | -0.76 | -0.71 | -0.13 | -0.45 | -0.95 | -0.34 | -0.89 | -0.3 | -0.94 | -0.77 | -0.38 | -0.52 | 0.02 | -0.38 | -0.11 | -0.89 | -0.1075 | -0.14 | 0.1725 | 0.598908 | 0.715519 | 0.624042 | 1 | 0.999631 | 0.855702 |
| Bcas1_Q80YN3_S525s_2 | -0.39 | 1.53 | 1.55 | 0.86 | 0.77 | 1.04 | 0.24 | 0.45 | 0.35 | 0.9 | 0.41 | 1.46 | 1.34 | 1.16 | 1.53 | 2.43 | -0.2625 | -0.1075 | 0.7275 | 0.175064 | 0.576502 | 0.828834 | 1 | 0.999631 | 0.642116 |
| Bcas1_Q80YN3_S601s_1 | 0.69 | 1.2 | 1.24 | 1.32 | 1.54 | 1.6 | 0.73 | 1.25 | 1.54 | 0.85 | 1.59 | 1.45 | -0.14 | 1.73 | 2.62 | 1.59 | 0.1675 | 0.245 | 0.3375 | 0.545071 | 0.58497 | 0.422189 | 1 | 0.999631 | 0.833658 |
| Bcas1_Q80YN3_S615s_1 | -0.76 | -0.05 | -0.09 | -0.63 | -0.34 | 0.55 | -0.94 | -0.56 | -0.52 | -0.26 | -0.12 | 0.1 | 0.01 | 0.35 | -0.45 | -0.19 | 0.06 | 0.1825 | 0.3125 | 0.34937 | 0.874203 | 0.547083 | 1 | 0.999631 | 0.736792 |
| Bcas1_Q80YN3_T379t_1 | -2.02 | -1.64 | -1.38 | -1.08 | -3.47 | -0.76 | -0.92 | -1.23 | -2 | -0.77 | -0.62 | -2.35 | -1.66 | -0.56 | -2.45 | -1.61 | -0.065 | 0.095 | -0.04 | 0.927469 | 0.913587 | 0.837752 | 1 | 0.999631 | 0.976306 |
| Bcas1_Q80YN3_T523t_1 | -1.58 | -0.94 | -0.64 | -1.06 | -0.99 | -0.59 | -1.42 | -1.28 | -0.75 | -1.29 | -1.08 | -1.05 | -2.07 | -0.49 | -0.84 | -0.96 | -0.015 | 0.0125 | -0.035 | 0.932195 | 0.962337 | 0.966835 | 1 | 0.999631 | 0.978188 |
| Bcas1_Q80YN3_T523t_2 | -0.39 | 1.53 | 1.55 | 0.86 | 0.77 | 1.04 | 0.24 | 0.45 | 0.35 | 0.9 | 0.41 | 1.46 | 1.34 | 1.16 | 1.53 | 2.43 | -0.2625 | -0.1075 | 0.7275 | 0.175064 | 0.576502 | 0.828834 | 1 | 0.999631 | 0.642116 |
| Bcas1_Q80YN3_T591t_1 | -0.91 | -1.45 | -0.56 | -0.87 | -1.32 | -1.56 | -1.24 | -0.95 | -1.14 | -1.15 | -1.25 | -1.29 | -0.44 | -0.94 | -0.51 | -1.3 | -0.32 | -0.26 | 0.15 | 0.659574 | 0.289642 | 0.368487 | 1 | 0.999631 | 0.880252 |
| Bcas3_Q0VF62_S503s_1 | -0.28 | -0.62 | -1.38 | -0.99 | -0.22 | -1.33 | -0.74 | -1.02 | -1.21 | -1.08 | -0.52 | -0.92 | -0.15 | -0.2 | -0.59 | -1.04 | -0.01 | -0.115 | 0.3225 | 0.383413 | 0.977738 | 0.729388 | 1 | 0.999631 | 0.754581 |
| Bcas3_Q0VF62_S518s_1 | 1.16 | 0.31 | 0.31 | 0.13 | 0.28 | -0.43 | -0.56 | 0.57 | 0.58 | 0.72 | 0.1 | -0.33 | 0.69 | 0.75 | -0.29 | -0.09 | -0.5125 | -0.21 | -0.2125 | 0.584375 | 0.19234 | 0.567248 | 1 | 0.999631 | 0.849121 |
| Bcas3_Q0VF62_S550s_1 | -0.5 | -1.11 | -0.26 | -0.6 | -0.73 | -0.74 | -0.99 | -0.56 | -0.52 | -0.37 | -0.87 | -0.66 | -0.08 | -0.16 | -0.61 | -0.16 | -0.1375 | 0.0125 | 0.365 | 0.255745 | 0.626086 | 0.965842 | 1 | 0.999631 | 0.690775 |
| Bcas3_Q0VF62_S854s_1 | -0.74 | -0.07 | -0.25 | -0.47 | 0 | -0.12 | -0.22 | -0.31 | 0.06 | -0.18 | -0.21 | -0.24 | -0.32 | -0.19 | -0.29 | -0.46 | 0.22 | 0.24 | 0.0675 | 0.8124 | 0.409629 | 0.384271 | 1 | 0.999631 | 0.937041 |
| Bcas3_Q0VF62_S862s_1 | 0.14 | 1.7 | 1.04 | 0.44 | 0.99 | 1.62 | 0.17 | 0.82 | 1.23 | 0.46 | 0.83 | 1.51 | 1.69 | 1.78 | 0.91 | 0.62 | 0.07 | 0.1775 | 0.42 | 0.35987 | 0.874385 | 0.672871 | 1 | 0.999631 | 0.741182 |
| Bcas3_Q0VF62_S871s_1 | -0.58 | -0.02 | -1.48 | -0.8 | 0.26 | -0.12 | -0.49 | -1.91 | -0.64 | -0.91 | -0.64 | -1.42 | 0.41 | -0.16 | -2.08 | -3.23 | 0.155 | -0.1825 | -0.545 | 0.491989 | 0.766613 | 0.630475 | 1 | 0.999631 | 0.809399 |
| Bckdha_Q3U3J1_S338s_1 | 0.01 | -3.57 | -3.23 | -3.83 | -2.69 | -1.86 | -0.86 | -3.17 | -2.58 | -1.3 | -3.8 | -4.92 | -2.28 | -3.42 | -3.65 | -5.27 | 0.51 | -0.495 | -1 | 0.299062 | 0.570735 | 0.631109 | 1 | 0.999631 | 0.712318 |
| Bckdha_Q3U3J1_S348s_1 | -1.09 | -0.18 | 0.19 | -0.4 | -0.34 | -0.24 | -0.61 | -0.31 | -0.29 | -0.62 | -0.22 | 0.03 | 0.59 | 0.3 | 0.18 | 0.15 | -0.005 | 0.095 | 0.675 | 0.075301 | 0.987731 | 0.782536 | 1 | 0.999631 | 0.541044 |
| Bckdha_Q3U3J1_T339t_1 | 0.44 | 0.01 | 0.07 | 0.36 | -0.24 | 0.58 | -0.44 | 0.75 | 0 | 0.79 | -0.26 | 0.75 | 0.33 | 0 | -0.17 | -0.67 | -0.0575 | 0.1 | -0.3475 | 0.29022 | 0.867777 | 0.76551 | 1 | 0.999631 | 0.706164 |
| Bckdk_D3Z7R0_S31s_1 | -0.96 | -1.36 | -1.11 | -1.34 | -2.34 | -1.25 | -0.73 | -1.75 | -1.45 | -0.58 | -1.48 | -1.52 | -0.39 | -0.91 | -1.39 | -1 | -0.325 | -0.065 | 0.27 | 0.400809 | 0.394289 | 0.834963 | 1 | 0.999631 | 0.763209 |
| Bcl11a_Q5STS9_S205s_1 | -0.5 | -0.26 | -0.29 | -0.68 | -0.46 | 0.06 | -0.44 | 0.41 | -0.21 | 0.16 | 0.15 | -0.97 | -0.07 | -0.05 | 0.12 | 0.43 | 0.325 | 0.215 | 0.54 | 0.080254 | 0.289646 | 0.523084 | 1 | 0.999631 | 0.545579 |
| Bcl11a_Q5STS9_S328s_1 | -0.79 | 0.72 | -0.12 | -0.43 | 0.22 | 0.57 | 0.52 | -0.54 | 0.47 | -0.4 | 0.93 | -0.39 | 0.79 | 0.74 | 0.38 | -0.06 | 0.3475 | 0.3075 | 0.6175 | 0.146755 | 0.406937 | 0.501225 | 1 | 0.999631 | 0.618703 |
| Bcl11a_Q5STS9_S431s_1 | 0.2 | 1.46 | 0.99 | -0.35 | 0.14 | 0.94 | 0.08 | -0.04 | 1.1 | 1.51 | 0.84 | 1.07 | 0.77 | 1.16 | 1.31 | 0.19 | -0.295 | 0.555 | 0.2825 | 0.548308 | 0.513609 | 0.213986 | 1 | 0.999631 | 0.834633 |
| Bcl11a_Q5STS9_S432s_1 | 2.27 | 0.87 | 2.17 | 1.22 | 0.13 | 1.35 | 1.43 | 0.89 | 1.36 | 2.05 | 1.48 | 1.07 | 0.75 | 1.24 | 1.33 | 1.33 | -0.6825 | -0.1425 | -0.47 | 0.255633 | 0.148103 | 0.729426 | 1 | 0.999631 | 0.690775 |
| Bcl11a_Q5STS9_S608s_1 | -0.15 | 0.65 | 0.77 | 0.72 | -0.08 | -0.02 | 0.04 | 0.71 | -0.01 | 0.06 | 0.81 | 0.49 | 0.46 | 0.28 | 0.02 | 0.55 | -0.335 | -0.16 | -0.17 | 0.603478 | 0.320022 | 0.637246 | 1 | 0.999631 | 0.857824 |
| Bcl11a_Q5STS9_S718s_1 | 0.41 | -0.19 | 0.43 | 0.56 | -0.63 | -0.09 | 0.09 | 0.57 | -0.76 | 0.28 | -0.68 | -0.15 | 0.48 | -0.3 | 0.61 | 0.68 | -0.3175 | -0.63 | 0.065 | 0.850047 | 0.356738 | 0.089212 | 1 | 0.999631 | 0.949525 |
| Bcl11a_Q5STS9_S86s_1 | -0.76 | 0.16 | 0.1 | -0.12 | -0.79 | -0.14 | -0.5 | 0.06 | -0.44 | -0.06 | -0.21 | 0.07 | 0.52 | 0.09 | 0.25 | 0.66 | -0.1875 | -0.005 | 0.535 | 0.123708 | 0.569019 | 0.987002 | 1 | 0.999631 | 0.594905 |
| Bcl11b_Q99PV8_S309s_1 | -0.47 | 0.42 | -0.64 | -0.2 | -0.56 | -0.19 | -0.8 | -0.42 | -0.28 | -0.02 | 0.31 | 0.38 | 0.72 | -0.42 | -0.55 | 0.3 | -0.27 | 0.32 | 0.235 | 0.563081 | 0.40217 | 0.344897 | 1 | 0.999631 | 0.841038 |
| Bcl11b_Q99PV8_S690s_1 | 0.32 | 0.27 | -0.53 | -0.42 | -0.32 | -0.52 | -0.27 | -0.3 | 0.02 | 0.85 | -0.51 | -0.05 | 0.55 | 0.43 | 0.04 | 0.25 | -0.2625 | 0.1675 | 0.4075 | 0.230627 | 0.386093 | 0.66247 | 1 | 0.999631 | 0.681463 |
| Bcl11b_Q99PV8_T344t_1 | -0.63 | -0.44 | -0.46 | -0.93 | -0.51 | -0.07 | -0.99 | -0.87 | -0.34 | -0.41 | -0.66 | 0.1 | 0.33 | -0.59 | -0.86 | -0.29 | 0.005 | 0.2875 | 0.2625 | 0.450478 | 0.986666 | 0.327012 | 1 | 0.999631 | 0.787759 |
| Bcl11b_Q99PV8_T672t_1 | -0.15 | 0.26 | 0.01 | -0.29 | -0.23 | -0.55 | -0.67 | -0.18 | 0.16 | -0.15 | -0.24 | 0 | 0.88 | 0.2 | 0.14 | -0.23 | -0.365 | -0.015 | 0.29 | 0.391592 | 0.190943 | 0.954813 | 1 | 0.999631 | 0.758901 |
| Bcl2l13_P59017_S35s_1 | -0.63 | 0.68 | -0.31 | -0.24 | -0.2 | -0.07 | -0.49 | -0.24 | -0.6 | -0.81 | -0.69 | -0.43 | 0.41 | -0.44 | -0.97 | 0.02 | -0.125 | -0.5075 | -0.12 | 0.77644 | 0.708921 | 0.157448 | 1 | 0.999631 | 0.921191 |
| Bcl2l13_P59017_S377s_1 | -0.29 | -1.37 | -0.52 | -0.84 | -0.87 | -0.57 | -0.97 | -0.76 | -1.1 | -0.2 | -1.02 | -0.7 | -0.4 | -0.78 | -1.1 | -0.34 | -0.0375 | 0 | 0.1 | 0.775739 | 0.902681 | 1 | 1 | 1 | 0.920998 |
| Bcl2l13_P59017_S387s_1 | -2.91 | -7.48 | -4.95 | -6.49 | -5.17 | -4.52 | 0.7 | -5.87 | -4.21 | -2.22 | -2.35 | -2.91 | -6.4 | -7.33 | -4.94 | -7.44 | 1.7425 | 2.535 | -1.07 | 0.292398 | 0.269479 | 0.022437 | 1 | 0.999631 | 0.706671 |
| Bcl7a_Q9CXE2_S207s_1 | 0.73 | 1.29 | 0.56 | 0.34 | 0.82 | 1.07 | 0.56 | 0.06 | 1.32 | 1.3 | -0.16 | 0.63 | 0.33 | 1.03 | 0.79 | 0.46 | -0.1025 | 0.0425 | -0.0775 | 0.814927 | 0.759876 | 0.917699 | 1 | 0.999631 | 0.937235 |
| Bcl9_Q9D219_S291s_1 | -0.64 | 0.85 | -1.74 | -0.94 | 0.59 | 0.21 | -0.56 | -0.57 | -0.19 | -0.78 | -0.32 | -0.16 | 1.03 | 0.27 | -0.45 | -0.58 | 0.535 | 0.255 | 0.685 | 0.268717 | 0.352101 | 0.629013 | 1 | 0.999631 | 0.696662 |
| Bcl9_Q9D219_S686s_1 | 0.87 | 1.96 | 1.66 | 1.36 | 1.51 | 1.41 | 0.96 | 2.16 | 1.48 | 0.86 | 1.85 | 2.37 | 1.28 | 1.65 | 2.58 | 2.34 | 0.0475 | 0.1775 | 0.5 | 0.232882 | 0.895597 | 0.661957 | 1 | 0.999631 | 0.682826 |
| Bcl9l_Q67FY2_S1014s_1 | 1.65 | 1.9 | 1.35 | 0.92 | 1.3 | 1.68 | 0.99 | 1.52 | 1.98 | 3.19 | 1.36 | 1.6 | 0.25 | 2.66 | 1.49 | 1.92 | -0.0825 | 0.5775 | 0.125 | 0.809416 | 0.79174 | 0.218564 | 1 | 0.999631 | 0.936582 |
| Bcl9l_Q67FY2_S116s_1 | 1.94 | 1.85 | 1.74 | 1.35 | 1.3 | 1.26 | 1.73 | 0.94 | 2.58 | 2.18 | 1.75 | 1.94 | 0.99 | 1.81 | 1.2 | 1.65 | -0.4125 | 0.3925 | -0.3075 | 0.344168 | 0.169754 | 0.209105 | 1 | 0.999631 | 0.733869 |
| Bclaf1_F8WI22_S177s_1 | -1.1 | -1.52 | -1.06 | -1.66 | -0.98 | -1.76 | -1.11 | -1.47 | -1.27 | -1.54 | -1.14 | -1.23 | -0.39 | -1.22 | -0.95 | -1.25 | 0.005 | 0.04 | 0.3825 | 0.258183 | 0.986572 | 0.884609 | 1 | 0.999631 | 0.692624 |
| Bclaf1_F8WI22_S221s_1 | -1.45 | -1.39 | -0.23 | -0.45 | -1.22 | -1.79 | -1.12 | -0.14 | -1.34 | -2.33 | -0.4 | -0.32 | -2.95 | -0.33 | -1.36 | 0.79 | -0.1875 | -0.2175 | -0.0825 | 0.912454 | 0.678054 | 0.682769 | 1 | 0.999631 | 0.972191 |
| Bclaf1_F8WI22_S261s_2 | -1.12 | 1.5 | 1.22 | 0.56 | 0.22 | 0.96 | 0.59 | 1.36 | 0.47 | 0.1 | 0.61 | -0.02 | 0.99 | 1.32 | 0.79 | 1.1 | 0.2425 | -0.25 | 0.51 | 0.368575 | 0.676308 | 0.656576 | 1 | 0.999631 | 0.745968 |
| Bclaf1_F8WI22_S261s_3 | -0.82 | 1.79 | 2.67 | 1.56 | 1.16 | 1.14 | 1.6 | 2.76 | 1.36 | -0.28 | 1.52 | 3.91 | 2.7 | 1.56 | 0.99 | 0.48 | 0.365 | 0.3275 | 0.1325 | 0.864149 | 0.622422 | 0.740265 | 1 | 0.999631 | 0.955131 |
| Bclaf1_F8WI22_S263s_3 | 1.48 | 0.56 | 2.37 | 1.2 | 1.05 | 0.32 | 1.51 | 2.22 | 0.07 | 0.49 | 2.36 | 3.28 | 2.29 | 2.47 | 2.32 | 1.31 | -0.1275 | 0.1475 | 0.695 | 0.150575 | 0.803129 | 0.844162 | 1 | 0.999631 | 0.623175 |
| Bclaf1_F8WI22_S267s_1 | -1.8 | -5.4 | -3.97 | -4.19 | -4.08 | -5.83 | -2.98 | -4.47 | -4.89 | -4.13 | -4.14 | -4.67 | -2.62 | -5.71 | -5.26 | -5.42 | -0.5 | -0.6175 | -0.9125 | 0.320578 | 0.550344 | 0.380938 | 1 | 0.999631 | 0.723684 |
| Bclaf1_F8WI22_S267s_2 | -1.22 | 1.11 | 1.29 | 0.62 | 0.5 | 0.53 | 0.71 | 1.41 | 0.8 | -0.33 | 0.74 | -0.01 | 1.33 | 0.74 | 0.96 | 1.07 | 0.3375 | -0.15 | 0.575 | 0.307265 | 0.550542 | 0.797706 | 1 | 0.999631 | 0.717856 |
| Bclaf1_F8WI22_S267s_3 | -0.82 | 1.79 | 2.24 | 1.03 | 1.16 | 1.14 | 1.48 | 1.96 | 1.36 | -0.28 | 2.67 | 2.98 | 2.7 | 1.56 | 2.76 | 1.61 | 0.375 | 0.6225 | 1.0975 | 0.125765 | 0.554708 | 0.482207 | 1 | 0.999631 | 0.596745 |
| Bclaf1_F8WI22_S284s_2 | -0.33 | -0.42 | 0.37 | -0.18 | -0.18 | -0.04 | -0.64 | -0.46 | 0.02 | 0.18 | -0.44 | 0.23 | 0.71 | -0.05 | 0.15 | 0.44 | -0.19 | 0.1375 | 0.4525 | 0.182701 | 0.520416 | 0.654037 | 1 | 0.999631 | 0.643861 |
| Bclaf1_F8WI22_S289s_1 | 0.22 | -0.24 | 0.34 | 0.19 | -0.94 | -0.83 | -0.51 | -0.04 | -1.46 | -1.35 | -0.59 | -0.55 | 0.32 | -0.88 | 0.37 | 0.09 | -0.7075 | -1.115 | -0.1525 | 0.67689 | 0.03827 | 0.006586 | 1 | 0.999631 | 0.886865 |
| Bclaf1_F8WI22_S289s_2 | -1.03 | -1.09 | 0.37 | -0.18 | -0.64 | -0.76 | -0.64 | -0.46 | -0.84 | -0.95 | -0.44 | 0.23 | -0.41 | -0.84 | 0.15 | 0.44 | -0.1425 | -0.0175 | 0.3175 | 0.487619 | 0.704061 | 0.968089 | 1 | 0.999631 | 0.805946 |
| Bclaf1_F8WI22_S395s_1 | -1.49 | -2.17 | 1.57 | -1.46 | -0.71 | -2.39 | 3.25 | 0.4 | -0.72 | -2.01 | 2.19 | 3.73 | -0.29 | -2.52 | 1.43 | 0.22 | 1.025 | 1.685 | 0.5975 | 0.555379 | 0.415606 | 0.224115 | 1 | 0.999631 | 0.838109 |
| Bclaf1_F8WI22_S494s_1 | -0.46 | -0.35 | 0.05 | -0.39 | -1.15 | -0.87 | -0.5 | -0.07 | -1 | -0.3 | -0.28 | -0.58 | -0.72 | -0.88 | 0.12 | -0.02 | -0.36 | -0.2525 | -0.0875 | 0.796978 | 0.267255 | 0.39314 | 1 | 0.999631 | 0.929975 |
| Bclaf1_F8WI22_S510s_1 | -0.89 | -0.09 | 0.19 | -0.44 | -0.22 | -0.15 | -0.51 | -0.05 | -0.1 | -0.31 | -0.1 | 0.07 | 0.92 | 0.61 | 0.36 | 0.54 | 0.075 | 0.1975 | 0.915 | 0.018994 | 0.808424 | 0.53076 | 1 | 0.999631 | 0.391024 |
| Bclaf1_F8WI22_S646s_1 | 0.71 | -0.37 | 0.13 | -0.14 | -0.21 | 0.21 | 0.38 | -0.53 | 0.53 | 1.29 | -0.04 | -0.73 | 0.27 | -0.36 | -0.12 | -0.53 | -0.12 | 0.18 | -0.2675 | 0.449797 | 0.727625 | 0.703254 | 1 | 0.999631 | 0.78723 |
| Bclaf1_F8WI22_S656s_1 | -0.22 | 0.06 | -0.07 | -0.55 | -0.08 | -0.07 | -0.68 | 0 | -0.1 | 0.2 | -0.1 | 0.01 | 0.84 | 0.58 | 0.16 | 0.61 | -0.0125 | 0.1975 | 0.7425 | 0.030425 | 0.964819 | 0.463988 | 1 | 0.999631 | 0.438141 |
| Bclaf1_F8WI22_S759s_1 | -0.04 | -1.3 | 1.25 | -0.01 | -0.57 | -0.85 | 2.23 | -0.11 | -0.27 | -2.28 | 0.39 | 0.87 | -1.42 | -3.36 | 0.83 | -0.95 | 0.2 | -0.2975 | -1.2 | 0.188689 | 0.794203 | 0.698044 | 1 | 0.999631 | 0.648183 |
| Bclaf1_F8WI22_T256t_2 | -1.4 | 0.37 | 1.91 | 1.13 | 0.91 | -0.32 | 1.63 | 1.86 | 1.27 | -1.18 | 1.76 | 0.05 | 1.83 | -0.45 | 2.27 | 0.73 | 0.5175 | -0.0275 | 0.5925 | 0.471801 | 0.49882 | 0.974 | 1 | 0.999631 | 0.798564 |
| Bclaf1_F8WI22_T256t_3 | -4.43 | 2.22 | 2.37 | 1.2 | 1.33 | 1.5 | 1.51 | 2.22 | 1.94 | -0.34 | 2.36 | 3.28 | 3.27 | 1.66 | 2.32 | 1.31 | 1.3 | 1.47 | 1.8 | 0.218667 | 0.3573 | 0.344817 | 1 | 0.999631 | 0.669859 |
| Bclaf1_F8WI22_T305t_1 | -0.03 | -0.55 | -0.66 | -1.56 | -0.5 | -0.34 | -0.34 | -0.71 | -0.91 | 0.16 | -0.81 | -0.8 | 0.52 | 0.45 | 0.3 | -0.4 | 0.2275 | 0.11 | 0.9175 | 0.043282 | 0.526968 | 0.789751 | 1 | 0.999631 | 0.475443 |
| Bclaf1_F8WI22_T339t_1 | -0.53 | 0.57 | 0.79 | 0.49 | -0.75 | 0.56 | -0.36 | -0.03 | 0.85 | 0.77 | 0.29 | 0.29 | 0.4 | -0.16 | -0.26 | 0.05 | -0.475 | 0.22 | -0.3225 | 0.393321 | 0.25882 | 0.547385 | 1 | 0.999631 | 0.759134 |
| Bcor_Q8CGN4_S1391s_1 | 2.55 | 0.64 | 1.68 | 1.43 | 0.05 | 0.96 | 1.78 | 0.52 | 0.45 | 1.92 | 0.18 | -0.54 | 0.15 | -0.48 | 0.72 | -0.22 | -0.7475 | -1.0725 | -1.5325 | 0.007624 | 0.161112 | 0.095534 | 1 | 0.999631 | 0.358173 |
| Bcr_Q6PAJ1_S356s_1 | 0.77 | 0.32 | -0.23 | 0.51 | 0.86 | 0.6 | 0.18 | 1.1 | 0.47 | 0.39 | 0.69 | 0 | -0.14 | 0.91 | 0.54 | 0.58 | 0.3425 | 0.045 | 0.13 | 0.716862 | 0.313697 | 0.887306 | 1 | 0.999631 | 0.903251 |
| Bcr_Q6PAJ1_S384s_1 | -0.68 | -0.35 | -0.45 | -1.2 | -0.94 | -0.7 | -1.43 | -1.73 | -0.91 | -0.39 | -1.99 | -0.76 | 2.1 | -0.13 | 0.23 | -1.53 | -0.53 | -0.3425 | 0.8375 | 0.240119 | 0.139993 | 0.404933 | 1 | 0.999631 | 0.683719 |
| Bcr_Q6PAJ1_S697s_1 | 2.09 | 0.64 | 0.67 | 1.04 | 0.72 | 1.38 | 1.64 | 0.76 | 1.25 | 1.07 | 1 | 0.67 | -1.01 | 0.31 | 0.92 | -0.32 | 0.015 | -0.1125 | -1.135 | 0.046963 | 0.97074 | 0.768463 | 1 | 0.999631 | 0.480102 |
| Begain_F8WIG2_S16s_1 | 3.82 | 0.99 | 0.51 | 1.99 | 1.52 | 1.14 | 1.82 | 0.86 | 1.2 | 0.82 | 0.68 | 0.39 | -1.32 | -0.68 | 0.21 | -0.29 | -0.4925 | -1.055 | -2.3475 | 0.007455 | 0.473046 | 0.140854 | 1 | 0.999631 | 0.358173 |
| Begain_F8WIG2_S265s_1 | -0.63 | -0.61 | -0.59 | -0.95 | -0.99 | -1.02 | -1.18 | -0.8 | -1.16 | -0.87 | -0.96 | -0.52 | -0.46 | -0.73 | -1.16 | -0.53 | -0.3025 | -0.1825 | -0.025 | 0.932113 | 0.236304 | 0.50352 | 1 | 0.999631 | 0.978188 |
| Begain_F8WIG2_S460s_1 | 1.76 | -1.36 | -1.69 | 0.31 | 0.82 | -1.14 | 1.39 | -0.36 | 0.16 | -0.6 | -0.15 | -2.84 | -3.6 | -1.25 | -2.15 | -2.17 | 0.4225 | -0.6125 | -2.0475 | 0.028657 | 0.622481 | 0.506338 | 1 | 0.999631 | 0.427571 |
| Begain_F8WIG2_S482s_1 | -1.56 | 1.06 | -1.03 | -1.13 | 0.84 | 1.54 | -0.92 | -2.14 | 1.52 | -1.72 | -1.5 | -0.89 | 1.51 | -0.62 | -1.14 | -1.6 | 0.495 | 0.0175 | 0.2025 | 0.797561 | 0.578921 | 0.983133 | 1 | 0.999631 | 0.930124 |
| Begain_F8WIG2_S492s_1 | -0.02 | -0.21 | -0.24 | -0.32 | -0.35 | 0.09 | -0.62 | -0.41 | 0.15 | -0.11 | -0.35 | -0.29 | 0.4 | -0.03 | -0.35 | -0.12 | -0.125 | 0.0475 | 0.1725 | 0.555059 | 0.636637 | 0.854095 | 1 | 0.999631 | 0.838109 |
| Begain_F8WIG2_S529s_1 | 0.52 | -0.64 | -0.99 | 0.25 | 0.37 | -0.03 | 0 | 0.18 | -0.05 | -0.45 | 0.34 | -0.71 | -1.1 | -0.32 | 0.03 | -0.26 | 0.345 | -0.0025 | -0.1975 | 0.652788 | 0.377045 | 0.995317 | 1 | 0.999631 | 0.877763 |
| Begain_F8WIG2_S579s_1 | -0.64 | -0.59 | -0.18 | -1.26 | -1.33 | -0.7 | -1.1 | -0.96 | -1.8 | -0.66 | -1.61 | -2.99 | 0.21 | -0.74 | -1.11 | -0.49 | -0.355 | -1.0975 | 0.135 | 0.728502 | 0.271998 | 0.050444 | 1 | 0.999631 | 0.905923 |
| Begain_F8WIG2_S589s_1 | 0.49 | 0.99 | 0.46 | 0.06 | 0.46 | 1.03 | -0.18 | 0.33 | 0.28 | 0.71 | 0.5 | 0.25 | 0.56 | 1.17 | 0.36 | 1.1 | -0.09 | -0.065 | 0.2975 | 0.39183 | 0.794535 | 0.827075 | 1 | 0.999631 | 0.758901 |
| Begain_F8WIG2_Y495y_1 | -0.71 | -0.77 | -1.05 | -0.4 | -0.57 | -1.23 | -0.6 | -0.45 | -0.76 | -0.85 | -0.62 | -0.05 | -1.33 | -0.96 | -0.69 | -0.71 | 0.02 | 0.1625 | -0.19 | 0.534224 | 0.945186 | 0.590875 | 1 | 0.999631 | 0.828493 |
| Begain_F8WIG2_Y614y_1 | -0.2 | -1.55 | -1.25 | -1.11 | 0.59 | -0.05 | 0 | -2.11 | -0.4 | -2.35 | -0.58 | -0.68 | -0.05 | 1.26 | -2.4 | -0.81 | 0.635 | 0.025 | 0.5275 | 0.470977 | 0.300929 | 0.960736 | 1 | 0.999631 | 0.797962 |
| Bet1_O35623_S50s_1 | 0.4 | 0.98 | 0.24 | 0.77 | 1.53 | 0.51 | 0.35 | 0.34 | 1.21 | -0.11 | 0.28 | -0.61 | -1 | 1.48 | -0.28 | 0.87 | 0.085 | -0.405 | -0.33 | 0.548007 | 0.811906 | 0.349916 | 1 | 0.999631 | 0.834507 |
| Bicd1_G3UXK5_S546s_1 | 0.08 | -0.44 | -1.85 | 0.53 | 0.34 | -0.56 | -0.25 | 0.17 | 0.07 | -0.35 | -0.18 | -0.05 | -0.1 | -0.07 | 0.02 | -0.32 | 0.345 | 0.2925 | 0.3025 | 0.548328 | 0.508941 | 0.560275 | 1 | 0.999631 | 0.834633 |
| Bicd2_D3Z390_S504s_1 | -0.2 | 0.07 | 0.2 | -0.01 | -0.11 | 0.14 | -0.19 | 0.17 | 0.24 | 0.38 | 0.33 | 0.61 | 0.61 | 0.75 | 0.61 | 0.72 | -0.0125 | 0.375 | 0.6575 | 0.030219 | 0.959745 | 0.162213 | 1 | 0.999631 | 0.437261 |
| Bin1_Q6P1B9_S245s_1 | -0.2 | 1.51 | 1.12 | -1.11 | 2.01 | 0.37 | 2.21 | -0.17 | 2.5 | -0.64 | 1.52 | 2.88 | 2.97 | -0.22 | -0.08 | -0.27 | 0.775 | 1.235 | 0.27 | 0.754226 | 0.311104 | 0.176244 | 1 | 0.999631 | 0.913377 |
| Bin1_Q6P1B9_S265s_1 | -0.72 | -0.79 | -0.77 | -0.96 | -0.95 | -0.57 | -1.46 | -0.95 | -0.63 | -0.51 | -1.33 | -0.98 | 0.41 | -0.2 | -1.28 | -0.67 | -0.1725 | -0.0525 | 0.375 | 0.349314 | 0.535775 | 0.853548 | 1 | 0.999631 | 0.736792 |
| Bin1_Q6P1B9_S265s_2 | -0.5 | 0.08 | 0.27 | -0.58 | -0.23 | -0.47 | -0.35 | -0.11 | -0.21 | -0.57 | 0 | 0.18 | 0.91 | -0.24 | -0.29 | 0.17 | -0.1075 | 0.0325 | 0.32 | 0.411718 | 0.714738 | 0.919745 | 1 | 0.999631 | 0.767937 |
| Bin1_Q6P1B9_S267s_1 | -0.47 | -0.51 | -1.62 | -1.77 | -0.56 | -0.56 | -1.92 | -1.55 | -0.56 | -0.82 | -1.89 | -1.54 | 0.25 | -0.21 | -1.93 | -1.43 | -0.055 | -0.11 | 0.2625 | 0.646772 | 0.906743 | 0.809302 | 1 | 0.999631 | 0.874255 |
| Bin1_Q6P1B9_S267s_2 | -0.37 | -0.64 | 0.23 | -0.49 | -0.18 | -0.99 | -0.29 | -0.22 | -0.71 | -0.71 | 0.08 | -0.01 | 0.49 | -1.39 | -0.63 | 0.16 | -0.1025 | -0.02 | -0.025 | 0.956393 | 0.748423 | 0.952515 | 1 | 0.999631 | 0.984803 |
| Bin1_Q6P1B9_S273s_1 | 0.01 | 0.97 | 0.28 | -0.48 | 0 | 0.07 | -0.79 | -0.81 | -0.18 | 0.6 | -0.01 | 0.03 | 1.35 | 0.87 | -0.07 | 0.23 | -0.5775 | -0.085 | 0.4 | 0.375792 | 0.165513 | 0.819753 | 1 | 0.999631 | 0.748307 |
| Bin1_Q6P1B9_S273s_2 | -0.47 | 0.41 | 0.32 | -0.71 | -0.14 | -0.1 | -0.43 | 0.03 | -0.13 | -0.37 | -0.11 | 0.4 | 0.92 | 0.18 | 0.1 | 0.19 | -0.0475 | 0.06 | 0.46 | 0.241063 | 0.887239 | 0.866988 | 1 | 0.999631 | 0.684338 |
| Bin1_Q6P1B9_S293s_1 | -0.8 | -0.87 | -1.09 | -1.07 | -0.64 | -0.82 | -0.97 | -2.7 | -0.94 | -1.02 | -0.85 | -0.88 | -0.74 | -1.25 | -1.2 | -0.95 | -0.325 | 0.035 | -0.0775 | 0.781031 | 0.487855 | 0.886083 | 1 | 0.999631 | 0.921543 |
| Bin2_S4R2J8_S393s_1 | 0.7 | 0.84 | 1.26 | 0.77 | -0.29 | 0.7 | 0.41 | 1.36 | 0.27 | 1.39 | 0.59 | 1.49 | 0.16 | 1.69 | 1.75 | 1.63 | -0.3475 | 0.0425 | 0.415 | 0.33307 | 0.369461 | 0.905663 | 1 | 0.999631 | 0.728079 |
| Bin2_S4R2J8_S415s_1 | 0.01 | 0.22 | 0.24 | -1.28 | 0.16 | -0.01 | -0.54 | -0.08 | 0.79 | -0.91 | 0.27 | -4.49 | 0.39 | 0.01 | -1.96 | -1.61 | 0.085 | -0.8825 | -0.59 | 0.35356 | 0.830867 | 0.417992 | 1 | 0.999631 | 0.73949 |
| Bin2_S4R2J8_S415s_2 | 0.48 | 0.25 | -0.07 | 0.9 | -0.72 | -0.55 | -1.12 | -0.35 | 0.33 | 0.99 | 0.32 | -0.01 | 0.01 | 0.47 | -2.1 | 0.81 | -1.075 | 0.0175 | -0.5925 | 0.350543 | 0.006345 | 0.958775 | 1 | 0.999631 | 0.737333 |
| Bin2_S4R2J8_S428s_1 | 1.06 | 1.41 | 0.65 | 0.69 | 0.39 | 0.97 | 0.63 | 0.42 | 1.17 | 1.78 | 1 | 0.45 | 2.15 | 1.34 | 1.19 | 0.89 | -0.35 | 0.1475 | 0.44 | 0.248495 | 0.248966 | 0.683179 | 1 | 0.999631 | 0.685751 |
| Bin2_S4R2J8_S448s_1 | 1.25 | 0.49 | 1.96 | 1.44 | 1.03 | 1.27 | 1.27 | 1.13 | 1.05 | 1.44 | 1.05 | 0.86 | -1.38 | 0.84 | 1.9 | 0.46 | -0.11 | -0.185 | -0.83 | 0.232586 | 0.749148 | 0.610489 | 1 | 0.999631 | 0.682545 |
| Bin2_S4R2J8_T392t_1 | 1.05 | 1.01 | 1.29 | 1.1 | -0.16 | 0.79 | 0.52 | 1.5 | -0.05 | 1.65 | 0.2 | 1.68 | -0.31 | 1.77 | 1.88 | 1.72 | -0.45 | -0.2425 | 0.1525 | 0.763204 | 0.240026 | 0.596887 | 1 | 0.999631 | 0.915608 |
| Birc6_S4R1L5_S2216s_1 | 1.61 | 0.05 | 1.28 | 1.06 | -0.87 | 0.67 | 1.97 | -0.14 | -0.11 | 2.18 | 0.29 | -0.05 | -0.13 | -0.8 | 0.3 | 0.34 | -0.5925 | -0.4225 | -1.0725 | 0.030989 | 0.354156 | 0.475225 | 1 | 0.999631 | 0.44065 |
| Birc6_S4R1L5_S552s_2 | -0.2 | 0.47 | 0.14 | 0.01 | 0.37 | 0 | 0.26 | 0.11 | 0.67 | 0.83 | 0.88 | 0.79 | 0.17 | 0.55 | -0.67 | 0.32 | 0.08 | 0.6875 | -0.0125 | 0.971852 | 0.761586 | 0.025457 | 1 | 0.999631 | 0.989788 |
| Birc6_S4R1L5_S556s_1 | 0.21 | -1 | 0.13 | -0.36 | -1.41 | 0.54 | 0.1 | -0.26 | -0.75 | 0.4 | -1.15 | -1.18 | -0.27 | -0.57 | -1.1 | 0.3 | -0.0025 | -0.415 | -0.155 | 0.711673 | 0.99581 | 0.370464 | 1 | 0.999631 | 0.900631 |
| Birc6_S4R1L5_S556s_2 | -0.2 | 0.47 | 0.14 | 0.01 | 0.37 | 0 | 0.26 | 0.11 | 0.67 | 0.83 | 0.88 | 0.79 | 0.17 | 0.55 | -0.67 | 0.32 | 0.08 | 0.6875 | -0.0125 | 0.971852 | 0.761586 | 0.025457 | 1 | 0.999631 | 0.989788 |
| Bles03_Q8VD62_S59s_1 | -0.72 | 2.37 | -0.61 | 0.5 | 0.37 | 0.73 | -0.23 | -1.12 | 1.6 | 1.34 | -0.35 | -1.22 | 0.81 | 1.89 | -0.24 | -0.03 | -0.4475 | -0.0425 | 0.2225 | 0.769597 | 0.541227 | 0.960549 | 1 | 0.999631 | 0.918554 |
| Bloc1s4_Q8VED2_S11s_1 | -0.03 | -1.17 | -0.67 | -0.79 | 0.32 | -0.05 | -0.26 | 0.4 | -0.59 | 1.56 | -0.78 | -0.6 | 0.14 | 1.38 | -1.94 | -0.17 | 0.7675 | 0.5625 | 0.5175 | 0.433753 | 0.038572 | 0.326805 | 1 | 0.999631 | 0.779567 |
| Bmp2k_Q91Z96_S1087s_1 | 0.04 | 0.16 | -0.41 | -0.68 | 0.45 | 1.05 | -0.05 | -0.09 | 0.32 | -0.75 | -0.37 | -0.04 | 0.43 | 1 | -0.15 | 0.02 | 0.5625 | 0.0125 | 0.5475 | 0.158264 | 0.138539 | 0.970983 | 1 | 0.999631 | 0.628615 |
| Bmp2k_Q91Z96_T644t_1 | 0.37 | -1.29 | 0.78 | 0.62 | -1.2 | -1.4 | 0.96 | 0.5 | -0.73 | -0.48 | 1.32 | 0.3 | -1.13 | -1.54 | 0.23 | 0.89 | -0.405 | -0.0175 | -0.5075 | 0.451851 | 0.553668 | 0.97687 | 1 | 0.999631 | 0.788148 |
| Bmpr2_O35607_S515s_1 | -0.51 | -0.64 | -0.55 | -0.74 | 0.12 | -0.3 | -0.76 | -0.73 | -0.31 | -0.27 | -0.63 | -0.44 | -0.14 | -0.09 | -0.44 | -0.6 | 0.1925 | 0.1975 | 0.2925 | 0.303755 | 0.508442 | 0.433598 | 1 | 0.999631 | 0.715879 |
| Bmpr2_O35607_S586s_1 | 0.72 | -0.59 | -3.01 | 1.12 | 1.77 | 0.1 | 1.51 | 0.63 | 0.68 | -0.66 | 0.25 | 0.7 | -1.34 | -0.37 | -0.48 | 0.03 | 1.4425 | 0.6825 | -0.1 | 0.905738 | 0.123775 | 0.434456 | 1 | 0.999631 | 0.969317 |
| Bmpr2_O35607_S680s_1 | -3.79 | -0.43 | -0.14 | 0.52 | 0.25 | -0.19 | 0.36 | 0.69 | 0.11 | -0.27 | 1.49 | 0.68 | 0.93 | -0.16 | 0.48 | 0.45 | 1.2375 | 1.4625 | 1.385 | 0.129819 | 0.16925 | 0.129381 | 1 | 0.999631 | 0.601246 |
| Bmpr2_O35607_S862s_1 | -0.98 | -0.97 | -0.87 | -1.34 | -1.27 | -0.56 | -1.49 | -1.71 | -1.47 | -1.05 | -0.93 | -0.63 | 0.18 | -1.03 | -1.7 | -0.67 | -0.2175 | 0.02 | 0.235 | 0.577795 | 0.500166 | 0.944828 | 1 | 0.999631 | 0.84622 |
| Bmpr2_O35607_S863s_1 | -0.81 | -0.89 | -1.07 | -1.49 | -1.04 | -1.13 | -1.27 | -1.45 | -0.99 | -0.7 | -0.99 | -0.47 | -0.27 | -0.73 | -1.54 | -0.75 | -0.1575 | 0.2775 | 0.2425 | 0.501477 | 0.562107 | 0.345257 | 1 | 0.999631 | 0.814092 |
| Bmpr2_O35607_T517t_1 | -0.7 | -0.51 | -0.59 | -0.28 | -0.6 | -1.03 | -0.73 | -0.01 | -0.81 | -0.54 | -0.48 | 0.51 | -0.25 | -0.59 | -0.24 | -0.38 | -0.0725 | 0.19 | 0.155 | 0.572922 | 0.807428 | 0.584577 | 1 | 0.999631 | 0.844402 |

| | | | | | | | | | | | | | | | | | | | | | | | | |
|---|---|---|---|---|---|---|---|---|---|---|---|---|---|---|---|---|---|---|---|---|---|---|---|---|
| Bnip3_O55003_S79s_1 | 0.25 | 0.2 | -0.21 | -0.26 | -0.12 | 0.14 | -0.57 | -0.54 | -0.04 | 0.25 | -0.26 | -0.09 | 0.1 | 0.54 | 0.17 | -0.29 | -0.2675 | -0.03 | 0.135 | 0.664713 | 0.366499 | 0.912949 | 1 | 0.999631 | 0.882317 |
| Bnip3_O55003_S88s_1 | -1.54 | -0.09 | -0.06 | -0.52 | -0.51 | -0.22 | -0.73 | -0.65 | -0.29 | -0.4 | -0.44 | 0.12 | 0.47 | 0.15 | -0.09 | 0.29 | 0.025 | 0.3 | 0.7575 | 0.077904 | 0.947206 | 0.445663 | 1 | 0.999631 | 0.543121 |
| Bnip3_O55003_T66t_1 | 2.62 | 0.46 | 0.56 | 0.68 | -0.54 | 0.03 | 2.06 | -0.05 | -0.35 | 0.84 | -0.41 | -0.2 | -2.18 | -1.43 | -0.89 | -0.93 | -0.705 | -1.11 | -2.4375 | 0.001335 | 0.317234 | 0.067414 | 1 | 0.999631 | 0.351308 |
| Bnip3l_Q91Z78_S165s_1 | -0.45 | 1.03 | 0.46 | 0.45 | 0.55 | 1.39 | 0.12 | 0.88 | 1.31 | 1.09 | 0.77 | 0.8 | 0.94 | 1.35 | 1.05 | 0.69 | 0.3625 | 0.62 | 0.635 | 0.114253 | 0.383989 | 0.112489 | 1 | 0.999631 | 0.579001 |
| Bnip3l_Q91Z78_S167s_1 | 0.02 | 2.64 | 2.35 | 1.2 | 0.99 | 2.37 | 0.35 | 2.3 | 2.59 | 2.5 | 1.44 | 1.87 | 2.95 | 2.73 | 1.74 | 1.81 | -0.05 | 0.5475 | 0.755 | 0.235448 | 0.942175 | 0.371287 | 1 | 0.999631 | 0.683301 |
| Bod1l_E9Q6J5_S1989s_1 | -0.16 | 0.66 | 0.49 | -0.15 | 0.38 | -0.68 | -0.42 | -0.56 | 0.42 | -0.81 | 0.23 | 0.08 | 0.12 | 0.79 | 0.87 | -1.22 | -0.53 | -0.23 | -0.07 | 0.889616 | 0.153328 | 0.540863 | 1 | 0.999631 | 0.963505 |
| Bod1l_E9Q6J5_S2898s_1 | 0.77 | 0.43 | 1.4 | -0.2 | 1.04 | 0.39 | 0.54 | 0.69 | 2.02 | 1.5 | 0.62 | 1.36 | 1.73 | 1.54 | 1.66 | 0.56 | 0.065 | 0.775 | 0.7725 | 0.101303 | 0.863043 | 0.101741 | 1 | 0.999631 | 0.564448 |
| Bod1l_E9Q6J5_T2890t_1 | 0.33 | 3.17 | 1.67 | 1.43 | 0.99 | 2.43 | 0.53 | 1.88 | 1.65 | 2.41 | 2.54 | 1.66 | 2.14 | 3.22 | 0.86 | 2.81 | -0.1925 | 0.415 | 0.6075 | 0.389985 | 0.766617 | 0.479494 | 1 | 0.999631 | 0.758901 |
| Bola1_Q9D8S9_S81s_1 | -0.2 | 0.1 | 1.2 | 0.83 | 0.64 | 0.91 | -0.16 | 0.34 | 0.97 | 1.12 | 0.64 | 1.1 | -0.58 | 2.32 | 1.38 | 1.39 | -0.05 | 0.475 | 0.645 | 0.314841 | 0.900465 | 0.217854 | 1 | 0.999631 | 0.720305 |
| Bop1_P97452_S559s_1 | 0.24 | -0.68 | -0.13 | -0.96 | -0.73 | -3.01 | -0.78 | -0.63 | -0.03 | -1.6 | 0.05 | 1.11 | -0.08 | -0.95 | -0.86 | -1.05 | -0.905 | 0.265 | -0.3525 | 0.368416 | 0.141208 | 0.643918 | 1 | 0.999631 | 0.745968 |
| Bptf_E9Q6A7_S228s_1 | 0.11 | 0.33 | 0.47 | -0.07 | 0.08 | 0.1 | -0.16 | -0.02 | 0.02 | 0.55 | 0.61 | 0.68 | 0.55 | 0.72 | 0.16 | 0.63 | -0.21 | 0.255 | 0.305 | 0.307626 | 0.41252 | 0.380056 | 1 | 0.999631 | 0.717856 |
| Bptf_E9Q6A7_S2353s_1 | 0.49 | 0.4 | 0.97 | 0.32 | 0.23 | 0.19 | -0.24 | 0.41 | 0.17 | 1.02 | 0.41 | 0.83 | 0.55 | 1.03 | 0.63 | 0.63 | -0.3975 | 0.0625 | 0.165 | 0.578562 | 0.179143 | 0.840277 | 1 | 0.999631 | 0.84622 |
| Braf_P28028_S348s_1 | 0.29 | 0.74 | 1.34 | 0.92 | 0.67 | 0.82 | 0.22 | 1.16 | 0.41 | 0.74 | 0.53 | 1.29 | 0.77 | 0.71 | 1.23 | 1.38 | -0.105 | -0.08 | 0.2 | 0.559236 | 0.752361 | 0.813314 | 1 | 0.999631 | 0.839222 |
| Braf_P28028_S714s_1 | 0 | -0.12 | 0.6 | 0.06 | -1.06 | -0.16 | -0.2 | 0.2 | -0.72 | 0.06 | 0.11 | -0.06 | -0.48 | -0.32 | 0.39 | 0.36 | -0.44 | -0.2875 | -0.1475 | 0.666784 | 0.220364 | 0.370818 | 1 | 0.999631 | 0.88264 |
| Braf_P28028_S735s_1 | -0.21 | 0.59 | 0.24 | -0.31 | -0.16 | 0.24 | -0.72 | 0.01 | 0.29 | -0.37 | -0.31 | -0.11 | 0.23 | -0.21 | 0.12 | 0.36 | -0.235 | -0.2025 | 0.0475 | 0.882886 | 0.485039 | 0.528293 | 1 | 0.999631 | 0.963505 |
| Braf_P28028_T384t_1 | -0.02 | 0.29 | 0.21 | 0.68 | -0.21 | -0.11 | 0.12 | 0.81 | 0.23 | -0.05 | 0.04 | 0.36 | 0.25 | 0.31 | 0.54 | 0.62 | -0.1375 | -0.145 | 0.14 | 0.631475 | 0.668862 | 0.601126 | 1 | 0.999631 | 0.867623 |
| Braf_P28028_T790t_1 | 1.07 | 1.34 | -0.2 | 1.77 | 1.64 | 1.53 | 1.3 | 0.45 | 1.09 | 0.43 | 1.29 | 0.58 | -1.66 | 1.4 | 0.56 | 1.61 | 0.235 | -0.1475 | -0.5175 | 0.497863 | 0.624092 | 0.748076 | 1 | 0.999631 | 0.811908 |
| Brap_Q99MP8_S86s_1 | 0.03 | 1.49 | 1.59 | 1.4 | 0.9 | 0.59 | 0.87 | 0.26 | 2.01 | 0.57 | 0.68 | 2.83 | 1.35 | 0.61 | 1.36 | 1.53 | -0.4725 | 0.395 | 0.085 | 0.843225 | 0.255844 | 0.514575 | 1 | 0.999631 | 0.94821 |
| Brap_Q99MP8_S86s_2 | 0.05 | 0.69 | 0.6 | 1.52 | 0.94 | 0.61 | 1.07 | 0.79 | 1.04 | 0.59 | 2.36 | 1.97 | 1.07 | 0.47 | 0.73 | 1.58 | 0.1375 | 0.775 | 0.2475 | 0.546847 | 0.694603 | 0.135412 | 1 | 0.999631 | 0.834213 |
| Brd1_G5E8P1_T885t_1 | 0.01 | 2.73 | 3.45 | -0.1 | 2.97 | 0.56 | 5.6 | 0.96 | 5.58 | 0.7 | 4.05 | 5.19 | 5.11 | -1.35 | 2.49 | -0.79 | 1 | 2.3575 | -0.1575 | 0.915325 | 0.433535 | 0.080115 | 1 | 0.999631 | 0.97286 |
| Brd3_Q8K2F0_S262s_1 | -0.4 | 0.2 | 0.58 | 0.13 | 0.07 | -0.3 | -0.17 | -0.08 | -0.27 | 0.08 | 0.02 | 0.15 | 0.49 | 0.23 | -0.2 | 0.59 | -0.2475 | -0.1325 | 0.15 | 0.65707 | 0.400826 | 0.659219 | 1 | 0.999631 | 0.879129 |
| Brd4_Q3UH70_S603s_1 | -0.69 | 0.41 | 1.12 | 0.62 | 0.43 | 0.35 | -0.05 | 0.67 | 0.2 | 0.31 | 0.81 | 0.93 | 0.58 | 1.01 | 0.67 | 0.81 | -0.015 | 0.1975 | 0.4025 | 0.339731 | 0.970799 | 0.643237 | 1 | 0.999631 | 0.732291 |
| Brd8_D3YZC7_S547s_1 | 1.11 | 0.7 | 1.78 | 1.74 | 0.31 | 1.01 | 2.26 | 0.38 | 0.59 | 1.31 | 0.93 | 0.77 | 0.51 | -0.03 | 0.86 | -1.17 | -0.3425 | -0.4325 | -1.29 | 0.024514 | 0.491142 | 0.22885 | 1 | 0.999631 | 0.417261 |
| Brd8_D3YZC7_T264t_1 | 0.48 | 2.03 | 1.79 | 0.95 | 1.2 | 1.65 | -0.03 | 0.3 | 1.5 | 0.7 | 1.44 | 1.33 | 2.15 | 2.31 | 1.64 | 1.54 | -0.5325 | -0.07 | 0.5975 | 0.17718 | 0.300049 | 0.865206 | 1 | 0.999631 | 0.642116 |
| Brd9_Q3UQU0_S565s_1 | 2.06 | 0.83 | 1.8 | 0.33 | 0.17 | 0.91 | 1.26 | 1.08 | 0.57 | 1.64 | 0.33 | 0.57 | -0.72 | 0.03 | 0.22 | 0.12 | -0.4 | -0.4775 | -1.3425 | 0.013782 | 0.388387 | 0.333353 | 1 | 0.999631 | 0.385092 |
| Brf1_G3X8S2_S358s_1 | 1.06 | 2.49 | 2.96 | 1.75 | 1.48 | 2.27 | 3.14 | 1.27 | 1.93 | 0.88 | 1.93 | 1.65 | 1.39 | 0.55 | 1.73 | 0.54 | -0.025 | -0.4675 | -1.0125 | 0.063738 | 0.963536 | 0.332191 | 1 | 0.999631 | 0.513532 |
| Brf1_G3X8S2_S552s_1 | 1.14 | 1.34 | 0.48 | 1.16 | 1.02 | 0.17 | 0.18 | 1.45 | 1.38 | 0.91 | 1.1 | 1.7 | 1.19 | 1.25 | 1 | 0.56 | -0.325 | 0.2425 | -0.03 | 0.926232 | 0.404477 | 0.45164 | 1 | 0.999631 | 0.97588 |
| Brsk1_Q5RJI5_S384s_1 | 1.7 | 0.27 | -0.08 | 1.42 | 1.48 | 1.26 | 1.49 | 1.57 | 0.39 | 0.15 | 0.89 | -0.37 | -1.48 | 1.21 | 0.14 | 0.47 | 0.6225 | -0.5625 | -0.7425 | 0.263867 | 0.170126 | 0.262448 | 1 | 0.999631 | 0.693392 |
| Brsk1_Q5RJI5_S399s_1 | -0.21 | -0.33 | -0.92 | -0.1 | 0.03 | -0.13 | -0.25 | 0.05 | -0.11 | -0.3 | -0.54 | -0.24 | 0.41 | -0.16 | -0.54 | -0.16 | 0.315 | 0.0925 | 0.2775 | 0.417483 | 0.274555 | 0.750203 | 1 | 0.999631 | 0.771469 |
| Brsk1_Q5RJI5_S427s_1 | 1.26 | -0.51 | -0.45 | 0.03 | 0.61 | 0.75 | 0.65 | 0.48 | 0.32 | -0.01 | -0.01 | -0.47 | -0.08 | 0.06 | 0.78 | -0.85 | 0.54 | -0.125 | -0.105 | 0.835375 | 0.211387 | 0.775524 | 1 | 0.999631 | 0.944699 |
| Brsk1_Q5RJI5_S432s_1 | 0.86 | -0.32 | 0.47 | -0.06 | 0.49 | 0.53 | 0.41 | 0.75 | 0.49 | -0.76 | 0.97 | 1.08 | 0.49 | 1.03 | 0.64 | 0.87 | 0.3075 | 0.2075 | 0.52 | 0.158457 | 0.350608 | 0.665879 | 1 | 0.999631 | 0.628615 |
| Brsk1_Q5RJI5_S450s_1 | 0.43 | 0.12 | -0.82 | 0.15 | 0.99 | 0.39 | -0.08 | 0.27 | 0.25 | -0.26 | -0.11 | -0.32 | -1.14 | 0.6 | 0.11 | -0.1 | 0.4225 | -0.08 | -0.1025 | 0.821394 | 0.270237 | 0.816196 | 1 | 0.999631 | 0.938464 |
| Brsk1_Q5RJI5_S499s_1 | 0.92 | 2.3 | 1.78 | 0.57 | 0.76 | 1.48 | -0.02 | 1.13 | 1.41 | 2.01 | 1.04 | 1.02 | 0.66 | 2.53 | 1.2 | 0.85 | -0.555 | -0.0225 | -0.0825 | 0.878615 | 0.266368 | 0.960033 | 1 | 0.999631 | 0.96244 |
| Brsk1_Q5RJI5_S508s_1 | 0.74 | 0.37 | -0.61 | 0.75 | 1.28 | 1.08 | 0.45 | 1.16 | 0.83 | 0.33 | 0.92 | 0.01 | -1.06 | 1.16 | 0.65 | 0.94 | 0.68 | 0.21 | 0.11 | 0.842919 | 0.100106 | 0.600658 | 1 | 0.999631 | 0.94811 |
| Brsk1_Q5RJI5_S519s_1 | -0.73 | -0.69 | -1.46 | -0.35 | 0.09 | 0.4 | -0.19 | 0.37 | -0.06 | -0.28 | -0.41 | -0.72 | -0.23 | 0.59 | -0.4 | -0.32 | 0.975 | 0.44 | 0.7175 | 0.076254 | 0.011779 | 0.198394 | 1 | 0.999631 | 0.542251 |
| Brsk1_Q5RJI5_S519s_2 | -0.25 | -0.06 | -1.11 | 0.58 | 0.98 | 0.9 | 0.52 | 0.53 | 0.23 | -0.17 | 1.69 | 0.56 | 0.01 | -0.34 | 1.33 | 0.87 | 0.9425 | 0.7875 | 0.6775 | 0.193851 | 0.031422 | 0.141217 | 1 | 0.999631 | 0.650692 |
| Brsk1_Q5RJI5_S527s_1 | -0.57 | -0.65 | 0.2 | -0.57 | -0.8 | -0.33 | -0.82 | -0.21 | -0.84 | -0.5 | -0.22 | 0.2 | -0.07 | 0.52 | 0.34 | 0.56 | -0.1425 | 0.0575 | 0.735 | 0.044089 | 0.647486 | 0.866407 | 1 | 0.999631 | 0.475443 |
| Brsk1_Q5RJI5_S751s_1 | 1.04 | 0.58 | 0.11 | 1.16 | 0.64 | 1.19 | 0.91 | 1.33 | 1.04 | 0.7 | 1.03 | 0.15 | -5.05 | 1.18 | 0.81 | 1.17 | 0.295 | 0.0075 | -1.195 | 0.370105 | 0.377764 | 0.983096 | 1 | 0.999631 | 0.746338 |
| Brsk1_Q5RJI5_T535t_1 | -0.53 | -0.05 | -0.28 | 0.1 | 0.21 | -0.19 | -0.21 | 0.44 | -0.2 | -0.52 | 0.34 | -0.06 | 0.05 | -0.2 | 0.31 | 0.76 | 0.2525 | 0.08 | 0.42 | 0.215874 | 0.386778 | 0.790515 | 1 | 0.999631 | 0.668602 |
| Brsk1_Q5RJI5_T739t_1 | 1.9 | -1.31 | 1.5 | 1.21 | 0.24 | 0.57 | 1.26 | 0.42 | -0.39 | 1.33 | -0.76 | 0.05 | -1.13 | 0.24 | 0.85 | -0.01 | -0.2025 | -0.7675 | -0.8375 | 0.270928 | 0.7644 | 0.32342 | 1 | 0.999631 | 0.698674 |
| Brsk2_G3UWV4_S368s_1 | 1.16 | 0.43 | 0.31 | 1.26 | 1.31 | 1.65 | 1.24 | 1.57 | 0.87 | 0.55 | 1.54 | 0.55 | -1.25 | 1.42 | 0.81 | 1.29 | 0.6525 | 0.0875 | -0.2225 | 0.714304 | 0.061725 | 0.811798 | 1 | 0.999631 | 0.901694 |
| Brsk2_G3UWV4_S383s_1 | -0.25 | 0.02 | -0.17 | 0.51 | 0.41 | -0.58 | 0.18 | 0.77 | 0.19 | -0.76 | 0.39 | 0.07 | -0.3 | 0.11 | -0.41 | 0.39 | 0.1675 | -0.055 | -0.08 | 0.80729 | 0.641253 | 0.873844 | 1 | 0.999631 | 0.935604 |
| Brsk2_G3UWV4_S394s_1 | 0.17 | -0.12 | -0.77 | -0.4 | -0.23 | -0.52 | -0.88 | -0.23 | -0.52 | -0.54 | -0.26 | -0.07 | -0.23 | 0.39 | 0.37 | 0.27 | -0.185 | -0.0675 | 0.48 | 0.162849 | 0.553072 | 0.823957 | 1 | 0.999631 | 0.631442 |
| Brsk2_G3UWV4_S417s_1 | -0.44 | 1.39 | 0.56 | 0.45 | 1.19 | 0.66 | 0.31 | 0.22 | 0.63 | -0.26 | 0.51 | 1.1 | 0.66 | 0.97 | 0.53 | 0.59 | 0.105 | 0.005 | 0.1975 | 0.629579 | 0.805967 | 0.991265 | 1 | 0.999631 | 0.866951 |
| Brsk2_G3UWV4_S458s_1 | 1.47 | -0.08 | -0.38 | 0.58 | 1.2 | 0.72 | 1.09 | 0.85 | 0.69 | 0.56 | 0.94 | -0.07 | -1.78 | 0.72 | 0.38 | 0.09 | 0.5675 | 0.1325 | -0.545 | 0.393635 | 0.197745 | 0.770225 | 1 | 0.999631 | 0.759138 |
| Brsk2_G3UWV4_S462s_1 | 1.08 | 0.64 | -0.67 | 0.53 | 1.52 | 1.02 | 0.16 | 0.02 | 1.16 | 0.53 | 0.25 | -0.07 | -1.17 | 1.31 | 0.05 | 0.32 | 0.285 | 0.0725 | -0.2675 | 0.646847 | 0.562526 | 0.871537 | 1 | 0.999631 | 0.874255 |
| Brsk2_G3UWV4_S478s_1 | -0.59 | 0.06 | -0.6 | -0.8 | -0.59 | -0.84 | -0.4 | 0.34 | -1.28 | -0.7 | -0.04 | 0.37 | 0.13 | -0.39 | 0.28 | -0.17 | 0.11 | 0.07 | 0.445 | 0.187519 | 0.751529 | 0.86594 | 1 | 0.999631 | 0.646474 |
| Brsk2_G3UWV4_S512s_1 | 0.1 | 0.82 | 0.35 | 0.62 | 1.87 | 2.09 | 0.64 | 1.63 | 1.29 | 0.19 | 1.19 | 1.06 | -0.34 | 2.02 | 1.69 | 0.97 | 1.085 | 0.46 | 0.6125 | 0.253482 | 0.014997 | 0.198864 | 1 | 0.999631 | 0.688735 |
| Brsk2_G3UWV4_S536s_1 | -0.91 | -0.07 | 0.11 | -0.41 | 0.21 | -0.12 | -0.79 | -0.62 | 0.03 | -0.49 | -0.72 | -0.35 | 0.69 | 0.02 | -0.1 | 0.06 | -0.01 | -0.0625 | 0.4875 | 0.180426 | 0.977192 | 0.84881 | 1 | 0.999631 | 0.64376 |
| Brsk2_G3UWV4_T454t_1 | 1.91 | 0.87 | 1.56 | 1.85 | 0.96 | 2.08 | 1.83 | 1.54 | 0.51 | 1.55 | 1.11 | 0.25 | -1.63 | 1.29 | 0.89 | 1.25 | 0.055 | -0.6925 | -1.0975 | 0.120958 | 0.879131 | 0.103422 | 1 | 0.999631 | 0.588397 |
| Brsk2_G3UWV4_T466t_1 | 0.74 | 1.24 | 1.31 | 1.11 | 0.17 | 2.33 | 0.77 | 1.78 | 1.28 | 0.37 | 1.29 | 0.4 | -0.45 | 0.75 | -0.02 | 1.87 | 0.1625 | -0.265 | -0.5625 | 0.277717 | 0.734181 | 0.439017 | 1 | 0.999631 | 0.700874 |
| Brsk2_G3UWV4_T470t_1 | 1.89 | 1.34 | 0.7 | 1.96 | 1.19 | 1.79 | 2.3 | 1.6 | 1.2 | 2.15 | 1.09 | -0.29 | -0.48 | 1.12 | 0.39 | 0.82 | 0.2475 | -0.435 | -1.01 | 0.046494 | 0.523629 | 0.429174 | 1 | 0.999631 | 0.479949 |
| Brsk2_G3UWV4_T486t_1 | -0.43 | 0.3 | 0.23 | -0.84 | -0.48 | -0.1 | -1.11 | -0.17 | -0.07 | -0.29 | -0.29 | 0.29 | 0.87 | 0.35 | -0.12 | -0.05 | -0.28 | 0.095 | 0.4475 | 0.264294 | 0.460647 | 0.784433 | 1 | 0.999631 | 0.6935 |
| Brwd1_Q921C3_S1751s_1 | -0.33 | -0.43 | -0.87 | -0.74 | -1.08 | -0.52 | -0.71 | -1.33 | -0.35 | -0.34 | -0.96 | -1.44 | -1.17 | -1.07 | -0.65 | -1.37 | -0.3175 | -0.18 | -0.4725 | 0.14064 | 0.291893 | 0.599107 | 1 | 0.999631 | 0.612258 |
| Brwd1_Q921C3_S1830s_1 | -0.45 | 1.44 | -2.42 | -2.06 | -0.48 | -0.55 | -2.48 | -2.7 | 0.15 | 0.15 | -3.03 | -1.64 | 1.44 | 0.25 | -3.95 | -3.22 | -0.68 | -0.22 | -0.4975 | 0.709347 | 0.467302 | 0.827821 | 1 | 0.999631 | 0.899175 |
| Brwd1_Q921C3_S1857s_1 | -1.02 | -2.15 | -0.37 | -0.55 | -2.08 | -2.35 | -0.77 | -0.96 | -1.94 | -0.83 | -0.77 | 0.86 | -0.71 | -1.75 | 1.15 | 0.55 | -0.5175 | 0.3525 | 0.8325 | 0.23812 | 0.334183 | 0.580969 | 1 | 0.999631 | 0.683301 |
| Brwd1_Q921C3_S2035s_1 | -1.53 | -1.79 | 0 | 0.44 | -2.14 | 0.39 | 0.59 | -0.53 | -1.77 | -1.48 | 0.15 | 0.39 | -1.68 | -1.74 | 0.28 | -0.91 | 0.2975 | 0.0425 | -0.2925 | 0.654777 | 0.685851 | 0.951283 | 1 | 0.999631 | 0.878941 |
| Bscl2_Q9Z2E9_S280s_1 | 0.41 | 2.88 | 5.49 | 1.36 | 3 | 3.22 | 5.49 | 2.88 | 3.72 | 1.06 | 5.85 | 6.79 | 4.98 | 0.17 | 4.29 | 0.6 | 1.1125 | 1.82 | -0.025 | 0.985718 | 0.319788 | 0.224544 | 1 | 0.999631 | 0.995133 |
| Bsg_K3W4Q8_S201s_1 | -0.44 | 1.72 | 1.42 | 1.38 | -0.78 | 2.05 | 2.49 | -0.17 | 0.29 | 0.92 | -0.18 | 1.18 | 0.51 | -0.1 | 1.19 | 0.28 | -0.1225 | -0.4675 | -0.55 | 0.311934 | 0.881736 | 0.396347 | 1 | 0.999631 | 0.719361 |
| Bsg_K3W4Q8_T203t_1 | 0.61 | -4.96 | 0.48 | -0.45 | -4.6 | -6.52 | -0.46 | 0.15 | -7.15 | -2.07 | -0.11 | 0.27 | -6.51 | -7.13 | 0.89 | 0.69 | -1.7775 | -1.185 | -1.935 | 0.375244 | 0.325502 | 0.520653 | 1 | 0.999631 | 0.748001 |
| Bsn_O88737_S105s_1 | -0.54 | -0.99 | 0.23 | -0.05 | -1.14 | -0.7 | 0.53 | -0.08 | -1.23 | -1.72 | 0.32 | -0.6 | -1.29 | -1.98 | -0.42 | -1.3 | -0.01 | -0.47 | -0.91 | 0.055474 | 0.981888 | 0.351164 | 1 | 0.999631 | 0.494338 |
| Bsn_O88737_S1100s_1 | -1.04 | -0.45 | -0.99 | -2.52 | -1.56 | -0.55 | -2.21 | -1.8 | -1.42 | -0.92 | -1.78 | -1.55 | 0.53 | -0.97 | -1.19 | -1.31 | -0.28 | -0.1675 | 0.515 | 0.375323 | 0.596823 | 0.719519 | 1 | 0.999631 | 0.748013 |
| Bsn_O88737_S1108s_1 | -0.76 | 0.03 | -0.8 | -2.05 | -0.15 | 0.09 | -1.89 | -1.63 | -0.85 | -0.41 | -1.27 | -1.3 | 0.44 | 0.17 | -1.43 | -0.9 | 0 | -0.0625 | 0.465 | 0.421461 | 1 | 0.892807 | 1 | 0.999631 | 0.772817 |
| Bsn_O88737_S1114s_1 | -1.59 | -0.36 | -1.78 | -3.06 | -0.94 | -1.74 | -2.67 | -3 | -1.6 | -1.03 | -2.14 | -2.12 | -0.67 | -0.95 | -2.43 | -1.97 | -0.39 | -0.025 | 0.1925 | 0.758545 | 0.549879 | 0.964608 | 1 | 0.999631 | 0.914449 |
| Bsn_O88737_S1137s_1 | -0.62 | 0.79 | -0.46 | -0.12 | 0.58 | 0.09 | -0.36 | -0.58 | 0.45 | 0.52 | -0.05 | -0.2 | 0.7 | 0.82 | -0.37 | -0.35 | 0.035 | 0.2825 | 0.3025 | 0.507084 | 0.931536 | 0.467458 | 1 | 0.999631 | 0.81564 |
| Bsn_O88737_S1173s_1 | -1.57 | -1.42 | -1.52 | -0.9 | -0.35 | -1.51 | -0.94 | -1.01 | -1.04 | -1.82 | -1.3 | -1.03 | -1.13 | -1.31 | -1.19 | -1.19 | 0.4 | 0.055 | 0.1475 | 0.607843 | 0.239115 | 0.858778 | 1 | 0.999631 | 0.858527 |
| Bsn_O88737_S121s_1 | 1.3 | -0.4 | 0.5 | 0.71 | -0.26 | 0.32 | 1.2 | 0.88 | 0.16 | 0.96 | 0.45 | -0.16 | -1.8 | -1 | 0.44 | -0.34 | 0.0075 | -0.175 | -1.2025 | 0.050785 | 0.986929 | 0.682461 | 1 | 0.999631 | 0.489558 |
| Bsn_O88737_S1236s_1 | -0.96 | -0.29 | 1.72 | 0.42 | 0.54 | -0.77 | 0.92 | -0.32 | -0.32 | -1.3 | 0.3 | 0.01 | -1.12 | -2.17 | -0.34 | -0.3 | -0.13 | -0.55 | -1.205 | 0.08722 | 0.83438 | 0.377145 | 1 | 0.999631 | 0.553514 |
| Bsn_O88737_S1369s_1 | -0.44 | -0.91 | 0.53 | -0.27 | 1.26 | -0.36 | 1.94 | 1.27 | 0.53 | -2.82 | 1.99 | 1.65 | 0.28 | -1.64 | 0.94 | -0.09 | 1.3 | 0.61 | 0.145 | 0.800862 | 0.032566 | 0.539218 | 1 | 0.999631 | 0.931985 |
| Bsn_O88737_S1370s_1 | -0.79 | 1.17 | 0.13 | -0.58 | 3.09 | 0.82 | 1.26 | 0.43 | 2.72 | -1.21 | 0.4 | 0.98 | 0.97 | 0.08 | 0.35 | 0.14 | 1.4175 | 0.74 | 0.4025 | 0.405378 | 0.053211 | 0.371623 | 1 | 0.999631 | 0.765965 |
| Bsn_O88737_S138s_2 | 2.76 | 0.34 | -0.01 | 1.19 | 1.75 | 0.53 | 2.21 | 0.07 | 0.52 | 0.42 | -0.14 | -0.26 | -0.07 | -0.25 | -0.14 | -1.16 | 0.07 | -0.935 | -1.475 | 0.033213 | 0.920726 | 0.138171 | 1 | 0.999631 | 0.44294 |
| Bsn_O88737_S1407s_1 | 0.77 | -0.77 | -1.36 | -0.03 | 1.38 | 0.02 | 0.44 | 0.05 | -0.3 | -1.8 | 0.16 | -1.51 | -2.19 | -0.62 | -0.9 | -1.28 | 0.82 | -0.515 | -0.9 | 0.118338 | 0.138939 | 0.400355 | 1 | 0.999631 | 0.585395 |
| Bsn_O88737_S142s_1 | 1.45 | -0.65 | 0.82 | 0.48 | 1.05 | 0.71 | 0.82 | -0.62 | 0.56 | 0.2 | -0.03 | -0.94 | -2.42 | -0.05 | -0.24 | -0.13 | -0.035 | -0.5775 | -1.235 | 0.080615 | 0.947774 | 0.276211 | 1 | 0.999631 | 0.545579 |
| Bsn_O88737_S142s_2 | 2.76 | 0.34 | -0.01 | 1.19 | 1.75 | 0.53 | 2.21 | 0.07 | 0.52 | 0.42 | -0.14 | -0.26 | -0.07 | -0.25 | -0.14 | -1.16 | 0.07 | -0.935 | -1.475 | 0.033213 | 0.920726 | 0.138171 | 1 | 0.999631 | 0.44294 |
| Bsn_O88737_S1460s_1 | 0.04 | -1.2 | -1.24 | -0.32 | 0.03 | -0.71 | -0.91 | -0.79 | -0.96 | -0.99 | -0.88 | -1.16 | -0.41 | -0.2 | -0.5 | -1.1 | 0.085 | -0.3175 | 0.1275 | 0.749771 | 0.828486 | 0.3869 | 1 | 0.999631 | 0.912137 |
| Bsn_O88737_S1482s_1 | -1.59 | -0.54 | -0.7 | -1.67 | -1.18 | -1.37 | -1.46 | -1.46 | -1.12 | -0.86 | -1.34 | -0.86 | 0.33 | -0.96 | -0.75 | -0.78 | -0.2425 | 0.08 | 0.585 | 0.19043 | 0.478704 | 0.820893 | 1 | 0.999631 | 0.648674 |
| Bsn_O88737_S151s_1 | 1.49 | -0.93 | -0.6 | 1.53 | 2.17 | 0.4 | 1.19 | 1.18 | -0.08 | -0.6 | -0.17 | -0.46 | -0.77 | 0.19 | 0.7 | -0.39 | 0.8625 | -0.7 | -0.44 | 0.508672 | 0.217746 | 0.267814 | 1 | 0.999631 | 0.816932 |
| Bsn_O88737_S1650s_1 | 1.53 | -0.88 | -0.28 | 0.67 | 1.14 | 0.29 | 0.54 | 0.79 | -0.56 | -0.55 | 0.35 | 0.33 | 0.14 | -0.08 | 0.5 | -0.56 | 0.43 | -0.3675 | -0.26 | 0.630746 | 0.417051 | 0.50687 | 1 | 0.999631 | 0.867509 |
| Bsn_O88737_S1990s_1 | -1.09 | 0.26 | -0.23 | 0.22 | 0.09 | -0.16 | -0.6 | -0.42 | -0.62 | -1.25 | -0.75 | -0.26 | -0.53 | -0.39 | 0.16 | -0.05 | -0.0625 | -0.51 | 0.0075 | 0.984424 | 0.865031 | 0.212467 | 1 | 0.999631 | 0.995133 |
| Bsn_O88737_S2046s_1 | -0.1 | -0.08 | -0.45 | 0.22 | 0.42 | 0.11 | -0.04 | 0.29 | -0.12 | -0.42 | -0.14 | 0.11 | -0.29 | 0.22 | 0.16 | 0.24 | 0.2975 | -0.04 | 0.185 | 0.536569 | 0.280279 | 0.884912 | 1 | 0.999631 | 0.830094 |
| Bsn_O88737_S241s_1 | 1.45 | -2.19 | -2.55 | 1.34 | 1.26 | -0.68 | 1.26 | 0.65 | -0.78 | -0.64 | -1.06 | -2.22 | -3.49 | -1.56 | -1.38 | -0.66 | 1.11 | -0.6875 | -1.285 | 0.243697 | 0.289734 | 0.493432 | 1 | 0.999631 | 0.685637 |
| Bsn_O88737_S245s_1 | 0.52 | -1.3 | -1.82 | 0.59 | 0.97 | 0.25 | 0.72 | 0.74 | 0.04 | -0.81 | 0.38 | -0.9 | -2.9 | 0 | -0.23 | -0.27 | 1.1725 | 0.18 | -0.3475 | 0.667422 | 0.067035 | 0.77499 | 1 | 0.999631 | 0.88264 |
| Bsn_O88737_S2578s_1 | -0.3 | 1.19 | 1.7 | 0.94 | 0.56 | 1.3 | 0.52 | 1.07 | 0.83 | 0.43 | 1.88 | 1.1 | 1.55 | 0.58 | 2.01 | 1.41 | -0.02 | 0.1775 | 0.505 | 0.323172 | 0.964543 | 0.721894 | 1 | 0.999631 | 0.724485 |
| Bsn_O88737_S2659s_1 | 1.22 | -0.05 | -1.16 | -0.7 | -0.34 | -0.85 | -0.15 | -1.89 | -0.92 | -1.08 | -2.43 | -0.99 | -1.91 | -1.98 | -2.91 | -1.38 | -0.635 | -1.1825 | -1.8725 | 0.007217 | 0.294518 | 0.064478 | 1 | 0.999631 | 0.358173 |
| Bsn_O88737_S2805s_1 | 1.14 | -0.45 | 0.48 | 0.52 | 1.2 | 0.01 | 0.85 | 0.91 | -0.5 | -0.66 | 0.04 | -0.6 | 0.01 | 0.88 | -0.35 | -0.96 | 0.32 | -0.8525 | -0.5275 | 0.29429 | 0.44686 | 0.048327 | 1 | 0.999631 | 0.708293 |
| Bsn_O88737_S2808s_1 | 0.98 | 2.46 | 0.17 | -0.72 | 1.08 | 0.5 | -0.1 | 0.57 | -1.12 | -0.5 | -0.26 | -1.56 | -1.07 | -0.4 | -0.78 | 0.27 | -0.21 | -1.5825 | -1.2175 | 0.088542 | 0.743931 | 0.035967 | 1 | 0.999631 | 0.553858 |
| Bsn_O88737_S2811s_1 | -0.63 | -1.11 | -0.22 | -0.26 | 0.76 | 0.15 | -0.27 | 0.18 | 0.07 | -0.45 | -0.4 | -0.44 | 0 | -1.11 | -0.16 | 0.02 | 0.76 | 0.25 | 0.2425 | 0.524754 | 0.04382 | 0.427545 | 1 | 0.999631 | 0.822727 |
| Bsn_O88737_S2822s_1 | 0.85 | -0.2 | -1.39 | -0.17 | 1.98 | -0.41 | 1.09 | -0.59 | 0.57 | -2.2 | -0.15 | -0.66 | -0.66 | -1.47 | -0.82 | -1.68 | 0.745 | -0.3825 | -0.93 | 0.086428 | 0.289488 | 0.569658 | 1 | 0.999631 | 0.55299 |
| Bsn_O88737_S284s_1 | 1.27 | 0.47 | -0.3 | 0.68 | 1.07 | 0.77 | 0.28 | 1.28 | 0.01 | -0.82 | 0.23 | -1.02 | -1.09 | 0.1 | 0.16 | 0.04 | 0.32 | -0.93 | -0.7275 | 0.12448 | 0.427316 | 0.058057 | 1 | 0.999631 | 0.596063 |
| Bsn_O88737_S2858s_1 | -1.14 | -0.64 | -1.25 | -2.38 | -1.43 | -1.23 | -1.15 | -1.53 | -0.99 | -1.2 | -1.54 | -2 | -0.13 | -0.9 | -1.69 | -1.33 | 0.0175 | -0.08 | 0.34 | 0.486561 | 0.963954 | 0.851872 | 1 | 0.999631 | 0.804772 |
| Bsn_O88737_S2860s_1 | -1.28 | -1.03 | -0.26 | -1.2 | -1.46 | -1.15 | -1.5 | -1.08 | -1.16 | -0.91 | -1.1 | 0.03 | 0.25 | -0.64 | -0.54 | -0.04 | -0.355 | 0.1575 | 0.7 | 0.077058 | 0.270156 | 0.681981 | 1 | 0.999631 | 0.542683 |
| Bsn_O88737_S2866s_1 | 0.19 | 0.15 | 0.88 | 0.67 | 0.15 | 0.92 | 0.38 | 0.25 | 0.22 | 0.56 | 0.04 | 0.27 | 0.13 | 0.89 | 0.42 | 0.77 | -0.0475 | -0.2 | 0.08 | 0.806594 | 0.876841 | 0.500059 | 1 | 0.999631 | 0.935434 |
| Bsn_O88737_S2908s_1 | -1.27 | -0.79 | -0.67 | -1.17 | -0.99 | -1.16 | -0.84 | -1.18 | -1.01 | -1.24 | -0.86 | -0.69 | 0.13 | -1.07 | -0.92 | -0.66 | -0.0675 | 0.025 | 0.345 | 0.344755 | 0.798836 | 0.929199 | 1 | 0.999631 | 0.734195 |
| Bsn_O88737_S301s_1 | 1.16 | -0.38 | -1.84 | 1.06 | 2.01 | 0.05 | 0.98 | 0.7 | 0.05 | -0.44 | 0.81 | -1.24 | -1.18 | 0.08 | -0.51 | -0.51 | 0.935 | -0.205 | -0.53 | 0.437398 | 0.21411 | 0.780008 | 1 | 0.999631 | 0.781057 |
| Bsn_O88737_S3022s_1 | -1.35 | -0.1 | -0.25 | -0.06 | 0.15 | -0.09 | -0.42 | -0.25 | -0.29 | -1.04 | -0.11 | -0.26 | -0.44 | -0.02 | -0.23 | 0.07 | 0.2875 | 0.015 | 0.285 | 0.447758 | 0.426666 | 0.969169 | 1 | 0.999631 | 0.786228 |
| Bsn_O88737_S3158s_1 | 0.9 | -0.18 | -0.21 | 1.08 | 1.16 | 0.79 | 0.46 | 0.65 | 0.28 | -0.47 | 0.51 | -0.1 | 0.45 | 0.12 | -0.31 | 0.14 | 0.3675 | -0.3425 | -0.2975 | 0.464916 | 0.352062 | 0.415181 | 1 | 0.999631 | 0.794369 |
| Bsn_O88737_S3218s_1 | -0.71 | -1.45 | -1.82 | -1.36 | -1.87 | -1.22 | -1.07 | -1.64 | -2.44 | -1.76 | -0.98 | -1.75 | -2.93 | -2.02 | -2.64 | -1.04 | -0.115 | -0.3975 | -0.8225 | 0.103534 | 0.731536 | 0.324959 | 1 | 0.999631 | 0.567302 |
| Bsn_O88737_S3258s_1 | -0.07 | -0.79 | 0.44 | 0.4 | -1.49 | -1.23 | 0.33 | -0.01 | -1.19 | -0.21 | -1.11 | -1.06 | -1.33 | -2.02 | -1.39 | -1.11 | -0.595 | -0.8875 | -1.4575 | 0.003228 | 0.250209 | 0.041942 | 1 | 0.999631 | 0.351308 |
| Bsn_O88737_S3301s_1 | -0.79 | -0.58 | -0.46 | -0.2 | -0.71 | -0.83 | -0.49 | -0.58 | -0.83 | -0.54 | -0.59 | -0.55 | -0.92 | -1.16 | -1.28 | -0.54 | -0.145 | -0.12 | -0.4675 | 0.147833 | 0.573725 | 0.649294 | 1 | 0.999631 | 0.620569 |
| Bsn_O88737_S3382s_1 | 0.41 | 1.12 | 1.08 | 0.75 | 0.59 | 1.46 | 0.22 | 1.01 | 0.51 | 0.71 | 0.75 | 0.72 | 0.75 | 1.26 | 0.99 | 1.54 | -0.02 | -0.1675 | 0.295 | 0.36794 | 0.953886 | 0.548142 | 1 | 0.999631 | 0.745968 |
| Bsn_O88737_S3460s_1 | 0.3 | -0.62 | -0.77 | 1.61 | 1.8 | 1.39 | 1.87 | 0.76 | 1.07 | -0.85 | 0.62 | -0.13 | -1.91 | -1.16 | 0.43 | -1.52 | 1.325 | 0.0475 | -1.17 | 0.105082 | 0.03578 | 0.939411 | 1 | 0.999631 | 0.568213 |
| Bsn_O88737_S3466s_1 | 3.14 | -0.56 | -0.76 | 1.39 | 0.71 | 0.28 | 1.41 | -0.75 | -0.48 | 0.92 | -0.58 | -1.62 | -2.61 | -0.75 | -0.83 | -0.91 | -0.39 | -1.2425 | -2.0775 | 0.037583 | 0.661405 | 0.196415 | 1 | 0.999631 | 0.458256 |
| Bsn_O88737_S3512s_1 | -1.55 | -0.51 | -2.18 | -2.92 | -1.85 | -1.55 | -2.67 | -3.08 | -1.34 | -0.62 | -2.47 | -2.66 | 0.4 | -1.16 | -3.04 | -2.23 | -0.4975 | 0.0175 | 0.2825 | 0.720446 | 0.390044 | 0.977952 | 1 | 0.999631 | 0.904869 |
| Bsn_O88737_S3512s_2 | -0.55 | 0.14 | -0.5 | -0.61 | 0.3 | -0.4 | -0.7 | -0.05 | -0.26 | -0.78 | -0.54 | 0.02 | 0.11 | -0.26 | -0.61 | -0.13 | 0.1675 | -0.01 | 0.1575 | 0.621342 | 0.607253 | 0.974407 | 1 | 0.999631 | 0.861888 |
| Bsn_O88737_S3516s_1 | -7.15 | -9.04 | -4.12 | -4.56 | -9.66 | -7.32 | -3.92 | -5.37 | -8.71 | -6.51 | -3.63 | -4.1 | -3.22 | -6.37 | -5.33 | -5.5 | -0.35 | 0.48 | 1.1125 | 0.337812 | 0.808122 | 0.732768 | 1 | 0.999631 | 0.730926 |
| Bsn_O88737_S3516s_2 | -1.79 | -1.97 | -0.17 | -1.5 | -2.67 | -2.19 | -1.16 | -1.16 | -2.52 | -2.28 | 0.24 | 0.33 | 0.16 | -0.97 | -1.35 | -1.34 | -0.4375 | 0.3 | 0.4825 | 0.358611 | 0.407003 | 0.698761 | 1 | 0.999631 | 0.741182 |
| Bsn_O88737_S3519s_1 | -1.51 | -1.23 | -1.29 | -1.63 | -1.35 | -0.3 | -1.7 | -1.04 | -1.05 | -0.71 | -1.51 | -1.5 | 0.03 | 0.57 | -0.83 | -1.15 | 0.3175 | 0.2225 | 1.07 | 0.026742 | 0.367882 | 0.458163 | 1 | 0.999631 | 0.422285 |
| Bsn_O88737_S3519s_2 | -0.86 | -0.42 | -0.48 | -1.07 | -0.33 | -0.77 | -1.09 | -1.17 | -0.74 | -1.15 | 0.01 | 0.16 | -0.08 | -0.56 | -1.11 | -1.38 | -0.1325 | 0.2775 | -0.075 | 0.839827 | 0.665368 | 0.463476 | 1 | 0.999631 | 0.946365 |
| Bsn_O88737_S3545s_1 | -1.09 | -1.23 | -0.79 | -0.54 | -0.06 | -0.5 | -0.3 | -0.48 | -0.28 | -1.52 | -1.26 | -0.61 | -1.6 | -0.31 | -0.99 | -0.45 | 0.5775 | -0.005 | 0.075 | 0.840558 | 0.057506 | 0.988874 | 1 | 0.999631 | 0.946805 |
| Bsn_O88737_S3730s_1 | 2.47 | 1.48 | 0.16 | 0.49 | 0.51 | 2.54 | 0.96 | 1.66 | 0.3 | 1.87 | -0.04 | -0.34 | -0.77 | 0.85 | 1.97 | -1.02 | 0.2675 | -0.7025 | -0.8925 | 0.262002 | 0.665378 | 0.291746 | 1 | 0.999631 | 0.692758 |
| Bsn_O88737_S435s_1 | 2.4 | -1.31 | 0.04 | 0.32 | -1.58 | 2.5 | 3.38 | -1.91 | -1.92 | 1.45 | -1.18 | -0.45 | -1.92 | -1.58 | -0.07 | -0.71 | 0.235 | -0.8875 | -1.4325 | 0.085605 | 0.859969 | 0.34446 | 1 | 0.999631 | 0.551124 |
| Bsn_O88737_S572s_1 | 2.36 | -0.79 | 0.15 | 0.83 | -3.59 | 0.51 | 3.06 | -1.06 | -5.07 | 0.55 | -2.68 | -1.81 | -2.5 | -1.77 | 0.13 | -1.2 | -0.9075 | -2.89 | -1.9725 | 0.024925 | 0.49522 | 0.029023 | 1 | 0.999631 | 0.418003 |
| Bsn_O88737_S817s_1 | 0.26 | -0.16 | -0.24 | -0.81 | 0.56 | 1.29 | -0.24 | -1.83 | 0.24 | 0.74 | 0.12 | -0.3 | 1.11 | 1.65 | -0.99 | -0.57 | 0.1825 | 0.4375 | 0.5375 | 0.39023 | 0.772893 | 0.227354 | 1 | 0.999631 | 0.758901 |
| Bsn_O88737_S90s_1 | 2.89 | -0.86 | -0.86 | 2.1 | 1.21 | 1.88 | 3.38 | 1.29 | 0.22 | -0.07 | 0.62 | -0.65 | -3.58 | 0.26 | 0.81 | 0.01 | 1.1225 | -0.7875 | -1.4425 | 0.24173 | 0.255519 | 0.384019 | 1 | 0.999631 | 0.684831 |
| Bsn_O88737_T2595t_1 | -0.19 | -0.03 | 4.85 | 3.41 | 0.03 | -1.05 | 3.17 | 3.39 | -0.19 | -0.78 | 5.88 | 5.96 | -0.35 | -1.51 | 3.17 | 4.18 | -0.625 | 0.7075 | -0.6375 | 0.683464 | 0.663229 | 0.709863 | 1 | 0.999631 | 0.889602 |
| Bsn_O88737_T3254t_1 | 0.35 | 3.02 | 3.25 | 1.32 | 2.87 | 2.87 | 3.42 | 2.07 | 2.5 | -1.4 | 3.29 | 2.42 | 1.36 | 0.13 | 2.13 | 0.54 | 0.8225 | -0.2825 | -0.945 | 0.213268 | 0.235407 | 0.794658 | 1 | 0.999631 | 0.666787 |
| Bsn_O88737_T3896t_1 | 0.25 | 0.16 | -0.42 | 0.07 | 0.66 | -0.08 | 0.03 | 0.12 | 0.2 | -0.35 | -0.38 | 0.67 | 0.26 | 0.7 | 0.9 | 0.87 | 0.1675 | 0.02 | 0.6675 | 0.051309 | 0.569571 | 0.952806 | 1 | 0.999631 | 0.489777 |
| Bsn_O88737_T55t_1 | 0.55 | -0.78 | -0.83 | 0.73 | 0.63 | -0.48 | 0.61 | 0.58 | -0.98 | -1.2 | 0.19 | -1.51 | 0.21 | -0.46 | -0.24 | -0.31 | 0.4175 | -0.7925 | -0.1175 | 0.7917 | 0.388931 | 0.15433 | 1 | 0.999631 | 0.926882 |
| Btbd10_E9Q6L8_S44s_1 | 1.49 | 0.11 | 0.95 | 0.61 | -0.18 | 0.07 | 0.61 | 0.11 | -0.39 | 1.06 | 0.37 | -0.16 | -0.74 | -0.21 | 0.12 | -0.46 | -0.6375 | -0.57 | -1.1125 | 0.014031 | 0.100244 | 0.207546 | 1 | 0.999631 | 0.385092 |
| Btbd10_E9Q6L8_S71s_1 | 0.05 | -1.24 | 0.35 | 0.42 | -0.65 | -1.1 | 0.53 | -0.72 | -0.17 | 0.66 | -0.59 | -0.33 | -0.69 | -1.94 | 0.27 | -1.14 | -0.38 | -0.0025 | -0.77 | 0.188541 | 0.447526 | 0.995654 | 1 | 0.999631 | 0.647893 |
| Btbd9_Q8C726_T578t_1 | -0.56 | 1.33 | -0.44 | 0.46 | 1.19 | 1.57 | -0.63 | 0.93 | 1.09 | -0.76 | -0.55 | -0.2 | 1.1 | 0.17 | -0.81 | 0.19 | 0.5675 | -0.3025 | -0.035 | 0.949047 | 0.349779 | 0.591155 | 1 | 0.999631 | 0.983587 |
| Bzw1_Q9CQC6_S411s_1 | -0.21 | 0.09 | 1.91 | 0.49 | -1.23 | 0.09 | 0.79 | -0.46 | -0.08 | 0.16 | 0.24 | 0.98 | -0.99 | -0.84 | 0.64 | -0.3 | -0.7725 | -0.245 | -0.9425 | 0.113225 | 0.201606 | 0.623557 | 1 | 0.999631 | 0.577412 |
| Bzw1_Q9CQC6_S413s_1 | 0.07 | -0.5 | 2.75 | 0.34 | -0.02 | 1.34 | 0.03 | 1.21 | 1.61 | 1.69 | 0.41 | 1.19 | 0.52 | 1.5 | 1.04 | -0.47 | -0.025 | 0.56 | -0.0175 | 0.980937 | 0.971941 | 0.425593 | 1 | 0.999631 | 0.994608 |
| C0HKD9_S94s_1 | 0.37 | 1.56 | 2.13 | 1.24 | 0.48 | 1.89 | 1.05 | 1.63 | 1.24 | 1.09 | 1.17 | 0.89 | 1.15 | 1.63 | 1.72 | 1.65 | -0.14 | 0.17 | 0.5525 | 0.606027 | 0.892622 | 0.564797 | 1 | 0.999631 | 0.85817 |
| C2cd2_E9Q3C1_S436s_1 | -0.26 | 0.38 | 0.33 | -0.4 | -0.42 | 0.02 | -0.4 | 0.29 | 0.14 | 0.43 | 0.16 | 0 | 0.79 | 0.6 | 0.39 | 0.48 | 0.415 | 0.525 | 0.625 | 0.099743 | 0.658562 | 0.571268 | 1 | 0.999631 | 0.562805 |
| C2cd2l_Q80X80_S374s_1 | -2.38 | -0.21 | -0.65 | -1.98 | -0.37 | -0.98 | -1.03 | -1.18 | -0.4 | -2.39 | -0.21 | -0.12 | 0.57 | -1.17 | -0.29 | -1.83 | -0.1625 | -0.2125 | 0.235 | 0.354648 | 0.42577 | 0.441352 | 1 | 0.999631 | 0.739951 |
| C2cd2l_Q80X80_S396s_1 | -0.88 | -1.47 | -1.97 | -1.66 | -1.58 | -1.82 | -1.34 | -1.89 | -1.8 | -1.17 | -2.07 | -1.79 | -0.95 | -1.28 | -1.6 | -1.21 | -0.175 | -0.085 | 0.355 | 0.488528 | 0.606918 | 0.538807 | 1 | 0.999631 | 0.806721 |
| C2cd2l_Q80X80_S411s_1 | -0.64 | -1.03 | -0.75 | -0.97 | -0.92 | -1.32 | -1 | -0.85 | -1.16 | -0.82 | -1.13 | -0.62 | -0.26 | -0.58 | -0.55 | -0.58 | -0.57 | 0.0375 | -0.335 | 0.21232 | 0.495493 | 0.752969 | 1 | 0.999631 | 0.666473 |
| C2cd2l_Q80X80_S468s_1 | -1.47 | -5.51 | -4.92 | -5.78 | -2.99 | -5.36 | -5.56 | -6.05 | -3.9 | -2.02 | -4.75 | -6.86 | -3.24 | -4.5 | -5.69 | -5.59 | -0.34 | 0.01 | -0.39 | 0.734562 | 0.586138 | 0.975297 | 1 | 0.999631 | 0.907462 |
| C2cd2l_Q80X80_S48s_2 | -0.69 | -0.61 | -0.91 | 0.19 | -0.42 | -1.67 | -0.38 | -0.91 | -0.96 | -0.81 | -0.42 | 0.21 | -0.2 | -1.04 | -1.65 | -0.69 | -0.72 | -0.3125 | 0.0025 | 0.35058 | 0.395517 | 0.978878 | 1 | 0.999631 | 0.737333 |
| C2cd2l_Q80X80_S59s_1 | -0.73 | -1.76 | -0.31 | -1.9 | -1.69 | -1.88 | -1.73 | -2.28 | -1.73 | -0.59 | -2.04 | -1.59 | 1.14 | -1.04 | -1.92 | -2.87 | -0.2625 | -0.0475 | 0.585 | 0.997559 | 0.104802 | 0.519379 | 1 | 0.999631 | 0.99922 |
| C2cd2l_Q80X80_S613s_1 | -0.72 | 0.12 | -0.16 | -1.51 | 0.29 | -0.25 | -1.35 | -2.01 | -0.78 | -0.95 | -0.65 | -0.08 | 1.23 | -0.13 | -0.41 | -0.62 | -0.4775 | -0.5475 | 1.1275 | 0.276368 | 0.650045 | 0.908296 | 1 | 0.999631 | 0.700187 |
| C2cd2l_Q80X80_S619s_1 | -1.16 | 0.02 | 1.2 | -0.61 | -0.71 | -0.77 | -1.04 | 0.06 | -0.78 | -0.59 | 1.02 | -2.39 | 2.11 | -0.08 | 0.5 | 1.43 | -0.065 | 0.73 | 1.405 | 0.099334 | 0.368779 | 0.477539 | 1 | 0.999631 | 0.562298 |
| C2cd2l_Q80X80_S623s_1 | -1.83 | -0.52 | 0.34 | -0.72 | -1.02 | -0.9 | -0.79 | -0.28 | -0.64 | -0.25 | 0.37 | 0.71 | 1.55 | -0.05 | 0.57 | 0.82 | 0.4825 | -0.06 | 0.295 | 0.021652 | 0.886803 | 0.174328 | 1 | 0.999631 | 0.405544 |
| C2cd2l_Q80X80_S660s_1 | -2.02 | -0.79 | -0.4 | -0.03 | -0.5 | -1.14 | -0.3 | 0.63 | -1.25 | -1.66 | 0.27 | -0.84 | -0.76 | -0.95 | -0.21 | -0.14 | 0.265 | -0.09 | 0.4075 | 0.532317 | 0.368766 | 0.91377 | 1 | 0.999631 | 0.8277 |
| C2cd2l_Q80X80_S662s_1 | -1.68 | -0.89 | -0.89 | 0.36 | -0.72 | -1.2 | -0.19 | 0.07 | -1.05 | -1.45 | -0.57 | -0.39 | -0.2 | -0.84 | -0.25 | -0.18 | 0.675 | 0.34 | 0.4525 | 0.374852 | 0.584447 | 0.847847 | 1 | 0.999631 | 0.74795 |
| C2cd2l_Q80X80_T388t_1 | -0.07 | 0.13 | 0.51 | 0.4 | 0.88 | 1.87 | 0.12 | 0.8 | 1.2 | 0.67 | 0.3 | 0.16 | 0.36 | 1.1 | 1.03 | 0.29 | -0.55 | 0.005 | -0.1125 | 0.188111 | 0.109581 | 0.307666 | 1 | 0.999631 | 0.647379 |
| C2cd2l_Q80X80_T417t_1 | -1.6 | -2.26 | -3.18 | -3.81 | -2.73 | -3.12 | -3.58 | -3.62 | -2.66 | -1.92 | -2.55 | -3.7 | -0.98 | -2.27 | -3.65 | -4.4 | 0.11 | -0.2725 | 0.335 | 0.886249 | 0.285979 | 0.992924 | 1 | 0.999631 | 0.963505 |
| C2cd2l_Q80X80_T417t_2 | -1.21 | -2.75 | -1.14 | 0.5 | -1.13 | -3.24 | -0.62 | 0.83 | -2.07 | -2.69 | -0.7 | -0.23 | -1.71 | -2.17 | 0.54 | 0.08 | 0.21 | -0.0475 | 0.0075 | 0.682534 | 0.90546 | 0.725274 | 1 | 0.999631 | 0.889581 |
| C2cd2l_Q80X80_T428t_1 | 0.52 | -1.61 | -0.15 | 0.63 | -0.22 | -0.66 | 0.47 | 0.64 | 0.16 | -0.23 | -0.54 | -0.19 | -0.76 | -0.09 | 0.13 | 0.14 | 0.21 | -0.085 | -0.0325 | 0.988598 | 0.702967 | 0.925148 | 1 | 0.999631 | 0.995409 |
| C2cd2l_Q80X80_T428t_2 | -1.21 | -2.75 | -0.97 | 0.65 | -1.13 | -3.24 | 0.21 | 0.72 | -2.07 | -2.69 | 0.38 | -0.24 | -1.71 | -2.17 | -0.5 | -0.03 | 1.1375 | 0.1775 | 0.28 | 0.965549 | 0.82856 | 0.92283 | 1 | 0.999631 | 0.988053 |
| C2cd4cC2CD4 family_Q5HZI2_S153s_1 | -0.44 | 0.83 | 0.08 | 0.75 | 1.68 | 0.52 | 1.46 | 2.11 | 0.8 | 0 | 1.57 | -0.44 | -0.09 | 0.65 | 0.04 | 1.74 | 0.915 | -0.685 | -0.75 | 0.574019 | 0.025597 | 0.726742 | 1 | 0.999631 | 0.84468 |
| C2cd4cC2CD4 family_Q5HZI2_S175s_1 | 1.37 | 0.04 | -1.33 | -0.79 | 1.58 | 0.3 | 0.49 | 0.58 | 0.31 | -0.68 | -1.29 | -1.79 | -0.68 | -0.28 | -1.48 | -1.27 | 1.0725 | -1.2025 | -1.755 | 0.224647 | 0.146175 | 0.316512 | 1 | 0.999631 | 0.67471 |
| C2cd4cC2CD4 family_Q5HZI2_S203s_1 | 2.43 | -2 | -2.19 | 0.96 | 2.73 | -1.32 | 1.39 | 0.69 | -0.27 | -3.69 | -0.07 | -1.58 | -2.21 | -1.58 | -1.22 | -2.81 | 1.0375 | -0.71 | -1.3 | 0.1081 | 0.384315 | 0.331252 | 1 | 0.999631 | 0.569498 |
| C2cd4cC2CD4 family_Q5HZI2_S205s_1 | 0.99 | -2 | -0.8 | 1.32 | 1.67 | -0.82 | 1.87 | 0.94 | -0.72 | -1.9 | 0.38 | -1.09 | -2.2 | -1.03 | -0.91 | -1.55 | -0.47 | 0.3525 | 0.6725 | 0.101057 | 0.246112 | 0.385352 | 1 | 0.999631 | 0.564448 |
| C2cd4cC2CD4 family_Q5HZI2_S259s_1 | 0.43 | 0.63 | 1.8 | 1.46 | 0.02 | 0.42 | 1.21 | 0.79 | 1.82 | 1.37 | -0.13 | 2.67 | 1.72 | 0.68 | 2.38 | 2.23 | 0.6525 | -0.51 | -0.835 | 0.189025 | 0.271913 | 0.56682 | 1 | 0.999631 | 0.648465 |
| C2cd4cC2CD4 family_Q5HZI2_T282t_1 | 1.81 | -1.41 | -1.67 | 1.06 | 1.66 | -0.93 | 1.09 | 0.58 | -0.14 | -1.16 | 0.3 | -1.25 | -0.94 | -1.06 | -1.11 | -0.44 | -0.0275 | -0.005 | 0.075 | 0.29649 | 0.472993 | 0.544857 | 1 | 0.999631 | 0.709528 |
| C2cd5_E9Q026_S295s_1 | -1.08 | -0.67 | -0.86 | -0.93 | -0.65 | -1.64 | -0.91 | -0.45 | -0.71 | -2.24 | -0.3 | -0.31 | -1.62 | -0.23 | -0.26 | -1.13 | -0.23 | 0.27 | 0.475 | 0.845563 | 0.931693 | 0.991254 | 1 | 0.999631 | 0.948565 |
| C2cd5_Q7TPS5_S284s_1 | -0.7 | 1.29 | 0.6 | 0.52 | 0.49 | 0.88 | -0.08 | -0.5 | 1.22 | 0.16 | 1.08 | 0.33 | 2.17 | 1.02 | 0.26 | 0.16 | 0.335 | -0.535 | -1.2175 | 0.415188 | 0.63747 | 0.571859 | 1 | 0.999631 | 0.769452 |
| C2cd5_Q7TPS5_T330t_1 | 1.41 | 1.01 | 0.49 | 1.68 | 1 | 1.87 | 1.42 | 1.64 | 0.17 | 1.09 | 0.88 | 0.31 | -1.89 | 1.27 | 0.12 | 0.22 | -0.1175 | -0.3325 | 0.1025 | 0.080552 | 0.348186 | 0.169387 | 1 | 0.999631 | 0.545579 |
| C2cd5_Q7TPS5_T809t_1 | -0.5 | -0.91 | -1.2 | -0.74 | -0.58 | -1.01 | -0.73 | -1.5 | -0.74 | -1.33 | -1.1 | -1.51 | -0.61 | -0.79 | -0.69 | -0.85 | -0.0875 | 0.1625 | 0.1525 | 0.719661 | 0.703241 | 0.282011 | 1 | 0.999631 | 0.904782 |
| C530008M17Rik_E9Q5L4_S1076s_1 | -0.36 | 0.06 | -0.89 | -1.19 | -0.1 | -0.84 | -0.67 | -1.12 | 0 | -0.72 | -0.5 | -0.51 | 0.05 | -0.32 | -0.46 | -1.04 | -0.3025 | -0.1875 | 0.2575 | 0.696041 | 0.813124 | 0.647748 | 1 | 0.999631 | 0.895683 |
| C530008M17Rik_E9Q5L4_S1100s_1 | -0.41 | -0.55 | 0.55 | -0.51 | -0.96 | -0.78 | -0.24 | -0.15 | -0.88 | -0.19 | -0.22 | -0.38 | -0.47 | -0.43 | 0.57 | 0.44 | 0.42 | 0.3825 | 0.8275 | 0.527594 | 0.4035 | 0.592113 | 1 | 0.999631 | 0.824554 |
| C530008M17Rik_E9Q5L4_S1259s_1 | 0.37 | 0.46 | -0.82 | 0.57 | 0.67 | 0.43 | 0.45 | 0.71 | 0.74 | 0 | 0.49 | 0.88 | 0.66 | 1.04 | 1.34 | 0.85 | -0.4875 | -0.625 | -1.6175 | 0.054471 | 0.257205 | 0.342862 | 1 | 0.999631 | 0.491994 |
| C530008M17Rik_E9Q5L4_S571s_1 | 0.48 | -0.97 | 0.85 | 0.86 | -1.27 | -0.59 | 0.87 | 0.26 | -0.58 | 0.43 | -0.73 | -0.4 | -1.27 | -2.02 | -1.13 | -0.83 | 0.0725 | 0.425 | 0.845 | 0.007279 | 0.410437 | 0.216586 | 1 | 0.999631 | 0.358173 |
| C530008M17Rik_E9Q5L4_S730s_1 | -0.42 | 0.69 | -1.2 | -1.23 | -1.32 | 0.83 | -1.23 | -0.15 | 0.83 | 0.23 | -0.53 | -0.99 | -0.35 | 0.48 | 0.37 | 0.72 | 0.6275 | 0.04 | 0.7125 | 0.10811 | 0.905584 | 0.453752 | 1 | 0.999631 | 0.569498 |
| C530008M17Rik_E9Q5L4_S867s_1 | -0.28 | -0.06 | 0.38 | -0.46 | 0.17 | 0.93 | -0.01 | 1 | 0.22 | -0.75 | 0.1 | 0.17 | -0.02 | 0.41 | 0.98 | 1.06 | 0.0775 | 0.0525 | 0.625 | 0.072019 | 0.095187 | 0.905981 | 1 | 0.999631 | 0.532005 |
| C530008M17Rik_E9Q5L4_T1033t_1 | -0.71 | -0.14 | 0.08 | -0.74 | -0.01 | 0.03 | -0.39 | -0.83 | -0.65 | 0.31 | -0.82 | -0.14 | 0.28 | 0.77 | -0.32 | 0.26 | 0.3875 | 0.1125 | -0.0825 | 0.103361 | 0.813975 | 0.884397 | 1 | 0.999631 | 0.566655 |
| C78339_Q3URQ4_T70t_1 | 2.34 | -2.74 | 0.96 | 0.7 | 0.16 | 1.41 | 0.77 | 0.47 | -1.73 | 3.11 | 0.49 | -0.16 | -1.43 | 1.82 | 0.34 | 0.2 | 0.2825 | -0.16 | 0.0125 | 0.939003 | 0.687308 | 0.928945 | 1 | 0.999631 | 0.980404 |
| Cables1_Q3U148_T376t_1 | -0.4 | -1.22 | 0.27 | 0.93 | -0.51 | -0.16 | -0.05 | 1.43 | -0.3 | -0.53 | -0.25 | 0.02 | -0.08 | -2.03 | 0.59 | 1.15 | -0.7025 | -0.3925 | -0.08 | 0.98638 | 0.624025 | 0.729167 | 1 | 0.999631 | 0.995135 |
| Cables2_Q8K3M5_S103s_1 | -0.24 | -0.56 | 0.53 | 0.26 | -0.55 | -0.81 | -0.67 | -0.79 | -0.63 | -0.76 | -0.11 | -0.08 | -0.46 | -0.33 | 0.29 | 0.17 | -0.115 | 0.1075 | 0.23 | 0.823166 | 0.043505 | 0.270427 | 1 | 0.999631 | 0.938864 |
| Cabp1_D3Z1M4_S101s_1 | -0.08 | 0.84 | 0.49 | 0.53 | 0.56 | 0.44 | -0.29 | 0.61 | 0.25 | 0.52 | 0.54 | 0.9 | 0.8 | 0.73 | 0.39 | 0.78 | -0.87 | -0.9675 | -1.03 | 0.463436 | 0.725998 | 0.725375 | 1 | 0.999631 | 0.793615 |
| Cachd1_Z4YK18_S1169s_1 | -1.55 | -5.96 | -4.4 | -3.56 | -4.51 | -5.1 | -2.7 | -6.64 | -6.21 | -4.2 | -4.16 | -4.77 | -4.55 | -4.48 | -5.52 | -5.04 | -0.0625 | 0.0075 | 0.25 | 0.230576 | 0.41725 | 0.297624 | 1 | 0.999631 | 0.681463 |
| Cacna1a_E9Q1R5_S1899s_1 | -1.4 | -1.13 | -1.12 | -1.08 | -1.17 | -1.45 | -1.36 | -1 | -1.36 | -1.34 | -1.18 | -0.82 | -0.42 | -1.1 | -1.17 | -1.04 | 0.11 | -0.3175 | -0.16 | 0.409005 | 0.801516 | 0.977325 | 1 | 0.999631 | 0.766996 |
| Cacna1a_E9Q1R5_S1934s_1 | 0.67 | 0.52 | -0.2 | 1.12 | 1.02 | 0.08 | 0.38 | 1.07 | 0.41 | -0.12 | 0.36 | 0.19 | -0.51 | 0.68 | 0.53 | 0.77 | -0.1875 | -0.35 | 0.455 | 0.702917 | 0.772529 | 0.36523 | 1 | 0.999631 | 0.896857 |
| Cacna1a_E9Q1R5_S1951s_1 | -0.93 | -0.83 | 0.57 | -0.33 | -0.91 | -0.81 | -0.31 | -0.24 | -0.86 | -2.5 | 0.18 | 0.26 | 0.58 | -0.37 | -0.25 | 0.34 | -0.315 | -0.4175 | 0.0525 | 0.297329 | 0.633628 | 0.595491 | 1 | 0.999631 | 0.710672 |
| Cacna1a_E9Q1R5_S1969s_1 | 0.49 | -0.44 | -0.48 | 0.4 | -0.29 | -0.48 | -0.21 | -0.31 | -0.73 | -0.26 | -0.39 | -0.32 | -1.03 | 0.33 | 0.2 | 0.68 | -0.2125 | -0.1675 | 0.23 | 0.907671 | 0.334116 | 0.228614 | 1 | 0.999631 | 0.96989 |
| Cacna1a_E9Q1R5_S1981s_1 | -0.33 | -0.3 | 0.07 | -0.54 | -0.61 | -0.47 | -0.45 | -0.42 | -0.57 | -0.3 | -0.2 | -0.7 | -0.38 | 0.19 | -0.43 | 0.44 | -0.3 | -1.1025 | -1.095 | 0.486443 | 0.407473 | 0.546566 | 1 | 0.999631 | 0.804724 |
| Cacna1a_E9Q1R5_S2173s_1 | -0.75 | -0.75 | 0.57 | -0.47 | -1.07 | -0.72 | -0.41 | -0.4 | -2.11 | -1.95 | -1.54 | -0.21 | 0.22 | -3.01 | -1.31 | -1.68 | 0.1425 | -0.0775 | 0.44 | 0.120164 | 0.425536 | 0.051111 | 1 | 0.999631 | 0.587703 |

| | | | | | | | | | | | | | | | | | | | | | | | | | |
|---|---|---|---|---|---|---|---|---|---|---|---|---|---|---|---|---|---|---|---|---|---|---|---|---|---|
| Cacna1a_E9Q1R5_S2205s_1 | 0.1 | -0.7 | -0.12 | -0.04 | -0.02 | 0.4 | -0.57 | 0 | -0.37 | -0.43 | -0.45 | 0.18 | -0.54 | 0.89 | 0.3 | 0.35 | 0.0075 | 0.1 | 0.285 | 0.263362 | 0.654017 | 0.798987 | 1 | 0.999631 | 0.693034 |
| Cacna1a_E9Q1R5_S2226s_1 | -0.82 | -0.88 | -0.39 | -0.44 | -1.17 | -0.52 | -0.37 | -0.44 | -1.05 | -0.32 | -0.65 | -0.11 | -0.22 | -0.63 | -0.23 | -0.31 | -0.1225 | -0.2425 | 0.13 | 0.331019 | 0.979545 | 0.746969 | 1 | 0.999631 | 0.727302 |
| Cacna1a_E9Q1R5_S2256s_1 | -0.11 | 0.47 | 0.34 | -0.52 | -0.25 | 0.69 | -0.44 | -0.31 | -0.19 | 0.21 | -0.19 | -0.62 | 0.64 | -0.02 | -0.28 | 0.36 | -0.0175 | 0.0625 | 0.5375 | 0.715947 | 0.737526 | 0.472265 | 1 | 0.999631 | 0.902765 |
| Cacna1a_E9Q1R5_S705s_1 | -0.41 | -1.3 | -1.7 | -1.92 | -0.91 | -0.96 | -1.72 | -1.81 | -1.3 | -0.73 | -1.71 | -1.34 | 0.2 | -0.77 | -1.34 | -1.27 | -0.36 | 0.0575 | 0.2875 | 0.273704 | 0.96598 | 0.876267 | 1 | 0.999631 | 0.700187 |
| Cacna1a_E9Q1R5_S745s_1 | -0.18 | 0.71 | 1.66 | 1.2 | -0.08 | 0.51 | 0.45 | 1.07 | 0.73 | 0.59 | 0.76 | 1.54 | 1.22 | 0.47 | 1.06 | 1.79 | -0.2225 | -0.05 | 0.255 | 0.543562 | 0.426306 | 0.89659 | 1 | 0.999631 | 0.832881 |
| Cacna1a_E9Q1R5_S820s_1 | -0.6 | -0.67 | -0.67 | -0.88 | -0.94 | -0.46 | -1.34 | -0.97 | -0.97 | -0.23 | -0.78 | -1.04 | 0.04 | -0.28 | -1.15 | -0.41 | 0.22 | 0.14 | 0.345 | 0.44871 | 0.428074 | 0.86066 | 1 | 0.999631 | 0.786796 |
| Cacna1a_E9Q1R5_T1888t_1 | -0.78 | -0.17 | -0.66 | -0.79 | -0.09 | -0.23 | -0.78 | -0.42 | -0.21 | -0.29 | -0.99 | -0.35 | 0.01 | -0.05 | -0.48 | -0.5 | 0.0425 | -0.125 | 0.595 | 0.270241 | 0.447808 | 0.646804 | 1 | 0.999631 | 0.698438 |
| Cacna1a_E9Q1R5_T986t_1 | -1.35 | -0.72 | -1.5 | -1.47 | -0.61 | -1.28 | -1.64 | -1.34 | -1.71 | -1.51 | -1.34 | -0.98 | -0.27 | -1.21 | -0.89 | -0.29 | 0.2125 | 0.13 | 0.615 | 0.114455 | 0.896615 | 0.686432 | 1 | 0.999631 | 0.579001 |
| Cacna1b_O55017_S1907s_1 | -0.18 | 0.4 | 0.3 | 0.77 | 0.57 | 0.59 | -0.04 | 1.02 | 0.44 | -0.75 | 1.54 | 0.58 | 0.56 | 0.09 | 1.41 | 1.69 | 0.5675 | 0.2775 | 0.2475 | 0.17301 | 0.527295 | 0.789469 | 1 | 0.999631 | 0.642116 |
| Cacna1b_O55017_S1929s_1 | -0.01 | 0.41 | -0.79 | 0.03 | 0.94 | 0.52 | 0.05 | 0.4 | 0.55 | 0.02 | 0.03 | 0.15 | -0.13 | 0.5 | 0.1 | 0.16 | 0.1475 | 0.5375 | 0.44 | 0.4787 | 0.123717 | 0.411781 | 1 | 0.999631 | 0.801051 |
| Cacna1b_O55017_S1951s_1 | -0.83 | -0.07 | -0.62 | -0.79 | -0.08 | -0.58 | -0.03 | -1.03 | 0.02 | -0.38 | -0.15 | 0.35 | 0.67 | -0.46 | -0.33 | -0.43 | 0.3575 | -0.1775 | -0.035 | 0.248091 | 0.659262 | 0.102849 | 1 | 0.999631 | 0.685751 |
| Cacna1b_O55017_S2007s_1 | -0.02 | -1.05 | -1.56 | -0.22 | 0.03 | -0.75 | -0.12 | -0.58 | -0.74 | -1 | -0.69 | -1.13 | -1.34 | -0.44 | -0.37 | -0.84 | 0.3125 | -0.4375 | 0.37 | 0.935259 | 0.388649 | 0.651796 | 1 | 0.999631 | 0.979194 |
| Cacna1b_O55017_S2062s_1 | -0.09 | -0.21 | -0.74 | -0.42 | 0.38 | -0.24 | 0.02 | -0.37 | -0.66 | -1.18 | -0.49 | -0.88 | 0.59 | -0.14 | 0.1 | -0.53 | 0.1825 | 0.1275 | -0.115 | 0.290745 | 0.296002 | 0.156129 | 1 | 0.999631 | 0.706164 |
| Cacna1b_O55017_S2221s_1 | 1.3 | 1.8 | 0.3 | 1.15 | 1.94 | 1.78 | 0.55 | 1.01 | 2.03 | 0.99 | 0.91 | 1.13 | 0.32 | 1.66 | 0.97 | 1.14 | -0.06 | 0.0175 | 0.2 | 0.787774 | 0.679513 | 0.757527 | 1 | 0.999631 | 0.924613 |
| Cacna1b_O55017_S2244s_1 | -0.01 | -0.14 | -0.49 | -0.1 | 0.03 | -0.18 | -0.53 | -0.3 | -0.16 | 0.08 | -0.33 | -0.26 | 0.21 | 0.25 | -0.5 | 0.1 | 0.775 | 0.7625 | 0.545 | 0.515754 | 0.818357 | 0.946619 | 1 | 0.999631 | 0.819449 |
| Cacna1b_O55017_S424s_1 | -0.69 | 1.28 | 0.49 | 0.4 | 1.92 | 1.03 | 0.58 | 1.05 | 2.06 | -0.79 | 1.4 | 1.86 | 0.55 | 1.3 | 1.13 | 0.68 | 0.5175 | 0.3025 | 0.725 | 0.237788 | 0.123494 | 0.283238 | 1 | 0.999631 | 0.683301 |
| Cacna1b_O55017_S446s_1 | -0.01 | 0.71 | 0 | -0.77 | 1.35 | 0.57 | -0.12 | 0.2 | 1.64 | -0.6 | -0.51 | 0.61 | 0.66 | 1.39 | 0.73 | 0.05 | -0.2725 | 0.24 | 0.625 | 0.109202 | 0.246376 | 0.593098 | 1 | 0.999631 | 0.572424 |
| Cacna1b_O55017_S745s_1 | -1.04 | -0.91 | -0.17 | -0.29 | -1.57 | -1.39 | -0.4 | -0.14 | -0.82 | -0.95 | 0.17 | 0.15 | -0.31 | -0.81 | 0.52 | 0.69 | 0.18 | 0.0825 | -0.04 | 0.164297 | 0.515976 | 0.542543 | 1 | 0.999631 | 0.632578 |
| Cacna1b_O55017_S770s_1 | -0.32 | -2.26 | -1.03 | -0.97 | -1.68 | -0.74 | -0.83 | -0.61 | -1.7 | -0.64 | -0.96 | -0.95 | -0.27 | -1.66 | -1.18 | -1.63 | 0.0375 | -0.49 | -0.4575 | 0.935867 | 0.693172 | 0.855579 | 1 | 0.999631 | 0.97939 |
| Cacna1b_O55017_S773s_2 | 0.86 | 0.17 | 0.44 | -0.07 | -0.04 | 0.4 | 0.54 | 0.65 | -0.23 | 0.55 | -0.47 | -0.41 | -0.15 | -0.07 | 0.48 | -0.69 | -0.1 | 0.285 | 0.7675 | 0.223599 | 0.902892 | 0.181806 | 1 | 0.999631 | 0.67471 |
| Cacna1b_O55017_S783s_1 | -0.6 | 0.64 | 0.19 | -0.05 | 0.24 | 0.03 | -0.64 | 0.15 | 0.55 | 0.41 | -0.31 | 0.67 | 1.27 | 0.65 | 0.47 | 0.86 | 0.6225 | -0.025 | 1.09 | 0.055027 | 0.77767 | 0.445808 | 1 | 0.999631 | 0.492066 |
| Cacna1b_O55017_S875s_1 | -4.51 | -4.73 | -3.91 | -3.42 | -5.85 | -4.56 | -3.19 | -0.48 | -7.5 | -4.34 | -1.17 | -3.66 | -4.58 | -5.78 | -0.16 | -1.69 | -0.71 | -0.81 | -1.7275 | 0.344194 | 0.545652 | 0.982583 | 1 | 0.999631 | 0.733869 |
| Cacna1b_O55017_S915s_1 | 0.17 | -3.12 | -1.32 | -1.66 | -3.45 | -2.5 | -0.66 | -2.16 | -3.62 | -0.43 | -2.42 | -2.7 | -2.79 | -4.46 | -2.19 | -3.4 | 0.4225 | -0.4225 | -0.9475 | 0.037483 | 0.372474 | 0.343081 | 1 | 0.999631 | 0.458129 |
| Cacna1b_O55017_T1935t_1 | 1.11 | 1.1 | -0.07 | 2.2 | 2.08 | 1.06 | 1.81 | 1.08 | 1.23 | -0.51 | 1.67 | 0.26 | -1.08 | 0.54 | 0.08 | 1.01 | -0.8225 | -1.0175 | -2.4675 | 0.131958 | 0.405303 | 0.495409 | 1 | 0.999631 | 0.603456 |
| Cacna1c_F7C376_S1825s_1 | 2.32 | -1.33 | 0.56 | -0.49 | -1.67 | -0.7 | 0.95 | -0.81 | -1.61 | 0.29 | -1.11 | -0.58 | -1.89 | -3.84 | -1.21 | -1.87 | 0.9875 | 0.0275 | -0.405 | 0.01412 | 0.33655 | 0.211766 | 1 | 0.999631 | 0.385092 |
| Cacna1c_F7C376_S2208s_1 | 1.15 | -0.88 | -1.88 | 0.74 | 1.21 | -0.05 | 1.06 | 0.86 | 0.24 | -0.37 | 0.19 | -0.82 | -1.32 | 0.13 | -0.8 | -0.5 | -0.145 | -0.1925 | 0.21 | 0.557213 | 0.166921 | 0.967297 | 1 | 0.999631 | 0.838305 |
| Cacna1c_F7C376_S840s_1 | -0.99 | -1.13 | -1.45 | -1.54 | -1.35 | -1.52 | -1.47 | -1.35 | -1.43 | -0.93 | -1.64 | -1.88 | -0.17 | -1.14 | -1.46 | -1.5 | -0.07 | 0.0475 | 0.26 | 0.578836 | 0.569833 | 0.537273 | 1 | 0.999631 | 0.84629 |
| Cacna1c_F7C376_S863s_1 | -0.09 | 0.48 | 0.23 | -0.15 | -0.47 | 0.83 | -0.16 | -0.01 | 0.15 | -0.22 | -0.07 | 0.8 | 0.84 | 0.66 | -0.35 | 0.36 | -0.285 | -0.185 | -0.0975 | 0.469213 | 0.839682 | 0.883664 | 1 | 0.999631 | 0.796658 |
| Cacna1c_F7C376_S863s_2 | -0.28 | 1.01 | 0.21 | -0.33 | -0.15 | 0.3 | -0.76 | 0.08 | -0.18 | 0.7 | -0.35 | -0.3 | 0.16 | 0.47 | -0.76 | 0.35 | -0.285 | -0.185 | -0.0975 | 0.819626 | 0.474527 | 0.650506 | 1 | 0.999631 | 0.938303 |
| Cacna1c_F7C376_T867t_2 | -0.28 | 1.01 | 0.21 | -0.33 | -0.15 | 0.3 | -0.76 | 0.08 | -0.18 | 0.7 | -0.35 | -0.3 | 0.16 | 0.47 | -0.76 | 0.35 | -0.1175 | -0.775 | -1.675 | 0.819626 | 0.474527 | 0.650506 | 1 | 0.999631 | 0.938303 |
| Cacna1d_Q99246_S1684s_1 | 0.76 | -0.27 | 2.04 | 1.53 | 0.36 | 0.85 | 1.35 | 1.03 | -0.31 | 1.06 | 0.21 | 0 | -2.73 | -0.4 | -0.22 | 0.71 | -0.125 | -0.1525 | 0.0875 | 0.05287 | 0.817067 | 0.174734 | 1 | 0.999631 | 0.489777 |
| Cacna1e_Q61290_S1051s_1 | -0.21 | -0.5 | -0.94 | -0.69 | -0.41 | -0.7 | -0.98 | -0.75 | -0.74 | -0.74 | -0.87 | -0.6 | -0.3 | -0.5 | -0.62 | -0.57 | -0.1725 | -0.255 | -0.295 | 0.763034 | 0.65398 | 0.577585 | 1 | 0.999631 | 0.915608 |
| Cacna1e_Q61290_S1099s_1 | -0.8 | -0.08 | 0.15 | -0.06 | -0.39 | -0.86 | -0.13 | -0.1 | -0.93 | -0.77 | -0.09 | -0.02 | -0.3 | -1.2 | -0.12 | -0.35 | -0.3175 | -0.92 | -1.3775 | 0.426282 | 0.592574 | 0.472973 | 1 | 0.999631 | 0.774781 |
| Cacna1e_Q61290_S15s_1 | 1.92 | -0.44 | 2.9 | 0.94 | -0.99 | 1.33 | 3.55 | 0.16 | -1.06 | 1.87 | 0.81 | 0.02 | -0.42 | -0.5 | 0.44 | 0.29 | 2.075 | 3.2725 | 1.0475 | 0.063253 | 0.759049 | 0.281545 | 1 | 0.999631 | 0.512978 |
| Cacna1e_Q61290_S2054s_1 | -1 | 0.75 | 0.97 | -0.41 | 3.37 | 0.14 | 3.68 | 1.42 | 3.92 | -0.88 | 4.99 | 5.37 | 2.36 | -0.17 | 1.6 | 0.71 | 0.14 | 0.1575 | 0.07 | 0.1298 | 0.030738 | 0.028747 | 1 | 0.999631 | 0.601246 |
| Cacna1e_Q61290_S2073s_1 | -0.65 | -0.96 | -0.95 | -0.37 | -0.55 | -1.22 | -0.44 | -0.16 | -0.77 | -0.81 | -0.27 | -0.45 | -0.9 | -0.83 | -0.65 | -0.27 | 0.4025 | 0.7275 | 0.28 | 0.816983 | 0.659206 | 0.582103 | 1 | 0.999631 | 0.937777 |
| Cacna1e_Q61290_S2097s_1 | -1.29 | -0.97 | -1.18 | -0.65 | -0.59 | -0.92 | -0.63 | -0.34 | -1.28 | -1.33 | 0.96 | 0.47 | -0.27 | -0.79 | -0.95 | -0.96 | 0.3975 | -0.015 | 0.47 | 0.375681 | 0.163312 | 0.21499 | 1 | 0.999631 | 0.748307 |
| Cacna1e_Q61290_S2099s_1 | 0 | 0.37 | -1.1 | 0.34 | 1.22 | 0.51 | 0.16 | -0.69 | 0.49 | -0.62 | -0.12 | -0.2 | 0.58 | 0.52 | -0.03 | 0.42 | -0.565 | -0.73 | -1.4675 | 0.253615 | 0.429147 | 0.971344 | 1 | 0.999631 | 0.688735 |
| Cacna1e_Q61290_S20s_1 | 0.68 | -1.49 | 1.17 | -0.68 | -2.6 | -0.64 | 2.25 | -1.59 | -1.5 | 0.6 | -2 | -0.34 | -1.48 | -1.93 | -0.98 | -1.8 | 0.1275 | 0.2425 | 0.4975 | 0.030484 | 0.589459 | 0.341215 | 1 | 0.999631 | 0.438141 |
| Cacna1e_Q61290_S737s_1 | -0.85 | -0.74 | -0.74 | -0.88 | -0.55 | -0.55 | -0.89 | -0.71 | -0.61 | -0.59 | -0.79 | -0.25 | 0.09 | -0.34 | -0.58 | -0.39 | 0.2875 | 0.135 | 0.2575 | 0.10086 | 0.594196 | 0.352758 | 1 | 0.999631 | 0.564448 |
| Cacna1e_Q61290_S740s_1 | -2.05 | -1.86 | -0.43 | -0.99 | -1.58 | -1.77 | -0.55 | -0.28 | -1.77 | -1.37 | -0.49 | -1.16 | -0.95 | -1.92 | -0.72 | -0.71 | 0.2225 | 1.415 | 1.0325 | 0.58423 | 0.56648 | 0.766478 | 1 | 0.999631 | 0.849121 |
| Cacna1e_Q61290_S793s_1 | 0 | 1.39 | 4.27 | 1.71 | 2.35 | 0.68 | 1.99 | 3.24 | 3.6 | -0.03 | 4.27 | 5.19 | 3.78 | 0.17 | 4.3 | 3.25 | 0.905 | 2.035 | 1.215 | 0.35591 | 0.804342 | 0.270003 | 1 | 0.999631 | 0.74034 |
| Cacna1e_Q61290_S794s_1 | -0.44 | 3 | 2.74 | 0.26 | 3.55 | 1.23 | 2.88 | 1.52 | 5.03 | -0.06 | 3.9 | 4.83 | 5.82 | 0.64 | 2.6 | 1.36 | 0.29 | -0.295 | 0.05 | 0.330121 | 0.322979 | 0.130037 | 1 | 0.999631 | 0.726892 |
| Cacna1e_Q61290_S826s_1 | -0.15 | 0.03 | 0.57 | 0.16 | 0.31 | 0.18 | 0.39 | 0.89 | -0.38 | -0.43 | 0.46 | -0.22 | -0.12 | 0.57 | 0.1 | 0.26 | 0.89 | -0.2025 | -1.8025 | 0.870867 | 0.329923 | 0.364258 | 1 | 0.999631 | 0.958735 |
| Cacna1e_Q61290_S852s_1 | 1.73 | -2.19 | -2.2 | 1.42 | 1.43 | -1.36 | 1.55 | 0.7 | 0.22 | -1.07 | -0.12 | -1.08 | -2.75 | -2.34 | -0.52 | -2.84 | 0.425 | 0.4475 | -0.025 | 0.106932 | 0.425454 | 0.83715 | 1 | 0.999631 | 0.568797 |
| Cacna1e_Q61290_S856s_1 | -2.58 | -3.84 | -4.69 | -3.5 | -1.96 | -3.48 | -3.6 | -3.87 | -3.08 | -2.56 | -3.49 | -3.69 | -2.45 | -4.24 | -3.81 | -4.21 | 0.8525 | -0.03 | -0.535 | 0.964432 | 0.455111 | 0.364331 | 1 | 0.999631 | 0.987331 |
| Cacna1e_Q61290_S873s_1 | 1.11 | -0.72 | -1.45 | 1.28 | 1.41 | -0.01 | 1.06 | 1.17 | 0.64 | -0.84 | 0.8 | -0.5 | -1.23 | -0.32 | -0.27 | -0.1 | 0.01 | -0.36 | -0.125 | 0.417465 | 0.218732 | 0.965918 | 1 | 0.999631 | 0.771469 |
| Cacna1e_Q61290_S948s_1 | -1.4 | -1.87 | -0.59 | 0.14 | -1.93 | -1.69 | -0.12 | 0.06 | -2.1 | -2.3 | -0.47 | -0.29 | -1.98 | -1.98 | -0.32 | 0.06 | 0.4025 | -0.1875 | -0.375 | 0.842886 | 0.98699 | 0.567582 | 1 | 0.999631 | 0.94811 |
| Cacna1e_Q61290_S988s_1 | 1.39 | 1.12 | 0.49 | 1.49 | 1.67 | 1.42 | 1.14 | 1.87 | 0.97 | 0.32 | 1.47 | 0.98 | -0.54 | 0.8 | 0.95 | 1.78 | 0.07 | -0.4175 | -0.92 | 0.464782 | 0.229755 | 0.604413 | 1 | 0.999631 | 0.794369 |
| Cacna1e_Q61290_T2225t_1 | 1.34 | 1.58 | 0.74 | 1.59 | 1.97 | 0.89 | 1.23 | 1.44 | 0.62 | 0.32 | 1.53 | 1.11 | -1.07 | 0.8 | 0.49 | 1.35 | 0.2075 | -0.0775 | 0.0875 | 0.103703 | 0.835972 | 0.268145 | 1 | 0.999631 | 0.567918 |
| Cacna1e_Q61290_T29t_1 | 0.21 | -0.16 | -0.42 | 0.23 | 0.23 | 0.02 | -0.09 | 0.53 | 0.26 | -0.42 | -0.03 | -0.26 | -0.49 | -0.05 | 0.26 | 0.49 | 0.3275 | 0.77 | 0.57 | 0.793521 | 0.472033 | 0.793459 | 1 | 0.999631 | 0.928057 |
| Cacna1e_Q61290_T441t_1 | -1.3 | 0.65 | 1.56 | -0.05 | -0.28 | 0.31 | 1.04 | 1.1 | 0.36 | -0.74 | 2.17 | 2.15 | 1.23 | -0.46 | 1.84 | 0.53 | -0.4 | -0.9425 | -0.1275 | 0.418502 | 0.598372 | 0.358543 | 1 | 0.999631 | 0.771707 |
| Cacna1g_Q5SUG2_S1063s_1 | -1.74 | -4.25 | -0.38 | -2.63 | -3.51 | -4.01 | -0.76 | -2.32 | -5.13 | -3.36 | -1.1 | -3.18 | -3.05 | -3.44 | -1.35 | -1.67 | -0.0625 | 0.4975 | -0.4425 | 0.878247 | 0.671154 | 0.357 | 1 | 0.999631 | 0.962347 |
| Cacna1g_Q5SUG2_S2068s_1 | 0.3 | 1.75 | 0.39 | -0.39 | 1.64 | 0 | 1.15 | -0.99 | 2.64 | 0.12 | -0.14 | 1.42 | 2.62 | 0.18 | -1.23 | -1.29 | 0.41 | 0.335 | 0.725 | 0.617188 | 0.924531 | 0.481325 | 1 | 0.999631 | 0.861376 |
| Cacna1g_Q5SUG2_S467s_1 | -1.05 | -1.21 | -1.27 | -1.32 | -0.66 | -0.62 | -1.14 | -0.79 | -1.1 | -0.61 | -1.08 | -0.72 | 0.21 | -0.3 | -1.4 | -0.46 | 0.735 | 0.0525 | -0.325 | 0.079222 | 0.126921 | 0.219707 | 1 | 0.999631 | 0.545579 |
| Cacna1g_Q5SUG2_S518s_1 | 0.81 | -1.03 | -1.38 | -0.2 | -0.01 | 0.13 | 0.43 | 0.59 | -0.02 | -0.02 | -0.22 | -1.33 | -1.2 | -0.86 | -0.19 | -0.85 | -0.6775 | -0.27 | -1.56 | 0.525441 | 0.149299 | 0.922468 | 1 | 0.999631 | 0.823288 |
| Cacna1h_E9Q6P9_S1203s_1 | 1.72 | 0.59 | 1.79 | 1.26 | -0.07 | 0.82 | 0.83 | 1.07 | 1.05 | 1.52 | 2.2 | -0.49 | -0.58 | -0.37 | 0.34 | -0.27 | -0.1 | 0.0625 | 0.535 | 0.001965 | 0.102946 | 0.644057 | 1 | 0.999631 | 0.351308 |
| Cacna1h_E9Q6P9_S2195s_1 | 0.01 | 0.27 | 0.25 | -0.31 | 0.13 | -0.17 | -0.48 | 0.34 | 0.37 | -0.17 | -0.01 | 0.28 | 1.23 | 0.12 | 0.56 | 0.45 | -0.1775 | -0.3575 | -0.405 | 0.135893 | 0.733615 | 0.823964 | 1 | 0.999631 | 0.608661 |
| Cacna1h_E9Q6P9_T1172t_1 | 0.2 | -1.07 | -0.89 | 0.02 | 0.28 | -1.29 | -0.64 | -0.8 | -1.27 | -0.03 | -0.99 | -0.88 | -0.67 | -0.61 | -1.23 | -0.85 | 0.865 | 0.535 | 1.285 | 0.303986 | 0.690401 | 0.403396 | 1 | 0.999631 | 0.715879 |
| Cacna1h_E9Q6P9_T751t_1 | -0.14 | 0.47 | 1.47 | 0.3 | 1.26 | 1.39 | 0.91 | 2 | 1.36 | 0.79 | 0.74 | 1.35 | 2.28 | 1.53 | 1.75 | 1.68 | -0.385 | -0.1325 | 0.055 | 0.008858 | 0.05754 | 0.196962 | 1 | 0.999631 | 0.365244 |
| Cacna1i_E9Q7P2_S1017s_1 | 0.35 | 1.17 | 0 | 0.13 | 0.49 | 0.69 | -0.56 | -0.51 | 0.76 | 0.7 | -0.34 | 0 | 0.59 | 0.92 | -0.4 | 0.76 | -0.26 | -0.065 | -0.7775 | 0.894102 | 0.366681 | 0.73608 | 1 | 0.999631 | 0.965259 |
| Cacna1i_E9Q7P2_S1744s_1 | 0.99 | -0.47 | 0.67 | 0.49 | 0 | -0.19 | 0.71 | 0.12 | -0.21 | 0.38 | 0.77 | 0.48 | -0.71 | -0.66 | 0.16 | -0.22 | 0.0725 | 0.3275 | -0.5325 | 0.075305 | 0.501213 | 0.86822 | 1 | 0.999631 | 0.541044 |
| Cacna1i_E9Q7P2_S1745s_1 | 0.4 | -0.84 | -0.66 | -0.4 | 0.76 | -1.11 | -0.38 | -0.48 | 0.22 | 0.36 | -0.54 | -0.23 | -1.45 | -1.04 | -0.44 | -0.7 | 0.63 | -0.325 | -0.87 | 0.186593 | 0.874024 | 0.386551 | 1 | 0.999631 | 0.646474 |
| Cacna1i_E9Q7P2_S2037s_1 | 1.89 | -0.84 | -1.43 | 0.13 | 1.12 | -0.05 | 1.08 | 0.12 | 0.16 | 0.01 | -0.39 | -1.33 | -1.08 | -0.47 | -1.46 | -0.72 | 0.1275 | -0.085 | -0.5475 | 0.216981 | 0.377428 | 0.649057 | 1 | 0.999631 | 0.669465 |
| Cacna1i_E9Q7P2_S2157s_1 | 1.18 | 0.56 | -0.48 | 1.06 | 0.92 | 0.47 | 1.02 | 0.42 | 0.41 | -0.41 | 1.29 | 0.69 | -0.41 | -0.04 | -0.75 | 1.33 | 0.335 | -0.9575 | -0.8725 | 0.33281 | 0.755401 | 0.863026 | 1 | 0.999631 | 0.728079 |
| Cacna1i_E9Q7P2_S895s_1 | 1.37 | -0.4 | 0.06 | 0.43 | 1.8 | 0.02 | 0.81 | 0.17 | -0.22 | -0.88 | -0.61 | -0.66 | -0.7 | 0.07 | -0.32 | -1.08 | 0.2625 | 0.165 | -0.13 | 0.075789 | 0.519295 | 0.039017 | 1 | 0.999631 | 0.541044 |
| Cacna1i_E9Q7P2_T469t_1 | -0.01 | 1 | 1.13 | 1.38 | 0.61 | 1.21 | 1.28 | 1.45 | 1.23 | 0.38 | 1.32 | 1.23 | 0.71 | 0.17 | 1.02 | 1.08 | 0.1625 | 0.3425 | 0.5175 | 0.743286 | 0.487224 | 0.675688 | 1 | 0.999631 | 0.90972 |
| Cacnb1_A2A543_S464s_1 | -0.16 | -0.25 | -1.35 | -0.4 | -0.25 | 0.32 | -1.06 | -0.52 | -0.1 | 0.88 | -0.98 | -0.59 | 0.14 | 0.37 | -0.42 | -0.18 | 0.06 | 0.3325 | 0.7025 | 0.181762 | 0.686446 | 0.472596 | 1 | 0.999631 | 0.643861 |
| Cacnb1_Q8R3Z5_S186s_1 | 0.19 | 0.9 | -0.18 | 0.76 | -0.45 | 0.75 | 0.69 | 0.92 | 0.88 | 1.2 | 0.44 | 0.48 | 0.75 | 1.73 | 1.12 | 0.88 | -0.1475 | -1.1075 | -0.8775 | 0.083716 | 0.881912 | 0.352415 | 1 | 0.999631 | 0.549732 |
| Cacnb1_Q8R3Z5_S436s_1 | 0.01 | -0.01 | -1.81 | -2.58 | -0.17 | -0.97 | -1.41 | -2.43 | -0.76 | -1.47 | -2.66 | -3.93 | -1.04 | -1.16 | -1.9 | -3.8 | 0.63 | 0.3425 | 0.2725 | 0.285321 | 0.83571 | 0.205114 | 1 | 0.999631 | 0.705851 |
| Cacnb1_Q8R3Z5_S53s_2 | 0.27 | 0.31 | 0 | 1.04 | 1.59 | 0.52 | 0.46 | 1.57 | 1.32 | 0.43 | 0.42 | 0.82 | -0.19 | 1.39 | 0.34 | 1.17 | -0.1825 | 0.8175 | 1.4 | 0.535237 | 0.133192 | 0.337545 | 1 | 0.999631 | 0.829114 |
| Cacnb1_Q8R3Z5_S68s_1 | -1.67 | 0.05 | -0.04 | -1.34 | -0.97 | -1.11 | -0.46 | -1.19 | 0.72 | -0.3 | -0.34 | 0.19 | 2.53 | -0.27 | 0.45 | -0.11 | -0.6575 | -0.3475 | -0.515 | 0.067065 | 0.68864 | 0.116751 | 1 | 0.999631 | 0.520345 |
| Cacnb1_Q8R3Z5_T205t_1 | -0.3 | -1.35 | 0.17 | -0.55 | -1.43 | -1.1 | -0.89 | -1.24 | -1.31 | -0.18 | -0.97 | -0.96 | -1.14 | -1.03 | -0.91 | -1.01 | 0.19 | 0.1675 | 0.835 | 0.18144 | 0.093361 | 0.402543 | 1 | 0.999631 | 0.643843 |
| Cacnb1_Q8R3Z5_T217t_1 | -2.36 | -0.17 | -0.13 | -0.11 | -0.63 | -0.6 | -0.71 | -0.07 | -0.73 | -1.5 | -0.29 | 0.42 | 0.72 | -0.33 | 0.29 | -0.11 | -0.425 | -0.59 | -0.3875 | 0.157412 | 0.721366 | 0.787753 | 1 | 0.999631 | 0.628054 |
| Cacnb1_Q8R3Z5_T499t_1 | -0.07 | 0.44 | 0.92 | 0.17 | 1.06 | 0.77 | -0.48 | -1.59 | 1.08 | 0.16 | -0.38 | -1.76 | 0.31 | 0.07 | -0.54 | 0.07 | 0.115 | 0.095 | 0.555 | 0.273662 | 0.474212 | 0.321221 | 1 | 0.999631 | 0.700187 |
| Cacnb1_Q8R3Z5_T540t_1 | -1.36 | -0.39 | -0.5 | -0.91 | 0.07 | -0.61 | -1.13 | -1.03 | -0.08 | -0.94 | -0.83 | -0.93 | 1.17 | -0.73 | -0.64 | -0.74 | -0.3525 | 0.1475 | 0.045 | 0.279713 | 0.756044 | 0.783094 | 1 | 0.999631 | 0.701922 |
| Cacnb2_Q8CC27_S485s_1 | -1.07 | -0.51 | -0.57 | -0.13 | -1.03 | -1.33 | -0.7 | -0.63 | -0.62 | -0.78 | -0.56 | 0.27 | -0.77 | -0.7 | -0.64 | 0.01 | -0.11 | 0.09 | -0.4675 | 0.892876 | 0.269223 | 0.671319 | 1 | 0.999631 | 0.964596 |
| Cacnb2_Q8CC27_S508s_1 | -0.17 | -0.38 | -0.25 | 0.71 | -0.65 | -1.01 | 0.05 | 1.08 | -0.2 | -1 | 0.21 | 1.26 | -0.42 | -0.55 | -0.57 | -0.42 | -0.07 | -0.065 | -0.16 | 0.175042 | 0.823187 | 0.858101 | 1 | 0.999631 | 0.642116 |
| Cacnb2_Q8CC27_S510s_1 | -0.17 | -0.38 | -1.88 | -2.32 | -0.65 | -1.01 | -1.45 | -1.92 | -0.2 | -1 | -1.86 | -1.95 | -0.42 | -0.55 | -1.76 | -2.66 | -0.06 | -0.26 | 0.4025 | 0.812308 | 0.899116 | 0.915523 | 1 | 0.999631 | 0.937041 |
| Cacnb2_Q8CC27_S552s_1 | -0.72 | -1.84 | -0.22 | -1.17 | -2.02 | -0.51 | -1.13 | -0.53 | -1.38 | -0.52 | -1.38 | -1.71 | -0.44 | -0.28 | -0.66 | -0.96 | -0.325 | -0.565 | -1.565 | 0.325944 | 0.898542 | 0.54753 | 1 | 0.999631 | 0.724678 |
| Cacnb2_Q8CC27_S570s_1 | 0.38 | -0.42 | -0.18 | 0.08 | -0.77 | -0.35 | 0.06 | -0.38 | -0.43 | -0.11 | -1.08 | -0.78 | -2.4 | -0.73 | -1.48 | -1.79 | -0.13 | 0.435 | -0.3475 | 0.003134 | 0.297412 | 0.108454 | 1 | 0.999631 | 0.351308 |
| Cacnb2_Q8CC27_S69s_1 | 1.35 | 0.85 | 0.93 | 0.52 | 0.29 | 1.56 | 0.88 | 0.4 | 1.08 | 1.82 | 1.08 | 1.41 | 0.62 | 0.4 | 0.48 | 0.76 | 0.03 | 0.635 | 1.2 | 0.259377 | 0.71853 | 0.184182 | 1 | 0.999631 | 0.692624 |
| Cacnb2_Q8CC27_S83s_1 | -2.64 | 2.35 | 1.3 | -2.55 | 0.87 | 1.26 | -0.91 | -2.64 | 1.75 | 0.17 | -0.81 | -0.11 | 4.58 | -0.19 | -0.57 | -0.56 | -0.11 | -0.255 | -0.17 | 0.435621 | 0.9821 | 0.599449 | 1 | 0.999631 | 0.780319 |
| Cacnb2_Q8CC27_T549t_1 | 0.51 | 0.41 | 3.33 | 0.95 | -0.16 | -0.09 | 1.9 | 3.11 | -0.48 | 0.37 | 2.32 | 1.97 | -0.54 | -1.46 | 2.89 | 3.63 | -0.2625 | -0.025 | 0.2975 | 0.888121 | 0.903827 | 0.760281 | 1 | 0.999631 | 0.963505 |
| Cacnb3_D3Z3Z3_S151s_1 | -0.54 | -0.84 | -0.3 | -0.81 | -1.12 | -0.79 | -0.79 | -0.84 | -1.17 | -0.33 | -0.44 | -0.65 | 0.14 | -0.58 | -0.79 | -0.07 | 0.7475 | -0.3875 | 0.7625 | 0.373652 | 0.322083 | 0.933753 | 1 | 0.999631 | 0.747035 |
| Cacnb3_D3Z3Z3_S31s_1 | -2.31 | -1.73 | -1.71 | -0.83 | 0.44 | -1.67 | -1.85 | -0.51 | -1.12 | -4.82 | -0.57 | -1.62 | 0.61 | -0.59 | -1.84 | -1.71 | -0.055 | 0.245 | 0.4475 | 0.217074 | 0.205048 | 0.658448 | 1 | 0.999631 | 0.669465 |
| Cacnb3_D3Z3Z3_S392s_1 | -0.38 | -0.53 | -0.7 | -0.96 | -0.91 | -0.45 | -0.73 | -0.7 | -0.27 | 0.02 | -0.67 | -0.67 | 0.29 | -0.1 | -0.59 | -0.38 | 0.0975 | 0.1825 | 0.735 | 0.177474 | 0.832953 | 0.411156 | 1 | 0.999631 | 0.642116 |
| Cacnb3_D3Z3Z3_S402s_1 | -1.68 | -0.92 | -0.12 | -0.69 | -1.07 | -0.79 | -0.68 | -0.48 | -1.06 | -1.44 | 0.1 | -0.28 | -0.04 | -0.95 | 0.49 | 0.03 | 0.525 | 0.7125 | 0.84 | 0.121596 | 0.790264 | 0.695073 | 1 | 0.999631 | 0.590097 |
| Cacnb3_D3Z3Z3_S406s_1 | -1.17 | -0.83 | -0.95 | -1.13 | -0.88 | 0.57 | -1.04 | -0.63 | 0.2 | 0.19 | -0.55 | -1.07 | -0.02 | -0.13 | 0.01 | -0.58 | -0.315 | -0.3 | -0.52 | 0.013719 | 0.193516 | 0.068687 | 1 | 0.999631 | 0.385092 |
| Cacnb4_Q8R0S4_S16s_1 | 0.52 | 0.41 | -0.13 | -0.48 | -0.72 | 0.11 | -0.57 | 0.24 | -1.04 | 0.65 | -0.08 | -0.41 | -1.56 | 0.2 | -0.39 | -0.01 | 0.225 | 0.2 | 0.0825 | 0.268632 | 0.390782 | 0.48544 | 1 | 0.999631 | 0.696662 |
| Cacnb4_Q8R0S4_S38s_1 | -0.64 | 0.06 | -1.1 | -0.23 | 0.11 | -0.26 | -0.89 | 0.03 | -0.43 | -0.44 | -0.24 | 0 | -0.95 | -0.03 | -0.66 | 0.06 | 0.3275 | 0.1775 | -0.12 | 0.830325 | 0.538331 | 0.544792 | 1 | 0.999631 | 0.942193 |
| Cacnb4_Q8R0S4_S423s_1 | 0.03 | -0.2 | -0.94 | -0.9 | 0.25 | 0.23 | -0.14 | -1.04 | 0.54 | -1.38 | -0.49 | 0.03 | -0.24 | -0.28 | -1.33 | -0.64 | 0.235 | 0.04 | -1.1275 | 0.755901 | 0.416232 | 0.70283 | 1 | 0.999631 | 0.913733 |
| Cacnb4_Q8R0S4_S443s_2 | 0.44 | 1.05 | 0.66 | 1.9 | 1.4 | 1.06 | 0.78 | 1.75 | 1.16 | 0.93 | 1.32 | 0.8 | -0.97 | 0.18 | 0.38 | -0.05 | -0.445 | 0.495 | -0.2425 | 0.026822 | 0.552526 | 0.913795 | 1 | 0.999631 | 0.422285 |
| Cacnb4_Q8R0S4_S447s_1 | 0.59 | 0.29 | 1.42 | -0.48 | -0.43 | 1 | -0.37 | -0.16 | 0.88 | 2.25 | 0.39 | 0.28 | 1.05 | -0.04 | 0.45 | -0.61 | 0.255 | 0.13 | -0.965 | 0.632524 | 0.37172 | 0.382095 | 1 | 0.999631 | 0.868271 |
| Cacnb4_Q8R0S4_S447s_2 | 0.86 | 1.13 | 0.66 | 1.9 | 1.61 | 1.43 | 0.78 | 1.75 | 1.44 | 1.51 | 1.32 | 0.8 | -0.24 | 0.6 | 0.38 | -0.05 | -0.3625 | -0.8625 | -1.55 | 0.025422 | 0.492093 | 0.713557 | 1 | 0.999631 | 0.422114 |
| Cacnb4_Q8R0S4_S507s_1 | 0.34 | -1.64 | 1.14 | 0.37 | -1.29 | -1.16 | 1.86 | -0.65 | -2.19 | 0.45 | -0.82 | -0.68 | -1.61 | -1.92 | -0.41 | -2.05 | -0.13 | 0.435 | -0.3475 | 0.032156 | 0.661455 | 0.24487 | 1 | 0.999631 | 0.442395 |
| Cacnb4_Q8R0S4_S50s_1 | 1.35 | 0.85 | 0.93 | 0.52 | 0.29 | 1.56 | 0.88 | 0.4 | 1.08 | 1.82 | 1.08 | 1.41 | 0.62 | 0.4 | 0.48 | 0.76 | 0.22 | 0.58 | 1.0425 | 0.259377 | 0.71853 | 0.184182 | 1 | 0.999631 | 0.692624 |
| Cacnb4_Q8R0S4_S64s_1 | -2.64 | 2.35 | -0.65 | -3.64 | 0.87 | 1.26 | -1.95 | -3.88 | 1.75 | 0.17 | -2.05 | -2.13 | 4.58 | -0.19 | -2.23 | -2.57 | -1.47 | -1.8375 | -1.435 | 0.558125 | 0.885033 | 0.675517 | 1 | 0.999631 | 0.838323 |
| Cacng2_O88602_S249s_1 | 1.74 | -1.89 | -0.88 | -2.03 | -3.76 | -1.42 | 0.32 | -4.08 | -4.15 | 0.82 | -3.55 | -3.53 | -0.84 | -2.33 | -3.63 | -2 | 1.2175 | 0.62 | 1.0625 | 0.134773 | 0.224073 | 0.160014 | 1 | 0.999631 | 0.606296 |
| Cacng2_O88602_S253s_1 | -2.4 | 1.58 | 1.02 | -0.44 | 1.42 | 2.06 | 0.37 | 0.78 | 1.69 | -1.26 | 1.02 | 0.79 | 2.35 | 0.75 | 0.86 | 0.05 | -0.4325 | -0.795 | -0.985 | 0.241994 | 0.168785 | 0.517292 | 1 | 0.999631 | 0.684914 |
| Cacng2_O88602_Y288y_1 | 0.74 | 0.65 | 0.49 | 1.06 | 0.1 | 0.38 | 0.43 | 0.3 | 0.15 | 0.12 | 0.11 | -0.62 | -0.44 | -0.69 | -0.4 | 0.53 | 1.6125 | 1.5425 | 1.755 | 0.017197 | 0.113624 | 0.022872 | 1 | 0.999631 | 0.391024 |
| Cacng3_Q9JJV5_S248s_1 | -4.62 | 2.65 | 2.37 | -0.47 | 2.27 | 1.63 | 1.06 | 1.42 | 2.17 | -2.01 | 3.03 | 2.91 | 3.21 | 0.86 | 1.93 | 0.95 | 0.705 | -0.6675 | -0.7075 | 0.2558 | 0.281492 | 0.38851 | 1 | 0.999631 | 0.690775 |
| Cacng3_Q9JJV5_S302s_1 | 2.24 | 0.42 | 0.55 | 1.54 | 1.89 | 2.1 | 1.99 | 1.59 | 1.09 | 0.2 | 1.46 | -0.67 | 0.32 | 0.23 | 0.85 | 0.52 | -0.555 | -0.1425 | -1.375 | 0.139972 | 0.12927 | 0.272414 | 1 | 0.999631 | 0.611718 |
| Cacng4_Q9JJV4_S228s_1 | 0.12 | -0.99 | 0.12 | -0.56 | -1.12 | -1.15 | 0.02 | -1.28 | -0.33 | -0.45 | -0.57 | -0.53 | -0.6 | -2.32 | -0.92 | -2.97 | -1.21 | -1.5625 | -1.6575 | 0.035801 | 0.195266 | 0.667419 | 1 | 0.999631 | 0.453811 |
| Cacng4_Q9JJV4_S253s_1 | 1.3 | -2.18 | -1.49 | -1.18 | -3.17 | -2.59 | 0.3 | -2.93 | -4.96 | 0.43 | -2.88 | -2.39 | -1.03 | -3.66 | -2.32 | -3.17 | 1.0325 | 1.19 | 1.66 | 0.068549 | 0.224843 | 0.196494 | 1 | 0.999631 | 0.523794 |
| Cacng4_Q9JJV4_S259s_1 | -2.78 | 1.39 | 1.14 | -0.54 | 1.13 | 0.68 | 0.21 | 1.32 | 1.38 | -1.48 | 1.93 | 2.14 | 2.66 | 0.79 | 1.86 | 0.54 | -0.3 | 0.0975 | 0.5375 | 0.09812 | 0.248871 | 0.293193 | 1 | 0.999631 | 0.561078 |
| Cacng8_S4R2F1_S251s_1 | -2.39 | -0.89 | -0.66 | -1.02 | -1.57 | -1.62 | -1.4 | -1.57 | -1.22 | -1.39 | -0.97 | -0.99 | 0.27 | -0.98 | -0.8 | -1.3 | 0.165 | 0.3875 | 0.86 | 0.29408 | 0.458252 | 0.81246 | 1 | 0.999631 | 0.708293 |
| Cacng8_S4R2F1_S254s_1 | -1.85 | -1.27 | -1.1 | -1.47 | -1.29 | -0.97 | -1.51 | -1.26 | -1.4 | -0.96 | -0.77 | -1.01 | 0.01 | -0.98 | -0.63 | -0.65 | 0.4475 | -0.335 | -0.1375 | 0.025545 | 0.557221 | 0.205455 | 1 | 0.999631 | 0.422114 |
| Cacng8_S4R2F1_S257s_1 | 0.5 | -0.95 | -0.76 | 0.2 | 1.14 | -0.83 | 0.56 | -0.09 | -0.02 | -1.34 | -0.48 | -0.51 | -0.79 | 0.21 | -0.35 | -0.63 | 0.5175 | 0.1375 | 0.5025 | 0.748173 | 0.394417 | 0.456538 | 1 | 0.999631 | 0.911931 |
| Cacng8_S4R2F1_S290s_1 | -1.1 | 0.23 | 0.02 | -0.73 | 0.51 | -0.06 | -0.27 | 0.31 | -0.11 | -1.44 | 0.28 | 0.24 | 0.55 | 0.3 | -0.22 | -0.2 | 0.935 | 0.4275 | -0.1175 | 0.220933 | 0.188865 | 0.778332 | 1 | 0.999631 | 0.671424 |
| Cacng8_S4R2F1_S357s_1 | -0.17 | -0.08 | -1.77 | 0.16 | 1.19 | 0.01 | 0.47 | 0.21 | 0.29 | -0.78 | 0.25 | 0.09 | -1.51 | 0 | -0.37 | -0.45 | -0.09 | -1.3025 | -2.0525 | 0.820595 | 0.077358 | 0.388236 | 1 | 0.999631 | 0.938379 |
| Cacng8_S4R2F1_T415t_1 | 1.77 | 0.77 | 1.76 | 2.33 | 0.69 | 2 | 2.01 | 1.57 | -0.17 | 0.91 | 1.07 | -0.39 | -2.04 | 0.17 | -0.03 | 0.32 | -0.23 | 0.445 | 0.07 | 0.005529 | 0.837094 | 0.019901 | 1 | 0.999631 | 0.351308 |
| Cactin_Q9CS00_S148s_1 | -0.4 | 0.52 | -1.16 | -0.24 | -0.03 | -0.66 | -1.3 | -0.21 | 0.25 | -0.52 | 0.41 | 0.36 | 0.54 | -0.78 | -0.96 | 0.2 | 0.0875 | -0.1075 | -0.235 | 0.885503 | 0.599974 | 0.296939 | 1 | 0.999631 | 0.963505 |
| Cactin_Q9CS00_S96s_1 | 1.08 | 1.13 | 1.21 | 0.98 | 1.38 | 1.14 | 1.12 | 1.11 | 1.33 | 1.05 | 0.98 | 0.61 | -0.02 | 1.53 | 0.51 | 1.44 | 0.4525 | 0.5225 | 0.4475 | 0.561361 | 0.711057 | 0.689875 | 1 | 0.999631 | 0.840316 |
| Cad_E9QAI5_T1770t_1 | -1.11 | -1.5 | -2.03 | -1.23 | -1.12 | -1.44 | -1 | -0.5 | -1.32 | -0.77 | -0.79 | -0.9 | -1.54 | -0.58 | -1.33 | -0.63 | -0.385 | -0.4275 | 0.655 | 0.23766 | 0.187162 | 0.116016 | 1 | 0.999631 | 0.683301 |
| Cadm2_G3UZM4_S423s_1 | -3.55 | -2.1 | 0.28 | 1.15 | -2.85 | -2.98 | -0.39 | 0.46 | -2.42 | -3.12 | -0.11 | -0.28 | -1.74 | -1.5 | 0.85 | 0.79 | -0.0975 | -0.185 | 0.45 | 0.553667 | 0.74533 | 0.70639 | 1 | 0.999631 | 0.837943 |
| Cadm4_Q8R464_S361s_1 | -1.07 | -1.44 | -1.28 | -1.64 | -1.88 | -1.28 | -1.51 | -1.15 | -1.9 | -0.76 | -1.57 | -1.94 | -0.54 | -1.07 | -1.21 | -0.81 | 1.075 | 0.6075 | 0.3625 | 0.15457 | 0.729222 | 0.592243 | 1 | 0.999631 | 0.625677 |
| Cadps_Q80TJ1_S375s_1 | -1.32 | -0.51 | -1.71 | -1.11 | 0.78 | -0.15 | 0.61 | -1.59 | -0.27 | -1.33 | 0 | -0.62 | 0.57 | -0.73 | -1.66 | -1.38 | -0.1475 | -0.4525 | 0.3825 | 0.494719 | 0.073713 | 0.148638 | 1 | 0.999631 | 0.810404 |
| Cadps_Q80TJ1_S486s_1 | -0.66 | -2.21 | -0.55 | -2.36 | -1.81 | -2.25 | -0.31 | -2 | -2.54 | -1.48 | -2.76 | -0.81 | -1.57 | -1.32 | -1.19 | -0.17 | 1.32 | 0.1075 | -0.2525 | 0.483342 | 0.803418 | 0.462796 | 1 | 0.999631 | 0.803286 |
| Cadps_Q80TJ1_S89s_1 | 1.2 | 1.07 | -1.99 | 1.42 | 2.58 | 1.4 | 2.25 | 0.75 | 2.09 | -0.91 | 1.24 | -0.29 | 1.06 | 0.71 | -0.18 | -0.9 | -0.225 | -0.2875 | 0.2225 | 0.753595 | 0.121032 | 0.907066 | 1 | 0.999631 | 0.913377 |
| Cadps_Q80TJ1_S96s_1 | -0.17 | -0.56 | 1.84 | 0.22 | -0.23 | 0.5 | -0.32 | 0.48 | -0.28 | 0.32 | -0.58 | 0.72 | -0.31 | 0.56 | 0.85 | 1.12 | -0.135 | -0.305 | 0.115 | 0.694789 | 0.671921 | 0.608264 | 1 | 0.999631 | 0.895302 |
| Cadps2_S4R1C6_S161s_1 | -0.81 | -0.5 | -0.66 | -0.74 | -0.74 | -0.75 | -1.12 | -0.64 | -0.81 | -0.86 | -1.08 | -1.18 | -0.34 | -0.62 | -0.19 | -1.1 | -0.12 | 0.2175 | 0.5475 | 0.709364 | 0.589815 | 0.244607 | 1 | 0.999631 | 0.899175 |
| Cadps2_S4R1C6_S39s_1 | -0.76 | -0.84 | 0.77 | 0.07 | -0.65 | 0.49 | -0.98 | -0.1 | -0.02 | 0.55 | -1.04 | 0.62 | -0.94 | 0.97 | 0.64 | 0.76 | 0.085 | 0.355 | -0.3575 | 0.325178 | 0.80041 | 0.670998 | 1 | 0.999631 | 0.72463 |
| Cadps2_S4R1C6_T811t_1 | -0.33 | -0.67 | 0.15 | -0.22 | -0.71 | -0.51 | 0.49 | 0 | -0.61 | -0.28 | 0.29 | 0.95 | -0.52 | -1.06 | -0.31 | -0.61 | -0.0475 | -0.165 | -0.1175 | 0.275631 | 0.807349 | 0.380295 | 1 | 0.999631 | 0.700187 |
| Calcoco1_E9Q7U2_S563s_1 | -0.27 | -0.21 | -0.81 | 0.04 | -0.39 | -0.41 | -0.76 | 0.12 | -0.44 | -0.61 | -0.62 | -0.24 | -0.61 | -0.73 | -0.19 | -0.19 | 0.695 | 0.3475 | 0.2025 | 0.710469 | 0.878163 | 0.570074 | 1 | 0.999631 | 0.899851 |
| Calml3_P0DP28_S82s_1 | -0.07 | 3.76 | 3.03 | 1.47 | 2.83 | 3.56 | 2.36 | 2.22 | 3.47 | 0.95 | 2.97 | 2.19 | 3.06 | 2.65 | 2.08 | 1.21 | -0.6925 | -0.1525 | 0.44 | 0.804817 | 0.389035 | 0.694793 | 1 | 0.999631 | 0.934231 |
| Camk1_Q91YS8_S324s_1 | -1.53 | -0.82 | 0.5 | -0.09 | -0.35 | -1.1 | -1.02 | -2.24 | -0.93 | -1.04 | -0.38 | -0.2 | 0.65 | -0.16 | -1.06 | 0.39 | 0.8475 | -0.225 | 0.49 | 0.425142 | 0.221578 | 0.746013 | 1 | 0.999631 | 0.774293 |
| Camk1_Q91YS8_T177t_1 | -1.23 | 0.17 | -2.05 | -2.64 | -0.41 | 1.89 | -1.5 | -2.34 | -0.97 | -0.68 | -2.21 | -2.79 | -0.8 | 2.53 | -3.38 | -2.14 | 0.465 | -0.4325 | 0.2925 | 0.6828 | 0.382003 | 0.749068 | 1 | 0.999631 | 0.889602 |
| Camk1_Q91YS8_T339t_1 | -0.72 | -0.11 | -0.89 | -0.63 | -0.04 | 0.06 | -0.45 | -0.06 | -1.05 | -1.69 | -0.92 | -0.42 | -0.31 | -0.32 | -0.09 | -0.46 | 0.2175 | 0.0725 | 0.2675 | 0.335221 | 0.123308 | 0.235156 | 1 | 0.999631 | 0.730219 |
| Camk1d_Q8BW96_S319s_1 | -0.9 | -0.59 | -1.2 | -0.8 | -0.09 | -0.96 | -0.76 | -0.81 | -0.53 | -1.13 | -1.1 | -0.44 | -0.42 | -0.31 | -1.11 | -0.58 | -0.15 | 0.0975 | -0.6325 | 0.398644 | 0.469943 | 0.80884 | 1 | 0.999631 | 0.761726 |
| Camk1d_Q8BW96_S349s_1 | 0.83 | -0.32 | 0.23 | 0.85 | -0.42 | 0.19 | 0.72 | 0.5 | 0.31 | 1.06 | 0.45 | 0.16 | -0.35 | -0.8 | 0.46 | -0.25 | 0.7975 | -0.175 | 0.2175 | 0.140725 | 0.697426 | 0.792195 | 1 | 0.999631 | 0.612258 |
| Camk1d_Q8BW96_T172t_1 | -1.39 | -0.42 | 0.39 | -0.11 | -1.55 | 1.43 | -0.14 | 1.92 | -1.74 | -1.13 | 1.61 | -0.97 | -1.33 | 1.42 | -0.43 | -0.32 | -0.2775 | 2.4525 | 2.0825 | 0.726838 | 0.311549 | 0.812042 | 1 | 0.999631 | 0.905774 |
| Camk2a_F8WIS9_S272s_1 | -0.24 | 5.45 | 5.74 | 0.29 | 3.35 | 2.14 | 2.57 | 2.07 | 7.31 | -0.3 | 5.59 | 8.45 | 11.45 | 1.64 | 3.87 | 2.61 | -0.7825 | -0.5075 | -0.4675 | 0.374056 | 0.842056 | 0.267371 | 1 | 0.999631 | 0.747035 |
| Camk2a_F8WIS9_S275s_1 | 0.21 | 1.11 | 2.89 | 0.38 | -0.1 | 0.34 | 0.61 | 0.61 | 0.23 | 0.73 | 1.09 | 0.51 | 0.75 | 0.35 | 0.82 | 0.8 | 0.2225 | -0.5925 | 0.5725 | 0.42293 | 0.196672 | 0.395174 | 1 | 0.999631 | 0.773293 |
| Camk2a_F8WIS9_S275s_2 | 0.11 | 2.18 | 4.17 | -0.03 | 3.46 | 0.74 | 0.02 | 3.1 | 0.7 | 1.61 | 0.46 | 1.29 | 5 | 1.54 | 1.16 | 1.02 | -1.5475 | -0.6725 | -0.185 | 0.62433 | 0.842519 | 0.512041 | 1 | 0.999631 | 0.86356 |
| Camk2a_F8WIS9_S279s_1 | 0.27 | 1.66 | 3.85 | -0.55 | -1.54 | 0.38 | 0.16 | 0.04 | -0.26 | 1.18 | 1.27 | 0.35 | 2.3 | 0.72 | 0.3 | 1.17 | 0.2225 | -0.5925 | 0.5725 | 0.838596 | 0.115531 | 0.459028 | 1 | 0.999631 | 0.946365 |
| Camk2a_F8WIS9_S279s_2 | 0.11 | 2.18 | 4.17 | -0.03 | 3.46 | 0.74 | 0.02 | 3.1 | 0.7 | 1.61 | 0.46 | 1.29 | 5 | 1.54 | 1.16 | 1.02 | -0.435 | 0.0875 | 0.085 | 0.62433 | 0.842519 | 0.512041 | 1 | 0.999631 | 0.86356 |
| Camk2a_F8WIS9_S481s_1 | 0.23 | -0.12 | -0.01 | 0.21 | -0.89 | -0.11 | 0.09 | -0.52 | 0.09 | -0.35 | 1.53 | -0.61 | 0.7 | -0.25 | 0.11 | 0.09 | -0.52 | 0.4375 | -0.8325 | 0.783871 | 0.167758 | 0.852004 | 1 | 0.999631 | 0.922628 |
| Camk2a_F8WIS9_S485s_1 | -0.57 | -1.5 | -0.7 | -0.22 | -1.11 | -2.76 | -1.12 | -0.08 | -1.65 | -0.71 | 0.12 | 1 | -1.41 | -1.62 | -2.28 | -1.01 | -0.8875 | -0.32 | 1.255 | 0.061081 | 0.36686 | 0.45488 | 1 | 0.999631 | 0.509355 |
| Camk2a_F8WIS9_T253t_1 | -0.55 | 2.16 | 2.74 | 0.81 | 0.22 | 1.33 | -0.29 | 0.35 | 0.96 | 0.92 | 0.76 | 1.24 | 4.7 | 2.11 | 1.07 | 2.3 | -0.685 | -0.57 | -0.6925 | 0.190826 | 0.232797 | 0.632125 | 1 | 0.999631 | 0.648674 |
| Camk2a_F8WIS9_T286t_1 | 0.97 | 2.18 | 3.09 | 1.34 | 0.51 | 1.96 | 1.22 | 1.15 | 0.82 | 1.75 | 2.03 | 0.7 | 1.34 | 1.51 | 0.28 | 1.68 | 0.32 | 0.7725 | 0.3475 | 0.213245 | 0.206674 | 0.303511 | 1 | 0.999631 | 0.666787 |
| Camk2a_P11798_S330s_1 | -0.03 | 1 | -1.67 | -1.93 | 0.03 | 0.92 | -0.76 | -1.54 | 1.89 | 0.67 | -0.77 | -1.33 | 1.54 | 0.07 | -1.43 | -1.42 | -0.0975 | -0.1675 | -0.0125 | 0.687069 | 0.676805 | 0.387433 | 1 | 0.999631 | 0.891291 |
| Camk2a_P11798_S331s_1 | -1.64 | -0.85 | 1.05 | -0.27 | -0.87 | -1.47 | 0.08 | 0.16 | -1.73 | -1.27 | 0.43 | 0.19 | -0.67 | -1.8 | 0.12 | 0.59 | -0.055 | 0.0325 | -0.27 | 0.985444 | 0.874967 | 0.809337 | 1 | 0.999631 | 0.995133 |
| Camk2a_P11798_S333s_2 | -1.44 | -1.13 | -1.97 | -2.1 | -1.12 | -0.94 | -1.62 | -3.18 | -1.04 | -0.91 | -1.78 | -2.78 | -0.49 | -1.58 | -3.02 | -2.63 | 0.06 | 0.155 | 0.0725 | 0.634805 | 0.915408 | 0.944768 | 1 | 0.999631 | 0.868886 |
| Camk2a_P11798_T334t_1 | -0.53 | 0.4 | -0.1 | -0.13 | -0.28 | 0.37 | 0.21 | -0.42 | -0.19 | 0.09 | 0.26 | 0.1 | 0.04 | -0.57 | 0.23 | 0.23 | 0.145 | 0.005 | -0.0525 | 0.829601 | 0.850457 | 0.600774 | 1 | 0.999631 | 0.942058 |
| Camk2a_P11798_T334t_2 | -2.56 | -2.68 | 0.25 | -0.01 | -2.06 | -2.17 | -0.01 | -0.18 | -2.25 | -2.16 | -0.28 | -0.29 | -1.66 | -3.11 | 0.09 | -0.53 | -0.4725 | 0.2525 | 0.12 | 0.954072 | 0.865437 | 0.995276 | 1 | 0.999631 | 0.984441 |
| Camk2a_P11798_T336t_1 | -1.32 | 0.78 | 0.76 | -0.37 | -0.71 | -0.08 | -0.61 | -0.64 | 0.25 | 0.14 | 0.18 | 0.29 | 1.39 | -0.51 | -0.36 | -0.19 | 0.2675 | 0.1325 | -0.0875 | 0.843806 | 0.350601 | 0.604716 | 1 | 0.999631 | 0.948343 |
| Camk2a_P11798_T336t_2 | -1.85 | -1.33 | -0.13 | -0.33 | -0.5 | -1.16 | -0.24 | -0.67 | -1.11 | -0.9 | -0.56 | -0.54 | -1 | -1.73 | -0.39 | -0.87 | -0.005 | 0.475 | 0.1175 | 0.855312 | 0.547283 | 0.758698 | 1 | 0.999631 | 0.951817 |
| Camk2a_P11798_T337t_1 | -1.4 | -0.57 | -0.22 | -0.6 | -1.27 | -0.44 | -0.19 | -0.91 | -0.62 | -0.6 | 0.29 | 0.04 | 0.24 | -1.48 | -0.31 | -0.77 | 0.005 | 0.0025 | -0.0675 | 0.791226 | 0.989119 | 0.216409 | 1 | 0.999631 | 0.926539 |
| Camk2a_P11798_T337t_2 | -1.61 | -1.09 | 0.15 | -0.09 | -1.16 | -1.01 | -0.12 | -0.33 | -1.02 | -1 | -0.29 | -0.32 | -0.42 | -1.69 | -0.05 | -0.75 | -0.215 | -0.4525 | 0.0125 | 0.89578 | 0.991417 | 0.995589 | 1 | 0.999631 | 0.966139 |
| Camk2b_P28652_S331s_1 | 0.42 | -0.11 | 0.53 | 0.32 | 0.65 | -0.14 | -0.25 | 0.04 | -0.79 | -0.19 | -0.39 | 0.72 | 0.58 | 0.01 | 0.15 | 0.47 | 0.235 | -0.1275 | -0.435 | 0.966713 | 0.485693 | 0.244607 | 1 | 0.999631 | 0.988351 |
| Camk2b_P28652_S363s_1 | 0.61 | 0.33 | 0.11 | 1.18 | 0.76 | 0.96 | 0.81 | 0.64 | 0.79 | 0.51 | 0.54 | -0.12 | 0.04 | 0.63 | -0.18 | 0 | -0.2075 | -0.2175 | -0.32 | 0.230677 | 0.443895 | 0.712109 | 1 | 0.999631 | 0.681463 |
| Camk2b_P28652_S367s_1 | -0.59 | -0.31 | 0.6 | 0.02 | -0.65 | -0.25 | 0.36 | -0.57 | -0.63 | -0.47 | 0.36 | -0.41 | -0.11 | -0.95 | 0.24 | -0.74 | 0.1225 | -0.2425 | 0.2025 | 0.431558 | 0.57265 | 0.557107 | 1 | 0.999631 | 0.778356 |
| Camk2b_P28652_S371s_1 | -0.16 | 0.64 | 0.98 | 1.43 | 0.17 | 0.84 | 0.81 | 1.56 | -0.16 | 0.27 | 0.32 | 1.49 | 0.2 | 0.64 | 0.96 | 1.9 | -0.3825 | -0.3225 | -0.4025 | 0.673688 | 0.776628 | 0.608923 | 1 | 0.999631 | 0.885515 |
| Camk2b_P28652_T287t_1 | 0.48 | 1.1 | 1.78 | 0.63 | 0.09 | 1.03 | 0.7 | 0.64 | 0.4 | 1.25 | 0.79 | 0.26 | 0.59 | 1.04 | -0.06 | 0.81 | -0.3475 | -1.9075 | -0.94 | 0.327402 | 0.314841 | 0.412395 | 1 | 0.999631 | 0.724678 |
| Camk2b_P28652_T382t_1 | -0.25 | 1.15 | 1.34 | 0.4 | -1.14 | 1.81 | 0.72 | -0.14 | -1.68 | -0.75 | 1.48 | -4.04 | -0.05 | 0.85 | -2.17 | 0.25 | 0.045 | 0.7 | 0.2775 | 0.18262 | 0.594627 | 0.090092 | 1 | 0.999631 | 0.643861 |
| Camk2b_P28652_T400t_1 | -2 | -0.6 | -0.08 | -1.05 | -0.99 | -0.93 | -0.26 | -1.37 | -0.12 | -0.69 | 0.04 | -0.16 | 0.53 | -1.37 | -0.84 | -0.94 | -0.135 | 0.25 | 0.1525 | 0.610319 | 0.920596 | 0.131692 | 1 | 0.999631 | 0.859048 |
| Camk2b_P28652_T400t_2 | -0.98 | -0.29 | -0.54 | -1.1 | -1.1 | -0.19 | -0.69 | -1.47 | -0.41 | -0.03 | -0.91 | -0.56 | 0.33 | -0.86 | -0.74 | -1.03 | -0.13 | 0.0475 | 0.15 | 0.697278 | 0.707275 | 0.443857 | 1 | 0.999631 | 0.895731 |
| Camk2b_P28652_T401t_2 | -1.3 | -0.88 | -0.42 | -1.03 | -1.37 | -0.68 | -0.78 | -1.32 | -0.84 | -0.47 | -0.97 | -1.16 | 0.01 | -1.17 | -0.53 | -1.34 | 0.06 | 0.485 | 0.17 | 0.701662 | 0.677415 | 0.876593 | 1 | 0.999631 | 0.896835 |
| Camk2b_Q5SVI1_S427s_1 | -1.01 | -0.55 | -1.16 | -0.63 | -1.49 | -0.87 | -0.61 | -0.14 | -0.4 | -0.64 | 0.27 | -0.64 | -1.17 | -0.7 | -0.77 | -0.03 | 0.045 | 0.7 | 0.2775 | 0.621222 | 0.863112 | 0.153145 | 1 | 0.999631 | 0.861888 |
| Camk2b_Q5SVJ0_T400t_1 | -2 | -0.6 | -0.08 | -1.05 | -0.99 | -0.93 | -0.26 | -1.37 | -0.12 | -0.69 | 0.04 | -0.16 | 0.53 | -1.37 | -0.84 | -0.94 | -0.135 | 0.25 | 0.1525 | 0.610319 | 0.920596 | 0.131692 | 1 | 0.999631 | 0.859048 |
| Camk2b_Q5SVJ0_T400t_2 | -0.98 | -0.29 | -0.54 | -1.1 | -1.1 | -0.19 | -0.69 | -1.47 | -0.41 | -0.03 | -0.91 | -0.56 | 0.33 | -0.86 | -0.74 | -1.03 | -0.13 | 0.0475 | 0.15 | 0.697278 | 0.707275 | 0.443857 | 1 | 0.999631 | 0.895731 |
| Camk2b_Q5SVJ0_T401t_2 | -1.3 | -0.88 | -0.42 | -1.03 | -1.37 | -0.68 | -0.78 | -1.32 | -0.84 | -0.47 | -0.97 | -1.16 | 0.01 | -1.17 | -0.53 | -1.34 | -0.225 | -0.0875 | 0.415 | 0.701662 | 0.677415 | 0.876593 | 1 | 0.999631 | 0.896835 |
| Camk2d_E9Q1T1_S235s_1 | -0.56 | -0.56 | -0.82 | -0.93 | -0.8 | -0.84 | -1.15 | -0.98 | -1 | -1.38 | -0.7 | -0.14 | 0.1 | -0.33 | -0.76 | -0.22 | -1.1775 | 0.1925 | 0.865 | 0.191542 | 0.374417 | 0.791231 | 1 | 0.999631 | 0.648674 |
| Camk2d_E9Q1T1_S276s_1 | -0.39 | 0.98 | 1.26 | -1.03 | -1.28 | -0.82 | -0.79 | -1 | 0.04 | 1.02 | -0.45 | 0.98 | 3.5 | -0.05 | 0.11 | 0.72 | -2.045 | -0.3975 | 1.015 | 0.326239 | 0.044308 | 0.748055 | 1 | 0.999631 | 0.724678 |
| Camk2d_E9Q1T1_S276s_2 | -0.8 | 3.5 | 3.91 | -0.52 | -1.85 | 1.08 | -0.94 | -0.38 | 1.03 | 1.86 | 1.04 | 0.57 | 5.62 | 1.6 | 1.23 | 1.7 | -1.855 | -0.31 | 0.2325 | 0.466301 | 0.113076 | 0.720994 | 1 | 0.999631 | 0.795267 |
| Camk2d_E9Q1T1_S280s_1 | -0.41 | 1.6 | 2.5 | -1.33 | -3.04 | -0.25 | -0.9 | -0.87 | -0.54 | 1.06 | 0.28 | 0.32 | 3.22 | 0.16 | -1.03 | 0.94 | -2.045 | -0.3975 | 1.015 | 0.828171 | 0.069513 | 0.707699 | 1 | 0.999631 | 0.941315 |
| Camk2d_E9Q1T1_S280s_2 | -0.8 | 3.5 | 3.91 | -0.52 | -1.85 | 1.08 | -0.94 | -0.38 | 1.03 | 1.86 | 1.04 | 0.57 | 5.62 | 1.6 | 1.23 | 1.7 | 0.165 | 0.1225 | 0.265 | 0.466301 | 0.113076 | 0.720994 | 1 | 0.999631 | 0.795267 |
| Camk2d_E9Q1T1_S338s_1 | 0.42 | -0.21 | 0.42 | 0.62 | 0.27 | -0.38 | 1.2 | 0.82 | 0.31 | -0.11 | 1.15 | 0.39 | 0.09 | -0.14 | 1.27 | 1.09 | -0.535 | -0.4175 | 0.4075 | 0.525755 | 0.677238 | 0.730512 | 1 | 0.999631 | 0.82353 |

| | | | | | | | | | | | | | | | | | | | | | | | | | |
|---|---|---|---|---|---|---|---|---|---|---|---|---|---|---|---|---|---|---|---|---|---|---|---|---|---|
| Camk2d_E9Q1T1_S524s_1 | -0.49 | 0.33 | -0.45 | 0.07 | -0.52 | -1.03 | -0.81 | -0.32 | 0.17 | -1.04 | -0.53 | -0.81 | -0.01 | 0.82 | -0.1 | 0.38 | 0.0025 | 0.5575 | -0.105 | 0.259295 | 0.109045 | 0.265904 | 1 | 0.999631 | 0.692624 |
| Camk2d_E9Q1T1_S528s_1 | -0.81 | -0.5 | 0.87 | 0.47 | -1.05 | -0.72 | 0.16 | 1.65 | -0.61 | -0.47 | 1.84 | 1.5 | -1.53 | -1.27 | 0.59 | 1.82 | -0.6875 | -0.5725 | -0.7 | 0.892396 | 0.996905 | 0.417485 | 1 | 0.999631 | 0.964421 |
| Camk2d_E9Q1T1_T287t_1 | 0.97 | 2.18 | 3.11 | 1.35 | 0.51 | 1.96 | 1.23 | 1.16 | 0.82 | 1.75 | 2.04 | 0.71 | 1.34 | 1.51 | 0.28 | 1.68 | -0.1025 | -0.4825 | -0.61 | 0.210283 | 0.206788 | 0.303303 | 1 | 0.999631 | 0.66441 |
| Camk2g_Q923T9_S334s_1 | 1.23 | -0.39 | -0.38 | 0.35 | 0.2 | -0.93 | 0.93 | 0.2 | -0.39 | -0.44 | 0.62 | -0.91 | -0.12 | -0.19 | -1.52 | 0.2 | 0.115 | -0.1375 | 0.255 | 0.251691 | 0.839882 | 0.328634 | 1 | 0.999631 | 0.687531 |
| Camk2g_Q923T9_S349s_1 | -0.02 | 0.71 | 0.19 | 0.18 | 0.05 | 0.77 | 0.28 | 0.42 | 0.22 | -0.25 | 0.22 | 0.32 | 0.27 | 0.59 | 0.34 | 0.88 | -0.5675 | -0.515 | -0.59 | 0.413787 | 0.692042 | 0.636111 | 1 | 0.999631 | 0.768655 |
| Camk2g_Q923T9_T287t_1 | 1.14 | 2.06 | 2.56 | 1.45 | 0.51 | 1.82 | 1.23 | 1.38 | 0.61 | 1.69 | 2.03 | 0.82 | 1.04 | 1.09 | 0.92 | 1.8 | -0.0875 | -0.9775 | -0.7775 | 0.161515 | 0.193114 | 0.27351 | 1 | 0.999631 | 0.631442 |
| Camk4_P08414_S337s_1 | -0.12 | -1.72 | -2.83 | -1.03 | -1.82 | -1.82 | -1.07 | -1.34 | -1.88 | -2.48 | -1.65 | -3.6 | -0.82 | -2.98 | -3.57 | -1.44 | 1.1975 | -0.4125 | 0.32 | 0.316594 | 0.873866 | 0.154334 | 1 | 0.999631 | 0.720734 |
| Camk4_P08414_T196t_1 | -1.18 | 0.74 | 0.49 | -0.14 | -0.42 | 2.84 | 0.04 | 2.24 | -1.05 | -1.11 | 2.37 | -1.95 | -0.75 | 2.93 | -0.48 | -0.51 | 0.1 | 0.12 | -0.0325 | 0.706618 | 0.155426 | 0.652251 | 1 | 0.999631 | 0.898087 |
| Camkk1_Q8VBY2_S458s_1 | -0.84 | 0.72 | 0.32 | -0.33 | 0.38 | 0.21 | -0.06 | -0.26 | 0.14 | -0.6 | 0.65 | 0.16 | 0.29 | 0.02 | -0.23 | -0.34 | 0.1625 | 0.32 | 0.3725 | 0.934895 | 0.795079 | 0.780545 | 1 | 0.999631 | 0.979037 |
| Camkk1_Q8VBY2_S475s_1 | -1.31 | 1.13 | 0.33 | 0.63 | 0.39 | 0.37 | 0.14 | 0.53 | 0.59 | 0 | 0.58 | 0.89 | 0.56 | 0.2 | 0.81 | 0.7 | 0.2 | 0.1475 | 0.225 | 0.476333 | 0.746011 | 0.545775 | 1 | 0.999631 | 0.799882 |
| Camkk1_Q8VBY2_S492s_1 | -1.81 | 0.62 | -0.37 | -0.48 | 0.23 | -0.99 | -0.36 | -0.12 | 0.6 | -2.1 | -0.35 | 0.4 | 0.58 | -0.48 | -0.84 | -0.4 | -0.8475 | -1.145 | -0.6175 | 0.680826 | 0.702005 | 0.834118 | 1 | 0.999631 | 0.888714 |
| Camkk1_Q8VBY2_S82s_1 | 0.26 | 0.48 | -0.04 | 0.63 | -0.52 | -1.15 | -0.18 | -0.21 | -0.88 | -2.83 | 0.06 | 0.4 | -1.2 | 0.65 | -0.09 | -0.5 | 0.125 | -1.5875 | -2.65 | 0.167858 | 0.022439 | 0.109269 | 1 | 0.999631 | 0.637399 |
| Camkk1_Q8VBY2_T108t_1 | 1.89 | 0.99 | 0.69 | 2.74 | 2.01 | 0.85 | 2.6 | 1.35 | -0.13 | -0.1 | 2.18 | -1.99 | -4.13 | 0.54 | -0.35 | -0.35 | 0.2275 | -0.13 | -0.11 | 0.020392 | 0.820693 | 0.086202 | 1 | 0.999631 | 0.393617 |
| Camkk2_Q8C078_S129s_1 | 0.46 | 0.71 | -1.11 | 0.44 | 0.67 | 0.55 | 0.17 | 0.02 | 0.64 | -0.18 | -0.54 | 0.06 | -0.81 | 0.88 | -0.01 | 0 | -0.03 | -0.2475 | -0.0675 | 0.83037 | 0.602667 | 0.781727 | 1 | 0.999631 | 0.942193 |
| Camkk2_Q8C078_S133s_1 | -0.28 | 0.21 | -0.49 | 0.99 | -0.21 | 0.1 | -0.22 | 0.64 | -0.19 | -0.86 | -0.03 | 0.52 | -0.15 | -0.38 | 0.46 | 0.23 | 0.485 | 0.51 | 0.305 | 0.866628 | 0.93911 | 0.570553 | 1 | 0.999631 | 0.956432 |
| Camkk2_Q8C078_S495s_1 | -1.19 | -0.06 | -0.28 | -0.79 | 0.11 | 0 | -0.16 | -0.33 | 0.08 | -0.96 | 0.5 | 0.1 | 0.45 | -0.43 | -0.62 | -0.5 | 0.35 | 0.2325 | 0.4475 | 0.436624 | 0.152928 | 0.232794 | 1 | 0.999631 | 0.780748 |
| Camkk2_Q8C078_S511s_1 | 0.42 | 1.41 | 0.65 | 0.38 | 0.72 | 1.35 | 0.92 | 1.27 | 0.71 | 0.47 | 1.2 | 1.41 | 0.94 | 1.28 | 0.8 | 1.63 | 0.315 | 0.0875 | -1.65 | 0.226957 | 0.297152 | 0.519934 | 1 | 0.999631 | 0.678092 |
| Camkk2_Q8C078_S522s_1 | 1.82 | -2.11 | -0.27 | 0.17 | -0.91 | 0.31 | 2.5 | -1.03 | -1.74 | 2.1 | 0.57 | -0.97 | -2.73 | -2.38 | -0.89 | -0.99 | -0.0275 | 0.1875 | 0.4825 | 0.065901 | 0.750784 | 0.931102 | 1 | 0.999631 | 0.516933 |
| Camkk2_Q8C078_S572s_1 | 0.28 | 0.95 | 0.67 | 0.48 | 0.45 | 0.85 | 0.08 | 0.89 | 0.64 | 0.55 | 0.57 | 1.37 | 0.88 | 1.45 | 0.86 | 1.12 | 0.46 | -0.02 | -0.405 | 0.133414 | 0.927053 | 0.549515 | 1 | 0.999631 | 0.60492 |
| Camkk2_Q8C078_S99s_1 | 0.87 | -0.3 | -0.96 | 0.87 | 1.28 | -0.48 | 0.96 | 0.56 | 0.23 | -0.25 | 0.54 | -0.12 | -1.15 | 0.62 | -0.45 | -0.16 | 0.435 | 0.48 | 0.06 | 0.463021 | 0.408014 | 0.966076 | 1 | 0.999631 | 0.793564 |
| Camkv_Q3UHL1_S387s_1 | -1.47 | 0.65 | 0.59 | 0.5 | 0.13 | 0.48 | 1.15 | 0.25 | 1.02 | -0.75 | 1.14 | 0.78 | 0.54 | -0.65 | 0.43 | 0.19 | -0.0975 | 0.31 | -0.4775 | 0.911748 | 0.412749 | 0.441182 | 1 | 0.999631 | 0.971848 |
| Camkv_Q3UHL1_S392s_1 | -0.1 | -0.24 | -0.04 | 0.05 | -0.26 | -0.34 | 0.43 | -0.55 | 0.24 | -0.01 | 0.33 | 0.35 | -0.44 | -0.87 | -0.25 | -0.68 | -0.3575 | -0.205 | -1.4425 | 0.114405 | 0.738488 | 0.233541 | 1 | 0.999631 | 0.579001 |
| Camkv_Q3UHL1_S436s_1 | 3 | 0.71 | 0.72 | 1.75 | 0.29 | 1.2 | 2.53 | 0.73 | 1.31 | 2.22 | 1.27 | 0.56 | -0.78 | -0.07 | 0.65 | 0.61 | -0.8025 | -0.0525 | -1.2925 | 0.031416 | 0.585088 | 0.727214 | 1 | 0.999631 | 0.441911 |
| Camkv_Q3UHL1_S436s_2 | 0.77 | 1.93 | 2.17 | 1.21 | 0.51 | -0.19 | 1.98 | 0.57 | 1.59 | 1.63 | 1.44 | 1.21 | 1.55 | -0.7 | -0.88 | 0.94 | 0.0075 | -0.815 | -0.92 | 0.059252 | 0.145416 | 0.885947 | 1 | 0.999631 | 0.504286 |
| Camkv_Q3UHL1_S506s_1 | 3.61 | -0.23 | -2.4 | -0.48 | 1.51 | -0.15 | 0.8 | -1.63 | 0.14 | -0.97 | -0.67 | -1.26 | -1.29 | -1.05 | -0.68 | -0.16 | 0.3075 | -0.2175 | 0.32 | 0.407633 | 0.995089 | 0.469853 | 1 | 0.999631 | 0.766996 |
| Camkv_Q3UHL1_T430t_1 | -0.81 | -0.35 | -0.93 | -0.65 | -0.36 | -0.11 | -0.78 | -0.26 | -0.55 | -0.87 | -1.39 | -0.8 | -0.26 | 0.19 | -0.7 | -0.69 | -0.38 | -0.8775 | -1.8 | 0.336588 | 0.281236 | 0.46931 | 1 | 0.999631 | 0.730499 |
| Camkv_Q3UHL1_T438t_1 | 2.42 | -0.35 | 0.25 | 0.36 | -0.38 | 0.39 | 1.49 | -0.34 | -0.61 | 1.16 | -0.24 | -1.14 | -1.74 | -1.03 | -0.64 | -1.11 | -0.5475 | 0.1075 | -0.7575 | 0.011598 | 0.570322 | 0.225597 | 1 | 0.999631 | 0.385092 |
| Camkv_Q3UHL1_T438t_2 | 0.54 | 0.84 | -0.83 | -0.55 | -0.19 | -0.77 | -0.53 | -0.7 | 0.75 | 1.09 | -0.6 | -0.81 | 0.6 | -1.29 | -1.62 | -0.72 | -0.705 | -0.73 | -0.9225 | 0.213851 | 0.214604 | 0.851755 | 1 | 0.999631 | 0.667209 |
| Camkv_Q3UHL1_T441t_1 | 1.59 | -0.28 | 1.07 | -0.19 | -1.52 | 0.42 | 0.95 | -0.48 | -1.2 | 1.53 | -0.32 | -0.74 | 0.94 | -1.06 | -0.88 | -0.5 | -0.9125 | -0.0225 | -1.8375 | 0.143277 | 0.285217 | 0.296146 | 1 | 0.999631 | 0.614578 |
| Camkv_Q3UHL1_T441t_2 | 1.01 | -0.12 | 2.12 | 1.18 | -0.66 | -1.24 | 1.92 | 0.52 | 0.04 | 1.56 | 1.36 | 1.14 | -0.53 | -2.47 | -1 | 0.84 | -0.1575 | -0.0325 | 0.215 | 0.027933 | 0.233748 | 0.9663 | 1 | 0.999631 | 0.423977 |
| Camkv_Q3UHL1_T446t_1 | -0.31 | -0.3 | -0.66 | -0.87 | -0.46 | -0.53 | -1.05 | -0.73 | -0.44 | -0.35 | -0.8 | -0.68 | 0.09 | -0.12 | -0.89 | -0.36 | 0.0575 | -0.0225 | 0.3425 | 0.51725 | 0.572632 | 0.90627 | 1 | 0.999631 | 0.819449 |
| Camkv_Q3UHL1_T446t_2 | -1.46 | -1.78 | -1.46 | -0.9 | -1.39 | -2.17 | -0.98 | -0.83 | -1.66 | -2.26 | -0.74 | -1.03 | -0.96 | -1.17 | -1.41 | -0.69 | 0.1925 | 0.04 | 0.2625 | 0.300165 | 0.87648 | 0.95491 | 1 | 0.999631 | 0.713499 |
| Camkv_Q3UHL1_T470t_1 | -0.32 | -0.4 | -0.38 | -0.54 | -0.01 | -0.38 | -0.51 | 0.03 | -0.09 | -0.39 | -0.52 | -0.48 | -0.53 | 0.28 | -0.34 | 0 | 0.4125 | 0.1425 | 0.18 | 0.385528 | 0.457008 | 0.873563 | 1 | 0.999631 | 0.756126 |
| Camsap1_A0A0A0MQE5_S1069s_1 | -0.68 | -0.83 | -0.56 | 0.01 | 0.6 | -0.43 | -0.18 | -0.4 | 0.07 | -1.24 | -0.52 | 0.2 | 0.29 | -0.86 | -0.75 | -0.02 | 0.4425 | -0.46 | -0.88 | 0.632573 | 0.241389 | 0.717366 | 1 | 0.999631 | 0.868271 |
| Camsap1_A0A0A0MQE5_S1133s_1 | 0.46 | -0.93 | 0.33 | 0.42 | 0.36 | -0.4 | -0.17 | 2.26 | -0.23 | -0.55 | -0.09 | -0.69 | -1.39 | -0.54 | -0.55 | -0.76 | 0.145 | 0.3775 | -0.145 | 0.053206 | 0.481625 | 0.247468 | 1 | 0.999631 | 0.490801 |
| Camsap1_A0A0A0MQE5_S1407s_1 | -0.09 | 1.11 | -0.23 | 0.82 | 1.34 | -0.33 | 0.36 | 0.82 | 1.91 | -0.9 | 1.2 | 0.91 | 0.71 | -0.66 | 0.13 | 0.85 | -0.235 | 0.0875 | 0.125 | 0.757158 | 0.755217 | 0.546192 | 1 | 0.999631 | 0.914382 |
| Camsap1_A0A0A0MQE5_S550s_1 | -1.25 | -1.02 | 0.45 | -1.04 | -1.6 | -0.78 | -0.97 | -0.45 | -0.52 | -0.61 | -1.08 | -0.3 | 0.08 | -1.44 | -0.42 | -0.58 | 0.7625 | 0.945 | 0.4175 | 0.79749 | 0.600899 | 0.83725 | 1 | 0.999631 | 0.930124 |
| Camsap1_A0A0A0MQE5_S550s_2 | -0.53 | -0.16 | -0.72 | -0.62 | -0.02 | 0.58 | -0.03 | 0.49 | 0.56 | 0.39 | 0.65 | 0.15 | 0.87 | -0.16 | -1.47 | 0.4 | 0.715 | -0.0225 | 0.28 | 0.411064 | 0.021223 | 0.005874 | 1 | 0.999631 | 0.767937 |
| Camsap1_A0A0A0MQE5_S553s_1 | 1.43 | -1.31 | -0.77 | 1.51 | 1.09 | 0.6 | 0.86 | 1.17 | -0.02 | 0.19 | 0.52 | 0.08 | -0.39 | 0.96 | 0.47 | 0.94 | 0.7625 | 0.945 | 0.4175 | 0.69301 | 0.295656 | 0.973003 | 1 | 0.999631 | 0.894912 |
| Camsap1_A0A0A0MQE5_S553s_2 | -0.53 | -0.16 | -0.72 | -0.62 | -0.02 | 0.58 | -0.03 | 0.49 | 0.56 | 0.39 | 0.65 | 0.15 | 0.87 | -0.16 | -1.47 | 0.4 | 0.32 | 0.0675 | -0.605 | 0.411064 | 0.021223 | 0.005874 | 1 | 0.999631 | 0.767937 |
| Camsap1_A0A0A0MQE5_S626s_1 | 1.88 | -0.13 | -1.05 | 1.3 | 1.34 | 0.06 | 0.86 | 1.02 | 0.7 | 0.5 | 0.9 | 0.17 | -0.69 | 0.38 | -0.11 | 0 | 0.485 | -0.005 | -0.0775 | 0.351286 | 0.62157 | 0.913348 | 1 | 0.999631 | 0.738069 |
| Camsap1_A0A0A0MQE5_S718s_1 | -0.63 | -0.76 | -1.39 | -1.64 | -0.17 | -0.59 | -0.66 | -1.06 | -1.24 | -1.2 | -0.76 | -1.24 | -0.81 | -0.91 | -1.27 | -1.74 | 0.5375 | 0.1425 | 0.1 | 0.83279 | 0.175463 | 0.98774 | 1 | 0.999631 | 0.94321 |
| Camsap1_A0A0A0MQE5_T559t_1 | 0.97 | -0.09 | -0.64 | 1.13 | 1.4 | 0.28 | 0.7 | 1.14 | 0.67 | -0.01 | 1.24 | 0.04 | -0.45 | 0.83 | 0.57 | 0.82 | 0.2275 | 0.1125 | 0.2375 | 0.841496 | 0.268146 | 0.772826 | 1 | 0.999631 | 0.947343 |
| Camsap1_V9GXD7_S104s_1 | 0.63 | 0.57 | 1.25 | 1.16 | 0.82 | 1.51 | 0.91 | 1.28 | 0.56 | 0.82 | 1.3 | 1.38 | 0.17 | 1.01 | 1.66 | 1.72 | -0.1325 | 0.0225 | 0.1475 | 0.571928 | 0.456487 | 0.727187 | 1 | 0.999631 | 0.844031 |
| Camsap2_H7BX08_S1009s_1 | 0.52 | 1.5 | 0.51 | 0.5 | 0.64 | 0.65 | 0.18 | 1.03 | 0.94 | 0.55 | 0.66 | 0.97 | 0.58 | 0.84 | 0.89 | 1.31 | 0.045 | -0.115 | -0.2575 | 0.673371 | 0.696529 | 0.944695 | 1 | 0.999631 | 0.885515 |
| Camsap2_H7BX08_S1302s_1 | 0.27 | 0.09 | 0.73 | 0.64 | 0.68 | 0.09 | 0.46 | 0.68 | 0.21 | -0.22 | 0.83 | 0.45 | -0.84 | 0.37 | 0.47 | 0.7 | -0.0725 | 0.31 | 0.5325 | 0.525001 | 0.873874 | 0.723229 | 1 | 0.999631 | 0.822727 |
| Camsap2_H7BX08_S1308s_1 | -0.13 | 0.95 | 0.3 | -0.14 | 0.45 | 0.43 | -0.4 | 0.21 | 0.49 | 0.56 | 0.52 | 0.65 | 0.72 | 1.03 | 0.56 | 0.8 | 0.55 | 0.09 | -0.215 | 0.140392 | 0.836985 | 0.34343 | 1 | 0.999631 | 0.611934 |
| Camsap2_H7BX08_S408s_1 | 0.07 | 1.06 | -0.15 | 0.72 | 0.96 | 1.24 | 0.57 | 1.13 | 0.54 | -0.2 | 0.89 | 0.83 | -1.48 | 1.29 | 0.18 | 0.85 | -0.365 | -0.0125 | 0.5675 | 0.724507 | 0.137608 | 0.818647 | 1 | 0.999631 | 0.905674 |
| Camsap2_H7BX08_S589s_1 | -0.61 | -0.23 | -0.23 | -0.42 | -0.96 | -0.8 | -0.88 | -0.31 | -0.21 | -0.78 | -0.3 | -0.25 | 0.7 | -0.46 | 0.43 | 0.11 | -0.4225 | -0.06 | -0.01 | 0.114288 | 0.193612 | 0.96312 | 1 | 0.999631 | 0.579001 |
| Camsap2_H7BX08_S590s_1 | -0.29 | -0.02 | -0.08 | -0.44 | -0.51 | -0.84 | -0.68 | -0.49 | -0.35 | -0.23 | -0.39 | -0.1 | 0.42 | -0.43 | -0.43 | -0.43 | 0.035 | -0.9525 | -1.4 | 0.974929 | 0.114754 | 0.81352 | 1 | 0.999631 | 0.991108 |
| Camsap2_H7BX08_S674s_1 | 2.04 | -0.48 | -0.51 | 1.5 | 0.16 | 0.16 | 1.6 | 0.77 | -0.63 | -0.06 | -0.29 | -0.28 | -2.04 | -1.26 | 0.33 | -0.08 | 0.6975 | 0.3975 | -0.3575 | 0.087092 | 0.957921 | 0.143404 | 1 | 0.999631 | 0.553514 |
| Camsap2_H7BX08_S853s_1 | 1.34 | 0.34 | -0.5 | 1.89 | 1.5 | 0.53 | 1.66 | 2.17 | 1.08 | 0.52 | 1.96 | 1.1 | -1.41 | 0.92 | 0.87 | 1.26 | -0.175 | -0.075 | 0.605 | 0.619823 | 0.243216 | 0.484515 | 1 | 0.999631 | 0.861888 |
| Camsap2_H7BX08_S922s_1 | -1.28 | 0.41 | -0.24 | -0.98 | -0.49 | -0.4 | -0.96 | -0.94 | -0.51 | -1.02 | -0.48 | -0.38 | 0.24 | 0.25 | 0.33 | -0.49 | 0.1 | 0.0425 | -0.1425 | 0.183358 | 0.669474 | 0.855825 | 1 | 0.999631 | 0.644372 |
| Camsap3_Q80VC9_S1095s_1 | 1.59 | 0.98 | 0.29 | 1.07 | 1.21 | 1.23 | 0.32 | 1.57 | 1.29 | 1.07 | 0.97 | 0.77 | 0.21 | 1.28 | 0.63 | 1.24 | 0.4775 | 0.76 | 0.285 | 0.720347 | 0.796663 | 0.899372 | 1 | 0.999631 | 0.904858 |
| Camsap3_Q80VC9_S345s_1 | 0.01 | 1.18 | 0.1 | 0.62 | 1.54 | 1.01 | 1.12 | 0.15 | 1.88 | 0.18 | 1.64 | 1.25 | 0.73 | 0.69 | 1.63 | 0 | 2.0825 | 1.83 | 1.5775 | 0.51734 | 0.251587 | 0.118158 | 1 | 0.999631 | 0.819449 |
| Camsap3_Q80VC9_S346s_1 | -4.61 | -1.76 | 1.37 | 1.24 | 0.63 | 0.16 | 1.37 | 2.41 | 0.67 | -1.09 | 2.71 | 1.27 | -0.85 | -0.68 | 2.44 | 1.64 | -0.5125 | 0.3125 | 0.59 | 0.270229 | 0.128278 | 0.206603 | 1 | 0.999631 | 0.698438 |
| Camsap3_Q80VC9_S359s_1 | -1.63 | -0.94 | 0.03 | -1.43 | -2.04 | -2.13 | -1.17 | -0.68 | -0.74 | -0.58 | -0.74 | -0.66 | -1.29 | -0.38 | 0.55 | -0.49 | 0.5525 | 0.1025 | 0.4825 | 0.258857 | 0.302428 | 0.430505 | 1 | 0.999631 | 0.692624 |
| Camsap3_Q80VC9_S380s_1 | -1.45 | -0.28 | 1.45 | -0.8 | 0.99 | -0.8 | 0.28 | 0.66 | -0.05 | 0.05 | -0.28 | -0.39 | 0.46 | -0.4 | 1.48 | -0.69 | 0.105 | -0.16 | -0.4175 | 0.496162 | 0.407142 | 0.85913 | 1 | 0.999631 | 0.811209 |
| Camsap3_Q80VC9_S443s_1 | 1.17 | 0.37 | -0.54 | 0.92 | 1.1 | 0.13 | 0.62 | 0.49 | 0.62 | 0.32 | 0.33 | 0.01 | -0.74 | 0.53 | -0.09 | 0.55 | -0.095 | 0.06 | 0.54 | 0.389022 | 0.804196 | 0.695726 | 1 | 0.999631 | 0.758181 |
| Camsap3_Q80VC9_S560s_1 | -1.08 | -0.79 | -0.88 | -1.06 | -0.89 | -1.1 | -1.21 | -0.99 | -1.17 | -0.89 | -1.21 | -0.3 | 0.25 | -0.71 | -0.5 | -0.69 | -0.035 | 0.0325 | 0.52 | 0.115517 | 0.693572 | 0.840921 | 1 | 0.999631 | 0.580709 |
| Camsap3_Q80VC9_S598s_1 | -0.71 | -0.29 | -0.52 | -0.55 | -0.29 | -0.61 | -0.92 | -0.39 | -0.43 | -0.64 | -0.55 | -0.32 | -0.39 | 0.38 | -0.09 | 0.11 | 0.1 | 0.1525 | -0.1975 | 0.102857 | 0.894499 | 0.897517 | 1 | 0.999631 | 0.565572 |
| Camsap3_Q80VC9_S779s_1 | 0.9 | 0.06 | -0.58 | -0.17 | 0.41 | 0.04 | 0.06 | 0.1 | 0.59 | 0.34 | 0.17 | -0.28 | -0.35 | 0.15 | -0.33 | -0.05 | 0.2075 | 0.71 | 0.6025 | 0.600219 | 0.776642 | 0.691312 | 1 | 0.999631 | 0.856618 |
| Camsap3_Q80VC9_S785s_1 | 1.33 | 0.46 | 0.2 | 0.36 | 1.55 | 0.31 | 0.03 | 1.29 | 1.66 | 1.39 | 0.45 | 1.69 | 1.25 | -0.25 | 1.97 | 1.79 | 0.33 | 0.85 | 0.905 | 0.27167 | 0.636666 | 0.099788 | 1 | 0.999631 | 0.699288 |
| Camsap3_Q80VC9_S824s_1 | -0.04 | -0.26 | -0.26 | 0.07 | -0.24 | -0.36 | 0.09 | 1.34 | -0.17 | 0.05 | 1.03 | 2 | 0.29 | 0.2 | 1.28 | 1.36 | -0.125 | 0.09 | 0.5275 | 0.031057 | 0.419563 | 0.103835 | 1 | 0.999631 | 0.44065 |
| Camsap3_Q80VC9_T842t_1 | 0.11 | 1.02 | 0.92 | 0.37 | 0.3 | 1.15 | -0.06 | 0.53 | 0.73 | 0.87 | 0.27 | 0.91 | 1.21 | 1.39 | 0.74 | 1.19 | -0.265 | -0.195 | -0.0975 | 0.13474 | 0.728531 | 0.779843 | 1 | 0.999631 | 0.606296 |
| Camta1_A2A891_S1228s_1 | -0.4 | 0.54 | 0.1 | -0.18 | -0.73 | -0.87 | 0.06 | 0.54 | -0.84 | 0.36 | -0.46 | 0.22 | -1.29 | 0.49 | 0.31 | 0.16 | -0.1175 | 0.2975 | 0.1525 | 0.829709 | 0.508642 | 0.604621 | 1 | 0.999631 | 0.942058 |
| Cand1_Q6ZQ38_S1219s_1 | 0.02 | 1.69 | 1.98 | 2.02 | 0.28 | 0.72 | 1.6 | 2.64 | 1.58 | 0.36 | 3.04 | 1.92 | 1.59 | 1.66 | 2.23 | 0.84 | -0.44 | 0.19 | 0.0625 | 0.770878 | 0.85255 | 0.650841 | 1 | 0.999631 | 0.919413 |
| Cand1_Q6ZQ38_S1220s_1 | 0.12 | -1.17 | 0.17 | -0.36 | -1.48 | -0.47 | -1.01 | -0.04 | 0.23 | -1.63 | 0.77 | 0.15 | 0.49 | -1.66 | 0.31 | -0.13 | 0.4725 | -0.29 | -0.4875 | 0.90798 | 0.322377 | 0.735227 | 1 | 0.999631 | 0.96989 |
| Cand1_Q6ZQ38_S1226s_1 | -0.5 | -1.06 | -0.52 | -1.21 | -0.43 | -0.46 | 0.61 | -1.12 | -0.94 | -1.36 | -0.88 | -1.27 | -1.09 | -1.26 | -2.18 | -0.71 | 0.0225 | -0.1525 | 0.0075 | 0.231013 | 0.259418 | 0.341502 | 1 | 0.999631 | 0.681463 |
| Cand1_Q6ZQ38_S1230s_1 | -0.85 | -1.27 | -1.8 | -2.12 | -1.12 | -1.25 | -1.33 | -2.25 | -1.4 | -1.13 | -2.27 | -1.85 | -0.85 | -1.7 | -1.53 | -1.93 | -0.0475 | 0.2725 | 0.4 | 0.984768 | 0.953984 | 0.69885 | 1 | 0.999631 | 0.995133 |
| Cand1_Q6ZQ38_S558s_1 | -0.66 | -0.43 | -0.87 | -0.88 | -0.64 | -0.3 | -1.37 | -0.72 | -1.17 | -0.63 | -0.42 | 0.47 | 0.2 | -0.31 | -0.67 | -0.46 | 0.0225 | -0.1525 | 0.0075 | 0.214644 | 0.876573 | 0.479037 | 1 | 0.999631 | 0.66742 |
| Cand1_Q6ZQ38_T1229t_1 | -0.85 | -1.27 | -1.8 | -2.12 | -1.12 | -1.25 | -1.33 | -2.25 | -1.4 | -1.13 | -2.27 | -1.85 | -0.85 | -1.7 | -1.53 | -1.93 | 0.13 | 0.525 | 0.115 | 0.984768 | 0.953984 | 0.69885 | 1 | 0.999631 | 0.995133 |
| Canx_P35564_S563s_1 | -0.45 | 0.95 | 0.39 | -0.2 | -0.06 | 1.39 | -0.39 | 0.27 | 0.82 | 1.78 | 0.04 | 0.15 | 0.43 | 0.56 | 0.09 | 0.07 | -0.1 | 0.28 | 0.4475 | 0.759937 | 0.7839 | 0.294984 | 1 | 0.999631 | 0.914614 |
| Canx_P35564_S582s_1 | -0.66 | -1.21 | -0.79 | -0.29 | -0.75 | -1.36 | -0.98 | -0.26 | -0.59 | -0.57 | -0.31 | -0.36 | 0.24 | -0.62 | -0.64 | -0.14 | -0.185 | -0.705 | -0.6075 | 0.214011 | 0.766269 | 0.345477 | 1 | 0.999631 | 0.667209 |
| Cap1_P40124_S307s_1 | -0.61 | -3.99 | 1.64 | 1 | -1.86 | -2.85 | 1.87 | 0.14 | -2.26 | -2.21 | -0.44 | 0.13 | -2.32 | -3.03 | 1.1 | -0.14 | 0.32 | -0.2525 | -0.43 | 0.64997 | 0.894334 | 0.559849 | 1 | 0.999631 | 0.875355 |
| Cap1_P40124_S309s_1 | 0.76 | -0.95 | 1.62 | 0.35 | 0.76 | 0.53 | 1.62 | 0.15 | 0.8 | 0.34 | -0.77 | 0.4 | -0.53 | 0.5 | 0.97 | -0.88 | -0.175 | -0.0725 | 0.0325 | 0.494938 | 0.574415 | 0.664511 | 1 | 0.999631 | 0.810518 |
| Cap1_P40124_S34s_1 | -0.61 | -0.5 | -0.75 | -0.7 | -0.84 | -0.86 | -0.63 | -0.93 | -0.88 | -0.19 | -1.17 | -0.61 | -0.4 | -0.4 | -1.05 | -0.58 | -0.8175 | -0.5225 | -1.2875 | 0.909876 | 0.465872 | 0.806533 | 1 | 0.999631 | 0.970597 |
| Cap2_D3YTR7_S145s_1 | 2.67 | 1.33 | 2.19 | 2.01 | 0.11 | 1.96 | 2.39 | 0.47 | 1.13 | 2.84 | 1.46 | 0.68 | 0.33 | 0.75 | 0.98 | 0.99 | -0.305 | 0.3125 | -0.075 | 0.005115 | 0.171934 | 0.32035 | 1 | 0.999631 | 0.351308 |
| Cap2_D3YTR7_S149s_1 | -0.12 | 0.59 | 0.6 | 0.47 | -0.36 | 0.58 | -0.26 | 0.36 | 0.8 | 0.89 | 0.34 | 0.76 | -0.31 | -0.1 | 0.73 | 0.92 | 0.8625 | 1.945 | 0.66 | 0.844604 | 0.364858 | 0.300897 | 1 | 0.999631 | 0.94856 |
| Cap2_D3YTR7_S199s_1 | 0.98 | 2.28 | 2.36 | -0.37 | 3.62 | 1.11 | 2.91 | 1.06 | 5.23 | 1.05 | 3.54 | 3.21 | 5.19 | 1.01 | 1 | 0.69 | 0.935 | 1.9475 | 0.8175 | 0.540127 | 0.293996 | 0.059981 | 1 | 0.999631 | 0.831295 |
| Cap2_D3YTR7_T198t_1 | -0.39 | 3.12 | 3.11 | 0.25 | 3.67 | 1.33 | 3.34 | 1.49 | 4.93 | 0.51 | 3.86 | 4.58 | 4.97 | 1.24 | 2.24 | 0.91 | -0.2125 | 0.3275 | 0.435 | 0.468136 | 0.341236 | 0.122639 | 1 | 0.999631 | 0.795565 |
| Capn5_O08688_T44t_1 | 0.42 | 0.2 | 0.18 | 0.55 | -0.15 | 0.34 | 0.1 | 0.21 | 1.15 | 0.97 | 0.26 | 0.28 | 1.45 | 0.03 | 0.82 | 0.79 | 0.2275 | 0.3 | 0.525 | 0.24005 | 0.409771 | 0.30933 | 1 | 0.999631 | 0.683719 |
| Capza2_P47754_T63t_1 | -0.39 | 1.5 | 0.71 | 0.4 | 1.15 | 1.61 | -0.02 | 0.39 | 0.52 | 0.8 | 0.66 | 1.44 | 1.49 | 1.21 | 0.67 | 0.95 | 0.38 | 0.205 | 0.5775 | 0.244729 | 0.652168 | 0.497798 | 1 | 0.999631 | 0.685751 |
| Capzb_P47757_S263s_1 | 1.34 | 1.1 | 0.42 | 0.75 | 1.88 | 1.27 | 0.37 | 1.61 | 1.19 | 1.14 | 0.56 | 1.54 | 0.9 | 1.77 | 1.75 | 1.5 | 0.6375 | -0.03 | 1.1025 | 0.119157 | 0.345542 | 0.544371 | 1 | 0.999631 | 0.586721 |
| Carhsp1_Q9CR86_S31s_1 | -1.37 | 1.98 | 1.41 | 0.05 | 1.75 | 1.56 | 0.46 | 0.85 | 1.64 | -0.75 | 0.34 | 0.72 | 2.56 | 2.01 | 1.22 | 0.69 | 0.0175 | -0.1725 | -0.235 | 0.164504 | 0.381086 | 0.969615 | 1 | 0.999631 | 0.632897 |
| Carhsp1_Q9CR86_S42s_1 | 0.6 | 1.03 | -0.67 | 0.27 | 0.83 | 0.52 | -0.33 | 0.28 | 0.81 | 0.72 | -0.32 | -0.67 | 0.08 | 1.11 | -0.36 | -0.54 | -0.2075 | -0.435 | -0.2525 | 0.637466 | 0.967432 | 0.727162 | 1 | 0.999631 | 0.870011 |
| Carhsp1_Q9CR86_S53s_1 | 0.73 | 1.32 | 1.34 | 1.19 | 1.02 | 1.01 | 0.21 | 1.51 | 0.47 | 1.2 | 0.36 | 0.81 | -0.39 | 1.67 | 1.15 | 1.14 | -1.1525 | -0.7775 | -0.775 | 0.586847 | 0.545236 | 0.178558 | 1 | 0.999631 | 0.849744 |
| Cars_Q9ER72_S19s_1 | -0.39 | -1.3 | -0.52 | -1.3 | -1.88 | -3.01 | -1.16 | -2.07 | -2.34 | -1.46 | -1.79 | -1.03 | -2.38 | -2.24 | -1.19 | -0.8 | -0.3775 | -0.1075 | 0.0975 | 0.113494 | 0.02468 | 0.068949 | 1 | 0.999631 | 0.57792 |
| Cars_Q9ER72_S307s_1 | -0.76 | 1.2 | 1.34 | 0.18 | 0.38 | 0.08 | -0.18 | 0.17 | -0.14 | -0.15 | 0.67 | 1.15 | 0.87 | 0.25 | 0.6 | 0.63 | 0.1025 | 0.6525 | 1.0975 | 0.84197 | 0.437318 | 0.843355 | 1 | 0.999631 | 0.947766 |
| Cartpt_P56388_S48s_1 | -1.46 | -0.84 | -0.45 | -1.21 | -0.26 | -0.94 | -0.86 | -1.49 | 0.06 | -1.95 | -0.39 | 0.93 | 0.58 | -0.27 | 0.77 | -0.65 | -0.38 | 0.3425 | 0.8325 | 0.023824 | 0.775532 | 0.282481 | 1 | 0.999631 | 0.415644 |
| Casc3_Q8K3W3_S262s_1 | -0.74 | 0.85 | -0.81 | -0.17 | 0.37 | -1.83 | -0.96 | 0.03 | -0.19 | -1.01 | 1.02 | 0.68 | 0.86 | 0.5 | 1.02 | 0.08 | 0.0725 | -0.095 | 0.745 | 0.08269 | 0.511913 | 0.538969 | 1 | 0.999631 | 0.548633 |
| Casc3_Q8K3W3_S44s_1 | -1.39 | -2.82 | -0.75 | -0.65 | -3.06 | -2.58 | -0.3 | 0.62 | -2.23 | -2.34 | -0.29 | -1.13 | -1.79 | -0.89 | 0.78 | -0.73 | 0.0575 | 1.0625 | 0.2825 | 0.271367 | 0.934421 | 0.880176 | 1 | 0.999631 | 0.699267 |
| Casc4_E9PZJ6_S231s_1 | -0.13 | 1.09 | 0.99 | 0 | 0.72 | 0.55 | 1.35 | -0.44 | 2.23 | -0.06 | 0.86 | 3.17 | 2.71 | 0.12 | -0.48 | 0.73 | 0.0575 | 1.0625 | 0.2825 | 0.678804 | 0.902144 | 0.152105 | 1 | 0.999631 | 0.887183 |
| Casc4_E9PZJ6_S232s_1 | -0.13 | 1.09 | 0.99 | 0 | 0.72 | 0.55 | 1.35 | -0.44 | 2.23 | -0.06 | 0.86 | 3.17 | 2.71 | 0.12 | -0.48 | 0.73 | -0.11 | -0.1 | 0.1225 | 0.678804 | 0.902144 | 0.152105 | 1 | 0.999631 | 0.887183 |
| Cask_O70589_S571s_1 | -0.83 | -0.24 | -0.36 | -0.69 | -0.51 | -0.57 | -0.84 | -0.64 | -0.42 | -0.73 | -0.87 | -0.5 | 0.46 | -0.67 | -0.71 | -0.71 | 0.18 | 0.01 | -1.2025 | 0.738511 | 0.673958 | 0.717346 | 1 | 0.999631 | 0.90804 |
| Cask_O70589_S582s_1 | 0.21 | -0.53 | 0.37 | -1.28 | -1.35 | 0.56 | 2.64 | -2.36 | -0.74 | 0.59 | -0.61 | -0.43 | 0.11 | -2.42 | -1.27 | -2.46 | -0.155 | -0.3325 | -0.165 | 0.085659 | 0.857616 | 0.982969 | 1 | 0.999631 | 0.551124 |
| Cask_O70589_S583s_1 | -0.48 | -1.18 | 0.06 | -0.38 | -0.81 | -1.16 | -0.25 | -0.38 | -0.94 | -0.8 | -1.36 | -0.21 | -0.84 | -1.36 | -0.68 | 0.24 | -0.035 | 0.1025 | 0.5 | 0.701688 | 0.664839 | 0.385055 | 1 | 0.999631 | 0.896835 |
| Cask_O70589_T182t_1 | -0.37 | -0.96 | -0.2 | -0.32 | -0.49 | -1.22 | -0.39 | 0.11 | -1.1 | -0.99 | 0.61 | 0.04 | 1.32 | -0.72 | -0.09 | -0.36 | 0.3425 | -0.1375 | -0.08 | 0.300186 | 0.920556 | 0.816365 | 1 | 0.999631 | 0.713499 |
| Caskin1_Q6P9K8_S1012s_1 | 0 | -1.46 | -1.31 | -0.35 | -0.15 | -0.98 | -0.24 | -0.38 | -0.98 | -0.74 | -0.87 | -1.08 | -0.48 | -1.25 | -1.18 | -0.53 | -0.105 | -0.045 | 0.375 | 0.850396 | 0.407362 | 0.721671 | 1 | 0.999631 | 0.949525 |
| Caskin1_Q6P9K8_S1188s_1 | -0.24 | 0.41 | 0.98 | -0.05 | 0.01 | 0.24 | -0.12 | 0.55 | -0.08 | 0.15 | 0.19 | 0.66 | 0.37 | 0.49 | 0.44 | 1.3 | -0.0125 | 0.2325 | 0.1775 | 0.339438 | 0.759624 | 0.89804 | 1 | 0.999631 | 0.732185 |
| Caskin1_Q6P9K8_S1233s_1 | -0.47 | -2.24 | 0 | -0.15 | -0.83 | -0.58 | -0.98 | -0.52 | -0.35 | -1.07 | -0.37 | -0.14 | -0.81 | -1.29 | 0.25 | -0.3 | 0.245 | 0.3375 | 0.38 | 0.754833 | 0.979898 | 0.657312 | 1 | 0.999631 | 0.913582 |
| Caskin1_Q6P9K8_S1288s_1 | 0.21 | -0.73 | -0.16 | -0.94 | -0.29 | -0.01 | -0.39 | 0.05 | -0.38 | 0.08 | 1.32 | -1.29 | -0.2 | 0.51 | -0.56 | 0.15 | -0.41 | 0.42 | -0.005 | 0.333983 | 0.460495 | 0.54879 | 1 | 0.999631 | 0.72834 |
| Caskin1_Q6P9K8_S1330s_1 | -1.6 | 0.32 | -1.27 | -1.99 | 0.72 | -4.08 | -1.04 | -1.78 | -0.13 | -0.4 | -1.01 | -1.32 | 0.72 | -3.3 | -0.7 | -1.28 | 0.0325 | -0.0875 | 0.36 | 0.995277 | 0.671318 | 0.441212 | 1 | 0.999631 | 0.997705 |
| Caskin1_Q6P9K8_S358s_1 | -0.56 | -0.01 | -0.25 | -0.74 | -0.09 | -0.2 | -0.45 | -0.69 | -0.29 | -1 | -0.3 | -0.32 | -0.1 | 0.06 | -0.19 | 0.11 | 0.73 | -0.3725 | -0.46 | 0.236516 | 0.909443 | 0.776388 | 1 | 0.999631 | 0.683301 |
| Caskin1_Q6P9K8_S390s_1 | 1.86 | -0.68 | -0.68 | 1.39 | 1.61 | 0.27 | 1.32 | 1.61 | -0.22 | -0.99 | 1.35 | 0.26 | -1.43 | -0.14 | 1.02 | 0.6 | 0.1525 | -0.705 | -0.8475 | 0.546234 | 0.286043 | 0.615162 | 1 | 0.999631 | 0.834213 |
| Caskin1_Q6P9K8_S423s_1 | 1.3 | 0.7 | -0.7 | 0.88 | 1.1 | 0.38 | 0.96 | 0.35 | 0.24 | -0.48 | -0.08 | -0.32 | -0.08 | -0.42 | -0.81 | 0.1 | -0.405 | -0.0325 | 0.065 | 0.094903 | 0.739087 | 0.142639 | 1 | 0.999631 | 0.558128 |
| Caskin1_Q6P9K8_S432s_1 | -0.14 | 0.42 | 0.39 | -0.51 | -0.36 | 0.44 | -0.73 | -0.81 | 0.17 | 0.29 | -0.47 | 0.04 | 0.44 | 0.61 | -0.03 | -0.6 | 0.035 | -0.13 | 0.205 | 0.86591 | 0.298245 | 0.921747 | 1 | 0.999631 | 0.955982 |
| Caskin1_Q6P9K8_S579s_1 | -0.65 | -0.23 | -0.51 | -0.3 | -0.03 | -0.25 | -0.64 | -0.63 | -0.09 | -0.1 | -1.06 | -0.96 | 0.03 | -0.55 | -0.54 | 0.19 | 0.1425 | 0.4125 | 2.36E-16 | 0.512339 | 0.896906 | 0.696825 | 1 | 0.999631 | 0.818048 |
| Caskin1_Q6P9K8_S720s_1 | 0.4 | -0.75 | -0.29 | 0.29 | -0.58 | 1.21 | -0.25 | -0.16 | 0.47 | 0.65 | -0.05 | 0.23 | -0.67 | 0.42 | -0.15 | 0.05 | 0.7225 | -0.5775 | -0.7675 | 1 | 0.756448 | 0.253508 | 1 | 0.999631 | 1 |
| Caskin1_Q6P9K8_S750s_1 | 0.92 | -0.21 | -1.24 | 0.88 | 0.96 | 0.59 | 0.77 | 0.92 | -0.18 | -0.89 | -0.26 | -0.63 | -1.75 | 0.12 | -0.86 | -0.23 | 0.5575 | -0.7625 | -1.0875 | 0.21972 | 0.163916 | 0.273595 | 1 | 0.999631 | 0.670699 |
| Caskin1_Q6P9K8_S757s_1 | 2.23 | -1.31 | -0.45 | 1.09 | 1.94 | -0.4 | 1.65 | 0.6 | 0.05 | -1.25 | 0.03 | -0.32 | -1.46 | -0.57 | -0.2 | -0.56 | -0.6175 | -0.3825 | -2.005 | 0.161042 | 0.506461 | 0.32115 | 1 | 0.999631 | 0.631211 |
| Caskin1_Q6P9K8_S820s_1 | 2.11 | 1.11 | 2.27 | 2.21 | 0.58 | 1.2 | 2.42 | 1.03 | 0.49 | 2.17 | 1.92 | 1.59 | -0.8 | -0.85 | -0.03 | 1.36 | -0.26 | -0.7425 | -1.7075 | 0.004083 | 0.200148 | 0.405733 | 1 | 0.999631 | 0.351308 |
| Caskin1_Q6P9K8_S822s_1 | 2 | 0.07 | 1.09 | 1.6 | 0.47 | 1.47 | 1.73 | 0.05 | 0.66 | 1.96 | -0.15 | -0.68 | -1.87 | -0.32 | 0.28 | -0.16 | -0.77 | -0.3875 | 0.465 | 0.013468 | 0.62807 | 0.263047 | 1 | 0.999631 | 0.385092 |
| Caskin1_Q6P9K8_S918s_1 | 1.16 | 0.81 | 0.66 | 2.07 | 0.14 | 0.78 | -0.91 | 1.61 | -0.14 | 2.54 | 0.65 | 0.1 | 0.38 | 1.93 | 1.57 | 2.68 | 0.12 | -1.0625 | -2.025 | 0.39636 | 0.194071 | 0.536228 | 1 | 0.999631 | 0.760744 |
| Caskin1_Q6P9K8_S965s_1 | 2.85 | -1.47 | -0.49 | 1.1 | 0.59 | 0.36 | 1.52 | 0 | 0.35 | 0.41 | -1.31 | -1.71 | -2.93 | -0.42 | -1.15 | -1.61 | 0.235 | 1.2725 | 0.8575 | 0.051339 | 0.889988 | 0.278853 | 1 | 0.999631 | 0.489777 |
| Caskin1_Q6P9K8_T1197t_1 | -0.16 | 1.73 | 1.59 | -0.49 | 1.52 | 0.81 | 1.15 | 0.13 | 2.45 | -0.33 | 2.61 | 3.03 | 3.14 | 0.68 | 2.05 | 0.23 | 0.2075 | -0.3625 | -0.305 | 0.280807 | 0.69057 | 0.152958 | 1 | 0.999631 | 0.701922 |
| Caskin1_Q6P9K8_T670t_1 | 0.25 | 0.26 | 0.37 | 0.55 | 0.53 | 0.72 | 0.26 | 0.75 | 0.15 | -0.15 | 0.01 | -0.03 | -0.56 | 0.81 | 0.12 | -0.16 | 0.0725 | -0.01 | 0.2525 | 0.395825 | 0.417483 | 0.165472 | 1 | 0.999631 | 0.76043 |
| Caskin1_Q6P9K8_T708t_1 | -0.16 | -0.11 | -0.06 | -0.26 | -0.02 | 0.12 | -0.45 | 0.05 | -0.29 | -0.13 | -0.18 | -0.03 | -0.16 | 0.11 | 0.08 | 0.39 | -0.0075 | -0.8175 | -1.8875 | 0.365086 | 0.774169 | 0.966967 | 1 | 0.999631 | 0.745324 |
| Caskin1_Q6P9K8_T745t_1 | 1.86 | -0.02 | -0.98 | 0.02 | 1.01 | 0.17 | -0.07 | -0.26 | 0.43 | -1.35 | -0.27 | -1.2 | -5.82 | 0.22 | -0.52 | -0.55 | 1.305 | -0.2225 | -1.6825 | 0.162735 | 0.989915 | 0.229296 | 1 | 0.999631 | 0.631442 |
| Caskin1_Q6P9K8_T759t_1 | 2.74 | 0.19 | -1.17 | 1.43 | 2.71 | 1 | 2.54 | 2.16 | 1.32 | -0.38 | 1.1 | 0.26 | -3.56 | -0.2 | -0.01 | 0.23 | 0.31 | -0.1375 | -0.1875 | 0.131369 | 0.129245 | 0.783936 | 1 | 0.999631 | 0.603456 |
| Caskin1_Q6P9K8_T996t_1 | 2.09 | 0.9 | 0.67 | 2 | 2.37 | 1.39 | 1.51 | 1.63 | 1.56 | 0.43 | 1.3 | 1.82 | 0.47 | 1.72 | 1.53 | 1.19 | 0.1175 | -0.0325 | -0.7775 | 0.681692 | 0.469927 | 0.766342 | 1 | 0.999631 | 0.88924 |
| Caskin1_Q6P9K8_Y457y_1 | 0.21 | -1.44 | -2.56 | -0.95 | -0.09 | -1.42 | -1.22 | -1.54 | -1.38 | -0.88 | -1.68 | -0.93 | -1.5 | -0.72 | -4 | -1.63 | -0.155 | -0.085 | -0.3325 | 0.340301 | 0.844655 | 0.953586 | 1 | 0.999631 | 0.733032 |
| Caskin2_Q8VHK1_S694s_1 | -0.38 | -0.93 | -0.35 | -0.44 | -1.07 | -0.96 | -0.45 | -0.24 | -1.64 | -0.6 | -0.32 | 0.12 | -0.66 | -1.85 | -0.4 | -0.52 | 0.3375 | 0.455 | 0.4125 | 0.40415 | 0.610763 | 0.834677 | 1 | 0.999631 | 0.765215 |
| Casp3_P70677_S26s_1 | -0.16 | -0.24 | -0.63 | -0.02 | 0.48 | -0.09 | -0.1 | 0.01 | 0.07 | -0.02 | 0.36 | 0.36 | 0.21 | 0.45 | -0.03 | -0.03 | -0.22 | -0.1125 | 0.2025 | 0.178572 | 0.239004 | 0.118563 | 1 | 0.999631 | 0.642903 |
| Cast_Q921U7_S117s_1 | -2.01 | -0.67 | -2.39 | -1.76 | -1.98 | -1.22 | -2.12 | -2.39 | -1.63 | -1.19 | -2.65 | -1.81 | -1.02 | -0.82 | -1.87 | -2.31 | -0.23 | -0.2825 | 0.205 | 0.681394 | 0.616017 | 0.80896 | 1 | 0.999631 | 0.889229 |
| Cbfa2t2_O70374_S235s_1 | 0.43 | 0.66 | 1.72 | 0.67 | 0.3 | 0.49 | 0.64 | 1.13 | 0.45 | 0.75 | 0.23 | 0.92 | 0.32 | 0.8 | 1.79 | 1.39 | 0.5875 | 1.1525 | -0.7225 | 0.641559 | 0.52924 | 0.440384 | 1 | 0.999631 | 0.871335 |
| Cbfa2t2_O70374_S380s_1 | 0.34 | 2.25 | 1.45 | -0.41 | 2.55 | 0.33 | 3.1 | 0 | 3.63 | 0.15 | 0.91 | 3.55 | 1.78 | -0.03 | -0.33 | -0.68 | -0.3125 | -0.3525 | -0.39 | 0.322746 | 0.494129 | 0.234346 | 1 | 0.999631 | 0.724485 |
| Cbfa2t2_O70374_S522s_1 | 0.67 | 0.02 | 0.9 | -0.17 | -1.4 | 0.24 | 0.82 | 0.51 | -0.61 | 1.09 | 0.65 | -1.12 | -0.77 | -0.65 | 1.16 | 0.12 | 0.1 | 0.8325 | 1.2375 | 0.436576 | 0.550098 | 0.517771 | 1 | 0.999631 | 0.780748 |
| Cbl_P22682_S617s_1 | -1.5 | -0.44 | -0.48 | -0.09 | -1.06 | -1.11 | -0.19 | 0.25 | -0.24 | -0.45 | -0.08 | 1.59 | -0.08 | -0.39 | 3.53 | -0.62 | 0.8025 | 0.7525 | 1.03 | 0.183441 | 0.82021 | 0.1358 | 1 | 0.999631 | 0.644372 |
| Cbl_P22682_S861s_1 | -1.25 | -0.83 | -1.08 | -1.11 | 0.22 | -0.22 | -0.65 | -0.41 | 0.08 | -0.12 | -0.46 | -0.76 | 0.29 | 0.73 | -0.28 | -0.89 | -0.9825 | -1.35 | -1.87 | 0.023405 | 0.016748 | 0.02565 | 1 | 0.999631 | 0.41509 |
| Cbx3_D3Z313_S93s_1 | -0.05 | -2 | 1.2 | 1.47 | -1.89 | -1.83 | 1.02 | -0.61 | -2.47 | -1.1 | 0.25 | -1.46 | -2.93 | -2.42 | -0.04 | -1.47 | -0.7875 | -0.17 | -0.8875 | 0.054994 | 0.292327 | 0.135564 | 1 | 0.999631 | 0.492066 |
| Cbx3_D3Z313_S95s_1 | 1.49 | 1.28 | 2.51 | 2.59 | 0.22 | 1.31 | 2.13 | 1.06 | 1.34 | 2.06 | 1.23 | 2.56 | 0.91 | 0.66 | 1.19 | 1.56 | -0.7425 | 0.0225 | -0.595 | 0.051852 | 0.132142 | 0.708035 | 1 | 0.999631 | 0.489777 |
| Cbx5_Q61686_S110s_1 | 0.89 | 0.61 | 1.04 | 0.73 | -0.63 | 0.61 | 0.75 | -0.43 | 0.42 | 1.07 | 0.82 | 1.05 | 0.81 | -0.59 | 0.34 | 0.33 | -0.265 | 0.0525 | 0.3925 | 0.120956 | 0.074352 | 0.935393 | 1 | 0.999631 | 0.588397 |
| Cbx5_Q61686_S92s_1 | -1.03 | 0.16 | 0.07 | -0.1 | -0.55 | -0.56 | -0.5 | -0.35 | -0.44 | -0.29 | 0.01 | 0.03 | 0.82 | -0.4 | -0.18 | 0.43 | -0.0925 | -0.6125 | -0.1225 | 0.349818 | 0.421082 | 0.877813 | 1 | 0.999631 | 0.736956 |
| Cby1_Q9D1C2_S20s_1 | -0.22 | -0.15 | -0.63 | 0.24 | -0.45 | -0.41 | 0.05 | -0.32 | -0.83 | -0.15 | -0.95 | -1.28 | -1.16 | 0.23 | -0.48 | 0.16 | 0.56 | -0.055 | 0.6675 | 0.757803 | 0.747338 | 0.097711 | 1 | 0.999631 | 0.914449 |
| Cby1_Q9D1C2_S9s_1 | -1.31 | -0.51 | -0.2 | 0.09 | 0.42 | 0.25 | -0.18 | -0.18 | -0.38 | -1.21 | -1.29 | 0.73 | 0.5 | 1.16 | -0.59 | -0.33 | 0.1525 | 0.065 | 0.42 | 0.189498 | 0.144871 | 0.916276 | 1 | 0.999631 | 0.648674 |
| Cc2d1a_E9PX94_S267s_1 | -1.01 | -0.29 | -0.71 | -0.03 | -0.76 | -0.96 | -0.18 | 0.47 | -0.41 | -0.42 | -0.9 | -0.05 | 0.05 | -0.23 | -0.13 | -0.05 | 0.6525 | 0.7775 | 0.8925 | 0.20199 | 0.700345 | 0.844235 | 1 | 0.999631 | 0.658952 |
| Cc2d1a_E9PX94_S390s_1 | -1.32 | 1.44 | 1.47 | -0.26 | 1.36 | 1.36 | 0.82 | 0.4 | 1.78 | 0.37 | 1.03 | 1.26 | 1.97 | 0.73 | 1.53 | 0.67 | -0.1225 | -0.2675 | 0.0675 | 0.204213 | 0.325091 | 0.260286 | 1 | 0.999631 | 0.659781 |
| Ccar2_Q8VDP4_S298s_1 | -2.56 | -2.35 | 1.03 | 0.52 | -1.73 | -3 | 0.22 | 0.66 | -2.31 | -2.5 | 0.32 | 0.06 | -2.68 | -2.62 | 1.33 | 0.88 | 0.25 | 0.405 | 0.775 | 0.955678 | 0.910302 | 0.796611 | 1 | 0.999631 | 0.984649 |
| Ccar2_Q8VDP4_S612s_1 | -1.26 | 1.12 | 3.51 | 1.44 | 0.46 | 1.09 | 2.02 | 2.24 | 0.5 | -0.34 | 3.34 | 2.93 | 0.88 | 1.07 | 2.72 | 3.24 | -1.14 | -1.325 | -1.9625 | 0.435209 | 0.785446 | 0.723296 | 1 | 0.999631 | 0.779854 |
| Ccar2_Q8VDP4_S625s_1 | 2.46 | 1.83 | 2.54 | 1.22 | 0.59 | 1.2 | 1.89 | -0.19 | -0.01 | 1.76 | 0.65 | 0.35 | 0.09 | 0.09 | 0.13 | -0.11 | -0.5675 | -1.555 | 0.575 | 0.000297 | 0.044991 | 0.018143 | 1 | 0.999631 | 0.351308 |
| Ccdc115_Q8VE99_T90t_1 | -0.29 | 1.3 | 1.59 | 1.47 | 0.01 | -0.47 | -0.06 | 2.32 | 0.41 | -0.42 | 0.4 | -2.54 | 2.03 | 0.88 | 2.59 | 0.87 | 0.01 | -0.2475 | -1.58 | 0.324 | 0.414904 | 0.055771 | 1 | 0.999631 | 0.724485 |
| Ccdc120_A2AEV7_S253s_1 | 1.47 | 0.41 | 1.62 | 0.89 | 0.88 | 1.57 | 2.15 | -0.17 | 1.15 | 1.3 | 0.72 | 0.23 | -0.83 | -0.78 | 0.35 | -0.67 | -0.175 | 0.1825 | 0.4475 | 0.003154 | 0.984886 | 0.52637 | 1 | 0.999631 | 0.351308 |
| Ccdc120_A2AEV7_S283s_1 | 1.39 | 0.87 | -0.09 | 0.87 | 0.2 | 0.55 | 0.76 | 0.83 | 0.86 | 2.03 | 0.81 | 0.07 | 3.21 | 0.49 | 0.36 | 0.77 | -0.37 | 0.055 | 0.1075 | 0.504223 | 0.630918 | 0.708603 | 1 | 0.999631 | 0.814292 |
| Ccdc132_Q8CI71_S15s_1 | -0.5 | -0.5 | 0.54 | 0.07 | -0.66 | -0.89 | -0.31 | -0.01 | -0.68 | -0.07 | 0.17 | 0.41 | 0.11 | -0.66 | 0.15 | 0.44 | -0.0575 | -0.8425 | -0.6025 | 0.777775 | 0.301929 | 0.882021 | 1 | 0.999631 | 0.921382 |
| Ccdc132_Q8CI71_S494s_1 | 0.29 | -0.77 | 0.14 | -0.13 | -1.04 | 0.65 | 0.07 | -0.38 | -1.08 | -0.95 | -0.6 | -1.21 | -0.98 | -0.34 | -0.44 | -1.12 | -1.2025 | -0.7575 | -0.615 | 0.114976 | 0.8916 | 0.025402 | 1 | 0.999631 | 0.579336 |
| Ccdc132_Q8CI71_S498s_1 | -0.39 | -1.33 | -0.35 | -2.27 | -2.65 | -1.67 | -0.98 | -3.85 | -2.2 | 0.27 | -2.21 | -3.23 | 0.14 | -1.8 | -2.66 | -2.48 | -0.005 | 0.1525 | 0.6075 | 0.387439 | 0.104092 | 0.337124 | 1 | 0.999631 | 0.756827 |
| Ccdc136_D3Z165_T102t_1 | 0.6 | 1.5 | -0.02 | 0.34 | 0.94 | 0.96 | -0.2 | 0.7 | 1.09 | 1.07 | 0.3 | 0.57 | 2.18 | 1.15 | 0.43 | 1.09 | -0.005 | -0.26 | -0.955 | 0.219225 | 0.99049 | 0.699306 | 1 | 0.999631 | 0.669859 |
| Ccdc177_Q3UHB8_S209s_1 | 0.83 | -0.4 | -0.43 | 0.17 | -0.23 | -0.15 | 0.28 | 0.25 | -0.18 | 0.4 | -0.76 | -0.33 | -1.49 | -0.65 | -0.62 | -0.89 | -0.9075 | -0.5875 | -0.9275 | 0.031764 | 0.988678 | 0.515955 | 1 | 0.999631 | 0.442395 |
| Ccdc177_Q3UHB8_S209s_2 | 0.15 | -0.06 | 0.31 | -0.43 | -0.97 | -1.79 | -0.39 | -0.51 | -0.82 | 0.09 | -0.44 | -1.21 | -0.94 | -0.95 | -0.83 | -1.02 | -0.48 | -0.4225 | -0.5825 | 0.008578 | 0.033245 | 0.123214 | 1 | 0.999631 | 0.365244 |
| Ccdc177_Q3UHB8_S211s_1 | 0.29 | -0.3 | 0.31 | -0.32 | -0.52 | -0.4 | -0.48 | -0.54 | -0.51 | 0.32 | -0.73 | -0.79 | -0.16 | -0.57 | -0.85 | -0.77 | -0.3875 | -0.3925 | -0.85 | 0.091775 | 0.100881 | 0.245566 | 1 | 0.999631 | 0.557085 |
| Ccdc177_Q3UHB8_S211s_2 | 0.26 | -0.52 | -0.1 | -0.32 | -0.73 | -0.63 | -0.46 | -0.41 | -0.57 | 0.06 | -0.55 | -1.19 | -0.95 | -1.13 | -0.83 | -1.17 | 0.225 | 0.305 | 0.315 | 0.015529 | 0.176975 | 0.273452 | 1 | 0.999631 | 0.388832 |
| Ccdc177_Q3UHB8_S273s_1 | -0.41 | 0 | 0.44 | 0.69 | 0.62 | 0.28 | 0.28 | 0.44 | 0.5 | 0.6 | 0.62 | 0.22 | 0.64 | 0.6 | 0.31 | 0.43 | 0.6225 | 0.4325 | -0.02 | 0.350197 | 0.475161 | 0.351761 | 1 | 0.999631 | 0.737144 |
| Ccdc177_Q3UHB8_S281s_1 | -0.06 | -0.85 | -0.17 | -0.82 | 0.82 | 0.62 | -0.05 | -0.8 | 0.25 | 0.69 | 0.01 | -1.12 | 0.42 | -0.37 | -1.53 | -0.5 | 0.1425 | -0.0075 | 4.91E-17 | 0.964554 | 0.160029 | 0.333726 | 1 | 0.999631 | 0.987331 |
| Ccdc177_Q3UHB8_S303s_1 | -0.48 | -0.47 | -0.92 | -0.83 | -0.47 | -0.56 | -0.51 | -0.59 | -0.76 | -0.39 | -0.83 | -0.75 | -0.46 | -0.69 | -0.81 | -0.74 | -0.9375 | -0.6975 | -0.345 | 1 | 0.567139 | 0.977603 | 1 | 0.999631 | 1 |
| Ccdc177_Q3UHB8_S310s_1 | -1.48 | -1.96 | -2.91 | -4.08 | -3.83 | -3.12 | -2.99 | -4.24 | -3.24 | -2.51 | -4.61 | -2.86 | -3.64 | -2.95 | -1.97 | -3.25 | -0.4275 | -0.2875 | -1.615 | 0.576573 | 0.133835 | 0.304934 | 1 | 0.999631 | 0.846204 |
| Ccdc177_Q3UHB8_S423s_1 | 0.12 | -2.34 | -2.64 | -1.08 | -2.04 | -4.03 | 0.06 | -1.64 | -3.19 | -1.59 | -0.57 | -1.74 | -1.57 | -3.32 | -1.96 | -5.55 | -0.3025 | -0.5425 | -0.49 | 0.111314 | 0.640151 | 0.697051 | 1 | 0.999631 | 0.577068 |
| Ccdc177_Q3UHB8_S90s_1 | -0.02 | -0.33 | 0.16 | 0.03 | -0.25 | -0.35 | -0.55 | -0.22 | -0.85 | -0.4 | -0.6 | -0.48 | -1.07 | -0.18 | -0.63 | -0.24 | -0.26 | -0.135 | -0.735 | 0.145221 | 0.243646 | 0.062789 | 1 | 0.999631 | 0.616182 |
| Ccdc177_Q3UHB8_T208t_1 | 1.48 | -1.03 | 0.55 | 0.5 | -0.16 | 0.55 | 0.12 | -0.05 | 0.52 | 1.63 | -0.24 | -0.95 | -0.73 | -0.01 | -0.12 | -0.58 | -0.2725 | -0.34 | -0.82 | 0.178151 | 0.610396 | 0.842179 | 1 | 0.999631 | 0.642627 |
| Ccdc177_Q3UHB8_T208t_2 | 0.29 | -0.63 | -0.26 | -0.29 | -0.69 | -0.43 | -0.49 | -0.37 | -0.52 | 0.05 | -0.6 | -1.18 | -0.95 | -1.16 | -0.83 | -1.23 | 0.1225 | 0.88 | 0.9925 | 0.022154 | 0.345815 | 0.347283 | 1 | 0.999631 | 0.405544 |
| Ccdc25_Q78PG9_S204s_1 | -0.27 | -0.63 | -0.07 | -0.92 | -0.17 | -1.24 | 0.2 | -0.19 | 0.28 | -0.44 | 1.01 | 0.78 | 0.19 | 0.46 | 0.99 | 0.44 | -0.245 | 0.2475 | 0.0575 | 0.011997 | 0.746156 | 0.045061 | 1 | 0.999631 | 0.385092 |
| Ccdc6_D3YZP9_S233s_1 | -0.48 | 0.71 | 1.03 | 0.47 | -0.17 | 0.53 | -0.52 | 0.91 | 0.5 | 0.66 | 0.83 | 0.73 | -0.84 | 0.85 | 0.69 | 1.26 | -0.1175 | 0.1925 | 0.4425 | 0.913426 | 0.585313 | 0.501285 | 1 | 0.999631 | 0.972551 |
| Ccdc6_D3YZP9_S237s_1 | -0.42 | -0.26 | -0.87 | -0.87 | -0.03 | -1.04 | -1 | -0.82 | -0.28 | -0.21 | -0.84 | -0.32 | 0.01 | -0.5 | -0.11 | -0.05 | 0.5625 | 0.385 | 0.365 | 0.162612 | 0.719743 | 0.517972 | 1 | 0.999631 | 0.631442 |
| Ccdc6_D3YZP9_S316s_1 | 0.36 | 0.58 | 0.79 | 1.44 | 0.49 | 0.86 | 1.66 | 2.41 | 0.53 | 0.93 | 2.36 | 0.89 | 0.57 | 1.03 | 1.86 | 1.17 | 0.17 | 0.0775 | 0.1075 | 0.356203 | 0.245339 | 0.407176 | 1 | 0.999631 | 0.74034 |
| Ccdc6_D3YZP9_S345s_1 | 0.46 | 0.67 | 0.49 | 0.76 | 0.91 | 0.5 | 0.66 | 0.99 | 0.42 | 0.2 | 0.89 | 1.18 | 0.4 | 0.25 | 1.11 | 1.05 | 0.1575 | -0.5 | 0.175 | 0.735649 | 0.504727 | 0.799716 | 1 | 0.999631 | 0.90759 |
| Ccdc6_D3YZP9_S360s_1 | 0.1 | -0.31 | 0.09 | 0.24 | 0.19 | 0.11 | 0.46 | -0.01 | -0.1 | -0.47 | -0.63 | -0.68 | 0.02 | 0.59 | -0.11 | 0.32 | 0.6175 | 0.4125 | 0.9275 | 0.564368 | 0.548518 | 0.096234 | 1 | 0.999631 | 0.84139 |
| Ccdc6_D3YZP9_S45s_1 | -0.26 | 1.51 | 1.05 | 0.52 | 0.83 | 1.5 | 0.77 | 2.19 | 1.2 | 0.7 | 1.02 | 1.55 | 1.12 | 1.55 | 1.87 | 1.99 | 0.015 | 0.39 | -0.965 | 0.054419 | 0.217295 | 0.340657 | 1 | 0.999631 | 0.491994 |
| Ccdc85a_E0CX32_S216s_1 | 0.04 | 0.82 | 2.92 | 0.51 | 0.62 | 0.73 | 1.79 | 1.21 | 1.07 | 0.73 | 2.11 | 1.94 | 0.16 | -0.72 | 1.23 | -0.24 | -0.8025 | -1.31 | -1.41 | 0.175917 | 0.980689 | 0.5503 | 1 | 0.999631 | 0.642116 |
| Ccdc85a_E0CX32_S240s_1 | 1.4 | -0.79 | 0.19 | -0.07 | -2.12 | -0.25 | 2.09 | -2.2 | -2.45 | 0.32 | -1.35 | -1.03 | -1.63 | -1.49 | -0.43 | -1.36 | -0.87 | -0.4175 | -0.145 | 0.018071 | 0.40962 | 0.069917 | 1 | 0.999631 | 0.391024 |

| | | | | | | | | | | | | | | | | | | | | | | | | | |
|---|---|---|---|---|---|---|---|---|---|---|---|---|---|---|---|---|---|---|---|---|---|---|---|---|---|
| Ccdc85a_E0CX32_S250s_1 | -2.06 | -5.37 | 2.22 | 1.83 | -3.67 | -5.8 | 1.66 | 0.95 | -4.49 | -3.34 | 1.4 | 1.38 | -2.79 | -4.41 | 1.62 | 1.62 | 0.225 | -0.05 | -0.4325 | 0.941344 | 0.685794 | 0.834902 | 1 | 0.999631 | 0.981286 |
| Ccdc85a_E0CX32_S273s_1 | 2.6 | 2.76 | 2.31 | 1.62 | 1.49 | 2.65 | 4.05 | 2 | 1.89 | 2.5 | 2.31 | 2.39 | 2.08 | 1.8 | 2.63 | 1.05 | -0.9525 | -0.65 | -1.77 | 0.321973 | 0.687514 | 0.88111 | 1 | 0.999631 | 0.724228 |
| Ccdc85a_E0CX32_S284s_1 | 1.96 | -1.7 | 0.4 | 1.04 | -2.66 | -0.9 | 3.17 | -1.72 | -2.02 | 1.84 | 0.16 | -0.88 | -1.9 | -2.24 | 0.15 | -1.39 | 0.415 | 1.6025 | 1.4175 | 0.052128 | 0.462696 | 0.510733 | 1 | 0.999631 | 0.489777 |
| Ccdc85a_E0CX32_S308s_1 | 0.21 | 1.43 | 0.83 | -1.12 | 2.24 | 0.06 | 1.34 | -0.63 | 3.4 | -0.22 | 1.47 | 3.11 | 4.13 | 0.66 | 1.8 | 0.43 | 0.13 | 0.3925 | 0.525 | 0.127426 | 0.578593 | 0.090577 | 1 | 0.999631 | 0.59788 |
| Ccdc88a_Q5SNZ0_S1332s_1 | -1.04 | 0.04 | 0 | -0.71 | -0.35 | -0.33 | -0.58 | 0.07 | 0.13 | -0.79 | -0.28 | 0.8 | 0.66 | -0.09 | 0.05 | -0.23 | 0.64 | 0.5075 | 0.2475 | 0.180118 | 0.700265 | 0.370965 | 1 | 0.999631 | 0.64376 |
| Ccdc88a_Q5SNZ0_S1362s_1 | 1.43 | 1.16 | -0.47 | 2 | 2.07 | 1.62 | 1.34 | 1.65 | 1.3 | 0.91 | 2.2 | 1.74 | -0.23 | 1.58 | 1.62 | 2.14 | 0.1025 | 0.1425 | -0.735 | 0.709709 | 0.231217 | 0.370262 | 1 | 0.999631 | 0.899334 |
| Ccdc88a_Q5SNZ0_S1505s_1 | 1.8 | -0.74 | 0.47 | 1.26 | 0.55 | 1.16 | 1.4 | 0.09 | 0.85 | 1.39 | 0.41 | 0.71 | -0.86 | 0.36 | 0.44 | -0.09 | -0.2675 | 0.13 | 0.3575 | 0.221404 | 0.85746 | 0.794354 | 1 | 0.999631 | 0.671697 |
| Ccdc88a_Q5SNZ0_S1594s_1 | -0.34 | 0.1 | -0.19 | -0.06 | -0.22 | -0.38 | -0.45 | -0.51 | -0.22 | 0.12 | 0.08 | 0.05 | 0.58 | 0.31 | -0.47 | 0.52 | 0.33 | -0.4175 | -0.8875 | 0.295721 | 0.289148 | 0.613443 | 1 | 0.999631 | 0.708981 |
| Ccdc88a_Q5SNZ0_S1726s_1 | 1 | 0.32 | -0.84 | -0.11 | 0.55 | 0.67 | 0.13 | 0.34 | -0.37 | -0.32 | 0.62 | -1.23 | -0.01 | 0.41 | -2.63 | -0.95 | 0.07 | -0.1075 | 0.54 | 0.217936 | 0.424041 | 0.421086 | 1 | 0.999631 | 0.669859 |
| Ccdc88c_Q6VGS5_S1563s_1 | -1.13 | -1.05 | 0.23 | -0.92 | -1.07 | -0.95 | -0.43 | -0.14 | -0.23 | -0.55 | -0.51 | -2.01 | -0.54 | -0.26 | 0.69 | -0.6 | -0.47 | -0.38 | -1.2775 | 0.237931 | 0.858899 | 0.826163 | 1 | 0.999631 | 0.683301 |
| Ccdc88c_Q6VGS5_S1675s_1 | 1.67 | -0.71 | -1.17 | -0.68 | -1.24 | -0.44 | 0.04 | -1.13 | -0.66 | 1.45 | -1.76 | -1.44 | -2.1 | -0.92 | -1.24 | -1.74 | -0.07 | -0.02 | 0.08 | 0.064228 | 0.46532 | 0.653917 | 1 | 0.999631 | 0.514389 |
| Ccdc88c_Q6VGS5_S227s_1 | -0.6 | 0.15 | -0.02 | 0.32 | -0.63 | -0.24 | -0.22 | 0.66 | -0.15 | -0.95 | 0 | 0.87 | 0.54 | 0.39 | 0.08 | -0.84 | -0.14 | -0.1725 | 0.345 | 0.839685 | 0.846372 | 0.9624 | 1 | 0.999631 | 0.946365 |
| Ccdc92_Q8VDN4_S194s_1 | -0.21 | 1.08 | 0.17 | 0.24 | 0.19 | 0.69 | -0.46 | 0.3 | 0.23 | 0.27 | -0.03 | 0.12 | 0.71 | 1.19 | 0.29 | 0.47 | -0.23 | -0.7625 | -2.1275 | 0.367529 | 0.71116 | 0.605202 | 1 | 0.999631 | 0.745968 |
| Ccdc92_Q8VDN4_S261s_1 | 2.73 | 0.52 | 0.57 | 1.38 | 0.34 | 1.53 | 2.75 | -0.34 | 0.57 | 2.54 | -0.11 | -0.85 | -2.8 | -0.09 | -0.09 | -0.33 | 0.2475 | 0.4075 | 0.92 | 0.015292 | 0.759486 | 0.343262 | 1 | 0.999631 | 0.388832 |
| Ccdc97_Q9DBT3_S210s_1 | -0.89 | -0.59 | -0.64 | -0.65 | -0.51 | -0.16 | -1.11 | 0 | -0.52 | 0.46 | -0.1 | -0.98 | 0.54 | -0.35 | 1.19 | -0.47 | -0.2675 | -0.475 | -1.0575 | 0.047306 | 0.432576 | 0.263661 | 1 | 0.999631 | 0.480102 |
| Cckbr_P56481_S257s_1 | -0.21 | -0.72 | 0.47 | -0.2 | -0.47 | -0.46 | -0.19 | -0.61 | -1.38 | 0.36 | -0.14 | -1.4 | -1.71 | -1.6 | -0.66 | -0.92 | 0.22 | 0.3825 | 0.3475 | 0.019506 | 0.401366 | 0.340266 | 1 | 0.999631 | 0.391668 |
| Cckbr_P56481_S443s_1 | -0.78 | -0.38 | -0.29 | -0.58 | -0.36 | -0.62 | -0.61 | 0.44 | -0.31 | -0.56 | -0.17 | 0.54 | -0.38 | -0.01 | -0.1 | -0.15 | -0.0325 | -0.005 | 0.52 | 0.228384 | 0.497675 | 0.249351 | 1 | 0.999631 | 0.679038 |
| Cckbr_P56481_T433t_1 | -0.9 | -1.04 | -0.96 | -0.87 | -1.28 | -0.74 | -1.06 | -0.82 | -1.71 | -0.46 | -0.91 | -0.71 | -0.02 | -0.63 | -0.36 | -0.68 | 0.27 | 0.5775 | 0.485 | 0.091821 | 0.89639 | 0.987798 | 1 | 0.999631 | 0.557085 |
| Ccm2_Q8K2Y9_S164s_1 | -0.12 | -0.48 | -0.25 | -2.52 | 0.67 | -0.56 | -0.91 | -1.49 | 0.52 | 0.28 | -0.9 | -0.96 | 1.29 | 0.23 | -1.39 | -1.56 | -0.1525 | -0.0125 | 0.0775 | 0.534969 | 0.678222 | 0.362745 | 1 | 0.999631 | 0.828951 |
| Ccm2_Q8K2Y9_S393s_1 | 0.45 | 1.25 | 0.95 | 0.85 | 0.97 | 0.7 | 0.12 | 1.1 | 0.8 | 0.75 | 0.79 | 1.11 | 0.21 | 1.35 | 1.16 | 1.09 | -0.6775 | -0.9525 | -2.085 | 0.827574 | 0.636252 | 0.964492 | 1 | 0.999631 | 0.941202 |
| Ccm2_Q8K2Y9_S417s_1 | 1.99 | -1.72 | -0.44 | -0.77 | -2.17 | -0.83 | 1.97 | -2.62 | -2.31 | 1.78 | -1.89 | -2.33 | -1.61 | -1.95 | -2.01 | -3.71 | 0.165 | 0.2825 | 0.4875 | 0.024817 | 0.546585 | 0.392547 | 1 | 0.999631 | 0.418003 |
| Ccm2_Q8K2Y9_S425s_1 | -0.5 | 0.62 | 0.16 | -0.34 | -0.17 | -0.02 | 0.17 | 0.62 | 0.09 | 0.34 | -0.11 | 0.75 | 0.56 | 0.81 | 0.27 | 0.25 | -0.17 | 0.0075 | -0.19 | 0.180918 | 0.630491 | 0.43052 | 1 | 0.999631 | 0.64376 |
| Ccnk_Q3U3M5_S329s_1 | 1.02 | 0.85 | 1.15 | 0.41 | 0.47 | 1.08 | 0.72 | 0.48 | 0.35 | 1.46 | 0.65 | 1 | 0.2 | 0.73 | 0.74 | 1 | 0.0925 | 0.435 | 0.6675 | 0.553916 | 0.559334 | 0.982186 | 1 | 0.999631 | 0.837952 |
| Ccnk_Q3U3M5_S341s_1 | 0.2 | 0.13 | 1.71 | 0.61 | 0.52 | 0.12 | 0.91 | 1.47 | 0.62 | 0.45 | 1.56 | 1.76 | 0.91 | 0.53 | 1.94 | 1.94 | -0.3575 | 0.3575 | 0.485 | 0.196714 | 0.836909 | 0.369303 | 1 | 0.999631 | 0.653373 |
| Ccnl1_Q52KE7_S358s_1 | 0.48 | 1.9 | 1.94 | 1.81 | 0.76 | 1.27 | 1.03 | 1.64 | 1.93 | 1.12 | 2.24 | 2.27 | 1.5 | 1.87 | 2.16 | 2.54 | -0.2975 | 0.07 | -0.195 | 0.270609 | 0.383712 | 0.422897 | 1 | 0.999631 | 0.698674 |
| Ccnl1_Q52KE7_S380s_1 | 1.23 | 0.93 | 1.79 | 1.24 | 0.47 | 1.03 | 1.07 | 1.43 | 1.17 | 1.81 | 1.4 | 1.09 | 0.79 | 0.95 | 1.37 | 1.3 | -0.4575 | 0.1025 | 0.1 | 0.540002 | 0.359599 | 0.821095 | 1 | 0.999631 | 0.831295 |
| Ccnl2_Q9JJA7_S366s_1 | 0.12 | 1.61 | 0.98 | 0.67 | 0.76 | 0.56 | -0.17 | 0.4 | 1.09 | 0.6 | 0.49 | 1.61 | 1.14 | 1.71 | 0.12 | 0.81 | 1.23 | 2.6375 | 1.0875 | 0.825185 | 0.248917 | 0.803032 | 1 | 0.999631 | 0.940309 |
| Ccnt2_Q7TQK0_S549s_1 | 0.96 | 4.51 | 2.42 | -0.74 | 4.62 | 3.57 | 2.9 | 0.98 | 7.13 | 2.76 | 3.43 | 4.38 | 7.42 | 2.95 | 1.13 | 0 | -0.255 | -0.0225 | 0.29 | 0.515584 | 0.301524 | 0.059133 | 1 | 0.999631 | 0.819449 |
| Ccnt2_Q7TQK0_S596s_1 | 0.69 | 1.07 | 0.73 | 0.48 | 0.46 | 0.58 | 0.14 | 0.77 | 1.01 | 0.68 | 1.01 | 0.18 | 0.82 | 1.25 | 0.47 | 1.59 | -1.2075 | -0.365 | -1.7375 | 0.401787 | 0.358234 | 0.940933 | 1 | 0.999631 | 0.763221 |
| Ccnt2_Q7TQK0_S657s_1 | 2.87 | 0.87 | 1.58 | 1.69 | 0.81 | -0.48 | 1.76 | 0.09 | 0.88 | 1.87 | 0.99 | 1.81 | 0.33 | -0.29 | 0.55 | -0.53 | 0.7675 | -0.845 | -0.8325 | 0.004223 | 0.060239 | 0.447645 | 1 | 0.999631 | 0.351308 |
| Ccny_Q8BGU5_S25s_1 | 0.31 | -0.4 | -1.46 | -1.24 | -0.36 | 0.87 | -0.22 | -0.01 | -1.98 | -1.7 | -0.84 | -1.65 | -2.66 | 0.48 | -1.32 | -2.62 | -0.4175 | -0.0875 | 0.5 | 0.277238 | 0.126069 | 0.092036 | 1 | 0.999631 | 0.700628 |
| Ccny_Q8BGU5_S324s_1 | -0.72 | -0.4 | -0.07 | -0.89 | -1.06 | -1.06 | -1.23 | -0.4 | -1.43 | -0.7 | -0.21 | -0.09 | 0.2 | -0.28 | -0.13 | 0.13 | -0.2425 | -0.0075 | 0.37 | 0.12885 | 0.201276 | 0.816868 | 1 | 0.999631 | 0.59942 |
| Ccny_Q8BGU5_S326s_1 | -0.72 | -0.78 | -0.59 | -1.01 | -0.97 | -0.91 | -1.35 | -0.84 | -1.02 | -0.61 | -0.84 | -0.66 | 0.06 | -0.54 | -0.76 | -0.38 | -0.1825 | 0.09 | 0.1375 | 0.235767 | 0.354338 | 0.976786 | 1 | 0.999631 | 0.683301 |
| Ccny_Q8BGU5_S33s_1 | -0.41 | 0.63 | 0.07 | -0.6 | 0.03 | 0.03 | -0.7 | -0.4 | 0.14 | 0.31 | -0.49 | 0.09 | 0.7 | -0.05 | -0.64 | 0.23 | -0.5825 | 0.1425 | -0.055 | 0.738316 | 0.608942 | 0.802243 | 1 | 0.999631 | 0.908027 |
| Ccny_Q8BGU5_S45s_1 | 0.59 | -0.16 | 1.15 | -0.25 | -0.35 | 0.42 | -0.62 | -0.45 | 0.2 | 1.65 | -0.35 | 0.4 | 0.76 | -0.39 | 0.56 | 0.18 | 0.405 | -0.4025 | -0.52 | 0.897409 | 0.173134 | 0.779085 | 1 | 0.999631 | 0.966758 |
| Ccny_Q8BGU5_S83s_1 | 0.63 | -0.32 | -0.79 | 1.12 | 0.91 | -0.12 | 1 | 0.47 | -0.38 | -0.73 | 0.13 | 0.01 | -0.79 | -0.25 | -0.76 | 0.36 | 0.215 | -0.295 | 0.455 | 0.305526 | 0.406752 | 0.394723 | 1 | 0.999631 | 0.716537 |
| Ccny_Q8BGU5_S99s_1 | -0.63 | 0.76 | 0.5 | -0.45 | -0.08 | 0.96 | -0.24 | 0.4 | 0.29 | -0.59 | -0.22 | -0.48 | 0.87 | 1.11 | 0.13 | -0.11 | 0.175 | 0.1825 | 0.4275 | 0.324484 | 0.618752 | 0.473286 | 1 | 0.999631 | 0.72463 |
| Ccny_Q8BGU5_T37t_1 | -0.27 | 1.77 | 0.67 | -0.08 | 0.5 | 1.75 | 0.44 | 0.1 | 0.95 | 0.93 | 0.42 | 0.52 | 0.37 | 1.37 | 0.92 | 1.14 | -0.17 | 0.39 | 0.4575 | 0.393444 | 0.746756 | 0.697125 | 1 | 0.999631 | 0.759134 |
| Ccnyl1_E9Q226_S274s_1 | -0.57 | -0.63 | -0.73 | -0.73 | -0.62 | -0.59 | -1.53 | -0.6 | -0.52 | -0.14 | -0.24 | -0.2 | 0.3 | -0.44 | -0.52 | -0.17 | -0.255 | 0.175 | 0.4825 | 0.147566 | 0.57283 | 0.138745 | 1 | 0.999631 | 0.620119 |
| Ccnyl1_E9Q226_S276s_1 | -0.82 | -0.22 | -0.26 | -0.69 | -0.59 | -0.76 | -1.18 | -0.48 | -0.49 | -0.42 | -0.35 | -0.03 | 0.22 | -0.27 | -0.22 | 0.21 | 0.1025 | -0.1725 | 0.2475 | 0.135154 | 0.386817 | 0.53577 | 1 | 0.999631 | 0.607096 |
| Ccnyl1_E9Q226_S52s_1 | -1.12 | -0.02 | 0.05 | -0.04 | -0.43 | -0.15 | -0.52 | 0.38 | -0.87 | -0.05 | -0.49 | -0.41 | 0.05 | 0.3 | -0.59 | 0.1 | -0.8775 | -0.805 | -1.795 | 0.516841 | 0.779746 | 0.633595 | 1 | 0.999631 | 0.819449 |
| Ccp110_Q7TSH4_S176s_1 | 1.59 | -0.9 | 0.92 | 0.12 | -1.97 | 0.14 | 1.62 | -1.57 | -1.51 | 1.43 | -0.73 | -0.68 | -1.55 | -1.9 | -0.9 | -1.1 | 0.7625 | -0.4975 | -0.16 | 0.007555 | 0.318751 | 0.28748 | 1 | 0.999631 | 0.358173 |
| Ccs_Q9WU84_S267s_1 | 0.98 | 0.51 | -0.26 | 0.48 | 1.3 | 1.87 | 0.31 | 1.28 | 0.06 | 0.09 | -0.57 | 0.14 | 0.18 | 0.86 | -0.58 | 0.61 | -0.88 | -0.2 | -0.9625 | 0.704048 | 0.089038 | 0.174078 | 1 | 0.999631 | 0.897149 |
| Ccsap_Q8QZT2_S129s_1 | 0.91 | 2.33 | 1.77 | 0.74 | 1.58 | 0.5 | 0.79 | -0.64 | 1.75 | 1.33 | 1.05 | 0.82 | 1.44 | 0.97 | -1.49 | 0.98 | 0.92 | 2.1525 | -1.12 | 0.177822 | 0.130576 | 0.639687 | 1 | 0.999631 | 0.642116 |
| Ccsap_Q8QZT2_S222s_1 | 0.09 | 2.22 | 2.74 | -0.15 | 2.69 | 0.38 | 5.54 | -0.03 | 5.06 | 0.3 | 3.26 | 4.89 | 3.18 | -1.32 | -0.68 | -0.76 | -0.0525 | 0.0025 | -0.425 | 0.31304 | 0.470631 | 0.083144 | 1 | 0.999631 | 0.719861 |
| Ccser1_Q8C0C4_S375s_1 | -0.71 | -1.01 | -0.98 | -0.97 | -0.16 | -1.05 | -1.23 | -1.44 | -0.75 | -0.86 | -1.18 | -0.87 | -0.91 | -0.92 | -2.23 | -1.31 | 0.185 | 0.0575 | -0.0125 | 0.260222 | 0.873901 | 0.992136 | 1 | 0.999631 | 0.692624 |
| Ccser2_Q3UHI0_S222s_1 | 0.02 | 0.25 | -1.94 | -0.12 | 0.14 | -0.12 | -0.34 | -0.73 | -0.1 | -0.01 | -0.62 | -0.83 | -1.25 | 0.2 | -0.79 | 0 | -0.6425 | -0.13 | 0.5475 | 0.982228 | 0.71325 | 0.910277 | 1 | 0.999631 | 0.994846 |
| Cct2_P80314_S3s_1 | 0.06 | 1.11 | 0.6 | 0.2 | 0.52 | 0.4 | -0.32 | -1.2 | 0.68 | 0.15 | 0.18 | 0.44 | 0.96 | 1.96 | 0.39 | 0.85 | 2.2175 | 3.06 | 1.99 | 0.211487 | 0.172594 | 0.688778 | 1 | 0.999631 | 0.665474 |
| Ccz1_Q8C1Y8_S264s_1 | -0.3 | 0.5 | 0.89 | -0.23 | 4.71 | 2.11 | 1.96 | 0.95 | 5.83 | 0.4 | 2.64 | 4.23 | 5.06 | 1.86 | 2.52 | -0.62 | 0.04 | -0.07 | -1.3825 | 0.076395 | 0.013765 | 0.013581 | 1 | 0.999631 | 0.542525 |
| Cd200_Q80VX2_S270s_1 | 1.59 | -2.79 | -1.15 | -0.54 | -1.37 | -1.33 | 0.91 | -0.94 | -0.79 | -0.25 | -1.1 | -1.03 | -2.21 | -3.23 | -1.09 | -1.89 | -0.245 | 0.31 | 0.555 | 0.136168 | 0.964968 | 0.931208 | 1 | 0.999631 | 0.609095 |
| Cd248_Q91V98_S754s_1 | 0.82 | 1.41 | 1.31 | 0.8 | 0.97 | 1.41 | 0.69 | 0.29 | 1.38 | 2.06 | 0.91 | 1.23 | 1.16 | 1.76 | 1.77 | 1.87 | 0.075 | 0.215 | 0.12 | 0.102846 | 0.461435 | 0.369283 | 1 | 0.999631 | 0.565572 |
| Cd2ap_Q9JLQ0_S404s_1 | 0.59 | -0.79 | -0.75 | 1.13 | -0.02 | -0.75 | 0.49 | 0.76 | 0 | 0.2 | 0.46 | 0.38 | -1.15 | 0.13 | 1.15 | 0.53 | 0.1225 | 0.04 | -2E-16 | 0.84665 | 0.88952 | 0.652855 | 1 | 0.999631 | 0.948838 |
| Cd2ap_Q9JLQ0_S510s_1 | 0.21 | -0.17 | -0.55 | -0.27 | 0.5 | -0.17 | -0.3 | -0.32 | -0.06 | 0.51 | -0.73 | -0.34 | 0.49 | -0.36 | -0.33 | -0.58 | 0.0925 | -0.1725 | 0.4325 | 1 | 0.690926 | 0.907945 | 1 | 0.999631 | 1 |
| Cd2ap_Q9JLQ0_S580s_1 | -0.21 | 2.18 | 0.67 | 0.5 | 1.93 | 1.47 | 0.07 | 0.04 | 2.09 | 0.06 | 0.14 | 0.16 | 2 | 1.74 | 0.78 | 0.35 | -0.2025 | -0.5325 | -0.07 | 0.464974 | 0.882549 | 0.786234 | 1 | 0.999631 | 0.794369 |
| Cd34_Q64314_S343s_1 | -0.66 | -0.23 | 0.09 | 0.44 | -0.33 | -0.4 | -0.33 | -0.11 | -1.09 | -0.13 | -1.74 | 0.47 | -0.22 | -0.24 | 0.21 | -0.39 | -0.18 | -0.0375 | 0.475 | 0.835019 | 0.50899 | 0.313418 | 1 | 0.999631 | 0.944416 |
| Cd44_Q80X37_S544s_1 | 0.35 | 1.01 | 1.54 | 0.28 | 0.01 | 0.85 | 0.41 | 1.19 | 0.35 | 0.25 | 1.33 | 1.1 | 1.24 | 1.26 | 0.74 | 1.84 | 0.085 | -0.1125 | -0.02 | 0.250344 | 0.65308 | 0.926977 | 1 | 0.999631 | 0.686417 |
| Cdc23_Q8BGZ4_S460s_1 | -0.58 | -0.06 | -0.41 | 0.17 | 0.29 | -0.37 | -0.42 | -0.04 | 0.09 | -1.14 | 0.3 | -0.58 | -0.41 | -0.26 | -0.21 | -0.08 | 0.155 | -0.2975 | 0.27 | 0.945915 | 0.776947 | 0.771931 | 1 | 0.999631 | 0.982217 |
| Cdc23_Q8BGZ4_S470s_1 | -0.07 | 0.22 | -1.01 | -0.68 | 0.55 | 0.05 | -0.87 | -0.65 | 0.48 | 0.43 | -2.32 | -1.32 | 0.01 | 0.72 | -0.19 | -1 | 0.2175 | 0.2625 | 0.8075 | 0.55225 | 0.715439 | 0.656499 | 1 | 0.999631 | 0.836594 |
| Cdc23_Q8BGZ4_T478t_1 | -1.05 | 0.16 | 0.14 | -0.58 | 0.11 | -0.51 | -0.35 | 0.29 | -0.59 | -0.3 | 0.36 | 0.25 | 1.09 | 0.83 | -0.17 | 0.15 | -0.2025 | -0.1025 | -0.4175 | 0.081845 | 0.558924 | 0.504741 | 1 | 0.999631 | 0.54792 |
| Cdc27_Q8R568_S371s_1 | -0.2 | 0.2 | -0.04 | 0.48 | 0.03 | -0.59 | -0.21 | 0.4 | 0.08 | 0.01 | -0.3 | 0.24 | -1.25 | -0.1 | 0.12 | 0 | -0.3775 | -0.1825 | 0.025 | 0.291705 | 0.518737 | 0.716779 | 1 | 0.999631 | 0.706164 |
| Cdc42bpa_F6Q5A5_S234s_1 | 0.33 | 2.05 | 1.76 | 0.79 | 0.43 | 1.36 | 0.8 | 0.83 | 1.88 | 0.41 | 0.81 | 1.1 | 1.21 | 1.48 | 1.72 | 0.62 | 0.365 | 0.4625 | 0.185 | 0.956789 | 0.396349 | 0.708726 | 1 | 0.999631 | 0.984803 |
| Cdc42bpa_F6Q5A5_S945s_1 | 0.99 | 0.64 | 0.14 | 1.28 | 0.86 | 1.37 | 0.89 | 1.39 | 1.57 | 0.88 | 1.18 | 1.27 | -0.31 | 1.41 | 1.41 | 1.28 | 0.7225 | 0.4825 | 0.1975 | 0.697751 | 0.281595 | 0.186989 | 1 | 0.999631 | 0.895731 |
| Cdc42bpa_F6Q5A5_S967s_1 | -0.13 | -0.79 | -0.87 | -0.37 | 0.09 | 0.11 | 0.13 | 0.4 | 0.3 | -0.2 | -0.11 | -0.22 | -0.77 | -0.02 | -0.21 | -0.37 | -1.14 | -0.385 | -1.5425 | 0.541211 | 0.024281 | 0.126013 | 1 | 0.999631 | 0.83207 |
| Cdc42bpb_Q7TT50_S1692s_1 | 0.58 | -0.16 | 1.02 | -0.85 | -4.64 | 0.79 | 0.11 | -0.23 | -1.85 | -0.62 | 0.33 | 1.19 | 0.54 | -4.73 | 0.16 | -1.55 | -0.1925 | -0.135 | 0.41 | 0.175162 | 0.319495 | 0.579065 | 1 | 0.999631 | 0.642116 |
| Cdc42ep1_Q91W92_S121s_1 | -0.06 | 1.54 | 0.83 | 0.26 | 0.87 | 0.73 | -0.07 | 0.27 | 1.08 | 0.28 | -0.27 | 0.94 | 1.19 | 1.24 | 0.96 | 0.82 | -0.2475 | -0.1425 | -0.295 | 0.310583 | 0.641708 | 0.768611 | 1 | 0.999631 | 0.71924 |
| Cdc42ep2_Q8JZX9_S137s_1 | -0.75 | 0.48 | 0.12 | 0.54 | -0.99 | -0.74 | 0.1 | 1.03 | -0.86 | -1.28 | 0.33 | 1.63 | -1.39 | -0.85 | 0.72 | 0.73 | 0.14 | -0.2875 | -0.2175 | 0.607918 | 0.628815 | 0.825691 | 1 | 0.999631 | 0.858527 |
| Cdc42ep4_Q9JM96_S116s_1 | -0.38 | 0.09 | 1.47 | 0.8 | 0.59 | 0.11 | 1 | 0.84 | 0.24 | -0.16 | 0.56 | 0.19 | -0.97 | 0.45 | 0.52 | 1.11 | -0.365 | -0.07 | 0.035 | 0.696683 | 0.749883 | 0.509284 | 1 | 0.999631 | 0.895731 |
| Cdc42ep4_Q9JM96_S140s_1 | -0.23 | -0.19 | -0.66 | -1.25 | -1.12 | -0.25 | -1.7 | -0.72 | -0.54 | 0.03 | -0.9 | -1.2 | 0.12 | -0.57 | -0.83 | -0.91 | 0.21 | 0.44 | 0.4625 | 0.926144 | 0.370899 | 0.854821 | 1 | 0.999631 | 0.97588 |
| Cdc42ep4_Q9JM96_S64s_1 | -0.65 | 0.6 | 0.29 | -0.11 | 0.33 | 0.13 | 0.01 | 0.5 | 0.52 | -0.65 | 0.79 | 1.23 | -0.14 | 0.36 | 1.09 | 0.67 | -0.345 | -0.9975 | -0.1525 | 0.262888 | 0.530555 | 0.358851 | 1 | 0.999631 | 0.693034 |
| Cdc5l_Q6A068_T385t_1 | -0.26 | 0.25 | 0.3 | 0.67 | -0.57 | 0.04 | -0.35 | 0.46 | -0.32 | -1.46 | -0.79 | -0.46 | -0.81 | -0.16 | -0.56 | 1.88 | -0.2 | -0.11 | -0.195 | 0.795339 | 0.316471 | 0.017546 | 1 | 0.999631 | 0.929228 |
| Cdh10_P70408_T701t_1 | 0.49 | 0.13 | -0.35 | -0.2 | -0.11 | -0.04 | -0.31 | -0.27 | -0.07 | 0.21 | -0.42 | -0.09 | -0.25 | 0.15 | -0.52 | -0.09 | -0.2525 | 0.055 | -0.235 | 0.544311 | 0.480054 | 0.716182 | 1 | 0.999631 | 0.833226 |
| Cdh12_Q5RJH3_S649s_1 | 0.32 | 1.23 | 1.1 | 0.68 | 0.44 | 0.98 | 0.61 | 0.29 | 1.04 | 0.66 | 1.13 | 0.72 | 0.94 | 0.86 | 0.07 | 0.52 | -0.6025 | -0.3875 | -0.36 | 0.502763 | 0.423703 | 0.857999 | 1 | 0.999631 | 0.814092 |
| Cdh22_Q9WTP5_S705s_1 | 0.21 | -0.22 | -0.16 | -0.27 | -0.55 | -0.52 | -0.72 | -1.06 | -0.82 | -0.11 | -0.91 | -0.15 | 0.19 | -0.35 | -1.47 | -0.25 | 0.92 | -0.57 | -1.1275 | 0.373859 | 0.043359 | 0.227968 | 1 | 0.999631 | 0.747035 |
| Cdip1_F6X7Q6_T17t_1 | 1.52 | -0.82 | -1.77 | 0.93 | 1.65 | 0.08 | 1.04 | 0.77 | -0.12 | -1.29 | -0.05 | -0.96 | -3.33 | 0.07 | -0.66 | -0.73 | 0.195 | 0.3525 | 0.855 | 0.237042 | 0.226882 | 0.441577 | 1 | 0.999631 | 0.683301 |
| Cdk10_Q3UMM4_T196t_1 | -1.02 | 0.09 | -0.22 | -0.02 | 0.25 | 0.43 | -0.52 | -0.55 | -0.14 | 0.11 | -0.06 | 0.33 | 0.67 | 0.8 | 0.27 | 0.51 | -0.05 | 0.4775 | -0.27 | 0.028728 | 0.604828 | 0.295114 | 1 | 0.999631 | 0.427571 |
| Cdk11b_P24788_S47s_1 | 0.5 | 1.19 | 1.49 | 0.92 | 0.45 | 1.27 | 0.47 | 1.71 | 1.52 | 1.16 | 1.9 | 1.43 | -0.08 | 1.05 | 0.89 | 1.16 | -0.28 | 0.0725 | 0.085 | 0.489548 | 0.896838 | 0.158525 | 1 | 0.999631 | 0.807494 |
| Cdk11b_P24788_S578s_1 | 0.06 | 0.51 | 0.76 | 0.07 | 0.15 | 0.1 | -0.18 | 0.21 | 0.21 | 0.49 | 0.37 | 0.62 | 0.55 | 0.45 | 0.33 | 0.41 | 0.0875 | 0.8625 | 0.3575 | 0.772307 | 0.32538 | 0.799273 | 1 | 0.999631 | 0.919649 |
| Cdk11b_P24788_S65s_1 | -0.3 | -1.15 | -3.48 | -1.41 | -0.02 | -2.53 | -2.11 | -1.33 | -1.07 | -0.98 | -0.74 | -0.1 | 0.02 | -1.54 | -1.59 | -1.8 | -0.7 | -0.2575 | -1.6625 | 0.613761 | 0.908781 | 0.198504 | 1 | 0.999631 | 0.860866 |
| Cdk11b_P24788_S72s_1 | 2.18 | 1.45 | 2.01 | 1.88 | 0.39 | 2.29 | 2.18 | -0.14 | 1.33 | 2.97 | 1.12 | 1.07 | 0.03 | 0.23 | 1.16 | -0.55 | 0.135 | 0.2625 | 0.445 | 0.002154 | 0.245597 | 0.582301 | 1 | 0.999631 | 0.351308 |
| Cdk12_Q14AX6_S1079s_1 | -0.31 | 1.19 | 1.96 | 0.86 | 0.86 | 1.52 | 0.92 | 0.94 | 1.28 | -0.09 | 1.52 | 2.04 | 1.01 | 1.21 | 1.38 | 1.88 | 0.5275 | 0.54 | -0.07 | 0.373395 | 0.775381 | 0.661753 | 1 | 0.999631 | 0.747035 |
| Cdk13_Q69ZA1_S326s_1 | 1.44 | 5.24 | 6.21 | 2.5 | 4.17 | 4.38 | 5.06 | 3.89 | 4.7 | 1.26 | 6.06 | 5.53 | 4.51 | 3.28 | 4.8 | 2.52 | -0.81 | -0.385 | -1.6325 | 0.947249 | 0.596784 | 0.685568 | 1 | 0.999631 | 0.982895 |
| Cdk13_Q69ZA1_T1087t_1 | 2.16 | 0.66 | 2.07 | 1.35 | -0.01 | 1.04 | 1.47 | 0.5 | 1.18 | 2.24 | 0.43 | 0.85 | -0.11 | -0.84 | 0.8 | -0.14 | 0.125 | 0.015 | 0.19 | 0.006025 | 0.101578 | 0.445151 | 1 | 0.999631 | 0.351545 |
| Cdk13_Q69ZA1_T588t_1 | -0.09 | -0.18 | 0.96 | 0.64 | 0.41 | 0.01 | 0.56 | 0.85 | -0.75 | -0.26 | 1.38 | 1.02 | -0.37 | 0.43 | 1.11 | 0.92 | 0.1175 | -0.08 | 0.1125 | 0.665012 | 0.725549 | 0.977774 | 1 | 0.999631 | 0.882433 |
| Cdk13_Q69ZA1_T871t_1 | -1.02 | 1.92 | 1.08 | -0.13 | 0.27 | 1.32 | 0.47 | 0.26 | 0.91 | -0.82 | 1.8 | -0.36 | 1.26 | 1.33 | 0.48 | -0.77 | 0.46 | -0.24 | -0.555 | 0.875362 | 0.850863 | 0.917894 | 1 | 0.999631 | 0.960522 |
| Cdk14_D3YV23_S32s_1 | -0.81 | 1.1 | 0 | 0.16 | 1.02 | 0.77 | 0.8 | -0.3 | 0.33 | -1.19 | 0.8 | -0.45 | -0.36 | 0.19 | -1.1 | -0.5 | 0.7175 | -0.1875 | -1.2625 | 0.250158 | 0.338651 | 0.661459 | 1 | 0.999631 | 0.686417 |
| Cdk16_Q04735_S103s_1 | 1.39 | 0.07 | -0.94 | 1.74 | 1.85 | 0.34 | 1.42 | 1.52 | 1.1 | -0.6 | 0.6 | 0.41 | -2.93 | 0.22 | -0.06 | -0.02 | -0.1975 | 0.105 | 0.19 | 0.155741 | 0.269052 | 0.770694 | 1 | 0.999631 | 0.626452 |
| Cdk16_Q04735_S43s_1 | 1.04 | 1.17 | -0.51 | 0.39 | -0.4 | 1.67 | -0.51 | 0.54 | 1.7 | 1.34 | -0.01 | -0.52 | 0.25 | 1.55 | 0 | 1.05 | 0.345 | -0.105 | 0.085 | 0.705931 | 0.733183 | 0.860471 | 1 | 0.999631 | 0.897883 |
| Cdk16_Q04735_S60s_1 | -0.09 | 0.54 | 0 | -0.03 | 0.5 | 0.67 | 0.12 | 0.51 | 0.58 | 0.23 | -0.06 | -0.75 | 0.32 | -0.26 | -0.04 | 0.74 | 0.2225 | -0.3375 | -0.6175 | 0.799168 | 0.227692 | 0.767365 | 1 | 0.999631 | 0.930937 |
| Cdk16_Q04735_S75s_1 | 0.96 | 0.82 | 0.22 | 0.9 | 1.4 | 0.67 | 0.73 | 0.99 | 0.76 | 0.4 | 0.21 | 0.18 | -1 | 0.78 | 0.25 | 0.4 | 0.2675 | -0.0075 | -0.46 | 0.172946 | 0.465795 | 0.271229 | 1 | 0.999631 | 0.642116 |
| Cdk16_Q04735_S84s_1 | 0.52 | -0.14 | -0.45 | 0.76 | 0.7 | 0.01 | 0.18 | 0.87 | 0.32 | -0.64 | 0.31 | 0.67 | -1.28 | 0.45 | -0.61 | 0.29 | 0.7675 | 0.415 | 0.685 | 0.348798 | 0.473369 | 0.985285 | 1 | 0.999631 | 0.736697 |
| Cdk17_Q8K0D0_S38s_1 | -0.76 | -0.49 | -0.51 | -0.02 | 0.17 | -0.5 | 0.31 | 1.31 | 0.46 | -0.26 | -0.53 | 0.21 | 0.25 | 0.03 | 0.34 | 0.34 | 0.2625 | -0.6625 | -0.7925 | 0.037274 | 0.083395 | 0.223216 | 1 | 0.999631 | 0.457763 |
| Cdk17_Q8K0D0_S53s_1 | 1.63 | -0.13 | -0.18 | 1.4 | 0.79 | 0.04 | 1.75 | 1.19 | -0.64 | -0.93 | 1.45 | 0.19 | -0.54 | 0.1 | -1.23 | 1.22 | 0.095 | 0.2025 | 0.29 | 0.233902 | 0.636902 | 0.320302 | 1 | 0.999631 | 0.683077 |
| Cdk17_Q8K0D0_S81s_1 | -0.04 | 0.63 | -1.26 | -0.46 | 0.32 | -0.6 | -0.61 | 0.14 | 0.07 | 0.14 | -0.47 | -0.06 | 0.15 | 0.77 | -0.62 | -0.27 | 0.68 | -0.22 | -0.215 | 0.550477 | 0.832595 | 0.632768 | 1 | 0.999631 | 0.836038 |
| Cdk17_Q8K0D0_S96s_1 | 0.64 | 0.17 | -1.2 | 1.12 | 1.07 | 0.48 | 0.89 | 1.01 | 0.27 | -0.92 | 0.54 | -0.04 | -1.08 | 0.8 | 0.22 | -0.07 | 0.4825 | 0.355 | 0.04 | 0.713242 | 0.185304 | 0.689665 | 1 | 0.999631 | 0.900989 |
| Cdk18_Q04899_S109s_1 | 0.13 | 0.09 | -0.84 | 0.16 | 0.86 | 0.42 | -0.04 | 0.23 | 0.47 | 0.13 | -0.05 | 0.41 | -1.66 | 0.61 | 0.43 | 0.32 | 0.8475 | 0.46 | 0.1725 | 0.941441 | 0.179722 | 0.291468 | 1 | 0.999631 | 0.981286 |
| Cdk18_Q04899_S66s_1 | -0.05 | -0.46 | -0.28 | 1.41 | 1.21 | 0.92 | 0.55 | 1.33 | 0.44 | 0.2 | 1.11 | 0.71 | -0.89 | 0.38 | 1.04 | 0.78 | 0.9275 | -0.12 | -0.5425 | 0.758348 | 0.082278 | 0.327807 | 1 | 0.999631 | 0.914449 |
| Cdk5r1_P61809_T138t_1 | 1.34 | 0.19 | -0.71 | 1.26 | 1.59 | 1.4 | 1.26 | 1.54 | 0.64 | -0.04 | 1.17 | -0.17 | -2.68 | 1.11 | 0.59 | 0.89 | 0.5675 | 0.51 | 0.07 | 0.54046 | 0.072449 | 0.823693 | 1 | 0.999631 | 0.831391 |
| Cdkl5_Q3UTQ8_S306s_1 | -0.39 | -1 | -0.86 | -0.79 | 0.44 | -0.82 | 0.23 | -0.62 | 0.26 | -1.34 | -0.22 | 0.3 | -0.72 | -1.32 | -0.08 | -0.64 | 0.3475 | -1.0275 | -0.36 | 0.839682 | 0.139205 | 0.233693 | 1 | 0.999631 | 0.946365 |
| Cdkl5_Q3UTQ8_S377s_1 | 1.3 | -0.71 | 0.46 | 1.76 | 1.14 | -0.4 | 1.41 | 2.05 | -0.25 | -1.58 | 0.23 | 0.3 | -0.55 | -0.65 | 0.94 | 1.63 | 1.1775 | 0.415 | 0.2725 | 0.606171 | 0.605498 | 0.126961 | 1 | 0.999631 | 0.85817 |
| Cdkl5_Q3UTQ8_S407s_1 | -1.02 | 0.91 | -0.16 | 0.53 | 2.07 | 1.04 | 0.64 | 1.22 | 0.71 | -0.74 | 1.08 | 0.87 | -0.85 | 0.54 | 0.76 | 0.9 | 0.07 | 0.1225 | -0.575 | 0.619443 | 0.035352 | 0.456728 | 1 | 0.999631 | 0.861888 |
| Cdkl5_Q3UTQ8_S497s_1 | 1.15 | 1.02 | 2.24 | 2.3 | 0.86 | 1.18 | 2.3 | 2.65 | 1.33 | 0.81 | 2.77 | 2.29 | -1.09 | 1.02 | 2.42 | 2.06 | 0.3425 | 0.085 | -0.5375 | 0.453093 | 0.891481 | 0.816134 | 1 | 0.999631 | 0.788717 |
| Cdkl5_Q3UTQ8_S681s_1 | 0.61 | 2.47 | 2.14 | 1.77 | 1.02 | 2.34 | 2.56 | 2.44 | 1.88 | 1.4 | 2.13 | 1.92 | 0.44 | 1.78 | 0.58 | 2.04 | 0.2575 | -0.015 | -0.6275 | 0.330806 | 0.503846 | 0.844059 | 1 | 0.999631 | 0.727302 |
| Cdkl5_Q3UTQ8_S691s_1 | 0.28 | -0.12 | 0 | -0.26 | 0.51 | 0.36 | -0.08 | 0.14 | -0.1 | 0.44 | -0.28 | -0.22 | -0.7 | -0.03 | -1.31 | -0.57 | 0.1125 | 0.035 | 0.2975 | 0.094415 | 0.347981 | 0.958473 | 1 | 0.999631 | 0.558128 |
| Cdkl5_Q3UTQ8_S761s_1 | 0.29 | 1.07 | -0.46 | 0.85 | 0.78 | 1.08 | -0.25 | 0.59 | 0.6 | 0.85 | -0.07 | 0.51 | 0.49 | 1.54 | 0.74 | 0.17 | 1.2175 | 0.4875 | 0.355 | 0.512245 | 0.79599 | 0.930844 | 1 | 0.999631 | 0.818025 |
| Cdkl5_Q3UTQ8_Y171y_1 | -0.41 | -4.19 | -4.73 | -1.22 | -1.71 | -1.39 | -1.8 | -0.78 | -1.71 | -1.33 | -3.18 | -2.38 | -2.06 | -2.08 | -2.77 | -2.22 | 0.295 | 0.8125 | -0.035 | 0.703944 | 0.217558 | 0.624357 | 1 | 0.999631 | 0.897149 |
| Cdkn1b_P46414_S10s_1 | -0.7 | 0.88 | 0.64 | -0.25 | 0.89 | -0.14 | 0.95 | 0.05 | 1.4 | -0.26 | 1.03 | 1.65 | 0.63 | -0.15 | -0.05 | 0 | -0.4325 | 0.28 | 0.89 | 0.934041 | 0.517047 | 0.147536 | 1 | 0.999631 | 0.978764 |
| Cdkn1b_P46414_S178s_1 | -0.53 | 1.04 | 0.28 | 1.04 | 0 | -0.04 | -0.15 | 0.29 | 0.98 | -0.46 | 0.45 | 1.98 | 1.16 | 0.6 | 2.53 | 1.1 | -0.7475 | -0.7825 | -1.7475 | 0.114872 | 0.287114 | 0.631269 | 1 | 0.999631 | 0.579336 |
| Cdkn2aip_Q8BI72_S169s_1 | 3.03 | 1.31 | 1.99 | 1.81 | 0.64 | 1.47 | 2.74 | 0.3 | 1.28 | 2.1 | 0.81 | 0.82 | -0.11 | 0.1 | 1.11 | 0.05 | -0.0325 | -0.135 | 0.1525 | 0.003087 | 0.225184 | 0.113482 | 1 | 0.999631 | 0.351308 |
| Cds1_P98191_S56s_1 | -0.75 | 0.34 | -0.23 | -0.51 | -0.14 | -0.25 | -0.63 | -0.26 | -0.51 | 0.08 | -0.59 | -0.67 | 0.56 | 0.15 | -0.58 | -0.67 | -0.5725 | -0.3675 | -1.4675 | 0.704664 | 0.916922 | 0.690142 | 1 | 0.999631 | 0.897611 |
| Cds2_Q6PBC0_S20s_1 | 0.34 | -0.35 | 1.32 | 1.51 | -0.58 | -0.44 | 0.97 | 0.58 | -0.26 | 0.64 | 0.48 | 0.49 | -1.39 | -1.21 | 0.28 | -0.73 | -0.6375 | -0.615 | -1.3025 | 0.019909 | 0.29825 | 0.437361 | 1 | 0.999631 | 0.391668 |
| Cds2_Q6PBC0_S32s_1 | 1.11 | -0.31 | 0.51 | 0.56 | -0.81 | -0.26 | 0.44 | -0.05 | -0.47 | 0.97 | -0.48 | -0.61 | -0.69 | -0.92 | -0.4 | -1.33 | -0.155 | 0.045 | -1.185 | 0.006598 | 0.132223 | 0.203528 | 1 | 0.999631 | 0.351545 |
| Cds2_Q6PBC0_S32s_2 | 1.13 | 0.56 | 0.51 | 0.35 | 0.44 | 1.21 | 0.36 | -0.08 | 1.41 | 1.76 | -0.47 | 0.03 | -0.28 | -0.87 | -0.5 | -0.54 | -0.16 | 0.035 | -1.2675 | 0.003007 | 0.657551 | 0.931805 | 1 | 0.999631 | 0.351308 |
| Cds2_Q6PBC0_S36s_2 | 1.13 | 0.56 | 1.03 | 0.8 | 0.44 | 1.21 | 0.88 | 0.35 | 1.41 | 1.76 | -0.46 | 0.95 | -0.28 | -0.87 | -0.08 | -0.32 | 0.995 | 0.8975 | -0.6025 | 0.002014 | 0.596591 | 0.942154 | 1 | 0.999631 | 0.351308 |
| Cdv3_Q4VAA2_S51s_1 | 0.2 | 0 | 0.88 | -0.48 | 1.38 | 0.57 | 2.05 | 0.58 | 1.94 | -0.3 | 1.02 | 1.53 | -0.04 | -0.86 | 0.82 | -1.73 | 0.445 | -0.485 | -1.1675 | 0.302819 | 0.045559 | 0.113409 | 1 | 0.999631 | 0.714978 |
| Cdv3_Q4VAA2_S58s_1 | 1.28 | -1.85 | -0.26 | 0.8 | 0.5 | -0.13 | 1.1 | 0.28 | -1.1 | -0.01 | -0.43 | -0.43 | -1.26 | -1.83 | -0.61 | -1 | -0.0475 | 0.1375 | 0.57 | 0.101876 | 0.504567 | 0.465517 | 1 | 0.999631 | 0.564448 |
| Cecr6_Q99MX7_S521s_1 | -0.42 | 0.12 | -0.08 | -0.38 | -0.42 | -0.05 | -0.43 | -0.05 | -0.33 | 0.13 | 0.32 | -0.33 | 0.66 | 0.1 | 0.03 | 0.73 | 0.0125 | 0.0575 | 0.415 | 0.093207 | 0.858064 | 0.640049 | 1 | 0.999631 | 0.558128 |
| Cecr6_Q99MX7_S527s_1 | -1.13 | -0.43 | -0.31 | -1.33 | -0.36 | -0.79 | -0.79 | -1.21 | -0.81 | -0.49 | -0.96 | -0.71 | 0.25 | -0.62 | -0.55 | -0.62 | 0.2 | 0.1075 | 0.43 | 0.279746 | 0.970743 | 0.860125 | 1 | 0.999631 | 0.701922 |
| Cecr6_Q99MX7_S548s_1 | -1.93 | -0.81 | -1.22 | -1.57 | -0.84 | -1.47 | -1.19 | -1.23 | -0.91 | -1.6 | -1.13 | -1.46 | -0.2 | -1.42 | -1.2 | -0.99 | -1.0175 | 0.0025 | -0.94 | 0.283836 | 0.536555 | 0.748842 | 1 | 0.999631 | 0.704942 |
| Celf4_Q7TSY6_S164s_1 | 1.99 | 0.58 | 1.46 | 1.66 | -0.9 | 0.83 | 1.62 | 0.07 | 0.57 | 2.49 | 1.42 | 1.22 | 1.4 | 0.3 | 0.7 | -0.47 | -0.095 | 0.02 | 0.0975 | 0.073687 | 0.095845 | 0.995839 | 1 | 0.999631 | 0.539601 |
| Celsr2_F7BHW1_S637s_1 | -0.14 | -0.75 | 0.05 | 0.46 | 0.15 | -0.56 | -0.35 | 0 | -0.97 | -0.4 | 0.34 | 0.73 | -0.34 | -0.53 | 0.48 | 0.4 | 0.4275 | -0.2325 | 0.1475 | 0.802521 | 0.778004 | 0.964319 | 1 | 0.999631 | 0.932677 |
| Celsr2_F7BHW1_S808s_1 | 0.87 | -0.23 | -0.92 | -0.33 | 0.62 | -0.22 | 0.23 | 0.47 | -0.65 | -0.1 | -0.64 | -0.15 | 0.46 | -0.79 | -0.06 | 0.37 | -0.745 | -0.4625 | -2.0475 | 0.750516 | 0.315751 | 0.574036 | 1 | 0.999631 | 0.912627 |
| Cend1_Q9JKC6_S109s_1 | 1.89 | 0.02 | 2.02 | 1.54 | -0.11 | 0.9 | 1.12 | 0.58 | 0.57 | 2.55 | 0.07 | 0.43 | -0.83 | -1.07 | 0.15 | -0.97 | -1.3575 | -1.405 | -1.6525 | 0.002482 | 0.159163 | 0.483157 | 1 | 0.999631 | 0.351308 |
| Cend1_Q9JKC6_S10s_1 | 1.69 | -0.22 | 0.65 | 0.39 | -3.05 | -0.49 | 1.24 | -0.62 | -0.88 | -0.5 | -0.75 | -0.98 | -2.13 | -0.95 | -0.24 | -0.78 | -0.105 | -0.215 | 0.4475 | 0.010297 | 0.131176 | 0.007014 | 1 | 0.999631 | 0.383871 |
| Cend1_Q9JKC6_S90s_1 | -0.14 | -0.94 | -0.13 | -0.37 | -0.31 | 0.06 | -0.89 | -0.86 | -0.4 | 0.1 | -1.61 | -0.53 | 0.15 | 0.78 | -0.27 | -0.45 | -0.1475 | 0.1375 | -0.755 | 0.248001 | 0.754861 | 0.605415 | 1 | 0.999631 | 0.685751 |
| Cenpa_D6RJ71_S15s_1 | 0.46 | -1.18 | -0.26 | -0.58 | -0.82 | -0.41 | -0.23 | -0.69 | -0.31 | 0.7 | -0.7 | -0.7 | -1.42 | -1.07 | -0.5 | -1.59 | -0.4875 | -1.0075 | -0.8125 | 0.101667 | 0.69898 | 0.766333 | 1 | 0.999631 | 0.564448 |
| Cenpv_Q9CXS4_S18s_1 | 0.71 | -0.31 | 0.77 | 0.78 | -0.65 | -0.14 | 0.43 | 0.36 | -0.68 | -0.9 | 0.21 | -0.71 | -0.93 | -0.8 | 0.76 | -0.33 | 0.0475 | -1.3525 | -1.84 | 0.102538 | 0.218786 | 0.024299 | 1 | 0.999631 | 0.565479 |
| Cenpv_Q9CXS4_S6s_1 | 2.67 | 1.49 | 1.21 | 2.35 | 1.41 | 1.7 | 2.45 | 2.35 | 1.01 | 0.65 | 1.31 | -0.66 | -1.4 | 0.96 | 0.56 | 0.24 | -0.11 | -0.6725 | -1.5775 | 0.00889 | 0.910465 | 0.02552 | 1 | 0.999631 | 0.365244 |
| Cep128_F7CIB0_S536s_1 | 2.64 | 0.65 | 0.76 | 2.34 | 1.09 | 1.62 | 2.76 | 0.48 | 0.64 | 1.12 | 0.67 | 1.27 | -1.6 | 0 | 1.49 | 0.19 | 0.2675 | -0.69 | -0.8525 | 0.050719 | 0.862629 | 0.209471 | 1 | 0.999631 | 0.489558 |
| Cep131_B1AXI9_S47s_1 | 1.04 | 1.29 | 0.32 | 0.7 | 0.32 | 1.85 | 1.03 | 1.22 | 0.19 | 0.05 | 1.66 | -1.31 | -1.8 | 1.74 | -0.42 | 0.42 | 0.25 | 0.1125 | -0.6775 | 0.23273 | 0.496377 | 0.257711 | 1 | 0.999631 | 0.682707 |
| Cep131_B1AXI9_S78s_1 | 0.59 | 0.6 | -0.62 | -0.06 | 0.39 | 0.54 | 0.36 | 0.22 | 0.55 | 0.21 | -0.25 | 0.45 | -0.67 | 0.84 | -1.68 | -0.69 | 0.0175 | 0.165 | 0.035 | 0.239505 | 0.464983 | 0.761768 | 1 | 0.999631 | 0.683719 |
| Cep170_H7BX26_S1048s_1 | 0.57 | 0.59 | -0.37 | 0.63 | 0.55 | 0.82 | 0.18 | -0.06 | 1.2 | 1.3 | -0.52 | 0.1 | 0.5 | 1.25 | -0.44 | 0.25 | 0.395 | -0.0225 | -1.0675 | 0.935377 | 0.959358 | 0.732694 | 1 | 0.999631 | 0.979217 |
| Cep170_H7BX26_S1155s_1 | 1.57 | 0.62 | 0.27 | 1.76 | 1.43 | 0.83 | 2.33 | 1.21 | 1.43 | 0.52 | 1.83 | 0.35 | -1.72 | 0.24 | 0.5 | 0.93 | -0.8875 | -0.2375 | -2.13 | 0.111062 | 0.401454 | 0.963002 | 1 | 0.999631 | 0.577068 |
| Cep170_H7BX26_S1157s_1 | 1.61 | 1.48 | 1.28 | 1.75 | 1.15 | -0.45 | 2.8 | -0.93 | 1.53 | 1.13 | 1.35 | 1.16 | -1.74 | -0.5 | -1.35 | 1.19 | -0.395 | -0.515 | -1.3775 | 0.005051 | 0.253031 | 0.373305 | 1 | 0.999631 | 0.351308 |
| Cep170_H7BX26_S1215s_1 | 2.03 | -1.33 | 0.06 | 0.81 | -0.03 | -1.62 | 1.87 | -0.23 | -0.98 | 0.26 | -0.24 | 0.47 | -2.88 | -1.38 | -0.56 | 0.88 | 0.2675 | 0.195 | -0.255 | 0.15221 | 0.651927 | 0.462095 | 1 | 0.999631 | 0.625198 |
| Cep170_H7BX26_S1260s_1 | 0.77 | 0.71 | -1.06 | -0.26 | 0.85 | -0.34 | 0.32 | 0.4 | 0.29 | 0.16 | 0.06 | 0.43 | -0.62 | 0.27 | -0.49 | -0.02 | 1.485 | 0.16 | -0.7425 | 0.591657 | 0.576734 | 0.658427 | 1 | 0.999631 | 0.852627 |
| Cep170_H7BX26_S1296s_1 | 0.97 | 0.35 | -1.17 | 0.92 | 2.49 | 1.31 | 1.14 | 2.07 | 0.68 | 0.21 | 1.1 | -0.28 | -2.58 | 0.58 | 0.9 | -0.8 | -0.2775 | 0.0025 | 0.3725 | 0.372638 | 0.020425 | 0.767706 | 1 | 0.999631 | 0.747035 |
| Cep170_H7BX26_S1511s_1 | -0.26 | 0.04 | -1.16 | -0.83 | -0.38 | -0.85 | -0.59 | -1.5 | -0.6 | 0.15 | -1.09 | -0.66 | -0.19 | 0.47 | -0.38 | -0.62 | -0.2775 | 0.0025 | 0.3725 | 0.349646 | 0.469781 | 0.994891 | 1 | 0.999631 | 0.736894 |
| Cep170_H7BX26_S1512s_1 | -0.26 | 0.04 | -1.16 | -0.83 | -0.38 | -0.85 | -0.59 | -1.5 | -0.6 | 0.15 | -1.09 | -0.66 | -0.19 | 0.47 | -0.38 | -0.62 | 0.0875 | 0.115 | 0.4 | 0.349646 | 0.469781 | 0.994891 | 1 | 0.999631 | 0.736894 |
| Cep170_H7BX26_S1512s_2 | -0.49 | 1.13 | -0.77 | -2.39 | -0.98 | 0.02 | -0.86 | -0.35 | 0.07 | -0.55 | -0.79 | -0.79 | 0.22 | 0.97 | -0.49 | -1.62 | -0.3725 | 0.26 | 0.0675 | 0.615578 | 0.896517 | 0.86381 | 1 | 0.999631 | 0.861203 |
| Cep170_H7BX26_S1519s_1 | -0.64 | -1.26 | 0.24 | 0.66 | -1.34 | -1.07 | 0.06 | -0.14 | -1.44 | -0.46 | 0.7 | 1.24 | -1.4 | -0.25 | 0.23 | 0.69 | 0.0875 | 0.115 | 0.4 | 0.906281 | 0.474688 | 0.695841 | 1 | 0.999631 | 0.969391 |
| Cep170_H7BX26_S1519s_2 | -0.49 | 1.13 | -0.77 | -2.39 | -0.98 | 0.02 | -0.86 | -0.35 | 0.07 | -0.55 | -0.79 | -0.79 | 0.22 | 0.97 | -0.49 | -1.62 | -0.55 | 0.48 | -1.555 | 0.615578 | 0.896517 | 0.86381 | 1 | 0.999631 | 0.861203 |
| Cep170_H7BX26_S378s_1 | 1.69 | -2.56 | -3.72 | -4.08 | -1.54 | -1.99 | -1.14 | -6.2 | -0.64 | 0.92 | -4.09 | -2.94 | -0.41 | -0.99 | -5.92 | -7.57 | 0.1525 | 0.3925 | 0.33 | 0.410349 | 0.714923 | 0.746186 | 1 | 0.999631 | 0.767937 |
| Cep170_H7BX26_S443s_1 | -0.04 | 0.86 | 0.93 | 0.81 | 0.79 | 0.85 | 0.35 | 1.18 | 0.95 | 0.81 | 0.93 | 1.44 | -0.29 | 1.35 | 1.25 | 1.57 | -0.0025 | 0.4825 | 0.29 | 0.489856 | 0.64339 | 0.243694 | 1 | 0.999631 | 0.807612 |
| Cep170_H7BX26_S829s_1 | 0.54 | 2.02 | -0.24 | -0.38 | 0.63 | 1.79 | -0.57 | 0.08 | 1.57 | 2.06 | 0.04 | 0.2 | 1.11 | 1.8 | 0 | 0.19 | 0.3325 | 0.3175 | -0.285 | 0.644369 | 0.996985 | 0.47531 | 1 | 0.999631 | 0.872436 |
| Cep170_H7BX26_S914s_1 | 0.47 | 0.84 | 0.4 | 1.82 | 1.36 | 0.22 | 1.67 | 1.61 | 1.05 | 0.02 | 2.09 | 1.64 | -0.67 | 0.36 | 1.58 | 1.12 | 0.705 | 0.235 | 0.1475 | 0.606434 | 0.469845 | 0.545551 | 1 | 0.999631 | 0.858185 |
| Cep170_H7BX26_S914s_2 | -0.33 | 2.32 | 1.2 | 1.72 | 2.41 | 1.55 | 1.46 | 2.31 | 2.27 | 0.15 | 1.47 | 1.96 | 1.3 | 1.87 | 1.16 | 1.17 | 0.8475 | 0.8975 | -0.12 | 0.788954 | 0.231372 | 0.722409 | 1 | 0.999631 | 0.925513 |
| Cep170_H7BX26_S950s_1 | 0.58 | 1.66 | 0.61 | 1.64 | 2.38 | 1.68 | 2.08 | 1.74 | 2.44 | 0.43 | 2.64 | 2.57 | -0.65 | 1.46 | 1.88 | 1.32 | 0.9375 | 0.6575 | -0.8025 | 0.837983 | 0.040735 | 0.136325 | 1 | 0.999631 | 0.946137 |
| Cep170_H7BX26_S985s_1 | -0.03 | 1.35 | 1.06 | 1.42 | 1.64 | 2.13 | 2.41 | 1.37 | 2.54 | 0.19 | 2.17 | 1.53 | -0.64 | 0.7 | 0.53 | 0 | 0.37 | 0.12 | -0.7625 | 0.09907 | 0.04286 | 0.263428 | 1 | 0.999631 | 0.562298 |
| Cep170_H7BX26_T394t_1 | 0.38 | 1.32 | 0.66 | 2.09 | 1.38 | 1.54 | 0.84 | 2.17 | 1.32 | 0.92 | 2.13 | 0.56 | -1.12 | 0.7 | 1.3 | 0.52 | -0.0075 | -0.0125 | -0.02 | 0.214384 | 0.421697 | 0.805233 | 1 | 0.999631 | 0.66742 |
| Cep170_H7BX26_T912t_1 | 0 | 1.55 | 0.76 | 0.51 | 0.86 | 0.92 | 0.42 | 0.59 | 1.21 | 0.53 | 0.63 | 0.4 | 0.51 | 1.03 | 0.41 | 0.79 | 0.705 | 0.235 | 0.1475 | 0.958566 | 0.983582 | 0.974249 | 1 | 0.999631 | 0.98564 |
| Cep170_H7BX26_T912t_2 | -0.33 | 2.32 | 1.2 | 1.72 | 2.41 | 1.55 | 1.46 | 2.31 | 2.27 | 0.15 | 1.47 | 1.96 | 1.3 | 1.87 | 1.16 | 1.17 | -0.23 | 0.245 | 0.185 | 0.788954 | 0.231372 | 0.722409 | 1 | 0.999631 | 0.925513 |
| Cep170b_Q80U49_S1032s_1 | -0.4 | 0.84 | 0.73 | 0.1 | -0.08 | 0.33 | -0.4 | 0.5 | 0.25 | 0.53 | 0.46 | 1.01 | 0.21 | 0.38 | 0.21 | 1.21 | 0.1925 | -0.0025 | -0.465 | 0.645296 | 0.539733 | 0.50449 | 1 | 0.999631 | 0.873083 |
| Cep170b_Q80U49_S1101s_1 | 1.51 | 1.16 | 0.64 | 1.05 | 1.19 | 1.49 | 1.5 | 0.95 | 1.33 | 1.1 | 1.49 | 0.43 | -0.49 | 1.17 | 0.12 | 1.7 | 0.0925 | 0.015 | -0.825 | 0.366521 | 0.514944 | 0.994126 | 1 | 0.999631 | 0.745796 |
| Cep170b_Q80U49_S1122s_1 | 0.64 | 0.41 | 0.26 | 1.45 | 0.47 | 0.01 | 1.55 | 1.1 | 0.17 | 0.02 | 1.98 | 0.65 | -1.21 | -0.13 | -0.54 | 1.34 | -0.14 | -0.415 | -0.2675 | 0.161478 | 0.82769 | 0.975741 | 1 | 0.999631 | 0.631442 |
| Cep170b_Q80U49_S1166s_1 | 0.55 | -0.61 | 0.3 | -1.17 | 0.14 | 0.11 | -0.3 | -1.44 | -1.78 | 0.51 | -0.28 | -1.04 | -0.92 | -0.16 | -1.23 | 0.31 | 0.305 | 0.085 | 0.125 | 0.600348 | 0.783267 | 0.481217 | 1 | 0.999631 | 0.856683 |
| Cep170b_Q80U49_S1347s_1 | -0.01 | -0.48 | 0.03 | 0.03 | 0.25 | -0.06 | -0.1 | 0.7 | -0.13 | -0.79 | 0.51 | 0.32 | -0.56 | -0.14 | 0.32 | 0.45 | 0.225 | 0.205 | -0.215 | 0.708287 | 0.310888 | 0.809523 | 1 | 0.999631 | 0.898901 |
| Cep170b_Q80U49_S1533s_1 | 0.59 | -0.52 | -1.32 | 0.39 | -0.21 | 0.13 | -0.19 | 0.31 | 0.15 | -0.71 | -0.39 | 0.91 | -0.97 | -0.57 | -0.87 | 0.69 | 0.6175 | 0.215 | 0.13 | 0.693949 | 0.615066 | 0.69914 | 1 | 0.999631 | 0.895131 |
| Cep170b_Q80U49_S371s_1 | 0.16 | 0.3 | -0.43 | 0.56 | 0.53 | 0.41 | 0.54 | 1.58 | 0.47 | 0.7 | 0.78 | -0.5 | -0.52 | 0.34 | 0.81 | 0.48 | 0.695 | 0.545 | 0.2575 | 0.736531 | 0.115651 | 0.577296 | 1 | 0.999631 | 0.90765 |
| Cep170b_Q80U49_S478s_1 | 1.21 | 0.46 | 1.03 | 0.82 | 1.18 | 2.33 | 1.4 | 1.39 | 1.37 | 1.29 | 1.51 | 1.53 | -1.1 | 2.09 | 1.65 | 1.91 | 0.82 | 0.215 | 0.5175 | 0.707312 | 0.063714 | 0.071382 | 1 | 0.999631 | 0.898523 |
| Cep170b_Q80U49_S534s_1 | -0.03 | -0.81 | -1.68 | -0.5 | -0.11 | -0.11 | 0.02 | 0.46 | -0.62 | -0.93 | 0.63 | -1.24 | -0.88 | -0.25 | -0.63 | 0.81 | 0.0625 | -0.0125 | -0.97 | 0.305723 | 0.055472 | 0.67329 | 1 | 0.999631 | 0.716639 |
| Cep170b_Q80U49_S819s_1 | 1.47 | 0.15 | 0.56 | 0.53 | 0.36 | 0.86 | 1.35 | 0.39 | 1.19 | 1.03 | 0.37 | 0.07 | 0.05 | -0.63 | -0.19 | -0.4 | 0.1925 | -0.8375 | -0.63 | 0.021855 | 0.868914 | 0.974997 | 1 | 0.999631 | 0.405544 |
| Cep170b_Q80U49_S840s_1 | 1.49 | -0.96 | -0.21 | 1.18 | 1.4 | -1.2 | 1.26 | 0.81 | 0.18 | -1.41 | -0.3 | -0.32 | -1.39 | 0.32 | 0.02 | 0.03 | 0.465 | -0.77 | -0.715 | 0.328069 | 0.793625 | 0.190101 | 1 | 0.999631 | 0.72534 |
| Cep170b_Q80U49_S843s_1 | 1.29 | -1.17 | -0.82 | 1.32 | 1.37 | -1.38 | 1.49 | 1 | -0.21 | -1.64 | 0.13 | -0.74 | -1.72 | -0.1 | -0.21 | -0.21 | 0.115 | 0.2925 | 0.76 | 0.311372 | 0.576901 | 0.279194 | 1 | 0.999631 | 0.71924 |
| Cep170b_Q80U49_S965s_1 | -0.96 | -0.9 | -0.55 | -1.51 | -1.28 | -0.77 | -0.92 | -0.49 | -1.19 | -0.42 | -1.19 | 0.05 | -0.14 | -0.28 | 0.29 | -0.75 | -0.075 | -0.02 | 0.21 | 0.0512 | 0.713035 | 0.453612 | 1 | 0.999631 | 0.489558 |
| Cep170b_Q80U49_S981s_1 | -0.19 | 0.32 | -0.35 | -0.9 | -0.03 | -0.09 | -0.86 | -0.44 | -0.05 | 0.17 | -0.35 | -0.97 | 0.78 | 0.2 | -0.75 | -0.51 | -0.49 | -0.1225 | -1.0625 | 0.631173 | 0.828853 | 0.957617 | 1 | 0.999631 | 0.867571 |
| Cep250_E9Q5A8_S2322s_1 | 1.25 | 0.24 | 0.5 | 0.02 | -0.13 | 0 | 0.47 | -0.29 | -0.41 | 1.35 | 0.27 | 0.31 | -0.13 | -0.59 | -1.05 | -0.47 | -0.7175 | -0.7675 | -0.255 | 0.015673 | 0.17831 | 0.783152 | 1 | 0.999631 | 0.388832 |
| Cep41_H9KUZ8_S286s_1 | 0.95 | -0.69 | -0.72 | 0.7 | -1.8 | 0.55 | -1.09 | -0.29 | -1.28 | 2.1 | -1.75 | -1.9 | -2.3 | 1.01 | -0.2 | 0.71 | 0.27 | 0.87 | 0.5225 | 0.738533 | 0.255448 | 0.406724 | 1 | 0.999631 | 0.90804 |
| Cep68_Q3TRP8_S419s_1 | -0.88 | 0.18 | -0.12 | 0.01 | 0.14 | 0.28 | -0.4 | 0.25 | 0.76 | 1.08 | 0.24 | 0.59 | -0.28 | 1.06 | 0.49 | 0.01 | -0.545 | -0.9725 | -0.2925 | 0.210754 | 0.416695 | 0.026556 | 1 | 0.999631 | 0.665026 |
| Cep97_Q9CZ62_S449s_1 | -0.04 | 0.83 | 1.08 | 1.44 | 0.66 | -0.21 | 0.49 | 0.19 | 0.21 | -1.09 | 0.45 | -0.15 | -0.98 | 0.87 | 1.02 | 1.23 | 0.4175 | 1.4125 | 0.38 | 0.601574 | 0.175112 | 0.054297 | 1 | 0.999631 | 0.856765 |

| | | | | | | | | | | | | | | | | | | | | | | | | | |
|---|---|---|---|---|---|---|---|---|---|---|---|---|---|---|---|---|---|---|---|---|---|---|---|---|---|
| Cep97_Q9CZ62_S725s_1 | 1.6 | 2.53 | 1.34 | 0.92 | 2.99 | 0.99 | 2.91 | 1.17 | 4.89 | -0.34 | 3.34 | 4.15 | 4.82 | 1.43 | 2.1 | -0.44 | -0.1175 | 0.175 | 0.81 | 0.698373 | 0.478508 | 0.196551 | 1 | 0.999631 | 0.895731 |
| Cetn2_B1AUQ7_S11s_1 | 0.68 | 1.02 | 1.01 | 0.71 | 0.74 | 1.35 | 0.26 | 0.6 | 1.02 | 1.42 | 0.47 | 1.21 | 1.65 | 1.74 | 1.34 | 1.93 | -0.43 | -0.8025 | -1.175 | 0.015993 | 0.700944 | 0.563207 | 1 | 0.999631 | 0.388832 |
| Cfl1_P18760_S24s_1 | 1.17 | 0.08 | 0.84 | 0.73 | -0.44 | 0.68 | 1.35 | -0.49 | -0.69 | 0.24 | 0.23 | -0.17 | -0.47 | -0.73 | -0.39 | -0.29 | 0.565 | -0.0225 | -0.9075 | 0.004497 | 0.378406 | 0.044136 | 1 | 0.999631 | 0.351308 |
| Cfl1_P18760_S3s_1 | 1.52 | 1.13 | -0.59 | 1.87 | 2.41 | 0.52 | 1.42 | 1.84 | 1.29 | 0.41 | 1.65 | 0.49 | -1.95 | 1.32 | 0.53 | 0.4 | -0.3675 | -0.0475 | -0.4225 | 0.260859 | 0.36441 | 0.968687 | 1 | 0.999631 | 0.69271 |
| Cfl1_P18760_T25t_1 | 1.13 | -0.38 | 0.88 | 0.6 | -0.65 | 0.13 | 1.25 | 0.03 | 0.04 | 1.34 | 0.46 | 0.2 | -0.13 | -0.73 | 1 | 0.4 | 0.4925 | -0.0575 | -0.54 | 0.389493 | 0.456009 | 0.913281 | 1 | 0.999631 | 0.758666 |
| Cfl2_P45591_S3s_1 | 1.3 | 1.03 | -0.73 | 1.64 | 2.05 | 0.71 | 1.12 | 1.33 | 1.18 | 0.36 | 1.09 | 0.38 | -1.26 | 1.41 | 0.49 | 0.44 | -0.4725 | -0.7175 | -1.6175 | 0.436616 | 0.380011 | 0.914259 | 1 | 0.999631 | 0.780748 |
| Cgn_D3YUW7_S208s_1 | 2.17 | 0.51 | 1.23 | 1.43 | 0.46 | 0.45 | 1.85 | 0.69 | 0.75 | 1.76 | 0.01 | -0.05 | -0.46 | -0.48 | 0.3 | -0.49 | -0.0575 | 0.0625 | 0.4225 | 0.002689 | 0.316494 | 0.182916 | 1 | 0.999631 | 0.351308 |
| Cgnl1_D3Z3H4_S203s_1 | -1.15 | -1.24 | -0.93 | -2.18 | -1.64 | -1.4 | -1.22 | -1.47 | -1 | -0.88 | -2.1 | -1.27 | -0.13 | -1.33 | -1.14 | -1.21 | -0.265 | -0.08 | 0.3125 | 0.315951 | 0.861892 | 0.876229 | 1 | 0.999631 | 0.720734 |
| Cgnl1_D3Z3H4_S257s_1 | -0.06 | -0.42 | 0.36 | 0.08 | -0.23 | 0.15 | -0.55 | -0.47 | -0.48 | 0.16 | -0.15 | 0.11 | 0.8 | -0.13 | 0.14 | 0.4 | -0.605 | -0.0125 | -0.98 | 0.354733 | 0.377687 | 0.788198 | 1 | 0.999631 | 0.739951 |
| Champ1_Q8K327_S110s_1 | -1.01 | -2.39 | 0.58 | 0.02 | -2.8 | -2.85 | 0.3 | 0.13 | -2.77 | -1.22 | 0.39 | 0.75 | -3.59 | -3.19 | 0.59 | -0.53 | -0.215 | -0.1825 | -0.485 | 0.353088 | 0.526125 | 0.988946 | 1 | 0.999631 | 0.73949 |
| Champ1_Q8K327_S204s_1 | 0.51 | -0.25 | 0.03 | -0.04 | 0.28 | 0.01 | -0.46 | -0.44 | 0.06 | 0.63 | -0.53 | -0.64 | -0.33 | -0.15 | -0.72 | -0.49 | -0.735 | -0.5675 | -1.2625 | 0.13316 | 0.483158 | 0.619218 | 1 | 0.999631 | 0.60492 |
| Champ1_Q8K327_S265s_1 | 1.83 | 0.46 | 1.27 | 1.25 | -0.52 | 0.69 | 1.55 | 0.15 | 0.53 | 1.48 | 0.03 | 0.5 | 0.14 | -0.47 | 0.67 | -0.58 | 0.2775 | 0.3225 | 0.48 | 0.01219 | 0.157166 | 0.196329 | 1 | 0.999631 | 0.385092 |
| Champ1_Q8K327_S405s_1 | 0.17 | 0.91 | -1.06 | -0.59 | 0.33 | 0.55 | 0.1 | -0.44 | 0.03 | -0.36 | 0.23 | 0.82 | 1.45 | -0.12 | -0.69 | 0.71 | -0.3775 | 0.0575 | 0.205 | 0.420081 | 0.551786 | 0.50534 | 1 | 0.999631 | 0.772641 |
| Champ1_Q8K327_S416s_1 | 0.87 | 0.69 | -0.14 | -0.27 | 0.16 | 0 | -0.37 | -0.15 | 0.43 | 1.03 | -0.08 | 0 | 0.31 | 1.2 | 0.24 | 0.22 | -0.14 | -0.12 | 0.3825 | 0.608908 | 0.286137 | 0.884992 | 1 | 0.999631 | 0.858851 |
| Champ1_Q8K327_S425s_1 | -1.55 | -1.2 | 0.19 | -0.57 | -1.09 | -1.15 | -0.79 | -0.66 | -1.77 | -1.06 | -0.2 | -0.58 | 0.49 | -1.48 | -0.48 | -0.13 | 0.1375 | 0.45 | 0.4925 | 0.475457 | 0.728977 | 0.805849 | 1 | 0.999631 | 0.799882 |
| Champ1_Q8K327_S532s_1 | 0.69 | 1.43 | 0.13 | 1.36 | 0.86 | 1.11 | 0.69 | 1.5 | 1.25 | 1.09 | 1.89 | 1.18 | 0.53 | 1.32 | 1.68 | 2.05 | 0.0275 | 0.025 | 0.1875 | 0.285562 | 0.712076 | 0.253077 | 1 | 0.999631 | 0.705851 |
| Chat_Q8BQV2_S366s_1 | -0.01 | 0.18 | 0.15 | 0.07 | 0.12 | -0.23 | -0.07 | 0.68 | -0.05 | 0.14 | -0.34 | 0.74 | 0.8 | 0.12 | 0.3 | -0.08 | 0.0625 | 0.0275 | -0.1825 | 0.535521 | 0.9223 | 0.934524 | 1 | 0.999631 | 0.829362 |
| Chchd2_Q9D1L0_S45s_1 | 0.11 | -0.43 | -0.48 | -0.4 | 0.06 | -0.69 | -0.31 | -0.01 | -0.3 | 0.12 | -0.31 | -0.6 | -1.07 | -0.19 | -0.53 | -0.14 | -0.335 | -0.315 | -0.1175 | 0.582213 | 0.829779 | 0.924198 | 1 | 0.999631 | 0.847594 |
| Chchd2_Q9D1L0_T13t_1 | 0.63 | -0.11 | 0.55 | 0.18 | -0.51 | 0.23 | 0.19 | 0 | -0.05 | 0.83 | -0.35 | -0.44 | 0.35 | 0.18 | 0.28 | -0.03 | -0.48 | 0.115 | -1.1675 | 0.69482 | 0.283783 | 0.399215 | 1 | 0.999631 | 0.895302 |
| Chchd3_Q9D9P1_S112s_1 | -0.14 | -2.54 | -0.93 | -0.92 | -3.11 | -1.59 | -0.29 | -1.46 | -2.25 | 0.6 | -0.66 | -1.76 | -1.49 | -2.96 | -2.59 | -2.16 | -0.1875 | 0.07 | -0.555 | 0.057126 | 0.486748 | 0.872838 | 1 | 0.999631 | 0.49679 |
| Chchd6_E9Q4M4_S164s_1 | 0.48 | -0.29 | -0.22 | -0.45 | -0.69 | -0.05 | 0 | -0.49 | 0.07 | 0.63 | -0.21 | -0.69 | -0.48 | -0.24 | -0.76 | -1.22 | -1.6025 | -1.55 | -0.84 | 0.137114 | 0.55706 | 0.850376 | 1 | 0.999631 | 0.610369 |
| Chchd6_E9Q4M4_S172s_1 | 0.72 | 0.42 | 1.33 | 0.22 | -1.4 | -1.82 | 0.84 | -1.34 | -2.47 | -1.03 | 0.88 | -0.89 | -0.23 | -1.64 | 0.39 | 0.81 | -0.475 | -0.055 | -0.035 | 0.148038 | 0.020235 | 0.037351 | 1 | 0.999631 | 0.621175 |
| Chd1_P40201_S1690s_1 | 0.25 | 0.09 | 0.47 | -0.09 | -0.73 | 0.65 | 0.25 | -1.35 | 0.32 | 0.69 | -0.41 | -0.1 | 0.9 | 0.53 | -0.8 | -0.05 | 0.01 | 0.1425 | -0.14 | 0.931721 | 0.309417 | 0.865432 | 1 | 0.999631 | 0.978006 |
| Chd2_E9PZM4_S1365s_1 | -2.15 | -1.63 | -0.59 | -1.45 | -1.29 | -2.23 | -1.01 | -1.25 | -0.81 | -2.79 | -1.11 | -0.54 | -0.41 | -2.25 | -1.94 | -1.78 | -0.7425 | -0.0775 | -0.4625 | 0.779159 | 0.980893 | 0.797829 | 1 | 0.999631 | 0.921416 |
| Chd3_B1AR17_S142s_1 | 0.78 | -0.01 | 0.83 | 0.26 | 0.16 | -0.96 | 0.1 | -0.41 | 0.05 | 1.04 | 0.24 | 0.22 | 0.47 | 0.25 | 0.7 | -1.41 | -0.1225 | -0.895 | -0.4025 | 0.365108 | 0.06058 | 0.821944 | 1 | 0.999631 | 0.745324 |
| Chd3_B1AR17_S1651s_1 | 0.55 | -0.46 | 1.12 | 0.67 | 0.13 | 0.2 | 0.27 | 0.79 | 0.03 | -0.2 | -1.98 | 0.45 | -1.61 | -0.17 | 0.75 | 1.3 | 0.09 | 0.295 | -0.8625 | 0.536817 | 0.747339 | 0.145842 | 1 | 0.999631 | 0.830094 |
| Chd4_E9QAS4_S1329s_1 | 0.54 | 1.6 | 2.82 | 1.81 | 1.04 | 2.52 | 2.51 | 1.06 | 2.21 | 1.19 | 2.69 | 1.86 | 0.56 | 0.11 | 1.68 | 0.97 | -0.255 | -0.1075 | -0.575 | 0.132398 | 0.87535 | 0.579162 | 1 | 0.999631 | 0.60434 |
| Chd4_E9QAS4_S1515s_1 | 1.42 | 0.81 | 1.06 | 0.48 | 0.6 | 1.14 | 0.49 | 0.52 | 1.02 | 1.49 | 0.27 | 0.56 | 0.22 | 0.7 | 0.55 | 0 | -0.455 | -0.1225 | 0.1275 | 0.10457 | 0.415569 | 0.768128 | 1 | 0.999631 | 0.568213 |
| Chd5_E9PYL1_S1633s_1 | 1.06 | 2.02 | 1.78 | 1.04 | 0.76 | 1.77 | 0.66 | 0.89 | 1.45 | 1.65 | 1.12 | 1.19 | 1.22 | 2 | 1.11 | 2.08 | -0.2425 | -0.415 | 0.6175 | 0.743004 | 0.241421 | 0.711672 | 1 | 0.999631 | 0.90972 |
| Chd6_A3KFM7_S2688s_1 | -0.16 | 0.76 | 0.87 | -0.1 | 0.15 | 0.11 | 0.22 | -0.08 | -0.59 | 0.8 | -1.24 | 0.74 | 0.69 | 1.14 | 1.39 | 0.62 | -0.075 | 0.56 | 0.8525 | 0.120607 | 0.463032 | 0.445414 | 1 | 0.999631 | 0.588344 |
| Chd8_F7AL76_S659s_1 | -0.06 | 0.33 | 1.58 | 1 | 0.35 | 0.26 | 0.88 | 1.06 | 0.84 | 0.34 | 1.72 | 2.19 | 1.41 | 1.05 | 1.67 | 2.13 | -0.145 | 0.0825 | 0.21 | 0.073301 | 0.855449 | 0.296299 | 1 | 0.999631 | 0.537539 |
| Chd9_Q8BYH8_S2078s_1 | 0.54 | -0.24 | 0.07 | 0.18 | 0.09 | 0.03 | 0.3 | -0.45 | 0.21 | 0.92 | 0.16 | -0.41 | 1.14 | 0.25 | 0.53 | -0.53 | -0.955 | -0.93 | -1.74 | 0.605654 | 0.623976 | 0.815592 | 1 | 0.999631 | 0.85817 |
| Chd9_Q8BYH8_S2734s_1 | 2.02 | -0.41 | -0.45 | -0.21 | -1.25 | -0.56 | 0.66 | -1.72 | -1.27 | 1.55 | -1.23 | -1.82 | -1.02 | -0.57 | -3.42 | -1 | 0.1375 | 0.19 | 0.48 | 0.044589 | 0.193033 | 0.285967 | 1 | 0.999631 | 0.475443 |
| Chga_P26339_S282s_1 | 0.34 | 2.29 | 3.78 | 1.31 | 0.77 | 2.59 | 2.83 | 2.08 | 1.96 | 0.92 | 2.62 | 2.98 | 2.39 | 2.08 | 3.5 | 1.67 | 0.4 | 0.7025 | 0.8125 | 0.517201 | 0.856396 | 0.80288 | 1 | 0.999631 | 0.819449 |
| Chga_P26339_S350s_1 | -0.08 | 1.09 | 0.74 | 0.01 | 0.98 | 1.53 | 0.14 | 0.71 | 1.81 | 2.08 | 0.03 | 0.65 | 1.49 | 2 | 0.98 | 0.54 | 0.2875 | 0.3575 | -0.2425 | 0.084167 | 0.337777 | 0.2019 | 1 | 0.999631 | 0.549732 |
| Chga_P26339_S414s_1 | 1.09 | 0.14 | 1.67 | 0.84 | 0.24 | 1.64 | 2.55 | 0.46 | 1.29 | 1.82 | 1.3 | 0.76 | 0.52 | 0.28 | 1.33 | 0.64 | -0.125 | -0.1125 | 0.2075 | 0.55619 | 0.6161 | 0.378646 | 1 | 0.999631 | 0.838305 |
| Chgb_P16014_S100s_1 | -0.26 | -0.48 | -1 | -0.94 | -0.51 | -0.75 | -1.04 | -0.88 | -0.47 | -1.16 | -0.55 | -0.95 | 0.8 | -0.58 | -0.83 | -1.24 | -0.135 | -0.1625 | -0.7475 | 0.659105 | 0.663999 | 0.717715 | 1 | 0.999631 | 0.880034 |
| Chgb_P16014_S129s_1 | 0.09 | -1.34 | -0.29 | -0.67 | -1.15 | -0.7 | 0.48 | -1.38 | -1.13 | -0.21 | -0.82 | -0.7 | -0.97 | -1.42 | -1.15 | -1.66 | 0.4375 | 0.8175 | 1.0325 | 0.07145 | 0.7813 | 0.67139 | 1 | 0.999631 | 0.531592 |
| Chgb_P16014_S220s_1 | -0.85 | -0.63 | 0.3 | -0.55 | -0.34 | 0.34 | 0.12 | -0.1 | 0.34 | 0.41 | 0.28 | 0.51 | 0.99 | 0.28 | 0.95 | 0.18 | -0.545 | -0.22 | -1.4175 | 0.018392 | 0.207506 | 0.026525 | 1 | 0.999631 | 0.391024 |
| Chgb_P16014_S234s_2 | 0.34 | -0.36 | -0.09 | -0.07 | -0.81 | -1.06 | 0.73 | -1.22 | -0.29 | 1.17 | -0.25 | -1.69 | 0.29 | -2.55 | -1.96 | -1.63 | -0.2225 | -0.415 | -1.625 | 0.032369 | 0.247523 | 0.693413 | 1 | 0.999631 | 0.442395 |
| Chgb_P16014_S237s_1 | 0.79 | -0.81 | 0.32 | -0.38 | -1.18 | -0.09 | 1.27 | -0.97 | -1.09 | 0.83 | -0.54 | -0.94 | -0.8 | -1.7 | -1.3 | -2.78 | -0.565 | -0.2525 | -1.44 | 0.010325 | 0.710529 | 0.439337 | 1 | 0.999631 | 0.383871 |
| Chgb_P16014_S237s_2 | 0.27 | -0.34 | -0.08 | -0.08 | -1 | -1.04 | 0.77 | -1.22 | -0.28 | 0.92 | -0.31 | -1.57 | 0.17 | -2.63 | -2.03 | -1.5 | 0.345 | 0.59 | 0.515 | 0.027656 | 0.240456 | 0.614003 | 1 | 0.999631 | 0.423977 |
| Chgb_P16014_S300s_1 | 0.28 | -0.89 | -3.74 | -3.87 | -0.15 | -0.45 | -2.75 | -3.49 | -0.51 | 0.81 | -2.64 | -3.52 | -0.15 | 0.84 | -3.54 | -3.31 | -0.3575 | 0.07 | 0.3875 | 0.688824 | 0.761937 | 0.632179 | 1 | 0.999631 | 0.892096 |
| Chgb_P16014_S301s_1 | -1.66 | -3.18 | 0.86 | 0.88 | -2.18 | -3.48 | 0.49 | 0.64 | -2.79 | -1.61 | 0.53 | 1.05 | -2.05 | -2.06 | 1.12 | 1.44 | 0.035 | 0.0175 | 0.385 | 0.742411 | 0.768975 | 0.951606 | 1 | 0.999631 | 0.909615 |
| Chgb_P16014_S308s_1 | -0.79 | -2.03 | -0.91 | -1.22 | -1.98 | -1.42 | -0.53 | -0.88 | -1.82 | -0.91 | -1.21 | -0.94 | -1.26 | -1.34 | -0.45 | -0.36 | 0.5025 | 0.2075 | 1.4175 | 0.351434 | 0.933372 | 0.962744 | 1 | 0.999631 | 0.738077 |
| Chgb_P16014_S308s_2 | -0.26 | 0.5 | 0.35 | 0.3 | 0.01 | 1.49 | 1.34 | 0.06 | 0.57 | 1.6 | -0.73 | 0.28 | 1.97 | 2.07 | 1.54 | 0.98 | -0.135 | 0.245 | -0.07 | 0.002377 | 0.255857 | 0.670368 | 1 | 0.999631 | 0.351308 |
| Chgb_P16014_S342s_1 | 0.43 | 0.63 | 1.22 | 0.6 | 0.51 | 0.76 | 0.47 | 0.6 | 1.06 | 1.47 | 0.67 | 0.66 | 0.63 | 0.51 | 0.83 | 0.63 | 0.1175 | -0.485 | -1.8625 | 0.813188 | 0.623008 | 0.449365 | 1 | 0.999631 | 0.937041 |
| Chgb_P16014_S365s_1 | 0.79 | -5.36 | -2.49 | -2.68 | -3.71 | -2.54 | 1.25 | -4.27 | -5.66 | 0.41 | -2.8 | -3.63 | -3.59 | -4.56 | -3.52 | -5.52 | -0.325 | -0.355 | -1.715 | 0.126436 | 0.937503 | 0.749374 | 1 | 0.999631 | 0.596745 |
| Chgb_P16014_S378s_1 | 1.71 | -1.14 | 1.27 | 0.19 | -1.55 | 0.87 | 2.81 | -1.4 | -0.38 | 2.44 | -0.29 | -1.16 | -0.63 | -1.68 | -0.91 | -1.61 | 0.02 | 0.845 | -0.22 | 0.018871 | 0.755818 | 0.688003 | 1 | 0.999631 | 0.391024 |
| Chgb_P16014_S390s_1 | 0.78 | 2.39 | 1.4 | 0.16 | 2.99 | 1.32 | 1.13 | -0.63 | 2.98 | 2.34 | 1.42 | 1.37 | 5.5 | 1.88 | -3.1 | -0.43 | -0.045 | 0.2525 | -0.7975 | 0.889142 | 0.979276 | 0.158419 | 1 | 0.999631 | 0.963505 |
| Chgb_P16014_S408s_1 | 0.73 | -0.15 | 1.44 | -0.74 | -0.12 | 0.09 | 1.88 | -0.75 | 1.31 | 0.67 | 0.49 | -0.18 | 1.11 | -1.78 | -0.23 | -1.01 | -0.185 | -0.205 | 0.43 | 0.26669 | 0.945601 | 0.636859 | 1 | 0.999631 | 0.695664 |
| Chgb_P16014_S428s_1 | 0.55 | 0.87 | 1.78 | 0.87 | 0.42 | 0.71 | 0.9 | 1.3 | 0.66 | 0.81 | 0.84 | 0.94 | 1.02 | 1.76 | 1.39 | 1.62 | 0.065 | 0.4625 | 0.84 | 0.249768 | 0.601166 | 0.534083 | 1 | 0.999631 | 0.686359 |
| Chgb_P16014_S99s_1 | -0.77 | -1.22 | -1.25 | -1.1 | -0.83 | -1.6 | -1.1 | -0.55 | -0.98 | -1.16 | -0.18 | -0.17 | 0.29 | -0.44 | -0.21 | -0.62 | -0.1775 | -0.3225 | -0.0525 | 0.021885 | 0.832279 | 0.18678 | 1 | 0.999631 | 0.405544 |
| Chic1_Q8CBW7_S80s_1 | 1.03 | 1.19 | -0.14 | -0.58 | -0.31 | 0.33 | 0.09 | 0.68 | -0.66 | 0.09 | -0.12 | 0.9 | 0.09 | 0.8 | 0.28 | 0.12 | -0.145 | -0.1625 | 0.1575 | 0.909021 | 0.701776 | 0.532472 | 1 | 0.999631 | 0.970293 |
| Chm_Q3UR39_S622s_1 | -0.36 | -0.22 | -1.14 | -0.91 | -0.14 | -0.11 | -1.76 | -1.2 | -0.39 | -0.07 | -1.27 | -1.55 | 0.06 | 0 | -1.4 | -0.66 | -0.06 | -0.3025 | 0.6225 | 0.708154 | 0.746603 | 0.698712 | 1 | 0.999631 | 0.898901 |
| Chmp7_Q8R1T1_S232s_1 | -1.23 | -0.77 | -0.79 | -0.09 | -1.13 | -0.58 | -0.89 | -0.52 | -1.53 | -0.82 | -1.04 | -0.7 | -0.05 | -0.18 | -0.02 | -0.14 | -0.5775 | 0.0975 | 0.4 | 0.076015 | 0.852077 | 0.386731 | 1 | 0.999631 | 0.542138 |
| Chrm1_P12657_S244s_1 | 0.84 | 0.05 | 3.17 | 0.85 | -0.12 | 0.78 | 0.97 | 0.97 | 0.01 | 1.02 | 3.16 | 1.11 | 1.37 | 0.18 | 3.09 | 1.87 | -0.26 | 0.8825 | 0.585 | 0.614546 | 0.381427 | 0.905987 | 1 | 0.999631 | 0.861203 |
| Chrm1_P12657_S251s_1 | 0.78 | 1.07 | 1.2 | 0.81 | 0.41 | 1.28 | 0.14 | 0.99 | 0.94 | 1.77 | 1.89 | 2.79 | 1.53 | 0.89 | 1.94 | 1.84 | -0.185 | 0.615 | 0.3575 | 0.10119 | 0.431488 | 0.050649 | 1 | 0.999631 | 0.564448 |
| Chrm1_P12657_S254s_1 | 0.56 | 0.69 | 0.47 | 0.55 | 0.05 | 0.76 | 0.26 | 0.46 | 0.38 | 1.39 | 0.91 | 2.05 | 1.44 | 0.21 | 0.96 | 1.09 | -0.445 | -0.54 | -2.105 | 0.298222 | 0.484735 | 0.130171 | 1 | 0.999631 | 0.711311 |
| Chrm1_P12657_S451s_1 | 2.14 | -0.31 | 1.61 | 1.85 | -0.59 | 0.91 | 2.65 | 0.54 | -0.07 | 1.79 | 0.62 | 0.79 | -0.67 | -0.93 | -0.34 | -1.19 | 0.03 | -0.105 | 0.045 | 0.003053 | 0.56354 | 0.387889 | 1 | 0.999631 | 0.351308 |
| Chrm2_Q9ERZ4_S232s_1 | -0.58 | -0.51 | -0.45 | -0.08 | -0.85 | -0.56 | -0.41 | 0.32 | -1.11 | -1.18 | -0.03 | 0.28 | -0.55 | -0.6 | -0.17 | -0.12 | 0.2225 | -0.0775 | -0.2025 | 0.876241 | 0.925387 | 0.793512 | 1 | 0.999631 | 0.961058 |
| Chrm3_Q9ERZ3_S288s_1 | -0.27 | -0.34 | -2.27 | -2.59 | -0.35 | -1.17 | -1.23 | -1.83 | -0.51 | -0.97 | -2.42 | -1.88 | -0.77 | -0.91 | -2.64 | -1.96 | -0.1675 | -0.145 | -0.1125 | 0.765465 | 0.719817 | 0.908406 | 1 | 0.999631 | 0.916326 |
| Chrm3_Q9ERZ3_S384s_1 | 0.1 | 0.35 | -0.4 | -0.44 | -0.02 | -0.16 | -0.66 | -0.22 | -0.58 | -0.39 | -0.33 | 0.33 | 0.27 | 0.37 | -1.59 | 0.11 | -0.13 | -0.215 | -1.485 | 0.81685 | 0.580069 | 0.661337 | 1 | 0.999631 | 0.937777 |
| Chrm3_Q9ERZ3_S472s_1 | 1.07 | 0.77 | 1.24 | 0.75 | 0.09 | 1.46 | 1.64 | 0.12 | 1.11 | 0.73 | 1 | 0.13 | -0.28 | -0.75 | 0.04 | -1.12 | 0.19 | -0.3475 | -1.01 | 0.001477 | 0.761774 | 0.498749 | 1 | 0.999631 | 0.351308 |
| Chrm3_Q9ERZ3_S577s_1 | 0.35 | 0.2 | -0.55 | 0.79 | 0.57 | 1 | -0.18 | 0.16 | -0.21 | -0.36 | 0.27 | -0.3 | -1.67 | 0.35 | -1.87 | -0.06 | -0.4775 | 0.145 | -0.005 | 0.103891 | 0.626275 | 0.335703 | 1 | 0.999631 | 0.568213 |
| Chrm4_P32211_S262s_1 | -0.22 | -0.74 | -1.62 | -0.56 | -1.22 | -2.45 | -0.87 | -0.51 | -0.73 | -0.98 | -0.46 | -0.39 | -1.12 | -1.29 | -0.64 | -0.11 | 0.18 | -0.02 | 0.1375 | 0.990383 | 0.339638 | 0.688651 | 1 | 0.999631 | 0.995933 |
| Chrm4_P32211_T320t_1 | -0.43 | 0.4 | -0.18 | -0.77 | 1.19 | 0.02 | -0.45 | -1.02 | 0.83 | -0.6 | -0.63 | -0.66 | -0.55 | -0.02 | -0.21 | 0.35 | 0.22 | -0.0675 | -0.15 | 0.703361 | 0.718296 | 0.963472 | 1 | 0.999631 | 0.897068 |
| Chrna4_O70174_S468s_1 | -0.01 | -0.6 | -0.92 | -0.22 | -0.26 | -0.33 | -0.59 | 0.31 | -0.32 | -0.16 | -0.26 | -1.28 | -0.92 | -0.36 | -0.85 | -0.22 | 0.29 | 0.0675 | 0.075 | 0.65661 | 0.501023 | 0.852007 | 1 | 0.999631 | 0.879056 |
| Chrna4_O70174_S491s_1 | -0.17 | -0.13 | 0.19 | 0.73 | 0.65 | 0.07 | 0.04 | 1.02 | 0.36 | 0.17 | 0.12 | 0.24 | 0.35 | 0.26 | -0.55 | 0.86 | -0.055 | -0.0025 | 0.74 | 0.847127 | 0.411769 | 0.819249 | 1 | 0.999631 | 0.948838 |
| Chrna4_O70174_S543s_1 | -0.46 | 0.36 | 0.64 | -0.84 | 0.01 | 0.23 | -0.85 | 0.09 | -0.42 | -0.1 | 0.21 | 0 | 1.38 | 0.36 | 0.64 | 0.28 | 0.2625 | -0.055 | 0.745 | 0.112092 | 0.895611 | 0.994853 | 1 | 0.999631 | 0.577068 |
| Chrnb2_Q9ERK7_S395s_1 | -0.25 | -0.66 | 0.03 | 0.1 | -0.54 | 0.04 | -0.08 | 0.85 | -0.8 | -0.75 | 0.27 | 0.28 | 0.64 | 0.32 | 0.87 | 0.37 | -0.485 | -0.265 | -1.0925 | 0.034281 | 0.472516 | 0.883118 | 1 | 0.999631 | 0.446273 |
| Chst15_Q91XQ5_S185s_1 | 0.11 | -4.87 | -2.16 | -1.77 | -2.65 | -3.81 | -0.79 | -3.38 | -3.21 | -0.64 | -2.3 | -3.6 | -2.7 | -4 | -2.76 | -3.6 | -1.4225 | -0.475 | -0.725 | 0.254431 | 0.646008 | 0.800971 | 1 | 0.999631 | 0.689457 |
| Cic_Q924A2_S1343s_1 | -0.15 | 0.94 | 2.75 | 0.12 | -1.5 | 0.81 | -1.03 | -0.31 | 0.21 | 1.8 | -1.13 | 0.88 | 0.96 | -0.22 | 1.44 | -1.42 | 0.02 | -0.1125 | 0.515 | 0.373854 | 0.074754 | 0.551357 | 1 | 0.999631 | 0.747035 |
| Cic_Q924A2_S1809s_1 | 0.23 | 0.3 | -0.53 | -0.79 | 0.1 | 0.27 | -0.54 | -0.54 | 0.24 | -0.78 | -0.27 | -0.43 | 0.79 | 0.41 | 0.54 | -0.47 | -0.4275 | 0.045 | 0.3025 | 0.22449 | 0.956387 | 0.762682 | 1 | 0.999631 | 0.67471 |
| Cic_Q924A2_S2148s_1 | 0.74 | 1.75 | 1.89 | 1.32 | 1.06 | 0.91 | 0.63 | 1.39 | 1.46 | 1 | 1.54 | 1.88 | 1.54 | 1.86 | 1.72 | 1.79 | -0.185 | -0.3725 | 0.0875 | 0.378095 | 0.226344 | 0.898538 | 1 | 0.999631 | 0.750904 |
| Cic_Q924A2_S2195s_1 | 0.48 | 0.2 | 0.32 | 0.02 | 0.2 | 0.27 | -0.2 | 0.01 | -0.18 | 0.38 | -0.27 | -0.4 | 0.43 | -0.24 | 0.01 | 1.17 | -0.115 | 0.34 | 0.5925 | 0.812113 | 0.475012 | 0.21405 | 1 | 0.999631 | 0.936949 |
| Cisd3_Z4YKM2_S71s_1 | -0.27 | -0.57 | -0.03 | -0.75 | -0.46 | -0.32 | -0.85 | -0.45 | -0.69 | 0.15 | -0.11 | 0.39 | 0.87 | 0.25 | -0.73 | 0.36 | 0.0275 | -0.105 | -0.0575 | 0.157767 | 0.68125 | 0.31965 | 1 | 0.999631 | 0.628615 |
| Cit_E9QL53_S1348s_1 | -0.08 | -0.99 | -1.45 | -0.93 | -0.91 | -0.59 | -1.18 | -0.66 | -1.48 | -0.37 | -0.79 | -1.23 | -1.13 | -0.49 | -1 | -1.06 | 0.0425 | 0.085 | -0.4075 | 0.875192 | 0.936721 | 0.789187 | 1 | 0.999631 | 0.960439 |
| Cit_E9QL53_S1458s_1 | -0.19 | -0.66 | 0.31 | 0.6 | -0.16 | -0.58 | 0.52 | 0.45 | -0.23 | 0 | 0.05 | 0.58 | -1.25 | -0.63 | 0.26 | 0.05 | -0.125 | -0.1125 | 0.81 | 0.368712 | 0.913251 | 0.813378 | 1 | 0.999631 | 0.745981 |
| Cit_E9QL53_S1608s_1 | -0.31 | 0.27 | 0.28 | -0.03 | -0.72 | 0.17 | -0.01 | 0.27 | 0.24 | -0.6 | -0.48 | 0.6 | 1.33 | 0.89 | 0.21 | 1.02 | -0.455 | 0 | -0.2025 | 0.035888 | 0.692878 | 0.752412 | 1 | 0.999631 | 0.453815 |
| Cit_E9QL53_S1883s_1 | 0.15 | -0.33 | -0.01 | -0.31 | -0.75 | -0.67 | -0.4 | -0.5 | -0.17 | 0.25 | -0.09 | -0.49 | -0.22 | -0.52 | -0.41 | -0.16 | 0.2175 | -0.26 | -0.8925 | 0.475905 | 0.098943 | 1 | 1 | 1 | 0.799882 |
| Cit_E9QL53_S1966s_1 | 1.47 | -0.68 | -1.71 | 0.13 | 0.09 | -0.61 | 0.62 | -0.02 | -0.21 | 0.16 | -0.64 | -1.14 | -0.88 | -1.17 | -1.05 | -1.26 | 0.8625 | 0.3475 | -0.0975 | 0.167989 | 0.735175 | 0.691877 | 1 | 0.999631 | 0.637399 |
| Cit_E9QL53_S479s_1 | 0.46 | 0 | -0.93 | 1.2 | 1.48 | 0.58 | 0.81 | 1.31 | 0.31 | -0.16 | 1.1 | 0.87 | -1.73 | 0.48 | 0.63 | 0.96 | 2.0425 | 2.9875 | 0.8575 | 0.885249 | 0.091344 | 0.493335 | 1 | 0.999631 | 0.963505 |
| Cit_E9QL53_S698s_1 | -1.14 | 3.57 | 3.97 | -1.22 | 4.12 | 2.32 | 6.82 | 0.09 | 6.09 | -0.31 | 4.68 | 6.67 | 6.77 | -0.44 | 2.81 | -0.53 | 0.0025 | -0.05 | 0.2575 | 0.648373 | 0.249372 | 0.12356 | 1 | 0.999631 | 0.874615 |
| Ciz1_F7AT58_S192s_1 | -0.01 | -0.19 | 0.23 | 0.98 | -0.12 | 0.39 | -0.06 | 0.81 | 0.47 | -0.25 | -0.36 | 0.95 | 0.6 | 0.84 | -0.12 | 0.72 | 0.195 | 0.345 | 0.825 | 0.497717 | 0.994459 | 0.902976 | 1 | 0.999631 | 0.811908 |
| Ckap4_Q8BMK4_S367s_1 | -1.05 | -0.06 | -0.09 | -0.32 | -0.23 | -0.33 | -0.35 | 0.17 | 0.12 | -0.15 | -0.56 | 0.45 | 0.96 | 0.24 | 0.3 | 0.28 | -0.315 | -0.735 | -1.8375 | 0.036255 | 0.537894 | 0.339812 | 1 | 0.999631 | 0.453963 |
| Ckap4_Q8BMK4_S68s_1 | 1.14 | 0.31 | 1.4 | 0.94 | 0.23 | 0.8 | 0.96 | 0.54 | 0.53 | 1.4 | -0.37 | -0.71 | -0.65 | -1.13 | -0.93 | -0.85 | -0.0175 | 0.155 | 0.19 | 0.000236 | 0.34587 | 0.165326 | 1 | 0.999631 | 0.351308 |
| Ckap5_Z4YL78_S816s_1 | -0.04 | -0.8 | -0.08 | -0.57 | -0.71 | -0.69 | -0.23 | 0.07 | -0.44 | 0.06 | -0.35 | -0.14 | -0.18 | -0.3 | 0.12 | -0.37 | -0.3225 | -0.235 | 0.185 | 0.543589 | 0.955851 | 0.604064 | 1 | 0.999631 | 0.832881 |
| Ckb_Q04447_S199s_1 | -0.25 | 0.21 | 0.36 | -0.11 | -0.37 | -0.26 | -0.4 | -0.05 | -0.25 | -0.34 | -0.28 | 0.14 | 0.23 | 0.4 | 0.05 | 0.27 | -0.23 | -0.0675 | 0.3325 | 0.521291 | 0.238193 | 0.406639 | 1 | 0.999631 | 0.820259 |
| Ckb_Q04447_T322t_1 | -0.53 | 0.31 | 0.1 | 0.02 | -0.01 | -0.24 | -0.76 | -0.01 | -0.73 | 0 | 0.27 | 0.09 | 0.17 | 0.58 | 0.13 | 0.35 | -0.375 | -0.2875 | 0.315 | 0.291214 | 0.46219 | 0.839628 | 1 | 0.999631 | 0.706164 |
| Ckb_Q04447_T35t_1 | -0.25 | 0.8 | 0.45 | -0.07 | -0.02 | 0.05 | -0.68 | 0.08 | 0.08 | -0.05 | -0.36 | 0.11 | 0.74 | 1.01 | 0.03 | 0.41 | -0.0325 | -0.28 | 0.4725 | 0.39848 | 0.281972 | 0.381638 | 1 | 0.999631 | 0.761726 |
| Ckmt1_P30275_S319s_1 | -0.43 | 0.49 | 0.84 | 0.34 | 0.04 | 0.67 | -0.13 | 0.53 | 0.12 | -0.49 | 0.63 | -0.14 | 0.47 | 0.94 | 0.99 | 0.73 | -0.0825 | -0.0625 | -1.7175 | 0.197543 | 0.927126 | 0.467052 | 1 | 0.999631 | 0.654636 |
| Clasp1_E9Q6L0_S676s_1 | 2.72 | -0.73 | -0.18 | 1.08 | 0.65 | 0.39 | 1.5 | 0.02 | 1.26 | 0.59 | 0.3 | 0.49 | -3.02 | -0.82 | 0.38 | -0.52 | -0.1175 | -0.6475 | -0.0525 | 0.082245 | 0.909892 | 0.929429 | 1 | 0.999631 | 0.54792 |
| Clasp1_E9QKH0_S1111s_2 | -0.17 | -0.52 | -1.36 | -0.58 | -0.16 | -0.68 | -0.64 | -1.62 | -0.17 | 0.49 | -1.93 | -3.61 | 0.07 | -0.05 | -1.91 | -0.95 | 0.5075 | 0.3725 | -0.295 | 0.916561 | 0.769158 | 0.445921 | 1 | 0.999631 | 0.973495 |
| Clasp1_E9QNY9_S1118s_1 | -1.53 | 0.17 | 0.49 | -1.78 | 0.28 | -0.01 | 0.11 | -1 | 0.49 | -2.23 | 0.12 | 0.46 | -0.38 | -0.97 | -0.69 | -1.79 | 0.4525 | 0.1575 | 0.0875 | 0.622799 | 0.396296 | 0.629829 | 1 | 0.999631 | 0.862913 |
| Clasp1_E9QNY9_S1184s_1 | -1.03 | -1.43 | -0.11 | 0.84 | -0.07 | -1.39 | 0.97 | 0.57 | 0.01 | -1.04 | -0.26 | 0.19 | -2.72 | -0.65 | 0.52 | 1.47 | 0.33 | -0.2625 | -0.7725 | 0.921847 | 0.489455 | 0.769001 | 1 | 0.999631 | 0.975032 |
| Clasp1_E9QNY9_S1211s_1 | 0.63 | 1.06 | 2.28 | 1.5 | 0.91 | 1.91 | 2.29 | 1.68 | 1.41 | 0.5 | 1.43 | 1.08 | -0.15 | 0.58 | 1.93 | 0.02 | -0.5775 | -0.645 | -0.84 | 0.180276 | 0.463486 | 0.534261 | 1 | 0.999631 | 0.64376 |
| Clasp1_E9QNY9_S572s_1 | 1.81 | -1.07 | 0.13 | -0.06 | -1.67 | -0.68 | 1.99 | -1.14 | -1.94 | 1.08 | -0.3 | -0.61 | -0.43 | -1.08 | -0.52 | -0.52 | 0.665 | 0.345 | 0.015 | 0.161655 | 0.514208 | 0.404679 | 1 | 0.999631 | 0.631442 |
| Clasp1_E9QNY9_S594s_1 | 0.97 | -0.59 | -1 | 0.42 | 0.45 | 0.62 | 0.61 | 0.78 | 0.58 | 0.12 | 0.66 | -0.18 | -0.57 | 0.28 | 0.58 | -0.43 | 0.125 | 0.0825 | -0.165 | 0.976321 | 0.157415 | 0.474273 | 1 | 0.999631 | 0.991938 |
| Clasp1_E9QNY9_S598s_1 | 1.04 | 0.91 | 0.86 | 1.33 | 1.21 | 1.02 | 0.9 | 1.51 | 1.26 | 0.99 | 0.93 | 1.29 | 0.03 | 1.56 | 0.87 | 1.02 | 0.3425 | 0.21 | 0.07 | 0.6606 | 0.64074 | 0.753495 | 1 | 0.999631 | 0.880315 |
| Clasp1_E9QNY9_S600s_1 | 0.89 | 0.8 | 0.67 | 0.65 | 1.13 | 0.95 | 0.69 | 1.61 | 1.11 | 0.94 | 0.74 | 1.06 | -0.1 | 1.45 | 0.83 | 1.11 | 0.1625 | -0.01 | 1.0075 | 0.852413 | 0.24219 | 0.408128 | 1 | 0.999631 | 0.950222 |
| Clasp1_E9QNY9_S620s_1 | -0.42 | 0.01 | 0.23 | -1.61 | -1.08 | 0.27 | -0.07 | -0.26 | -0.47 | -0.08 | -0.12 | -1.16 | 2.12 | 0 | 0.09 | 0.03 | 0.275 | -0.1275 | -0.4925 | 0.119091 | 0.733213 | 0.982854 | 1 | 0.999631 | 0.586721 |
| Clasp1_E9QNY9_S646s_1 | 1.23 | 0 | 0.31 | 0.75 | 0.83 | 0.1 | 1.34 | 1.12 | -0.06 | 0.19 | 0.87 | 0.78 | -1.11 | 0.44 | 0.88 | 0.11 | 1.1225 | 0.2225 | -0.29 | 0.32474 | 0.485835 | 0.734601 | 1 | 0.999631 | 0.72463 |
| Clasp1_E9QNY9_S646s_2 | 1.25 | -2.7 | -0.66 | 0.77 | 1.27 | 0.75 | 0.96 | 0.17 | 0.55 | -0.7 | 0.32 | -0.62 | -0.84 | 0.1 | -0.64 | -1.12 | 1.1075 | 0.55 | -0.0875 | 0.719392 | 0.182641 | 0.787643 | 1 | 0.999631 | 0.904782 |
| Clasp1_E9QNY9_S647s_2 | 1.25 | -2.7 | -0.48 | 1.53 | 1.27 | 0.75 | 1.56 | 0.45 | 0.55 | -0.7 | 1.58 | 0.37 | -0.84 | 0.1 | 0.33 | -0.34 | 0.2375 | -1.1375 | -2.49 | 0.919914 | 0.223645 | 0.560495 | 1 | 0.999631 | 0.974901 |
| Clasp1_E9QNY9_S685s_1 | 3.04 | -1.06 | -0.97 | 1.41 | 0.07 | 1.14 | 2.09 | 0.07 | -0.51 | 0.35 | -0.03 | -1.94 | -3.37 | -1.32 | -1.39 | -1.46 | 0.3075 | 0.4725 | 0.0625 | 0.024225 | 0.80346 | 0.254854 | 1 | 0.999631 | 0.415644 |
| Clasp1_E9QNY9_S779s_1 | 0.22 | 0.83 | -0.73 | 0.27 | 0.44 | 0.83 | 0.13 | 0.42 | 0.58 | -0.17 | 0.95 | 1.12 | -1.2 | 1.53 | -0.39 | 0.9 | 0.5275 | 1.015 | 0.74 | 0.920782 | 0.416522 | 0.288412 | 1 | 0.999631 | 0.975032 |
| Clasp2_F7DCH5_S1294s_1 | -1.04 | 0.88 | -2.15 | -1.68 | -0.35 | 0.56 | -1.16 | -0.93 | 0.74 | -0.4 | -0.66 | 0.39 | 0.36 | -0.25 | -0.15 | -0.99 | 0.055 | 0.435 | 0.74 | 0.269585 | 0.446425 | 0.150944 | 1 | 0.999631 | 0.697839 |
| Clasp2_F7DCH5_S1307s_1 | 0.08 | -0.14 | -0.59 | 0.94 | 0.35 | -1.43 | 1.38 | 0.21 | -1.19 | 0.16 | 1.14 | 1.92 | 0.12 | -0.48 | 1.55 | 2.06 | 0.67 | 0.7225 | 0.315 | 0.245333 | 0.926885 | 0.518571 | 1 | 0.999631 | 0.685751 |
| Clasp2_F7DCH5_S14s_1 | -0.37 | 1.99 | 0.22 | -0.83 | 2.11 | 1.01 | 0.78 | -0.21 | 2.45 | -0.83 | 0.65 | 1.63 | 2.42 | 0.2 | 0.37 | -0.72 | -0.0925 | -0.15 | -0.4325 | 0.691296 | 0.344444 | 0.388725 | 1 | 0.999631 | 0.893997 |
| Clasp2_F7DCH5_S370s_1 | 0.65 | -0.24 | -0.67 | 0.46 | -0.61 | 0.63 | 0.25 | -0.44 | 0.2 | 0.08 | 0.45 | -1.13 | -0.06 | 0.01 | -0.33 | -1.15 | 0.1575 | 0.24 | -0.0025 | 0.316188 | 0.825467 | 0.742686 | 1 | 0.999631 | 0.720734 |
| Clasp2_F7DCH5_S374s_1 | 0.64 | 0.36 | 0.63 | 0.69 | 0.91 | 0.56 | 0.46 | 1.02 | 0.64 | 0.62 | 1.07 | 0.95 | -0.14 | 0.45 | 1 | 1 | 0.1 | 0.105 | -0.0375 | 0.994197 | 0.548047 | 0.367526 | 1 | 0.999631 | 0.997705 |
| Clasp2_F7DCH5_S376s_1 | 0.69 | 0.13 | 0.54 | 0.61 | 0.7 | 0.49 | 0.37 | 0.81 | 0.45 | 0.27 | 0.89 | 0.78 | 0.35 | 0.02 | 0.67 | 0.78 | 0.11 | 0.0175 | -0.4475 | 0.903071 | 0.703644 | 0.711914 | 1 | 0.999631 | 0.968183 |
| Clasp2_F7DCH5_S413s_1 | -0.79 | 0.09 | -0.09 | 0 | -0.52 | -1 | 0.73 | 0.44 | -0.22 | -1.17 | 0.22 | 0.45 | -1.76 | -0.65 | -0.09 | -0.08 | -0.0725 | -0.5375 | -1.8925 | 0.326725 | 0.80271 | 0.966402 | 1 | 0.999631 | 0.724678 |
| Clasp2_F7DCH5_S426s_1 | 4.09 | -1.2 | 0.65 | 1.29 | 0.99 | 0.85 | 2.29 | 0.41 | 0.53 | 1.64 | 0.41 | 0.1 | -0.64 | -0.3 | -0.45 | -1.35 | 0.535 | -0.6175 | -1.9425 | 0.072574 | 0.942435 | 0.590205 | 1 | 0.999631 | 0.533932 |
| Clasp2_F7DCH5_S461s_1 | 2.54 | 0 | 0.06 | 1.86 | 1.61 | 1.5 | 2.05 | 1.44 | 1.16 | 0.87 | 0.9 | -0.94 | -2.89 | 0.55 | -0.13 | -0.84 | 0.11 | -0.2725 | -1.75 | 0.042713 | 0.378146 | 0.39648 | 1 | 0.999631 | 0.475443 |
| Clasp2_F7DCH5_S465s_1 | 2.59 | -0.69 | -0.66 | 1.38 | 1.01 | 0.33 | 1.33 | 0.39 | 0.77 | 0.42 | 0.05 | 0.29 | -2.17 | -1.26 | 0.07 | -1.02 | 0.1425 | -0.9025 | -1.2825 | 0.052296 | 0.881933 | 0.708756 | 1 | 0.999631 | 0.489777 |
| Clasp2_F7DCH5_S531s_1 | 2.8 | -0.34 | 0.61 | 1.41 | 0.87 | 1.77 | 1.87 | 0.54 | 1.15 | 1.59 | -1.15 | -0.72 | -0.95 | 0.79 | 0.87 | -1.36 | 0.075 | -0.72 | -1.9425 | 0.120631 | 0.829329 | 0.29211 | 1 | 0.999631 | 0.588344 |
| Clasp2_F7DCH5_S535s_1 | 2.68 | -2.3 | -1.74 | 0.56 | 0.21 | -0.84 | 1.27 | -1.14 | -0.4 | -0.22 | -1.64 | -1.42 | -2.8 | -1.39 | -1.76 | -2.62 | 0.49 | 0.72 | 0.1525 | 0.080251 | 0.944857 | 0.490788 | 1 | 0.999631 | 0.545579 |
| Clasp2_F7DCH5_S591s_1 | 0.28 | 0.64 | 0.38 | 1.21 | 1.08 | 1.28 | 0.85 | 1.26 | 1.65 | 1.19 | 1.29 | 1.26 | -0.28 | 1 | 1.17 | 1.23 | -0.2325 | 0.1175 | 0.2125 | 0.720044 | 0.123291 | 0.03764 | 1 | 0.999631 | 0.904809 |
| Clasp2_F7DCH5_S635s_1 | -1.68 | -2.48 | -1.33 | -1.82 | -2.38 | -2.53 | -1.38 | -1.95 | -2.34 | -1.26 | -2.49 | -0.75 | -1.12 | -2.68 | -1.01 | -1.65 | 0.015 | -0.1975 | -0.07 | 0.638524 | 0.534834 | 0.802396 | 1 | 0.999631 | 0.870831 |
| Clasp2_F7DCH5_S641s_1 | -2.19 | -0.26 | -0.99 | -0.79 | -1.72 | -1.41 | -0.76 | -0.28 | -1.69 | -0.93 | -1.93 | -0.47 | -1.79 | -0.8 | -0.14 | -1.78 | 0.0725 | -0.0925 | -0.37 | 0.896205 | 0.975571 | 0.694817 | 1 | 0.999631 | 0.966232 |
| Clasp2_F7DCH5_S905s_1 | -0.77 | 0.77 | 0.32 | 0.23 | 0.14 | 0.62 | -0.09 | 0.17 | 0.3 | -0.36 | -0.13 | 0.37 | -0.64 | 0.05 | 0.16 | -0.5 | 0.515 | 0.29 | -0.5525 | 0.369083 | 0.846204 | 0.811325 | 1 | 0.999631 | 0.746189 |
| Clasp2_F7DCH5_S968s_1 | 0.29 | -0.02 | -1.61 | 0.64 | 0.99 | -0.07 | 0.34 | 0.1 | 0.68 | -0.34 | 0.13 | -0.01 | -0.94 | -0.16 | -0.89 | -0.92 | -0.255 | -0.2025 | -0.1525 | 0.290598 | 0.327088 | 0.572829 | 1 | 0.999631 | 0.706164 |
| Clasp2_F7DCH5_T564t_1 | 0.23 | 0.2 | -0.44 | -0.52 | 0.35 | -0.79 | -0.75 | -0.36 | -0.07 | 0.06 | -0.57 | -0.76 | 0.31 | -0.37 | -0.4 | -0.68 | 0.6725 | 0.6025 | 0.26 | 0.663643 | 0.481454 | 0.545869 | 1 | 0.999631 | 0.881991 |
| Clasp2_Q8BRT1_S8s_1 | -0.37 | 1.99 | -1.05 | -1.83 | 2.11 | 1.01 | -0.26 | -1.43 | 2.45 | -0.83 | -0.55 | 0.08 | 2.42 | 0.2 | -0.98 | -1.86 | -0.1775 | -0.4825 | -0.4275 | 0.805759 | 0.493811 | 0.53527 | 1 | 0.999631 | 0.934783 |
| Clasrp_Q8CFC7_S294s_1 | 1.21 | -0.38 | -0.45 | -0.22 | 0.06 | -0.74 | 0.33 | -0.2 | -0.38 | -0.31 | -0.4 | -0.68 | -0.51 | -0.39 | -0.56 | -0.09 | -0.8 | -0.115 | -0.1925 | 0.321948 | 0.688896 | 0.256038 | 1 | 0.999631 | 0.724228 |
| Clasrp_Q8CFC7_S541s_1 | -0.8 | -1.08 | -0.61 | -1.65 | -1.61 | -2.52 | -1.44 | -1.77 | -1.53 | -1.06 | -1.16 | -0.85 | -0.63 | -1.4 | -2.53 | -0.35 | -0.1725 | 0.07 | 0.6175 | 0.707296 | 0.045311 | 0.72343 | 1 | 0.999631 | 0.898523 |
| Clcn6_O35454_S684s_1 | -0.47 | -0.69 | 0.03 | 0.08 | -0.93 | -1.09 | 0.02 | 0.26 | -0.83 | -0.25 | 0.11 | 0.2 | -0.16 | -0.19 | 1.08 | 0.69 | -0.045 | -0.1775 | 0.675 | 0.141601 | 0.662676 | 0.838665 | 1 | 0.999631 | 0.612726 |
| Cldn10_Q9Z0S6_S147s_1 | 0.69 | -0.17 | -0.39 | 0.2 | -0.82 | -0.32 | 0.34 | 0.95 | -0.66 | 0.52 | -0.77 | 0.53 | -0.04 | 0.43 | 1.46 | 1.18 | -0.5175 | -0.15 | 0.26 | 0.137288 | 0.918551 | 0.680369 | 1 | 0.999631 | 0.610369 |
| Cldn11_Q60771_S194s_1 | -1.93 | -1.47 | -0.98 | -1.24 | -1.57 | -1.41 | -1.45 | -3.26 | -1.4 | -1.23 | -2.04 | -1.55 | -1.11 | -1.7 | -0.55 | -1.22 | -0.525 | -0.5775 | -0.2475 | 0.476306 | 0.284569 | 0.644657 | 1 | 0.999631 | 0.799882 |
| Cldn11_Q60771_S196s_1 | -1.77 | -1.82 | -1.55 | -2.24 | -2.36 | -2.08 | -1.55 | -3.49 | -2.92 | -2.12 | -2.39 | -2.26 | -1.86 | -2.64 | -1.5 | -2.37 | -0.8125 | -1.315 | 0.52 | 0.485376 | 0.23749 | 0.079946 | 1 | 0.999631 | 0.804287 |
| Cldn11_Q60771_S197s_1 | -1.19 | -2.04 | 0.72 | 2.7 | -2.43 | -1.71 | -1.39 | 2.47 | -1.94 | -1.62 | -0.55 | -0.96 | -2.3 | -1.75 | 2.39 | 3.93 | -0.4875 | -0.655 | 0.0925 | 0.739586 | 0.535424 | 0.189697 | 1 | 0.999631 | 0.908631 |
| Cldn11_Q60771_S198s_1 | -1.19 | -2.31 | -1.67 | -1.2 | -2.68 | -2.29 | -1.8 | -1.55 | -2.22 | -1.27 | -3.03 | -2.47 | -1.64 | -2.12 | -1.55 | -0.69 | 0.1725 | -0.1325 | -0.73 | 0.82341 | 0.218171 | 0.164442 | 1 | 0.999631 | 0.938864 |
| Cldn12_Q9ET43_S228s_1 | -0.25 | -0.61 | -1.52 | 0.92 | 0.41 | -1.98 | 0.55 | 0.25 | -1.08 | -0.25 | -0.05 | -0.61 | -1.48 | -0.82 | -1.67 | -0.41 | 0.2425 | -1.1725 | -1.145 | 0.200198 | 0.803804 | 0.798872 | 1 | 0.999631 | 0.658104 |
| Cldn5_O54942_T207t_1 | -0.47 | -0.8 | 0.11 | -0.84 | -2.57 | 1.34 | -0.21 | 0.41 | -4.85 | 0.65 | 0.49 | -2.98 | -1.59 | -0.33 | -2.08 | -2.58 | -0.0975 | 0.055 | 0.22 | 0.044127 | 0.749153 | 0.330152 | 1 | 0.999631 | 0.475443 |
| Clec16a_Z4YLG3_S406s_1 | -1.37 | 1.15 | -0.45 | 0 | -0.34 | 0.15 | -0.62 | -0.25 | 1.11 | -1.56 | -0.1 | 0.1 | 0.7 | 0.1 | -0.32 | -0.27 | -0.725 | -0.865 | -1.28 | 0.683429 | 0.848528 | 0.935395 | 1 | 0.999631 | 0.889602 |
| Clec2l_P0C7M9_S29s_1 | 1.14 | -1.13 | 0.06 | -0.27 | -1.3 | -0.59 | -0.03 | -1.18 | -1.34 | 0.58 | -1.59 | -1.31 | -1.09 | -1.57 | -1.07 | -1.59 | 0.175 | 0.24 | 0.525 | 0.021745 | 0.181165 | 0.186409 | 1 | 0.999631 | 0.405544 |
| Clec2l_P0C7M9_S47s_1 | -1.01 | -0.57 | -0.29 | -0.29 | -0.41 | -0.3 | -0.45 | -0.3 | -0.34 | -0.55 | -0.4 | 0.09 | 0.02 | -0.35 | 0.41 | -0.14 | 0.355 | 0.7475 | 0.6825 | 0.123293 | 0.519165 | 0.427617 | 1 | 0.999631 | 0.594105 |
| Clec2l_P0C7M9_S52s_1 | -2.2 | -0.47 | -0.62 | -1.42 | -0.32 | -0.75 | -0.83 | -1.39 | -0.06 | -1.28 | -0.42 | 0.04 | 0.08 | -0.61 | -0.64 | -0.81 | 0.7125 | 0.2275 | -0.1375 | 0.149025 | 0.430754 | 0.143288 | 1 | 0.999631 | 0.621356 |
| Clint1_Q5SUH6_S227s_1 | 0.65 | 4.22 | 4.99 | 1.71 | 2.64 | 4.57 | 4.7 | 2.51 | 3.94 | 1.34 | 3.5 | 3.7 | 4.2 | 1.72 | 3.53 | 1.57 | -0.3 | -0.4025 | -0.575 | 0.894506 | 0.48974 | 0.824222 | 1 | 0.999631 | 0.965376 |
| Clint1_Q5SUH6_S299s_1 | 1.09 | 0.45 | 1.67 | 0.77 | 0.06 | 1.03 | 0.9 | 0.79 | 0.35 | 1.52 | 0.12 | 0.38 | 0.17 | 0.3 | 0.28 | 0.93 | 0.61 | 0.3825 | 0.2475 | 0.134831 | 0.415499 | 0.343399 | 1 | 0.999631 | 0.606296 |
| Clint1_Q5SUH6_S457s_1 | -0.43 | -0.24 | -1.42 | -0.89 | 0.66 | -0.51 | -0.41 | -0.28 | 0.08 | -0.66 | -0.6 | -0.27 | -0.78 | 0.18 | -0.09 | -1.3 | -0.78 | -0.3125 | -1.3775 | 0.570822 | 0.138722 | 0.291957 | 1 | 0.999631 | 0.84338 |
| Clint1_Q5SUH6_T308t_1 | 1.61 | 0.82 | 1.76 | 0.94 | -0.7 | 1.26 | 0.92 | 0.53 | 0.79 | 2.49 | 0.08 | 0.52 | -1.13 | 0.16 | 0.38 | 0.21 | 0.9625 | -0.3425 | -1.445 | 0.008828 | 0.119326 | 0.564712 | 1 | 0.999631 | 0.365244 |
| Clint1_Q5SUH7_S457s_1 | 1.85 | -1.57 | -2.61 | 0.21 | 1.7 | -1.19 | 1.85 | -0.63 | -0.12 | -0.71 | -0.7 | -1.96 | -2.39 | -1.45 | -1.99 | -2.07 | -0.185 | 0.3625 | 0.435 | 0.119889 | 0.382228 | 0.710003 | 1 | 0.999631 | 0.587703 |
| Clip1_D3Z3M7_S1116s_1 | -0.15 | 1.35 | -0.05 | 0.42 | 0.78 | 0.1 | -0.44 | 0.39 | 1.33 | -0.05 | 0.56 | 1.18 | 0.38 | 1.39 | 0.91 | 0.63 | 0.0425 | -0.36 | -0.47 | 0.312789 | 0.663156 | 0.433618 | 1 | 0.999631 | 0.719861 |
| Clip1_D3Z3M7_S194s_1 | 0.68 | 1.03 | 0.07 | 1.01 | 0.93 | 0.91 | 0.46 | 0.66 | 0.3 | -0.58 | 1.5 | 0.13 | -0.28 | 0.42 | 0.11 | 0.66 | 0.7675 | -0.2025 | -0.125 | 0.206048 | 0.890035 | 0.450902 | 1 | 0.999631 | 0.6605 |
| Clip1_D3Z3M7_S347s_1 | 0.9 | 0.2 | -0.38 | 0.71 | 1.64 | 0.74 | 1.07 | 1.05 | 0.67 | -0.07 | 0.28 | -0.26 | -0.64 | 0.76 | 0.45 | 0.36 | 0.2675 | 0.05 | -0.02 | 0.770144 | 0.057964 | 0.593441 | 1 | 0.999631 | 0.918998 |
| Clip2_Q9Z0H8_S196s_1 | 0.37 | 0.46 | 0.24 | 0.76 | 0.47 | 1.21 | 0.31 | 0.91 | 0.7 | 0.55 | 0.55 | 0.23 | -0.26 | 0.99 | 0.73 | 0.29 | -0.7025 | -0.0025 | 0.1 | 0.954578 | 0.379633 | 0.850607 | 1 | 0.999631 | 0.984441 |
| Clip2_Q9Z0H8_S203s_1 | -1.59 | -0.93 | 0.31 | -1.88 | -1.61 | -2.2 | -1.31 | -1.78 | -1.54 | -0.87 | -1.31 | -0.38 | 0.61 | -1.97 | -1.21 | -1.12 | -0.28 | 0.085 | 0.2025 | 0.878612 | 0.174625 | 0.996138 | 1 | 0.999631 | 0.96244 |
| Clip2_Q9Z0H8_S208s_1 | -0.45 | 0.5 | 0.12 | -1.11 | -0.5 | -0.56 | -0.52 | -0.48 | -0.06 | 0.51 | -0.57 | -0.48 | 0.03 | -0.33 | 0.18 | -0.01 | 0.2975 | 0.12 | 0.115 | 0.610028 | 0.455948 | 0.842864 | 1 | 0.999631 | 0.859048 |
| Clip2_Q9Z0H8_S212s_1 | 0.75 | -0.21 | 0.28 | 0.61 | 1.13 | 0.49 | 0.08 | 0.92 | 0.97 | 0.19 | 0.44 | 0.31 | 0.35 | 0.52 | 0.63 | 0.39 | 0.6925 | 0.0575 | 0.6475 | 0.71424 | 0.400634 | 0.71513 | 1 | 0.999631 | 0.901694 |
| Clip2_Q9Z0H8_S39s_1 | -0.84 | -0.73 | -0.5 | 0.04 | 1.33 | 0.17 | -0.54 | -0.22 | -1.22 | 0.17 | -0.85 | 0.1 | 0.97 | 0.66 | 0.06 | -1.13 | -0.355 | -0.2725 | -0.9775 | 0.20337 | 0.140101 | 0.887624 | 1 | 0.999631 | 0.659781 |
| Clip2_Q9Z0H8_S43s_1 | 1.96 | 2.07 | 1.88 | 1.05 | 0.78 | 2.42 | 1.5 | 0.84 | 1.45 | 2.66 | 1.05 | 0.71 | 1.27 | 0.81 | 0.56 | 0.41 | -0.075 | 0.2775 | 0.03 | 0.018715 | 0.424307 | 0.56468 | 1 | 0.999631 | 0.391024 |
| Clip2_Q9Z0H8_S50s_1 | 0.01 | 0.11 | -0.64 | -0.11 | -0.06 | -0.21 | -0.27 | -0.39 | 0.35 | 0.11 | -0.11 | 0.13 | 0.34 | -0.33 | -0.33 | -0.19 | 0.05 | 0.7725 | 0.0825 | 0.924588 | 0.782517 | 0.340986 | 1 | 0.999631 | 0.975422 |
| Clip2_Q9Z0H8_S924s_1 | 1.1 | 2.46 | 1.13 | -0.04 | 2.22 | 0.26 | 2.11 | 0.26 | 3.79 | 0.65 | 1.25 | 2.05 | 2.32 | 1.28 | 0.94 | 0.44 | 0.25 | -0.1675 | -0.285 | 0.889115 | 0.940308 | 0.318173 | 1 | 0.999631 | 0.963505 |
| Clip2_Q9Z0H8_S974s_1 | 1.04 | 1.15 | 0.32 | 1.4 | 1.78 | 1.17 | 0.99 | 0.97 | 0.84 | 0.4 | 1.57 | 0.43 | -0.08 | 1.33 | 0.52 | 1 | -0.0225 | 0.1925 | 0.51 | 0.487664 | 0.464209 | 0.660818 | 1 | 0.999631 | 0.805946 |
| Clip3_B9EHT4_S399s_1 | -0.21 | 0.2 | -0.81 | -1.03 | -0.08 | 0.36 | -1.19 | -1.03 | 0.06 | 0.54 | -0.87 | -0.81 | 0.87 | 0.02 | -0.29 | -0.41 | 0.32 | -0.435 | -0.78 | 0.239687 | 0.960172 | 0.66263 | 1 | 0.999631 | 0.683719 |
| Clip4_Q8CI96_S580s_1 | 1.32 | 1.68 | 1.08 | 1.78 | 1.01 | 2.3 | 1.32 | 2.51 | 1.44 | 0.86 | 1.83 | -0.01 | -0.01 | 1.61 | 0.27 | 0.87 | 0.0375 | -0.135 | -0.035 | 0.082035 | 0.43507 | 0.32553 | 1 | 0.999631 | 0.54792 |
| Clip4_Q8CI96_S608s_1 | 0.82 | 1.11 | 0.74 | 1.14 | 0.8 | 0.64 | 0.88 | 1.64 | 0.67 | -0.17 | 1.37 | 1.4 | 0.57 | 1.33 | -0.07 | 1.84 | -0.65 | -0.29 | -1.695 | 0.93612 | 0.901803 | 0.734529 | 1 | 0.999631 | 0.97939 |
| Clk2_D3Z7L4_S17s_1 | 1.27 | 0.63 | 0.59 | 0.43 | -0.87 | 0.74 | 1.02 | -0.57 | 0.01 | 2.02 | 0.06 | -0.33 | -0.95 | -0.81 | -0.93 | -1.17 | -0.3325 | -0.0325 | -0.6225 | 0.000234 | 0.195574 | 0.58599 | 1 | 0.999631 | 0.351308 |
| Clk2_D3Z7L4_S50s_1 | 0.4 | 0.72 | 1.22 | 0.49 | -0.1 | 0.33 | 0.68 | 0.59 | 1.04 | 1.12 | 0.34 | 0.2 | -0.75 | 0.21 | 0.96 | -0.08 | -0.6025 | -0.05 | 0.1925 | 0.156296 | 0.297161 | 0.92443 | 1 | 0.999631 | 0.626452 |
| Clk3_O35492_S157s_1 | 0.36 | 1.13 | 0.85 | 0.41 | 0.71 | -0.08 | -0.21 | -0.08 | 0.5 | 0.6 | 0.86 | 0.59 | 0.9 | 1.31 | 0.86 | 0.45 | -0.2775 | -0.2725 | 0.08 | 0.562019 | 0.088119 | 0.862153 | 1 | 0.999631 | 0.84072 |
| Clmn_Q8C5W0_S358s_1 | -0.73 | -0.99 | -1.3 | -0.97 | -1.36 | -1.71 | -1.06 | -0.97 | -1.67 | -1.27 | -1.27 | -0.87 | -1.02 | -0.75 | -1.34 | -0.56 | 0.1125 | 0.1925 | 0.695 | 0.793519 | 0.338644 | 0.356347 | 1 | 0.999631 | 0.928057 |
| Clmn_Q8C5W0_S419s_1 | 0.62 | 1.63 | 0.58 | 0.58 | 0.78 | 1.91 | 0.53 | 0.64 | 1.12 | 1.35 | 0.22 | 1.49 | 1.28 | 1.96 | 1.15 | 1.8 | -0.135 | -0.2275 | 0.35 | 0.083583 | 0.784489 | 0.631143 | 1 | 0.999631 | 0.549732 |
| Cln3_D3YVA8_S14s_1 | 0.58 | 0.61 | 0.05 | -0.5 | 0.12 | 0.11 | -0.05 | 0.02 | 0.36 | 0.25 | -0.79 | 0.01 | 0.46 | 1.12 | 0.29 | 0.27 | -0.18 | 0.2025 | 0.2625 | 0.357078 | 0.670558 | 0.560521 | 1 | 0.999631 | 0.740804 |
| Clns1a_Q923F1_S3s_1 | -0.46 | 0.39 | 0 | 0.13 | -0.09 | -0.44 | -0.11 | -0.02 | 0.13 | 0 | 0.29 | 0.45 | 0 | 0.6 | 0.09 | 0.42 | -0.0775 | 0.1575 | 0.015 | 0.413175 | 0.526508 | 0.489891 | 1 | 0.999631 | 0.768474 |
| Clstn3_D3Z601_S841s_1 | 0.66 | 0.96 | 0.69 | 0.71 | 0.59 | 0.88 | 0.7 | 0.54 | 1.02 | 1.09 | 0.89 | 0.65 | 1.49 | 0.63 | 0.62 | 0.34 | 0.465 | 1.12 | 1.58 | 0.963864 | 0.748566 | 0.540654 | 1 | 0.999631 | 0.987331 |
| Clstn3_D3Z601_S914s_1 | -0.92 | -1.76 | -0.88 | -1.71 | -1.45 | -0.75 | -1.26 | 0.05 | -0.45 | -0.33 | -0.32 | 0.31 | 1.02 | -0.43 | 0.05 | 0.41 | -0.035 | -0.0025 | 0.445 | 0.00301 | 0.275395 | 0.008029 | 1 | 0.999631 | 0.351308 |

| | | | | | | | | | | | | | | | | | | | | | | | | | |
|---|---|---|---|---|---|---|---|---|---|---|---|---|---|---|---|---|---|---|---|---|---|---|---|---|---|
| Clta_B1AWD8_S105s_1 | 0.07 | 0.91 | -0.06 | -0.25 | 0.36 | 0.78 | -0.5 | -0.11 | 0.7 | 0.61 | -0.32 | -0.33 | 1.2 | 0.73 | 0.01 | 0.51 | -0.4725 | -0.315 | 0.225 | 0.266977 | 0.928068 | 0.994961 | 1 | 0.999631 | 0.695664 |
| Cltc_Q68FD5_S576s_1 | -0.59 | 0.16 | 0.96 | 0.77 | -0.9 | -0.2 | 0.18 | 0.33 | -0.48 | -0.59 | 0.62 | 0.49 | 0.38 | -0.11 | 1.1 | 0.83 | 0.265 | 0.0575 | 0.59 | 0.612663 | 0.29201 | 0.498127 | 1 | 0.999631 | 0.860335 |
| Cltc_Q68FD5_S67s_1 | -1.36 | -0.27 | -0.98 | -1.02 | -0.17 | -0.61 | -1.05 | -0.74 | -0.91 | -0.58 | -1.15 | -0.76 | 0.22 | -0.25 | -0.55 | -0.69 | 0.15 | 0.74 | 0.0325 | 0.122218 | 0.433315 | 0.857012 | 1 | 0.999631 | 0.590804 |
| Cltc_Q68FD5_T235t_1 | -1.18 | 0.34 | 0.3 | -1.79 | 0.47 | 0.37 | 0.39 | -2.96 | 0.47 | -2.02 | 1.11 | 1.07 | 1.1 | -0.54 | -1.45 | -1.31 | 0.7575 | -1.8575 | 0.8 | 0.963099 | 0.862571 | 0.367157 | 1 | 0.999631 | 0.987129 |
| Cltc_Q68FD5_T302t_1 | -2.92 | -1.55 | -1.91 | -2.89 | -1.67 | -0.53 | -2.09 | -1.95 | -5.11 | -3.38 | -2.57 | -5.64 | -1.52 | -0.41 | -2.25 | -1.89 | -0.3975 | -0.12 | -0.0875 | 0.136298 | 0.133033 | 0.024801 | 1 | 0.999631 | 0.60941 |
| Cluh_Z4YLI8_S1287s_1 | 0.94 | 0.23 | 0.27 | 0.06 | -0.37 | 0.31 | -0.1 | 0.07 | 0.13 | 0.61 | -0.07 | 0.35 | 0.64 | 0.29 | 0.29 | -0.07 | 2.3975 | 3.69 | 0.3425 | 0.786536 | 0.208618 | 0.699803 | 1 | 0.999631 | 0.923689 |
| Clvs1_Q9D4C9_S34s_1 | -1.75 | 3.94 | 5.55 | 1.8 | 5.94 | 4.82 | 5.4 | 2.97 | 8.5 | 1.29 | 6.67 | 7.84 | 6.54 | -0.24 | 4.49 | 0.12 | -0.0675 | -0.465 | -1.395 | 0.858 | 0.122217 | 0.079975 | 1 | 0.999631 | 0.952251 |
| Clvs1_Q9D4C9_S41s_1 | 3.76 | 0.63 | 0.19 | 1.41 | 1.99 | 0.97 | 1.77 | 0.99 | 1.46 | 1.51 | 0.6 | 0.56 | -0.91 | 0.4 | 0.98 | -0.06 | -0.155 | -0.4025 | -0.4875 | 0.097363 | 0.926989 | 0.534275 | 1 | 0.999631 | 0.559562 |
| Clvs1_Q9D4C9_S50s_1 | 0.76 | 1.21 | 1.26 | 1.8 | 0.82 | 1.03 | 0.9 | 1.66 | 0.85 | 0.26 | 1.6 | 0.71 | -0.19 | 1.27 | 0.87 | 1.13 | 0.5275 | 0.0125 | -0.2125 | 0.252896 | 0.638731 | 0.298188 | 1 | 0.999631 | 0.688432 |
| Clvs2_Q8BG92_S302s_1 | 0.49 | 1.83 | 0.36 | 1.89 | 1.4 | 2.1 | 1.1 | 2.08 | 1.27 | 0.2 | 2.37 | 0.78 | -0.1 | 2.1 | 1.14 | 0.58 | 0.17 | -0.155 | 0.26 | 0.712035 | 0.27194 | 0.982494 | 1 | 0.999631 | 0.9008 |
| Cmc2_F8WHQ0_S47s_1 | 0.4 | -0.26 | -1.06 | -0.44 | 0.27 | -0.33 | 0.07 | -0.69 | -1.41 | 0.32 | -0.15 | -0.74 | 0.02 | -0.07 | 0.03 | -0.3 | 1.305 | 1.355 | 0.725 | 0.47597 | 0.6573 | 0.739588 | 1 | 0.999631 | 0.799882 |
| Cmip_Q9D486_S349s_1 | -1.36 | 3.17 | 3.92 | 0.39 | 3.63 | 2.44 | 2.99 | 2.28 | 4.09 | -1.13 | 4.22 | 4.36 | 4.3 | 1.05 | 2.88 | 0.79 | -0.3575 | -0.0325 | -0.86 | 0.564827 | 0.246267 | 0.38968 | 1 | 0.999631 | 0.84139 |
| Cmpk1_Q9DBP5_S147s_1 | 0.97 | 1.07 | 1.3 | 0.26 | -0.04 | 1.34 | 0.79 | 0.08 | 1.07 | 1.04 | 0.61 | 0.75 | 0.63 | -0.15 | 0.12 | -0.44 | -1.0625 | -0.9975 | -1.4125 | 0.037675 | 0.379516 | 0.917667 | 1 | 0.999631 | 0.458331 |
| Cmtm4_Q8CJ61_S194s_1 | -1.77 | -5.78 | -3.38 | -3.46 | -6 | -5.79 | -2.61 | -4.24 | -7.38 | -3.17 | -3.53 | -4.3 | -5.26 | -6.13 | -4.25 | -4.4 | 0.5925 | -0.0025 | 1.1875 | 0.106183 | 0.294449 | 0.369582 | 1 | 0.999631 | 0.568797 |
| Cmtm4_Q8CJ61_S199s_1 | -1.03 | 1.14 | -0.1 | -0.22 | 1.59 | 1.28 | -0.26 | -0.45 | 0.94 | -0.52 | -0.33 | -0.31 | 1.72 | 1.09 | 1.12 | 0.61 | -0.405 | -0.4575 | -1.1225 | 0.032441 | 0.349965 | 0.996191 | 1 | 0.999631 | 0.442395 |
| Cmtm6_Q9CZ69_S8s_1 | -0.8 | 0.98 | 1.53 | -1.07 | -2.01 | 0.47 | 0.32 | 0.24 | -0.8 | 0.3 | -0.64 | -0.05 | 0.88 | -1.11 | -3.6 | -0.02 | -0.2225 | 0.4925 | -1.1025 | 0.275078 | 0.599598 | 0.471996 | 1 | 0.999631 | 0.700187 |
| Cmtm7_Q9ESD6_S2s_1 | 1.64 | -1.16 | -0.89 | -0.61 | -1.21 | -1.32 | 1.17 | -0.55 | 1.44 | -1.7 | 0.85 | 0.36 | 0.91 | -2.41 | -3.9 | -0.03 | 0.91 | -0.19 | -0.7825 | 0.321369 | 0.768884 | 0.552 | 1 | 0.999631 | 0.72408 |
| Cnbp_P53996_T165t_1 | -0.04 | 1.55 | 0.86 | 1.52 | 0.98 | 2.15 | 1.58 | 2.82 | 1.6 | 1.29 | 1.2 | -0.96 | -3.12 | 1.78 | 1.8 | 0.3 | 0.285 | 0.92 | 0.8225 | 0.457386 | 0.097477 | 0.762927 | 1 | 0.999631 | 0.790722 |
| Cnksr2_Q80YA9_S12s_1 | -1.28 | -0.13 | -1.79 | -0.66 | -1.27 | -0.48 | -0.64 | -0.33 | -0.32 | -1.02 | 0.7 | 0.46 | 0.45 | -0.23 | -0.77 | -0.02 | 0.1025 | 0.1725 | -0.3 | 0.087764 | 0.497218 | 0.093155 | 1 | 0.999631 | 0.553787 |
| Cnksr2_Q80YA9_S325s_1 | -2.07 | -3.82 | -2.21 | -2.11 | -2.15 | -3.02 | -1.74 | -2.89 | -2.98 | -2.13 | -2.23 | -2.18 | -2.9 | -2.3 | -3.47 | -2.74 | 0.565 | -0.305 | -0.855 | 0.532266 | 0.834756 | 0.706877 | 1 | 0.999631 | 0.8277 |
| Cnksr2_Q80YA9_S325s_2 | 0.39 | -2.51 | -0.53 | -0.34 | 0.45 | -2.46 | 0.5 | 0.78 | -1.5 | -2.96 | 0.11 | 0.14 | -3.19 | -2.12 | -0.83 | -0.27 | -3.9E-17 | 0.27 | 0.47 | 0.292646 | 0.513009 | 0.718876 | 1 | 0.999631 | 0.706671 |
| Cnksr2_Q80YA9_S390s_1 | -0.38 | -0.27 | -0.89 | -0.83 | -0.33 | -0.31 | -0.97 | -0.76 | -0.24 | -0.19 | -0.16 | -0.7 | 0.2 | 0.11 | -0.67 | -0.13 | 0.265 | -1.1625 | -1.5975 | 0.172383 | 1 | 0.361124 | 1 | 0.999631 | 0.642116 |
| Cnksr2_Q80YA9_S430s_1 | 2.55 | -1.75 | -0.3 | 1.82 | 1.19 | -1.03 | 2.76 | 0.46 | -1.64 | -1.51 | 1.07 | -0.25 | -1.5 | -1.09 | -0.38 | -1.1 | 1.2775 | -0.5275 | -0.4325 | 0.091878 | 0.806375 | 0.266631 | 1 | 0.999631 | 0.557085 |
| Cnksr2_Q80YA9_S467s_1 | 2.03 | -1.69 | -1.6 | 1.19 | 1.96 | 0.3 | 1.84 | 0.94 | 0.21 | -2.4 | 0.75 | -0.74 | 0.52 | -0.29 | -2.19 | 0.16 | 0.5475 | 0.1925 | 0.645 | 0.656554 | 0.175551 | 0.606442 | 1 | 0.999631 | 0.879056 |
| Cnksr2_Q80YA9_S503s_1 | -0.79 | -0.83 | -1.01 | -1.01 | -0.36 | -0.63 | -0.23 | -0.23 | 0.05 | 0.01 | -0.83 | -2.1 | 0.19 | -0.23 | -0.49 | -0.53 | 0.2875 | 0.09 | 0.285 | 0.048356 | 0.045954 | 0.692522 | 1 | 0.999631 | 0.48343 |
| Cnksr2_Q80YA9_S505s_1 | -0.41 | -0.8 | -1.55 | -0.75 | -0.29 | -0.74 | -0.43 | -0.9 | -0.67 | -0.24 | -0.85 | -1.39 | -0.73 | -0.13 | -1.06 | -0.45 | 0.19 | 0.39 | 0.9325 | 0.437015 | 0.384279 | 0.806637 | 1 | 0.999631 | 0.781057 |
| Cnksr2_Q80YA9_S702s_1 | -1.18 | -0.18 | -0.27 | -0.51 | -0.44 | 0.03 | -0.7 | -0.27 | -0.12 | -0.39 | -0.3 | 0.23 | 0.8 | 0.38 | 0.09 | 0.32 | 0.2425 | 0.2975 | 0.195 | 0.018664 | 0.557013 | 0.244237 | 1 | 0.999631 | 0.391024 |
| Cnksr2_Q80YA9_S756s_1 | 0.55 | 1.04 | -0.33 | 0.18 | 0.51 | 1.47 | 0.07 | 0.36 | 1.3 | 0.89 | 0.58 | -0.14 | 0.18 | 1.51 | 0.29 | 0.24 | -0.1525 | 0.2075 | 0.435 | 0.656434 | 0.563916 | 0.486938 | 1 | 0.999631 | 0.879056 |
| Cnksr2_Q80YA9_S906s_1 | -0.69 | -0.34 | -0.5 | -0.57 | -0.62 | -0.6 | -1.06 | -0.43 | -0.38 | -0.63 | -0.3 | 0.04 | 0.11 | 0.13 | -0.56 | -0.04 | 0.64 | -0.215 | -0.6725 | 0.159767 | 0.55996 | 0.447119 | 1 | 0.999631 | 0.629764 |
| Cnksr2_Q80YA9_T339t_2 | 0.34 | -2.11 | -0.53 | -0.34 | 0.46 | -1.82 | 0.5 | 0.78 | -1.32 | -2.43 | 0.11 | 0.14 | -2.51 | -1.72 | -0.83 | -0.27 | 0.7975 | -0.0225 | -1.085 | 0.309462 | 0.373755 | 0.764816 | 1 | 0.999631 | 0.718845 |
| Cnksr2_Q80YA9_T809t_1 | 2.29 | 1.28 | 0.22 | 2.15 | 2.6 | 2.79 | 1.98 | 1.76 | 2.82 | 2.03 | 1.73 | -0.73 | -2.03 | 1.94 | 0.87 | 0.82 | 0.49 | -0.3175 | -0.825 | 0.218423 | 0.136697 | 0.97734 | 1 | 0.999631 | 0.669859 |
| Cnnm1_Q0GA42_S850s_1 | 1.16 | -0.38 | -1.42 | 0.65 | 1.39 | -0.21 | 0.56 | 0.23 | 0.18 | -0.6 | 0.28 | -1.12 | -1.47 | 0.05 | -0.98 | -0.89 | -0.8875 | -0.995 | -1.14 | 0.18813 | 0.422738 | 0.600809 | 1 | 0.999631 | 0.647379 |
| Cnnm1_Q0GA42_T824t_1 | 1.31 | 0.42 | 1.76 | 1.27 | -0.34 | 0.18 | 1.25 | 0.12 | -0.04 | 0.93 | 0.03 | -0.14 | -0.46 | -0.59 | 0.93 | 0.32 | 0.595 | 0.6825 | 0.9675 | 0.028036 | 0.062303 | 0.027561 | 1 | 0.999631 | 0.423977 |
| Cnnm3_Q32NY4_S667s_1 | -1.56 | -0.7 | -0.36 | -0.63 | -0.21 | -0.33 | -0.73 | 0.4 | 0.23 | -0.66 | 0.15 | -0.24 | 0.5 | 0.49 | -0.18 | -0.19 | 0.6425 | 0.41 | 0.22 | 0.023793 | 0.130868 | 0.084291 | 1 | 0.999631 | 0.415644 |
| Cnnm4_Q69ZF7_S663s_1 | 0.86 | 0.21 | -0.42 | 0.96 | 1.8 | 0.58 | 0.75 | 1.05 | 2.18 | 0.17 | 0.83 | 0.07 | 0.58 | 0.7 | -0.16 | 1.37 | -0.38 | 0.5125 | -0.235 | 0.625475 | 0.14659 | 0.455836 | 1 | 0.999631 | 0.86386 |
| Cnnm4_Q69ZF7_S766s_1 | 0.23 | 0.89 | 0.59 | -0.14 | 0.22 | 0.62 | 0.13 | -0.92 | 0.6 | 1.35 | 1.35 | 0.32 | 0.78 | -0.54 | -0.4 | 0.79 | -1.02 | 0.0275 | -0.01 | 0.590191 | 0.354787 | 0.189359 | 1 | 0.999631 | 0.851598 |
| Cnot10_Q8BH15_S520s_1 | 0 | 0.61 | -0.33 | 0.19 | -0.51 | -1.6 | -0.61 | -0.89 | -0.02 | 0.48 | -0.48 | 0.6 | 0.42 | 0.2 | 0.36 | -0.55 | 0.1525 | 0.3875 | -0.0275 | 0.977374 | 0.014007 | 0.937913 | 1 | 0.999631 | 0.992174 |
| Cnot2_E9Q8D5_S165s_1 | 0.48 | 0.68 | 1.32 | 0.14 | 0.74 | 0.52 | 1.1 | 0.87 | 1.19 | 0.51 | 1.38 | 1.09 | 0.51 | 0.42 | 0.99 | 0.59 | -0.4275 | -0.265 | -0.7425 | 0.935844 | 0.638171 | 0.28395 | 1 | 0.999631 | 0.97939 |
| Cnot3_F6YBV1_S256s_1 | 1.77 | 0.83 | 1.7 | 1.54 | 0.04 | 1.2 | 1.96 | 0.93 | 1.73 | 1.4 | 0.6 | 1.05 | 0.29 | 0.79 | 1.11 | 0.68 | -0.265 | -1.92 | -1.515 | 0.050084 | 0.342665 | 0.466028 | 1 | 0.999631 | 0.488436 |
| Cnp_P16330_S22s_1 | 0.29 | 0.16 | 1.23 | 0.87 | -2.58 | 1.59 | 3.35 | -0.87 | -2.33 | 0.57 | -2.59 | -0.78 | -1.29 | -1.05 | 0.45 | -1.62 | -0.175 | 0.3625 | 0.7275 | 0.011945 | 0.816409 | 0.019007 | 1 | 0.999631 | 0.385092 |
| Cnp_P16330_S382s_1 | -1.37 | -0.62 | -0.3 | -1.06 | -1.28 | -0.45 | -1.07 | -1.25 | -0.37 | -0.88 | -0.8 | 0.15 | -0.21 | -0.02 | 0.14 | -0.35 | -0.2 | -0.3025 | 0.3325 | 0.049553 | 0.6093 | 0.333045 | 1 | 0.999631 | 0.487521 |
| Cnp_P16330_S408s_1 | -0.34 | 0.24 | 0.69 | -0.08 | -0.61 | 0.51 | -0.04 | -0.15 | -0.21 | 0.13 | -0.52 | -0.1 | 0.36 | 0.4 | 0.8 | 0.28 | 0.0075 | -0.2075 | -0.9725 | 0.322204 | 0.5702 | 0.355464 | 1 | 0.999631 | 0.72427 |
| Cnp_P16330_S4s_1 | 1.66 | -0.86 | -1.13 | 0.75 | -0.72 | -1.18 | 1.32 | 1.03 | -0.4 | 1.07 | -0.62 | -0.46 | -1.24 | -1.04 | -0.6 | -0.59 | 0.1325 | -0.1325 | -0.0025 | 0.13972 | 0.992472 | 0.763573 | 1 | 0.999631 | 0.611718 |
| Cnr1_P47746_S317s_1 | 0.76 | 1.47 | 1.23 | 1.55 | 0.92 | 1.83 | 0.86 | 1.93 | 1.18 | 0.93 | 1.52 | 0.85 | 0.46 | 1.74 | 1.53 | 1.27 | 2.095 | 2.9525 | 0.305 | 0.994646 | 0.714196 | 0.665323 | 1 | 0.999631 | 0.997705 |
| Cnr1_P47746_S442s_1 | 1.62 | 1.48 | 3.57 | -0.67 | 4.13 | 3.6 | 5.09 | 1.56 | 6.52 | 1.26 | 4.36 | 5.67 | 5.81 | -0.5 | 1.76 | 0.15 | -0.235 | 0.67 | 0.6175 | 0.827258 | 0.056021 | 0.03595 | 1 | 0.999631 | 0.941153 |
| Cnst_Q8CBC4_S279s_1 | 0.03 | -0.13 | -1.42 | -0.82 | -0.45 | -1.17 | -1.15 | -0.51 | -0.13 | -0.11 | 0.03 | 0.55 | 0.16 | 0.57 | -0.28 | -0.32 | 0.0375 | 0.3075 | 0.13 | 0.156471 | 0.554847 | 0.109113 | 1 | 0.999631 | 0.626488 |
| Cntnap1_O54991_S1384s_1 | -1.51 | -0.37 | -1.06 | -2.58 | -0.71 | -1.4 | -1.45 | -1.81 | -1.01 | -1.07 | -1.49 | -0.72 | -0.52 | -0.9 | -1.4 | -2.18 | -0.5375 | -0.59 | -0.8 | 0.81204 | 0.93871 | 0.520021 | 1 | 0.999631 | 0.936949 |
| Coasy_Q9DBL7_S177s_1 | 1.9 | 0 | 2.03 | 1.35 | 0.13 | 0.78 | 1.93 | 0.29 | 0.76 | 1.8 | -0.22 | 0.58 | -0.91 | 0.73 | 1.49 | 0.77 | 0.885 | -0.11 | -0.485 | 0.217843 | 0.351973 | 0.316442 | 1 | 0.999631 | 0.669859 |
| Cobl_G3UWY3_S1121s_1 | 1.32 | 0.1 | -1.29 | 1.3 | 1.66 | 0.92 | 1.3 | 1.09 | 0.4 | -0.24 | 0.45 | 0.38 | -1.75 | 0.34 | 0.33 | 0.57 | 0.025 | -0.0575 | -0.055 | 0.50925 | 0.151054 | 0.851015 | 1 | 0.999631 | 0.816932 |
| Cobl_G3UWY3_S669s_1 | 0.13 | 0.02 | 0.06 | 0.17 | -0.25 | -0.07 | 0.03 | 0.77 | -0.49 | -0.3 | 0.44 | 0.5 | -0.47 | -0.14 | 0.16 | 0.61 | -0.2 | -0.01 | 0.355 | 0.862552 | 0.932307 | 0.856121 | 1 | 0.999631 | 0.954762 |
| Cobl_G3UWY3_S819s_1 | -0.55 | 0.57 | 0 | -0.05 | 0.1 | 0.24 | -0.42 | -0.75 | -0.49 | 0.3 | 0.23 | -0.11 | 0.62 | 0.67 | -0.21 | 0.31 | 0.485 | -0.3575 | -0.69 | 0.332819 | 0.573577 | 0.976416 | 1 | 0.999631 | 0.728079 |
| Cobl_Q5NBX1_S269s_1 | 1.19 | -0.68 | 0.19 | 0.55 | 0.72 | 0.55 | 1.17 | 0.75 | 0.18 | -0.53 | 0.04 | 0.13 | -1.18 | 0.08 | -0.06 | -0.35 | -0.325 | -0.27 | -0.245 | 0.165394 | 0.255678 | 0.408882 | 1 | 0.999631 | 0.633733 |
| Cobll1_B1AZ15_S234s_1 | 0.65 | 1.86 | 2.02 | 0.59 | 0.34 | 1.73 | 0.6 | 1.15 | 0.99 | 1.35 | 1.16 | 0.54 | 0.64 | 1.12 | 1.05 | 1.33 | 1.63 | 2.3775 | 1.895 | 0.564474 | 0.493755 | 0.527119 | 1 | 0.999631 | 0.84139 |
| Cobll1_B1AZ15_S281s_1 | 0.8 | 7.86 | 5.72 | 1.58 | 8.22 | 5.9 | 5 | 3.36 | 9.16 | -0.07 | 7.05 | 9.33 | 8.74 | 4.33 | 6.82 | 3.65 | 0.0925 | 0.2725 | 0.66 | 0.28419 | 0.339041 | 0.3235 | 1 | 0.999631 | 0.705132 |
| Cog3_E9QL65_S212s_1 | -0.48 | -0.13 | -0.22 | -0.62 | -0.28 | -0.48 | -0.39 | 0.07 | 0.1 | -0.42 | -0.29 | 0.25 | -0.2 | 0.38 | 0.22 | 0.79 | 0.17 | 1.3175 | 0.105 | 0.061 | 0.727321 | 0.351689 | 1 | 0.999631 | 0.509355 |
| Coil_E9Q284_S149s_1 | -1.42 | -3.41 | -4.06 | -3.76 | -2.27 | -3.34 | -2.56 | -3.8 | -2.84 | -2.62 | -0.83 | -1.09 | -1.19 | -2.37 | -3.37 | -5.3 | -0.3225 | 0.205 | -0.195 | 0.907904 | 0.784723 | 0.085251 | 1 | 0.999631 | 0.96989 |
| Coil_E9Q284_S174s_1 | 0.06 | -0.77 | -0.39 | -0.92 | -1.37 | -0.3 | -0.45 | -1.19 | 0.09 | 0.4 | -0.86 | -0.83 | -0.14 | -0.98 | -0.75 | -0.93 | -0.1775 | -0.1175 | -0.325 | 0.579847 | 0.38702 | 0.611212 | 1 | 0.999631 | 0.84667 |
| Col15a1_O35206_S125s_1 | 0.07 | 0.35 | 0.04 | 0.22 | 0.34 | -0.1 | 0.24 | -0.51 | 0.08 | -0.27 | 0.28 | 0.12 | -0.3 | -0.3 | -0.24 | 0.22 | -0.525 | 0.3975 | 0.2875 | 0.263114 | 0.535034 | 0.654005 | 1 | 0.999631 | 0.693034 |
| Col4a3bp_Q9EQG9_S123s_1 | -2.04 | -0.41 | -0.76 | -2.22 | -1.77 | -2.41 | -0.84 | -2.51 | -0.98 | -1.91 | -1.06 | 0.11 | 0.81 | -2.42 | -0.59 | -2.08 | 0.225 | 0.99 | 0.3275 | 0.706863 | 0.34824 | 0.48788 | 1 | 0.999631 | 0.898287 |
| Col4a3bp_Q9EQG9_S123s_2 | -0.59 | 0.44 | 0.07 | -1.29 | 0.04 | 0.03 | 0.31 | -0.85 | 0.62 | 0.18 | 0.8 | 0.99 | 0.63 | 0.01 | 0.62 | -1.32 | 0.79 | 0.835 | 0.5125 | 0.557675 | 0.614365 | 0.038686 | 1 | 0.999631 | 0.838305 |
| Col4a3bp_Q9EQG9_S126s_1 | -2.68 | -0.22 | 0.5 | -1.32 | 0.06 | -0.19 | 0.18 | -0.61 | 0.89 | -2.61 | 0.42 | 0.92 | 0.81 | -1.36 | 0.19 | -1.31 | 0.225 | 0.99 | 0.3275 | 0.511303 | 0.235466 | 0.388097 | 1 | 0.999631 | 0.817687 |
| Col4a3bp_Q9EQG9_S126s_2 | -0.59 | 0.44 | 0.07 | -1.29 | 0.04 | 0.03 | 0.31 | -0.85 | 0.62 | 0.18 | 0.8 | 0.99 | 0.63 | 0.01 | 0.62 | -1.32 | -0.4025 | -0.0475 | 0.39 | 0.557675 | 0.614365 | 0.038686 | 1 | 0.999631 | 0.838305 |
| Comt_O88587_S218s_1 | -0.27 | 0.16 | 1.44 | 0.85 | -0.37 | -0.31 | 0.31 | 0.94 | -0.25 | 0.21 | 0.93 | 1.1 | 0.85 | 0.39 | 1.04 | 1.46 | -0.3925 | -0.0075 | 0.905 | 0.385921 | 0.396242 | 0.920076 | 1 | 0.999631 | 0.756141 |
| Copa_F8WHL2_S173s_1 | -0.27 | -0.1 | 0.07 | -0.11 | -0.58 | -0.74 | -0.46 | -0.2 | -0.12 | -0.93 | 0.64 | -0.03 | 1.13 | 0.58 | 0.75 | 0.75 | -0.27 | -0.4425 | 0.3 | 0.008012 | 0.142619 | 0.983431 | 1 | 0.999631 | 0.364291 |
| Copb2_O55029_S860s_1 | 0.7 | -0.81 | 0.54 | -0.14 | -1.02 | 0.32 | 0.36 | -0.45 | -0.22 | 0.32 | -1.49 | -0.09 | -1.09 | 1.44 | 0.28 | 0.86 | 0.1275 | 0.3675 | 0.16 | 0.612461 | 0.560554 | 0.382615 | 1 | 0.999631 | 0.860287 |
| Copg1_Q9QZE5_T584t_1 | -0.48 | 0.08 | -0.47 | 0.08 | -0.09 | -0.01 | -0.2 | 0.02 | 0.34 | -0.33 | 0.13 | 0.54 | 0.13 | 0.25 | -0.71 | 0.18 | -0.17 | 0.39 | -0.3825 | 0.640713 | 0.633004 | 0.25602 | 1 | 0.999631 | 0.871305 |
| Cops3_O88543_S410s_1 | -1.13 | -1.16 | -0.71 | -0.45 | -0.97 | -1.68 | -0.58 | -0.9 | -0.49 | -0.38 | -0.69 | -0.33 | -1.28 | -2.31 | -0.82 | -0.57 | -0.3325 | 0.0425 | -0.6425 | 0.382973 | 0.608777 | 0.189638 | 1 | 0.999631 | 0.754387 |
| Cops3_O88543_S423s_1 | 0.48 | -0.15 | 0.57 | 0.24 | 0.58 | -0.62 | 0.31 | -0.46 | 0.66 | -0.13 | 0.11 | 0.67 | -0.88 | -0.84 | 0.3 | -0.01 | 0.4075 | 0.0175 | 0.51 | 0.11197 | 0.363819 | 0.893748 | 1 | 0.999631 | 0.577068 |
| Cops5_O35864_S284s_1 | -0.01 | -0.82 | -1.3 | -0.66 | 0.7 | -0.04 | -0.94 | -0.88 | -0.19 | -0.96 | -0.9 | -0.67 | 0.68 | -0.63 | -0.73 | -0.07 | -0.4125 | -0.45 | 0.01 | 0.251029 | 0.379741 | 0.960646 | 1 | 0.999631 | 0.686822 |
| Coro1a_O89053_S426s_1 | 0.46 | 0.01 | -0.18 | -0.36 | 0.21 | -0.44 | -0.76 | -0.73 | -0.54 | -0.29 | -0.4 | -0.64 | 0.34 | -0.05 | -0.66 | 0.34 | -0.31 | -0.1275 | -0.5125 | 0.977257 | 0.228068 | 0.137676 | 1 | 0.999631 | 0.992174 |
| Coro1a_O89053_T418t_1 | 1.44 | 0.77 | 0.5 | 1.26 | 1.25 | -0.69 | 1.47 | 0.7 | 0.91 | 0.45 | 1.31 | 0.79 | 0.08 | 0.6 | 0.12 | 1.12 | -0.2125 | -0.09 | -0.08 | 0.186452 | 0.538215 | 0.701438 | 1 | 0.999631 | 0.646474 |
| Coro1a_O89053_T424t_1 | 0.35 | 0.65 | 0.47 | 1.19 | -0.39 | 0.09 | 0.9 | 1.21 | -0.06 | 0.65 | 0.84 | 0.87 | -0.71 | 0.82 | 0.92 | 1.31 | -0.305 | -0.1825 | 0.43 | 0.865182 | 0.607385 | 0.78827 | 1 | 0.999631 | 0.955708 |
| Coro7_Q9D2V7_S459s_1 | -1 | -0.64 | 0.04 | -0.28 | -0.66 | -0.83 | -0.65 | -0.96 | -0.21 | -1.72 | -1.01 | 0.33 | 0.99 | -0.78 | -0.5 | 0.13 | -0.5125 | -0.37 | -0.9875 | 0.351596 | 0.322364 | 0.705918 | 1 | 0.999631 | 0.738113 |
| Cotl1_Q9CQI6_S141s_1 | 0.12 | -2.18 | -0.79 | -0.53 | -1.75 | -1.75 | -0.2 | -1.73 | -1.67 | -0.74 | -1.08 | -1.37 | -1.59 | -2.98 | -1.29 | -1.47 | -0.09 | 0.25 | -1.145 | 0.108419 | 0.375171 | 0.463628 | 1 | 0.999631 | 0.570434 |
| Cox6a1_Q9DCW5_S28s_1 | -0.15 | -1.05 | 0.02 | -0.38 | -0.81 | -0.68 | 0.37 | -0.8 | -0.51 | 0.93 | -0.55 | -0.43 | -1.96 | -2.15 | -0.46 | -1.57 | -0.03 | 0.1175 | -0.005 | 0.026186 | 0.813771 | 0.562516 | 1 | 0.999631 | 0.422285 |
| Cpeb2_E9Q5X2_S514s_1 | -0.12 | 0.29 | 0.01 | 0.16 | 0.38 | -0.14 | 0.18 | -0.2 | 0.01 | 0.18 | 0.6 | 0.02 | -0.25 | 0.39 | -0.15 | 0.33 | 0.355 | 0.22 | 0.3425 | 0.986539 | 0.90934 | 0.667067 | 1 | 0.999631 | 0.995165 |
| Cpeb2_E9Q5X2_S517s_1 | -0.14 | -0.24 | -1.54 | -0.68 | 0.43 | 0.05 | -0.62 | -1.04 | -0.13 | 0.09 | -0.61 | -1.07 | 0.34 | 0.51 | -0.89 | -1.19 | 0.215 | -0.0225 | 0.5 | 0.505743 | 0.432805 | 0.599407 | 1 | 0.999631 | 0.814836 |
| Cpeb3_D3Z1R6_S197s_1 | 1.2 | 0.51 | 0.5 | 0.63 | 0.94 | 1.33 | 0.62 | 0.81 | 0.78 | 1.22 | 0.23 | 0.52 | 1.43 | 1.76 | 0.51 | 1.14 | 0.425 | -0.2025 | 0.06 | 0.187808 | 0.469281 | 0.944631 | 1 | 0.999631 | 0.647111 |
| Cpeb3_D3Z1R6_S291s_1 | 0.37 | -0.58 | -0.39 | 0.21 | 0 | 0.69 | 0.37 | 0.25 | -0.01 | -0.68 | -0.47 | -0.04 | -0.61 | 0.94 | -0.52 | 0.04 | 0.085 | -0.385 | 0.15 | 0.889335 | 0.203501 | 0.546666 | 1 | 0.999631 | 0.963505 |
| Cpeb3_D3Z1R6_S413s_1 | -0.65 | 0.64 | -0.74 | -0.24 | 0.01 | -0.11 | -0.01 | -0.54 | -0.03 | -0.55 | -1.4 | -0.55 | 0.48 | 0.37 | -1.24 | 0 | 0.37 | -0.03 | 0.3225 | 0.758802 | 0.814843 | 0.375852 | 1 | 0.999631 | 0.914449 |
| Cpeb4_Q5SU48_S97s_1 | 0.17 | 0.19 | -0.69 | 0.09 | 0.42 | 0.57 | -0.04 | 0.29 | -0.14 | -0.38 | 0.08 | 0.08 | 0.21 | 0.48 | -0.13 | 0.49 | -0.1175 | 0.08 | 0.6275 | 0.340817 | 0.244743 | 0.922249 | 1 | 0.999631 | 0.733032 |
| Cpeb4_Q5SU48_S99s_1 | -0.36 | -0.9 | 0.21 | -0.44 | -0.83 | -0.54 | -0.6 | 0.01 | -0.74 | -0.98 | 0.24 | 0.31 | -0.37 | -0.25 | 0.6 | 1.04 | 0.255 | 0.07 | 0.5625 | 0.158942 | 0.72273 | 0.84535 | 1 | 0.999631 | 0.629092 |
| Cplx1_P63040_S93s_1 | -0.55 | -0.08 | 0.76 | 0.18 | 0.02 | 0.86 | 0 | 0.45 | 0.21 | -0.04 | 0.1 | 0.32 | 0.06 | 0.88 | 0.61 | 1.01 | -0.2475 | 0.0225 | -0.115 | 0.161868 | 0.488136 | 0.833847 | 1 | 0.999631 | 0.631442 |
| Cplx2_P84086_T95t_1 | 1.22 | 1.37 | 0.33 | -0.1 | 0.57 | 1.11 | -0.04 | 0.19 | 0.72 | 2.02 | 0.02 | 0.15 | 0.11 | 1.38 | 0.4 | 0.47 | -0.025 | 0.0025 | -0.2 | 0.797326 | 0.565159 | 0.966598 | 1 | 0.999631 | 0.930124 |
| Cplx3_Q8R1B5_T113t_1 | -0.54 | 0.84 | 1.58 | 1.08 | 0.66 | 0.81 | 0.23 | 1.16 | 0.44 | 0.38 | 0.86 | 1.29 | 1.06 | 0.38 | 0.41 | 0.31 | -0.56 | -0.3525 | -0.69 | 0.674572 | 0.95753 | 0.995844 | 1 | 0.999631 | 0.885515 |
| Cpne1_B7ZCP4_T469t_1 | 1.25 | 0.08 | 0.9 | 0.83 | -0.31 | 0.15 | 0.96 | 0.02 | -0.14 | 1.12 | 0.21 | 0.46 | -0.04 | -0.19 | 0.32 | 0.21 | -0.0225 | -0.41 | 0.43 | 0.064469 | 0.162506 | 0.368301 | 1 | 0.999631 | 0.514485 |
| Cpne5_Q8JZW4_T138t_1 | -0.38 | -0.77 | -0.16 | 0.04 | -0.16 | -0.56 | -0.64 | 0 | -0.28 | -0.67 | -1.95 | -0.01 | 0.12 | -0.04 | 0.18 | 0.19 | -0.155 | -0.295 | 0.305 | 0.16652 | 0.939668 | 0.376119 | 1 | 0.999631 | 0.63587 |
| Cpne6_Q3UYN2_S330s_1 | -0.09 | -0.73 | -0.2 | 0.11 | -0.81 | -0.56 | -0.43 | 0.27 | -0.63 | 0.17 | -0.78 | -0.85 | 0.06 | -0.38 | 0.17 | 0.46 | -0.0075 | 0.22 | -0.3175 | 0.361618 | 0.64288 | 0.396263 | 1 | 0.999631 | 0.741948 |
| Cpne6_Q3UYN2_S553s_1 | 0.65 | -0.2 | -1.43 | -0.75 | -0.03 | -0.29 | -0.37 | -1.07 | 0.04 | 1.14 | -1.24 | -0.79 | -1.17 | 0.17 | -0.93 | -1.07 | 0.32 | 0.5725 | 0.575 | 0.538877 | 0.987251 | 0.7231 | 1 | 0.999631 | 0.830899 |
| Cpne6_Q3UYN2_S553s_2 | -1.23 | -0.77 | -0.26 | 0.1 | -0.46 | 0.61 | -0.33 | -0.7 | 0.52 | -0.05 | -0.44 | 0.1 | -0.33 | 1.03 | 0.18 | -0.74 | -0.325 | -0.2825 | 0.38 | 0.239142 | 0.441671 | 0.151786 | 1 | 0.999631 | 0.683719 |
| Cpne6_Q3UYN2_S555s_1 | -1.22 | -1.89 | -0.43 | -1.66 | -1.7 | -1.4 | -1.65 | -1.75 | -1.81 | -1.11 | -2.08 | -1.33 | -0.22 | -1.06 | -1.22 | -1.18 | -0.0275 | 0.1075 | 0.1525 | 0.370145 | 0.372541 | 0.487702 | 1 | 0.999631 | 0.746338 |
| Cpne6_Q3UYN2_S555s_2 | -0.44 | -0.82 | 0.39 | -0.39 | -0.38 | -0.41 | -0.29 | -0.29 | -0.4 | -0.33 | -0.19 | 0.09 | -0.34 | 0.09 | 0 | -0.4 | -0.6025 | -0.07 | -0.325 | 0.658786 | 0.929394 | 0.744416 | 1 | 0.999631 | 0.87993 |
| Cpne6_Q3UYN2_S555s_3 | 0.83 | 0.47 | 0.74 | 1.22 | -0.66 | 0.17 | 0.83 | 0.51 | 0.11 | 0.56 | 1.23 | 1.08 | 0.07 | 0.33 | 0.68 | 0.88 | -0.17 | 0.36 | 0.7075 | 0.323648 | 0.130602 | 0.838267 | 1 | 0.999631 | 0.724485 |
| Cpne6_Q3UYN2_S68s_1 | -0.5 | 0.04 | 0.32 | 0.11 | -0.55 | -0.06 | -0.18 | 0.08 | 0.18 | 0.15 | 0.21 | 0.87 | 0.79 | 0.91 | 0.56 | 0.54 | -0.12 | -0.1725 | 0.195 | 0.037896 | 0.56291 | 0.265071 | 1 | 0.999631 | 0.459387 |
| Cpne6_Q3UYN2_T547t_1 | -0.22 | -0.8 | 0.52 | 0.24 | -0.26 | -0.24 | -0.12 | -0.12 | -0.43 | -0.03 | -0.35 | -0.14 | -0.71 | 0.34 | 0.47 | 0.42 | 0.01 | 0.17 | 0.235 | 0.642299 | 0.718127 | 0.619472 | 1 | 0.999631 | 0.871335 |
| Cpne6_Q3UYN2_T547t_2 | -0.48 | -0.73 | 0.53 | -0.42 | -0.34 | -0.24 | -0.21 | -0.27 | -0.27 | -0.15 | -0.12 | 0.12 | -0.22 | 0.23 | 0.15 | -0.32 | -0.6025 | -0.07 | -0.325 | 0.51699 | 0.975326 | 0.615541 | 1 | 0.999631 | 0.819449 |
| Cpne6_Q3UYN2_T547t_3 | 0.83 | 0.47 | 0.74 | 1.22 | -0.66 | 0.17 | 0.83 | 0.51 | 0.11 | 0.56 | 1.23 | 1.08 | 0.07 | 0.33 | 0.68 | 0.88 | 0.09 | -0.015 | -0.87 | 0.323648 | 0.130602 | 0.838267 | 1 | 0.999631 | 0.724485 |
| Cpne6_Q3UYN2_T551t_2 | -0.6 | -1.5 | 0.49 | 0.73 | -0.77 | -0.88 | 0.01 | 1.12 | -0.61 | -1.75 | 0.13 | 1.29 | -1.35 | -0.15 | -1.58 | -1.28 | -0.6025 | -0.07 | -0.325 | 0.14461 | 0.884848 | 0.983534 | 1 | 0.999631 | 0.616037 |
| Cpne6_Q3UYN2_T551t_3 | 0.83 | 0.47 | 0.74 | 1.22 | -0.66 | 0.17 | 0.83 | 0.51 | 0.11 | 0.56 | 1.23 | 1.08 | 0.07 | 0.33 | 0.68 | 0.88 | 1.0225 | 1.4275 | 0.3175 | 0.323648 | 0.130602 | 0.838267 | 1 | 0.999631 | 0.724485 |
| Cpox_P36552_S99s_1 | -2.36 | -10.01 | -4.74 | -5.14 | -5.56 | -5.96 | -3.2 | -3.44 | -5.65 | -3.05 | -2.9 | -4.94 | -5.28 | -5.56 | -5.09 | -5.05 | -0.4725 | -0.0375 | -0.2475 | 0.814799 | 0.497373 | 0.348011 | 1 | 0.999631 | 0.937235 |
| Cpsf7_Q8BTV2_S48s_1 | 0.48 | 1.25 | 2.08 | 1.06 | 0.46 | 0.24 | 1 | 1.28 | 0.61 | 0.88 | 1.66 | 1.57 | 0.59 | 0.24 | 1.13 | 1.92 | -0.385 | 0.1525 | -0.1175 | 0.60801 | 0.264659 | 0.929144 | 1 | 0.999631 | 0.858527 |
| Cpsf7_Q8BTV2_T194t_1 | 0.24 | 1.68 | 1.82 | 0.75 | -0.16 | 1.37 | 0.64 | 1.1 | 0.59 | 1.11 | 1.79 | 1.61 | 1.27 | 0.44 | 0.27 | 2.04 | -0.1 | 0.82 | -0.27 | 0.822881 | 0.429418 | 0.737597 | 1 | 0.999631 | 0.938864 |
| Cracr2b_E9Q2F1_S224s_1 | 0.41 | 1.47 | 0.59 | -0.17 | 1.86 | 0.56 | 0.27 | -0.79 | 1.79 | 0.84 | 0.94 | 2.01 | 0.48 | -0.57 | 0.72 | 0.59 | -0.4 | -0.0225 | -0.0575 | 0.551626 | 0.863826 | 0.08973 | 1 | 0.999631 | 0.836275 |
| Crebbp_F8VPR5_S2350s_1 | 0.61 | 1.4 | 1.07 | 1 | -0.48 | 1.37 | 0.66 | 0.93 | 0.86 | 1.55 | 0.73 | 0.85 | 0.96 | 1.19 | 0.76 | 0.94 | -0.315 | -0.32 | -1.59 | 0.845973 | 0.35459 | 0.942577 | 1 | 0.999631 | 0.948565 |
| Creg2_Q8BGC9_S109s_2 | 0.54 | 0.1 | -0.24 | 0.05 | -0.83 | -1.06 | 0.6 | 0.48 | -0.7 | 0.37 | -0.46 | -0.04 | -1.72 | -2.41 | -0.54 | -1.24 | -0.315 | -0.32 | -1.59 | 0.00411 | 0.486261 | 0.350779 | 1 | 0.999631 | 0.351308 |
| Creg2_Q8BGC9_S111s_2 | 0.54 | 0.1 | -0.24 | 0.05 | -0.83 | -1.06 | 0.6 | 0.48 | -0.7 | 0.37 | -0.46 | -0.04 | -1.72 | -2.41 | -0.54 | -1.24 | 0.6575 | 1.035 | 0.6625 | 0.00411 | 0.486261 | 0.350779 | 1 | 0.999631 | 0.351308 |
| Crhr1_P35347_S386s_1 | -1.79 | -0.56 | -1.34 | -1.64 | 0.09 | -1.06 | -0.6 | -1.13 | 0.36 | -2.54 | -0.02 | 1.01 | 0.45 | -1.26 | -0.6 | -1.27 | 0.03 | 0.44 | 0.315 | 0.185546 | 0.122834 | 0.178573 | 1 | 0.999631 | 0.646224 |
| Crhr1_P35347_S396s_1 | 0.23 | 0.74 | -0.09 | 0.33 | 0.04 | 0.86 | 0 | 0.43 | 0.73 | 1.11 | 0.71 | 0.42 | 0.35 | 0.66 | 0.74 | 0.72 | -0.1075 | 0.5025 | 0.5775 | 0.307286 | 0.924138 | 0.164232 | 1 | 0.999631 | 0.717856 |
| Crhr1_P35347_S400s_1 | -1.63 | -1.43 | 0.1 | -1.08 | -2.25 | -1.03 | -0.58 | -0.61 | -1.52 | -0.57 | -0.26 | 0.32 | 0.15 | -1.08 | -0.29 | -0.51 | 0.0325 | 0.5825 | 0.4625 | 0.226959 | 0.833992 | 0.340546 | 1 | 0.999631 | 0.678092 |
| Crhr1_P35347_S405s_1 | -0.22 | 0.49 | 1.11 | 0.17 | 0.15 | 0.73 | 0.11 | 0.69 | 1.01 | 1.17 | 0.4 | 1.3 | 1.16 | 0.05 | 0.62 | 1.57 | 0.805 | -0.3575 | -0.61 | 0.305065 | 0.926874 | 0.140729 | 1 | 0.999631 | 0.716239 |
| Crip2_Q9DCT8_S115s_1 | 1.06 | 0.37 | -0.72 | 1.44 | 1.52 | 1.14 | 1.16 | 1.55 | 0.34 | -0.57 | 1.53 | -0.58 | -0.59 | 0.98 | -0.77 | 0.09 | -0.125 | -0.205 | -0.2275 | 0.298914 | 0.108756 | 0.569409 | 1 | 0.999631 | 0.712131 |
| Crk_Q8JZR2_S40s_1 | 2.12 | 2.59 | 0.87 | 0.28 | 3.24 | 1.29 | 0.19 | 0.64 | 3.09 | 0.29 | 0.51 | 1.15 | 1.34 | 2.49 | 0.64 | 0.48 | -0.1175 | -0.125 | 0.3275 | 0.720946 | 0.86848 | 0.781125 | 1 | 0.999631 | 0.905276 |
| Crkl_P47941_S107s_1 | -0.85 | -0.34 | -0.87 | -0.25 | -1.18 | -0.79 | -0.65 | -0.16 | -0.39 | -0.6 | -1.21 | -0.61 | -0.11 | -0.61 | -0.51 | 0.23 | -0.1025 | 0.1925 | 0.1675 | 0.331369 | 0.711846 | 0.687242 | 1 | 0.999631 | 0.727302 |
| Crmp1_P97427_S495s_1 | -0.57 | 0.45 | 0.49 | -0.36 | 0.07 | 0.05 | -0.04 | -0.48 | 0.02 | 0.14 | 0.5 | 0.12 | 0.32 | -0.11 | -0.05 | 0.52 | -0.35 | -0.145 | 0.18 | 0.644487 | 0.762376 | 0.573376 | 1 | 0.999631 | 0.87245 |
| Crmp1_P97427_S540s_1 | 0.29 | 1.06 | 2.43 | 2.16 | 0.47 | 0.43 | 0.88 | 2.76 | 0.2 | -0.17 | 3.25 | 2.08 | 1.47 | 0.98 | 2.05 | 2.16 | 0.5025 | 0.095 | 0.365 | 0.735846 | 0.598972 | 0.860909 | 1 | 0.999631 | 0.90759 |
| Crmp1_P97427_S542s_1 | 1.73 | 1.09 | 0.27 | 1.41 | 2.58 | 1.79 | 0.61 | 1.53 | 1.5 | 1.49 | 1.26 | 0.63 | 2.14 | 1.8 | 0.73 | 1.29 | 0.29 | -0.505 | -0.83 | 0.417109 | 0.313819 | 0.808149 | 1 | 0.999631 | 0.771432 |
| Crmp1_P97427_S570s_1 | 1.7 | 1.21 | 0.27 | 1.79 | 1.98 | 1.78 | 1 | 1.37 | 1.32 | 0.37 | 0.89 | 0.37 | -0.58 | 1.48 | 0.36 | 0.39 | -0.435 | -0.3475 | -0.82 | 0.132506 | 0.485914 | 0.247693 | 1 | 0.999631 | 0.60434 |
| Crmp1_P97427_S8s_1 | 0.47 | -1.32 | -0.95 | -0.21 | -1.56 | -1.24 | -0.21 | -0.74 | -1.46 | -0.08 | -0.95 | -0.91 | -2.65 | -1.16 | -1.22 | -0.26 | 0.035 | -0.24 | -0.055 | 0.180398 | 0.367511 | 0.468485 | 1 | 0.999631 | 0.64376 |
| Crmp1_P97427_T509t_1 | 0.26 | 1.24 | 0.99 | 0.62 | 0.5 | 1.7 | 0.12 | 0.93 | 0.01 | 0.6 | 1.03 | 0.51 | 0.09 | 1.58 | 0.42 | 0.8 | -0.865 | -0.95 | -1.865 | 0.892298 | 0.93087 | 0.489883 | 1 | 0.999631 | 0.964417 |
| Crmp1_Q6P1J1_T27t_1 | 2.66 | -0.16 | 2.5 | 1.32 | -0.26 | 1.18 | 1.77 | 0.17 | 1.02 | 1.7 | -0.53 | 0.33 | -0.93 | -0.95 | 0.39 | 0.35 | -0.715 | -0.5025 | -0.8975 | 0.018546 | 0.239731 | 0.206169 | 1 | 0.999631 | 0.391024 |
| Crocc_Q8CJ40_S1043s_1 | 0.99 | 1 | 2.1 | 0.71 | -0.5 | 0.51 | 1.47 | 0.46 | 0.16 | 0.19 | 1.4 | 1.04 | -0.95 | 0.26 | 1.38 | 0.52 | -0.675 | -0.6475 | -1.3675 | 0.117828 | 0.159403 | 0.264375 | 1 | 0.999631 | 0.585087 |
| Crocc_Q8CJ40_S1289s_1 | 2.42 | -0.64 | 0.83 | 0.65 | -0.51 | -0.63 | 2.47 | -0.77 | -1.08 | 1.7 | 0.28 | -0.23 | -1.24 | -0.92 | 0.29 | -0.34 | 0.17 | 0.215 | 0.3025 | 0.05495 | 0.443851 | 0.400839 | 1 | 0.999631 | 0.492066 |
| Crocc_Q8CJ40_S1304s_1 | 0.08 | 0.65 | -1.37 | -0.89 | -0.04 | 0.25 | -0.89 | -0.17 | -0.22 | 0.05 | -0.51 | 0.01 | 0.13 | 0.26 | -0.37 | -0.34 | -0.0925 | -0.25 | -0.085 | 0.527404 | 0.728415 | 0.643425 | 1 | 0.999631 | 0.824554 |
| Crocc_Q8CJ40_S1319s_1 | -0.33 | -0.16 | 0.21 | -0.52 | -0.47 | -0.07 | 0.01 | -0.64 | -0.86 | -0.58 | -0.49 | 0.13 | 0.42 | -0.17 | -0.89 | -0.5 | 0.5 | 0.5975 | 0.6 | 0.815712 | 0.751106 | 0.441892 | 1 | 0.999631 | 0.937576 |
| Crocc_Q8CJ40_S1404s_1 | 0.35 | 0.73 | 0.04 | -0.43 | 1.27 | 1.17 | 0.04 | 0.21 | 0.89 | 0.54 | 0.58 | 1.07 | 1.28 | 1.49 | 0.44 | -0.12 | -0.1475 | 0.13 | 0.315 | 0.200263 | 0.235151 | 0.094387 | 1 | 0.999631 | 0.658107 |
| Crocc_Q8CJ40_S337s_2 | 0.09 | -0.21 | 1.58 | 1.07 | 0.19 | -0.11 | 0.48 | 1.38 | 0.52 | 0.38 | 0.83 | 1.32 | 0.78 | 0.79 | 1.5 | 0.72 | 0.4875 | 0.6525 | 0.7725 | 0.493024 | 0.766257 | 0.775757 | 1 | 0.999631 | 0.80955 |
| Crocc_Q8CJ40_S337s_3 | -1.66 | 0.49 | 0.6 | 0.47 | 0.71 | 0.39 | 0.26 | 0.49 | 0.64 | 0.17 | 0.89 | 0.81 | 0.3 | 0.85 | 1.26 | 0.58 | -0.1475 | 0.13 | 0.315 | 0.177222 | 0.35582 | 0.238162 | 1 | 0.999631 | 0.642116 |
| Crocc_Q8CJ40_T330t_2 | 0.09 | -0.21 | 1.58 | 1.07 | 0.19 | -0.11 | 0.48 | 1.38 | 0.52 | 0.38 | 0.83 | 1.32 | 0.78 | 0.79 | 1.5 | 0.72 | 0.4875 | 0.6525 | 0.7725 | 0.493024 | 0.766257 | 0.775757 | 1 | 0.999631 | 0.80955 |
| Crocc_Q8CJ40_T330t_3 | -1.66 | 0.49 | 0.6 | 0.47 | 0.71 | 0.39 | 0.26 | 0.49 | 0.64 | 0.17 | 0.89 | 0.81 | 0.3 | 0.85 | 1.26 | 0.58 | 0.08 | 0.235 | 0.05 | 0.177222 | 0.35582 | 0.238162 | 1 | 0.999631 | 0.642116 |
| Cry2_Q9R194_S557s_1 | 0.28 | 1.18 | 1.28 | 1.29 | 0.95 | 1.33 | 0.89 | 1.18 | 0.89 | 0.9 | 1.23 | 1.95 | 1.24 | 1.03 | 1.44 | 0.52 | -0.2475 | -0.0425 | 0.135 | 0.890171 | 0.800033 | 0.533771 | 1 | 0.999631 | 0.9636 |
| Cs_Q9CZU6_S226s_1 | -0.42 | -0.85 | -0.71 | -0.95 | -1.24 | -1.12 | -0.77 | -0.79 | -1.65 | -0.87 | -0.47 | -0.11 | -0.03 | -1.01 | -1.02 | -0.33 | 0.9425 | 0.255 | 0.085 | 0.692055 | 0.360373 | 0.909686 | 1 | 0.999631 | 0.894017 |
| Csdc2_Q91YQ3_S48s_1 | 0.15 | -0.04 | 1.27 | 2.2 | 1.63 | 1.85 | 1.6 | 2.27 | 0.87 | -0.26 | 2.29 | 1.7 | -1.02 | 1.34 | 2.41 | 1.19 | 0.255 | -0.27 | 0.8525 | 0.912915 | 0.088701 | 0.70843 | 1 | 0.999631 | 0.97231 |
| Csde1_Q91W50_S123s_1 | -1.11 | 0.29 | 0.19 | -0.84 | 0.24 | 0.46 | -0.47 | -0.68 | -0.65 | -0.84 | -0.49 | -0.57 | 0.59 | 0.82 | 0.12 | 0.41 | 0.35 | -0.11 | -0.13 | 0.058149 | 0.563701 | 0.487401 | 1 | 0.999631 | 0.499755 |
| Csnk1a1_E9Q2U6_T349t_1 | 0.2 | -0.69 | -1.94 | -1.86 | -0.02 | 0.12 | -1.28 | -1.71 | -0.94 | -0.3 | -1.47 | -2.02 | 0.71 | -0.19 | -3.15 | -2.18 | 0.555 | -0.415 | -0.84 | 0.88297 | 0.573189 | 0.849027 | 1 | 0.999631 | 0.963505 |
| Csnk1d_Q9DC28_S331s_1 | 2.27 | 0.16 | 0.51 | 1.52 | 1.63 | 0.95 | 1.81 | 2.29 | 0.94 | 0.41 | 1.22 | 0.23 | -0.9 | 1.16 | 0.57 | 0.27 | 0.735 | 0.695 | 0.565 | 0.177406 | 0.297422 | 0.419124 | 1 | 0.999631 | 0.642116 |
| Csnk1d_Q9DC28_T347t_1 | -0.4 | -1.66 | -1.84 | -0.18 | -0.85 | -0.32 | -0.47 | 0.5 | -0.18 | -0.81 | -0.44 | 0.13 | -0.29 | -0.58 | -0.93 | -0.02 | 0.055 | -0.345 | -0.85 | 0.23918 | 0.152342 | 0.15277 | 1 | 0.999631 | 0.683719 |
| Csnk1e_Q9JMK2_S363s_1 | 0.24 | 0.12 | 1.13 | 0.85 | -0.25 | 0.91 | 1.51 | 0.39 | -0.03 | 0.5 | 0.2 | 0.29 | -0.59 | -0.76 | 0.68 | -0.39 | 0.455 | -0.02 | 0.185 | 0.065339 | 0.899487 | 0.300151 | 1 | 0.999631 | 0.516298 |
| Csnk1e_Q9JMK2_S389s_1 | 0.9 | -0.72 | -0.96 | 0.08 | 0.58 | 0.54 | 0.07 | -0.07 | 0.01 | -0.33 | -0.07 | -0.39 | 0.23 | -0.38 | 0.03 | 0.16 | 0.305 | -0.29 | -0.34 | 0.678519 | 0.315314 | 0.962998 | 1 | 0.999631 | 0.887183 |
| Csnk1g1_Q8BTH8_S361s_1 | 1.15 | 0.59 | -1.48 | 0.84 | 1.47 | -0.2 | 0.86 | 0.19 | 0.65 | -0.13 | 0.05 | -0.63 | -0.47 | 0.52 | 0.03 | -0.34 | -0.12 | -0.505 | -1.0375 | 0.563323 | 0.629601 | 0.62797 | 1 | 0.999631 | 0.841038 |
| Csnk1g1_Q8BTH8_T214t_1 | 1.14 | 1.03 | 1.3 | 0.45 | 0.51 | 1.71 | 0.84 | 0.38 | 1.14 | 1.19 | 0.26 | -0.69 | 0.1 | 1.15 | -0.43 | -1.05 | 0.115 | 0.1175 | 0.3775 | 0.054647 | 0.746451 | 0.293243 | 1 | 0.999631 | 0.492066 |
| Csnk2b_G3UXG7_S158s_1 | -3.14 | -4.33 | -2.57 | -2.77 | -2.68 | -4.79 | -2.19 | -2.69 | -3.51 | -3.85 | -2.54 | -2.44 | -2.11 | -3.85 | -2.32 | -3.02 | 0.0425 | 0.15 | 0.32 | 0.477795 | 0.854826 | 0.814491 | 1 | 0.999631 | 0.800237 |
| Cspg5_Q71M36_S467s_1 | -1.47 | -0.02 | 0.17 | -1.17 | -0.78 | -0.41 | -0.56 | -0.57 | -1.11 | -0.88 | 0.08 | 0.02 | 0.02 | -0.83 | -0.34 | -0.06 | 0.155 | -0.45 | 0.2525 | 0.483292 | 0.918254 | 0.758904 | 1 | 0.999631 | 0.803286 |
| Cspg5_Q71M36_S467s_1_1 | -1.56 | -0.06 | -0.45 | -1.69 | -0.92 | -0.66 | -0.65 | -0.91 | -1.45 | -1.13 | -1.51 | -1.47 | -0.05 | -0.35 | -1.46 | -0.89 | -0.2 | 0.035 | 0.4825 | 0.609519 | 0.706868 | 0.296596 | 1 | 0.999631 | 0.858923 |
| Cspg5_Q71M36_S475s_1 | -0.91 | -0.94 | -0.62 | -0.98 | -0.79 | -1.35 | -1.06 | -1.05 | -1.09 | -0.14 | -1.49 | -0.59 | 0.76 | -1.28 | -0.59 | -0.41 | -0.41 | -0.09 | 0.21 | 0.284434 | 0.439372 | 0.919507 | 1 | 0.999631 | 0.705468 |
| Cspg5_Q71M36_S475s_1_1 | -1.25 | -0.43 | -0.68 | -0.81 | -1.13 | -1.39 | -1.23 | -1.06 | -0.96 | -0.8 | -1.08 | -0.69 | -0.53 | -0.56 | -0.99 | -0.25 | 0.2025 | 0.015 | 0.0325 | 0.511917 | 0.157543 | 0.752056 | 1 | 0.999631 | 0.817687 |
| Cspg5_Q71M36_S541s_1 | -0.18 | -1.03 | -0.6 | -0.25 | 0.04 | -0.73 | -0.38 | -0.18 | -0.79 | -0.88 | -0.2 | -0.13 | -1 | -0.05 | -0.6 | -0.28 | 0.095 | 0.0275 | 0.1725 | 0.924868 | 0.517887 | 0.96357 | 1 | 0.999631 | 0.975422 |
| Cspg5_Q71M36_S543s_1 | -1.12 | -0.66 | -1.08 | -0.66 | -0.49 | -0.75 | -0.78 | -1.12 | -0.75 | -1.17 | -0.83 | -0.66 | -0.82 | -0.44 | -0.9 | -0.67 | -0.49 | 0.1675 | -0.085 | 0.551538 | 0.727328 | 0.919997 | 1 | 0.999631 | 0.836275 |
| Cspp1_B2RX88_S806s_1 | 0.88 | 0.89 | 0.12 | 0.39 | 0.33 | 0.02 | 0.11 | -0.14 | 1.37 | 1.59 | -0.42 | 0.41 | 0.31 | 0.79 | 0.09 | 0.75 | 0.0725 | 0.11 | 0.3325 | 0.796747 | 0.113233 | 0.728033 | 1 | 0.999631 | 0.929952 |
| Csrp1_P97315_S192s_1 | -0.82 | 0.25 | -0.33 | -0.38 | 0.12 | -0.05 | -0.7 | -0.36 | -0.44 | -0.48 | -0.29 | 0.37 | 0.39 | -0.07 | -0.25 | -0.02 | -0.3275 | -0.1825 | -0.4 | 0.327651 | 0.824839 | 0.747226 | 1 | 0.999631 | 0.725013 |
| Csrp1_P97315_S81s_1 | 0.12 | 0.67 | 0.59 | 1.4 | 0.05 | 0.31 | 0.79 | 0.32 | -0.28 | 0.15 | 1.2 | 0.98 | -1.15 | 0.25 | 0.69 | 1.39 | -0.43 | -0.155 | -0.1375 | 0.476795 | 0.348363 | 0.675635 | 1 | 0.999631 | 0.799947 |
| Cstf2_F6ZKC7_S145s_1 | -0.13 | 0.3 | 0.67 | 0.21 | -0.78 | 0.16 | -0.09 | 0.04 | 0.24 | -0.81 | 0.16 | 0.84 | -0.58 | -0.29 | 0.47 | 0.9 | -0.2625 | -0.5725 | -1.1075 | 0.732535 | 0.196101 | 0.695117 | 1 | 0.999631 | 0.907231 |
| Cstf3_Q99LI7_S691s_1 | 0.96 | 0.74 | 2.4 | 1.86 | 0.37 | 1.33 | 2.74 | 0.47 | 0.45 | 1.72 | 0.54 | 0.96 | 0.01 | -0.42 | 1.41 | 0.53 | -0.035 | 0.4525 | 0.3 | 0.056226 | 0.66629 | 0.240241 | 1 | 0.999631 | 0.495761 |
| Ctage5_H3BK44_S496s_1 | -0.07 | 0.94 | 0.45 | 0.51 | 0.65 | 0.27 | 0.08 | 0.69 | 1.17 | 0.46 | 1.12 | 0.89 | 0.45 | 0.91 | 0.7 | 0.97 | 0.375 | 0.065 | -0.79 | 0.36052 | 0.910034 | 0.18105 | 1 | 0.999631 | 0.741182 |
| Ctage5_H3BK44_S543s_1 | 0.96 | 0.5 | -0.81 | 1.36 | 1.77 | -0.31 | 1.25 | 0.8 | 1.15 | -0.4 | 1.36 | 0.16 | -1.04 | 0.28 | -0.46 | 0.07 | 0.3075 | 0.4675 | -0.06 | 0.154908 | 0.527004 | 0.910195 | 1 | 0.999631 | 0.6258 |
| Ctage5_H3BK44_S593s_1 | 0.26 | -0.21 | 0.89 | 0.24 | 0.49 | 0.86 | 0.08 | 0.98 | 0.97 | 0.77 | 0.75 | 0.56 | 0.46 | 0.35 | -0.68 | 0.81 | -0.0475 | -0.0475 | 0.1175 | 0.883773 | 0.375329 | 0.154026 | 1 | 0.999631 | 0.963505 |
| Ctage5_H3BK44_S636s_1 | -1.64 | 0.28 | -0.67 | -0.71 | -1.09 | -0.15 | -0.7 | -0.99 | -0.6 | -1.06 | -1.07 | -0.2 | -0.21 | -0.53 | -0.99 | -0.54 | -0.07 | -0.3575 | -0.2725 | 0.785533 | 0.913013 | 0.91388 | 1 | 0.999631 | 0.923063 |
| Ctdp1_Q7TSG2_S502s_1 | 0.03 | -0.21 | -1.56 | -0.6 | -0.1 | -0.71 | -1.18 | -0.63 | -0.68 | -0.34 | -1.52 | -1.23 | -1.13 | -0.37 | -0.99 | -0.94 | -0.0225 | -0.54 | -0.7625 | 0.508881 | 0.865345 | 0.420869 | 1 | 0.999631 | 0.816932 |
| Ctdspl2_Q8BG15_S9s_1 | 0.88 | 0.29 | -0.9 | -0.05 | 0.63 | -0.27 | 0.31 | -0.54 | -0.39 | -0.27 | 0.04 | -1.32 | -0.73 | -0.4 | -1.39 | -0.31 | -0.0225 | 0.16 | 0.7975 | 0.111504 | 0.959551 | 0.258211 | 1 | 0.999631 | 0.577068 |
| Ctif_Q6PEE2_S18s_1 | -0.22 | 1.68 | 1.09 | 0.58 | 0.83 | 1.07 | 0.08 | 1.06 | 1.13 | 0.84 | 0.92 | 0.88 | 1.8 | 1.79 | 1.31 | 1.42 | 0.0525 | 0.3775 | 0.3925 | 0.088014 | 0.960067 | 0.700071 | 1 | 0.999631 | 0.553858 |
| Ctnna2_E0CXB9_S275s_1 | -0.94 | -0.92 | -0.76 | -0.37 | -0.47 | -0.71 | -0.67 | -0.93 | -0.65 | -0.68 | -0.13 | -0.02 | 0.43 | -0.62 | -0.84 | -0.39 | -0.895 | 0.04 | 0.0075 | 0.288179 | 0.841945 | 0.222354 | 1 | 0.999631 | 0.706035 |
| Ctnna2_E0CXB9_S667s_1 | -2.1 | -3.9 | -4.45 | -5.68 | -3.92 | -3.93 | -4.88 | -6.98 | -3.31 | -2.58 | -4.71 | -5.37 | -0.4 | -3.87 | -6.17 | -5.66 | -0.405 | -0.54 | 0.6625 | 0.995277 | 0.330975 | 0.962648 | 1 | 0.999631 | 0.997705 |
| Ctnna2_E0CXB9_S667s_2 | -1.66 | -1.03 | 0.31 | -0.97 | -1.01 | -1.45 | -0.94 | -1.57 | -1.2 | -1.54 | -3.55 | 0.78 | -0.32 | -0.47 | 0.68 | -0.59 | 0.2775 | 0.3175 | 0.7525 | 0.195001 | 0.360591 | 0.531363 | 1 | 0.999631 | 0.652768 |
| Ctnna2_E0CXB9_S914s_1 | -1.2 | -0.59 | -0.09 | -1.12 | -0.42 | -0.43 | -0.79 | -0.25 | -0.62 | -0.49 | -0.26 | -0.36 | -0.17 | 0.28 | 0.26 | -0.36 | 1.07 | 1.9775 | 0.8775 | 0.056777 | 0.403856 | 0.341542 | 1 | 0.999631 | 0.496275 |
| Ctnna2_E0CXB9_S952s_1 | -1.96 | -0.25 | 0.2 | -1.37 | 0.87 | -0.71 | 0.98 | -0.24 | 1.69 | -1.42 | 1.57 | 2.69 | 2.52 | -1.25 | 0.25 | -1.39 | -0.35 | -0.1025 | 0.1875 | 0.337957 | 0.093621 | 0.04704 | 1 | 0.999631 | 0.862054 |
| Ctnna2_E0CXB9_T274t_1 | -1.14 | -0.64 | -0.04 | -0.4 | -0.46 | -0.6 | -0.7 | -1.86 | -0.93 | -0.65 | -0.81 | -0.24 | -1.1 | -0.32 | -0.32 | 0.09 | -0.51 | -1.5675 | 0.6725 | 0.625199 | 0.391655 | 0.755623 | 1 | 0.999631 | 0.863768 |
| Ctnna2_E0CXB9_T666t_2 | -1.58 | -1.03 | 0.91 | -0.55 | -0.81 | -1.26 | -0.55 | -1.67 | -1.01 | -1.7 | -7.34 | 1.53 | -0.21 | -0.36 | 1.33 | -0.32 | 0.32 | 0.4075 | 0.735 | 0.285649 | 0.35791 | 0.353806 | 1 | 0.999631 | 0.705851 |
| Ctnna2_E0CXB9_T670t_1 | -2.63 | -2.21 | -2.32 | -2.07 | -2.65 | -1.09 | -1.82 | -2.39 | -1.71 | -1.79 | -2.59 | -1.51 | -0.82 | -2.08 | -1.97 | -1.42 | -0.105 | 0.58 | 0.91 | 0.065078 | 0.407937 | 0.226265 | 1 | 0.999631 | 0.515898 |
| Ctnna2_E0CXB9_T670t_2 | -1.7 | -1.12 | -2.3 | -1.6 | -1.09 | -1.82 | -2.01 | -2.22 | -1.44 | -1.54 | -0.39 | -1.03 | -0.12 | -0.75 | -1.35 | -0.86 | -0.1425 | 0.67 | 0.365 | 0.036662 | 0.774382 | 0.149564 | 1 | 0.999631 | 0.454456 |
| Ctnnb1_E9Q6A9_S191s_1 | -1.66 | 0.17 | -1.18 | -1.95 | -0.68 | -1.34 | -1.53 | -1.64 | 0.36 | -0.35 | -1.56 | -0.39 | -0.36 | -0.42 | -1.32 | -1.06 | 0.12 | 0.2925 | 0.3525 | 0.47377 | 0.770242 | 0.254504 | 1 | 0.999631 | 0.799779 |

| | | | | | | | | | | | | | | | | | | | | | | | | | |
|---|---|---|---|---|---|---|---|---|---|---|---|---|---|---|---|---|---|---|---|---|---|---|---|---|---|
| Ctnnb1_Q02248_S552s_1 | -1.24 | 0.12 | 0.11 | 0.01 | -0.21 | -0.15 | -0.55 | 0.39 | -0.21 | -0.44 | 0.31 | 0.51 | 0.15 | -0.13 | 0.02 | 0.37 | -0.07 | 0.11 | 0.135 | 0.367139 | 0.759797 | 0.478528 | 1 | 0.999631 | 0.745796 |
| Ctnnb1_Q02248_S675s_1 | -0.42 | -0.03 | -0.41 | -0.99 | 0.09 | 0.12 | -1.67 | -0.67 | -0.21 | -0.19 | -0.3 | -0.71 | -0.68 | -0.01 | -0.39 | -0.23 | 0.1825 | 0.5425 | 0.3575 | 0.677401 | 0.876229 | 0.718696 | 1 | 0.999631 | 0.887183 |
| Ctnnb1_Q02248_T551t_1 | -1.3 | -0.39 | 0.75 | 0.75 | -0.47 | -0.33 | 0.06 | 1.28 | -0.5 | -0.77 | 1.53 | 1.72 | -0.36 | -0.48 | 1.02 | 1.06 | 0.0625 | 0.5525 | 0.7975 | 0.551632 | 0.752813 | 0.463354 | 1 | 0.999631 | 0.836275 |
| Ctnnbip1_Q9JJN6_S10s_1 | -0.99 | -0.83 | -0.81 | -0.86 | -0.93 | -0.54 | -0.83 | -0.94 | -0.87 | -0.73 | -0.02 | 0.34 | 0.19 | -0.17 | -0.66 | 0.34 | -0.255 | 0.2625 | 0.6025 | 0.027672 | 0.795713 | 0.126485 | 1 | 0.999631 | 0.423977 |
| Ctnnbl1_Q9CWL8_S545s_1 | -0.19 | 0.56 | 1.08 | 1 | 0.61 | 0.03 | 0.2 | 0.59 | 0.39 | 0.1 | 1.66 | 1.35 | 1.07 | 0.5 | 2.08 | 1.21 | -0.325 | -0.805 | -1.1775 | 0.192313 | 0.475626 | 0.572147 | 1 | 0.999631 | 0.648866 |
| Ctnnd1_P30999_S269s_1 | 2.58 | 1.28 | 0.03 | 1.7 | 2.12 | 0.67 | 1.44 | 0.06 | 1.08 | 0.8 | 0.29 | 0.2 | -0.22 | 0 | 0.16 | 0.94 | -0.3625 | -0.58 | -1.695 | 0.053356 | 0.604937 | 0.15401 | 1 | 0.999631 | 0.490856 |
| Ctnnd1_P30999_S312s_1 | 1.62 | 0.48 | 0.48 | 0.11 | -0.05 | 0.31 | 0.76 | 0.22 | -0.07 | 1.25 | -0.64 | -0.17 | -0.82 | -2.34 | 0.08 | -1.01 | 0.11 | 0.1875 | 0.2 | 0.011458 | 0.353328 | 0.25921 | 1 | 0.999631 | 0.385092 |
| Ctnnd1_P30999_S346s_1 | 0.44 | -0.06 | 0.24 | 0.47 | 0.45 | 0.91 | 0.04 | 0.13 | 0.78 | 0.9 | 0.18 | -0.02 | 0.43 | 0.27 | 0.27 | 0.92 | -0.31 | 0.205 | 0.2675 | 0.511219 | 0.711933 | 0.558086 | 1 | 0.999631 | 0.817687 |
| Ctnnd1_P30999_S349s_1 | -0.16 | 0.28 | 0.17 | 0.15 | -0.1 | -0.19 | -0.32 | -0.19 | 0.17 | 0.55 | 0.25 | 0.29 | -0.04 | 0.12 | 0.14 | 1.29 | -0.07 | 0.2525 | 0.3325 | 0.471433 | 0.219797 | 0.432812 | 1 | 0.999631 | 0.798132 |
| Ctnnd1_P30999_S47s_1 | -0.12 | -0.1 | -0.09 | -0.59 | -0.36 | 0.04 | -0.56 | -0.3 | -0.24 | 0.06 | -0.04 | 0.33 | -0.31 | 0.93 | -0.06 | -0.13 | -0.25 | -0.5425 | -1.75 | 0.362188 | 0.794543 | 0.368195 | 1 | 0.999631 | 0.742225 |
| Ctnnd1_P30999_T310t_1 | 1.61 | 0.18 | 0.04 | 0.29 | 0.13 | 0.74 | 0.61 | -0.36 | 0.5 | 1.07 | -0.62 | -1 | -0.73 | -0.93 | -1.27 | -1.95 | 0.455 | 0.135 | -0.04 | 0.002931 | 0.566521 | 0.342666 | 1 | 0.999631 | 0.351308 |
| Ctnnd2_B7ZNF6_S1054s_1 | 0.99 | 0.74 | 0.67 | 1.19 | 0.48 | 2.34 | 0.62 | 1.97 | 1.16 | 1.09 | 1.05 | 0.83 | -0.39 | 2.16 | 1.09 | 0.57 | -0.1475 | -0.545 | -0.6625 | 0.938181 | 0.339439 | 0.608544 | 1 | 0.999631 | 0.979905 |
| Ctnnd2_B7ZNF6_S1056s_1 | -0.62 | -0.68 | -0.88 | -1.24 | -1.01 | -0.63 | -1.35 | -1.02 | -1.67 | -0.25 | -1.41 | -2.27 | -0.88 | -0.86 | -2.05 | -2.28 | -0.11 | 0.875 | 0.945 | 0.13561 | 0.603659 | 0.235194 | 1 | 0.999631 | 0.607928 |
| Ctnnd2_B7ZNF6_S1061s_1 | -2.22 | -5.36 | -2.64 | -3.22 | -3.77 | -4.08 | -1.94 | -4.09 | -4.14 | -3.2 | -1.08 | -1.52 | -2.83 | -4.81 | -1.05 | -0.97 | -0.63 | -0.3025 | -1.7625 | 0.346241 | 0.884982 | 0.328298 | 1 | 0.999631 | 0.735175 |
| Ctnnd2_B7ZNF6_S1068s_1 | 1.41 | 0.07 | 1.02 | 1.19 | -0.75 | 0.36 | 0.78 | 0.78 | 0.62 | 1.68 | 0.05 | 0.13 | -1.06 | -0.16 | -1.26 | -0.88 | -0.5525 | -0.9225 | -2.7325 | 0.001358 | 0.183966 | 0.51862 | 1 | 0.999631 | 0.351308 |
| Ctnnd2_B7ZNF6_S1068s_2 | 2.15 | -0.04 | 1.3 | 1.18 | 0.19 | 0.48 | 1.32 | 0.39 | -0.33 | 1.4 | 0.07 | -0.24 | -2.51 | -0.44 | -1.42 | -1.97 | -0.4725 | -0.3725 | -1.2925 | 0.00084 | 0.272357 | 0.122791 | 1 | 0.999631 | 0.351308 |
| Ctnnd2_B7ZNF6_S1072s_1 | 0.45 | -1.62 | 0.04 | -0.68 | -2 | -0.73 | 0.31 | -1.28 | -1.55 | -0.17 | -0.45 | -1.13 | -1.43 | -2.3 | -1.5 | -1.75 | -0.5925 | -0.9525 | -2.705 | 0.021347 | 0.439463 | 0.478833 | 1 | 0.999631 | 0.403597 |
| Ctnnd2_B7ZNF6_S1072s_2 | 1.46 | -0.82 | 1.3 | 1.18 | -0.73 | -0.23 | 1.32 | 0.39 | -1.03 | 0.51 | 0.07 | -0.24 | -2.71 | -1.6 | -1.42 | -1.97 | 0.03 | -0.105 | -0.2 | 0.000729 | 0.353943 | 0.122772 | 1 | 0.999631 | 0.351308 |
| Ctnnd2_B7ZNF6_S1123s_1 | -1.75 | -0.61 | -0.18 | 0.07 | -0.85 | -0.87 | -0.18 | -0.45 | -1.37 | -0.94 | -0.56 | -0.02 | -1.49 | -0.61 | -0.69 | -0.48 | -0.0275 | -0.4875 | -0.3575 | 0.663899 | 0.944131 | 0.825452 | 1 | 0.999631 | 0.881993 |
| Ctnnd2_B7ZNF6_S1216s_1 | -0.79 | -0.17 | 0.51 | -0.63 | -0.29 | -0.04 | -0.19 | -0.67 | -0.91 | -1.11 | -1.16 | 0.15 | -1.55 | 1.09 | -1.48 | -0.57 | 0.5225 | -0.295 | -1.3675 | 0.565183 | 0.937344 | 0.267887 | 1 | 0.999631 | 0.84139 |
| Ctnnd2_B7ZNF6_S200s_1 | 1.2 | -0.42 | 0.83 | 1.14 | 0.5 | 0.79 | 1.92 | 1.63 | 0.14 | -0.2 | 1.32 | 0.31 | -0.91 | -0.48 | -1 | -0.33 | -0.4525 | 0.2475 | 0.15 | 0.008506 | 0.291331 | 0.542618 | 1 | 0.999631 | 0.365244 |
| Ctnnd2_B7ZNF6_S263s_1 | -0.14 | -0.1 | -0.02 | -0.45 | -0.86 | -0.68 | -0.65 | -0.33 | -0.17 | 0.52 | -0.05 | -0.02 | 0.99 | 0.19 | -0.87 | -0.42 | 0.0475 | -0.0375 | -0.2425 | 0.724615 | 0.102035 | 0.384662 | 1 | 0.999631 | 0.905674 |
| Ctnnd2_B7ZNF6_S263s_2 | -0.19 | -0.64 | -0.33 | -0.69 | -0.29 | -0.48 | -0.63 | -0.26 | -0.53 | -0.33 | -0.6 | -0.54 | -0.9 | -0.04 | -0.91 | -0.97 | -0.185 | 0.14 | -0.055 | 0.46644 | 0.853681 | 0.885356 | 1 | 0.999631 | 0.795267 |
| Ctnnd2_B7ZNF6_S272s_1 | -0.6 | -0.91 | -1.15 | -1.31 | -1.46 | -0.59 | -1.05 | -1.61 | -1.12 | -0.38 | -0.98 | -0.93 | -0.05 | -0.63 | -2.69 | -0.82 | 0.0475 | -0.0375 | -0.2425 | 0.920386 | 0.569429 | 0.642866 | 1 | 0.999631 | 0.975032 |
| Ctnnd2_B7ZNF6_S272s_2 | -0.19 | -0.64 | -0.33 | -0.69 | -0.29 | -0.48 | -0.63 | -0.26 | -0.53 | -0.33 | -0.6 | -0.54 | -0.9 | -0.04 | -0.91 | -0.97 | 0.135 | -0.045 | -0.6625 | 0.46644 | 0.853681 | 0.885356 | 1 | 0.999631 | 0.795267 |
| Ctnnd2_B7ZNF6_S281s_1 | 0.65 | -0.02 | -1.37 | -0.69 | 0.23 | -0.5 | -0.31 | -0.31 | -0.11 | -0.13 | -0.65 | -0.72 | -0.62 | -0.4 | -1.97 | -1.09 | 0.2 | 0.625 | 0.0675 | 0.225577 | 0.763176 | 0.920988 | 1 | 0.999631 | 0.676382 |
| Ctnnd2_B7ZNF6_S311s_1 | -2.76 | -0.19 | -0.25 | -1.5 | -1 | -0.69 | -0.77 | -1.44 | -0.59 | -1.39 | -0.25 | 0.03 | -0.25 | -0.78 | -1.39 | -2.01 | 0.3125 | 0.18 | -0.0725 | 0.916589 | 0.728479 | 0.32511 | 1 | 0.999631 | 0.973495 |
| Ctnnd2_B7ZNF6_S323s_1 | -1.32 | 0.32 | -0.01 | -1.31 | -0.27 | -0.34 | 0.07 | -0.53 | -0.15 | -1.15 | 0.36 | -0.66 | 0.13 | -0.39 | -0.89 | -1.46 | 0.225 | 0.15 | -0.0975 | 0.888781 | 0.480262 | 0.724247 | 1 | 0.999631 | 0.963505 |
| Ctnnd2_B7ZNF6_S32s_1 | -1.28 | -0.82 | -1.31 | -1.44 | -0.76 | -1.07 | -1.02 | -1.1 | -1.35 | -1.43 | -0.87 | -0.6 | -0.92 | -1 | -1.85 | -1.47 | -0.005 | -0.14 | -0.1125 | 0.767907 | 0.397208 | 0.628758 | 1 | 0.999631 | 0.917335 |
| Ctnnd2_B7ZNF6_S394s_1 | 0.6 | -1.04 | -1.28 | -0.9 | -0.77 | -0.3 | -1.19 | -0.38 | -0.46 | -0.39 | -1.16 | -1.17 | -0.93 | 0.05 | -1.36 | -0.83 | 0.145 | 0.1075 | -0.2725 | 0.821082 | 0.991217 | 0.762653 | 1 | 0.999631 | 0.938464 |
| Ctnnd2_B7ZNF6_S411s_1 | -0.79 | -0.21 | -1 | -0.65 | -0.43 | -0.41 | -0.78 | -0.45 | -0.52 | -0.46 | -0.75 | -0.49 | -1.02 | -0.26 | -1.4 | -1.06 | 0.0125 | -0.6075 | -0.915 | 0.443536 | 0.600855 | 0.700594 | 1 | 0.999631 | 0.782783 |
| Ctnnd2_B7ZNF6_S434s_1 | 0.26 | -0.46 | -2.97 | -1.77 | 0.23 | -0.98 | -1.76 | -2.38 | -0.91 | -0.59 | -2.96 | -2.91 | -1.6 | -1.53 | -2.91 | -2.56 | 0.105 | -0.7625 | -1.45 | 0.21298 | 0.987439 | 0.473441 | 1 | 0.999631 | 0.666787 |
| Ctnnd2_B7ZNF6_S436s_1 | 0.91 | -0.38 | -1.29 | -0.19 | 0.8 | -0.48 | 0.04 | -0.89 | -0.92 | -1.19 | -0.96 | -0.93 | -2.18 | -0.56 | -2.46 | -1.55 | 0.255 | -0.0025 | -0.1025 | 0.027271 | 0.844564 | 0.113807 | 1 | 0.999631 | 0.423171 |
| Ctnnd2_B7ZNF6_S457s_1 | -0.51 | -0.15 | -1.09 | -0.62 | 0.08 | -0.21 | -0.57 | -0.65 | -0.3 | -0.66 | -0.71 | -0.71 | -0.14 | -0.15 | -1.32 | -1.17 | 0.5325 | -0.0675 | -0.2625 | 0.797198 | 0.421052 | 0.993276 | 1 | 0.999631 | 0.930124 |
| Ctnnd2_B7ZNF6_S470s_1 | 0.25 | 0.24 | -0.27 | 0.49 | 1.24 | 0.56 | 0.26 | 0.78 | 0.31 | -0.46 | -0.23 | 0.82 | -0.47 | 0.44 | -0.3 | -0.01 | 0.0875 | 0.04 | -0.03 | 0.433396 | 0.114226 | 0.851633 | 1 | 0.999631 | 0.779548 |
| Ctnnd2_B7ZNF6_S486s_1 | 0.1 | -0.01 | -1.05 | -0.39 | -0.31 | 0.2 | -0.29 | -0.6 | 0.06 | -0.26 | -0.06 | -0.93 | 0.22 | 0.08 | -0.94 | -0.83 | -0.0775 | 0.1725 | -0.235 | 0.942199 | 0.798295 | 0.913645 | 1 | 0.999631 | 0.981629 |
| Ctnnd2_B7ZNF6_S528s_1 | -2.97 | -6.52 | -1.82 | -2.26 | -4.94 | -4.49 | -1.93 | -2.52 | -5.28 | -3.51 | -1.67 | -2.42 | -4.59 | -4.62 | -2.6 | -2.7 | -0.325 | -0.2625 | 0.085 | 0.819889 | 0.944303 | 0.879408 | 1 | 0.999631 | 0.938303 |
| Ctnnd2_B7ZNF6_S530s_1 | -1.04 | -1.61 | -0.65 | -1.49 | -1.23 | -1.84 | -1.56 | -1.46 | -1.3 | -1.58 | -1.61 | -1.35 | -0.33 | -2.11 | -0.96 | -1.05 | -0.525 | -0.57 | -0.24 | 0.845238 | 0.307597 | 0.399128 | 1 | 0.999631 | 0.948565 |
| Ctnnd2_B7ZNF6_S533s_1 | -1.97 | -2.86 | -1.18 | -2.68 | -2.85 | -3.08 | -2.41 | -2.45 | -3.2 | -2.59 | -2.57 | -2.61 | -2.24 | -2.55 | -2.71 | -2.15 | 0.52 | 0.135 | -0.1575 | 0.569556 | 0.223878 | 0.192833 | 1 | 0.999631 | 0.84338 |
| Ctnnd2_B7ZNF6_S7s_1 | 0.3 | -0.83 | -1.3 | 0.15 | 0.1 | -1.16 | 0.88 | 0.58 | -0.84 | -1.42 | 0.03 | 1.09 | -0.97 | -0.53 | 0.25 | -1.06 | -0.0325 | -0.1075 | 0.1675 | 0.741195 | 0.351554 | 0.824401 | 1 | 0.999631 | 0.909055 |
| Ctnnd2_B7ZNF6_T1052t_1 | -0.63 | -1.87 | -0.25 | -0.29 | -2.21 | -0.99 | -0.51 | 0.54 | -1.66 | -1.04 | -0.36 | -0.41 | -1.26 | -1.28 | -0.29 | 0.46 | -0.04 | 0.015 | -0.665 | 0.753224 | 0.957917 | 0.819567 | 1 | 0.999631 | 0.913377 |
| Ctnnd2_B7ZNF6_T468t_1 | 0.84 | 1.19 | -0.47 | -0.09 | 0.32 | 0.49 | 0.11 | 0.39 | 0.56 | 0.66 | -0.01 | 0.32 | 0.25 | -0.32 | -0.56 | -0.56 | 0.145 | 1.12 | 0.26 | 0.151527 | 0.920426 | 0.97146 | 1 | 0.999631 | 0.62505 |
| Ctps1_P70698_S562s_1 | -1.49 | -8.04 | -3.79 | -3.94 | -4.45 | -5.84 | -3.44 | -2.95 | -5.74 | -3.25 | -1.52 | -2.27 | -3.22 | -4.72 | -3.51 | -4.77 | 0.0375 | 0.4625 | 0.0275 | 0.828832 | 0.909823 | 0.431541 | 1 | 0.999631 | 0.941583 |
| Ctr9_Q62018_S925s_1 | 0.33 | 1.25 | 1.36 | 1.31 | 0.56 | 1.81 | 0.97 | 1.06 | 1.95 | 1.89 | 0.86 | 1.4 | 0.81 | 1.16 | 1.48 | 0.91 | 0.0075 | -0.005 | 0.2225 | 0.936764 | 0.920115 | 0.238031 | 1 | 0.999631 | 0.97939 |
| Cttn_Q921L6_S380s_1 | -0.26 | -0.57 | -0.6 | -0.71 | -0.29 | 0.05 | -1.04 | -0.83 | -0.3 | -0.5 | -0.8 | -0.56 | 0.14 | -0.31 | -0.51 | -0.57 | -0.0425 | -0.035 | 0.2675 | 0.460519 | 0.981115 | 0.984799 | 1 | 0.999631 | 0.791962 |
| Cttn_Q921L6_S380s_2 | -0.75 | -0.27 | -0.42 | -0.87 | -0.49 | -0.39 | -1.1 | -0.5 | -0.68 | -0.6 | -0.73 | -0.44 | -0.19 | -0.45 | -0.49 | -0.11 | -0.04 | 0.015 | 0.1675 | 0.365973 | 0.882548 | 0.895856 | 1 | 0.999631 | 0.745796 |
| Cttn_Q921L6_S381s_1 | -0.35 | -0.32 | -0.81 | -1.07 | -0.27 | -0.09 | -1.21 | -1.14 | -0.47 | -0.14 | -1.06 | -0.82 | -0.05 | -0.18 | -0.79 | -0.86 | 0.66 | -0.1425 | -0.3375 | 0.625085 | 0.912492 | 0.963337 | 1 | 0.999631 | 0.863768 |
| Cttn_Q921L6_T286t_1 | 0.41 | -1.38 | -1.77 | 1.37 | 0.81 | -0.74 | 1.08 | 0.12 | -0.65 | -1.97 | 0.87 | -0.19 | -0.97 | -0.68 | -1.33 | 0.26 | -0.61 | 0.065 | 0.6525 | 0.641466 | 0.385002 | 0.863487 | 1 | 0.999631 | 0.871335 |
| Cttn_Q921L6_T364t_1 | -1.86 | -0.82 | -0.21 | -1.96 | -2.12 | -1.3 | -2.3 | -1.57 | -2.06 | -0.72 | -1.37 | -0.44 | 0.98 | -1.22 | -1.29 | -0.71 | -0.01 | -0.0475 | 0.29 | 0.302146 | 0.207244 | 0.90071 | 1 | 0.999631 | 0.714978 |
| Cttn_Q921L6_T364t_2 | -0.73 | -0.21 | -0.47 | -1.07 | -0.49 | -0.38 | -1.11 | -0.54 | -0.66 | -0.6 | -0.8 | -0.61 | -0.17 | -0.43 | -0.52 | -0.2 | -0.7675 | -0.75 | -1.615 | 0.351366 | 0.973823 | 0.866738 | 1 | 0.999631 | 0.738077 |
| Cttnbp2_B9EJA2_S1529s_1 | 1.73 | -1.24 | 0.53 | 0.32 | -0.74 | -0.59 | 0.38 | -0.78 | -1.04 | 1.04 | -0.59 | -1.07 | -3.23 | -0.97 | 0.12 | -1.04 | 0.5125 | -0.0325 | -0.1575 | 0.069606 | 0.22301 | 0.299223 | 1 | 0.999631 | 0.526681 |
| Cttnbp2_B9EJA2_S367s_1 | -1.27 | -1.6 | -2.27 | -2.46 | -0.97 | -1.62 | -1.98 | -0.98 | -2.05 | -1.49 | -2.58 | -1.61 | -1.61 | -1.59 | -3.39 | -1.64 | -0.0025 | -0.0275 | 0.155 | 0.754107 | 0.20414 | 0.933568 | 1 | 0.999631 | 0.913377 |
| Cttnbp2_B9EJA2_S465s_1 | -0.81 | -0.02 | -0.09 | -0.26 | 0.03 | -0.59 | -0.32 | -0.31 | -0.5 | -0.62 | 0.47 | -0.64 | 0.32 | -0.95 | 0.3 | -0.23 | 0.3875 | -0.325 | -0.7925 | 0.686564 | 0.993129 | 0.938398 | 1 | 0.999631 | 0.890911 |
| Cttnbp2_B9EJA2_S472s_1 | 0.74 | 0.88 | 1.23 | 1.31 | 1.08 | 1.69 | 1.94 | 1 | 1.5 | 0.13 | 1.1 | 0.13 | -0.71 | 0.88 | 1.41 | -0.59 | -0.045 | -0.32 | -0.15 | 0.149187 | 0.240492 | 0.413811 | 1 | 0.999631 | 0.621356 |
| Cttnbp2_B9EJA2_T15t_1 | 1.01 | -0.22 | 0.04 | 0.83 | 1.11 | 0.16 | 0.01 | 0.2 | 0.45 | 0.27 | -0.17 | -0.17 | 0.14 | -0.03 | 0.9 | 0.05 | 0.2525 | 0.6475 | 0.6825 | 0.704964 | 0.909406 | 0.392708 | 1 | 0.999631 | 0.897793 |
| Cttnbp2nl_Q99LJ0_S477s_1 | -1.05 | -1.18 | -0.61 | -0.22 | -0.54 | -0.87 | -0.77 | 0.13 | -0.6 | -1.13 | -0.01 | 1.27 | -0.66 | -0.88 | 0.61 | 0.6 | 0.1125 | -0.2175 | 0.04 | 0.154734 | 0.471361 | 0.236175 | 1 | 0.999631 | 0.6258 |
| Cttnbp2nl_Q99LJ0_S481s_1 | -0.9 | -1.07 | -0.24 | -0.81 | -0.28 | -1.21 | -0.6 | -0.48 | -0.84 | -1.12 | -1.19 | -0.74 | -0.23 | -0.63 | -0.84 | -1.16 | -0.2775 | -0.0025 | -0.055 | 0.904882 | 0.724579 | 0.464476 | 1 | 0.999631 | 0.968908 |
| Ctu1_Q99J10_S378s_1 | 0.23 | -0.99 | 1.29 | 0.15 | -0.48 | 0.02 | -0.17 | 0.2 | -0.66 | 0.3 | 0.1 | 0.93 | -0.72 | -0.04 | 0.8 | 0.42 | -0.1725 | 0.65 | 0.68 | 0.917903 | 0.555373 | 0.996246 | 1 | 0.999631 | 0.97388 |
| Cul3_F6UY44_S80s_1 | 0.27 | 0.78 | 0.33 | 0.72 | 0.26 | 0.7 | -0.06 | 0.51 | 1.28 | 1.4 | 1.16 | 0.86 | 1.25 | 1.83 | 1.05 | 0.69 | -0.015 | -0.08 | 0.3875 | 0.067732 | 0.549935 | 0.038437 | 1 | 0.999631 | 0.521762 |
| Cul4b_J3QJX0_S126s_1 | 0.11 | 0.63 | -0.25 | -0.54 | -0.06 | 0.83 | -0.39 | -0.49 | 0.31 | 0.05 | -0.55 | -0.18 | 1.37 | 0.24 | -0.38 | 0.27 | -0.62 | -0.3225 | -0.7975 | 0.392437 | 0.969868 | 0.819425 | 1 | 0.999631 | 0.758901 |
| Cul4b_J3QJX0_T32t_1 | 0.6 | 1.12 | 1.34 | 0.42 | 0.75 | 0.19 | 0.09 | -0.03 | 1.3 | -0.02 | 0.08 | 0.83 | -0.22 | 0.1 | 0.12 | 0.29 | -0.6075 | -0.0675 | 0.3375 | 0.030494 | 0.079028 | 0.423384 | 1 | 0.999631 | 0.438141 |
| Cul9_E9QP09_S1983s_1 | -0.38 | 1.23 | -0.63 | -0.48 | -0.58 | 0.47 | -1.35 | -1.23 | 0.3 | 0.3 | -0.47 | -0.66 | 1.11 | 0.01 | 0 | -0.03 | -0.215 | 0.245 | 0.365 | 0.501369 | 0.286753 | 0.888602 | 1 | 0.999631 | 0.814092 |
| Cul9_E9QP09_S2440s_1 | -0.47 | -0.75 | -0.19 | -0.59 | -0.6 | -0.87 | -0.95 | -0.44 | -0.68 | 0.05 | -0.33 | -0.06 | -0.04 | -0.24 | -0.22 | -0.04 | -0.2925 | 0.34 | 0.415 | 0.205474 | 0.425879 | 0.404969 | 1 | 0.999631 | 0.6605 |
| Cul9_E9QP09_S584s_1 | -0.31 | -0.02 | 0.39 | -0.59 | -0.35 | -0.14 | -0.94 | -0.27 | 0.02 | 0.39 | -0.07 | 0.49 | 0.51 | 0.1 | 0.11 | 0.41 | 0.7025 | 1.4475 | -0.525 | 0.212397 | 0.373273 | 0.294383 | 1 | 0.999631 | 0.666508 |
| Cul9_E9QP09_S592s_1 | 0.7 | 0.79 | 3.01 | 0.25 | 2.05 | 0.29 | 4.39 | 0.83 | 3.08 | 0.41 | 2.93 | 4.12 | 1.94 | -0.41 | 1.91 | -0.79 | -0.665 | -0.095 | -1.015 | 0.532533 | 0.466118 | 0.121997 | 1 | 0.999631 | 0.8277 |
| Cwc22_B1AYU7_S39s_1 | 1.37 | -1.16 | 0.59 | 1 | -0.31 | -0.76 | 0.96 | -0.75 | 0.76 | 1.31 | -0.46 | -0.19 | 0.5 | -0.8 | -0.17 | -1.79 | -0.245 | 0.2375 | 0.0475 | 0.149199 | 0.299119 | 0.879797 | 1 | 0.999631 | 0.621356 |
| Cwc22_B1AYU7_S61s_1 | 0.61 | 0.94 | 1.23 | 0.71 | 0.22 | 1.03 | 0.54 | 0.72 | 0.96 | 1.26 | 0.93 | 1.29 | 0.51 | 1.07 | 0.76 | 1.34 | -0.4425 | 0.115 | -0.325 | 0.880281 | 0.407143 | 0.394459 | 1 | 0.999631 | 0.963151 |
| Cwc22_B1AYU7_S824s_1 | 0.95 | 0.57 | 1.22 | 1.06 | -0.21 | 0.99 | 0.86 | 0.39 | 1.07 | 1.55 | 1.11 | 0.53 | 0.97 | 0.25 | 1.04 | 0.24 | 0.1575 | 0.2125 | 0.775 | 0.339493 | 0.21305 | 0.715379 | 1 | 0.999631 | 0.732185 |
| Cx3cl1_O35188_S262s_1 | -1.2 | 0.11 | -0.09 | -0.96 | -0.43 | -0.15 | -0.81 | -0.12 | 0.06 | -1.22 | 0.06 | -0.19 | -0.17 | 0.08 | 0.28 | 0.77 | -0.145 | -0.39 | 0.36 | 0.077253 | 0.675632 | 0.63 | 1 | 0.999631 | 0.542683 |
| Cx3cl1_O35188_S384s_1 | -0.02 | 1.13 | -0.28 | 0.1 | 0.3 | 0.14 | 0.04 | -0.13 | 0.36 | -0.35 | -0.04 | -0.6 | 0.6 | 0.41 | 0.45 | 0.91 | -0.2275 | 0.6325 | 0.7825 | 0.344309 | 0.680783 | 0.329868 | 1 | 0.999631 | 0.733953 |
| Cxadr_P97792_S332s_1 | -0.66 | -0.96 | -0.46 | -0.32 | -1.14 | -0.94 | -0.57 | -0.66 | 0.14 | 0.22 | -0.11 | -0.12 | 0.66 | -0.21 | -0.46 | 0.74 | -0.015 | 0.1275 | -0.145 | 0.058826 | 0.418326 | 0.039926 | 1 | 0.999631 | 0.501907 |
| Cyb561_Q60720_S246s_2 | -0.83 | -1.69 | -0.76 | -0.77 | -1.36 | -1.11 | -0.91 | -0.73 | -0.64 | -0.59 | -1.07 | -1.24 | -2.39 | -1.45 | -0.46 | -0.33 | -0.015 | 0.1275 | -0.145 | 0.77505 | 0.96201 | 0.699965 | 1 | 0.999631 | 0.920998 |
| Cyb561_Q60720_S248s_2 | -0.83 | -1.69 | -0.76 | -0.77 | -1.36 | -1.11 | -0.91 | -0.73 | -0.64 | -0.59 | -1.07 | -1.24 | -2.39 | -1.45 | -0.46 | -0.33 | -0.005 | -0.0525 | 0.3075 | 0.77505 | 0.96201 | 0.699965 | 1 | 0.999631 | 0.920998 |
| Cyb5b_Q9CQX2_S33s_1 | -1.14 | -0.98 | -1.57 | -1.13 | -0.97 | -1.16 | -1.09 | -1.62 | -1.08 | -1.45 | -1.25 | -1.25 | -0.48 | -0.86 | -0.99 | -1.26 | -0.225 | -0.3675 | -0.7 | 0.326636 | 0.98554 | 0.843135 | 1 | 0.999631 | 0.724678 |
| Cyb5b_Q9CQX2_T3t_1 | 0.89 | -0.98 | -0.95 | -1 | -0.97 | -0.58 | -0.29 | -1.1 | -1.04 | 0.36 | -1.54 | -1.29 | -0.78 | -1.25 | -1.07 | -1.74 | -0.425 | -0.085 | -0.2825 | 0.175081 | 0.638747 | 0.52989 | 1 | 0.999631 | 0.642116 |
| Cyfip1_Q7TMB8_S1233s_1 | 0.27 | 0 | -0.97 | -1.37 | -1.31 | -0.41 | -0.71 | -1.34 | -0.2 | -0.07 | -1.3 | -0.84 | 0.09 | -1.05 | -1.58 | -0.66 | -0.1525 | -0.2575 | 0.33 | 0.576379 | 0.345997 | 0.855907 | 1 | 0.999631 | 0.846204 |
| Cyfip1_Q7TMB8_S872s_1 | -0.8 | -0.59 | -0.6 | -0.68 | -0.68 | -0.48 | -0.89 | -1.23 | -1.6 | -0.41 | -1.16 | -0.53 | -0.02 | -0.58 | -1.11 | 0.36 | 0.15 | -0.075 | 0.0125 | 0.381092 | 0.568891 | 0.446705 | 1 | 0.999631 | 0.752619 |
| Cyfip1_Q7TMB8_T1234t_1 | -0.46 | -0.73 | -0.89 | -0.97 | -1.05 | -0.56 | 0.03 | -0.87 | -0.69 | -0.61 | -1.27 | -0.78 | -0.39 | -0.84 | -1.08 | -0.69 | -0.9425 | -1.065 | -1.69 | 0.96621 | 0.635086 | 0.790184 | 1 | 0.999631 | 0.988334 |
| Cyld_Z4YJJ5_S344s_1 | 2.52 | 0.22 | 1.25 | 0.96 | -0.23 | 0.33 | 1.34 | -0.26 | -0.06 | 1.46 | 0.4 | -1.11 | -0.32 | -1.03 | -0.13 | -0.33 | -0.2375 | -0.0575 | -0.01 | 0.006496 | 0.115253 | 0.123655 | 1 | 0.999631 | 0.351545 |
| Cyld_Z4YJJ5_S397s_1 | -1.53 | -0.94 | -0.61 | -0.89 | -1.46 | -1.2 | -1.01 | -1.25 | -1.48 | -1.02 | -1.12 | -0.58 | -0.35 | -1.41 | -1.07 | -1.18 | -0.15 | -0.0375 | 0.375 | 0.977436 | 0.417869 | 0.859233 | 1 | 0.999631 | 0.992174 |
| Cyld_Z4YJJ5_S398s_1 | -0.9 | 0.01 | -0.01 | -0.29 | -0.25 | -0.35 | -0.64 | -0.55 | -0.19 | -0.57 | -0.18 | -0.4 | 0.14 | 0.24 | 0.01 | -0.08 | 0.2975 | 0.035 | 0.03 | 0.249037 | 0.615049 | 0.901875 | 1 | 0.999631 | 0.686327 |
| Cys1_Q8R4T1_S116s_1 | 0.54 | 0.09 | -1.06 | 0.03 | 0.45 | 0.78 | 0.5 | -0.94 | 0.28 | 0.32 | 0.62 | -1.48 | 1.29 | 0.77 | -0.72 | -1.62 | -0.1175 | -0.8625 | -1.455 | 0.964267 | 0.544098 | 0.948698 | 1 | 0.999631 | 0.987331 |
| Cys1_Q8R4T1_T55t_1 | 2.54 | -1.16 | -0.93 | -0.09 | 0.01 | 0.55 | 0.6 | -1.27 | -0.1 | 0.73 | -1.97 | -1.75 | -1.37 | -0.51 | -1.36 | -2.22 | 0.28 | -0.3775 | -0.08 | 0.093227 | 0.887394 | 0.363904 | 1 | 0.999631 | 0.558128 |
| D17Wsu92e_B2KF52_S215s_1 | 0.21 | 0.13 | -0.86 | 0.6 | 1.29 | -0.17 | 0.05 | 0.03 | 0.6 | -0.79 | -0.75 | -0.49 | -0.49 | 0.76 | 0.11 | -0.62 | 0.4425 | 0.215 | 0.5 | 0.857229 | 0.531555 | 0.404357 | 1 | 0.999631 | 0.952021 |
| D17Wsu92e_B2KF52_S265s_1 | -0.86 | 0.17 | -0.54 | -1.09 | 0.07 | 0.3 | -0.66 | -0.26 | 0.13 | -0.83 | -0.61 | -0.15 | 0.58 | -0.1 | -0.06 | -0.74 | -0.505 | -0.0675 | 0.3625 | 0.235828 | 0.244014 | 0.569617 | 1 | 0.999631 | 0.683301 |
| D2Wsu81e_Q3UHX9_S338s_1 | -0.02 | -0.74 | 0.37 | 0.01 | -0.22 | -0.95 | -0.37 | -0.86 | -0.06 | 0.18 | -1.11 | 0.34 | 0.51 | -0.19 | 0.31 | 0.44 | 0.085 | 0.0825 | 0.47 | 0.305206 | 0.152981 | 0.868899 | 1 | 0.999631 | 0.716239 |
| D430041D05Rik_F6ZGR6_S1044s_1 | -0.31 | -0.49 | 0.46 | -0.01 | -0.5 | -0.55 | -0.16 | 1.2 | -0.76 | -0.04 | 0.97 | -0.19 | -0.07 | -0.2 | -0.03 | 1.83 | 0.1625 | 0.3375 | 0.4925 | 0.361128 | 0.848717 | 0.844079 | 1 | 0.999631 | 0.74139 |
| D430041D05Rik_F6ZGR6_S1070s_1 | -0.39 | 1.2 | 0.25 | 0.66 | 0.87 | 0.81 | -0.1 | 0.79 | 1 | 0.16 | 1 | 0.91 | 0.36 | 0.45 | 1.42 | 1.46 | 0.0175 | 0.12 | 0.26 | 0.28771 | 0.691703 | 0.411479 | 1 | 0.999631 | 0.705972 |
| D430041D05Rik_F6ZGR6_S1088s_1 | -0.75 | -0.28 | -0.57 | -0.85 | -0.42 | -0.69 | -0.68 | -0.59 | -0.4 | -0.78 | -0.54 | -0.25 | 0.59 | -0.38 | -1.18 | -0.44 | 0.2375 | -0.3775 | -0.395 | 0.52552 | 0.945045 | 0.66243 | 1 | 0.999631 | 0.823288 |
| D430041D05Rik_F6ZGR6_S1104s_1 | 1.19 | 1.13 | 0.28 | 1.21 | 1.37 | 1.11 | 0.73 | 1.55 | 0.54 | 0.27 | 1.21 | 0.28 | -0.41 | 0.67 | 0.59 | 1.38 | -0.1975 | -1.015 | -0.9475 | 0.375753 | 0.476239 | 0.299077 | 1 | 0.999631 | 0.748307 |
| D430041D05Rik_F6ZGR6_S1108s_1 | 1.66 | 1.52 | -0.05 | 0.22 | 0.66 | 1.13 | 1.16 | -0.39 | 0.18 | 0.03 | -0.37 | -0.55 | -0.85 | 0.77 | -0.26 | -0.1 | -0.445 | -0.3925 | -1.185 | 0.093574 | 0.708573 | 0.048721 | 1 | 0.999631 | 0.558128 |
| D430041D05Rik_F6ZGR6_S1226s_1 | -1.25 | -4.14 | -2.13 | -1.85 | -3.72 | -2.96 | -0.61 | -3.86 | -3.66 | -2.7 | -2.1 | -2.48 | -3.97 | -4.33 | -2.78 | -3.03 | -0.165 | -1.0275 | -1.61 | 0.093842 | 0.60343 | 0.543328 | 1 | 0.999631 | 0.558128 |
| D430041D05Rik_F6ZGR6_S726s_1 | 1.3 | 1.66 | -0.02 | 0.21 | 0.96 | 0.55 | 0.45 | 0.53 | -0.02 | -0.51 | -0.35 | -0.08 | -1.93 | 0.87 | -0.89 | -1.34 | 0.0275 | -0.31 | -0.6325 | 0.031353 | 0.695816 | 0.034846 | 1 | 0.999631 | 0.441911 |
| D430041D05Rik_F6ZGR6_S762s_1 | 0.21 | -0.25 | -1.29 | -0.85 | -0.47 | 0.07 | 0.08 | -1.75 | -0.49 | 0.16 | -1.32 | -1.77 | 0.23 | -1.03 | -1.67 | -2.24 | 0.1 | -0.04 | 0.4225 | 0.286196 | 0.956678 | 0.547893 | 1 | 0.999631 | 0.705972 |
| D430041D05Rik_F6ZGR6_S785s_1 | -0.8 | -0.51 | -0.78 | -0.86 | -0.62 | -0.43 | -0.85 | -0.65 | -0.81 | -0.91 | -0.72 | -0.67 | 0.14 | -0.38 | -0.69 | -0.33 | -0.26 | -0.2725 | -0.3075 | 0.176989 | 0.68497 | 0.871591 | 1 | 0.999631 | 0.642116 |
| D430041D05Rik_F6ZGR6_S927s_1 | -1.27 | 0.32 | -0.15 | -0.16 | -0.78 | -0.56 | -0.76 | -0.2 | -0.53 | -0.56 | -0.22 | -1.04 | -0.82 | -0.8 | -0.7 | -0.17 | 0.0225 | -0.5425 | 0.5975 | 0.445378 | 0.496751 | 0.493739 | 1 | 0.999631 | 0.784262 |
| D430041D05Rik_F6ZGR6_S993s_2 | -0.5 | 1.51 | 1.09 | -0.59 | 0.35 | 0.66 | 0.45 | 0.14 | -0.1 | -0.18 | -0.26 | -0.12 | 1.33 | 0.32 | 2.01 | 0.24 | -0.12 | 0.5975 | 0.6575 | 0.348056 | 0.964976 | 0.302409 | 1 | 0.999631 | 0.736697 |
| D430041D05Rik_F6ZGR6_S994s_1 | -0.66 | 0.34 | -0.49 | -2.14 | -0.27 | -0.39 | -0.56 | -2.21 | 0.06 | -0.85 | 0.59 | -0.36 | 1.03 | 0.02 | -0.15 | -1.22 | 0.615 | 0.4075 | 0.4425 | 0.306871 | 0.846181 | 0.296623 | 1 | 0.999631 | 0.717667 |
| D430041D05Rik_F6ZGR6_S994s_2 | -0.5 | 1.51 | 0.12 | -0.05 | 0.35 | 0.66 | 0.72 | 1.81 | -0.1 | -0.18 | 1.64 | 1.35 | 1.33 | 0.32 | -0.37 | 1.57 | -0.69 | -0.61 | -1.965 | 0.449003 | 0.240383 | 0.494212 | 1 | 0.999631 | 0.786796 |
| D430042O09Rik_F6XQI1_T1128t_1 | 2.98 | 0.51 | 2.02 | 1.49 | 0.1 | 1.6 | 1.9 | 0.64 | 0.84 | 3.3 | 0.23 | 0.19 | -0.74 | 0.17 | 0.02 | -0.31 | 0.16 | 0.01 | 0.2525 | 0.00361 | 0.266662 | 0.446847 | 1 | 0.999631 | 0.351308 |
| D630045J12Rik_D3YTS3_S1197s_1 | 0.6 | 0.45 | -0.03 | 0.98 | 0.89 | 0.83 | 0.15 | 0.77 | 0.47 | 0.34 | 0.41 | 0.82 | -0.43 | 1 | 1.11 | 1.33 | 0.9825 | 1.0425 | 1.305 | 0.577838 | 0.618312 | 0.973889 | 1 | 0.999631 | 0.84622 |
| D630045J12Rik_D3YTS3_S1303s_1 | -1.46 | -1.8 | -0.6 | -2.1 | -0.12 | -0.05 | -0.69 | -1.17 | -0.44 | -0.19 | -0.31 | -0.85 | -0.35 | 0.82 | -0.74 | -0.47 | 0.35 | -0.255 | 0.165 | 0.017568 | 0.037582 | 0.019293 | 1 | 0.999631 | 0.391024 |
| D630045J12Rik_D3YTS3_S1429s_1 | -0.14 | 0.28 | 0.19 | -0.04 | 0.43 | 0.42 | 0.16 | 0.68 | -0.37 | -0.09 | 0.41 | -0.68 | 0.39 | 0.45 | -0.41 | 0.52 | -0.06 | -0.105 | 0.265 | 0.609467 | 0.192466 | 0.426048 | 1 | 0.999631 | 0.858923 |
| D630045J12Rik_D3YTS3_S1587s_1 | -0.15 | 0.16 | -0.25 | -0.43 | 0.29 | -0.39 | -0.68 | -0.13 | -0.03 | -0.72 | -0.02 | -0.32 | 0.7 | -0.15 | -0.16 | 0 | 0.16 | -0.02 | 0.025 | 0.417329 | 0.842429 | 0.718959 | 1 | 0.999631 | 0.771463 |
| D630045J12Rik_D3YTS3_S1799s_1 | -0.12 | -0.53 | 0.45 | -0.21 | -0.11 | 0.66 | 0.14 | -0.46 | 0.31 | -0.27 | 0.06 | -0.59 | 0.9 | -0.51 | -0.78 | 0.08 | 0.6325 | 0.2625 | 0.8 | 0.953797 | 0.643859 | 0.952085 | 1 | 0.999631 | 0.984441 |
| Dab2ip_F6RK07_S1062s_1 | 0.46 | 0.36 | -0.77 | -0.22 | 1 | 1.43 | -0.55 | 0.48 | 0.99 | 0.51 | -0.22 | -0.4 | 0.93 | 1.38 | 0.49 | 0.23 | 0.6025 | -0.365 | -0.2975 | 0.070777 | 0.212312 | 0.545734 | 1 | 0.999631 | 0.530826 |
| Dab2ip_F6RK07_S641s_1 | 1.07 | 0.42 | -1.21 | 1.03 | 1.9 | -0.08 | 0.93 | 0.97 | 0.53 | -0.61 | 0.25 | -0.32 | -0.11 | 0.47 | -0.33 | 0.09 | 0.1525 | 0.135 | 0.6825 | 0.575638 | 0.331588 | 0.51195 | 1 | 0.999631 | 0.846085 |
| Dab2ip_F6RK07_S657s_1 | -0.83 | -0.22 | -0.4 | -0.53 | -0.33 | -0.35 | -0.61 | -0.08 | -0.3 | -0.89 | -0.2 | -0.05 | 0.6 | 0.02 | -0.11 | 0.24 | 0.775 | 0.0675 | -0.6175 | 0.044966 | 0.56925 | 0.654716 | 1 | 0.999631 | 0.475465 |
| Dab2ip_F6RK07_S789s_1 | 0.73 | 0.54 | -0.36 | 1.28 | 1.48 | 1.35 | 1.34 | 1.12 | 0.74 | 0.53 | 0.78 | 0.41 | -1.38 | 0.74 | 0.16 | 0.2 | 0.015 | 0.1775 | 0.39 | 0.264121 | 0.058237 | 0.857581 | 1 | 0.999631 | 0.6935 |
| Dab2ip_F6RK07_S872s_1 | -0.22 | 0.22 | -0.21 | -0.22 | -0.15 | 0.09 | -0.4 | 0.09 | 0.06 | 0.29 | 0.08 | -0.15 | 0.43 | 0.29 | -0.14 | 0.55 | 0.655 | -0.2475 | -0.4375 | 0.208152 | 0.954354 | 0.504505 | 1 | 0.999631 | 0.661649 |
| Dab2ip_F6RK07_S889s_1 | 0.37 | -2.83 | -1.69 | -0.4 | -0.02 | -1.08 | -0.24 | -0.59 | -1.25 | -1.54 | -1.04 | -1.71 | -1.77 | -1.65 | -1.22 | -1.66 | -1.1625 | -0.6225 | -0.4825 | 0.502082 | 0.335101 | 0.703995 | 1 | 0.999631 | 0.814092 |
| Dact1_D3Z5V0_S492s_1 | 0.04 | -0.49 | 0.35 | 0.72 | -1.39 | -2.5 | 0.3 | -0.44 | -1.28 | -0.5 | 0.64 | -0.73 | -0.47 | -0.79 | 0.33 | -0.38 | 0.7675 | 0.875 | 1.35 | 0.226217 | 0.075187 | 0.200701 | 1 | 0.999631 | 0.677133 |
| Dact3_Q0PHV7_S10s_1 | -3.74 | -0.06 | -1.58 | -3.15 | -0.87 | -0.41 | -2.1 | -2.08 | -0.77 | -1.23 | -1.46 | -1.57 | 1.59 | -0.72 | -1.74 | -2.26 | 0.21 | 0.4625 | 0.685 | 0.203728 | 0.356588 | 0.259302 | 1 | 0.999631 | 0.659781 |
| Dact3_Q0PHV7_S165s_1 | -0.43 | -0.04 | 0.57 | 0.86 | 0.09 | 0.35 | 0.55 | 0.81 | -0.05 | -0.03 | 1.31 | 1.58 | 0.23 | 0.73 | 1.78 | 0.96 | 0.82 | -0.925 | 0.3225 | 0.142211 | 0.558132 | 0.362145 | 1 | 0.999631 | 0.612726 |
| Dact3_Q0PHV7_S217s_1 | 1.64 | 0.56 | -1.76 | -0.74 | 2.14 | 2 | -0.14 | -1.02 | 0.26 | -0.49 | -1.83 | -1.94 | 0.84 | 1.07 | -0.73 | -0.19 | -0.125 | -0.0025 | 0.22 | 0.669307 | 0.389853 | 0.266097 | 1 | 0.999631 | 0.883599 |
| Dact3_Q0PHV7_S237s_1 | 0.05 | -0.04 | 0.99 | 0.45 | 0.06 | 0.36 | 0.09 | 0.44 | 0 | 0.95 | -0.07 | 0.56 | -0.3 | 0.24 | 1.41 | 0.98 | -0.445 | -0.4175 | -0.935 | 0.624858 | 0.687147 | 0.994554 | 1 | 0.999631 | 0.863768 |
| Dact3_Q0PHV7_S268s_1 | 1.87 | -0.57 | 0.49 | 0.41 | -1.21 | 0.33 | 1.62 | -0.32 | -0.89 | 1.66 | 0.25 | -0.49 | -0.53 | -0.29 | 0.33 | -1.05 | -0.12 | -0.0175 | 0.64 | 0.107799 | 0.52354 | 0.539476 | 1 | 0.999631 | 0.569416 |
| Dact3_Q0PHV7_S290s_1 | -0.32 | -2.84 | -0.35 | -0.57 | -1.32 | -1.07 | -0.95 | -1.22 | -1.15 | -0.13 | -2.06 | -0.81 | -0.67 | -0.15 | -0.03 | -0.67 | -0.395 | 0.0775 | -0.8425 | 0.285429 | 0.83091 | 0.978677 | 1 | 0.999631 | 0.705851 |
| Dact3_Q0PHV7_S370s_1 | 1.63 | 0.55 | 0.01 | 1.29 | 0.83 | -0.71 | 0.31 | 1.47 | 0.84 | 1.38 | 0.84 | 0.73 | -0.49 | 0.24 | 0.49 | -0.13 | 0.085 | 0.19 | -0.345 | 0.074307 | 0.471629 | 0.847784 | 1 | 0.999631 | 0.539897 |
| Dact3_Q0PHV7_S370s_2 | 0.97 | 1.52 | 1.85 | 1.63 | 1.16 | 2.15 | 1.18 | 1.82 | 1.6 | 2.35 | 1.51 | 1.27 | 0.72 | 1.08 | 1.5 | 1.29 | -0.285 | 0.355 | 0.7725 | 0.305329 | 0.803923 | 0.582316 | 1 | 0.999631 | 0.716239 |
| Dact3_Q0PHV7_S374s_1 | -2.85 | -7.12 | -3.81 | -5.14 | -6.3 | -5.31 | -3.86 | -4.59 | -7.27 | -3.14 | -2.91 | -4.18 | -2.28 | -5.31 | -3.67 | -4.57 | 0.085 | 0.19 | -0.345 | 0.433128 | 0.756482 | 0.761879 | 1 | 0.999631 | 0.779401 |
| Dact3_Q0PHV7_S374s_2 | 0.97 | 1.52 | 1.85 | 1.63 | 1.16 | 2.15 | 1.18 | 1.82 | 1.6 | 2.35 | 1.51 | 1.27 | 0.72 | 1.08 | 1.5 | 1.29 | 0.0925 | 0.1925 | 0.355 | 0.305329 | 0.803923 | 0.582316 | 1 | 0.999631 | 0.716239 |
| Dact3_Q0PHV7_S409s_1 | 0.46 | 1.44 | 2.12 | 1.88 | 1.45 | 1.24 | 1.27 | 2.31 | 1.69 | 1.18 | 2.34 | 1.46 | 0.58 | 1.85 | 2.27 | 2.62 | -0.0425 | 0.3 | 0.2275 | 0.51518 | 0.83154 | 0.661811 | 1 | 0.999631 | 0.819449 |
| Dact3_Q0PHV7_S6s_1 | 0.74 | 1.37 | 1.03 | 0.97 | 1.36 | 0.82 | 0.56 | 1.2 | 1.29 | 1.04 | 1.47 | 1.51 | 0.65 | 1.34 | 0.79 | 2.24 | 0.4175 | -0.7025 | -0.15 | 0.577161 | 0.884485 | 0.288394 | 1 | 0.999631 | 0.84622 |
| Dag1_Q62165_T788t_1 | 0.5 | 0.01 | -1.11 | -0.26 | 0.37 | 1.14 | -0.71 | 0.01 | -0.51 | -0.23 | 0.04 | -2.97 | -0.26 | 1.26 | -0.95 | -1.51 | -0.93 | -1.345 | -1.7975 | 0.809536 | 0.397564 | 0.320742 | 1 | 0.999631 | 0.936616 |
| Dagla_Q6WQJ1_S1032s_1 | 0.5 | -2.42 | -1.53 | -0.77 | -3.35 | -2.28 | 0.3 | -2.61 | -2.83 | -0.8 | -2.93 | -3.04 | -2.99 | -3.01 | -2.96 | -2.45 | 0.3025 | -0.6825 | -0.6625 | 0.010754 | 0.301225 | 0.088574 | 1 | 0.999631 | 0.383871 |
| Dagla_Q6WQJ1_S744s_1 | 0.29 | -0.14 | -1.97 | 0.14 | 0.75 | -0.93 | -0.31 | 0.02 | -0.36 | -1.15 | -1.09 | -1.81 | -2.28 | -0.31 | -1.37 | -0.37 | 0.08 | -0.495 | -0.3075 | 0.30926 | 0.60063 | 0.238782 | 1 | 0.999631 | 0.718845 |
| Dagla_Q6WQJ1_S954s_1 | 0.58 | 0.92 | -0.62 | 0.52 | 0.69 | 0.57 | 0.07 | 0.39 | -0.01 | -0.44 | 0.24 | -0.37 | -0.46 | 0.14 | -0.35 | 0.84 | -0.2725 | -0.385 | -0.73 | 0.49701 | 0.831793 | 0.222439 | 1 | 0.999631 | 0.811348 |
| Dap_Q91XC8_S2s_1 | 1.75 | 0.31 | 1.15 | 0.55 | -0.01 | 0.95 | 1.48 | 0.25 | 0.46 | 1.05 | 0.52 | 0.19 | 0.07 | 0.04 | 0.89 | -0.16 | -0.2725 | -0.385 | -0.73 | 0.102197 | 0.549722 | 0.333456 | 1 | 0.999631 | 0.564448 |
| Dap_Q91XC8_S3s_1 | 1.75 | 0.31 | 1.15 | 0.55 | -0.01 | 0.95 | 1.48 | 0.25 | 0.46 | 1.05 | 0.52 | 0.19 | 0.07 | 0.04 | 0.89 | -0.16 | 0.405 | 0.6475 | -0.25 | 0.102197 | 0.549722 | 0.333456 | 1 | 0.999631 | 0.564448 |
| Dapk1_Q80YE7_S326s_1 | -1.27 | 0.97 | 1.46 | -0.46 | 0.02 | 0.77 | 1.97 | -0.44 | 0.82 | 0.96 | 0.61 | 0.9 | 1.05 | 0.36 | -1.1 | -0.61 | 0.23 | 0.31 | 0.66 | 0.723023 | 0.578968 | 0.28183 | 1 | 0.999631 | 0.905674 |
| Dapk2_Q8VDF3_S349s_1 | -1.01 | 0.76 | -0.49 | -0.14 | 0.21 | 0.83 | -0.29 | -0.71 | 0.68 | 0 | 0 | -0.32 | 0.5 | 0.63 | 0.52 | 0.11 | -0.155 | 0.0075 | 0.58 | 0.130078 | 0.629638 | 0.47502 | 1 | 0.999631 | 0.601275 |
| Dars_Q922B2_S249s_1 | -0.37 | 0.47 | 0.3 | -0.5 | -0.4 | -0.11 | -0.2 | -0.01 | -0.46 | -0.47 | 0.55 | 0.31 | -0.03 | 0.17 | 0.78 | 1.3 | -0.3 | -0.44 | 0.5725 | 0.175619 | 0.619167 | 0.984205 | 1 | 0.999631 | 0.642116 |
| Daxx_O35613_S515s_1 | 0.44 | 0.9 | 0.12 | 0.31 | -0.04 | 0.3 | -0.2 | 0.51 | -0.16 | 0.7 | -0.39 | -0.14 | 1.63 | 0.63 | 0.36 | 1.44 | 0.25 | 0.77 | 0.4925 | 0.15817 | 0.325348 | 0.213126 | 1 | 0.999631 | 0.628615 |
| Dbi_M0QWU8_S2s_1 | -0.97 | -1.05 | 3.21 | 1.97 | 0.9 | -0.01 | 1.27 | 2 | 0.3 | 1.92 | 1.88 | 2.14 | 0.88 | -0.3 | 1.31 | 3.24 | -0.0175 | 0.0875 | 0.6575 | 0.656843 | 0.800514 | 0.445838 | 1 | 0.999631 | 0.879056 |
| Dbi_M0QWU8_Y29y_1 | -0.44 | -0.1 | -0.06 | -0.43 | -0.11 | 0.11 | -0.56 | -0.54 | 0.14 | -0.39 | -0.59 | 0.16 | 0.74 | 0.67 | 0.35 | -0.16 | -0.125 | 0.36 | 0.5275 | 0.060268 | 0.94976 | 0.767706 | 1 | 0.999631 | 0.505962 |
| Dbn1_F7CPL2_S78s_1 | 0.28 | 0.05 | -0.06 | 0.22 | -0.06 | 0.19 | -0.02 | -0.12 | 0.51 | 0.69 | 0.38 | 0.35 | 0.83 | 0.8 | 0.41 | 0.56 | 0.025 | 0.2475 | 0.255 | 0.07857 | 0.607193 | 0.17541 | 1 | 0.999631 | 0.544927 |
| Dbn1_F7CPL2_S79s_1 | -0.03 | -0.24 | -0.14 | -0.04 | -0.22 | 0.04 | -0.21 | 0.04 | -0.04 | 0.23 | 0.16 | 0.19 | -0.26 | 0.29 | 0.17 | 0.37 | -0.255 | 0.0725 | -0.2325 | 0.375907 | 0.916122 | 0.322767 | 1 | 0.999631 | 0.748307 |
| Dbn1_Q9QXS6_S241s_1 | 0.16 | 1.22 | 0.09 | 0.34 | 0.65 | -0.15 | 0.42 | -0.13 | 0.86 | -0.09 | 0.3 | 1.03 | 1.33 | -0.29 | 0.02 | -0.18 | -0.4825 | -0.49 | -1.3975 | 0.611132 | 0.479139 | 0.851029 | 1 | 0.999631 | 0.859461 |
| Dbn1_Q9QXS6_S344s_1 | 0.05 | -1.18 | -0.22 | -0.46 | -1.19 | -1.88 | 0.14 | -0.81 | -0.75 | -0.33 | -1.68 | -1.01 | -2.02 | -2.23 | -1.32 | -1.83 | 0.0825 | -0.0025 | 0.13 | 0.003474 | 0.321128 | 0.238562 | 1 | 0.999631 | 0.351308 |
| Dbn1_Q9QXS6_S385s_1 | 0.23 | 0.16 | -1.1 | -1.01 | 0.05 | 0.22 | -0.77 | -0.89 | -0.04 | -0.64 | -0.45 | -0.6 | 0.19 | 0.27 | -0.78 | -0.88 | 0.415 | 0.28 | 0.465 | 0.780645 | 0.852817 | 0.995001 | 1 | 0.999631 | 0.921426 |
| Dbn1_Q9QXS6_S387s_1 | -1.38 | -0.92 | -1.14 | -1.06 | -0.96 | -0.98 | -0.52 | -0.38 | -1.28 | -1.21 | -0.59 | -0.3 | -0.42 | -0.58 | -0.64 | -1 | 0.0125 | 0.1425 | 0.5 | 0.125994 | 0.149769 | 0.388996 | 1 | 0.999631 | 0.596745 |
| Dbn1_Q9QXS6_S658s_1 | -0.69 | -0.75 | -0.48 | -0.55 | -0.4 | -0.48 | -0.77 | -0.77 | -0.46 | -0.66 | -0.69 | -0.09 | 0.22 | -0.13 | -0.52 | -0.04 | -0.2825 | 0.0525 | -0.1925 | 0.106881 | 0.959313 | 0.595712 | 1 | 0.999631 | 0.568797 |
| Dbn1_Q9QXS6_T394t_1 | -0.28 | 0.18 | -1.05 | -0.46 | -0.88 | -0.6 | -0.93 | -0.33 | -0.29 | -0.06 | -0.4 | -0.65 | 0.01 | -1.25 | -0.88 | -0.26 | 0.95 | 0.6 | -0.5375 | 0.636408 | 0.402186 | 0.87463 | 1 | 0.999631 | 0.869568 |
| Dbndd1_H3BK17_S6s_1 | 1.23 | -0.79 | -0.85 | 1.03 | 1.65 | 1.15 | 1.01 | 0.61 | 1.32 | 1.34 | 0.31 | 0.05 | -2.09 | 0.91 | -0.04 | -0.31 | 0.3725 | -0.055 | 0.21 | 0.47465 | 0.111085 | 0.329384 | 1 | 0.999631 | 0.799827 |
| Dbnl_Q62418_S141s_1 | 0.47 | -0.69 | -1.21 | -1.72 | -0.16 | -0.36 | -0.41 | -0.73 | -0.33 | -0.11 | -1.73 | -1.2 | -0.09 | -0.49 | -0.71 | -1.02 | -0.255 | -0.78 | -1.5725 | 0.669491 | 0.429113 | 0.921253 | 1 | 0.999631 | 0.883626 |
| Dbnl_Q62418_S235s_1 | 2.08 | 0.5 | 1.13 | 1.23 | 0.77 | 1.08 | 1.19 | 0.88 | 0.78 | 1.72 | -0.08 | -0.6 | -1.03 | -0.54 | 0.65 | -0.43 | 0.425 | -0.67 | -0.8225 | 0.007111 | 0.486107 | 0.183856 | 1 | 0.999631 | 0.358173 |
| Dbnl_Q62418_S277s_1 | 2.13 | 0.74 | 0.46 | 1.48 | 1.42 | 1.98 | 1.54 | 1.57 | 0.93 | 0.32 | 1.39 | -0.51 | -1.2 | 1.86 | -0.04 | 0.9 | 0.4825 | -0.605 | -0.6525 | 0.239809 | 0.302641 | 0.219662 | 1 | 0.999631 | 0.683719 |
| Dbnl_Q62418_T299t_1 | 1.91 | 1.53 | 0.77 | 2.15 | 1.48 | 2.69 | 2 | 2.12 | 1.37 | 0.53 | 2.44 | -0.4 | 0.41 | 2.27 | 0.36 | 0.71 | -0.3875 | 0.21 | 0.3025 | 0.223707 | 0.24192 | 0.336638 | 1 | 0.999631 | 0.67471 |
| Dbp_Q60925_S82s_1 | -0.62 | -0.3 | 0.17 | -0.31 | -0.3 | -0.81 | -0.92 | -0.58 | -0.34 | -0.34 | -0.25 | 0.71 | 0.13 | -0.14 | -0.16 | 0.32 | -1.01 | 0.53 | -2.1575 | 0.329677 | 0.198198 | 0.547057 | 1 | 0.999631 | 0.726674 |
| Dbr1_Q923B1_S505s_1 | 3.41 | 1.37 | 2.78 | 1.45 | 1.21 | 1.44 | 2.33 | -0.01 | 4.57 | 3.11 | 2.03 | 1.42 | 1.37 | -0.07 | 0.95 | -1.87 | -0.33 | -0.8575 | -0.615 | 0.017636 | 0.131011 | 0.48812 | 1 | 0.999631 | 0.391024 |
| Dbr1_Q923B1_S520s_1 | 0.91 | 2.35 | 1.24 | 1.51 | 1.38 | 0.91 | 0.95 | 1.45 | 0.59 | 1.43 | 0.36 | 0.2 | 0.38 | 1.04 | 1.58 | 0.55 | -0.265 | -0.125 | -0.3275 | 0.167115 | 0.372066 | 0.063273 | 1 | 0.999631 | 0.636719 |
| Dcaf10_A2AKB9_S100s_1 | 1.2 | 0.66 | 1.37 | 1.65 | 0.68 | 1.05 | 1.06 | 1.03 | 1.13 | 1.21 | 1.51 | 0.53 | 0.4 | 0.93 | 1.58 | 0.66 | -0.0825 | -0.335 | 0.445 | 0.386526 | 0.379334 | 0.713054 | 1 | 0.999631 | 0.756141 |
| Dcaf6_Q9DC22_S584s_1 | -0.52 | 0.21 | 0.91 | 0.56 | -0.25 | -0.14 | 0.02 | 1.2 | -0.63 | -0.06 | 0.4 | 0.11 | 0.6 | 0.31 | 1.29 | 0.74 | 0.355 | 0.2075 | -1.125 | 0.275837 | 0.851688 | 0.401078 | 1 | 0.999631 | 0.700187 |
| Dcaf8_Q8N7N5_S100s_1 | -0.92 | 0.15 | 1.01 | 0.17 | -0.1 | 0.17 | 1.59 | 0.17 | 0.75 | -0.02 | 0.15 | 0.36 | -0.09 | -2.23 | -0.34 | -1.43 | 0.8075 | 0.6025 | -0.2425 | 0.077344 | 0.494202 | 0.630329 | 1 | 0.999631 | 0.542683 |
| Dcbld2_Q91ZV3_S582s_1 | -0.28 | 0.14 | -0.73 | -0.42 | 0.84 | 0.11 | 0.51 | 0.48 | 0.54 | -0.51 | 0.66 | 0.43 | -0.03 | -0.02 | -1.13 | -1.08 | 0.165 | 0.65 | 0.425 | 0.538448 | 0.021055 | 0.116608 | 1 | 0.999631 | 0.830863 |
| Dcbld2_Q91ZV3_T738t_1 | 0.04 | 1.03 | -0.16 | -0.58 | 0.28 | 0.43 | -0.06 | 0.34 | 0.63 | -0.2 | 1.19 | 1.31 | 1.14 | 0.64 | 0.11 | 0.14 | 0.3125 | -0.1925 | -0.645 | 0.333465 | 0.660629 | 0.187409 | 1 | 0.999631 | 0.728079 |
| Dcc_P70211_T1135t_1 | 1.52 | 0.06 | -1.25 | 1.7 | 0.98 | 0.75 | 0.44 | 1.11 | 0.19 | 0.83 | 0.25 | -0.01 | -2.68 | 0.73 | 0.16 | 1.24 | 0.2175 | 0.2475 | 0.27 | 0.505237 | 0.623844 | 0.765232 | 1 | 0.999631 | 0.814727 |
| Dclk1_Q1EDH1_S25s_1 | 0.52 | 1.4 | -0.13 | 0.4 | 0.99 | 1.16 | 0.25 | 0.66 | 1.46 | 0.75 | 0.64 | 0.33 | 0.45 | 1.43 | 0.77 | 0.62 | -0.005 | 0.055 | 0.1225 | 0.509417 | 0.576244 | 0.546546 | 1 | 0.999631 | 0.816932 |
| Dclk1_Q1EDH1_S30s_1 | 0.4 | -0.22 | 0.12 | 0.51 | 0.61 | -0.33 | 0.17 | 0.34 | -0.1 | 0.22 | 0.39 | 0.52 | 0.2 | -0.23 | 0.61 | 0.72 | 0.3 | 0.28 | 0.53 | 0.717661 | 0.987157 | 0.851404 | 1 | 0.999631 | 0.903591 |
| Dclk1_Q1EDH1_S33s_1 | 0.68 | 0.63 | 0.53 | 0.91 | 1.11 | 1.22 | 0.47 | 1.15 | 1.34 | 0.49 | 0.95 | 1.09 | 1.28 | 0.88 | 1.54 | 1.17 | -0.155 | -0.43 | -0.615 | 0.087507 | 0.293113 | 0.340003 | 1 | 0.999631 | 0.553616 |
| Dclk1_Q1EDH1_S40s_1 | 1.5 | -1.71 | -0.73 | 0.22 | -0.38 | -1.87 | 0.51 | 0.4 | -0.72 | -0.08 | -0.74 | -0.9 | -1.75 | -1.26 | 0.22 | -0.39 | 0.065 | 0.38 | 0.24 | 0.401264 | 0.840265 | 0.505876 | 1 | 0.999631 | 0.763221 |
| Dclk1_Q1EDH1_S414s_1 | 0.27 | 0.24 | 0.98 | 0.65 | 0.22 | 0.18 | 1.07 | 0.93 | 0.49 | 0.6 | 1.41 | 1.16 | 0.77 | 0.23 | 0.97 | 1.13 | 0.2425 | -0.0075 | -0.06 | 0.477771 | 0.844681 | 0.26954 | 1 | 0.999631 | 0.800237 |
| Dclk1_Q1EDH1_S45s_1 | 0.54 | -0.84 | -1.31 | -0.86 | -0.76 | -0.39 | -0.04 | -0.31 | -0.23 | -0.04 | -1.13 | -1.1 | -1.05 | -0.62 | -0.83 | -0.21 | 0.2575 | 0.38 | -0.125 | 0.891896 | 0.56911 | 0.98737 | 1 | 0.999631 | 0.964276 |
| Dclk1_Q1EDH1_S45s_2 | 0.8 | 0.03 | -0.15 | 0.79 | 1.08 | -0.32 | 0.64 | 1.1 | 0.56 | 0.9 | 0.41 | 1.12 | -0.43 | 0.35 | 0.91 | 0.14 | 0.7 | 0.14 | -0.3075 | 0.75399 | 0.538038 | 0.281599 | 1 | 0.999631 | 0.913377 |
| Dclk1_Q1EDH1_S46s_1 | 0.65 | -1.19 | -1.63 | -0.07 | -0.1 | 0.23 | 0.49 | -0.06 | -1.08 | 0.52 | -0.5 | -0.62 | -1.54 | -1.34 | -0.53 | -0.06 | 0.7225 | 0.315 | 1.57E-16 | 0.595451 | 0.186224 | 0.806199 | 1 | 0.999631 | 0.854508 |
| Dclk1_Q1EDH1_S46s_2 | 0.64 | 0.14 | -1.56 | -0.15 | 0.6 | 0.57 | 0.39 | 0.4 | 0.32 | 0.48 | -0.25 | -0.22 | -0.8 | 0.26 | 0.06 | -0.45 | 0.5025 | 0.2225 | -0.1575 | 1 | 0.137812 | 0.520834 | 1 | 0.999631 | 1 |
| Dclk1_Q1EDH1_S48s_2 | 0.63 | 0.22 | -0.99 | 0.21 | 0.57 | 0.37 | 0.48 | 0.66 | 0.34 | 0.31 | 0 | 0.31 | -0.99 | 0.28 | 0.38 | -0.23 | -0.03 | 0.09 | -0.0925 | 0.734239 | 0.198225 | 0.562413 | 1 | 0.999631 | 0.907462 |
| Dclk1_Q9JLM8_S307s_1 | 0.24 | 0.31 | 0.16 | 0.15 | -0.04 | 0.37 | -0.19 | 0.6 | -0.29 | 0.55 | 0.34 | 0.62 | -0.16 | 0.26 | 0.09 | 0.3 | -0.5925 | -0.01 | 0.7325 | 0.732514 | 0.912717 | 0.759343 | 1 | 0.999631 | 0.907231 |

| | | | | | | | | | | | | | | | | | | | | | | | | | |
|---|---|---|---|---|---|---|---|---|---|---|---|---|---|---|---|---|---|---|---|---|---|---|---|---|---|
| Dclk1_Q9JLM8_S307s_2 | -0.49 | -0.2 | 0.4 | -0.14 | -1.43 | -0.38 | -0.76 | -0.23 | -0.4 | 0.2 | -0.04 | -0.23 | 0.43 | 0.68 | 0.2 | 1.19 | -0.5925 | -0.01 | 0.7325 | 0.055218 | 0.118075 | 0.973493 | 1 | 0.999631 | 0.492913 |
| Dclk1_Q9JLM8_S323s_2 | -0.49 | -0.2 | 0.4 | -0.14 | -1.43 | -0.38 | -0.76 | -0.23 | -0.4 | 0.2 | -0.04 | -0.23 | 0.43 | 0.68 | 0.2 | 1.19 | 0.0025 | 0.355 | 0.65 | 0.055218 | 0.118075 | 0.973493 | 1 | 0.999631 | 0.492913 |
| Dclk1_Q9JLM8_S32s_1 | 0.27 | 0.12 | -0.07 | -0.27 | -0.02 | -0.14 | -0.07 | 0.29 | 0.28 | 0.73 | 0.02 | 0.44 | -0.13 | 0.99 | 0.78 | 1.01 | -0.4325 | -0.1 | 0.12 | 0.087715 | 0.992273 | 0.228691 | 1 | 0.999631 | 0.553787 |
| Dclk1_Q9JLM8_S363s_1 | 0.41 | 0.62 | 1.44 | -0.25 | 0.25 | -0.2 | -0.09 | 0.53 | -0.02 | 0.15 | 1.1 | 0.59 | 0.45 | 0.06 | 0.89 | 1.3 | -0.005 | -0.075 | -0.025 | 0.786413 | 0.28686 | 0.815741 | 1 | 0.999631 | 0.923689 |
| Dclk1_Q9JLM8_S364s_1 | -0.38 | -0.45 | -0.17 | -0.1 | -0.57 | -0.91 | 0.14 | 0.22 | -0.86 | -0.9 | 0.14 | 0.22 | -0.79 | -0.58 | -0.19 | 0.36 | 0.08 | 0.2025 | 0.3325 | 0.940524 | 0.987872 | 0.83241 | 1 | 0.999631 | 0.981146 |
| Dclk1_Q9JLM8_S392s_1 | -0.06 | 0.15 | -0.85 | -0.45 | 0.09 | -0.09 | -0.4 | -0.49 | 0.01 | -0.2 | 0.02 | -0.23 | 0.25 | 0.12 | -0.61 | 0.36 | 0.1225 | 0.0575 | 0.635 | 0.366879 | 0.798016 | 0.509843 | 1 | 0.999631 | 0.745796 |
| Dclk1_Q9JLM8_S736s_1 | -1.19 | -0.35 | 0.09 | -0.34 | -0.36 | -0.44 | -0.44 | -0.06 | -0.19 | -0.91 | -0.25 | -0.21 | 0.32 | 0.14 | -0.03 | 0.32 | 0.17 | 0.335 | 0.425 | 0.088432 | 0.708336 | 0.871429 | 1 | 0.999631 | 0.553858 |
| Dclk1_Q9JLM8_S742s_1 | -1.04 | 0.31 | -0.47 | -0.76 | -0.23 | 0.24 | -1.27 | -0.02 | -0.11 | -0.41 | -0.13 | 0.03 | 0.72 | 0.33 | -0.56 | -0.75 | -0.1625 | 0.36 | 0.74 | 0.360002 | 0.694829 | 0.345721 | 1 | 0.999631 | 0.741182 |
| Dclk1_Q9JLM8_S77s_1 | -0.01 | 0.57 | 0.43 | -0.22 | 0.14 | 0.4 | -0.49 | 0.07 | 0.54 | 0.75 | 0.26 | 0.66 | 1.52 | 0.71 | 0.57 | 0.93 | 0.9225 | 0.7325 | -0.4775 | 0.052363 | 0.608174 | 0.240231 | 1 | 0.999631 | 0.489777 |
| Dclk2_Q6PGN3_S343s_1 | -0.41 | 0.05 | 0.28 | -0.08 | 1.46 | 0.17 | 1.14 | 0.76 | 0.88 | -0.98 | 1.57 | 1.3 | -1.14 | -1.12 | 0.83 | -0.64 | 0.4375 | 0.29 | -0.295 | 0.32664 | 0.022116 | 0.203455 | 1 | 0.999631 | 0.724678 |
| Dclk2_Q6PGN3_S346s_1 | -0.35 | -0.33 | -0.48 | -1.21 | -0.03 | 0.2 | 0.22 | -1.01 | 0.16 | -1.6 | 0.44 | -0.21 | 0.32 | -0.94 | -1.05 | -1.88 | 0.6 | 0.08 | 0.1525 | 0.54584 | 0.259286 | 0.548476 | 1 | 0.999631 | 0.833922 |
| Dcp1a_Q91YD3_S542s_1 | 0.94 | 1.47 | -0.33 | 0.57 | 1.95 | 1.59 | 0.46 | 1.05 | 1.59 | 0.87 | 0.31 | 0.2 | 0.37 | 1.65 | 0.86 | 0.38 | -0.09 | -0.12 | -0.615 | 0.747536 | 0.225523 | 0.866574 | 1 | 0.999631 | 0.911931 |
| Dcps_Q9DAR7_T21t_1 | 1.26 | 2.26 | 1.71 | 1.86 | 1.5 | 2.03 | 1.51 | 1.69 | 1.64 | 2.23 | 0.89 | 1.85 | 0.71 | 1.19 | 2.18 | 0.55 | -0.0575 | -1.39 | -2.8625 | 0.174114 | 0.766173 | 0.749214 | 1 | 0.999631 | 0.642116 |
| Dctn1_E9Q586_S2s_1 | 4.58 | -0.97 | -1.13 | 2 | 0.44 | 1.31 | 2.14 | 0.36 | 0.32 | -0.51 | 0 | -0.89 | -4.11 | -0.89 | -0.4 | -1.57 | 0.6575 | 0.0525 | -0.035 | 0.054744 | 0.962168 | 0.25878 | 1 | 0.999631 | 0.492066 |
| Dctn4_Q8CBY8_S196s_1 | 0.65 | 0.31 | -0.95 | 1.26 | 1.1 | 0.64 | 0.98 | 1.18 | 0.28 | 0.02 | 1.4 | -0.22 | -0.21 | 0.44 | 0.72 | 0.18 | 0.4175 | 1.07 | 0.9675 | 0.942847 | 0.176211 | 0.923293 | 1 | 0.999631 | 0.981629 |
| Dcx_O88809_S306s_1 | 0.47 | 0.06 | -0.42 | -0.2 | 0.44 | 0.88 | 0.05 | 0.21 | 2.23 | 2.35 | 0.06 | -0.45 | 1.09 | 1.57 | 0.78 | 0.34 | -0.3975 | 0.5675 | 0.825 | 0.023172 | 0.205065 | 0.135441 | 1 | 0.999631 | 0.414424 |
| Dcx_O88809_S332s_1 | -1.12 | -0.89 | -0.7 | -1.62 | -2.02 | -1.14 | -1.57 | -1.19 | -0.33 | -1.14 | -0.79 | 0.2 | 0.76 | -1.01 | -0.56 | -0.22 | 0.1575 | 0.4675 | 0.1775 | 0.08033 | 0.243156 | 0.154705 | 1 | 0.999631 | 0.545579 |
| Dcx_O88809_S332s_2 | 0.36 | -0.22 | -0.96 | 0.49 | 0.26 | -0.24 | -0.14 | 0.42 | 0.45 | 0.94 | -0.37 | 0.52 | -0.68 | 0.55 | 0.01 | 0.5 | 0.49 | 0.2525 | 0.395 | 0.687962 | 0.679711 | 0.291233 | 1 | 0.999631 | 0.891733 |
| Dcx_O88809_S339s_1 | 0.72 | -0.13 | -0.69 | 0.86 | 0.48 | 0.47 | 0.4 | 1.37 | 0.34 | 0.46 | 0.4 | 0.57 | 0.33 | 0.72 | 0.45 | 0.84 | 0.215 | 0.3975 | 0.595 | 0.34196 | 0.266762 | 0.519373 | 1 | 0.999631 | 0.73347 |
| Dda1_D3Z6F8_S30s_1 | 0.15 | 2.39 | 2.1 | 1.13 | 1.86 | 1.66 | 1.26 | 1.85 | 2.27 | 1.04 | 1.95 | 2.1 | 2.26 | 2.21 | 2.15 | 1.53 | -0.4575 | 0.13 | 0.07 | 0.259966 | 0.665645 | 0.466423 | 1 | 0.999631 | 0.692624 |
| Ddah1_Q9CWS0_S33s_1 | 0.17 | -0.09 | 0.07 | 0.38 | -0.31 | -0.38 | -0.28 | -0.33 | 0.43 | -0.12 | 0.23 | 0.51 | 0.31 | -0.27 | 0.19 | 0.58 | 0.27 | 0.3675 | 0.6225 | 0.817948 | 0.081961 | 0.638217 | 1 | 0.999631 | 0.938303 |
| Ddhd1_F8WHP5_S707s_1 | -1.19 | 0.24 | -0.26 | -0.46 | 0.16 | 0.74 | -0.97 | -0.52 | 0.6 | -0.15 | -0.47 | -0.18 | 0.73 | 0.47 | 0.19 | -0.57 | -0.1025 | 0.2775 | 0.375 | 0.161896 | 0.560448 | 0.358644 | 1 | 0.999631 | 0.631442 |
| Ddhd1_F8WHP5_T710t_1 | -1.63 | 0.21 | 0.53 | -0.55 | -0.68 | -0.57 | -0.83 | 0.23 | -0.33 | -0.81 | 0.27 | 0.54 | 0.44 | 0.05 | -0.47 | 0.04 | 0.1575 | 0.14 | 0.79 | 0.455338 | 0.83809 | 0.603421 | 1 | 0.999631 | 0.790041 |
| Ddhd2_Q80Y98_S447s_1 | -0.36 | 1.33 | 1.15 | 0.6 | 1.2 | 0.95 | -0.02 | 1.22 | 0.8 | 0.76 | 0.58 | 1.14 | 1.58 | 0.89 | 1.8 | 1.61 | -0.31 | -0.455 | 0.07 | 0.093639 | 0.732638 | 0.731337 | 1 | 0.999631 | 0.558128 |
| Ddi2_A2ADY9_S121s_1 | -1.05 | -0.49 | -0.09 | -0.59 | -1 | -0.99 | -0.76 | -0.71 | -1.69 | -0.39 | -0.55 | -1.41 | -0.82 | -0.6 | -0.12 | -0.4 | 0.3025 | 0.11 | 0.3275 | 0.829711 | 0.293948 | 0.261903 | 1 | 0.999631 | 0.942058 |
| Ddi2_A2ADY9_S128s_1 | -0.4 | -0.35 | -1.14 | -0.85 | -0.23 | -0.2 | -0.67 | -0.43 | -0.67 | -0.73 | -0.77 | -0.13 | -0.24 | 0.58 | -1.06 | -0.71 | 0.11 | 0.1225 | -0.07 | 0.43947 | 0.310432 | 0.722769 | 1 | 0.999631 | 0.781983 |
| Ddn_Q80TS7_S138s_1 | -0.29 | -0.54 | -0.21 | -0.05 | -0.11 | -0.21 | -0.15 | -0.18 | -0.51 | -0.05 | 0.12 | -0.16 | 0.05 | -0.19 | -1.06 | -0.17 | -0.195 | 0.0025 | -0.035 | 0.834594 | 0.650976 | 0.655663 | 1 | 0.999631 | 0.944144 |
| Ddn_Q80TS7_S354s_1 | -0.68 | -0.72 | -0.63 | -1.11 | -0.89 | -1.22 | -1.07 | -0.74 | -0.83 | -0.72 | -0.92 | -0.66 | -1.05 | -1.27 | -0.72 | -0.24 | 0.5725 | 0.115 | 0.0825 | 0.9146 | 0.457899 | 0.99222 | 1 | 0.999631 | 0.972695 |
| Ddn_Q80TS7_S388s_1 | 0.02 | 0.31 | 0.55 | 0.35 | 1.04 | 0.52 | 0.38 | 1.58 | 0.79 | -0.47 | 0.39 | 0.98 | -0.07 | 1.23 | -0.91 | 1.31 | -0.015 | -0.0975 | -0.84 | 0.873473 | 0.110694 | 0.755391 | 1 | 0.999631 | 0.959788 |
| Ddn_Q80TS7_S524s_1 | 0.98 | 1.98 | 0.35 | 1.33 | 1.75 | 1.27 | 1.36 | 0.2 | 1.77 | 0.62 | 0.75 | 1.11 | -0.47 | 1.1 | -0.21 | 0.86 | -0.055 | -0.0425 | -0.4075 | 0.114339 | 0.97379 | 0.81959 | 1 | 0.999631 | 0.579001 |
| Ddn_Q80TS7_S77s_1 | -0.18 | -0.42 | -0.31 | -0.41 | -0.3 | -0.48 | -0.41 | -0.35 | -1.03 | -0.07 | -0.11 | -0.28 | -0.61 | -1.13 | -0.95 | -0.26 | -1.095 | -1.0725 | -2 | 0.198559 | 0.813321 | 0.888673 | 1 | 0.999631 | 0.656516 |
| Ddx17_Q3U741_S522s_1 | 2.17 | 0.66 | 1.56 | 1.06 | 0.4 | -0.68 | 1.91 | -0.56 | -0.21 | 1.37 | 0.4 | -0.4 | -0.65 | -0.86 | -0.85 | -0.19 | -0.2475 | 0.6675 | 0.205 | 0.000462 | 0.099365 | 0.051137 | 1 | 0.999631 | 0.351308 |
| Ddx20_Q9JJY4_S472s_1 | 0.45 | -0.58 | 1.29 | -1.08 | -0.84 | -0.42 | 1.02 | -0.67 | -0.87 | 0.09 | 2.06 | 1.47 | -0.21 | 0.12 | 0.88 | 0.11 | -0.2 | 0.1175 | -0.365 | 0.705329 | 0.686834 | 0.38352 | 1 | 0.999631 | 0.89781 |
| Ddx20_Q9JJY4_S496s_1 | 1.02 | 0.42 | 1.35 | 0.89 | -0.24 | 0.75 | 1.03 | 1.34 | 0.91 | 1.44 | 0.79 | 1.01 | 0.93 | 0.44 | 0.43 | 0.42 | -0.445 | -0.6325 | -1.2575 | 0.265238 | 0.617339 | 0.703954 | 1 | 0.999631 | 0.694315 |
| Ddx21_Q9JIK5_S243s_1 | 0.17 | -0.09 | -0.04 | -0.39 | -0.8 | -0.08 | 0.32 | -1.57 | -0.33 | 0.39 | -1.09 | -1.85 | -0.33 | -1.33 | -1.45 | -2.27 | -0.5275 | -0.4325 | -0.9725 | 0.013438 | 0.30986 | 0.20588 | 1 | 0.999631 | 0.385092 |
| Ddx23_D3Z0M9_S108s_1 | 1.48 | -0.2 | 1.22 | 0.58 | -0.18 | -0.33 | 0.8 | 0.68 | -0.17 | 0.55 | 0.79 | 0.18 | -0.23 | -0.31 | -0.16 | -0.11 | -0.0225 | -0.8025 | -0.2325 | 0.033303 | 0.264019 | 0.326899 | 1 | 0.999631 | 0.44294 |
| Ddx24_F8WJA0_S126s_1 | -1.04 | 0.2 | 0.74 | 0.35 | 0.24 | -0.01 | 0.21 | -0.28 | -1.08 | -1.54 | -0.37 | 0.03 | -0.42 | -1.19 | 0.42 | 0.51 | -0.8475 | -1.685 | -1.53 | 0.658267 | 0.955677 | 0.130074 | 1 | 0.999631 | 0.879726 |
| Ddx27_H3BLJ6_S133s_1 | 2.17 | 1.66 | 1.88 | 1.15 | 0.13 | 3.03 | 0.75 | -0.44 | 1.2 | 2.51 | -1.71 | -1.88 | 0.78 | 0.09 | 0.42 | -0.55 | 2.385 | 0.5725 | -1.995 | 0.002612 | 0.243869 | 0.104253 | 1 | 0.999631 | 0.351308 |
| Ddx3x_Q62167_S612s_1 | -1.04 | 0.13 | -1.04 | -2.75 | 2.03 | 1.44 | 2.58 | -1.21 | 2.59 | -6.66 | 1.1 | 0.56 | -2.05 | -2.48 | -1.71 | -6.44 | -0.38 | 0.27 | 0.1425 | 0.085297 | 0.022154 | 0.753387 | 1 | 0.999631 | 0.551124 |
| Ddx51_Q6P9R1_S77s_1 | -0.1 | -0.53 | -0.06 | -0.22 | -0.78 | -0.77 | -0.07 | -0.81 | -0.11 | 0.3 | -0.21 | 0.19 | 0.26 | -0.51 | 0.32 | -0.41 | 0.5575 | 0.4475 | -0.095 | 0.660476 | 0.202961 | 0.331057 | 1 | 0.999631 | 0.880315 |
| Ddx54_Q8K4L0_S774s_1 | -0.38 | -1.21 | -0.29 | -1.4 | -0.52 | -0.15 | -0.09 | -0.29 | -0.11 | 0.45 | -0.72 | -1.11 | -0.05 | -1.41 | -0.24 | -1.96 | -0.3025 | 0.12 | -0.385 | 0.853096 | 0.12193 | 0.32249 | 1 | 0.999631 | 0.950673 |
| Ddx59_Q9DBN9_S150s_1 | 0.98 | 2.69 | 1.76 | 0.98 | 1.75 | 1.66 | 1.08 | 0.71 | 2.29 | 1.56 | 1.42 | 1.62 | 0.32 | 2.11 | 1.3 | 1.14 | 0.5125 | 0.4325 | -0.4925 | 0.463904 | 0.512864 | 0.787159 | 1 | 0.999631 | 0.793747 |
| Dek_E9Q8Y1_S33s_1 | -0.93 | 2.2 | 2.1 | 1.63 | 1.31 | 2.49 | 1.34 | 1.91 | 1.97 | 0.95 | 1.69 | 2.12 | 1.24 | 0.37 | 1.33 | 0.09 | -0.2575 | 0.2825 | 0.42 | 0.492101 | 0.468971 | 0.538701 | 1 | 0.999631 | 0.809399 |
| Dek_E9Q8Y1_S55s_1 | 0.78 | 1.78 | 1.19 | 1.53 | 1.09 | 1.15 | 0.52 | 1.49 | 0.94 | 1.74 | 1.78 | 1.95 | 1.26 | 1.85 | 1.99 | 1.86 | -0.04 | 0.235 | 0.8675 | 0.232461 | 0.448382 | 0.429212 | 1 | 0.999631 | 0.682449 |
| Dennd1a_A2ALU3_S86s_1 | -0.18 | 0.34 | 0.95 | 0.04 | -0.07 | 0.65 | -0.23 | 0.64 | 0.22 | 1.08 | 0.22 | 0.57 | 1.34 | 0.81 | 1.26 | 1.21 | -0.09 | -0.2525 | 0.17 | 0.026329 | 0.91167 | 0.512641 | 1 | 0.999631 | 0.422285 |
| Dennd1a_Q8K382_S473s_1 | 0.34 | -0.66 | 0.15 | -0.16 | -0.43 | -0.29 | -0.1 | 0.13 | -0.5 | 0.15 | 0.02 | -1.01 | -0.45 | 0.71 | -0.36 | 0.45 | 0.6275 | -0.0475 | 0.0875 | 0.665822 | 0.769898 | 0.501065 | 1 | 0.999631 | 0.88264 |
| Dennd1a_Q8K382_T707t_1 | 0.87 | -1.21 | -1.6 | 0.2 | 0.85 | -0.18 | 0.14 | -0.04 | -0.32 | -0.39 | -0.95 | -0.27 | -0.96 | -0.15 | -0.44 | 0.16 | -0.565 | -0.245 | -0.72 | 0.879618 | 0.286435 | 0.932017 | 1 | 0.999631 | 0.962995 |
| Dennd4a_E9Q8V6_S1053s_1 | -0.03 | 0.53 | -0.39 | 0.22 | -0.28 | -0.31 | -0.33 | -1.01 | -0.27 | 0.64 | -0.27 | -0.75 | -0.34 | -0.28 | -1.29 | -0.64 | -0.085 | -0.1075 | -0.24 | 0.065868 | 0.097229 | 0.517923 | 1 | 0.999631 | 0.516933 |
| Dennd4a_E9Q8V6_S1251s_1 | 1.04 | 0.82 | -0.05 | 0.86 | 0.77 | 0.64 | 0.31 | 0.61 | 0.52 | 0.71 | 0.53 | 0.48 | -0.03 | 0.8 | 0.6 | 0.34 | -0.225 | 0.04 | -0.2375 | 0.504791 | 0.787211 | 0.732511 | 1 | 0.999631 | 0.814292 |
| Dennd4a_E9Q8V6_S1252s_1 | 0.92 | 0.71 | 0.66 | 0.7 | 0.67 | 0.39 | 0.41 | 0.62 | 0.83 | 0.74 | 0.87 | 0.71 | -0.13 | 0.85 | 0.69 | 0.63 | -0.925 | -0.765 | -1.2225 | 0.457191 | 0.35692 | 0.868593 | 1 | 0.999631 | 0.790722 |
| Dennd4a_E9Q8V6_S1512s_1 | 1.54 | 0.11 | -1.17 | -0.24 | -1.31 | -0.93 | -0.21 | -1.01 | -0.78 | 0.24 | -1.37 | -0.91 | -1.51 | -0.73 | -1.29 | -1.12 | -0.3725 | -0.04 | 0.48 | 0.046131 | 0.121689 | 0.22104 | 1 | 0.999631 | 0.47959 |
| Dennd4a_E9Q8V6_S732s_1 | -0.57 | -0.07 | 1.09 | 1.37 | 0.47 | 0.11 | -0.11 | -0.14 | 0.37 | -0.03 | -0.03 | 1.35 | 0.83 | 0.16 | 2.34 | 0.41 | -0.72 | -0.4225 | -0.79 | 0.439246 | 0.428991 | 0.939665 | 1 | 0.999631 | 0.781983 |
| Dennd4b_Q3U1Y4_S736s_1 | 1.42 | 0.4 | 1.24 | 0.2 | -0.47 | 0.66 | 0.55 | -0.36 | 0.59 | 1.63 | 0.07 | -0.72 | 0.39 | -0.11 | 0.06 | -0.24 | -0.05 | -0.055 | 0.3175 | 0.056165 | 0.110886 | 0.440037 | 1 | 0.999631 | 0.495761 |
| Dennd4b_Q3U1Y4_S930s_1 | -0.34 | -0.13 | 0.4 | -0.5 | 0.14 | -0.24 | -0.38 | -0.29 | -0.66 | -0.15 | -0.08 | 0.1 | 0.21 | 0.04 | 0.18 | 0.27 | 0.0775 | 0.0075 | -1.07 | 0.309251 | 0.865248 | 0.862222 | 1 | 0.999631 | 0.718845 |
| Dennd4b_Q3U1Y4_S951s_1 | 1.39 | 1.09 | 0.66 | 1.44 | 1.04 | 1.53 | 1.22 | 1.1 | 1.45 | 1.73 | 0.95 | 0.48 | -0.57 | 0.67 | 0.37 | -0.17 | -0.61 | -0.8975 | -1.6325 | 0.015263 | 0.786377 | 0.983436 | 1 | 0.999631 | 0.388832 |
| Dennd4b_Q3U1Y4_S961s_1 | 1.16 | 0.47 | 0.69 | 0.62 | -0.05 | -0.14 | 0.67 | 0.02 | -0.59 | 0.66 | -0.36 | -0.36 | -1.23 | -0.96 | -0.25 | -1.15 | 0.44 | -0.0175 | -0.665 | 0.000666 | 0.065704 | 0.028 | 1 | 0.999631 | 0.351308 |
| Dennd4c_E9Q449_S1323s_1 | 1.06 | -0.84 | -0.42 | -0.01 | 0.28 | 0.56 | 0.13 | 0.58 | 0.1 | 0.81 | -0.25 | -0.94 | -0.97 | -0.31 | -0.69 | -0.9 | -0.0325 | 0.1425 | 0.4325 | 0.152069 | 0.307461 | 0.972852 | 1 | 0.999631 | 0.625198 |
| Dennd5a_Q6PAL8_S193s_1 | -0.5 | -0.05 | -0.35 | -0.44 | -0.46 | -0.14 | -0.73 | -0.14 | -0.05 | 0.32 | -0.54 | -0.5 | 0.3 | 0.21 | -0.01 | -0.11 | -0.3175 | -0.0025 | 0.13 | 0.143175 | 0.903556 | 0.638555 | 1 | 0.999631 | 0.614578 |
| Dennd5b_A2RSQ0_S178s_1 | -0.17 | 0.5 | 0.13 | 0.19 | -0.12 | -0.09 | -0.18 | -0.23 | 0.11 | 0.02 | 0.33 | 0.18 | 0.26 | 0.51 | 0.02 | 0.38 | -0.275 | 0.4 | -0.0625 | 0.656493 | 0.230958 | 0.99252 | 1 | 0.999631 | 0.879056 |
| Dennd6a_Q8BH65_S16s_1 | 0.82 | 1.46 | 1.28 | 1.67 | 1.07 | 0.85 | 1.17 | 1.04 | 1.91 | 2.08 | 1.47 | 1.37 | 0.93 | 1.48 | 1.75 | 0.82 | -0.11 | 1.1275 | 0.9125 | 0.856808 | 0.333835 | 0.221543 | 1 | 0.999631 | 0.951933 |
| Denr_E9Q0G1_S73s_1 | 0.63 | 1.42 | 1.34 | 0.14 | 0.07 | 0.51 | 0.87 | 1.64 | 1.74 | 1.55 | 2.11 | 2.64 | 1.45 | 1.15 | 2.21 | 2.37 | 0.4225 | 0.345 | 0.9 | 0.056609 | 0.802337 | 0.017533 | 1 | 0.999631 | 0.496238 |
| Denr_E9Q0G1_S9s_1 | 0.2 | 2.14 | -1.32 | -0.27 | 0.52 | 0.95 | 0.29 | 0.68 | 0.54 | 1.04 | 0.63 | -0.08 | 2.81 | 0.89 | -0.06 | 0.71 | -0.0475 | 0.03 | -0.385 | 0.289043 | 0.524943 | 0.613963 | 1 | 0.999631 | 0.706046 |
| Depdc5_P61460_S1496s_1 | 1.54 | 0.86 | 0.02 | 0.44 | 0.91 | 0.91 | 0.25 | 0.6 | 1.15 | 0.89 | 0.72 | 0.22 | -0.6 | 1.33 | 0.34 | 0.25 | -0.39 | -0.275 | 0.035 | 0.441095 | 0.899369 | 0.93926 | 1 | 0.999631 | 0.781983 |
| Dfnb31_Q80VW5_S246s_1 | -0.76 | -0.56 | -0.74 | -0.91 | -1.53 | -1.61 | -0.72 | -0.67 | -1.69 | -1.51 | -0.14 | -0.73 | -0.73 | -1.29 | -0.84 | 0.03 | -0.4025 | -0.05 | 0.515 | 0.91908 | 0.234195 | 0.481618 | 1 | 0.999631 | 0.974421 |
| Dfnb31_Q80VW5_S685s_1 | -0.06 | 0.37 | 0.56 | -0.21 | -1.05 | 0.25 | -0.3 | 0.15 | -0.13 | 1.04 | -0.13 | -0.32 | 0.81 | 1.53 | -0.37 | 0.75 | 0.1275 | -0.08 | 0.0525 | 0.255306 | 0.287748 | 0.895657 | 1 | 0.999631 | 0.6904 |
| Dgkb_Q6NS52_S418s_1 | -0.48 | 0.04 | -0.02 | 0.2 | -0.06 | 0.05 | -0.15 | 0.41 | 0.06 | -0.52 | -0.1 | -0.02 | -0.73 | 0.46 | 0.01 | 0.21 | 0.7675 | 0.0475 | 0.02 | 0.881155 | 0.646009 | 0.77973 | 1 | 0.999631 | 0.963364 |
| Dgkb_Q6NS52_S94s_1 | 0.27 | -0.85 | -0.38 | 0.98 | 0.77 | -0.26 | 1.18 | 1.4 | 0.06 | -0.88 | 1.15 | -0.12 | -0.95 | 0.1 | 0.56 | 0.39 | 0.855 | 0.175 | -0.4 | 0.968076 | 0.153252 | 0.929591 | 1 | 0.999631 | 0.988771 |
| Dgkb_Q6NS52_S95s_1 | 1.15 | -0.31 | -1.06 | 1.09 | 1.47 | 0.74 | 1.2 | 0.88 | 1.04 | 0.09 | 0.64 | -0.2 | -1.63 | 0.74 | 0.17 | -0.01 | 0.3125 | -0.125 | -0.105 | 0.551133 | 0.128372 | 0.756173 | 1 | 0.999631 | 0.836275 |
| Dgkb_Q6NS52_T117t_1 | 0.54 | -0.57 | -0.66 | 0.2 | 0.23 | 0.05 | -0.03 | 0.51 | -0.47 | 0.04 | -0.11 | -0.45 | -0.87 | 0.03 | -0.12 | 0.05 | 1.38 | 0.2825 | -1.5925 | 0.789732 | 0.37877 | 0.725727 | 1 | 0.999631 | 0.925636 |
| Dgkd_E9PUQ8_S44s_1 | 2.67 | 0.69 | -1.01 | 2.69 | 2.89 | 2.4 | 2.57 | 2.7 | 2.6 | 0.78 | 2.45 | 0.34 | -4.25 | 1.25 | 0.91 | 0.76 | -0.25 | -0.44 | 0.3325 | 0.250023 | 0.10339 | 0.75921 | 1 | 0.999631 | 0.686417 |
| Dgkg_F2Z409_S98s_1 | -0.81 | -0.58 | -1.79 | -1.44 | -1.26 | -0.19 | -1.54 | -2.63 | -1.34 | -1.23 | -2.18 | -1.63 | -0.58 | 0.04 | -1.4 | -1.35 | -0.72 | -0.9325 | -1.5275 | 0.460475 | 0.639268 | 0.258327 | 1 | 0.999631 | 0.791962 |
| Dgkg_F2Z409_T87t_1 | 2.45 | 0.62 | 2.48 | 1.61 | -0.31 | 1.47 | 2.12 | 1 | 0.63 | 2.26 | 0.38 | 0.16 | 0.02 | -0.14 | 1.01 | 0.16 | -0.2325 | -0.02 | 0.3425 | 0.010373 | 0.254845 | 0.139555 | 1 | 0.999631 | 0.383871 |
| Dgkh_D3YXJ0_S1067s_1 | -0.61 | -0.63 | -1.41 | -1.78 | -1.01 | -1.08 | -1.55 | -1.72 | -0.65 | -0.7 | -1.83 | -1.33 | -0.47 | -0.33 | -1.02 | -1.24 | 0.0675 | -0.0175 | 0.4025 | 0.392048 | 0.525401 | 0.961221 | 1 | 0.999631 | 0.758901 |
| Dgkh_D3YXJ0_S677s_1 | -0.91 | -1.25 | -1.17 | -1.67 | -1.19 | -1.3 | -0.84 | -1.4 | -1.54 | -0.9 | -1.53 | -1.1 | -0.36 | -0.95 | -1.06 | -1.02 | -0.1 | 0.035 | -0.005 | 0.22105 | 0.80988 | 0.953513 | 1 | 0.999631 | 0.671424 |
| Dgki_D3YWQ0_S705s_1 | 0.07 | 0.79 | 0.06 | 0.45 | 0.32 | 0.29 | -0.34 | 0.7 | 0.19 | 0.05 | 0.45 | 0.82 | 0.19 | 0.4 | 0.1 | 0.66 | -0.465 | -0.4275 | -0.0775 | 0.987091 | 0.757437 | 0.910176 | 1 | 0.999631 | 0.995165 |
| Dgki_D3YWQ0_S906s_1 | -0.64 | -0.98 | -0.86 | -1.06 | -1.37 | -1.56 | -1.06 | -1.41 | -1.51 | -1.21 | -1.51 | -1.02 | -0.84 | -1.08 | -1.12 | -0.81 | 0.07 | -0.43 | -0.115 | 0.776802 | 0.091712 | 0.133006 | 1 | 0.999631 | 0.921293 |
| Dgki_D3YWQ0_T704t_1 | 0.03 | 0.42 | -1.14 | 0.04 | 0.34 | -0.4 | -0.49 | 0.18 | -0.24 | -0.34 | 0.02 | -1.81 | 0.87 | 0.68 | -3.26 | 0.6 | 0.35 | 0.4125 | 0.7825 | 0.898924 | 0.861267 | 0.403632 | 1 | 0.999631 | 0.967185 |
| Dgkq_D6RI30_S17s_1 | 0.64 | 0.8 | -0.43 | 0.44 | 0.79 | 0.82 | 0.39 | 0.85 | 0.34 | 0.75 | 0.87 | 1.14 | 0.46 | 1.51 | 1.23 | 1.38 | 0.105 | 0.375 | 0.695 | 0.068613 | 0.308461 | 0.263705 | 1 | 0.999631 | 0.523794 |
| Dgkq_D6RI30_S22s_1 | -0.06 | -0.67 | -0.38 | -0.56 | -0.07 | -0.36 | -0.61 | -0.21 | -0.27 | 0.03 | 0.05 | 0.02 | 0.01 | -0.11 | 0.7 | 0.51 | -0.1225 | -0.485 | -1.7275 | 0.051149 | 0.697749 | 0.182741 | 1 | 0.999631 | 0.489558 |
| Dgkq_D6RI30_S39s_1 | 2.5 | -0.37 | 0.08 | 0.52 | -0.24 | 1.43 | 1.14 | -0.09 | 0.23 | 1.42 | -0.37 | -0.49 | -0.82 | -1.6 | -0.42 | -1.34 | 0.805 | -0.4125 | -1.055 | 0.018153 | 0.856278 | 0.485484 | 1 | 0.999631 | 0.391024 |
| Dgkz_Q80UP3_S706s_1 | 2.23 | -0.43 | -1.16 | 1.59 | 1.25 | 1.08 | 1.64 | 1.48 | 0.11 | 0.65 | 0.43 | -0.61 | -3.15 | 0.74 | 0.45 | -0.03 | -0.4625 | -0.12 | -0.205 | 0.31876 | 0.278854 | 0.586104 | 1 | 0.999631 | 0.722239 |
| Dhrs13_B0QZT2_S291s_1 | 0.47 | 1.02 | 0.64 | 0.18 | 0.2 | 0.03 | 0.14 | 0.09 | 0.64 | 1.07 | -0.06 | 0.18 | 0.4 | 0.8 | 0.34 | -0.05 | -0.2075 | -0.425 | -0.5775 | 0.532705 | 0.112237 | 0.730102 | 1 | 0.999631 | 0.8277 |
| Dhrs13_B0QZT2_S307s_1 | -0.02 | -1.26 | 0.5 | 0.1 | -0.49 | -0.68 | -0.07 | -0.27 | 0.14 | -0.94 | -0.7 | -0.88 | -0.16 | -0.41 | -1.44 | -0.98 | -0.0975 | -0.0475 | 0.3075 | 0.235186 | 0.610446 | 0.353702 | 1 | 0.999631 | 0.683301 |
| Dhx29_Q8BN72_S198s_1 | -0.64 | 0 | 0.41 | -0.49 | -0.27 | -0.28 | -0.79 | 0.23 | 0.3 | -0.39 | -0.93 | 0.11 | 0.43 | -0.07 | 0.08 | 0.07 | 0.3475 | 0.7275 | 1.0175 | 0.366621 | 0.779558 | 0.901737 | 1 | 0.999631 | 0.745796 |
| Dhx30_Q99PU8_S226s_1 | -1.34 | -0.34 | 0.31 | 0.48 | 0.01 | 0.06 | -0.05 | 0.48 | -0.02 | 0.49 | 0.59 | 0.96 | 0.95 | 0.38 | 0.66 | 1.19 | -0.565 | 0.3225 | 0.3725 | 0.043491 | 0.420585 | 0.130727 | 1 | 0.999631 | 0.475443 |
| Dhx32_Q8BZS9_S12s_1 | 0.48 | 1.16 | 0.37 | 0.66 | 0.05 | -0.13 | 0 | 0.49 | 0.8 | 1.36 | 0.59 | 1.21 | 1.43 | 1.07 | 0.76 | 0.9 | -0.055 | 0.0375 | 0.545 | 0.254166 | 0.076635 | 0.317912 | 1 | 0.999631 | 0.689137 |
| Dhx9_O70133_S136s_1 | -1.04 | -0.26 | 0.24 | -0.27 | -0.1 | -0.41 | -0.6 | -0.44 | -0.41 | -0.28 | -0.46 | -0.03 | 0.22 | 0.27 | 0.06 | 0.3 | 0.0775 | 0.325 | 0.785 | 0.129655 | 0.866528 | 0.910032 | 1 | 0.999631 | 0.601246 |
| Dhx9_O70133_S137s_1 | -0.52 | -0.73 | 0.08 | -0.28 | -0.59 | -0.47 | -0.37 | 0.29 | -0.43 | -0.37 | 0.31 | 0.34 | 0.88 | -0.07 | 0.42 | 0.46 | -0.11 | 0.2925 | 0.44 | 0.03716 | 0.805537 | 0.332877 | 1 | 0.999631 | 0.457023 |
| Diap1_E9PV41_S22s_1 | 0.11 | -0.26 | -0.22 | 0.07 | -1.06 | 0.55 | -0.22 | -0.01 | 0 | 0.22 | -0.01 | 0.66 | -0.19 | 0.22 | 0.63 | 0.8 | 0.5475 | 0.7375 | 1.0125 | 0.192937 | 0.764951 | 0.310991 | 1 | 0.999631 | 0.649499 |
| Dido1_Q8C9B9_S1726s_1 | -0.59 | 2.9 | 1.85 | 0.83 | 2.29 | 1.61 | 1.43 | 1.85 | 2.58 | -0.76 | 2.73 | 3.39 | 2.92 | 1.87 | 2.42 | 1.83 | 0.7125 | -0.94 | -0.245 | 0.169333 | 0.42929 | 0.479185 | 1 | 0.999631 | 0.638979 |
| Dip2b_Q3UH60_S50s_1 | 0.83 | -1.12 | 0.07 | 0.51 | 1.3 | 0.62 | 0.61 | 0.61 | 0.11 | -0.31 | -0.93 | -2.34 | -0.33 | 1.02 | -0.14 | -1.24 | -1.0025 | -0.185 | 0.52 | 0.673658 | 0.134411 | 0.153276 | 1 | 0.999631 | 0.885515 |
| Dip2b_Q3UH60_S782s_1 | -0.56 | -0.69 | -0.91 | -1.04 | -1.33 | -1.69 | -1.44 | -2.75 | -0.72 | -1.09 | -0.02 | -2.11 | -0.45 | 0.02 | -0.35 | -0.34 | 0.125 | -0.31 | -0.2375 | 0.089363 | 0.019343 | 0.67741 | 1 | 0.999631 | 0.555695 |
| Dip2b_Q3UH60_T70t_1 | -0.52 | 0.47 | -0.36 | 0.07 | 0.32 | -0.17 | 0.01 | 0 | -0.93 | -0.87 | 0.93 | -0.71 | -1.16 | -0.57 | 0.5 | -0.06 | 1.445 | -0.5925 | -1.9 | 0.581971 | 0.682744 | 0.521054 | 1 | 0.999631 | 0.847567 |
| Dip2c_B7ZMZ7_T123t_1 | 2.79 | -0.74 | -1.95 | 2.06 | 2.65 | 1.02 | 2.01 | 2.26 | 0.89 | -1.04 | 1.49 | -1.55 | -3.93 | -0.21 | -1.1 | -0.2 | -0.2525 | -0.195 | -1.1575 | 0.139875 | 0.176028 | 0.609711 | 1 | 0.999631 | 0.611718 |
| Diras2_Q5PR73_S126s_1 | 1.19 | 0.27 | 0.22 | -0.07 | -0.66 | 0.41 | 1.72 | -0.87 | -0.02 | 1.41 | -0.14 | -0.42 | 0.21 | -0.89 | -0.82 | -1.52 | 0.3825 | -0.5175 | -0.23 | 0.025867 | 0.670904 | 0.682121 | 1 | 0.999631 | 0.422248 |
| Diras2_Q5PR73_S175s_1 | -0.87 | 0.26 | 0.92 | 0.72 | -0.15 | 0.38 | 0.3 | 2.03 | -0.22 | -1.01 | 1.76 | -1.57 | -0.14 | 0.24 | -0.51 | 0.52 | -0.2225 | -0.9725 | -0.0775 | 0.615779 | 0.506132 | 0.487436 | 1 | 0.999631 | 0.861203 |
| Diras2_Q5PR73_S35s_1 | -1.43 | -0.58 | -0.45 | -1.1 | -2.29 | -0.26 | -0.93 | -0.97 | -2.43 | -1.72 | -0.14 | -3.16 | -0.86 | 0.12 | -2.19 | -0.94 | -0.515 | -0.3375 | -0.955 | 0.877375 | 0.63215 | 0.140878 | 1 | 0.999631 | 0.9617 |
| Dis3l2_Q8CI75_S864s_1 | 1.02 | -0.2 | -0.35 | -0.59 | -0.39 | -0.29 | 0.14 | -1.64 | 0.04 | 1 | -1.47 | -1.04 | 0.23 | -0.82 | -1.38 | -1.97 | -0.0475 | 0.1225 | 0.35 | 0.106438 | 0.3108 | 0.578286 | 1 | 0.999631 | 0.568797 |
| Disp2_Q8CIP5_S1119s_1 | -0.58 | -0.46 | -0.35 | -0.83 | -0.8 | -0.33 | -0.58 | -0.7 | -0.1 | -0.63 | -0.49 | -0.51 | 0.09 | 0.19 | -0.88 | -0.22 | 0.005 | 0.3625 | 0.3625 | 0.307477 | 0.852751 | 0.649074 | 1 | 0.999631 | 0.717856 |
| Disp2_Q8CIP5_S1121s_1 | 0.58 | 0.17 | -0.37 | 0.04 | 0.44 | 0.18 | 0.18 | -0.36 | 0.72 | 0.54 | 0.26 | 0.35 | 0.17 | 0.93 | 0.41 | 0.36 | 0.4325 | -0.0225 | -0.04 | 0.285768 | 0.987199 | 0.242861 | 1 | 0.999631 | 0.705851 |
| Disp2_Q8CIP5_S1317s_1 | 0.12 | 0.5 | 0.12 | -0.22 | 1.18 | 1.16 | -0.11 | 0.02 | 0.56 | 0.35 | -0.17 | -0.31 | 0.36 | 0.14 | -0.05 | -0.09 | -0.12 | 0.3475 | 0.1525 | 0.891693 | 0.283128 | 0.943274 | 1 | 0.999631 | 0.964276 |
| Disp2_Q8CIP5_S1345s_1 | 0.53 | 0.04 | -0.65 | -1 | 0.17 | -0.12 | -0.3 | -1.31 | 0.81 | 0.53 | -0.95 | -0.08 | 0.21 | 0.32 | -0.05 | -0.95 | -0.565 | -0.0325 | -0.9475 | 0.734113 | 0.791457 | 0.487674 | 1 | 0.999631 | 0.907462 |
| Disp2_Q8CIP5_S6s_1 | 2.19 | -0.52 | 0.26 | 0.48 | -1.01 | 0.29 | 0.85 | 0.02 | 0.29 | 1.83 | 0.21 | -0.05 | -0.69 | -0.31 | -0.36 | -0.02 | -0.31 | 0.325 | 0.33 | 0.107725 | 0.373662 | 0.959591 | 1 | 0.999631 | 0.569416 |
| Dixdc1_Q80Y83_S13s_1 | 0.37 | 0.51 | 0.53 | -0.13 | 0.08 | 0.65 | -0.81 | 0.12 | 0.95 | 1.59 | -0.69 | 0.73 | 1.57 | 0.55 | 0.17 | 0.31 | -0.34 | -0.9975 | -1.6875 | 0.400585 | 0.401258 | 0.506721 | 1 | 0.999631 | 0.763209 |
| Dixdc1_Q80Y83_S335s_1 | 1.59 | -0.33 | -0.72 | 0.78 | -0.75 | -0.03 | 1.09 | -0.35 | -0.41 | -0.52 | -0.66 | -1.08 | -1.29 | -1.51 | -0.64 | -1.99 | -0.6225 | -0.395 | -1.2075 | 0.011667 | 0.57131 | 0.077204 | 1 | 0.999631 | 0.385092 |
| Dixdc1_Q80Y83_S50s_1 | 0.04 | -1.4 | -0.46 | -1.21 | -2.06 | -1.08 | -0.43 | -1.95 | -1.38 | -0.05 | -1.68 | -1.5 | -0.8 | -2.71 | -2 | -2.35 | -0.535 | -0.53 | -0.8925 | 0.036222 | 0.216965 | 0.419491 | 1 | 0.999631 | 0.453963 |
| Dkc1_Q9ESX5_S508s_1 | -0.16 | -2.26 | -1.25 | -1.21 | -1.78 | -1.67 | -1.42 | -2.15 | -2 | -1.08 | -2.12 | -1.8 | -2.18 | -1.93 | -1.86 | -2.48 | 0.085 | -0.055 | 0.045 | 0.070891 | 0.24431 | 0.276809 | 1 | 0.999631 | 0.531292 |
| Dkk3_Q9QUN9_S295s_1 | -0.27 | -0.88 | -0.34 | -0.23 | -0.55 | -0.41 | -0.29 | -0.13 | -0.79 | 0.04 | -0.45 | -0.74 | -0.41 | -0.12 | -1.02 | 0.01 | 0.6275 | 0.5475 | 0.405 | 0.894681 | 0.752715 | 0.859434 | 1 | 0.999631 | 0.965377 |
| Dlc1_Q9R0Z9_S584s_1 | 0.45 | -0.54 | 0.15 | 1.3 | 0.79 | 1.23 | 0.83 | 1.02 | 1.06 | 1.24 | 0.93 | 0.32 | -0.22 | 0.75 | 1 | 1.45 | -0.175 | -0.23 | 0.47 | 0.424059 | 0.139884 | 0.222269 | 1 | 0.999631 | 0.773835 |
| Dlg1_D3Z3B8_S149s_1 | -0.74 | -1.98 | -1.2 | -1.3 | -1.01 | -1.7 | -0.98 | -2.23 | -1.4 | -0.89 | -2.18 | -1.67 | -0.89 | -0.72 | -0.89 | -0.84 | -0.2175 | -0.0425 | 0.3625 | 0.177783 | 0.662543 | 0.557894 | 1 | 0.999631 | 0.642116 |
| Dlg1_D3Z3B8_S40s_1 | -1.1 | -0.59 | -0.09 | 0 | -1.27 | -1.06 | -0.51 | 0.19 | -0.9 | -0.98 | -0.07 | 0 | -0.18 | -0.45 | 0.28 | 0.02 | -0.175 | 0.2775 | 0.9775 | 0.31712 | 0.6005 | 0.912002 | 1 | 0.999631 | 0.720971 |
| Dlg1_D3Z3B8_S41s_1 | -0.68 | 0.19 | 1.2 | 0.29 | -0.16 | 0.45 | -0.38 | 0.39 | -0.28 | 0.61 | 0.59 | 1.19 | 1.1 | 0.7 | 1.52 | 1.59 | 0.3 | -0.2425 | 0.3675 | 0.047292 | 0.683819 | 0.560295 | 1 | 0.999631 | 0.480102 |
| Dlg1_D3Z3B8_S487s_1 | -1.03 | -0.34 | -0.38 | -1.31 | -0.59 | -0.33 | 0.09 | -1.03 | 0 | -1.54 | 0.54 | -3.03 | -0.19 | 0.07 | -0.52 | -0.95 | 0.4475 | -0.24 | -0.085 | 0.332393 | 0.413337 | 0.744764 | 1 | 0.999631 | 0.728079 |
| Dlg1_D3Z3B8_S492s_1 | -0.63 | 0.17 | -0.1 | -0.27 | 0.54 | 0.28 | 0.48 | -0.34 | 0.07 | -1.31 | -0.06 | -0.49 | -0.28 | -0.26 | -0.05 | -0.58 | 0.0575 | 0.355 | 0.125 | 0.779059 | 0.176283 | 0.529023 | 1 | 0.999631 | 0.921416 |
| Dlg1_E9Q9H0_S102s_1 | -0.76 | -0.84 | -0.26 | -0.63 | -1.31 | -0.54 | -0.44 | 0.03 | -0.35 | -0.91 | -0.1 | 0.29 | -0.2 | -0.45 | -1.3 | -0.04 | 0.1925 | 0.1325 | 0.205 | 0.728377 | 0.865957 | 0.300572 | 1 | 0.999631 | 0.905923 |
| Dlg1_E9Q9H0_S122s_1 | -2.26 | -0.57 | -0.32 | -1.44 | -0.62 | -0.9 | -0.89 | -1.41 | -1.19 | -1.71 | -0.92 | -0.24 | -0.33 | -1.27 | -0.97 | -1.2 | 0.15 | 0.1475 | 0.205 | 0.669158 | 0.673005 | 0.794297 | 1 | 0.999631 | 0.883599 |
| Dlg2_E9Q2L2_S28s_1 | -1.71 | -0.55 | -0.2 | -2.05 | -0.89 | -0.64 | -0.96 | -1.42 | -0.62 | -2.11 | -0.75 | -0.44 | 0.3 | -1.41 | -1.07 | -1.51 | -0.22 | -0.1975 | 0.1325 | 0.717411 | 0.742943 | 0.786942 | 1 | 0.999631 | 0.903387 |
| Dlg2_E9Q2L2_S318s_1 | 0.32 | 1.22 | 0.92 | 0.41 | 0.5 | 1.13 | -0.22 | 0.58 | 0.64 | 0.61 | 0.36 | 0.47 | 1.46 | 1.05 | -0.11 | 1 | -0.025 | 0.0025 | 0.19 | 0.749688 | 0.554501 | 0.514636 | 1 | 0.999631 | 0.912137 |
| Dlg2_E9Q2L2_S323s_1 | 0.33 | 1.19 | 0.07 | 0.33 | 0.61 | 0.68 | -0.21 | 0.74 | 0.47 | 0.47 | 0.6 | 0.39 | 0.64 | 0.34 | 0.27 | 1.43 | 0.1 | -0.2225 | -0.07 | 0.628781 | 0.944075 | 0.993627 | 1 | 0.999631 | 0.866321 |
| Dlg2_E9Q2L2_S328s_1 | -0.53 | -1.7 | -1.85 | -1.29 | -0.61 | -1.71 | -1.47 | -1.18 | -1.39 | -1.81 | -1.58 | -1.48 | -0.96 | -1.24 | -1.61 | -1.84 | 0.94 | 0.875 | -0.1475 | 0.856266 | 0.797016 | 0.528097 | 1 | 0.999631 | 0.951817 |
| Dlg2_E9Q2L2_S360s_1 | -1.27 | -1.24 | -0.96 | -2.25 | 0.04 | -0.68 | 0.81 | -2.13 | 1.19 | -2.15 | -0.62 | -0.64 | 0.59 | -1.92 | -1.98 | -3 | 1.1225 | 1.66 | 0.7375 | 0.837534 | 0.151445 | 0.207379 | 1 | 0.999631 | 0.946036 |
| Dlg2_E9Q2L2_S365s_1 | -0.13 | 1.36 | 1.33 | -0.66 | 2.18 | 1.14 | 2.32 | 0.75 | 2.81 | 0.39 | 2.44 | 2.9 | 3.51 | 0.75 | 0.1 | 0.49 | 0.6575 | -0.0525 | -0.3825 | 0.373513 | 0.078965 | 0.036701 | 1 | 0.999631 | 0.747035 |
| Dlg2_E9Q2L2_S406s_1 | 1.38 | 0.17 | -1.19 | 0.72 | 1.76 | 0.88 | 0.38 | 0.69 | 0.44 | 0.15 | 0.5 | -0.22 | 0.07 | -0.12 | -0.89 | 0.49 | -0.055 | -0.1925 | 9.81E-17 | 0.506795 | 0.262892 | 0.921387 | 1 | 0.999631 | 0.815431 |
| Dlg2_E9Q2L2_S414s_1 | 0.01 | -0.16 | -0.55 | 0.11 | -0.05 | -0.14 | -0.54 | -0.08 | -0.41 | -0.3 | -0.3 | -0.35 | -0.36 | 0.02 | -0.45 | 0.2 | -0.1775 | -0.1425 | 0.265 | 1 | 0.840542 | 0.473999 | 1 | 0.999631 | 1 |
| Dlg2_E9Q2L2_S45s_1 | -0.01 | 0.6 | 0.94 | 0.03 | 0.35 | 0.77 | -0.47 | 0.2 | 0.18 | 0.79 | -0.43 | 0.45 | 0.86 | 0.42 | 0.32 | 1.02 | 0.0175 | -0.255 | -0.4625 | 0.450609 | 0.629687 | 0.702773 | 1 | 0.999631 | 0.787797 |
| Dlg2_E9Q2L2_S528s_1 | -0.09 | -2.78 | -0.76 | -2.06 | -1.77 | -1.56 | -0.69 | -1.6 | -2.02 | -1.42 | -1.28 | -1.99 | -1.52 | -2.11 | -1.88 | -2.03 | 0.7325 | 0.245 | -0.3025 | 0.428317 | 0.976476 | 0.664352 | 1 | 0.999631 | 0.776069 |
| Dlg2_E9Q2L2_S530s_1 | -1.5 | -0.75 | -1.25 | -1.53 | -0.27 | -0.7 | -0.23 | -0.9 | -0.69 | -0.9 | -0.66 | -1.8 | -1.63 | -1.44 | -2.15 | -1.02 | -0.7375 | -0.2725 | -1.3725 | 0.398808 | 0.034683 | 0.498603 | 1 | 0.999631 | 0.761726 |
| Dlg2_E9Q2L2_S656s_1 | 1.92 | 1.51 | 2.51 | 1.78 | 0.58 | 1.67 | 2.2 | 0.32 | 1.2 | 2.86 | 1.54 | 1.03 | 0.79 | -0.6 | 1.14 | 0.9 | -0.015 | -0.0525 | 0.18 | 0.011073 | 0.140607 | 0.55272 | 1 | 0.999631 | 0.385092 |
| Dlg2_E9Q2L2_S65s_1 | -2.02 | -0.95 | -1.55 | -1.7 | -1.2 | -1.26 | -1.59 | -2.23 | -1.39 | -1.99 | -1.8 | -1.25 | -0.11 | -1.5 | -1.94 | -1.95 | -0.18 | -0.1075 | 0.4575 | 0.706426 | 0.966092 | 0.874783 | 1 | 0.999631 | 0.898066 |
| Dlg2_E9Q2L2_T317t_1 | 0.32 | 1.22 | 1.44 | 0.77 | 0.53 | 1.01 | 0.3 | 1.19 | 0.97 | 0.7 | 0.81 | 0.84 | 1.49 | 1.14 | 1.12 | 1.83 | -0.3375 | -0.3375 | 0.1325 | 0.215727 | 0.611036 | 0.734997 | 1 | 0.999631 | 0.668551 |
| Dlg3_Q52KF7_S139s_1 | -0.31 | 0.21 | 0.68 | 0.14 | -0.07 | 0.45 | -0.45 | -0.56 | -0.16 | -0.28 | -0.35 | 0.16 | 0.13 | 0.68 | 0.01 | 0.43 | 0.4125 | -0.265 | -0.3425 | 0.687537 | 0.333734 | 0.283115 | 1 | 0.999631 | 0.891556 |
| Dlg3_Q52KF7_S368s_1 | 0.18 | -1.36 | -1.59 | -0.42 | 0.33 | -1.05 | -0.37 | -0.45 | -0.85 | -1.45 | -0.69 | -1.26 | -1.65 | -0.95 | -0.88 | -1.08 | -0.08 | 0.015 | 0.165 | 0.451367 | 0.394872 | 0.553202 | 1 | 0.999631 | 0.788148 |
| Dlg3_Q52KF7_S491s_1 | 0.18 | -1.25 | -0.72 | -1.11 | -0.58 | -0.6 | -0.39 | -1.65 | -0.79 | -0.32 | -1.14 | -0.59 | -0.34 | -0.25 | -1 | -0.65 | -0.395 | -0.315 | -0.3775 | 0.675486 | 0.850487 | 0.968889 | 1 | 0.999631 | 0.885853 |
| Dlg3_Q52KF7_S493s_1 | 0.08 | -0.13 | 0.58 | -0.46 | -0.79 | -0.88 | 0.45 | -0.29 | -0.63 | -0.21 | 0.05 | -0.4 | -0.17 | -0.81 | -0.3 | -0.16 | 0.305 | -0.085 | -0.3925 | 0.276144 | 0.319117 | 0.338199 | 1 | 0.999631 | 0.700187 |
| Dlg3_Q52KF7_S495s_1 | 0.09 | -0.22 | -0.63 | -0.5 | 0.27 | 0.15 | 0.11 | -0.57 | 0.17 | -0.46 | -0.38 | -0.93 | -0.73 | -0.16 | -0.91 | -1.03 | 0.3875 | -0.13 | -0.17 | 0.247325 | 0.331966 | 0.796465 | 1 | 0.999631 | 0.685751 |
| Dlg3_Q52KF7_T365t_1 | 0.03 | -1.15 | -1.14 | -0.35 | 0.34 | -1.01 | -0.27 | -0.12 | -0.28 | -1.22 | -0.55 | -1.08 | -0.99 | -0.99 | -0.62 | -0.69 | -0.1325 | 0.0675 | 0.0375 | 0.638773 | 0.353475 | 0.737567 | 1 | 0.999631 | 0.870943 |
| Dlg4_Q62108_S139s_1 | -0.08 | -0.58 | -0.09 | -0.29 | -0.68 | -0.34 | -0.38 | -0.17 | -0.19 | -0.2 | -0.2 | -0.18 | 0.14 | -0.59 | -0.39 | -0.05 | -0.0825 | -0.1625 | -0.5575 | 0.901616 | 0.614494 | 0.79093 | 1 | 0.999631 | 0.968183 |
| Dlg4_Q62108_S292s_1 | 0.14 | -1.32 | -0.83 | -0.77 | -0.95 | -0.29 | -0.73 | -1.14 | -1 | 0.12 | -1.05 | -1.5 | -0.8 | -1.51 | -1.33 | -1.37 | -0.435 | 0.0925 | 0.4625 | 0.163464 | 0.824747 | 0.719241 | 1 | 0.999631 | 0.631937 |
| Dlg4_Q62108_S412s_1 | -1.15 | -2.23 | -1.04 | -2.02 | -2.54 | -2.18 | -1.84 | -1.62 | -2.46 | -1.57 | -0.76 | -1.28 | -0.44 | -1.71 | -1.65 | -0.79 | -0.355 | 0.0175 | 0.4 | 0.307082 | 0.265492 | 0.83912 | 1 | 0.999631 | 0.717856 |
| Dlg4_Q62108_S415s_1 | -1.28 | -0.87 | -0.58 | -1.3 | -1.6 | -1.09 | -1.5 | -1.26 | -1.55 | -0.8 | -0.96 | -0.65 | -0.15 | -0.72 | -0.57 | -0.99 | 0.4025 | 0.0375 | -0.045 | 0.236673 | 0.231601 | 0.956528 | 1 | 0.999631 | 0.683301 |
| Dlg4_Q62108_S419s_1 | -0.43 | 0.43 | 0.04 | -0.13 | 0.57 | 0.55 | -0.17 | 0.57 | 0.23 | -0.19 | 0.11 | -0.09 | -0.31 | 0.14 | -0.25 | 0.15 | 1.1475 | 1.1925 | 0.365 | 0.884851 | 0.215454 | 0.896923 | 1 | 0.999631 | 0.963505 |
| Dlg4_Q62108_S507s_1 | -0.46 | 0.92 | -1.37 | 1.91 | 0.95 | 1.12 | 1.16 | 2.36 | 1.15 | -0.32 | 2.23 | 2.71 | 0.09 | -0.14 | 2.31 | 0.2 | 0.1475 | -0.14 | -0.3125 | 0.651463 | 0.127391 | 0.188734 | 1 | 0.999631 | 0.876488 |
| Dlg4_Q62108_S510s_1 | -0.31 | 0.97 | 1.18 | 0.71 | 0.3 | 1.27 | 0.97 | 0.6 | 0.41 | 0.16 | 0.83 | 0.59 | 0.21 | 0.13 | 0.29 | 0.67 | -0.0975 | 0.0225 | 0.13 | 0.424411 | 0.71152 | 0.714069 | 1 | 0.999631 | 0.773927 |
| Dlg4_Q62108_S603s_1 | -0.28 | -1.04 | -0.65 | -0.68 | -0.99 | -0.28 | -1.11 | -0.66 | -0.85 | 0.14 | -0.88 | -0.97 | -0.86 | 0.08 | -0.88 | -0.47 | -0.7175 | -0.8125 | 0.105 | 0.702461 | 0.748149 | 0.948075 | 1 | 0.999631 | 0.896857 |
| Dlg4_Q62108_S70s_1 | -1.07 | -0.71 | -0.15 | -0.46 | -1.14 | -1.31 | -1.19 | -1.62 | -1.71 | -1.32 | -1.56 | -1.05 | -0.09 | -1.09 | -0.51 | -0.28 | -0.01 | 0.1775 | 0.1275 | 0.764103 | 0.032153 | 0.024322 | 1 | 0.999631 | 0.915775 |
| Dlg5_E9Q9I2_S109s_1 | -0.64 | 0.12 | -0.52 | -0.53 | -0.57 | -0.1 | -0.51 | -0.43 | -0.54 | -0.67 | 0.37 | -0.02 | -0.59 | -0.4 | 0.25 | -0.32 | 0.435 | 0.3225 | -0.1 | 0.697515 | 0.971669 | 0.60515 | 1 | 0.999631 | 0.895731 |
| Dlg5_E9Q9I2_S1209s_1 | -0.68 | -0.47 | 0.69 | 0.84 | 0.26 | 0.49 | 0.46 | 0.91 | 0.26 | -0.45 | 1.05 | 0.81 | -0.92 | -1.03 | 0.27 | 1.66 | 0.38 | 0.945 | 0.9325 | 0.879838 | 0.305985 | 0.515001 | 1 | 0.999631 | 0.962995 |
| Dlg5_E9Q9I2_S818s_1 | -1.67 | -0.1 | 0.66 | 0.85 | -0.22 | -0.31 | 0.54 | 1.25 | -0.29 | 0.6 | 1.84 | 1.37 | 0.95 | -0.1 | 1.32 | 1.3 | 0.0525 | 0.2275 | 0.32 | 0.145606 | 0.538784 | 0.176214 | 1 | 0.999631 | 0.616968 |
| Dlgap1_Q9D415_S169s_1 | -0.17 | -0.66 | -1.04 | -1.28 | -0.89 | -0.34 | -0.82 | -0.89 | -0.48 | -0.07 | -0.52 | -1.17 | 0.23 | -0.47 | -0.86 | -0.77 | -0.3925 | -0.165 | 0.045 | 0.410022 | 0.870669 | 0.535523 | 1 | 0.999631 | 0.767806 |
| Dlgap1_Q9D415_S389s_1 | -0.01 | 0.52 | -0.08 | -0.43 | -0.19 | -0.27 | -0.6 | -0.51 | -0.4 | 0.24 | -0.25 | -0.25 | 0.17 | -0.04 | -0.27 | 0.32 | -0.3525 | -1.66 | -1.9375 | 0.887761 | 0.197264 | 0.596129 | 1 | 0.999631 | 0.963505 |
| Dlgap1_Q9D415_S403s_1 | 1.72 | -1.48 | 0.3 | -0.4 | -1.23 | 0.01 | 1.25 | -1.3 | -3.09 | -0.51 | -0.89 | -2.01 | -1.71 | -2.8 | -0.89 | -2.21 | 0.5325 | -0.425 | -0.1775 | 0.01805 | 0.656312 | 0.056615 | 1 | 0.999631 | 0.391024 |
| Dlgap1_Q9D415_S418s_1 | 1.44 | 0.23 | 0.01 | 1.01 | 1.53 | 1.25 | 1.13 | 0.91 | 0.32 | -0.15 | 0.76 | 0.06 | 0.19 | 0.59 | 0.29 | 0.91 | 0.0675 | -0.0025 | 0.095 | 0.656459 | 0.176991 | 0.302213 | 1 | 0.999631 | 0.879056 |
| Dlgap1_Q9D415_S421s_1 | 0.42 | 0.06 | 0.81 | 0.73 | 0.44 | 0.48 | 0.4 | 0.97 | 0.27 | -0.08 | 0.89 | 0.93 | 0.65 | -0.1 | 0.89 | 0.96 | 0.48 | 0.0525 | 0.415 | 0.787587 | 0.815509 | 0.994179 | 1 | 0.999631 | 0.9245 |
| Dlgap1_Q9D415_S437s_1 | -0.98 | -1.15 | -1.03 | -1.18 | -0.46 | -0.19 | -1.07 | -0.7 | -0.87 | -0.88 | -0.94 | -1.44 | -0.17 | -1.09 | -1.01 | -0.41 | 0.1475 | 0.055 | 0.425 | 0.209982 | 0.10803 | 0.842441 | 1 | 0.999631 | 0.66441 |
| Dlgap1_Q9D415_S509s_1 | 0.1 | -0.5 | -1.17 | -0.42 | -0.05 | -0.02 | -0.48 | -0.85 | -0.6 | 0.31 | -0.79 | -0.69 | 0.22 | 0.17 | -0.38 | -0.3 | 0.1 | -0.1875 | -0.495 | 0.249985 | 0.678046 | 0.886071 | 1 | 0.999631 | 0.686417 |
| Dlgap1_Q9D415_S516s_1 | 0.03 | 0.64 | 0.59 | 0.3 | -0.73 | 1.12 | 0.58 | 0.99 | -0.07 | 0.42 | 0.68 | -0.22 | -1.01 | 0.25 | 0.26 | 0.08 | 0.67 | 0.51 | 0.895 | 0.206884 | 0.81846 | 0.556381 | 1 | 0.999631 | 0.660763 |
| Dlgap1_Q9D415_S52s_1 | -0.92 | -1 | -1.14 | -2.11 | 0 | -0.54 | -0.66 | -1.29 | -0.66 | -0.7 | -1.01 | -0.76 | 1.19 | -0.25 | -1.9 | -0.63 | -0.1925 | -0.485 | -1.635 | 0.174936 | 0.111409 | 0.152117 | 1 | 0.999631 | 0.642116 |
| Dlgap1_Q9D415_S937s_1 | 2.24 | 0.88 | 0.37 | 0.62 | 0.65 | 1.59 | 1.2 | -0.1 | 0.7 | 1.58 | 0.08 | -0.19 | -0.89 | -0.02 | -0.79 | -0.73 | 0.295 | -0.1175 | 0.1225 | 0.004871 | 0.710267 | 0.374838 | 1 | 0.999631 | 0.351308 |
| Dlgap1_Q9D415_S963s_1 | -0.5 | 0.33 | -0.67 | -0.79 | 0.36 | 0.88 | -0.74 | -0.95 | -0.3 | -0.42 | -0.91 | -0.47 | 0.23 | -0.06 | -1.05 | -0.26 | 0.0425 | -0.2925 | 0.2725 | 0.758823 | 0.54239 | 0.72635 | 1 | 0.999631 | 0.914449 |
| Dlgap1_Q9D415_T436t_1 | -0.65 | -0.19 | -1.24 | -1.42 | -0.34 | -0.43 | -0.99 | -1.57 | -1.02 | -0.52 | -1.78 | -1.35 | -0.1 | 0.13 | -1.55 | -0.89 | -0.1675 | -0.935 | -3.185 | 0.563207 | 0.91601 | 0.470679 | 1 | 0.999631 | 0.841038 |
| Dlgap2_Q0VF59_S1041s_1 | 2.54 | -2.52 | -0.78 | 0.06 | -2.53 | 0.55 | 3.56 | -2.95 | -2.31 | 2.21 | -1.91 | -2.43 | -3.83 | -3.63 | -1.85 | -4.13 | 0.1325 | 0.2125 | 0.28 | 0.009296 | 0.914652 | 0.480311 | 1 | 0.999631 | 0.367426 |
| Dlgap2_Q0VF59_S1048s_1 | -0.5 | 0.24 | -0.38 | -0.11 | 0.3 | 0.09 | -0.42 | -0.19 | 0.01 | -0.34 | 0.06 | 0.37 | 0.22 | 0.21 | -0.15 | 0.09 | 0.445 | 0.905 | -0.475 | 0.355772 | 0.654865 | 0.481345 | 1 | 0.999631 | 0.74034 |
| Dlgap2_Q0VF59_S249s_1 | 0.76 | 0.31 | 0.22 | 0.14 | 1.34 | 0.26 | 4.89 | -3.28 | 2.52 | -0.36 | 1.07 | 1.82 | 1.58 | -0.36 | -1.59 | -0.1 | 0.3525 | -0.1325 | -0.66 | 0.440858 | 0.756525 | 0.142426 | 1 | 0.999631 | 0.781983 |
| Dlgap2_Q0VF59_S390s_1 | 2.38 | 2.28 | 0.12 | 2.36 | 2.97 | 1.4 | 2.09 | 2.09 | 2.24 | 1.5 | 2.08 | 0.79 | 0.16 | 1.26 | 1.17 | 1.91 | -1.1175 | -0.0275 | -1.13 | 0.288905 | 0.549038 | 0.822318 | 1 | 0.999631 | 0.706046 |
| Dlgap2_Q0VF59_S454s_2 | -1.47 | -0.32 | 1.03 | -0.62 | -2.18 | -0.32 | -0.68 | -2.67 | 0.29 | 0.18 | -1.01 | -0.95 | -0.81 | -1.8 | -1.71 | -1.58 | -0.745 | -0.88 | -1.515 | 0.055857 | 0.126679 | 0.96186 | 1 | 0.999631 | 0.495701 |
| Dlgap2_Q0VF59_S457s_1 | 0.37 | -0.41 | 0.41 | -0.63 | -0.84 | -0.23 | -0.39 | -1.78 | -1.21 | 0.96 | -1.48 | -2.05 | -1.1 | -1.37 | -1.19 | -2.66 | -0.245 | -0.1725 | 0.24 | 0.006826 | 0.10835 | 0.191018 | 1 | 0.999631 | 0.358173 |
| Dlgap2_Q0VF59_S461s_1 | 0.47 | 1.65 | 2.13 | 1.33 | 0.89 | 1.33 | 0.82 | 1.56 | 0.88 | 0.96 | 1.37 | 1.68 | 1.58 | 1.15 | 1.41 | 2.4 | -0.04 | -0.3375 | -0.4825 | 0.591034 | 0.541456 | 0.672092 | 1 | 0.999631 | 0.852188 |
| Dlgap2_Q0VF59_S668s_1 | 0.42 | -0.46 | -0.95 | 0.05 | 0.31 | -0.7 | -0.22 | -0.49 | -0.42 | -0.63 | -0.64 | -0.6 | -0.93 | -0.47 | -1.34 | -0.13 | -0.2525 | -0.3425 | -1.045 | 0.261585 | 0.91666 | 0.340783 | 1 | 0.999631 | 0.69271 |
| Dlgap2_Q0VF59_S746s_1 | 0.23 | -0.77 | -0.08 | -0.3 | -0.96 | -0.62 | 0.38 | -0.73 | -0.84 | 0.36 | -0.78 | -1.03 | -0.81 | -1.68 | -1.05 | -1.56 | -1.2E-16 | -0.055 | 0.1925 | 0.012986 | 0.508857 | 0.394467 | 1 | 0.999631 | 0.385092 |

| | | | | | | | | | | | | | | | | | | | | | | | | | |
|---|---|---|---|---|---|---|---|---|---|---|---|---|---|---|---|---|---|---|---|---|---|---|---|---|---|
| Dlgap2_Q0VF59_S777s_1 | -0.27 | 0.73 | -0.17 | 0.11 | 0.48 | 0.33 | -0.4 | -0.01 | 0.36 | -0.16 | -0.14 | 0.12 | 0.39 | 0.32 | -0.16 | 0.62 | 0.0925 | -0.4575 | -0.6225 | 0.576454 | 1 | 0.862542 | 1 | 0.999631 | 0.846204 |
| Dlgap2_Q0VF59_S807s_1 | 1.66 | 0.66 | -1.68 | 0.86 | 1.8 | -0.76 | 1.54 | -0.71 | 0.03 | -1.73 | 2.14 | -0.77 | 0.63 | -0.71 | -1.62 | 0.71 | 0.8275 | 0.69 | 0.59 | 0.441038 | 0.915082 | 0.630908 | 1 | 0.999631 | 0.781983 |
| Dlgap2_Q0VF59_S812s_1 | -1.5 | -1.37 | -1.26 | -1.57 | 0.3 | -0.98 | -0.29 | -1.42 | 0.39 | -2.04 | -0.52 | -0.77 | 1.34 | -1.84 | -1.41 | -1.43 | -0.3225 | -0.11 | 0.085 | 0.379994 | 0.058001 | 0.176745 | 1 | 0.999631 | 0.752193 |
| Dlgap2_Q0VF59_S984s_1 | -0.37 | -0.62 | -0.55 | -0.79 | -1.14 | -0.75 | -0.95 | -0.78 | -1.13 | -0.43 | -0.58 | -0.63 | -0.34 | -0.63 | -0.88 | -0.14 | -0.03 | 0.2025 | 0.3125 | 0.774151 | 0.214493 | 0.691854 | 1 | 0.999631 | 0.920998 |
| Dlgap2_Q0VF59_T460t_1 | 0.78 | 2.52 | 2.43 | 1.48 | 1.26 | 3.03 | 1.07 | 1.73 | 2.13 | 1.86 | 1.93 | 2.1 | 1.7 | 2.1 | 1.96 | 2.7 | -0.385 | -0.285 | -0.4075 | 0.501777 | 0.956825 | 0.633183 | 1 | 0.999631 | 0.814092 |
| Dlgap2_Q0VF59_T744t_1 | 0.5 | -2.4 | -0.46 | -0.36 | -1.03 | -1.27 | -0.52 | -1.44 | -1.2 | -0.59 | -0.86 | -1.21 | -0.72 | -1.6 | -0.57 | -1.46 | 0.1975 | 0.565 | 0.6625 | 0.505544 | 0.515034 | 0.624191 | 1 | 0.999631 | 0.814836 |
| Dlgap2_Q0VF59_T816t_1 | 0.12 | 0.17 | 0.2 | 0.28 | 0.52 | 0.16 | 0.87 | 0.01 | 0.61 | 0.59 | 0.35 | 1.48 | 1.31 | -0.25 | 1.38 | 0.98 | 1.61 | 2.625 | -0.2825 | 0.12385 | 0.483876 | 0.097116 | 1 | 0.999631 | 0.595021 |
| Dlgap3_B1AS06_S185s_1 | 0.27 | 1 | -0.01 | -0.84 | 2.44 | 1.37 | 3.94 | -0.89 | 3.62 | 0.26 | 2.57 | 4.47 | 1.03 | 0.67 | -0.84 | -1.57 | -0.195 | 0.36 | -0.535 | 0.665059 | 0.110504 | 0.011937 | 1 | 0.999631 | 0.882433 |
| Dlgap3_B1AS06_S407s_2 | 0.64 | 0 | 1.95 | -0.06 | 0.28 | -0.94 | 0.83 | 1.58 | 0.29 | 0.6 | 1.49 | 1.59 | 0.95 | -2.43 | 1.01 | 0.86 | -1.04 | -0.795 | -0.385 | 0.526589 | 0.758635 | 0.502571 | 1 | 0.999631 | 0.824261 |
| Dlgap3_B1AS06_S410s_1 | 0.79 | -1.24 | 0 | -0.74 | -1.41 | -1.6 | -0.4 | -1.94 | -2.95 | 0.9 | -0.6 | -1.72 | -0.69 | -0.27 | -1.15 | -0.62 | -0.195 | 0.36 | -0.535 | 0.419986 | 0.067299 | 0.342304 | 1 | 0.999631 | 0.772605 |
| Dlgap3_B1AS06_S410s_2 | 0.64 | 0 | 1.95 | -0.06 | 0.28 | -0.94 | 0.83 | 1.58 | 0.29 | 0.6 | 1.49 | 1.59 | 0.95 | -2.43 | 1.01 | 0.86 | -0.33 | -0.38 | -1.9625 | 0.526589 | 0.758635 | 0.502571 | 1 | 0.999631 | 0.824261 |
| Dlgap3_B1AS06_S429s_1 | 2.86 | 0.95 | 0.91 | 1.69 | 0.54 | 1.5 | 2.76 | 0.29 | 0.67 | 2.42 | 1.3 | 0.5 | -0.67 | -0.94 | -0.16 | 0.33 | -0.425 | -0.3275 | 0.05 | 0.003121 | 0.611545 | 0.515599 | 1 | 0.999631 | 0.351308 |
| Dlgap3_B1AS06_S559s_1 | 0.01 | -0.91 | -0.79 | -0.6 | -0.93 | -1.07 | -0.89 | -1.1 | -0.69 | -0.59 | -1.45 | -0.87 | 0.02 | -0.24 | -1.04 | -0.83 | -0.03 | -0.3025 | 0.1 | 0.891446 | 0.160415 | 0.336066 | 1 | 0.999631 | 0.964253 |
| Dlgap3_B1AS06_S58s_1 | -0.73 | -0.68 | -0.71 | -1.08 | -0.4 | -0.95 | -1.07 | -0.9 | -0.91 | -1.31 | -1.02 | -1.17 | 0.07 | -0.93 | -0.8 | -1.14 | 1.3025 | 2.4375 | -0.8825 | 0.771663 | 0.911094 | 0.258198 | 1 | 0.999631 | 0.919413 |
| Dlgap3_B1AS06_S712s_1 | 1.59 | 2.06 | 2.43 | 0.06 | 3.79 | 1.32 | 4.74 | 1.5 | 4.86 | 1.1 | 4.68 | 5.25 | 2.63 | 0.12 | 1.17 | -1.31 | -0.165 | -0.3325 | 0.365 | 0.312993 | 0.154879 | 0.026849 | 1 | 0.999631 | 0.719861 |
| Dlgap3_B1AS06_S753s_1 | -0.93 | -0.27 | -0.45 | -0.68 | -0.58 | -0.63 | -0.92 | -0.86 | -0.72 | -1.12 | -1.41 | -0.41 | -0.09 | -0.37 | -0.26 | -0.15 | 1.3575 | -0.13 | -1.1325 | 0.218652 | 0.537174 | 0.314223 | 1 | 0.999631 | 0.669859 |
| Dlgap3_B1AS06_S771s_1 | -0.34 | 0.81 | -1.05 | 1.61 | 1.67 | 1.01 | 2.15 | 1.63 | 0.81 | -1.17 | 1.57 | -0.7 | -4.07 | 0.31 | -0.69 | 0.95 | -0.2275 | 0.2025 | -0.9325 | 0.306438 | 0.038808 | 0.865575 | 1 | 0.999631 | 0.717667 |
| Dlgap3_B1AS06_S779s_1 | 1.73 | 0.89 | 1.4 | 2.13 | 0.77 | 1.74 | 1.57 | 1.16 | 1.73 | 1.71 | 1.44 | 2.08 | -0.43 | 0.6 | 1 | 1.25 | -0.4275 | 0.09 | -0.74 | 0.06186 | 0.534906 | 0.554348 | 1 | 0.999631 | 0.509355 |
| Dlgap3_B1AS06_S783s_1 | 0.04 | -0.6 | -0.52 | -0.45 | -1.06 | -0.07 | -0.83 | -1.28 | -0.16 | 1.01 | -1.15 | -0.87 | -1.14 | -0.59 | -1.33 | -1.43 | -0.21 | -0.0375 | -0.2425 | 0.040357 | 0.223951 | 0.851876 | 1 | 0.999631 | 0.472911 |
| Dlgap3_B1AS06_S919s_1 | 0.51 | 0.28 | 0.02 | 0.52 | -0.46 | 0.37 | 0.42 | 0.16 | 0.27 | 0.49 | 0.13 | 0.29 | 0.16 | 0.04 | 0.02 | 0.14 | -0.38 | -0.05 | 0.3025 | 0.384925 | 0.487342 | 0.886154 | 1 | 0.999631 | 0.75538 |
| Dlgap3_B1AS06_S922s_1 | 0.2 | 1.12 | 0.98 | 0.11 | 0.18 | 1 | -0.26 | -0.03 | 1.01 | 1.1 | 0.11 | -0.01 | 0.97 | 0.24 | 1.27 | 1.14 | -1.18 | -0.9825 | -1.715 | 0.436369 | 0.339767 | 0.901076 | 1 | 0.999631 | 0.780748 |
| Dlgap4_B1AZP2_S678s_1 | 1.36 | -0.69 | 0.12 | 0.09 | -0.9 | -1.91 | 0.45 | -1.48 | -1.7 | 0.33 | -0.84 | -0.84 | -0.25 | -2.32 | -2.46 | -0.95 | -0.0725 | -0.365 | -1.345 | 0.018634 | 0.074086 | 0.099502 | 1 | 0.999631 | 0.391024 |
| Dlgap4_B1AZP2_S681s_1 | 1.27 | 0.2 | 0.91 | 1.04 | 0.63 | 0.57 | 1.28 | 0.65 | -0.01 | 0.1 | 0.91 | 0.96 | -1.33 | -0.5 | 0.44 | -0.57 | 0.065 | -0.2625 | -0.87 | 0.010808 | 0.824928 | 0.339435 | 1 | 0.999631 | 0.383923 |
| Dlgap4_B7ZNS2_S156s_1 | 0.45 | -0.75 | -0.88 | -0.2 | -0.37 | -0.3 | 0.16 | -0.61 | -0.41 | -0.37 | -0.65 | -1 | -1.04 | -0.9 | -1.29 | -1.63 | 0.645 | -0.4925 | -0.4725 | 0.041836 | 0.85851 | 0.479092 | 1 | 0.999631 | 0.475443 |
| Dlgap4_B7ZNS2_S35s_1 | 1.17 | 0.24 | -1.91 | 0.67 | 1.86 | -0.4 | 0.93 | 0.36 | 0.15 | -1.3 | -0.04 | -0.61 | -0.59 | -0.43 | -0.12 | -0.58 | -0.5525 | -0.2925 | -0.54 | 0.453864 | 0.386761 | 0.470816 | 1 | 0.999631 | 0.789094 |
| Dlgap4_B7ZNS2_S377s_1 | 1.15 | 0.18 | 1.46 | -0.13 | -0.41 | 1.19 | 0.17 | -0.5 | 0.32 | 1.8 | -0.29 | -0.34 | 0.88 | -0.26 | 0.28 | -0.4 | -0.4225 | 0.5125 | -0.165 | 0.266292 | 0.291716 | 0.613782 | 1 | 0.999631 | 0.695664 |
| Dlgap4_B7ZNS2_S405s_1 | 0.49 | 0.67 | 1.01 | 0.78 | 0.15 | -0.12 | 0.79 | 0.44 | 0.97 | 0.96 | 1.65 | 1.42 | 0.9 | -0.28 | 0.14 | 1.53 | -0.1775 | -0.185 | -0.57 | 0.699766 | 0.171391 | 0.101511 | 1 | 0.999631 | 0.896612 |
| Dlgap4_B7ZNS2_S405s_2 | 0.93 | 1.46 | -0.7 | 1.39 | 1.68 | -0.8 | 0.81 | 0.68 | 1.2 | -0.11 | 1.05 | 0.2 | 0.52 | 0.03 | -0.47 | 0.72 | 0.1225 | -0.1875 | 0.145 | 0.300493 | 0.783059 | 0.738302 | 1 | 0.999631 | 0.71369 |
| Dlgap4_B7ZNS2_S410s_1 | 0.28 | -1.21 | 0.1 | 0.7 | 0.23 | -1.53 | 0.85 | 0.81 | -0.46 | -1.42 | 1.06 | -0.06 | 0.28 | -1.01 | -0.02 | 1.2 | -0.1775 | -0.185 | -0.57 | 0.798353 | 0.844213 | 0.755009 | 1 | 0.999631 | 0.930748 |
| Dlgap4_B7ZNS2_S410s_2 | 0.93 | 1.46 | -0.7 | 1.39 | 1.68 | -0.8 | 0.81 | 0.68 | 1.2 | -0.11 | 1.05 | 0.2 | 0.52 | 0.03 | -0.47 | 0.72 | 0.06 | 0.4975 | 0.4875 | 0.300493 | 0.783059 | 0.738302 | 1 | 0.999631 | 0.71369 |
| Dlgap4_B7ZNS2_S415s_1 | -3.3 | -2.04 | -3.42 | -1.97 | -1.92 | -3.73 | -2.7 | -2.14 | -1.63 | -3.32 | -1.99 | -1.8 | -1.74 | -2.05 | -2.35 | -2.64 | -0.4175 | -0.1275 | 0.34 | 0.283798 | 0.908505 | 0.347932 | 1 | 0.999631 | 0.704942 |
| Dlgap4_B7ZNS2_S517s_1 | -1.37 | -0.07 | -0.74 | -0.94 | -1.78 | -1.02 | -0.94 | -1.05 | -1.63 | -0.89 | -0.91 | -0.2 | 0.23 | -1.6 | 0.05 | -0.44 | 0.2525 | -0.055 | 0.06 | 0.483607 | 0.261162 | 0.754665 | 1 | 0.999631 | 0.803422 |
| Dlgap4_B7ZNS2_S649s_1 | 1.42 | 2.79 | 0.5 | 0.66 | 2.26 | 2.27 | 0.5 | 1.35 | 2.49 | 0 | 0.96 | 1.7 | 0.01 | 3.71 | 1 | 0.89 | -0.07 | 0.1725 | 0.33 | 0.942326 | 0.678588 | 0.934271 | 1 | 0.999631 | 0.981629 |
| Dlgap4_B7ZNS2_S715s_1 | -1.01 | -0.82 | -1.81 | -1.15 | -1.24 | -0.83 | -1.24 | -1.76 | -1.49 | 0.07 | -1.47 | -1.21 | -0.43 | -0.55 | -1.31 | -1.18 | -0.03 | 0.1025 | 0.4475 | 0.368407 | 0.831684 | 0.688301 | 1 | 0.999631 | 0.745968 |
| Dlgap4_B7ZNS2_S956s_1 | -0.6 | -0.04 | -0.44 | -0.51 | 0.05 | 0.14 | -1.35 | -0.55 | 0.08 | -0.22 | -0.01 | -1.03 | 0.43 | 0.24 | -0.85 | 0.38 | 0.005 | -0.0675 | 0.1025 | 0.243929 | 0.936873 | 0.758142 | 1 | 0.999631 | 0.685637 |
| Dlgap4_B7ZNS2_T543t_1 | 0.02 | -0.37 | -0.43 | -0.3 | -0.21 | -0.08 | -0.43 | -0.34 | -0.42 | -0.1 | -0.33 | -0.5 | -0.26 | 0.08 | -0.36 | -0.13 | 0.46 | 0.625 | 0.41 | 0.713429 | 0.983966 | 0.794961 | 1 | 0.999631 | 0.900989 |
| Dlgap4_I1E4X0_S228s_1 | -0.37 | 0.49 | -0.99 | -0.78 | 1.15 | 0.51 | -0.5 | -0.97 | 0.64 | 1.09 | -0.36 | -0.52 | 0.88 | 0.32 | -1.04 | -0.17 | -1.2625 | -1.115 | -1.5625 | 0.420344 | 0.400014 | 0.218484 | 1 | 0.999631 | 0.77267 |
| Dmap1_Q9JI44_S455s_1 | 2.34 | 0.35 | 1.88 | 2.18 | -2.79 | 0.9 | 2.91 | 0.68 | -0.6 | 1.42 | 1.26 | 0.21 | -0.58 | -1 | 1.19 | 0.89 | -0.1375 | -0.05 | -0.28 | 0.031713 | 0.262212 | 0.087803 | 1 | 0.999631 | 0.442395 |
| Dmd_P11531_S3606s_1 | -0.21 | -1.1 | -0.78 | -0.01 | -0.78 | -0.9 | -0.69 | -0.28 | -0.78 | -0.68 | -0.23 | -0.61 | -0.4 | -1.35 | -0.64 | -0.83 | 0.27 | 0.085 | -0.1075 | 0.452712 | 0.676369 | 0.879803 | 1 | 0.999631 | 0.788431 |
| Dmtn_Q9WV69_S11s_1 | 0.54 | 0.01 | -0.43 | 0.63 | 0.86 | 0.09 | 0.19 | 0.69 | 0.03 | 0.27 | 0.86 | -0.07 | -0.97 | 0.03 | 0.28 | 0.98 | -0.015 | -0.9775 | -2.11 | 0.817447 | 0.439268 | 0.812362 | 1 | 0.999631 | 0.938024 |
| Dmtn_Q9WV69_S152s_1 | 2.81 | 0.06 | 0.55 | 3.05 | 0.47 | 1.71 | 2.89 | 1.34 | 0.58 | 0.43 | 1.6 | -0.05 | -2.11 | 0.06 | 0.06 | 0.02 | -0.6425 | -0.565 | -1.0525 | 0.025315 | 0.985018 | 0.209052 | 1 | 0.999631 | 0.422114 |
| Dmtn_Q9WV69_S156s_1 | 1.15 | -0.02 | 1.24 | 0.29 | -0.76 | 0.35 | 0.91 | -0.41 | -0.21 | 0.71 | -0.04 | -0.06 | -0.53 | -0.16 | -0.39 | -0.47 | 0.0425 | 0.74 | 0.5225 | 0.015766 | 0.190446 | 0.171603 | 1 | 0.999631 | 0.388832 |
| Dmtn_Q9WV69_S21s_1 | -1.27 | 0.8 | 2.3 | 0.58 | 1.13 | -0.12 | 0.08 | 1.49 | 1.21 | -0.17 | 1.98 | 2.35 | 0.71 | 0.1 | 1.87 | 1.82 | 0.28 | 4.91E-16 | 0.845 | 0.490106 | 0.953628 | 0.369856 | 1 | 0.999631 | 0.807727 |
| Dmtn_Q9WV69_S226s_1 | -2.47 | -0.55 | -0.66 | -1.04 | -0.52 | -1.23 | -1.14 | -0.71 | -1.31 | -1.31 | -1.26 | -0.84 | -0.06 | -0.1 | -0.73 | -0.45 | -0.0925 | 0.2925 | 0.5675 | 0.092478 | 0.543277 | 1 | 1 | 1 | 0.557085 |
| Dmtn_Q9WV69_S26s_1 | -0.59 | 0.19 | 0.12 | 0.1 | 0.32 | -0.42 | -0.06 | -0.39 | 0.22 | -0.74 | 1.11 | 0.4 | 0.51 | 0.43 | 0.56 | 0.59 | 0.7475 | -0.4975 | -0.0975 | 0.079469 | 0.764452 | 0.49601 | 1 | 0.999631 | 0.545579 |
| Dmtn_Q9WV69_S279s_1 | -0.18 | -0.05 | -0.64 | 0.33 | -0.42 | 1.62 | 0.52 | 0.73 | -0.36 | -0.47 | -0.06 | -1.64 | -1.24 | 1.38 | -0.11 | -0.96 | 0.145 | -0.1725 | 0.13 | 0.864807 | 0.120544 | 0.241939 | 1 | 0.999631 | 0.9555 |
| Dmtn_Q9WV69_S287s_1 | 0.91 | 1.5 | 1.11 | 0.66 | 1.47 | 1.24 | 0.68 | 1.37 | 1.1 | 1.05 | 0.31 | 1.03 | 1.42 | 1.37 | 0.95 | 0.96 | -0.4875 | -0.18 | 0.1125 | 0.677312 | 0.638973 | 0.591409 | 1 | 0.999631 | 0.887183 |
| Dmtn_Q9WV69_S289s_1 | -0.7 | -0.52 | -0.31 | -0.42 | -1.54 | -1.11 | -0.62 | -0.63 | -1.63 | -0.98 | -0.09 | 0.03 | -0.33 | -0.85 | -0.25 | -0.07 | 0.2525 | -0.5175 | -0.62 | 0.70561 | 0.127542 | 0.661313 | 1 | 0.999631 | 0.89781 |
| Dmtn_Q9WV69_S303s_1 | 0.8 | 0.64 | -0.4 | 1.02 | 1.09 | 0.48 | 0.81 | 0.69 | 0.45 | -0.2 | 0.07 | -0.33 | -1.02 | 0.59 | -0.29 | 0.3 | 0.33 | 0.0375 | 0.215 | 0.204128 | 0.491219 | 0.19522 | 1 | 0.999631 | 0.659781 |
| Dmtn_Q9WV69_S307s_1 | -0.48 | -0.41 | -0.68 | -0.54 | 0.2 | -0.02 | -0.56 | -0.41 | -0.27 | -1.16 | -0.27 | -0.26 | -0.71 | -0.09 | -0.66 | 0.21 | -0.53 | 0.0775 | 0.67 | 0.503005 | 0.244525 | 0.901669 | 1 | 0.999631 | 0.814092 |
| Dmtn_Q9WV69_S307s_1_1 | -1.36 | -0.33 | -0.05 | -0.93 | -1.16 | -1.47 | -0.89 | -1.27 | -0.96 | -0.29 | -0.99 | -0.12 | 0.59 | -0.24 | -0.27 | -0.07 | -0.07 | -0.1575 | 0.0525 | 0.108416 | 0.151985 | 0.842025 | 1 | 0.999631 | 0.570434 |
| Dmtn_Q9WV69_S315s_1 | -1.11 | -1.3 | -0.86 | -1.02 | -1.65 | -1.52 | -0.84 | -0.56 | -1.58 | -1.6 | -0.7 | -1.04 | -0.52 | -1.37 | -0.93 | -1.26 | 0.14 | 0.0525 | 0.4125 | 0.864701 | 0.82903 | 0.610749 | 1 | 0.999631 | 0.955486 |
| Dmtn_Q9WV69_S333s_1 | 0.43 | 0.85 | 0.21 | 0.88 | 0.55 | 1 | 0.56 | 0.82 | 0.61 | 0.22 | 0.4 | 1.35 | 0.17 | 1.56 | 1.12 | 1.17 | 0.24 | -0.1975 | -0.735 | 0.287909 | 0.618036 | 0.877873 | 1 | 0.999631 | 0.705972 |
| Dmtn_Q9WV69_S87s_1 | 0.79 | -0.27 | 0.02 | 0.57 | 0.71 | 0.31 | 0.73 | 0.32 | 0.37 | 0.04 | 0.17 | -0.26 | -0.89 | -0.19 | -0.21 | -0.54 | -0.385 | 0.46 | 1.0375 | 0.058646 | 0.458843 | 0.553042 | 1 | 0.999631 | 0.501503 |
| Dmtn_Q9WV69_S92s_1 | -1.39 | 0.27 | 0.32 | -1.04 | -1.25 | -0.41 | -1.32 | -0.4 | -0.38 | 0.01 | -0.05 | 0.42 | 1.25 | 0.51 | -0.05 | 0.6 | 0.28 | -0.1375 | -0.03 | 0.05888 | 0.432696 | 0.329383 | 1 | 0.999631 | 0.501953 |
| Dmtn_Q9WV69_S92s_2 | -0.38 | 0.36 | -0.01 | 0.56 | 0.55 | 0.14 | 0.21 | 0.75 | 0.22 | -0.33 | -0.06 | 0.15 | -0.59 | 0.57 | 0.14 | 0.29 | 0.34 | 0.22 | 0.23 | 0.934912 | 0.374943 | 0.658468 | 1 | 0.999631 | 0.979037 |
| Dmtn_Q9WV69_S96s_2 | -0.36 | 0.25 | -0.38 | 0.04 | 0.56 | 0.14 | 0.01 | 0.2 | 0.31 | -0.54 | 0.42 | 0.24 | -0.48 | 0.58 | -0.19 | 0.56 | 0.04 | -0.4175 | -0.075 | 0.526705 | 0.239542 | 0.502923 | 1 | 0.999631 | 0.824261 |
| Dmwd_E9Q6J8_S111s_1 | -0.01 | 0.61 | -1.17 | -1.79 | 0.1 | 0.23 | -1.17 | -1.36 | 0.26 | -0.04 | -1.7 | -2.55 | 0.01 | 0.38 | -1.24 | -1.81 | -0.035 | 0.03 | 0.3125 | 0.910725 | 0.948183 | 0.586552 | 1 | 0.999631 | 0.971199 |
| Dmwd_E9Q6J8_S487s_1 | 0.36 | 0.3 | -0.18 | -0.62 | 0.26 | 0.27 | -0.55 | -0.26 | 0.48 | 0.47 | -0.54 | -0.43 | 0.4 | 0.51 | 0.19 | 0.01 | 1.645 | 2.78 | 0.675 | 0.353983 | 0.918114 | 0.937322 | 1 | 0.999631 | 0.73969 |
| Dmwd_E9Q6J8_S537s_1 | -1.17 | 2.25 | 2.41 | 0.89 | 3.59 | 1.21 | 4.73 | 1.43 | 5.19 | -0.13 | 4.66 | 5.78 | 4.21 | 0.08 | 2.28 | 0.51 | -0.2925 | -0.1075 | 0.4875 | 0.531647 | 0.131882 | 0.063676 | 1 | 0.999631 | 0.827186 |
| Dmwd_E9Q6J8_S634s_1 | -0.27 | 0.6 | 0.24 | -0.43 | -0.15 | -0.84 | -0.18 | 0.14 | -0.41 | 0.13 | 0.16 | -0.17 | 0.88 | 0.31 | 0.55 | 0.35 | -0.24 | -0.4 | -0.1675 | 0.170301 | 0.406744 | 0.74274 | 1 | 0.999631 | 0.64073 |
| Dmwd_E9Q6J8_T453t_1 | 0.77 | 1.69 | 1.52 | 1.43 | 0.92 | 1.6 | 0.65 | 1.28 | 0.71 | 0.78 | 1.17 | 1.15 | 0.5 | 1.41 | 2.03 | 0.8 | -0.0025 | -0.0625 | 0.1425 | 0.685604 | 0.473747 | 0.2112 | 1 | 0.999631 | 0.890608 |
| Dmwd_E9Q6J8_T457t_1 | 0.43 | 0.16 | -0.53 | 0.17 | 0.02 | 0.21 | -0.2 | 0.19 | 0.05 | 0.38 | -0.35 | -0.1 | -0.15 | 0.21 | 0.44 | 0.3 | -0.205 | 0.175 | 0.5025 | 0.659802 | 0.993217 | 0.843995 | 1 | 0.999631 | 0.880252 |
| Dmwd_E9Q6J8_T461t_1 | -0.82 | -0.51 | -0.22 | -0.18 | -0.4 | -1.14 | -0.2 | -0.81 | -0.73 | 0.14 | 0.09 | -0.53 | 1.11 | -0.21 | -1.01 | 0.39 | -0.0675 | 0.085 | 0.65 | 0.295286 | 0.515271 | 0.590198 | 1 | 0.999631 | 0.708769 |
| Dmwd_Q08274_S659s_1 | -0.77 | -0.61 | -0.56 | -0.14 | -0.47 | -0.86 | -0.56 | -0.46 | -0.46 | -0.93 | -0.24 | -0.11 | -0.14 | -0.34 | 0.56 | 0.44 | 0.56 | 0.65 | 0.8425 | 0.072613 | 0.798257 | 0.777625 | 1 | 0.999631 | 0.533932 |
| Dmxl1_Q6PNC0_S1255s_1 | -0.93 | -1.41 | 0.76 | -1.28 | -0.49 | 0.75 | -1.08 | 0.2 | -0.1 | 0.03 | 0.22 | -0.41 | 0.05 | 0.44 | 0.26 | -0.24 | -0.2075 | 0.065 | -0.3575 | 0.116521 | 0.348228 | 0.208846 | 1 | 0.999631 | 0.583887 |
| Dmxl1_Q6PNC0_S1286s_1 | 1.04 | 1.15 | 0.48 | 1.03 | 0.71 | 1.26 | -0.21 | 1.11 | 0.84 | 1.2 | 0.74 | 1.18 | 0.18 | 0.84 | 0.83 | 0.42 | -0.3025 | -0.005 | 0.3425 | 0.268722 | 0.585619 | 0.818986 | 1 | 0.999631 | 0.696662 |
| Dmxl1_Q6PNC0_S1900s_1 | -0.2 | 0.86 | 0.86 | 0.22 | -0.34 | 0.45 | 0.2 | 0.22 | 0.58 | 0.33 | -0.01 | 0.82 | 1.55 | 0.33 | 0.33 | 0.9 | -0.535 | -0.8425 | -0.44 | 0.410911 | 0.387255 | 0.988672 | 1 | 0.999631 | 0.767937 |
| Dmxl1_Q6PNC0_S206s_1 | 1.41 | 0.09 | 2.29 | 1.13 | -0.24 | 1.26 | 1.2 | 0.56 | 0.62 | 1.84 | -0.74 | -0.17 | 0.25 | 1.31 | 1.19 | 0.41 | -0.2525 | -0.12 | -0.09 | 0.391521 | 0.325504 | 0.213339 | 1 | 0.999631 | 0.758901 |
| Dmxl1_Q6PNC0_S2372s_1 | 0.03 | -0.18 | -0.6 | 0.14 | -0.21 | -0.51 | -0.33 | -0.57 | -0.67 | -0.44 | 0.03 | -0.01 | -0.09 | -0.11 | -0.43 | -0.34 | -0.2825 | -0.3375 | 0.175 | 0.761189 | 0.36505 | 0.696279 | 1 | 0.999631 | 0.91472 |
| Dmxl1_Q6PNC0_S916s_1 | -0.59 | -0.7 | -1.28 | -1.62 | -1.07 | -0.97 | -1.4 | -1.88 | -1.43 | -1.08 | -1.42 | -1.61 | -0.75 | -0.79 | -0.69 | -1.26 | 0.1825 | 0.02 | 0.08 | 0.610051 | 0.425714 | 0.311644 | 1 | 0.999631 | 0.859048 |
| Dmxl1_Q6PNC0_S922s_1 | -0.13 | -0.13 | -1.15 | -0.31 | -0.05 | -0.21 | -0.36 | -0.37 | -0.26 | -0.05 | -0.61 | -0.72 | -0.64 | 0.05 | -0.44 | -0.37 | -0.22 | -0.43 | -0.575 | 0.816697 | 0.559809 | 0.952584 | 1 | 0.999631 | 0.937777 |
| Dmxl1_Q6PNC0_T661t_1 | 1.44 | -0.42 | 0.7 | 0.23 | -0.27 | 0.85 | 0.75 | -0.26 | 0.08 | 0.83 | -0.28 | -0.4 | 0.21 | 0.3 | -0.57 | -0.29 | -0.265 | -0.05 | 0.325 | 0.215147 | 0.644287 | 0.36637 | 1 | 0.999631 | 0.668192 |
| Dmxl2_Q8BPN8_S1141s_1 | -0.44 | 0.01 | -0.16 | -0.72 | -0.36 | -0.56 | -0.98 | -0.47 | -0.2 | -0.45 | -0.61 | -0.25 | 0.33 | -0.04 | 0 | -0.3 | -0.26 | -0.13 | 0.35 | 0.301284 | 0.364603 | 0.859051 | 1 | 0.999631 | 0.714243 |
| Dmxl2_Q8BPN8_S1144s_1 | -1.09 | -1.36 | -0.81 | -0.57 | -1.54 | -1.13 | -1.39 | -0.81 | -0.92 | -1.92 | -1.03 | -0.48 | -0.01 | -1.43 | -0.13 | -0.86 | -0.24 | -0.1125 | 0.3125 | 0.389626 | 0.393429 | 0.728109 | 1 | 0.999631 | 0.758766 |
| Dmxl2_Q8BPN8_S1152s_1 | -0.77 | 0.84 | 1.03 | 0.5 | 0.13 | 0.4 | -0.36 | 0.47 | 0.3 | -0.15 | 0.36 | 0.64 | 0.8 | 0.73 | 0.45 | 0.87 | -0.0025 | -0.1425 | 0.57 | 0.469273 | 0.585034 | 0.795845 | 1 | 0.999631 | 0.796658 |
| Dmxl2_Q8BPN8_S1288s_1 | -0.92 | -0.84 | -0.8 | -0.98 | -1.11 | -0.96 | -1.1 | -0.38 | -0.64 | -1.02 | -1.3 | -1.15 | 0.73 | -0.36 | -0.46 | -1.17 | 0.0625 | 0.2 | -0.05 | 0.18746 | 0.992601 | 0.593293 | 1 | 0.999631 | 0.646474 |
| Dmxl2_Q8BPN8_S1399s_1 | 1.06 | 1.78 | 1.35 | 1.96 | 1 | 1.74 | 1.7 | 1.96 | 1.4 | 1.24 | 2.57 | 1.74 | 0.84 | 1.59 | 1.56 | 1.96 | -0.4525 | -0.1675 | -0.1375 | 0.889614 | 0.850593 | 0.602585 | 1 | 0.999631 | 0.963505 |
| Dmxl2_Q8BPN8_S1403s_1 | -0.33 | -0.09 | -0.49 | -0.1 | -0.48 | -0.73 | -0.76 | -0.85 | -0.3 | 0 | -1.48 | 0.1 | 0.76 | -0.98 | -0.87 | -0.47 | -0.21 | -0.09 | 0.375 | 0.74531 | 0.09353 | 0.670629 | 1 | 0.999631 | 0.910585 |
| Dmxl2_Q8BPN8_S1856s_1 | -1.06 | -0.81 | 0.09 | -1.12 | -0.65 | -0.7 | -1.15 | -1.24 | -0.98 | -0.61 | -0.8 | -0.87 | -0.21 | -0.72 | -0.5 | 0.03 | 0.26 | 0.695 | -0.115 | 0.321713 | 0.55047 | 0.790011 | 1 | 0.999631 | 0.724092 |
| Dmxl2_Q8BPN8_S1925s_1 | 0.99 | 2.23 | 1.37 | 0.89 | 0.89 | 2.61 | 1.14 | 1.88 | 1.9 | 2.8 | 2.11 | 1.45 | 0.92 | 1.59 | 1.2 | 1.31 | -0.2125 | -0.3525 | -1.6825 | 0.759285 | 0.583464 | 0.121643 | 1 | 0.999631 | 0.914614 |
| Dmxl2_Q8BPN8_S2010s_1 | 1.62 | -0.94 | -0.12 | -0.51 | -1.35 | -0.74 | 1.25 | 0.04 | -1.46 | 0.87 | -0.33 | -0.44 | -0.91 | -2.87 | -0.83 | -2.07 | -0.0525 | 0.1375 | -0.5025 | 0.028197 | 0.762458 | 0.596418 | 1 | 0.999631 | 0.424192 |
| Dmxl2_Q8BPN8_S221s_1 | -1.46 | -0.98 | -2.4 | -2.04 | -1.55 | -1.96 | -1.12 | -2.46 | -1.38 | -1.95 | -1.78 | -1.22 | -0.9 | -1.22 | -4.24 | -2.53 | 0.0525 | -0.015 | 0 | 0.493485 | 0.900543 | 0.717729 | 1 | 0.999631 | 0.80955 |
| Dmxl2_Q8BPN8_S2395s_1 | -0.77 | -0.89 | -1.03 | -0.64 | -0.2 | -1.24 | -0.69 | -0.99 | -0.79 | -0.56 | -0.92 | -1.12 | -0.7 | -0.55 | -0.93 | -1.15 | -0.2725 | 0.0175 | -0.0425 | 1 | 0.861516 | 0.95461 | 1 | 0.999631 | 1 |
| Dmxl2_Q8BPN8_S2626s_1 | -0.9 | -0.98 | -0.19 | -0.74 | -1.52 | -1.05 | -0.83 | -0.5 | -0.71 | -1.07 | -0.44 | -0.52 | -0.65 | -1.29 | -0.54 | -0.5 | 0.0825 | 0.4275 | 0.1425 | 0.897519 | 0.408828 | 0.953685 | 1 | 0.999631 | 0.966758 |
| Dmxl2_Q8BPN8_S2697s_1 | -0.7 | 0.66 | 0.86 | 0.15 | -0.07 | 0.88 | 0.39 | 0.1 | 0.7 | 0.4 | 0.9 | 0.68 | 1.06 | 0.06 | 0.17 | 0.25 | -0.1675 | 0.2375 | 0.3575 | 0.73877 | 0.839302 | 0.280564 | 1 | 0.999631 | 0.90804 |
| Dmxl2_Q8BPN8_S2698s_1 | -0.29 | 0.74 | 0.78 | 0.12 | -0.04 | 0.46 | 0.27 | -0.01 | 0.57 | 0.49 | 0.64 | 0.6 | 1.22 | 0.37 | 0.47 | 0.72 | -0.0675 | 0.065 | -0.7125 | 0.340602 | 0.611207 | 0.464201 | 1 | 0.999631 | 0.733032 |
| Dmxl2_Q8BPN8_S423s_1 | -0.54 | 0.56 | 0.95 | -0.22 | -0.34 | 1 | 0.37 | -0.55 | 0.15 | 0.37 | 0.26 | 0.23 | -0.19 | -1.12 | -0.09 | -0.7 | 0.4425 | 0.2375 | 0.27 | 0.119962 | 0.885699 | 0.861519 | 1 | 0.999631 | 0.587703 |
| Dmxl2_Q8BPN8_S423s_2 | -1.61 | 0.08 | -0.65 | -0.31 | 0.27 | -0.37 | -0.54 | -0.08 | 0.43 | -1.4 | -0.27 | -0.3 | 0.04 | 0.38 | -0.42 | -1.41 | 0.1475 | 0.1525 | -0.5125 | 0.596288 | 0.289646 | 0.634628 | 1 | 0.999631 | 0.854508 |
| Dmxl2_Q8BPN8_S434s_1 | -0.67 | -0.82 | 0.15 | -0.26 | -0.63 | -0.79 | 0.74 | -0.33 | -0.44 | -0.66 | 0.02 | 0.09 | -0.24 | -1.49 | -0.57 | -1.35 | 0.4425 | 0.2375 | 0.27 | 0.217091 | 0.718696 | 0.649052 | 1 | 0.999631 | 0.669465 |
| Dmxl2_Q8BPN8_S434s_2 | -1.61 | 0.08 | -0.65 | -0.31 | 0.27 | -0.37 | -0.54 | -0.08 | 0.43 | -1.4 | -0.27 | -0.3 | 0.04 | 0.38 | -0.42 | -1.41 | -0.1275 | 0.085 | 0.435 | 0.596288 | 0.289646 | 0.634628 | 1 | 0.999631 | 0.854508 |
| Dmxl2_Q8BPN8_S451s_1 | -2.05 | -2.02 | -0.84 | -1.16 | -1.35 | -2.96 | -1.49 | -0.78 | -1.84 | -2.55 | -0.5 | -0.84 | -0.41 | -1.77 | -1.43 | -0.72 | -0.76 | -1.1175 | -1.8975 | 0.336076 | 0.805407 | 0.871206 | 1 | 0.999631 | 0.730466 |
| Dmxl2_Q8BPN8_S462s_1 | 1.4 | -1.42 | -0.12 | 0.22 | -2.29 | -0.62 | 0.89 | -0.94 | -1.82 | 0.35 | -0.77 | -2.15 | -1.05 | -3.05 | -1.4 | -2.01 | -0.52 | -0.5825 | -2.055 | 0.01467 | 0.333566 | 0.144054 | 1 | 0.999631 | 0.386791 |
| Dmxl2_Q8BPN8_S463s_1 | 1.94 | -1.46 | -0.1 | 0.49 | -1.09 | 0.04 | 0.93 | -1.09 | -1.23 | 1.01 | -0.94 | -0.3 | -1.82 | -2.69 | -1.18 | -1.66 | 0.95 | 1.32 | 0.875 | 0.012727 | 0.49541 | 0.451041 | 1 | 0.999631 | 0.385092 |
| Dmxl2_Q8BPN8_S481s_1 | -1.59 | 2.48 | 1.52 | -0.85 | 2.19 | 0.93 | 1.59 | 0.65 | 2.93 | -1.24 | 2.32 | 2.83 | 3.28 | -0.06 | 1.38 | 0.46 | -0.04 | 0.0625 | 1.11 | 0.405594 | 0.298134 | 0.280738 | 1 | 0.999631 | 0.765965 |
| Dnaja1_P63037_S112s_1 | -0.61 | 2.6 | 0.22 | -0.15 | -0.19 | 1.93 | -0.54 | 0.7 | -0.79 | 2 | 1 | 0.1 | 1.08 | 1.26 | 1.65 | 2.51 | -0.245 | 0.11 | -0.1375 | 0.133968 | 0.959547 | 0.93899 | 1 | 0.999631 | 0.60492 |
| Dnaja1_P63037_S335s_1 | -2.46 | -3.23 | -2.22 | -2.7 | -2.64 | -3.57 | -2.1 | -3.28 | -3.14 | -3.13 | -2.27 | -1.63 | -3.64 | -3.61 | -2.02 | -1.89 | 0.2775 | 0.355 | -0.3425 | 0.78558 | 0.542812 | 0.796465 | 1 | 0.999631 | 0.923063 |
| Dnajb1_Q9QYJ3_S16s_1 | 0.94 | 0.03 | 0.12 | 0.02 | 1.12 | 0.67 | 0.31 | 0.12 | 0.43 | 1.27 | 0.69 | 0.14 | 0.73 | -0.43 | 0.06 | -0.62 | -0.54 | -0.175 | -0.525 | 0.399739 | 0.429 | 0.336946 | 1 | 0.999631 | 0.762224 |
| Dnajb2_Q9QYI5_S311s_1 | 0.41 | -1.11 | 1.62 | -0.35 | -1.16 | -1.09 | 0.01 | 0.65 | -2.06 | -0.24 | 0.93 | 1.24 | -1.24 | -1.66 | 1.14 | 0.23 | -0.18 | -0.0475 | -0.2175 | 0.497304 | 0.41569 | 0.832819 | 1 | 0.999631 | 0.811649 |
| Dnajb2_Q9QYI5_S78s_1 | -0.26 | -1.19 | -0.03 | -0.35 | -1.05 | -1.03 | -0.34 | -0.13 | -1.4 | -0.38 | -0.17 | -0.07 | -1.22 | -1.19 | -0.24 | -0.05 | 0.5725 | -0.6125 | -1.4125 | 0.602776 | 0.625547 | 0.90686 | 1 | 0.999631 | 0.857278 |
| Dnajb4_Q9D832_S148s_1 | 1.91 | 0.54 | 0.9 | 1.9 | 1.61 | 2.05 | 2.43 | 1.45 | 1 | 0.08 | 1.25 | 0.47 | -1.34 | 0.12 | 1.06 | -0.24 | 0.22 | 0.5775 | 0.9825 | 0.028755 | 0.187259 | 0.181207 | 1 | 0.999631 | 0.427571 |
| Dnajc2_Q3TRX6_S47s_1 | -0.03 | 2.22 | 0.45 | 1.74 | 1.88 | 0.23 | 1.13 | 2.02 | 1.5 | 0.04 | 2.2 | 2.95 | 2.01 | 2.01 | 2.33 | 1.96 | -0.3125 | -0.385 | 0.125 | 0.078039 | 0.716821 | 0.432186 | 1 | 0.999631 | 0.543121 |
| Dnajc5_P60904_S10s_1 | -1.19 | -1.61 | -1.64 | -1.79 | -1.83 | -1.83 | -1.71 | -2.11 | -1.9 | -1.46 | -2.37 | -2.04 | -0.74 | -1.56 | -2.04 | -1.39 | -0.2975 | -0.32 | 0.1575 | 0.723942 | 0.246817 | 0.222465 | 1 | 0.999631 | 0.905674 |
| Dnajc5_P60904_S12s_1 | -1.08 | -1.76 | -1.96 | -2.02 | -1.96 | -1.79 | -1.72 | -2.54 | -2.14 | -1.71 | -2.56 | -1.69 | -0.98 | -1.65 | -2.07 | -1.49 | -0.055 | -0.18 | 0.205 | 0.664014 | 0.374457 | 0.361223 | 1 | 0.999631 | 0.881993 |
| Dnajc5_P60904_S15s_1 | -0.54 | -0.21 | -0.97 | -1.88 | -1.38 | -0.2 | -1.55 | -0.69 | -1.92 | -0.47 | -0.68 | -1.25 | -0.81 | 0.09 | -1.6 | -0.46 | -0.5675 | -0.15 | -0.05 | 0.676014 | 0.904614 | 0.702637 | 1 | 0.999631 | 0.886277 |
| Dnajc5_P60904_T11t_1 | -0.75 | -1.76 | -1.05 | -1.32 | -2.19 | -2.51 | -1.35 | -1.1 | -2.04 | -0.89 | -1.45 | -1.1 | -1.75 | -1.4 | -0.93 | -1 | -0.05 | -0.1325 | -0.0775 | 0.885294 | 0.179568 | 0.680092 | 1 | 0.999631 | 0.963505 |
| Dnajc6_B1B0B7_S38s_1 | -0.52 | -0.64 | -0.5 | -0.73 | -0.41 | -0.27 | -0.91 | -1 | -0.84 | -0.91 | -0.75 | -0.42 | -0.63 | -0.61 | -1.4 | -0.06 | -0.0975 | -0.06 | 0.355 | 0.821463 | 0.855809 | 0.6066 | 1 | 0.999631 | 0.938464 |
| Dnajc6_B1B0B7_S39s_1 | -0.85 | -0.78 | -0.67 | -0.92 | -0.9 | -0.94 | -1.06 | -0.71 | -1.03 | -0.86 | -0.71 | -0.86 | -0.22 | -0.59 | -0.64 | -0.35 | -0.0025 | 0.135 | -0.045 | 0.208984 | 0.683097 | 0.806751 | 1 | 0.999631 | 0.662575 |
| Dnajc6_B1B0B7_T403t_1 | -1.03 | -0.69 | -0.56 | -1.51 | -0.52 | -0.86 | -0.68 | -1.74 | -0.93 | -1.08 | -0.09 | -1.15 | 0.56 | -0.28 | -2.09 | -2.16 | 1.16 | -0.4075 | -1.0425 | 0.943725 | 0.994543 | 0.707276 | 1 | 0.999631 | 0.981629 |
| Dnajc6_Q80TZ3_S591s_2 | 2.55 | -0.28 | -1.51 | 1.98 | 2.86 | 0.51 | 2.57 | 1.44 | 1.18 | -1.13 | 1.25 | -0.19 | -3.13 | 1.13 | 0.2 | 0.37 | 1.16 | -0.4075 | -1.0425 | 0.370665 | 0.23837 | 0.673546 | 1 | 0.999631 | 0.746338 |
| Dnajc6_Q80TZ3_S595s_2 | 2.55 | -0.28 | -1.51 | 1.98 | 2.86 | 0.51 | 2.57 | 1.44 | 1.18 | -1.13 | 1.25 | -0.19 | -3.13 | 1.13 | 0.2 | 0.37 | 0.6775 | -0.5075 | -0.5675 | 0.370665 | 0.23837 | 0.673546 | 1 | 0.999631 | 0.746338 |
| Dnajc6_Q80TZ3_S734s_1 | 1.61 | -1.61 | -0.31 | 1.18 | 1.71 | 0.49 | 0.8 | 0.58 | 0.17 | -0.51 | -0.5 | -0.32 | -2.6 | 0.64 | 0.28 | 0.28 | 0.69 | -0.3925 | -0.1475 | 0.535721 | 0.342691 | 0.457709 | 1 | 0.999631 | 0.829362 |
| Dnajc6_Q80TZ3_S739s_1 | 1.88 | 0.01 | -1.09 | -0.92 | 1.34 | 0.06 | 0.73 | 0.51 | -0.16 | -0.65 | -0.28 | -0.6 | 0.46 | -0.02 | -0.05 | -1.1 | 0.91 | -0.2225 | 0.075 | 0.826959 | 0.304726 | 0.534591 | 1 | 0.999631 | 0.940917 |
| Dnajc6_Q80TZ3_S769s_1 | -0.9 | -0.67 | 0.23 | -0.33 | 0.05 | 0.65 | 1.05 | 0.22 | -0.28 | -1.03 | -0.31 | -0.94 | -0.4 | -0.79 | -0.14 | -0.04 | 0.7775 | -0.1675 | 0.3575 | 0.831586 | 0.027684 | 0.533558 | 1 | 0.999631 | 0.942722 |
| Dnajc6_Q80TZ3_S770s_1 | 0.11 | -1.29 | -0.39 | -0.01 | 0.53 | 0.28 | 0.98 | -0.26 | -0.17 | -0.85 | -0.75 | -0.48 | -1.19 | 0.98 | 0.69 | -0.63 | 0.685 | 0.51 | -0.5875 | 0.529729 | 0.082124 | 0.657447 | 1 | 0.999631 | 0.826087 |
| Dnajc6_Q80TZ3_T590t_1 | 1.91 | -0.53 | 0.2 | -1.43 | 2.17 | 0.49 | -0.1 | 0.33 | 0.74 | -0.19 | 0.66 | 0.98 | -2.48 | 1.4 | -0.1 | -1.02 | -1.1 | 0.995 | 0.895 | 0.530528 | 0.377364 | 0.45558 | 1 | 0.999631 | 0.826702 |
| Dnajc8_E9PZT4_S35s_1 | -0.42 | 3.3 | 3.09 | 2.02 | -0.29 | 1.31 | 0.48 | 2.09 | 2.6 | 2.64 | 2.85 | 3.88 | 2.59 | 2.2 | 3.21 | 3.57 | -0.145 | -0.275 | 0.2 | 0.273902 | 0.222844 | 0.22963 | 1 | 0.999631 | 0.700187 |
| Dnal1_D3YYU7_S56s_1 | 0.41 | 1.48 | 1.26 | 1.05 | 1.07 | 1.08 | 0.55 | 0.92 | 1.53 | 1.1 | 0.71 | -0.24 | 0.94 | 2.24 | 0.71 | 1.11 | -0.0225 | 0.16 | 0.235 | 0.638032 | 0.646033 | 0.534608 | 1 | 0.999631 | 0.87033 |
| Dnm1_P39053_S512s_1 | -0.7 | -0.66 | 0.18 | -0.47 | -0.19 | -0.75 | -0.44 | -0.36 | -0.77 | -0.3 | 0.07 | -0.01 | -0.09 | -0.83 | 0.14 | 0.07 | -0.595 | -0.3325 | -0.5075 | 0.512982 | 0.939971 | 0.630313 | 1 | 0.999631 | 0.818435 |
| Dnm1_P39053_S61s_1 | 0.47 | 0.33 | 1.04 | 0.5 | -0.17 | -0.12 | 0.64 | -0.39 | -0.73 | 0.63 | 0.81 | 0.3 | -0.54 | 0.04 | 0.27 | 0.54 | 0.3075 | -0.095 | 0.3425 | 0.159865 | 0.088386 | 0.406955 | 1 | 0.999631 | 0.629764 |
| Dnm1_P39053_S644s_1 | 0.6 | -1.59 | -0.98 | -2.04 | -0.91 | -0.07 | -0.77 | -1.03 | -1.43 | -0.98 | -0.6 | -1.38 | -0.92 | 0.52 | -1.86 | -0.38 | 1.5825 | -1.57 | -2.8975 | 0.616081 | 0.587352 | 0.865602 | 1 | 0.999631 | 0.861203 |
| Dnm1_P39053_S774s_1 | 3.69 | -2.28 | -2.69 | 2.54 | 3.24 | -0.37 | 2.98 | 1.74 | 0.79 | -4.23 | 0.87 | -2.45 | -4.55 | -1.75 | -1.72 | -2.31 | 1.2925 | -2.5125 | -3.9575 | 0.074695 | 0.321195 | 0.379517 | 1 | 0.999631 | 0.540402 |
| Dnm1_P39053_S774s_2 | 4.5 | -3.25 | -2.07 | 2.12 | 3.48 | -1.43 | 3.6 | 0.82 | -0.08 | -4.63 | -0.23 | -3.81 | -4.81 | -3.27 | -3.04 | -3.41 | -0.3925 | -0.4875 | -0.7075 | 0.02705 | 0.486401 | 0.191256 | 1 | 0.999631 | 0.422893 |
| Dnm1_P39053_S777s_1 | 0.58 | 1.36 | 1.08 | 0.4 | -0.27 | 1.87 | 0.05 | 0.2 | 0.53 | 1.51 | 0.2 | -0.77 | 0.64 | 0.74 | -0.45 | -0.34 | 0.625 | -0.81 | -1.87 | 0.10548 | 0.436377 | 0.336903 | 1 | 0.999631 | 0.568524 |
| Dnm1_P39053_S778s_1 | 2.82 | -0.61 | 0.11 | 1.69 | 1.87 | 0.79 | 2.22 | 1.63 | 0.81 | -0.17 | 0.84 | -0.71 | -1.96 | -0.8 | -0.16 | -0.55 | 1.395 | -2.865 | -4.07 | 0.031271 | 0.402144 | 0.301479 | 1 | 0.999631 | 0.441911 |
| Dnm1_P39053_S778s_2 | 4.61 | -3.52 | -2.65 | 2.28 | 3.63 | -1.51 | 3.57 | 0.61 | 0.1 | -6.62 | -0.29 | -3.93 | -5.59 | -3.39 | -3.03 | -3.55 | 1.345 | 0.19 | -0.53 | 0.0356 | 0.481371 | 0.19962 | 1 | 0.999631 | 0.453811 |
| Dnm1_P39053_S795s_1 | 1.95 | -0.81 | -2 | 1.91 | 1.88 | 1.23 | 1.4 | 1.92 | 0.85 | -0.24 | 1.6 | -0.4 | -2.9 | 1.12 | 0.68 | 0.03 | 0.0925 | -0.965 | -1.2075 | 0.643714 | 0.147641 | 0.841907 | 1 | 0.999631 | 0.872211 |
| Dnm1_P39053_S822s_1 | 1.62 | -0.48 | -1.23 | 0.42 | 0.93 | -1.09 | 0.9 | -0.04 | -0.15 | -1.09 | -1.06 | -1.23 | -0.64 | -0.49 | -1.76 | -1.61 | 0.105 | -0.295 | -0.255 | 0.078339 | 0.893211 | 0.133479 | 1 | 0.999631 | 0.544068 |
| Dnm1_P39053_S847s_1 | 1.09 | 1.1 | 1.49 | 1.39 | 0.99 | 1.33 | 1.95 | 1.22 | 1.07 | 1.22 | 1.07 | 0.53 | 0.14 | 1.36 | 1.17 | 1.38 | 0.705 | 0.0875 | -0.64 | 0.485458 | 0.723208 | 0.306227 | 1 | 0.999631 | 0.804287 |
| Dnm1_P39053_S851s_1 | 1.01 | 0.48 | -0.61 | 1.32 | 1.83 | 1.06 | 0.75 | 1.38 | 1.23 | 0.24 | 0.99 | 0.09 | -2.77 | 1.05 | 0.56 | 0.8 | 1.155 | -0.18 | -1.1725 | 0.463146 | 0.150903 | 0.856933 | 1 | 0.999631 | 0.793564 |
| Dnm1_P39053_S853s_1 | 2.23 | -0.92 | -1.4 | 1.11 | 2.84 | 1.15 | 0.89 | 0.76 | 0.69 | -0.77 | 0.49 | -0.11 | -4.95 | 0.76 | 0.28 | 0.24 | 1.63 | -0.4425 | -1.215 | 0.392754 | 0.196657 | 0.822719 | 1 | 0.999631 | 0.758901 |
| Dnm1_P39053_S857s_1 | 2.11 | -1.2 | -1.79 | 1.52 | 3.23 | 0.94 | 1.59 | 1.4 | 0.77 | -1.5 | 0.87 | -1.27 | -3.23 | 0.41 | -0.59 | -0.81 | -0.7375 | 0.2175 | 0.76 | 0.266482 | 0.108514 | 0.660591 | 1 | 0.999631 | 0.695664 |
| Dnm1_P39053_T404t_1 | -0.01 | -0.25 | -1.21 | -0.76 | -1.35 | -1.22 | -1.15 | -1.46 | 0.6 | 0.53 | -1.86 | -0.63 | 0.58 | 0.28 | 0.12 | -0.17 | 1.125 | -1.96 | -3.475 | 0.056999 | 0.042908 | 0.710736 | 1 | 0.999631 | 0.496676 |
| Dnm1_P39053_T776t_2 | 3.97 | -2.36 | -1.22 | 1.84 | 2.82 | -1.11 | 3.7 | 1.32 | -0.86 | -1.48 | 0.05 | -3.32 | -2.79 | -2.82 | -2.82 | -3.24 | 0.7675 | -0.015 | -0.3075 | 0.016472 | 0.463862 | 0.175234 | 1 | 0.999631 | 0.389192 |
| Dnm1l_Q8K1M6_S532s_1 | 1.35 | 0.35 | -1.48 | 0.98 | 1.56 | 0.6 | 0.73 | 1.38 | 0.77 | -0.04 | 0.59 | -0.18 | -1.4 | 0.66 | 0.44 | 0.27 | 0.3 | 0.0925 | -0.2075 | 0.660375 | 0.224487 | 0.980325 | 1 | 0.999631 | 0.880315 |
| Dnm1l_Q8K1M6_S535s_1 | -0.08 | -1.8 | -0.63 | 1.22 | -0.62 | -1.69 | 0.81 | 1.41 | -1.51 | -1.47 | 1.18 | 0.88 | -2.56 | -1.35 | 0.76 | 1.03 | 0.7625 | 0.32 | 0.36 | 0.820936 | 0.714942 | 0.912362 | 1 | 0.999631 | 0.938464 |
| Dnm1l_Q8K1M6_S622s_1 | -0.05 | 0.26 | -0.42 | 1.06 | 1.18 | 1.01 | 0.7 | 1.01 | 0.56 | -0.04 | 0.84 | 0.77 | -0.65 | 0.99 | 0.84 | 1.11 | -0.94 | 0.1675 | 0.59 | 0.474781 | 0.054773 | 0.420373 | 1 | 0.999631 | 0.799827 |
| Dnm3_E9QLL2_S763s_1 | -1.78 | -0.66 | 0.07 | -1.48 | -2.76 | -1.41 | -1.96 | -1.48 | -1.4 | -0.13 | -1 | -0.65 | 0.33 | -0.8 | -0.64 | -0.38 | 0.81 | -0.4725 | -1.4 | 0.232397 | 0.07996 | 0.726625 | 1 | 0.999631 | 0.682449 |
| Dnm3_E9QLL2_S763s_2 | 1.71 | -0.36 | -1.5 | 1.49 | 1.63 | 0.62 | 1.28 | 1.05 | 0.49 | -0.02 | 0.08 | -1.1 | -2.31 | -0.17 | -0.5 | -1.28 | 0.0325 | -0.3675 | -1.1225 | 0.100275 | 0.266798 | 0.52885 | 1 | 0.999631 | 0.563325 |
| Dnm3_E9QLL2_S767s_1 | 0.78 | -0.6 | -0.47 | 0.87 | 0.26 | -0.59 | 0.85 | 0.19 | -0.4 | 0.05 | 0 | -0.54 | -0.82 | -1.52 | -0.69 | -0.88 | 0.7425 | -0.4925 | -1.295 | 0.026742 | 0.944777 | 0.394028 | 1 | 0.999631 | 0.422285 |
| Dnm3_E9QLL2_S767s_2 | 1.46 | -0.48 | -2.04 | 1.27 | 1.29 | 0.14 | 1.02 | 0.73 | 0.4 | -0.25 | -0.46 | -1.45 | -2.27 | -0.4 | -0.89 | -1.41 | 0.9425 | -0.43 | -1.525 | 0.128423 | 0.337337 | 0.54067 | 1 | 0.999631 | 0.599259 |
| Dnm3_E9QLL2_T769t_2 | 2.09 | -0.16 | -0.88 | 1.76 | 2.17 | 1.37 | 1.6 | 1.44 | 0.66 | 0.38 | 0.72 | -0.67 | -2.38 | 0.22 | -0.02 | -1.11 | -0.34 | 0.7475 | 0.1325 | 0.085541 | 0.178368 | 0.545945 | 1 | 0.999631 | 0.551124 |
| Dnmt3a_G3UXY2_S28s_1 | 0.2 | -1.29 | -1.24 | -1.48 | -1.19 | -0.95 | -1.51 | -1.52 | 0.31 | 0.63 | -0.96 | -0.8 | -0.06 | -0.11 | -1.51 | -1.6 | 0.1525 | 0.0225 | 0.32 | 0.805711 | 0.416534 | 0.173543 | 1 | 0.999631 | 0.934783 |
| Dnmt3a_G3UXY2_T46t_1 | 0.87 | -0.12 | 0.37 | -0.33 | 0.23 | 0.54 | 0.33 | 0.3 | 0.99 | 0.82 | -0.89 | -0.04 | 0.99 | 0.52 | 0.44 | 0.12 | -0.1325 | 0.035 | 0.465 | 0.392948 | 0.638088 | 0.96311 | 1 | 0.999631 | 0.758901 |
| Dnpep_Q8BPW9_T201t_1 | 0.75 | 1.76 | 1.43 | 1.11 | 1.23 | 1.29 | 0.36 | 1.64 | 1.2 | 1.16 | 1.11 | 1.72 | 2.11 | 1.52 | 1.55 | 1.73 | 0.7325 | -0.1 | -0.025 | 0.179672 | 0.719158 | 0.912614 | 1 | 0.999631 | 0.64376 |
| Doc2a_H3BJT3_S156s_1 | -0.16 | -0.57 | -1.55 | -0.16 | 0.56 | -0.22 | -0.08 | 0.23 | -0.34 | -1.02 | -0.3 | -1.18 | -1.2 | 0.04 | -0.61 | -0.77 | 0.3125 | 0.7675 | 0.725 | 0.953246 | 0.080322 | 0.806509 | 1 | 0.999631 | 0.984325 |
| Dock1_Q8BUR4_S1858s_1 | 0.02 | 0 | -0.85 | 0.23 | 0.05 | -0.3 | -0.34 | 1.24 | 0.3 | 0.16 | 1.2 | 0.81 | 0.88 | 0.39 | 0.61 | 0.42 | -0.6925 | -0.15 | -1.1475 | 0.051673 | 0.475111 | 0.059749 | 1 | 0.999631 | 0.489777 |
| Dock11_A2AF67_S1095s_1 | 1.5 | -1.46 | 3.53 | 3 | 0.02 | -0.07 | 1.26 | 2.59 | -0.41 | 0.69 | 3.13 | 2.56 | -1.72 | -1.59 | 3.09 | 2.2 | -0.32 | -0.02 | -0.0725 | 0.425079 | 0.532628 | 0.899567 | 1 | 0.999631 | 0.774293 |
| Dock3_Q8CIQ7_S1655s_1 | -0.05 | -0.77 | -0.3 | -0.1 | -0.61 | -0.95 | -0.44 | -0.5 | -0.5 | -0.48 | -0.04 | -0.28 | -0.84 | -0.05 | -0.58 | -0.04 | -0.025 | 0.3475 | 0.42 | 0.826739 | 0.27044 | 0.944249 | 1 | 0.999631 | 0.940834 |
| Dock3_Q8CIQ7_S1763s_1 | 0.64 | 1 | 0.75 | 0.8 | 0.81 | 1.12 | 0.49 | 0.67 | 1.3 | 1.48 | 0.73 | 1.07 | 1.01 | 1.48 | 1.25 | 1.13 | 0.02 | -0.1675 | -1.5075 | 0.148501 | 0.923133 | 0.230342 | 1 | 0.999631 | 0.62132 |
| Dock3_Q8CIQ7_S1765s_1 | 0.51 | 0.27 | -5.28 | -5.15 | 0.21 | 0 | -4.82 | -4.96 | 1.09 | 0.5 | -4.71 | -7.2 | -1 | 0.71 | -6.75 | -8.64 | 0.2825 | -1.29 | -1.255 | 0.517001 | 0.99128 | 0.938842 | 1 | 0.999631 | 0.819449 |
| Dock3_Q8CIQ7_T200t_1 | -0.54 | 1.13 | 1.9 | 0.7 | -1.84 | 3.47 | 0.55 | 2.14 | -1.78 | -0.62 | 3.1 | -2.67 | -3.28 | 2.92 | -0.69 | -0.78 | 0.2725 | -0.3 | 0.465 | 0.295698 | 0.792366 | 0.28657 | 1 | 0.999631 | 0.708981 |
| Dock4_P59764_S1608s_1 | -1.17 | 0 | -0.25 | -0.83 | 0.09 | -0.34 | -0.48 | -0.43 | -0.82 | -1.63 | -1.12 | 0.12 | 0.65 | -0.4 | -0.28 | -0.36 | -0.24 | 0.0125 | -0.12 | 0.254641 | 0.424165 | 0.507043 | 1 | 0.999631 | 0.689511 |
| Dock4_P59764_S1629s_1 | 0.15 | -0.05 | -0.07 | -0.14 | -0.31 | -0.31 | -0.06 | -0.39 | -0.12 | 0.33 | -0.07 | -0.2 | -0.26 | -0.06 | -0.2 | -0.07 | 0.3225 | 0.0625 | -0.02 | 0.648934 | 0.328635 | 0.961603 | 1 | 0.999631 | 0.874615 |
| Dock4_P59764_S1709s_1 | 0.47 | -0.13 | -0.57 | 0.4 | 0.55 | 0.22 | 0.17 | 0.52 | 0.05 | 0.09 | 0.65 | -0.37 | -0.72 | 0.72 | -0.16 | 0.25 | 1.8025 | 0.305 | -0.9275 | 0.961071 | 0.319389 | 0.860937 | 1 | 0.999631 | 0.986594 |
| Dock4_P59764_S1726s_1 | -0.06 | -0.36 | -1.07 | 0.01 | 1.9 | 0.85 | 2.05 | 0.93 | 0.96 | -1.34 | 1.15 | -1.03 | -2.17 | -0.45 | -0.51 | -2.06 | -0.1525 | 0.1475 | 0.145 | 0.092183 | 0.001263 | 0.629821 | 1 | 0.999631 | 0.557085 |
| Dock4_P59764_T1869t_1 | 0.24 | 0.89 | 0.68 | 0.91 | 0.65 | 0.55 | 0.27 | 0.64 | 0.6 | 0.58 | 1.15 | 0.98 | -0.18 | 1.19 | 0.92 | 1.37 | -0.3275 | -0.78 | 0.7725 | 0.719894 | 0.576283 | 0.617147 | 1 | 0.999631 | 0.904808 |
| Dock5_B2RY04_S1832s_1 | -0.48 | 0.08 | -0.72 | -0.25 | -0.32 | -0.98 | -0.93 | -0.45 | -1.79 | -1.4 | -1.4 | 0.1 | 0.2 | 0.44 | 0.92 | 0.16 | 0.0575 | 0.7 | 0.76 | 0.035611 | 0.291534 | 0.10444 | 1 | 0.999631 | 0.453811 |
| Dock6_Q8VDR9_S178s_1 | -0.46 | 1.57 | 1.23 | -0.2 | 0.7 | 1.29 | -0.16 | 0.54 | 1.42 | 1.46 | 1.09 | 0.97 | 1.03 | 1.07 | 1.66 | 1.42 | -0.26 | -0.0175 | 0.075 | 0.154923 | 0.915379 | 0.180101 | 1 | 0.999631 | 0.6258 |
| Dock7_Q8R1A4_S1382s_1 | 0.2 | 1.05 | 1.19 | 1.01 | 0.4 | 0.79 | 0.71 | 0.51 | 0.99 | 0.88 | 0.77 | 0.74 | 0.54 | 1.08 | 0.76 | 1.37 | 0.4075 | -0.8725 | -0.28 | 0.829304 | 0.399308 | 0.954061 | 1 | 0.999631 | 0.942014 |
| Dock7_Q8R1A4_S1428s_1 | 0.06 | -0.44 | -0.62 | -0.67 | 0.64 | 0 | -0.38 | -0.3 | 0.23 | -2.32 | -2.21 | -0.86 | -0.67 | 0.04 | -1.07 | -1.09 | -0.1575 | 0.3275 | -0.6025 | 0.445412 | 0.232823 | 0.153808 | 1 | 0.999631 | 0.784262 |
| Dock7_Q8R1A4_S180s_1 | 0.95 | 1.18 | 0.01 | 0.41 | 0.83 | 0.73 | 0.33 | 0.03 | 0.76 | 2.06 | 0.7 | 0.34 | -0.82 | 1.03 | -0.03 | -0.04 | -0.24 | -0.1325 | -0.6925 | 0.208218 | 0.655295 | 0.473872 | 1 | 0.999631 | 0.661649 |
| Dock7_Q8R1A4_S182s_1 | 1.88 | 0.55 | -0.27 | 1.63 | 1.01 | 0.22 | 1.04 | 0.56 | 1.01 | 0.22 | 1.16 | 0.87 | -0.77 | 0.89 | -0.14 | 1.04 | 0.215 | 0.525 | 0.2225 | 0.264891 | 0.634063 | 0.794865 | 1 | 0.999631 | 0.693761 |
| Dock7_Q8R1A4_S30s_1 | -0.41 | -0.7 | 0.27 | -0.18 | 0.67 | 1.23 | -0.88 | -1.18 | 0.66 | 0.08 | 0.29 | 0.05 | -1.52 | 1.89 | -0.41 | -0.09 | -0.105 | -0.2825 | -0.11 | 0.737127 | 0.705216 | 0.118451 | 1 | 0.999631 | 0.907787 |
| Dock7_Q8R1A4_S864s_1 | -0.56 | -0.21 | -0.71 | -0.62 | -1.42 | -0.13 | -0.34 | -0.63 | -1.17 | -0.51 | -0.29 | -1.26 | -0.02 | -0.85 | -0.33 | -1.34 | -0.4325 | -0.0675 | -0.015 | 0.760554 | 0.757112 | 0.390134 | 1 | 0.999631 | 0.914614 |
| Dock7_Q8R1A4_S900s_1 | 1.46 | 0.15 | -1.98 | 0.63 | -1.2 | 0.88 | -0.76 | -0.39 | -1.82 | 3.23 | -0.15 | -1.27 | -2.09 | 3.43 | -2.33 | 1.19 | -0.1725 | -0.03 | 0.145 | 0.990924 | 0.569623 | 0.953418 | 1 | 0.999631 | 0.99628 |
| Dock7_Q8R1A4_S929s_1 | -0.19 | -0.55 | 0.4 | 1.22 | -0.53 | -0.66 | 0.91 | 0.47 | -0.21 | -1.02 | 0.87 | 1.12 | -0.03 | -0.4 | 0.56 | 1.33 | 0.02 | 0.685 | -0.055 | 0.77774 | 0.734739 | 0.9585 | 1 | 0.999631 | 0.921382 |
| Dock7_Q8R1A4_T214t_1 | 0.59 | 1.22 | 1.07 | 1.13 | 0.48 | 1.16 | 1.18 | 1.27 | 1.18 | 1.86 | 1.59 | 2.12 | 1.3 | 0.53 | 1.17 | 0.79 | 0.4025 | 0.41 | 0.6 | 0.86123 | 0.946181 | 0.04975 | 1 | 0.999631 | 0.954096 |
| Dock9_E9QMR2_S1191s_1 | -0.64 | -1.46 | -1.23 | -1.87 | -1.27 | -0.62 | -0.89 | -0.81 | -1.29 | -1.3 | -0.37 | -0.6 | -0.28 | -1.28 | -0.64 | -0.6 | 0.37 | -0.5025 | 0.035 | 0.130905 | 0.242526 | 0.289763 | 1 | 0.999631 | 0.602409 |

| | | | | | | | | | | | | | | | | | | | | | | | | | | |
|---|---|---|---|---|---|---|---|---|---|---|---|---|---|---|---|---|---|---|---|---|---|---|---|---|---|---|
| Dock9_E9QMR2_S924s_1 | -1.84 | -1.78 | -1.87 | -2.98 | -2.52 | -0.4 | -2.27 | -1.8 | -2.59 | -2.04 | -1.34 | -4.51 | -1.27 | -0.86 | -2.97 | -3.23 | -0.21 | -0.2275 | -0.3325 | 0.953644 | 0.479002 | 0.454995 | | 1 | 0.999631 | 0.984441 |
| Dopey2_Q3UHQ6_S601s_1 | 0.05 | -1.38 | 0.52 | -0.16 | -1.24 | -0.64 | 0.14 | -0.07 | -0.88 | 0.05 | -0.84 | -0.21 | -1.68 | -1.53 | 0.61 | 0.3 | 0.565 | 0.02 | 0.45 | 0.610642 | 0.664809 | 0.619673 | | 1 | 0.999631 | 0.859267 |
| Dos_E9PW07_S136s_1 | 0.6 | -0.14 | -0.03 | 0.23 | 1.86 | 0.08 | 0.8 | 0.18 | -0.34 | -1.1 | 1.13 | 1.05 | -0.33 | 0.79 | 1.04 | 0.96 | -0.0375 | 0.0075 | 0.205 | 0.264359 | 0.211047 | 0.969976 | | 1 | 0.999631 | 0.6935 |
| Dos_E9PW07_S282s_1 | -0.7 | -0.49 | 0.52 | -0.11 | -0.1 | -0.32 | -0.76 | 0.25 | -0.92 | -0.28 | -0.28 | 0.73 | -0.93 | -0.31 | 0.33 | 0.95 | -0.1275 | -0.1425 | -0.6925 | 0.666885 | 0.91772 | 0.986137 | | 1 | 0.999631 | 0.88264 |
| Dos_E9PW07_S320s_1 | 1.86 | 0.2 | 0.25 | 0.48 | -0.05 | 1.35 | 0.54 | 0.44 | 1.13 | 1.87 | -0.3 | -0.48 | -0.35 | 0.62 | -0.07 | -0.18 | 0.05 | 0.165 | 0.53 | 0.144346 | 0.784753 | 0.819514 | | 1 | 0.999631 | 0.616037 |
| Dos_E9PW07_S485s_1 | -1.23 | 0.6 | -0.56 | -0.84 | -0.18 | 0.04 | -0.85 | -0.84 | -0.2 | -1.09 | -0.32 | 0.24 | 0.55 | 0.26 | -0.48 | -0.24 | -0.1775 | 0.7075 | 0.7975 | 0.26053 | 0.90995 | 0.724204 | | 1 | 0.999631 | 0.692624 |
| Dos_E9PW07_S527s_1 | -1.8 | -3.85 | -2.7 | -2.57 | -2.7 | -2.95 | -3.2 | -2.78 | -2.26 | -1.71 | -2.61 | -1.51 | -1.02 | -2.35 | -2.46 | -1.9 | 0.28 | 0.1075 | 0.1275 | 0.140667 | 0.68044 | 0.159128 | | 1 | 0.999631 | 0.612258 |
| Dos_E9PW07_S544s_1 | 1.11 | 1.34 | 0.16 | 1.29 | 1.76 | 0.9 | 1.18 | 1.18 | 1.81 | 0.86 | 0.72 | 0.94 | 0.43 | 1.61 | 0.93 | 1.44 | 0.145 | 0.3 | 0.2725 | 0.75349 | 0.43907 | 0.782175 | | 1 | 0.999631 | 0.913377 |
| Dos_E9PW07_S617s_1 | 0.44 | 0.32 | 0.21 | 0.19 | 0.35 | 0.78 | 0.02 | 0.59 | 1 | 0.83 | 0.13 | 0.4 | 0.6 | 0.42 | 0.77 | 0.46 | 0.2825 | 0.4625 | 1.1575 | 0.319623 | 0.591565 | 0.316942 | | 1 | 0.999631 | 0.723075 |
| Dos_E9PW07_T600t_1 | -0.5 | -0.93 | -0.93 | -1.77 | -0.68 | 0.27 | -1.3 | -1.29 | -0.64 | 0.45 | -1.18 | -0.91 | 0.66 | 1.16 | -0.65 | -0.67 | 0.1625 | -0.06 | 0.095 | 0.043494 | 0.527526 | 0.308017 | | 1 | 0.999631 | 0.475443 |
| Dos_E9PW07_T698t_1 | -0.6 | 1.38 | -0.26 | -0.58 | 0.11 | -0.41 | -0.26 | 1.15 | -0.73 | -0.12 | 0.11 | 0.44 | 0.04 | -0.59 | 0.18 | 0.69 | 0.15 | 0.5225 | 0.7825 | 0.853037 | 0.764459 | 0.905145 | | 1 | 0.999631 | 0.950673 |
| Dpf1_D3YYG6_T3t_1 | -0.66 | -1.48 | 0.04 | -0.96 | -0.08 | -1.4 | -0.25 | -0.73 | -0.38 | -0.84 | 0.8 | -0.55 | -0.55 | -1.39 | 1.15 | 0.86 | -0.36 | -0.2075 | -0.045 | 0.221766 | 0.725956 | 0.277706 | | 1 | 0.999631 | 0.672397 |
| Dpf2_F8WIP7_T204t_1 | 1.09 | 2.15 | 2.31 | 1.63 | 0.96 | 1.88 | 1.28 | 1.62 | 1.11 | 1.85 | 1.41 | 1.98 | 1.02 | 1.7 | 2.07 | 2.21 | -0.305 | 0.185 | 0.27 | 0.91148 | 0.334291 | 0.5773 | | 1 | 0.999631 | 0.971796 |
| Dpp10_E9QN98_S28s_1 | -0.3 | -0.76 | 0.08 | -0.76 | -0.72 | -0.55 | -0.58 | -1.11 | -0.47 | 0.17 | -0.42 | -0.28 | -0.05 | -0.52 | 0.06 | -0.15 | -0.145 | -0.8625 | -1.5475 | 0.409094 | 0.325531 | 0.559417 | | 1 | 0.999631 | 0.766996 |
| Dpp10_E9QN98_T11t_1 | 0.55 | -1.85 | 0.28 | -0.24 | -0.92 | -1.14 | 0.44 | -0.22 | -2.46 | 0.58 | -0.13 | -2.7 | -1.71 | -2.25 | -1.01 | -2.48 | -0.1025 | -0.265 | -0.22 | 0.021782 | 0.804678 | 0.328046 | | 1 | 0.999631 | 0.405544 |
| Dpp6_Q80VM5_S13s_1 | -0.56 | -1.18 | 1.02 | -0.13 | -1.52 | -1.35 | 0.85 | 0.76 | -2.62 | 0.06 | 0.72 | -0.07 | -1.69 | -0.99 | 0.33 | 0.62 | 0.8225 | 0.2875 | 0.0475 | 0.733312 | 0.884278 | 0.73002 | | 1 | 0.999631 | 0.907462 |
| Dpp8_Q80YA7_S568s_1 | 0.92 | 0.69 | 0.19 | -0.1 | 0.43 | 1.54 | 1.13 | 1.89 | 0.75 | 1.53 | 0.22 | 0.35 | 0.89 | 2.27 | -0.29 | -0.98 | -0.1825 | 0.075 | 0.3375 | 0.943378 | 0.061148 | 0.469247 | | 1 | 0.999631 | 0.981629 |
| Dpy19l1_A6X919_S22s_1 | -0.2 | 0.48 | -0.11 | -0.77 | -0.17 | -0.45 | -0.64 | -0.07 | -1 | 0.47 | 0.08 | 0.15 | 0.13 | -0.33 | 0.06 | 0.89 | 0.085 | 0.0525 | -0.2175 | 0.397063 | 0.582143 | 0.856627 | | 1 | 0.999631 | 0.761093 |
| Dpy19l1_A6X919_S23s_1 | -0.69 | -0.59 | 0.41 | 0.01 | -0.33 | -0.44 | -0.9 | 1.15 | -0.99 | -0.48 | 1.02 | -0.2 | -0.25 | -0.19 | -2.09 | 0.8 | 0.1125 | -0.2025 | -0.57 | 0.716984 | 0.861095 | 0.912772 | | 1 | 0.999631 | 0.903294 |
| Dpy19l1_A6X919_S46s_1 | -0.24 | -2.42 | -1.6 | -0.79 | -0.46 | -2.9 | -0.91 | -0.33 | -3.13 | -0.13 | -0.7 | -1.9 | -1.25 | -1.54 | -2.49 | -2.05 | -0.665 | -0.925 | -1.585 | 0.287877 | 0.868168 | 0.780628 | | 1 | 0.999631 | 0.705972 |
| Dpysl2_O08553_S517s_2 | 2.9 | 1.73 | 1.45 | 1.34 | 0.74 | 1.82 | 1.85 | 0.35 | 1.14 | 2 | 0.14 | 0.44 | -0.44 | 0.47 | 0.55 | 0.5 | -0.6975 | -0.8275 | -0.82 | 0.004336 | 0.197532 | 0.097506 | | 1 | 0.999631 | 0.351308 |
| Dpysl2_O08553_S518s_1 | 1.84 | 0.43 | 1.14 | 0.68 | -0.75 | 0.92 | 1.28 | -0.15 | -0.1 | 1.02 | -0.05 | -0.09 | 0.03 | 0.17 | 0.35 | 0.26 | -0.415 | -0.7575 | -1.9 | 0.044772 | 0.202276 | 0.072257 | | 1 | 0.999631 | 0.475443 |
| Dpysl2_O08553_S518s_2 | 3.08 | 1.16 | 1.88 | 1.93 | 0.96 | 2.39 | 2.32 | 0.72 | 0.86 | 2.81 | 0.45 | 0.9 | -1.05 | 0.16 | 0.78 | 0.56 | 0.715 | 0.145 | 0.105 | 0.005 | 0.453374 | 0.227321 | | 1 | 0.999631 | 0.351308 |
| Dpysl2_O08553_S540s_1 | 1.73 | 1.5 | 0.6 | 0.97 | 3.12 | 2.13 | 1.12 | 1.29 | 2.25 | 0.85 | 1.56 | 0.72 | 1.79 | 2.08 | 0.3 | 1.05 | 0.77 | -0.0975 | 0.4975 | 0.821765 | 0.170109 | 0.738875 | | 1 | 0.999631 | 0.938464 |
| Dpysl2_O08553_S542s_1 | 1.25 | 1.3 | 0.63 | 2.16 | 2.93 | 1.79 | 1.23 | 2.47 | 0.89 | 0.46 | 1.82 | 1.78 | 1.37 | 2.51 | 1.75 | 1.7 | 0.1975 | -0.58 | -0.945 | 0.246029 | 0.123675 | 0.829014 | | 1 | 0.999631 | 0.685751 |
| Dpysl2_O08553_S570s_1 | 1.36 | 1.3 | 0.92 | 2 | 1.91 | 1.63 | 1.17 | 1.66 | 0.68 | 0.27 | 1.53 | 0.78 | -0.03 | 0.89 | 0.28 | 0.66 | -0.7775 | -1.025 | -2.005 | 0.022371 | 0.541084 | 0.142642 | | 1 | 0.999631 | 0.405775 |
| Dpysl2_O08553_S62s_1 | 3.3 | 2.64 | 2.42 | 2.39 | 1.93 | 3.18 | 1.9 | 0.63 | 1.91 | 3.62 | 0.48 | 0.64 | 0.28 | 1.91 | 0.59 | -0.05 | -0.215 | -0.015 | 0.2725 | 0.001621 | 0.159251 | 0.153542 | | 1 | 0.999631 | 0.351308 |
| Dpysl2_O08553_T509t_1 | 0.26 | 0.99 | 0.58 | 0.27 | 0.37 | 0.57 | -0.08 | 0.38 | 0.58 | 0.58 | 0.19 | 0.69 | 0.58 | 0.97 | 0.86 | 0.78 | -0.8675 | -1.02 | -1.67 | 0.371638 | 0.46658 | 0.95872 | | 1 | 0.999631 | 0.746522 |
| Dpysl2_O08553_T512t_1 | 1.97 | 1.17 | 3.3 | 1.89 | -0.17 | 1.59 | 2.34 | 1.1 | 0.86 | 1.7 | 2.49 | -0.8 | 0.36 | 0.4 | 0.47 | 0.42 | -0.605 | -0.7075 | -1.0825 | 0.003648 | 0.183477 | 0.188039 | | 1 | 0.999631 | 0.351308 |
| Dpysl2_O08553_T514t_1 | 1.79 | 1.16 | 1.57 | 1.25 | -0.01 | 1.15 | 1.47 | 0.74 | 0.49 | 1.69 | 0.37 | 0.39 | -0.22 | 0.24 | 0.85 | 0.57 | -0.4325 | -0.7475 | -1.67 | 0.00894 | 0.126218 | 0.084268 | | 1 | 0.999631 | 0.365396 |
| Dpysl2_O08553_T514t_2 | 2.92 | 1.3 | 1.83 | 1.68 | 0.89 | 2.04 | 2.27 | 0.8 | 0.97 | 2.41 | 0.5 | 0.86 | -0.73 | 0.19 | 0.77 | 0.82 | -0.0425 | 0.56 | 0.615 | 0.006003 | 0.384611 | 0.168538 | | 1 | 0.999631 | 0.351545 |
| Dpysl3_E9PWE8_S101s_1 | 0.64 | 0.71 | 1.01 | 0.13 | 0.38 | 1.31 | 0.12 | 0.51 | 1.41 | 2.1 | 0.09 | 1.13 | 1.72 | 1.62 | 0.49 | 1.12 | -0.3 | 0.7425 | 1.14 | 0.124975 | 0.90229 | 0.230135 | | 1 | 0.999631 | 0.596063 |
| Dpysl3_E9PWE8_S585s_1 | -0.17 | 1.68 | 0.15 | 0.31 | 0.59 | 0.67 | -0.61 | 0.12 | 1.31 | 1.58 | 0.39 | 1.66 | 2.57 | 1.8 | 1.22 | 0.94 | -0.065 | 0.03 | 0.17 | 0.048397 | 0.533865 | 0.146719 | | 1 | 0.999631 | 0.48343 |
| Dpysl3_E9PWE8_S631s_2 | -0.2 | 0.56 | -0.8 | 0.57 | 0.56 | -0.7 | -0.61 | 0.62 | -0.17 | -0.94 | 0.68 | 0.68 | 0.41 | 0.03 | 0.7 | -0.33 | -0.355 | 0.02 | -0.19 | 0.683608 | 0.889276 | 0.950965 | | 1 | 0.999631 | 0.889602 |
| Dpysl3_E9PWE8_S631s_3 | 1.38 | 2.87 | 0.75 | 2.44 | 2.58 | 0.33 | 0.43 | 2.68 | 2.43 | 0.62 | 2.28 | 2.19 | 0.8 | 2.57 | 1.51 | 1.8 | -0.3 | 0.36 | 0.3125 | 0.735923 | 0.62202 | 0.972848 | | 1 | 0.999631 | 0.90759 |
| Dpysl3_E9PWE8_S635s_1 | 0.04 | -0.56 | -0.29 | -0.53 | -0.77 | -0.71 | -0.65 | -0.41 | -0.31 | 0.61 | -0.18 | -0.02 | 0.17 | -0.18 | -0.13 | 0.05 | -0.045 | 0.2125 | -0.1125 | 0.289949 | 0.268828 | 0.265743 | | 1 | 0.999631 | 0.706164 |
| Dpysl3_E9PWE8_S652s_1 | 0.31 | 0.49 | 0.36 | 1.17 | 0.85 | 0.11 | 0.3 | 0.89 | 1.4 | 0.03 | 1.24 | 0.51 | 0.51 | 0.77 | 0.41 | 0.19 | 0.5325 | 0.405 | 0.385 | 0.724358 | 0.889704 | 0.591451 | | 1 | 0.999631 | 0.905674 |
| Dpysl3_E9PWE8_S654s_1 | 1.5 | 0.87 | -0.16 | 1.27 | 2.65 | 1.39 | 0.36 | 1.21 | 1.54 | 1.28 | 1.36 | 0.92 | 1.09 | 1.99 | 1 | 0.94 | -0.33 | 0.11 | 0.1 | 0.395228 | 0.344518 | 0.326112 | | 1 | 0.999631 | 0.759996 |
| Dpysl3_E9PWE8_T621t_1 | 0.5 | -0.17 | -0.07 | 0.21 | -0.02 | -0.13 | -0.51 | -0.19 | -0.32 | 0.57 | 0.01 | 0.65 | 0.33 | -0.07 | 0.28 | 0.33 | -0.1275 | 0.425 | 0.7125 | 0.734038 | 0.244525 | 0.738815 | | 1 | 0.999631 | 0.907462 |
| Dpysl3_E9PWE8_T622t_1 | -0.61 | 0.25 | 0.33 | -0.06 | 0 | -0.13 | -0.53 | 0.06 | 0.34 | 0.26 | 0.14 | 0.87 | 0.72 | 0.65 | 0.61 | 0.78 | -0.7325 | -0.47 | -1.635 | 0.041489 | 0.680025 | 0.209716 | | 1 | 0.999631 | 0.474979 |
| Dpysl3_E9PWE8_T627t_1 | 2.94 | 0.18 | 1.31 | 1.38 | -0.43 | 0.63 | 3.14 | -0.46 | 0.41 | 2.83 | 0.34 | 0.35 | -0.62 | -0.7 | 0.4 | 0.19 | -0.76 | -0.205 | -2.0675 | 0.017176 | 0.415031 | 0.530021 | | 1 | 0.999631 | 0.391024 |
| Dpysl3_E9PWE8_T627t_2 | 3.25 | 1.58 | 0.79 | 0.87 | 0.72 | 1.19 | 2.09 | -0.55 | 1.53 | 3.4 | 0.78 | -0.04 | 0.42 | -0.25 | -1.64 | -0.31 | -0.355 | 0.02 | -0.19 | 0.008799 | 0.293489 | 0.801598 | | 1 | 0.999631 | 0.365244 |
| Dpysl3_E9PWE8_T627t_3 | 1.38 | 2.87 | 0.75 | 2.44 | 2.58 | 0.33 | 0.43 | 2.68 | 2.43 | 0.62 | 2.28 | 2.19 | 0.8 | 2.57 | 1.51 | 1.8 | -0.325 | -0.31 | -1.26 | 0.735923 | 0.62202 | 0.972848 | | 1 | 0.999631 | 0.90759 |
| Dpysl3_E9PWE8_T632t_1 | 3.13 | 1.97 | 1.99 | 1.15 | 0.83 | 1.92 | 2.67 | 1.52 | 1.58 | 2.83 | 1.75 | 0.84 | 0.79 | -0.06 | 1.39 | 1.08 | -0.4775 | 0.23 | -0.945 | 0.026831 | 0.536019 | 0.567985 | | 1 | 0.999631 | 0.422285 |
| Dpysl3_E9PWE8_T632t_2 | 2.69 | 1.3 | -0.25 | 0.76 | 0.67 | 0.71 | 0.41 | 0.8 | 1.09 | 2.61 | 0.97 | 0.75 | 0.41 | -0.16 | 0.68 | -0.21 | -0.13 | 0.5025 | 0.7275 | 0.135306 | 0.406677 | 0.731107 | | 1 | 0.999631 | 0.607096 |
| Dpysl3_E9PWE8_T637t_1 | -0.38 | -0.69 | -0.55 | -1.06 | -0.83 | -0.74 | -0.71 | -0.92 | -0.21 | 0.41 | -0.57 | -0.3 | 0.35 | 0.1 | 0.06 | -0.28 | -0.4775 | -0.335 | -0.3775 | 0.033339 | 0.61789 | 0.135551 | | 1 | 0.999631 | 0.44294 |
| Dpysl4_Q3TMU8_S570s_1 | 0.98 | 2.17 | 1.49 | 1.53 | 1.37 | 1.16 | 0.78 | 0.95 | 0.92 | 1.39 | 1.38 | 1.14 | 0.92 | 1.08 | 0.84 | 1.82 | -0.1275 | 0.2625 | 0.1925 | 0.323552 | 0.161048 | 0.315617 | | 1 | 0.999631 | 0.724485 |
| Dpysl4_Q3TMU8_S572s_1 | -0.11 | 0.04 | -0.11 | 0.03 | -0.22 | -0.35 | -0.02 | -0.07 | 0.06 | 0.36 | 0.01 | 0.47 | -0.01 | 0.24 | 0.15 | 0.24 | -0.35 | 0.135 | 0.015 | 0.466899 | 0.592998 | 0.317329 | | 1 | 0.999631 | 0.795267 |
| Dpysl5_Q9EQF6_T509t_1 | 1.12 | 1.42 | 1.2 | 1.07 | 1.02 | 0.68 | 0.49 | 1.22 | 1.23 | 1.39 | 1.33 | 1.4 | 0.73 | 1.21 | 1.18 | 1.75 | -0.2375 | 0.25 | -0.1475 | 0.961762 | 0.217614 | 0.585454 | | 1 | 0.999631 | 0.986594 |
| Dpysl5_Q9EQF6_T509t_2 | 0.34 | 1.98 | 1.54 | 1.81 | 1.41 | 0.95 | 0.78 | 1.58 | 1.63 | 1.41 | 1.52 | 2.11 | 0.71 | 1.12 | 1.31 | 1.94 | 0.075 | 0.4575 | 0.205 | 0.743285 | 0.569384 | 0.545296 | | 1 | 0.999631 | 0.90972 |
| Dpysl5_Q9EQF6_T514t_1 | -0.14 | 1.15 | 0.79 | 1.14 | 1.18 | 0.34 | 0.76 | 0.96 | 1.3 | 0.72 | 0.86 | 1.89 | 0.55 | 0.73 | 1.22 | 1.26 | -0.2375 | 0.25 | -0.1475 | 0.596766 | 0.839585 | 0.280091 | | 1 | 0.999631 | 0.854617 |
| Dpysl5_Q9EQF6_T514t_2 | 0.34 | 1.98 | 1.54 | 1.81 | 1.41 | 0.95 | 0.78 | 1.58 | 1.63 | 1.41 | 1.52 | 2.11 | 0.71 | 1.12 | 1.31 | 1.94 | -0.385 | -0.515 | -0.61 | 0.743285 | 0.569384 | 0.545296 | | 1 | 0.999631 | 0.90972 |
| Drd1_Q61616_S441s_1 | 1.33 | -2.34 | -1.45 | 0 | -1.25 | -1.72 | 0.37 | -1.4 | -0.25 | -0.14 | -1.52 | -2.61 | -0.09 | -1.83 | -1.37 | -1.61 | -0.2525 | -0.0825 | -0.745 | 0.443577 | 0.637977 | 0.557074 | | 1 | 0.999631 | 0.782783 |
| Drd1_Q61616_T446t_1 | 0.06 | -2.09 | -2.11 | -1.09 | -1.36 | -2.21 | -1.43 | -1.24 | -1.66 | -0.06 | -2.15 | -1.69 | -2.14 | -1.41 | -2.35 | -2.31 | 0.4275 | 0.565 | 0.6025 | 0.17868 | 0.629015 | 0.8946 | | 1 | 0.999631 | 0.642903 |
| Drd5_Q8BLD9_S385s_1 | -1.03 | -1.49 | -1.07 | -1.69 | -0.37 | -0.98 | -0.79 | -1.43 | 0 | -2.31 | -0.85 | 0.14 | -0.06 | -0.8 | -1.12 | -0.89 | -0.4775 | -0.7975 | -0.9225 | 0.103241 | 0.20278 | 0.312764 | | 1 | 0.999631 | 0.566608 |
| Drosha_Q5HZJ0_T272t_1 | 0.43 | -0.36 | 1.81 | 1.48 | -1.64 | 0.42 | 2.15 | 0.52 | -1.45 | 0.87 | 0.4 | 0.35 | -1.66 | -0.22 | 1.21 | 0.34 | -0.3175 | 0.105 | 0.31 | 0.203498 | 0.556352 | 0.233119 | | 1 | 0.999631 | 0.659781 |
| Dst_S4R1P5_S106s_1 | -0.7 | -0.36 | 1.22 | 0.15 | -0.17 | -0.53 | -0.71 | 0.45 | -0.29 | 0.3 | 0.21 | 0.51 | 0.25 | 0.12 | 0.19 | 0.99 | -0.6425 | -0.64 | -0.9675 | 0.505242 | 0.503108 | 0.813652 | | 1 | 0.999631 | 0.814727 |
| Dst_S4R1P5_S140s_1 | 0.32 | -1.92 | -0.7 | -1.19 | -2.92 | -1.83 | -0.2 | -1.11 | -2.84 | -0.05 | -1.28 | -1.88 | -1.6 | -2.07 | -1.49 | -2.2 | 0.005 | -0.16 | -0.1275 | 0.069551 | 0.34359 | 0.35196 | | 1 | 0.999631 | 0.526652 |
| Dst_S4R1P5_S4072s_1 | -1.55 | -1.3 | -1.59 | -1.08 | -1.52 | -1.75 | -1.04 | -1.19 | -1.75 | -1.81 | -1.37 | -1.23 | -1.65 | -1.59 | -1.45 | -1.34 | 0.62 | 0.205 | 0.5875 | 0.648345 | 0.985762 | 0.572429 | | 1 | 0.999631 | 0.874615 |
| Dst_S4R1P5_S5775s_1 | -0.4 | -0.24 | -2.95 | -0.94 | -0.23 | -0.39 | -0.85 | -0.58 | -1.02 | -0.27 | -1.56 | -0.86 | -0.16 | -0.56 | 0.01 | -1.47 | -0.0425 | 0.89 | 0.715 | 0.36647 | 0.299701 | 0.739529 | | 1 | 0.999631 | 0.745796 |
| Dst_S4R1P5_S77s_1 | -2.38 | -6.55 | -4.17 | -3.16 | -3.7 | -6.08 | -3.49 | -3.16 | -4.97 | -4.16 | -1.71 | -1.86 | -2.84 | -4.07 | -3.14 | -3.35 | 0.0825 | -0.3075 | -0.3075 | 0.39246 | 0.964989 | 0.406621 | | 1 | 0.999631 | 0.758901 |
| Dstn_Q9R0P5_S3s_1 | -0.23 | 0.05 | -0.12 | 1.54 | -0.09 | -0.16 | 1.41 | 0.41 | -0.08 | -0.85 | 1.02 | -0.08 | 0.2 | -0.55 | 1.78 | -1.42 | -0.225 | 0.19 | -0.13 | 0.663777 | 0.872096 | 0.564114 | | 1 | 0.999631 | 0.881991 |
| Dtd1_Q9DD18_S197s_1 | -0.79 | -2.32 | -0.73 | -0.12 | -1.76 | -2.37 | -0.42 | -0.31 | -1.71 | -1.05 | -0.28 | -0.16 | -1.24 | -2.02 | -0.25 | -0.97 | 0.0075 | 0.095 | -0.63 | 0.814028 | 0.717588 | 0.729833 | | 1 | 0.999631 | 0.937041 |
| Dtd1_Q9DD18_S205s_1 | 0.31 | 0.85 | 1.51 | 0.87 | 0.73 | 1.11 | 1.23 | 0.5 | 0.97 | 0.97 | 0.85 | 1.13 | -0.51 | 0.24 | 1.02 | 0.27 | -0.4075 | -0.5825 | -1.7775 | 0.150874 | 0.982157 | 0.763594 | | 1 | 0.999631 | 0.623359 |
| Dtna_Q8CFR5_S500s_1 | 2.34 | -1.19 | 0.8 | 0.28 | -1.78 | 0.85 | 3.06 | -1.53 | -1.16 | 2.5 | -0.55 | -0.89 | -1.2 | -1.31 | -0.25 | -2.12 | 0.1975 | 0.1975 | -0.525 | 0.032192 | 0.724637 | 0.551493 | | 1 | 0.999631 | 0.442395 |
| Dtna_Q9D2N4_S306s_1 | 1.98 | 1.46 | 1.87 | 1.34 | 0.91 | 2.38 | 2.2 | 1.95 | 1.84 | 2.55 | 1.62 | 1.43 | 0.28 | 1.81 | 2.25 | 0.21 | -0.1075 | -0.2525 | -1.3925 | 0.322821 | 0.603811 | 0.561548 | | 1 | 0.999631 | 0.724485 |
| Dtna_Q9D2N4_S308s_1 | 2.77 | 1.17 | 1.58 | 2.01 | 0.75 | 2.04 | 2.57 | 1.74 | 2.17 | 2.27 | 1.26 | 0.82 | -1.28 | 1.06 | 1.78 | 0.4 | 0.155 | 0.3275 | 0.0825 | 0.057415 | 0.824572 | 0.596633 | | 1 | 0.999631 | 0.49679 |
| Dtna_Q9D2N4_T501t_1 | 0.8 | 0.06 | -1.72 | -0.75 | 0.8 | -0.28 | -1 | -0.51 | 0 | 1.1 | -0.71 | -0.69 | 0.54 | -0.16 | -0.79 | -0.87 | 0.0525 | 0.165 | 0.18 | 0.88702 | 0.795662 | 0.601673 | | 1 | 0.999631 | 0.963505 |
| Dtnb_G3UXK0_S364s_1 | 0.61 | 0.65 | 0.51 | 0.56 | 0.62 | 0.83 | 0.49 | 0.6 | 0.65 | 0.97 | 0.61 | 0.76 | 0.19 | 0.97 | 0.69 | 1.2 | -0.9575 | -0.675 | -1.59 | 0.566847 | 0.82312 | 0.506604 | | 1 | 0.999631 | 0.842178 |
| Dtnb_G3UXK0_S501s_2 | 3.33 | -1.12 | 0.81 | 0.43 | -0.94 | -0.18 | 0.77 | -0.03 | -0.82 | 2.62 | -0.6 | -0.45 | -1.21 | -1.06 | 0.23 | -0.87 | -0.37 | -0.675 | -1.015 | 0.08496 | 0.279753 | 0.52835 | | 1 | 0.999631 | 0.550457 |
| Dtnb_G3UXK0_T504t_2 | 0.99 | -0.59 | 0.44 | 0.9 | -0.22 | -0.15 | 0.78 | -0.15 | -0.18 | 0.25 | -0.32 | -0.71 | -0.93 | -1.57 | 0.52 | -0.34 | -0.57 | -1.015 | -1.135 | 0.082906 | 0.395716 | 0.130419 | | 1 | 0.999631 | 0.549443 |
| Dtnb_G3UXK0_T507t_2 | 1.82 | -1.13 | 0.91 | 0.43 | -1.19 | -0.11 | 0.91 | 0.14 | -1.7 | 0.54 | -0.61 | -0.26 | -1.12 | -1.05 | 0.43 | -0.77 | -0.665 | -0.9 | -1.6675 | 0.101838 | 0.404685 | 0.163956 | | 1 | 0.999631 | 0.564448 |
| Dtnb_G3UZJ3_S494s_1 | 2.44 | 1.84 | 0.02 | -0.16 | 0.94 | 0.66 | 0.33 | -0.45 | 1.31 | 1.87 | -1.75 | -0.89 | -0.34 | 0.47 | -1.04 | -1.62 | -0.8275 | -0.235 | -0.2075 | 0.036438 | 0.313513 | 0.349634 | | 1 | 0.999631 | 0.453963 |
| Dtx4_Q6PDK8_S273s_1 | 0.03 | -0.06 | 0.98 | -0.59 | -0.94 | -1.85 | -0.3 | 0.14 | -0.73 | 0.3 | 0.44 | -0.59 | -0.21 | -0.28 | -0.18 | 0.2 | -0.05 | -0.015 | 0.0125 | 0.588044 | 0.126712 | 0.595238 | | 1 | 0.999631 | 0.850634 |
| Dtx4_Q6PDK8_T72t_1 | 0.55 | 0.78 | 0.73 | 0.55 | -0.58 | 1.62 | 0.08 | 1.29 | 0.51 | 0.8 | 1.59 | -0.35 | 0.16 | 1.52 | 0.5 | 0.48 | -0.5075 | 0.1725 | 1.0725 | 0.971889 | 0.91845 | 0.970929 | | 1 | 0.999631 | 0.989788 |
| Dync1h1_Q9JHU4_S1228s_1 | -0.18 | 0.63 | 0.73 | 0.12 | -0.72 | -0.12 | -0.34 | 0.45 | 0.28 | 0.45 | 0.52 | 0.74 | 1.82 | 1.26 | 1.28 | 1.23 | 0.31 | 0.205 | 0.645 | 0.008255 | 0.172969 | 0.57608 | | 1 | 0.999631 | 0.365244 |
| Dync1h1_Q9JHU4_S4366s_1 | -0.78 | 0.63 | 0.37 | -0.51 | 0.4 | 0.23 | 0.25 | 0.07 | 0.34 | -0.53 | 0.24 | 0.48 | 1.03 | 0.54 | 0.49 | 0.23 | -0.015 | 0.265 | 0.735 | 0.131839 | 0.406369 | 0.622526 | | 1 | 0.999631 | 0.603456 |
| Dync1i1_O88485_S142s_1 | -1.3 | -0.13 | -0.38 | -0.63 | -0.35 | -0.5 | -0.78 | -0.87 | -0.47 | -0.5 | -0.42 | 0.01 | 0.42 | 0.16 | -0.05 | -0.03 | -0.5475 | -0.345 | 0.2475 | 0.052109 | 0.963077 | 0.430584 | | 1 | 0.999631 | 0.489777 |
| Dync1i1_O88485_S50s_1 | -0.27 | -0.16 | 1.07 | 0.51 | -0.38 | -0.27 | -0.09 | -0.3 | -1.1 | 0.36 | -0.08 | 0.59 | 0.38 | -0.09 | 0.09 | 1.76 | 0.0225 | -0.13 | 0.2575 | 0.623191 | 0.140242 | 0.47019 | | 1 | 0.999631 | 0.862913 |
| Dync1i1_O88485_S80s_1 | -0.22 | 0.41 | 0.09 | -0.08 | 0.45 | 0.09 | -0.21 | -0.04 | 0.05 | -0.33 | -0.12 | 0.08 | 0.07 | 0.49 | 0.16 | 0.51 | -0.03 | -0.1175 | 0.0375 | 0.388308 | 0.935499 | 0.634966 | | 1 | 0.999631 | 0.757772 |
| Dync1i2_Q3TPJ8_S81s_1 | -0.25 | 0.33 | -0.36 | -0.05 | 0.12 | -0.09 | -0.43 | -0.05 | -0.09 | -0.75 | 0.09 | -0.05 | -0.21 | 0.01 | -0.12 | 0.14 | -0.36 | 0.0075 | 0.255 | 0.897052 | 0.913412 | 0.704926 | | 1 | 0.999631 | 0.966612 |
| Dync1i2_Q3TPJ8_S95s_2 | -0.22 | -1.3 | 0.46 | -0.46 | -1.46 | -1.69 | -0.15 | 0.34 | -1.51 | 0.16 | 0.4 | -0.54 | -1.06 | -0.62 | 0.6 | 0.58 | -0.6325 | -0.8575 | -0.45 | 0.62926 | 0.525547 | 0.988612 | | 1 | 0.999631 | 0.866631 |
| Dync1i2_Q3TPJ8_S98s_1 | 0.7 | 0.67 | 0.81 | 1.56 | -0.75 | 0.55 | 0.21 | 1.2 | -0.96 | -0.26 | 1.31 | 0.22 | 0.85 | 0.39 | 0.29 | 0.41 | -0.545 | -0.085 | 0.2275 | 0.185908 | 0.176765 | 0.108747 | | 1 | 0.999631 | 0.646224 |
| Dync1i2_Q3TPJ8_S98s_2 | -0.22 | -1.3 | -0.02 | -0.95 | -1.46 | -1.69 | -0.75 | -0.77 | -1.51 | 0.16 | -0.48 | -1 | -1.06 | -0.62 | 0.11 | -0.01 | -0.0125 | -0.1325 | 0.5625 | 0.590967 | 0.186943 | 0.852287 | | 1 | 0.999631 | 0.852188 |
| Dync1i2_Q3TPJ8_T174t_1 | -0.76 | 0.66 | 0.39 | -0.42 | 0.16 | 0.05 | -0.35 | -0.04 | -0.68 | -0.28 | -0.16 | 0.46 | 1.11 | -0.12 | 0.6 | 0.53 | -0.745 | -0.8625 | -0.2525 | 0.208455 | 0.973021 | 0.749818 | | 1 | 0.999631 | 0.662061 |
| Dync1i2_Q3TPJ8_T89t_1 | 0.83 | 0.44 | 0.49 | 0.7 | -0.33 | -0.29 | 0.51 | -0.41 | -0.42 | 0.28 | -0.12 | -0.73 | -0.48 | 0.47 | 0.23 | 1.23 | 0.07 | 0.1625 | 0.395 | 0.524597 | 0.029809 | 0.017261 | | 1 | 0.999631 | 0.822727 |
| Dync1li1_Q8R1Q8_S207s_1 | -0.98 | -0.21 | -0.72 | -0.91 | -0.27 | -0.47 | -1.01 | -0.79 | -0.13 | -0.98 | -0.51 | -0.55 | 0.16 | -0.12 | -0.8 | -0.48 | -0.115 | -0.47 | -1.325 | 0.259941 | 0.81646 | 0.604856 | | 1 | 0.999631 | 0.692624 |
| Dync1li1_Q8R1Q8_S405s_1 | 2.71 | 0.84 | 2.05 | 1.43 | 1.35 | 2.85 | 1.54 | 0.83 | 1.49 | 2.9 | 0.49 | 0.27 | -0.87 | 1.47 | 0.84 | 0.29 | 1.1275 | 0.725 | 0.4525 | 0.044217 | 0.831945 | 0.475368 | | 1 | 0.999631 | 0.475443 |
| Dync1li1_Q8R1Q8_S421s_1 | 0.59 | 0.13 | -0.38 | 0.5 | 1.24 | 1.75 | 1.08 | 1.28 | 1.07 | 0.59 | 1.17 | 0.91 | -0.62 | 1.62 | 1.15 | 0.5 | -0.4225 | -0.095 | 0.4325 | 0.382664 | 0.00494 | 0.042608 | | 1 | 0.999631 | 0.754123 |
| Dync1li1_Q8R1Q8_S516s_1 | -0.8 | -1.06 | 0.09 | -0.96 | -1.01 | -1.33 | -0.85 | -1.23 | -1.62 | -0.75 | -0.36 | -0.38 | 0.57 | -0.93 | -0.5 | -0.14 | -0.4475 | 0.0175 | 0.1625 | 0.32053 | 0.216658 | 0.814716 | | 1 | 0.999631 | 0.723684 |
| Dync1li1_Q8R1Q8_T512t_1 | -0.08 | 0.14 | 0.5 | -0.03 | -0.57 | -0.2 | -0.33 | -0.16 | -0.35 | 0.02 | 0.21 | 0.72 | 0.39 | 0.25 | 0.35 | 0.19 | 0.0975 | 0.26 | 0.37 | 0.563008 | 0.112876 | 0.956211 | | 1 | 0.999631 | 0.841038 |
| Dync1li2_Q6PDL0_S194s_1 | -1.58 | -0.43 | -0.26 | -1.05 | -0.6 | -0.69 | -0.69 | -0.95 | -0.46 | -1.24 | -0.44 | -0.14 | -0.05 | -0.85 | -0.42 | -0.52 | 0.205 | 0.535 | 0.6675 | 0.342465 | 0.777259 | 0.516982 | | 1 | 0.999631 | 0.73348 |
| Dync1li2_Q6PDL0_S407s_1 | 0.27 | 0.33 | 1.28 | 1.24 | 0.41 | 1.79 | 0.73 | 1.01 | 0.83 | 1.3 | 0.88 | 2.25 | -0.43 | 2.55 | 2.54 | 1.13 | -0.3275 | -0.0475 | 0.11 | 0.337402 | 0.6163 | 0.232213 | | 1 | 0.999631 | 0.730499 |
| Dync1li2_Q6PDL0_S487s_1 | 0.14 | 0.41 | -0.32 | 0.17 | -0.04 | -0.66 | 0.21 | -0.42 | -0.17 | 0.54 | 0.15 | -0.31 | 0.34 | 0.29 | 0.04 | 0.17 | -0.4725 | -0.955 | -1.465 | 0.704008 | 0.298204 | 0.878267 | | 1 | 0.999631 | 0.897149 |
| Dync1li2_Q6PDL0_T376t_1 | 2.93 | 0.97 | 2.21 | 1.29 | 0.72 | 2.53 | 1.75 | 0.51 | 0.57 | 2.61 | 0.15 | 0.25 | -0.87 | 1.11 | 1.29 | 0.01 | -0.565 | 0.3425 | -1.5075 | 0.035234 | 0.429129 | 0.168313 | | 1 | 0.999631 | 0.452817 |
| Dyrk1a_F6U6X3_S491s_1 | 1.73 | -0.51 | -0.07 | -0.3 | -1.72 | -1.16 | 2.25 | -0.78 | -0.17 | 1.73 | 0.48 | 0.18 | -0.79 | -1.32 | -1.19 | -1.88 | 0.17 | 0.3 | -0.1125 | 0.016095 | 0.529515 | 0.571805 | | 1 | 0.999631 | 0.388832 |
| Dyrk1a_F6U6X3_S720s_1 | -0.77 | -1.95 | -0.34 | -1.33 | -1.27 | -1.54 | -0.64 | -0.26 | -1.75 | -0.95 | -0.58 | 0.09 | -2.54 | -1.97 | 0.49 | -0.82 | 0.24 | 0.16 | 0.565 | 0.867397 | 0.701501 | 0.547293 | | 1 | 0.999631 | 0.9567 |
| Dyrk1a_F6U6X3_Y283y_1 | -2.22 | -0.28 | -0.12 | -0.76 | -0.57 | -0.49 | -0.84 | -0.52 | -0.51 | -1.31 | -0.64 | -0.28 | 0.35 | -0.53 | -0.39 | -0.55 | -0.095 | -0.17 | 0.615 | 0.275061 | 0.607317 | 0.749357 | | 1 | 0.999631 | 0.700187 |
| Dyrk4_Q8BI55_Y344y_1 | -0.41 | 0.3 | 0.38 | 0.15 | -0.25 | 0.16 | -0.49 | 0.62 | 0.1 | -0.39 | -0.35 | 0.38 | 1.16 | 0.4 | 0.6 | 0.72 | 0.2575 | -0.195 | 0.165 | 0.079749 | 0.779032 | 0.595757 | | 1 | 0.999631 | 0.545579 |
| Dzank1_E9Q4Z6_S769s_1 | -0.91 | 0.02 | -1.12 | -0.76 | -0.25 | -0.6 | -0.63 | -0.26 | -0.87 | -1.78 | -0.67 | -0.23 | -0.34 | -0.51 | -0.35 | -0.91 | -0.3325 | 0.08 | 0.6025 | 0.633449 | 0.427036 | 0.640688 | | 1 | 0.999631 | 0.868569 |
| Dzip3_F6Z1P8_S98s_1 | -0.35 | 0.42 | 0.4 | 0.7 | -0.24 | -0.37 | 0.42 | 0.03 | 0.07 | 0.72 | -0.28 | 0.98 | 0.57 | 0.6 | 1.62 | 0.79 | -1.0075 | -1.17 | -0.525 | 0.130769 | 0.323318 | 0.835831 | | 1 | 0.999631 | 0.602177 |
| E030010N08Rik_F6XLV1_T56t_1 | -0.31 | 0.96 | 0.3 | 0.52 | -1.74 | 0.06 | -0.54 | -0.34 | -0.48 | -1.03 | -0.89 | -0.81 | 0.11 | -0.09 | -0.58 | -0.07 | -0.5275 | -0.8 | -2.2325 | 0.162283 | 0.047486 | 0.005871 | | 1 | 0.999631 | 0.631442 |
| E9Q0B3_S233s_1 | 2.5 | -0.92 | 0.79 | 1.68 | -0.43 | 0.26 | 2.12 | -0.01 | -0.68 | 1.13 | 0.22 | 0.18 | -2.51 | -1.86 | -0.16 | -0.35 | -0.35 | -0.3025 | 0.48 | 0.019073 | 0.517731 | 0.286952 | | 1 | 0.999631 | 0.391024 |
| Ebag9_Q9D0V7_S36s_1 | -0.07 | 0.59 | 0.69 | 0.59 | 0.4 | -0.09 | 0.03 | 0.06 | 0.01 | 0.07 | 0.37 | 0.14 | 1.21 | 0.74 | 0.94 | 0.83 | 0.4025 | 0.31 | 1.3475 | 0.137427 | 0.234343 | 0.301705 | | 1 | 0.999631 | 0.610456 |
| Ece2_E9Q1S5_S27s_1 | -1.72 | -1.32 | 0.02 | -1.77 | -0.43 | -0.69 | -1.14 | -0.92 | -0.31 | -0.34 | -1.95 | -0.95 | 0.84 | -0.14 | 0.32 | -0.42 | 0.4 | 0.3275 | 0.0575 | 0.018329 | 0.365788 | 0.562278 | | 1 | 0.999631 | 0.391024 |
| Edc3_Q8K2D3_S131s_1 | -0.55 | 0.48 | -0.48 | -1.3 | 0.21 | -0.23 | 0.19 | -0.42 | -0.19 | -0.47 | -0.01 | 0.13 | -0.06 | -0.28 | 0.03 | -1.31 | -0.0125 | -0.1625 | 0.2875 | 0.902304 | 0.331242 | 0.42093 | | 1 | 0.999631 | 0.968183 |
| Edc4_G5E896_S6s_1 | 0.1 | 1.22 | -0.29 | 0.16 | 0.6 | 0.41 | -0.24 | 0.37 | 0.35 | -0.56 | 0.14 | 0.61 | 0.43 | 0.94 | 0.28 | 0.69 | 0.7575 | 0.445 | 1.0625 | 0.463574 | 0.973945 | 0.696285 | | 1 | 0.999631 | 0.793701 |
| Edc4_G5E896_S727s_1 | 0.17 | -0.27 | -1.35 | 0.77 | 1.5 | -0.58 | 0.21 | 1.22 | 0.63 | -0.25 | 0.49 | 0.23 | 1.55 | 1.45 | 1.51 | -0.94 | -0.73 | -0.2825 | 0.1 | 0.139189 | 0.220746 | 0.356127 | | 1 | 0.999631 | 0.611718 |
| Eef1a1_P10126_T176t_1 | -0.06 | 0.15 | 0.64 | 0.04 | -1.01 | -0.03 | -1.02 | -0.09 | 0.03 | -0.52 | 0.07 | 0.06 | 0.1 | -0.18 | 0.67 | 0.58 | -0.105 | 0.585 | 0.5025 | 0.761522 | 0.058524 | 0.348099 | | 1 | 0.999631 | 0.914725 |
| Eef1b2_O70251_S82s_1 | 0.73 | 1.19 | 0.78 | 0.02 | 1.76 | 1.2 | -0.42 | -0.24 | 1.4 | 0.96 | 1.26 | 1.44 | 1.15 | 1.63 | 1 | 0.95 | -0.315 | -0.1725 | 0.19 | 0.170009 | 0.846352 | 0.095254 | | 1 | 0.999631 | 0.64073 |
| Eef1b2_O70251_S83s_1 | 0.69 | 2.23 | 0.87 | 0.57 | 0.38 | 1.68 | 0.1 | 0.94 | 0.38 | 1.67 | 0.66 | 0.96 | 1.8 | 1.53 | 1.03 | 0.76 | -0.06 | 0.44 | 0.205 | 0.674201 | 0.524382 | 0.709347 | | 1 | 0.999631 | 0.885515 |
| Eef1d_D3YZT9_S152s_1 | 0.06 | 0.91 | 0.74 | 0.59 | 0.44 | 0.53 | 0.39 | 0.7 | 1.05 | 1.19 | 0.55 | 1.27 | 0.78 | 0.85 | 0.88 | 0.61 | 0.0875 | 0.3775 | 0.065 | 0.499137 | 0.829599 | 0.179436 | | 1 | 0.999631 | 0.813279 |
| Eef1d_D3YZT9_S71s_1 | -1.13 | -1.48 | -1.91 | -0.93 | -1.47 | -1.43 | -0.95 | -1.25 | -1.56 | -1.62 | -0.66 | -0.1 | -1.52 | -1.19 | -1.08 | -1.4 | -0.615 | -0.02 | -1.0325 | 0.83917 | 0.774336 | 0.386262 | | 1 | 0.999631 | 0.946365 |
| Eef1d_D3YZT9_T166t_1 | 2.48 | 1.22 | 2.67 | 2.61 | 0.65 | 1.39 | 2.65 | 1.83 | 1.49 | 2.55 | 2.42 | 2.44 | 0.87 | 0.46 | 2.08 | 1.44 | 0.14 | -0.155 | 0.7725 | 0.052936 | 0.242533 | 0.962409 | | 1 | 0.999631 | 0.489777 |
| Eef2_P58252_T57t_1 | -0.68 | 2.19 | -0.45 | 1.68 | 0.77 | 2.08 | 0.69 | -0.24 | -0.08 | -1.37 | 2.46 | 1.11 | -1.51 | 3.04 | 2.3 | 2 | 0.755 | 2.2275 | -0.445 | 0.473714 | 0.854792 | 0.870436 | | 1 | 0.999631 | 0.799779 |
| Eef2k_F6ZEU2_S42s_1 | 1.31 | -0.43 | 1.3 | -1.21 | 1.63 | 0.9 | 3.09 | -1.63 | 3.8 | -0.39 | 2.74 | 3.73 | 3.38 | -1.63 | -0.96 | -1.6 | -0.4325 | 0.3575 | 0.5675 | 0.700622 | 0.460782 | 0.049342 | | 1 | 0.999631 | 0.896835 |
| Eefsec_Q9JHW4_T532t_1 | 0.63 | 1.49 | 1.47 | 1.35 | 0.42 | 1.17 | 0.41 | 1.21 | 1.48 | 1.6 | 1.25 | 2.04 | 0.58 | 2.67 | 1.97 | 1.99 | 0.1125 | 0.365 | -0.495 | 0.247564 | 0.221197 | 0.281841 | | 1 | 0.999631 | 0.685751 |
| Eepd1_Q3TGW2_S106s_2 | -0.68 | -2.64 | -1.4 | -0.8 | -2.23 | -2.69 | 0.38 | -0.53 | -2.95 | -0.66 | -0.2 | -0.25 | -3.87 | -2.98 | -0.08 | -0.57 | -0.2875 | 0.185 | 0.0975 | 0.577834 | 0.87994 | 0.607036 | | 1 | 0.999631 | 0.84622 |
| Eepd1_Q3TGW2_S110s_1 | -0.77 | -1.31 | -1.51 | -2.67 | -2.05 | -2.15 | -1.3 | -1.91 | -1.71 | -0.45 | -2.16 | -1.2 | -0.59 | -1.97 | -1.66 | -1.65 | -0.1075 | 0.94 | -0.3225 | 0.840713 | 0.512221 | 0.718484 | | 1 | 0.999631 | 0.946805 |
| Eepd1_Q3TGW2_S16s_1 | 0.79 | 1.72 | 4.86 | 3.44 | 1.45 | 1.37 | 3.46 | 4.1 | 2.93 | 0.37 | 6.44 | 4.83 | 2.23 | 0.17 | 3.52 | 3.6 | 0.2375 | 0.01 | -0.04 | 0.754826 | 0.912873 | 0.493096 | | 1 | 0.999631 | 0.913582 |
| Eepd1_Q3TGW2_S25s_1 | 0.37 | 0.94 | 0.17 | 1.11 | 0.76 | 1.19 | 0.64 | 0.95 | 0.92 | 0.43 | 0.82 | 0.46 | -0.48 | 1.01 | 1.01 | 0.89 | 0.6025 | 0.1225 | -0.505 | 0.926487 | 0.450464 | 0.974885 | | 1 | 0.999631 | 0.976003 |
| Efhd2_Q8C845_S74s_1 | 0.77 | 1 | -0.75 | 1.11 | 1.76 | 0.77 | 0.85 | 1.16 | 1.05 | -0.08 | 1.22 | 0.43 | -1.02 | 0.74 | 0.2 | 0.19 | 0.61 | -0.4525 | -0.3875 | 0.356714 | 0.216314 | 0.806292 | | 1 | 0.999631 | 0.740499 |
| Efhd2_Q8C845_S76s_1 | 1.32 | 0.73 | -0.36 | 0.52 | 1.49 | 1.98 | 0.37 | 0.81 | 0.09 | 0.38 | 0.4 | -0.47 | -0.86 | 1.11 | 0.16 | 0.25 | 0.145 | -0.2025 | 0.0475 | 0.451996 | 0.217977 | 0.285901 | | 1 | 0.999631 | 0.788148 |
| Efnb1_P52795_S280s_1 | -0.43 | 0.25 | -1.07 | 0.18 | -0.64 | 0.22 | -0.24 | 0.17 | -0.29 | -1.25 | 0.15 | -0.49 | 0.14 | -0.68 | -0.48 | 0.14 | -0.1775 | 0.205 | 0.18 | 0.90515 | 0.70433 | 0.636197 | | 1 | 0.999631 | 0.969056 |
| Efnb1_P52795_S286s_1 | -0.74 | -0.44 | -0.53 | -0.76 | -0.72 | -1.28 | -0.64 | -0.54 | -0.59 | -0.82 | -0.29 | 0.05 | -0.38 | -1.1 | -0.08 | -0.19 | 0.0675 | -0.0625 | -0.6925 | 0.580254 | 0.519817 | 0.485719 | | 1 | 0.999631 | 0.846774 |
| Efnb1_P52795_S337s_1 | 0.38 | -0.3 | -0.43 | -0.44 | 0.03 | -0.51 | 0.25 | -0.29 | -0.6 | -0.44 | 0.03 | -0.03 | -0.72 | -0.62 | -1.68 | -0.54 | 0.2525 | 1.485 | 0.4975 | 0.086435 | 0.828628 | 0.841985 | | 1 | 0.999631 | 0.55299 |
| Efnb3_O35393_S268s_1 | -0.9 | -0.28 | 2.59 | 1.36 | 2.58 | -0.44 | 2.1 | -0.46 | 3.57 | 0.08 | 1.75 | 3.31 | 3.9 | -0.2 | 1.51 | -0.45 | -0.03 | 0.035 | 0.425 | 0.648761 | 0.796165 | 0.152331 | | 1 | 0.999631 | 0.874615 |
| Efnb3_O35393_S274s_1 | 0.06 | -0.78 | 0.91 | -0.67 | -0.88 | -0.28 | 0.32 | 0.24 | 0.11 | -0.67 | 0.35 | -0.13 | 0.3 | -0.5 | 0.41 | 1.01 | -0.0425 | -0.035 | 0.26 | 0.388936 | 0.947928 | 0.936826 | | 1 | 0.999631 | 0.758158 |
| Efnb3_O35393_S332s_1 | -0.77 | -1.05 | -0.62 | -1.19 | -0.68 | -1.12 | -0.86 | -1.14 | -0.54 | -0.96 | -1.27 | -1 | -0.36 | -0.65 | -0.81 | -0.77 | -0.14 | 0.0375 | 0.545 | 0.376615 | 0.873351 | 0.903082 | | 1 | 0.999631 | 0.748839 |
| Efr3a_Q8BG67_S220s_1 | 0.54 | 1.86 | 1.64 | 0.58 | 0.51 | 2.11 | 0.21 | 1.23 | 0.94 | 0.65 | 1.36 | 1.82 | 1.84 | 1.53 | 1.85 | 1.58 | -0.6 | -0.4325 | -0.1675 | 0.181447 | 0.784023 | 0.9304 | | 1 | 0.999631 | 0.643843 |
| Efr3b_Q6ZQ18_S214s_1 | -0.2 | -0.21 | -0.16 | 0.03 | -1 | -0.75 | -0.52 | -0.67 | -1.02 | -0.26 | -0.71 | -0.28 | -0.36 | -0.45 | -0.46 | 0.06 | -0.5075 | -0.2525 | -0.1225 | 0.549462 | 0.032624 | 0.151341 | | 1 | 0.999631 | 0.835338 |
| Efr3b_Q6ZQ18_S216s_1 | -0.4 | -0.24 | -0.21 | -0.04 | -0.95 | -0.26 | -0.59 | -1.12 | -0.68 | 0.01 | -1.06 | -0.17 | 0.15 | -0.15 | -1.14 | -0.24 | -0.03 | -0.01 | 0.4525 | 0.724777 | 0.098118 | 0.433138 | | 1 | 0.999631 | 0.905674 |
| Eftud1_Q8C0D5_S474s_1 | -0.73 | -0.09 | -1.3 | -0.59 | -0.31 | -0.3 | -1.15 | -1.07 | -0.48 | 0.02 | -1.29 | -1 | 0.44 | 0.46 | -0.77 | -1.03 | -0.2025 | 0.1675 | -0.07 | 0.337393 | 0.934092 | 0.979831 | | 1 | 0.999631 | 0.730499 |
| Eftud1_Q8C0D5_S59s_1 | -1.11 | 0.48 | -0.52 | -0.53 | -0.53 | -1.21 | -0.8 | 0.05 | 0.16 | -1.63 | 0.41 | 0.05 | -0.07 | -0.73 | -0.25 | -0.91 | 1.095 | 0.41 | -0.165 | 0.863032 | 0.630914 | 0.753604 | | 1 | 0.999631 | 0.954983 |
| Ehbp1_Q69ZW3_S1022s_1 | 1.31 | -0.12 | -1.99 | 1.1 | 1.95 | 0.67 | 1.79 | 0.27 | 1.05 | -0.36 | 1.1 | 0.15 | -0.65 | 0.19 | -0.21 | 0.31 | -0.515 | -0.135 | -1.395 | 0.8134 | 0.170708 | 0.582615 | | 1 | 0.999631 | 0.937041 |
| Ehbp1_Q69ZW3_S1038s_1 | 2.37 | -0.08 | -0.11 | 1.48 | 0.67 | -0.39 | 0.69 | 0.63 | 0.58 | 2.34 | 0.37 | -0.17 | -1.08 | -0.34 | 0.35 | -0.85 | 0.1875 | -0.065 | -0.5625 | 0.045876 | 0.400704 | 0.852278 | | 1 | 0.999631 | 0.477883 |
| Ehbp1_Q69ZW3_S335s_1 | 0.78 | 3.32 | 4.31 | 1.64 | 1.65 | 3.64 | 3.51 | 2 | 2.76 | 1.25 | 3 | 2.78 | 2.26 | 1.58 | 2.52 | 1.44 | 0.54 | -0.0825 | -0.175 | 0.45338 | 0.820397 | 0.934051 | | 1 | 0.999631 | 0.788717 |
| Ehbp1_Q69ZW3_S588s_1 | -0.19 | -0.56 | -1.05 | -0.91 | 0.51 | -0.37 | 0.12 | -0.81 | -0.17 | -0.62 | -1.45 | -0.8 | -1.07 | -0.52 | -0.87 | -0.95 | 0.2725 | -0.1775 | -0.4775 | 0.581059 | 0.163335 | 0.819189 | | 1 | 0.999631 | 0.847114 |
| Ehbp1_Q69ZW3_S764s_1 | 0.5 | -0.46 | 0.71 | 1.48 | 1 | 0.21 | 1.55 | 0.56 | 0.68 | 0.32 | -0.08 | 0.6 | -0.87 | 0.16 | 0.78 | 0.25 | 0.44 | 0.6125 | 0.8025 | 0.354588 | 0.565447 | 0.682864 | | 1 | 0.999631 | 0.739951 |
| Ehbp1_Q69ZW3_S765s_1 | 0.1 | 0.01 | -0.22 | 0.26 | 0.71 | -0.29 | 0.41 | 1.08 | 0.21 | -0.34 | 1.43 | 1.3 | -0.11 | 1.36 | 0.83 | 1.28 | -0.4925 | -0.175 | -0.2875 | 0.059699 | 0.217122 | 0.182808 | | 1 | 0.999631 | 0.50521 |
| Ehbp1l1_G5E8Y6_S502s_1 | 1.29 | 1.04 | -0.07 | 0.36 | 1.17 | 0.26 | -0.25 | -0.53 | 0.34 | 1.65 | -0.38 | 0.31 | 0.63 | 0.58 | 0.47 | -0.21 | 1.2775 | 1.615 | 0.0175 | 0.472662 | 0.304062 | 0.726639 | | 1 | 0.999631 | 0.799014 |
| Ehbp1l1_G5E8Y6_S518s_1 | -1.11 | 1.05 | 0.89 | 0.04 | 2.11 | 0.58 | 3.05 | 0.24 | 2.79 | -0.05 | 1.08 | 3.51 | 1.66 | -0.92 | 1.48 | -1.28 | 0.0425 | -0.055 | 0.76 | 0.982563 | 0.103541 | 0.076021 | | 1 | 0.999631 | 0.994846 |
| Ehd1_Q9WVK4_S456s_1 | -1.23 | 0.9 | 0.34 | -0.55 | 0.2 | -0.29 | -0.34 | 0.06 | 0.02 | -0.32 | -0.52 | 0.06 | 0.9 | 1.06 | 0.52 | 0.02 | 0.0475 | -0.035 | 0.6125 | 0.153091 | 0.927514 | 0.907548 | | 1 | 0.999631 | 0.625198 |
| Ehd3_Q9QXY6_S456s_1 | -1.43 | -0.52 | -0.12 | -0.62 | -0.58 | -0.4 | -0.87 | -0.65 | -0.83 | -1.28 | -0.53 | -0.19 | 0.3 | -0.29 | -0.21 | -0.04 | -0.1925 | 0.075 | 0.47 | 0.109944 | 0.886081 | 0.926731 | | 1 | 0.999631 | 0.573646 |
| Ehd3_Q9QXY6_S479s_1 | -0.11 | 0.25 | 0.89 | -0.57 | -0.2 | 0.45 | -0.38 | -0.18 | -0.16 | 0.36 | 0.31 | 0.25 | 0.44 | 0.79 | 0.34 | 0.77 | -0.67 | -0.44 | 0.3575 | 0.223896 | 0.609149 | 0.836 | | 1 | 0.999631 | 0.67471 |
| Ei24_Q61070_S330s_1 | -1.28 | -0.61 | 0.45 | -0.98 | -2.12 | -0.85 | -1.25 | -0.88 | -2.11 | -0.23 | -0.86 | -0.98 | 0.1 | -0.94 | -0.27 | 0.12 | 0.555 | 0.2225 | 0.15 | 0.434661 | 0.168158 | 0.399805 | | 1 | 0.999631 | 0.779717 |
| Eif2b4_D3Z780_S63s_1 | 1.22 | -0.67 | 0.11 | 0.91 | 0.96 | -0.04 | 0.68 | 2.19 | 0.55 | -0.08 | 1.41 | 0.58 | -0.47 | 0.91 | 1.8 | -0.07 | -0.3775 | 0.105 | 0.2875 | 0.803916 | 0.344896 | 0.655904 | | 1 | 0.999631 | 0.93396 |
| Eif2b5_Q8CHW4_S540s_1 | -0.09 | 0.49 | 0.55 | 0.2 | 0.06 | -0.33 | -0.13 | 0.04 | 0 | 0.15 | 0.63 | 0.79 | 0.75 | 0.38 | 0.37 | 0.8 | -0.1975 | -0.0075 | 0.295 | 0.344964 | 0.180626 | 0.734071 | | 1 | 0.999631 | 0.734195 |
| Eif2s2_Q99L45_S2s_1 | 0.06 | 0.25 | 0.43 | 0.18 | 0.15 | -0.06 | -0.24 | 0.28 | 0.05 | -0.04 | 0.23 | 0.65 | 0.28 | 0.6 | 0.46 | 0.76 | -0.1375 | 0.7375 | -0.8125 | 0.297752 | 0.443284 | 0.978225 | | 1 | 0.999631 | 0.711185 |
| Eif3a_P23116_S1159s_1 | 2.31 | 1.2 | 0.99 | 2.04 | 0.16 | 2.21 | 0.87 | 2.75 | 1.34 | 3.4 | 2.46 | 2.29 | -1.28 | 0.31 | 2.46 | 1.8 | 0.1575 | 0.325 | -0.02 | 0.311025 | 0.82155 | 0.164555 | | 1 | 0.999631 | 0.71924 |
| Eif3a_P23116_S1300s_1 | 1.23 | 1.42 | 0.91 | 1.79 | 1.32 | 1.66 | 1.09 | 1.91 | 1.43 | 1.84 | 1.49 | 1.89 | -0.98 | 2.28 | 1.91 | 2.06 | -0.7175 | 0.5775 | -0.86 | 0.977231 | 0.615914 | 0.289638 | | 1 | 0.999631 | 0.992174 |
| Eif3a_P23116_T574t_1 | 0.91 | 2.37 | 1.98 | 2.22 | 0.55 | 1.25 | 1.3 | 1.51 | 2.79 | 2.49 | 2.43 | 2.08 | 0.06 | 0.94 | 1.41 | 1.63 | -0.2625 | -0.1375 | 0.025 | 0.091482 | 0.095099 | 0.153909 | | 1 | 0.999631 | 0.557085 |
| Eif3a_P23116_T982t_1 | 0.33 | 1.34 | 0.33 | 0.11 | 0.71 | 0.23 | 0.18 | -0.06 | 0.82 | 0.82 | 0.41 | -0.49 | 0.86 | 0.49 | 0.26 | 0.6 | -0.2125 | -0.1375 | 0.0075 | 0.943918 | 0.458954 | 0.742885 | | 1 | 0.999631 | 0.981629 |
| Eif3b_Q8JZQ9_S111s_1 | 0.25 | 1.03 | 0.98 | -0.23 | 0.25 | -0.22 | 0.67 | 0.48 | 0.19 | -0.39 | 1.06 | 0.62 | -0.41 | 1.13 | 0.54 | 0.8 | -0.535 | 0.205 | -0.365 | 0.986653 | 0.574234 | 0.750738 | | 1 | 0.999631 | 0.995165 |
| Eif3b_Q8JZQ9_S120s_1 | 0.68 | -0.48 | 0.34 | -0.33 | -1.09 | -1.01 | -0.14 | 0.31 | -0.36 | 1.63 | -0.11 | -0.13 | -1.17 | -0.16 | -0.15 | 0.23 | -0.005 | 0.0625 | 0.185 | 0.394258 | 0.232028 | 0.687187 | | 1 | 0.999631 | 0.759303 |
| Eif3b_Q8JZQ9_S120s_2 | -0.47 | 0.56 | 0.57 | 0.11 | 0.08 | 0.19 | 0.14 | 0.34 | 0.11 | 0.4 | 0.1 | 0.41 | 0.03 | 0.8 | 0.24 | 0.44 | -0.005 | 0.0625 | 0.185 | 0.601022 | 0.987048 | 0.845178 | | 1 | 0.999631 | 0.856689 |
| Eif3b_Q8JZQ9_S123s_2 | -0.47 | 0.56 | 0.57 | 0.11 | 0.08 | 0.19 | 0.14 | 0.34 | 0.11 | 0.4 | 0.1 | 0.41 | 0.03 | 0.8 | 0.24 | 0.44 | -0.685 | -0.1275 | 0.355 | 0.601022 | 0.987048 | 0.845178 | | 1 | 0.999631 | 0.856689 |
| Eif3b_Q8JZQ9_S75s_1 | 0.12 | 1.22 | 1.06 | 0.5 | -0.38 | 0.04 | 0.05 | 0.45 | -0.01 | 0.18 | 0.97 | 1.25 | 0.72 | 0.76 | 1.42 | 1.42 | -0.2675 | 0.0875 | 0.165 | 0.344377 | 0.067873 | 0.753805 | | 1 | 0.999631 | 0.733953 |
| Eif3b_Q8JZQ9_S75s_2 | -0.02 | 0.55 | 1.31 | 0.73 | -0.13 | 0.14 | 0.27 | 1.22 | 0.14 | 0.63 | 0.92 | 1.23 | 0.25 | 0.72 | 1.03 | 1.23 | -0.2675 | 0.0875 | 0.165 | 0.667328 | 0.512895 | 0.818319 | | 1 | 0.999631 | 0.88264 |
| Eif3b_Q8JZQ9_S79s_2 | -0.02 | 0.55 | 1.31 | 0.73 | -0.13 | 0.14 | 0.27 | 1.22 | 0.14 | 0.63 | 0.92 | 1.23 | 0.25 | 0.72 | 1.03 | 1.23 | -0.2475 | 0.035 | -0.235 | 0.667328 | 0.512895 | 0.818319 | | 1 | 0.999631 | 0.88264 |
| Eif3b_Q8JZQ9_S88s_1 | 0.41 | 0.55 | 1.75 | 1.39 | 0.18 | 1.05 | 1.2 | 0.68 | 0.3 | 1.01 | 1.23 | 1.7 | 0.82 | -0.08 | 1.28 | 1.14 | -0.075 | 0.2375 | -0.1575 | 0.600931 | 0.540329 | 0.935641 | | 1 | 0.999631 | 0.856689 |
| Eif3b_Q8JZQ9_S90s_1 | 0.8 | 0.86 | 0.48 | 0.54 | 0.71 | 0.59 | 0.6 | 0.48 | 0.87 | 1.37 | 0.8 | 0.59 | 0.11 | 0.84 | 0.64 | 0.46 | -0.215 | 0.0925 | -0.5225 | 0.595455 | 0.7572 | 0.410777 | | 1 | 0.999631 | 0.854508 |
| Eif3c_Q8R1B4_S166s_1 | 1.06 | 1.39 | 2.42 | 1.43 | 0.92 | 1.63 | 1.27 | 1.62 | 1.66 | 1.78 | 1.5 | 1.73 | 0.85 | 0.83 | 1.6 | 0.93 | -0.0025 | 0.33 | 0.6475 | 0.191717 | 0.555737 | 0.787836 | | 1 | 0.999631 | 0.648674 |
| Eif3c_Q8R1B4_S39s_1 | -0.24 | 0.82 | 0.47 | 0.44 | 0.61 | 0.36 | -0.21 | 0.72 | 0.78 | 0.41 | 0.68 | 0.94 | 1.15 | 1.21 | 0.87 | 0.85 | -0.02 | -0.06 | 0.8375 | 0.067455 | 0.994115 | 0.306103 | | 1 | 0.999631 | 0.521594 |
| Eif3e_P60229_S203s_1 | -0.29 | 0.52 | -0.34 | 0.69 | 0.74 | 0.31 | -0.85 | 0.3 | -1.68 | 0.41 | 1.05 | 0.56 | 1.48 | 0.67 | 1.18 | 0.6 | 0.02 | -0.1425 | 0.23 | 0.04777 | 0.962586 | 0.920517 | | 1 | 0.999631 | 0.481942 |
| Eif3f_Q9DCH4_S262s_1 | 0.96 | 0.53 | 1.84 | 0.98 | 0.97 | 0.74 | 0.83 | 1.85 | 0.24 | 1.63 | 1.56 | 0.31 | 0.65 | 0.89 | 1.91 | 1.78 | 0.115 | 0.255 | 0.515 | 0.593323 | 0.958608 | 0.756228 | | 1 | 0.999631 | 0.853305 |
| Eif3g_Q9Z1D1_S42s_1 | 0.07 | 0.99 | 0.35 | 0.01 | 0.85 | 0.68 | -0.24 | 0.59 | 0.57 | 0.88 | 0.74 | 0.25 | 0.94 | 1.15 | 0.36 | 1.03 | 0.1125 | 0.33 | 0.1825 | 0.158515 | 0.747357 | 0.434696 | | 1 | 0.999631 | 0.628615 |
| Eif3j1_Q3UGC7_T112t_1 | -0.51 | -0.07 | -0.16 | -0.06 | -0.15 | -0.08 | -0.08 | -0.04 | -0.31 | -0.36 | 0.33 | 0.86 | -0.33 | -0.26 | 0.15 | 0.37 | -0.4075 | -0.4675 | -1.7275 | 0.548817 | 0.645243 | 0.355164 | | 1 | 0.999631 | 0.835034 |
| Eif4a2_Q8BTU6_S2s_1 | 1.56 | -0.99 | -1.76 | 0.33 | -1.77 | -0.5 | 1.59 | -1.81 | -1.35 | 0.67 | -1.17 | -0.88 | -2.98 | -1.47 | -2.62 | -0.7 | -0.76 | -0.05 | -1.08 | 0.049531 | 0.66439 | 0.544058 | | 1 | 0.999631 | 0.487521 |

| | | | | | | | | | | | | | | | | | | | | | | | | | | |
|---|---|---|---|---|---|---|---|---|---|---|---|---|---|---|---|---|---|---|---|---|---|---|---|---|---|---|
| Eif4a3_Q91VC3_S12s_1 | 2.53 | 1.3 | 0.38 | 1.12 | 0.65 | 0.48 | 0.87 | 0.29 | 1.33 | 1.97 | 0.8 | 1.03 | 0.18 | 0.57 | 0.84 | -0.58 | 0.3675 | -0.12 | -0.96 | 0.058396 | 0.114591 | 0.918611 | | 1 | 0.999631 | 0.500325 |
| Eif4b_Q8BGD9_S207s_1 | 1.13 | 1.03 | -0.91 | 1.55 | 1.54 | 0.73 | 1.11 | 0.89 | 1.08 | 0.2 | 1.37 | -0.33 | -1.88 | 0.71 | -0.65 | 0.78 | 0.5825 | 0.205 | -2.0375 | 0.212287 | 0.495188 | 0.844606 | | 1 | 0.999631 | 0.666473 |
| Eif4b_Q8BGD9_S263s_1 | -0.76 | -7.63 | -3.96 | 0.42 | -3.76 | -5.34 | 0.67 | -1.17 | -4.74 | -2.76 | -1.16 | -2.45 | -6.29 | -5.16 | -3.7 | -4.93 | 0.69 | 1.1125 | -0.8875 | 0.215668 | 0.759048 | 0.901337 | | 1 | 0.999631 | 0.668551 |
| Eif4b_Q8BGD9_S406s_1 | 1.5 | -4.54 | 0.58 | 0.01 | -0.72 | -1.27 | 2.28 | 0.02 | 0.52 | -0.78 | 1.65 | 0.61 | -2.19 | -2.36 | -1.68 | 0.23 | 0.14 | -0.32 | -2.225 | 0.480746 | 0.604308 | 0.374931 | | 1 | 0.999631 | 0.801848 |
| Eif4b_Q8BGD9_S409s_1 | 1.71 | 0.24 | 1.18 | 2.05 | -0.5 | 2.38 | 3.53 | 0.33 | 0.32 | 2.42 | 0.94 | 0.22 | -2.87 | -0.77 | 0.62 | -0.7 | -0.6175 | -0.9375 | -1.4425 | 0.011251 | 0.872362 | 0.588936 | | 1 | 0.999631 | 0.385092 |
| Eif4b_Q8BGD9_S422s_2 | 1.41 | 0.6 | 1.79 | 0.93 | 0.19 | 0.53 | 1.31 | 0.23 | 0.18 | 1.35 | -0.18 | -0.37 | 0.16 | -0.82 | -0.08 | -0.3 | -0.8375 | -0.04 | -1.6575 | 0.003074 | 0.129971 | 0.06315 | | 1 | 0.999631 | 0.351308 |
| Eif4b_Q8BGD9_S424s_1 | 1.77 | -0.23 | 0.51 | 0.14 | -0.63 | -0.63 | 1.15 | -1.05 | 0.03 | 2.09 | 0.03 | -0.12 | -0.02 | -1.34 | -2.22 | -0.86 | -0.1775 | 0.0225 | 0.2475 | 0.01613 | 0.179964 | 0.948519 | | 1 | 0.999631 | 0.388832 |
| Eif4b_Q8BGD9_S445s_1 | 0.51 | 1.97 | 1.4 | 1.48 | 1.06 | 1.47 | 0.76 | 1.36 | 1.58 | 1.19 | 1.38 | 1.3 | 1.56 | 1.84 | 1.25 | 1.7 | 0.1525 | 0.1525 | -0.7725 | 0.509788 | 0.62845 | 0.949079 | | 1 | 0.999631 | 0.817176 |
| Eif4b_Q8BGD9_S495s_2 | -0.11 | -0.95 | 0.58 | 0.43 | -0.24 | 0.75 | 0.54 | -0.49 | -0.41 | 0.61 | 0.47 | -0.11 | 0.32 | -1.74 | -0.32 | -1.4 | -0.325 | -0.3975 | -0.8875 | 0.180501 | 0.731759 | 0.719438 | | 1 | 0.999631 | 0.64376 |
| Eif4b_Q8BGD9_S497s_1 | 0.88 | 1.12 | 1.22 | 1.53 | 0.16 | 1.16 | 1.12 | 1.01 | 0.68 | 1.21 | 0.51 | 0.76 | 0.09 | 0.04 | 0.95 | 0.12 | -0.065 | 0.31 | -1.605 | 0.021096 | 0.323063 | 0.189897 | | 1 | 0.999631 | 0.400544 |
| Eif4b_Q8BGD9_S497s_2 | 0.32 | 0.63 | 0.64 | 1.31 | 0.63 | 0.11 | 0.87 | 1.03 | 0.6 | 0.85 | 1.51 | 1.18 | -3.29 | -0.21 | 0.61 | -0.63 | -0.1175 | -0.335 | -1.1175 | 0.057603 | 0.844187 | 0.366938 | | 1 | 0.999631 | 0.497148 |
| Eif4b_Q8BGD9_S498s_2 | 0.79 | -0.21 | 0.81 | 0.49 | -0.06 | 0.85 | 0.79 | -0.17 | 0.51 | 1.07 | -0.53 | -0.51 | -0.73 | -0.61 | 0.13 | -1.38 | -0.67 | -0.1975 | -0.4825 | 0.019838 | 0.755269 | 0.465015 | | 1 | 0.999631 | 0.391668 |
| Eif4b_Q8BGD9_S504s_1 | 0.27 | -1.58 | 0.11 | -1.59 | -2.35 | -0.93 | -0.52 | -1.67 | -0.88 | 0.07 | -1.72 | -1.05 | 0.35 | -1.7 | -0.99 | -2.38 | -0.2125 | -0.505 | -1.015 | 0.490242 | 0.274048 | 0.733997 | | 1 | 0.999631 | 0.807727 |
| Eif4b_Q8BGD9_S504s_2 | 0.63 | -0.24 | 0.79 | 0.37 | -0.21 | 0.57 | 0.74 | -0.4 | 0.17 | 0.73 | -0.67 | -0.7 | -0.38 | -0.65 | 0 | -1.48 | -0.2925 | -0.1875 | 0.5275 | 0.029736 | 0.57581 | 0.245636 | | 1 | 0.999631 | 0.433947 |
| Eif4b_Q8BGD9_S93s_1 | 0.49 | 1.32 | 1.53 | 1.42 | 1.04 | 0.87 | 0.49 | 1.19 | 1.12 | 0.12 | 1.24 | 1.53 | 1.91 | 1.39 | 1.61 | 1.96 | -0.885 | -1.11 | -1.27 | 0.142542 | 0.37903 | 0.641033 | | 1 | 0.999631 | 0.613207 |
| Eif4b_Q8BGD9_T420t_2 | 1.4 | 0.18 | 1.5 | 0.58 | -0.31 | -0.16 | 0.99 | -0.4 | -0.63 | 1.44 | -0.79 | -0.8 | 0.67 | -1.1 | -0.54 | -0.45 | -0.1525 | 0.155 | -0.7925 | 0.022349 | 0.069529 | 0.080589 | | 1 | 0.999631 | 0.405775 |
| Eif4b_Q8BGD9_Y211y_1 | 0.18 | 0.01 | -0.13 | 0.81 | 0.37 | -0.28 | 0.1 | 0.07 | 0.22 | 0.24 | 1.09 | -0.06 | -3.06 | -0.66 | 1.69 | -0.27 | 0.4225 | 0.915 | 0.77 | 0.368609 | 0.619671 | 0.666621 | | 1 | 0.999631 | 0.745968 |
| Eif4e2_M0QWD1_S2s_1 | -0.21 | -1.05 | -3.42 | -0.56 | -0.39 | -1.73 | -0.96 | -0.47 | -0.01 | -0.99 | -0.53 | -0.05 | -0.16 | -0.06 | -0.3 | -1.64 | 0.2425 | 0.4325 | 0.8575 | 0.298163 | 0.547963 | 0.199079 | | 1 | 0.999631 | 0.711311 |
| Eif4enif1_Q9EST3_S539s_1 | -0.52 | 0.91 | 0.98 | 0.4 | 0.99 | 0.93 | 0.14 | 0.68 | 1.1 | 0.71 | 0.65 | 1.04 | 1.38 | 1.36 | 1.06 | 1.4 | 0.2975 | -0.26 | 0.0075 | 0.047712 | 0.548643 | 0.276418 | | 1 | 0.999631 | 0.481835 |
| Eif4enif1_Q9EST3_S562s_1 | 0.88 | 1.2 | -0.44 | 0.96 | 1.01 | 0.78 | 0.53 | 1.47 | 0.51 | -0.07 | 0.73 | 0.39 | -0.35 | 1.48 | 0.96 | 0.54 | 0.02 | 0.3825 | 0.55 | 0.988239 | 0.481696 | 0.533136 | | 1 | 0.999631 | 0.995409 |
| Eif4enif1_Q9EST3_S77s_1 | 0.16 | 1.1 | 0.61 | 0.66 | 0.43 | 0.72 | 0.37 | 1.09 | 0.81 | 0.8 | 1.29 | 1.16 | 1.12 | 1.44 | 0.85 | 1.32 | -0.0975 | 0.09 | 0.2875 | 0.107309 | 0.948358 | 0.225568 | | 1 | 0.999631 | 0.569416 |
| Eif4enif1_Q9EST3_S925s_1 | 1.18 | 1.5 | 0.94 | 1.08 | 1.15 | 1.35 | 0.85 | 0.96 | 1.3 | 1.11 | 1.37 | 1.28 | 1.01 | 2.08 | 1.23 | 1.53 | 0.81 | -0.7825 | -1.1375 | 0.396178 | 0.711767 | 0.728926 | | 1 | 0.999631 | 0.760566 |
| Eif4g1_E9PVC6_S1149s_1 | -0.31 | 1 | -1.81 | -0.7 | 2.45 | 0.18 | 0.78 | -1.99 | 2.31 | -0.77 | -2.64 | -3.85 | 0.36 | 0.63 | -4.28 | -3.08 | 0.015 | 0.325 | -0.21 | 0.336418 | 0.396756 | 0.535256 | | 1 | 0.999631 | 0.730499 |
| Eif4g1_E9PVC6_S1164s_1 | 0.41 | -0.08 | 0.93 | 1.4 | 0.95 | -0.59 | 1.32 | 1.04 | 0.77 | 0.2 | 1.55 | 1.44 | -0.44 | -0.06 | 1.25 | 1.07 | -0.75 | -0.135 | -0.34 | 0.676896 | 0.976161 | 0.469557 | | 1 | 0.999631 | 0.886865 |
| Eif4g1_E9PVC6_S1184s_1 | 0.7 | 1.62 | 0.78 | 0.21 | 0.65 | 0.15 | -0.21 | -0.28 | 0.67 | 1.01 | 0.38 | 0.71 | 0.44 | 0.77 | 0.38 | 0.36 | -0.275 | -0.3225 | -1.0775 | 0.354418 | 0.070587 | 0.705133 | | 1 | 0.999631 | 0.739951 |
| Eif4g1_E9PVC6_S1550s_1 | 1.18 | -0.12 | 0.36 | 0.32 | -0.75 | 0.17 | 2.7 | -1.48 | -0.73 | 2.02 | -0.6 | -0.24 | -0.49 | -0.89 | -0.68 | -0.51 | -0.055 | 0.7 | -0.275 | 0.01036 | 0.740357 | 0.611833 | | 1 | 0.999631 | 0.383871 |
| Eif4g1_E9PVC6_T1166t_1 | 1.39 | 1.92 | 1.37 | 1.9 | 1.3 | 1.97 | 1.57 | 1.52 | 2.57 | 2.37 | 2.32 | 2.12 | 0.47 | 1.59 | 1.91 | 1.51 | 0.415 | -0.02 | 0.8375 | 0.478031 | 0.846618 | 0.029057 | | 1 | 0.999631 | 0.800501 |
| Eif4g1_E9PVC6_T165t_1 | 0.26 | -0.7 | -0.62 | -0.46 | 0.12 | 0.81 | -0.39 | -0.4 | -0.1 | -0.55 | -0.02 | -0.93 | 1.17 | 0.16 | 0.21 | 0.29 | -0.685 | -0.3325 | 0.3475 | 0.043152 | 0.284649 | 0.953898 | | 1 | 0.999631 | 0.475443 |
| Eif4g1_E9PVC6_T167t_1 | -0.56 | 0.68 | 0.98 | 0.3 | -0.39 | -0.39 | -0.55 | -0.01 | -0.25 | 0.25 | -1.08 | 1.15 | 0.29 | 1.06 | 0.26 | 1.18 | 0.15 | 0.2825 | 0.615 | 0.421646 | 0.089528 | 0.538354 | | 1 | 0.999631 | 0.772942 |
| Eif4g2_F7CBP1_T469t_1 | -0.39 | 0.24 | 0.89 | 0.52 | 0.22 | 0.63 | 0.23 | 0.78 | 0.2 | 0.61 | 0.4 | 1.18 | 0.66 | 1.02 | 1.27 | 0.77 | -0.3575 | -0.045 | 0.315 | 0.107803 | 0.660718 | 0.452639 | | 1 | 0.999631 | 0.569416 |
| Eif4g3_Z4YKC4_S237s_1 | -0.21 | -0.96 | -0.49 | -0.08 | 0.65 | -2.2 | -0.92 | -0.7 | 0.13 | -1.79 | -0.22 | -0.04 | -0.39 | -0.09 | 0.23 | -0.23 | -0.04 | 0.3025 | 0.445 | 0.336044 | 0.529205 | 0.923328 | | 1 | 0.999631 | 0.730466 |
| Eif4g3_Z4YKC4_S274s_1 | 0.29 | 1.22 | 0.93 | 0.77 | 0.89 | 0.92 | 0.18 | 1.06 | 0.9 | 1.01 | 1.14 | 1.37 | 0.99 | 1.42 | 1.03 | 1.55 | 0.47 | -0.205 | -0.0625 | 0.1873 | 0.901466 | 0.322891 | | 1 | 0.999631 | 0.646474 |
| Eif4g3_Z4YKC4_S288s_1 | 0.53 | -0.25 | -0.35 | 0.76 | 0.89 | 0.11 | 0.62 | 0.95 | 0.35 | -0.9 | 0.14 | 0.28 | -0.89 | 0.08 | 0.79 | 0.46 | -0.5925 | -0.28 | -0.28 | 0.890541 | 0.212468 | 0.619867 | | 1 | 0.999631 | 0.963768 |
| Eif4g3_Z4YKC4_S479s_1 | 0.83 | 0.45 | 0.63 | 0.79 | 0.04 | 0.11 | 0.2 | -0.02 | 0.54 | 0.71 | -0.22 | 0.55 | 0.06 | 0.89 | 0.07 | 0.56 | 0.3225 | 0.57 | 1.055 | 0.379184 | 0.031554 | 0.363604 | | 1 | 0.999631 | 0.751637 |
| Eif4g3_Z4YKC4_T175t_1 | -0.71 | 2.46 | 2.36 | 1.26 | 1.54 | 2.04 | 1.14 | 1.94 | 1.98 | 0.84 | 2.33 | 2.5 | 2.01 | 2.67 | 2.3 | 2.61 | 0.295 | 0.45 | 0.855 | 0.139436 | 0.636551 | 0.443127 | | 1 | 0.999631 | 0.611718 |
| Eif5b_Q05D44_S137s_2 | -1.17 | 0.67 | 0.29 | 0.65 | 0.97 | 0.29 | -0.22 | 0.58 | 0.58 | -0.11 | 0.53 | 1.24 | 0.9 | 0.95 | 1.18 | 0.83 | 0.295 | 0.45 | 0.855 | 0.077931 | 0.540289 | 0.370599 | | 1 | 0.999631 | 0.543121 |
| Eif5b_Q05D44_S139s_2 | -1.17 | 0.67 | 0.29 | 0.65 | 0.97 | 0.29 | -0.22 | 0.58 | 0.58 | -0.11 | 0.53 | 1.24 | 0.9 | 0.95 | 1.18 | 0.83 | 0.295 | 0.16 | -0.69 | 0.077931 | 0.540289 | 0.370599 | | 1 | 0.999631 | 0.543121 |
| Eif5b_Q05D44_S165s_1 | 0.73 | 0.16 | 1.22 | 0.35 | 0.94 | 0.47 | 3.17 | -0.94 | 1.3 | 0.86 | 0 | 0.94 | 1.67 | -1.05 | 0.15 | -1.07 | -0.8675 | -0.6475 | -1.8425 | 0.283328 | 0.704112 | 0.676894 | | 1 | 0.999631 | 0.704903 |
| Elac2_B1ATP7_S191s_1 | 3.25 | 1.49 | 2.22 | 1.82 | 0.01 | 1.93 | 2.77 | 0.6 | 1.25 | 3.1 | 1 | 0.84 | -0.05 | 0.39 | 1.04 | 0.03 | -0.425 | -0.87 | -0.97 | 0.002214 | 0.205639 | 0.289746 | | 1 | 0.999631 | 0.351308 |
| Elavl2_B1AXZ0_S119s_1 | 1.18 | -1.01 | 1.84 | 0.62 | -0.27 | 0.29 | 0.95 | -0.04 | -0.22 | 0.64 | -1.02 | -0.25 | -0.97 | -0.02 | -0.04 | -0.22 | -0.5825 | -0.2 | 0.0175 | 0.125892 | 0.48476 | 0.188996 | | 1 | 0.999631 | 0.596745 |
| Elavl2_B1AXZ0_S221s_1 | 0.03 | 0.67 | 0.27 | -0.02 | -0.82 | -0.08 | -0.62 | 0.14 | 0.13 | -0.53 | 0.38 | 0.17 | -0.03 | 0.41 | 0.39 | 0.25 | -0.0875 | 0.0375 | 0.07 | 0.952995 | 0.094813 | 0.530301 | | 1 | 0.999631 | 0.984317 |
| Elavl4_Q61701_S36s_1 | -0.41 | -0.14 | 0.47 | 0.1 | -0.31 | -0.37 | 0.17 | 0.18 | -0.25 | -0.21 | 0.23 | 0.4 | -0.33 | -0.05 | 0.44 | 0.24 | 0.34 | 0.68 | 1.0325 | 0.830857 | 0.771741 | 0.904408 | | 1 | 0.999631 | 0.942418 |
| Elfn1_Q8C8T7_S773s_1 | -0.69 | -1.3 | 0.14 | -0.79 | -0.46 | 0.11 | -0.77 | -0.16 | -0.02 | 0.31 | -0.87 | 0.66 | -0.37 | 0.92 | 1.16 | -0.22 | -1.09 | -0.16 | 0.005 | 0.052505 | 0.370595 | 0.145143 | | 1 | 0.999631 | 0.489777 |
| Elfn1_Q8C8T7_T517t_1 | -0.17 | -1.71 | 1.15 | 0.06 | -3.05 | -0.71 | -0.51 | -0.76 | -0.73 | -0.75 | 0.08 | 0.09 | -0.66 | -0.53 | -0.08 | 0.62 | 0.8225 | 0.21 | -0.075 | 0.993328 | 0.163588 | 0.783731 | | 1 | 0.999631 | 0.997705 |
| Elfn2_Q68FM6_S451s_1 | 0.61 | -0.25 | -0.88 | 0.21 | 1.3 | 1.15 | 0.04 | 0.49 | 0.39 | 0.07 | 0.22 | -0.15 | -0.96 | 0.28 | 0.32 | -0.25 | 0.795 | 0.3675 | 0.235 | 0.864936 | 0.078774 | 0.571899 | | 1 | 0.999631 | 0.955539 |
| Elfn2_Q68FM6_S471s_1 | -0.94 | 0.42 | -0.6 | 0.83 | 0.83 | 1.06 | 0.27 | 0.73 | 0.81 | 0.27 | 0.15 | -0.05 | -0.59 | 0.76 | 0.19 | 0.29 | 1.3425 | 0.1875 | -0.92 | 0.63053 | 0.094322 | 0.420367 | | 1 | 0.999631 | 0.867509 |
| Elfn2_Q68FM6_S639s_1 | 1.55 | -0.37 | -0.91 | 1.74 | 2.23 | 1.35 | 1.99 | 1.81 | 1.5 | -0.15 | 1.26 | 0.15 | -1.66 | 0.39 | 0.21 | -0.61 | 0.0975 | -1.0775 | -1.9825 | 0.221293 | 0.054091 | 0.788383 | | 1 | 0.999631 | 0.671697 |
| Elfn2_Q68FM6_S647s_1 | 1.08 | -2.06 | -3.13 | -1.23 | -0.39 | -2 | -0.36 | -2.2 | -1.91 | -1.34 | -3.44 | -2.96 | -4.94 | -1.76 | -3.74 | -2.83 | 0.16 | -0.0875 | -0.3075 | 0.061961 | 0.912447 | 0.240871 | | 1 | 0.999631 | 0.509355 |
| Elfn2_Q68FM6_S671s_1 | 0.72 | -0.44 | -0.34 | 0.75 | 0.67 | -0.36 | 0.18 | 0.84 | 0.04 | -0.3 | 0.54 | 0.06 | -0.65 | -0.24 | 0.16 | 0.19 | 0.4225 | 0.2225 | 0.0775 | 0.452525 | 0.703518 | 0.821255 | | 1 | 0.999631 | 0.788364 |
| Elfn2_Q68FM6_S750s_1 | -0.1 | -0.23 | -0.07 | 0.1 | 0.44 | 0.05 | 0.27 | 0.63 | 0.11 | -0.36 | 0.68 | 0.16 | -0.59 | -0.31 | 0.45 | 0.46 | 0.1325 | -0.1225 | -0.325 | 0.820133 | 0.121392 | 0.464495 | | 1 | 0.999631 | 0.938303 |
| Elfn2_Q68FM6_T493t_1 | 0.53 | -0.41 | 0.37 | 0.76 | 0.36 | 0.61 | -0.01 | 0.82 | -0.21 | 0.33 | 0.43 | 0.21 | -0.97 | 0.27 | 0.33 | 0.32 | -0.21 | -1.045 | 0.4525 | 0.447076 | 0.700555 | 0.717358 | | 1 | 0.999631 | 0.785579 |
| Elfn2_Q68FM6_T655t_1 | -1.63 | -1.75 | -1.32 | -1.01 | -1.85 | -1.67 | -2.29 | -0.74 | -1.77 | -1.9 | -2.81 | -3.41 | -1.01 | -1.27 | -1.6 | -0.02 | 0.195 | 0.005 | 0.2075 | 0.275921 | 0.583199 | 0.031983 | | 1 | 0.999631 | 0.700187 |
| Elmo1_Q8BPU7_S344s_1 | -0.16 | -0.4 | -0.32 | -0.12 | -0.2 | 0.25 | -0.31 | 0.04 | -0.34 | -0.43 | -0.02 | -0.19 | -0.01 | -0.25 | 0.05 | 0.04 | -0.15 | 0.28 | 0.66 | 0.442806 | 0.45102 | 0.98419 | | 1 | 0.999631 | 0.782367 |
| Elmsan1_E9Q2I4_S456s_1 | -0.23 | 0.12 | 0.75 | -0.05 | -0.23 | 0.46 | -0.54 | 0.3 | 0.34 | 0.24 | 0.15 | 0.98 | -0.22 | 1 | 1.57 | 0.88 | -0.075 | -0.1225 | 0.2725 | 0.152993 | 0.66643 | 0.410564 | | 1 | 0.999631 | 0.625198 |
| Emb_P21995_S312s_1 | -0.93 | -0.93 | 0.24 | 0.34 | -0.48 | -0.91 | 0.48 | -0.67 | -1.64 | -1.19 | -0.16 | 1.22 | 0.31 | -1.78 | 1.52 | -0.24 | 0.055 | 0.1075 | 0.3125 | 0.691828 | 0.867692 | 0.851155 | | 1 | 0.999631 | 0.894017 |
| Emc4_Q9CZX9_S36s_1 | -1.24 | -0.38 | -0.74 | -0.76 | -0.52 | -1.08 | -0.87 | -0.43 | -0.65 | -1.4 | -0.14 | -0.5 | -0.05 | -0.53 | -0.89 | -0.4 | -0.39 | -1.3625 | -2.5125 | 0.348565 | 0.853433 | 0.762273 | | 1 | 0.999631 | 0.736697 |
| Eml1_D3Z4J9_S115s_1 | 2.64 | -4.95 | -2.75 | -2.41 | -3.91 | -3.33 | 3.04 | -4.83 | -7.09 | 1.62 | -4.19 | -3.26 | -4.05 | -5.34 | -4.04 | -4.09 | -0.1375 | -0.12 | 0.2625 | 0.09242 | 0.847614 | 0.510233 | | 1 | 0.999631 | 0.557085 |
| Eml1_D3Z4J9_S154s_1 | -0.31 | 0.42 | 1.01 | 0.55 | 0.25 | 0.32 | -0.3 | 0.85 | 0.24 | 0.55 | -0.11 | 0.51 | 0.58 | 1.29 | 0.19 | 0.66 | -0.63 | -0.6925 | -1.235 | 0.50326 | 0.715506 | 0.733377 | | 1 | 0.999631 | 0.814092 |
| Eml3_Q8VC03_S157s_1 | 2.34 | 0.11 | 1.31 | 0.73 | -0.3 | 0.52 | 1.59 | 0.16 | 0.02 | 1.8 | -0.05 | -0.05 | -0.25 | -0.73 | 0.58 | -0.05 | 0.1525 | 0.3025 | -0.195 | 0.035735 | 0.282541 | 0.265231 | | 1 | 0.999631 | 0.453811 |
| Eml3_Q8VC03_S177s_1 | 1.77 | 3.05 | 0.48 | 0.26 | 2.08 | 2.7 | 0.6 | 0.79 | 3.19 | 2.21 | 0.68 | 0.69 | 1.48 | 1.74 | 1 | 0.56 | 0.1025 | 0.0975 | 0.3075 | 0.756951 | 0.833164 | 0.700679 | | 1 | 0.999631 | 0.914348 |
| Eml4_F8WJ93_S146s_1 | 0.3 | -0.3 | -0.06 | 0.57 | 0.29 | 0.22 | 0.11 | 0.3 | 0.4 | 0.34 | -0.05 | 0.21 | 0.49 | 0.37 | 0.43 | 0.45 | -0.0975 | 0.32 | 0.3675 | 0.318398 | 0.714413 | 0.742736 | | 1 | 0.999631 | 0.72215 |
| Eml4_F8WJ93_S908s_1 | -0.84 | -0.56 | -0.61 | -0.79 | -0.59 | -1.13 | -0.82 | -0.65 | -0.39 | -0.71 | -0.22 | -0.2 | 0.29 | -0.61 | -0.24 | -0.77 | 0.25 | -0.2075 | 0.31 | 0.272797 | 0.701684 | 0.238299 | | 1 | 0.999631 | 0.700187 |
| Enah_J3QNM3_S129s_1 | -0.34 | -0.1 | 0.43 | 0.71 | 0.5 | 0.05 | 0.85 | 0.3 | -0.8 | -0.75 | 1.29 | 0.13 | 1.61 | -0.38 | -0.28 | 0.99 | 0.06 | -0.4975 | 0.4225 | 0.551145 | 0.459589 | 0.68743 | | 1 | 0.999631 | 0.836275 |
| Enah_J3QNM3_S240s_1 | -0.18 | -0.28 | -0.94 | -0.74 | -0.04 | -0.97 | -0.63 | -0.26 | -0.43 | -1.13 | -1.17 | -1.4 | 0.38 | 0.37 | 0.29 | -1.49 | 0.075 | 0.3275 | 0.1275 | 0.388516 | 0.851872 | 0.154825 | | 1 | 0.999631 | 0.757772 |
| Enah_J3QNM3_S339s_1 | -0.18 | -0.68 | -1.34 | -0.28 | -0.56 | -0.82 | -0.25 | -0.55 | -0.1 | 0.07 | -0.4 | -0.74 | -0.62 | -0.63 | -0.5 | -0.22 | 0.38 | -0.3175 | -0.5025 | 0.710831 | 0.820062 | 0.366267 | | 1 | 0.999631 | 0.900199 |
| Enah_J3QNM3_S339s_2 | 0.17 | -0.15 | -0.46 | 0.9 | 0.4 | 0.29 | 1.07 | 0.22 | 0.24 | -0.77 | -0.07 | -0.21 | -2.56 | -0.45 | 0.45 | 1.01 | -0.055 | -0.2275 | 0.02 | 0.500192 | 0.317453 | 0.413919 | | 1 | 0.999631 | 0.813379 |
| Enah_J3QNM3_S340s_1 | 0.29 | 0.41 | 0.33 | 0.17 | 0.4 | 1.22 | -0.08 | -0.56 | 0.48 | 0.5 | -0.58 | -0.11 | 0.05 | 1.05 | 0.08 | 0.1 | 0.38 | -0.3175 | -0.5025 | 0.951063 | 0.888118 | 0.485904 | | 1 | 0.999631 | 0.984079 |
| Enah_J3QNM3_S340s_2 | 0.17 | -0.15 | -0.46 | 0.9 | 0.4 | 0.29 | 1.07 | 0.22 | 0.24 | -0.77 | -0.07 | -0.21 | -2.56 | -0.45 | 0.45 | 1.01 | -0.165 | 0.0475 | 0.3475 | 0.500192 | 0.317453 | 0.413919 | | 1 | 0.999631 | 0.813379 |
| Enah_J3QNM3_S727s_1 | 0.82 | 1.17 | 0.51 | 0.42 | 0.67 | 0.74 | 0.13 | 0.72 | 0.6 | 0.81 | 0.69 | 1.01 | 1.18 | 1.02 | 1.05 | 1.06 | -0.3675 | 0.485 | -0.82 | 0.250159 | 0.576325 | 0.867204 | | 1 | 0.999631 | 0.686417 |
| Enah_J3QNM3_T708t_1 | 1.34 | 0.53 | 0.6 | 0.61 | 0.54 | 0.35 | 0.75 | -0.03 | 1.36 | 1.77 | 0.38 | 1.51 | -1.8 | 0.35 | 0.83 | 0.42 | -0.2925 | -0.3925 | -0.6375 | 0.174598 | 0.251627 | 0.222066 | | 1 | 0.999631 | 0.642116 |
| Eno1_P17182_S263s_1 | 1.16 | -0.4 | 0.66 | -0.03 | -1.02 | 0.2 | 1.77 | -0.73 | -0.68 | 1.53 | -0.53 | -0.5 | 0.23 | -1.04 | 0.14 | -0.49 | 0.01 | -0.015 | 0.52 | 0.182321 | 0.650551 | 0.503091 | | 1 | 0.999631 | 0.643861 |
| Eno2_P17183_S79s_1 | -0.82 | -0.05 | 1.11 | -0.12 | 0.32 | -0.67 | -0.71 | 1.22 | -0.75 | -0.43 | 0.56 | 0.68 | 1.1 | -0.06 | 0.39 | 0.77 | -0.17 | -0.0325 | -0.2325 | 0.278162 | 0.98568 | 0.97634 | | 1 | 0.999631 | 0.700874 |
| Enox1_Q8BHR2_S403s_1 | 0.38 | 1.25 | 2.52 | 1.31 | 0.8 | 0.82 | 1.17 | 1.99 | 1.1 | 1.22 | 1.46 | 1.55 | 0.23 | 1.42 | 1.54 | 1.34 | 0.7475 | 0.3575 | -0.015 | 0.64926 | 0.729447 | 0.941746 | | 1 | 0.999631 | 0.874768 |
| Ensa_P60840_S43s_1 | 1.28 | 1.59 | -0.84 | 1.29 | 1.49 | 2.42 | 0.9 | 1.5 | 1.51 | 1.37 | 1.36 | 0.51 | -1.26 | 2.18 | 1.33 | 1.01 | -0.035 | -0.0075 | -0.3375 | 0.985122 | 0.220035 | 0.527306 | | 1 | 0.999631 | 0.995133 |
| Ep300_B2RWS6_S500s_1 | 0.74 | 0.49 | -0.35 | -0.73 | 0.19 | 0.76 | 0.3 | -1.24 | 0.76 | 0.15 | 0.03 | -0.82 | -0.16 | 0.28 | -0.29 | -1.03 | -0.7325 | -0.4075 | -1.0775 | 0.452565 | 0.945801 | 0.987028 | | 1 | 0.999631 | 0.788364 |
| Ep400_Q8CHI8_S721s_1 | 1.15 | 0.95 | 2.21 | 1.82 | 0.66 | 0.18 | 1.4 | 0.96 | 0.25 | 0.96 | 1.35 | 1.94 | -1.27 | -0.02 | 1.65 | 1.46 | -0.3125 | 0.36 | -0.2875 | 0.129616 | 0.088772 | 0.376337 | | 1 | 0.999631 | 0.601246 |
| Ep400_Q8CHI8_S728s_1 | 1.18 | 1.36 | 1.6 | 1.69 | 0.84 | 0.92 | 1.1 | 1.72 | 1.54 | 1.94 | 1.67 | 2.12 | 0.42 | 1.26 | 1.45 | 1.55 | 0.1725 | 0.6625 | 0.4325 | 0.411751 | 0.306064 | 0.212768 | | 1 | 0.999631 | 0.767937 |
| Ep400_Q8CHI8_S923s_1 | -0.24 | 1.39 | 0.41 | -0.39 | 1.4 | 0.06 | 0.23 | 0.17 | 2.03 | 1.47 | -0.3 | 0.62 | 2.03 | 0.01 | 0.23 | 0.63 | 0.1725 | 0.6625 | 0.4325 | 0.448908 | 0.722361 | 0.280373 | | 1 | 0.999631 | 0.786796 |
| Ep400_Q8CHI8_T922t_1 | -0.24 | 1.39 | 0.41 | -0.39 | 1.4 | 0.06 | 0.23 | 0.17 | 2.03 | 1.47 | -0.3 | 0.62 | 2.03 | 0.01 | 0.23 | 0.63 | -0.1625 | 0.52 | 0.7725 | 0.448908 | 0.722361 | 0.280373 | | 1 | 0.999631 | 0.786796 |
| Epb4.1_P48193_S182s_1 | 0.48 | 1.84 | 0.65 | 0.93 | 0.81 | 1.59 | 0.27 | 0.58 | 1.64 | 1.13 | 2.04 | 1.17 | 2.22 | 1.64 | 1.44 | 1.69 | -0.61 | -0.115 | -0.1275 | 0.065762 | 0.694728 | 0.202315 | | 1 | 0.999631 | 0.516933 |
| Epb4.1_P48193_S543s_1 | -0.66 | -0.1 | 0.14 | 0.45 | -0.48 | -1.4 | -0.44 | -0.29 | -0.52 | 0.09 | 0.02 | -0.22 | 0.08 | -1.13 | -0.12 | 0.49 | 0.03 | -0.1925 | -0.1225 | 0.765431 | 0.119728 | 0.72558 | | 1 | 0.999631 | 0.916326 |
| Epb4.1_P48193_S556s_1 | -0.44 | -0.84 | -0.06 | 0.35 | 0.07 | -0.5 | -0.38 | -0.06 | -0.32 | -1.23 | 0.07 | -0.28 | -0.37 | 0.06 | -0.23 | -0.94 | 0.68 | 0.3525 | -0.515 | 0.742452 | 0.927411 | 0.626077 | | 1 | 0.999631 | 0.909615 |
| Epb4.1_P48193_S86s_1 | 0.73 | 0.32 | -0.35 | 1.71 | 2.06 | 0.68 | 1.08 | 1.31 | 1.28 | 0.23 | 1.14 | 1.17 | -1.78 | 0.72 | 0.28 | 1.13 | 1.0475 | 0.5075 | 0.355 | 0.462062 | 0.186652 | 0.465873 | | 1 | 0.999631 | 0.792786 |
| Epb4.1_P48193_S87s_1 | -0.01 | 0.63 | -0.87 | 0.87 | 2.33 | 1.43 | 0.75 | 0.3 | 1.48 | -0.87 | 0.71 | 1.33 | -0.16 | 1.18 | 0.59 | 0.43 | -0.5975 | -0.47 | -0.8375 | 0.454616 | 0.077728 | 0.409393 | | 1 | 0.999631 | 0.789731 |
| Epb4.1_P48193_S97s_1 | 0.1 | -0.66 | -0.63 | 0.3 | -1.03 | -1.78 | 0.33 | -0.8 | -0.61 | -0.66 | -0.27 | -1.23 | -0.79 | -1.13 | -0.57 | -1.75 | -0.3575 | -0.58 | -0.025 | 0.052655 | 0.229043 | 0.205026 | | 1 | 0.999631 | 0.489777 |
| Epb4.1_P48193_T62t_1 | -1.13 | -1.12 | 0.89 | 0.31 | -1.29 | -1.97 | 0.29 | 0.49 | -1.53 | -2.36 | -0.08 | 0.6 | -0.36 | -1.88 | 0.67 | 0.42 | 0.3625 | 0.635 | 0.405 | 0.970838 | 0.611118 | 0.445027 | | 1 | 0.999631 | 0.989618 |
| Epb4.1l1_A2AUK5_S441s_1 | 1.01 | 1.62 | -0.78 | -0.96 | 2.13 | 1.64 | -0.86 | -0.57 | 2.05 | 1.89 | -0.65 | 0.14 | 2.05 | 1.52 | -0.17 | -0.89 | 0.9375 | 0.7625 | 0.38 | 0.624277 | 0.676155 | 0.440786 | | 1 | 0.999631 | 0.86356 |
| Epb4.1l1_A2AUK5_S441s_2 | -0.77 | -0.12 | -1.39 | 0.95 | 0.48 | -0.23 | 0.69 | 1.48 | 0.39 | -0.18 | 1.11 | 0.4 | 0 | 0.59 | -0.76 | 0.36 | 0.3575 | -0.4275 | -0.59 | 0.489007 | 0.118482 | 0.171935 | | 1 | 0.999631 | 0.807201 |
| Epb4.1l1_E9PV14_S1077s_1 | 0.51 | 1.53 | 2.12 | 1.89 | 0.96 | 2.46 | 2.85 | 1.21 | 1.52 | -0.46 | 1.2 | 2.08 | 0.44 | 0.53 | 2.11 | 0.61 | -0.105 | -0.19 | 0.4175 | 0.261944 | 0.511352 | 0.478262 | | 1 | 0.999631 | 0.692758 |
| Epb4.1l1_E9PV14_S1228s_1 | -0.29 | -0.47 | -0.35 | -0.59 | -0.71 | -0.22 | -0.76 | -0.43 | -0.67 | -0.4 | -1.02 | -0.37 | 0.02 | -0.02 | 0 | -0.03 | -0.39 | -0.18 | -1.6375 | 0.131942 | 0.681932 | 0.490701 | | 1 | 0.999631 | 0.603456 |
| Epb4.1l1_E9PV14_S1255s_1 | 1.22 | -0.89 | -0.42 | 0.09 | -0.72 | -0.26 | 0.11 | -0.69 | -0.2 | 1.23 | -0.8 | -0.95 | -2.1 | -2.03 | -1.15 | -1.27 | 1.2875 | 1.02 | 1.5625 | 0.007628 | 0.416072 | 0.768639 | | 1 | 0.999631 | 0.358173 |
| Epb4.1l1_E9PV14_S1325s_1 | -1.65 | 3.03 | 2.58 | 0.33 | 3.39 | 2.24 | 1.4 | 2.41 | 3.18 | -1.42 | 3.03 | 3.58 | 3.84 | 1.86 | 3.13 | 1.71 | -0.4 | -0.19 | 0.5775 | 0.149645 | 0.215869 | 0.460912 | | 1 | 0.999631 | 0.6218 |
| Epb4.1l1_E9PV14_S430s_1 | -0.7 | 1.57 | 1.76 | 0.22 | 0.26 | 1.56 | 0.1 | -0.67 | 1.1 | 0.62 | -0.47 | 0.84 | 1.09 | 1.72 | 1.11 | 1.24 | 0.4775 | 0.59 | -3.9E-17 | 0.31343 | 0.549785 | 0.757503 | | 1 | 0.999631 | 0.719861 |
| Epb4.1l1_E9PV14_S440s_1 | 0.5 | -0.13 | -1.26 | -1.26 | 0.59 | 0.51 | -0.94 | -0.4 | 0.74 | 0.89 | -1.65 | 0.23 | -0.16 | -0.15 | -0.3 | -1.54 | -0.2875 | 0.28 | -0.19 | 1 | 0.377155 | 0.378476 | | 1 | 0.999631 | 1 |
| Epb4.1l1_E9PV14_S713s_1 | 0.16 | -1.58 | 0.62 | 1.67 | -0.14 | -1.31 | 0.64 | 0.53 | 0.34 | -1.21 | 1.29 | 1.57 | -1.06 | -0.98 | 1.5 | 0.65 | 0.7675 | -0.16 | -0.52 | 0.813614 | 0.689351 | 0.73003 | | 1 | 0.999631 | 0.937041 |
| Epb4.1l1_E9PV14_T1217t_1 | 1.65 | -0.75 | -0.5 | 0.98 | 0.72 | 1.22 | 1.4 | 1.11 | 0.87 | 0.33 | 0.22 | -0.68 | -1.48 | 0.36 | 0.58 | -0.16 | -1.335 | -0.855 | -2.2325 | 0.439703 | 0.183081 | 0.79085 | | 1 | 0.999631 | 0.781983 |
| Epb4.1l1_E9PV14_T1257t_1 | 1.23 | 1.84 | 0.96 | 0.52 | -0.01 | -0.31 | 0.5 | -0.97 | 0.63 | 1.34 | -0.61 | -0.23 | -0.84 | -1.25 | -0.7 | -1.59 | -0.23 | 0.055 | 0.4125 | 0.000161 | 0.008669 | 0.107868 | | 1 | 0.999631 | 0.351308 |
| Epb4.1l1_Q9Z2H5_S378s_1 | -0.51 | -0.58 | -0.19 | -0.6 | -0.78 | -0.68 | -0.87 | -0.47 | -0.72 | -0.19 | -0.55 | -0.2 | 0.12 | -0.05 | -0.34 | 0.04 | -0.305 | 0.1225 | 0.5075 | 0.160757 | 0.36802 | 0.839322 | | 1 | 0.999631 | 0.630575 |
| Epb4.1l1_Q9Z2H5_S430s_1 | -0.54 | 0.14 | 1.34 | 0.27 | -0.49 | 0.01 | -0.09 | 0.56 | -0.24 | -0.04 | 0.76 | 1.22 | 0.57 | 0.31 | 1.16 | 1.2 | -0.3775 | -0.5225 | -1.5025 | 0.272197 | 0.488753 | 0.803952 | | 1 | 0.999631 | 0.699819 |
| Epb4.1l1_Q9Z2H5_S461s_1 | 1.46 | -0.84 | 0.53 | -0.33 | -1.76 | 0.06 | 2.74 | -1.73 | -1.29 | 1.74 | -0.91 | -0.81 | -0.71 | -1.67 | -1.49 | -1.32 | 0.2875 | 0.2525 | 0.485 | 0.014804 | 0.70927 | 0.495548 | | 1 | 0.999631 | 0.388832 |
| Epb4.1l1_Q9Z2H5_S461s_2 | -1.9 | 0.3 | -0.52 | -1.29 | -0.75 | -0.06 | -0.64 | -0.81 | -0.61 | -0.26 | -1.1 | -0.43 | 1.29 | -0.18 | -1.61 | -0.97 | -0.72 | -0.265 | -0.77 | 0.492207 | 0.553047 | 0.606867 | | 1 | 0.999631 | 0.809399 |
| Epb4.1l1_Q9Z2H5_S466s_1 | 0.43 | -0.43 | 1.53 | -0.3 | -1.83 | 0.54 | 0.27 | -0.63 | -0.78 | 1.42 | -0.16 | -0.31 | 0 | -1.39 | -0.21 | -0.25 | 0.09 | -0.02 | 0.5325 | 0.160249 | 0.26734 | 0.659106 | | 1 | 0.999631 | 0.629764 |
| Epb4.1l1_Q9Z2H5_S466s_2 | -1.54 | 0.55 | 0.44 | -0.26 | -0.61 | 0.31 | -0.03 | -0.12 | -0.28 | -0.17 | -0.62 | 0.18 | 1.21 | -0.03 | -0.34 | 0.48 | 1.5925 | 1.985 | 1.015 | 0.342756 | 0.853553 | 0.96714 | | 1 | 0.999631 | 0.73348 |
| Epb4.1l1_Q9Z2H5_S510s_1 | 0.17 | 2.16 | 1.69 | 0.8 | 3.83 | 1.83 | 3.57 | 1.96 | 4.35 | 0.75 | 3.61 | 4.05 | 3.74 | 1.43 | 2.11 | 1.6 | -0.065 | -0.15 | -0.065 | 0.127724 | 0.026462 | 0.034818 | | 1 | 0.999631 | 0.598268 |
| Epb4.1l1_Q9Z2H5_S544s_1 | 0.4 | 0.43 | 0.65 | 0.69 | 0.36 | 0.22 | 0.2 | 1.13 | 0.44 | 0.47 | 0.14 | 0.52 | 0.13 | 0.46 | 0.39 | 0.93 | -0.295 | -0.2425 | 0.0475 | 0.825743 | 0.827208 | 0.557068 | | 1 | 0.999631 | 0.940475 |
| Epb4.1l1_Q9Z2H5_S544s_2 | -0.01 | 0.57 | -0.38 | -0.38 | -0.28 | 0.27 | -1.19 | -0.18 | -0.32 | 0.27 | -0.18 | -0.94 | 0.82 | -0.17 | -0.42 | -0.24 | -0.175 | 0.045 | 0 | 0.902758 | 0.454863 | 0.512188 | | 1 | 0.999631 | 0.968183 |
| Epb4.1l1_Q9Z2H5_S546s_1 | 0.27 | 0.39 | 0.2 | 0.16 | -0.06 | 0.57 | -0.46 | 0.27 | 0.37 | 0.66 | -0.02 | 0.19 | -0.26 | 0.84 | 0.25 | 0.19 | -0.295 | -0.2425 | 0.0475 | 1 | 0.556059 | 0.866206 | | 1 | 0.999631 | 1 |
| Epb4.1l1_Q9Z2H5_S546s_2 | -0.01 | 0.57 | -0.38 | -0.38 | -0.28 | 0.27 | -1.19 | -0.18 | -0.32 | 0.27 | -0.18 | -0.94 | 0.82 | -0.17 | -0.42 | -0.24 | 0.36 | 0.1525 | -0.0725 | 0.902758 | 0.454863 | 0.512188 | | 1 | 0.999631 | 0.968183 |
| Epb4.1l1_Q9Z2H5_S639s_1 | 1.03 | 1.58 | 0.13 | 1.7 | 1.87 | 1.49 | 0.48 | 2.04 | 1.68 | 0.93 | 1.12 | 1.32 | -0.46 | 2.03 | 1.18 | 1.4 | -0.2475 | -0.195 | 0.37 | 0.901532 | 0.45561 | 0.706045 | | 1 | 0.999631 | 0.968183 |
| Epb4.1l1_Q9Z2H5_S666s_1 | -0.49 | -0.46 | 0.22 | -0.06 | -0.7 | -0.7 | -0.51 | 0.13 | -0.57 | -0.07 | -0.55 | -0.38 | 0.45 | -0.35 | 0.14 | 0.45 | -0.47 | -0.2775 | 0.21 | 0.276383 | 0.43692 | 0.50772 | | 1 | 0.999631 | 0.700187 |
| Epb4.1l1_Q9Z2H5_S671s_1 | -0.67 | -0.3 | 0.5 | -0.17 | -0.91 | -0.86 | -0.72 | -0.03 | -1.33 | 0 | 0.1 | -0.52 | -0.11 | -0.31 | 0.24 | 0.38 | -0.3875 | -0.0125 | 0.365 | 0.550977 | 0.198189 | 0.507863 | | 1 | 0.999631 | 0.836275 |
| Epb4.1l1_Q9Z2H5_S677s_1 | -0.14 | 0.52 | 0.25 | -0.38 | -0.23 | 0.12 | -0.56 | -0.63 | -0.11 | 0.45 | -0.27 | 0.13 | 1.05 | 0.56 | 0.15 | -0.05 | -0.2725 | -0.2775 | -0.245 | 0.327172 | 0.237587 | 0.968475 | | 1 | 0.999631 | 0.724678 |
| Epb4.1l1_Q9Z2H5_S684s_1 | 0.15 | 0.51 | 0.77 | 0.75 | 0.27 | 0.26 | -0.11 | 0.67 | 0.21 | 0.18 | 0.1 | 0.58 | -0.44 | 0.73 | 0.17 | 0.74 | 1.9425 | 1.9925 | 1.2725 | 0.504708 | 0.355919 | 0.331874 | | 1 | 0.999631 | 0.814292 |
| Epb4.1l1_Q9Z2H5_S69s_1 | -0.82 | 4.88 | 3.35 | 1.27 | 5.44 | 3.63 | 3.71 | 3.67 | 6.08 | -0.37 | 5.35 | 5.59 | 4.74 | 2.84 | 4.52 | 1.67 | 0.49 | 0.4225 | 1.0425 | 0.309376 | 0.109808 | 0.24891 | | 1 | 0.999631 | 0.718845 |
| Epb4.1l1_Q9Z2H5_S75s_1 | -0.64 | 3.29 | 2.06 | 1.79 | 2.08 | 2.67 | 1.36 | 2.35 | 2.82 | 0.26 | 2.43 | 2.68 | 2.67 | 2.72 | 2.93 | 2.35 | 1.565 | -0.005 | -0.28 | 0.176351 | 0.52457 | 0.635628 | | 1 | 0.999631 | 0.642116 |
| Epb4.1l1_Q9Z2H5_S782s_1 | 1.29 | -1.13 | -2.49 | 0.8 | 1.93 | 0.8 | 0.95 | 1.05 | 0.48 | -0.84 | 0.2 | -1.39 | -2.05 | 0.41 | -0.62 | -0.39 | -0.5 | -0.445 | -1.2275 | 0.749772 | 0.073725 | 0.995327 | | 1 | 0.999631 | 0.912137 |
| Epb4.1l1_Q9Z2H5_S868s_1 | 1.38 | -1.23 | -0.27 | 0.05 | -1.04 | -0.57 | 0.13 | -0.59 | -0.73 | 0.65 | -0.82 | -0.95 | -1.57 | -1.58 | -0.65 | -1.18 | -0.0175 | -0.31 | 0.4825 | 0.044521 | 0.368582 | 0.463474 | | 1 | 0.999631 | 0.475443 |
| Epb4.1l1_Q9Z2H5_T30t_1 | -0.45 | -2.66 | -3.31 | -1.19 | -2.22 | -2.34 | -1.35 | -1.77 | -1.7 | -0.93 | -3.4 | -2.82 | -2.11 | -1.31 | -1.46 | -0.8 | 2.36E-16 | 0.0925 | 0.2325 | 0.456827 | 0.977593 | 0.683778 | | 1 | 0.999631 | 0.790722 |
| Epb4.1l1_Q9Z2H5_T475t_1 | -0.56 | -1.34 | 0.06 | -0.28 | -0.91 | -0.94 | -0.37 | 0.1 | -0.89 | -0.46 | -0.28 | -0.12 | -0.84 | -0.68 | 0.06 | 0.27 | -0.0175 | -0.2725 | -0.015 | 0.581214 | 1 | 0.802427 | | 1 | 0.999631 | 0.847114 |
| Epb4.1l1_Q9Z2H5_T475t_2 | -0.4 | 0.89 | 1.06 | 0.4 | 0.05 | 1.1 | 0.37 | 0.36 | 0.72 | -0.22 | -0.26 | 0.62 | -0.2 | 0.3 | 0.45 | 1.34 | 0.19 | -0.075 | -0.0725 | 0.973674 | 0.965083 | 0.523561 | | 1 | 0.999631 | 0.990719 |
| Epb4.1l1_Q9Z2H5_T489t_1 | 0.41 | 0.05 | -0.84 | -0.59 | 0.34 | 0.2 | -0.45 | -0.3 | 0.03 | 0.09 | -0.68 | -0.71 | -0.76 | 0.4 | -0.6 | -0.3 | -0.0725 | -0.145 | 0.1125 | 0.858286 | 0.605675 | 0.844357 | | 1 | 0.999631 | 0.952251 |
| Epb4.1l1_Q9Z2H5_T685t_1 | 0.46 | 0.19 | -0.02 | 0.13 | 0.07 | 0.19 | -0.16 | 0.37 | -0.09 | 0.13 | 0.01 | 0.13 | -0.15 | 0.62 | 0.16 | 0.58 | -0.1375 | 0.22 | 0.3075 | 0.714528 | 0.778961 | 0.570031 | | 1 | 0.999631 | 0.901866 |
| Epb4.1l2_Q80UE4_S116s_1 | 0.29 | 1.37 | 1.85 | 1.09 | 0.49 | 1.6 | 0.78 | 1.18 | 1.25 | 1.74 | 1.07 | 1.42 | 1.58 | 1.23 | 1.14 | 1.88 | 0.5025 | 0.6425 | 1.12 | 0.443783 | 0.736381 | 0.565171 | | 1 | 0.999631 | 0.782802 |
| Epb4.1l2_Q80UE4_S379s_1 | -0.48 | 0 | -1.95 | -2.49 | 0.25 | -0.77 | -1.11 | -1.28 | 0.16 | -0.3 | -0.71 | -1.5 | 0.18 | 0.52 | -0.24 | -0.9 | -0.955 | 0.305 | 0.625 | 0.088383 | 0.421728 | 0.31574 | | 1 | 0.999631 | 0.553858 |
| Epb4.1l2_Q80UE4_S38s_1 | -7.28 | -5.2 | -0.63 | -0.43 | -7.76 | -7.82 | -0.87 | -0.91 | -6.53 | -4.32 | -0.87 | -0.6 | -4.2 | -5.43 | -1.08 | -0.33 | -0.335 | 0.1 | 0.3875 | 0.722056 | 0.666783 | 0.871152 | | 1 | 0.999631 | 0.905674 |
| Epb4.1l2_Q80UE4_S492s_1 | -0.04 | 0.41 | -0.24 | -0.62 | 0.03 | 0.03 | -0.82 | -1.07 | 0.83 | 0.36 | -0.61 | -0.67 | 0.67 | 0.22 | -0.02 | 0.19 | 0.0825 | 0.0125 | 0.3575 | 0.259051 | 0.380561 | 0.815153 | | 1 | 0.999631 | 0.692624 |
| Epb4.1l2_Q80UE4_S543s_1 | -0.61 | 0.06 | 0.25 | -0.2 | 0.05 | 0.25 | -0.53 | 0.06 | -0.21 | -0.35 | -0.19 | 0.3 | 0.59 | 0.07 | -0.26 | 0.53 | -0.1975 | -0.085 | -0.2175 | 0.306296 | 0.789084 | 0.967338 | | 1 | 0.999631 | 0.717521 |
| Epb4.1l2_Q80UE4_S628s_1 | 0.96 | -0.45 | -0.55 | -0.05 | -0.8 | -0.17 | -0.04 | 0.13 | -0.31 | 1.01 | -0.72 | -0.41 | -0.27 | -0.42 | 0.28 | -0.55 | 0.4175 | 1.045 | 0.295 | 0.597687 | 0.62618 | 0.862579 | | 1 | 0.999631 | 0.855033 |
| Epb4.1l2_Q80UE4_S86s_1 | -0.06 | 1.09 | 0.28 | -0.99 | 1.46 | 0.4 | 0.66 | -0.53 | 1.93 | -0.3 | 1.05 | 1.82 | 1.48 | 0.41 | 0.14 | -0.53 | -0.015 | 7.85E-17 | 0.3275 | 0.598648 | 0.451845 | 0.111257 | | 1 | 0.999631 | 0.85557 |
| Epb4.1l3_D0VYV6_S451s_1 | -0.68 | 0.05 | 0.12 | -0.29 | -0.07 | -0.48 | -0.46 | 0.15 | -0.23 | -0.48 | -0.13 | 0.04 | 0.16 | -0.03 | 0.19 | 0.19 | 0.5775 | -0.1625 | 0.85 | 0.28768 | 0.960273 | 1 | | 1 | 1 | 0.705972 |
| Epb4.1l4b_B1AXV1_S114s_1 | -0.07 | -0.72 | -0.62 | -1.69 | 0.08 | -0.92 | 0.12 | -0.07 | -0.16 | -0.45 | -2.14 | -1 | -0.07 | -0.26 | -0.35 | 0.98 | 0.245 | -0.04 | -0.05 | 0.084949 | 0.185099 | 0.754407 | | 1 | 0.999631 | 0.550457 |
| Epb41l3_Q9WV92_S428s_1 | 1.25 | 1.64 | 0.04 | 1.03 | 1.23 | 2.01 | 0.93 | 0.77 | 1.72 | 1.49 | 0.42 | 0.17 | 1.03 | 1.45 | 0.43 | 0.85 | -0.2525 | -0.1775 | 0.02 | 0.904379 | 0.572086 | 0.934913 | | 1 | 0.999631 | 0.96853 |
| Epb41l3_Q9WV92_S451s_1 | -0.36 | -0.41 | -0.04 | -0.52 | -0.62 | -0.69 | -0.65 | -0.38 | -0.84 | -0.59 | -0.27 | -0.34 | -0.06 | -0.55 | -0.45 | -0.19 | 0.6125 | -0.2475 | -0.385 | 0.943732 | 0.322762 | 0.519416 | | 1 | 0.999631 | 0.981629 |
| Epb41l3_Q9WV92_S456s_1 | -0.32 | -0.57 | -1.26 | 0.07 | 0.6 | 0.27 | -0.35 | -0.15 | -0.7 | -0.77 | -0.75 | -0.85 | -2.16 | 0.18 | -0.94 | -0.7 | 0.525 | 0.0075 | 0.1625 | 0.470078 | 0.122332 | 0.462802 | | 1 | 0.999631 | 0.79732 |
| Epb41l3_Q9WV92_S470s_1 | -0.77 | 1.22 | 0.25 | 0.54 | 1.39 | 0.96 | 0.39 | 0.6 | 0.9 | -0.98 | 0.87 | 0.48 | 0.01 | 1.2 | 0.57 | 0.11 | 0.1525 | -0.1625 | 0.2075 | 0.735726 | 0.262073 | 0.989294 | | 1 | 0.999631 | 0.90759 |
| Epb41l3_Q9WV92_S486s_1 | -0.12 | -0.71 | -0.69 | 0.82 | -0.38 | -0.86 | 0.41 | 0.74 | -0.99 | -1.14 | 0.33 | 0.45 | -1.23 | -0.16 | 0.79 | 0.73 | 0.5625 | -0.1225 | -0.4475 | 0.707946 | 0.753069 | 0.755091 | | 1 | 0.999631 | 0.898881 |
| Epb41l3_Q9WV92_S601s_1 | 1.22 | -0.49 | -1.21 | 0.52 | 1.04 | 0.29 | 0.73 | 0.23 | 0.27 | -0.38 | 0.11 | -0.45 | -2.11 | 0.35 | -0.11 | 0.12 | 0.12 | -0.0475 | 0.4 | 0.522627 | 0.301228 | 0.817032 | | 1 | 0.999631 | 0.821224 |
| Epb41l3_Q9WV92_S63s_1 | -0.13 | -0.44 | 0.14 | -0.27 | 0.67 | -0.38 | -0.39 | -0.12 | 0.12 | -0.22 | -0.5 | -0.29 | 0.13 | 0.45 | -0.08 | 0.4 | -0.415 | -0.285 | 0.1975 | 0.190911 | 0.711666 | 0.864215 | | 1 | 0.999631 | 0.648674 |
| Epb41l3_Q9WV92_S655s_1 | -0.2 | -0.69 | -0.15 | -0.61 | -1.01 | -0.49 | -0.95 | -0.86 | -0.56 | -1.09 | -1.7 | 0.56 | -0.76 | 0.03 | -0.09 | -0.04 | 0.14 | -0.355 | -0.1725 | 0.53678 | 0.149975 | 0.555769 | | 1 | 0.999631 | 0.830094 |
| Epb41l3_Q9WV92_S804s_1 | -0.04 | 0.09 | -0.22 | 0.27 | 0.38 | -0.16 | 0.11 | 0.33 | 0.12 | -0.66 | -0.37 | -0.41 | -0.79 | 0.27 | -0.04 | -0.03 | 0.3675 | -0.0375 | -0.2425 | 0.59928 | 0.596581 | 0.231534 | | 1 | 0.999631 | 0.855995 |
| Epb41l3_Q9WV92_S96s_1 | 0.5 | 0.28 | -0.61 | 0.82 | 0.97 | 0.56 | 0 | 0.93 | 0.46 | 0.12 | 0.05 | 0.21 | -1.82 | 0.88 | 0.6 | 0.36 | -0.16 | 0.175 | 0.2075 | 0.698268 | 0.356575 | 0.915911 | | 1 | 0.999631 | 0.895731 |
| Epb41l3_Q9WV92_T409t_1 | 0.93 | 2.06 | 1.93 | 1.53 | 1.11 | 1.83 | 0.94 | 1.93 | 1.87 | 1.43 | 1.93 | 1.92 | -0.52 | 2.75 | 2.43 | 2.62 | -0.59 | -0.015 | 0.09 | 0.775287 | 0.669518 | 0.600506 | | 1 | 0.999631 | 0.920998 |
| Epb41l3_Q9WV92_T493t_1 | -0.44 | 0.56 | 0.44 | 0.21 | -0.34 | -0.31 | -0.61 | -0.33 | -0.12 | 0.43 | 0.21 | 0.19 | 0.65 | 0.43 | -0.09 | 0.14 | -0.4175 | -0.075 | 0.1425 | 0.791795 | 0.072082 | 0.961892 | | 1 | 0.999631 | 0.926887 |
| Epb41l3_Q9WV92_T495t_1 | -0.75 | -1.04 | 0.5 | -0.25 | -1.11 | -1.36 | -0.62 | -0.12 | -1.1 | -1.13 | -0.16 | 0.55 | -0.44 | -1.06 | 0.01 | 0.52 | 0.1625 | -0.03 | 0.0925 | 0.760739 | 0.34118 | 0.880713 | | 1 | 0.999631 | 0.914614 |
| Epb41l3_Q9WV92_T518t_1 | 0.47 | 0.5 | 0.78 | 0.62 | 0.6 | 0.9 | 0.43 | 1.09 | 0.22 | 0.56 | 0.73 | 0.74 | -0.89 | 1.24 | 1.11 | 1.28 | 0.0425 | -0.05 | 0.135 | 0.855021 | 0.542019 | 0.908921 | | 1 | 0.999631 | 0.951781 |
| Epb41l3_Q9WV92_T64t_1 | 0.2 | 0.24 | 0.63 | 0.32 | 0.75 | 0.6 | 0 | 0.21 | 0.41 | 0.4 | 0.04 | 0.34 | -0.15 | 0.84 | 0.46 | 0.78 | -0.3675 | -0.32 | 0.3425 | 0.679208 | 0.879268 | 0.846885 | | 1 | 0.999631 | 0.887281 |
| Epb41l3_Q9WV92_T687t_1 | 0.08 | -0.34 | 1.09 | 0.49 | -0.34 | -0.41 | 0.09 | 0.51 | 0.2 | -0.47 | -1.22 | 1.53 | 0.56 | 0.48 | 0.56 | 1.09 | 0.365 | 0.58 | 0.45 | 0.371555 | 0.35 | 0.596443 | | 1 | 0.999631 | 0.746522 |
| Epb41l3_Q9WV92_T725t_1 | -0.15 | 2.32 | 1.77 | 0.08 | 1.54 | 1.58 | 1.01 | 1.35 | 2.26 | -0.03 | 1.98 | 2.13 | 1.85 | 1.46 | 1.81 | 0.7 | -0.305 | -0.07 | -0.1875 | 0.465094 | 0.52691 | 0.431578 | | 1 | 0.999631 | 0.794442 |
| Epb41l3_Q9WV92_T923t_1 | 0.03 | -0.36 | 0.29 | -0.03 | -0.6 | -0.36 | -0.32 | -0.01 | -0.44 | 0.31 | 0.1 | -0.32 | -0.07 | -0.36 | -0.05 | -0.34 | -0.3825 | -1.0125 | -1.31 | 0.515639 | 0.276828 | 0.814776 | | 1 | 0.999631 | 0.819449 |
| Epb41l4a_P52963_S402s_1 | 0.71 | -1.43 | 0.99 | 0.22 | -1.68 | -0.89 | 1.21 | 0.32 | -1.3 | -0.08 | -1.36 | -0.82 | -1.84 | -1.84 | 0.34 | -1.41 | -0.78 | 0.58 | 0.575 | 0.074255 | 0.60637 | 0.099728 | | 1 | 0.999631 | 0.539897 |
| Epb41l4a_P52963_S541s_1 | -1.62 | -1.59 | -0.58 | -1.86 | -2.97 | -1.86 | -2.72 | -1.22 | -1.09 | 0.05 | -1.01 | -1.28 | 0.68 | -1.18 | -1.5 | -1.35 | -0.78 | 0.58 | 0.575 | 0.305084 | 0.1213 | 0.186434 | | 1 | 0.999631 | 0.716239 |
| Epb41l4a_P52963_S543s_1 | -1.62 | -1.59 | -0.58 | -1.86 | -2.97 | -1.86 | -2.72 | -1.22 | -1.09 | 0.05 | -1.01 | -1.28 | 0.68 | -1.18 | -1.5 | -1.35 | 0.1175 | -0.4875 | 0.1025 | 0.305084 | 0.1213 | 0.186434 | | 1 | 0.999631 | 0.716239 |
| Epha5_Q6PFV6_Y649y_1 | -0.38 | -2.41 | -1.67 | -0.79 | -0.89 | -1.76 | -0.83 | -1.3 | -2 | -1.74 | -1.37 | -2.09 | -1.11 | -1.17 | -1.4 | -1.16 | -0.1825 | -0.41 | 0.455 | 0.821953 | 0.805606 | 0.310456 | | 1 | 0.999631 | 0.938464 |
| Epha7_Q61772_S946s_1 | -1.27 | -0.45 | 0.4 | 0.18 | -0.05 | -0.97 | -0.89 | 0.04 | -0.85 | -1.16 | -0.79 | 0.02 | 0.7 | -0.78 | 0.28 | 0.48 | 0.1425 | -0.495 | -0.065 | 0.357033 | 0.683712 | 0.367409 | | 1 | 0.999631 | 0.740804 |
| Epn1_Q80VP1_S418s_1 | 0.2 | 0.35 | -1.27 | 0.34 | 0.28 | 0.22 | -0.33 | 0.02 | -0.61 | -0.84 | -0.52 | -0.39 | -0.54 | 0.61 | -0.49 | -0.22 | 0.13 | -0.305 | -0.545 | 0.889137 | 0.731708 | 0.247095 | | 1 | 0.999631 | 0.963505 |
| Epn1_Q80VP1_S419s_1 | 0.72 | 0.85 | -1.07 | -0.17 | 0.79 | 0.19 | -0.14 | 0.01 | 0.56 | -0.94 | 0.05 | -0.56 | -0.86 | 0.2 | -0.9 | -0.29 | 0.1325 | -0.15 | 0.305 | 0.287089 | 0.781503 | 0.561933 | | 1 | 0.999631 | 0.705972 |
| Epn1_Q80VP1_S434s_1 | -0.17 | -0.03 | -1.22 | 0.96 | 0.4 | -0.24 | -0.27 | 0.18 | -0.38 | -1.11 | 0.21 | 0.22 | 0.11 | -0.14 | -0.04 | 0.83 | -0.0375 | 0.035 | 0.49 | 0.531163 | 0.772109 | 0.770756 | | 1 | 0.999631 | 0.826936 |
| Epn1_Q80VP1_S453s_1 | -0.91 | -0.15 | -0.35 | -0.57 | -0.6 | -0.37 | -0.8 | -0.36 | -0.46 | -1.12 | -0.3 | 0.04 | 0.24 | -0.11 | -0.15 | 0 | -0.01 | -0.115 | 0.465 | 0.121788 | 0.892497 | 0.917729 | | 1 | 0.999631 | 0.590509 |
| Epn1_Q80VP1_T463t_1 | -0.67 | -0.59 | -0.17 | -0.44 | -0.37 | -0.69 | -0.68 | -0.17 | -0.65 | -1.14 | -0.53 | -0.01 | 0.21 | -0.65 | 0.29 | 0.14 | -0.01 | -0.085 | 0.0675 | 0.172773 | 0.969936 | 0.71837 | | 1 | 0.999631 | 0.642116 |
| Epn1_Q80VP1_T469t_1 | -0.23 | -0.09 | -1.48 | -0.8 | 0.11 | -0.59 | -1.05 | -1.11 | -0.6 | -0.83 | -0.83 | -0.68 | -0.7 | -0.13 | -0.64 | -0.86 | 0.225 | -0.195 | -0.26 | 0.861226 | 0.980971 | 0.811799 | | 1 | 0.999631 | 0.954096 |
| Epn1_Q80VP1_T493t_1 | 1.31 | 0.13 | 0.26 | 0.61 | 0.99 | 1.31 | 0.55 | 0.36 | 0.81 | 0.52 | -0.09 | 0.29 | -1.04 | 1.34 | 0.57 | 0.4 | -0.1725 | -0.19 | 0.1775 | 0.625161 | 0.539139 | 0.590082 | | 1 | 0.999631 | 0.863768 |
| Epn2_Q5NCM5_S173s_1 | 0.16 | 0.89 | 0.5 | 0.29 | 0.46 | 0.42 | -0.31 | 0.58 | 0.33 | 0.41 | 0.05 | 0.29 | 0.68 | 1.01 | 0.29 | 0.57 | -0.5175 | 0.13 | 0.3075 | 0.571351 | 0.58273 | 0.498947 | | 1 | 0.999631 | 0.843666 |
| Epn2_Q5NCM5_S192s_2 | -1.35 | -0.16 | -0.5 | -0.64 | -2.25 | -0.3 | -1.14 | -1.03 | -0.67 | -0.06 | -0.93 | -0.47 | -0.61 | -0.12 | -0.28 | -0.41 | -0.2825 | -0.23 | 0.275 | 0.373119 | 0.272774 | 0.710975 | | 1 | 0.999631 | 0.747035 |
| Epn2_Q5NCM5_S195s_1 | 0.27 | -1.35 | -0.6 | -0.16 | -1.38 | -0.8 | -0.18 | -0.61 | -0.88 | -0.34 | -0.98 | -0.56 | -0.44 | -0.04 | 0.17 | -0.43 | -1.195 | -0.105 | 0.575 | 0.497938 | 0.507574 | 0.560831 | | 1 | 0.999631 | 0.811908 |
| Epn2_Q5NCM5_S199s_1 | -1.33 | -1.06 | -0.56 | -1.98 | -3.59 | -1.57 | -2.12 | -2.43 | -2.29 | -0.22 | -2.13 | -0.71 | -0.18 | -1.3 | -0.96 | -0.19 | -0.295 | 0.0225 | 0.3825 | 0.1929 | 0.033016 | 0.848634 | | 1 | 0.999631 | 0.649499 |
| Epn2_Q5NCM5_S199s_2 | -1.28 | -0.48 | -0.5 | -0.64 | -1.42 | -0.49 | -1.14 | -1.03 | -0.75 | -0.66 | -0.93 | -0.47 | -0.5 | -0.18 | -0.28 | -0.41 | 0.4675 | -0.14 | -1.1625 | 0.225717 | 0.366635 | 0.938889 | | 1 | 0.999631 | 0.676382 |
| Epn2_Q5NCM5_S478s_1 | 1.47 | 0.01 | -0.87 | 1.74 | 1.4 | 0.29 | 1.43 | 1.1 | 0.42 | 0.19 | 1.37 | -0.19 | -2.39 | 0.33 | -0.34 | 0.1 | -0.0875 | 0.085 | 0.5225 | 0.152353 | 0.447517 | 0.824943 | | 1 | 0.999631 | 0.625198 |
| Epn2_Q5NCM5_S482s_1 | -0.93 | -0.71 | -1.81 | -1.72 | -0.75 | -1.21 | -1.66 | -1.9 | -0.85 | -0.98 | -1.69 | -1.31 | -0.09 | -0.4 | -1.3 | -1.29 | 0.2625 | -0.175 | -0.0425 | 0.237931 | 0.820947 | 0.816005 | | 1 | 0.999631 | 0.683301 |
| Epn2_Q5NCM5_T504t_1 | 1.21 | 1.37 | 0.32 | 1.4 | 1.85 | 1.51 | 0.88 | 1.11 | 1.53 | 0.72 | 0.74 | 0.61 | -0.14 | 2.09 | 1.2 | 0.98 | -0.565 | -0.3725 | -0.9875 | 0.932459 | 0.469961 | 0.631807 | | 1 | 0.999631 | 0.978264 |

| | | | | | | | | | | | | | | | | | | | | | | | | | |
|---|---|---|---|---|---|---|---|---|---|---|---|---|---|---|---|---|---|---|---|---|---|---|---|---|---|
| Epn2_Q5NCM6_S192s_1 | 0.12 | -2.23 | 0 | -0.75 | -2.16 | -1.97 | -0.18 | -0.81 | -1.79 | -1.08 | -1.08 | -0.4 | -2.14 | -2.81 | -0.67 | -1.19 | -0.2975 | -0.3125 | -0.555 | 0.15062 | 0.38879 | 0.513014 | 1 | 0.999631 | 0.623175 |
| Epn2_Q5NCM6_S192s_2 | 0.96 | -0.41 | -0.46 | -0.68 | -0.75 | -0.12 | -0.34 | -0.57 | -0.89 | 0.71 | -0.92 | -0.74 | -0.48 | -0.95 | -0.68 | -0.7 | 0.07 | -0.0475 | 0.3625 | 0.192926 | 0.465397 | 0.544669 | 1 | 0.999631 | 0.649499 |
| Epn2_Q5NCM6_S195s_1 | 0.5 | -0.44 | -2 | -2.16 | -0.74 | -0.32 | -1.1 | -1.66 | -0.48 | 0.75 | -2.98 | -1.58 | 0.4 | -0.38 | -1.7 | -0.97 | -0.28 | -0.25 | -0.515 | 0.60288 | 0.911163 | 0.957271 | 1 | 0.999631 | 0.857306 |
| Epn2_Q5NCM6_S195s_2 | 0.9 | -0.55 | -0.46 | -0.68 | -0.72 | -0.28 | -0.34 | -0.57 | -0.83 | 0.7 | -0.92 | -0.74 | -0.57 | -0.9 | -0.68 | -0.7 | -0.825 | -0.9175 | 0.2 | 0.216633 | 0.479751 | 0.622032 | 1 | 0.999631 | 0.669465 |
| Epn3_Q91W69_S257s_1 | 1.77 | -0.26 | 2.23 | 1.05 | -0.24 | 0.75 | 0.08 | 0.9 | -0.01 | 2.42 | -1.16 | -0.13 | 1.47 | 1.69 | 1.13 | 1.3 | -0.645 | 0.5325 | 0.69 | 0.703835 | 0.160935 | 0.277735 | 1 | 0.999631 | 0.897149 |
| Eprs_Q8CGC7_S944s_1 | 0.3 | 2.55 | 0.5 | 1.15 | 1.1 | 0.23 | 0.32 | 0.27 | 2.6 | 1.31 | 0.83 | 1.89 | 2.78 | 1.88 | 0.95 | 1.65 | 0.1875 | 0.185 | 0.49 | 0.252411 | 0.226901 | 0.371285 | 1 | 0.999631 | 0.688149 |
| Eps15_F6W2Q5_S324s_1 | -1.25 | -1.83 | -0.64 | -0.81 | -1.23 | -1.25 | -0.79 | -0.51 | 0.05 | -2.62 | -0.8 | -0.42 | -0.89 | -1.44 | 0.04 | -0.28 | -0.23 | -0.13 | 0.2525 | 0.271069 | 0.594854 | 0.753033 | 1 | 0.999631 | 0.698674 |
| Eps15l1_Q60902_S229s_1 | -0.82 | -0.88 | -0.83 | -1.26 | -1.08 | -1.16 | -1.29 | -1.18 | -1.16 | -1.07 | -1.42 | -0.66 | -0.1 | -1.18 | -0.89 | -0.61 | 0.025 | 0.215 | 0.375 | 0.448723 | 0.358722 | 0.647775 | 1 | 0.999631 | 0.786796 |
| Eps15l1_Q60902_S353s_1 | -0.6 | 0 | -1 | -0.62 | -0.37 | -0.61 | -0.6 | -0.54 | -0.13 | -0.68 | -0.41 | -0.14 | 0.5 | -0.23 | -0.93 | -0.06 | 0.265 | 0.3325 | -0.225 | 0.347875 | 0.930719 | 0.494949 | 1 | 0.999631 | 0.736498 |
| Eps15l1_Q60902_T364t_1 | -0.24 | -2.03 | 0.94 | 0.61 | -0.63 | 0.33 | 0.14 | 0.5 | -0.36 | -0.47 | 0.8 | 0.64 | -2.24 | -0.92 | 0.39 | 1.15 | -0.1525 | -0.355 | 0.215 | 0.794831 | 0.678138 | 0.619446 | 1 | 0.999631 | 0.928846 |
| Eps8_Q08509_S624s_1 | -1.1 | -0.39 | 0.2 | -0.1 | -0.06 | -0.51 | -0.82 | -0.61 | -0.79 | -1.81 | -0.57 | 0.36 | 0.08 | -0.16 | 0.23 | -0.68 | 0.5425 | -0.0075 | 0.765 | 0.573816 | 0.664703 | 0.482381 | 1 | 0.999631 | 0.84468 |
| Eps8l1_E9Q4X5_S300s_1 | -1.16 | -2.69 | -2.79 | 0.64 | -0.77 | -0.2 | -0.86 | -2 | -2.69 | -2.39 | -1.58 | 0.63 | -0.37 | -0.32 | -2.47 | 0.22 | 0.07 | -0.4975 | 0.455 | 0.386153 | 0.491208 | 0.993714 | 1 | 0.999631 | 0.756141 |
| Eps8l1_E9Q4X5_S685s_1 | -0.14 | -1.44 | -1.65 | 1.84 | -0.73 | -0.09 | -0.49 | 0.2 | -3.18 | -0.9 | -0.09 | 0.79 | -0.62 | 0.11 | -0.11 | 1.05 | -0.785 | -1.1525 | -2.1325 | 0.556181 | 0.923502 | 0.624226 | 1 | 0.999631 | 0.838305 |
| Eps8l2_Q99K30_S242s_1 | 2.21 | 1.17 | 2.38 | 1.83 | 0.63 | 1.68 | 1.71 | 0.43 | 1.02 | 2.21 | -0.71 | 0.46 | 0.07 | -0.04 | 0.23 | -1.2 | -0.6425 | -1.01 | -2.3 | 0.000588 | 0.090132 | 0.081098 | 1 | 0.999631 | 0.351308 |
| Erbb2ip_Q80TH2_S1013s_1 | 2.54 | 0.04 | -0.36 | 1.12 | 0.36 | -0.09 | 1.07 | -0.57 | 0 | 0.81 | -0.82 | -0.69 | -2.03 | -1.71 | -0.92 | -1.2 | -0.155 | 0.01 | -0.3075 | 0.004204 | 0.340227 | 0.155534 | 1 | 0.999631 | 0.351308 |
| Erbb2ip_Q80TH2_S1125s_1 | 1.58 | 0.95 | 0.63 | 1.71 | 1.48 | 0.23 | 0.92 | 1.62 | 1.85 | 0.54 | 1.11 | 1.41 | 0.14 | 1.28 | 0.98 | 1.24 | 0.5 | 1.2525 | -0.895 | 0.444308 | 0.704924 | 0.979642 | 1 | 0.999631 | 0.783263 |
| Erbb2ip_Q80TH2_S1209s_1 | 2.31 | 3.02 | 0.47 | -0.82 | 2.57 | 2.17 | 1.98 | 0.26 | 4.83 | 1.05 | 1.19 | 2.92 | 2.32 | -0.26 | -0.48 | -0.18 | -0.42 | -0.1025 | -0.2525 | 0.35235 | 0.571719 | 0.259231 | 1 | 0.999631 | 0.73909 |
| Erbb2ip_Q80TH2_S849s_1 | 0.18 | -0.05 | -0.22 | -0.32 | -0.04 | -1.11 | -0.41 | -0.53 | -0.03 | 0.06 | -0.65 | -0.2 | -0.58 | -0.14 | -0.61 | -0.09 | -0.2625 | 0.05 | -0.155 | 0.397563 | 0.190937 | 0.719221 | 1 | 0.999631 | 0.761181 |
| Erbb2ip_Q80TH2_S854s_1 | -0.03 | -0.97 | -0.77 | -0.57 | -0.81 | -0.88 | -0.93 | -0.77 | -0.61 | -0.08 | -0.99 | -0.46 | -0.68 | -0.75 | -0.64 | -0.89 | -0.1575 | -0.165 | 0.11 | 0.615815 | 0.365175 | 0.879248 | 1 | 0.999631 | 0.861203 |
| Erbb4_Q61527_S997s_1 | -0.19 | -0.51 | 0.54 | 0.07 | -0.77 | -0.03 | 0.16 | -0.08 | -0.42 | -0.07 | 0.24 | -0.5 | -0.14 | -0.28 | -0.21 | 0.98 | -0.5225 | -0.4525 | 0.1875 | 0.782416 | 0.642589 | 0.620525 | 1 | 0.999631 | 0.922325 |
| Erbb4_Q61527_T699t_1 | -0.27 | -0.13 | -0.37 | -0.6 | -0.9 | -0.66 | -1.11 | -0.79 | -0.91 | -0.57 | -0.87 | -0.83 | 0.39 | -0.4 | -0.07 | -0.54 | -0.3475 | -0.34 | -1.5775 | 0.556818 | 0.062293 | 0.103144 | 1 | 0.999631 | 0.838305 |
| Erc1_V9GWT6_S17s_1 | 2.33 | -0.43 | 1.54 | 1.98 | -0.16 | 1.22 | 2.14 | 0.83 | 0.51 | 2.02 | 1.03 | 0.5 | -2.21 | -0.35 | 1.15 | 0.52 | 0.3725 | -0.1725 | -1.1025 | 0.081447 | 0.616963 | 0.598989 | 1 | 0.999631 | 0.546723 |
| Erc1_V9GWT6_S21s_1 | 2.01 | -1.56 | 0.96 | 0.88 | 1.04 | -0.86 | 1.94 | 1.66 | 0.83 | 0.52 | 0.47 | -0.22 | -1.32 | -1.33 | 0.66 | -0.13 | 0.015 | -0.1375 | -0.815 | 0.181779 | 0.664498 | 0.805853 | 1 | 0.999631 | 0.643861 |
| Erc1_V9GWT6_S37s_1 | 1.51 | 0.22 | 0.32 | 1.22 | 0.46 | 1.42 | 2.06 | -0.61 | 1.22 | 1.35 | 0.46 | -0.31 | -0.93 | 0.46 | 0.35 | 0.13 | -0.275 | 0.25 | -0.605 | 0.095257 | 0.980078 | 0.775257 | 1 | 0.999631 | 0.558128 |
| Erc1_V9GWT6_S415s_1 | 1.19 | 1.2 | 1.98 | 1.56 | 0.63 | 2.07 | 1.69 | 0.44 | 1.84 | 2.8 | 1.28 | 1.01 | 0.46 | 0.71 | 0.57 | 1.77 | -0.275 | 0.25 | -0.605 | 0.141237 | 0.527845 | 0.56862 | 1 | 0.999631 | 0.61245 |
| Erc1_V9GWT6_S416s_1 | 1.19 | 1.2 | 1.98 | 1.56 | 0.63 | 2.07 | 1.69 | 0.44 | 1.84 | 2.8 | 1.28 | 1.01 | 0.46 | 0.71 | 0.57 | 1.77 | -0.155 | -0.2225 | -1.3325 | 0.141237 | 0.527845 | 0.56862 | 1 | 0.999631 | 0.61245 |
| Erc1_V9GWT6_S702s_1 | 0.97 | -0.92 | -0.74 | -0.14 | -0.89 | -0.08 | -0.01 | -0.47 | -0.83 | 0.87 | -0.71 | -1.05 | -1.88 | -1.52 | -1.07 | -1.69 | -0.28 | 0.045 | 0.1525 | 0.014393 | 0.733828 | 0.694321 | 1 | 0.999631 | 0.385092 |
| Erc1_V9GWT6_S937s_1 | -1.53 | -1.78 | -0.97 | -2.42 | -1.72 | -2.49 | -1.95 | -1.66 | -1.9 | -2.18 | -0.97 | -1.47 | -0.84 | -1.62 | -1.35 | -2.28 | 0.2875 | 0.2725 | 0.2275 | 0.72475 | 0.458797 | 0.912245 | 1 | 0.999631 | 0.905674 |
| Erc2_Q3UHT7_S14s_1 | 0.23 | -0.35 | 0.61 | 0.16 | 0.39 | 0.46 | 0.14 | 0.81 | 0.1 | 0.55 | 1.07 | 0.02 | 0.25 | -0.04 | 0.35 | 1 | -0.22 | 0.18 | -0.6 | 0.521813 | 0.353082 | 0.444559 | 1 | 0.999631 | 0.820623 |
| Erc2_Q3UHT7_S329s_1 | -0.07 | -0.42 | 0.34 | 0.08 | -0.47 | -0.09 | 0.35 | -0.74 | -0.14 | 0.67 | 0.19 | -0.07 | 0.11 | -1.55 | -0.34 | -0.69 | -0.1875 | 0.17 | 0.4325 | 0.162064 | 0.507258 | 0.564961 | 1 | 0.999631 | 0.631442 |
| Erc2_Q3UHT7_S966s_1 | -0.46 | -0.57 | -0.6 | -1.61 | -0.7 | -1.06 | -0.88 | -1.35 | -0.15 | -0.6 | -0.84 | -0.97 | 0.3 | -0.92 | -0.34 | -0.55 | -0.315 | -0.0025 | 0.005 | 0.290495 | 0.582316 | 0.636633 | 1 | 0.999631 | 0.706164 |
| Ercc5_E9QM61_S341s_1 | 1.37 | 2.21 | 1.65 | 1.2 | 1.64 | 1.47 | 0.7 | 1.36 | 1.82 | 1.76 | 1.39 | 1.45 | 1.49 | 2.04 | 1.31 | 1.61 | -0.3475 | -0.0075 | -0.005 | 0.988172 | 0.362852 | 0.993604 | 1 | 0.999631 | 0.995409 |
| Ercc5_E9QM61_S384s_1 | 0.8 | 1.08 | 2.05 | 1.34 | 0.88 | 0.76 | 0.92 | 1.32 | 0.8 | 0.71 | 1.59 | 2.14 | 0.87 | 0.98 | 1.57 | 1.83 | -0.1525 | -0.7325 | -0.625 | 0.989663 | 0.3112 | 0.986121 | 1 | 0.999631 | 0.995892 |
| Erg_E9PY05_S222s_1 | 1.62 | 0.3 | 1.7 | 0.99 | 0.32 | 0.58 | 1.97 | 1.13 | 0.04 | 0.53 | 0.71 | 0.4 | 1.07 | -1.16 | 1.31 | 0.89 | -0.31 | -0.3075 | -0.735 | 0.30965 | 0.744497 | 0.077199 | 1 | 0.999631 | 0.718889 |
| Erg_E9PY05_S88s_1 | 0.49 | 1.2 | 3.17 | 0.99 | 1.2 | 0.87 | 1.85 | 0.69 | 0.73 | 1.01 | 1.31 | 1.57 | 0.26 | 0.25 | 1.31 | 1.09 | 0.55 | 0.235 | -0.71 | 0.234103 | 0.597138 | 0.590437 | 1 | 0.999631 | 0.683077 |
| Ermn_Q5EBJ4_S211s_1 | -3.93 | -6.75 | -2.47 | -2.46 | -3.61 | -7.4 | -0.76 | -1.64 | -5.34 | -5.14 | -1.43 | -2.76 | -3.6 | -6.42 | -4.44 | -3.99 | -0.015 | 0.1175 | 0.445 | 0.489753 | 0.717386 | 0.843047 | 1 | 0.999631 | 0.807612 |
| Ermn_Q5EBJ4_S223s_1 | -1.07 | -0.48 | -0.24 | -0.75 | -0.35 | -0.79 | -0.78 | -0.68 | -0.41 | -0.52 | -0.89 | -0.25 | 0.27 | -0.41 | -0.16 | -0.46 | -0.135 | -0.05 | 0.3875 | 0.190255 | 0.957807 | 0.6967 | 1 | 0.999631 | 0.648674 |
| Ermn_Q5EBJ4_S227s_1 | -0.34 | -0.31 | -0.57 | -0.95 | -0.58 | -0.41 | -1.12 | -0.6 | -0.81 | -0.87 | -0.45 | -0.24 | 0.78 | -0.06 | -0.72 | -0.62 | 0.2075 | -0.3925 | 0.1075 | 0.343345 | 0.640222 | 0.864581 | 1 | 0.999631 | 0.73348 |
| Ermn_Q5EBJ4_S230s_1 | -0.72 | -0.18 | -0.32 | -0.63 | 0.01 | 0.23 | -0.51 | -0.75 | -0.63 | -1.02 | -0.97 | -0.8 | 0.04 | -0.6 | -0.54 | -0.32 | 0.205 | 0.075 | 0.46 | 0.720349 | 0.512102 | 0.165957 | 1 | 0.999631 | 0.904858 |
| Ermn_Q5EBJ4_S9s_1 | -0.29 | -0.56 | 0.69 | 0.22 | 0.23 | -0.21 | 0.17 | 0.69 | 0.39 | 0.38 | 0.41 | -0.82 | 0.19 | -0.14 | 0.78 | 1.07 | 0.0725 | -0.1325 | 0.295 | 0.27673 | 0.569948 | 0.856944 | 1 | 0.999631 | 0.700187 |
| Ermn_Q5EBJ4_T234t_1 | -0.89 | -0.03 | 0.33 | -0.25 | 0.11 | -0.19 | -0.69 | 0.22 | -0.58 | -0.35 | -0.22 | -0.22 | 0.1 | -0.24 | 0.41 | 0.07 | 0.045 | -2.095 | -0.7975 | 0.40496 | 0.837866 | 0.685108 | 1 | 0.999631 | 0.765844 |
| Ermn_Q5EBJ4_T257t_1 | -0.48 | 2.79 | 3.04 | 1.17 | -1.09 | 3.85 | 0.86 | 3.08 | -0.49 | 0 | 2.45 | -3.82 | -1.12 | 3.67 | 0.59 | 0.19 | 0.315 | 0.165 | 0.185 | 0.476812 | 0.969611 | 0.132385 | 1 | 0.999631 | 0.799947 |
| Esyt2_Q3TZZ7_S667s_1 | -1.33 | -1.36 | -1.33 | -1.03 | -1.2 | -1.07 | -0.66 | -0.86 | -1.04 | -1.37 | -0.89 | -1.09 | -1.65 | -1.03 | -0.77 | -0.86 | -0.2425 | -0.18 | 0.15 | 0.552454 | 0.235094 | 0.526213 | 1 | 0.999631 | 0.836655 |
| Etl4_E9QAU4_S1100s_1 | 0.21 | 0.4 | 0.62 | 0.07 | -0.27 | 0.58 | -0.26 | 0.28 | -0.04 | 0.45 | 0.16 | 0.01 | 0.39 | 0.56 | 0.19 | 0.76 | 0.6825 | -0.3075 | -0.0975 | 0.607437 | 0.430547 | 0.511638 | 1 | 0.999631 | 0.858527 |
| Etl4_E9QAU4_S1123s_1 | -0.42 | -1.1 | -0.49 | -0.14 | 0.54 | -0.54 | 0.48 | 0.1 | -0.4 | -1.97 | -0.32 | -0.69 | -0.02 | -1.35 | -0.01 | -1.16 | -0.2725 | -0.55 | -0.855 | 0.818327 | 0.075164 | 0.48239 | 1 | 0.999631 | 0.938303 |
| Etl4_E9QAU4_S1138s_1 | 1.69 | 0.43 | 1.23 | 1.32 | 1.41 | 0.25 | 1.63 | 0.29 | 1.25 | -0.05 | 0.87 | 0.4 | 0.79 | -0.59 | -0.08 | 1.13 | 7.85E-17 | -0.0675 | 0.2975 | 0.091329 | 0.538528 | 0.190478 | 1 | 0.999631 | 0.557085 |
| Etl4_E9QAU4_S1164s_1 | -0.74 | -0.44 | 0.07 | 0.13 | -0.6 | -0.63 | -0.28 | 0.53 | -0.48 | -1.02 | 0.08 | 0.17 | -0.15 | -0.26 | 0.44 | 0.18 | 0.065 | 0.3925 | 0.2425 | 0.382168 | 1 | 0.856202 | 1 | 0.999631 | 0.753668 |
| Etl4_E9QAU4_S1698s_1 | -0.69 | -0.01 | -1.07 | -1.61 | -0.22 | 0.1 | -1.35 | -1.65 | 0.53 | -0.62 | -1.01 | -0.71 | 0.8 | 0.04 | -1.78 | -1.47 | -0.075 | -0.355 | -1.045 | 0.701792 | 0.897853 | 0.405675 | 1 | 0.999631 | 0.896856 |
| Etl4_E9QAU4_S1813s_1 | 1.76 | 0.43 | 0.33 | 1.31 | 0.85 | 1.18 | 1.89 | -0.39 | 0.7 | 0.58 | 0.71 | 0.42 | -0.62 | -0.55 | 0.64 | 0.18 | 0.74 | 0.2075 | 0.0775 | 0.04208 | 0.889932 | 0.3576 | 1 | 0.999631 | 0.475443 |
| Etl4_E9QAU4_S1847s_1 | 0.74 | -0.42 | -0.17 | 1.03 | 0.94 | 1.32 | 1.18 | 0.7 | 0.71 | 0.23 | 0.92 | 0.15 | 0.03 | 1.12 | 0.03 | 0.31 | 0.5125 | -0.075 | 0.1125 | 0.859716 | 0.079621 | 0.611408 | 1 | 0.999631 | 0.953272 |
| Etl4_E9QAU4_S1859s_1 | 0.04 | -0.7 | -0.88 | 0.05 | -0.13 | 0.17 | 0.19 | 0.33 | -0.6 | -0.8 | 0.49 | -0.88 | -0.5 | 0 | -0.31 | -0.23 | 0.3375 | 0.0475 | 0.2325 | 0.737342 | 0.127416 | 0.854513 | 1 | 0.999631 | 0.907787 |
| Etl4_E9QAU4_S1894s_1 | -1.13 | -1.02 | -0.37 | -0.6 | -0.2 | -0.72 | -0.51 | -0.34 | -0.75 | -1.27 | -0.57 | -0.34 | -0.28 | -0.45 | -0.4 | -1.06 | 0.0375 | 0.13 | 0.5075 | 0.482102 | 0.255198 | 0.883174 | 1 | 0.999631 | 0.802327 |
| Etl4_E9QAU4_S1919s_1 | -0.4 | -0.53 | -0.47 | -0.8 | -0.62 | -0.63 | -0.57 | -0.23 | -0.77 | -0.21 | -0.47 | -0.23 | -0.06 | 0.12 | -0.57 | 0.34 | -0.5325 | -0.8475 | -0.7 | 0.12375 | 0.880795 | 0.63089 | 1 | 0.999631 | 0.594905 |
| Etl4_E9QAU4_S1950s_1 | -0.15 | -0.26 | 1.58 | 1.56 | -0.31 | -0.1 | 1.27 | -0.26 | -0.45 | 0 | -0.28 | 0.07 | 0.15 | -0.07 | -0.37 | 0.22 | 0.0225 | -0.22 | -0.3975 | 0.187216 | 0.367958 | 0.115058 | 1 | 0.999631 | 0.646474 |
| Etl4_E9QAU4_S364s_1 | -0.97 | -1 | -0.35 | -0.86 | -0.47 | -1.12 | -1.02 | -0.48 | -0.43 | -1.51 | -1.05 | -1.07 | -0.88 | -1.71 | -1.07 | -1.11 | -0.8425 | -0.155 | 0.395 | 0.231282 | 0.939396 | 0.503319 | 1 | 0.999631 | 0.681463 |
| Etl4_E9QAU4_S410s_1 | 0.36 | 1.31 | 2.4 | -0.25 | 0.39 | 0.44 | -0.55 | 0.17 | 0.43 | 0.17 | 0.94 | 1.66 | 2.24 | 0.78 | 0.29 | 2.09 | 0.5075 | -0.015 | -0.1475 | 0.560139 | 0.160902 | 0.798076 | 1 | 0.999631 | 0.839805 |
| Etl4_E9QAU4_S412s_1 | 0.69 | -0.58 | -0.87 | -0.28 | 0.8 | 0.48 | -0.37 | 0.08 | -0.1 | -0.7 | 0.35 | -0.65 | -0.82 | -0.33 | -0.32 | -0.16 | -0.5925 | -0.995 | -1.7125 | 0.709634 | 0.244053 | 0.971662 | 1 | 0.999631 | 0.899334 |
| Etl4_E9QAU4_T521t_1 | 1.44 | -1.37 | -0.5 | 0.22 | -0.82 | -1.17 | 0.43 | -1.02 | -1.09 | 0.26 | -1.54 | -1.82 | -1.17 | -1.65 | -1.98 | -2.26 | 0.7175 | 1.63 | 0.67 | 0.014291 | 0.355587 | 0.162514 | 1 | 0.999631 | 0.385092 |
| Etl4_E9QAU4_T918t_1 | 0.15 | 1.11 | 3.46 | 1.15 | 2.37 | 1.17 | 2.25 | 2.95 | 3.68 | 0.07 | 3.9 | 4.74 | 5.66 | -0.59 | 3.14 | 0.34 | -0.2225 | 0.3825 | 0.28 | 0.617287 | 0.321031 | 0.153457 | 1 | 0.999631 | 0.861376 |
| Etv6_E9Q8J8_S126s_1 | 1.28 | 2.44 | 2.01 | 1.76 | 1.63 | 1.84 | 1.2 | 1.93 | 2.46 | 2.25 | 1.91 | 2.4 | 2.05 | 2.47 | 2.18 | 1.91 | -0.02 | 0.3125 | -0.265 | 0.415046 | 0.507709 | 0.258684 | 1 | 0.999631 | 0.769328 |
| Eva1a_D3Z511_S118s_1 | -0.11 | 0.09 | -2.56 | -0.52 | 0.28 | -2.06 | -0.4 | -1 | -0.42 | -0.99 | -0.7 | 0.26 | -1.08 | -0.62 | -1.98 | -0.48 | -0.29 | -0.4 | 0.0025 | 0.674589 | 0.977025 | 0.607695 | 1 | 0.999631 | 0.885515 |
| Evc2_Q8K1G2_S331s_1 | 0.2 | -1.4 | -0.39 | -0.72 | -1.29 | -0.4 | -1 | -0.78 | -0.73 | -0.44 | -1.18 | -1.56 | -1.31 | 0.25 | -0.75 | -0.49 | 0.01 | 0.345 | 0.0125 | 0.995657 | 0.464909 | 0.351472 | 1 | 0.999631 | 0.997705 |
| Evi5l_H3BJ67_S579s_1 | 0.9 | 1.37 | 0.01 | 1.01 | 1.67 | 0.57 | 0.37 | 0.72 | 1.83 | 0.97 | 0.92 | 0.95 | 1.36 | 0.89 | 0.61 | 0.48 | -0.2875 | 0.3025 | 0.54 | 0.973937 | 0.980458 | 0.379171 | 1 | 0.999631 | 0.990749 |
| Evi5l_H3BJ67_S638s_1 | -0.32 | -0.27 | -0.25 | -0.79 | -0.29 | -0.79 | -0.96 | -0.74 | -0.24 | -0.21 | -0.08 | 0.11 | 0.66 | -0.23 | -0.31 | 0.41 | -0.1975 | -0.0525 | -0.07 | 0.133506 | 0.312835 | 0.272606 | 1 | 0.999631 | 0.60492 |
| Evi5l_H3BJX1_S394s_1 | -0.7 | -0.88 | -0.34 | -1.11 | -1.95 | -0.89 | -0.99 | 0.01 | -0.96 | -0.38 | -1.64 | -0.26 | -1.02 | -0.95 | -0.72 | -0.62 | -0.825 | 0.0475 | 0.6225 | 0.814015 | 0.644569 | 0.889629 | 1 | 0.999631 | 0.937041 |
| Evl_E9PVP4_S117s_1 | -1.17 | -0.16 | -0.62 | -0.17 | -1.32 | -0.76 | -1.99 | -1.35 | -1 | -0.9 | 0.39 | -0.42 | 0.26 | -0.34 | 0.2 | 0.25 | -0.5375 | -0.6925 | -1.2125 | 0.09325 | 0.045648 | 0.906918 | 1 | 0.999631 | 0.558128 |
| Evl_E9PVP4_S137s_1 | -1.78 | -6.77 | -4.26 | -3.84 | -4.79 | -5.83 | -3.67 | -4.51 | -7.13 | -4.07 | -3.58 | -4.64 | -4.62 | -6.1 | -5.18 | -5.6 | -0.0725 | 0.5925 | -0.07 | 0.208134 | 0.579665 | 0.536964 | 1 | 0.999631 | 0.661649 |
| Evl_E9PVP4_S231s_1 | 0.86 | -0.32 | 0.96 | 0.78 | 0.49 | 0.12 | 0.76 | 0.62 | 1.46 | 1.72 | 0.21 | 1.26 | -0.52 | 0.46 | 1.89 | 0.17 | 0.0675 | 0.2075 | -0.1425 | 0.89832 | 0.838368 | 0.200073 | 1 | 0.999631 | 0.966961 |
| Evl_E9PVP4_S291s_1 | 0.35 | -0.26 | 0.53 | 1.36 | 0.27 | 0.27 | 0.5 | 1.21 | 0.26 | 0.6 | 0.61 | 1.34 | -0.17 | 0.49 | -0.23 | 1.32 | 0.3 | 0.51 | 0.8675 | 0.766317 | 0.867313 | 0.616332 | 1 | 0.999631 | 0.916961 |
| Evl_E9PVP4_S316s_1 | -0.74 | -1.16 | -0.74 | -0.67 | 0.17 | -1.01 | -0.76 | -0.51 | -0.67 | -0.72 | -0.24 | 0.36 | 0.15 | -0.28 | 0.09 | 0.2 | 1.4675 | -0.2775 | -0.5975 | 0.011516 | 0.364473 | 0.143744 | 1 | 0.999631 | 0.385092 |
| Exoc1_Q5PPR2_S448s_1 | 0.13 | -1.31 | -1.17 | 1.32 | 1.42 | -0.11 | 2.06 | 1.47 | -0.86 | -2.03 | 1.14 | -0.39 | -3.43 | -1.21 | 0.96 | 0.26 | 0.7025 | 0.1075 | -0.225 | 0.547576 | 0.054807 | 0.72667 | 1 | 0.999631 | 0.834277 |
| Exoc1_Q5PPR2_S455s_1 | 0.37 | -1.22 | -2.05 | 0.11 | 0.02 | -0.76 | 0.13 | 0.63 | -1.66 | -0.81 | -0.08 | 0.19 | -0.67 | -1.21 | -0.77 | -1.04 | 1.095 | 0.2725 | -0.1525 | 0.680321 | 0.243856 | 0.8657 | 1 | 0.999631 | 0.888282 |
| Exoc1_Q5PPR2_S458s_1 | 1.33 | -0.19 | -1.6 | 0.9 | 1.45 | 1.23 | 0.84 | 1.3 | 1.62 | -0.28 | 0.85 | -0.66 | -0.91 | 0.84 | 0.32 | -0.42 | 0.825 | -0.47 | -1.27 | 0.821949 | 0.093833 | 0.713589 | 1 | 0.999631 | 0.938464 |
| Exoc1_Q5PPR2_S469s_1 | 2.5 | -0.93 | -0.7 | 1.09 | 1.86 | 0.35 | 1.83 | 1.22 | 0.33 | -1.08 | 1.12 | -0.29 | -2.28 | 0.24 | 0.04 | -1.12 | -0.0275 | -0.415 | -1.1775 | 0.163076 | 0.300238 | 0.568556 | 1 | 0.999631 | 0.631442 |
| Exoc1_Q6P1Y9_S470s_1 | 1.63 | -1.22 | -0.27 | 0.96 | -0.57 | -0.29 | 1.37 | 0.48 | -1.37 | 0.72 | -0.06 | 0.15 | -1.56 | -1.35 | 0.01 | -0.71 | 0.3025 | -0.4175 | -1.5825 | 0.095282 | 0.967851 | 0.551296 | 1 | 0.999631 | 0.558128 |
| Exoc2_Q9D4H1_S432s_1 | 2.92 | -0.22 | -0.7 | 2.09 | 1.14 | 0.83 | 2.56 | 0.77 | 0.86 | 0.98 | 0.71 | -0.13 | -1.01 | -0.44 | -0.21 | -0.58 | 1.3025 | 0.47 | -0.66 | 0.066097 | 0.721378 | 0.604888 | 1 | 0.999631 | 0.516933 |
| Exoc7_O35250_S283s_1 | 1.88 | 0.45 | -0.39 | 2.25 | 2.26 | 2.57 | 2.4 | 2.17 | 2.04 | 0.75 | 2.43 | 0.85 | -1.94 | 1.62 | 1.29 | 0.58 | 0.92 | 0.425 | -1.865 | 0.457974 | 0.042518 | 0.488277 | 1 | 0.999631 | 0.790722 |
| Exoc8_Q6PGF7_S28s_1 | 3.19 | 0.43 | -4.97 | -0.27 | 1.64 | 1.53 | -0.92 | -0.19 | 1.29 | 1.31 | -1.2 | -1.32 | -2.21 | 0.5 | -2.74 | -4.63 | -0.22 | 0.3 | 0.065 | 0.280408 | 0.552428 | 0.786925 | 1 | 0.999631 | 0.701922 |
| Exosc5_Q9CRA8_S23s_1 | 0.8 | 1.09 | 1.32 | 0.84 | 0.48 | 0.69 | 0.92 | 1.08 | 1.06 | 1.32 | 1.55 | 1.32 | 0.14 | 1.46 | 1.59 | 1.12 | -0.05 | 0.165 | 0.62 | 0.865424 | 0.424064 | 0.280337 | 1 | 0.999631 | 0.955768 |
| Exph5_Q0VAV2_S1071s_1 | -0.6 | 0.36 | -0.54 | -1.49 | -0.35 | -1.61 | -0.59 | 0.08 | -0.61 | -1.43 | -0.16 | 0.59 | 0.96 | 0.2 | -0.21 | -0.74 | 0.3175 | -0.12 | 0.9075 | 0.232486 | 0.91879 | 0.755911 | 1 | 0.999631 | 0.682449 |
| Exph5_Q0VAV2_S1117s_1 | 0.04 | -0.64 | 1.31 | 1.22 | 1.51 | 0.25 | -0.6 | 2.04 | 1.75 | -0.65 | -0.14 | 0.49 | 1.62 | 0.71 | 2.78 | 0.45 | -0.3025 | -0.365 | -0.09 | 0.176383 | 0.641251 | 0.849727 | 1 | 0.999631 | 0.642116 |
| Exph5_Q0VAV2_S331s_1 | 0.12 | 0.79 | 0.9 | -0.31 | -0.45 | 0.84 | 0.2 | -0.3 | 0.18 | 0.37 | -0.3 | -0.21 | -0.21 | 0.46 | -0.05 | 0.94 | -0.2125 | -0.0475 | -0.43 | 0.825229 | 0.465722 | 0.324274 | 1 | 0.999631 | 0.940309 |
| Exph5_Q0VAV2_S915s_1 | 0.93 | 1.12 | 0.81 | 1.43 | 0.58 | 0.57 | 1.26 | 1.03 | 0 | 0.01 | 2.29 | 1.8 | -0.02 | 0.52 | 0.46 | 1.61 | 0.33 | 0.8025 | 0.2 | 0.292626 | 0.469846 | 0.932887 | 1 | 0.999631 | 0.706671 |
| Eya1_Q8C9D0_S280s_1 | -0.04 | 2.38 | 1.61 | 0 | 2.05 | 1.31 | 1.01 | 0.9 | 2.59 | 0.33 | 2.25 | 1.99 | 3.03 | 0.28 | 1.45 | -0.01 | -0.4075 | -0.3375 | 0.3425 | 0.802492 | 0.581287 | 0.267355 | 1 | 0.999631 | 0.932677 |
| Eya4_Q8BY78_S338s_1 | -0.74 | 0.26 | 1.18 | -0.15 | -0.54 | -0.36 | -0.61 | 0.43 | -0.39 | -0.56 | 0.23 | -0.08 | 0.24 | 0.48 | 0.32 | 0.88 | -0.1175 | -0.6 | -0.615 | 0.437691 | 0.378168 | 0.446529 | 1 | 0.999631 | 0.781154 |
| F11r_O88792_S285s_1 | -0.57 | -0.22 | 1.19 | 0.68 | -2.37 | 0.81 | 0.63 | 1.54 | -1.81 | -0.45 | 1.87 | -0.93 | -2.66 | 0.21 | 0.36 | 0.71 | -0.5475 | 0.0475 | -0.8675 | 0.431143 | 0.887386 | 0.44754 | 1 | 0.999631 | 0.778271 |
| Fadd_Q61160_T204t_1 | 0.19 | -0.53 | 0.92 | 0.65 | -0.45 | -0.67 | 0.21 | -0.05 | -0.64 | 0.62 | 0.58 | 0.86 | -0.71 | -1.54 | -0.21 | 0.22 | -0.6025 | -0.6375 | -1.6775 | 0.094851 | 0.17697 | 0.916462 | 1 | 0.999631 | 0.558128 |
| Faf1_P54731_T268t_1 | 2.07 | 0.04 | 1.09 | 0.4 | -0.21 | 0.89 | 0.87 | -0.36 | -0.22 | 2.05 | -0.23 | -0.55 | -0.23 | -0.39 | -2 | -0.49 | -0.4975 | 0.0825 | 0.6525 | 0.012899 | 0.263586 | 0.353308 | 1 | 0.999631 | 0.385092 |
| Fahd2_Q3TC72_S129s_1 | -1.36 | -0.13 | -0.31 | -0.89 | -2.26 | -0.64 | -1.35 | -0.43 | -0.59 | -0.7 | -1.23 | 0.16 | 0.07 | -0.34 | -0.09 | 0.28 | -0.1075 | -0.5 | -0.9525 | 0.093866 | 0.308903 | 0.840264 | 1 | 0.999631 | 0.558128 |
| Fam102a_Q78T81_S244s_1 | 1.17 | 0.67 | -0.76 | 0.9 | 0.43 | 0.69 | 0.47 | -0.04 | 0.15 | 0.26 | -0.04 | -0.39 | -2.37 | 0.25 | 0.71 | -0.42 | 0.2425 | 0.415 | -0.1725 | 0.200634 | 0.808728 | 0.278635 | 1 | 0.999631 | 0.658263 |
| Fam104a_A2A6P4_S62s_2 | 0.48 | 1.7 | 1.38 | 2.08 | 1.47 | 1.84 | 1.25 | 2.05 | 1.49 | 1.61 | 1.78 | 2.42 | 0.39 | 1.86 | 1.05 | 1.65 | 0.1225 | 0.0525 | 0.055 | 0.712521 | 0.541433 | 0.322508 | 1 | 0.999631 | 0.9008 |
| Fam107a_Q78TU8_T138t_1 | 0.39 | -0.48 | -0.59 | 0.35 | 0.2 | -0.32 | -0.15 | 0.43 | -0.21 | 0.03 | -0.09 | 0.15 | -0.6 | 0.29 | -0.21 | 0.41 | -0.15 | 0.11 | -0.2825 | 0.88625 | 0.722942 | 0.872654 | 1 | 0.999631 | 0.963505 |
| Fam110b_Q8C739_S172s_1 | 0.06 | -0.58 | -0.03 | 0.02 | -0.56 | -0.54 | 0.03 | -0.06 | -0.33 | 0.09 | -0.13 | 0.28 | -0.65 | -0.57 | -0.76 | 0.32 | 0.05 | -0.07 | 0.53 | 0.425457 | 0.606148 | 0.703675 | 1 | 0.999631 | 0.774624 |
| Fam110b_Q8C739_S234s_1 | -1.17 | -0.82 | 1.32 | 0.32 | -0.69 | -0.86 | 0.62 | 0.78 | -1.32 | -0.9 | 0.36 | 1.23 | -0.34 | -0.53 | 1.73 | 0.91 | -0.1325 | 0.1375 | 0.1275 | 0.450792 | 0.937319 | 0.922722 | 1 | 0.999631 | 0.787797 |
| Fam110b_Q8C739_S95s_1 | 0.45 | 0.26 | 0.51 | 0.28 | 0.29 | 0.54 | -0.36 | 0.5 | 0.35 | 1.13 | 0.13 | 0.44 | 0.26 | 0.8 | 0.39 | 0.56 | 0.2975 | 0.2075 | 0.2225 | 0.64614 | 0.648776 | 0.648929 | 1 | 0.999631 | 0.873763 |
| Fam114a1_Q9D281_T197t_1 | 0.19 | -1.01 | -0.61 | -0.34 | 0.43 | -0.71 | -0.25 | -0.05 | -0.58 | 0.07 | -0.24 | -0.19 | 0.56 | -0.56 | -0.59 | -0.29 | 0.38 | 0.01 | 0.3 | 0.576469 | 0.424329 | 0.53892 | 1 | 0.999631 | 0.846204 |
| Fam114a2_Q8VE88_S112s_1 | -0.69 | -0.69 | 0.06 | 0.47 | 0.38 | -0.09 | -0.16 | 0.54 | -0.39 | -0.75 | -0.07 | 0.4 | -0.26 | 0.06 | 0.17 | 0.38 | -0.215 | 0.2925 | 0.13 | 0.418453 | 0.305113 | 0.979694 | 1 | 0.999631 | 0.771707 |
| Fam117b_Q3U3E2_S10s_1 | -0.82 | 0.12 | -0.14 | -0.26 | -0.42 | -0.92 | -0.28 | -0.34 | -0.34 | -0.47 | 0.22 | 0.66 | -0.11 | -0.66 | 0.24 | -0.05 | -0.525 | -0.3125 | -1.18 | 0.702382 | 0.486857 | 0.42562 | 1 | 0.999631 | 0.896857 |
| Fam117b_Q3U3E2_S176s_1 | 2.33 | 1.09 | 1.49 | 1.28 | 0.2 | 1.58 | 1.69 | 0.62 | 1.62 | 2.4 | 0.37 | 0.55 | 0.4 | -0.25 | 0.93 | 0.39 | 0.3325 | -0.0525 | -0.225 | 0.012434 | 0.253179 | 0.548994 | 1 | 0.999631 | 0.385092 |
| Fam117b_Q3U3E2_S386s_1 | 0.03 | -0.32 | 0.44 | 0.45 | -0.11 | 0.08 | 0.95 | 1.01 | 0.17 | -0.11 | 0.37 | -0.04 | -1.34 | 0.13 | 0.13 | 0.78 | 0.1725 | 0.285 | -0.2475 | 0.636654 | 0.372888 | 0.858838 | 1 | 0.999631 | 0.869788 |
| Fam117b_Q3U3E2_S444s_1 | 1.53 | 0.97 | -1.02 | -1.05 | 1.4 | 1.19 | -0.42 | -1.05 | 2.34 | 1.43 | -0.69 | -1.51 | -1.08 | 2.57 | -1.04 | -1.01 | -0.1425 | 0.0225 | 0.3225 | 0.797524 | 0.826575 | 0.768907 | 1 | 0.999631 | 0.930124 |
| Fam117b_Q3U3E2_T451t_1 | 0.73 | 0.11 | 0.1 | 0.24 | -0.13 | 0.27 | 0.08 | 0.39 | 0.18 | -0.12 | 0.25 | 0.96 | 0.3 | 0.12 | 1.01 | 1.04 | 0.035 | -0.3475 | 0.1075 | 0.358466 | 0.605966 | 0.945115 | 1 | 0.999631 | 0.741182 |
| FAM120A_Q6A0A9_S382s_1 | -0.23 | -0.87 | -0.72 | 0.09 | -0.24 | -1.18 | -0.03 | -0.14 | -0.36 | -0.8 | -1.44 | -0.52 | -0.41 | 0.02 | -0.36 | -0.55 | 0.7675 | 0.9125 | 0.1775 | 0.743802 | 0.923681 | 0.345038 | 1 | 0.999631 | 0.909852 |
| Fam122a_Q9DB52_S144s_1 | 0.01 | 1.54 | -1.72 | 1.06 | 1.19 | 0.97 | 0.66 | 1.14 | 1.32 | 0.37 | 1.86 | 0.99 | 1.41 | 1.54 | 0.24 | -1.59 | -0.2775 | 0.03 | 0.245 | 0.84074 | 0.257243 | 0.212738 | 1 | 0.999631 | 0.946805 |
| Fam122a_Q9DB52_S267s_1 | -0.32 | 1.05 | 0.55 | -0.01 | 0.32 | 0.11 | -0.26 | -0.01 | 0.68 | -0.24 | 0.54 | 0.41 | 0.7 | 0.33 | 0.46 | 0.76 | 0.1575 | 0.17 | 0.25 | 0.50769 | 0.440288 | 0.937836 | 1 | 0.999631 | 0.816264 |
| Fam126a_Q6P9N1_S433s_1 | -0.47 | -0.77 | -1.04 | -0.01 | -0.7 | -0.54 | -0.16 | -0.26 | -0.18 | -1.41 | -0.3 | 0.28 | -0.41 | -0.67 | -0.32 | 0.11 | 0.5425 | 0.2425 | 0.0775 | 0.46774 | 0.61281 | 0.688876 | 1 | 0.999631 | 0.795565 |
| Fam126a_Q6P9N1_S453s_1 | -0.08 | -0.64 | -0.59 | -0.11 | 0.66 | -0.22 | 0.04 | 0.27 | -0.06 | -0.86 | 0.06 | 0.41 | -0.58 | -0.44 | -0.1 | 0.01 | 0.625 | -0.095 | -0.0975 | 0.799539 | 0.096847 | 0.491594 | 1 | 0.999631 | 0.931051 |
| Fam126b_Q8C729_S321s_1 | 1.4 | 1.2 | 0.37 | 2.17 | 1.85 | 2.37 | 1.04 | 2.38 | 1.3 | 0.97 | 1.4 | 1.09 | -1.23 | 2.21 | 1.99 | 1.78 | 0.1925 | 0.0825 | 0.3975 | 0.900341 | 0.199678 | 0.810784 | 1 | 0.999631 | 0.967793 |
| Fam126b_Q8C729_S346s_1 | -1 | -1.14 | -1.79 | -1.09 | -1.07 | -1.06 | -0.94 | -1.18 | -1.1 | -1.53 | -0.92 | -1.14 | -0.41 | -1.32 | -0.84 | -0.86 | 0.1225 | -0.07 | 0.145 | 0.247976 | 0.488718 | 0.782862 | 1 | 0.999631 | 0.685751 |
| Fam126b_Q8C729_S575s_1 | -0.01 | -0.72 | -0.06 | 0.1 | 0.63 | -0.04 | -0.27 | -0.52 | -0.18 | 0.02 | -0.25 | -0.56 | 0.83 | -1.02 | -0.17 | 0.25 | 0.3625 | -0.0675 | -0.13 | 0.740253 | 0.721357 | 0.814836 | 1 | 0.999631 | 0.90877 |
| Fam126b_Q8C729_S577s_1 | -0.58 | -0.49 | -0.06 | 0.1 | 0.64 | 0.57 | -0.27 | -0.52 | -0.45 | -0.04 | -0.25 | -0.56 | -0.26 | -1.37 | -0.17 | 0.25 | 0.5125 | 0.0175 | 0.19 | 0.748503 | 0.327174 | 0.815054 | 1 | 0.999631 | 0.911931 |
| Fam126b_Q8C729_T306t_1 | 0.34 | -0.54 | -1.12 | 0.45 | 0.58 | 0.1 | 0.18 | 0.32 | -0.1 | -0.27 | 0.09 | -0.52 | -0.56 | 0.32 | -0.06 | 0.19 | 0.0175 | 0.0825 | -0.9825 | 0.659777 | 0.214168 | 0.965534 | 1 | 0.999631 | 0.880252 |
| Fam126b_Q8C729_T30t_1 | 0.49 | 2 | -1.28 | 0.44 | -0.76 | 1.39 | -0.16 | 1.25 | 0.05 | 0.75 | 1.11 | 0.07 | 0.85 | 1.39 | -2.71 | -1.81 | -0.07 | 0.335 | -1.455 | 0.350114 | 0.981367 | 0.898567 | 1 | 0.999631 | 0.737144 |
| Fam129a_Q3UW53_S595s_1 | -0.75 | 0.59 | 1.45 | -1.25 | 0.53 | 0.11 | 1.15 | -2.03 | 1.33 | -1.12 | 0.26 | 0.91 | 0.62 | -2.06 | -0.79 | -3.55 | -0.37 | -1.1175 | -1.06 | 0.141487 | 0.930927 | 0.645667 | 1 | 0.999631 | 0.612532 |
| Fam131b_E9Q8P8_S130s_1 | 1.32 | -0.67 | -0.18 | 0.64 | 0.84 | -0.82 | 0.47 | -0.86 | -0.44 | -0.86 | -0.49 | -1.57 | -1.51 | -0.79 | -0.25 | -0.58 | -0.0375 | -0.005 | 0.4375 | 0.05419 | 0.518534 | 0.043011 | 1 | 0.999631 | 0.491994 |
| Fam131b_E9Q8P8_S145s_1 | -2.16 | -1.62 | -0.22 | -0.65 | -1.63 | -1.52 | -1.01 | -0.64 | -2.26 | -2.19 | 0.06 | -0.28 | -0.72 | -1.53 | -0.33 | -0.32 | 0.61 | 0.4975 | 1.1 | 0.393668 | 0.936981 | 0.994106 | 1 | 0.999631 | 0.759138 |
| Fam131b_E9Q8P8_S325s_1 | -2.34 | 1.17 | 0.53 | -0.08 | 0.49 | 0.92 | -0.38 | 0.69 | 0.87 | -1.02 | 0.78 | 0.64 | 1.35 | 1.02 | 0.57 | 0.74 | 0.11 | -0.045 | 0.2375 | 0.137011 | 0.405211 | 0.527678 | 1 | 0.999631 | 0.610369 |
| Fam131b_E9Q8P8_S350s_1 | -1.01 | -1 | -0.68 | -0.91 | -0.82 | -0.45 | -1.08 | -0.81 | -1.52 | -0.82 | -1.08 | -0.36 | -0.67 | -1.03 | -0.54 | -0.41 | -0.0625 | 0.1775 | 0.52 | 0.411257 | 0.671485 | 0.886974 | 1 | 0.999631 | 0.767937 |
| Fam131b_E9Q8P8_S75s_1 | -0.8 | 0.08 | -0.39 | -0.67 | -0.32 | -0.47 | -0.75 | -0.49 | -0.08 | -0.3 | -0.41 | -0.28 | 0.71 | 0.03 | -0.41 | -0.03 | 0.2375 | -0.38 | 0.205 | 0.166477 | 0.828226 | 0.546204 | 1 | 0.999631 | 0.63587 |
| Fam131c_A2ADB2_S35s_1 | -0.71 | -0.35 | 1.32 | -0.8 | 0.08 | 0.24 | 0.47 | -0.38 | -0.46 | -1.28 | -0.54 | 0.22 | 0.14 | -0.3 | 0.11 | 0.33 | -0.03 | -0.1775 | -0.595 | 0.6786 | 0.63307 | 0.48837 | 1 | 0.999631 | 0.887183 |
| Fam133b_Q9CVI2_S82s_1 | 0.81 | 0.87 | -0.12 | 0.93 | 0.55 | 0.17 | 0.39 | 1.26 | 0.71 | 0.34 | 0.17 | 0.56 | 0.02 | 1.1 | -1.15 | 0.14 | 0.25 | 0.4725 | 0.2825 | 0.252259 | 0.934304 | 0.592126 | 1 | 0.999631 | 0.68813 |
| Fam134c_Q9CQV4_S26s_1 | -0.07 | 0.53 | -0.39 | -0.37 | 0.41 | 0.3 | -0.06 | 0.05 | 0.59 | 0 | 0.54 | 0.46 | 0.54 | 0.39 | -0.26 | 0.16 | -0.6125 | -0.5125 | 0.03 | 0.415612 | 0.416227 | 0.158226 | 1 | 0.999631 | 0.769842 |
| Fam13c_G3X9S1_S48s_1 | -0.45 | -1.24 | -1.17 | -1.16 | -1.76 | -1.6 | -1.95 | -1.16 | -1.86 | -0.9 | -1.44 | -1.87 | -0.72 | -1.25 | -1.09 | -0.84 | 0.5975 | -0.215 | 0.89 | 0.923457 | 0.070518 | 0.155314 | 1 | 0.999631 | 0.975032 |
| Fam149b1_Q6NSV7_S59s_1 | -0.49 | 0.11 | 1.19 | 0.44 | -0.33 | 0.78 | 1.02 | 2.17 | -0.82 | -2.37 | 1.37 | 2.21 | 1.14 | 0.13 | 2.08 | 1.46 | 0.4425 | 0.0875 | 0.0225 | 0.10577 | 0.305924 | 0.82133 | 1 | 0.999631 | 0.568797 |
| Fam160a2_F8WHJ1_S482s_1 | 1.15 | 0.17 | 1.39 | 0.64 | 1.35 | 1.53 | 0.96 | 1.28 | 0.34 | 0.16 | 1.18 | 2.02 | -1.03 | 1.2 | 1.65 | 1.62 | 0.0975 | -0.2875 | -0.85 | 0.971339 | 0.206914 | 0.856772 | 1 | 0.999631 | 0.989721 |
| Fam160a2_F8WHJ1_S529s_1 | 0.32 | -0.96 | -2.32 | -1.44 | -0.68 | -0.59 | -0.66 | -2.08 | -0.58 | -1.38 | -0.84 | -2.75 | -2.63 | -0.43 | -2.85 | -1.89 | 1.0675 | 0.0575 | -1.335 | 0.23594 | 0.869841 | 0.663645 | 1 | 0.999631 | 0.683301 |
| Fam160a2_F8WHJ1_S789s_1 | 1.76 | -0.23 | -0.37 | 2.69 | 1.73 | 1.75 | 2.46 | 2.18 | 1.56 | -0.43 | 2.34 | 0.61 | -2.04 | 1.57 | -1.31 | 0.29 | 0.3525 | -0.13 | 0.345 | 0.181172 | 0.143948 | 0.945467 | 1 | 0.999631 | 0.643763 |
| Fam160a2_F8WHJ1_S834s_1 | -0.5 | 0.69 | 0.2 | -0.02 | 1.18 | 1.04 | -0.37 | -0.07 | 0.95 | -0.08 | -0.11 | -0.91 | 0.34 | 1.08 | -0.14 | 0.47 | -0.5825 | -0.95 | -0.615 | 0.3805 | 0.437868 | 0.771391 | 1 | 0.999631 | 0.752468 |
| Fam160b1_Q8CDM8_S552s_1 | 2.03 | 1.35 | 2.07 | 1.57 | -0.71 | 2.09 | 3.36 | -0.05 | -0.13 | 1.86 | 0.29 | 1.2 | 0.95 | 0.59 | 2.04 | 0.98 | -0.5825 | -0.95 | -0.615 | 0.137777 | 0.490808 | 0.065523 | 1 | 0.999631 | 0.610682 |
| Fam160b1_Q8CDM8_T551t_1 | 2.03 | 1.35 | 2.07 | 1.57 | -0.71 | 2.09 | 3.36 | -0.05 | -0.13 | 1.86 | 0.29 | 1.2 | 0.95 | 0.59 | 2.04 | 0.98 | 0.295 | -0.3075 | -0.1175 | 0.137777 | 0.490808 | 0.065523 | 1 | 0.999631 | 0.610682 |
| Fam163b_Q8BUM6_S79s_1 | 0.76 | -0.71 | -0.57 | 0.22 | 0.6 | 0.22 | 0.05 | 0.01 | -0.44 | -0.52 | 0.31 | -0.88 | -0.81 | 0.51 | -1.25 | 0.78 | -0.555 | -0.92 | -0.6225 | 0.833619 | 0.448209 | 0.477477 | 1 | 0.999631 | 0.943778 |
| Fam169a_Q5XG69_S359s_1 | 1.6 | 2.17 | 2.15 | 2.27 | 0.77 | 1.48 | 2.04 | 1.68 | 1.85 | 1.67 | 0.82 | 0.17 | 1.61 | 1.48 | 1.49 | 1.12 | 1.2525 | 1.6575 | 0.985 | 0.057767 | 0.12842 | 0.051246 | 1 | 0.999631 | 0.497861 |
| Fam169a_Q5XG69_S512s_1 | -0.28 | 5.2 | 3.55 | 0.66 | 4.75 | 3.48 | 3.74 | 2.17 | 6.02 | 0.09 | 4.3 | 5.35 | 6.35 | 1.79 | 3.48 | 1.45 | -0.2125 | -0.275 | -0.7025 | 0.493951 | 0.30092 | 0.303156 | 1 | 0.999631 | 0.81007 |
| Fam169a_Q5XG69_T358t_1 | 1.97 | 1.69 | 2.51 | 2.21 | 1.32 | 1.95 | 2.42 | 1.84 | 1.76 | 2.28 | 2 | 1.24 | 0.8 | 1.08 | 2 | 1.69 | -0.4325 | -1.0225 | -0.7825 | 0.080836 | 0.521631 | 0.41616 | 1 | 0.999631 | 0.545579 |
| Fam171a1_A2ATL1_S346s_1 | -1.46 | -3.76 | -5.09 | -5.31 | -2.9 | -3.78 | -4.04 | -6.63 | -4.5 | -3.08 | -5.72 | -6.41 | -4.49 | -2.39 | -4.93 | -6.94 | 0.49 | 0.0925 | 0.695 | 0.479834 | 0.674469 | 0.316122 | 1 | 0.999631 | 0.801333 |
| Fam171a1_A2ATL1_S400s_1 | -0.6 | 0.71 | -0.14 | -0.21 | 1.1 | 0.46 | -0.29 | 0.45 | 0.59 | -0.09 | -0.21 | -0.16 | 0.97 | 0.86 | 0.3 | 0.41 | 0.3125 | 0.28 | 0.3125 | 0.081872 | 0.239224 | 0.799769 | 1 | 0.999631 | 0.54792 |
| Fam171a1_A2ATL1_S699s_1 | 0.4 | -0.08 | -2.73 | -1.39 | -0.24 | 0.03 | -1.1 | -1.24 | 0.32 | 0.01 | -1.54 | -1.47 | 1.27 | 0 | -2.24 | -1.58 | 0.8125 | 0.4625 | 1.0425 | 0.732235 | 0.64918 | 0.711614 | 1 | 0.999631 | 0.907231 |
| Fam171a1_A2ATL1_S724s_1 | -0.33 | 0.86 | -0.37 | -0.22 | 1.17 | 0.58 | 0.73 | 0.71 | 1.18 | -1.41 | 1.2 | 0.82 | 1.53 | 1.15 | 0.48 | 0.95 | 0.2825 | -0.4825 | -0.0525 | 0.022772 | 0.040229 | 0.465785 | 1 | 0.999631 | 0.409409 |
| Fam171a1_A2ATL1_T323t_1 | -1.54 | -1.19 | -0.01 | 0.29 | -0.83 | -1.31 | 0.29 | 0.53 | -1.16 | -2.41 | -0.21 | -0.6 | -1.53 | -1.56 | 0.3 | 0.13 | 0.2225 | 0.0825 | 0.29 | 0.931734 | 0.622691 | 0.426677 | 1 | 0.999631 | 0.978006 |
| Fam171a2_A2A699_S261s_1 | -0.17 | -0.29 | -0.73 | -0.03 | 0.22 | -0.44 | -0.27 | 0.16 | -0.11 | -0.27 | -0.32 | -0.19 | -0.26 | 0.34 | -0.1 | -0.04 | 0.0225 | -0.2175 | -0.795 | 0.347808 | 0.452444 | 0.759805 | 1 | 0.999631 | 0.736498 |
| Fam171a2_A2A699_S593s_1 | 1.5 | -0.02 | -0.32 | 0.43 | 0.34 | 0.78 | 0.66 | -0.1 | 0.64 | 1.16 | -0.03 | -1.05 | -0.42 | 0.04 | -0.47 | -0.74 | -0.1575 | -0.0275 | 0.9275 | 0.092364 | 0.958664 | 0.704373 | 1 | 0.999631 | 0.557085 |
| Fam171a2_A2A699_T378t_1 | -1.29 | -0.81 | 1.21 | 0.23 | -0.4 | -0.46 | -0.02 | -0.41 | -0.79 | -1.21 | -0.02 | 1.25 | 0.64 | -0.6 | 1.75 | 1.26 | 1.225 | 0.77 | 0.7275 | 0.189009 | 0.764576 | 0.96828 | 1 | 0.999631 | 0.648465 |
| Fam171b_Q14CH0_S410s_1 | 0.08 | 0.87 | -0.25 | 0.34 | 1.62 | 1.16 | 1.25 | 1.91 | 1.05 | -0.44 | 1.41 | 2.1 | 1.49 | -0.19 | 1.75 | 0.9 | -0.1625 | -0.29 | -0.8925 | 0.151228 | 0.004196 | 0.179444 | 1 | 0.999631 | 0.624065 |
| Fam171b_Q14CH0_S432s_1 | 0.01 | 0.47 | 0.39 | -0.14 | -0.18 | 0.57 | 0.4 | -0.71 | 0.03 | 0.23 | -0.33 | -0.36 | 0.22 | -1.17 | -0.27 | -1.62 | 1.4675 | 1.635 | 0.945 | 0.068049 | 0.647629 | 0.331259 | 1 | 0.999631 | 0.522238 |
| Fam171b_Q14CH0_S497s_1 | -1.53 | 1.56 | -0.44 | -0.88 | 2.83 | 0.46 | 1.11 | 0.18 | 3.71 | -1.03 | 1.33 | 1.24 | 2.97 | 0.45 | -0.65 | -0.28 | 0.7375 | -0.0575 | -0.0425 | 0.310985 | 0.085161 | 0.131347 | 1 | 0.999631 | 0.71924 |
| Fam171b_Q14CH0_S698s_1 | -1.33 | -0.84 | -0.92 | -0.01 | 0 | -0.24 | 0.34 | -0.25 | -0.4 | -1.55 | -0.55 | -0.83 | -1.28 | -1.5 | -0.32 | -0.17 | -0.1275 | 0.05 | 0.7425 | 0.922766 | 0.053808 | 0.883467 | 1 | 0.999631 | 0.975032 |
| Fam171b_Q14CH0_S788s_1 | -1.26 | -1.22 | -0.41 | -0.88 | -1.17 | -1.55 | -0.92 | -0.64 | -1.21 | -1.48 | -0.59 | -0.29 | 0.53 | -0.87 | -0.31 | -0.15 | -0.0975 | 0.45 | 0.155 | 0.07642 | 0.693461 | 0.891527 | 1 | 0.999631 | 0.542525 |
| Fam171b_Q14CH0_T787t_1 | -0.52 | -0.39 | -3.72 | -3.01 | -1.04 | -0.67 | -3.4 | -2.92 | -0.29 | -0.74 | -1.77 | -3.04 | 0.28 | -0.66 | -3.08 | -3.56 | 0.0725 | 0.3175 | 0.625 | 0.885135 | 0.917318 | 0.62355 | 1 | 0.999631 | 0.963505 |
| Fam177a1_Q8BR63_S151s_1 | -1.49 | -2.48 | -0.29 | -0.66 | -1.58 | -1.36 | -1.15 | -0.54 | -0.94 | -0.8 | -1.18 | -0.73 | -0.54 | -0.83 | -0.23 | -0.82 | 0.23 | -0.0225 | 0.2825 | 0.220288 | 0.885012 | 0.510908 | 1 | 0.999631 | 0.670699 |
| Fam184a_F6RSY8_S671s_1 | -0.19 | 1 | -0.94 | -1.04 | 1.43 | 0.63 | -1.05 | -1.26 | -0.07 | -0.67 | -0.29 | -0.23 | 0.21 | 0.09 | -0.36 | 0.02 | 0.4075 | 0.15 | -0.08 | 0.555327 | 0.746814 | 0.961858 | 1 | 0.999631 | 0.838109 |
| Fam189a1_Q6A044_S455s_1 | 0.53 | -0.65 | -0.07 | 0.49 | 0.4 | 0.36 | 0.76 | 0.41 | 0.45 | 0.16 | 0.8 | -0.51 | -0.89 | 0.25 | 0.27 | 0.35 | 0.05 | 0.3575 | 0.57 | 0.848769 | 0.238927 | 0.710971 | 1 | 0.999631 | 0.949525 |
| Fam189a2_E9QP68_S586s_1 | -0.39 | -0.17 | 0.34 | -0.51 | -0.21 | 0.01 | -0.53 | 0.2 | -0.31 | 0.41 | 0.27 | 0.33 | 0.83 | -0.03 | 0.52 | 0.23 | -0.165 | -0.19 | 0.145 | 0.112127 | 0.869472 | 0.271129 | 1 | 0.999631 | 0.577068 |
| Fam195b_Q3UGS4_S21s_1 | 0.22 | 0.23 | 0.49 | 0.38 | 0.08 | -0.11 | -0.01 | 0.7 | -0.35 | -0.32 | 0.51 | 0.72 | -0.59 | 0.43 | 0.84 | 1.22 | 0.2775 | 0.255 | 0.165 | 0.725933 | 0.555792 | 0.573067 | 1 | 0.999631 | 0.90568 |
| Fam196a_E9PZW7_S177s_1 | -0.45 | -1.63 | -0.04 | 0.34 | -0.82 | -0.46 | 0.07 | 0.54 | -0.89 | -0.39 | -0.24 | 0.76 | -0.65 | -0.62 | 0.88 | -0.73 | -0.155 | 0.18 | 0.1575 | 0.759959 | 0.574683 | 0.623646 | 1 | 0.999631 | 0.914614 |
| Fam207a_P58468_S38s_1 | -0.14 | 0.66 | -0.47 | -0.89 | -0.28 | -0.24 | -0.52 | -0.42 | 0.69 | 0.06 | -0.29 | -0.58 | 0.28 | 0.22 | -0.8 | 0.09 | 0.3775 | 0.215 | 0.03 | 0.711663 | 0.667119 | 0.674939 | 1 | 0.999631 | 0.900631 |
| Fam208a_Q69ZR9_S1108s_1 | -0.18 | 0.44 | -0.44 | -0.46 | 0.6 | 0.23 | -0.22 | 0.26 | 0.51 | -0.98 | -0.1 | 0.79 | -0.21 | 0.57 | -0.78 | -0.1 | -0.495 | -0.1025 | -0.8725 | 0.937416 | 0.253049 | 0.627005 | 1 | 0.999631 | 0.979547 |
| Fam208a_Q69ZR9_S339s_1 | 0.95 | 1.02 | 1.76 | 1.49 | 0.37 | 0.89 | 1.23 | 0.75 | 1.37 | 0.6 | 1.94 | 0.9 | -0.6 | 0.22 | 1.32 | 0.79 | 0.1875 | -0.0125 | -0.3875 | 0.077532 | 0.139571 | 0.785033 | 1 | 0.999631 | 0.543121 |
| Fam208a_Q69ZR9_S962s_1 | -0.28 | 1.16 | -0.42 | -0.66 | 0.67 | 0.85 | 0.22 | -1.19 | 1.61 | -1.5 | -0.68 | 0.32 | 1.21 | 0.42 | -1.33 | -2.05 | -0.255 | 0.3425 | 0.4875 | 0.609909 | 0.740335 | 0.98574 | 1 | 0.999631 | 0.859048 |
| Fam21_Q6PGL7_S747s_1 | 0.23 | 0.21 | 1.07 | 0.03 | 0.27 | 0.19 | -0.21 | 0.27 | 0.42 | 0.36 | 0.71 | 1.42 | 1.68 | 0.09 | 1.09 | 0.63 | 0.4025 | 0.4325 | 1.14 | 0.264882 | 0.42307 | 0.360934 | 1 | 0.999631 | 0.693761 |
| Fam21_Q6PGL7_S981s_1 | 0.09 | 1.58 | -0.79 | 0.39 | 1.42 | 0.95 | 0.17 | 0.34 | 1.36 | 0.93 | -0.1 | 0.81 | 1.4 | 1.66 | 1.69 | 1.08 | 0.1625 | 1.5875 | 0.3925 | 0.040019 | 0.451117 | 0.429614 | 1 | 0.999631 | 0.472195 |
| Fam212b_Q80VY2_S262s_1 | -0.64 | -0.48 | 4.24 | 2.04 | -0.66 | -0.51 | 3.81 | 3.17 | 0.52 | -0.58 | 5.16 | 6.41 | 0.62 | 0.18 | 3.81 | 2.12 | -0.485 | -0.52 | -0.05 | 0.744585 | 0.908001 | 0.375519 | 1 | 0.999631 | 0.910374 |
| Fam217b_A2AJW5_S106s_1 | -0.58 | -1.8 | 0.05 | -1.3 | -1.74 | -1.68 | -1.2 | -0.95 | -0.65 | -1.79 | -1.24 | -2.03 | -0.59 | -0.76 | -0.27 | -2.21 | 0.1525 | 0.3575 | 0.05 | 0.927496 | 0.283063 | 0.299993 | 1 | 0.999631 | 0.976306 |
| Fam219a_Q9D772_S44s_1 | -0.3 | 1.12 | 0.84 | 1.05 | 0.77 | 0.99 | 0.85 | 0.71 | 1.19 | 0.32 | 0.9 | 1.73 | 0.25 | 0.69 | 1.26 | 0.71 | -0.885 | -0.3025 | -0.5975 | 0.902774 | 0.673407 | 0.423286 | 1 | 0.999631 | 0.968183 |
| Fam222a_Q6PGH4_S313s_1 | 1.21 | 2.37 | 1.16 | 1.43 | 1.05 | 0.5 | 0.55 | 0.53 | 1.88 | 2 | 0.3 | 0.78 | 0.95 | 1.41 | 0.83 | 0.59 | -0.145 | -0.6725 | -0.1175 | 0.131799 | 0.026298 | 0.534689 | 1 | 0.999631 | 0.603456 |
| Fam49b_Q921M7_S117s_1 | -0.82 | -0.94 | 0.14 | -0.41 | -1.08 | -1.14 | -1.09 | 0.7 | -1.18 | -1.66 | -0.72 | -1.16 | -3.55 | -0.39 | 0.15 | 1.29 | 0.175 | 0.2975 | 0.355 | 0.897866 | 0.765134 | 0.078958 | 1 | 0.999631 | 0.966758 |
| Fam53b_Q3U0R8_S168s_1 | 0.18 | 1.16 | 0.37 | 0.75 | 0.89 | 1.03 | 0.28 | 0.96 | 0.57 | 0.32 | 1.48 | 1.28 | 1.09 | 0.8 | 0.86 | 1.13 | 0.47 | -0.735 | -0.565 | 0.279209 | 0.590766 | 0.43702 | 1 | 0.999631 | 0.70145 |
| Fam53b_Q3U0R8_S180s_1 | 1.19 | 0.01 | -0.73 | -1.1 | 0.76 | 0.8 | -0.56 | 0.25 | -0.32 | 0.55 | -2.87 | -0.93 | -1.58 | 0.68 | -0.33 | -1.66 | 0.22 | 0.14 | 0.17 | 0.408813 | 0.399929 | 0.356392 | 1 | 0.999631 | 0.766996 |
| Fam53c_Q8BXQ8_S122s_1 | 0.62 | 1.97 | 0.42 | 1.7 | 1.5 | 1.32 | 0.83 | 1.94 | 1.27 | 0.73 | 1.38 | 1.89 | 0.61 | 1.89 | 1.32 | 1.57 | 0.0775 | 0.4325 | 0.445 | 0.715208 | 0.617497 | 0.754061 | 1 | 0.999631 | 0.902377 |
| Fam53c_Q8BXQ8_S162s_1 | -0.93 | 0.64 | 0.64 | -0.47 | 0.23 | -0.02 | 0.03 | -0.05 | 0.21 | -0.31 | 0.92 | 0.79 | 1.08 | 0.18 | 0.63 | -0.23 | 0.0825 | 0.305 | 0.225 | 0.360766 | 0.848304 | 0.369924 | 1 | 0.999631 | 0.741241 |

| | | | | | | | | | | | | | | | | | | | | | | | | | | |
|---|---|---|---|---|---|---|---|---|---|---|---|---|---|---|---|---|---|---|---|---|---|---|---|---|---|---|
| Fam53c_Q8BXQ8_S273s_1 | 1.27 | 1.62 | 1.09 | 1.55 | 1.45 | 1.47 | 0.9 | 2.04 | 2.07 | 1.89 | 1.68 | 1.11 | 0.88 | 2.09 | 1.72 | 1.74 | -0.45 | 0.2875 | -0.2175 | 0.519629 | 0.79362 | 0.337423 | | 1 | 0.999631 | 0.819456 |
| Fam57b_Q7TNV1_S215s_1 | 0.43 | -0.58 | 0.35 | 0.27 | -0.67 | -0.65 | -0.18 | 0.17 | 0.09 | 0.46 | 0.59 | 0.48 | -0.3 | -0.36 | -0.21 | 0.47 | -0.17 | -0.115 | -0.5275 | 0.54526 | 0.209853 | 0.377944 | | 1 | 0.999631 | 0.833783 |
| Fam57b_Q7TNV1_S215s_2 | 0.29 | -1.46 | -0.16 | -0.07 | -0.76 | -0.72 | -0.13 | -0.47 | -0.66 | 0.6 | -0.87 | -0.93 | -0.63 | -0.82 | -0.84 | -1.22 | -0.17 | -0.115 | -0.5275 | 0.224675 | 0.679175 | 0.817675 | | 1 | 0.999631 | 0.67471 |
| Fam57b_Q7TNV1_S219s_2 | 0.29 | -1.46 | -0.16 | -0.07 | -0.76 | -0.72 | -0.13 | -0.47 | -0.66 | 0.6 | -0.87 | -0.93 | -0.63 | -0.82 | -0.84 | -1.22 | 0.17 | 0.1975 | 0.305 | 0.224675 | 0.679175 | 0.817675 | | 1 | 0.999631 | 0.67471 |
| Fam63a_Q76LS9_S103s_1 | -0.43 | -1.34 | -2.12 | -1.88 | -0.88 | -0.65 | -1.52 | -2.04 | -0.59 | -0.5 | -2.05 | -1.84 | -0.36 | -0.35 | -2.26 | -1.58 | -0.2525 | -0.23 | -0.2525 | 0.588044 | 0.716937 | 0.70474 | | 1 | 0.999631 | 0.850634 |
| Fam63b_Q6PDI6_S21s_1 | 0.4 | 0.28 | 0.39 | -0.02 | -0.23 | 0.09 | -0.09 | 0.27 | 0.12 | 0.4 | -0.24 | -0.15 | 0.41 | -0.14 | -0.02 | -0.21 | 0.375 | -0.1525 | 0.08 | 0.393238 | 0.337531 | 0.413833 | | 1 | 0.999631 | 0.759134 |
| Fam63b_Q6PDI6_S82s_1 | 0.66 | 1.83 | -0.29 | 1.35 | 1.73 | 0.63 | 1.45 | 1.24 | 1.01 | 0.4 | 1.19 | 0.34 | 1.47 | 1.04 | 0.18 | 1.18 | 0.065 | 0.1775 | 0.7 | 0.875512 | 0.448461 | 0.753922 | | 1 | 0.999631 | 0.960584 |
| Fam65a_Q68FE6_S347s_1 | -0.99 | 0.04 | -0.02 | -0.16 | 0.06 | -0.4 | -0.53 | 0 | -0.38 | -0.64 | -0.26 | 0.86 | -0.1 | 0.69 | 0.87 | 0.21 | -0.33 | -0.1675 | -0.31 | 0.08213 | 0.841552 | 0.66992 | | 1 | 0.999631 | 0.54792 |
| Fam65a_Q68FE6_S755s_2 | -0.14 | 0.59 | -0.4 | -0.7 | -0.6 | -1.53 | -0.41 | 0.57 | -1.22 | -0.71 | 0.25 | 0.36 | -1.14 | -0.45 | 0.29 | -0.59 | 0.38 | 1.6125 | 0.1075 | 0.464611 | 0.500123 | 0.715826 | | 1 | 0.999631 | 0.794281 |
| Fam65a_Q68FE6_S875s_1 | 1.32 | 3.13 | 3.03 | 0.51 | 3.53 | 1.54 | 2.69 | 1.75 | 3.93 | 0.44 | 4.6 | 5.47 | 3.9 | 0.8 | 2.72 | 1 | -0.145 | 0.4575 | 0.44 | 0.899272 | 0.590663 | 0.165164 | | 1 | 0.999631 | 0.967236 |
| Fam65b_Q80U16_S441s_1 | 0.47 | 3.01 | 2.73 | 0.24 | 1.24 | 2.32 | 1.01 | 1.3 | 2.74 | 1.34 | 1.94 | 2.26 | 3.46 | 1.22 | 2.81 | 0.72 | -0.2625 | 0.035 | -0.2475 | 0.605089 | 0.834925 | 0.518324 | | 1 | 0.999631 | 0.85817 |
| Fam73b_Q8BK03_S276s_1 | 0.68 | 0.75 | -0.13 | 0.25 | -0.13 | 0.98 | -0.78 | 0.43 | 1.07 | 1.3 | -0.9 | 0.22 | 0.34 | 0.6 | -0.61 | 0.23 | -0.74 | -0.6725 | -0.9725 | 0.511954 | 0.539579 | 0.945103 | | 1 | 0.999631 | 0.817687 |
| Fam76b_D6RG49_S193s_1 | 1.21 | 1.73 | 0.6 | 0.56 | 1.05 | 0.53 | -0.48 | 0.04 | 1.71 | 0.11 | -0.6 | 0.19 | 0.23 | 0.5 | 0.13 | -0.65 | -0.84 | -0.71 | -1.8025 | 0.032159 | 0.105993 | 0.219701 | | 1 | 0.999631 | 0.442395 |
| Fam83h_Q148V8_S1041s_1 | 1.37 | 0.53 | 0.07 | 0.23 | -0.28 | -0.13 | -0.08 | -0.67 | -0.29 | 1.3 | -1.06 | -0.59 | -1.39 | -1.69 | -0.83 | -1.1 | -0.2475 | 0.705 | 0.72 | 0.000781 | 0.034819 | 0.213512 | | 1 | 0.999631 | 0.351308 |
| Fam83h_Q148V8_S948s_1 | 0.31 | 0.42 | -0.05 | -0.32 | -0.88 | -0.59 | 0.12 | 0.72 | 0.68 | 0.76 | 1.33 | 0.41 | 0.75 | 0.51 | 1.05 | 0.93 | 0.1325 | 0.3 | 0.305 | 0.037439 | 0.541703 | 0.048336 | | 1 | 0.999631 | 0.458129 |
| Fam84a_Q9D650_S67s_1 | 0.34 | 1.09 | -0.29 | 0.14 | 1.05 | 0.86 | -0.04 | -0.06 | 2.2 | -0.28 | -0.67 | 1.23 | 2.45 | 0.89 | -0.22 | -0.62 | -0.195 | -0.44 | -0.75 | 0.648551 | 0.747456 | 0.647749 | | 1 | 0.999631 | 0.874615 |
| Fam91a1_Q3UVG3_S671s_1 | 0.8 | -0.05 | 0.83 | 0.5 | 0.19 | 0.13 | 0.84 | 0.14 | -0.56 | 1.17 | 0.16 | -0.45 | 0.5 | -0.45 | -0.6 | -0.37 | 0.14 | -0.265 | -0.2675 | 0.063543 | 0.542544 | 0.33104 | | 1 | 0.999631 | 0.513165 |
| Farp1_F8VPU2_S20s_1 | -0.69 | -0.78 | -0.62 | -0.27 | -0.58 | -0.12 | -0.85 | -0.25 | -0.94 | -1.39 | -0.33 | -0.76 | -1.18 | -0.66 | -0.67 | -0.92 | 0.745 | 0.2725 | -0.0925 | 0.364488 | 0.619479 | 0.404864 | | 1 | 0.999631 | 0.744849 |
| Farp1_F8VPU2_S373s_1 | 0.79 | 0.52 | -0.58 | 0.61 | 0.95 | 1.5 | 0.52 | 1.35 | 0.87 | 0.55 | 0.93 | 0.08 | -0.12 | 1.27 | -0.12 | -0.06 | -0.345 | -0.115 | 0.385 | 0.839887 | 0.080135 | 0.485453 | | 1 | 0.999631 | 0.946365 |
| Farp1_F8VPU2_S389s_1 | -0.76 | 0.08 | 0.27 | 0.49 | -0.62 | -0.14 | -0.72 | 0.18 | -0.2 | -0.28 | 0.01 | 0.09 | 1.13 | -0.29 | 0.39 | 0.39 | -0.1925 | 0.17 | 0.4425 | 0.363879 | 0.356838 | 0.732261 | | 1 | 0.999631 | 0.743941 |
| Farp1_F8VPU2_S403s_1 | -1.02 | -0.87 | 0.17 | -1.38 | -0.98 | -1.19 | -0.99 | -0.71 | -0.63 | -0.75 | -0.77 | -0.27 | -0.04 | -0.53 | -0.23 | -0.53 | -0.28 | 0.17 | 0.3075 | 0.268049 | 0.602438 | 0.65313 | | 1 | 0.999631 | 0.696016 |
| Farp1_F8VPU2_S427s_1 | 0.01 | 0.51 | 0.24 | -0.25 | -0.21 | 0.23 | -0.41 | -0.22 | 0.62 | 0.48 | -0.25 | 0.34 | 0.25 | 0.57 | 0.24 | 0.68 | 0.545 | 0.32 | 0.815 | 0.320513 | 0.340471 | 0.591657 | | 1 | 0.999631 | 0.723684 |
| Farp1_F8VPU2_S875s_1 | -0.61 | -0.48 | -2.79 | -3.22 | 0.03 | -0.02 | -2.27 | -2.66 | -0.24 | -0.19 | -2.01 | -3.38 | 0.25 | 0.11 | -1.58 | -2.62 | -0.1125 | 0.2225 | 0.3775 | 0.35542 | 0.53921 | 0.726477 | | 1 | 0.999631 | 0.74034 |
| Farp1_F8VPU2_S892s_1 | -0.61 | -0.41 | 0.43 | -0.03 | -0.56 | -0.55 | -0.2 | 0.24 | -0.19 | 0.35 | -0.27 | 0.38 | 0.2 | -0.08 | 0.39 | 0.38 | -0.0425 | -0.34 | -0.5225 | 0.267269 | 0.737629 | 0.512026 | | 1 | 0.999631 | 0.695893 |
| Farp1_F8VPU2_T24t_1 | 0.68 | 0.32 | -0.14 | 0.67 | 0.49 | 0.17 | 0.14 | 0.56 | 0.1 | 0.13 | 0.24 | -0.3 | -0.37 | 0.41 | -0.68 | 0.08 | -0.1625 | 0.2325 | 0.235 | 0.167732 | 0.884091 | 0.2755 | | 1 | 0.999631 | 0.637399 |
| Fasn_P19096_S1577s_1 | -0.02 | 0.56 | 0.29 | 0.68 | 0.08 | 0.4 | 0.3 | 0.08 | 0.4 | 0.58 | 0.42 | 1.04 | 0.7 | 0.58 | 0.47 | 0.7 | 0.275 | 0.25 | 0.4875 | 0.422617 | 0.54941 | 0.439288 | | 1 | 0.999631 | 0.773293 |
| Fat1_F2Z4A3_T153t_1 | -0.29 | 0.24 | -0.14 | -0.34 | -0.29 | 0.06 | -0.04 | 0.84 | -0.17 | -0.14 | 0.8 | -0.02 | 0.16 | 0.33 | 0.21 | 0.72 | -0.455 | -0.1275 | -0.3475 | 0.122362 | 0.403577 | 0.445415 | | 1 | 0.999631 | 0.590804 |
| Fbxl17_Q9QZN1_S42s_1 | 0.77 | 0.03 | -0.75 | -0.44 | -0.58 | -0.71 | -0.3 | -0.62 | 0.07 | 0.29 | -0.16 | -1.1 | 0.44 | -0.61 | -0.84 | -0.77 | 0.2125 | 0.5775 | 0.995 | 0.443365 | 0.230118 | 0.7743 | | 1 | 0.999631 | 0.782783 |
| Fbxl20_Q9CZV8_T403t_1 | -0.85 | 1.34 | 1.21 | -0.8 | 1.27 | 0.09 | 0.61 | -0.22 | 1.49 | -1.13 | 1.12 | 1.73 | 2.62 | 0.01 | 1.53 | 0.72 | 2.36E-16 | 0.7025 | 0.4925 | 0.192754 | 0.732166 | 0.46885 | | 1 | 0.999631 | 0.649499 |
| Fbxl4_V9GWU9_S45s_1 | -0.88 | -0.49 | -0.75 | 0.51 | -1.01 | -1.02 | -0.76 | 1.18 | -0.83 | -0.56 | 1.13 | 1.46 | 0.04 | -0.46 | 1.64 | -0.86 | 2.36E-16 | 0.7025 | 0.4925 | 0.405114 | 1 | 0.260223 | | 1 | 0.999631 | 0.765844 |
| Fbxl4_V9GWU9_T44t_1 | -0.88 | -0.49 | -0.75 | 0.51 | -1.01 | -1.02 | -0.76 | 1.18 | -0.83 | -0.56 | 1.13 | 1.46 | 0.04 | -0.46 | 1.64 | -0.86 | -0.745 | -0.0175 | -0.7975 | 0.405114 | 1 | 0.260223 | | 1 | 0.999631 | 0.765844 |
| Fbxo10_F6Q7Q7_S165s_1 | 1.73 | -0.29 | 0.69 | 0.98 | -0.94 | 0.02 | 1.06 | -0.01 | 0.44 | 1.74 | 0.16 | 0.7 | -0.07 | -0.51 | 0.84 | -0.34 | -0.5425 | -0.48 | -0.47 | 0.130794 | 0.188264 | 0.972653 | | 1 | 0.999631 | 0.602177 |
| Fbxo30_Q8BJL1_S573s_2 | -0.24 | -0.26 | -0.37 | -0.22 | -0.89 | -0.98 | -0.62 | -0.77 | -1.15 | -0.26 | -1.02 | -0.58 | -1.2 | -0.29 | -1.47 | -0.01 | -0.5425 | -0.48 | -0.47 | 0.240666 | 0.042307 | 0.124337 | | 1 | 0.999631 | 0.684338 |
| Fbxo30_Q8BJL1_T574t_2 | -0.24 | -0.26 | -0.37 | -0.22 | -0.89 | -0.98 | -0.62 | -0.77 | -1.15 | -0.26 | -1.02 | -0.58 | -1.2 | -0.29 | -1.47 | -0.01 | -0.135 | -0.555 | 0.0675 | 0.240666 | 0.042307 | 0.124337 | | 1 | 0.999631 | 0.684338 |
| Fbxo31_Q3TQF0_S33s_2 | 0.07 | 1.11 | 0.4 | 1.26 | 0.84 | 0.08 | -0.17 | 1.55 | 1.4 | 0.77 | -0.67 | -0.88 | 0.65 | 1.71 | -0.28 | 1.03 | -0.315 | 0.35 | 0.1275 | 0.889567 | 0.770445 | 0.34418 | | 1 | 0.999631 | 0.963505 |
| Fbxo31_Q3TQF0_T37t_1 | -1.04 | -0.52 | 0.64 | -0.18 | -0.99 | -0.96 | -0.41 | 0 | -0.51 | -0.26 | 0.11 | 0.96 | -0.18 | -0.87 | 0.29 | 0.17 | -0.135 | -0.555 | 0.0675 | 0.772984 | 0.460679 | 0.456752 | | 1 | 0.999631 | 0.92024 |
| Fbxo31_Q3TQF0_T37t_2 | 0.07 | 1.11 | 0.4 | 1.26 | 0.84 | 0.08 | -0.17 | 1.55 | 1.4 | 0.77 | -0.67 | -0.88 | 0.65 | 1.71 | -0.28 | 1.03 | 0.01 | 0.155 | 0.45 | 0.889567 | 0.770445 | 0.34418 | | 1 | 0.999631 | 0.963505 |
| Fbxo4_Q8CHQ0_S11s_1 | -0.38 | -0.71 | -0.15 | -0.56 | -0.83 | -0.51 | -0.68 | 0.26 | -0.68 | -0.66 | 0.01 | 0.15 | 0.37 | -0.29 | -0.24 | 0.16 | -1.075 | -0.9275 | -1.335 | 0.15933 | 0.974991 | 0.623579 | | 1 | 0.999631 | 0.629514 |
| Fbxo41_Q6NS60_S363s_1 | 1.6 | 0.27 | 1.29 | 0.38 | -0.22 | 0.45 | 0.92 | -1.91 | -0.52 | 0.88 | -0.05 | -0.48 | 0.09 | -0.65 | -0.15 | -1.09 | 0.2125 | 0.88 | -0.075 | 0.010575 | 0.112579 | 0.064596 | | 1 | 0.999631 | 0.383871 |
| Fbxo41_Q6NS60_S476s_1 | -0.09 | 0.1 | 0.17 | -0.52 | 1.11 | -0.99 | 1.09 | -0.7 | 2.06 | -1.44 | 1.14 | 1.42 | 3.44 | -1.66 | -1.06 | -1.36 | 0.935 | 1.2175 | 0.6625 | 0.942313 | 0.694653 | 0.22737 | | 1 | 0.999631 | 0.981629 |
| Fbxo41_Q6NS60_S476s_2 | -2.06 | -0.06 | -0.11 | -0.61 | 0.3 | -0.06 | 0.74 | -0.08 | 0.95 | -0.93 | 0.75 | 1.26 | 2.9 | -1.22 | -0.18 | -1.69 | -0.385 | -0.37 | -0.445 | 0.499892 | 0.075285 | 0.071827 | | 1 | 0.999631 | 0.813279 |
| Fbxo41_Q6NS60_S838s_1 | 2.48 | 0.95 | 0.88 | 0.75 | 0.21 | 1.22 | 1.57 | 0.52 | 0.4 | 2.11 | 0.99 | 0.08 | 1.15 | 0.92 | 1.29 | -0.08 | -0.4975 | 1.0825 | -0.215 | 0.376231 | 0.434896 | 0.512408 | | 1 | 0.999631 | 0.748365 |
| Fbxo41_Q6NS60_T477t_1 | 1.07 | -0.19 | 0.38 | -0.3 | -0.26 | -1.17 | 0.72 | -0.32 | 2.07 | -0.36 | 1.58 | 2 | 1.84 | -0.41 | -0.39 | -0.94 | 0.935 | 1.2175 | 0.6625 | 0.732108 | 0.307858 | 0.093859 | | 1 | 0.999631 | 0.907231 |
| Fbxo41_Q6NS60_T477t_2 | -2.06 | -0.06 | -0.11 | -0.61 | 0.3 | -0.06 | 0.74 | -0.08 | 0.95 | -0.93 | 0.75 | 1.26 | 2.9 | -1.22 | -0.18 | -1.69 | 0.3775 | 0.145 | -0.14 | 0.499892 | 0.075285 | 0.071827 | | 1 | 0.999631 | 0.813279 |
| Fbxo42_Q6PDJ6_S365s_1 | 1 | 0.38 | -0.52 | 0.98 | 1.15 | 0.43 | 0.76 | 1.01 | 0.82 | 0.04 | 1.27 | 0.29 | -0.64 | 0.74 | 0.45 | 0.73 | -0.2375 | -0.39 | -0.245 | 0.76758 | 0.352448 | 0.744573 | | 1 | 0.999631 | 0.917335 |
| Fbxo42_Q6PDJ6_S373s_1 | -1.1 | -0.13 | -0.26 | -0.36 | -0.08 | -1.37 | -0.68 | -0.67 | -0.62 | -0.94 | -0.88 | -0.97 | -0.37 | -0.51 | -0.8 | -1.15 | 0.285 | 0.1625 | -1.3925 | 0.479121 | 0.518432 | 0.219564 | | 1 | 0.999631 | 0.801052 |
| Fbxo42_Q6PDJ6_S429s_1 | 2 | -0.1 | -0.77 | 1.08 | 1.14 | 0.57 | 1.04 | 0.6 | 1.54 | 1.13 | 0.76 | -0.57 | -1.47 | -0.01 | -0.39 | -1.49 | -0.005 | 0.275 | -0.63 | 0.053759 | 0.622819 | 0.812506 | | 1 | 0.999631 | 0.4909 |
| Fbxo42_Q6PDJ6_S488s_1 | 1.31 | 0.2 | -0.55 | 0.51 | 0.71 | 0.58 | 0.29 | -0.13 | 1.33 | 1.66 | 0.15 | -0.57 | -0.7 | 0.27 | -0.11 | -0.51 | -0.3675 | -0.32 | 0.0875 | 0.176967 | 0.990548 | 0.642817 | | 1 | 0.999631 | 0.642116 |
| Fbxo42_Q6PDJ6_S578s_1 | -0.54 | 0.55 | 0.84 | 0.72 | 0.06 | -0.05 | -0.3 | 0.39 | -0.72 | 0.08 | 0.72 | 0.21 | 0.66 | -0.37 | 0.46 | 1.17 | -0.5875 | -0.295 | -1.48 | 0.845661 | 0.329844 | 0.466265 | | 1 | 0.999631 | 0.948565 |
| Fbxo9_Q8BK06_S22s_1 | 1.23 | 0.34 | 0.79 | 0.53 | -0.11 | 0.13 | 1.23 | -0.71 | 0.67 | 1.12 | -0.19 | 0.11 | -1.09 | -1.02 | 0.18 | -1.1 | -0.03 | 0.255 | 0.3475 | 0.003618 | 0.200473 | 0.438566 | | 1 | 0.999631 | 0.351308 |
| Fcgr1_P26151_S347s_1 | 1.25 | 0.94 | 0.85 | 0.43 | 1.02 | 0.99 | 0.36 | 0.98 | 0.55 | 1.54 | 1.36 | 1.04 | 0.91 | 1.13 | 1.31 | 1.51 | -0.4825 | 0.125 | 0.795 | 0.275095 | 0.91964 | 0.444938 | | 1 | 0.999631 | 0.700187 |
| Fcho1_Q8K285_S498s_1 | 0.03 | 0.05 | 1.83 | 0.2 | -0.75 | 0.68 | -0.19 | 0.44 | 0.32 | 1.24 | -0.19 | 1.24 | 1.02 | 1.39 | 1.4 | 1.48 | 0.605 | 2.4775 | 0.135 | 0.100117 | 0.352497 | 0.812336 | | 1 | 0.999631 | 0.562917 |
| Fcho2_Q3UQN2_S394s_1 | 0.12 | 1.45 | 1.67 | -0.11 | 3.08 | 0.39 | 2.93 | -0.85 | 4.96 | 0.82 | 3.28 | 3.98 | 2.53 | 0.17 | 1.45 | -0.48 | 0.06 | 0.005 | -0.5675 | 0.850357 | 0.517219 | 0.016339 | | 1 | 0.999631 | 0.949525 |
| Fcho2_Q3UQN2_S403s_1 | 0.48 | 0.43 | 0.42 | 1.03 | 0.32 | 0.46 | 1.25 | 0.57 | 0.66 | -0.03 | 1.23 | 0.52 | 0.03 | -0.23 | 0.11 | 0.18 | -0.2175 | -0.3075 | -0.015 | 0.074682 | 0.846231 | 0.988325 | | 1 | 0.999631 | 0.540402 |
| Fcho2_Q3UQN2_S487s_1 | -0.34 | 0.56 | 0.52 | 0.75 | 0.63 | 0.08 | -0.28 | 0.19 | -0.3 | -0.09 | 0.49 | 0.16 | 0.16 | 0.34 | 0.04 | 0.89 | 3.14E-16 | 0.1525 | 0.0975 | 0.966568 | 0.528155 | 0.377753 | | 1 | 0.999631 | 0.988351 |
| Fcho2_Q3UQN2_T682t_1 | 0.66 | 0.82 | 1.46 | 0.83 | -0.08 | 1.89 | 0.69 | 1.27 | 1.15 | 1.26 | 1.45 | 0.52 | 1.31 | 1.25 | 1.09 | 0.51 | 0.22 | 0.1975 | 0.3575 | 0.767347 | 1 | 0.640097 | | 1 | 0.999631 | 0.917335 |
| Fchsd2_Q3USJ8_S630s_1 | 0.24 | 0.65 | -0.07 | 0.71 | 0.85 | 0.46 | 0.18 | 0.92 | 0.82 | 0.74 | 0.97 | -0.21 | 0.38 | 0.95 | 0.83 | 0.8 | -0.14 | 0.1625 | 0.395 | 0.26889 | 0.481906 | 0.584665 | | 1 | 0.999631 | 0.69692 |
| Fdx1l_Q9CPW2_S51s_1 | -0.64 | -0.47 | -0.15 | -0.47 | -0.56 | -0.15 | -0.78 | -0.8 | -0.3 | 0.19 | -0.77 | -0.2 | 0.22 | -0.21 | -0.32 | 0.16 | -0.285 | 0.64 | 0.405 | 0.192859 | 0.609394 | 0.589976 | | 1 | 0.999631 | 0.649499 |
| Fer_Q3TZJ5_S411s_1 | -1.44 | -0.37 | 0.03 | -1 | -1.43 | -0.73 | -1.08 | -0.68 | -0.42 | 0.19 | -0.72 | 0.73 | 0.31 | -0.71 | -0.27 | -0.49 | -0.1825 | 0.2775 | 0.5025 | 0.336944 | 0.46278 | 0.177635 | | 1 | 0.999631 | 0.730499 |
| Fez1_Q8K0X8_S298s_1 | 0.55 | 1.65 | 1.7 | 1.38 | 0.82 | 1.5 | 0.97 | 1.26 | 1.72 | 1.37 | 1.52 | 1.78 | 1.24 | 2.03 | 1.82 | 2.2 | -0.0375 | 0.175 | 0.265 | 0.202466 | 0.595026 | 0.412895 | | 1 | 0.999631 | 0.659062 |
| Fez1_Q8K0X8_S316s_1 | -0.02 | -0.36 | -0.15 | 0.16 | 0.14 | -0.43 | 0.06 | -0.29 | -0.21 | -0.63 | 0.36 | 0.81 | -0.11 | -0.14 | 0.78 | 0.16 | 0.0525 | 0.075 | -0.2375 | 0.418211 | 0.889058 | 0.633535 | | 1 | 0.999631 | 0.771699 |
| Fez2_Q3TN06_S200s_1 | 0.42 | 0.67 | 0.49 | 0 | 0.52 | 0.89 | 0.4 | -0.02 | 0.64 | 0.87 | -0.24 | 0.61 | 0.02 | 0.37 | 0.54 | -0.3 | -0.0675 | -0.34 | -0.6825 | 0.463041 | 0.860619 | 0.821652 | | 1 | 0.999631 | 0.793564 |
| Fgd6_Q69ZL1_S1167s_1 | 0.64 | 0.62 | 0.39 | 0.47 | -0.07 | 0.94 | 0.86 | 0.12 | 0.64 | 0.39 | 0.28 | -0.55 | 0.46 | -0.39 | 0.44 | -1.12 | 0.3725 | -0.0375 | 0.025 | 0.115861 | 0.830281 | 0.306098 | | 1 | 0.999631 | 0.581581 |
| Fgd6_Q69ZL1_S583s_1 | 0.81 | 1.28 | 0.61 | 1.47 | 1.08 | 1.48 | 0.88 | 2.22 | 0.7 | 1.14 | 1.32 | 0.86 | -0.79 | 1.75 | 1.33 | 1.98 | -0.2825 | -0.0775 | 0.4475 | 0.966988 | 0.331734 | 0.903865 | | 1 | 0.999631 | 0.988373 |
| Fgf12_P61329_S212s_1 | -0.23 | 1.05 | 0.61 | 0.05 | -0.26 | 0.58 | -0.38 | 0.41 | 0.33 | 0.3 | 0.28 | 0.26 | 0.9 | 1.37 | 0.42 | 0.58 | 0.08 | 0.1825 | -0.4725 | 0.263597 | 0.468645 | 0.81729 | | 1 | 0.999631 | 0.693034 |
| Fgf12_P61329_S226s_1 | 0.43 | -0.27 | -0.66 | -1.55 | -0.1 | 0.26 | -0.23 | -1.66 | 0.52 | 0.83 | -1.98 | -0.69 | -0.44 | -0.61 | -0.97 | -1.92 | -0.0475 | 0.185 | -0.545 | 0.360274 | 0.882875 | 0.78888 | | 1 | 0.999631 | 0.741182 |
| Fgf12_P61329_S227s_1 | -0.11 | 0.32 | 0.26 | 0.68 | -0.02 | 0.61 | 1.14 | -0.77 | 0.58 | 1.24 | 0.22 | -0.15 | -0.12 | -0.46 | 0.14 | -0.59 | -0.5925 | 0.26 | -0.95 | 0.110082 | 0.912608 | 0.616224 | | 1 | 0.999631 | 0.57368 |
| Fgf12_P61329_T229t_1 | 1.27 | -0.37 | 1.37 | 1.33 | -0.4 | 0.34 | 1.11 | 0.18 | 1.08 | 2.1 | 0.48 | 0.98 | -0.9 | -0.36 | 0.77 | 0.29 | 0.5775 | -0.1475 | -0.9425 | 0.095953 | 0.24856 | 0.613892 | | 1 | 0.999631 | 0.558577 |
| Fhod3_Q76LL6_S345s_1 | 0.76 | -0.52 | 0.13 | 1.84 | 1.74 | -0.04 | 1.77 | 1.05 | 0.6 | -0.63 | 1.21 | 0.44 | -0.86 | 0.2 | -0.32 | -0.58 | 0.385 | 0.4275 | 0.505 | 0.09458 | 0.343789 | 0.799173 | | 1 | 0.999631 | 0.558128 |
| Fkbp15_Q6P9Q6_S1091s_1 | -0.07 | 1.21 | 0.64 | 0.63 | 1.53 | 0.81 | 0.48 | 1.13 | 1.31 | 0.84 | 0.77 | 1.2 | 0.87 | 1.62 | 1.32 | 0.62 | -0.1375 | -0.0275 | 0.315 | 0.20453 | 0.306555 | 0.227086 | | 1 | 0.999631 | 0.660045 |
| Fkbp1a_P26883_S9s_1 | -0.73 | -0.21 | 0.37 | -0.09 | -0.4 | -0.2 | -0.45 | -0.16 | -0.72 | -1.08 | 0.44 | 0.59 | 0.01 | 0.05 | 0.26 | 0.28 | 0.3 | -0.2825 | 0.2475 | 0.337057 | 0.648555 | 0.952362 | | 1 | 0.999631 | 0.730499 |
| Fkbp8_O35465_S286s_1 | -0.5 | -0.34 | -1.13 | -0.3 | -0.12 | -0.74 | 0.33 | -0.54 | -0.42 | -1.2 | -0.27 | -1.51 | -0.83 | -0.26 | -0.11 | -0.08 | -0.52 | -0.11 | -0.0925 | 0.462312 | 0.388044 | 0.462719 | | 1 | 0.999631 | 0.792786 |
| Flad1_Q8R123_S468s_1 | 0.78 | 1.11 | 2.63 | 1.66 | 0.42 | 1.07 | 0.83 | 1.78 | 0.92 | 1.38 | 1.72 | 1.72 | 0.41 | 1.03 | 2.32 | 2.05 | -0.71 | -0.46 | -1.2975 | 0.868217 | 0.285641 | 0.803383 | | 1 | 0.999631 | 0.95689 |
| Flcn_Q8QZS3_S571s_1 | 2.44 | 0.31 | 1.69 | 1.07 | -0.44 | 0.81 | 2.2 | 0.1 | 0.33 | 2.26 | 0.78 | 0.3 | 0.09 | -0.3 | 0.78 | -0.25 | 1.185 | 1.81 | 0.3125 | 0.024211 | 0.290594 | 0.443238 | | 1 | 0.999631 | 0.415644 |
| Flcn_Q8QZS3_S62s_1 | -0.26 | 1.52 | 1.29 | -1.25 | 2.66 | 0.44 | 3.02 | -0.08 | 3.92 | -0.53 | 1.83 | 3.32 | 3.59 | -0.39 | 0.27 | -0.92 | 1.87 | 3.1275 | 0.675 | 0.762155 | 0.200497 | 0.102642 | | 1 | 0.999631 | 0.915029 |
| Flcn_Q8QZS3_S62s_2 | -0.59 | -1.58 | 1.06 | -1.76 | 2.65 | 0.24 | 3.45 | -1.73 | 3.1 | 0.62 | 2.09 | 3.83 | 1.81 | -2.11 | 0.79 | -0.66 | 1.87 | 3.1275 | 0.675 | 0.470218 | 0.128055 | 0.003744 | | 1 | 0.999631 | 0.79732 |
| Flcn_Q8QZS3_S73s_2 | -0.59 | -1.58 | 1.06 | -1.76 | 2.65 | 0.24 | 3.45 | -1.73 | 3.1 | 0.62 | 2.09 | 3.83 | 1.81 | -2.11 | 0.79 | -0.66 | 0.08 | 0.315 | 0.2425 | 0.470218 | 0.128055 | 0.003744 | | 1 | 0.999631 | 0.79732 |
| Flcn_Q8QZS3_T313t_1 | 1.2 | 0.8 | 0.77 | 1.37 | 1.15 | 0.68 | 1.15 | 1.48 | 0.89 | 1.19 | 1.94 | 1.38 | 0.79 | 1.11 | 1.19 | 2.02 | -0.3975 | -1.1525 | -1.485 | 0.49956 | 0.784226 | 0.342256 | | 1 | 0.999631 | 0.813279 |
| Flii_Q9JJ28_S436s_1 | 1.98 | 1.77 | 1.23 | 1.17 | 0.46 | 1.76 | 1.59 | 0.75 | 0.25 | 1.36 | 0.26 | -0.33 | -0.73 | 0.34 | 0.23 | 0.37 | 0.075 | 0.165 | 0.6 | 0.002436 | 0.31585 | 0.018266 | | 1 | 0.999631 | 0.351308 |
| Flna_B7FAV1_S1435s_1 | -0.4 | -0.11 | 1.06 | 0.57 | 0.33 | 0.56 | -0.16 | 0.69 | -0.05 | 0.21 | 0.37 | 1.25 | 0.74 | 0.32 | 1.34 | 1.12 | 1.03 | 1.725 | 0.3025 | 0.170156 | 0.847038 | 0.703674 | | 1 | 0.999631 | 0.64073 |
| Flna_B7FAV1_S2088s_1 | -0.18 | 0.56 | 0.99 | 0.18 | 2.56 | 2.55 | 0.59 | -0.03 | 1.07 | 2.93 | 1.23 | 3.22 | -0.38 | 1.11 | 0.87 | 1.16 | -0.105 | -1.645 | -0.925 | 0.499838 | 0.131642 | 0.011919 | | 1 | 0.999631 | 0.813279 |
| Flrt3_Q8BGT1_S644s_1 | 1.06 | 0.83 | 0.18 | -0.27 | -0.43 | 1.2 | 0.49 | 0.12 | -0.23 | 0.54 | -0.18 | -4.91 | 0.65 | -0.56 | -1.04 | -0.95 | -0.5975 | -0.9825 | 0.505 | 0.077839 | 0.813158 | 0.16011 | | 1 | 0.999631 | 0.543121 |
| Flt3_Q3UEW6_S988s_1 | 0.41 | 0.37 | 0.23 | -0.19 | 1 | -0.79 | -1.17 | -0.61 | 0.02 | -0.74 | -1.61 | -0.78 | 1.88 | 0.24 | 0.66 | 0.06 | -0.375 | 0.2675 | -0.535 | 0.264318 | 0.226361 | 0.026333 | | 1 | 0.999631 | 0.6935 |
| Fmn1_E9Q7P6_S168s_1 | 0.1 | 0.15 | -0.13 | -0.65 | -0.82 | -0.58 | -0.71 | 0.08 | -0.04 | 0.28 | 0.37 | -0.07 | -0.24 | -0.56 | -1.39 | -0.48 | 0.45 | -0.1 | -0.6475 | 0.159575 | 0.258784 | 0.375239 | | 1 | 0.999631 | 0.629764 |
| Fmn1_E9Q7P6_S588s_1 | 1.06 | -0.57 | -0.64 | 1.03 | 0.45 | 0.29 | 0.83 | 1.11 | 0.27 | -0.42 | 0.67 | -0.04 | -1.97 | -0.13 | -0.05 | 0.44 | 0.385 | 0.7175 | -0.7625 | 0.325163 | 0.363172 | 0.842135 | | 1 | 0.999631 | 0.72463 |
| Fmn2_Q9JL04_S16s_1 | 0.48 | -0.26 | -0.97 | -1.9 | 0.42 | -0.62 | 0.47 | -1.38 | 0.21 | -0.37 | -0.24 | 0.62 | 1.19 | -2.73 | -1.37 | -2.79 | 0.8875 | 0.655 | 0.0925 | 0.412132 | 0.531352 | 0.190331 | | 1 | 0.999631 | 0.767962 |
| Fmn2_Q9JL04_S373s_1 | 0.37 | 1.08 | -1.29 | 1.45 | 1.47 | 1.71 | 0.5 | 1.48 | 1.22 | -1.15 | 1.81 | 2.35 | -0.3 | 0.08 | 1.46 | 0.74 | -0.2825 | -0.495 | -0.48 | 0.886194 | 0.162156 | 0.451462 | | 1 | 0.999631 | 0.963505 |
| Fmn2_Q9JL04_S405s_1 | 0.64 | 0.07 | -2.51 | -1.59 | -1.06 | -1.07 | -0.54 | -1.85 | -1.33 | -0.35 | -1.81 | -1.88 | -0.16 | -2.05 | -1.93 | -1.17 | -0.5875 | -0.1175 | 0.0825 | 0.524277 | 0.682851 | 0.496271 | | 1 | 0.999631 | 0.822727 |
| Fmn2_Q9JL04_S409s_1 | -0.78 | -1.31 | -1.35 | -2.51 | -2.91 | -0.96 | -1.84 | -2.59 | -1.3 | -0.29 | -2 | -2.83 | -1.26 | -1.4 | -1.29 | -1.67 | -0.2775 | 0.0425 | -0.045 | 0.836852 | 0.279373 | 0.843123 | | 1 | 0.999631 | 0.945549 |
| Fmn2_Q9JL04_S409s_2 | 0.23 | -0.74 | -0.74 | -1.82 | -1 | -0.76 | -1.11 | -1.31 | -1.14 | 0.47 | -1.31 | -0.92 | -0.5 | -1.14 | -0.82 | -0.79 | -0.1925 | 0.0525 | -0.0025 | 0.918773 | 0.52054 | 0.937493 | | 1 | 0.999631 | 0.974321 |
| Fmn2_Q9JL04_S413s_2 | -0.25 | -0.77 | -0.74 | -1.82 | -1.08 | -0.85 | -1.11 | -1.31 | -1.47 | 0.33 | -1.31 | -0.92 | -0.58 | -1.4 | -0.82 | -0.79 | -0.115 | 0.1525 | 0.3075 | 0.994999 | 0.600359 | 0.916037 | | 1 | 0.999631 | 0.997705 |
| Fmn2_Q9JL04_S493s_1 | -0.39 | -0.65 | -1.07 | -0.84 | -0.83 | -0.76 | -0.88 | -0.94 | -0.73 | 0.02 | -1.18 | -0.45 | -0.32 | -0.23 | -0.52 | -0.65 | -0.055 | 0.13 | 0.175 | 0.304687 | 0.656995 | 0.652774 | | 1 | 0.999631 | 0.716239 |
| Fmn2_Q9JL04_S614s_1 | -0.06 | 0.02 | 0.31 | -0.07 | 0.02 | 0.03 | -0.21 | 0.14 | 0.38 | -0.02 | 0.15 | 0.21 | 0.58 | 0.41 | -0.17 | 0.08 | 0.0075 | 0.32 | 0.29 | 0.560659 | 0.822758 | 0.61345 | | 1 | 0.999631 | 0.839977 |
| Fmn2_Q9JL04_S629s_1 | -0.69 | -1.12 | -0.36 | -0.59 | -1.23 | -1.04 | -0.68 | 0.22 | -0.85 | -0.15 | -0.07 | -0.41 | -0.45 | -0.57 | -0.68 | 0.1 | 0.36 | 0.2975 | -0.175 | 0.373962 | 0.984027 | 0.311344 | | 1 | 0.999631 | 0.747035 |
| Fmn2_Q9JL04_S69s_1 | 2.15 | 1.92 | -0.06 | 1.37 | 2.3 | 1.78 | 1.37 | 1.37 | 2.24 | 1.13 | 1.6 | 1.6 | 0.35 | 0.99 | 1.83 | 1.51 | 0.125 | 0.29 | -0.825 | 0.75118 | 0.483665 | 0.566161 | | 1 | 0.999631 | 0.912766 |
| Fmn2_Q9JL04_S720s_2 | 0.82 | 0.91 | 0.34 | 1.44 | 1.08 | 1.03 | 1.21 | 0.69 | 1.62 | 0.63 | 1.37 | 1.05 | -1.34 | 0.53 | 0.67 | 0.35 | -0.1875 | 0.4875 | 0.305 | 0.122055 | 0.685966 | 0.415789 | | 1 | 0.999631 | 0.590723 |
| Fmn2_Q9JL04_S724s_1 | -0.55 | -1.19 | -1.17 | -1.34 | -1.41 | -1 | -0.96 | -1.63 | -0.6 | -0.17 | -0.78 | -0.75 | -0.09 | -1.18 | -0.88 | -0.88 | 0.125 | 0.29 | -0.825 | 0.392143 | 0.53746 | 0.129055 | | 1 | 0.999631 | 0.758901 |
| Fmn2_Q9JL04_S724s_2 | 0.82 | 0.91 | 0.34 | 1.44 | 1.08 | 1.03 | 1.21 | 0.69 | 1.62 | 0.63 | 1.37 | 1.05 | -1.34 | 0.53 | 0.67 | 0.35 | 0.4375 | 0.5475 | -0.16 | 0.122055 | 0.685966 | 0.415789 | | 1 | 0.999631 | 0.590723 |
| Fmnl1_Q9JL26_S995s_1 | -2.7 | -5.57 | -3.08 | -4.29 | -2.88 | -4.31 | -3.34 | -3.36 | -4.12 | -2.82 | -3.45 | -3.06 | -3.06 | -4.5 | -4.2 | -4.52 | -0.1025 | -0.18 | 0.155 | 0.807992 | 0.49943 | 0.40063 | | 1 | 0.999631 | 0.936206 |
| Fmnl1_Q9JL26_T483t_1 | -0.69 | 0.49 | -0.31 | -0.61 | -0.02 | -0.66 | -0.59 | -0.26 | -0.74 | -0.45 | -0.52 | -0.13 | -0.09 | -0.15 | -0.28 | 0.02 | 0.4025 | 0.2225 | 0.2925 | 0.651013 | 0.764709 | 0.601033 | | 1 | 0.999631 | 0.876297 |
| Fmnl2_E9PXE6_S183s_1 | -0.22 | -1.67 | -0.5 | -0.22 | -0.02 | -0.27 | -0.62 | -0.09 | -0.54 | -0.43 | -0.45 | -0.3 | -1.22 | -0.2 | -0.01 | -0.01 | 0.2175 | 0.3675 | -0.3525 | 0.520234 | 0.307227 | 0.555845 | | 1 | 0.999631 | 0.819528 |
| Fmnl2_E9PXE6_S981s_1 | -2.74 | -6.17 | -2.73 | -3.37 | -2.82 | -5.22 | -2.64 | -3.46 | -3.68 | -4.34 | -2.92 | -2.6 | -3.82 | -4.38 | -5.38 | -2.84 | -0.53 | 0.0525 | 0.0075 | 0.67822 | 0.803859 | 0.646518 | | 1 | 0.999631 | 0.887183 |
| Fnbp1_Q80TY0_S217s_1 | -0.35 | -0.68 | 0.45 | -0.56 | -0.92 | -1.04 | -0.82 | -0.48 | -0.7 | -0.06 | -0.18 | 0.01 | -0.19 | -0.58 | 0.03 | -0.37 | -0.2675 | 0.1725 | -0.205 | 0.98269 | 0.12794 | 0.878631 | | 1 | 0.999631 | 0.994846 |
| Fnbp1_Q80TY0_S220s_1 | 0.45 | -1.02 | -0.02 | -0.63 | -0.72 | -0.18 | -0.57 | -0.82 | -0.43 | 0.63 | -0.42 | -0.31 | -0.66 | -0.83 | -0.07 | -0.48 | -0.0775 | -2E-16 | -0.775 | 0.60383 | 0.477837 | 0.680968 | | 1 | 0.999631 | 0.858061 |
| Fnbp1_Q80TY0_S402s_1 | 0.43 | 1.83 | 0.46 | 0.16 | 1.18 | 0.44 | 0.82 | 0.13 | 0.94 | 0.79 | 0.37 | 0.78 | 0.85 | -0.27 | -0.35 | -0.45 | -0.725 | -0.1925 | 0.1375 | 0.125075 | 0.857373 | 1 | | 1 | 1 | 0.596063 |
| Fnbp1_Q80TY0_S422s_1 | 0.53 | 1.77 | 2.14 | 0.41 | -0.24 | 2.19 | 0.3 | -0.3 | 1.49 | 2.31 | -0.04 | 0.32 | 1.95 | 1.19 | 0.83 | 1.43 | -0.8175 | -0.0025 | 0.2275 | 0.775212 | 0.280714 | 0.759435 | | 1 | 0.999631 | 0.920998 |
| Fnbp1_Q80TY0_S451s_1 | -0.92 | -2.31 | -0.63 | -2.48 | -3.65 | -2.24 | -1.69 | -2.03 | -2.6 | -0.63 | -1.61 | -1.51 | -0.51 | -1.65 | -1.39 | -1.88 | 0.72 | 0.305 | 0.03 | 0.667315 | 0.181486 | 0.996504 | | 1 | 0.999631 | 0.88264 |
| Fnbp1_Q80TY0_S451s_2 | -1.27 | -2.1 | -1.64 | -1.15 | -0.99 | -0.51 | -1.17 | -0.61 | -0.58 | -0.83 | -1.93 | -1.6 | -2.15 | -1.22 | -0.99 | -1.68 | 0.72 | 0.305 | 0.03 | 0.936091 | 0.043223 | 0.449126 | | 1 | 0.999631 | 0.97939 |
| Fnbp1_Q80TY0_S455s_2 | -1.27 | -2.1 | -1.64 | -1.15 | -0.99 | -0.51 | -1.17 | -0.61 | -0.58 | -0.83 | -1.93 | -1.6 | -2.15 | -1.22 | -0.99 | -1.68 | 0.015 | 0.04 | -1.1175 | 0.936091 | 0.043223 | 0.449126 | | 1 | 0.999631 | 0.97939 |
| Fnbp1l_E9PUK3_S153s_1 | -0.41 | -0.9 | -0.12 | -1.37 | -0.99 | -0.71 | 1.12 | -2.16 | -0.66 | -1.03 | -0.23 | -0.72 | -0.6 | -2.43 | -1.11 | -3.13 | 0.0075 | 0.24 | 0.0725 | 0.082322 | 0.98167 | 0.910551 | | 1 | 0.999631 | 0.54792 |
| Fnbp1l_E9PUK3_S295s_1 | -0.16 | 0.3 | -0.19 | 0.22 | 0.38 | -0.23 | -0.24 | 0.29 | 0.24 | 0.3 | 0.17 | 0.42 | -0.9 | 0.48 | 0.31 | 0.57 | 1.2075 | 1.6525 | 0.6725 | 0.853811 | 0.978985 | 0.369119 | | 1 | 0.999631 | 0.950951 |
| Fnbp1l_E9PUK3_S430s_1 | -1.13 | 0.33 | 0.78 | -0.22 | 2.21 | -1.22 | 3.38 | 0.22 | 2.86 | -1.72 | 2.04 | 3.19 | 2.38 | -0.42 | 0.34 | 0.15 | 0.065 | 0.5925 | 0.5125 | 0.320744 | 0.223918 | 0.135278 | | 1 | 0.999631 | 0.723691 |
| Fnbp1l_E9PUK3_S443s_1 | 0.09 | 1.02 | -0.28 | -0.5 | 0.7 | 1.32 | -0.71 | -0.72 | 0.86 | 1.44 | -0.22 | 0.62 | 1.44 | 1.25 | 0.15 | -0.46 | 0.0175 | 0.2725 | 0.6275 | 0.344844 | 0.907696 | 0.223042 | | 1 | 0.999631 | 0.734195 |
| Fnbp4_Q6ZQ03_S487s_1 | 0.2 | 0.74 | 2.27 | 1.92 | 0.6 | 1.19 | 1.27 | 2.14 | 1.32 | 0.82 | 1.65 | 2.43 | 1.55 | 0.97 | 2.55 | 2.57 | -0.645 | 0.115 | 0.5425 | 0.290832 | 0.973945 | 0.622113 | | 1 | 0.999631 | 0.706164 |
| Fndc3a_E0CXY0_S202s_1 | 0.63 | 0 | 0.26 | -0.75 | -0.48 | -0.87 | -0.72 | -0.37 | 0.14 | 1.15 | -0.67 | -0.02 | 1.56 | 0.84 | -0.45 | 0.36 | 0.395 | 0.3075 | 1.095 | 0.286736 | 0.08632 | 0.803902 | | 1 | 0.999631 | 0.705972 |
| Fndc3a_E0CXY0_S213s_1 | -0.36 | -0.6 | -0.06 | 0.24 | -0.13 | -0.69 | 0.12 | 1.5 | -0.86 | -0.1 | 0.33 | 1.08 | 1.29 | 0.92 | 0.73 | 0.66 | -0.865 | -0.6325 | -1.38 | 0.005827 | 0.414192 | 0.490205 | | 1 | 0.999631 | 0.351545 |
| Fndc4_Q3TR08_S209s_1 | 1.84 | -0.75 | 0.91 | 0.33 | -1.23 | -0.45 | 1.15 | -0.6 | -1.08 | 1.11 | 0.21 | -0.44 | -0.67 | -1.4 | -0.37 | -0.75 | -0.775 | -0.2075 | -0.9775 | 0.027863 | 0.210957 | 0.339124 | | 1 | 0.999631 | 0.423977 |
| Fnip1_F6RJ64_S237s_1 | 0.35 | 0.78 | 1.4 | 2.19 | 0.21 | -1.15 | 0.8 | 1.76 | 0.91 | 0.08 | 1.13 | 1.77 | -2.31 | -0.25 | 1.86 | 1.51 | -0.0025 | 0.065 | 0.365 | 0.289044 | 0.251072 | 0.680942 | | 1 | 0.999631 | 0.706046 |
| Foxk1_P42128_S199s_1 | 0.69 | 1.08 | 1.07 | 0.78 | 1.04 | 1.3 | 0.37 | 0.9 | 0.9 | 1.08 | 0.82 | 1.08 | 1.01 | 1.43 | 1.19 | 1.45 | -0.295 | 0.1625 | 0.2125 | 0.212745 | 0.993136 | 0.799039 | | 1 | 0.999631 | 0.66658 |
| Foxk1_P42128_S431s_1 | 0.34 | 0.82 | 1.15 | 0.88 | 0.38 | 0.41 | 0.58 | 0.64 | 0.53 | 0.38 | 1.47 | 1.46 | 0.3 | 0.9 | 1.1 | 1.74 | 0.0475 | 0.135 | 0.06 | 0.577609 | 0.291693 | 0.659677 | | 1 | 0.999631 | 0.84622 |
| Foxk1_P42128_S87s_1 | 0.75 | 0.78 | 1.23 | 1.5 | 0.93 | 0.84 | 1.03 | 1.65 | 1.21 | 0.98 | 1.28 | 1.33 | 1.21 | 1.07 | 0.94 | 1.28 | -0.0275 | 0.3225 | 0.7125 | 0.84219 | 0.878967 | 0.639864 | | 1 | 0.999631 | 0.947766 |
| Foxk2_Q3UCQ1_S389s_1 | 0.18 | 1.4 | 1.96 | 1.62 | 0.32 | 1.06 | 1.32 | 2.35 | 0.48 | 1.17 | 2.19 | 2.61 | 1.02 | 1.51 | 2.71 | 2.77 | -0.5625 | -0.475 | -0.99 | 0.210948 | 0.958478 | 0.574729 | | 1 | 0.999631 | 0.665076 |
| Fra10ac1_Q8BP78_S243s_1 | 2.56 | 1.62 | 2.05 | 2.31 | 1.18 | 1.7 | 2.09 | 1.32 | 2.15 | 3 | 0.78 | 0.71 | 0.97 | 0.76 | 2.17 | 0.68 | 0.2675 | 0.3875 | 0.0475 | 0.036199 | 0.111821 | 0.395915 | | 1 | 0.999631 | 0.453963 |
| Frmd3_Q8BHD4_S63s_1 | -1.39 | -0.89 | -0.44 | -0.3 | -1.05 | -0.48 | -0.36 | -0.06 | -0.66 | -1.49 | -0.21 | 0.89 | -1.28 | -1.04 | -0.19 | -0.32 | 0.6275 | -0.0025 | -1.1325 | 0.903555 | 0.452639 | 0.462738 | | 1 | 0.999631 | 0.968183 |
| Frmd4a_Q3TB04_S372s_1 | 0.44 | -2.02 | -2.53 | -0.78 | -0.71 | -0.84 | 0.56 | -1.39 | -1.01 | -0.93 | -1.15 | -1.81 | -4.58 | -1.39 | -2.3 | -1.15 | -0.1625 | 0.88 | 1.1 | 0.220219 | 0.375687 | 0.99682 | | 1 | 0.99985 | 0.670699 |
| Frmd4a_Q3TB04_S385s_1 | -1.59 | -0.31 | -0.13 | -1.78 | -1.31 | -0.59 | -1.25 | -1.31 | -0.07 | -0.14 | -0.04 | -0.04 | 1.4 | -0.35 | -0.13 | -0.33 | -0.4175 | -0.3425 | -0.5775 | 0.071368 | 0.7167 | 0.063095 | | 1 | 0.999631 | 0.531405 |
| Frmd4a_Q3TB04_S402s_1 | 1.33 | 0.54 | 0.32 | 0.48 | -0.05 | 0.79 | 0.46 | -0.2 | 0.59 | 1.48 | -0.68 | -0.09 | 0.87 | 0.15 | -0.21 | -0.45 | -0.125 | 0.3825 | -0.6 | 0.164577 | 0.251071 | 0.49187 | | 1 | 0.999631 | 0.632941 |
| Frmd4a_Q3TB04_S48s_1 | 1.51 | -1.42 | -0.64 | -0.23 | -0.46 | 0.23 | 0 | -1.05 | 0.27 | 1.03 | 0.15 | -0.7 | -0.94 | -0.24 | -0.91 | -1.09 | -0.1475 | 0.2675 | 0.0725 | 0.32387 | 0.838498 | 0.555523 | | 1 | 0.999631 | 0.724485 |
| Frmd4a_Q3TB04_S491s_1 | 0.23 | 0.82 | 0.76 | 0.71 | 0.64 | 0.07 | 0.35 | 0.87 | 1.01 | 0.61 | 1.23 | 0.74 | 0.89 | 0.51 | 0.71 | 0.7 | 0.395 | 0.45 | 0.26 | 0.799092 | 0.61455 | 0.359356 | | 1 | 0.999631 | 0.930937 |
| Frmd4a_Q3TB04_S514s_1 | 0.24 | -0.02 | -0.89 | -1.09 | -0.4 | 0.55 | -0.2 | -0.13 | -0.23 | 0.85 | -0.66 | 0.08 | 0.25 | -0.23 | -0.41 | -0.33 | 0.2975 | 0.07 | 0.595 | 0.507712 | 0.327066 | 0.327306 | | 1 | 0.999631 | 0.816264 |
| Frmd4a_Q3TB04_S65s_1 | -1.05 | -0.48 | -0.35 | -0.3 | -0.43 | -0.21 | -0.24 | -0.11 | -0.97 | -0.63 | -0.42 | 0.12 | 0.22 | -0.03 | -0.26 | 0.27 | -0.3775 | -0.2975 | -0.1225 | 0.077025 | 0.2917 | 0.834587 | | 1 | 0.999631 | 0.542683 |
| Frmd5_F6XM46_S270s_1 | 1.3 | 1.35 | 1.02 | 0.16 | 1.33 | 0.83 | 0.8 | -0.64 | 0.29 | 1.98 | -0.01 | 0.38 | 1.95 | 1.54 | -0.54 | 0.39 | -0.3625 | -0.1325 | -0.265 | 0.832123 | 0.438811 | 0.55477 | | 1 | 0.999631 | 0.94321 |
| Frmd8_Q3UFK8_S21s_1 | 0.75 | -0.92 | 0.32 | -0.28 | -1.07 | -0.27 | 0 | -0.24 | -0.4 | 0.09 | 0.02 | -0.37 | 0.04 | -0.92 | 0.55 | -0.86 | -0.4725 | 0.1325 | -0.485 | 0.59229 | 0.403299 | 0.740059 | | 1 | 0.999631 | 0.852797 |
| Frmd8_Q3UFK8_S24s_1 | 1.41 | 1.13 | 1.56 | 0.76 | 0.32 | 1.25 | 0.7 | 0.7 | 1.49 | 2.11 | 0.75 | 1.04 | 0.79 | 0.25 | 1.26 | 0.62 | 0.5925 | 0.16 | -0.2875 | 0.174615 | 0.154425 | 0.722007 | | 1 | 0.999631 | 0.642116 |
| Frs2_Q8C180_S428s_1 | 1.4 | -0.16 | -0.59 | 0.9 | 1.52 | 0.32 | 1.16 | 0.92 | 0.86 | 0.3 | 0.69 | 0.34 | -1.14 | 0.49 | 0.82 | 0.23 | 1.7325 | 1.345 | 0.46 | 0.621631 | 0.24797 | 0.731646 | | 1 | 0.999631 | 0.861925 |
| Fry_E9Q8I9_S1936s_1 | -1.04 | 1.77 | -0.38 | 0.08 | 2.98 | 1.11 | 2.21 | 1.06 | 3.16 | -0.84 | 1.7 | 1.79 | 1.63 | 0.63 | 0.5 | -0.49 | -0.5325 | -0.4775 | -0.995 | 0.4926 | 0.026568 | 0.155785 | | 1 | 0.999631 | 0.80955 |
| Fry_E9Q8I9_S1940s_1 | 0.93 | -0.14 | 0.57 | 0.67 | -0.42 | -0.56 | 0.61 | 0.27 | -0.57 | 0.35 | 0.1 | 0.24 | -0.9 | -0.62 | 0.11 | -0.54 | 0.3025 | -0.1675 | 0.1075 | 0.019046 | 0.178823 | 0.191789 | | 1 | 0.999631 | 0.391024 |
| Fry_E9Q8I9_S1955s_1 | -0.75 | 0.5 | -0.42 | -0.49 | 0.31 | 0.25 | -0.62 | 0.11 | -0.14 | -0.15 | -0.67 | -0.87 | -0.16 | -0.14 | -0.04 | -0.39 | -0.18 | 0.2625 | 0.4025 | 0.755326 | 0.418465 | 0.645106 | | 1 | 0.999631 | 0.913663 |
| Fry_E9Q8I9_S2459s_1 | -0.04 | 1 | 0.6 | 0.33 | 0.31 | 0.58 | 0.11 | 0.17 | 0.82 | 0.9 | 0.32 | 0.9 | 1.31 | 0.87 | 0.73 | 0.59 | 0.5725 | 0.0525 | -0.4275 | 0.249444 | 0.556559 | 0.420116 | | 1 | 0.999631 | 0.686359 |
| Fry_E9Q8I9_S2515s_1 | 2.06 | 0.7 | -0.2 | 1.79 | 2.11 | 0.86 | 1.37 | 2.3 | 1.19 | 0.49 | 1.98 | 0.9 | -0.84 | 1.39 | 1.41 | 0.68 | -0.385 | 0.4125 | 0.445 | 0.523407 | 0.323677 | 0.925393 | | 1 | 0.999631 | 0.821972 |
| Fry_E9Q8I9_S2815s_1 | 0.72 | 1.18 | 1.24 | 1.1 | -0.08 | 0.92 | 1.26 | 0.6 | 1.56 | 1.67 | 1.45 | 1.21 | 2.37 | 1.04 | 1.52 | 1.09 | -0.7325 | -0.635 | 0.64 | 0.248026 | 0.276654 | 0.146325 | | 1 | 0.999631 | 0.685751 |
| Fry_E9Q8I9_S3019s_1 | 0.12 | 0.02 | 1 | 1.05 | 0.17 | -0.49 | 0.28 | -0.7 | -0.37 | -0.21 | -0.31 | 0.54 | 0.93 | 1.24 | 0.63 | 1.95 | -0.1125 | 0.25 | 1.035 | 0.144343 | 0.07867 | 0.114201 | | 1 | 0.999631 | 0.616037 |
| Fry_E9Q8I9_T1965t_1 | -1.7 | 0.51 | 0.06 | -0.54 | -0.37 | -0.95 | -0.89 | 0.09 | 0.18 | -0.67 | -0.32 | 0.14 | -0.02 | -0.33 | 1.78 | 1.04 | 0.1675 | 0.1375 | 0.1625 | 0.118123 | 0.823085 | 0.615376 | | 1 | 0.999631 | 0.585087 |
| Fryl_F8VQ05_T1960t_1 | -0.16 | 1.04 | 1 | 1.09 | 0.95 | 0.85 | 0.52 | 1.32 | 0.69 | 0.22 | 1.37 | 1.24 | 0.07 | 1.01 | 0.94 | 1.6 | -0.36 | 0.2225 | -0.06 | 0.712558 | 0.647714 | 0.737525 | | 1 | 0.999631 | 0.9008 |
| Fscn1_Q61553_T403t_1 | 0.51 | 0.89 | 0.76 | 0.43 | -0.03 | 0.75 | 0.33 | 0.1 | 0.94 | 0.99 | 0.78 | 0.77 | 0.93 | 0.64 | 0.29 | 0.49 | 0.2925 | 0.645 | 0.26 | 0.836618 | 0.220654 | 0.393405 | | 1 | 0.999631 | 0.945389 |
| Fsd1_Q7TPM6_S300s_1 | 1.66 | 1.63 | 1.68 | 2.05 | 1.95 | 1.93 | 1.65 | 2.66 | 2.34 | 2.17 | 2.83 | 2.26 | 0.44 | 2.11 | 2.82 | 2.69 | -0.3875 | -0.16 | -0.445 | 0.622099 | 0.34179 | 0.040295 | | 1 | 0.999631 | 0.862299 |
| Fsd1_Q7TPM6_S469s_1 | 0.62 | -0.28 | -0.23 | 0.78 | -0.7 | -0.57 | 0.67 | -0.06 | -0.63 | 0.1 | 0.47 | 0.31 | -0.68 | -0.8 | 0.64 | -0.05 | -0.8 | -0.3475 | -0.91 | 0.321772 | 0.361402 | 0.680859 | | 1 | 0.999631 | 0.724092 |
| Fsd1_Q7TPM6_S477s_1 | 2.02 | -0.78 | -0.8 | -0.89 | -1.52 | -0.58 | 0.2 | -1.75 | -2 | 2.15 | -0.67 | -1.32 | 0.02 | -1.5 | -1.4 | -1.21 | -0.2875 | 0.32 | -0.735 | 0.215237 | 0.29453 | 0.728455 | | 1 | 0.999631 | 0.668192 |
| Fsd1_Q7TPM6_S477s_2 | 0.03 | -1.29 | -0.35 | -0.52 | -1.68 | -1.42 | 0.02 | -0.2 | -0.34 | 0.54 | -0.36 | -0.69 | -0.07 | -1.28 | -2.36 | -1.36 | -0.95 | -0.7525 | -0.5375 | 0.176201 | 0.554566 | 0.42773 | | 1 | 0.999631 | 0.642116 |
| Fsd1l_E0CXZ1_S314s_1 | 2.03 | 1.74 | 2.39 | 1.29 | -0.37 | 1.82 | 2.16 | 0.04 | 0.36 | 2.47 | 0.7 | 0.91 | 2.07 | 0.31 | 1.57 | 1.35 | 0.5675 | 0.0625 | -0.3525 | 0.240347 | 0.141587 | 0.153029 | | 1 | 0.999631 | 0.683935 |
| Fsd1l_E0CXZ1_S319s_1 | 1.88 | 0.5 | -0.48 | 0.71 | 2.18 | 1.92 | 0.37 | 0.41 | 1.06 | 1.59 | 0.07 | 0.14 | -0.76 | 1.2 | 0.32 | 0.44 | 0.485 | -0.3475 | -1.28 | 0.546512 | 0.367167 | 0.911344 | | 1 | 0.999631 | 0.834213 |
| Fth1_P09528_S5s_1 | 1.69 | -0.56 | -1.03 | -0.36 | 0.17 | 0.21 | 3.16 | -1.86 | 0.14 | 1.33 | -1.76 | -1.36 | -0.2 | -0.83 | -2.12 | -2.23 | -0.56 | -1.51 | -1.06 | 0.088981 | 0.638602 | 0.671401 | | 1 | 0.999631 | 0.554372 |
| Ftsj3_Q9DBE9_S576s_1 | 0.15 | 0.2 | -0.7 | 0.21 | -0.66 | -0.5 | -1.17 | -0.05 | -0.56 | -1.12 | -1.88 | -2.62 | -1.44 | -1.06 | -0.81 | -1.07 | 0.4775 | 0.0475 | 0.72 | 0.0088 | 0.133626 | 0.010261 | | 1 | 0.999631 | 0.365244 |
| Fubp1_Q3TUE1_S628s_1 | -1.43 | 0.72 | -1.37 | -1.69 | 0.82 | 0.63 | -1.35 | -1.96 | 1.09 | -1.14 | -1.66 | -1.87 | 1.82 | 0.8 | -1.61 | -1.9 | 0.135 | 0.4 | 0.415 | 0.440743 | 0.545826 | 0.95104 | | 1 | 0.999631 | 0.781983 |
| Fxr1_Q61584_S409s_1 | -0.39 | -0.62 | -0.71 | 1.01 | -0.66 | -0.51 | 0.08 | 0.92 | 0.49 | -0.3 | -0.79 | 1.49 | 0.28 | 0.38 | -0.43 | 0.72 | -0.5225 | 0.3025 | -0.3275 | 0.379272 | 0.788886 | 0.499746 | | 1 | 0.999631 | 0.751637 |
| Fxr2_Q6P5B5_S477s_1 | 0.61 | 0.13 | -0.91 | 1.07 | 0.1 | -1.35 | -0.41 | 0.47 | 0.15 | 0.14 | 0.68 | 1.14 | 0.41 | 0.18 | 0.33 | -1.33 | -0.305 | 0.23 | 0.2675 | 0.555188 | 0.340821 | 0.525 | | 1 | 0.999631 | 0.838109 |
| Fxr2_Q6P5B5_S603s_1 | -1.28 | -1.54 | -1.1 | -0.5 | -1.46 | -1.78 | -1.16 | -1.24 | -1.1 | -1.88 | -0.39 | -0.13 | -0.87 | -0.94 | -1.03 | -0.51 | -0.495 | -0.9125 | -1.0525 | 0.420173 | 0.342791 | 0.607023 | | 1 | 0.999631 | 0.77267 |
| Fxyd7_P59648_S60s_1 | 2.22 | 1.78 | 1.01 | 0.66 | -0.39 | 2.58 | 1.94 | -0.44 | 0.16 | 3.52 | -1.34 | -0.32 | 1.4 | 0.85 | -0.13 | -0.66 | 0.08 | -0.2725 | -0.3575 | 0.077909 | 0.516779 | 0.354265 | | 1 | 0.999631 | 0.543121 |
| Fxyd7_P59648_S72s_1 | -0.91 | -0.24 | -0.43 | -0.81 | -0.32 | -0.41 | -0.94 | -0.4 | -0.98 | -0.85 | -0.06 | -1.59 | -0.24 | -0.68 | -1.82 | -1.08 | 0.1 | 0.3775 | 0.525 | 0.377731 | 0.780699 | 0.474923 | | 1 | 0.999631 | 0.750463 |
| Fxyd7_P59648_S73s_1 | -1.42 | -0.51 | -2.13 | -2.2 | -1.46 | 0.32 | -2.62 | -2.1 | -1.59 | -0.94 | -1.01 | -1.21 | -0.68 | -0.46 | -1.79 | -1.23 | -0.5425 | -1.0625 | -1.06 | 0.288578 | 0.881087 | 0.381339 | | 1 | 0.999631 | 0.706046 |
| Fxyd7_P59648_T62t_1 | 1.28 | 0.82 | 1.09 | 1.01 | -0.57 | 1.52 | 2 | -0.92 | -0.29 | 1.32 | -1.05 | -0.03 | 0.04 | 0.54 | 0.12 | -0.74 | 0.9225 | 1.21 | 0.8675 | 0.010988 | 0.418829 | 0.049593 | | 1 | 0.999631 | 0.385092 |
| Fyn_D3YZ57_S21s_1 | -1.67 | 0.89 | 1.66 | -0.13 | 1.1 | 0.66 | 1.14 | 1.54 | 1.82 | -1.41 | 2.48 | 2.7 | 2.06 | -0.46 | 2.12 | 0.5 | -0.4125 | 0.415 | 0.4575 | 0.309594 | 0.184575 | 0.257627 | | 1 | 0.999631 | 0.718889 |
| Fyn_D3YZ57_Y185y_1 | -1.15 | -0.74 | -1.22 | -0.82 | -1.28 | -1.33 | -1.52 | -1.45 | -0.79 | -0.58 | -1.03 | 0.13 | 0.2 | -1.16 | -0.68 | -0.46 | -0.4175 | -0.3975 | -0.6225 | 0.220077 | 0.12442 | 0.227026 | | 1 | 0.999631 | 0.670699 |
| Fyttd1_Q91Z49_S23s_1 | 1.83 | 1.27 | 2.19 | 1.79 | 0.97 | 0.89 | 2.1 | 1.45 | 1.01 | 2.06 | 1.29 | 1.13 | 1.39 | 1.26 | 1.71 | 0.23 | 0.4325 | 0.4375 | 0.8875 | 0.140737 | 0.263219 | 0.266034 | | 1 | 0.999631 | 0.612258 |
| Fzd3_Q61086_S588s_1 | -1.35 | 0 | -0.56 | -0.9 | -0.44 | 0.03 | -0.22 | -0.45 | 0.07 | -0.09 | -0.88 | -0.16 | 0.56 | 0.12 | 0.2 | -0.14 | -0.535 | -0.2525 | -0.195 | 0.033068 | 0.223855 | 0.263442 | | 1 | 0.999631 | 0.44294 |
| Fzd3_Q61086_S665s_1 | 0.32 | -1.03 | -1.25 | -0.9 | -1.38 | -0.86 | -1.63 | -1.13 | -0.51 | 0.09 | -1.71 | -1.74 | -1.07 | -1.28 | -1.29 | 0 | 1.1 | 0.3725 | 0.365 | 0.673955 | 0.197233 | 0.639359 | | 1 | 0.999631 | 0.885515 |
| Fzd3_Q61086_T542t_1 | -0.99 | 1.84 | 1.18 | 1.22 | 2.54 | 1.75 | 1.53 | 1.83 | 2.09 | -0.82 | 1.89 | 1.58 | 0.24 | 1.27 | 1.64 | 1.56 | -0.08 | 0.4975 | 0.9225 | 0.565786 | 0.089175 | 0.645177 | | 1 | 0.999631 | 0.841575 |
| G3bp1_P97855_T268t_1 | -0.35 | 0.31 | 0.6 | 0.09 | 0.04 | -0.12 | -0.28 | 0.69 | 0.5 | 0.05 | 0.76 | 1.33 | 0.65 | 1.06 | 1.05 | 1.58 | -0.0225 | 0.1225 | 0.14 | 0.020584 | 0.809825 | 0.193541 | | 1 | 0.999631 | 0.396554 |
| G3bp2_P97379_S225s_1 | -0.53 | -0.05 | 0.13 | 0.01 | 0.36 | -0.32 | -0.55 | -0.02 | 0.35 | -0.51 | 0.28 | -0.07 | -0.8 | 0.3 | 0.24 | 0.38 | -0.0725 | 0.0875 | 0.3925 | 0.699932 | 0.940984 | 0.694822 | | 1 | 0.999631 | 0.896612 |

| | | | | | | | | | | | | | | | | | | | | | | | | | |
|---|---|---|---|---|---|---|---|---|---|---|---|---|---|---|---|---|---|---|---|---|---|---|---|---|---|
| G3bp2_P97379_T227t_1 | 0.16 | 1.09 | 0.92 | 0.62 | 0.81 | 0.74 | 0.17 | 0.78 | 0.73 | 0.54 | 0.83 | 1.04 | 1.02 | 1.26 | 0.95 | 1.13 | 0.5325 | 0.415 | 0.03 | 0.223406 | 0.815112 | 0.772892 | 1 | 0.999631 | 0.67471 |
| Gab2_Q3ZB57_S211s_1 | 1.05 | 0.53 | 0.52 | 0.87 | 1.04 | 1.51 | 1.31 | 1.24 | 1.74 | 1.09 | 1.42 | 0.38 | 0.18 | 0.66 | 1.42 | 0.83 | -0.11 | -0.0575 | -0.01 | 0.931071 | 0.066945 | 0.259815 | 1 | 0.999631 | 0.977912 |
| Gab2_Q3ZB57_S402s_1 | 0.79 | 1.43 | 0.56 | 0.78 | 0.99 | 0.59 | 0.49 | 1.05 | 0.8 | 0.81 | 0.74 | 0.98 | 0.7 | 1.34 | 0.97 | 0.51 | -0.03 | -0.2175 | -0.175 | 0.975959 | 0.713469 | 0.840443 | 1 | 0.999631 | 0.991761 |
| Gab2_Q3ZB57_T388t_1 | 0.88 | 1.12 | 0.98 | 1.43 | 1.01 | 1.57 | 0.27 | 1.44 | 0.92 | 1.03 | 0.65 | 0.94 | 0 | 1.53 | 1.53 | 0.65 | 0.0725 | -0.1625 | -1.8225 | 0.670713 | 0.931133 | 0.418722 | 1 | 0.999631 | 0.884315 |
| Gabarapl2_P60521_S10s_1 | 1.53 | -1.87 | -0.55 | -0.76 | -1.2 | -0.92 | 1.99 | -1.23 | -1.46 | 1.07 | -1.23 | -0.68 | -0.97 | -2.1 | -2.66 | -3.21 | -0.4275 | -0.4925 | -0.82 | 0.033497 | 0.936563 | 0.839062 | 1 | 0.999631 | 0.443263 |
| Gabbr1_Q9WV18_S877s_1 | 0.45 | -0.81 | -0.14 | -1.51 | -1.59 | -0.39 | 1.88 | -3.62 | -2.27 | 0.18 | -0.54 | -1.35 | -0.09 | -1.89 | -1.12 | -2.19 | -0.1 | -0.195 | 0.2125 | 0.178899 | 0.685344 | 0.430219 | 1 | 0.999631 | 0.643093 |
| Gabbr1_Q9WV18_S909s_1 | 0.16 | -0.6 | -0.77 | -0.44 | -0.11 | -0.48 | -1 | -0.46 | -0.41 | -0.45 | -1.16 | -0.41 | 0.51 | -1.01 | -0.18 | -0.12 | 0.57 | 0.04 | -0.2825 | 0.595532 | 0.755614 | 0.555469 | 1 | 0.999631 | 0.854508 |
| Gabbr1_Q9WV18_S949s_1 | 1.18 | -0.59 | -0.85 | 0.72 | 0.95 | 0.2 | 0.65 | 0.94 | 0.29 | -0.02 | 0.37 | -0.02 | -1.19 | 0.42 | 0.13 | -0.03 | -0.0325 | -0.5125 | 0.23 | 0.617352 | 0.264749 | 0.93399 | 1 | 0.999631 | 0.861376 |
| Gabbr1_Q9WV18_T872t_1 | -0.64 | -0.44 | 0.28 | -0.84 | -0.9 | -0.17 | -0.75 | 0.05 | -1.57 | -0.9 | -0.16 | -1.06 | -0.21 | -0.2 | 0.1 | -0.41 | 0.3975 | 0.7775 | 0.73 | 0.497703 | 0.927675 | 0.214476 | 1 | 0.999631 | 0.811908 |
| Gabbr1_Q9WV18_T929t_1 | -1.61 | -0.63 | -0.36 | -1.62 | -0.33 | -0.73 | -0.55 | -1.02 | -0.08 | -1.32 | -0.08 | 0.37 | 1.3 | -0.83 | -0.75 | -1.02 | -0.5875 | -0.065 | -0.3975 | 0.229213 | 0.3035 | 0.124458 | 1 | 0.999631 | 0.679776 |
| Gabbr2_Q80T41_S770s_1 | -3.97 | -4.49 | -5.32 | -5.65 | -5.5 | -5.14 | -4.37 | -6.77 | -5.73 | -3.97 | -3.73 | -6.26 | -4.84 | -4.79 | -5.22 | -6.17 | 0.0575 | -0.14 | -0.0025 | 0.41939 | 0.319648 | 0.921725 | 1 | 0.999631 | 0.772152 |
| Gabbr2_Q80T41_S775s_1 | 0.05 | -0.6 | -1.83 | -1.08 | -0.46 | -0.36 | -0.73 | -1.68 | -0.97 | -0.31 | -1.45 | -1.29 | -0.69 | -0.84 | -1.18 | -0.76 | 0.035 | 0.4375 | 0.14 | 0.995279 | 0.903292 | 0.760241 | 1 | 0.999631 | 0.997705 |
| Gabbr2_Q80T41_S778s_2 | 0.04 | 0.11 | -0.1 | 0.79 | 0.44 | -0.49 | 0.2 | 0.83 | 0.84 | 0.22 | 0.45 | 1.08 | 0.68 | 0.92 | -1.01 | 0.81 | -0.175 | 0.015 | -0.26 | 0.771885 | 0.923137 | 0.204532 | 1 | 0.999631 | 0.919413 |
| Gabbr2_Q80T41_S783s_1 | 0.92 | 0.6 | -1.75 | -0.6 | 0.2 | -0.21 | -0.21 | -1.31 | 0.72 | 0.2 | -0.43 | -1.26 | 1.06 | -0.43 | -0.82 | -1.68 | -0.4125 | 0.4775 | 0.2825 | 0.724579 | 0.778154 | 0.98203 | 1 | 0.999631 | 0.905674 |
| Gabbr2_Q80T41_S792s_1 | 0.45 | -0.86 | 0.19 | -0.37 | -1.21 | 0.66 | -0.67 | -1.02 | 0.41 | 0.23 | -0.25 | 0.93 | 0.05 | 0.28 | 0.43 | -0.22 | 0.3175 | -0.0175 | 0.4 | 0.450673 | 0.404204 | 0.245297 | 1 | 0.999631 | 0.787797 |
| Gabbr2_Q80T41_S883s_1 | -1.92 | -1.09 | -1 | -1.83 | -1.1 | -1.03 | -1.39 | -1.05 | -1.5 | -2.1 | -1.1 | -1.21 | -0.51 | -1.23 | -1.29 | -1.21 | 0.605 | -1.1125 | -1.1975 | 0.27678 | 0.319549 | 0.961353 | 1 | 0.999631 | 0.700187 |
| Gabbr2_Q80T41_T762t_1 | 1.47 | 0.94 | 1.05 | 2.08 | 1.35 | 2.32 | 1.8 | 2.49 | -0.31 | -0.19 | 2.63 | -1.04 | -2 | 1.49 | 0.41 | 0.85 | 1.3925 | 1.4575 | 0.765 | 0.116727 | 0.135037 | 0.160903 | 1 | 0.999631 | 0.584493 |
| Gabra2_P26048_T393t_1 | 1.36 | -1.58 | 0.24 | -0.11 | 0.79 | 1.26 | 2.39 | 1.04 | 0.73 | 2.38 | 1.39 | 1.24 | -0.55 | 0.48 | 1.85 | 1.19 | -0.44 | -0.58 | -0.0175 | 0.292037 | 0.048752 | 0.041287 | 1 | 0.999631 | 0.706164 |
| Gabra3_B1AVY2_S474s_1 | -0.69 | -0.07 | -0.48 | -0.9 | -0.61 | -1.42 | -1.08 | -0.79 | -1.02 | -0.97 | -1.21 | -1.26 | -0.07 | -0.77 | -0.77 | -0.6 | -1.0475 | -0.65 | -0.24 | 0.956763 | 0.174792 | 0.06446 | 1 | 0.999631 | 0.984803 |
| Gabra3_B1AVY2_T468t_1 | -1.06 | -0.83 | -0.68 | -0.26 | -2.11 | -1.82 | -1.43 | -1.66 | -1.71 | -1.12 | -2.41 | -0.19 | -0.9 | -1.2 | -0.49 | -1.2 | -0.1775 | -0.32 | -0.3875 | 0.46069 | 0.004816 | 0.196453 | 1 | 0.999631 | 0.791962 |
| Gabra4_Q3USX9_T449t_1 | 0.3 | -0.13 | 0.02 | 0.55 | 0.05 | 0.47 | -0.13 | -0.36 | -0.72 | 0.41 | -0.33 | 0.1 | -0.63 | -0.4 | 0.54 | -0.32 | -0.09 | -0.4525 | -0.03 | 0.286649 | 0.553727 | 0.353905 | 1 | 0.999631 | 0.705972 |
| Gabrd_P22933_S390s_1 | -0.93 | -0.48 | -1.07 | -0.59 | -0.54 | -0.78 | -1.52 | -0.59 | -0.81 | -0.49 | -1.99 | -1.59 | -0.57 | -0.68 | -1.06 | -0.88 | 0.095 | -0.065 | 0.325 | 0.918278 | 0.776468 | 0.262454 | 1 | 0.999631 | 0.974076 |
| Gabrg2_P22723_S393s_1 | 0.53 | 0.47 | 0.66 | 0.51 | 1 | 0.75 | 0.3 | 0.5 | 0.82 | 0.77 | 0.24 | 0.08 | 0.57 | 0.83 | 0.75 | 1.32 | 0.825 | -0.0775 | -0.0375 | 0.275133 | 0.716769 | 0.81895 | 1 | 0.999631 | 0.700187 |
| Gabrg2_P22723_T395t_1 | -0.62 | -0.24 | 0.16 | 0.19 | -1.73 | 2.53 | 0.51 | 1.48 | -0.74 | -0.59 | 1.79 | -1.28 | -1.34 | 1.63 | -0.53 | -0.42 | -0.905 | -0.085 | 0.0325 | 0.950277 | 0.321627 | 0.90329 | 1 | 0.999631 | 0.984079 |
| Gad1_P48318_S527s_1 | -0.52 | -2.22 | 0.72 | -0.04 | -2.56 | -3.3 | -0.13 | 0.31 | -1.85 | -1.4 | 0.13 | 0.72 | -1.51 | -1.64 | 0.73 | 0.49 | 0.16 | -0.7875 | -0.2275 | 0.966602 | 0.346369 | 0.911911 | 1 | 0.999631 | 0.988351 |
| Gad1_P48318_S55s_1 | 0.28 | 1.15 | 0.3 | 0.33 | 0.82 | 1.37 | 0.91 | -0.4 | 0.72 | -0.26 | -1.21 | -0.34 | 1 | 1.01 | 0.35 | -1.21 | 0.6425 | -0.4925 | 0.0775 | 0.669087 | 0.70848 | 0.100074 | 1 | 0.999631 | 0.883599 |
| Gad1_P48318_S77s_1 | 0.51 | -1.68 | -1.75 | 0.01 | 0.47 | 0.1 | -0.35 | -0.56 | -0.36 | -1.45 | -1.47 | -1.6 | -0.26 | 0.05 | -1.11 | -1.28 | -0.42 | 0.165 | 0.555 | 0.897804 | 0.274756 | 0.413559 | 1 | 0.999631 | 0.966758 |
| Gak_F6SLK1_S112s_1 | 0.9 | 1.6 | 1.81 | 1.55 | 0.7 | 1.83 | 0.63 | 1.02 | 1.34 | 1.67 | 1.77 | 1.74 | 2.04 | 1.78 | 1.77 | 2.49 | -0.2975 | -0.515 | -1.15 | 0.118101 | 0.261846 | 0.582921 | 1 | 0.999631 | 0.585087 |
| Gap43_P06837_S122s_1 | 1.1 | -1.6 | -0.27 | -0.78 | -1.05 | -0.5 | 0.55 | -1.74 | -1.43 | 0.35 | -1.63 | -0.9 | -1.05 | -2 | -0.84 | -2.26 | -0.3725 | -0.285 | -0.9325 | 0.081532 | 0.65532 | 0.434631 | 1 | 0.999631 | 0.546935 |
| Gap43_P06837_S128s_1 | 1.89 | -0.89 | -0.47 | -0.29 | -1.17 | 0.42 | 0.9 | -1.4 | 0 | 1.04 | -1.21 | -0.73 | -0.12 | -0.99 | -0.56 | -1.82 | -0.8025 | -1.2 | -1.405 | 0.171593 | 0.618483 | 0.687052 | 1 | 0.999631 | 0.641657 |
| Gap43_P06837_S133s_1 | 1.57 | -0.82 | 0.07 | 0.15 | -0.9 | -0.34 | 1.08 | -2.08 | -1.51 | 0.83 | -1.87 | -1.28 | -0.86 | -1.84 | -0.54 | -1.41 | -0.28 | -0.62 | -0.88 | 0.024001 | 0.283342 | 0.110576 | 1 | 0.999631 | 0.415644 |
| Gap43_P06837_S142s_1 | 0.2 | -2.05 | -2.31 | -1.78 | -2.26 | -2.02 | 0.08 | -2.86 | -2.73 | -0.49 | -2.57 | -2.63 | -1.56 | -2.31 | -2.7 | -2.89 | -0.2425 | 0.1825 | 0.525 | 0.157688 | 0.711026 | 0.385802 | 1 | 0.999631 | 0.628615 |
| Gap43_P06837_S192s_1 | 0.27 | 0.4 | 0.31 | 0.11 | 0.37 | -0.22 | -0.45 | 0.42 | 0.47 | 0.5 | 0.09 | 0.76 | 1.47 | 0.55 | 0.69 | 0.48 | -0.2925 | 0.1675 | 0.5075 | 0.124522 | 0.416777 | 0.498553 | 1 | 0.999631 | 0.596063 |
| Gap43_P06837_S193s_1 | 0.33 | 0.4 | 0.34 | 0.11 | 0.29 | -0.24 | -0.44 | 0.4 | 0.44 | 0.47 | 0.13 | 0.81 | 1.49 | 0.54 | 0.68 | 0.5 | -0.1625 | -0.0525 | 0.1375 | 0.138556 | 0.321753 | 0.534942 | 1 | 0.999631 | 0.611124 |
| Gap43_P06837_S86s_1 | -0.41 | -0.51 | -1.24 | -1.3 | -1.3 | -0.77 | -0.78 | -1.26 | -1 | -1.41 | -0.71 | -0.55 | -0.13 | -1.23 | -0.54 | -1.01 | -0.095 | 0.46 | 0.475 | 0.716678 | 0.616934 | 0.878761 | 1 | 0.999631 | 0.903179 |
| Gap43_P06837_S96s_1 | -0.32 | -0.58 | -0.04 | -0.08 | -0.65 | -0.57 | -0.07 | -0.11 | 0.11 | 0.14 | 0.12 | 0.45 | 0.6 | -0.5 | 0.61 | 0.17 | 0.01 | 0.1825 | 0.0375 | 0.192945 | 0.733843 | 0.108326 | 1 | 0.999631 | 0.649499 |
| Gap43_P06837_S96s_2 | -0.08 | -0.42 | 0.02 | 0.1 | -0.18 | -0.14 | -0.1 | 0.08 | -0.31 | 0.1 | 0.2 | 0.36 | 0.29 | -0.74 | 0.73 | -0.51 | 0.6575 | 0.79 | 0.8225 | 0.923562 | 0.968028 | 0.519411 | 1 | 0.999631 | 0.975032 |
| Gap43_P06837_T171t_1 | -2.32 | -1.01 | -1.4 | -0.95 | -0.23 | -1.56 | -0.9 | -0.36 | 0.21 | -2.4 | -0.12 | -0.21 | 0.67 | -1.57 | -0.72 | -0.77 | 0.4075 | 0.6875 | 0.49 | 0.143402 | 0.149722 | 0.217454 | 1 | 0.999631 | 0.614578 |
| Gap43_P06837_T172t_1 | -1.85 | -0.74 | -1.17 | -2.01 | -0.27 | -1.2 | -1.04 | -1.63 | -0.18 | -1.67 | -0.87 | -0.3 | 0.45 | -1.37 | -1.38 | -1.51 | 0.09 | 0.3125 | 0.245 | 0.360101 | 0.33345 | 0.146023 | 1 | 0.999631 | 0.741182 |
| Gap43_P06837_T95t_1 | -0.51 | -0.93 | -2.74 | -3.16 | -0.7 | -1.09 | -2.38 | -2.81 | -0.64 | -0.34 | -2.28 | -2.83 | -0.03 | -0.76 | -2.47 | -3.1 | -0.845 | -0.7125 | -0.5 | 0.771475 | 0.901634 | 0.692257 | 1 | 0.999631 | 0.919413 |
| Gap43_P06837_T95t_2 | -2.3 | -4.74 | 1.08 | -0.26 | -3.01 | -4.59 | -0.12 | -1.88 | -2.82 | -3.96 | -1.38 | -0.91 | -3.73 | -3.05 | -1.22 | -0.22 | -0.08 | -0.195 | 0.21 | 0.694887 | 0.533565 | 0.567991 | 1 | 0.999631 | 0.895302 |
| Gapdh_P16858_S208s_1 | 0.04 | 0.95 | 0.37 | 0.18 | -0.21 | 0.44 | -0.27 | 1.26 | 0.86 | 0.57 | -0.97 | 0.3 | 0.91 | 0.86 | -0.2 | 0.81 | -0.1725 | 0.0175 | 0.2775 | 0.576825 | 0.845048 | 0.661866 | 1 | 0.999631 | 0.84622 |
| Gapdh_P16858_T209t_1 | -0.08 | 1.31 | 0.8 | 0.27 | 0.28 | 0.69 | -0.13 | 0.77 | 0.85 | 0.36 | 0.66 | 0.5 | 0.95 | 1.27 | 0.74 | 0.45 | 0.05 | -0.16 | 0.27 | 0.475838 | 0.652616 | 0.960892 | 1 | 0.999631 | 0.799882 |
| Gapvd1_Q6PAR5_S1041s_1 | -0.52 | -0.73 | -0.36 | 0.08 | -0.52 | 0.15 | -0.53 | -0.43 | -0.9 | 0.02 | -0.88 | -0.41 | 0.19 | -0.11 | -0.42 | -0.11 | 0.175 | 0.2875 | 0.38 | 0.390435 | 0.867495 | 0.630187 | 1 | 0.999631 | 0.758901 |
| Gapvd1_Q6PAR5_S1041s_2 | 0.1 | 1.01 | 1.39 | 0.82 | 0.89 | 1.4 | 0.63 | 1.1 | 0.96 | 1.09 | 0.83 | 1.59 | 0.44 | 1.55 | 0.76 | 2.09 | -0.22 | -0.0575 | 0.16 | 0.414833 | 0.61626 | 0.425161 | 1 | 0.999631 | 0.769212 |
| Gapvd1_Q6PAR5_S1045s_1 | -0.96 | 0.45 | -0.47 | -0.03 | -0.44 | -0.66 | -0.81 | 0.02 | -1.23 | -0.16 | -0.27 | 0.42 | -0.26 | 0.15 | 0.01 | -0.27 | 0.175 | 0.2875 | 0.38 | 0.662633 | 0.555804 | 0.897817 | 1 | 0.999631 | 0.881311 |
| Gapvd1_Q6PAR5_S1045s_2 | 0.1 | 1.01 | 1.39 | 0.82 | 0.89 | 1.4 | 0.63 | 1.1 | 0.96 | 1.09 | 0.83 | 1.59 | 0.44 | 1.55 | 0.76 | 2.09 | 1.08 | 1.1725 | -0.1175 | 0.414833 | 0.61626 | 0.425161 | 1 | 0.999631 | 0.769212 |
| Gapvd1_Q6PAR5_S881s_1 | 0.48 | 1 | 0.47 | 0.47 | 2.14 | 1.07 | 2.51 | 1.02 | 1.98 | 0.47 | 2.09 | 2.57 | 0.07 | 0.47 | 0.7 | 0.71 | -0.0375 | 0.1225 | 0.2075 | 0.698569 | 0.021975 | 0.027724 | 1 | 0.999631 | 0.895842 |
| Gapvd1_Q6PAR5_S893s_1 | -0.38 | 0.15 | -0.66 | -0.15 | -0.23 | -0.19 | -0.02 | -0.75 | 0.03 | 0.13 | -0.26 | -0.45 | 0.57 | 0.47 | -0.55 | -0.7 | 0.1325 | 0.15 | 0.5575 | 0.605567 | 0.899803 | 0.680953 | 1 | 0.999631 | 0.85817 |
| Gapvd1_Q6PAR5_T741t_1 | -0.73 | -0.71 | 0.5 | 0.43 | -0.25 | 0.16 | 0.05 | 0.06 | -0.6 | -0.08 | -0.16 | 0.93 | -1.05 | 0.84 | 1.08 | 0.85 | -0.0625 | 0.03 | 0.235 | 0.33101 | 0.722035 | 0.744254 | 1 | 0.999631 | 0.727302 |
| Garem_Q3UFT3_S493s_1 | -0.28 | 0.53 | -0.4 | -0.02 | 0.32 | 0.18 | -0.73 | -0.19 | 0.18 | 0.07 | -0.01 | -0.29 | 0.32 | 0.63 | -0.46 | 0.28 | 0.045 | 0.36 | 0.4975 | 0.518388 | 0.856181 | 0.921134 | 1 | 0.999631 | 0.819449 |
| Garem_Q3UFT3_S698s_1 | -0.64 | -0.12 | -1.28 | -0.81 | -0.68 | 0.01 | -0.88 | -1.12 | 0.34 | 0.03 | -1.37 | -0.41 | -0.01 | -0.38 | -0.31 | -0.16 | -0.12 | -0.29 | 0.3425 | 0.152626 | 0.901359 | 0.420127 | 1 | 0.999631 | 0.625198 |
| Garem_Q3UFT3_S721s_1 | -0.03 | 0.78 | 0.71 | 0.24 | 0.18 | 0.47 | -0.23 | 0.8 | 0.14 | -0.37 | 0.14 | 0.63 | 0.43 | 1.41 | 0.58 | 0.65 | 0.0475 | 0.215 | 0.545 | 0.339338 | 0.71855 | 0.392115 | 1 | 0.999631 | 0.732185 |
| Garem_Q3UFT3_T738t_1 | -0.66 | -0.32 | -0.13 | -0.25 | -0.4 | -0.27 | -0.64 | 0.14 | -0.48 | -0.38 | 0.22 | 0.14 | 0.34 | -0.3 | 0.49 | 0.29 | -0.215 | -0.4625 | 0.5575 | 0.099126 | 0.865353 | 0.470365 | 1 | 0.999631 | 0.562298 |
| Gareml_Q6PAJ3_S509s_1 | -0.14 | -0.36 | -0.77 | -1.04 | -0.39 | -0.09 | -1.23 | -1.46 | -0.68 | -0.19 | -1.95 | -1.34 | 0.91 | 0.04 | -0.22 | -0.81 | 0.2125 | 0.715 | 1.2225 | 0.206986 | 0.58703 | 0.299478 | 1 | 0.999631 | 0.660822 |
| Gareml_Q6PAJ3_T507t_1 | -1.37 | -1.36 | -1.23 | -0.15 | 0.19 | -2.08 | -0.54 | -0.83 | -0.63 | -0.9 | -0.43 | 0.71 | 1.04 | -1.46 | 1.37 | -0.17 | 0.2725 | 0.225 | 0.705 | 0.079041 | 0.682746 | 0.137376 | 1 | 0.999631 | 0.545579 |
| Garnl3_A2AUQ6_S26s_1 | 0 | 0.14 | -0.12 | -0.36 | 0.36 | 0.55 | -0.3 | 0.14 | 0.02 | -0.31 | 0.28 | 0.57 | 1.04 | 0.39 | 0.41 | 0.64 | 0.975 | 0.0625 | 0.2525 | 0.035836 | 0.352925 | 0.453653 | 1 | 0.999631 | 0.453811 |
| Garnl3_B1AW31_S26s_1 | 0.82 | 0.17 | 0.24 | 1.63 | 2.04 | 1.17 | 1.92 | 1.63 | 1.62 | -0.58 | 1.25 | 0.82 | -0.86 | 1.96 | 0.84 | 1.93 | 0.2425 | 0.355 | 0.755 | 0.704982 | 0.031754 | 0.908783 | 1 | 0.999631 | 0.897793 |
| Garnl3_Q3V0G7_S6s_1 | -0.39 | -0.17 | 0.45 | 0.75 | -0.6 | 0.19 | 0.66 | 1.36 | 0.04 | -0.11 | 0.97 | 1.16 | 0.23 | 0.65 | 1.53 | 1.25 | 0.745 | 0.155 | 0.1625 | 0.0914 | 0.606386 | 0.407038 | 1 | 0.999631 | 0.557085 |
| Garnl3_Q3V0G7_S844s_3 | 0.22 | -0.01 | -0.02 | 1.37 | 1.4 | 0.81 | 0.4 | 1.93 | 0.68 | 0.75 | 0.19 | 0.56 | -0.7 | 1.03 | 1.15 | 0.73 | -0.525 | 0.0675 | 0.0875 | 0.751857 | 0.125478 | 0.682582 | 1 | 0.999631 | 0.912945 |
| Garnl3_Q3V0G7_S848s_1 | 0.03 | -0.49 | 0.65 | -0.16 | -1.75 | -0.56 | 0.65 | -0.41 | -0.4 | 0.71 | 0.15 | -0.16 | 0.82 | -0.2 | -0.26 | 0.02 | -0.285 | 0.5425 | 0.6075 | 0.818566 | 0.316875 | 0.854394 | 1 | 0.999631 | 0.938303 |
| Garnl3_Q3V0G7_S848s_2 | -0.52 | -0.47 | -0.11 | -1.04 | -1.2 | -0.46 | -0.95 | -0.67 | -0.35 | 1.1 | -0.41 | -0.31 | 1.26 | -0.39 | -0.38 | -0.2 | 0.745 | 0.155 | 0.1625 | 0.191744 | 0.365169 | 0.213221 | 1 | 0.999631 | 0.648674 |
| Garnl3_Q3V0G7_S848s_3 | 0.22 | -0.01 | -0.02 | 1.37 | 1.4 | 0.81 | 0.4 | 1.93 | 0.68 | 0.75 | 0.19 | 0.56 | -0.7 | 1.03 | 1.15 | 0.73 | -0.045 | -0.1625 | -0.0075 | 0.751857 | 0.125478 | 0.682582 | 1 | 0.999631 | 0.912945 |
| Garnl3_Q3V0G7_T851t_1 | 0.34 | 0.4 | 0.22 | 0.03 | 0.35 | 0.25 | 0.34 | -0.13 | 0.09 | 0.76 | -0.02 | -0.49 | 0.92 | -0.24 | -0.19 | 0.47 | -0.285 | 0.5425 | 0.6075 | 0.982819 | 0.859342 | 0.620049 | 1 | 0.999631 | 0.99485 |
| Garnl3_Q3V0G7_T851t_2 | -0.52 | -0.47 | -0.11 | -1.04 | -1.2 | -0.46 | -0.95 | -0.67 | -0.35 | 1.1 | -0.41 | -0.31 | 1.26 | -0.39 | -0.38 | -0.2 | 0.745 | 0.155 | 0.1625 | 0.191744 | 0.365169 | 0.213221 | 1 | 0.999631 | 0.648674 |
| Garnl3_Q3V0G7_T851t_3 | 0.22 | -0.01 | -0.02 | 1.37 | 1.4 | 0.81 | 0.4 | 1.93 | 0.68 | 0.75 | 0.19 | 0.56 | -0.7 | 1.03 | 1.15 | 0.73 | -0.475 | -0.3275 | 0.315 | 0.751857 | 0.125478 | 0.682582 | 1 | 0.999631 | 0.912945 |
| Garnl3_Q3V0G7_T901t_1 | -1.66 | -0.16 | -0.13 | -0.53 | -0.98 | -1.92 | -0.46 | -1.02 | -2.46 | -0.85 | 0.34 | -0.82 | 0.26 | -0.65 | -0.79 | -0.04 | -0.4425 | -0.1025 | -1.2425 | 0.479807 | 0.308234 | 0.596939 | 1 | 0.999631 | 0.801333 |
| Gars_Q9CZD3_S643s_1 | 1.22 | 0.69 | 2.07 | 1.04 | -0.01 | 1.68 | 1.15 | 0.43 | 0.89 | 2.06 | 0.51 | 1.15 | -0.35 | 0.29 | 0.18 | -0.07 | -0.7525 | -1.2425 | -1.2375 | 0.006758 | 0.345586 | 0.814867 | 1 | 0.999631 | 0.358173 |
| Gas2_P11862_S283s_1 | 0.26 | -1.85 | 1.62 | 1.55 | -2.61 | -1.73 | 2.7 | 0.21 | -2.5 | -1.09 | -0.44 | 0.64 | -2.4 | -2.24 | 0.64 | 0.63 | -0.01 | 0.005 | 0.0975 | 0.236904 | 0.541084 | 0.191989 | 1 | 0.999631 | 0.683301 |
| Gas2l1_Q8JZP9_S292s_1 | 0.28 | 0.21 | 0.59 | -0.15 | 0.18 | 0.71 | -0.07 | 0.07 | 0.04 | 0.57 | 0.2 | 0.14 | -0.05 | 0.52 | 0.36 | 0.49 | -0.09 | -0.53 | -1.275 | 0.748123 | 0.972962 | 0.98593 | 1 | 0.999631 | 0.911931 |
| Gas2l1_Q8JZP9_S297s_1 | 2.41 | 0.96 | 2.86 | 1.81 | 0.58 | 3.29 | 2.23 | 1.58 | 1.34 | 2.87 | 0.74 | 0.97 | -1.23 | 1.75 | 1.41 | 1.01 | 0.07 | -0.0875 | -0.17 | 0.092407 | 0.886255 | 0.370808 | 1 | 0.999631 | 0.557085 |
| Gas2l1_Q8JZP9_S306s_1 | 0.37 | -0.72 | -0.61 | -0.05 | 0.11 | 0.01 | -0.5 | -0.35 | -0.3 | 0.08 | -0.66 | -0.48 | -0.6 | -0.29 | -0.19 | -0.61 | -0.0225 | 0.095 | 0.3675 | 0.619821 | 0.833245 | 0.798801 | 1 | 0.999631 | 0.861888 |
| Gas2l1_Q8JZP9_S352s_1 | -0.1 | -0.37 | -0.45 | -0.53 | -0.45 | -0.25 | -0.31 | -0.53 | -0.34 | -0.19 | -0.08 | -0.46 | 0.06 | 0.03 | -0.34 | 0.27 | -0.35 | -0.6575 | -1.26 | 0.216678 | 0.926684 | 0.712449 | 1 | 0.999631 | 0.669465 |
| Gas2l1_Q8JZP9_S352s_2 | 1.09 | 0.5 | -0.22 | -0.64 | 0 | -0.47 | 0.11 | -0.31 | -0.77 | 0.94 | -1.14 | -0.93 | -2.28 | 0.72 | -1.06 | -1.69 | 0.8475 | -0.32 | -0.8275 | 0.085179 | 0.399504 | 0.261255 | 1 | 0.999631 | 0.550808 |
| Gas2l1_Q8JZP9_S482s_1 | 2.29 | -0.12 | -0.63 | 1.37 | 2.29 | 1.12 | 1.5 | 1.39 | 0.87 | 0.09 | 0.63 | 0.04 | -1.53 | 0.78 | 0.23 | 0.12 | 0.625 | -0.7625 | -1.435 | 0.276737 | 0.207017 | 0.615895 | 1 | 0.999631 | 0.700187 |
| Gas7_B1ATI9_S312s_1 | 2.46 | -0.04 | -1.15 | 1.79 | 2.84 | -0.73 | 2.82 | 0.63 | 0.47 | -0.71 | 0.85 | -0.6 | -1.82 | -0.32 | -0.07 | -0.47 | 0.255 | -0.6425 | -0.5625 | 0.096953 | 0.549959 | 0.353917 | 1 | 0.999631 | 0.55886 |
| Gas7_B1ATI9_S47s_1 | 0.85 | -0.47 | 0.41 | 0.29 | 0.65 | 0.79 | 0.32 | 0.34 | 0.33 | 0.06 | -0.57 | -1.31 | -0.73 | 0.05 | -0.46 | -0.03 | 0.8375 | -0.075 | -0.0975 | 0.1532 | 0.454514 | 0.174501 | 1 | 0.999631 | 0.625198 |
| Gas7_B1ATI9_S53s_1 | 0.25 | -0.23 | -0.38 | 0.34 | 0.94 | 0.77 | 1.01 | 0.61 | 0.14 | -0.55 | 0.21 | -0.12 | -0.77 | 0.46 | 0.3 | -0.4 | 0.3275 | -0.1025 | -0.39 | 0.796577 | 0.012959 | 0.810387 | 1 | 0.999631 | 0.929952 |
| Gas7_B1ATI9_S88s_1 | 1.22 | 0.67 | -0.37 | 1.09 | 0.76 | 1.06 | 0.81 | 1.29 | 1.1 | 0.13 | 1.07 | -0.1 | 0.03 | 0.41 | 0.5 | 0.11 | 0.7675 | 0.3675 | -0.565 | 0.343156 | 0.409437 | 0.824965 | 1 | 0.999631 | 0.73348 |
| Gas7_B1ATI9_S92s_1 | 1.27 | 0.09 | -1.38 | 0.92 | 1.2 | 1.06 | 1.19 | 0.52 | 1.16 | 0.29 | 0.75 | 0.17 | -1.32 | 0.4 | -0.24 | -0.2 | -0.1525 | -0.11 | -0.8125 | 0.375168 | 0.191164 | 0.530055 | 1 | 0.999631 | 0.748001 |
| Gas7_B1ATI9_S98s_1 | -1.89 | -4.46 | -0.56 | 0.1 | -3.9 | -3.43 | 0.21 | -0.3 | -4.17 | -2.98 | -0.06 | -0.04 | -4.89 | -3.95 | -0.3 | -0.92 | -0.0125 | 0.0625 | -0.4925 | 0.527942 | 0.902329 | 0.92938 | 1 | 0.999631 | 0.824683 |
| Gas7_B1ATI9_S99s_1 | 1.16 | 1.26 | 0.98 | 1.33 | 0.8 | 1.55 | 1.2 | 1.13 | 1.73 | 1.65 | 1.02 | 0.58 | 0 | 0.84 | 1.07 | 0.85 | -0.405 | 0.1225 | -0.29 | 0.153308 | 0.962663 | 0.851216 | 1 | 0.999631 | 0.625198 |
| Gatad2b_Q8VHR5_S335s_1 | -0.01 | 0.21 | 0.76 | 0.29 | -0.47 | 0.35 | -0.67 | 0.42 | -0.15 | 0.27 | 0.54 | 1.08 | 0.15 | 0.24 | -1.59 | 1.29 | -0.165 | -0.07 | 0.295 | 0.613155 | 0.265626 | 0.724058 | 1 | 0.999631 | 0.860866 |
| Gatad2b_Q8VHR5_S339s_1 | -0.27 | 0.18 | 0.16 | 0.17 | -0.22 | -0.52 | 0.2 | 0.12 | 0.1 | -0.38 | -0.04 | 0.28 | 0.29 | -0.06 | 0.33 | 0.86 | 0.1225 | 0.3 | 0.9075 | 0.354952 | 0.559335 | 0.801221 | 1 | 0.999631 | 0.740147 |
| Gatad2b_Q8VHR5_S487s_1 | -1.22 | -0.43 | 0.16 | -0.02 | -0.79 | 0.13 | -0.08 | -0.28 | 0.53 | -1.33 | 0.37 | 0.12 | 0.29 | 0.17 | 0.92 | 0.74 | -0.51 | 0.33 | 0.5625 | 0.038241 | 0.746863 | 0.550984 | 1 | 0.999631 | 0.459475 |
| Gatad2b_Q8VHR5_T121t_1 | 0.25 | 1.08 | 0.03 | 0.19 | 0.15 | 0.35 | -0.65 | -0.34 | 0.7 | 0.14 | 0.42 | 1.61 | 0.53 | 1.18 | 0.48 | 1.61 | 0.0325 | 0.2975 | 0.16 | 0.170879 | 0.172553 | 0.424452 | 1 | 0.999631 | 0.641032 |
| Gba2_Q69ZF3_S893s_1 | 0.54 | 0.62 | 0.3 | 0.67 | 0.1 | 0.87 | 0.55 | 0.74 | 0.92 | 0.56 | 0.94 | 0.9 | 0.31 | 1.08 | 0.32 | 1.06 | 0.1275 | 0.1975 | 0.455 | 0.61722 | 0.905705 | 0.261957 | 1 | 0.999631 | 0.861376 |
| Gbf1_Q6DFZ1_S1318s_1 | 0.34 | 1.75 | 0.51 | 0.24 | 1.28 | 1.35 | -0.04 | 0.76 | 1.95 | 1 | 0.08 | 0.6 | 1.44 | 2.2 | 0.55 | 0.47 | -0.2675 | -0.1475 | 0.7775 | 0.384094 | 0.780603 | 0.694572 | 1 | 0.999631 | 0.754721 |
| Gbf1_Q6DFZ1_S1786s_1 | 0.02 | 0.79 | 0.35 | 0.11 | -0.53 | 1.32 | -0.3 | -0.29 | 1.17 | 0.1 | 0.03 | -0.62 | 1.02 | 0.86 | 1.09 | 1.41 | -0.065 | 0.1925 | 0.1 | 0.027301 | 0.552423 | 0.722698 | 1 | 0.999631 | 0.423171 |
| Gbf1_Q6DFZ1_S1791s_1 | 1.25 | 0.16 | 1.04 | 0.73 | -0.19 | 0.48 | 0.89 | 1.74 | 0.95 | 1.02 | 1.21 | 0.77 | 0.55 | 0.8 | 1.38 | 0.85 | -0.31 | 0.0875 | -1.3025 | 0.775988 | 0.885539 | 0.546744 | 1 | 0.999631 | 0.920998 |
| Gcfc2_S4R198_S169s_1 | 0.21 | -0.37 | 1.11 | 0.95 | -0.01 | -0.72 | 0.63 | 0.76 | 0.73 | 0.38 | 0.52 | 0.62 | -1.7 | -0.91 | 0.14 | -0.84 | 0.1375 | -0.0675 | 0.3475 | 0.022735 | 0.507684 | 0.815944 | 1 | 0.999631 | 0.409409 |
| Gdap1l1_Q8VE33_S139s_1 | -0.86 | -0.21 | 0.04 | -1.2 | 0.78 | -0.42 | -0.98 | -1.06 | -0.29 | -0.83 | -0.55 | -0.83 | 0.6 | 0.05 | -1.25 | -0.24 | 0.2075 | -0.155 | 0.215 | 0.46744 | 0.776645 | 0.847982 | 1 | 0.999631 | 0.795527 |
| Gdap1l1_Q8VE33_S37s_1 | 0.42 | -0.22 | 0.21 | 0.06 | 0.04 | 1.02 | 0.18 | 0.06 | -0.02 | 0.18 | -0.48 | 0.17 | 0.47 | 0.78 | -0.25 | 0.33 | 0.2075 | -0.155 | 0.215 | 0.518045 | 0.519583 | 0.596297 | 1 | 0.999631 | 0.819449 |
| Gdap1l1_Q8VE33_S38s_1 | 0.42 | -0.22 | 0.21 | 0.06 | 0.04 | 1.02 | 0.18 | 0.06 | -0.02 | 0.18 | -0.48 | 0.17 | 0.47 | 0.78 | -0.25 | 0.33 | -0.3925 | -0.5675 | 0.16 | 0.518045 | 0.519583 | 0.596297 | 1 | 0.999631 | 0.819449 |
| Gdi1_P50396_S121s_1 | -0.66 | 0.18 | -0.29 | -0.15 | -0.73 | -0.54 | -1.01 | -0.21 | 0.06 | -0.33 | -2.02 | -0.9 | 0.52 | 0.81 | -0.84 | -0.77 | 0.175 | 0.12 | -0.38 | 0.727485 | 0.214616 | 0.244446 | 1 | 0.999631 | 0.905923 |
| Gemin5_E9PUU4_S1410s_2 | 1.36 | 1.76 | 1.17 | 1.22 | 1.65 | 1.84 | 1.13 | 1.59 | 1.63 | 1.77 | 1.61 | 0.98 | -0.66 | 1.88 | 1.15 | 1.62 | -0.5 | 0.2825 | 0.8325 | 0.493278 | 0.538031 | 0.688968 | 1 | 0.999631 | 0.80955 |
| Gemin5_E9PUU4_S1414s_1 | -0.56 | 0.55 | 0.92 | -0.74 | -0.74 | -0.01 | -0.87 | -0.21 | -0.02 | 0.99 | -0.34 | 0.67 | 2.36 | 0.79 | -0.16 | 0.51 | 0.23 | 0.4225 | 0.2775 | 0.193353 | 0.276419 | 0.565727 | 1 | 0.999631 | 0.650018 |
| Gemin5_Q8BX17_S1410s_2 | 0.66 | 1.63 | 0.22 | -0.46 | 1.5 | 1.17 | 0.38 | -0.08 | 0.8 | 1.39 | 0.67 | 0.88 | 0.21 | 1.35 | 0.35 | 1.25 | 0.23 | 0.4225 | 0.2775 | 0.585323 | 0.663258 | 0.364105 | 1 | 0.999631 | 0.849461 |
| Gemin5_Q8BX17_S1414s_2 | 0.66 | 1.63 | 0.22 | -0.46 | 1.5 | 1.17 | 0.38 | -0.08 | 0.8 | 1.39 | 0.67 | 0.88 | 0.21 | 1.35 | 0.35 | 1.25 | -0.1775 | -0.085 | -1.775 | 0.585323 | 0.663258 | 0.364105 | 1 | 0.999631 | 0.849461 |
| Gemin7_D3Z684_S3s_1 | -2.02 | 0.25 | -0.31 | -0.26 | -0.98 | -1.79 | -0.22 | -0.06 | -1.03 | -0.67 | -0.33 | -0.65 | -0.05 | -3.18 | -2.76 | -3.45 | -0.02 | 0.0075 | 0.495 | 0.049142 | 0.758758 | 0.862575 | 1 | 0.999631 | 0.487138 |
| Gfap_P03995_S12s_1 | 0.05 | 0.21 | 0.8 | -0.16 | -0.37 | 1.21 | -0.01 | -0.01 | -0.13 | 0.45 | 0.75 | -0.14 | 1.08 | 0.79 | 0.9 | 0.11 | -0.2025 | -0.9775 | -1.5325 | 0.180542 | 0.960547 | 0.982623 | 1 | 0.999631 | 0.64376 |
| Gfap_P03995_S390s_1 | 2.6 | 0.32 | -0.35 | 0.66 | 0.79 | 1.23 | 1.15 | -0.75 | 0.37 | 0.41 | -0.26 | -1.2 | -1.55 | -0.1 | -0.5 | -0.75 | -0.3175 | 0.0625 | 0.84 | 0.03396 | 0.770804 | 0.162249 | 1 | 0.999631 | 0.445973 |
| Gfap_P03995_T40t_1 | -0.67 | -0.34 | 0.11 | -0.47 | -0.37 | -0.68 | -1.29 | -0.3 | -0.15 | -0.04 | -0.47 | -0.46 | 0.71 | 0.26 | 0.61 | 0.41 | 0.065 | 0.0275 | 0.7075 | 0.017479 | 0.340376 | 0.827825 | 1 | 0.999631 | 0.391024 |
| Gfpt1_P47856_S243s_1 | -0.23 | 1.75 | 0.28 | 0.22 | 0.73 | 0.97 | -0.01 | 0.59 | 0.74 | 0.29 | 0.11 | 0.99 | 1.11 | 1.53 | 1.24 | 0.97 | -0.1475 | -0.1025 | 0.2925 | 0.136916 | 0.887374 | 0.952496 | 1 | 0.999631 | 0.610369 |
| Gga3_S4R2D2_S159s_1 | -0.07 | -0.12 | 0.14 | 0.17 | 0.01 | -0.05 | -0.45 | 0.02 | -0.17 | 0.18 | -0.32 | 0.02 | 0.64 | 0.26 | 0.15 | 0.24 | -0.1025 | 0.07 | 0.4125 | 0.30308 | 0.561426 | 0.69348 | 1 | 0.999631 | 0.714978 |
| Ggt7_Q99JP7_S83s_1 | -0.37 | -0.29 | -0.13 | -1.23 | -0.47 | -0.21 | -1.35 | -0.4 | -0.52 | 0.15 | -0.83 | -0.54 | 0.47 | 0.09 | -0.36 | -0.57 | 0.5125 | 0.2825 | -0.1675 | 0.289043 | 0.782903 | 0.844689 | 1 | 0.999631 | 0.706046 |
| Gigyf1_Q99MR1_S137s_1 | 0.52 | 0.64 | 0.13 | 1.52 | 1.32 | 0.9 | 1.11 | 1.53 | 0.93 | 0.28 | 1.66 | 1.07 | -1.02 | 1.07 | 1.1 | 0.99 | 0.43 | 0.4875 | -0.295 | 0.762628 | 0.167585 | 0.500236 | 1 | 0.999631 | 0.915369 |
| Gigyf1_Q99MR1_S157s_1 | 0.04 | -0.27 | -0.57 | 0.84 | 0.8 | -0.76 | 1.1 | 0.62 | 0.78 | -0.72 | 0.99 | 0.94 | -0.83 | -0.16 | 0.46 | -0.61 | 0.0725 | 0.175 | 0.0625 | 0.49429 | 0.384202 | 0.329243 | 1 | 0.999631 | 0.81007 |
| Gigyf1_Q99MR1_S24s_1 | -0.69 | 0.91 | 0.36 | -0.13 | 0.26 | 0.42 | -0.14 | 0.2 | 0.85 | -0.79 | 0.25 | 0.84 | -0.41 | 0.84 | 0.13 | 0.14 | -5.9E-17 | -0.12 | 0.385 | 0.885355 | 0.847347 | 0.722332 | 1 | 0.999631 | 0.963505 |
| Gigyf1_Q99MR1_S413s_1 | -0.31 | -0.52 | -1.16 | -0.19 | -0.02 | -0.41 | -0.53 | -1.22 | -0.86 | -0.07 | -1.56 | -0.17 | 0.22 | -0.32 | -0.67 | 0.13 | -0.0975 | 0.3075 | 0.47 | 0.29107 | 1 | 0.772306 | 1 | 0.999631 | 0.706164 |
| Gigyf1_Q99MR1_S540s_1 | -0.63 | -0.46 | -0.48 | 0 | -0.51 | -0.73 | -0.63 | -0.09 | -0.3 | -0.19 | -0.46 | 0.61 | 0.34 | -0.62 | 0.24 | 0.35 | 0.09 | -0.065 | 0.075 | 0.185111 | 0.727081 | 0.359893 | 1 | 0.999631 | 0.646072 |
| Gigyf1_Q99MR1_S755s_1 | -0.42 | 0.22 | 0.66 | -0.38 | -0.31 | 0.11 | -0.21 | 0.85 | -0.08 | -0.28 | -0.05 | 0.23 | -1 | -0.37 | 1.18 | 0.57 | 0.09 | -0.065 | 0.075 | 0.885172 | 0.813567 | 0.8443 | 1 | 0.999631 | 0.963505 |
| Gigyf1_Q99MR1_S756s_1 | -0.42 | 0.22 | 0.66 | -0.38 | -0.31 | 0.11 | -0.21 | 0.85 | -0.08 | -0.28 | -0.05 | 0.23 | -1 | -0.37 | 1.18 | 0.57 | 0.555 | 0.655 | 0.595 | 0.885172 | 0.813567 | 0.8443 | 1 | 0.999631 | 0.963505 |
| Gigyf2_G3UWR2_S161s_1 | 0.37 | 0.38 | 0.68 | 1.11 | 0.8 | 1.39 | 0.89 | 1.68 | 1.04 | 1.2 | 1.5 | 1.42 | 0.3 | 1.24 | 1.68 | 1.7 | 0.805 | 0.5375 | -0.02 | 0.15653 | 0.107428 | 0.044681 | 1 | 0.999631 | 0.626488 |
| Gigyf2_G3UWR2_S202s_1 | 1.48 | 1.65 | 0.35 | 1.94 | 2.35 | 2.5 | 1.61 | 2.18 | 2.54 | 1.5 | 2.19 | 1.34 | -0.76 | 2.66 | 1.89 | 1.55 | 0.28 | 0.415 | 0.9725 | 0.977791 | 0.069315 | 0.243484 | 1 | 0.999631 | 0.992341 |
| Gigyf2_G3UWR2_S237s_1 | -0.43 | 1.09 | 0.84 | 0.41 | 0.58 | 1.43 | 0.37 | 0.65 | 1.14 | 0.75 | 0.7 | 0.98 | 1.06 | 1.72 | 1.53 | 1.49 | 0.15 | 0.8 | 1.785 | 0.02997 | 0.49722 | 0.283104 | 1 | 0.999631 | 0.435495 |
| Gigyf2_G3UWR2_T25t_1 | -1.27 | 1.17 | -0.28 | -0.82 | -0.37 | 0.21 | 0.05 | -0.49 | 0.14 | 0.42 | 0.05 | 1.39 | 1.86 | 0.54 | 2.04 | 1.5 | -0.1525 | -0.05 | 0.4375 | 0.010712 | 0.77197 | 0.178763 | 1 | 0.999631 | 0.383871 |
| Git1_Q5F258_S361s_1 | -0.89 | -0.2 | -0.56 | -1.25 | -0.33 | -0.7 | -1.41 | -1.07 | -0.65 | -0.77 | -0.9 | -0.78 | 0.36 | -0.34 | -0.67 | -0.5 | -0.1925 | 0.0125 | 0.32 | 0.248028 | 0.666187 | 0.869151 | 1 | 0.999631 | 0.685751 |
| Git1_Q5F258_S362s_1 | -0.84 | -0.48 | -0.46 | -0.78 | -0.81 | -0.82 | -0.95 | -0.75 | -0.78 | -0.74 | -0.52 | -0.47 | 0.1 | -0.52 | -0.56 | -0.3 | -0.1 | -0.6325 | -0.325 | 0.291631 | 0.435728 | 0.961202 | 1 | 0.999631 | 0.706164 |
| Git1_Q5F258_S561s_1 | 0.45 | 0.05 | -0.39 | -0.03 | -0.34 | 0.35 | -0.5 | 0.17 | 0.26 | -0.29 | -1.36 | -1.06 | 0.03 | -0.18 | -0.38 | -0.69 | -0.235 | 0.0275 | 0.0125 | 0.318438 | 0.752426 | 0.149276 | 1 | 0.999631 | 0.72215 |
| Git2_F7BIK4_S360s_1 | 1.32 | 0.72 | 0.56 | 0.8 | 0.51 | 0.41 | 0.61 | 0.93 | 0.95 | 1.48 | 0.34 | 0.74 | 0.75 | 0.7 | 1.41 | 0.59 | 0.9 | -0.3525 | -0.385 | 0.969366 | 0.410622 | 0.934834 | 1 | 0.999631 | 0.989089 |
| Gja1_P23242_S255s_1 | -1.4 | -2.21 | -3.99 | -2.37 | -1.53 | -1.41 | -1.38 | -2.05 | -1.75 | -2.23 | -3.84 | -3.56 | -3.06 | -2.04 | -3.83 | -2.58 | -0.6275 | -0.305 | 0.375 | 0.527379 | 0.110471 | 0.598801 | 1 | 0.999631 | 0.824554 |
| Gja1_P23242_S306s_1 | -0.06 | 0.15 | -0.91 | -0.08 | 0.06 | -0.65 | -0.03 | -2.79 | -0.11 | 0.05 | -1.1 | -0.96 | 0.24 | 0.27 | -0.32 | 0.41 | -0.5925 | -0.29 | -0.0725 | 0.291994 | 0.333417 | 0.442713 | 1 | 0.999631 | 0.706164 |
| Gja1_P23242_S314s_1 | -1.24 | -1.5 | -0.49 | -0.59 | -1.49 | -2.47 | -1.12 | -1.11 | -2.06 | -1.7 | -0.8 | -0.42 | -0.84 | -2.16 | -0.63 | -0.48 | -0.365 | -0.125 | 0.3625 | 0.873064 | 0.166865 | 0.520989 | 1 | 0.999631 | 0.959648 |
| Gja1_P23242_S328s_1 | -0.9 | -1.03 | -0.93 | -1.31 | -1.37 | -1.18 | -1.52 | -1.56 | -1.21 | -0.68 | -1.89 | -0.89 | -0.38 | -0.85 | -0.83 | -0.66 | -0.19 | 0.4 | -0.115 | 0.214848 | 0.165972 | 0.707074 | 1 | 0.999631 | 0.667764 |
| Gja1_P23242_S364s_1 | -1.02 | -0.81 | 0.35 | 0.27 | -0.53 | -1.45 | -0.02 | 0.03 | -0.64 | -0.73 | 0.24 | 1.52 | -1.01 | -1.34 | 0.72 | -0.04 | -0.305 | 0.2175 | -0.31 | 0.833988 | 0.68804 | 0.495664 | 1 | 0.999631 | 0.943876 |
| Gja1_P23242_S365s_1 | -0.48 | 0.55 | 1.25 | 0.96 | 0.76 | -0.8 | 0.08 | 1.02 | 0.39 | -0.38 | 1.2 | 1.94 | -0.11 | -0.69 | 1.32 | 0.52 | -0.03 | 0.3775 | 0.7275 | 0.5671 | 0.558422 | 0.70691 | 1 | 0.999631 | 0.842252 |
| Gja1_P23242_S373s_1 | -1.13 | -1.31 | -0.26 | -1.03 | -1.16 | -0.77 | -1.31 | -0.61 | -0.85 | -0.76 | -0.69 | 0.08 | 0.07 | -0.8 | 0.19 | -0.28 | -0.525 | -0.5875 | -0.445 | 0.070332 | 0.926832 | 0.299374 | 1 | 0.999631 | 0.530212 |
| Gjc2_Q8BQU6_T190t_1 | 1.02 | 1.04 | 1.79 | 0.84 | 0.5 | 0.46 | 0.23 | 1.4 | 0.7 | 0.13 | 1.29 | 0.22 | 0.75 | 0.42 | 0.42 | 1.32 | -0.5975 | -1.2575 | -0.38 | 0.226821 | 0.165112 | 0.134717 | 1 | 0.999631 | 0.678076 |
| Gjc3_Q921C1_S249s_1 | -1.3 | -0.99 | 0.18 | -1.43 | -2.99 | -0.75 | -1.07 | -1.12 | -3.59 | -1.52 | -1.34 | -2.12 | -1.46 | -1.16 | -2.03 | -0.41 | -0.1375 | 0.42 | 0.0475 | 0.438811 | 0.310033 | 0.051545 | 1 | 0.999631 | 0.781983 |
| Glcci1_E9QKK4_S137s_1 | 1.07 | 1.19 | 0.77 | 1.05 | 0.16 | 1.23 | 0.96 | 1.18 | 1.74 | 1.74 | 1.35 | 0.93 | 1.07 | 0.9 | 1.36 | 0.94 | 0.1375 | -0.115 | -0.725 | 0.864259 | 0.663849 | 0.175917 | 1 | 0.999631 | 0.955131 |
| Glcci1_E9QKK4_S140s_1 | 1.62 | 1.26 | 1.44 | 1.28 | 1.34 | 2.25 | 1.2 | 1.36 | 1.85 | 1.92 | 1.03 | 0.34 | 0.2 | 1.18 | 1.28 | 0.04 | 0.03 | 0.355 | 0.635 | 0.076101 | 0.658622 | 0.772495 | 1 | 0.999631 | 0.542138 |
| Glcci1_E9QKK4_S215s_1 | -0.01 | 0.78 | 1.08 | 0.78 | -0.11 | 0.68 | 0.98 | 1.2 | 0.68 | 0.06 | 1.52 | 1.79 | 0.34 | 0.97 | 1.83 | 2.03 | -0.7025 | -0.3 | -0.8275 | 0.182847 | 0.937364 | 0.438372 | 1 | 0.999631 | 0.643861 |
| Glcci1_E9QKK4_S26s_1 | -0.08 | 0.7 | -0.1 | -1.17 | -0.71 | -0.59 | -0.8 | -1.36 | 0.35 | -0.2 | -0.7 | -1.3 | 0.22 | -1.71 | -0.3 | -2.17 | -0.07 | 0.1075 | 0.0425 | 0.202012 | 0.116988 | 0.548668 | 1 | 0.999631 | 0.658952 |
| Glcci1_E9QKK4_S336s_1 | -0.27 | -0.54 | 0.34 | -0.29 | -0.62 | -0.14 | -0.05 | -0.23 | 0.13 | -0.41 | -0.66 | 0.61 | -0.37 | 0.09 | -0.02 | -0.29 | 0.9225 | 0.585 | 0.42 | 0.890996 | 0.811742 | 0.770734 | 1 | 0.999631 | 0.96413 |
| Glcci1_E9QKK4_S389s_1 | 0.38 | -0.4 | -0.76 | 1.29 | 0.68 | 0.58 | 1.48 | 1.46 | 0.54 | -1.21 | 1.58 | 1.94 | -0.08 | 0.41 | 1.36 | 0.5 | 0.005 | 0.115 | 0.23 | 0.423614 | 0.082851 | 0.438543 | 1 | 0.999631 | 0.773835 |
| Glod4_F6ZTG3_S109s_1 | 0.01 | 0.38 | 0.06 | 0.36 | 0.22 | 0.1 | 0.18 | 0.33 | 0.25 | -0.2 | 0.12 | 1.1 | 0.7 | -0.21 | 1 | 0.24 | -0.0175 | -0.0825 | 0.03 | 0.508867 | 0.983581 | 0.7356 | 1 | 0.999631 | 0.816932 |
| Glrb_P48168_T391t_1 | -0.6 | -0.7 | -0.47 | -0.49 | -1.06 | -0.49 | -0.39 | -0.39 | -0.95 | -0.6 | -1.18 | 0.14 | -1.35 | -0.07 | -0.53 | -0.19 | -0.865 | -0.87 | -1.525 | 0.931735 | 0.947574 | 0.808013 | 1 | 0.999631 | 0.978006 |
| Gltscr1l_Q8CHH5_S930s_1 | 2.29 | 0.52 | 2.19 | 1.58 | -0.35 | 0.11 | 1.87 | 1.49 | 0.96 | 2.02 | -0.25 | 0.37 | -0.15 | -1.16 | 1.12 | 0.67 | -0.235 | -0.12 | 0.57 | 0.025561 | 0.17537 | 0.156353 | 1 | 0.999631 | 0.422114 |
| Gltscr2_Q8BK35_S305s_1 | -0.58 | 0.07 | 0.08 | -0.82 | -0.53 | -1.47 | -0.46 | 0.27 | -0.22 | -1.6 | -1.15 | 1.24 | -0.23 | -1.13 | 0.56 | 1.83 | 0.1 | -0.0825 | 0.1325 | 0.359057 | 0.578707 | 0.843327 | 1 | 0.999631 | 0.741182 |
| Glud1_P26443_S450s_1 | -1.05 | -0.96 | 1.15 | -0.19 | -0.78 | -0.64 | 0.38 | 0.39 | -1.13 | -1.4 | 0.79 | 0.36 | 0.05 | -1.2 | 0.69 | -0.06 | -0.1575 | -0.33 | 0.05 | 0.821959 | 0.855804 | 0.901334 | 1 | 0.999631 | 0.938464 |
| Glul_P15105_S19s_1 | 0.1 | 0.89 | 0.53 | -0.36 | 0.82 | 0.33 | 0.21 | -0.83 | -0.3 | 0.62 | 0.02 | -0.5 | 0.43 | 0.72 | -0.74 | 0.95 | -0.535 | -0.1175 | -0.27 | 0.913002 | 0.715676 | 0.400003 | 1 | 0.999631 | 0.97231 |
| Glul_P15105_S343s_1 | 0.2 | 0.79 | -0.06 | -0.17 | 0.38 | 0.11 | -1.14 | -0.73 | 0.62 | 0.6 | -0.67 | -0.26 | 0.73 | 0.56 | -1.05 | -0.56 | 0.745 | 0.405 | 0.56 | 0.570149 | 0.215556 | 0.768856 | 1 | 0.999631 | 0.84338 |
| Gm13889_Q3UPL5_S210s_1 | 0.54 | 1.18 | 0.34 | 1.11 | 1.64 | 1.65 | 0.77 | 2.09 | 1.63 | 1.09 | 1.57 | 0.5 | -0.02 | 2.55 | 2 | 0.88 | -0.395 | 0.3275 | 0.54 | 0.333746 | 0.065635 | 0.282823 | 1 | 0.999631 | 0.728079 |
| Gm15800_E9Q2E4_S1496s_1 | -0.55 | -0.76 | -0.72 | -1.18 | -1.05 | -1.96 | -0.76 | -1.02 | -0.29 | -0.43 | -0.56 | -0.62 | 1.67 | -1.69 | -0.37 | -0.66 | 0.0475 | 0.195 | 0.2875 | 0.410234 | 0.253822 | 0.238564 | 1 | 0.999631 | 0.767924 |
| Gm15800_E9Q2E4_S1498s_1 | 0.14 | -0.86 | -1.34 | -1.36 | -0.57 | -0.43 | -0.82 | -1.41 | -0.09 | -0.61 | -1.31 | -0.63 | 0.27 | 0.29 | -2.24 | -0.59 | -0.06 | 0.295 | 0.4525 | 0.64662 | 0.908121 | 0.652081 | 1 | 0.999631 | 0.874181 |
| Gm15800_E9Q2E4_S1562s_1 | -0.72 | -0.02 | 0.21 | -0.76 | -0.48 | -0.35 | -0.64 | -0.06 | 0.03 | -0.84 | 0.14 | 0.56 | 0.73 | -0.63 | 0.06 | 0.36 | 0.43 | 0.4325 | 0.5275 | 0.275772 | 0.852026 | 0.463929 | 1 | 0.999631 | 0.700187 |
| Gm15800_E9Q2E4_S1916s_1 | -1.94 | -0.08 | -1.02 | -1.59 | -0.25 | -0.83 | -0.17 | -1.66 | 0.07 | -2.07 | -1.13 | 0.23 | 0.89 | -1.11 | -1.27 | -1.03 | -0.2075 | -0.0025 | 0.3125 | 0.384851 | 0.398295 | 0.487009 | 1 | 0.999631 | 0.75538 |
| Gm15800_E9Q2E4_T2503t_1 | -0.17 | 0.21 | -0.03 | -0.45 | -0.07 | -0.15 | -0.52 | -0.53 | -0.23 | 0.21 | -0.29 | -0.14 | 0.63 | 0 | -0.02 | 0.2 | -0.01 | -0.1025 | 0.21 | 0.318117 | 0.453574 | 0.992809 | 1 | 0.999631 | 0.721765 |
| Gm20521_D3Z5F7_S177s_1 | -0.65 | 0.44 | 0.31 | -0.56 | 0.27 | 0.01 | -0.28 | -0.5 | -0.13 | -0.45 | -0.28 | -0.01 | 0.47 | -0.2 | -0.1 | 0.21 | 0.5575 | -0.0525 | 0.0325 | 0.570919 | 0.977594 | 0.76578 | 1 | 0.999631 | 0.84338 |
| Gm27021_M0QWE2_T124t_1 | -0.72 | -0.11 | -0.63 | -0.99 | -0.18 | 0.21 | -0.32 | 0.07 | 0.1 | -1.07 | -0.62 | -1.07 | 0.84 | -0.12 | -1.82 | -1.22 | -0.2025 | -0.24 | -0.03 | 0.95448 | 0.079539 | 0.885066 | 1 | 0.999631 | 0.984441 |
| Gm27029_Q3THG9_S88s_1 | -0.68 | -0.66 | -0.38 | -0.83 | -1.27 | -0.61 | -0.73 | -0.75 | -1.45 | -0.23 | -0.97 | -0.86 | -0.65 | -0.76 | -1.09 | -0.17 | 0.0725 | -0.05 | 0.71 | 0.922422 | 0.45734 | 0.465806 | 1 | 0.999631 | 0.975032 |
| Gm28036_F6YJB6_S336s_1 | -0.31 | 1.57 | 1.63 | 0.37 | 0.54 | 1.97 | 0.57 | 0.47 | 1.23 | 0.7 | 0.24 | 0.89 | 0.01 | 2.44 | 2.18 | 1.47 | -0.185 | 0.23 | 0.4175 | 0.288589 | 0.894627 | 0.91911 | 1 | 0.999631 | 0.706046 |
| Gm7694_J3QNH8_S221s_1 | 0.14 | -0.17 | -0.45 | -0.17 | -0.12 | -0.4 | -0.24 | -0.63 | 0.25 | 0.47 | -0.32 | -0.13 | 0.15 | 0.32 | 0.21 | 0.34 | -0.0375 | 0.505 | -0.1575 | 0.15204 | 0.488516 | 0.444393 | 1 | 0.999631 | 0.625198 |
| Gm9774_Q9JKV1_S211s_1 | 0.58 | 1.58 | 1.89 | 0.91 | 1.15 | 1.22 | 0.92 | 1.52 | 1.69 | 1.24 | 1.99 | 2.06 | 0.83 | 0.51 | 1.35 | 1.64 | 0.3125 | -0.265 | -0.515 | 0.702703 | 0.915268 | 0.201435 | 1 | 0.999631 | 0.896857 |
| Gm9774_Q9JKV1_S213s_1 | 1.36 | 1.67 | 1.47 | 1.2 | 1.56 | 2.59 | 1.27 | 1.53 | 1.22 | 1.8 | 1.95 | -0.33 | -0.71 | 2.28 | 0.89 | 1.18 | 0.21 | -0.015 | -0.67 | 0.380651 | 0.371271 | 0.600434 | 1 | 0.999631 | 0.752619 |
| Gm9774_Q9JKV1_S213s_2 | 1.91 | 0.81 | 0.91 | 2.04 | 1.55 | 2.23 | 1.41 | 1.32 | 1.91 | 2.5 | 0.9 | 0.3 | -0.65 | 1.87 | 0.93 | 0.84 | 0.6125 | 0.465 | 0.265 | 0.253849 | 0.594614 | 0.978137 | 1 | 0.999631 | 0.688747 |
| Gm996_A2AJA9_S109s_1 | -3.57 | -1.17 | -1.21 | -4.12 | -0.84 | -1.84 | -1.34 | -3.6 | -0.91 | -5.4 | -1.29 | -0.61 | -0.09 | -3.39 | -1.48 | -4.05 | -0.14 | -0.23 | 0.425 | 0.794509 | 0.47811 | 0.692134 | 1 | 0.999631 | 0.928784 |
| Gm996_A2AJA9_S286s_1 | -0.75 | -2.05 | -0.51 | -1.23 | -1.21 | -1.14 | -1.27 | -1.48 | -1.26 | -1.19 | -2.22 | -0.79 | -0.57 | -0.63 | -0.75 | -0.89 | 0.35 | 0.7975 | -0.1125 | 0.281283 | 0.704084 | 0.609717 | 1 | 0.999631 | 0.702476 |
| Gm996_A2AJA9_S435s_1 | -1.46 | 0.1 | -0.2 | -1.52 | 0.19 | -0.9 | -0.05 | -0.92 | 0.35 | -1.54 | 0.44 | 0.86 | -0.24 | -1.25 | -0.58 | -1.46 | -0.425 | -0.3675 | 0.225 | 0.818385 | 0.47382 | 0.215776 | 1 | 0.999631 | 0.938303 |
| Gm996_A2AJA9_S466s_1 | -0.51 | -0.38 | -0.28 | -0.79 | -1.66 | -1.14 | -0.92 | 0.06 | -1.5 | -0.63 | -0.67 | -0.63 | 0.38 | -0.65 | 0.07 | -0.86 | -0.0675 | -0.3525 | -1.0975 | 0.537987 | 0.287388 | 0.252377 | 1 | 0.999631 | 0.830863 |
| Gm996_A2AJA9_S509s_1 | 0.64 | 0.72 | 0.46 | 0.31 | 0.83 | 0.89 | 0.95 | -0.81 | 0.71 | 0.64 | -0.13 | -0.5 | 0.45 | -0.79 | -0.55 | -1.37 | -0.3375 | -0.1925 | 0.4475 | 0.021574 | 0.8747 | 0.326208 | 1 | 0.999631 | 0.405125 |
| Gm996_A2AJA9_S560s_1 | -1.48 | -1.17 | -0.41 | -1.16 | -1.85 | -1.23 | -1.48 | -1.01 | -1.57 | -1.19 | -1.68 | -0.55 | -0.15 | -1.44 | 0.19 | -1.03 | 0.4725 | 0.425 | -0.205 | 0.325601 | 0.321197 | 0.604969 | 1 | 0.999631 | 0.724678 |
| Gm996_A2AJA9_S697s_1 | 0.55 | 1.15 | 1.48 | -0.23 | 0.79 | 1.32 | 2.49 | 0.24 | 0.76 | 0.41 | 1.99 | 1.49 | 1.98 | -0.46 | 0.81 | -0.2 | -0.125 | -0.0475 | 0.295 | 0.736883 | 0.406087 | 0.397953 | 1 | 0.999631 | 0.907709 |
| Gm996_A2AJA9_S761s_1 | -1.42 | -1.91 | -2.44 | -2.19 | -2.45 | -2.55 | -1.78 | -1.68 | -2.86 | -1.41 | -2.48 | -1.4 | -0.92 | -2.72 | -1.25 | -1.89 | -0.3425 | 0.0025 | 0.2825 | 0.517823 | 0.718052 | 0.91227 | 1 | 0.999631 | 0.819449 |
| Gm996_A2AJA9_S821s_1 | -0.94 | -1.18 | -0.97 | -1.46 | -1.5 | -1.36 | -1.61 | -1.45 | -1.46 | -0.85 | -1.1 | -1.13 | 0.01 | -1.24 | -1.36 | -0.83 | 0.065 | 0.4125 | 0.335 | 0.455658 | 0.193097 | 0.992762 | 1 | 0.999631 | 0.790041 |
| Gmps_Q3THK7_S332s_1 | -0.25 | 0.52 | 0.72 | -0.16 | 0.43 | 0.23 | 0.19 | 0.24 | 0.62 | 0.17 | 0.66 | 1.03 | 1.03 | 0.4 | 0.42 | 0.32 | -0.15 | 0.09 | -0.165 | 0.34982 | 0.831997 | 0.247142 | 1 | 0.999631 | 0.736956 |
| Gnao1_P18872_S29s_1 | -0.18 | -1.26 | -2.18 | -2.07 | -0.94 | -0.98 | -1.66 | -2.71 | -0.72 | -0.46 | -1.91 | -2.24 | -0.59 | -1.14 | -2.44 | -2.18 | -0.5975 | -0.9375 | -0.875 | 0.777235 | 0.791231 | 0.87722 | 1 | 0.999631 | 0.921382 |
| Gnas_Z4YKV1_S337s_1 | 1.22 | -0.79 | -0.11 | -0.22 | -1.32 | -0.32 | 0.16 | -0.81 | -1.64 | 0.34 | -1.05 | -1.3 | -0.3 | -0.84 | -0.87 | -1.39 | 0.915 | -0.9375 | 2.0125 | 0.087009 | 0.248445 | 0.119663 | 1 | 0.999631 | 0.553514 |

| | | | | | | | | | | | | | | | | | | | | | | | | | |
|---|---|---|---|---|---|---|---|---|---|---|---|---|---|---|---|---|---|---|---|---|---|---|---|---|---|
| Gnaz_O70443_S27s_1 | -3.25 | 1 | -1.43 | -0.65 | 1.34 | 1.05 | -0.11 | -2.95 | -3.27 | -4.52 | -1.45 | 1.16 | 1.45 | 0.24 | 1.47 | 0.56 | -0.6825 | -0.425 | -1.0225 | 0.030992 | 0.425142 | 0.475129 | 1 | 0.999631 | 0.44065 |
| Gnb5_P62881_S63s_1 | 1.68 | 1.44 | 1.39 | 1.03 | 0.39 | 1.6 | 0.67 | 0.15 | 1.21 | 2.08 | 0.37 | 0.18 | 0.78 | 0.59 | 0.19 | -0.11 | -0.0875 | 0.2425 | 0.32 | 0.00928 | 0.086537 | 0.354079 | 1 | 0.999631 | 0.367426 |
| Gng12_Q9DAS9_S10s_1 | -0.47 | 0.09 | -0.04 | -0.05 | -0.66 | 0.18 | -0.05 | -0.29 | 0.12 | 0.03 | 0.71 | -0.36 | 0.44 | 0.36 | 0 | 0.01 | 0.31 | 0.19 | 0.44 | 0.283402 | 0.762673 | 0.449258 | 1 | 0.999631 | 0.704903 |
| Gng12_Q9DAS9_S10s_2 | -1.48 | 1.57 | 1.05 | 0.32 | 0.69 | 0.99 | 0.22 | 0.8 | 1.09 | -0.66 | 0.63 | 1.16 | 1.23 | 1.06 | 0.5 | 0.43 | 0.31 | 0.19 | 0.44 | 0.489134 | 0.617689 | 0.786864 | 1 | 0.999631 | 0.807201 |
| Gng12_Q9DAS9_S7s_2 | -1.48 | 1.57 | 1.05 | 0.32 | 0.69 | 0.99 | 0.22 | 0.8 | 1.09 | -0.66 | 0.63 | 1.16 | 1.23 | 1.06 | 0.5 | 0.43 | -0.1525 | -0.455 | 0.005 | 0.489134 | 0.617689 | 0.786864 | 1 | 0.999631 | 0.807201 |
| Gng12_Q9DAS9_T8t_1 | -0.5 | 0.17 | 0.64 | -0.06 | -1.34 | 0.88 | -0.2 | 0.3 | -0.59 | -0.33 | 0.54 | -1.19 | -0.36 | 0.39 | -0.26 | 0.5 | -0.5425 | -0.4125 | -0.02 | 0.989151 | 0.758858 | 0.304401 | 1 | 0.999631 | 0.995574 |
| Gng2_P63213_T52t_1 | -0.32 | -0.03 | 0.13 | -0.57 | -0.59 | -0.63 | -1.06 | -0.68 | -0.75 | -0.33 | -0.81 | -0.55 | 0.33 | -0.21 | -1.06 | 0.07 | -0.1675 | 0.16 | 0.8525 | 0.957701 | 0.072973 | 0.167753 | 1 | 0.999631 | 0.985344 |
| Gng3_P63216_S12s_1 | -1.37 | 0.15 | -0.25 | -0.02 | -0.47 | -0.08 | -0.92 | -0.69 | -0.13 | 0.24 | -0.98 | 0.02 | 1.51 | -0.17 | -0.06 | 0.64 | -0.1575 | -0.1525 | 0.4325 | 0.110197 | 0.671654 | 0.712224 | 1 | 0.999631 | 0.573987 |
| Gng3_P63216_T10t_1 | -0.81 | 0.48 | 0.35 | -0.46 | -0.11 | 0.14 | -0.38 | -0.72 | -0.02 | 0.34 | -0.44 | -0.93 | 0.87 | 0.13 | -0.15 | 0.44 | -0.005 | 0.1075 | 0.555 | 0.298297 | 0.677844 | 0.716817 | 1 | 0.999631 | 0.711324 |
| Gng3_P63216_T5t_1 | -1.33 | -0.9 | -1.07 | -1 | -0.91 | -1.02 | -1.36 | -1.03 | -1.14 | -0.91 | -0.99 | -0.83 | 0.07 | -0.89 | -0.89 | -0.37 | -0.485 | -0.0925 | -0.075 | 0.113179 | 0.984143 | 0.672296 | 1 | 0.999631 | 0.577412 |
| Gng3_P63216_T5t_2 | 0.52 | 0.25 | -0.54 | -0.7 | -0.99 | -0.11 | -0.76 | -0.55 | -0.25 | 0.41 | -0.49 | -0.51 | 0.12 | -0.03 | -1.26 | 0.4 | -0.2925 | -0.275 | -0.9525 | 0.871728 | 0.209821 | 0.810708 | 1 | 0.999631 | 0.959119 |
| Gng4_P50153_S12s_1 | -0.42 | -0.74 | 0.91 | 0.28 | -0.4 | -1.41 | -0.1 | 0.77 | -0.47 | 0.29 | -0.77 | -0.12 | 0.35 | -1.22 | -1.14 | -1.77 | -0.4825 | 0.3025 | 0.0775 | 0.105056 | 0.588308 | 0.528859 | 1 | 0.999631 | 0.568213 |
| Gng4_P50153_S12s_2 | 1.2 | 0.28 | 0.44 | 0.86 | 0.83 | 0.16 | -0.42 | 0.28 | 1.26 | 1.8 | -0.03 | 0.96 | 1.1 | 0.75 | 0.96 | 0.28 | -0.235 | -0.6175 | 0.04 | 0.819764 | 0.196379 | 0.491709 | 1 | 0.999631 | 0.938303 |
| Gng4_P50153_S9s_1 | 0.4 | -0.83 | -2.1 | -0.49 | -1.51 | -1.01 | -0.37 | -1.07 | -0.54 | -0.3 | -1.56 | -3.09 | -0.11 | -0.44 | -1.14 | -1.17 | -0.42 | 0.23 | 0.035 | 0.941172 | 0.656955 | 0.404155 | 1 | 0.999631 | 0.981286 |
| Gng4_P50153_S9s_2 | 0.53 | 0.6 | 0.44 | 0.86 | 0.49 | 0.4 | -0.42 | 0.28 | 1 | 1.42 | -0.03 | 0.96 | 0.62 | 0.71 | 0.96 | 0.28 | -0.65 | -0.3925 | 0.0775 | 0.903514 | 0.175331 | 0.522712 | 1 | 0.999631 | 0.968183 |
| Gnl1_P36916_S51s_1 | -0.43 | -0.24 | 0.29 | -0.32 | -1.37 | -0.77 | -0.61 | -0.55 | -1.01 | -0.23 | -0.45 | -0.58 | 0.13 | -0.52 | -0.14 | 0.14 | 0.5475 | -0.14 | -0.605 | 0.804713 | 0.05582 | 0.214928 | 1 | 0.999631 | 0.934231 |
| Gnl1_P36916_S51s_2 | -0.29 | 0.09 | -0.84 | 0.88 | 0.55 | 0.2 | 0.67 | 0.61 | -0.57 | 0.35 | -0.16 | -0.34 | -1.27 | 0.22 | -0.91 | -0.62 | 1.2875 | -0.165 | -0.3225 | 0.218066 | 0.178797 | 0.737041 | 1 | 0.999631 | 0.669859 |
| Gnl1_P36916_S68s_1 | 0.93 | -0.13 | 0.14 | 2.71 | 2.49 | 1.18 | 2.34 | 2.79 | 1.27 | -0.48 | 2.51 | -0.31 | -0.96 | 1.07 | 1.16 | 1.09 | -0.245 | -0.3525 | -1.135 | 0.658249 | 0.072635 | 0.843234 | 1 | 0.999631 | 0.879726 |
| Gnl1_P36916_S89s_1 | -0.45 | 3.11 | 3.09 | 2.06 | 0.83 | 2.46 | 2.27 | 1.27 | 1.62 | 0.37 | 2.41 | 2 | -0.59 | 0.29 | 2.4 | 1.17 | 0.625 | -0.0475 | -0.8 | 0.229332 | 0.761798 | 0.671667 | 1 | 0.999631 | 0.679824 |
| Gnl1_P36916_T48t_2 | -0.55 | -1.32 | -1.09 | 1.03 | 0.03 | -0.61 | 0.58 | 0.57 | -1.05 | -0.37 | -0.2 | -0.5 | -3.31 | -0.39 | -0.68 | -0.75 | 0.0725 | -0.04 | -0.545 | 0.304228 | 0.272438 | 0.92785 | 1 | 0.999631 | 0.716239 |
| Gnl1_P36916_T50t_2 | -0.29 | 0.09 | -1.62 | 1.2 | 0.55 | 0.2 | 0.02 | -1.1 | -0.57 | 0.35 | -0.14 | -0.42 | -1.27 | 0.22 | -0.29 | -1.46 | 0.1025 | 0.105 | -0.17 | 0.400428 | 0.905983 | 0.943724 | 1 | 0.999631 | 0.763209 |
| Gnl3_Q8CI11_S472s_1 | -0.3 | -0.42 | -0.22 | -1.23 | -0.41 | -0.13 | -0.56 | -0.66 | -0.42 | -0.02 | -0.25 | -1.06 | -0.52 | -0.27 | -0.74 | -1.32 | 0.0125 | -0.1725 | 0.6125 | 0.645491 | 0.743795 | 0.769209 | 1 | 0.999631 | 0.873231 |
| Gnpat_Q05DV2_S12s_1 | -0.48 | 0.05 | -0.58 | 0.14 | -0.21 | 0.33 | -1.66 | 0.72 | 0.06 | 0.27 | -0.59 | -1.3 | 0.99 | 0.3 | -0.22 | 0.51 | 1.395 | 2.285 | 0.285 | 0.112935 | 0.980585 | 0.673364 | 1 | 0.999631 | 0.577408 |
| Golga3_E9QP99_S1479s_1 | 0.76 | 1.57 | 1.69 | -0.16 | 3.11 | 1.27 | 3.6 | 1.46 | 4.36 | 1.12 | 3.26 | 4.26 | 2.59 | 0.27 | 1.65 | 0.49 | -0.345 | 0.0025 | -0.08 | 0.648934 | 0.053632 | 0.013101 | 1 | 0.999631 | 0.874615 |
| Golga3_E9QP99_S285s_1 | 0.03 | -1.68 | -2.01 | -2.04 | -1.86 | -2.19 | -2 | -1.03 | -1.46 | -1.11 | -1.9 | -1.22 | -0.72 | -1.63 | -1.26 | -2.41 | -0.7625 | -0.27 | 0.11 | 0.886629 | 0.508559 | 0.995977 | 1 | 0.999631 | 0.963505 |
| Golga3_E9QP99_S312s_1 | -0.09 | -1.2 | -0.43 | -0.5 | -1.98 | -0.27 | -1.08 | -1.94 | -1.05 | -0.67 | -1.31 | -0.27 | -1.23 | -0.29 | -0.1 | -0.16 | 0.1425 | 0.3725 | -0.055 | 0.775681 | 0.116347 | 0.45875 | 1 | 0.999631 | 0.920998 |
| Golga3_E9QP99_T1391t_1 | 0.21 | 0.49 | 0.96 | 0.31 | -0.1 | 1.27 | 0.99 | 0.38 | 0.77 | 0.42 | 1.05 | 1.22 | -0.44 | 0.65 | 1.04 | 0.5 | -0.71 | -0.8225 | -0.8275 | 0.886986 | 0.700104 | 0.246563 | 1 | 0.999631 | 0.963505 |
| GOLGA7B_Q9D428_S14s_1 | 0.73 | -0.22 | -0.38 | 0.3 | -0.97 | -1 | 0.3 | -0.74 | -1.88 | -0.37 | -0.01 | -0.6 | -0.91 | -1.29 | 0.17 | -0.85 | -0.525 | -0.2325 | -0.545 | 0.070672 | 0.101728 | 0.101921 | 1 | 0.999631 | 0.530826 |
| GOLGA7B_Q9D428_S16s_1 | 2.25 | 0.85 | 1.86 | 1.9 | 0.47 | 1.71 | 1.77 | 0.81 | 0.9 | 2.31 | 0.89 | 1.83 | 0.6 | 1.18 | 1.95 | 0.95 | -0.0025 | 0.565 | 0.7025 | 0.219795 | 0.244315 | 0.610859 | 1 | 0.999631 | 0.670699 |
| Golim4_D3YVW2_S568s_1 | -0.15 | 1.4 | 2.36 | 1.19 | 0.89 | 1.56 | 1.32 | 1.02 | 1.78 | 1.19 | 1.99 | 2.1 | 0.65 | 1.38 | 3 | 2.58 | -0.6975 | -0.5575 | -1.5575 | 0.30692 | 0.996038 | 0.293496 | 1 | 0.999631 | 0.717667 |
| Golph3_Q9CRA5_S35s_1 | 1.77 | -0.03 | 1.31 | 1.04 | -0.41 | 0.34 | 2.05 | -0.68 | 0.51 | 1.01 | -0.2 | 0.54 | -0.27 | -0.92 | 0.55 | -1.5 | -0.4025 | -0.2625 | -0.47 | 0.015877 | 0.295406 | 0.232425 | 1 | 0.999631 | 0.388832 |
| Gon4l_E9Q507_S1394s_1 | 1.06 | 0.42 | 1.15 | 1.22 | -0.14 | 0.57 | 0.95 | 0.86 | -0.31 | 0.88 | 1.25 | 0.98 | -0.62 | -0.22 | 1.09 | 1.72 | -0.2825 | 0.0025 | -0.495 | 0.39402 | 0.256347 | 0.519097 | 1 | 0.999631 | 0.759303 |
| Gorab_Q8BRM2_S335s_1 | 0.32 | -0.62 | 0.03 | 0.08 | -0.92 | 0.08 | -0.44 | -0.04 | 0.39 | 0.81 | -0.89 | -0.49 | 0.03 | -0.4 | -0.93 | -0.87 | -0.0075 | -0.04 | 0.21 | 0.184292 | 0.412613 | 0.995423 | 1 | 0.999631 | 0.644768 |
| Gorasp2_Q99JX3_S350s_1 | -0.42 | -0.11 | -0.71 | -0.43 | -0.36 | -0.52 | -0.57 | -0.25 | -0.71 | -0.65 | -0.29 | -0.18 | -0.03 | -0.38 | -0.38 | -0.04 | 0.5425 | 0.275 | 0.1 | 0.467951 | 0.976541 | 0.885754 | 1 | 0.999631 | 0.795565 |
| Gorasp2_Q99JX3_T154t_1 | 0.76 | -0.1 | -0.52 | 1.38 | 1.33 | 1.01 | 1.02 | 0.33 | 1.47 | 0.59 | 0.6 | -0.04 | -1.29 | 2.05 | -0.01 | 1.17 | 1.1625 | 0.0575 | -0.475 | 0.892516 | 0.253285 | 0.585477 | 1 | 0.999631 | 0.964449 |
| Gorasp2_Q99JX3_T157t_1 | 1.48 | -0.13 | -1.51 | 0.8 | 2.57 | 1.04 | 1.15 | 0.53 | 0.91 | 0.69 | -0.43 | -0.3 | -2.44 | 1.4 | -0.13 | -0.09 | -0.06 | 0.2125 | 0.74 | 0.593393 | 0.117754 | 0.930166 | 1 | 0.999631 | 0.853305 |
| Gpalpp1_Q69ZC8_S104s_1 | -0.32 | 1.78 | 1.74 | 0.57 | 0.94 | 1.38 | 0.36 | 0.85 | 1.05 | 0.99 | 0.91 | 1.67 | 1.55 | 2.21 | 1.27 | 1.7 | 0.1125 | 0.3725 | 0.6175 | 0.172578 | 0.906386 | 0.675691 | 1 | 0.999631 | 0.642116 |
| Gpatch8_A2A6A1_S1018s_1 | -0.02 | 1.28 | -0.19 | -0.29 | 0.42 | 1.17 | -0.58 | 0.22 | 1.32 | 0.58 | 0.2 | 0.17 | 1.86 | 1.13 | 0.02 | 0.24 | -0.1775 | -0.02 | 0.1275 | 0.258235 | 0.816614 | 0.412497 | 1 | 0.999631 | 0.692624 |
| Gpatch8_A2A6A1_S1039s_1 | 0.46 | -0.54 | -0.64 | -0.63 | -1.01 | -0.63 | -0.39 | -0.03 | -0.82 | -0.09 | -0.14 | -0.38 | 0.54 | -0.78 | -0.48 | -0.12 | -0.3225 | -0.15 | -0.715 | 0.75567 | 0.624556 | 0.954698 | 1 | 0.999631 | 0.913663 |
| Gpatch8_A2A6A1_S1179s_1 | 0.78 | 0.01 | 0.52 | 0.49 | -0.37 | 0.15 | 0.48 | 0.25 | 0.21 | 1.37 | -0.07 | -0.31 | -0.51 | -1.03 | 0.53 | -0.05 | -0.3375 | -0.5475 | -1.24 | 0.094934 | 0.300428 | 0.716688 | 1 | 0.999631 | 0.558128 |
| Gpatch8_A2A6A1_S1179s_2 | 0.08 | -0.53 | 1.01 | 0.69 | -0.85 | -0.07 | 0.7 | 0.12 | -0.23 | 0.44 | -0.3 | -0.85 | -1.05 | -1.78 | 0 | -0.88 | -0.3375 | -0.5475 | -1.24 | 0.026536 | 0.461183 | 0.22287 | 1 | 0.999631 | 0.422285 |
| Gpatch8_A2A6A1_S1185s_2 | 0.08 | -0.53 | 1.01 | 0.69 | -0.85 | -0.07 | 0.7 | 0.12 | -0.23 | 0.44 | -0.3 | -0.85 | -1.05 | -1.78 | 0 | -0.88 | -0.9825 | -0.95 | -1.565 | 0.026536 | 0.461183 | 0.22287 | 1 | 0.999631 | 0.422285 |
| Gpatch8_A2A6A1_S451s_1 | 2.35 | 0.1 | 1.25 | 0.75 | -0.42 | 0.03 | 1.14 | -0.23 | -0.39 | 1.98 | -0.16 | -0.78 | -0.43 | -0.71 | -0.09 | -0.58 | 0.1375 | 0.43 | 0.04 | 0.008107 | 0.0951 | 0.193327 | 1 | 0.999631 | 0.365244 |
| Gpatch8_A2A6A1_S648s_1 | -0.07 | 0.09 | 0.03 | -0.29 | 0.13 | -0.11 | -0.13 | 0.42 | 0.63 | 0.83 | -0.4 | 0.42 | 0.59 | 0.02 | -0.15 | -0.54 | -0.355 | -0.055 | 0.0275 | 0.902412 | 0.59867 | 0.216278 | 1 | 0.999631 | 0.968183 |
| Gpatch8_A2A6A1_S735s_1 | 0.35 | 0.32 | 0.35 | -0.17 | -0.23 | -0.22 | -0.2 | 0.08 | 0.07 | 0.53 | 0.05 | -0.02 | 0.58 | 0.19 | 0.09 | 0.1 | -0.0825 | 0.2425 | -0.625 | 0.92463 | 0.188918 | 0.843414 | 1 | 0.999631 | 0.975422 |
| Gpatch8_A2A6A1_S753s_1 | -0.05 | -0.48 | -0.71 | -0.68 | -0.2 | -0.61 | -1.01 | -0.43 | -0.72 | 0.6 | 0.08 | -0.91 | -0.64 | -0.96 | -1.76 | -1.06 | -0.33 | 0.0175 | -0.6875 | 0.092826 | 0.780527 | 0.545434 | 1 | 0.999631 | 0.557387 |
| Gpbp1_Q6NXH3_S114s_1 | 2.56 | 2.28 | 1.38 | 0.98 | 0.65 | 2.16 | 0.86 | 2.21 | 1.98 | 2.54 | 1.98 | 0.77 | -1.08 | 1.8 | 1.7 | 2.03 | 0.5525 | -0.0175 | -0.955 | 0.354786 | 0.528385 | 0.972039 | 1 | 0.999631 | 0.739951 |
| Gpbp1l1_Q6NZP2_T353t_1 | 1.3 | 0.45 | -0.15 | 0.66 | 1.55 | 0.23 | 1.86 | 0.83 | 0.91 | 0.07 | 1.02 | 0.19 | -1.27 | 0.42 | -0.02 | -0.69 | -0.135 | -0.105 | -0.96 | 0.064469 | 0.243017 | 0.964953 | 1 | 0.999631 | 0.514485 |
| Gpd2_Q64521_S415s_1 | 0.54 | 0.56 | 0.18 | -0.88 | -0.15 | 0.71 | 0.46 | -1.16 | 0.2 | 0.01 | 0.18 | -0.41 | 0.02 | -0.48 | -1.74 | -1.24 | -0.435 | -0.66 | -1.6975 | 0.07706 | 0.788289 | 0.785766 | 1 | 0.999631 | 0.542683 |
| Gpd2_Q64521_T413t_1 | 0.57 | -0.69 | 1.15 | 0.45 | -0.45 | -0.6 | 1.06 | -0.27 | -0.37 | 0.24 | 0.09 | -1.12 | -2.57 | -1.11 | -0.17 | -1.46 | 0.1225 | 0.425 | 0.075 | 0.013905 | 0.399481 | 0.185481 | 1 | 0.999631 | 0.385092 |
| Gphn_Q8BUV3_S262s_2 | -0.07 | 0.13 | -0.79 | -0.88 | 0.08 | -0.23 | -0.13 | -0.84 | 0 | 0.3 | 0.02 | -0.23 | 0.15 | 0.1 | -1.02 | -0.54 | -0.2225 | -0.165 | 0.135 | 0.851647 | 0.727415 | 0.216211 | 1 | 0.999631 | 0.950099 |
| Gphn_Q8BUV3_S268s_2 | 0.41 | -1.2 | -0.4 | -0.98 | -0.89 | -0.7 | -0.19 | -1.28 | -1 | -1.02 | -0.77 | -0.04 | 0.06 | 0.1 | -0.97 | -0.82 | 0.06 | 0.24 | 0.385 | 0.766823 | 0.599344 | 0.700221 | 1 | 0.999631 | 0.916961 |
| Gphn_Q8BUV3_S270s_1 | -1.25 | 0.07 | -0.26 | -0.82 | -0.72 | -0.39 | -0.36 | -0.55 | -0.39 | -0.23 | -0.3 | -0.38 | 0 | 0.01 | -0.29 | -0.44 | -0.575 | -0.44 | 0.46 | 0.301524 | 0.859853 | 0.486751 | 1 | 0.999631 | 0.714243 |
| Gphn_Q8BUV3_S270s_2 | -0.16 | -0.32 | 0.38 | -0.27 | -0.29 | -1.19 | -0.87 | -0.32 | -0.76 | -0.29 | -1.39 | 0.31 | 0.21 | -0.17 | 0.84 | 0.59 | -0.2325 | 0.09 | 0.5275 | 0.195655 | 0.097235 | 0.292099 | 1 | 0.999631 | 0.653056 |
| Gphn_Q8BUV3_S280s_1 | 0.47 | -0.12 | -0.24 | 0.09 | -0.4 | 0.11 | -0.17 | -0.27 | 0.19 | 0.38 | -0.23 | 0.22 | 1.32 | 0.6 | 0.36 | 0.03 | -0.31 | 0.7475 | 1.1675 | 0.167929 | 0.407831 | 0.756387 | 1 | 0.999631 | 0.637399 |
| Gphn_Q8BUV3_S283s_1 | 0.13 | 1.03 | 0.91 | 0.09 | 0.02 | 0.81 | 0.15 | -0.06 | 1.49 | 1.15 | 0.21 | 2.3 | 2.48 | 1.47 | 1.25 | 1.63 | -0.435 | 0.0175 | -0.75 | 0.013371 | 0.384015 | 0.143179 | 1 | 0.999631 | 0.385092 |
| Gphn_Q8BUV3_S338s_1 | 0.24 | -0.4 | 1.58 | 0.72 | -0.73 | -0.52 | 2.69 | -1.04 | -0.22 | -0.01 | 0.93 | 1.51 | 0.34 | -1.06 | -1.9 | 1.76 | 0.47 | -0.04 | -0.06 | 0.354178 | 0.606626 | 0.974172 | 1 | 0.999631 | 0.73989 |
| Gphn_Q8BUV3_T286t_1 | 0.47 | 0.35 | -0.7 | 0.14 | 0.57 | 0.51 | 0.5 | 0.56 | 0.06 | 0.23 | -0.28 | 0.09 | -0.18 | 0.06 | -0.1 | 0.24 | -0.2875 | -0.165 | 0.0775 | 0.861055 | 0.159526 | 0.904769 | 1 | 0.999631 | 0.954096 |
| Gpi_P06745_T109t_1 | -3.24 | -6.66 | -1.46 | -3.5 | -4.38 | -5.42 | -2.4 | -3.81 | -4.93 | -4.14 | -2.68 | -3.77 | -3.66 | -5.73 | -2.26 | -2.9 | 1.6425 | 3.1075 | 1.14 | 0.944755 | 0.788893 | 0.871003 | 1 | 0.999631 | 0.982141 |
| Gpkow_Q56A08_S348s_1 | -0.67 | 3.89 | 2.79 | -0.3 | 5.08 | 1.54 | 4.68 | 0.98 | 7.19 | -0.38 | 4.79 | 6.54 | 6.76 | 0.42 | 3.17 | -0.08 | -0.2425 | -0.1825 | 0.1775 | 0.487068 | 0.230488 | 0.099243 | 1 | 0.999631 | 0.805351 |
| Gpm6a_P35802_S256s_1 | -0.55 | -1.14 | -1.82 | -1.2 | -1.17 | -0.93 | -0.81 | -2.77 | -1.34 | -0.02 | -2.21 | -1.87 | -0.12 | -0.92 | -1.63 | -1.33 | -0.3 | -0.755 | -0.61 | 0.679165 | 0.625304 | 0.723615 | 1 | 0.999631 | 0.887281 |
| Gpm6a_P35802_S267s_1 | -0.31 | -1.1 | -1.32 | -0.98 | -2.11 | -0.25 | -1.01 | -1.54 | -2.13 | -0.11 | -1.91 | -2.58 | -2.05 | -0.63 | -2.19 | -1.28 | -0.24 | -0.28 | 0.575 | 0.178409 | 0.500301 | 0.186304 | 1 | 0.999631 | 0.642903 |
| Gpm6a_P35802_S267s_2 | -2.19 | -1.54 | 1.04 | 0.57 | -2.3 | -1.28 | 0.2 | 0.3 | -1.79 | -1.69 | 0.11 | 0.13 | -1.56 | -0.92 | 0.35 | 2.31 | -0.38 | -0.5025 | 0.1825 | 0.565903 | 0.783477 | 0.737676 | 1 | 0.999631 | 0.841575 |
| Gpm6a_P35802_T268t_2 | -2.45 | -2.78 | 1.04 | 0.57 | -2.94 | -2.7 | 0.2 | 0.3 | -3.07 | -2.8 | 0.11 | 0.13 | -2.26 | -3.29 | 0.35 | 2.31 | -0.325 | 0.14 | -0.47 | 0.892966 | 0.739368 | 0.661135 | 1 | 0.999631 | 0.964596 |
| Gpm6b_P35803_S16s_1 | 0.15 | -0.71 | 0.3 | 0.18 | -0.88 | -0.39 | -0.04 | -0.07 | -0.49 | 0.6 | -0.78 | 1.15 | -1.44 | -0.24 | 0.02 | -0.3 | -0.2525 | -0.1875 | -0.065 | 0.273126 | 0.349591 | 0.774218 | 1 | 0.999631 | 0.700187 |
| Gpm6b_P35803_S278s_1 | -0.66 | -0.24 | -0.32 | -0.49 | -0.93 | -0.26 | -0.79 | -0.74 | -0.63 | -0.76 | -0.38 | -0.69 | -0.43 | -0.27 | -0.72 | -0.55 | -0.315 | 0.6375 | -0.3725 | 0.814395 | 0.357322 | 0.471603 | 1 | 0.999631 | 0.937111 |
| Gpm6b_P35803_S280s_1 | -4.63 | -6.01 | -5.7 | -6.31 | -5.78 | -4.91 | -5.37 | -7.85 | -5.68 | -3.66 | -4.5 | -6.26 | -5.23 | -5.64 | -6.71 | -6.56 | -0.685 | -0.3825 | -1.215 | 0.456168 | 0.636812 | 0.320351 | 1 | 0.999631 | 0.790326 |
| Gpm6b_P35803_T13t_1 | 1.07 | 0.47 | 0.81 | 0.56 | 0.29 | -0.53 | -0.19 | 0.6 | 0.06 | 1.01 | 0.09 | 0.22 | 0 | 0.5 | -2.2 | -0.25 | -0.475 | 0.2525 | -0.3525 | 0.052367 | 0.05955 | 0.250821 | 1 | 0.999631 | 0.489777 |
| Gpm6b_P35803_T288t_1 | -1.99 | -2.87 | -2.03 | -2.22 | -3.26 | -2.58 | -2.24 | -2.93 | -2.31 | -1.62 | -1.89 | -2.28 | -2.55 | -2.39 | -2.74 | -2.84 | -0.2575 | -1.4425 | 0.1925 | 0.278966 | 0.180553 | 0.439343 | 1 | 0.999631 | 0.701431 |
| Gpr101_Q80T62_S241s_1 | -0.14 | -2.26 | 0.25 | -0.6 | -1.38 | -0.79 | -0.99 | -0.62 | -1.58 | -1.92 | -2.21 | -2.81 | 0.41 | -0.38 | -0.74 | -1.27 | -0.0325 | 0.225 | 0.3525 | 0.747877 | 0.630423 | 0.027061 | 1 | 0.999631 | 0.911931 |
| Gpr101_Q80T62_S378s_1 | 0.79 | -0.8 | -0.16 | -0.58 | -0.69 | 0.05 | -0.33 | 0.09 | 0.16 | 0.43 | -1.27 | 0.83 | -0.22 | 0.05 | 0.83 | 0 | 0.09 | 0.445 | 0.2625 | 0.419324 | 0.935295 | 0.676659 | 1 | 0.999631 | 0.772152 |
| Gpr101_Q80T62_T509t_1 | -0.16 | -2.1 | 0.19 | -0.3 | -1.33 | -0.8 | -0.06 | 0.18 | -0.64 | -0.09 | 0.27 | -0.13 | -1.13 | -0.47 | 0.14 | 0.14 | -0.4025 | 0.11 | 0.2925 | 0.637075 | 0.873371 | 0.396292 | 1 | 0.999631 | 0.870011 |
| Gpr137c_E9Q343_S351s_1 | -0.07 | 0.87 | 1.22 | -0.16 | 0.09 | 0.03 | 0.2 | -0.07 | 0.19 | 0.24 | 1.1 | 0.77 | 0.36 | 0.27 | 1.14 | 1.26 | -0.5925 | -0.35 | -1.11 | 0.506027 | 0.288528 | 0.790279 | 1 | 0.999631 | 0.814836 |
| Gpr153_Q8K0Z9_S229s_1 | 1.2 | -0.25 | 0.12 | -0.38 | -0.37 | 0.07 | -0.52 | -0.86 | -0.61 | 1.63 | -0.33 | -1.4 | -1.55 | -0.09 | -1.26 | -0.85 | -0.555 | -0.49 | -1.315 | 0.037053 | 0.169068 | 0.599072 | 1 | 0.999631 | 0.456881 |
| Gpr155_A2AWR3_S835s_1 | 1.2 | -0.54 | 0.77 | 0.58 | -0.55 | -0.38 | 0.58 | 0.14 | -0.64 | 0.82 | 0.03 | -0.16 | -0.91 | -1.12 | -0.37 | -0.85 | -0.4875 | -0.17 | -0.8025 | 0.009775 | 0.225845 | 0.306353 | 1 | 0.999631 | 0.374514 |
| Gpr155_A2AWR3_S835s_2 | 1.45 | -0.27 | 0.52 | 0.54 | -0.09 | -0.24 | 0.56 | 0.06 | 0.07 | 1.18 | 0.36 | -0.05 | 0.24 | -0.83 | -0.12 | -0.26 | -0.165 | -0.3025 | -0.945 | 0.083814 | 0.238493 | 0.701469 | 1 | 0.999631 | 0.549732 |
| Gpr155_A2AWR3_S839s_1 | 1.09 | -0.34 | 0.56 | -0.13 | -0.28 | 0.21 | 0.34 | 0.25 | 0.04 | 0.69 | -0.04 | -0.72 | -0.72 | -0.77 | -0.55 | -0.56 | -0.4875 | -0.17 | -0.8025 | 0.027661 | 0.659603 | 0.491315 | 1 | 0.999631 | 0.423977 |
| Gpr155_A2AWR3_S839s_2 | 1.45 | -0.27 | 0.52 | 0.54 | -0.09 | -0.24 | 0.56 | 0.06 | 0.07 | 1.18 | 0.36 | -0.05 | 0.24 | -0.83 | -0.12 | -0.26 | -0.2475 | -0.0125 | 0.55 | 0.083814 | 0.238493 | 0.701469 | 1 | 0.999631 | 0.549732 |
| Gpr155_A2AWR3_S860s_1 | -0.52 | -0.86 | 0.43 | 0.05 | -0.66 | -0.74 | -0.5 | 0.01 | -0.75 | 0.3 | 0.24 | -0.74 | -0.23 | -0.34 | 1.06 | 0.81 | 0.5675 | -0.1925 | 0.265 | 0.244354 | 0.495082 | 0.976014 | 1 | 0.999631 | 0.685751 |
| Gpr158_Q8C419_S1059s_1 | -0.52 | 0 | -2.06 | -0.62 | 0.59 | -0.71 | -0.47 | -0.34 | -0.54 | -2.04 | -0.8 | -0.59 | -0.58 | 0.03 | -0.51 | -1.08 | 0.2575 | -1.2875 | -2.1575 | 0.58503 | 0.267279 | 0.71667 | 1 | 0.999631 | 0.849277 |
| Gpr158_Q8C419_S239s_1 | 2.49 | -3.53 | -2.27 | 0.04 | -1.59 | -1.2 | 3.03 | -2.48 | -3 | 0.25 | -2.49 | -3.18 | -2.15 | -2.32 | -3.11 | -4.32 | 0.6575 | -0.3925 | 0.8325 | 0.096309 | 0.866452 | 0.342246 | 1 | 0.999631 | 0.55886 |
| Gpr158_Q8C419_S249s_1 | -1.02 | -3.21 | -2.87 | -1.38 | -0.57 | -1.95 | -1.14 | -2.19 | -1.96 | -2.54 | -1.64 | -3.91 | 0.55 | -0.46 | -2.05 | -3.19 | 0.0925 | -0.1875 | 0.3075 | 0.343346 | 0.284226 | 0.556654 | 1 | 0.999631 | 0.73348 |
| Gpr158_Q8C419_S692s_1 | -0.64 | -0.88 | -1.19 | -1.07 | -0.99 | -0.75 | -1.02 | -0.65 | -0.72 | -1.26 | -1.98 | -0.57 | -0.63 | -0.27 | -1 | -0.65 | 1.05 | -0.215 | -1.5575 | 0.316757 | 0.720935 | 0.613659 | 1 | 0.999631 | 0.720734 |
| Gpr158_Q8C419_S819s_1 | 1.88 | 0.59 | 0.87 | 2.38 | 2.99 | 1.09 | 3.61 | 2.23 | 1.82 | -0.56 | 2.49 | 1.11 | -0.74 | 0.11 | 0.83 | -0.71 | 0.29 | -0.5275 | -1.58 | 0.013961 | 0.113961 | 0.75719 | 1 | 0.999631 | 0.385092 |
| Gpr158_Q8C419_S881s_1 | 2.36 | -0.03 | -1.02 | 1.78 | 0.75 | 0.88 | 1.95 | 0.67 | 0.76 | 0.87 | 0.31 | -0.96 | -2.5 | -0.18 | -0.09 | -0.46 | 0.72 | 0.02 | -0.3925 | 0.08448 | 0.69568 | 0.505494 | 1 | 0.999631 | 0.549801 |
| Gpr158_Q8C419_S902s_1 | 2.14 | 0.43 | -0.6 | 1.97 | 2.41 | 1.75 | 1.47 | 1.19 | 1.74 | -0.1 | 1.36 | 1.02 | 0 | 1.4 | 0.3 | 0.67 | 0.285 | 0.0175 | -0.1375 | 0.547581 | 0.270376 | 0.976611 | 1 | 0.999631 | 0.834277 |
| Gpr158_Q8C419_S944s_1 | -0.38 | -1 | -1.97 | -0.62 | -0.3 | -0.87 | -0.84 | -0.82 | -0.79 | -0.91 | -1.25 | -0.95 | -0.99 | -1.07 | -1.43 | -1.03 | 0.3675 | 0.8025 | 0.455 | 0.726975 | 0.466859 | 0.963589 | 1 | 0.999631 | 0.905774 |
| Gpr158_Q8C419_S946s_1 | -0.42 | -0.07 | -2.13 | -0.41 | -0.08 | -0.72 | -0.38 | -0.38 | -0.47 | 0.43 | -0.22 | 0.44 | 0.38 | -0.99 | -0.19 | -0.41 | -0.055 | -0.06 | -0.14 | 0.387353 | 0.434517 | 0.128835 | 1 | 0.999631 | 0.756827 |
| Gpr158_Q8C419_S976s_1 | -0.23 | 0.16 | -0.12 | 0.65 | 0 | 0.01 | 0.14 | 0.09 | 0.27 | -0.67 | 0.38 | 0.24 | -0.29 | -0.31 | -0.17 | 0.67 | 0.3875 | -0.295 | -0.8375 | 0.696059 | 0.844447 | 0.864363 | 1 | 0.999631 | 0.895683 |
| Gpr162_Q3UN16_S255s_1 | 0.44 | -0.98 | -0.16 | -0.57 | 0.11 | 1.84 | 0.05 | -1.72 | -0.5 | 0.47 | -1.66 | -0.76 | 0.31 | -1.23 | -1.59 | -2.11 | -0.3125 | -0.425 | -1.015 | 0.156399 | 0.581234 | 0.561939 | 1 | 0.999631 | 0.626452 |
| Gpr162_Q3UN16_S256s_1 | 0.09 | -1.84 | 0.24 | -0.22 | -1.14 | -0.68 | -0.28 | -0.88 | -1.53 | -0.14 | -0.21 | -1.55 | -1.3 | -1.63 | -2.04 | -0.82 | 0.14 | -0.435 | -0.4175 | 0.072747 | 0.523097 | 0.462925 | 1 | 0.999631 | 0.534241 |
| Gpr162_Q3UN16_S413s_1 | 0.95 | -0.24 | -0.9 | 0.33 | 0.5 | -0.3 | 0.44 | 0.06 | -0.49 | -0.32 | -0.35 | -0.44 | -0.97 | -0.25 | -0.28 | -0.03 | 0.9875 | -1.5125 | -1.3725 | 0.359673 | 0.744697 | 0.299381 | 1 | 0.999631 | 0.741182 |
| Gpr162_Q3UN16_S534s_1 | 1.88 | -3.04 | -1.76 | 1.04 | 1.79 | -1.41 | 1.55 | 0.14 | -1.3 | -3.1 | -1.85 | -1.68 | -3.1 | -1.05 | -1.49 | -1.73 | 0.46 | 1.02 | 0.8675 | 0.213094 | 0.408541 | 0.172963 | 1 | 0.999631 | 0.666787 |
| Gpr162_Q3UN16_S542s_1 | -2.51 | -1.97 | -2.77 | -1.52 | -2.73 | -0.6 | -2.42 | -1.18 | -1.52 | -2.03 | -0.5 | -0.64 | -0.92 | -2 | -1.66 | -0.72 | -0.41 | -0.2275 | -0.57 | 0.063414 | 0.39733 | 0.0446 | 1 | 0.999631 | 0.512978 |
| Gpr162_Q3UN16_S561s_1 | 0.26 | -2.26 | 0.84 | 1.25 | -0.9 | -3.04 | 1.4 | 0.99 | -1.36 | -0.62 | 0.85 | 0.31 | -1.05 | -1.73 | 0.48 | 0.11 | -0.2025 | 0.9375 | 0.2025 | 0.490642 | 0.710381 | 0.779897 | 1 | 0.999631 | 0.807996 |
| Gpr162_Q3UN16_S565s_1 | 1.06 | 0.11 | 0.83 | 0 | 0.77 | 1.02 | -0.5 | -0.1 | 1.87 | 3.08 | 0.04 | 0.76 | 1.2 | 0.5 | 1 | 0.11 | 0.46 | 0.0125 | 0.7425 | 0.605915 | 0.642633 | 0.167919 | 1 | 0.999631 | 0.85817 |
| Gpr162_Q3UN16_T587t_1 | -1.19 | -0.78 | 0.1 | -0.22 | -0.37 | 0.72 | -0.44 | -0.16 | -0.89 | -0.6 | -0.49 | -0.06 | 0.1 | 1.4 | -0.03 | -0.59 | -0.0475 | 0.215 | 0.5425 | 0.153415 | 0.264421 | 0.972674 | 1 | 0.999631 | 0.625198 |
| Gpr17_Q6NS65_S323s_1 | -0.73 | -0.61 | -0.43 | -0.73 | -0.65 | -0.27 | -1.08 | -0.69 | -0.33 | -0.3 | -0.5 | -0.51 | 0.68 | -0.07 | -0.64 | -0.3 | -0.3125 | 0.07 | -0.545 | 0.141925 | 0.86093 | 0.392584 | 1 | 0.999631 | 0.612726 |
| Gpr17_Q6NS65_S332s_1 | -0.42 | -1.79 | -0.91 | -1.82 | -2.49 | -0.85 | -1.03 | -1.82 | -1.22 | -0.29 | -1.33 | -1.82 | -0.73 | -2.41 | -1.67 | -2.31 | 0.1325 | 0.1825 | 0.65 | 0.287454 | 0.522629 | 0.878426 | 1 | 0.999631 | 0.705972 |
| Gpr26_Q8BZA7_S329s_1 | -1.02 | -0.12 | -1.07 | -1.51 | -0.04 | -0.93 | -1.22 | -1 | -0.47 | 0.07 | -1.62 | -0.97 | -0.5 | 1.08 | -0.44 | -1.26 | 0.7325 | -0.3975 | -1.0225 | 0.242053 | 0.738815 | 0.688351 | 1 | 0.999631 | 0.684914 |
| Gpr34_Q9R1K6_S341s_1 | 1.41 | -0.21 | -0.13 | 0.31 | -0.19 | 2.24 | 1.22 | 1.04 | 0.03 | 1.01 | 0.37 | -1.62 | -1.44 | 0.74 | -0.83 | -1.18 | -0.27 | -0.115 | 0.3425 | 0.09599 | 0.216867 | 0.520261 | 1 | 0.999631 | 0.558577 |
| Gpr37l1_Q99JG2_S471s_1 | -0.25 | 0.09 | 0.81 | 0.13 | 0.05 | 0.01 | -0.37 | 0.01 | 0.32 | -0.18 | -0.29 | 0.47 | 0.87 | 0.42 | 0.47 | 0.39 | -0.4525 | -0.3725 | 0.405 | 0.307422 | 0.383492 | 0.733561 | 1 | 0.999631 | 0.717856 |
| Gpr56_Q8K209_S681s_1 | 0.14 | -0.94 | -0.96 | -0.62 | -1.2 | -1.13 | -0.89 | -0.97 | -0.93 | -0.48 | -1.29 | -1.17 | 0.02 | -0.52 | -0.14 | -0.12 | -0.6375 | -0.205 | -0.8325 | 0.255786 | 0.175573 | 0.303669 | 1 | 0.999631 | 0.690775 |
| Gpr61_Q8C010_S258s_1 | 1.62 | -0.95 | -0.03 | -0.16 | -0.45 | -0.52 | 0.18 | -1.28 | -0.59 | 1.45 | -0.46 | -0.74 | -0.64 | -1.01 | -0.53 | -0.67 | 0.2 | 0.02 | 0.3625 | 0.132962 | 0.274247 | 0.759419 | 1 | 0.999631 | 0.60492 |
| Gprc5b_D2DFA9_S337s_1 | -0.16 | 0.06 | -0.43 | 0.29 | 0.2 | 0.21 | -0.06 | 0.21 | -0.27 | 0.03 | 0.42 | -0.34 | 0.09 | 0.42 | 0.63 | 0.07 | 0.0975 | 0.2525 | 0.295 | 0.251523 | 0.458061 | 0.947529 | 1 | 0.999631 | 0.687255 |
| Gprin1_Q3UNH4_S166s_1 | -0.56 | 0.68 | 0.77 | 0.64 | -0.18 | 0.5 | 0.44 | 1.16 | 0.35 | 0.19 | 0.71 | 1.29 | 0.72 | 0.5 | 0.59 | 0.9 | 0.29 | 0.215 | 0.1775 | 0.432856 | 0.814551 | 0.539656 | 1 | 0.999631 | 0.779247 |
| Gprin1_Q3UNH4_S182s_1 | 0.19 | -0.44 | -0.64 | -0.19 | 0.45 | -0.11 | -0.28 | 0.02 | 0.15 | -0.07 | -0.21 | -0.09 | -0.31 | 0.32 | -0.35 | -0.03 | 0.375 | 0.135 | -0.14 | 0.581979 | 0.345996 | 0.457728 | 1 | 0.999631 | 0.847567 |
| Gprin1_Q3UNH4_S190s_1 | 0.84 | -0.73 | -1.48 | -0.65 | -0.28 | 0.14 | -0.26 | -0.12 | -0.45 | 0.27 | -0.63 | -0.67 | -0.15 | -0.84 | -0.79 | -0.8 | 0.02 | 0.2175 | 0.515 | 0.776939 | 0.434335 | 0.789115 | 1 | 0.999631 | 0.921303 |
| Gprin1_Q3UNH4_S219s_1 | -0.56 | -1.02 | -0.4 | -0.84 | -0.71 | -1.23 | -0.56 | -0.24 | -1.01 | -0.66 | -0.01 | -0.27 | -0.17 | -0.72 | -0.11 | 0.24 | 0.515 | 0.155 | 0.08 | 0.13417 | 0.947953 | 0.50145 | 1 | 0.999631 | 0.60492 |
| Gprin1_Q3UNH4_S369s_1 | -0.19 | -0.7 | -0.55 | -0.2 | 0.42 | 0.2 | -0.18 | -0.02 | 0 | -0.48 | 0.1 | -0.64 | -0.76 | -0.01 | -0.44 | -0.11 | -1.42 | 0.075 | -0.3575 | 0.795629 | 0.083256 | 0.60618 | 1 | 0.999631 | 0.929355 |
| Gprin1_Q3UNH4_S433s_1 | 0.7 | -0.95 | 1.42 | -1.21 | -1.13 | -1.34 | -0.7 | -2.55 | -1.29 | 0.5 | 0.7 | 0.35 | -1.5 | -1.47 | 0.85 | 0.65 | 0.86 | 0.5525 | 0.61 | 0.653303 | 0.056289 | 0.914448 | 1 | 0.999631 | 0.87811 |
| Gprin1_Q3UNH4_S446s_1 | 0.76 | -0.63 | -1.52 | 0.61 | 1.18 | 0.28 | 0.79 | 0.41 | 0.89 | -0.62 | 0.66 | 0.5 | -0.28 | 0.62 | 0.42 | 0.9 | 0.3025 | 0.47 | -0.2375 | 0.28579 | 0.131197 | 0.35463 | 1 | 0.999631 | 0.705851 |
| Gprin1_Q3UNH4_S468s_1 | 0.44 | -1.54 | -1.17 | 0.21 | -0.5 | -0.51 | 0.51 | -0.35 | -0.43 | 0.02 | 0.44 | -0.21 | -1.2 | -1.17 | -0.65 | 0.01 | 0.005 | 0.0775 | 0.485 | 0.657814 | 0.558544 | 0.358566 | 1 | 0.999631 | 0.879549 |
| Gprin1_Q3UNH4_S495s_1 | -0.5 | -0.77 | -1.05 | -1 | -0.77 | -0.8 | -1.11 | -0.62 | -0.72 | -0.4 | -0.92 | -0.97 | 0.08 | -0.25 | -0.67 | -0.54 | 0.77 | 0.14 | 0.1975 | 0.137117 | 0.984834 | 0.781414 | 1 | 0.999631 | 0.610369 |
| Gprin1_Q3UNH4_S519s_1 | 0.54 | -0.6 | -1.47 | 0.07 | 0.48 | 0.45 | 0.31 | 0.38 | 0.14 | -0.24 | -0.16 | -0.64 | -0.76 | 0.31 | -0.19 | -0.03 | 0.6425 | 0.3375 | -0.3125 | 0.680172 | 0.098273 | 0.758472 | 1 | 0.999631 | 0.888282 |
| Gprin1_Q3UNH4_S53s_1 | -0.7 | -1.18 | -1.14 | -1.59 | -0.05 | -0.98 | -0.56 | -0.45 | -0.61 | -1 | -0.34 | -1.31 | -1.16 | -1.79 | -1.24 | -1.67 | 0.7625 | 0.495 | 0.48 | 0.342814 | 0.0651 | 0.322178 | 1 | 0.999631 | 0.73348 |
| Gprin1_Q3UNH4_S576s_1 | -0.26 | -0.6 | -2.63 | -1.49 | 0.28 | -0.01 | -1.22 | -0.98 | 0.07 | -0.48 | -1.18 | -1.41 | -0.21 | 0.12 | -1.59 | -1.38 | 0.225 | 0.2 | 1.1125 | 0.442155 | 0.211586 | 0.399718 | 1 | 0.999631 | 0.782298 |
| Gprin1_Q3UNH4_S57s_1 | -0.87 | -0.41 | -0.47 | -1.52 | -0.08 | 0.04 | -1.38 | -0.95 | 0.03 | -0.38 | -0.81 | -1.31 | -0.07 | 0.34 | 1.06 | -0.15 | -0.815 | -0.525 | -1.525 | 0.018241 | 0.596905 | 0.61751 | 1 | 0.999631 | 0.391024 |
| Gprin1_Q3UNH4_S618s_2 | 1.18 | -1.31 | 0.26 | -0.1 | -1.1 | -1.43 | 0.75 | -1.45 | -0.87 | 1.12 | -1.27 | -1.05 | -0.81 | -2.7 | -1.32 | -1.24 | -0.0375 | -0.6875 | -1.4225 | 0.027467 | 0.233105 | 0.444356 | 1 | 0.999631 | 0.423809 |
| Gprin1_Q3UNH4_S620s_1 | 2.64 | -0.61 | -0.22 | 0.91 | -0.03 | 0.89 | 2 | -0.29 | 0.19 | 1.25 | -0.19 | -1.28 | -1.37 | -0.31 | -0.51 | -0.78 | -0.595 | -0.6575 | -1.845 | 0.059204 | 0.961734 | 0.392585 | 1 | 0.999631 | 0.504286 |
| Gprin1_Q3UNH4_S620s_2 | 1.81 | -0.78 | 0.35 | 0.17 | -0.57 | -0.64 | 0.84 | -0.46 | -0.48 | 1.26 | -0.76 | -1.1 | -0.97 | -1.46 | -2.44 | -0.96 | -0.42 | -0.975 | -1.45 | 0.009719 | 0.319577 | 0.339871 | 1 | 0.999631 | 0.374514 |
| Gprin1_Q3UNH4_S622s_1 | 2.14 | -0.42 | 0.36 | 0.15 | -0.24 | 0.74 | 0.81 | -0.76 | -0.23 | 0.99 | -0.7 | -1.73 | -1.24 | -0.83 | -0.17 | -1.33 | -0.6875 | -0.6025 | -1.69 | 0.026507 | 0.494177 | 0.187137 | 1 | 0.999631 | 0.422285 |
| Gprin1_Q3UNH4_S622s_2 | 1.54 | -1.02 | 0.31 | 0.06 | -0.81 | -0.99 | 0.8 | -0.86 | -0.66 | 1.19 | -0.97 | -1.08 | -0.89 | -2 | -1.9 | -1.08 | -0.0025 | 0.065 | -0.1 | 0.011562 | 0.274685 | 0.378221 | 1 | 0.999631 | 0.385092 |
| Gprin1_Q3UNH4_S649s_1 | -1.11 | -3.48 | -1.77 | -2.6 | -1.76 | -2.62 | -2.15 | -2.44 | -2.31 | -2.68 | -1.68 | -2.03 | -2.19 | -2.82 | -1.57 | -2.78 | 0.445 | 0.4425 | 0.53 | 0.855881 | 0.996086 | 0.900729 | 1 | 0.999631 | 0.951817 |
| Gprin1_Q3UNH4_S668s_1 | 0.64 | 0.73 | -1.22 | -1.78 | 0.79 | 1.59 | -0.8 | -1.43 | 1.02 | 1.63 | -1.21 | -1.3 | 0.58 | 0.94 | -1.13 | 0.1 | 0.315 | 0.0275 | 0.2575 | 0.453183 | 0.592354 | 0.610916 | 1 | 0.999631 | 0.788717 |
| Gprin1_Q3UNH4_S675s_1 | -0.05 | -1.15 | -2.37 | -0.67 | -0.15 | -1.19 | -0.79 | -0.85 | -0.94 | -0.92 | -1.14 | -1.13 | -0.55 | -0.47 | -0.85 | -1.34 | -0.0275 | 0.1175 | 0.2475 | 0.612422 | 0.535161 | 0.953912 | 1 | 0.999631 | 0.860287 |
| Gprin1_Q3UNH4_S691s_1 | -0.15 | -0.33 | -1.16 | -0.89 | -0.48 | -0.58 | -1.04 | -0.54 | -0.44 | 0.02 | -0.69 | -0.95 | 0.04 | -0.42 | -0.76 | -0.4 | -0.0625 | 0.2075 | 0.62 | 0.481467 | 0.931061 | 0.738952 | 1 | 0.999631 | 0.802053 |
| Gprin1_Q3UNH4_S714s_1 | -0.29 | -0.04 | -0.39 | -0.3 | -0.19 | -0.19 | -0.54 | -0.35 | -0.11 | 0.04 | -0.1 | -0.02 | 0.84 | 0.17 | 0.02 | 0.43 | 0.8425 | -0.1975 | -0.5325 | 0.061612 | 0.798292 | 0.405577 | 1 | 0.999631 | 0.509355 |
| Gprin1_Q3UNH4_S764s_1 | 1 | -1.17 | -1.49 | 0.31 | 0.79 | 0.05 | 0.72 | 0.46 | -0.45 | -0.51 | 0.18 | -1.36 | -1.91 | -0.23 | -0.67 | -0.67 | 0.4925 | -0.5025 | -0.385 | 0.405208 | 0.158149 | 0.746975 | 1 | 0.999631 | 0.765844 |
| Gprin1_Q3UNH4_S764s_2 | 0.96 | -0.64 | -2.33 | 0.35 | 0.8 | -0.31 | 0.11 | -0.29 | -0.39 | -0.93 | -0.43 | -1.92 | -0.4 | 0.11 | -1.93 | -0.98 | -0.4475 | 0.04 | 0.68 | 0.605311 | 0.47401 | 0.486358 | 1 | 0.999631 | 0.85817 |
| Gprin1_Q3UNH4_S771s_1 | -1.35 | -1.02 | -0.37 | -1.26 | -2.09 | -1.35 | -1.42 | -0.93 | -1.56 | -0.43 | -0.92 | -0.93 | 0.69 | -1.36 | -0.69 | 0.08 | 0.4925 | -0.5025 | -0.385 | 0.180994 | 0.224938 | 0.910469 | 1 | 0.999631 | 0.64376 |
| Gprin1_Q3UNH4_S771s_2 | 0.96 | -0.64 | -2.33 | 0.35 | 0.8 | -0.31 | 0.11 | -0.29 | -0.39 | -0.93 | -0.43 | -1.92 | -0.4 | 0.11 | -1.93 | -0.98 | 0.7375 | -0.2425 | -0.2625 | 0.605311 | 0.47401 | 0.486358 | 1 | 0.999631 | 0.85817 |
| Gprin1_Q3UNH4_S817s_1 | 0.75 | 0.16 | -1.06 | 0.99 | 1.22 | 0.97 | 0.4 | 1.2 | 0.14 | -0.58 | 0.48 | -0.17 | -2.32 | 0.95 | 0.66 | 0.5 | 0.9625 | -0.1025 | 0.0075 | 0.73609 | 0.142411 | 0.621166 | 1 | 0.999631 | 0.90759 |
| Gprin1_Q3UNH4_S917s_1 | 1.61 | -1.27 | -2.2 | 0.34 | 1.84 | -0.44 | 0.44 | 0.49 | -0.3 | -0.98 | -0.11 | -0.54 | -0.25 | -0.77 | -0.69 | 0.22 | -0.565 | 0.1025 | -0.4525 | 0.992193 | 0.269343 | 0.893203 | 1 | 0.999631 | 0.997164 |
| Gprin1_Q3UNH4_T256t_1 | 1.93 | 0.53 | -0.58 | -0.19 | -0.15 | 0.12 | 0.19 | -0.73 | 1.07 | 1.74 | -0.17 | -0.54 | 0.08 | 0.82 | -0.28 | -0.74 | 0.0175 | 0.015 | 0.4 | 0.449081 | 0.3125 | 0.880955 | 1 | 0.999631 | 0.786796 |
| Gprin1_Q3UNH4_T795t_1 | -0.78 | 0.01 | -0.17 | -0.56 | -0.06 | -0.33 | -0.56 | -0.48 | -0.11 | -0.53 | -0.57 | -0.23 | 0.4 | 0.01 | -0.33 | 0.02 | 0.0175 | -0.0675 | 0.645 | 0.227777 | 0.951201 | 0.959303 | 1 | 0.999631 | 0.678949 |
| Gprin3_Q8BWS5_S327s_1 | -0.51 | -0.12 | -0.68 | -0.68 | -0.09 | -0.43 | -0.9 | -0.5 | -0.28 | -0.53 | -0.94 | -0.51 | 0.7 | 0.68 | -0.38 | -0.41 | 1.2725 | 0.0775 | -0.8275 | 0.111857 | 0.95132 | 0.811992 | 1 | 0.999631 | 0.577068 |
| Gprin3_Q8BWS5_S572s_1 | 2.51 | 0.4 | -2.69 | 0.37 | 2.33 | 0.05 | 2.13 | 1.17 | 1.36 | 0.64 | -0.2 | -0.9 | -2.86 | 0.23 | 0.56 | -0.65 | -0.4225 | -0.1575 | 0.0325 | 0.466192 | 0.232495 | 0.939168 | 1 | 0.999631 | 0.795267 |
| Gps1_G3UXW9_S275s_1 | -0.29 | 0.56 | 0.13 | -0.05 | -0.01 | -0.1 | -0.76 | -0.47 | -0.5 | 0.05 | -0.27 | 0.44 | 0.18 | 0.43 | 0.19 | -0.32 | -0.26 | 0.3 | 0.185 | 0.919347 | 0.190018 | 0.63151 | 1 | 0.999631 | 0.974502 |
| Gps1_G3UXW9_S4s_1 | 1.57 | -0.14 | 0.73 | 0.26 | 0.34 | 0.23 | 0.4 | 0.41 | 0.45 | 0.91 | 0.93 | 1.33 | 0.45 | 1.42 | 0.43 | 0.86 | -0.4475 | 0.06 | 0.18 | 0.674586 | 0.501611 | 0.475752 | 1 | 0.999631 | 0.885515 |
| Gps1_G3UXW9_S509s_1 | 0.04 | 0.22 | 0.22 | -0.09 | -0.65 | -0.24 | -0.05 | -0.46 | 0.25 | 0.48 | -0.08 | -0.02 | 0.54 | -0.36 | 1.21 | -0.28 | 0.05 | -0.5375 | -0.55 | 0.655951 | 0.108054 | 0.821616 | 1 | 0.999631 | 0.879056 |
| Gps1_G3UXW9_S518s_1 | 0.66 | 0.39 | -0.55 | -0.11 | 0.47 | 0.33 | 0.08 | -0.29 | -0.56 | -0.53 | -0.22 | -0.45 | -0.13 | -1.31 | 0.08 | -0.45 | 1.0075 | 0.475 | 0.1425 | 0.210281 | 0.885381 | 0.128679 | 1 | 0.999631 | 0.66441 |
| Gpsm1_F6SX81_S293s_1 | 0.91 | 0.3 | -0.12 | 0.64 | 1.74 | 1.51 | 1.03 | 1.48 | 1.67 | 0.55 | 0.83 | 0.58 | -0.08 | 1.08 | 0.8 | 0.5 | 0.355 | 0.52 | 0.3275 | 0.703962 | 0.009576 | 0.21974 | 1 | 0.999631 | 0.897149 |
| Gpsm1_F6SX81_S315s_1 | -0.22 | 1.29 | -0.67 | -0.03 | 0.87 | 0.68 | -0.25 | 0.49 | 1.02 | 0.69 | 0.04 | 0.7 | 0.17 | 0.99 | 0.44 | 0.08 | 0.1375 | 0.0425 | 0.1775 | 0.484029 | 0.452919 | 0.271806 | 1 | 0.999631 | 0.803422 |
| Gpsm1_F6SX81_S367s_1 | -1.03 | -0.39 | -0.47 | -0.45 | -0.32 | -0.81 | -0.77 | 0.11 | -0.56 | -0.75 | -0.41 | -0.45 | 0.01 | -0.65 | -0.04 | -0.95 | -0.315 | 0.1625 | 0.3275 | 0.606133 | 0.663658 | 0.876397 | 1 | 0.999631 | 0.85817 |
| Gpsm1_F6SX81_S477s_1 | -0.48 | -0.37 | -0.32 | 0.06 | -0.39 | -0.49 | -0.9 | -0.59 | -0.61 | -0.2 | -0.06 | 0.41 | -0.08 | 0.01 | 0.16 | 0.11 | -0.385 | -0.095 | -0.1075 | 0.25127 | 0.248217 | 0.601694 | 1 | 0.999631 | 0.687113 |
| Gpsm3_G3UZQ1_S38s_1 | 0.47 | 0.21 | 0.69 | -0.06 | -0.31 | -0.24 | 0.5 | -0.18 | -0.38 | 0.19 | 0.96 | 0.16 | 0.38 | 0.03 | 0.32 | 0.15 | -0.5475 | 0.1225 | 0.3275 | 0.715748 | 0.227697 | 0.789597 | 1 | 0.999631 | 0.902706 |
| Gramd1a_F8WJ74_S11s_1 | -0.28 | 0.53 | 0.16 | -0.44 | -0.72 | -0.28 | -0.61 | -0.61 | 0.22 | 0.68 | -0.47 | 0.03 | 1.03 | 0.47 | -0.37 | 0.15 | -0.065 | 0.1075 | -0.19 | 0.414261 | 0.09399 | 0.732939 | 1 | 0.999631 | 0.768655 |
| Gramd1a_F8WJ74_S11s_2 | 0.83 | -0.27 | -0.09 | 0.5 | -0.2 | -0.18 | 0.22 | 0.87 | 0.7 | 0.4 | 0.33 | -0.03 | -0.74 | 0.42 | 0.21 | 0.32 | 0.2575 | 0.205 | 0.805 | 0.633661 | 0.862093 | 0.753182 | 1 | 0.999631 | 0.868569 |
| Gramd1a_F8WJ74_S269s_1 | -1.21 | 0.2 | -0.54 | -1.05 | -0.03 | -0.29 | -0.83 | -0.42 | -0.63 | -0.6 | -0.39 | -0.16 | 1 | -0.07 | -0.09 | -0.22 | -0.07 | 0.0675 | 0.4425 | 0.086987 | 0.497244 | 0.578025 | 1 | 0.999631 | 0.553514 |
| Gramd1a_F8WJ74_T279t_1 | 0.03 | -0.74 | 0.05 | 0.15 | -0.66 | -0.8 | 0.11 | 0.56 | -0.31 | -0.23 | 0.23 | 0.07 | -0.23 | 0.32 | 0.36 | 0.81 | 0.23 | 0.33 | 0.7575 | 0.227219 | 0.857822 | 0.827494 | 1 | 0.999631 | 0.678384 |
| Gramd1b_Q80TI0_S165s_1 | 0.12 | 0 | -0.59 | 0.19 | 0.31 | 0.25 | -0.2 | 0.28 | 0.48 | 0.19 | 0.25 | 0.12 | 0.74 | 0.8 | 0.75 | 0.46 | 0.3325 | 0.575 | 0.4175 | 0.028001 | 0.432589 | 0.26417 | 1 | 0.999631 | 0.423977 |
| Gramd1b_Q80TI0_S174s_1 | 0.48 | -0.32 | -0.18 | 0.27 | 0.49 | 0.5 | 0.2 | 0.39 | 0.75 | 0.81 | 0.53 | 0.46 | -0.11 | 1.07 | 0.59 | 0.37 | -0.1975 | -0.5675 | -1.37 | 0.261469 | 0.253673 | 0.072215 | 1 | 0.999631 | 0.69271 |
| Gramd1b_Q80TI0_S194s_1 | -3.43 | -6.18 | -3.44 | -3.02 | -4.3 | -5.16 | -1.57 | -5.83 | -6.53 | -4.08 | -3.24 | -4.49 | -5.51 | -7.02 | -5.56 | -3.46 | 0.6775 | 0.6125 | 0.505 | 0.146965 | 0.846256 | 0.522792 | 1 | 0.999631 | 0.618703 |
| Gramd1b_Q80TI0_S197s_1 | -1.32 | -3.61 | -0.83 | -0.52 | -1.49 | -1.36 | -0.51 | -0.21 | -0.66 | -2.49 | -0.02 | -0.66 | -1.92 | -1.01 | -0.38 | -0.95 | 0.2 | 0.4075 | 0.395 | 0.466616 | 0.332553 | 0.436585 | 1 | 0.999631 | 0.795267 |
| Gramd1b_Q80TI0_S418s_1 | -0.32 | -0.5 | -0.88 | 0.91 | -0.69 | -0.41 | -0.07 | 1.18 | -0.61 | -0.49 | 0.99 | 0.95 | -0.08 | 0.35 | -0.18 | 0.7 | 0.0475 | 0.6825 | 1.045 | 0.379895 | 0.70382 | 0.460633 | 1 | 0.999631 | 0.752193 |
| Gramd1b_Q80TI0_S694s_1 | -2.22 | -1.54 | -0.67 | -0.61 | -1.29 | -2.2 | -0.65 | -0.71 | -1.23 | -1.59 | -0.05 | 0.56 | 0.19 | -1.32 | 0.45 | -0.18 | -0.43 | -0.1125 | -0.105 | 0.067126 | 0.923425 | 0.25636 | 1 | 0.999631 | 0.520345 |
| Gramd1b_Q80TI0_T200t_1 | 0.58 | 0.11 | 0.92 | 0.77 | -0.46 | 0.22 | 0.32 | 0.58 | -0.2 | 0.71 | 0.5 | 0.92 | -0.19 | 0.28 | 1.15 | 0.72 | 0.1475 | 0.105 | 0.3125 | 0.780488 | 0.208292 | 0.743788 | 1 | 0.999631 | 0.921416 |
| Grasp_Q9JJA9_S384s_1 | -0.71 | 1.04 | 0.58 | -0.18 | 0.86 | 0.51 | -0.14 | 0.09 | 0.72 | -0.91 | 0.6 | 0.74 | 1.76 | 0.35 | -0.02 | -0.11 | 0.1475 | 0.29 | 0.28 | 0.568715 | 0.736545 | 0.841101 | 1 | 0.999631 | 0.843185 |
| Grasp_Q9JJA9_S93s_1 | -0.24 | -0.57 | -0.77 | -0.71 | -0.72 | -0.18 | -0.42 | -0.38 | -0.73 | -0.13 | -0.25 | -0.02 | -0.32 | -0.25 | -0.61 | 0.01 | -0.1625 | 0.1375 | 0.06 | 0.348434 | 0.578487 | 0.324281 | 1 | 0.999631 | 0.736697 |
| Grasp_Q9JJA9_T76t_1 | 0.57 | 2.48 | 1.84 | 1.28 | 1.79 | 1.65 | 0.92 | 1.16 | 1.9 | 2.17 | 1.9 | 0.75 | 0.49 | 2.14 | 2.13 | 1.65 | 0.26 | -0.775 | -1.9425 | 0.909586 | 0.714 | 0.779785 | 1 | 0.999631 | 0.970597 |
| Grb10_B1ATS7_S50s_1 | 3.06 | -0.26 | 1.1 | 2.28 | 0.81 | 1.36 | 2.81 | 2.24 | 0.82 | 1.32 | 0.84 | 0.1 | -2.91 | 0.28 | 1.41 | -0.37 | -0.1675 | 0.29 | 0.5975 | 0.075351 | 0.728541 | 0.273862 | 1 | 0.999631 | 0.541044 |
| Grb10_B1ATS7_S96s_1 | -1.13 | -0.07 | -0.64 | -0.85 | -0.68 | -0.65 | -0.87 | -1.16 | 0.05 | -0.34 | -0.89 | -0.35 | 0.72 | 0.02 | 0.02 | -1.06 | 0.97 | 0.02 | -0.21 | 0.19145 | 0.590738 | 0.40418 | 1 | 0.999631 | 0.648674 |
| Grb14_Q9JLM9_S364s_1 | 0.82 | -1.1 | -0.23 | -0.2 | 0.93 | -0.17 | 1.78 | 0.63 | -0.2 | -0.91 | 0.52 | -0.04 | 0.45 | -1.09 | 0.04 | -0.95 | 0.005 | -0.1075 | -0.355 | 0.684683 | 0.087983 | 0.966258 | 1 | 0.999631 | 0.889888 |
| Gria2_G5E8H1_Y876y_1 | 0.95 | 0.98 | -1.41 | -1.39 | 0.96 | 0.38 | -1.03 | -1.16 | 0.3 | 0.6 | -1.5 | -0.7 | 0.41 | 0.96 | -2.53 | -1.13 | 0.085 | -0.075 | 0.6525 | 0.693867 | 0.994713 | 0.884242 | 1 | 0.999631 | 0.895131 |
| Gria2_P23819_S899s_1 | -1.02 | -0.26 | 0.44 | 0.15 | -0.58 | -0.11 | 0.18 | 0.16 | -0.56 | -0.76 | 0.48 | -0.15 | -0.02 | 0.54 | 0.68 | 0.72 | 0.2425 | -0.1275 | 0.3325 | 0.118455 | 0.822015 | 0.858783 | 1 | 0.999631 | 0.585447 |
| Gria2_P23819_T912t_1 | -1.23 | -0.62 | -0.72 | -0.44 | -0.3 | -0.44 | -0.75 | -0.55 | -0.69 | -0.84 | -1 | -0.99 | 0 | -0.32 | -0.98 | -0.38 | 0.0475 | -0.325 | -0.6325 | 0.333251 | 0.392335 | 0.651416 | 1 | 0.999631 | 0.728079 |
| Gria3_B0QZW1_Y871y_1 | -3.47 | -3.03 | -1.76 | -2.65 | -1.97 | -3.58 | -2.02 | -3.15 | -2.96 | -2.73 | -3.15 | -3.37 | -2.21 | -4.25 | -3.63 | -3.35 | -0.2575 | -0.165 | -0.285 | 0.248172 | 0.925484 | 0.42509 | 1 | 0.999631 | 0.685751 |
| Gria3_B0QZW1_Y881y_1 | -0.07 | -0.05 | -0.51 | -0.1 | -0.21 | -0.15 | -0.7 | -0.7 | -0.11 | -0.46 | -0.5 | -0.32 | -0.12 | -0.29 | -0.8 | -0.66 | 0.23 | 0.2 | 0.5825 | 0.352288 | 0.358684 | 0.533824 | 1 | 0.999631 | 0.73909 |

| | | | | | | | | | | | | | | | | | | | | | | | | | |
|---|---|---|---|---|---|---|---|---|---|---|---|---|---|---|---|---|---|---|---|---|---|---|---|---|---|
| Gria4_G5E863_T875t_1 | -0.75 | -0.39 | -0.17 | -0.24 | -0.14 | 0.41 | -0.79 | -0.11 | -0.48 | -0.12 | -0.22 | 0.07 | 0.62 | -0.37 | 0.34 | 0.19 | 0.2225 | -0.1625 | -0.165 | 0.096678 | 0.482804 | 0.474073 | 1 | 0.999631 | 0.55886 |
| Grid1_Q61627_S882s_1 | 1.21 | 0.15 | -0.31 | 0.01 | 0.91 | 0.62 | 0.02 | 0.4 | 0.46 | 0.21 | -0.14 | -0.12 | -0.35 | 0.9 | -0.38 | 0.23 | -0.5875 | -0.265 | -1.5225 | 0.712697 | 0.570104 | 0.671107 | 1 | 0.999631 | 0.9008 |
| Grid2ip_Q0QWG9_S45s_1 | 2 | -1.55 | 0.92 | 0.75 | -1.3 | 0.09 | 1.32 | -0.34 | -0.2 | 2.02 | -0.51 | -0.25 | -0.57 | -1.61 | -0.35 | -1.44 | -0.24 | -0.2075 | 0.2025 | 0.055559 | 0.472186 | 0.750505 | 1 | 0.999631 | 0.494662 |
| Grik5_Q61626_S961s_1 | -0.41 | -0.72 | -0.48 | -0.91 | -1.05 | -0.61 | -1.06 | -0.76 | -1.41 | -0.33 | -0.92 | -0.69 | 0 | -0.65 | -0.48 | -0.58 | -0.1925 | -0.235 | -0.4625 | 0.49989 | 0.370419 | 0.516347 | 1 | 0.999631 | 0.813279 |
| Grik5_Q61626_Y891y_1 | -0.27 | -1.21 | 0.04 | -0.44 | -1.03 | -1.79 | -0.2 | 0.37 | -1.77 | -1.42 | -0.08 | 0.45 | -1.42 | -1.53 | 0.09 | -0.87 | -0.0275 | -0.1625 | -0.575 | 0.3206 | 0.705099 | 0.670283 | 1 | 0.999631 | 0.723684 |
| Grin2a_P35436_S1025s_1 | -1.15 | -1.72 | -3.75 | -3.13 | -1.21 | -2.12 | -2.79 | -3.74 | -2.31 | -1.78 | -3.22 | -3.09 | -1.89 | -2.37 | -3.37 | -4.42 | -0.025 | -0.6325 | -1.2075 | 0.436505 | 0.969161 | 0.795511 | 1 | 0.999631 | 0.780748 |
| Grin2a_P35436_S1198s_1 | 0.43 | -0.48 | 1.4 | 0.44 | -0.3 | 0 | 3.82 | -1.83 | 0.42 | -0.84 | -0.06 | -0.26 | 1.47 | -2.66 | -0.81 | -1.04 | -0.73 | -0.7725 | -1.7325 | 0.159378 | 0.981526 | 0.185298 | 1 | 0.999631 | 0.629514 |
| Grin2a_P35436_S1204s_1 | 0.05 | -1.65 | -0.58 | -1.31 | -2.52 | -2.18 | 0.29 | -2 | -2.24 | -0.26 | -1.61 | -2.47 | -0.51 | -3.98 | -2.07 | -3.86 | -0.415 | -0.1 | -0.5025 | 0.050147 | 0.286296 | 0.199443 | 1 | 0.999631 | 0.488453 |
| Grin2a_P35436_S1459s_1 | 0.97 | -0.19 | -1.15 | -1.41 | -0.3 | -0.32 | -1.49 | -1.33 | -0.87 | 0.43 | -1.12 | -0.62 | -0.84 | -0.71 | -1.61 | -0.63 | -0.0725 | 0.0425 | 0.3575 | 0.367441 | 0.473801 | 0.864089 | 1 | 0.999631 | 0.745967 |
| Grin2a_P35436_S882s_1 | -1.41 | -1.15 | -1.84 | -1.99 | -1.3 | -1.76 | -1.92 | -1.7 | -1.85 | -1.53 | -1.76 | -1.08 | -0.04 | -1.03 | -2.17 | -1.72 | -0.2175 | -0.0375 | 0.215 | 0.46806 | 0.808165 | 0.894007 | 1 | 0.999631 | 0.795565 |
| Grin2a_P35436_S890s_1 | -0.6 | 0.16 | 0.17 | -0.75 | -0.28 | -0.17 | -0.62 | -0.82 | -0.34 | -0.43 | -0.22 | -0.18 | 0.48 | 0.05 | -0.75 | 0.06 | 0.02 | -0.2125 | -0.2075 | 0.581221 | 0.513351 | 0.905113 | 1 | 0.999631 | 0.847114 |
| Grin2a_P35436_S929s_1 | 0.59 | 1.75 | 0.24 | 0.61 | 1.25 | 1.37 | 0.03 | 0.62 | 1 | 1.01 | 0.15 | 0.18 | 0.78 | 0.58 | 0 | 1 | -0.955 | -0.905 | -1.99 | 0.61567 | 0.963689 | 0.610449 | 1 | 0.999631 | 0.861203 |
| Grin2b_G3X9V4_S1116s_1 | 2.11 | -0.63 | 1.01 | 1.13 | -0.86 | -0.54 | 1.59 | -0.39 | -0.34 | 1.51 | -0.85 | -0.32 | -0.41 | -2.24 | -0.3 | -1.39 | 0.135 | 0.1625 | 0.8175 | 0.011651 | 0.195231 | 0.207732 | 1 | 0.999631 | 0.385092 |
| Grin2b_G3X9V4_S1143s_1 | -1.1 | 0.21 | -0.79 | -1.52 | -0.18 | -0.07 | -1.18 | -1.23 | -0.19 | -0.63 | -0.94 | -0.79 | 0.75 | 0.24 | -0.42 | -0.5 | -0.0375 | -0.4725 | -1.39 | 0.102887 | 0.770846 | 0.693049 | 1 | 0.999631 | 0.565572 |
| Grin2b_G3X9V4_S1166s_1 | 2.36 | 0.72 | 1.41 | 1.85 | 1.02 | 0.99 | 2.52 | 1.66 | 1.12 | 1.3 | 1.26 | 0.77 | -0.71 | 0.34 | 0.82 | 0.33 | -0.205 | -0.7525 | -1.4775 | 0.012988 | 0.937021 | 0.240125 | 1 | 0.999631 | 0.385092 |
| Grin2b_G3X9V4_S1284s_1 | 1.72 | 0.07 | 0.4 | 0.75 | 0.33 | 0.24 | 1.03 | 0.52 | -0.12 | 0.9 | -0.18 | -0.67 | -1.29 | -0.96 | -0.18 | -0.54 | 1.15 | 1.655 | 0.2325 | 0.006585 | 0.611931 | 0.133398 | 1 | 0.999631 | 0.351545 |
| Grin2b_G3X9V4_S1303s_1 | -0.32 | 1.87 | 1.66 | -0.22 | 1.64 | 1.05 | 3.12 | 1.78 | 2.13 | 0.32 | 2.66 | 4.5 | 3.37 | 0.38 | -0.26 | 0.43 | 0.1275 | -0.4725 | 0.015 | 0.788909 | 0.103965 | 0.092807 | 1 | 0.999631 | 0.925513 |
| Grin2b_G3X9V4_S882s_1 | -0.98 | -0.36 | -0.82 | -1.41 | -0.27 | -0.42 | -1.19 | -1.18 | -1.17 | -0.98 | -0.26 | -3.05 | -0.36 | 0.26 | -2.05 | -1.36 | 0.35 | 0.07 | -0.66 | 0.977213 | 0.719385 | 0.423818 | 1 | 0.999631 | 0.992174 |
| Grin2b_G3X9V4_S882s_2 | -0.46 | -1.6 | 3.33 | 1.07 | -1.07 | -1.12 | 1.74 | 4.19 | -1.05 | -0.12 | 1.71 | 2.08 | 0.34 | -1.33 | -1.59 | 2.28 | -0.23 | 0.0825 | 0.425 | 0.578357 | 0.804211 | 0.949966 | 1 | 0.999631 | 0.84622 |
| Grin2b_G3X9V4_S886s_1 | -0.41 | -0.81 | -0.89 | -0.92 | -0.54 | -1.33 | -0.96 | -1.12 | -0.71 | -0.63 | -0.94 | -0.42 | 0.42 | -0.91 | -0.53 | -0.31 | 0.35 | 0.07 | -0.66 | 0.250064 | 0.424634 | 0.760341 | 1 | 0.999631 | 0.686417 |
| Grin2b_G3X9V4_S886s_2 | -0.46 | -1.6 | 3.33 | 1.07 | -1.07 | -1.12 | 1.74 | 4.19 | -1.05 | -0.12 | 1.71 | 2.08 | 0.34 | -1.33 | -1.59 | 2.28 | 0.09 | 0.2 | 0.625 | 0.578357 | 0.804211 | 0.949966 | 1 | 0.999631 | 0.84622 |
| Grin2b_G3X9V4_S917s_1 | -1.65 | -1.1 | -0.81 | -1.27 | -1.15 | -1.2 | -1.13 | -0.99 | -1.28 | -1.67 | -0.86 | -0.22 | -0.12 | -1.2 | -0.5 | -0.51 | -0.015 | -0.095 | -0.51 | 0.094865 | 0.74093 | 0.600379 | 1 | 0.999631 | 0.558128 |
| Grin2b_G3X9V4_S929s_1 | 1.23 | 0.92 | 0.98 | 1.49 | 0.96 | 1.19 | 1.19 | 1.22 | 1.04 | 1.17 | 1.41 | 0.62 | -0.63 | 0.77 | 0.9 | 1.54 | 0.59 | -7.9E-17 | -0.12 | 0.289584 | 0.953155 | 0.746569 | 1 | 0.999631 | 0.706164 |
| Grin2b_G3X9V4_S929s_2 | 0.07 | 0.79 | -0.31 | 1.15 | 1.57 | 0.65 | 0.73 | 1.11 | 0.43 | 0.36 | 0.37 | 0.54 | -0.7 | 0.61 | 0.65 | 0.66 | -0.3025 | -1.065 | -1.0575 | 0.796351 | 0.161617 | 1 | 1 | 1 | 0.929895 |
| Grin2b_G3X9V4_S930s_1 | 0.99 | 0.87 | 0.73 | 0.87 | 0.86 | 0.53 | 0.48 | 0.38 | -0.16 | -0.46 | -0.07 | -0.11 | -0.82 | -0.23 | -0.33 | 0.61 | 0.4225 | -0.04 | -0.1475 | 0.012937 | 0.23697 | 0.001724 | 1 | 0.999631 | 0.385092 |
| Grin2b_G3X9V4_S930s_2 | -0.04 | 0.48 | -0.51 | 0.78 | 1 | 0.19 | 0.45 | 0.76 | 0.16 | 0.06 | 0.2 | 0.13 | -1.13 | 0.52 | -0.03 | 0.76 | 0.2525 | -0.3425 | -1.6225 | 0.763705 | 0.257123 | 0.904974 | 1 | 0.999631 | 0.915694 |
| Grin2b_G3X9V4_S940s_1 | 1.53 | 0.41 | 0.27 | 1.14 | 1.18 | 1.11 | 1.61 | 0.46 | 0.9 | 0.75 | 0.53 | -0.2 | -1.29 | -0.61 | -0.4 | -0.84 | 2.015 | 2.865 | 0.415 | 0.001793 | 0.522517 | 0.399978 | 1 | 0.999631 | 0.351308 |
| Grin2b_G3X9V4_Y1039y_1 | -0.41 | 2.86 | 2.08 | -0.22 | 4.73 | 1.63 | 3.8 | 2.21 | 4.62 | 0.31 | 4.91 | 5.93 | 3.59 | 0.59 | 1.29 | 0.5 | -0.655 | -0.5275 | -0.2675 | 0.659694 | 0.054353 | 0.045661 | 1 | 0.999631 | 0.880252 |
| Grin2b_G3X9V4_Y1049y_1 | -0.61 | -1.06 | 0.27 | -0.15 | -1.63 | -0.85 | -0.7 | -0.99 | -0.93 | -1.36 | -0.56 | -0.81 | -0.79 | -1.03 | -0.34 | -0.46 | -0.0925 | -0.2375 | -0.0475 | 0.47589 | 0.102164 | 0.169413 | 1 | 0.999631 | 0.799882 |
| Grin2b_G3X9V4_Y1070y_1 | -0.81 | -3.03 | -2.28 | -1.3 | -1.5 | -2.52 | -1.46 | -2.31 | -2.71 | -1.55 | -1.99 | -2.12 | -1.77 | -2.13 | -1.61 | -2.1 | 0.5225 | 0.5 | 0.29 | 0.923232 | 0.860239 | 0.648379 | 1 | 0.999631 | 0.975032 |
| Grin2b_G3X9V4_Y1225y_1 | 0.99 | 1.31 | 0.88 | -0.01 | 1.7 | 0.28 | 2.18 | 1.1 | 2.24 | -0.54 | 1.69 | 1.78 | 2.27 | 0.43 | 1.04 | 0.59 | -0.3075 | -0.54 | -0.37 | 0.554643 | 0.285334 | 0.427702 | 1 | 0.999631 | 0.838109 |
| Grin2b_G3X9V4_Y932y_1 | -0.57 | -1.35 | -2.27 | -1.8 | -1.69 | -1.29 | -1.9 | -2.34 | -2.08 | -1.07 | -2.38 | -2.62 | -1.03 | -1.48 | -2.07 | -2.89 | 0.13 | -0.135 | -0.415 | 0.476919 | 0.469323 | 0.274733 | 1 | 0.999631 | 0.799947 |
| Grin2b_G3X9V4_Y932y_2 | -0.16 | 0.29 | -0.52 | 0.76 | 0.49 | -0.24 | 0.31 | 0.33 | -0.06 | -0.41 | 0.03 | 0.27 | -1.64 | 0.2 | -0.25 | 0.4 | 0.0025 | -0.225 | 0.715 | 0.424291 | 0.710774 | 0.70085 | 1 | 0.999631 | 0.773927 |
| Grip1_H7BX90_S380s_1 | -0.67 | -1.4 | 0.01 | 0.05 | -1.26 | -0.15 | -0.83 | 0.24 | -1.17 | -1.35 | -0.26 | -0.13 | 0.23 | -0.18 | 0.58 | 0.22 | -0.2725 | -0.0725 | 0.585 | 0.097929 | 0.995659 | 0.621007 | 1 | 0.999631 | 0.560722 |
| Grip1_H7BX90_S43s_1 | -1.01 | -1.36 | 0 | -0.81 | -1.37 | -1.44 | -1.04 | -0.42 | -1.63 | -1.02 | -0.51 | -0.31 | 0.53 | -1.45 | -0.09 | 0.17 | -0.29 | -1.0725 | -0.55 | 0.257223 | 0.482374 | 0.862003 | 1 | 0.999631 | 0.691981 |
| Grip1_H7BX90_S709s_1 | 0.58 | -0.27 | 1.31 | -0.03 | -0.74 | -0.12 | 1.99 | -0.7 | -1.61 | -0.63 | -0.47 | 0.01 | 0.74 | -1.46 | 0.5 | -0.39 | 0.0875 | 0.2925 | 0.09 | 0.34213 | 0.65922 | 0.044372 | 1 | 0.999631 | 0.73347 |
| Grip1_H7BX90_S794s_1 | -1.6 | -3.13 | 0.17 | -0.45 | -2.31 | -2.05 | 0.16 | -0.46 | -2.26 | -1.79 | 0.19 | 0.02 | -1.86 | -1.91 | -0.49 | -0.39 | -0.005 | 0.0725 | -0.4775 | 0.902739 | 0.915095 | 0.727157 | 1 | 0.999631 | 0.968183 |
| Grip1_H7BX90_S885s_1 | 1.54 | 0.14 | 0.51 | 0.99 | 0.33 | 1.18 | 1.21 | 0.44 | 1.11 | 1.27 | 0.71 | 0.38 | -0.16 | 0.46 | 0.7 | 0.27 | 0.2975 | -0.0025 | 0.2425 | 0.234347 | 0.989805 | 0.850211 | 1 | 0.999631 | 0.683223 |
| Grip1_H7BX90_S928s_1 | 0.12 | 0.27 | -1.01 | -0.28 | 0.27 | 0.1 | -0.29 | 0.21 | -0.23 | -0.19 | -0.15 | -0.34 | 0.12 | 0.26 | -0.78 | 0.47 | 0.3425 | -0.415 | -0.3425 | 0.561294 | 0.397762 | 0.994073 | 1 | 0.999631 | 0.840316 |
| Grip1_H7BX90_T904t_1 | -0.38 | 1 | 0.29 | 0.26 | 0.02 | 1.61 | 0.33 | 0.58 | 0.13 | 0.07 | 0.76 | -1.45 | -2.32 | 1.84 | -0.19 | 0.47 | -0.2875 | -0.2575 | 0.1625 | 0.667939 | 0.438754 | 0.428552 | 1 | 0.999631 | 0.883098 |
| Grip2_E0CXE0_S10s_1 | -0.53 | 0.09 | 0.83 | 0.86 | -0.51 | -0.54 | 0.1 | 1.05 | -0.69 | -0.4 | -0.33 | 1.64 | 0.5 | -0.12 | 1.77 | -0.25 | -0.115 | 0.0375 | 0.3975 | 0.761175 | 0.548341 | 0.657081 | 1 | 0.999631 | 0.91472 |
| Grip2_E0CXE0_S773s_1 | 0.49 | 1.28 | 0.45 | -0.02 | 0.92 | 1.35 | -0.43 | -0.1 | 0.96 | 1.2 | 0.03 | 0.16 | 1.5 | 1.26 | 0.37 | 0.66 | 0.06 | 0.925 | -0.2425 | 0.33245 | 0.808364 | 0.926276 | 1 | 0.999631 | 0.728079 |
| Grip2_E0CXE0_S774s_1 | -0.47 | -0.61 | -5.51 | -5.14 | -0.5 | -0.39 | -4.84 | -5.76 | -0.62 | -0.34 | -2.92 | -4.15 | 0.1 | -0.63 | -7.19 | -4.98 | -0.2575 | -0.325 | -0.6925 | 0.896244 | 0.971377 | 0.51777 | 1 | 0.999631 | 0.966232 |
| Grip2_E0CXE0_S810s_1 | -0.2 | -0.27 | -1.03 | -1.13 | -0.95 | 0.41 | -1.12 | -2 | -0.2 | -1.34 | -1.34 | -1.05 | -1.34 | -1.2 | -2.53 | -0.33 | -0.245 | 0.095 | 0.645 | 0.1828 | 0.620158 | 0.406842 | 1 | 0.999631 | 0.643861 |
| Grip2_E0CXS4_S43s_1 | -0.88 | -0.91 | -0.49 | -1.14 | -1.36 | -0.88 | -1.21 | -0.95 | -0.92 | -0.67 | -0.81 | -0.64 | 0.64 | -0.21 | -0.93 | -0.34 | -0.105 | -0.065 | 0.395 | 0.117309 | 0.372856 | 0.721658 | 1 | 0.999631 | 0.585087 |
| Grip2_E0CXS4_S996s_1 | -1.42 | -2.44 | -0.08 | -0.95 | -2.28 | -1.86 | -0.64 | -0.53 | -2.04 | -2.01 | -0.54 | -0.56 | -1.49 | -1.23 | -0.82 | 0.23 | 0.0075 | 0.6425 | -0.805 | 0.494947 | 0.860036 | 0.912768 | 1 | 0.999631 | 0.810518 |
| Gripap1_Q8VD04_S237s_1 | 0.09 | 2.03 | 1.21 | 0.54 | 1.06 | 0.66 | 1.56 | 0.62 | 2.37 | 0.2 | 1.59 | 2.28 | 0.71 | -0.22 | 0.79 | -0.63 | -0.2925 | 0.1275 | 0.4125 | 0.144098 | 0.986901 | 0.297211 | 1 | 0.999631 | 0.616037 |
| Gripap1_Q8VD04_S657s_1 | -0.49 | 0.35 | 0.41 | -0.67 | -0.54 | -0.09 | -0.78 | -0.16 | -0.05 | -0.22 | -0.02 | 0.4 | 0.59 | 0.36 | -0.04 | 0.34 | -0.4525 | -0.205 | 0.6025 | 0.26694 | 0.414432 | 0.715719 | 1 | 0.999631 | 0.695664 |
| Grm1_P97772_S969s_1 | -0.86 | -0.88 | -0.22 | -0.38 | -1.04 | -1.67 | -0.48 | -0.96 | -0.56 | -0.71 | -0.55 | -1.34 | 0.75 | -1.25 | -0.25 | 0.82 | 0.025 | 0.2625 | 0.3325 | 0.241969 | 0.197449 | 0.519632 | 1 | 0.999631 | 0.684914 |
| Grm1_P97772_T988t_1 | -0.09 | -0.16 | -1.36 | -1.77 | -0.33 | -0.94 | -1.05 | -0.96 | -0.83 | 0.29 | -1.12 | -0.67 | -0.35 | -0.63 | -1.22 | 0.15 | 0.225 | -0.345 | -0.0375 | 0.504472 | 0.954893 | 0.599816 | 1 | 0.999631 | 0.814292 |
| Grm2_Q14BI2_S843s_1 | -0.75 | -0.13 | -1.09 | -0.88 | -0.86 | -0.33 | -0.8 | 0.04 | -1.2 | -2.37 | -0.29 | -0.37 | -0.31 | -1.32 | -0.61 | -0.76 | -0.29 | 0.17 | 0.41 | 0.915044 | 0.506182 | 0.494773 | 1 | 0.999631 | 0.97286 |
| Grm2_Q14BI2_S871s_1 | -0.67 | -0.52 | -1.01 | -0.89 | -1.19 | -1.29 | -1.24 | -0.53 | -1.14 | 0.17 | -0.22 | -1.22 | -0.03 | -0.27 | -0.7 | -0.45 | -0.11 | -0.03 | 0.3575 | 0.183826 | 0.323057 | 0.657554 | 1 | 0.999631 | 0.644372 |
| Grm3_Q9QYS2_S845s_1 | -0.7 | 0.18 | -0.63 | -0.46 | -0.31 | -0.4 | -0.73 | -0.61 | -0.15 | -0.54 | -0.33 | -0.71 | 0.43 | 0.03 | -0.36 | -0.28 | -0.6075 | -0.2725 | -0.165 | 0.302655 | 0.707413 | 0.92183 | 1 | 0.999631 | 0.714978 |
| Grm3_Q9QYS2_S878s_1 | -0.49 | -0.77 | -0.15 | -0.85 | -0.49 | -1.22 | -1.54 | -1.44 | -0.78 | -0.05 | -1.46 | -1.06 | -0.19 | -0.32 | -1.62 | -0.79 | 0.6325 | 1.145 | 0.875 | 0.673666 | 0.088625 | 0.463654 | 1 | 0.999631 | 0.885515 |
| Grm5_E9QMC2_S1005s_1 | -0.67 | 0.9 | 1.35 | -0.64 | 1.11 | 1.25 | 0.74 | 0.37 | 1.82 | 0.82 | 0.94 | 1.94 | 1.98 | 0.37 | 2.1 | -0.01 | -0.3275 | -0.2125 | 0.2175 | 0.212615 | 0.241045 | 0.061956 | 1 | 0.999631 | 0.66658 |
| Grm5_E9QMC2_S1141s_1 | -0.65 | -0.12 | 0 | -1.04 | -0.46 | -0.65 | -1.25 | -0.76 | -0.55 | -0.93 | -0.64 | -0.54 | 0.67 | -0.09 | -0.88 | -0.64 | 0.4075 | 0.2625 | 0.6 | 0.614468 | 0.338528 | 0.509868 | 1 | 0.999631 | 0.861203 |
| Grm5_E9QMC2_S1153s_1 | -2.54 | -1.15 | -0.37 | -0.58 | -0.18 | -1.09 | -0.68 | -1.06 | -0.94 | -1.71 | -0.64 | -0.3 | 0.24 | -1.09 | -1.02 | -0.37 | 0.7875 | -0.08 | -0.5325 | 0.283308 | 0.422922 | 0.625757 | 1 | 0.999631 | 0.704903 |
| Grm5_E9QMC2_S839s_1 | 1.18 | 1.09 | -0.97 | 1.93 | 1.91 | 1.75 | 1.04 | 1.68 | 1.61 | -0.3 | 1.64 | -0.04 | -1.7 | 1.25 | 0.82 | 0.73 | 0.7575 | 0.2425 | -0.095 | 0.508328 | 0.202303 | 0.911449 | 1 | 0.999631 | 0.816932 |
| Grm5_E9QMC2_S860s_1 | 0.99 | 0.35 | 0.3 | 1.54 | 1.85 | 1.08 | 1.34 | 1.94 | 1.18 | -0.05 | 1.75 | 1.27 | -0.89 | 0.93 | 1.39 | 1.37 | 1.8175 | 0.3975 | -0.08 | 0.867112 | 0.066949 | 0.606511 | 1 | 0.999631 | 0.956562 |
| Grm5_E9QMC2_S901s_1 | 1.23 | 0.29 | -0.43 | 2.31 | 3.15 | 1.55 | 2.78 | 3.19 | 1.61 | -0.69 | 2.81 | 1.26 | -0.57 | 1.12 | 1.64 | 0.89 | 1.2325 | -1.0075 | -1.0625 | 0.905845 | 0.01608 | 0.63036 | 1 | 0.999631 | 0.969329 |
| Grm5_E9QMC2_T840t_1 | 0.87 | 0.12 | -1.37 | 1.46 | 1.21 | 1.5 | 1.31 | 1.99 | -0.3 | -1.57 | 1.56 | -2.64 | -1.88 | 1.12 | -1.19 | -1.22 | 0.47 | -0.5625 | -2.2925 | 0.196077 | 0.05573 | 0.30101 | 1 | 0.999631 | 0.653373 |
| Grm5_Q3UVX5_S870s_1 | 3.22 | 0.11 | -1.02 | 2.19 | 2.23 | 0.61 | 2.27 | 1.27 | 1.12 | 0.42 | 1.01 | -0.3 | -3.14 | -0.69 | 0.04 | -0.88 | 0.06 | 0.295 | 0.42 | 0.043888 | 0.605462 | 0.529782 | 1 | 0.999631 | 0.475443 |
| Grm7_Q68ED2_S900s_1 | -0.9 | -0.65 | -0.24 | -0.99 | -0.5 | -0.73 | -0.9 | -0.41 | -0.09 | 0.04 | -0.97 | -0.58 | 0.04 | -0.25 | -0.9 | 0.01 | -0.005 | 0.1875 | 0.53 | 0.235237 | 0.831176 | 0.38868 | 1 | 0.999631 | 0.683301 |
| Gsg1l_D3Z7H4_S274s_1 | -0.77 | -0.27 | -0.15 | -0.41 | -0.56 | -0.29 | -0.6 | -0.17 | -0.43 | -0.22 | -0.03 | -0.17 | 0.69 | -0.34 | -0.12 | 0.29 | 0.26 | 0.6525 | 1.0075 | 0.136418 | 0.985022 | 0.49168 | 1 | 0.999631 | 0.609678 |
| Gsk3a_D3Z7E5_S39s_1 | -1.89 | 0.89 | 0.56 | -1.07 | 0.16 | 0.23 | -0.64 | -0.22 | 0.04 | -0.4 | 0.26 | 1.2 | 0.96 | 0.29 | 0.91 | 0.36 | -0.285 | -0.1225 | 0.405 | 0.128109 | 0.676794 | 0.339455 | 1 | 0.999631 | 0.598425 |
| Gsk3a_D3Z7E5_Y279y_1 | 0.42 | 1.38 | 1.42 | 0.86 | 0.7 | 0.73 | 0.43 | 1.08 | 0.97 | 1.18 | 1.34 | 0.1 | 1.42 | 1.31 | 1.57 | 1.4 | 0.725 | 0.095 | 0.015 | 0.228869 | 0.383315 | 0.750378 | 1 | 0.999631 | 0.679434 |
| Gsk3b_E9QAQ5_S25s_1 | 0.84 | 0.67 | -0.68 | 1 | 1.47 | 0.88 | 0.92 | 1.46 | 1.25 | -0.11 | 0.51 | 0.56 | 0.22 | 0.8 | 0.79 | 0.08 | -0.13 | -0.045 | 0.195 | 0.972454 | 0.106212 | 0.837024 | 1 | 0.999631 | 0.990167 |
| Gsk3b_E9QAQ5_S402s_1 | -0.44 | -0.21 | 0.03 | -0.4 | -0.51 | -0.28 | -0.43 | -0.32 | -0.6 | -0.06 | -0.35 | -0.19 | 0.12 | -0.12 | -0.25 | 0.01 | 0.7825 | -0.59 | -0.8775 | 0.486879 | 0.600854 | 0.867334 | 1 | 0.999631 | 0.805169 |
| Gsk3b_E9QAQ5_S9s_1 | 1.82 | 0.5 | 1.13 | 2.48 | 1.27 | 2.38 | 2.85 | 2.56 | 0.89 | 0.45 | 2.34 | -0.11 | -0.99 | 1.61 | 0.61 | 1.19 | -0.32 | -0.6125 | -0.55 | 0.191955 | 0.150513 | 0.348574 | 1 | 0.999631 | 0.648674 |
| Gsk3b_E9QAQ5_T433t_1 | 0.76 | -3.48 | -1.03 | -1.28 | -1.37 | -2.77 | 0.23 | -2.4 | -3.51 | -0.66 | -1.89 | -1.42 | -0.05 | -3.25 | -1.52 | -2.41 | -0.15 | -0.0525 | 0.23 | 0.565506 | 0.736138 | 0.509637 | 1 | 0.999631 | 0.84139 |
| Gsn_P13020_S307s_1 | -0.42 | 0.1 | -0.1 | 0.01 | -0.03 | -0.49 | -0.38 | -0.11 | -0.31 | 0.56 | -0.6 | -0.27 | 0.32 | 0.24 | -0.26 | 0.21 | 0.735 | 0.1175 | 0.0125 | 0.437285 | 0.56889 | 0.87276 | 1 | 0.999631 | 0.781057 |
| Gspt2_Q149F3_S195s_1 | 1.82 | 0.63 | -0.71 | 1.36 | 2.28 | 1.23 | 1.29 | 1.24 | 1.92 | 0.01 | 1.24 | 0.4 | 0.29 | 1.61 | 0.73 | 0.52 | 0.0225 | 0.375 | -0.105 | 0.9826 | 0.207867 | 0.852332 | 1 | 0.999631 | 0.994846 |
| Gsto1_O09131_S139s_1 | -0.19 | 0.21 | 0.67 | -0.15 | 0.05 | 0.58 | 0.49 | -0.49 | 0.84 | 0.33 | 0.36 | 0.51 | 0.33 | -0.06 | 0.19 | -0.34 | -0.5975 | -0.615 | -1.1675 | 0.747815 | 0.948267 | 0.236054 | 1 | 0.999631 | 0.911931 |
| Gtf2f1_Q3THK3_T331t_1 | 2.49 | 1.85 | 1.94 | 2.2 | 0.74 | 1.92 | 2 | 1.43 | 1.15 | 2.54 | 1.69 | 0.64 | 0.25 | 1.23 | 1.85 | 0.48 | -0.5375 | -0.63 | 0.085 | 0.016344 | 0.114085 | 0.175274 | 1 | 0.999631 | 0.388832 |
| Gtf2i_Q3UHU8_S783s_1 | -0.04 | 0.7 | 0.94 | 0.92 | 0.14 | -0.07 | -0.35 | 0.65 | -0.16 | 0.29 | -0.08 | -0.05 | 0.51 | 1.22 | 0.32 | 0.81 | 0.0075 | 0.375 | 0.5575 | 0.811128 | 0.142607 | 0.069011 | 1 | 0.999631 | 0.936949 |
| Gtf3c1_Q8K284_S1794s_1 | -1.08 | 0.74 | 0.22 | -0.08 | 0.22 | -0.22 | -0.44 | 0.27 | 0.89 | -0.32 | -0.02 | 0.75 | 1.43 | 0.78 | -0.02 | -0.16 | -0.3625 | -0.2175 | 0.06 | 0.287196 | 0.985634 | 0.430143 | 1 | 0.999631 | 0.705972 |
| Gtf3c4_Q8BMQ2_S17s_1 | 0.8 | -0.18 | 0.42 | 0.83 | -0.62 | -0.36 | 0.83 | 0.57 | -0.13 | 0.15 | 0.22 | 0.76 | 0.82 | 0.08 | 1.03 | 0.18 | 0.4875 | 0.715 | 1.0725 | 0.872173 | 0.396602 | 0.530651 | 1 | 0.999631 | 0.959391 |
| Gtpbp1_O08582_S24s_1 | -0.61 | 0.53 | 0.02 | -0.28 | 1.07 | 0.73 | -0.75 | 0.56 | 0.83 | 0.49 | 0.57 | 0.63 | 1.67 | 1.45 | 0.27 | 0.56 | -0.1375 | 0.2175 | 0.5875 | 0.02867 | 0.294117 | 0.04445 | 1 | 0.999631 | 0.427571 |
| Gtpbp1_O08582_S580s_1 | 0.03 | 0.88 | 0.14 | -0.35 | 0.39 | 0.38 | -0.35 | -0.27 | 0.61 | 0.39 | 0.15 | 0.42 | 1.31 | 0.75 | 0.47 | 0.52 | -0.2025 | -0.0625 | 0.06 | 0.13258 | 0.698588 | 0.512047 | 1 | 0.999631 | 0.604411 |
| Gtpbp1_O08582_S69s_1 | -0.51 | 0.14 | -0.48 | -0.7 | -0.47 | -0.71 | -0.83 | -0.35 | -1.21 | -0.53 | -0.02 | -0.04 | -0.14 | -0.21 | -0.66 | -0.3 | -0.0025 | 0.245 | 0.015 | 0.846555 | 0.486407 | 0.863829 | 1 | 0.999631 | 0.948838 |
| Gtpbp1_O08582_S8s_1 | -0.6 | -0.1 | 0.34 | -0.44 | -0.3 | -0.54 | -0.44 | 0.47 | 0.06 | -0.72 | 0.37 | 0.47 | -0.15 | -0.25 | -0.19 | -0.15 | 0.0475 | -0.13 | 0.0375 | 0.960987 | 0.994181 | 0.51134 | 1 | 0.999631 | 0.986594 |
| Gucy1a2_F8VQK3_S41s_1 | 0.06 | -0.44 | -0.4 | -0.58 | -0.7 | -0.19 | 0.27 | -0.55 | -0.59 | 0.06 | -0.63 | -0.72 | -0.21 | -0.19 | -0.56 | -0.25 | -0.47 | 0.0225 | 0.4425 | 0.896188 | 0.878619 | 0.667612 | 1 | 0.999631 | 0.966232 |
| Gys1_Q9Z1E4_S645s_1 | -1.12 | -1.68 | -0.37 | -1.16 | -1.53 | -1.63 | -1.35 | -1.7 | -1.26 | -0.87 | -1.57 | -0.54 | 0.07 | -1.13 | -0.68 | -0.82 | 0.725 | 0.355 | 0.0225 | 0.280286 | 0.170407 | 0.952194 | 1 | 0.999631 | 0.701922 |
| Gys1_Q9Z1E4_S653s_2 | -1.1 | 1.9 | 3.96 | 2.86 | 1.84 | 2.67 | 2.94 | 3.07 | 1.53 | 2.06 | 2.4 | 3.05 | 2.64 | 1.32 | 2.17 | 1.58 | 0.725 | 0.355 | 0.0225 | 0.981386 | 0.459713 | 0.717627 | 1 | 0.999631 | 0.994608 |
| Gys1_Q9Z1E4_S657s_2 | -1.1 | 1.9 | 3.96 | 2.86 | 1.84 | 2.67 | 2.94 | 3.07 | 1.53 | 2.06 | 2.4 | 3.05 | 2.64 | 1.32 | 2.17 | 1.58 | -0.1875 | -1.175 | -2.2725 | 0.981386 | 0.459713 | 0.717627 | 1 | 0.999631 | 0.994608 |
| Gys1_Q9Z1E4_S698s_1 | 2.84 | -0.71 | -0.3 | 0.41 | -1.39 | 0.03 | 3.88 | -1.03 | -1.57 | 1.47 | -1.87 | -0.49 | -2.19 | -2.11 | -0.61 | -1.94 | 0.185 | -0.35 | 0.115 | 0.013499 | 0.879017 | 0.234803 | 1 | 0.999631 | 0.385092 |
| H1f0_P10922_S92s_1 | -1.2 | -5.62 | -4.51 | -3.22 | -2.75 | -4.2 | -1.51 | -5.35 | -3.93 | -3.87 | -3.02 | -5.13 | -1.16 | -3.97 | -5.15 | -3.81 | -0.4175 | -0.4725 | -0.9225 | 0.915118 | 0.864558 | 0.700433 | 1 | 0.999631 | 0.97286 |
| H2afx_P27661_S140s_1 | 0.51 | -2.52 | -2.19 | -1.35 | -2.41 | -1.51 | -0.91 | -2.39 | -1.82 | -0.11 | -2.47 | -3.04 | -0.62 | -1.84 | -3.12 | -3.66 | -0.0575 | -0.56 | 0.015 | 0.283975 | 0.545288 | 0.565063 | 1 | 0.999631 | 0.704942 |
| H2afy_Q9QZQ8_T178t_1 | -0.87 | -0.85 | -1.4 | -1.25 | -0.8 | -1.06 | -1.5 | -1.24 | -0.92 | -1.27 | -1.79 | -2.63 | -0.47 | -1.09 | -1.58 | -1.17 | 0.215 | 0.795 | 1.13 | 0.964365 | 0.83825 | 0.18939 | 1 | 0.999631 | 0.987331 |
| Habp4_E9QKB2_S109s_1 | -0.18 | 1.9 | 1.47 | 0.07 | 1.5 | 0.88 | 0.63 | 1.11 | 2.03 | 0.47 | 1.4 | 2.54 | 3.35 | 1.18 | 1.96 | 1.29 | 0.2075 | 0.695 | 0.4325 | 0.103313 | 0.67375 | 0.216825 | 1 | 0.999631 | 0.566655 |
| Habp4_E9QKB2_S75s_1 | 1.1 | 1.24 | 1.17 | 1.66 | 0.93 | 1.55 | 1.26 | 2.26 | 1.81 | 1.69 | 2.27 | 2.18 | 0.24 | 2 | 2.1 | 2.56 | -0.0625 | 0.0775 | 0.435 | 0.398125 | 0.549188 | 0.032023 | 1 | 0.999631 | 0.761726 |
| Habp4_E9QKB2_S7s_1 | -0.41 | 0.62 | 0.45 | 0.04 | 0.53 | -0.08 | -0.15 | 0.15 | 0.4 | 0.12 | 0.06 | 0.43 | 1.02 | 0.36 | 0.5 | 0.56 | 0.2025 | 0.26 | 0.5725 | 0.217008 | 0.84639 | 0.804876 | 1 | 0.999631 | 0.669465 |
| Hacd3_Q8K2C9_S114s_1 | -1.52 | -0.38 | -0.23 | -0.56 | -0.37 | -0.28 | -0.88 | -0.35 | -0.36 | -0.68 | -0.32 | -0.29 | 0.26 | -0.33 | -0.33 | 0 | -0.035 | 0.5525 | 0.8275 | 0.142956 | 0.566826 | 0.459012 | 1 | 0.999631 | 0.614053 |
| Hars2_G5E823_S66s_1 | 0.13 | 1.39 | 0.77 | -0.91 | 1.22 | 0.33 | -0.03 | -0.28 | 1.09 | -0.33 | 1.26 | 1.57 | 1.46 | 0.72 | 1.23 | 1.28 | -0.2225 | -0.77 | 0.3 | 0.119179 | 0.948597 | 0.361102 | 1 | 0.999631 | 0.586721 |
| Hbbt1_E9Q223_S88s_1 | 0.22 | 1.35 | -1.01 | 1.85 | 0.52 | 0.27 | 0.83 | -0.1 | 0.06 | 0.22 | -0.08 | -0.87 | 1.49 | 0.51 | 0.9 | 0.71 | -1.2775 | -0.6125 | -1.0525 | 0.624486 | 0.712346 | 0.231354 | 1 | 0.999631 | 0.86366 |
| Hcar1_E9PZR8_S331s_1 | 2.23 | 0.62 | 1.53 | 1.65 | 1.27 | 0.58 | -0.5 | -0.43 | 0.82 | 2.23 | 0.55 | -0.02 | -0.45 | 0.59 | 1.63 | 0.05 | -0.0675 | -0.33 | 0.2475 | 0.06821 | 0.028548 | 0.274587 | 1 | 0.999631 | 0.522688 |
| Hcfc1_Q61191_S3s_1 | -1.03 | -3.09 | 0.71 | -0.07 | -0.33 | -3.74 | 0.15 | 0.17 | -1.98 | -1.89 | -0.2 | -0.73 | 0.24 | -2.6 | 0.52 | -0.65 | -0.2275 | -0.4325 | -0.4375 | 0.790361 | 0.949695 | 0.686865 | 1 | 0.999631 | 0.926161 |
| Hcls1_P49710_S333s_1 | 0.09 | 1.35 | 0.84 | 0.6 | 0.48 | 0.83 | -0.06 | 0.72 | 0.11 | -0.14 | 0.08 | 1.1 | 0.95 | -0.47 | -0.11 | 0.76 | 1.98 | 1.6 | -0.49 | 0.32778 | 0.525941 | 0.289325 | 1 | 0.999631 | 0.725014 |
| Hcn1_O88704_S53s_1 | -1.75 | -0.26 | 0.62 | 0.41 | 3.18 | 0.18 | 3.9 | -0.32 | 2.7 | -1.8 | 2.3 | 2.22 | 1.32 | -1.11 | -0.53 | -2.62 | 1.3075 | 0.6975 | 0.4775 | 0.56521 | 0.076389 | 0.141565 | 1 | 0.999631 | 0.84139 |
| Hcn1_O88704_S69s_1 | -2.26 | -2.38 | 0.55 | -1.68 | 0.34 | 0.22 | -0.14 | -0.96 | -0.98 | -1.06 | -0.97 | 0.03 | -1.74 | 0.05 | -1.07 | -1.1 | -0.215 | -0.85 | -0.8025 | 0.494201 | 0.072182 | 0.301587 | 1 | 0.999631 | 0.81007 |
| Hcn1_O88704_S810s_1 | 0.13 | 0.19 | 0.19 | 0.74 | 0.26 | 0.05 | 0.24 | -0.16 | -0.78 | -0.19 | -1.15 | -0.03 | -1.52 | -0.19 | -0.59 | 0.34 | 0.1625 | -0.035 | -0.3125 | 0.084619 | 0.430536 | 0.030422 | 1 | 0.999631 | 0.549982 |
| Hcn1_O88704_S867s_1 | 1.04 | 0.75 | 0.54 | 0.91 | 1.03 | 1.34 | 0.65 | 0.87 | 0.62 | 0.58 | 0.9 | 1 | -0.47 | 0.76 | 0.23 | 1.47 | -0.28 | -0.46 | -0.7525 | 0.475302 | 0.553113 | 0.895208 | 1 | 0.999631 | 0.799882 |
| Hcn1_O88704_S9s_1 | 0.93 | 0.98 | 0.5 | 0.26 | 0.58 | 0.8 | 0.78 | -0.61 | 1.23 | 1.1 | -0.18 | -1.32 | 0.45 | 0.27 | -0.07 | -0.99 | -0.265 | 0.075 | 0.2825 | 0.079208 | 0.476556 | 0.430867 | 1 | 0.999631 | 0.545579 |
| Hcn1_O88704_T39t_1 | -0.73 | -0.75 | -0.91 | -1.11 | -1.02 | -0.75 | -1.33 | -1.46 | -1.05 | -0.41 | -0.7 | -1.04 | 0.46 | -0.51 | -1.49 | -0.83 | 0.1325 | 0.27 | 0.33 | 0.511777 | 0.342247 | 0.787088 | 1 | 0.999631 | 0.817687 |
| Hcn2_O88703_S119s_1 | -0.56 | -0.55 | -0.88 | -1.38 | -0.7 | -0.77 | -0.99 | -0.38 | -0.75 | 0.09 | -0.63 | -1 | 0.02 | -0.74 | -0.84 | -0.49 | -0.2975 | -0.165 | 0.45 | 0.342735 | 0.657461 | 0.442685 | 1 | 0.999631 | 0.73348 |
| Hcn2_O88703_S131s_1 | -0.6 | -0.35 | -0.11 | -0.56 | -1.08 | -0.82 | -0.83 | -0.08 | -0.12 | -0.95 | -0.52 | -0.69 | 0.54 | -0.38 | 0.11 | -0.09 | -0.17 | -0.4025 | -0.0175 | 0.173079 | 0.339619 | 0.574791 | 1 | 0.999631 | 0.642116 |
| Hcn2_O88703_S134s_1 | 0.18 | -0.38 | -0.62 | -0.71 | -0.28 | -0.41 | -0.67 | -0.85 | -0.77 | -0.8 | -1.04 | -0.53 | 0.44 | -0.37 | -0.84 | -0.83 | 0.6325 | 0.725 | 1.2275 | 0.964256 | 0.574501 | 0.201864 | 1 | 0.999631 | 0.987331 |
| Hcn2_O88703_S67s_1 | -1.83 | -1.61 | -0.81 | -0.83 | -0.38 | -1.21 | -0.56 | -0.4 | -0.51 | -1.88 | -0.18 | 0.39 | 0.78 | -0.46 | 0.6 | -1.09 | 0.265 | -0.165 | 0.275 | 0.030676 | 0.099341 | 0.182755 | 1 | 0.999631 | 0.438906 |
| Hcn2_O88703_S726s_1 | -0.44 | -1.64 | 0.09 | 0.13 | 0.23 | -0.41 | -0.39 | -0.23 | -0.42 | -1.02 | -1.17 | 0.09 | -1.38 | 0.29 | 0.37 | -0.04 | 0.275 | 0.2875 | 0.39 | 0.613641 | 0.542081 | 0.733274 | 1 | 0.999631 | 0.860866 |
| Hcn2_O88703_S750s_1 | -0.75 | 0.33 | -0.68 | -0.62 | -0.09 | -0.02 | -0.53 | 0.02 | -0.1 | -0.34 | -0.23 | 0.1 | 0.75 | 0.09 | -0.51 | -0.49 | -0.1625 | -0.0975 | 0.185 | 0.353263 | 0.409999 | 0.388405 | 1 | 0.999631 | 0.73949 |
| Hcn2_O88703_S771s_1 | -0.59 | -0.58 | -0.65 | -0.79 | -0.74 | -0.52 | -1.17 | -0.83 | -0.88 | -0.37 | -1.12 | -0.63 | -0.28 | -0.34 | -0.58 | -0.67 | -0.45 | -0.2775 | -0.005 | 0.49592 | 0.529082 | 0.722392 | 1 | 0.999631 | 0.811036 |
| Hcn2_O88703_S795s_1 | -0.64 | -0.35 | -0.16 | -0.86 | -1.04 | -0.97 | -0.8 | -1 | -1.3 | -0.3 | -0.75 | -0.77 | 0.18 | -0.9 | -1.1 | -0.21 | -0.345 | -0.9225 | -1.5075 | 0.989361 | 0.11265 | 0.392167 | 1 | 0.999631 | 0.995688 |
| Hcn2_O88703_S818s_1 | 1.24 | -2.92 | 0.6 | -0.07 | -1.74 | -0.65 | 1.49 | -1.63 | -2.23 | 0.16 | -1.52 | -1.25 | -2.32 | -1.7 | -1.13 | -2.03 | -0.5425 | -0.6425 | -1.21 | 0.092725 | 0.735186 | 0.323141 | 1 | 0.999631 | 0.557333 |
| Hcn2_O88703_S834s_1 | 1.32 | -0.37 | 0.03 | 0.04 | -0.7 | -0.15 | 0.18 | -0.48 | -0.57 | 1.22 | -1.12 | -1.08 | -0.7 | -1.41 | -0.61 | -1.1 | 0.0425 | -0.125 | 0.18 | 0.015951 | 0.209818 | 0.300908 | 1 | 0.999631 | 0.388832 |
| Hcn2_O88703_S840s_1 | -0.04 | 0.16 | -1.17 | -1.33 | -0.54 | 0.19 | -0.94 | -0.92 | -0.58 | -0.32 | -1.09 | -0.89 | 0.11 | -0.34 | -0.52 | -0.91 | 0.1175 | 0.0975 | 0.72 | 0.683616 | 0.924515 | 0.766819 | 1 | 0.999631 | 0.889602 |
| Hcn2_O88703_S842s_1 | -1.33 | 0.16 | 0.32 | -0.1 | -0.23 | -0.01 | -0.49 | 0.25 | -0.49 | -0.9 | 0.17 | 0.66 | 0.34 | 0.38 | 0.45 | 0.76 | -0.335 | -0.015 | -0.08 | 0.100953 | 0.773296 | 0.841397 | 1 | 0.999631 | 0.564448 |
| Hcn2_O88703_S90s_1 | 0.57 | -0.44 | -0.28 | -0.56 | -0.67 | -0.47 | -0.74 | -0.17 | -0.54 | 0.68 | -0.46 | -0.45 | 0.01 | -0.2 | -0.32 | -0.52 | 0.06 | -0.1725 | -0.4 | 0.815266 | 0.320885 | 0.970068 | 1 | 0.999631 | 0.937414 |
| Hcn2_O88703_T70t_1 | 0.65 | -1.95 | 0.35 | 0.25 | -1.52 | 0.09 | 0.73 | 0.24 | -0.81 | 0.19 | -0.42 | -0.35 | -0.89 | -0.8 | -0.15 | -0.46 | -0.0875 | -1.05 | -0.2675 | 0.490052 | 0.930093 | 0.766362 | 1 | 0.999631 | 0.807727 |
| Hcn2_O88703_T763t_1 | 1.49 | 0.4 | -0.95 | -0.85 | 1.47 | 0.37 | -1.25 | -0.85 | -0.03 | -2.33 | -0.96 | -0.79 | 0.18 | 0.07 | -0.7 | -0.53 | 0.3875 | 0.38 | 0.3425 | 0.640192 | 0.906192 | 0.142114 | 1 | 0.999631 | 0.870995 |
| Hcn3_O88705_S33s_1 | 0.5 | -1.11 | -0.8 | -2.07 | 0.61 | 0.33 | -1.4 | -1.47 | -0.38 | 0.98 | -2.05 | -0.51 | 0.56 | 0.42 | -1.62 | -1.47 | -0.0025 | 0.145 | 0.2075 | 0.627859 | 0.572216 | 0.601979 | 1 | 0.999631 | 0.865919 |
| Hcn3_O88705_S633s_1 | -0.63 | -0.17 | -0.58 | -0.61 | -0.41 | -0.53 | -0.73 | -0.33 | -0.74 | -0.05 | -0.76 | 0.14 | -0.11 | -0.13 | -0.67 | -0.25 | -0.4625 | 0.4225 | 0.71 | 0.480253 | 0.992144 | 0.650342 | 1 | 0.999631 | 0.801597 |
| Hcn3_O88705_S648s_1 | -1.01 | -0.77 | -0.96 | -0.08 | -1.11 | -1.2 | -1.21 | -1.15 | -0.69 | -0.29 | -0.57 | 0.42 | 0.7 | -0.12 | 0.09 | -0.65 | 0.345 | 0.2625 | 0.1675 | 0.089411 | 0.133388 | 0.259152 | 1 | 0.999631 | 0.555695 |
| Hcn3_O88705_S654s_1 | 0.39 | 0.27 | -0.19 | 0.22 | 0.8 | 0.66 | -0.11 | 0.72 | 0.54 | -0.4 | 0.54 | 1.06 | 0.45 | 0.17 | 0.22 | 0.52 | 0.06 | -0.07 | 0.465 | 0.557738 | 0.274085 | 0.47399 | 1 | 0.999631 | 0.838305 |
| Hcn3_O88705_S720s_1 | -0.36 | -0.94 | 0.01 | 0.17 | -0.58 | -0.73 | -0.11 | 0.54 | -0.46 | -0.1 | -0.72 | -0.12 | -0.98 | 0.21 | 1.23 | 0.28 | -0.3775 | -0.115 | -1.9475 | 0.358721 | 0.876861 | 0.83499 | 1 | 0.999631 | 0.741182 |
| Hcn3_O88705_S733s_1 | 1.69 | -3.62 | -0.33 | -0.81 | -3.36 | -0.54 | 2.56 | -3.24 | -1.91 | 1.96 | -1.79 | -1.79 | -2.33 | -2.83 | -1.69 | -4.01 | 0.58 | -0.015 | -0.3675 | 0.081222 | 0.801963 | 0.925933 | 1 | 0.999631 | 0.54635 |
| Hcn3_O88705_T16t_1 | -0.17 | -0.69 | 0.33 | -0.69 | 0.55 | 0.4 | 0.95 | -0.8 | 0.05 | -0.05 | -1.34 | 0.06 | -0.04 | -0.65 | -0.46 | -1.54 | -0.16 | 0.0075 | 0.58 | 0.38602 | 0.205882 | 0.971628 | 1 | 0.999631 | 0.756141 |
| Hcn4_B2RY58_S1026s_1 | -0.59 | -1.47 | -0.51 | -0.51 | -1.13 | -1.23 | -0.67 | -0.69 | -1.04 | -0.28 | -1.23 | -0.5 | 0.04 | -0.47 | -0.03 | -0.3 | -0.375 | 0.0375 | 0.66 | 0.106042 | 0.621715 | 0.983265 | 1 | 0.999631 | 0.568797 |
| Hcn4_B2RY58_S918s_1 | 0.48 | -0.33 | -0.98 | 0.4 | -0.01 | -1.45 | 0.18 | -0.65 | 0.28 | -0.66 | -0.73 | 0.83 | 1.19 | -0.11 | 1.94 | -0.81 | 1.19 | 0.1825 | 0.4875 | 0.314878 | 0.439864 | 0.938684 | 1 | 0.999631 | 0.720305 |
| Hcn4_B2RY58_S936s_1 | -0.28 | -1.28 | -3.1 | -1.4 | -0.06 | -0.77 | -0.5 | 0.03 | -0.77 | -1.46 | -2.3 | -0.8 | -1.02 | -0.57 | -1.05 | -1.47 | 0.22 | 0.48 | 0.9025 | 0.398445 | 0.057025 | 0.769486 | 1 | 0.999631 | 0.761726 |
| Hcn4_B2RY58_T1071t_1 | -0.98 | -1.81 | -0.89 | -1.27 | -1.28 | -1.18 | -1.14 | -0.47 | -1.55 | 0.09 | -1.52 | -0.05 | 0.41 | -0.14 | -1.31 | -0.3 | 0.175 | 0.2775 | 0.625 | 0.05599 | 0.501625 | 0.327 | 1 | 0.999631 | 0.495761 |
| Hdac4_Q6NZM9_S465s_1 | -1.02 | 0.34 | 0.81 | 0.21 | 0.3 | 0.45 | 0 | 0.29 | 0.46 | -0.06 | 0.47 | 0.58 | 0.39 | 0.82 | 0.73 | 0.9 | -0.18 | -0.0725 | 0.205 | 0.158925 | 0.666267 | 0.513104 | 1 | 0.999631 | 0.629092 |
| Hdac4_Q6NZM9_S629s_1 | -0.45 | 0.13 | -0.16 | -0.49 | -0.33 | -0.44 | -0.73 | -0.19 | -0.35 | -0.35 | -0.24 | -0.32 | 0.35 | 0.26 | -0.5 | -0.26 | -0.5925 | -0.0125 | 0.1525 | 0.534785 | 0.51457 | 0.784478 | 1 | 0.999631 | 0.828799 |
| Hdac4_Q6NZM9_S630s_1 | 0.33 | 1.16 | 0.36 | 0.21 | -0.2 | 1.03 | -0.51 | -0.63 | 0.18 | 0.78 | 0.24 | 0.81 | 1.03 | 1.4 | 0.31 | -0.07 | 0.2675 | 0.365 | 0.435 | 0.713925 | 0.189627 | 0.96965 | 1 | 0.999631 | 0.901439 |
| Hdac4_Q6NZM9_T463t_1 | -1.21 | 0.32 | -0.05 | -0.23 | 0.13 | 0.06 | -0.3 | 0.01 | 0.5 | 0.07 | -0.23 | -0.05 | -0.21 | 0.78 | 0.14 | -0.14 | 0.06 | -0.005 | -0.29 | 0.306811 | 0.466968 | 0.351705 | 1 | 0.999631 | 0.717667 |
| Hdac5_Q3UJF1_S652s_1 | -0.14 | 0.56 | -0.46 | -0.23 | -1.04 | 0.98 | 0.47 | -0.44 | 0.07 | -0.45 | -0.28 | 0.37 | -0.87 | 0.22 | -0.85 | 0.07 | 0.2425 | -0.475 | -0.02 | 0.467266 | 0.899922 | 0.988077 | 1 | 0.999631 | 0.795438 |
| Hdac5_Q3UJF1_S662s_1 | 0.06 | -0.52 | -0.7 | -0.24 | -0.47 | 0.17 | 0.24 | -0.37 | -1.21 | -0.89 | -0.49 | -0.71 | -0.54 | -0.63 | -0.46 | 0.15 | 0.5975 | 0.26 | 0.57 | 0.950601 | 0.434828 | 0.138473 | 1 | 0.999631 | 0.984079 |
| Hdac5_Q3UJF1_T488t_1 | -0.55 | 1.6 | 0.74 | 0.6 | 1.51 | 1.85 | 0.43 | 0.99 | 1.15 | 0.14 | 1.19 | 0.95 | -0.25 | 2.32 | 1.1 | 1.5 | -0.1175 | 0.12 | 0.21 | 0.374068 | 0.254157 | 0.593663 | 1 | 0.999631 | 0.747035 |
| Hdac6_Q9Z2V5_S1148s_1 | -0.24 | -0.38 | -0.07 | -0.14 | -0.22 | -0.66 | -0.21 | -0.21 | 0.03 | -0.31 | -0.26 | 0.19 | 0.71 | -0.55 | 0 | -0.15 | -0.015 | 0.34 | 0.3125 | 0.53853 | 0.640891 | 0.647424 | 1 | 0.999631 | 0.830863 |
| Hdac6_Q9Z2V5_S43s_1 | 0.57 | 1.48 | 1.18 | 0.82 | 0.97 | 1.8 | 0.35 | 0.87 | 1.46 | 1.42 | 0.7 | 1.83 | -0.02 | 2.58 | 1.81 | 0.93 | 0.24 | 0.02 | 0.28 | 0.575195 | 0.968152 | 0.342415 | 1 | 0.999631 | 0.846045 |
| Hdac6_Q9Z2V5_S59s_1 | -0.02 | 0.49 | -0.29 | 0.28 | 0.76 | -0.16 | 0.58 | 0.24 | 0.3 | -0.53 | 0.14 | 0.63 | -0.09 | 0.43 | 0.97 | 0.27 | 0.5425 | 0.3175 | -0.055 | 0.421317 | 0.454411 | 0.953339 | 1 | 0.999631 | 0.772817 |
| Hdac6_Q9Z2V5_S909s_1 | 1.2 | 0.05 | -1.24 | 1.5 | 1.42 | 0.18 | 1.22 | 0.86 | 1.09 | -0.38 | 1.25 | 0.82 | -0.59 | 0.54 | 0.53 | 0.81 | -0.135 | 0.1425 | 0.115 | 0.930249 | 0.385778 | 0.627903 | 1 | 0.999631 | 0.977851 |
| Hdac9_Q99N13_S218s_1 | 0.59 | 1.07 | 1.15 | 0.8 | 0.57 | 1.22 | 0.35 | 0.93 | 0.95 | 1.46 | 0.79 | 0.98 | 0.72 | 1.31 | 1.01 | 1.03 | -0.7425 | -1.01 | -1.6425 | 0.695096 | 0.650992 | 0.619096 | 1 | 0.999631 | 0.895345 |
| Hdgf_E0CXA0_S101s_1 | 1.81 | 0.37 | 2.63 | 1.97 | -0.85 | 0.86 | 3.4 | 0.4 | -0.23 | 1.84 | 0.95 | 0.18 | -0.42 | -0.74 | 1.05 | 0.32 | -0.32 | 0.1925 | 0.08 | 0.015735 | 0.406114 | 0.117506 | 1 | 0.999631 | 0.388832 |
| Hdgf_E0CXA0_S133s_1 | 0.72 | 1.99 | 0.98 | 0.68 | 1.11 | 1.23 | 0.12 | 0.63 | 1.4 | 1.58 | 0.85 | 1.31 | 1.24 | 2.29 | 1.17 | -0.01 | -0.2475 | -0.3025 | 0.22 | 0.879555 | 0.432967 | 0.606127 | 1 | 0.999631 | 0.962995 |
| Hdgf_E0CXA0_S174s_1 | -0.34 | 3.74 | 3.11 | 1.72 | 1.98 | 2.77 | 0.75 | 1.74 | 1.89 | 1.59 | 1.32 | 2.22 | 2.37 | 2.7 | 2 | 2.04 | 0.5425 | 0.605 | -0.125 | 0.783686 | 0.77474 | 0.709773 | 1 | 0.999631 | 0.922628 |
| Hdgfrp2_Q3UMU9_S576s_1 | 1.07 | 2.43 | 0.82 | 0.53 | 2.47 | 3.16 | 0.57 | 0.82 | 2.39 | 2.32 | 1.41 | 1.15 | 1.37 | 2.37 | 0.2 | 0.41 | -0.3775 | 0.1775 | -0.49 | 0.833625 | 0.428432 | 0.243503 | 1 | 0.999631 | 0.943778 |
| Hdgfrp2_Q3UMU9_S628s_2 | 0.43 | 0.98 | 2 | 1.45 | 0.02 | 1.05 | 0.97 | 1.31 | 1.3 | 1.82 | 1.25 | 1.2 | 0.14 | 1.06 | 0.98 | 0.72 | -0.3475 | -0.0125 | -0.66 | 0.251165 | 0.389337 | 0.645832 | 1 | 0.999631 | 0.68701 |
| Hdgfrp2_Q3UMU9_S635s_1 | 1.01 | 0.84 | 0.5 | 0.77 | 0.25 | 0.63 | 0.82 | 0.03 | 0.11 | 1.48 | 0.84 | 0.64 | -0.51 | 0.25 | 0.61 | 0.13 | -0.35 | -0.005 | -0.46 | 0.068861 | 0.240522 | 0.971053 | 1 | 0.999631 | 0.523934 |
| Hdgfrp2_Q3UMU9_S635s_2 | 0.52 | 1.21 | 2 | 1.45 | 0.48 | 1.02 | 0.97 | 1.31 | 1.1 | 1.61 | 1.25 | 1.2 | 0.75 | 0.89 | 0.98 | 0.72 | -0.9125 | -0.8375 | -1.045 | 0.222079 | 0.355837 | 0.988899 | 1 | 0.999631 | 0.672638 |
| Hdgfrp2_Q3UMU9_S659s_1 | 2.18 | 1.47 | 1.63 | 1.48 | 1.16 | 0.96 | 1.04 | -0.05 | 2.05 | 0.88 | 1.06 | -0.58 | 1.26 | 0.97 | 0.47 | -0.12 | -0.8025 | -0.275 | -1.38 | 0.019422 | 0.025764 | 0.13859 | 1 | 0.999631 | 0.391668 |
| Hdgfrp2_Q3UMU9_S661s_1 | 2.03 | 1.49 | 1.67 | 1.75 | 0.83 | 0.95 | 1.71 | 0.24 | 1.06 | 2.2 | 1.29 | 1.29 | -0.07 | 1.11 | 0.58 | -0.2 | -0.7325 | -0.3375 | -1.2175 | 0.003613 | 0.042819 | 0.412312 | 1 | 0.999631 | 0.351308 |
| Hdlbp_Q8VDJ3_S944s_1 | 2.45 | 0.96 | 1.77 | 1.18 | 0.06 | 1.15 | 2.06 | 0.16 | 0.85 | 2.32 | 1.09 | 0.75 | 0.24 | 0.19 | 0.87 | 0.19 | 0.25 | 0.255 | 0.535 | 0.011332 | 0.190636 | 0.483183 | 1 | 0.999631 | 0.385092 |
| Heatr5b_Q8C547_S1737s_1 | -1.14 | 0.51 | 0.83 | -0.81 | -0.03 | 0.17 | 0.33 | -0.08 | 0.3 | -0.33 | 0.6 | -0.16 | 0.57 | 0.46 | -0.44 | 0.94 | 0.3125 | 0.515 | 0.14 | 0.326822 | 0.597489 | 0.613225 | 1 | 0.999631 | 0.724678 |
| Heatr5b_Q8C547_S1739s_1 | -1.77 | 0.22 | 0.12 | -0.35 | 0.21 | 0.13 | -0.63 | -0.24 | 0.18 | -0.16 | -0.6 | 0.86 | -0.35 | -0.35 | 0.19 | -0.71 | -0.705 | -0.765 | -0.1375 | 0.771145 | 0.513977 | 0.333892 | 1 | 0.999631 | 0.919413 |
| Hebp1_Q9R257_S111s_1 | -0.56 | 0.1 | -0.03 | -0.17 | -0.88 | -0.62 | -1.08 | -0.9 | -0.88 | -0.6 | -1.48 | -0.76 | -0.11 | -0.43 | -0.18 | -0.49 | -0.4225 | -0.1575 | -2.1425 | 0.637322 | 0.023532 | 0.030819 | 1 | 0.999631 | 0.870011 |
| Hectd1_F8WIE5_S357s_1 | 2.11 | 0.76 | 1.48 | 1.13 | 0.21 | 1.52 | 1.44 | 0.62 | 1.54 | 2.45 | 0.51 | 0.35 | -1.83 | -0.83 | 0.21 | -0.64 | -0.3775 | -0.405 | -1.1475 | 0.001386 | 0.331412 | 0.766849 | 1 | 0.999631 | 0.351308 |
| Hectd1_F8WIE5_S358s_1 | 1.59 | 0.07 | 2.29 | 1.81 | -0.43 | 1.15 | 2.72 | 0.81 | -0.05 | 2.17 | 1.11 | 0.91 | -0.3 | -0.54 | 1.67 | 0.34 | 0.65 | -0.2925 | -0.9675 | 0.090006 | 0.598662 | 0.506185 | 1 | 0.999631 | 0.557085 |
| Hecw1_Q8K4P8_S1071s_1 | 0.99 | 0.13 | 0.01 | 1.16 | 1.58 | 1.08 | 1.22 | 1.01 | 0.46 | -0.59 | 1.07 | 0.18 | -0.63 | 0.02 | -0.73 | -0.24 | -0.455 | -0.4275 | -0.8875 | 0.026575 | 0.087385 | 0.516127 | 1 | 0.999631 | 0.422285 |
| Hecw1_Q8K4P8_S1199s_1 | 1.07 | 1.14 | 1.31 | 0.14 | 1.34 | 0.02 | -0.18 | 0.66 | -1.07 | 0.55 | 1.05 | 1.42 | 0.6 | 0.79 | -1.56 | 0.28 | 1.265 | 0.595 | 1.9675 | 0.134662 | 0.30043 | 0.453567 | 1 | 0.999631 | 0.606296 |
| Hecw1_Q8K4P8_S396s_1 | -4.08 | -0.7 | -0.14 | -1.06 | -0.76 | -0.75 | -0.17 | 0.76 | -1.94 | -2.1 | 0.6 | -0.16 | 0.7 | -0.09 | 0.23 | 1.05 | 0.1875 | 0.2525 | -0.695 | 0.031561 | 0.150614 | 0.538193 | 1 | 0.999631 | 0.442292 |
| Hecw1_Q8K4P8_S718s_1 | 1.17 | 1.43 | -0.17 | 1.11 | 0.82 | 1.56 | 1.37 | 0.54 | 2.01 | 0.7 | 1.22 | 0.62 | -1.17 | 0.72 | 0.91 | 0.3 | 0.6325 | 0.0725 | 0.0125 | 0.225631 | 0.659758 | 0.590433 | 1 | 0.999631 | 0.676382 |
| Hecw1_Q8K4P8_S786s_1 | 0.61 | -0.2 | 0.12 | 0.32 | 0.35 | 0.09 | 0.55 | 2.39 | -0.27 | -0.13 | 0.53 | 1.01 | -0.47 | 0.16 | 0.44 | 0.77 | 0.33 | -0.2525 | -0.1125 | 0.972426 | 0.236194 | 0.844928 | 1 | 0.999631 | 0.990167 |
| Hecw1_Q8K4P8_S869s_1 | 0.01 | 0.04 | 0.11 | 0.85 | -0.12 | 0.39 | 0.78 | 1.28 | -0.39 | -0.11 | 0.15 | 0.35 | 0.24 | 0.14 | -1 | 1.18 | 0.105 | 0.15 | 0.2625 | 0.813523 | 0.387768 | 0.434765 | 1 | 0.999631 | 0.937041 |
| Hecw1_Q8K4P8_S937s_1 | -0.06 | 0.29 | 0.08 | -0.04 | 0.2 | 0.3 | -0.26 | 0.45 | 0.09 | 0.33 | 0.48 | -0.03 | 0.43 | 0.61 | 0.18 | 0.1 | -0.1825 | 0.155 | -1.45 | 0.358687 | 0.696051 | 0.570791 | 1 | 0.999631 | 0.741182 |
| Hecw2_Q6I6G8_S1046s_1 | 2.35 | -0.56 | -1.12 | 1.61 | 0.45 | -1.46 | 2.1 | 0.46 | 1.67 | 0.05 | 0.46 | 0.72 | 0.04 | -1.43 | -0.78 | -1.35 | 0.66 | 1.7375 | 0.97 | 0.089946 | 0.848564 | 0.845221 | 1 | 0.999631 | 0.557085 |
| Hecw2_Q6I6G8_S1048s_1 | 1.09 | 0.87 | 3.28 | -1.07 | 1.45 | 1.12 | 3.72 | 0.52 | 3.25 | 1.68 | 1.84 | 4.35 | 5.79 | -0.13 | 3.04 | -0.65 | 0.16 | -0.4325 | -0.835 | 0.512617 | 0.50357 | 0.091072 | 1 | 0.999631 | 0.818235 |
| Hecw2_Q6I6G8_S1173s_1 | 0.8 | -0.03 | -0.34 | 1.14 | 0.61 | -0.36 | 0.67 | 1.29 | -0.06 | -0.58 | 0.38 | 0.1 | -3.06 | 0.45 | -0.07 | 0.91 | 0.29 | -0.115 | -0.89 | 0.328876 | 0.731289 | 0.304822 | 1 | 0.999631 | 0.726182 |
| Hecw2_Q6I6G8_S1173s_2 | 1.47 | 0.05 | 0.1 | 1.5 | 1.15 | 0.42 | 1.29 | 1.42 | 0.36 | 0.86 | 0.87 | 0.57 | -0.81 | 0.17 | 0.55 | -0.35 | 0.0925 | -0.1825 | -0.9225 | 0.092114 | 0.523025 | 0.787566 | 1 | 0.999631 | 0.557085 |
| Hecw2_Q6I6G8_S1176s_2 | 1.73 | 0.41 | 0.3 | 1.44 | 1.48 | 0.3 | 1.21 | 1.26 | 0.53 | 0.88 | 0.92 | 0.82 | -0.23 | 0.31 | 0.5 | -0.39 | 0.63 | -0.175 | -0.69 | 0.052965 | 0.831671 | 0.640753 | 1 | 0.999631 | 0.489777 |

| | | | | | | | | | | | | | | | | | | | | | | | | | |
|---|---|---|---|---|---|---|---|---|---|---|---|---|---|---|---|---|---|---|---|---|---|---|---|---|---|
| Hecw2_Q6I6G8_S1181s_2 | 1.15 | -0.37 | -0.73 | 1.25 | 0.77 | 0.54 | 1.27 | 1.24 | 0.18 | 0.83 | 0.21 | -0.62 | -1.53 | 0.01 | 0.2 | -0.14 | -0.2025 | 0.215 | -0.3825 | 0.259272 | 0.232151 | 0.750016 | 1 | 0.999631 | 0.692624 |
| Hecw2_Q6I6G8_S406s_1 | -2.51 | -2.56 | -3.28 | -3.45 | -2.73 | -3.22 | -2.7 | -3.96 | -2.69 | -2.02 | -2.58 | -3.65 | -2.29 | -3.9 | -2.78 | -4.36 | 0.115 | 0.2525 | 0.34 | 0.460989 | 0.60608 | 0.610894 | 1 | 0.999631 | 0.792192 |
| Hecw2_Q6I6G8_S48s_1 | 0.76 | 0.03 | 0.45 | 1.21 | 0.89 | 0.32 | 0.69 | 1.01 | 0.84 | 0.18 | 1.34 | 1.1 | -0.03 | 0.67 | 1.76 | 1.41 | 0.07 | 0.0325 | 0.4975 | 0.469118 | 0.729427 | 0.507876 | 1 | 0.999631 | 0.796658 |
| Hecw2_Q6I6G8_S596s_1 | -1.22 | 1.23 | 0.09 | 0.12 | 0.56 | 0.38 | -0.35 | -0.09 | 0.64 | -0.09 | 0.15 | -0.35 | 1.62 | 0.36 | 0.08 | 0.15 | 0.085 | -0.0525 | -0.1175 | 0.391718 | 0.890188 | 0.949366 | 1 | 0.999631 | 0.758901 |
| Hecw2_Q6I6G8_S858s_1 | 1.4 | 0.93 | 0.31 | 0.43 | 1.35 | 0.99 | 0.44 | 0.63 | 1.15 | 0.76 | 0.23 | 0.72 | -0.01 | 1.08 | 0.46 | 1.07 | -0.055 | 0.295 | 0.5025 | 0.764799 | 0.80834 | 0.881214 | 1 | 0.999631 | 0.916147 |
| Hepacam_Q640R3_S280s_1 | -1.3 | -0.79 | -0.06 | -0.4 | -1.11 | -0.59 | -0.72 | -0.35 | -0.51 | -0.24 | -0.42 | -0.2 | 0.18 | -0.63 | 0.09 | -0.18 | -0.0175 | 0.475 | -0.09 | 0.192002 | 0.872727 | 0.380506 | 1 | 0.999631 | 0.648674 |
| Hepacam_Q640R3_S301s_2 | -2 | -2.47 | -2.37 | -2.52 | -2.84 | -2.52 | -1.15 | -2.92 | -2.3 | -2.77 | -0.94 | -1.45 | -1.41 | -3.43 | -1.73 | -3.15 | -0.2025 | -0.155 | 0.295 | 0.856207 | 0.966956 | 0.283245 | 1 | 0.999631 | 0.951817 |
| Hepacam_Q640R3_S320s_1 | -0.51 | 0.89 | -0.06 | -0.98 | -0.14 | -1.1 | -0.53 | 0.3 | -0.23 | -1.72 | 0.65 | 0.02 | 1.82 | -0.64 | 0.09 | -0.75 | 0.4375 | 0.775 | 0.615 | 0.647901 | 0.669996 | 0.791774 | 1 | 0.999631 | 0.874615 |
| Hepacam_Q640R3_S321s_1 | -0.21 | 3.76 | 4.3 | 0.81 | 2.38 | 3.16 | 2.27 | 2.6 | 3.64 | 0.25 | 3.93 | 3.94 | 3.51 | 1.77 | 3.43 | 2.41 | -0.255 | -0.005 | 0.1525 | 0.545477 | 0.651946 | 0.528698 | 1 | 0.999631 | 0.833826 |
| Hepacam_Q640R3_S345s_1 | -0.41 | -0.62 | -0.76 | -0.49 | -0.67 | -1.02 | -0.94 | -0.67 | -0.52 | -0.57 | -0.72 | -0.49 | -0.34 | -0.16 | -0.65 | -0.52 | -0.045 | 0.115 | -0.3525 | 0.583579 | 0.314777 | 0.983837 | 1 | 0.999631 | 0.848843 |
| Hepacam_Q640R3_S352s_1 | 0.87 | 0.98 | 0.19 | 0.93 | 1.4 | 0.49 | -0.03 | 0.93 | 0.88 | 0.62 | 0.68 | 1.25 | -1.47 | 0.93 | 0.84 | 1.26 | -1.1625 | -0.605 | -1.9475 | 0.55791 | 0.904166 | 0.707315 | 1 | 0.999631 | 0.838323 |
| Hepacam_Q640R3_S366s_2 | 2.92 | -0.4 | 0.29 | 1.85 | 0.45 | -0.25 | 1.45 | -1.64 | 0.53 | 2.07 | -0.49 | 0.13 | -1.92 | -0.56 | -0.05 | -0.6 | -1.1625 | -0.605 | -1.9475 | 0.024597 | 0.198317 | 0.463536 | 1 | 0.999631 | 0.417284 |
| Hepacam_Q640R3_S370s_2 | 2.92 | -0.4 | 0.29 | 1.85 | 0.45 | -0.25 | 1.45 | -1.64 | 0.53 | 2.07 | -0.49 | 0.13 | -1.92 | -0.56 | -0.05 | -0.6 | -0.395 | -0.2375 | 0.02 | 0.024597 | 0.198317 | 0.463536 | 1 | 0.999631 | 0.417284 |
| Hepacam_Q640R3_T297t_1 | -0.54 | 0.43 | 0.9 | 0.16 | -0.3 | 0.3 | -0.49 | -0.14 | -0.38 | 0.14 | 0.12 | 0.12 | -0.55 | 0.56 | 0.6 | 0.42 | -0.0175 | 0.475 | -0.09 | 0.961928 | 0.294873 | 0.514069 | 1 | 0.999631 | 0.986594 |
| Hepacam_Q640R3_T297t_2 | -2 | -2.47 | -2.37 | -2.52 | -2.84 | -2.52 | -1.15 | -2.92 | -2.3 | -2.77 | -0.94 | -1.45 | -1.41 | -3.43 | -1.73 | -3.15 | 1.0925 | 1.4225 | 0.245 | 0.856207 | 0.966956 | 0.283245 | 1 | 0.999631 | 0.951817 |
| Hepacam_Q640R3_T378t_1 | 0.92 | -0.83 | 0.32 | -1.29 | 2.21 | 0.24 | 2.45 | -1.41 | 3.87 | 0.02 | -0.21 | 1.13 | 3.42 | -0.69 | -1.46 | -1.17 | -0.755 | -0.5175 | -0.5825 | 0.818189 | 0.243345 | 0.147778 | 1 | 0.999631 | 0.938303 |
| Herc1_E9PZP8_S133s_1 | -0.36 | -2.45 | -1.74 | -2.02 | -3.82 | -3.49 | -1.05 | -1.23 | -2.66 | -1.2 | -2.46 | -2.32 | -2.59 | -2.46 | -1.61 | -2.24 | 0.32 | -0.69 | -2.105 | 0.247356 | 0.329461 | 0.335243 | 1 | 0.999631 | 0.685751 |
| Herc1_E9PZP8_S1406s_1 | 2.23 | 0.17 | 0.88 | 1.18 | 1.3 | 0.74 | 2.89 | 0.81 | 0.52 | 0.35 | 1.23 | -0.4 | -2.15 | -0.78 | -0.13 | -0.9 | 0.085 | 0.1925 | 0.285 | 0.003507 | 0.593854 | 0.1988 | 1 | 0.999631 | 0.351308 |
| Herc1_E9PZP8_S1428s_1 | -0.39 | -1.64 | -1.17 | -1.19 | -0.31 | -0.83 | -1.47 | -1.44 | -0.43 | -0.2 | -1.76 | -1.23 | 0.27 | -0.83 | -1.19 | -1.5 | 0.0875 | 0.29 | -0.2575 | 0.53925 | 0.827013 | 0.662911 | 1 | 0.999631 | 0.830899 |
| Herc1_E9PZP8_S1491s_1 | 0.82 | 0.79 | 0.55 | 1.06 | 0.7 | 0.82 | 1 | 1.05 | 1.08 | 0.77 | 1.38 | 1.15 | -0.36 | 0.78 | 0.91 | 0.86 | -0.2425 | 0.0025 | -0.1725 | 0.487758 | 0.72822 | 0.300159 | 1 | 0.999631 | 0.80597 |
| Herc1_E9PZP8_S2718s_1 | 1.27 | 1.13 | 1.62 | 0.82 | 0.84 | 1.24 | 1.06 | 0.73 | 1.41 | 1.28 | 1.62 | 0.54 | 0.18 | 1.37 | 1.47 | 1.13 | -0.5925 | -0.0725 | 0.28 | 0.648587 | 0.397922 | 0.994055 | 1 | 0.999631 | 0.874615 |
| Herc1_E9PZP8_S2718s_2 | 0.4 | -0.16 | 0.84 | 0.88 | -1.09 | -0.17 | 0.58 | 0.27 | -0.62 | 0.43 | 0.99 | 0.87 | 0.24 | 0.03 | 0.98 | 1.83 | -0.2325 | 0.035 | 0.5075 | 0.551794 | 0.188882 | 0.867912 | 1 | 0.999631 | 0.836275 |
| Herc1_E9PZP8_S2721s_1 | 0.57 | 0.89 | 1.41 | -0.45 | 0.12 | 1.63 | -0.19 | -0.07 | 0.72 | 2.06 | 0.43 | -0.65 | 2.02 | 1.39 | 0.37 | 0.67 | -0.2325 | 0.14 | -0.1575 | 0.334258 | 0.664266 | 0.954786 | 1 | 0.999631 | 0.728785 |
| Herc1_E9PZP8_S2736s_1 | 0.54 | 0.43 | 0.03 | -0.51 | -0.23 | 0.7 | -0.63 | -0.28 | 1.03 | 0.8 | -0.52 | -0.26 | 0.85 | -0.19 | -0.6 | -0.2 | -0.0625 | -0.555 | 0.3 | 0.701508 | 0.547348 | 0.753218 | 1 | 0.999631 | 0.896835 |
| Herc1_E9PZP8_S2745s_1 | -0.89 | -1.22 | -0.35 | -1.05 | -0.68 | -1.45 | -0.84 | -0.79 | -0.75 | -1.91 | -1.39 | -1.68 | -0.66 | -0.63 | -1.09 | 0.07 | 0.375 | 0.11 | 0.9275 | 0.409389 | 0.840591 | 0.138824 | 1 | 0.999631 | 0.766996 |
| Herc1_E9PZP8_S4855s_1 | -2.14 | -0.91 | 0.11 | -1.13 | -0.54 | -1.4 | -0.04 | -0.59 | -0.54 | -1.04 | -1.66 | -0.39 | -0.34 | -1.18 | 0.79 | 0.37 | -0.025 | 0.335 | 0.6225 | 0.133699 | 0.464585 | 0.829993 | 1 | 0.999631 | 0.60492 |
| Herc1_E9PZP8_T2699t_1 | -1.07 | 0.67 | -1.25 | -0.12 | -0.5 | -0.35 | -0.8 | -0.22 | -0.33 | -0.47 | 0.39 | -0.02 | 0.5 | 0.03 | 0.44 | -0.25 | -0.3075 | -0.3725 | -0.42 | 0.205546 | 0.95547 | 0.481073 | 1 | 0.999631 | 0.6605 |
| Herpud2_Q9JJC9_S90s_1 | 1.1 | -0.56 | 0.38 | -0.21 | -0.81 | -0.26 | 0.64 | -0.09 | -0.61 | 0.19 | -0.01 | -0.35 | 0.24 | -0.43 | -0.23 | -0.55 | -0.0025 | -0.0975 | 0.265 | 0.327277 | 0.503091 | 0.376093 | 1 | 0.999631 | 0.724678 |
| Hgs_Q3UMA3_S240s_1 | -1.23 | 0.32 | -0.18 | -0.66 | 0 | -0.48 | -0.73 | -0.55 | -0.31 | -1.22 | -0.6 | -0.01 | 0.37 | -0.25 | -0.61 | -0.2 | 0.065 | 0.475 | 0.8575 | 0.52063 | 0.994743 | 0.817586 | 1 | 0.999631 | 0.819896 |
| Hgsnat_Q3UDW8_S206s_1 | -0.9 | -0.39 | -0.1 | -1.03 | -0.59 | -0.31 | -0.95 | -0.31 | -0.21 | -0.39 | -0.3 | 0.38 | 0.71 | 0.11 | 0.21 | -0.02 | 0.03 | 0.4725 | -0.4975 | 0.027278 | 0.837163 | 0.173148 | 1 | 0.999631 | 0.423171 |
| Hid1_Q8R1F6_S592s_1 | 0.8 | 1.24 | 0.92 | 0.56 | 0.45 | 1.75 | 0.83 | 0.61 | 1.84 | 2.18 | 0.64 | 0.75 | 0.33 | 0.39 | 0.72 | 0.09 | 0.61 | 0.5525 | 0.82 | 0.120183 | 0.93196 | 0.273048 | 1 | 0.999631 | 0.587703 |
| Hid1_Q8R1F6_S669s_1 | -0.34 | 0.2 | 0.37 | 0.62 | 0.7 | 1.02 | 0.23 | 1.34 | 1 | 0.22 | 1.22 | 0.62 | -0.27 | 2.25 | 1.33 | 0.82 | -0.3075 | 0.3175 | -1.025 | 0.144933 | 0.101492 | 0.131605 | 1 | 0.999631 | 0.616037 |
| Hid1_Q8R1F6_T590t_1 | 1.83 | 1.86 | 1.49 | 1.34 | 0.7 | 2.41 | 1.21 | 0.97 | 2.47 | 3.23 | 0.75 | 1.34 | 0.41 | 1.1 | 1.01 | -0.1 | 0.24 | 0.385 | 1.2925 | 0.015933 | 0.451133 | 0.555198 | 1 | 0.999631 | 0.388832 |
| Hint1_B0R1E3_S45s_1 | -2.76 | 2.25 | 1.42 | 0.53 | 1.06 | 0.8 | 0.39 | 0.15 | 1.24 | -0.37 | 1.32 | 0.79 | 1.26 | 2.05 | 1.75 | 1.55 | -0.5125 | 0.7675 | -0.295 | 0.195441 | 0.80322 | 0.702722 | 1 | 0.999631 | 0.653056 |
| Hip1r_Q9JKY5_S19s_1 | 0.7 | -0.15 | 0.06 | 1.26 | -0.06 | -0.85 | 0.76 | -0.03 | 0.82 | 0.97 | 1.41 | 1.74 | -2.93 | 0.67 | 1 | 1.95 | -0.1325 | 0.1125 | 0.36 | 0.75832 | 0.266146 | 0.077519 | 1 | 0.999631 | 0.914449 |
| Hipk2_Q9QZR5_Y354y_1 | 0.07 | 0.66 | 0.34 | 0.26 | 0.41 | 0.17 | 0.22 | 0 | 0.46 | 0.64 | 0.1 | 0.58 | 0.85 | 0.93 | 0.69 | 0.3 | -0.52 | -0.125 | -0.54 | 0.242617 | 0.609546 | 0.684089 | 1 | 0.999631 | 0.685099 |
| Hipk3_Q9ERH7_Y359y_1 | 0.34 | -0.79 | 0.38 | -0.65 | -0.48 | -1.66 | -0.04 | -0.62 | -1.86 | 0.57 | -0.27 | 0.34 | -0.66 | -2.26 | 0.42 | -0.38 | -0.0725 | 0.03 | -0.12 | 0.369606 | 0.263829 | 0.829056 | 1 | 0.999631 | 0.746338 |
| Hirip3_Q8BLH7_S575s_1 | 1.32 | 1.81 | 1 | 1.53 | 1.64 | 1.16 | 1.14 | 1.43 | 1.88 | 1.11 | 1.2 | 1.59 | 0.91 | 1.7 | 1.44 | 1.13 | -0.63 | -0.7925 | -0.9325 | 0.711458 | 0.799303 | 0.923499 | 1 | 0.999631 | 0.900631 |
| Hirip3_Q8BLH7_S80s_1 | 2.02 | -0.19 | 2.54 | 2.33 | 0.11 | 0.71 | 2.04 | 1.32 | 1.04 | 1.25 | 1.18 | 0.06 | 0.86 | -0.53 | 1.48 | 1.16 | 0.5425 | 0.32 | 0.3125 | 0.194422 | 0.35895 | 0.223438 | 1 | 0.999631 | 0.65187 |
| Hist1h1a_P43275_S2s_1 | -2.12 | -2.76 | -1.43 | -2.4 | -1.71 | -1.73 | -1.71 | -1.39 | -2.41 | -0.89 | -2.3 | -1.83 | -1.56 | -1.48 | -2.61 | -1.81 | 0.5425 | 0.32 | 0.3125 | 0.446218 | 0.127406 | 0.474262 | 1 | 0.999631 | 0.784601 |
| Hist1h1a_P43275_T4t_1 | -2.12 | -2.76 | -1.43 | -2.4 | -1.71 | -1.73 | -1.71 | -1.39 | -2.41 | -0.89 | -2.3 | -1.83 | -1.56 | -1.48 | -2.61 | -1.81 | -0.1075 | 0.08 | -0.1775 | 0.446218 | 0.127406 | 0.474262 | 1 | 0.999631 | 0.784601 |
| Hist1h1b_P43276_S18s_1 | -1.78 | -2 | -3.65 | -3.71 | -2.69 | -2.14 | -2.82 | -3.92 | -2.41 | -1.51 | -2.92 | -3.98 | -1.28 | -2.03 | -4.35 | -4.19 | -0.2475 | 0.1 | 0.3175 | 0.826407 | 0.853082 | 0.902989 | 1 | 0.999631 | 0.940813 |
| Hist1h1b_P43276_S2s_1 | -1.25 | -0.03 | -0.01 | -0.27 | -0.46 | -0.72 | -0.63 | -0.74 | -0.27 | -0.4 | -0.2 | -0.29 | 0.07 | -0.52 | -0.23 | 0.39 | -0.2475 | 0.1 | 0.3175 | 0.417974 | 0.468679 | 0.769485 | 1 | 0.999631 | 0.771539 |
| Hist1h1b_P43276_T4t_1 | -1.25 | -0.03 | -0.01 | -0.27 | -0.46 | -0.72 | -0.63 | -0.74 | -0.27 | -0.4 | -0.2 | -0.29 | 0.07 | -0.52 | -0.23 | 0.39 | 0.17 | -0.555 | -0.6525 | 0.417974 | 0.468679 | 0.769485 | 1 | 0.999631 | 0.771539 |
| Hist1h1c_P15864_S36s_1 | 0.34 | 0.27 | 0.67 | 1.11 | 1.32 | -0.64 | 0.62 | 1.77 | -0.54 | -0.81 | 0.69 | 0.83 | -1.44 | 0.64 | -0.78 | 1.36 | -0.7325 | -1.37 | -0.8825 | 0.297656 | 0.744049 | 0.237538 | 1 | 0.999631 | 0.711122 |
| Hist1h1c_P15864_T146t_1 | 1.34 | 0.81 | 2.53 | 1.74 | -1.48 | 1.81 | 2.38 | 0.78 | -0.6 | 1.1 | 0.51 | -0.07 | 0.36 | 0.2 | 2.19 | 0.14 | -0.05 | -0.0875 | 0.1275 | 0.141352 | 0.374711 | 0.018509 | 1 | 0.999631 | 0.61245 |
| Hist1h1d_P43277_S2s_1 | -0.99 | -0.79 | -0.98 | -0.79 | -1.07 | -1.11 | -1.12 | -0.45 | -1.5 | -0.71 | -0.79 | -0.9 | -0.45 | -1.03 | -0.87 | -0.69 | -0.0875 | -0.1 | 0.165 | 0.647895 | 0.8518 | 0.757104 | 1 | 0.999631 | 0.874615 |
| Hist1h1d_P43277_T18t_1 | -1.43 | -1.36 | -0.84 | -1.13 | -1.08 | -1.63 | -0.92 | -1.48 | -1.58 | -0.78 | -1 | -1.8 | -0.98 | -1.48 | -0.44 | -1.2 | -0.07 | -0.2225 | 0.095 | 0.619877 | 0.761094 | 0.760774 | 1 | 0.999631 | 0.861888 |
| Hist1h1d_P43277_T4t_1 | -0.99 | -0.79 | -1.2 | -0.79 | -1.07 | -1.11 | -1.2 | -0.67 | -1.5 | -0.71 | -1.06 | -1.39 | -0.45 | -1.03 | -0.82 | -1.09 | -0.785 | -0.5375 | -0.2925 | 0.745301 | 0.787393 | 0.449609 | 1 | 0.999631 | 0.910585 |
| Hist1h1e_P43274_S2s_1 | -1.03 | 0.51 | -0.69 | -1.58 | -1.2 | -1.28 | -1.49 | -1.96 | -0.22 | -0.49 | -1.67 | -2.56 | 0.28 | -0.59 | -1.88 | -1.77 | -0.0225 | -0.3825 | 0.355 | 0.636195 | 0.110154 | 0.404265 | 1 | 0.999631 | 0.869397 |
| Hist1h1e_P43274_T18t_1 | -0.99 | -1.38 | -1.26 | -0.35 | -1.36 | -1.51 | -0.74 | -0.46 | -1.43 | -2.22 | -0.86 | -1 | 0.16 | -1.71 | -0.36 | -0.65 | -0.785 | -0.5375 | -0.2925 | 0.441059 | 0.950407 | 0.346885 | 1 | 0.999631 | 0.781983 |
| Hist1h1e_P43274_T4t_1 | -1.03 | 0.51 | -0.69 | -1.58 | -1.2 | -1.28 | -1.49 | -1.96 | -0.22 | -0.49 | -1.67 | -2.56 | 0.28 | -0.59 | -1.88 | -1.77 | -0.0475 | 0.0075 | 0.135 | 0.636195 | 0.110154 | 0.404265 | 1 | 0.999631 | 0.869397 |
| Hist1h4a_P62806_S48s_1 | -0.61 | -0.32 | -0.48 | -0.61 | -0.43 | -0.27 | -0.7 | -0.81 | -0.78 | -0.37 | -0.47 | -0.37 | -0.06 | -0.63 | -0.54 | -0.25 | -0.0725 | 0.2025 | 0.69 | 0.633773 | 0.852052 | 0.976508 | 1 | 0.999631 | 0.868569 |
| Hivep2_Q3UHF7_S1077s_1 | -0.27 | 0.69 | 0.08 | -0.07 | 0.49 | 0.29 | -0.45 | -0.19 | 0.45 | 0.46 | -0.12 | 0.45 | 1.33 | 0.94 | 0.38 | 0.54 | 0.175 | 0.81 | 0.24 | 0.073797 | 0.829391 | 0.52502 | 1 | 0.999631 | 0.539631 |
| Hivep2_Q3UHF7_S169s_1 | 0.75 | 0.39 | 1.34 | 1.21 | 0.7 | 0.52 | 1.36 | 1.81 | 1.68 | 1.83 | 2.33 | 1.09 | 0.66 | 0.72 | 1.5 | 1.77 | 0.2 | -0.5425 | -1.4925 | 0.538304 | 0.648568 | 0.048381 | 1 | 0.999631 | 0.830863 |
| Hivep2_Q3UHF7_S1921s_1 | 1.59 | -0.78 | -0.1 | 0.79 | 0.49 | -0.24 | 1.32 | 0.73 | -0.88 | -0.09 | 0.61 | -0.31 | -1.7 | -1.81 | -0.42 | -0.54 | -0.8975 | -0.14 | -1.9375 | 0.026606 | 0.720688 | 0.341854 | 1 | 0.999631 | 0.422285 |
| Hivep2_Q3UHF7_S2085s_1 | 1.99 | 1.16 | 2.08 | 1.12 | 0.41 | 0.53 | 2.1 | -0.28 | 1.25 | 2.08 | 1.51 | 0.95 | 0.27 | -0.66 | -1.65 | 0.64 | -0.4375 | 0.385 | -0.075 | 0.004637 | 0.111873 | 0.711189 | 1 | 0.999631 | 0.351308 |
| Hivep2_Q3UHF7_S2295s_1 | 0.57 | 0.53 | 1.36 | 0.39 | -0.49 | 0.83 | 0.28 | 0.48 | 0.74 | 1.23 | 0.89 | 1.53 | 0.74 | 0.48 | 0.67 | 0.66 | 0.065 | 0.4825 | 0.695 | 0.811862 | 0.257181 | 0.263589 | 1 | 0.999631 | 0.936949 |
| Hivep2_Q3UHF7_S314s_1 | -0.07 | -0.91 | -0.07 | -0.22 | -1.06 | 0.14 | -0.3 | 0.21 | 0.21 | 0.24 | -0.06 | 0.27 | 1.02 | -0.26 | 0.5 | 0.25 | -0.14 | 0.2325 | -0.415 | 0.087009 | 0.861209 | 0.126758 | 1 | 0.999631 | 0.553514 |
| Hivep2_Q3UHF7_S395s_1 | 0.63 | 0.36 | 0.79 | 0.83 | 0.54 | 0.56 | 0.89 | 0.06 | 0.48 | 0.83 | 1.55 | 0.68 | 0.12 | 0.39 | -0.97 | 1.41 | 0.0925 | 0.055 | 0.2225 | 0.400785 | 0.620943 | 0.470525 | 1 | 0.999631 | 0.763209 |
| Hivep2_Q3UHF7_S41s_1 | 0.35 | 0.55 | 0.29 | 0.49 | 0.64 | 0.24 | 0.82 | 0.35 | 0.47 | -0.15 | 1.4 | 0.18 | 0.15 | 0.84 | -0.01 | 1.59 | -0.055 | -0.935 | -0.205 | 0.577965 | 0.719035 | 0.881054 | 1 | 0.999631 | 0.84622 |
| Hivep2_Q3UHF7_S865s_1 | 0.98 | 0.35 | 2.41 | 1.58 | 0.93 | 1.81 | 0.89 | 1.47 | 0.93 | 0.63 | 0.28 | -0.26 | 0.64 | 1.34 | 0.96 | 1.56 | -0.7625 | 0.06 | -0.9725 | 0.665985 | 0.906622 | 0.07896 | 1 | 0.999631 | 0.88264 |
| Hivep2_Q3UHF7_T2294t_1 | 1.58 | -0.11 | 1.79 | 0.47 | 1.02 | -0.41 | 0.48 | -0.41 | 0.8 | 1.75 | 0.8 | 0.62 | 0.38 | 0.03 | -0.71 | 0.14 | -0.135 | 0.095 | -0.1875 | 0.071307 | 0.172979 | 0.903393 | 1 | 0.999631 | 0.531405 |
| Hmga1_Q3TE85_S101s_1 | -0.57 | -0.15 | -0.94 | 0.07 | -0.87 | 0.1 | -0.69 | -0.67 | -0.1 | -0.45 | -0.01 | -0.65 | -0.86 | -0.76 | -0.32 | -0.4 | -0.2575 | -0.195 | -0.5125 | 0.575675 | 0.695695 | 0.770649 | 1 | 0.999631 | 0.846085 |
| Hmga1_Q3TE85_S102s_1 | -0.9 | -0.27 | -0.01 | -0.21 | -1.19 | 0.32 | -0.52 | -1.03 | -0.06 | -0.75 | -0.38 | -0.98 | -0.39 | -1.24 | -0.92 | -0.89 | -0.1175 | 0.0475 | 0.245 | 0.146031 | 0.520551 | 0.559424 | 1 | 0.999631 | 0.617311 |
| Hmga1_Q3TE85_S36s_1 | 0.15 | 1.03 | 0.87 | 0.9 | 0.79 | 0.32 | 0.15 | 1.22 | 0.55 | 0.42 | 0.94 | 1.23 | 1.12 | 0.83 | 0.54 | 1.44 | -0.295 | 1.0875 | 1.23 | 0.478728 | 0.734061 | 0.884282 | 1 | 0.999631 | 0.801051 |
| Hmgcr_F2Z470_S384s_1 | -0.84 | 0.27 | 0.7 | -1.3 | 0.6 | -1.21 | -0.63 | -1.11 | 0.82 | 0.09 | 1.33 | 0.94 | 1.04 | 0.93 | 0.3 | 1.48 | -0.58 | -0.97 | -1.27 | 0.032413 | 0.607069 | 0.053934 | 1 | 0.999631 | 0.442395 |
| Hmgn3_Q9DCB1_S6s_1 | 2.22 | 0.47 | 2.41 | 1.48 | -0.57 | 1.17 | 3.47 | 0.19 | -0.69 | 1.66 | 0.6 | 1.13 | -0.01 | -0.33 | 1.5 | 0.34 | 0.28 | -0.3775 | -2.1225 | 0.041508 | 0.501572 | 0.135039 | 1 | 0.999631 | 0.474979 |
| Hn1_P97825_S70s_1 | 1.37 | -1.74 | -2.89 | -1.11 | 0.02 | -1.8 | 0.03 | -1.5 | -1.24 | -0.21 | -1.94 | -2.49 | -1.86 | -2.51 | -3.42 | -5.07 | -0.6675 | -0.3125 | -0.6025 | 0.051031 | 0.752595 | 0.673438 | 1 | 0.999631 | 0.489558 |
| Hn1_P97825_S79s_1 | 2.77 | 1.23 | 1.91 | 1.96 | -0.15 | 1.6 | 2.49 | 1.26 | 1.16 | 2.62 | 2 | 0.84 | 0.55 | 0.99 | 2.34 | 1.58 | 0.045 | -0.1725 | 0.655 | 0.231466 | 0.263119 | 0.527337 | 1 | 0.999631 | 0.681606 |
| Hn1_P97825_S87s_1 | -0.53 | 0.64 | 0.21 | 0.32 | 0.31 | 0.25 | -0.14 | 0.4 | 0.13 | -0.23 | 0.15 | -0.1 | 1.04 | 0.69 | 0.7 | 0.83 | 0.12 | 0.7325 | 1.055 | 0.072185 | 0.88867 | 0.594918 | 1 | 0.999631 | 0.532251 |
| Hnrnpa1_Q5EBP8_S257s_1 | -2.61 | -1.64 | -0.56 | -1.97 | -2.12 | -1.51 | -1.37 | -1.3 | -1.66 | -1.15 | -0.68 | -0.36 | -0.67 | -1.45 | -0.24 | -0.2 | 0.045 | 0.3375 | 0.135 | 0.055953 | 0.790543 | 0.158476 | 1 | 0.999631 | 0.495761 |
| Hnrnpa1_Q5EBP8_S369s_1 | 1.55 | -2.62 | 0.28 | -1.42 | -0.59 | -0.55 | 0.08 | -0.97 | -0.6 | 0.44 | -0.21 | -0.49 | 0.46 | -0.73 | -0.72 | -0.68 | -0.115 | 0.2525 | 0.6675 | 0.871833 | 0.956354 | 0.685335 | 1 | 0.999631 | 0.959119 |
| Hnrnpa1_Q5EBP8_S6s_1 | -0.01 | 0.24 | 0.65 | 0.41 | -0.11 | 0.26 | -0.09 | 0.77 | 0.17 | 0.63 | 0.79 | 0.71 | 0.79 | 1.07 | 0.85 | 1.25 | 0.17 | 0.5475 | 1.225 | 0.042176 | 0.708314 | 0.387936 | 1 | 0.999631 | 0.475443 |
| Hnrnpa2b1_O88569_S259s_1 | -1.4 | 0.16 | -0.63 | -0.83 | -0.46 | -0.4 | -0.68 | -0.48 | -0.38 | -0.08 | 0.06 | -0.11 | 0.7 | 0.25 | 0.92 | 0.33 | -0.0525 | 0.0625 | 0.1 | 0.009768 | 0.63381 | 0.157403 | 1 | 0.999631 | 0.374514 |
| Hnrnpa2b1_O88569_S344s_1 | -0.1 | 0.43 | -0.03 | -0.32 | 0.62 | -0.51 | 0.03 | -0.37 | 0.78 | -0.66 | -0.21 | 0.32 | 0.61 | 0.67 | -0.59 | -0.31 | -0.775 | -0.4575 | -0.2425 | 0.796858 | 0.875773 | 0.867739 | 1 | 0.999631 | 0.929952 |
| Hnrnpa3_Q8BG05_S359s_1 | 0.41 | -0.4 | -0.52 | -0.9 | -1.68 | -0.96 | -0.47 | -1.4 | -0.79 | -0.45 | -1.22 | -0.78 | 0.02 | -0.46 | -1.13 | -0.81 | 0.3125 | -0.2225 | -0.74 | 0.544326 | 0.071275 | 0.216351 | 1 | 0.999631 | 0.833226 |
| Hnrnpa3_Q8BG05_S376s_1 | 0.22 | -1.03 | -0.26 | -2.43 | -0.01 | -0.95 | -0.75 | -0.54 | -1.57 | -0.05 | -1.79 | -0.98 | -1.13 | -1.65 | -0.92 | -2.76 | 0.2025 | 0.2475 | 0.7925 | 0.263419 | 0.580206 | 0.724669 | 1 | 0.999631 | 0.693034 |
| Hnrnpc_Q9Z204_S218s_1 | -0.84 | -0.04 | -0.04 | -1.41 | 0.48 | -0.26 | -0.82 | -0.92 | 0.14 | -0.45 | -0.75 | -0.28 | 1.2 | 0.77 | -0.74 | -0.39 | -0.255 | -0.5175 | -0.92 | 0.160652 | 0.653169 | 0.536519 | 1 | 0.999631 | 0.630408 |
| Hnrnpc_Q9Z204_S228s_1 | -0.01 | -1.1 | 0.62 | 0.61 | -0.42 | -0.82 | 0.56 | -0.22 | -0.71 | 0.02 | -0.43 | -0.83 | -0.05 | -1.42 | -0.58 | -1.51 | -0.9925 | -1.035 | -1.485 | 0.095035 | 0.592822 | 0.258075 | 1 | 0.999631 | 0.558128 |
| Hnrnpc_Q9Z204_S236s_1 | 2.18 | 0.71 | 1.88 | 2.04 | -0.06 | 0.84 | 1.49 | 0.57 | -0.16 | 1.77 | 0.51 | 0.55 | 0.18 | -0.28 | 0.8 | 0.17 | -0.27 | -0.0125 | -1.245 | 0.004967 | 0.049112 | 0.061067 | 1 | 0.999631 | 0.351308 |
| Hnrnpc_Q9Z204_S293s_1 | 0.75 | 2.43 | 2.77 | 1.69 | 1.56 | 2.47 | 2.11 | 0.42 | 2.78 | 2.87 | 0.76 | 1.18 | 1.41 | 0.14 | 0.88 | 0.23 | -0.2025 | -0.085 | -1.2925 | 0.032915 | 0.640994 | 0.984276 | 1 | 0.999631 | 0.44294 |
| Hnrnpd_Q60668_S80s_1 | 1.76 | 3.25 | 3.52 | 1.76 | 1.8 | 2.65 | 3.6 | 1.43 | 2.51 | 1.71 | 2.92 | 2.81 | 1.69 | 1.16 | 2.64 | -0.37 | 0.3975 | 0.9025 | -1.625 | 0.088396 | 0.740045 | 0.868606 | 1 | 0.999631 | 0.553858 |
| Hnrnpd_Q60668_S80s_2 | 0.98 | 3.85 | 3.1 | -0.79 | 2.62 | 2.47 | 3.2 | 0.44 | 3.99 | 2.33 | 1.81 | 2.62 | 3.49 | -0.61 | 0.48 | -2.72 | 0.325 | -0.0925 | -1.1875 | 0.263015 | 0.703267 | 0.374244 | 1 | 0.999631 | 0.693034 |
| Hnrnpd_Q60668_S82s_1 | 0.69 | 1.44 | 0.24 | -0.7 | 0.69 | 1.48 | 1.25 | -0.45 | 1.11 | 1.19 | -0.38 | -0.62 | 0.82 | -0.72 | -1.06 | -2.12 | 0.4875 | 0.4925 | -0.84 | 0.102213 | 0.569494 | 0.876945 | 1 | 0.999631 | 0.564448 |
| Hnrnpd_Q60668_S83s_1 | 0.75 | 3.19 | 3.58 | 0.62 | 2.01 | 2.65 | 3.89 | 1.54 | 2.95 | 0.99 | 2.93 | 3.24 | 2.55 | -0.37 | 2.32 | 0.28 | 0.3975 | 0.9025 | -1.625 | 0.372201 | 0.553457 | 0.55297 | 1 | 0.999631 | 0.746884 |
| Hnrnpd_Q60668_S83s_2 | 0.98 | 3.85 | 3.1 | -0.79 | 2.62 | 2.47 | 3.2 | 0.44 | 3.99 | 2.33 | 1.81 | 2.62 | 3.49 | -0.61 | 0.48 | -2.72 | 0.195 | -0.1425 | 0.1175 | 0.263015 | 0.703267 | 0.374244 | 1 | 0.999631 | 0.693034 |
| Hnrnpdl_D3YTQ3_S241s_1 | -0.23 | 0.08 | -1.15 | -0.98 | -0.34 | 0.68 | -0.96 | -0.88 | -1.25 | -0.78 | -0.51 | -0.31 | -0.64 | -0.76 | -0.15 | -0.26 | 0.7025 | 1.7725 | 0.475 | 0.752128 | 0.672753 | 0.708706 | 1 | 0.999631 | 0.913038 |
| Hnrnph1_Q8C2Q7_S104s_1 | -0.02 | 0.54 | 0.47 | -0.98 | 0.7 | -0.27 | 2.38 | 0.01 | 2.14 | -0.44 | 1.66 | 3.74 | 2.01 | -1.12 | 1.44 | -0.42 | -0.195 | -1.4 | -0.86 | 0.517009 | 0.273638 | 0.052037 | 1 | 0.999631 | 0.819449 |
| Hnrnph2_P70333_S4s_1 | 0.52 | -2.43 | -0.08 | -0.54 | -1.87 | -1.25 | 0.79 | -0.98 | -2.2 | -0.57 | -3.31 | -2.05 | -1.55 | -2.9 | -0.39 | -1.13 | 0.42 | 1.17 | 0.22 | 0.255105 | 0.79536 | 0.087211 | 1 | 0.999631 | 0.690222 |
| Hnrnpk_P61979_S284s_1 | 0.5 | 2.79 | 2.19 | 0.42 | 2.33 | 1.83 | 2.07 | 1.35 | 3.77 | 0.69 | 2.74 | 3.38 | 3.02 | 1.18 | 1.95 | 0.63 | -0.7425 | -0.415 | -0.275 | 0.754362 | 0.47273 | 0.166071 | 1 | 0.999631 | 0.913377 |
| Hnrnpl_G5E924_S174s_1 | 1.03 | 0.67 | 0.32 | 0.1 | 0.49 | -0.82 | 0.18 | -0.7 | 0.26 | 0.28 | 0.07 | -0.15 | 1.1 | -0.18 | 0.38 | -0.28 | -0.2475 | 0.365 | -0.11 | 0.497717 | 0.081935 | 0.190528 | 1 | 0.999631 | 0.811908 |
| Hnrnpl_G5E924_S90s_1 | 0.4 | 1.24 | 0.26 | 0.28 | 0.66 | -0.14 | 0.4 | 0.27 | 0.73 | 0.85 | 0.47 | 1.59 | 0.62 | 1.1 | 0.34 | -0.32 | 0.0525 | -0.245 | 0.1425 | 0.784216 | 0.458781 | 0.330588 | 1 | 0.999631 | 0.92266 |
| Hnrnpll_Q921F4_S37s_1 | -0.16 | 0.01 | 0.53 | 0.16 | 0.18 | -0.07 | 0.28 | 0.36 | -1.33 | -0.26 | 0.63 | 0.52 | 0.55 | -0.71 | 0.65 | 0.62 | 1.06 | 0.6675 | -0.455 | 0.716155 | 0.844968 | 0.598669 | 1 | 0.999631 | 0.902805 |
| Hnrnpm_B8JK33_S413s_1 | 0.38 | 0.31 | -0.93 | 1.97 | 1.7 | 0.86 | 2.13 | 1.28 | 1.43 | 0.74 | 0.97 | 1.26 | -1.06 | -0.07 | 0.75 | 0.29 | 0.98 | 0.235 | -0.73 | 0.481085 | 0.098478 | 0.255298 | 1 | 0.999631 | 0.801848 |
| Hnrnpm_B8JK33_S488s_1 | -0.37 | 0.63 | -0.94 | 2.05 | 2.12 | 0.41 | 1.61 | 1.15 | 1.43 | 0.1 | -0.18 | 0.96 | -1.32 | 0.82 | -1.32 | 0.27 | -0.5075 | -0.1375 | -0.43 | 0.343026 | 0.164484 | 0.728292 | 1 | 0.999631 | 0.73348 |
| Hnrnpm_B8JK33_S535s_1 | 1.19 | 0.59 | 1.04 | 1.05 | 0.52 | 0.62 | 0.72 | -0.02 | 0.49 | 0.93 | 1.04 | 0.86 | 0.2 | 0.6 | 0.79 | 0.56 | -0.3275 | -1.1275 | -1.405 | 0.165402 | 0.100718 | 0.622106 | 1 | 0.999631 | 0.633733 |
| Hnrnpm_B8JK33_S578s_1 | 3.71 | 0.15 | -0.62 | 1.18 | 0.98 | -0.27 | 1.73 | 0.67 | 0.17 | -0.81 | 0.86 | -0.31 | -1.08 | 0.44 | -1.63 | 1.07 | -0.29 | -0.915 | -0.1575 | 0.170762 | 0.714367 | 0.218959 | 1 | 0.999631 | 0.641032 |
| Hnrnpm_B8JK33_S597s_1 | 0.45 | -0.36 | -0.78 | 0.7 | 0.45 | -0.57 | 0.11 | -1.14 | -0.66 | -1.75 | 0.45 | -1.69 | -0.05 | -0.43 | -0.21 | 0.07 | 0.115 | 0.3625 | 1.0625 | 0.688338 | 0.542377 | 0.134417 | 1 | 0.999631 | 0.891814 |
| Hnrnpu_Q8VEK3_S247s_1 | -0.92 | 0.54 | 0.42 | -0.31 | 0.1 | 0.13 | 0.05 | -0.09 | 0.13 | -0.39 | 0.52 | 0.92 | 1.87 | 0.36 | 0.98 | 0.77 | 0.08 | 0.21 | -0.1075 | 0.041053 | 0.753253 | 0.415487 | 1 | 0.999631 | 0.474979 |
| Hnrnpul2_Q00PI9_S159s_1 | -0.23 | 0.42 | -1.61 | -1.44 | 0.16 | 0.09 | -1.35 | -1.44 | 0.62 | 0.5 | -1.46 | -1.68 | 0.62 | -0.3 | -1.52 | -2.09 | 0.3425 | 0.2825 | -0.15 | 0.876611 | 0.892795 | 0.76458 | 1 | 0.999631 | 0.961275 |
| Hnrnpul2_Q00PI9_S159s_2 | -0.78 | 1.07 | 0.18 | -0.45 | 0.96 | 0.75 | -0.22 | -0.1 | 1.23 | 0.2 | -0.08 | -0.2 | 0.68 | -0.44 | -0.38 | -0.44 | -0.3875 | 0.0675 | -0.4225 | 0.754179 | 0.479706 | 0.57179 | 1 | 0.999631 | 0.913377 |
| Hnrnpul2_Q00PI9_S166s_1 | 0.23 | 0.28 | 0.93 | 0.23 | -0.43 | 0.18 | 0.23 | 0.14 | 0.52 | 0.87 | 0.22 | 0.33 | 0.55 | -0.79 | 0.29 | -0.07 | 0.3375 | 0.3475 | -0.145 | 0.277092 | 0.211381 | 0.821817 | 1 | 0.999631 | 0.700555 |
| Hnrnpul2_Q00PI9_S166s_2 | -0.75 | 0.83 | 0.18 | -0.45 | 0.83 | 0.65 | -0.22 | -0.1 | 1.09 | 0.39 | -0.08 | -0.2 | 0.55 | -0.5 | -0.38 | -0.44 | -0.94 | 0.0825 | -1.07 | 0.739836 | 0.440553 | 0.445862 | 1 | 0.999631 | 0.908631 |
| Hnrnpul2_Q00PI9_S183s_1 | 2.07 | 1.18 | 1.86 | -0.27 | -0.43 | 0.74 | 0.98 | -0.21 | 1.26 | 2.27 | 1.05 | 0.59 | 0.6 | -0.4 | 0.44 | -0.08 | -0.7825 | -0.75 | -2.0675 | 0.071006 | 0.126938 | 0.887421 | 1 | 0.999631 | 0.531307 |
| Hnrnpul2_Q00PI9_S191s_1 | 1.94 | -1.54 | 0.46 | 0.26 | -1.79 | -0.73 | 2.7 | -2.19 | -1.74 | 1.46 | -0.55 | -1.05 | -1.25 | -2.52 | -1.14 | -2.24 | -1.0775 | -1.04 | -0.7825 | 0.013913 | 0.493796 | 0.394962 | 1 | 0.999631 | 0.385092 |
| Hnrnpul2_Q00PI9_S224s_1 | 1.41 | 1.63 | 2.91 | 1.76 | -0.07 | 1.93 | 0.88 | 0.66 | 0.52 | 1.75 | 0.42 | 0.86 | 0.78 | 0.78 | 1.86 | 1.16 | -1.0825 | -1.1375 | -0.935 | 0.092843 | 0.053338 | 0.039519 | 1 | 0.999631 | 0.557387 |
| Hnrnpul2_Q00PI9_S226s_1 | 1.42 | 1.88 | 3 | 1.93 | 0.01 | 2.27 | 0.97 | 0.65 | 0.55 | 1.77 | 0.43 | 0.93 | 0.55 | 0.72 | 1.95 | 1.27 | 0.1475 | 0.425 | 0.17 | 0.063376 | 0.067474 | 0.027192 | 1 | 0.999631 | 0.512978 |
| Homer1_Q9Z2Y3_S286s_1 | -1.26 | -2.47 | -2.51 | -1.89 | -1.4 | -2.63 | -1.86 | -1.65 | -1.52 | -1.31 | -2.18 | -1.42 | -1.18 | -1.18 | -1.66 | -3.43 | -0.4275 | -0.2425 | -0.3725 | 0.763468 | 0.713307 | 0.275604 | 1 | 0.999631 | 0.915649 |
| Homer3_Q99JP6_S159s_1 | 0.86 | -0.26 | 0.22 | 0.58 | -0.04 | -0.75 | 0.19 | 0.29 | 0.31 | 0.1 | -0.16 | 0.18 | 0.52 | -0.05 | -0.53 | -0.03 | -0.3975 | 0.65 | -0.205 | 0.323976 | 0.253075 | 0.45624 | 1 | 0.999631 | 0.724485 |
| Hook3_Q8BUK6_S238s_1 | -1.74 | -3.33 | -3.63 | -1.79 | -3.41 | -3.96 | -2.23 | -2.48 | -3.31 | -2.36 | -1.23 | -0.99 | -1.8 | -2.76 | -3.54 | -3.21 | 0.39 | -0.215 | -0.31 | 0.722517 | 0.50026 | 0.335795 | 1 | 0.999631 | 0.905674 |
| Hook3_Q8BUK6_S707s_1 | 1.74 | 0.15 | 0.04 | 0.76 | 1.04 | 1.21 | 0.82 | 1.18 | 0.43 | 1.09 | 0.49 | -0.18 | -0.93 | 0.76 | 0.8 | 0.82 | 0.22 | -0.5425 | -0.545 | 0.570871 | 0.344812 | 0.639619 | 1 | 0.999631 | 0.84338 |
| Hopx_Q8R1H0_S61s_1 | 0.32 | 0.34 | 1.37 | 0.86 | -0.19 | 1.88 | 0.28 | 1.8 | -0.19 | 0.28 | 1.12 | -0.49 | -1.3 | 0.2 | 0.8 | 1.01 | -0.645 | -0.1125 | 0.03 | 0.326091 | 0.683596 | 0.226434 | 1 | 0.999631 | 0.724678 |
| Hp1bp3_Z4YKA3_S475s_1 | -0.18 | 0.34 | 0.03 | 0.39 | -0.67 | -0.46 | -0.18 | -0.69 | -0.53 | -0.17 | 0.74 | 0.09 | 0.22 | 0.2 | 0.09 | 0.19 | -0.5825 | 0.035 | -0.44 | 0.914049 | 0.036402 | 0.743619 | 1 | 0.999631 | 0.972551 |
| Hp1bp3_Z4YKA3_T13t_1 | -0.64 | -2.1 | -0.67 | -0.9 | -1.75 | -3.08 | -0.78 | -1.03 | -1.74 | -1.67 | -0.65 | -0.11 | -1.49 | -2.55 | -0.68 | -1.35 | -0.125 | 0.13 | 0.1975 | 0.386275 | 0.317725 | 0.94418 | 1 | 0.999631 | 0.756141 |
| Hpcal1_P62748_S175s_1 | 0.15 | 0.63 | 0.4 | 0.24 | 0.17 | 0.24 | -0.02 | 0.53 | 0.18 | 0.93 | 0.51 | 0.32 | 0.84 | 0.43 | 0.13 | 0.81 | 0.06 | 0.1225 | -0.1125 | 0.517991 | 0.632895 | 0.650227 | 1 | 0.999631 | 0.819449 |
| Hpcal1_P62748_T144t_1 | -0.01 | -1.45 | -0.54 | -0.82 | -1.41 | -0.29 | -0.23 | -0.65 | -0.47 | -0.22 | -0.43 | -1.21 | -0.9 | -1.32 | -0.25 | -0.8 | -0.0025 | -0.285 | -0.325 | 0.777529 | 0.882492 | 0.752601 | 1 | 0.999631 | 0.921382 |
| Hpcal4_F2Z3Z1_T146t_1 | 0.32 | -0.68 | -0.37 | -0.48 | -0.34 | -0.5 | 0.12 | -0.5 | -0.56 | -0.32 | -1.12 | -0.35 | 0.43 | -0.87 | -1.2 | -0.87 | 0.4175 | 0.195 | 0.45 | 0.456207 | 0.993646 | 0.403519 | 1 | 0.999631 | 0.790326 |
| Hrh1_P70174_S230s_1 | -1.23 | 0.24 | -0.56 | -0.62 | -0.12 | 0.52 | -0.63 | -0.27 | -0.41 | -0.16 | -1.03 | 0.21 | -1.52 | 0.28 | 0.3 | 0.57 | -0.325 | -0.4375 | -0.8225 | 0.40682 | 0.302475 | 0.634249 | 1 | 0.999631 | 0.766722 |
| Hrh1_P70174_S264s_1 | 0.97 | 0.27 | 0.06 | 0.72 | 0.58 | -0.68 | 0.38 | 0.44 | -0.1 | -0.05 | 0.17 | 0.25 | -1.73 | 0.12 | -0.06 | 0.4 | 0.0575 | 0.37 | 0.3375 | 0.124438 | 0.39334 | 0.167581 | 1 | 0.999631 | 0.596063 |
| Hrh3_E9Q292_S302s_1 | 0.54 | -1.03 | -0.8 | -1.46 | -0.75 | -1.82 | -1.01 | 1.06 | -0.57 | 0.63 | -0.47 | -0.86 | 0.61 | -1.26 | -0.56 | -0.19 | -0.02 | 0.755 | 0.9425 | 0.538005 | 0.930902 | 0.474703 | 1 | 0.999631 | 0.830863 |
| Hrh3_E9Q292_S332s_1 | -1.48 | -5.32 | -6.25 | -3.22 | -3.2 | -4.37 | -3.49 | -5.29 | -4.02 | -2.37 | -2.87 | -3.99 | -2.07 | -4.01 | -2.67 | -3.75 | -0.18 | 0.1925 | 0.395 | 0.354742 | 0.98412 | 0.453172 | 1 | 0.999631 | 0.739951 |
| Hrh3_E9Q292_S343s_1 | -0.7 | -0.47 | 0.33 | -0.27 | -0.92 | -0.09 | -0.58 | -0.24 | -0.54 | -0.31 | 0.31 | 0.2 | 0.96 | -0.21 | 0.33 | -0.61 | 0.345 | 0.4625 | 1.43 | 0.357477 | 0.58791 | 0.578079 | 1 | 0.999631 | 0.740986 |
| Hrh3_E9Q292_S439s_1 | -0.75 | -0.58 | -1.23 | -0.53 | -0.57 | 0.25 | -0.77 | -0.62 | -0.21 | 0.05 | -1.02 | -0.06 | 1.04 | 0.38 | 0.85 | 0.36 | 0.2775 | 0.605 | 0.6125 | 0.001136 | 0.302392 | 0.191938 | 1 | 0.999631 | 0.351308 |
| Hsd17b4_P51660_S308s_1 | 0.11 | -0.29 | 0.41 | -0.79 | -0.23 | 0.72 | 0 | 0.06 | 0.78 | 0.22 | 0.68 | 0.18 | -0.05 | 1.18 | 0.47 | 0.29 | 0.245 | 0.4775 | 0.1525 | 0.143723 | 0.44324 | 0.104179 | 1 | 0.999631 | 0.615182 |
| Hsp90aa1_P07901_S231s_1 | 0.45 | 1.94 | 2.88 | 2.87 | 1.8 | 1.59 | 2.85 | 2.88 | 1.6 | 0.97 | 3.27 | 4.21 | 2.21 | 0.85 | 4.43 | 1.26 | 0.4375 | 0.5425 | 0.8475 | 0.857924 | 0.684944 | 0.565109 | 1 | 0.999631 | 0.952251 |
| Hsp90aa1_P07901_S263s_1 | -1.71 | 1.68 | 1.72 | 0.39 | 1.16 | 1.15 | 0.75 | 0.77 | 1.33 | -0.47 | 1.28 | 2.11 | 2.06 | 1.17 | 1.25 | 0.99 | -0.1275 | 0.2925 | 0.1275 | 0.267406 | 0.545829 | 0.527214 | 1 | 0.999631 | 0.695909 |
| Hsp90aa1_P07901_S454s_1 | -0.78 | -0.47 | -0.36 | -0.61 | -0.8 | -0.64 | -0.13 | -1.16 | -0.54 | 0.2 | -0.19 | -0.52 | -0.6 | -0.35 | -0.65 | -0.11 | 0.215 | 0.3775 | 0.045 | 0.654899 | 0.669588 | 0.320259 | 1 | 0.999631 | 0.878984 |
| Hsp90ab1_P11499_S226s_1 | 0.3 | 1.35 | 1.29 | 1.14 | 1.33 | 1.3 | 0.72 | 1.59 | 1.48 | 1.46 | 1.1 | 1.55 | 1.36 | 0.96 | 1.63 | 0.31 | 0.615 | 0.4375 | 0.2325 | 0.910509 | 0.531834 | 0.258428 | 1 | 0.999631 | 0.971071 |
| Hsp90ab1_P11499_S255s_1 | -1.04 | 1.2 | -1.51 | -2.62 | 1.08 | 0.9 | -1.28 | -2.21 | 1.33 | -0.11 | -1.51 | -1.93 | 1.66 | 0.52 | -2.63 | -2.59 | -0.1725 | 0.3375 | 0.32 | 0.8398 | 0.535858 | 0.64519 | 1 | 0.999631 | 0.946365 |
| Hsp90ab1_P11499_S445s_1 | 1.27 | 0.06 | 0.48 | 0.22 | 0.21 | 0.48 | 0.49 | 0.16 | 0.51 | 1.02 | 0.71 | 1.14 | 0.52 | 1.09 | 0.99 | 0.71 | -0.005 | 0.4425 | 0.495 | 0.375165 | 0.599494 | 0.341809 | 1 | 0.999631 | 0.748001 |
| Hsp90ab1_P11499_T446t_1 | -0.23 | -1.27 | 0.3 | -0.2 | -0.85 | -0.73 | 0.35 | -0.19 | -0.43 | 0.16 | 0.39 | 0.25 | 0.2 | -0.44 | 0.73 | 0.09 | -1.1825 | -0.49 | -1.3675 | 0.256381 | 0.990581 | 0.275321 | 1 | 0.999631 | 0.691343 |
| Hsp90b1_P08113_T44t_1 | 0.94 | 0.74 | 3.77 | 0.18 | -0.36 | 1.11 | 0.05 | 0.1 | -0.41 | 0.52 | 2.58 | 0.98 | -0.23 | 0.17 | 0.56 | -0.34 | -0.0825 | -0.12 | -0.0025 | 0.085628 | 0.142198 | 0.583253 | 1 | 0.999631 | 0.551124 |
| Hspa12a_Q8K0U4_S23s_1 | 0.66 | 0.83 | 1.06 | 0.95 | 0.63 | 0.83 | 0.56 | 1.15 | 0.42 | 0.6 | 1.18 | 0.82 | 0.3 | 1.32 | 0.66 | 1.21 | -0.265 | -0.025 | -0.3175 | 0.993932 | 0.752569 | 0.670553 | 1 | 0.999631 | 0.997705 |
| Hspa12a_Q8K0U4_S24s_1 | 0.85 | 1.44 | 1.94 | 0.92 | 0.97 | 0.96 | 0.63 | 1.53 | 0.68 | 0.96 | 2.01 | 1.4 | -0.31 | 0.96 | 1.62 | 1.61 | -0.03 | -0.1225 | -0.525 | 0.528437 | 0.45159 | 0.94995 | 1 | 0.999631 | 0.825257 |
| Hspa12a_Q8K0U4_S24s_2 | 1.36 | 0.03 | 0.68 | 0.38 | 1.25 | 0.16 | 0.27 | 0.65 | 0.5 | 0.04 | 0.57 | 0.85 | -0.87 | 0.27 | 0.14 | 0.81 | 0.08 | 0.2175 | 0.3725 | 0.259348 | 0.937912 | 0.735071 | 1 | 0.999631 | 0.692624 |
| Hspa12a_Q8K0U4_S631s_1 | -0.27 | 0.32 | -0.17 | -0.34 | 0.28 | -0.09 | -0.68 | 0.35 | 0.26 | -0.14 | 0 | 0.29 | 0.59 | 0.43 | 0.03 | -0.02 | -0.39 | 0.1975 | -0.0125 | 0.243777 | 0.805221 | 0.44385 | 1 | 0.999631 | 0.685637 |
| Hspa12b_Q9CZJ2_S276s_1 | -0.26 | 0.72 | 0.91 | 0.61 | 0.66 | 0.08 | -0.17 | -0.15 | 0.53 | 0.66 | 0.63 | 0.95 | 0.42 | 0.35 | 0.5 | 0.66 | 0.2375 | 0.5625 | 0.4675 | 0.970311 | 0.283144 | 0.550959 | 1 | 0.999631 | 0.989586 |
| Hspa12b_Q9CZJ2_S433s_1 | -0.12 | 0.83 | 0.56 | 0.31 | 0.49 | 0.63 | 0.5 | 0.91 | 1.22 | 0.76 | 0.73 | 1.12 | 0.98 | 0.52 | 0.76 | 1.19 | 0.0075 | -0.05 | 0.3975 | 0.172997 | 0.425965 | 0.091881 | 1 | 0.999631 | 0.642116 |
| Hspa12b_Q9CZJ2_S44s_1 | -0.95 | -0.02 | -0.46 | -0.96 | 0.02 | -0.3 | -0.87 | -1.21 | -0.18 | -1.05 | -1.09 | -0.27 | 0.52 | 0.3 | -1.34 | -0.28 | -0.08 | 0.6475 | 0.2575 | 0.401737 | 0.983964 | 0.890612 | 1 | 0.999631 | 0.763221 |
| Hspa12b_Q9CZJ2_T42t_1 | -0.87 | -1.31 | -0.81 | -1.04 | -1.03 | -1.43 | -0.65 | -1.24 | -0.26 | -1.45 | 0.32 | -0.05 | -0.46 | -1.12 | -0.44 | -0.98 | -0.6975 | -0.04 | -0.2425 | 0.409252 | 0.776397 | 0.136438 | 1 | 0.999631 | 0.766996 |
| Hspa1a_Q61696_T636t_1 | -1 | 0.06 | 0.46 | 0.93 | -0.31 | -1.55 | -0.1 | -0.38 | -0.39 | -0.15 | 0.13 | 0.7 | -0.38 | -1.13 | -0.11 | 1.1 | -0.0375 | 0.42 | 0.32 | 0.67233 | 0.180154 | 0.930749 | 1 | 0.999631 | 0.885053 |
| Hspa4_Q3U2G2_S503s_1 | -0.64 | 1.75 | 1.64 | 0.91 | 0.27 | 1.9 | 0.96 | 0.38 | 1.38 | 0.92 | 1.39 | 1.65 | 0.94 | 1.1 | 1.19 | 1.71 | -0.085 | 0.03 | 0.24 | 0.555884 | 0.950195 | 0.438506 | 1 | 0.999631 | 0.838305 |
| Hspa4_Q3U2G2_S506s_1 | -1.02 | 0.93 | 0.94 | 0.55 | 0.14 | 0.16 | 0.85 | -0.09 | -0.01 | -0.18 | 0.82 | 0.89 | 0.79 | 0.11 | 0.89 | 0.57 | -0.13 | 0.035 | -0.745 | 0.62061 | 0.859736 | 0.953135 | 1 | 0.999631 | 0.861888 |
| Hspa4_Q3U2G2_S548s_1 | -0.03 | 0.28 | 1.43 | 1.25 | 1.04 | -0.2 | 1.2 | 0.37 | 0.82 | 0.66 | 1.01 | 0.58 | -0.55 | -0.55 | 0.79 | 0.26 | -0.41 | 0.4175 | 0.5725 | 0.139899 | 0.77862 | 0.928178 | 1 | 0.999631 | 0.611718 |
| Hspa4_Q3U2G2_S694s_1 | -0.97 | -0.24 | -0.88 | -0.8 | -1.29 | -1.48 | -0.68 | -1.08 | -0.23 | -0.51 | -0.23 | -0.25 | 1.16 | -0.69 | -1.03 | -0.04 | -0.4325 | -0.44 | -0.5375 | 0.260463 | 0.193484 | 0.156349 | 1 | 0.999631 | 0.692624 |
| Hspa4_Q3U2G2_T468t_1 | 1.19 | 0.01 | 1.08 | -0.13 | -0.9 | 0.38 | 0.65 | 0.29 | -0.1 | 0.32 | -0.07 | 0.24 | 0.2 | -0.22 | 0.78 | -0.76 | -0.34 | 0.1625 | 0.3225 | 0.266382 | 0.364891 | 0.267778 | 1 | 0.999631 | 0.695664 |
| Hspa4_Q3U2G2_T540t_1 | 0.88 | 1.51 | 1.26 | 1.48 | 0.77 | 1.13 | 0.66 | 1.21 | 1.45 | 1.31 | 1.32 | 1.7 | 1.48 | 1.56 | 1.56 | 1.82 | 0.8275 | 1.1525 | 0.8275 | 0.27714 | 0.242107 | 0.55801 | 1 | 0.999631 | 0.700555 |
| Hspa4l_P48722_S53s_1 | -1.59 | 1.02 | 0.17 | -1.89 | 0.96 | 0.5 | 0.37 | -0.81 | 1.45 | -1.26 | 0.67 | 1.46 | 2.05 | -0.12 | -0.23 | -0.68 | 0.6175 | 0.8675 | 0.815 | 0.318928 | 0.255999 | 0.186486 | 1 | 0.999631 | 0.722245 |
| Hspa4l_P48722_S805s_1 | -0.36 | -0.77 | -3.03 | -1.33 | -0.26 | -0.38 | -0.83 | -1.55 | -0.31 | -0.44 | -1.03 | -0.24 | 0.73 | 0.03 | -0.75 | -2.24 | -0.1575 | -0.2225 | -0.7525 | 0.297835 | 0.312152 | 0.148999 | 1 | 0.999631 | 0.711216 |
| Hspa4l_P48722_S811s_1 | -0.57 | -0.92 | -1.05 | -0.62 | -1.23 | -0.76 | -0.25 | -1.55 | -1.05 | -0.51 | -1.3 | -1.19 | -1.29 | -1.64 | -1.41 | -1.83 | 0.915 | 1.095 | 0.35 | 0.024459 | 0.645835 | 0.454691 | 1 | 0.999631 | 0.417027 |
| Hspa4l_P48722_T524t_1 | -1.18 | 0.24 | 1.52 | -1.08 | 0.32 | 0.41 | 2.65 | -0.22 | 1.34 | -1.06 | 0.75 | 2.85 | 2.81 | -0.8 | -0.11 | -1 | -0.655 | -0.365 | 0.155 | 0.708958 | 0.261776 | 0.239619 | 1 | 0.999631 | 0.898972 |
| Hspa4l_P48722_T572t_1 | 0.51 | 1.36 | 0.97 | 1.29 | 0.5 | 0.6 | -0.31 | 0.72 | 0.44 | 0.68 | 0.94 | 0.61 | 0.5 | 1.6 | 0.87 | 1.78 | 0.375 | 0.2475 | 0.145 | 0.691622 | 0.077777 | 0.239071 | 1 | 0.999631 | 0.894017 |
| Hspa4l_P48722_T797t_1 | -4.18 | -2.65 | 0.13 | -0.83 | -2.97 | -1.44 | -0.37 | -1.25 | -2.21 | -2.26 | -0.95 | -1.12 | -2.9 | -2.38 | -0.59 | -1.08 | 0.2475 | 0.51 | 0.6275 | 0.878002 | 0.693828 | 0.780625 | 1 | 0.999631 | 0.962284 |
| Hsph1_E9Q0U7_S733s_1 | -1.06 | -0.29 | -0.31 | -1.29 | 0.71 | -0.38 | -0.81 | -1.48 | -0.62 | -0.61 | -0.05 | 0.37 | 0.96 | 0.28 | -1.51 | -0.17 | -0.075 | 0.19 | 0.2425 | 0.26404 | 0.61917 | 0.195831 | 1 | 0.999631 | 0.693486 |
| Hsph1_E9Q0U7_S769s_1 | -2.15 | -4.05 | -3.83 | -2.54 | -2.25 | -4.57 | -3.14 | -2.91 | -3.6 | -3.16 | -2.64 | -2.41 | -1.76 | -3.03 | -3.24 | -3.57 | 0.155 | 0.0775 | 0.45 | 0.67017 | 0.902184 | 0.710122 | 1 | 0.999631 | 0.884054 |
| Hsph1_E9Q0U7_T445t_1 | -0.19 | 0.46 | 0.14 | -0.17 | 0.17 | 0.32 | 0.16 | 0.21 | -0.82 | -0.11 | 0.77 | 0.71 | 0.22 | 0.77 | 0.33 | 0.72 | -0.09 | 0.2525 | 0.73 | 0.16291 | 0.55648 | 0.851284 | 1 | 0.999631 | 0.631442 |
| Hsph1_E9Q0U7_T462t_1 | -0.32 | 0.71 | 0.28 | 0.12 | 0.35 | 0.58 | -0.34 | -0.16 | 0.56 | 0.42 | -0.18 | 1 | 1.33 | 1.06 | 0.49 | 0.83 | -0.165 | -1.29 | -1.77 | 0.054436 | 0.790545 | 0.486278 | 1 | 0.999631 | 0.491994 |
| Htatsf1_Q8BGC0_S388s_1 | 1.02 | 2.1 | 1 | 1.44 | 0.91 | 0.72 | 2.92 | 0.35 | -0.02 | -0.77 | 1.49 | -0.3 | -2.02 | 0.07 | -1.72 | 2.15 | 0.22 | -0.065 | -0.56 | 0.062802 | 0.77442 | 0.030723 | 1 | 0.999631 | 0.512978 |
| Htr2a_P35363_S280s_1 | 0.63 | 0.23 | -2.16 | 0 | 1.16 | -0.45 | -0.46 | -0.67 | 0.68 | -0.74 | -0.84 | -0.66 | -1.78 | -0.17 | -0.94 | -0.65 | 0.275 | 1.285 | 1.5075 | 0.390841 | 0.743936 | 0.920112 | 1 | 0.999631 | 0.758901 |
| Htr2c_P34968_S408s_1 | -0.02 | -1.15 | -0.81 | -0.53 | -0.16 | -0.38 | -0.53 | -0.34 | 0.76 | 0.71 | 0.02 | 1.14 | 0.15 | 0.88 | 1.34 | 1.15 | 0.2525 | 0.115 | -0.17 | 0.002822 | 0.381605 | 0.005417 | 1 | 0.999631 | 0.351308 |
| Htt_G3X9H5_S1180s_1 | 0.56 | -0.02 | 0.39 | 0.74 | 0.58 | 0.79 | 0.6 | 0.71 | 0.4 | 0.56 | 0.62 | 0.55 | -0.79 | 0.54 | 0.65 | 0.59 | 1.3875 | 2.4725 | -0.3325 | 0.675517 | 0.354754 | 0.675869 | 1 | 0.999631 | 0.885853 |
| Htt_G3X9H5_S1842s_1 | 0.06 | 3.93 | 4.12 | 0.05 | 3.75 | 2.92 | 6 | 1.04 | 6.43 | 0.25 | 4.97 | 6.4 | 6.26 | -0.63 | 2.63 | -1.43 | -0.1025 | -0.2075 | 0.565 | 0.849451 | 0.304085 | 0.141108 | 1 | 0.999631 | 0.949525 |
| Htt_G3X9H5_S1854s_1 | -0.06 | -1.23 | 1 | 0.18 | 0.08 | -1.01 | 0.28 | 0.13 | -1.6 | -0.01 | 0.54 | 0.13 | 1.5 | -1.2 | 0.9 | 0.95 | 0.37 | 0.505 | 0.2425 | 0.408745 | 0.840989 | 0.729881 | 1 | 0.999631 | 0.766996 |
| Htt_G3X9H5_S638s_1 | -1.02 | 0.45 | -0.56 | -0.87 | -0.1 | 0.84 | -0.47 | -0.79 | -0.01 | 0.17 | -0.49 | 0.35 | 0.05 | -0.22 | 0.23 | -1.09 | 0.225 | -0.16 | -0.0775 | 0.587283 | 0.432122 | 0.218282 | 1 | 0.999631 | 0.850015 |
| Htt_G3X9H5_T2923t_1 | -0.38 | -0.29 | 0.34 | -0.42 | -0.11 | 0.22 | -0.24 | 0.28 | -0.63 | -0.33 | -0.02 | -0.41 | -0.83 | -0.16 | -0.08 | 0.01 | -0.7025 | -0.4525 | -1.6175 | 0.815986 | 0.444945 | 0.593016 | 1 | 0.999631 | 0.93761 |
| Huwe1_Q7TMY8_S1298s_1 | 0.48 | -0.12 | 1.34 | 1.67 | -0.95 | -0.49 | 1.36 | 0.64 | -0.69 | 0.91 | 0.09 | 1.25 | -0.93 | -1.94 | 0.37 | -0.6 | 0.8275 | 1.1325 | 0.9475 | 0.017489 | 0.259668 | 0.420413 | 1 | 0.999631 | 0.391024 |
| Huwe1_Q7TMY8_S1368s_1 | -0.92 | 2.26 | 1.42 | -0.19 | 2.15 | 1.66 | 1.01 | 1.06 | 2.56 | 0.08 | 2.03 | 2.43 | 2.44 | 1.07 | 2.14 | 0.71 | -0.105 | 0.2775 | 0.28 | 0.2182 | 0.247217 | 0.18574 | 1 | 0.999631 | 0.669859 |
| Huwe1_Q7TMY8_S1395s_1 | -0.26 | -1.16 | -0.66 | -0.8 | -0.82 | -1.2 | -0.72 | -0.56 | -0.75 | -0.31 | -0.66 | -0.05 | -0.2 | -0.67 | -0.24 | -0.65 | -0.055 | 0.2125 | 0.3575 | 0.383093 | 0.723918 | 0.384117 | 1 | 0.999631 | 0.754387 |

| | | | | | | | | | | | | | | | | | | | | | | | | | |
|---|---|---|---|---|---|---|---|---|---|---|---|---|---|---|---|---|---|---|---|---|---|---|---|---|---|
| Huwe1_Q7TMY8_S1907s_1 | -0.41 | 0.6 | 0.43 | 0.07 | 0.26 | 0.1 | -0.11 | 0.22 | 0.32 | 0.15 | 0.74 | 0.33 | 0.7 | 0.49 | 0.09 | 0.84 | -0.135 | -0.2425 | 0.13 | 0.308308 | 0.854926 | 0.508467 | 1 | 0.999631 | 0.718521 |
| Huwe1_Q7TMY8_S2917s_1 | -0.86 | -0.91 | 0.22 | -1.21 | -0.91 | -1.29 | -0.36 | -0.74 | -0.34 | -2.35 | -0.36 | -0.68 | -0.21 | -1.31 | -0.88 | 0.16 | -0.41 | -0.7475 | -1.2275 | 0.774203 | 0.723697 | 0.65124 | 1 | 0.999631 | 0.920998 |
| Huwe1_Q7TMY8_S2918s_1 | -0.35 | -0.65 | 1.11 | 0.39 | -0.47 | -0.56 | 0.37 | -0.48 | -1.2 | -0.11 | -0.45 | -0.73 | -0.68 | -1.99 | -0.77 | -0.97 | 0.1325 | 0.2525 | 0.9225 | 0.027205 | 0.362005 | 0.120518 | 1 | 0.999631 | 0.423171 |
| Huwe1_Q7TMY8_S3374s_1 | -0.04 | 0.03 | 0.9 | 0.15 | 0.35 | 0.66 | -0.23 | 0.79 | 0.55 | -0.32 | 0.34 | 1.48 | 1.3 | 1.25 | 1.34 | 0.84 | 0.0325 | 0.0575 | 0.6325 | 0.016326 | 0.702515 | 0.560504 | 1 | 0.999631 | 0.388832 |
| Huwe1_Q7TMY8_S3663s_1 | -1.08 | -0.71 | -0.35 | -1.66 | -0.91 | -0.78 | -0.96 | -1.02 | -0.91 | -0.89 | -0.89 | -0.88 | -0.33 | -0.49 | -0.16 | -0.29 | -0.175 | 0.19 | 0.4075 | 0.092662 | 0.920875 | 0.862177 | 1 | 0.999631 | 0.557279 |
| Huwe1_Q7TMY8_S3758s_1 | -1.59 | 1.39 | -0.07 | -0.65 | -0.17 | -0.06 | 0.01 | -1.4 | 0.24 | -0.52 | -0.28 | 0.4 | 0.79 | 0.87 | -0.5 | -0.45 | 0.0625 | 0.19 | 0.815 | 0.537135 | 0.782857 | 0.752568 | 1 | 0.999631 | 0.830294 |
| Huwe1_Q7TMY8_S3818s_1 | 0.02 | -1.17 | 0.55 | -0.26 | -0.28 | -0.58 | 0.11 | 0.14 | -0.33 | -0.08 | -0.13 | 0.44 | 0.03 | 0.5 | 1.06 | 0.81 | 0.0475 | -0.2875 | 0.145 | 0.082572 | 0.876454 | 0.641156 | 1 | 0.999631 | 0.548531 |
| Huwe1_Q7TMY8_S3922s_1 | -0.56 | -0.68 | -0.94 | -0.42 | -0.26 | -1.29 | -0.82 | -0.04 | -1.32 | -1.25 | -0.5 | -0.68 | -0.75 | -0.83 | -0.42 | -0.02 | 0.25 | 0.2575 | 0.765 | 0.641148 | 0.888428 | 0.356619 | 1 | 0.999631 | 0.871335 |
| Huwe1_Q7TMY8_T3927t_1 | -0.57 | 0.81 | -0.67 | -1.76 | 0.2 | -0.38 | -0.58 | -0.43 | -0.2 | -0.78 | -0.33 | 0.15 | -0.21 | 0.79 | -0.1 | 0.39 | -0.73 | -0.5575 | 0.16 | 0.176998 | 0.629595 | 0.625951 | 1 | 0.999631 | 0.642116 |
| Huwe1_Q7TMY8_T3930t_1 | -0.29 | -0.08 | -0.9 | -1.52 | -1.49 | -1.76 | -0.8 | -1.66 | -1.7 | -1.17 | -1.47 | -0.68 | -0.57 | 0.46 | -1.6 | -0.44 | 0.75 | -0.16 | 0.5825 | 0.752851 | 0.09023 | 0.188453 | 1 | 0.999631 | 0.913377 |
| Ibtk_Q6ZPR6_S945s_1 | -0.61 | 2.73 | 2.27 | 1 | 2.16 | 2.34 | 1.23 | 2.66 | 2.08 | -0.1 | 0.84 | 1.93 | 1.53 | 2.01 | 2.12 | 2.06 | 0.3 | 0.7425 | 0.66 | 0.398967 | 0.307679 | 0.840593 | 1 | 0.999631 | 0.761767 |
| Ica1l_Q3TY65_S294s_1 | -0.25 | 1.52 | -4.06 | -2.34 | 0.27 | 1.12 | -2.71 | -2.61 | 1.14 | 1.13 | -2.34 | -2.09 | 1.88 | 1.56 | -3.06 | -2.87 | 0.2 | 0.1925 | -2E-17 | 0.666971 | 0.821713 | 0.578652 | 1 | 0.999631 | 0.88264 |
| Ice1_E9Q286_T1607t_1 | -1.12 | -1.07 | 0.06 | -1.02 | -0.27 | -0.31 | -0.89 | -0.88 | -0.62 | -1.67 | 0.08 | -0.17 | -0.93 | -0.92 | -1.53 | 0.23 | 0.7725 | 0.5975 | 0.115 | 1 | 0.578196 | 0.679948 | 1 | 0.999631 | 1 |
| Ifngr1_P15261_S296s_1 | 0.9 | 0.6 | 0.66 | 0.72 | 1.93 | 1.66 | 1.23 | 1.15 | 1.67 | -0.09 | 1.99 | 1.7 | -0.24 | 1.47 | 1.67 | 0.44 | -0.455 | -0.36 | -0.05 | 0.799044 | 0.018638 | 0.218395 | 1 | 0.999631 | 0.930937 |
| Ifngr1_P15261_T373t_1 | -0.45 | 0.38 | 0.13 | 0.05 | -1 | -0.16 | -0.64 | 0.09 | -1.31 | 0.07 | -0.27 | 0.18 | -0.61 | 0.01 | 0.32 | 0.19 | 0.5425 | 1.27 | 1.215 | 0.882089 | 0.197381 | 0.373778 | 1 | 0.999631 | 0.963505 |
| Ifngr2_Q63953_T331t_1 | -2.84 | 0.19 | -0.35 | -1.79 | -0.36 | -0.88 | -0.65 | -0.73 | 0.55 | -1.02 | 0.17 | 0.59 | 1.58 | -0.11 | -0.61 | -0.79 | 0.055 | 0.0975 | 0.3325 | 0.136709 | 0.39564 | 0.094288 | 1 | 0.999631 | 0.61018 |
| Ift46_Q9DB07_T283t_1 | -1.04 | -0.45 | -0.01 | -0.23 | 0.02 | -0.48 | -0.62 | -0.43 | -0.37 | -0.41 | -0.36 | -0.2 | 0 | -0.15 | 0.15 | -0.4 | -0.5625 | -0.2 | -0.545 | 0.322398 | 0.860839 | 0.746832 | 1 | 0.999631 | 0.724386 |
| Ift74_B1AWG1_S300s_1 | 1.02 | -0.48 | 0.67 | -0.28 | -1.1 | -0.37 | 0.17 | -0.02 | -0.45 | 0.68 | -0.1 | 0 | -0.62 | -0.19 | -0.14 | -0.3 | 0.16 | 0.165 | 0.2925 | 0.19562 | 0.225262 | 0.64498 | 1 | 0.999631 | 0.653056 |
| Igbp1b_Q9QZ29_S282s_1 | -2.27 | -2.94 | -2.49 | -0.92 | -1.2 | -2.49 | -1.84 | -2.45 | -2.28 | -2.17 | -1.73 | -1.78 | -1.98 | -3.04 | -1.21 | -1.22 | 0.6775 | -0.3475 | -0.325 | 0.607653 | 0.748349 | 0.713473 | 1 | 0.999631 | 0.858527 |
| Igsf5_D3YXH0_S284s_1 | 2.5 | 1.08 | -0.43 | 1.5 | 2.45 | 1.35 | 2.1 | 1.46 | 1.81 | 0.15 | 1.22 | 0.08 | 0.71 | 0.99 | 1.53 | 0.12 | -0.2575 | 0.045 | 0.03 | 0.598588 | 0.274238 | 0.60327 | 1 | 0.999631 | 0.85557 |
| Igsf9b_E9PZ19_S1207s_1 | -0.39 | -0.14 | -1.09 | -0.72 | -1.17 | -0.72 | -0.87 | -0.61 | -0.98 | -0.3 | -0.9 | 0.02 | -0.25 | -0.62 | -0.55 | -0.8 | -0.265 | 0.03 | 0.665 | 0.92516 | 0.401812 | 0.898716 | 1 | 0.999631 | 0.975509 |
| Igsf9b_E9PZ19_S1215s_1 | -0.56 | -0.36 | -0.19 | -0.71 | -1.4 | -0.24 | -0.62 | -0.62 | -0.63 | -0.28 | -0.51 | -0.28 | 1.13 | -0.12 | -0.13 | -0.04 | -0.9575 | -0.7525 | -0.5925 | 0.096751 | 0.413632 | 0.909311 | 1 | 0.999631 | 0.55886 |
| Igsf9b_E9PZ19_S1258s_1 | 0.92 | -2.55 | -0.27 | -2.28 | -3.29 | -0.96 | -0.58 | -3.18 | -4.12 | 0.58 | -1.93 | -1.72 | -0.06 | -2.81 | -1.61 | -2.07 | -0.145 | 0.44 | 0.845 | 0.503236 | 0.324434 | 0.495716 | 1 | 0.999631 | 0.814092 |
| Igsf9b_E9PZ19_S775s_1 | -0.78 | 0.03 | -0.79 | 0.15 | -0.85 | -0.31 | -0.39 | -0.42 | 0.46 | -0.06 | 0.15 | -0.18 | 1.03 | 0.39 | -0.1 | 0.67 | -0.08 | 0.3175 | 1.0125 | 0.048353 | 0.657605 | 0.212122 | 1 | 0.999631 | 0.48343 |
| Igsf9b_E9PZ19_S783s_1 | -0.92 | -1.77 | -0.82 | -1.65 | -1.96 | -0.54 | -1.96 | -1.02 | -1.84 | 0.68 | -1.15 | -1.58 | 0.71 | 0.04 | -1.46 | -0.4 | 0.4425 | 0.445 | 0.955 | 0.064459 | 0.850574 | 0.580326 | 1 | 0.999631 | 0.514485 |
| Ikbkap_Q7TT37_T1285t_1 | -1.15 | 0.01 | 0.16 | -0.03 | -0.21 | -0.1 | -0.17 | 1.24 | 0.31 | -0.57 | 0.78 | 0.25 | 0.57 | 0.32 | 1.1 | 0.82 | -0.3175 | -0.0175 | 0.1325 | 0.028954 | 0.335748 | 0.300474 | 1 | 0.999631 | 0.429864 |
| Ikbkb_Q5D0E0_S672s_1 | 0.26 | 0.15 | -0.2 | 0.1 | -0.4 | -0.04 | -0.5 | -0.02 | -0.08 | 0.72 | -0.14 | -0.26 | 0.7 | 0.36 | 0.16 | -0.38 | -0.1225 | -0.405 | 0.44 | 0.684658 | 0.242234 | 0.955063 | 1 | 0.999631 | 0.889888 |
| Ikbkg_Q8VC91_S379s_1 | -0.78 | -0.58 | -0.62 | -1.25 | -1.4 | -0.7 | -1.16 | -0.46 | -1.23 | -0.8 | -0.6 | -2.22 | -0.05 | -0.32 | -0.79 | -0.31 | 0.8375 | -0.39 | -1.905 | 0.17826 | 0.698212 | 0.328025 | 1 | 0.999631 | 0.642792 |
| Ildr2_B5TVM2_S287s_1 | 3.38 | 0.33 | -1.54 | 2.35 | 3.01 | 1.24 | 2.43 | 1.19 | 1.86 | 0.7 | 0.96 | -0.56 | -2.39 | 0.02 | -0.26 | -0.47 | 0.5975 | 0.42 | 0.44 | 0.091859 | 0.418029 | 0.707265 | 1 | 0.999631 | 0.557085 |
| Ildr2_B5TVM2_S384s_1 | 0.13 | 0.61 | -0.6 | 0.37 | 1.26 | 0.72 | -0.11 | 1.03 | 1.23 | 0.2 | 0.51 | 0.25 | 0.67 | 0.96 | 0.14 | 0.5 | -0.225 | 0.0375 | -0.275 | 0.241191 | 0.159674 | 0.28109 | 1 | 0.999631 | 0.684359 |
| Ildr2_B5TVM2_S435s_1 | -0.09 | -0.65 | 0.06 | 0.04 | -0.43 | -0.55 | -0.38 | -0.18 | -0.4 | -0.07 | 0 | -0.02 | -0.46 | -0.48 | -0.36 | -0.44 | -0.06 | -0.0125 | 0.285 | 0.353388 | 0.4179 | 0.894878 | 1 | 0.999631 | 0.73949 |
| Ildr2_B5TVM2_S457s_1 | -0.83 | 0.09 | -0.6 | -0.59 | -0.24 | -0.37 | -0.9 | -0.66 | -0.33 | -0.83 | -0.58 | -0.24 | 0.07 | -0.1 | -0.45 | -0.31 | -0.4 | -0.4725 | -0.2075 | 0.377799 | 0.844606 | 0.967617 | 1 | 0.999631 | 0.750463 |
| Ildr2_B5TVM2_T438t_1 | 0.6 | -0.17 | 0.35 | 1.79 | -0.42 | -0.18 | 0.89 | 0.68 | 0.02 | 0.39 | 0.44 | -0.17 | -0.48 | -0.45 | 1.61 | 1.06 | 0.0225 | -0.195 | -0.905 | 0.735164 | 0.425156 | 0.294279 | 1 | 0.999631 | 0.90759 |
| Ilf3_Q9Z1X4_S382s_1 | -1.7 | -4.7 | -4.45 | -2.11 | -1.78 | -6.09 | -2.49 | -2.51 | -3.64 | -2.22 | -3.78 | -4.1 | -4.9 | -3.71 | -2.89 | -5.08 | -0.3 | 0.0225 | 0.2375 | 0.280622 | 0.983164 | 0.801749 | 1 | 0.999631 | 0.701922 |
| Ilf3_Q9Z1X4_T67t_1 | -0.78 | 0.4 | 1.3 | 0.3 | 0.09 | -1.01 | 0.55 | 0.39 | -0.02 | -0.28 | 0.47 | 1.14 | 1.02 | 0.31 | -0.17 | 1.01 | 0.0275 | -0.345 | -0.3225 | 0.633212 | 0.562999 | 0.96413 | 1 | 0.999631 | 0.868569 |
| Ina_P46660_S30s_1 | 0.49 | 0.08 | 0.27 | 0.39 | 0.12 | 0.6 | 0.38 | 0.24 | 0.37 | -0.61 | -0.21 | 0.3 | 0.6 | -0.3 | -0.13 | -0.23 | -0.6325 | -0.26 | -0.73 | 0.318621 | 0.913222 | 0.284963 | 1 | 0.999631 | 0.722158 |
| Ina_P46660_S335s_1 | 0.67 | -0.84 | 0.08 | -0.19 | -1.35 | -0.16 | -0.12 | -1.18 | -0.39 | 0.83 | -0.56 | -1.2 | -0.41 | -0.95 | -0.42 | -1.42 | 0.725 | -0.555 | -0.7925 | 0.101398 | 0.173431 | 0.605649 | 1 | 0.999631 | 0.564448 |
| Ina_P46660_S37s_1 | 0.97 | -0.91 | -2.07 | 0.63 | 0.45 | -0.64 | 0.34 | 1.37 | -0.2 | -0.77 | -1.2 | -1.43 | -1.89 | -0.38 | -1.4 | -0.88 | -0.445 | -0.7425 | -0.49 | 0.267411 | 0.327208 | 0.419993 | 1 | 0.999631 | 0.695909 |
| Inf2_E9QLA5_S1170s_1 | 0.08 | 0.68 | -0.97 | -0.46 | 0.88 | -0.51 | -0.74 | -2.08 | 0.64 | -0.62 | -3.19 | -0.47 | -0.05 | 0.07 | -1.31 | -1.34 | 0.44 | 0.19 | -0.77 | 0.340568 | 0.485579 | 0.351904 | 1 | 0.999631 | 0.733032 |
| Inf2_E9QLA5_S1172s_1 | -0.03 | -1.55 | -0.28 | 0.23 | -0.25 | -0.2 | -0.26 | 0.84 | -0.3 | -1.03 | 0.02 | 0.44 | -1 | -1.7 | -0.84 | -1.17 | 0.125 | 0.0825 | 0.195 | 0.104576 | 0.349575 | 0.694227 | 1 | 0.999631 | 0.568213 |
| Inf2_E9QLA5_S1184s_1 | 0.92 | 0.28 | 0.74 | 1.16 | 0.29 | 0.18 | 0.94 | 2.19 | 0.5 | 0.42 | 1.05 | 1.46 | -0.02 | 0.57 | 1.55 | 1.78 | 0.215 | 0.21 | -0.06 | 0.670703 | 0.791956 | 0.812725 | 1 | 0.999631 | 0.884315 |
| Inpp4a_E9Q9A0_S186s_1 | 2 | 3.69 | -0.58 | -0.1 | 3.44 | 3 | 0.11 | -0.68 | 3.71 | 2.93 | -0.36 | -0.43 | 3.48 | 2.7 | -0.7 | -0.71 | 0.6525 | -0.22 | -0.045 | 0.961764 | 0.859624 | 0.866893 | 1 | 0.999631 | 0.986594 |
| Inpp4a_E9Q9A0_S516s_1 | 0.23 | 0.22 | -2.22 | -0.55 | 1.89 | -0.23 | -0.37 | -1 | 0.67 | -0.93 | -1.08 | -1.86 | -0.08 | 0.33 | -0.98 | -1.77 | 0.6525 | -0.22 | -0.045 | 0.945974 | 0.393375 | 0.753043 | 1 | 0.999631 | 0.982217 |
| Inpp4a_E9Q9A0_S517s_1 | 0.23 | 0.22 | -2.22 | -0.55 | 1.89 | -0.23 | -0.37 | -1 | 0.67 | -0.93 | -1.08 | -1.86 | -0.08 | 0.33 | -0.98 | -1.77 | -0.1175 | 0.36 | 0.4075 | 0.945974 | 0.393375 | 0.753043 | 1 | 0.999631 | 0.982217 |
| Inpp4a_E9Q9A0_S916s_1 | -0.82 | 1.79 | 1.51 | 0.59 | 0.51 | 0.81 | 0.35 | 0.93 | 1.24 | 0.06 | 1.46 | 1.75 | 1.19 | 1.35 | 1.08 | 1.08 | 0.5925 | -0.8525 | 0.515 | 0.465289 | 0.831719 | 0.570001 | 1 | 0.999631 | 0.794642 |
| Inpp4b_E9Q2C4_S324s_1 | -1.95 | 3.09 | 0.28 | 0.2 | 2.31 | 1.14 | -0.11 | 0.65 | -0.48 | -1.87 | -0.03 | 0.59 | 2.53 | 0.49 | 0.5 | 0.16 | -0.3575 | -0.45 | -0.815 | 0.608123 | 0.553478 | 0.40306 | 1 | 0.999631 | 0.858527 |
| Inpp4b_E9Q2C4_T316t_1 | 0.37 | -1.57 | -0.31 | -0.11 | -0.96 | -0.97 | -0.26 | -0.86 | -0.98 | 0.43 | -1.91 | -0.96 | -1.69 | -1.02 | -1.15 | -1.02 | 1.0275 | 2.4275 | 1.4475 | 0.091144 | 0.420997 | 0.446755 | 1 | 0.999631 | 0.557085 |
| Inpp5b_F2Z437_S156s_1 | -0.21 | 2.33 | 2.4 | -0.09 | 4.16 | 1.04 | 2.88 | 0.46 | 6.76 | -2.13 | 4.56 | 4.95 | 6.64 | 1.09 | 1.7 | 0.79 | -0.14 | 0.09 | 0.31 | 0.284746 | 0.30087 | 0.189846 | 1 | 0.999631 | 0.705486 |
| Inpp5d_Q9ES52_S709s_1 | 0.09 | 0.08 | 0.42 | -0.53 | 0.13 | -0.24 | -0.32 | -0.07 | -0.18 | 0.18 | 0.15 | 0.27 | 0.09 | 0.22 | -0.04 | 1.03 | 0.1775 | 0.3025 | 0.2675 | 0.399549 | 0.63335 | 0.763589 | 1 | 0.999631 | 0.762046 |
| Inpp5e_Q9JII1_S103s_1 | 0.23 | 0.21 | -0.08 | 0.17 | 0.25 | 0.1 | 0.01 | 0.88 | 0.71 | 0.41 | -0.04 | 0.66 | -0.02 | 0.34 | 0.63 | 0.65 | -0.225 | 0.02 | 0.37 | 0.368487 | 0.537866 | 0.297227 | 1 | 0.999631 | 0.745968 |
| Inpp5f_Q8CDA1_S1103s_1 | 0.56 | -1.07 | -0.81 | 0.39 | -0.44 | -1.09 | -0.05 | -0.25 | -0.22 | -0.15 | -0.88 | 0.4 | -0.21 | -0.41 | 1.13 | 0.04 | -0.4075 | -0.1325 | -0.1025 | 0.474922 | 0.622074 | 0.966192 | 1 | 0.999631 | 0.799827 |
| Inpp5f_Q8CDA1_S879s_1 | -0.4 | 1.1 | 0.41 | -0.27 | -0.46 | 0.17 | -0.64 | 0.14 | 0.04 | 0.33 | -0.36 | 0.3 | 0.11 | 0.38 | -0.18 | 0.12 | -0.115 | -0.005 | 0.3875 | 0.794311 | 0.327444 | 0.73814 | 1 | 0.999631 | 0.928662 |
| Inpp5f_Q8CDA1_S908s_1 | -0.51 | -0.32 | -1.06 | -0.71 | -1.02 | -0.89 | -0.59 | -0.56 | -0.58 | -0.34 | -0.72 | -0.98 | 0.07 | -0.24 | -0.68 | -0.2 | 0.1525 | 0.0125 | 0.095 | 0.233419 | 0.680244 | 0.986308 | 1 | 0.999631 | 0.683077 |
| Inpp5f_Q8CDA1_S936s_1 | 0.86 | 0.65 | 0.66 | 0.58 | 0.8 | 1.44 | 0.32 | 0.8 | 0.65 | 0.92 | 0.56 | 0.67 | 0.38 | 0.86 | 1.1 | 0.79 | -0.1775 | -0.3475 | -0.665 | 0.740852 | 0.613909 | 0.959857 | 1 | 0.999631 | 0.909055 |
| Inpp5f_Q8CDA1_S941s_1 | 1.24 | -0.57 | -0.59 | -0.24 | -0.12 | -0.17 | -0.21 | -0.37 | 0 | 0.7 | -1.27 | -0.98 | -0.66 | -0.1 | -0.99 | -1.07 | 0.185 | -0.23 | -0.1775 | 0.182854 | 0.680648 | 0.550084 | 1 | 0.999631 | 0.643861 |
| Inpp5f_Q8CDA1_S941s_2 | 1.58 | 1.54 | 0.24 | -0.45 | 0.67 | 1.94 | 0.06 | 0.98 | 1.48 | 1.44 | -0.89 | -0.04 | 0.41 | 0.54 | 1.09 | 0.16 | 0.185 | -0.23 | -0.1775 | 0.729159 | 0.749558 | 0.737877 | 1 | 0.999631 | 0.905923 |
| Inpp5f_Q8CDA1_S946s_2 | 1.58 | 1.54 | 0.24 | -0.45 | 0.67 | 1.94 | 0.06 | 0.98 | 1.48 | 1.44 | -0.89 | -0.04 | 0.41 | 0.54 | 1.09 | 0.16 | 0.265 | -0.1175 | -0.345 | 0.729159 | 0.749558 | 0.737877 | 1 | 0.999631 | 0.905923 |
| Inpp5j_P59644_S266s_1 | 1.1 | -0.12 | 0.41 | 0.41 | 0.59 | 0.56 | 0.75 | 0.96 | 0.17 | 0.46 | 0.68 | 0.02 | -1.48 | 0.5 | 0.7 | 0.7 | 0.0925 | -0.3125 | -0.87 | 0.532067 | 0.411484 | 0.72843 | 1 | 0.999631 | 0.827588 |
| Inpp5j_P59644_S691s_1 | 1.96 | -0.45 | 0.06 | 0.41 | 0.2 | 0.63 | 1.52 | 0 | 0.84 | 0.98 | -0.5 | -0.59 | 0.38 | -0.53 | -0.16 | -1.19 | 0.3375 | 0.26 | 0.4175 | 0.148579 | 0.870008 | 0.609569 | 1 | 0.999631 | 0.62132 |
| Inpp5j_P59644_S69s_1 | 1.32 | -0.19 | -0.23 | 0.25 | 0.49 | 0.71 | 0.39 | 0.91 | 0.4 | 0.8 | 0.37 | 0.62 | -0.25 | 1.08 | 0.95 | 1.04 | -0.5575 | 0.4475 | 1.0575 | 0.385688 | 0.394626 | 0.511327 | 1 | 0.999631 | 0.756141 |
| Ints1_K3W4P2_S131s_1 | 1.46 | 1.58 | 1.62 | 0.68 | 0.15 | 0.89 | 0.94 | 1.13 | 1.84 | 1.62 | 1.68 | 1.99 | 4.32 | 1.61 | 1.72 | 1.92 | -0.1425 | 0.5525 | 0.5025 | 0.111479 | 0.127797 | 0.167132 | 1 | 0.999631 | 0.577068 |
| Ints1_K3W4P2_S1354s_1 | -0.87 | 0.43 | 1.14 | 0.18 | -0.4 | -0.15 | 0.1 | 0.76 | 0.59 | 0.03 | 1.1 | 1.37 | 0.17 | 0.34 | 1.12 | 1.26 | 0.0375 | -0.145 | 0.235 | 0.311741 | 0.759325 | 0.273762 | 1 | 0.999631 | 0.71924 |
| Ints10_E9Q361_S231s_1 | -0.05 | -0.64 | -0.58 | -0.46 | -0.33 | -0.74 | -0.22 | -0.29 | -0.56 | -0.76 | -0.47 | -0.52 | 0.04 | -0.13 | -0.44 | -0.26 | 0.0925 | 0.58 | 0.7675 | 0.42389 | 0.889423 | 0.587143 | 1 | 0.999631 | 0.773835 |
| Ints12_Q9D168_S127s_1 | 0.19 | 0.46 | 0.34 | 0.25 | -0.31 | 0.46 | 0.59 | 0.87 | 1.08 | 0.9 | 1.07 | 0.51 | 1.17 | 0.8 | 1.37 | 0.97 | 0.22 | 0.2275 | 0.3375 | 0.018669 | 0.767628 | 0.050124 | 1 | 0.999631 | 0.391024 |
| Ip6k1_H3BIW9_S40s_1 | -0.5 | -0.46 | -0.49 | -0.13 | 0.11 | -0.24 | -0.18 | -0.39 | -0.15 | -0.33 | -0.28 | 0.09 | -0.12 | 0.19 | -0.12 | -0.18 | 0.5425 | 0.075 | -1.08 | 0.234699 | 0.394256 | 0.388332 | 1 | 0.999631 | 0.683301 |
| Ipcef1_Q5DU31_S192s_1 | 1.75 | -0.49 | -2.72 | 1.1 | 1.31 | -0.49 | 0.72 | 0.27 | 0.06 | -0.27 | 0.15 | 0 | -1.34 | -0.67 | -1.78 | -0.89 | 0.3475 | 1.46 | 1.0625 | 0.240098 | 0.559083 | 0.931303 | 1 | 0.999631 | 0.683719 |
| Ipo11_Q8K2V6_S343s_1 | -0.41 | 0.38 | 1.55 | 0.05 | -0.55 | 1.72 | -0.14 | 1.93 | 1.65 | 0.92 | 2.38 | 2.46 | 1.99 | 0.57 | 1.86 | 1.4 | 0.375 | -0.0125 | -0.325 | 0.057364 | 0.608874 | 0.017693 | 1 | 0.999631 | 0.49679 |
| Iqgap2_Q3UQ44_S16s_1 | 1.23 | 1.97 | 0.6 | 1.69 | 1.74 | 2.12 | 1.51 | 1.62 | 1.82 | 1.22 | 2.07 | 0.33 | -0.7 | 2.56 | 0.46 | 1.87 | -0.495 | 0.1675 | -0.175 | 0.643345 | 0.304191 | 0.978854 | 1 | 0.999631 | 0.871827 |
| Iqgap2_Q3UQ44_S396s_1 | -0.15 | -0.16 | -1.27 | -1.46 | -0.99 | -2.38 | -0.21 | -1.44 | -0.73 | -1.05 | 0.41 | -1 | -1.17 | -0.96 | -1.45 | -0.16 | 0.205 | 0.09 | 0.0275 | 0.69676 | 0.36168 | 0.723781 | 1 | 0.999631 | 0.895731 |
| Iqsec1_Q8R0S2_S104s_1 | 0.25 | 0.49 | -0.79 | 0.17 | 0.5 | 0.57 | -0.1 | -0.03 | 0.39 | 0 | -0.44 | 0.53 | 0.07 | 0.89 | -1.48 | 0.75 | 0.1025 | -0.895 | -1.0525 | 0.961156 | 0.568846 | 0.812357 | 1 | 0.999631 | 0.986594 |
| Iqsec1_Q8R0S2_S159s_1 | 2.09 | -0.72 | -0.64 | 0.39 | 0.14 | 0.38 | 2.85 | -1.84 | -0.89 | 0.38 | -0.13 | -1.82 | -0.86 | -0.87 | -0.22 | -1.14 | 0.2725 | 0.05 | -0.24 | 0.113033 | 0.918313 | 0.232499 | 1 | 0.999631 | 0.577408 |
| Iqsec1_Q8R0S2_S179s_1 | -0.9 | -0.06 | -0.52 | -0.41 | 0.29 | -0.51 | 0.1 | -0.68 | -0.08 | -1.62 | 0.1 | -0.09 | -0.67 | -0.77 | -0.99 | -0.42 | 0.1025 | -0.245 | -0.18 | 0.441535 | 0.419305 | 0.90836 | 1 | 0.999631 | 0.781983 |
| Iqsec1_Q8R0S2_S197s_1 | 0.38 | 0.4 | -0.11 | 0.22 | 0.88 | 0.19 | 0.32 | -0.09 | 0.1 | -0.53 | 0.45 | -0.11 | 0.53 | 0.2 | -0.45 | -0.11 | 0.13 | -0.1625 | -0.0675 | 0.579184 | 0.732558 | 0.432252 | 1 | 0.999631 | 0.846484 |
| Iqsec1_Q8R0S2_S362s_1 | 0.54 | 1.44 | -0.65 | 0.04 | 0.6 | 1 | -0.02 | 0.31 | 0.16 | 0.76 | 0.35 | -0.55 | 0.03 | 0.93 | -0.33 | 0.47 | -0.15 | -0.255 | -0.9125 | 0.891836 | 0.781413 | 0.74234 | 1 | 0.999631 | 0.964276 |
| Iqsec1_Q8R0S2_S415s_1 | 1.39 | 0.16 | 0.45 | 0.09 | 0.19 | 0.85 | 0.83 | -0.38 | 0.91 | 0.98 | -0.29 | -0.53 | 0.17 | -0.96 | -0.12 | -0.65 | 0.52 | 0.4275 | 0.83 | 0.047481 | 0.71968 | 0.595249 | 1 | 0.999631 | 0.481399 |
| Iqsec1_Q8R0S2_S513s_1 | -1.36 | 0.24 | -0.1 | -0.52 | 0.24 | 0.12 | -0.33 | 0.31 | 0.13 | -1.09 | 0.43 | 0.5 | 0.75 | 0.3 | 0.26 | 0.27 | -0.1025 | -0.0325 | 0.4125 | 0.057063 | 0.197846 | 0.387327 | 1 | 0.999631 | 0.49679 |
| Iqsec1_Q8R0S2_S795s_1 | -0.44 | -0.01 | 0.29 | -0.55 | -0.66 | 0.13 | -0.37 | -0.22 | -0.45 | -0.26 | 0.01 | -0.14 | 0.71 | 0.02 | -0.25 | 0.46 | 0.0275 | -0.1125 | -0.04 | 0.253748 | 0.74145 | 0.912767 | 1 | 0.999631 | 0.688735 |
| Iqsec1_Q8R0S2_S923s_1 | 1.23 | 1.26 | 1.03 | 1.1 | 1.47 | 0.88 | 0.92 | 1.46 | 0.86 | 0.72 | 1.14 | 1.45 | 1.01 | 1.07 | 0.59 | 1.79 | -0.1075 | -0.2175 | -0.1475 | 0.903299 | 0.91807 | 0.683086 | 1 | 0.999631 | 0.968183 |
| Iqsec1_Q8R0S2_S937s_1 | 0.15 | 0.08 | -0.49 | 0.29 | 0.31 | -0.74 | -0.03 | 0.06 | 0.03 | -0.89 | -0.01 | 0.03 | -0.37 | 0.22 | -0.41 | 0 | -0.1 | -0.1625 | -1.15 | 0.642517 | 0.742596 | 0.519521 | 1 | 0.999631 | 0.871335 |
| Iqsec2_E9QAD8_S111s_1 | 1.58 | -0.13 | 1.68 | 0.72 | 0.95 | 0.14 | 1.83 | 0.53 | -0.17 | 0.37 | 1.73 | 1.27 | -0.64 | -1.13 | 0.45 | 0.57 | 0.5575 | -0.035 | -0.4925 | 0.059754 | 0.846905 | 0.770855 | 1 | 0.999631 | 0.50521 |
| Iqsec2_E9QAD8_S1143s_1 | 1.78 | 0.89 | -0.45 | 1.79 | 2.11 | 1.12 | 1.35 | 1.66 | 1.37 | 0.51 | 1.43 | 0.56 | -0.71 | 0.87 | 0.43 | 1.45 | -0.07 | 0.07 | 0.715 | 0.441869 | 0.305013 | 0.948573 | 1 | 0.999631 | 0.78215 |
| Iqsec2_E9QAD8_S1158s_1 | -1.05 | -0.09 | 0.12 | -0.97 | -0.48 | -0.32 | -1.11 | -0.36 | -0.45 | -0.19 | -0.55 | -0.52 | 0.26 | 0.08 | 0.1 | 0.43 | -0.055 | 0.39 | 0.4375 | 0.070437 | 0.849952 | 0.841422 | 1 | 0.999631 | 0.530609 |
| Iqsec2_E9QAD8_S1161s_1 | -0.73 | -0.26 | -3.44 | -2.84 | -1.49 | -0.13 | -2.39 | -3.48 | -0.68 | -0.81 | -2.23 | -1.99 | 0.15 | 0.1 | -3.59 | -2.18 | 0.2025 | 0.22 | -0.175 | 0.671099 | 0.951827 | 0.615128 | 1 | 0.999631 | 0.884528 |
| Iqsec2_E9QAD8_S15s_1 | 1.15 | -0.19 | -1.02 | -1.06 | 0.02 | 1.11 | -0.3 | -1.14 | 0.05 | 1.41 | -1.04 | -0.66 | -0.33 | 0.65 | -1.23 | -0.91 | -0.3575 | -0.63 | -0.17 | 0.771786 | 0.746304 | 0.742937 | 1 | 0.999631 | 0.919413 |
| Iqsec2_E9QAD8_S214s_1 | -0.89 | -2.7 | -1.14 | -1.48 | -2.4 | -2.58 | -1.11 | -1.55 | -3 | -1.74 | -2.16 | -1.83 | -1.5 | -2.3 | -1.8 | -1.29 | 0.1525 | 0.2975 | 0.475 | 0.708984 | 0.479346 | 0.205474 | 1 | 0.999631 | 0.898972 |
| Iqsec2_E9QAD8_S246s_1 | -0.2 | 0.22 | -0.95 | -0.8 | -0.34 | -0.19 | -0.28 | -0.31 | -0.35 | -0.48 | -0.17 | 0.46 | -0.04 | -0.21 | -0.05 | 0.47 | -0.08 | 0.01 | 0.1825 | 0.205942 | 0.636299 | 0.428986 | 1 | 0.999631 | 0.6605 |
| Iqsec2_E9QAD8_S262s_1 | -0.93 | -0.73 | -1.08 | -0.44 | -0.85 | -0.85 | 0.05 | -1.85 | -0.89 | -0.95 | -0.91 | -0.39 | -0.22 | -0.36 | -2.07 | 0.2 | 0.305 | 0.0975 | 0.0975 | 0.714617 | 0.845856 | 0.971888 | 1 | 0.999631 | 0.901866 |
| Iqsec2_E9QAD8_S393s_1 | -1.85 | -0.58 | -0.87 | -1.19 | -0.38 | -0.9 | -0.71 | -1.28 | -0.77 | -2.05 | -0.49 | -0.79 | -0.6 | -1.08 | -1.39 | -1.03 | -0.1125 | -0.2625 | 0.0725 | 0.789189 | 0.401642 | 0.824058 | 1 | 0.999631 | 0.925513 |
| Iqsec2_E9QAD8_S412s_1 | -0.45 | -0.97 | -0.19 | 0.07 | -0.39 | -1.06 | -0.22 | -0.32 | -0.92 | -1.06 | -0.68 | 0.07 | -0.53 | -0.67 | -0.04 | -0.01 | -0.0275 | 0.25 | 0.3525 | 0.832278 | 0.735702 | 0.478832 | 1 | 0.999631 | 0.94321 |
| Iqsec2_E9QAD8_S528s_1 | -0.81 | -0.26 | 0.21 | -0.28 | -0.42 | -0.53 | -0.22 | -0.08 | -0.27 | -0.72 | 0.43 | 0.42 | 0.43 | -0.61 | 0.07 | 0.38 | -0.0225 | -1.17 | -2.8925 | 0.344934 | 0.926183 | 0.509765 | 1 | 0.999631 | 0.734195 |
| Iqsec2_E9QAD8_S567s_1 | 1.9 | -2.82 | -1.5 | 0.01 | -1.04 | -0.66 | 2.09 | -2.89 | -2.13 | -1.3 | -1.52 | -2.14 | -5.41 | -3.7 | -2.34 | -2.53 | 0.75 | -0.2175 | -0.475 | 0.019375 | 0.985408 | 0.214511 | 1 | 0.999631 | 0.391668 |
| Iqsec2_E9QAD8_S627s_1 | 1.09 | -2.93 | -4.35 | -1.63 | -0.76 | -0.71 | -0.87 | -2.48 | -1.52 | -0.49 | -3.16 | -3.52 | -1.96 | -2.18 | -2.02 | -3.56 | -0.7 | -0.72 | -0.3475 | 0.648993 | 0.484085 | 0.851722 | 1 | 0.999631 | 0.874615 |
| Iqsec2_E9QAD8_S741s_1 | -0.53 | -0.9 | 0.86 | -0.59 | -0.86 | -1.57 | -0.44 | -1.09 | -1.99 | -1.2 | 0.11 | -0.96 | -0.12 | -1.47 | -0.7 | -0.26 | -0.1075 | -0.065 | 0.205 | 0.476163 | 0.138847 | 0.205484 | 1 | 0.999631 | 0.799882 |
| Iqsec2_E9QAD8_S744s_1 | -0.84 | -0.48 | 0.01 | -0.44 | -0.15 | -0.91 | -0.31 | -0.81 | 0.44 | -1.77 | -0.73 | 0.05 | 1.4 | -1.35 | -0.4 | -0.58 | 0.0475 | 0.21 | 0.025 | 0.716048 | 0.729426 | 0.895069 | 1 | 0.999631 | 0.902781 |
| Iqsec2_E9QAD8_S82s_1 | -2.3 | -6.41 | -4.11 | -4.3 | -4.29 | -4.76 | -3.25 | -4.63 | -4.87 | -3.91 | -3.43 | -4.07 | -3.24 | -4.69 | -3.84 | -5.25 | 0.685 | 0.1325 | -0.345 | 0.975852 | 0.952216 | 0.789281 | 1 | 0.999631 | 0.991751 |
| Iqsec2_E9QAD8_Y89y_1 | 0.11 | -1.27 | -1.96 | -0.81 | -0.12 | 0.09 | -0.88 | -0.28 | -0.12 | -0.11 | -1.92 | -1.25 | -1.67 | -0.79 | -0.8 | -2.05 | 0.2025 | 0.195 | 0.5925 | 0.504594 | 0.160938 | 0.816928 | 1 | 0.999631 | 0.814292 |
| Iqsec3_Q3TES0_S1068s_1 | -0.75 | -0.01 | -0.8 | -1.16 | -0.52 | -0.73 | -0.97 | 0.31 | -0.49 | -0.65 | -0.44 | -0.36 | 0.34 | -0.55 | 0.03 | -0.17 | -0.675 | -0.8225 | -1.2875 | 0.120873 | 0.597843 | 0.538138 | 1 | 0.999631 | 0.588397 |
| Iqsec3_Q3TES0_S1152s_1 | 1.71 | -0.28 | -0.01 | 0.52 | -0.77 | -0.11 | 1.03 | -0.91 | -0.52 | 0.66 | -0.71 | -0.78 | -1.2 | -0.47 | -0.28 | -1.26 | 0.8775 | 0.5675 | 0.9625 | 0.023517 | 0.253438 | 0.138545 | 1 | 0.999631 | 0.415644 |
| Iqsec3_Q3TES0_S132s_1 | 0.15 | -0.44 | -1.24 | -0.82 | 0.75 | 0.62 | -0.41 | 0.2 | 0.14 | -0.26 | -0.35 | 0.39 | 1.02 | 0.64 | 0.45 | -0.61 | -0.0925 | 0.1775 | 0.735 | 0.056353 | 0.049674 | 0.148622 | 1 | 0.999631 | 0.496118 |
| Iqsec3_Q3TES0_S256s_1 | -0.59 | -0.29 | 0.32 | -0.43 | -0.55 | -0.63 | -0.35 | 0.17 | -0.75 | -0.6 | 0.78 | 0.29 | 0.62 | -0.12 | 1.19 | 0.26 | 0.395 | 0.05 | 0.2175 | 0.076276 | 0.771323 | 0.673248 | 1 | 0.999631 | 0.542251 |
| Iqsec3_Q3TES0_S646s_1 | 0.37 | -0.01 | -0.78 | 0.08 | 0.71 | 0.27 | -0.09 | 0.35 | 0.6 | -0.18 | -0.4 | -0.16 | 0.49 | 0.49 | 0.02 | -0.47 | 0.0975 | -0.48 | -0.065 | 0.565532 | 0.254413 | 0.889866 | 1 | 0.999631 | 0.84139 |
| Iqsec3_Q3TES0_T357t_1 | 0.83 | -0.25 | -0.35 | 0.11 | 1.12 | -0.14 | 0.06 | -0.31 | 0.07 | -0.17 | -0.23 | -1.25 | 1 | 0.17 | -0.47 | -0.62 | -0.2375 | 0.0725 | 0.355 | 0.885873 | 0.814683 | 0.254396 | 1 | 0.999631 | 0.963505 |
| Iqsec3_Q3TES0_T653t_1 | -1.19 | -0.09 | -0.4 | -0.54 | -1.28 | -0.73 | -0.74 | -0.42 | -0.97 | -0.19 | -0.55 | -0.22 | 0.17 | -0.52 | 0.05 | -0.5 | 0.3575 | -0.5175 | -2.01 | 0.324081 | 0.480807 | 0.831212 | 1 | 0.999631 | 0.724507 |
| Ireb2_Q811J3_S178s_1 | 2.73 | -0.31 | -1.59 | 1.89 | 0.94 | 0.34 | 1.85 | 1.02 | 0.48 | 0.09 | 0.39 | -0.31 | -3.05 | -0.2 | -0.94 | -1.13 | -0.3875 | -0.1025 | -0.51 | 0.066583 | 0.692053 | 0.558917 | 1 | 0.999631 | 0.518767 |
| Irf2bp1_Q8R3Y8_S365s_1 | 2.13 | 1.17 | 1.87 | 1.07 | 1.11 | 1.29 | 1.64 | 0.65 | 1.6 | 2.88 | 0.75 | 0.6 | 1.29 | 1.23 | 1.34 | 0.34 | -0.4125 | 0.0475 | -0.4475 | 0.205699 | 0.293042 | 0.850235 | 1 | 0.999631 | 0.6605 |
| Irf2bp1_Q8R3Y8_S434s_1 | 0.99 | 1.11 | 1.53 | 1.02 | 0.06 | 0.41 | 1.1 | 1.43 | 0.47 | 1.36 | 1.57 | 1.44 | 0.85 | 1.18 | 0.34 | 0.49 | 0.0675 | -0.0625 | -0.03 | 0.17674 | 0.269147 | 0.885976 | 1 | 0.999631 | 0.642116 |
| Irf2bp2_E9Q1P8_S226s_1 | 1.18 | 1.85 | 0.44 | 1.3 | 1.17 | 1.84 | 0.91 | 1.12 | 1.78 | 1.09 | 0.77 | 0.88 | 0.58 | 2.17 | 0.24 | 1.66 | -0.325 | 0.17 | 0.0875 | 0.953123 | 0.855684 | 0.871641 | 1 | 0.999631 | 0.984317 |
| Irf2bp2_E9Q1P8_S343s_1 | 0.54 | 1.48 | 2.18 | 1.4 | 0.88 | 0.9 | 0.84 | 1.68 | 1.11 | 1.08 | 1.93 | 2.16 | 1.01 | 1.48 | 1.58 | 1.88 | -0.19 | 0.2825 | 0.05 | 0.828444 | 0.421813 | 0.69624 | 1 | 0.999631 | 0.941315 |
| Irf2bp2_E9Q1P8_S440s_1 | -0.41 | 2.1 | 1.74 | 0.31 | 0.75 | 0.53 | 0.8 | 0.9 | 1.46 | 0.39 | 0.99 | 2.03 | 0.16 | 1.19 | 1.6 | 0.99 | 0.0325 | -0.39 | -1.21 | 0.934083 | 0.729595 | 0.651281 | 1 | 0.999631 | 0.978764 |
| Irf2bp2_E9Q1P8_S443s_1 | 0.34 | 0.78 | 1.87 | 0.85 | 1.02 | 0.31 | 1.49 | 1.15 | 0.73 | -0.3 | 0.92 | 0.93 | -1.29 | -0.07 | 0.96 | -0.6 | -0.0225 | 0.2725 | 0.2675 | 0.045145 | 0.936538 | 0.379697 | 1 | 0.999631 | 0.476115 |
| Irf2bpl_Q8K3X4_S195s_1 | 1.03 | 2.02 | 0.33 | 0.87 | 0.7 | 1.9 | 0.81 | 0.75 | 1.67 | 2.22 | 0.58 | 0.87 | 1.2 | 2.11 | 0.86 | 1.15 | -0.185 | -0.0575 | -1.3925 | 0.551741 | 0.959378 | 0.581497 | 1 | 0.999631 | 0.836275 |
| Irf2bpl_Q8K3X4_S313s_1 | 2.37 | 3.04 | 2.16 | 2.36 | 2.5 | 2.6 | 2.23 | 1.86 | 3.3 | 3.23 | 2.06 | 1.11 | 0.55 | 0.89 | 1.75 | 1.17 | -0.7275 | -0.0125 | 0.09 | 0.00323 | 0.553385 | 0.91226 | 1 | 0.999631 | 0.351308 |
| Irf2bpl_Q8K3X4_S526s_1 | 0.87 | 1.51 | 2.17 | 1.3 | 0.93 | 0.83 | 0.28 | 0.9 | 1.85 | 1.41 | 1.04 | 1.5 | 1.77 | 2.05 | 0.9 | 1.49 | -0.5325 | -0.4925 | 0.0225 | 0.819274 | 0.057168 | 0.971841 | 1 | 0.999631 | 0.938303 |
| Irf2bpl_Q8K3X4_S636s_2 | 0.44 | 0.79 | 1.63 | 0.6 | 0.27 | -0.49 | 0.67 | 0.88 | -0.05 | -0.27 | 0.74 | 1.07 | 0.43 | 0.89 | 1.31 | 0.92 | 0.1225 | 0.745 | 0.095 | 0.950936 | 0.207999 | 0.256218 | 1 | 0.999631 | 0.984079 |
| Irf2bpl_Q8K3X4_S638s_1 | 0.22 | 1.39 | 0.83 | -0.35 | 0.7 | 0.7 | 0.73 | 0.45 | 1.4 | 1.07 | 0.79 | 1.81 | 1.45 | 0.37 | 0.14 | 0.51 | -0.45 | -0.4875 | 0.115 | 0.838433 | 0.75423 | 0.110802 | 1 | 0.999631 | 0.946365 |
| Irf2bpl_Q8K3X4_S638s_2 | 0.44 | 0.79 | 1.28 | 0.27 | 0.27 | -0.49 | 0.46 | 0.74 | -0.05 | -0.27 | 0.19 | 0.96 | 0.43 | 0.89 | 0.99 | 0.93 | -0.335 | -0.1225 | 0.165 | 0.728643 | 0.237074 | 0.212459 | 1 | 0.999631 | 0.905923 |
| Irf2bpl_Q8K3X4_S641s_1 | -0.33 | 2.06 | 0.45 | -0.24 | -0.09 | 1.57 | 0.62 | -1.5 | 1.27 | -1.32 | 0.57 | 0.93 | 1.21 | 0.81 | 1.46 | -0.88 | 0.77 | 0.3275 | 0.2225 | 0.808558 | 0.655652 | 0.863278 | 1 | 0.999631 | 0.93635 |
| Irs1_Q543V3_S1072s_1 | 0.51 | 0.1 | -0.38 | 1.31 | 0.84 | 0.93 | 1.78 | 1.07 | 0.32 | 0.66 | 1.49 | 0.38 | 0.3 | 1.06 | 0.23 | 0.84 | 2.115 | 1.825 | -0.0775 | 0.601673 | 0.088281 | 0.465632 | 1 | 0.999631 | 0.856765 |
| Irs1_Q543V3_S1094s_1 | -0.12 | 2.84 | 0.64 | -0.09 | 5.24 | 2.38 | 3.47 | 0.64 | 5.45 | 0.07 | 2.14 | 2.91 | 5.25 | 0.9 | -1.63 | -1.56 | -0.4775 | -0.395 | -1.48 | 0.958101 | 0.062073 | 0.129637 | 1 | 0.999631 | 0.985497 |
| Irs1_Q543V3_S325s_1 | 2.17 | 0.1 | 1.23 | 1.13 | -0.33 | 1.2 | 1.96 | -0.11 | 0.41 | 1.81 | 0.37 | 0.46 | -0.8 | -0.7 | 1.13 | -0.92 | 0.1175 | -0.06 | 0.615 | 0.029014 | 0.44818 | 0.451386 | 1 | 0.999631 | 0.429864 |
| Irs1_Q543V3_S336s_1 | -2.18 | -0.01 | 0.49 | 0.37 | -0.31 | -1.53 | 0.52 | 0.46 | -0.68 | -0.79 | -0.45 | 0.35 | -0.22 | -1.39 | 1.31 | 1.43 | 0.6775 | 0.6425 | 0.9175 | 0.449033 | 0.865961 | 0.922033 | 1 | 0.999631 | 0.786796 |
| Irs1_Q543V3_S369s_1 | -0.25 | 1.66 | 0.4 | 1.42 | 1.69 | 2.31 | 0.44 | 1.5 | 1.99 | 1.79 | 0.77 | 1.25 | 1.14 | 2.13 | 1.96 | 1.67 | 0.4175 | -0.0575 | 0.445 | 0.080697 | 0.232048 | 0.216768 | 1 | 0.999631 | 0.545579 |
| Irs1_Q543V3_T448t_1 | 0.23 | -0.74 | -0.51 | -0.51 | 0.39 | 0.38 | -0.68 | 0.05 | 0.11 | 0.58 | -1.09 | -1.36 | 0.57 | -0.47 | 0.15 | 0 | 0.62 | 1.155 | -0.01 | 0.228247 | 0.256491 | 0.906331 | 1 | 0.999631 | 0.679038 |
| Irs2_P81122_S1165s_1 | 1.01 | -0.83 | 0.17 | 0.69 | 1.37 | 0.19 | 1.46 | 0.5 | 1.66 | 0.68 | 1.34 | 1.98 | 0.61 | -0.26 | 0.94 | -0.29 | 0.26 | 0.175 | 0.975 | 0.983706 | 0.219657 | 0.032815 | 1 | 0.999631 | 0.995133 |
| Irs2_P81122_S1190s_1 | -0.61 | -1.06 | -0.73 | -0.21 | -0.09 | -0.64 | -0.67 | -0.17 | -0.64 | -0.78 | 0.7 | -1.19 | 1.67 | -0.85 | 0.28 | 0.19 | 0.8325 | -0.09 | -0.6875 | 0.083605 | 0.391948 | 0.692232 | 1 | 0.999631 | 0.549732 |
| Irs2_P81122_S303s_1 | 0.31 | -0.6 | -1.19 | -0.11 | 1.08 | 0.14 | 0.96 | -0.44 | 0.15 | -0.93 | 0.18 | -1.35 | -0.78 | -0.8 | -1.46 | -1.3 | 0.1225 | -0.27 | -1.485 | 0.105413 | 0.096836 | 0.852094 | 1 | 0.999631 | 0.568461 |
| Irs2_P81122_S362s_1 | 2.68 | 1.03 | 1.32 | 3 | 1.83 | 2.42 | 2.83 | 1.44 | 2.05 | 2.1 | 1.94 | 0.86 | -1.24 | 0.14 | 2.02 | 1.17 | 0.325 | -0.015 | 0.2575 | 0.0717 | 0.818504 | 0.614452 | 1 | 0.999631 | 0.532005 |
| Irs2_P81122_S388s_1 | 0.05 | -0.16 | 0.09 | 1.05 | 0.63 | 0.87 | 0.29 | 0.54 | 0.79 | 0.76 | 0.13 | -0.71 | -0.09 | 1.44 | 0.54 | 0.17 | 0.06 | -0.0625 | 0.3275 | 0.5577 | 0.342719 | 0.972726 | 1 | 0.999631 | 0.838305 |
| Irs2_P81122_S573s_1 | 0.63 | 1.06 | 1.16 | 0.76 | 0.92 | 1.11 | 0.46 | 1.36 | 0.75 | 0.76 | 0.92 | 0.93 | 1.14 | 1.19 | 1.07 | 1.52 | 0.1 | -0.145 | -0.5225 | 0.268065 | 0.838748 | 0.810147 | 1 | 0.999631 | 0.696016 |
| Irs2_P81122_S604s_1 | 1.7 | 0.91 | 0.3 | 0.69 | 0.99 | 1.6 | 1.21 | 0.2 | 1.26 | 1.33 | 0.46 | -0.03 | 0.84 | 0.62 | 0.33 | -0.28 | -0.07 | -0.83 | 0.3075 | 0.215284 | 0.809747 | 0.740743 | 1 | 0.999631 | 0.668192 |
| Irs2_P81122_S66s_1 | 0.16 | 0.14 | 0.05 | -0.04 | -0.39 | 0.31 | -0.67 | 0.78 | 0.46 | 0.29 | -0.97 | -2.79 | -0.33 | -0.11 | 0.93 | 1.05 | 0.2925 | 0.1725 | 0.1825 | 0.435084 | 0.844961 | 0.236066 | 1 | 0.999631 | 0.779781 |
| Irs2_P81122_S675s_1 | -0.58 | -0.3 | -0.27 | -0.97 | -0.48 | -0.12 | -0.07 | -0.28 | -0.47 | -0.5 | 0.19 | -0.65 | -0.63 | -0.84 | 0.44 | -0.36 | 0.0225 | -0.14 | -0.2375 | 0.622153 | 0.300816 | 0.584208 | 1 | 0.999631 | 0.862299 |
| Irs2_P81122_S723s_1 | 0.63 | 0.34 | -0.28 | 0.52 | 0.24 | -0.46 | 1.03 | 0.49 | -0.18 | 0.61 | 0.33 | -0.11 | -0.03 | -0.3 | -0.25 | 0.84 | 0.1225 | -0.175 | 0.415 | 0.529702 | 0.953041 | 0.671881 | 1 | 0.999631 | 0.826087 |
| Irs2_P81122_S723s_2 | -0.6 | 1.11 | -0.48 | -0.24 | 0.61 | 0.38 | -0.29 | -0.42 | 0.22 | 0.39 | -0.53 | -0.99 | 0.54 | 0.12 | 0.4 | 0.39 | -0.445 | 0.24 | 0.545 | 0.333724 | 0.786405 | 0.720361 | 1 | 0.999631 | 0.728079 |
| Irs2_P81122_S727s_1 | -1.69 | -0.44 | 1.81 | -0.34 | -1.5 | -0.69 | -0.34 | 0.09 | -0.46 | 0.16 | 0.02 | 0.58 | 0.57 | -0.6 | 0.66 | 0.89 | 0.1225 | -0.175 | 0.415 | 0.448196 | 0.533644 | 0.723777 | 1 | 0.999631 | 0.786727 |
| Irs2_P81122_S727s_2 | -0.6 | 1.11 | -0.48 | -0.24 | 0.61 | 0.38 | -0.29 | -0.42 | 0.22 | 0.39 | -0.53 | -0.99 | 0.54 | 0.12 | 0.4 | 0.39 | -0.485 | 0.2975 | 0.2975 | 0.333724 | 0.786405 | 0.720361 | 1 | 0.999631 | 0.728079 |
| Irs2_P81122_S728s_1 | 0.01 | -0.04 | 1.17 | -0.46 | -0.93 | -0.28 | 0.08 | -0.13 | -0.09 | 0.68 | 1.09 | 0.19 | 0.94 | 0.26 | -0.11 | 0.78 | -0.0575 | 0.2625 | -0.0825 | 0.496335 | 0.255796 | 0.498543 | 1 | 0.999631 | 0.811248 |
| Irs2_P81122_S924s_1 | -0.34 | -1.05 | -0.22 | -0.3 | -0.37 | -0.67 | -0.64 | -0.46 | -0.69 | -0.53 | 0.36 | 0 | 0.16 | -1.87 | -0.9 | 0.37 | 0.26 | 0.175 | 0.975 | 0.874298 | 0.83927 | 0.458624 | 1 | 0.999631 | 0.960237 |
| Irs2_P81122_T1189t_1 | -0.61 | -1.06 | -0.73 | -0.21 | -0.09 | -0.64 | -0.67 | -0.17 | -0.64 | -0.78 | 0.7 | -1.19 | 1.67 | -0.85 | 0.28 | 0.19 | 0.4275 | 0.17 | -0.0325 | 0.083605 | 0.391948 | 0.692232 | 1 | 0.999631 | 0.549732 |
| Irs2_P81122_T347t_1 | -0.09 | -0.15 | 0.81 | 1.3 | 0.55 | 1.27 | 0.43 | 1.33 | 0.16 | 0.61 | 0.8 | 0.98 | -1.88 | 1.28 | 1.82 | 0.52 | 0.2725 | 0.2 | -0.195 | 0.966652 | 0.322764 | 0.676128 | 1 | 0.999631 | 0.988351 |
| Irs2_P81122_T517t_1 | -0.41 | 0.73 | 0.7 | -0.38 | 0.42 | 0.52 | 0.29 | 0.5 | 0.19 | 0.24 | 0.59 | 0.42 | -1.5 | 0.66 | 0.28 | 0.42 | -0.06 | -0.215 | 0.0525 | 0.723014 | 0.446828 | 0.585736 | 1 | 0.999631 | 0.905674 |
| Irs2_P81122_T524t_1 | 0.22 | 0.09 | 1.35 | 1 | 0.47 | 0.95 | 0.87 | 0.13 | 0.73 | 0.71 | 0.98 | -0.62 | 1.2 | 0.35 | 0.06 | 1.26 | -0.1925 | 0.0525 | 0.365 | 0.903827 | 0.87277 | 0.642079 | 1 | 0.999631 | 0.968183 |
| Iscu_D3Z7W0_S15s_1 | 0.13 | 0.71 | 0.59 | 0.23 | 0.14 | 0.49 | -0.17 | 0.43 | 0.2 | 0.6 | 0.59 | 0.48 | 0.8 | 0.74 | 0.68 | 0.9 | -0.0075 | 0.5575 | 0.5925 | 0.213916 | 0.502471 | 0.847715 | 1 | 0.999631 | 0.667209 |
| Iscu_D3Z7W0_S30s_1 | 0.39 | 0.87 | 1.11 | 0.67 | 0.44 | 0.87 | 0.7 | 1 | 1.19 | 1.3 | 1.69 | 1.09 | 1.08 | 1.3 | 1.42 | 1.61 | -0.2075 | -0.1125 | -0.4375 | 0.06989 | 0.978465 | 0.077965 | 1 | 0.999631 | 0.52844 |
| Islr2_E9Q9N7_S694s_1 | 0.03 | -0.08 | -0.31 | -0.31 | -0.44 | -0.52 | 0.06 | -0.6 | -0.41 | -0.79 | 0.57 | -0.49 | 0.62 | -1.23 | -1.66 | -0.15 | -0.22 | 0.1925 | 0.4175 | 0.392394 | 0.444649 | 0.746758 | 1 | 0.999631 | 0.758901 |
| Islr2_E9Q9N7_S698s_1 | -0.15 | 0.43 | 0.84 | -0.28 | 0.27 | -0.13 | -0.36 | 0.18 | 0.28 | 0.53 | 0.26 | 0.54 | 1.07 | 0.44 | 0.68 | 0.32 | -0.2375 | -0.3325 | 0.0375 | 0.261508 | 0.517103 | 0.558978 | 1 | 0.999631 | 0.69271 |
| Islr2_E9Q9N7_S739s_1 | -0.02 | -0.44 | 0.04 | -0.53 | -0.29 | -0.71 | -0.74 | -0.16 | -0.9 | -0.79 | -0.97 | 0.38 | -0.12 | -0.7 | -0.44 | 0.46 | 0.1175 | 0.515 | 0.8975 | 0.914084 | 0.411284 | 0.385216 | 1 | 0.999631 | 0.972551 |
| Isyna1_Q9JHU9_S524s_1 | -0.09 | -0.63 | -0.8 | -0.13 | 0.18 | -0.4 | -0.61 | -0.35 | 0.37 | -0.08 | -0.65 | 0.77 | 0.16 | 0.59 | 0.86 | 0.33 | 0.3075 | 0.5975 | 0.9725 | 0.017172 | 0.700521 | 0.193275 | 1 | 0.999631 | 0.391024 |
| Isyna1_Q9JHU9_T523t_1 | 0.44 | -0.54 | 0.13 | 0.53 | 0.75 | 0.93 | -0.05 | 0.16 | 1.05 | 0.99 | -0.17 | 1.08 | 0.8 | 2.34 | 0.91 | 0.4 | 0.21 | 0.1825 | 0.0675 | 0.064039 | 0.402392 | 0.158799 | 1 | 0.999631 | 0.513532 |
| Itfg2_Q91WI7_S219s_1 | -0.75 | 0.86 | 1.21 | 0.57 | 0.48 | 0.86 | 0.52 | 0.87 | 1.08 | -0.03 | 0.88 | 0.69 | 0.42 | 0.8 | 0.55 | 0.39 | -1.1825 | -0.6725 | -1.245 | 0.878343 | 0.628857 | 0.701064 | 1 | 0.999631 | 0.962349 |
| Itfg2_Q91WI7_T440t_1 | 1.64 | -2.61 | -1.49 | -1.54 | -2.24 | -2.5 | -1.26 | -2.73 | -2.58 | 0.59 | -2.42 | -2.28 | -2.08 | -2.49 | -2.6 | -1.81 | -0.585 | -0.6 | -1.2375 | 0.149138 | 0.183796 | 0.51663 | 1 | 0.999631 | 0.621356 |
| Itga3_Q62470_S1013s_1 | 1.49 | -1.26 | -0.37 | -0.91 | -1.75 | -0.51 | 0.48 | -1.61 | -1.94 | 1.08 | -1.03 | -1.56 | -1.54 | -1.44 | -1.55 | -1.47 | 0.21 | 0.1625 | -0.145 | 0.05034 | 0.418865 | 0.45955 | 1 | 0.999631 | 0.488571 |
| Itga6_F6VSK8_S658s_1 | 0.87 | 1.13 | -1.81 | -1.9 | 1.02 | 1.59 | -1.35 | -2.13 | 1.38 | 0.79 | -1.54 | -1.69 | 0.52 | 1.33 | -1.93 | -2.21 | -0.105 | 0.8375 | 0.6125 | 0.888125 | 0.84134 | 0.869247 | 1 | 0.999631 | 0.963505 |
| Itgav_F6RWM8_S57s_1 | -0.35 | -0.91 | -1.29 | 1.06 | -1.39 | -0.47 | 0.22 | -0.27 | 0.11 | -0.11 | 0.7 | 1.16 | -0.09 | -0.56 | 1.06 | 0.55 | 0.1625 | 0.245 | 0.3125 | 0.301616 | 0.851901 | 0.149813 | 1 | 0.999631 | 0.714243 |
| Itm2c_Q91VK4_T39t_1 | -1.08 | -0.68 | -0.52 | -0.85 | -0.48 | -0.84 | -0.75 | -0.41 | -0.53 | -1.06 | -0.47 | -0.09 | -0.11 | -0.64 | -0.63 | -0.5 | -0.3625 | -0.3475 | -0.5925 | 0.297246 | 0.538539 | 0.429485 | 1 | 0.999631 | 0.710639 |
| Itpka_Q8R071_S185s_1 | -0.99 | 0.68 | 0.54 | 1.11 | -0.05 | -0.83 | 0.7 | 0.07 | -0.15 | -0.86 | 0.71 | 0.25 | -2.09 | -0.29 | 0.5 | 0.85 | 1.5075 | -1.705 | -3.4675 | 0.411223 | 0.488465 | 0.516205 | 1 | 0.999631 | 0.767937 |
| Itpka_Q8R071_S20s_1 | 3.41 | -1.68 | -1.4 | 2.49 | 3.94 | 0.27 | 2.85 | 1.79 | 0.61 | -2.06 | 1.42 | -3.97 | -6.46 | -0.51 | -1.66 | -2.42 | -0.26 | -0.3325 | -0.295 | 0.045559 | 0.262898 | 0.281643 | 1 | 0.999631 | 0.477252 |
| Itpkb_B2RXC2_S269s_1 | 0.71 | -0.57 | -0.12 | -0.16 | -0.56 | -0.33 | 0.26 | -0.55 | -0.87 | 0 | -0.1 | -0.5 | 0.03 | -0.27 | -0.73 | -0.35 | -0.2125 | -0.2275 | -0.335 | 0.420959 | 0.471561 | 0.372637 | 1 | 0.999631 | 0.772817 |
| Itpr1_P11881_S1573s_1 | 0.3 | 0.95 | 1.02 | 0.84 | 0.34 | 0.6 | 0.6 | 0.72 | 0.32 | 0.6 | 1 | 0.28 | 0.14 | 0.79 | 0.25 | 0.59 | -0.2425 | -0.275 | 0.32 | 0.298552 | 0.442099 | 0.46221 | 1 | 0.999631 | 0.711767 |
| Itpr1_P11881_S1700s_1 | -1.33 | -1.27 | 0.07 | 0.11 | -1 | -1.9 | -0.43 | -0.06 | -1.52 | -1.92 | -0.16 | 0.08 | -0.38 | -1.42 | 0.14 | 0.52 | 0.195 | 0.2975 | 0.18 | 0.559156 | 0.646162 | 0.639065 | 1 | 0.999631 | 0.839222 |
| Itpr3_P70227_S934s_1 | -1.09 | -0.02 | -0.91 | -0.74 | -0.43 | 0.06 | -0.77 | -0.84 | -1.07 | -0.51 | -0.21 | 0.22 | -0.62 | 0.4 | -1.24 | -0.58 | -0.4325 | 0.03 | 0.3425 | 0.672212 | 0.574365 | 0.441611 | 1 | 0.999631 | 0.885035 |
| Itsn1_E9Q0N0_S1135s_1 | -0.2 | 0.06 | 0.1 | -0.19 | -0.65 | -0.43 | -0.54 | -0.34 | -0.27 | 0.21 | 0 | -0.05 | 0.29 | 0.27 | 0.26 | 0.32 | 0.0525 | -0.03 | -0.065 | 0.211677 | 0.098625 | 0.907213 | 1 | 0.999631 | 0.665474 |
| Itsn1_E9Q0N0_S203s_1 | -0.15 | 0.43 | -0.17 | 0.39 | 0.34 | 0.01 | 0.04 | 0.32 | 0.13 | -0.49 | 0.53 | 0.21 | -0.54 | 0.12 | 0.15 | 0.51 | 0.205 | 0.16 | 0.19 | 0.848206 | 0.848276 | 0.926465 | 1 | 0.999631 | 0.949393 |
| Itsn1_E9Q0N0_S315s_1 | -1.18 | -0.64 | -0.82 | 0.11 | -0.14 | -0.83 | -0.39 | -0.35 | -0.1 | -0.78 | -0.35 | -0.66 | -0.8 | -0.48 | -0.2 | -0.29 | 1.2275 | 0.3675 | -0.2375 | 0.595657 | 0.552558 | 0.65005 | 1 | 0.999631 | 0.854508 |
| Itsn1_E9Q0N0_S324s_1 | -0.96 | 1.89 | -1.07 | 0.35 | 1.79 | 1.34 | 1.03 | 0.96 | 1.48 | -1.57 | 1.53 | 0.24 | -1.89 | 1.41 | 0.24 | -0.5 | -0.095 | 0.005 | 0.2275 | 0.779579 | 0.080473 | 0.675666 | 1 | 0.999631 | 0.921416 |
| Itsn1_E9Q0N0_S335s_1 | -0.69 | -0.46 | -0.22 | -0.19 | -0.54 | -0.66 | -0.29 | -0.45 | -0.86 | -0.3 | -0.21 | -0.17 | -0.21 | -0.62 | -0.17 | 0.35 | -0.485 | 0.21 | -0.67 | 0.47883 | 0.709592 | 0.98611 | 1 | 0.999631 | 0.801051 |
| Itsn1_E9Q0N0_S558s_1 | 1.9 | 1.74 | 1.67 | 2.04 | 1.06 | 1.84 | 1.77 | 0.74 | 2.58 | 2.25 | 1.83 | 1.53 | 0.46 | 0.46 | 2.58 | 1.17 | -0.485 | 0.4575 | -0.7675 | 0.191387 | 0.159924 | 0.505432 | 1 | 0.999631 | 0.648674 |
| Itsn1_E9Q0N0_S623s_1 | 1.12 | -1.27 | -1.7 | 0.3 | -1.06 | -1.75 | -1.04 | 0.36 | -0.1 | 1.59 | -0.79 | -0.42 | -1.04 | -0.67 | -0.83 | -2.08 | 0.3675 | -0.0175 | 0.04 | 0.259475 | 0.496065 | 0.54402 | 1 | 0.999631 | 0.692624 |
| Itsn1_E9Q0N0_S902s_1 | 0.27 | 0.32 | -0.53 | 0.26 | 0.93 | 0.58 | 0.13 | 0.15 | 0.28 | -0.25 | 0.02 | 0.2 | -0.69 | 0.94 | -0.06 | 0.29 | -0.14 | -0.25 | -0.3325 | 0.922889 | 0.273648 | 0.954374 | 1 | 0.999631 | 0.975032 |

| | | | | | | | | | | | | | | | | | | | | | | | | | |
|---|---|---|---|---|---|---|---|---|---|---|---|---|---|---|---|---|---|---|---|---|---|---|---|---|---|
| Itsn1_E9Q0N0_S976s_1 | -0.21 | 0.29 | -0.04 | 0.16 | 0.07 | 0.08 | -0.44 | -0.07 | -0.22 | -0.11 | 0.13 | -0.6 | -0.79 | -0.35 | -0.18 | 0.19 | -0.105 | -0.1325 | -0.5025 | 0.308407 | 0.598421 | 0.38576 | 1 | 0.999631 | 0.718521 |
| Itsn1_E9Q0N0_S984s_1 | 0.28 | 0.13 | -0.23 | 0.99 | 0.32 | 0.72 | -0.29 | 0 | -0.29 | 0.58 | 0.28 | 0.07 | -2.81 | 1.21 | 0.44 | 0.32 | -0.125 | 0.0275 | 0.185 | 0.536813 | 0.770568 | 0.707767 | 1 | 0.999631 | 0.830094 |
| Itsn1_E9Q0N0_T1142t_1 | -0.28 | -0.36 | -0.58 | -0.75 | -0.7 | -0.6 | -0.59 | -0.58 | -0.46 | -0.42 | -0.64 | -0.34 | -0.52 | 0.14 | -0.59 | -0.26 | 0.5125 | 0.07 | 0.46 | 0.542887 | 0.610425 | 0.914612 | 1 | 0.999631 | 0.832881 |
| Itsn2_E9QNG1_S110s_1 | -0.83 | 0.11 | -0.63 | -0.27 | 0.59 | 0.6 | -0.38 | -0.38 | 0 | -0.49 | -0.35 | -0.5 | -0.09 | 0.2 | 0 | 0.11 | -0.4275 | 0.435 | -0.8025 | 0.160545 | 0.186075 | 0.820023 | 1 | 0.999631 | 0.630397 |
| Iws1_Q8C1D8_S261s_1 | 0.93 | 0.9 | 2.75 | 2.33 | 0.81 | 1.12 | 1.64 | 1.63 | 1.65 | 2.19 | 2.7 | 2.11 | 0.67 | 0.7 | 0.82 | 1.51 | -0.7225 | -0.405 | -1.6525 | 0.129391 | 0.391586 | 0.391292 | 1 | 0.999631 | 0.600575 |
| Iws1_Q8C1D8_S295s_1 | 1.5 | 1.45 | 2.68 | 2.21 | 0.37 | 1.23 | 2 | 1.35 | 1.41 | 1.95 | 1.45 | 1.41 | -0.19 | 0.17 | 1.4 | -0.15 | -0.065 | -0.65 | -1.035 | 0.00519 | 0.122556 | 0.274662 | 1 | 0.999631 | 0.351308 |
| Iws1_Q8C1D8_S667s_1 | 2.1 | 2.29 | -0.06 | 2.07 | 1.38 | 1.33 | 1.65 | 1.78 | 1.44 | 0.97 | 2.12 | -0.73 | 1.41 | -0.44 | 0.02 | 1.27 | 0.3025 | -0.105 | 0.375 | 0.133264 | 0.901317 | 0.383087 | 1 | 0.999631 | 0.60492 |
| Jag2_Q9QYE5_S1193s_1 | 0.64 | 1.57 | 0.87 | -0.11 | 1.81 | 2.31 | 0.53 | -0.47 | 1.93 | 1.34 | -0.39 | -0.33 | 0.84 | 1.67 | 0.75 | 1.21 | -0.7575 | -0.5025 | -0.7825 | 0.380374 | 0.639364 | 0.865238 | 1 | 0.999631 | 0.752364 |
| Jag2_Q9QYE5_S1199s_1 | 0.93 | -0.3 | 1.26 | 0.45 | -1.15 | -0.09 | 1.33 | -0.78 | -0.72 | 1.11 | -0.03 | -0.03 | -0.51 | -0.26 | 0.4 | -0.42 | 0.59 | 0.36 | 0.6175 | 0.082268 | 0.2145 | 0.315161 | 1 | 0.999631 | 0.54792 |
| Jag2_Q9QYE5_T1164t_1 | -1.08 | 0.61 | 1.03 | 0.17 | 0.89 | 0.88 | 0.43 | 0.89 | 0.85 | -0.98 | 0.95 | 1.35 | 0.87 | 0.83 | 1.09 | 0.41 | 0.2175 | 0.745 | 0.4575 | 0.2074 | 0.213049 | 0.567929 | 1 | 0.999631 | 0.661377 |
| Jak2_Q62120_S523s_1 | 0 | -0.07 | -1.74 | -1.52 | -0.71 | -0.17 | -0.89 | -0.69 | 0.55 | 0.53 | -0.85 | -0.58 | -0.1 | 0.18 | -0.91 | -0.67 | -0.2525 | -1.0525 | -1.515 | 0.373511 | 0.642711 | 0.195013 | 1 | 0.999631 | 0.747035 |
| Jak2_Q62120_Y570y_1 | 0.48 | 0.2 | -0.58 | 0.44 | 0.25 | -0.65 | 0.1 | -0.17 | -1.31 | -1.08 | -0.6 | -0.68 | -2.52 | 0.1 | -2.08 | -1.02 | -0.4975 | -0.29 | 0.3625 | 0.024818 | 0.472817 | 0.011308 | 1 | 0.999631 | 0.418003 |
| Jakmip1_G3UYZ9_T285t_1 | -0.16 | -0.26 | -0.54 | -0.06 | -0.63 | -0.57 | -1.39 | -0.42 | -0.83 | 0.5 | -0.91 | -0.94 | -0.46 | 0.79 | -0.18 | 0.28 | 0.1 | 0.18 | -0.79 | 0.313775 | 0.123871 | 0.457163 | 1 | 0.999631 | 0.719939 |
| Jakmip2_D3YXK0_S259s_1 | -0.67 | -0.36 | 0.96 | 0.64 | -0.64 | -0.3 | 1.12 | 0.79 | 0 | 0.09 | 0.27 | 0.93 | -0.75 | -1.74 | -0.61 | 0.51 | -0.015 | 0.325 | -0.8875 | 0.179404 | 0.850813 | 0.682931 | 1 | 0.999631 | 0.64376 |
| Jakmip2_D3YXK0_S260s_1 | -1.31 | -1.63 | -0.55 | -0.72 | -1.33 | -1.65 | -0.09 | -1.2 | -1.08 | -1.14 | -0.5 | -0.19 | -1.73 | -2.56 | -0.87 | -2.6 | 0.26 | 0.805 | 0.3325 | 0.083475 | 0.971414 | 0.388662 | 1 | 0.999631 | 0.549732 |
| Jakmip2_D3YXK0_S278s_1 | 0.31 | -0.98 | -2.29 | -1.88 | -0.7 | -0.54 | -1.3 | -1.26 | -0.43 | 0.71 | -0.97 | -0.93 | -0.36 | -0.3 | -1.21 | -1.64 | -0.0875 | -0.14 | 0.145 | 0.584425 | 0.64196 | 0.220543 | 1 | 0.999631 | 0.849121 |
| Jakmip3_E9Q7K7_S384s_1 | -0.12 | -0.64 | 1.43 | 0.05 | -0.65 | 0.01 | -0.12 | 1.13 | -0.45 | 0.3 | 0.36 | -0.05 | 0.1 | -0.09 | 0.16 | 1.13 | -0.65 | -0.2875 | -0.19 | 0.771307 | 0.869917 | 0.763776 | 1 | 0.999631 | 0.919413 |
| Jazf1_Q80ZQ5_T98t_1 | 1.32 | 1.08 | 0.76 | 1.15 | 0.41 | 0.67 | 0.89 | -0.26 | 1.08 | 0.91 | 0.86 | 0.31 | 0.89 | 0.61 | 0.83 | 1.22 | 0.05 | 0.035 | 0.05 | 0.518141 | 0.067297 | 0.33414 | 1 | 0.999631 | 0.819449 |
| Jcad_Q5DTX6_S351s_1 | -0.05 | 0.21 | 0.11 | 0.39 | -0.18 | 0.45 | 0.25 | 0.34 | -0.29 | 0.1 | 0.64 | 0.35 | -0.07 | 0.19 | 0.03 | 0.71 | -0.0025 | 0.015 | 0.3925 | 0.868086 | 0.850215 | 0.906105 | 1 | 0.999631 | 0.95689 |
| Jcad_Q5DTX6_S405s_1 | -1.08 | -0.06 | 0.02 | -0.61 | -0.2 | -0.66 | -0.75 | -0.13 | -0.37 | -0.47 | -0.78 | -0.05 | 0.14 | -0.44 | -0.17 | 0.31 | -0.1925 | -0.345 | -0.5875 | 0.287552 | 0.994094 | 0.965004 | 1 | 0.999631 | 0.705972 |
| Jcad_Q5DTX6_S498s_1 | 0.09 | 0.54 | 0.39 | -0.87 | -0.11 | -0.15 | -0.32 | -0.04 | -0.03 | 0.03 | -0.44 | -0.79 | -0.2 | -0.88 | -0.85 | -0.27 | -0.285 | 0.0775 | -1.23 | 0.157993 | 0.586205 | 0.38424 | 1 | 0.999631 | 0.628615 |
| Jcad_Q5DTX6_S543s_1 | 1.47 | -1.17 | -0.02 | -0.13 | -0.88 | -0.55 | 0.63 | -0.19 | -0.07 | 1.3 | -0.24 | -0.53 | -1.94 | -0.91 | -0.25 | -1.67 | 0.125 | -0.0425 | -0.345 | 0.065079 | 0.622746 | 0.899724 | 1 | 0.999631 | 0.515898 |
| Jcad_Q5DTX6_S655s_1 | 0.01 | -0.68 | -0.28 | 0.96 | 0.1 | -0.13 | 0.16 | 0.38 | -0.51 | -0.24 | 0.97 | -0.38 | -0.59 | -0.35 | -0.2 | -0.23 | 0.335 | 0.225 | 0.02 | 0.386273 | 0.741878 | 0.928095 | 1 | 0.999631 | 0.756141 |
| Jcad_Q5DTX6_S888s_1 | -0.69 | -0.86 | -0.71 | -0.92 | -0.06 | -0.2 | -0.97 | -0.61 | -0.53 | -0.78 | -0.65 | -0.32 | -0.54 | 0.11 | -1.47 | -1.2 | 0.225 | -0.4475 | 0.0625 | 0.958967 | 0.261116 | 0.383747 | 1 | 0.999631 | 0.985853 |
| Jcad_Q5DTX6_S941s_1 | 1.69 | -0.61 | 0.78 | 0.78 | 0.38 | 0.35 | 0.99 | 1.82 | 0.56 | 0 | 0.81 | -0.52 | -0.04 | 1.09 | 0.75 | 1.09 | -0.2375 | 0.04 | 0.4225 | 0.903516 | 0.6783 | 0.402834 | 1 | 0.999631 | 0.968183 |
| Jcad_Q5DTX6_T1001t_1 | -0.01 | 0.08 | -0.02 | 0.57 | 0.07 | -0.24 | -0.09 | -0.07 | 0.48 | -0.61 | 0.74 | 0.17 | 0.46 | 0.92 | 0.77 | 0.16 | 0.01 | 0.525 | 0.4825 | 0.195662 | 0.370994 | 0.911113 | 1 | 0.999631 | 0.653056 |
| Jmjd1c_G3UZM1_S1970s_1 | -0.05 | -0.16 | 1.29 | 0.09 | -0.19 | 0.29 | 0.31 | 0.8 | 0.35 | 0.75 | 1.12 | 1.05 | 0.31 | -0.11 | 1.66 | 1.24 | -0.2725 | 0.2725 | 0.6125 | 0.350648 | 0.979924 | 0.203793 | 1 | 0.999631 | 0.737333 |
| Jmjd1c_G3UZM1_S617s_1 | 0.62 | 0.39 | 0.73 | 0.46 | 0.29 | 0.2 | 0 | 0.62 | 1.31 | 0.95 | 0.51 | 0.52 | 1.27 | 1.25 | 0.94 | 1.19 | 0.67 | 0.325 | 0.6 | 0.043042 | 0.305581 | 0.361129 | 1 | 0.999631 | 0.475443 |
| Jmy_Q9QXM1_S108s_1 | 0.39 | 0.9 | 0.11 | 0.8 | 1.44 | 1.48 | 0.65 | 1.31 | 1.64 | 0.59 | 0.33 | 0.94 | 0.6 | 1.87 | 1.15 | 0.98 | -0.4575 | -0.6 | -1.6 | 0.126508 | 0.057251 | 0.386122 | 1 | 0.999631 | 0.596745 |
| Jmy_Q9QXM1_S75s_1 | 2.88 | 0.81 | 1.56 | 1.32 | 0.12 | 1.59 | 2.26 | 0.77 | 0.82 | 2.33 | 0.8 | 0.22 | -1.11 | 0.38 | 1.05 | -0.15 | 0.6325 | 0.7175 | 0.3325 | 0.019294 | 0.440613 | 0.313757 | 1 | 0.999631 | 0.391668 |
| Jph1_Q9ET80_S157s_1 | -1.28 | -2.37 | -3.15 | -2.04 | -1.06 | -2.53 | -1.38 | -1.34 | -1.36 | -1.18 | -1.64 | -1.79 | -2.07 | -1.4 | -2.35 | -1.69 | 0.31 | -0.7 | 0.8225 | 0.459085 | 0.207735 | 0.109759 | 1 | 0.999631 | 0.79152 |
| Jph1_Q9ET80_S186s_1 | -2.16 | -2.72 | -0.85 | -0.18 | -2.57 | -1.35 | -1.39 | 0.64 | -2.91 | -2.42 | -1.47 | -1.91 | -1.1 | -2.41 | 0.28 | 0.61 | 0.48 | 0.025 | 0.165 | 0.311663 | 0.690279 | 0.262318 | 1 | 0.999631 | 0.71924 |
| Jph1_Q9ET80_S216s_1 | -0.54 | 0.76 | 0.36 | 0.41 | 0.6 | 1.26 | 0.44 | 0.61 | 0.56 | 0.02 | 0.68 | -0.17 | -0.53 | 0.73 | 0.1 | 1.35 | -0.2025 | 0.6625 | 0.75 | 0.730324 | 0.199431 | 0.946354 | 1 | 0.999631 | 0.906591 |
| Jph1_Q9ET80_S448s_1 | -0.03 | 1.06 | 0.75 | -0.24 | -0.21 | 0.82 | -0.15 | 0.27 | 0.53 | 1.24 | 0.92 | 1.5 | 0.93 | 0.38 | 1.45 | 1.78 | -0.475 | -0.89 | -1.725 | 0.112195 | 0.611814 | 0.114801 | 1 | 0.999631 | 0.577068 |
| Jph1_Q9ET80_S490s_1 | 2.58 | -0.29 | 0.79 | 1.32 | -0.37 | 0.69 | 1.57 | 0.61 | -0.23 | 1.44 | 0.34 | -0.71 | -1.07 | -1.01 | -0.18 | -0.24 | 0.1075 | -0.215 | -0.08 | 0.014147 | 0.465004 | 0.20799 | 1 | 0.999631 | 0.385092 |
| Jph3_Q9ET77_S166s_1 | 0.09 | -0.26 | 0.14 | 0.09 | 0.12 | -0.19 | 0.26 | 0.3 | -0.35 | -0.19 | 0.03 | -0.29 | -0.67 | 0 | 0.06 | 0.35 | 0.185 | 0.5 | 0.6225 | 0.802174 | 0.675676 | 0.411115 | 1 | 0.999631 | 0.932677 |
| Jph3_Q9ET77_S188s_1 | -0.06 | -1.18 | 0.06 | -0.28 | 0.84 | -1.14 | -0.33 | -0.09 | 0.95 | 1.38 | 0.09 | -1.88 | -0.52 | 1.42 | 0.4 | -0.27 | -0.3625 | -0.3825 | 0.065 | 0.227988 | 0.695664 | 0.475952 | 1 | 0.999631 | 0.679038 |
| Jph3_Q9ET77_S420s_1 | -0.71 | -0.25 | -0.93 | -1.13 | -1.16 | -1.23 | -1.19 | -0.89 | -1.14 | -0.91 | -1.28 | -1.22 | -0.23 | -1.24 | -0.79 | -0.5 | -0.2825 | 0.24 | -0.1275 | 0.85134 | 0.21933 | 0.208745 | 1 | 0.999631 | 0.950061 |
| Jph3_Q9ET77_S506s_1 | 0.96 | 0.88 | 1.39 | 1.11 | 0.48 | 1.29 | 0.94 | 0.5 | 1.31 | 1.47 | 1.22 | 1.3 | 0.96 | 0.69 | 1.01 | 1.17 | 0.105 | -0.6475 | -0.1875 | 0.653303 | 0.347103 | 0.360712 | 1 | 0.999631 | 0.87811 |
| Jph3_Q9ET77_S578s_1 | -0.85 | -2.63 | -1.15 | -0.58 | -1.61 | -1.52 | -1.19 | -0.47 | -2.3 | -0.6 | -3.15 | -1.75 | -1.73 | -0.96 | -2.41 | -0.86 | -0.6175 | 0.17 | 0.655 | 0.731369 | 0.831809 | 0.321266 | 1 | 0.999631 | 0.906897 |
| Jph3_Q9ET77_T162t_1 | -0.17 | 0.14 | 0.53 | 0.02 | -0.52 | -0.32 | -0.62 | -0.49 | 0.77 | 0.68 | -0.61 | 0.36 | 1.35 | 0.26 | 0.64 | 0.89 | 0.555 | -0.175 | -0.115 | 0.076759 | 0.038223 | 0.651147 | 1 | 0.999631 | 0.542673 |
| Jph4_Q80WT0_T569t_1 | 0.29 | -0.77 | -1.98 | -0.3 | -0.03 | -0.57 | -0.57 | 0.63 | -0.01 | -1.77 | -0.99 | -0.69 | -0.51 | -0.7 | -1.47 | -0.54 | -0.425 | 0.535 | 0.6425 | 0.8206 | 0.300276 | 0.754302 | 1 | 0.999631 | 0.938379 |
| Jun_P05627_S73s_1 | 0.05 | 0.49 | 1.02 | 1.45 | -0.4 | -0.03 | 0.67 | 1.07 | 0.71 | 0.5 | 1.84 | 2.1 | 1.07 | 0.12 | 1.52 | 2.87 | 0.0025 | 0.5325 | 0.7525 | 0.292889 | 0.34572 | 0.2832 | 1 | 0.999631 | 0.706829 |
| Kalrn_F6QYT9_S1586s_1 | -2.42 | -2.93 | -1.73 | -1.65 | -1.17 | -3.36 | -1.52 | -2.67 | -2.33 | -2.7 | -0.72 | -0.85 | -1.99 | -1.76 | -1.01 | -0.96 | 0.5125 | 0.585 | -0.845 | 0.094653 | 0.996327 | 0.342713 | 1 | 0.999631 | 0.558128 |
| Kalrn_F6QYT9_S1785s_1 | -0.42 | 0.52 | 0.96 | -0.68 | 0.82 | 0.35 | 1.49 | -0.23 | 1.88 | 0.74 | -0.69 | 0.79 | 0.92 | -1.73 | -0.86 | -1.33 | -0.7375 | -0.1275 | -0.145 | 0.201546 | 0.317462 | 0.338965 | 1 | 0.999631 | 0.658952 |
| Kalrn_F6QYT9_S465s_1 | -0.43 | -0.76 | -0.38 | -0.86 | -1.35 | -1.79 | -1.07 | -1.17 | -0.84 | -0.58 | -0.83 | -0.69 | -0.25 | -1.02 | -1.19 | -0.55 | -0.5475 | -0.045 | -0.6225 | 0.656809 | 0.023781 | 0.626121 | 1 | 0.999631 | 0.879056 |
| Kank1_E9Q238_S887s_1 | 0.81 | -0.23 | 0.43 | 0.3 | -0.07 | -0.32 | 0 | -0.49 | 0.15 | 0.76 | 0.14 | 0.08 | -0.5 | -0.26 | -0.13 | -0.29 | -0.2725 | -0.095 | -0.5275 | 0.072231 | 0.095919 | 0.889624 | 1 | 0.999631 | 0.532251 |
| Kank3_Q9Z1P7_S166s_1 | -0.24 | 0.43 | 0.44 | -0.43 | -0.31 | 0.28 | -0.38 | -0.48 | -0.06 | -0.12 | 0.17 | -0.17 | 0.22 | 0.24 | -2.04 | -0.33 | -0.2725 | -0.095 | -0.5275 | 0.343956 | 0.412887 | 0.75781 | 1 | 0.999631 | 0.733852 |
| Kank3_Q9Z1P7_S167s_1 | -0.24 | 0.43 | 0.44 | -0.43 | -0.31 | 0.28 | -0.38 | -0.48 | -0.06 | -0.12 | 0.17 | -0.17 | 0.22 | 0.24 | -2.04 | -0.33 | -0.19 | 0.07 | -0.235 | 0.343956 | 0.412887 | 0.75781 | 1 | 0.999631 | 0.733852 |
| Kank3_Q9Z1P7_S290s_1 | -0.43 | -0.09 | -0.1 | -1.01 | -0.12 | -0.88 | -0.79 | -0.6 | 0.03 | -0.31 | -0.8 | -0.27 | 0.06 | -0.77 | -0.68 | -1.18 | -0.975 | -0.47 | -1.27 | 0.53568 | 0.55805 | 0.831547 | 1 | 0.999631 | 0.829362 |
| Kank3_Q9Z1P7_T292t_1 | 2.24 | 0.13 | 2.41 | 1.91 | -0.55 | 1.48 | 2.3 | -0.44 | 0.43 | 2.69 | 1.28 | 0.41 | -0.35 | 0.17 | 0.97 | 0.82 | 0.1125 | -0.075 | 0.5525 | 0.044307 | 0.226156 | 0.489449 | 1 | 0.999631 | 0.475443 |
| Kank4_Q6P9J5_S74s_1 | -0.66 | 0.85 | 0.22 | 0.65 | 0.7 | 0.11 | 0.09 | 0.61 | 0.03 | -0.37 | 0.11 | 0.99 | 1.06 | 0.76 | 0.57 | 0.88 | -0.28 | -0.035 | -0.86 | 0.172306 | 0.769726 | 0.863813 | 1 | 0.999631 | 0.642116 |
| Kat7_Q5SVQ0_S124s_1 | 0.93 | 1.14 | 1.8 | 0.34 | 0.5 | 1.01 | 1.35 | 0.23 | 1.39 | 1.18 | 0.39 | 1.11 | 1.26 | 0.27 | 0.27 | -1.03 | 0.0325 | 0.42 | 0.135 | 0.125885 | 0.487107 | 0.928286 | 1 | 0.999631 | 0.596745 |
| Kat7_Q5SVQ0_T88t_1 | -0.18 | 1.39 | 1.1 | 0.04 | 0.79 | 1.15 | 0.09 | 0.45 | 1.46 | 0.47 | 0.88 | 1.22 | 0.46 | 1.11 | 0.4 | 0.92 | -0.42 | -0.1475 | 0.23 | 0.754888 | 0.940742 | 0.349969 | 1 | 0.999631 | 0.913582 |
| Kat8_Q9D1P2_S42s_1 | -0.14 | -0.12 | -0.16 | -0.06 | -0.22 | -0.72 | -0.85 | -0.37 | -0.33 | 0.14 | -0.58 | -0.3 | 0.74 | 0.08 | 0.16 | -0.54 | -0.425 | -0.5225 | -1.4125 | 0.495693 | 0.127321 | 0.582951 | 1 | 0.999631 | 0.810961 |
| Katnb1_Q8BG40_S360s_1 | 0.99 | -1 | 0.58 | 0.25 | -1.04 | -0.3 | 1.1 | -0.64 | -0.84 | 0.77 | -0.41 | -0.79 | -0.4 | -1.81 | -0.7 | -1.92 | 0.545 | -0.6075 | -0.825 | 0.024091 | 0.466996 | 0.338687 | 1 | 0.999631 | 0.415644 |
| Kazn_B7ZCA7_S210s_1 | 1.08 | -0.07 | -0.05 | 0.56 | 1.52 | -0.01 | 1.05 | 1.14 | 0.51 | -1.36 | 0.91 | -0.97 | -0.52 | 0.35 | -1.74 | 0.13 | 0.6225 | 0.0025 | -0.1 | 0.133526 | 0.217248 | 0.300113 | 1 | 0.999631 | 0.60492 |
| Kazn_B7ZCA7_S231s_1 | 1.8 | 0.87 | 0.22 | 1.56 | 1.84 | 1.54 | 1.55 | 2.01 | 1.47 | 1.11 | 0.95 | 0.93 | -0.06 | 1.62 | 1.21 | 1.28 | -0.3375 | 0.3725 | -0.1775 | 0.839433 | 0.130865 | 0.994927 | 1 | 0.999631 | 0.946365 |
| Kazn_Q69ZS8_S8s_1 | -0.04 | -0.41 | 0.62 | 0.44 | -0.68 | -0.34 | 0.16 | 0.12 | 0.31 | 0.39 | 0.07 | 1.33 | -0.45 | -0.72 | 1.12 | -0.05 | -0.05 | -0.23 | 0.0325 | 0.701655 | 0.335895 | 0.342859 | 1 | 0.999631 | 0.896835 |
| Kbtbd11_Q8BNW9_S107s_1 | -0.14 | -1.12 | 1.56 | 0.65 | -0.06 | -0.52 | -0.07 | 1.4 | -0.92 | 0 | 0.72 | 0.23 | -0.58 | -0.36 | 0.75 | 1.27 | -0.4725 | 0.1275 | -1.0925 | 0.959975 | 0.937066 | 0.705057 | 1 | 0.999631 | 0.986196 |
| Kbtbd11_Q8BNW9_S316s_1 | 1.05 | 0.27 | -0.13 | -0.58 | -0.47 | 0.1 | 0.3 | -1.21 | 0.24 | 0.82 | 0.25 | -0.19 | -0.33 | -0.49 | -0.92 | -2.02 | -0.4825 | -0.9125 | -1.785 | 0.048428 | 0.321289 | 0.756664 | 1 | 0.999631 | 0.48343 |
| Kbtbd11_Q8BNW9_S320s_1 | 0.45 | -0.52 | 1.24 | 0.27 | -0.58 | -0.18 | 0.73 | -0.46 | -0.72 | 0.6 | -1.14 | -0.95 | -0.77 | -2.19 | -0.72 | -2.02 | -1.255 | 0.055 | -0.275 | 0.005425 | 0.299418 | 0.096048 | 1 | 0.999631 | 0.351308 |
| Kbtbd11_Q8BNW9_S62s_1 | 0.02 | -2.54 | -0.27 | -0.76 | -1.75 | -1.2 | -0.84 | -4.78 | -1.22 | -1.03 | -0.52 | -0.56 | -0.87 | -2.73 | -1.09 | 0.04 | -0.4675 | -0.075 | -0.175 | 0.702913 | 0.194543 | 0.920949 | 1 | 0.999631 | 0.896857 |
| Kbtbd11_Q8BNW9_S70s_1 | -0.26 | 0.09 | 0.43 | 0.03 | -0.48 | -0.66 | -0.24 | -0.2 | -0.33 | 0.08 | -0.17 | 0.41 | 0.21 | -0.22 | -0.29 | -0.11 | -0.4125 | 0.29 | 0.49 | 0.555048 | 0.108306 | 0.799744 | 1 | 0.999631 | 0.838109 |
| Kbtbd11_Q8BNW9_S70s_2 | -0.97 | 0.13 | 0.48 | -0.87 | -0.51 | -0.73 | -0.97 | -0.67 | 0.1 | 0.1 | -0.03 | -0.24 | 0.59 | -0.54 | 0.2 | 0.48 | -0.2975 | 0.55 | 0.68 | 0.284636 | 0.298536 | 0.461707 | 1 | 0.999631 | 0.705486 |
| Kbtbd11_Q8BNW9_S73s_2 | -1.32 | 0.19 | 1.02 | -0.42 | -0.67 | -0.86 | -0.11 | -0.08 | -0.18 | 0.02 | 0.61 | 1.22 | 0.88 | -0.57 | 0.59 | 1.29 | -0.0925 | -0.7825 | -0.87 | 0.259771 | 0.554492 | 0.325546 | 1 | 0.999631 | 0.692624 |
| Kcna1_P16388_S13s_1 | 0.49 | -1.08 | -0.47 | -1.32 | -0.39 | -0.82 | -0.43 | -1.11 | -1.53 | -0.97 | -1.16 | -1.85 | -2.46 | -0.77 | -1.6 | -1.03 | -0.71 | -0.7025 | 0.0425 | 0.11865 | 0.829776 | 0.102205 | 1 | 0.999631 | 0.585512 |
| Kcna1_P16388_S23s_1 | -0.08 | -1.16 | -0.83 | 0.11 | -1.28 | -1.12 | -1.45 | -0.95 | -1.26 | -0.33 | -2.04 | -1.14 | -1.29 | -1.09 | -0.04 | 0.63 | -0.075 | -0.4675 | -0.375 | 0.934201 | 0.065927 | 0.144479 | 1 | 0.999631 | 0.978787 |
| Kcna1_P16388_S5s_1 | 0.01 | -0.87 | -1.07 | -1.26 | -1.05 | -1.3 | -0.7 | -0.44 | -1.19 | -0.67 | -1.88 | -1.32 | -0.49 | -1.18 | -1.68 | -1.34 | 0.1525 | -0.23 | 0.5575 | 0.359721 | 0.835741 | 0.250039 | 1 | 0.999631 | 0.741182 |
| Kcna2_P63141_S434s_1 | -1.35 | -1.04 | -0.37 | -0.37 | -0.9 | -0.96 | -0.61 | -0.05 | -0.84 | -2.02 | -0.45 | -0.74 | -0.36 | -0.38 | 0.28 | -0.44 | -0.055 | -0.2175 | 0.33 | 0.138152 | 0.665436 | 0.591783 | 1 | 0.999631 | 0.611124 |
| Kcna2_P63141_S440s_1 | -0.44 | 1.93 | -0.06 | -0.25 | -0.16 | 1.1 | -0.55 | 0.57 | 0.22 | 0.78 | -0.57 | -0.12 | 0.25 | 1.5 | 0.12 | 0.63 | -0.305 | -0.215 | 0.505 | 0.572835 | 0.926775 | 0.703715 | 1 | 0.999631 | 0.844402 |
| Kcna2_P63141_S441s_1 | 0.15 | 0.31 | 0.21 | -0.38 | -0.55 | 0.45 | -0.57 | -0.26 | -0.24 | 0.5 | -0.51 | -0.32 | 0.85 | 1.09 | -0.13 | 0.5 | -0.1975 | -0.085 | 0.0525 | 0.179791 | 0.3623 | 0.513796 | 1 | 0.999631 | 0.64376 |
| Kcna2_P63141_S449s_1 | -0.31 | -1.77 | -1.07 | -1.51 | -1.63 | -1.39 | -0.08 | -2.35 | -3.07 | -0.38 | -0.04 | -1.51 | 0.25 | -2.03 | -1 | -1.67 | -0.2275 | 0.0075 | -0.905 | 0.924286 | 0.709567 | 0.899705 | 1 | 0.999631 | 0.975391 |
| Kcna4_Q61423_S79s_1 | 0.59 | -2.06 | -1.19 | 0.24 | -1.14 | -1.89 | 0.63 | -0.93 | -0.74 | 0.34 | -1.2 | -0.79 | -1.93 | -1.45 | -0.7 | -1.96 | -0.295 | -0.3875 | -0.9175 | 0.166987 | 0.752029 | 0.99054 | 1 | 0.999631 | 0.636719 |
| Kcna4_Q61423_S82s_1 | 0.38 | -3.89 | -2.97 | -0.93 | -1.46 | -3.74 | -1.48 | -1.91 | -1.69 | -2.27 | -2.47 | -2.53 | -2.2 | -3.31 | -2.12 | -3.45 | 0.085 | -0.1975 | -0.62 | 0.315359 | 0.758036 | 0.65404 | 1 | 0.999631 | 0.72069 |
| Kcna5_Q61762_S29s_1 | 0.98 | -1.26 | -0.92 | -0.71 | -0.6 | -1.13 | -0.31 | 0.47 | -0.03 | -0.22 | -2.4 | -0.05 | -2.25 | -0.37 | -0.15 | -1.62 | -0.43 | 0.06 | -0.725 | 0.342718 | 0.877311 | 0.772212 | 1 | 0.999631 | 0.73348 |
| Kcna5_Q61762_S535s_1 | 0.86 | -1.02 | -0.58 | 0.12 | -0.79 | -0.8 | 0.05 | -0.8 | 0.26 | 1.27 | -1.54 | -0.37 | -0.78 | -1.11 | 0.39 | -2.02 | 0.805 | 0.03 | 0.2775 | 0.237902 | 0.349024 | 0.925987 | 1 | 0.999631 | 0.683301 |
| Kcna6_Q61923_S511s_1 | 0.56 | 0.33 | 0 | 1.11 | 1.47 | 0.94 | 2.25 | 0.56 | 0.85 | 0.9 | 0.13 | 0.24 | 0.64 | 0.96 | 0.48 | 1.03 | 0.025 | 0.015 | 0.0775 | 0.417237 | 0.084302 | 0.931304 | 1 | 0.999631 | 0.771432 |
| Kcna6_Q61923_S6s_1 | 0.25 | -0.02 | -0.41 | -0.27 | 0.16 | -0.02 | -0.55 | 0.06 | -0.06 | -0.12 | -0.2 | -0.01 | -0.46 | 0.11 | -0.22 | 0.43 | -0.0975 | 0.4425 | 0.92 | 0.810496 | 0.93085 | 0.955069 | 1 | 0.999631 | 0.936878 |
| Kcnab2_E0CXI4_S111s_1 | -0.18 | 1.21 | 1.1 | 0.01 | 0.16 | 1.23 | -0.05 | 0.41 | 1.06 | 0.83 | 0.68 | 1.34 | 1.49 | 1.04 | 2.09 | 1.2 | -0.0975 | 0.4425 | 0.92 | 0.05657 | 0.826094 | 0.284877 | 1 | 0.999631 | 0.496238 |
| Kcnab2_E0CXI4_S112s_1 | -0.18 | 1.21 | 1.1 | 0.01 | 0.16 | 1.23 | -0.05 | 0.41 | 1.06 | 0.83 | 0.68 | 1.34 | 1.49 | 1.04 | 2.09 | 1.2 | 0.4775 | -0.1925 | -0.28 | 0.05657 | 0.826094 | 0.284877 | 1 | 0.999631 | 0.496238 |
| Kcnab2_E0CXI4_S20s_1 | 1.13 | 0.12 | -0.19 | 1.04 | 1.45 | 0.53 | 0.86 | 1.17 | 0.5 | 0.13 | 0.75 | -0.05 | -0.66 | 0.93 | 0.4 | 0.31 | 0.15 | 0.2725 | 0.11 | 0.547703 | 0.242046 | 0.62554 | 1 | 0.999631 | 0.834277 |
| Kcnab2_E0CXI4_S9s_1 | 0.21 | 0.87 | 0.77 | 0.69 | 0.63 | 0.94 | 0.48 | 1.09 | 0.68 | 1.02 | 1.07 | 0.86 | 0.13 | 0.82 | 0.88 | 1.15 | 0.8 | -0.7825 | -1.205 | 0.741757 | 0.597189 | 0.333867 | 1 | 0.999631 | 0.909308 |
| Kcnab2_E0CXI4_Y25y_1 | 0.55 | -0.76 | -2.85 | 0.78 | 1.72 | -0.98 | 0.96 | -0.78 | -0.22 | -2.25 | -1.09 | -1.85 | -2.6 | -1.02 | -1.51 | -1.97 | -0.16 | -0.005 | 0.345 | 0.147604 | 0.393366 | 0.355629 | 1 | 0.999631 | 0.620119 |
| Kcnb1_Q03717_S15s_1 | -0.94 | -0.07 | -0.41 | -1.08 | -0.65 | -0.4 | -1.23 | -0.86 | -0.48 | -0.51 | -0.96 | -0.57 | 0.09 | -0.19 | -0.35 | -0.67 | 0.1025 | 0.3925 | 0.88 | 0.329378 | 0.632534 | 0.98751 | 1 | 0.999631 | 0.726484 |
| Kcnb1_Q03717_S444s_1 | -2.62 | -2.48 | -1.25 | -0.7 | -2.8 | -1.54 | -1.03 | -1.27 | -2.24 | -2.2 | -0.16 | -0.88 | -0.45 | -2.11 | -1.53 | 0.56 | 0.5175 | 0.375 | 0.0625 | 0.210926 | 0.854838 | 0.536197 | 1 | 0.999631 | 0.665076 |
| Kcnb1_Q03717_S484s_1 | -1.12 | -0.38 | -0.01 | -1.26 | 0.11 | -0.33 | 0.09 | -0.57 | 0.12 | -1.41 | -0.11 | 0.13 | 0.38 | -1.1 | -0.79 | -1.01 | -1.9325 | -6.655 | -2.655 | 0.890071 | 0.176629 | 0.424235 | 1 | 0.999631 | 0.9636 |
| Kcnb1_Q03717_S511s_1 | 1.51 | -2.78 | -0.7 | -1.68 | -5.17 | -0.8 | 1.33 | -6.74 | -20.8 | 0.44 | -4.82 | -5.09 | -2.41 | -4.99 | -3.53 | -3.34 | 0.9325 | -0.0725 | -0.4475 | 0.014489 | 0.288269 | 0.115981 | 1 | 0.999631 | 0.385092 |
| Kcnb1_Q03717_S541s_1 | 0.78 | -0.41 | -0.91 | 1.51 | 1.29 | 0.31 | 1.72 | 1.38 | 0.64 | -1.16 | 1.13 | 0.07 | -1.33 | 0.24 | 0.08 | 0.19 | 0.655 | -0.7025 | -1.73 | 0.46731 | 0.128164 | 0.912942 | 1 | 0.999631 | 0.795438 |
| Kcnb1_Q03717_S633s_1 | 2.65 | -1.53 | -1.28 | 1.64 | 1.19 | 0.45 | 2.21 | 0.25 | 0.24 | -0.57 | 0.24 | -1.24 | -2.95 | -0.92 | -0.66 | -0.91 | 0.6425 | -0.06 | -0.0775 | 0.109225 | 0.508425 | 0.469545 | 1 | 0.999631 | 0.572424 |
| Kcnb1_Q03717_S655s_1 | -0.53 | -0.76 | -1.62 | -0.18 | 0.32 | -0.13 | -0.38 | -0.33 | -0.78 | -1.1 | -0.55 | -0.9 | -1.31 | -0.36 | -1.28 | -0.45 | -0.14 | 0.055 | 0.1975 | 0.852231 | 0.104106 | 0.867791 | 1 | 0.999631 | 0.950222 |
| Kcnb2_A6H8H5_S481s_2 | -1.28 | -1.43 | -0.78 | -1.01 | -0.93 | -2.27 | -0.78 | -1.08 | -0.99 | -0.65 | -1.11 | -1.53 | -0.94 | -1.23 | -0.86 | -0.68 | -0.5925 | -0.3225 | -0.3075 | 0.509096 | 0.714969 | 0.856595 | 1 | 0.999631 | 0.816932 |
| Kcnb2_A6H8H5_S488s_1 | 0.6 | 0.16 | 1.35 | -0.13 | -0.77 | -0.01 | 0.91 | -0.52 | -0.98 | 0.5 | 0.57 | 0.6 | -0.36 | 0.35 | 0.17 | 0.59 | -0.14 | 0.055 | 0.1975 | 0.452022 | 0.225647 | 0.507852 | 1 | 0.999631 | 0.788148 |
| Kcnb2_A6H8H5_S488s_2 | -1.28 | -1.43 | -0.78 | -1.01 | -0.93 | -2.27 | -0.78 | -1.08 | -0.99 | -0.65 | -1.11 | -1.53 | -0.94 | -1.23 | -0.86 | -0.68 | 0.29 | 0.6525 | -0.25 | 0.509096 | 0.714969 | 0.856595 | 1 | 0.999631 | 0.816932 |
| Kcnb2_A6H8H5_S851s_1 | 0.56 | 0.72 | -0.84 | -1.01 | 0.63 | 1.55 | -0.52 | -1.07 | 0.65 | 2.58 | -0.67 | -0.52 | 0.33 | 0.07 | -1.29 | -0.68 | 0.11 | -0.2175 | -0.265 | 0.648183 | 0.662286 | 0.407328 | 1 | 0.999631 | 0.874615 |
| Kcnb2_A6H8H5_T725t_1 | 0.81 | 0 | 0.03 | 0.53 | 0.35 | 0.72 | 0.13 | 0.61 | 0.09 | 0.18 | 0.51 | -0.28 | -0.46 | 0.42 | -0.13 | 0.48 | 0.9625 | 1.31 | 1.395 | 0.460763 | 0.715197 | 0.49853 | 1 | 0.999631 | 0.791962 |
| Kcnc1_P15388_S468s_1 | -1.69 | 3.13 | 0.2 | -0.68 | 3.54 | 2.07 | 0.34 | -1.14 | 4.58 | -0.65 | 0.9 | 1.37 | 5.5 | 1.42 | 0.62 | -1 | -0.0525 | -0.335 | 0.035 | 0.348539 | 0.444735 | 0.323321 | 1 | 0.999631 | 0.736697 |
| Kcnc1_P15388_T527t_1 | -0.58 | -0.43 | -0.22 | -0.45 | -0.3 | -0.41 | -0.83 | -0.35 | -0.75 | -0.94 | -0.43 | -0.9 | -0.26 | -0.93 | 0.1 | -0.45 | 1.295 | 1.285 | 0.56 | 0.911569 | 0.836959 | 0.21959 | 1 | 0.999631 | 0.971796 |
| Kcnc2_Q14B80_S508s_1 | -0.26 | 1.99 | 2.74 | 0.56 | 2.79 | 1.56 | 2.29 | 3.57 | 2.83 | -1.52 | 3.87 | 4.99 | 3.71 | -0.07 | 2.93 | 0.7 | 0.875 | 1.7325 | 0.8175 | 0.564309 | 0.091798 | 0.35124 | 1 | 0.999631 | 0.84139 |
| Kcnc2_Q14B80_S509s_1 | -0.72 | 0.7 | 1.25 | -1.6 | 1.2 | 0.4 | 1.09 | 0.44 | 2.53 | -1.26 | 2.22 | 3.07 | 1.04 | 0.42 | 1.13 | 0.31 | 0.0725 | -0.1625 | 0.15 | 0.207184 | 0.177877 | 0.113499 | 1 | 0.999631 | 0.660895 |
| Kcnc2_Q14B80_S557s_1 | -1.07 | -0.12 | -0.49 | -0.52 | -0.67 | -0.27 | -0.87 | -0.1 | -0.98 | -0.86 | -0.81 | -0.2 | -0.18 | -0.52 | -0.51 | -0.39 | 1 | -0.1175 | -0.315 | 0.627897 | 0.81828 | 0.615046 | 1 | 0.999631 | 0.865919 |
| Kcnc2_Q14B80_S604s_1 | 0.48 | -1.03 | -2.65 | 0.06 | 1.27 | -0.75 | 0.46 | -0.12 | -0.43 | -1.03 | -0.03 | -2.12 | -1.58 | -0.62 | -0.96 | -1.24 | 0.3875 | 0.0525 | 0.67 | 0.63116 | 0.188236 | 0.873401 | 1 | 0.999631 | 0.867571 |
| Kcnc2_Q14B80_S619s_1 | 0.01 | 0.93 | -0.08 | 0.12 | 0.69 | 1.28 | 0.09 | 0.47 | 0.64 | 0.24 | 0.2 | 0.11 | 0.28 | 1.51 | 0.85 | 1.02 | 0.7 | -0.3125 | -0.3425 | 0.102132 | 0.299804 | 0.869578 | 1 | 0.999631 | 0.564448 |
| Kcnc3_E9QKI2_S732s_1 | 0.87 | 0.16 | -0.58 | -0.14 | 1.29 | 1.57 | 0.18 | 0.07 | 1.06 | -0.38 | -0.57 | -1.05 | -0.47 | 1.08 | -0.73 | -0.94 | -0.0975 | -0.4575 | -0.4 | 0.515103 | 0.156923 | 0.547678 | 1 | 0.999631 | 0.819449 |
| Kcnc3_E9QKI2_T751t_1 | 0.04 | -0.33 | -0.33 | 0.19 | 0.06 | -0.13 | -0.5 | -0.25 | -0.59 | -0.59 | -0.42 | -0.66 | -0.55 | -0.58 | -0.28 | -0.62 | -0.5275 | 0.395 | 0.41 | 0.17961 | 0.718311 | 0.106612 | 1 | 0.999631 | 0.64376 |
| Kcnd1_Q03719_S547s_1 | -0.21 | -0.75 | 0.27 | 0.23 | -0.86 | -0.97 | -0.33 | -0.41 | -0.47 | 0.2 | 0.42 | 0.97 | 0.33 | -0.98 | 0.79 | 1.04 | 0.2125 | -0.2975 | -0.7125 | 0.411682 | 0.132815 | 0.33101 | 1 | 0.999631 | 0.767937 |
| Kcnd1_Q03719_S550s_1 | 0.12 | -0.91 | -0.16 | -0.93 | 0.26 | -0.32 | 0.06 | -1.03 | -0.98 | -0.5 | -0.35 | -1.24 | 0.29 | -1.31 | -1.76 | -1.95 | -0.1475 | -0.07 | 0.3225 | 0.205152 | 0.593829 | 0.425208 | 1 | 0.999631 | 0.6605 |
| Kcnd1_Q03719_S555s_1 | -0.06 | -0.13 | -0.17 | -0.49 | -0.58 | -0.31 | -0.57 | 0.02 | -0.65 | -0.18 | -0.39 | 0.09 | 0.53 | -0.06 | -0.54 | 0.51 | -0.0625 | -0.83 | -3.275 | 0.353058 | 0.583196 | 0.803048 | 1 | 0.999631 | 0.73949 |
| Kcnd1_Q03719_S568s_1 | 3.21 | -1.37 | -0.07 | 0.44 | 0.62 | -0.23 | 2.18 | -0.61 | -0.32 | 1.66 | -0.48 | -1.97 | -5.69 | -1.47 | -1.74 | -1.99 | -0.655 | 1.0475 | 0.87 | 0.017974 | 0.949439 | 0.436728 | 1 | 0.999631 | 0.391024 |
| Kcnd2_Q9Z0V2_S546s_1 | 0 | 0.17 | -1.82 | -3.66 | -1.54 | -1.12 | -1.55 | -3.72 | -0.14 | 0.67 | -0.84 | -0.81 | 0.55 | -2.23 | 0.34 | -0.49 | -0.7825 | -0.085 | 0.1575 | 0.366948 | 0.485834 | 0.233567 | 1 | 0.999631 | 0.745796 |
| Kcnd2_Q9Z0V2_S546s_2 | -0.48 | -1.56 | 0.77 | -0.76 | -0.77 | -1.68 | -1.23 | -1.48 | -1.3 | -1.34 | 0.17 | 0.1 | -1.33 | -1.54 | 0.54 | 0.93 | -0.2925 | -0.16 | -0.03 | 0.823392 | 0.135515 | 0.884652 | 1 | 0.999631 | 0.938864 |
| Kcnd2_Q9Z0V2_S548s_1 | 0.33 | -0.75 | -0.38 | -0.48 | -0.96 | -0.21 | -0.44 | -0.84 | -0.55 | 0.63 | -0.77 | -1.23 | -0.05 | -0.5 | -0.74 | -0.11 | -0.345 | -0.205 | 0.125 | 0.930535 | 0.385522 | 0.722695 | 1 | 0.999631 | 0.977897 |
| Kcnd2_Q9Z0V2_S548s_2 | -0.79 | -1.43 | -0.71 | -1.01 | -1.17 | -1.64 | -1.28 | -1.23 | -1.3 | -1.28 | -1.12 | -1.06 | -0.78 | -1.45 | -0.89 | -0.32 | -0.42 | -0.01 | 0.3925 | 0.717409 | 0.232046 | 0.462811 | 1 | 0.999631 | 0.903387 |
| Kcnd2_Q9Z0V2_S552s_1 | -0.7 | -0.31 | 0.07 | -0.58 | -0.75 | -0.73 | -1.16 | -0.56 | -0.64 | -0.18 | -0.72 | -0.02 | 0.51 | -0.37 | -0.19 | 0.1 | -0.345 | -0.205 | 0.125 | 0.251975 | 0.16638 | 0.974222 | 1 | 0.999631 | 0.687756 |
| Kcnd2_Q9Z0V2_S552s_2 | -0.79 | -1.43 | -0.71 | -1.01 | -1.17 | -1.64 | -1.28 | -1.23 | -1.3 | -1.28 | -1.12 | -1.06 | -0.78 | -1.45 | -0.89 | -0.32 | -0.06 | -0.295 | -1.2525 | 0.717409 | 0.232046 | 0.462811 | 1 | 0.999631 | 0.903387 |
| Kcnd2_Q9Z0V2_S572s_1 | 1.93 | 0.46 | 0.26 | 1.13 | 0.76 | 0.92 | 1.52 | 0.34 | 0.94 | 1.5 | 0.44 | -0.28 | -1.23 | 0 | -0.03 | 0.03 | -0.13 | -0.1175 | -0.1125 | 0.023347 | 0.891064 | 0.562812 | 1 | 0.999631 | 0.41509 |
| Kcnd3_Q9Z0V1_S550s_1 | -0.84 | -0.68 | -1.24 | -0.91 | -0.97 | -1.15 | -1 | -1.07 | -1.04 | -1.41 | -1.09 | -0.6 | -1.4 | -1.06 | -1.02 | -0.64 | -0.4725 | -0.1025 | -0.26 | 0.708928 | 0.60328 | 0.686545 | 1 | 0.999631 | 0.898972 |
| Kcnd3_Q9Z0V1_S566s_1 | -0.77 | -1.35 | -1.12 | -1.17 | -1.7 | -1.33 | -1.35 | -1.92 | -1.17 | -1.44 | -1.5 | -0.71 | -0.97 | -2.18 | -1.04 | -1.26 | 0.1775 | -0.7075 | -0.82 | 0.47225 | 0.110078 | 0.7303 | 1 | 0.999631 | 0.79887 |
| Kcnh1_Q3UHC9_S831s_1 | 1.89 | -0.34 | -2.03 | 1.2 | 1.01 | -0.49 | 0.91 | 0 | 0.53 | -1.04 | -0.43 | -1.17 | -1.02 | -0.85 | -0.43 | -0.26 | 0.5125 | 0.69 | 0.015 | 0.305779 | 0.829207 | 0.405508 | 1 | 0.999631 | 0.716639 |
| Kcnh1_Q3UHC9_S872s_1 | -1.08 | -7.16 | -2.28 | -2.07 | -1.61 | -3.31 | -3.03 | -2.59 | -3.66 | -1.52 | -2.09 | -2.56 | -3.46 | -2.9 | -2.33 | -3.84 | 0.17 | 0.16 | 0.4875 | 0.98991 | 0.672021 | 0.577452 | 1 | 0.999631 | 0.995892 |
| Kcnh1_Q3UHC9_S877s_1 | 0.53 | 0.44 | 0.86 | 1.15 | 0.58 | 0.86 | 1.31 | 0.91 | 0.38 | 0.33 | 1.77 | 1.14 | -0.08 | 0.6 | 2 | 2.41 | -0.1725 | 0.37 | 0.8975 | 0.394505 | 0.563309 | 0.686065 | 1 | 0.999631 | 0.759319 |
| Kcnh1_Q3UHC9_S951s_1 | 0.17 | 1.19 | 0.41 | -0.74 | 0.32 | 1.38 | -0.76 | -0.6 | 1.2 | 1.7 | -0.19 | -0.2 | 2.85 | 1.2 | 0.33 | 0.24 | -0.09 | 0.1175 | 0.7825 | 0.188218 | 0.764965 | 0.525202 | 1 | 0.999631 | 0.647434 |
| Kcnh1_Q3UHC9_S951s_2 | -0.39 | 0.47 | 1.62 | 1.6 | 0.03 | 0.81 | 0.91 | 1.19 | 1.19 | 1.41 | 0.93 | 0.24 | 2.68 | 0.26 | 1.22 | 2.27 | 0.0225 | -0.5175 | -0.14 | 0.248978 | 0.859743 | 0.819934 | 1 | 0.999631 | 0.686327 |
| Kcnh2_Q53Z09_S1140s_1 | -0.82 | -0.95 | -1.12 | -1.56 | -0.52 | -0.92 | -0.75 | -2.17 | -0.14 | -1.81 | -3.49 | -1.08 | 0.3 | -0.02 | -2.5 | -2.79 | -0.0225 | -0.2825 | -0.06 | 0.847369 | 0.955663 | 0.435562 | 1 | 0.999631 | 0.948838 |
| Kcnh2_Q53Z09_S241s_1 | 0.19 | 0.27 | -0.26 | -0.47 | 0.13 | -0.01 | -0.09 | -0.39 | 0.08 | -0.63 | -1.23 | 0.38 | -0.34 | 0.07 | 0.19 | -0.43 | 0.0525 | 0.3225 | 0.8175 | 0.850643 | 0.937073 | 0.496895 | 1 | 0.999631 | 0.949698 |
| Kcnh2_Q53Z09_S286s_1 | -1.24 | -1.05 | 0.84 | -0.14 | -0.89 | -0.82 | 0.04 | 0.29 | -0.71 | -0.13 | 0.24 | 0.3 | 0.53 | -0.57 | 0.8 | 0.92 | -0.205 | -0.3925 | -0.1075 | 0.157229 | 0.919917 | 0.525649 | 1 | 0.999631 | 0.627565 |
| Kcnh7_Q9ER47_S1135s_1 | 0.35 | -0.24 | -0.58 | 0.26 | 0 | -0.53 | -0.42 | -0.08 | -0.81 | -0.38 | -0.51 | -0.08 | -0.51 | -0.21 | 0.12 | -0.04 | -0.1275 | -0.5425 | 0.1625 | 0.745375 | 0.512355 | 0.24264 | 1 | 0.999631 | 0.910585 |
| Kcnh7_Q9ER47_S1147s_1 | -0.52 | -1.5 | -0.44 | -0.36 | -0.71 | -0.89 | -0.86 | -0.87 | -0.94 | -0.88 | -1.47 | -1.7 | -0.43 | -0.22 | -1.84 | 0.32 | -0.2725 | -0.015 | 0.115 | 0.749146 | 0.690431 | 0.159926 | 1 | 0.999631 | 0.912137 |
| Kcnh7_Q9ER47_S174s_1 | -0.41 | 0.26 | -0.12 | -0.07 | 0.33 | -0.69 | -0.67 | -0.4 | 0.13 | 0.21 | -0.37 | -0.37 | -0.33 | 0.07 | -0.1 | 0.48 | -0.2525 | -0.865 | -0.925 | 0.712823 | 0.406244 | 0.959027 | 1 | 0.999631 | 0.900825 |
| Kcnh7_Q9ER47_S209s_1 | 1.24 | -1.41 | 0.09 | 0.24 | -0.18 | 0.02 | 0.25 | -0.94 | -0.58 | 0.02 | -1.53 | -1.21 | -0.48 | -0.74 | -1.21 | -1.11 | -1.1875 | -0.9175 | -1.2975 | 0.108762 | 0.650116 | 0.165677 | 1 | 0.999631 | 0.571276 |
| Kcnh7_Q9ER47_S350s_1 | 1.24 | -0.03 | 0.61 | 0.34 | -2.2 | -0.16 | 0.32 | -0.55 | -1.24 | 0.54 | -0.49 | -0.32 | 0.04 | -1.21 | -0.71 | -1.15 | -1.08 | 0.4975 | -1.42 | 0.009645 | 0.055988 | 0.063228 | 1 | 0.999631 | 0.374514 |
| Kcnh7_Q9ER47_S905s_1 | 1.96 | -3.22 | -1.54 | -0.55 | -2.92 | -2.76 | -0.77 | -1.22 | -2.68 | 2.6 | -0.71 | -0.57 | -2.16 | -2.53 | -3.36 | -0.98 | -0.185 | 0.1975 | -0.375 | 0.183863 | 0.312786 | 0.705551 | 1 | 0.999631 | 0.644372 |
| Kcnh7_Q9ER47_S912s_1 | -0.59 | -1.45 | -0.51 | -1.95 | -1.36 | -2.05 | -0.47 | -1.36 | -1.85 | -1.38 | -0.89 | 0.41 | -1.12 | -2.41 | -1.46 | -1.01 | -0.6425 | -0.895 | -1.945 | 0.426889 | 0.686632 | 0.722067 | 1 | 0.999631 | 0.775206 |
| Kcnh7_Q9ER47_T882t_1 | 1.49 | -1.01 | 1.5 | 1.63 | -0.95 | 0.33 | 1.24 | 0.42 | -1.08 | 1.39 | -0.16 | -0.12 | -1.04 | -1.94 | -0.59 | -0.6 | 0.2125 | 0.3475 | 0.5725 | 0.011906 | 0.364878 | 0.235897 | 1 | 0.999631 | 0.385092 |
| Kcnip2_Q3YAB2_S19s_1 | -1.17 | 0.38 | 0.03 | -0.51 | 0.16 | -0.46 | -0.26 | 0.14 | 0.37 | -1.23 | 0.43 | 0.55 | 1.02 | -0.01 | 0.18 | -0.17 | -0.6125 | -0.825 | -1.2875 | 0.208236 | 0.581998 | 0.500874 | 1 | 0.999631 | 0.661649 |
| Kcnip2_Q3YAB2_S9s_1 | 0.94 | -0.78 | -0.1 | -0.11 | -1.45 | -0.32 | -0.18 | -0.55 | -1.26 | 0.38 | -1.12 | -1.35 | -1.04 | -1.67 | -1.07 | -1.42 | -0.31 | -0.4675 | -0.0975 | 0.009423 | 0.188044 | 0.130742 | 1 | 0.999631 | 0.369343 |
| Kcnip4_Q6PHZ8_S8s_1 | -0.97 | -1.05 | -0.95 | -0.9 | -1.55 | -1.49 | -1.12 | -0.95 | -1.34 | -2.37 | -0.75 | -1.28 | -0.9 | -1.16 | -1.13 | -1.07 | -0.79 | 0.42 | 0.6925 | 0.708269 | 0.246171 | 0.223954 | 1 | 0.999631 | 0.898901 |
| Kcnj10_Q9JM63_S338s_1 | -0.41 | -1.55 | -1.04 | -0.91 | -1.94 | -1.23 | -1.94 | -1.96 | -0.83 | 0.06 | -1.26 | -0.2 | -0.45 | 0.19 | -0.18 | -0.7 | 0.295 | 0.3525 | 0.495 | 0.074882 | 0.037386 | 0.302212 | 1 | 0.999631 | 0.540556 |
| Kcnj12_B1ATI1_S502s_1 | -0.19 | 2.23 | 2.05 | 0.99 | 2.1 | 1.86 | 1.18 | 1.12 | 2.4 | 0.12 | 1.96 | 2.01 | 2.59 | 1.41 | 1.85 | 1.21 | -0.2375 | -0.2575 | 0.4 | 0.405216 | 0.599538 | 0.604797 | 1 | 0.999631 | 0.765844 |
| Kcnj12_B1ATI1_T456t_1 | -0.3 | 0.1 | 0.19 | -0.13 | -0.06 | 0.08 | -0.68 | -0.43 | -0.18 | 0.26 | -0.55 | -0.7 | 0.81 | 0.23 | 0.05 | 0.37 | -0.265 | -1.405 | 0.185 | 0.204013 | 0.411057 | 0.413847 | 1 | 0.999631 | 0.659781 |
| Kcnj13_P86046_T327t_1 | 0.18 | 0.61 | 0.79 | 0.11 | -0.56 | 0.89 | 0.13 | 0.17 | -0.44 | -0.24 | -3.48 | 0.23 | 0.18 | 0.9 | 0.81 | 0.54 | -0.435 | -0.3125 | 0.325 | 0.563131 | 0.470177 | 0.089139 | 1 | 0.999631 | 0.841038 |
| Kcnj16_Q9Z307_S374s_1 | -0.54 | 0.02 | 0.33 | -0.22 | 0.02 | -0.31 | -1.23 | -0.63 | -0.41 | -0.93 | -0.12 | -0.2 | 0.2 | -0.03 | 0.19 | 0.53 | -0.2375 | -0.4375 | -0.01 | 0.309149 | 0.235074 | 0.340058 | 1 | 0.999631 | 0.718845 |
| Kcnj3_P63250_S368s_1 | -1.13 | 0.32 | -0.47 | -0.63 | -0.97 | -0.58 | -0.86 | -0.45 | -0.92 | -1.3 | -0.7 | -0.74 | -0.68 | -0.5 | -0.62 | -0.15 | 0.0125 | -0.175 | 0.4075 | 0.978286 | 0.505066 | 0.24526 | 1 | 0.999631 | 0.992746 |
| Kcnj3_P63250_S369s_1 | -1.12 | -0.27 | -0.71 | -0.97 | -0.5 | -0.68 | -1.03 | -0.81 | -0.73 | -1.36 | -0.56 | -1.12 | -0.65 | -0.33 | -0.18 | -0.28 | -0.505 | -2.0525 | -2.375 | 0.205915 | 0.965514 | 0.587306 | 1 | 0.999631 | 0.6605 |
| Kcnj3_P63250_S421s_1 | 3.78 | -0.16 | 0.82 | 1.95 | -0.71 | 1.39 | 3.56 | 0.13 | -0.86 | -0.56 | -0.39 | -0.01 | -2.58 | -0.81 | 0.06 | 0.22 | 0.7125 | -0.4175 | -0.4725 | 0.024905 | 0.6408 | 0.021667 | 1 | 0.999631 | 0.418003 |
| Kcnj3_P63250_S442s_1 | 1.17 | -0.45 | -1.72 | 1.17 | 1.16 | 0.3 | 1.07 | 0.49 | -0.28 | -1.16 | 0.55 | -0.61 | -1.15 | 0.26 | -0.66 | -0.17 | -0.0375 | -0.46 | -0.4525 | 0.492819 | 0.290681 | 0.554808 | 1 | 0.999631 | 0.80955 |
| Kcnj3_P63250_S447s_1 | 0 | -0.11 | -0.28 | -1.39 | -0.3 | -0.54 | -0.84 | -0.25 | -0.92 | -0.34 | -1.25 | -1.11 | -0.46 | 0 | -1.45 | -1.68 | 0.37 | 0.2 | -0.9225 | 0.368926 | 0.918644 | 0.259237 | 1 | 0.999631 | 0.74602 |
| Kcnj4_Q8R435_T344t_1 | -0.51 | -0.38 | -1.86 | -0.48 | -0.99 | -0.27 | 0.23 | -0.72 | -1.83 | -0.17 | -0.59 | 0.16 | -3.34 | -0.92 | -0.96 | -1.7 | -0.1575 | -0.175 | 0.9325 | 0.151012 | 0.401841 | 0.703918 | 1 | 0.999631 | 0.623678 |
| Kcnj6_P48542_S7s_1 | -0.32 | -1.1 | -0.42 | -0.68 | -0.58 | -1.81 | -0.56 | -0.2 | -0.34 | -1.41 | -1.36 | -0.11 | 0.08 | -0.28 | 1.21 | 0.2 | -0.0475 | -0.4025 | -0.3125 | 0.036222 | 0.693208 | 0.659352 | 1 | 0.999631 | 0.453963 |
| Kcnj6_Q8C4T8_S386s_1 | 0.49 | 1.32 | -0.43 | 1.47 | 1.4 | -0.04 | 0.04 | 1.26 | 1.11 | -0.11 | 0.08 | 0.16 | -0.67 | 0.57 | 0.54 | 1.16 | 0.1525 | 0.285 | 0.515 | 0.568529 | 0.92954 | 0.421161 | 1 | 0.999631 | 0.843185 |
| Kcnk1_O08581_S326s_1 | -1.02 | -0.16 | -1.07 | -1.14 | -0.52 | -0.43 | -1 | -0.83 | -0.47 | -0.29 | -0.9 | -0.59 | 0.57 | -0.4 | -1.02 | -0.48 | 0.16 | 0.1925 | -0.3525 | 0.235801 | 0.631931 | 0.385971 | 1 | 0.999631 | 0.683301 |
| Kcnk12_Q76M80_S5s_1 | 0.8 | -1.07 | -0.54 | 0.58 | 0.01 | 0.19 | 0.39 | -0.18 | 0.13 | 0.44 | 0.47 | -0.5 | -0.52 | -0.6 | -0.48 | -0.04 | -0.5675 | 0.3375 | 0.025 | 0.44976 | 0.721947 | 0.68978 | 1 | 0.999631 | 0.78723 |
| Kcnk2_Q6P6P9_S336s_1 | 0.01 | 0.89 | 1.1 | 0 | 0.35 | -0.23 | -0.37 | -0.02 | 0.68 | -0.1 | 1.13 | 1.64 | 1.11 | 0.27 | 0.5 | 0.22 | 0.2025 | 0.685 | 0.9025 | 0.948366 | 0.13419 | 0.466819 | 1 | 0.999631 | 0.983277 |
| Kcnk2_Q6P6P9_S354s_1 | 0.02 | 0.45 | 0.26 | 0.58 | 0.38 | 0.27 | 0.95 | 0.52 | 1.23 | 0.4 | 0.88 | 1.54 | 0.56 | 0.81 | 1.81 | 1.74 | -0.045 | -0.0275 | 0.41 | 0.035804 | 0.470991 | 0.059508 | 1 | 0.999631 | 0.453811 |
| Kcnk4_O88454_S324s_1 | -0.47 | 0.06 | -0.24 | -0.27 | -0.35 | 0.49 | -0.99 | -0.25 | -0.34 | 0.22 | -0.9 | -0.01 | 0.98 | 0.48 | -0.77 | 0.03 | -0.1525 | 0.125 | 0.2525 | 0.326842 | 0.897942 | 0.932212 | 1 | 0.999631 | 0.724678 |
| Kcnma1_J3QN27_S582s_1 | -0.96 | -0.72 | -0.64 | -1.18 | -0.81 | -1.22 | -1.03 | -1.05 | -0.94 | -0.91 | -0.6 | -0.55 | 0.03 | -0.81 | -1.1 | -0.61 | 0.1025 | -0.1125 | -0.05 | 0.460069 | 0.55718 | 0.644799 | 1 | 0.999631 | 0.791962 |
| Kcnma1_J3QN27_S809s_1 | 0.87 | 1.17 | 0.29 | 1.37 | 1.4 | 1.21 | 0.29 | 1.21 | 0.93 | 0.51 | 0.72 | 1.09 | -0.01 | 1.12 | 0.93 | 1.46 | 0.2125 | 0.195 | 0.4375 | 0.903148 | 0.778876 | 0.72891 | 1 | 0.999631 | 0.968183 |
| Kcnma1_J3QN27_T580t_1 | -0.17 | -0.85 | -0.98 | -1.7 | -0.26 | -0.63 | -0.68 | -1.28 | 0.08 | -0.66 | -1.03 | -1.31 | 0.71 | 0.22 | -1.34 | -1.54 | 0.0125 | -0.205 | 0.255 | 0.463184 | 0.586532 | 0.653881 | 1 | 0.999631 | 0.793564 |
| Kcnma1_Q08460_S672s_1 | -1.43 | -0.96 | -0.89 | -0.76 | -0.57 | -1.06 | -0.92 | -1.44 | -0.81 | -1.74 | -1.13 | -1.18 | -1.16 | -0.2 | -0.92 | -0.74 | -1.4225 | -0.905 | -0.815 | 0.443058 | 0.966419 | 0.513116 | 1 | 0.999631 | 0.782541 |
| Kcnma1_Q08460_S782s_1 | 0.73 | -1.31 | 0.65 | -1 | -2.53 | -0.8 | -0.02 | -3.27 | -2.53 | 1.2 | -2.59 | -0.63 | 0.42 | -1.91 | -0.68 | -2.02 | 0.145 | -0.3175 | 0.32 | 0.25978 | 0.102559 | 0.333734 | 1 | 0.999631 | 0.692624 |
| Kcnma1_Q08460_S966s_1 | -0.69 | 0.25 | -0.65 | -1.17 | -0.07 | -0.05 | -0.85 | -0.71 | -0.8 | -1.05 | -1.23 | -0.45 | 1 | -0.48 | -0.37 | -1.13 | -0.4275 | -0.475 | -0.2325 | 0.53479 | 0.701982 | 0.399046 | 1 | 0.999631 | 0.828799 |
| Kcnma1_Q08460_S977s_1 | -0.83 | -0.91 | -0.63 | -1.1 | -1.14 | -1.16 | -1.03 | -1.85 | -1.32 | -0.56 | -2.16 | -1.33 | -0.2 | -1.26 | -2.12 | -0.82 | -0.41 | -0.085 | 0.0325 | 0.587704 | 0.158361 | 0.217818 | 1 | 0.999631 | 0.850383 |
| Kcnma1_Q08460_S978s_1 | -0.54 | -0.99 | -0.9 | -1.29 | -1.15 | -1.28 | -1.34 | -1.59 | -1.29 | -0.49 | -1.32 | -0.96 | -0.13 | -1.18 | -1.54 | -0.74 | -0.1375 | -0.0425 | 0.35 | 0.931478 | 0.153687 | 0.785982 | 1 | 0.999631 | 0.978006 |
| Kcnma1_Q08460_S978s_2 | -0.31 | 0.46 | 0.62 | 0.47 | 0.12 | 0.19 | -0.18 | 0.56 | -0.17 | 0.08 | 0.64 | 0.52 | 0.57 | 0.58 | 0.5 | 0.99 | -0.0875 | -0.005 | 0.4025 | 0.289638 | 0.661453 | 0.898356 | 1 | 0.999631 | 0.706164 |
| Kcnma1_Q08460_S982s_2 | -0.31 | 0.46 | 0.69 | 0.31 | 0.12 | 0.19 | -0.2 | 0.69 | -0.17 | 0.08 | 0.57 | 0.65 | 0.57 | 0.58 | 0.57 | 1.04 | -0.5725 | -0.2825 | -0.055 | 0.231835 | 0.788721 | 0.988174 | 1 | 0.999631 | 0.682192 |
| Kcnma1_Q08460_T1088t_1 | -0.43 | -0.66 | -0.54 | -0.66 | -1.26 | -0.86 | -1.38 | -1.08 | -1.17 | -0.3 | -1.07 | -0.88 | 0.34 | -0.9 | -1.14 | -0.81 | -0.045 | 0.705 | 0.7975 | 0.883209 | 0.041887 | 0.340928 | 1 | 0.999631 | 0.963505 |
| Kcnma1_Q08460_T734t_1 | 0.68 | -0.77 | -0.72 | -0.69 | -0.76 | 0.27 | -0.48 | -0.71 | 0.64 | 0.86 | -0.44 | 0.26 | 0.97 | 0.93 | -0.19 | -0.02 | 0.0825 | -0.27 | 0.2175 | 0.107983 | 0.914733 | 0.138105 | 1 | 0.999631 | 0.569416 |
| Kcnma1_Q08460_T970t_1 | -0.69 | -0.25 | -0.98 | -0.85 | -0.28 | -0.43 | -0.9 | -0.83 | -0.89 | -1.09 | -0.94 | -0.93 | 0.68 | -1 | -0.44 | -1.14 | -0.08 | -0.235 | -0.315 | 0.626631 | 0.776747 | 0.333844 | 1 | 0.999631 | 0.864779 |
| Kcnn2_P58390_S152s_1 | 0.33 | -0.69 | -1.33 | -0.37 | -0.46 | -0.74 | -0.26 | -0.92 | -0.52 | -0.35 | -0.81 | -1.32 | -0.91 | -0.88 | -0.89 | -0.64 | -0.435 | -0.69 | -1.985 | 0.42077 | 0.835748 | 0.571805 | 1 | 0.999631 | 0.772817 |

| | | | | | | | | | | | | | | | | | | | | | | | | | |
|---|---|---|---|---|---|---|---|---|---|---|---|---|---|---|---|---|---|---|---|---|---|---|---|---|---|
| Kcnn2_P58390_S827s_2 | 1.49 | -0.7 | 0.18 | 0.03 | -0.41 | 0.05 | 0.39 | -0.77 | -0.05 | 0.52 | -0.49 | -1.74 | -1.04 | -1.87 | -2.24 | -1.79 | -0.8 | -0.8575 | -1.075 | 0.00255 | 0.386219 | 0.268913 | 1 | 0.999631 | 0.351308 |
| Kcnn2_P58390_S838s_1 | 1.14 | -0.65 | -0.83 | -1.64 | -1.45 | -0.75 | -0.46 | -2.52 | -1 | -0.95 | -2.03 | -1.43 | -0.55 | -1.41 | -1.4 | -2.92 | -0.8 | -0.8575 | -1.075 | 0.138431 | 0.245316 | 0.164691 | 1 | 0.999631 | 0.611124 |
| Kcnn2_P58390_S839s_1 | 1.14 | -0.65 | -0.83 | -1.64 | -1.45 | -0.75 | -0.46 | -2.52 | -1 | -0.95 | -2.03 | -1.43 | -0.55 | -1.41 | -1.4 | -2.92 | -0.1675 | -0.365 | 0.715 | 0.138431 | 0.245316 | 0.164691 | 1 | 0.999631 | 0.611124 |
| Kcnq2_B7ZBV3_S238s_1 | -1.16 | -0.51 | 1.22 | 0.87 | -0.58 | -0.08 | -0.21 | 0.62 | -0.89 | -0.79 | -0.67 | 1.31 | -0.12 | 0.46 | 1.48 | 1.46 | 0.255 | -0.2425 | -0.1925 | 0.266435 | 0.766548 | 0.598016 | 1 | 0.999631 | 0.695664 |
| Kcnq2_B7ZBV3_S302s_1 | 0.3 | 0.45 | -0.21 | 0.59 | 0.59 | -0.16 | 0.64 | 1.08 | 0.57 | 0.02 | -0.33 | -0.1 | 0.2 | 0.32 | -0.77 | 0.61 | -0.64 | -0.665 | -0.935 | 0.616257 | 0.464723 | 0.454179 | 1 | 0.999631 | 0.861203 |
| Kcnq2_B7ZBV3_S313s_1 | 1.56 | 0.66 | 0.45 | 0.49 | -0.39 | 0.85 | 0.57 | -0.43 | 0.15 | 1.62 | -0.35 | -0.92 | -0.15 | -0.03 | 0.12 | -0.52 | 0.0275 | 0.335 | 0.3975 | 0.02209 | 0.148162 | 0.250777 | 1 | 0.999631 | 0.405544 |
| Kcnq2_B7ZBV3_S322s_1 | -0.56 | -1.19 | -0.43 | -0.8 | -1.17 | -1 | -0.6 | -0.1 | -0.48 | -0.35 | -0.3 | -0.51 | -0.12 | -0.43 | -0.69 | -0.15 | -0.9675 | -0.1425 | 0.855 | 0.217513 | 0.933798 | 0.243408 | 1 | 0.999631 | 0.669718 |
| Kcnq2_B7ZBV3_S332s_1 | -0.93 | -0.93 | 0.78 | -1.44 | -2.07 | -0.89 | -2.07 | -1.36 | -1.02 | -0.47 | -1.66 | 0.06 | 0.93 | -1 | 0.72 | 0.25 | -0.335 | -0.21 | 0.2725 | 0.171797 | 0.089599 | 0.800088 | 1 | 0.999631 | 0.641657 |
| Kcnq2_B7ZBV3_S529s_1 | -0.34 | -0.76 | -0.15 | -0.6 | -0.99 | -0.63 | -0.78 | -0.79 | -0.95 | -0.34 | -0.91 | -0.49 | -0.07 | -0.28 | -0.21 | -0.2 | 1.27 | 1.66 | -0.005 | 0.341822 | 0.216986 | 0.474907 | 1 | 0.999631 | 0.733397 |
| Kcnq2_B7ZBV3_S585s_1 | -0.14 | 1.13 | 0.48 | 0.33 | 3.68 | 1.43 | 0.82 | 0.95 | 5.1 | 0.41 | 1.81 | 1.12 | 2.88 | -0.65 | -0.48 | 0.03 | -0.125 | -0.985 | -0.145 | 0.994737 | 0.071679 | 0.098705 | 1 | 0.999631 | 0.997705 |
| Kcnq2_B7ZBV8_S579s_1 | -1.2 | -1.31 | -0.39 | -0.86 | -1.67 | -1.5 | -0.81 | -0.28 | -2.04 | -1.78 | -1.84 | -2.04 | -1.17 | -1.75 | -0.55 | -0.87 | -0.515 | -0.98 | -1.5925 | 0.69681 | 0.74935 | 0.007589 | 1 | 0.999631 | 0.895731 |
| Kcnq3_Q8K3F6_S666s_1 | 1.38 | -1.04 | 0.26 | 1.14 | -1.18 | -0.84 | 1.95 | -0.25 | -1.43 | -0.59 | 0.08 | -0.24 | -2.16 | -2.24 | 0.21 | -0.44 | -0.1 | -0.2425 | 0.3425 | 0.050025 | 0.513927 | 0.117756 | 1 | 0.999631 | 0.488436 |
| Kcnq3_Q8K3F6_T580t_1 | -0.43 | -0.25 | -0.27 | -0.43 | -0.19 | -0.4 | -0.68 | -0.51 | -0.56 | -0.4 | -0.68 | -0.71 | 0.16 | 0.15 | -0.36 | 0.04 | 0.2425 | 0.325 | 0.8225 | 0.233153 | 0.684179 | 0.335726 | 1 | 0.999631 | 0.683035 |
| Kcnq3_Q8K3F6_T82t_1 | -0.33 | -1.95 | 0 | 0.1 | -0.2 | -0.73 | -0.47 | 0.19 | -0.06 | 0 | -0.58 | -0.24 | 0.08 | -0.04 | 0.5 | 0.57 | -0.275 | -1.3525 | -2.235 | 0.113989 | 0.620775 | 0.500173 | 1 | 0.999631 | 0.579001 |
| Kcnq5_E9Q9F1_S461s_1 | 2.92 | -0.38 | -0.19 | 1.63 | 0.16 | 0.4 | 2.37 | -0.05 | -0.59 | 0.71 | -0.58 | -0.97 | -2.22 | -1.13 | -0.95 | -0.66 | 0.5125 | 0.21 | 0.49 | 0.013191 | 0.74332 | 0.10196 | 1 | 0.999631 | 0.385092 |
| Kcnq5_E9Q9F1_S467s_2 | -1.25 | -1.38 | -1.31 | -0.35 | -0.75 | -1.71 | -0.15 | 0.37 | -1.68 | -1.16 | 0.38 | -0.99 | 0 | -1.3 | -0.64 | -0.39 | -0.42 | 0.14 | 0.225 | 0.231109 | 0.301297 | 0.664042 | 1 | 0.999631 | 0.681463 |
| Kcnq5_E9Q9F1_S477s_1 | -0.71 | -0.42 | -0.21 | -0.8 | -1.12 | -0.46 | -0.94 | -1.3 | -0.76 | -0.31 | -0.22 | -0.29 | -0.22 | -0.55 | 0 | -0.47 | 0.5125 | 0.21 | 0.49 | 0.453589 | 0.175207 | 0.619398 | 1 | 0.999631 | 0.78875 |
| Kcnq5_E9Q9F1_S477s_2 | -1.25 | -1.38 | -1.31 | -0.35 | -0.75 | -1.71 | -0.15 | 0.37 | -1.68 | -1.16 | 0.38 | -0.99 | 0 | -1.3 | -0.64 | -0.39 | 0.785 | -0.7725 | 0.4625 | 0.231109 | 0.301297 | 0.664042 | 1 | 0.999631 | 0.681463 |
| Kcnq5_E9Q9F1_S79s_1 | 0.5 | 1.44 | -1.49 | 1.63 | 1.87 | 1.39 | -0.02 | 1.98 | -0.5 | -0.25 | 0.28 | -0.54 | 1.56 | 1.31 | -0.59 | 1.65 | -0.22 | -0.2675 | -0.115 | 0.55532 | 0.307179 | 0.260748 | 1 | 0.999631 | 0.838109 |
| Kcns2_O35174_S441s_1 | -0.1 | 0.29 | 0.53 | 0.53 | 0.01 | -0.3 | 0.14 | 0.52 | -0.19 | -0.03 | 0.26 | 0.14 | -0.03 | 0.28 | -0.16 | 0.7 | 0.0225 | 0.2775 | -0.295 | 0.722404 | 0.460792 | 0.347805 | 1 | 0.999631 | 0.905674 |
| Kcnt1_Q6ZPR4_S1028s_1 | -0.81 | -2.01 | 1.36 | -0.13 | -1.5 | -1.65 | 0.61 | 1.04 | -2.23 | -2.34 | 1.16 | 2.93 | -2.19 | -1.55 | 0.72 | 0.25 | 0.2475 | 0.205 | 0.2725 | 0.731577 | 0.979126 | 0.826275 | 1 | 0.999631 | 0.906935 |
| Kcnt1_Q6ZPR4_S34s_1 | -0.26 | 0.1 | -0.53 | -0.13 | 0.53 | -0.26 | -0.09 | -0.01 | -0.26 | -0.24 | 0.29 | 0.21 | 0.01 | -0.01 | 0.09 | 0.18 | -0.265 | -0.4125 | -0.0575 | 0.339611 | 0.40018 | 0.479721 | 1 | 0.999631 | 0.732185 |
| Kcnt1_Q6ZPR4_S44s_1 | 0.57 | -0.23 | -0.79 | -0.74 | -0.31 | -0.31 | -0.42 | -1.21 | -1.38 | 0.66 | -1 | -1.12 | 0.06 | -0.53 | -0.2 | -0.75 | -0.3175 | 0.27 | 0.0575 | 0.883287 | 0.503505 | 0.439795 | 1 | 0.999631 | 0.963505 |
| Kcnt1_Q6ZPR4_S47s_1 | -0.66 | -1.32 | -0.48 | -0.37 | -1.85 | -0.85 | -1.05 | -0.35 | -0.78 | 0.25 | -0.57 | -0.65 | -0.36 | -1.56 | -0.29 | -0.39 | 0.06 | -0.1825 | -0.62 | 0.884832 | 0.420888 | 0.451431 | 1 | 0.999631 | 0.963505 |
| Kcnt2_D3Z592_S1053s_1 | 0.78 | -0.37 | -3.76 | -4.46 | 0.35 | -0.85 | -2.65 | -4.42 | -0.72 | -1.08 | -2.96 | -3.78 | -0.9 | -0.19 | -5.7 | -3.5 | 0.4275 | 0.1375 | 0.5 | 0.681706 | 0.965781 | 0.884664 | 1 | 0.999631 | 0.88924 |
| Kcnv1_Q8BZN2_S179s_1 | -0.35 | 1.33 | 0.69 | 0.24 | 1.74 | 1.4 | 0.31 | 0.17 | 1.51 | -1 | 0.97 | 0.98 | 2.3 | 0.51 | 0.66 | 0.44 | -0.7525 | -0.0375 | -0.625 | 0.359259 | 0.398334 | 0.818502 | 1 | 0.999631 | 0.741182 |
| Kctd1_E9Q6T9_S364s_1 | 1.52 | 2.3 | 1.82 | 1.55 | 1.07 | 0.94 | 1.23 | 0.94 | 1.92 | 1.9 | 1.77 | 1.45 | 0.86 | 1.48 | 0.71 | 1.64 | 0.3025 | -0.0975 | 0.4325 | 0.0975 | 0.020763 | 0.898249 | 1 | 0.999631 | 0.560034 |
| Kctd1_E9Q6T9_S616s_1 | -0.12 | 0.6 | 0.24 | -0.19 | 0.46 | 0.75 | 0.06 | 0.47 | 0.54 | -0.13 | -0.36 | 0.09 | 0.72 | 0.23 | 0.62 | 0.69 | 0.1075 | 0.24 | 0.485 | 0.183251 | 0.321281 | 0.762683 | 1 | 0.999631 | 0.644372 |
| Kctd12_Q6WVG3_S187s_1 | -0.84 | -0.55 | -0.91 | -0.94 | -0.64 | -0.7 | -0.89 | -0.58 | -0.45 | -0.7 | -0.8 | -0.33 | -0.34 | -0.07 | -0.58 | -0.31 | 0.98 | 0.6875 | 0.2925 | 0.10519 | 0.661212 | 0.371183 | 1 | 0.999631 | 0.568213 |
| Kctd12_Q6WVG3_S189s_1 | -1.68 | -1.54 | 0.65 | -2.03 | 0 | -0.35 | -0.28 | -0.05 | -0.9 | -1.32 | 1.1 | -0.73 | -0.75 | -1.03 | -1.14 | -0.51 | 0.005 | -0.1425 | 0.1825 | 0.613309 | 0.106124 | 0.351122 | 1 | 0.999631 | 0.860866 |
| Kctd12_Q6WVG3_S200s_1 | -0.55 | 0.13 | -0.16 | -0.73 | -0.23 | -0.7 | -0.49 | 0.13 | -0.47 | -0.77 | -0.23 | -0.41 | 0.2 | -0.26 | -0.41 | -0.11 | 0.09 | -0.01 | 0.245 | 0.569595 | 0.987325 | 0.636645 | 1 | 0.999631 | 0.84338 |
| Kctd12_Q6WVG3_S202s_1 | -1.55 | -2.45 | -0.28 | -1.04 | -1.78 | -1.76 | -0.81 | -0.61 | -1.63 | -1.81 | -1.57 | -0.35 | -2.24 | -1.56 | -0.03 | -0.51 | -0.0975 | -0.385 | 0.115 | 0.690785 | 0.860641 | 0.984882 | 1 | 0.999631 | 0.893731 |
| Kctd12_Q6WVG3_T198t_1 | 0.01 | -1.72 | -0.28 | -0.34 | -0.94 | -0.93 | -0.45 | -0.4 | -1.16 | -2.03 | -0.37 | -0.31 | -0.26 | -0.48 | -0.3 | -0.83 | -0.1025 | -0.08 | 0.6325 | 0.784969 | 0.813416 | 0.468859 | 1 | 0.999631 | 0.922909 |
| Kctd16_Q5DTY9_S130s_1 | -0.62 | 0.37 | 0.93 | 0.5 | -0.72 | 0.64 | 0.17 | 0.68 | -0.36 | 0.59 | 0.2 | 0.43 | 0.8 | 0.54 | 0.66 | 1.71 | -0.0825 | -0.155 | 0.1825 | 0.163967 | 0.81878 | 0.8416 | 1 | 0.999631 | 0.632208 |
| Kctd16_Q5DTY9_S137s_1 | -0.72 | -0.31 | -0.42 | -0.37 | -0.74 | -0.23 | -0.88 | -0.3 | -0.65 | -0.59 | -0.74 | -0.46 | -0.09 | -0.16 | -0.96 | 0.12 | -0.1975 | -0.145 | 0.475 | 0.581701 | 0.763102 | 0.541841 | 1 | 0.999631 | 0.847567 |
| Kctd16_Q5DTY9_S143s_1 | -0.9 | -0.38 | -0.07 | -0.88 | -0.75 | -0.09 | -1.01 | -1.17 | -0.55 | 0.03 | -1.39 | -0.9 | 1.27 | 0.43 | -1.45 | -0.58 | -0.2675 | -0.2025 | -0.55 | 0.417598 | 0.56986 | 0.705157 | 1 | 0.999631 | 0.771539 |
| Kctd16_Q5DTY9_S173s_1 | 0.77 | -0.04 | 0.46 | 0.38 | -1.33 | 0.84 | 0.98 | 0.01 | -0.27 | 0.86 | 0.12 | 0.05 | 0 | -0.33 | 0.25 | -0.55 | -0.0325 | -0.025 | 0.7225 | 0.112938 | 0.607199 | 0.553009 | 1 | 0.999631 | 0.577408 |
| Kctd16_Q5DTY9_S389s_1 | -0.64 | -0.07 | -0.45 | -0.4 | -0.2 | -0.29 | -0.7 | -0.5 | -0.44 | -0.12 | -1.27 | 0.17 | 0.41 | 0.21 | -0.12 | 0.83 | -0.0375 | 0.075 | 0.0425 | 0.042959 | 0.90179 | 0.945185 | 1 | 0.999631 | 0.475443 |
| Kctd16_Q5DTY9_T340t_1 | -0.34 | 1.62 | 1.73 | 1.18 | 0.28 | 1.74 | 0.94 | 1.08 | 0.86 | 0.72 | 1.76 | 1.15 | 0.38 | 1.25 | 0.85 | 1.88 | -0.1375 | -0.525 | -0.4725 | 0.936887 | 0.94277 | 0.881293 | 1 | 0.999631 | 0.97939 |
| Kctd17_F7B6A4_S166s_1 | 0.36 | 0.05 | 0.05 | 0.81 | 0.13 | 0.29 | -0.3 | 0.6 | -0.16 | 0.27 | -0.15 | -0.79 | -1.61 | -0.03 | 0.23 | 0.79 | -0.3875 | 0.02 | -0.8475 | 0.3705 | 0.661713 | 0.139008 | 1 | 0.999631 | 0.746338 |
| Kctd3_Q8BFX3_S712s_1 | 1.76 | -0.45 | 0.08 | 0.59 | -0.46 | 0.19 | 0.94 | -0.24 | 0.4 | 1.68 | 0.31 | -0.33 | -0.68 | -0.42 | -0.09 | -0.22 | 0.22 | 2.035 | 0.87 | 0.099815 | 0.465333 | 0.972461 | 1 | 0.999631 | 0.562904 |
| Kctd4_Q9D7X1_S20s_1 | 0.97 | 0.99 | 0.18 | -0.95 | 2.43 | 2.21 | 2.71 | -5.28 | 4.96 | 0.58 | 1.86 | 1.93 | 4 | 0.34 | 1.54 | -1.21 | 0.3825 | 0.5475 | 0.97 | 0.401068 | 0.895882 | 0.044965 | 1 | 0.999631 | 0.763221 |
| Kctd8_Q50H33_S413s_1 | 0.33 | 0.44 | 0.07 | 0.31 | 1.28 | 0.66 | 0 | 0.74 | 0.84 | 0.81 | 0.56 | 1.13 | 1.23 | 1.43 | 1.42 | 0.95 | 0.0125 | -0.0875 | 0.035 | 0.005481 | 0.251052 | 0.060027 | 1 | 0.999631 | 0.351308 |
| Kdm1a_Q6ZQ88_S850s_1 | -0.41 | 0.18 | -0.07 | -0.63 | -0.12 | -0.33 | -0.42 | -0.01 | 0.07 | -0.53 | -0.48 | -0.34 | 0.57 | -0.11 | -0.48 | -0.77 | -0.685 | 0.23 | -0.59 | 0.926191 | 0.964608 | 0.771463 | 1 | 0.999631 | 0.97588 |
| Kdm2a_F6YRW4_S422s_1 | 1.23 | 0.68 | 1.74 | 1.45 | 0.05 | 1.07 | 0.78 | 0.46 | 0.75 | 2.5 | 1.39 | 1.38 | 0.32 | 0.3 | 1.27 | 0.85 | -0.63 | -1.3875 | -2.29 | 0.132174 | 0.071279 | 0.59343 | 1 | 0.999631 | 0.603903 |
| Kdm2a_F6YRW4_S739s_1 | 2.29 | -0.42 | 1.03 | 1.08 | -0.97 | 0.53 | 1.93 | -0.03 | -0.8 | 1.35 | -1.2 | -0.92 | -1.59 | -1.67 | -1.24 | -0.68 | -0.63 | -1.3875 | -2.29 | 0.002036 | 0.393993 | 0.077508 | 1 | 0.999631 | 0.351308 |
| Kdm2a_F6YRW4_S740s_1 | 2.29 | -0.42 | 1.03 | 1.08 | -0.97 | 0.53 | 1.93 | -0.03 | -0.8 | 1.35 | -1.2 | -0.92 | -1.59 | -1.67 | -1.24 | -0.68 | -0.03 | 0.3625 | -0.2475 | 0.002036 | 0.393993 | 0.077508 | 1 | 0.999631 | 0.351308 |
| Kdm5b_Q80Y84_S1405s_1 | -0.64 | 0.08 | 0.93 | 0.41 | -0.25 | -0.17 | 0.66 | 0.42 | 0.5 | -0.5 | 1.24 | 0.99 | 0.16 | -0.69 | 0.23 | 0.09 | -0.355 | 0.29 | 0.22 | 0.550494 | 0.940304 | 0.460913 | 1 | 0.999631 | 0.836038 |
| Kdm5c_P41230_S856s_1 | -0.78 | 0.03 | 1.62 | 1.23 | -1.82 | 0.64 | 0.49 | 1.37 | -0.65 | 0.38 | 1.29 | 2.24 | 0.59 | 0.19 | 0.64 | 1.56 | -0.4475 | -1.1975 | -1.485 | 0.702074 | 0.648052 | 0.693646 | 1 | 0.999631 | 0.896857 |
| Keap1_Q9Z2X8_T43t_1 | 1.09 | -1.74 | -0.82 | -0.58 | -1.32 | -1.93 | -0.5 | -0.09 | -2.98 | -0.13 | -1.57 | -2.16 | -2.96 | -2.35 | -2.74 | 0.06 | -0.3375 | 0.2025 | 0.56 | 0.086515 | 0.492253 | 0.132147 | 1 | 0.999631 | 0.553155 |
| Khdrbs1_Q60749_S38s_1 | 0.54 | 1.5 | 1.97 | 0.65 | 0.1 | 1.9 | 0.08 | 1.23 | 1.33 | 1.86 | 1.34 | 0.94 | 1.3 | 1.98 | 1.36 | 2.26 | -0.345 | 0.025 | 0.24 | 0.209006 | 0.523292 | 0.617538 | 1 | 0.999631 | 0.662575 |
| Khdrbs1_Q60749_S58s_1 | 0.32 | 1.15 | 0.92 | 0.48 | 0.35 | 0.49 | 0.07 | 0.58 | 0.65 | 0.76 | 0.61 | 0.95 | 1.16 | 0.97 | 0.7 | 1 | -0.35 | 0.1 | 0.0575 | 0.44481 | 0.254673 | 0.931566 | 1 | 0.999631 | 0.783608 |
| Khsrp_Q3U0V1_S182s_1 | 0.6 | 1.81 | 1.86 | 1.2 | 1.14 | 1.13 | 0.62 | 1.18 | 1.37 | 1.49 | 1.49 | 1.52 | 1.27 | 1.59 | 1.28 | 1.56 | -0.405 | -0.55 | -0.745 | 0.872859 | 0.333745 | 0.770602 | 1 | 0.999631 | 0.959629 |
| Khsrp_Q3U0V1_S481s_1 | 1.49 | 1.79 | 1.9 | 1.6 | 1.35 | 1.15 | 1.46 | 1.2 | 1.2 | 1.64 | 0.73 | 1.01 | 0.2 | 0.83 | 1.89 | 0.88 | -0.4625 | 0.1425 | 0.45 | 0.080608 | 0.123726 | 0.086236 | 1 | 0.999631 | 0.545579 |
| Kiaa0195_Q7TSH8_S445s_1 | -0.3 | -1.15 | -1.46 | -1.79 | -0.85 | -1.76 | -1.93 | -2.01 | -1.42 | 0.01 | -1.22 | -1.5 | 0.74 | -1.02 | -1.48 | -1.14 | 0.5425 | 0.585 | 0.1275 | 0.421715 | 0.280721 | 0.758324 | 1 | 0.999631 | 0.772942 |
| Kiaa1045_Q80TL4_S18s_1 | 1 | 1.87 | -0.09 | 1.72 | 1.62 | 2.59 | 0.75 | 1.71 | 2.51 | 1.32 | 1.33 | 1.68 | -0.1 | 2.5 | 1.41 | 1.2 | -0.0825 | -0.2875 | -0.3075 | 0.839273 | 0.326909 | 0.259521 | 1 | 0.999631 | 0.946365 |
| Kiaa1045_Q80TL4_S309s_1 | 0.3 | 0.1 | -0.41 | 0.11 | -0.07 | 0.27 | -0.27 | -0.16 | -0.51 | 0.68 | -0.43 | -0.79 | -0.38 | 0.39 | -0.9 | -0.24 | 0.0825 | 0.115 | -0.2425 | 0.398783 | 0.765876 | 0.455956 | 1 | 0.999631 | 0.761726 |
| Kiaa1045_Q80TL4_S309s_2 | -0.07 | 0.72 | -0.59 | 0.96 | 1.08 | -0.66 | 0.47 | 0.46 | 0.31 | 0.19 | 1.04 | -0.06 | -2.06 | 0.7 | 0.82 | 0.59 | 0.135 | 0.1175 | 0.3975 | 0.72671 | 0.864077 | 0.788599 | 1 | 0.999631 | 0.905774 |
| Kiaa1045_Q80TL4_S347s_1 | -0.67 | -0.12 | -0.96 | -0.66 | -0.96 | -0.35 | -0.01 | -0.55 | -0.73 | -0.73 | -0.22 | -0.26 | -0.55 | -0.16 | -0.17 | 0.06 | 0.1675 | -0.0825 | 0.0775 | 0.218958 | 0.670331 | 0.69716 | 1 | 0.999631 | 0.669859 |
| Kiaa1045_Q80TL4_S348s_1 | -0.67 | -0.12 | -0.58 | -1.97 | -0.96 | -0.35 | -0.77 | -0.59 | -0.73 | -0.73 | -1.65 | -0.56 | -0.55 | -0.16 | -1.23 | -1.09 | 0.4775 | -0.2025 | -0.5775 | 0.866644 | 0.687952 | 0.8566 | 1 | 0.999631 | 0.956432 |
| Kiaa1045_Q80TL4_S43s_1 | 0.92 | -0.19 | -0.87 | 0.51 | 1 | 0.15 | 0.27 | 0.86 | 0.21 | -0.16 | 0.27 | -0.76 | -1.17 | 0.36 | -0.62 | -0.51 | 0.1725 | -0.0975 | -0.595 | 0.253887 | 0.289389 | 0.654833 | 1 | 0.999631 | 0.688747 |
| Kiaa1045_Q80TL4_S51s_1 | 0.57 | 2.14 | -0.6 | -0.08 | 0.35 | 1.85 | 0.27 | 0.25 | 1.68 | 0.53 | 0.58 | -1.15 | -0.73 | 0.91 | -0.12 | -0.41 | -0.485 | 0.0375 | -0.505 | 0.353035 | 0.786758 | 0.894759 | 1 | 0.999631 | 0.73949 |
| Kiaa1045_Q80TL4_S73s_1 | 0.43 | -0.64 | 0.86 | 1.04 | 0.2 | -0.63 | 0.18 | 0 | -0.18 | -0.14 | 0.61 | 1.55 | 0.22 | -0.36 | -0.28 | 0.09 | -0.12 | 0.0775 | 0.4925 | 0.243488 | 0.26473 | 0.942472 | 1 | 0.999631 | 0.685637 |
| Kiaa1045_Q80TL4_T100t_1 | 0.27 | 1.84 | 1.57 | 0.62 | 0.35 | 2.04 | 0.28 | 1.15 | 0.95 | 1.65 | 0.72 | 1.29 | 1.19 | 1.9 | 1.23 | 1.95 | -0.105 | 0.275 | 0.2075 | 0.272696 | 0.81682 | 0.855915 | 1 | 0.999631 | 0.700187 |
| Kiaa1045_Q80TL4_T47t_1 | 0.4 | 1.18 | 1.42 | 0.93 | 0.72 | 0.88 | 0.75 | 1.16 | 0.53 | 0.57 | 2.1 | 1.83 | -0.04 | 0.78 | 2.09 | 1.93 | 0.6625 | -0.245 | -0.4975 | 0.690315 | 0.72867 | 0.549915 | 1 | 0.999631 | 0.893349 |
| Kiaa1107_Q80TK0_S1051s_1 | 0.59 | -0.78 | -0.32 | 0.69 | 1.5 | -0.03 | 0.96 | 0.4 | 0.03 | -1.11 | 0.29 | -0.01 | -1.5 | -0.15 | -0.06 | -0.1 | -0.6275 | -0.4175 | -1.11 | 0.317169 | 0.177604 | 0.597145 | 1 | 0.999631 | 0.720971 |
| Kiaa1107_Q80TK0_S353s_1 | 1.04 | 0.94 | 0.8 | -0.14 | -0.49 | 0.61 | 0.56 | -0.55 | 0.25 | 1.77 | -0.15 | -0.9 | 0.98 | -0.88 | -0.82 | -1.08 | 0.515 | 0.13 | 0.64 | 0.055296 | 0.155145 | 0.472849 | 1 | 0.999631 | 0.493177 |
| Kiaa1107_Q80TK0_S481s_1 | 0.3 | -0.76 | 0.33 | 0.17 | 0.76 | 0.9 | 0.23 | 0.21 | -0.04 | 0.72 | -0.13 | 0.01 | 0.25 | 1.08 | 0.67 | 0.6 | -0.075 | 0.33 | 0.86 | 0.099528 | 0.159748 | 0.717726 | 1 | 0.999631 | 0.562298 |
| Kiaa1107_Q80TK0_S695s_1 | 0.31 | -0.94 | 0.94 | 0.18 | -0.98 | 0.42 | 0.03 | 0.72 | -0.25 | 0.11 | 0.67 | 1.28 | 1.37 | 0.53 | 0.67 | 1.36 | -0.19 | 0.0525 | -0.655 | 0.079573 | 0.881671 | 0.50644 | 1 | 0.999631 | 0.545579 |
| Kiaa1107_Q80TK0_S714s_1 | -0.08 | -1.33 | 0.53 | -0.87 | -0.63 | -1.37 | 1.02 | -1.53 | -0.26 | -1.12 | -0.01 | -0.15 | -0.49 | -2.03 | -0.19 | -1.66 | 0.3425 | 0.1975 | -1.1325 | 0.26048 | 0.76674 | 0.910541 | 1 | 0.999631 | 0.692624 |
| Kiaa1107_Q80TK0_S714s_2 | 0.08 | -1.08 | 0.34 | -0.59 | -0.77 | -0.1 | 0.87 | 0.12 | -0.64 | 0.02 | 0.02 | 0.14 | -2.65 | -1.25 | -0.92 | -0.96 | 0.3425 | 0.1975 | -1.1325 | 0.043543 | 0.454419 | 0.611795 | 1 | 0.999631 | 0.475443 |
| Kiaa1107_Q80TK0_S722s_2 | 0.08 | -1.08 | 0.34 | -0.59 | -0.77 | -0.1 | 0.87 | 0.12 | -0.64 | 0.02 | 0.02 | 0.14 | -2.65 | -1.25 | -0.92 | -0.96 | 0.01 | 0.2075 | -0.0325 | 0.043543 | 0.454419 | 0.611795 | 1 | 0.999631 | 0.475443 |
| Kiaa1107_Q80TK0_S783s_2 | -1.21 | 0.22 | 0.28 | 0.21 | -0.33 | -0.19 | -0.09 | 0.15 | -0.56 | -0.72 | 0.74 | 0.87 | -0.28 | -0.48 | -0.1 | 0.23 | 0.0725 | 0.2475 | -0.1525 | 0.936859 | 0.979332 | 0.690974 | 1 | 0.999631 | 0.97939 |
| Kiaa1107_Q80TK0_S785s_1 | -0.38 | 0.16 | -0.15 | -0.57 | -0.13 | -0.19 | -0.14 | -0.19 | 0.18 | -0.27 | 0.21 | -0.07 | -0.46 | -0.08 | -0.19 | -0.82 | 0.01 | 0.2075 | -0.0325 | 0.630902 | 0.781773 | 0.394821 | 1 | 0.999631 | 0.867552 |
| Kiaa1107_Q80TK0_S785s_2 | -1.21 | 0.22 | 0.28 | 0.21 | -0.33 | -0.19 | -0.09 | 0.15 | -0.56 | -0.72 | 0.74 | 0.87 | -0.28 | -0.48 | -0.1 | 0.23 | -0.4375 | 0.3875 | -0.08 | 0.936859 | 0.979332 | 0.690974 | 1 | 0.999631 | 0.97939 |
| Kiaa1107_Q80TK0_S828s_1 | 0.55 | 3.42 | 1 | 0.45 | 0.55 | 3.01 | 0.12 | -0.01 | 1.49 | 2.98 | 1.51 | 0.99 | 4.08 | 2.44 | -1.67 | 0.25 | -0.1925 | -0.6625 | 0.0475 | 0.947501 | 0.614542 | 0.59669 | 1 | 0.999631 | 0.983056 |
| Kiaa1109_A2AAE1_S1294s_1 | -0.18 | -0.31 | -0.18 | -0.82 | -0.76 | 0.42 | -1.3 | -0.62 | -0.33 | -1.79 | -0.85 | -1.17 | -0.16 | -0.03 | -0.43 | -0.68 | -0.0025 | 0.195 | -0.76 | 0.877111 | 0.629223 | 0.097702 | 1 | 0.999631 | 0.961513 |
| Kiaa1109_A2AAE1_S1418s_1 | -0.15 | -1.28 | -0.66 | -0.7 | -0.45 | -0.68 | -0.74 | -0.93 | -0.49 | -0.01 | -0.87 | -0.64 | -1.55 | -0.89 | -1.96 | -1.43 | -0.535 | -0.49 | -0.4475 | 0.060082 | 0.993519 | 0.569243 | 1 | 0.999631 | 0.505461 |
| Kiaa1109_A2AAE1_S1981s_1 | -0.26 | 0.1 | -0.69 | -0.89 | -1.18 | -0.08 | -1.08 | -1.54 | -1.2 | -1.55 | -0.69 | -0.26 | -0.97 | -0.03 | -1.37 | -1.16 | -0.445 | -0.39 | -1.2075 | 0.274234 | 0.192862 | 0.218461 | 1 | 0.999631 | 0.700187 |
| Kiaa1109_A2AAE1_S2350s_1 | 0.47 | -1.74 | 0.1 | -0.49 | -1.27 | -1.01 | -0.1 | -1.06 | -0.97 | 0.25 | -1.56 | -0.94 | -1.51 | -1.67 | -1.38 | -1.93 | 0.02 | -0.17 | -0.18 | 0.029465 | 0.394172 | 0.496518 | 1 | 0.999631 | 0.433091 |
| Kiaa1109_A2AAE1_S2368s_1 | 0 | -0.17 | -0.54 | -0.37 | 0.05 | -0.15 | -0.57 | -0.33 | -0.36 | -0.34 | -0.51 | -0.55 | -0.79 | 0.27 | -0.62 | -0.66 | -0.5475 | -0.3725 | 0.22 | 0.596114 | 0.940815 | 0.515097 | 1 | 0.999631 | 0.854508 |
| Kiaa1109_A2AAE1_S2809s_1 | -0.4 | -0.01 | -0.34 | -0.43 | -1.34 | -0.15 | -0.99 | -0.89 | -0.36 | -0.14 | -1.35 | -0.82 | -0.17 | 0.87 | -0.88 | -0.12 | 0.6575 | 0.175 | 0.5825 | 0.583228 | 0.109713 | 0.280945 | 1 | 0.999631 | 0.848468 |
| Kiaa1109_A2AAE1_S903s_1 | -1.04 | -4 | -0.06 | -0.31 | -0.94 | -1.61 | -0.23 | 0 | -2.61 | -0.77 | -0.04 | -1.29 | -2.91 | -0.95 | -0.07 | 0.85 | -0.6475 | -0.3725 | -2.9425 | 0.575823 | 0.446104 | 0.848681 | 1 | 0.999631 | 0.846085 |
| Kiaa1109_A2AAE1_S925s_1 | 3.05 | 1.52 | 2.46 | 2.39 | 1.07 | 1.84 | 2.38 | 1.54 | 2 | 2.67 | 1.51 | 1.75 | -2.29 | 0.48 | -0.15 | -0.39 | 0.5075 | -0.1625 | -1.5725 | 0.000734 | 0.143213 | 0.374349 | 1 | 0.999631 | 0.351308 |
| Kiaa1109_A2AAE1_S936s_1 | 1.58 | 1.44 | -0.02 | 2.07 | 2.7 | 2.05 | 0.95 | 1.4 | 1.85 | 0.17 | 1.63 | 0.77 | -2.69 | 1.72 | 0.13 | -0.38 | 0.175 | 0.0175 | -1.7625 | 0.097819 | 0.36131 | 0.768426 | 1 | 0.999631 | 0.560722 |
| Kiaa1109_F6YAT5_S213s_1 | 1.11 | 0.18 | -0.04 | 0.98 | 0.77 | 0.8 | 0.61 | 0.75 | 0.68 | 1.18 | 0.38 | 0.06 | -3.13 | -0.2 | -0.52 | -0.97 | -0.5 | -0.995 | -2.6075 | 0.02009 | 0.598979 | 0.964149 | 1 | 0.999631 | 0.391668 |
| Kiaa1109_F6YAT5_S475s_1 | 1.94 | -1.18 | 0.14 | -0.15 | -1.37 | -0.11 | 1.27 | -1.04 | -1.56 | 1.33 | -1.5 | -1.5 | -1.93 | -1.43 | -2.92 | -3.4 | 0.2375 | 0.3525 | 0.43 | 0.003827 | 0.520362 | 0.25505 | 1 | 0.999631 | 0.351308 |
| Kiaa1109_F6YAT5_S889s_1 | -1.81 | -0.98 | -2.76 | -1.64 | -1.16 | -1.52 | -1.95 | -1.61 | -1.26 | -0.32 | -1.88 | -2.32 | -0.92 | -0.93 | -2.21 | -1.41 | -0.295 | -0.2725 | -0.4975 | 0.368362 | 0.560254 | 0.511584 | 1 | 0.999631 | 0.745968 |
| Kiaa1109_F6YAT5_T577t_1 | -0.66 | -0.52 | -0.26 | -0.32 | -0.93 | -0.48 | -0.94 | -0.59 | -0.88 | -0.8 | -0.21 | -0.96 | -0.88 | -0.29 | -1.3 | -1.28 | 0.0625 | -0.8825 | -1.1975 | 0.152981 | 0.268996 | 0.351034 | 1 | 0.999631 | 0.625198 |
| Kiaa1522_A2A7S8_S227s_1 | -0.21 | -2.67 | -1.74 | -1.56 | -1.79 | -1.64 | -0.21 | -2.29 | -2.57 | -2.88 | -1.98 | -2.28 | -2.68 | -3.43 | -2.34 | -2.52 | 0.1975 | -0.18 | 0.655 | 0.043768 | 0.9182 | 0.10959 | 1 | 0.999631 | 0.475443 |
| Kiaa1522_A2A7S8_S264s_1 | -0.59 | -1.77 | -0.3 | -0.49 | 0.73 | -0.24 | -1.77 | -1.08 | -0.81 | -0.34 | -0.59 | -2.13 | 0.2 | -1.02 | -0.09 | 0.38 | -0.2075 | -0.21 | 0.825 | 0.170447 | 0.732715 | 0.716718 | 1 | 0.999631 | 0.64073 |
| Kiaa1522_A2A7S8_S289s_1 | -0.6 | -0.66 | -0.23 | -1 | -1.38 | -0.75 | -0.66 | -0.53 | -1.78 | -0.49 | -0.32 | -0.74 | 1.1 | -0.23 | -0.37 | 0.31 | -0.015 | -0.3325 | 0.315 | 0.059475 | 0.501572 | 0.587049 | 1 | 0.999631 | 0.50521 |
| Kiaa1522_A2A7S8_S289s_2 | 0.02 | -0.15 | -0.18 | -0.56 | 0.03 | -0.14 | -0.56 | -0.26 | 0.01 | -0.72 | -0.15 | -1.34 | -0.16 | 0.32 | -0.3 | 0.53 | 0.16 | -0.455 | 0.4325 | 0.333086 | 0.955422 | 0.369165 | 1 | 0.999631 | 0.728079 |
| Kiaa1522_A2A7S8_S294s_2 | 0.02 | -0.15 | -0.16 | 0.16 | 0.03 | -0.14 | 0.19 | 0.43 | 0.01 | -0.72 | -0.07 | -1.17 | -0.16 | 0.32 | 0.25 | 1.19 | 0.345 | -0.495 | -0.945 | 0.235491 | 0.535188 | 0.198547 | 1 | 0.999631 | 0.683301 |
| Kiaa1522_A2A7S8_S306s_1 | 1.37 | 0.58 | 0.32 | 0.91 | 0.7 | 1.69 | 1.83 | 0.34 | 0.36 | 0.44 | 0.2 | 0.2 | -1.53 | 0.4 | 0.43 | 0.1 | 1.485 | -0.755 | -0.5425 | 0.081402 | 0.424864 | 0.129685 | 1 | 0.999631 | 0.546723 |
| Kiaa1522_A2A7S8_S332s_1 | 1 | 1.08 | -0.34 | 1.51 | 1.36 | 3.25 | 1.9 | 2.68 | 0.68 | -0.6 | 0.94 | -0.79 | -3.76 | 2.94 | 0.9 | 1 | 0.6825 | -0.315 | -0.2425 | 0.665333 | 0.018637 | 0.191828 | 1 | 0.999631 | 0.882566 |
| Kiaa1522_A2A7S8_S430s_1 | 0.67 | -1.29 | -2.78 | -1.71 | 0.13 | -0.44 | -0.57 | -1.5 | -1.45 | -1.67 | -1.52 | -1.73 | -2.14 | -0.3 | -2.51 | -1.13 | 0.1375 | 0.03 | 0.2775 | 0.752792 | 0.344437 | 0.630404 | 1 | 0.999631 | 0.913377 |
| Kiaa1522_A2A7S8_S796s_1 | -0.36 | -0.17 | -2.08 | -0.37 | -0.21 | -0.94 | -0.74 | -0.54 | -0.86 | -1.68 | 0.48 | -0.8 | 0.22 | -0.26 | -0.85 | -0.98 | -0.1175 | -0.0825 | -0.565 | 0.584049 | 0.763283 | 0.958661 | 1 | 0.999631 | 0.849058 |
| Kiaa1522_A2A7S8_S890s_1 | 0.54 | -1.36 | 0.46 | -0.01 | -0.89 | -0.41 | 0.46 | 0 | -1.35 | 0.17 | 0.04 | 0.44 | -1.3 | -1.54 | 0.46 | -0.25 | -0.6125 | -0.055 | -1.105 | 0.347261 | 0.812164 | 0.880914 | 1 | 0.999631 | 0.736373 |
| Kiaa1598_Q8K2Q9_S3s_1 | 0.47 | -0.23 | 0.44 | 1.43 | -0.16 | -0.59 | 0.22 | 0.19 | 0.09 | 1.56 | 0.13 | 0.11 | -0.55 | -0.94 | -0.38 | -0.44 | 0.5825 | 0.38 | 0.4325 | 0.017103 | 0.145982 | 0.908455 | 1 | 0.999631 | 0.391024 |
| Kiaa1671_Q8BRV5_S166s_1 | 1.33 | 1.92 | 1.4 | 1.56 | 2.15 | 2.56 | 1.24 | 2.59 | 2.15 | 1.45 | 2.38 | 1.75 | 0.76 | 2.53 | 2.22 | 2.43 | -0.0725 | -0.19 | -1.145 | 0.335943 | 0.132968 | 0.240834 | 1 | 0.999631 | 0.730466 |
| Kiaa1671_Q8BRV5_S175s_1 | 0.74 | 1.81 | 1.1 | 1.31 | 0.62 | 1.94 | 0.9 | 1.21 | 1.57 | 1.26 | 1.6 | -0.23 | -0.72 | 0.75 | 0.56 | -0.21 | -0.295 | -0.2625 | -0.9525 | 0.02023 | 0.847424 | 0.687781 | 1 | 0.999631 | 0.392698 |
| Kiaa1671_Q8BRV5_S205s_1 | -0.45 | -1.99 | -0.92 | -0.92 | -1.61 | -1.67 | -0.58 | -1.6 | -1.9 | -0.71 | -1.1 | -1.62 | -1.48 | -2.67 | -1.69 | -2.25 | -0.5575 | -0.5225 | -0.6425 | 0.048369 | 0.483951 | 0.538776 | 1 | 0.999631 | 0.48343 |
| Kiaa1671_Q8BRV5_S277s_1 | -1.45 | -3.46 | 0.33 | 1.63 | -3.78 | -3.43 | 1.18 | 0.85 | -4.31 | -2.52 | 1.35 | 0.44 | -2.19 | -3.57 | 0.86 | -0.62 | 0.2225 | -0.2925 | -0.965 | 0.605867 | 0.705716 | 0.720847 | 1 | 0.999631 | 0.85817 |
| Kiaa1671_Q8BRV5_S302s_1 | 1.36 | 0.73 | 0.91 | 1.57 | 1.53 | 0.84 | 1.89 | 1.2 | 0.33 | 0.54 | 1.97 | 0.56 | -0.25 | 0.88 | -0.49 | 0.57 | 1.0575 | -1.6275 | -2.5625 | 0.034985 | 0.511785 | 0.497096 | 1 | 0.999631 | 0.450183 |
| Kiaa1671_Q8BRV5_T215t_1 | 4.1 | -1.78 | -2.55 | 0.88 | 2.8 | -0.01 | 2.07 | 0.02 | 0.24 | -3.15 | -0.82 | -2.13 | -3.2 | -1.64 | -2.52 | -2.24 | 0.09 | 0.4825 | 0.8175 | 0.071981 | 0.461456 | 0.270824 | 1 | 0.999631 | 0.532005 |
| Kidins220_E9Q9B7_S1578s_1 | -0.36 | 0 | -1.29 | -1.25 | -0.12 | -0.01 | -1.21 | -1.2 | 0.51 | 0.31 | -0.62 | -1.17 | 1.12 | 0.59 | -0.53 | -0.81 | 0.0725 | -0.4925 | -0.435 | 0.144207 | 0.840513 | 0.335264 | 1 | 0.999631 | 0.616037 |
| Kidins220_E9Q9B7_S1704s_1 | 0.97 | 0.6 | -0.08 | 1.25 | 0.94 | 0.86 | 0.21 | 1.02 | 0.03 | 0.1 | 0.15 | 0.49 | -0.14 | 0.52 | 0.2 | 0.42 | 0.2375 | 0.53 | 0.7525 | 0.253343 | 0.842119 | 0.177458 | 1 | 0.999631 | 0.688735 |
| Kidins220_E9Q9B7_S885s_1 | -1.33 | -0.68 | 0.13 | -0.95 | -0.64 | 0.03 | -0.39 | -0.88 | 0.04 | -0.62 | -0.41 | 0.28 | 0.09 | -0.27 | 0.82 | -0.46 | -0.3475 | -0.015 | -0.49 | 0.10314 | 0.535328 | 0.194309 | 1 | 0.999631 | 0.566498 |
| Kif13b_E9Q4K7_S1380s_1 | 1.2 | 1.1 | 1.48 | 1.64 | 0.73 | 0.56 | 1.28 | 1.46 | 1.17 | 1.16 | 1.29 | 1.74 | -0.56 | 0.67 | 1.87 | 1.48 | 0.2975 | 0.1725 | 0.235 | 0.3578 | 0.273186 | 0.957384 | 1 | 0.999631 | 0.741099 |
| Kif13b_E9Q4K7_S1794s_1 | -0.39 | 0.21 | -0.05 | 0.44 | 0.8 | 0.09 | -0.21 | 0.72 | -0.08 | -0.2 | 0.48 | 0.7 | -0.03 | 0.38 | 0.62 | 0.18 | 0.42 | 0.1475 | -0.0725 | 0.461759 | 0.389716 | 0.605587 | 1 | 0.999631 | 0.792786 |
| Kif13b_E9Q4K7_S1795s_1 | -0.62 | -0.19 | -0.05 | 0.11 | 0.52 | -0.13 | -0.09 | 0.63 | -0.65 | -0.24 | 0.48 | 0.25 | -1.31 | 0.07 | 0.46 | -0.26 | 0.98 | 0.985 | 0.46 | 0.863824 | 0.195429 | 0.66743 | 1 | 0.999631 | 0.955131 |
| Kif1a_Q6TA13_S1102s_1 | -1.13 | -1.1 | -0.74 | -0.65 | 0.73 | 0.64 | -0.93 | -0.14 | 0.19 | 0.21 | -0.28 | 0.2 | -1.89 | 0.19 | -0.18 | 0.1 | 0.2225 | 0.29 | 0.24 | 0.355637 | 0.035223 | 0.004955 | 1 | 0.999631 | 0.74034 |
| Kif1a_Q6TA13_S1345s_1 | -0.12 | -0.67 | -1.1 | -0.9 | 0.02 | -0.5 | -0.72 | -0.7 | -0.1 | -0.29 | -0.72 | -0.52 | -0.18 | 0.37 | -0.84 | -1.18 | 0.1825 | 0.4025 | 0.94 | 0.569862 | 0.4931 | 0.36769 | 1 | 0.999631 | 0.84338 |
| Kif1a_Q6TA13_S1378s_1 | -1.45 | 0.49 | 0.19 | -0.2 | 0.42 | 0.03 | -0.41 | -0.28 | 0.2 | -0.08 | 0.14 | 0.38 | 0.77 | 0.75 | 0.81 | 0.46 | 0.13 | 0.01 | 0.3025 | 0.053982 | 0.685765 | 0.365282 | 1 | 0.999631 | 0.491994 |
| Kif1a_Q6TA13_S1438s_1 | -0.42 | 0.59 | -2.27 | -3.72 | 0.04 | 0.18 | -1.99 | -3.53 | 0.07 | -0.74 | -2.49 | -2.62 | 0.84 | -0.43 | -2.48 | -2.54 | 0.1375 | -0.19 | 0.225 | 0.779115 | 0.907367 | 0.992065 | 1 | 0.999631 | 0.921416 |
| Kif1a_Q6TA13_S1485s_1 | -0.2 | 0.73 | 0.3 | 0.56 | 0.23 | 0.98 | 0.26 | 0.47 | 0.74 | 0 | 0.16 | -0.27 | 0.61 | 1.25 | 0.73 | -0.3 | -0.1475 | -0.16 | 0.05 | 0.579802 | 0.665865 | 0.57931 | 1 | 0.999631 | 0.84667 |
| Kif1a_Q6TA13_S1503s_1 | -0.89 | -0.62 | 0.03 | -0.44 | -0.3 | -1.24 | -0.65 | -0.32 | -0.4 | -1.16 | -0.4 | -0.6 | -0.28 | -0.84 | -0.22 | -0.38 | -0.11 | 0.0725 | -0.34 | 0.87632 | 0.658733 | 0.621516 | 1 | 0.999631 | 0.961058 |
| Kif1a_Q6TA13_S1556s_1 | 0.5 | -0.55 | 0.26 | 0.16 | -0.22 | -0.27 | 0.3 | 0.12 | 0.02 | 0.65 | 0.21 | -0.22 | -0.45 | -0.31 | 0 | -0.23 | 0.38 | -0.375 | 0.3875 | 0.307818 | 0.727864 | 0.83027 | 1 | 0.999631 | 0.718124 |
| Kif1a_Q6TA13_S411s_1 | -0.05 | -0.06 | -0.52 | -0.1 | 0.32 | -0.13 | 0.04 | 0.56 | -0.01 | -0.82 | -0.79 | -0.61 | 0.09 | 0.16 | 0.14 | 0.43 | 1.005 | -0.15 | -0.6125 | 0.184052 | 0.189414 | 0.227248 | 1 | 0.999631 | 0.644602 |
| Kif1a_Q6TA13_S425s_1 | 1.66 | -1.32 | -2 | 0.91 | 1.27 | 0.41 | 1.26 | 0.33 | 0.26 | -0.65 | 0.29 | -1.25 | -1.15 | 0.03 | -0.78 | -1.3 | -0.4775 | 0.015 | -0.01 | 0.453493 | 0.226571 | 0.856982 | 1 | 0.999631 | 0.788717 |
| Kif1a_Q6TA13_S428s_1 | 0.61 | 0.64 | 0.89 | -0.48 | -0.47 | 0.25 | -0.09 | 0.06 | 0.39 | 0.53 | 0.22 | 0.58 | -0.93 | 0.7 | 1.46 | 0.39 | 0.3275 | -0.0725 | -0.0625 | 0.985349 | 0.208673 | 0.966063 | 1 | 0.999631 | 0.995133 |
| Kif1a_Q6TA13_S941s_1 | -1.04 | 0.68 | -0.37 | -0.46 | 0.58 | -0.21 | -0.31 | 0.06 | -0.67 | -0.88 | 0.05 | 0.02 | -1.14 | -0.55 | -0.42 | 0.67 | -0.2475 | 0.2725 | -0.5475 | 0.90019 | 0.43211 | 0.865904 | 1 | 0.999631 | 0.967733 |
| Kif1a_Q6TA13_S946s_1 | 0.41 | -0.05 | 0.46 | 0.53 | -0.37 | 0.26 | 0.45 | 0.02 | 0.46 | 0.93 | 0.48 | 0.57 | -0.12 | -0.32 | -0.16 | -0.24 | 0.1975 | 0.1225 | 0.0175 | 0.072806 | 0.404192 | 0.333991 | 1 | 0.999631 | 0.534296 |
| Kif1a_Q6TA13_T1511t_1 | -1.15 | -1.06 | 0.04 | -0.29 | -0.38 | -1.01 | -0.2 | -0.08 | -0.75 | -0.77 | -0.19 | -0.26 | 0.04 | -0.78 | -1.2 | -0.45 | 0.4225 | -0.2325 | 0.79 | 0.965962 | 0.599922 | 0.735775 | 1 | 0.999631 | 0.988179 |
| Kif1b_Q60575_S1165s_1 | -3.19 | -1.18 | -0.97 | -2.68 | -1.36 | -2.36 | -1.23 | -1.38 | -1.77 | -2.74 | -2.01 | -2.43 | -0.84 | -1.73 | -1.79 | -0.5 | 0.2125 | 0.2725 | -0.5775 | 0.196417 | 0.454571 | 0.672148 | 1 | 0.999631 | 0.653373 |
| Kif1b_Q60575_S1408s_1 | 0.52 | -0.28 | -0.33 | -0.19 | 0.25 | -0.31 | 0.56 | 0.07 | 0.43 | -0.14 | 0.17 | 0.35 | -0.68 | -0.56 | -0.19 | -1.16 | 0.92 | 0.6875 | 1.025 | 0.11722 | 0.509494 | 0.38373 | 1 | 0.999631 | 0.585087 |
| Kif1b_Q60575_S1567s_1 | -2.68 | 1.06 | 1.21 | -0.24 | 1.53 | 0.35 | 0.63 | 0.52 | 1.5 | -0.35 | 0.39 | 0.56 | 2.61 | 0.63 | 0.34 | -0.13 | 0.09 | 0.0025 | 0.3325 | 0.286122 | 0.276585 | 0.428284 | 1 | 0.999631 | 0.705972 |
| Kif1b_Q60575_S1613s_1 | -0.84 | 0.22 | -0.46 | -0.37 | -0.06 | -0.46 | -0.47 | -0.1 | -0.3 | -0.44 | -0.52 | -0.18 | 0.28 | 0.03 | -0.22 | -0.21 | 0.57 | -0.145 | -0.7875 | 0.321333 | 0.76831 | 0.993439 | 1 | 0.999631 | 0.72408 |
| Kif1b_Q60575_T1406t_1 | -0.71 | 0.19 | 0.47 | -0.18 | 0.44 | 0.29 | 1.36 | -0.04 | 0.48 | -0.87 | 0.26 | -0.68 | -0.81 | -1.39 | -0.32 | -0.86 | -0.315 | -0.35 | -1.1275 | 0.058231 | 0.174932 | 0.732747 | 1 | 0.999631 | 0.499755 |
| Kif1c_E9Q9B0_S1043s_1 | 2.09 | -0.25 | 0.68 | 0.69 | -0.15 | 0.52 | 1.35 | 0.23 | -0.3 | 2.27 | -0.04 | -0.12 | -0.64 | -0.54 | 0.18 | -0.3 | -0.33 | -0.025 | 0.2075 | 0.043265 | 0.558669 | 0.615403 | 1 | 0.999631 | 0.475443 |
| Kif1c_E9Q9B0_T1092t_1 | 0.6 | 0.65 | 0.51 | -0.04 | 0.14 | 0.11 | -0.4 | 0.55 | 0.31 | 0.57 | 0.24 | 0.5 | 0.68 | 0.57 | 0.59 | 0.71 | 0.6725 | -0.2925 | -0.585 | 0.475939 | 0.298825 | 0.928178 | 1 | 0.999631 | 0.799882 |
| Kif21a_E9Q0J5_S1241s_1 | 2.56 | -0.03 | -0.43 | 2.02 | 2.51 | -0.26 | 2.4 | 2.16 | 1.12 | -0.05 | 1.39 | 0.49 | -0.09 | 0.81 | 0.88 | 0.18 | -0.225 | -0.075 | 0.31 | 0.405877 | 0.441724 | 0.684518 | 1 | 0.999631 | 0.765969 |
| Kif21a_E9Q0J5_S258s_1 | -0.18 | -0.7 | -0.53 | -0.54 | -0.83 | -0.6 | -0.63 | -0.79 | -0.89 | -0.66 | -0.51 | -0.19 | -0.28 | -0.19 | -0.19 | -0.05 | 0.075 | 0.345 | 0.88 | 0.270992 | 0.375063 | 0.789188 | 1 | 0.999631 | 0.698674 |
| Kif21a_E9Q0J5_S524s_1 | 0.14 | 1.21 | 1.21 | 1.27 | 0.9 | 1.3 | 0.71 | 1.22 | 1.22 | 0.76 | 1.14 | 2.09 | 1.39 | 1.63 | 2.15 | 2.18 | -0.01 | -0.175 | -0.19 | 0.038162 | 0.825786 | 0.401001 | 1 | 0.999631 | 0.459475 |
| Kif21a_E9Q0J5_S855s_1 | 0.68 | 1.1 | 0.49 | 0.58 | 0.92 | 1.57 | 0 | 0.32 | 1.39 | 0.73 | 0.31 | -0.28 | -0.98 | 2.2 | 0.4 | 0.47 | 0.025 | 0.4375 | 0.005 | 0.753946 | 0.97918 | 0.657276 | 1 | 0.999631 | 0.913377 |
| Kif21b_Q9QXL1_S1150s_1 | 0.2 | -0.31 | -0.32 | -0.23 | 0.13 | -0.54 | -0.24 | 0.09 | 0.29 | 0.44 | -0.04 | 0.4 | -0.26 | -0.34 | -0.12 | 0.08 | 0.1725 | 0.665 | 0.4725 | 0.985933 | 0.928935 | 0.132014 | 1 | 0.999631 | 0.995133 |
| Kif21b_Q9QXL1_S1198s_1 | 0.62 | 1.48 | 0.52 | 0.97 | 1.23 | 1.32 | 0.11 | 1.62 | 1.7 | 1.68 | 1.12 | 1.75 | 0.91 | 1.45 | 1.18 | 1.94 | 0.175 | 0.5375 | 0.3175 | 0.208004 | 0.667508 | 0.061782 | 1 | 0.999631 | 0.661649 |
| Kif21b_Q9QXL1_S1217s_1 | 0.48 | -0.08 | -0.18 | 0.17 | 0.26 | 0.27 | 0.05 | 0.51 | 0.25 | 1.02 | 0.74 | 0.53 | 0 | 0.21 | 0.53 | 0.92 | -0.0075 | -0.2875 | 0.045 | 0.342215 | 0.519436 | 0.096049 | 1 | 0.999631 | 0.73347 |
| Kif21b_Q9QXL1_S222s_1 | 0.15 | -0.11 | -0.09 | 0.42 | -0.31 | -1.13 | 1.14 | 0.64 | -0.22 | 0.17 | 0.35 | -1.08 | 0.4 | -0.54 | 1.1 | -0.41 | -0.8175 | -0.7475 | -0.765 | 0.914012 | 0.987766 | 0.44374 | 1 | 0.999631 | 0.972551 |
| Kif21b_Q9QXL1_S223s_1 | 1.43 | 0.35 | 0.75 | 0.71 | -1.34 | -0.62 | 0.95 | 0.98 | -0.39 | 0.02 | 0.4 | 0.22 | -0.04 | -0.49 | 0.5 | 0.21 | 1.375 | 2.69 | 1.0425 | 0.054891 | 0.172559 | 0.045793 | 1 | 0.999631 | 0.492066 |
| Kif21b_Q9QXL1_S969s_1 | -0.61 | 2.15 | 0.89 | -0.96 | 2.95 | 1.02 | 2.59 | 0.41 | 5.18 | 0.32 | 2.72 | 4.01 | 5.37 | 0.23 | 1.11 | -1.07 | 0.6225 | 0.71 | 1.0075 | 0.43748 | 0.118619 | 0.031366 | 1 | 0.999631 | 0.781057 |
| Kif21b_Q9QXL1_T1621t_1 | -1.42 | 0.58 | -0.48 | -1.22 | -0.63 | 0.51 | -0.23 | 0.3 | 0.89 | -2.48 | 0.5 | 1.39 | 0.59 | 0.95 | 0.93 | -0.98 | 0.13 | -0.165 | -0.5475 | 0.111711 | 0.224094 | 0.415738 | 1 | 0.999631 | 0.577068 |
| Kif2a_F8VQ42_S111s_1 | 0.94 | 0.41 | -0.11 | 0.27 | 0.89 | 0.47 | 0.37 | 0.3 | -0.28 | 0.13 | 0.77 | 0.23 | 0.53 | 0.68 | -0.76 | -1.13 | -0.5275 | 0.1425 | 0.6125 | 0.276683 | 0.675999 | 0.636632 | 1 | 0.999631 | 0.700187 |
| Kif2a_F8VQ42_S123s_1 | -0.81 | -1.09 | -0.71 | -0.72 | -1.86 | -1.46 | -1.02 | -1.1 | -1.04 | -0.85 | -0.56 | -0.31 | 0 | -0.8 | 0.12 | -0.2 | 0.12 | 0.1925 | 0.6025 | 0.07407 | 0.09039 | 0.613515 | 1 | 0.999631 | 0.539897 |
| Kif2a_F8VQ42_S190s_1 | -0.39 | 1.74 | -0.24 | -0.19 | 0.49 | 1.06 | -0.21 | 0.06 | 0.55 | 0.15 | 0.69 | 0.3 | 0.74 | 0.85 | 0.63 | 1.11 | -0.7025 | -1.6525 | -2.205 | 0.241568 | 0.82173 | 0.697999 | 1 | 0.999631 | 0.684831 |
| Kif2a_F8VQ42_S410s_1 | 1.12 | -1.06 | -0.22 | 0.25 | -2.32 | -0.6 | 1.92 | -1.72 | -2.47 | -0.05 | -2.32 | -1.68 | -2.93 | -2.17 | -1.88 | -1.75 | -0.3175 | -0.325 | 0.065 | 0.001368 | 0.443271 | 0.027128 | 1 | 0.999631 | 0.351308 |
| Kif2a_F8VQ42_S568s_1 | 0.03 | 0.3 | 0.39 | -0.52 | -0.24 | -0.28 | -0.25 | -0.3 | -0.13 | -0.42 | -0.39 | -0.16 | 0.39 | 0 | -0.59 | 0.66 | -0.02 | 0.0775 | 0.405 | 0.863375 | 0.27832 | 0.289481 | 1 | 0.999631 | 0.955107 |
| Kif2a_F8VQ42_S569s_1 | -0.58 | 0.12 | -0.27 | 0.27 | 0.04 | -0.19 | -0.26 | -0.13 | -0.12 | 0.02 | -0.46 | 0.41 | 0.42 | 0.08 | 0.04 | 0.62 | 0.415 | -0.04 | -0.185 | 0.224903 | 0.943421 | 0.809793 | 1 | 0.999631 | 0.67471 |
| Kif2a_P28740_S100s_1 | 1.2 | -0.49 | -1.2 | 0.01 | 0.23 | 0.69 | 0.16 | 0.1 | 0.56 | -0.25 | -0.31 | -0.64 | -0.29 | 0.63 | -1.02 | -0.54 | 0.48 | -0.09 | -0.3275 | 0.744239 | 0.407922 | 0.939684 | 1 | 0.999631 | 0.910233 |
| Kif3a_B1AQZ2_S713s_1 | 1.07 | 0.06 | 0.17 | 1.08 | 1.97 | 0.68 | 0.65 | 1 | 0.79 | -0.34 | 0.64 | 0.93 | -1.54 | 0.95 | 0.63 | 1.03 | -0.285 | -0.285 | 0.0875 | 0.592874 | 0.263025 | 0.825762 | 1 | 0.999631 | 0.853138 |
| Kif3b_Q61771_S681s_1 | -0.8 | 1.04 | 0.79 | -0.04 | 0.13 | -0.62 | -0.11 | 0.45 | -0.22 | -0.42 | 0.43 | 0.06 | -0.57 | 1.35 | 0.22 | 0.34 | 1.14 | 0.2125 | -0.5275 | 0.870789 | 0.53707 | 0.529942 | 1 | 0.999631 | 0.958735 |
| Kif3c_O35066_S768s_1 | 1.43 | 1.65 | 0.37 | 0.36 | 2.27 | 2.63 | 1.5 | 1.97 | 2.52 | 0.66 | 1.43 | 0.05 | -2.24 | 2.73 | 0.58 | 0.63 | 0.0275 | -0.2375 | 0.0025 | 0.571604 | 0.01977 | 0.715023 | 1 | 0.999631 | 0.843675 |
| Kif5a_P33175_S931s_1 | -0.27 | -1.49 | -0.51 | 0.27 | -0.16 | -0.68 | -1.02 | -0.03 | -0.32 | -0.47 | -1.49 | -0.67 | -1.16 | -0.01 | -0.04 | -0.78 | 0.12 | 0.1 | 0.4675 | 0.995654 | 0.948629 | 0.595852 | 1 | 0.999631 | 0.997705 |
| Kif5b_Q61768_S933s_1 | -0.6 | -1.09 | -0.1 | -0.35 | -0.43 | -0.46 | -0.22 | -0.55 | -0.38 | -0.06 | -1.1 | -0.2 | -0.35 | -0.08 | 0.28 | -0.12 | 0.3475 | 0.0075 | 0.635 | 0.173141 | 0.682522 | 0.776326 | 1 | 0.999631 | 0.642116 |
| Kif5b_Q61768_S945s_1 | 0.03 | 0.03 | -0.48 | 0.17 | 0.89 | 1 | -0.03 | -0.72 | 0.08 | -0.16 | -0.45 | 0.31 | 1.07 | 0.09 | 0.82 | 0.31 | 0.1525 | 0.165 | 0.725 | 0.08257 | 0.421933 | 0.9798 | 1 | 0.999631 | 0.548531 |
| Kif5c_P28738_S934s_1 | -0.86 | -1.72 | -1.14 | -1.32 | -1.06 | -0.79 | -0.84 | -1.74 | -0.96 | 0.05 | -2.21 | -1.26 | -0.49 | -0.55 | -0.13 | -0.97 | 0.4275 | 0.3125 | 0.3775 | 0.047223 | 0.642609 | 0.73132 | 1 | 0.999631 | 0.480102 |
| Kif5c_P28738_T396t_1 | -0.78 | -0.17 | -0.29 | -0.39 | 0.34 | 0.11 | -0.42 | 0.05 | -0.87 | 0.32 | -0.26 | 0.43 | 0.16 | 1.05 | -0.69 | -0.64 | -0.2625 | -0.0775 | 0.16 | 0.395492 | 0.155732 | 0.395263 | 1 | 0.999631 | 0.760219 |
| Kif7_J3QNW9_S662s_1 | 0.44 | -0.2 | 0.42 | 0.22 | -0.03 | -0.24 | -0.03 | 0.13 | -0.03 | 0.18 | -0.09 | 0.51 | 0.66 | -0.04 | 0.37 | 0.53 | -0.235 | -0.1 | -0.315 | 0.606258 | 0.334762 | 0.788548 | 1 | 0.999631 | 0.858173 |
| Kifap3_P70188_S102s_1 | 0.27 | 0.69 | 0.63 | 1.05 | 0.14 | 0.09 | 0.79 | 0.68 | 0.64 | 0.52 | 0.71 | 0.37 | 0.55 | 0.16 | 0.6 | 0.07 | -0.01 | 0.5775 | 0.6575 | 0.317456 | 0.44428 | 0.718918 | 1 | 0.999631 | 0.721033 |
| Kifap3_P70188_S60s_1 | -0.58 | 0.73 | -0.23 | -0.06 | -0.22 | 0.43 | -0.32 | -0.07 | 0.4 | 0.2 | 1.16 | 0.41 | 0.39 | 0.65 | 0.41 | 1.04 | 0.08 | 0.365 | -0.0575 | 0.094397 | 0.97729 | 0.146253 | 1 | 0.999631 | 0.558128 |
| Kifc2_G3X8Q6_S643s_1 | 0.9 | 0.55 | 0.9 | 0.28 | 0.09 | 0.65 | 0.83 | 1.38 | 0.15 | 0.84 | 1.74 | 1.36 | -0.56 | 0.47 | 1.29 | 1.2 | -0.4925 | -0.3675 | 0.265 | 0.898682 | 0.814329 | 0.362944 | 1 | 0.999631 | 0.967127 |
| Kit_P05532_S185s_1 | -0.28 | -0.12 | -0.03 | -0.23 | -1.42 | -0.57 | -0.49 | -0.15 | -1.25 | -0.98 | -0.23 | 0.33 | 0.64 | -0.74 | 0.26 | 0.24 | -0.9025 | -0.9475 | -1.375 | 0.460778 | 0.149925 | 0.350329 | 1 | 0.999631 | 0.791962 |
| Kiz_Q3UXL4_S454s_1 | 2 | 0.74 | 1.66 | 0.32 | -0.39 | 0.53 | 1.09 | -0.12 | 0.54 | 1.59 | -0.58 | -0.62 | -1.59 | 0.07 | 1.1 | -0.36 | 0.4975 | 0.9975 | 1.215 | 0.046773 | 0.087717 | 0.137293 | 1 | 0.999631 | 0.480102 |
| Klc1_Q5UE59_S162s_1 | -1.29 | 1.66 | 1.33 | -0.32 | 0.82 | 1.5 | 0.71 | 0.34 | 1.82 | 0.63 | 1.21 | 1.71 | 2.84 | 1.58 | 1.15 | 0.67 | -0.0625 | 0.2775 | 0.6 | 0.123363 | 0.45587 | 0.15949 | 1 | 0.999631 | 0.594166 |
| Klc1_Q5UE59_S460s_1 | -0.42 | 0.37 | -0.27 | -0.58 | -0.21 | 0.07 | -0.73 | -0.28 | 0.11 | 0.07 | -0.27 | 0.3 | 0.9 | 0.43 | 0.03 | 0.14 | -0.205 | -0.1225 | -0.0275 | 0.106623 | 0.843529 | 0.378654 | 1 | 0.999631 | 0.568797 |
| Klc1_Q5UE59_S521s_1 | -0.61 | -0.87 | 0.32 | -0.17 | -0.71 | -1.03 | -0.02 | -0.39 | -0.56 | -0.94 | -0.21 | -0.11 | -0.36 | -1 | -0.5 | 0.42 | 0.1125 | 0.3075 | 0.165 | 0.94665 | 0.574194 | 0.732091 | 1 | 0.999631 | 0.982771 |
| Klc1_Q5UE59_S524s_1 | -1.32 | -0.2 | -0.63 | -0.93 | -0.35 | -0.45 | -0.45 | -1.38 | -0.09 | -0.5 | -0.66 | -0.6 | -0.14 | -0.37 | -1.14 | -0.77 | -0.6225 | 0.19 | -0.9675 | 0.655412 | 0.756085 | 0.355785 | 1 | 0.999631 | 0.879056 |
| Klc1_Q5UE59_S534s_1 | -0.08 | 0.27 | 0.56 | 0.06 | -0.32 | -0.36 | 0.1 | -1.1 | 0.21 | 0.87 | 0.33 | 0.16 | -1.81 | -1.14 | -0.21 | 0.1 | 0.4625 | 0.675 | -0.21 | 0.055692 | 0.082375 | 0.524404 | 1 | 0.999631 | 0.495418 |

| | | | | | | | | | | | | | | | | | | | | | | | | | |
|---|---|---|---|---|---|---|---|---|---|---|---|---|---|---|---|---|---|---|---|---|---|---|---|---|---|
| Klc2_D3YXZ3_S151s_1 | 1.69 | 1.87 | 0.75 | 0.1 | 1.87 | 3.59 | 0.14 | 0.66 | 2.99 | 3.04 | 0.48 | 0.6 | 0.88 | 1.98 | 0.57 | 0.14 | 0.2725 | 0.1025 | 0.36 | 0.69591 | 0.548651 | 0.367451 | 1 | 0.999631 | 0.895683 |
| Klc2_D3YXZ3_S443s_1 | -0.04 | 0.05 | -0.54 | 0.46 | 0.29 | 0.57 | -0.44 | 0.6 | -0.33 | 0.5 | -0.18 | 0.35 | -1.4 | 1.46 | 0.64 | 0.67 | 1.0075 | 0.245 | -0.5475 | 0.545374 | 0.440881 | 0.76056 | 1 | 0.999631 | 0.833826 |
| Klc2_D3YXZ3_S551s_1 | 0.48 | 0.13 | -1.47 | 0.72 | 1.87 | 0.72 | 0.85 | 0.45 | 1.14 | -0.38 | 0.63 | -0.55 | -1.97 | 0.93 | -0.6 | -0.69 | -0.475 | -0.075 | 0.465 | 0.432643 | 0.085732 | 0.676257 | 1 | 0.999631 | 0.779247 |
| Klc2_D3YXZ3_S552s_1 | 0.32 | -0.1 | 0.7 | 0.84 | 1.13 | 0.18 | -2.4 | 0.95 | 1.18 | -0.57 | 0.34 | 0.51 | 1.07 | 0.91 | 1.33 | 0.31 | -0.065 | 0.445 | 0.5275 | 0.210298 | 0.525534 | 0.858542 | 1 | 0.999631 | 0.66441 |
| Klc2_D3YXZ3_S577s_1 | 0.94 | 1.4 | 2.35 | 2.9 | 1.49 | 1.64 | 1.22 | 2.98 | 1.81 | 1.31 | 2.99 | 3.26 | 0.72 | 2.11 | 3.44 | 3.43 | 0.5325 | -0.16 | -0.035 | 0.456242 | 0.905047 | 0.456897 | 1 | 0.999631 | 0.790326 |
| Klc2_D3YXZ3_S584s_1 | -0.68 | -0.06 | -2.35 | -0.58 | 0.63 | -0.41 | -1.08 | -0.68 | 0.19 | -1.03 | -1.44 | -2.03 | 0.25 | -0.74 | -1.82 | -1.5 | -0.165 | -0.4125 | -0.0175 | 0.954513 | 0.355819 | 0.796326 | 1 | 0.999631 | 0.984441 |
| Klc2_D3YXZ3_S605s_1 | 1.59 | 0.9 | 0.47 | 0.7 | 0.92 | 1 | 0.27 | 0.81 | 0.6 | 0.63 | 0.38 | 0.4 | 0.56 | 1.5 | 0.57 | 0.96 | 0.0625 | 0.0575 | 0.1975 | 0.962282 | 0.621622 | 0.209616 | 1 | 0.999631 | 0.986642 |
| Klc2_D3YXZ3_S606s_1 | -0.04 | 0.19 | -0.37 | -0.09 | 0.06 | 0.38 | -0.46 | -0.04 | 0.1 | 0.01 | -0.49 | 0.3 | -0.54 | 0.48 | 0.24 | 0.3 | -0.0725 | 0.425 | 0.1375 | 0.551449 | 0.826307 | 0.842788 | 1 | 0.999631 | 0.836275 |
| Klc4_Q9DBS5_S590s_1 | -0.08 | 0.6 | 0.25 | 0.39 | 0.33 | 0.51 | -0.14 | 0.17 | 0.58 | 0.24 | 1.04 | 1 | 0.8 | 0.36 | -0.07 | 0.62 | -0.375 | -0.4725 | -0.145 | 0.668827 | 0.795855 | 0.188044 | 1 | 0.999631 | 0.883535 |
| Klc4_Q9DBS5_S611s_1 | -0.22 | 0.32 | -0.01 | -0.16 | -0.35 | -0.37 | -0.52 | -0.33 | -0.67 | -0.8 | -0.1 | -0.39 | 0.15 | 0.01 | -0.64 | -0.17 | 0.3675 | 0.465 | 0.83 | 0.638862 | 0.156529 | 0.123857 | 1 | 0.999631 | 0.870943 |
| Klf16_P58334_S103s_1 | -0.82 | 0.31 | -1.58 | -0.23 | 0.09 | -0.21 | -1.03 | 0.3 | 0.29 | 0.53 | -1.1 | -0.18 | 0.56 | 1.39 | -0.75 | -0.2 | -0.15 | -0.0125 | 0.125 | 0.167081 | 0.446205 | 0.373683 | 1 | 0.999631 | 0.636719 |
| Klf3_Q60980_S249s_1 | 1.09 | 1.9 | 1.08 | 0.75 | 0.86 | 1.81 | 0.38 | 1.17 | 1.88 | 1.78 | 1.15 | -0.04 | 1.51 | 2.28 | 0.76 | 0.77 | 0.0325 | 0.03 | -0.0775 | 0.77684 | 0.704234 | 0.979356 | 1 | 0.999631 | 0.921293 |
| Klhdc7a_A2APT9_S361s_1 | -0.06 | 0.72 | 0.42 | 0.2 | 0.49 | 0.44 | -0.13 | 0.61 | 0.16 | 0.05 | 0.36 | 0.83 | -0.38 | 0.57 | 0.14 | 0.64 | -0.07 | -0.385 | -0.5225 | 0.823403 | 0.913152 | 0.922396 | 1 | 0.999631 | 0.938864 |
| Klhl22_D3YZH5_S18s_1 | 0.02 | -0.5 | -1.05 | 0.02 | -0.82 | -0.27 | -0.15 | -0.55 | -0.95 | 0.2 | -0.09 | -2.21 | -0.6 | 0.42 | -1.65 | -1.77 | -0.305 | 0.31 | -0.0525 | 0.342027 | 0.834361 | 0.492257 | 1 | 0.999631 | 0.73347 |
| Klhl26_Q8BGY4_S13s_1 | 0.91 | 0.01 | 0.37 | -0.48 | 0.48 | -1.07 | -0.21 | 0.39 | 0.68 | 2.02 | -1.04 | 0.39 | 0.61 | 1.25 | -0.77 | -0.49 | -0.25 | -0.115 | -0.29 | 0.920188 | 0.50052 | 0.622841 | 1 | 0.999631 | 0.97499 |
| Klhl9_Q6ZPT1_S603s_1 | 0.27 | 0.28 | 0.16 | 0.15 | 0.14 | -0.02 | -0.5 | 0.24 | 0.17 | 0.19 | -0.46 | 0.5 | 0.62 | -0.1 | -1.36 | 0.54 | -0.3975 | 0.47 | 0.3225 | 0.528525 | 0.35776 | 0.692986 | 1 | 0.999631 | 0.825257 |
| Kmt2a_P55200_S2539s_1 | -0.9 | -1.62 | -1.16 | -1.07 | -1.67 | -1.44 | -1.03 | -2.2 | -1.39 | -0.92 | -0.12 | -0.44 | -0.62 | -0.45 | -0.52 | -1.87 | -0.365 | 0.0225 | -0.055 | 0.424055 | 0.246168 | 0.205199 | 1 | 0.999631 | 0.773835 |
| Kmt2a_P55200_S2931s_1 | 0.84 | -0.87 | 1.14 | 0.42 | -0.44 | -0.44 | 0.55 | 0.4 | -0.69 | 0.68 | 1.16 | 0.47 | -0.06 | -0.37 | 0.87 | 0.87 | -0.22 | -0.5825 | -0.2325 | 0.915253 | 0.460645 | 0.967261 | 1 | 0.999631 | 0.97286 |
| Kmt2a_P55200_S3029s_1 | -0.56 | 0.41 | -0.01 | -0.18 | -0.76 | -0.88 | 0.34 | 0.08 | -0.75 | -1.29 | -0.33 | -0.3 | -1.22 | -0.08 | 0.77 | -0.74 | -0.365 | 0.135 | 0.09 | 0.620866 | 0.564084 | 0.118323 | 1 | 0.999631 | 0.861888 |
| Kmt2b_F6W623_S158s_1 | 0.65 | 1.3 | 1.43 | 0.87 | 0.93 | 0.46 | 0.15 | 1.25 | 1.03 | 0.9 | 1.45 | 1.41 | 1.64 | 0.45 | 1.05 | 1.47 | 0.0325 | -0.2325 | 0.005 | 0.806305 | 0.297637 | 0.656739 | 1 | 0.999631 | 0.935204 |
| Kmt2d_Q6PDK2_S2318s_1 | -0.18 | -0.69 | 1 | 0.42 | 0.32 | -0.49 | 0.17 | 0.68 | -0.98 | -0.77 | 0.39 | 0.98 | 0.11 | 0.24 | 0.19 | 0.03 | -0.4325 | 0.0275 | -0.2925 | 0.989913 | 0.939934 | 0.674152 | 1 | 0.999631 | 0.995892 |
| Kmt2d_Q6PDK2_S3618s_1 | 0.09 | 2.72 | 1.03 | -0.05 | 0.63 | 1.39 | -0.82 | 0.86 | 2.39 | 0.84 | -0.7 | 1.37 | 0.9 | 0.91 | 1.46 | -0.65 | -0.3275 | -0.145 | 0.12 | 0.675055 | 0.541815 | 0.972416 | 1 | 0.999631 | 0.885588 |
| Kmt2d_Q6PDK2_S4410s_1 | 0.63 | 1.65 | 1.83 | 1.41 | 0.84 | 1.18 | 0.7 | 1.49 | 1.1 | 0.99 | 1.13 | 1.72 | 0.84 | 1.84 | 1.39 | 1.93 | -0.4625 | -0.1225 | 0.3425 | 0.760156 | 0.358623 | 0.680183 | 1 | 0.999631 | 0.914614 |
| Kmt2d_Q6PDK2_S4789s_1 | 0.97 | 2.07 | 2.28 | 1.8 | 1.32 | 1.05 | 1.15 | 1.75 | 1.26 | 1.08 | 2.06 | 2.23 | 1.39 | 2.14 | 2.54 | 2.42 | -0.935 | -0.74 | -1.3725 | 0.408404 | 0.210915 | 0.766703 | 1 | 0.999631 | 0.766996 |
| Kmt2e_Q3UG20_S263s_1 | 2.46 | 1.64 | 2.76 | 2.63 | 0.94 | 1.25 | 2.1 | 1.46 | 0.81 | 2.89 | 1.88 | 0.95 | 0.91 | 0.84 | 1.69 | 0.56 | 0.0775 | 0.0575 | 0.1075 | 0.004752 | 0.028488 | 0.171308 | 1 | 0.999631 | 0.351308 |
| Kndc1_Q0KK55_S794s_1 | -0.81 | -0.95 | 0.03 | -0.27 | -0.75 | -0.08 | -0.54 | -0.32 | -0.29 | -0.46 | -1.06 | 0.04 | -0.43 | 0.01 | -0.47 | -0.68 | -0.4075 | -0.7325 | -1.64 | 0.751085 | 0.808458 | 0.872768 | 1 | 0.999631 | 0.912766 |
| Kndc1_Q0KK55_S843s_1 | 2.34 | 0.83 | 0.92 | 0.44 | 0.23 | 0.59 | 1.54 | 0.54 | -0.03 | 1.12 | 0.18 | 0.33 | -1.27 | -0.37 | 0.48 | -0.87 | -0.8225 | -0.57 | 0.63 | 0.010567 | 0.402474 | 0.142835 | 1 | 0.999631 | 0.383871 |
| Kndc1_Q0KK55_S951s_1 | -2.26 | -0.67 | 0.25 | -0.75 | -3.11 | 0.27 | -1.03 | -2.85 | -0.74 | -0.96 | -1.85 | -2.16 | 0.39 | -0.51 | 0 | -0.79 | -0.32 | -0.2725 | -0.175 | 0.262987 | 0.333846 | 0.331889 | 1 | 0.999631 | 0.693034 |
| Kndc1_Q0KK55_S951s_2 | -0.87 | -0.09 | 0.24 | 0.49 | -1.08 | -0.4 | 0.46 | -0.49 | -0.21 | -1.19 | 0.37 | -0.29 | -1.12 | -0.07 | -0.01 | 0.27 | 0.075 | -0.65 | -0.5325 | 0.686603 | 0.460163 | 0.534952 | 1 | 0.999631 | 0.890911 |
| Kndc1_Q0KK55_S967s_1 | -0.66 | -0.31 | -0.82 | -0.42 | -1.8 | -0.95 | -0.1 | 0.94 | -1.19 | -1.58 | -1.6 | -0.44 | -1.04 | -0.8 | -1.52 | -0.98 | -0.32 | -0.2725 | -0.175 | 0.099275 | 0.891209 | 0.080502 | 1 | 0.999631 | 0.562298 |
| Kndc1_Q0KK55_S967s_2 | -0.87 | -0.09 | 0.24 | 0.49 | -1.08 | -0.4 | 0.46 | -0.49 | -0.21 | -1.19 | 0.37 | -0.29 | -1.12 | -0.07 | -0.01 | 0.27 | -0.0875 | 0.2475 | 0.085 | 0.686603 | 0.460163 | 0.534952 | 1 | 0.999631 | 0.890911 |
| Kpna6_O35345_S6s_1 | 1.2 | 0.94 | 1.05 | 0.94 | 0.57 | 0.88 | 1.2 | 1.13 | 0.98 | 1.57 | 1.33 | 1.24 | 0.68 | 1.09 | 1.22 | 1.48 | -0.4525 | 0.1975 | -0.55 | 0.772181 | 0.73732 | 0.35447 | 1 | 0.999631 | 0.919605 |
| Kri1_V9GXJ9_S24s_1 | 0.36 | 2.35 | -0.09 | -0.27 | 0.23 | -0.6 | 0.21 | 0.7 | 0.17 | 0.72 | 1.02 | 1.23 | 1.14 | -0.76 | 0.47 | -0.7 | -0.51 | -0.13 | -0.595 | 0.42483 | 0.455231 | 0.738092 | 1 | 0.999631 | 0.774276 |
| Krit1_Q6S5J6_S22s_1 | 1.77 | 1.4 | 1.17 | 0.99 | 0.67 | 0.89 | 1.23 | 0.5 | 1.51 | 2.06 | 0.52 | 0.72 | 0.99 | 0.78 | 1.04 | 0.14 | -0.69 | -2.02 | -0.5975 | 0.100463 | 0.11017 | 0.748579 | 1 | 0.999631 | 0.56376 |
| Krt8_P11679_S24s_1 | 0.73 | 1.68 | 3.01 | 0.73 | 0.97 | 0.59 | 1.29 | 0.54 | 1.2 | -0.65 | -4.04 | 1.56 | 0.7 | 0.48 | 1.04 | 1.54 | -1.5625 | 0.2975 | 0.64 | 0.289477 | 0.209694 | 0.114632 | 1 | 0.999631 | 0.706164 |
| Krtap9-1_Q64526_S4s_1 | 0.08 | -1.96 | 2.55 | 1 | -1.08 | -0.94 | -0.64 | -1.92 | -1.68 | 0.61 | 1.53 | 2.4 | -1.69 | 4.34 | 0.62 | 0.96 | 0.6325 | -0.25 | -0.53 | 0.628309 | 0.091805 | 0.788239 | 1 | 0.999631 | 0.865998 |
| Ksr1_Q5SXE4_S320s_1 | 1.07 | 0.24 | -0.98 | 1.42 | 1.49 | 0.5 | 1.31 | 0.98 | 0.39 | -0.38 | 0.93 | -0.19 | -1.03 | 0.46 | -0.37 | 0.57 | 0.095 | -0.1575 | -0.235 | 0.383579 | 0.252265 | 0.658456 | 1 | 0.999631 | 0.754618 |
| Ksr1_Q5SXE4_S392s_1 | 0.69 | 0.49 | 0.09 | 0.68 | 0.43 | 0.47 | 0.59 | 0.84 | 0.35 | -0.1 | 1.13 | -0.06 | -0.28 | 0.43 | 0.12 | 0.74 | -0.2125 | 0.555 | 0.805 | 0.484228 | 0.721367 | 0.658349 | 1 | 0.999631 | 0.803422 |
| Ksr2_M0QW59_T273t_1 | -0.66 | 0.5 | -0.46 | 0.35 | -0.52 | 0.41 | 0.12 | -1.13 | 1.32 | 0.53 | -0.15 | 0.25 | 1.61 | 0.66 | 0.48 | 0.2 | 1.265 | 0.27 | -1.0125 | 0.084963 | 0.629329 | 0.21223 | 1 | 0.999631 | 0.550457 |
| Ksr2_M0QW59_T291t_1 | 1.74 | -1.61 | -1.41 | 1.11 | 1.55 | 0.61 | 1.33 | 1.4 | 0.29 | 0.05 | 1.17 | -0.6 | -2.7 | -0.25 | -0.79 | -0.48 | -0.105 | -0.5975 | 0.0425 | 0.268266 | 0.126521 | 0.741644 | 1 | 0.999631 | 0.696361 |
| Kti12_Q9D1R2_T156t_1 | 0 | -0.18 | -0.43 | -0.71 | 0.01 | 0.1 | -1.36 | -0.49 | -0.31 | -0.49 | -0.95 | -1.96 | -0.61 | 1.06 | -0.52 | -1.08 | 0.165 | 0.455 | 0.5425 | 0.929033 | 0.783299 | 0.166809 | 1 | 0.999631 | 0.977175 |
| Ktn1_Q61595_S124s_1 | 0.02 | -0.09 | 1 | 0.37 | 0.16 | 0.16 | 0.4 | 1.24 | 0.49 | 0.45 | 1.39 | 0.79 | 0.44 | 0.68 | 1.37 | 0.98 | -0.6175 | 0.0575 | -1.055 | 0.158618 | 0.659088 | 0.226175 | 1 | 0.999631 | 0.628696 |
| Ktn1_Q61595_S75s_1 | 1.77 | 0.42 | 2.39 | 0.4 | -0.47 | 0.66 | 1.38 | 0.94 | 0.58 | 2.66 | 1.02 | 0.95 | -0.03 | -0.22 | 0.26 | 0.75 | -0.725 | 0.1625 | -0.5125 | 0.063024 | 0.300109 | 0.92566 | 1 | 0.999631 | 0.512978 |
| L1cam_Q6PGJ3_S1165s_1 | 1.17 | 2.23 | 2.38 | 0.91 | 0.43 | 2.09 | 0.89 | 0.38 | 1.51 | 3 | 1.35 | 1.48 | 0.74 | 1.3 | 1.76 | 0.84 | -0.8575 | -0.1575 | -0.915 | 0.261885 | 0.175541 | 0.749509 | 1 | 0.999631 | 0.692758 |
| L1cam_Q6PGJ3_S1180s_1 | -0.12 | -1.34 | 0.09 | 0.29 | -1.56 | -0.57 | -0.32 | -2.06 | -1.28 | 0.04 | 0.24 | -0.71 | -0.06 | -1.26 | -0.37 | -3.05 | 0.1875 | 0.2775 | 0.1675 | 0.199627 | 0.119098 | 0.745866 | 1 | 0.999631 | 0.657804 |
| L1cam_Q6PGJ3_S1180s_2 | -1.11 | 0.2 | -0.05 | -0.11 | -0.28 | -0.09 | -0.02 | 0.07 | 0.11 | -0.19 | 0.29 | -0.17 | 0.9 | -1.04 | -0.12 | -0.14 | -0.355 | 0.1075 | -0.3275 | 0.726379 | 0.579661 | 0.435781 | 1 | 0.999631 | 0.90568 |
| L1cam_Q6PGJ3_S1183s_1 | 0.53 | -1.24 | 0.23 | -0.55 | -0.77 | -0.26 | -0.25 | -1.17 | -0.57 | 0.37 | -0.05 | -0.35 | -0.52 | -1.51 | 0.25 | -0.56 | 0.1875 | 0.2775 | 0.1675 | 0.525876 | 0.43105 | 0.808027 | 1 | 0.999631 | 0.823594 |
| L1cam_Q6PGJ3_S1183s_2 | -1.11 | 0.2 | -0.05 | -0.11 | -0.28 | -0.09 | -0.02 | 0.07 | 0.11 | -0.19 | 0.29 | -0.17 | 0.9 | -1.04 | -0.12 | -0.14 | -0.27 | -0.1675 | 0.215 | 0.726379 | 0.579661 | 0.435781 | 1 | 0.999631 | 0.90568 |
| L1cam_Q6PGJ3_S1250s_1 | -0.84 | 0.28 | -0.08 | -0.63 | 0.17 | -0.07 | -1.2 | -1.25 | -0.23 | -0.47 | -0.7 | -0.54 | 0.48 | 0.21 | -0.53 | -0.57 | 0.8625 | 1.4925 | 1.1525 | 0.588902 | 0.542672 | 0.611533 | 1 | 0.999631 | 0.851154 |
| L1cam_Q6PGJ3_T1174t_1 | -0.72 | 5.2 | 6.7 | 2.89 | 3.82 | 4.62 | 4.38 | 4.7 | 5.06 | 0.46 | 6.88 | 7.64 | 4.86 | 3.15 | 6.27 | 4.4 | 0.08 | 0.7875 | 1.06 | 0.437538 | 0.536877 | 0.446534 | 1 | 0.999631 | 0.781057 |
| Lactb2_Q99KR3_T279t_1 | -0.09 | 0.42 | 0.46 | 0.8 | 0.45 | 0.13 | 0.29 | 1.04 | 0.95 | 0.6 | 1.43 | 1.76 | 1.76 | 1.11 | 1.67 | 1.29 | 0.115 | 0.19 | -0.8325 | 0.007572 | 0.8022 | 0.046961 | 1 | 0.999631 | 0.358173 |
| Lad1_P57016_T19t_1 | 1.02 | 1.94 | 1.28 | 2.18 | 2.03 | 1.72 | 1.3 | 1.83 | 1.45 | 1.84 | 2.39 | 1.5 | -0.01 | 1.57 | 1.07 | 0.46 | 0.145 | 0.1775 | 0.34 | 0.084058 | 0.739526 | 0.61335 | 1 | 0.999631 | 0.549732 |
| Lamtor1_Q9CQ22_T28t_1 | -0.37 | -0.1 | -0.95 | -1.57 | -0.03 | -0.02 | -1.18 | -1.18 | -0.4 | 0.21 | -1.24 | -0.85 | 0.31 | 0.25 | -1.36 | -0.83 | 0.255 | -0.0375 | 0.0425 | 0.503464 | 0.748147 | 0.690822 | 1 | 0.999631 | 0.814092 |
| Lancl2_F6RJV6_S302s_1 | -0.54 | 0.11 | -0.18 | -1.38 | -0.08 | -0.31 | 0.01 | -0.59 | -0.07 | -0.82 | -1.44 | 0.19 | -0.1 | 0.15 | 0.13 | -2 | -0.135 | -0.615 | -0.5425 | 0.939916 | 0.494606 | 0.93681 | 1 | 0.999631 | 0.981146 |
| Larp1_Z4YJT3_S1023s_1 | 1.98 | 0.87 | 0.96 | -0.51 | 1.14 | 0.48 | 1.05 | 0.09 | 1.58 | 0.62 | 0.07 | -1.43 | 1.45 | -0.01 | 0.52 | -0.83 | -0.3325 | 0.16 | -1.0175 | 0.399346 | 0.797704 | 0.402012 | 1 | 0.999631 | 0.762042 |
| Larp1_Z4YJT3_S494s_1 | 1.44 | 0.64 | 1.48 | 1.04 | 0.38 | 0.79 | 1.39 | 0.71 | 1.14 | 1.86 | 1.28 | 0.96 | 0.38 | -0.51 | 0.72 | -0.06 | -0.3775 | -0.4225 | -1.47 | 0.019786 | 0.326341 | 0.629307 | 1 | 0.999631 | 0.391668 |
| Larp1_Z4YJT3_S494s_2 | 2.09 | 0.7 | 1.54 | 1.9 | 0.7 | 0.89 | 1.97 | 1.16 | 0.74 | 1.89 | 1.31 | 0.6 | -0.86 | 0.13 | 0.75 | 0.33 | -0.53 | -0.0025 | -0.62 | 0.008619 | 0.372652 | 0.334906 | 1 | 0.999631 | 0.365244 |
| Larp1_Z4YJT3_S498s_1 | 1.16 | -0.41 | 0.36 | 0.03 | -0.49 | -0.5 | 0.52 | -0.51 | 0.17 | 1.47 | -0.45 | -0.06 | 0.17 | -1.12 | 0.09 | -0.48 | -0.3775 | -0.4225 | -1.47 | 0.186379 | 0.222017 | 0.996036 | 1 | 0.999631 | 0.646474 |
| Larp1_Z4YJT3_S498s_2 | 2.09 | 0.7 | 1.54 | 1.9 | 0.7 | 0.89 | 1.97 | 1.16 | 0.74 | 1.89 | 1.31 | 0.6 | -0.86 | 0.13 | 0.75 | 0.33 | -0.2425 | 0.11 | 0.3575 | 0.008619 | 0.372652 | 0.334906 | 1 | 0.999631 | 0.365244 |
| Larp1_Z4YJT3_S68s_1 | 0.77 | 0.75 | 2.02 | 0.67 | -0.17 | 1.97 | 0.51 | 0.93 | 0.54 | 1.78 | 1 | 1.33 | 0.73 | 1.36 | 1.88 | 1.67 | -0.1775 | 0.2675 | 0.3375 | 0.405231 | 0.639188 | 0.793434 | 1 | 0.999631 | 0.765844 |
| Larp1_Z4YJT3_S68s_2 | 0.18 | 1.58 | 1.91 | 0.95 | 0.56 | 1.72 | 0.55 | 1.08 | 1.02 | 1.78 | 1.15 | 1.74 | 0.54 | 2.02 | 1.61 | 1.8 | 0.3025 | -0.075 | 0.0625 | 0.493113 | 0.696714 | 0.536873 | 1 | 0.999631 | 0.80955 |
| Larp1_Z4YJT3_S743s_1 | 0.42 | 1.3 | 0.22 | 0.7 | 1.43 | 1.31 | -0.11 | 1.22 | 1.17 | 0.46 | 0.29 | 0.42 | 0.19 | 1.65 | 0.7 | 0.35 | -0.455 | -0.0475 | 0.2425 | 0.880948 | 0.4816 | 0.829417 | 1 | 0.999631 | 0.963364 |
| Larp1_Z4YJT3_S751s_1 | -0.38 | 0.21 | -0.07 | 0.25 | -0.27 | -0.67 | -0.63 | -0.24 | -0.39 | -0.11 | 0.08 | 0.24 | -0.23 | 0.25 | 0.64 | 0.32 | -0.185 | 0.1575 | 0.1725 | 0.451791 | 0.120954 | 0.868853 | 1 | 0.999631 | 0.788148 |
| Larp1_Z4YJT3_S81s_1 | 0.1 | 0.5 | 0.14 | 0.11 | -0.09 | 0.19 | -0.17 | 0.18 | 0.1 | 0.65 | 0.33 | 0.4 | 0.54 | 0.43 | 0.31 | 0.26 | -0.1775 | 0.2675 | 0.3375 | 0.528746 | 0.468619 | 0.55665 | 1 | 0.999631 | 0.825257 |
| Larp1_Z4YJT3_S81s_2 | 0.18 | 1.58 | 1.91 | 0.95 | 0.56 | 1.72 | 0.55 | 1.08 | 1.02 | 1.78 | 1.15 | 1.74 | 0.54 | 2.02 | 1.61 | 1.8 | -0.2725 | 0.2175 | 0.6775 | 0.493113 | 0.696714 | 0.536873 | 1 | 0.999631 | 0.80955 |
| Larp1_Z4YJT3_S828s_1 | -0.69 | 0.2 | 0.6 | -0.63 | -0.26 | -0.18 | -0.92 | -0.25 | 0.07 | -0.11 | -0.21 | 0.6 | 1.15 | 0.6 | 0.21 | 0.23 | 0.66 | 0.34 | -0.1775 | 0.119546 | 0.475336 | 0.574826 | 1 | 0.999631 | 0.587113 |
| Larp1_Z4YJT3_T503t_1 | 1.67 | 0.3 | 0.05 | 1.56 | 1.84 | 1.03 | 1.65 | 1.7 | 1.46 | 1 | 1.63 | 0.85 | -0.71 | 1.37 | 1.16 | 1.05 | 0.02 | 0.2975 | 0.63 | 0.761794 | 0.159543 | 0.456511 | 1 | 0.999631 | 0.914904 |
| Larp1_Z4YJT3_T626t_1 | -1.08 | 0.48 | 0.68 | 0.07 | 0.54 | -0.18 | -0.37 | 0.24 | 0.62 | 0.05 | 0.09 | 0.58 | 1.09 | 0.55 | 0.6 | 0.43 | -0.2425 | 0.445 | 0.62 | 0.163904 | 0.963219 | 0.488693 | 1 | 0.999631 | 0.632208 |
| Larp1b_Q3TQS1_S60s_1 | 0.22 | 0.14 | -0.31 | -1.47 | -0.5 | -0.48 | -1.04 | -0.37 | -0.07 | -0.51 | 1.46 | -0.52 | 1.2 | -0.41 | 0.4 | -0.13 | -0.305 | -0.1175 | 0.175 | 0.235909 | 0.562708 | 0.436257 | 1 | 0.999631 | 0.683301 |
| Larp4_E9Q066_S578s_1 | -0.88 | 0.59 | 0.93 | 1.15 | 0.15 | -0.03 | -0.18 | 0.63 | 0.19 | -0.34 | 0.24 | 1.23 | 0.38 | 0.08 | 0.93 | 1.1 | 0.04 | -0.6175 | -0.3475 | 0.724343 | 0.518992 | 0.823473 | 1 | 0.999631 | 0.905674 |
| Larp4b_Q6A0A2_S517s_1 | -0.55 | -1.12 | 0.1 | 0.48 | -0.28 | -0.44 | -0.49 | 0.28 | -1.43 | -1.29 | -0.43 | -0.41 | -2.01 | -0.77 | 0.09 | 0.21 | 0.065 | -0.03 | 0.2675 | 0.548017 | 0.920229 | 0.183742 | 1 | 0.999631 | 0.834507 |
| Larp4b_Q6A0A2_S519s_1 | 0.43 | 0.73 | 0.68 | 0.77 | 0.91 | 0.83 | 0.35 | 0.78 | 0.56 | 0.41 | 0.62 | 0.9 | 0.61 | 0.95 | 0.72 | 1.4 | -0.6925 | -0.8775 | -0.9925 | 0.380348 | 0.800374 | 0.907342 | 1 | 0.999631 | 0.752364 |
| Larp4b_Q6A0A2_S637s_1 | 2.09 | 0.72 | 1.77 | 1.17 | -0.04 | 0.59 | 1.35 | 1.08 | 0.34 | 1.66 | 0.41 | -0.17 | 0.16 | 0.58 | 0.91 | 0.13 | 0.3775 | -0.275 | -1.515 | 0.026788 | 0.128326 | 0.090335 | 1 | 0.999631 | 0.422285 |
| Larp4b_Q6A0A2_S655s_1 | 0.59 | -0.73 | 0.47 | 0.19 | 0.84 | -0.59 | 1.8 | -0.02 | 0.22 | -0.21 | -0.36 | -0.23 | -1.11 | -1.41 | -1.31 | -1.71 | -0.9 | -1.035 | -1.14 | 0.002012 | 0.498804 | 0.449991 | 1 | 0.999631 | 0.351308 |
| Larp4b_Q6A0A2_S655s_2 | 2.06 | 1.19 | 1.33 | 1.39 | -0.25 | 0.35 | 1.64 | 0.63 | 1.23 | 1.22 | -0.36 | -0.26 | -0.31 | 0.52 | 0.49 | 0.71 | -0.45 | 0.2375 | -0.15 | 0.008452 | 0.060007 | 0.048999 | 1 | 0.999631 | 0.365244 |
| Larp4b_Q6A0A2_T651t_1 | 0.43 | -1.94 | -1.52 | -1.14 | -1.34 | -1.94 | -1.4 | -1.29 | -1.49 | -0.3 | -0.71 | -0.72 | -0.34 | -1.81 | -0.96 | -1.66 | -0.9 | -1.035 | -1.14 | 0.792751 | 0.382202 | 0.658676 | 1 | 0.999631 | 0.927581 |
| Larp4b_Q6A0A2_T651t_2 | 2.06 | 1.19 | 1.33 | 1.39 | -0.25 | 0.35 | 1.64 | 0.63 | 1.23 | 1.22 | -0.36 | -0.26 | -0.31 | 0.52 | 0.49 | 0.71 | -0.43 | -0.07 | 0.3225 | 0.008452 | 0.060007 | 0.048999 | 1 | 0.999631 | 0.365244 |
| Larp7_Q05CL8_S253s_2 | 0.47 | 2.85 | 1.41 | 1.37 | 2.49 | 0.19 | 0.98 | 0.72 | 0.92 | 1.83 | 2.16 | 0.91 | 1.83 | 1.92 | 2.31 | 1.33 | -0.43 | -0.07 | 0.3225 | 0.528833 | 0.497148 | 0.897685 | 1 | 0.999631 | 0.825257 |
| Larp7_Q05CL8_S256s_2 | 0.47 | 2.85 | 1.41 | 1.37 | 2.49 | 0.19 | 0.98 | 0.72 | 0.92 | 1.83 | 2.16 | 0.91 | 1.83 | 1.92 | 2.31 | 1.33 | -0.9925 | -0.6575 | -1.43 | 0.528833 | 0.497148 | 0.897685 | 1 | 0.999631 | 0.825257 |
| Las1l_A2BE28_S20s_1 | 2.62 | 0.85 | 2.93 | 1.89 | 0.39 | 1.01 | 2.09 | 0.83 | 0.61 | 1.93 | 1.58 | 1.54 | -0.09 | 0.12 | 1.75 | 0.79 | -0.21 | 0.005 | 0.0475 | 0.02944 | 0.091084 | 0.217909 | 1 | 0.999631 | 0.433091 |
| Las1l_A2BE28_S238s_1 | 1.17 | 0.99 | 1.29 | 1.15 | 1.16 | 1.23 | 0.46 | 0.91 | 1.15 | 1.62 | 0.86 | 0.99 | 0.64 | 1.75 | 1.1 | 1.3 | -0.3025 | -0.525 | -0.875 | 0.882293 | 0.449337 | 0.985608 | 1 | 0.999631 | 0.963505 |
| Lats1_Q8BYR2_S204s_1 | 1.11 | -0.51 | 1.68 | 0.93 | -0.09 | 0.63 | 1.07 | 0.39 | 0.31 | 1.32 | 0.47 | -0.99 | -1.42 | -0.44 | 1.03 | 0.54 | 0.025 | -0.1775 | -1.0125 | 0.194519 | 0.543923 | 0.396093 | 1 | 0.999631 | 0.65187 |
| Lbr_Q3U9G9_S101s_1 | 1.58 | -0.15 | 0.89 | 1.22 | 1.19 | 0.76 | 1.16 | 0.53 | 0.72 | 1.17 | 0.39 | 0.55 | -1.1 | -0.29 | 0.77 | 0.11 | -0.83 | -0.7275 | -1.205 | 0.072026 | 0.950973 | 0.669951 | 1 | 0.999631 | 0.532005 |
| Lbr_Q3U9G9_S67s_1 | 1.88 | 0.53 | 1.62 | 0.94 | 0.01 | 0.72 | 1.2 | -0.28 | 0.44 | 1.72 | 0.19 | -0.29 | 0.83 | -0.8 | 0.08 | 0.04 | -0.265 | -0.365 | -1.17 | 0.022257 | 0.086122 | 0.169673 | 1 | 0.999631 | 0.405775 |
| Lbr_Q3U9G9_S67s_2 | 1.88 | 0.12 | 1.03 | 1.24 | 1.28 | 0.27 | 1.47 | 0.19 | 0.8 | 1.03 | 0.57 | 0.41 | -0.38 | -0.27 | 0.37 | -0.13 | -0.265 | -0.365 | -1.17 | 0.017122 | 0.576192 | 0.372692 | 1 | 0.999631 | 0.391024 |
| Lbr_Q3U9G9_S71s_2 | 1.88 | 0.12 | 1.03 | 1.24 | 1.28 | 0.27 | 1.47 | 0.19 | 0.8 | 1.03 | 0.57 | 0.41 | -0.38 | -0.27 | 0.37 | -0.13 | -0.45 | -0.1125 | -0.11 | 0.017122 | 0.576192 | 0.372692 | 1 | 0.999631 | 0.391024 |
| Lca5_G5E887_S7s_1 | 0.29 | 0.48 | 0.87 | 0.66 | -0.23 | 0.45 | -0.31 | 0.59 | 0.34 | 1.04 | -0.3 | 0.77 | -0.65 | 0.68 | 1.32 | 0.51 | -0.2175 | -0.205 | 0.0275 | 0.800484 | 0.17406 | 0.751206 | 1 | 0.999631 | 0.931939 |
| Lcp1_Q61233_S5s_1 | 0.82 | 0.45 | 0.2 | -0.7 | -0.33 | 0.89 | -0.47 | -0.19 | -0.2 | 1.19 | -0.95 | -0.09 | 0.21 | 0.75 | 0.4 | -0.48 | -0.195 | -0.3075 | -1.245 | 0.948386 | 0.621108 | 0.692616 | 1 | 0.999631 | 0.983277 |
| Ldb1_D3Z1C5_S266s_1 | 1.97 | 1.34 | 2.21 | 1.68 | 0.86 | 2.21 | 2.26 | 1.09 | 2.04 | 2.12 | 0.7 | 1.11 | 0.1 | 0.71 | 1.67 | -0.26 | 0.36 | 1.1575 | -0.6675 | 0.020206 | 0.637454 | 0.456165 | 1 | 0.999631 | 0.392698 |
| Ldb3_Q9JKS4_S179s_1 | -2.63 | -3.17 | 0.45 | -0.2 | -1.78 | -3 | 0.69 | -0.02 | -2.94 | -1.05 | 0.81 | 2.26 | -3.2 | -3.23 | 0.07 | -1.86 | -0.0175 | 1.9075 | -1.4425 | 0.513194 | 0.732064 | 0.358003 | 1 | 0.999631 | 0.818518 |
| Lect1_G5E8Z6_S224s_3 | 1.21 | 0.47 | 1.58 | -0.2 | 1.83 | -0.9 | 4.25 | -2.19 | 3.34 | 0.04 | 2.95 | 4.36 | 3.21 | -3.79 | 0.08 | -2.21 | -0.0175 | 1.9075 | -1.4425 | 0.290691 | 0.988991 | 0.052078 | 1 | 0.999631 | 0.706164 |
| Lect1_G5E8Z6_T228t_3 | 1.21 | 0.47 | 1.58 | -0.2 | 1.83 | -0.9 | 4.25 | -2.19 | 3.34 | 0.04 | 2.95 | 4.36 | 3.21 | -3.79 | 0.08 | -2.21 | -0.0175 | 1.9075 | -1.4425 | 0.290691 | 0.988991 | 0.052078 | 1 | 0.999631 | 0.706164 |
| Lect1_G5E8Z6_T229t_3 | 1.21 | 0.47 | 1.58 | -0.2 | 1.83 | -0.9 | 4.25 | -2.19 | 3.34 | 0.04 | 2.95 | 4.36 | 3.21 | -3.79 | 0.08 | -2.21 | 0.405 | -0.5375 | -2.0475 | 0.290691 | 0.988991 | 0.052078 | 1 | 0.999631 | 0.706164 |
| Lemd3_E9QP59_S331s_1 | 2.35 | -0.59 | -1.56 | 0.95 | 1.16 | 0.02 | 1.38 | 0.21 | 0.39 | -0.9 | 0.9 | -1.39 | -3.77 | -1.25 | -1.12 | -0.9 | -0.3625 | -0.4675 | -0.8025 | 0.050727 | 0.617781 | 0.54637 | 1 | 0.999631 | 0.489558 |
| Leng8_D3YWS8_S352s_1 | 1.36 | 0.05 | 1.52 | 1.5 | 0.75 | 0.52 | 1.17 | 0.54 | -0.36 | 1.62 | 0.32 | 0.98 | 0.28 | 0.3 | 0.19 | 0.45 | -0.7775 | -0.7 | -1.4575 | 0.061876 | 0.366797 | 0.379201 | 1 | 0.999631 | 0.509355 |
| Leo1_Q5XJE5_S162s_1 | 2.17 | 1.59 | 2.6 | 1.84 | 0.12 | 2.02 | 2.44 | 0.51 | 0.73 | 2.65 | 1.21 | 0.81 | 0.01 | 0.66 | 1.35 | 0.35 | -0.79 | -0.455 | -1.13 | 0.003661 | 0.183162 | 0.165598 | 1 | 0.999631 | 0.351308 |
| Leo1_Q5XJE5_S198s_1 | 1.88 | 1.17 | 2.81 | 2 | 0.32 | 1.27 | 2.27 | 0.84 | 0.95 | 2.31 | 1.5 | 1.28 | 0.26 | 0.42 | 1.14 | 1.52 | -0.57 | -0.19 | -1.325 | 0.028485 | 0.137968 | 0.314062 | 1 | 0.999631 | 0.42728 |
| Leo1_Q5XJE5_S206s_1 | 2.14 | 1.68 | 2.36 | 2.09 | 0.72 | 2.34 | 1.81 | 1.12 | 2.08 | 2.85 | 1.01 | 1.57 | 1.05 | 0.52 | 1.28 | 0.12 | 0.2175 | -0.0175 | -1.17 | 0.003559 | 0.170389 | 0.652243 | 1 | 0.999631 | 0.351308 |
| Leo1_Q5XJE5_T189t_1 | -0.06 | 0.37 | 1.1 | 1.33 | -0.01 | 1.33 | 1.76 | 0.53 | -0.03 | 2.04 | 0.26 | 0.4 | -1.58 | -0.54 | 0.99 | -0.81 | 0.3675 | 0.6975 | 0.5175 | 0.065833 | 0.6542 | 0.973611 | 1 | 0.999631 | 0.516933 |
| Letmd1_Q924L1_S35s_1 | -1.03 | -0.59 | 0.14 | 0.12 | 0.45 | 0.12 | -0.15 | -0.31 | 1.66 | -0.34 | 0.18 | -0.07 | 1.7 | -0.74 | -0.16 | -0.09 | -0.08 | 0.01 | 0.3525 | 0.363551 | 0.316202 | 0.186998 | 1 | 0.999631 | 0.743747 |
| Lgalsl_Q8VED9_S25s_1 | -0.58 | 0.26 | -0.24 | -0.38 | -0.11 | -0.27 | -0.55 | -0.33 | -0.02 | -0.18 | -0.34 | -0.36 | 0.35 | 0.24 | -0.3 | 0.18 | -0.5825 | -0.2775 | 0.0425 | 0.280736 | 0.776306 | 0.972098 | 1 | 0.999631 | 0.701922 |
| Lgalsl_Q8VED9_S4s_1 | 0.41 | 1.03 | 2.43 | 2.01 | 0.82 | 0.34 | 0.97 | 1.42 | 0.75 | 0.53 | 1.7 | 1.79 | 0.91 | 0.98 | 1.8 | 2.36 | 0.075 | 0.05 | 0.34 | 0.937074 | 0.245676 | 0.60008 | 1 | 0.999631 | 0.97939 |
| Lima1_Q9ERG0_S200s_1 | -1.01 | -2.34 | -2.33 | -1.58 | -1.54 | -1.5 | -1.35 | -2.57 | -1.37 | -0.78 | -1.9 | -3.01 | -0.21 | -1.32 | -2.1 | -2.27 | 0.13 | 0.5375 | 0.49 | 0.528669 | 0.858819 | 0.925403 | 1 | 0.999631 | 0.825257 |
| Lima1_Q9ERG0_S212s_1 | -0.89 | -0.61 | -1.3 | -1.88 | -1.29 | -0.5 | -1 | -1.37 | -0.17 | -0.89 | -1.24 | -0.23 | -0.19 | -0.37 | -0.98 | -1.18 | -0.4325 | -0.66 | 0.3375 | 0.23034 | 0.720021 | 0.19398 | 1 | 0.999631 | 0.681463 |
| Lima1_Q9ERG0_S328s_1 | -1.35 | -2.27 | -0.69 | -0.59 | -1.43 | -3.39 | -0.89 | -0.92 | -2.64 | -3.53 | -0.84 | -0.53 | -0.27 | -2.51 | -0.4 | -0.37 | -0.25 | -0.07 | -0.4925 | 0.580178 | 0.499631 | 0.372745 | 1 | 0.999631 | 0.846774 |
| Limch1_D3Z589_S10s_1 | -0.14 | 0.06 | 0.12 | 0.6 | -0.45 | 0.04 | 0.49 | -0.44 | 0.23 | -0.04 | 0.53 | -0.36 | -0.04 | 0.01 | -1.06 | -0.24 | 0.7825 | 0.07 | 0.125 | 0.181385 | 0.443689 | 0.822644 | 1 | 0.999631 | 0.643843 |
| Limch1_D3Z589_S144s_1 | -0.99 | 0.48 | -1.51 | -0.75 | 0.45 | 0.19 | -0.03 | -0.25 | 0.05 | -0.81 | -0.23 | -1.5 | -2.36 | 0.21 | 0.22 | -0.34 | 0.16 | -0.0025 | -0.275 | 0.850743 | 0.098241 | 0.891285 | 1 | 0.999631 | 0.949706 |
| Limch1_D3Z589_S218s_1 | 1 | 0.86 | -0.03 | 0.71 | 1.36 | 0.72 | 0.49 | 0.61 | 1.65 | -0.26 | 0.7 | 0.44 | -0.33 | 1.29 | 0.12 | 0.36 | -0.325 | 0.1575 | 0.415 | 0.520265 | 0.637126 | 0.99555 | 1 | 0.999631 | 0.819528 |
| Limch1_D3Z589_S364s_1 | 0.11 | -0.15 | 0.76 | 0.11 | 0.7 | 0.14 | -0.19 | -1.12 | 0.39 | 0.21 | -0.1 | 0.96 | 0.38 | 0.91 | -0.39 | 1.59 | -0.065 | -0.305 | -0.255 | 0.374256 | 0.449042 | 0.645467 | 1 | 0.999631 | 0.747198 |
| Limch1_D3Z589_S560s_1 | -0.58 | -0.8 | -0.38 | -0.17 | -0.71 | -1.11 | -0.38 | 0.01 | -0.92 | -0.85 | -0.61 | -0.77 | -1.07 | -0.71 | -0.75 | -0.42 | -0.045 | 0.185 | 0.5 | 0.401146 | 0.839652 | 0.268554 | 1 | 0.999631 | 0.763221 |
| Limch1_D3Z589_S58s_1 | -0.1 | -0.28 | -0.31 | -0.76 | -0.44 | -0.13 | -0.96 | -0.1 | -0.14 | 0.33 | -0.58 | -0.32 | 0.34 | 0.23 | -0.02 | 0 | -0.035 | 0.3925 | 0.3225 | 0.107025 | 0.882261 | 0.551065 | 1 | 0.999631 | 0.568797 |
| Limch1_D3Z589_S598s_1 | 0.68 | 1.03 | 1.3 | 0.77 | 0.01 | 0.8 | 0.86 | 1.97 | 1.02 | 1.51 | 1.25 | 1.57 | 0.84 | 1.12 | 1.31 | 1.8 | -0.3825 | -0.33 | -0.5925 | 0.332766 | 0.933792 | 0.186559 | 1 | 0.999631 | 0.728079 |
| Limch1_D3Z589_S72s_1 | 1.1 | -0.57 | -1.11 | -1.18 | -0.91 | -0.31 | -0.87 | -1.2 | -0.95 | 1.07 | -1.75 | -1.45 | -0.4 | -0.47 | -1.62 | -1.64 | -0.33 | -0.4275 | 0.4875 | 0.321262 | 0.470149 | 0.654229 | 1 | 0.999631 | 0.72408 |
| Limch1_D3Z589_S74s_1 | -0.83 | -2.17 | 0.64 | -1.07 | -2.2 | -2.45 | -0.02 | -0.08 | -2.51 | -1.61 | -0.06 | -0.96 | -1.15 | -1.47 | 1.32 | -0.18 | -0.005 | -0.2025 | -0.66 | 0.519614 | 0.66884 | 0.540679 | 1 | 0.999631 | 0.819456 |
| Limch1_D3Z589_S814s_1 | 0.56 | -0.06 | -0.22 | -0.33 | 0.17 | -0.09 | 0.12 | -0.27 | -0.33 | -0.19 | -0.86 | 0.52 | -0.99 | -0.35 | -0.25 | -1.1 | 0.165 | 0.315 | 0.0375 | 0.083978 | 0.98635 | 0.5899 | 1 | 0.999631 | 0.549732 |
| Limd1_Q9QXD8_S659s_1 | 1.15 | 0.78 | -1.51 | -0.85 | 1.95 | 0.13 | -0.79 | -1.06 | 0.66 | 0.59 | -0.81 | 0.39 | 1.67 | -0.07 | -0.56 | -1.32 | 0.65 | 1.3125 | 0.4725 | 0.961911 | 0.839284 | 0.630847 | 1 | 0.999631 | 0.986594 |
| Lin7a_Q8JZS0_S233s_1 | -1.29 | -0.05 | -3.67 | -1.39 | -0.05 | -0.71 | -0.7 | -2.34 | 0.56 | -0.22 | -0.25 | -1.24 | 1.13 | -0.55 | -2.01 | -3.08 | 0.0975 | 0.3775 | 0.605 | 0.642358 | 0.416703 | 0.102219 | 1 | 0.999631 | 0.871335 |
| Lin7b_O88951_S115s_1 | -0.64 | -0.15 | 0.25 | -0.34 | -0.39 | -0.03 | -0.46 | 0.39 | -0.29 | -0.12 | 0.32 | 0.72 | 0.82 | 0.17 | 0.59 | -0.04 | -1.1925 | -1.0325 | -1.5 | 0.097131 | 0.760086 | 0.282125 | 1 | 0.999631 | 0.55886 |
| Lin7b_O88951_S199s_1 | 0.29 | -1.57 | -1.46 | -0.66 | -2.1 | -3.36 | 0.24 | -2.95 | -1.53 | -0.5 | -3.63 | -1.87 | -1.72 | -5.07 | -1.21 | -1.4 | -0.1475 | 0.0175 | 0.1275 | 0.109816 | 0.158039 | 0.161043 | 1 | 0.999631 | 0.573646 |
| Lingo1_Q9D1T0_S596s_1 | -0.06 | -0.03 | -0.21 | -0.17 | -0.27 | -0.11 | -0.48 | -0.2 | -0.25 | 0.09 | -0.27 | 0.03 | 0.21 | 0 | -0.17 | 0 | -0.215 | 0.2425 | 0.73 | 0.631508 | 0.539181 | 0.944135 | 1 | 0.999631 | 0.867623 |
| Lingo1_Q9D1T0_S602s_1 | -0.96 | -0.59 | -0.46 | -1.36 | -0.98 | -1.01 | -1.07 | -1.17 | -0.68 | -0.3 | -0.88 | -0.54 | 0.99 | -0.52 | -0.49 | -0.43 | 0.07 | -0.06 | 0.2675 | 0.11297 | 0.455926 | 0.436765 | 1 | 0.999631 | 0.577408 |
| Lipe_E9Q4M2_S963s_1 | -0.23 | 0.14 | -0.27 | -0.06 | 0.23 | -0.07 | -0.21 | -0.09 | 0.02 | -0.24 | -0.35 | -0.09 | 0.02 | 0.28 | -0.02 | 0.37 | 0.07 | -0.06 | 0.2675 | 0.345697 | 0.780831 | 0.815225 | 1 | 0.999631 | 0.735045 |
| Lipe_E9Q4M2_S964s_1 | -0.23 | 0.14 | -0.27 | -0.06 | 0.23 | -0.07 | -0.21 | -0.09 | 0.02 | -0.24 | -0.35 | -0.09 | 0.02 | 0.28 | -0.02 | 0.37 | 0.1825 | 0.3425 | 0.955 | 0.345697 | 0.780831 | 0.815225 | 1 | 0.999631 | 0.735045 |
| Llgl1_Q80Y17_S986s_1 | -2.31 | -1.77 | -1.72 | -2.15 | -1.83 | -1.95 | -1.71 | -1.73 | -1.87 | -1.84 | -1.72 | -1.15 | -0.25 | -1.39 | -1.12 | -1.37 | -0.105 | 0.05 | 0.2525 | 0.021583 | 0.488082 | 0.268353 | 1 | 0.999631 | 0.405125 |
| Lmf1_Q3U3R4_S19s_1 | -0.14 | -0.71 | -0.8 | 0.14 | -1.05 | -0.48 | -0.54 | 0.14 | -0.53 | 0.13 | -0.52 | -0.39 | -0.37 | 0.19 | -0.44 | 0.12 | 0.125 | 0.7975 | 0.5075 | 0.466776 | 0.769355 | 0.878879 | 1 | 0.999631 | 0.795267 |
| Lmf1_Q3U3R4_S7s_1 | 1.18 | -0.32 | 0.44 | 0.1 | 0.81 | -1.22 | 1.24 | 1.07 | 0.22 | 1.72 | 1.74 | 0.91 | 2.02 | -1.82 | 1.57 | 1.66 | -0.0525 | -0.095 | -0.23 | 0.543323 | 0.832878 | 0.113262 | 1 | 0.999631 | 0.832881 |
| Lmna_P48678_S22s_1 | 0.14 | -0.35 | -0.24 | -0.52 | -0.33 | -0.61 | -0.17 | -0.07 | -0.41 | -0.22 | -0.14 | -0.58 | -0.28 | -0.18 | -1.14 | -0.29 | -0.22 | 0.095 | 0.13 | 0.496537 | 0.847228 | 0.730283 | 1 | 0.999631 | 0.811248 |
| Lmna_P48678_S390s_1 | 0.34 | 0.48 | 0.03 | -0.16 | 0.01 | 0.48 | 0.02 | -0.7 | -0.19 | 0.97 | -0.07 | 0.36 | 0.2 | 0.25 | 0.16 | 0.6 | -0.395 | 0.08 | -0.065 | 0.658487 | 0.506115 | 0.782247 | 1 | 0.999631 | 0.879726 |
| Lmna_P48678_S392s_1 | 0.29 | 0.14 | -0.09 | -0.32 | -0.4 | -0.23 | -0.41 | -0.52 | -0.4 | 0.38 | 0.34 | 0.02 | 0.21 | -0.26 | -0.08 | -0.11 | -0.095 | 5.89E-17 | -0.3925 | 0.821743 | 0.147357 | 0.790318 | 1 | 0.999631 | 0.938464 |
| Lmna_P48678_S395s_1 | 1.41 | 1.89 | 1.71 | 1.51 | 1.1 | 1.76 | 1.14 | 2.14 | 1.51 | 1.71 | 1.73 | 1.57 | 0.11 | 1.88 | 1.37 | 1.59 | -0.5575 | -1.2275 | -1.47 | 0.359392 | 0.767648 | 1 | 1 | 1 | 0.741182 |
| Lmna_P48678_S407s_1 | 1.94 | -0.85 | 0.7 | -0.33 | -0.71 | -0.15 | 1.22 | -1.13 | -1.8 | 0.36 | -0.88 | -1.13 | -0.98 | -1.78 | -0.73 | -0.93 | -1.2975 | 0.4125 | -1.16 | 0.032057 | 0.438403 | 0.097059 | 1 | 0.999631 | 0.442395 |
| Lmna_P48678_S429s_1 | 0.99 | 0.72 | 0.82 | -0.13 | -1.52 | 0.15 | 0.13 | -1.55 | 0.27 | 3.2 | 0.71 | -0.13 | 0.43 | -1.34 | -1.39 | 0.06 | -0.38 | -0.3175 | 0.2275 | 0.042599 | 0.027381 | 0.560694 | 1 | 0.999631 | 0.475443 |
| Lmna_P48678_T19t_1 | 0.19 | 0.09 | -0.48 | -1.53 | -0.15 | -0.36 | -0.52 | -2.22 | -0.62 | -0.31 | -0.99 | -1.08 | 0.5 | 0.02 | -0.64 | -0.7 | -0.1075 | -0.43 | -2.96 | 0.634 | 0.505597 | 0.468029 | 1 | 0.999631 | 0.868569 |
| Lmna_P48678_T3t_1 | 2.96 | -0.95 | -1.02 | -0.91 | -0.34 | -0.07 | 1.52 | -1.46 | -0.13 | 2.96 | -2 | -2.47 | -1.53 | -4.45 | -3.15 | -2.63 | -0.2 | -0.59 | -0.0975 | 0.012576 | 0.913947 | 0.749213 | 1 | 0.999631 | 0.385092 |
| Lmnb1_P14733_S17s_1 | 0.13 | 0.62 | -0.18 | 0.18 | 0.16 | -0.12 | 0 | -0.09 | -0.05 | -0.26 | -0.76 | -0.54 | 0.7 | 0.26 | -0.91 | 0.31 | -0.0825 | -0.11 | 0.36 | 0.810209 | 0.464356 | 0.074695 | 1 | 0.999631 | 0.936878 |
| Lmnb1_P14733_S24s_1 | 0.01 | 0 | -1.27 | -0.99 | -0.13 | -0.5 | -0.78 | -1.17 | -0.77 | 0.09 | -0.98 | -1.03 | 0.23 | -0.35 | -0.24 | -0.45 | -0.2925 | -0.2725 | -0.5875 | 0.370376 | 0.837475 | 0.795546 | 1 | 0.999631 | 0.746338 |
| Lmnb1_P14733_S394s_1 | 0.4 | -0.2 | -0.11 | -0.23 | -0.62 | -0.17 | -0.15 | -0.37 | -0.56 | 0.59 | -0.54 | -0.72 | -0.58 | -0.56 | -0.59 | -0.76 | -0.2425 | -0.495 | -0.0425 | 0.061197 | 0.298121 | 0.462902 | 1 | 0.999631 | 0.509355 |
| Lmnb1_P14733_T21t_1 | -0.16 | -0.27 | -0.23 | -0.37 | -0.1 | -0.54 | -0.54 | -0.82 | -1.12 | -0.37 | -0.32 | -1.2 | 0.12 | -0.77 | -0.22 | -0.33 | -0.225 | 0.4925 | 0.3675 | 0.886462 | 0.362385 | 0.133121 | 1 | 0.999631 | 0.963505 |
| Lmnb2_P21619_S397s_1 | -1.12 | 0.74 | 0.27 | 0.47 | -0.39 | -0.64 | -0.26 | 0.75 | 0.31 | 0.27 | 0.24 | 1.51 | -0.32 | 0.63 | 0.57 | 0.95 | -0.5625 | 0.395 | -0.3975 | 0.452087 | 0.645277 | 0.330622 | 1 | 0.999631 | 0.788148 |
| Lmnb2_P21619_S403s_1 | 0.04 | -1.66 | 0.35 | 1.38 | -2.19 | 0.04 | 0.21 | -0.2 | -1.75 | 0.32 | 2.04 | 1.08 | -0.97 | -0.13 | -0.46 | 0.08 | -0.66 | -0.2875 | -0.9575 | 0.519065 | 0.454661 | 0.65873 | 1 | 0.999631 | 0.819456 |
| Lmnb2_P21619_S427s_1 | 0.31 | -1.18 | -0.24 | -0.52 | -1.11 | -1.55 | -0.74 | -0.87 | -1.19 | -0.17 | -0.6 | -0.82 | -1.23 | -1.71 | -0.94 | -1.58 | -0.125 | 0.29 | 0.415 | 0.030736 | 0.102416 | 0.469886 | 1 | 0.999631 | 0.439142 |
| Lmnb2_P21619_T494t_1 | -1.08 | -0.67 | -0.94 | -0.66 | -0.84 | -0.99 | -1.28 | -0.74 | -0.62 | -0.1 | -1.01 | -0.46 | 0.29 | -0.71 | -0.64 | -0.63 | -1.255 | -0.525 | -0.9425 | 0.229609 | 0.633733 | 0.339075 | 1 | 0.999631 | 0.680148 |
| Lmo7_E9PYI7_S1073s_1 | 0.09 | -0.84 | 0.36 | -0.44 | -1.81 | -1.15 | -0.89 | -2 | -1.47 | 0.54 | -0.99 | -1.01 | 0.12 | -1.98 | -1.11 | -1.63 | -0.315 | -0.8575 | -0.33 | 0.087385 | 0.00854 | 0.299137 | 1 | 0.999631 | 0.553616 |
| Lmo7_E9PYI7_S122s_1 | -0.24 | 0.96 | -0.74 | -0.99 | 0.01 | -0.22 | -1.01 | -1.05 | -1.15 | -1.29 | -0.92 | -1.08 | 0.57 | -0.99 | -1.07 | -0.84 | 0.2725 | -0.1925 | -0.0925 | 0.546786 | 0.519273 | 0.073946 | 1 | 0.999631 | 0.834213 |
| Lmo7_E9PYI7_S1269s_1 | -0.53 | -1.11 | -0.7 | 0.17 | -0.06 | -0.98 | 0.24 | -0.28 | -0.58 | -1.43 | -0.63 | -0.3 | -2.91 | -0.16 | 0.59 | -0.06 | 0.05 | -0.215 | 0.54 | 0.898338 | 0.483646 | 0.616545 | 1 | 0.999631 | 0.966961 |
| Lmo7_E9PYI7_S127s_1 | -0.65 | -0.32 | -1.19 | 0.25 | -0.42 | -0.54 | -0.82 | 0.07 | -0.54 | -0.74 | -0.69 | -0.8 | 0.47 | 0.02 | -0.02 | -0.22 | 0.075 | -0.135 | -0.585 | 0.171367 | 0.892987 | 0.540101 | 1 | 0.999631 | 0.641657 |
| Lmo7_E9PYI7_S1300s_1 | 0.95 | 0.66 | -0.34 | 0.65 | 0.53 | 1.5 | -0.04 | 0.23 | 0.85 | 0.43 | -0.18 | 0.28 | -0.46 | -0.02 | -0.25 | 0.31 | 0.4875 | -0.8775 | -2.4675 | 0.137062 | 0.861627 | 0.72101 | 1 | 0.999631 | 0.610369 |
| Lmo7_E9PYI7_S1369s_1 | 2.62 | -1.3 | -1.89 | 0.97 | 0.47 | 0.28 | 1.74 | -0.14 | -0.47 | 0.08 | -0.5 | -2.22 | -3.98 | -1.81 | -1.51 | -2.17 | -0.345 | -0.7825 | -1.65 | 0.032284 | 0.615116 | 0.388987 | 1 | 0.999631 | 0.442395 |
| Lmo7_E9PYI7_S1369s_2 | 1.73 | -0.73 | 0 | 0.69 | -0.24 | -0.34 | 1.11 | -0.22 | -0.46 | 0.21 | -0.3 | -0.89 | -2.03 | -1.27 | -0.79 | -0.82 | 0.845 | 1.1875 | 0.1775 | 0.0132 | 0.549681 | 0.164859 | 1 | 0.999631 | 0.385092 |
| Lmo7_E9PYI7_S1376s_1 | 0.01 | -0.37 | 0.23 | -0.57 | 1.83 | -0.81 | 1.86 | -0.2 | 2.53 | -0.7 | 0.66 | 1.56 | 1.87 | -1.32 | 0.38 | -0.92 | 0.78 | -0.3575 | -0.685 | 0.789496 | 0.206105 | 0.088577 | 1 | 0.999631 | 0.925593 |
| Lmo7_E9PYI7_S1391s_1 | 0.09 | -1.02 | -1.81 | -0.88 | 0.62 | 0.42 | -0.83 | -0.71 | -0.37 | -2.12 | -1.41 | -1.15 | -1.68 | -1.2 | -1.9 | -1.58 | -0.535 | -0.34 | -0.6525 | 0.131905 | 0.146908 | 0.483353 | 1 | 0.999631 | 0.603456 |
| Lmo7_E9PYI7_S206s_1 | 0.26 | -0.74 | -0.17 | -0.43 | -1.57 | -0.74 | -0.12 | -0.79 | -0.92 | 0.05 | -0.8 | -0.77 | -0.59 | -0.91 | -1.35 | -0.84 | 0.04 | -0.145 | -0.2775 | 0.074544 | 0.179851 | 0.340378 | 1 | 0.999631 | 0.540075 |
| Lmo7_E9PYI7_S253s_1 | 0.02 | 0.51 | 0.03 | 0.9 | 0.68 | 0.73 | 0.07 | 0.14 | 0.35 | -0.17 | 0.37 | 0.33 | -0.14 | 0.23 | 0.05 | 0.21 | 0.3875 | -0.0175 | -0.3125 | 0.389399 | 0.900892 | 0.645383 | 1 | 0.999631 | 0.758626 |
| Lmo7_E9PYI7_S466s_1 | 0.95 | 1.09 | -0.32 | 1.61 | 1.98 | 0.42 | 1.77 | 0.71 | 1.15 | 0.24 | 1.13 | 0.74 | -0.21 | 0.7 | 0.81 | 0.78 | 0.0975 | 0.5325 | 0.5125 | 0.509162 | 0.464213 | 0.969108 | 1 | 0.999631 | 0.816932 |
| Lmo7_E9PYI7_S580s_1 | -0.2 | -0.05 | -0.37 | -0.22 | 0.93 | -0.51 | 0.35 | -1.22 | -0.06 | 0.05 | 0.61 | 0.69 | 0.5 | 0.14 | -0.1 | 0.67 | 0.2075 | -0.0575 | -0.4775 | 0.107956 | 0.831703 | 0.090272 | 1 | 0.999631 | 0.569416 |
| Lmo7_E9PYI7_S589s_1 | 1.1 | -0.91 | -2.21 | -0.64 | 0.32 | -0.36 | -0.81 | -0.98 | -0.35 | -0.93 | -0.34 | -1.27 | -1.68 | 0.02 | -1.73 | -1.18 | -0.135 | 0.185 | -0.0675 | 0.501985 | 0.753538 | 0.929077 | 1 | 0.999631 | 0.814092 |
| Lmo7_E9PYI7_S787s_1 | -1.28 | -2 | -1.11 | -1.71 | -1.2 | -2.41 | -1.29 | -1.74 | -1.68 | -1.67 | -1.32 | -0.69 | -1.61 | -2.15 | -1.25 | -1.36 | -0.345 | -0.7825 | -1.65 | 0.845206 | 0.711733 | 0.598006 | 1 | 0.999631 | 0.948565 |
| Lmo7_E9PYI7_T1365t_2 | 1.73 | -0.73 | 0 | 0.69 | -0.24 | -0.34 | 1.11 | -0.22 | -0.46 | 0.21 | -0.3 | -0.89 | -2.03 | -1.27 | -0.79 | -0.82 | 0.1025 | 0.34 | 0.43 | 0.0132 | 0.549681 | 0.164859 | 1 | 0.999631 | 0.385092 |
| Lmtk2_Q3TYD6_S1104s_1 | -0.53 | -0.86 | -1.19 | -1.26 | -0.47 | -0.96 | -1.34 | -0.66 | -0.98 | -0.49 | -0.69 | -0.32 | 0.06 | -0.56 | -0.88 | -0.74 | 0.4525 | 0.2925 | -0.09 | 0.219178 | 0.740835 | 0.270258 | 1 | 0.999631 | 0.669859 |
| Lmtk2_Q3TYD6_S1359s_1 | 0.12 | -1.04 | -0.45 | 0.03 | 0.19 | -0.22 | 0 | 0.5 | -0.15 | 0.06 | -0.26 | 0.18 | -2 | 0.77 | -0.91 | 0.44 | 0.0475 | 0.1575 | 0.3175 | 0.885504 | 0.20527 | 0.390691 | 1 | 0.999631 | 0.963505 |
| Lmtk2_Q3TYD6_S1418s_1 | 0.23 | 0.06 | 0.3 | 0.44 | 0.41 | 0.34 | -0.16 | 0.63 | 0.14 | 0.29 | 0.39 | 0.84 | -0.49 | 0.98 | 0.92 | 0.89 | 0.67 | -0.53 | 0.01 | 0.42665 | 0.861991 | 0.569258 | 1 | 0.999631 | 0.775108 |
| Lmtk2_Q3TYD6_S611s_1 | 0.78 | 0.34 | -0.73 | 1.67 | 1.67 | 1.25 | 0.56 | 1.26 | 0.72 | -0.02 | -0.05 | -0.71 | -1.05 | 1.57 | 0.81 | 0.77 | 0.32 | 0.0225 | 0.375 | 0.988035 | 0.210902 | 0.337376 | 1 | 0.999631 | 0.995409 |
| Lmtk2_Q3TYD6_S742s_1 | -0.54 | -0.62 | -0.26 | -0.02 | 0.15 | 0.02 | -0.35 | 0.02 | -0.55 | -0.38 | -0.34 | -0.08 | -0.18 | 0.3 | -0.13 | 0.07 | -0.245 | -0.095 | 0.37 | 0.21834 | 0.250984 | 0.934389 | 1 | 0.999631 | 0.669859 |
| Lmtk3_Q5XJV6_S232s_1 | -0.21 | -0.45 | -0.34 | -0.68 | -0.75 | -0.55 | -0.76 | -0.6 | -0.78 | -0.36 | -0.41 | -0.51 | 0.33 | -0.13 | -0.52 | 0.12 | 0.28 | -0.0475 | -0.075 | 0.244582 | 0.329302 | 0.716693 | 1 | 0.999631 | 0.685751 |
| Lmtk3_Q5XJV6_S531s_1 | 0.38 | -0.56 | -0.68 | -0.22 | 0.55 | -0.06 | 0.07 | -0.52 | -0.14 | -0.56 | -0.23 | -0.34 | -0.4 | -0.13 | -0.55 | -0.3 | 0.1025 | -0.0575 | 0.5825 | 0.819563 | 0.43406 | 0.880693 | 1 | 0.999631 | 0.938303 |
| Lmtk3_Q5XJV6_S947s_1 | -1.24 | -0.84 | 0.46 | -0.3 | -0.79 | -1.13 | -0.04 | 0.45 | -0.74 | -1.32 | 0.73 | -0.82 | 0.68 | -0.53 | -0.39 | 0.65 | -0.2775 | -0.1325 | 0.16 | 0.24108 | 0.832609 | 0.914473 | 1 | 0.999631 | 0.684338 |
| Lmtk3_Q5XJV6_S962s_1 | -0.21 | -0.29 | -0.41 | -1.03 | -0.69 | -0.65 | -1.03 | -0.68 | -1.08 | -0.21 | -0.83 | -0.35 | 0.06 | -0.41 | -0.66 | -0.29 | 0.25 | 0.2 | 0.61 | 0.620487 | 0.340797 | 0.688182 | 1 | 0.999631 | 0.861888 |
| Lmtk3_Q5XJV6_S977s_1 | -0.92 | -0.26 | -0.31 | -0.51 | -0.35 | -0.24 | -0.57 | 0.16 | -0.3 | -0.99 | -0.08 | 0.17 | 0.22 | -0.2 | 0.09 | 0.33 | 0.2125 | -0.44 | 0.095 | 0.063268 | 0.394681 | 0.557463 | 1 | 0.999631 | 0.512978 |
| Lnp_Q7TQ95_S194s_1 | 0.3 | 1.11 | -2.57 | 0.89 | 0.77 | -0.86 | 0.78 | -0.11 | 0.08 | -0.16 | -0.94 | -1.01 | 1.33 | -1.62 | 0.96 | -0.56 | -0.06 | -0.315 | -0.1975 | 0.919314 | 0.795233 | 0.578953 | 1 | 0.999631 | 0.974502 |

| | | | | | | | | | | | | | | | | | | | | | | | | | |
|---|---|---|---|---|---|---|---|---|---|---|---|---|---|---|---|---|---|---|---|---|---|---|---|---|---|
| Lnp_Q7TQ95_S222s_1 | -0.29 | -0.44 | -1.46 | -1.53 | -0.83 | -1.64 | -0.93 | -0.56 | -1.34 | -1.28 | -0.89 | -1.47 | -0.69 | -1.36 | -1.87 | -0.59 | -0.105 | 0.1125 | 0.3825 | 0.659208 | 0.881762 | 0.410341 | 1 | 0.999631 | 0.880034 |
| Lnp_Q7TQ95_S411s_1 | -0.31 | 0.3 | -0.34 | -0.12 | -0.08 | -0.22 | -0.32 | -0.27 | -0.24 | 0.03 | -0.08 | 0.27 | 0.66 | 0.27 | -0.33 | 0.46 | -0.17 | 0.185 | 0.3625 | 0.265335 | 0.688166 | 0.688707 | 1 | 0.999631 | 0.694352 |
| Loh12cr1_E9Q2W8_S76s_1 | -0.11 | -0.27 | 0.12 | 0.04 | -0.46 | -0.01 | -0.53 | 0.1 | 0.06 | 0.08 | 0.04 | 0.34 | 0.6 | 0.22 | -0.27 | 0.68 | -0.7075 | -0.78 | -1.1225 | 0.270922 | 0.534812 | 0.469991 | 1 | 0.999631 | 0.698674 |
| Lpar1_P61793_S323s_1 | 1.87 | 1.03 | 1.91 | 1.25 | 0.43 | 1.17 | 1.31 | 0.32 | 0.76 | 2.12 | -0.13 | 0.19 | 0.62 | 0.08 | 0.94 | -0.07 | 0.1 | 0.18 | 0.3875 | 0.011363 | 0.073249 | 0.151831 | 1 | 0.999631 | 0.385092 |
| Lpgat1_E9QL80_S24s_1 | -0.36 | -0.72 | 0.33 | -0.39 | -0.79 | 0.12 | -0.11 | 0.04 | -0.28 | -0.32 | -0.1 | 0.28 | 0.34 | -0.33 | 0.79 | -0.39 | -0.2575 | -0.1 | -1.1075 | 0.331854 | 0.768141 | 0.576416 | 1 | 0.999631 | 0.727673 |
| Lpgat1_E9QL80_S272s_1 | 0.72 | -0.81 | 0.36 | -0.65 | -1.38 | 0.14 | 1.54 | -1.71 | -0.64 | 0.16 | -0.14 | -0.16 | 0.23 | -1.98 | -0.65 | -2.41 | 0.0175 | -0.15 | -0.295 | 0.108784 | 0.727383 | 0.809969 | 1 | 0.999631 | 0.571276 |
| Lpin1_E9QKQ5_S328s_1 | 1.15 | 0.24 | 0.88 | 0.78 | 0.7 | 0.94 | 0.55 | 0.93 | 0.73 | 0.93 | 0.35 | 0.44 | 0.27 | 0.86 | 0.64 | 0.1 | -0.0725 | -0.05 | 0.45 | 0.382191 | 0.951388 | 0.624761 | 1 | 0.999631 | 0.753668 |
| Lpin2_E9PWN0_S144s_1 | 0.47 | 1.05 | 0.6 | 0.65 | 0.68 | 1.08 | 0.12 | 0.6 | 0.96 | 0.55 | 0.56 | 0.5 | 0.88 | 1.73 | 0.62 | 1.34 | -0.1975 | -0.0725 | -0.31 | 0.206331 | 0.807957 | 0.854127 | 1 | 0.999631 | 0.6605 |
| Lpin2_E9PWN0_T329t_1 | -0.62 | -0.87 | 0.87 | -0.32 | -1.11 | -0.89 | 0.09 | 0.18 | -0.55 | 0.11 | -0.3 | -0.49 | -1.12 | -0.84 | 0.39 | -0.61 | 0.23 | -0.2 | 0.6175 | 0.530034 | 0.682554 | 0.862123 | 1 | 0.999631 | 0.826436 |
| Lppr2_Q8VCY8_S312s_1 | -0.8 | -0.16 | -0.65 | -0.7 | -0.04 | -0.01 | -0.83 | -0.51 | -0.49 | -0.74 | -1.02 | -0.86 | 0.95 | 0.21 | -1.05 | 0.05 | 0.9175 | -0.085 | -0.165 | 0.18207 | 0.455659 | 0.479864 | 1 | 0.999631 | 0.643861 |
| Lppr2_Q8VCY8_S330s_1 | 1.13 | 0.26 | -0.21 | 0.47 | 1.88 | 1.44 | 0.78 | 1.22 | 1.64 | -0.6 | 0.41 | -0.14 | 0.59 | 0.51 | -0.46 | 0.35 | 1.61 | 2.415 | 0.195 | 0.677521 | 0.032966 | 0.870916 | 1 | 0.999631 | 0.887183 |
| Lppr2_Q8VCY8_S330s_1_1 | 0.79 | 1.57 | 2.87 | -1.35 | 3.02 | 2.23 | 5.64 | -0.57 | 4.14 | 0.59 | 3.85 | 4.96 | 2.82 | -1.05 | 2.09 | 0.8 | 0.5675 | 0.2825 | -0.205 | 0.851932 | 0.241621 | 0.053826 | 1 | 0.999631 | 0.950099 |
| Lppr2_Q8VCY8_S364s_1 | 0.08 | 0.02 | 0.3 | 0.87 | 0.82 | 0.83 | 0.87 | 1.02 | 0.69 | 0.37 | 0.79 | 0.55 | -0.39 | 0.07 | 0.63 | 0.14 | 0.145 | -0.1125 | 0.1825 | 0.556508 | 0.066549 | 0.350191 | 1 | 0.999631 | 0.838305 |
| Lppr2_Q8VCY8_S398s_1 | -0.66 | 0.02 | -1.25 | 0.68 | 0.36 | -0.47 | -0.59 | 0.07 | -0.63 | -0.55 | -0.33 | -0.15 | -0.33 | -0.15 | -0.92 | 0.92 | 0.725 | 0.405 | 0.145 | 0.732435 | 0.751087 | 0.794161 | 1 | 0.999631 | 0.907231 |
| Lppr3_Q7TPB0_S311s_1 | -0.44 | -1.44 | 0.28 | -0.33 | -0.84 | 0.17 | 1.73 | -0.09 | -0.5 | -0.24 | 0.61 | -0.18 | -0.51 | -0.44 | 0.51 | -0.91 | 0.37 | 0.435 | 0.3325 | 0.753341 | 0.235323 | 0.356666 | 1 | 0.999631 | 0.913377 |
| Lppr3_Q7TPB0_S346s_1 | -0.07 | 0.42 | -0.03 | 0.09 | 0.91 | 0.59 | 0.06 | 0.33 | 1.04 | 0.22 | 0.16 | 0.73 | 0.16 | 0.64 | 0.52 | 0.42 | 0.155 | 0.1075 | 0.5225 | 0.256398 | 0.21734 | 0.17953 | 1 | 0.999631 | 0.691343 |
| Lppr3_Q7TPB0_S351s_1 | -0.43 | -0.13 | -0.78 | -0.73 | -0.14 | -0.26 | -0.97 | -0.08 | -0.29 | -0.28 | -1.15 | 0.08 | -0.05 | 0.55 | -0.31 | -0.17 | 1.0375 | -0.315 | -0.585 | 0.129025 | 0.619464 | 0.755313 | 1 | 0.999631 | 0.59942 |
| Lppr3_Q7TPB0_S382s_1 | 1.31 | -1.21 | -1.79 | 0.83 | 1.73 | 0.2 | 0.52 | 0.84 | -0.19 | -0.94 | -0.75 | -0.24 | -1.58 | -0.17 | -1.35 | -0.1 | 0.44 | 0.305 | 0.0225 | 0.441178 | 0.176375 | 0.652124 | 1 | 0.999631 | 0.781983 |
| Lppr3_Q7TPB0_S426s_1 | 0.04 | -0.78 | -1.55 | -0.3 | 0.47 | -0.46 | -0.29 | -0.55 | -0.15 | -0.05 | -0.73 | -0.44 | -0.83 | -0.3 | -0.99 | -0.38 | 0.1975 | 0.195 | 0.55 | 0.955671 | 0.302312 | 0.44498 | 1 | 0.999631 | 0.984649 |
| Lppr3_Q7TPB0_S505s_1 | -0.86 | -1.02 | -0.88 | -0.98 | -0.44 | -0.93 | -1.04 | -0.54 | -0.64 | -0.74 | -0.9 | -0.68 | -0.19 | -0.46 | -0.31 | -0.58 | -0.095 | -0.1325 | -0.4775 | 0.060876 | 0.451894 | 0.427791 | 1 | 0.999631 | 0.509205 |
| Lppr3_Q7TPB0_S567s_1 | 0.2 | -3.64 | -0.47 | 0.08 | -2.42 | -1.73 | 0.18 | -0.24 | -1.94 | -0.57 | -1.05 | -0.8 | -1.78 | -2.55 | 0.13 | -1.54 | -0.01 | 0.06 | -0.385 | 0.605476 | 0.919781 | 0.873819 | 1 | 0.999631 | 0.85817 |
| Lppr3_Q7TPB0_S575s_1 | 0.92 | 0.41 | 0.52 | 1.01 | 0.93 | 0.06 | 0.51 | 1.32 | 0.75 | 1.24 | 0.57 | 0.54 | 0.39 | 0.77 | 0.16 | 0 | 0.045 | 0.2825 | 0.4525 | 0.236925 | 0.976679 | 0.840194 | 1 | 0.999631 | 0.683301 |
| Lppr3_Q7TPB0_S641s_1 | -0.02 | 0.13 | -0.25 | -0.01 | 0.19 | -0.35 | -0.05 | 0.24 | 0.23 | 0.04 | 0.23 | 0.48 | 0.74 | 0.43 | 0.27 | 0.22 | -0.385 | 0.535 | 0.75 | 0.128911 | 0.862911 | 0.28379 | 1 | 0.999631 | 0.59942 |
| Lppr3_Q7TPB0_T342t_1 | -0.15 | 0.92 | 0.92 | 0.08 | -0.3 | 0.3 | -0.21 | 0.44 | 1.33 | 0.86 | 0.66 | 1.06 | 1.65 | 0.87 | 0.96 | 1.29 | 0.305 | 0.1 | 0.2175 | 0.067141 | 0.298081 | 0.152084 | 1 | 0.999631 | 0.520345 |
| Lppr3_Q7TPB0_T374t_1 | 0.15 | 0.18 | -0.38 | 0.22 | 0.7 | 0.53 | -0.51 | 0.67 | 0.68 | 0.18 | -0.45 | 0.16 | 0.31 | 0.59 | -0.09 | 0.23 | 0.005 | 0.34 | 0.495 | 0.477579 | 0.393432 | 0.759589 | 1 | 0.999631 | 0.800237 |
| Lppr4_Q7TME0_S347s_1 | -0.93 | -0.35 | -1 | -0.64 | -0.64 | -0.42 | -1.09 | -0.75 | -0.12 | -0.18 | -0.6 | -0.66 | 0.48 | -0.5 | -0.72 | -0.2 | 0.965 | 2.585 | -1.3525 | 0.183645 | 0.985876 | 0.258312 | 1 | 0.999631 | 0.644372 |
| Lppr4_Q7TME0_S37s_1 | -0.9 | -7.17 | -1.68 | -0.97 | -4.01 | -2.8 | 1.79 | -1.84 | 2.19 | -1.02 | -0.63 | -0.92 | -3.95 | -5.32 | -5.05 | -1.81 | 0.7025 | 0.4525 | 0.66 | 0.356504 | 0.563617 | 0.097486 | 1 | 0.999631 | 0.740467 |
| Lppr4_Q7TME0_S386s_1 | -0.47 | 1.03 | 0.61 | 1.06 | 1.59 | 1.09 | 0.57 | 1.79 | 1.1 | -0.39 | 1.98 | 1.35 | 0.55 | 1.07 | 1.63 | 1.62 | 0.6425 | -0.775 | -1.2125 | 0.158929 | 0.13347 | 0.433069 | 1 | 0.999631 | 0.629092 |
| Lppr4_Q7TME0_S435s_1 | 2.48 | -0.66 | -0.88 | 0.51 | 2.23 | 0.75 | 0.69 | 0.35 | 0.67 | -1.28 | -0.05 | -0.99 | -2.36 | 0.18 | -0.35 | -0.87 | 0.5425 | -0.375 | -0.72 | 0.161922 | 0.41098 | 0.334408 | 1 | 0.999631 | 0.631442 |
| Lppr4_Q7TME0_S439s_1 | 1.32 | -0.46 | -0.69 | 0.94 | 1 | 0.18 | 1.01 | 1.09 | 0.02 | 0.24 | 0.14 | -0.79 | -1.68 | -0.3 | -0.09 | 0.3 | 0.02 | -0.1125 | 0.195 | 0.248739 | 0.300502 | 0.4764 | 1 | 0.999631 | 0.685877 |
| Lppr4_Q7TME0_S474s_1 | -0.05 | -0.64 | -0.67 | -0.57 | -0.37 | -0.31 | -0.79 | -0.38 | -0.7 | -0.59 | -0.57 | -0.52 | -0.22 | -0.19 | -0.55 | -0.19 | -0.17 | -0.3725 | 0.1625 | 0.506257 | 0.941417 | 0.673979 | 1 | 0.999631 | 0.815079 |
| Lppr4_Q7TME0_S549s_1 | -0.94 | -2.11 | -3.13 | -1.5 | -1.82 | -1.82 | -2.35 | -2.37 | -2.67 | -1.43 | -2.95 | -2.12 | -1.48 | -1.87 | -2.01 | -1.67 | 0.05 | 0.4375 | 1.15 | 0.731338 | 0.71875 | 0.493379 | 1 | 0.999631 | 0.906897 |
| Lppr4_Q7TME0_S594s_1 | -1.46 | -1.44 | 0.26 | -1.36 | -1.49 | -0.01 | -0.3 | -2 | -1.22 | -0.13 | -0.79 | -0.11 | 0.32 | 0.76 | -0.3 | -0.18 | 0.8575 | -0.1725 | -1.04 | 0.033563 | 0.930851 | 0.372634 | 1 | 0.999631 | 0.443263 |
| Lppr4_Q7TME0_S608s_1 | 0.4 | -0.11 | -0.49 | -0.04 | 1.22 | -0.6 | 2.15 | 0.42 | 0.73 | -1.39 | 0.6 | -0.87 | -0.24 | -2.4 | -0.89 | -0.87 | 0.7425 | 0.005 | 0.1175 | 0.051984 | 0.149523 | 0.742904 | 1 | 0.999631 | 0.489777 |
| Lppr4_Q7TME0_S649s_1 | 0.12 | -0.66 | -0.16 | -0.16 | 1.22 | 0.75 | 0.06 | 0.08 | 0.22 | -1.56 | 0.19 | 0.31 | -0.67 | 0.31 | 0.52 | -0.55 | 0.77 | 0.895 | -0.04 | 0.756647 | 0.058012 | 0.991407 | 1 | 0.999631 | 0.914089 |
| Lppr4_Q7TME0_S685s_1 | -0.87 | 1.32 | 1.23 | 1.23 | 0.89 | 1.95 | 1.81 | 1.34 | 2.24 | 0.44 | 1.8 | 2.01 | 0.04 | 0.73 | 1.63 | 0.35 | -0.1725 | -0.7675 | -1.93 | 0.945179 | 0.175449 | 0.164089 | 1 | 0.999631 | 0.982141 |
| Lppr4_Q7TME0_S746s_1 | 2.11 | -0.64 | -0.71 | 0.5 | -0.4 | 0 | 1.73 | -0.76 | -0.97 | 0.94 | -0.53 | -1.25 | -2.26 | -1.6 | -1.33 | -1.27 | 0.6225 | -0.1325 | -0.125 | 0.011275 | 0.819286 | 0.305411 | 1 | 0.999631 | 0.385092 |
| Lppr4_Q7TME0_T417t_1 | 0.34 | 0.34 | -0.64 | 0.71 | 0.8 | 0.93 | 0.4 | 1.11 | 0.15 | -0.25 | 0.87 | -0.55 | -0.28 | 0.63 | 0.08 | -0.18 | -0.0375 | -0.1775 | 0.5 | 0.746763 | 0.103073 | 0.754998 | 1 | 0.999631 | 0.91173 |
| Lppr4_Q7TME0_T548t_1 | -0.9 | -1.99 | -1.39 | -1.29 | -1.14 | -1.47 | -1.39 | -1.72 | -1.91 | -1.09 | -1.97 | -1.31 | -0.53 | -0.61 | -1.22 | -1.21 | -0.0775 | -0.0025 | -0.2225 | 0.174858 | 0.903751 | 0.616361 | 1 | 0.999631 | 0.642116 |
| Lppr5_Q8BJ52_S302s_1 | -0.47 | -1.02 | -0.72 | -1.07 | -0.73 | -1.06 | -0.94 | -0.86 | -1.19 | -0.52 | -0.69 | -0.89 | -0.85 | -1.43 | -1.15 | -0.74 | 0.3925 | 0.3325 | -0.7725 | 0.474349 | 0.766553 | 0.99308 | 1 | 0.999631 | 0.799827 |
| Lrba_E9Q3Y4_S994s_1 | -0.04 | 1.7 | 2.37 | 0.34 | 1.14 | 1.5 | 2.51 | 0.79 | 2.13 | -0.12 | 1.56 | 2.13 | 1.24 | -0.6 | 1.02 | -0.38 | -0.3225 | 0.3325 | 0.655 | 0.25853 | 0.524571 | 0.633271 | 1 | 0.999631 | 0.692624 |
| Lrch1_P62046_S522s_1 | -1.21 | -0.67 | -0.91 | -0.9 | -1.72 | -0.4 | -2.14 | -0.72 | -1.07 | -0.53 | -0.5 | -0.26 | -0.12 | -0.29 | -0.51 | -0.15 | -0.42 | -0.51 | -0.2175 | 0.038367 | 0.450008 | 0.268026 | 1 | 0.999631 | 0.460131 |
| Lrch3_F6YZL6_S212s_1 | -0.18 | -1.02 | 1.05 | 0.62 | -0.65 | -1.49 | -0.2 | 1.13 | -0.4 | -0.93 | -0.28 | 0.04 | 0.13 | -0.9 | 0.72 | -0.35 | 0.56 | -0.2825 | -0.0625 | 0.686172 | 0.515209 | 0.302202 | 1 | 0.999631 | 0.890692 |
| Lrfn1_Q2WF71_S669s_1 | -0.02 | -0.58 | -1.75 | -0.06 | 0.59 | -0.76 | -0.2 | 0.2 | -0.68 | -1.42 | -1.14 | -0.3 | -1.13 | -0.42 | 0.11 | -1.22 | 0.26 | -0.0775 | 0.325 | 0.898894 | 0.253054 | 0.543443 | 1 | 0.999631 | 0.967185 |
| Lrfn1_Q2WF71_S713s_1 | -1.26 | 0.06 | -1.04 | -2.15 | 0.56 | 0.06 | -1.32 | -2.65 | -1 | -0.7 | -1.53 | -1.47 | -0.46 | -0.11 | -1.71 | -0.81 | -0.0825 | -0.03 | 0.2225 | 0.546753 | 0.729554 | 0.870833 | 1 | 0.999631 | 0.834213 |
| Lrfn1_Q2WF71_S726s_1 | -0.22 | -0.04 | -0.33 | 0.18 | -0.08 | -0.26 | -0.62 | 0.22 | -0.41 | -0.04 | 0.05 | -0.13 | -0.04 | 0.34 | 0.25 | -0.07 | 0.3175 | 0.2225 | 0.3575 | 0.439124 | 0.772317 | 0.910115 | 1 | 0.999631 | 0.781983 |
| Lrfn2_Q80TG9_S742s_1 | -0.69 | 0.21 | -0.77 | -0.31 | 0.32 | -0.22 | -0.16 | -0.23 | 0.2 | -0.12 | -0.76 | 0.01 | 1.02 | -0.25 | -0.5 | -0.4 | -0.045 | 0.125 | -0.27 | 0.412507 | 0.323349 | 0.525603 | 1 | 0.999631 | 0.768114 |
| Lrfn4_Q80XU8_S575s_1 | 0.1 | -0.47 | 0.15 | -0.69 | -0.51 | -0.42 | -0.01 | -0.15 | -0.23 | -0.04 | 0.01 | -0.15 | 0.15 | -1 | -0.23 | -0.91 | 0.1425 | 0.25 | 0.6325 | 0.484636 | 0.881154 | 0.673934 | 1 | 0.999631 | 0.803422 |
| Lrfn4_Q80XU8_S585s_1 | -0.18 | 0.66 | 0.54 | 0.04 | 0.46 | 0.3 | -0.25 | 1.12 | 0.47 | 0.32 | 0.12 | 1.15 | 0.84 | 0.98 | 0.9 | 0.87 | 0.645 | 0.8275 | 0.865 | 0.059766 | 0.698178 | 0.472743 | 1 | 0.999631 | 0.50521 |
| Lrfn4_Q80XU8_S611s_1 | 0.69 | 2.31 | 1.95 | 1.61 | 2.55 | 2.5 | 1.68 | 2.41 | 3.37 | 1.43 | 1.93 | 3.14 | 2.94 | 2.21 | 3.14 | 1.73 | -0.4225 | -0.6175 | -1.3575 | 0.088836 | 0.135403 | 0.15066 | 1 | 0.999631 | 0.553932 |
| Lrfn4_Q80XU8_S627s_1 | -0.41 | -3.89 | -1.62 | -1.04 | -3.22 | -3.05 | 0.25 | -2.63 | -3.38 | -1.9 | -1.74 | -2.41 | -3.2 | -3.63 | -2.52 | -3.04 | 1.1 | 2.085 | 0.375 | 0.07706 | 0.660928 | 0.416071 | 1 | 0.999631 | 0.542683 |
| Lrp1_Q91ZX7_S4521s_1 | 0.17 | 0.44 | 1.34 | 0.66 | 1.78 | 1.15 | 3.47 | 0.61 | 3.46 | 0 | 3.71 | 3.78 | 2.26 | -0.07 | 1.72 | 0.2 | 0.4225 | 0.49 | 0.97 | 0.517166 | 0.093461 | 0.029424 | 1 | 0.999631 | 0.819449 |
| Lrp4_Q8VI56_S1830s_1 | 0.27 | -0.74 | 0.52 | -0.26 | 0.87 | 0.44 | -0.21 | 0.38 | 1.04 | 1.02 | -0.1 | -0.21 | 0.37 | 0.86 | 1.35 | 1.09 | -0.1525 | 0.3075 | 0.33 | 0.027934 | 0.275209 | 0.280228 | 1 | 0.999631 | 0.423977 |
| Lrpap1_P55302_S53s_1 | 0.5 | -1.05 | -0.08 | -0.39 | -0.7 | -0.59 | 0.12 | -0.46 | -0.05 | 0.83 | -0.2 | -0.37 | -0.28 | -0.53 | 0.83 | 0.28 | -0.0425 | 0.005 | -0.45 | 0.464155 | 0.69155 | 0.472282 | 1 | 0.999631 | 0.793831 |
| Lrrc16a_D3Z030_S1099s_1 | -0.75 | -3.26 | -0.93 | -0.78 | -2.3 | -2.21 | -0.65 | -0.73 | -2.99 | -1.05 | -0.46 | -1.2 | -2.73 | -2.76 | -1.08 | -0.95 | -0.9075 | -1.29 | -0.61 | 0.524842 | 0.949804 | 0.994502 | 1 | 0.999631 | 0.822727 |
| Lrrc16a_D3Z030_S1149s_1 | 2.35 | -3.2 | -2.99 | -1.55 | -3.84 | -2.51 | 2.96 | -5.63 | -7.16 | 1.85 | -3.36 | -1.88 | -1.72 | -3.03 | -1.81 | -1.27 | -0.2675 | 0.0025 | -1.0175 | 0.594181 | 0.635418 | 0.505851 | 1 | 0.999631 | 0.85392 |
| Lrrc16b_Q3UFQ8_T131t_1 | 1.25 | -0.59 | -1.42 | -0.81 | -0.81 | -0.26 | -0.57 | -1 | 0.28 | 1.54 | -1.6 | -1.78 | -0.67 | -0.83 | -1.44 | -2.7 | 0.25 | 0.745 | 0.475 | 0.146176 | 0.627489 | 0.997667 | 1 | 1 | 0.61767 |
| Lrrc4b_P0C192_S698s_1 | -1.69 | -3.61 | -5.92 | -1.9 | -3.24 | -3.12 | -3.17 | -2.59 | -3.86 | -2.39 | -2.13 | -1.76 | -2.42 | -2.83 | -2.27 | -3.7 | 0.95 | 1.615 | 1.0175 | 0.59577 | 0.771642 | 0.434875 | 1 | 0.999631 | 0.854508 |
| Lrrc4b_P0C192_S700s_1 | -5.24 | -13.59 | -4.12 | -5.88 | -5.63 | -9.09 | -3.99 | -6.32 | -7.34 | -4.59 | -3.63 | -6.81 | -8.27 | -6.98 | -3.64 | -5.87 | -0.9575 | -0.3775 | -0.3325 | 0.609255 | 0.641487 | 0.421357 | 1 | 0.999631 | 0.858851 |
| Lrrc4c_Q8C031_S631s_1 | -1.78 | -3.08 | -1.69 | -2.18 | -4.3 | -4.64 | -1.36 | -2.26 | -3.86 | -2.46 | -1.9 | -2.02 | -4.23 | -2.6 | -1.57 | -1.66 | -0.2775 | 0.1025 | 0.1475 | 0.598468 | 0.221308 | 0.472044 | 1 | 0.999631 | 0.855552 |
| Lrrc7_E9Q6L9_S1123s_1 | -0.11 | 0.67 | 0.62 | 0.49 | 0.2 | -0.12 | -0.16 | 0.64 | 0.12 | 0.29 | 0.58 | 1.09 | 0.36 | 0.09 | 0.64 | 1.17 | -0.035 | 0.025 | -0.385 | 0.67465 | 0.383691 | 0.756824 | 1 | 0.999631 | 0.885515 |
| Lrrc7_E9Q6L9_S1137s_1 | 0.72 | 0.47 | -0.1 | 0.66 | 0.53 | 0.22 | 0.14 | 0.72 | 0.58 | 0.43 | 0.55 | 0.29 | -0.99 | 0.38 | 0.34 | 0.48 | -0.3225 | -0.12 | -0.2775 | 0.361924 | 0.906019 | 0.930602 | 1 | 0.999631 | 0.742006 |
| Lrrc7_E9Q6L9_S1162s_1 | 0.76 | 0.36 | -0.17 | -0.17 | 0.46 | -0.17 | -0.48 | -0.32 | 0.04 | 0.24 | -0.01 | 0.03 | 0.03 | -0.13 | -0.86 | 0.63 | -0.46 | -0.065 | 0.2125 | 0.496582 | 0.355073 | 0.694874 | 1 | 0.999631 | 0.811248 |
| Lrrc7_E9Q6L9_S1211s_1 | 0.46 | 1.17 | 1.71 | 1.07 | 0.65 | 0.65 | 0.54 | 0.73 | 0.68 | 0.92 | 0.91 | 1.64 | 0.87 | 0.52 | 1.74 | 2.13 | 0.1425 | -0.18 | -0.445 | 0.639505 | 0.163847 | 0.85758 | 1 | 0.999631 | 0.870995 |
| Lrrc7_E9Q6L9_S1395s_1 | 0.17 | -0.44 | -0.17 | 0.45 | 0.45 | -0.37 | 0.27 | 0.23 | -0.19 | -1.4 | 0.18 | 0.7 | -1.84 | -0.85 | 0.45 | 0.47 | 0.5075 | 0.415 | 0.5525 | 0.42675 | 0.652817 | 0.702964 | 1 | 0.999631 | 0.775151 |
| Lrrc7_E9Q6L9_S1444s_1 | -1.24 | 0.62 | 0.43 | -0.21 | 0.61 | -0.07 | 0.53 | 0.56 | 0.71 | -1.5 | 1.16 | 0.89 | 0.98 | 0.08 | 0.48 | 0.27 | 0.1325 | -0.0575 | 0.1425 | 0.244907 | 0.263259 | 0.536799 | 1 | 0.999631 | 0.685751 |
| Lrrc7_E9Q6L9_S713s_1 | 0.88 | 0.14 | -0.43 | 1.21 | 1.89 | 0.03 | 0.63 | -0.22 | 0.22 | -0.12 | 0.95 | 0.52 | 1.69 | -0.43 | 0.17 | 0.94 | 1.13 | 0.2175 | -0.645 | 0.795559 | 0.809469 | 0.89402 | 1 | 0.999631 | 0.929355 |
| Lrrc7_E9Q6L9_S855s_1 | -0.14 | -0.92 | -1.78 | 0.66 | 1.18 | 0.02 | 0.6 | 0.54 | 0.15 | -0.65 | 0.14 | -0.95 | -2.71 | -0.46 | -1.02 | -0.57 | 0.2275 | -0.06 | 0.3275 | 0.340855 | 0.056106 | 0.693358 | 1 | 0.999631 | 0.733032 |
| Lrrc7_E9Q6L9_S874s_1 | -0.18 | 0.16 | -0.63 | -0.02 | 0.05 | 0.42 | -0.33 | 0.1 | 0.02 | 0.09 | -1.79 | 0.77 | -1.35 | 0.57 | 0.62 | 0.8 | -0.4425 | 0.1675 | 0.33 | 0.523475 | 0.44838 | 0.910577 | 1 | 0.999631 | 0.821972 |
| Lrrc7_E9Q6L9_S954s_1 | -0.62 | 0.05 | 0.04 | -0.62 | -0.49 | -0.76 | -1.03 | -0.64 | -0.24 | -0.02 | -0.28 | 0.06 | 0.47 | -0.34 | -0.39 | 0.43 | -0.3375 | -0.3925 | 0.0775 | 0.365158 | 0.153887 | 0.570248 | 1 | 0.999631 | 0.745324 |
| Lrrc7_E9Q6L9_T870t_1 | -2.68 | -5.52 | -0.86 | -0.2 | -4.46 | -5.51 | -0.64 | 0 | -5.82 | -4.54 | -0.26 | -0.21 | -3.49 | -5.14 | -0.49 | 0.17 | -0.58 | -0.23 | -0.66 | 0.957394 | 0.826343 | 0.805569 | 1 | 0.999631 | 0.985227 |
| Lrrc75a_Q7TSF4_S15s_1 | 0.93 | -0.03 | -0.27 | -0.43 | -0.37 | -0.64 | -0.29 | -0.82 | -0.14 | 0.57 | -0.19 | -0.96 | -0.37 | -0.97 | -0.74 | -0.36 | -0.4125 | -0.29 | 0.15 | 0.104392 | 0.126536 | 0.598832 | 1 | 0.999631 | 0.568213 |
| Lrrc75a_Q7TSF4_S322s_1 | -0.17 | 0.03 | 1.13 | 0.18 | 0.18 | -0.35 | 0.06 | -0.37 | -0.19 | 0.57 | -1 | 0.63 | 0.86 | 0.4 | 0.1 | 0.41 | -0.09 | 0.22 | 0.7025 | 0.686749 | 0.255905 | 0.536779 | 1 | 0.999631 | 0.890988 |
| Lrrc8b_Q5DU41_S215s_1 | -1.25 | -0.26 | -0.96 | -1.05 | -0.1 | -1.09 | -1.46 | -1.23 | -0.27 | -0.6 | -1.11 | -0.66 | 1.17 | -0.64 | -0.99 | -0.25 | -0.3 | -0.2575 | -0.035 | 0.18055 | 0.813785 | 0.50878 | 1 | 0.999631 | 0.64376 |
| Lrrfip1_E9Q9T1_S236s_1 | 0.35 | -0.79 | -0.28 | -0.71 | -0.83 | -0.52 | -0.64 | -0.64 | -0.34 | -0.26 | -0.81 | -1.05 | -0.61 | 0.24 | -1.55 | 0.35 | -0.565 | -0.255 | -0.9525 | 0.943395 | 0.357091 | 0.477061 | 1 | 0.999631 | 0.981629 |
| Lrrfip1_E9Q9T1_S88s_1 | -1.42 | -3.69 | -0.16 | 1.96 | -4.31 | -4.05 | 2.79 | 0 | -2.6 | -1.81 | 0.78 | -0.7 | -2.15 | -3.22 | -0.65 | -1.1 | -0.2375 | -0.215 | -1.3 | 0.402232 | 0.748076 | 0.830511 | 1 | 0.999631 | 0.763561 |
| Lrrfip1_E9Q9T1_S90s_1 | 0.64 | -0.8 | -0.03 | -0.83 | -1.38 | 0.57 | 0.52 | -1.68 | -0.73 | 1.52 | -1.04 | -1.63 | -1.06 | -1.28 | -1.65 | -2.23 | -0.1475 | 0.1075 | -0.0175 | 0.013347 | 0.705188 | 0.755751 | 1 | 0.999631 | 0.385092 |
| Lrrfip2_E9QN52_S103s_1 | -0.15 | 0.77 | 0.52 | -0.04 | -0.25 | 1.09 | -0.01 | -0.32 | 0.51 | 0.03 | 0.44 | 0.55 | -0.29 | 0.33 | 0.68 | 0.31 | 0.17 | 0.3 | 0.1175 | 0.96048 | 0.712831 | 0.733375 | 1 | 0.999631 | 0.986517 |
| Lrrfip2_E9QN52_S103s_2 | -0.19 | 0.01 | -0.02 | 0.77 | 0.57 | 0.09 | -0.02 | 0.61 | -0.29 | 0.25 | 1.18 | 0.63 | 0.01 | 0.55 | 0 | 0.48 | -0.0375 | 0.09 | 0.295 | 0.72413 | 0.595357 | 0.451672 | 1 | 0.999631 | 0.905674 |
| Lrrfip2_E9QN52_S111s_1 | -0.31 | 0.1 | -1.14 | -0.8 | -0.28 | -0.85 | -0.86 | -0.31 | -0.39 | -0.41 | -0.78 | -0.21 | 0.1 | -0.35 | -0.66 | -0.06 | 0.1375 | -0.2225 | 0.2475 | 0.427547 | 0.91402 | 0.792546 | 1 | 0.999631 | 0.775223 |
| Lrrfip2_E9QN52_S95s_1 | -0.19 | 0.7 | -0.6 | -0.31 | 0 | 1.64 | 0.37 | -1.86 | -0.54 | -0.36 | 0.31 | -0.7 | 1.04 | -0.85 | 0.25 | 0.15 | 0.745 | 0.4925 | -0.1725 | 0.600108 | 0.841582 | 0.561471 | 1 | 0.999631 | 0.856618 |
| Lrrfip2_E9QN52_T99t_1 | -0.23 | 0.06 | -0.56 | 0.52 | 1.23 | 1.41 | 0.89 | -0.76 | 1.18 | -0.52 | 0.49 | 0.61 | -2.58 | 0.45 | 0.68 | 0.55 | 0.09 | -0.9575 | -0.8475 | 0.811527 | 0.166732 | 0.261787 | 1 | 0.999631 | 0.936949 |
| Lrrtm2_Q8BGA3_S458s_1 | 2.28 | 1.31 | 1.25 | 1.43 | 1.17 | 1.88 | 1.97 | 1.61 | 0.88 | 1.16 | 0.55 | -0.15 | 0.38 | 0.79 | 1.03 | 0.68 | -0.0875 | 0.0075 | -0.1925 | 0.029851 | 0.789695 | 0.031825 | 1 | 0.999631 | 0.434385 |
| Lrrtm4_E9Q8Y9_S462s_1 | 1.31 | 1.21 | 0.64 | 1.76 | 1.22 | 1.45 | 1.2 | 0.7 | 1.35 | 0.82 | 1.32 | 1.46 | 0.42 | 1.34 | 1.07 | 1.32 | -0.38 | -0.0775 | 0.0825 | 0.598192 | 0.787569 | 0.981624 | 1 | 0.999631 | 0.855276 |
| Lsm11_Q8BUV6_S155s_1 | -1.09 | -0.06 | -1.58 | -0.43 | -1.9 | -0.32 | -1.24 | -1.22 | -1.09 | -1.23 | -0.77 | -0.38 | -0.18 | -0.67 | -0.14 | -1.84 | -0.3325 | -1.13 | -1.13 | 0.868749 | 0.409743 | 0.846445 | 1 | 0.999631 | 0.957311 |
| Lsm11_Q8BUV6_S15s_1 | 0.86 | -1.04 | 1.35 | -0.53 | -1.33 | -0.19 | 2.65 | -1.82 | -2.54 | 0.64 | -1.54 | -0.44 | -0.48 | -1.42 | -0.84 | -1.14 | 0.05 | 0.2225 | -0.365 | 0.065357 | 0.737287 | 0.17117 | 1 | 0.999631 | 0.516298 |
| Lsm11_Q8BUV6_S15s_2 | -0.14 | -0.87 | -0.82 | -0.54 | -0.7 | -0.25 | -0.55 | -0.67 | -0.49 | 0.08 | -0.09 | -0.98 | -0.78 | -1.37 | -1.29 | -0.39 | -0.6875 | 0.0225 | -0.35 | 0.304452 | 0.857732 | 0.51373 | 1 | 0.999631 | 0.716239 |
| Lsm11_Q8BUV6_S18s_1 | 0.14 | -0.47 | 0.27 | -0.72 | -1.63 | -1.03 | -0.28 | -0.59 | -0.92 | 0.15 | 0.07 | 0.01 | -0.95 | -1.1 | -0.28 | 0.15 | -0.2075 | 0.135 | 0.19 | 0.391894 | 0.1008 | 0.951755 | 1 | 0.999631 | 0.758901 |
| Lsm11_Q8BUV6_S21s_1 | -0.13 | -0.52 | 0.37 | -0.77 | -0.78 | -0.61 | 0.03 | -0.52 | -1.29 | -0.1 | 0.91 | -0.03 | -0.48 | -0.49 | 0.55 | 0.13 | 0.02 | 0.1925 | -0.2725 | 0.625349 | 0.545009 | 0.783561 | 1 | 0.999631 | 0.863802 |
| Lsm11_Q8BUV6_S21s_2 | -0.14 | -0.91 | -0.82 | -0.54 | -0.73 | -0.38 | -0.55 | -0.67 | -0.67 | 0.1 | -0.09 | -0.98 | -0.63 | -1.19 | -1.29 | -0.39 | -0.3425 | -0.2825 | 0.365 | 0.432746 | 0.942104 | 0.581276 | 1 | 0.999631 | 0.779247 |
| Lsm11_Q8BUV6_S285s_1 | -2.8 | -2.19 | -2.23 | -3.77 | -2.66 | -2.44 | -2.77 | -4.49 | -3.73 | -2.19 | -2.17 | -4.03 | -0.23 | -2.34 | -4.11 | -2.85 | -0.1625 | 0.055 | -0.585 | 0.640161 | 0.537054 | 0.620873 | 1 | 0.999631 | 0.870995 |
| Lsm11_Q8BUV6_T105t_1 | 0.67 | 0.34 | -0.5 | 0.09 | 0.26 | -0.43 | 0.13 | -0.01 | 0.7 | 0.6 | 0.34 | -0.82 | -0.21 | -0.63 | -1.13 | 0.23 | 0.2125 | 0.365 | 0.7725 | 0.169281 | 0.624679 | 0.897795 | 1 | 0.999631 | 0.638979 |
| Lsm12_Q9D0R8_T75t_1 | -0.27 | 2.03 | 1.54 | 0.86 | 1.42 | 1.74 | 0.52 | 1.33 | 1.64 | 1.09 | 1.27 | 1.62 | 1.71 | 2.07 | 1.47 | 2 | -0.0525 | 0.225 | 0.53 | 0.142979 | 0.684735 | 0.464679 | 1 | 0.999631 | 0.614053 |
| Lsm14a_Q8K2F8_S182s_1 | 0.65 | 1.59 | 1.25 | 1.02 | 0.96 | 1.49 | 0.62 | 1.23 | 1.21 | 1.2 | 1.4 | 1.6 | 1.14 | 1.91 | 1.6 | 1.98 | -0.04 | 0.2125 | 0.5825 | 0.142364 | 0.869578 | 0.45646 | 1 | 0.999631 | 0.612726 |
| Lsm14a_Q8K2F8_S183s_1 | 0.5 | 1.51 | 1.08 | 0.96 | 1.06 | 1.28 | 0.4 | 1.15 | 1.17 | 1.12 | 1.23 | 1.38 | 1.23 | 1.96 | 1.55 | 1.64 | -0.03 | -0.135 | 0.2625 | 0.10179 | 0.902949 | 0.478299 | 1 | 0.999631 | 0.564448 |
| Lsm14a_Q8K2F8_S192s_1 | -0.32 | 0.71 | 0.1 | -0.2 | 0.32 | 0.14 | -0.22 | -0.07 | 0.27 | -0.11 | -0.15 | -0.26 | 0.7 | 0.47 | -0.05 | 0.22 | 0.45 | 0.2525 | 0.165 | 0.451591 | 0.923427 | 0.6728 | 1 | 0.999631 | 0.788148 |
| Lsm14a_Q8K2F8_S216s_1 | 0.85 | 0.45 | -0.49 | 0.94 | 1.18 | 0.85 | 0.5 | 1.02 | 0.68 | 0.43 | 0.95 | 0.7 | -0.13 | 1.23 | 0.7 | 0.61 | 0.1825 | -0.4125 | -0.315 | 0.705921 | 0.247123 | 0.501268 | 1 | 0.999631 | 0.897883 |
| Lsr_D3YYU9_S258s_1 | 0.08 | 0.49 | 0.01 | -0.14 | -0.34 | 0.69 | 0.26 | 0.56 | -0.8 | -0.75 | -0.17 | 0.51 | -1.08 | -0.19 | 0.55 | -0.1 | 1.3375 | 3.35 | 0.76 | 0.42799 | 0.567447 | 0.274564 | 1 | 0.999631 | 0.77566 |
| Ltbp4_D3Z598_S1491s_1 | -0.9 | -1.37 | -0.84 | 0.24 | 2.11 | 0.83 | 0.02 | -0.48 | 1.8 | 2.22 | 2.74 | 3.77 | -0.84 | 0.18 | 0.29 | 0.54 | 1.0025 | 0.3325 | -1.5875 | 0.117928 | 0.046397 | 8.63E-05 | 1 | 0.441561 | 0.585087 |
| Ltn1_Q14B86_S18s_1 | 2.25 | -0.13 | -0.79 | 2.65 | 1.92 | 1.57 | 2.33 | 2.17 | 2.1 | 1.52 | 2.09 | -0.4 | -3.77 | 0.62 | 1.35 | -0.57 | 0.185 | 0.48 | 0.6875 | 0.205708 | 0.209451 | 0.713649 | 1 | 0.999631 | 0.6605 |
| Luc7l3_Q5SUF2_S425s_1 | -0.37 | 0.77 | 0.82 | 0.62 | 1.09 | 0.32 | 0 | 1.17 | 0.86 | 0.72 | 0.78 | 1.4 | 0.91 | 1.87 | 0.81 | 1 | -0.12 | -0.2025 | -1.095 | 0.107653 | 0.648767 | 0.198497 | 1 | 0.999631 | 0.569416 |
| Lurap1_Q9D6I9_S118s_1 | -0.27 | -1.38 | -1.28 | -1.32 | -1.39 | -0.9 | -1.35 | -1.09 | -0.72 | -0.6 | -1.82 | -1.92 | -1.84 | -2.4 | -3.07 | -1.32 | -0.51 | 0.37 | 0.02 | 0.03485 | 0.717086 | 0.644296 | 1 | 0.999631 | 0.44933 |
| Lurap1_Q9D6I9_S126s_1 | -0.66 | -0.56 | 0.2 | -0.46 | -1.67 | -0.84 | -0.79 | -0.22 | -0.27 | 0.68 | -0.28 | -0.13 | -1.43 | -0.5 | 0.4 | 0.13 | -0.8525 | 0.1325 | 0.88 | 0.964484 | 0.192891 | 0.296513 | 1 | 0.999631 | 0.987331 |
| Lurap1l_Q8K2P1_S72s_1 | -0.58 | -1.26 | -1.75 | -1.59 | -1.86 | -2.41 | -1.61 | -2.71 | -1.87 | -0.78 | -0.57 | -1.43 | -0.48 | 0.18 | -0.17 | -1.19 | -0.655 | -0.74 | -0.825 | 0.053331 | 0.044384 | 0.744283 | 1 | 0.999631 | 0.490856 |
| Luzp1_Q8R4U7_S395s_1 | 0.71 | -1.44 | 1.56 | 1.03 | -1.82 | -1.03 | 1.29 | 0.8 | -2.19 | -0.15 | 0.85 | 0.39 | -1.18 | -1.27 | 0.95 | 0.06 | 0.865 | -0.3075 | -0.995 | 0.282989 | 0.452698 | 0.378829 | 1 | 0.999631 | 0.704545 |
| Luzp1_Q8R4U7_S612s_1 | 0.73 | 1.02 | 0.01 | 2.38 | 2.63 | 1.04 | 2.76 | 1.17 | 1.46 | -1.7 | 1.74 | 1.41 | -0.96 | 0.26 | 1.32 | -0.46 | 0.9075 | -0.2875 | -0.445 | 0.138914 | 0.178373 | 0.712698 | 1 | 0.999631 | 0.611718 |
| Luzp1_Q8R4U7_S746s_1 | 1.64 | 1.22 | 0.63 | 1.48 | 1.96 | 2.67 | 1.28 | 2.69 | 1.37 | 1.47 | 0.26 | 0.72 | -2.31 | 2.79 | 1.54 | 1.17 | -0.37 | -0.1525 | -0.015 | 0.64326 | 0.046294 | 0.458773 | 1 | 0.999631 | 0.871827 |
| Luzp2_Q8BGY3_S279s_1 | -0.35 | 0.18 | 0.04 | -0.44 | -0.43 | -0.87 | -0.67 | -0.08 | -0.19 | -0.41 | -0.21 | -0.37 | -0.21 | -0.46 | 0 | 0.04 | 0.185 | -0.28 | 0.34 | 0.959779 | 0.22775 | 0.575298 | 1 | 0.999631 | 0.986095 |
| Lysmd1_Q9D0E3_S23s_1 | 0.22 | 0.72 | -0.04 | 0.74 | 0.97 | 0.66 | 0.11 | 0.64 | 1.29 | -1 | 0.42 | -0.19 | 0.84 | 0.91 | 0.76 | 0.49 | 1.555 | 1.965 | 1.3075 | 0.285661 | 0.559701 | 0.574948 | 1 | 0.999631 | 0.705851 |
| Lysmd2_G3X915_S24s_1 | -2 | 2.77 | 2.5 | -1.05 | 3.22 | 1.23 | 2.66 | 1.33 | 4.07 | -1.84 | 3.5 | 4.35 | 4.73 | 0.33 | 1.96 | 0.43 | 0.19 | -0.125 | 0.095 | 0.341645 | 0.188125 | 0.242371 | 1 | 0.999631 | 0.73317 |
| Lysmd2_G3X915_S31s_1 | -0.13 | 0.09 | -0.16 | 0.38 | 0.37 | 0.2 | 0.01 | 0.36 | -0.06 | -0.34 | 0.11 | -0.03 | 0.35 | -0.11 | 0.06 | 0.26 | -0.02 | 0.3225 | 0.5225 | 0.741018 | 0.468469 | 0.643459 | 1 | 0.999631 | 0.909055 |
| Lyst_G5E8Q0_S2098s_1 | -0.38 | 0.37 | 0.7 | 0.08 | 0.41 | 0.17 | -0.36 | 0.47 | 0.43 | 0.5 | 0.33 | 0.8 | 0.69 | 0.3 | 1.04 | 0.83 | -0.4475 | 0.02 | 0.12 | 0.14868 | 0.952369 | 0.318545 | 1 | 0.999631 | 0.62132 |
| Lyst_G5E8Q0_S2117s_1 | -0.08 | 1.01 | -0.15 | -0.43 | 0.19 | -0.2 | -0.68 | -0.75 | 0.82 | 0.29 | -0.43 | -0.25 | 0.75 | -0.04 | 0.04 | 0.08 | 0.125 | 0.365 | 0.0375 | 0.760837 | 0.271348 | 0.962478 | 1 | 0.999631 | 0.914614 |
| Lyst_G5E8Q0_S2202s_1 | -0.46 | -0.31 | -1.94 | -0.71 | -0.06 | -0.66 | -1.46 | -0.74 | -0.13 | -0.39 | -0.4 | -1.04 | 0.18 | -0.31 | -1.76 | -1.38 | -0.405 | -1.015 | -0.515 | 0.945161 | 0.782526 | 0.395873 | 1 | 0.999631 | 0.982141 |
| Lyst_G5E8Q0_S2559s_1 | 0.05 | -2.41 | -1.26 | -1.67 | -2.52 | -2.14 | -0.9 | -1.35 | -2.88 | -1.45 | -2.02 | -3 | -2.16 | -2.83 | -1.31 | -1.05 | -0.1375 | -0.4025 | -0.7075 | 0.398983 | 0.487917 | 0.105462 | 1 | 0.999631 | 0.761767 |
| Lyst_G5E8Q0_S2559s_2 | -0.02 | -0.43 | -0.51 | -1.9 | -1.41 | -0.38 | -0.56 | -1.06 | -0.46 | 0.11 | -2.15 | -1.97 | -1.2 | -0.88 | -2.14 | -1.47 | 0.075 | -0.11 | -0.6525 | 0.160257 | 0.762662 | 0.524605 | 1 | 0.999631 | 0.629764 |
| Lyst_G5E8Q0_S2563s_1 | -0.05 | -3.36 | -1.44 | -2.03 | -2.68 | -1.86 | -0.78 | -1.26 | -2.34 | -0.26 | -2.95 | -1.77 | -4.87 | -1.79 | -1.86 | -0.97 | -0.1375 | -0.4025 | -0.7075 | 0.494063 | 0.915401 | 0.888895 | 1 | 0.999631 | 0.81007 |
| Lyst_G5E8Q0_S2563s_2 | -0.02 | -0.43 | -0.51 | -1.9 | -1.41 | -0.38 | -0.56 | -1.06 | -0.46 | 0.11 | -2.15 | -1.97 | -1.2 | -0.88 | -2.14 | -1.47 | -0.0325 | -0.06 | -0.1925 | 0.160257 | 0.762662 | 0.524605 | 1 | 0.999631 | 0.629764 |
| Lyst_G5E8Q0_S2613s_1 | 0.08 | -1.3 | -0.86 | -0.35 | -0.92 | -0.99 | -0.25 | -0.4 | -1.13 | -0.26 | -0.72 | -0.56 | -1.08 | -1.02 | -0.48 | -0.62 | 0.0425 | -0.5725 | -0.185 | 0.609605 | 0.930111 | 0.873006 | 1 | 0.999631 | 0.858923 |
| Lyst_G5E8Q0_S669s_1 | -0.35 | -0.71 | -1.04 | -0.04 | 0.18 | -0.72 | -0.95 | -0.48 | -0.68 | -0.63 | -2.08 | -1.04 | -0.31 | -0.9 | -0.89 | -0.78 | 0.3425 | 0.56 | 0.8275 | 0.579098 | 0.904299 | 0.183476 | 1 | 0.999631 | 0.846484 |
| Lyst_G5E8Q0_S866s_1 | -0.24 | 1.13 | 1.19 | -0.06 | 0.33 | 1.45 | 0.51 | 1.1 | 0.69 | 0.78 | 1.5 | 1.29 | 1.35 | 1.98 | 1.71 | 0.29 | 0.09 | -0.53 | -1.625 | 0.126147 | 0.449675 | 0.210996 | 1 | 0.999631 | 0.596745 |
| Lzts3_A2AHG0_T561t_1 | 1.62 | -1.14 | -0.66 | 1.91 | -0.26 | -0.23 | 2.27 | 0.31 | -0.63 | 0.09 | 0.62 | -0.47 | -2.02 | -2.45 | -0.44 | 0.14 | -1.1075 | -0.2675 | -1.705 | 0.083121 | 0.915812 | 0.476043 | 1 | 0.999631 | 0.549732 |
| Lzts3_A2AHG0_T563t_1 | 2.2 | -1.27 | 0.93 | 0.03 | -1.35 | -0.36 | -0.63 | -0.2 | -1.06 | 2.85 | -0.64 | -0.33 | -2.48 | -1.17 | -0.2 | -1.08 | -0.125 | -0.2375 | 0.0975 | 0.046212 | 0.131349 | 0.788886 | 1 | 0.999631 | 0.479949 |
| Macf1_Q9QXZ0_S3889s_1 | -0.83 | -1.99 | -0.25 | -0.32 | -1.43 | -1.19 | -0.79 | -0.48 | -1.99 | -0.96 | -1.19 | -0.2 | -1.5 | -1.1 | -0.13 | -0.27 | -0.08 | 0.1175 | 0.425 | 0.844856 | 0.777351 | 0.645566 | 1 | 0.999631 | 0.948565 |
| Macf1_Q9QXZ0_S4458s_1 | -0.6 | -0.27 | -0.89 | -1.34 | -0.47 | -0.56 | -1.1 | -1.29 | -0.41 | -0.26 | -1.03 | -0.93 | 0.5 | -0.14 | -0.96 | -0.8 | 0.5575 | 1.095 | 1.33 | 0.322635 | 0.813536 | 0.730953 | 1 | 0.999631 | 0.724485 |
| Macf1_Q9QXZ0_S4483s_1 | -0.09 | -0.91 | -2.87 | -2.93 | -0.58 | -0.96 | -2.04 | -0.99 | -0.5 | -0.2 | -1.15 | -0.57 | 0.51 | -0.76 | -0.84 | -0.39 | -0.0775 | 0.01 | 0.65 | 0.078044 | 0.427238 | 0.123676 | 1 | 0.999631 | 0.543121 |
| Macf1_Q9QXZ0_S4611s_1 | -0.97 | -0.88 | -1.14 | -1.13 | -1.05 | -0.85 | -0.74 | -1.79 | -0.73 | -0.87 | -2.2 | -0.28 | 0.27 | -0.73 | -0.59 | -0.47 | -0.6225 | -0.49 | 0.3575 | 0.063592 | 0.799179 | 0.981027 | 1 | 0.999631 | 0.513165 |
| Macf1_Q9QXZ0_S5551s_1 | -0.33 | -0.13 | -0.56 | 0.34 | -0.91 | -0.96 | -1.14 | -0.16 | -0.58 | -0.38 | -1.28 | -0.4 | 0.09 | 0.25 | -0.17 | 0.58 | 0.3225 | 0.2275 | 0.0325 | 0.287018 | 0.083994 | 0.165693 | 1 | 0.999631 | 0.705972 |
| Macf1_Q9QXZ0_S57s_1 | -1.44 | -1.59 | -1.08 | -1.16 | -0.78 | -1.8 | -0.65 | -0.75 | -0.92 | -1.86 | -0.94 | -0.64 | -1.23 | -1.64 | -0.63 | -1.64 | 0.2325 | 0.5075 | 1.055 | 0.922837 | 0.346025 | 0.50599 | 1 | 0.999631 | 0.975032 |
| Macf1_Q9QXZ0_S5988s_1 | -1.02 | 1.76 | 0.86 | -0.34 | 1.57 | 0.17 | 0.47 | -0.02 | 1.58 | -1.65 | 1.24 | 2.12 | 2.59 | 0.75 | 1.59 | 0.55 | 0.1425 | -0.24 | -0.2125 | 0.148395 | 0.716657 | 0.579716 | 1 | 0.999631 | 0.62132 |
| Macf1_Q9QXZ0_S6923s_1 | 0.35 | 0.64 | -0.5 | 0.22 | 0.82 | -0.09 | 0.14 | 0.41 | 0.23 | -0.5 | -0.3 | 0.32 | -0.07 | -0.08 | -0.07 | 0.08 | 0.175 | 0.13 | 0.2775 | 0.517043 | 0.679965 | 0.500695 | 1 | 0.999631 | 0.819449 |
| Macf1_Q9QXZ0_S7244s_1 | 0.89 | 1.53 | 1.4 | 1.12 | 1 | 1.8 | 0.94 | 1.9 | 1.26 | 1.39 | 1.35 | 1.46 | 1.32 | 1.8 | 1.14 | 1.79 | -0.0575 | 0.375 | 0.6975 | 0.383539 | 0.601171 | 0.626815 | 1 | 0.999631 | 0.754618 |
| Macf1_Q9QXZ0_S907s_1 | -0.12 | -1.1 | -0.34 | -0.24 | -0.53 | 0.3 | -0.93 | -0.87 | -0.75 | 0.02 | -0.04 | 0.47 | 0.69 | 0.56 | -0.45 | 0.19 | -0.4675 | -0.25 | -0.6425 | 0.088059 | 0.878193 | 0.318513 | 1 | 0.999631 | 0.553858 |
| Macrod2_Q3UYG8_S257s_1 | 1.44 | 1.96 | 3.17 | 1.85 | 1.14 | 2.34 | 2.08 | 0.99 | 1.76 | 2.3 | 1.37 | 1.99 | 0.89 | 1.02 | 2.21 | 1.73 | -0.2825 | 0.0175 | 0.365 | 0.195265 | 0.33877 | 0.557945 | 1 | 0.999631 | 0.652992 |
| Macrod2_Q3UYG8_S266s_1 | 0.72 | 1.42 | 1.39 | 0.98 | 0.39 | 1.29 | 0.6 | 1.1 | 0.93 | 1.46 | 1.05 | 1.14 | 1.35 | 1.69 | 1.14 | 1.79 | 0.4525 | 0.2475 | 0.1075 | 0.262824 | 0.38524 | 0.95181 | 1 | 0.999631 | 0.693034 |
| Macrod2_Q3UYG8_S283s_1 | -1.28 | 0.83 | 2.2 | -1.3 | 0.61 | 0.36 | 1.71 | -0.42 | 1.29 | -1.79 | 0.37 | 1.57 | 1.99 | -2.41 | 0.95 | 0.35 | -0.0625 | -0.2275 | 0.2125 | 0.920798 | 0.591905 | 0.802775 | 1 | 0.999631 | 0.975032 |
| Macrod2_Q3UYG8_S284s_1 | -1.28 | 0.83 | 2.2 | -1.3 | 0.61 | 0.36 | 1.71 | -0.42 | 1.29 | -1.79 | 0.37 | 1.57 | 1.99 | -2.41 | 0.95 | 0.35 | 0.4525 | 0.2475 | 0.1075 | 0.920798 | 0.591905 | 0.802775 | 1 | 0.999631 | 0.975032 |
| Madd_E9QN47_S1038s_1 | -0.5 | -0.85 | -0.78 | -1.34 | -0.9 | -0.63 | -1.01 | -1.04 | -1.11 | -0.06 | -0.82 | -0.7 | -0.02 | -0.44 | -1.01 | -0.5 | -0.0275 | 0.195 | 0.375 | 0.280023 | 0.921637 | 0.561211 | 1 | 0.999631 | 0.701922 |
| Madd_E9QN47_S1089s_1 | -1.98 | -2.35 | -3.35 | -2.23 | -1.13 | -2.8 | -2.66 | -2.47 | -1.39 | -2.51 | -2.82 | -2.33 | -0.44 | -2.15 | -1.99 | -3.42 | 0.2125 | 0.215 | 0.4775 | 0.445926 | 0.65027 | 0.619167 | 1 | 0.999631 | 0.784402 |
| Madd_E9QN47_S1218s_1 | -0.67 | -0.52 | -0.78 | -0.71 | -0.68 | -0.84 | -1.06 | -1.07 | -0.65 | -0.75 | -1.05 | -0.83 | 0.14 | -0.35 | -0.58 | -0.44 | -0.2425 | -0.15 | 0.3625 | 0.227255 | 0.33163 | 0.551428 | 1 | 0.999631 | 0.678384 |
| Madd_E9QN47_S1247s_1 | 0.99 | 0.2 | -0.37 | -0.42 | 0.15 | 0.61 | 0.26 | 0.04 | 0.69 | 1.32 | 1.33 | 1.29 | -0.21 | -0.63 | 1.58 | 0.5 | 0.165 | 1.0575 | 0.21 | 0.700909 | 0.656748 | 0.019429 | 1 | 0.999631 | 0.896835 |
| Madd_E9QN47_S1480s_1 | -0.04 | 0.1 | 0.32 | -0.1 | -0.42 | 0.44 | 0.73 | -0.18 | 0.29 | -0.17 | 0.23 | 0.22 | 0.94 | 0.1 | 0.56 | 0.14 | 0.0725 | 0.0725 | 0.365 | 0.257706 | 0.824298 | 0.782676 | 1 | 0.999631 | 0.692624 |
| Madd_E9QN47_S1627s_1 | -0.1 | -0.71 | -0.26 | -0.62 | -1.11 | -0.95 | 0.08 | -0.47 | -0.31 | -0.25 | 0.22 | 0.5 | 0.13 | -0.83 | -0.32 | -0.01 | -0.19 | 0.4625 | 0.165 | 0.619935 | 0.579143 | 0.158574 | 1 | 0.999631 | 0.861888 |
| Madd_E9QN47_S817s_1 | 0.33 | -1.9 | 0.53 | -0.93 | -2.46 | -0.33 | -0.37 | -1.26 | -1.18 | 1.33 | -1.35 | -1.55 | 0.14 | -1.25 | -0.09 | -1.56 | -0.6125 | -0.195 | -0.1975 | 0.757123 | 0.373871 | 0.802374 | 1 | 0.999631 | 0.914382 |
| Madd_E9QN47_S819s_1 | 1.57 | -1.09 | 0 | 0.66 | -0.59 | -0.22 | 1.08 | -0.28 | -0.71 | 0.65 | -0.55 | -1.07 | -1.74 | -1.79 | -0.68 | -1.49 | -0.2875 | -0.705 | -1.71 | 0.012435 | 0.636776 | 0.265463 | 1 | 0.999631 | 0.385092 |
| Madd_E9QN47_S828s_1 | -0.42 | 0.82 | -2.04 | -1.41 | 0.23 | -0.58 | -0.67 | -0.8 | -0.25 | -0.19 | -1.32 | -1 | 0.61 | 0.32 | -0.99 | -0.59 | 0.3075 | 0.0725 | 0.6 | 0.368908 | 0.611533 | 0.906785 | 1 | 0.999631 | 0.74602 |
| Madd_E9QN47_S829s_1 | -0.54 | -0.44 | -1.06 | -0.83 | -0.02 | -0.51 | -0.8 | -0.95 | -0.73 | -0.91 | -0.35 | -1 | 0.14 | -0.3 | -0.73 | -0.52 | 0.1475 | -0.03 | 0.365 | 0.267842 | 0.632324 | 0.917223 | 1 | 0.999631 | 0.696016 |
| Madd_E9QN47_S881s_1 | 1.68 | 1.14 | -0.18 | 0.69 | 2.13 | 2.43 | 0.61 | 1.57 | 2.03 | 0.62 | 0.45 | 1.1 | 1.6 | 1.82 | 1.36 | 0.68 | 0.8525 | 0.2175 | 0.5325 | 0.26286 | 0.126395 | 0.666168 | 1 | 0.999631 | 0.693034 |
| Madd_E9QN47_S909s_1 | 2.16 | -1.13 | 0.09 | -0.04 | -1.34 | -0.28 | 1.15 | -1.19 | -2.06 | 1.44 | -0.79 | -2.04 | -2.1 | -1.64 | -0.81 | -1.57 | -0.685 | -1.1325 | -1.8 | 0.019962 | 0.390749 | 0.241605 | 1 | 0.999631 | 0.391668 |
| Madd_E9QN47_S968s_1 | 2.1 | 0.15 | 0.82 | 1.2 | -0.93 | 0.8 | 1.51 | -0.76 | -0.23 | 1.33 | 0.26 | 0.39 | -0.61 | -1.1 | -0.29 | -1 | -0.9125 | -0.63 | -1.8175 | 0.002225 | 0.179437 | 0.223405 | 1 | 0.999631 | 0.351308 |
| Madd_E9QN47_T1620t_1 | -0.48 | -0.78 | -0.17 | -0.81 | -0.52 | -0.41 | -0.34 | -0.25 | -0.07 | -0.12 | -0.27 | 0.09 | -0.4 | -0.23 | 0.7 | -0.4 | 0.18 | 0.4675 | 0.4775 | 0.200178 | 0.497975 | 0.11177 | 1 | 0.999631 | 0.658104 |
| Madd_E9QN47_T1620t_2 | -1.58 | -1.22 | -1.27 | -0.05 | -1.76 | -1.16 | -0.46 | -0.5 | -1.13 | -1.12 | 0.12 | -0.17 | -0.35 | -0.69 | 0.58 | 0.34 | 0.06 | 0.455 | 1 | 0.046314 | 0.892235 | 0.3307 | 1 | 0.999631 | 0.479949 |
| Mag_P20917_S547s_1 | -0.34 | -0.16 | -0.07 | -0.56 | -0.82 | -0.56 | -1.1 | -1.43 | -1.01 | -0.52 | -1.66 | -0.8 | -0.57 | -0.19 | -0.32 | -0.53 | -0.695 | -0.715 | -0.12 | 0.668542 | 0.034895 | 0.048482 | 1 | 0.999631 | 0.883335 |
| Mag_P20917_S547s_1_1 | -0.34 | -0.16 | -0.07 | -0.56 | -0.82 | -0.56 | -1.1 | -1.43 | -1.01 | -0.52 | -1.66 | -0.8 | -0.57 | -0.19 | -0.32 | -0.53 | -0.695 | -0.715 | -0.12 | 0.668542 | 0.034895 | 0.048482 | 1 | 0.999631 | 0.883335 |
| Mag_P20917_S565s_1 | 2.45 | 2.32 | 0.59 | 1.03 | 0.49 | 2.6 | -0.05 | 0 | 1.02 | 2.28 | 0.6 | 1.37 | -0.07 | 1.17 | 1.78 | 0.76 | -0.8375 | -0.28 | -0.6875 | 0.237896 | 0.24317 | 0.609266 | 1 | 0.999631 | 0.683301 |
| Maged2_Q9ER67_S84s_1 | -0.46 | 1.04 | -0.56 | -0.46 | -0.01 | -0.24 | -1.01 | -1.49 | -0.53 | -0.18 | 0.36 | -0.33 | 1.05 | 0.27 | 0.86 | -0.05 | -0.5775 | -0.06 | 0.6425 | 0.181254 | 0.252316 | 0.888638 | 1 | 0.999631 | 0.643828 |
| Maged2_Q9ER67_T121t_1 | 0.38 | -0.28 | 0.76 | 0.27 | -0.21 | -0.14 | -0.14 | 0.85 | 0.12 | 0.24 | 0.36 | -0.46 | 0.36 | 0.15 | 0.61 | 1.73 | -0.1925 | -0.2175 | 0.43 | 0.323264 | 0.593114 | 0.517369 | 1 | 0.999631 | 0.724485 |
| Magi1_Q4VBG2_S732s_1 | 2.06 | -2.4 | -1.99 | -0.37 | -2.21 | -1.58 | 0.83 | -2.36 | -2.43 | 0.49 | -2.79 | -2.68 | -3.75 | -4.01 | -3.68 | -3.43 | -0.655 | -1.1775 | -3.0425 | 0.005869 | 0.545852 | 0.299175 | 1 | 0.999631 | 0.351545 |
| Magi1_Q6RHR9_T1235t_1 | -1.21 | -0.45 | -0.47 | -0.56 | -0.15 | -0.72 | -1.31 | -0.85 | -0.59 | -0.97 | -0.65 | -1 | -0.05 | -0.06 | -0.81 | -0.86 | -0.085 | -0.13 | 0.2275 | 0.517683 | 0.800865 | 0.658647 | 1 | 0.999631 | 0.819449 |
| Magi2_Q9WVQ1_S522s_1 | -1.06 | -0.23 | -0.87 | -0.12 | -0.12 | -0.11 | -0.46 | -0.34 | -0.18 | -0.72 | -0.03 | -0.2 | -0.29 | 0.02 | -0.65 | -0.17 | 0.3125 | 0.2875 | 0.2975 | 0.388894 | 0.320581 | 0.392677 | 1 | 0.999631 | 0.758158 |
| Magi2_Q9WVQ1_S539s_1 | -0.44 | -0.06 | -0.22 | -0.18 | -0.02 | -0.01 | -0.75 | 0.2 | -0.12 | -0.5 | 0.34 | 0.54 | 0.15 | 0 | -0.01 | -0.04 | 0.08 | 0.29 | 0.25 | 0.355897 | 0.784553 | 0.364205 | 1 | 0.999631 | 0.74034 |
| Magi2_Q9WVQ1_S563s_1 | -0.51 | -0.32 | -0.85 | -0.15 | -0.45 | -0.1 | -0.49 | -0.04 | -0.38 | -0.48 | 0.2 | -0.13 | -0.34 | -0.39 | -0.96 | -0.03 | 0.1875 | 0.26 | 0.0275 | 0.93238 | 0.501747 | 0.387495 | 1 | 0.999631 | 0.978264 |
| Magi2_Q9WVQ1_S56s_1 | 0.59 | 0.19 | 0.08 | 1.93 | 1.24 | 0.55 | 1.7 | 0.89 | 1.08 | 0.34 | 0.74 | 1.19 | -1.25 | 0.27 | 0.92 | 0.14 | 0.3975 | 0.14 | -0.6775 | 0.255044 | 0.405395 | 0.758972 | 1 | 0.999631 | 0.690222 |
| Magi2_Q9WVQ1_S711s_2 | 0.9 | -0.08 | -0.81 | -1.71 | -0.63 | 0.01 | -1.05 | -2.95 | -0.75 | 0.89 | -2.11 | -2.23 | -2.86 | -1.66 | -2.56 | -2.91 | -0.73 | -0.625 | -2.0725 | 0.004623 | 0.337575 | 0.443637 | 1 | 0.999631 | 0.351308 |
| Magi2_Q9WVQ1_S714s_1 | 1.27 | -1.58 | -1.4 | 0 | -0.4 | -0.3 | 0.66 | 0.02 | -0.82 | 0.03 | -0.74 | -0.41 | -0.61 | -3.02 | -1.24 | -0.78 | 0.4225 | -0.0575 | -0.985 | 0.214095 | 0.510576 | 0.926977 | 1 | 0.999631 | 0.667209 |
| Magi2_Q9WVQ1_S720s_1 | 1.01 | -1.88 | -1.49 | -0.96 | -2.15 | -2.67 | -0.31 | -2.27 | -3.73 | -0.09 | -2.33 | -2.4 | -2.18 | -3.92 | -1.76 | -1.58 | -1.02 | -1.3075 | -1.53 | 0.06029 | 0.184366 | 0.153719 | 1 | 0.999631 | 0.505962 |
| Magi2_Q9WVQ1_S720s_2 | 0.09 | -1.67 | -0.46 | 0.07 | -2.14 | -1.04 | 0.94 | -0.34 | -1.71 | 0.21 | -1.5 | -1.37 | -2.63 | -2.45 | -1.53 | -0.87 | -0.1525 | -0.6 | -1.3775 | 0.027858 | 0.82263 | 0.296603 | 1 | 0.999631 | 0.423977 |
| Magi2_Q9WVQ1_S720s_3 | -0.54 | -2.02 | -0.54 | -1.57 | -2.26 | -1.36 | -0.43 | -1.57 | -2.66 | -1.73 | -1 | -2.73 | -1.17 | -3.22 | -2.65 | -2.16 | -0.2375 | -0.8625 | -1.1325 | 0.057336 | 0.635605 | 0.123187 | 1 | 0.999631 | 0.49679 |
| Magi2_Q9WVQ1_S721s_1 | -0.05 | -2.26 | -1.67 | -0.99 | -2.3 | -1.71 | -0.22 | -2.46 | -2.69 | -0.92 | -2.83 | -3.33 | -1.87 | -3.16 | -1.62 | -1.97 | -0.43 | -1.2 | -0.9125 | 0.118854 | 0.497278 | 0.085184 | 1 | 0.999631 | 0.585965 |

| | | | | | | | | | | | | | | | | | | | | | | | | | |
|---|---|---|---|---|---|---|---|---|---|---|---|---|---|---|---|---|---|---|---|---|---|---|---|---|---|
| Magi2_Q9WVQ1_S721s_2 | -0.65 | -1.82 | -0.81 | -1.71 | -2.81 | -1.43 | -1.05 | -2.95 | -1.68 | 0.04 | -2.11 | -2.23 | -2.19 | -2.57 | -2.56 | -2.91 | -0.8125 | -0.2475 | -1.31 | 0.005463 | 0.145992 | 0.659519 | 1 | 0.999631 | 0.351308 |
| Magi2_Q9WVQ1_S721s_3 | -0.54 | -2.02 | -0.54 | -1.57 | -2.26 | -1.36 | -0.43 | -1.57 | -2.66 | -1.73 | -1 | -2.73 | -1.17 | -3.22 | -2.65 | -2.16 | -0.2375 | -0.8625 | -1.1325 | 0.057336 | 0.635605 | 0.123187 | 1 | 0.999631 | 0.49679 |
| Magi2_Q9WVQ1_S850s_1 | -0.3 | 0 | -0.47 | -0.91 | -0.05 | -0.39 | -0.91 | -0.86 | -0.53 | -0.26 | -0.69 | -0.45 | 0.51 | 0 | -0.64 | 0.12 | -0.1325 | -0.0625 | 0.4175 | 0.258988 | 0.683969 | 0.83118 | 1 | 0.999631 | 0.692624 |
| Magi2_Q9WVQ1_Y198y_1 | -1 | -0.91 | 0.5 | -0.35 | -0.09 | 0.16 | -0.87 | -0.41 | -1.27 | 1.18 | 0.2 | 0.38 | 0.01 | -0.36 | -0.09 | 0.32 | 0.1375 | 0.5625 | 0.41 | 0.316003 | 0.738014 | 0.334195 | 1 | 0.999631 | 0.720734 |
| Magi3_G5E8T6_S236s_1 | -0.6 | 0.88 | -0.63 | 0 | 0.15 | 1.44 | -0.21 | 0.94 | 1.23 | -0.19 | -1.33 | 0.68 | 0.77 | 0.7 | 0.17 | -0.41 | 0.6675 | 0.185 | 0.395 | 0.387006 | 0.191116 | 0.758971 | 1 | 0.999631 | 0.756701 |
| Magi3_G5E8T6_S237s_1 | -0.85 | 0.89 | 1.54 | 1.4 | -0.67 | 1.4 | 1.28 | 1.66 | 0.98 | 0.76 | 1.72 | 2.13 | 1.09 | 1.28 | 0.81 | 1.07 | 0.1725 | 0.6525 | 0.3175 | 0.54969 | 0.800188 | 0.27848 | 1 | 0.999631 | 0.835438 |
| Magi3_G5E8T6_S568s_1 | -0.09 | -0.59 | -0.99 | -0.22 | -0.14 | 0.45 | -1.49 | -1.15 | 0.01 | -0.65 | -1.54 | -0.22 | 0.09 | 0.08 | -1.82 | -0.71 | -0.11 | -0.1275 | -0.1175 | 0.806789 | 0.814635 | 0.75433 | 1 | 0.999631 | 0.935523 |
| Magi3_G5E8T6_S662s_1 | 2.11 | 0.3 | 1.18 | 1.18 | -0.96 | 0.84 | 1.64 | -0.38 | -0.65 | 1.94 | -0.1 | 0.07 | 0.17 | -0.16 | 0.7 | 0.2 | -0.9075 | -0.8775 | -0.965 | 0.042403 | 0.168077 | 0.173864 | 1 | 0.999631 | 0.475443 |
| Magi3_G5E8T6_S833s_1 | 0.3 | -1 | -0.55 | -1.17 | -0.94 | -0.76 | -0.94 | -0.45 | -0.59 | -0.02 | -0.95 | 0 | -0.16 | -0.55 | -0.91 | -0.3 | -0.1675 | 0.215 | 0.125 | 0.752059 | 0.650658 | 0.60248 | 1 | 0.999631 | 0.913038 |
| Magi3_G5E8T6_S954s_1 | -0.24 | -2.04 | -0.83 | -1.19 | -0.77 | -2.06 | -0.28 | -1.52 | -1.29 | -0.72 | -1.26 | -1.14 | -1.06 | -1.9 | -0.47 | -1.42 | -0.0825 | -0.0275 | -0.1375 | 0.771034 | 0.871199 | 0.946185 | 1 | 0.999631 | 0.919413 |
| Mal2_Q8BI08_S2s_1 | -0.69 | 0.32 | -0.6 | -1.5 | -0.75 | 0.16 | -0.78 | -1.47 | 0.37 | -0.32 | -1.43 | -0.97 | -0.77 | -0.01 | -0.37 | -1.01 | -0.0925 | 0.03 | 0.0775 | 0.859385 | 0.84575 | 0.953066 | 1 | 0.999631 | 0.953128 |
| Maoa_Q64133_S388s_1 | -0.78 | 0.18 | -1.26 | -1.17 | -1.03 | -0.51 | -0.72 | -0.86 | -0.22 | -1.28 | -1.11 | -0.48 | 0.73 | -0.04 | -0.63 | -0.4 | -0.0225 | -0.015 | 0.6725 | 0.15362 | 0.951107 | 0.9714 | 1 | 0.999631 | 0.625198 |
| Map1a_Q9QYR6_S1008s_2 | -2.32 | -0.54 | -0.15 | -2.71 | -2.34 | -1.54 | -2.65 | -2.61 | -1.89 | -0.88 | -2.26 | -0.42 | 0.68 | -1.54 | -1.49 | -0.91 | -0.855 | 0.0675 | 0.615 | 0.404684 | 0.187901 | 0.921464 | 1 | 0.999631 | 0.765844 |
| Map1a_Q9QYR6_S1052s_1 | -0.09 | 0.98 | 1.18 | 0.67 | -0.02 | 1.6 | 0.07 | 0.4 | 1.04 | 1.33 | 0.15 | 1.22 | 0.68 | 0.97 | 0.77 | 1.31 | -0.1725 | 0.25 | 0.2475 | 0.498144 | 0.702592 | 0.535736 | 1 | 0.999631 | 0.811985 |
| Map1a_Q9QYR6_S1062s_1 | -0.51 | 0.15 | 0.18 | -0.21 | 0.04 | -0.39 | -0.25 | 0.13 | -0.21 | -0.54 | -0.19 | -0.05 | -0.02 | 0.09 | 0.45 | -0.06 | -0.02 | -0.15 | 0.2125 | 0.488465 | 0.943543 | 0.601349 | 1 | 0.999631 | 0.806721 |
| Map1a_Q9QYR6_S1190s_1 | -0.93 | -0.3 | 0.15 | 0.03 | -0.54 | 0.12 | -0.34 | 0.22 | -0.53 | -0.97 | 0.2 | 0.55 | -0.29 | 0.38 | 0.46 | 0.59 | 0.1275 | 0.075 | 0.5475 | 0.15205 | 0.707465 | 0.859294 | 1 | 0.999631 | 0.625198 |
| Map1a_Q9QYR6_S1196s_1 | 0.22 | 1.72 | 1.98 | 0.8 | 0.65 | 0.76 | 0.02 | 1.2 | 0.75 | 0.68 | 0.97 | 1.42 | 2.44 | 1.39 | 1.63 | 1.2 | -0.5225 | -0.225 | 0.485 | 0.323019 | 0.269317 | 0.609169 | 1 | 0.999631 | 0.724485 |
| Map1a_Q9QYR6_S1205s_1 | -0.26 | 0.45 | 0.86 | 0.37 | -0.03 | 0.01 | -0.2 | 0.26 | -0.04 | -0.01 | 0.5 | 0.69 | 0.34 | 0.28 | 0.48 | 0.73 | -0.345 | -0.07 | 0.1025 | 0.755042 | 0.277604 | 0.837137 | 1 | 0.999631 | 0.913659 |
| Map1a_Q9QYR6_S1305s_1 | 1.97 | 2.7 | 2.61 | 2.14 | 2.69 | 2.21 | 0.51 | 1.89 | 1.91 | 2.19 | 1.97 | 2.03 | 2.67 | 2.64 | 2.42 | 2.8 | -0.53 | -0.33 | 0.2775 | 0.365819 | 0.281022 | 0.259358 | 1 | 0.999631 | 0.745796 |
| Map1a_Q9QYR6_S1306s_1 | 1.78 | 2.73 | 3.19 | 1.92 | 1.64 | 2.25 | 1.36 | 2.86 | 1.6 | 2.03 | 3.04 | 1.58 | 1.63 | 3.28 | 2.29 | 3.5 | -0.3775 | -0.3425 | 0.27 | 0.606146 | 0.415688 | 0.467372 | 1 | 0.999631 | 0.85817 |
| Map1a_Q9QYR6_S1310s_1 | -0.5 | 0.24 | 0.52 | 0.43 | -0.11 | 0.02 | -0.16 | 0.26 | -0.27 | -0.31 | 0.07 | -0.09 | 0.27 | 0.79 | 0.64 | 0.95 | -0.17 | -0.3225 | 0.49 | 0.170548 | 0.583009 | 0.316315 | 1 | 0.999631 | 0.64073 |
| Map1a_Q9QYR6_S1360s_1 | -0.49 | 1.02 | 0.82 | 0.77 | 0.3 | 0.96 | 0.19 | 0.85 | 0.41 | -0.06 | 0.45 | 1.51 | 0.63 | 0.61 | 0.32 | 0.62 | 0.045 | 0.0475 | 0.015 | 0.968926 | 0.910229 | 0.918403 | 1 | 0.999631 | 0.988839 |
| Map1a_Q9QYR6_S1370s_1 | 0.17 | 2.91 | 1.62 | 2.11 | 2.93 | 4.57 | 4.43 | 2.67 | 2.51 | -0.4 | 4.32 | 1.45 | -1.65 | 1.49 | 1.44 | -1 | 1.9475 | 0.2675 | -1.6325 | 0.083008 | 0.015605 | 0.786788 | 1 | 0.999631 | 0.549628 |
| Map1a_Q9QYR6_S1580s_1 | -0.75 | 0.23 | -0.17 | -0.16 | 0.25 | -0.54 | -0.57 | -0.22 | 0.61 | 0.2 | -0.33 | 0.1 | 0.66 | 0.16 | -0.19 | -0.7 | -0.0575 | 0.3575 | 0.195 | 0.614092 | 0.858882 | 0.294988 | 1 | 0.999631 | 0.860996 |
| Map1a_Q9QYR6_S1606s_1 | -0.76 | -0.11 | -0.4 | -1.06 | -0.69 | -0.35 | -1.4 | -1.03 | -0.62 | -0.68 | -0.85 | -0.89 | 0.01 | -0.21 | -0.59 | -0.85 | -0.285 | -0.1775 | 0.1725 | 0.61854 | 0.4118 | 0.553448 | 1 | 0.999631 | 0.861888 |
| Map1a_Q9QYR6_S1648s_1 | 1.09 | 1.12 | 0.39 | 0.46 | 0.66 | 0.94 | 0.09 | 0.64 | 0.49 | 0.89 | 0.17 | 0.45 | 0.19 | 0.48 | 0.68 | 0.47 | -0.1825 | -0.265 | -0.31 | 0.333442 | 0.567061 | 0.405053 | 1 | 0.999631 | 0.728079 |
| Map1a_Q9QYR6_S1747s_1 | -0.49 | 0.12 | -0.18 | 0.29 | 0.23 | -0.41 | 0.02 | 0.1 | -0.24 | -0.51 | 0.24 | 0.29 | -0.21 | 0.04 | 0.02 | 0.32 | 0.05 | 0.01 | 0.1075 | 0.724389 | 0.863702 | 0.974963 | 1 | 0.999631 | 0.905674 |
| Map1a_Q9QYR6_S1768s_1 | -0.07 | 0.68 | 0.96 | 0.17 | 0.23 | 0.56 | -0.03 | 0.71 | 0.34 | -0.17 | 0.59 | 0.94 | 0.63 | 1 | 0.3 | 0.7 | -0.0675 | -0.01 | 0.2225 | 0.517696 | 0.837665 | 0.977985 | 1 | 0.999631 | 0.819449 |
| Map1a_Q9QYR6_S1772s_1 | -0.61 | 0.49 | -0.03 | -1.31 | -0.59 | -0.25 | -1.04 | -0.83 | -0.86 | -0.52 | -0.72 | -0.65 | 2.06 | -0.08 | 0.08 | 0.41 | -0.3125 | -0.3225 | 0.9825 | 0.112099 | 0.462158 | 0.432495 | 1 | 0.999631 | 0.577068 |
| Map1a_Q9QYR6_S1789s_1 | 0.29 | 0.4 | 0.51 | 0.43 | 0.28 | 0.31 | 0 | 0.56 | 0.12 | 0.35 | 0.3 | 0.32 | 0.27 | 0.71 | 0.48 | 0.65 | -0.12 | -0.135 | 0.12 | 0.657398 | 0.630505 | 0.579499 | 1 | 0.999631 | 0.879224 |
| Map1a_Q9QYR6_S1993s_1 | -0.22 | 0.06 | -0.2 | -0.31 | -0.36 | -0.4 | -0.26 | -0.64 | -0.84 | -0.64 | -0.77 | -0.44 | 0.19 | -0.26 | -0.48 | -0.23 | -0.2475 | -0.505 | -0.0275 | 0.923476 | 0.32489 | 0.071512 | 1 | 0.999631 | 0.975032 |
| Map1a_Q9QYR6_S2029s_1 | -0.03 | 2.3 | 1.5 | 0.99 | 0.71 | 0.99 | 0.2 | 0.54 | 1.68 | -0.22 | 0.53 | 0.41 | 2.25 | 1.59 | 0.89 | 0.97 | -0.58 | -0.59 | 0.235 | 0.666498 | 0.250783 | 0.319771 | 1 | 0.999631 | 0.88264 |
| Map1a_Q9QYR6_S2048s_1 | -0.7 | -0.35 | -0.26 | -0.43 | -0.66 | -0.54 | -0.83 | -0.52 | -0.62 | -0.72 | -0.62 | -0.49 | -0.08 | -0.17 | -0.2 | -0.21 | -0.2025 | -0.1775 | 0.27 | 0.324488 | 0.41965 | 0.485001 | 1 | 0.999631 | 0.72463 |
| Map1a_Q9QYR6_S2156s_1 | -0.64 | -0.49 | -1.77 | -0.1 | -0.44 | -0.86 | -0.48 | -0.33 | -1.16 | -1.8 | -0.69 | -0.79 | -0.5 | 0.08 | -0.53 | -0.57 | 0.2225 | -0.36 | 0.37 | 0.376161 | 0.568477 | 0.416513 | 1 | 0.999631 | 0.748365 |
| Map1a_Q9QYR6_S2162s_1 | 0.37 | 0.49 | 0.01 | 1.5 | 1.33 | -0.28 | 1.31 | 1.15 | -0.32 | -1.16 | 1.05 | -0.18 | 0.25 | -0.31 | 0.58 | 0.59 | 0.285 | -0.745 | -0.315 | 0.443297 | 0.553777 | 0.175816 | 1 | 0.999631 | 0.782783 |
| Map1a_Q9QYR6_S2199s_1 | -0.06 | 0.7 | -0.29 | 0.26 | 0.48 | 0.46 | -0.22 | 0.09 | 0.05 | -0.36 | 0.04 | 0.08 | -0.21 | 0.6 | -0.11 | 0.28 | 0.05 | -0.2 | -0.0125 | 0.971087 | 0.87579 | 0.520658 | 1 | 0.999631 | 0.989676 |
| Map1a_Q9QYR6_S2209s_1 | 0.35 | -0.62 | -0.45 | 0.68 | 0.39 | -0.65 | 0.5 | 0.33 | -0.52 | -0.96 | 0.18 | -0.45 | -0.82 | -0.63 | 0.06 | 0.05 | 0.1525 | -0.4275 | -0.325 | 0.432489 | 0.71109 | 0.302385 | 1 | 0.999631 | 0.779247 |
| Map1a_Q9QYR6_S2226s_1 | 0.02 | 0.05 | -1.04 | -0.27 | 0.88 | 0.35 | -0.3 | 0.16 | 0.56 | -0.75 | -0.67 | -0.38 | 0.02 | 0.78 | 0.38 | -0.13 | 0.5825 | -2.4E-16 | 0.5725 | 0.14368 | 0.139822 | 1 | 1 | 1 | 0.615182 |
| Map1a_Q9QYR6_S2230s_1 | -0.63 | -0.78 | 0.73 | -0.41 | -0.29 | -0.38 | -1.3 | 0.82 | -0.33 | -0.18 | 0.27 | 0.77 | -0.05 | 0.15 | 1.15 | 0.45 | -0.015 | 0.405 | 0.6975 | 0.133978 | 0.976731 | 0.35179 | 1 | 0.999631 | 0.60492 |
| Map1a_Q9QYR6_S2591s_1 | 3.5 | 2.51 | 2.53 | 2.08 | 0.12 | 2.49 | 3.17 | 0.56 | 1.83 | 3.16 | 1.43 | 1.04 | 0.19 | 0.55 | 1.48 | 0.86 | -1.07 | -0.79 | -1.885 | 0.001145 | 0.152174 | 0.150335 | 1 | 0.999631 | 0.351308 |
| Map1a_Q9QYR6_S2630s_1 | 1.36 | 1.25 | 1.26 | 0.72 | 0.49 | 1.16 | 1.25 | 0.7 | 1.33 | 1.94 | 0.93 | 0.77 | 0.38 | 0.35 | 1.2 | 1.08 | -0.2475 | 0.095 | -0.395 | 0.256693 | 0.414263 | 0.781587 | 1 | 0.999631 | 0.691567 |
| Map1a_Q9QYR6_S2637s_1 | -0.98 | 0.09 | -2.91 | -2.28 | 0.68 | -0.16 | -1.73 | -2.76 | 0.3 | 0.1 | -1.87 | -1.99 | -1.06 | -0.26 | -2.48 | -2.22 | 0.5275 | 0.655 | 0.015 | 0.983909 | 0.555276 | 0.421062 | 1 | 0.999631 | 0.995133 |
| Map1a_Q9QYR6_S384s_1 | 0.66 | 0.48 | 0.08 | 0.79 | 0.62 | -0.21 | 0.02 | 1.09 | 0.73 | 0.63 | 0.55 | 0.64 | 0.51 | 0.45 | 0.5 | -0.2 | -0.1225 | 0.135 | -0.1875 | 0.558142 | 0.732497 | 0.618791 | 1 | 0.999631 | 0.838323 |
| Map1a_Q9QYR6_S500s_1 | 1.54 | 1.33 | 2.98 | 1.07 | 0.75 | 2.83 | 1.31 | 1.54 | 0.53 | 2.39 | 0.26 | 0.67 | -3.01 | 2.18 | 2.37 | 0.26 | -0.1225 | -0.7675 | -1.28 | 0.269287 | 0.827192 | 0.213329 | 1 | 0.999631 | 0.697419 |
| Map1a_Q9QYR6_S611s_1 | 1.46 | 1.01 | 0.25 | 0.13 | 0.77 | 0.9 | 0.36 | -0.49 | 1.01 | 1.6 | -0.1 | -0.34 | 0.17 | 0.1 | -0.17 | -0.67 | -0.3275 | -0.17 | -0.855 | 0.052481 | 0.458505 | 0.745536 | 1 | 0.999631 | 0.489777 |
| Map1a_Q9QYR6_S644s_1 | 1.53 | 1.66 | 2.56 | 2.59 | 0.99 | 1.81 | 2.02 | 1.46 | 1.67 | 2.19 | 2.18 | 1.93 | 1.38 | 0.67 | 1.95 | 1.73 | -0.515 | -0.0925 | -0.6525 | 0.13927 | 0.193569 | 0.791417 | 1 | 0.999631 | 0.611718 |
| Map1a_Q9QYR6_S667s_1 | -0.1 | 1.83 | 0.09 | -0.56 | 1.1 | 1.31 | -0.64 | -0.36 | 1.15 | 1.1 | -0.23 | -0.25 | 1.73 | 1.78 | -0.36 | 0.05 | 0.0375 | 0.1275 | 0.485 | 0.482032 | 0.953609 | 0.831008 | 1 | 0.999631 | 0.802327 |
| Map1a_Q9QYR6_S678s_1 | 1.85 | 1.54 | 0.47 | 2.3 | 2.43 | 2.36 | 1.87 | 2.1 | 1.68 | 1.29 | 2 | 1.09 | -1.96 | 1.71 | 1.65 | 1.28 | 0.65 | -0.025 | -0.87 | 0.311833 | 0.136984 | 0.954221 | 1 | 0.999631 | 0.719291 |
| Map1a_Q9QYR6_S852s_1 | -0.7 | 0.81 | 0.18 | 0.22 | -0.08 | 0.14 | -0.09 | 0.16 | -0.07 | -1.14 | 0.13 | 0.78 | -0.02 | -0.18 | 0.47 | 0.23 | -0.095 | -0.2025 | -0.0025 | 0.994735 | 0.785488 | 0.677049 | 1 | 0.999631 | 0.997705 |
| Map1a_Q9QYR6_S895s_1 | -0.53 | 0.33 | 0.47 | -0.09 | 0.02 | -0.35 | -0.39 | -0.31 | -0.12 | -0.26 | -0.42 | 0.5 | 0.55 | 0.23 | -0.03 | 0.22 | -0.3025 | -0.12 | 0.1975 | 0.552414 | 0.332611 | 0.728473 | 1 | 0.999631 | 0.836655 |
| Map1a_Q9QYR6_S908s_1 | -0.06 | 1.09 | 0.62 | 0.22 | 0.52 | 0.72 | -0.27 | 0.17 | 0.39 | 0.68 | -0.26 | 0.74 | 0.95 | 1.04 | 0.35 | 0.38 | -0.1825 | -0.08 | 0.2125 | 0.558241 | 0.611246 | 0.827913 | 1 | 0.999631 | 0.838323 |
| Map1a_Q9QYR6_S981s_1 | 0.29 | 0.66 | 1.52 | 0.65 | 0.43 | 0.38 | 0.41 | 0.63 | 0.43 | 0.87 | 0.76 | 0.86 | 0.76 | 0.75 | 0.82 | 1.07 | -0.3175 | -0.05 | 0.07 | 0.836173 | 0.32985 | 0.880213 | 1 | 0.999631 | 0.945157 |
| Map1a_Q9QYR6_S991s_1 | 0.08 | 0.24 | -0.06 | -0.14 | -0.32 | -0.28 | -0.22 | -0.61 | -0.49 | 0.19 | -0.6 | 0 | 0.18 | 0.26 | -0.39 | 0.16 | -0.3875 | -0.255 | 0.0225 | 0.938183 | 0.140611 | 0.391918 | 1 | 0.999631 | 0.979905 |
| Map1a_Q9QYR6_S991s_2 | -2.32 | -0.54 | -0.15 | -2.71 | -2.34 | -1.54 | -2.65 | -2.61 | -1.89 | -0.88 | -2.26 | -0.42 | 0.68 | -1.54 | -1.49 | -0.91 | -0.855 | 0.0675 | 0.615 | 0.404684 | 0.187901 | 0.921464 | 1 | 0.999631 | 0.765844 |
| Map1a_Q9QYR6_T1712t_1 | 0.42 | 0.3 | -1.66 | 1.64 | 1.24 | -0.01 | 1.29 | 1.24 | -0.51 | -0.46 | 0.7 | -0.14 | -0.87 | -0.25 | -0.02 | 1.27 | 0.765 | -0.2775 | -0.1425 | 0.84348 | 0.26961 | 0.676185 | 1 | 0.999631 | 0.94821 |
| Map1a_Q9QYR6_T1806t_1 | 1.42 | -0.69 | -1.22 | 2.14 | 3.72 | 0.17 | 2.6 | 2.04 | 0.36 | -1.88 | 0.83 | -1.21 | -3.65 | -0.23 | -0.53 | -0.32 | 1.72 | -0.8875 | -1.595 | 0.130765 | 0.094523 | 0.335589 | 1 | 0.999631 | 0.602177 |
| Map1a_Q9QYR6_T1928t_1 | 2.4 | -0.85 | -1.56 | 0.99 | 1.72 | -1.08 | 3.06 | 0.47 | 0.17 | -0.91 | 0.8 | -1.46 | -2.44 | -2.12 | -1.78 | 0.05 | 0.7975 | -0.595 | -1.8175 | 0.069131 | 0.466263 | 0.512415 | 1 | 0.999631 | 0.524251 |
| Map1a_Q9QYR6_T2031t_1 | -0.05 | 1.14 | 0.87 | 0.33 | 0.57 | 0.71 | 0.05 | 0.47 | 0.25 | 0.18 | 0.18 | -0.01 | 0.95 | 0.97 | 0.42 | 0.71 | -0.1225 | -0.4225 | 0.19 | 0.59219 | 0.718322 | 0.216504 | 1 | 0.999631 | 0.852774 |
| Map1a_Q9QYR6_T2182t_1 | 2.29 | -0.09 | -1.41 | 1.54 | 2.79 | 0.72 | 1.92 | 1.21 | 0.67 | -1.47 | 0.86 | -0.95 | -1.88 | 0.3 | -0.83 | -0.19 | 1.0775 | -0.805 | -1.2325 | 0.159372 | 0.210282 | 0.372004 | 1 | 0.999631 | 0.629514 |
| Map1a_Q9QYR6_T633t_1 | -0.47 | 0.37 | 0.64 | 0.13 | 0.13 | 0.22 | -0.08 | -0.06 | 0.15 | -0.05 | 0 | 0.26 | 0.89 | 0.38 | 0.88 | -0.02 | -0.115 | -0.0775 | 0.365 | 0.33278 | 0.707939 | 0.804327 | 1 | 0.999631 | 0.728079 |
| Map1b_P14873_S1013s_1 | 0.46 | 0.58 | 1.67 | 1.53 | 0.47 | 0.8 | 1.28 | 0.8 | 0.71 | 1.66 | 0.66 | 0.88 | 0.67 | 0.01 | 0.98 | 0.08 | -0.2225 | -0.0825 | -0.625 | 0.150646 | 0.554159 | 0.837471 | 1 | 0.999631 | 0.623175 |
| Map1b_P14873_S1073s_1 | 0.9 | 0.76 | 0.81 | 0.5 | 0.34 | 0.82 | 0.98 | 0.38 | 0.96 | 1.37 | 0.37 | 0.51 | 0.97 | 0.84 | 0.6 | 0.37 | -0.1125 | 0.06 | -0.0475 | 0.867795 | 0.679947 | 0.846689 | 1 | 0.999631 | 0.956838 |
| Map1b_P14873_S1151s_1 | -1.22 | 0.16 | -0.19 | -0.8 | -0.26 | -0.16 | -0.78 | -0.23 | -0.11 | -1.02 | -0.34 | 0.18 | 0.61 | 0.27 | 0.08 | 0.03 | 0.155 | 0.19 | 0.76 | 0.065578 | 0.669947 | 0.643837 | 1 | 0.999631 | 0.516933 |
| Map1b_P14873_S1204s_1 | -2.62 | -2.18 | -2.26 | -3.04 | -1.9 | -2.49 | -2.05 | -3.09 | -2.1 | -2.16 | -2.38 | -2.22 | -1.24 | -2.56 | -2.6 | -2.93 | 0.1425 | 0.31 | 0.1925 | 0.656114 | 0.690806 | 0.301017 | 1 | 0.999631 | 0.879056 |
| Map1b_P14873_S1233s_1 | 1.29 | 1.43 | -0.14 | 0.69 | 1.5 | 1.39 | 0.7 | 0.46 | 1.37 | 1.09 | 0.13 | 0.15 | 0.33 | 1.6 | 0.6 | 0.01 | 0.195 | -0.1325 | -0.1825 | 0.705309 | 0.652381 | 0.775994 | 1 | 0.999631 | 0.89781 |
| Map1b_P14873_S1247s_1 | -1.02 | 3.1 | 1.66 | 0.79 | 2.04 | 3.03 | 1.56 | 1.95 | 3.28 | 0.14 | 2.35 | 2.3 | 2.13 | 2.31 | 2.34 | 1.53 | 1.0125 | 0.885 | 0.945 | 0.238693 | 0.225057 | 0.359372 | 1 | 0.999631 | 0.68341 |
| Map1b_P14873_S1255s_1 | -0.96 | 0.18 | -0.11 | -0.76 | -0.38 | 0.87 | -1.67 | 0.04 | -0.17 | 0.12 | 1.7 | 1.08 | 0.4 | 0.33 | 0.91 | 2 | 0.1275 | 1.095 | 1.3225 | 0.016064 | 0.815377 | 0.045588 | 1 | 0.999631 | 0.388832 |
| Map1b_P14873_S1260s_1 | -0.3 | 1.06 | 0.48 | 0.21 | 0.69 | 0.55 | 0.05 | 0.18 | 0.64 | 0.1 | 0.32 | 0.57 | 0.98 | 0.59 | 0.3 | 0.85 | 0.005 | 0.045 | 0.3175 | 0.395134 | 0.988596 | 0.897318 | 1 | 0.999631 | 0.759958 |
| Map1b_P14873_S1307s_1 | -0.17 | 0.49 | -0.29 | -0.73 | 0.66 | 0.32 | -0.82 | -0.64 | 0.6 | 0.72 | -0.69 | -0.13 | 1.29 | 0.72 | -0.22 | -0.23 | 0.055 | 0.3 | 0.565 | 0.227721 | 0.898534 | 0.480316 | 1 | 0.999631 | 0.678949 |
| Map1b_P14873_S1317s_1 | -0.74 | -1.85 | -1.66 | -1.58 | -1.57 | -1.56 | -1.23 | -2.14 | -2.16 | -1.13 | -2.18 | -2.14 | -1.35 | -1.56 | -1.87 | -1.71 | -0.1675 | -0.445 | -0.165 | 0.626589 | 0.627587 | 0.257455 | 1 | 0.999631 | 0.864779 |
| Map1b_P14873_S1334s_1 | -0.66 | -1.47 | -1.8 | -2.09 | -1.33 | -1.79 | -1.73 | -2.18 | -2.12 | -1.46 | -2.35 | -1.91 | -0.3 | -1.26 | -1.91 | -2.03 | -0.2525 | -0.455 | 0.13 | 0.789264 | 0.502415 | 0.252382 | 1 | 0.999631 | 0.925513 |
| Map1b_P14873_S1373s_1 | -0.49 | -0.32 | 0.36 | -0.28 | -0.28 | -0.43 | -0.35 | -0.15 | -0.38 | -0.25 | 0.02 | 0.48 | 0.48 | -0.08 | 0.52 | 0.42 | -0.12 | 0.15 | 0.5175 | 0.12766 | 0.667995 | 0.644489 | 1 | 0.999631 | 0.598244 |
| Map1b_P14873_S1382s_1 | -0.33 | 0.87 | 0.34 | 0.02 | 0.09 | -0.31 | -0.27 | 0.2 | -0.19 | -0.48 | -0.74 | -0.83 | 1.04 | 0.32 | 1.02 | 1.04 | -0.2975 | -0.785 | 0.63 | 0.10474 | 0.374679 | 0.040767 | 1 | 0.999631 | 0.568213 |
| Map1b_P14873_S1384s_1 | -0.53 | 0.45 | 0.08 | -0.1 | -0.03 | -0.05 | -0.44 | -0.03 | -0.01 | -0.32 | -0.1 | 0.16 | 0.78 | 0.38 | 0.22 | 0.54 | -0.1125 | -0.0425 | 0.505 | 0.137597 | 0.703614 | 0.887917 | 1 | 0.999631 | 0.610681 |
| Map1b_P14873_S1391s_1 | -1.8 | -1.4 | -0.82 | -1.13 | -1.31 | -1.9 | -1.1 | -0.84 | -1.68 | -2.36 | -0.63 | -0.77 | -0.46 | -1.41 | -0.51 | -0.6 | -1.6E-16 | -0.0725 | 0.5425 | 0.151599 | 1 | 0.87155 | 1 | 0.999631 | 0.625093 |
| Map1b_P14873_S1395s_1 | -1.19 | 0.38 | 0.48 | -0.78 | -0.92 | -0.27 | -1.01 | -0.59 | -0.35 | -0.14 | -0.44 | 0.25 | 1.11 | 0.04 | 0.06 | 0.26 | -0.42 | 0.1075 | 0.645 | 0.195725 | 0.350213 | 0.80713 | 1 | 0.999631 | 0.653056 |
| Map1b_P14873_S1422s_1 | 2.21 | -0.81 | -1.87 | 1.61 | 2.68 | -0.2 | 2.16 | 0.81 | 0.85 | -0.2 | 0.62 | -1.05 | -2.23 | -0.31 | -1.12 | -1.14 | 1.0775 | -0.23 | -1.485 | 0.123948 | 0.298874 | 0.803164 | 1 | 0.999631 | 0.595021 |
| Map1b_P14873_S1438s_1 | -1.21 | -1.65 | 0.48 | 0.49 | -1.53 | -1.61 | -0.02 | 0.49 | -1.66 | -1.48 | 0.48 | 0.93 | -1.94 | -1.56 | 0.75 | 0.97 | -0.195 | 0.04 | 0.0275 | 0.973218 | 0.776178 | 0.958306 | 1 | 0.999631 | 0.990577 |
| Map1b_P14873_S1497s_1 | -0.06 | -0.08 | -0.18 | 0.1 | 0.29 | -0.1 | -0.2 | 0.19 | -0.05 | -0.02 | -0.15 | -0.02 | -0.1 | 0.3 | 0.03 | 0.09 | 0.1 | -0.005 | 0.135 | 0.615871 | 0.690668 | 0.983436 | 1 | 0.999631 | 0.861203 |
| Map1b_P14873_S1523s_1 | 0.65 | -0.02 | 0.74 | 0.33 | -0.59 | 0.09 | 0.17 | 1.28 | 0.3 | 0.35 | 1.27 | 0.53 | 0.89 | 0.27 | 0.37 | 1.53 | -0.1875 | 0.1875 | 0.34 | 0.375594 | 0.656603 | 0.576803 | 1 | 0.999631 | 0.748307 |
| Map1b_P14873_S1616s_1 | 0.79 | -0.31 | 1.69 | 0.95 | -0.49 | 0.13 | 0.58 | -0.11 | 0.26 | 0.44 | -0.25 | 1 | 0.05 | 0.82 | 1.24 | 1.12 | -0.7525 | -0.4175 | 0.0275 | 0.954131 | 0.121359 | 0.384766 | 1 | 0.999631 | 0.984441 |
| Map1b_P14873_S1627s_1 | 2.8 | -0.3 | -0.46 | 0.98 | 1.05 | 0.47 | 1.61 | 0.46 | 0.86 | 1.33 | -0.22 | -0.82 | -4.13 | -0.42 | -1.26 | -2.25 | 0.1425 | -0.4675 | -2.77 | 0.013979 | 0.840732 | 0.557996 | 1 | 0.999631 | 0.385092 |
| Map1b_P14873_S1627s_2 | 0.94 | 1.47 | 0.52 | 2.07 | 1.18 | 0.82 | 1.8 | 1.4 | 1.74 | 1.33 | 1.09 | 0.95 | -3.14 | 0.02 | 0.61 | -0.69 | 0.05 | 0.0275 | -2.05 | 0.023412 | 0.900034 | 0.944178 | 1 | 0.999631 | 0.41509 |
| Map1b_P14873_S1649s_1 | -0.37 | -0.17 | 0.36 | -0.73 | -0.45 | -0.49 | -0.46 | -0.07 | -0.26 | -0.55 | -0.08 | -0.3 | 0.44 | 0.07 | -0.2 | 0.35 | -0.14 | -0.07 | 0.3925 | 0.26106 | 0.649352 | 0.822952 | 1 | 0.999631 | 0.69271 |
| Map1b_P14873_S1775s_1 | -1.29 | -0.54 | 0.38 | -0.95 | -0.81 | -0.81 | -1.16 | -0.7 | -0.57 | -0.32 | -0.35 | -0.19 | 1 | -0.29 | 0 | -0.16 | -0.27 | 0.2425 | 0.7375 | 0.132464 | 0.489196 | 0.535817 | 1 | 0.999631 | 0.60434 |
| Map1b_P14873_S1781s_1 | -1.88 | -0.25 | 0.14 | -1.72 | -2.1 | -0.62 | -2.27 | -0.89 | -0.55 | 0.02 | -0.64 | 0.28 | 0.49 | -0.13 | 0.02 | 0.3 | -0.5425 | 0.705 | 1.0975 | 0.051583 | 0.374039 | 0.198439 | 1 | 0.999631 | 0.489777 |
| Map1b_P14873_S1781s_2 | -0.67 | -1.54 | -1.34 | -0.36 | -0.45 | -0.82 | -0.8 | -0.09 | -1.79 | -0.34 | -0.41 | -0.94 | 0.27 | -0.26 | -0.21 | -0.49 | 0.4375 | 0.1075 | 0.805 | 0.048924 | 0.234843 | 0.804012 | 1 | 0.999631 | 0.48544 |
| Map1b_P14873_S1789s_1 | -0.21 | -0.62 | -0.21 | -0.79 | -0.12 | -0.87 | -0.96 | -0.44 | -0.51 | -0.29 | -0.7 | -0.12 | -0.03 | -0.33 | -0.05 | -0.03 | -0.14 | 0.0525 | 0.3475 | 0.244726 | 0.647114 | 0.853998 | 1 | 0.999631 | 0.685751 |
| Map1b_P14873_S1789s_2 | -0.95 | -0.37 | -0.45 | -0.38 | -0.41 | -0.25 | -0.66 | -0.08 | -0.73 | -0.43 | -0.67 | -0.74 | 0.12 | -0.48 | -0.23 | -0.65 | 0.1875 | -0.105 | 0.2275 | 0.470027 | 0.498604 | 0.696927 | 1 | 0.999631 | 0.79732 |
| Map1b_P14873_S1814s_1 | -0.34 | -0.49 | -0.69 | -0.49 | -0.63 | -0.69 | -0.54 | -0.65 | -0.64 | -0.28 | -0.79 | -0.43 | -0.28 | -0.6 | -0.39 | -0.23 | -0.125 | -0.0325 | 0.1275 | 0.637846 | 0.597449 | 0.900435 | 1 | 0.999631 | 0.87033 |
| Map1b_P14873_S1877s_1 | -1.34 | 0.25 | 3.07 | 1.24 | 0.11 | -0.05 | 1.7 | 1.27 | 0.42 | -0.74 | 1.96 | 3.03 | 0.48 | 0.34 | 1.46 | 0.71 | -0.0475 | 0.3625 | -0.0575 | 0.944747 | 0.957051 | 0.735144 | 1 | 0.999631 | 0.982141 |
| Map1b_P14873_S1911s_1 | 0.69 | 0.29 | -1.54 | -0.45 | 1.34 | 0.57 | -0.38 | -0.58 | 0.83 | 0.36 | -1.03 | -1.05 | -0.86 | 1.06 | -1.01 | -1.01 | 0.49 | 0.03 | -0.2025 | 0.750207 | 0.421046 | 0.9615 | 1 | 0.999631 | 0.91245 |
| Map1b_P14873_S2030s_1 | -0.92 | -0.79 | -0.13 | -0.38 | -0.8 | -0.88 | -0.56 | -0.39 | -0.6 | -0.45 | -0.54 | 0.1 | -0.04 | -0.4 | 0.16 | 0.03 | -0.1025 | 0.1825 | 0.4925 | 0.136822 | 0.722525 | 0.560001 | 1 | 0.999631 | 0.610369 |
| Map1b_P14873_S2054s_1 | 0.7 | -1.81 | -1.65 | 1.08 | -0.26 | -0.99 | 0.57 | 0.35 | -0.59 | -1.41 | 1.08 | 0.55 | -3.86 | -0.52 | -1.17 | 0.23 | 0.3375 | 0.3275 | -0.91 | 0.373781 | 0.648625 | 0.693196 | 1 | 0.999631 | 0.747035 |
| Map1b_P14873_S2122s_1 | 0.2 | -1.14 | 0.63 | -0.07 | -0.47 | -0.13 | -0.48 | -0.69 | -0.97 | 0.56 | -0.72 | -0.52 | -0.95 | -0.07 | 0.91 | -0.25 | -0.3475 | -0.3175 | 0.005 | 0.992189 | 0.393466 | 0.51671 | 1 | 0.999631 | 0.997164 |
| Map1b_P14873_S2267s_1 | 0.64 | 0.48 | 0.58 | 0.79 | 0.54 | 1 | 0.37 | 1.04 | 0.87 | 1.17 | 0.47 | 1.1 | 0.13 | 1.27 | 1.07 | 1.21 | 0.115 | 0.28 | 0.2975 | 0.390745 | 0.67226 | 0.321251 | 1 | 0.999631 | 0.758901 |
| Map1b_P14873_S2410s_1 | -1.4 | 0.34 | -0.51 | 0.18 | -0.27 | -0.15 | -0.1 | -0.64 | -0.5 | -0.13 | -0.53 | 0.02 | 0.89 | -0.07 | 0.46 | 0.31 | 0.0575 | 0.0625 | 0.745 | 0.117944 | 0.889315 | 0.882179 | 1 | 0.999631 | 0.585087 |
| Map1b_P14873_S25s_1 | -0.74 | 0.34 | -0.19 | -0.29 | -0.05 | 0 | -0.59 | -0.14 | 0.26 | -0.08 | -0.29 | 0.14 | 0.86 | 0.39 | -0.48 | -0.07 | 0.025 | 0.2275 | 0.395 | 0.327416 | 0.936236 | 0.477307 | 1 | 0.999631 | 0.724678 |
| Map1b_P14873_S336s_1 | -1.06 | -1.47 | 0.6 | -0.55 | -1.68 | -0.87 | 0.12 | -0.19 | -1.13 | -0.54 | 0.07 | 0.1 | -0.72 | -1.11 | -0.1 | 1.01 | -0.035 | 0.245 | 0.39 | 0.512858 | 0.94928 | 0.63001 | 1 | 0.999631 | 0.818365 |
| Map1b_P14873_S339s_1 | -0.7 | 0.69 | 0.18 | -0.4 | -0.37 | -0.86 | -0.05 | -0.12 | 0.29 | -0.29 | 0.26 | 0 | 0.58 | 0.34 | -0.03 | 0.6 | -0.2925 | 0.1225 | 0.43 | 0.272361 | 0.442942 | 0.738879 | 1 | 0.999631 | 0.699889 |
| Map1b_P14873_S341s_1 | -1.4 | 0.08 | 0.01 | -0.12 | 0.05 | 0.15 | -0.05 | 0.45 | -0.24 | -0.16 | 0.56 | 0.86 | 0.84 | -0.37 | 0.7 | 0.38 | 0.5075 | 0.6125 | 0.745 | 0.117399 | 0.202189 | 0.184079 | 1 | 0.999631 | 0.585087 |
| Map1b_P14873_S343s_1 | 0.18 | 0.48 | 0.68 | -0.04 | 0.04 | 0.13 | 0.16 | 0.15 | 0.38 | 0.1 | 0.12 | 0.11 | 0.7 | 0.38 | 0.48 | 0.53 | -0.205 | -0.1475 | 0.1975 | 0.502463 | 0.442884 | 0.595255 | 1 | 0.999631 | 0.814092 |
| Map1b_P14873_S601s_1 | 0.83 | 0.62 | 0.99 | 1.04 | 0.02 | 1.44 | 1.81 | 0.67 | 1.07 | 0.84 | 1.17 | 0.84 | 0.07 | -0.11 | 1.02 | 0.22 | 0.115 | 0.11 | -0.57 | 0.113488 | 0.779707 | 0.67032 | 1 | 0.999631 | 0.57792 |
| Map1b_P14873_S614s_1 | 1.81 | 1.21 | 0.73 | 0.1 | -0.47 | 1.54 | 1.69 | -1.23 | 0.64 | 1.6 | -0.38 | -0.26 | 1.11 | 0.4 | 0.39 | -0.23 | -0.58 | -0.5625 | -0.545 | 0.245721 | 0.427434 | 0.315177 | 1 | 0.999631 | 0.685751 |
| Map1b_P14873_S934s_1 | -0.97 | 0.58 | 1.06 | 0 | 0.68 | 0.42 | 0.22 | 0.59 | 0.74 | -0.35 | 0.85 | 1.71 | 1.25 | 0.74 | 1.02 | 0.9 | 0.31 | 0.57 | 0.81 | 0.095592 | 0.484234 | 0.322796 | 1 | 0.999631 | 0.558128 |
| Map1b_P14873_S960s_1 | 0.15 | 0.83 | 1.11 | 0.4 | 0.4 | 0.74 | 0.33 | 0.57 | 0.61 | 1.4 | 0.88 | 0.95 | 0.28 | 0.39 | 1.15 | 0.87 | -0.1125 | 0.3375 | 0.05 | 0.887044 | 0.706822 | 0.31382 | 1 | 0.999631 | 0.963505 |
| Map1b_P14873_S989s_1 | 1.87 | 1.07 | 0.97 | 1.14 | 0.62 | 1.39 | 1.08 | 0.92 | 1.63 | 2.42 | 1.21 | 0.75 | 0.37 | 0.29 | 1.02 | 0.24 | -0.26 | 0.24 | -0.7825 | 0.041097 | 0.415382 | 0.566768 | 1 | 0.999631 | 0.474979 |
| Map1b_P14873_S989s_2 | 0.71 | 1.13 | 2.37 | 1.75 | 0.7 | 1.79 | 1.8 | 1.12 | 1.53 | 2.19 | 1.27 | 1.06 | 0.2 | -0.08 | 0.65 | 1.09 | -0.1375 | 0.0225 | -1.025 | 0.041372 | 0.755085 | 0.958734 | 1 | 0.999631 | 0.474979 |
| Map1b_P14873_S992s_1 | 1.46 | 1.59 | 2.12 | 1.27 | 0.86 | 1.78 | 1.47 | 0.87 | 1.53 | 2.62 | 1.38 | 1.5 | 0.81 | 0.82 | 1.4 | 0.95 | -0.365 | 0.1475 | -0.615 | 0.075457 | 0.287184 | 0.691003 | 1 | 0.999631 | 0.541044 |
| Map1b_P14873_S992s_2 | 0.71 | 1.13 | 2.37 | 1.75 | 0.7 | 1.79 | 1.8 | 1.12 | 1.53 | 2.19 | 1.27 | 1.06 | 0.2 | -0.08 | 0.65 | 1.09 | -0.1375 | 0.0225 | -1.025 | 0.041372 | 0.755085 | 0.958734 | 1 | 0.999631 | 0.474979 |
| Map1b_P14873_T1202t_1 | -0.83 | 0.53 | 0.27 | -0.47 | 0.52 | -0.07 | -0.09 | -0.34 | 0.31 | -0.47 | 0.11 | 1.07 | 0.93 | 0.17 | 0.56 | -0.09 | 0.13 | 0.38 | 0.5175 | 0.222934 | 0.732301 | 0.400058 | 1 | 0.999631 | 0.673942 |
| Map1b_P14873_T1499t_1 | -0.37 | -0.52 | 0.04 | 0.41 | -0.06 | -0.38 | -0.09 | 0.49 | -0.46 | -0.33 | 0.12 | 0.35 | -0.34 | -0.33 | 0.43 | 0.34 | 0.1 | 0.03 | 0.135 | 0.702122 | 0.757544 | 0.928188 | 1 | 0.999631 | 0.896857 |
| Map1b_P14873_T1629t_1 | 2.96 | -0.38 | -0.42 | 1.43 | 1.06 | 0.51 | 2.22 | 0.68 | 1.21 | 1.13 | 0.32 | -0.86 | -2.87 | -0.79 | -0.59 | -1.32 | 0.22 | -0.4475 | -2.29 | 0.019458 | 0.779746 | 0.59061 | 1 | 0.999631 | 0.391668 |
| Map1b_P14873_T1629t_2 | 0.94 | 1.47 | 0.52 | 2.07 | 1.18 | 0.82 | 1.8 | 1.4 | 1.74 | 1.33 | 1.09 | 0.95 | -3.14 | 0.02 | 0.61 | -0.69 | 0.05 | 0.0275 | -2.05 | 0.023412 | 0.900034 | 0.944178 | 1 | 0.999631 | 0.41509 |
| Map1b_P14873_T1784t_1 | 2.07 | -1.08 | -1.26 | 0.77 | 1.81 | -1.18 | 1.23 | -0.11 | 0.26 | -1.73 | -0.68 | -1.13 | -1.41 | -1.06 | -1.4 | -0.85 | 0.3125 | -0.945 | -1.305 | 0.091454 | 0.729288 | 0.249123 | 1 | 0.999631 | 0.557085 |
| Map1b_P14873_T1784t_2 | -0.67 | -1.54 | -1.34 | -0.36 | -0.45 | -0.82 | -0.8 | -0.09 | -1.79 | -0.34 | -0.41 | -0.94 | 0.27 | -0.26 | -0.21 | -0.49 | 0.4375 | 0.1075 | 0.805 | 0.048924 | 0.234843 | 0.804012 | 1 | 0.999631 | 0.48544 |
| Map1b_P14873_T1925t_1 | -0.39 | 0.03 | -0.46 | -0.81 | -0.51 | -1.83 | -0.79 | -0.19 | 0.3 | -0.03 | -0.58 | -0.56 | 0.14 | 0.58 | -0.46 | -0.52 | -0.4225 | 0.19 | 0.3425 | 0.355025 | 0.30444 | 0.566343 | 1 | 0.999631 | 0.740148 |
| Map1b_P14873_T1928t_1 | 0.08 | -0.04 | -0.48 | -0.26 | 0.15 | -0.05 | -0.48 | 0.24 | 0.06 | 0.29 | -0.39 | 0.32 | 0.09 | 0.26 | 0.21 | 0.06 | 0.14 | 0.245 | 0.33 | 0.249767 | 0.621553 | 0.408284 | 1 | 0.999631 | 0.686359 |
| Map1b_P14873_T1945t_1 | -0.35 | 0.33 | -0.44 | -0.53 | 0.12 | -0.27 | -0.83 | -0.95 | -0.1 | 0.13 | -0.52 | -0.63 | 0.47 | 0.46 | -0.46 | 0.53 | -0.235 | -0.0325 | 0.4975 | 0.186287 | 0.504426 | 0.919691 | 1 | 0.999631 | 0.646474 |
| Map1b_P14873_T1962t_1 | 0.73 | -0.48 | -0.64 | 1.05 | 0.89 | -0.46 | 0.69 | 0.68 | 0.36 | -0.64 | 0.52 | 0.16 | -1.93 | 0.47 | 0.34 | 0.07 | 0.285 | -0.065 | -0.4275 | 0.505724 | 0.566701 | 0.891711 | 1 | 0.999631 | 0.814836 |
| Map1b_P14873_T2047t_1 | 1.62 | -0.9 | -1.5 | 1.11 | 2.59 | 0.35 | 1.65 | 0.53 | 1.09 | -0.66 | 0 | -2.52 | -2.01 | -0.86 | -1.04 | -1.43 | 1.1975 | -0.605 | -1.4175 | 0.068605 | 0.159391 | 0.519676 | 1 | 0.999631 | 0.523794 |
| Map1b_P14873_T2301t_1 | 0.96 | 0.51 | 1.17 | 0.92 | 0.35 | 0.89 | 0.98 | 0.76 | 0.76 | 1.3 | 0.33 | 0.56 | 0.4 | 0.6 | 1.35 | 0.17 | -0.145 | -0.1525 | -0.26 | 0.462342 | 0.605352 | 0.628712 | 1 | 0.999631 | 0.792786 |
| Map1b_P14873_T527t_1 | -0.96 | 1.2 | 1.24 | 0.88 | 0.73 | 0.79 | 0.49 | 1.02 | 1 | 0.03 | 1.32 | 1.49 | 0.9 | 0.86 | 1.27 | 1.41 | 0.1675 | 0.37 | 0.52 | 0.323709 | 0.738482 | 0.519052 | 1 | 0.999631 | 0.724485 |
| Map1b_P14873_T945t_1 | 0.15 | 1.05 | 0.87 | 0.71 | 0.49 | 1.01 | 0.71 | 0.55 | 1.15 | 0.86 | 1.08 | 1.43 | 1.14 | 0.64 | 1.56 | 1.07 | -0.005 | 0.435 | 0.4075 | 0.245175 | 0.986439 | 0.172038 | 1 | 0.999631 | 0.685751 |
| Map1s_Q8C052_S458s_1 | 0.19 | 1.84 | 1.84 | 1.16 | 0.38 | 1.86 | 1.05 | 2.03 | 0.32 | 0.45 | 2.14 | -0.34 | -0.47 | 1.88 | -0.07 | 0.92 | 0.0725 | -0.615 | -0.6925 | 0.263318 | 0.886828 | 0.317929 | 1 | 0.999631 | 0.693034 |
| Map1s_Q8C052_S462s_1 | 0.84 | 0.09 | 0.32 | 0.37 | -0.31 | 0.11 | 0.63 | -0.39 | 0.45 | 1.06 | 0.4 | 0.58 | 0.3 | -0.77 | 0.51 | -0.09 | -0.395 | 0.2175 | -0.4175 | 0.271446 | 0.243615 | 0.470278 | 1 | 0.999631 | 0.699288 |
| Map1s_Q8C052_S462s_2 | 1.33 | 1.09 | 1.62 | 1.17 | 0.75 | 1.11 | 1.12 | 1.73 | 1.09 | 2.25 | 0.62 | 0.93 | -0.85 | 0.56 | 0.6 | -0.36 | -0.125 | -0.08 | -1.315 | 0.00767 | 0.676853 | 0.838075 | 1 | 0.999631 | 0.358173 |
| Map1s_Q8C052_S465s_1 | 0.66 | 0.47 | 2.19 | 0.54 | -0.07 | 0.86 | 0.6 | 0.19 | 0.89 | 1.41 | 0.93 | 0.47 | 0.21 | -0.23 | -0.13 | 0.58 | -0.57 | -0.04 | -0.8575 | 0.08005 | 0.220468 | 0.928469 | 1 | 0.999631 | 0.545579 |
| Map1s_Q8C052_S465s_2 | 1.33 | 1.09 | 1.62 | 1.17 | 0.75 | 1.11 | 1.12 | 1.73 | 1.09 | 2.25 | 0.62 | 0.93 | -0.85 | 0.56 | 0.6 | -0.36 | -0.125 | -0.08 | -1.315 | 0.00767 | 0.676853 | 0.838075 | 1 | 0.999631 | 0.358173 |
| Map1s_Q8C052_S684s_1 | 1.43 | 0.55 | -0.14 | 1.48 | 1.4 | 0.42 | 1.06 | 0.93 | 0.5 | 0.52 | 0.92 | 0.28 | -0.58 | 0.51 | 0.11 | 0.32 | 0.1225 | -0.275 | -0.74 | 0.125878 | 0.775844 | 0.512715 | 1 | 0.999631 | 0.596745 |
| Map2_P20357_S1013s_1 | -2.79 | -6.28 | -2.23 | -2.68 | -6.93 | -2.79 | -0.93 | -2.85 | -13.52 | -3.47 | -1.68 | -4.47 | -5.82 | -4.05 | -3.02 | -4.39 | 0.12 | -2.29 | -0.825 | 0.391969 | 0.928532 | 0.345173 | 1 | 0.999631 | 0.758901 |
| Map2_P20357_S1023s_1 | 0.27 | 1.41 | -0.02 | -0.55 | 0.92 | 1.36 | -0.39 | 0.68 | 1.29 | 1.37 | 0.5 | 0.55 | 1.74 | 1.29 | 0.28 | 0.51 | 0.365 | 0.65 | 0.6775 | 0.203654 | 0.486231 | 0.181386 | 1 | 0.999631 | 0.659781 |
| Map2_P20357_S1161s_1 | -0.17 | 0.9 | 0.85 | 0.39 | 0.48 | 0.64 | 0.07 | 0.89 | 0.4 | 0.46 | 0.73 | 1.12 | 0.8 | 1.05 | 0.97 | 1.24 | 0.0275 | 0.185 | 0.5225 | 0.141803 | 0.935224 | 0.59152 | 1 | 0.999631 | 0.612726 |
| Map2_P20357_S1352s_1 | -1.51 | 1.12 | 5.1 | 1.48 | 1 | 0.02 | 4.07 | 3.91 | 1.65 | -2.4 | 6.34 | 7.39 | 1.93 | -0.7 | 5.87 | 3.05 | 0.7025 | 1.6975 | 0.99 | 0.542707 | 0.628684 | 0.452606 | 1 | 0.999631 | 0.832881 |
| Map2_P20357_S1485s_1 | 2.29 | 0.65 | 1.85 | 1.46 | 0.89 | 1.45 | 1.54 | 1.15 | 1.12 | 2.13 | 0.48 | 0.53 | -0.41 | 0.56 | 1.08 | 0.04 | -0.305 | -0.4975 | -1.245 | 0.022061 | 0.439447 | 0.326434 | 1 | 0.999631 | 0.405544 |
| Map2_P20357_S1539s_1 | -1.65 | -2.59 | -0.28 | 0.14 | -0.86 | -3.17 | -0.35 | -0.08 | -1.73 | -2.11 | -0.02 | 1.03 | -0.53 | -2.65 | 0.65 | -0.38 | -0.02 | 0.3875 | 0.3675 | 0.652939 | 0.980566 | 0.64833 | 1 | 0.999631 | 0.877851 |
| Map2_P20357_S1560s_1 | 1.12 | 0.7 | -0.59 | 1.78 | 1.89 | 0.48 | 1.05 | 1.36 | 0.69 | -0.18 | 1.22 | 1.14 | -1.33 | 1.58 | 1.08 | 0.98 | 0.4425 | -0.035 | -0.175 | 0.808889 | 0.416985 | 0.949249 | 1 | 0.999631 | 0.93635 |
| Map2_P20357_S1625s_1 | 0.37 | 0.4 | 1.88 | 0.53 | 0.64 | 0.86 | 0.1 | 0.98 | 0.3 | 0.31 | -0.01 | 1.37 | -0.26 | 0.05 | 1.98 | 0.94 | -0.15 | -0.3025 | -0.1175 | 0.83724 | 0.716326 | 0.515399 | 1 | 0.999631 | 0.945882 |
| Map2_P20357_S1635s_1 | -0.74 | -0.96 | -0.91 | 1.87 | 0.33 | -0.03 | 1.78 | 1.85 | 0.4 | -1.42 | 1.85 | 0.65 | -0.28 | -0.15 | 1.98 | 1.3 | 1.1675 | 0.555 | 0.8975 | 0.261645 | 0.138839 | 0.514443 | 1 | 0.999631 | 0.69271 |
| Map2_P20357_S1654s_2 | 1.57 | -0.05 | -1.08 | 0.94 | 1.78 | 1.4 | 1.73 | 2.39 | 1.22 | 0.56 | 1.37 | 0.34 | -1.69 | 0.6 | 0.2 | 0.82 | 1.48 | 0.5275 | -0.3625 | 0.61664 | 0.024058 | 0.373223 | 1 | 0.999631 | 0.861203 |
| Map2_P20357_S1743s_1 | 1.95 | 0.89 | -1.14 | 2.35 | 2.01 | 0.47 | 2.17 | 0.89 | 1.29 | 0.51 | 2.13 | 0.98 | -2.53 | 0.55 | -0.69 | 0.32 | 0.3725 | 0.215 | -1.6 | 0.10119 | 0.632782 | 0.775368 | 1 | 0.999631 | 0.564448 |
| Map2_P20357_S1783s_1 | 0.28 | 2.04 | 1.74 | -0.36 | 0.26 | 0.68 | 3.11 | 0.31 | 0.49 | 0.55 | 1.47 | 3.07 | 1.45 | -0.38 | 0.83 | 0.45 | 0.165 | 0.47 | -0.3375 | 0.591735 | 0.832424 | 0.528019 | 1 | 0.999631 | 0.852627 |
| Map2_P20357_S1791s_1 | 1.51 | 0.93 | 2.09 | 1.29 | -0.25 | 1.14 | 1.5 | 0.77 | 0.24 | 1.82 | 0.9 | 0.94 | -0.07 | 0.53 | 1.4 | 0.72 | -0.665 | -0.48 | -0.81 | 0.070578 | 0.152259 | 0.261232 | 1 | 0.999631 | 0.530826 |
| Map2_P20357_S238s_1 | 1.96 | 0.86 | -1.04 | 1.47 | 3.5 | 0.97 | 1.27 | 1.02 | 1.03 | -1.5 | 0.63 | -0.25 | -2.9 | 1.06 | 0.11 | 0.45 | 0.8775 | -0.835 | -1.1325 | 0.247895 | 0.278209 | 0.288439 | 1 | 0.999631 | 0.685751 |
| Map2_P20357_S283s_1 | 2.3 | -1.61 | -2.51 | 1.4 | 2.76 | -0.6 | 2.37 | 1.24 | -0.3 | -2.27 | 0.36 | -1.91 | -2.25 | -0.92 | -1.31 | -1.32 | 1.5475 | -0.925 | -1.345 | 0.206445 | 0.209922 | 0.422109 | 1 | 0.999631 | 0.6605 |
| Map2_P20357_S28s_1 | -1.53 | -2.3 | -2 | -2.07 | -3.04 | -2.06 | -1.21 | -2.87 | -3.63 | -1.74 | -3.19 | -3.09 | -3.54 | -2.5 | -2.68 | -1.58 | -0.32 | -0.9375 | -0.6 | 0.191328 | 0.471929 | 0.054088 | 1 | 0.999631 | 0.648674 |
| Map2_P20357_S42s_1 | 0.96 | 0.45 | 0.8 | 1.85 | 1.46 | 0.57 | 2.63 | 1.23 | 0.78 | 0.33 | 1.22 | 1.26 | -0.12 | 0.06 | 0.23 | -0.01 | 0.4575 | -0.1175 | -0.975 | 0.020002 | 0.363582 | 0.76222 | 1 | 0.999631 | 0.391668 |
| Map2_P20357_S447s_1 | 0.45 | 0.81 | 0.41 | 0.54 | 0.66 | 0.66 | 0.24 | 0.43 | 0.71 | 0.3 | 0.8 | 0.92 | 0.84 | 1.15 | 0.2 | 1.24 | -0.055 | 0.13 | 0.305 | 0.362969 | 0.827655 | 0.633244 | 1 | 0.999631 | 0.74343 |
| Map2_P20357_S472s_1 | 0.05 | 0.42 | -1.01 | 1.14 | 0.72 | 0.12 | 0.78 | 0.38 | 0.36 | -0.43 | 0.67 | -0.1 | -0.53 | 0.4 | 0.25 | 0.98 | 0.35 | -0.025 | 0.125 | 0.80897 | 0.449785 | 0.959045 | 1 | 0.999631 | 0.93635 |
| Map2_P20357_S476s_1 | 0.66 | 0.86 | -0.24 | 0.87 | 1.6 | 1.41 | 0.83 | 0.72 | 0.9 | -0.03 | 0.94 | 0.5 | -0.05 | 1.43 | 0.29 | 0.9 | 0.6025 | 0.04 | 0.105 | 0.806983 | 0.121288 | 0.914541 | 1 | 0.999631 | 0.935523 |
| Map2_P20357_S599s_1 | 1.01 | 2.29 | 1.12 | 0.95 | 1.68 | 2.81 | 0.87 | 1.01 | 2.15 | 1.69 | 0.79 | 0.83 | 1.29 | 1.74 | 0.87 | 2.03 | 0.25 | 0.0225 | 0.14 | 0.739705 | 0.625156 | 0.960232 | 1 | 0.999631 | 0.908631 |
| Map2_P20357_S608s_1 | -0.15 | 0.93 | 1 | 0.19 | 0.5 | 0.59 | 0.13 | 0.77 | 0.29 | 0.15 | 0.68 | 0.86 | 0.91 | 1.17 | 1.14 | 0.87 | 0.005 | 0.0025 | 0.53 | 0.1524 | 0.988418 | 0.994455 | 1 | 0.999631 | 0.625198 |
| Map2_P20357_S63s_1 | 1.54 | 0.96 | 0.52 | 0.48 | 0.74 | 1.43 | 1.83 | -0.28 | 1.46 | 0.93 | 0.36 | -0.67 | -0.08 | 0.02 | -0.23 | -0.19 | 0.055 | -0.355 | -0.995 | 0.011873 | 0.911081 | 0.478058 | 1 | 0.999631 | 0.385092 |
| Map2_P20357_S654s_1 | -1.16 | -1.08 | -0.53 | -1.11 | -0.41 | -0.37 | -0.39 | -1.16 | -0.17 | -1.41 | -1.56 | -1.34 | -0.46 | -0.43 | -0.56 | -0.4 | 0.3875 | -0.15 | 0.5075 | 0.097098 | 0.220723 | 0.691376 | 1 | 0.999631 | 0.55886 |
| Map2_P20357_S655s_1 | -0.15 | -0.22 | -0.35 | -0.54 | -0.54 | -0.56 | -0.29 | -0.43 | -0.6 | -0.28 | -0.75 | -0.28 | -0.03 | -0.28 | -0.13 | 0.23 | -0.14 | -0.1625 | 0.2625 | 0.356175 | 0.5668 | 0.542895 | 1 | 0.999631 | 0.74034 |
| Map2_P20357_S823s_1 | 3.37 | -2.32 | -4.15 | 2.94 | 3.93 | -2.54 | 3.87 | 1.21 | -0.44 | -10.38 | 0.98 | -3.29 | -3.1 | -1.88 | -3.48 | -2.38 | 1.6575 | -3.2425 | -2.67 | 0.120717 | 0.425537 | 0.240838 | 1 | 0.999631 | 0.588344 |
| Map2_P20357_S825s_1 | 2.1 | -0.48 | -3.83 | 2.78 | 2.78 | -0.03 | 3.69 | 1.14 | 1.1 | -1.04 | 0.91 | -2.81 | -2.6 | 0.9 | -3.12 | -2.12 | 1.7525 | -0.6025 | -1.8775 | 0.22038 | 0.246104 | 0.688129 | 1 | 0.999631 | 0.670752 |
| Map2_P20357_S938s_1 | 1.47 | 1.46 | 1.33 | 1.07 | 0.8 | 1.12 | 1.07 | 0.63 | 0.88 | 1.38 | -0.23 | -0.09 | 0.63 | 0.55 | 0.64 | 0.2 | -0.4275 | -0.8475 | -0.8275 | 0.013134 | 0.121308 | 0.060811 | 1 | 0.999631 | 0.385092 |
| Map2_P20357_T1160t_1 | -0.33 | 0.68 | 1.1 | 1.4 | 0.34 | 0.75 | 0.6 | 1.67 | 0.39 | -0.02 | 2.39 | 2.18 | 0.68 | 1.03 | 1.87 | 1.63 | 0.1275 | 0.5225 | 0.59 | 0.218932 | 0.780645 | 0.428199 | 1 | 0.999631 | 0.669859 |
| Map2_P20357_T1260t_1 | -0.2 | 0.4 | -1.33 | 1.28 | 2.2 | 1.98 | 1.9 | 1.82 | 0.65 | -1.94 | 1.52 | -0.39 | -1.33 | 1.02 | -0.77 | 0.13 | 1.9375 | -0.0775 | -0.275 | 0.6833 | 0.003772 | 0.923728 | 1 | 0.999631 | 0.889602 |
| Map2_P20357_T1358t_1 | -1.64 | 1.05 | 1.29 | -0.11 | 1 | 1.08 | 0.28 | 0.65 | 0.41 | -0.69 | 0.64 | 0.91 | 1.66 | 1.31 | 1.4 | 0.81 | 0.605 | 0.17 | 1.1475 | 0.091181 | 0.344067 | 0.80168 | 1 | 0.999631 | 0.557085 |
| Map2_P20357_T1599t_2 | -0.95 | 1.28 | 2.5 | 1.85 | 1.2 | 1.08 | 0.03 | 2.03 | -0.63 | 0.88 | 1.25 | 0.21 | -0.95 | 0.27 | 2.39 | 2.18 | -0.085 | -0.7425 | -0.1975 | 0.833723 | 0.90975 | 0.338092 | 1 | 0.999631 | 0.943785 |
| Map2_P20357_T1609t_1 | 0.14 | 1.55 | 1.58 | 0.76 | 0.81 | 1.14 | 0.45 | 1.22 | 1.19 | 0.73 | 1.22 | 1.85 | 1.49 | 1.75 | 1.6 | 1.96 | -0.1025 | 0.24 | 0.6925 | 0.100342 | 0.795282 | 0.570238 | 1 | 0.999631 | 0.563392 |
| Map2_P20357_T1609t_2 | -0.64 | 0.12 | -0.14 | -0.38 | 0.38 | -0.17 | -0.07 | -0.12 | -0.17 | 0.19 | 0.04 | -0.46 | -0.57 | 0.33 | 0.01 | 0.09 | 0.265 | 0.16 | 0.225 | 0.49693 | 0.361517 | 0.591165 | 1 | 0.999631 | 0.811348 |
| Map2_P20357_T1617t_1 | 1.69 | 0.84 | 0.16 | 1.3 | 1.81 | 1.54 | 1.27 | 1.5 | 1.57 | 1.4 | 1.42 | 0.87 | -1.43 | 1.75 | 1.1 | 0.96 | 0.5325 | 0.3175 | -0.4025 | 0.560334 | 0.169323 | 0.415978 | 1 | 0.999631 | 0.839886 |
| Map2_P20357_T1617t_2 | 1.41 | 0.92 | -0.25 | 0.65 | 1.53 | 1.54 | 0.81 | 1.02 | 1 | 1.33 | 0.94 | 0.67 | -2.7 | 1.18 | 0.61 | 0.78 | 0.5425 | 0.3025 | -0.715 | 0.400924 | 0.194896 | 0.446008 | 1 | 0.999631 | 0.763209 |
| Map2_P20357_T1620t_1 | 1.54 | 1.46 | 1.76 | 1.27 | 1.26 | 2.98 | 1.19 | 1.99 | 1.47 | 1.94 | 1.78 | 1.67 | -1.79 | 2.43 | 1.77 | 1.93 | 0.3475 | 0.2075 | -0.4225 | 0.618861 | 0.419832 | 0.437418 | 1 | 0.999631 | 0.861888 |
| Map2_P20357_T1623t_1 | 1.19 | 0.77 | 1.39 | 1.04 | 1.16 | 2.54 | 0.9 | 1.28 | 1.07 | 1.27 | 1.09 | 0.69 | -1.34 | 1.79 | 1.19 | 1.45 | 0.3725 | -0.0675 | -0.325 | 0.620126 | 0.356689 | 0.808415 | 1 | 0.999631 | 0.861888 |
| Map2_P20357_T1623t_2 | 1.41 | 0.92 | -0.27 | -0.17 | 1.53 | 1.54 | 0.5 | 0.52 | 1 | 1.33 | 0.66 | 0.06 | -2.7 | 1.18 | 0.09 | 0.54 | 0.55 | 0.29 | -0.695 | 0.408366 | 0.269334 | 0.54578 | 1 | 0.999631 | 0.766996 |

| | | | | | | | | | | | | | | | | | | | | | | | | | |
|---|---|---|---|---|---|---|---|---|---|---|---|---|---|---|---|---|---|---|---|---|---|---|---|---|---|
| Map2_P20357_T1650t_1 | 0.58 | 0.88 | 1.01 | 1.65 | 1.43 | -0.5 | 0.12 | 1.42 | 1.54 | 1.39 | 1.78 | 2.88 | 2.56 | 1.77 | 1.89 | 1.36 | -0.4125 | 0.8675 | 0.865 | 0.040936 | 0.417662 | 0.058544 | 1 | 0.999631 | 0.474979 |
| Map2_P20357_T1650t_2 | 1.57 | -0.05 | -1.08 | 0.94 | 1.78 | 1.4 | 1.73 | 2.39 | 1.22 | 0.56 | 1.37 | 0.34 | -1.69 | 0.6 | 0.2 | 0.82 | 1.48 | 0.5275 | -0.3625 | 0.61664 | 0.024058 | 0.373223 | 1 | 0.999631 | 0.861203 |
| Map2k1_P31938_S231s_1 | -0.7 | -0.14 | 0.02 | -0.71 | 0.24 | -0.21 | -0.18 | 0.31 | -0.09 | -0.27 | 0.16 | 0.29 | -0.64 | -0.09 | 0.65 | -0.49 | 0.4225 | 0.405 | 0.24 | 0.532689 | 0.178251 | 0.200238 | 1 | 0.999631 | 0.8277 |
| Map2k1_P31938_T286t_1 | -0.67 | -0.18 | 0.11 | 0.67 | 0.41 | 0.57 | 0.14 | 1.17 | 0.66 | 0.33 | 0.72 | 1.07 | 0.39 | 1.06 | 0.69 | 0.28 | 0.59 | 0.7125 | 0.6225 | 0.118029 | 0.137055 | 0.068469 | 1 | 0.999631 | 0.585087 |
| Map2k2_Q91YS7_S295s_1 | 0.53 | -0.33 | 0.68 | 1.15 | 0.4 | 0.22 | 1.09 | 1.51 | -0.51 | -1.08 | 0.79 | 0.89 | -0.73 | -0.25 | 1.46 | 0.93 | 0.2975 | -0.485 | -0.155 | 0.779301 | 0.489427 | 0.376856 | 1 | 0.999631 | 0.921416 |
| Map2k2_Q91YS7_S306s_1 | 1.91 | -0.74 | 0.34 | 0.43 | 0.38 | 1.18 | 0.92 | 0.62 | 0.38 | 0.45 | -0.11 | 0.66 | -1.15 | 0.61 | -0.19 | -0.88 | 0.29 | -0.14 | -0.8875 | 0.167896 | 0.586303 | 0.792191 | 1 | 0.999631 | 0.637399 |
| Map2k2_Q91YS7_T394t_1 | 1.71 | -0.86 | 1.94 | 1.57 | -0.74 | -0.65 | 1.66 | 0.63 | -0.89 | 0.51 | 0.44 | 0.01 | -1.2 | -1.64 | 0.64 | -0.29 | -0.865 | -1.0725 | -1.7125 | 0.038221 | 0.272687 | 0.126362 | 1 | 0.999631 | 0.459475 |
| Map2k4_P47809_S392s_1 | -0.97 | -0.47 | -0.68 | -1.11 | -0.36 | -0.55 | -1.01 | -0.68 | -0.59 | -1 | -0.57 | -0.36 | 0.26 | -0.7 | -0.41 | -0.65 | 0.1575 | 0.1775 | 0.4325 | 0.214641 | 0.576749 | 0.539473 | 1 | 0.999631 | 0.66742 |
| Map2k4_P47809_S88s_1 | -0.72 | -0.33 | 0.06 | -0.04 | -0.16 | -0.17 | -0.19 | 0.09 | -0.12 | -0.4 | 0.07 | 0.33 | 0.17 | 0.4 | 0.57 | 0.37 | 0.15 | 0.2275 | 0.635 | 0.056184 | 0.587437 | 0.462291 | 1 | 0.999631 | 0.495761 |
| Map3k1_P53349_S915s_1 | 0.72 | -1.14 | -0.44 | -0.4 | 0 | -1.23 | -0.29 | 0.12 | -0.69 | -0.3 | -0.43 | -0.61 | -1.22 | -0.7 | -1.1 | -0.51 | -0.035 | -0.1925 | -0.5675 | 0.205207 | 0.940425 | 0.63622 | 1 | 0.999631 | 0.6605 |
| Map3k10_D3YXM8_S502s_1 | -0.56 | -0.48 | -0.1 | -0.72 | -0.74 | -0.54 | -0.87 | -0.81 | -0.68 | -0.23 | -0.63 | -0.26 | 0.25 | 0.1 | -0.41 | -0.05 | -0.275 | 0.015 | 0.4375 | 0.166308 | 0.300991 | 0.956866 | 1 | 0.999631 | 0.63587 |
| Map3k10_D3YXM8_S714s_1 | 2.82 | -0.74 | -3.79 | -0.22 | 2.65 | 1.43 | -0.05 | -0.06 | 0.68 | 0.09 | -0.78 | -1.77 | -4.43 | 0.68 | -2.29 | -2.39 | 1.475 | 0.0375 | -1.625 | 0.275065 | 0.265931 | 0.975926 | 1 | 0.999631 | 0.700187 |
| Map3k10_D3YXM8_S818s_1 | 0.45 | 0.33 | -0.41 | 0.54 | 0.82 | 0.94 | 0.37 | 0.17 | 1.26 | 0.48 | 0.32 | -0.14 | 0.06 | 0.63 | -0.33 | 0.23 | 0.3475 | 0.2525 | -0.08 | 0.819763 | 0.302093 | 0.515117 | 1 | 0.999631 | 0.938303 |
| Map3k10_D3YXM8_T552t_1 | 0.71 | 1.37 | 0.92 | 0.84 | 1.11 | 0.92 | 0.24 | 1.51 | 1.06 | 0.82 | 1.14 | 1.83 | 1.2 | 2.21 | 1.14 | 1.66 | -0.015 | 0.2525 | 0.5925 | 0.111356 | 0.964564 | 0.437301 | 1 | 0.999631 | 0.577068 |
| Map3k12_Q60700_S569s_1 | 2.15 | -0.5 | 0.25 | 0.84 | 0.02 | 0.47 | 1.05 | -0.02 | 0.09 | 1 | -0.13 | 0.03 | -1.38 | -0.71 | -0.05 | -0.57 | -0.305 | -0.4375 | -1.3625 | 0.036566 | 0.589171 | 0.447382 | 1 | 0.999631 | 0.453963 |
| Map3k12_Q60700_T43t_1 | -1.52 | 3.5 | 1.97 | -0.32 | 3.22 | 2.32 | 1.67 | 2.18 | 3.47 | -1.48 | 3.39 | 4.02 | 4.3 | 2.48 | 3.15 | 1.06 | 1.44 | 1.4425 | 1.84 | 0.123435 | 0.175562 | 0.333897 | 1 | 0.999631 | 0.594232 |
| Map3k13_Q1HKZ5_S38s_1 | 0.99 | 1 | 1.09 | 0.92 | 1.79 | 0.92 | 0.76 | 1.01 | 1.48 | 0.43 | 1.18 | 1.17 | -0.24 | 1.34 | 0.42 | 1.27 | 0.12 | 0.065 | -0.3025 | 0.457486 | 0.687547 | 0.829308 | 1 | 0.999631 | 0.790722 |
| Map3k13_Q1HKZ5_S680s_1 | 1.14 | 0.7 | -0.07 | -0.18 | 0.77 | 1.65 | 0.33 | -0.22 | 1.54 | 1.15 | -0.14 | 0.72 | 0.3 | 0.51 | -0.03 | -0.18 | 0.235 | 0.42 | -0.2475 | 0.524943 | 0.625958 | 0.376559 | 1 | 0.999631 | 0.822727 |
| Map3k13_Q1HKZ5_S914s_1 | 0.93 | 1.29 | 0.06 | 1.05 | 1.33 | 1.36 | 0.59 | 1.27 | 1.47 | 1.2 | 1.09 | 1.66 | 0.11 | 1.51 | 1.48 | 1.24 | 0.305 | 0.5225 | 0.2525 | 0.562876 | 0.396561 | 0.150074 | 1 | 0.999631 | 0.841038 |
| Map3k2_G5E8L8_S163s_1 | -1.97 | -1.58 | -1.34 | -1.53 | -0.98 | -1.31 | -1.17 | -1.11 | -1.6 | -2.17 | -1.22 | -0.78 | -0.26 | -1.28 | -1.44 | -1.6 | 0.4625 | 0.1625 | 0.46 | 0.23395 | 0.097421 | 0.651208 | 1 | 0.999631 | 0.683077 |
| Map3k4_O08648_S116s_1 | 1.55 | 1.11 | 1.31 | 1.03 | 0.45 | 1.41 | 0.84 | 0.87 | 1.39 | 1.96 | 0.05 | 0.79 | -0.65 | 1.32 | 1.37 | -0.05 | -0.3575 | -0.2025 | -0.7525 | 0.152781 | 0.244516 | 0.636672 | 1 | 0.999631 | 0.625198 |
| Map3k4_O08648_S424s_1 | -1.46 | -1.04 | -1.72 | -0.71 | -0.28 | -2.08 | -0.9 | -1.37 | -0.64 | -1.41 | -1.38 | -0.6 | -0.37 | -0.63 | -0.64 | -0.65 | 0.075 | 0.225 | 0.66 | 0.060822 | 0.862143 | 0.528817 | 1 | 0.999631 | 0.509205 |
| Map3k4_O08648_S454s_1 | -1.39 | -1.22 | -1.59 | -1.84 | -2.17 | -1.09 | -1.79 | -3.2 | -1.16 | -0.67 | -1.07 | -1.76 | -0.42 | -0.31 | -1.65 | -1.26 | -0.5525 | 0.345 | 0.6 | 0.142276 | 0.234022 | 0.296938 | 1 | 0.999631 | 0.612726 |
| Map3k7_Q923A8_S389s_1 | 1.95 | -0.06 | -0.4 | 0.36 | -0.14 | -0.22 | 0.46 | -0.07 | -0.25 | 0.35 | 0.61 | -0.73 | -0.54 | -0.09 | -0.68 | 0.53 | -0.455 | -0.4675 | -0.6575 | 0.24671 | 0.379257 | 0.408934 | 1 | 0.999631 | 0.685751 |
| Map3k9_Q3U1V8_S540s_1 | -1.49 | -0.43 | -0.17 | -1.37 | -1.55 | -1.09 | -1.58 | -1.23 | -0.97 | -0.34 | -0.31 | -0.01 | 0.68 | -0.51 | -0.43 | -0.08 | -0.4975 | 0.4575 | 0.78 | 0.097016 | 0.2004 | 0.269597 | 1 | 0.999631 | 0.55886 |
| Map4_E9PZ43_S298s_1 | 1.25 | 0.73 | 0.12 | 1 | 0.41 | 0.3 | 1.42 | 0.27 | 1.13 | 0.84 | 1.16 | -0.28 | -0.78 | -0.22 | 0.26 | 1.4 | -0.175 | -0.0625 | -0.61 | 0.240897 | 0.647033 | 0.88169 | 1 | 0.999631 | 0.684338 |
| Map4_E9PZ43_S366s_1 | 0.18 | 0.57 | -0.62 | 0.11 | 1.34 | -0.56 | 0.43 | 0.49 | 0.26 | -0.21 | 0.01 | 0.17 | -0.66 | 0.05 | -0.63 | 0.88 | 0.365 | -0.0025 | -0.15 | 0.734906 | 0.422039 | 0.993854 | 1 | 0.999631 | 0.90759 |
| Map4_E9PZ43_S514s_1 | 1.36 | 0.77 | 1.23 | 0.66 | 0.46 | 1.14 | 0.65 | 0.65 | 0.83 | 1.66 | 0.6 | 0.6 | 0.36 | 0.6 | 0.77 | 0.49 | -0.28 | -0.0825 | -0.45 | 0.154996 | 0.351368 | 0.811687 | 1 | 0.999631 | 0.6258 |
| Map4_E9PZ43_S632s_1 | 0.72 | 0.41 | -0.12 | 0.36 | 0.9 | 0.34 | 0.27 | 0.96 | 0.73 | 0.69 | 0.51 | 0.68 | -0.51 | 0.9 | 0.57 | 0.77 | 0.275 | 0.31 | 0.09 | 0.819152 | 0.381856 | 0.281984 | 1 | 0.999631 | 0.938303 |
| Map4_E9PZ43_S645s_1 | 0.84 | 0.41 | 0.46 | 1.42 | 0.84 | 1.34 | 1.52 | 2.02 | 1.58 | 1.37 | 2 | 1.89 | -0.13 | 1.78 | 2.32 | 2.04 | 0.6475 | 0.9275 | 0.72 | 0.216486 | 0.097424 | 0.017125 | 1 | 0.999631 | 0.669465 |
| Map4_E9PZ43_S675s_1 | 1.92 | 0.69 | 0.02 | 1.91 | 1.14 | 0.97 | 1.89 | 1 | 1.43 | 1.97 | 1.41 | 1.42 | -0.98 | 0.28 | 0.56 | 1.02 | 0.115 | 0.4225 | -0.915 | 0.140326 | 0.814025 | 0.381866 | 1 | 0.999631 | 0.611934 |
| Map4_E9PZ43_S723s_1 | 1.46 | 0.33 | -0.18 | 1.74 | 1.38 | 0.89 | 1.71 | 1.23 | 0.72 | -0.17 | 1.27 | 0.16 | -1.65 | 0.58 | -0.5 | 0.66 | 0.465 | -0.3425 | -1.065 | 0.119408 | 0.330558 | 0.515821 | 1 | 0.999631 | 0.586997 |
| Map4_E9PZ43_S855s_1 | 0.56 | 0.23 | -0.26 | 0.9 | 0.45 | -0.41 | 0.57 | 1.27 | 0.11 | -0.04 | 0.42 | 0.49 | -0.51 | 0.65 | 0.16 | 0.38 | 0.1125 | -0.1125 | -0.1875 | 0.627247 | 0.788998 | 0.733647 | 1 | 0.999631 | 0.865372 |
| Map4_E9PZ43_S93s_1 | 1.36 | -2.55 | -3.14 | 0.69 | 1.61 | -1.4 | 0.68 | -0.01 | -0.6 | -1.6 | -1.36 | -1.81 | -1.94 | -1.21 | -1.52 | -2.16 | 1.13 | -0.4325 | -0.7975 | 0.426319 | 0.322225 | 0.667847 | 1 | 0.999631 | 0.774781 |
| Map4_E9PZ43_T505t_1 | 0.23 | 1.71 | 1.82 | 0.56 | 0.55 | 1.77 | 0.44 | 1.33 | 1.16 | 1.45 | 1.2 | 1.4 | 1.26 | 1.71 | 1.28 | 2.3 | -0.0575 | 0.2225 | 0.5575 | 0.246343 | 0.905477 | 0.593837 | 1 | 0.999631 | 0.685751 |
| Map4_E9PZ43_T694t_1 | 0.23 | 0.91 | 0.59 | 0.17 | 0.41 | 0.56 | 0.08 | 0.48 | 0.54 | 0.89 | 0.31 | 0.52 | 0.83 | 1.11 | 0.56 | 0.87 | -0.0925 | 0.09 | 0.3675 | 0.245991 | 0.743078 | 0.759139 | 1 | 0.999631 | 0.685751 |
| Map4_P27546_S506s_1 | -0.12 | 0.05 | -0.17 | -0.78 | -0.33 | 0.06 | -0.54 | -0.46 | -0.32 | -0.14 | -0.1 | -0.07 | -0.29 | 0.52 | 0.09 | -0.37 | -0.0625 | 0.0975 | 0.2425 | 0.480095 | 0.831414 | 0.731288 | 1 | 0.999631 | 0.801463 |
| Map4_P27546_S517s_1 | -1.25 | 1.28 | 1.35 | -0.51 | 1.12 | 0.77 | 0.45 | 0.59 | 1.58 | -1.07 | 1.21 | 1.88 | 2.45 | 1.02 | 1.36 | 0.45 | 0.515 | 0.6825 | 1.1025 | 0.133505 | 0.401986 | 0.413416 | 1 | 0.999631 | 0.60492 |
| Map4k2_D3YUH9_S284s_1 | -0.2 | 0.26 | -0.25 | 0.2 | 0.31 | 0.84 | 0.17 | -0.33 | 0.27 | 0.73 | -0.28 | 1.03 | 0.52 | 1.39 | 0.92 | 1.13 | 0.245 | 0.435 | 0.9875 | 0.009926 | 0.453048 | 0.23581 | 1 | 0.999631 | 0.376086 |
| Map4k4_E9PVG7_S661s_1 | -0.55 | 0.48 | 0.67 | 0.39 | 0.48 | 0.43 | 0.67 | 0.85 | 0.73 | 0 | 1.2 | 0.76 | 0.52 | 0.23 | 0.37 | 0.38 | 0.36 | 0.425 | 0.1275 | 0.710192 | 0.290439 | 0.287744 | 1 | 0.999631 | 0.899723 |
| Map4k4_E9PVG7_S676s_1 | 0.18 | -2.26 | -1.94 | -1.1 | -0.97 | -2.01 | -0.91 | -2.25 | -0.99 | -0.6 | -1.14 | -1.22 | -1.53 | -2.51 | -1.61 | -2.71 | -0.255 | 0.2925 | -0.81 | 0.179743 | 0.664818 | 0.58116 | 1 | 0.999631 | 0.64376 |
| Map4k4_E9PVG7_S684s_1 | 0.41 | 0.64 | 0.29 | 0.03 | -0.72 | 0.05 | 0.7 | -0.08 | 0.74 | 0.87 | 0.46 | 0.39 | 1.13 | 0.65 | 1.26 | 0.91 | -0.355 | 0.2725 | 0.645 | 0.050652 | 0.321039 | 0.33362 | 1 | 0.999631 | 0.489558 |
| Map4k4_E9PVG7_S761s_1 | 2 | -0.32 | 1.87 | 1.16 | -0.84 | 1.12 | 1.84 | -0.02 | -0.43 | 2.03 | 0.65 | 0.12 | -1.05 | -0.01 | 0.78 | -0.36 | -0.6525 | -0.585 | -1.3375 | 0.04592 | 0.365987 | 0.39289 | 1 | 0.999631 | 0.477883 |
| Map4k4_E9PVG7_S763s_1 | 1.96 | -0.53 | 0.68 | 0.48 | -0.69 | 0.58 | 1.53 | -0.43 | 0.09 | 1.41 | 0.14 | -0.27 | -1.18 | -0.26 | 0.32 | -0.59 | -0.4 | -0.305 | -1.075 | 0.077323 | 0.538748 | 0.600096 | 1 | 0.999631 | 0.542683 |
| Map4k4_E9PVG7_S849s_1 | -1.03 | -1.06 | -0.03 | -0.06 | -0.31 | -0.56 | -0.37 | 0.14 | -1.46 | 0.02 | -0.35 | 0.11 | -0.62 | -0.97 | 0.76 | 0.25 | 0.27 | 0.125 | 0.4 | 0.410496 | 0.450764 | 0.782572 | 1 | 0.999631 | 0.767937 |
| Map4k4_E9PVG7_S850s_1 | 0.33 | 0.61 | -0.37 | -1.32 | -0.84 | 0.08 | -0.65 | -0.06 | 0.34 | 0.06 | -0.15 | -0.44 | 0.97 | 0.67 | -0.06 | -0.54 | -0.18 | 0.14 | 0.4475 | 0.398792 | 0.698888 | 0.756852 | 1 | 0.999631 | 0.761726 |
| Map4k4_E9PVG7_S891s_1 | 1.31 | 0.98 | -0.06 | 0.71 | 1.38 | 1.79 | 0.51 | 0.76 | 0.87 | 1.61 | 0.63 | 0.44 | 0.43 | 0.17 | 1.71 | 0.02 | 0.375 | 0.1525 | -0.1525 | 0.747387 | 0.372998 | 0.704413 | 1 | 0.999631 | 0.911931 |
| Map4k4_E9PVG7_S949s_1 | 0.78 | 0.09 | 0.5 | 0.39 | 0.53 | 0.76 | 0.78 | 1.61 | -0.12 | 0.01 | 0.61 | -0.14 | -0.7 | 1.11 | 0.48 | 1.52 | 0.48 | -0.35 | 0.1625 | 0.739398 | 0.159931 | 0.262929 | 1 | 0.999631 | 0.908594 |
| Map4k5_E9Q1T3_S268s_1 | 0.15 | 0.7 | 0.46 | 1.15 | 0.52 | 1.07 | 0.02 | 1.18 | 0.92 | 0.59 | 1.23 | 1.31 | 0.23 | 0.84 | 0.97 | 1.1 | 0.0825 | 0.3975 | 0.17 | 0.625163 | 0.82038 | 0.238174 | 1 | 0.999631 | 0.863768 |
| Map4k5_E9Q1T3_S366s_1 | 0.07 | 0.67 | -0.53 | -0.27 | 1.24 | 0.81 | -0.3 | -0.46 | 0.27 | 0.1 | -0.35 | -0.03 | -0.14 | 1.72 | -0.46 | -0.28 | 0.3375 | 0.0125 | 0.225 | 0.67503 | 0.476953 | 0.970443 | 1 | 0.999631 | 0.885588 |
| Map6_Q7TSJ2_S207s_1 | -0.35 | 0.39 | 0.86 | 0.67 | 0.6 | 0.2 | 1.25 | 0.93 | 0.44 | 0.01 | 1.55 | 0.77 | -0.08 | -0.42 | 0.61 | 1.73 | 0.3525 | 0.3 | 0.0675 | 0.895773 | 0.3496 | 0.483518 | 1 | 0.999631 | 0.966139 |
| Map6_Q7TSJ2_S294s_1 | -0.67 | -2.87 | -0.48 | -0.95 | 0.31 | -1.98 | 2.44 | -1.59 | -1.14 | -0.79 | -1.55 | -0.89 | -0.41 | -0.69 | -1.06 | -2 | 1.0375 | 0.15 | 0.2025 | 0.733971 | 0.310099 | 0.780301 | 1 | 0.999631 | 0.907462 |
| Map6_Q7TSJ2_S386s_1 | 1.01 | 1.46 | 0.83 | 0.41 | 0.85 | 1.19 | 0.71 | 0.76 | 1.2 | 0.54 | 0.98 | 1.08 | 0.41 | 0.79 | 0.94 | 0.55 | -0.05 | 0.0225 | -0.255 | 0.440914 | 0.869049 | 0.943945 | 1 | 0.999631 | 0.781983 |
| Map6_Q7TSJ2_S424s_1 | 0.28 | 0.42 | 0.43 | -0.12 | -0.33 | -0.43 | 0.11 | 0.06 | -0.83 | 0.22 | -0.2 | 0.24 | 0.21 | 0.05 | 0.38 | 0.02 | -0.4 | -0.395 | -0.0875 | 0.758076 | 0.167773 | 0.252285 | 1 | 0.999631 | 0.914449 |
| Map6_Q7TSJ2_S632s_1 | -0.02 | -0.21 | 0.13 | 1.4 | -0.13 | -0.03 | 0.93 | 0.52 | -0.43 | -0.35 | 0.22 | 0.65 | -0.04 | -0.09 | 0.72 | 0.47 | -0.0025 | -0.3025 | -0.06 | 0.887854 | 0.995381 | 0.497239 | 1 | 0.999631 | 0.963505 |
| Map6_Q7TSJ2_S687s_1 | 0.26 | -0.05 | -0.26 | 0.65 | 0.47 | 0.19 | 0.15 | 0.44 | 0.28 | -0.19 | -0.16 | 0.26 | -0.65 | 0.72 | 0.38 | 0.22 | 0.1625 | -0.1025 | 0.0175 | 0.963764 | 0.575813 | 0.738714 | 1 | 0.999631 | 0.987331 |
| Map6_Q7TSJ2_S832s_1 | -0.23 | 0.14 | 0.92 | 0.39 | 0.74 | 0.45 | 0.27 | 0.93 | -0.02 | -0.7 | 0.7 | 0.75 | 0.85 | 0.29 | 1.13 | 0.73 | 0.2925 | -0.1225 | 0.445 | 0.226042 | 0.380362 | 0.771851 | 1 | 0.999631 | 0.676934 |
| Map6_Q7TSJ2_S86s_1 | 2.31 | 0.95 | -0.63 | 1.36 | 1.64 | 0.61 | 1.51 | 0.24 | 0.98 | 0.27 | 0.35 | -0.09 | -0.17 | 0.1 | -0.74 | 0.44 | 0.0025 | -0.62 | -1.09 | 0.094776 | 0.996827 | 0.311307 | 1 | 0.999631 | 0.558128 |
| Map6_Q7TSJ2_S888s_1 | -1.15 | -0.1 | 0.25 | 0.19 | 0.18 | -0.25 | -0.02 | 0.31 | 0.09 | -1.03 | 0.35 | 0.11 | 0.83 | 0.53 | 0.56 | 0.25 | 0.2575 | 0.0825 | 0.745 | 0.074361 | 0.488723 | 0.852142 | 1 | 0.999631 | 0.539897 |
| Map6_Q7TSJ2_S905s_1 | -0.86 | -0.22 | -0.23 | -0.49 | -0.54 | -0.7 | -0.66 | -0.56 | -0.58 | -0.64 | -0.47 | -0.32 | 0.14 | -0.33 | -0.37 | -0.17 | -0.165 | -0.0525 | 0.2675 | 0.379568 | 0.530295 | 0.847298 | 1 | 0.999631 | 0.751933 |
| Map6_Q7TSJ2_T634t_1 | 0.77 | 1.95 | 0.39 | 0.18 | -0.3 | 0.22 | 0.13 | 0.31 | 1.63 | 0.5 | 0.39 | 0.49 | 0.55 | 1.28 | 0.34 | -0.13 | -0.7325 | -0.07 | -0.3125 | 0.518424 | 0.102787 | 0.882639 | 1 | 0.999631 | 0.819449 |
| Map6d1_Q14BB9_S154s_1 | -0.36 | 0.55 | 0.49 | 0.09 | -0.51 | -0.24 | 0.3 | 0.88 | 0.32 | -0.26 | 1.21 | 0.59 | 0.16 | 0.59 | 0.62 | 1.07 | -0.085 | 0.2725 | 0.4175 | 0.241702 | 0.82503 | 0.488821 | 1 | 0.999631 | 0.684831 |
| Map6d1_Q14BB9_S53s_1 | 1.64 | 2.33 | 3.26 | 2.1 | 0.38 | 2.72 | 1.44 | 2.21 | 1.39 | 2.27 | 2.55 | 0.14 | 0.51 | 0.99 | 0.26 | 1.19 | -0.645 | -0.745 | -1.595 | 0.003224 | 0.266346 | 0.223098 | 1 | 0.999631 | 0.351308 |
| Map7d1_A8Y5P4_S118s_1 | -0.38 | -1.04 | -0.07 | -0.77 | -0.03 | 0.12 | -0.39 | -0.89 | -0.15 | 0.03 | -0.41 | -0.49 | 1.21 | -0.02 | -0.58 | -0.29 | 0.2675 | 0.31 | 0.645 | 0.171626 | 0.443031 | 0.332022 | 1 | 0.999631 | 0.641657 |
| Map7d1_A8Y5P4_S256s_1 | 0.68 | 0.33 | 0.75 | 0.63 | 0.45 | 0.2 | -0.02 | -0.27 | 0.58 | -0.18 | 0.38 | -0.56 | 0.33 | 0.72 | 0.5 | 0.17 | -0.5075 | -0.5425 | -0.1675 | 0.556511 | 0.085543 | 0.124012 | 1 | 0.999631 | 0.838305 |
| Map7d1_A8Y5P4_S275s_1 | 0.06 | 1.41 | -0.2 | 0.82 | 1.18 | 1.02 | 0.53 | 0.93 | 1.68 | 0.03 | 0.76 | 1.4 | 0.53 | 1.1 | 0.56 | 0.27 | 0.3925 | 0.445 | 0.0925 | 0.825574 | 0.335739 | 0.377919 | 1 | 0.999631 | 0.940387 |
| Map7d1_A8Y5P4_S78s_1 | 0.4 | -0.53 | -0.73 | 0.66 | -0.09 | -0.24 | -0.36 | 1.03 | 0.09 | -0.18 | -0.56 | 0.37 | -0.43 | 0.85 | 0.63 | 0.66 | 0.135 | -0.02 | 0.4775 | 0.300559 | 0.765331 | 0.960577 | 1 | 0.999631 | 0.71369 |
| Map7d1_A8Y5P4_S95s_2 | 1.95 | 1.46 | -0.82 | 2.08 | 1.98 | 0.49 | 2.12 | 1.76 | 1.39 | 1.6 | 1.04 | 0.52 | -0.14 | 1.4 | 1.25 | 1.31 | 0.42 | -0.03 | -0.2125 | 0.756465 | 0.543621 | 0.962846 | 1 | 0.999631 | 0.914089 |
| Map7d1_A8Y5P4_T99t_1 | -0.66 | 0.69 | 0.84 | -0.3 | -0.6 | 0.83 | -0.16 | 0.38 | 0.47 | 1.11 | 0.7 | 0.93 | 1.41 | 0.79 | 0.74 | 1.23 | -0.03 | 0.66 | 0.9 | 0.053047 | 0.948215 | 0.126978 | 1 | 0.999631 | 0.489777 |
| Map7d1_A8Y5P4_T99t_2 | 1.95 | 1.46 | -0.82 | 2.08 | 1.98 | 0.49 | 2.12 | 1.76 | 1.39 | 1.6 | 1.04 | 0.52 | -0.14 | 1.4 | 1.25 | 1.31 | 0.42 | -0.03 | -0.2125 | 0.756465 | 0.543621 | 0.962846 | 1 | 0.999631 | 0.914089 |
| Map7d1_F6WF36_S146s_1 | 1.18 | 2.07 | 2.03 | 1.21 | 0.96 | 2.13 | 1.03 | 1.56 | 2.09 | 2.02 | 1.8 | 1.7 | 1.22 | 2.27 | 1.8 | 2.35 | -0.2025 | 0.28 | 0.2875 | 0.466452 | 0.597323 | 0.393734 | 1 | 0.999631 | 0.795267 |
| Map7d1_F6WF36_S189s_1 | 1.58 | 1.54 | -0.17 | 1.69 | 2.41 | 1.23 | 1.46 | 1.26 | 1.72 | 0.79 | 1.3 | 0.52 | -1.37 | 1.81 | 0.95 | 0.99 | 0.43 | -0.0775 | -0.565 | 0.440265 | 0.392915 | 0.875223 | 1 | 0.999631 | 0.781983 |
| Map7d1_F6WF36_S205s_1 | -0.83 | 0.62 | 0.87 | 0.1 | 0.83 | 0.88 | 0.13 | 0.16 | 0.87 | -0.33 | 0.09 | 0.45 | 0.76 | 0.92 | 0.96 | 0.46 | 0.31 | 0.08 | 0.585 | 0.176173 | 0.469927 | 0.857978 | 1 | 0.999631 | 0.642116 |
| Map7d1_F6WF36_S214s_1 | 0.43 | 0.3 | 0.55 | -0.07 | -0.03 | 0.69 | 0.8 | 0.56 | -0.24 | 1.14 | 0.33 | 0.35 | -1.77 | 1.03 | 0.69 | 1.05 | 0.2025 | 0.0925 | -0.0525 | 0.933013 | 0.498395 | 0.792824 | 1 | 0.999631 | 0.978327 |
| Map7d1_F6WF36_S264s_1 | 1.19 | 0.61 | 1.37 | 1.1 | 0.26 | 0.47 | 1.22 | 1.29 | 1.27 | 1.43 | -0.04 | 0.35 | 0.41 | 1.27 | 0.8 | 0.23 | -0.2575 | -0.315 | -0.39 | 0.272748 | 0.457687 | 0.441085 | 1 | 0.999631 | 0.700187 |
| Map7d1_F6WF36_S289s_1 | -1 | -1.28 | -0.57 | -0.93 | -0.76 | -0.99 | -1.27 | -1.21 | -0.99 | -0.99 | -0.59 | -0.43 | 0.13 | -0.41 | -0.59 | -0.25 | -0.1125 | 0.195 | 0.665 | 0.052502 | 0.682722 | 0.506457 | 1 | 0.999631 | 0.489777 |
| Map7d1_F6WF36_S293s_1 | -1.42 | 0.14 | 0.71 | -0.23 | -0.46 | -0.41 | -1 | -0.08 | -0.47 | 0.12 | -0.59 | 0.18 | 0.83 | 0.07 | 1.3 | 0 | -0.2875 | 0.01 | 0.75 | 0.1703 | 0.542093 | 0.983156 | 1 | 0.999631 | 0.64073 |
| Map7d1_F6WF36_S60s_1 | 1 | 1.22 | 1.18 | 1.55 | 1.42 | 1.3 | 1.47 | 1.61 | 1.37 | 1.01 | 1.16 | 1.54 | 0.38 | 1.73 | 1.73 | 1.9 | 0.2125 | 0.0325 | 0.1975 | 0.621537 | 0.406003 | 0.904677 | 1 | 0.999631 | 0.861912 |
| Map7d2_A2AG50_S131s_1 | 1.13 | 0.96 | 1.03 | 0.9 | 1.36 | 1.23 | 1.14 | 1.17 | 1 | 0.54 | 1.26 | 1.17 | 0.05 | 1.67 | 1.11 | 0.78 | 0.22 | -0.0125 | -0.1025 | 0.787061 | 0.356524 | 0.963492 | 1 | 0.999631 | 0.923988 |
| Map7d2_A2AG50_S162s_1 | 2.74 | 0.26 | -0.33 | 2.2 | 2.14 | 2.62 | 2.22 | 1.37 | 1.8 | 1.41 | 0.6 | -0.47 | -2.56 | 1.18 | 0.85 | -0.81 | 0.87 | -0.3825 | -1.5525 | 0.13417 | 0.230269 | 0.628919 | 1 | 0.999631 | 0.60492 |
| Map7d2_A2AG50_S173s_1 | 0.59 | 0.26 | 1 | 1.03 | 1.12 | 0.82 | 0.69 | 0.79 | 1.17 | 0.85 | 0.56 | 1.31 | -0.39 | 1.34 | 1.55 | 0.91 | 0.135 | 0.2525 | 0.1325 | 0.775799 | 0.63563 | 0.428355 | 1 | 0.999631 | 0.920998 |
| Map7d2_A2AG50_S515s_1 | 0.1 | 0.43 | -0.68 | 0.17 | 0.38 | 0.58 | -0.34 | 1.05 | 0.3 | -1.36 | 0.32 | 0.79 | 0.85 | 0.66 | 0.03 | 0.64 | 0.4125 | 0.0075 | 0.54 | 0.149066 | 0.299577 | 0.988046 | 1 | 0.999631 | 0.621356 |
| Map7d2_A2AG50_S567s_1 | 1.1 | -1.74 | -2.69 | 0.46 | 1.39 | -1.23 | 1.23 | 0.21 | -0.35 | -1.91 | -1.09 | -1.2 | -2.17 | -1.16 | -1.04 | -1.89 | 1.1175 | -0.42 | -0.8475 | 0.312101 | 0.248585 | 0.615832 | 1 | 0.999631 | 0.719585 |
| Map7d2_A2AG50_S572s_1 | -0.31 | 0.98 | -0.86 | -0.86 | 0.68 | 0.38 | -0.16 | -0.43 | 0.59 | -0.59 | -0.16 | -0.32 | 0.66 | 0.29 | 0.2 | -0.54 | 0.38 | 0.1425 | 0.415 | 0.401444 | 0.432994 | 0.767686 | 1 | 0.999631 | 0.763221 |
| Map7d2_Q8C0A6_S34s_1 | 2.01 | 1.08 | 0.82 | 2.47 | 2.64 | 2.67 | 1.9 | 2.43 | 2.01 | 1.09 | 2.25 | 1.37 | -7.4 | 2.3 | 1.84 | 1.68 | 0.815 | 0.085 | -1.99 | 0.327049 | 0.077825 | 0.853436 | 1 | 0.999631 | 0.724678 |
| Map9_Q3TRR0_S257s_1 | 1.49 | 0.69 | 0.32 | 1.42 | -0.46 | 0.61 | 1.74 | 0.37 | 0.51 | 1.23 | 1.92 | 2.39 | -0.07 | -0.65 | 1.93 | -0.42 | -0.415 | 0.5325 | -0.7825 | 0.210042 | 0.416754 | 0.28272 | 1 | 0.999631 | 0.66441 |
| Map9_Q3TRR0_S260s_1 | 1.39 | -0.18 | 0.61 | 0.3 | 0.48 | 0.43 | 0.88 | 0.07 | 0.97 | 1.65 | -0.26 | 0.16 | -0.73 | -0.37 | -0.17 | -0.28 | -0.065 | 0.1 | -0.9175 | 0.035658 | 0.864739 | 0.843994 | 1 | 0.999631 | 0.453811 |
| Mapk10_Q3TQZ7_T221t_1 | 0.04 | 0.03 | 0.19 | 0.01 | 0.47 | 0.4 | -0.13 | 0.45 | -1.01 | -0.38 | 0.64 | 0.29 | 0.88 | -0.68 | -0.34 | -0.04 | 0.23 | -0.1825 | -0.1125 | 0.765311 | 0.382235 | 0.638913 | 1 | 0.999631 | 0.916326 |
| Mapk10_Q3TQZ7_Y223y_1 | -1.38 | -1.35 | -1.21 | -1.52 | -1.58 | -1.74 | -1.47 | -1.6 | -1.52 | -1.66 | -1.57 | -1.27 | -0.63 | -1.08 | -1.33 | -1.25 | -0.2325 | -0.14 | 0.2925 | 0.325139 | 0.337838 | 0.579372 | 1 | 0.999631 | 0.72463 |
| Mapk14_B2KF34_Y182y_1 | 1.12 | -1.03 | -0.95 | -1.05 | 0.7 | -0.8 | 2.24 | -1.22 | 1.2 | 1.56 | 2.02 | 3.13 | 1.79 | -0.9 | 0.44 | 1.13 | 0.7075 | 2.455 | 1.0925 | 0.139979 | 0.401699 | 0.002643 | 1 | 0.999631 | 0.611718 |
| Mapk3_D3Z3G6_T203t_1 | -1.18 | 0.46 | 0.63 | -0.35 | -0.56 | 0.78 | -0.44 | 1.29 | -0.54 | -0.43 | 0.26 | -1.47 | -0.49 | 1.07 | 0.18 | 0.11 | 0.3775 | -0.435 | 0.3275 | 0.519052 | 0.507985 | 0.407265 | 1 | 0.999631 | 0.819456 |
| Mapk4_Q6P5G0_S382s_1 | 0.61 | 1.22 | 0.64 | -0.51 | 0.82 | 0.26 | -0.38 | -0.05 | -0.07 | 0.58 | 0.37 | 0.83 | 1.17 | 1.07 | -0.09 | 0.68 | -0.3275 | -0.0625 | 0.2175 | 0.635921 | 0.456538 | 0.880151 | 1 | 0.999631 | 0.869397 |
| Mapk4_Q6P5G0_S430s_1 | 0.43 | 1.14 | 0.56 | 0.85 | 0.89 | 0.5 | 0.6 | 0.21 | 0.36 | 0.24 | 0.89 | 0.97 | 0.87 | 0.82 | -0.99 | 1.45 | -0.195 | -0.13 | -0.2075 | 0.691714 | 0.501275 | 0.676469 | 1 | 0.999631 | 0.894017 |
| Mapk4_Q6P5G0_S504s_1 | 0.42 | 0.44 | 0.88 | 0.63 | 0.17 | 0.15 | 0.24 | 0.86 | 0.28 | 0.69 | 0.97 | 0.83 | 0.56 | 0.44 | 0.52 | 1.14 | -0.2375 | 0.1 | 0.0725 | 0.808817 | 0.406661 | 0.721959 | 1 | 0.999631 | 0.93635 |
| Mapk8_D3Z1Z4_S301s_1 | -0.25 | -1.41 | 0 | -0.51 | -0.07 | 0.24 | -0.46 | -0.22 | -0.1 | -0.71 | 0.19 | -1.3 | -1.34 | 0.53 | -1.27 | -0.95 | 0.415 | 0.0625 | -0.215 | 0.673932 | 0.269071 | 0.888309 | 1 | 0.999631 | 0.885515 |
| Mapk8ip1_Q9WVI9_S14s_1 | -1.33 | -0.78 | -1.26 | -1.54 | -0.63 | -1.18 | -1.05 | -1.55 | -0.74 | -0.88 | -1.53 | -1 | -0.54 | -1.32 | -1.36 | -1.11 | 0.125 | 0.19 | 0.145 | 0.657755 | 0.68457 | 0.53923 | 1 | 0.999631 | 0.879549 |
| Mapk8ip1_Q9WVI9_S189s_1 | 0.94 | 1.05 | 0.73 | 0.01 | 5.51 | 2.54 | 2.27 | 0.78 | 6.77 | 0.53 | 1.83 | 2.93 | 4.56 | 0.37 | 1.62 | -0.69 | 2.0925 | 2.3325 | 0.7825 | 0.43717 | 0.037178 | 0.070277 | 1 | 0.999631 | 0.781057 |
| Mapk8ip1_Q9WVI9_S231s_1 | 1.68 | -1.92 | -2.04 | -1.14 | -2.23 | -0.57 | 0.34 | -2.02 | -2.02 | 0.82 | -1.94 | -2.2 | -2.87 | -1.91 | -1.26 | -3.49 | -0.265 | -0.48 | -1.5275 | 0.101697 | 0.773064 | 0.624805 | 1 | 0.999631 | 0.564448 |
| Mapk8ip1_Q9WVI9_S351s_1 | 0.16 | 0.03 | -0.42 | 0.53 | -0.21 | -0.13 | -0.21 | 1.32 | -0.19 | 0.86 | 0.42 | 0.51 | -0.01 | -0.12 | 1.05 | 1.15 | 0.1175 | 0.325 | 0.4425 | 0.294406 | 0.78 | 0.350073 | 1 | 0.999631 | 0.708293 |
| Mapk8ip2_Q9ERE9_S128s_1 | -0.3 | -0.48 | -0.9 | -0.66 | -0.06 | -0.89 | -0.88 | -0.68 | -0.18 | -0.73 | -0.44 | -0.86 | 0.1 | 0.26 | -0.59 | -0.53 | -0.0425 | 0.0325 | 0.395 | 0.245408 | 0.88662 | 0.90997 | 1 | 0.999631 | 0.685751 |
| Mapk8ip2_Q9ERE9_S175s_1 | 0.85 | 0.22 | -0.79 | 1.55 | 0.09 | 0.34 | 0.96 | 1.36 | 0.88 | 0.06 | 0.89 | 0.83 | -0.91 | 0.29 | 0.71 | 1.17 | 0.23 | 0.2075 | -0.1425 | 0.814505 | 0.666463 | 0.683403 | 1 | 0.999631 | 0.937111 |
| Mapk8ip2_Q9ERE9_S248s_1 | 0.57 | -0.11 | 0.05 | 0.28 | 0.33 | 0.39 | 0.2 | 0.02 | 0.55 | 0.34 | -0.05 | -0.3 | -0.12 | 0.25 | 1.05 | 0.46 | 0.0375 | -0.0625 | 0.2125 | 0.54311 | 0.887888 | 0.84 | 1 | 0.999631 | 0.832881 |
| Mapk8ip2_Q9ERE9_S249s_1 | 0.57 | -0.11 | 0.05 | 0.28 | 0.33 | 0.39 | 0.2 | 0.02 | 0.55 | 0.34 | -0.05 | -0.3 | -0.12 | 0.25 | 1.05 | 0.46 | 0.0375 | -0.0625 | 0.2125 | 0.54311 | 0.887888 | 0.84 | 1 | 0.999631 | 0.832881 |
| Mapk8ip2_Q9ERE9_S249s_2 | 0.48 | 0.71 | -0.43 | 0 | -0.4 | -0.16 | -0.04 | 0.29 | 0.21 | -0.1 | 0.48 | 0.84 | 0.2 | 0.22 | -1.71 | -0.8 | -0.2675 | 0.1675 | -0.7125 | 0.178731 | 0.428358 | 0.641468 | 1 | 0.999631 | 0.642903 |
| Mapk8ip2_Q9ERE9_S370s_1 | -0.42 | -0.62 | -0.67 | -0.15 | -0.04 | -0.17 | -0.23 | 0.14 | 0.24 | -0.36 | -0.26 | -0.24 | -1.09 | 0.18 | -0.13 | -0.06 | 0.39 | 0.31 | 0.19 | 0.596322 | 0.150477 | 0.280946 | 1 | 0.999631 | 0.854508 |
| Mapk8ip2_Q9ERE9_S380s_1 | 1.59 | -0.06 | -0.34 | 1.27 | 1.89 | 1.38 | 1.05 | 0.9 | 1.41 | 0.74 | -0.38 | -0.14 | -0.99 | 1.29 | 0.75 | 0.59 | 0.69 | -0.2075 | -0.205 | 0.741472 | 0.184932 | 0.72078 | 1 | 0.999631 | 0.909067 |
| Mapk8ip3_K3W4S4_S611s_1 | 0.58 | 0.04 | 0.77 | 0.84 | 0.22 | 0.21 | 0.7 | 0.93 | 1.06 | 0.46 | 1.72 | 1.78 | 0.71 | 0.43 | 1.51 | 1.32 | -0.0425 | 0.6975 | 0.435 | 0.245413 | 0.89095 | 0.09237 | 1 | 0.999631 | 0.685751 |
| Mapk8ip3_K3W4S4_S685s_1 | 0.46 | -0.12 | -0.77 | -0.47 | -0.26 | 0.26 | 0.11 | -0.66 | 0.5 | 0.83 | -0.33 | -0.44 | 0.06 | -0.51 | -1.01 | -1.69 | 0.0875 | 0.365 | -0.5625 | 0.233135 | 0.807601 | 0.38814 | 1 | 0.999631 | 0.683035 |
| Mapk8ip3_K3W4S4_S763s_1 | -0.22 | -2.86 | 0.91 | -1.06 | -2.33 | -1.4 | 2.45 | -5.71 | 0.07 | -0.44 | -0.47 | 0.27 | 0.37 | -1.95 | -2.15 | -1.63 | -0.94 | 0.665 | -0.5325 | 0.535924 | 0.553606 | 0.368819 | 1 | 0.999631 | 0.829551 |
| Mapk8ip3_K3W4S4_T1199t_1 | 2.83 | 1.17 | 2.06 | 1.54 | 0.09 | 1.86 | 2.11 | 0.25 | 0.48 | 3.42 | 0.26 | 0.6 | -0.21 | 0.38 | 0.89 | 0.41 | -0.8225 | -0.71 | -1.5325 | 0.005041 | 0.17845 | 0.345443 | 1 | 0.999631 | 0.351308 |
| Mapkap1_Q8BKH7_S318s_1 | 0.25 | 1.84 | 1.86 | 1.24 | 1.39 | 1.84 | 1.42 | 1.71 | 2.27 | 1.37 | 1.83 | 2.43 | 1.88 | 1.61 | 1.54 | 2.42 | 0.2925 | 0.6775 | 0.565 | 0.212314 | 0.469184 | 0.148509 | 1 | 0.999631 | 0.666473 |
| Mapkbp1_Q6NS57_S756s_1 | 2.37 | -0.09 | 0.62 | 1.52 | 0.3 | 0.61 | 1.67 | 0.52 | 0.91 | 2.32 | 0.46 | -0.11 | -0.07 | -0.45 | -0.1 | -0.37 | -0.33 | -0.21 | -1.3525 | 0.023837 | 0.561156 | 0.7536 | 1 | 0.999631 | 0.415644 |
| Mapre2_Q8R001_S199s_1 | 0.66 | -0.27 | 0.7 | -0.24 | -1.37 | 0.47 | 1.47 | -0.53 | -0.18 | 0.45 | -1.19 | 0.07 | -2.18 | -0.18 | -0.35 | -0.63 | -0.2025 | -0.425 | -1.0475 | 0.06159 | 0.738814 | 0.34456 | 1 | 0.999631 | 0.509355 |
| Mapre2_Q8R001_S222s_1 | 1.56 | -0.15 | -0.21 | 0.33 | 0.47 | 1.98 | 0.09 | 0.33 | 1.88 | 2.2 | 1.01 | 1.39 | -1.4 | 1.02 | 1.82 | 0.83 | 0.335 | 1.2375 | 0.185 | 0.79475 | 0.542976 | 0.024204 | 1 | 0.999631 | 0.928846 |
| Mapre2_Q8R001_S9s_1 | -1.88 | -1.46 | -1.04 | -1.25 | -2.18 | -3.21 | -1.01 | -1.03 | -2.02 | -2.07 | -1.6 | -0.75 | -2.51 | -2.97 | -0.75 | -1.15 | -0.45 | -0.2025 | -0.4375 | 0.415624 | 0.39433 | 0.594503 | 1 | 0.999631 | 0.769842 |
| Mapre3_D3Z6G3_S147s_1 | 1.97 | 0.79 | 1.19 | 0.78 | -0.66 | 1.53 | 2.08 | -0.28 | 0.57 | 1.71 | -0.23 | -0.06 | 1.17 | 0.38 | 0.7 | 0.08 | -0.515 | -0.685 | -0.6 | 0.149296 | 0.435148 | 0.188274 | 1 | 0.999631 | 0.621356 |
| Mapre3_Q6PER3_S162s_1 | 1.97 | 0.79 | 1.19 | 0.78 | -0.66 | 1.53 | 2.08 | -0.28 | 0.57 | 1.71 | -0.23 | -0.06 | 1.17 | 0.38 | 0.7 | 0.08 | -0.515 | -0.685 | -0.6 | 0.149296 | 0.435148 | 0.188274 | 1 | 0.999631 | 0.621356 |
| Mapt_B1AQW2_T42t_1 | -0.15 | -1.16 | -2.05 | -0.94 | -1.51 | -1.31 | -0.7 | -1.66 | -1.38 | -0.4 | -0.78 | -0.54 | -0.93 | -1.76 | -1.95 | -0.12 | -0.22 | 0.3 | -0.115 | 0.830514 | 0.614653 | 0.501871 | 1 | 0.999631 | 0.942238 |
| Mapt_B1AQW2_T54t_1 | -0.72 | -0.74 | -0.56 | -0.92 | -1.45 | -1.29 | -0.42 | -1.95 | -0.82 | -0.08 | -0.14 | -1.26 | -0.71 | -1.52 | -0.46 | -0.99 | -0.5425 | 0.16 | -0.185 | 0.567896 | 0.148926 | 0.638769 | 1 | 0.999631 | 0.843071 |
| Mapt_P10637_S167s_2 | -0.54 | -1.02 | -0.95 | -1.14 | -1.05 | -1.38 | -1.15 | -1.3 | -1.23 | -0.62 | -1.1 | -0.59 | 0 | -0.57 | -1.06 | -0.5 | -0.3075 | 0.0275 | 0.38 | 0.263504 | 0.250898 | 0.925104 | 1 | 0.999631 | 0.693034 |
| Mapt_P10637_S180s_1 | 0.81 | -0.53 | -0.03 | 0.22 | -0.72 | -0.88 | -0.27 | 0.36 | -0.24 | 0.82 | 0.07 | 0.37 | -1.22 | -0.36 | 0.86 | -0.38 | -0.495 | 0.1375 | -0.3925 | 0.432485 | 0.232804 | 0.718609 | 1 | 0.999631 | 0.779247 |
| Mapt_P10637_S180s_2 | 0.59 | -0.36 | 0.08 | -0.3 | -0.05 | -0.64 | -0.58 | -0.38 | 0.43 | 0.74 | -0.75 | -0.22 | -0.52 | -0.27 | -0.3 | -0.17 | -0.415 | 0.0475 | -0.3175 | 0.329307 | 0.202279 | 0.90718 | 1 | 0.999631 | 0.726484 |
| Mapt_P10637_S184s_2 | 1.14 | 0.6 | 0.57 | 1.12 | 2.53 | 1.78 | 0.51 | 1.69 | 1.17 | 0.41 | 0.55 | 1.16 | -0.6 | 0.59 | 1.39 | -0.34 | 0.77 | -0.035 | -0.5975 | 0.223779 | 0.102208 | 0.91194 | 1 | 0.999631 | 0.67471 |
| Mapt_P10637_S187s_1 | 0.01 | 0.23 | 0.19 | 0.27 | -0.13 | -0.1 | -0.08 | -0.43 | -0.19 | 0.27 | -0.68 | -0.34 | 0.62 | 0.13 | 0.08 | -0.29 | -0.36 | -0.41 | -0.04 | 0.894115 | 0.157917 | 0.180379 | 1 | 0.999631 | 0.965259 |
| Mapt_P10637_S187s_2 | -0.39 | -0.34 | 1.08 | 0.08 | -0.74 | 0.03 | -0.79 | 0.09 | -0.62 | 0.84 | -0.65 | 0.96 | 0.57 | 0.64 | 0.77 | -0.29 | -0.46 | 0.025 | 0.315 | 0.467223 | 0.282609 | 0.96194 | 1 | 0.999631 | 0.795438 |
| Mapt_P10637_S187s_3 | -0.24 | 0.34 | 1.14 | 1.11 | -0.33 | 0.67 | 0.39 | 1.34 | 0.11 | 0.13 | 1.08 | 1.5 | 0.66 | 0.26 | 1.93 | 1.05 | -0.07 | 0.1175 | 0.3875 | 0.423068 | 0.87902 | 0.801282 | 1 | 0.999631 | 0.773407 |
| Mapt_P10637_S188s_1 | -0.56 | -0.7 | -0.38 | -1.24 | -1.07 | -0.8 | -0.86 | -1 | -1.25 | -0.54 | -0.77 | -0.51 | -0.31 | -1.08 | -1.02 | -0.72 | -0.2125 | -0.0475 | -0.0625 | 0.849697 | 0.45231 | 0.880251 | 1 | 0.999631 | 0.949525 |
| Mapt_P10637_S188s_2 | -0.35 | -0.61 | -0.13 | -0.94 | -0.79 | -0.67 | -0.78 | -0.63 | -0.65 | -0.36 | -0.22 | -0.21 | 0.21 | -0.57 | -0.2 | -0.29 | -0.21 | 0.1475 | 0.295 | 0.367871 | 0.444951 | 0.612389 | 1 | 0.999631 | 0.745968 |
| Mapt_P10637_S188s_3 | -0.21 | 0.52 | 0.86 | 0.98 | -0.2 | 0.77 | 0.36 | 1.17 | 0.23 | 0.23 | 1.03 | 1.25 | 0.65 | 0.32 | 1.63 | 0.94 | -0.0125 | 0.1475 | 0.3475 | 0.401939 | 0.975088 | 0.708453 | 1 | 0.999631 | 0.763288 |
| Mapt_P10637_S191s_1 | 0.33 | -0.13 | -0.42 | -0.66 | -0.79 | -0.66 | -0.67 | -0.63 | -0.25 | 0.57 | -0.47 | -0.84 | 0.45 | -0.4 | -0.35 | -0.23 | -0.4675 | -0.0275 | 0.0875 | 0.802034 | 0.129699 | 0.943141 | 1 | 0.999631 | 0.932677 |
| Mapt_P10637_S191s_2 | -0.34 | -0.61 | -0.99 | -1.66 | -0.85 | -0.56 | -1.48 | -1.6 | -0.47 | -0.37 | -0.49 | -0.7 | 0.28 | -0.52 | -1.07 | -0.8 | -0.2225 | 0.3925 | 0.3725 | 0.38636 | 0.57052 | 0.26516 | 1 | 0.999631 | 0.756141 |
| Mapt_P10637_S191s_3 | -0.39 | 0.39 | 0.64 | 0.85 | -0.43 | 0.49 | 0.17 | 1.04 | 0.07 | 0.14 | 0.8 | 1.36 | 0.82 | 0.16 | 1.65 | 1.06 | -0.055 | 0.22 | 0.55 | 0.212584 | 0.89318 | 0.59741 | 1 | 0.999631 | 0.66658 |
| Mapt_P10637_S197s_1 | 1.06 | -0.36 | 0.15 | -0.25 | 1.82 | 0.16 | 0.36 | -1.09 | 0.96 | 0.78 | -0.18 | 0.29 | 1.71 | 0.67 | -0.4 | -0.5 | 0.1625 | 0.3125 | 0.22 | 0.69789 | 0.790397 | 0.460321 | 1 | 0.999631 | 0.895731 |
| Mapt_P10637_S224s_2 | 3.01 | -0.63 | 1.24 | 1.36 | 1.03 | 0.39 | 1.91 | 1.3 | 0.52 | 0.93 | 1.58 | 1.45 | -0.75 | 0.16 | 1.26 | 0.49 | -0.0875 | -0.125 | -0.955 | 0.221392 | 0.902338 | 0.857799 | 1 | 0.999631 | 0.671697 |
| Mapt_P10637_S313s_1 | 0.65 | 0.45 | 1.2 | 2.04 | 2.6 | 2.98 | 2.76 | 3.24 | 1.95 | 1.46 | 2.46 | 0.14 | 0.83 | 2.53 | 2.96 | 1.75 | 1.81 | 0.4175 | 0.9325 | 0.113012 | 0.000986 | 0.465716 | 1 | 0.999631 | 0.577408 |
| Mapt_P10637_S345s_1 | 0.76 | 0.38 | -0.84 | 1.26 | 1.06 | 0.04 | 0.53 | 0.76 | 0.36 | -0.18 | 0.32 | 0.72 | -0.87 | 0.41 | -0.01 | 0.61 | 0.2075 | -0.085 | -0.355 | 0.502955 | 0.66222 | 0.856088 | 1 | 0.999631 | 0.814092 |
| Mapt_P10637_S35s_1 | -0.43 | -0.36 | 1.19 | 1.11 | -0.24 | -0.44 | 0.16 | 1.14 | -0.48 | -0.46 | 0.58 | 0.71 | -0.3 | 0.08 | 0.87 | 0.98 | -0.2225 | -0.29 | 0.03 | 0.953522 | 0.6737 | 0.578616 | 1 | 0.999631 | 0.984441 |
| Mapt_P10637_S385s_1 | 0.39 | 1.24 | 0.88 | 0.26 | -0.21 | 1 | 0.18 | 0.14 | 0.55 | 1.45 | 0.92 | 0.5 | 0.37 | 0.99 | 0.64 | 1.16 | -0.415 | 0.1625 | 0.0975 | 0.778925 | 0.270114 | 0.646627 | 1 | 0.999631 | 0.921416 |
| Mapt_P10637_S385s_2 | 0.69 | 1.59 | 0.37 | 0.64 | 0.62 | 1.87 | 0.57 | 0.6 | 1.2 | 1.54 | 0.77 | 0.11 | 0.02 | 1.57 | 0.44 | 0.44 | 0.0925 | 0.0825 | -0.205 | 0.637364 | 0.822951 | 0.841767 | 1 | 0.999631 | 0.870011 |
| Mapt_P10637_S385s_3 | -0.33 | 1.2 | 0.81 | 1.41 | 0.72 | 1.33 | 0.89 | 1.33 | 0.81 | 1.03 | 1.27 | 0.79 | -0.11 | 0.74 | 1.02 | 1.28 | 0.295 | 0.2025 | -0.04 | 0.933284 | 0.483653 | 0.624487 | 1 | 0.999631 | 0.978327 |
| Mapt_P10637_S389s_1 | -1.61 | -0.05 | 0.71 | -1.74 | -2.89 | -1.8 | -1.04 | -1.95 | -1.29 | 0.18 | -1.66 | -0.05 | 1.54 | -1.3 | 0.14 | -0.13 | -1.2475 | -0.0325 | 0.735 | 0.330894 | 0.074124 | 0.961445 | 1 | 0.999631 | 0.727302 |
| Mapt_P10637_S389s_2 | -2.6 | 0.57 | 0.99 | -1.34 | -3.29 | -0.68 | -1.65 | -0.87 | 0.04 | 0.79 | -0.5 | 0.54 | 1.61 | -0.06 | 0.32 | 0.31 | -1.0275 | 0.8125 | 1.14 | 0.174933 | 0.265314 | 0.311439 | 1 | 0.999631 | 0.642116 |
| Mapt_P10637_S389s_3 | -0.33 | 1.2 | 0.81 | 1.41 | 0.72 | 1.33 | 0.89 | 1.33 | 0.81 | 1.03 | 1.27 | 0.79 | -0.11 | 0.74 | 1.02 | 1.28 | 0.295 | 0.2025 | -0.04 | 0.933284 | 0.483653 | 0.624487 | 1 | 0.999631 | 0.978327 |
| Mapt_P10637_S393s_1 | -0.45 | -0.51 | -0.4 | -0.71 | -0.69 | -0.75 | -0.68 | -0.6 | -0.62 | -0.37 | -0.16 | -0.44 | 0.5 | -0.4 | -0.55 | -0.03 | -0.1625 | 0.12 | 0.3975 | 0.237083 | 0.49317 | 0.638744 | 1 | 0.999631 | 0.683301 |
| Mapt_P10637_S393s_2 | 0.03 | 1.06 | 0.88 | -0.43 | -0.39 | 1.09 | -0.61 | -0.27 | 0.76 | 1.16 | 0.08 | 0.5 | 0.81 | 0.96 | 0.42 | 0.45 | -0.43 | 0.24 | 0.275 | 0.497821 | 0.391248 | 0.572572 | 1 | 0.999631 | 0.811908 |
| Mapt_P10637_S393s_3 | -0.33 | 1.2 | 0.81 | 1.41 | 0.72 | 1.33 | 0.89 | 1.33 | 0.81 | 1.03 | 1.27 | 0.79 | -0.11 | 0.74 | 1.02 | 1.28 | 0.295 | 0.2025 | -0.04 | 0.933284 | 0.483653 | 0.624487 | 1 | 0.999631 | 0.978327 |
| Mapt_P10637_S402s_1 | -0.65 | -0.16 | 1.84 | -0.21 | 1.18 | 0.33 | 1.87 | 0.6 | 2.44 | -2.72 | 2.14 | 2.57 | 4.12 | -0.93 | 1.04 | 0.98 | 0.79 | 0.9025 | 1.0975 | 0.292418 | 0.202079 | 0.45666 | 1 | 0.999631 | 0.706671 |
| Mapt_P10637_S405s_1 | -0.6 | 2.31 | 0.88 | -0.54 | 2.98 | 0.68 | 1.56 | 0.4 | 3.38 | -0.91 | 1.36 | 1.56 | 3.37 | 0.49 | 1.17 | -0.18 | 0.8925 | 0.835 | 0.7 | 0.439807 | 0.273776 | 0.397262 | 1 | 0.999631 | 0.781983 |
| Mapt_P10637_S57s_1 | -0.42 | 0.43 | 0.4 | -0.13 | -0.07 | 0.04 | -0.39 | 0.03 | -0.44 | -0.67 | -0.2 | 0.03 | 0.78 | -0.12 | -0.02 | 0.19 | -0.1675 | -0.39 | 0.1375 | 0.693923 | 0.57556 | 0.23858 | 1 | 0.999631 | 0.895131 |
| Mapt_P10637_T170t_1 | 0.15 | -0.89 | -0.17 | -0.56 | -0.97 | -0.69 | -0.4 | -0.68 | -0.54 | 0.28 | -0.65 | -0.69 | -0.61 | -0.88 | -0.48 | -0.67 | -0.3175 | -0.0325 | -0.2925 | 0.374627 | 0.319287 | 0.927439 | 1 | 0.999631 | 0.747792 |
| Mapt_P10637_T170t_2 | -0.15 | -0.82 | -0.41 | -0.71 | -0.74 | -1.15 | -0.86 | -0.83 | -0.67 | -0.17 | -0.93 | -0.41 | -0.14 | -0.48 | -0.67 | -0.34 | -0.3725 | -0.0225 | 0.115 | 0.700046 | 0.187663 | 0.940125 | 1 | 0.999631 | 0.896612 |
| Mapt_P10637_T194t_1 | 0.88 | -0.58 | 0.17 | 0.64 | 0.95 | 1.38 | 0.62 | 0.91 | 0.08 | -0.89 | 0.78 | -0.05 | -0.6 | 0.27 | 0.65 | 0.85 | 0.6875 | -0.2975 | 0.015 | 0.973516 | 0.090908 | 0.520617 | 1 | 0.999631 | 0.990676 |
| Mapt_P10637_T194t_2 | 0.17 | -0.4 | 0.42 | -0.25 | -0.07 | 0.57 | -0.17 | 0.46 | -0.12 | -0.01 | 0.11 | -0.13 | 0.01 | 0.15 | 0.59 | 0.09 | 0.2125 | -0.0225 | 0.225 | 0.482493 | 0.506045 | 0.937375 | 1 | 0.999631 | 0.802587 |
| Mapt_P10637_T194t_3 | -0.35 | -0.49 | 0.65 | 0.92 | -0.92 | 0.22 | 0.61 | 1.03 | -0.37 | -0.3 | 1.33 | 0.44 | 0.68 | -0.03 | 0.89 | 0.44 | 0.0525 | 0.0925 | 0.3125 | 0.462096 | 0.918129 | 0.854107 | 1 | 0.999631 | 0.792786 |
| Mapt_P10637_T201t_1 | 1.77 | 0.09 | -0.49 | 1.84 | 2.3 | 1.96 | 1.45 | 1.86 | 1.08 | 0.3 | 1.29 | 0.14 | -2.45 | 1.37 | 0.99 | 0.61 | 1.09 | -0.1 | -0.6725 | 0.465456 | 0.077412 | 0.867095 | 1 | 0.999631 | 0.794794 |
| Mapt_P10637_T206t_1 | -0.05 | 0.76 | 1.38 | -0.16 | -0.43 | 0.19 | -0.81 | 0.47 | 0.63 | 1.5 | 0.62 | 1.75 | 1.03 | 1.31 | 1.58 | 1.54 | -0.6275 | 0.6425 | 0.8825 | 0.051002 | 0.184432 | 0.180555 | 1 | 0.999631 | 0.489558 |
| Mapt_P10637_T220t_1 | -3.46 | -9.24 | -5.55 | -5.74 | -7.28 | -7.6 | -4.05 | -6.01 | -7.76 | -4.47 | -4.29 | -6.02 | -7.34 | -9.94 | -6.61 | -5.57 | -0.2375 | 0.3625 | -1.3675 | 0.299963 | 0.846976 | 0.769557 | 1 | 0.999631 | 0.713477 |
| Mapt_P10637_T220t_2 | 3.01 | -0.52 | 1.24 | 1.36 | 0.93 | 0.47 | 1.91 | 1.3 | 0.54 | 0.83 | 1.58 | 1.45 | -0.88 | 0.07 | 1.26 | 0.49 | -0.12 | -0.1725 | -1.0375 | 0.185169 | 0.862567 | 0.800333 | 1 | 0.999631 | 0.646072 |
| Mapt_P10637_T252t_1 | 1.66 | 0.12 | -0.06 | 1.76 | 1.95 | 0.38 | 2.01 | 2.36 | 0.99 | -0.18 | 2.28 | 1.29 | -1.88 | 0.43 | 1.53 | 1.44 | 0.805 | 0.225 | -0.49 | 0.549077 | 0.196766 | 0.72346 | 1 | 0.999631 | 0.835034 |
| Mapt_P10637_T392t_1 | -1.46 | -0.24 | -0.4 | -0.64 | -2.74 | -1.7 | -0.72 | -0.88 | -1.19 | 0.05 | -0.52 | -0.34 | 1.23 | -0.86 | -0.68 | -0.5 | -0.825 | 0.185 | 0.4825 | 0.366786 | 0.125089 | 0.638285 | 1 | 0.999631 | 0.745796 |
| Mapt_P10637_T392t_2 | 0.05 | 1.84 | -4.92 | -4.36 | -0.11 | 1.43 | -3.12 | -3.51 | 1.02 | 1.6 | -5.37 | -5.99 | 0.18 | 1.44 | -4.66 | -5.1 | 0.52 | -0.3375 | -0.1875 | 0.923733 | 0.763598 | 0.878589 | 1 | 0.999631 | 0.975109 |
| Mapt_P10637_T403t_1 | -1.14 | 1.93 | 2.18 | 0.62 | 2.8 | 1.32 | 2.85 | 0.83 | 3.11 | -0.69 | 1.97 | 3.52 | 2.75 | 0.63 | 1.01 | 0.42 | 1.0525 | 1.08 | 0.305 | 0.706403 | 0.209658 | 0.31593 | 1 | 0.999631 | 0.898066 |
| March11_Q8CBH7_S115s_1 | 0.7 | -0.29 | -0.05 | -0.09 | -0.59 | -0.14 | -0.21 | -0.18 | -0.03 | 0.53 | -0.15 | -0.17 | -0.28 | -0.74 | -0.31 | -0.3 | -0.3475 | -0.0225 | -0.475 | 0.164482 | 0.266878 | 0.945106 | 1 | 0.999631 | 0.632897 |
| Marcks_P26645_S112s_1 | -0.23 | 0.12 | -0.09 | -0.24 | 0.11 | -0.26 | -0.66 | -0.25 | -0.16 | 0.24 | -0.25 | -0.29 | 1.1 | 0.37 | 0.29 | 0.1 | -0.155 | -0.005 | 0.575 | 0.095657 | 0.569622 | 0.984942 | 1 | 0.999631 | 0.558128 |
| Marcks_P26645_S113s_1 | -0.21 | 1.18 | 0.02 | 0.53 | 0.11 | 0.29 | -0.59 | -0.11 | -0.02 | -0.5 | -0.43 | -0.21 | 1.37 | 0.38 | 0.55 | 0.01 | -0.455 | -0.67 | 0.1975 | 0.64751 | 0.245567 | 0.087592 | 1 | 0.999631 | 0.874615 |
| Marcks_P26645_S122s_1 | -2.43 | -1.35 | -0.01 | -0.76 | -3.38 | -3.08 | -0.95 | 0.89 | -1.54 | -0.27 | 0.42 | 1.24 | -1.3 | -1.43 | -0.28 | 0.96 | -0.4925 | 1.1 | 0.625 | 0.363413 | 0.612804 | 0.13745 | 1 | 0.999631 | 0.743747 |
| Marcks_P26645_S138s_1 | -0.55 | -0.61 | -0.6 | -0.8 | -0.42 | -0.62 | -0.54 | -0.64 | -0.36 | -0.45 | -0.24 | -0.12 | 0.18 | -0.67 | -0.47 | -0.28 | 0.085 | 0.3475 | 0.33 | 0.28357 | 0.717267 | 0.180697 | 1 | 0.999631 | 0.704942 |
| Marcks_P26645_S138s_2 | 0.1 | 0.13 | 0.08 | 0.38 | 0.8 | 0.6 | 0.45 | 0.21 | 0.42 | 0.23 | 0.39 | 0.01 | -0.45 | 0.32 | 0.22 | -0.1 | 0.3425 | 0.09 | -0.175 | 0.559391 | 0.200279 | 0.72411 | 1 | 0.999631 | 0.839333 |
| Marcks_P26645_S140s_1 | -0.09 | 0.96 | 0.23 | -0.32 | 0.58 | 0.85 | -0.18 | -0.02 | 0.08 | 0.19 | -0.27 | 0.3 | -0.03 | -0.69 | 0.05 | 0.59 | 0.1125 | -0.12 | -0.215 | 0.597671 | 0.769107 | 0.729681 | 1 | 0.999631 | 0.855033 |

| | | | | | | | | | | | | | | | | | | | | | | | | | |
|---|---|---|---|---|---|---|---|---|---|---|---|---|---|---|---|---|---|---|---|---|---|---|---|---|---|
| Marcks_P26645_S141s_2 | -0.21 | 0.02 | -0.05 | -0.48 | 0.29 | 0.43 | -0.09 | -0.93 | -0.25 | -0.45 | -0.36 | 0.14 | -0.18 | 0 | -0.37 | -0.8 | 0.105 | -0.05 | -0.1575 | 0.607509 | 0.766329 | 0.855591 | 1 | 0.999631 | 0.858527 |
| Marcks_P26645_S160s_1 | 0.94 | 1.3 | 0.93 | 1.22 | 0.53 | 1.51 | 0.63 | 1.21 | 2.06 | 1.4 | 1.65 | 1.57 | 0.98 | 1.61 | 1.11 | 2.54 | -0.1275 | 0.5725 | 0.4625 | 0.257311 | 0.681008 | 0.059828 | 1 | 0.999631 | 0.691981 |
| Marcks_P26645_S163s_1 | 1.09 | 1.47 | 0.69 | 1.46 | 1.75 | 1.89 | 0.9 | 2.07 | 1.76 | 1.01 | 1.45 | 1.72 | 0.63 | 2.19 | 1.86 | 1.91 | 0.475 | 0.3075 | 0.47 | 0.269081 | 0.194109 | 0.34251 | 1 | 0.999631 | 0.697063 |
| Marcks_P26645_S171s_1 | 0.45 | 1.07 | 1.11 | 0.9 | 0.35 | 0.85 | 0.65 | 1.02 | 0.44 | 0.77 | 0.81 | 0.72 | 0.69 | 0.3 | 0.69 | 0.67 | -0.165 | -0.1975 | -0.295 | 0.327709 | 0.566075 | 0.479528 | 1 | 0.999631 | 0.725013 |
| Marcks_P26645_S26s_1 | 0.27 | -0.55 | 0.33 | 0.25 | -0.16 | 0.12 | -0.06 | -0.12 | 0.06 | 0.85 | -0.03 | 0.53 | 0.29 | 0.26 | 0.46 | -0.18 | -0.13 | 0.2775 | 0.1325 | 0.686107 | 0.655341 | 0.42205 | 1 | 0.999631 | 0.890692 |
| Marcks_P26645_S27s_1 | 0.11 | -0.17 | 0.11 | -0.34 | -0.38 | -0.08 | -0.32 | 0.08 | 0.34 | 0.51 | 0.34 | 0.54 | 1.02 | 0.25 | 0.53 | 0.7 | -0.1025 | 0.505 | 0.6975 | 0.039902 | 0.694145 | 0.072976 | 1 | 0.999631 | 0.471899 |
| Marcks_P26645_S29s_1 | 0.89 | 1.18 | 0.24 | -0.41 | 0.89 | 1.05 | -0.66 | -0.21 | 0.98 | 1.19 | -0.84 | 0.1 | 1.73 | 1.13 | -0.08 | -0.01 | -0.2075 | -0.1175 | 0.2175 | 0.684213 | 0.685558 | 0.828619 | 1 | 0.999631 | 0.889649 |
| Marcks_P26645_S46s_1 | -0.26 | -0.83 | -0.4 | -0.63 | -0.42 | -0.61 | -0.42 | -0.46 | -0.56 | -0.1 | -0.35 | -0.03 | 0.03 | -0.28 | -0.12 | 0.01 | 0.0525 | 0.27 | 0.44 | 0.140059 | 0.834861 | 0.340941 | 1 | 0.999631 | 0.611718 |
| Marcks_P26645_T143t_1 | 0.21 | 0.13 | 0.6 | 0.12 | 0.35 | 0.2 | 0.48 | 0.54 | 0.73 | 0.26 | 0.81 | 1.03 | 0.17 | 0.56 | 0.89 | 0.71 | 0.1275 | 0.4425 | 0.3175 | 0.301614 | 0.616141 | 0.146917 | 1 | 0.999631 | 0.714243 |
| Marcks_P26645_T143t_2 | 0.13 | 0.15 | 0.4 | 0.54 | 0.95 | 0.59 | 0.53 | 0.2 | 0.51 | 0.34 | 0.39 | -0.14 | -0.52 | 0.34 | 0.27 | -0.12 | 0.2625 | -0.03 | -0.3125 | 0.330414 | 0.347325 | 0.91344 | 1 | 0.999631 | 0.727302 |
| Marcks_P26645_T79t_1 | 0.71 | 1.29 | 1.21 | 1.24 | 0.87 | 1.05 | 0.75 | 1.15 | 1.38 | 1.24 | 0.76 | 1.62 | 1.58 | 1.33 | 1.63 | 1.64 | -0.1575 | 0.1375 | 0.4325 | 0.150247 | 0.553165 | 0.64975 | 1 | 0.999631 | 0.622782 |
| Marcksl1_P28667_S101s_1 | 0.01 | 0.57 | 0.63 | 0.38 | 0.09 | 1.04 | 0.12 | 0.12 | 1.49 | 1.15 | 1.42 | 1.14 | 1.01 | 0.96 | 1.25 | 1.86 | -0.055 | 0.9025 | 0.8725 | 0.021805 | 0.863383 | 0.007719 | 1 | 0.999631 | 0.405544 |
| Marcksl1_P28667_S104s_1 | 0.87 | 1.23 | 0.84 | 1.1 | 1.55 | 1.5 | 0.68 | 1.78 | 1.77 | 1.57 | 1.63 | 2.03 | 1.08 | 1.69 | 1.98 | 2.53 | 0.3675 | 0.74 | 0.81 | 0.046652 | 0.255849 | 0.017162 | 1 | 0.999631 | 0.480102 |
| Marcksl1_P28667_S120s_1 | -1.52 | -0.9 | -1.12 | -1.66 | -1.24 | -1.18 | -1.5 | -1.25 | -0.18 | -0.19 | -1.09 | -0.91 | 0.38 | -0.68 | -0.86 | -0.44 | 0.0075 | 0.7075 | 0.9 | 0.032517 | 0.978266 | 0.061193 | 1 | 0.999631 | 0.442395 |
| Marcksl1_P28667_S132s_1 | -1.44 | -1.49 | -0.47 | -0.45 | -1.06 | -0.51 | -0.2 | -0.09 | -0.29 | -0.44 | 0.1 | 0.37 | -0.14 | -0.93 | 0.21 | 0.35 | 0.4975 | 0.8975 | 0.835 | 0.070772 | 0.207797 | 0.034231 | 1 | 0.999631 | 0.530826 |
| Marcksl1_P28667_S135s_1 | -1.76 | -1.14 | -1.19 | -2.42 | -1.84 | -1.15 | -1.76 | -2.01 | -0.61 | -0.27 | -2.29 | -0.6 | 1.15 | -0.15 | -1.42 | -2.36 | -0.0625 | 0.685 | 0.9325 | 0.217006 | 0.866229 | 0.20323 | 1 | 0.999631 | 0.669465 |
| Marcksl1_P28667_S151s_1 | -0.25 | -0.95 | 1.1 | 0.53 | -0.75 | -0.49 | 0.4 | 0.26 | -0.42 | 0.46 | 0.22 | 0.49 | -0.36 | -1.16 | 0.61 | 0.41 | -0.2525 | 0.08 | -0.2325 | 0.678515 | 0.613415 | 0.866694 | 1 | 0.999631 | 0.887183 |
| Marcksl1_P28667_S151s_2 | 0.13 | -1.23 | -0.85 | -1.2 | -0.61 | -0.68 | -0.68 | -0.49 | -0.32 | 0.43 | -0.47 | -1.46 | -0.94 | -1.75 | -1.8 | -1.78 | 0.1725 | 0.3325 | -0.78 | 0.076731 | 0.624336 | 0.495242 | 1 | 0.999631 | 0.542673 |
| Marcksl1_P28667_S22s_1 | -0.47 | -0.55 | -0.19 | -0.2 | -0.4 | -0.33 | -0.09 | -0.12 | 0.12 | 0.49 | 0.39 | 0.59 | 0.76 | -0.21 | 0.33 | 0.75 | 0.1175 | 0.75 | 0.76 | 0.038743 | 0.636076 | 0.015915 | 1 | 0.999631 | 0.462703 |
| Marcksl1_P28667_S36s_1 | -0.16 | -1.16 | -1.09 | -1.27 | -0.62 | -0.75 | -0.34 | -1.15 | -0.12 | -0.09 | -0.11 | -0.47 | -0.36 | -0.47 | -0.75 | -0.29 | 0.205 | 0.7225 | 0.4525 | 0.203603 | 0.55143 | 0.048566 | 1 | 0.999631 | 0.659781 |
| Marcksl1_P28667_T122t_1 | -1.78 | -1.64 | -1.74 | -1.43 | -1.41 | -2.34 | -1.76 | -1.54 | -0.93 | -1.13 | -1.41 | -0.74 | 0.18 | -1.01 | -2.25 | -1.45 | -0.115 | 0.595 | 0.515 | 0.310469 | 0.693639 | 0.050314 | 1 | 0.999631 | 0.71924 |
| Marcksl1_P28667_T148t_1 | 0.05 | -0.88 | -0.7 | -0.44 | -0.75 | -0.31 | -0.44 | -0.61 | 0.26 | 0.65 | -0.64 | -0.64 | -0.02 | -0.82 | -0.9 | -0.71 | -0.035 | 0.4 | -0.12 | 0.729238 | 0.904858 | 0.327066 | 1 | 0.999631 | 0.905923 |
| Marcksl1_P28667_T148t_2 | 0.13 | -1.23 | -0.85 | -1.2 | -0.61 | -0.68 | -0.68 | -0.49 | -0.32 | 0.43 | -0.47 | -1.46 | -0.94 | -1.75 | -1.8 | -1.78 | 0.1725 | 0.3325 | -0.78 | 0.076731 | 0.624336 | 0.495242 | 1 | 0.999631 | 0.542673 |
| Marcksl1_P28667_T41t_1 | -0.51 | -1.2 | -1.39 | -1.28 | -1.37 | -1.56 | -0.42 | -1.27 | -0.83 | -0.78 | -0.62 | -0.81 | -0.7 | -1.05 | -0.78 | -0.81 | -0.06 | 0.335 | 0.26 | 0.40787 | 0.864011 | 0.265275 | 1 | 0.999631 | 0.766996 |
| Marcksl1_P28667_T85t_1 | 0.11 | -1.16 | -0.96 | -1.05 | -0.61 | -0.73 | -1.38 | -0.77 | 0.28 | 0.79 | -1.09 | -1.21 | -0.16 | -0.98 | -1.24 | -1.07 | -0.1075 | 0.4575 | -0.0975 | 0.808889 | 0.767687 | 0.404843 | 1 | 0.999631 | 0.93635 |
| Mark1_Q8VHJ5_S382s_1 | -0.19 | 0.52 | 0.02 | 0.13 | 0.33 | 0.08 | -0.34 | -0.04 | 0.06 | 0.04 | 0.02 | 0.27 | 0.53 | 0.42 | 0.08 | 0.32 | -0.1125 | -0.0225 | 0.2175 | 0.464048 | 0.691607 | 0.933554 | 1 | 0.999631 | 0.793831 |
| Mark1_Q8VHJ5_S403s_1 | 1.11 | 0.16 | -0.64 | 1.21 | 1.01 | 1.2 | 0.98 | 1.36 | 0.85 | 0.67 | 0.66 | -0.07 | -1.57 | 0.99 | 0.26 | 0.97 | 0.6775 | 0.0675 | -0.2975 | 0.655929 | 0.143445 | 0.884932 | 1 | 0.999631 | 0.879056 |
| Mark1_Q8VHJ5_S414s_1 | 1.86 | 0.86 | 2.63 | 2.34 | 0.22 | 1.41 | 2.97 | 1.4 | 0.65 | 1.4 | 1.25 | 1.79 | 0.29 | 0.61 | 2.51 | 2.18 | -0.4225 | -0.65 | -0.525 | 0.401713 | 0.49816 | 0.168747 | 1 | 0.999631 | 0.763221 |
| Mark1_Q8VHJ5_S423s_1 | -1.23 | -0.67 | 0.47 | 0.16 | -0.22 | -0.5 | -0.43 | 0.11 | -0.55 | -0.76 | -0.32 | -0.55 | -0.19 | -0.01 | -0.16 | -0.16 | 0.0575 | -0.2275 | 0.1875 | 0.648393 | 0.888678 | 0.579422 | 1 | 0.999631 | 0.874615 |
| Mark1_Q8VHJ5_S444s_1 | 1.46 | 0.68 | 1.38 | 2.06 | 1.46 | 0.8 | 1.61 | 1.42 | 0.78 | 0.81 | 1.17 | 1.13 | 0.24 | -0.09 | 0.86 | 0.5 | -0.0725 | -0.4225 | -1.0175 | 0.022118 | 0.839916 | 0.237594 | 1 | 0.999631 | 0.405544 |
| Mark1_Q8VHJ5_S468s_1 | 1.49 | 0.23 | -0.94 | 0.12 | 1.07 | 0.33 | 0.55 | -0.05 | 1.35 | -0.37 | 0.14 | -0.76 | -0.09 | -0.18 | -0.96 | -0.82 | 0.25 | -0.135 | -0.7375 | 0.17456 | 0.627515 | 0.826032 | 1 | 0.999631 | 0.642116 |
| Mark1_Q8VHJ5_S46s_1 | 0.03 | -1 | 0.1 | 1.23 | 0.98 | 0.17 | 1.06 | 1.17 | 0.07 | -1.13 | 0.94 | 0.63 | -1.02 | -0.14 | 1.13 | 0.33 | 0.755 | 0.0375 | -0.015 | 0.979568 | 0.141549 | 0.94922 | 1 | 0.999631 | 0.993554 |
| Mark1_Q8VHJ5_S488s_1 | 0.71 | -1.36 | 0.01 | 0.27 | -0.3 | -0.25 | 0.39 | 0.69 | -0.05 | -0.64 | 0.29 | 0.01 | -2.23 | 0.16 | 0.16 | 0.26 | 0.225 | -0.005 | -0.32 | 0.635719 | 0.642026 | 0.991518 | 1 | 0.999631 | 0.869397 |
| Mark1_Q8VHJ5_S588s_1 | 1.92 | -0.23 | -1.75 | 1.28 | 0.93 | 0.48 | 1.6 | 1.11 | 0.81 | 0.4 | 0.65 | -0.43 | -1.57 | 0.36 | -0.01 | -0.42 | 0.725 | 0.0525 | -0.715 | 0.382577 | 0.344881 | 0.945086 | 1 | 0.999631 | 0.754097 |
| Mark1_Q8VHJ5_S590s_1 | 0.36 | -0.65 | -0.82 | -0.15 | -0.79 | -0.48 | 0.49 | 0.01 | -1.25 | -0.37 | 0.68 | -0.77 | -1.46 | -0.07 | -0.89 | -0.69 | 0.1225 | -0.1125 | -0.4625 | 0.274973 | 0.756258 | 0.811821 | 1 | 0.999631 | 0.700187 |
| Mark1_Q8VHJ5_S593s_1 | 0.74 | 0.39 | 0.62 | -0.02 | 0.38 | 0.6 | 0.31 | 0.22 | -0.23 | 0.72 | -0.17 | 0.18 | -0.3 | 0.29 | -0.64 | 0.17 | -0.055 | -0.3075 | -0.5525 | 0.126678 | 0.840894 | 0.360239 | 1 | 0.999631 | 0.597196 |
| Mark1_Q8VHJ5_S633s_1 | -0.1 | -1.76 | -1.08 | -0.59 | -0.63 | -1.64 | -0.14 | -0.81 | -0.76 | -1.22 | -0.78 | -0.53 | -1.4 | -1.15 | -0.92 | -1.07 | 0.0775 | 0.06 | -0.2525 | 0.527287 | 0.864675 | 0.879706 | 1 | 0.999631 | 0.824554 |
| Mark1_Q8VHJ5_S649s_1 | 1.12 | 0.5 | 1.15 | 0.71 | 1.86 | 0.49 | 1.81 | 1.65 | 1.2 | 0.95 | 1.44 | 1.76 | -0.48 | 1.13 | 1.63 | 1.1 | 0.5825 | 0.4675 | -0.025 | 0.957862 | 0.145299 | 0.149976 | 1 | 0.999631 | 0.985412 |
| Mark1_Q8VHJ5_S666s_1 | 2.03 | 1.84 | 0.38 | 1.17 | 1.33 | 1.54 | 1.99 | -0.16 | 1.95 | 0.51 | 0.5 | 0.3 | -0.29 | -1.06 | -0.55 | -1.37 | -0.18 | -0.54 | -2.1725 | 0.000668 | 0.743253 | 0.299826 | 1 | 0.999631 | 0.351308 |
| Mark2_Q05512_S390s_1 | 0.79 | -1.33 | -0.46 | 1.11 | 1.99 | 0.78 | 1.49 | 1.03 | 1.26 | -0.24 | 0.09 | -0.05 | -1.89 | 0.53 | 0.58 | -0.21 | 1.295 | 0.2375 | -0.275 | 0.700979 | 0.043983 | 0.693356 | 1 | 0.999631 | 0.896835 |
| Mark2_Q05512_S408s_1 | 0.11 | 0.3 | -0.6 | -0.02 | 0.63 | -0.37 | -0.58 | -0.14 | 0.28 | -0.21 | 0.29 | -0.19 | -0.27 | 0.09 | 0.26 | -0.18 | -0.0625 | 0.095 | 0.0275 | 0.930675 | 0.860042 | 0.7584 | 1 | 0.999631 | 0.977897 |
| Mark2_Q05512_S409s_1 | 0.11 | 0.3 | -0.6 | -0.02 | 0.63 | -0.37 | -0.58 | -0.14 | 0.28 | -0.21 | 0.29 | -0.19 | -0.27 | 0.09 | 0.26 | -0.18 | -0.0625 | 0.095 | 0.0275 | 0.930675 | 0.860042 | 0.7584 | 1 | 0.999631 | 0.977897 |
| Mark2_Q05512_S40s_1 | 1.03 | -0.78 | -1.57 | 0.96 | 1.52 | -0.21 | 0.44 | 1.23 | -0.12 | -0.33 | -0.08 | -0.99 | -1.57 | -0.57 | -0.58 | -0.75 | 0.835 | -0.29 | -0.7775 | 0.230826 | 0.233147 | 0.640123 | 1 | 0.999631 | 0.681463 |
| Mark2_Q05512_S453s_1 | 2.93 | -1.63 | -2.99 | 1.97 | 3.3 | -1.18 | 2.72 | 1.48 | 0.1 | -3.54 | 0.78 | -1.75 | -2.75 | -1.32 | -1.46 | -1.49 | 1.51 | -1.1725 | -1.825 | 0.160271 | 0.317268 | 0.430208 | 1 | 0.999631 | 0.629764 |
| Mark2_Q05512_S483s_1 | 0.96 | 0.43 | -0.45 | 1.27 | 1.52 | 1.16 | 1.01 | 1.51 | 0.67 | 0.27 | 0.97 | -0.05 | -1.91 | 0.68 | 0.28 | 0.42 | 0.7475 | -0.0875 | -0.685 | 0.296501 | 0.087292 | 0.840657 | 1 | 0.999631 | 0.709528 |
| Mark2_Q05512_S566s_1 | 1.67 | -0.27 | -1.19 | 0.68 | 1.09 | 0.35 | 1.3 | 0.41 | -0.37 | 0.22 | -0.5 | -0.44 | -1.44 | 0.06 | -2.43 | -0.31 | 0.565 | -0.495 | -1.2525 | 0.112966 | 0.355475 | 0.405398 | 1 | 0.999631 | 0.577408 |
| Mark2_Q05512_S616s_1 | 0.06 | -0.13 | -0.03 | 0.77 | 0.95 | 0.23 | 0.88 | 1.29 | -0.08 | -0.3 | 0.57 | 0.84 | 0.34 | -1.05 | -0.01 | 0.33 | 0.67 | 0.09 | -0.265 | 0.51931 | 0.071372 | 0.806237 | 1 | 0.999631 | 0.819456 |
| Mark3_Q03141_S215s_1 | -0.63 | 0.15 | 1.55 | 0.82 | -0.51 | 1.14 | 0.58 | 1.8 | -0.66 | -0.81 | 1.45 | -0.83 | 0.1 | 1.13 | 0.37 | 1.15 | 0.28 | -0.685 | 0.215 | 0.674827 | 0.647227 | 0.307448 | 1 | 0.999631 | 0.885515 |
| Mark3_Q03141_S419s_1 | -0.69 | 0.11 | -0.03 | -0.34 | 0.04 | -0.36 | -0.36 | -0.37 | -0.16 | -0.43 | -0.06 | 0.08 | 0.24 | 0.21 | 0.27 | 0.23 | -0.025 | 0.095 | 0.475 | 0.128989 | 0.929524 | 0.745415 | 1 | 0.999631 | 0.59942 |
| Mark3_Q03141_S42s_1 | 0.79 | -0.96 | -2.31 | -0.79 | 1.63 | 0.03 | -0.14 | -2.01 | 0.21 | -0.62 | -1.57 | -2.01 | -0.44 | 0.12 | -2.5 | -1.6 | 0.695 | -0.18 | -0.2875 | 0.705418 | 0.422024 | 0.801502 | 1 | 0.999631 | 0.89781 |
| Mark3_Q03141_S469s_1 | 1.21 | -0.27 | -0.57 | 1.55 | 1.46 | 0.71 | 1.52 | 1.24 | 0.64 | 0.06 | 1.32 | 0.25 | -0.32 | 0.88 | 0.63 | 0.35 | 0.7525 | 0.0875 | -0.095 | 0.862252 | 0.17002 | 0.874126 | 1 | 0.999631 | 0.954601 |
| Mark3_Q03141_S489s_1 | 0.69 | -0.91 | -0.55 | 1.37 | 1.8 | -0.58 | 0.91 | 1.18 | 0.46 | 0.11 | 0 | -0.15 | -2.13 | -0.15 | -0.56 | -1.72 | 0.6775 | -0.045 | -1.29 | 0.066506 | 0.314198 | 0.930344 | 1 | 0.999631 | 0.518767 |
| Mark3_Q03141_S499s_1 | -0.11 | -0.16 | -0.16 | 0.77 | 0.47 | 0.55 | 0.29 | -0.22 | 0.06 | 1.65 | -0.53 | -0.67 | -2.46 | 0.32 | 0.01 | 0.94 | 0.1875 | 0.0425 | -0.3825 | 0.583891 | 0.571737 | 0.937074 | 1 | 0.999631 | 0.848949 |
| Mark3_Q03141_S543s_1 | 1.04 | 0.04 | -0.51 | 0.55 | 0.42 | 0.83 | 0.7 | 0.66 | 0.29 | 0.65 | -0.03 | -0.2 | -0.8 | 0.68 | 0.78 | 0.02 | 0.3725 | -0.1025 | -0.11 | 0.818809 | 0.320929 | 0.79615 | 1 | 0.999631 | 0.938303 |
| Mark3_Q03141_S563s_1 | 1.01 | 3.08 | 3.02 | 2.66 | 1.61 | 4.19 | 3.29 | 4.34 | 0.93 | 0.31 | 4.4 | -1.23 | -2.06 | 3.47 | 1.59 | -0.01 | 0.915 | -1.34 | -1.695 | 0.140766 | 0.212502 | 0.24308 | 1 | 0.999631 | 0.612258 |
| Mark3_Q03141_S583s_1 | 0.15 | -1.66 | -2.76 | -0.94 | 0.17 | -0.99 | -0.37 | -1.03 | -1.18 | -1.69 | -1.19 | -2.39 | -2.43 | -1.19 | -2.33 | -1.81 | 0.7475 | -0.31 | -0.6375 | 0.311554 | 0.237868 | 0.615223 | 1 | 0.999631 | 0.71924 |
| Mark3_Q03141_S619s_1 | -1.73 | -4.71 | -6.03 | -6.12 | -3.15 | -4.18 | -5.24 | -8.1 | -4.61 | -2.1 | -5.56 | -5.92 | -4.59 | -4.74 | -6.42 | -6.93 | -0.52 | 0.1 | -1.0225 | 0.32436 | 0.680986 | 0.93053 | 1 | 0.999631 | 0.72463 |
| Mark3_Q03141_T211t_1 | -1.97 | 0.65 | 0.34 | -0.5 | -2.1 | 2.81 | -0.17 | 1.11 | -2.28 | -1.32 | 0.79 | -2.19 | -0.14 | 2.84 | -0.7 | -0.7 | 0.7825 | -0.88 | 0.695 | 0.441942 | 0.451318 | 0.292293 | 1 | 0.999631 | 0.78215 |
| Mark4_Q8CIP4_S438s_1 | -1.08 | -2.2 | 0.41 | 0.82 | -1.51 | -1.37 | 0.09 | 1.05 | -1.49 | -1.15 | 0.46 | 0.86 | -1.7 | -1.5 | 0.21 | 0.76 | 0.0775 | 0.1825 | -0.045 | 0.955512 | 0.923585 | 0.818715 | 1 | 0.999631 | 0.984649 |
| Mark4_Q8CIP4_S458s_1 | 3.21 | -0.16 | 1.68 | 1.64 | 1.54 | 1.05 | 2.97 | 1.7 | 0.69 | 1.12 | 0.83 | -0.16 | 0.57 | 0.36 | 0.8 | -0.26 | 0.2225 | -0.9725 | -1.225 | 0.084083 | 0.753795 | 0.166604 | 1 | 0.999631 | 0.549732 |
| Mark4_Q8CIP4_S45s_1 | 0.42 | -0.44 | 0.18 | 1.74 | 1.2 | 0.46 | 1.51 | 2.11 | 0.43 | -0.39 | 0.48 | 0.17 | 0.26 | 0.5 | 0.62 | 1.08 | 0.845 | -0.3025 | 0.14 | 0.769593 | 0.135179 | 0.53307 | 1 | 0.999631 | 0.918554 |
| Mark4_Q8CIP4_S464s_1 | 2.99 | 0.64 | 0.78 | 2.11 | 1.18 | 1.71 | 2.68 | 1.88 | 0.46 | 1.78 | 1.54 | 0.86 | -2.05 | 0 | 0.96 | 0.35 | 0.2325 | -0.47 | -1.815 | 0.034823 | 0.690895 | 0.429298 | 1 | 0.999631 | 0.44933 |
| Mark4_Q8CIP4_S567s_1 | 0.71 | 1.29 | 0.9 | 1.24 | 2.3 | 2.95 | 2.3 | 2.95 | 1.25 | -0.43 | 2.92 | -0.09 | 0.3 | 2.45 | 2.26 | -0.42 | 1.59 | -0.1225 | 0.1125 | 0.862983 | 0.000368 | 0.8591 | 1 | 0.999631 | 0.954983 |
| Mark4_Q8CIP4_S594s_1 | -1.01 | -0.19 | -0.38 | -0.4 | 0.64 | 0.11 | -0.15 | -0.39 | -0.76 | -1.07 | 0.05 | -0.43 | 0.23 | -0.27 | -0.88 | -0.41 | 0.5475 | -0.0575 | 0.1625 | 0.642501 | 0.118685 | 0.866653 | 1 | 0.999631 | 0.871335 |
| Mark4_Q8CIP4_S654s_1 | 1.05 | -0.15 | -1.13 | 0.02 | 1.93 | 0.06 | 2.24 | 0.11 | 1.57 | -0.79 | 1.71 | 0.2 | 0.02 | 0.35 | -1.11 | -1.53 | 1.1375 | 0.725 | -0.515 | 0.384835 | 0.106825 | 0.291861 | 1 | 0.999631 | 0.75538 |
| Mast1_Q9R1L5_S1138s_1 | 3.64 | -2.21 | -2.04 | 1.83 | 1.67 | 1.6 | 3.15 | 0.93 | 0.79 | -0.27 | 0.7 | -1.77 | -3.06 | -1.08 | -0.47 | -2.61 | 1.5325 | -0.4425 | -2.11 | 0.135778 | 0.255135 | 0.741324 | 1 | 0.999631 | 0.608415 |
| Mast1_Q9R1L5_S1244s_1 | -1.58 | 2.54 | 1.12 | -1.3 | 3.11 | 2.12 | 1.2 | 0.35 | 3.52 | -0.49 | 0.43 | 2.34 | 3.49 | 1.24 | 0.44 | -0.76 | 1.5 | 1.255 | 0.9075 | 0.430619 | 0.154342 | 0.290883 | 1 | 0.999631 | 0.778235 |
| Mast1_Q9R1L5_S1414s_1 | 0.83 | -0.45 | -1.56 | 0.38 | -0.14 | -0.99 | 0.39 | -0.1 | 0.26 | 0.24 | 0.36 | -1.27 | -0.86 | -0.11 | -0.58 | 0.21 | -0.01 | 0.0975 | -0.135 | 0.802577 | 0.985458 | 0.870154 | 1 | 0.999631 | 0.932677 |
| Mast1_Q9R1L5_S346s_1 | -0.2 | -0.69 | -0.97 | -1.09 | -1.95 | -0.56 | -0.51 | -0.42 | -0.03 | -0.31 | -0.32 | -2.14 | -1.24 | -2.66 | -0.84 | -1.01 | -0.1225 | 0.0375 | -0.7 | 0.148196 | 0.767297 | 0.939849 | 1 | 0.999631 | 0.62132 |
| Mast1_Q9R1L5_S718s_1 | 0.03 | -1.5 | -1.32 | -0.19 | -0.18 | 0.11 | -0.52 | -1.01 | -1.45 | -0.73 | -0.87 | -1.25 | -0.03 | -1.4 | -1.1 | -1.11 | 0.345 | -0.33 | -0.165 | 0.730721 | 0.443988 | 0.443942 | 1 | 0.999631 | 0.906863 |
| Mast1_Q9R1L5_S791s_1 | -0.56 | 0.28 | -0.15 | -0.66 | -0.27 | -2.19 | -0.1 | -0.75 | -0.24 | -0.01 | 0.58 | 0.9 | 0.22 | 0.48 | 0.84 | 0.23 | -0.555 | 0.58 | 0.715 | 0.052822 | 0.274247 | 0.138855 | 1 | 0.999631 | 0.489777 |
| Mast1_Q9R1L5_S895s_1 | 1.38 | 1.96 | 2.36 | -1.07 | 3.33 | 1.96 | 2.6 | 1.26 | 3.59 | 1.84 | 3.75 | 4.64 | 2.09 | 2.03 | -0.58 | 1.61 | 1.13 | 2.2975 | 0.13 | 0.879969 | 0.168531 | 0.020729 | 1 | 0.999631 | 0.962995 |
| Mast1_Q9R1L5_S90s_1 | 0 | 0.61 | -0.17 | -0.42 | 0.5 | -0.08 | -0.35 | -0.09 | -0.03 | -0.02 | -0.45 | 1.09 | 0.86 | 0.14 | 0.91 | 0.18 | -0.01 | 0.1425 | 0.5175 | 0.168166 | 0.975574 | 0.72615 | 1 | 0.999631 | 0.637399 |
| Mast1_Q9R1L5_S968s_1 | 0.13 | -0.17 | 0.1 | -0.28 | -0.17 | 0.07 | -0.1 | -0.66 | 0.29 | 0.77 | -0.34 | 0.17 | -0.32 | 0.51 | 0.9 | 0.29 | -0.16 | 0.2775 | 0.4 | 0.25585 | 0.562758 | 0.386401 | 1 | 0.999631 | 0.690775 |
| Mast1_Q9R1L5_T1371t_1 | 0.58 | -0.14 | 0.54 | 0.18 | 0.11 | 0.16 | 0.87 | -0.47 | 0.06 | 0.76 | -0.38 | -0.77 | -0.07 | 0.05 | 0.11 | -0.56 | -0.1225 | -0.3725 | -0.4075 | 0.215819 | 0.727807 | 0.348334 | 1 | 0.999631 | 0.668602 |
| Mast1_Q9R1L5_T351t_1 | -0.18 | 0.56 | -0.49 | -0.32 | 0.62 | 0.8 | -0.15 | -0.31 | 0.73 | 0.89 | -0.36 | -1.34 | 0.89 | 0.18 | -0.25 | -0.74 | 0.3475 | 0.0875 | 0.1275 | 0.76517 | 0.36584 | 0.869277 | 1 | 0.999631 | 0.916326 |
| Mast2_B1AST8_S1177s_1 | 0.98 | 0.86 | 1.74 | 0.81 | 0.99 | 0.54 | 0.72 | 0.44 | 1.24 | 1.1 | 1.66 | 0.34 | 0.85 | 0.63 | 0.7 | 1.82 | -0.425 | -0.0125 | -0.0975 | 0.80036 | 0.186933 | 0.973411 | 1 | 0.999631 | 0.931901 |
| Mast3_Q3U214_S107s_1 | -0.04 | 0.73 | -0.52 | -0.16 | 0.57 | 0.4 | -0.46 | 0.39 | 0.54 | -0.16 | 0.01 | 0.37 | 0.6 | 0.88 | 0.1 | 0.31 | 0.2225 | 0.1875 | 0.47 | 0.213392 | 0.550834 | 0.593215 | 1 | 0.999631 | 0.666971 |
| Mast3_Q3U214_S156s_1 | 1.13 | 0.33 | -0.32 | 1.5 | 1.49 | 0.94 | 1.23 | 0.77 | 0.69 | 0.45 | 0.88 | -0.01 | -1.13 | 0.21 | -0.5 | -0.34 | 0.4475 | -0.1575 | -1.1 | 0.041659 | 0.311681 | 0.723692 | 1 | 0.999631 | 0.475443 |
| Mast3_Q3U214_S159s_1 | 0.59 | 1.03 | 1.2 | 1.08 | 0.05 | 0.98 | 0.8 | 0.07 | 0.31 | 1.15 | 1.32 | -0.38 | -0.25 | -0.19 | -1.15 | 0.3 | -0.5 | -0.375 | -1.2975 | 0.0055 | 0.145071 | 0.381841 | 1 | 0.999631 | 0.351308 |
| Mast3_Q3U214_S168s_1 | 0.71 | -0.74 | -0.42 | 0.19 | 0.05 | 0.37 | -0.11 | 0.46 | 0.17 | -0.5 | 0.14 | -1.66 | -0.52 | -0.72 | 0.49 | -0.36 | 0.2575 | -0.3975 | -0.2125 | 0.621352 | 0.490156 | 0.440171 | 1 | 0.999631 | 0.861888 |
| Mast3_Q3U214_S61s_1 | 0.93 | -0.07 | -0.26 | 1.41 | 1.21 | 1.17 | 1.02 | 0.7 | 0.33 | 0.29 | 0.22 | -0.14 | -0.86 | -0.04 | -0.07 | 0.38 | 0.5225 | -0.3275 | -0.65 | 0.185522 | 0.226396 | 0.440876 | 1 | 0.999631 | 0.646224 |
| Mast3_Q3U214_S803s_1 | -0.74 | 1.47 | -0.08 | 0.97 | 0.36 | -0.79 | 0.53 | 0.26 | 0.53 | 0.05 | 0.36 | 0.04 | -0.09 | -0.2 | -0.67 | 0.29 | -0.315 | -0.16 | -0.5725 | 0.278225 | 0.560979 | 0.744823 | 1 | 0.999631 | 0.700874 |
| Mast3_Q3U214_S942s_1 | 1.53 | 0.22 | 0.48 | 0.66 | 0.04 | -0.14 | 1.57 | -0.76 | 0.61 | 1.58 | 0.63 | -0.11 | 0.74 | -0.77 | -1.14 | -0.6 | -0.545 | -0.045 | -1.165 | 0.034352 | 0.315724 | 0.918913 | 1 | 0.999631 | 0.446273 |
| Mast3_Q3U214_S947s_1 | 2.85 | 0.1 | -0.83 | 1.55 | 2.21 | 0.82 | 2.47 | 0.16 | 1.71 | 0.9 | 0.79 | -0.9 | -1.33 | -0.91 | -1.25 | -0.85 | 0.4975 | -0.2925 | -2.0025 | 0.018506 | 0.56258 | 0.732334 | 1 | 0.999631 | 0.391024 |
| Mast4_E9PWX8_S1176s_1 | 1.29 | 1.05 | 0.49 | 1.03 | 1.67 | 1.22 | 0.26 | 0.92 | 2.09 | 0.71 | 1.65 | 0.53 | 1.16 | 0.71 | 1.2 | 0.93 | 0.0525 | 0.28 | 0.035 | 0.908375 | 0.884938 | 0.505504 | 1 | 0.999631 | 0.969941 |
| Mast4_E9PWX8_S2161s_1 | 3.32 | -0.6 | 0.26 | 1.76 | 2.44 | 0.74 | 2.93 | 0.4 | 1.75 | 1.4 | 0.2 | -0.81 | -5.1 | -0.9 | -0.21 | -2.03 | 0.4425 | -0.55 | -3.245 | 0.018601 | 0.632297 | 0.547536 | 1 | 0.999631 | 0.391024 |
| Mat2b_Q99LB6_S271s_1 | 0.04 | -0.42 | 1.49 | 0.9 | 0.05 | 0.84 | 0.14 | 0.77 | 0.74 | 1.28 | -0.27 | 1.12 | 0.01 | 2 | 0.97 | 1.18 | -0.0525 | 0.215 | 0.5375 | 0.341002 | 0.908455 | 0.679546 | 1 | 0.999631 | 0.733032 |
| Matr3_Q8K310_S188s_1 | 1.36 | 1.35 | 1.19 | 1.48 | 0.72 | 1.08 | 1.37 | 0.38 | 1.21 | 2.02 | 0.96 | 0.55 | 0.24 | 0.15 | -0.31 | 0.69 | -0.4575 | -0.16 | -1.1525 | 0.003702 | 0.141771 | 0.651899 | 1 | 0.999631 | 0.351308 |
| Matr3_Q8K310_S211s_1 | -0.07 | -1.12 | -0.3 | 0.23 | -1.16 | -0.88 | 0.18 | -0.89 | -2.21 | -0.15 | 0.57 | -1.3 | -2.44 | -1.92 | -4.14 | -2.88 | -0.3725 | -0.4575 | -2.53 | 0.000711 | 0.377468 | 0.464378 | 1 | 0.999631 | 0.351308 |
| Matr3_Q8K310_S4s_1 | -0.22 | 1.06 | 0.83 | 0.89 | 0.96 | 0.73 | 2.11 | 0.72 | 1.06 | -0.08 | 2.01 | 0.95 | -0.03 | 0.49 | -0.02 | 0.61 | 0.49 | 0.345 | -0.3775 | 0.327022 | 0.272961 | 0.489248 | 1 | 0.999631 | 0.724678 |
| Matr3_Q8K310_S598s_1 | 1.53 | 0.31 | 1.09 | 0.65 | -0.45 | 0.37 | 0.91 | -0.1 | -0.21 | 1.28 | 0.07 | -0.05 | -0.17 | -0.57 | 0.32 | -0.33 | -0.7125 | -0.6225 | -1.0825 | 0.013914 | 0.100259 | 0.171082 | 1 | 0.999631 | 0.385092 |
| Matr3_Q8K310_S619s_1 | -0.88 | -3.36 | 1.68 | 1.3 | -3.81 | -3.17 | 2.56 | -1.15 | -4.62 | -1.96 | -0.04 | 0.49 | -2.69 | -4.06 | 0.88 | 0.06 | -1.0775 | -1.2175 | -1.1375 | 0.417818 | 0.49604 | 0.392663 | 1 | 0.999631 | 0.771539 |
| Matr3_Q8K310_S746s_1 | -0.2 | 2.87 | 2.54 | 1.47 | 1.4 | 2.28 | 1.3 | 1.08 | 2.41 | 1.88 | 1.74 | 1.96 | 2.54 | 2.77 | 1.4 | 2.44 | -0.155 | 0.3275 | 0.6175 | 0.369735 | 0.814259 | 0.609592 | 1 | 0.999631 | 0.746338 |
| Matr3_Q8K310_S9s_1 | -0.97 | 1.11 | -0.05 | -0.14 | 0.6 | 0.89 | 1.12 | 0.18 | 0.89 | -0.95 | 1.24 | 1.15 | 0.9 | 0.45 | 0 | 0 | 0.71 | 0.595 | 0.35 | 0.462272 | 0.14303 | 0.341012 | 1 | 0.999631 | 0.792786 |
| Matr3_Q8K310_T150t_1 | -0.93 | 2.27 | 0.91 | 0.44 | -0.41 | 0.53 | -0.65 | 0.09 | 1.61 | 1.63 | 0.37 | 1.84 | 1.87 | 1.23 | 0.38 | 1.82 | -0.7825 | 0.69 | 0.6525 | 0.339596 | 0.237485 | 0.312577 | 1 | 0.999631 | 0.732185 |
| Mb21d2_D3Z742_S433s_1 | 1.06 | 0.75 | -0.71 | 1.09 | 1.4 | 1.07 | 0.26 | -0.11 | 0.79 | 0.35 | -0.47 | 0.04 | -0.65 | 1.04 | -2.01 | 0.74 | 0.1075 | -0.37 | -0.7675 | 0.303212 | 0.834233 | 0.449275 | 1 | 0.999631 | 0.715044 |
| Mb21d2_D3Z742_S433s_2 | 0.29 | -0.02 | -0.74 | -0.55 | 0.12 | 0.35 | -0.32 | -1.02 | 0.18 | 0.84 | -0.41 | -1.14 | -0.07 | 0.27 | -1.45 | -0.67 | 0.0375 | 0.1225 | -0.225 | 0.616236 | 0.923656 | 0.793815 | 1 | 0.999631 | 0.861203 |
| Mb21d2_D3Z742_S439s_1 | -0.62 | -1.16 | -0.74 | -1.58 | -1.29 | -1.01 | -1.7 | -1.5 | -1.1 | -0.47 | -0.87 | -1.53 | 0.17 | -0.7 | -1.46 | -0.54 | -0.35 | 0.0325 | 0.3925 | 0.355702 | 0.282243 | 0.925986 | 1 | 0.999631 | 0.74034 |
| Mb21d2_D3Z742_S439s_2 | 0.14 | 0 | -0.08 | -0.93 | -0.01 | 0.12 | -0.18 | -0.52 | 0.02 | 0.63 | 0 | -0.38 | -0.04 | 0.24 | -1.17 | -0.36 | 0.07 | 0.285 | -0.115 | 0.778666 | 0.82903 | 0.430564 | 1 | 0.999631 | 0.921416 |
| Mb21d2_D3Z742_S439s_3 | -0.59 | -0.81 | -2.34 | 0.35 | 1.35 | -1.02 | -0.6 | 0.19 | 0.43 | -0.87 | 1.27 | 0.31 | 0.36 | 0.23 | 0.24 | -0.23 | 0.8275 | 1.1325 | 0.9975 | 0.08718 | 0.240457 | 0.102054 | 1 | 0.999631 | 0.553514 |
| Mb21d2_D3Z742_S442s_2 | -0.32 | 0.04 | 0.59 | -1.5 | -0.38 | -0.55 | -0.35 | -0.7 | -0.43 | 0.02 | 0.05 | -0.14 | 0.04 | 0.16 | -0.53 | -0.19 | -0.1975 | 0.1725 | 0.1675 | 0.71741 | 0.653899 | 0.701077 | 1 | 0.999631 | 0.903387 |
| Mbp_P04370_S139s_1 | -1.04 | -0.57 | -0.82 | -1.33 | -0.75 | -0.61 | -1.57 | -1.03 | -0.66 | -0.68 | -1.3 | -1.09 | 0.15 | -0.1 | -0.43 | -0.73 | -0.05 | 0.0075 | 0.6625 | 0.066007 | 0.874612 | 0.980109 | 1 | 0.999631 | 0.516933 |
| Mbp_P04370_S145s_1 | -4.06 | -1.82 | -1.93 | -2.81 | -1.24 | -2.44 | -2.24 | -2.56 | -1.36 | -4.02 | -1.61 | -1.65 | -0.2 | -1.97 | -1.84 | -3.31 | 0.535 | 0.495 | 0.825 | 0.270799 | 0.34271 | 0.496766 | 1 | 0.999631 | 0.698674 |
| Mbp_P04370_S151s_1 | -1.78 | -2.99 | -2.02 | -2.3 | -3.9 | -0.88 | -1.81 | -2.3 | -4.09 | -2.49 | -2 | -3.97 | -1.06 | -1.06 | -2.56 | -3.53 | 0.05 | -0.865 | 0.22 | 0.715326 | 0.935209 | 0.136875 | 1 | 0.999631 | 0.902377 |
| Mbp_P04370_S172s_1 | -1.01 | 0.43 | -1.1 | -0.85 | -0.06 | -0.08 | -1.01 | -1.46 | 0.14 | -0.73 | -0.52 | -0.67 | 0.23 | 0.33 | -1.01 | -0.87 | -0.02 | 0.1875 | 0.3025 | 0.538137 | 0.966377 | 0.653336 | 1 | 0.999631 | 0.830863 |
| Mbp_P04370_S176s_1 | -0.49 | -0.65 | -0.32 | -0.42 | 0.17 | -0.63 | -0.86 | -0.43 | -0.3 | -0.18 | -0.7 | -0.83 | 1.28 | -0.31 | -0.16 | -0.3 | 0.0325 | -0.0325 | 0.5975 | 0.167499 | 0.912812 | 0.905736 | 1 | 0.999631 | 0.637399 |
| Mbp_P04370_S201s_1 | -0.41 | -0.53 | -0.5 | -0.68 | -0.57 | -0.62 | -0.87 | -1.65 | -0.88 | 0.04 | -0.89 | -0.46 | 0.06 | -0.16 | -0.76 | -1.44 | -0.3975 | -0.0175 | -0.045 | 0.905305 | 0.220176 | 0.953718 | 1 | 0.999631 | 0.969056 |
| Mbp_P04370_S243s_1 | 0.16 | -0.27 | 0.34 | 0.29 | -1.04 | 0.2 | 0.31 | -0.82 | -0.11 | 1.14 | -0.75 | -0.49 | -0.17 | -0.97 | 0.42 | -1.48 | -0.4675 | -0.1825 | -0.68 | 0.149183 | 0.241708 | 0.677686 | 1 | 0.999631 | 0.621356 |
| Mbp_P04370_S245s_1 | 0.67 | -0.4 | 0.53 | 0.2 | -0.62 | 0.13 | 0.23 | -0.9 | -0.67 | 1.12 | -0.88 | -0.21 | -0.62 | -1.3 | 0.02 | -1.38 | -0.54 | -0.41 | -1.07 | 0.026802 | 0.177098 | 0.407882 | 1 | 0.999631 | 0.422285 |
| Mbp_P04370_S66s_1 | -0.11 | -1.06 | -0.81 | -1.33 | -1.32 | -0.95 | -0.68 | -1.23 | -0.94 | 0.05 | -0.75 | -0.84 | -0.44 | -1.2 | -0.86 | -0.92 | -0.2175 | 0.2075 | -0.0275 | 0.938573 | 0.522533 | 0.580467 | 1 | 0.999631 | 0.980156 |
| Mbp_P04370_S70s_1 | 0.08 | -1.94 | -1.71 | -2.45 | -2.95 | -1.77 | -1.92 | -1.92 | -2.86 | -0.32 | -1.4 | -3.61 | -1.67 | -1.66 | -6.87 | -4.43 | -0.635 | -0.5425 | -2.1525 | 0.087906 | 0.274267 | 0.506169 | 1 | 0.999631 | 0.553787 |
| Mbp_P04370_T122t_1 | 0.91 | -1.54 | -1.43 | -1.08 | -2.14 | -1.46 | -0.58 | -2.22 | -2.23 | -0.15 | -2.05 | -1.54 | -1.12 | -1.09 | -1.03 | -1.31 | -0.815 | -0.7075 | -0.3525 | 0.517989 | 0.208208 | 0.302047 | 1 | 0.999631 | 0.819449 |
| Mbp_P04370_T149t_1 | -0.29 | -2 | -1.27 | -1.16 | -2 | -0.84 | -0.74 | -1.99 | -2.26 | -0.91 | -1.45 | -2.07 | -1.46 | -1.21 | -2.06 | -1.75 | -0.2125 | -0.4925 | -0.44 | 0.300615 | 0.653327 | 0.294473 | 1 | 0.999631 | 0.71369 |
| Mbp_P04370_T152t_1 | 0.41 | -1.9 | -1.69 | -2.53 | -2.55 | -0.83 | -1.86 | -2.48 | -2.77 | -0.31 | -1.57 | -4.69 | -1.34 | -1.11 | -3.33 | -3.05 | -0.5025 | -0.9075 | -0.78 | 0.31337 | 0.459295 | 0.363887 | 1 | 0.999631 | 0.719861 |
| Mbp_P04370_T167t_1 | -1.41 | 0.73 | 1.16 | -1.03 | 1.6 | 0.45 | 0.76 | -0.67 | 2.97 | -0.46 | 1.45 | 2.48 | 3.6 | -0.5 | -0.14 | -0.37 | 0.6725 | 1.7475 | 0.785 | 0.441336 | 0.348181 | 0.066988 | 1 | 0.999631 | 0.781983 |
| Mbp_P04370_T174t_1 | 0.06 | 0.09 | -0.14 | -0.72 | -0.14 | -0.04 | -0.96 | -0.34 | 0.12 | 0.08 | -0.31 | -0.44 | -0.15 | -0.11 | -0.07 | -0.15 | -0.1925 | 0.04 | 0.0575 | 0.846912 | 0.5563 | 0.895822 | 1 | 0.999631 | 0.948838 |
| Mbp_P04370_T211t_1 | -0.38 | -0.53 | -0.12 | -1.57 | -1.12 | -1.23 | 2.08 | -0.68 | -1.19 | -0.15 | -2.91 | -1.53 | -0.91 | -0.95 | 0.18 | -0.87 | 0.4125 | -0.795 | 0.0125 | 0.976702 | 0.581234 | 0.202604 | 1 | 0.999631 | 0.992016 |
| Mbp_P04370_T229t_1 | 1.5 | -1.21 | -0.58 | -0.97 | -2.09 | -0.62 | 0.03 | -1.84 | -2.46 | 0.36 | -2.04 | -1.63 | -1.02 | -1.66 | -1.17 | -1.78 | -0.815 | -1.1275 | -1.0925 | 0.088633 | 0.264397 | 0.168036 | 1 | 0.999631 | 0.553858 |
| Mbp_P04370_Y146y_1 | -0.11 | 1.58 | -1.49 | -1.1 | 0.21 | 0.41 | -1.07 | -1.93 | -0.19 | 0.15 | -1.26 | -2.36 | 1.29 | 1.25 | -1.52 | -1.46 | -0.315 | -0.635 | 0.17 | 0.851001 | 0.683958 | 0.425145 | 1 | 0.999631 | 0.94989 |
| Mbp_P04370_Y169y_1 | -1.04 | -1.82 | -3.45 | -1.22 | -1.26 | -1.44 | -1.41 | -2.28 | -0.95 | -1.26 | -2.38 | -2.5 | -2.21 | -0.37 | -2.74 | -2.16 | 0.285 | 0.11 | 0.0125 | 0.985155 | 0.604654 | 0.857298 | 1 | 0.999631 | 0.995133 |
| Mcf2l_E9PXE2_S1146s_1 | 1.47 | 0.25 | 0.52 | 1.51 | 0.61 | 1.77 | 1.66 | 1.2 | 1.36 | 2.06 | 1.12 | -0.09 | 0.08 | 1.46 | 1.61 | -0.71 | 0.3725 | 0.175 | -0.3275 | 0.582098 | 0.380208 | 0.736583 | 1 | 0.999631 | 0.847567 |
| Mcf2l_E9PXE2_S558s_1 | 0.49 | -1.03 | -1.04 | -0.56 | 0.16 | -0.73 | 0.51 | -1.32 | -0.37 | -0.55 | -0.49 | -1 | -1.78 | -2.37 | -1.36 | -2.39 | 0.19 | -0.0675 | -1.44 | 0.00801 | 0.711855 | 0.865391 | 1 | 0.999631 | 0.364291 |
| Mcf2l_E9PXE2_S936s_1 | 0.01 | 0.27 | -0.12 | 0.52 | -0.09 | -0.05 | -0.2 | 0.52 | -1.39 | 0.52 | 0.88 | 0.97 | 0.05 | 0.22 | 1.2 | 0.29 | -0.125 | 0.075 | 0.27 | 0.448909 | 0.666031 | 0.888129 | 1 | 0.999631 | 0.786796 |
| Mcf2l_E9PXE2_S946s_1 | -1.99 | -2.78 | -2.37 | -2.83 | -2.21 | -2.8 | -2.31 | -2.89 | -2.24 | -1.25 | -2.53 | -2.8 | -0.46 | -3.1 | -1.96 | -2.45 | -0.06 | 0.2875 | 0.5 | 0.376056 | 0.848201 | 0.481002 | 1 | 0.999631 | 0.748307 |
| Mchr1_Q8JZL2_S325s_1 | -0.89 | 0.93 | 0.91 | 1.34 | 0.18 | 1.29 | 0.49 | 0.74 | 0.28 | -0.82 | 1.65 | -0.23 | 1.36 | 1.75 | 0.11 | 0.22 | 0.1025 | -0.3525 | 0.2875 | 0.627791 | 0.841536 | 0.591258 | 1 | 0.999631 | 0.865919 |
| Mcm2_P97310_S40s_1 | 0.27 | 0.98 | 0.59 | 0.11 | 0.45 | 0.23 | 0.73 | 0.17 | 0.8 | 0.82 | -0.07 | 0.04 | 0.8 | 0.86 | 1.46 | 0.54 | -0.0925 | -0.09 | 0.4275 | 0.226241 | 0.75539 | 0.795881 | 1 | 0.999631 | 0.677133 |
| Mcmbp_H9KV04_S154s_1 | 0.45 | 0.18 | 1.31 | -0.21 | 0.06 | 0.52 | -0.13 | 0.81 | 0.45 | 0.88 | 0.3 | 0.72 | 0.2 | 0.84 | 1.3 | 0.67 | -0.1175 | 0.155 | 0.32 | 0.444466 | 0.765877 | 0.679364 | 1 | 0.999631 | 0.78327 |
| Mcoln1_Q99J21_S547s_1 | -0.74 | 0.08 | -0.03 | -0.26 | -0.04 | -0.05 | -0.56 | -0.19 | -0.25 | -0.01 | 0.02 | 0.37 | 0.56 | 0.2 | 0.1 | 0.28 | 0.0275 | 0.27 | 0.5225 | 0.111602 | 0.92447 | 0.3773 | 1 | 0.999631 | 0.577068 |
| Mcoln1_Q99J21_T34t_1 | -0.78 | -0.8 | -0.91 | -1.45 | -0.23 | -0.78 | -0.99 | -0.72 | -0.83 | -0.57 | -0.9 | -0.59 | 0.05 | -0.73 | -0.99 | -1.07 | 0.305 | 0.2625 | 0.3 | 0.405833 | 0.31276 | 0.356557 | 1 | 0.999631 | 0.765969 |
| Mctp1_E9PV86_S56s_1 | -0.55 | -0.39 | -0.87 | -1.1 | -0.33 | -0.2 | -1.35 | -1.08 | -0.19 | 0.06 | -0.78 | -0.23 | 0.94 | -0.48 | -0.64 | -0.44 | -0.0125 | 0.4425 | 0.5725 | 0.189703 | 0.971605 | 0.172796 | 1 | 0.999631 | 0.648674 |
| Mctp1_E9PV86_S941s_1 | -0.18 | -0.52 | 0.43 | 0.25 | -0.25 | -0.38 | 0.04 | 0.68 | -0.49 | 0.12 | 0.24 | 1.09 | -0.27 | 0.01 | 0.95 | 1.12 | 0.0275 | 0.245 | 0.4575 | 0.289146 | 0.937104 | 0.545376 | 1 | 0.999631 | 0.706127 |
| Mctp1_E9PV86_T145t_1 | -0.38 | -1.11 | -0.7 | -0.28 | -0.63 | -0.64 | -0.2 | -0.26 | -0.5 | -0.27 | -0.32 | -0.45 | -0.68 | -0.27 | -0.17 | -0.2 | 0.185 | 0.2325 | 0.2875 | 0.368189 | 0.529802 | 0.42369 | 1 | 0.999631 | 0.745968 |
| Mctp1_E9PYR7_S174s_1 | -1.15 | -0.64 | -0.41 | 0.01 | -1.44 | -0.89 | -0.16 | 0.42 | -1.13 | -0.72 | 1.01 | 0.85 | -1.17 | -0.47 | 0.45 | 0.35 | 0.03 | 0.55 | 0.3375 | 0.459641 | 0.947517 | 0.330842 | 1 | 0.999631 | 0.791604 |
| Mdc1_E9QK89_T624t_1 | 3.26 | 0.63 | 1.45 | 1.63 | 1.05 | 1 | 1.99 | 0.76 | 1.1 | 2.54 | 0.8 | 0.4 | -0.27 | 0.7 | 1.24 | -0.13 | -0.5425 | -0.5325 | -1.3575 | 0.043725 | 0.345607 | 0.420267 | 1 | 0.999631 | 0.475443 |
| Mdh1_P14152_S241s_1 | -0.09 | -2.51 | -0.84 | -1.39 | -2.51 | -2.32 | -0.71 | -2.03 | -2.11 | -1.39 | -0.84 | -1.67 | -1.89 | -2.05 | -2.18 | -0.69 | -0.685 | -0.295 | -0.495 | 0.391966 | 0.262858 | 0.584762 | 1 | 0.999631 | 0.758901 |
| Mdh1_P14152_S242s_1 | -0.35 | 1.07 | 0.21 | 0.32 | 0.21 | 1.06 | 0.12 | -0.22 | 1.59 | 1.36 | 0.36 | 1.33 | 1.4 | 0.96 | 0.57 | 0.38 | -0.02 | 0.8475 | 0.515 | 0.21272 | 0.960323 | 0.06134 | 1 | 0.999631 | 0.66658 |
| Mdn1_J3QMC5_S4822s_1 | -0.06 | 2.29 | 1.36 | 1.86 | 1.14 | 1.59 | 0.94 | 1.94 | 2.22 | 0.45 | 2.36 | 3.1 | 1.72 | 2.17 | 1.14 | 1.96 | 0.04 | 0.67 | 0.385 | 0.46968 | 0.938477 | 0.336214 | 1 | 0.999631 | 0.796991 |
| Mecp2_Q9Z2D6_S13s_1 | -0.68 | 1.75 | 2.21 | 1.43 | -0.21 | 1.31 | 1.46 | 0.89 | 0.5 | -0.55 | 1.17 | 1.76 | 1.35 | 1.01 | 1.49 | 1.95 | -0.315 | -0.4575 | 0.2725 | 0.655358 | 0.635807 | 0.528459 | 1 | 0.999631 | 0.879056 |
| Mecp2_Q9Z2D6_S229s_1 | -0.34 | -0.79 | 0.08 | 0.06 | -1.25 | -1.01 | -0.64 | 0.1 | -1.63 | -0.51 | 0.35 | 0.71 | -0.07 | -1.31 | 0.22 | -0.26 | -0.4525 | -0.0225 | -0.1075 | 0.793399 | 0.245524 | 0.965738 | 1 | 0.999631 | 0.928057 |
| Mecp2_Q9Z2D6_S421s_1 | 0.07 | 0.85 | -0.31 | -0.23 | 0.16 | 1.14 | -0.22 | -0.55 | 0.97 | 0.71 | -0.95 | -0.75 | 0.66 | 0.89 | -0.18 | -0.58 | 0.0375 | -0.1 | 0.1025 | 0.815773 | 0.931971 | 0.848645 | 1 | 0.999631 | 0.937576 |
| Mecp2_Q9Z2D6_S80s_1 | -0.91 | -1.12 | 0.05 | -0.3 | -1.04 | -0.99 | -0.81 | -0.86 | -1.26 | -1.13 | -0.11 | -1.55 | -0.49 | -0.63 | -0.54 | -0.45 | -0.355 | -0.4425 | 0.0425 | 0.900185 | 0.28578 | 0.304271 | 1 | 0.999631 | 0.967733 |
| Med1_Q925J9_T1036t_1 | 1.21 | 2.95 | 1.24 | -0.19 | 2.12 | 2.55 | 0.15 | 0.55 | 2.29 | 1.24 | 0.83 | 0.7 | 2.15 | 2.63 | 0.59 | 1.37 | 0.04 | -0.0375 | 0.3825 | 0.585002 | 0.958115 | 0.954721 | 1 | 0.999631 | 0.849277 |
| Med15_E9Q7C1_T564t_1 | -1.16 | 0.66 | 1.02 | 0.96 | 0.36 | 0.31 | 0.38 | 0.76 | -0.41 | 0.59 | 0.85 | 1.24 | 2.22 | 0.75 | 1.48 | 1.12 | 0.0825 | 0.1975 | 1.0225 | 0.091926 | 0.867539 | 0.731384 | 1 | 0.999631 | 0.557085 |
| Med26_Q7TN02_S435s_1 | -0.23 | 1.09 | 1.49 | 0.63 | 0.69 | 0.57 | 0.06 | 0.62 | 0.88 | 0.43 | 1.82 | 0.98 | 2.18 | 1.12 | 1.88 | 1.42 | -0.26 | 0.2825 | 0.905 | 0.063232 | 0.520825 | 0.541273 | 1 | 0.999631 | 0.512978 |
| Med7_Q9CZB6_S195s_1 | 0.68 | 0.95 | 1.04 | -0.07 | 1.63 | 1.97 | 0.12 | -0.18 | 0.44 | 2.04 | 0.35 | 0.84 | 0.01 | 1.68 | 0.92 | 0.71 | 0.235 | 0.2675 | 0.18 | 0.678833 | 0.66759 | 0.558744 | 1 | 0.999631 | 0.887183 |
| Mef2a_Q60929_S255s_1 | 0.32 | 0.91 | 1.22 | 1.29 | 0.92 | 0.16 | 0.2 | 1.29 | 0.81 | 0.9 | 0.68 | 1.89 | 0.47 | 0.92 | 1.95 | 1.39 | -0.2925 | 0.135 | 0.2475 | 0.546928 | 0.43948 | 0.721574 | 1 | 0.999631 | 0.834213 |
| Mef2a_Q60929_S400s_1 | 1.23 | 2.09 | 2.53 | 1.23 | 1.21 | 1.72 | 1.11 | 2.09 | 1.87 | 1.52 | 1.68 | 2.59 | 1.7 | 2.4 | 1.8 | 1.81 | -0.2375 | 0.145 | 0.1575 | 0.688203 | 0.557168 | 0.723904 | 1 | 0.999631 | 0.891814 |
| Mef2c_Q8CFN5_S222s_1 | 0.03 | 1.12 | 1.76 | 1.25 | 0.91 | 1.03 | 1.06 | 1.5 | 1.25 | 0.75 | 1.8 | 2.1 | 1.1 | 1.48 | 1.55 | 2.16 | 0.085 | 0.435 | 0.5325 | 0.236053 | 0.828993 | 0.354699 | 1 | 0.999631 | 0.683301 |
| Mef2c_Q8CFN5_S240s_1 | 0.47 | 0.69 | 0.9 | 1.09 | 1.06 | 0.36 | 0.75 | 1.19 | 0.6 | 0.61 | 1.23 | 0.95 | 1.22 | 0.77 | 0.88 | 1.77 | 0.0525 | 0.06 | 0.3725 | 0.278556 | 0.85883 | 0.835597 | 1 | 0.999631 | 0.70098 |
| Mef2c_Q8CFN5_S396s_1 | 1.04 | 1.98 | 1.82 | 1.56 | 1.19 | 1.34 | 0.92 | 1.65 | 1.32 | 1.45 | 2.01 | 1.85 | 1.19 | 1.79 | 1.47 | 2.18 | -0.325 | 0.0575 | 0.0575 | 0.870204 | 0.309544 | 0.857915 | 1 | 0.999631 | 0.958461 |
| Mef2c_Q8CFN5_S446s_1 | -0.56 | 2.97 | 4.55 | 2.49 | 2.02 | 2.36 | 2.86 | 3.78 | 2.47 | 0.14 | 4.6 | 3.86 | 1.46 | 2.31 | 4.78 | 3.8 | 0.3925 | 0.405 | 0.725 | 0.516804 | 0.689203 | 0.744563 | 1 | 0.999631 | 0.819449 |
| Mef2d_E9QKT0_S250s_1 | 1.01 | -1.7 | -0.96 | 0.08 | -0.79 | -1.07 | 0.16 | -0.48 | -1.16 | -1.1 | -0.87 | -1.25 | -1.31 | -1.25 | -0.88 | -0.5 | -0.1525 | -0.7025 | -0.5925 | 0.314532 | 0.795732 | 0.224231 | 1 | 0.999631 | 0.720156 |
| Mef2d_E9QKT0_S429s_1 | 2.18 | 1.55 | 1.11 | 1.61 | 1.67 | 1.69 | 1.58 | 1.39 | 1.55 | 1.88 | 1.4 | 0.94 | 0.18 | 1.33 | 1.19 | 0.77 | -0.03 | -0.17 | -0.745 | 0.071659 | 0.919286 | 0.618858 | 1 | 0.999631 | 0.532005 |
| Mef2d_E9QKT0_S435s_1 | 0.93 | 0.12 | -0.28 | -0.38 | 0.9 | 0.22 | -0.06 | 0.23 | 0.69 | -0.3 | -0.19 | -0.29 | 0.16 | -0.01 | -0.8 | -0.03 | 0.225 | -0.12 | -0.2675 | 0.50266 | 0.553509 | 0.762254 | 1 | 0.999631 | 0.814092 |
| Men1_F7CJP6_S41s_1 | -0.71 | -0.62 | -0.43 | -1.74 | 0.14 | -0.81 | -0.95 | -0.91 | -0.05 | -0.19 | -1.39 | -1.39 | 1.24 | 0.15 | -1.18 | -0.62 | 0.2425 | 0.12 | 0.7725 | 0.186829 | 0.545683 | 0.793868 | 1 | 0.999631 | 0.646474 |
| Mepce_Q8K3A9_S192s_1 | 1.02 | 2.25 | 3 | 1.36 | 0.63 | 1.97 | 1.97 | 1.31 | 2.78 | 2.28 | 2.45 | 2.72 | 1.51 | 1.31 | 2.41 | 2.68 | -0.4375 | 0.65 | 0.07 | 0.894115 | 0.402453 | 0.173386 | 1 | 0.999631 | 0.965259 |
| Mepce_Q8K3A9_T188t_1 | 0.56 | 1.3 | 1.77 | 0.93 | -0.12 | 0.99 | 1.18 | 0.41 | 0.98 | 1.7 | 1.57 | 1.46 | 0.48 | 0.77 | 1.35 | 1.28 | -0.525 | 0.2875 | -0.17 | 0.650189 | 0.207023 | 0.413264 | 1 | 0.999631 | 0.875535 |
| Met_F8VQL0_S988s_1 | -1.6 | 0.41 | -0.67 | -1.22 | -0.09 | -0.29 | -1.29 | -0.84 | 0.53 | 0.3 | -0.41 | -0.88 | -1.63 | 0.07 | -0.89 | -0.05 | 0.1425 | 0.655 | 0.145 | 0.792171 | 0.769772 | 0.221197 | 1 | 0.999631 | 0.927114 |
| Mettl22_D3Z0D7_S83s_1 | 0.67 | 0.85 | 0.63 | 0.4 | 0.4 | 0.36 | 0.7 | 0.46 | 0.6 | 1.14 | 0.99 | 0.95 | 0.42 | 0.81 | 0.59 | 0.71 | -0.1575 | 0.2825 | -0.005 | 0.98543 | 0.527842 | 0.300802 | 1 | 0.999631 | 0.995133 |
| Mex3c_Q05A36_S530s_1 | 2.18 | 0.69 | 1.79 | 1.91 | 1.53 | 2.24 | 1.9 | 1.33 | 1.25 | 2.37 | 1.2 | 0.43 | 0.69 | -0.27 | 1.38 | 0.64 | 0.1075 | -0.33 | -1.0325 | 0.047122 | 0.784411 | 0.507679 | 1 | 0.999631 | 0.480102 |
| Mff_Q6PCP5_S129s_1 | 1.72 | 0.94 | 0.32 | 1.32 | 1.58 | 1.49 | 1.65 | 1.59 | 1.75 | 1.08 | 1.61 | 1.58 | 0.1 | 0.34 | 0.91 | 0.87 | 0.5025 | 0.43 | -0.52 | 0.20166 | 0.158734 | 0.253962 | 1 | 0.999631 | 0.658952 |
| Mff_Q6PCP5_S129s_2 | 1.06 | -1.45 | -2.91 | 1.02 | 1.93 | 0.39 | 1.2 | 1.16 | 0.51 | -1.65 | 0.37 | -1.67 | -2.66 | -0.86 | -1.14 | -2.53 | 1.74 | -0.04 | -1.2275 | 0.206532 | 0.07456 | 0.967838 | 1 | 0.999631 | 0.6605 |
| Mff_Q6PCP5_S131s_1 | 0.91 | 0.8 | 0.57 | 0.4 | 0.11 | 1.09 | -0.26 | 0.82 | 1.27 | 1.51 | 1.06 | 0.95 | 0.5 | 0.32 | 1.24 | 0.48 | -0.23 | 0.5275 | -0.035 | 0.9127 | 0.524237 | 0.078353 | 1 | 0.999631 | 0.972191 |
| Mff_Q6PCP5_S131s_2 | 1.06 | -1.45 | -2.91 | 1.02 | 1.93 | 0.39 | 1.2 | 1.16 | 0.51 | -1.65 | 0.37 | -1.67 | -2.66 | -0.86 | -1.14 | -2.53 | 1.74 | -0.04 | -1.2275 | 0.206532 | 0.07456 | 0.967838 | 1 | 0.999631 | 0.6605 |
| Mff_Q6PCP5_S146s_1 | -0.73 | 3.03 | 1.24 | 0.67 | 2.41 | 2.2 | 2.15 | 2.59 | 1.48 | -1.29 | 3.16 | 1.38 | 2.05 | 0.78 | 1.22 | 0.52 | 1.285 | 0.13 | 0.09 | 0.903789 | 0.089391 | 0.90031 | 1 | 0.999631 | 0.968183 |
| Mff_Q6PCP5_S146s_1_1 | -1.72 | 0.94 | 1.04 | 0.22 | 1.95 | 1 | 1.45 | 1.76 | 1.65 | -1.61 | 1.85 | 2.18 | 0.96 | 0.48 | 0.9 | 1.09 | 1.42 | 0.8975 | 0.7375 | 0.234464 | 0.039253 | 0.353758 | 1 | 0.999631 | 0.683301 |
| Mff_Q6PCP5_S151s_1 | -2.85 | -1.24 | 0.27 | -1.11 | -3.3 | -2.52 | -0.7 | -0.82 | -2.66 | -0.98 | -1.24 | -0.28 | 2.06 | -2.03 | -0.19 | -0.75 | -0.6025 | -0.0575 | 1.005 | 0.288345 | 0.45312 | 0.936244 | 1 | 0.999631 | 0.706035 |
| Mff_Q6PCP5_S151s_2 | -2.76 | -1.56 | -1.7 | -2.74 | -1.62 | -1.68 | -2.82 | -1.82 | -1.32 | -2 | -1.68 | -1.86 | -0.25 | -1.53 | -1.69 | -1.79 | 0.205 | 0.475 | 0.875 | 0.088605 | 0.630632 | 0.229234 | 1 | 0.999631 | 0.553858 |

| | | | | | | | | | | | | | | | | | | | | | | | | | |
|---|---|---|---|---|---|---|---|---|---|---|---|---|---|---|---|---|---|---|---|---|---|---|---|---|---|
| Mff_Q6PCP5_T149t_2 | -2.57 | -1.22 | -1.7 | -2.74 | -1.14 | -1.31 | -2.82 | -1.82 | -1.08 | -2.03 | -1.68 | -1.86 | 0.01 | -1.28 | -1.69 | -1.79 | 0.285 | 0.395 | 0.87 | 0.118023 | 0.565433 | 0.35757 | 1 | 0.999631 | 0.585087 |
| Mfsd5_Q921Y4_S440s_1 | -0.09 | 1.45 | -0.3 | -0.12 | 0.42 | -0.55 | -0.24 | 0.3 | -0.36 | -0.67 | -0.86 | 0.94 | -2.17 | -0.93 | -0.92 | 0.89 | -0.2525 | -0.4725 | -1.0175 | 0.151819 | 0.578675 | 0.388051 | 1 | 0.999631 | 0.625198 |
| Mfsd6_D3Z183_S646s_1 | -0.63 | -0.04 | 0.08 | -0.22 | -0.53 | -0.81 | -0.55 | -0.45 | -0.62 | -1.04 | -0.58 | -0.08 | 0.45 | -0.3 | 0.15 | 0.24 | -0.3825 | -0.3775 | 0.3375 | 0.294833 | 0.176052 | 0.247727 | 1 | 0.999631 | 0.708705 |
| Mfsd6_D3Z183_S647s_1 | -0.61 | -0.03 | -0.42 | -0.22 | -0.23 | -0.39 | -0.56 | -0.62 | -0.47 | -0.2 | -0.3 | -0.54 | 0.13 | -0.04 | -0.33 | -0.15 | -0.13 | -0.0575 | 0.2225 | 0.442795 | 0.618368 | 0.828337 | 1 | 0.999631 | 0.782367 |
| Mfsd6_D3Z183_S736s_1 | 2.13 | -1.13 | -0.14 | -0.1 | -1.41 | 0.01 | 0.81 | -1.58 | -1.18 | 0.75 | -1.37 | -0.91 | -0.46 | -0.97 | -0.99 | -1.51 | -0.7325 | -0.8675 | -1.1725 | 0.094782 | 0.361866 | 0.261386 | 1 | 0.999631 | 0.558128 |
| Mfsd8_Q8BH31_S18s_1 | 0.01 | 0.38 | 1.33 | 0.8 | 0.22 | 0.58 | 0.51 | 0.74 | 0.39 | 0.98 | 0.77 | 0.96 | 1.01 | 1.02 | 0.58 | 1.52 | -0.1175 | 0.145 | 0.4025 | 0.302789 | 0.729827 | 0.682079 | 1 | 0.999631 | 0.714978 |
| Mga_H7BX50_S287s_1 | 0.04 | -1.31 | 0.27 | -1.03 | -1.64 | -0.86 | 0.44 | -0.64 | -1.33 | -0.63 | -1.57 | -1.25 | -2.41 | -3.37 | 0.11 | -2.41 | -0.1675 | -0.6875 | -1.5125 | 0.064023 | 0.754633 | 0.138434 | 1 | 0.999631 | 0.513532 |
| Mgea5_Q9EQQ9_T709t_1 | 0.11 | 0.69 | -0.06 | 0.58 | 0.36 | 0.49 | -0.09 | 0.13 | 0.46 | 0.18 | 0.05 | 0.87 | 0.72 | 1.25 | 0.35 | 0.92 | -0.1075 | 0.06 | 0.48 | 0.170859 | 0.713341 | 0.850199 | 1 | 0.999631 | 0.641032 |
| Mgll_E9Q3B9_S205s_1 | -0.6 | 0.66 | 0.72 | 0.28 | -0.06 | 0.48 | -0.29 | 0.21 | 0.11 | -0.63 | 0.05 | 0.87 | 0.88 | 0.65 | 0.94 | 0.65 | -0.18 | -0.165 | 0.515 | 0.176472 | 0.625888 | 0.702745 | 1 | 0.999631 | 0.642116 |
| Mgll_E9Q3B9_S22s_1 | 1.28 | 0.28 | 0.51 | 0.3 | -0.01 | 0.38 | 0.51 | 0.26 | 0.12 | 0.78 | 0.08 | -0.21 | -0.69 | -0.48 | -0.51 | -0.45 | -0.3075 | -0.4 | -1.125 | 0.005487 | 0.338137 | 0.272599 | 1 | 0.999631 | 0.351308 |
| Mgrn1_Q9D074_S106s_1 | 1.51 | 1.77 | 2.56 | 1.01 | 0.34 | 2.05 | 1.14 | 1.09 | 1.62 | 2.6 | 0.58 | 1.41 | 0.45 | 0.87 | 1.19 | 1.17 | -0.5575 | -0.16 | -0.7925 | 0.067693 | 0.241749 | 0.749186 | 1 | 0.999631 | 0.521762 |
| Mgrn1_Q9D074_S428s_1 | 0.3 | -1.02 | 0.06 | 0.05 | -0.88 | -0.45 | 0.4 | -0.39 | -0.61 | 0.36 | -0.31 | -0.21 | -0.57 | -0.52 | -0.14 | -0.2 | -0.1775 | -0.04 | -0.205 | 0.57485 | 0.659756 | 0.916138 | 1 | 0.999631 | 0.845659 |
| Mgrn1_Q9D074_S437s_1 | -0.26 | -0.35 | 0.34 | 1.71 | 0 | 0.15 | 1.04 | 2.13 | 0.49 | 0.68 | 1.66 | 2.49 | -1.37 | 0.98 | 1.58 | 2.31 | 0.47 | 0.97 | 0.515 | 0.527887 | 0.45156 | 0.132963 | 1 | 0.999631 | 0.824683 |
| Mia3_Q8BI84_S1733s_1 | -0.86 | -0.28 | -0.23 | 0.3 | 0.04 | 0.2 | 0.83 | 0.3 | 0.17 | -0.71 | 1.45 | 0.84 | -0.7 | 0.61 | 0.33 | 0.87 | 0.61 | 0.705 | 0.545 | 0.221321 | 0.091154 | 0.176823 | 1 | 0.999631 | 0.671697 |
| Mia3_Q8BI84_S1915s_1 | 1.06 | -0.4 | 1.36 | 0.72 | 0.21 | 1.24 | 0.52 | 0.56 | 0.53 | 1.9 | -0.19 | 0.05 | -1.01 | 0.96 | 0.62 | 0.47 | -0.0525 | -0.1125 | -0.425 | 0.440777 | 0.903334 | 0.840488 | 1 | 0.999631 | 0.781983 |
| Mia3_Q8BI84_S820s_1 | -0.22 | 0.27 | 0.13 | -2.15 | -0.1 | -0.23 | -0.35 | -0.46 | 0.26 | -0.81 | -0.76 | 0.64 | 0.44 | 0.39 | 0.3 | 0.71 | 0.2075 | 0.325 | 0.9525 | 0.098747 | 0.69404 | 0.596211 | 1 | 0.999631 | 0.562298 |
| Mib2_Q8R516_S251s_1 | -0.01 | -0.46 | -0.5 | -0.13 | -0.8 | -0.57 | -0.83 | -0.14 | -0.3 | 0.15 | -0.09 | -0.18 | -0.89 | -0.14 | -0.45 | -0.03 | -0.31 | 0.17 | -0.1025 | 0.745803 | 0.284974 | 0.529757 | 1 | 0.999631 | 0.910993 |
| Mical1_E9PUI4_S777s_1 | 0.03 | -0.08 | 0.03 | 0.03 | -0.17 | -0.25 | -0.64 | -0.53 | -0.43 | 0.13 | -0.5 | -0.12 | 0.56 | -0.17 | -0.18 | 0.42 | -0.4 | -0.2325 | 0.155 | 0.608782 | 0.12731 | 0.390691 | 1 | 0.999631 | 0.858851 |
| Mical2_Q8BML1_S965s_1 | -0.34 | -0.89 | 0.8 | -0.03 | -0.71 | 0.56 | -0.16 | -0.53 | -0.88 | -0.82 | -0.32 | 0.17 | -1.24 | 0.27 | -0.55 | -0.05 | -0.095 | -0.3475 | -0.2775 | 0.559969 | 0.82891 | 0.426563 | 1 | 0.999631 | 0.839805 |
| Mical3_Q8CJ19_S1131s_1 | 0.83 | 1.72 | 0.7 | -0.25 | 0.86 | 2.2 | -0.56 | -0.61 | 2.18 | 2.07 | -0.76 | -0.15 | 1.31 | 0.87 | -0.69 | -0.17 | -0.2775 | 0.085 | -0.42 | 0.463911 | 0.689506 | 0.910402 | 1 | 0.999631 | 0.793747 |
| Mical3_Q8CJ19_S1167s_1 | 0.64 | 0.1 | 0.05 | 0.7 | 0.27 | 0.67 | -0.66 | 0.24 | 0.37 | 0.22 | 0.13 | 0.11 | 0.14 | 0.51 | -0.72 | 0.7 | -0.2425 | -0.165 | -0.215 | 0.583748 | 0.500706 | 0.558969 | 1 | 0.999631 | 0.848861 |
| Mical3_Q8CJ19_S1187s_1 | -0.88 | -0.47 | -0.93 | -1.15 | -0.59 | -0.86 | -1.36 | -1.22 | -0.75 | -0.73 | -1.29 | -0.98 | -0.2 | -0.48 | -1.24 | -0.92 | -0.15 | -0.08 | 0.1475 | 0.663778 | 0.611623 | 0.778961 | 1 | 0.999631 | 0.881991 |
| Mical3_Q8CJ19_S1645s_1 | -1.56 | -2.89 | -2.7 | -2.7 | -1.01 | -3.42 | -1.67 | -3.14 | -2.74 | -2.85 | -2.72 | -2.93 | -2.72 | -2.68 | -3.6 | -3.26 | 0.1525 | -0.3475 | -0.6025 | 0.155202 | 0.796931 | 0.331287 | 1 | 0.999631 | 0.62632 |
| Mical3_Q8CJ19_S1691s_1 | 0.33 | 0.53 | 0.14 | 1.63 | 0.76 | 1.03 | 0.63 | 1.67 | 0.49 | 0.09 | 1.37 | 0.8 | -1.59 | 0.89 | 1.78 | 1.43 | 0.365 | 0.03 | -0.03 | 0.967303 | 0.380407 | 0.944206 | 1 | 0.999631 | 0.988463 |
| Mical3_Q8CJ19_S1692s_1 | -0.02 | 0.53 | 0.14 | 1.63 | 0.41 | 0.82 | 0.63 | 1.67 | 0.28 | -0.17 | 1.37 | 0.8 | -1.3 | 0.71 | 1.78 | 1.43 | 0.3125 | 1.57E-16 | 0.085 | 0.902567 | 0.490842 | 1 | 1 | 1 | 0.968183 |
| Mical3_Q8CJ19_S1695s_1 | -0.15 | -0.37 | -0.31 | 0.76 | -0.39 | -0.17 | -0.06 | 0.35 | -0.73 | -1.6 | 0.52 | 0.62 | -2.12 | -0.23 | 0.46 | 0.12 | -0.05 | -0.28 | -0.425 | 0.471157 | 0.883219 | 0.612963 | 1 | 0.999631 | 0.797981 |
| Mical3_Q8CJ19_S1716s_1 | 0.85 | 2.33 | 1.01 | 0.99 | 1.7 | 1.95 | 0.26 | 1.41 | 1.47 | 1.49 | 1.33 | 1.55 | 2.09 | 2.62 | 1.56 | 1.18 | 0.035 | 0.165 | 0.5675 | 0.237069 | 0.942096 | 0.661317 | 1 | 0.999631 | 0.683301 |
| Mical3_Q8CJ19_S1736s_1 | 1.4 | 1.16 | 0.75 | 1.93 | 1.96 | 2.95 | 1.32 | 2.02 | 2.37 | 1.28 | 1.92 | 1.52 | 0.11 | 2.11 | 2.39 | 2.1 | 0.7525 | 0.4625 | 0.3675 | 0.503567 | 0.094367 | 0.230879 | 1 | 0.999631 | 0.814092 |
| Mical3_Q8CJ19_S1769s_1 | 2.05 | -0.53 | -1.7 | 0.31 | -0.3 | 0.55 | 0.44 | 0.04 | 1 | 1.32 | -1.18 | 0.85 | -0.3 | 0.74 | 0.83 | -0.16 | 0.15 | 0.465 | 0.245 | 0.741384 | 0.83431 | 0.586607 | 1 | 0.999631 | 0.909067 |
| Mical3_Q8CJ19_S685s_1 | 0.46 | -0.66 | 0.04 | 0.08 | -0.37 | -0.52 | 0.27 | -0.14 | -0.13 | 0.35 | -0.17 | -0.61 | -0.71 | -1.41 | -0.3 | -1.01 | -0.17 | -0.12 | -0.8375 | 0.045015 | 0.609718 | 0.729669 | 1 | 0.999631 | 0.475465 |
| Mical3_Q8CJ19_S687s_1 | 0.17 | -1.56 | -0.6 | -0.39 | -0.26 | -0.98 | 0.08 | -0.92 | -0.78 | -0.57 | -0.12 | -0.75 | -1.36 | -1.72 | -1.25 | -1.87 | 0.075 | 0.04 | -0.955 | 0.038762 | 0.86269 | 0.920781 | 1 | 0.999631 | 0.462703 |
| Mical3_Q8CJ19_S867s_1 | 0.51 | 0.02 | 0.09 | 0.55 | 0.82 | 0.17 | 0.29 | 0.19 | 0.18 | -0.05 | 0.6 | 0.38 | -1.13 | -0.33 | 0.45 | 0.25 | 0.075 | -0.015 | -0.4825 | 0.248406 | 0.79178 | 0.958153 | 1 | 0.999631 | 0.685751 |
| Mical3_Q8CJ19_T684t_1 | -0.17 | -0.69 | -0.4 | -0.35 | -1.13 | -0.92 | -0.21 | -0.92 | -0.82 | 0.07 | -0.71 | -1.34 | -1.88 | -1.65 | -0.8 | -1.67 | -0.3925 | -0.2975 | -1.0975 | 0.007611 | 0.204276 | 0.404111 | 1 | 0.999631 | 0.358173 |
| Micall1_Q8BGT6_S628s_1 | 0.55 | 1.54 | 0.13 | 0.39 | 0.76 | 1.5 | 0.29 | 0.81 | 2.03 | 1.08 | -0.4 | 0.11 | 0.41 | 0.52 | -0.13 | 0.64 | 0.1875 | 0.0525 | -0.2925 | 0.454293 | 0.641425 | 0.926587 | 1 | 0.999631 | 0.789679 |
| Micall1_Q8BGT6_S647s_1 | 0.83 | -0.66 | -0.29 | -0.63 | -0.67 | -0.12 | -0.46 | -0.41 | 1.63 | 1.27 | -0.51 | 0.3 | 0.36 | -0.43 | -0.06 | -0.15 | -0.2275 | 0.86 | 0.1175 | 0.772836 | 0.553633 | 0.143708 | 1 | 0.999631 | 0.920172 |
| Micall2_E9PZD2_S67s_1 | 0.37 | 0.26 | 0.35 | -0.19 | -0.34 | 0.29 | 0 | -0.17 | 0.94 | 0.85 | -0.49 | 0.22 | -0.64 | 0.35 | -0.02 | -1.08 | -0.2525 | 0.1825 | -0.545 | 0.173696 | 0.368912 | 0.632172 | 1 | 0.999631 | 0.642116 |
| Mid1_B1AV01_S96s_1 | 0.09 | -0.43 | -2.25 | -3.53 | -1.39 | -0.87 | -2.19 | -2.12 | 0.53 | 0.68 | -1 | -0.23 | -0.5 | -0.81 | -1.7 | -2.52 | -0.1125 | 1.525 | 0.1475 | 0.858297 | 0.885322 | 0.083525 | 1 | 0.999631 | 0.952251 |
| Mid1ip1_Q9CQ20_S71s_1 | -0.68 | -1.19 | -0.68 | -0.78 | -1.21 | -1.31 | -0.77 | -0.46 | -1.19 | -1.33 | -0.57 | -1.18 | 0.49 | -0.56 | -0.29 | -0.83 | -0.105 | -0.235 | 0.535 | 0.159298 | 0.724699 | 0.429267 | 1 | 0.999631 | 0.629514 |
| Mid2_B1AVF6_S96s_1 | -1.42 | -5.25 | 0.08 | 0.02 | -3.07 | -5.66 | 0.12 | 0.5 | -5.21 | -2.92 | 0.45 | 0.2 | -7.03 | -4.28 | 0.36 | 0.06 | -0.385 | -0.2275 | -1.08 | 0.556646 | 0.812526 | 0.884342 | 1 | 0.999631 | 0.838305 |
| Mief1_Q8BGV8_S59s_1 | -0.39 | -0.76 | -0.84 | -0.91 | -1.39 | -1.11 | -0.77 | -1.35 | -0.98 | -0.28 | -1.24 | -1.03 | -0.14 | -0.88 | -1.13 | -0.76 | -0.43 | -0.1575 | -0.0025 | 0.993797 | 0.138871 | 0.611342 | 1 | 0.999631 | 0.997705 |
| Mief1_Q8BGV8_S94s_1 | -0.2 | 0.36 | 0.65 | 0.48 | -0.11 | 0.07 | 0.04 | 0.61 | -0.28 | -0.03 | 0.86 | 1.01 | 0.27 | 0.4 | 0.41 | 0.75 | -0.17 | 0.0675 | 0.135 | 0.662095 | 0.577749 | 0.861772 | 1 | 0.999631 | 0.881128 |
| Mier1_Q5UAK0_S195s_1 | 0.88 | 0.51 | 1.25 | 0.56 | 1.37 | 0.23 | 0.39 | 1.07 | -0.23 | 0.8 | 0.54 | 0.6 | 0.85 | 0.84 | 0.76 | 0.83 | -0.035 | -0.3725 | 0.02 | 0.945136 | 0.920342 | 0.279736 | 1 | 0.999631 | 0.982141 |
| Mink1_Q9JM52_S537s_1 | 1.76 | 1.14 | 1.18 | 1.54 | 0.95 | 2.61 | 1.94 | 0.18 | 1.9 | 1.86 | 0.8 | -0.28 | -0.78 | 1.49 | 0.35 | 0.73 | 0.015 | -0.335 | -0.9575 | 0.070136 | 0.976835 | 0.514568 | 1 | 0.999631 | 0.529516 |
| Mink1_Q9JM52_S559s_1 | -0.17 | 0.44 | -0.02 | -0.37 | 0.07 | 0.21 | -0.45 | -0.14 | -0.12 | -0.06 | -0.33 | -0.17 | 0.52 | 0.83 | -0.46 | -0.03 | -0.0475 | -0.14 | 0.245 | 0.517549 | 0.871255 | 0.618705 | 1 | 0.999631 | 0.819449 |
| Mink1_Q9JM52_S603s_1 | 1.21 | 0.84 | 0.04 | 1.4 | 1.2 | 1.33 | 1.12 | 1.56 | 1.08 | 0.68 | 1.38 | 0.4 | -1.92 | 0.88 | 0.55 | 0.37 | 0.43 | 0.0125 | -0.9025 | 0.177648 | 0.233833 | 0.974303 | 1 | 0.999631 | 0.642116 |
| Mink1_Q9JM52_S644s_1 | -0.99 | -0.42 | -0.26 | -0.25 | 0.02 | -0.49 | -0.6 | 0.04 | -0.26 | -0.34 | 0.23 | 0.07 | 0.33 | 0 | 0.11 | 0.38 | 0.2225 | 0.405 | 0.685 | 0.043016 | 0.468698 | 0.195746 | 1 | 0.999631 | 0.475443 |
| Mink1_Q9JM52_S699s_1 | 2.19 | -0.19 | -1.02 | 2.14 | 1.37 | 1.01 | 2.24 | 1.41 | 0.62 | 0.48 | 1.43 | 0.27 | -2.69 | -0.38 | -0.21 | -0.11 | 0.7275 | -0.08 | -1.6275 | 0.089893 | 0.346141 | 0.915618 | 1 | 0.999631 | 0.557085 |
| Mink1_Q9JM52_S729s_1 | -0.55 | -1.09 | -1.09 | -0.9 | -1.02 | -1.01 | -0.81 | -0.66 | -1.5 | -0.89 | -1.33 | -0.88 | -2.08 | -0.85 | -0.79 | -0.78 | 0.0325 | -0.2425 | -0.2175 | 0.569984 | 0.900353 | 0.410376 | 1 | 0.999631 | 0.84338 |
| Mink1_Q9JM52_S745s_1 | 0.08 | -0.44 | -0.39 | -0.18 | -0.19 | 0.51 | -0.09 | -0.11 | 0.11 | -0.76 | -0.39 | -0.21 | -0.41 | 0.55 | -0.02 | 0.9 | 0.2625 | -0.08 | 0.4875 | 0.199201 | 0.361022 | 0.787501 | 1 | 0.999631 | 0.657167 |
| Mink1_Q9JM52_S769s_1 | 1.92 | 0.81 | -0.53 | 2.18 | 2.38 | 0.83 | 1.78 | 2.24 | 2 | 0.96 | 1.78 | 1.79 | -1.06 | 0.87 | 1.71 | 0.52 | 0.7125 | 0.5375 | -0.585 | 0.439551 | 0.278562 | 0.380946 | 1 | 0.999631 | 0.781983 |
| Mink1_Q9JM52_S894s_1 | -0.67 | -0.96 | -1.22 | -0.73 | -0.57 | -1.13 | -0.77 | -0.58 | -1.2 | -0.69 | -0.62 | -0.51 | -0.75 | -0.7 | -0.63 | -0.71 | 0.1325 | 0.14 | 0.1975 | 0.478246 | 0.627633 | 0.627624 | 1 | 0.999631 | 0.800612 |
| Mink1_Q9JM52_S903s_1 | 1.79 | 0.22 | 0.33 | -0.06 | -0.72 | 0.13 | 0.66 | -0.59 | -0.76 | 0.81 | -1 | -0.85 | -0.17 | -0.05 | -0.46 | -0.85 | -0.7 | -1.02 | -0.9525 | 0.056687 | 0.178901 | 0.088256 | 1 | 0.999631 | 0.496275 |
| Mink1_Q9JM52_T675t_1 | 2.29 | 1.42 | -0.02 | 1.31 | 3.32 | 0.95 | 1.79 | 2.43 | 2.71 | 0.47 | 1.83 | -0.08 | -2.3 | 2.16 | 0.74 | 0.38 | 0.8725 | -0.0175 | -1.005 | 0.280392 | 0.182361 | 0.980219 | 1 | 0.999631 | 0.701922 |
| Mios_Q8VE19_S766s_1 | -0.25 | 0.97 | -0.2 | -0.53 | 0.71 | 0.32 | -0.46 | -0.25 | 1.18 | 0.44 | 0.21 | 0.5 | 1.89 | 0.74 | -0.02 | -0.06 | 0.0825 | 0.585 | 0.64 | 0.245232 | 0.844815 | 0.169587 | 1 | 0.999631 | 0.685751 |
| Misp_Q9D279_S416s_3 | 1 | -1.41 | 0.43 | 1 | -1.77 | -1.87 | 0.73 | -0.34 | -1.77 | -3 | -0.64 | -0.67 | -2.56 | -1.3 | 0.39 | 1.02 | -1.0675 | -1.775 | -0.8675 | 0.325839 | 0.172872 | 0.030082 | 1 | 0.999631 | 0.724678 |
| Misp_Q9D279_S419s_3 | 1 | -1.41 | 0.43 | 1 | -1.77 | -1.87 | 0.73 | -0.34 | -1.77 | -3 | -0.64 | -0.67 | -2.56 | -1.3 | 0.39 | 1.02 | -1.0675 | -1.775 | -0.8675 | 0.325839 | 0.172872 | 0.030082 | 1 | 0.999631 | 0.724678 |
| Misp_Q9D279_T420t_3 | 1 | -1.41 | 0.43 | 1 | -1.77 | -1.87 | 0.73 | -0.34 | -1.77 | -3 | -0.64 | -0.67 | -2.56 | -1.3 | 0.39 | 1.02 | -1.0675 | -1.775 | -0.8675 | 0.325839 | 0.172872 | 0.030082 | 1 | 0.999631 | 0.724678 |
| Misp_Q9D279_T423t_3 | 1 | -1.41 | 0.43 | 1 | -1.77 | -1.87 | 0.73 | -0.34 | -1.77 | -3 | -0.64 | -0.67 | -2.56 | -1.3 | 0.39 | 1.02 | -1.0675 | -1.775 | -0.8675 | 0.325839 | 0.172872 | 0.030082 | 1 | 0.999631 | 0.724678 |
| Mkl1_D3YUI2_S621s_1 | 0.91 | -1.32 | -1.07 | 0.56 | 1.36 | -1.47 | 0.66 | 0.39 | -0.59 | -1.19 | 0.3 | -0.75 | -1.3 | -0.75 | -0.7 | -0.44 | 0.465 | -0.3275 | -0.5675 | 0.316291 | 0.527417 | 0.581458 | 1 | 0.999631 | 0.720734 |
| Mkl1_D3YUI2_S855s_1 | -1.27 | -1.29 | 0.28 | 0.95 | -0.23 | -1.25 | 0.15 | 1.2 | -1.45 | -1.43 | 1.18 | 1.27 | -0.17 | -1.63 | 1.06 | 1.8 | 0.3 | 0.225 | 0.5975 | 0.470611 | 0.657514 | 0.78764 | 1 | 0.999631 | 0.797474 |
| Mkl1_D3YUI2_T503t_1 | -1.36 | 0.37 | 0.07 | -1 | -0.83 | -0.61 | -1.41 | -0.13 | -0.25 | -0.02 | -0.52 | -0.22 | 0.9 | 0.26 | -0.51 | 0.25 | -0.265 | 0.2275 | 0.705 | 0.170746 | 0.575912 | 0.597657 | 1 | 0.999631 | 0.641032 |
| Mkl2_Q5DTZ3_S207s_1 | -0.18 | 0.73 | -0.11 | -1.24 | 0.01 | -0.14 | -0.47 | -0.5 | 0.08 | -1.13 | -0.39 | -0.17 | 0.43 | 0.65 | 0.25 | -0.6 | -0.075 | -0.2025 | 0.3825 | 0.428828 | 0.857903 | 0.66516 | 1 | 0.999631 | 0.776582 |
| Mkl2_Q5DTZ3_S209s_1 | -1 | -0.03 | 0.22 | -0.4 | -0.11 | -0.56 | -0.54 | -0.26 | -0.53 | -1.08 | -0.1 | -0.23 | 0.23 | -0.39 | -0.39 | -0.23 | -0.065 | -0.1825 | 0.1075 | 0.762956 | 0.84341 | 0.624368 | 1 | 0.999631 | 0.915608 |
| Mlc1_E9QP87_S189s_1 | -0.32 | -0.09 | 0.71 | -0.59 | 0 | -1.43 | -0.34 | -0.85 | -0.92 | -0.65 | -0.66 | -0.49 | -0.02 | 0.09 | 0.68 | -0.67 | -0.5825 | -0.6075 | 0.0925 | 0.822268 | 0.183773 | 0.098586 | 1 | 0.999631 | 0.938608 |
| Mlc1_E9QP87_T41t_1 | -0.2 | -0.95 | 3.26 | 2.39 | -0.63 | -0.51 | 1.29 | 1.54 | -0.13 | 0.26 | 3.76 | 4.21 | -0.14 | -1.08 | 2.35 | 2.11 | -0.7025 | 0.9 | -0.315 | 0.778202 | 0.487796 | 0.493735 | 1 | 0.999631 | 0.921416 |
| Mlf2_Q99KX1_S237s_1 | 0.12 | 0.82 | 0.59 | 0.03 | 0.25 | 0.18 | -0.29 | 0.04 | 0.19 | 0.41 | 0.15 | 0.33 | 0.74 | 0.71 | 0.17 | 0.4 | -0.345 | -0.12 | 0.115 | 0.718295 | 0.255953 | 0.67737 | 1 | 0.999631 | 0.903971 |
| Mlf2_Q99KX1_S32s_1 | -1.1 | 0.14 | -0.2 | -0.46 | -0.15 | -0.12 | -1.07 | -0.73 | 0.14 | 0.16 | -0.53 | 0.01 | 1.03 | 0.25 | -0.48 | -0.49 | -0.1125 | 0.35 | 0.4825 | 0.294951 | 0.760807 | 0.328016 | 1 | 0.999631 | 0.708769 |
| Mlh3_Q68FG1_S1283s_2 | -0.15 | -0.15 | 0.28 | -0.53 | -0.24 | -0.32 | -0.39 | 0.21 | -0.46 | 0.9 | -0.63 | 0.14 | -0.45 | -1.06 | -0.75 | -0.58 | -0.0475 | 0.125 | -0.5725 | 0.087043 | 0.86881 | 0.753505 | 1 | 0.999631 | 0.553514 |
| Mlip_V9GWW6_S61s_1 | -0.2 | -1.33 | -0.94 | -0.8 | -0.89 | -1.23 | -1.15 | -0.79 | -2.02 | -0.92 | -2 | -1.47 | -0.85 | -1.17 | -1.09 | -0.81 | -0.1975 | -0.785 | -0.1625 | 0.621437 | 0.529784 | 0.059581 | 1 | 0.999631 | 0.86189 |
| Mllt4_E9Q9C3_S1090s_1 | -0.42 | 0.69 | 0.34 | -0.21 | -0.14 | 0.24 | -0.55 | -0.04 | -0.24 | 0.05 | -0.57 | -0.08 | 0.53 | 0.84 | 0.21 | 0.6 | -0.2225 | -0.31 | 0.445 | 0.217134 | 0.515305 | 0.3663 | 1 | 0.999631 | 0.669465 |
| Mllt4_E9Q9C3_S1114s_1 | -0.77 | -0.27 | -0.73 | -0.09 | 0.46 | -0.1 | -0.01 | -0.49 | -0.12 | -1.04 | -0.67 | -0.66 | -0.18 | 0.01 | -0.02 | -0.33 | 0.43 | -0.1575 | 0.335 | 0.274452 | 0.189351 | 0.620626 | 1 | 0.999631 | 0.700187 |
| Mllt4_E9Q9C3_S1179s_1 | 0.7 | 0.79 | 0.59 | 0.06 | 0.76 | 1.11 | 0.04 | 0.29 | 0.55 | 0.69 | 0.56 | 0.46 | 0.02 | 1.57 | 0.55 | 0.35 | 0.015 | 0.03 | 0.0875 | 0.825943 | 0.96378 | 0.912909 | 1 | 0.999631 | 0.940599 |
| Mllt4_E9Q9C3_S1180s_2 | -0.11 | -1.04 | -0.15 | -0.3 | -0.32 | -0.59 | -0.33 | -0.15 | -1.22 | -0.98 | -0.63 | -1.01 | -0.78 | -0.24 | -0.36 | -0.82 | 0.0525 | -0.56 | -0.15 | 0.654092 | 0.860703 | 0.099191 | 1 | 0.999631 | 0.878477 |
| Mllt4_E9Q9C3_S1189s_1 | -0.12 | 1.19 | 0.03 | -0.47 | 0.86 | 0.97 | -0.42 | 0.06 | 1.13 | 0.25 | 0.08 | 0.15 | 1.72 | 0.68 | 0.05 | 0.2 | 0.21 | 0.245 | 0.505 | 0.32474 | 0.654696 | 0.574868 | 1 | 0.999631 | 0.72463 |
| Mllt4_E9Q9C3_S1189s_2 | -0.11 | -1.04 | -0.15 | -0.3 | -0.32 | -0.59 | -0.33 | -0.15 | -1.22 | -0.98 | -0.63 | -1.01 | -0.78 | -0.24 | -0.36 | -0.82 | 0.0525 | -0.56 | -0.15 | 0.654092 | 0.860703 | 0.099191 | 1 | 0.999631 | 0.878477 |
| Mllt4_E9Q9C3_S1218s_1 | -1.35 | -0.42 | -0.8 | -1.3 | -0.81 | -0.76 | -1.31 | -1.14 | -0.8 | -0.72 | -1.12 | -0.79 | 0.31 | -0.19 | -0.65 | -0.59 | -0.0375 | 0.11 | 0.6875 | 0.081091 | 0.904109 | 0.721439 | 1 | 0.999631 | 0.546121 |
| Mllt4_E9Q9C3_S1282s_1 | 0.22 | -0.21 | -1.33 | 0.07 | 0.07 | -0.31 | 0.09 | -0.04 | 0.8 | 0 | 0.03 | -0.15 | -0.82 | -0.8 | -0.9 | -1.54 | 0.265 | 0.4825 | -0.7025 | 0.111417 | 0.488191 | 0.262981 | 1 | 0.999631 | 0.577068 |
| Mllt4_E9Q9C3_S1461s_1 | -0.02 | 0.23 | 1.28 | 0.47 | 0.58 | 1.83 | 0.04 | 0.85 | 0.49 | 0.5 | -0.37 | 0.2 | 0.08 | 2.17 | 0.99 | 0.68 | 0.335 | -0.285 | 0.49 | 0.338972 | 0.46546 | 0.452386 | 1 | 0.999631 | 0.732185 |
| Mllt4_E9Q9C3_S216s_1 | 2.58 | 0.68 | 0.34 | 1.28 | 1.4 | 1.26 | 0.86 | 1.24 | 1.43 | 0.56 | 1.21 | 1.2 | -1.69 | 1.01 | 1.26 | 0.71 | -0.03 | -0.12 | -0.8975 | 0.242554 | 0.950125 | 0.810645 | 1 | 0.999631 | 0.685099 |
| Mllt4_E9Q9C3_S246s_1 | -1 | -0.3 | -0.82 | -1.34 | 0.28 | 0.41 | -0.08 | -1.29 | 1.05 | -0.54 | -0.46 | -0.06 | 0.31 | -0.59 | -1.16 | -1.51 | 0.695 | 0.8625 | 0.1275 | 0.777742 | 0.133854 | 0.067076 | 1 | 0.999631 | 0.921382 |
| Mllt4_E9Q9C3_S2s_1 | 1.67 | 0.74 | 2.1 | 1.44 | -0.62 | 1.63 | 2.15 | 0.06 | 1.12 | 2.42 | 1.07 | 1.14 | 0.53 | -0.35 | 0.75 | 0.5 | -0.6825 | -0.05 | -1.13 | 0.016532 | 0.297805 | 0.907862 | 1 | 0.999631 | 0.389691 |
| Mllt4_E9Q9C3_S391s_1 | -0.13 | 0.27 | -1.42 | -1.19 | -0.32 | 0.34 | -2.18 | -2.03 | 0.36 | -0.44 | -1.55 | -1.7 | 1.25 | 0.07 | -2.45 | -1.58 | -0.43 | -0.215 | -0.06 | 0.940482 | 0.522535 | 0.712584 | 1 | 0.999631 | 0.981146 |
| Mllt4_E9Q9C3_S562s_1 | 2.29 | -1.26 | 0.81 | 0.24 | -0.31 | 0.51 | 1.07 | -0.23 | -0.37 | 1.29 | 0.08 | -0.32 | -2.01 | -1.02 | -0.29 | -0.6 | -0.26 | -0.35 | -1.5 | 0.06192 | 0.714935 | 0.635589 | 1 | 0.999631 | 0.509355 |
| Mllt4_E9Q9C3_S655s_1 | -1.06 | -2.37 | -1.23 | -1.67 | -2.25 | -2.02 | -0.84 | -1.46 | -2.11 | -0.96 | -1.78 | -1.12 | -1.98 | -2 | -1.08 | -0.93 | -0.06 | 0.09 | 0.085 | 0.840246 | 0.887378 | 0.825447 | 1 | 0.999631 | 0.946665 |
| Mllt4_E9Q9C3_T214t_1 | 0.86 | 0.63 | 0.26 | 1.18 | 2.38 | 1.44 | 1.17 | 2.39 | 1.7 | 0.74 | 1.15 | 0.93 | -0.98 | 1.72 | 1.14 | 1.94 | 1.1125 | 0.3975 | 0.2225 | 0.723437 | 0.015085 | 0.25123 | 1 | 0.999631 | 0.905674 |
| Mllt6_B1AR09_S273s_1 | -0.29 | -0.17 | 0.48 | 0.25 | 0.28 | -0.06 | -0.16 | 0.43 | -0.08 | -0.62 | 0.46 | 0.96 | 0.3 | 0.53 | 0.93 | 1.03 | 0.055 | 0.1125 | 0.63 | 0.076862 | 0.852015 | 0.777989 | 1 | 0.999631 | 0.542683 |
| Mmrn2_A6H6E2_S403s_1 | 0.3 | 0.59 | 0.93 | 0.48 | -0.14 | 1.27 | -0.09 | 0.61 | -0.35 | 1.21 | 0.49 | 0.42 | 1.18 | 1.21 | 0.97 | 1.64 | -0.1625 | -0.1325 | 0.675 | 0.04475 | 0.665934 | 0.72235 | 1 | 0.999631 | 0.475443 |
| Mobp_Q9D2P8_S107s_1 | 2.11 | 0.8 | 1.23 | 0.82 | -0.17 | 1.13 | 0.53 | -0.04 | 0.7 | 1.93 | -0.42 | -0.14 | 0.34 | 0.49 | 0.32 | 0.1 | -0.8775 | -0.7225 | -0.9275 | 0.026945 | 0.060941 | 0.218959 | 1 | 0.999631 | 0.422772 |
| Mobp_Q9D2P8_S74s_1 | 2.04 | 0.38 | 0.74 | 0.16 | -1.46 | 0.21 | 0.54 | -1.03 | -0.09 | 1.19 | -1.1 | -1.03 | 0.32 | 0.25 | -0.15 | -1 | -1.265 | -1.0875 | -0.975 | 0.073742 | 0.051462 | 0.103006 | 1 | 0.999631 | 0.539615 |
| Mobp_Q9D2P8_S85s_1 | -1.69 | 0.16 | 0.02 | -1.84 | 1.23 | 0.07 | 0.14 | -1.58 | 3.15 | -1.36 | 1.03 | 1.67 | 4.23 | -0.46 | -0.65 | -1.55 | 0.8025 | 1.96 | 1.23 | 0.315364 | 0.26771 | 0.059947 | 1 | 0.999631 | 0.72069 |
| Mobp_Q9D2P8_S98s_1 | -0.4 | 0.09 | 0.68 | 0.07 | -0.2 | 0.68 | -0.63 | -0.24 | 0.65 | 0.93 | -0.29 | 0.82 | 1.17 | 0.98 | 0.96 | 0.65 | -0.2075 | 0.4175 | 0.83 | 0.026617 | 0.579818 | 0.28567 | 1 | 0.999631 | 0.422285 |
| Mon1a_Q6PDG8_S56s_1 | -0.87 | -0.1 | 0.29 | -0.1 | -0.21 | -0.4 | -0.63 | 0.05 | -0.01 | -0.5 | -0.01 | 0.37 | 0.34 | 0.13 | 0.33 | 0.42 | -0.1025 | 0.1575 | 0.5 | 0.149401 | 0.753932 | 0.648965 | 1 | 0.999631 | 0.621541 |
| Mon2_B9EKJ3_S1175s_1 | -1.08 | 0.77 | 0.66 | -0.52 | 0.32 | -0.07 | 0.09 | -0.19 | 0.57 | -1.5 | 0.87 | 1.38 | 0.95 | -0.22 | 0.69 | -0.38 | 0.08 | 0.3725 | 0.3025 | 0.569172 | 0.858913 | 0.593804 | 1 | 0.999631 | 0.84338 |
| Mon2_B9EKJ3_S1213s_1 | -0.28 | 1.63 | 0.8 | 0.5 | 0.95 | 1.78 | 1.35 | 0.01 | 1.75 | 0.02 | 0.5 | 0.51 | 0.25 | 0.82 | 0.61 | 0.28 | 0.36 | 0.0325 | -0.1725 | 0.687623 | 0.48577 | 0.949177 | 1 | 0.999631 | 0.891556 |
| Mon2_B9EKJ3_S534s_1 | 0.2 | 0.72 | 1.18 | 0.76 | -0.13 | 0.73 | 0.54 | 0.38 | 0.18 | 0.86 | 0.38 | 0.37 | 0.13 | 0.65 | 0.68 | 0.71 | -0.335 | -0.2675 | -0.1725 | 0.596506 | 0.309746 | 0.401734 | 1 | 0.999631 | 0.854508 |
| Mon2_Q80TL7_S534s_1 | 0.22 | 1.57 | 1.69 | 1.03 | 0.93 | 1.43 | 0.68 | 0.88 | 1.41 | 1.35 | 1.05 | 0.93 | 1.19 | 1.42 | 1.38 | 0.85 | -0.1475 | 0.0575 | 0.0825 | 0.832368 | 0.701058 | 0.878983 | 1 | 0.999631 | 0.94321 |
| Morc2a_Q69ZX6_S614s_1 | 0.23 | 0.6 | 0.8 | 0.36 | 0.27 | 0.54 | 0.39 | 0.85 | 0.35 | 0.34 | 0.59 | 1.38 | 0.86 | 0.99 | 0.62 | 1.78 | 0.015 | 0.1675 | 0.565 | 0.123916 | 0.955702 | 0.613007 | 1 | 0.999631 | 0.595021 |
| Morc2a_Q69ZX6_S775s_1 | 1.04 | 1.31 | 0.55 | 0.26 | 0.87 | 0.47 | -0.08 | 0.27 | 0.97 | 0.47 | 0.86 | 1.74 | -0.05 | 0.41 | 0.94 | -0.19 | -0.4075 | 0.22 | -0.5125 | 0.200696 | 0.251534 | 0.564753 | 1 | 0.999631 | 0.658263 |
| Morc2a_Q69ZX6_S854s_1 | 0.34 | -0.32 | -0.46 | 0.11 | -0.86 | -0.17 | -0.36 | -0.58 | -0.04 | 0.04 | -0.05 | 0.39 | -0.24 | 0.18 | 0.32 | -0.05 | -0.41 | 0.1675 | 0.135 | 0.668064 | 0.193588 | 0.572632 | 1 | 0.999631 | 0.883098 |
| Mpdz_D3YUX2_S230s_1 | 0.47 | -0.04 | 0.15 | 0.31 | 0.05 | -0.18 | -0.23 | -0.46 | 0.4 | 0.24 | -0.01 | 0.15 | 0.08 | 0.52 | -0.27 | 0.04 | -0.4275 | -0.0275 | -0.13 | 0.666963 | 0.122758 | 0.916295 | 1 | 0.999631 | 0.88264 |
| Mpdz_D3YUX2_S231s_1 | 0.53 | -0.01 | -0.82 | -0.04 | -0.02 | -0.37 | -0.55 | -0.7 | 0.21 | 0.27 | -0.26 | -0.18 | -0.3 | 0.55 | -0.1 | 0 | -0.325 | 0.095 | 0.1225 | 0.743119 | 0.358565 | 0.785507 | 1 | 0.999631 | 0.90972 |
| Mphosph8_Q3TYA6_S25s_1 | 0.01 | 0.24 | 0.43 | -0.03 | -0.9 | -0.26 | 0.19 | 0.18 | -1.33 | 0.92 | -0.2 | -0.18 | -1.71 | 0.24 | 0.09 | -0.61 | -0.36 | -0.36 | -0.66 | 0.168413 | 0.281685 | 0.440701 | 1 | 0.999631 | 0.637627 |
| Mpp1_B7ZCM1_S57s_1 | 0.16 | -0.65 | -1.41 | 0.38 | 0.31 | 0.37 | -0.02 | 0.2 | -0.25 | -0.9 | -0.12 | 0.07 | -1.61 | -0.05 | 0.16 | -0.14 | 0.595 | 0.08 | -0.03 | 0.955522 | 0.17431 | 0.859083 | 1 | 0.999631 | 0.984649 |
| Mpp1_B7ZCM1_S93s_1 | -0.52 | 0.12 | -0.3 | 0.74 | -0.13 | -0.57 | 0.15 | 0.51 | -0.08 | 0.09 | 0.22 | 0.04 | -0.43 | -0.44 | 0.69 | 0.14 | -0.02 | 0.0575 | -0.02 | 0.96079 | 0.957426 | 0.863219 | 1 | 0.999631 | 0.986594 |
| Mpp2_Q9WV34_S121s_1 | -1.15 | -0.24 | -1.21 | -1.58 | -0.9 | -0.64 | -1.73 | -1.11 | -0.55 | -0.81 | -0.16 | -0.98 | 0.41 | -0.65 | -2.47 | -0.77 | -0.05 | 0.42 | 0.175 | 0.771369 | 0.895471 | 0.267606 | 1 | 0.999631 | 0.919413 |
| Mpp2_Q9WV34_S42s_1 | -0.21 | -0.54 | -1.64 | -0.2 | -0.34 | -0.66 | -0.75 | -0.55 | -0.71 | -0.75 | -0.58 | -0.78 | -2.34 | -0.41 | -1.08 | -0.31 | 0.0725 | -0.0575 | -0.3875 | 0.479191 | 0.844666 | 0.876605 | 1 | 0.999631 | 0.801052 |
| Mpp2_Q9WV34_T117t_1 | -1.41 | -1 | -0.62 | -1.13 | -1.53 | -0.86 | -1.22 | -1.13 | -1.27 | -1.46 | -1.08 | -0.9 | 0.49 | -0.54 | -0.94 | -0.66 | -0.145 | -0.1375 | 0.6275 | 0.127418 | 0.616867 | 0.637162 | 1 | 0.999631 | 0.59788 |
| Mpp3_Q6XE40_S307s_1 | -0.68 | -0.02 | -0.38 | -0.5 | -0.57 | -0.14 | -0.9 | -0.2 | -0.79 | -0.56 | -0.04 | 0.01 | -0.19 | -0.06 | 0.12 | 0.32 | -0.0575 | 0.05 | 0.4425 | 0.153984 | 0.844794 | 0.871512 | 1 | 0.999631 | 0.625198 |
| Mpp3_Q6XE40_S354s_1 | 0.83 | 0.01 | 1.24 | 0.2 | -1.02 | 0.25 | 0.51 | 0.2 | -0.55 | -0.27 | -0.06 | -0.52 | -0.4 | -0.71 | 0.6 | -0.72 | -0.585 | -0.92 | -0.8775 | 0.064261 | 0.199103 | 0.023182 | 1 | 0.999631 | 0.514389 |
| Mpp6_Q9JLB0_S199s_1 | -1.14 | -0.34 | -0.68 | -1.26 | 0.28 | -0.62 | -1.12 | -1.98 | -0.42 | -1.37 | -0.92 | -0.02 | -0.58 | -0.07 | -0.91 | 0.07 | -0.005 | 0.1725 | 0.4825 | 0.200887 | 0.991826 | 0.654187 | 1 | 0.999631 | 0.65842 |
| Mprip_F6RND9_S350s_1 | -0.74 | -0.2 | -0.46 | -0.92 | 0.2 | -0.1 | -0.59 | -0.16 | -0.17 | -0.72 | -0.65 | -0.23 | 0.03 | 0.53 | -0.71 | -0.47 | 0.4175 | 0.1375 | 0.425 | 0.260048 | 0.178439 | 0.641854 | 1 | 0.999631 | 0.692624 |
| Mprip_F6RND9_S89s_2 | 0.06 | 0.89 | 1.11 | -0.09 | 0.89 | 0.34 | 0.05 | 0.46 | 0.77 | 0.84 | -0.25 | 0.49 | 1.44 | 0.69 | -0.25 | -0.09 | -0.0575 | -0.03 | -0.045 | 0.924847 | 0.875106 | 0.940259 | 1 | 0.999631 | 0.975422 |
| Mprip_F6RND9_S93s_2 | 0.06 | 0.89 | 1.11 | -0.09 | 0.89 | 0.34 | 0.05 | 0.46 | 0.77 | 0.84 | -0.25 | 0.49 | 1.44 | 0.69 | -0.25 | -0.09 | -0.0575 | -0.03 | -0.045 | 0.924847 | 0.875106 | 0.940259 | 1 | 0.999631 | 0.975422 |
| Mprip_F6RND9_S962s_1 | -0.68 | 1.02 | 0.42 | -0.42 | 0.14 | 1 | -0.67 | -0.02 | 0.61 | 0.94 | -0.44 | -0.37 | 0.9 | 0.55 | -1.09 | 0.36 | 0.0275 | 0.1 | 0.095 | 0.862008 | 0.955206 | 0.840448 | 1 | 0.999631 | 0.954601 |
| Mprip_F6RND9_S976s_1 | 0.5 | -0.2 | -0.27 | 1.17 | 2.03 | 1.45 | 0.56 | 1.22 | 1.71 | -0.03 | 0.73 | 1.13 | -0.28 | 0.65 | 0.56 | 0.57 | 1.015 | 0.585 | 0.075 | 0.857484 | 0.04224 | 0.239882 | 1 | 0.999631 | 0.95214 |
| Mprip_F6RND9_S978s_1 | 0.73 | 1.02 | 0.03 | 1.11 | 1.39 | 1.43 | 0.66 | 1.26 | 1.48 | 0.23 | 1.15 | 1.11 | 0.12 | 1.22 | 0.73 | 1.15 | 0.4625 | 0.27 | 0.0825 | 0.830383 | 0.193424 | 0.486629 | 1 | 0.999631 | 0.942193 |
| Mprip_Q5SWZ5_S998s_1 | 1.64 | 0.8 | 0.12 | 2.2 | 2.28 | 1.35 | 1.89 | 2.2 | 0.82 | -0.39 | 2.2 | 1.36 | -1.51 | 0.86 | 1.28 | 0.96 | 0.74 | -0.1925 | -0.7925 | 0.273156 | 0.145783 | 0.764044 | 1 | 0.999631 | 0.700187 |
| Msantd1_Q8BIL2_S161s_1 | -0.9 | -0.61 | -0.19 | -0.67 | -0.39 | -0.85 | -0.48 | -0.89 | -0.37 | -0.61 | -0.86 | -0.06 | 0.17 | 0.01 | -0.39 | 0.17 | -0.06 | 0.1175 | 0.5825 | 0.077436 | 0.829371 | 0.697453 | 1 | 0.999631 | 0.542951 |
| Msh6_P54276_S63s_1 | 1.36 | 1.03 | 1.87 | 1.37 | 0.59 | 0.83 | 1.01 | 0.96 | 1.05 | 1.62 | 0.44 | 1.13 | 0.82 | 0.79 | 1.09 | 0.8 | -0.56 | -0.3475 | -0.5325 | 0.097363 | 0.068703 | 0.322789 | 1 | 0.999631 | 0.559562 |
| Msi2_Q920Q6_S279s_1 | 0.33 | -0.1 | 0.42 | -0.4 | -0.25 | -0.28 | -0.19 | 0.63 | -0.11 | -0.01 | -0.3 | 0.43 | 0.48 | 0.98 | -0.16 | 0.89 | -0.085 | -0.06 | 0.485 | 0.206567 | 0.797566 | 0.84754 | 1 | 0.999631 | 0.6605 |
| Msi2_Q920Q6_S6s_1 | -0.41 | -0.61 | 0.01 | -0.28 | -0.92 | -0.96 | -0.33 | -0.84 | -0.49 | 0.21 | -0.31 | 0.41 | -0.43 | -0.9 | -0.15 | 0.31 | -0.44 | 0.2775 | 0.03 | 0.930783 | 0.13819 | 0.386051 | 1 | 0.999631 | 0.97791 |
| Msl3_Q9WVG9_S345s_1 | -0.79 | 0.03 | -0.31 | -0.74 | -0.08 | -0.82 | -0.19 | -0.42 | 0.09 | -0.46 | -0.33 | -0.35 | 0.95 | -0.02 | -0.24 | 0.14 | 0.075 | 0.19 | 0.66 | 0.097069 | 0.808538 | 0.534027 | 1 | 0.999631 | 0.55886 |
| Msrb3_D3YUC9_S105s_1 | 2.61 | 0.67 | 0.87 | 1.5 | 0.46 | 1.04 | 1.74 | 0.24 | 0.09 | 2.12 | 0.51 | 0.78 | 0.31 | -0.78 | 0.57 | -0.66 | -0.5425 | -0.5375 | -1.5525 | 0.013215 | 0.304662 | 0.356688 | 1 | 0.999631 | 0.385092 |
| Mt3_P28184_S33s_1 | 0.22 | 0.7 | 0.32 | 0.02 | 0.14 | 0.53 | 0.98 | 0.21 | 0.93 | -0.21 | -0.07 | -0.39 | 0.23 | -0.53 | -0.31 | -0.47 | 0.15 | -0.25 | -0.585 | 0.086427 | 0.620459 | 0.494064 | 1 | 0.999631 | 0.55299 |
| Mta1_F8WHY8_S432s_1 | -0.31 | 1.45 | 1.02 | -0.28 | 1 | 0.97 | 0.18 | 0.67 | 1.22 | -0.31 | 0.67 | 1.41 | 1.59 | 1.16 | 1.03 | 0.53 | 0.235 | 0.2775 | 0.6075 | 0.228555 | 0.616984 | 0.615913 | 1 | 0.999631 | 0.679038 |
| Mta1_F8WHY8_S505s_1 | -0.57 | 0.49 | 1 | 0.07 | 0.31 | 0.28 | -0.13 | -0.36 | 0.64 | 0.39 | -0.46 | 0.16 | 0.65 | 0.99 | 0.85 | 0.73 | -0.2225 | -0.065 | 0.5575 | 0.162586 | 0.564079 | 0.875078 | 1 | 0.999631 | 0.631442 |
| Mta1_F8WHY8_S622s_1 | 0.4 | 1.84 | 1.63 | 1.34 | 0.44 | 1.26 | 0.22 | 1.43 | 1.26 | 1.11 | 1.5 | 1.69 | 0.32 | 1.97 | 1.38 | 2.39 | -0.465 | 0.0875 | 0.2125 | 0.683942 | 0.292922 | 0.813296 | 1 | 0.999631 | 0.889602 |
| Mta3_H3BIY7_S176s_1 | -0.57 | 2.64 | 3.5 | -0.77 | 3.01 | 0.01 | 3.54 | -0.28 | 4.75 | -0.02 | 5.17 | 5.91 | 3.99 | 0.04 | 4.02 | -0.14 | 0.37 | 2.7525 | 0.7775 | 0.567898 | 0.769108 | 0.087032 | 1 | 0.999631 | 0.843071 |
| Mta3_H3BIY7_S207s_1 | -0.71 | 0.48 | -0.14 | 0.15 | -0.53 | -0.13 | 0.49 | 0.77 | -0.13 | -0.6 | 1.34 | 1.85 | 0.74 | 0.08 | 1.6 | 1.66 | 0.205 | 0.67 | 1.075 | 0.036234 | 0.605821 | 0.266606 | 1 | 0.999631 | 0.453963 |
| Mta3_H3BIY7_S85s_1 | 1.02 | 1.17 | 2.22 | 1.63 | 0.78 | 1.25 | 1.22 | 1.47 | 1.42 | 1.48 | 1.87 | 1.58 | 0.5 | 1.27 | 1.54 | 1.47 | -0.33 | 0.0775 | -0.315 | 0.427399 | 0.344948 | 0.817687 | 1 | 0.999631 | 0.77521 |
| Mtch1_Q791T5_S364s_1 | -0.23 | 0.01 | 0.2 | -0.49 | -0.47 | 0.04 | -0.73 | 0.34 | -0.44 | 0.33 | 0.62 | 0.32 | 0.66 | 0.92 | 0.28 | 0.48 | -0.0775 | 0.335 | 0.7125 | 0.038078 | 0.813057 | 0.318353 | 1 | 0.999631 | 0.459475 |
| Mtcl1_H3BL83_S1095s_1 | 1.07 | 1.37 | 0.69 | 2.14 | 1.31 | 1.34 | 1.69 | 1.68 | 0.93 | 0.98 | 1.96 | 2.1 | -1.68 | 0.87 | 0.14 | 1.93 | 0.1875 | 0.175 | -1.0025 | 0.186924 | 0.597403 | 0.688629 | 1 | 0.999631 | 0.646474 |
| Mtcl1_H3BL83_S1369s_2 | 2.22 | 0.42 | 0.94 | 2.11 | 0.4 | 1.42 | 2.02 | 1.96 | 0.8 | 1.28 | 1.04 | -0.05 | -3.11 | -0.18 | 0.49 | -0.29 | 0.0275 | -0.655 | -2.195 | 0.018973 | 0.95898 | 0.211496 | 1 | 0.999631 | 0.391024 |
| Mtcl1_H3BL83_T1366t_2 | 2.22 | 0.42 | 0.94 | 2.11 | 0.4 | 1.42 | 2.02 | 1.96 | 0.8 | 1.28 | 1.04 | -0.05 | -3.11 | -0.18 | 0.49 | -0.29 | 0.0275 | -0.655 | -2.195 | 0.018973 | 0.95898 | 0.211496 | 1 | 0.999631 | 0.391024 |
| Mtcl1_Q3UHU5_S255s_1 | 0.05 | -0.87 | -0.81 | 0.45 | 0.11 | -0.4 | -0.09 | -0.81 | -1.17 | -0.29 | 0.04 | -0.16 | -1.49 | -0.06 | -1.13 | 0.8 | -0.0025 | -0.1 | -0.175 | 0.75803 | 0.994889 | 0.81327 | 1 | 0.999631 | 0.914449 |
| Mtcl1_Q3UHU5_S312s_1 | 0.84 | 0.36 | -0.84 | 0.77 | 1.15 | 0.62 | 0.54 | 0.67 | 0.4 | -0.03 | 0.14 | 0.03 | -1.71 | 0.85 | -0.56 | 0.7 | 0.4625 | -0.1475 | -0.4625 | 0.477786 | 0.277384 | 0.718949 | 1 | 0.999631 | 0.800237 |
| Mtdh_Q80WJ7_S297s_1 | 0.18 | 1.15 | -0.22 | 1.38 | 2.28 | 0.61 | 1.62 | 0.62 | 1.11 | 0.07 | 1.55 | 0.53 | -0.75 | 1.1 | -0.52 | 1.15 | 0.66 | 0.1925 | -0.3775 | 0.522947 | 0.223605 | 0.690261 | 1 | 0.999631 | 0.821522 |
| Mtdh_Q80WJ7_S423s_1 | 0.75 | 2.12 | 0.93 | 0.56 | 1.54 | 1.15 | 0.08 | 0.67 | 1.49 | 1.55 | 0.52 | 1.01 | 2.38 | 2.16 | 0.87 | 1.16 | -0.23 | 0.0525 | 0.5525 | 0.276635 | 0.615162 | 0.901976 | 1 | 0.999631 | 0.700187 |
| Mtdh_Q80WJ7_S565s_1 | -0.55 | 0.52 | 0.51 | 0 | 0.28 | 0.09 | 0 | 0.14 | 0.51 | -0.01 | 0.79 | 0.63 | 0.4 | 0.54 | 0.3 | 0.64 | 0.0075 | 0.36 | 0.35 | 0.309416 | 0.980898 | 0.314666 | 1 | 0.999631 | 0.718845 |
| Mtfr1l_Q9CWE0_S100s_1 | -0.24 | 1.59 | 1.3 | 0.27 | 0.98 | 1.35 | -0.15 | 1.19 | 1.13 | 0.61 | 0.9 | 1.18 | 0.51 | 1.54 | 1.38 | 1.42 | 0.1125 | 0.225 | 0.4825 | 0.325249 | 0.82574 | 0.613886 | 1 | 0.999631 | 0.72463 |
| Mtfr1l_Q9CWE0_S107s_1 | 0.44 | -0.79 | -1.46 | -1.17 | -1.54 | -1.91 | -0.4 | -1.57 | -2.26 | -0.84 | -0.64 | -0.62 | -1.64 | -1.52 | -2.97 | -1.76 | -0.61 | -0.345 | -1.2275 | 0.034639 | 0.240056 | 0.523693 | 1 | 0.999631 | 0.448552 |
| Mtfr1l_Q9CWE0_S235s_1 | -0.52 | 0.69 | 0.45 | 0.45 | 0.6 | 0.59 | -0.29 | 0.55 | 0.58 | -0.08 | 0.18 | 0.78 | 0.55 | 0.69 | 0.41 | 0.85 | 0.095 | 0.0975 | 0.3575 | 0.314093 | 0.795517 | 0.788374 | 1 | 0.999631 | 0.719959 |
| Mtif2_Q5M6W9_S83s_1 | 0.83 | -2.75 | -0.78 | -0.66 | -2.79 | -1.99 | -1.14 | -1.94 | -2.85 | -0.14 | -1.58 | -1.81 | -1.43 | -2.24 | -1.01 | -0.58 | -1.125 | -0.755 | -0.475 | 0.514081 | 0.138983 | 0.361864 | 1 | 0.999631 | 0.81893 |
| Mtmr1_I7HJQ9_S659s_1 | -0.16 | -1.2 | -0.45 | -0.59 | -1.84 | -1.52 | -0.11 | -0.97 | -1.76 | -0.56 | -0.53 | -0.24 | -0.79 | -1 | -0.79 | -0.89 | -0.51 | -0.1725 | -0.2675 | 0.403388 | 0.252724 | 0.675128 | 1 | 0.999631 | 0.764348 |
| Mtmr1_I7HJQ9_T665t_2 | 1.78 | 0.72 | -2.14 | 0.1 | 1.53 | 1.96 | 0.76 | 0.83 | -0.16 | 0.78 | 0.38 | -1.11 | -1.26 | 0.64 | -1.52 | 0.15 | 1.155 | -0.1425 | -0.6125 | 0.477236 | 0.156103 | 0.860274 | 1 | 0.999631 | 0.800086 |
| Mtmr2_Q9Z2D1_S21s_1 | -0.23 | -1.26 | -2.34 | -1.85 | -1.5 | -1.54 | -1.1 | -3.47 | -1.91 | -2.56 | -1.97 | -3.18 | -0.17 | -3.65 | -2.05 | -2.33 | -0.4825 | -0.985 | -0.63 | 0.407139 | 0.449062 | 0.078832 | 1 | 0.999631 | 0.766996 |
| Mtmr3_M0QW74_S633s_1 | 0.35 | 0.05 | 0.6 | 0.82 | -0.39 | -0.11 | 0.55 | 0.5 | 0.15 | 0.39 | 0.75 | 0 | -0.08 | -0.54 | 0.26 | 0.05 | -0.3175 | -0.1325 | -0.5325 | 0.120176 | 0.344265 | 0.665216 | 1 | 0.999631 | 0.587703 |
| Mtmr4_Q91XS1_S572s_1 | -0.35 | -0.52 | -0.49 | -0.48 | -0.15 | -0.23 | -0.1 | 0.05 | -0.19 | 0.22 | 0.39 | -0.27 | 0.11 | 0.41 | 0.58 | 0.11 | 0.3525 | 0.4975 | 0.7625 | 0.017808 | 0.153843 | 0.091791 | 1 | 0.999631 | 0.391024 |
| Mtmr6_Q8VE11_S585s_1 | 2.23 | -0.28 | -0.94 | 0.75 | 1.7 | -0.25 | 1.28 | 0.59 | 0.87 | 0.62 | -0.75 | -0.91 | 0.06 | -0.29 | -0.86 | -1.47 | 0.39 | -0.4825 | -1.08 | 0.137558 | 0.588946 | 0.515775 | 1 | 0.999631 | 0.610681 |
| Mtmr7_Q9Z2C9_S593s_1 | 0.38 | 1.32 | 0.41 | 0.02 | 0.05 | -0.23 | -0.3 | 0.07 | 0.01 | -0.73 | 0.22 | 1.34 | 0.34 | 0.43 | 0.03 | 0.33 | -0.635 | -0.3225 | -0.25 | 0.479222 | 0.080707 | 0.513045 | 1 | 0.999631 | 0.801052 |
| Mtmr7_Q9Z2C9_S595s_1 | -0.2 | 0.53 | -0.26 | -0.42 | 0.11 | -0.37 | -0.69 | 0.14 | -0.05 | -0.34 | -0.05 | 0.22 | 0.54 | 0.25 | -0.03 | -0.01 | -0.115 | 0.0325 | 0.275 | 0.408565 | 0.729209 | 0.916104 | 1 | 0.999631 | 0.766996 |
| Mtmr7_Q9Z2C9_S634s_1 | 0.88 | -0.86 | -2.05 | -2.2 | -0.89 | 0.08 | -0.69 | -3.24 | 0.07 | 1.11 | -2.41 | -3.58 | -0.57 | -0.67 | -2.79 | -3.7 | -0.1275 | -0.145 | -0.875 | 0.347724 | 0.884201 | 0.896312 | 1 | 0.999631 | 0.736498 |
| Mtmr7_Q9Z2C9_T578t_1 | -0.6 | -0.7 | -0.09 | -0.82 | -1.32 | -1.15 | -0.86 | 0.28 | -1.51 | -0.24 | -0.44 | -0.11 | -0.93 | -0.31 | 0.55 | 0.58 | -0.21 | -0.0225 | 0.525 | 0.224379 | 0.601304 | 0.952694 | 1 | 0.999631 | 0.67471 |
| Mtor_Q9JLN9_S567s_1 | -0.31 | 0.57 | -0.35 | -0.53 | 0.07 | 0.37 | -0.52 | -0.55 | -0.94 | -0.71 | -0.99 | -0.61 | -0.9 | -0.44 | -0.74 | -0.37 | -0.0025 | -0.6575 | -0.4575 | 0.199703 | 0.994438 | 0.064238 | 1 | 0.999631 | 0.657804 |
| Mtpn_P62774_T31t_1 | -0.05 | 0.2 | 0.23 | 1.12 | 0.22 | -0.37 | 0.44 | 0.55 | 0.79 | -0.01 | 1.68 | 1.72 | 0.71 | 0.6 | 0.66 | 0.53 | -0.165 | 0.67 | 0.25 | 0.457459 | 0.643631 | 0.175672 | 1 | 0.999631 | 0.790722 |
| Mtss1l_D3YWC9_S199s_1 | -0.91 | -0.81 | -1.12 | -0.61 | -1.41 | -1.43 | -0.83 | -0.72 | -1.35 | -2.13 | -0.37 | -0.17 | 0.69 | -1.76 | -2.21 | -1.33 | -0.235 | -0.1425 | -0.29 | 0.626648 | 0.423587 | 0.755183 | 1 | 0.999631 | 0.864779 |
| Mtss1l_D3YWC9_S329s_1 | 0.37 | 0.34 | -0.31 | -0.59 | 0 | 0.15 | -0.22 | -0.16 | 0.66 | 0.39 | 0.1 | 0.2 | 1.03 | 1.43 | -0.04 | 0.34 | -0.01 | 0.385 | 0.7375 | 0.10447 | 0.974183 | 0.253744 | 1 | 0.999631 | 0.568213 |
| Mtss1l_D3YWC9_S357s_1 | 0.82 | 0.5 | 2.75 | 1.57 | 0.62 | 0.29 | 1.69 | 1.35 | 1.79 | 1.14 | 3.46 | 3.93 | 0.01 | 1.24 | 3.98 | 1.89 | -0.4225 | 1.17 | 0.37 | 0.661225 | 0.446673 | 0.136187 | 1 | 0.999631 | 0.880543 |
| Mtss1l_D3YWC9_S502s_1 | 0.58 | -0.62 | 0.68 | 0.03 | -0.33 | 0.04 | 0.15 | 0.26 | 0.08 | 0.66 | 0.47 | 0.24 | 0.33 | 0.52 | 0.7 | 0.31 | -0.1375 | 0.195 | 0.2975 | 0.4186 | 0.697775 | 0.589636 | 1 | 0.999631 | 0.77172 |
| Mtss1l_D3YWC9_S540s_1 | -0.64 | -0.23 | -0.42 | -0.13 | -0.24 | -0.19 | -0.48 | -0.49 | -0.37 | -0.76 | 0 | -0.44 | -0.11 | -0.11 | -0.12 | 0 | 0.005 | -0.0375 | 0.27 | 0.33213 | 0.984224 | 0.895062 | 1 | 0.999631 | 0.728025 |
| Mtss1l_D3YWC9_T195t_1 | -0.55 | -0.04 | -1.11 | -1.06 | -0.86 | -1 | -0.77 | -0.55 | -0.64 | -0.46 | -0.33 | -1.15 | -0.31 | -0.7 | -0.67 | -0.35 | -0.105 | 0.045 | 0.1825 | 0.591992 | 0.741592 | 0.897287 | 1 | 0.999631 | 0.852728 |
| Mtss1l_F7D291_S157s_1 | 1.25 | 0.71 | 0.37 | 0.94 | 1.26 | 1.68 | 0.96 | 0.09 | 1.07 | 1.44 | 0.58 | 0.05 | -1.59 | -0.27 | 0.23 | -0.81 | 0.18 | -0.0325 | -1.4275 | 0.008002 | 0.647955 | 0.931388 | 1 | 0.999631 | 0.364291 |
| Mtus2_Q3UHD3_S109s_1 | 1.64 | 0.3 | -0.87 | -0.04 | 1.4 | 0.76 | 0.33 | -0.14 | 0.74 | 0.1 | -0.73 | -0.64 | -2.3 | 0.73 | -1.13 | -0.89 | 0.33 | -0.39 | -1.155 | 0.131831 | 0.561289 | 0.502722 | 1 | 0.999631 | 0.603456 |
| Mtus2_Q3UHD3_S1324s_1 | -1.25 | -0.28 | -0.4 | -0.23 | 0.38 | 0 | 1.01 | 0.67 | 0.28 | -1.04 | -0.77 | 0.79 | -0.23 | -0.41 | -0.04 | -0.31 | 1.055 | 0.355 | 0.2925 | 0.382031 | 0.012591 | 0.46202 | 1 | 0.999631 | 0.753668 |
| Mtus2_Q3UHD3_S1340s_2 | 4.12 | -2.09 | -1.1 | 1.98 | 1.38 | -0.8 | 3.38 | -0.23 | 0.1 | -0.85 | -0.06 | -2.44 | -3.83 | -2.44 | -2.57 | -2.09 | 0.205 | -1.54 | -3.46 | 0.018363 | 0.887305 | 0.258141 | 1 | 0.999631 | 0.391024 |
| Mtus2_Q3UHD3_S1344s_1 | 2.95 | -1.87 | -0.82 | 0.98 | 0.88 | -0.03 | 3 | -0.24 | -0.38 | -0.81 | 0.21 | -2.21 | -2.68 | -2.12 | -2.36 | -2.4 | 0.5925 | -1.1075 | -2.7 | 0.013788 | 0.594768 | 0.290861 | 1 | 0.999631 | 0.385092 |
| Mtus2_Q3UHD3_S1344s_2 | 4.12 | -2.09 | -1.1 | 1.98 | 1.38 | -0.8 | 3.38 | -0.23 | 0.1 | -0.85 | -0.06 | -2.44 | -3.83 | -2.44 | -2.57 | -2.09 | 0.205 | -1.54 | -3.46 | 0.018363 | 0.887305 | 0.258141 | 1 | 0.999631 | 0.391024 |
| Mtus2_Q3UHD3_S141s_1 | 1.31 | -0.61 | 0.56 | 1.25 | 1.26 | 0.78 | 0.91 | 0.29 | 0.56 | 0.05 | -0.16 | 0.6 | -0.78 | 0.6 | 0.03 | 0.19 | 0.1825 | -0.365 | -0.6175 | 0.241033 | 0.697097 | 0.443399 | 1 | 0.999631 | 0.684338 |
| Mtus2_Q3UHD3_S332s_1 | 0.4 | -0.21 | 0.04 | 0.17 | -0.66 | 0.15 | -0.09 | 0 | 0.43 | 0.96 | 0.14 | 0.2 | -0.77 | -0.08 | -0.41 | -0.6 | -0.25 | 0.3325 | -0.565 | 0.084049 | 0.39783 | 0.285126 | 1 | 0.999631 | 0.549732 |

| | | | | | | | | | | | | | | | | | | | | | | | | | |
|---|---|---|---|---|---|---|---|---|---|---|---|---|---|---|---|---|---|---|---|---|---|---|---|---|---|
| Mtus2_Q3UHD3_S435s_2 | 0.36 | -1.18 | -1.55 | -1.63 | -1.43 | -2.14 | -0.17 | -0.47 | -0.8 | -0.4 | -1.2 | -0.47 | -1.54 | -2.17 | -1.22 | -2.04 | -0.0525 | 0.2825 | -0.7425 | 0.155601 | 0.928659 | 0.558863 | 1 | 0.999631 | 0.626444 |
| Mtus2_Q3UHD3_S951s_1 | 0.72 | -0.93 | -1.59 | -0.06 | 1.12 | 0.21 | 0.45 | -0.08 | 0.16 | -0.72 | -0.58 | -0.74 | -1.59 | -0.21 | -0.72 | -0.62 | 0.89 | -0.005 | -0.32 | 0.559141 | 0.114368 | 0.992248 | 1 | 0.999631 | 0.839222 |
| Mtus2_Q3UHD3_T1331t_1 | -1.47 | -2.1 | -2.86 | -0.24 | 0.6 | -1.02 | 0.4 | -0.47 | -0.1 | -3.33 | -0.19 | -0.47 | -1.59 | -1.46 | -1.76 | -2.16 | 1.545 | 0.645 | -0.075 | 0.889083 | 0.027483 | 0.445574 | 1 | 0.999631 | 0.963505 |
| Mtus2_Q3UHD3_T433t_2 | 0.36 | -1.18 | -1.55 | -1.63 | -1.43 | -2.14 | -0.17 | -0.47 | -0.8 | -0.4 | -1.2 | -0.47 | -1.54 | -2.17 | -1.22 | -2.04 | -0.0525 | 0.2825 | -0.7425 | 0.155601 | 0.928659 | 0.558863 | 1 | 0.999631 | 0.626444 |
| Mtx3_D3YTP3_S249s_1 | 0.23 | -0.29 | 0.42 | 0.14 | -0.15 | 0.52 | -0.34 | 0.1 | -0.11 | 0.12 | -1.03 | 0 | -0.55 | 0.53 | 0.33 | 0.17 | -0.0925 | -0.38 | -0.005 | 0.988337 | 0.759097 | 0.285992 | 1 | 0.999631 | 0.995409 |
| Mtx3_D3YTP3_S284s_1 | 0 | -0.84 | -0.65 | -0.57 | -0.32 | -0.72 | -0.7 | -0.47 | -0.58 | -0.41 | -0.34 | -0.83 | -0.72 | -0.22 | -0.44 | -0.77 | -0.0375 | -0.025 | -0.0225 | 0.9426 | 0.894549 | 0.932092 | 1 | 0.999631 | 0.981629 |
| Mtx3_D3YTP3_S311s_1 | -0.37 | -0.3 | -0.13 | 0.4 | -0.62 | -0.47 | -1.09 | 0.21 | -0.29 | -0.53 | -0.54 | -0.88 | -0.92 | -0.19 | 0.05 | 0.26 | -0.3925 | -0.46 | -0.1 | 0.781564 | 0.277686 | 0.141767 | 1 | 0.999631 | 0.921707 |
| Mvb12a_Q78HU3_S168s_1 | 1.22 | 1.28 | 0.48 | 1.09 | 1.97 | 1.19 | 0.4 | 0.94 | 1.33 | 0.65 | 1.08 | 1.05 | -0.39 | 1.53 | 0.41 | 1.1 | 0.1075 | 0.01 | -0.355 | 0.442384 | 0.780558 | 0.973733 | 1 | 0.999631 | 0.782367 |
| Mvb12b_Q6KAU4_S307s_1 | -0.06 | -0.31 | -0.2 | -0.35 | -0.41 | -0.25 | -0.47 | -0.11 | -0.28 | 0.17 | -0.07 | -0.06 | 0.2 | 0.03 | -0.22 | 0.29 | -0.08 | 0.17 | 0.305 | 0.283408 | 0.741336 | 0.50643 | 1 | 0.999631 | 0.704903 |
| Mxra7_Q9D1T2_S118s_1 | 0.09 | 0.1 | -0.25 | -0.16 | -0.73 | -0.03 | -0.58 | 0.29 | 0 | -0.13 | -0.32 | 0.74 | 0.58 | 0.94 | 0.35 | 0.34 | -0.2075 | 0.1275 | 0.6075 | 0.057449 | 0.507504 | 0.685143 | 1 | 0.999631 | 0.49679 |
| Mxra7_Q9D1T2_S79s_1 | 1.02 | 1.86 | -1.36 | -0.04 | -0.34 | 1.9 | -0.58 | -0.11 | 1.76 | 1.9 | -1.55 | -0.34 | 2.18 | 0.62 | 0.55 | -0.04 | -0.1525 | 0.0725 | 0.4575 | 0.540689 | 0.846144 | 0.938744 | 1 | 0.999631 | 0.831517 |
| Mybbp1a_Q7TPV4_S1164s_1 | -1.39 | -0.7 | -1.33 | -0.36 | -1.12 | -0.31 | -0.67 | -0.82 | -1.04 | 0.17 | -0.13 | -0.27 | -0.33 | -0.23 | -0.27 | -0.27 | 0.215 | 0.6275 | 0.67 | 0.063379 | 0.528751 | 0.123921 | 1 | 0.999631 | 0.512978 |
| Mybbp1a_Q7TPV4_S1253s_1 | -1 | -1.19 | -0.68 | -1.29 | -1.5 | -1.1 | -1.17 | -2.64 | -1.28 | -0.82 | -1.31 | -1.16 | -1.34 | -0.84 | -0.4 | 0.12 | -0.5625 | -0.1025 | 0.425 | 0.275467 | 0.171581 | 0.71172 | 1 | 0.999631 | 0.700187 |
| Mybbp1a_Q7TPV4_S722s_1 | 1.4 | 1.92 | 3.46 | 1.68 | 0.88 | 2.41 | 2.04 | 1.53 | 1.56 | 2.65 | 0.97 | 1.48 | 1.41 | 0.56 | 1.7 | 0.56 | -0.4 | -0.45 | -1.0575 | 0.063995 | 0.453859 | 0.413589 | 1 | 0.999631 | 0.513532 |
| Mycbp2_F6SMY7_S2298s_1 | -0.89 | -0.84 | -0.43 | -0.08 | -1.12 | -1.52 | -0.37 | -0.84 | -1.15 | -1.08 | -0.53 | -0.48 | -0.68 | -1.07 | -0.34 | -0.08 | -0.4025 | -0.25 | 0.0175 | 0.959705 | 0.256215 | 0.441691 | 1 | 0.999631 | 0.986095 |
| Mycbp2_F6SMY7_S2801s_1 | 2.95 | -0.33 | 1.35 | 1.57 | -0.12 | 0.88 | 2.6 | 0.29 | -0.58 | 2.22 | 0.45 | -0.33 | -2.1 | -0.14 | 0.61 | -0.6 | -0.4725 | -0.945 | -1.9425 | 0.028731 | 0.552114 | 0.260242 | 1 | 0.999631 | 0.427571 |
| Mycbp2_F6SMY7_S3893s_1 | -0.82 | -0.49 | -0.97 | -1.23 | -0.25 | -0.65 | -1.12 | -1.19 | -0.36 | -0.95 | -0.92 | -1.19 | -0.14 | -0.66 | -1.05 | -0.72 | 0.075 | 0.0225 | 0.235 | 0.473533 | 0.813697 | 0.941234 | 1 | 0.999631 | 0.799776 |
| Mycbp2_F6SMY7_T1589t_1 | 1.09 | 0.14 | 0.57 | 0.46 | 0.31 | 0.76 | -0.06 | 0.27 | -0.37 | 2.23 | 0.39 | -0.38 | 1.13 | 1.04 | -0.92 | -2.08 | -0.245 | -0.0975 | -0.7725 | 0.293955 | 0.440799 | 0.868689 | 1 | 0.999631 | 0.708293 |
| Mycbp2_F6SMY7_T3882t_1 | -1.7 | 0.03 | -0.52 | -0.7 | -0.78 | -1.02 | -1.19 | -0.74 | -0.78 | -0.65 | 0.41 | -0.21 | 0.41 | -1.91 | 0.2 | -0.88 | -0.21 | 0.415 | 0.1775 | 0.764156 | 0.589701 | 0.360824 | 1 | 0.999631 | 0.915775 |
| Myef2_Q8C854_S16s_1 | 0.25 | 0.17 | 2.12 | 0.87 | -0.4 | 0.8 | 0.58 | 0.91 | 0.14 | 0.63 | 0.18 | 1.21 | -0.87 | 0.45 | 0.96 | 0.38 | -0.38 | -0.3125 | -0.6225 | 0.275985 | 0.458915 | 0.528891 | 1 | 0.999631 | 0.700187 |
| Myef2_Q8C854_S403s_1 | -0.11 | 0.14 | 0.21 | 1.28 | 0.7 | 0.23 | 1.3 | 0.56 | 0.65 | 0.6 | 0 | 0.58 | -0.84 | -0.26 | 0.49 | 0.17 | 0.3175 | 0.0775 | -0.49 | 0.270316 | 0.423181 | 0.83455 | 1 | 0.999631 | 0.698438 |
| Myh10_Q5SV64_S1970s_1 | 1.64 | 0.42 | -1.21 | 1.76 | 2.61 | 0.91 | 1.73 | 1.7 | 1.28 | 0.02 | 1.34 | 1.08 | -1.05 | 1.42 | 0.62 | 0.9 | 1.085 | 0.2775 | -0.18 | 0.814065 | 0.137764 | 0.682825 | 1 | 0.999631 | 0.937041 |
| Myh10_Q5SV64_S1996s_1 | -0.29 | 2.65 | 1.64 | 0 | 2.21 | 2.69 | 1.06 | -0.32 | 2.5 | 0.8 | 0.76 | 1.08 | 1.25 | 0.99 | 0.76 | -0.75 | 0.41 | 0.285 | -0.4375 | 0.552193 | 0.627472 | 0.692045 | 1 | 0.999631 | 0.836594 |
| Myh10_Q5SV64_S2006s_1 | -1.29 | -0.07 | 1.12 | -1.24 | 0.4 | -0.79 | 0.94 | -0.57 | 1.17 | -1.54 | 1.24 | 1.46 | 1 | -1.87 | 0.27 | -1.75 | 0.365 | 0.9525 | -0.2175 | 0.786481 | 0.56525 | 0.251505 | 1 | 0.999631 | 0.923689 |
| Myh10_Q5SV64_T1995t_1 | -0.29 | 2.65 | 1.64 | 0 | 2.21 | 2.69 | 1.06 | -0.32 | 2.5 | 0.8 | 0.76 | 1.08 | 1.25 | 0.99 | 0.76 | -0.75 | 0.41 | 0.285 | -0.4375 | 0.552193 | 0.627472 | 0.692045 | 1 | 0.999631 | 0.836594 |
| Myh11_E9QPE7_S1954s_1 | -0.55 | -0.31 | -0.25 | 0.62 | 3.7 | 0.47 | 0.02 | -0.84 | 1.13 | 3.73 | 3.68 | 5.19 | -0.84 | 0.24 | 0.98 | 0.3 | 0.96 | 3.555 | 0.2925 | 0.522569 | 0.294474 | 0.001095 | 1 | 0.999631 | 0.821224 |
| Myh9_Q8VDD5_S1943s_1 | 1.48 | 0.71 | -3.43 | -3.78 | 0.23 | 1.16 | -2.18 | -4.53 | 1.44 | 2.46 | -3.35 | -4.81 | 0.41 | 0.49 | -4.46 | -5.29 | -0.075 | 0.19 | -0.9575 | 0.582112 | 0.962228 | 0.920226 | 1 | 0.999631 | 0.847567 |
| Myl12a_Q6ZWQ9_S20s_1 | 1.62 | 0.91 | 0.85 | 1.2 | 1.15 | 0.95 | 1.24 | 1.39 | 0.66 | 0.6 | 1.02 | 0.88 | -0.11 | 1.16 | 0.77 | 1.44 | 0.0375 | -0.355 | -0.33 | 0.421085 | 0.893378 | 0.237185 | 1 | 0.999631 | 0.772817 |
| Myo18a_E9PUR2_S279s_1 | 0.93 | 1.18 | 0.27 | -0.14 | -0.49 | 0.58 | -0.17 | -0.31 | -0.9 | 1.84 | -0.2 | -0.41 | -0.38 | -0.91 | 0.05 | 0.51 | -0.6575 | -0.4775 | -0.7425 | 0.111523 | 0.117445 | 0.443372 | 1 | 0.999631 | 0.577068 |
| Myo18a_E9PUR2_S52s_1 | -0.23 | -0.79 | -0.66 | 1.1 | 0.21 | -0.75 | 0.18 | 0.3 | -0.77 | -0.04 | -0.96 | 0.34 | -0.7 | 0.06 | 0.49 | 0.7 | 0.13 | -0.2125 | 0.2825 | 0.579923 | 0.783731 | 0.673066 | 1 | 0.999631 | 0.84667 |
| Myo18a_E9QA74_S1595s_1 | 1.03 | 1.23 | 1.16 | 1.09 | -0.15 | 2.28 | 0.97 | 0.84 | 1.63 | 2.65 | 1.48 | 1.3 | 2.52 | 1.71 | 1.68 | 2.32 | -0.1425 | 0.6375 | 0.93 | 0.012626 | 0.764884 | 0.090878 | 1 | 0.999631 | 0.385092 |
| Myo18a_E9QA74_S472s_1 | 2 | 1.41 | 0.29 | 2.23 | 2.05 | 1.24 | 1.99 | 0.98 | 1.91 | 0.48 | 1.45 | 1.24 | -0.27 | 0.24 | 0.39 | 0.35 | 0.0825 | -0.2125 | -1.305 | 0.015846 | 0.864175 | 0.6717 | 1 | 0.999631 | 0.388832 |
| Myo18a_Q9JMH9_S164s_1 | 0.11 | -0.32 | 1.18 | 1.07 | 1.52 | 1.05 | 1.45 | -0.94 | -0.19 | -0.97 | 0.86 | -0.12 | -2.06 | 0.62 | 0.71 | 1.09 | 0.26 | -0.615 | -0.42 | 0.561358 | 0.674592 | 0.235562 | 1 | 0.999631 | 0.840316 |
| Myo18a_Q9JMH9_S340s_1 | -1.92 | -4.17 | -1.15 | -1.44 | -4.51 | -5.28 | -1.52 | -1.37 | -5.75 | -3.38 | -1.52 | -1.49 | -3.57 | -4.93 | -1.32 | -1.01 | -1 | -0.865 | -0.5375 | 0.590492 | 0.351097 | 0.418187 | 1 | 0.999631 | 0.851647 |
| Myo5a_D3YZ62_S1425s_1 | -0.52 | -0.88 | -0.73 | -0.6 | -0.82 | -1.06 | -0.95 | -0.5 | -0.88 | -0.8 | -0.88 | -0.47 | -0.61 | -0.6 | -0.63 | -0.25 | -0.15 | -0.075 | 0.16 | 0.560724 | 0.561239 | 0.770822 | 1 | 0.999631 | 0.839977 |
| Myo5a_D3YZ62_S1625s_1 | -0.05 | -0.33 | -0.65 | 0.27 | 0.36 | -0.26 | -0.02 | 0.28 | -0.51 | -0.71 | -0.44 | -0.02 | -1.05 | -0.13 | -0.29 | 0.21 | 0.28 | -0.23 | -0.125 | 0.737595 | 0.366512 | 0.465943 | 1 | 0.999631 | 0.907795 |
| Myo5a_D3YZ62_S600s_1 | -0.4 | -0.55 | -0.43 | -0.81 | -0.82 | -0.89 | -0.72 | -0.86 | -0.61 | -0.35 | -0.6 | -0.5 | 0.35 | -0.29 | -0.53 | -0.44 | -0.275 | 0.0325 | 0.32 | 0.31776 | 0.269767 | 0.897508 | 1 | 0.999631 | 0.721252 |
| Myo5a_D3YZ62_T1623t_1 | 0.1 | -0.73 | -0.33 | 0.44 | 0.19 | -1.14 | 0.05 | 0.2 | 0.33 | -1.03 | 0.26 | 0.11 | -0.24 | -0.81 | -0.3 | 0.4 | -0.045 | 0.0475 | -0.1075 | 0.781604 | 0.912801 | 0.908654 | 1 | 0.999631 | 0.921707 |
| Myo6_E9Q3L1_S1123s_1 | -0.47 | -0.24 | -0.56 | -0.37 | 0.11 | -0.26 | -0.83 | 0.24 | -0.18 | -0.73 | -0.45 | 0.29 | 0.47 | -0.34 | -1.18 | -0.5 | 0.225 | 0.1425 | 0.0225 | 0.952914 | 0.469803 | 0.639009 | 1 | 0.999631 | 0.984317 |
| Myo9a_D3Z3A8_S1259s_1 | 0.38 | 0.95 | 0.69 | 0.31 | -0.22 | 0.08 | 0.11 | 0.66 | 0.09 | 0.02 | 0.69 | 0.69 | 0.19 | 0.31 | 1.21 | 0.66 | -0.425 | -0.21 | 0.01 | 0.976423 | 0.177529 | 0.498504 | 1 | 0.999631 | 0.991938 |
| Myo9a_D3Z3A8_S1300s_1 | 0.47 | 0.07 | -1.43 | -0.62 | -0.05 | 0.7 | -0.85 | -1.4 | 0.35 | 0.65 | -0.14 | -0.25 | -0.18 | 0.44 | -0.25 | -0.72 | -0.0225 | 0.53 | 0.2 | 0.671865 | 0.968276 | 0.262287 | 1 | 0.999631 | 0.884779 |
| Myo9b_E9QKV6_S1113s_1 | -0.83 | 0.06 | -0.12 | -1.16 | 0.3 | -0.51 | -0.67 | -1.21 | -0.03 | 0.59 | -0.2 | -0.49 | 1.24 | 0.6 | -0.01 | -0.78 | -0.01 | 0.48 | 0.775 | 0.142835 | 0.981031 | 0.23354 | 1 | 0.999631 | 0.613952 |
| Myo9b_E9QKV6_S1264s_1 | -0.92 | -0.51 | -0.88 | -1.51 | -0.34 | -1.14 | 0.1 | -1.2 | 0.06 | -1.41 | -1.16 | 0.38 | 0.68 | -0.61 | -0.88 | -1.39 | 0.31 | 0.4225 | 0.405 | 0.402711 | 0.431849 | 0.380376 | 1 | 0.999631 | 0.764046 |
| Myo9b_E9QKV6_S1335s_1 | -4.4 | -0.88 | -0.77 | -2.05 | -1.22 | -2.09 | -1.57 | -1.83 | -1.5 | -1.54 | -1.01 | -0.97 | 0.23 | -1.08 | -0.86 | -2.55 | 0.3475 | 0.77 | 0.96 | 0.289997 | 0.648075 | 0.322032 | 1 | 0.999631 | 0.706164 |
| Myof_E9Q390_S174s_1 | 0.37 | 0.42 | 1.06 | 0.95 | 0.62 | 0.03 | 0.35 | 0.27 | 0.99 | 0.61 | -0.07 | 0.79 | -0.81 | 0.73 | 0.44 | 0.59 | -0.3825 | -0.12 | -0.4625 | 0.278564 | 0.205339 | 0.72315 | 1 | 0.999631 | 0.70098 |
| Mypop_Q8R4U1_S182s_1 | -0.11 | 0.05 | 0.84 | -0.18 | -0.95 | -0.32 | -0.62 | 0.31 | -0.24 | -0.45 | 0.73 | 0.71 | 0.84 | 0.51 | 1.17 | 0.55 | -0.545 | 0.0375 | 0.6175 | 0.096067 | 0.166907 | 0.925478 | 1 | 0.999631 | 0.558708 |
| Myrip_Q8K3I4_S540s_1 | 0.72 | 0.58 | 0.77 | 0.8 | 0.69 | 0.41 | 0.11 | 1.04 | 0.27 | 0.5 | 0.99 | 0.55 | -0.13 | 0.46 | 0.62 | 0.63 | -0.155 | -0.14 | -0.3225 | 0.29135 | 0.586505 | 0.605652 | 1 | 0.999631 | 0.706164 |
| Myrip_Q8K3I4_S618s_1 | 0.08 | 0.46 | -1.13 | -1.51 | -0.56 | -0.13 | -2.35 | -1.81 | -0.78 | 0.04 | -1.99 | -1.95 | -0.73 | 0.01 | -0.91 | -0.33 | -0.6875 | -0.645 | 0.035 | 0.943565 | 0.290393 | 0.310444 | 1 | 0.999631 | 0.981629 |
| Myt1l_P97500_S391s_1 | 0.68 | 1.62 | 0.12 | -0.06 | 0.58 | -0.11 | 0.4 | 0.86 | 0.32 | 0.95 | 0.89 | 0.74 | 0.98 | 0.88 | 0.82 | 1.59 | -0.1575 | 0.135 | 0.4775 | 0.279064 | 0.710935 | 0.743089 | 1 | 0.999631 | 0.701431 |
| Myt1l_P97500_S470s_1 | 1.64 | 0.14 | 1.69 | 2.19 | -0.35 | 0.53 | 1.77 | 1.01 | 0.44 | 1.55 | 1.23 | 1.71 | -0.52 | -0.89 | 1.25 | 1.07 | -0.675 | -0.1825 | -1.1875 | 0.08491 | 0.254899 | 0.715133 | 1 | 0.999631 | 0.550457 |
| Myt1l_P97500_S685s_1 | -0.92 | 0.06 | -1.15 | -0.68 | -0.58 | -0.54 | -1.4 | -0.26 | -0.41 | -0.18 | -0.82 | -0.02 | -0.47 | -0.2 | 0.18 | -0.68 | -0.0225 | 0.315 | 0.38 | 0.313377 | 0.952144 | 0.381538 | 1 | 0.999631 | 0.719861 |
| Myt1l_P97500_T372t_1 | -0.08 | 0.71 | 0.96 | 0.14 | 0.27 | 0.37 | 0.33 | -0.03 | 0.9 | 0.12 | 0.75 | 0.03 | 0.73 | 0.39 | -0.98 | 0.16 | -0.1975 | 0.0175 | -0.3575 | 0.428747 | 0.531248 | 0.961199 | 1 | 0.999631 | 0.776573 |
| N4bp1_Q6A037_S560s_1 | -0.26 | -0.38 | -0.38 | -0.12 | -0.21 | -0.32 | -0.94 | -0.97 | -0.25 | -0.2 | -1.93 | -0.42 | 0.15 | 0.38 | 0.77 | -0.25 | -0.325 | -0.415 | 0.5475 | 0.104378 | 0.271513 | 0.336337 | 1 | 0.999631 | 0.568213 |
| Nab2_Q61127_S147s_1 | 2.79 | 0.63 | 2.09 | 2.34 | -0.38 | 0.79 | 2.01 | 0.88 | 0.03 | 1.89 | 0.71 | 1.42 | -0.61 | -0.37 | 1.77 | 0.16 | -1.1375 | -0.95 | -1.725 | 0.02107 | 0.086688 | 0.120558 | 1 | 0.999631 | 0.400544 |
| Nab2_Q61127_S6s_1 | 1.02 | 1.24 | 1.87 | 1.19 | 0.4 | 0.06 | 0.83 | 1.29 | 0.08 | 0.95 | 1.21 | 1.33 | 0.65 | 0.52 | 1.75 | 1.6 | -0.685 | -0.4375 | -0.2 | 0.614828 | 0.076456 | 0.251097 | 1 | 0.999631 | 0.861203 |
| Nab2_Q61127_T434t_1 | 0.31 | 1.55 | 2.42 | 0.78 | 1.07 | 0.72 | 0.99 | 1.87 | 0.39 | 1.04 | 1.82 | 1.66 | 0.22 | 1.43 | 2.01 | 2.34 | -0.1025 | -0.0375 | 0.235 | 0.695616 | 0.835385 | 0.943417 | 1 | 0.999631 | 0.895676 |
| Nacad_Q5SWP3_S1077s_1 | 0.3 | 1.56 | 0.09 | 1.11 | 1.99 | 2.8 | 1.46 | 0.69 | 2.9 | 0.63 | 2.07 | 1.5 | -0.75 | 2.64 | 1.59 | 0.73 | 0.97 | 1.01 | 0.2875 | 0.684238 | 0.087799 | 0.089617 | 1 | 0.999631 | 0.889649 |
| Nacad_Q5SWP3_S1214s_1 | 0.19 | -1.32 | -2.57 | -2.01 | -1.8 | -1.65 | -0.86 | -1.55 | -1.31 | -1.41 | -1.39 | -0.56 | -1.31 | -0.68 | -2.87 | -1.57 | -0.0375 | 0.26 | -0.18 | 0.789171 | 0.947966 | 0.654587 | 1 | 0.999631 | 0.925513 |
| Nacad_Q5SWP3_S1268s_1 | -0.17 | 3.91 | 3.44 | 0.82 | 2.61 | 2.19 | 2.13 | 2.36 | 4 | 0.59 | 3.79 | 4.11 | 3.35 | 2.06 | 3.68 | 1.65 | 0.3225 | 1.1225 | 0.685 | 0.47691 | 0.710561 | 0.329331 | 1 | 0.999631 | 0.799947 |
| Nadk_P58058_S46s_1 | 1.18 | -0.19 | -0.33 | 0.21 | 1.13 | -0.28 | -0.03 | 0.1 | 0.58 | -1.35 | 0.44 | -0.3 | 0.23 | 1.4 | -1.05 | -1.34 | 0.0125 | -0.375 | -0.4075 | 0.531046 | 0.977635 | 0.479462 | 1 | 0.999631 | 0.82688 |
| Nadk_P58058_S48s_1 | 0.55 | 0.21 | -0.34 | 0.57 | 0.37 | -0.08 | 0.5 | -0.14 | 0.3 | 0.15 | 0.3 | -0.4 | -0.16 | 0.73 | -0.48 | 0.14 | -0.085 | -0.16 | -0.19 | 0.613733 | 0.788532 | 0.625065 | 1 | 0.999631 | 0.860866 |
| Nans_Q99J77_S275s_1 | 1.55 | 0.65 | 1.03 | 0.92 | 0.15 | 1.07 | 1.3 | 0.14 | 1.4 | 1.94 | 0.8 | 0.96 | 1.28 | 0.79 | 1.06 | 0.77 | -0.3725 | 0.2375 | -0.0625 | 0.842225 | 0.332096 | 0.507493 | 1 | 0.999631 | 0.947766 |
| Napa_Q9DB05_S26s_1 | -2.95 | 0.39 | -1.7 | -2.7 | 0.88 | -0.9 | 0.44 | -2.24 | 0.75 | -4.04 | -0.5 | 0.49 | 0.26 | -2.49 | -0.98 | -2.98 | 1.285 | 0.915 | 0.1925 | 0.832795 | 0.174545 | 0.434 | 1 | 0.999631 | 0.94321 |
| Napepld_Q8BH82_S27s_1 | 1.41 | 0.27 | 0.27 | 1.07 | -0.2 | 0.92 | 1.43 | 0.63 | 0.24 | 1.18 | 0.33 | -0.83 | -1.96 | -0.16 | -1.12 | -0.72 | -0.06 | -0.525 | -1.745 | 0.003759 | 0.890546 | 0.291457 | 1 | 0.999631 | 0.351308 |
| Napg_Q9CWZ7_S284s_1 | 1.01 | 0.22 | 0.96 | -0.52 | -1.15 | 0.63 | -0.72 | 0.05 | -0.29 | 0.96 | 0.79 | 0.33 | -0.03 | 0.27 | 0.2 | 0.33 | -0.715 | 0.03 | -0.225 | 0.573034 | 0.176735 | 0.946614 | 1 | 0.999631 | 0.844445 |
| Napg_Q9CWZ7_T287t_1 | 2.6 | 2.04 | 1.41 | 1.06 | 0.6 | 2.73 | 1.34 | 0.82 | 1.2 | 2.43 | 0.26 | 0.36 | 0.39 | 1.55 | 0.98 | 0.82 | -0.405 | -0.715 | -0.8425 | 0.071985 | 0.460716 | 0.221604 | 1 | 0.999631 | 0.532005 |
| Nasp_B1AU75_S712s_1 | 2.97 | 1.62 | 3.49 | 2.07 | 0.5 | 1.73 | 2.47 | 1.13 | 1.47 | 2.91 | 1.77 | 1 | 0.52 | 0.6 | 1.11 | 0.8 | -1.08 | -0.75 | -1.78 | 0.002446 | 0.073175 | 0.190466 | 1 | 0.999631 | 0.351308 |
| Nav1_Q8CH77_S1010s_1 | -0.71 | -0.54 | -0.75 | -0.57 | -0.26 | -1.33 | -0.75 | -0.44 | -0.42 | -0.82 | -0.42 | 0.12 | -0.13 | -0.08 | -0.38 | -0.04 | -0.0525 | 0.2575 | 0.485 | 0.091458 | 0.861904 | 0.381383 | 1 | 0.999631 | 0.557085 |
| Nav1_Q8CH77_S1251s_1 | -0.72 | -0.05 | -0.24 | -0.58 | 0.08 | -0.04 | -0.79 | -0.19 | -0.55 | -0.58 | -0.19 | 0.06 | 0.85 | -0.84 | 0.23 | 0.09 | 0.1625 | 0.0825 | 0.48 | 0.251417 | 0.597315 | 0.781127 | 1 | 0.999631 | 0.687149 |
| Nav1_Q8CH77_S291s_1 | 0.52 | 1.61 | 0.8 | 0.67 | 1.07 | -0.11 | -0.28 | -0.5 | 0.32 | -0.58 | 0.99 | 0.65 | -0.27 | 1.05 | -0.24 | 0.98 | -0.855 | -0.555 | -0.52 | 0.256955 | 0.066594 | 0.20654 | 1 | 0.999631 | 0.691567 |
| Nav1_Q8CH77_S454s_1 | 1.48 | -0.8 | -1.31 | 1.63 | 1.71 | 0.35 | 1.01 | 1.49 | 0.64 | 0.51 | 1.23 | 0.41 | -2.38 | 1.4 | 1.18 | 0.53 | 0.89 | 0.4475 | -0.0675 | 0.945674 | 0.236255 | 0.525653 | 1 | 0.999631 | 0.982217 |
| Nav1_Q8CH77_S455s_1 | 1.74 | -0.15 | -0.49 | 1.51 | 2.33 | -0.03 | 1.66 | 1.72 | 1.8 | 0.07 | 1.27 | 1.36 | 0.65 | 0.6 | 0.69 | 0.87 | 0.7675 | 0.4725 | 0.05 | 0.925498 | 0.272533 | 0.449382 | 1 | 0.999631 | 0.975764 |
| Nav1_Q8CH77_S510s_1 | 3.04 | 1.46 | 0.97 | 1.47 | 0.72 | 1.28 | 2.61 | 0.77 | 0.86 | 1.48 | 0.69 | 1.13 | -0.95 | 0.04 | 1.21 | -0.24 | -0.39 | -0.695 | -1.72 | 0.013769 | 0.501657 | 0.159988 | 1 | 0.999631 | 0.385092 |
| Nav1_Q8CH77_S544s_1 | 0.21 | -0.22 | -1.56 | -0.45 | 0.2 | 0.57 | -0.31 | 0.02 | 0.25 | 0.21 | -0.56 | -0.54 | -0.29 | 0.07 | -1.21 | -0.61 | 0.625 | 0.345 | -0.005 | 0.991311 | 0.156982 | 0.436723 | 1 | 0.999631 | 0.996474 |
| Nav1_Q8CH77_S762s_1 | -0.18 | 1.12 | -0.6 | -0.17 | 0.82 | 0.75 | -0.4 | 0.42 | 0.07 | 0.45 | -0.22 | -0.09 | 1.23 | 1.21 | -0.26 | 0.25 | 0.355 | 0.01 | 0.565 | 0.275686 | 0.438103 | 0.980457 | 1 | 0.999631 | 0.700187 |
| Nav1_Q8CH77_S810s_1 | -1.58 | -0.67 | -0.67 | -1.21 | -1.13 | -1.24 | -0.07 | -0.8 | -0.94 | -1.47 | -0.23 | -0.33 | -0.77 | -1.22 | -0.86 | -0.58 | 0.2225 | 0.29 | 0.175 | 0.600581 | 0.546841 | 0.457538 | 1 | 0.999631 | 0.856688 |
| Nav1_Q8CH77_S93s_1 | 0.43 | 0.93 | 0.73 | 1.08 | 0.98 | 0.94 | 0.65 | 1.12 | 1.47 | 0.37 | 1.25 | 1.63 | 1.19 | 1.07 | 0.96 | 1.43 | 0.13 | 0.3875 | 0.37 | 0.222615 | 0.628754 | 0.286538 | 1 | 0.999631 | 0.673692 |
| Nav1_Q8CH77_S952s_1 | 0.51 | -1.08 | -1.17 | 0.75 | 0.35 | -0.93 | 0.51 | 0.44 | 0.38 | -0.5 | 0.52 | 0.21 | -1.18 | -0.47 | -0.08 | -0.02 | 0.34 | 0.4 | -0.19 | 0.724736 | 0.548321 | 0.451227 | 1 | 0.999631 | 0.905674 |
| Nav1_Q8CH77_S998s_1 | 0.93 | -0.35 | -1.21 | 1.12 | 0.72 | -0.02 | 0.82 | 0.84 | 0.33 | 0.21 | 0.41 | -0.36 | -1.61 | 0.24 | 0.48 | 0.2 | 0.4675 | 0.025 | -0.295 | 0.654745 | 0.397304 | 0.962923 | 1 | 0.999631 | 0.878941 |
| Nav1_Q8CH77_T360t_1 | -0.05 | 2.22 | 1.27 | 0.35 | 1.66 | 2.46 | 1.54 | 1.98 | 1.69 | 0.28 | 2.26 | 1.36 | 0.5 | 1.45 | 1.78 | -0.3 | 0.9625 | 0.45 | -0.09 | 0.885523 | 0.083785 | 0.458219 | 1 | 0.999631 | 0.963505 |
| Nav2_V9GXT3_S1240s_1 | -0.64 | 0.26 | 0.66 | -1.31 | 0.81 | -0.06 | 0.54 | 0.92 | 1.63 | -0.8 | 1.6 | 2 | 0.61 | -0.64 | 0.61 | 0.31 | 0.81 | 1.365 | 0.48 | 0.356019 | 0.110733 | 0.074202 | 1 | 0.999631 | 0.74034 |
| Nav3_E9QMF5_S1539s_1 | 0.59 | 0.09 | 0.68 | 0.28 | -0.56 | 1.09 | 0.8 | 1.16 | 0.83 | 1.29 | 1.26 | 1.64 | 0.25 | -0.84 | 0.9 | 0.68 | 0.2125 | 0.845 | -0.1625 | 0.701419 | 0.615317 | 0.01627 | 1 | 0.999631 | 0.896835 |
| Nav3_E9QMF5_S1848s_1 | 0.52 | -2.8 | 0.91 | -0.02 | -1.38 | -1.14 | 0.11 | 0.91 | -1.5 | 0.44 | -0.4 | -0.67 | -1.72 | -0.97 | -1.14 | -1.21 | -0.0275 | -0.185 | -0.9125 | 0.241991 | 0.974632 | 0.820474 | 1 | 0.999631 | 0.684914 |
| Nav3_E9QMF5_S275s_1 | 0.01 | -0.25 | -0.26 | 0.43 | 0.17 | 0.25 | 0.37 | -0.02 | 0.51 | -0.11 | 0.54 | 0.44 | 0.03 | -0.25 | -0.18 | 0.79 | 0.21 | 0.3625 | 0.115 | 0.741155 | 0.446755 | 0.244224 | 1 | 0.999631 | 0.909055 |
| Nav3_E9QMF5_S354s_1 | 0.94 | -0.55 | 0.4 | -1.14 | 1.11 | 0.96 | 0.66 | 1.45 | 0.92 | 0.22 | 0.81 | 1.04 | -0.2 | 0.68 | 0.57 | 0.39 | 1.1325 | 0.835 | 0.4475 | 0.370275 | 0.035332 | 0.107501 | 1 | 0.999631 | 0.746338 |
| Nbas_E9Q411_S472s_1 | -0.59 | 0.45 | -0.21 | -0.15 | -0.16 | -0.01 | -0.53 | 0.61 | 0.07 | 0.09 | -0.5 | -0.02 | -0.47 | -0.49 | -0.38 | 0.2 | 0.1025 | 0.035 | -0.16 | 0.637212 | 0.769838 | 0.912167 | 1 | 0.999631 | 0.870011 |
| Nbas_E9Q411_S474s_1 | -0.07 | -0.48 | 0.63 | -0.21 | -0.08 | -0.01 | -0.08 | -0.38 | -1.81 | 0.59 | 0.24 | -0.03 | -0.21 | -0.91 | 0.6 | 0.32 | -0.105 | -0.22 | -0.0175 | 0.966807 | 0.733509 | 0.686584 | 1 | 0.999631 | 0.988351 |
| Nbea_Q9EPN1_S1311s_1 | 0.56 | 0.66 | 0.35 | 0.44 | 0.4 | 0.8 | 0.27 | 0.52 | 0.72 | 1.02 | 0.4 | 0.04 | 0.45 | 1.05 | 0.33 | 0.62 | -0.005 | 0.0425 | 0.11 | 0.705897 | 0.984098 | 0.88685 | 1 | 0.999631 | 0.897883 |
| Nbea_Q9EPN1_S1519s_1 | -0.49 | -0.21 | 0.36 | -0.29 | -0.24 | -0.15 | -0.31 | 0.24 | -0.17 | 0.13 | 0.22 | 0.47 | 0.43 | 0.24 | 0.48 | 0.81 | 0.0425 | 0.32 | 0.6475 | 0.059617 | 0.883765 | 0.301868 | 1 | 0.999631 | 0.50521 |
| Nbea_Q9EPN1_S1596s_1 | -1.23 | -2.46 | -1.01 | -0.37 | -1.61 | -0.99 | -0.49 | -0.79 | -1.88 | -1.67 | -1.21 | -0.95 | -1.52 | -1.19 | -1.01 | -0.51 | 0.2975 | -0.16 | 0.21 | 0.659198 | 0.533744 | 0.734007 | 1 | 0.999631 | 0.880034 |
| Nbea_Q9EPN1_S1724s_1 | 1.09 | 0.34 | 1.11 | 1.61 | 1.59 | 1.11 | 0.24 | 1.14 | 1.2 | -0.7 | -0.22 | 1.26 | -0.08 | 1.36 | 0.57 | 0.66 | -0.0175 | -0.6525 | -0.41 | 0.331898 | 0.964411 | 0.234056 | 1 | 0.999631 | 0.727673 |
| Nbea_Q9EPN1_S2565s_1 | 0 | -0.56 | -0.88 | -0.09 | 0.06 | -0.57 | 0.27 | -0.02 | -0.25 | -0.47 | 0.15 | -1.52 | -0.87 | -0.17 | 0.07 | 0.16 | 0.3175 | -0.14 | 0.18 | 0.619877 | 0.334824 | 0.737481 | 1 | 0.999631 | 0.861888 |
| Ncam1_A0A0A6YY91_S1019s_1 | -0.4 | 0.08 | -0.43 | -0.15 | 0.16 | 0.2 | -0.6 | -0.02 | -0.13 | -1.21 | -0.12 | -0.16 | 0 | -0.1 | 0.05 | 0.22 | 0.16 | -0.18 | 0.2675 | 0.347312 | 0.585147 | 0.598731 | 1 | 0.999631 | 0.736373 |
| Ncam1_P13595_S1005s_1 | -0.47 | -1.14 | -1.14 | -0.9 | -0.94 | -1.06 | -0.93 | -1.02 | -1.14 | -0.94 | -0.93 | -0.82 | -0.03 | -0.62 | -0.68 | -0.66 | -0.075 | -0.045 | 0.415 | 0.204759 | 0.775565 | 0.869989 | 1 | 0.999631 | 0.660366 |
| Ncam1_P13595_S1016s_1 | 0.12 | 0.2 | -1.07 | -1.33 | -0.21 | 0.45 | -0.65 | -0.61 | 0.77 | 0.45 | -1.6 | -0.73 | -0.39 | 1.24 | -0.57 | -0.91 | 0.265 | 0.2425 | 0.3625 | 0.53062 | 0.563021 | 0.693301 | 1 | 0.999631 | 0.82672 |
| Ncam1_P13595_S774s_1 | -6.75 | -7.16 | -2.92 | -4.39 | -4.99 | -7.56 | -2.03 | -3 | -5.06 | -6.47 | -0.73 | -0.64 | -3.23 | -7.25 | -1.65 | -3.87 | 0.91 | 2.08 | 1.305 | 0.329988 | 0.503372 | 0.195054 | 1 | 0.999631 | 0.726756 |
| Ncam1_P13595_S817s_1 | -0.63 | -2.17 | -1.93 | -2.25 | -1.65 | -2.05 | -1.25 | -1.95 | -2.09 | -1.05 | -2.11 | -2.14 | -2.63 | -2.63 | -2.1 | -2.68 | 0.02 | -0.1025 | -0.765 | 0.090761 | 0.961614 | 0.820857 | 1 | 0.999631 | 0.557085 |
| Ncam1_P13595_S887s_1 | -0.3 | -1.14 | -0.78 | -0.82 | -0.97 | -0.64 | -0.74 | -0.61 | -0.46 | -0.35 | -0.56 | -0.59 | -0.08 | -1.05 | -0.7 | -0.1 | 0.02 | 0.27 | 0.2775 | 0.436294 | 0.942294 | 0.345947 | 1 | 0.999631 | 0.780748 |
| Ncam1_P13595_S946s_1 | -0.34 | -0.42 | -0.8 | -0.3 | -0.09 | -0.58 | -0.33 | 0.28 | -0.48 | -0.33 | -0.57 | -0.29 | -0.41 | -0.54 | -0.27 | -0.07 | 0.285 | 0.0475 | 0.1425 | 0.617096 | 0.336141 | 0.854626 | 1 | 0.999631 | 0.861376 |
| Ncam1_P13595_S946s_2 | 0.58 | 0.59 | -0.99 | -0.2 | 0.27 | 0.6 | -0.76 | -0.26 | 0.28 | 0.76 | -0.81 | 0.42 | -2.41 | -0.92 | -0.05 | -0.35 | -0.0325 | 0.1675 | -0.9275 | 0.139877 | 0.943774 | 0.73064 | 1 | 0.999631 | 0.611718 |
| Ncam1_P13595_S958s_1 | -0.66 | -0.22 | -0.53 | -0.66 | -0.64 | -0.72 | -0.9 | -0.17 | -0.55 | -0.71 | -0.79 | -0.22 | 0.53 | -0.12 | -0.56 | -0.28 | -0.09 | -0.05 | 0.41 | 0.230466 | 0.743467 | 0.8539 | 1 | 0.999631 | 0.681463 |
| Ncam1_P13595_S958s_2 | 0.58 | 0.59 | -0.99 | -0.2 | 0.27 | 0.6 | -0.76 | -0.26 | 0.28 | 0.76 | -0.81 | 0.42 | -2.41 | -0.92 | -0.05 | -0.35 | -0.0325 | 0.1675 | -0.9275 | 0.139877 | 0.943774 | 0.73064 | 1 | 0.999631 | 0.611718 |
| Ncam1_P13595_T1030t_1 | 0.02 | 1.21 | -0.65 | -0.05 | 0.32 | 1 | -0.37 | -0.02 | 0.77 | 0.69 | -0.33 | 0.04 | -0.03 | 1.33 | 0.01 | 0.31 | 0.1 | 0.16 | 0.2725 | 0.577918 | 0.829914 | 0.727782 | 1 | 0.999631 | 0.84622 |
| Ncam1_P13595_T783t_1 | 0.98 | 2.55 | -0.63 | -1.07 | 1.09 | 2.04 | -0.67 | -0.97 | 2.48 | 2.48 | -0.65 | -0.83 | 2.64 | 2.61 | -1.15 | -0.46 | -0.085 | 0.4125 | 0.4525 | 0.682455 | 0.928182 | 0.700764 | 1 | 0.999631 | 0.889581 |
| Ncam1_P13595_T783t_1_1 | 0.98 | 2.55 | -0.24 | -0.74 | 1.09 | 2.04 | -0.34 | -0.65 | 2.48 | 2.48 | -0.28 | -0.41 | 2.64 | 2.61 | -0.71 | 0 | -0.1025 | 0.43 | 0.4975 | 0.612366 | 0.902801 | 0.652395 | 1 | 0.999631 | 0.860287 |
| Ncam1_P13595_T788t_1 | 1.57 | -1.88 | -0.24 | -0.43 | -1.99 | -0.92 | 1.02 | -1.44 | -1.82 | 0.87 | -1.45 | -1.43 | -1.97 | -2.61 | -0.89 | -1.75 | -0.5875 | -0.7125 | -1.56 | 0.047252 | 0.489171 | 0.395123 | 1 | 0.999631 | 0.480102 |
| Ncam1_P13595_T788t_1_1 | 1.57 | -1.88 | -0.24 | -0.43 | -1.99 | -0.92 | 1.02 | -1.44 | -1.82 | 0.87 | -1.45 | -1.43 | -1.97 | -2.61 | -0.89 | -1.75 | -0.5875 | -0.7125 | -1.56 | 0.047252 | 0.489171 | 0.395123 | 1 | 0.999631 | 0.480102 |
| Ncam1_P13595_T803t_1 | -1.78 | -2.7 | 3.46 | -0.27 | -1.05 | -1.56 | 3.37 | 1.28 | -0.81 | -1.17 | 4.58 | 6.62 | -1.55 | -1.53 | 2.64 | 0.21 | 0.8325 | 2.6275 | 0.265 | 0.851127 | 0.581947 | 0.210974 | 1 | 0.999631 | 0.949927 |
| Ncam2_O35136_S765s_1 | -3.75 | -1.58 | -1.09 | -2.37 | 1.32 | -1.41 | -1.43 | -1.53 | 1.26 | -1.8 | 1.64 | 1.7 | 3.1 | -0.94 | 0.43 | -2.08 | 1.435 | 2.8975 | 2.325 | 0.051649 | 0.094791 | 0.008407 | 1 | 0.999631 | 0.489777 |
| Ncam2_O35136_S786s_1 | -1.28 | -1.4 | -0.09 | -0.08 | -0.83 | -1.49 | -0.39 | -0.04 | -1.08 | -1.45 | 1 | 0.74 | 0.51 | -0.97 | 1.04 | 1.26 | 0.025 | 0.515 | 1.1725 | 0.063063 | 0.956597 | 0.43512 | 1 | 0.999631 | 0.512978 |
| Ncbp1_Q3UYV9_S7s_1 | 0.71 | 3.61 | 1.35 | 0.32 | 4.12 | 1.14 | 5.03 | 0.78 | 5.74 | -0.54 | 4.06 | 2.2 | 4.66 | -2.55 | -1.92 | -2.04 | 1.27 | 1.3675 | -1.96 | 0.22846 | 0.26762 | 0.310218 | 1 | 0.999631 | 0.679038 |
| Ncdn_Q9Z0E0_S566s_1 | -0.31 | 0.4 | 0.47 | -0.03 | 0.04 | 0.22 | -0.41 | 0.18 | 0.94 | -0.34 | -0.05 | 0.31 | 0.88 | 0.95 | 0.35 | 0.62 | -0.125 | 0.0825 | 0.5675 | 0.097081 | 0.676637 | 0.8204 | 1 | 0.999631 | 0.55886 |
| Ncf2_O70145_S332s_1 | 0.54 | 0.97 | 0.33 | -0.2 | -0.38 | 0.81 | 0.06 | -0.07 | 0.88 | 1.7 | -0.02 | 0.39 | 1.23 | 0.66 | 0.63 | 1.25 | -0.305 | 0.3275 | 0.5325 | 0.153938 | 0.41735 | 0.461738 | 1 | 0.999631 | 0.625198 |
| Nckap1_P28660_S337s_1 | 0.49 | -1.1 | -1.25 | -1.31 | -1.24 | -0.76 | -0.84 | -1.53 | -0.85 | 0.55 | -1.22 | -0.82 | -0.91 | -1.27 | -1.53 | -1.74 | -0.3 | 0.2075 | -0.57 | 0.23361 | 0.509999 | 0.701506 | 1 | 0.999631 | 0.683077 |
| Nckap5l_Q6GQX2_S1284s_1 | 0.7 | -0.81 | -0.5 | 0.25 | 0.32 | -0.04 | 0.55 | 0.24 | 0.28 | 0.23 | 0.56 | -0.47 | -0.46 | 0.01 | -0.17 | -0.63 | 0.3575 | 0.24 | -0.2225 | 0.579591 | 0.357447 | 0.565819 | 1 | 0.999631 | 0.84667 |
| Nckipsd_Q9ESJ4_S122s_1 | -0.39 | -2.79 | -1.41 | -1.31 | -2.71 | -2.47 | -1.36 | -1.68 | -3.01 | -2.34 | -2.21 | -2.26 | -3.07 | -2.35 | -1.46 | -1.46 | -0.58 | -0.98 | -0.61 | 0.30578 | 0.299382 | 0.075134 | 1 | 0.999631 | 0.716639 |
| Nckipsd_Q9ESJ4_T174t_1 | 0.3 | 0.49 | 0.03 | 0.13 | 0.3 | 0.63 | -0.14 | 0.15 | 0.19 | 0.14 | 0.18 | -0.22 | -0.2 | 1.18 | 0.22 | 0.08 | -0.0025 | -0.165 | 0.0825 | 0.820589 | 0.992753 | 0.533891 | 1 | 0.999631 | 0.938379 |
| Ncl_P09405_S303s_1 | 0.36 | 1.79 | 0.66 | 0.64 | 1 | 1.26 | 0.12 | 0.33 | 0.68 | 0.47 | 0.24 | 1.79 | 1.28 | 1.82 | 0.38 | 1.09 | -0.185 | -0.0675 | 0.28 | 0.52757 | 0.656775 | 0.882323 | 1 | 0.999631 | 0.824554 |
| Ncl_P09405_S403s_1 | -0.05 | 1.87 | 1.41 | 0.73 | 0.59 | 0.96 | -0.11 | 0.61 | 1.45 | 0.91 | 1.01 | 2.35 | 1.67 | 1.63 | 1.19 | 2.25 | -0.4775 | 0.44 | 0.695 | 0.157099 | 0.309664 | 0.391927 | 1 | 0.999631 | 0.627294 |
| Ncoa1_P70365_S570s_1 | 0.01 | 1.81 | 1.62 | -0.5 | 0.61 | 1.6 | 1.91 | -0.26 | 2.48 | 0.63 | 0.57 | 0.85 | 0.93 | 0.09 | 0.09 | -1.34 | 0.23 | 0.3975 | -0.7925 | 0.25098 | 0.733715 | 0.549783 | 1 | 0.999631 | 0.686822 |
| Ncoa2_Q61026_S493s_1 | 0.67 | 2.09 | 1.69 | 1.26 | 1.67 | 2.11 | 0.96 | 1.28 | 1.79 | 1.66 | 1.07 | 1.15 | 1.55 | 2.44 | 1.56 | 1.27 | 0.0775 | -0.01 | 0.2775 | 0.50679 | 0.845605 | 0.978831 | 1 | 0.999631 | 0.815431 |
| Ncoa2_Q61026_S493s_2 | 1.51 | 0.93 | 0.77 | 0.07 | 0.49 | 0.5 | 0.56 | -0.02 | 0.82 | 1.05 | 0.29 | -0.14 | 0.22 | 0.98 | -0.57 | 0.25 | -0.4375 | -0.315 | -0.6 | 0.191603 | 0.234021 | 0.447226 | 1 | 0.999631 | 0.648674 |
| Ncoa2_Q61026_S499s_2 | 1.51 | 0.93 | 0.77 | 0.07 | 0.49 | 0.5 | 0.56 | -0.02 | 0.82 | 1.05 | 0.29 | -0.14 | 0.22 | 0.98 | -0.57 | 0.25 | -0.4375 | -0.315 | -0.6 | 0.191603 | 0.234021 | 0.447226 | 1 | 0.999631 | 0.648674 |
| Ncoa2_Q61026_S565s_1 | -0.53 | 0.78 | -0.99 | -0.25 | 0.69 | 0.28 | -0.58 | -0.3 | -0.48 | -0.42 | -0.49 | 0.14 | 1.75 | 1.01 | -0.01 | 0.31 | 0.27 | -0.065 | 1.0125 | 0.072652 | 0.555715 | 0.874484 | 1 | 0.999631 | 0.533932 |
| Ncoa5_B7ZC24_S126s_1 | 1.12 | -1.27 | 0.84 | 0.77 | -1.54 | -0.2 | 1.63 | -1.2 | -0.42 | 0.87 | 0.14 | -1.17 | -0.74 | -1.33 | -0.22 | -0.79 | -0.6925 | -0.51 | -1.135 | 0.063144 | 0.388572 | 0.427599 | 1 | 0.999631 | 0.512978 |
| Ncoa5_B7ZC24_S162s_1 | 1.34 | 1.31 | 1.53 | 1.32 | 0.96 | 1.37 | 1.34 | 1.26 | 0.63 | 1.65 | 0.38 | 1.16 | 0.01 | 0.48 | 1.4 | 0.54 | -0.1425 | -0.42 | -0.7675 | 0.049992 | 0.560741 | 0.230229 | 1 | 0.999631 | 0.488436 |
| Ncoa5_B7ZC24_S34s_1 | -0.97 | -3.73 | -3.63 | -2.49 | -1.56 | -4.19 | -3.07 | -1.91 | -4.16 | -1.96 | -2.01 | -3.42 | -2.53 | -2.62 | -2.66 | -6.3 | 0.0225 | -0.1825 | -0.8225 | 0.40306 | 0.976637 | 0.806361 | 1 | 0.999631 | 0.764348 |
| Ncoa5_B7ZC24_S378s_1 | -0.59 | -0.79 | -1.02 | -0.57 | -1.2 | -1.96 | -1.16 | -1.12 | -1.91 | -0.83 | -0.99 | -1.35 | 0.1 | -0.62 | -0.56 | 0.39 | -0.6175 | -0.5275 | 0.57 | 0.115074 | 0.058853 | 0.12444 | 1 | 0.999631 | 0.579336 |
| Ncoa5_B7ZC24_S6s_1 | 1.26 | -1.39 | -1.42 | 0.15 | -0.45 | -0.65 | 0.35 | -0.25 | -0.43 | 0.68 | -0.28 | -0.17 | -1.28 | -0.73 | -0.48 | -0.33 | 0.1 | 0.3 | -0.355 | 0.569046 | 0.871059 | 0.635344 | 1 | 0.999631 | 0.84338 |
| Ncoa5_B7ZC24_S9s_1 | 0.52 | -0.17 | 0.81 | -0.15 | -0.5 | -0.18 | 1.12 | 0.31 | 0.28 | 0.55 | 0.41 | -0.35 | -0.75 | -1.17 | 0.43 | -1.16 | -0.065 | -0.03 | -0.915 | 0.06474 | 0.8781 | 0.932243 | 1 | 0.999631 | 0.515799 |
| Ncoa5_Q91W39_S377s_1 | 0.51 | -0.16 | -0.06 | -0.17 | 0.68 | 0.04 | -0.24 | -0.06 | -0.12 | 0.57 | 0.07 | -0.09 | 0.87 | 0.36 | -0.02 | 0.18 | 0.075 | 0.0775 | 0.3175 | 0.343222 | 0.809926 | 0.797654 | 1 | 0.999631 | 0.73348 |
| Ncoa7_B9EIW5_S342s_1 | -0.29 | -2.56 | -1.52 | -0.84 | -1.89 | -2.08 | -0.23 | -1.77 | -1.74 | -0.57 | -1.15 | -1.96 | -2.02 | -3.04 | -1.79 | -3.01 | -0.19 | -0.0525 | -1.1625 | 0.05611 | 0.74696 | 0.922489 | 1 | 0.999631 | 0.495761 |
| Ncor1_Q5RIM6_S1123s_1 | 0.62 | 1.58 | 1.22 | 0.6 | 0.55 | 1.05 | 0.82 | 0.94 | 1.07 | 1.49 | 0.72 | 0.23 | 1.23 | 1.09 | 0.33 | 0.82 | -0.165 | -0.1275 | -0.1375 | 0.703836 | 0.602398 | 0.738293 | 1 | 0.999631 | 0.897149 |
| Ncor1_Q5RIM6_S1334s_1 | 0.89 | 0.69 | 0.93 | 0.74 | 0.78 | 1.27 | 0.56 | 1.33 | 1.45 | 1.5 | 1.17 | 1.05 | 0.72 | 1.54 | 0.46 | 1.21 | 0.1725 | 0.48 | 0.17 | 0.604841 | 0.540287 | 0.085741 | 1 | 0.999631 | 0.85817 |
| Ncor1_Q5RIM6_S2117s_1 | -0.08 | -0.76 | 0.31 | 0.61 | 1.07 | 0.35 | -0.01 | 0.25 | -0.68 | 1.2 | 1 | 0.21 | -0.39 | 1.37 | -1.44 | 0.58 | 0.395 | 0.4125 | 0.01 | 0.986981 | 0.319701 | 0.413374 | 1 | 0.999631 | 0.995165 |
| Ncor1_Q5RIM6_S2135s_1 | 0.45 | 0.56 | 0.13 | -0.21 | 0.43 | 0.36 | 0.21 | 0.71 | 0.39 | 0.39 | 0.51 | 0.79 | 0.72 | 1.06 | 1.15 | 0.74 | 0.195 | 0.2875 | 0.685 | 0.045291 | 0.494642 | 0.329175 | 1 | 0.999631 | 0.476658 |
| Ncor2_E9PY55_S1818s_1 | -0.12 | -0.64 | 0.26 | -0.33 | -0.25 | -0.84 | 0.49 | 0.12 | -0.18 | 0.14 | -1.15 | -0.31 | -0.54 | -0.97 | -0.14 | -1.38 | 0.0875 | -0.1675 | -0.55 | 0.160017 | 0.809629 | 0.647483 | 1 | 0.999631 | 0.629764 |
| Ncor2_E9PY55_S2211s_1 | 0.66 | 1.06 | 1.81 | 1.22 | 0.85 | 0.9 | 0.83 | 1.29 | 1.58 | 0.5 | 1.71 | 2.02 | 1.19 | 1.54 | 1.73 | 1.64 | -0.22 | 0.265 | 0.3375 | 0.326966 | 0.489464 | 0.526424 | 1 | 0.999631 | 0.724678 |
| Ncor2_E9PY55_T1349t_1 | 0.88 | 1.82 | 1.84 | 1.25 | 1.04 | 1.05 | 0.84 | 0.71 | 1.3 | 1.72 | 1.07 | 1.02 | 1.07 | 1.33 | 0.98 | 1.32 | -0.5375 | -0.17 | -0.2725 | 0.412589 | 0.103711 | 0.612036 | 1 | 0.999631 | 0.768114 |
| Ncor2_E9PY55_T1405t_1 | 1.16 | 1.79 | 1.84 | 1.72 | 1.34 | 0.82 | 1.22 | 2.3 | 1.54 | 1.12 | 2.23 | 2.23 | 1.02 | 1.94 | 2.03 | 1.49 | -0.2075 | 0.1525 | -0.0075 | 0.982557 | 0.577739 | 0.667238 | 1 | 0.999631 | 0.994846 |
| Ndel1_Q9ERR1_S194s_1 | -0.89 | -0.2 | -0.4 | -0.44 | -0.75 | -1.25 | -0.35 | -0.28 | -0.41 | -0.34 | -0.57 | -0.55 | 0.42 | -0.59 | -0.84 | -0.73 | -0.175 | 0.015 | 0.0475 | 0.897325 | 0.58348 | 0.955396 | 1 | 0.999631 | 0.966758 |
| Ndel1_Q9ERR1_S194s_2 | -0.58 | -0.13 | -0.31 | 0.02 | -0.41 | -0.63 | -0.76 | -0.25 | -0.48 | -0.06 | -0.15 | -0.44 | 0.14 | -0.27 | 0.11 | -0.29 | -0.2625 | -0.0325 | 0.1725 | 0.557733 | 0.338587 | 0.905063 | 1 | 0.999631 | 0.838305 |
| Ndel1_Q9ERR1_S198s_1 | -2.25 | -2.1 | -1.51 | -2.95 | -3.19 | -2.56 | -2.52 | -2.58 | -2.81 | -1.68 | -2.12 | -2.6 | -0.31 | -1.82 | -1.65 | -1.82 | -0.51 | -0.1 | 0.8025 | 0.107693 | 0.178626 | 0.803288 | 1 | 0.999631 | 0.569416 |
| Ndel1_Q9ERR1_S198s_2 | -0.67 | -0.17 | -0.31 | 0.02 | -0.5 | -0.75 | -0.76 | -0.25 | -0.58 | -0.02 | -0.15 | -0.44 | 0.15 | -0.28 | 0.11 | -0.29 | -0.2825 | -0.015 | 0.205 | 0.496418 | 0.318735 | 0.958061 | 1 | 0.999631 | 0.811248 |
| Ndel1_Q9ERR1_S215s_1 | 0.02 | 0.59 | -0.01 | -0.33 | 0.54 | 0.51 | -0.69 | -0.02 | 0.27 | -0.07 | -0.54 | -0.01 | 0.13 | -0.04 | 0.25 | 0.06 | 0.0175 | -0.155 | 0.0325 | 0.914541 | 0.961888 | 0.626678 | 1 | 0.999631 | 0.972695 |
| Ndel1_Q9ERR1_S231s_1 | -0.1 | -0.24 | -0.63 | -0.47 | -0.39 | -0.3 | -0.66 | -0.26 | -0.51 | -0.16 | 0.06 | -0.69 | 0.16 | -0.1 | -0.3 | -0.27 | -0.0425 | 0.035 | 0.2325 | 0.423312 | 0.86886 | 0.904258 | 1 | 0.999631 | 0.77354 |
| Ndel1_Q9ERR1_S336s_1 | -0.03 | 0.66 | 0.05 | -0.49 | -0.65 | 0.47 | -0.46 | -0.35 | 0.38 | -0.25 | -0.98 | -0.83 | -0.36 | 0.67 | 0.6 | 0.55 | -0.295 | -0.4675 | 0.3175 | 0.408464 | 0.42537 | 0.261144 | 1 | 0.999631 | 0.766996 |
| Ndel1_Q9ERR1_T219t_1 | -0.14 | -0.09 | -3.28 | -1.55 | 0.03 | -0.64 | -1.18 | -0.19 | -1.09 | -0.13 | -0.53 | -0.02 | 0.66 | -0.65 | -1.61 | -1.04 | 0.77 | 0.8225 | 0.605 | 0.445551 | 0.290444 | 0.259344 | 1 | 0.999631 | 0.784372 |
| Ndfip2_M0QWK1_S12s_1 | -0.73 | 0.09 | 0.55 | -0.62 | -0.18 | -0.3 | -1.43 | -0.09 | -0.25 | 0.05 | -1.11 | -0.58 | 0.41 | -0.02 | 0.39 | -0.47 | -0.3225 | -0.295 | 0.255 | 0.522678 | 0.458172 | 0.470386 | 1 | 0.999631 | 0.821224 |
| Ndfip2_M0QWK1_T216t_1 | 0.99 | -0.41 | 0.87 | 0.31 | -0.72 | 0.28 | 0.9 | -0.78 | -0.75 | 0.48 | -0.99 | -1.11 | -1.07 | -1.36 | 0.09 | -1.55 | -0.52 | -1.0325 | -1.4125 | 0.013204 | 0.301512 | 0.049833 | 1 | 0.999631 | 0.385092 |
| Ndrg1_Q62433_S330s_1 | -0.37 | -0.82 | 0.41 | -0.26 | -0.13 | -0.16 | -0.41 | -0.92 | -0.14 | -0.24 | -0.4 | -0.31 | 0.59 | 0.17 | 0.64 | 0.05 | -0.145 | -0.0125 | 0.6225 | 0.100147 | 0.675642 | 0.968792 | 1 | 0.999631 | 0.562917 |
| Ndrg1_Q62433_S330s_2 | -1.4 | 0.14 | 1.42 | 0.21 | -1.29 | -0.34 | 0.49 | 0.62 | -1.18 | -1.08 | 0.03 | 1.27 | 0.39 | -0.7 | 0.96 | 1.2 | -0.2225 | -0.3325 | 0.37 | 0.568645 | 0.732449 | 0.646826 | 1 | 0.999631 | 0.843185 |
| Ndrg1_Q62433_S333s_2 | -1.4 | 0.14 | 1.01 | 0.19 | -1.29 | -0.34 | -0.27 | -0.88 | -1.18 | -1.08 | 0.23 | 0.36 | 0.39 | -0.7 | 0.58 | 0.45 | -0.68 | -0.4025 | 0.195 | 0.720938 | 0.20975 | 0.50307 | 1 | 0.999631 | 0.905276 |
| Ndrg1_Q62433_T328t_2 | -1.61 | -1.6 | 1.19 | 0.2 | -1.7 | -1.39 | 0.09 | -0.16 | -1.27 | -1.01 | 0.15 | 0.77 | 0.27 | -0.23 | 0.75 | 0.79 | -0.335 | 0.115 | 0.85 | 0.215558 | 0.6461 | 0.87757 | 1 | 0.999631 | 0.668551 |
| Ndrg1_Q62433_T375t_1 | 0.24 | -0.04 | -0.61 | -0.01 | -0.08 | 0.12 | -0.46 | 0.42 | 0.01 | 0.61 | -0.31 | 0.14 | 0.21 | 0.08 | 0.19 | 0.93 | 0.105 | 0.2175 | 0.4575 | 0.192243 | 0.736752 | 0.503221 | 1 | 0.999631 | 0.648844 |
| Ndrg2_Q9QYG0_S314s_2 | 0.2 | -0.76 | -0.31 | -0.78 | -1.05 | -0.74 | -0.64 | 0.16 | -0.05 | -0.34 | -1.16 | -0.2 | -0.96 | 0.41 | 0.1 | 0.65 | -0.155 | -0.025 | 0.4625 | 0.298016 | 0.673676 | 0.945777 | 1 | 0.999631 | 0.711311 |
| Ndrg2_Q9QYG0_S318s_1 | -1.15 | -0.58 | -0.01 | -1.11 | -0.88 | -0.65 | -0.65 | -0.58 | -0.68 | -0.26 | -0.22 | 0.38 | 0.51 | -0.45 | -0.34 | -0.36 | 0.0225 | 0.5175 | 0.5525 | 0.171603 | 0.944287 | 0.185762 | 1 | 0.999631 | 0.641657 |
| Ndrg2_Q9QYG0_S318s_2 | -0.11 | 0.16 | -0.28 | -0.14 | -0.54 | 0.04 | -0.78 | 0.56 | -0.12 | 0.06 | -0.28 | 0.14 | 0.41 | 0.33 | 0.27 | 0.22 | -0.0875 | 0.0425 | 0.4 | 0.15692 | 0.800584 | 0.86972 | 1 | 0.999631 | 0.626839 |
| Ndrg2_Q9QYG0_S324s_1 | -1.08 | -0.43 | 0.15 | -1.36 | -1.05 | -0.45 | -1.42 | -0.73 | -0.58 | -0.05 | -0.67 | 0.47 | 0.68 | -0.49 | -0.48 | -0.02 | -0.2325 | 0.4725 | 0.6025 | 0.191548 | 0.56574 | 0.286229 | 1 | 0.999631 | 0.648674 |
| Ndrg2_Q9QYG0_S324s_2 | -0.71 | 0.05 | 0.03 | -0.47 | -0.22 | 0.08 | -0.66 | 0.37 | 0 | 0.14 | -0.32 | 0.28 | 1.02 | 0.1 | -0.21 | 0.24 | 0.1675 | 0.3 | 0.5625 | 0.148509 | 0.614722 | 0.333652 | 1 | 0.999631 | 0.62132 |
| Ndrg2_Q9QYG0_S336s_1 | -0.04 | 0.49 | 0.53 | -0.38 | 0.08 | 0.35 | -0.37 | 0.59 | 0.02 | 0.44 | -0.36 | 0.4 | -0.29 | 0.5 | -0.16 | 0.3 | 0.0125 | -0.025 | -0.0625 | 0.857235 | 0.970426 | 0.940709 | 1 | 0.999631 | 0.952021 |
| Ndrg2_Q9QYG0_S336s_2 | 0.26 | -0.07 | -0.65 | -0.07 | -0.92 | -0.05 | -1.14 | -0.21 | -0.53 | 0.83 | -0.83 | -0.37 | -0.07 | 0.96 | -0.18 | -0.27 | -0.4475 | -0.0925 | 0.2425 | 0.527357 | 0.22478 | 0.824053 | 1 | 0.999631 | 0.824554 |
| Ndrg2_Q9QYG0_S338s_1 | -0.72 | -0.37 | -0.27 | -0.59 | -1.08 | -1.18 | -0.75 | 0.09 | -1.17 | -0.14 | -1.39 | -0.14 | -0.55 | -0.37 | -0.08 | -0.57 | -0.2425 | -0.2225 | 0.095 | 0.738064 | 0.482476 | 0.554828 | 1 | 0.999631 | 0.907936 |
| Ndrg2_Q9QYG0_S339s_1 | -0.61 | 0.04 | -0.26 | -1.07 | -0.76 | -0.56 | -1.04 | -1.03 | -0.86 | -0.34 | -0.78 | -0.64 | 1.39 | -0.77 | -0.75 | -0.3 | -0.3725 | -0.18 | 0.3675 | 0.49345 | 0.255869 | 0.579746 | 1 | 0.999631 | 0.80955 |
| Ndrg2_Q9QYG0_T316t_1 | -0.04 | -0.02 | 0.89 | 0.07 | 0.39 | -0.71 | -0.52 | 0.99 | 0.22 | 0.51 | -0.09 | 1.64 | 0.38 | 0.31 | 1.16 | 1.46 | -0.1875 | 0.345 | 0.6025 | 0.14689 | 0.673443 | 0.435708 | 1 | 0.999631 | 0.618703 |
| Ndrg2_Q9QYG0_T316t_2 | -0.64 | 0.04 | -0.02 | -0.43 | -0.23 | -0.11 | -0.66 | 0.45 | -0.26 | 0.04 | -0.34 | 0.35 | 0.46 | 0.14 | -0.03 | 0.33 | 0.125 | 0.21 | 0.4875 | 0.12868 | 0.701881 | 0.491846 | 1 | 0.999631 | 0.599259 |
| Ndrg2_Q9QYG0_T334t_2 | -0.01 | -0.28 | 0.54 | -0.6 | -0.9 | -0.39 | -0.24 | 1.71 | -0.49 | 0.56 | -0.31 | -0.73 | 0.19 | 0.33 | 0.06 | 0.31 | 0.1325 | -0.155 | 0.31 | 0.353625 | 0.815354 | 0.69093 | 1 | 0.999631 | 0.73949 |
| Ndrg3_Q9QYF9_S335s_1 | -0.16 | 1.2 | 1.08 | 0.94 | 1.33 | 0.28 | 1.18 | -0.15 | 1.62 | 0.13 | 1.84 | 2.96 | 2 | 0.06 | 1.23 | 1.12 | -0.105 | 0.8725 | 0.3375 | 0.495453 | 0.81796 | 0.169749 | 1 | 0.999631 | 0.810828 |
| Ndrg3_Q9QYF9_S352s_1 | -0.64 | 0.2 | 0.19 | -0.55 | -0.32 | -1.01 | -0.77 | -0.55 | -1.01 | 0.26 | -0.36 | -0.02 | 0.02 | -1.22 | -0.07 | 0.31 | -0.4625 | -0.0825 | -0.04 | 0.924127 | 0.171143 | 0.827888 | 1 | 0.999631 | 0.975324 |

| | | | | | | | | | | | | | | | | | | | | | | | | | |
|---|---|---|---|---|---|---|---|---|---|---|---|---|---|---|---|---|---|---|---|---|---|---|---|---|---|
| Ndrg3_Q9QYF9_S361s_1 | -0.53 | 0.25 | 0.35 | -0.04 | -0.22 | -0.33 | -0.4 | -0.13 | -0.23 | -0.17 | -0.19 | 0.39 | 0.88 | -0.33 | 0.52 | 0.45 | -0.2775 | -0.0575 | 0.3725 | 0.323134 | 0.340351 | 0.853829 | 1 | 0.999631 | 0.724485 |
| Ndrg3_Q9QYF9_S374s_1 | 1.04 | 2.13 | 1.35 | -0.29 | 3.35 | 1.09 | 2.77 | 0.21 | 4.55 | 0.67 | 2.15 | 2.77 | 4.22 | 0.5 | 1.31 | -0.67 | 0.7975 | 1.4775 | 0.2825 | 0.775817 | 0.317482 | 0.100087 | 1 | 0.999631 | 0.920998 |
| Ndrg3_Q9QYF9_T355t_1 | -0.9 | -0.25 | 0.03 | -0.25 | -0.54 | 0 | -0.76 | -0.42 | -0.38 | -0.44 | -0.43 | 0.26 | -0.05 | 0.1 | 0.1 | 0.37 | -0.0875 | 0.095 | 0.4725 | 0.150158 | 0.777652 | 0.766852 | 1 | 0.999631 | 0.622782 |
| Ndrg4_Q8BTG7_S298s_1 | 0.1 | 0.3 | 0.66 | -0.08 | -0.27 | 0.33 | 0 | 0.58 | 0.1 | 0.3 | 0.64 | 0.65 | 0.82 | 0.71 | 0.88 | 1.4 | -0.085 | 0.1775 | 0.7075 | 0.043709 | 0.780237 | 0.547277 | 1 | 0.999631 | 0.475443 |
| Ndrg4_Q8BTG7_S298s_2 | -1.25 | 1.05 | -0.25 | 1.05 | 0.53 | 0.95 | 0.29 | 0.41 | 0.2 | 0.13 | 1.25 | 0.6 | 0.37 | 1.24 | -0.39 | -0.75 | 0.395 | 0.395 | -0.0325 | 0.959476 | 0.464786 | 0.490895 | 1 | 0.999631 | 0.986095 |
| Ndrg4_Q8BTG7_S306s_1 | -0.71 | 1.06 | 1.64 | -0.57 | 0.76 | 0.84 | 0.98 | 0.41 | 1.56 | -1.1 | 1.54 | 0.9 | 1.26 | 0.24 | 1.35 | -0.56 | 0.3925 | 0.37 | 0.2175 | 0.743597 | 0.48156 | 0.627592 | 1 | 0.999631 | 0.90976 |
| Ndrg4_Q8BTG7_S306s_2 | -1.25 | 1.05 | -0.25 | 1.05 | 0.53 | 0.95 | 0.29 | 0.41 | 0.2 | 0.13 | 1.25 | 0.6 | 0.37 | 1.24 | -0.39 | -0.75 | 0.395 | 0.395 | -0.0325 | 0.959476 | 0.464786 | 0.490895 | 1 | 0.999631 | 0.986095 |
| Ndrg4_Q8BTG7_S317s_1 | -0.09 | -0.13 | 0.63 | 0 | -0.95 | -0.16 | -0.08 | 1.02 | -0.33 | 0.5 | -0.14 | 1.58 | 1.01 | 0.46 | 1.32 | 1.13 | -0.145 | 0.3 | 0.8775 | 0.022357 | 0.738297 | 0.514759 | 1 | 0.999631 | 0.405775 |
| Ndrg4_Q8BTG7_S317s_2 | 0.09 | 1.22 | 0.45 | 0.18 | 0.45 | 0.56 | 0.21 | 0.83 | 0.76 | 1.55 | 0.8 | -0.14 | 0.88 | 1.11 | 1 | 0.47 | 0.0275 | 0.2575 | 0.38 | 0.291905 | 0.933354 | 0.552927 | 1 | 0.999631 | 0.706164 |
| Ndrg4_Q8BTG7_S337s_1 | -1.3 | -1.14 | 0.35 | -0.96 | -0.81 | 0.34 | -0.66 | 0.35 | -0.23 | 0.76 | -2.19 | 0.11 | 0.19 | 0.75 | 0.92 | 0.92 | 0.5675 | 0.375 | 1.4575 | 0.006174 | 0.243528 | 0.574496 | 1 | 0.999631 | 0.351545 |
| Ndrg4_Q8BTG7_T315t_2 | 0.09 | 1.22 | 0.45 | 0.18 | 0.45 | 0.56 | 0.21 | 0.83 | 0.76 | 1.55 | 0.8 | -0.14 | 0.88 | 1.11 | 1 | 0.47 | 0.0275 | 0.2575 | 0.38 | 0.291905 | 0.933354 | 0.552927 | 1 | 0.999631 | 0.706164 |
| Ndufb10_Q9DCS9_S21s_1 | -0.41 | 0.37 | -0.62 | -0.42 | -0.09 | -0.61 | -0.89 | -1.19 | -0.53 | -0.46 | -0.08 | -0.12 | -0.04 | -0.21 | -0.35 | -0.04 | -0.425 | -0.0275 | 0.11 | 0.72952 | 0.242364 | 0.929864 | 1 | 0.999631 | 0.905923 |
| Ndufs5_Q99LY9_T93t_1 | -0.04 | -0.01 | 0.72 | 0.36 | -1.3 | -0.14 | 0.62 | -0.3 | -0.52 | 0.79 | -0.22 | -0.73 | -1.51 | -0.78 | 0.68 | -0.46 | -0.5375 | -0.4275 | -0.775 | 0.128033 | 0.226986 | 0.294981 | 1 | 0.999631 | 0.598425 |
| Ndufs8_Q8K3J1_S108s_1 | 1.28 | -0.25 | 0.35 | 0.37 | -0.77 | -0.1 | 0.45 | -0.65 | 0.67 | 1.45 | 0.36 | 0.26 | -0.2 | -0.5 | -0.04 | -0.86 | -0.705 | 0.2475 | -0.8375 | 0.054744 | 0.116803 | 0.558199 | 1 | 0.999631 | 0.492066 |
| Ndufv3_Q3U422_S465s_1 | -1.07 | -2.1 | -1.92 | -1.33 | -1.46 | -2.91 | -1.9 | -1.52 | -1.46 | -1.04 | -1.2 | -0.63 | -0.2 | -1.91 | -0.89 | -1.3 | -0.3425 | 0.5225 | 0.53 | 0.243834 | 0.415195 | 0.150579 | 1 | 0.999631 | 0.685637 |
| Nedd4_P46935_S309s_1 | 0.58 | -0.05 | -0.15 | 0.45 | 0.5 | 0.28 | 0.4 | 0.85 | 0.39 | 0.43 | 0.07 | 0.05 | 0.14 | -0.07 | -0.06 | 0.33 | 0.3 | 0.0275 | -0.1225 | 0.688528 | 0.314722 | 0.924829 | 1 | 0.999631 | 0.891939 |
| Nedd4_P46935_T287t_1 | -0.41 | 0.05 | -0.05 | -0.12 | 0.39 | 0.07 | 0.08 | 0.03 | 0.06 | 0.33 | -0.29 | -0.26 | 0.99 | 0.48 | 0.57 | 0.18 | 0.275 | 0.0925 | 0.6875 | 0.04193 | 0.286947 | 0.739299 | 1 | 0.999631 | 0.475443 |
| Nedd4l_Q8CFI0_S504s_1 | -0.92 | 1.55 | -0.65 | -0.01 | 1.07 | 1.15 | -0.44 | 0.34 | 1.28 | -0.03 | -0.3 | 0.36 | 0.47 | 1.44 | -0.3 | -0.24 | 0.5375 | 0.335 | 0.35 | 0.575884 | 0.38178 | 0.576462 | 1 | 0.999631 | 0.846085 |
| Nedd4l_Q8CFI0_S508s_1 | -1.34 | -2.16 | -1.19 | -1.03 | -0.97 | -1.88 | -0.79 | -1.26 | -2.15 | -2.28 | -0.92 | -0.85 | -1.6 | -1.82 | -1.45 | -0.86 | 0.205 | -0.12 | -0.0025 | 0.994583 | 0.579358 | 0.790339 | 1 | 0.999631 | 0.997705 |
| Nefh_P19246_S122s_1 | -1.27 | -2.13 | -0.29 | -0.46 | -1.11 | -1.32 | -1.16 | -1.5 | -0.56 | -1.14 | -0.47 | -0.15 | 1.32 | -1 | 0.42 | -0.02 | -0.235 | 0.4575 | 1.2175 | 0.060875 | 0.58337 | 0.330911 | 1 | 0.999631 | 0.509205 |
| Nefh_P19246_S345s_1 | 1.07 | -0.85 | 0 | -0.54 | -0.13 | 0.35 | 0.14 | -0.41 | -0.65 | 1.18 | -1.39 | -0.48 | 0.07 | -0.99 | -0.53 | -1.31 | 0.0675 | -0.255 | -0.61 | 0.237313 | 0.878188 | 0.682992 | 1 | 0.999631 | 0.683301 |
| Nefh_P19246_S416s_1 | -1.81 | -0.51 | -1.42 | -1.68 | -0.75 | -1 | -1.35 | -1.75 | -0.34 | -2.19 | -1.61 | -1 | -0.17 | -1.01 | -0.58 | -1.91 | 0.1425 | 0.07 | 0.4375 | 0.358894 | 0.707756 | 0.88297 | 1 | 0.999631 | 0.741182 |
| Nefh_P19246_S500s_1 | 0.36 | 1.82 | 2.16 | 1.31 | 0.29 | 1.37 | 0.1 | 1.3 | 0.25 | 1.52 | 0.77 | 1.52 | 1.42 | 1.32 | 1.57 | 2.16 | -0.6475 | -0.3975 | 0.205 | 0.642087 | 0.202711 | 0.416093 | 1 | 0.999631 | 0.871335 |
| Nefh_P19246_S535s_1 | -0.74 | -4.53 | -4.09 | -2.61 | -3.28 | -4.26 | -2.39 | -4.89 | -4.11 | -2.11 | -4.06 | -5.51 | -1.64 | -3.08 | -4.96 | -5.74 | -0.7125 | -0.955 | -0.8625 | 0.429258 | 0.426975 | 0.331008 | 1 | 0.999631 | 0.776658 |
| Nefh_P19246_S54s_1 | 0.53 | -0.87 | -0.47 | -0.03 | 0.4 | 0.14 | -0.22 | -0.44 | -0.48 | 0.54 | -0.71 | -1.99 | -1.39 | 0.31 | -1.62 | -0.81 | 0.18 | -0.45 | -0.6675 | 0.203989 | 0.629962 | 0.425912 | 1 | 0.999631 | 0.659781 |
| Nefh_P19246_S577s_2 | -0.99 | -1.75 | -0.93 | -0.71 | -0.64 | -2.68 | -1.48 | -0.94 | -0.76 | -1.23 | -1.15 | -0.85 | 0.62 | -1.08 | -1.46 | -0.96 | -0.34 | 0.0975 | 0.375 | 0.453404 | 0.482666 | 0.758188 | 1 | 0.999631 | 0.788717 |
| Nefh_P19246_S583s_2 | -0.99 | -1.75 | -0.93 | -0.71 | -0.64 | -2.68 | -1.48 | -0.94 | -0.76 | -1.23 | -1.15 | -0.85 | 0.62 | -1.08 | -1.46 | -0.96 | -0.34 | 0.0975 | 0.375 | 0.453404 | 0.482666 | 0.758188 | 1 | 0.999631 | 0.788717 |
| Nefh_P19246_S607s_1 | -1.77 | -8.01 | -4.22 | -3.91 | -4.34 | -6.32 | -4 | -5.75 | -6.61 | -2.48 | -3.07 | -4.5 | -5.89 | -6.73 | -4.4 | -6.74 | -0.625 | 0.3125 | -1.4625 | 0.23828 | 0.606132 | 0.817738 | 1 | 0.999631 | 0.683378 |
| Nefh_P19246_S61s_1 | -0.53 | -0.36 | -0.62 | -0.83 | -0.8 | 0.07 | -0.57 | -0.31 | 0.1 | 0.06 | -0.73 | -0.02 | 0.89 | 0.55 | 0.02 | -0.03 | 0.1825 | 0.4375 | 0.9425 | 0.014207 | 0.52789 | 0.16429 | 1 | 0.999631 | 0.385092 |
| Nefh_P19246_S637s_1 | -1.7 | -5.41 | -3.43 | -2.64 | -3.58 | -4.19 | -2.92 | -2.38 | -4.79 | -3.05 | -1.92 | -2.29 | -4.45 | -4.5 | -2.53 | -2.97 | 0.0275 | 0.2825 | -0.3175 | 0.697874 | 0.971559 | 0.74947 | 1 | 0.999631 | 0.895731 |
| Nefh_P19246_S727s_1 | -1.45 | -3.47 | -2.89 | -2.84 | -2.88 | -3.5 | -2.67 | -3.61 | -2.88 | -1.83 | -1.94 | -3.09 | -2.07 | -2.19 | -2.44 | -2.85 | -0.5025 | 0.2275 | 0.275 | 0.55149 | 0.29515 | 0.654843 | 1 | 0.999631 | 0.836275 |
| Nefh_P19246_S769s_1 | -1.87 | -4.08 | -2.26 | -1.94 | -2.36 | -3.56 | -2.27 | -1.76 | -3.29 | -2.36 | -1.55 | -1.84 | -1.66 | -3.19 | -0.79 | -1.92 | 0.05 | 0.2775 | 0.6475 | 0.329092 | 0.931915 | 0.640147 | 1 | 0.999631 | 0.726344 |
| Nefh_P19246_S795s_1 | -1.93 | -2.49 | -3.31 | -2.25 | -2.44 | -3.08 | -2.48 | -2.62 | -3.05 | -2.31 | -2.43 | -2.41 | -2.12 | -4 | -2.49 | -2.6 | -0.16 | -0.055 | -0.3075 | 0.53358 | 0.654314 | 0.881261 | 1 | 0.999631 | 0.828143 |
| Nefh_P19246_S834s_1 | -1.09 | -1.6 | -2.87 | -2.6 | -1.25 | -1.64 | -2.51 | -3.15 | -1.49 | -1.18 | -2.51 | -2.57 | -0.34 | -0.7 | -2.05 | -2.27 | -0.0975 | 0.1025 | 0.7 | 0.248435 | 0.859118 | 0.842408 | 1 | 0.999631 | 0.685751 |
| Nefh_P19246_S834s_2 | -0.37 | 1.08 | 0.62 | 0.65 | 0.79 | 0.52 | 0.01 | 0.3 | 1.21 | 1.12 | -0.28 | 1.36 | 1.56 | 0.74 | 2.01 | 1.55 | -0.09 | 0.3575 | 0.97 | 0.040315 | 0.807008 | 0.455665 | 1 | 0.999631 | 0.472911 |
| Nefh_P19246_S867s_1 | -0.55 | -4.97 | -4.74 | -4.59 | -3.8 | -5.07 | -3.79 | -6.76 | -5.92 | -2.52 | -4.03 | -5.05 | -3.04 | -4.63 | -3.41 | -6.41 | -1.1425 | -0.6675 | -0.66 | 0.554211 | 0.307615 | 0.549193 | 1 | 0.999631 | 0.837952 |
| Nefh_P19246_S888s_1 | 2.14 | 0.96 | 0.04 | 0.71 | 1.84 | 0.51 | -0.1 | -0.09 | 3 | 3.18 | -1.14 | 1.19 | 3.42 | -0.19 | 0.91 | -0.46 | -0.4225 | 0.595 | -0.0425 | 0.960296 | 0.469168 | 0.535197 | 1 | 0.999631 | 0.986428 |
| Nefh_P19246_S947s_1 | -0.61 | 0.19 | 0.18 | -0.13 | -0.17 | 0.03 | -0.58 | -0.15 | -0.1 | -0.1 | -0.24 | 0.09 | 0.7 | 0.26 | 0.02 | 0.8 | -0.125 | 0.005 | 0.5375 | 0.130736 | 0.673121 | 0.986134 | 1 | 0.999631 | 0.602177 |
| Nefh_P19246_S965s_1 | 0.01 | 0.42 | -1.83 | -1.1 | -0.85 | -0.52 | -1.55 | -1.85 | 0.66 | 0.55 | -1.62 | -0.77 | 0.72 | -0.64 | -0.98 | -1.55 | -0.5675 | 0.33 | 0.0125 | 0.984286 | 0.315571 | 0.62583 | 1 | 0.999631 | 0.995133 |
| Nefh_P19246_T839t_1 | 0.09 | -0.69 | -0.7 | 0.08 | -0.22 | -0.01 | -0.45 | -0.85 | 0 | 0.11 | -1.19 | 0.28 | 0.26 | -0.12 | -0.79 | -0.14 | -0.0775 | 0.105 | 0.1075 | 0.766783 | 0.81432 | 0.798557 | 1 | 0.999631 | 0.916961 |
| Nefh_P19246_T839t_2 | -0.37 | 1.08 | 0.62 | 0.65 | 0.79 | 0.52 | 0.01 | 0.3 | 1.21 | 1.12 | -0.28 | 1.36 | 1.56 | 0.74 | 2.01 | 1.55 | -0.09 | 0.3575 | 0.97 | 0.040315 | 0.807008 | 0.455665 | 1 | 0.999631 | 0.472911 |
| Nefl_P08551_S473s_1 | -0.57 | 3.76 | 3.52 | 0.95 | 2.87 | 3.03 | 1.87 | 2.9 | 3.3 | 0.51 | 3.19 | 4.07 | 4.03 | 2.65 | 3.7 | 1.97 | 0.7525 | 0.8525 | 1.1725 | 0.249786 | 0.427083 | 0.452741 | 1 | 0.999631 | 0.686359 |
| Nefl_P08551_S532s_1 | 1.72 | 1.48 | 1.96 | 1.75 | 0.91 | 1.78 | 1.09 | 1.44 | 1.59 | 2.46 | 0.92 | 1.41 | 1.13 | 0.89 | 1.59 | 0.54 | -0.4225 | -0.1325 | -0.69 | 0.054206 | 0.166457 | 0.71799 | 1 | 0.999631 | 0.491994 |
| Nefl_P08551_S56s_1 | -1.05 | -0.38 | -0.1 | -0.97 | -0.39 | -0.37 | -0.37 | -0.03 | -0.18 | -1.13 | -0.29 | -0.53 | 1.06 | -0.58 | -0.5 | -0.41 | 0.335 | 0.0925 | 0.5175 | 0.26802 | 0.287379 | 0.792568 | 1 | 0.999631 | 0.696016 |
| Nefm_P08553_S28s_1 | 0.77 | -0.7 | -0.14 | 0.54 | -0.06 | -0.48 | 0.36 | 0.02 | 0.02 | -0.42 | 0.3 | -0.22 | 0.02 | 0.36 | -0.14 | 0.53 | -0.1575 | -0.1975 | 0.075 | 0.849373 | 0.684735 | 0.612171 | 1 | 0.999631 | 0.949525 |
| Nefm_P08553_S30s_1 | -0.12 | -0.99 | -1.04 | -1.25 | -0.91 | -0.14 | -1.25 | -0.52 | -1.23 | -0.15 | -1.08 | -0.85 | -0.86 | -0.48 | -0.46 | -0.29 | 0.145 | 0.0225 | 0.3275 | 0.348855 | 0.6936 | 0.951748 | 1 | 0.999631 | 0.736697 |
| Nefm_P08553_S428s_1 | 1.67 | 0.5 | -1.41 | 1.33 | 1.46 | 0.11 | 1.52 | 0.91 | 0.04 | -0.28 | 0.98 | -0.87 | 0.74 | -0.16 | -0.87 | -0.34 | 0.4775 | -0.555 | -0.68 | 0.33147 | 0.48677 | 0.437724 | 1 | 0.999631 | 0.727302 |
| Nefm_P08553_S44s_1 | 0.16 | -0.65 | -0.57 | -0.35 | -0.33 | 0.37 | -0.42 | 0.35 | 0.53 | 0.33 | 0.02 | 0.24 | 0.67 | 0.54 | 0.62 | 0.26 | 0.345 | 0.6325 | 0.875 | 0.01543 | 0.302842 | 0.053445 | 1 | 0.999631 | 0.388832 |
| Nefm_P08553_S610s_1 | 0.98 | 1.64 | 1.94 | 1.13 | 0.45 | 2.5 | 1.12 | 1.48 | 1.41 | 2.29 | 1.36 | 1.25 | 0.8 | 2.07 | 1.93 | 1.79 | -0.035 | 0.155 | 0.225 | 0.569993 | 0.939592 | 0.668557 | 1 | 0.999631 | 0.84338 |
| Nefm_P08553_S645s_1 | 1.77 | 0.17 | 3.15 | 2.58 | -0.85 | 0.71 | 3.1 | 0.35 | 0.05 | 1.43 | 0.91 | 1.36 | 0.5 | -0.24 | 2.31 | 1.43 | -1.09 | -0.98 | -0.9175 | 0.238586 | 0.248383 | 0.154516 | 1 | 0.999631 | 0.683403 |
| Nefm_P08553_S669s_1 | 0.36 | -2.73 | 0.69 | -1.13 | -2.61 | -0.47 | 1.37 | -1.74 | -3.42 | -0.22 | -0.7 | -0.13 | -1.11 | -2.98 | 0.86 | -0.12 | -0.16 | -0.415 | -0.135 | 0.889402 | 0.873237 | 0.665515 | 1 | 0.999631 | 0.963505 |
| Nefm_P08553_S715s_1 | 0.5 | -1.53 | -0.32 | -1.02 | -2.19 | -0.47 | 0.23 | -2.12 | -1.21 | 1.12 | -1.53 | -1.67 | -1.62 | -2.12 | -1.42 | -2.99 | -0.545 | -0.23 | -1.445 | 0.019898 | 0.421481 | 0.74383 | 1 | 0.999631 | 0.391668 |
| Nefm_P08553_S715s_2 | -0.46 | -0.93 | -0.77 | -0.52 | -1.02 | -0.18 | -0.47 | -0.91 | -0.13 | 0.81 | -1.78 | -1.78 | -0.66 | -1.44 | -1.31 | -2.99 | 0.025 | -0.05 | -0.93 | 0.080926 | 0.931957 | 0.932853 | 1 | 0.999631 | 0.545727 |
| Nefm_P08553_S715s_3 | -0.73 | 0.07 | 1.37 | 0.62 | -0.53 | 0.04 | 0.61 | 0.29 | -0.11 | 0.67 | 0.3 | -0.14 | 0 | -0.89 | -0.17 | -0.76 | -0.23 | -0.1525 | -0.7875 | 0.124844 | 0.632636 | 0.744774 | 1 | 0.999631 | 0.596063 |
| Nefm_P08553_S720s_1 | -0.33 | -1.18 | -0.8 | -0.32 | -1.62 | -0.82 | -0.95 | -1.13 | -1.06 | 0.61 | -1.66 | -1.43 | 0.02 | -2.62 | -1.65 | -2.9 | -0.4725 | -0.2275 | -1.13 | 0.094876 | 0.162199 | 0.6631 | 1 | 0.999631 | 0.558128 |
| Nefm_P08553_S720s_3 | -0.73 | 0.07 | 1.37 | 0.62 | -0.53 | 0.04 | 0.61 | 0.29 | -0.11 | 0.67 | 0.3 | -0.14 | 0 | -0.89 | -0.17 | -0.76 | -0.23 | -0.1525 | -0.7875 | 0.124844 | 0.632636 | 0.744774 | 1 | 0.999631 | 0.596063 |
| Nefm_P08553_S723s_1 | 1.52 | 0.08 | -0.54 | -0.63 | -0.84 | 1.13 | -0.46 | -0.86 | 0.49 | 2.16 | -1.34 | -1.14 | -0.03 | -0.87 | -0.07 | -3.05 | -0.365 | -0.065 | -1.1125 | 0.165007 | 0.557177 | 0.937932 | 1 | 0.999631 | 0.633483 |
| Nefm_P08553_S723s_2 | -0.46 | -0.93 | -0.36 | -0.14 | -1.02 | -0.18 | -0.11 | -0.57 | -0.13 | 0.81 | -1.5 | -1.38 | -0.66 | -1.44 | -0.78 | -2.56 | 0.0025 | -0.0775 | -0.8875 | 0.077797 | 0.99371 | 0.884786 | 1 | 0.999631 | 0.543121 |
| Nefm_P08553_S723s_3 | -0.73 | 0.07 | 1.37 | 0.62 | -0.53 | 0.04 | 0.61 | 0.29 | -0.11 | 0.67 | 0.3 | -0.14 | 0 | -0.89 | -0.17 | -0.76 | -0.23 | -0.1525 | -0.7875 | 0.124844 | 0.632636 | 0.744774 | 1 | 0.999631 | 0.596063 |
| Nefm_P08553_S769s_1 | -0.97 | -0.91 | -1.09 | -1.27 | -0.89 | -1.11 | -1.37 | -0.85 | -0.82 | -0.73 | -0.76 | -0.56 | 0.27 | -0.02 | -0.4 | -0.38 | 0.005 | 0.3425 | 0.9275 | 0.009509 | 0.984373 | 0.189259 | 1 | 0.999631 | 0.371283 |
| Nefm_P08553_T565t_1 | -1.85 | 2.36 | 4 | 1.68 | 2.16 | 2.08 | 2 | 2.99 | 2.41 | -0.86 | 3.58 | 4.58 | 3.49 | 2.58 | 3.87 | 3.08 | 0.76 | 0.88 | 1.7075 | 0.135957 | 0.485288 | 0.548613 | 1 | 0.999631 | 0.608682 |
| Nek1_B2RXQ4_S972s_1 | 0.43 | 2.74 | 1.91 | 1.52 | 1.85 | 2.07 | 0.56 | 2.04 | 2.38 | 1.87 | 1.4 | 1.39 | 2.43 | 2.81 | 2.08 | 1.99 | -0.02 | 0.11 | 0.6775 | 0.191691 | 0.970982 | 0.827738 | 1 | 0.999631 | 0.648674 |
| Nek4_F8WGA9_S423s_1 | 0.14 | 2.77 | 2.3 | 2.15 | 1.18 | 1.99 | 1.47 | 2.17 | 2.01 | 0.72 | 2.26 | 2.37 | 1.52 | 2.18 | 2.77 | 3.2 | -0.1375 | -1.6E-16 | 0.5775 | 0.363612 | 0.809484 | 1 | 1 | 1 | 0.743747 |
| Nelfa_Q8BG30_S363s_1 | -0.51 | 0.97 | 0.81 | 0.28 | 0.33 | 0.3 | 0.23 | 0.08 | 0.4 | -0.3 | -0.32 | 0.88 | 0.79 | 1.1 | 0.17 | 0.85 | -0.1525 | -0.2225 | 0.34 | 0.412711 | 0.6742 | 0.614222 | 1 | 0.999631 | 0.768201 |
| Nelfb_Q8C4Y3_S557s_1 | 0.11 | 0.81 | 0.54 | -0.07 | 0.36 | 0.32 | -0.17 | -0.04 | 0.32 | 0.33 | 0.13 | 0.43 | 0.62 | 0.83 | 0.3 | 0.71 | -0.23 | -0.045 | 0.2675 | 0.407214 | 0.452503 | 0.87791 | 1 | 0.999631 | 0.766996 |
| Nelfb_Q8C4Y3_T559t_1 | 0.18 | 1.06 | 0.87 | 0.35 | 0.49 | 0.43 | 0.07 | 0.74 | 0.64 | 0.08 | 0.88 | 1.09 | 0.71 | 1.14 | 1.09 | 1.1 | -0.1825 | 0.0575 | 0.395 | 0.231811 | 0.556395 | 0.867386 | 1 | 0.999631 | 0.682192 |
| Nelfe_P19426_S115s_1 | 0.5 | 1.21 | 0.61 | 1.02 | 1.2 | 0.85 | 0.99 | 0.66 | 0.84 | 1.13 | 0.86 | 1.13 | 0.89 | 1.43 | 1.02 | 1.34 | 0.09 | 0.155 | 0.335 | 0.291042 | 0.750153 | 0.584945 | 1 | 0.999631 | 0.706164 |
| Nelfe_P19426_S181s_1 | 2.56 | 1.27 | 1.78 | 1.45 | 0.46 | 0.72 | 1.83 | 0.39 | 0.89 | 1.85 | 0.97 | 0.27 | 0.04 | 0.35 | 0.46 | -0.44 | -0.915 | -0.77 | -1.6625 | 0.001437 | 0.056594 | 0.099316 | 1 | 0.999631 | 0.351308 |
| Nelfe_P19426_S49s_1 | -0.02 | 0.44 | 0.25 | -0.02 | -0.18 | 0.73 | -0.27 | -0.2 | 0.85 | 0.22 | 0.29 | 0.81 | 1.28 | 1.03 | -0.2 | 0.35 | -0.1425 | 0.38 | 0.4525 | 0.257925 | 0.652151 | 0.208299 | 1 | 0.999631 | 0.692624 |
| Neo1_Q7TQG5_S1182s_1 | -0.35 | -0.75 | -0.56 | -0.84 | -0.31 | -1.17 | -0.86 | -0.53 | -0.74 | -0.84 | -1.12 | -0.05 | -0.5 | -0.63 | -0.23 | 0.11 | -0.0925 | -0.0625 | 0.3125 | 0.312729 | 0.749927 | 0.842866 | 1 | 0.999631 | 0.719861 |
| Neo1_Q7TQG5_S1198s_2 | 4.05 | -0.15 | 1.31 | 0.79 | 1.11 | 0.31 | 1.7 | 1.8 | 1.05 | 1.97 | 0.63 | 2.15 | 0.94 | 0.45 | 0.46 | 0.57 | -0.27 | -0.05 | -0.895 | 0.275084 | 0.74817 | 0.953026 | 1 | 0.999631 | 0.700187 |
| Neo1_Q7TQG5_S1224s_1 | 1.34 | -1.84 | 0.5 | -0.11 | 2.06 | 1.06 | 1.7 | -0.04 | 1.94 | 1.12 | 1.31 | 1.87 | 2.7 | -1.02 | 0.16 | -0.27 | 1.2225 | 1.5875 | 0.42 | 0.64435 | 0.114253 | 0.030004 | 1 | 0.999631 | 0.872436 |
| Neo1_Q7TQG5_S1405s_1 | 0.46 | -1 | -1.17 | -0.3 | 0.12 | -0.52 | -0.24 | -0.73 | -0.37 | -1.06 | -0.51 | -1.21 | -0.92 | -0.77 | -0.64 | -0.49 | 0.16 | -0.285 | -0.2025 | 0.618982 | 0.699775 | 0.508267 | 1 | 0.999631 | 0.861888 |
| Neo1_Q7TQG5_T1202t_1 | -0.75 | -0.43 | -1.05 | -0.31 | -0.13 | -0.37 | -0.46 | 0.14 | -0.3 | -0.77 | -1.11 | -0.26 | -0.97 | -0.09 | -0.36 | -0.01 | 0.43 | 0.025 | 0.2775 | 0.421833 | 0.158263 | 0.937928 | 1 | 0.999631 | 0.772942 |
| Neo1_Q7TQG5_T1202t_2 | 4.05 | -0.15 | 1.31 | 0.79 | 1.11 | 0.31 | 1.7 | 1.8 | 1.05 | 1.97 | 0.63 | 2.15 | 0.94 | 0.45 | 0.46 | 0.57 | -0.27 | -0.05 | -0.895 | 0.275084 | 0.74817 | 0.953026 | 1 | 0.999631 | 0.700187 |
| Nes_Q6P5H2_S1793s_1 | 0.71 | 1.17 | -0.24 | -0.41 | 0.46 | -0.12 | -0.11 | -1.06 | 0.56 | 1.41 | -0.28 | -0.02 | 0.47 | 1.33 | 1.11 | 0.64 | -0.515 | 0.11 | 0.58 | 0.202122 | 0.28815 | 0.827342 | 1 | 0.999631 | 0.658952 |
| Neto1_Q8R4I7_S520s_1 | 0.73 | 1.41 | 0.82 | 1.12 | 1.59 | 1.47 | 0.9 | 2.26 | 0.22 | 1.07 | 1.13 | 0.68 | -1.1 | 1.37 | 1.11 | 2.08 | 0.535 | -0.245 | -0.155 | 0.807037 | 0.148709 | 0.451313 | 1 | 0.999631 | 0.935523 |
| Neurl4_H9H9T3_S247s_1 | 0.02 | -0.39 | -0.39 | -0.71 | -0.48 | -0.31 | -0.78 | -0.57 | -0.05 | -0.2 | -0.09 | 0.09 | -0.23 | 0.47 | 0.13 | 0.18 | -0.1675 | 0.305 | 0.505 | 0.122708 | 0.539825 | 0.275745 | 1 | 0.999631 | 0.591845 |
| Neurl4_H9H9T3_S647s_1 | -1.14 | -0.95 | 1.13 | -0.15 | 0.06 | -1.29 | -0.09 | -0.07 | -0.99 | 0.45 | 0.17 | 0.8 | 0.43 | -0.13 | 2.18 | 1.19 | -0.07 | 0.385 | 1.195 | 0.088486 | 0.899455 | 0.518617 | 1 | 0.999631 | 0.553858 |
| Neurl4_H9H9T3_S652s_1 | -1.59 | -2.15 | -1.59 | -1.2 | -2.6 | -1.95 | -1.47 | -2.37 | -2.03 | -2.31 | -1 | -1.19 | -1.2 | -1.76 | -0.69 | -1.44 | -0.465 | 2.36E-16 | 0.36 | 0.321138 | 0.201192 | 1 | 1 | 1 | 0.72408 |
| Nexn_Q7TPW1_S16s_1 | 0.73 | 0.2 | 0.39 | 0.23 | -0.29 | -0.24 | -0.25 | 0.28 | 0.11 | 1.25 | -0.17 | 0.09 | 0.94 | 0.78 | 0.38 | 0.9 | -0.5125 | -0.0675 | 0.3625 | 0.233529 | 0.084129 | 0.854415 | 1 | 0.999631 | 0.683077 |
| Nexn_Q7TPW1_S296s_1 | -0.44 | -1.28 | -0.95 | -0.44 | -0.8 | -1.74 | -1.03 | 0.02 | -1.37 | -0.17 | -0.83 | 0.1 | -0.74 | -0.8 | -0.9 | -0.5 | -0.11 | 0.21 | 0.0425 | 0.892073 | 0.791212 | 0.603664 | 1 | 0.999631 | 0.964276 |
| Nf1_Q04690_S2169s_1 | -0.32 | -0.66 | -0.47 | -0.79 | -0.52 | -0.55 | -0.95 | -0.42 | -0.7 | 0.22 | -0.6 | -0.72 | 0.39 | -0.13 | -0.33 | -0.13 | -0.05 | 0.11 | 0.51 | 0.109412 | 0.847838 | 0.725815 | 1 | 0.999631 | 0.572573 |
| Nf1_Q04690_S2477s_1 | 0.17 | 0.2 | 0.36 | -0.07 | -0.48 | 0.39 | -0.23 | 0.16 | -0.24 | 0.38 | -0.31 | 0.23 | 0.12 | 0.35 | 0.27 | 0.73 | -0.205 | -0.15 | 0.2025 | 0.4832 | 0.482031 | 0.60061 | 1 | 0.999631 | 0.803286 |
| Nf1_Q04690_S2496s_1 | 0.15 | 0.24 | -0.13 | -0.12 | 0.95 | 0.07 | -0.08 | -0.61 | 0.48 | -0.27 | -0.32 | 0.24 | -1.19 | 0.25 | 0.45 | 0 | 0.0475 | -0.0025 | -0.1575 | 0.696467 | 0.895184 | 0.993269 | 1 | 0.999631 | 0.895731 |
| Nf1_Q04690_S2502s_1 | 1.21 | -0.28 | 0.09 | 0.03 | -0.09 | -0.44 | -0.4 | -0.01 | -0.25 | -0.06 | -0.24 | -0.12 | 0.46 | -0.09 | -0.67 | 0.48 | -0.4975 | -0.43 | -0.2175 | 0.616984 | 0.193799 | 0.252026 | 1 | 0.999631 | 0.861376 |
| Nf1_Q04690_S2504s_1 | -0.6 | 0.02 | 0.25 | -0.07 | 0.07 | -0.1 | -0.22 | -0.26 | -0.2 | 0.05 | 0.12 | 0.25 | 0.37 | -0.13 | -0.44 | 0.48 | -0.0275 | 0.155 | 0.17 | 0.622547 | 0.921138 | 0.595215 | 1 | 0.999631 | 0.862727 |
| Nf1_Q04690_S2524s_1 | -1.84 | -0.26 | -0.65 | -1.16 | 0.36 | -0.38 | 1.16 | -0.45 | 1.19 | -1.49 | 0.06 | 0.55 | 0.84 | -0.68 | -0.94 | -0.34 | 1.15 | 1.055 | 0.6975 | 0.183432 | 0.036168 | 0.10653 | 1 | 0.999631 | 0.644372 |
| Nf1_Q04690_S2578s_1 | -0.64 | -0.17 | 0.2 | -0.3 | 0.18 | -0.41 | -0.53 | -0.2 | -0.13 | -0.68 | -0.22 | -0.09 | 0.29 | -0.14 | 0.38 | 0.32 | -0.0125 | -0.0525 | 0.44 | 0.173991 | 0.966509 | 0.86034 | 1 | 0.999631 | 0.642116 |
| Nf1_Q04690_S2783s_1 | 0.26 | 0.45 | 0.52 | 0.04 | 0.1 | 0.13 | 0.37 | 0.58 | 0.54 | 0.63 | -0.59 | -0.81 | -0.05 | 0.83 | -0.63 | -0.03 | -0.0225 | -0.375 | -0.2875 | 0.438801 | 0.931104 | 0.361126 | 1 | 0.999631 | 0.781983 |
| Nf1_Q04690_S2798s_1 | 1.96 | 4.11 | 4.03 | 1.1 | 4.53 | 3.62 | 6.4 | 1.76 | 6.27 | 0.39 | 3.77 | 6.33 | 5.01 | 1 | 2.71 | 1.42 | 1.2775 | 1.39 | -0.265 | 0.792155 | 0.242425 | 0.318843 | 1 | 0.999631 | 0.927114 |
| Nf1_Q04690_S659s_1 | 1.83 | -0.48 | -1.59 | 1.47 | 2.29 | 0.76 | 1.27 | 1.21 | 0.91 | -0.04 | 0.35 | -0.86 | -1.48 | 0.38 | -0.43 | -0.41 | 1.075 | -0.2175 | -0.7925 | 0.324399 | 0.182084 | 0.78185 | 1 | 0.999631 | 0.72463 |
| Nf1_Q04690_S866s_1 | -2.46 | -0.77 | -0.69 | -2.77 | -0.43 | -1.5 | -0.31 | -2.42 | -0.31 | -3.06 | -0.21 | -0.03 | 0.2 | -2.61 | -0.71 | -3.32 | 0.5075 | 0.77 | 0.0625 | 0.941478 | 0.448497 | 0.345317 | 1 | 0.999631 | 0.981286 |
| Nf1_Q04690_S878s_1 | -0.09 | 0.36 | -1.18 | 0.94 | 1.82 | 0.39 | 0.28 | 0.44 | 0.45 | -0.22 | -0.12 | 0.04 | 1.73 | 0.76 | 0.01 | -0.06 | 0.725 | 0.03 | 0.6025 | 0.301079 | 0.195324 | 0.947919 | 1 | 0.999631 | 0.714243 |
| Nf1_Q04690_T2545t_1 | -1.66 | -0.05 | 0.14 | -0.17 | 0.07 | -0.21 | -0.65 | -0.2 | 0.08 | -0.34 | -0.65 | 0.45 | -0.02 | -0.03 | 0.41 | -0.12 | 0.1875 | 0.32 | 0.495 | 0.271819 | 0.66486 | 0.495891 | 1 | 0.999631 | 0.699374 |
| Nf1_Q04690_T2546t_1 | -1.12 | -0.15 | 0.22 | -0.19 | -0.08 | -0.27 | -0.59 | -0.26 | -0.09 | -0.15 | -0.48 | 0.36 | 0.07 | -0.14 | 0.27 | -0.14 | 0.01 | 0.22 | 0.325 | 0.370575 | 0.976498 | 0.549732 | 1 | 0.999631 | 0.746338 |
| Nfat5_Q80TR0_S117s_1 | 0.34 | 0.64 | -0.09 | 0.35 | 0.05 | 0.52 | -0.31 | 0.35 | 0.37 | -0.18 | 0.68 | 0.86 | 0.09 | 0.49 | 0.74 | 0.35 | -0.1575 | 0.1225 | 0.1075 | 0.724157 | 0.601682 | 0.709053 | 1 | 0.999631 | 0.905674 |
| Nfic_P70255_S194s_1 | 1.47 | 1.28 | 2.89 | 1.73 | 0.33 | 2.08 | 3.85 | 0.41 | 0.69 | 2.18 | 0.78 | 1.06 | 0.74 | -0.18 | 1.3 | 0.14 | -0.175 | -0.665 | -1.3425 | 0.017005 | 0.825312 | 0.186632 | 1 | 0.999631 | 0.391024 |
| Nfic_P70255_S305s_1 | -0.93 | 0.16 | -0.81 | -3.26 | -0.35 | 0.41 | -1.46 | -0.89 | -0.14 | -1.6 | -0.11 | 0.35 | 0 | 0.14 | 1.8 | 0.96 | 0.6375 | 0.835 | 1.935 | 0.02372 | 0.391609 | 0.277166 | 1 | 0.999631 | 0.415644 |
| Nfic_P70255_S339s_1 | -0.43 | 0.63 | 0.54 | 0.31 | -0.01 | -0.07 | -0.5 | 0.59 | 0.01 | 0.56 | 0.41 | 0.46 | 0.02 | 0.16 | 0.39 | 0.2 | -0.26 | 0.0975 | -0.07 | 0.831081 | 0.470421 | 0.764564 | 1 | 0.999631 | 0.942552 |
| Nfix_D3YZ00_S301s_1 | -0.29 | -0.04 | 0.29 | -0.69 | -0.29 | -0.49 | -0.54 | -0.28 | -0.24 | -0.37 | -0.39 | -0.15 | 0.2 | 0.28 | -0.57 | 0.11 | -0.2175 | -0.105 | 0.1875 | 0.589645 | 0.459203 | 0.722806 | 1 | 0.999631 | 0.851414 |
| Nfx1_B1AY10_S197s_1 | 0.96 | 2.15 | -0.05 | 0.69 | 1.85 | 2.91 | 1.36 | 0.94 | 1.57 | -0.16 | -0.04 | 0.24 | 1.71 | 0.75 | -0.21 | 0.22 | 0.8275 | -0.535 | -0.32 | 0.576709 | 0.168503 | 0.350923 | 1 | 0.999631 | 0.846204 |
| Ngef_E9QK62_S606s_1 | 0.85 | 0.78 | -0.29 | 1.1 | 1.45 | 0.7 | 0.27 | 1.15 | 0.65 | 0.18 | 0.28 | 0.92 | -0.36 | 1.14 | 0.3 | 0.92 | 0.2825 | -0.1025 | -0.11 | 0.808615 | 0.490332 | 0.785508 | 1 | 0.999631 | 0.93635 |
| Ngef_E9QK62_S696s_1 | 0.67 | -2.06 | -1.74 | 1.61 | 1.21 | -2.4 | 1.16 | -0.19 | 0.03 | -0.65 | -0.06 | -0.27 | -2.76 | -2.29 | -0.74 | -2.36 | 0.325 | 0.1425 | -1.6575 | 0.080374 | 0.759549 | 0.858879 | 1 | 0.999631 | 0.545579 |
| Nhp2l1_Q9D0T1_S122s_1 | -0.73 | 0.8 | -1.63 | -1.37 | 1.18 | -0.13 | -0.95 | -2.24 | 0.51 | 0.88 | -1.63 | -0.78 | 1.21 | 0.58 | -2.8 | -0.14 | 0.1975 | 0.4775 | 0.445 | 0.62063 | 0.802243 | 0.504966 | 1 | 0.999631 | 0.861888 |
| Nhs_B9EJX2_S715s_1 | 0.23 | -0.1 | -0.03 | 0.03 | -0.22 | 0.19 | 0.01 | 0.34 | 0.6 | 0.2 | -0.15 | 0.54 | 1.27 | 0.84 | 0.74 | 0.16 | 0.0475 | 0.265 | 0.72 | 0.045692 | 0.851744 | 0.358767 | 1 | 0.999631 | 0.477252 |
| Nhsl1_Q8CAF4_S1075s_1 | -0.05 | -0.01 | 0.19 | 0.16 | 1 | 0.97 | -0.46 | 0.74 | 0.89 | 0.7 | 0.11 | 0.49 | 0.7 | 0.97 | 0.87 | 0.44 | 0.49 | 0.475 | 0.6725 | 0.032563 | 0.205401 | 0.111955 | 1 | 0.999631 | 0.442395 |
| Nhsl1_Q8CAF4_S1360s_1 | 1.89 | -2.67 | -1.15 | -0.16 | -2.08 | -1.6 | 3.56 | -2.98 | -4 | 0.84 | -1.99 | -2.23 | -3.52 | -4.49 | -1.44 | -2.51 | -0.2525 | -1.3225 | -2.4675 | 0.030066 | 0.865318 | 0.280445 | 1 | 0.999631 | 0.436111 |
| Nhsl1_Q8CAF4_S535s_1 | 0.3 | -0.76 | 0.26 | -1.19 | -2.7 | -0.72 | -0.28 | -1.21 | -0.49 | 0.94 | -1.53 | -1.21 | -1.08 | -2.92 | -1.36 | -2.54 | -0.88 | -0.225 | -1.6275 | 0.012668 | 0.156036 | 0.710336 | 1 | 0.999631 | 0.385092 |
| Nhsl2_B1AXH1_S1145s_1 | -0.96 | -0.02 | 1.52 | -0.97 | -2.11 | -0.61 | 0.13 | 1.44 | -0.99 | -0.24 | 1.35 | 1.52 | -0.9 | -0.01 | 2.47 | 2.15 | -0.18 | 0.5175 | 1.035 | 0.252456 | 0.826957 | 0.495096 | 1 | 0.999631 | 0.688149 |
| Nhsl2_B1AXH1_S1208s_1 | 0.92 | 1.23 | 0.14 | 1.54 | 1.64 | 0.79 | 1.36 | 1.57 | 0.95 | -0.13 | 1.28 | 0.65 | -0.44 | 0.98 | -0.26 | 0.71 | 0.3825 | -0.27 | -0.71 | 0.144176 | 0.319422 | 0.530861 | 1 | 0.999631 | 0.616037 |
| Nhsl2_B1AXH1_S690s_1 | 0.01 | -0.22 | -0.57 | 0.19 | 0.45 | -0.32 | 0.18 | 0.69 | -0.67 | -0.49 | 0.07 | 0.39 | -0.54 | -0.04 | -0.46 | -0.12 | 0.3975 | -0.0275 | -0.1425 | 0.642074 | 0.232697 | 0.935581 | 1 | 0.999631 | 0.871335 |
| Nhsl2_B1AXH1_T284t_1 | 0.48 | 0.81 | 1.36 | 1.57 | 1.57 | 1.95 | 1.4 | 2.16 | 0.93 | 0.16 | 2.13 | 0.53 | -1.14 | 1.83 | 0.88 | 0.71 | 0.715 | -0.1175 | -0.485 | 0.43289 | 0.058111 | 0.805344 | 1 | 0.999631 | 0.779247 |
| Nicn1_Q9CQM0_S142s_1 | -0.07 | -0.23 | -0.21 | -0.68 | -0.25 | -0.11 | -0.56 | -0.55 | 0.05 | 0.29 | -0.28 | -0.09 | 0.35 | 0.36 | 0.04 | 0.35 | -0.07 | 0.29 | 0.5725 | 0.066771 | 0.794217 | 0.311128 | 1 | 0.999631 | 0.519446 |
| Nifk_Q91VE6_S256s_1 | 1.82 | -0.49 | -0.01 | 0.66 | -0.9 | 0.61 | 0.57 | -0.84 | -0.44 | 0.77 | -1.26 | -1.47 | 0.01 | 0.07 | -0.74 | -0.95 | -0.635 | -1.095 | -0.8975 | 0.114691 | 0.298733 | 0.113327 | 1 | 0.999631 | 0.579117 |
| Nipbl_Q6KCD5_S2652s_1 | -0.72 | -0.53 | -0.51 | -1.16 | -0.87 | -1.02 | -1.14 | -1.58 | -0.62 | -1.01 | -1.31 | -0.55 | -0.07 | -0.71 | -1.37 | -0.58 | -0.4225 | -0.1425 | 0.0475 | 0.894496 | 0.165359 | 0.64205 | 1 | 0.999631 | 0.965376 |
| Nipbl_Q6KCD5_T599t_1 | -0.06 | -0.85 | -1.05 | -0.45 | -1.42 | -1.39 | -0.62 | -0.66 | -1.64 | 0.02 | -0.53 | -1.03 | -0.8 | -0.46 | -0.62 | 0.34 | -0.42 | -0.1925 | 0.2175 | 0.564275 | 0.240708 | 0.648426 | 1 | 0.999631 | 0.84139 |
| Nisch_Q80TM9_S1128s_1 | -0.21 | 1.33 | 0.45 | -0.3 | 1.23 | 1.15 | -0.17 | 0.46 | 1.98 | 1.51 | 0.25 | 0.49 | 2.07 | 1.42 | 0.13 | 0.37 | 0.35 | 0.74 | 0.68 | 0.233976 | 0.467546 | 0.179264 | 1 | 0.999631 | 0.683077 |
| Nisch_Q80TM9_S492s_1 | -0.98 | -0.3 | 0.27 | -0.33 | -0.28 | -0.87 | -0.51 | -0.68 | -0.48 | -0.76 | -0.3 | 0.02 | -0.18 | -0.44 | 0.25 | -0.23 | -0.25 | -0.045 | 0.185 | 0.600148 | 0.452354 | 0.896302 | 1 | 0.999631 | 0.856618 |
| Nkd2_Q8VE28_S259s_1 | -0.51 | -1.22 | -0.52 | -0.75 | -0.54 | -0.88 | -0.34 | -1.34 | -1.08 | -0.16 | 0.4 | -0.81 | -0.07 | -0.31 | -0.84 | -0.15 | -0.025 | 0.3375 | 0.4075 | 0.224047 | 0.938011 | 0.395502 | 1 | 0.999631 | 0.67471 |
| Nktr_P30415_S1139s_1 | 0.53 | 0.68 | 1.76 | 0.75 | -0.96 | -0.35 | 0.02 | 1.02 | 0.44 | -0.1 | -0.81 | 2.36 | 0.64 | 0.6 | 2.1 | 1.88 | -0.9975 | -0.4575 | 0.375 | 0.437293 | 0.059204 | 0.493395 | 1 | 0.999631 | 0.781057 |
| Nktr_P30415_S468s_1 | 0.7 | -0.64 | -0.45 | -0.79 | -1.07 | -1.06 | -1.16 | -0.96 | -1.04 | -0.62 | -1.17 | 0.31 | -0.3 | 0.43 | -0.85 | -0.63 | -0.7675 | -0.335 | -0.0425 | 0.923347 | 0.057454 | 0.475165 | 1 | 0.999631 | 0.975032 |
| Nlgn2_Q69ZK9_S714s_1 | -0.88 | -0.75 | -0.97 | -1.21 | -0.85 | -1 | -1.41 | -1.17 | -0.93 | -0.68 | -1.11 | -0.78 | 0.13 | -0.56 | -1.08 | -0.55 | -0.155 | 0.0775 | 0.4375 | 0.21167 | 0.554535 | 0.766327 | 1 | 0.999631 | 0.665474 |
| Nlgn3_Q8BYM5_S722s_1 | -1.59 | -4.91 | -4.27 | -3.27 | -3.19 | -4.44 | -2.99 | -4 | -3.85 | -2.6 | -2.4 | -3.05 | -2.42 | -3.73 | -3.76 | -4.14 | -0.145 | 0.535 | -0.0025 | 0.997227 | 0.837501 | 0.454571 | 1 | 0.999631 | 0.998985 |
| Nlgn4l_B0F2B4_S880s_1 | 0.79 | -1.65 | -1.26 | 0.09 | -0.06 | -0.31 | 0.31 | -0.45 | -0.26 | -0.02 | -0.4 | -0.86 | -1.25 | -1.74 | -0.25 | -1.32 | 0.38 | 0.1225 | -0.6325 | 0.302437 | 0.492407 | 0.824833 | 1 | 0.999631 | 0.714978 |
| Nlk_O54949_T298t_1 | 0.27 | -1.81 | -2.14 | -0.6 | -0.1 | 0.03 | -0.22 | -1 | -0.32 | 0.57 | -0.99 | -1.29 | 0.4 | -0.27 | -3.62 | -0.59 | 0.7475 | 0.5625 | 0.05 | 0.955882 | 0.196983 | 0.379343 | 1 | 0.999631 | 0.984741 |
| Nmt1_O70310_S47s_1 | 0.85 | 1.46 | 1.08 | 0.82 | 1.06 | 0.96 | 0.8 | 1.64 | 1.29 | 1.36 | 1.67 | 1.57 | 0.76 | 1.71 | 1.68 | 1.7 | 0.0625 | 0.42 | 0.41 | 0.246866 | 0.834515 | 0.150701 | 1 | 0.999631 | 0.685751 |
| Nmt2_O70311_S38s_1 | 1.02 | 1.45 | 1.16 | 1 | 1.04 | 1.12 | 0.55 | 0.73 | 0.88 | 1.61 | 1.06 | 1.25 | 0.75 | 1.23 | 0.61 | 0.84 | -0.2975 | 0.0425 | -0.3 | 0.313382 | 0.279341 | 0.880256 | 1 | 0.999631 | 0.719861 |
| Nmt2_O70311_S66s_1 | 0.42 | 0.86 | 1.37 | 0.41 | -0.1 | 1.29 | 0.94 | 1.01 | 0.86 | 1.61 | 1.33 | 1.28 | 1.48 | 0.16 | 2.3 | 0.46 | 0.02 | 0.505 | 0.335 | 0.518055 | 0.958981 | 0.147805 | 1 | 0.999631 | 0.819449 |
| No66_Q9JJF3_S105s_1 | 0.55 | 1.27 | 0.96 | 1.19 | 0.5 | 0.49 | 1.06 | 1.15 | 0.64 | 0.51 | 1.98 | 1.24 | 1.17 | 1.22 | 1.77 | 2.75 | -0.1925 | 0.1 | 0.735 | 0.10172 | 0.527551 | 0.797767 | 1 | 0.999631 | 0.564448 |
| Noc2l_J3QK52_S60s_1 | -0.25 | -1.85 | -1.62 | -1.43 | -1.49 | -2.84 | -1.21 | -2.73 | -1.14 | -1.29 | -2.3 | -1.65 | -1.02 | -3.33 | -3.33 | -2.7 | -0.78 | -0.3075 | -1.3075 | 0.049401 | 0.151562 | 0.487022 | 1 | 0.999631 | 0.487521 |
| Noc2l_J3QK52_T59t_1 | 2.53 | 0.32 | -0.26 | -2.01 | -0.09 | 0.52 | -0.84 | -1.17 | -0.07 | 1.18 | -1.3 | -2.13 | -0.08 | -0.86 | -1.57 | -1.7 | -0.54 | -0.725 | -1.1975 | 0.190288 | 0.541734 | 0.483401 | 1 | 0.999631 | 0.648674 |
| Nol3_Q9D1X0_S142s_1 | -0.97 | -0.1 | 0.96 | 1.74 | 0.21 | 0.04 | 0.51 | 2.31 | 1.13 | 1.57 | 2.86 | 2.44 | 1.38 | -0.52 | 2.31 | 1.84 | 0.36 | 1.5925 | 0.845 | 0.278081 | 0.609636 | 0.031211 | 1 | 0.999631 | 0.700874 |
| Nol3_Q9D1X0_T114t_1 | -1.67 | 0.76 | 0.74 | 0.19 | -0.8 | -0.4 | 0.43 | 0.72 | 0.12 | -0.37 | 1.1 | -0.64 | -0.41 | -0.02 | 0.83 | 0.9 | -0.0175 | 0.0475 | 0.32 | 0.596218 | 0.977009 | 0.939299 | 1 | 0.999631 | 0.854508 |
| Nol4_F6XSA1_S135s_1 | 0.97 | 1.25 | 0.99 | 0.4 | 0.76 | 0.54 | 0.53 | 0.11 | 0.55 | 1.15 | -0.74 | 0.55 | -0.11 | 1.13 | -0.41 | 0.67 | -0.4175 | -0.525 | -0.5825 | 0.179918 | 0.176955 | 0.244998 | 1 | 0.999631 | 0.64376 |
| Nol4l_Q3UH85_S17s_1 | 0.65 | 0.24 | 1.18 | 1.11 | -0.13 | 0.16 | 0.45 | 1.24 | 0.3 | 1.15 | 1.42 | 1.63 | 0.67 | 1.3 | 1.48 | 1.77 | -0.365 | 0.33 | 0.51 | 0.184167 | 0.349353 | 0.400467 | 1 | 0.999631 | 0.644602 |
| Nol9_Q3TZX8_S88s_1 | -0.67 | 0.58 | -0.68 | -0.51 | 0.12 | -0.53 | -0.88 | -0.52 | 0.14 | -0.39 | -0.47 | 0.38 | 0.25 | -0.04 | -0.05 | 0.24 | -0.1325 | 0.235 | 0.42 | 0.262624 | 0.728397 | 0.545864 | 1 | 0.999631 | 0.693034 |
| Nolc1_E9Q5C9_S563s_1 | -2.29 | -0.98 | -0.41 | -0.39 | -1.33 | -1.61 | -0.92 | -0.54 | -1 | -1.62 | -0.44 | -0.85 | -0.57 | -1.37 | -0.19 | -0.91 | -0.0825 | 0.04 | 0.2575 | 0.603385 | 0.862887 | 0.934382 | 1 | 0.999631 | 0.857824 |
| Nolc1_E9Q5C9_S626s_1 | -6.5 | -5.63 | -3.14 | -5.07 | -5.41 | -6.25 | -3.82 | -5.91 | -5.75 | -2.31 | -2.92 | -3.09 | -1.37 | -5.07 | -4.65 | -3.83 | -0.2625 | 1.5675 | 1.355 | 0.170547 | 0.737263 | 0.108886 | 1 | 0.999631 | 0.64073 |
| Nono_Q99K48_T452t_1 | -0.51 | 1.58 | 1.43 | 1.03 | 1.06 | 1.02 | 0.01 | 0.52 | 1.02 | 0.61 | 0.92 | 1.51 | 1.7 | 1.8 | 0.98 | 1.47 | -0.23 | 0.1325 | 0.605 | 0.237685 | 0.649638 | 0.787092 | 1 | 0.999631 | 0.683301 |
| Nop56_Q9D6Z1_S536s_1 | -0.05 | 0.57 | -0.12 | -0.43 | -0.1 | 0.01 | -0.73 | -0.87 | 0.02 | -0.35 | -0.23 | -0.11 | -0.39 | 0.71 | -0.32 | -0.56 | -0.415 | -0.16 | -0.1325 | 0.733072 | 0.240182 | 0.596679 | 1 | 0.999631 | 0.907434 |
| Nop58_Q6DFW4_S509s_1 | -0.62 | 0.18 | 0.21 | -0.07 | 0.23 | 0.59 | -0.63 | -0.17 | 0.43 | -0.26 | 0.13 | 0.47 | 0.21 | 0.65 | 0.4 | 0.49 | 0.08 | 0.2675 | 0.5125 | 0.12066 | 0.820796 | 0.408291 | 1 | 0.999631 | 0.588344 |
| Nop58_Q6DFW4_S521s_1 | -0.3 | 0.56 | -0.28 | -0.81 | 0.49 | 0.09 | -0.62 | -0.05 | -0.02 | 0.15 | -0.94 | 0.22 | 1.73 | 0.39 | -0.19 | -0.14 | 0.185 | 0.06 | 0.655 | 0.214393 | 0.627513 | 0.881397 | 1 | 0.999631 | 0.66742 |
| Nos1_S4R255_S1412s_1 | -0.16 | -0.35 | -0.55 | 0.86 | -0.03 | -0.21 | -0.66 | 0.5 | -0.41 | -1.46 | -0.52 | 0.24 | -1.12 | 0.34 | -0.14 | 0.02 | -0.05 | -0.4875 | -0.175 | 0.695281 | 0.900328 | 0.301997 | 1 | 0.999631 | 0.895471 |
| Nos1_S4R255_S287s_1 | -1.88 | -2.79 | -1.5 | -1.82 | -2.24 | -3 | -1.63 | -1.73 | -2.52 | -2.3 | -1.43 | -1.42 | -1.13 | -2.89 | -1.63 | -1.42 | -0.1525 | 0.08 | 0.23 | 0.625002 | 0.714566 | 0.844642 | 1 | 0.999631 | 0.863768 |
| Nos1_S4R255_S847s_1 | 0.61 | 0.78 | -0.26 | 0.65 | 1.15 | 0.45 | 0.22 | 1.13 | 0.37 | 0.17 | 0.23 | 0.98 | -0.08 | 0.8 | 0.17 | 0.45 | 0.2925 | -0.0075 | -0.11 | 0.757966 | 0.42428 | 0.98261 | 1 | 0.999631 | 0.914449 |
| Nos1_S4R255_S857s_1 | 0.05 | -1.42 | 0.56 | 0.05 | -1.53 | -1.16 | 0.33 | 0.16 | -1.86 | -0.41 | 0.2 | -0.03 | -0.85 | -1.71 | -0.02 | 0.03 | -0.36 | -0.335 | -0.4475 | 0.424698 | 0.535828 | 0.565328 | 1 | 0.999631 | 0.774174 |
| Nos1_S4R255_T116t_1 | -0.54 | -0.28 | -0.37 | -0.01 | -0.37 | -0.95 | -1.41 | -0.64 | -0.55 | -0.42 | -0.71 | -1.33 | -0.12 | 0.28 | -0.68 | -0.29 | -0.5425 | -0.4525 | 0.0975 | 0.757813 | 0.101845 | 0.158784 | 1 | 0.999631 | 0.914449 |
| Nos1_S4R255_T128t_1 | 0.15 | 0.65 | 0.56 | 0.46 | 0.07 | 0.72 | 0.12 | 0.3 | 0.64 | 0.63 | 0.51 | 0.58 | 1.1 | 1.19 | 1.12 | 1.05 | -0.1525 | 0.135 | 0.66 | 0.032423 | 0.579028 | 0.596177 | 1 | 0.999631 | 0.442395 |
| Nos1ap_Q9D3A8_S465s_1 | 0.28 | -0.97 | -1.62 | -0.26 | -1.26 | -1.58 | -0.54 | -1.82 | -0.81 | -0.79 | -1.01 | -1.19 | -2.04 | -3.84 | -0.03 | -0.69 | -0.6575 | -0.3075 | -1.0075 | 0.23506 | 0.187451 | 0.476407 | 1 | 0.999631 | 0.683301 |
| Nostrin_Q6WKZ7_S429s_1 | 0 | -0.29 | 1.44 | 0.47 | -0.43 | 0.38 | -0.27 | -0.34 | -0.12 | -0.14 | -0.3 | 0.03 | 0.4 | 0.6 | 0.35 | 1.18 | -0.57 | -0.5375 | 0.2275 | 0.600602 | 0.194012 | 0.198032 | 1 | 0.999631 | 0.856688 |
| Npdc1_Q64322_S236s_2 | 1.5 | -0.27 | 0.37 | 0.44 | 0.39 | 0.69 | 0.67 | 0.34 | 0.16 | 1.26 | -0.5 | -0.8 | -1.14 | -0.53 | -0.35 | -0.51 | 0.0125 | -0.48 | -1.1425 | 0.020033 | 0.974259 | 0.387418 | 1 | 0.999631 | 0.391668 |
| Npdc1_Q64322_S243s_1 | 1.09 | -1.33 | 0.29 | -0.2 | -0.97 | -0.4 | -0.12 | -0.32 | -0.5 | 0.83 | -0.86 | -0.96 | -1.05 | -0.94 | -0.88 | -1.5 | -0.415 | -0.335 | -1.055 | 0.058217 | 0.418285 | 0.577064 | 1 | 0.999631 | 0.499755 |
| Npdc1_Q64322_T239t_1 | 1.3 | -0.92 | 0.14 | 0.67 | 0.05 | 0.06 | 1.08 | 0.96 | 0.06 | 0.8 | 0.11 | 0.08 | -1.17 | -1.07 | -0.18 | -0.59 | 0.24 | -0.035 | -1.05 | 0.058458 | 0.639858 | 0.942101 | 1 | 0.999631 | 0.500444 |
| Npdc1_Q64322_T239t_2 | 1.5 | -0.27 | 0.37 | 0.44 | 0.39 | 0.69 | 0.67 | 0.34 | 0.16 | 1.26 | -0.5 | -0.8 | -1.14 | -0.53 | -0.35 | -0.51 | 0.0125 | -0.48 | -1.1425 | 0.020033 | 0.974259 | 0.387418 | 1 | 0.999631 | 0.391668 |
| Npepps_E9Q039_S746s_1 | -0.66 | -0.11 | 0.03 | -0.55 | -0.56 | 0.33 | -0.42 | -0.91 | 0.07 | 0.04 | -0.29 | 0.3 | 0.64 | 0.77 | 0.49 | 0.38 | -0.0675 | 0.3525 | 0.8925 | 0.01241 | 0.844147 | 0.244348 | 1 | 0.999631 | 0.385092 |
| Npy1r_Q04573_S367s_1 | 0.14 | 0.46 | -0.43 | -0.39 | 0.22 | 1.4 | -0.75 | -0.58 | 0.92 | 1.19 | -0.71 | -0.59 | 1.78 | 1.68 | -0.88 | 0.23 | 0.1275 | 0.2575 | 0.7575 | 0.232294 | 0.799556 | 0.615941 | 1 | 0.999631 | 0.682449 |
| Nr2f1_Q32NY6_T48t_1 | -0.94 | 0.64 | 0.77 | -0.33 | -0.19 | -0.26 | 0.24 | 0.52 | 0.41 | 0.35 | 0.35 | 1.03 | 0.41 | 0.94 | 0.45 | 1.21 | 0.0425 | 0.5 | 0.7175 | 0.134549 | 0.922363 | 0.268512 | 1 | 0.999631 | 0.606095 |
| Nr3c1_E9Q310_T168t_1 | 1.4 | 2.68 | 1.02 | 1.43 | 2.32 | 1.61 | 1.46 | 2.12 | 2.23 | 0.97 | 0.39 | -0.08 | 1.8 | 1.12 | 0.68 | 0.84 | 0.245 | -0.755 | -0.5225 | 0.253178 | 0.557202 | 0.204157 | 1 | 0.999631 | 0.688735 |
| Nr3c2_D6RIL1_S299s_1 | 0.77 | 1.32 | 1.51 | 0.36 | 0.72 | 0.98 | 0.55 | 1.76 | 0.61 | 0.12 | 1.36 | 0.9 | 1.05 | 2.07 | 1.51 | 1.15 | 0.0125 | -0.2425 | 0.455 | 0.252314 | 0.974107 | 0.535693 | 1 | 0.999631 | 0.68813 |
| Nrbp1_D3YUV1_S2s_1 | -0.45 | 0.08 | 0.25 | 0.1 | -0.31 | -0.1 | 0.35 | -0.17 | -0.77 | -0.28 | -0.03 | 0.38 | 0.34 | -0.22 | 0.51 | 0.09 | -0.0525 | -0.17 | 0.185 | 0.556924 | 0.854036 | 0.613485 | 1 | 0.999631 | 0.838305 |
| Nrbp1_D3YUV1_S430s_1 | -0.66 | -0.58 | -0.59 | -1.08 | -0.47 | -0.72 | -0.86 | -0.86 | -0.74 | -0.46 | -0.84 | -0.72 | -0.09 | -0.22 | -0.68 | -0.75 | 3.93E-17 | 0.0375 | 0.2925 | 0.347449 | 1 | 0.886879 | 1 | 0.999631 | 0.736498 |
| Nrbp1_D3YUV1_T439t_1 | 0.68 | 1.38 | 0.57 | 1.14 | 0.99 | 1.29 | 0.73 | 0.83 | 1.21 | 0.54 | 0.66 | 0.87 | 0.18 | 0.86 | 1.37 | 1.25 | 0.0175 | -0.1225 | -0.0275 | 0.940917 | 0.952587 | 0.693028 | 1 | 0.999631 | 0.981286 |
| Nrbp1_D3YUV1_T441t_1 | 0.48 | -0.21 | 1.39 | 1.09 | -0.56 | -0.12 | 1.44 | 0.88 | -1.02 | 0.17 | 1.46 | 1.69 | -1.74 | -1.06 | 1.14 | 1.66 | -0.2775 | -0.1125 | -0.6875 | 0.391032 | 0.605511 | 0.862299 | 1 | 0.999631 | 0.758901 |
| Nrbp2_Q91V36_T409t_1 | -1.18 | 0.4 | -0.53 | 0.32 | -1.7 | -0.19 | -0.05 | 0.83 | -0.32 | -0.66 | 0.45 | 0.48 | 0.67 | 0.87 | 0.3 | 0.35 | -0.03 | 0.235 | 0.795 | 0.079294 | 0.959106 | 0.610839 | 1 | 0.999631 | 0.545579 |
| Nrcam_Q810U4_S1178s_1 | -0.97 | 1.2 | 2.51 | 1.54 | 0.4 | 1.03 | 0.87 | 1.37 | 0.6 | 0.62 | 1.23 | 2.26 | 1.15 | 1.23 | 1.77 | 2.76 | -0.1525 | 0.1075 | 0.6575 | 0.375179 | 0.821753 | 0.88381 | 1 | 0.999631 | 0.748001 |
| Nrcam_Q810U4_S1179s_1 | -0.33 | 0.57 | -2.09 | -1.28 | -0.45 | 0.59 | -1.8 | -2.8 | -0.01 | 0.14 | -1.93 | -2.24 | 0.19 | 0.51 | -2.9 | -1.6 | -0.3325 | -0.2275 | -0.1675 | 0.844367 | 0.686537 | 0.762305 | 1 | 0.999631 | 0.94856 |
| Nrcam_Q810U4_S1182s_1 | -1.9 | -2.59 | -1.12 | -1.58 | -1.57 | -3.37 | -1.63 | -1.64 | -2.07 | -1.99 | -1.26 | -0.96 | -0.08 | -2.08 | -1.12 | -0.83 | -0.255 | 0.2275 | 0.77 | 0.143722 | 0.614332 | 0.58821 | 1 | 0.999631 | 0.615182 |
| Nrcam_Q810U4_T1173t_1 | 0.28 | 2.25 | 1.35 | 0.26 | 0.39 | 2.52 | -0.12 | 0.29 | 1.59 | 1.48 | 0.17 | 0.65 | 1.85 | 1.42 | 0.76 | 1.34 | -0.265 | -0.0625 | 0.3075 | 0.545836 | 0.696561 | 0.908643 | 1 | 0.999631 | 0.833922 |

| | | | | | | | | | | | | | | | | | | | | | | | | | |
|---|---|---|---|---|---|---|---|---|---|---|---|---|---|---|---|---|---|---|---|---|---|---|---|---|---|
| Nrd1_Q8BHG1_S85s_1 | 0.46 | -0.06 | 0.58 | 0.57 | -0.28 | -0.29 | 0.65 | -0.35 | 0.41 | 1.15 | 0.89 | 0.25 | 1.06 | -0.65 | 0.17 | -0.59 | -0.455 | 0.2875 | -0.39 | 0.379248 | 0.185654 | 0.377292 | 1 | 0.999631 | 0.751637 |
| Nrg2_D3YZR2_S464s_1 | 1.02 | 0.15 | -0.49 | -0.7 | -0.11 | 0.8 | -0.04 | -0.72 | 1.16 | -0.68 | -0.66 | -1.31 | -0.69 | 0.49 | -0.54 | -1.7 | -0.0125 | -0.3675 | -0.605 | 0.286851 | 0.978893 | 0.543917 | 1 | 0.999631 | 0.705972 |
| Nrg2_D3YZR2_S470s_1 | 0.84 | -0.27 | 0.77 | 0.25 | -1.29 | -0.29 | 0.82 | -0.53 | -1.13 | -0.42 | 0.04 | -0.41 | -1.86 | -0.61 | -0.44 | -0.4 | -0.72 | -0.8775 | -1.225 | 0.017637 | 0.157353 | 0.040226 | 1 | 0.999631 | 0.391024 |
| Nrgn_P60761_S76s_1 | 2.13 | -2.48 | 0.07 | -0.21 | -2.35 | -0.79 | 3.02 | -2.28 | -5.41 | 0.81 | -1.77 | -2.67 | -1.31 | -4.26 | -2.42 | -1.83 | -0.4775 | -2.1375 | -2.3325 | 0.035814 | 0.721868 | 0.139579 | 1 | 0.999631 | 0.453811 |
| Nrxn1_E0CY11_S1450s_1 | -1.66 | 0.77 | -0.29 | -2.06 | -0.28 | -0.91 | 0.05 | -0.74 | 0.31 | -2.19 | 0.74 | 0.97 | 1.24 | -1.94 | -0.78 | -2.12 | 0.34 | 0.7675 | -0.09 | 0.917735 | 0.583926 | 0.378006 | 1 | 0.999631 | 0.97388 |
| Nrxn1_E0CY11_S1466s_1 | -0.22 | -1.03 | 0.02 | -0.54 | -0.11 | -0.42 | -0.46 | 0.23 | -1.6 | -0.74 | -0.58 | -0.61 | -0.35 | -0.83 | -0.71 | -0.02 | 0.2525 | -0.44 | -0.035 | 0.920185 | 0.442508 | 0.243618 | 1 | 0.999631 | 0.97499 |
| Nsd1_E9QAE4_S2232s_1 | -0.9 | -1.26 | -1.42 | -0.92 | -1.42 | -1.69 | -1.18 | -0.96 | -1.71 | -1.45 | -1.13 | -0.54 | -0.42 | -0.95 | -0.75 | -1.24 | -0.1875 | -0.0825 | 0.285 | 0.367683 | 0.511154 | 0.804566 | 1 | 0.999631 | 0.745968 |
| Nsd1_E9QAE4_S2370s_1 | 0.24 | 0.07 | -0.32 | -1.49 | -1.17 | -0.05 | -0.4 | -0.79 | -0.32 | 0.35 | -1.18 | -0.97 | -0.04 | -0.43 | -0.49 | -0.35 | -0.2275 | -0.155 | 0.0475 | 0.909311 | 0.611465 | 0.754863 | 1 | 0.999631 | 0.970501 |
| Nsf_P46460_S207s_1 | -0.12 | 1 | -0.25 | -0.19 | -0.06 | 0.59 | -0.26 | -0.19 | 0.37 | -0.05 | -0.05 | -0.38 | 0.67 | 0.39 | 0.56 | 0.3 | -0.09 | -0.1375 | 0.37 | 0.316603 | 0.809379 | 0.707541 | 1 | 0.999631 | 0.720734 |
| Nsf_P46460_S298s_1 | 0.01 | 0.72 | 1.27 | 0.31 | -0.06 | -0.25 | -0.15 | 0.82 | 0.19 | 1.04 | 0.7 | 0.54 | 0.98 | -0.02 | 0.37 | 1.22 | -0.4875 | 0.04 | 0.06 | 0.883946 | 0.220027 | 0.911212 | 1 | 0.999631 | 0.963505 |
| Nsf_P46460_S31s_1 | 0.31 | 0.35 | 0.26 | -1.03 | -1.16 | -1.47 | 1.07 | -0.78 | -0.49 | -0.81 | 0.31 | 0.11 | -0.82 | -1.05 | -0.95 | -0.34 | -0.5575 | -0.1925 | -0.7625 | 0.078097 | 0.361768 | 0.652264 | 1 | 0.999631 | 0.543121 |
| Nsfl1c_Q9CZ44_S114s_1 | -0.42 | 0.7 | 0.22 | -0.09 | -0.03 | 0.55 | -0.29 | 0.31 | 0.26 | 0.41 | -0.02 | 0.52 | 0.59 | 1.11 | 0.52 | 0.61 | 0.0325 | 0.19 | 0.605 | 0.09903 | 0.923349 | 0.559729 | 1 | 0.999631 | 0.562298 |
| Nsfl1c_Q9CZ44_S140s_1 | 2.7 | -1.22 | 0.32 | 0.89 | -2.37 | 0.76 | 2.39 | 0.24 | -2.23 | 2 | 0.46 | 0.04 | -1.64 | -1.16 | 0.06 | -0.14 | -0.4175 | -0.605 | -1.3925 | 0.102632 | 0.703573 | 0.559416 | 1 | 0.999631 | 0.565572 |
| Nsfl1c_Q9CZ44_S176s_1 | -0.66 | 1.62 | 1.46 | -0.49 | 2.78 | 0.43 | 3.44 | 0.17 | 3.87 | -0.35 | 2.83 | 4.11 | 3.86 | -0.22 | 1.7 | -0.67 | 1.2225 | 2.1325 | 0.685 | 0.509135 | 0.190725 | 0.062049 | 1 | 0.999631 | 0.816932 |
| Nsmf_Q99NF2_S148s_1 | 0.79 | 0.32 | 0.37 | 0.42 | 0.75 | 0.25 | -0.33 | 1.05 | 0.39 | -0.18 | 0.64 | 0.35 | -0.81 | 0.14 | 0.52 | -0.17 | -0.045 | -0.175 | -0.555 | 0.140878 | 0.897655 | 0.549322 | 1 | 0.999631 | 0.61245 |
| Nsmf_Q99NF2_S153s_1 | 0.51 | 1.77 | 0.3 | 0.31 | 0.38 | 1.61 | 0.23 | 0.63 | 1.23 | 1.43 | -0.3 | -0.64 | 1.21 | 2.23 | 0.65 | 0.36 | -0.01 | -0.2925 | 0.39 | 0.455128 | 0.982365 | 0.616457 | 1 | 0.999631 | 0.790041 |
| Nsmf_Q99NF2_S158s_1 | 0.04 | 0.32 | 0.88 | 0.18 | -0.01 | 0.87 | 0.3 | 1.25 | 0.96 | 1.1 | 1.25 | 0.9 | 1.22 | 1.48 | 1.21 | 0.91 | 0.2475 | 0.6975 | 0.85 | 0.019607 | 0.497814 | 0.034121 | 1 | 0.999631 | 0.391668 |
| Nsmf_Q99NF2_S292s_1 | -0.8 | -0.12 | 0.27 | -0.67 | -0.69 | -0.07 | -0.4 | 0.1 | -0.42 | -0.34 | 0.73 | 0.65 | 0.61 | 0.47 | 1.34 | 0.56 | 0.065 | 0.485 | 1.075 | 0.01374 | 0.848247 | 0.250594 | 1 | 0.999631 | 0.385092 |
| Nsmf_Q99NF2_S36s_1 | -0.6 | -0.3 | -1 | -0.82 | 0.49 | -1.03 | -1.25 | -1.08 | -1.14 | -0.77 | -1.25 | -1.31 | 0.27 | 0.35 | -0.96 | 0.39 | -0.0375 | -0.4375 | 0.6925 | 0.099319 | 0.929785 | 0.148307 | 1 | 0.999631 | 0.562298 |
| Nt5c_Q9JM14_T102t_1 | -0.86 | 1.99 | 3.21 | 2.31 | 1.34 | 2.26 | 1.15 | 2.6 | 1.32 | -0.67 | 2.21 | 3.67 | 2.35 | 2.62 | 3.86 | 2.76 | 0.175 | -0.03 | 1.235 | 0.153812 | 0.832057 | 0.977954 | 1 | 0.999631 | 0.625198 |
| Ntrk3_Q6VNS1_S561s_1 | -0.21 | 0.05 | 0.91 | -0.25 | -0.72 | 0.03 | -0.11 | -0.12 | -0.59 | 0.14 | 0.31 | 1.12 | 0.8 | 0.13 | 1.01 | 0.79 | -0.355 | 0.12 | 0.5575 | 0.156718 | 0.320902 | 0.784642 | 1 | 0.999631 | 0.626749 |
| Ntsr1_O88319_S407s_1 | -0.15 | -2.63 | -2.77 | -1.51 | -1.84 | -2.27 | -0.85 | -1.64 | -2.33 | -1.34 | -2.27 | -2.01 | -2.89 | -1.95 | -2.54 | -1.81 | 0.115 | -0.2225 | -0.5325 | 0.384563 | 0.850401 | 0.708038 | 1 | 0.999631 | 0.75538 |
| Nuak1_Q641K5_T212t_1 | -0.43 | 1.44 | -0.76 | 0.43 | 0.22 | 2.24 | 0.4 | 0.62 | -0.38 | -0.84 | 0.23 | -0.38 | 0.16 | 1.97 | 0.58 | 0.57 | 0.7 | -0.5125 | 0.65 | 0.277908 | 0.267462 | 0.326916 | 1 | 0.999631 | 0.700874 |
| Nucb1_Q02819_S368s_1 | -0.65 | 0.59 | 1.79 | 0.46 | -0.01 | -0.13 | 0.13 | 0.46 | 0.17 | -0.08 | 1.12 | 1.53 | 0.12 | 0.56 | 0.88 | 1.21 | -0.435 | 0.1375 | 0.145 | 0.780752 | 0.381202 | 0.811721 | 1 | 0.999631 | 0.921446 |
| Nucb1_Q02819_S456s_1 | -1.51 | -1.59 | 0.09 | -1.21 | -1.18 | -1.55 | -0.81 | -0.25 | -0.66 | -0.92 | -0.58 | -0.38 | -0.32 | -0.67 | -0.32 | 0.17 | 0.1075 | 0.42 | 0.77 | 0.100019 | 0.815147 | 0.322209 | 1 | 0.999631 | 0.562917 |
| Nucb1_Q02819_S85s_1 | -1.27 | -0.62 | -0.74 | -0.7 | -0.65 | -1.37 | -0.83 | -0.69 | -0.14 | -0.67 | -0.34 | -0.36 | 0.89 | 0.01 | -0.53 | -0.48 | -0.0525 | 0.455 | 0.805 | 0.062089 | 0.857431 | 0.129257 | 1 | 0.999631 | 0.509355 |
| Nucks1_Q80XU3_S181s_1 | 0.77 | 1.68 | -1.2 | -0.07 | 0.62 | 1.39 | -0.74 | 0.17 | 1.52 | 2.01 | -1.33 | -0.17 | 1.63 | 2.02 | -0.79 | 0.28 | 0.065 | 0.2125 | 0.49 | 0.532706 | 0.923046 | 0.804744 | 1 | 0.999631 | 0.8277 |
| Nucks1_Q80XU3_S214s_1 | 0.51 | 1.88 | 2.82 | 1.38 | 1.07 | 2.06 | 1.63 | 1.5 | 1.48 | 1.48 | 1.78 | 1.26 | -0.45 | 0.75 | 1.88 | 0.56 | -0.0825 | -0.1475 | -0.9625 | 0.141325 | 0.867043 | 0.756969 | 1 | 0.999631 | 0.61245 |
| Nucks1_Q80XU3_S58s_2 | -0.71 | 1.85 | 3.12 | 2 | 1.24 | -0.03 | 2.55 | 0.92 | 1.17 | 1.1 | 1.68 | 1.99 | 0.87 | -1.02 | 0.17 | 1.33 | -0.395 | -0.08 | -1.2275 | 0.16229 | 0.641128 | 0.913894 | 1 | 0.999631 | 0.631442 |
| Nucks1_Q80XU3_S61s_2 | -0.71 | 1.85 | 3.12 | 2 | 1.24 | -0.03 | 2.55 | 0.92 | 1.17 | 1.1 | 1.68 | 1.99 | 0.87 | -1.02 | 0.17 | 1.33 | -0.395 | -0.08 | -1.2275 | 0.16229 | 0.641128 | 0.913894 | 1 | 0.999631 | 0.631442 |
| Nucks1_Q80XU3_S79s_1 | 0.32 | -1.72 | 3.72 | 1.62 | 1.17 | -2.38 | 4.66 | 1.13 | -0.02 | 0.37 | 2.75 | 3.35 | 1.22 | -0.78 | 2.15 | 0.69 | 0.16 | 0.6275 | -0.165 | 0.880866 | 0.917989 | 0.60773 | 1 | 0.999631 | 0.963364 |
| Nudc_O35685_S136s_1 | -5.53 | -0.58 | -4.86 | -5.59 | 0.68 | -0.65 | -2.88 | -6.35 | 0.37 | -2.33 | -0.9 | -6.68 | 2 | -0.05 | -5.19 | -4.04 | 1.84 | 1.755 | 2.32 | 0.199849 | 0.280367 | 0.302479 | 1 | 0.999631 | 0.657804 |
| Nudc_O35685_S139s_1 | 0.59 | 1.73 | 1.58 | 0.62 | 0.9 | 2.03 | 1.47 | 0.63 | 1.77 | 1.68 | 0.97 | 1.07 | 1.82 | 1.44 | 0.75 | 0.99 | 0.1275 | 0.2425 | 0.12 | 0.768428 | 0.766281 | 0.534494 | 1 | 0.999631 | 0.9177 |
| Nudc_O35685_T108t_1 | 0.18 | 0.02 | 0.58 | 0.75 | -0.03 | -0.15 | 0.36 | 0.25 | 0 | 0.35 | 0.86 | 0.16 | -0.09 | 0.19 | 0.74 | 0.75 | -0.275 | -0.04 | 0.015 | 0.964461 | 0.345469 | 0.89909 | 1 | 0.999631 | 0.987331 |
| Nudt11_P0C028_T150t_1 | -1.79 | -1.33 | -0.63 | -1.33 | -0.67 | -1.35 | -0.98 | 0.61 | -0.86 | -1.39 | -3.37 | -1.53 | -0.27 | -0.56 | -0.22 | -0.65 | 0.6725 | -0.5175 | 0.845 | 0.02746 | 0.172093 | 0.36038 | 1 | 0.999631 | 0.423809 |
| Nufip2_Q5F2E7_S213s_1 | 0.06 | 0.44 | 1.12 | 0.88 | -0.39 | 0.51 | 0.94 | 0.8 | 0.97 | 0.8 | 0.88 | 1.65 | 0.82 | -0.52 | 1.01 | 1.1 | -0.16 | 0.45 | -0.0225 | 0.95976 | 0.682246 | 0.214173 | 1 | 0.999631 | 0.986095 |
| Nufip2_Q5F2E7_S376s_1 | 1.94 | 1.98 | 0.8 | 2.52 | 0.91 | 3.34 | 1.38 | 2.09 | 1.83 | 1.66 | 1.42 | 2.21 | 1.1 | 2.53 | 1.4 | 2.63 | 0.12 | -0.03 | 0.105 | 0.835846 | 0.836661 | 0.94122 | 1 | 0.999631 | 0.945038 |
| Nufip2_Q5F2E7_S626s_1 | -0.28 | 1.17 | 1.51 | 1.02 | 0.55 | 0.55 | 0.67 | 1 | 0.64 | -0.64 | 1.64 | 1.31 | -0.2 | 1.04 | 1.92 | 1.5 | -0.1625 | -0.1175 | 0.21 | 0.707932 | 0.690985 | 0.840718 | 1 | 0.999631 | 0.898881 |
| Numa1_E9Q7G0_S1739s_1 | -0.27 | -0.52 | 0.37 | 0.23 | -1.45 | -0.3 | -1.53 | 1.18 | -0.73 | -0.68 | -2.05 | 0.9 | -0.07 | -0.52 | 1.57 | 0.62 | -0.4775 | -0.5925 | 0.4475 | 0.366559 | 0.435898 | 0.324101 | 1 | 0.999631 | 0.745796 |
| Numa1_E9Q7G0_S398s_1 | 0.41 | -0.17 | 0.17 | 0.77 | -0.66 | 0.05 | 0 | 0.29 | -0.27 | -0.22 | 0 | 0.36 | -0.13 | -0.57 | -0.03 | -1.19 | -0.375 | -0.3275 | -0.775 | 0.059614 | 0.268204 | 0.30677 | 1 | 0.999631 | 0.50521 |
| Numa1_E9Q7G0_T1982t_1 | -1.03 | -1.66 | -0.09 | -0.92 | -2.34 | -2.66 | -0.41 | -1.98 | -1.91 | -0.61 | -1.36 | -2.28 | -1.85 | -1.93 | -2.78 | -1.88 | -0.9225 | -0.615 | -1.185 | 0.015231 | 0.115973 | 0.210577 | 1 | 0.999631 | 0.388832 |
| Numb_Q9QZS3_S230s_1 | -1.09 | -1.09 | -1.43 | -1.41 | -1.83 | -1.93 | -1.9 | -1.08 | -1.13 | -1.54 | -1.46 | -0.61 | 0.89 | -0.45 | -0.59 | -1.18 | -0.43 | 0.07 | 0.9225 | 0.062086 | 0.165028 | 0.81815 | 1 | 0.999631 | 0.509355 |
| Numb_Q9QZS3_S351s_1 | 0.21 | -0.8 | -1.55 | 1.47 | 0.25 | -0.1 | 0.81 | -0.08 | 0.06 | -0.3 | -0.79 | -1.43 | 0.02 | 0.16 | 1.11 | 0.28 | 0.3875 | -0.4475 | 0.56 | 0.384148 | 0.535155 | 0.499947 | 1 | 0.999631 | 0.754721 |
| Numb_Q9QZS3_T378t_1 | 3.33 | -0.13 | 0.19 | 2 | 0.98 | 0.96 | 2.81 | 0.39 | 0.26 | 0.74 | 0.59 | -0.57 | -1.59 | -0.75 | -0.07 | -0.82 | -0.0625 | -1.0925 | -2.155 | 0.016741 | 0.94067 | 0.173165 | 1 | 0.999631 | 0.391024 |
| Numbl_O08919_S263s_1 | 0.18 | 0.28 | 0.54 | 0.38 | 0.21 | 0.31 | -0.81 | -0.5 | -0.21 | 0.19 | 0.22 | -0.51 | 0.39 | 0.57 | -0.57 | 0.44 | -0.5425 | -0.4225 | -0.1375 | 0.686155 | 0.121231 | 0.159354 | 1 | 0.999631 | 0.890692 |
| Numbl_O08919_S270s_1 | -0.34 | -0.32 | 0.67 | -0.88 | -0.07 | -0.41 | -0.66 | -0.45 | -0.85 | -0.36 | -1.6 | 0.44 | 1.84 | -0.22 | 0.22 | 0.18 | -0.18 | -0.375 | 0.7225 | 0.190805 | 0.624882 | 0.466037 | 1 | 0.999631 | 0.648674 |
| Numbl_O08919_S411s_1 | 1.92 | -0.65 | 0.06 | 1 | -0.5 | 0.35 | 1.64 | 0.15 | -0.92 | 0.86 | 0.2 | -0.73 | -1.64 | -1.15 | -0.37 | -0.8 | -0.1725 | -0.73 | -1.5725 | 0.019268 | 0.788179 | 0.263382 | 1 | 0.999631 | 0.391668 |
| Numbl_O08919_T279t_1 | 0.13 | -0.01 | -0.45 | 0.16 | 1.01 | 0.92 | 0.22 | 0.9 | 0.55 | 0.13 | -0.2 | -0.16 | -0.73 | 1.14 | 0.59 | 0.55 | 0.805 | 0.1225 | 0.43 | 0.328989 | 0.02068 | 0.683909 | 1 | 0.999631 | 0.726274 |
| Numbl_O08919_T409t_1 | 2.24 | -1.31 | 0.05 | 0.81 | -1.1 | 0.25 | 1.19 | 0.34 | -1.06 | 0.49 | -0.5 | -1.13 | -3.82 | -2.14 | -0.75 | -1.83 | -0.2775 | -0.9975 | -2.5825 | 0.01155 | 0.719457 | 0.196009 | 1 | 0.999631 | 0.385092 |
| Nup107_E9Q4V9_S11s_1 | -1.74 | -0.68 | -0.32 | -1.41 | -0.77 | -0.71 | -0.96 | -1 | -0.86 | -0.71 | -1.12 | -0.09 | 0.6 | -0.13 | -0.75 | -1.41 | 0.1775 | 0.3425 | 0.615 | 0.247328 | 0.622394 | 0.404925 | 1 | 0.999631 | 0.685751 |
| Nup160_Q9Z0W3_S1123s_1 | -1.12 | -0.24 | 0.07 | -0.92 | -0.67 | -0.77 | -1.15 | -1.02 | -0.54 | -0.42 | 0.02 | -1.14 | 0.42 | -0.14 | 0.04 | -0.57 | -0.35 | 0.0325 | 0.49 | 0.219098 | 0.31386 | 0.933034 | 1 | 0.999631 | 0.669859 |
| Nup188_Q6ZQH8_S1719s_1 | -0.16 | 0.1 | 0.57 | -0.58 | -0.36 | 0.18 | -0.43 | 0.04 | 0.21 | -0.99 | -0.55 | -0.3 | 0.49 | 0.08 | 1.26 | 0.66 | -0.125 | -0.39 | 0.64 | 0.116062 | 0.702999 | 0.310092 | 1 | 0.999631 | 0.582016 |
| Nup205_B9EJ54_S1165s_1 | -0.95 | -0.52 | -0.7 | -1.15 | -1.36 | -1.28 | -1 | -1.49 | -0.84 | -0.67 | -1.52 | -1.28 | -0.44 | -0.8 | -1.05 | -0.6 | -0.4525 | -0.2475 | 0.1075 | 0.719669 | 0.115806 | 0.430296 | 1 | 0.999631 | 0.904782 |
| Nup210_Q9QY81_S1848s_1 | -1.17 | -1.87 | -1.89 | -1.98 | -2.52 | -2.12 | -2.07 | -3.14 | -2.43 | -1.18 | -2.96 | -2.7 | -0.8 | -2.83 | -2.9 | -1.24 | -0.735 | -0.59 | -0.215 | 0.689336 | 0.054784 | 0.195442 | 1 | 0.999631 | 0.892416 |
| Nup35_Q8R4R6_S258s_1 | -0.5 | -0.6 | 0.21 | -0.22 | -0.73 | -0.69 | -0.5 | -0.25 | -0.47 | -1.07 | -0.27 | -0.33 | 0.53 | -1.1 | -0.28 | -0.04 | -0.265 | -0.2575 | 0.055 | 0.891982 | 0.36619 | 0.426899 | 1 | 0.999631 | 0.964276 |
| Nup35_Q8R4R6_T265t_1 | -0.55 | -0.58 | -0.18 | -0.73 | -1.3 | -0.36 | -0.97 | -0.71 | -0.94 | -0.56 | -0.49 | -0.1 | 0.26 | -0.12 | -0.34 | 0.09 | -0.325 | -0.0125 | 0.4825 | 0.12219 | 0.288598 | 0.96583 | 1 | 0.999631 | 0.590804 |
| Nup35_Q8R4R6_T274t_1 | -0.24 | -0.98 | -0.94 | -1.06 | -0.57 | -1.04 | -0.9 | -1.88 | -1.22 | -0.68 | -1.36 | -0.52 | 0.43 | -1.19 | -1.85 | -1.68 | -0.2925 | -0.14 | -0.2675 | 0.611037 | 0.425559 | 0.673239 | 1 | 0.999631 | 0.859461 |
| Nup50_Q9JIH2_S322s_1 | 2.14 | 0.22 | 0.67 | 1.01 | -0.28 | 0.24 | 1.57 | -0.83 | -0.04 | 1.88 | -0.6 | -0.61 | 0.37 | -0.86 | 0.19 | -1.12 | -0.835 | -0.8525 | -1.365 | 0.024381 | 0.182289 | 0.207656 | 1 | 0.999631 | 0.416717 |
| Nup88_Q8CEC0_S699s_1 | -0.41 | -0.71 | -0.16 | -0.62 | -0.66 | -1 | -0.85 | -0.8 | -0.27 | -0.44 | -0.48 | 0.03 | -0.23 | 0.2 | -0.41 | 0.39 | -0.3525 | 0.185 | 0.4625 | 0.161714 | 0.187602 | 0.504273 | 1 | 0.999631 | 0.631442 |
| Nup93_Q8BJ71_S644s_1 | 0.33 | -1.58 | -0.68 | -0.45 | -0.66 | -1.88 | -1.08 | -0.87 | -1.61 | -0.46 | -0.66 | -1.64 | -2.16 | -1.95 | -1.2 | -1.49 | -0.5275 | -0.4975 | -1.105 | 0.031385 | 0.264977 | 0.31442 | 1 | 0.999631 | 0.441911 |
| Nwd2_Q6P5U7_S1662s_1 | 1.12 | 0.74 | 1.92 | 1.54 | -0.28 | 1.84 | 2.7 | 0.42 | 0.16 | 1.44 | 0.82 | -0.67 | -0.98 | 0.1 | 0.71 | -0.11 | -0.16 | -0.8925 | -1.4 | 0.009001 | 0.804119 | 0.096156 | 1 | 0.999631 | 0.365396 |
| Nwd2_Q6P5U7_S792s_1 | -0.09 | 0.68 | 1.5 | 1.19 | -0.11 | 0.64 | 1.01 | 0.64 | 0.29 | 0.64 | 1.6 | 1.79 | 0.35 | 0.78 | 2.08 | 2.59 | -0.275 | 0.26 | 0.63 | 0.292754 | 0.513999 | 0.592639 | 1 | 0.999631 | 0.706671 |
| Nyap1_E9PXJ0_T705t_1 | -0.01 | 1.12 | 0.76 | -0.14 | -1.16 | 0.5 | -0.62 | 0.24 | -0.09 | 0.34 | 0.11 | -0.25 | 1.29 | 1.98 | 1.03 | -0.18 | -0.6925 | -0.405 | 0.5975 | 0.262445 | 0.161333 | 0.278396 | 1 | 0.999631 | 0.693034 |
| Nyap2_Q8BM65_S184s_1 | -0.1 | 0.22 | 0.19 | -0.1 | 0.44 | -0.07 | -0.44 | -0.39 | -0.05 | -0.14 | 0.35 | 0.85 | 0.26 | 0.66 | 0.35 | 0.48 | -0.1675 | 0.2 | 0.385 | 0.18082 | 0.568563 | 0.523406 | 1 | 0.999631 | 0.64376 |
| Nyap2_Q8BM65_S238s_1 | 0.18 | 0.86 | 0.49 | 0.45 | 0.7 | 0.83 | 0.5 | 0.5 | 0.42 | 0.08 | 1.34 | 0.43 | 0.23 | 1.03 | 0.45 | 1.52 | 0.1375 | 0.0725 | 0.3125 | 0.404139 | 0.603437 | 0.834026 | 1 | 0.999631 | 0.765215 |
| Nyap2_Q8BM65_S732s_1 | -0.25 | 0.06 | -0.76 | -0.31 | -0.06 | -0.73 | -0.73 | -0.74 | -0.38 | -0.1 | -0.1 | -0.25 | 0.59 | 0.11 | -1.3 | 0.01 | -0.25 | 0.1075 | 0.1675 | 0.705156 | 0.414651 | 0.701646 | 1 | 0.999631 | 0.89781 |
| Nyap2_Q8BM65_Y300y_1 | 0.05 | 1.29 | 1.04 | 0.74 | 0.45 | 1.02 | 0.31 | 0.63 | 0.79 | 1.2 | 0.7 | 1.2 | 1.79 | 1.08 | 1.19 | 1.11 | -0.1775 | 0.1925 | 0.5125 | 0.17988 | 0.606847 | 0.577497 | 1 | 0.999631 | 0.64376 |
| O35375-5_O35375_S869s_1 | -0.87 | -2 | -0.58 | -0.43 | -1 | -0.65 | -0.07 | -0.29 | -1.11 | -1.74 | -0.58 | -0.66 | -0.8 | -0.9 | -1.27 | -1.53 | 0.4675 | -0.0525 | -0.155 | 0.70714 | 0.270573 | 0.904833 | 1 | 0.999631 | 0.898523 |
| Ociad1_Q9CRD0_T146t_1 | -0.86 | -0.8 | -0.88 | -1.35 | -0.48 | -0.4 | -0.78 | -0.62 | -1.17 | -1.29 | -0.72 | -0.7 | -0.64 | -0.65 | -0.96 | -0.88 | 0.4025 | 0.0025 | 0.19 | 0.506365 | 0.143748 | 0.993049 | 1 | 0.999631 | 0.815124 |
| Ocln_Q61146_S368s_1 | 0.01 | 0.39 | -0.43 | -0.01 | 1.19 | 0.85 | 0.24 | -0.71 | 0.98 | 1.07 | -0.08 | -0.31 | 1.17 | 0.92 | 1.66 | -1.03 | 0.4025 | 0.425 | 0.69 | 0.241231 | 0.36887 | 0.306956 | 1 | 0.999631 | 0.684359 |
| Ogfr_Q99PG2_S403s_1 | -0.62 | 0.18 | -0.13 | -0.47 | 0.3 | -0.03 | -0.61 | -0.1 | -0.27 | -0.88 | -0.34 | 0.23 | -0.09 | 0.18 | 0.15 | 0.56 | 0.15 | -0.055 | 0.46 | 0.164702 | 0.633473 | 0.870264 | 1 | 0.999631 | 0.633184 |
| Ogfrl1_Q8VE52_S406s_1 | 0.61 | -0.02 | 1.1 | 0.61 | -0.69 | -0.73 | 0.59 | -0.35 | 0.81 | 0.98 | 0.31 | 0.06 | -0.27 | -1.62 | -1.42 | -0.49 | -0.87 | -0.035 | -1.525 | 0.004361 | 0.047777 | 0.920724 | 1 | 0.999631 | 0.351308 |
| Ogfrl1_Q8VE52_S65s_1 | 0.62 | 1.31 | -0.25 | 0.11 | 0.84 | 1.77 | 0.21 | 0.41 | 1.21 | 1.52 | 0.49 | 0.25 | 1.65 | 1.69 | 0.15 | 0.79 | 0.36 | 0.42 | 0.6225 | 0.218897 | 0.444853 | 0.356215 | 1 | 0.999631 | 0.669859 |
| Opalin_Q7M750_S100s_1 | -0.33 | -0.47 | -0.3 | -0.72 | -0.85 | -1.01 | -0.85 | -0.84 | -0.97 | -1.02 | -1.4 | -1.16 | -0.44 | -0.67 | 0.02 | -0.87 | -0.4325 | -0.6825 | -0.035 | 0.90979 | 0.098554 | 0.024444 | 1 | 0.999631 | 0.970597 |
| Opalin_Q7M750_S90s_1 | -0.48 | -1.07 | -0.81 | -0.88 | -1.48 | -0.84 | -1.19 | -0.98 | -1.42 | -0.56 | -1.81 | -1.09 | -1.56 | -1.4 | -0.74 | -2.54 | -0.3125 | -0.41 | -0.75 | 0.091677 | 0.270055 | 0.243791 | 1 | 0.999631 | 0.557085 |
| Ophn1_B1AXS4_T52t_1 | -0.28 | -0.54 | -0.11 | -0.51 | -0.91 | -0.6 | -0.4 | -0.12 | -0.09 | 0.48 | 0.16 | 0.17 | 0.4 | -0.16 | 0.15 | 0.3 | -0.1475 | 0.54 | 0.5325 | 0.086345 | 0.598567 | 0.06768 | 1 | 0.999631 | 0.55299 |
| Oprl1_B0R0C0_S329s_1 | 1.07 | -0.79 | -0.37 | 0.1 | -0.19 | 0.57 | 0.73 | -0.76 | 0.21 | 0.62 | -0.53 | -0.7 | -1.03 | -1.2 | -0.54 | -2.19 | 0.085 | -0.1025 | -1.2425 | 0.031458 | 0.864092 | 0.832538 | 1 | 0.999631 | 0.441911 |
| Oprm1_E9Q2X5_S311s_1 | -0.51 | 0.22 | 0.77 | 0.71 | -0.53 | 1.42 | 1.05 | 0.96 | -0.19 | -1.67 | 1.34 | 0.1 | -0.51 | -2.11 | 0.54 | -0.86 | 0.4275 | -0.4025 | -1.0325 | 0.095679 | 0.394025 | 0.5214 | 1 | 0.999631 | 0.558128 |
| Oprm1_E9Q2X5_T306t_1 | 0.01 | -2.91 | -4.75 | -2.19 | -0.78 | -2.1 | -1.4 | -2.4 | -2.73 | -2.81 | -2.17 | -1.88 | -2.83 | -2.84 | -5.58 | -3.95 | 0.79 | 0.0625 | -1.34 | 0.203589 | 0.393566 | 0.94326 | 1 | 0.999631 | 0.659781 |
| Osbp_Q3B7Z2_S236s_1 | -0.08 | 0.69 | 1.48 | 1.03 | 0.68 | -0.88 | -0.26 | 0.68 | 0.96 | -0.24 | 0.23 | 1.39 | -1.03 | 0.56 | 1.58 | 1.15 | -0.725 | -0.195 | -0.215 | 0.721147 | 0.153281 | 0.68169 | 1 | 0.999631 | 0.905416 |
| Osbp_Q3B7Z2_S238s_1 | -0.97 | -0.73 | -0.63 | -0.89 | -0.42 | -1.12 | -1.01 | -0.69 | -1.39 | -0.68 | -1.25 | -0.46 | -0.4 | -0.82 | -0.57 | -0.16 | -0.005 | -0.14 | 0.3175 | 0.281676 | 0.985181 | 0.649416 | 1 | 0.999631 | 0.702855 |
| Osbp_Q3B7Z2_S380s_1 | -0.68 | -0.01 | 0.25 | -0.6 | 0.33 | -0.01 | -0.41 | -1.62 | 0.68 | 0.54 | -1.01 | 0.97 | 1.32 | 0.37 | 0.14 | 0 | -0.1675 | 0.555 | 0.7175 | 0.096172 | 0.717532 | 0.263464 | 1 | 0.999631 | 0.55886 |
| Osbp_Q3B7Z2_S383s_1 | -0.85 | 0.26 | -0.6 | -0.65 | -0.44 | -0.14 | -0.37 | 0.29 | -0.09 | -0.44 | 0.81 | -0.45 | 0.93 | -0.48 | -0.03 | -0.21 | 0.295 | 0.4175 | 0.5125 | 0.231084 | 0.387696 | 0.308687 | 1 | 0.999631 | 0.681463 |
| Osbp_Q3B7Z2_S384s_1 | -0.59 | 0.28 | -1.12 | -0.73 | -0.43 | -0.41 | -0.6 | -0.77 | -0.12 | -2.29 | -0.99 | -1.21 | -0.49 | -0.75 | -1.72 | -1.66 | -0.0125 | -0.6125 | -0.615 | 0.180084 | 0.970773 | 0.24436 | 1 | 0.999631 | 0.64376 |
| Osbp2_Q5QNQ6_S759s_1 | 0.67 | -1.31 | -0.37 | -1.53 | -0.98 | -1.11 | 4.42 | -2.95 | 1.74 | 0.51 | -1.89 | 1.18 | -0.7 | -1.44 | -2.12 | -1.85 | 0.48 | 1.02 | -0.8925 | 0.128086 | 0.735174 | 0.235677 | 1 | 0.999631 | 0.598425 |
| Osbpl10_S4R1M9_S28s_1 | -0.03 | 1.05 | 1.19 | -0.6 | -0.68 | 0.52 | 0 | -0.06 | 0.23 | 1.06 | 0.28 | 0.58 | 0.64 | 1.59 | 1.6 | -0.07 | -0.4575 | 0.135 | 0.5375 | 0.340769 | 0.344934 | 0.769098 | 1 | 0.999631 | 0.733032 |
| Osbpl10_S4R1M9_S58s_1 | 0.68 | 0.84 | 0.85 | 0.56 | 0.3 | 1.2 | 0.2 | 1.18 | 1.14 | 1.14 | 0.53 | 1.43 | 0.27 | 1.3 | 1.09 | 1.14 | -0.0125 | 0.3275 | 0.2175 | 0.504813 | 0.969324 | 0.273312 | 1 | 0.999631 | 0.814292 |
| Osbpl10_S4R1M9_S58s_2 | 0.96 | -0.03 | 1.05 | 0.97 | 0.05 | 0.21 | 0.49 | 1.32 | 0.31 | 0.81 | 0.81 | 0.89 | -2.67 | -0.15 | 1.12 | 0.78 | -0.22 | -0.0325 | -0.9675 | 0.233979 | 0.576098 | 0.923067 | 1 | 0.999631 | 0.683077 |
| Osbpl10_S4R1M9_S66s_1 | 0.01 | -0.95 | 0.25 | -0.41 | -0.77 | -1.21 | -0.07 | -0.48 | -0.4 | -0.52 | -0.47 | -0.48 | -1.01 | -1.12 | -0.81 | -0.22 | -0.3575 | -0.1925 | -0.515 | 0.187443 | 0.349797 | 0.553919 | 1 | 0.999631 | 0.646474 |
| Osbpl10_S4R1M9_S66s_2 | 0.96 | -0.03 | 0.76 | 0.8 | 0.05 | 0.21 | 0.47 | 1.07 | 0.31 | 0.81 | 0.63 | 0.41 | -2.67 | -0.15 | 0.81 | 0.5 | -0.1725 | -0.0825 | -1 | 0.188282 | 0.620787 | 0.792548 | 1 | 0.999631 | 0.647439 |
| Osbpl3_D3Z194_S250s_1 | 1.22 | 0.99 | -0.15 | 1.06 | 1.93 | 1.29 | 0.62 | 0.66 | 0.94 | 0.51 | 0.07 | 0.6 | -0.52 | 0.74 | 0.47 | 0.88 | 0.345 | -0.25 | -0.3875 | 0.39415 | 0.431989 | 0.517852 | 1 | 0.999631 | 0.759303 |
| Osbpl3_D3Z194_S400s_1 | -0.64 | -0.3 | -0.61 | -0.15 | -0.57 | -0.64 | -0.73 | 0.29 | -0.41 | -0.22 | -0.32 | -0.28 | -0.37 | -0.25 | -0.38 | -0.08 | 0.0125 | 0.1175 | 0.155 | 0.580651 | 0.968411 | 0.649535 | 1 | 0.999631 | 0.846995 |
| Osbpl3_D3Z194_S478s_1 | -0.72 | 0.8 | -0.29 | 0.13 | 0.51 | -0.34 | -0.47 | 0.63 | -0.27 | -0.71 | 0.24 | -0.23 | 0.69 | 0.61 | -0.38 | -0.4 | 0.1025 | -0.2225 | 0.15 | 0.735874 | 0.80935 | 0.574573 | 1 | 0.999631 | 0.90759 |
| Osbpl3_D3Z194_T477t_1 | -0.72 | 0.8 | -0.29 | 0.13 | 0.51 | -0.34 | -0.47 | 0.63 | -0.27 | -0.71 | 0.24 | -0.23 | 0.69 | 0.61 | -0.38 | -0.4 | 0.1025 | -0.2225 | 0.15 | 0.735874 | 0.80935 | 0.574573 | 1 | 0.999631 | 0.90759 |
| Osbpl5_Q9ER64_S311s_1 | 1.5 | -0.2 | 0.66 | 0.76 | -0.41 | 0.16 | 0.8 | -0.37 | -0.09 | 1.79 | -0.93 | -1.33 | -1.07 | -1.56 | -0.65 | -1.36 | -0.635 | -0.82 | -1.84 | 0.001266 | 0.169446 | 0.253493 | 1 | 0.999631 | 0.351308 |
| Osbpl6_F8WIZ7_S290s_1 | -0.85 | 0.56 | -0.4 | -0.3 | 0.58 | -0.04 | -0.51 | -0.07 | 0.2 | -0.32 | -1.18 | 0.04 | -0.79 | -0.02 | 0.09 | -0.4 | 0.2375 | -0.0675 | -0.0325 | 0.933127 | 0.538663 | 0.874328 | 1 | 0.999631 | 0.978327 |
| Osbpl6_F8WIZ7_S9s_1 | 2.59 | 0.81 | 1.27 | 2.28 | 0.55 | 1.35 | 2.04 | 2.09 | 1.52 | 1.38 | 0.92 | 0.04 | -1.73 | 0.5 | 0.81 | 0.12 | -0.23 | -0.7725 | -1.8125 | 0.016304 | 0.656366 | 0.150656 | 1 | 0.999631 | 0.388832 |
| Osbpl6_F8WIZ7_T288t_1 | -0.04 | -1.03 | -0.44 | 0.26 | -0.88 | -0.53 | -0.65 | 1.03 | -0.73 | -1.27 | -0.43 | 0.66 | -1.79 | -0.35 | -0.77 | 0.24 | 0.055 | -0.13 | -0.355 | 0.476233 | 0.910158 | 0.784512 | 1 | 0.999631 | 0.799882 |
| Osbpl8_B9EJ86_S342s_1 | -0.73 | 2.45 | 2.06 | -0.09 | 0.48 | 1.98 | 1 | 0.43 | 1.32 | 1.01 | 2.15 | 1.32 | 2.34 | 1.63 | 2.4 | 2.17 | 0.05 | 0.5275 | 1.2125 | 0.112453 | 0.947311 | 0.474973 | 1 | 0.999631 | 0.577306 |
| Osbpl8_B9EJ86_S65s_1 | -0.93 | -0.6 | -1.23 | -0.64 | -0.14 | -0.56 | -0.93 | -1.05 | 0.07 | -3.5 | -1.06 | -0.3 | -0.82 | -0.34 | -0.97 | -0.69 | 0.18 | -0.3475 | 0.145 | 0.633149 | 0.562957 | 0.633027 | 1 | 0.999631 | 0.868569 |
| Osbpl8_B9EJ86_T39t_1 | -0.47 | 0.21 | -0.45 | 0.02 | 0.23 | 0.57 | -0.3 | 0.42 | 0.02 | -0.7 | 0.06 | -0.09 | 0.53 | 0.9 | 0.9 | 0.04 | 0.4025 | -0.005 | 0.765 | 0.042496 | 0.214393 | 0.987197 | 1 | 0.999631 | 0.475443 |
| Otud5_Q3U2S4_S165s_1 | 0.34 | 1.29 | 0.95 | 1.02 | -0.67 | 0.89 | 0.58 | 0.38 | 1.24 | 1.26 | 0.97 | -0.89 | 0.02 | 1.04 | 0.62 | 0.63 | -0.605 | -0.255 | -0.3225 | 0.365646 | 0.152298 | 0.625784 | 1 | 0.999631 | 0.745725 |
| Otud5_Q3U2S4_S64s_1 | 0.12 | 1.42 | 1.27 | -0.33 | 1.48 | -0.41 | 0.57 | 0.38 | 0.44 | 1.1 | 1.85 | 1.49 | 2.35 | -0.19 | 0.69 | 1.86 | -0.115 | 0.6 | 0.5575 | 0.396013 | 0.829702 | 0.246779 | 1 | 0.999631 | 0.760506 |
| Otud7a_Q8R554_S593s_1 | 1.61 | -0.36 | 0.67 | -0.19 | -0.92 | -0.76 | 0.85 | -0.35 | -0.93 | 0.51 | -0.01 | 0.12 | -0.27 | -0.23 | 0.38 | 0.14 | -0.7275 | -0.51 | -0.4275 | 0.37253 | 0.209224 | 0.333812 | 1 | 0.999631 | 0.747035 |
| Otud7a_Q8R554_S711s_1 | 0.41 | -0.05 | 0.6 | 0.59 | 0.41 | 0.34 | 0.25 | 0.53 | 0.08 | 0.61 | 0.61 | 1.68 | 0.23 | 1.13 | 0.44 | 1.13 | -0.005 | 0.3575 | 0.345 | 0.32605 | 0.984855 | 0.366811 | 1 | 0.999631 | 0.724678 |
| Otud7a_Q8R554_S749s_1 | 1.51 | 0.74 | 1.1 | 1.48 | 1.01 | 1.01 | 2.25 | 0.73 | 1.47 | 0.91 | 1.45 | 0.52 | -0.86 | -0.36 | 0.92 | -0.98 | 0.0425 | -0.12 | -1.5275 | 0.007722 | 0.913789 | 0.723963 | 1 | 0.999631 | 0.358173 |
| Otud7a_Q8R554_S771s_1 | 1.08 | -0.03 | 0.54 | 0.35 | 0 | 0.32 | 0.17 | -0.06 | 0.14 | 0.9 | -0.39 | -0.55 | -0.02 | 0.16 | 0.13 | -0.17 | -0.3775 | -0.46 | -0.46 | 0.176383 | 0.234911 | 0.276487 | 1 | 0.999631 | 0.642116 |
| Otud7a_Q8R554_T804t_1 | -1.47 | 2.11 | 1.65 | 0.61 | 0.51 | 1.84 | 0.35 | 1.79 | 0.44 | -0.03 | 1.36 | 0.16 | 1.11 | 1.85 | 0.5 | 0.65 | 0.3975 | -0.2425 | 0.3025 | 0.687169 | 0.612703 | 0.748129 | 1 | 0.999631 | 0.891307 |
| Otud7a_Q8R554_T829t_1 | -0.65 | -0.12 | -0.07 | -0.67 | -0.94 | -0.11 | -0.71 | -0.08 | -1.03 | -0.49 | -0.32 | -1.72 | 0.01 | 0.01 | -1.34 | -1.49 | -0.0825 | -0.5125 | -0.325 | 0.471222 | 0.796364 | 0.19659 | 1 | 0.999631 | 0.797981 |
| Oxr1_Q4KMM3_S104s_1 | -0.27 | 1.04 | -0.2 | 0.09 | 0.02 | -0.17 | -0.7 | -0.36 | -0.38 | -0.08 | -0.05 | -0.24 | -0.58 | 0.33 | -0.31 | -0.21 | -0.4675 | -0.3525 | -0.3575 | 0.367821 | 0.21585 | 0.327459 | 1 | 0.999631 | 0.745968 |
| Oxr1_Q4KMM3_S13s_1 | 0.9 | 0.55 | 0.55 | 0.77 | 1.23 | 0.69 | 1.05 | 0.42 | 0.42 | -1.38 | 0.86 | 0.48 | -0.73 | 0.45 | -0.11 | 1.24 | 0.155 | -0.5975 | -0.48 | 0.283971 | 0.584327 | 0.23787 | 1 | 0.999631 | 0.704942 |
| Oxr1_Q4KMM3_S15s_1 | 1.36 | 1.1 | 0.57 | 1.05 | 1.82 | 1.36 | 0.81 | 0.59 | 0.74 | 0.19 | 0.52 | 0.77 | -0.8 | 0.61 | 0.35 | 1.2 | 0.125 | -0.465 | -0.68 | 0.153381 | 0.722556 | 0.137966 | 1 | 0.999631 | 0.625198 |
| Oxr1_Q4KMM3_S202s_1 | -0.16 | -0.24 | -0.08 | 0.87 | 0.14 | -0.53 | 0.13 | 0.59 | -0.31 | -0.36 | 0.94 | 1.03 | -0.41 | -0.27 | 0.6 | 1.22 | -0.015 | 0.2275 | 0.1875 | 0.683572 | 0.967386 | 0.61698 | 1 | 0.999631 | 0.889602 |
| Oxr1_Q4KMM3_S323s_1 | -0.12 | 0.15 | 1.2 | -0.98 | -0.61 | 0.61 | 0.95 | -0.6 | -0.9 | 0.21 | -0.48 | -1.04 | 0 | -0.66 | 0.76 | -1.28 | 0.025 | -0.615 | -0.3575 | 0.539453 | 0.964016 | 0.239047 | 1 | 0.999631 | 0.830899 |
| Oxr1_Q4KMM3_S488s_1 | 3.27 | 0.05 | 4.7 | -1.45 | -1.42 | -0.15 | 3.11 | -1.79 | 5.47 | 3.19 | 4.91 | 2 | -0.61 | -0.29 | 0.49 | -1.19 | -1.705 | 2.25 | -2.0425 | 0.121929 | 0.28031 | 0.128774 | 1 | 0.999631 | 0.590723 |
| Oxr1_Q4KMM3_S518s_1 | -0.08 | -0.13 | -0.62 | 0.2 | 1.04 | 2.47 | 0.29 | 0.91 | 0.75 | -0.54 | 0.33 | 0.02 | -0.77 | 1.64 | 0.89 | -0.19 | 1.335 | 0.2975 | 0.55 | 0.314289 | 0.01667 | 0.412213 | 1 | 0.999631 | 0.720084 |
| Oxr1_Q4KMM3_S568s_1 | 0.5 | 0.42 | 0.24 | 0.36 | 1.56 | 1.5 | -0.14 | 0.87 | 1.24 | 0.72 | 0.82 | 0.42 | 0.13 | -0.18 | 0.26 | 0.02 | 0.5675 | 0.42 | -0.3225 | 0.248096 | 0.179966 | 0.154096 | 1 | 0.999631 | 0.685751 |
| Oxtr_P97926_S360s_1 | -0.1 | -1.02 | -0.37 | -0.53 | -0.84 | 0.1 | -0.21 | -0.01 | 0.19 | 0.95 | -1.02 | -0.05 | 0.28 | -0.1 | -0.3 | -0.11 | 0.265 | 0.5225 | 0.4475 | 0.178052 | 0.427231 | 0.256328 | 1 | 0.999631 | 0.642493 |
| P0C913_S28s_1 | -2.37 | -1.8 | -2.98 | -4.25 | -1.77 | -1.12 | -2.82 | -3.56 | -1.25 | -1.94 | -2.26 | -3.76 | -1.13 | -1.81 | -2.92 | -3.96 | 0.5325 | 0.5475 | 0.395 | 0.58578 | 0.435367 | 0.420914 | 1 | 0.999631 | 0.849524 |
| P2ry12_Q9CPV9_S342s_1 | 0.11 | 0.4 | 0.3 | 0.49 | 0.1 | -0.16 | 0.1 | 0.03 | 0.14 | 0.17 | 0.01 | 0.07 | 0.58 | 0.32 | -0.03 | 0.35 | -0.3075 | -0.2275 | -0.02 | 0.9435 | 0.221837 | 0.366442 | 1 | 0.999631 | 0.981629 |
| P32507-2_P32507_S414s_1 | 0.78 | 0.53 | 0.74 | 0.36 | 0.43 | 0.44 | 0.59 | 0.45 | 1.17 | 0.8 | 0.75 | 0.08 | 1.05 | 0.82 | 0.82 | 1.54 | -0.125 | 0.0975 | 0.455 | 0.153089 | 0.60793 | 0.755521 | 1 | 0.999631 | 0.625198 |
| Pabpc4_V9GXG3_S96s_1 | -0.08 | 1.48 | 0.38 | 0.6 | 0.96 | 1.21 | -0.13 | 0.76 | 1.54 | 0.38 | -0.07 | 0.56 | 0.41 | 1.36 | 1.42 | 0.49 | 0.105 | 0.0075 | 0.325 | 0.459134 | 0.807293 | 0.986956 | 1 | 0.999631 | 0.79152 |
| Pacrgl_Q9D3X5_S47s_1 | -0.44 | 0.3 | 1.72 | 0.76 | 0.7 | 0.51 | 0.72 | 1.44 | 0.71 | 0.51 | 1.7 | 1.41 | 0.31 | 1.1 | 1.61 | 1.28 | 0.2575 | 0.4975 | 0.49 | 0.343971 | 0.58841 | 0.336444 | 1 | 0.999631 | 0.733852 |
| Pacs1_Q8K212_S23s_1 | -1.55 | -2.2 | -2.04 | -0.94 | -2.04 | -2.05 | -0.86 | -1.09 | -2.53 | -1.8 | -0.12 | -1.55 | -1.44 | -1.98 | -0.96 | -1.24 | 0.1725 | 0.1825 | 0.2775 | 0.479904 | 0.681313 | 0.735506 | 1 | 0.999631 | 0.801333 |
| Pacs1_Q8K212_S28s_1 | -0.59 | -0.59 | -1.04 | -1.09 | -0.42 | -0.94 | -1.09 | -0.61 | -0.32 | 0.03 | -0.96 | -0.94 | 0.02 | -0.17 | -0.65 | -0.76 | 0.0625 | 0.28 | 0.4375 | 0.189558 | 0.825521 | 0.406533 | 1 | 0.999631 | 0.648674 |
| Pacs1_Q8K212_S493s_1 | -0.73 | -3.44 | -1.4 | 0.12 | -1.99 | -2.22 | 1.46 | -0.43 | -2.4 | -1.23 | -0.54 | -1.86 | -3.03 | -3.66 | -1.59 | -1.16 | 0.5675 | -0.145 | -0.9975 | 0.249157 | 0.566462 | 0.848266 | 1 | 0.999631 | 0.686359 |
| Pacs1_Q8K212_S526s_1 | 2.33 | -3.89 | -2.55 | -0.79 | -2.98 | -1.21 | 4 | -4.61 | -3.65 | 1.71 | -3.21 | -3.21 | -5.52 | -5.93 | -4.55 | -4.78 | 0.025 | -0.865 | -3.97 | 0.00657 | 0.989718 | 0.584814 | 1 | 0.999631 | 0.351545 |
| Pacs1_Q8K212_S532s_1 | -0.17 | 0.16 | -1.12 | -2.97 | -0.5 | 0.39 | -2.12 | -3.12 | -0.02 | -0.47 | -2.39 | -1.65 | -0.02 | 0.38 | -0.95 | -1.67 | -0.3125 | -0.1075 | 0.46 | 0.538135 | 0.73299 | 0.890454 | 1 | 0.999631 | 0.830863 |
| Pacs1_Q8K212_T44t_1 | -0.4 | -0.06 | -0.18 | -0.1 | -0.78 | -0.19 | -0.29 | -0.01 | 0.04 | 0.16 | -0.28 | 0.67 | 0.8 | 0.5 | 0.12 | 0.31 | -0.1325 | 0.3325 | 0.6175 | 0.053508 | 0.62782 | 0.273905 | 1 | 0.999631 | 0.4909 |
| Pacs2_E9Q7E9_S349s_1 | 0.18 | 0.95 | 0.25 | 1.24 | 0.91 | -0.25 | 1.02 | 0.84 | 0.31 | 0.23 | -0.28 | 1.22 | -0.73 | 0.77 | 1 | -0.19 | -0.025 | -0.285 | -0.4425 | 0.359805 | 0.950023 | 0.495898 | 1 | 0.999631 | 0.741182 |
| Pacs2_E9Q7E9_S417s_1 | 0.62 | 1.26 | 0.43 | 0.96 | 1.31 | 1.36 | 0.28 | 1.2 | 1.13 | 0.43 | 1.12 | 0.36 | -0.2 | 1.88 | 0.8 | 0.68 | 0.22 | -0.0575 | -0.0275 | 0.952194 | 0.528993 | 0.862235 | 1 | 0.999631 | 0.984317 |
| Pacs2_E9Q7E9_S454s_1 | 1.04 | -1.11 | 0.17 | 0.35 | -1.06 | -0.39 | 0.89 | -0.74 | -0.64 | 0.7 | 0.02 | -0.45 | -0.8 | -1.33 | -0.17 | -0.94 | -0.4375 | -0.205 | -0.9225 | 0.084332 | 0.446475 | 0.688062 | 1 | 0.999631 | 0.549732 |
| Pacsin1_Q61644_S326s_1 | -0.53 | -0.85 | -1.94 | -0.22 | 1.23 | 0.21 | -0.11 | 0.58 | -0.37 | -2.19 | 0.19 | -0.29 | 0.34 | -0.85 | -0.85 | -0.63 | 1.3625 | 0.22 | 0.3875 | 0.411298 | 0.013009 | 0.709257 | 1 | 0.999631 | 0.767937 |
| Pacsin1_Q61644_S343s_1 | -1.67 | -1.22 | -0.95 | -1.29 | -0.94 | -1.64 | -1.22 | -0.7 | -1.02 | -2.37 | -0.83 | -0.13 | 0.04 | -1.86 | -0.44 | -0.58 | 0.1575 | 0.195 | 0.5725 | 0.2099 | 0.611036 | 0.681755 | 1 | 0.999631 | 0.66441 |
| Pacsin1_Q61644_S343s_2 | 1.67 | 1.69 | 1.88 | 1.92 | 1.31 | 2.4 | 1.68 | 1.03 | 1.89 | 2.74 | 1.31 | 0.94 | -1.25 | 0.06 | 0.49 | 1.07 | -0.185 | -0.07 | -1.6975 | 0.005313 | 0.588202 | 0.863343 | 1 | 0.999631 | 0.351308 |
| Pacsin1_Q61644_S345s_2 | -0.93 | -0.53 | 0.69 | 1.33 | -0.53 | -1.52 | 0.55 | 1.34 | -0.87 | -1.93 | 1.33 | 1.73 | -0.55 | -1.41 | 0.73 | 1.53 | -0.18 | -0.075 | -0.065 | 0.92983 | 0.802741 | 0.932749 | 1 | 0.999631 | 0.97754 |
| Pacsin1_Q61644_S346s_1 | 0.58 | -0.76 | -0.05 | -0.02 | -1.34 | -0.49 | 0.06 | -1.25 | -0.87 | 0.53 | -0.86 | -0.56 | -0.93 | -1.53 | -0.65 | -1.77 | -0.6925 | -0.3775 | -1.1575 | 0.015157 | 0.127317 | 0.389238 | 1 | 0.999631 | 0.388832 |
| Pacsin1_Q61644_S427s_1 | -0.02 | 0.52 | -0.19 | -0.51 | 0.82 | 0.2 | -0.11 | 0.01 | -0.06 | -0.07 | -0.18 | 0.24 | 0.06 | 0.64 | -1.4 | 0.13 | 0.28 | 0.0325 | -0.0925 | 0.846013 | 0.413463 | 0.915147 | 1 | 0.999631 | 0.948565 |
| Pacsin1_Q61644_S76s_1 | 0.6 | 1.43 | 1.09 | 0.46 | 0.23 | 0.39 | 0.06 | 0.55 | 1.28 | 1.74 | 1.38 | 1.54 | 1.43 | 1.1 | 0.79 | 1.48 | -0.5875 | 0.59 | 0.305 | 0.380089 | 0.079048 | 0.082862 | 1 | 0.999631 | 0.752237 |
| Pacsin1_Q61644_T181t_1 | 0.46 | -1.33 | 0.79 | 0.61 | -0.9 | -0.78 | 0.98 | -0.11 | -0.73 | -0.38 | 0.34 | 0.26 | -1.05 | -1.69 | 0.56 | -0.29 | -0.335 | -0.26 | -0.75 | 0.247622 | 0.57475 | 0.620066 | 1 | 0.999631 | 0.685751 |
| Pacsin3_F6W8Y8_S44s_1 | -1.05 | -0.5 | -0.09 | -0.93 | -0.59 | -1.68 | -0.05 | -0.1 | -0.05 | -1.13 | 0.31 | 0.49 | 0.36 | -0.81 | 0.02 | 0.53 | 0.0375 | 0.5475 | 0.6675 | 0.116002 | 0.930343 | 0.217561 | 1 | 0.999631 | 0.582001 |
| Pacsin3_F6W8Y8_S79s_1 | -2.26 | -1.39 | 2.62 | 0.46 | -1.77 | -2.68 | 0.13 | 0.05 | -1.34 | -2.05 | -0.59 | 1.36 | -1.46 | -1.46 | 3.14 | 1.04 | -0.925 | -0.5125 | 0.4575 | 0.72701 | 0.409339 | 0.649617 | 1 | 0.999631 | 0.905774 |
| Pafah1b2_Q61206_S64s_1 | -1.71 | 0.01 | 0.31 | -0.46 | -0.32 | -0.23 | -0.74 | 0.01 | -0.43 | -0.26 | -0.68 | 0.15 | 0.35 | 0.35 | -0.38 | 0.6 | 0.1425 | 0.1575 | 0.6925 | 0.170429 | 0.754234 | 0.734835 | 1 | 0.999631 | 0.64073 |
| Pag1_Q3U1F9_S379s_1 | -0.2 | -0.65 | 0.03 | 0.42 | -0.35 | -0.48 | 0.03 | 0.2 | -0.96 | -0.47 | -0.13 | 0.51 | -0.41 | -0.1 | 0.59 | 0.71 | -0.05 | -0.1625 | 0.2975 | 0.445798 | 0.876373 | 0.682449 | 1 | 0.999631 | 0.784402 |
| Pag1_Q3U1F9_Y224y_1 | 0.07 | -0.53 | -0.38 | -0.43 | -0.72 | -0.52 | -0.52 | -1.09 | -0.33 | -0.71 | -0.79 | -0.93 | 0.7 | -0.01 | -0.59 | -0.64 | -0.395 | -0.3725 | 0.1825 | 0.630947 | 0.173647 | 0.205311 | 1 | 0.999631 | 0.867552 |
| Pagr1a_Q99L02_S147s_2 | -0.29 | 0.85 | 2.77 | 0.33 | 0.06 | 1.45 | -0.31 | 0.67 | 0.51 | 1.05 | -0.02 | 0.52 | 2.1 | 1.86 | 1.4 | 1.82 | -0.4475 | -0.4 | 0.88 | 0.173548 | 0.514641 | 0.52922 | 1 | 0.999631 | 0.642116 |
| Pagr1a_Q99L02_T137t_2 | -0.29 | 0.85 | 2.77 | 0.33 | 0.06 | 1.45 | -0.31 | 0.67 | 0.51 | 1.05 | -0.02 | 0.52 | 2.1 | 1.86 | 1.4 | 1.82 | -0.4475 | -0.4 | 0.88 | 0.173548 | 0.514641 | 0.52922 | 1 | 0.999631 | 0.642116 |
| Paics_Q9DCL9_S107s_1 | -1.61 | 0.22 | -0.35 | -1.4 | -0.79 | -0.28 | -0.6 | -0.69 | 0.26 | -1.47 | -0.2 | 1.05 | 0.67 | -0.47 | -1.13 | -0.51 | 0.195 | 0.695 | 0.425 | 0.436236 | 0.656835 | 0.277571 | 1 | 0.999631 | 0.780748 |
| Paip1_G3UZS6_S52s_1 | -0.36 | 0.09 | 0.23 | 1.08 | -0.5 | 0.62 | -0.03 | 1.04 | 0.66 | 0.45 | -1.04 | -0.07 | -0.04 | -0.97 | -1.65 | -0.07 | 0.0225 | -0.26 | -0.9425 | 0.072267 | 0.959382 | 0.581741 | 1 | 0.999631 | 0.532251 |
| Pak1_G5E884_S144s_1 | -1.36 | 0.65 | 0.98 | -0.18 | -1.16 | 1.3 | -0.05 | -0.04 | -1.15 | -0.54 | 0.77 | -1.89 | -0.17 | 1.17 | -1.31 | -0.27 | -0.01 | -0.725 | -0.1675 | 0.797753 | 0.987665 | 0.304501 | 1 | 0.999631 | 0.930242 |
| Pak1_G5E884_S222s_1 | -1.13 | 0.09 | 0.02 | -0.39 | -0.51 | -0.42 | -0.7 | -0.47 | -0.38 | -0.51 | 0.12 | 0.16 | 0.4 | 0.01 | -0.04 | 0.4 | -0.1725 | 0.2 | 0.545 | 0.149566 | 0.602303 | 0.582395 | 1 | 0.999631 | 0.621721 |
| Pak1_G5E884_S28s_1 | 1.8 | 0.15 | 0.14 | 1.02 | -0.4 | -0.23 | 1.98 | 1.32 | -0.25 | 0.38 | 2.36 | -0.29 | -1.51 | -1.46 | 0.87 | 0.76 | -0.11 | -0.2275 | -1.1125 | 0.130061 | 0.862143 | 0.732068 | 1 | 0.999631 | 0.601275 |
| Pak1_G5E884_T146t_1 | -1.65 | 0.7 | 0.58 | -0.51 | -1.14 | 1.3 | -0.62 | -0.46 | -1.11 | -0.63 | 0.28 | -1.73 | -0.01 | 1.18 | -0.89 | -0.28 | -0.01 | -0.5775 | 0.22 | 0.728164 | 0.988218 | 0.368059 | 1 | 0.999631 | 0.905923 |
| Pak1_G5E884_T211t_1 | 1.69 | -0.72 | -2.74 | 1.8 | 2.53 | 0.76 | 1.63 | 1.42 | 0.25 | -0.92 | 0.76 | -0.56 | -2.99 | 0.54 | -0.53 | -0.41 | 1.5775 | -0.125 | -0.855 | 0.452596 | 0.133774 | 0.899809 | 1 | 0.999631 | 0.788364 |
| Pak1_G5E884_T228t_1 | -1.17 | -0.67 | -0.23 | -1.28 | -1.22 | -1.49 | -0.79 | -0.59 | -0.46 | -1.18 | -0.11 | -0.49 | 0.56 | -0.76 | -0.09 | 0.11 | -0.185 | 0.2775 | 0.7925 | 0.067356 | 0.59693 | 0.450345 | 1 | 0.999631 | 0.521449 |
| Pak1_G5E884_T229t_1 | -1.06 | -1.18 | -0.51 | -0.51 | -0.47 | -0.75 | -0.89 | -0.94 | -0.85 | -0.92 | -0.93 | -0.33 | 0.6 | -0.31 | -0.59 | -0.21 | 0.0525 | 0.0575 | 0.6875 | 0.080462 | 0.853375 | 0.849299 | 1 | 0.999631 | 0.545579 |
| Pak1ip1_Q9DCE5_S361s_1 | 2.32 | 1.35 | 1.99 | 1.57 | 0.27 | 2.25 | 2.08 | 0.75 | 1.1 | 2.7 | 1.2 | 0.95 | 1.19 | 0.45 | 1.7 | 1.33 | -0.47 | -0.32 | -0.64 | 0.114291 | 0.359475 | 0.484637 | 1 | 0.999631 | 0.579001 |
| Pak2_Q8CIN4_S141s_1 | 0.4 | 1.05 | 0.98 | 0.76 | 0.7 | 1.02 | 0.19 | 0.76 | 0.9 | 0.66 | 0.74 | 0.99 | 0.69 | 1.37 | 1.35 | 1.3 | -0.13 | 0.025 | 0.38 | 0.240204 | 0.660518 | 0.926691 | 1 | 0.999631 | 0.683719 |
| Pak2_Q8CIN4_T143t_1 | -0.93 | -0.12 | -0.69 | -1.8 | -0.22 | -0.53 | -1.2 | -1.27 | -0.76 | -0.53 | -0.39 | -1.11 | 0.03 | 0.16 | -1.21 | -0.82 | 0.08 | 0.1875 | 0.425 | 0.377035 | 0.851228 | 0.638374 | 1 | 0.999631 | 0.749236 |
| Pak3_Q61036_S4s_1 | -0.7 | -0.68 | -0.15 | -0.36 | -0.56 | -0.72 | -0.53 | -0.33 | -0.56 | -1.24 | -0.74 | -0.82 | 0.28 | -0.29 | -0.19 | -0.61 | -0.0625 | -0.3675 | 0.27 | 0.400706 | 0.810272 | 0.218654 | 1 | 0.999631 | 0.763209 |
| Pak4_Q8BTW9_S104s_1 | 0.67 | -0.21 | 0.4 | 0.19 | -0.03 | -0.49 | 0.51 | -0.6 | -0.37 | 0.79 | -0.07 | -0.14 | -0.3 | -0.19 | 0.1 | -0.24 | -0.415 | -0.21 | -0.42 | 0.189634 | 0.247641 | 0.554865 | 1 | 0.999631 | 0.648674 |
| Pak4_Q8BTW9_S181s_1 | -0.17 | 0.53 | 1.07 | 1.06 | 0.94 | 0.13 | 0.13 | 0.88 | 0.42 | 0.06 | 1.26 | 0.96 | -0.11 | 0.9 | 1.32 | 1.43 | -0.1025 | 0.0525 | 0.2625 | 0.566365 | 0.788714 | 0.897244 | 1 | 0.999631 | 0.84194 |
| Pak4_Q8BTW9_S266s_1 | -0.23 | 0.83 | 0.63 | -0.25 | -0.36 | 0.9 | -0.63 | 0.03 | 0.11 | 0.52 | -0.76 | -0.99 | 0.46 | 0.94 | -0.84 | 0.22 | -0.26 | -0.525 | -0.05 | 0.914066 | 0.548847 | 0.2581 | 1 | 0.999631 | 0.972551 |
| Pak4_Q8BTW9_S476s_1 | -0.52 | 0.07 | 0.22 | -0.18 | -0.09 | -0.66 | -0.45 | -0.22 | -0.2 | -0.68 | -0.26 | 0.18 | 0.05 | -0.25 | 0.34 | 0.09 | -0.2525 | -0.1375 | 0.16 | 0.600602 | 0.382684 | 0.657049 | 1 | 0.999631 | 0.856688 |
| Pak6_Q3ULB5_S348s_2 | 2.51 | -0.83 | -0.02 | 1.7 | -0.04 | -0.51 | 2.78 | 0.03 | -0.36 | 1.4 | 0.2 | -0.66 | -1.76 | -2.77 | -1.76 | -0.74 | -0.275 | -0.695 | -2.5975 | 0.006464 | 0.766733 | 0.38542 | 1 | 0.999631 | 0.351545 |
| Pak6_Q3ULB5_S352s_1 | 1.42 | 0.15 | 1.21 | 0.14 | -0.98 | 1.49 | 1.14 | -0.01 | -0.38 | 2.11 | 0.46 | 0.03 | 0.11 | -0.26 | 0.29 | 0.49 | -0.32 | -0.175 | -0.5725 | 0.174198 | 0.59368 | 0.766466 | 1 | 0.999631 | 0.642116 |
| Pak6_Q3ULB5_S352s_2 | 2.51 | -0.83 | -0.02 | 1.7 | -0.04 | -0.51 | 2.78 | 0.03 | -0.36 | 1.4 | 0.2 | -0.66 | -1.76 | -2.77 | -1.76 | -0.74 | -0.275 | -0.695 | -2.5975 | 0.006464 | 0.766733 | 0.38542 | 1 | 0.999631 | 0.351545 |
| Pak6_Q3ULB5_S561s_1 | -0.89 | -0.12 | -0.21 | -0.33 | -0.52 | -0.5 | -0.68 | -0.44 | -0.61 | -0.71 | -0.31 | -0.01 | 0.29 | -0.17 | -0.2 | 0.01 | -0.1475 | -0.0225 | 0.37 | 0.243416 | 0.589406 | 0.941272 | 1 | 0.999631 | 0.685637 |
| Pak7_Q8C015_S104s_1 | -0.17 | -0.34 | -0.07 | -0.68 | -1.13 | -0.49 | -0.55 | -0.82 | -0.71 | -0.2 | 0.05 | -0.08 | 0.02 | -0.41 | -0.74 | 0.09 | -0.4325 | 0.08 | 0.055 | 0.863424 | 0.145869 | 0.786361 | 1 | 0.999631 | 0.955107 |

| | | | | | | | | | | | | | | | | | | | | | | | | | | |
|---|---|---|---|---|---|---|---|---|---|---|---|---|---|---|---|---|---|---|---|---|---|---|---|---|---|---|
| Pak7_Q8C015_S104s_2 | -0.6 | -0.52 | -0.13 | -0.46 | -1.07 | -0.69 | -0.43 | -0.75 | -0.94 | -1.15 | -0.75 | -1.31 | -0.8 | -0.41 | -0.69 | -0.47 | -0.3075 | -0.61 | -0.165 | 0.556705 | 0.263279 | 0.044938 | | 1 | 0.999631 | 0.838305 |
| Pak7_Q8C015_S277s_1 | 0.68 | -1.69 | -1.32 | -0.77 | -1.34 | -1 | -0.47 | -1.28 | -1.98 | 0.25 | -1.05 | -1.68 | -3.17 | -1.18 | -1.7 | -1.62 | -0.2475 | -0.34 | -1.1425 | 0.087083 | 0.634597 | 0.601378 | | 1 | 0.999631 | 0.553514 |
| Pak7_Q8C015_S288s_1 | -0.16 | 0.68 | -0.04 | 0.1 | 0.28 | -0.88 | 0.09 | -0.1 | 0.47 | -0.47 | 0.09 | 0.39 | -0.06 | 0.09 | -0.89 | -1.29 | -0.2975 | -0.025 | -0.6825 | 0.113963 | 0.399958 | 0.94012 | | 1 | 0.999631 | 0.579001 |
| Pak7_Q8C015_S290s_1 | -0.1 | -0.23 | 0.21 | 0.76 | 0.16 | -0.45 | -0.12 | 0.46 | -0.13 | -0.91 | 0.91 | 1.07 | -0.53 | -0.54 | 1.07 | 0.61 | -0.1475 | 0.075 | -0.0075 | 0.986941 | 0.65946 | 0.878527 | | 1 | 0.999631 | 0.995165 |
| Pak7_Q8C015_S341s_1 | -0.34 | -0.53 | -1.03 | -1.4 | -0.48 | -0.26 | -0.93 | -0.8 | -0.25 | -0.6 | -0.36 | -1.72 | 0.35 | -0.78 | -0.26 | -1.13 | 0.2075 | 0.0925 | 0.37 | 0.384873 | 0.530799 | 0.82511 | | 1 | 0.999631 | 0.75538 |
| Pak7_Q8C015_T107t_1 | 0.55 | -0.23 | -0.1 | 0.17 | 0.17 | 0.33 | 0.06 | 0.21 | -0.04 | 0.45 | 0.47 | 0.01 | -1.71 | -0.22 | 0.26 | -0.23 | 0.095 | 0.125 | -0.5725 | 0.22856 | 0.72719 | 0.676511 | | 1 | 0.999631 | 0.679038 |
| Pak7_Q8C015_T107t_2 | -0.6 | -0.52 | -0.13 | -0.46 | -1.07 | -0.69 | -0.43 | -0.75 | -0.94 | -1.15 | -0.75 | -1.31 | -0.8 | -0.41 | -0.69 | -0.47 | -0.3075 | -0.61 | -0.165 | 0.556705 | 0.263279 | 0.044938 | | 1 | 0.999631 | 0.838305 |
| Pakap_F7AA26_S1043s_1 | 0.24 | -1.12 | -0.89 | 0.24 | 0.95 | -0.41 | 0.4 | 0.7 | -0.49 | -0.99 | 0.69 | -0.17 | -1.44 | -0.76 | 0.65 | 0.43 | 0.7925 | 0.1425 | 0.1025 | 0.856489 | 0.104177 | 0.768944 | | 1 | 0.999631 | 0.951817 |
| Pakap_F7AA26_S239s_1 | -0.41 | -0.98 | -0.93 | -1.31 | -1.36 | -0.97 | -1.11 | -0.94 | -1.23 | -0.38 | -2.09 | -1.23 | -0.15 | -0.46 | -1.27 | -0.85 | -0.1875 | -0.325 | 0.225 | 0.533151 | 0.515831 | 0.430716 | | 1 | 0.999631 | 0.827798 |
| Pald1_P70261_S347s_1 | -1.2 | 0.81 | 1.25 | 1.24 | 0.18 | 1.16 | -0.1 | 1.01 | 0.83 | -0.12 | 0.42 | 1.54 | 1.62 | 1.92 | 0.61 | 1.2 | 0.0375 | 0.1425 | 0.8125 | 0.191757 | 0.949895 | 0.817537 | | 1 | 0.999631 | 0.648674 |
| Palm_Q9Z0P4_S116s_1 | 2.17 | 1.21 | 0 | -0.67 | 0.65 | 2.02 | -0.61 | -0.58 | 0.7 | 2.19 | 0.09 | 0.1 | -0.31 | 0.04 | -0.08 | -0.28 | -0.3075 | 0.0925 | -0.835 | 0.174648 | 0.693272 | 0.896667 | | 1 | 0.999631 | 0.642116 |
| Palm_Q9Z0P4_S116s_2 | -0.13 | 0.36 | 1.42 | 1.26 | -0.12 | 0.24 | 0.9 | 1.72 | 0.05 | -0.13 | 1.07 | 1.81 | 0.44 | 0.13 | 1.36 | 1.88 | -0.0425 | -0.0275 | 0.225 | 0.665565 | 0.933616 | 0.959643 | | 1 | 0.999631 | 0.88264 |
| Palm_Q9Z0P4_S116s_3 | 0.22 | 1.84 | 0.88 | 0.72 | 0.87 | 1.76 | 0.63 | 1.41 | 1.09 | 1 | 1.27 | 1.24 | 1.63 | 1.74 | 1.22 | 1.36 | 0.2525 | 0.235 | 0.5725 | 0.164044 | 0.551841 | 0.531359 | | 1 | 0.999631 | 0.632208 |
| Palm_Q9Z0P4_S122s_1 | -0.14 | 0.45 | -0.39 | -1.11 | -0.29 | 0.01 | -0.68 | -0.95 | 0.28 | -0.39 | -0.39 | -0.15 | 1.03 | 0.2 | -0.95 | -0.73 | -0.18 | 0.135 | 0.185 | 0.725665 | 0.648921 | 0.723952 | | 1 | 0.999631 | 0.90568 |
| Palm_Q9Z0P4_S122s_2 | -0.52 | 0.18 | -1.64 | -2.19 | -0.5 | -0.02 | -1.61 | -2.08 | -0.32 | -0.37 | -1.56 | -1.7 | 0.69 | -0.01 | -2.03 | -2.08 | -0.01 | 0.055 | 0.185 | 0.811502 | 0.987558 | 0.926301 | | 1 | 0.999631 | 0.936949 |
| Palm_Q9Z0P4_S122s_3 | 0.22 | 1.84 | 0.88 | 0.72 | 0.87 | 1.76 | 0.63 | 1.41 | 1.09 | 1 | 1.27 | 1.24 | 1.63 | 1.74 | 1.22 | 1.36 | 0.2525 | 0.235 | 0.5725 | 0.164044 | 0.551841 | 0.531359 | | 1 | 0.999631 | 0.632208 |
| Palm_Q9Z0P4_S124s_1 | -1.03 | -1.12 | -2.27 | -2.92 | -1.55 | -1.49 | -1.99 | -3.34 | -0.82 | -1.16 | -2.01 | -2.2 | 0.68 | -1.3 | -2.44 | -2.87 | -0.2575 | 0.2875 | 0.3525 | 0.660483 | 0.654676 | 0.589746 | | 1 | 0.999631 | 0.880315 |
| Palm_Q9Z0P4_S124s_2 | -1.04 | -0.76 | -1.33 | -1.77 | -1.22 | -0.97 | -1.28 | -1.56 | -1.17 | -1.23 | -1.29 | -1.3 | 0.06 | -0.89 | -1.58 | -1.59 | -0.0325 | -0.0225 | 0.225 | 0.616067 | 0.915325 | 0.939568 | | 1 | 0.999631 | 0.861203 |
| Palm_Q9Z0P4_S124s_3 | 0.22 | 1.84 | 0.88 | 0.72 | 0.87 | 1.76 | 0.63 | 1.41 | 1.09 | 1 | 1.27 | 1.24 | 1.63 | 1.74 | 1.22 | 1.36 | 0.2525 | 0.235 | 0.5725 | 0.164044 | 0.551841 | 0.531359 | | 1 | 0.999631 | 0.632208 |
| Palm_Q9Z0P4_S157s_1 | 0.04 | -0.74 | -0.52 | -0.25 | -0.61 | -0.23 | 0.06 | -0.19 | -0.5 | -0.22 | -0.48 | -0.99 | -1.51 | -0.75 | -0.76 | -0.93 | 0.125 | -0.18 | -0.62 | 0.080096 | 0.667776 | 0.558566 | | 1 | 0.999631 | 0.545579 |
| Palm_Q9Z0P4_S157s_2 | -1.15 | -0.76 | -1.44 | -0.53 | -0.48 | -0.83 | -0.84 | -0.72 | -0.98 | -0.79 | -1.19 | -1 | -1.05 | -0.72 | -1.04 | -1.03 | 0.2525 | -0.02 | 0.01 | 0.974306 | 0.393986 | 0.946302 | | 1 | 0.999631 | 0.990847 |
| Palm_Q9Z0P4_S161s_1 | 1.01 | -0.66 | -1.49 | 0.77 | 1.43 | 1.03 | 0.84 | 0.45 | 1.11 | -0.07 | 0.02 | -0.68 | -0.92 | 0.48 | 0.08 | -1.37 | 1.03 | 0.1875 | -0.34 | 0.606923 | 0.096794 | 0.767447 | | 1 | 0.999631 | 0.858527 |
| Palm_Q9Z0P4_S161s_2 | -3.2 | -0.58 | -1.37 | -0.63 | -0.75 | -0.85 | -1.31 | -0.75 | -0.96 | -0.83 | -0.76 | -0.7 | -0.38 | -2.22 | -0.64 | -1.3 | 0.53 | 0.6325 | 0.31 | 0.640149 | 0.364404 | 0.279981 | | 1 | 0.999631 | 0.870995 |
| Palm_Q9Z0P4_S200s_1 | -0.13 | 1.26 | 1.03 | -0.54 | 0.27 | 0.45 | 0.25 | 0.39 | -0.04 | 0.29 | 0.49 | 1.38 | 1.1 | 0.61 | 0.44 | 0.63 | -0.065 | 0.125 | 0.29 | 0.529294 | 0.880285 | 0.804407 | | 1 | 0.999631 | 0.825786 |
| Palm_Q9Z0P4_S204s_1 | 0.4 | 0.77 | 1.31 | 1.79 | 0.53 | 0.73 | 0.68 | 0.96 | 0.34 | 0.26 | 1.08 | 1 | 0.53 | -0.75 | 0.85 | 0.69 | -0.3425 | -0.3975 | -0.7375 | 0.13825 | 0.337466 | 0.321582 | | 1 | 0.999631 | 0.611124 |
| Palm_Q9Z0P4_S301s_1 | -0.47 | -1.01 | -1.56 | -0.79 | -0.56 | -0.8 | -0.66 | -0.45 | -0.5 | -1.26 | -0.89 | -0.86 | -0.46 | -0.48 | -0.92 | -0.52 | 0.34 | 0.08 | 0.3625 | 0.285438 | 0.276771 | 0.80838 | | 1 | 0.999631 | 0.705851 |
| Palm_Q9Z0P4_S321s_1 | -1.09 | -0.34 | -0.47 | -1.28 | -0.64 | -0.99 | -1.01 | -0.99 | -0.86 | -1.03 | -0.69 | -0.67 | 0.07 | -0.6 | -0.77 | -0.69 | -0.1125 | -0.0175 | 0.2975 | 0.409324 | 0.713472 | 0.955145 | | 1 | 0.999631 | 0.766996 |
| Palm_Q9Z0P4_T141t_1 | -0.53 | -0.38 | -0.32 | -0.4 | -0.53 | -0.74 | -0.68 | -0.58 | -0.9 | -1.3 | -0.72 | -0.58 | -0.08 | -0.44 | -0.56 | 0.01 | -0.225 | -0.4675 | 0.14 | 0.619832 | 0.34444 | 0.109378 | | 1 | 0.999631 | 0.861888 |
| Palm_Q9Z0P4_T141t_2 | -0.45 | -0.11 | -0.11 | -0.11 | -0.48 | -0.07 | -0.59 | -0.4 | -0.67 | -0.95 | 0.07 | -0.13 | -0.11 | -0.46 | -0.19 | 0.39 | -0.19 | -0.225 | 0.1025 | 0.734228 | 0.461709 | 0.480406 | | 1 | 0.999631 | 0.907462 |
| Palm_Q9Z0P4_T145t_1 | -1.4 | -0.56 | -0.56 | -1.05 | -0.9 | -0.91 | -0.79 | -0.86 | -0.88 | -0.82 | -1.21 | -0.47 | 0.4 | -0.67 | -0.93 | -0.31 | 0.0275 | 0.0475 | 0.515 | 0.202594 | 0.922814 | 0.880813 | | 1 | 0.999631 | 0.659204 |
| Palm_Q9Z0P4_T145t_2 | -0.45 | -0.11 | -0.11 | -0.11 | -0.48 | -0.07 | -0.59 | -0.4 | -0.67 | -0.95 | 0.07 | -0.13 | -0.11 | -0.46 | -0.19 | 0.39 | -0.19 | -0.225 | 0.1025 | 0.734228 | 0.461709 | 0.480406 | | 1 | 0.999631 | 0.907462 |
| Palm_Q9Z0P4_T153t_2 | -0.98 | -0.9 | -1.37 | -0.44 | -0.5 | -0.91 | -0.94 | -0.7 | -1.1 | -0.89 | -1.21 | -0.81 | -1.17 | -0.7 | -0.88 | -1.15 | 0.16 | -0.08 | -0.0525 | 0.866678 | 0.582924 | 0.785813 | | 1 | 0.999631 | 0.956432 |
| Palm_Q9Z0P4_T291t_1 | 0.13 | 0.2 | -0.63 | 0.56 | 1.65 | -0.08 | 0.45 | 0.74 | 1.51 | -0.55 | 0.59 | -0.33 | 1.39 | 0.17 | -0.47 | -0.48 | 0.625 | 0.24 | 0.0875 | 0.85778 | 0.170127 | 0.636449 | | 1 | 0.999631 | 0.952251 |
| Palm_Q9Z0P4_T319t_1 | -1.74 | -1.14 | -0.28 | -1.13 | -1.21 | -1.03 | -0.87 | -0.37 | -1.13 | -1.71 | -0.49 | -0.71 | 0.06 | -1.17 | -0.28 | -0.57 | 0.2025 | 0.0625 | 0.5825 | 0.180904 | 0.58625 | 0.878925 | | 1 | 0.999631 | 0.64376 |
| Palm2_Q8BR92_S225s_1 | 0.88 | 1.74 | -0.92 | 0.69 | 1.64 | 2.21 | 0.25 | 1.28 | 0.28 | 1.13 | -0.47 | 0.58 | -1.08 | 2.39 | -1.76 | 1.15 | 0.7475 | -0.2175 | -0.4225 | 0.659609 | 0.246877 | 0.713874 | | 1 | 0.999631 | 0.880252 |
| Palm2_Q8BR92_S355s_1 | 0.51 | 1.66 | 2.05 | 1.26 | 0.59 | 1.06 | 0.89 | 1.19 | 0.61 | 1.08 | 1.05 | 1.81 | 1.03 | 0.41 | 1.88 | 2.98 | -0.4375 | -0.2325 | 0.205 | 0.729468 | 0.256039 | 0.580191 | | 1 | 0.999631 | 0.905923 |
| Palm3_A2TJV2_S270s_1 | -0.35 | 0.64 | 0.05 | -1.4 | -0.22 | -0.81 | -0.97 | -0.15 | -0.21 | -0.2 | 0.08 | -1.22 | 0.98 | -0.17 | 0.03 | 0.65 | -0.2725 | -0.1225 | 0.6375 | 0.211488 | 0.55543 | 0.803385 | | 1 | 0.999631 | 0.665474 |
| Palm3_A2TJV2_S524s_1 | 0.77 | 3.2 | 2.52 | 0.57 | 3.13 | 2.7 | 2.31 | 1.56 | 3.99 | 0.49 | 2.71 | 2.91 | 4.47 | 1.1 | 1.82 | 0.28 | 0.66 | 0.76 | 0.1525 | 0.87328 | 0.323958 | 0.38417 | | 1 | 0.999631 | 0.959782 |
| Palm3_A2TJV2_S565s_1 | -1.25 | -0.73 | 0.12 | 0.47 | -0.27 | -0.19 | -0.14 | 0.59 | 0.17 | 0.13 | 0.05 | 0.93 | 0.64 | 0.05 | 0.56 | 0.96 | 0.345 | 0.6675 | 0.9 | 0.063433 | 0.431769 | 0.151634 | | 1 | 0.999631 | 0.512978 |
| Palm3_A2TJV2_S601s_1 | -1.67 | -1.64 | -0.74 | -1.63 | -1.58 | -1.41 | -0.31 | -1.69 | -0.31 | -1.98 | -1.09 | -0.95 | -0.28 | -2.6 | -1.01 | -2.19 | 0.1725 | 0.3375 | -0.1 | 0.853493 | 0.66481 | 0.426279 | | 1 | 0.999631 | 0.950701 |
| Palm3_A2TJV2_S620s_1 | -1.18 | 0.12 | -1.56 | -1.7 | -0.32 | -0.24 | -1.32 | -1.47 | -0.2 | -0.97 | -1.64 | -1.96 | 0.14 | -0.07 | -1.67 | -0.69 | 0.2425 | -0.1125 | 0.5075 | 0.360258 | 0.625991 | 0.832467 | | 1 | 0.999631 | 0.741182 |
| Palm3_A2TJV2_S87s_1 | -0.62 | 1.98 | 2.28 | 0.73 | 1.7 | 2.15 | 1.18 | 1.71 | 2.46 | 0.53 | 1.79 | 1.61 | 2.24 | 1.22 | 1.62 | 0.73 | 0.5925 | 0.505 | 0.36 | 0.587238 | 0.351917 | 0.470646 | | 1 | 0.999631 | 0.850015 |
| Palmd_Q3UVT7_S364s_1 | 2.38 | -0.18 | -0.66 | 1.07 | 0.42 | 1.41 | 1.89 | 0.55 | 0.91 | 1.8 | 0.48 | -0.96 | -2.13 | 0.13 | 0.63 | -0.82 | 0.415 | -0.095 | -1.2 | 0.154587 | 0.545927 | 0.903566 | | 1 | 0.999631 | 0.625677 |
| Palmd_Q3UVT7_S370s_1 | 0.08 | 0.47 | 0.43 | 0.87 | 0.9 | 0.97 | 1.21 | 1.02 | 0.66 | 0.77 | 1.26 | 0.01 | -1.02 | 0.67 | 0.53 | 1.24 | 0.5625 | 0.2125 | -0.1075 | 0.826786 | 0.059292 | 0.542682 | | 1 | 0.999631 | 0.940834 |
| Palmd_Q3UVT7_S385s_1 | 0.01 | 0.36 | 0.04 | 0.57 | 0.22 | -0.08 | 0.55 | 0.37 | 0.25 | 0.53 | 0.49 | 0.25 | -0.03 | 0.47 | 0.36 | 0.91 | 0.02 | 0.135 | 0.1825 | 0.570855 | 0.942065 | 0.616241 | | 1 | 0.999631 | 0.84338 |
| Pank2_V9GXP9_S43s_1 | 1.69 | 1.57 | 0.63 | 1.39 | 1.26 | 1.63 | 1.17 | 1.35 | 1.43 | 1.51 | 1.23 | 0.91 | 0.88 | 1.66 | 0.63 | 1.39 | 0.0325 | -0.05 | -0.18 | 0.632519 | 0.916967 | 0.878562 | | 1 | 0.999631 | 0.868271 |
| Pank2_V9GXP9_S44s_1 | 1.45 | 1.71 | 0.5 | 0.94 | 1.23 | 2.1 | 1.16 | 0.86 | 1.49 | 1.57 | 1.19 | 0.36 | -0.08 | 1.98 | 0.34 | 0.97 | 0.1875 | 0.0025 | -0.3475 | 0.492678 | 0.63124 | 0.994995 | | 1 | 0.999631 | 0.80955 |
| Pank4_Q80YV4_S63s_1 | -0.54 | 0.62 | 0.19 | -0.33 | 0.73 | 0.14 | 1.03 | -0.46 | 1.55 | -0.22 | 0.47 | 0.93 | 1.52 | -0.34 | -0.23 | -0.75 | 0.375 | 0.6975 | 0.065 | 0.902707 | 0.379637 | 0.143487 | | 1 | 0.999631 | 0.968183 |
| Papd5_E9QJT6_S97s_1 | 0.7 | 1.08 | -0.37 | -0.58 | 0.78 | 0 | -0.24 | -1.2 | 0.4 | 0.71 | -0.38 | -0.51 | 1.52 | -1.18 | 0.66 | -0.34 | -0.3725 | -0.1525 | -0.0425 | 0.947131 | 0.488431 | 0.751605 | | 1 | 0.999631 | 0.982895 |
| Papola_F6VTN8_S234s_1 | 0.97 | 0.23 | 1.71 | 1.51 | 0.25 | 0.46 | 1.19 | 1.17 | 0.91 | 1.82 | 1.77 | 1.77 | 1.08 | 0.59 | 1.63 | 1.43 | -0.3375 | 0.4625 | 0.0775 | 0.852519 | 0.418819 | 0.271668 | | 1 | 0.999631 | 0.950236 |
| Paqr9_Q6TCG2_S33s_1 | 2.17 | 1.33 | 1.23 | 0.87 | 0.42 | 0.99 | 1.41 | 0.47 | 0.02 | 1.3 | -0.29 | 0.1 | 0.3 | 0.02 | 0.45 | 0.1 | -0.5775 | -1.1175 | -1.1825 | 0.006545 | 0.14872 | 0.028065 | | 1 | 0.999631 | 0.351545 |
| Pard3_B7ZNY3_S1203s_1 | 0.78 | 0.82 | -0.3 | 0.7 | 0.67 | 0.87 | 0.43 | 0.51 | 1.1 | 0.16 | -0.32 | 0.3 | -0.66 | 0.91 | -0.02 | -0.48 | 0.12 | -0.19 | -0.5625 | 0.223545 | 0.715068 | 0.642709 | | 1 | 0.999631 | 0.67471 |
| Pard3_B7ZNY3_S383s_1 | -0.85 | -0.41 | -1.18 | -1.26 | -0.58 | -1 | -0.93 | -1.76 | -0.07 | -0.4 | -2.02 | -1.15 | -0.23 | 0.29 | -1.5 | -0.98 | -0.1425 | 0.015 | 0.32 | 0.475956 | 0.681833 | 0.974047 | | 1 | 0.999631 | 0.799882 |
| Pard3_B7ZNY3_S849s_1 | -0.22 | 0.58 | 1.38 | 0.79 | 1.27 | 0.73 | 0.79 | 1.48 | 1.47 | 0.24 | 1.34 | 1.86 | 0.88 | 0.41 | 1.45 | 0.72 | 0.435 | 0.595 | 0.2325 | 0.576656 | 0.277979 | 0.220483 | | 1 | 0.999631 | 0.846204 |
| Pard3_B7ZNY3_S854s_1 | 0.4 | -0.19 | 1.06 | 0.71 | -0.18 | 0.51 | -0.01 | 0.25 | 0.3 | 1.12 | -0.12 | 0.35 | 0.85 | -0.43 | 0.79 | 0.2 | -0.3525 | -0.0825 | -0.1425 | 0.732285 | 0.313908 | 0.831779 | | 1 | 0.999631 | 0.907231 |
| Pard3b_Q5SV54_S338s_1 | -0.1 | 0.2 | 0.31 | -0.27 | -0.61 | -0.72 | -0.43 | -0.16 | -0.6 | 0.49 | -1.11 | -1.08 | -0.78 | 0.43 | 1.01 | -0.02 | -0.515 | -0.61 | 0.125 | 0.76422 | 0.082461 | 0.15673 | | 1 | 0.999631 | 0.915775 |
| Pard3b_Q5SV54_S746s_1 | -0.55 | -0.01 | -0.16 | -0.1 | 0.08 | -1.06 | -0.29 | -0.73 | -0.17 | -0.38 | -0.68 | 0.65 | 0.09 | -0.27 | 0.55 | 0.88 | -0.295 | 0.06 | 0.5175 | 0.153618 | 0.371239 | 0.863404 | | 1 | 0.999631 | 0.625198 |
| Parg_G3X8U8_S297s_1 | 0.11 | 1.3 | 0.47 | -0.19 | 0.72 | 0.26 | 0.03 | -0.51 | 0.82 | 1.34 | 0.52 | 0.59 | 0.78 | 0.55 | 0.66 | 0.12 | -0.2975 | 0.395 | 0.105 | 0.785641 | 0.475189 | 0.323541 | | 1 | 0.999631 | 0.923063 |
| Parn_Q8VDG3_S556s_1 | 0.78 | 1.17 | 0.38 | 0.28 | 0.5 | 0.51 | 0.21 | 0.27 | 1.2 | 1.24 | 0.1 | 0.3 | 0.88 | 0.99 | 0.53 | 0.21 | -0.28 | 0.0575 | -1.6E-16 | 1 | 0.345822 | 0.880232 | | 1 | 0.999631 | 1 |
| Pbrm1_D3Z1N4_S1468s_1 | 1.05 | -0.59 | 0.04 | 0.67 | -1.71 | -1.02 | 0.96 | -0.34 | -1.01 | 0.76 | -0.26 | 0.16 | -0.01 | -1.31 | -0.82 | -0.79 | -0.82 | -0.38 | -1.025 | 0.042632 | 0.198161 | 0.448261 | | 1 | 0.999631 | 0.475443 |
| Pbx2_Q3UR63_S248s_1 | 1.3 | 0.17 | 1.77 | 1.38 | 0.21 | 1.06 | 1.89 | 0.77 | 1.51 | 0.49 | 1.77 | 1.78 | -0.11 | 1.24 | 1.59 | 1.32 | -0.1725 | 0.2325 | -0.145 | 0.769244 | 0.713483 | 0.608604 | | 1 | 0.999631 | 0.918358 |
| Pcbp1_P60335_S189s_1 | -0.68 | -1.11 | -0.45 | -0.5 | -0.66 | -1.22 | -0.58 | -0.25 | -1.04 | -1.08 | -0.34 | -0.12 | -0.32 | -0.71 | -0.66 | -0.19 | 0.0075 | 0.04 | 0.215 | 0.481121 | 0.980553 | 0.904951 | | 1 | 0.999631 | 0.801848 |
| Pcbp1_P60335_S190s_1 | -0.41 | 0.05 | -0.45 | -0.5 | -0.14 | -0.32 | -0.6 | -0.4 | -0.3 | -0.6 | -0.5 | -0.37 | -0.02 | -0.2 | -0.43 | 0.05 | -0.0375 | -0.115 | 0.1775 | 0.542291 | 0.886059 | 0.664118 | | 1 | 0.999631 | 0.832854 |
| Pcbp2_B2M1R7_S185s_1 | 0.24 | -0.43 | 1.39 | 0.86 | 0.32 | 0.31 | 0.86 | 0.35 | 0.13 | -0.32 | 0.17 | 1.13 | -0.73 | 1.17 | 0.8 | 0.5 | -0.055 | -0.2375 | -0.08 | 0.88087 | 0.894162 | 0.621565 | | 1 | 0.999631 | 0.963364 |
| Pcdh1_E0CXD4_S343s_1 | 0.85 | -0.56 | -1.25 | 1.32 | 1.17 | -0.01 | 0.51 | 0.78 | 0.05 | 0.05 | 0.42 | -0.26 | -2.15 | -0.44 | -0.29 | 0.94 | 0.5225 | -0.025 | -0.575 | 0.458927 | 0.384157 | 0.964771 | | 1 | 0.999631 | 0.79152 |
| Pcdh1_F7BJK1_S1010s_1 | 0.62 | -0.52 | -0.65 | -0.03 | 0.17 | -0.71 | 0.01 | -0.52 | -0.19 | 0.02 | -0.26 | -0.29 | -2.36 | -0.81 | -0.52 | 0.05 | -0.1175 | -0.035 | -0.765 | 0.185029 | 0.753268 | 0.918214 | | 1 | 0.999631 | 0.646072 |
| Pcdh1_F7BJK1_S1010s_2 | -0.79 | -1.06 | -0.64 | -0.99 | -1.23 | -1.11 | -0.81 | -0.93 | -0.93 | -0.21 | -0.74 | -0.32 | -0.06 | -0.69 | -0.37 | -0.71 | -0.15 | 0.32 | 0.4125 | 0.182463 | 0.55489 | 0.279025 | | 1 | 0.999631 | 0.643861 |
| Pcdh1_F7BJK1_S1012s_1 | -0.62 | -1.37 | -0.41 | -1.21 | -1.49 | -1.24 | -1.32 | -1.06 | -1.41 | -0.39 | -0.82 | -0.57 | -0.13 | -0.83 | -0.58 | -0.4 | -0.375 | 0.105 | 0.4175 | 0.236447 | 0.238054 | 0.768248 | | 1 | 0.999631 | 0.683301 |
| Pcdh1_F7BJK1_S1012s_2 | -0.79 | -1.06 | -0.64 | -0.99 | -1.23 | -1.11 | -0.81 | -0.93 | -0.93 | -0.21 | -0.74 | -0.32 | -0.06 | -0.69 | -0.37 | -0.71 | -0.15 | 0.32 | 0.4125 | 0.182463 | 0.55489 | 0.279025 | | 1 | 0.999631 | 0.643861 |
| Pcdh1_F7BJK1_S741s_1 | 0.4 | -0.91 | -0.01 | 0.7 | 0.87 | 1.58 | 0.78 | 1.69 | -0.8 | 0.04 | 2.05 | -0.43 | -2.1 | 0.84 | 1.41 | -0.16 | 1.185 | 0.17 | -0.0475 | 0.94907 | 0.017083 | 0.794524 | | 1 | 0.999631 | 0.983587 |
| Pcdh1_F7BJK1_S788s_1 | -0.13 | -0.14 | -0.09 | -0.16 | 0.33 | 0.17 | -0.38 | -0.13 | 0.17 | -0.24 | -0.1 | -0.32 | -0.63 | 0.17 | -0.39 | 0.11 | 0.1275 | 0.0075 | -0.055 | 0.854731 | 0.62802 | 0.976202 | | 1 | 0.999631 | 0.951562 |
| Pcdh1_F7BJK1_S801s_1 | -0.06 | 1.41 | 1.11 | 0.49 | 2.43 | 0.63 | 1.76 | 0.96 | 2.5 | -1.12 | 2.07 | 2.77 | 2.13 | 0 | 1.04 | 0.94 | 0.7075 | 0.8175 | 0.29 | 0.577501 | 0.171956 | 0.342609 | | 1 | 0.999631 | 0.84622 |
| Pcdh1_F7BJK1_S812s_1 | -0.19 | -0.44 | -1.35 | 0.03 | -0.02 | -0.07 | -0.23 | -0.64 | -0.52 | -0.32 | -0.07 | -0.58 | 0.17 | 0.23 | -1.05 | -0.36 | 0.2475 | 0.115 | 0.235 | 0.589072 | 0.495528 | 0.74882 | | 1 | 0.999631 | 0.851233 |
| Pcdh1_F7BJK1_S823s_1 | -0.71 | -0.98 | -0.55 | 0.03 | -0.66 | -0.86 | -0.51 | 0.13 | -0.64 | -0.99 | -0.05 | -0.06 | -0.79 | -0.66 | 0.33 | 0.45 | 0.0775 | 0.1175 | 0.385 | 0.356294 | 0.818973 | 0.739257 | | 1 | 0.999631 | 0.74038 |
| Pcdh1_F7BJK1_S823s_2 | 0.66 | -0.81 | -0.25 | 0.65 | 1.65 | -0.18 | 0.94 | 1.31 | -0.18 | -1.31 | 0.47 | -0.34 | -0.42 | -0.46 | 0.02 | -0.15 | 0.8675 | -0.4025 | -0.315 | 0.440148 | 0.109235 | 0.420845 | | 1 | 0.999631 | 0.781983 |
| Pcdh1_F7BJK1_S978s_1 | -2.61 | -1.38 | -0.59 | -0.41 | -0.79 | -1.09 | 0.18 | -0.23 | -1.01 | -1.5 | -0.33 | -0.43 | -1.07 | -1.53 | 0.39 | -0.18 | 0.765 | 0.43 | 0.65 | 0.295131 | 0.172809 | 0.427182 | | 1 | 0.999631 | 0.708769 |
| Pcdh1_F7BJK1_S979s_1 | -1.4 | 0.4 | -0.35 | -0.03 | 0.26 | 0.38 | -0.23 | 0.37 | 0.46 | -0.62 | -0.31 | 0.48 | 0.56 | 0.04 | -0.46 | 0.14 | 0.54 | 0.3475 | 0.415 | 0.357285 | 0.2082 | 0.457437 | | 1 | 0.999631 | 0.740933 |
| Pcdh10_Q925I8_S841s_1 | 0.41 | -1.08 | -1.59 | -0.03 | -0.14 | -0.9 | 0.02 | 0.13 | -0.53 | -1.62 | -0.53 | 0.33 | -0.58 | -0.2 | -0.02 | -0.02 | 0.35 | -0.015 | 0.3675 | 0.441476 | 0.478709 | 0.978713 | | 1 | 0.999631 | 0.781983 |
| Pcdh10_Q925I8_S898s_1 | 0.94 | 0.22 | -1.36 | -0.13 | 0.56 | -0.24 | -0.32 | -0.52 | -1.11 | -1.14 | -1.97 | -0.32 | -2.44 | 0.13 | -0.01 | -0.41 | -0.0475 | -1.0525 | -0.6 | 0.388679 | 0.924559 | 0.078101 | | 1 | 0.999631 | 0.757945 |
| Pcdh11x_Q2TJH7_S333s_1 | 0.37 | -0.17 | 0.33 | 0.25 | -0.26 | -0.49 | 0.17 | 0.3 | 0.33 | 1.07 | 0.32 | 0.81 | 0.24 | -0.27 | 0.37 | 0.04 | -0.265 | 0.4375 | -0.1 | 0.736776 | 0.374635 | 0.167063 | | 1 | 0.999631 | 0.907709 |
| Pcdh17_E9PXF0_S1024s_1 | 1.02 | -0.51 | -1.76 | -0.28 | 0.5 | 0.23 | -0.03 | 0.13 | 0.75 | -0.16 | 0.38 | -0.48 | -0.57 | -0.57 | -0.19 | -0.38 | 0.59 | 0.505 | -0.045 | 0.933465 | 0.286015 | 0.393202 | | 1 | 0.999631 | 0.978417 |
| Pcdh17_E9PXF0_S1036s_1 | -0.61 | -0.03 | -0.11 | -0.47 | -0.33 | -0.29 | -0.44 | 0.23 | -0.39 | -0.24 | 0.55 | 0.38 | -0.33 | -0.1 | 0.74 | 0.41 | 0.0975 | 0.38 | 0.485 | 0.178467 | 0.73097 | 0.25931 | | 1 | 0.999631 | 0.642903 |
| Pcdh17_E9PXF0_S1109s_1 | 0.74 | -0.2 | -0.6 | -0.04 | 0.26 | -0.64 | -0.18 | -0.23 | 0.13 | 0.02 | 0.13 | 0.22 | 0.04 | 0.28 | -0.15 | -0.34 | -0.1725 | 0.15 | -0.0175 | 0.961228 | 0.63347 | 0.654593 | | 1 | 0.999631 | 0.986594 |
| Pcdh17_E9PXF0_S1110s_1 | 0.88 | 0.04 | -0.66 | -0.68 | 0.51 | -0.47 | 0.06 | -1.98 | 0.05 | 0.41 | -1.35 | -0.83 | 0.29 | 0.24 | -0.66 | -0.67 | -0.365 | -0.325 | -0.095 | 0.834098 | 0.542262 | 0.531381 | | 1 | 0.999631 | 0.943897 |
| Pcdh18_Q8VHR0_S781s_2 | -0.61 | -1.34 | -0.95 | -0.5 | -1.37 | -0.97 | -0.3 | -0.2 | -0.48 | -0.07 | -0.38 | -0.03 | -0.82 | -0.6 | -1.06 | -1.19 | 0.14 | 0.61 | -0.0675 | 0.831325 | 0.698651 | 0.064239 | | 1 | 0.999631 | 0.942635 |
| Pcdh7_E9Q2S0_S1000s_2 | 0.22 | -1.1 | -1.12 | 0.34 | 0.2 | -0.3 | 0.01 | 0.79 | -0.79 | -0.39 | -0.28 | 0.06 | -1.75 | -0.21 | -0.18 | 0.53 | 0.59 | 0.065 | 0.0125 | 0.982678 | 0.20819 | 0.881332 | | 1 | 0.999631 | 0.994846 |
| Pcdh7_E9Q2S0_S1011s_1 | -1.23 | -1.8 | -1.08 | -1.21 | -1.76 | -1.82 | -1.33 | -1.17 | -1.79 | -1.36 | -1.31 | -0.92 | -1.06 | -1.43 | -0.96 | -0.76 | -0.19 | -0.015 | 0.2775 | 0.378528 | 0.522911 | 0.961174 | | 1 | 0.999631 | 0.75162 |
| Pcdh7_E9Q2S0_S1011s_2 | 0.22 | -1.1 | -1.26 | 0.23 | 0.2 | -0.3 | 0 | 0.53 | -0.79 | -0.39 | -0.35 | -0.23 | -1.75 | -0.21 | -0.33 | 0.39 | 0.585 | 0.0375 | 0.0025 | 0.996455 | 0.197988 | 0.929758 | | 1 | 0.999631 | 0.9984 |
| Pcdh7_E9Q2S0_S1235s_1 | 0.54 | -0.46 | -0.39 | 0.38 | 0.68 | 0.52 | 0.26 | 0.69 | 0.03 | -0.19 | 0.17 | 0.75 | -1.07 | 0.58 | 0.52 | 0.75 | 0.52 | 0.1725 | 0.1775 | 0.713396 | 0.131515 | 0.633844 | | 1 | 0.999631 | 0.900989 |
| Pcdh7_E9Q2S0_S974s_1 | -0.83 | -0.45 | -0.22 | -0.4 | -0.42 | -0.37 | 0.03 | -0.28 | -0.23 | -0.9 | -0.22 | -0.24 | -0.69 | -0.58 | -0.24 | -0.55 | 0.215 | 0.0775 | -0.04 | 0.888847 | 0.42288 | 0.791985 | | 1 | 0.999631 | 0.963505 |
| Pcdh7_E9Q2S0_S989s_1 | -1.87 | -0.83 | -1.25 | -2.52 | -0.85 | -0.78 | -1.49 | -2.34 | -0.7 | -1.46 | -0.95 | -1.11 | -0.01 | -1.74 | -1.13 | -2.7 | 0.2525 | 0.5625 | 0.2225 | 0.716701 | 0.607094 | 0.191338 | | 1 | 0.999631 | 0.903179 |
| Pcdh8_Q7TSK3_S798s_1 | -0.3 | -0.51 | -0.07 | -0.71 | -0.29 | -0.39 | -0.59 | -0.25 | -0.47 | 1.05 | -0.21 | -0.89 | -0.23 | 0.66 | 0.03 | 0.04 | 0.0175 | 0.2675 | 0.5225 | 0.124786 | 0.946504 | 0.543778 | | 1 | 0.999631 | 0.596063 |
| Pcdh8_Q7TSK3_S802s_1 | -0.68 | -1.74 | -1.56 | -2.27 | -2.98 | -2.51 | -1.59 | -2.28 | -1.79 | -0.57 | -2.17 | -0.56 | 0.3 | -1.26 | -0.99 | -0.68 | -0.7775 | 0.29 | 0.905 | 0.076179 | 0.096115 | 0.567608 | | 1 | 0.999631 | 0.542251 |
| Pcdh8_Q7TSK3_S904s_1 | 0.64 | -0.47 | 0.3 | 0.21 | 3.82 | 2.93 | 2.42 | 0.68 | 2.87 | 1 | 1.44 | 2.15 | 2.36 | 0.13 | 1.54 | 0.27 | 2.2925 | 1.695 | 0.905 | 0.119715 | 0.004497 | 0.004225 | | 1 | 0.999631 | 0.587662 |
| Pcdh9_F8VPK8_S1109s_1 | 0.74 | 1.83 | 0.05 | -0.35 | 0.86 | 1.43 | -0.63 | -0.69 | -0.6 | 0.72 | -0.4 | 0.4 | 1.5 | 0.74 | -0.63 | 0 | -0.325 | -0.5375 | -0.165 | 0.784874 | 0.614287 | 0.32713 | | 1 | 0.999631 | 0.922904 |
| Pcdh9_F8VPK8_S1156s_1 | 1.54 | -1.4 | -1.55 | 0.9 | 1.75 | -0.39 | 1.43 | 0.76 | 0 | -1.01 | 0.7 | -1.02 | -1.03 | -0.29 | -0.39 | -0.44 | 1.015 | -0.205 | -0.41 | 0.568749 | 0.225777 | 0.794408 | | 1 | 0.999631 | 0.843185 |
| Pcdh9_F8VPK8_S1231s_1 | -0.8 | -0.58 | -0.8 | -1.71 | -0.85 | -0.81 | -1.05 | -1.81 | 0.04 | -1.23 | -1.83 | -1.48 | -0.61 | -1.47 | -1.56 | -2.31 | -0.1575 | -0.1525 | -0.515 | 0.253552 | 0.666658 | 0.743141 | | 1 | 0.999631 | 0.688735 |
| Pcdh9_F8VPK8_S932s_1 | -0.83 | -0.53 | -0.61 | -0.69 | -0.55 | -0.39 | -0.74 | -0.59 | -0.26 | -0.52 | -0.5 | -0.16 | 0.27 | -0.26 | -0.36 | -0.31 | 0.0975 | 0.305 | 0.5 | 0.104609 | 0.685175 | 0.244158 | | 1 | 0.999631 | 0.568213 |
| Pcdha9_Q91Y11_S789s_1 | 0.38 | 0.63 | 0.11 | 0.77 | 0.25 | -0.38 | 0.57 | 0.43 | 0.86 | 0.61 | 0.26 | -0.06 | -1.95 | -0.03 | 0.59 | -0.14 | -0.255 | -0.055 | -0.855 | 0.131421 | 0.419385 | 0.860585 | | 1 | 0.999631 | 0.603456 |
| Pcdha9_Q91Y11_S922s_1 | 1.11 | 0.93 | -0.48 | 0.68 | 1.87 | 0.51 | 0.16 | 0.26 | 1.14 | 0.63 | -0.46 | 0.02 | 0.63 | 0.76 | 0.02 | 0.3 | 0.14 | -0.2275 | -0.1325 | 0.748369 | 0.780051 | 0.637459 | | 1 | 0.999631 | 0.911931 |
| Pcdhga12_Q91XY7_S136s_1 | -1.08 | -0.32 | -0.46 | 0.22 | -0.28 | -0.4 | 0.4 | -0.04 | -0.29 | -1.01 | 0.06 | -0.04 | -0.46 | 0.44 | 0.54 | 0.52 | 0.33 | 0.09 | 0.67 | 0.109914 | 0.356765 | 0.813654 | | 1 | 0.999631 | 0.573646 |
| Pcdhgc3_Q91XX1_S780s_1 | 0.96 | 1.03 | 0.45 | 0.68 | 0.65 | 0.19 | 0.29 | 0.87 | 1.47 | 1.25 | 0.72 | 1.25 | 0.54 | 0.32 | 0.48 | 1.22 | -0.28 | 0.3925 | -0.14 | 0.664576 | 0.33682 | 0.199768 | | 1 | 0.999631 | 0.882317 |
| Pcdhgc5_Q91XW9_S746s_1 | -0.7 | -1.41 | -0.11 | -0.73 | -1.09 | -0.39 | -0.86 | -0.58 | -0.34 | -0.22 | -0.5 | -0.35 | 0.56 | -0.07 | -0.09 | -0.31 | 0.0075 | 0.385 | 0.76 | 0.062043 | 0.98246 | 0.255823 | | 1 | 0.999631 | 0.509355 |
| Pcif1_P59114_S116s_1 | 0.9 | 2.45 | 0.58 | -0.13 | 1.38 | 0.75 | -0.05 | 0.25 | 1.4 | 1.84 | -0.08 | 0.11 | 0.5 | 1.53 | 0.36 | 0.44 | -0.3675 | -0.1325 | -0.2425 | 0.668589 | 0.524998 | 0.838276 | | 1 | 0.999631 | 0.883335 |
| Pclo_Q9QYX7_S1116s_1 | -0.66 | -0.04 | -1.61 | -1.43 | -0.5 | -0.36 | -1.44 | -2.84 | 0.17 | -0.06 | -1.79 | -0.89 | 0.5 | -0.21 | -1.67 | -1.57 | -0.35 | 0.2925 | 0.1975 | 0.737499 | 0.569172 | 0.587609 | | 1 | 0.999631 | 0.907787 |
| Pclo_Q9QYX7_S1292s_1 | -0.83 | -0.39 | -0.44 | -0.95 | -0.64 | -1.06 | -1.13 | -1.11 | -0.91 | -0.76 | -1.06 | -1.08 | -0.08 | -0.47 | -0.87 | -0.55 | -0.3325 | -0.3 | 0.16 | 0.607133 | 0.239502 | 0.28124 | | 1 | 0.999631 | 0.858527 |
| Pclo_Q9QYX7_S131s_1 | 1.59 | 1.02 | 0.66 | 2.28 | 1.95 | 2.21 | 1.06 | 2.1 | 1.29 | 0.46 | 1.32 | 0.73 | -0.15 | 1.93 | 1.43 | 1.15 | 0.4425 | -0.4375 | -0.2975 | 0.579459 | 0.318787 | 0.308605 | | 1 | 0.999631 | 0.84667 |
| Pclo_Q9QYX7_S1332s_1 | -0.54 | 0.34 | -0.41 | -0.1 | 0.5 | -0.16 | -0.46 | -0.36 | -0.53 | -0.15 | -0.23 | 0.13 | 0.99 | 0.08 | 0.12 | -0.19 | 0.0575 | -0.0175 | 0.4275 | 0.260772 | 0.862357 | 0.954606 | | 1 | 0.999631 | 0.69271 |
| Pclo_Q9QYX7_S1341s_1 | -1.43 | -0.6 | -0.51 | -0.99 | -0.85 | -0.82 | -1.06 | -0.42 | -1.05 | -1.12 | -0.73 | -0.45 | 0.03 | -0.34 | -0.84 | -0.61 | 0.095 | 0.045 | 0.4425 | 0.217006 | 0.757225 | 0.888229 | | 1 | 0.999631 | 0.669465 |
| Pclo_Q9QYX7_S1378s_1 | -0.03 | -0.71 | 0.03 | -0.01 | -0.94 | -0.45 | 0.05 | 0.21 | -0.33 | 0.18 | 0.01 | 0.38 | 0.47 | -0.77 | 0.21 | -0.03 | -0.1025 | 0.24 | 0.15 | 0.683081 | 0.767618 | 0.438283 | | 1 | 0.999631 | 0.889602 |
| Pclo_Q9QYX7_S1439s_1 | -0.57 | 0.41 | -0.62 | -0.3 | 0.11 | 0.14 | -0.74 | 0 | 0.16 | -0.38 | 0.07 | -0.14 | 0.42 | 0.58 | -0.14 | 0.1 | 0.1475 | 0.1975 | 0.51 | 0.163538 | 0.671883 | 0.545102 | | 1 | 0.999631 | 0.631937 |
| Pclo_Q9QYX7_S1444s_1 | -0.8 | -0.82 | -0.71 | -0.5 | -0.92 | -1.73 | -0.49 | -1.43 | -0.87 | -1.14 | -0.98 | -1.3 | -0.59 | -1.51 | -0.09 | -0.84 | -0.435 | -0.365 | -0.05 | 0.888459 | 0.204146 | 0.174516 | | 1 | 0.999631 | 0.963505 |
| Pclo_Q9QYX7_S1444s_2 | -0.92 | -1.21 | -0.46 | -2.34 | -2.58 | -2.03 | -1.78 | -1.5 | -2.45 | -1.02 | -1.38 | -2.95 | -0.38 | -1.25 | -0.74 | -1.08 | -0.74 | -0.7175 | 0.37 | 0.414307 | 0.122379 | 0.217719 | | 1 | 0.999631 | 0.768655 |
| Pclo_Q9QYX7_S1457s_1 | -3.82 | -3.52 | 0.09 | -2.52 | -4.18 | -3.06 | -1.35 | -1.71 | -3.41 | -1.99 | -0.25 | -1.54 | -0.29 | -2.2 | 0.09 | -2.44 | -0.1325 | 0.645 | 1.2325 | 0.211358 | 0.888975 | 0.504303 | | 1 | 0.999631 | 0.665474 |
| Pclo_Q9QYX7_S1457s_2 | -0.92 | -1.21 | -0.46 | -2.34 | -2.58 | -2.03 | -1.78 | -1.5 | -2.45 | -1.02 | -1.38 | -2.95 | -0.38 | -1.25 | -0.74 | -1.08 | -0.74 | -0.7175 | 0.37 | 0.414307 | 0.122379 | 0.217719 | | 1 | 0.999631 | 0.768655 |
| Pclo_Q9QYX7_S1484s_1 | -1.11 | 0.1 | -3.1 | -3.81 | -0.2 | -0.54 | -3.05 | -2.11 | 0.03 | -0.27 | -1.62 | -2.13 | 0.37 | 0.09 | -2.35 | -2.94 | 0.505 | 0.9825 | 0.7725 | 0.467592 | 0.60354 | 0.291641 | | 1 | 0.999631 | 0.795565 |
| Pclo_Q9QYX7_S1550s_1 | -0.04 | 0.29 | -2.17 | -1.84 | -0.4 | 0.65 | -1.66 | -2.32 | -0.06 | 0.23 | -2.14 | -2.13 | -0.11 | -0.6 | -3.09 | -2.9 | 0.0075 | -0.085 | -0.735 | 0.39996 | 0.992449 | 0.913934 | | 1 | 0.999631 | 0.762503 |
| Pclo_Q9QYX7_S1551s_1 | -0.72 | -3.44 | 0.15 | -0.24 | -1.84 | -2.38 | -0.27 | 0.02 | -2.18 | -1.6 | -0.39 | 0.01 | -2.65 | -3.04 | 0 | -0.27 | -0.055 | 0.0225 | -0.4275 | 0.66076 | 0.949504 | 0.978584 | | 1 | 0.999631 | 0.880315 |
| Pclo_Q9QYX7_S1553s_1 | -0.76 | 0.59 | 1.51 | 0.11 | -0.55 | -0.09 | 0.24 | 0.07 | -0.04 | -0.25 | 0.98 | 1.73 | -0.17 | -0.94 | 0.64 | 0.66 | -0.445 | 0.2425 | -0.315 | 0.578599 | 0.362524 | 0.688177 | | 1 | 0.999631 | 0.84622 |
| Pclo_Q9QYX7_S1563s_1 | -1.59 | 0.44 | -0.52 | -1.71 | 1.22 | 0.08 | -0.04 | -0.21 | 2.06 | -2.52 | 0.19 | 1.35 | 2.48 | -0.55 | -0.87 | -1.51 | 1.1075 | 1.115 | 0.7325 | 0.414219 | 0.06834 | 0.268981 | | 1 | 0.999631 | 0.768655 |
| Pclo_Q9QYX7_S1585s_1 | -0.46 | -1.05 | -0.77 | 0.27 | -0.11 | -1.15 | -0.57 | 0.05 | -1.02 | -0.81 | -0.2 | -0.46 | -1.43 | -0.72 | -0.39 | -0.39 | 0.0575 | -0.12 | -0.23 | 0.568318 | 0.884841 | 0.744188 | | 1 | 0.999631 | 0.843185 |
| Pclo_Q9QYX7_S1766s_1 | -0.76 | 0.03 | -0.8 | -2.05 | -0.15 | 0.09 | -1.89 | -1.63 | -0.85 | -0.41 | -1.27 | -1.3 | 0.44 | 0.17 | -1.43 | -0.9 | 0 | -0.0625 | 0.465 | 0.421461 | 1 | 0.892807 | | 1 | 0.999631 | 0.772817 |
| Pclo_Q9QYX7_S1772s_1 | -1.59 | -0.36 | -1.78 | -3.06 | -0.94 | -1.74 | -2.67 | -3 | -1.6 | -1.03 | -2.14 | -2.12 | -0.67 | -0.95 | -2.43 | -1.97 | -0.39 | -0.025 | 0.1925 | 0.758545 | 0.549879 | 0.964608 | | 1 | 0.999631 | 0.914449 |
| Pclo_Q9QYX7_S1829s_1 | -0.72 | -0.18 | -1.02 | -0.46 | 0.09 | -0.32 | -0.47 | -0.11 | -0.61 | -0.46 | -0.46 | -0.66 | -0.19 | -0.31 | -0.55 | -0.27 | 0.3925 | 0.0475 | 0.265 | 0.387668 | 0.196069 | 0.866156 | | 1 | 0.999631 | 0.757129 |
| Pclo_Q9QYX7_S212s_1 | -0.27 | -0.99 | -1.15 | -0.32 | 0.03 | 0.11 | -0.89 | -0.08 | -0.21 | -0.45 | -1.29 | -0.62 | -0.52 | 0.01 | -0.17 | -0.47 | 0.475 | 0.04 | 0.395 | 0.250546 | 0.19765 | 0.911032 | | 1 | 0.999631 | 0.686448 |
| Pclo_Q9QYX7_S2288s_1 | 1.03 | 0.6 | 0.01 | 0.96 | 1.12 | 1.42 | 0.62 | 1.67 | 0.31 | 0.02 | 0.93 | 0.08 | -0.87 | 1.09 | 0.75 | 0.66 | 0.5575 | -0.315 | -0.2425 | 0.615709 | 0.137757 | 0.379775 | | 1 | 0.999631 | 0.861203 |
| Pclo_Q9QYX7_S2495s_1 | -0.74 | -0.81 | -2.47 | -0.21 | -0.66 | -0.63 | -0.65 | 0.89 | -1.31 | -1.14 | -1.34 | -1.7 | -1.44 | -0.59 | -0.08 | -0.7 | 0.795 | -0.315 | 0.355 | 0.50809 | 0.183288 | 0.518593 | | 1 | 0.999631 | 0.816745 |
| Pclo_Q9QYX7_S258s_1 | 0.22 | -1.01 | 0.93 | 0.38 | -0.02 | 0.52 | 0.27 | 0.76 | 0.11 | 0.22 | 0.29 | 0.98 | 0.28 | 0.05 | 1.17 | 0.62 | 0.2525 | 0.27 | 0.4 | 0.401175 | 0.563144 | 0.549369 | | 1 | 0.999631 | 0.763221 |
| Pclo_Q9QYX7_S3358s_1 | 1.81 | 0.81 | 0.47 | 0.6 | 1.23 | 1.03 | 1.09 | 0.97 | 0.7 | 0.85 | 1.54 | 0.53 | 0.21 | 0.49 | 0.85 | 0.61 | 0.1575 | -0.0175 | -0.3825 | 0.318127 | 0.64745 | 0.964383 | | 1 | 0.999631 | 0.721765 |
| Pclo_Q9QYX7_S3514s_1 | -0.89 | -2.23 | -2.93 | -3.14 | -1.87 | -2.24 | -2.61 | -3.41 | -1.89 | -1.81 | -2.58 | -3.1 | -1.14 | -2.09 | -2.63 | -2.69 | -0.235 | -0.0475 | 0.16 | 0.780245 | 0.673031 | 0.931042 | | 1 | 0.999631 | 0.921416 |
| Pclo_Q9QYX7_S3545s_1 | 0.38 | -0.39 | -0.7 | 0.05 | 0.92 | 0.03 | -0.08 | 0.51 | -0.18 | -0.76 | 0.12 | -0.83 | -1.21 | -0.08 | -0.35 | 0.32 | 0.51 | -0.2475 | -0.165 | 0.693381 | 0.175171 | 0.500079 | | 1 | 0.999631 | 0.895102 |
| Pclo_Q9QYX7_S3561s_1 | 0.57 | 0.56 | -1.31 | 0.93 | 1.23 | 0.17 | 0.94 | 0.72 | 0.34 | -0.41 | -0.21 | -0.91 | 0.33 | 0.61 | -0.15 | -0.31 | 0.5775 | -0.485 | -0.0675 | 0.896571 | 0.278554 | 0.371946 | | 1 | 0.999631 | 0.966483 |
| Pclo_Q9QYX7_S3598s_1 | 1.49 | -1.23 | -1.08 | 0.69 | -0.02 | -0.95 | 0.27 | -0.37 | -1.2 | -0.59 | -0.72 | -1.88 | -2.46 | -1.52 | -1.07 | -1.04 | -0.235 | -1.065 | -1.49 | 0.046429 | 0.715416 | 0.128926 | | 1 | 0.999631 | 0.479949 |
| Pclo_Q9QYX7_S3608s_1 | -2.19 | -1.26 | -0.88 | -2.13 | -2.69 | -2.44 | -2.07 | -2.26 | -2.29 | -1.15 | -1.49 | -1.18 | -0.06 | -1.65 | -1.69 | -0.79 | -0.75 | 0.0875 | 0.5675 | 0.262266 | 0.065796 | 0.83572 | | 1 | 0.999631 | 0.69292 |
| Pclo_Q9QYX7_S3610s_1 | -1.63 | -1.24 | -0.69 | -2.23 | -2.63 | -2.21 | -2.65 | -2.55 | -2.19 | -0.87 | -1.84 | -1.03 | -0.08 | -1.57 | -1.58 | -0.95 | -1.0625 | -0.035 | 0.4025 | 0.400927 | 0.014391 | 0.93746 | | 1 | 0.999631 | 0.763209 |
| Pclo_Q9QYX7_S3610s_2 | -2.64 | -1.03 | -0.96 | -0.21 | -0.53 | -1.3 | -0.36 | -0.3 | -1.16 | -1.23 | -0.57 | -0.6 | -1.09 | -1.83 | -0.11 | 0.22 | 0.5875 | 0.32 | 0.5075 | 0.425595 | 0.275668 | 0.535009 | | 1 | 0.999631 | 0.774624 |
| Pclo_Q9QYX7_S3616s_1 | -0.44 | -0.51 | 0.73 | -0.16 | -0.95 | -0.2 | -0.14 | -0.38 | -0.61 | 0.11 | -0.69 | -0.61 | -0.06 | -0.02 | 0.36 | -0.08 | -0.3225 | -0.355 | 0.145 | 0.685087 | 0.383428 | 0.348362 | | 1 | 0.999631 | 0.890187 |
| Pclo_Q9QYX7_S3616s_2 | -3.35 | -1.78 | -0.96 | -0.21 | -1.31 | -1.49 | -0.36 | -0.3 | -2.35 | -2.11 | -0.57 | -0.6 | -1.55 | -2.39 | -0.11 | 0.22 | 0.71 | 0.1675 | 0.6175 | 0.443936 | 0.29735 | 0.818845 | | 1 | 0.999631 | 0.782877 |
| Pclo_Q9QYX7_S3763s_1 | -0.5 | 2.18 | 1.78 | 1.36 | 1.75 | 1.61 | 1.15 | 1.58 | 2.01 | -0.04 | 1.73 | 2.07 | 2.67 | 1.72 | 1.7 | 1.98 | 0.3175 | 0.2375 | 0.8125 | 0.184065 | 0.570845 | 0.731783 | | 1 | 0.999631 | 0.644602 |
| Pclo_Q9QYX7_S3777s_1 | 1.26 | 0.51 | 0.23 | -0.14 | -1.03 | 1.47 | -0.31 | 0.69 | 0.19 | 1.42 | 0.6 | -0.17 | -1.83 | 0.63 | 0.6 | 0.14 | -0.26 | 0.045 | -0.58 | 0.342815 | 0.649597 | 0.919484 | | 1 | 0.999631 | 0.73348 |
| Pclo_Q9QYX7_S4016s_1 | 0.23 | -1.52 | -0.72 | -0.06 | 0.26 | -1.06 | -0.34 | -0.8 | -1.23 | -1.11 | -0.8 | -1.14 | -0.87 | -1.55 | -1.48 | -0.86 | 0.0325 | -0.5525 | -0.6725 | 0.146949 | 0.944103 | 0.197284 | | 1 | 0.999631 | 0.618703 |
| Pclo_Q9QYX7_S4066s_1 | -1.08 | -2.08 | -1.02 | -1.48 | -1.45 | -1.9 | -0.9 | -0.57 | -2.37 | -1.85 | -0.97 | -1.06 | -2.3 | -2.25 | -1.34 | -1.07 | 0.21 | -0.1475 | -0.325 | 0.439891 | 0.593676 | 0.72486 | | 1 | 0.999631 | 0.781983 |
| Pclo_Q9QYX7_S4132s_1 | 0.5 | 0.61 | 0.3 | 0.33 | 0.46 | 0.33 | -0.25 | 0.49 | 0.2 | 0.36 | 0.5 | -0.03 | 0.54 | 0.3 | 0.21 | 0.83 | -0.1775 | -0.1775 | 0.035 | 0.902189 | 0.522589 | 0.500844 | | 1 | 0.999631 | 0.968183 |
| Pclo_Q9QYX7_S4325s_1 | -0.07 | 0.3 | 1.01 | 0.12 | 0.36 | 0.21 | 0.2 | 0.78 | 0.98 | -0.54 | 0.72 | 2.12 | 0.77 | -0.19 | 1.45 | 0.23 | 0.0475 | 0.48 | 0.225 | 0.605487 | 0.882003 | 0.393192 | | 1 | 0.999631 | 0.85817 |
| Pclo_Q9QYX7_S4544s_1 | 2.14 | -0.22 | -0.07 | 1.04 | 0.55 | 0.85 | 1.41 | 0.44 | -0.01 | -0.04 | -0.07 | -1 | -1.55 | -0.38 | -0.35 | -0.44 | 0.09 | -1.0025 | -1.4025 | 0.032185 | 0.868503 | 0.095934 | | 1 | 0.999631 | 0.442395 |
| Pclo_Q9QYX7_S4598s_1 | -0.43 | -2.04 | -1.35 | -1.48 | -1.64 | -1.65 | -1.25 | -1.39 | -1.97 | -0.89 | -2.49 | -1.69 | -1.18 | -2.06 | -2.59 | -1.96 | -0.1575 | -0.435 | -0.6225 | 0.182461 | 0.669691 | 0.355605 | | 1 | 0.999631 | 0.643861 |
| Pclo_Q9QYX7_S4706s_1 | -1 | -0.25 | -1.01 | -0.49 | -0.86 | -0.35 | -0.93 | -0.26 | -1.56 | -0.77 | -0.48 | -0.99 | -0.21 | -0.4 | -1.56 | -0.73 | 0.0875 | -0.2625 | -0.0375 | 0.922585 | 0.77856 | 0.449336 | | 1 | 0.999631 | 0.975032 |
| Pclo_Q9QYX7_S4814s_1 | 2.08 | -0.23 | -0.75 | 0.04 | -0.14 | 0.89 | 0.82 | -0.42 | -0.88 | 1.43 | -1.27 | -0.61 | -1.23 | -1.36 | -0.84 | -1.93 | 0.0025 | -0.6175 | -1.625 | 0.020883 | 0.996837 | 0.426379 | | 1 | 0.999631 | 0.399638 |
| Pclo_Q9QYX7_S5063s_1 | -0.29 | -2.07 | 0.26 | -0.78 | -0.99 | -0.25 | -0.19 | -0.7 | -0.57 | -0.28 | -0.41 | 0.71 | -0.01 | -0.18 | 0.78 | 0.75 | 0.1875 | 0.5825 | 1.055 | 0.067987 | 0.708263 | 0.292509 | | 1 | 0.999631 | 0.522149 |
| Pclo_Q9QYX7_S599s_1 | 0.94 | -0.07 | -2.22 | 0.81 | 1.66 | 0.17 | 1.19 | 0.12 | 0.36 | -0.3 | 0.03 | -0.83 | 0.23 | -0.13 | -1.06 | -1 | 0.92 | -0.05 | -0.355 | 0.617362 | 0.224619 | 0.94216 | | 1 | 0.999631 | 0.861376 |
| Pclo_Q9QYX7_S625s_1 | -0.74 | -1.07 | 0.29 | 0.05 | -1 | -1.01 | 0.12 | -0.19 | -0.55 | -0.45 | 0.69 | 0.82 | 0.03 | -0.84 | 0.85 | 0.88 | -0.1525 | 0.495 | 0.5975 | 0.247341 | 0.72034 | 0.305936 | | 1 | 0.999631 | 0.685751 |
| Pclo_Q9QYX7_S709s_1 | -0.28 | 0.19 | -0.3 | -0.04 | -0.14 | -0.02 | -0.43 | -0.48 | -0.21 | 0.18 | 0.13 | -0.08 | 0.37 | -0.15 | -0.61 | 0.15 | -0.16 | 0.1125 | 0.0475 | 0.882857 | 0.545917 | 0.672421 | | 1 | 0.999631 | 0.963505 |
| Pclo_Q9QYX7_S709s_3 | -0.15 | 0.2 | 0.75 | 0.65 | 0.79 | 0.09 | -0.04 | 1.44 | 0.41 | 0.32 | 0.59 | 1.7 | 0.31 | 0.93 | 0.88 | 1.21 | 0.2075 | 0.3925 | 0.47 | 0.191985 | 0.60962 | 0.334457 | | 1 | 0.999631 | 0.648674 |
| Pclo_Q9QYX7_S75s_1 | 1.01 | -0.52 | -0.6 | 0.15 | -0.06 | -0.01 | -0.04 | -0.28 | -0.53 | 0.34 | -0.26 | -0.72 | -0.71 | 0.08 | -0.19 | -0.24 | -0.1075 | -0.3025 | -0.275 | 0.51802 | 0.78205 | 0.493232 | | 1 | 0.999631 | 0.819449 |
| Pclo_Q9QYX7_S769s_1 | -1.7 | -4.88 | -5.06 | -3.55 | -3.93 | -5.46 | -2.9 | -6.41 | -4.43 | -3.1 | -4.36 | -5.5 | -4.54 | -5.05 | -5.94 | -8.46 | -0.8775 | -0.55 | -2.2 | 0.048654 | 0.36667 | 0.500012 | | 1 | 0.999631 | 0.484645 |
| Pclo_Q9QYX7_S813s_1 | 0.21 | 0.26 | -0.22 | 0.55 | -0.17 | 0.61 | 0.18 | 0.71 | 0.21 | 0.5 | 0.17 | 0.29 | -0.48 | 0.22 | 0.58 | 0.49 | 0.1325 | 0.0925 | 0.0025 | 0.99425 | 0.672171 | 0.738579 | | 1 | 0.999631 | 0.997705 |
| Pclo_Q9QYX7_S844s_1 | 1.42 | 0.57 | 0.65 | 1.93 | 2.07 | 1.09 | 0.98 | 1.41 | 1.04 | 0.11 | 0.92 | 1.8 | 0.44 | 0.73 | 1.43 | 1.18 | 0.245 | -0.175 | -0.1975 | 0.633156 | 0.551842 | 0.705428 | | 1 | 0.999631 | 0.868569 |
| Pclo_Q9QYX7_T1120t_1 | -0.85 | -0.97 | -0.47 | -0.35 | -0.84 | -0.85 | -0.71 | -0.23 | -0.99 | -0.65 | -0.69 | -0.62 | -0.03 | -0.76 | -0.46 | -0.35 | 0.0025 | -0.0775 | 0.26 | 0.40712 | 0.99299 | 0.778288 | | 1 | 0.999631 | 0.766996 |
| Pclo_Q9QYX7_T2998t_1 | -0.68 | 0.57 | -0.73 | -0.4 | 0.95 | 0.23 | -0.98 | -0.57 | 0.39 | -0.45 | -0.59 | -0.92 | -1.23 | 0.24 | -0.5 | -0.36 | 0.2175 | -0.0825 | -0.1525 | 0.725838 | 0.660831 | 0.84292 | | 1 | 0.999631 | 0.90568 |
| Pclo_Q9QYX7_T3376t_1 | -0.79 | 0.04 | -0.23 | -0.38 | 0 | -0.96 | -0.14 | -0.51 | -0.14 | -0.88 | 0.51 | -0.52 | -0.49 | -1.48 | -0.52 | -0.14 | -0.0625 | 0.0825 | -0.3175 | 0.406099 | 0.84631 | 0.824133 | | 1 | 0.999631 | 0.766128 |
| Pclo_Q9QYX7_T815t_1 | 0.18 | -0.26 | -1.18 | -0.7 | 0.16 | 0.09 | -0.63 | -1.1 | 0.3 | 0.62 | -0.65 | -0.45 | -0.3 | 0.66 | -2.3 | -1.42 | 0.12 | 0.445 | -0.35 | 0.586251 | 0.773803 | 0.30621 | | 1 | 0.999631 | 0.849723 |
| Pclo_Q9QYX7_T860t_1 | 1.7 | -0.61 | -0.86 | 0.57 | 1.14 | -0.42 | 0.71 | -0.33 | -0.2 | -0.84 | -0.34 | -1 | 0.02 | -0.15 | -1.15 | -0.34 | 0.075 | -0.795 | -0.605 | 0.317645 | 0.905402 | 0.185148 | | 1 | 0.999631 | 0.721152 |
| Pcm1_Q9R0L6_S1471s_1 | 1.84 | 1.1 | 1.84 | 0.43 | 2.04 | 0.66 | 3.76 | -0.69 | 3.19 | 0.52 | 1.41 | 2.34 | 2.21 | -1.09 | 1.57 | -0.67 | 0.14 | 0.5625 | -0.7975 | 0.315234 | 0.873096 | 0.366208 | | 1 | 0.999631 | 0.72069 |
| Pcm1_Q9R0L6_S65s_1 | -0.17 | 0.35 | -0.27 | -0.72 | -0.27 | -0.13 | -0.41 | -1.01 | 0.45 | 0.14 | -0.31 | -0.41 | 1.1 | 0.63 | -0.17 | -0.73 | -0.2525 | 0.17 | 0.41 | 0.381071 | 0.454563 | 0.621019 | | 1 | 0.999631 | 0.752619 |
| Pcm1_Q9R0L6_S996s_1 | 0.09 | 1.31 | 1.58 | 0.94 | 0.78 | 0.43 | 0.98 | 1.43 | 1.12 | 0.43 | 1.24 | 1.91 | 1.5 | 1.09 | 1.94 | 1.73 | -0.075 | 0.195 | 0.585 | 0.163489 | 0.848696 | 0.658716 | | 1 | 0.999631 | 0.631937 |
| Pcnp_D3YWA4_S80s_1 | 0.51 | 0.38 | 0.7 | 0.96 | 1.03 | 0.49 | 0.56 | 0.47 | 1.19 | 0.83 | 0.51 | 0.47 | 0.75 | 1.04 | 0.99 | 2.25 | -1.6E-16 | 0.1125 | 0.62 | 0.135264 | 1 | 0.70173 | | 1 | 0.999631 | 0.607096 |
| Pcnx_Q9QYC1_S574s_1 | 1.05 | 0.26 | 0.2 | -0.12 | -0.82 | 0.07 | 0.37 | -0.52 | 0.18 | 0.98 | -0.13 | -0.42 | -0.5 | -0.51 | -0.65 | -0.77 | -0.5725 | -0.195 | -0.955 | 0.014998 | 0.156085 | 0.629894 | | 1 | 0.999631 | 0.388832 |
| Pcnxl2_Q5DU28_S660s_1 | -0.74 | -0.98 | 1.64 | 1.31 | -0.75 | -0.02 | 1.09 | 0.33 | 0.08 | 0.61 | 0.26 | 0.52 | -0.34 | -0.56 | 1.16 | 0.53 | -0.145 | 0.06 | -0.11 | 0.874784 | 0.833763 | 0.923387 | | 1 | 0.999631 | 0.960237 |
| Pcnxl3_Q8VI59_S1955s_1 | -0.22 | -0.66 | -1.22 | -1.33 | -0.78 | -0.23 | -1.42 | -1.17 | -0.86 | -1.01 | -1.42 | -1.33 | 0.52 | 0.09 | -1.07 | -1.15 | -0.0425 | -0.2975 | 0.455 | 0.352694 | 0.910812 | 0.387634 | | 1 | 0.999631 | 0.73949 |

| | | | | | | | | | | | | | | | | | | | | | | | | | |
|---|---|---|---|---|---|---|---|---|---|---|---|---|---|---|---|---|---|---|---|---|---|---|---|---|---|
| Pcnxl3_Q8VI59_S287s_1 | 1.03 | -0.83 | -1.45 | -0.31 | -0.87 | -1.23 | -0.39 | -0.59 | -0.28 | -0.24 | -0.96 | -0.4 | -1.64 | -1.54 | -1.77 | -1.85 | -0.38 | -0.08 | -1.31 | 0.025563 | 0.470101 | 0.877613 | 1 | 0.999631 | 0.422114 |
| Pcnxl3_Q8VI59_S297s_1 | -0.19 | -0.55 | 0.94 | -0.93 | -1.5 | -0.77 | -0.9 | -0.53 | -0.08 | 0.4 | -0.17 | -0.57 | 0.19 | 0.28 | -0.09 | -0.52 | -0.7425 | 0.0775 | 0.1475 | 0.74006 | 0.116994 | 0.86134 | 1 | 0.999631 | 0.908664 |
| Pcnxl3_Q8VI59_S96s_1 | -0.94 | -0.69 | -0.45 | -1.46 | -0.96 | -1.46 | -1.56 | -1.21 | -1 | -1.17 | -1.01 | -0.68 | -0.12 | -1.86 | -0.9 | -0.49 | -0.4125 | -0.08 | 0.0425 | 0.922546 | 0.203706 | 0.795492 | 1 | 0.999631 | 0.975032 |
| Pcnxl3_Q8VI59_T199t_1 | -0.87 | -0.58 | 0.03 | -0.11 | 0.05 | -0.67 | -0.79 | 0.14 | -0.39 | -0.51 | -0.39 | 0.12 | 0.27 | -0.1 | -0.14 | 0.66 | 0.065 | 0.09 | 0.555 | 0.129146 | 0.851951 | 0.775334 | 1 | 0.999631 | 0.599711 |
| Pcnxl3_Q8VI59_T370t_1 | -1.47 | -0.86 | -1.65 | -2.3 | -0.28 | -1.59 | -0.29 | -3.22 | -0.51 | -1.74 | -0.76 | 0.56 | 1.17 | -1.66 | -2.26 | -2.8 | 0.225 | 0.9575 | 0.1825 | 0.822086 | 0.73867 | 0.092124 | 1 | 0.999631 | 0.938505 |
| Pcnxl3_Q8VI59_T97t_1 | 0.15 | -0.12 | -0.4 | -1.44 | -0.25 | -0.81 | -1.72 | -1.12 | -0.8 | -0.16 | -0.76 | -0.45 | 0.48 | -0.15 | -0.83 | -0.26 | -0.5225 | -0.09 | 0.2625 | 0.556069 | 0.261142 | 0.819363 | 1 | 0.999631 | 0.838305 |
| Pcp4l1_Q6W8Q3_S16s_1 | -0.29 | -1.45 | -0.23 | 1.52 | -0.29 | -1.37 | 0.02 | 0.25 | 0.63 | 0.77 | -0.05 | 0.53 | -1.86 | -0.97 | 0.88 | -2.19 | -0.235 | 0.5825 | -0.9225 | 0.266779 | 0.712349 | 0.331712 | 1 | 0.999631 | 0.695664 |
| Pcsk2_P21661_S526s_2 | -0.81 | -0.17 | -0.83 | -0.49 | -1.48 | -0.68 | -1.88 | 1.07 | -1.2 | -0.71 | -0.58 | -0.3 | -1.14 | -0.72 | 2.03 | 0.12 | -0.1675 | -0.1225 | 0.6475 | 0.329439 | 0.782552 | 0.694709 | 1 | 0.999631 | 0.726484 |
| Pcsk2_P21661_T525t_2 | -0.81 | -0.17 | -0.83 | -0.49 | -1.48 | -0.68 | -1.88 | 1.07 | -1.2 | -0.71 | -0.58 | -0.3 | -1.14 | -0.72 | 2.03 | 0.12 | -0.1675 | -0.1225 | 0.6475 | 0.329439 | 0.782552 | 0.694709 | 1 | 0.999631 | 0.726484 |
| Pcyt1a_P49586_S319s_1 | 2.86 | 0.07 | 2.55 | 1.14 | -0.56 | 0.87 | 1.39 | 0.65 | 0.15 | 2.35 | 0.48 | -0.21 | -0.34 | -0.39 | 0.42 | -0.34 | -1.0675 | -0.9625 | -1.8175 | 0.01341 | 0.141321 | 0.22535 | 1 | 0.999631 | 0.385092 |
| Pcyt1a_P49586_S331s_1 | 1.06 | 1.95 | 3.65 | 2.52 | 1.14 | 1.82 | 2.03 | 2.69 | 2.3 | 1.78 | 3.41 | 3.53 | 0.67 | 2.18 | 3.68 | 3.52 | -0.375 | 0.46 | 0.2175 | 0.779219 | 0.517645 | 0.467913 | 1 | 0.999631 | 0.921416 |
| Pcyt1a_P49586_S343s_1 | 1.01 | 1.7 | 1.56 | 1.39 | 1.34 | 0.89 | 0.99 | 1.27 | 1.85 | 1.58 | 1.6 | 1.7 | 1.42 | 2.05 | 1.57 | 1.91 | -0.2925 | 0.2675 | 0.3225 | 0.306733 | 0.298793 | 0.335616 | 1 | 0.999631 | 0.717667 |
| Pcyt1a_P49586_S343s_2 | 0.72 | 1.17 | 1.32 | 1.43 | 0.81 | 1.47 | 1.2 | 1.26 | 1.7 | 1.76 | 0.99 | 1.15 | 2.08 | 1.97 | 0.85 | 1.66 | 0.025 | 0.24 | 0.48 | 0.208261 | 0.929997 | 0.45097 | 1 | 0.999631 | 0.661649 |
| Pcyt1b_Q811Q9_S289s_1 | 0.92 | 0.79 | 1.26 | -0.14 | -0.31 | 0.61 | 0.24 | -0.22 | 0.91 | 1.96 | -0.07 | 0 | 0.37 | 0.17 | 0.19 | 0.42 | -0.6275 | -0.0075 | -0.42 | 0.256895 | 0.124021 | 0.988616 | 1 | 0.999631 | 0.691567 |
| Pcyt1b_Q811Q9_S289s_2 | 1.62 | 0.55 | 0.95 | 1.5 | 0.61 | 0.61 | 1.84 | 0.72 | 0.85 | 1.9 | 0.7 | 0.49 | -0.08 | -0.35 | 0.1 | 0.24 | -0.21 | -0.17 | -1.1775 | 0.006 | 0.598158 | 0.678796 | 1 | 0.999631 | 0.351545 |
| Pcyt1b_Q811Q9_S293s_1 | 0.03 | -0.15 | 0.71 | -0.16 | -0.72 | -0.84 | -0.3 | -0.25 | 0.04 | 0.74 | -0.26 | 0.58 | 0.26 | -1.43 | 0.13 | -0.26 | -0.635 | 0.1675 | -0.4325 | 0.337472 | 0.063406 | 0.634042 | 1 | 0.999631 | 0.730499 |
| Pcyt1b_Q811Q9_S293s_2 | 1.62 | 0.55 | 0.95 | 1.5 | 0.61 | 0.61 | 1.84 | 0.72 | 0.85 | 1.9 | 0.7 | 0.49 | -0.08 | -0.35 | 0.1 | 0.24 | -0.21 | -0.17 | -1.1775 | 0.006 | 0.598158 | 0.678796 | 1 | 0.999631 | 0.351545 |
| Pcyt1b_Q811Q9_S316s_1 | 1.13 | 0.66 | 2.48 | 1.02 | 0.01 | 1.23 | 0.69 | 0.49 | 1.01 | 2.8 | 0.5 | 1.06 | 1.17 | 0.83 | 1.36 | 0.88 | -0.7175 | 0.02 | -0.2625 | 0.543 | 0.138594 | 0.97277 | 1 | 0.999631 | 0.832881 |
| Pdap1_Q3UHX2_S176s_1 | 0.47 | 0.4 | -2.59 | 1.72 | 2.59 | 2.05 | 1.38 | 1.55 | 1.03 | -0.94 | 1.35 | -1.95 | -2.32 | 1.26 | -1.22 | -0.88 | 1.8925 | -0.1275 | -0.79 | 0.441544 | 0.043685 | 0.902562 | 1 | 0.999631 | 0.781983 |
| Pdap1_Q3UHX2_S19s_1 | -0.36 | -0.59 | 0.04 | -0.24 | -0.08 | 0.2 | -0.64 | 0.94 | -0.16 | -0.19 | -0.43 | -0.25 | 0.56 | -1.05 | -0.47 | 0.4 | 0.3925 | 0.03 | 0.1475 | 0.723622 | 0.305032 | 0.909445 | 1 | 0.999631 | 0.905674 |
| Pdap1_Q3UHX2_S48s_1 | 1.65 | 0.35 | 1.8 | 1 | 0.94 | 1.27 | 1.3 | 1.07 | 0.2 | 1.9 | 0.52 | 0.13 | 1.41 | -0.4 | 0.5 | -0.21 | -0.055 | -0.5125 | -0.875 | 0.106991 | 0.880015 | 0.321033 | 1 | 0.999631 | 0.568797 |
| Pdap1_Q3UHX2_S63s_1 | -4.4 | -5.6 | -1 | -2.98 | -3.52 | -3.9 | -2.61 | -1.57 | -7.71 | -3.57 | -1.65 | -1.3 | -4.58 | -6.12 | -1.22 | -1.27 | 0.595 | -0.0625 | 0.1975 | 0.881693 | 0.540678 | 0.966889 | 1 | 0.999631 | 0.963505 |
| Pdap1_Q3UHX2_T18t_1 | 0.3 | 0.41 | -1.78 | 2.98 | 1.2 | 0.79 | 1.3 | 2.62 | 0.41 | -0.73 | 2.7 | -1.13 | -2.22 | 1.53 | 0.62 | 0.63 | 1 | -0.165 | -0.3375 | 0.755276 | 0.287232 | 0.882433 | 1 | 0.999631 | 0.913663 |
| Pdcd4_Q61823_S71s_1 | -0.33 | 2.58 | 2.16 | 0.46 | 1.38 | 2.33 | 0.77 | 1.08 | 1.73 | 0.47 | 1.7 | 1.79 | 3.61 | 2.41 | 2.48 | 2.31 | 0.1725 | 0.205 | 1.485 | 0.048608 | 0.80038 | 0.763153 | 1 | 0.999631 | 0.484645 |
| Pdcd4_Q61823_S76s_1 | 0.65 | 0.64 | 0.56 | 1.73 | 0.88 | 1.29 | 1.06 | 1.75 | 1.75 | 0.64 | 1.55 | 1.43 | -0.32 | 0.67 | 1.08 | 1.23 | 0.35 | 0.4475 | -0.23 | 0.609242 | 0.343304 | 0.265843 | 1 | 0.999631 | 0.858851 |
| Pdcd5_D3Z7Q5_S198s_1 | -1.29 | 0.2 | -0.17 | 0.01 | -0.43 | 0.23 | -0.42 | 0.1 | -0.81 | -0.23 | -0.07 | 0.28 | -0.11 | -0.45 | -0.1 | -0.3 | 0.1825 | 0.105 | 0.0725 | 0.849004 | 0.638738 | 0.79881 | 1 | 0.999631 | 0.949525 |
| Pdcl_Q9DBX2_S296s_1 | -1.02 | -0.18 | 0.46 | -0.17 | -0.22 | -0.2 | -0.15 | 0.13 | -0.32 | -0.25 | 0.19 | 0.65 | -0.27 | -0.3 | 0.05 | 0.16 | 0.1175 | 0.295 | 0.1375 | 0.709936 | 0.734751 | 0.459883 | 1 | 0.999631 | 0.89951 |
| Pde10a_F8WHK3_T706t_1 | -2.29 | -4.02 | -4.67 | -2.86 | -2.79 | -3.28 | -2.39 | -2.87 | -4.19 | -3.06 | -2.46 | -3.43 | -3.07 | -3.33 | -5.45 | -2.78 | 0.6275 | 0.175 | -0.1975 | 0.783695 | 0.252327 | 0.768546 | 1 | 0.999631 | 0.922628 |
| Pde11a_P0C1Q2_S163s_1 | -1.39 | -1.01 | -1.61 | -0.56 | -2.07 | -1.34 | -1.1 | -2.22 | -1.06 | -0.28 | -2.19 | -1.22 | -1.01 | -1.51 | -1.11 | -1.28 | -0.54 | -0.045 | -0.085 | 0.796595 | 0.171112 | 0.919794 | 1 | 0.999631 | 0.929952 |
| Pde11a_P0C1Q2_T136t_1 | -0.14 | 1.28 | 1.3 | 0.92 | 0.73 | 1.38 | 1.29 | 0.92 | 1.27 | 0.25 | 1.4 | 0.88 | -0.06 | 0.85 | 0.5 | 0.66 | 0.24 | 0.11 | -0.3525 | 0.398673 | 0.535323 | 0.796556 | 1 | 0.999631 | 0.761726 |
| Pde1b_Q01065_S7s_1 | -0.65 | 0.11 | 0.12 | -0.39 | -0.36 | -0.72 | -0.81 | -0.59 | -0.37 | -0.59 | -0.2 | 0.11 | 0.52 | 0.43 | -0.04 | 0.06 | -0.4175 | -0.06 | 0.445 | 0.184358 | 0.169636 | 0.846258 | 1 | 0.999631 | 0.64478 |
| Pde1c_Q8CDV2_S440s_1 | 0.07 | -0.69 | -0.25 | 0.32 | 0.22 | -1.32 | 0.06 | 0.24 | 0.31 | 0.5 | -0.82 | 0.51 | 0.54 | 0.44 | 0.91 | 0.37 | -0.0625 | 0.2625 | 0.7025 | 0.053032 | 0.88364 | 0.515176 | 1 | 0.999631 | 0.489777 |
| Pde1c_Q8CDV2_S441s_1 | 0.07 | -0.69 | -0.25 | 0.32 | 0.22 | -1.32 | 0.06 | 0.24 | 0.31 | 0.5 | -0.82 | 0.51 | 0.54 | 0.44 | 0.91 | 0.37 | -0.0625 | 0.2625 | 0.7025 | 0.053032 | 0.88364 | 0.515176 | 1 | 0.999631 | 0.489777 |
| Pde2a_Q5PR72_S116s_1 | -0.1 | 1.4 | 1.52 | 0.59 | 1.09 | 1.15 | 0.02 | 1.05 | 1.94 | 1.6 | 0.29 | 1.44 | 2.17 | 1.72 | 2.63 | 1.5 | -0.025 | 0.465 | 1.1525 | 0.027077 | 0.955613 | 0.359518 | 1 | 0.999631 | 0.422893 |
| Pde2a_Q5PR72_S13s_1 | 1.37 | -2.21 | -2.84 | 0.15 | 0.75 | 0.54 | 1.44 | 0.34 | 0.46 | -0.72 | -1.33 | -0.57 | -2.37 | -1.3 | -0.16 | -2.78 | 1.65 | 0.3425 | -0.77 | 0.441316 | 0.086166 | 0.709679 | 1 | 0.999631 | 0.781983 |
| Pde2a_Q5PR72_S204s_1 | 0.43 | 1.03 | -3.15 | -3.5 | 0.27 | 0.46 | -2.33 | -2.87 | 0.12 | 0.72 | -3.23 | -3.72 | 0.95 | 1.2 | -3.24 | -2.76 | 0.18 | -0.23 | 0.335 | 0.811523 | 0.885067 | 0.868958 | 1 | 0.999631 | 0.936949 |
| Pde4a_O89084_S13s_1 | 1.37 | 1.57 | 0.38 | 1.71 | 1.91 | 0.71 | 1.18 | 1.11 | 1.25 | 0.34 | 1.49 | 0.8 | -1.06 | 0.87 | -0.79 | 1.95 | -0.03 | -0.2875 | -1.015 | 0.161931 | 0.939682 | 0.48275 | 1 | 0.999631 | 0.631442 |
| Pde4a_O89084_S147s_1 | 0.26 | -0.81 | 0.01 | -0.72 | -1.49 | -1.38 | -0.54 | -1.25 | -0.6 | 0.35 | -0.48 | 0.01 | -0.5 | -0.61 | -0.44 | -0.36 | -0.85 | 0.135 | -0.1625 | 0.632308 | 0.039031 | 0.717255 | 1 | 0.999631 | 0.868256 |
| Pde4a_O89084_S204s_1 | 1.14 | 0.67 | -1.77 | 0.83 | 1.09 | 1.54 | 0.77 | 2.03 | 1.27 | 1.3 | 1.14 | 1.41 | -1.48 | 1.1 | 1.8 | 0.77 | 1.14 | 1.0625 | 0.33 | 0.697215 | 0.102904 | 0.106806 | 1 | 0.999631 | 0.895731 |
| Pde4a_O89084_S333s_1 | -1.86 | -3.55 | -1.83 | -1.44 | -1.55 | -2.89 | -1.18 | -3.15 | -1.8 | -2.26 | -1.36 | -1.86 | -1.58 | -2.43 | -1.69 | -1.84 | -0.0225 | 0.35 | 0.285 | 0.563216 | 0.97053 | 0.475236 | 1 | 0.999631 | 0.841038 |
| Pde4a_O89084_S49s_1 | 2.62 | -0.2 | 0.13 | 2.22 | 0.85 | 0.61 | 2.38 | 0.01 | 0.41 | 1.31 | 0.74 | -0.42 | -2.05 | -0.88 | -0.32 | -1.12 | -0.23 | -0.6825 | -2.285 | 0.008726 | 0.764937 | 0.350324 | 1 | 0.999631 | 0.365244 |
| Pde4a_O89084_S786s_1 | -0.56 | -0.3 | -0.87 | -0.39 | -0.16 | -0.61 | -0.13 | -0.58 | -0.9 | -0.96 | -0.38 | 0.46 | -1.02 | -0.88 | -0.68 | -0.28 | 0.16 | 0.085 | -0.185 | 0.547191 | 0.559135 | 0.821274 | 1 | 0.999631 | 0.834277 |
| Pde4b_Q8VBU5_S139s_1 | -2.91 | 0.18 | -1.18 | -0.05 | 0.18 | -1.5 | -0.39 | -0.35 | -0.89 | -1.96 | -1.51 | -1.42 | -1.17 | -1.08 | -0.42 | -0.21 | 0.475 | -0.455 | 0.27 | 0.686121 | 0.501591 | 0.497754 | 1 | 0.999631 | 0.890692 |
| Pde4b_Q8VBU5_S275s_1 | -1.61 | -0.15 | 0.29 | -0.54 | 0.22 | -0.92 | -0.23 | -0.29 | -0.19 | -1.88 | 0.31 | 0.63 | 0.74 | -0.57 | 0.04 | -0.08 | 0.1975 | 0.22 | 0.535 | 0.276421 | 0.663863 | 0.725505 | 1 | 0.999631 | 0.700187 |
| Pde4b_Q8VBU5_S294s_1 | 2.64 | 0.64 | 0.96 | 1.61 | 1.65 | 1.05 | 1.96 | 1.81 | 0.35 | 0.4 | 2.48 | 1.37 | -1.58 | 2 | 1.09 | 1.3 | 0.155 | -0.3125 | -0.76 | 0.345751 | 0.738685 | 0.609302 | 1 | 0.999631 | 0.735045 |
| Pde4b_Q8VBU5_S304s_1 | 0.2 | 0.95 | 0.57 | -0.05 | 1.92 | 1.39 | -0.34 | 0.05 | 1.37 | 1.42 | 0.39 | 0.86 | 1.08 | 1.26 | 0.28 | 0.41 | 0.3375 | 0.5925 | 0.34 | 0.368084 | 0.532277 | 0.124036 | 1 | 0.999631 | 0.745968 |
| Pde4d_E9PVU7_S105s_1 | 1.3 | -0.2 | 0.51 | 1 | -0.57 | 0.45 | 1.11 | -0.3 | -0.06 | 1.49 | 0.39 | -0.06 | -0.46 | -0.92 | 0.42 | -0.32 | -0.48 | -0.2125 | -0.9725 | 0.046363 | 0.326621 | 0.655049 | 1 | 0.999631 | 0.479949 |
| Pde4d_E9PVU7_S137s_1 | -1.37 | 1.06 | -0.48 | 0.15 | 0.21 | 0.06 | -0.33 | 0.26 | 0.46 | 0.17 | 0.05 | 0.27 | -0.02 | 0.68 | -0.64 | 0.18 | 0.21 | 0.3975 | 0.21 | 0.699528 | 0.674006 | 0.429534 | 1 | 0.999631 | 0.896612 |
| Pde4d_E9PVU7_S14s_1 | 0.67 | -0.12 | -1.28 | -1.04 | 0.21 | 0.56 | -0.84 | -1.31 | 0.84 | 0.89 | -1.22 | -0.93 | 0.55 | 0.14 | -1.31 | -1.21 | 0.0975 | 0.3375 | -0.015 | 0.979819 | 0.864297 | 0.604533 | 1 | 0.999631 | 0.993709 |
| Pde4d_E9PVU7_S185s_1 | 0.53 | 1.39 | 0.59 | 0.8 | 1.09 | 1.53 | 0.99 | 0.72 | 1.19 | 1.12 | 1.67 | 0.64 | 1.73 | 1.23 | -0.05 | 1.94 | 0.255 | 0.3275 | 0.385 | 0.426016 | 0.42228 | 0.342057 | 1 | 0.999631 | 0.774781 |
| Pde4d_E9PVU7_S257s_1 | 1.84 | 0.07 | 0.12 | 0.95 | 0.8 | 1.18 | 1.61 | 0.28 | -0.4 | 0.08 | 1.74 | -0.83 | -0.84 | 0.49 | 1.15 | 2.06 | 0.2225 | -0.5975 | -0.03 | 0.963781 | 0.643314 | 0.356261 | 1 | 0.999631 | 0.987331 |
| Pde4d_E9PVU7_S343s_1 | -0.54 | 0.63 | 0.32 | -0.8 | 1.74 | 0.18 | 0.25 | 0.05 | 3.17 | -0.41 | -0.38 | 1.2 | 3.6 | -0.21 | -0.77 | -0.36 | 0.6525 | 0.9925 | 0.6625 | 0.479039 | 0.20574 | 0.23407 | 1 | 0.999631 | 0.801052 |
| Pde7b_E9Q0W7_S478s_1 | -0.39 | -0.34 | 0.5 | -0.1 | -0.53 | -0.39 | -0.62 | 0.39 | -0.84 | -1.57 | 0.13 | 0.87 | -0.82 | 0.34 | 0.42 | 0.27 | -0.205 | -0.27 | 0.135 | 0.72858 | 0.552654 | 0.615969 | 1 | 0.999631 | 0.905923 |
| Pde8a_O88502_S452s_1 | -0.46 | -0.6 | -0.39 | -0.72 | -0.5 | -0.79 | -1 | -1.85 | -0.95 | -1.22 | -0.74 | -0.28 | -0.24 | -1 | -0.29 | -0.31 | -0.4925 | -0.255 | 0.0825 | 0.783886 | 0.166226 | 0.395064 | 1 | 0.999631 | 0.922628 |
| Pde8b_E9PUL4_S304s_1 | -1.47 | -1.44 | -2.27 | -1.75 | -1.39 | -1.53 | -1.06 | -2.2 | -1.67 | -1.91 | -1.15 | -1.53 | -0.78 | -1.39 | -2.36 | -1.85 | 0.1875 | 0.1675 | 0.1375 | 0.736348 | 0.585795 | 0.59629 | 1 | 0.999631 | 0.907605 |
| Pde8b_E9PUL4_S381s_1 | -0.28 | -1.11 | -0.16 | -0.55 | -0.91 | -1.08 | -0.07 | -0.3 | -0.54 | -0.02 | 0.48 | -0.44 | -1.39 | -1.06 | -1.4 | 0.16 | -0.065 | 0.395 | -0.3975 | 0.368761 | 0.852649 | 0.277901 | 1 | 0.999631 | 0.745981 |
| Pde8b_E9PUL4_S385s_1 | 0.01 | -0.54 | -0.96 | -1.03 | -0.73 | -0.81 | -0.7 | -1.36 | -0.57 | -0.78 | -0.16 | -1.08 | -0.42 | -0.62 | -2.17 | -0.84 | -0.27 | -0.0175 | -0.3825 | 0.411692 | 0.418596 | 0.959859 | 1 | 0.999631 | 0.767937 |
| Pde8b_E9PUL4_S470s_1 | 0.33 | -0.76 | -1.47 | -0.15 | 0.3 | -0.77 | -0.53 | -0.72 | -0.12 | -0.73 | -0.68 | -0.8 | -0.43 | -0.71 | -0.79 | -0.31 | 0.0825 | -0.07 | -0.0475 | 0.90977 | 0.853672 | 0.868256 | 1 | 0.999631 | 0.970597 |
| Pde9a_F6ZAU1_S177s_1 | 0.85 | -0.87 | 1.53 | 0.28 | 0.23 | -1.58 | 1.8 | -0.06 | 0.14 | -0.01 | 0.99 | 0.93 | -0.63 | -2.24 | 2.12 | -1.93 | -0.35 | 0.065 | -1.1175 | 0.259556 | 0.644254 | 0.902871 | 1 | 0.999631 | 0.692624 |
| Pdha1_P35486_S232s_1 | -0.48 | -0.76 | -0.31 | 0 | 1.03 | -1.04 | 1.44 | -0.02 | -0.13 | -1.63 | 0.41 | -0.12 | -3.54 | -2.25 | -1.4 | -2.06 | 0.74 | 0.02 | -1.925 | 0.002036 | 0.187865 | 0.964993 | 1 | 0.999631 | 0.351308 |
| Pdha1_P35486_S293s_1 | -1.52 | -2.03 | -2.23 | -3.33 | -0.58 | -2.22 | 1.12 | -3.48 | -0.17 | -3.23 | -1.19 | -1.22 | 0.59 | -3.22 | -3.64 | -4.19 | 0.9875 | 0.825 | -0.3375 | 0.732613 | 0.299829 | 0.235608 | 1 | 0.999631 | 0.907231 |
| Pdha1_P35486_S293s_2 | -0.9 | -1.07 | -1.17 | -1.95 | 0.18 | -0.11 | 1.5 | -1.02 | -0.88 | -2.02 | -1 | -0.99 | -0.84 | -2.6 | -1.61 | -2.94 | 1.41 | 0.05 | -0.725 | 0.174338 | 0.022505 | 0.894647 | 1 | 0.999631 | 0.642116 |
| Pdha1_P35486_S295s_1 | 0.39 | -2.37 | -0.19 | -1.82 | -2.34 | -1.99 | 1.53 | -1.57 | -0.9 | -2.51 | -0.16 | -0.85 | -0.27 | -3.4 | -1.44 | -1.68 | -0.095 | -0.1075 | -0.7 | 0.391697 | 0.920421 | 0.882665 | 1 | 0.999631 | 0.758901 |
| Pdha1_P35486_S300s_1 | -0.88 | -0.1 | 0.04 | -0.6 | -0.06 | -1.15 | 1.4 | -0.33 | 0.07 | -1.08 | 0.3 | 0.73 | 0.24 | -1.19 | -0.3 | -1 | 0.35 | 0.39 | -0.1775 | 0.667161 | 0.513913 | 0.383607 | 1 | 0.999631 | 0.88264 |
| Pdha1_P35486_S300s_2 | -0.9 | -1.07 | -1.17 | -1.95 | 0.18 | -0.11 | 1.5 | -1.02 | -0.88 | -2.02 | -1 | -0.99 | -0.84 | -2.6 | -1.61 | -2.94 | 1.41 | 0.05 | -0.725 | 0.174338 | 0.022505 | 0.894647 | 1 | 0.999631 | 0.642116 |
| Pdha1_P35486_Y301y_1 | -1.43 | 0.12 | -0.03 | -0.63 | -0.17 | -1.71 | -0.06 | -0.12 | -0.21 | -1.29 | 0.8 | 1.54 | 1.35 | -1.56 | -1.06 | -1.56 | -0.0225 | 0.7025 | -0.215 | 0.756648 | 0.964004 | 0.287243 | 1 | 0.999631 | 0.914089 |
| Pdlim1_S4R1V0_S185s_1 | 0.58 | -1.62 | -0.46 | 0.39 | 0.72 | 0.7 | 0.65 | 0.3 | -0.13 | 1 | -0.44 | 0.06 | -1.04 | 0.53 | 0.11 | 0.85 | 0.87 | 0.4 | 0.39 | 0.515691 | 0.096845 | 0.470943 | 1 | 0.999631 | 0.819449 |
| Pdlim4_Q5SWV3_S111s_1 | -0.2 | 1.15 | -0.29 | -0.95 | 0.27 | 0.11 | 0.14 | -0.62 | 1.17 | 0.46 | 0.32 | -1.06 | 0.65 | 0.13 | -0.95 | -1.29 | 0.0475 | 0.295 | -0.2925 | 0.616641 | 0.918334 | 0.617329 | 1 | 0.999631 | 0.861203 |
| Pdlim4_Q5SWV3_S118s_1 | 0.51 | 0.92 | 0.74 | 0.45 | 0.25 | 0.79 | -0.09 | 1.06 | 0.8 | 0.8 | 0.18 | 1.06 | -0.22 | 1.31 | 0.66 | 0.97 | -0.1525 | 0.055 | 0.025 | 0.947661 | 0.64122 | 0.852651 | 1 | 0.999631 | 0.983122 |
| Pdlim5_D9J2Z9_S119s_1 | 1.62 | 1.14 | 1.33 | 1.62 | 0.4 | 1.82 | 2.54 | 0.33 | 0.96 | 2.19 | 1.28 | 2.01 | 0.49 | 0.63 | 1.26 | 0.6 | -0.155 | 0.1825 | -0.6825 | 0.047264 | 0.765024 | 0.607473 | 1 | 0.999631 | 0.480102 |
| Pdlim5_D9J2Z9_S223s_1 | -1.86 | -0.36 | -0.46 | -0.82 | -0.17 | -0.91 | -1.03 | -0.77 | -0.48 | -0.72 | -0.46 | -0.62 | 0.42 | -0.49 | -0.88 | -0.87 | 0.155 | 0.305 | 0.42 | 0.367212 | 0.697734 | 0.42281 | 1 | 0.999631 | 0.745797 |
| Pdlim5_D9J2Z9_S250s_1 | 0.82 | 0.54 | 0.62 | 0.98 | 1.61 | 0.4 | 0.41 | 0.41 | 1.97 | 0.93 | 1.13 | 1.53 | -0.29 | 1.19 | 1.31 | 1.01 | -0.0325 | 0.65 | 0.065 | 0.872422 | 0.925447 | 0.0621 | 1 | 0.999631 | 0.959468 |
| Pdpk1_F2Z3Z9_S117s_1 | 0.13 | 0.89 | 0.17 | -0.43 | -0.1 | 0.46 | -0.56 | -0.27 | -0.03 | 0.1 | -0.29 | -0.19 | 0.5 | 0.72 | 0.02 | -0.03 | -0.3075 | -0.2925 | 0.1125 | 0.761412 | 0.409564 | 0.390466 | 1 | 0.999631 | 0.914725 |
| Pds5b_F8WHU5_S1162s_2 | 0.64 | -0.47 | 0 | 0.72 | -0.95 | -0.51 | 0.13 | 0.2 | -1.99 | 0.57 | 0.15 | -1.07 | -0.34 | -1.11 | -0.16 | 0.11 | -0.505 | -0.8075 | -0.5975 | 0.16303 | 0.223826 | 0.193658 | 1 | 0.999631 | 0.631442 |
| Pds5b_F8WHU5_S1166s_1 | 0.3 | -0.38 | 0.46 | 0.03 | -0.18 | -0.08 | -0.01 | -0.83 | 0.3 | 0.19 | -0.52 | 0.02 | 0.01 | 0.15 | -0.11 | 0.18 | -0.3775 | -0.105 | -0.045 | 0.881001 | 0.248176 | 0.742542 | 1 | 0.999631 | 0.963364 |
| Pds5b_F8WHU5_S1166s_2 | 0.64 | -0.47 | 0 | 0.72 | -0.95 | -0.51 | 0.13 | 0.2 | -1.99 | 0.57 | 0.15 | -1.07 | -0.34 | -1.11 | -0.16 | 0.11 | -0.505 | -0.8075 | -0.5975 | 0.16303 | 0.223826 | 0.193658 | 1 | 0.999631 | 0.631442 |
| Pds5b_F8WHU5_S1359s_1 | 0.36 | 1.65 | 0.37 | 0.34 | 1.34 | 1.63 | -0.24 | 0.12 | 1.79 | 1.8 | 0.13 | -0.17 | 2.88 | 3.08 | 0.24 | 0.16 | 0.0325 | 0.2075 | 0.91 | 0.248407 | 0.950051 | 0.716285 | 1 | 0.999631 | 0.685751 |
| Pds5b_F8WHU5_S1384s_1 | 0.55 | 0.12 | 1.3 | 0.91 | 0.34 | -0.09 | 0.88 | 0.32 | -0.24 | 0.56 | 1.23 | 0.93 | -0.94 | -0.14 | 1.16 | 1.21 | -0.3575 | -0.1 | -0.3975 | 0.470287 | 0.320672 | 0.808498 | 1 | 0.999631 | 0.79732 |
| Pds5b_F8WHU5_S1386s_1 | 1.18 | 1.18 | 0.33 | 0.4 | 1.02 | 0.98 | 0.01 | 0.6 | 1.49 | 2.39 | -0.16 | -0.15 | 1.07 | 0.87 | -0.1 | -0.64 | -0.12 | 0.12 | -0.4725 | 0.319357 | 0.737643 | 0.844675 | 1 | 0.999631 | 0.722792 |
| Pds5b_F8WHU5_T1255t_1 | -1.45 | 1.03 | 0.87 | -0.37 | 1.12 | 0.74 | 0.05 | 0.22 | 0.79 | -0.71 | 0.62 | 1.31 | 1.1 | 0.49 | 0.55 | 0.58 | 0.5125 | 0.4825 | 0.66 | 0.251817 | 0.382242 | 0.464657 | 1 | 0.999631 | 0.68769 |
| Pdxdc1_Q99K01_T687t_1 | 0.38 | 1.45 | 0.37 | 0.96 | 1.73 | 1.31 | 0.6 | 1.28 | 1.24 | 0.42 | 1.26 | 1.21 | -0.39 | 1.79 | 1.19 | 1.15 | 0.44 | 0.2425 | 0.145 | 0.77544 | 0.250249 | 0.50835 | 1 | 0.999631 | 0.920998 |
| Pdzd2_E9Q1M1_S1061s_1 | -0.46 | -0.33 | -1.1 | -0.91 | -0.48 | 0 | -0.69 | -1.71 | 0.08 | -0.74 | -0.7 | -0.35 | -0.11 | -0.4 | -0.57 | 0.12 | -0.02 | 0.2725 | 0.46 | 0.173183 | 0.960637 | 0.406189 | 1 | 0.999631 | 0.642116 |
| Pdzd2_E9Q1M1_S323s_1 | 0.06 | 0.29 | 0.15 | -0.02 | -0.34 | -0.13 | -0.1 | -0.64 | 0.13 | -0.06 | 0.32 | 0.7 | 0.2 | -0.09 | -0.3 | 0.72 | -0.4225 | 0.1525 | 0.0125 | 0.968534 | 0.121523 | 0.584414 | 1 | 0.999631 | 0.988771 |
| Pdzd2_E9Q1M1_S677s_1 | 0.18 | 0.18 | -0.64 | -0.04 | -0.33 | -0.61 | -1.06 | 0.1 | 0.26 | -0.71 | 0.1 | -0.38 | -1.74 | 0.19 | -0.71 | 0.5 | -0.395 | -0.1025 | -0.36 | 0.487917 | 0.267443 | 0.763615 | 1 | 0.999631 | 0.806103 |
| Pdzd2_E9Q1M1_S947s_1 | -0.09 | 0.48 | 0.16 | 0.79 | 0.05 | -0.03 | 0.18 | 0.62 | 0.39 | 0.3 | 1.2 | 1.04 | 0.51 | -0.03 | 0.86 | 1.46 | -0.13 | 0.3975 | 0.365 | 0.365103 | 0.667945 | 0.26128 | 1 | 0.999631 | 0.745324 |
| Pdzd4_Q9QY39_S278s_1 | -0.36 | -0.77 | 1.36 | 0.64 | -0.54 | -0.67 | 0.12 | 0.5 | -1.09 | -1.62 | 0.86 | 1.44 | 0.13 | -0.48 | -0.91 | -0.1 | -0.365 | -0.32 | -0.5575 | 0.287576 | 0.486009 | 0.682003 | 1 | 0.999631 | 0.705972 |
| Pdzd8_B9EJ80_S515s_1 | 0.69 | 1.03 | 0.87 | 1 | 0.69 | 1.06 | 0.94 | 1.31 | 0.91 | 1.39 | 1.34 | 1.62 | 0.92 | 1.12 | 1.34 | 1.78 | 0.1025 | 0.4175 | 0.3925 | 0.214068 | 0.692462 | 0.149896 | 1 | 0.999631 | 0.667209 |
| Pdzd8_B9EJ80_S532s_1 | 1.02 | -0.73 | -0.67 | 0.97 | 1.17 | 0.19 | 0.49 | 0.64 | 0.14 | -0.31 | -0.23 | 0.36 | -1.6 | 0.82 | 0.22 | 0.05 | 0.475 | -0.1575 | -0.275 | 0.669267 | 0.351961 | 0.748642 | 1 | 0.999631 | 0.883599 |
| Pdzrn4_E9Q5P6_S520s_1 | 0.54 | -1.09 | -0.87 | -1.22 | -1.02 | -0.68 | -0.24 | -1.05 | -1.39 | 0.11 | -1.55 | -0.36 | -0.5 | -0.58 | -0.75 | -1.4 | -0.0875 | -0.1375 | -0.1475 | 0.745022 | 0.841417 | 0.796651 | 1 | 0.999631 | 0.910583 |
| Pea15_Q62048_S94s_1 | -0.15 | -0.11 | 0.33 | -0.29 | -0.11 | 0.07 | -0.29 | -0.19 | 0.22 | 0.33 | 0.37 | 0.15 | 0.37 | 0.56 | 0.27 | -0.04 | -0.075 | 0.3225 | 0.345 | 0.259583 | 0.773129 | 0.239251 | 1 | 0.999631 | 0.692624 |
| Peak1_Q69Z38_S1363s_1 | -0.26 | 1.58 | 1.42 | 1 | 1.38 | 1.34 | 0.49 | 1.5 | 1.25 | 0.11 | 1.56 | 1.01 | 0.37 | 0.78 | 1.2 | 0.65 | 0.2425 | 0.0475 | -0.185 | 0.682102 | 0.599096 | 0.923516 | 1 | 0.999631 | 0.889471 |
| Peak1_Q69Z38_S282s_1 | -0.39 | 0.04 | -0.13 | -0.88 | -0.3 | -0.21 | -1.14 | -0.41 | -0.26 | 0.17 | -0.34 | -0.54 | 0.43 | 0.23 | -0.26 | -0.53 | -0.175 | 0.0975 | 0.3075 | 0.392912 | 0.600776 | 0.756828 | 1 | 0.999631 | 0.758901 |
| Peak1_Q69Z38_T1141t_1 | -0.49 | -0.42 | -0.42 | -0.99 | -0.18 | -0.67 | -0.81 | -0.18 | -0.31 | -0.83 | -0.6 | -0.45 | -0.24 | -0.2 | -0.51 | -0.49 | 0.12 | 0.0325 | 0.22 | 0.448795 | 0.678049 | 0.90669 | 1 | 0.999631 | 0.786796 |
| Peak1_Q69Z38_T851t_1 | -0.8 | -1.15 | -1.02 | -0.12 | -1.7 | -1.24 | -1.56 | -0.59 | -2.29 | -0.71 | -0.72 | -0.58 | -1.71 | -0.63 | -0.56 | -1.58 | -0.5 | -0.3025 | -0.3475 | 0.398216 | 0.186963 | 0.511583 | 1 | 0.999631 | 0.761726 |
| Pebp1_P70296_S113s_1 | -0.71 | 0.91 | 0.56 | -0.19 | -0.16 | -0.27 | 0.04 | -0.33 | 0.14 | -0.6 | -0.52 | 0.19 | -0.18 | 0.56 | -1.03 | 0.17 | -0.3225 | -0.34 | -0.2625 | 0.589487 | 0.411196 | 0.429936 | 1 | 0.999631 | 0.851414 |
| Pebp1_P70296_S185s_1 | -1.66 | 0.37 | -0.17 | -0.85 | 0.44 | -0.3 | -0.74 | -0.72 | 0.41 | -0.39 | -0.15 | 0.25 | 0.63 | -0.37 | -0.55 | -0.62 | 0.2475 | 0.6075 | 0.35 | 0.491859 | 0.614842 | 0.208982 | 1 | 0.999631 | 0.809348 |
| Pebp1_P70296_S52s_1 | -0.74 | 0.61 | 0.2 | -0.41 | 0.23 | 0.19 | -0.75 | 0.16 | 0.1 | 0.06 | -0.09 | 1.11 | 0.76 | 0.85 | 0.23 | 0.37 | 0.0425 | 0.38 | 0.6375 | 0.114326 | 0.913583 | 0.370115 | 1 | 0.999631 | 0.579001 |
| Pebp1_P70296_T44t_1 | -0.78 | 0.35 | 1.65 | 1.05 | -0.06 | -0.73 | 1.09 | 2.06 | -0.37 | -0.76 | 1.62 | 2.16 | 1.04 | -0.22 | 2.02 | 3.44 | 0.0225 | 0.095 | 1.0025 | 0.235309 | 0.974829 | 0.903408 | 1 | 0.999631 | 0.683301 |
| Peg10_Q7TN75_S30s_1 | 0.09 | 0.85 | 1.65 | 0.62 | 0.74 | 0.73 | 1.01 | 0.87 | 0.91 | 0.66 | 0.6 | 0.97 | 1.57 | 0.86 | 0.94 | -0.01 | 0.035 | -0.0175 | 0.0375 | 0.934324 | 0.921742 | 0.961875 | 1 | 0.999631 | 0.978794 |
| Pelo_Q80X73_S55s_1 | 1.12 | 1.41 | 0.89 | 1.19 | 0.06 | 1.07 | 0.96 | 1.47 | 1.02 | 1.66 | 0.69 | 1.19 | -1.87 | 0.28 | 1.87 | -0.96 | -0.2625 | -0.0125 | -1.3225 | 0.091175 | 0.456421 | 0.967026 | 1 | 0.999631 | 0.557085 |
| Penk_P22005_S252s_1 | -0.26 | -0.37 | -1.45 | -0.84 | 0.3 | -0.7 | -1.42 | -0.72 | 0.22 | -0.34 | -1.2 | -0.38 | 0.81 | 0.12 | 0.12 | -0.31 | 0.095 | 0.305 | 0.915 | 0.037344 | 0.827346 | 0.460953 | 1 | 0.999631 | 0.458065 |
| Pex1_Q5BL07_S1142s_1 | 1.15 | 0.04 | 1.01 | 0.26 | 0.18 | 0.41 | 1.26 | -0.25 | 0.1 | 0.78 | 0.21 | 0.09 | -1.26 | 0.1 | 0.51 | 0.08 | -0.215 | -0.32 | -0.7575 | 0.127326 | 0.608144 | 0.377901 | 1 | 0.999631 | 0.59788 |
| Pex1_Q5BL07_S1172s_1 | -0.99 | -0.09 | -1.25 | -2.18 | 0.41 | -0.73 | -1.12 | -1.39 | -0.05 | -1.27 | -1.65 | -0.6 | -0.12 | -0.02 | -0.97 | -0.54 | 0.42 | 0.235 | 0.715 | 0.152274 | 0.443134 | 0.654629 | 1 | 0.999631 | 0.625198 |
| Pex14_Q9R0A0_S370s_1 | 1.44 | -1.1 | 0.14 | -0.12 | -1.02 | -0.67 | 0.8 | -0.45 | -1.54 | 0.45 | -0.18 | -1.94 | -2.19 | -3.31 | -1.17 | -2.32 | -0.425 | -0.8925 | -2.3375 | 0.003479 | 0.480213 | 0.21324 | 1 | 0.999631 | 0.351308 |
| Pex14_Q9R0A0_T48t_1 | -2.61 | -1.01 | -1.11 | -1.21 | -2.27 | -2.14 | -1.26 | -1.75 | -1.22 | -2.22 | -0.43 | -2.62 | -1.12 | -1.08 | -2.52 | -2.78 | -0.37 | -0.1375 | -0.39 | 0.482002 | 0.400499 | 0.810116 | 1 | 0.999631 | 0.802327 |
| Pex16_Q91XC9_S181s_1 | -0.5 | -0.19 | -0.37 | -0.74 | -1.48 | -0.23 | -0.04 | -0.32 | -1.4 | -0.32 | 0.03 | -0.85 | -0.15 | -0.66 | -0.43 | -0.78 | -0.0675 | -0.185 | -0.055 | 0.851787 | 0.853842 | 0.613778 | 1 | 0.999631 | 0.950099 |
| Pex19_Q8VCI5_S35s_1 | -0.12 | 0.08 | -0.48 | -0.75 | -0.4 | 0.42 | -0.82 | -0.46 | -0.41 | 0.77 | -0.45 | -0.08 | 0.95 | 0.5 | 0.49 | 0.44 | 0.0025 | 0.275 | 0.9125 | 0.014137 | 0.994296 | 0.460776 | 1 | 0.999631 | 0.385092 |
| Pex5l_F8SLQ1_S240s_1 | 0.61 | 0.81 | -0.36 | -0.26 | 0.38 | 0.39 | -0.04 | 0.13 | 0.52 | 0.63 | 0.25 | 0.33 | -0.1 | 0.88 | 1.46 | 0.72 | 0.015 | 0.2325 | 0.54 | 0.239243 | 0.96547 | 0.511094 | 1 | 0.999631 | 0.683719 |
| Pex5l_F8SLQ1_S277s_1 | 0.24 | 0.68 | -0.46 | 1.62 | 1.49 | 1.23 | 0.78 | 1.98 | 1.39 | -0.14 | 0.74 | -0.01 | -1.43 | 1.48 | 0.81 | 0.71 | 0.85 | -0.025 | -0.1275 | 0.851801 | 0.099721 | 0.962084 | 1 | 0.999631 | 0.950099 |
| Pex5l_F8SLQ1_S285s_1 | -1.01 | 0.99 | 0.42 | -0.77 | 2.34 | 0.29 | 2.34 | -0.35 | 2.68 | 2.01 | 0.74 | 2.6 | 3.44 | 1.6 | -1.07 | 0.02 | 1.2475 | 2.1 | 1.09 | 0.261559 | 0.11774 | 0.005568 | 1 | 0.999631 | 0.69271 |
| Pex5l_F8SLQ1_S461s_1 | 0.86 | 0.13 | 0.44 | 0.24 | -0.45 | 0.74 | 0.47 | -0.29 | 0.81 | 1.35 | 0.14 | 0.03 | 0.46 | -0.01 | -0.03 | -0.01 | -0.3 | 0.165 | -0.315 | 0.311656 | 0.408569 | 0.660198 | 1 | 0.999631 | 0.71924 |
| Pex5l_F8SLQ1_S471s_1 | -0.4 | -0.27 | 1.26 | 0.94 | -0.42 | -0.19 | 0.18 | 1.27 | -0.24 | -0.12 | 1.3 | 1.68 | 0.49 | 0.35 | 1.04 | 1.91 | -0.1725 | 0.2725 | 0.565 | 0.291544 | 0.741667 | 0.645474 | 1 | 0.999631 | 0.706164 |
| Pfkfb2_B2Z892_S447s_1 | 0.52 | 0.65 | 1.27 | 0.56 | 1.37 | 0.56 | -0.12 | 1.05 | 1.09 | 1.06 | 1.52 | 1.53 | 0.33 | 1.19 | 1.89 | 1.37 | -0.035 | 0.55 | 0.445 | 0.276211 | 0.926863 | 0.089084 | 1 | 0.999631 | 0.700187 |
| Pfkfb2_B2Z892_S464s_1 | 1.03 | 0.87 | 1.01 | 2.05 | 1.85 | 2.56 | 1.62 | 2.09 | 2.2 | 1.5 | 1.8 | 1.24 | -1.15 | 1.36 | 1.56 | 1.26 | 0.79 | 0.445 | -0.4825 | 0.448776 | 0.050793 | 0.246424 | 1 | 0.999631 | 0.786796 |
| Pfkfb2_B2Z892_T449t_1 | 0.13 | 1.42 | 0.46 | -0.29 | 1.32 | 1.12 | -0.64 | -0.2 | 1.48 | -0.13 | 0.06 | 0.05 | 1.09 | 1.55 | 0.13 | -0.2 | -0.03 | -0.065 | 0.2125 | 0.682517 | 0.956819 | 0.895742 | 1 | 0.999631 | 0.889581 |
| Pfkfb4_Q6DTY7_S176s_1 | 0.55 | 0.71 | -0.13 | 0.6 | 0.25 | 0.77 | 0.3 | 0.88 | 0.89 | -0.19 | 0.94 | 0.23 | -0.02 | 1.08 | 0.34 | 0.43 | 0.1175 | 0.035 | 0.025 | 0.943497 | 0.702697 | 0.923264 | 1 | 0.999631 | 0.981629 |
| Pfkl_P12382_S775s_1 | -0.33 | -0.29 | -8.62 | -10.79 | -0.64 | -0.63 | -8.68 | -14.57 | -0.51 | -0.16 | -6.39 | -12.51 | -0.79 | -0.43 | -12.14 | -11.46 | -1.1225 | 0.115 | -1.1975 | 0.73434 | 0.759938 | 0.972767 | 1 | 0.999631 | 0.907462 |
| Pfkl_P12382_T304t_1 | 0.12 | 0.02 | 0.08 | -0.4 | -0.78 | -0.55 | -0.43 | -0.22 | -0.5 | -0.01 | 0.24 | -0.29 | 0.16 | 0.01 | -0.07 | 0.34 | -0.45 | -0.095 | 0.155 | 0.586037 | 0.114789 | 0.742719 | 1 | 0.999631 | 0.849534 |
| Pfkm_P47857_S377s_1 | -1.58 | 0.72 | -0.97 | -0.54 | 0.41 | -0.24 | -0.99 | -0.66 | 0.66 | -2.16 | -1.33 | -0.01 | 1.38 | -1.36 | 0.5 | -0.91 | 0.2225 | -0.1175 | 0.495 | 0.488977 | 0.675738 | 0.868841 | 1 | 0.999631 | 0.807201 |
| Pfkm_P47857_S667s_1 | -0.64 | -0.06 | -0.44 | -0.51 | 0.12 | -0.61 | -1 | -1.15 | 0.12 | -0.25 | -0.12 | -0.74 | 0.65 | 0.08 | -1.21 | -0.19 | -0.2475 | 0.165 | 0.245 | 0.564207 | 0.476874 | 0.582805 | 1 | 0.999631 | 0.84139 |
| Pfkp_Q8C605_S2s_1 | -0.62 | -0.33 | -0.38 | -0.57 | -0.47 | -0.45 | -0.96 | -0.53 | -0.58 | 0.27 | -0.65 | -0.67 | -0.1 | -0.15 | -0.86 | -0.23 | -0.1275 | 0.0675 | 0.14 | 0.640862 | 0.617896 | 0.826233 | 1 | 0.999631 | 0.871335 |
| Pgam1_Q9DBJ1_S14s_1 | 0.77 | 1.9 | 1.25 | 1.51 | 1.7 | 1.1 | 0.81 | 1.09 | 2.26 | 1.21 | 2.7 | 1.56 | 1.27 | 0.86 | 0.27 | 1.54 | -0.1825 | 0.575 | -0.3725 | 0.35401 | 0.592369 | 0.188992 | 1 | 0.999631 | 0.73969 |
| Pgam1_Q9DBJ1_Y92y_1 | -0.91 | 0.41 | 0.97 | 1.2 | 2.38 | -0.35 | 2.35 | 0.64 | 2.12 | -2.6 | 2.6 | 3.6 | 2.85 | -1.32 | 1.68 | 0.01 | 0.8375 | 1.0125 | 0.3875 | 0.664666 | 0.264352 | 0.424502 | 1 | 0.999631 | 0.882317 |
| Pgbd5_D3YZI9_S519s_1 | -1.39 | -0.29 | -1.81 | -1.57 | -0.28 | -1.06 | -1.68 | -1.34 | -0.18 | -0.55 | -2.19 | -1.75 | 0.95 | -1.02 | -1.99 | -0.9 | 0.175 | 0.0975 | 0.525 | 0.414139 | 0.690647 | 0.857512 | 1 | 0.999631 | 0.768655 |
| Pgbd5_D3YZI9_S520s_1 | -0.71 | -0.7 | -0.85 | -0.68 | -0.78 | -0.81 | -0.73 | -0.5 | -0.91 | -0.76 | -0.71 | -0.77 | -0.31 | -0.54 | -0.74 | -0.44 | 0.03 | -0.0525 | 0.2275 | 0.402505 | 0.898571 | 0.826678 | 1 | 0.999631 | 0.763938 |
| Pgk1_P09411_S203s_1 | -1.26 | -0.15 | 0.01 | -0.66 | -0.48 | -0.46 | -0.7 | -0.36 | -0.31 | -0.38 | -0.27 | 0.18 | 0.29 | 0.15 | -0.16 | 0.26 | 0.015 | 0.32 | 0.65 | 0.092521 | 0.96416 | 0.373351 | 1 | 0.999631 | 0.557085 |
| Pgls_F6X8L5_S43s_1 | -1.58 | 0.09 | -0.58 | -0.54 | -0.39 | -0.25 | -0.51 | -0.52 | -0.77 | -1.2 | -0.48 | -0.44 | -0.09 | -0.31 | -1.02 | -0.36 | 0.235 | -0.07 | 0.2075 | 0.619293 | 0.529225 | 0.86096 | 1 | 0.999631 | 0.861888 |
| Pgm2_Q7TSV4_S173s_1 | -2.25 | 1.64 | 1.29 | -1.35 | 0.09 | 0.44 | 0.84 | -1.17 | 0.64 | -2.66 | 1.39 | -1.97 | 1.46 | -0.93 | -0.08 | -2.3 | 0.2175 | -0.4825 | -0.295 | 0.780933 | 0.811931 | 0.68356 | 1 | 0.999631 | 0.921543 |
| Pgm2_Q9D0F9_S117s_1 | -1.49 | -1.26 | -1.45 | -1.64 | -1.11 | -1.92 | -1.52 | -1.87 | -0.64 | -0.96 | -1.2 | 0.42 | 0.23 | -0.53 | 0.72 | -0.49 | -0.145 | 0.865 | 1.4425 | 0.002423 | 0.609784 | 0.046036 | 1 | 0.999631 | 0.351308 |
| Pgm2_Q9D0F9_T507t_1 | -0.58 | 0.44 | 0.4 | -0.11 | -0.64 | -0.47 | -0.19 | -1.56 | 0.32 | -0.31 | 0.97 | 0.34 | 0.75 | -0.11 | 0.89 | -0.33 | -0.7525 | 0.2925 | 0.2625 | 0.524616 | 0.078474 | 0.446832 | 1 | 0.999631 | 0.822727 |
| Pgm2l1_Q8CAA7_S175s_1 | -0.9 | -1.02 | -0.32 | -0.88 | -0.64 | -0.55 | -0.72 | -0.43 | -0.95 | -0.91 | -0.09 | -0.46 | -0.68 | -0.67 | -0.05 | -0.5 | 0.195 | 0.1775 | 0.305 | 0.336613 | 0.469552 | 0.579789 | 1 | 0.999631 | 0.730499 |
| Pgm3_Q8BWW3_S64s_1 | -0.61 | -0.31 | 0.11 | -0.14 | -0.8 | -0.51 | -0.48 | 0 | -0.94 | -0.89 | -0.39 | -0.75 | -0.29 | -0.11 | -0.74 | -0.5 | -0.21 | -0.505 | -0.1725 | 0.573633 | 0.479763 | 0.102345 | 1 | 0.999631 | 0.844597 |
| Pgrmc1_O55022_S181s_1 | -0.61 | -0.07 | 0.61 | 0.06 | -0.59 | 0.15 | -0.45 | -0.06 | -0.3 | 0.15 | -0.24 | 0.24 | 0.3 | 0.21 | 0.49 | 0.34 | -0.235 | -0.035 | 0.3375 | 0.320299 | 0.493677 | 0.916579 | 1 | 0.999631 | 0.723684 |
| Pgrmc2_Q80UU9_S84s_1 | -0.04 | -0.03 | -0.6 | -0.11 | -0.37 | 0.05 | -0.26 | -0.13 | -0.23 | -0.14 | -0.24 | -0.06 | 0.22 | 0.79 | 0 | 0.01 | 0.0175 | 0.0275 | 0.45 | 0.176929 | 0.947044 | 0.916744 | 1 | 0.999631 | 0.642116 |
| Phactr1_Q2M3X8_S230s_1 | 0.74 | 1.52 | 1.15 | 0.8 | 0.5 | 1.84 | 0.55 | 0.97 | 0.84 | 1.54 | 0.92 | 0.93 | 0.19 | 1.88 | 1.15 | 1.63 | -0.0875 | 0.005 | 0.16 | 0.707463 | 0.815507 | 0.987119 | 1 | 0.999631 | 0.898603 |
| Phactr1_Q2M3X8_S498s_1 | 1.41 | 0.77 | 0.04 | 1.24 | 2.26 | 1.1 | 0.9 | 1.56 | 0.85 | 0.31 | 1.45 | 1.1 | 1.39 | 1.23 | 1.08 | 0.63 | 0.59 | 0.0625 | 0.2175 | 0.572532 | 0.18607 | 0.876135 | 1 | 0.999631 | 0.844402 |
| Phactr1_Q2M3X8_S60s_1 | 2.22 | -0.53 | -0.17 | 1.63 | 2.67 | 0.07 | 2.25 | 1.41 | 0.83 | -0.2 | 1.58 | 1.61 | 0.83 | 0.89 | 0.94 | 1.44 | 0.8125 | 0.1675 | 0.2375 | 0.70303 | 0.307718 | 0.812842 | 1 | 0.999631 | 0.896857 |
| Phactr1_Q2M3X8_T97t_1 | 1.05 | -0.36 | -1.17 | 0.49 | 1.19 | 0.62 | 1.09 | 1.03 | 0.33 | -0.51 | 0.99 | -0.34 | -0.06 | 0.38 | 0.36 | -0.54 | 0.98 | 0.115 | 0.0325 | 0.948899 | 0.063498 | 0.834805 | 1 | 0.999631 | 0.983587 |
| Phactr2_B1AVN9_S204s_1 | 0.17 | -0.48 | 0.2 | 1.03 | 0.53 | -0.17 | 0.21 | 1.12 | 1.45 | 0.17 | 0.04 | 0.13 | -1.57 | 1.69 | 0.69 | -0.44 | 0.1925 | 0.2175 | -0.1375 | 0.840846 | 0.64198 | 0.629462 | 1 | 0.999631 | 0.946819 |
| Phactr2_B1AVN9_S444s_1 | 1.34 | 1.21 | 1.47 | 1.88 | 2.04 | 0.88 | 1.22 | 2.04 | 1.77 | 1.41 | 1.24 | 1.91 | 0.97 | 1.85 | 0.92 | 1.94 | 0.07 | 0.1075 | -0.055 | 0.878711 | 0.84356 | 0.715707 | 1 | 0.999631 | 0.962442 |
| Phactr2_B1AVN9_T25t_1 | 0.08 | 1.57 | 1.17 | 0.74 | 1.17 | 1.13 | 0.55 | 0.9 | 1.55 | -0.11 | 1.25 | 2.25 | 0.99 | 1.93 | 1.48 | 1.38 | 0.0475 | 0.345 | 0.555 | 0.184426 | 0.897363 | 0.532011 | 1 | 0.999631 | 0.644799 |
| Phactr3_Q8BYK5_S189s_1 | -0.88 | 0.21 | 0.49 | -1.09 | -0.7 | -0.84 | -0.11 | 0.09 | -0.33 | -0.15 | 0.43 | -0.05 | 0.48 | -0.57 | 0.83 | 1.11 | -0.0725 | 0.2925 | 0.78 | 0.149866 | 0.86908 | 0.49713 | 1 | 0.999631 | 0.622212 |
| Phactr3_Q8BYK5_T18t_1 | 0.14 | 0.56 | 0.85 | 0.69 | 0.59 | 0.67 | -0.24 | 0.13 | 0.67 | 1.01 | 0.33 | 0.79 | 1.42 | 1.66 | 0.83 | 1.3 | -0.2725 | 0.14 | 0.7425 | 0.038584 | 0.394489 | 0.633615 | 1 | 0.999631 | 0.461663 |
| Phactr3_Q8BYK5_T195t_1 | 0.37 | 1.49 | 0.84 | 1.23 | 0.15 | 0.37 | 0.14 | 0.7 | 2.24 | -0.6 | 0.21 | 0.79 | -0.43 | 1.86 | 2.16 | 1.26 | -0.6425 | -0.3225 | 0.23 | 0.691341 | 0.070644 | 0.587972 | 1 | 0.999631 | 0.893997 |
| Phactr4_Q501J7_S147s_1 | 1 | 0.41 | 1.11 | 1.05 | 0.72 | 0.9 | 0.43 | 0.45 | 1.26 | 1.26 | 0.51 | 0.78 | -1.09 | 0.73 | 0.69 | 0.36 | -0.2675 | 0.06 | -0.72 | 0.137287 | 0.350463 | 0.847737 | 1 | 0.999631 | 0.610369 |
| Phactr4_Q501J7_S430s_1 | -0.53 | 1.04 | 1.27 | 0.8 | 0.79 | 0.73 | 0.05 | 0.64 | 0.64 | 0.56 | 0.77 | 1.53 | 0.32 | 1.45 | 1.86 | 1.3 | -0.0925 | 0.23 | 0.5875 | 0.254719 | 0.829638 | 0.61265 | 1 | 0.999631 | 0.68954 |
| Phb2_O35129_S267s_1 | 1.53 | -0.84 | -0.03 | -0.01 | -1.08 | -0.04 | 0.18 | -0.9 | -0.73 | 0.72 | -0.71 | -0.65 | -0.18 | -0.61 | -0.84 | -1.02 | -0.6225 | -0.505 | -0.825 | 0.125433 | 0.264978 | 0.379098 | 1 | 0.999631 | 0.596551 |
| Phc3_F8WIN6_S281s_1 | 0.99 | 0.56 | 0.38 | 0.09 | 0.12 | 0.54 | 0.15 | -0.09 | 0.33 | 1.16 | 0.16 | -0.06 | 0.21 | 0.29 | 0.02 | -0.36 | -0.325 | -0.1075 | -0.465 | 0.168827 | 0.287643 | 0.765446 | 1 | 0.999631 | 0.637839 |
| Phf10_K4DI61_S326s_1 | -0.88 | 1.69 | 0.75 | -0.12 | 0.13 | -0.76 | -0.32 | 0.52 | -0.27 | 0.43 | 0.69 | -2.32 | 0.6 | 0.55 | -0.29 | -0.29 | -0.4675 | -0.7275 | -0.2175 | 0.699985 | 0.417117 | 0.358511 | 1 | 0.999631 | 0.896612 |
| Phf2_Q9WTU0_S1057s_1 | 1.41 | 0.25 | 0.89 | 1.16 | -0.13 | 0.7 | 1.08 | 0.03 | -0.4 | 1.24 | -0.09 | -0.76 | -0.62 | -0.99 | -0.44 | 0.19 | -0.5075 | -0.93 | -1.3925 | 0.004427 | 0.210896 | 0.078259 | 1 | 0.999631 | 0.351308 |
| Phf2_Q9WTU0_S1060s_1 | 1.47 | -1.01 | 0.92 | 0.04 | -1.34 | 0.15 | 0.1 | -0.45 | -0.27 | 1.47 | -1.52 | -0.87 | -1.08 | -1.18 | -0.3 | -1.64 | -0.74 | -0.6525 | -1.405 | 0.029795 | 0.2243 | 0.389666 | 1 | 0.999631 | 0.434193 |
| Phf20_Q8BLG0_S894s_1 | 0.42 | 0.41 | 1.18 | 0.66 | 0.52 | 0.13 | 0.24 | -0.05 | 0.17 | 0.1 | 1.8 | 1 | 1.09 | 0.06 | 1.18 | 0.69 | -0.4575 | 0.1 | 0.0875 | 0.808771 | 0.137364 | 0.818655 | 1 | 0.999631 | 0.93635 |
| Phf3_B2RQG2_S377s_1 | 0.05 | 1.12 | 0.21 | 0.64 | 0.38 | 0.61 | 0.2 | 0.2 | 0.85 | 0.4 | -0.01 | -0.13 | 0.94 | 1.04 | 0.06 | 0.57 | -0.1575 | -0.2275 | 0.1475 | 0.691049 | 0.616328 | 0.531779 | 1 | 0.999631 | 0.893959 |
| Phf3_B2RQG2_S660s_1 | 0.17 | 1.05 | 1.3 | 0.56 | 0.44 | 0.56 | 0.75 | 0.14 | 0.83 | -0.15 | 0.29 | 1.32 | 0.76 | 0.85 | 0.56 | 1.86 | -0.2975 | -0.1975 | 0.2375 | 0.561731 | 0.372736 | 0.634841 | 1 | 0.999631 | 0.840624 |
| Phf6_E9QAG2_S203s_1 | 2.54 | 1.51 | 3.19 | 2.46 | 0.12 | 1.88 | 2.96 | 1.64 | 0.52 | 1.8 | 1.78 | 1.64 | 0.04 | 0.99 | 2.05 | 1.4 | -0.775 | -0.99 | -1.305 | 0.028141 | 0.225001 | 0.050983 | 1 | 0.999631 | 0.423977 |
| Phip_F8VQ93_S1783s_1 | -2.3 | -0.53 | -1.02 | -0.74 | -0.83 | -1.06 | -0.47 | -1.53 | -1.15 | -0.11 | -0.45 | 0.27 | 0.68 | -1.03 | 0.23 | 0.1 | 0.175 | 0.7875 | 1.1425 | 0.046487 | 0.693261 | 0.124891 | 1 | 0.999631 | 0.479949 |
| Phkb_Q7TSH2_S19s_1 | 0.09 | 0.95 | 0.26 | 0.92 | 1.16 | 0.65 | 0.38 | 0.92 | 0.7 | -0.11 | 0.99 | 0.69 | -0.12 | 1.13 | 0.57 | 0.3 | 0.2225 | 0.0125 | -0.085 | 0.823045 | 0.497762 | 0.972162 | 1 | 0.999631 | 0.938864 |
| Phldb1_D3Z4N0_S382s_1 | 0.15 | 0.91 | 0.67 | 0.33 | 0.41 | -0.01 | -0.2 | 0.6 | 0.52 | 0.28 | 1.07 | 1.19 | 0.57 | -0.03 | 1.53 | 0.69 | -0.315 | 0.25 | 0.175 | 0.657049 | 0.319614 | 0.45455 | 1 | 0.999631 | 0.879129 |
| Phldb1_D3Z4N0_S445s_1 | 0.17 | 0.2 | 0.07 | -0.19 | -0.24 | -0.4 | -0.56 | 0.26 | -0.36 | 0.17 | -0.44 | -0.08 | 0.13 | 0.04 | 0.18 | 0.32 | -0.2975 | -0.24 | 0.105 | 0.696885 | 0.302226 | 0.387799 | 1 | 0.999631 | 0.895731 |
| Phldb1_D3Z4N0_S463s_1 | 0.39 | 0.26 | 0.82 | 0.85 | 0.48 | 0.12 | -0.38 | 0.45 | 0.53 | 0.2 | 0.01 | 1.04 | 0.8 | 0.08 | 1.11 | 1.24 | -0.4125 | -0.135 | 0.2275 | 0.524685 | 0.200035 | 0.679961 | 1 | 0.999631 | 0.822727 |
| Phldb1_D3Z4N0_S472s_1 | 0.35 | 0.26 | -0.05 | 0.41 | 0.23 | -0.43 | -0.65 | 0.78 | 0 | -0.2 | 0.24 | -0.06 | 0.22 | 0.42 | -0.3 | 0.56 | -0.26 | -0.2475 | -0.0175 | 0.954836 | 0.479675 | 0.355036 | 1 | 0.999631 | 0.984441 |
| Phldb1_D3Z4N0_S491s_1 | 1.84 | 1.09 | 0.78 | 0.8 | 0.88 | 0.18 | 0.08 | 1.2 | 1.51 | 1.21 | -0.44 | -0.42 | 0.03 | -0.36 | 0.92 | -0.18 | -0.5425 | -0.6625 | -1.025 | 0.026567 | 0.176375 | 0.236062 | 1 | 0.999631 | 0.422285 |
| Phldb1_D3Z4N0_S51s_1 | 1.41 | 2.13 | 1.87 | 0.37 | 0.63 | 0.57 | 0.08 | 1.74 | 1.38 | 0.62 | 0.76 | 1.26 | 4.46 | 1.62 | 1.87 | 0.28 | -0.69 | -0.44 | 0.6125 | 0.46682 | 0.182939 | 0.319379 | 1 | 0.999631 | 0.795267 |
| Phldb1_D3Z4N0_S541s_1 | -0.15 | 0.77 | 0.46 | 0.21 | -0.05 | 0.43 | -0.56 | 0.84 | 0.08 | 0.18 | 0.42 | 0.44 | 0.68 | 0.91 | -0.16 | -0.01 | -0.1575 | -0.0425 | 0.0325 | 0.929724 | 0.676131 | 0.885461 | 1 | 0.999631 | 0.97754 |
| Phldb1_D3Z4N0_S950s_1 | -0.88 | -1.42 | 0.07 | -0.77 | -1.38 | -1.4 | -0.89 | -0.67 | -1.49 | -0.7 | -0.47 | -0.04 | -0.44 | -1.16 | -0.46 | -0.25 | -0.335 | 0.075 | 0.1725 | 0.663416 | 0.380708 | 0.862019 | 1 | 0.999631 | 0.881968 |
| Phldb2_D3Z069_S456s_1 | 0.78 | 1.37 | 1.65 | 1.06 | 0.93 | 1.39 | 1.11 | 0.87 | 1.31 | 1.69 | 1.28 | 1.88 | 0.77 | 1.51 | 1.82 | 2 | -0.14 | 0.325 | 0.31 | 0.413011 | 0.633327 | 0.305801 | 1 | 0.999631 | 0.768474 |
| Phlpp1_Q8CHE4_S304s_1 | -0.5 | 0.05 | -0.34 | -0.05 | 0.23 | -0.5 | -0.97 | -1.58 | -0.24 | -0.86 | -0.95 | -0.85 | -0.5 | 0.78 | -0.16 | -0.68 | -0.495 | -0.515 | 0.07 | 0.855119 | 0.239889 | 0.102172 | 1 | 0.999631 | 0.951787 |
| Phlpp2_J3QM82_S1214s_1 | 0.06 | 0.32 | 0.26 | 0.14 | 0.14 | -0.15 | -0.31 | 0.13 | 0.27 | 0.67 | 0.3 | 1.04 | 0.35 | 0.52 | 1.16 | 0.83 | -0.2425 | 0.375 | 0.52 | 0.104492 | 0.34154 | 0.206385 | 1 | 0.999631 | 0.568213 |
| Phlpp2_J3QM82_S1242s_1 | -0.45 | 0.19 | 0.3 | 1.14 | 0.25 | -0.72 | 0.63 | 1.14 | -0.2 | -0.53 | 1.14 | 1.78 | -0.37 | 0.34 | 1.07 | 1.32 | 0.03 | 0.2525 | 0.295 | 0.54744 | 0.950472 | 0.666532 | 1 | 0.999631 | 0.834277 |

| | | | | | | | | | | | | | | | | | | | | | | | | | |
|---|---|---|---|---|---|---|---|---|---|---|---|---|---|---|---|---|---|---|---|---|---|---|---|---|---|
| Phospho2_Q9D9M5_T203t_1 | 0.02 | -0.1 | 3.05 | 1.84 | 0.78 | 1.49 | 2.33 | 1.91 | 0.56 | 1.52 | 2.71 | 0.98 | -0.73 | 1.6 | 1.26 | 2.81 | 0.425 | 0.24 | 0.0325 | 0.97147 | 0.563445 | 0.759727 | 1 | 0.999631 | 0.989755 |
| Phrf1_A6H619_S622s_1 | -0.13 | 0.6 | 0.13 | -0.06 | 0.22 | 0.22 | 0.14 | -0.28 | 0.27 | -0.02 | 0.47 | -0.01 | 0.37 | 0.76 | -0.21 | -0.23 | -0.06 | 0.0425 | 0.0375 | 0.914411 | 0.831816 | 0.883215 | 1 | 0.999631 | 0.972695 |
| Phrf1_F7ANA1_S247s_1 | 0.88 | -0.11 | 0.14 | -0.12 | -0.61 | -1.17 | 0.01 | -0.77 | -0.72 | 0.2 | -0.72 | -0.96 | -0.17 | -0.65 | -0.52 | -0.36 | -0.8325 | -0.7475 | -0.6225 | 0.083616 | 0.042013 | 0.069363 | 1 | 0.999631 | 0.549732 |
| Phrf1_F7ANA1_S277s_1 | 0.73 | 0.81 | 1.31 | -0.03 | 0.04 | 0.42 | 0.53 | 0.02 | 0.76 | 1.16 | 0.51 | 0.4 | 1.23 | 0.64 | 0.38 | 0.06 | -0.4525 | 0.0025 | -0.1275 | 0.748711 | 0.204486 | 0.994426 | 1 | 0.999631 | 0.911931 |
| Phyhipl_Q8BGT8_T116t_1 | -0.19 | 2.73 | 1.73 | 0.69 | 0.64 | 2.69 | 0.1 | 2.19 | 1.38 | 0.22 | 2.23 | -1.3 | 2.39 | 1.8 | -0.77 | 1.8 | 0.165 | -0.6075 | 0.065 | 0.937049 | 0.831162 | 0.488409 | 1 | 0.999631 | 0.97939 |
| Pi4k2a_Q2TBE6_S47s_1 | -0.1 | 0.67 | -0.51 | -0.82 | -0.6 | 0.06 | -1.08 | -1 | -0.38 | -0.04 | -1.17 | -0.5 | 0.43 | 0.6 | -0.88 | -0.54 | -0.465 | -0.3325 | 0.0925 | 0.845127 | 0.276438 | 0.423902 | 1 | 0.999631 | 0.948565 |
| Pi4k2a_Q2TBE6_S47s_2 | 0.38 | 0.68 | 0.93 | 0.3 | 0.39 | 0.37 | 0.22 | 0.73 | 0.46 | 1.22 | 0.22 | 0.43 | 0.32 | 0.8 | 1.04 | 0.46 | -0.145 | 0.01 | 0.0825 | 0.790985 | 0.595568 | 0.975143 | 1 | 0.999631 | 0.926419 |
| Pi4k2a_Q2TBE6_S51s_2 | 0.38 | 0.68 | 0.93 | 0.3 | 0.39 | 0.37 | 0.22 | 0.73 | 0.46 | 1.22 | 0.22 | 0.43 | 0.32 | 0.8 | 1.04 | 0.46 | -0.145 | 0.01 | 0.0825 | 0.790985 | 0.595568 | 0.975143 | 1 | 0.999631 | 0.926419 |
| Pi4k2a_Q2TBE6_S9s_1 | -0.42 | 1.29 | 0.52 | -0.25 | 0.88 | 0.25 | -0.08 | 0.42 | -0.07 | -0.31 | -0.05 | 0.29 | 0.78 | 0.4 | 0 | -0.08 | 0.0825 | -0.32 | -0.01 | 0.981944 | 0.848451 | 0.449383 | 1 | 0.999631 | 0.994846 |
| Pi4ka_E9Q3L2_S1427s_1 | -2.64 | -1 | 0.1 | -0.68 | -0.98 | -0.74 | -1.22 | -1.14 | -3.6 | -1.37 | -0.97 | -1.27 | -1.14 | -1.59 | -0.78 | -0.97 | 0.035 | -0.7475 | -0.065 | 0.907225 | 0.948279 | 0.325377 | 1 | 0.999631 | 0.96989 |
| Pi4ka_E9Q3L2_S259s_1 | 0.45 | -1.07 | -0.25 | -0.5 | 0.17 | 1 | -0.39 | 0.12 | 0.53 | 0.41 | 0.14 | -0.35 | 0.15 | 0.09 | 0.29 | -0.48 | 0.5675 | 0.525 | 0.355 | 0.37139 | 0.199321 | 0.197662 | 1 | 0.999631 | 0.746522 |
| Pi4ka_E9Q3L2_S263s_2 | -0.76 | -0.4 | -0.24 | -0.3 | -0.53 | -0.9 | -0.26 | 0.3 | -0.98 | -0.5 | -0.07 | -0.23 | -0.69 | -1.26 | 0.06 | -0.15 | 0.0775 | -0.02 | -0.085 | 0.815735 | 0.810905 | 0.947457 | 1 | 0.999631 | 0.937576 |
| Pi4ka_E9Q3L2_S268s_1 | -1.39 | -0.86 | -1.05 | -1.29 | -1.12 | -0.75 | -1.25 | -0.9 | -1.3 | -0.86 | -1.16 | -0.87 | -0.55 | -0.59 | -0.74 | -1.03 | 0.1425 | 0.1 | 0.42 | 0.165088 | 0.591672 | 0.712831 | 1 | 0.999631 | 0.633483 |
| Pi4ka_E9Q3L2_S268s_2 | -0.76 | -0.4 | -0.24 | -0.3 | -0.53 | -0.9 | -0.26 | 0.3 | -0.98 | -0.5 | -0.07 | -0.23 | -0.69 | -1.26 | 0.06 | -0.15 | 0.0775 | -0.02 | -0.085 | 0.815735 | 0.810905 | 0.947457 | 1 | 0.999631 | 0.937576 |
| Pi4kb_E9Q8A3_S523s_1 | 0.73 | 0.57 | 0 | 0.8 | 1.06 | 0.56 | 0.54 | 0.77 | 1.23 | -0.1 | 0.61 | 0.73 | -0.2 | 0.79 | 0.32 | 0.92 | 0.2075 | 0.0925 | -0.0675 | 0.851823 | 0.480074 | 0.798164 | 1 | 0.999631 | 0.950099 |
| Pianp_Q6P1B3_S32s_1 | 1.52 | 0.37 | 1.39 | 0.94 | 0.19 | 0.74 | 0.95 | 0.44 | 0.38 | 1.27 | -0.07 | -1.14 | -0.24 | 0.41 | 0.73 | 0.26 | -0.475 | -0.945 | -0.765 | 0.062074 | 0.187067 | 0.098635 | 1 | 0.999631 | 0.509355 |
| Pias1_Q3U556_S485s_1 | 0.25 | 0.62 | 1.67 | 0.64 | -0.66 | -0.37 | 0.1 | 0.08 | 0.28 | 0.13 | 0.35 | 1.37 | -0.24 | 1.43 | 0.79 | 1.4 | -1.0075 | -0.2625 | 0.05 | 0.917176 | 0.021303 | 0.535855 | 1 | 0.999631 | 0.973513 |
| Pik3c3_E9Q824_S282s_1 | -0.37 | -0.16 | -0.54 | -1.64 | -0.89 | -0.24 | -0.98 | -0.6 | -0.76 | -0.06 | -0.74 | -0.45 | 0.22 | -0.17 | -0.68 | -1.16 | 7.85E-17 | 0.175 | 0.23 | 0.609028 | 1 | 0.65279 | 1 | 0.999631 | 0.858851 |
| Pik3cd_B0QZL5_S516s_1 | -0.66 | -0.88 | -0.14 | -0.21 | -0.72 | -0.17 | 0.14 | -0.03 | -0.12 | -0.21 | 0.88 | 0.61 | 0.35 | -0.03 | 0.63 | 0.59 | 0.2775 | 0.7625 | 0.8575 | 0.021104 | 0.383269 | 0.055513 | 1 | 0.999631 | 0.400544 |
| Pikfyve_D3Z5N5_S318s_1 | 1.16 | 0.38 | 0.18 | 0.18 | 1.28 | -0.06 | 0.47 | 0.68 | 0.66 | 0.37 | 0.26 | 0.29 | 0.52 | 0.48 | 0.73 | 0.88 | 0.1175 | -0.08 | 0.1775 | 0.590489 | 0.755774 | 0.79934 | 1 | 0.999631 | 0.851647 |
| Pip4k2b_Q80XI4_S131s_1 | 1.06 | -0.95 | 0.28 | 0.57 | -0.93 | -0.37 | 0.59 | 0.21 | -0.89 | 1.08 | -0.03 | 0.01 | -0.86 | -1.39 | 0.25 | -0.53 | -0.365 | -0.1975 | -0.8725 | 0.117039 | 0.477448 | 0.718303 | 1 | 0.999631 | 0.585087 |
| Pip4k2c_Q91XU3_S129s_1 | 1.51 | -0.74 | 0.97 | 0.31 | -0.68 | -0.2 | 0.32 | -0.72 | -1.02 | -0.04 | -0.65 | 0.44 | -1.63 | -1.21 | -0.65 | -1.64 | -0.8325 | -0.83 | -1.795 | 0.00538 | 0.12498 | 0.14915 | 1 | 0.999631 | 0.351308 |
| Pip5k1a_F8WI18_S457s_1 | -0.4 | -1.15 | -0.64 | -0.49 | -0.16 | -1.55 | 0.17 | -0.28 | -1.44 | -0.8 | -0.18 | -0.2 | -0.24 | -0.89 | -0.61 | -1.06 | 0.215 | 0.015 | -0.03 | 0.926275 | 0.604184 | 0.967634 | 1 | 0.999631 | 0.97588 |
| Pip5k1c_F8WHW6_S475s_1 | -0.58 | -1.23 | -0.86 | -0.25 | -0.02 | -1.4 | -0.4 | -0.73 | -1.1 | -1.95 | -0.81 | -0.6 | -1.53 | -1.31 | -0.35 | -0.18 | 0.0925 | -0.385 | -0.1125 | 0.785973 | 0.805359 | 0.327853 | 1 | 0.999631 | 0.923346 |
| Pip5k1c_F8WHW6_S538s_1 | 2.68 | 1.01 | 1.5 | 1.58 | 0.2 | 1.7 | 2.29 | 0.75 | 0.44 | 1.94 | 1.13 | 0.32 | -1.5 | 0 | 0.2 | 0.43 | -0.4575 | -0.735 | -1.91 | 0.004535 | 0.405997 | 0.15682 | 1 | 0.999631 | 0.351308 |
| Pip5k1c_F8WHW6_S540s_1 | 1.89 | 0.11 | 1.01 | 1.22 | -0.54 | 0.26 | 1.56 | 0.23 | -0.43 | 1.37 | 0.08 | 0.16 | -1.28 | -0.72 | 0.2 | -0.74 | -0.68 | -0.7625 | -1.6925 | 0.004618 | 0.217177 | 0.15234 | 1 | 0.999631 | 0.351308 |
| Pip5k1c_F8WHW6_S554s_1 | -0.08 | 0.7 | -0.23 | 0.06 | 0.79 | 0.31 | -0.25 | 0.19 | 0.37 | -0.28 | -0.02 | -0.23 | -0.57 | 0.23 | -0.23 | -0.07 | 0.1475 | -0.1525 | -0.2725 | 0.422734 | 0.66069 | 0.630837 | 1 | 0.999631 | 0.773293 |
| Pip5k1c_F8WHW6_S671s_1 | 1.08 | 0.07 | -2.6 | 0.03 | 1.87 | 0.72 | 0.69 | -0.65 | 0.4 | -1.18 | -1.11 | -1.79 | -1.66 | 0.29 | -0.31 | -0.89 | 1.0125 | -0.565 | -0.2875 | 0.712588 | 0.23538 | 0.487034 | 1 | 0.999631 | 0.9008 |
| Pip5k1c_F8WHW6_S681s_1 | 0.21 | -2.32 | 0.35 | -0.38 | -2.85 | -0.57 | 1.69 | -1.94 | -0.8 | 0.69 | -0.34 | -0.9 | -0.16 | -2.06 | -0.17 | -1.69 | -0.3825 | 0.1975 | -0.485 | 0.495253 | 0.703254 | 0.759477 | 1 | 0.999631 | 0.810677 |
| Pip5k1c_F8WHW6_T661t_1 | 0.46 | -0.79 | -2.04 | -0.04 | 0.98 | 0.2 | 0.28 | 0.39 | 1.04 | -0.94 | -0.86 | -2.05 | 0.88 | -0.98 | -1.75 | -0.67 | 1.065 | -0.1 | -0.0275 | 0.968123 | 0.068031 | 0.892936 | 1 | 0.999631 | 0.988771 |
| Pitpnc1_Q8K4R4_S274s_1 | 0.14 | 0.75 | 0.41 | 0.97 | 0.84 | 0.95 | 0.57 | 1.36 | 0.51 | 0.07 | 0.92 | 0.43 | 0.34 | 0.92 | 0.45 | 1.13 | 0.3625 | -0.085 | 0.1425 | 0.670459 | 0.252158 | 0.788082 | 1 | 0.999631 | 0.884249 |
| Pitpnc1_Q8K4R4_S299s_1 | -0.88 | -0.82 | -2.12 | -0.72 | -0.03 | -1.38 | -1.24 | -0.72 | -0.26 | -0.37 | -0.67 | -0.35 | 0.66 | 0.33 | -0.58 | -0.78 | 0.2925 | 0.7225 | 1.0425 | 0.047685 | 0.509654 | 0.074893 | 1 | 0.999631 | 0.481835 |
| Pitpnm1_O35954_S326s_1 | -0.73 | -1.25 | -1.61 | -2.26 | -1.64 | -1.49 | -0.67 | -1.18 | -1.64 | -1.47 | -1.6 | -0.71 | -0.8 | -1.53 | -1.51 | -0.52 | 0.2175 | 0.1075 | 0.3725 | 0.38741 | 0.583337 | 0.788628 | 1 | 0.999631 | 0.756827 |
| Pitpnm1_O35954_S593s_1 | 0.29 | 0.16 | -0.04 | -0.23 | 0.19 | 0.06 | -0.36 | 0.07 | 0.19 | 0.31 | 0.28 | 0.09 | 0.32 | 0.37 | 0.22 | 0.18 | -0.055 | 0.1725 | 0.2275 | 0.413399 | 0.835469 | 0.506614 | 1 | 0.999631 | 0.768474 |
| Pitpnm1_O35954_S621s_1 | 1.78 | 0.75 | -3.78 | 1.39 | 1.64 | 0.21 | 1.89 | 0.87 | 1.03 | -0.53 | 0.42 | -1.6 | -1.52 | -0.13 | -1.32 | -1.54 | 1.1175 | -0.205 | -1.1625 | 0.317247 | 0.343083 | 0.865236 | 1 | 0.999631 | 0.720971 |
| Pitpnm1_O35954_S664s_1 | 0.74 | -0.77 | -0.6 | -0.44 | 0.08 | 0.41 | -0.12 | -0.97 | 0.25 | 0.09 | -0.15 | -0.44 | 0.78 | 0 | 0.74 | -0.37 | 0.1175 | 0.205 | 0.555 | 0.231179 | 0.789528 | 0.60243 | 1 | 0.999631 | 0.681463 |
| Pitpnm1_O35954_S895s_1 | -0.3 | 0.01 | -1.99 | -1.37 | 0.06 | -0.61 | -0.76 | -2.21 | 0.12 | 0.13 | -0.9 | -1.05 | 0.74 | 0 | -0.6 | -0.33 | 0.0325 | 0.4875 | 0.865 | 0.119478 | 0.95692 | 0.366579 | 1 | 0.999631 | 0.587062 |
| Pitpnm1_O35954_T270t_1 | 0.01 | 1.16 | 0.16 | 0.39 | 1.18 | 1.12 | 1.02 | -0.28 | 1.61 | 0.62 | 0.98 | 0.82 | 0.28 | 0.43 | 0.85 | -0.39 | 0.33 | 0.5775 | -0.1375 | 0.726345 | 0.445387 | 0.136877 | 1 | 0.999631 | 0.90568 |
| Pitpnm2_Q6ZPQ6_S1209s_1 | 1.25 | -0.03 | -0.88 | 1.56 | 1.61 | 0.57 | 1.36 | 0.83 | 0.11 | -0.76 | 0.82 | 0.11 | -1.24 | 0.28 | 0.09 | 0.43 | 0.6175 | -0.405 | -0.585 | 0.356626 | 0.287842 | 0.502032 | 1 | 0.999631 | 0.740467 |
| Pitpnm2_Q6ZPQ6_S1230s_1 | 1.73 | -0.01 | 0.49 | 1.24 | 0.45 | 1.11 | 2.14 | 1.69 | 0.48 | -0.49 | 1.17 | -1.55 | -2.5 | 0.47 | -1.71 | 0.58 | 0.485 | -0.96 | -1.6525 | 0.051527 | 0.343579 | 0.155727 | 1 | 0.999631 | 0.489777 |
| Pitpnm2_Q6ZPQ6_S1234s_1 | 1.17 | -0.1 | -1.33 | 0.54 | -0.7 | -1.07 | 0.77 | -0.83 | -0.83 | -0.1 | 0.09 | -0.99 | -1.13 | -0.7 | -3.84 | -1.06 | -0.5275 | -0.5275 | -1.7525 | 0.046884 | 0.39666 | 0.351333 | 1 | 0.999631 | 0.480102 |
| Pitpnm2_Q6ZPQ6_S302s_1 | 0.24 | -1.33 | -0.46 | -0.22 | -0.51 | -0.5 | 0.38 | -0.75 | -0.16 | -0.3 | -0.28 | -0.13 | -0.58 | -1.49 | 0.23 | -1.34 | 0.0975 | 0.225 | -0.3525 | 0.481872 | 0.813024 | 0.539996 | 1 | 0.999631 | 0.802327 |
| Pitpnm2_Q6ZPQ6_S365s_1 | -0.31 | 0.58 | -1.18 | -0.51 | 0.69 | 0.09 | 0.1 | -0.84 | 0.43 | -0.21 | 0.02 | -1.4 | 0.21 | 0.55 | -1.49 | -0.31 | 0.365 | 0.065 | 0.095 | 0.859958 | 0.436571 | 0.897712 | 1 | 0.999631 | 0.953438 |
| Pitpnm2_Q6ZPQ6_S397s_1 | 0.62 | 1.35 | -0.41 | 0.7 | 0.6 | 0.31 | -0.11 | 0.37 | 0.07 | -0.7 | -0.01 | 0.19 | 0.89 | 0.93 | -0.87 | 0.33 | -0.2725 | -0.6775 | -0.245 | 0.64176 | 0.498801 | 0.130461 | 1 | 0.999631 | 0.871335 |
| Pitpnm3_Q3UHE1_S279s_1 | -1.48 | -0.97 | -0.22 | -1.98 | -1.75 | -1.24 | -1.68 | -0.79 | -1.15 | -0.71 | -1.37 | -0.34 | 0.47 | -0.52 | -0.33 | -0.4 | -0.2025 | 0.27 | 0.9675 | 0.049658 | 0.638712 | 0.540605 | 1 | 0.999631 | 0.487521 |
| Pitpnm3_Q3UHE1_S305s_1 | 0.88 | 0.17 | -0.78 | 0.8 | 1.61 | 0.72 | 0.73 | 0.17 | 0.9 | -0.09 | 0.37 | -0.62 | -0.82 | 1.18 | 0.38 | 0.05 | 0.54 | -0.1275 | -0.07 | 0.894952 | 0.262062 | 0.791435 | 1 | 0.999631 | 0.965552 |
| Pitpnm3_Q3UHE1_S479s_1 | -0.44 | -0.27 | -0.53 | -0.23 | -0.74 | -0.2 | -0.63 | -0.3 | -0.51 | -0.44 | -0.35 | -0.18 | -0.49 | -0.07 | -0.4 | -0.33 | -0.1 | -0.0025 | 0.045 | 0.868109 | 0.698176 | 0.991991 | 1 | 0.999631 | 0.95689 |
| Pitpnm3_Q3UHE1_S488s_1 | 0.17 | 0.37 | -0.65 | -0.54 | -0.29 | 0.23 | -0.63 | -0.14 | -0.19 | 0.75 | -1.01 | -0.68 | 1.12 | 0.76 | -0.68 | -0.69 | -0.045 | -0.12 | 0.29 | 0.573954 | 0.895711 | 0.790272 | 1 | 0.999631 | 0.84468 |
| Pitpnm3_Q3UHE1_S891s_1 | 0.09 | 1.15 | 0.44 | 1.33 | 1.12 | 0.78 | 0.42 | 1.16 | 0.97 | 0.18 | 1.01 | 1.24 | 0.44 | 1.79 | 1.23 | 1.41 | 0.1175 | 0.0975 | 0.465 | 0.284463 | 0.746213 | 0.8028 | 1 | 0.999631 | 0.705468 |
| Pitpnm3_Q3UHE1_S912s_1 | -0.85 | 1.6 | 1.08 | 0.04 | -0.84 | 2.26 | -0.02 | 1.05 | -0.67 | -0.19 | 1.09 | -1.44 | -0.26 | 2.17 | -0.42 | -0.36 | 0.145 | -0.77 | -0.185 | 0.801233 | 0.848588 | 0.274122 | 1 | 0.999631 | 0.932069 |
| Pitpnm3_Q3UHE1_S93s_1 | -0.95 | -1.75 | 0.32 | 0.09 | -1.33 | -0.82 | -0.16 | 0.12 | -0.76 | -1.71 | 0.31 | 0.69 | -0.32 | -0.73 | -0.36 | 0.52 | 0.025 | 0.205 | 0.35 | 0.504601 | 0.962751 | 0.753424 | 1 | 0.999631 | 0.814292 |
| Pja1_O55176_S137s_1 | 0.93 | 0.78 | 2.11 | 1.08 | -0.01 | 1.3 | 1.02 | 1.34 | 0.69 | 1.78 | 1.2 | 1.46 | 0.48 | 1.15 | 2.03 | 1.35 | -0.3125 | 0.0575 | 0.0275 | 0.950305 | 0.472321 | 0.884033 | 1 | 0.999631 | 0.984079 |
| Pja1_O55176_S32s_1 | 1.9 | -0.06 | 0.46 | 1.62 | 2.04 | 0.51 | 1.75 | 1.66 | 1.1 | 0.43 | 1.81 | 0.58 | -1.51 | 1.28 | 0.5 | 0.92 | 0.51 | -7.9E-17 | -0.6825 | 0.335849 | 0.349105 | 1 | 1 | 1 | 0.730466 |
| Pja2_Q80U04_S320s_1 | 0.38 | 0.23 | -0.74 | 0.75 | 0.87 | 0.51 | 0.35 | 0.72 | 0.64 | 0.11 | 0.95 | 0.29 | -0.58 | 0.8 | 0.18 | 0.53 | 0.4575 | 0.3425 | 0.0775 | 0.86016 | 0.224258 | 0.386244 | 1 | 0.999631 | 0.953558 |
| Pkd2_O35245_S754s_1 | -1.36 | -2.22 | -0.45 | -0.05 | -2.1 | -2 | -0.97 | -0.73 | -0.51 | -0.97 | -0.83 | -0.13 | -0.54 | -2.94 | -1.18 | -2.09 | -0.43 | 0.41 | -0.6675 | 0.312449 | 0.441128 | 0.415615 | 1 | 0.999631 | 0.719861 |
| Pkib_F7D7A9_S24s_1 | 0.86 | 0.13 | 0.64 | 0.05 | 0.48 | 0.49 | 0.3 | -0.56 | -0.04 | 0.53 | -0.23 | -0.42 | -0.09 | -0.31 | -0.36 | -0.45 | -0.2425 | -0.46 | -0.7225 | 0.037945 | 0.491548 | 0.189603 | 1 | 0.999631 | 0.459387 |
| Pkn1_P70268_S540s_1 | -0.86 | 1.21 | -0.17 | 0.27 | 1.34 | 0.96 | -0.66 | -0.59 | -0.42 | 0.57 | 0.13 | 0.4 | 0.01 | 0.93 | 1.09 | 0.66 | 0.15 | 0.0575 | 0.56 | 0.260359 | 0.805634 | 0.902308 | 1 | 0.999631 | 0.692624 |
| Pkn1_P70268_S920s_1 | -0.9 | 0.07 | -0.45 | -0.42 | -0.44 | -0.48 | -0.66 | -0.5 | -0.14 | -0.51 | 0.22 | 0.54 | 0.29 | -0.27 | -0.15 | 0.04 | -0.095 | 0.4525 | 0.4025 | 0.224509 | 0.737397 | 0.208715 | 1 | 0.999631 | 0.67471 |
| Pkn2_G3UXH4_S534s_1 | 0.46 | 1.28 | 0.61 | -0.17 | -0.07 | 0.09 | -0.29 | 0.34 | 0.62 | 0.19 | 0.6 | 0.28 | 1.25 | 0.64 | 0.48 | 0.35 | -0.5275 | -0.1225 | 0.135 | 0.729134 | 0.158519 | 0.729987 | 1 | 0.999631 | 0.905923 |
| Pkp2_Q9CQ73_S82s_1 | -0.51 | -0.21 | -0.15 | -0.06 | -0.3 | -0.12 | 0.32 | -0.24 | -0.24 | -0.7 | 0.26 | -0.1 | -0.57 | 0.17 | 0.12 | 0.63 | 0.1475 | 0.0375 | 0.32 | 0.351155 | 0.583392 | 0.900061 | 1 | 0.999631 | 0.738069 |
| Pkp4_Q68FH0_S1036s_1 | -0.97 | -1.25 | -0.08 | -1.29 | -0.23 | -0.91 | -1.04 | -1.5 | -1.7 | -1.46 | -0.94 | -0.74 | -0.12 | -0.35 | 0 | -1.47 | -0.0225 | -0.3125 | 0.4125 | 0.361038 | 0.954261 | 0.420382 | 1 | 0.999631 | 0.741378 |
| Pkp4_Q68FH0_S1090s_1 | -1.2 | -1.97 | -0.48 | -1.49 | -1.73 | -1.86 | -1.39 | -1.88 | -1.91 | -1.37 | -0.77 | -1.01 | -0.34 | -1.64 | -1.02 | -0.29 | -0.43 | 0.02 | 0.4625 | 0.313944 | 0.246387 | 0.960805 | 1 | 0.999631 | 0.719939 |
| Pkp4_Q68FH0_S1090s_2 | -1.06 | -1.61 | -1.47 | -1.89 | -1.28 | -1.91 | -1.79 | -1.33 | -1.75 | -1.5 | -1.73 | -1.6 | -0.8 | -1.27 | -1.38 | -1.04 | -0.07 | -0.1375 | 0.385 | 0.232176 | 0.815065 | 0.625325 | 1 | 0.999631 | 0.682449 |
| Pkp4_Q68FH0_S1099s_1 | 0.02 | -1.75 | -0.79 | -1.09 | -0.74 | -1.13 | -1.64 | -1.57 | -1.17 | -0.77 | -2.44 | -1.2 | -0.39 | -0.48 | -0.78 | -0.96 | -0.3675 | -0.4925 | 0.25 | 0.545097 | 0.390182 | 0.32926 | 1 | 0.999631 | 0.833658 |
| Pkp4_Q68FH0_S1133s_1 | -0.78 | -0.66 | -0.76 | -0.84 | -1.02 | -0.79 | -1.3 | -1.06 | -0.88 | -0.61 | -0.77 | -0.61 | -0.03 | -0.57 | -0.84 | -0.24 | -0.2825 | 0.0425 | 0.34 | 0.265524 | 0.263649 | 0.861232 | 1 | 0.999631 | 0.694352 |
| Pkp4_Q68FH0_S1136s_1 | -0.88 | 0.39 | 0.09 | 0.02 | -0.55 | 0.22 | -0.21 | 0.06 | 0.33 | 0.22 | -0.17 | 0.45 | 1.11 | 0.31 | 0.42 | 0.59 | -0.025 | 0.3025 | 0.7025 | 0.081049 | 0.943092 | 0.392094 | 1 | 0.999631 | 0.546121 |
| Pkp4_Q68FH0_S213s_1 | 2.43 | -0.04 | -0.17 | 1.54 | 1.03 | 1 | 1.46 | 0.6 | 0.51 | 0.81 | 0.04 | -0.55 | -2 | -0.06 | -0.29 | -0.53 | 0.0825 | -0.7375 | -1.66 | 0.033216 | 0.889252 | 0.258905 | 1 | 0.999631 | 0.44294 |
| Pkp4_Q68FH0_S220s_1 | 0.26 | -0.43 | -0.69 | -0.96 | -0.41 | 0.29 | -1.26 | -1.37 | -1.07 | 0.35 | -1.58 | -0.87 | 0.22 | -0.92 | -2.2 | -1.32 | -0.2325 | -0.3375 | -0.6 | 0.274995 | 0.61009 | 0.4783 | 1 | 0.999631 | 0.700187 |
| Pkp4_Q68FH0_S246s_1 | -0.67 | 0.44 | -1.1 | -0.82 | -0.29 | 0.01 | -1.35 | -1.24 | 0.26 | -1.08 | -0.66 | -1.07 | 0.24 | 0.16 | -0.1 | -0.61 | -0.18 | -0.1 | 0.46 | 0.275416 | 0.696414 | 0.825039 | 1 | 0.999631 | 0.700187 |
| Pkp4_Q68FH0_S272s_1 | 2.73 | -0.4 | 0.32 | -0.72 | 1.17 | 1.52 | 0.08 | -0.31 | 0.29 | 1.57 | -0.14 | -0.21 | -1.9 | -0.64 | 0.09 | -1.08 | 0.1325 | -0.105 | -1.365 | 0.102085 | 0.86534 | 0.892427 | 1 | 0.999631 | 0.564448 |
| Pkp4_Q68FH0_S280s_1 | -1.22 | -3.17 | -0.79 | -0.77 | -2.41 | -2.85 | -1.05 | -1.53 | -2.08 | -1.55 | -1.88 | -1.14 | -1.04 | -2.01 | -1.14 | -1.25 | -0.4725 | -0.175 | 0.1275 | 0.8216 | 0.459631 | 0.755005 | 1 | 0.999631 | 0.938464 |
| Pkp4_Q68FH0_S313s_1 | -0.61 | -0.61 | -0.6 | -0.72 | -0.78 | -0.76 | -1.26 | -0.53 | -1 | -0.41 | -0.51 | -0.48 | -0.01 | -0.4 | -0.71 | -0.31 | -0.1975 | 0.035 | 0.2775 | 0.336055 | 0.455282 | 0.893479 | 1 | 0.999631 | 0.730466 |
| Pkp4_Q68FH0_S326s_2 | -0.76 | 1.81 | -2.27 | -1.92 | 1.23 | 0.71 | -2.2 | -1.32 | 1.39 | 0.4 | -1.29 | -1.53 | 1.77 | 1.62 | -0.93 | -1.27 | 0.39 | 0.5275 | 1.0825 | 0.314702 | 0.712841 | 0.600067 | 1 | 0.999631 | 0.720305 |
| Pkp4_Q68FH0_S336s_1 | -0.32 | -0.25 | -0.9 | -1.21 | -0.75 | -0.41 | -1.59 | -1.02 | -0.61 | 0.1 | -0.22 | -0.42 | 0.79 | -0.13 | -0.6 | -0.86 | -0.2725 | 0.3825 | 0.47 | 0.295554 | 0.459078 | 0.262628 | 1 | 0.999631 | 0.708945 |
| Pkp4_Q68FH0_S336s_2 | 0.01 | 0.2 | -0.98 | -0.97 | -1.82 | 0.01 | -0.31 | 0.88 | -0.29 | -0.34 | 0.3 | -0.46 | -0.81 | -0.43 | -0.45 | 0.87 | 0.125 | 0.2375 | 0.23 | 0.628938 | 0.830784 | 0.535633 | 1 | 0.999631 | 0.86642 |
| Pkp4_Q68FH0_S405s_1 | -0.66 | -2.26 | -1.75 | -1.76 | -2.32 | -2.62 | -1.64 | -2.26 | -2.26 | -1.67 | -2.19 | -1.73 | -2.32 | -2.13 | -1.87 | -1.52 | -0.6025 | -0.355 | -0.3525 | 0.390432 | 0.15505 | 0.371436 | 1 | 0.999631 | 0.758901 |
| Pkp4_Q68FH0_S421s_1 | -0.32 | -0.38 | -0.3 | -0.65 | -0.82 | -0.32 | -0.96 | -0.33 | -0.42 | -0.2 | -0.42 | -0.35 | 0.3 | -0.38 | -0.29 | 0.12 | -0.195 | 0.065 | 0.35 | 0.252044 | 0.481132 | 0.793536 | 1 | 0.999631 | 0.687761 |
| Pkp4_Q68FH0_S426s_1 | -0.9 | -1.68 | -1.69 | -1.65 | -2.01 | -1.58 | -1.55 | -1.79 | -2.38 | -0.57 | -2.16 | -2.49 | -1.23 | -1.43 | -2.18 | -1.61 | -0.2525 | -0.42 | -0.1325 | 0.700821 | 0.395727 | 0.383322 | 1 | 0.999631 | 0.896835 |
| Pkp4_Q68FH0_S437s_1 | -0.23 | -0.67 | -0.96 | -1 | -1.8 | -1.43 | -0.83 | -0.88 | -1.32 | -0.67 | -1.74 | -1.46 | -1.27 | -0.83 | -1.93 | -0.99 | -0.52 | -0.5825 | -0.54 | 0.150589 | 0.141471 | 0.107914 | 1 | 0.999631 | 0.623175 |
| Pkp4_Q68FH0_S446s_1 | 1.12 | 1.19 | 0.19 | 0.95 | 1.27 | 1.19 | 0.81 | 0.95 | 1.37 | 0.07 | 0.61 | 0.65 | 0.08 | 1.31 | 1.15 | 1.29 | 0.1925 | -0.1875 | 0.095 | 0.811804 | 0.537512 | 0.620426 | 1 | 0.999631 | 0.936949 |
| Pkp4_Q68FH0_S460s_1 | 1.61 | 1.5 | 0.2 | 0.78 | 1.18 | 1.35 | 0.52 | 0.52 | 1.19 | 1.29 | 0.66 | -0.05 | -0.13 | 0.85 | 0.18 | 0.93 | -0.13 | -0.25 | -0.565 | 0.207039 | 0.745652 | 0.576816 | 1 | 0.999631 | 0.660822 |
| Pkp4_Q68FH0_S75s_1 | -0.11 | 0.03 | 0.76 | 0.86 | -0.02 | -0.08 | -0.13 | 0.97 | -0.5 | -0.86 | -0.54 | 0.35 | 0.12 | 0.21 | -0.23 | 0.04 | -0.2 | -0.7725 | -0.35 | 0.309691 | 0.597804 | 0.065974 | 1 | 0.999631 | 0.718889 |
| Pkp4_Q68FH0_T1012t_1 | -1.06 | -0.12 | -0.07 | -0.66 | -0.08 | -0.07 | -1.22 | -0.42 | -0.33 | -0.79 | -0.29 | -0.01 | 1.16 | 0.09 | -0.04 | -0.53 | 0.03 | 0.1225 | 0.6475 | 0.157959 | 0.936124 | 0.715216 | 1 | 0.999631 | 0.628615 |
| Pkp4_Q68FH0_T1016t_1 | -1.07 | 1.41 | 1.61 | 0.26 | 0.56 | 1.29 | 0.22 | 1.74 | 0.45 | -0.95 | 1.55 | 1.59 | 1.98 | 1.26 | 1.65 | 1.73 | 0.4 | 0.1075 | 1.1025 | 0.082575 | 0.532011 | 0.887235 | 1 | 0.999631 | 0.548531 |
| Pkp4_Q68FH0_T282t_1 | 0.38 | -0.55 | 0.75 | -0.17 | -0.59 | 0.23 | -0.54 | -0.26 | 0.12 | 1.15 | -2.38 | -1.46 | -0.25 | 0.16 | 0.06 | -0.39 | -0.3925 | -0.745 | -0.2075 | 0.570992 | 0.297076 | 0.329196 | 1 | 0.999631 | 0.84338 |
| Plcb1_Q9Z1B3_S1197s_1 | 0.7 | 1.36 | 1.36 | 1.58 | 0.86 | 1.14 | 1.46 | 0.99 | 1.54 | 1.36 | 1.9 | 1.75 | 0.9 | 0.79 | 1.21 | 1.54 | -0.1375 | 0.3875 | -0.14 | 0.67188 | 0.644387 | 0.21676 | 1 | 0.999631 | 0.884779 |
| Plcb1_Q9Z1B3_S1200s_1 | 0.38 | 0.35 | 0.78 | 0.85 | 0.4 | 0.21 | 0.69 | 0.57 | 0.32 | -0.11 | 0.42 | 0.11 | -0.92 | -0.04 | 0.27 | 0.35 | -0.1225 | -0.405 | -0.675 | 0.088561 | 0.646123 | 0.166383 | 1 | 0.999631 | 0.553858 |
| Plcb1_Q9Z1B3_S848s_1 | 0.08 | 0.01 | 0.27 | 0.34 | -0.24 | -0.54 | -0.27 | -0.11 | -0.2 | 0.09 | 0.56 | -0.07 | -0.22 | -0.78 | 0.11 | -0.01 | -0.465 | -0.08 | -0.4 | 0.213572 | 0.083945 | 0.775597 | 1 | 0.999631 | 0.667146 |
| Plcb1_Q9Z1B3_S981s_1 | -0.38 | -0.83 | -2.54 | -1.41 | -0.89 | -0.97 | -1.09 | -1.17 | -1.26 | -0.2 | -2.07 | -1.71 | -0.67 | -0.89 | -1.82 | -2.48 | 0.26 | -0.02 | -0.175 | 0.762232 | 0.57002 | 0.971936 | 1 | 0.999631 | 0.915029 |
| Plcb1_Q9Z1B3_S982s_1 | -0.16 | -0.3 | -0.49 | -0.54 | -0.69 | -0.77 | -0.76 | -0.75 | -1.06 | -0.55 | -1.06 | -0.6 | -0.18 | -0.28 | -0.76 | -0.21 | -0.37 | -0.445 | 0.015 | 0.958247 | 0.143422 | 0.126815 | 1 | 0.999631 | 0.98551 |
| Plcb1_Q9Z1B3_T1164t_1 | -0.37 | -0.97 | -0.45 | -0.55 | -0.89 | -1.01 | -0.72 | -0.54 | -1.27 | -0.28 | -0.96 | -0.53 | -0.97 | -0.4 | -0.83 | -0.27 | -0.205 | -0.175 | -0.0325 | 0.916251 | 0.447706 | 0.585621 | 1 | 0.999631 | 0.973495 |
| Plcb3_P51432_S1105s_1 | 0.06 | 3.29 | 2.02 | 1.24 | 1.7 | 3.36 | 2.26 | 1.48 | 2.43 | 0.33 | 1.7 | 1.99 | 1.98 | 2.07 | 1.86 | 1.46 | 0.5475 | -0.04 | 0.19 | 0.761556 | 0.444954 | 0.955873 | 1 | 0.999631 | 0.914725 |
| Plcg1_Q62077_S1248s_1 | 1.86 | 0.16 | 0.13 | 2.05 | 1.31 | 0.67 | 2.23 | 1 | 1.38 | 0.79 | 1.82 | 1.08 | -1.42 | -0.15 | 0.49 | 0.28 | 0.2525 | 0.2175 | -1.25 | 0.064837 | 0.657953 | 0.683215 | 1 | 0.999631 | 0.515898 |
| Plcg1_Q62077_S1263s_1 | -0.44 | -1.19 | 0.24 | 0.71 | -0.14 | -1.96 | 0.51 | -0.39 | -1.73 | -1.92 | -2.66 | 0.56 | -1.25 | -0.76 | -0.11 | 0.59 | -0.325 | -1.2675 | -0.2125 | 0.694055 | 0.592502 | 0.102753 | 1 | 0.999631 | 0.895131 |
| Plch2_H3BIW6_S1024s_1 | 0.39 | -0.32 | -0.85 | -0.34 | -0.19 | -0.6 | -0.21 | -0.59 | -0.62 | -0.12 | -0.67 | -0.55 | -0.24 | -0.6 | -1.64 | -0.61 | -0.1175 | -0.21 | -0.4925 | 0.249575 | 0.717893 | 0.533524 | 1 | 0.999631 | 0.686359 |
| Plch2_H3BIW6_S1044s_1 | 0.66 | 0.49 | -0.42 | -0.01 | 0.81 | 1.2 | -0.29 | -0.09 | 1.32 | -0.87 | 0.12 | -0.23 | 0.81 | 1.39 | -1.09 | -0.6 | 0.2275 | -0.095 | -0.0525 | 0.927868 | 0.596057 | 0.848029 | 1 | 0.999631 | 0.976452 |
| Plch2_H3BIW6_S558s_1 | 0.08 | 0.66 | -0.55 | -0.97 | 0.93 | 0.3 | -0.22 | -0.25 | 0.85 | -1.54 | -0.57 | 0.78 | -0.37 | -0.61 | 0.2 | -1.69 | 0.385 | 0.075 | -0.4225 | 0.413972 | 0.391469 | 0.902798 | 1 | 0.999631 | 0.768655 |
| Plch2_H3BIW6_S559s_1 | -1.29 | 0.72 | -0.8 | -0.9 | 2.44 | 1.28 | 0.22 | -0.61 | 2.07 | -2.41 | 0.9 | 1.06 | 0.17 | -0.26 | 1.23 | -1.19 | 1.4 | 0.9725 | 0.555 | 0.371715 | 0.070593 | 0.308619 | 1 | 0.999631 | 0.746522 |
| Plch2_H3BIW6_S575s_1 | -0.44 | -0.57 | -0.39 | -0.61 | -0.68 | -0.58 | -0.91 | -0.87 | -0.62 | -0.63 | -0.92 | -0.27 | -0.18 | -0.59 | -0.66 | -0.42 | -0.2575 | -0.1075 | 0.04 | 0.883147 | 0.296028 | 0.684246 | 1 | 0.999631 | 0.963505 |
| Plch2_H3BIW6_S585s_1 | -0.22 | -1.91 | -0.44 | -1.79 | -0.09 | -0.79 | -1.72 | -1.93 | -1.21 | -0.84 | -1.41 | 0.13 | -1.22 | 0.66 | -1.34 | -0.41 | -0.0425 | 0.2575 | 0.5125 | 0.390738 | 0.93961 | 0.624945 | 1 | 0.999631 | 0.758901 |
| Plch2_H3BIW6_S886s_1 | 0.18 | -0.05 | -0.35 | -0.83 | -0.27 | -0.36 | -0.67 | -0.71 | -0.4 | 0.21 | -0.75 | -0.72 | 0.68 | -0.67 | -0.44 | -0.38 | -0.24 | -0.1525 | 0.06 | 0.879942 | 0.43755 | 0.665044 | 1 | 0.999631 | 0.962995 |
| Plch2_H3BIW6_S911s_1 | 2.67 | 1.34 | 2.02 | 1.87 | -0.68 | 2.31 | 2.75 | 0.37 | 0.44 | 1.91 | -0.01 | -0.4 | 0.53 | 0.81 | 1.37 | 0.34 | -0.7875 | -1.49 | -1.2125 | 0.009875 | 0.30811 | 0.018674 | 1 | 0.999631 | 0.375526 |
| Plch2_H3BIW6_S947s_1 | -0.86 | -0.4 | -0.71 | -0.81 | -1.3 | -0.54 | -1.64 | -0.71 | 0.05 | -1.13 | -0.8 | -1.79 | -0.25 | -0.54 | -0.81 | -0.37 | -0.3525 | -0.2225 | 0.2025 | 0.484319 | 0.289439 | 0.586469 | 1 | 0.999631 | 0.803422 |
| Plcl1_Q3USB7_S48s_1 | 0.08 | 1.05 | -0.73 | -0.21 | 0.89 | 1 | -0.55 | -1.19 | 1.36 | 1.57 | -1.02 | -2.23 | 1.05 | 0.74 | 0.04 | -0.03 | -0.01 | -0.1275 | 0.4025 | 0.385959 | 0.986569 | 0.883545 | 1 | 0.999631 | 0.756141 |
| Plcl1_Q3USB7_S570s_1 | 0.74 | -0.45 | 0.06 | 0.42 | -0.55 | -0.11 | 0.69 | 0.03 | 0.87 | 0.22 | -0.27 | 0.4 | -1.05 | -1.78 | 0.11 | -0.88 | -0.1775 | 0.1125 | -1.0925 | 0.036461 | 0.639583 | 0.763477 | 1 | 0.999631 | 0.453963 |
| Plcl1_Q3USB7_S77s_1 | 0.58 | -0.31 | -1.16 | -0.33 | 0.9 | -0.86 | -1.07 | -0.13 | 0.54 | -0.49 | -1.71 | -0.59 | 0.35 | -0.23 | -0.71 | -0.46 | 0.015 | -0.2575 | 0.0425 | 0.921187 | 0.977278 | 0.635764 | 1 | 0.999631 | 0.975032 |
| Plcl1_Q3USB7_S78s_1 | 0.44 | -0.84 | -1.16 | -0.33 | 0.7 | -1.29 | -1.07 | -0.13 | 0.27 | -0.67 | -1.71 | -0.59 | 0.16 | -0.65 | -0.71 | -0.46 | 0.025 | -0.2025 | 0.0575 | 0.890224 | 0.962477 | 0.689958 | 1 | 0.999631 | 0.9636 |
| Plcl1_Q3USB7_S96s_1 | -0.04 | 0.71 | 0.37 | 0.91 | 1.37 | 1.02 | 1 | 1.56 | 0.63 | 0.72 | 1.25 | 1.48 | 0.99 | 1.48 | 1.17 | 0.84 | 0.75 | 0.5325 | 0.6325 | 0.076478 | 0.032758 | 0.140025 | 1 | 0.999631 | 0.542546 |
| Plcl2_Q8K394_S1114s_1 | -0.27 | 0.23 | 0.03 | 0 | 0 | -0.06 | -0.29 | 0.08 | 0.05 | -0.03 | 0.36 | 0.18 | 0.21 | 0.16 | 0.15 | 0.36 | -0.065 | 0.1425 | 0.2225 | 0.419224 | 0.795384 | 0.586292 | 1 | 0.999631 | 0.772152 |
| Plcl2_Q8K394_S124s_1 | 1.34 | 0.65 | -0.63 | 1.75 | 1.32 | 0.91 | 1.26 | 1.18 | 0.77 | -0.16 | 1.03 | 1.06 | 0.01 | 0.7 | 0.87 | 0.69 | 0.39 | -0.1025 | -0.21 | 0.689373 | 0.439877 | 0.852356 | 1 | 0.999631 | 0.892416 |
| Plcl2_Q8K394_S18s_1 | 0.71 | -1.09 | -1.21 | 0.69 | 2.35 | 0.8 | 1.72 | 1.03 | 0.43 | -0.81 | 1.18 | -0.43 | -2.92 | 0.1 | -0.62 | -0.43 | 1.7 | 0.3175 | -0.7425 | 0.335494 | 0.01463 | 0.616045 | 1 | 0.999631 | 0.730235 |
| Plcl2_Q8K394_T585t_1 | -2.16 | -0.45 | -2.63 | -3.47 | -0.65 | -0.84 | -2.29 | -3.08 | -0.79 | -1.16 | -2.63 | -3.23 | -0.28 | -0.78 | -3.08 | -3.75 | 0.4625 | 0.225 | 0.205 | 0.822613 | 0.54561 | 0.767816 | 1 | 0.999631 | 0.938864 |
| Plcl2_Q8K394_T85t_1 | 1.45 | -0.5 | -3.26 | 1.84 | 2.34 | 0.1 | 2.46 | 1.78 | 0.77 | -1.62 | 1.44 | -0.99 | -2.59 | 0.6 | -0.15 | -0.73 | 1.7875 | 0.0175 | -0.6 | 0.602749 | 0.129151 | 0.988099 | 1 | 0.999631 | 0.857278 |
| Plec_Q9QXS1_S1324s_1 | -0.37 | -0.44 | -0.43 | -0.57 | -0.78 | -0.42 | -1.03 | -0.86 | -0.92 | -0.26 | -0.64 | -0.59 | -0.15 | -0.38 | -0.38 | -0.73 | -0.32 | -0.15 | 0.0425 | 0.876996 | 0.224038 | 0.571492 | 1 | 0.999631 | 0.96149 |
| Plec_Q9QXS1_S4274s_1 | 0.75 | -0.3 | -0.82 | -0.19 | 0.85 | -0.09 | -0.39 | 0.02 | 0.09 | -0.43 | -0.3 | -0.76 | -1.38 | 0.12 | -0.98 | -1.03 | 0.2375 | -0.21 | -0.6775 | 0.160588 | 0.572924 | 0.591949 | 1 | 0.999631 | 0.630397 |
| Plec_Q9QXS1_S4530s_1 | -0.15 | 0.2 | -1.11 | -0.69 | -0.21 | -0.66 | -0.65 | -0.59 | -0.31 | -0.95 | -0.5 | -0.31 | 0.44 | -0.17 | -0.56 | -0.49 | -0.09 | -0.08 | 0.2425 | 0.543529 | 0.792838 | 0.824133 | 1 | 0.999631 | 0.832881 |
| Plec_Q9QXS1_S609s_1 | -1.61 | -0.21 | -0.56 | -1.17 | -1.19 | -0.31 | -0.78 | -0.82 | -0.74 | -1.44 | -0.97 | -0.53 | 0.37 | -0.58 | -0.8 | -0.12 | 0.1125 | -0.0325 | 0.605 | 0.171178 | 0.765117 | 0.932992 | 1 | 0.999631 | 0.641448 |
| Plekha1_Q8BUL6_S308s_1 | 1.52 | 1.51 | 0.92 | 1.66 | 1.55 | 2.02 | 1.24 | 1.78 | 2.18 | 1.58 | 1.54 | 1.37 | -0.37 | 1.95 | 1.7 | 1.55 | 0.245 | 0.265 | -0.195 | 0.711199 | 0.419854 | 0.401061 | 1 | 0.999631 | 0.900552 |
| Plekha1_Q8BUL6_S372s_1 | 0.31 | -0.2 | -0.04 | 0.96 | -0.56 | -1.21 | 0.46 | 0.47 | 0.98 | -1.18 | 1.24 | 0.52 | -0.78 | 0.09 | -0.98 | 1.01 | -0.4675 | 0.1325 | -0.4225 | 0.407351 | 0.327204 | 0.811891 | 1 | 0.999631 | 0.766996 |
| Plekha2_F8WIK5_S312s_1 | 0.57 | 0.28 | -0.7 | 0.94 | 1.61 | -0.13 | 0.54 | 0.46 | 0.67 | 0.01 | 0.77 | 0.29 | -0.09 | 0.33 | 0.5 | 0.91 | 0.3475 | 0.1625 | 0.14 | 0.739873 | 0.47388 | 0.688166 | 1 | 0.999631 | 0.908631 |
| Plekha5_E9Q6H8_S1020s_1 | 0.09 | 0.25 | -0.24 | 0.1 | 0.21 | 0.5 | -0.31 | 0.2 | 0.23 | 0.06 | -0.44 | -0.24 | 0.42 | 0.55 | -0.01 | 0.25 | 0.1 | -0.1475 | 0.2525 | 0.386522 | 0.721225 | 0.600515 | 1 | 0.999631 | 0.756141 |
| Plekha5_E9Q6H8_S141s_1 | -0.33 | -0.03 | -1.42 | -1.88 | -1.42 | -1.23 | -1.42 | -1.18 | -0.77 | -0.14 | -1 | -1.03 | -0.24 | -0.56 | -2.1 | -0.78 | -0.3975 | 0.18 | -0.005 | 0.992823 | 0.370241 | 0.702135 | 1 | 0.999631 | 0.997601 |
| Plekha5_E9Q6H8_S417s_1 | -0.82 | 0.18 | -0.46 | 1 | 1.33 | 0.42 | 0.26 | 0.8 | 0.38 | 0.49 | 0.82 | 0.91 | 0.32 | 0.78 | 0.5 | 0.89 | 0.7275 | 0.675 | 0.6475 | 0.153969 | 0.129933 | 0.133996 | 1 | 0.999631 | 0.625198 |
| Plekha5_E9Q6H8_S484s_1 | 0.21 | -1.52 | -1.46 | -1.82 | -1.28 | -0.5 | -1.42 | -1.81 | -0.55 | 1.08 | -2.8 | -1.74 | -1.14 | -0.71 | -2.5 | -1.11 | -0.105 | 0.145 | -0.2175 | 0.698076 | 0.834167 | 0.861584 | 1 | 0.999631 | 0.895731 |
| Plekha5_E9Q6H8_S532s_1 | 0.79 | 2.16 | 4.24 | 1 | 2.84 | 1.86 | 4.89 | 1.93 | 5.68 | 1.14 | 5.79 | 6.86 | 5.84 | 0.79 | 2.37 | 0.33 | 0.8325 | 2.82 | 0.285 | 0.819462 | 0.374196 | 0.049303 | 1 | 0.999631 | 0.938303 |
| Plekha5_E9Q6H8_S56s_1 | -1.2 | -1.38 | -0.77 | -0.77 | -1.02 | -1.9 | -0.56 | -0.49 | -1.43 | -1.86 | 0.14 | -0.15 | -1.17 | -0.93 | -0.4 | -0.65 | 0.0375 | 0.205 | 0.2425 | 0.449211 | 0.920358 | 0.674903 | 1 | 0.999631 | 0.786889 |
| Plekha5_E9Q6H8_S574s_1 | -0.62 | -0.46 | -1.14 | -1.38 | -1.12 | -0.91 | -1.2 | -1.31 | -0.69 | -0.23 | -0.84 | -0.87 | 0.47 | -0.62 | -1.29 | -0.76 | -0.235 | 0.2425 | 0.35 | 0.428016 | 0.436871 | 0.456536 | 1 | 0.999631 | 0.77566 |
| Plekha6_Q7TQG1_S336s_1 | -0.13 | -0.41 | -0.99 | -0.63 | -0.55 | 0.12 | -0.87 | -0.07 | -1 | -0.57 | -1.03 | -1.29 | -0.28 | -0.24 | -0.79 | -0.56 | 0.1975 | -0.4325 | 0.0725 | 0.817049 | 0.553762 | 0.179529 | 1 | 0.999631 | 0.937777 |
| Plekha6_Q7TQG1_S426s_1 | -0.95 | -0.49 | -0.42 | -0.34 | -0.28 | -0.13 | -0.75 | 0.14 | -0.33 | -0.26 | -0.13 | -0.16 | 0.02 | 0.28 | -0.12 | -0.08 | 0.295 | 0.33 | 0.575 | 0.068763 | 0.333132 | 0.230048 | 1 | 0.999631 | 0.523794 |
| Plekha6_Q7TQG1_S573s_1 | 1.95 | -1.79 | -0.56 | -0.29 | -2.56 | -0.81 | -0.19 | -0.7 | -2.16 | -0.06 | -1.16 | -1.32 | -1.44 | -1.86 | -0.93 | -1.71 | -0.8925 | -1.0025 | -1.3125 | 0.08976 | 0.288805 | 0.220369 | 1 | 0.999631 | 0.556851 |
| Plekha6_Q7TQG1_S574s_1 | 0.17 | -0.94 | 0.41 | -0.44 | -1.12 | -0.23 | 0.69 | -1.21 | -0.97 | 0.23 | -0.78 | -2.02 | -1.28 | -1.38 | -0.89 | -1.41 | -0.2675 | -0.685 | -1.04 | 0.017252 | 0.598404 | 0.207484 | 1 | 0.999631 | 0.391024 |
| Plekha6_Q7TQG1_S816s_1 | -0.5 | -0.82 | -0.36 | -0.45 | -0.32 | -0.95 | -1.18 | -1.54 | -1.2 | -1.17 | -1.3 | -1.83 | -0.49 | -0.48 | -1.67 | 0 | -0.465 | -0.8425 | -0.1275 | 0.747968 | 0.171088 | 0.012769 | 1 | 0.999631 | 0.911931 |
| Plekha6_Q7TQG1_S864s_1 | -0.62 | -1.12 | 1.24 | 0.1 | -0.93 | -0.99 | 0.09 | 0.58 | -0.6 | -0.35 | 0.47 | 0.79 | 0 | -0.05 | 0.78 | 1.15 | -0.2125 | 0.1775 | 0.57 | 0.313709 | 0.716192 | 0.752506 | 1 | 0.999631 | 0.719939 |
| Plekha6_Q7TQG1_S901s_1 | -0.38 | -0.43 | -0.12 | -0.55 | -0.4 | -0.33 | -0.76 | -0.26 | -0.64 | -0.46 | -0.69 | -0.2 | 0.2 | 0.06 | -0.39 | 0.04 | -0.0675 | -0.1275 | 0.3475 | 0.240158 | 0.791854 | 0.630364 | 1 | 0.999631 | 0.683719 |
| Plekha6_Q7TQG1_S967s_1 | -0.94 | -1.23 | -1.54 | -0.68 | -0.9 | -1.09 | -1.07 | -0.6 | -1.28 | -1.24 | -0.79 | -1.38 | -2.78 | -1.15 | -0.94 | -0.16 | 0.1825 | -0.075 | -0.16 | 0.768348 | 0.532647 | 0.803857 | 1 | 0.999631 | 0.9177 |
| Plekha6_Q7TQG1_T1045t_1 | 0.92 | 0.03 | 0.21 | 0.52 | 0.36 | -0.51 | 0.73 | 0.45 | -0.54 | 0 | 0.4 | -0.17 | -0.78 | -0.09 | 0.49 | 0.25 | -0.1625 | -0.4975 | -0.4525 | 0.246898 | 0.650353 | 0.153751 | 1 | 0.999631 | 0.685751 |
| Plekha7_Q3UIL6_S373s_1 | -1.35 | -1.26 | -0.65 | -1.65 | -1.36 | -0.88 | -1.13 | -1.28 | -0.98 | -1.32 | -1.13 | -0.24 | -0.43 | -1.64 | -1.18 | -1.48 | 0.065 | 0.31 | 0.045 | 0.905239 | 0.827903 | 0.389905 | 1 | 0.999631 | 0.969056 |
| Plekha7_Q3UIL6_S668s_1 | -1.17 | -0.01 | -0.66 | -1.28 | -1.02 | -0.43 | -1.65 | 0.43 | -0.62 | -1.1 | 0.26 | 0 | 0.47 | -0.12 | 0.16 | 0.43 | 0.1125 | 0.415 | 1.015 | 0.018392 | 0.820954 | 0.34004 | 1 | 0.999631 | 0.391024 |
| Plekhf2_Q91WB4_S16s_1 | 0.06 | 1.25 | 0.48 | 0.43 | 1.19 | 0.83 | -0.37 | 1.26 | 0.3 | 0.86 | 0.37 | -0.92 | 1.2 | 1.06 | 0.34 | 0.76 | 0.1725 | -0.4025 | 0.285 | 0.438825 | 0.696454 | 0.377583 | 1 | 0.999631 | 0.781983 |
| Plekhf2_Q91WB4_S226s_1 | -0.15 | 0.72 | 0.43 | 0.91 | 1.14 | 1.26 | 0.31 | 1.12 | 0.97 | 0.32 | 0.82 | 0.83 | 0.56 | 1.5 | 1.11 | 0.67 | 0.48 | 0.2575 | 0.4825 | 0.204413 | 0.18942 | 0.437881 | 1 | 0.999631 | 0.659876 |
| Plekhg1_Q3UX37_S107s_1 | -0.42 | -0.7 | -1.14 | -1.82 | -1.57 | -0.63 | -1.4 | -1.06 | -1.22 | -0.07 | -1.21 | -1.94 | -0.52 | -0.44 | -1.31 | -1.34 | -0.145 | -0.09 | 0.1175 | 0.774645 | 0.704907 | 0.849553 | 1 | 0.999631 | 0.920998 |
| Plekhg1_Q3UX37_S613s_1 | 1.06 | 1.31 | 0.92 | 1.19 | 1.11 | 0.72 | 0.92 | 0.17 | 1.16 | 1.51 | 1.16 | 1.36 | 0.51 | 0.83 | 1.21 | 0.97 | -0.39 | 0.1775 | -0.24 | 0.415272 | 0.200459 | 0.492055 | 1 | 0.999631 | 0.769468 |
| Plekhg5_B1AS67_S925s_1 | 0.63 | 1.57 | 1.76 | 1.23 | 1.26 | 1.2 | 0.79 | 1.12 | 1.51 | 1.21 | 2 | 1.37 | 0.56 | 1.48 | 1.97 | 2.19 | -0.205 | 0.225 | 0.2525 | 0.570541 | 0.524207 | 0.517755 | 1 | 0.999631 | 0.84338 |
| Pllp_Q9DCU2_S13s_1 | -0.52 | -0.33 | 2.16 | -0.63 | -0.76 | -0.84 | 1.03 | 1.65 | -0.56 | -0.34 | 2.3 | 2.03 | 0.01 | -0.55 | 2.81 | 2.56 | 0.1 | 0.6875 | 1.0375 | 0.282987 | 0.900603 | 0.441618 | 1 | 0.999631 | 0.704545 |
| Pllp_Q9DCU2_S14s_1 | -0.46 | 0.03 | 0.8 | -0.4 | -0.28 | -0.74 | -0.77 | -0.22 | -0.3 | -0.15 | -0.26 | -0.47 | 0.12 | -0.15 | 0.16 | 0.02 | -0.495 | -0.2875 | 0.045 | 0.898612 | 0.18253 | 0.409527 | 1 | 0.999631 | 0.967127 |
| Pllp_Q9DCU2_S182s_1 | 0.45 | -1.37 | -0.44 | -0.17 | -0.54 | -0.88 | -0.53 | -0.18 | -0.35 | 0.19 | -0.27 | 0.21 | -2.97 | -0.15 | 0.03 | -0.37 | -0.15 | 0.3275 | -0.4825 | 0.501346 | 0.712627 | 0.43532 | 1 | 0.999631 | 0.814092 |
| Pln_P61014_S16s_1 | 0.48 | -0.13 | -0.47 | -0.06 | 2.6 | -0.96 | -0.2 | -0.94 | -0.23 | 2.61 | 0.17 | 2.49 | -1.44 | 0.35 | -0.04 | -0.63 | 0.17 | 1.305 | -0.395 | 0.379209 | 0.823082 | 0.083874 | 1 | 0.999631 | 0.751637 |
| Plp1_P60202_S114s_1 | -0.01 | 0.27 | 0 | -0.09 | -0.67 | 0.44 | -0.66 | -0.79 | 0.34 | -0.15 | -0.23 | 0.22 | 0.86 | 0.46 | 0.22 | -0.04 | -0.4625 | 0.0025 | 0.3325 | 0.291379 | 0.190229 | 0.992603 | 1 | 0.999631 | 0.706164 |
| Plp1_P60202_T118t_1 | -0.03 | -1.16 | -0.45 | -0.77 | -1.47 | 0.11 | -1.05 | -1.26 | -1.78 | -0.3 | 0.31 | -2.03 | -0.91 | -0.83 | -0.4 | -0.86 | -0.315 | -0.3475 | -0.1475 | 0.66199 | 0.461884 | 0.544112 | 1 | 0.999631 | 0.881128 |
| Pls3_B1AX58_S549s_1 | -0.44 | 0.43 | 1.03 | 0.07 | 0.41 | 0.61 | -0.12 | -0.18 | 0.54 | -0.25 | 0.6 | 0.75 | 0.04 | 0.12 | 0.81 | 0.98 | -0.0925 | 0.1375 | 0.215 | 0.601978 | 0.807566 | 0.729227 | 1 | 0.999631 | 0.856857 |
| Plxdc2_B1AY86_S456s_1 | -1.08 | -0.57 | -0.37 | -0.44 | -0.97 | -0.71 | -0.63 | -0.13 | -0.32 | -0.28 | 0 | 0.08 | -0.07 | -0.45 | 0.08 | 0.32 | 0.005 | 0.485 | 0.585 | 0.087808 | 0.986716 | 0.111227 | 1 | 0.999631 | 0.553787 |
| Plxna1_P70206_S1630s_1 | 0.1 | -0.87 | -0.01 | -0.19 | -0.48 | -0.78 | -0.16 | -0.14 | -0.31 | -0.46 | 0.23 | 0.45 | -0.95 | -0.9 | -0.62 | -0.29 | -0.1475 | 0.22 | -0.4475 | 0.201553 | 0.642426 | 0.530963 | 1 | 0.999631 | 0.658952 |
| Plxna2_P70207_S1630s_1 | -0.57 | -0.86 | -0.83 | -1.2 | -1.23 | -1.09 | -1.13 | -1.39 | -1.03 | -0.4 | -0.83 | -1.28 | -0.06 | -0.78 | -1.31 | -0.59 | -0.345 | -0.02 | 0.18 | 0.607041 | 0.199065 | 0.947072 | 1 | 0.999631 | 0.858527 |
| Plxna2_P70207_S1633s_1 | 1.15 | -1.21 | -1.25 | 0 | 0.73 | -0.18 | 0.42 | -0.07 | -0.39 | -0.96 | -0.37 | -1.16 | -1.25 | -0.08 | -0.71 | -1.13 | 0.5525 | -0.3925 | -0.465 | 0.429328 | 0.33489 | 0.488534 | 1 | 0.999631 | 0.776658 |
| Plxna3_P70208_S1531s_1 | 0.66 | -0.92 | 0.18 | 0.3 | 0.29 | -0.88 | -0.29 | 0.15 | -0.08 | 0.29 | -0.38 | 0.03 | -2.41 | -0.98 | 0.16 | 0.16 | -0.2375 | -0.09 | -0.8225 | 0.211665 | 0.578905 | 0.816051 | 1 | 0.999631 | 0.665474 |
| Plxnc1_Q9QZC2_S984s_1 | -1.21 | -0.86 | -0.16 | -0.94 | -0.12 | -0.65 | -0.5 | 0.03 | -0.15 | -0.83 | -0.24 | -0.06 | 0.75 | 0.45 | 0.6 | -0.49 | 0.4825 | 0.4725 | 1.12 | 0.015278 | 0.156593 | 0.17847 | 1 | 0.999631 | 0.388832 |
| Plxnd1_Q3UH93_S1384s_1 | -0.25 | 1.04 | 0.51 | 0.47 | 0.77 | 0.34 | -0.96 | 0.87 | 0.11 | 0.96 | -1.96 | -0.1 | 1.47 | 0.76 | 1.57 | 0.64 | -0.1875 | -0.69 | 0.6675 | 0.109528 | 0.692976 | 0.273303 | 1 | 0.999631 | 0.572838 |
| Pml_F7BTZ2_S9s_1 | 0.37 | 0.37 | 0.83 | 0.56 | 0.39 | -0.38 | -0.01 | 0.18 | 0.28 | 0.25 | 0.18 | 0.8 | 0.13 | 0.69 | 0.58 | 0.39 | -0.4875 | -0.155 | -0.085 | 0.767943 | 0.105285 | 0.580693 | 1 | 0.999631 | 0.917335 |
| Pml_F7BTZ2_T473t_1 | 0.62 | 0.49 | -1.66 | -1.18 | -0.19 | -0.48 | -0.86 | -1.51 | -0.45 | 0.88 | -1.57 | -1.31 | -2.41 | 0.42 | -1.55 | -0.61 | -0.3275 | -0.18 | -0.605 | 0.421438 | 0.579011 | 0.800329 | 1 | 0.999631 | 0.772817 |
| Pmm1_Q91W01_S182s_1 | -0.68 | -2.04 | -1.1 | -0.73 | -1.98 | -1.38 | -0.87 | -0.66 | -1.07 | -1.38 | -0.5 | -0.16 | -0.36 | -1.74 | -0.19 | -0.75 | -0.085 | 0.36 | 0.3775 | 0.422221 | 0.841507 | 0.401865 | 1 | 0.999631 | 0.773241 |
| Pmpcb_Q9CXT8_T45t_1 | -0.08 | 0.57 | -0.68 | -0.6 | -0.27 | 0.09 | -0.92 | -0.3 | 0.4 | 0.29 | -0.29 | 0.61 | 1.42 | 0.25 | -0.04 | -0.88 | -0.1525 | 0.45 | 0.385 | 0.469108 | 0.684068 | 0.245431 | 1 | 0.999631 | 0.796658 |
| Pnck_Q9QYK9_S316s_1 | 0.65 | -0.95 | 0.48 | -0.44 | -0.6 | -0.12 | -0.2 | -0.31 | 0.42 | 0.12 | 0.12 | 0.16 | -1.25 | -1.65 | 1.02 | 0.31 | -0.2425 | 0.27 | -0.3275 | 0.622083 | 0.547113 | 0.504187 | 1 | 0.999631 | 0.862299 |
| Pnisr_A2AJT4_S211s_1 | 0.18 | 0.83 | 0.87 | 0.36 | 0.2 | 0.24 | -0.27 | 0.2 | 0.58 | 0.69 | 0.29 | 0.44 | 1.13 | 0.82 | 0.56 | 1.12 | -0.4675 | -0.06 | 0.3475 | 0.279973 | 0.126239 | 0.832984 | 1 | 0.999631 | 0.701922 |
| Pnisr_A2AJT4_S601s_1 | -0.44 | -3.91 | -5.6 | -4.54 | -0.62 | -4.13 | -3.14 | -6.88 | -2.75 | -0.39 | -3 | -4.37 | -2.64 | -4.44 | -6.14 | -7.48 | -0.07 | 0.995 | -1.5525 | 0.247211 | 0.961472 | 0.411747 | 1 | 0.999631 | 0.685751 |
| Pnn_Q3TUQ5_S100s_1 | 1.62 | 1.97 | 2.9 | 2.52 | 1.22 | 2.08 | 2.17 | 1.29 | 2.03 | 2.94 | 1.87 | 2.15 | 1.64 | 0.91 | 2.03 | 0.97 | -0.5625 | -0.005 | -0.865 | 0.057588 | 0.17094 | 0.989725 | 1 | 0.999631 | 0.497148 |
| Pnn_Q3TUQ5_S347s_1 | 1.45 | 1.82 | 2.39 | 1.9 | 1.66 | 1.33 | 1.07 | 1.96 | 2.31 | 2.1 | 1.82 | 2.23 | 2.29 | 1.8 | 2.22 | 1.92 | -0.385 | 0.225 | 0.1675 | 0.597349 | 0.248267 | 0.458107 | 1 | 0.999631 | 0.855033 |
| Pnn_Q3TUQ5_S443s_1 | -3.96 | -2.27 | -3.02 | -4.18 | -2.45 | -2.05 | -2.96 | -4.33 | -2.94 | -3.35 | -3.05 | -4.93 | -1.73 | -1.43 | -4.01 | -3.46 | 0.41 | -0.21 | 0.7 | 0.321694 | 0.499975 | 0.720332 | 1 | 0.999631 | 0.724092 |
| Pnn_Q3TUQ5_S66s_1 | -0.33 | 0.91 | 1.17 | 0.54 | -0.18 | 0.11 | -0.33 | -0.08 | 0.36 | 0.74 | 0.96 | 0.88 | 1.19 | 0.66 | 0.07 | 1.03 | -0.6925 | 0.1625 | 0.165 | 0.697465 | 0.080613 | 0.667944 | 1 | 0.999631 | 0.895731 |
| Pnn_Q3TUQ5_S699s_1 | 1.88 | -0.04 | 0.16 | 0.48 | -0.33 | 0.03 | 0.96 | -0.68 | 0.2 | 1.33 | 0.4 | -0.67 | 0.15 | -1.2 | -0.97 | -0.97 | -0.625 | -0.305 | -1.3675 | 0.020714 | 0.246683 | 0.583923 | 1 | 0.999631 | 0.397568 |

| | | | | | | | | | | | | | | | | | | | | | | | | | | |
|---|---|---|---|---|---|---|---|---|---|---|---|---|---|---|---|---|---|---|---|---|---|---|---|---|---|---|
| Pnpla6_Q3TRM4_S306s_1 | 0.17 | 0.18 | -0.33 | -0.87 | -1.3 | 0.35 | -0.8 | -0.05 | 0.67 | 0.5 | -0.64 | -2.04 | -0.35 | 0.52 | -0.18 | -2.3 | -0.2375 | -0.165 | -0.365 | 0.544438 | 0.588719 | 0.787608 | | 1 | 0.999631 | 0.833274 |
| Pnpo_Q91XF0_S164s_1 | 0.33 | 1.52 | 1.2 | 0.5 | 0.6 | 0.87 | 0.4 | 0.96 | 0.56 | 0.58 | 1.42 | 1.33 | 0.83 | 0.97 | 1.65 | 0.98 | -0.18 | 0.085 | 0.22 | 0.561997 | 0.60265 | 0.825663 | | 1 | 0.999631 | 0.84072 |
| Pnpo_Q91XF0_S165s_1 | 0.42 | 0.84 | 0.83 | -0.18 | 0.53 | 0.82 | -0.1 | 0.34 | 0.82 | 0.84 | 0.16 | 0.17 | 1.3 | 1.51 | 0.69 | 0.66 | -0.08 | 0.02 | 0.5625 | 0.148456 | 0.815151 | 0.954119 | | 1 | 0.999631 | 0.62132 |
| Pnpo_Q91XF0_S241s_1 | -0.39 | -0.43 | 0.96 | -0.29 | -0.56 | -0.62 | 0.09 | 0.62 | -0.73 | -0.46 | 0.22 | 0.63 | 0.2 | -0.41 | 1.95 | 1.28 | -0.08 | -0.0475 | 0.7925 | 0.189859 | 0.853981 | 0.915547 | | 1 | 0.999631 | 0.648674 |
| Pogz_D3YUW8_S1265s_1 | 0.24 | 0.38 | -0.23 | -0.8 | -0.37 | -1.27 | -0.78 | -1.26 | 0.01 | 0.11 | -1.39 | -0.76 | 0.08 | -0.23 | -0.66 | -0.4 | -0.8175 | -0.405 | -0.2 | 0.580408 | 0.046119 | 0.366199 | | 1 | 0.999631 | 0.846774 |
| Poldip3_F6VR84_S127s_1 | 0.17 | 0.41 | 1.5 | 0.55 | 0.8 | 1.07 | -0.44 | 0.67 | 1.02 | 1.14 | 0.61 | 0.17 | 1.02 | 2.15 | 0.93 | 1.37 | -0.1325 | 0.0775 | 0.71 | 0.113079 | 0.759923 | 0.840467 | | 1 | 0.999631 | 0.577408 |
| Pole4_E0CYA9_T11t_1 | 0.82 | 0.02 | -0.22 | 0.66 | 0.84 | -0.15 | 2.35 | -0.08 | -0.91 | 1.19 | -1.17 | 1.4 | -1.46 | -0.98 | -0.27 | -0.11 | 0.42 | -0.1925 | -1.025 | 0.031491 | 0.472298 | 0.767269 | | 1 | 0.999631 | 0.441911 |
| Poll_Q9QXE2_S175s_1 | 0.44 | 0.17 | 0.72 | 0.27 | 0.36 | 0.08 | -0.04 | 0.23 | 0.51 | 0.39 | 0.14 | 0.63 | 1.21 | 0.77 | 0.3 | 0.69 | -0.2425 | 0.0175 | 0.3425 | 0.288512 | 0.357708 | 0.948264 | | 1 | 0.999631 | 0.706046 |
| Polr2a_P08775_S1913s_1 | 0.38 | 0.73 | 0.81 | 0.36 | 0.67 | 1.17 | 0.33 | 0.07 | 1.23 | -0.13 | 0.26 | 0.31 | 1.05 | 0.8 | 1.41 | 1.15 | -0.01 | -0.1525 | 0.5325 | 0.091119 | 0.974712 | 0.66452 | | 1 | 0.999631 | 0.557085 |
| Polr2a_P08775_S1920s_1 | 0.19 | -0.34 | 0.35 | 0.58 | -0.15 | -0.37 | -0.26 | 0.24 | -0.1 | 0.09 | 0.78 | 0.69 | -0.09 | -0.49 | -0.29 | 0.38 | -0.33 | 0.17 | -0.3175 | 0.357799 | 0.286221 | 0.618198 | | 1 | 0.999631 | 0.741099 |
| Polr3c_F6QK97_S78s_1 | -0.19 | 1.58 | 1.82 | -0.63 | 1.09 | 1.1 | 1.98 | -1.84 | 0.83 | -0.8 | 0.03 | 0.71 | 0.26 | -0.46 | 0.65 | -1.33 | -0.0625 | -0.4525 | -0.865 | 0.21867 | 0.94449 | 0.491764 | | 1 | 0.999631 | 0.669859 |
| Polr3e_Q9CZT4_S136s_1 | -0.88 | 0.14 | 1.25 | 0.8 | 0.54 | -0.1 | -0.06 | 0.79 | 0.82 | 0.43 | 0.82 | 0.52 | 1 | -0.37 | 1.58 | 1.41 | -0.035 | 0.32 | 0.5775 | 0.336747 | 0.942374 | 0.492815 | | 1 | 0.999631 | 0.730499 |
| Pom121_Q8K3Z9_S159s_1 | 0.22 | -0.18 | -0.78 | 0.21 | -0.57 | -0.62 | -0.24 | -0.29 | -0.74 | -0.02 | 0.12 | -0.58 | -0.21 | -0.62 | -0.57 | 0.51 | -0.2975 | -0.1725 | -0.09 | 0.815073 | 0.348426 | 0.626678 | | 1 | 0.999631 | 0.937297 |
| Pom121_Q8K3Z9_T421t_1 | 1.46 | -0.83 | 0.7 | 0.27 | -0.57 | 0.24 | 0.72 | -0.52 | -1.05 | 1.62 | -0.04 | -0.71 | -1.08 | -2.03 | -0.27 | -0.88 | -0.4325 | -0.445 | -1.465 | 0.023736 | 0.421292 | 0.518118 | | 1 | 0.999631 | 0.415644 |
| Ppap2a_Q61469_T279t_1 | 0.46 | -1.74 | -0.43 | -0.21 | 0.72 | 0.87 | 0.93 | 0.5 | 0.53 | -0.88 | -0.32 | -0.29 | 0.74 | -0.71 | 0.36 | -1.86 | 1.235 | 0.24 | 0.1125 | 0.86569 | 0.020554 | 0.641705 | | 1 | 0.999631 | 0.955857 |
| Ppap2b_Q99JY8_S19s_1 | -0.6 | -1.55 | 0.08 | -0.73 | -1.69 | -1.04 | 0.12 | -0.19 | -1.25 | 0 | -0.19 | -0.61 | -0.62 | -0.97 | 0.34 | -0.67 | 4.42E-16 | 0.1875 | 0.22 | 0.620417 | 1 | 0.666 | | 1 | 0.999631 | 0.861888 |
| Ppap2c_G3XA61_S209s_1 | -1.41 | -0.45 | -0.12 | -0.9 | 0.57 | 0.18 | -0.64 | 0.34 | -0.68 | 0.19 | -0.72 | 0.65 | 1.16 | 0.61 | 0.35 | 0.41 | 0.8325 | 0.58 | 1.3525 | 0.004537 | 0.056523 | 0.203418 | | 1 | 0.999631 | 0.351308 |
| Ppapdc2_Q9D4F2_S24s_1 | 0.35 | -1.56 | -1.77 | -1.14 | -1.29 | -0.59 | -0.72 | -1.77 | -1.26 | -0.05 | -2.71 | -2.09 | -1.18 | -0.49 | -1.04 | -1.53 | -0.0625 | -0.4975 | -0.03 | 0.952295 | 0.902933 | 0.464253 | | 1 | 0.999631 | 0.984317 |
| Ppapdc2_Q9D4F2_S60s_1 | 1.99 | 0.27 | -1.35 | -1.17 | -0.16 | 0.99 | 0.13 | -2.86 | 0.5 | 1.76 | -1.92 | -2.38 | -0.91 | -0.02 | -2.82 | -2.8 | -0.41 | -0.445 | -1.5725 | 0.105158 | 0.676594 | 0.681061 | | 1 | 0.999631 | 0.568213 |
| Ppapdc2_Q9D4F2_S67s_1 | 0.38 | -0.51 | -0.91 | 0.43 | 0.34 | 0.01 | -0.15 | -0.42 | 0.01 | -0.33 | 0.35 | -0.27 | -1.16 | 0.2 | -0.69 | 0.14 | 0.0975 | 0.0925 | -0.225 | 0.628371 | 0.798305 | 0.810737 | | 1 | 0.999631 | 0.865998 |
| Ppapdc3_Q91WB2_S43s_1 | 1.25 | 1.33 | -0.79 | 0.21 | 0.7 | 0.62 | -0.37 | -2.58 | 1.12 | 1.44 | -1.23 | -1.18 | -1 | 0.26 | -0.97 | -0.12 | -0.9075 | -0.4625 | -0.9575 | 0.10588 | 0.271556 | 0.552966 | | 1 | 0.999631 | 0.568797 |
| Ppapdc3_Q91WB2_S62s_1 | -0.2 | 0.73 | 0.41 | 0.28 | 0.19 | 0.05 | -0.07 | 0.48 | -0.07 | -0.36 | -0.06 | 0.02 | 0.63 | 0.79 | 0.45 | 0.3 | -0.1425 | -0.4225 | 0.2375 | 0.453492 | 0.630276 | 0.171992 | | 1 | 0.999631 | 0.788717 |
| Ppfia1_B2RXQ2_S150s_1 | -0.08 | 0.64 | -0.69 | -1.19 | 0.39 | 0.09 | -1.13 | -2.48 | -0.35 | -0.3 | -1.05 | -0.18 | 1.34 | -0.5 | 0.23 | -0.55 | -0.4525 | -0.14 | 0.46 | 0.41205 | 0.509925 | 0.749448 | | 1 | 0.999631 | 0.767948 |
| Ppfia1_B2RXQ2_S239s_1 | -1.22 | -1.49 | -0.03 | -1.12 | -2.01 | -1.62 | -0.68 | -0.63 | -1.45 | -1.03 | -0.32 | -0.18 | -1 | -1.34 | -0.09 | -0.13 | -0.27 | 0.22 | 0.325 | 0.474827 | 0.555498 | 0.616243 | | 1 | 0.999631 | 0.799827 |
| Ppfia1_B2RXQ2_S732s_1 | 3.08 | 1.03 | 0.04 | 0.94 | 1.37 | 0.01 | 0.79 | 0.87 | 1.52 | 2.79 | -0.35 | -0.73 | -2.42 | 1 | -0.81 | -1.21 | -0.5125 | -0.465 | -2.1325 | 0.02624 | 0.423513 | 0.610804 | | 1 | 0.999631 | 0.422285 |
| Ppfia1_B2RXQ2_S775s_1 | -0.03 | -0.28 | 1.02 | -0.06 | -0.69 | -0.39 | -0.09 | 0.13 | -0.06 | 0.58 | -0.16 | 0.41 | 0.03 | -0.32 | 0.18 | 0.13 | -0.4225 | 0.03 | -0.1575 | 0.664244 | 0.262025 | 0.935284 | | 1 | 0.999631 | 0.882037 |
| Ppfia1_B2RXQ2_S797s_1 | 1.09 | 0.25 | 0.29 | 1.14 | 0.57 | -0.29 | 0.75 | 0.21 | 0.18 | 0.32 | 0.69 | 0.57 | -1.05 | -0.4 | -0.25 | 0.31 | -0.3825 | -0.2525 | -1.04 | 0.023937 | 0.301486 | 0.445559 | | 1 | 0.999631 | 0.415644 |
| Ppfia2_F7CYX4_S236s_1 | 1.91 | -1.02 | -2.29 | -1.51 | -1.25 | -0.56 | -1.57 | -2.18 | -0.95 | -0.18 | -2.28 | -1.92 | -0.33 | 0.19 | -1.72 | -1.35 | -0.6625 | -0.605 | -0.075 | 0.931985 | 0.442766 | 0.505808 | | 1 | 0.999631 | 0.978168 |
| Ppfia2_F7CYX4_S689s_1 | 0.07 | 0.16 | -0.21 | -0.19 | 0.59 | 0.1 | -0.66 | -0.27 | 0.06 | -0.04 | -0.73 | 0.03 | -0.93 | 0.43 | -0.36 | 0.19 | -0.0175 | -0.1275 | -0.125 | 0.731987 | 0.957171 | 0.664669 | | 1 | 0.999631 | 0.907231 |
| Ppfia2_F7CYX4_S774s_1 | -0.4 | -0.21 | 0.49 | -0.33 | -0.37 | -0.17 | -0.23 | -0.22 | -0.01 | 0.43 | 0.12 | 0.68 | 0.69 | 0.13 | -0.06 | 0.2 | -0.135 | 0.4175 | 0.3525 | 0.302192 | 0.637856 | 0.209541 | | 1 | 0.999631 | 0.714978 |
| Ppfia3_P60469_S1164s_1 | 0.23 | -0.2 | -1.19 | 0.25 | 0.54 | 0.66 | -0.09 | 0.03 | 0.05 | -1.08 | 0.02 | -1.37 | -1.3 | 0.34 | -1.19 | -0.87 | 0.5125 | -0.3675 | -0.5275 | 0.294268 | 0.211606 | 0.450697 | | 1 | 0.999631 | 0.708293 |
| Ppfia3_P60469_S142s_1 | -0.76 | -0.15 | -0.3 | -0.58 | -0.34 | -0.46 | -0.88 | -0.24 | -0.23 | -0.51 | -0.53 | 0.19 | 0.45 | 0.03 | 0 | 0.06 | -0.0325 | 0.1775 | 0.5825 | 0.068714 | 0.907195 | 0.553037 | | 1 | 0.999631 | 0.523794 |
| Ppfia3_P60469_S17s_1 | 0.41 | -0.01 | -0.06 | 0.76 | 0.73 | 0.03 | 0.44 | 0.66 | 0.21 | 0.19 | 0.28 | 0.03 | 0.2 | 0.19 | 0.21 | 0.55 | 0.19 | -0.0975 | 0.0125 | 0.967611 | 0.539795 | 0.735698 | | 1 | 0.999631 | 0.988581 |
| Ppfia3_P60469_S508s_1 | 0.9 | 2.12 | 1.51 | 1.49 | 1.53 | 1.64 | 0.43 | 1.43 | 1.79 | 1.62 | 1.21 | 1.86 | 0.55 | 2.17 | 1.75 | 1.95 | -0.2475 | 0.115 | 0.1 | 0.821102 | 0.524959 | 0.73313 | | 1 | 0.999631 | 0.938464 |
| Ppfia3_P60469_S537s_1 | 1.06 | -0.32 | -1.29 | -0.23 | 0.97 | 0.47 | -0.28 | -0.64 | -0.17 | 0.17 | -0.63 | -0.8 | -0.95 | 1.15 | -0.37 | -0.34 | 0.325 | -0.1625 | 0.0675 | 0.910395 | 0.560465 | 0.746863 | | 1 | 0.999631 | 0.97105 |
| Ppfia3_P60469_S645s_1 | -0.91 | 0.38 | 0.11 | -1.41 | 0.39 | -0.02 | -0.06 | 0.26 | 0.46 | -1.52 | 0.5 | 1.29 | -0.25 | -0.49 | -1.07 | -1.31 | 0.6 | 0.64 | -0.3225 | 0.503295 | 0.183139 | 0.34275 | | 1 | 0.999631 | 0.814092 |
| Ppfia3_P60469_S683s_1 | 1.25 | -0.38 | -1.31 | 0.78 | 1.46 | 0.23 | 0.47 | 0.78 | 0.25 | -0.15 | 0.11 | -0.97 | -1.9 | 0.31 | -0.25 | -0.27 | 0.65 | -0.275 | -0.6125 | 0.370259 | 0.277109 | 0.639925 | | 1 | 0.999631 | 0.746338 |
| Ppfia3_P60469_S69s_1 | -0.92 | -1.67 | -1.61 | -0.49 | -1.7 | -2.87 | -0.65 | -0.96 | -1.6 | -0.94 | -0.63 | -0.44 | -0.09 | -1.59 | -1.35 | -1.06 | -0.3725 | 0.27 | 0.15 | 0.733132 | 0.485629 | 0.5003 | | 1 | 0.999631 | 0.907434 |
| Ppfia3_P60469_S75s_1 | 1.34 | 1.03 | 0.43 | 1.28 | 1.56 | 1.75 | 0.45 | 1.31 | 1.61 | 0.63 | -0.29 | 0.52 | -0.06 | 1.85 | 0.68 | 0.85 | 0.2475 | -0.4025 | -0.19 | 0.671134 | 0.510627 | 0.368463 | | 1 | 0.999631 | 0.884528 |
| Ppfia3_P60469_T714t_1 | -0.61 | -0.48 | -0.89 | -1.18 | -0.51 | -0.56 | -1.24 | -0.98 | -0.78 | -0.06 | -0.92 | -0.71 | 0.33 | -0.28 | -0.62 | -0.38 | -0.0325 | 0.1725 | 0.5525 | 0.11756 | 0.913203 | 0.583437 | | 1 | 0.999631 | 0.585087 |
| Ppfia3_P60469_T714t_2 | 0.46 | 0.35 | -0.45 | 1.35 | 0.22 | -1.53 | 0.86 | 3.5 | -0.44 | -1.25 | 2.32 | 0.07 | -0.65 | -0.82 | 0.02 | 0.77 | 0.335 | -0.2525 | -0.5975 | 0.24582 | 0.726132 | 0.736945 | | 1 | 0.999631 | 0.685751 |
| Ppfia3_P60469_T718t_2 | 0.46 | 0.35 | -0.45 | 1.35 | 0.22 | -1.53 | 0.86 | 3.5 | -0.44 | -1.25 | 2.32 | 0.07 | -0.65 | -0.82 | 0.02 | 0.77 | 0.335 | -0.2525 | -0.5975 | 0.24582 | 0.726132 | 0.736945 | | 1 | 0.999631 | 0.685751 |
| Ppfia4_B8QI36_S140s_1 | -0.88 | -0.25 | -0.2 | -0.57 | -0.7 | -0.74 | -0.56 | -0.36 | -0.83 | -0.26 | -0.64 | 0.18 | -0.22 | -0.41 | 0.26 | 0.14 | -0.115 | 0.0875 | 0.4175 | 0.201921 | 0.672592 | 0.789587 | | 1 | 0.999631 | 0.658952 |
| Ppfia4_B8QI36_S641s_1 | -0.58 | 0.37 | -0.51 | -0.58 | -0.47 | -0.73 | -0.79 | -0.35 | -0.47 | -1.2 | -0.58 | -0.79 | -1.04 | -0.53 | -0.23 | -1.14 | -0.26 | -0.435 | -0.41 | 0.275229 | 0.410398 | 0.211437 | | 1 | 0.999631 | 0.700187 |
| Ppfia4_B8QI36_S701s_1 | 0.07 | -0.05 | 1.02 | -0.06 | -0.55 | -0.12 | -0.09 | 0.13 | 0.16 | 0.83 | -0.16 | 0.41 | 0.27 | -0.07 | 0.18 | 0.13 | -0.4025 | 0.065 | -0.1175 | 0.728235 | 0.247039 | 0.858406 | | 1 | 0.999631 | 0.905923 |
| Ppfibp1_F6S1C4_S128s_1 | 0.76 | 1.68 | 1.29 | 1.34 | 1.32 | 1.28 | 0.91 | 0.99 | 2.06 | 1.91 | 1.36 | 1.9 | 1.73 | 1.63 | 2.07 | 2.39 | -0.1425 | 0.54 | 0.6875 | 0.059989 | 0.623833 | 0.107427 | | 1 | 0.999631 | 0.505461 |
| Ppfibp1_F6S1C4_S48s_1 | -0.38 | 0 | 0.8 | -0.13 | -0.26 | -0.08 | 0.71 | 0.14 | 0.99 | 0.45 | 0.72 | 0.67 | 0.32 | -0.6 | 0.54 | -0.02 | 0.055 | 0.635 | -0.0125 | 0.974251 | 0.877351 | 0.078964 | | 1 | 0.999631 | 0.990847 |
| Ppfibp1_F6YZ95_S99s_1 | 0.38 | 1.16 | 1.07 | 0.29 | 0.45 | 0.72 | 0.03 | 0.19 | 0.73 | 1.13 | 0.3 | 1.18 | 0.7 | 0.87 | 0.43 | 0.82 | -0.3775 | 0.11 | -0.02 | 0.950943 | 0.255922 | 0.75153 | | 1 | 0.999631 | 0.984079 |
| Ppig_A2AR02_S413s_1 | 2.09 | 0.21 | 2.63 | -0.67 | 2.81 | -0.14 | 4.53 | 0.29 | 5.09 | 3.17 | 3.58 | 4.48 | 3.29 | -0.36 | 0.94 | -1.05 | 0.8075 | 3.015 | -0.36 | 0.73217 | 0.488357 | 0.003275 | | 1 | 0.999631 | 0.907231 |
| Ppig_A2AR02_S714s_1 | 1.81 | 0.65 | 1.51 | 0.98 | -0.53 | 0.92 | 2.4 | -1.01 | 0.21 | 1.34 | 0.39 | 0.27 | 0.42 | 0.02 | 0.89 | 0.32 | -0.7925 | -0.685 | -0.825 | 0.043655 | 0.284808 | 0.103359 | | 1 | 0.999631 | 0.475443 |
| Ppig_A2AR02_T356t_1 | -1.27 | -0.56 | 0.74 | -0.38 | -0.57 | -0.31 | -0.32 | -0.26 | 0.19 | 0.21 | 0.47 | -0.16 | 0.55 | 0.98 | -0.23 | -0.13 | 0.0025 | 0.545 | 0.66 | 0.197009 | 0.995228 | 0.228045 | | 1 | 0.999631 | 0.653716 |
| Ppip5k1_F6Q5C6_S140s_1 | -0.09 | 0.22 | 0.54 | 0.23 | 1.51 | 0.4 | -0.28 | 0.53 | 0.46 | 0.81 | 0.5 | 1.12 | 0.86 | 1.78 | 1 | 0.27 | 0.315 | 0.4975 | 0.7525 | 0.068321 | 0.434675 | 0.109322 | | 1 | 0.999631 | 0.523149 |
| Ppip5k1_F6Q5C6_S160s_1 | -0.07 | -0.38 | -0.52 | -0.17 | -0.07 | 0.07 | 0.18 | 0.41 | -0.11 | 0.05 | -0.29 | 0.14 | -0.54 | 0.43 | 0.01 | 0.07 | 0.4325 | 0.2325 | 0.2775 | 0.386554 | 0.115143 | 0.384466 | | 1 | 0.999631 | 0.756141 |
| Ppip5k2_E9Q9J4_S1005s_1 | -1.36 | -1.33 | 0.71 | -0.4 | -0.62 | -1.47 | -0.29 | 0.1 | -0.96 | -1.16 | 1.17 | -0.96 | -0.33 | 0.05 | 0.87 | -0.77 | 0.025 | 0.1175 | 0.55 | 0.335344 | 0.963342 | 0.858893 | | 1 | 0.999631 | 0.730219 |
| Ppm1a_Q9EQE2_S135s_1 | -0.45 | 1.18 | -0.6 | 0.1 | 0.44 | 0.35 | -0.4 | -0.73 | 0.1 | -0.24 | -0.99 | 0.48 | 0.85 | 1.02 | -0.03 | -0.07 | -0.1425 | -0.22 | 0.385 | 0.430681 | 0.762583 | 0.653408 | | 1 | 0.999631 | 0.778235 |
| Ppm1a_Q9EQE2_S216s_1 | 0.14 | 0.96 | 1.16 | 2.64 | 1.36 | 0.01 | 2.36 | 2.92 | -0.18 | 0.61 | 2.82 | 2.53 | -0.19 | 0.24 | 1.43 | 2.49 | 0.4375 | 0.22 | -0.2325 | 0.742796 | 0.550507 | 0.780534 | | 1 | 0.999631 | 0.90972 |
| Ppm1e_Q80TL0_S395s_1 | 0.48 | 0.84 | 0.52 | 1.32 | 0.41 | 1.5 | 1.28 | 1.53 | 0.64 | 0.42 | 1.75 | 0.34 | -1.35 | 1.2 | 1.19 | 1 | 0.39 | -0.0025 | -0.28 | 0.639557 | 0.285751 | 0.994946 | | 1 | 0.999631 | 0.870995 |
| Ppm1e_Q80TL0_S532s_1 | -0.03 | 1.12 | 0.93 | 0.53 | 1.05 | 0.74 | 0.09 | 1.08 | 1.12 | 0.88 | 0.67 | 1.45 | 0.91 | 1.55 | 1.34 | 1.15 | 0.1025 | 0.3925 | 0.6 | 0.108455 | 0.778693 | 0.272996 | | 1 | 0.999631 | 0.570434 |
| Ppm1e_Q80TL0_S545s_1 | 1 | 1.39 | 0.83 | 1.4 | 1.58 | 1.67 | 1.21 | 1.24 | 1.54 | 1.36 | 1.6 | 1.33 | -0.8 | 1.25 | 0.74 | 1.72 | 0.27 | 0.3025 | -0.4275 | 0.428732 | 0.335696 | 0.276971 | | 1 | 0.999631 | 0.776573 |
| Ppm1h_Q3UYC0_S118s_1 | 0.34 | -0.7 | -1.82 | -0.93 | -0.38 | -0.88 | -0.71 | -1.86 | -0.82 | -0.51 | -1.04 | -1.39 | -0.22 | -0.94 | -1.52 | -1.71 | -0.18 | -0.1625 | -0.32 | 0.545614 | 0.724646 | 0.728045 | | 1 | 0.999631 | 0.833826 |
| Ppm1h_Q3UYC0_S123s_1 | -1.31 | -2.29 | -1.08 | -1.03 | -1.11 | -2.51 | -0.61 | -0.88 | -2.52 | -1.08 | -0.86 | -0.28 | -1.98 | -1.92 | -0.58 | -0.1 | 0.15 | 0.2425 | 0.2825 | 0.59405 | 0.758273 | 0.645342 | | 1 | 0.999631 | 0.85389 |
| Ppm1h_Q3UYC0_S197s_1 | -0.98 | 1.17 | 1.74 | 0.78 | 0.61 | 0.91 | 0.79 | 1 | 0.52 | -0.46 | 1.65 | 2.41 | 1.59 | 0.47 | 1.52 | 1.82 | 0.15 | 0.3525 | 0.6725 | 0.278352 | 0.783355 | 0.644375 | | 1 | 0.999631 | 0.70098 |
| Ppm1h_Q3UYC0_S220s_1 | 1 | 0.72 | -0.23 | 0.59 | 1.29 | 1.03 | 0.48 | 0.5 | 0.72 | 0.8 | 0.67 | 0.29 | -0.91 | 1.3 | -0.15 | 0.99 | 0.305 | 0.1 | -0.2125 | 0.694321 | 0.402172 | 0.766235 | | 1 | 0.999631 | 0.895302 |
| Ppm1h_Q3UYC0_S220s_2 | 1.14 | 0.33 | -1.41 | 0.16 | 0.79 | -0.45 | -0.38 | -0.09 | 0.1 | 0.54 | 0.29 | -1.34 | -1.37 | 0.58 | 0.74 | 0.61 | -0.0875 | -0.1575 | 0.085 | 0.897068 | 0.874465 | 0.798587 | | 1 | 0.999631 | 0.966612 |
| Ppm1h_Q3UYC0_T109t_1 | 0.92 | 1.6 | 0.83 | 0.54 | 0.03 | 2.32 | 1.2 | 0.16 | 0.89 | 1.43 | 0.59 | -0.12 | 1.2 | 1.97 | -0.79 | 0.56 | -0.045 | -0.275 | -0.2375 | 0.681144 | 0.932775 | 0.5013 | | 1 | 0.999631 | 0.889016 |
| Ppm1h_Q3UYC0_T113t_1 | 1 | 0.7 | 0.94 | 1.28 | 1.38 | 1.38 | 0.19 | 1.13 | 1.11 | 1.17 | 1.33 | 0.67 | 0.06 | 1.07 | 1.3 | 1.06 | 0.04 | 0.09 | -0.1075 | 0.762256 | 0.906662 | 0.749181 | | 1 | 0.999631 | 0.915029 |
| Ppm1h_Q3UYC0_T223t_1 | 0.67 | 0.95 | 0.79 | 0.63 | 0.51 | 1.15 | 0.2 | 0.84 | 0.98 | 1.52 | 0.86 | 1.11 | 0.78 | 1.67 | 1.13 | 1.25 | -0.085 | 0.3575 | 0.4475 | 0.15999 | 0.769482 | 0.206447 | | 1 | 0.999631 | 0.629764 |
| Ppm1h_Q3UYC0_T223t_2 | 1.14 | 0.33 | -1.41 | 0.16 | 0.79 | -0.45 | -0.38 | -0.09 | 0.1 | 0.54 | 0.29 | -1.34 | -1.37 | 0.58 | 0.74 | 0.61 | -0.0875 | -0.1575 | 0.085 | 0.897068 | 0.874465 | 0.798587 | | 1 | 0.999631 | 0.966612 |
| Ppm1h_Q3UYC0_T361t_1 | -0.12 | 0.14 | 1.13 | 0.78 | 0 | -0.22 | 0.26 | 1.2 | 0.11 | -0.48 | 1.71 | 0.53 | 0.32 | 0.52 | 1.3 | 0.45 | -0.1725 | -0.015 | 0.165 | 0.674776 | 0.68249 | 0.976662 | | 1 | 0.999631 | 0.885515 |
| Ppm1j_Q149T7_S92s_1 | -0.01 | -1.16 | -1.06 | -0.7 | -0.47 | -1.45 | -0.05 | -0.67 | -0.3 | -0.39 | -0.76 | -0.81 | -0.68 | -1.14 | -0.63 | -0.76 | 0.0725 | 0.1675 | -0.07 | 0.839524 | 0.855299 | 0.621886 | | 1 | 0.999631 | 0.946365 |
| Ppm1k_Q8BXN7_S248s_1 | -1.61 | -0.32 | -1.43 | -1.39 | -0.29 | -0.62 | -0.93 | -0.66 | -0.6 | -1.77 | -1.55 | -1.04 | -0.3 | -0.76 | -0.1 | -1.17 | 0.5625 | -0.0525 | 0.605 | 0.154877 | 0.132489 | 0.896559 | | 1 | 0.999631 | 0.6258 |
| Ppm1m_Q8BU27_S42s_1 | 2.58 | -0.61 | 0.9 | 0.71 | -0.94 | 0.34 | 1.1 | -0.72 | -0.37 | 1.08 | -0.43 | -0.66 | -0.32 | 0.12 | 0.44 | -1.66 | -0.95 | -0.99 | -1.25 | 0.102291 | 0.204136 | 0.170197 | | 1 | 0.999631 | 0.564448 |
| Ppme1_Q8BVQ5_S243s_1 | -1.94 | -0.94 | -0.28 | -2.93 | -0.42 | -1.31 | -0.22 | -2.9 | -0.71 | -3.33 | -0.07 | 0.11 | -0.62 | -3.3 | -1.65 | -3.66 | 0.31 | 0.5225 | -0.785 | 0.339105 | 0.676748 | 0.547112 | | 1 | 0.999631 | 0.732185 |
| Ppp1ca_P62137_T320t_1 | -0.24 | 0.88 | 0.83 | 0.45 | 0.71 | 1.42 | -0.12 | 0.82 | 0.61 | 0.26 | 0.42 | 0.54 | 0.31 | 1.76 | 0.68 | 0.36 | 0.2275 | -0.0225 | 0.2975 | 0.49668 | 0.581295 | 0.944816 | | 1 | 0.999631 | 0.811278 |
| Ppp1cb_P62141_T316t_1 | -2.69 | 0.57 | 0.49 | -1.49 | 0.13 | 0.18 | -0.28 | -0.51 | 0.02 | -3.56 | 0.23 | 0.05 | 0.91 | 0.09 | -0.15 | -1.74 | 0.66 | -0.035 | 0.5575 | 0.511952 | 0.368055 | 0.973258 | | 1 | 0.999631 | 0.817687 |
| Ppp1r10_Q80W00_S313s_1 | 1.21 | 2.53 | 2.68 | 1.89 | 1.58 | 1.95 | 1.72 | 1.73 | 2.11 | 1.67 | 2.4 | 1.46 | 2.19 | 1.92 | 1.69 | 2.6 | -0.3325 | -0.1675 | 0.0225 | 0.956031 | 0.373514 | 0.681682 | | 1 | 0.999631 | 0.984741 |
| Ppp1r11_G3UZ30_T103t_1 | -0.88 | 0.25 | 0.11 | -0.58 | -0.59 | -0.08 | -0.58 | 0.19 | -0.42 | 0.03 | -0.19 | 0.09 | 0.46 | -0.28 | -0.59 | 0.13 | 0.01 | 0.1525 | 0.205 | 0.599243 | 0.977681 | 0.655656 | | 1 | 0.999631 | 0.855995 |
| Ppp1r12a_Q9DBR7_S299s_1 | -0.28 | 0.22 | 0.15 | -0.04 | 0.35 | -0.02 | -0.23 | 0.35 | -0.1 | -0.01 | -0.24 | 0.38 | -0.05 | 0.37 | 0.21 | 0.51 | 0.1 | -0.005 | 0.2475 | 0.398698 | 0.714021 | 0.985532 | | 1 | 0.999631 | 0.761726 |
| Ppp1r12a_Q9DBR7_S336s_1 | 1.14 | 1.76 | 0.35 | 1.68 | 2.11 | 1.46 | 0.5 | 0.88 | 1.39 | 0.6 | 1.33 | 0.98 | 0.11 | 1.71 | 0.83 | 0.68 | 0.005 | -0.1575 | -0.4 | 0.392684 | 0.991307 | 0.687262 | | 1 | 0.999631 | 0.758901 |
| Ppp1r12a_Q9DBR7_S422s_1 | -3.98 | -9.09 | -7.69 | -5.85 | -6.21 | -11.03 | -8.42 | -5.82 | -15.18 | -7.37 | -4.88 | -6.72 | -6.3 | -8.19 | -6.13 | -9.59 | -1.2175 | -1.885 | -0.9 | 0.449374 | 0.390253 | 0.388241 | | 1 | 0.999631 | 0.786904 |
| Ppp1r12a_Q9DBR7_S445s_1 | 0.16 | 0.63 | -0.28 | 0.83 | 0.62 | 0.8 | 0.54 | 0.79 | 1.15 | 0.06 | 0.81 | 1.67 | -0.67 | 0.53 | 1.68 | 0.44 | 0.3525 | 0.5875 | 0.16 | 0.755631 | 0.27335 | 0.18552 | | 1 | 0.999631 | 0.913663 |
| Ppp1r12a_Q9DBR7_T596t_1 | 2.02 | 0.4 | 0.53 | 1.33 | 0.77 | 1.14 | 1.11 | 0.32 | 1.09 | 1.55 | 0.14 | -0.16 | 0.1 | 0.66 | -0.47 | -1.06 | -0.235 | -0.415 | -1.2625 | 0.029376 | 0.578208 | 0.430248 | | 1 | 0.999631 | 0.433027 |
| Ppp1r12b_Q8BG95_S29s_1 | -0.32 | 0.23 | -0.58 | 0.5 | 0.21 | -0.12 | 0.3 | 1.1 | -0.12 | -0.06 | 0.81 | 0.65 | -0.82 | 0.97 | 1.21 | 1.34 | 0.415 | 0.3625 | 0.7175 | 0.195206 | 0.28318 | 0.341006 | | 1 | 0.999631 | 0.652992 |
| Ppp1r12b_Q8BG95_S446s_1 | 1.94 | 0.3 | -0.14 | 0.75 | 2.13 | 0.19 | 1.22 | 0.4 | 1.16 | -0.41 | 0.49 | 0.6 | -0.73 | 0.16 | 0.29 | -0.09 | 0.2725 | -0.2525 | -0.805 | 0.121408 | 0.636085 | 0.628974 | | 1 | 0.999631 | 0.589748 |
| Ppp1r12b_Q8BG95_S729s_1 | 1.81 | 0.31 | 0.28 | 1.27 | 1.74 | 1.37 | 1.37 | 0.83 | 0.87 | 0.33 | 0.67 | 0 | -0.32 | 1.57 | 0.49 | 0.54 | 0.41 | -0.45 | -0.3475 | 0.502846 | 0.338095 | 0.302896 | | 1 | 0.999631 | 0.814092 |
| Ppp1r12b_Q8BG95_T444t_1 | 0.63 | -0.14 | -0.08 | 0.47 | 1.73 | -0.25 | 1.11 | 0.06 | 0.48 | -0.44 | 0.55 | -0.15 | -0.8 | -0.11 | 0.04 | -0.37 | 0.4425 | -0.11 | -0.53 | 0.137939 | 0.361997 | 0.753303 | | 1 | 0.999631 | 0.611124 |
| Ppp1r12c_F6XWD4_S324s_1 | -1.01 | 0.15 | 0.7 | -0.24 | 0.4 | -0.39 | 0.38 | -0.14 | 0.57 | -1.5 | 0.2 | 1.1 | 0.18 | -0.35 | 1.42 | 0.48 | 0.1625 | 0.1925 | 0.5325 | 0.297517 | 0.692581 | 0.751465 | | 1 | 0.999631 | 0.710954 |
| Ppp1r12c_F6XWD4_S569s_1 | -0.12 | 0.71 | -0.5 | -0.34 | 0.02 | 0.15 | -0.54 | -0.5 | 0.63 | 0.65 | 0.01 | -0.01 | 0.96 | 0.64 | -0.16 | -0.28 | -0.155 | 0.3825 | 0.3525 | 0.408694 | 0.660088 | 0.302212 | | 1 | 0.999631 | 0.766996 |
| Ppp1r13b_Q62415_S335s_1 | -0.07 | 0.32 | 0.63 | 0.31 | 0.99 | 0.11 | 0.39 | 0.8 | 0.36 | 0.2 | 0.46 | 0.58 | -4.65 | 0.54 | 1.51 | 1.13 | 0.275 | 0.1025 | -0.665 | 0.589831 | 0.376692 | 0.70708 | | 1 | 0.999631 | 0.851414 |
| Ppp1r14a_Q91VC7_S136s_1 | -0.36 | -0.03 | 0.45 | -0.19 | 0.02 | 0.04 | -0.39 | -0.12 | 0.06 | -0.18 | 0.5 | 0.82 | 0.79 | 0.48 | 0.68 | 0.42 | -0.08 | 0.3325 | 0.625 | 0.059855 | 0.776158 | 0.331339 | | 1 | 0.999631 | 0.50521 |
| Ppp1r14a_Q91VC7_S26s_1 | 0.28 | 1.13 | 1.22 | 0.57 | 0.41 | 1.11 | 0.05 | -0.63 | 1.2 | 1.31 | 0.45 | 1.43 | 0.55 | 1.34 | 0.53 | 1.23 | -0.565 | 0.2975 | 0.1125 | 0.755907 | 0.201383 | 0.408108 | | 1 | 0.999631 | 0.913733 |
| Ppp1r16a_Q923M0_S353s_1 | 1.72 | 1.8 | 1.54 | 2.37 | 2.24 | 2.6 | 1.59 | 2.48 | 2.64 | 1.86 | 2.06 | 2.86 | 0.6 | 2.49 | 2.94 | 2.65 | 0.37 | 0.4975 | 0.3125 | 0.557393 | 0.277524 | 0.167455 | | 1 | 0.999631 | 0.838305 |
| Ppp1r18_Q8BQ30_S146s_1 | 0.09 | -1.36 | -2.45 | -1.45 | -0.41 | -1.35 | -1.46 | -2.5 | -1.39 | 0.09 | -1.68 | -1.17 | -0.33 | -1.06 | -1.94 | -2.62 | -0.1375 | 0.255 | -0.195 | 0.764454 | 0.821399 | 0.669667 | | 1 | 0.999631 | 0.915948 |
| Ppp1r18_Q8BQ30_S194s_1 | -0.42 | 0.43 | 0.69 | -0.05 | 0.54 | 1.17 | -0.22 | -0.45 | 0.92 | 0.07 | -0.13 | 1.15 | 1.2 | 1.08 | 0.35 | 0.97 | 0.0975 | 0.34 | 0.7375 | 0.063793 | 0.822685 | 0.41352 | | 1 | 0.999631 | 0.513532 |
| Ppp1r18_Q8BQ30_S234s_1 | 2.62 | 0.83 | 1.02 | 1.11 | 0.36 | 0.9 | 1.79 | 0.4 | 1.13 | 2.4 | 0.3 | 0.4 | 0.49 | 0.67 | 0.05 | 0.22 | -0.5325 | -0.3375 | -1.0375 | 0.037169 | 0.29912 | 0.565779 | | 1 | 0.999631 | 0.457023 |
| Ppp1r1a_Q9ERT9_S47s_1 | -0.76 | -0.12 | -1.57 | -1.08 | 0.26 | -0.27 | -0.53 | -0.44 | -0.39 | -0.2 | 0.11 | -0.04 | -0.01 | -0.09 | -0.45 | 0.26 | 0.6375 | 0.7525 | 0.81 | 0.053015 | 0.109708 | 0.058048 | | 1 | 0.999631 | 0.489777 |
| Ppp1r1a_Q9ERT9_S67s_1 | 1.65 | 0.54 | -0.87 | 1.86 | 2.39 | 1.14 | 1.84 | 1.84 | 1.34 | 0 | 1.6 | 0.19 | -2.09 | 1.13 | 0.73 | 0.44 | 1.0075 | -0.0125 | -0.7425 | 0.382806 | 0.123065 | 0.98503 | | 1 | 0.999631 | 0.754257 |
| Ppp1r1a_Q9ERT9_S6s_1 | 1.55 | 1.03 | 0.32 | 1.83 | 1.39 | 0.29 | 1.2 | 1.37 | 1.56 | 1.16 | 1.61 | 1.84 | 0.41 | 1.02 | 1.65 | 0.54 | -0.12 | 0.36 | -0.2775 | 0.531338 | 0.774682 | 0.357312 | | 1 | 0.999631 | 0.827082 |
| Ppp1r1b_Q60829_S10s_1 | 0.36 | -1.76 | -0.95 | -0.9 | -1.32 | -1.67 | -0.5 | -0.69 | -0.94 | 0.12 | -1.67 | -0.86 | -0.94 | -0.81 | -0.33 | -1.23 | -0.2325 | -0.025 | -0.015 | 0.974381 | 0.634695 | 0.962581 | | 1 | 0.999631 | 0.990847 |
| Ppp1r1b_Q60829_S10s_2 | -0.03 | -1.14 | -1.65 | -0.64 | 0.18 | -0.97 | 0.02 | -0.81 | -0.18 | -0.24 | -1.57 | -0.99 | -2.05 | -0.19 | -1.15 | -1.51 | 0.47 | 0.12 | -0.36 | 0.477715 | 0.299658 | 0.796437 | | 1 | 0.999631 | 0.800237 |
| Ppp1r1b_Q60829_S135s_1 | 0.02 | 0.6 | 2.09 | 0.7 | 3.54 | 1.56 | 3.52 | 1.11 | 2.56 | -1.85 | 2.22 | 0.97 | 1.21 | -0.84 | 2.23 | -0.81 | 1.58 | 0.1225 | -0.405 | 0.601831 | 0.042314 | 0.897085 | | 1 | 0.999631 | 0.856768 |
| Ppp1r1b_Q60829_S156s_1 | 0.06 | -0.04 | 0.03 | 0.13 | 0.09 | -0.31 | -0.44 | 1.07 | 0.01 | 0.28 | -0.28 | -0.05 | 0.62 | 0.7 | 1.29 | 0.22 | 0.0575 | -0.055 | 0.6625 | 0.056807 | 0.874702 | 0.829543 | | 1 | 0.999631 | 0.496275 |
| Ppp1r1b_Q60829_S61s_1 | -0.29 | -0.17 | -0.74 | 0.23 | 0.39 | -0.8 | -0.17 | 0.3 | -0.35 | -0.88 | -0.52 | -0.08 | -0.69 | 0.07 | -0.35 | -0.31 | 0.1725 | -0.215 | -0.0775 | 0.813569 | 0.634703 | 0.506867 | | 1 | 0.999631 | 0.937041 |
| Ppp1r1b_Q60829_S9s_1 | 1.79 | -1.38 | -2.38 | 0.91 | 0.59 | 0.02 | 0.23 | 0.54 | -0.48 | -0.4 | -0.93 | -1.95 | -3.35 | -0.5 | -0.97 | -1.16 | 0.61 | -0.675 | -1.23 | 0.233365 | 0.479947 | 0.459729 | | 1 | 0.999631 | 0.683077 |
| Ppp1r1b_Q60829_S9s_2 | -0.34 | -1.04 | -1.65 | -0.64 | -0.32 | -1.02 | 0.02 | -0.81 | -0.22 | -0.17 | -1.57 | -0.99 | -1.99 | -0.23 | -1.15 | -1.51 | 0.385 | 0.18 | -0.3025 | 0.516766 | 0.325602 | 0.68066 | | 1 | 0.999631 | 0.819449 |
| Ppp1r1b_Q60829_T39t_1 | 0.5 | 0.41 | 0.41 | 1.37 | 1.11 | 0.47 | 0.47 | 2.27 | 1.31 | 0.71 | 0.85 | 1.08 | 0.08 | 1.52 | 1.87 | 0.98 | 0.4075 | 0.315 | 0.44 | 0.342508 | 0.389239 | 0.343711 | | 1 | 0.999631 | 0.73348 |
| Ppp1r2_D3Z3A0_S122s_1 | -0.75 | -0.51 | 1.52 | 0.71 | -0.39 | -1.14 | 0.17 | 1.9 | -0.33 | -0.73 | 0.71 | 1.95 | 0.51 | -0.74 | -0.25 | 0.77 | -0.1075 | 0.1575 | -0.17 | 0.770843 | 0.883993 | 0.825065 | | 1 | 0.999631 | 0.919413 |
| Ppp1r2_D3Z3A0_S122s_2 | -1.14 | -0.09 | 0.84 | 0.75 | 0.7 | -0.19 | 0.45 | 1.09 | 0.97 | -1.29 | 0.25 | 0.75 | 0.88 | -0.82 | 0.45 | 0.8 | 0.4225 | 0.08 | 0.2375 | 0.673207 | 0.406847 | 0.897578 | | 1 | 0.999631 | 0.885515 |
| Ppp1r2_D3Z3A0_S123s_1 | 0.07 | 0.09 | -0.55 | -0.54 | -0.53 | 0.24 | -0.03 | -0.19 | 0.38 | -0.81 | -0.65 | 0.2 | 1.6 | 0.16 | -0.78 | -1.41 | 0.105 | 0.0125 | 0.125 | 0.839054 | 0.729326 | 0.973327 | | 1 | 0.999631 | 0.946365 |
| Ppp1r2_D3Z3A0_S123s_2 | -1.14 | -0.09 | 0.84 | 0.75 | 0.7 | -0.19 | 0.45 | 1.09 | 0.97 | -1.29 | 0.25 | 0.75 | 0.88 | -0.82 | 0.45 | 0.8 | 0.4225 | 0.08 | 0.2375 | 0.673207 | 0.406847 | 0.897578 | | 1 | 0.999631 | 0.885515 |
| Ppp1r2_D3Z3A0_S23s_1 | 0.4 | 0.26 | 0.43 | 0.44 | 0.61 | -0.04 | -0.21 | 0.72 | 0.7 | 0.75 | 0.05 | 0.97 | 1.32 | 1.39 | 0.81 | 0.51 | -0.1125 | 0.235 | 0.625 | 0.066324 | 0.707812 | 0.424727 | | 1 | 0.999631 | 0.517932 |
| Ppp1r21_Q3TDD9_S101s_1 | 1.42 | -0.05 | 0.58 | 1.5 | 0.47 | 0.79 | 1.43 | -0.1 | 0.39 | 1.12 | 1.16 | -0.03 | -1.1 | -0.41 | 0.73 | 0.58 | -0.215 | -0.2025 | -0.9125 | 0.111138 | 0.646208 | 0.6595 | | 1 | 0.999631 | 0.577068 |
| Ppp1r21_Q3TDD9_T652t_1 | -0.9 | -2.94 | -0.56 | 0.34 | -1.93 | -2.07 | 0.2 | 0.06 | -2.37 | -1.49 | 0.62 | 0.6 | -2.15 | -1.15 | 0.67 | 0.84 | 0.08 | 0.355 | 0.5675 | 0.51726 | 0.92117 | 0.691534 | | 1 | 0.999631 | 0.819449 |
| Ppp1r3d_A2AJW4_S54s_2 | -0.46 | 0 | -0.56 | -0.53 | 0.18 | 0.08 | -0.84 | -0.44 | -0.04 | 0.48 | -0.28 | -0.4 | -0.18 | 0.46 | -0.58 | -0.32 | 0.1325 | 0.3275 | 0.2325 | 0.487221 | 0.679696 | 0.298997 | | 1 | 0.999631 | 0.805474 |
| Ppp1r3d_A2AJW4_S58s_1 | -0.94 | -2.3 | -0.14 | -2.25 | -1.83 | -1.46 | -1.72 | -2.9 | -1.7 | -0.35 | -1.65 | -0.89 | 1.83 | -1.38 | -0.93 | -0.83 | -0.57 | 0.26 | 1.08 | 0.190212 | 0.323781 | 0.649465 | | 1 | 0.999631 | 0.648674 |
| Ppp1r3d_A2AJW4_S58s_2 | -0.29 | 0.67 | 0.75 | 0.17 | 0.4 | 0.76 | 0.1 | 0.55 | 0.23 | 1.07 | 0.58 | 0.11 | -0.12 | 0.87 | 0.36 | 0.78 | 0.1275 | 0.1725 | 0.1475 | 0.69324 | 0.69505 | 0.632005 | | 1 | 0.999631 | 0.895096 |
| Ppp1r3d_A2AJW4_T57t_2 | -0.82 | -1.74 | -2.12 | -1.18 | -0.27 | -1.78 | -2 | -1.46 | -0.61 | -1.11 | -1.11 | -0.85 | -0.29 | -0.46 | -1.53 | -1.48 | 0.0875 | 0.545 | 0.525 | 0.250624 | 0.849695 | 0.144636 | | 1 | 0.999631 | 0.686448 |
| Ppp1r3e_H3BJB2_S16s_1 | 0.02 | 1.35 | 1.34 | 1.7 | 1.29 | 1.17 | 1.01 | 1.89 | 1.4 | -0.19 | 1.28 | 1.7 | -0.09 | 1.53 | 1.43 | 1.46 | 0.2375 | -0.055 | -0.02 | 0.96881 | 0.570479 | 0.916623 | | 1 | 0.999631 | 0.988819 |
| Ppp1r3g_Q9CW07_S79s_1 | 0.69 | 1.91 | 1.2 | 1.1 | 0.81 | 0.63 | 0.11 | 1.02 | 0.83 | 0.47 | 0.74 | 1.68 | 0.48 | 0.87 | 0.94 | 1.95 | -0.5825 | -0.295 | -0.165 | 0.693887 | 0.119979 | 0.449784 | | 1 | 0.999631 | 0.895131 |
| Ppp1r3g_Q9CW07_S81s_1 | 0.34 | 0.69 | 1.25 | 1.55 | 0.5 | 0.37 | 0.32 | 0.97 | 0.27 | 0.12 | 1.45 | 2.2 | 0.57 | 0.54 | 1.43 | 2.19 | -0.4175 | 0.0525 | 0.225 | 0.633615 | 0.241996 | 0.920873 | | 1 | 0.999631 | 0.868569 |
| Ppp1r7_Q3UM45_S12s_1 | 0.35 | -0.75 | -0.75 | -0.19 | -0.42 | -0.84 | -0.25 | -0.67 | -0.61 | -0.36 | -0.43 | -0.95 | -0.93 | -0.86 | -0.79 | -0.57 | -0.21 | -0.2525 | -0.4525 | 0.203411 | 0.53317 | 0.464537 | | 1 | 0.999631 | 0.659781 |
| Ppp1r7_Q3UM45_S24s_1 | 0.87 | -1.29 | 0.91 | 0.31 | -1.41 | -0.66 | 0.02 | -0.46 | -0.51 | 1.1 | -1.07 | -1.35 | -0.66 | -2.87 | -0.35 | -1.38 | -0.8275 | -0.6575 | -1.515 | 0.046605 | 0.153547 | 0.34095 | | 1 | 0.999631 | 0.480102 |
| Ppp1r7_Q3UM45_S24s_2 | -0.24 | 0.42 | 1.99 | 1.83 | -0.12 | 0.12 | 1.58 | 0.83 | 0.09 | 0.69 | 0.85 | 0.92 | -0.53 | -1.38 | 0.74 | 0.01 | -0.3975 | -0.3625 | -1.29 | 0.065128 | 0.512958 | 0.504117 | | 1 | 0.999631 | 0.515898 |
| Ppp1r7_Q3UM45_S27s_1 | 1.87 | 1.24 | 2.24 | 2.14 | 0.84 | 1.37 | 2.41 | 0.56 | 1.66 | 2.34 | 1.36 | 1.37 | 0.57 | 0.12 | 0.92 | 0.16 | -0.5775 | -0.19 | -1.43 | 0.002132 | 0.219114 | 0.596964 | | 1 | 0.999631 | 0.351308 |
| Ppp1r7_Q3UM45_S27s_2 | -0.24 | 0.42 | 1.99 | 1.83 | -0.12 | 0.12 | 1.58 | 0.83 | 0.09 | 0.69 | 0.85 | 0.92 | -0.53 | -1.38 | 0.74 | 0.01 | -0.3975 | -0.3625 | -1.29 | 0.065128 | 0.512958 | 0.504117 | | 1 | 0.999631 | 0.515898 |
| Ppp1r7_Q3UM45_S45s_1 | -1.02 | 0.69 | 0.52 | -0.53 | 0.59 | -0.79 | -0.2 | -0.62 | 2.18 | -1.24 | 0.05 | 0.9 | 3.14 | -0.42 | -0.14 | -0.02 | -0.17 | 0.5575 | 0.725 | 0.382029 | 0.727077 | 0.45455 | | 1 | 0.999631 | 0.753668 |
| Ppp1r9a_H3BKQ7_S160s_1 | -0.23 | -1.48 | -1.85 | -1.03 | -0.55 | -2.57 | -0.91 | -0.34 | -0.43 | -1.03 | 0.17 | 0.15 | -0.19 | -1.73 | -0.54 | -2.23 | 0.055 | 0.8625 | -0.025 | 0.963921 | 0.921976 | 0.076566 | | 1 | 0.999631 | 0.987331 |
| Ppp1r9a_H3BKQ7_S187s_1 | -0.92 | -0.01 | -0.89 | -1.22 | -0.93 | -1.2 | -1.48 | -1.68 | -2.27 | -0.61 | -0.58 | -1.2 | -0.08 | -0.57 | -2.45 | -0.31 | -0.5625 | -0.405 | -0.0925 | 0.868213 | 0.124442 | 0.389077 | | 1 | 0.999631 | 0.95689 |
| Ppp1r9a_H3BKQ7_S192s_1 | -0.87 | -1.13 | -1.1 | -0.49 | -0.38 | -0.52 | -0.7 | -0.54 | -1.17 | -1.22 | -0.62 | -1.49 | -0.92 | -1.2 | -0.92 | -0.43 | 0.3625 | -0.2275 | 0.03 | 0.923074 | 0.18934 | 0.463549 | | 1 | 0.999631 | 0.975032 |
| Ppp1r9a_H3BKQ7_S199s_1 | 0.13 | 0.23 | -0.38 | -0.34 | 0.1 | -0.33 | -0.34 | -0.07 | 0.34 | -0.34 | -0.72 | -0.97 | -0.27 | 0.27 | -0.26 | -0.17 | -0.07 | -0.3325 | -0.0175 | 0.954045 | 0.799923 | 0.365854 | | 1 | 0.999631 | 0.984441 |
| Ppp1r9a_H3BKQ7_S372s_1 | -1.73 | -1.26 | -2.77 | -2.43 | -1.1 | -1.73 | -2.63 | -3.09 | -1.65 | -1.59 | -2.62 | -2.64 | -0.75 | -1.33 | -2.56 | -3.1 | -0.09 | -0.0775 | 0.1125 | 0.847746 | 0.862863 | 0.860958 | | 1 | 0.999631 | 0.949053 |
| Ppp1r9a_H3BKQ7_S841s_1 | -0.24 | -0.95 | -0.73 | -0.81 | -0.93 | -0.72 | -0.47 | -0.3 | -0.62 | -0.38 | -0.81 | -0.1 | -0.77 | -0.9 | -0.45 | 0.28 | 0.0775 | 0.205 | 0.2225 | 0.537766 | 0.785471 | 0.495374 | | 1 | 0.999631 | 0.830863 |
| Ppp1r9a_H3BKQ7_T371t_1 | -1.61 | -0.84 | -0.5 | -1.26 | -1.04 | -1.11 | -1.24 | -0.91 | -1.27 | -1.43 | -1.52 | -2.25 | 0.23 | -0.57 | -1.84 | -0.88 | -0.0225 | -0.565 | 0.2875 | 0.551449 | 0.941789 | 0.139611 | | 1 | 0.999631 | 0.836275 |
| Ppp1r9b_Q6R891_S100s_1 | 0.91 | 0.51 | -0.39 | 1.51 | 1.43 | 1.08 | 0.91 | 1.58 | 0.73 | 0.32 | 0.86 | 0.96 | -1.91 | 0.97 | 1.06 | 1.27 | 0.615 | 0.0825 | -0.2875 | 0.702378 | 0.168114 | 0.845883 | | 1 | 0.999631 | 0.896857 |
| Ppp1r9b_Q6R891_S192s_1 | 1.48 | 1.17 | -0.14 | 1.74 | 1.86 | 1.53 | 0.92 | 1.46 | 1.57 | 1.3 | 0.87 | 0.41 | -1.12 | 2.17 | 1.42 | 0.89 | 0.38 | -0.025 | -0.2225 | 0.75852 | 0.403607 | 0.957693 | | 1 | 0.999631 | 0.914449 |
| Ppp1r9b_Q6R891_S205s_1 | 1.36 | 1.41 | -0.42 | 1.43 | 2.05 | 1.35 | 0.9 | 1.36 | 1.39 | 0.17 | 0.61 | 0.64 | 0.36 | 1.67 | 0.25 | 1.3 | 0.47 | -0.2425 | -0.05 | 0.925829 | 0.344693 | 0.626539 | | 1 | 0.999631 | 0.97588 |
| Ppp1r9b_Q6R891_S382s_1 | 1.24 | 1.01 | 1.21 | 1.98 | 0.55 | 1.96 | 0.6 | 1.67 | 0.66 | 0.88 | 0.44 | 1.06 | -0.22 | 1.4 | 1.3 | 1.79 | -0.165 | -0.6 | -0.2925 | 0.544044 | 0.693797 | 0.081502 | | 1 | 0.999631 | 0.833226 |
| Ppp1r9b_Q6R891_S99s_1 | 0.41 | -0.37 | 0.08 | 1.38 | 0.66 | 0.48 | 0.41 | 1.31 | 0.27 | 0.07 | 0.58 | 0.36 | -1.1 | 0.76 | 0.71 | 0.8 | 0.34 | -0.055 | -0.0825 | 0.881283 | 0.42628 | 0.890287 | | 1 | 0.999631 | 0.963364 |
| Ppp1r9b_Q6R891_T193t_1 | 1.81 | 1.54 | 0.27 | 1.67 | 2.08 | 1.82 | 1.04 | 1.88 | 1.89 | 1.71 | 1.54 | 1.37 | -1.57 | 2.25 | 1.16 | 1.5 | 0.3825 | 0.305 | -0.4875 | 0.54141 | 0.371625 | 0.441227 | | 1 | 0.999631 | 0.832213 |
| Ppp2r5a_Q6PD03_S41s_1 | 2.38 | 1.77 | 1.28 | 1.52 | 1.65 | 2.44 | 1.49 | 0.64 | 1.9 | 1.15 | 1.11 | 1.31 | -3.81 | 1.98 | 0.65 | 0.32 | -0.1825 | -0.37 | -1.9525 | 0.095804 | 0.672608 | 0.295073 | | 1 | 0.999631 | 0.558128 |
| Ppp2r5d_Q91V89_S44s_1 | -0.49 | 1.47 | 2.09 | 1.11 | 1.34 | 1.6 | 1.27 | 1.92 | 2.03 | 0.5 | 1.9 | 2.39 | 1.41 | 1.56 | 2.82 | 1.75 | 0.4875 | 0.66 | 0.84 | 0.171931 | 0.366244 | 0.304436 | | 1 | 0.999631 | 0.641657 |
| Ppp2r5d_Q91V89_S565s_1 | 1.44 | 1.74 | 1.27 | 2.02 | 2.3 | 1.7 | 1.7 | 1.72 | 2.12 | 1.03 | 2.06 | 2.27 | 0.81 | 2 | 1.99 | 2.14 | 0.2375 | 0.2525 | 0.1175 | 0.760595 | 0.424424 | 0.489878 | | 1 | 0.999631 | 0.914614 |
| Ppp2r5d_Q91V89_S80s_1 | 0.04 | -1.04 | -1.83 | -0.42 | 1.25 | -1.85 | 0.28 | -1.19 | 0.75 | -0.76 | 0.54 | 0.06 | -1.13 | 0 | -1.76 | -1.5 | 0.435 | 0.96 | -0.285 | 0.59192 | 0.546862 | 0.079051 | | 1 | 0.999631 | 0.852728 |
| Ppp2r5d_Q91V89_S81s_1 | 0.48 | 0.48 | 0.35 | 1.01 | 1.45 | -0.22 | 0.5 | 1.17 | 1.01 | 0.37 | 1.04 | 0.88 | -0.89 | 1.03 | 0.96 | 0.87 | 0.145 | 0.245 | -0.0875 | 0.85337 | 0.719985 | 0.414724 | | 1 | 0.999631 | 0.950701 |
| Ppp2r5e_Q3UZJ4_S32s_1 | 2.02 | 0.1 | -0.31 | 0.62 | 1.23 | 1.2 | 0.61 | -0.86 | 1.32 | 1.31 | -0.14 | -0.52 | -0.17 | 0.51 | 0.09 | -1.37 | -0.0625 | -0.115 | -0.8425 | 0.176904 | 0.921206 | 0.855586 | | 1 | 0.999631 | 0.642116 |
| Ppp2r5e_Q3UZJ4_T7t_1 | 0.23 | 0.96 | -1.49 | -1.38 | 0.84 | 0.64 | -1.2 | -2.3 | 0.86 | 1.46 | -2.32 | -1.73 | 1.2 | 1.45 | -1.66 | -2.04 | -0.085 | -0.0125 | 0.1575 | 0.867755 | 0.919433 | 0.989645 | | 1 | 0.999631 | 0.956838 |
| Ppp3ca_P63328_S462s_1 | 0.88 | 0.84 | 1.4 | 1.1 | 0.22 | 0.99 | 0.3 | 1.2 | 0.84 | 1.08 | 0.59 | 0.97 | 1.2 | 1.01 | 1.16 | 0.5 | -0.3775 | -0.185 | -0.0875 | 0.774785 | 0.259519 | 0.502685 | | 1 | 0.999631 | 0.920998 |
| Ppp3ca_P63328_S469s_1 | 0.3 | -0.04 | -0.45 | 0.55 | 0.32 | 0.36 | -0.14 | 0.11 | -0.21 | -0.41 | -0.19 | -0.32 | -1.13 | 0.18 | -0.02 | 0.12 | 0.0725 | -0.3725 | -0.3025 | 0.457225 | 0.812048 | 0.232488 | | 1 | 0.999631 | 0.790722 |
| Ppp3ca_P63328_T427t_1 | -0.46 | 1.65 | -0.52 | 1.31 | 1.6 | 1.58 | 0.13 | 1.34 | 0.86 | 1.2 | 0.45 | 0.66 | -4.94 | 1.3 | 1.09 | 0.49 | 0.6675 | 0.2975 | -1.01 | 0.457908 | 0.285792 | 0.592021 | | 1 | 0.999631 | 0.790722 |
| Ppp3cb_E0CZ78_S116s_1 | -0.25 | 0.15 | -0.38 | -0.52 | -0.38 | 0.45 | -1.07 | -0.46 | 0.01 | -0.09 | -1.01 | -0.67 | 0.66 | 0.49 | 0.08 | 0.3 | -0.115 | -0.19 | 0.6325 | 0.056264 | 0.75323 | 0.570835 | | 1 | 0.999631 | 0.495761 |
| Ppp3cb_E0CZ78_S471s_1 | -0.32 | -0.17 | 0.25 | -0.27 | -0.85 | -0.49 | -0.54 | -0.09 | -0.23 | 0.28 | 0.08 | 0.32 | 0.08 | 0.05 | 0.09 | 0.39 | -0.365 | 0.24 | 0.28 | 0.335019 | 0.215504 | 0.398988 | | 1 | 0.999631 | 0.730134 |
| Ppp3cb_E0CZ78_S478s_1 | 0.63 | 0.77 | 0.06 | 1.47 | 1.3 | 1.03 | 0.8 | 1.45 | 1.08 | 0.67 | 1.27 | 0.88 | -0.83 | 1.16 | 1.02 | 0.95 | 0.4125 | 0.2425 | -0.1575 | 0.763148 | 0.258268 | 0.498707 | | 1 | 0.999631 | 0.915608 |
| Ppp4r4_Q8C0Y0_S777s_1 | 0.01 | 0.7 | -0.06 | 0.75 | 0.64 | 0.99 | 0.46 | 0.7 | 1.32 | -0.05 | 1.07 | 0.76 | 0.54 | 0.88 | -0.15 | 1.35 | 0.3475 | 0.425 | 0.305 | 0.457691 | 0.268909 | 0.28781 | | 1 | 0.999631 | 0.790722 |
| Ppp6r1_Q7TSI3_S739s_1 | -0.44 | -0.44 | 0 | -0.21 | -0.9 | -0.21 | -0.16 | -0.23 | -0.55 | -0.81 | 0.28 | 0.17 | -0.43 | 0.15 | -0.25 | 0.01 | -0.1025 | 0.045 | 0.1425 | 0.623991 | 0.718156 | 0.893399 | | 1 | 0.999631 | 0.863524 |
| Ppp6r1_Q7TSI3_S817s_1 | 0.75 | 1.11 | -0.3 | 0.38 | 2.17 | 1.11 | 2.17 | -0.22 | 2.93 | 0.11 | 2.05 | 2.74 | 1.11 | -0.02 | -1.05 | -0.47 | 0.8225 | 1.4725 | -0.5925 | 0.269061 | 0.180385 | 0.042272 | | 1 | 0.999631 | 0.697063 |
| Ppp6r1_Q7TSI3_S826s_1 | -0.68 | -0.56 | -0.63 | -0.64 | -0.41 | -0.68 | -0.76 | -1.13 | -0.98 | -0.62 | -0.63 | -0.19 | 0.25 | -0.17 | -0.66 | -0.24 | -0.1175 | 0.0225 | 0.4225 | 0.176914 | 0.650872 | 0.933934 | | 1 | 0.999631 | 0.642116 |
| Ppp6r2_G3X9K4_S435s_1 | -0.65 | -0.51 | -0.55 | -0.27 | 0.14 | -0.32 | -0.24 | -0.08 | -0.58 | -0.12 | -0.23 | 0.02 | -1.04 | -0.39 | -0.51 | 0.47 | 0.37 | 0.2675 | 0.1275 | 0.729365 | 0.16217 | 0.328674 | | 1 | 0.999631 | 0.905923 |
| Ppp6r2_G3X9K4_S670s_1 | 0.05 | 0.51 | 0.09 | 0.2 | 0.16 | -0.08 | -0.07 | 0.23 | 0.52 | 0.25 | 0.42 | 0.53 | 0.75 | 0.03 | 0.33 | 0.59 | -0.1525 | 0.2175 | 0.2125 | 0.481187 | 0.546875 | 0.404821 | | 1 | 0.999631 | 0.801848 |
| Ppp6r2_G3X9K4_S745s_1 | -0.24 | 0.53 | 1.07 | 0.62 | 0.97 | 0.86 | 0.91 | 1.23 | 1.1 | 0.25 | 1.39 | 0.45 | -0.61 | 0.72 | 1.22 | 1.91 | 0.4975 | 0.3025 | 0.315 | 0.57344 | 0.15191 | 0.452151 | | 1 | 0.999631 | 0.844597 |
| Ppp6r2_G3X9K4_S746s_1 | 0.08 | 0.66 | 0.83 | 0.73 | 0.56 | 1.4 | 0.6 | 1.47 | 0.59 | 2.37 | 0.86 | 0.57 | -0.07 | 1.7 | 1.2 | 1.29 | 0.4325 | 0.5225 | 0.455 | 0.301449 | 0.218462 | 0.264413 | | 1 | 0.999631 | 0.714243 |
| Ppp6r2_G3X9K4_S760s_1 | 0.29 | 0.74 | 0.6 | 0 | -0.09 | 1.09 | 0.42 | -0.2 | 0.02 | 1.34 | 0.26 | 1.53 | 0.8 | 0.87 | 0.21 | 2.07 | -0.1025 | 0.38 | 0.58 | 0.199074 | 0.776712 | 0.373738 | | 1 | 0.999631 | 0.657167 |
| Ppp6r2_G3X9K4_S760s_2 | 0.86 | 0.35 | 1.02 | 1.47 | 1.23 | 1.07 | 1.74 | 1.46 | 0.3 | 1.02 | 1.24 | 1.29 | -0.37 | 0.54 | 1.28 | 2.21 | 0.45 | 0.0375 | -0.01 | 0.985549 | 0.182386 | 0.916733 | | 1 | 0.999631 | 0.995133 |
| Ppp6r2_G3X9K4_S820s_1 | -1.11 | -2.12 | -1.23 | -1.37 | -1.94 | -2.67 | -1.42 | -1.14 | -1.5 | -1.27 | -1.13 | -0.94 | -0.31 | -1.7 | -0.73 | -1.04 | -0.335 | 0.2475 | 0.5125 | 0.215709 | 0.418886 | 0.443051 | | 1 | 0.999631 | 0.668551 |
| Prcc_Q9EQC8_S157s_1 | 0.83 | 0.87 | 2.99 | 1.88 | 0.17 | 1.47 | 1.42 | 0.94 | -0.82 | 0.62 | 0.39 | 2.01 | -0.18 | 0.88 | -0.09 | 0.46 | -0.6425 | -1.0925 | -1.375 | 0.025728 | 0.25595 | 0.13739 | | 1 | 0.999631 | 0.422114 |
| Prex1_I7HPV9_S1021s_1 | -0.49 | -0.28 | -1.32 | -1.7 | -0.08 | 1.33 | -1.01 | -3.03 | 0.1 | 0.03 | -1.16 | 0.39 | 1.5 | 1.41 | -1.5 | -1.04 | 0.25 | 0.7875 | 1.04 | 0.189656 | 0.768187 | 0.115627 | | 1 | 0.999631 | 0.648674 |
| Prex1_I7HPV9_S144s_1 | -0.58 | 1.33 | -0.26 | 1.08 | 1.39 | 1.08 | 0.74 | 1.09 | 1.79 | -0.49 | 2.17 | 1.21 | -0.48 | 0.77 | 1.74 | 0.44 | 0.6825 | 0.7775 | 0.225 | 0.709408 | 0.170089 | 0.267157 | | 1 | 0.999631 | 0.899175 |
| Prex1_I7HPV9_S973s_1 | 0.27 | -0.59 | -0.74 | -0.64 | -1.61 | -0.98 | -0.18 | -0.71 | -1.91 | 0 | -1.08 | -1.5 | -0.19 | -1.36 | -0.67 | -0.35 | -0.445 | -0.6975 | -0.2175 | 0.57349 | 0.267967 | 0.153073 | | 1 | 0.999631 | 0.844597 |
| Prickle1_Q3U5C7_S592s_1 | -0.52 | 0.21 | 0.1 | 0.64 | 0.19 | 0.62 | -0.01 | 0.61 | 0.4 | 0.67 | 0.21 | 1.22 | 0.28 | 0.96 | 1.52 | 0.99 | 0.245 | 0.5175 | 0.83 | 0.051986 | 0.462493 | 0.171717 | | 1 | 0.999631 | 0.489777 |
| Prickle1_Q3U5C7_S595s_1 | -1.07 | 0.16 | 0.67 | -0.68 | -2.37 | -0.52 | -1.21 | 0.4 | -2.49 | -0.16 | -1.13 | 0.45 | -0.05 | -0.8 | 1.8 | 1.09 | -0.695 | -0.6025 | 0.74 | 0.258732 | 0.286756 | 0.381431 | | 1 | 0.999631 | 0.692624 |

| | | | | | | | | | | | | | | | | | | | | | | | | | |
|---|---|---|---|---|---|---|---|---|---|---|---|---|---|---|---|---|---|---|---|---|---|---|---|---|---|
| Prickle2_Q80Y24_S319s_1 | -2.05 | -1.47 | -1.28 | -1.2 | 0.14 | 0.04 | -1.02 | -1.37 | -0.67 | -0.57 | -1.1 | -1.17 | 0.06 | -0.71 | -1.58 | -1.12 | 0.9475 | 0.6225 | 0.6625 | 0.133131 | 0.045229 | 0.069158 | 1 | 0.999631 | 0.60492 |
| Prickle2_Q80Y24_S321s_2 | -1.3 | -1.85 | -0.52 | -1.27 | -0.95 | -0.52 | -0.64 | -0.31 | -1.44 | -1.33 | -0.71 | -0.1 | -1.23 | -0.9 | -0.77 | -1.43 | 0.63 | 0.34 | 0.1525 | 0.674061 | 0.088197 | 0.423746 | 1 | 0.999631 | 0.885515 |
| Prickle2_Q80Y24_S322s_1 | -0.71 | -0.1 | -0.98 | -0.62 | -1.47 | -0.98 | -0.39 | -0.71 | -0.92 | -0.71 | -1.54 | -0.04 | -0.01 | -0.31 | -1.74 | -1.49 | -0.285 | -0.2 | -0.285 | 0.539475 | 0.401246 | 0.602556 | 1 | 0.999631 | 0.830899 |
| Prickle2_Q80Y24_S322s_2 | -1.3 | -1.85 | -0.52 | -1.27 | -0.95 | -0.52 | -0.64 | -0.31 | -1.44 | -1.33 | -0.71 | -0.1 | -1.23 | -0.9 | -0.77 | -1.43 | 0.63 | 0.34 | 0.1525 | 0.674061 | 0.088197 | 0.423746 | 1 | 0.999631 | 0.885515 |
| Prickle2_Q80Y24_S396s_1 | 0.04 | 0.95 | 0.87 | 0.96 | 0.93 | -0.1 | 1.14 | 0.82 | 1.12 | 0.41 | 1.46 | 1.46 | 0.66 | 0.1 | -0.53 | 1.95 | -0.0075 | 0.4075 | -0.16 | 0.765878 | 0.983857 | 0.278694 | 1 | 0.999631 | 0.916581 |
| Prickle2_Q80Y24_S414s_1 | -0.5 | -0.51 | -0.7 | -0.37 | -0.52 | -0.97 | -0.79 | -0.32 | -0.08 | -0.34 | -0.97 | -0.97 | 0.47 | -0.07 | -0.6 | 0.02 | -0.13 | -0.07 | 0.475 | 0.156139 | 0.621609 | 0.81939 | 1 | 0.999631 | 0.626452 |
| Prickle2_Q80Y24_S463s_1 | 0.69 | 3.26 | 2.27 | -0.63 | 2.71 | 1.96 | 2.97 | 0.7 | 3.53 | 1.22 | 3.23 | 1.9 | 4.24 | -0.06 | 0.82 | -0.04 | 0.6875 | 1.0725 | -0.1575 | 0.88929 | 0.434987 | 0.244221 | 1 | 0.999631 | 0.963505 |
| Prickle2_Q80Y24_S544s_1 | -0.63 | -0.27 | -0.51 | -0.42 | -0.45 | -0.24 | -0.85 | -0.38 | -0.42 | -0.36 | -0.58 | -0.37 | -0.01 | -0.39 | -0.09 | 0.13 | -0.0225 | 0.025 | 0.3675 | 0.204171 | 0.930615 | 0.919273 | 1 | 0.999631 | 0.659781 |
| Prickle2_Q80Y24_S572s_1 | 0.82 | 0.81 | -0.27 | 2.14 | 2.02 | 0.5 | 1.3 | 1.74 | 1.11 | -0.02 | 1.63 | 0.53 | -1.19 | 1.27 | 0.97 | 1.18 | 0.515 | -0.0625 | -0.3175 | 0.643135 | 0.357148 | 0.911352 | 1 | 0.999631 | 0.871773 |
| Prickle2_Q80Y24_S667s_1 | 0.6 | -0.88 | -1.22 | -0.22 | 0.22 | -0.79 | -0.08 | -0.11 | -1 | -1.29 | -0.1 | -1.01 | -0.89 | -0.41 | -0.89 | -0.89 | 0.24 | -0.42 | -0.34 | 0.434602 | 0.589737 | 0.375442 | 1 | 0.999631 | 0.779717 |
| Prickle2_Q80Y24_S696s_1 | 0.09 | -0.13 | -0.49 | 0.46 | 0.52 | 0.02 | 0.31 | 0.6 | -0.02 | 0.19 | 0.38 | -0.06 | -0.39 | 0.81 | -0.07 | 0.45 | 0.38 | 0.14 | 0.2175 | 0.563758 | 0.224958 | 0.642484 | 1 | 0.999631 | 0.841319 |
| Prickle2_Q80Y24_S732s_1 | 0.08 | 0.71 | 0.2 | 0.4 | 1.04 | 0.53 | 0.5 | 0.37 | 0.94 | -0.27 | 0.81 | 0.61 | -0.2 | 0.31 | -0.01 | 0.58 | 0.2625 | 0.175 | -0.1775 | 0.572797 | 0.362264 | 0.616032 | 1 | 0.999631 | 0.844402 |
| Prickle2_Q80Y24_S807s_1 | 0.17 | -0.83 | -1.02 | -0.4 | 0.29 | -0.16 | -0.62 | -0.21 | -0.66 | 0.36 | -0.7 | -0.59 | -0.38 | -0.13 | -0.96 | 0.26 | 0.345 | 0.1225 | 0.2175 | 0.584563 | 0.338708 | 0.751124 | 1 | 0.999631 | 0.849201 |
| Prickle2_Q80Y24_T535t_1 | -0.16 | -0.33 | 0.11 | 0.02 | -0.67 | -0.23 | -0.19 | 0.43 | -0.08 | 0.43 | -0.03 | 0.38 | 0.08 | -0.57 | 0.19 | 0.51 | -0.075 | 0.265 | 0.1425 | 0.662658 | 0.806084 | 0.342837 | 1 | 0.999631 | 0.881311 |
| Prickle2_Q80Y24_T535t_2 | 1.04 | 0.39 | 0.23 | 1.55 | 0.26 | -0.01 | 1.04 | -0.35 | -0.11 | 1 | 0.42 | -0.45 | -1.56 | -0.49 | -0.29 | 1.08 | -0.5675 | -0.5875 | -1.1175 | 0.074876 | 0.198001 | 0.199209 | 1 | 0.999631 | 0.540556 |
| Prickle2_Q80Y24_T540t_2 | 1.04 | 0.39 | 0.23 | 1.55 | 0.26 | -0.01 | 1.04 | -0.35 | -0.11 | 1 | 0.42 | -0.45 | -1.56 | -0.49 | -0.29 | 1.08 | -0.5675 | -0.5875 | -1.1175 | 0.074876 | 0.198001 | 0.199209 | 1 | 0.999631 | 0.540556 |
| Prickle2_Q80Y24_T806t_1 | 0.32 | -0.71 | -0.96 | -0.26 | 0.35 | -0.15 | -0.79 | 0.22 | -0.8 | 0.4 | -1.2 | -0.65 | -0.34 | -0.26 | -0.43 | 0.27 | 0.31 | -0.16 | 0.2125 | 0.565826 | 0.433306 | 0.715701 | 1 | 0.999631 | 0.841575 |
| Prkaa1_Q3TUQ7_S487s_1 | 0.64 | -0.24 | 0.04 | 0.4 | 0.28 | 0 | 0.42 | 0.34 | -0.25 | -0.17 | 0.04 | 0.28 | -0.57 | -0.07 | 0.56 | -0.23 | 0.05 | -0.235 | -0.2875 | 0.429241 | 0.862196 | 0.443881 | 1 | 0.999631 | 0.776658 |
| Prkaa1_Q3TUQ7_T174t_1 | -1.47 | 0.63 | 1 | -0.94 | -0.66 | 1.67 | 0.5 | -0.3 | -0.38 | -1.41 | 0.46 | -1.95 | -0.78 | 0.58 | -1.16 | -1.3 | 0.4975 | -0.625 | -0.47 | 0.481434 | 0.482515 | 0.390961 | 1 | 0.999631 | 0.802053 |
| Prkaa1_Q3TUQ7_T481t_1 | 2.14 | -0.43 | -0.03 | 0.01 | -0.06 | -0.56 | 1.39 | -1.66 | -1.36 | 0.67 | -1.2 | -1.37 | -1.86 | -1.57 | -1.42 | -1.55 | -0.645 | -1.2375 | -2.0225 | 0.004032 | 0.401683 | 0.094979 | 1 | 0.999631 | 0.351308 |
| Prkaa2_Q8BRK8_S481s_1 | 1.59 | -1.26 | -0.2 | -0.33 | -2.22 | -0.45 | 2.17 | -1.56 | -2.41 | 0.87 | -0.9 | -1.49 | -0.86 | -2.11 | -1.32 | -1.42 | -0.465 | -0.9325 | -1.3775 | 0.0399 | 0.636237 | 0.261336 | 1 | 0.999631 | 0.471899 |
| Prkab1_Q9R078_S182s_1 | -1.98 | -0.88 | -0.02 | -0.75 | -0.33 | -2.29 | -0.42 | -0.2 | -0.34 | -2.77 | 0.28 | 0.19 | 0.39 | -1.86 | 0.52 | 0.17 | 0.0975 | 0.2475 | 0.7125 | 0.269876 | 0.866707 | 0.734872 | 1 | 0.999631 | 0.698414 |
| Prkab2_Q6PAM0_S183s_1 | -0.82 | -0.03 | -0.48 | -0.88 | -0.29 | -0.43 | -0.78 | -0.33 | -0.58 | -0.54 | -0.45 | -0.4 | 0.27 | 0.29 | -0.27 | -0.27 | 0.095 | 0.06 | 0.5575 | 0.112932 | 0.746628 | 0.834941 | 1 | 0.999631 | 0.577408 |
| Prkaca_P05132_T188t_1 | -1.42 | 0.79 | 1.21 | -0.19 | -1.97 | 2.01 | -0.5 | 0.75 | -1.65 | -1.06 | 0.55 | -2.74 | -0.38 | 2.86 | -1.39 | 0.59 | -0.025 | -1.3225 | 0.3225 | 0.728978 | 0.977689 | 0.119892 | 1 | 0.999631 | 0.905923 |
| Prkaca_P05132_T190t_1 | -1.65 | 1.14 | 1.31 | -0.36 | -2.09 | 2.61 | 0.09 | 1.67 | -1.75 | -0.97 | 1.68 | -4 | 0.21 | 3.01 | -0.94 | -0.97 | 0.46 | -1.37 | 0.2175 | 0.827406 | 0.667067 | 0.258206 | 1 | 0.999631 | 0.941202 |
| Prkag2_Q91WG5_S122s_1 | -0.56 | 2.44 | 1.22 | 1.83 | 2.12 | 1.89 | 1.03 | 1.78 | 1.9 | 1.17 | 2.5 | 1.07 | 0.97 | 2.48 | 1.26 | 1.39 | 0.4725 | 0.4275 | 0.2925 | 0.655118 | 0.452577 | 0.518677 | 1 | 0.999631 | 0.879047 |
| Prkag2_Q91WG5_S196s_1 | -0.95 | -0.66 | -0.31 | -1.06 | -0.76 | -0.74 | -0.72 | -0.79 | -0.84 | -0.94 | -0.83 | -0.79 | -0.44 | -1.21 | -0.99 | -0.69 | -0.0075 | -0.105 | -0.0875 | 0.785221 | 0.977476 | 0.703188 | 1 | 0.999631 | 0.923063 |
| Prkag2_Q91WG5_S87s_1 | 0.25 | -0.79 | -0.82 | 0.64 | 0.21 | 0.34 | 0.16 | -0.18 | 0.3 | -0.04 | -0.77 | -0.25 | -1.71 | 0.27 | -0.52 | -0.18 | 0.3125 | -0.01 | -0.355 | 0.50701 | 0.43444 | 0.981475 | 1 | 0.999631 | 0.81564 |
| Prkar1a_Q9DBC7_S5s_1 | 2.22 | 2.12 | -1.75 | -1.17 | 0.86 | 1.14 | 0.36 | -1.44 | -0.24 | 1.76 | -0.51 | 0.59 | -1.62 | 1.48 | -0.97 | -1.65 | -0.125 | 0.045 | -1.045 | 0.351556 | 0.903678 | 0.96447 | 1 | 0.999631 | 0.738113 |
| Prkar1a_Q9DBC7_S77s_1 | -0.95 | 0.32 | -0.16 | -0.13 | -0.3 | -0.06 | -0.66 | 0.23 | 0.64 | 0.12 | -0.19 | 0.43 | 0.56 | 0.42 | 0.48 | 0.17 | 0.0325 | 0.48 | 0.6375 | 0.085747 | 0.92645 | 0.198366 | 1 | 0.999631 | 0.551251 |
| Prkar1a_Q9DBC7_S83s_1 | -0.53 | 0.32 | 0.12 | -0.38 | -0.26 | -0.01 | -0.72 | -0.05 | -0.33 | -0.57 | -0.39 | 0.04 | 0.61 | 0.09 | 0.22 | -0.11 | -0.1425 | -0.195 | 0.32 | 0.34115 | 0.649985 | 0.530948 | 1 | 0.999631 | 0.733032 |
| Prkar1a_Q9DBC7_T75t_1 | -0.35 | -0.02 | 0.69 | -0.67 | 0.31 | -0.1 | -0.37 | -0.2 | -0.25 | 0.54 | -0.48 | 0.62 | 1.08 | 0.48 | 0.4 | 0.97 | -0.0025 | 0.195 | 0.82 | 0.050796 | 0.994341 | 0.636187 | 1 | 0.999631 | 0.489558 |
| Prkar1b_P12849_S3s_1 | 0.24 | 0.39 | -0.46 | -0.45 | 1 | 0.63 | -0.29 | -1.29 | 0.15 | -0.48 | -0.31 | -1.19 | 0.95 | 0.4 | 0.01 | -0.17 | 0.0825 | -0.3875 | 0.3675 | 0.338351 | 0.87373 | 0.320904 | 1 | 0.999631 | 0.731571 |
| Prkar1b_P12849_S83s_1 | -1.34 | -1.13 | -0.97 | -0.7 | -0.9 | -1.44 | -0.69 | -0.81 | -1.09 | -1.73 | -0.69 | -0.21 | -0.18 | -1.19 | -0.24 | -0.69 | 0.075 | 0.105 | 0.46 | 0.193717 | 0.79448 | 0.779289 | 1 | 0.999631 | 0.650601 |
| Prkar1b_P12849_S9s_1 | -0.57 | 0.15 | -0.43 | -0.39 | 0.44 | 0.04 | -0.54 | -1.83 | -0.23 | -0.76 | -0.47 | 0.01 | -0.07 | 0.21 | -0.3 | -0.02 | -0.1625 | -0.0525 | 0.265 | 0.383741 | 0.740655 | 0.862203 | 1 | 0.999631 | 0.754721 |
| Prkar2a_Q8K1M3_S381s_1 | -0.33 | -0.2 | -0.71 | 0.01 | 1.12 | 1.74 | -0.85 | -1.38 | 1.43 | 0.66 | -0.16 | -0.39 | 2.68 | 1.94 | -0.34 | 0.04 | 0.465 | 0.6925 | 1.3875 | 0.060044 | 0.500551 | 0.137941 | 1 | 0.999631 | 0.505461 |
| Prkar2a_Q8K1M3_S97s_1 | -0.56 | 1.29 | 2.06 | 1.13 | 0.63 | 0.64 | 0.44 | 1.05 | 0.52 | 0.21 | 0.94 | 0.78 | 1.37 | 1.22 | 0.4 | 1.35 | -0.29 | -0.3675 | 0.105 | 0.850036 | 0.583751 | 0.498063 | 1 | 0.999631 | 0.949525 |
| Prkar2b_P31324_S112s_1 | -0.15 | 1.31 | 1.19 | 0.79 | 0.41 | 0.86 | 0.24 | 1.2 | 0.68 | 0.64 | 0.89 | 0.69 | 0.54 | 1.04 | 0.91 | 1 | -0.1075 | -0.06 | 0.0875 | 0.819824 | 0.788516 | 0.869644 | 1 | 0.999631 | 0.938303 |
| Prkca_Q4VA93_S13s_1 | -1.73 | 0.39 | 1.06 | -0.62 | 0.23 | -0.28 | 0.21 | -0.28 | 0.88 | -0.11 | 0.19 | 0.46 | 1.14 | -1.35 | 0.98 | 0.27 | 0.195 | 0.58 | 0.485 | 0.513514 | 0.733078 | 0.337271 | 1 | 0.999631 | 0.818773 |
| Prkca_Q4VA93_S226s_1 | -0.64 | -0.37 | -0.45 | -0.27 | -0.25 | -0.55 | -0.55 | -0.51 | -0.72 | -1 | -0.69 | -0.27 | -0.36 | -0.41 | -0.4 | -0.16 | -0.0325 | -0.2375 | 0.1 | 0.708313 | 0.893393 | 0.395867 | 1 | 0.999631 | 0.898901 |
| Prkca_Q4VA93_S319s_1 | -0.49 | -1.29 | -1.87 | -1.17 | -1.76 | -1.67 | -1.59 | -1.51 | -1.79 | -1.23 | -1.65 | -1.8 | -1.11 | -1.57 | -1.79 | -1.91 | -0.4275 | -0.4125 | -0.39 | 0.310492 | 0.214645 | 0.257686 | 1 | 0.999631 | 0.71924 |
| Prkcb_P68404_S11s_1 | -0.41 | -1.26 | 0.01 | -0.74 | -0.61 | -1.07 | -0.25 | -0.61 | -0.54 | -0.49 | 0.78 | -0.87 | -0.75 | -0.24 | 0.28 | -0.43 | -0.035 | 0.32 | 0.315 | 0.415894 | 0.919689 | 0.477987 | 1 | 0.999631 | 0.770095 |
| Prkcb_P68404_S206s_1 | -1.45 | -3.28 | -4.45 | -3.21 | -1.89 | -3.73 | -3.68 | -2.76 | -2.34 | -2.26 | -2.67 | -2.11 | -0.8 | -2.27 | -3.13 | -3.31 | 0.0825 | 0.7525 | 0.72 | 0.343366 | 0.902353 | 0.212908 | 1 | 0.999631 | 0.73348 |
| Prkcb_P68404_S660s_1 | -1.33 | 1.79 | 1.38 | 0.25 | 0.64 | 1.2 | 0.06 | 0.76 | 0.8 | -0.97 | 1.04 | 0.47 | 1.05 | 1.01 | 0.51 | 0.25 | 0.1425 | -0.1875 | 0.1825 | 0.779292 | 0.828296 | 0.798949 | 1 | 0.999631 | 0.921416 |
| Prkcb_P68404_T641t_1 | -1.46 | 2.87 | 2.65 | -0.42 | 4.02 | 1.35 | 2.37 | 0.43 | 5.51 | -0.72 | 3.73 | 5.09 | 6.36 | 1.02 | 1.98 | 0.37 | 1.1325 | 2.4925 | 1.5225 | 0.310003 | 0.333453 | 0.12685 | 1 | 0.999631 | 0.71896 |
| Prkcb_P68404_T642t_1 | -1.07 | -1.11 | -1.8 | -1.16 | -1.12 | -0.69 | -2.16 | -1.67 | -1.47 | -1.08 | -2.69 | -2.98 | -0.55 | -0.27 | -2.15 | -0.62 | -0.125 | -0.77 | 0.3875 | 0.403107 | 0.741675 | 0.129186 | 1 | 0.999631 | 0.764348 |
| Prkcb_P68404_T644t_1 | -0.59 | -0.27 | -0.59 | -0.47 | -0.76 | -0.13 | -1.15 | -0.23 | -0.88 | -0.26 | -1.05 | -0.94 | 0.64 | -0.06 | -0.66 | -0.34 | -0.0875 | -0.3025 | 0.375 | 0.29409 | 0.776162 | 0.30262 | 1 | 0.999631 | 0.708293 |
| Prkcd_Q1MX43_S389s_1 | -1.42 | 0.29 | 0.76 | -0.39 | -2.24 | 1.8 | -0.28 | 1.11 | -2.24 | -1.26 | 0.78 | -2.72 | -0.51 | 1.99 | -0.47 | -0.47 | 0.2875 | -1.17 | 0.325 | 0.640176 | 0.743001 | 0.165343 | 1 | 0.999631 | 0.870995 |
| Prkcd_Q1MX43_S528s_1 | 0.03 | 0.79 | 2.22 | 0.68 | 0.36 | 1.78 | 0.22 | 1.08 | 0.85 | 0.8 | 0.35 | 0.88 | 0.09 | 2.06 | 1.45 | 1.43 | -0.07 | -0.21 | 0.3275 | 0.570267 | 0.896672 | 0.65241 | 1 | 0.999631 | 0.84338 |
| Prkcd_Q1MX43_S547s_1 | -0.83 | 1.94 | 1.81 | 0.57 | 1.02 | 1.52 | 0.36 | 1.48 | 1.24 | 0.21 | 1.25 | 1.53 | 1.6 | 1.37 | 1.7 | 1.42 | 0.2225 | 0.185 | 0.65 | 0.288978 | 0.723901 | 0.772409 | 1 | 0.999631 | 0.706046 |
| Prkcd_Q1MX43_T390t_1 | -0.71 | 0.47 | 1.13 | -0.3 | -0.95 | 1.41 | -0.25 | 1.4 | -1.67 | -0.57 | 1.09 | -2.71 | -0.31 | 1.55 | -0.75 | -0.97 | 0.255 | -1.1125 | -0.2675 | 0.674612 | 0.694051 | 0.185401 | 1 | 0.999631 | 0.885515 |
| Prkcd_Q1MX43_Y196y_1 | -1.65 | -2.33 | -1.7 | -1.08 | -1.54 | -2.48 | -1.34 | -2.13 | -2.43 | -2.58 | -0.5 | -0.63 | -0.75 | -1.35 | -1.84 | -0.7 | -0.1825 | 0.155 | 0.53 | 0.202737 | 0.633052 | 0.785149 | 1 | 0.999631 | 0.659215 |
| Prkce_P16054_S234s_1 | 0.5 | -0.26 | -0.67 | 0.21 | 0.72 | 0.47 | 0.23 | 1 | 0.31 | -0.05 | 0.16 | -0.92 | -0.68 | 0.23 | 0.34 | -0.77 | 0.66 | -0.07 | -0.165 | 0.68802 | 0.077234 | 0.858589 | 1 | 0.999631 | 0.891733 |
| Prkce_P16054_S329s_1 | -1 | -1.23 | -0.71 | -1.66 | -0.89 | -1.53 | -1.29 | -1.21 | -1.08 | -0.88 | -1.17 | -1.71 | -0.86 | -1.24 | -1.16 | -2.16 | -0.08 | -0.06 | -0.205 | 0.592943 | 0.791179 | 0.85319 | 1 | 0.999631 | 0.853138 |
| Prkce_P16054_S329s_2 | -1.42 | -0.05 | -0.4 | -1.08 | 0.18 | -0.71 | -0.52 | -0.39 | 0.09 | -1.11 | -0.74 | -0.48 | 0.27 | -0.39 | -0.14 | -0.24 | 0.3775 | 0.1775 | 0.6125 | 0.130129 | 0.333064 | 0.666071 | 1 | 0.999631 | 0.601275 |
| Prkce_P16054_S337s_1 | -2.23 | -3.82 | -2.26 | -2.79 | -3.17 | -3.34 | -2.25 | -2.93 | -3.28 | -2.94 | -2.29 | -3.17 | -2.62 | -3.32 | -2.87 | -3.1 | -0.1475 | -0.145 | -0.2025 | 0.628462 | 0.733777 | 0.737269 | 1 | 0.999631 | 0.865998 |
| Prkce_P16054_S337s_2 | -1.42 | -0.05 | -0.46 | -0.7 | 0.18 | -0.71 | -0.22 | -0.3 | 0.09 | -1.11 | -0.38 | 0.14 | 0.27 | -0.39 | 0.44 | -0.76 | 0.395 | 0.3425 | 0.5475 | 0.208927 | 0.291261 | 0.416752 | 1 | 0.999631 | 0.662575 |
| Prkce_P16054_S346s_1 | -0.2 | 1.65 | -0.16 | -0.39 | 0.32 | 0.6 | -0.55 | -0.17 | 0.79 | 1.23 | -0.53 | -0.26 | 0.89 | 1.04 | 0.23 | -0.06 | -0.175 | 0.0825 | 0.3 | 0.564604 | 0.730237 | 0.887254 | 1 | 0.999631 | 0.84139 |
| Prkce_P16054_S346s_2 | -0.6 | 0.33 | 0.23 | -0.33 | -0.06 | 0 | -0.34 | 0.13 | 0.03 | 0.39 | -0.32 | -0.27 | 0.42 | 0.25 | 0.11 | 0.33 | 0.025 | 0.05 | 0.37 | 0.261028 | 0.934396 | 0.879298 | 1 | 0.999631 | 0.69271 |
| Prkce_P16054_S350s_1 | -1.77 | -1.62 | -1.16 | -2.24 | -2.06 | -2.01 | -2.09 | -2.12 | -2.22 | -1.07 | -1.89 | -1.49 | 0.12 | -2.02 | -1.78 | -1.74 | -0.3725 | 0.03 | 0.3425 | 0.510891 | 0.222827 | 0.934367 | 1 | 0.999631 | 0.817687 |
| Prkce_P16054_S350s_2 | -0.6 | 0.33 | 0.23 | -0.33 | -0.06 | 0 | -0.34 | 0.13 | 0.03 | 0.39 | -0.32 | -0.27 | 0.42 | 0.25 | 0.11 | 0.33 | 0.025 | 0.05 | 0.37 | 0.261028 | 0.934396 | 0.879298 | 1 | 0.999631 | 0.69271 |
| Prkce_P16054_S368s_1 | 0.88 | 0.27 | -0.38 | 0.94 | 1.21 | 0.31 | 0.55 | 0.76 | 0.52 | 0.03 | 0.45 | 0.18 | -0.61 | 0.41 | 0.26 | 0.4 | 0.28 | -0.1325 | -0.3125 | 0.454355 | 0.465053 | 0.714937 | 1 | 0.999631 | 0.789679 |
| Prkce_P16054_S388s_1 | -1.34 | -0.34 | -0.4 | -0.85 | -0.88 | -0.96 | -0.79 | -0.8 | -0.5 | -1.12 | -0.48 | -0.46 | 0.56 | -0.66 | -0.5 | -0.72 | -0.125 | 0.0925 | 0.4025 | 0.329855 | 0.677459 | 0.781248 | 1 | 0.999631 | 0.726756 |
| Prkce_P16054_S62s_1 | -0.71 | -0.25 | -0.65 | -0.02 | -0.45 | -0.62 | -0.52 | -0.33 | 0.13 | -0.16 | -0.45 | -0.6 | 0.77 | -0.55 | 0.41 | -0.58 | -0.0725 | 0.1375 | 0.42 | 0.309966 | 0.787828 | 0.652349 | 1 | 0.999631 | 0.71896 |
| Prkce_P16054_S729s_1 | -0.21 | 1.54 | 1.43 | 0.35 | -0.7 | 1.59 | 0.33 | 0.89 | -0.51 | -0.04 | 1.1 | -2.14 | -0.16 | 1.6 | -0.69 | 0.22 | -0.25 | -1.175 | -0.535 | 0.376572 | 0.670002 | 0.121358 | 1 | 0.999631 | 0.748839 |
| Prkce_P16054_T228t_1 | 0.52 | -0.23 | 0.03 | -0.22 | 0.08 | -0.3 | -0.32 | -0.18 | 0.22 | 0.75 | -0.21 | 0.01 | -0.33 | -0.52 | 0.02 | -0.08 | -0.205 | 0.1675 | -0.2525 | 0.421907 | 0.470091 | 0.609463 | 1 | 0.999631 | 0.772942 |
| Prkce_P16054_T710t_1 | -0.38 | 2.76 | 2.64 | 0.83 | -0.81 | 4.01 | 0.7 | 2.31 | -0.57 | 0.52 | 1.96 | -2.9 | -0.5 | 3.85 | -0.27 | 0.28 | 0.09 | -1.71 | -0.6225 | 0.566939 | 0.934713 | 0.142372 | 1 | 0.999631 | 0.842178 |
| Prkcg_P63318_S132s_1 | 1.43 | 0.3 | 0.94 | 0.1 | 0.18 | 1.54 | 1.14 | 0.59 | 0.66 | 0.84 | 0.06 | -4.27 | -0.93 | 0.87 | -0.3 | -0.22 | 0.17 | -1.37 | -0.8375 | 0.100079 | 0.688095 | 0.221182 | 1 | 0.999631 | 0.562917 |
| Prkcg_P63318_S148s_1 | 0.32 | 0.38 | -0.42 | -0.56 | -0.57 | 0.06 | -0.38 | 1.17 | -0.32 | 1.05 | 0.39 | 0.57 | 1.09 | 0.28 | 0.86 | 1.34 | 0.14 | 0.4925 | 0.9625 | 0.025732 | 0.753949 | 0.227335 | 1 | 0.999631 | 0.422114 |
| Prkcg_P63318_S326s_1 | -0.53 | -0.44 | 0.29 | 0.16 | -0.26 | -0.48 | -0.13 | 0.38 | -0.49 | -0.63 | 0.35 | -0.12 | 0.46 | -0.05 | -0.09 | 0.71 | 0.0075 | -0.0925 | 0.3875 | 0.27808 | 0.981434 | 0.78851 | 1 | 0.999631 | 0.700874 |
| Prkcg_P63318_S328s_1 | -0.91 | -1.96 | -3.33 | -3.09 | -2.24 | -1.94 | -1.84 | -3.46 | -1.49 | -0.92 | -2.43 | -1.89 | -0.79 | -1.76 | -2.31 | -2.1 | -0.0475 | 0.64 | 0.5825 | 0.339375 | 0.937405 | 0.291218 | 1 | 0.999631 | 0.732185 |
| Prkcg_P63318_S330s_1 | -1.14 | -1 | -0.52 | -1.68 | -1.48 | -1.3 | -1.9 | -1.16 | -1.35 | -0.82 | -1.29 | -0.5 | -0.2 | -0.96 | -0.63 | -0.97 | -0.375 | 0.095 | 0.395 | 0.279898 | 0.271306 | 0.786958 | 1 | 0.999631 | 0.701922 |
| Prkcg_P63318_S373s_1 | -0.94 | -1.77 | -2 | -1.86 | -1.68 | -1.38 | -1.97 | -2.11 | -1.51 | -0.61 | -1.13 | -0.91 | 0.59 | -1.39 | -1.31 | -0.69 | -0.1425 | 0.6025 | 0.9425 | 0.081362 | 0.667279 | 0.106815 | 1 | 0.999631 | 0.546723 |
| Prkcg_P63318_S510s_1 | 0.02 | 0.17 | 0.14 | -0.49 | 0.12 | -0.16 | -0.25 | 0.01 | 0.34 | -0.3 | 0.38 | 0.2 | 0.06 | -0.14 | 0.51 | 0.41 | -0.03 | 0.195 | 0.25 | 0.427464 | 0.911023 | 0.517161 | 1 | 0.999631 | 0.77521 |
| Prkcg_P63318_S639s_1 | -0.2 | -0.89 | 0.1 | 0.46 | -0.82 | -0.43 | 0.87 | -0.83 | -0.7 | -0.2 | 0.67 | -0.35 | -1.36 | -1.1 | -0.34 | -0.96 | -0.17 | -0.0125 | -0.8075 | 0.060731 | 0.718459 | 0.975907 | 1 | 0.999631 | 0.509205 |
| Prkcg_P63318_S687s_1 | -0.31 | -0.58 | -0.22 | -1.26 | -0.63 | -0.41 | -1.13 | -0.74 | -0.5 | -0.06 | -0.8 | -0.3 | -0.19 | -0.06 | -0.22 | -0.24 | -0.135 | 0.1775 | 0.415 | 0.215456 | 0.678564 | 0.596615 | 1 | 0.999631 | 0.668522 |
| Prkcg_P63318_S690s_1 | -0.06 | -0.73 | -1.95 | -0.25 | -0.06 | -1.31 | -0.26 | -0.4 | -0.91 | -1.53 | -0.09 | -1.7 | -0.77 | -1.08 | -2 | -0.27 | 0.24 | -0.31 | -0.2825 | 0.594405 | 0.619444 | 0.558059 | 1 | 0.999631 | 0.85392 |
| Prkcg_P63318_T514t_1 | -1.71 | 0.85 | 0.44 | -0.96 | -1.78 | 1.82 | -0.19 | 0.22 | -1.69 | -1.76 | 0.71 | -2.25 | -0.91 | 1.32 | -1.5 | -1.57 | 0.3625 | -0.9025 | -0.32 | 0.686151 | 0.662958 | 0.267413 | 1 | 0.999631 | 0.890692 |
| Prkcg_P63318_T655t_1 | -0.52 | -0.53 | -0.26 | -0.86 | -0.71 | -0.54 | -1.05 | -0.29 | -0.57 | -0.21 | -0.27 | -0.46 | 0.11 | -0.34 | -0.47 | -0.22 | -0.105 | 0.165 | 0.3125 | 0.298635 | 0.709858 | 0.538383 | 1 | 0.999631 | 0.711798 |
| Prkch_P23298_S675s_1 | -0.64 | 0.1 | 0.07 | 0.86 | 0.1 | -0.58 | 0.25 | 0.85 | -0.77 | -0.46 | 0.48 | 0.92 | 0.67 | 0.22 | 1.23 | 0.07 | 0.0575 | -0.055 | 0.45 | 0.295261 | 0.891106 | 0.908725 | 1 | 0.999631 | 0.708769 |
| Prkci_Q62074_T411t_1 | -1.64 | -0.42 | -0.62 | -1.46 | -1.25 | -0.09 | -1.36 | -0.56 | -2.02 | -1.65 | -0.35 | -2.66 | -1.13 | 0.6 | -2.71 | -1.74 | 0.22 | -0.635 | -0.21 | 0.756562 | 0.603803 | 0.252602 | 1 | 0.999631 | 0.914089 |
| Prkcq_Q02111_T536t_1 | -1.86 | -0.37 | -0.06 | -1.61 | -1.25 | 1.54 | -1.23 | 0.2 | -1.57 | -1.93 | 0.43 | -4.15 | -0.03 | 1.49 | -2.14 | -1.52 | 0.79 | -0.83 | 0.425 | 0.600691 | 0.28133 | 0.368927 | 1 | 0.999631 | 0.856689 |
| Prkcz_Q02956_T227t_1 | -0.61 | 1.12 | -0.24 | -0.67 | -0.21 | 0.32 | -0.62 | -0.07 | -1.34 | -0.12 | -0.46 | -0.99 | -1.16 | 0.57 | -0.09 | -0.78 | -0.045 | -0.6275 | -0.265 | 0.620349 | 0.919609 | 0.210223 | 1 | 0.999631 | 0.861888 |
| Prkd1_Q62101_S255s_1 | 0.84 | 0.76 | 0.46 | 0.66 | 0.72 | 1.43 | 1.07 | 1.39 | 1.25 | 1.36 | 1.02 | 1.03 | -0.52 | 0.62 | 0.77 | 1.03 | 0.4725 | 0.485 | -0.205 | 0.597109 | 0.108428 | 0.081119 | 1 | 0.999631 | 0.854923 |
| Prkd2_Q8BZ03_S197s_1 | 0.67 | -0.48 | -1.02 | -1 | -0.34 | -1.26 | -1.03 | -1.21 | -0.24 | 0.04 | -0.99 | -0.82 | -0.23 | -0.25 | -1.67 | -1.43 | -0.5025 | -0.045 | -0.4375 | 0.40869 | 0.268076 | 0.920961 | 1 | 0.999631 | 0.766996 |
| Prkd3_Q5FWX6_S735s_1 | -0.15 | 1.17 | 0.87 | -0.02 | -0.23 | 0.6 | -0.34 | 0.12 | 0.51 | 0.52 | 0.13 | -0.09 | 1.02 | 0.74 | 0.33 | 0.73 | -0.43 | -0.2 | 0.2375 | 0.543548 | 0.291529 | 0.601781 | 1 | 0.999631 | 0.832881 |
| Prkg1_P0C605_T532t_1 | -0.63 | 0.33 | 0.08 | -0.26 | -1.94 | 1.33 | -0.58 | 0.76 | -1.5 | -0.77 | 0.6 | -1.81 | -0.48 | 1 | -0.94 | -1.77 | 0.0125 | -0.75 | -0.4275 | 0.45922 | 0.985209 | 0.18395 | 1 | 0.999631 | 0.79152 |
| Prkg2_E9QPH0_S431s_1 | -0.81 | -0.87 | -0.82 | -0.87 | -0.65 | -0.45 | -1.31 | -0.69 | -0.8 | -0.8 | -0.68 | -0.19 | 0.01 | -0.69 | -0.6 | -0.26 | 0.0675 | 0.225 | 0.4575 | 0.134136 | 0.805695 | 0.404173 | 1 | 0.999631 | 0.60492 |
| Prkg2_E9QPH0_S97s_1 | -1.07 | -1 | -0.04 | -0.68 | -0.33 | -0.18 | 0.66 | -0.45 | -0.36 | -2.27 | 0.49 | 1.22 | -0.6 | -1.16 | 0.02 | -0.54 | 0.6225 | 0.4675 | 0.1275 | 0.735027 | 0.112879 | 0.510348 | 1 | 0.999631 | 0.90759 |
| Prkg2_E9QPH0_T109t_1 | -0.57 | -0.49 | -0.14 | -0.11 | 0.09 | -0.4 | -0.09 | -0.19 | -0.07 | -0.6 | -0.08 | -0.2 | -1.14 | -0.31 | 0.23 | -0.23 | 0.18 | 0.09 | -0.035 | 0.922327 | 0.495229 | 0.744032 | 1 | 0.999631 | 0.975032 |
| Prkg2_E9QPH0_T580t_1 | -1.83 | -0.19 | -0.83 | -1.23 | -1.72 | 1.01 | -0.73 | -0.06 | -1.49 | -1.27 | 0.17 | -1.36 | -0.04 | 0.86 | -0.5 | -1.89 | 0.645 | 0.0325 | 0.6275 | 0.31554 | 0.300381 | 0.947528 | 1 | 0.999631 | 0.72069 |
| Prkra_Q9WTX2_S18s_1 | -0.34 | 0.48 | 0.94 | 0.39 | 0.35 | 0.06 | -0.35 | 1.04 | 0.53 | 0.06 | 0.87 | 1.01 | 0.79 | 0.68 | 1.19 | 1.25 | -0.0925 | 0.25 | 0.61 | 0.109421 | 0.81691 | 0.501799 | 1 | 0.999631 | 0.572573 |
| Prmt7_Q922X9_T351t_1 | -0.18 | 1.17 | 0.99 | 0.47 | 0.76 | 1 | 0.52 | 0.96 | 1.13 | 1.31 | 0.66 | 0.9 | 1.28 | 1.54 | 1.17 | 1.66 | 0.1975 | 0.3875 | 0.8 | 0.051109 | 0.576797 | 0.302533 | 1 | 0.999631 | 0.489558 |
| Prnp_P04925_S43s_1 | 0.27 | -0.54 | 0.58 | 0.31 | -0.57 | 0.47 | 0.43 | -0.25 | 0.11 | 0.43 | 0.18 | 0.27 | -0.7 | -0.75 | 0.56 | 0.56 | -0.135 | 0.0925 | -0.2375 | 0.595584 | 0.716571 | 0.769338 | 1 | 0.999631 | 0.854508 |
| Proser1_Q5PRE5_S596s_1 | 0.42 | 0.95 | 1.39 | 0.96 | 0.96 | 1.02 | 1.05 | 1.11 | 1.14 | 1 | 1.42 | 1.56 | 0.72 | 1.33 | 1.62 | 1.76 | 0.105 | 0.35 | 0.4275 | 0.247905 | 0.709672 | 0.270076 | 1 | 0.999631 | 0.685751 |
| Proser1_Q5PRE5_T610t_1 | 1.13 | 0.09 | 0.1 | -0.66 | -0.84 | -0.18 | 0.39 | -0.74 | -0.74 | -0.03 | -0.36 | -0.5 | -0.65 | 0.6 | -0.44 | 0.29 | -0.5075 | -0.5725 | -0.215 | 0.645114 | 0.274771 | 0.180659 | 1 | 0.999631 | 0.873067 |
| Prox1_P48437_S179s_1 | 1.22 | 0.52 | 0.19 | -0.28 | -0.6 | -0.23 | 0.82 | -0.34 | -2.22 | 0.14 | -0.37 | -0.01 | 1.14 | -1.36 | -0.1 | -0.11 | -0.5 | -1.0275 | -0.52 | 0.361 | 0.266094 | 0.10059 | 1 | 0.999631 | 0.741378 |
| Prox1_P48437_S199s_1 | 0.78 | 0.33 | 0.82 | -0.47 | -0.7 | -0.11 | -0.03 | 0.06 | -0.24 | 0.43 | 0.2 | -0.43 | 0.06 | 0.3 | 0.1 | 0.97 | -0.56 | -0.375 | -0.0075 | 0.984804 | 0.149566 | 0.337089 | 1 | 0.999631 | 0.995133 |
| Prox1_P48437_S220s_1 | -1.32 | 0.31 | 0.84 | -0.38 | -0.98 | 0.67 | -0.71 | 0.07 | -1.64 | -1.6 | 0.02 | -0.94 | 0.07 | 0.2 | 0.55 | 0.62 | -0.1 | -0.9025 | 0.4975 | 0.307632 | 0.855661 | 0.131718 | 1 | 0.999631 | 0.717856 |
| Prpf3_Q922U1_S619s_1 | 0.2 | 2.17 | 1.49 | 1.35 | 0.72 | 2.17 | 1.09 | 0.6 | 2.21 | 1.75 | 1.06 | 0.83 | 1.06 | 1.25 | 0.59 | 0.05 | -0.1575 | 0.16 | -0.565 | 0.252671 | 0.756566 | 0.745856 | 1 | 0.999631 | 0.688432 |
| Prpf38a_Q4FK66_S209s_1 | 0.53 | -1.34 | 0.84 | 0.2 | -1.31 | -1.42 | 0.17 | -0.17 | -1.62 | 0.09 | -0.27 | -0.74 | -1.61 | -1.7 | -0.46 | -0.67 | -0.74 | -0.6925 | -1.1675 | 0.052868 | 0.215887 | 0.236393 | 1 | 0.999631 | 0.489777 |
| Prpf40a_Q9R1C7_S884s_1 | 1.83 | 1.17 | 2.82 | 2.51 | 0.16 | 1.45 | 2.48 | 1.38 | 1.21 | 2.22 | 1.74 | 1.68 | 0.6 | 0.02 | 1.71 | 0.72 | -0.715 | -0.37 | -1.32 | 0.021198 | 0.214317 | 0.392214 | 1 | 0.999631 | 0.401577 |
| Prpf40b_F6XW22_S218s_1 | 1.07 | 2.51 | 1.59 | 1.23 | 1.48 | 1.87 | 0.53 | 1.19 | 2.08 | 1.57 | 1.38 | 1.31 | 2.03 | 2.48 | 1.23 | 2.16 | -0.3325 | -0.015 | 0.375 | 0.389733 | 0.439505 | 0.968939 | 1 | 0.999631 | 0.758766 |
| Prpf4b_Q61136_S21s_1 | 1.17 | 2.02 | 2.16 | 1.83 | 0.97 | 1.89 | 2.01 | 0.82 | 1.92 | 2.79 | 1.42 | 1.54 | 0.58 | 0.24 | 1.78 | 0.23 | -0.3725 | 0.1225 | -1.0875 | 0.029036 | 0.347968 | 0.756543 | 1 | 0.999631 | 0.429864 |
| Prpf4b_Q61136_S24s_2 | 0.22 | -0.22 | 2.43 | 1.95 | -0.81 | -0.22 | 1.85 | 1.22 | 0.46 | 1.8 | 1.59 | 1.63 | 0.2 | 1.11 | -0.18 | 1.16 | -0.585 | 0.275 | -0.5225 | 0.430787 | 0.46102 | 0.670724 | 1 | 0.999631 | 0.778235 |
| Prpf4b_Q61136_S258s_1 | 2 | 0.64 | 1.55 | 0.23 | -0.05 | 1.28 | 0.18 | -1.5 | 0.03 | 1.74 | 0.05 | -0.26 | 0.69 | -0.21 | 0.23 | -0.8 | -1.1275 | -0.715 | -1.1275 | 0.043223 | 0.098108 | 0.223557 | 1 | 0.999631 | 0.475443 |
| Prpf4b_Q61136_S278s_1 | 0.5 | 0.31 | 0.68 | -0.04 | -0.37 | 0.57 | 0.12 | -0.26 | 0.58 | 1 | 0.04 | -0.1 | 0.3 | 0.29 | -0.55 | -0.18 | -0.3475 | 0.0175 | -0.3975 | 0.245986 | 0.284708 | 0.959066 | 1 | 0.999631 | 0.685751 |
| Prpf4b_Q61136_S33s_1 | -0.03 | 0.24 | 0.16 | -0.33 | -1.22 | -0.49 | -0.06 | -0.22 | 0.63 | -0.65 | -0.29 | -0.15 | -0.99 | -1 | -2.26 | -0.52 | -0.5075 | -0.125 | -1.2025 | 0.014312 | 0.146088 | 0.715664 | 1 | 0.999631 | 0.385092 |
| Prpf4b_Q61136_S33s_2 | 0.22 | -0.22 | 2.43 | 1.95 | -0.81 | -0.22 | 1.85 | 1.22 | 0.46 | 1.8 | 1.59 | 1.63 | 0.2 | 1.11 | -0.18 | 1.16 | -0.585 | 0.275 | -0.5225 | 0.430787 | 0.46102 | 0.670724 | 1 | 0.999631 | 0.778235 |
| Prpf4b_Q61136_S341s_1 | 1.68 | -5.43 | -3.2 | -1.48 | -3.94 | -4.02 | 1.63 | -5.02 | -15.45 | 0.15 | -3.39 | -2.92 | -2.83 | -4.35 | -3.47 | -4.4 | -0.73 | -3.295 | -1.655 | 0.223237 | 0.685776 | 0.311165 | 1 | 0.999631 | 0.674656 |
| Prpf4b_Q61136_S431s_1 | -0.06 | 4.18 | 4.51 | 2.15 | 3.09 | 3.74 | 3.81 | 3.37 | 4.23 | 0.72 | 4.65 | 3.89 | 4.15 | 2.82 | 4.35 | 3.06 | 0.8075 | 0.6775 | 0.9 | 0.36105 | 0.391896 | 0.571442 | 1 | 0.999631 | 0.741378 |
| Prpf4b_Q61136_S94s_1 | 1.32 | 1.31 | 2.32 | 1.43 | 0.86 | 1.35 | 1.42 | 1.35 | 1.57 | 2.06 | 1.9 | 1.52 | 1.51 | 1.5 | 1.71 | 1.7 | -0.35 | 0.1675 | 0.01 | 0.97557 | 0.292236 | 0.612895 | 1 | 0.999631 | 0.991563 |
| Prpf4b_Q61136_Y849y_1 | -0.88 | -0.01 | -0.49 | 0.07 | 0.42 | 0.05 | -0.58 | 0.29 | -0.59 | 0.2 | 0.98 | -0.43 | 0.7 | 0.33 | 0.44 | -0.23 | 0.3725 | 0.3675 | 0.6375 | 0.094172 | 0.295977 | 0.393976 | 1 | 0.999631 | 0.558128 |
| Prpsap2_Q8R574_S198s_1 | -0.82 | -1.5 | 0.48 | 0.74 | -0.56 | -2.72 | 0.55 | 0.37 | -0.45 | -2.58 | 0.61 | 0.58 | -0.49 | -2.46 | 0.82 | -0.28 | -0.315 | -0.185 | -0.3275 | 0.667459 | 0.695886 | 0.818452 | 1 | 0.999631 | 0.88264 |
| Prpsap2_Q8R574_S219s_1 | -0.7 | -1.1 | -0.93 | -0.71 | -0.16 | -0.28 | -1.19 | -0.88 | -0.2 | 0.37 | -0.93 | -0.42 | 0.71 | -0.61 | 0.21 | -0.58 | 0.2325 | 0.565 | 0.7925 | 0.056713 | 0.46655 | 0.116264 | 1 | 0.999631 | 0.496275 |
| Prpsap2_Q8R574_S227s_1 | -0.46 | 0.16 | -0.99 | -0.78 | 0.09 | -0.94 | -1.11 | -0.86 | 0.93 | -0.58 | -0.98 | -0.85 | 0.53 | -0.29 | -1.26 | -0.55 | -0.1875 | 0.1475 | 0.125 | 0.779973 | 0.625052 | 0.761691 | 1 | 0.999631 | 0.921416 |
| Prr12_E9PYL2_S648s_1 | -1.64 | 1.24 | 1.52 | -1.45 | 0.28 | -0.13 | 0.17 | -0.48 | 1.72 | -2.39 | 0.7 | 2.01 | 1.04 | -0.44 | 1.03 | 0.75 | 0.0425 | 0.5925 | 0.6775 | 0.405594 | 0.955262 | 0.602315 | 1 | 0.999631 | 0.765965 |
| Prr18_Q6PAN7_T91t_1 | -0.06 | 0.02 | -0.91 | 0.05 | -0.22 | 0.72 | -0.49 | -0.48 | 0.41 | 0.77 | -0.59 | 0.13 | -0.1 | 0.26 | 0.11 | -2.17 | 0.1075 | 0.405 | -0.25 | 0.660782 | 0.777669 | 0.309474 | 1 | 0.999631 | 0.880315 |
| Prr36_E9PV26_S197s_1 | -0.14 | 0.27 | 0.37 | 0.35 | 0.27 | -0.07 | -0.18 | -0.67 | -0.13 | -0.11 | -0.48 | -0.74 | -0.15 | 0.39 | 0.37 | -0.54 | -0.375 | -0.5775 | -0.195 | 0.556651 | 0.223357 | 0.066127 | 1 | 0.999631 | 0.838305 |
| Prr36_E9PV26_S519s_1 | -0.09 | 0.28 | 0.24 | -0.35 | 0.05 | 0.1 | -0.22 | -0.1 | -0.04 | 0.29 | 0.15 | 0.61 | 0.56 | 0.48 | 0.17 | 0.57 | -0.0625 | 0.2325 | 0.425 | 0.167933 | 0.813395 | 0.429299 | 1 | 0.999631 | 0.637399 |
| Prr5_Q812A5_S232s_1 | -0.41 | -0.38 | 0.47 | -0.27 | 0.21 | -0.26 | 0.45 | 1.64 | 0.21 | -0.29 | 0.07 | 0.96 | 0.2 | 0.89 | 1.07 | 1.04 | 0.6575 | 0.385 | 0.9475 | 0.020309 | 0.160116 | 0.306283 | 1 | 0.999631 | 0.393478 |
| Prr5_Q812A5_S302s_1 | 1.66 | -1.71 | -0.71 | -0.58 | -0.1 | -0.14 | 0.71 | -0.09 | -0.61 | 0.42 | -0.67 | -0.17 | -2.21 | -0.03 | -0.8 | -1.18 | 0.43 | 0.0775 | -0.72 | 0.343634 | 0.519212 | 0.908544 | 1 | 0.999631 | 0.733852 |
| Prr7_Q3V0I2_S185s_1 | -2.76 | -6.18 | -5.89 | -3.31 | -2.59 | -3.79 | -4.6 | -8.06 | -8.34 | -4.4 | -4 | -6.83 | -3.16 | -3.3 | -6.79 | -2.67 | -0.225 | -1.3575 | 0.555 | 0.614752 | 0.85698 | 0.258191 | 1 | 0.999631 | 0.861203 |
| Prr7_Q3V0I2_S68s_1 | -1.12 | -0.93 | -0.89 | -1.12 | -0.59 | -1.5 | -1.16 | 0.1 | -1.28 | -1.38 | -0.81 | -1.67 | -0.28 | 0.15 | -0.43 | -1.22 | 0.2275 | -0.27 | 0.57 | 0.12704 | 0.545545 | 0.351412 | 1 | 0.999631 | 0.59788 |
| Prrc2a_G3UX48_S1049s_1 | 0.37 | -0.26 | -0.49 | -0.3 | 0.03 | -0.91 | -0.23 | -0.16 | -0.14 | 0.36 | -0.84 | -0.96 | 0.35 | 0.49 | -0.82 | -0.1 | -0.1475 | -0.225 | 0.15 | 0.69657 | 0.649803 | 0.560089 | 1 | 0.999631 | 0.895731 |
| Prrc2a_G3UX48_S111s_1 | 0.33 | 0.8 | -2.18 | 0.88 | 1.44 | 0.69 | 0.46 | 0.5 | 0.97 | -0.07 | 1.03 | -0.09 | -2.35 | 1.05 | 0 | 0.13 | 0.815 | 0.5025 | -0.25 | 0.777457 | 0.244463 | 0.4794 | 1 | 0.999631 | 0.921382 |
| Prrc2a_G3UX48_S1162s_1 | -0.59 | 0.63 | 0.57 | 0.43 | 0.69 | 0.74 | 0.38 | 0.73 | 0.17 | -0.49 | 0.87 | 0.58 | 0.5 | 0.62 | 0.59 | 0.83 | 0.375 | 0.0225 | 0.375 | 0.299796 | 0.280631 | 0.956846 | 1 | 0.999631 | 0.7134 |
| Prrc2a_G3UX48_S295s_1 | -1.28 | 0.56 | 1.53 | -0.73 | -0.09 | 0.67 | -0.11 | 0.36 | 0.53 | -0.39 | 0.57 | 1.11 | 2.14 | -0.1 | 1.27 | 0.42 | 0.1875 | 0.435 | 0.9125 | 0.217988 | 0.754307 | 0.500053 | 1 | 0.999631 | 0.669859 |
| Prrc2a_G3UX48_S323s_1 | -0.05 | -0.14 | -0.71 | 1.46 | -0.63 | -0.1 | -0.81 | 1.99 | -0.62 | -0.01 | -0.43 | -0.5 | -0.18 | -0.16 | 0.43 | 1.05 | -0.0275 | -0.53 | 0.145 | 0.779949 | 0.968709 | 0.272821 | 1 | 0.999631 | 0.921416 |
| Prrc2a_G3UX48_S399s_1 | 1.41 | 1.41 | 1.88 | 1.81 | 1.97 | 0.87 | 1.3 | 1.3 | 0.69 | 0.59 | 3.02 | 0.79 | -0.65 | 1.63 | 2.86 | 1.22 | -0.2675 | -0.355 | -0.3625 | 0.585947 | 0.401915 | 0.525409 | 1 | 0.999631 | 0.849524 |
| Prrc2a_G3UX48_S706s_1 | -0.24 | 0.77 | 0.88 | 0.7 | 0.06 | 0.86 | 0.55 | 0.91 | 0.35 | 0.46 | 0.78 | 0.71 | -0.07 | 1.32 | 0.51 | 1.46 | 0.0675 | 0.0475 | 0.2775 | 0.536054 | 0.847998 | 0.885539 | 1 | 0.999631 | 0.829627 |
| Prrc2a_G3UX48_S834s_1 | 0.39 | 0.06 | 0.18 | 1.21 | 1.14 | 0.24 | 0.58 | 1.32 | 0.41 | 0.59 | 1.13 | 0.56 | -1.89 | 1.16 | 1.08 | 1.64 | 0.36 | 0.2125 | 0.0375 | 0.959759 | 0.349398 | 0.542326 | 1 | 0.999631 | 0.986095 |
| Prrc2a_G3UX48_T480t_1 | 1.15 | 1.29 | -0.04 | 1.44 | 1.94 | 1.88 | 0.39 | 0.83 | 1.04 | 0.62 | 0.3 | 1.21 | 1.11 | 0.91 | 0.46 | 1.37 | 0.3 | -0.1675 | 0.0025 | 0.995115 | 0.540364 | 0.681025 | 1 | 0.999631 | 0.997705 |
| Prrc2a_G3UX48_T554t_1 | 0 | 0.44 | 1.45 | 0.42 | 0.04 | 0.35 | -0.15 | 0.81 | -0.12 | 0.17 | 0.54 | 0.77 | 0.88 | 0.97 | 0.51 | 1.09 | -0.315 | -0.2375 | 0.285 | 0.451924 | 0.42048 | 0.541501 | 1 | 0.999631 | 0.788148 |
| Prrc2a_G3UX48_T727t_1 | 0.65 | 1.25 | 1.17 | 0.74 | 0.61 | 0.97 | 0.45 | 0.84 | 0.79 | 1.16 | 0.48 | 1.09 | 0.65 | 2.07 | 1.31 | 0.87 | -0.235 | -0.0725 | 0.2725 | 0.481889 | 0.403463 | 0.806909 | 1 | 0.999631 | 0.802327 |
| Prrc2b_F8WHT3_S1171s_1 | 2.13 | 0.31 | -0.2 | 1.26 | 0.34 | 1.42 | 1.51 | 1.56 | 0.35 | 1.13 | 0.69 | -1.11 | -1.66 | 1.21 | -0.23 | 1.09 | 0.3325 | -0.61 | -0.7725 | 0.313269 | 0.545487 | 0.351496 | 1 | 0.999631 | 0.719861 |
| Prrc2b_F8WHT3_S1681s_1 | -0.82 | 1.73 | 1.59 | -1.35 | 1.03 | -0.39 | 0.78 | -0.4 | 1.42 | -1.45 | 0.54 | 1.63 | 0.17 | -1.1 | -0.51 | -1.69 | -0.0325 | 0.2475 | -1.07 | 0.191746 | 0.96653 | 0.789058 | 1 | 0.999631 | 0.648674 |
| Prrc2b_F8WHT3_S415s_1 | 1.18 | 2.01 | 0.75 | 1.18 | 1.46 | 2.13 | 0.71 | 2.32 | 1.37 | 0.7 | 1.26 | 0.63 | -1.23 | 2.04 | 0.53 | 1.87 | 0.375 | -0.29 | -0.4775 | 0.505801 | 0.401883 | 0.426845 | 1 | 0.999631 | 0.814836 |
| Prrc2b_F8WHT3_S621s_1 | 1.14 | 2.27 | 0.6 | 0.51 | 1.33 | 1.48 | 0.43 | 0.08 | 1.68 | 0.69 | 0.88 | 0.9 | 2.06 | 2.18 | 1.04 | 0.81 | -0.3 | -0.0925 | 0.3925 | 0.447411 | 0.549737 | 0.837574 | 1 | 0.999631 | 0.785754 |
| Prrc2c_S4R294_S1965s_1 | 0.6 | 0.13 | 0.63 | 0.1 | 0.89 | 0.13 | 0.05 | 1.28 | 0.51 | -0.13 | -0.22 | 0.99 | -0.22 | 0.3 | 0.07 | 0.44 | 0.2225 | -0.0775 | -0.2175 | 0.480917 | 0.537281 | 0.827508 | 1 | 0.999631 | 0.801848 |
| Prrc2c_S4R294_S2646s_1 | 2.31 | 0.69 | 1.88 | 1.25 | -0.4 | 1.6 | 3.41 | 0.33 | -0.59 | 2.19 | -0.08 | 1.11 | -1.37 | 0.99 | 1.2 | -0.19 | -0.2975 | -0.875 | -1.375 | 0.049344 | 0.708262 | 0.196452 | 1 | 0.999631 | 0.487521 |
| Prrc2c_S4R294_S761s_1 | -1.42 | -0.27 | -0.04 | 1 | 0.36 | -0.06 | 2.22 | 1.24 | -0.34 | -2.41 | 1.29 | -0.24 | -1.37 | -0.7 | 0 | 0.49 | 1.1225 | -0.2425 | -0.2125 | 0.718317 | 0.101974 | 0.761257 | 1 | 0.999631 | 0.903971 |
| Prrc2c_S4R294_S899s_1 | 3.54 | -2.24 | 0.79 | 1.77 | -1.29 | -1.14 | 2.23 | 1.29 | -1.37 | 3.24 | 1.15 | 0.24 | -4.87 | -2.41 | 0.23 | -0.97 | -0.6925 | -0.15 | -2.97 | 0.052622 | 0.589721 | 0.909406 | 1 | 0.999631 | 0.489777 |
| Prrc2c_S4R294_S901s_2 | 0.93 | 0.89 | 1.54 | 1.24 | 1.46 | 2.01 | 1.46 | 1.6 | 1.6 | 1.88 | 0.06 | 1.26 | -0.82 | 0.11 | 1.71 | -0.49 | 0.4825 | 0.05 | -1.0225 | 0.083918 | 0.110266 | 0.907024 | 1 | 0.999631 | 0.549732 |
| Prrc2c_S4R294_S904s_1 | 0.45 | -0.04 | 1.16 | 1.08 | -2.63 | 0.55 | -0.88 | -0.72 | -0.39 | 2.77 | -1.7 | -0.7 | -0.39 | -1.58 | 0.14 | -0.9 | -1.5825 | -0.6675 | -1.345 | 0.014075 | 0.030903 | 0.453586 | 1 | 0.999631 | 0.385092 |
| Prrc2c_S4R294_S904s_2 | 1.01 | 0.76 | 1.54 | 1.24 | 1.08 | 1.8 | 1.46 | 1.6 | 1.28 | 1.92 | 0.06 | 1.26 | -0.73 | 0.06 | 1.71 | -0.49 | 0.3475 | -0.0075 | -1 | 0.087147 | 0.254027 | 0.985875 | 1 | 0.999631 | 0.553514 |
| Prrc2c_S4R294_T2625t_1 | -0.39 | 0.79 | 1.13 | 0.5 | 1.06 | 0.85 | 0.73 | 0.6 | 0.59 | -0.35 | 1.26 | 1.93 | -0.29 | 1.08 | 1.65 | 0.82 | 0.3025 | 0.35 | 0.3075 | 0.541922 | 0.412632 | 0.524484 | 1 | 0.999631 | 0.832536 |
| Prrc2c_S4R294_T358t_1 | 0.65 | -1.09 | 0.86 | 1.03 | 0.26 | -0.1 | 1.94 | -0.06 | 0.62 | 0.38 | 0.11 | -0.12 | -0.87 | -1.2 | 0.58 | -0.65 | 0.1475 | -0.115 | -0.8975 | 0.142318 | 0.811819 | 0.815242 | 1 | 0.999631 | 0.612726 |
| Prrt2_E9PUL5_S100s_1 | 0.63 | -0.3 | -0.78 | -0.51 | -0.21 | -0.59 | -0.44 | -0.64 | -0.54 | -0.08 | -1.66 | -1.2 | -1.63 | -1.11 | -0.81 | -1.24 | -0.23 | -0.63 | -0.9575 | 0.029644 | 0.516093 | 0.187264 | 1 | 0.999631 | 0.43391 |
| Prrt2_E9PUL5_S102s_1 | 0.75 | -1.29 | -0.36 | -0.41 | -0.4 | -0.78 | -0.42 | 0.16 | -1.04 | -0.76 | -0.45 | -0.79 | -0.92 | -1.46 | -0.61 | -1.35 | -0.0325 | -0.4325 | -0.7575 | 0.121796 | 0.941907 | 0.330866 | 1 | 0.999631 | 0.590509 |
| Prrt2_E9PUL5_S102s_2 | 1.12 | -0.57 | -0.2 | 0.62 | 0.85 | -0.05 | 0.39 | 0.31 | -0.06 | 0.03 | -0.9 | -0.56 | -2.31 | -0.73 | -0.67 | -1.01 | 0.1325 | -0.615 | -1.4225 | 0.018834 | 0.753653 | 0.182198 | 1 | 0.999631 | 0.391024 |
| Prrt2_E9PUL5_S104s_1 | 1.62 | -0.14 | -0.26 | 0.82 | 0.92 | 0 | 0.45 | 0.12 | 0.59 | -0.83 | -0.56 | -0.37 | -1.18 | -0.46 | -0.36 | -0.33 | -0.1375 | -0.8025 | -1.0925 | 0.041154 | 0.768258 | 0.139163 | 1 | 0.999631 | 0.474979 |
| Prrt2_E9PUL5_S104s_2 | 0.97 | -0.96 | 0.36 | 1.29 | 0.38 | -0.38 | 1.24 | 1.01 | -0.19 | -0.24 | 0.11 | -0.79 | -2.49 | -0.86 | 0.05 | -0.78 | 0.1475 | -0.6925 | -1.435 | 0.049294 | 0.793368 | 0.190406 | 1 | 0.999631 | 0.487521 |
| Prrt2_E9PUL5_S208s_1 | -3.15 | -0.88 | -0.96 | -2.03 | -1.01 | -2.85 | -0.94 | -2.15 | -0.94 | -3.43 | -0.93 | -0.42 | 0.41 | -2.9 | -1.52 | -2.13 | 0.0175 | 0.325 | 0.22 | 0.776815 | 0.977933 | 0.670615 | 1 | 0.999631 | 0.921293 |
| Prrt2_E9PUL5_S244s_1 | 0.32 | 1.77 | 1.65 | -0.62 | 1.92 | 0.99 | 4 | 0.2 | 1.83 | -0.67 | 1.93 | 1.58 | 1.22 | -1.8 | 0.19 | -2.06 | 0.9975 | 0.3875 | -1.3925 | 0.123579 | 0.266629 | 0.604542 | 1 | 0.999631 | 0.594646 |
| Prrt2_E9PUL5_S254s_1 | 0.38 | 3.88 | 2.09 | 0.12 | 3.98 | 2.62 | 2.72 | 0.48 | 5.85 | -0.71 | 3.18 | 3.34 | 5.19 | 0.62 | 1.18 | 0.05 | 0.8325 | 1.2975 | 0.1425 | 0.907774 | 0.403264 | 0.355986 | 1 | 0.999631 | 0.96989 |
| Prrt2_E9PUL5_S6s_1 | 1.18 | -0.23 | -1.12 | -0.65 | 0.07 | 0.03 | -0.3 | -0.66 | 0.25 | -0.15 | -0.76 | -1.46 | -0.33 | -0.39 | -1.25 | -0.71 | -0.01 | -0.325 | -0.465 | 0.374102 | 0.983812 | 0.57236 | 1 | 0.999631 | 0.747035 |
| Prrt2_E9PUL5_S98s_1 | 0.76 | -1.28 | -0.45 | -0.19 | -0.23 | -0.6 | -0.27 | -0.18 | -0.65 | 0.08 | -0.6 | -0.81 | -1.15 | -0.96 | -0.59 | -0.83 | -0.03 | -0.205 | -0.5925 | 0.198418 | 0.943644 | 0.653025 | 1 | 0.999631 | 0.656262 |
| Prrt2_E9PUL5_S98s_2 | 0.89 | -0.75 | -0.03 | 0.83 | 0.27 | -0.29 | 0.67 | 0.53 | -0.36 | -0.13 | -0.58 | -0.62 | -2.11 | -0.88 | -0.45 | -0.94 | 0.06 | -0.6575 | -1.33 | 0.02323 | 0.889998 | 0.135355 | 1 | 0.999631 | 0.414737 |
| Prrt2_E9PUL5_T214t_1 | -0.15 | -1.07 | -0.61 | -0.87 | -1.17 | -0.85 | -0.85 | -0.89 | -1.16 | -0.23 | -0.87 | -1.09 | -0.46 | -1.15 | -1.27 | -1.22 | -0.265 | -0.1625 | -0.35 | 0.316641 | 0.367277 | 0.632168 | 1 | 0.999631 | 0.720734 |
| Prrt3_Q6PE13_S783s_1 | -1.72 | -0.84 | -0.33 | -2.8 | -3.83 | -1.92 | -1.79 | -2.54 | -1.68 | -1.36 | -1.53 | -0.94 | 1.71 | -1.72 | -2.48 | -1.06 | -1.0975 | 0.045 | 0.535 | 0.561581 | 0.111088 | 0.932046 | 1 | 0.999631 | 0.840522 |
| Prrt3_Q6PE13_S783s_2 | -0.8 | -0.34 | 0.03 | -0.29 | -0.53 | -0.36 | -0.47 | -0.22 | -0.49 | -0.72 | 0.1 | 0.22 | 0 | -0.07 | 0.29 | 0.56 | -0.045 | 0.1275 | 0.545 | 0.1065 | 0.869041 | 0.703483 | 1 | 0.999631 | 0.568797 |
| Prrt3_Q6PE13_S791s_2 | -0.18 | 1.04 | -0.24 | 0.15 | 0.78 | 0.43 | -0.2 | 0.36 | -0.06 | 0.43 | -0.32 | 0.18 | 0.37 | 1.08 | 0.54 | 0.45 | 0.15 | -0.135 | 0.4175 | 0.281334 | 0.689806 | 0.713058 | 1 | 0.999631 | 0.702476 |
| Prrt3_Q6PE13_S799s_1 | -1.58 | -0.89 | -0.6 | -2.41 | -1.87 | -1.63 | -1.9 | -1.92 | -1.56 | -0.96 | -1.3 | -0.88 | 0.65 | -1.28 | -1.1 | -0.76 | -0.46 | 0.195 | 0.7475 | 0.196913 | 0.276846 | 0.652843 | 1 | 0.999631 | 0.653609 |
| Prrt3_Q6PE13_S799s_2 | -0.43 | 0.13 | -0.12 | -0.04 | 0.12 | 0.1 | -0.31 | 0.11 | -0.1 | -0.36 | -0.13 | 0.2 | 0.23 | 0.25 | 0.43 | 0.5 | 0.12 | 0.0175 | 0.4675 | 0.115709 | 0.647541 | 0.948633 | 1 | 0.999631 | 0.5811 |
| Prrt3_Q6PE13_S806s_1 | 1.45 | 2.1 | -0.06 | 2.28 | 2.7 | 1.31 | 1.36 | 2.01 | 1.88 | 0.55 | 1.78 | 1.38 | -3.42 | 1.47 | 0.85 | 1.49 | 0.4025 | -0.045 | -1.345 | 0.239972 | 0.48503 | 0.93634 | 1 | 0.999631 | 0.683719 |
| Prrt3_Q6PE13_S833s_1 | 0.18 | 0.1 | -0.16 | -0.04 | 0.01 | 0.13 | -0.62 | -0.35 | -0.05 | -0.17 | -0.4 | -0.37 | -0.62 | 0.27 | -0.36 | -0.01 | -0.2275 | -0.2675 | -0.2 | 0.519063 | 0.415544 | 0.304296 | 1 | 0.999631 | 0.819456 |
| Prrt3_Q6PE13_S845s_1 | -0.4 | 0.21 | -0.03 | -0.39 | -0.18 | -0.08 | -0.73 | -0.46 | -0.24 | -0.03 | -0.27 | -0.19 | 0.16 | 0.11 | -0.47 | 0 | -0.21 | -0.03 | 0.1025 | 0.73796 | 0.467544 | 0.911205 | 1 | 0.999631 | 0.907918 |
| Prrt3_Q6PE13_S865s_1 | 0.87 | 1.78 | 1.24 | 1.25 | 1.21 | 1.97 | 0.6 | 1.71 | 1.32 | 1.04 | 1.17 | 1.23 | 0.41 | 2.06 | 1.84 | 1.39 | 0.0875 | -0.095 | 0.14 | 0.741186 | 0.814414 | 0.740576 | 1 | 0.999631 | 0.909055 |

| | | | | | | | | | | | | | | | | | | | | | | | | | |
|---|---|---|---|---|---|---|---|---|---|---|---|---|---|---|---|---|---|---|---|---|---|---|---|---|---|
| Prune2_Q52KR3_S2850s_1 | -0.61 | 0.23 | 0.05 | -0.64 | 0.09 | 0.51 | -0.89 | 0.12 | 0.58 | 0.69 | -0.88 | -0.77 | 1.02 | 0.65 | 0.22 | 0.13 | 0.2 | 0.1475 | 0.7475 | 0.058372 | 0.605304 | 0.751127 | 1 | 0.999631 | 0.500325 |
| Prune2_Q52KR3_S3070s_1 | -0.7 | -0.5 | 0.18 | -0.04 | 0.63 | -0.22 | -0.31 | -0.1 | -0.17 | -0.93 | 0.65 | 0.26 | 1.16 | -0.34 | 0.68 | 0.61 | 0.265 | 0.2175 | 0.7925 | 0.070232 | 0.435116 | 0.593734 | 1 | 0.999631 | 0.52985 |
| Psd_Q5DTT2_S1011s_1 | 4.32 | -0.54 | -0.74 | 2.46 | 1.42 | 1.08 | 3.38 | 0.76 | 0.33 | -1.09 | 0.42 | -1.49 | -5.7 | -2.02 | -0.91 | -1.25 | 0.285 | -1.8325 | -3.845 | 0.018459 | 0.80621 | 0.129799 | 1 | 0.999631 | 0.391024 |
| Psd_Q5DTT2_S126s_1 | -1.45 | 0.1 | -0.71 | -1.24 | 2.16 | -0.41 | 1.91 | -0.76 | 2.73 | -2.27 | 1.02 | 2.53 | 1.68 | -0.51 | -1.7 | -0.98 | 1.55 | 1.8275 | 0.4475 | 0.534195 | 0.058211 | 0.103804 | 1 | 0.999631 | 0.828493 |
| Psd_Q5DTT2_S17s_1 | -1.27 | 0.95 | 0.8 | -1.12 | 0.52 | 0.59 | -0.09 | 0.02 | 0.86 | -1.24 | 0.47 | 1.06 | 0.93 | 0.17 | 0.55 | -0.99 | 0.42 | 0.4475 | 0.325 | 0.621025 | 0.466525 | 0.531421 | 1 | 0.999631 | 0.861888 |
| Psd_Q5DTT2_S283s_1 | 1.89 | -0.72 | -2.32 | 2.94 | 2.86 | -0.41 | 3.04 | 2.56 | -0.13 | -2.1 | 1.47 | -0.35 | -1.58 | 0.21 | -0.35 | 0.27 | 1.565 | -0.725 | -0.81 | 0.461826 | 0.225072 | 0.550636 | 1 | 0.999631 | 0.792786 |
| Psd_Q5DTT2_S34s_1 | -0.52 | 0.04 | -0.42 | -0.42 | 0.11 | -0.22 | -0.52 | -0.33 | -0.44 | -0.45 | -0.29 | -0.31 | -0.09 | 0 | -0.25 | 0.13 | 0.09 | -0.0425 | 0.2775 | 0.33718 | 0.741421 | 0.869887 | 1 | 0.999631 | 0.730499 |
| Psd_Q5DTT2_S720s_1 | 0.32 | -0.08 | -0.78 | 0.65 | 0.5 | -0.17 | 0.39 | 0.65 | -0.35 | -0.42 | 0.4 | -0.12 | -1.22 | 0.2 | -0.08 | -0.24 | 0.315 | -0.15 | -0.3625 | 0.417862 | 0.407298 | 0.694884 | 1 | 0.999631 | 0.771539 |
| Psd_Q5DTT2_S736s_1 | 0.16 | 1.13 | -0.91 | -0.97 | 0.45 | 0.93 | -0.67 | -0.4 | 0.35 | -0.4 | -0.31 | -0.95 | 0.11 | 0.96 | -0.4 | -0.63 | 0.225 | -0.18 | 0.1575 | 0.780482 | 0.692602 | 0.734148 | 1 | 0.999631 | 0.921416 |
| Psd2_Q6P1I6_S473s_1 | -0.82 | -1.73 | 0.16 | 0.8 | -2 | -2.8 | 0.31 | 0.83 | -2.15 | -2.27 | 0.47 | 0.94 | -1.83 | -2.3 | 0.96 | 0.51 | -0.5175 | -0.355 | -0.2675 | 0.755244 | 0.568579 | 0.688408 | 1 | 0.999631 | 0.913663 |
| Psd3_E9PUC5_S337s_1 | -0.97 | 2.34 | 0.01 | -0.96 | 2.88 | 1.32 | 1.91 | -1.48 | 3.86 | -0.2 | 1.59 | 2.27 | 3.36 | 0.6 | -1.36 | 1 | 1.0525 | 1.775 | 0.795 | 0.459459 | 0.327211 | 0.098287 | 1 | 0.999631 | 0.79152 |
| Psd3_E9PUC5_S339s_1 | -0.46 | 1.36 | 0.46 | -0.37 | 1.89 | 0.59 | 0.83 | 0.52 | 2.05 | 0.2 | 0.44 | 1.1 | 2.49 | 1.08 | -0.34 | 0.1 | 0.71 | 0.7 | 0.585 | 0.395619 | 0.176154 | 0.222422 | 1 | 0.999631 | 0.760249 |
| Psd3_E9PUC5_S340s_2 | -1.33 | 0.58 | 0.74 | -0.02 | 1.24 | -0.06 | 1.47 | 0.2 | 2.4 | -0.43 | 1.06 | 2.01 | 2.36 | -0.72 | -0.56 | -0.32 | 0.72 | 1.2675 | 0.1975 | 0.79514 | 0.212959 | 0.095185 | 1 | 0.999631 | 0.929101 |
| Psd3_E9PUC5_S342s_1 | 0.29 | 0.11 | 1.06 | -0.64 | 0.02 | -0.3 | 1.41 | 0.21 | 0.28 | -0.61 | 1.19 | 2.38 | 1.85 | 0.13 | 1.31 | 0.01 | 0.13 | 0.605 | 0.62 | 0.262452 | 0.788215 | 0.366302 | 1 | 0.999631 | 0.693034 |
| Psd3_E9PUC5_S348s_1 | -0.86 | -0.12 | 0.37 | -1.35 | -0.26 | -0.36 | -0.3 | -0.56 | 0.22 | -0.21 | 0.23 | 1.2 | 2.05 | -0.54 | -0.39 | -0.53 | 0.12 | 0.85 | 0.6375 | 0.347854 | 0.761472 | 0.094926 | 1 | 0.999631 | 0.736498 |
| Psd3_E9PUC5_S348s_2 | -1.03 | 0.52 | 0.48 | -0.27 | 0.95 | -0.47 | 1.26 | 0.13 | 1.76 | -0.65 | 0.8 | 1.08 | 1.59 | -0.77 | 0.2 | -0.3 | 0.5425 | 0.8225 | 0.255 | 0.660178 | 0.295774 | 0.175019 | 1 | 0.999631 | 0.880315 |
| Psd3_E9PUC5_S44s_1 | -0.45 | 0.37 | 0.66 | 1.05 | 0.89 | -0.59 | 0.66 | 0.77 | 1.2 | -0.39 | 0.61 | 0.36 | -1.22 | 0.92 | 1.32 | 1.12 | 0.025 | 0.0375 | 0.1275 | 0.834325 | 0.955854 | 0.933437 | 1 | 0.999631 | 0.943944 |
| Psd3_E9PUC5_S76s_1 | 1.57 | 0.48 | 0.55 | -0.03 | -1.32 | 1.42 | 1.24 | -1.27 | -1.73 | 1.1 | -0.99 | -0.14 | -0.68 | -1.42 | -0.24 | -0.89 | -0.625 | -1.0825 | -1.45 | 0.006358 | 0.401441 | 0.109486 | 1 | 0.999631 | 0.351545 |
| Psd3_E9PUC5_S78s_1 | 1.85 | 0.9 | 1.16 | 0.82 | -0.25 | 1.05 | 2.16 | -0.62 | -0.7 | 0.52 | -0.17 | -0.08 | -1.69 | -0.61 | 0.11 | -0.96 | -0.5975 | -1.29 | -1.97 | 0.001241 | 0.339901 | 0.005768 | 1 | 0.999631 | 0.351308 |
| Psd3_E9PUC5_S98s_1 | 1.46 | 0.98 | -0.87 | 1.83 | 2.42 | 1.88 | 1.18 | 1.49 | 1.34 | 0.11 | 1.39 | 0.34 | -2.39 | 1.42 | 0.54 | 1.02 | 0.8925 | -0.055 | -0.7025 | 0.445279 | 0.156234 | 0.929343 | 1 | 0.999631 | 0.784262 |
| Psen2_Q3U4P5_S25s_1 | -0.18 | -0.29 | 0.08 | -0.52 | -0.45 | -0.66 | -0.4 | -0.53 | -0.28 | 0.31 | 0 | 0.34 | 0.87 | -0.13 | -0.53 | 0.02 | -0.2825 | 0.32 | 0.285 | 0.443005 | 0.279878 | 0.276348 | 1 | 0.999631 | 0.782541 |
| Psen2_Q3U4P5_S52s_1 | 0.08 | -0.68 | 1.67 | 1.77 | -0.04 | 0.57 | 1.41 | 1.59 | -0.03 | 1.71 | 0.94 | 1.13 | -0.92 | -1.02 | 1.31 | -0.35 | 0.1725 | 0.2275 | -0.955 | 0.20213 | 0.787208 | 0.720955 | 1 | 0.999631 | 0.658952 |
| Psip1_Q99JF8_S106s_1 | 0.78 | 1.88 | 1.93 | 1.55 | 1.33 | 1.67 | 1.16 | 1.59 | 1.83 | 1.65 | 1.7 | 1.76 | 1.57 | 1.52 | 1.55 | 1.27 | -0.0975 | 0.2 | -0.0575 | 0.865639 | 0.768702 | 0.541683 | 1 | 0.999631 | 0.955857 |
| Psip1_Q99JF8_S176s_1 | -1.02 | -1.64 | -0.02 | -1.49 | -0.17 | -1.91 | -0.57 | -0.13 | -2.46 | -2.34 | -0.24 | -0.53 | -1.36 | -1.78 | 0.25 | -1.02 | 0.3475 | -0.35 | 0.065 | 0.903197 | 0.505682 | 0.578171 | 1 | 0.999631 | 0.968183 |
| Psip1_Q99JF8_T122t_1 | 1.1 | 1.76 | 0.25 | -1.25 | 0.73 | 3.06 | 1.59 | -1.29 | -0.34 | 1.43 | -0.75 | -1.82 | 1.25 | -1.54 | -1.69 | -2.81 | 0.5575 | -0.835 | -1.6625 | 0.097944 | 0.565941 | 0.322705 | 1 | 0.999631 | 0.560722 |
| Psma2_P49722_S7s_1 | -0.16 | 0.55 | 1.23 | 0.08 | -0.23 | -0.3 | -0.89 | 1.82 | -0.14 | 0.81 | 0.66 | 0.21 | 1.58 | 0.03 | 2.4 | 0.82 | -0.325 | -0.04 | 0.7825 | 0.177653 | 0.592008 | 0.918561 | 1 | 0.999631 | 0.642116 |
| Psma3_E0CX62_S175s_1 | -0.52 | 0 | -0.28 | -0.53 | -0.34 | -0.17 | 0.23 | -0.72 | -0.01 | 0.08 | -0.72 | -0.89 | -0.24 | -1.05 | -0.79 | -1.77 | 0.0825 | -0.0525 | -0.63 | 0.120097 | 0.782202 | 0.87312 | 1 | 0.999631 | 0.587703 |
| Psma5_Q9Z2U1_S16s_1 | 2.2 | 3.9 | 0.07 | 0.44 | 3.45 | 3.15 | 0.13 | 0.29 | 3.29 | 3.43 | 0.45 | 0.37 | 3.43 | 3.6 | -0.38 | 0.49 | 0.1025 | 0.2325 | 0.1325 | 0.907462 | 0.924091 | 0.825767 | 1 | 0.999631 | 0.96989 |
| Psma5_Q9Z2U1_S56s_1 | -0.6 | -0.15 | 0.01 | -0.22 | -0.22 | -0.48 | -0.54 | -0.21 | -0.29 | -0.19 | 0.11 | 0.5 | 0.82 | 0.18 | 0.13 | -0.06 | -0.1225 | 0.2725 | 0.5075 | 0.133117 | 0.639678 | 0.371024 | 1 | 0.999631 | 0.60492 |
| Psmd1_Q3TXS7_T273t_1 | -0.32 | 0.12 | -0.86 | -1.21 | -0.49 | 0.07 | -0.95 | -1.16 | 0.15 | 0.15 | -0.78 | -0.96 | 0.94 | 0.46 | -0.9 | -0.54 | -0.065 | 0.2075 | 0.5575 | 0.278062 | 0.871958 | 0.623771 | 1 | 0.999631 | 0.700874 |
| Psmd1_Q3TXS7_T311t_1 | 0.97 | 0.36 | -2.55 | -1.46 | -0.61 | 0.11 | 0.39 | 0.28 | 0.7 | 0.92 | 1.16 | 1.74 | 0.43 | 0.36 | 1.29 | -1.93 | 0.7125 | 1.8 | 0.7075 | 0.447081 | 0.348122 | 0.034519 | 1 | 0.999631 | 0.785579 |
| Psmd11_G3UX67_S7s_1 | 0.57 | 0.94 | -0.38 | 0.93 | 0.77 | 0.64 | 0.41 | 0.67 | 1.03 | 0.51 | 0.77 | 0.78 | -0.55 | 0.91 | 0.75 | 0.82 | 0.1075 | 0.2575 | -0.0325 | 0.943559 | 0.758732 | 0.481435 | 1 | 0.999631 | 0.981629 |
| Psmd2_Q8VDM4_S361s_1 | -0.13 | -0.22 | -0.33 | -0.55 | -0.33 | -0.51 | -0.76 | -0.69 | -0.54 | 0.4 | -0.34 | -0.34 | -0.04 | -0.54 | -1.18 | 0.04 | -0.265 | 0.1025 | -0.1225 | 0.728074 | 0.305816 | 0.734294 | 1 | 0.999631 | 0.905923 |
| Psmd2_Q8VDM4_T20t_1 | -0.72 | -1.3 | -0.57 | -0.52 | -0.92 | -1.01 | -0.69 | -0.59 | -0.93 | -0.36 | -0.7 | -0.6 | -0.46 | -0.21 | -0.1 | -0.96 | -0.025 | 0.13 | 0.345 | 0.313887 | 0.929504 | 0.661358 | 1 | 0.999631 | 0.719939 |
| Psmd2_Q8VDM4_T9t_1 | -0.88 | 3.15 | 0.18 | -0.67 | 2.81 | 1.89 | 0.84 | 0.31 | 3.41 | -0.1 | 1.14 | 1.23 | 3.99 | 1.96 | 0.11 | -0.97 | 1.0175 | 0.975 | 0.8275 | 0.499789 | 0.292123 | 0.35084 | 1 | 0.999631 | 0.813279 |
| Psmd4_F7AI87_T65t_1 | 0.18 | 0.27 | -0.4 | -1.55 | 0.38 | -0.01 | -1.07 | -1.09 | -0.63 | 0.85 | -1.25 | -0.46 | 0.22 | 0.38 | 0.26 | -0.08 | -0.0725 | 0.0025 | 0.57 | 0.210515 | 0.889305 | 0.996444 | 1 | 0.999631 | 0.664726 |
| Psmd9_Q9CR00_S128s_1 | -0.28 | -0.01 | -0.87 | -0.65 | 0.4 | -0.89 | -1.02 | -1.22 | 0.36 | -0.93 | -0.22 | -0.28 | 0.35 | 0.13 | 0 | -1.26 | -0.23 | 0.185 | 0.2575 | 0.545555 | 0.580424 | 0.609135 | 1 | 0.999631 | 0.833826 |
| Psmd9_Q9CR00_S2s_1 | 1.97 | -0.6 | -0.8 | -0.51 | -0.16 | -0.93 | 0.41 | -0.58 | -1.06 | 1.08 | -0.9 | -0.97 | -0.47 | -1.41 | -0.89 | -1.59 | -0.33 | -0.4775 | -1.105 | 0.105339 | 0.608493 | 0.521748 | 1 | 0.999631 | 0.568363 |
| Psmf1_Q8BHL8_S153s_1 | -0.09 | -1.32 | -2.96 | -2.07 | -0.82 | -1.12 | -1.68 | -2.81 | -1.61 | -0.75 | -2.06 | -1.38 | -1.15 | -1.45 | -2.64 | -3.2 | 0.0025 | 0.16 | -0.5 | 0.475172 | 0.997006 | 0.791867 | 1 | 0.999631 | 0.799882 |
| Psmg1_Q9JK23_T18t_1 | -1.1 | -2.1 | 0.31 | 0.54 | -1.25 | -3.96 | 0.26 | 0.07 | -1.44 | -1.87 | 0.62 | 0.98 | -2.37 | -1.63 | -0.08 | 0.5 | -0.6325 | 0.16 | -0.3075 | 0.699646 | 0.52864 | 0.847227 | 1 | 0.999631 | 0.896612 |
| Pspc1_Q8R326_S477s_1 | -0.03 | 1.21 | 1.05 | 0.29 | 0.23 | 0.65 | -0.11 | 0.45 | 0.61 | 0.7 | 0.26 | 0.74 | 0.91 | 0.91 | 0.72 | 1.11 | -0.325 | -0.0525 | 0.2825 | 0.43854 | 0.379557 | 0.882067 | 1 | 0.999631 | 0.781983 |
| Ptbp1_F7DCW4_S91s_1 | 2.15 | 0.44 | 1.36 | 0.94 | 0.08 | 0.47 | 2.32 | -0.85 | -0.21 | 1.95 | 0.28 | -0.07 | -0.83 | -0.34 | 0.17 | -0.35 | -0.7175 | -0.735 | -1.56 | 0.004184 | 0.301143 | 0.214987 | 1 | 0.999631 | 0.351308 |
| Ptbp2_Q91Z31_S308s_1 | 0.01 | 1.41 | 0.9 | 1.03 | 0.78 | 0.75 | 0.51 | 1.08 | 1.18 | 0.94 | 0.69 | 1.33 | 1.1 | 1.29 | 1.46 | 2.11 | -0.0575 | 0.1975 | 0.6525 | 0.122901 | 0.868974 | 0.586844 | 1 | 0.999631 | 0.592497 |
| Ptdss1_Q99LH2_S442s_1 | 2 | 0.34 | 1.01 | 0.11 | -0.15 | 0.66 | 0.45 | 0.01 | 0.12 | 1.88 | 0.44 | 0.03 | 0.04 | -0.01 | 0.14 | 0.23 | -0.6225 | -0.2475 | -0.765 | 0.102075 | 0.1869 | 0.658746 | 1 | 0.999631 | 0.564448 |
| Ptdss2_G3UYQ3_S16s_1 | -1.65 | -0.06 | 0.28 | -0.67 | -0.05 | -0.17 | -0.59 | -0.36 | 0.22 | -0.9 | 0.19 | 0.95 | 0.79 | 0.28 | 0.03 | -0.3 | 0.2325 | 0.64 | 0.725 | 0.147209 | 0.592301 | 0.247005 | 1 | 0.999631 | 0.619223 |
| Ptgdr2_Q9Z2J6_S375s_1 | 1.79 | 1.2 | 1.51 | 1.08 | 1.16 | 1.75 | 0.86 | 0.31 | 1.91 | 2.74 | 1.41 | 0.65 | 0.66 | 1.25 | 0.38 | 1.35 | -0.375 | 0.2825 | -0.485 | 0.180367 | 0.314725 | 0.539588 | 1 | 0.999631 | 0.64376 |
| Ptgfrn_Q9WV91_S875s_1 | 0.76 | 0.16 | -0.37 | -0.42 | -0.18 | -0.19 | 0.28 | -0.54 | -0.86 | 0.31 | -0.11 | -0.65 | 0.61 | -0.1 | -0.62 | 0.11 | -0.19 | -0.36 | -0.0325 | 0.935167 | 0.59151 | 0.374307 | 1 | 0.999631 | 0.979194 |
| Ptk2_P34152_S910s_1 | -0.38 | -0.1 | -0.02 | -0.4 | -0.34 | -0.39 | -0.67 | -0.71 | -0.55 | -0.4 | -0.45 | -0.37 | -0.11 | -0.26 | -0.61 | 0.05 | -0.3025 | -0.2175 | -0.0075 | 0.979374 | 0.248459 | 0.394624 | 1 | 0.999631 | 0.993455 |
| Ptk2_P34152_Y576y_1 | -0.78 | -0.59 | -1.09 | -0.51 | -0.53 | -0.99 | -0.55 | -0.32 | -0.75 | -1.18 | -0.7 | -0.43 | -0.95 | -0.97 | -0.43 | -0.93 | 0.145 | -0.0225 | -0.0775 | 0.793264 | 0.602165 | 0.93789 | 1 | 0.999631 | 0.928057 |
| Ptk2b_E9Q2A6_S375s_1 | 0.26 | 0.57 | 0.63 | 0.02 | 0.01 | 0.09 | -0.06 | 0.34 | 0.14 | -0.09 | 0.18 | 0.39 | 0.99 | 0.83 | 0.91 | 0.65 | -0.275 | -0.215 | 0.475 | 0.120604 | 0.313395 | 0.442643 | 1 | 0.999631 | 0.588344 |
| Ptma_P26350_S2s_1 | -1.11 | -0.43 | 1.49 | 0.98 | -0.5 | -0.08 | 0.59 | 0.73 | -0.38 | -0.64 | 0.7 | 1.19 | 0.26 | -0.2 | 1.17 | 1.28 | -0.0475 | -0.015 | 0.395 | 0.53686 | 0.937306 | 0.982045 | 1 | 0.999631 | 0.830094 |
| Ptms_Q9D0J8_S5s_1 | 0.23 | 1.49 | 1.55 | 1.27 | 0.94 | 1.16 | 0.3 | 1.25 | 1.12 | 1.15 | 1.35 | 1.69 | 1.18 | 1.78 | 1.57 | 1.86 | -0.2225 | 0.1925 | 0.4625 | 0.240929 | 0.567325 | 0.600364 | 1 | 0.999631 | 0.684338 |
| Ptov1_Q91VU8_S53s_1 | 1.5 | 1.2 | 1.71 | 0.96 | 0.61 | 0.61 | 1 | 1.02 | 0.8 | 1.17 | 1.44 | 0.84 | -0.41 | 1.01 | 0.83 | 0.82 | -0.5325 | -0.28 | -0.78 | 0.07121 | 0.082961 | 0.361622 | 1 | 0.999631 | 0.531405 |
| Ptpdc1_Q6NZK8_S475s_1 | 0.64 | -0.19 | 0.44 | 0.18 | -0.42 | -0.06 | 0.11 | 1.11 | 0.15 | 0.62 | -0.38 | 0.99 | 0.36 | 1.12 | 2.19 | 1.09 | -0.0825 | 0.0775 | 0.9225 | 0.052546 | 0.830241 | 0.835313 | 1 | 0.999631 | 0.489777 |
| Ptpn1_P35821_S364s_1 | 0.47 | -1.49 | -0.82 | -2 | -1.34 | -3.62 | -1.5 | -1.49 | -1.69 | -1.5 | -1.47 | -1.83 | -1.79 | -3.62 | -2.14 | -2.01 | -1.0275 | -0.6625 | -1.43 | 0.03959 | 0.15325 | 0.214432 | 1 | 0.999631 | 0.470002 |
| Ptpn11_P35235_Y542y_1 | 0.15 | 2.54 | 1.17 | -0.67 | 3.89 | 0.89 | 3.55 | 0.62 | 3.79 | -0.99 | 1.61 | 2.79 | 2.12 | -0.13 | -0.12 | -1.78 | 1.44 | 1.0025 | -0.775 | 0.403855 | 0.154055 | 0.359551 | 1 | 0.999631 | 0.764939 |
| Ptpn11_P35235_Y62y_1 | -0.51 | 0.06 | -0.07 | -0.65 | 0.65 | 0.28 | -0.25 | -0.53 | 0.57 | -0.94 | -0.85 | 0.42 | 0.62 | -0.67 | -0.03 | -0.34 | 0.33 | 0.0925 | 0.1875 | 0.612434 | 0.352018 | 0.831581 | 1 | 0.999631 | 0.860287 |
| Ptpn12_P35831_S19s_1 | 0.47 | 0.49 | 1.1 | 0.69 | -0.13 | 0.83 | 0.58 | 0.8 | 0.49 | 1.63 | 1.09 | 1.69 | 1.07 | 1.74 | 1.62 | 1.52 | -0.1675 | 0.5375 | 0.8 | 0.023974 | 0.600069 | 0.151325 | 1 | 0.999631 | 0.415644 |
| Ptpn12_P35831_S331s_1 | 0.22 | 1.14 | 1.02 | 0.57 | 0.8 | 1.05 | 0.27 | 0.76 | 1.22 | 1.22 | 0.49 | 1.11 | 1.54 | 0.83 | 1 | 0.9 | -0.0175 | 0.2725 | 0.33 | 0.337186 | 0.955973 | 0.415133 | 1 | 0.999631 | 0.730499 |
| Ptpn12_P35831_S368s_1 | -1 | 1.69 | -0.02 | -0.18 | 0.3 | 0.92 | -0.51 | -0.23 | 0.4 | 0 | -0.45 | -0.34 | 0.65 | 1.42 | -0.13 | -0.93 | -0.0025 | -0.22 | 0.13 | 0.847463 | 0.996596 | 0.691424 | 1 | 0.999631 | 0.948839 |
| Ptpn12_P35831_S396s_1 | -0.26 | -0.59 | 0.77 | 0.63 | 0.87 | 0.25 | 0.26 | 0.24 | 0.16 | -0.15 | -0.05 | 1.47 | 0.15 | -0.07 | 0.95 | 1.7 | 0.2675 | 0.22 | 0.545 | 0.291302 | 0.48773 | 0.649627 | 1 | 0.999631 | 0.706164 |
| Ptpn12_P35831_S434s_1 | -0.32 | 0.82 | 0.41 | 1.15 | 1.34 | 0.67 | 0.58 | 1.06 | 1.01 | -0.06 | 1.01 | 1.35 | -0.3 | 0.84 | 1.01 | 1.39 | 0.3975 | 0.3125 | 0.22 | 0.64268 | 0.306499 | 0.480691 | 1 | 0.999631 | 0.871387 |
| Ptpn12_P35831_S632s_1 | 0.01 | -0.79 | -1.32 | -1.3 | -0.85 | -1.8 | -1.04 | 0.31 | -0.48 | -2.5 | -0.17 | -0.55 | -0.07 | -1.31 | 0.06 | -1.87 | 0.005 | -0.075 | 0.0525 | 0.921279 | 0.992048 | 0.894779 | 1 | 0.999631 | 0.975032 |
| Ptpn12_P35831_T519t_1 | -0.5 | -0.29 | -0.72 | -0.95 | -0.48 | -0.96 | -0.79 | -0.83 | -0.89 | -0.04 | -1.21 | -0.82 | 0.22 | -0.83 | -0.23 | -0.6 | -0.15 | -0.125 | 0.255 | 0.45635 | 0.579647 | 0.709874 | 1 | 0.999631 | 0.790326 |
| Ptpn13_G5E8B1_S2059s_1 | 0.08 | -0.27 | -0.87 | -0.72 | -0.26 | -0.92 | -0.46 | -1.24 | -0.62 | 0.28 | -0.47 | -0.33 | -0.28 | 0.25 | -0.44 | -0.48 | -0.275 | 0.16 | 0.2075 | 0.545061 | 0.430752 | 0.638903 | 1 | 0.999631 | 0.833658 |
| Ptpn2_Q06180_S268s_1 | 0.76 | 1.65 | 0.93 | 0.05 | 0.18 | 1.28 | -0.05 | 0.92 | 0.5 | 0.89 | 0.31 | 0.75 | -0.45 | 1.54 | 0.92 | 0.12 | -0.265 | -0.235 | -0.315 | 0.546928 | 0.551032 | 0.536996 | 1 | 0.999631 | 0.834213 |
| Ptpn2_Q06180_S298s_1 | 1.31 | 0.87 | -0.27 | 1.34 | 1.25 | 0.69 | 0.45 | 0.23 | 1.49 | 0.91 | 0.11 | 1.15 | 0.55 | 0.95 | 0.03 | 0.35 | -0.1575 | 0.1025 | -0.3425 | 0.43418 | 0.713897 | 0.825052 | 1 | 0.999631 | 0.779717 |
| Ptpn23_Q6PB44_T744t_1 | -0.34 | -0.81 | -0.53 | -0.67 | -0.76 | -1.16 | -0.81 | -1.23 | -1.46 | -0.82 | -0.55 | -0.6 | -1.8 | -1.14 | -0.75 | -0.66 | -0.4025 | -0.27 | -0.5 | 0.165654 | 0.14614 | 0.385097 | 1 | 0.999631 | 0.634459 |
| Ptpn3_A2ALK8_S372s_1 | -1.47 | -1.02 | -1.11 | -1.34 | -1.51 | -1.65 | -1.32 | -0.96 | -1.27 | -1.25 | -0.58 | -1.62 | -0.04 | -0.75 | -1.64 | -0.72 | -0.125 | 0.055 | 0.4475 | 0.256007 | 0.647393 | 0.858742 | 1 | 0.999631 | 0.691015 |
| Ptpn3_A2ALK8_S434s_1 | -1 | 1.64 | 3.55 | -0.77 | 1.75 | 1.06 | 2.93 | 1.53 | 2.81 | 0.17 | 3 | 3.16 | 2.69 | -0.46 | 1.46 | -1.45 | 0.9625 | 1.43 | -0.295 | 0.806918 | 0.34288 | 0.217006 | 1 | 0.999631 | 0.935523 |
| Ptpn3_A2ALK8_S459s_1 | -0.07 | 0.18 | -0.85 | -0.99 | -0.58 | -0.92 | -1.3 | -1.1 | -1.2 | -1.88 | -1.69 | -1.67 | -1.87 | -1.21 | -3.51 | -2.73 | -0.5425 | -1.1775 | -1.8975 | 0.005393 | 0.148039 | 0.007823 | 1 | 0.999631 | 0.351308 |
| Ptpn4_E9Q0S8_S392s_1 | 1.63 | 0.04 | -0.23 | -1.04 | 0.68 | -0.13 | -0.36 | -1.79 | -0.33 | 0.45 | -1.16 | -0.67 | -0.5 | -1.02 | -1.94 | -1.58 | -0.5 | -0.5275 | -1.36 | 0.040618 | 0.465402 | 0.386293 | 1 | 0.999631 | 0.474979 |
| Ptpn4_E9Q0S8_S403s_1 | -0.12 | 0.32 | -0.73 | 0.19 | 0.33 | 0.17 | 0.35 | -0.06 | 0.13 | -0.43 | 0.2 | -0.09 | -0.56 | 0.12 | 0.08 | -0.12 | 0.2825 | 0.0375 | -0.035 | 0.918795 | 0.370774 | 0.908813 | 1 | 0.999631 | 0.974321 |
| Ptpn5_P54830_S319s_1 | -0.1 | -0.69 | -0.1 | 0.17 | -0.5 | -0.93 | -0.42 | -0.54 | -0.1 | -0.33 | 0.16 | 0.91 | 1.2 | -0.44 | 0.15 | 0.06 | -0.4175 | 0.34 | 0.4225 | 0.314754 | 0.169448 | 0.354784 | 1 | 0.999631 | 0.720305 |
| Ptpn5_P54830_S39s_1 | -0.39 | -0.39 | 0.12 | -0.25 | -0.69 | -0.16 | -0.38 | -0.48 | -0.99 | -0.86 | -1.27 | -0.57 | -0.86 | 0.17 | 0.43 | 0.81 | -0.2 | -0.695 | 0.365 | 0.372989 | 0.454872 | 0.032062 | 1 | 0.999631 | 0.747035 |
| Ptpra_Q91V35_S176s_2 | 1.15 | -0.39 | 0.78 | 1.34 | -0.36 | 0.71 | 1.05 | 0.35 | 0.48 | 0.69 | -0.04 | -0.99 | 0.65 | -0.43 | 0.16 | -1.37 | -0.2825 | -0.685 | -0.9675 | 0.099522 | 0.550542 | 0.199436 | 1 | 0.999631 | 0.562298 |
| Ptpra_Q91V35_S180s_1 | 0.67 | -0.71 | -0.33 | 0.16 | -0.73 | 0 | 0.84 | -0.19 | -0.35 | 0.45 | -0.36 | -0.17 | -0.43 | -1.29 | -0.61 | -1.24 | 0.0325 | -0.055 | -0.84 | 0.056445 | 0.94014 | 0.884371 | 1 | 0.999631 | 0.496238 |
| Ptpra_Q91V35_S180s_2 | 1.15 | -0.39 | 0.78 | 1.34 | -0.36 | 0.71 | 1.05 | 0.35 | 0.48 | 0.69 | -0.04 | -0.99 | 0.65 | -0.43 | 0.16 | -1.37 | -0.2825 | -0.685 | -0.9675 | 0.099522 | 0.550542 | 0.199436 | 1 | 0.999631 | 0.562298 |
| Ptprc_S4R1M0_S801s_1 | 0.06 | 0.79 | 0.56 | 0.11 | 0.04 | -0.17 | -0.66 | 0.26 | 0.62 | -0.16 | -0.35 | 1.15 | 0.64 | 0.15 | 0.92 | 0.46 | -0.5125 | -0.065 | 0.1625 | 0.615443 | 0.128728 | 0.871614 | 1 | 0.999631 | 0.861203 |
| Ptprd_Q64487_S1296s_1 | 1.78 | -4.59 | -1.81 | -1.95 | -5.04 | -3.05 | 2.27 | -4.88 | -8.15 | 1.07 | -3.61 | -4.85 | -4.61 | -5.51 | -3.52 | -4.92 | -1.0325 | -2.2425 | -2.9975 | 0.02588 | 0.573247 | 0.268651 | 1 | 0.999631 | 0.422248 |
| Ptprk_B2RRF0_S856s_1 | -0.15 | 0.82 | 0.24 | 0.36 | -0.4 | 0.02 | -0.17 | 0.64 | 0.69 | -0.33 | 0.06 | 0.02 | 1.06 | 0.62 | 0.02 | 0.8 | -0.295 | -0.2075 | 0.3075 | 0.392922 | 0.390534 | 0.543471 | 1 | 0.999631 | 0.758901 |
| Ptprn_Q60673_S307s_1 | -0.98 | -0.97 | -0.26 | -0.61 | -0.72 | -0.92 | -0.73 | -0.49 | -0.36 | -0.41 | -0.49 | -0.48 | 0.33 | -0.47 | 0.17 | 0.16 | -0.01 | 0.27 | 0.7525 | 0.040168 | 0.97119 | 0.340557 | 1 | 0.999631 | 0.472868 |
| Ptprn2_Q3UU93_S184s_1 | -0.9 | -0.43 | -0.65 | -1 | -1.31 | -0.54 | -0.77 | -0.77 | -0.56 | -1.35 | -0.67 | -0.97 | -0.52 | -0.06 | -0.72 | -0.68 | -0.1025 | -0.1425 | 0.25 | 0.415912 | 0.719444 | 0.633651 | 1 | 0.999631 | 0.770095 |
| Ptprn2_Q3UU93_S185s_1 | -0.35 | -0.93 | -1.09 | -1.35 | -0.37 | -1.21 | -0.82 | -0.96 | -1.09 | -1 | -1.25 | -0.73 | -0.27 | -1.21 | -0.84 | -0.71 | 0.09 | -0.0875 | 0.1725 | 0.621097 | 0.780234 | 0.776441 | 1 | 0.999631 | 0.861888 |
| Ptprn2_Q3UU93_S339s_1 | -1.12 | -1.06 | -1.56 | -0.81 | -1.21 | -0.98 | -0.9 | -1.06 | -0.9 | -0.59 | -1.01 | -1.51 | -0.1 | -0.8 | -1.51 | -1.11 | 0.1 | 0.135 | 0.2575 | 0.49807 | 0.708173 | 0.666994 | 1 | 0.999631 | 0.811985 |
| Ptprn2_Q3UU93_S423s_1 | -1.17 | -0.74 | -0.4 | -1.39 | -0.43 | -0.97 | -0.19 | -1.05 | -0.86 | -0.83 | -0.73 | -0.49 | 0.26 | -0.65 | -0.34 | -0.36 | 0.265 | 0.1975 | 0.6525 | 0.086064 | 0.442313 | 0.524171 | 1 | 0.999631 | 0.552688 |
| Ptprn2_Q3UU93_S678s_1 | -0.11 | -0.37 | 0.17 | 0.1 | -0.17 | 0.07 | -0.07 | -0.15 | -0.72 | 0 | 0.27 | 0.12 | 0.29 | -0.1 | 0.33 | 0.13 | -0.0275 | -0.03 | 0.215 | 0.456491 | 0.912895 | 0.924001 | 1 | 0.999631 | 0.790438 |
| Ptprn2_Q3UU93_S690s_1 | -0.33 | 0.83 | -1.01 | -0.66 | 1 | 0.42 | -0.66 | -0.52 | 0.84 | -0.41 | -1.22 | -0.49 | 1.15 | -0.18 | 0 | 0.07 | 0.3525 | -0.0275 | 0.5525 | 0.269989 | 0.503789 | 0.959603 | 1 | 0.999631 | 0.698438 |
| Ptprr_Q62132_S27s_1 | -0.88 | -2.8 | -0.24 | -1.27 | -1.66 | -1.07 | -1.38 | -0.28 | -1.23 | 0.38 | 0.42 | 0.72 | 0.13 | -0.14 | 0.74 | 0.08 | 0.2 | 1.37 | 1.5 | 0.018156 | 0.724545 | 0.052698 | 1 | 0.999631 | 0.391024 |
| Ptpru_Q3V360_S781s_1 | -1.92 | -0.89 | -1.15 | -1.91 | -0.04 | -2.2 | -1.52 | -1.5 | -1.55 | -0.36 | -2.04 | -1.32 | -0.22 | -0.78 | -0.71 | -1.77 | 0.1525 | 0.15 | 0.5975 | 0.183837 | 0.758189 | 0.731988 | 1 | 0.999631 | 0.644372 |
| Ptpru_Q3V360_S791s_1 | 0.03 | 0 | 0.31 | -0.39 | -0.38 | 0.41 | -0.37 | -0.29 | 0.26 | 0.52 | -0.15 | 0.13 | 0.16 | 0.87 | 0.09 | -0.13 | -0.145 | 0.2025 | 0.26 | 0.440855 | 0.631824 | 0.48805 | 1 | 0.999631 | 0.781983 |
| Ptprz1_B9EKR1_S1457s_1 | -3.42 | 1.49 | 1.78 | -1.17 | 1.21 | 1.03 | 1.3 | -0.75 | 1.9 | -3.79 | 1.65 | 3.24 | 2.1 | -0.41 | 1.8 | -0.92 | 1.0275 | 1.08 | 0.9725 | 0.433438 | 0.372765 | 0.524253 | 1 | 0.999631 | 0.779548 |
| Ptprz1_B9EKR1_S647s_1 | 0.77 | 0.68 | 1.29 | 0.34 | -0.03 | 1.08 | 0.84 | 1.65 | 0.81 | 2.48 | 1.47 | 1.52 | 0.02 | 0.78 | 1.41 | 1.57 | 0.115 | 0.8 | 0.175 | 0.676639 | 0.775904 | 0.072973 | 1 | 0.999631 | 0.886862 |
| Ptprz1_B9EKR1_T1681t_1 | -0.79 | -0.27 | -0.25 | -0.97 | -0.51 | -1.01 | -0.97 | -1.29 | -0.72 | -0.56 | -0.83 | -0.91 | 0.31 | -0.16 | -0.45 | -0.38 | -0.375 | -0.185 | 0.4 | 0.239281 | 0.23572 | 0.523806 | 1 | 0.999631 | 0.683719 |
| Pum1_Q80U78_S709s_1 | 0.5 | 1.39 | 1.43 | 1.32 | 0.77 | 0.85 | 0.47 | 0.92 | 0.8 | 0.94 | 0.92 | 1.15 | 0.3 | 1.15 | 1.1 | 1.31 | -0.4075 | -0.2075 | -0.195 | 0.594392 | 0.199442 | 0.501286 | 1 | 0.999631 | 0.85392 |
| Pum1_Q80U78_S98s_1 | -0.29 | 2.27 | 2.2 | -0.78 | 2.61 | 1.2 | 2.39 | 1.46 | 3.66 | 0.47 | 3.02 | 3.46 | 4.07 | 0.31 | 2.33 | -0.66 | 1.065 | 1.8025 | 0.6625 | 0.560199 | 0.187356 | 0.081983 | 1 | 0.999631 | 0.839805 |
| Pum1_Q80U78_T112t_1 | 0.75 | 0.34 | 0.48 | 0.02 | 0.61 | 1.1 | 0.47 | 0.87 | 1.13 | 0 | 1.1 | 0.56 | 0.17 | 1.32 | 0.88 | 1.11 | 0.365 | 0.3 | 0.4725 | 0.19726 | 0.218161 | 0.397607 | 1 | 0.999631 | 0.654124 |
| Pum2_Q3UR91_S136s_1 | 0.32 | 0.91 | 0.74 | 0.76 | 0.59 | 0.47 | 0.74 | 0.91 | 0.78 | 0.95 | 1.02 | 1.11 | 0.37 | 1.4 | 1.39 | 1.22 | -0.005 | 0.2825 | 0.4125 | 0.243503 | 0.984735 | 0.299233 | 1 | 0.999631 | 0.685637 |
| Pum2_Q3UR91_S181s_1 | 0.48 | 0.77 | 0.84 | 0.85 | 0.45 | 0.77 | 0.46 | 0.69 | 0.88 | 1.38 | 0.79 | 0.93 | 1.1 | 1.21 | 1.37 | 1.08 | -0.1425 | 0.26 | 0.455 | 0.115369 | 0.566559 | 0.346024 | 1 | 0.999631 | 0.580533 |
| Pura_P42669_S181s_1 | -0.27 | 0.05 | 0.26 | 0.17 | -0.11 | -0.02 | -0.13 | 0.43 | -0.62 | -0.34 | 0.14 | 0.19 | -0.09 | -0.04 | 0.03 | 0.38 | -0.01 | -0.21 | 0.0175 | 0.951007 | 0.970305 | 0.491873 | 1 | 0.999631 | 0.984079 |
| Pura_P42669_T182t_1 | -0.22 | 0.36 | 0.25 | 0.23 | -0.16 | 0.06 | -0.13 | 0.41 | -0.71 | -0.38 | 0.11 | 0.14 | 0 | 0.01 | -0.08 | 0.3 | -0.11 | -0.365 | -0.0975 | 0.731515 | 0.687732 | 0.255793 | 1 | 0.999631 | 0.906935 |
| Purb_O35295_S316s_1 | -0.09 | 0.07 | 0.78 | 1.18 | -0.27 | 0.22 | 1.1 | 0.57 | 0.59 | 0.9 | 0.66 | 0.56 | -0.39 | -0.52 | 0.47 | -0.31 | -0.08 | 0.1925 | -0.6725 | 0.115506 | 0.846785 | 0.583908 | 1 | 0.999631 | 0.580709 |
| Purb_O35295_S6s_2 | -0.57 | -0.61 | 1.34 | -0.46 | -0.62 | -1.17 | -0.23 | -1.04 | -0.54 | 0.28 | 1.37 | 0.08 | -1.32 | -1.01 | -0.25 | 0.71 | -0.69 | 0.3725 | -0.3925 | 0.515502 | 0.180356 | 0.517148 | 1 | 0.999631 | 0.819449 |
| Purb_O35295_S8s_2 | -0.57 | -0.61 | 1.34 | -0.46 | -0.62 | -1.17 | -0.23 | -1.04 | -0.54 | 0.28 | 1.37 | 0.08 | -1.32 | -1.01 | -0.25 | 0.71 | -0.69 | 0.3725 | -0.3925 | 0.515502 | 0.180356 | 0.517148 | 1 | 0.999631 | 0.819449 |
| Purg_Q8R4E6_S342s_1 | 0.27 | 0.83 | 0.73 | 0.5 | 0.5 | 0.84 | 0.51 | 0.82 | 1.06 | 1.09 | 1.12 | 0.91 | 0.45 | 0.69 | 1.04 | 0.7 | 0.085 | 0.4625 | 0.1375 | 0.639102 | 0.74491 | 0.099966 | 1 | 0.999631 | 0.870995 |
| Pvr_Q8K094_S384s_1 | 0.95 | -1.09 | 0.71 | -0.19 | -0.54 | 0.33 | 0.71 | -0.53 | -2.46 | -0.06 | -0.19 | -0.77 | -0.97 | -1.48 | -0.87 | -0.69 | -0.1025 | -0.965 | -1.0975 | 0.042665 | 0.844059 | 0.160449 | 1 | 0.999631 | 0.475443 |
| Pvr_Q8K094_S393s_1 | 0.34 | -1.09 | -0.5 | 0.17 | 0.44 | -0.36 | 0.58 | -0.19 | -0.79 | -1.23 | 0.65 | 0.08 | -0.92 | -1.46 | 0.32 | -0.1 | 0.3875 | -0.0525 | -0.27 | 0.589452 | 0.349119 | 0.917356 | 1 | 0.999631 | 0.851414 |
| Pvrl1_Q9JKF6_S421s_1 | -1.46 | -1.04 | -1.54 | -2.71 | -1.09 | -1.61 | -1.86 | -1.77 | -1.56 | -1.11 | -2.29 | -0.97 | -0.45 | -1.88 | -1.5 | -3.26 | 0.105 | 0.205 | -0.085 | 0.890567 | 0.79369 | 0.653693 | 1 | 0.999631 | 0.963768 |
| Pvrl1_Q9JKF6_S433s_1 | -0.79 | -0.85 | 0.19 | 0.44 | -1.52 | -0.78 | -0.32 | 0.29 | -1.47 | -0.58 | 0.35 | 0.65 | -1.33 | -0.45 | 0.6 | 0.78 | -0.33 | -0.01 | 0.1525 | 0.782395 | 0.496612 | 0.98528 | 1 | 0.999631 | 0.922325 |
| Pvrl1_Q9JKF6_S434s_1 | -1.14 | -0.55 | -0.49 | -0.27 | -0.4 | -0.62 | -0.53 | -0.36 | -0.77 | -1.01 | -0.37 | -0.81 | 0.68 | -0.46 | -0.21 | 0.09 | 0.135 | -0.1275 | 0.6375 | 0.099739 | 0.629638 | 0.675067 | 1 | 0.999631 | 0.562805 |
| Pwp1_Q99LL5_S493s_1 | 1.54 | 0.03 | 0.09 | 0.75 | 0.14 | 0.12 | 0.85 | 0.73 | 0.38 | 1.01 | 0.4 | 0.59 | -0.47 | -0.01 | 0.91 | -0.04 | -0.1425 | -0.0075 | -0.505 | 0.281286 | 0.725004 | 0.984873 | 1 | 0.999631 | 0.702476 |
| Pxn_F8VQ28_S258s_1 | 1.42 | 1.7 | 1.46 | 1.28 | 1.83 | 1.26 | 0.71 | 1.01 | 1.54 | 1.81 | 0.78 | 0.87 | 0.46 | 1.69 | 1.19 | 0.68 | -0.2625 | -0.215 | -0.46 | 0.205493 | 0.404632 | 0.511475 | 1 | 0.999631 | 0.6605 |
| Pygo2_Q3V113_T264t_1 | 0.34 | 1.06 | -0.13 | -0.35 | 0.63 | 0.44 | -0.69 | -1.12 | 0.85 | 0.46 | -0.52 | -0.8 | 0.51 | 0.65 | -0.51 | -0.22 | -0.415 | -0.2325 | -0.1225 | 0.775324 | 0.41181 | 0.6306 | 1 | 0.999631 | 0.920998 |
| Q3TQI7_S261s_1 | 0.24 | 1.46 | 1.38 | 0.44 | 0.94 | 0.64 | 0.62 | 0.82 | 1.05 | 0.07 | 0.76 | 0.86 | 1.32 | 1.51 | 0.77 | 0.83 | -0.125 | -0.195 | 0.2275 | 0.565395 | 0.723298 | 0.623787 | 1 | 0.999631 | 0.84139 |
| Q3TYS2_S168s_1 | 1.4 | 0.28 | 1.7 | 1.38 | 0.01 | 0.06 | 0.67 | 1.12 | -0.12 | 1.33 | 1.27 | 0.96 | 0.34 | -0.06 | 1.5 | 0.44 | -0.725 | -0.33 | -0.635 | 0.182899 | 0.10165 | 0.469881 | 1 | 0.999631 | 0.643861 |
| Q3UPC7_S376s_1 | -0.14 | -0.74 | -0.12 | -0.67 | 0.42 | -0.84 | -0.9 | -1.53 | -0.68 | 0.26 | -0.17 | -0.89 | 0.2 | -0.03 | -0.26 | -0.42 | -0.295 | 0.0475 | 0.29 | 0.359529 | 0.500146 | 0.891435 | 1 | 0.999631 | 0.741182 |
| Q3UTZ3_S517s_1 | 1.37 | 0.01 | -0.91 | 0.03 | 0.12 | 0.47 | 1.11 | -0.59 | 0.18 | 0.88 | 0.71 | -0.63 | -1.08 | 1.09 | -0.28 | 0.12 | 0.1525 | 0.16 | -0.1625 | 0.784833 | 0.778501 | 0.767027 | 1 | 0.999631 | 0.922904 |
| Q3UTZ3_S533s_1 | 1.16 | 0.87 | 0.97 | 2 | 1.31 | 2.46 | 2.32 | 1.78 | 1.43 | 1.38 | 1.64 | 1.64 | -1.17 | 1.22 | 0.94 | 0.08 | 0.7175 | 0.2725 | -0.9825 | 0.100143 | 0.084457 | 0.408181 | 1 | 0.999631 | 0.562917 |
| Q3UTZ3_S546s_1 | 1.01 | 0.04 | 0.23 | 0.85 | 0.18 | -0.01 | 0.06 | 3.38 | 0.7 | 2.61 | 1.62 | 0.05 | 0.14 | 1.05 | 0.46 | 1.17 | 0.37 | 0.7125 | 0.1725 | 0.648954 | 0.62592 | 0.221896 | 1 | 0.999631 | 0.874615 |
| Q3UTZ3_T541t_1 | -0.1 | 0.45 | 0.13 | -0.14 | -0.31 | 0.16 | -0.58 | -0.19 | 0.22 | 0.54 | 0.25 | 0.27 | 1.14 | 0.8 | 0.66 | 0.94 | -0.315 | 0.235 | 0.8 | 0.018773 | 0.281394 | 0.389963 | 1 | 0.999631 | 0.391024 |
| Q3UY34_S139s_1 | 2.88 | 0.99 | 0.87 | 1.52 | 0.87 | 0.79 | 2.12 | 0.88 | 1.05 | 1.53 | 0.55 | -0.61 | -0.79 | -0.18 | -0.5 | -0.3 | -0.4 | -0.935 | -2.0075 | 0.001617 | 0.449086 | 0.139467 | 1 | 0.999631 | 0.351308 |
| Q6DFW0-2_Q6DFW0_S9s_1 | -0.42 | 0.8 | 0.24 | -0.16 | 0.24 | 0.27 | -0.5 | -0.06 | 0.11 | -0.32 | -0.33 | 0.17 | 1.07 | 0.47 | 0.47 | 0.64 | -0.1275 | -0.2075 | 0.5475 | 0.145775 | 0.716182 | 0.547949 | 1 | 0.999631 | 0.616968 |
| Q6NV72-2_Q6NV72_S535s_1 | -1.73 | -2.46 | -1.06 | -0.98 | -2.94 | -2.16 | -0.42 | -0.93 | -2.24 | -1.78 | -1.13 | -1.23 | -2.02 | -1.71 | -1.85 | -1.15 | -0.055 | -0.0375 | -0.125 | 0.761417 | 0.927399 | 0.930444 | 1 | 0.999631 | 0.914725 |
| Q7TNG5-3_Q7TNG5_S151s_1 | -0.14 | -0.48 | 0.12 | 0.09 | 0.15 | -0.25 | 0.04 | 0.18 | 0.92 | 0.6 | 0.12 | 0.09 | 1.38 | 0.16 | 0.44 | 0.68 | 0.1325 | 0.535 | 0.7675 | 0.050385 | 0.621157 | 0.109776 | 1 | 0.999631 | 0.488571 |
| Q7TNG5-3_Q7TNG5_S153s_1 | 0.23 | 0.99 | 0.3 | -0.23 | -0.17 | 0.09 | -0.12 | 0.4 | 0.29 | 0.24 | 0.47 | 0.34 | 0.55 | 1.2 | 0.32 | 1.12 | -0.2725 | 0.0125 | 0.475 | 0.221051 | 0.412765 | 0.968597 | 1 | 0.999631 | 0.671424 |
| Q7TNG5-3_Q7TNG5_S154s_1 | 0.1 | -0.03 | 1.21 | 0.59 | 0.08 | 0.07 | 0.5 | 1.05 | 1.2 | 0.74 | 0.92 | 1.19 | 1.23 | 0.66 | 1.32 | 1.21 | -0.0425 | 0.545 | 0.6375 | 0.104953 | 0.910515 | 0.138148 | 1 | 0.999631 | 0.568213 |
| Q7TNG5-3_Q7TNG5_S170s_1 | -0.16 | -2.28 | -1.91 | -1.28 | -1.36 | -1.61 | -1.41 | -0.62 | -0.93 | -0.23 | -0.75 | -2.62 | -1.07 | -0.69 | -1.96 | -2.04 | 0.1575 | 0.275 | -0.0325 | 0.951561 | 0.745549 | 0.663163 | 1 | 0.999631 | 0.984079 |
| Q8BP99_S139s_1 | -1.67 | 0.04 | 0.62 | 0.64 | -0.94 | 0.48 | 0.35 | 1.23 | -0.91 | -0.93 | 1.57 | -1.03 | -0.53 | 1.01 | 0.63 | 1.17 | 0.3725 | -0.2325 | 0.6625 | 0.288886 | 0.559714 | 0.753194 | 1 | 0.999631 | 0.706046 |
| Q8BY89-2_Q8BY89_T12t_1 | -0.94 | -0.94 | -0.74 | -1.17 | -1.05 | -0.88 | -0.91 | -1.15 | -1.08 | -1.24 | -0.84 | -0.95 | -0.64 | -0.84 | -0.84 | -1.05 | -0.05 | -0.08 | 0.105 | 0.701401 | 0.837068 | 0.755345 | 1 | 0.999631 | 0.896835 |
| Q8BZR9_S563s_1 | 0.67 | 0.93 | 3.55 | 0.69 | 0.78 | 0.49 | 2.19 | 3.3 | 0.51 | 0.76 | 3.69 | 3.51 | -0.51 | 1.07 | 2.93 | 3.6 | 0.23 | 0.6575 | 0.3125 | 0.753943 | 0.782581 | 0.498019 | 1 | 0.999631 | 0.913377 |
| Q8C3W1_S130s_1 | -0.5 | -1.74 | -0.45 | -0.29 | -0.51 | -0.32 | -0.47 | -0.86 | -1.18 | -2.28 | -1.46 | -1.29 | -1.07 | 0.04 | -0.82 | -0.35 | 0.205 | -0.8075 | 0.195 | 0.648891 | 0.583912 | 0.079499 | 1 | 0.999631 | 0.874615 |
| Q8C3W1_S176s_1 | 0.18 | -0.91 | -0.99 | -0.12 | -0.03 | -0.58 | -0.5 | 0.03 | -0.19 | -0.52 | -0.35 | -0.31 | -0.77 | 0.15 | -0.3 | 0.02 | 0.19 | 0.1175 | 0.235 | 0.547397 | 0.596015 | 0.731855 | 1 | 0.999631 | 0.834277 |
| Q8C3W1_S278s_1 | -0.03 | -0.2 | 0.24 | -0.6 | -0.8 | -0.71 | -0.87 | -0.47 | -0.67 | 0.45 | -0.37 | -0.66 | 0.13 | -0.79 | -0.06 | -0.32 | -0.565 | -0.165 | -0.1125 | 0.737721 | 0.066422 | 0.642702 | 1 | 0.999631 | 0.907836 |
| Q8C3W1_T274t_1 | 0.43 | -0.37 | -0.35 | -0.56 | 0.27 | 0.13 | -0.16 | 0.15 | 0.31 | -0.82 | 0.38 | 0.36 | 0.72 | -0.62 | -0.55 | -0.05 | 0.31 | 0.27 | 0.0875 | 0.827783 | 0.31567 | 0.489188 | 1 | 0.999631 | 0.941227 |
| Q8C3W1_T276t_1 | -0.72 | -1.56 | -0.05 | -0.54 | -0.78 | -0.85 | -0.78 | -1.02 | -1.21 | -0.99 | 0.14 | -0.09 | 0.45 | -1.38 | -0.82 | -0.66 | -0.14 | 0.18 | 0.115 | 0.811253 | 0.689632 | 0.689293 | 1 | 0.999631 | 0.936949 |
| Q8C5K5-2_Q8C5K5_T196t_1 | 2.01 | 0.36 | 1.76 | 1.34 | 0.31 | 0.11 | 2.47 | 0.99 | 0.41 | 0.47 | 0.59 | 0.34 | 1.94 | -0.59 | 0.95 | -0.34 | -0.3975 | -0.915 | -0.8775 | 0.1822 | 0.502742 | 0.0373 | 1 | 0.999631 | 0.643861 |
| Q8K207_S115s_1 | -3.09 | -3.1 | 0.55 | -0.4 | -1.93 | -3.03 | -0.1 | 0.61 | -2.49 | -2.29 | 0.42 | 0.39 | -2.02 | -2.18 | 0.16 | 0.36 | 0.3975 | 0.5175 | 0.59 | 0.555123 | 0.71211 | 0.628648 | 1 | 0.999631 | 0.838109 |
| Q8K207_S95s_1 | -0.37 | -0.34 | -0.77 | -0.5 | -0.56 | -0.54 | -0.7 | -0.59 | -0.58 | -0.73 | -0.37 | -0.19 | -0.26 | -0.39 | -0.61 | -0.59 | -0.1025 | 0.0275 | 0.0325 | 0.905983 | 0.673083 | 0.918082 | 1 | 0.999631 | 0.969331 |
| Q91Z58_S529s_1 | 0.74 | 1.79 | 1.44 | 1.17 | 0.95 | 1.25 | 0.83 | 1.39 | 1.36 | 1.43 | 1.49 | 1.37 | 0.96 | 1.46 | 1.07 | 1.55 | -0.18 | 0.1275 | -0.025 | 0.940375 | 0.56628 | 0.672154 | 1 | 0.999631 | 0.981146 |
| Q99K99_S268s_1 | 0.76 | -0.28 | -0.24 | -0.68 | -0.72 | 0.31 | 0.26 | -0.55 | -0.49 | 1.36 | -0.74 | -0.55 | 0.76 | -0.27 | -1.01 | 0.36 | -0.065 | 0.005 | 0.07 | 0.883905 | 0.873409 | 0.992592 | 1 | 0.999631 | 0.963505 |
| Q99M80-2_Q99M80_S1005s_1 | -0.62 | -0.48 | -0.53 | -0.35 | 0.06 | 0.13 | -0.22 | 0.3 | 0.53 | 0.03 | -0.08 | -0.08 | -0.09 | 0.04 | -0.35 | -0.24 | 0.5625 | 0.595 | 0.335 | 0.229077 | 0.043892 | 0.04832 | 1 | 0.999631 | 0.679656 |
| Q9CPS8_S40s_1 | -0.6 | -1.34 | -2.58 | -0.92 | -1.43 | -1.25 | -2.83 | -2.37 | -1.62 | -1.87 | -3.54 | -3.5 | -0.77 | -0.68 | -3.84 | -0.54 | -0.61 | -1.2725 | -0.0975 | 0.901841 | 0.26781 | 0.060865 | 1 | 0.999631 | 0.968183 |
| Q9CPS8_T38t_1 | -1.04 | -1.6 | -2.58 | -0.92 | -1.34 | -1.26 | -2.83 | -2.37 | -1.54 | -0.91 | -3.54 | -3.5 | -0.59 | -1.66 | -3.84 | -0.54 | -0.415 | -0.8375 | -0.1225 | 0.871015 | 0.420568 | 0.243282 | 1 | 0.999631 | 0.958735 |
| Q9CQF4_S106s_1 | -1.63 | -0.75 | 0.27 | -0.68 | -0.5 | -0.65 | -0.18 | 0.17 | -0.52 | -1.31 | 0.47 | 0.29 | 0.2 | -0.68 | -0.23 | 0.03 | 0.4075 | 0.43 | 0.5275 | 0.24561 | 0.347613 | 0.423238 | 1 | 0.999631 | 0.685751 |
| Q9CZL2_S38s_1 | 0.77 | 0.21 | 0.13 | 0.55 | 0.27 | 0.2 | 0.38 | 0.93 | 0.42 | 0.77 | 1.09 | 0.66 | -0.54 | 0 | 0.91 | 0.1 | 0.03 | 0.32 | -0.2975 | 0.43437 | 0.918281 | 0.28547 | 1 | 0.999631 | 0.779717 |
| Q9D2Q3_S423s_1 | -0.52 | 0.17 | 0.14 | 0.03 | -0.01 | -0.14 | -0.21 | 0.18 | -0.03 | -0.05 | 0.17 | 0.06 | 0.34 | 0.61 | 0.35 | 0.25 | 9.81E-18 | 0.0825 | 0.4325 | 0.162909 | 1 | 0.763593 | 1 | 0.999631 | 0.631442 |
| Q9D5Q8-3_Q9D5Q8_S157s_1 | -1.73 | -2.12 | -3.32 | -3.19 | -1.24 | -1.84 | -2.38 | -2.9 | -1.88 | -1.67 | -1.82 | -2.93 | -1.41 | -1.42 | -3.37 | -2.76 | 0.5 | 0.515 | 0.35 | 0.549054 | 0.32834 | 0.289416 | 1 | 0.999631 | 0.835034 |
| Q9D6W8_S173s_1 | -1.47 | -1.69 | -1.72 | -0.68 | -1.29 | -1.13 | -0.77 | -1.24 | -0.26 | -1.12 | -0.6 | -1.38 | -1.01 | -1.55 | -0.9 | -1 | 0.2825 | 0.55 | 0.275 | 0.433224 | 0.385539 | 0.165487 | 1 | 0.999631 | 0.779438 |
| Q9D6W8_S43s_1 | -1.12 | 0.05 | -1.01 | -1.74 | -1.04 | -0.45 | -1.35 | -1.28 | -2.1 | -0.36 | -0.72 | -0.68 | -0.91 | 0.29 | -0.79 | -1.26 | -0.075 | -0.01 | 0.2875 | 0.556094 | 0.858247 | 0.984246 | 1 | 0.999631 | 0.838305 |
| Q9D6W8_T201t_1 | -0.43 | -1.38 | -0.08 | 0.13 | -1.66 | 0.04 | -0.31 | 0.84 | -0.4 | 0.29 | 0.3 | -0.52 | -1.11 | 0.51 | -0.59 | -0.01 | 0.1675 | 0.3575 | 0.14 | 0.767774 | 0.767325 | 0.390881 | 1 | 0.999631 | 0.917335 |
| Q9D9H8_S44s_1 | 2.25 | -0.76 | -0.04 | -0.17 | -2.3 | -0.78 | 3.41 | -3.08 | -2.46 | 2.2 | -1.76 | -0.38 | -1.62 | -3.07 | -1.5 | -1.31 | -1.0075 | -0.92 | -2.195 | 0.009235 | 0.46297 | 0.392129 | 1 | 0.999631 | 0.367426 |
| Q9D9H8_S77s_1 | 1 | -1.27 | 1.14 | 0.36 | -1.75 | 0.27 | 1.84 | -1.24 | -1.57 | 1.41 | -0.65 | -1.77 | -1.51 | -2.26 | -0.81 | -1.15 | -0.5275 | -0.9525 | -1.74 | 0.012793 | 0.539713 | 0.252358 | 1 | 0.999631 | 0.385092 |
| Qars_D3Z158_S70s_1 | -0.32 | 0.61 | 1.33 | 0.94 | 1 | 0.84 | 0.93 | 1.23 | 0.71 | 0.28 | 1.39 | 1.61 | 1.64 | 0.75 | 1.03 | 1.35 | 0.36 | 0.3575 | 0.5525 | 0.204929 | 0.351264 | 0.440414 | 1 | 0.999631 | 0.6605 |
| R3hcc1_E9PUE1_S167s_1 | 0.44 | 1.76 | 0.79 | 1.13 | 1.2 | 2.3 | 0.58 | 0.54 | 1.18 | 2 | 0.3 | 0.65 | 0.49 | 1.04 | 0.48 | 1.4 | 0.125 | 0.0025 | -0.1775 | 0.650614 | 0.791924 | 0.995604 | 1 | 0.999631 | 0.875991 |
| R3hdm1_E9Q9Q2_S1007s_1 | -0.39 | 0.07 | 0.49 | 0.19 | 0.23 | -0.33 | -0.18 | 0.63 | 0.06 | 0.16 | 0.59 | 0.32 | 0.14 | 0.5 | 0.73 | 0.56 | -0.0025 | 0.1925 | 0.3925 | 0.227424 | 0.993888 | 0.520379 | 1 | 0.999631 | 0.678689 |
| R3hdm1_E9Q9Q2_S280s_1 | -2.71 | -3.44 | 0.37 | 0.8 | -1.74 | -5.08 | 1.4 | -0.27 | -2.31 | -4.21 | 0.57 | 0.44 | -1.8 | -4.07 | 0.38 | -0.83 | -0.1775 | -0.1325 | -0.335 | 0.781545 | 0.904465 | 0.921327 | 1 | 0.999631 | 0.921707 |
| R3hdm1_E9Q9Q2_S282s_1 | -2 | -1.04 | -3.51 | -1.37 | -0.85 | -3.64 | -2.09 | -1.03 | -1.31 | -5.6 | -1.07 | -1.26 | -0.56 | -0.46 | -1.82 | -3.28 | 0.0775 | -0.33 | 0.45 | 0.553683 | 0.916649 | 0.755163 | 1 | 0.999631 | 0.837943 |
| R3hdm1_E9Q9Q2_S347s_1 | 0.23 | -0.22 | 0.69 | -0.37 | -0.7 | 0 | -0.61 | 0.42 | -0.36 | 1.27 | -1.09 | 0.36 | -0.18 | -0.28 | 0.02 | -0.48 | -0.305 | -0.0375 | -0.3125 | 0.358387 | 0.422408 | 0.942826 | 1 | 0.999631 | 0.741182 |
| R3hdm1_E9Q9Q2_S381s_1 | -0.02 | 0.53 | 0.33 | 0.36 | 0.72 | 0.36 | 0.31 | 0.69 | 0.54 | 0.38 | 0.93 | 0.96 | 0.23 | 0.79 | 0.68 | 0.51 | 0.22 | 0.4025 | 0.2525 | 0.391893 | 0.408569 | 0.174045 | 1 | 0.999631 | 0.758901 |
| R3hdm2_E9Q9D6_S263s_1 | 1.42 | 1.55 | 1.45 | 1.81 | 0.48 | 1.93 | 2.61 | 0.33 | 1.33 | 2.76 | 1.38 | 1.12 | -1.55 | -0.6 | -0.6 | 0.29 | -0.22 | 0.09 | -2.1725 | 0.000337 | 0.675256 | 0.82142 | 1 | 0.999631 | 0.351308 |
| R3hdm2_E9Q9D6_S349s_1 | -0.13 | 1.39 | 0.88 | 0.17 | 0.84 | 0.4 | -0.01 | 0.43 | 0.75 | 0.46 | 0.95 | 1.02 | 1.19 | 1.62 | 0.88 | 1.12 | -0.1625 | 0.2175 | 0.625 | 0.142051 | 0.680213 | 0.575661 | 1 | 0.999631 | 0.612726 |
| R3hdm2_E9Q9D6_S361s_1 | 1.62 | 1.29 | 1.59 | 1.74 | -0.02 | 1.12 | 2.2 | 0.29 | -0.02 | 1.71 | 1.17 | 0.86 | 0.78 | 0.35 | 1.25 | 0.98 | -0.6625 | -0.63 | -0.72 | 0.038842 | 0.188471 | 0.131636 | 1 | 0.999631 | 0.463118 |
| R3hdm2_E9Q9D6_S857s_1 | 0.16 | 0.8 | 0.27 | 0.07 | 0.2 | 0.22 | -0.04 | -0.06 | 0.34 | 0.45 | 0.59 | 0.67 | -0.01 | 0.86 | 0.08 | 0.86 | -0.245 | 0.1875 | 0.1225 | 0.725645 | 0.376524 | 0.505714 | 1 | 0.999631 | 0.90568 |
| R3hdm2_E9Q9D6_T35t_1 | 0.15 | -2.98 | -3.34 | -0.35 | 0.24 | -3.68 | -0.96 | -0.56 | -1.87 | -1.97 | -1.72 | -1.48 | -1.25 | -2.31 | -0.79 | -4.33 | 0.39 | -0.13 | -0.54 | 0.597682 | 0.713089 | 0.86922 | 1 | 0.999631 | 0.855033 |
| R3hdm2_Q80TM6_S263s_1 | 0.66 | 0.4 | 0.67 | 1.59 | 0.32 | 0.48 | 2.22 | 0.14 | 0.55 | 1.27 | 1.23 | 0.91 | -2.07 | -1.05 | -1.03 | -0.54 | -0.04 | 0.16 | -2.0025 | 0.000831 | 0.937542 | 0.648766 | 1 | 0.999631 | 0.351308 |
| Rab11a_P62492_S190s_1 | -2.15 | -0.56 | -0.65 | -1.64 | -0.55 | -1.06 | -0.84 | -1 | -0.17 | -2.03 | -0.69 | 0.02 | 0.78 | -1.14 | -0.99 | -1.24 | 0.3875 | 0.5325 | 0.6025 | 0.303004 | 0.350998 | 0.351192 | 1 | 0.999631 | 0.714978 |
| Rab11b_P46638_T77t_1 | -0.6 | -0.12 | -0.33 | -0.49 | 0 | 0.38 | -0.59 | -0.72 | 0.67 | -0.41 | -1.49 | 0.17 | 0.94 | 0.29 | -0.13 | 0.23 | 0.1525 | 0.12 | 0.7175 | 0.048171 | 0.639074 | 0.795376 | 1 | 0.999631 | 0.48343 |
| Rab11fip2_G3XA57_S277s_1 | 0.79 | 0.96 | 0.31 | 1.26 | 1.13 | 0.89 | 0.29 | 1.01 | 1.1 | 0.5 | 0.55 | 0.83 | -0.24 | 1.12 | 0.19 | 1.15 | -1.2E-16 | -0.085 | -0.275 | 0.512448 | 1 | 0.784204 | 1 | 0.999631 | 0.818094 |
| Rab11fip5_Q8R361_S296s_1 | 0.9 | -0.34 | -0.19 | 0.7 | 1.67 | 1.88 | 1.47 | 1.3 | 0.66 | -0.35 | 0.41 | -0.79 | -2.43 | 0.02 | 0.57 | -0.29 | 1.3125 | -0.285 | -0.8 | 0.238015 | 0.004434 | 0.530211 | 1 | 0.999631 | 0.683301 |

| | | | | | | | | | | | | | | | | | | | | | | | | | |
|---|---|---|---|---|---|---|---|---|---|---|---|---|---|---|---|---|---|---|---|---|---|---|---|---|---|
| Rab11fip5_Q8R361_S307s_1 | 0.29 | -1.96 | -0.56 | 0.47 | -0.24 | -0.88 | 0.55 | 0.96 | -0.39 | -1.29 | 0.96 | 0.25 | -1.02 | -0.98 | 0.78 | 0.66 | 0.5375 | 0.3225 | 0.3 | 0.653656 | 0.39533 | 0.625754 | 1 | 0.999631 | 0.878344 |
| Rab11fip5_Q8R361_S480s_2 | 0.41 | -0.43 | -0.41 | 1.1 | 0.23 | 0.62 | 0.59 | 0.6 | 0.21 | -0.37 | 0.19 | -0.44 | -0.9 | 1.11 | 1 | -0.21 | 0.3425 | -0.27 | 0.0825 | 0.88355 | 0.387832 | 0.518315 | 1 | 0.999631 | 0.963505 |
| Rab11fip5_Q8R361_S486s_1 | 0.96 | -1.31 | -0.37 | -0.23 | -0.45 | -0.42 | 0.97 | 0.96 | -0.83 | -0.31 | 0.7 | -0.61 | -1.98 | -0.29 | -0.39 | 0.34 | 0.5025 | -0.025 | -0.3425 | 0.580747 | 0.382479 | 0.962786 | 1 | 0.999631 | 0.846995 |
| Rab11fip5_Q8R361_S530s_1 | 0.26 | 0.34 | 0.64 | 0.33 | 1.06 | 0.74 | 0.16 | 0.6 | 0.8 | 0.23 | 0.33 | 0.68 | 1.14 | 2.25 | 0.91 | 0.68 | 0.2475 | 0.1175 | 0.8525 | 0.049631 | 0.391669 | 0.665238 | 1 | 0.999631 | 0.487521 |
| Rab11fip5_Q8R361_S539s_1 | 0 | 0.43 | -0.67 | 1.02 | 0.58 | 0.5 | 0.53 | 0.81 | -0.08 | 0.23 | 0.92 | 0.39 | -0.07 | 0.61 | -0.14 | 0.87 | 0.41 | 0.17 | 0.1225 | 0.780547 | 0.292705 | 0.685044 | 1 | 0.999631 | 0.921416 |
| Rab11fip5_Q8R361_S640s_1 | -1.7 | 0.1 | -0.8 | -1.26 | -0.38 | -0.21 | -0.79 | -0.62 | -0.55 | -1.31 | -1.34 | -1.25 | -0.98 | -0.48 | -2.59 | -1.13 | 0.415 | -0.1975 | -0.38 | 0.49704 | 0.321344 | 0.646607 | 1 | 0.999631 | 0.811348 |
| Rab12_P35283_S15s_1 | 0.37 | 1 | 0.99 | 0.47 | 0.16 | 0.04 | -0.38 | 0.3 | -0.21 | -0.14 | -0.04 | -0.76 | 0.22 | 0.33 | 1.01 | 0.43 | -0.6775 | -0.995 | -0.21 | 0.520316 | 0.040522 | 0.008255 | 1 | 0.999631 | 0.819528 |
| Rab12_P35283_S20s_1 | 1.8 | 0.9 | -0.23 | 1.1 | 0.69 | -0.97 | 0.58 | -0.26 | -0.3 | -0.75 | -0.34 | -1.28 | -1.11 | -0.07 | -1.55 | -1.16 | -0.8825 | -1.56 | -1.865 | 0.003931 | 0.120853 | 0.007196 | 1 | 0.999631 | 0.351308 |
| Rab14_Q91V41_S204s_1 | 0.1 | 0.46 | 0.88 | -0.93 | -0.56 | -0.11 | 0.28 | 0.4 | 0.09 | 0.18 | 0.67 | 0 | 0.15 | -0.05 | 0.67 | 0.89 | -0.125 | 0.1075 | 0.2875 | 0.5232 | 0.773569 | 0.797595 | 1 | 0.999631 | 0.821793 |
| Rab1A_P62821_S194s_1 | 0.94 | 0.07 | 0.49 | 0.21 | -0.56 | 0.43 | 0.66 | 0 | 0.49 | 1.28 | 0.5 | 0.12 | 0.5 | 0.38 | 0.45 | -0.1 | -0.295 | 0.17 | -0.12 | 0.708415 | 0.415638 | 0.628973 | 1 | 0.999631 | 0.898919 |
| Rab1A_P62821_T195t_1 | 0.97 | 0.03 | 0.49 | 0.2 | -0.53 | 0.47 | 0.71 | 0.03 | 0.5 | 1.27 | 0.52 | 0.09 | 0.48 | 0.33 | 0.47 | -0.16 | -0.2525 | 0.1725 | -0.1425 | 0.666675 | 0.49266 | 0.630084 | 1 | 0.999631 | 0.88264 |
| Rab1b_Q9D1G1_S179s_1 | -1.75 | -0.63 | -1.2 | -1.03 | -0.8 | -0.83 | -0.85 | -1.15 | -0.53 | -0.29 | -1.6 | 0.26 | -0.07 | -0.98 | -0.26 | -0.69 | 0.245 | 0.6125 | 0.6525 | 0.094185 | 0.429894 | 0.191737 | 1 | 0.999631 | 0.558128 |
| Rab22a_P35285_S187s_1 | 1.53 | -0.98 | 0.35 | 0.42 | -1.02 | -0.58 | 0.8 | -0.51 | -1.48 | 0.77 | 0.15 | -0.36 | -1.37 | -1.91 | -0.53 | -1.1 | -0.6575 | -0.56 | -1.5575 | 0.016454 | 0.278051 | 0.386038 | 1 | 0.999631 | 0.389192 |
| Rab23_Q9D4I9_S197s_1 | 1.14 | 0.71 | 0.37 | 0.95 | 1.66 | 1.28 | 0.62 | 1.47 | 1.24 | 0.45 | 0.94 | 1.49 | 0.23 | 1.41 | 1.22 | 1.33 | 0.465 | 0.2375 | 0.255 | 0.492146 | 0.174925 | 0.47842 | 1 | 0.999631 | 0.809399 |
| Rab31_Q3TXV4_S36s_1 | -0.84 | -1.62 | -0.44 | -0.59 | -1.27 | -0.98 | -0.89 | -0.86 | -1.16 | -0.88 | -0.68 | -0.17 | -0.41 | -0.58 | -0.69 | -0.21 | -0.1275 | 0.15 | 0.4 | 0.261813 | 0.694947 | 0.682708 | 1 | 0.999631 | 0.692758 |
| Rab37_Q544E8_T4t_1 | -0.88 | 0.1 | -0.95 | -0.1 | -0.78 | -0.34 | -0.37 | 0.86 | -0.15 | -1.28 | -1.17 | 0.62 | 1.02 | -0.19 | -1.58 | 0.59 | 0.3 | -0.0375 | 0.4175 | 0.477072 | 0.494766 | 0.939887 | 1 | 0.999631 | 0.800073 |
| Rab39b_Q8BHC1_S201s_1 | -0.85 | -0.86 | -1.18 | -0.67 | -0.95 | -0.87 | -1.21 | -0.84 | -1.5 | -0.93 | -0.96 | -0.7 | 0.14 | -0.86 | -0.67 | 0.27 | -0.0775 | -0.1325 | 0.61 | 0.110656 | 0.759213 | 0.647231 | 1 | 0.999631 | 0.575791 |
| Rab3a_P63011_S188s_1 | -0.53 | -0.38 | -0.17 | -0.52 | -0.48 | -0.46 | -0.71 | -0.61 | -0.5 | -0.48 | -0.68 | -0.29 | 0.2 | -0.34 | -0.17 | 0.06 | -0.165 | -0.0875 | 0.3375 | 0.247265 | 0.499464 | 0.730998 | 1 | 0.999631 | 0.685751 |
| Rab3a_P63011_S190s_1 | -1.09 | -1.08 | -0.67 | -0.89 | -1.03 | -1.05 | -0.99 | -0.78 | -0.89 | -1 | -1.05 | -0.13 | -0.39 | -0.84 | -0.84 | -0.51 | -0.03 | 0.165 | 0.2875 | 0.322701 | 0.902827 | 0.593579 | 1 | 0.999631 | 0.724485 |
| Rab3gap2_E9QKE4_S449s_1 | 0.01 | -0.17 | -0.74 | 0.09 | 0.18 | -0.54 | -0.67 | -0.28 | -0.39 | -0.48 | -0.39 | 0.25 | -0.59 | 0.2 | 0.24 | -0.02 | -0.125 | -0.05 | 0.16 | 0.635917 | 0.693415 | 0.874004 | 1 | 0.999631 | 0.869397 |
| Rab3ip_Q68EF0_S218s_1 | -0.46 | -0.28 | 0.26 | -0.11 | -0.14 | -0.64 | -0.56 | -0.11 | -0.34 | -0.62 | -0.25 | 0.1 | 0.12 | 0.01 | 0.26 | 0.02 | -0.215 | -0.13 | 0.25 | 0.394086 | 0.4558 | 0.660729 | 1 | 0.999631 | 0.759303 |
| Rab3ip_Q68EF0_S240s_1 | -0.3 | 0.46 | -0.7 | 0.33 | 0.31 | 0.28 | 0.35 | 0.22 | 0.44 | -0.19 | -0.13 | -0.03 | -0.84 | 0.76 | 0.02 | 0.24 | 0.3425 | 0.075 | 0.0975 | 0.823094 | 0.300528 | 0.829436 | 1 | 0.999631 | 0.938864 |
| Rab40c_E9Q016_S249s_1 | 0.6 | 0.92 | 0.92 | 0.6 | 0.63 | 0.85 | 0.66 | 1.45 | 0.61 | 1.43 | 1.76 | 1.72 | 1.39 | 1.26 | 1.36 | 1.73 | 0.1375 | 0.62 | 0.675 | 0.03322 | 0.633311 | 0.087247 | 1 | 0.999631 | 0.44294 |
| Rab5a_Q9CQD1_S193s_1 | 1.4 | -1.49 | -0.09 | 0.26 | -0.41 | 0.39 | 0.72 | 0.25 | -0.71 | 0.14 | -0.52 | -1.04 | -0.8 | -2.02 | -0.39 | -1.48 | 0.2175 | -0.5525 | -1.1925 | 0.081982 | 0.709494 | 0.36011 | 1 | 0.999631 | 0.54792 |
| Rab5b_P61021_S205s_1 | 0.65 | 1.41 | -0.42 | 0.21 | 0.52 | 1.5 | -0.05 | 0.67 | 0.57 | 0.74 | 0.4 | -0.18 | 1.59 | 1.09 | 0.33 | 0.55 | 0.1975 | -0.08 | 0.4275 | 0.371728 | 0.679062 | 0.853087 | 1 | 0.999631 | 0.746522 |
| Rab8a_P55258_S181s_1 | -0.2 | -0.59 | 0.14 | -0.39 | -1.35 | -1.54 | 0 | -0.65 | -0.5 | -0.47 | 0.14 | -0.23 | -0.68 | -0.87 | -0.14 | -0.01 | -0.625 | -0.005 | -0.165 | 0.620888 | 0.135182 | 0.986478 | 1 | 0.999631 | 0.861888 |
| Rab8a_P55258_S185s_1 | 0.13 | -0.24 | 0.92 | -0.13 | -0.54 | 0.46 | 0.44 | 0.02 | 0.31 | 1.09 | 0.47 | 0.3 | 0.3 | -0.17 | 0.64 | 0.34 | -0.075 | 0.3725 | 0.1075 | 0.765812 | 0.839438 | 0.310319 | 1 | 0.999631 | 0.916581 |
| Rab8b_P61028_S182s_1 | -1.28 | -1.73 | 0.41 | -0.34 | -1.21 | -1.49 | 0.43 | -0.34 | -1.71 | -1.56 | 0.04 | 0.26 | -0.76 | -2.32 | 0.05 | -0.92 | 0.0825 | -0.0075 | -0.2525 | 0.685672 | 0.888255 | 0.990592 | 1 | 0.999631 | 0.890608 |
| Rab8b_P61028_S183s_1 | 0.87 | -0.17 | 0.2 | -0.27 | -0.35 | 0.2 | 0.34 | -0.25 | 0.82 | 0 | 0.01 | -0.16 | 0.81 | 0.09 | -0.78 | -0.61 | -0.1725 | 0.01 | -0.28 | 0.534497 | 0.616276 | 0.978321 | 1 | 0.999631 | 0.828596 |
| Rab9a_Q9R0M6_S179s_1 | -1.13 | -1.79 | -0.39 | -0.52 | -1.03 | -1.16 | -0.5 | -0.89 | -1.26 | -1.73 | -0.15 | 0.08 | -0.96 | -1 | -0.18 | 0.31 | 0.0625 | 0.1925 | 0.5 | 0.282309 | 0.8658 | 0.706708 | 1 | 0.999631 | 0.703708 |
| Rabep1_J3QJV7_S323s_1 | 0.73 | 0.37 | -0.36 | 1.16 | 1.03 | 0.38 | 0.72 | 0.87 | 0.64 | 0.6 | 0.84 | 0.66 | -0.77 | 0.9 | 0.42 | 0.67 | 0.275 | 0.21 | -0.17 | 0.722904 | 0.463002 | 0.562689 | 1 | 0.999631 | 0.905674 |
| Rabep1_J3QJV7_S326s_1 | -0.47 | -0.49 | 1.54 | 0.28 | -0.39 | -0.61 | -0.49 | 0.91 | -0.76 | -0.41 | 0.8 | 1.19 | -0.52 | 0.01 | 0.55 | 1.33 | -0.36 | -0.01 | 0.1275 | 0.823289 | 0.514444 | 0.986855 | 1 | 0.999631 | 0.938864 |
| Rabgap1_A2AWA9_S42s_1 | 1.2 | -0.47 | 0.62 | -0.17 | -0.54 | -0.27 | 0.18 | -1.19 | -0.2 | 0.38 | -1.02 | -1.17 | -0.56 | -0.93 | -0.79 | -1.86 | -0.75 | -0.7975 | -1.33 | 0.016198 | 0.12539 | 0.134151 | 1 | 0.999631 | 0.388832 |
| Rabgap1_A2AWA9_S486s_1 | 1.9 | 0.56 | 0.96 | 1.48 | 0.47 | 1.12 | 1.52 | 0.89 | 0.87 | 1.63 | 0.99 | 0.67 | -1.22 | 0.82 | 1.11 | 0.53 | -0.225 | -0.185 | -0.915 | 0.124302 | 0.557287 | 0.629312 | 1 | 0.999631 | 0.596063 |
| Rabgef1_Q9JM13_S124s_1 | 1.54 | 0.42 | 0.81 | 1.57 | 1.67 | 2.41 | 1.18 | 1.44 | 1.31 | 1.06 | 1.4 | 0.91 | -2.64 | 2.8 | 1.57 | 1.24 | 0.59 | 0.085 | -0.3425 | 0.740581 | 0.157692 | 0.806154 | 1 | 0.999631 | 0.908846 |
| Rabgef1_Q9JM13_S132s_1 | 1.73 | 0.8 | 1.29 | 1.53 | 0.52 | 1.17 | 1.58 | 0.79 | 1.27 | 2.08 | 1.42 | 0.93 | 0.08 | 0.15 | 0.71 | 0.58 | -0.3225 | 0.0875 | -0.9575 | 0.014493 | 0.355073 | 0.80373 | 1 | 0.999631 | 0.385092 |
| Rabgef1_Q9JM13_S390s_1 | -2.34 | 0.42 | 1.08 | -2.65 | 0.67 | -0.04 | 1.27 | -1.7 | 1.7 | -3.24 | 1.14 | 1.54 | 0.36 | -3.36 | 0.06 | -3.77 | 0.9225 | 1.1575 | -0.805 | 0.515595 | 0.360674 | 0.380996 | 1 | 0.999631 | 0.819449 |
| Rabl6_Q5U3K5_S377s_1 | -0.68 | -0.33 | -0.49 | -0.42 | -0.06 | -0.81 | -0.28 | 0.24 | -0.75 | -0.63 | -0.27 | -0.94 | -0.34 | -0.35 | -1 | -0.3 | 0.2525 | -0.1675 | -0.0175 | 0.952743 | 0.406078 | 0.53891 | 1 | 0.999631 | 0.984317 |
| Rabl6_Q5U3K5_S482s_1 | -0.75 | 0.49 | -1.92 | -1.13 | -0.14 | 0.13 | -1.05 | -0.5 | 0.56 | 0.2 | -0.41 | -0.04 | -0.19 | 0.38 | 0.15 | -0.47 | 0.4375 | 0.905 | 0.795 | 0.142339 | 0.411231 | 0.101782 | 1 | 0.999631 | 0.612726 |
| Rabl6_Q5U3K5_S502s_1 | 0.01 | -0.27 | -0.11 | -0.46 | -0.4 | -0.01 | -0.4 | -0.48 | 0.07 | 0.32 | 0.16 | 0.07 | -0.61 | -0.3 | -0.11 | -0.17 | -0.115 | 0.3625 | -0.09 | 0.75075 | 0.655881 | 0.176498 | 1 | 0.999631 | 0.912676 |
| Rad23a_Q3TN85_S123s_1 | -1.02 | -0.27 | -3.14 | -1.91 | -0.73 | -1.07 | -2.17 | -1.9 | -0.94 | -0.91 | -1.39 | -3.63 | -1.22 | -0.63 | -1.39 | -0.57 | 0.1175 | -0.1325 | 0.6325 | 0.301582 | 0.852404 | 0.866132 | 1 | 0.999631 | 0.714243 |
| Rad23a_Q3TN85_S125s_1 | -0.45 | -0.17 | -2.04 | -1.53 | 0.03 | -2.05 | -0.58 | -0.56 | -0.38 | -0.99 | -1.11 | -0.97 | -0.6 | 0.13 | 0.56 | 0.21 | 0.2575 | 0.185 | 1.1225 | 0.041327 | 0.653304 | 0.688108 | 1 | 0.999631 | 0.474979 |
| Rad23a_Q3TN85_S92s_1 | -1.72 | -1.02 | 0.01 | -0.65 | -0.92 | -0.88 | -0.46 | -0.87 | -0.75 | -1.25 | -0.41 | -0.6 | 0 | -0.12 | -0.05 | 0.03 | 0.0625 | 0.0925 | 0.81 | 0.061353 | 0.871716 | 0.821983 | 1 | 0.999631 | 0.509355 |
| Rad23b_P54728_S160s_1 | -0.16 | 0.4 | 0.42 | 0.6 | 0.96 | 0.29 | 0.1 | 0.4 | 0.93 | 0.1 | -0.26 | 0.91 | 1.42 | 1.31 | 0.89 | 0.59 | 0.1225 | 0.105 | 0.7375 | 0.045214 | 0.689903 | 0.776121 | 1 | 0.999631 | 0.476358 |
| Rad23b_P54728_S160s_2 | 0.07 | -0.29 | -0.76 | -0.78 | -0.25 | -0.26 | -0.17 | -0.79 | -0.39 | -0.81 | -0.11 | 0.02 | -0.84 | -0.4 | -0.81 | -0.16 | 0.0725 | 0.1175 | -0.1125 | 0.736375 | 0.813254 | 0.72111 | 1 | 0.999631 | 0.907605 |
| Rad23b_P54728_T155t_1 | 0 | -0.05 | -0.48 | -0.21 | 0.6 | -0.36 | -0.71 | -0.4 | -0.02 | -0.55 | -1.26 | -0.1 | 0.44 | 0.18 | -1.44 | -1.4 | -0.0325 | -0.2975 | -0.37 | 0.460371 | 0.923599 | 0.398991 | 1 | 0.999631 | 0.791962 |
| Rad23b_P54728_T155t_2 | 0.07 | -0.29 | -0.81 | -0.99 | -0.25 | -0.26 | -0.46 | -1.38 | -0.39 | -0.81 | -0.27 | -1.36 | -0.84 | -0.4 | -0.48 | -0.29 | -0.0825 | -0.2025 | 0.0025 | 0.994084 | 0.826792 | 0.589106 | 1 | 0.999631 | 0.997705 |
| Radil_D3YWJ9_S206s_1 | 1.27 | 0.79 | 2 | 0.75 | 0.32 | 1.39 | 1.03 | 0.43 | 0.49 | 2.23 | 1.04 | 1.17 | -0.26 | -0.38 | 0.65 | 0.13 | -0.41 | 0.03 | -1.1675 | 0.013935 | 0.310751 | 0.94739 | 1 | 0.999631 | 0.385092 |
| Raf1_Q99N57_S233s_1 | -0.83 | 0.16 | 0.33 | 0.38 | -0.38 | -0.34 | 0.19 | 0.79 | -0.41 | -0.49 | -0.23 | -0.94 | -0.13 | -0.03 | -0.12 | -1.98 | 0.055 | -0.5275 | -0.575 | 0.284697 | 0.89043 | 0.162184 | 1 | 0.999631 | 0.705486 |
| Rai1_Q61818_S899s_1 | -0.99 | -1.52 | 0.39 | -0.25 | -1.09 | -0.62 | -0.95 | -0.38 | -1.33 | -1.3 | -0.75 | -0.18 | -1.19 | -1.33 | -0.91 | -0.44 | -0.1675 | -0.2975 | -0.375 | 0.420714 | 0.702553 | 0.538775 | 1 | 0.999631 | 0.772817 |
| Rala_P63321_S11s_1 | -0.42 | -0.26 | 0.03 | -0.25 | -1.06 | 0.07 | -0.51 | 0.06 | 0.01 | 0.39 | -0.58 | -1.14 | 0.37 | 0.25 | -0.07 | -0.08 | -0.135 | -0.105 | 0.3425 | 0.241342 | 0.68198 | 0.779173 | 1 | 0.999631 | 0.684481 |
| Ralbp1_Q62172_S11s_1 | -1.7 | -2.01 | -1.04 | -1.25 | -1.62 | -1.97 | -1 | -1.53 | -1.71 | -1.1 | -0.25 | -0.71 | -0.95 | -1.28 | -1.09 | -0.87 | -0.03 | 0.5575 | 0.4525 | 0.178693 | 0.928595 | 0.179211 | 1 | 0.999631 | 0.642903 |
| Ralbp1_Q62172_S34s_1 | -1.45 | -0.17 | 0.11 | -2.14 | -1.27 | -1.71 | -1.97 | -1.88 | -1.27 | -0.39 | -0.8 | -0.72 | 1.37 | -0.48 | -0.63 | -0.94 | -0.795 | 0.1175 | 0.7425 | 0.28037 | 0.146857 | 0.823331 | 1 | 0.999631 | 0.701922 |
| Ralbp1_Q62172_S34s_2 | 0.06 | 1.41 | 2.01 | 1.71 | 1.23 | 1.7 | 1.38 | 1.94 | 2.27 | 0.7 | 1.73 | 1.42 | 0.29 | 1.22 | 1.85 | 1.63 | 0.265 | 0.2325 | -0.05 | 0.923496 | 0.555147 | 0.65031 | 1 | 0.999631 | 0.975032 |
| Ralbp1_Q62172_S463s_1 | -0.37 | 1.12 | 0.43 | 0.45 | 0.9 | 0.71 | 0.09 | 0.65 | 0.72 | 0.14 | 0.78 | 1.03 | 0.78 | 1.43 | 0.64 | 0.9 | 0.18 | 0.26 | 0.53 | 0.188448 | 0.628342 | 0.499449 | 1 | 0.999631 | 0.64779 |
| Ralbp1_Q62172_S601s_1 | 1.07 | -0.79 | -1.02 | 0.46 | 0.96 | 0.91 | 0.85 | 0.19 | -0.55 | 0.51 | -0.06 | -1.07 | -1.47 | -1.25 | 0.22 | -0.84 | 0.7975 | -0.2225 | -0.765 | 0.203707 | 0.134172 | 0.690951 | 1 | 0.999631 | 0.659781 |
| Ralgapa1_Q6GYP7_S1003s_1 | -0.08 | -0.57 | 0.01 | -0.45 | -0.54 | 0.16 | -0.49 | 0.13 | -0.47 | 0.48 | 0.21 | -0.19 | 0.07 | 0.54 | 0.51 | -0.06 | 0.0875 | 0.28 | 0.5375 | 0.103166 | 0.770932 | 0.385333 | 1 | 0.999631 | 0.566498 |
| Ralgapa1_Q6GYP7_S1003s_2 | -0.85 | -0.32 | -0.15 | -0.98 | -0.69 | 0.31 | -0.69 | 0.06 | -0.26 | -0.47 | -0.41 | -0.3 | -1.06 | -0.06 | 0.01 | 0.21 | 0.3225 | 0.215 | 0.35 | 0.370868 | 0.372792 | 0.467169 | 1 | 0.999631 | 0.746338 |
| Ralgapa1_Q6GYP7_S1477s_1 | 0.92 | 0.95 | 1.54 | 1.04 | 0.38 | 1.37 | 0.97 | 1.24 | 0.7 | 1.87 | 0.96 | 1.2 | 0.84 | 1.32 | 0.74 | 1.63 | -0.1225 | 0.07 | 0.02 | 0.951471 | 0.698195 | 0.835561 | 1 | 0.999631 | 0.984079 |
| Ralgapa1_Q6GYP7_S368s_1 | -0.58 | -0.68 | -0.38 | -0.91 | -2.51 | -0.62 | -1.12 | -1.19 | -0.8 | 0.71 | -0.74 | 0.33 | 1.09 | -0.48 | 0.16 | 0.35 | -0.7225 | 0.5125 | 0.9175 | 0.033426 | 0.107388 | 0.226064 | 1 | 0.999631 | 0.44294 |
| Ralgapa1_Q6GYP7_S371s_1 | 0.52 | 0.55 | 1.14 | -0.07 | -0.1 | 0.18 | 0.02 | 0.06 | 0.6 | 1.39 | 0.28 | 0.69 | 1.38 | 0.44 | 0.8 | 1.04 | -0.495 | 0.205 | 0.38 | 0.309884 | 0.133793 | 0.58108 | 1 | 0.999631 | 0.71896 |
| Ralgapa1_Q6GYP7_S739s_1 | 1.74 | 1.28 | 0.87 | 0.77 | -0.62 | 0.91 | 1.3 | -0.47 | 0.45 | 1.53 | 0.79 | 0.35 | 0.96 | -0.24 | 1.74 | 1.16 | -0.885 | -0.385 | -0.26 | 0.578165 | 0.101162 | 0.315522 | 1 | 0.999631 | 0.84622 |
| Ralgapa1_Q6GYP7_S859s_1 | 0.7 | 0.99 | -0.14 | -0.11 | 0.89 | 0.43 | 0.07 | 0.22 | 1.01 | 0.42 | 0.42 | 0.5 | 0.94 | 0.85 | -0.06 | 0.64 | 0.0425 | 0.2275 | 0.2325 | 0.557805 | 0.906164 | 0.52663 | 1 | 0.999631 | 0.838305 |
| Ralgapa1_Q6GYP7_S863s_1 | -0.49 | 0.27 | 0.11 | -0.26 | -0.64 | 0.11 | -0.57 | 0.22 | 0.13 | -0.31 | 0.93 | 0.67 | -0.23 | 0.38 | 0.38 | 0.71 | -0.1275 | 0.4475 | 0.4025 | 0.244028 | 0.697159 | 0.232642 | 1 | 0.999631 | 0.685637 |
| Ralgapa1_Q6GYP7_T1001t_2 | -0.85 | -0.32 | -0.15 | -0.98 | -0.69 | 0.31 | -0.69 | 0.06 | -0.26 | -0.47 | -0.41 | -0.3 | -1.06 | -0.06 | 0.01 | 0.21 | 0.3225 | 0.215 | 0.35 | 0.370868 | 0.372792 | 0.467169 | 1 | 0.999631 | 0.746338 |
| Ralgapb_F8WHN4_S1450s_1 | -1.25 | 1.07 | -0.84 | -1.45 | -0.93 | -1.28 | -1.23 | -1 | 0.1 | 0.18 | -1.27 | -0.09 | -0.74 | -0.31 | -0.39 | -0.83 | -0.4925 | 0.3475 | 0.05 | 0.927473 | 0.370283 | 0.570658 | 1 | 0.999631 | 0.976306 |
| Ralgapb_F8WHN4_S359s_1 | -0.91 | 0.09 | 0.02 | -0.54 | -0.43 | -0.2 | -0.4 | -0.29 | -0.09 | -0.14 | -0.01 | -0.05 | 1.13 | 0.03 | -0.48 | 0.34 | 0.005 | 0.2625 | 0.59 | 0.185245 | 0.986856 | 0.403835 | 1 | 0.999631 | 0.646114 |
| Ralgapb_F8WHN4_T363t_1 | 0.16 | -0.8 | -0.2 | -0.45 | -0.24 | -0.35 | -0.28 | -0.42 | -0.63 | -0.01 | -0.22 | -0.46 | -0.74 | -0.37 | -0.18 | -0.26 | 2.94E-17 | -0.0075 | -0.065 | 0.839274 | 1 | 0.98073 | 1 | 0.999631 | 0.946365 |
| Ralgapb_F8WHN4_T363t_2 | -0.21 | 0.82 | 0.62 | 0.5 | 0.31 | 0.86 | 0.07 | 0.62 | 0.61 | 0.19 | 0.82 | 0.54 | -0.05 | 1.05 | 0.59 | 0.85 | 0.0325 | 0.1075 | 0.1775 | 0.633726 | 0.920723 | 0.737158 | 1 | 0.999631 | 0.868569 |
| Ralgapb_F8WHN4_T379t_1 | -0.12 | -0.13 | 0.02 | -0.02 | 0 | -0.29 | -0.25 | 0.15 | -0.63 | 0.11 | 0.1 | 0.19 | -0.2 | -0.12 | 0.12 | 0.3 | -0.035 | 0.005 | 0.0875 | 0.748663 | 0.885878 | 0.986036 | 1 | 0.999631 | 0.911931 |
| Ralgapb_F8WHN4_T734t_1 | -0.86 | 1.12 | -0.27 | -2.52 | 0.56 | -0.37 | -0.42 | -2.23 | 0.94 | -1.57 | 0 | 0.04 | 1.23 | -0.95 | -1.43 | -2.68 | 0.0175 | 0.485 | -0.325 | 0.733545 | 0.98316 | 0.550179 | 1 | 0.999631 | 0.907462 |
| Ralgps1_A2AR50_S292s_1 | 1.21 | -1.19 | 0.23 | 0.75 | -0.16 | -0.29 | 1.15 | 0.34 | -0.44 | 0.43 | 0.66 | -0.12 | -1.63 | -0.69 | 0.41 | -0.04 | 0.01 | -0.1175 | -0.7375 | 0.250473 | 0.98577 | 0.827178 | 1 | 0.999631 | 0.686448 |
| Ralgps1_A2AR50_S298s_1 | -0.55 | -4.7 | -3.4 | -1.51 | -0.68 | -3.03 | -1.82 | -1.81 | -2.1 | -2.59 | -1.8 | -2.34 | -1.64 | -2.95 | -2.66 | -3.51 | 0.705 | 0.3325 | -0.15 | 0.863668 | 0.444304 | 0.689389 | 1 | 0.999631 | 0.955131 |
| Ralgps1_A2AR50_S358s_1 | 1.57 | -0.43 | -1.85 | 1.22 | 1.87 | -0.48 | 1.5 | 1.06 | -0.01 | -1.62 | 0.05 | -0.76 | -2.06 | 0.04 | -0.62 | -0.86 | 0.86 | -0.7125 | -1.0025 | 0.222887 | 0.309859 | 0.369526 | 1 | 0.999631 | 0.673942 |
| Ralgps1_A2AR50_T319t_1 | 0.54 | 0.71 | -0.63 | -0.14 | 0.52 | 1.24 | -0.64 | -0.07 | -0.34 | 0.52 | -0.64 | -0.45 | 1.03 | 0.01 | -1.54 | -0.19 | 0.1425 | -0.3475 | -0.2925 | 0.607606 | 0.767555 | 0.406125 | 1 | 0.999631 | 0.858527 |
| Ralgps1_A2AR50_T331t_1 | -0.61 | 0.33 | 0.3 | 1.27 | 0.56 | 0.5 | 0.36 | 1.37 | 0.17 | 0.24 | 0.65 | 1.53 | 0.18 | 1.08 | 1.94 | 1.06 | 0.375 | 0.325 | 0.7425 | 0.162048 | 0.399064 | 0.500359 | 1 | 0.999631 | 0.631442 |
| Ralgps1_A2AR50_T360t_1 | -0.19 | 0.59 | 2.82 | 0.37 | 0.22 | 0.4 | 0.55 | 0.16 | 0.7 | 0.69 | 2.78 | 2.63 | 0.57 | 1.08 | 2.61 | 2.86 | -0.565 | 0.8025 | 0.8825 | 0.263164 | 0.359777 | 0.314435 | 1 | 0.999631 | 0.693034 |
| Raly_Q64012_S119s_1 | -0.6 | -0.07 | -0.63 | -1.33 | -0.31 | 0.26 | -1.78 | -1.33 | -0.44 | 0.5 | -0.46 | -0.66 | 0.66 | -0.21 | -0.28 | -0.39 | -0.1325 | 0.3925 | 0.6025 | 0.141849 | 0.791042 | 0.322268 | 1 | 0.999631 | 0.612726 |
| Raly_Q64012_S254s_1 | -1.08 | -1.3 | 0.65 | -0.67 | -0.94 | -0.83 | -0.11 | 1.72 | -0.48 | -0.57 | -0.5 | 0.41 | -1.03 | -1.32 | 1.04 | -0.05 | 0.56 | 0.315 | 0.26 | 0.678058 | 0.412427 | 0.513067 | 1 | 0.999631 | 0.887183 |
| Raly_Q64012_T258t_1 | -0.35 | -0.15 | -0.16 | -0.53 | 0 | -0.38 | -0.23 | 0.93 | 0.02 | -0.75 | -0.03 | 0.21 | 0.14 | 0.37 | 0.39 | 0.77 | 0.3775 | 0.16 | 0.715 | 0.029102 | 0.284639 | 0.599858 | 1 | 0.999631 | 0.430216 |
| Ranbp2_Q9ERU9_S2113s_1 | 1.02 | -0.3 | 1.03 | 1.29 | 0.32 | 0.84 | 1.27 | 1.11 | 0.04 | 0.72 | 1.31 | 0.89 | -0.46 | 0.14 | 1.57 | 0.68 | 0.125 | -0.02 | -0.2775 | 0.601478 | 0.762769 | 0.963761 | 1 | 0.999631 | 0.856765 |
| Ranbp2_Q9ERU9_S781s_1 | 1.16 | 2.15 | 3.75 | 0.04 | 2.81 | 2.57 | 4.78 | 0.93 | 4.21 | 1.7 | 3.77 | 4.64 | 4.8 | 0.97 | 1.44 | -2.06 | 0.9975 | 1.805 | -0.4875 | 0.719843 | 0.312955 | 0.066179 | 1 | 0.999631 | 0.904808 |
| Ranbp2_Q9ERU9_S788s_1 | -0.36 | -0.51 | -0.55 | -0.14 | -1.22 | -1.23 | -0.37 | -0.45 | -0.37 | -0.56 | -0.06 | -0.32 | 0.24 | 0.47 | -0.52 | 0.06 | -0.4275 | 0.0625 | 0.4525 | 0.175341 | 0.188323 | 0.811428 | 1 | 0.999631 | 0.642116 |
| Ranbp2_Q9ERU9_T898t_1 | 0.1 | -1.03 | 0.34 | 0.19 | -0.01 | 0.5 | 0.51 | -0.28 | -0.42 | -0.69 | -1.77 | -0.82 | -0.9 | 0.27 | 0.58 | -0.7 | 0.28 | -0.825 | -0.0875 | 0.852105 | 0.470201 | 0.079427 | 1 | 0.999631 | 0.950189 |
| Ranbp3_Q9CT10_S277s_1 | 2.05 | 1.01 | 0.72 | 0.77 | -1.31 | 0.64 | 1.07 | 0.45 | 0.99 | 2.27 | -0.9 | -0.98 | -1.77 | 0.39 | 0.45 | -1.29 | -0.925 | -0.7925 | -1.6925 | 0.016279 | 0.12177 | 0.303569 | 1 | 0.999631 | 0.388832 |
| Rangap1_P46061_S405s_1 | 1.04 | 0.45 | 1.3 | 0.41 | 0.08 | -0.02 | 0.39 | 0.19 | 0.83 | 1.42 | 0.34 | 0.34 | 0.47 | -0.04 | -0.04 | -0.03 | -0.64 | -0.0675 | -0.71 | 0.052563 | 0.055529 | 0.854205 | 1 | 0.999631 | 0.489777 |
| Rangap1_P46061_S444s_1 | -0.07 | 0.71 | 0.5 | 0.11 | -0.25 | 0.35 | -0.45 | -0.1 | -0.6 | 0.33 | 0.41 | 0.52 | 0.76 | 1.02 | 0.06 | 1.29 | -0.425 | -0.1475 | 0.47 | 0.216902 | 0.185325 | 0.676154 | 1 | 0.999631 | 0.669465 |
| Rangap1_P46061_T411t_1 | 1.9 | 0.32 | 0.2 | 1.13 | 0.74 | 0.73 | 1.09 | 0.94 | 0.59 | 1.33 | 0.19 | -0.44 | -1.81 | -0.14 | 0.16 | -0.62 | -0.0125 | -0.47 | -1.49 | 0.0201 | 0.975447 | 0.368872 | 1 | 0.999631 | 0.391668 |
| Rap1gap_A2ALS5_S439s_1 | 0.54 | -0.45 | -0.21 | 0.51 | 0.8 | -0.25 | 0.7 | 0.45 | -0.14 | -0.24 | 0.47 | 0.3 | -0.68 | -0.22 | 0.13 | -0.09 | 0.3275 | 7.85E-17 | -0.3125 | 0.390313 | 0.381141 | 1 | 1 | 1 | 0.758901 |
| Rap1gap_A2ALS5_S441s_1 | 0.96 | -0.15 | -0.49 | 0.7 | 1.08 | 0.14 | 0.98 | 0.45 | 0.17 | -0.33 | 0.71 | -0.08 | -1.15 | -0.01 | -0.04 | 0.01 | 0.4075 | -0.1375 | -0.5525 | 0.234006 | 0.332255 | 0.740793 | 1 | 0.999631 | 0.683077 |
| Rap1gap_A2ALS5_S460s_1 | 0.6 | -2.91 | -2.54 | -0.51 | -0.61 | -2.22 | 0.09 | -0.4 | -1.02 | -1.06 | -0.68 | -1.46 | -2.76 | -1.09 | -0.51 | -1.19 | 0.555 | 0.285 | -0.0475 | 0.954648 | 0.516478 | 0.705046 | 1 | 0.999631 | 0.984441 |
| Rap1gap_A2ALS5_S484s_1 | 0.97 | -0.6 | -0.28 | 1.76 | 1.62 | 0.12 | 1.18 | 1.56 | 0.66 | -0.84 | 0.88 | 0.74 | -2.51 | 1.04 | 1.53 | 1.18 | 0.6575 | -0.1025 | -0.1525 | 0.870989 | 0.280186 | 0.86813 | 1 | 0.999631 | 0.958735 |
| Rap1gap_A2ALS5_S498s_1 | -0.36 | -0.47 | -0.21 | -0.53 | 0.07 | -0.87 | -0.27 | -0.51 | 0.1 | -0.59 | 0.05 | 0.01 | -0.16 | -0.66 | -0.59 | 0.22 | -0.0025 | 0.285 | 0.095 | 0.759882 | 0.993014 | 0.317454 | 1 | 0.999631 | 0.914614 |
| Rap1gap_A2ALS5_S499s_1 | -0.36 | -0.47 | -0.21 | -0.53 | 0.07 | -0.87 | -0.27 | -0.51 | 0.1 | -0.59 | 0.05 | 0.01 | -0.16 | -0.66 | -0.59 | 0.22 | -0.0025 | 0.285 | 0.095 | 0.759882 | 0.993014 | 0.317454 | 1 | 0.999631 | 0.914614 |
| Rap1gap_A2ALS5_S515s_1 | 2.33 | 0.54 | 1.11 | 0.9 | -0.33 | 0.52 | 1.17 | 0.05 | -0.14 | 1.46 | -0.63 | -0.38 | -0.31 | -0.26 | 0.27 | -0.66 | -0.8675 | -1.1425 | -1.46 | 0.007162 | 0.095128 | 0.066437 | 1 | 0.999631 | 0.358173 |
| Rap1gap_A2ALS5_S541s_1 | -0.55 | -0.39 | 1.7 | -0.15 | -1.75 | -0.91 | -0.09 | 0.45 | -1.08 | -0.37 | -0.03 | 1.94 | 1.08 | -0.52 | 1.53 | 1.34 | -0.7275 | -0.0375 | 0.705 | 0.280592 | 0.269292 | 0.959313 | 1 | 0.999631 | 0.701922 |
| Rap1gap_A2ALS5_S541s_2 | -0.08 | -0.02 | 0.19 | -0.75 | -0.71 | -0.36 | -0.66 | -0.53 | -0.25 | 0.54 | -0.71 | -0.15 | 0.4 | 0.43 | 0.02 | -0.52 | -0.4 | 0.0225 | 0.2475 | 0.490549 | 0.188925 | 0.950291 | 1 | 0.999631 | 0.807972 |
| Rap1gap_A2ALS5_S542s_1 | -2.9 | -0.95 | 0.9 | -1.36 | -2.84 | -1.74 | -1.09 | -1.08 | -1.57 | -0.43 | -0.69 | -0.21 | 1.33 | -1.15 | -0.2 | -0.73 | -0.61 | 0.3525 | 0.89 | 0.295868 | 0.438472 | 0.635696 | 1 | 0.999631 | 0.709072 |
| Rap1gap_A2ALS5_S542s_2 | -0.08 | -0.02 | 0.19 | -0.75 | -0.71 | -0.36 | -0.66 | -0.53 | -0.25 | 0.54 | -0.71 | -0.15 | 0.4 | 0.43 | 0.02 | -0.52 | -0.4 | 0.0225 | 0.2475 | 0.490549 | 0.188925 | 0.950291 | 1 | 0.999631 | 0.807972 |
| Rap1gap_A2ALS5_S566s_1 | 0.35 | -0.22 | 0.19 | 0.84 | 0.39 | -0.36 | 0.18 | 0.96 | -0.29 | -0.11 | 0.36 | 0.32 | -0.46 | 0.01 | 0.24 | 0.97 | 0.0025 | -0.22 | -0.1 | 0.800979 | 0.994589 | 0.505416 | 1 | 0.999631 | 0.931985 |
| Rap1gap_A2ALS5_T522t_1 | 1.14 | -1.35 | -1.08 | 1.23 | -0.62 | -0.52 | 1.56 | 0.26 | -0.65 | -0.42 | -0.17 | -1.09 | -1.86 | -1.93 | -0.14 | -1.37 | 0.185 | -0.5675 | -1.31 | 0.091403 | 0.806219 | 0.392041 | 1 | 0.999631 | 0.557085 |
| Rap1gap2_Q5SVL6_S26s_1 | -0.51 | 0.09 | -0.01 | -0.4 | -0.16 | -0.46 | -0.55 | -0.29 | -0.15 | -0.34 | -0.23 | 0.02 | 0.55 | 0.03 | -0.2 | 0.25 | -0.1575 | 0.0325 | 0.365 | 0.256212 | 0.558192 | 0.904852 | 1 | 0.999631 | 0.691343 |
| Rap1gap2_Q5SVL6_S399s_1 | -0.31 | 0.13 | -0.51 | -0.35 | -0.03 | -0.7 | -0.51 | -0.09 | -1.1 | -1.13 | 0.42 | -0.52 | 0.75 | -0.48 | -1.75 | -0.57 | -0.0725 | -0.3225 | -0.2525 | 0.618975 | 0.800762 | 0.428038 | 1 | 0.999631 | 0.861888 |
| Rap1gap2_Q5SVL6_S488s_1 | 1.27 | 0.11 | 0.53 | 0.57 | 2.42 | 0.18 | 5.43 | 0.49 | 1.96 | 0.26 | 3.16 | 2.87 | -0.2 | -0.66 | -1.24 | -0.05 | 1.51 | 1.4425 | -1.1575 | 0.013171 | 0.174876 | 0.042555 | 1 | 0.999631 | 0.385092 |
| Rap1gap2_Q5SVL6_S525s_1 | -0.09 | 1.1 | 1.63 | 0.45 | 0.6 | 1.35 | 0.41 | 1.07 | 0.33 | 1.17 | 0.6 | 1.8 | -0.21 | 2.03 | 2.2 | 1.06 | 0.085 | 0.2025 | 0.4975 | 0.421232 | 0.84187 | 0.673067 | 1 | 0.999631 | 0.772817 |
| Rap1gap2_Q5SVL6_S581s_1 | 0.81 | -0.93 | -1.14 | 0.22 | -1.04 | -0.46 | 0.43 | -0.93 | -0.4 | 0.46 | -1.18 | -1.34 | -1.49 | -2.12 | -0.85 | -0.81 | -0.24 | -0.355 | -1.0575 | 0.067978 | 0.652281 | 0.53889 | 1 | 0.999631 | 0.522149 |
| Rap1gap2_Q5SVL6_S594s_1 | -1.13 | -0.41 | -1.04 | -1.48 | -1.39 | -1.31 | -1.4 | -1.74 | -0.75 | -0.59 | -0.93 | -1 | 0.47 | -0.57 | -0.98 | -0.69 | -0.445 | 0.1975 | 0.5725 | 0.180987 | 0.164784 | 0.527616 | 1 | 0.999631 | 0.64376 |
| Rap1gap2_Q5SVL6_S660s_1 | 0.21 | -0.21 | -0.36 | -0.62 | 0.48 | 0.03 | 0.01 | 0.55 | -0.85 | -1.1 | 0.56 | -0.19 | 0.2 | -0.14 | -0.05 | -0.06 | 0.5125 | -0.15 | 0.2325 | 0.441995 | 0.105858 | 0.718949 | 1 | 0.999631 | 0.78215 |
| Rap1gds1_E9Q912_S548s_1 | -0.13 | 0.76 | 0.93 | 0.63 | 1 | 0.34 | 0.64 | 0.53 | -0.03 | 0.87 | 0.98 | 0.49 | 1.54 | 0.14 | 0.8 | 1.37 | 0.08 | 0.03 | 0.415 | 0.326031 | 0.802813 | 0.933547 | 1 | 0.999631 | 0.724678 |
| Rapgef1_Q3UHC1_S308s_1 | 0.82 | 0.77 | 2.12 | 0.81 | -0.92 | 0.59 | 0.81 | 0.58 | 0.6 | 1.16 | 0.44 | 0.57 | 0.13 | -0.33 | 0.45 | 0.53 | -0.865 | -0.4375 | -0.935 | 0.040731 | 0.101435 | 0.273267 | 1 | 0.999631 | 0.474979 |
| Rapgef1_Q3UHC1_S326s_1 | 0.67 | 0.49 | 0.61 | 0.99 | 1.43 | 0.24 | 0.8 | 0.76 | 0.46 | -0.16 | 0.79 | 0.61 | 1.76 | 0.44 | 0.9 | 1.13 | 0.1175 | -0.265 | 0.3675 | 0.308885 | 0.711269 | 0.393814 | 1 | 0.999631 | 0.718845 |
| Rapgef1_Q3UHC1_S375s_1 | -0.62 | 0.29 | 0.59 | 0.97 | 1.32 | 1.19 | 1.16 | 0.79 | 1.05 | -0.73 | 0.98 | 0.51 | -0.56 | 0.9 | 0.8 | 0.48 | 0.8075 | 0.145 | 0.0975 | 0.834752 | 0.053154 | 0.774258 | 1 | 0.999631 | 0.944218 |
| Rapgef1_Q3UHC1_S381s_1 | 0.05 | 0.11 | 0.29 | 0.54 | 0.17 | -0.01 | 0.4 | 0.37 | -0.48 | -0.14 | 0.14 | 0.58 | -0.17 | 0.08 | -0.8 | 0.55 | -0.015 | -0.2225 | -0.3325 | 0.360231 | 0.953309 | 0.48447 | 1 | 0.999631 | 0.741182 |
| Rapgef2_E9QNQ4_S1086s_1 | 2.1 | 0.24 | -0.75 | 1.33 | 2.64 | 0.32 | 1.42 | 0.95 | 0.76 | -1.16 | 0.62 | 0.42 | -1.53 | 0.33 | -0.21 | 0.44 | 0.6025 | -0.57 | -0.9725 | 0.177504 | 0.399957 | 0.41363 | 1 | 0.999631 | 0.642116 |
| Rapgef2_E9QNQ4_S1113s_1 | -1.53 | -0.08 | 0.51 | 0.25 | -2.31 | -2.35 | -0.28 | 0.53 | -0.7 | -0.3 | -0.42 | 0.18 | 0.57 | -1.4 | 0.64 | 0.56 | -0.89 | -0.0975 | 0.305 | 0.620529 | 0.25515 | 0.836884 | 1 | 0.999631 | 0.861888 |
| Rapgef2_E9QNQ4_S1220s_1 | -0.95 | -0.24 | -1.19 | -0.57 | -1.32 | -1.08 | -0.35 | -0.62 | -1.54 | -1.74 | -0.6 | -1.17 | -0.33 | -1.52 | -1.53 | -0.89 | -0.105 | -0.525 | -0.33 | 0.403313 | 0.757115 | 0.167164 | 1 | 0.999631 | 0.764348 |
| Rapgef2_E9QNQ4_S1235s_1 | 2.45 | 1.4 | 0.96 | 1.66 | 1.91 | 2.52 | 2.28 | 1.1 | 2.29 | 1.92 | 1.08 | 0.93 | 1 | 0.67 | 0.42 | 1.27 | 0.335 | -0.0625 | -0.7775 | 0.071101 | 0.445335 | 0.888508 | 1 | 0.999631 | 0.531307 |
| Rapgef2_E9QNQ4_S1313s_1 | 0.58 | -0.51 | -3.8 | -1.38 | 0.17 | -0.41 | -1.58 | -2.67 | -0.19 | -0.17 | -2.4 | -2.02 | -1.6 | -1.22 | -2.98 | -3.54 | 0.155 | 0.0825 | -1.0575 | 0.271718 | 0.872958 | 0.931129 | 1 | 0.999631 | 0.699288 |
| Rapgef2_E9QNQ4_S928s_1 | -0.06 | 0.6 | 0.51 | 0.04 | 0.12 | 0.37 | -0.08 | -0.01 | 0.18 | 0.28 | -0.24 | 0.42 | 0.32 | 0.61 | 0.48 | 0.63 | -0.1725 | -0.1125 | 0.2375 | 0.427184 | 0.53777 | 0.705865 | 1 | 0.999631 | 0.775206 |
| Rapgef2_E9QNQ4_S931s_1 | 0.07 | 0.41 | -0.7 | -0.09 | -0.26 | 0.35 | 0.05 | 0.28 | 0.22 | -0.14 | 0.2 | 0.14 | -1.12 | 0.79 | 0.41 | 0.32 | 0.1825 | 0.1825 | 0.1775 | 0.706312 | 0.570227 | 0.562513 | 1 | 0.999631 | 0.898066 |
| Rapgef2_E9QNQ4_S933s_1 | 0.24 | -0.54 | -1.23 | -0.49 | 0.14 | -0.01 | -0.33 | 0.23 | -0.19 | -0.37 | -0.47 | -0.45 | 0.02 | -0.23 | -0.23 | -0.38 | 0.5125 | 0.135 | 0.3 | 0.414084 | 0.168443 | 0.698764 | 1 | 0.999631 | 0.768655 |
| Rapgef2_E9QNQ4_S957s_1 | 0.33 | 1.49 | 0.98 | 0.59 | 0.57 | 1.51 | 0.41 | 0.77 | 1.68 | 1.21 | 0.55 | 0.74 | 1.26 | 1.35 | 0.92 | 1.18 | -0.0325 | 0.1975 | 0.33 | 0.33918 | 0.929825 | 0.605492 | 1 | 0.999631 | 0.732185 |
| Rapgef2_E9QNQ4_S958s_1 | 0.29 | 1.57 | 1.36 | 1.09 | 0.51 | 1.35 | 0.84 | 1.09 | 1.52 | 0.93 | 0.96 | 1.33 | 1.13 | 1.14 | 1.57 | 1.73 | -0.13 | 0.1075 | 0.315 | 0.397466 | 0.717045 | 0.760932 | 1 | 0.999631 | 0.761181 |
| Rapgef3_H3BK39_S489s_1 | -0.66 | 0.44 | 0.21 | -1.06 | 0.19 | -0.64 | -0.01 | -1.47 | 0.04 | -0.57 | 0.45 | 0.52 | 0.69 | -1.23 | -1.34 | -1.65 | -0.215 | 0.3775 | -0.615 | 0.307546 | 0.66001 | 0.392386 | 1 | 0.999631 | 0.717856 |
| Rapgef6_Q5NCJ4_S809s_1 | 0.78 | 1.27 | -0.15 | 1.16 | 0.9 | 0.75 | 0.61 | 0.62 | 1.58 | 0.82 | 1.39 | 0.99 | 0.64 | 1.38 | 1.57 | 1.29 | -0.045 | 0.43 | 0.455 | 0.274709 | 0.89929 | 0.281327 | 1 | 0.999631 | 0.700187 |
| Raph1_F2Z3U3_S1027s_1 | 0.41 | 0.54 | 0.48 | 0.33 | 0.36 | 0.16 | 0.17 | 0.25 | 0.19 | 0.66 | 0.47 | 0.8 | 0.34 | 1.18 | 0.72 | 0.49 | -0.205 | 0.09 | 0.2425 | 0.423367 | 0.386953 | 0.731924 | 1 | 0.999631 | 0.77354 |
| Raph1_F2Z3U3_S1169s_1 | -0.93 | -0.25 | -0.7 | -0.98 | -0.27 | -0.48 | -1.06 | -1.05 | -0.29 | -0.79 | -1.29 | -0.64 | -0.14 | -0.18 | -0.66 | -0.62 | -1.2E-16 | -0.0375 | 0.315 | 0.323155 | 1 | 0.90761 | 1 | 0.999631 | 0.724485 |
| Raph1_F2Z3U3_S192s_1 | 1.16 | 0.73 | -0.08 | 1.28 | 0.96 | 1.07 | 0.94 | 0.83 | 1.53 | 1.02 | 1.3 | 0.82 | 0.22 | 1.08 | 0.75 | 1.13 | 0.1775 | 0.395 | 0.0225 | 0.954858 | 0.608516 | 0.302343 | 1 | 0.999631 | 0.984441 |
| Raph1_F2Z3U3_S850s_1 | 0.2 | -0.03 | -0.8 | 1.13 | 1.09 | 0.81 | 0.94 | 0.95 | 0.8 | -0.35 | 1.33 | 0.98 | -0.02 | 0.71 | 0.54 | 0.91 | 0.8225 | 0.565 | 0.41 | 0.368177 | 0.065472 | 0.282519 | 1 | 0.999631 | 0.745968 |
| Raph1_F2Z3U3_S899s_1 | -0.88 | -0.14 | -0.19 | -0.65 | -0.56 | -0.82 | -0.69 | -0.52 | -0.78 | -0.87 | -0.06 | -0.26 | 0.1 | -0.24 | -0.15 | 0.02 | -0.1825 | -0.0275 | 0.3975 | 0.206145 | 0.514546 | 0.932244 | 1 | 0.999631 | 0.6605 |
| Rasa2_P58069_S554s_1 | -0.48 | 0.45 | 0.21 | -0.39 | -0.7 | -0.26 | -0.75 | -0.56 | -0.21 | -1.43 | -0.16 | 1.24 | 0.06 | 0.24 | -0.09 | 0.09 | -0.515 | -0.0875 | 0.1275 | 0.691458 | 0.119696 | 0.873145 | 1 | 0.999631 | 0.894017 |
| Rasa3_Q60790_S809s_1 | -0.33 | 1.23 | 0.43 | 0.17 | 0.14 | 0.59 | 0.4 | 0.2 | 0.83 | 0.31 | 0.85 | 0.82 | 1.04 | 0.76 | 0.65 | 0.66 | -0.0425 | 0.3275 | 0.4025 | 0.299509 | 0.906842 | 0.392882 | 1 | 0.999631 | 0.712884 |
| Rasa3_Q60790_S833s_1 | -2.3 | -1.01 | 1.08 | -0.97 | -1.73 | -1.42 | -1 | -2.42 | -1.37 | -1.48 | -0.48 | 1.26 | -0.55 | -1.19 | -0.38 | 0.72 | -0.8425 | 0.2825 | 0.45 | 0.530928 | 0.231174 | 0.73279 | 1 | 0.999631 | 0.826822 |
| Rasal1_Q9Z268_S194s_1 | 0.01 | -0.01 | 0.19 | 0.16 | 0.1 | -0.04 | 0.02 | -0.21 | -0.86 | 0.24 | -0.22 | -0.3 | -0.04 | -0.25 | -0.41 | 0.64 | -0.12 | -0.3725 | -0.1025 | 0.749255 | 0.613652 | 0.238881 | 1 | 0.999631 | 0.912137 |
| Rasal1_Q9Z268_T400t_1 | 0.73 | 0.23 | 1.9 | 0.61 | 0.24 | 0.71 | 0.29 | 1.16 | 0.75 | 1.63 | 0.5 | 0.58 | 0.18 | 0.15 | 1.64 | 0.89 | -0.2675 | -0.0025 | -0.1525 | 0.754654 | 0.52519 | 0.99546 | 1 | 0.999631 | 0.913582 |
| Rasal2_D3YUS5_S1068s_1 | -1.33 | -0.86 | -2.53 | -2.29 | -1.43 | -1.14 | -1.55 | -1.88 | -1.16 | -1.06 | -2.1 | -2.02 | -0.12 | -0.71 | -2.19 | -2.27 | 0.2525 | 0.1675 | 0.43 | 0.48407 | 0.550853 | 0.720173 | 1 | 0.999631 | 0.803422 |
| Rasal2_D3YUS5_S1068s_2 | -1.11 | -1 | -0.32 | -1.23 | -0.51 | -1.09 | -1.25 | -0.76 | -0.63 | -0.32 | -0.6 | -0.65 | 0.29 | -0.61 | -0.59 | -0.81 | 0.0125 | 0.365 | 0.485 | 0.203964 | 0.968284 | 0.237695 | 1 | 0.999631 | 0.659781 |
| Rasal2_D3YUS5_S1076s_1 | -0.94 | -0.33 | -0.48 | -0.4 | -0.76 | -1.29 | -0.31 | -0.77 | -1.12 | -0.95 | -0.32 | -0.55 | -0.35 | -1.03 | -0.19 | -0.03 | -0.245 | -0.1975 | 0.1375 | 0.679732 | 0.427355 | 0.51877 | 1 | 0.999631 | 0.887852 |
| Rasal2_D3YUS5_S1076s_2 | -1.11 | -1 | -0.32 | -1.23 | -0.51 | -1.09 | -1.25 | -0.76 | -0.63 | -0.32 | -0.6 | -0.65 | 0.29 | -0.61 | -0.59 | -0.81 | 0.0125 | 0.365 | 0.485 | 0.203964 | 0.968284 | 0.237695 | 1 | 0.999631 | 0.659781 |
| Rasal2_D3YUS5_S115s_1 | 0.38 | 1.22 | 0.51 | 0.32 | 0.68 | 0.79 | -0.04 | 0.15 | 0.86 | 0.32 | 0.54 | 0.65 | 1 | 1.27 | 0.43 | 1.07 | -0.2125 | -0.015 | 0.335 | 0.336928 | 0.524757 | 0.960971 | 1 | 0.999631 | 0.730499 |
| Rasal2_D3YUS5_S140s_1 | -0.35 | -3.07 | -4.95 | -0.93 | -0.45 | -2.3 | -1.19 | -1.75 | -1.62 | -1.64 | -1.6 | -2.97 | -2.37 | -1.8 | -2.39 | -3.65 | 0.9025 | 0.3675 | -0.2275 | 0.813236 | 0.362708 | 0.701895 | 1 | 0.999631 | 0.937041 |
| Rasal2_D3YUS5_S164s_1 | -0.01 | 0.32 | -0.37 | -0.52 | 0.87 | 0.96 | 0.7 | -0.29 | 0.78 | 0.03 | -0.22 | -0.63 | 0.41 | -0.6 | -1.32 | 0.34 | 0.705 | 0.135 | -0.1475 | 0.744556 | 0.078045 | 0.720096 | 1 | 0.999631 | 0.910374 |
| Rasal2_D3YUS5_S785s_1 | 0.43 | -0.35 | 1.61 | 1.77 | -0.74 | -0.24 | 1.96 | 1.77 | 0.22 | -0.86 | 2.04 | 2.31 | -0.72 | 0.48 | 1.61 | 2.23 | -0.1775 | 0.0625 | 0.035 | 0.961483 | 0.812839 | 0.937438 | 1 | 0.999631 | 0.986594 |
| Rasal2_D3YUS5_S822s_1 | 1.47 | 0.9 | 0.72 | 1.72 | 1.42 | 1.44 | 1.51 | 1.33 | 1.25 | 1.19 | 1.31 | 1 | 0.06 | 0.77 | 1.03 | 1.01 | 0.2225 | -0.015 | -0.485 | 0.209403 | 0.465711 | 0.961523 | 1 | 0.999631 | 0.663627 |
| Rasal2_D3YUS5_S925s_1 | -1.22 | -0.63 | -0.94 | 0.29 | -0.72 | -1.29 | -0.74 | -0.11 | -0.49 | -0.8 | -0.33 | -1.11 | -0.36 | -1.11 | -3.02 | -0.53 | -0.09 | -0.0575 | -0.63 | 0.326803 | 0.825454 | 0.882315 | 1 | 0.999631 | 0.724678 |
| Rasal2_D3YUS5_S986s_1 | 1.63 | 0.89 | 0.62 | 1.18 | -0.21 | 0.8 | 1.41 | 0.07 | 1.07 | 1.89 | 0.57 | 0.64 | 0.84 | -0.76 | 0.13 | -0.15 | -0.5625 | -0.0375 | -1.065 | 0.02568 | 0.201769 | 0.923298 | 1 | 0.999631 | 0.422114 |
| Rasal2_D3YUS5_T742t_1 | 0.52 | 0.15 | 0.32 | 0.41 | 1.11 | 0.52 | 0.56 | 0.35 | 0.86 | -1.51 | -0.88 | 1.14 | 0.31 | 0.39 | 1.12 | 0 | 0.285 | -0.4475 | 0.105 | 0.748366 | 0.309117 | 0.459715 | 1 | 0.999631 | 0.911931 |
| Rasgef1a_A6H650_T157t_1 | 0.15 | -1.61 | 0.75 | 0.62 | -0.95 | -0.23 | 0.96 | 0.04 | -1.57 | 0.44 | -0.29 | -0.2 | -1.35 | -2.18 | 0.74 | -0.91 | -0.0225 | -0.3825 | -0.9025 | 0.231284 | 0.970389 | 0.543286 | 1 | 0.999631 | 0.681463 |
| Rasgrf1_P27671_S485s_1 | 0.86 | 0.73 | 0.27 | 0.84 | 1.09 | 0.3 | 0.72 | 1.16 | 1.26 | 0.51 | 0.9 | 0.52 | 0.59 | 0.64 | 0.17 | 0.75 | 0.1425 | 0.1225 | -0.1375 | 0.645292 | 0.639183 | 0.685365 | 1 | 0.999631 | 0.873083 |
| Rasgrf1_P27671_S726s_1 | -0.81 | 0.54 | 0.58 | -0.27 | -0.12 | -0.11 | -0.66 | -0.34 | 0.11 | 0.01 | 0 | 0.43 | 0.43 | 0.36 | 0.03 | -0.04 | -0.3175 | 0.1275 | 0.185 | 0.635062 | 0.407016 | 0.734848 | 1 | 0.999631 | 0.869104 |
| Rasgrf1_P27671_S727s_1 | -0.64 | 1.08 | 0.4 | 0.34 | 0.33 | -0.07 | -0.23 | -0.06 | 0.33 | 0.38 | 0.22 | 0.99 | 0.78 | 1.14 | 0.7 | 0.82 | -0.3025 | 0.185 | 0.565 | 0.173743 | 0.439204 | 0.649039 | 1 | 0.999631 | 0.642116 |
| Rasgrf1_P27671_S737s_1 | 0.64 | 0.65 | 0.26 | 1.3 | 1.46 | 1.08 | 0.97 | 1.42 | 0.91 | -0.02 | 1.39 | 0.49 | -0.76 | 1 | 0.75 | 0.88 | 0.52 | -0.02 | -0.245 | 0.596812 | 0.114128 | 0.958858 | 1 | 0.999631 | 0.854617 |
| Rasgrf1_P27671_S837s_1 | -0.41 | 0.91 | 1.51 | -0.81 | -0.43 | 1.23 | -0.68 | 0.45 | 0.9 | 1.08 | 0.33 | 1.36 | 2.21 | 1.59 | 0.66 | 0.55 | -0.1575 | 0.6175 | 0.9525 | 0.142803 | 0.801777 | 0.273864 | 1 | 0.999631 | 0.613952 |
| Rasgrf2_P70392_S484s_1 | 1.17 | 1.15 | 1.08 | 2.25 | 1.33 | 1.48 | 2 | 2.58 | 1.44 | 0.56 | 2.83 | 1.91 | -0.23 | 1.82 | 2.77 | 1.63 | 0.435 | 0.2725 | 0.085 | 0.891202 | 0.293859 | 0.600678 | 1 | 0.999631 | 0.96415 |
| Rasgrf2_P70392_S726s_1 | -0.61 | 0.38 | 0.61 | -0.46 | 0.18 | -0.2 | -0.29 | -0.3 | 0.25 | 0.21 | 0.31 | 0.6 | 0.82 | 0.26 | 0.01 | 0.32 | -0.1325 | 0.3625 | 0.3725 | 0.341392 | 0.707081 | 0.317682 | 1 | 0.999631 | 0.73317 |
| Rasgrf2_P70392_S801s_1 | 0.89 | 0.23 | 1.07 | 0.51 | -0.45 | 1.14 | 0.81 | -0.06 | 0.7 | 1.38 | 0.38 | 0.57 | -1.19 | 0.04 | 0.88 | -0.05 | -0.315 | 0.0825 | -0.755 | 0.124568 | 0.453507 | 0.806303 | 1 | 0.999631 | 0.596063 |
| Rasgrf2_P70392_S801s_2 | -0.12 | -0.21 | -1.19 | -0.37 | 0.34 | 0.01 | -0.32 | -0.3 | -0.14 | -0.01 | -0.6 | -0.56 | -1.64 | -0.06 | -0.33 | -0.86 | 0.405 | 0.145 | -0.25 | 0.566561 | 0.239126 | 0.666565 | 1 | 0.999631 | 0.84194 |
| Rasgrf2_P70392_S805s_1 | 0.28 | -1.88 | -2.04 | -0.58 | -0.56 | -0.9 | -0.79 | -0.85 | -0.51 | -0.47 | -1.72 | -1.58 | -0.93 | -1.91 | -1.68 | -2.19 | 0.28 | -0.015 | -0.6225 | 0.28766 | 0.591561 | 0.979719 | 1 | 0.999631 | 0.705972 |
| Rasgrf2_P70392_S805s_2 | -0.7 | -0.04 | -1.01 | 0.98 | 0.35 | -0.22 | 1.18 | 1.23 | -0.18 | 0.14 | 0.47 | -0.37 | -1.68 | -0.28 | 0.28 | -0.25 | 0.8275 | 0.2075 | -0.29 | 0.608425 | 0.136751 | 0.655835 | 1 | 0.999631 | 0.858627 |
| Rasgrp1_G3UYC3_S562s_1 | -0.56 | 0.01 | 0.29 | -0.86 | 0.52 | -0.06 | -0.05 | 0.65 | -0.45 | -0.44 | -0.15 | -0.1 | -0.53 | -0.32 | 1.2 | 0.46 | 0.545 | -0.005 | 0.4825 | 0.31385 | 0.143872 | 0.987908 | 1 | 0.999631 | 0.719939 |
| Rasgrp1_G3UYC3_S759s_1 | -0.41 | -1.71 | 0.82 | -0.61 | -1.07 | -1.29 | 0.39 | 0.09 | -1.44 | 0 | 0.13 | 1.38 | 1.28 | -0.37 | -0.37 | -0.5 | 0.0075 | 0.495 | 0.4875 | 0.430952 | 0.990032 | 0.480163 | 1 | 0.999631 | 0.778271 |
| Rasgrp2_Q9QUG9_S585s_1 | 1.02 | 0.56 | -0.02 | 1.13 | 0.93 | -0.16 | 1.12 | 1.53 | 0.66 | 0.37 | 0.92 | 0.1 | 0.29 | 1.21 | 0.34 | 1.3 | 0.1825 | -0.16 | 0.1125 | 0.779465 | 0.676393 | 0.652374 | 1 | 0.999631 | 0.921416 |
| Rasgrp2_Q9QUG9_S586s_1 | 1.02 | 0.56 | 0.05 | 0.91 | 0.93 | -0.16 | 0.82 | 0.93 | 0.66 | 0.37 | 0.76 | 0.34 | 0.29 | 1.21 | 0.17 | 0.97 | -0.005 | -0.1025 | 0.025 | 0.946783 | 0.989043 | 0.741329 | 1 | 0.999631 | 0.98281 |
| Rasip1_Q3U0S6_S286s_1 | -0.15 | -0.81 | 0.41 | 0.52 | -0.99 | -0.96 | 0.4 | 0.2 | -0.86 | -0.3 | 0.53 | 0.03 | -0.59 | 0.16 | 1.09 | 0.71 | -0.33 | -0.1425 | 0.35 | 0.460134 | 0.479411 | 0.737349 | 1 | 0.999631 | 0.791962 |
| Rasip1_Q3U0S6_T960t_1 | 0.54 | 1.56 | 0.79 | 0.62 | 2.55 | 0.42 | 0.92 | 0.71 | 3.12 | 0.92 | 1.88 | 1.74 | 2.74 | 0.09 | 0.35 | 0.64 | 0.2725 | 1.0375 | 0.0775 | 0.895965 | 0.587674 | 0.05632 | 1 | 0.999631 | 0.966232 |
| Rassf5_D3Z6G2_S177s_1 | -0.08 | 0.82 | 0.02 | 0.53 | -0.51 | -0.05 | 0.34 | 0.73 | 0.45 | 0.68 | 0.23 | 0.56 | 0.75 | 0.25 | -0.11 | 1.26 | -0.195 | 0.1575 | 0.215 | 0.58829 | 0.593517 | 0.608171 | 1 | 0.999631 | 0.850634 |
| Raver1_Q9CW46_S14s_1 | -0.64 | 0.42 | 0.34 | 0.34 | 0.38 | 0.78 | -0.13 | 0.58 | -0.38 | 0.32 | 0 | 0.45 | 0.91 | 0.39 | -0.18 | 0.26 | 0.2875 | -0.0175 | 0.23 | 0.544959 | 0.418495 | 0.960245 | 1 | 0.999631 | 0.833658 |
| Raver1_Q9CW46_S576s_1 | 0.59 | 1.74 | 1.55 | 0.85 | 1.06 | 0.66 | 0.82 | 1.69 | 1.04 | 0.47 | 1.73 | 1.91 | 1.13 | 0.91 | 1.17 | 2 | -0.125 | 0.105 | 0.12 | 0.760509 | 0.738172 | 0.807066 | 1 | 0.999631 | 0.914614 |
| Raver1_Q9CW46_S626s_1 | -0.18 | 0.82 | 1.83 | 0.81 | 0.6 | 0.54 | 0.57 | 1.26 | 0.24 | 0.55 | 0.77 | 1.4 | 0.19 | 0.75 | 1.63 | 1.5 | -0.0775 | -0.08 | 0.1975 | 0.697458 | 0.858632 | 0.863095 | 1 | 0.999631 | 0.895731 |

| | | | | | | | | | | | | | | | | | | | | | | | | | |
|---|---|---|---|---|---|---|---|---|---|---|---|---|---|---|---|---|---|---|---|---|---|---|---|---|---|
| Rb1_P13405_S243s_1 | 0.64 | 1.65 | 0.46 | 0.22 | 0.81 | 1.14 | 0.12 | 0.77 | 1.19 | 0.79 | 0.87 | 0.8 | 0.97 | 1.49 | 0.97 | 0.37 | -0.0325 | 0.17 | 0.2075 | 0.613729 | 0.933406 | 0.639512 | 1 | 0.999631 | 0.860866 |
| Rb1cc1_E9PX56_S222s_1 | -0.49 | 0.77 | 1.04 | 0.08 | -0.57 | 0.81 | 0.49 | 0.09 | 0.75 | 0.11 | 0.71 | 1.17 | 0.81 | 0.88 | 1.55 | 0.71 | -0.145 | 0.335 | 0.6375 | 0.145014 | 0.743788 | 0.426894 | 1 | 0.999631 | 0.616069 |
| Rb1cc1_E9PX56_S257s_1 | -0.21 | -0.77 | -0.81 | -0.92 | 0.34 | -0.02 | -0.1 | -0.14 | -0.4 | -1.18 | -1.39 | -1.27 | -0.11 | -0.49 | -1.44 | -1.15 | 0.6975 | -0.3825 | -0.12 | 0.752207 | 0.029026 | 0.261052 | 1 | 0.999631 | 0.913038 |
| Rb1cc1_E9PX56_S261s_1 | 0.49 | 0.46 | 1.64 | 1.85 | 1.15 | 1 | 0.67 | 1.2 | 0.77 | 0.41 | -0.32 | 0.47 | -0.48 | 0.35 | 2.19 | -1.77 | -0.105 | -0.7775 | -1.0375 | 0.210211 | 0.790436 | 0.098 | 1 | 0.999631 | 0.66441 |
| Rb1cc1_E9PX56_S629s_1 | -1.95 | -1.3 | -1.16 | -2.45 | -2.08 | -0.79 | -2.56 | -2.04 | -1.62 | -2.14 | -1.49 | -1.07 | -0.21 | -1.01 | -2.36 | -1.71 | -0.1525 | 0.135 | 0.3925 | 0.457363 | 0.741928 | 0.729445 | 1 | 0.999631 | 0.790722 |
| Rb1cc1_E9PX56_S629s_2 | 0.2 | -1.25 | -1.82 | -0.83 | -0.82 | -0.68 | -0.38 | -0.97 | -0.11 | -0.23 | -1.26 | -1.16 | 0.22 | -0.45 | -1.78 | -0.77 | 0.2125 | 0.235 | 0.23 | 0.67896 | 0.626884 | 0.637879 | 1 | 0.999631 | 0.887183 |
| Rb1cc1_E9PX56_S635s_2 | 0.2 | -1.25 | -1.82 | -0.83 | -0.82 | -0.68 | -0.38 | -0.97 | -0.11 | -0.23 | -1.26 | -1.16 | 0.22 | -0.45 | -1.78 | -0.77 | 0.2125 | 0.235 | 0.23 | 0.67896 | 0.626884 | 0.637879 | 1 | 0.999631 | 0.887183 |
| Rb1cc1_E9PX56_S649s_1 | -1.42 | -0.77 | -0.34 | -0.41 | -0.42 | -1.41 | -0.47 | -0.41 | -0.34 | -1.18 | 1.28 | 0.08 | 0.69 | -2.05 | 0.13 | -0.34 | 0.0575 | 0.695 | 0.3425 | 0.562814 | 0.875613 | 0.210974 | 1 | 0.999631 | 0.841038 |
| Rbbp6_P97868_S1244s_1 | -1.4 | -0.94 | -1.32 | -1.05 | -0.79 | -1.38 | -1.21 | -1.34 | -0.78 | -1.06 | -1.66 | -0.88 | 0.2 | -0.91 | -0.42 | -1.26 | -0.0025 | 0.0825 | 0.58 | 0.144791 | 0.992554 | 0.78414 | 1 | 0.999631 | 0.616037 |
| Rbl2_Q64700_S37s_1 | 0.82 | 1.01 | 0.56 | 0.85 | 0.75 | 0.23 | 0.66 | 0.7 | 0.46 | 1.06 | 0.89 | 1.18 | 0.64 | 0.92 | 1.19 | 1.18 | -0.225 | 0.0875 | 0.1725 | 0.550252 | 0.394118 | 0.755213 | 1 | 0.999631 | 0.836038 |
| Rbl2_Q64700_S946s_1 | 1.34 | -2.24 | 1.19 | 0.5 | -1.22 | -0.96 | 2.78 | 0.42 | 0.6 | -0.37 | 0.81 | 3.42 | 0.47 | -0.86 | 0.59 | 0.98 | 0.0575 | 0.9175 | 0.0975 | 0.903527 | 0.956757 | 0.371145 | 1 | 0.999631 | 0.968183 |
| Rbl2_Q64700_S960s_1 | -0.11 | 3.32 | 3.25 | 1.7 | 3.47 | 2.85 | 2.53 | 1.92 | 3.24 | 0.58 | 3.48 | 2.7 | 2.63 | 2.18 | 2.93 | 1.02 | 0.6525 | 0.46 | 0.15 | 0.850236 | 0.401855 | 0.614313 | 1 | 0.999631 | 0.949525 |
| Rbm10_Q99KG3_S646s_1 | 0 | 1.2 | 1.63 | 0.74 | 0.72 | 0.58 | 0.52 | 1.16 | 0.9 | 0.74 | 1.29 | 1.35 | 1.04 | 1.33 | 1.23 | 1.33 | -0.1475 | 0.1775 | 0.34 | 0.389322 | 0.704293 | 0.654006 | 1 | 0.999631 | 0.758621 |
| Rbm14_E9QL13_S582s_1 | 0.48 | 0.74 | 1.14 | 0.72 | 0.11 | 0.33 | 0.26 | 0.94 | 0.47 | 1.3 | 0.73 | 0.91 | 0.86 | 1.13 | 1.09 | 1.42 | -0.36 | 0.0825 | 0.355 | 0.243942 | 0.240858 | 0.782841 | 1 | 0.999631 | 0.685637 |
| Rbm14_E9QL13_T206t_1 | 0.06 | 0.88 | 0.99 | 0.26 | 0.27 | 0.44 | -0.06 | 0.54 | 0.69 | 0.66 | 0.66 | 0.83 | 1.2 | 1.08 | 0.93 | 1.32 | -0.25 | 0.1625 | 0.585 | 0.094595 | 0.4353 | 0.596172 | 1 | 0.999631 | 0.558128 |
| Rbm15_Q0VBL3_S293s_1 | -0.21 | 1.03 | 1.47 | 1.24 | 0.55 | 0.59 | 0.36 | 0.92 | 0.84 | 0.93 | 2.06 | 2.51 | 0.59 | 0.81 | 1.63 | 1.8 | -0.2775 | 0.7025 | 0.325 | 0.494283 | 0.491149 | 0.201072 | 1 | 0.999631 | 0.81007 |
| Rbm15_Q0VBL3_S655s_1 | 1.5 | 1.77 | 2.01 | 0.79 | 1.96 | 1.1 | 1.99 | 0.58 | 3.11 | 1.77 | 1.86 | 2.05 | 3.33 | 0.04 | 0.44 | 0.19 | -0.11 | 0.68 | -0.5175 | 0.48234 | 0.796903 | 0.124602 | 1 | 0.999631 | 0.802587 |
| Rbm15_Q0VBL3_S674s_1 | 1.66 | 1.43 | 2.37 | 1.81 | 0.5 | 0.77 | 1.86 | 0.12 | 1.41 | 2.1 | 1.28 | 1.03 | -0.1 | 0.24 | -0.13 | 1.86 | -1.005 | -0.3625 | -1.35 | 0.019631 | 0.035816 | 0.308703 | 1 | 0.999631 | 0.391668 |
| Rbm15_Q0VBL3_S782s_1 | 1.09 | 1.2 | 0.22 | 0.02 | 1.07 | 1.31 | -0.15 | 0.29 | 2.18 | 0.62 | -0.71 | 0.23 | -1.02 | 0.86 | 1.28 | -1.18 | -0.0025 | -0.0525 | -0.6475 | 0.318653 | 0.995465 | 0.93144 | 1 | 0.999631 | 0.722158 |
| Rbm15_Q0VBL3_T567t_1 | 0.25 | 2.67 | 2.16 | -0.39 | 3.11 | 2.75 | 1.17 | 1.68 | 4 | 0.14 | 2.92 | 3.19 | 3.37 | 3.07 | 2.45 | 1.8 | 1.005 | 1.39 | 1.5 | 0.05981 | 0.205792 | 0.17291 | 1 | 0.999631 | 0.50521 |
| Rbm15b_Q6PHZ5_S107s_1 | 1.89 | -0.17 | 1.27 | 0.77 | -1.34 | 0.84 | 1.54 | 0.05 | 0.59 | 1.46 | 0.05 | 0.07 | -1.15 | -1.06 | 0.31 | -0.34 | -0.6675 | -0.3975 | -1.5 | 0.015679 | 0.333171 | 0.446954 | 1 | 0.999631 | 0.388832 |
| Rbm15b_Q6PHZ5_S111s_1 | -0.5 | 0.3 | 0.23 | 0.01 | 1.03 | 0.59 | -0.74 | -1.17 | 0.36 | -0.33 | -0.64 | -0.24 | 0.66 | 0.85 | -0.66 | 0.36 | -0.0825 | -0.2225 | 0.2925 | 0.474963 | 0.873023 | 0.504416 | 1 | 0.999631 | 0.799827 |
| Rbm15b_Q6PHZ5_S261s_1 | 0.25 | 1.36 | -1.47 | 0.38 | 0.24 | 0.26 | -1.69 | -2.7 | 0.54 | 0.09 | -2.19 | -0.55 | 0.65 | 0.55 | -0.14 | -3.48 | -1.1025 | -0.6575 | -0.735 | 0.457418 | 0.199953 | 0.3853 | 1 | 0.999631 | 0.790722 |
| Rbm15b_Q6PHZ5_S263s_1 | 0.2 | -1.53 | 1.29 | 1.14 | -1.44 | -3.56 | 0.61 | 1.11 | -1.29 | -3.47 | 1.39 | 1.19 | -1.68 | -1.51 | 1.13 | 1.71 | -1.095 | -0.82 | -0.3625 | 0.700111 | 0.319798 | 0.475266 | 1 | 0.999631 | 0.896612 |
| Rbm15b_Q6PHZ5_S606s_1 | 1.22 | -2.14 | -0.83 | -1.36 | -2.81 | -1.1 | 0.99 | -5.19 | -1.19 | 0.14 | 0.04 | -0.9 | -2.72 | -2.96 | -1.32 | -1.49 | -1.25 | 0.3 | -1.345 | 0.090616 | 0.338215 | 0.671548 | 1 | 0.999631 | 0.557085 |
| Rbm15b_Q6PHZ5_T234t_1 | -1.89 | -0.25 | -0.05 | -1.24 | -0.16 | -0.31 | -0.59 | -0.87 | -0.27 | -2.22 | -0.23 | 0.58 | 1.53 | -0.86 | -0.65 | -1.38 | 0.375 | 0.3225 | 0.5175 | 0.458311 | 0.408439 | 0.626868 | 1 | 0.999631 | 0.790722 |
| Rbm15b_Q6PHZ5_T529t_1 | -0.87 | 2.6 | 2.73 | 0.93 | 2.28 | 1.6 | 1.27 | 2.02 | 2.56 | -0.41 | 2.59 | 2.99 | 3.33 | 1.62 | 2.87 | 1.64 | 0.445 | 0.585 | 1.0175 | 0.235998 | 0.564814 | 0.561001 | 1 | 0.999631 | 0.683301 |
| Rbm17_Q8JZX4_S155s_1 | -0.44 | 0.71 | 0.8 | 0.38 | 0.46 | 0.33 | 0.3 | 0.38 | 0.55 | 0.5 | 0.4 | 0.63 | 0.78 | 0.54 | 0.81 | 0.62 | 0.005 | 0.1575 | 0.325 | 0.361518 | 0.987808 | 0.640225 | 1 | 0.999631 | 0.741948 |
| Rbm17_Q8JZX4_S222s_1 | -0.06 | 0.67 | 0.6 | -0.19 | 0.13 | 0.01 | -0.01 | 0.05 | 0.16 | 0.43 | -0.26 | -0.2 | 0.57 | 0.66 | 0.15 | 0.62 | -0.21 | -0.2225 | 0.245 | 0.460661 | 0.479222 | 0.502703 | 1 | 0.999631 | 0.791962 |
| Rbm17_Q8JZX4_T71t_1 | 0.69 | 2.1 | 1.94 | 0.98 | 1.15 | 1.22 | 0.47 | 1.39 | 1.51 | 1.67 | 1.37 | 1.84 | 1.58 | 2.68 | 1.17 | 2.09 | -0.37 | 0.17 | 0.4525 | 0.345833 | 0.371784 | 0.659112 | 1 | 0.999631 | 0.735066 |
| Rbm20_Q3UQS8_S1184s_1 | 1.6 | 0.76 | 1.86 | 1.77 | 1.1 | 1.35 | 1.63 | 0.66 | 1.35 | 2.06 | 1.26 | 0.47 | 0.33 | 0.63 | 0.74 | -0.37 | -0.3125 | -0.2125 | -1.165 | 0.01221 | 0.385676 | 0.612089 | 1 | 0.999631 | 0.385092 |
| Rbm25_B2RY56_S578s_1 | 0.38 | -0.22 | 2.2 | 1.59 | -0.15 | 0.56 | 1.55 | 1.1 | 0.47 | 0.86 | 0.92 | 1.28 | 1.51 | -0.59 | 1.48 | 0.31 | -0.2225 | -0.105 | -0.31 | 0.644849 | 0.711056 | 0.845065 | 1 | 0.999631 | 0.872824 |
| Rbm25_B2RY56_S672s_1 | -0.09 | 1.03 | 1.08 | 0.53 | -0.23 | 0.35 | 0.22 | -0.71 | 1.42 | 0.86 | 0.82 | 1.04 | 1.12 | 0.4 | 1.56 | 1.07 | -0.73 | 0.3975 | 0.4 | 0.320273 | 0.077783 | 0.267737 | 1 | 0.999631 | 0.723684 |
| Rbm26_E9PUF4_S518s_1 | 0.32 | 1.68 | 2.46 | 1.21 | 0.73 | 1.58 | 1.31 | 1.4 | 1.32 | 1.33 | 2.12 | 1.99 | 1.28 | 0.93 | 2 | 2.45 | -0.1625 | 0.2725 | 0.2475 | 0.642435 | 0.7264 | 0.570735 | 1 | 0.999631 | 0.871335 |
| Rbm26_E9PUF4_S616s_1 | 0.42 | 1.15 | 1.28 | 0.35 | 0.4 | 0.76 | 0.28 | 0.64 | -0.54 | 0.19 | 0.19 | -0.16 | -0.27 | 0.71 | 0.33 | 1.11 | -0.28 | -0.88 | -0.33 | 0.420773 | 0.385542 | 0.026363 | 1 | 0.999631 | 0.772817 |
| Rbm27_Q5SFM8_S120s_1 | 1.29 | 0.75 | 1.53 | 2.13 | 0.77 | 0.64 | 1.55 | 1.47 | 1.21 | 1.9 | 1.57 | 1.29 | 0.53 | 0.35 | 1.64 | 1.63 | -0.3175 | 0.0675 | -0.3875 | 0.397272 | 0.414804 | 0.851077 | 1 | 0.999631 | 0.761181 |
| Rbm28_Q8CGC6_S252s_1 | 0.56 | 0.48 | -0.1 | -0.32 | 0.25 | 0.8 | -0.7 | 0.54 | 0.05 | 0.29 | 0.48 | 0.93 | 1.76 | 0.72 | 0.43 | 0.61 | 0.0675 | 0.2825 | 0.725 | 0.091198 | 0.865185 | 0.407289 | 1 | 0.999631 | 0.557085 |
| Rbm33_D3Z5I9_S203s_1 | -2.07 | 2.85 | 4.99 | 2.25 | 2.58 | 2.14 | 2.81 | 3.08 | 2.75 | -0.69 | 4.94 | 5.48 | 1.83 | 2.51 | 5.06 | 0.9 | 0.6475 | 1.115 | 0.57 | 0.695569 | 0.612995 | 0.523093 | 1 | 0.999631 | 0.895676 |
| Rbm33_D3Z5I9_S992s_1 | 1.56 | 1.05 | 1.69 | 1.29 | 0.33 | 1.58 | 1.88 | 0.93 | 1.03 | 1.57 | 0.51 | 0.55 | 0.46 | 0.08 | 1.05 | 1.06 | -0.2175 | -0.4825 | -0.735 | 0.053521 | 0.575372 | 0.172355 | 1 | 0.999631 | 0.4909 |
| Rbm39_B7ZD61_S135s_1 | -0.44 | 1.6 | 1.45 | 0.06 | 1.3 | 1.39 | 0.4 | 1.13 | 1.46 | -0.22 | 1.42 | 1.89 | 1.91 | 1.41 | 1.79 | 1.14 | 0.3875 | 0.47 | 0.895 | 0.103757 | 0.459075 | 0.457014 | 1 | 0.999631 | 0.567918 |
| Rbm39_B7ZD61_S97s_1 | 1.11 | 1.46 | 1.7 | -0.13 | 0.66 | 1.43 | 0.35 | 0.57 | 1.46 | 1.63 | 0.97 | 0.83 | 2.19 | 1.83 | 0.97 | 1.64 | -0.2825 | 0.1875 | 0.6225 | 0.205925 | 0.536975 | 0.673729 | 1 | 0.999631 | 0.6605 |
| Rbm45_Q8BHN5_S199s_1 | 0.42 | 0.79 | 0.95 | 0.31 | 0.45 | -0.13 | -0.1 | 0.5 | 1.21 | 1.2 | 0.75 | 0.44 | 0.3 | 0.58 | 0.99 | 0.19 | -0.4375 | 0.2825 | -0.1025 | 0.748147 | 0.161196 | 0.371211 | 1 | 0.999631 | 0.911931 |
| Rbm5_S4R1N4_S59s_1 | 0.01 | 3.54 | 4.34 | 0.84 | 3.27 | 2.64 | 4.03 | 2.04 | 4.45 | 0.02 | 4.3 | 4.48 | 3.14 | 0.68 | 3.15 | 0.2 | 0.8125 | 1.13 | -0.39 | 0.725783 | 0.410952 | 0.391633 | 1 | 0.999631 | 0.90568 |
| Rbm8a_Q9CWZ3_S42s_1 | 1.55 | 0.22 | 1.24 | 1.59 | 0.72 | 1.22 | 1.33 | 1.63 | 0.81 | 1.98 | 1.82 | 1.14 | 0.17 | 0.53 | 1.27 | 1.2 | 0.075 | 0.2875 | -0.3575 | 0.410235 | 0.844951 | 0.503612 | 1 | 0.999631 | 0.767924 |
| Rbm8a_Q9CWZ3_S42s_1_1 | 0.81 | 0.57 | 1.06 | 0.6 | 0.17 | 0.51 | 0.6 | 0.14 | 1.11 | 1.36 | 0.68 | 0.87 | 0.43 | 0.08 | 0.5 | 0.58 | -0.405 | 0.245 | -0.3625 | 0.222831 | 0.147862 | 0.393661 | 1 | 0.999631 | 0.673942 |
| Rbm8a_Q9CWZ3_S56s_1 | -0.52 | 0.63 | -0.68 | 0.16 | 0.01 | -0.22 | -0.39 | 0.83 | -0.09 | -0.27 | 0.29 | 0.23 | 0.81 | 0.25 | 0.98 | 0.01 | 0.16 | 0.1425 | 0.615 | 0.150245 | 0.695971 | 0.696069 | 1 | 0.999631 | 0.622782 |
| Rbmxl1_Q91VM5_S205s_1 | -1.03 | 0.1 | 0.19 | -0.25 | -0.28 | -0.06 | -0.65 | -0.25 | -0.15 | -0.14 | -0.07 | -0.05 | 0.59 | 0.29 | 0.12 | 0.45 | -0.0625 | 0.145 | 0.61 | 0.106884 | 0.853874 | 0.662409 | 1 | 0.999631 | 0.568797 |
| Rbmxl1_Q91VM5_S246s_1 | 0.3 | 0.69 | 0.92 | -0.48 | 0.97 | 0.49 | 1.66 | 0.76 | 0.97 | -0.64 | 2.27 | 2.37 | 1.71 | -0.39 | 1.56 | -0.49 | 0.6125 | 0.885 | 0.24 | 0.695876 | 0.149544 | 0.216719 | 1 | 0.999631 | 0.895683 |
| Rbmxl1_Q91VM5_S323s_1 | 0.03 | -1.41 | -0.63 | -0.07 | -0.81 | -0.32 | -0.07 | -0.47 | -0.74 | -0.15 | -0.22 | -0.23 | -0.83 | -1.35 | -1.47 | -1.92 | 0.1025 | 0.185 | -0.8725 | 0.05795 | 0.786967 | 0.628099 | 1 | 0.999631 | 0.498591 |
| Rbmxl1_Q91VM5_S349s_1 | 0.05 | -0.09 | 0.41 | 0.02 | -0.65 | -0.26 | -0.47 | 0 | -0.38 | 0.54 | -0.02 | 0.31 | -0.32 | 0.25 | 0.19 | 0.25 | -0.4425 | 0.015 | -0.005 | 0.986339 | 0.125305 | 0.96038 | 1 | 0.999631 | 0.995135 |
| Rbmxl1_Q91VM5_S88s_1 | 0.06 | 0.83 | 0.48 | -0.11 | 0.21 | 0.4 | 0.27 | 0.15 | 1.03 | 0.68 | 0.55 | 0.87 | 0.53 | 1.78 | 1.12 | 1.31 | -0.0575 | 0.4675 | 0.87 | 0.039603 | 0.842899 | 0.151113 | 1 | 0.999631 | 0.470002 |
| Rbmxl1_Q91VM5_T163t_1 | 0.1 | -1.24 | 0.35 | -0.06 | -0.52 | -0.86 | 0.01 | -0.02 | -0.89 | 0.39 | 0.02 | -0.24 | -0.37 | -0.45 | 0.17 | 0.13 | -0.135 | 0.0325 | 0.0825 | 0.839865 | 0.74265 | 0.940998 | 1 | 0.999631 | 0.946365 |
| Rbsn_Q80Y56_S214s_1 | -0.87 | -0.73 | -0.2 | -1.23 | -1.64 | -1.48 | 0 | -0.04 | -2.68 | -1.66 | -0.43 | -0.9 | -0.7 | -1.54 | -1.69 | -0.09 | -0.0325 | -0.66 | -0.2475 | 0.573856 | 0.94494 | 0.212694 | 1 | 0.999631 | 0.84468 |
| Rbsn_Q80Y56_S214s_2 | -1.56 | 0.62 | 0.5 | -1.61 | -0.71 | -0.81 | 0.15 | -0.76 | -0.54 | -0.65 | -0.72 | 0.4 | 0.31 | -0.28 | 0.77 | 0.51 | -0.02 | 0.135 | 0.84 | 0.183137 | 0.973261 | 0.825217 | 1 | 0.999631 | 0.644372 |
| Rbsn_Q80Y56_S216s_2 | -1.86 | 0.57 | -1.48 | -1.33 | -0.71 | -0.9 | -0.87 | -2.46 | -0.56 | -0.81 | -2.73 | -1.6 | 0.33 | -0.22 | -1.41 | -0.61 | -0.21 | -0.4 | 0.5475 | 0.36995 | 0.732577 | 0.545917 | 1 | 0.999631 | 0.746338 |
| Rbsn_Q80Y56_S218s_1 | -1.08 | -2.74 | -0.64 | -1.1 | -2.32 | -1.14 | -0.18 | -0.25 | -2.59 | -0.66 | -0.65 | -0.52 | 1.68 | -0.39 | -0.49 | -1.77 | 0.4175 | 0.285 | 1.1475 | 0.147814 | 0.500995 | 0.644871 | 1 | 0.999631 | 0.620569 |
| Rcan1_Q9JHG6_S32s_1 | -1.36 | -0.64 | -0.98 | -0.96 | -0.93 | -1.17 | -1.07 | -0.89 | -0.96 | -1.48 | -0.62 | -0.71 | -0.34 | -0.73 | -0.96 | -0.6 | -0.03 | 0.0425 | 0.3275 | 0.291107 | 0.908984 | 0.890998 | 1 | 0.999631 | 0.706164 |
| Rcc1_Q6PFB2_S11s_1 | 0.1 | 0.44 | 0.36 | 0.39 | -0.19 | -0.17 | -0.32 | 0.23 | 0.18 | 0.29 | -0.23 | -0.01 | 0.44 | 0.27 | 0.03 | 0.61 | -0.435 | -0.265 | 0.015 | 0.957355 | 0.112209 | 0.323523 | 1 | 0.999631 | 0.985227 |
| Rcor1_Q8CFE3_S454s_1 | 1.25 | 0.86 | 1.53 | 1.28 | 0.49 | 0.82 | 1.29 | 0.96 | 0.51 | 1.51 | 0.93 | 0.65 | 0.73 | 0.83 | 1.59 | 1.14 | -0.34 | -0.33 | -0.1575 | 0.625239 | 0.256299 | 0.316616 | 1 | 0.999631 | 0.863768 |
| Rcsd1_Q3UZA1_S149s_1 | 1.1 | 1.14 | 1.63 | 1.72 | 0.31 | 1.51 | 1.02 | 1.01 | 1.21 | 1.76 | 0.92 | 1.39 | 0.9 | 0.8 | 1.23 | 0.47 | -0.435 | -0.0775 | -0.5475 | 0.105596 | 0.212603 | 0.801491 | 1 | 0.999631 | 0.568797 |
| Rcsd1_Q3UZA1_S303s_1 | -0.67 | -0.63 | 0.28 | -0.08 | -1.05 | -0.5 | -0.08 | -0.73 | -0.41 | -0.44 | -0.66 | 0.46 | -0.1 | -0.29 | -0.32 | 0.38 | -0.315 | 0.0125 | 0.1925 | 0.577597 | 0.366572 | 0.972763 | 1 | 0.999631 | 0.84622 |
| Rcsd1_Q3UZA1_S90s_1 | 0.04 | 0.38 | -0.48 | -1.71 | -1.01 | -0.63 | -2.09 | -0.94 | -1.68 | 0.06 | -1.96 | -0.54 | 0.66 | 0.11 | -1.03 | -0.23 | -0.725 | -0.5875 | 0.32 | 0.557588 | 0.183848 | 0.340977 | 1 | 0.999631 | 0.838305 |
| Reep1_Q8BGH4_S152s_1 | 0.96 | 0.43 | -0.41 | 0.2 | 0.82 | 0.29 | 0.35 | 0.28 | 0.47 | 0.24 | 0.21 | 0.22 | -0.25 | 0.63 | -0.13 | 0.61 | 0.14 | -0.01 | -0.08 | 0.839738 | 0.685509 | 0.97635 | 1 | 0.999631 | 0.946365 |
| Reep1_Q8BGH4_S192s_1 | 1.69 | -1.16 | -1.49 | -0.4 | -1.56 | -0.16 | 1.23 | -1.21 | -0.45 | 1.15 | -1.61 | -2.25 | -0.27 | -1.5 | -1.67 | -2.14 | -0.085 | -0.45 | -1.055 | 0.166238 | 0.91812 | 0.618535 | 1 | 0.999631 | 0.63587 |
| Reep2_Q8VCD6_S201s_1 | 2.66 | -0.63 | -0.6 | 0.87 | 0.35 | 0.76 | 1.09 | -0.03 | 1.02 | 1.08 | 0.22 | -0.12 | -1.65 | -0.14 | -1.01 | -1.01 | -0.0325 | -0.025 | -1.5275 | 0.061144 | 0.963928 | 0.972923 | 1 | 0.999631 | 0.509355 |
| Rell2_E9Q812_S104s_1 | -0.5 | -0.81 | -0.54 | -0.5 | -1.37 | -0.09 | -0.87 | -0.73 | -1.28 | -0.3 | -0.66 | -0.99 | 0.89 | -0.48 | -0.65 | -0.79 | -0.1775 | -0.22 | 0.33 | 0.431172 | 0.583485 | 0.470071 | 1 | 0.999631 | 0.778271 |
| Rell2_E9Q812_S96s_1 | -0.33 | -0.92 | -1.29 | -1.04 | -1.3 | -1.07 | -0.78 | -2.58 | -1.1 | -0.02 | -1.05 | -0.39 | -0.1 | -1.04 | -1.23 | -1.05 | -0.5375 | 0.255 | 0.04 | 0.913797 | 0.236044 | 0.488526 | 1 | 0.999631 | 0.972551 |
| Rem2_E9Q4D5_S27s_1 | -0.2 | -0.32 | 0.49 | 0.94 | -0.61 | -1.02 | 0 | 1.09 | -0.22 | -1.04 | -0.32 | 0.97 | 0.34 | 0.23 | 1.93 | 1.09 | -0.3625 | -0.38 | 0.67 | 0.183574 | 0.4831 | 0.44398 | 1 | 0.999631 | 0.644372 |
| Rem2_E9Q4D5_S296s_1 | -0.29 | 0.81 | 3.06 | -0.18 | 1.59 | 0.53 | 2.5 | 1.92 | 2.63 | -0.72 | 3.27 | 4.39 | 2.7 | 0.03 | 2.24 | -0.09 | 0.785 | 1.5425 | 0.37 | 0.687462 | 0.322093 | 0.202334 | 1 | 0.999631 | 0.891556 |
| Rem2_E9Q4D5_S69s_1 | 1.02 | 1.19 | 0.94 | 1.39 | 1.41 | 0.86 | 0.71 | 1.83 | 1.5 | 1.34 | 1.48 | 1.63 | 0.77 | 1.67 | 1.84 | 1.51 | 0.0675 | 0.3525 | 0.3125 | 0.354698 | 0.834312 | 0.186726 | 1 | 0.999631 | 0.739951 |
| Reps1_O54916_S509s_1 | 1.88 | 1.16 | 0.05 | 0.3 | 0.83 | 1.44 | 0.2 | 0.19 | 1.59 | 1.45 | 0.33 | 0.68 | 1.07 | 0.38 | 0.37 | -0.11 | -0.1825 | 0.165 | -0.42 | 0.384051 | 0.707834 | 0.738 | 1 | 0.999631 | 0.754721 |
| Reps1_O54916_S93s_1 | 0.16 | -0.32 | -0.01 | -0.4 | -0.45 | -0.06 | 0.54 | -0.46 | -0.08 | 0.2 | 0.17 | 0.58 | -0.33 | -0.69 | -0.41 | 0.03 | 0.035 | 0.36 | -0.2075 | 0.496954 | 0.912426 | 0.222687 | 1 | 0.999631 | 0.811348 |
| Reps2_B9EI38_S481s_1 | 1.93 | 1.75 | 0.78 | 0.63 | 2.8 | 1.8 | 0.26 | 0.63 | 0.99 | 2.2 | 0.49 | 0.59 | 1.83 | 0.93 | 0.55 | 0.7 | 0.1 | -0.205 | -0.27 | 0.544229 | 0.867964 | 0.676955 | 1 | 0.999631 | 0.833226 |
| Reps2_B9EI38_S538s_1 | 0.77 | 0.48 | 1.36 | 0.13 | 0.13 | -0.14 | 0.24 | -0.48 | 0.64 | 1 | -1.07 | 0.26 | 0.26 | -0.3 | 0 | -0.42 | -0.7475 | -0.4775 | -0.8 | 0.044809 | 0.049979 | 0.347544 | 1 | 0.999631 | 0.475443 |
| Reps2_B9EI38_S619s_1 | 0.21 | 0.74 | 0.25 | 1.43 | 1.02 | 0.56 | 0.84 | 1.42 | 0.27 | -0.88 | 0.4 | 0.15 | 0.26 | 0.28 | 0.67 | 0.86 | 0.3025 | -0.6725 | -0.14 | 0.703093 | 0.409831 | 0.128967 | 1 | 0.999631 | 0.896857 |
| Reps2_B9EI38_T467t_1 | 0.05 | 0.29 | 0.32 | -0.12 | 0.04 | 0.12 | -0.35 | 0.17 | -0.07 | -0.12 | -0.58 | -0.25 | -0.02 | -0.27 | 0.28 | 0.46 | -0.14 | -0.39 | -0.0225 | 0.940015 | 0.594622 | 0.168397 | 1 | 0.999631 | 0.981146 |
| Rer1_Q9CQU3_S2s_1 | 0.93 | 1.17 | 1.36 | 1.5 | 0.8 | 1.54 | 0.93 | 1.54 | 0.49 | 2.23 | 1.36 | 0.94 | 1.57 | 1.48 | 1.64 | 2.09 | -0.0375 | 0.015 | 0.455 | 0.146601 | 0.899627 | 0.970203 | 1 | 0.999631 | 0.618444 |
| Rftn2_Q8CHX7_S228s_1 | -0.24 | -0.41 | 0.25 | -0.73 | -0.89 | 0.07 | -0.22 | -0.76 | -0.08 | 0.11 | -0.09 | -1.07 | 0.32 | -0.24 | -0.83 | -0.55 | -0.1675 | -7.9E-17 | -0.0425 | 0.907502 | 0.624329 | 1 | 1 | 1 | 0.96989 |
| Rftn2_Q8CHX7_S429s_1 | 0.37 | 0.46 | -0.15 | -0.21 | 0.23 | 0.57 | -0.66 | 0.01 | 0.01 | 0.25 | -0.37 | 0.33 | 0.42 | 0.94 | -0.26 | 0.93 | -0.08 | -0.0625 | 0.39 | 0.309305 | 0.816465 | 0.837884 | 1 | 0.999631 | 0.718845 |
| Rftn2_Q8CHX7_T408t_1 | -1.77 | -2.97 | -1.49 | -1.8 | -1.9 | -2.31 | -2.09 | -2.45 | -1.94 | -1.65 | -2.07 | -1.68 | -0.78 | -1.64 | -1.53 | -2.09 | -0.18 | 0.1725 | 0.4975 | 0.266937 | 0.627707 | 0.643626 | 1 | 0.999631 | 0.695664 |
| Rfx3_G3UWP0_S155s_1 | -0.21 | 0.45 | 0.58 | -0.39 | 0.53 | 1.54 | -0.09 | -0.19 | 1.25 | 0.56 | 0.25 | 0.74 | 0.64 | 1.04 | 0.07 | 1.27 | 0.34 | 0.5925 | 0.6475 | 0.118617 | 0.455601 | 0.119414 | 1 | 0.999631 | 0.585512 |
| Rgcc_Q9DBX1_S97s_1 | -0.78 | -0.25 | 0.36 | -0.85 | -0.23 | -2.87 | -0.48 | -0.55 | 0.37 | -1.33 | -0.38 | -0.3 | 1.11 | -0.7 | -0.65 | -1.07 | -0.6525 | -0.03 | 0.0525 | 0.920975 | 0.299681 | 0.945917 | 1 | 0.999631 | 0.975032 |
| Rgl1_E9Q8N0_S518s_1 | 0.29 | -0.12 | -0.32 | 1.48 | 1.29 | 0.63 | 0.85 | 1.61 | 0.59 | -0.35 | 1.49 | 0.62 | -1.7 | 1.08 | 0.8 | 1.52 | 0.7625 | 0.255 | 0.0925 | 0.899132 | 0.11163 | 0.624549 | 1 | 0.999631 | 0.967236 |
| Rgl3_Q3UYI5_S707s_1 | -0.61 | -0.46 | -0.42 | -0.56 | -1.2 | -0.6 | -0.41 | -1.04 | -1.96 | 0.24 | -0.97 | -1.79 | 0.78 | -0.13 | -2.08 | -1.42 | -0.3 | -0.6075 | -0.2 | 0.734552 | 0.291597 | 0.227776 | 1 | 0.999631 | 0.907462 |
| Rgma_Q8C4C4_S26s_1 | -0.55 | -0.41 | 0.24 | -0.15 | -0.35 | -1.2 | -0.21 | 0.87 | -0.74 | 0.04 | 0.29 | 0.85 | 0.32 | -0.06 | 0.23 | 0.38 | -0.005 | 0.3275 | 0.435 | 0.171857 | 0.991027 | 0.409964 | 1 | 0.999631 | 0.641657 |
| Rgs12_E9Q652_S1125s_1 | 3.16 | 0.81 | 0.47 | 2.22 | 1.4 | 1.87 | 2.24 | 1.65 | 1.39 | 1.74 | 1 | -0.02 | -2.35 | -0.05 | 0.34 | 0.04 | 0.125 | -0.6375 | -2.17 | 0.017308 | 0.832185 | 0.344569 | 1 | 0.999631 | 0.391024 |
| Rgs12_E9Q652_S1130s_1 | 1.98 | 0.08 | 0.88 | -0.64 | -0.47 | 0.56 | 1.27 | -0.74 | -0.22 | 1.41 | -0.08 | -0.3 | -0.3 | -0.56 | 0.3 | -2.17 | -0.42 | -0.3725 | -1.2575 | 0.090951 | 0.523521 | 0.555741 | 1 | 0.999631 | 0.557085 |
| Rgs12_E9Q652_S171s_1 | 0.12 | 0.06 | 0.29 | 0.65 | 0.44 | -0.17 | 0.54 | 1.07 | -0.13 | 0.23 | 1.28 | 1.26 | -0.65 | 0.35 | 0.34 | 1.15 | 0.19 | 0.38 | 0.0175 | 0.965943 | 0.566738 | 0.350491 | 1 | 0.999631 | 0.988179 |
| Rgs12_E9Q652_S850s_1 | -0.56 | -0.25 | -0.16 | -0.63 | -0.61 | -0.38 | -0.61 | -0.73 | -0.4 | -0.05 | -0.31 | -0.29 | 0 | -0.44 | 0.16 | 0.21 | -0.1825 | 0.1375 | 0.3825 | 0.217075 | 0.476297 | 0.601131 | 1 | 0.999631 | 0.669465 |
| Rgs12_E9Q652_S879s_1 | -1.06 | -0.31 | 0.5 | 0.52 | -0.42 | -0.34 | -0.21 | 0.14 | -0.89 | -0.4 | 0.12 | 0.28 | -0.32 | -0.44 | 0.31 | 0.71 | -0.12 | -0.135 | 0.1525 | 0.740396 | 0.765133 | 0.765571 | 1 | 0.999631 | 0.90883 |
| Rgs14_P97492_S203s_1 | 0.62 | 0.62 | 0.13 | 0.13 | -0.13 | 0.29 | -0.11 | 0.07 | 0.02 | 0.85 | -0.02 | 0.05 | 0.14 | 0.67 | 0 | 0.34 | -0.345 | -0.15 | -0.0875 | 0.773736 | 0.216712 | 0.636271 | 1 | 0.999631 | 0.920922 |
| Rgs14_P97492_S218s_1 | 0.36 | 0.8 | 0.81 | 1.11 | 0.45 | 0.79 | 0.42 | 1.24 | 0.8 | 0.92 | 1.1 | 0.92 | 0.1 | 1.7 | 1.67 | 1.1 | -0.045 | 0.165 | 0.3725 | 0.383405 | 0.882624 | 0.549049 | 1 | 0.999631 | 0.754581 |
| Rgs14_P97492_S289s_1 | 1.69 | 1.06 | 1 | 1.49 | 1 | 1.37 | 1.47 | 0.62 | 1.69 | 1.77 | 1.31 | 0.92 | -0.48 | 0.58 | 0.6 | 0.54 | -0.195 | 0.1125 | -1 | 0.018484 | 0.534259 | 0.724328 | 1 | 0.999631 | 0.391024 |
| Rgs19_Q8BFU4_S58s_1 | 0.35 | -0.12 | -0.46 | 0.04 | 0.43 | -0.33 | -0.08 | 0.05 | -0.14 | -0.23 | -0.16 | 0.2 | -0.1 | -0.12 | -0.06 | 0.08 | 0.065 | -0.035 | -0.0025 | 0.993158 | 0.826841 | 0.902307 | 1 | 0.999631 | 0.997705 |
| Rgs20_E9Q720_S219s_1 | -2.56 | -9.59 | -3.97 | -3.02 | -5.51 | -5.9 | -1.87 | -5.79 | -5.65 | -6.86 | -2.38 | -3.09 | -3.94 | -7.47 | -5.14 | -6.62 | 0.0175 | 0.29 | -1.0075 | 0.51111 | 0.991289 | 0.86017 | 1 | 0.999631 | 0.817687 |
| Rgs20_Q9QZB1_S13s_1 | 0.06 | -2.6 | -1.95 | -0.03 | -2.32 | -2.29 | -1.13 | -0.08 | -2.29 | -1.01 | -1.3 | -1.58 | -1.55 | -1.76 | -3.11 | -2.36 | -0.325 | -0.415 | -1.065 | 0.13935 | 0.668592 | 0.530205 | 1 | 0.999631 | 0.611718 |
| Rgs3_Q9DC04_S287s_1 | -0.98 | -1.42 | -1.26 | -1.72 | -1.41 | -1.67 | -1.32 | -1.22 | -0.88 | -1.09 | -0.99 | -0.15 | -0.63 | -1.24 | -0.78 | -1.49 | -0.06 | 0.5675 | 0.31 | 0.355877 | 0.825576 | 0.102734 | 1 | 0.999631 | 0.74034 |
| Rgs3_Q9DC04_S288s_1 | -0.98 | -1.42 | -1.02 | -1.01 | -1.41 | -1.67 | -0.84 | -0.61 | -0.88 | -1.09 | -0.98 | -0.12 | -0.63 | -1.24 | -0.31 | -0.59 | -0.025 | 0.34 | 0.415 | 0.204616 | 0.937116 | 0.289042 | 1 | 0.999631 | 0.660113 |
| Rgs6_Q9Z2H2_S244s_1 | -1.29 | -1.53 | -1.42 | -1.53 | -1.52 | -1.54 | -1.51 | -2.12 | -1.88 | -1.68 | -1.66 | -1.82 | -1.72 | -1.63 | -2.11 | -1.52 | -0.23 | -0.3175 | -0.3025 | 0.292995 | 0.390007 | 0.212594 | 1 | 0.999631 | 0.70692 |
| Rgs7_Q80XD3_S229s_1 | 0.6 | 1.31 | 0.98 | 1.04 | 0.75 | 1.98 | 0.44 | 1.42 | 0.59 | 0.31 | 1.23 | -0.34 | -0.59 | 1.73 | 0.94 | 0.69 | 0.165 | -0.535 | -0.29 | 0.554671 | 0.669904 | 0.180553 | 1 | 0.999631 | 0.838109 |
| Rgs7_Q80XD3_S239s_1 | -1.32 | 0.61 | 1.98 | 0.48 | 1.08 | -0.64 | 1.16 | 0.22 | 1.65 | 0.83 | 1.04 | 2.36 | 2.82 | 1.38 | 0.4 | 1.41 | 0.0175 | 1.0325 | 1.065 | 0.171801 | 0.98019 | 0.151914 | 1 | 0.999631 | 0.641657 |
| Rgs7bp_Q8BQP9_S18s_1 | 1.08 | 0.18 | -0.14 | 1.39 | 1.6 | 0.48 | 1.54 | 1.3 | 0.31 | -0.72 | 0.87 | 0.93 | -0.64 | 0.82 | 0.61 | 0.48 | 0.6025 | -0.28 | -0.31 | 0.519139 | 0.187604 | 0.578333 | 1 | 0.999631 | 0.819456 |
| Rgs8_D3Z5I4_S46s_1 | -0.1 | -1.11 | -1.28 | -0.12 | -1.32 | -1.47 | -0.75 | 0.35 | -0.3 | 0.04 | -0.17 | -0.67 | 0.04 | -0.96 | -0.3 | -0.09 | -0.145 | 0.3775 | 0.325 | 0.431444 | 0.767409 | 0.325997 | 1 | 0.999631 | 0.778288 |
| Rgs8_D3Z5I4_T47t_1 | 1.44 | 0.31 | 0.7 | 1.34 | -0.05 | 0.58 | 0.49 | 0.95 | 1.19 | 1.88 | 0.33 | -0.22 | -0.83 | 0.72 | -0.39 | -0.21 | -0.455 | -0.1525 | -1.125 | 0.024251 | 0.227261 | 0.763712 | 1 | 0.999631 | 0.415644 |
| Rhbdd3_Q8BP97_S365s_1 | 2.47 | -1.2 | 0.37 | 0.46 | -0.96 | -0.47 | 1.74 | -0.29 | -2.1 | 1.49 | -0.76 | -0.88 | -1.46 | -1.05 | -1.28 | -1.37 | -0.52 | -1.0875 | -1.815 | 0.021383 | 0.537417 | 0.255094 | 1 | 0.999631 | 0.403597 |
| Rhbdf1_F2Z3W3_S11s_1 | 0.89 | -0.07 | 0.98 | 1.19 | 0.16 | 0.93 | 1.52 | 0.47 | 0.18 | 0.59 | 0.63 | 0.87 | -0.61 | -0.61 | 1.99 | 0.49 | 0.0225 | -0.18 | -0.4325 | 0.486363 | 0.955981 | 0.611548 | 1 | 0.999631 | 0.804724 |
| Rhbdf1_F2Z3W3_S61s_1 | 0.89 | -0.56 | 0.11 | -0.14 | -0.96 | -0.35 | -0.28 | -0.73 | -0.52 | 0.52 | -0.57 | -0.3 | -1.1 | -0.43 | -0.02 | -0.32 | -0.655 | -0.2925 | -0.5425 | 0.198713 | 0.097907 | 0.477048 | 1 | 0.999631 | 0.656813 |
| Rhbdf1_F2Z3W3_S8s_1 | 1.98 | -0.46 | 0.17 | 0.9 | 0.48 | 0.89 | 0.77 | 0.34 | -0.19 | 1.84 | 0.39 | -0.63 | -1.51 | -0.49 | 0.45 | -0.44 | -0.0275 | -0.295 | -1.145 | 0.080824 | 0.956473 | 0.662224 | 1 | 0.999631 | 0.545579 |
| Rhbdl3_P58873_S6s_1 | -1.24 | 0.16 | 0.83 | -0.92 | 0.58 | 0.03 | 0.52 | -0.48 | 0.99 | -0.42 | 0.81 | 1.43 | 1.93 | -0.41 | 0.85 | 0.26 | 0.455 | 0.995 | 0.95 | 0.150807 | 0.377396 | 0.106843 | 1 | 0.999631 | 0.623336 |
| Rhob_P62746_S123s_1 | -1.38 | 2.17 | 1.69 | 0.15 | 2.42 | 1.6 | 1.5 | 1.73 | 2.68 | -0.31 | 3.05 | 3.22 | 3.24 | 0.57 | 1.74 | 0.78 | 1.155 | 1.5025 | 0.925 | 0.302684 | 0.138529 | 0.155963 | 1 | 0.999631 | 0.714978 |
| Rhog_P84096_T125t_1 | 0.78 | -0.17 | 0.47 | 0.72 | -0.78 | 0.14 | 0 | -0.1 | -0.69 | 0.2 | 0.5 | -0.5 | -1.11 | -0.75 | 0.58 | -0.84 | -0.635 | -0.5725 | -0.98 | 0.04672 | 0.083279 | 0.154288 | 1 | 0.999631 | 0.480102 |
| Ric8a_Q3TIR3_S435s_1 | -0.52 | -0.53 | -0.14 | -1.45 | -0.35 | 0.04 | -1.4 | -0.56 | 0.36 | 0.37 | -1.02 | 0.07 | 1.24 | 0.83 | -0.25 | 0.2 | 0.0925 | 0.605 | 1.165 | 0.02216 | 0.822309 | 0.181844 | 1 | 0.999631 | 0.405544 |
| Ric8a_Q3TIR3_S435s_2 | -1.23 | -0.29 | 0.69 | -0.05 | 0.1 | -0.1 | -0.02 | 0.67 | 0.77 | -0.34 | 0.57 | -0.13 | 0.01 | 0.5 | 0.33 | 0.94 | 0.3825 | 0.4375 | 0.665 | 0.156054 | 0.379568 | 0.357232 | 1 | 0.999631 | 0.626452 |
| Ric8a_Q3TIR3_T440t_2 | -1.23 | -0.29 | 0.69 | -0.05 | 0.1 | -0.1 | -0.02 | 0.67 | 0.77 | -0.34 | 0.57 | -0.13 | 0.01 | 0.5 | 0.33 | 0.94 | 0.3825 | 0.4375 | 0.665 | 0.156054 | 0.379568 | 0.357232 | 1 | 0.999631 | 0.626452 |
| Rictor_Q6QI06_S1234s_1 | -0.91 | 0.78 | 0.27 | -0.52 | 0.33 | 0.24 | -0.03 | 0.46 | 0.09 | -1.15 | -0.71 | -0.69 | -0.15 | 0.28 | -0.75 | -0.26 | 0.345 | -0.52 | -0.125 | 0.776484 | 0.397265 | 0.26583 | 1 | 0.999631 | 0.921191 |
| Rictor_Q6QI06_S1591s_1 | 0.37 | 1.41 | 0.03 | 1.54 | 0.7 | 1.39 | 1.06 | 1.35 | 1.27 | 0.48 | 1.9 | 1.46 | 0.85 | 1.9 | 1.08 | 1.56 | 0.2875 | 0.44 | 0.51 | 0.267789 | 0.487374 | 0.354976 | 1 | 0.999631 | 0.696016 |
| Rilpl1_Q9JJC6_S349s_1 | 0.41 | -0.88 | -0.72 | -0.09 | 0.51 | 0.89 | 0.46 | 0.58 | 0.4 | 0.37 | 0.37 | 0.13 | 0.09 | 0.77 | -0.1 | 0.32 | 0.93 | 0.6375 | 0.59 | 0.148106 | 0.02131 | 0.089913 | 1 | 0.999631 | 0.621206 |
| Rilpl1_Q9JJC6_S7s_1 | -0.81 | -1.89 | 0.02 | 0.16 | 1.14 | -0.06 | 0.13 | 0.61 | 0.09 | -0.73 | 0.17 | 0.49 | -0.2 | 0.41 | 0.59 | 0.54 | 1.085 | 0.635 | 0.965 | 0.071665 | 0.054953 | 0.230688 | 1 | 0.999631 | 0.532005 |
| Rimbp2_D3YXR8_S279s_1 | -0.06 | -0.2 | -0.44 | 0.61 | 0.45 | -0.51 | 0.31 | 0.74 | -0.32 | -0.8 | 0.05 | 0.69 | -0.62 | 0.13 | 0.47 | 0.72 | 0.27 | -0.0725 | 0.1975 | 0.619332 | 0.470515 | 0.855895 | 1 | 0.999631 | 0.861888 |
| Rimbp2_D3YXR8_S978s_1 | 1.3 | 3.36 | 2.03 | 0.67 | 4.1 | 2.29 | 3.85 | 0.21 | 5.75 | 1.59 | 1.56 | 3.52 | 4.65 | -0.09 | 0.51 | -0.38 | 0.7725 | 1.265 | -0.6675 | 0.551755 | 0.410405 | 0.222331 | 1 | 0.999631 | 0.836275 |
| Rims1_E9PWP6_S1077s_1 | -0.75 | -1.2 | -0.15 | -0.98 | -1.26 | -1.07 | -0.65 | -0.82 | -0.99 | -0.67 | -0.67 | -0.46 | -0.71 | -0.67 | -0.74 | -0.55 | -0.18 | 0.0725 | 0.1025 | 0.746731 | 0.570772 | 0.818001 | 1 | 0.999631 | 0.91173 |
| Rims1_E9PWP6_S1105s_1 | 0.11 | -0.07 | -0.75 | -0.16 | -0.52 | 0.17 | -0.54 | -0.45 | -0.28 | 0.64 | -0.83 | -0.22 | 0.47 | 0.3 | -0.06 | -0.29 | -0.1175 | 0.045 | 0.3225 | 0.338396 | 0.704011 | 0.905426 | 1 | 0.999631 | 0.731571 |
| Rims1_E9PWP6_S1166s_1 | 1.58 | 0.68 | -0.74 | 1.82 | 0.65 | 1.17 | 1.79 | 0.25 | 0.19 | 0.34 | -0.08 | -1.19 | -2.26 | -1.02 | -1.09 | -1.07 | 0.13 | -1.02 | -2.195 | 0.004074 | 0.829501 | 0.120813 | 1 | 0.999631 | 0.351308 |
| Rims1_E9PWP6_S1201s_2 | -0.53 | -0.82 | 0.28 | -0.03 | -0.6 | -0.55 | -0.66 | -0.15 | -0.55 | 0 | -0.42 | -0.15 | -0.11 | -1.05 | -0.39 | -0.07 | -0.215 | -0.005 | -0.13 | 0.729276 | 0.507034 | 0.987877 | 1 | 0.999631 | 0.905923 |
| Rims1_E9PWP6_S1205s_1 | -0.29 | -0.78 | -0.12 | -0.89 | -1 | -0.28 | -0.88 | -0.8 | -1.42 | -0.02 | -0.13 | -0.36 | 0.06 | -1.13 | -0.89 | -1.12 | -0.22 | 0.0375 | -0.25 | 0.511689 | 0.475982 | 0.923076 | 1 | 0.999631 | 0.817687 |
| Rims1_E9PWP6_S1205s_2 | -0.53 | -0.82 | 0.17 | -0.12 | -0.6 | -0.55 | -0.69 | -0.22 | -0.55 | 0 | -0.55 | -0.25 | -0.11 | -1.05 | -0.59 | -0.17 | -0.19 | -0.0125 | -0.155 | 0.667455 | 0.534223 | 0.968568 | 1 | 0.999631 | 0.88264 |
| Rims1_E9PWP6_S1229s_1 | -0.47 | -0.17 | -1.37 | -1.14 | -0.46 | -0.71 | -0.64 | -0.51 | -0.72 | -0.91 | -0.29 | -0.02 | -0.49 | -0.9 | -0.86 | -0.62 | 0.2075 | 0.3025 | 0.07 | 0.842672 | 0.532193 | 0.424614 | 1 | 0.999631 | 0.948041 |
| Rims1_E9PWP6_S1259s_2 | -0.48 | -0.9 | -0.94 | 0.18 | 0.04 | -0.42 | 0.45 | -1.24 | -1.28 | -0.65 | 0.19 | -1.35 | -0.88 | -1.37 | -1.47 | -0.65 | 0.2425 | -0.2375 | -0.5575 | 0.153638 | 0.580796 | 0.590437 | 1 | 0.999631 | 0.625198 |
| Rims1_E9PWP6_S1260s_2 | -0.48 | -0.9 | -0.94 | 0.18 | 0.04 | -0.42 | 0.45 | -1.24 | -1.28 | -0.65 | 0.19 | -1.35 | -0.88 | -1.37 | -1.47 | -0.65 | 0.2425 | -0.2375 | -0.5575 | 0.153638 | 0.580796 | 0.590437 | 1 | 0.999631 | 0.625198 |
| Rims1_E9PWP6_S1288s_1 | -2.04 | -2.91 | -0.45 | -1.64 | -1.44 | -1.72 | -0.91 | -2.08 | -1.48 | 0.29 | -1.46 | -0.35 | 1.77 | -1.5 | -1.48 | -0.43 | 0.2225 | 1.01 | 1.35 | 0.117306 | 0.673355 | 0.122555 | 1 | 0.999631 | 0.585087 |
| Rims1_E9PWP6_S1397s_1 | 1.35 | -0.5 | -2.16 | -0.52 | 0.3 | 0.28 | 1 | -0.21 | 0.19 | 0.92 | -0.34 | -1.41 | -0.85 | -0.4 | -2.25 | -2.46 | 0.8 | 0.2975 | -1.0325 | 0.20042 | 0.253227 | 0.698603 | 1 | 0.999631 | 0.658263 |
| Rims1_E9PWP6_S1402s_1 | 1.49 | -1.21 | -0.76 | 0.47 | -0.03 | 0.06 | 1.26 | 0.04 | -0.48 | 0.26 | -0.33 | -0.35 | -2.26 | -1.49 | -1.4 | -0.07 | 0.335 | -0.2225 | -1.3025 | 0.078974 | 0.589639 | 0.702348 | 1 | 0.999631 | 0.545579 |
| Rims1_E9PWP6_S1418s_1 | 1.38 | 0.42 | -0.26 | 0.69 | 1.13 | -0.12 | 1.02 | -0.2 | 0.62 | 0.12 | 0.66 | -0.03 | -0.32 | -0.36 | -1 | 0.57 | -0.1 | -0.215 | -0.835 | 0.095011 | 0.83154 | 0.589887 | 1 | 0.999631 | 0.558128 |
| Rims1_E9PWP6_S1626s_1 | -0.66 | 0.12 | -1.09 | -0.82 | -0.38 | 0.15 | -0.65 | -1.14 | 0.06 | -0.03 | -1.23 | -0.99 | 0.88 | -0.47 | -0.73 | -0.19 | 0.1075 | 0.065 | 0.485 | 0.287005 | 0.780416 | 0.877508 | 1 | 0.999631 | 0.705972 |
| Rims1_E9PWP6_S1626s_2 | -0.57 | 0.2 | -2.93 | -2.03 | 0.06 | -0.07 | -2.09 | -2.63 | -0.15 | 0.13 | -1.23 | -2.55 | 0.01 | 0.01 | -1.54 | -2.65 | 0.15 | 0.3825 | 0.29 | 0.727982 | 0.860957 | 0.641116 | 1 | 0.999631 | 0.905923 |
| Rims1_E9PWP6_S1629s_1 | -0.14 | -0.68 | 1.37 | -0.68 | -0.24 | -0.24 | -0.04 | 0.39 | -0.54 | -0.08 | 0.38 | 0.24 | 0.43 | -0.48 | 0.09 | -0.29 | 3.93E-17 | 0.0325 | -0.03 | 0.952313 | 1 | 0.948031 | 1 | 0.999631 | 0.984317 |
| Rims1_E9PWP6_S1629s_2 | -0.57 | 0.2 | -2.93 | -2.03 | 0.06 | -0.07 | -2.09 | -2.63 | -0.15 | 0.13 | -1.23 | -2.55 | 0.01 | 0.01 | -1.54 | -2.65 | 0.15 | 0.3825 | 0.29 | 0.727982 | 0.860957 | 0.641116 | 1 | 0.999631 | 0.905923 |
| Rims1_E9PWP6_S1641s_1 | -0.7 | -0.39 | -0.74 | -0.73 | -1.49 | -1.28 | -0.97 | -0.69 | -0.75 | -0.63 | -2.48 | -0.56 | -0.34 | -1.38 | -0.79 | -0.89 | -0.4675 | -0.465 | -0.21 | 0.511551 | 0.117174 | 0.321708 | 1 | 0.999631 | 0.817687 |
| Rims1_E9PWP6_S211s_1 | -1.08 | -1.2 | -1.03 | -1.1 | -0.75 | -1.36 | -1.03 | -1.01 | -0.93 | -1.55 | -0.77 | -0.63 | -0.42 | -0.68 | -0.84 | -0.9 | 0.065 | 0.1325 | 0.3925 | 0.168749 | 0.795402 | 0.650531 | 1 | 0.999631 | 0.637839 |
| Rims1_E9PWP6_S241s_1 | 0.31 | -0.13 | 0.52 | 0.42 | 0.58 | -0.08 | 0.93 | -0.06 | 0.07 | -0.71 | -0.34 | 0.28 | -0.95 | -0.44 | 0.32 | 0.64 | 0.0625 | -0.455 | -0.3875 | 0.353506 | 0.84936 | 0.178687 | 1 | 0.999631 | 0.73949 |
| Rims1_E9PWP6_S252s_1 | -0.4 | -0.55 | -0.06 | 0.08 | -0.97 | -1.63 | -0.72 | -0.47 | -1.17 | -0.44 | -0.37 | -0.14 | -0.41 | -0.61 | -0.03 | 0.16 | -0.715 | -0.2975 | 0.01 | 0.974723 | 0.05256 | 0.368515 | 1 | 0.999631 | 0.990998 |
| Rims1_E9PWP6_S346s_1 | 0.32 | 1.07 | 1.68 | 0.74 | 0.87 | 0.86 | 2.35 | 0.51 | 2.02 | 0.01 | 0.98 | 1.52 | 0.64 | -1.43 | 0.18 | -0.55 | 0.195 | 0.18 | -1.2425 | 0.032758 | 0.682691 | 0.715626 | 1 | 0.999631 | 0.44294 |
| Rims1_E9PWP6_S443s_1 | -1.42 | -3.23 | -1.22 | -1.75 | -2.9 | -2.59 | -1.83 | -1.88 | -3.1 | -1.86 | -1.41 | -1.38 | -1.62 | -2.29 | -1.55 | -1.23 | -0.395 | -0.0325 | 0.2325 | 0.636164 | 0.43253 | 0.953746 | 1 | 0.999631 | 0.869397 |
| Rims1_E9PWP6_S443s_2 | -0.58 | -0.45 | -0.84 | -0.48 | -0.61 | -0.59 | -0.95 | -0.81 | -0.85 | -0.45 | -0.54 | -1.1 | -0.26 | -0.66 | -0.92 | -0.42 | -0.1525 | -0.1475 | 0.0225 | 0.938041 | 0.542689 | 0.595366 | 1 | 0.999631 | 0.979905 |
| Rims1_E9PWP6_S447s_2 | -0.58 | -0.45 | -0.84 | -0.48 | -0.61 | -0.59 | -0.95 | -0.81 | -0.85 | -0.45 | -0.54 | -1.1 | -0.26 | -0.66 | -0.92 | -0.42 | -0.1525 | -0.1475 | 0.0225 | 0.938041 | 0.542689 | 0.595366 | 1 | 0.999631 | 0.979905 |
| Rims1_E9PWP6_S514s_1 | 0.17 | 2.62 | 2.48 | 0.57 | 2.38 | 2.55 | 1.25 | 2.1 | 2.3 | -0.92 | 2.31 | 3.04 | 2.69 | 0.98 | 2.37 | 1.7 | 0.61 | 0.2225 | 0.475 | 0.478812 | 0.342303 | 0.813739 | 1 | 0.999631 | 0.801051 |
| Rims1_E9PWP6_S726s_1 | 0.16 | -1.04 | -1.67 | -1.12 | -0.79 | -0.48 | -0.82 | -1.42 | -0.94 | -0.51 | -1.28 | -1.47 | -0.83 | -0.64 | -0.96 | -1 | 0.04 | -0.1325 | 0.06 | 0.884104 | 0.925156 | 0.761672 | 1 | 0.999631 | 0.963505 |
| Rims1_E9PWP6_S741s_1 | -0.89 | -1.21 | -1.2 | -1.5 | -1.26 | -1.14 | -1.52 | -1.59 | -1.3 | -0.9 | -1.37 | -1.01 | -0.1 | -0.63 | -1.05 | -0.91 | -0.1775 | 0.055 | 0.5275 | 0.126993 | 0.506154 | 0.84068 | 1 | 0.999631 | 0.59788 |
| Rims1_E9PWP6_S744s_1 | -0.79 | -0.14 | -0.69 | -0.61 | -0.07 | -0.53 | -0.76 | -0.62 | -0.25 | -0.53 | -0.96 | -0.74 | -0.48 | -0.48 | -0.69 | -0.32 | 0.0625 | -0.0625 | 0.065 | 0.821019 | 0.826148 | 0.830864 | 1 | 0.999631 | 0.938464 |
| Rims1_E9PWP6_S894s_1 | -0.21 | -0.72 | -1.61 | -0.49 | -0.46 | -0.35 | -1.03 | -0.59 | -0.5 | -0.4 | -1.23 | -1.15 | 0.43 | -0.45 | -0.72 | -0.6 | 0.15 | -0.0625 | 0.4225 | 0.322128 | 0.678802 | 0.872349 | 1 | 0.999631 | 0.72427 |
| Rims1_E9PWP6_S966s_2 | -0.91 | -2.38 | -2.28 | -0.87 | -0.49 | -2.98 | -0.35 | -0.67 | -0.95 | -1.61 | -0.79 | -1.34 | -1.23 | -2.31 | -1.04 | -2.61 | 0.4875 | 0.4375 | -0.1875 | 0.726403 | 0.47086 | 0.341144 | 1 | 0.999631 | 0.90568 |
| Rims1_E9PWP6_S968s_2 | -0.91 | -2.38 | -2.28 | -0.87 | -0.49 | -2.98 | -0.35 | -0.67 | -0.95 | -1.61 | -0.79 | -1.34 | -1.23 | -2.31 | -1.04 | -2.61 | 0.4875 | 0.4375 | -0.1875 | 0.726403 | 0.47086 | 0.341144 | 1 | 0.999631 | 0.90568 |
| Rims1_E9PWP6_S990s_1 | -1.09 | -1.33 | -1.8 | -1.83 | -1.63 | -1.56 | -1.07 | -2.07 | -1.74 | -0.95 | -2.26 | -2.26 | -1.49 | -1.82 | -2.63 | -1.9 | -0.07 | -0.29 | -0.4475 | 0.225857 | 0.827451 | 0.453139 | 1 | 0.999631 | 0.676578 |
| Rims1_E9PWP6_T1203t_1 | 0.23 | -0.1 | -0.5 | -0.17 | -0.4 | -0.64 | -0.69 | -1.01 | -0.76 | -0.19 | -0.48 | -0.58 | -0.47 | 0.01 | -0.29 | -0.2 | -0.55 | -0.3675 | -0.1025 | 0.728017 | 0.072233 | 0.215499 | 1 | 0.999631 | 0.905923 |
| Rims1_E9PWP6_T326t_1 | 2.3 | 2.01 | -0.59 | 0.77 | 2.46 | 1.65 | 0.25 | 0.94 | 2.27 | 1.25 | 0.08 | -0.19 | 0.25 | 1.41 | -0.18 | -0.15 | 0.2025 | -0.27 | -0.79 | 0.25914 | 0.777981 | 0.725428 | 1 | 0.999631 | 0.692624 |
| Rims1_E9PWP6_T743t_1 | -0.12 | -0.67 | 0.44 | -0.28 | -0.85 | -0.8 | -0.97 | -0.41 | -1.17 | -0.98 | -1.96 | -0.54 | -0.36 | -0.83 | -0.02 | 0.43 | -0.6 | -1.005 | -0.0375 | 0.922405 | 0.079772 | 0.026823 | 1 | 0.999631 | 0.975032 |
| Rims2_Q0VF51_S1075s_1 | -0.09 | 0.74 | 0.4 | 0.32 | 0.91 | 0.83 | 0.73 | 0.13 | 0.64 | -0.33 | 0.02 | 0.44 | -0.26 | -0.22 | -0.12 | 0.31 | 0.3075 | -0.15 | -0.415 | 0.200146 | 0.326402 | 0.65003 | 1 | 0.999631 | 0.658104 |
| Rims2_Q0VF51_S1099s_1 | -0.48 | -2.44 | -2.36 | -0.91 | -1.54 | -2.24 | -1.13 | -1.7 | -2.06 | -1.48 | -1.58 | -2.25 | -1.34 | -2.69 | -1.66 | -1.88 | -0.105 | -0.295 | -0.345 | 0.527055 | 0.837812 | 0.562453 | 1 | 0.999631 | 0.824554 |
| Rims2_Q0VF51_S1115s_2 | 0.45 | 0.43 | 0.31 | -0.55 | 0.13 | 0.1 | 0.7 | 1.28 | 0.14 | -0.06 | 0.61 | -1.28 | -0.57 | -0.03 | -0.54 | 0.69 | 0.3925 | -0.3075 | -0.2725 | 0.5032 | 0.315925 | 0.505946 | 1 | 0.999631 | 0.814092 |
| Rims2_Q0VF51_S1163s_1 | 2.69 | -0.46 | 0.4 | 1.63 | 0.61 | 1.01 | 1.97 | 0.41 | 0.22 | 1.15 | 0.52 | -0.76 | -0.96 | -0.42 | 0.06 | -0.88 | -0.065 | -0.7825 | -1.615 | 0.031127 | 0.924406 | 0.285246 | 1 | 0.999631 | 0.441042 |
| Rims2_Q0VF51_S1535s_1 | -0.36 | -1.56 | -1.71 | -0.5 | -0.56 | -0.75 | -0.46 | -1.31 | -0.89 | -0.98 | -0.9 | -0.91 | 0.18 | -1.43 | -1.13 | -1.28 | 0.2625 | 0.1125 | 0.1175 | 0.811004 | 0.518349 | 0.764897 | 1 | 0.999631 | 0.936949 |
| Rims2_Q0VF51_S1535s_2 | -0.17 | -2.35 | 0.28 | -0.18 | -0.27 | -1.34 | 0.21 | -0.02 | -1.92 | 0.36 | -0.18 | 0.12 | -0.69 | -0.88 | 0.34 | 0.64 | 0.25 | 0.2 | 0.4575 | 0.474971 | 0.685593 | 0.775252 | 1 | 0.999631 | 0.799827 |
| Rims2_Q0VF51_S1538s_1 | -0.62 | -1.04 | -0.84 | -3.02 | -1.51 | -0.81 | -1.67 | -1.32 | -1.11 | -0.65 | -1.71 | -1.83 | 0.33 | -0.79 | -1.29 | -1.58 | 0.0525 | 0.055 | 0.5475 | 0.392456 | 0.922254 | 0.922882 | 1 | 0.999631 | 0.758901 |
| Rims2_Q0VF51_S1538s_2 | -0.17 | -2.35 | 0.28 | -0.18 | -0.27 | -1.34 | 0.21 | -0.02 | -1.92 | 0.36 | -0.18 | 0.12 | -0.69 | -0.88 | 0.34 | 0.64 | 0.25 | 0.2 | 0.4575 | 0.474971 | 0.685593 | 0.775252 | 1 | 0.999631 | 0.799827 |
| Rims2_Q0VF51_S236s_1 | 2.75 | 0.68 | -1 | 0.96 | -0.1 | 0.9 | 1.05 | 0.01 | 0.01 | 2.15 | -0.63 | -0.36 | -0.44 | -0.25 | -0.44 | -1.22 | -0.3825 | -0.555 | -1.435 | 0.065353 | 0.600575 | 0.526722 | 1 | 0.999631 | 0.516298 |
| Rims2_Q0VF51_S250s_1 | 2.45 | 0.51 | 1.02 | 1.45 | 0.7 | 2.27 | 2.5 | -0.17 | 0.76 | 1.81 | 0.46 | -0.63 | -2.2 | -0.73 | -1 | -0.85 | -0.0325 | -0.7575 | -2.5525 | 0.000539 | 0.961586 | 0.220925 | 1 | 0.999631 | 0.351308 |
| Rims2_Q0VF51_S269s_1 | 1.23 | -0.65 | -0.39 | 0.24 | -0.37 | 0.03 | 0.07 | -0.52 | -0.41 | 0.64 | -0.84 | -1.3 | -1.38 | -1.33 | -1.37 | -1.38 | -0.305 | -0.585 | -1.4725 | 0.006002 | 0.487467 | 0.29916 | 1 | 0.999631 | 0.351545 |
| Rims2_Q0VF51_S271s_1 | 1.98 | 0.69 | -1.87 | 0.86 | 0.57 | 0.13 | 1.06 | 0.51 | 1.09 | 0.54 | -1.46 | -2.05 | -1.94 | -0.85 | -1.81 | -0.64 | 0.1525 | -0.885 | -1.725 | 0.045673 | 0.835788 | 0.369213 | 1 | 0.999631 | 0.477252 |
| Rims2_Q0VF51_S335s_1 | 1.16 | 0.97 | 0.94 | 1.18 | 0.67 | 1.19 | 1.1 | 1.4 | 0.87 | 1.05 | 1.16 | 0.92 | -0.04 | 1.27 | 1.08 | 1.05 | 0.0275 | -0.0625 | -0.2225 | 0.536409 | 0.917201 | 0.800108 | 1 | 0.999631 | 0.83005 |

| | | | | | | | | | | | | | | | | | | | | | | | | | |
|---|---|---|---|---|---|---|---|---|---|---|---|---|---|---|---|---|---|---|---|---|---|---|---|---|---|
| Rims2_Q0VF51_S432s_1 | -0.68 | -1.1 | -0.11 | -0.96 | -1.02 | -0.83 | 0.13 | -0.79 | -0.61 | -0.76 | -0.26 | -0.23 | 0.05 | -1.26 | -0.72 | -1.18 | 0.085 | 0.2475 | -0.065 | 0.870092 | 0.814046 | 0.442376 | 1 | 0.999631 | 0.958441 |
| Rims2_Q0VF51_S439s_1 | -0.21 | -0.06 | -0.55 | -0.51 | -0.23 | 0.23 | -0.44 | -0.55 | -0.51 | 0.21 | -0.73 | -1.14 | -0.24 | -0.13 | -1.11 | -0.14 | 0.085 | -0.21 | -0.0725 | 0.828412 | 0.766116 | 0.549275 | 1 | 0.999631 | 0.941315 |
| Rims2_Q0VF51_S482s_1 | -0.83 | 2.26 | 1.49 | -0.46 | 1.91 | 1.66 | 0.99 | 1.38 | 1.68 | -1.91 | 2.42 | 2.74 | 2.32 | 0.88 | 2.16 | 1.11 | 0.87 | 0.6175 | 1.0025 | 0.192195 | 0.224211 | 0.584779 | 1 | 0.999631 | 0.648844 |
| Rims2_Q0VF51_S925s_1 | 0.33 | -3.46 | -1.53 | -0.57 | -1.69 | -1.93 | -0.77 | -0.78 | -1.04 | -0.19 | -0.84 | -2.2 | -2 | -2.22 | -2 | -2.18 | 0.015 | 0.24 | -0.7925 | 0.285459 | 0.984239 | 0.76502 | 1 | 0.999631 | 0.705851 |
| Rims2_Q0VF51_T1114t_1 | 0.07 | -0.85 | -0.01 | -0.52 | -0.69 | -0.15 | -0.29 | -0.21 | -1.34 | -0.05 | -0.46 | -0.62 | 0.56 | -0.31 | 0.12 | -0.22 | -0.0075 | -0.29 | 0.365 | 0.311385 | 0.980492 | 0.443526 | 1 | 0.999631 | 0.71924 |
| Rims3_Q80U57_S22s_1 | 0.01 | 0.02 | -1.1 | 0.28 | 0.46 | 0.09 | -0.41 | 0.58 | -0.38 | -0.78 | 0.74 | 0.03 | -0.65 | 0.23 | -0.28 | -0.26 | 0.3775 | 0.1 | -0.0425 | 0.912698 | 0.343659 | 0.820899 | 1 | 0.999631 | 0.972191 |
| Rims3_Q80U57_S297s_1 | -0.37 | 0.47 | -0.01 | -0.66 | 0.32 | 0.1 | -0.67 | -0.37 | -0.46 | -0.22 | -0.6 | 0.09 | 1.36 | 0.13 | 0 | -0.12 | -0.0125 | -0.155 | 0.485 | 0.274471 | 0.972011 | 0.645652 | 1 | 0.999631 | 0.700187 |
| Rims4_P60191_S257s_1 | -1.02 | -0.82 | -0.64 | -0.36 | -0.27 | -1.11 | -0.51 | 0 | -0.53 | -1.87 | -0.04 | 0.63 | -0.62 | -0.94 | -0.8 | 0.2 | 0.2375 | 0.2575 | 0.17 | 0.628259 | 0.466951 | 0.61928 | 1 | 0.999631 | 0.865998 |
| Rin1_Q921Q7_S198s_1 | 0.84 | -0.65 | -1.03 | 0.83 | 1.68 | 0.33 | 0.44 | 0.75 | 0.42 | -0.4 | -0.86 | 0 | -0.88 | -0.08 | -0.29 | 0.03 | 0.8025 | -0.2075 | -0.3025 | 0.55324 | 0.155944 | 0.69395 | 1 | 0.999631 | 0.837597 |
| Rin1_Q921Q7_S246s_1 | 0.66 | 0.8 | 0.14 | -2.35 | 0.34 | 0.29 | -0.98 | 0.05 | -0.48 | -0.67 | -0.68 | -0.52 | 0.9 | 1.15 | 0.42 | 0.55 | 0.1125 | -0.4 | 0.9425 | 0.182253 | 0.873083 | 0.548767 | 1 | 0.999631 | 0.643861 |
| Rin1_Q921Q7_S280s_1 | 2.53 | 0.54 | 0.47 | 2.47 | 1.38 | 1.12 | 2.52 | 1.05 | 1.36 | 1.32 | 1.82 | 1.26 | -1.66 | -0.15 | 0.22 | 0.86 | 0.015 | -0.0625 | -1.685 | 0.034048 | 0.980185 | 0.90908 | 1 | 0.999631 | 0.446273 |
| Rin1_Q921Q7_S340s_1 | -0.13 | 1.21 | 0.75 | 0.42 | 0.61 | 0.85 | 0.08 | 0.7 | 1.1 | 0.21 | 0.75 | 0.63 | 1.15 | 1.54 | 1.22 | 0.89 | -0.0025 | 0.11 | 0.6375 | 0.101704 | 0.994371 | 0.764285 | 1 | 0.999631 | 0.564448 |
| Rin1_Q921Q7_S446s_1 | 0.13 | -0.22 | 0.06 | 0.48 | -0.17 | 0.45 | -0.16 | 0.33 | 0.01 | 0.56 | 0.47 | 0.71 | -0.16 | 0.86 | -0.87 | 0.16 | 1.57E-16 | 0.325 | -0.115 | 0.777906 | 1 | 0.281903 | 1 | 0.999631 | 0.921382 |
| Rin1_Q921Q7_S590s_1 | -0.21 | 0.12 | -0.86 | -0.07 | 0.2 | -0.1 | -0.42 | 0 | -0.29 | -0.2 | -0.39 | -0.75 | -0.38 | 0.28 | 0.03 | -0.04 | 0.175 | -0.1525 | 0.2275 | 0.493442 | 0.571493 | 0.62611 | 1 | 0.999631 | 0.80955 |
| Rin1_Q921Q7_Y35y_1 | -0.26 | -0.58 | -1.36 | -0.01 | -0.25 | -0.44 | -0.49 | -0.16 | -1.23 | -1.11 | -0.13 | -0.75 | -0.67 | -0.33 | -0.96 | -0.75 | 0.2175 | -0.2525 | -0.125 | 0.733701 | 0.525808 | 0.529343 | 1 | 0.999631 | 0.907462 |
| Ring1_O35730_S219s_1 | -0.7 | 0.57 | 0.19 | -0.06 | -0.03 | -0.18 | 0.1 | 0.14 | 0.4 | -1.01 | -0.91 | 0.84 | 0.09 | 0.17 | 1.07 | 1.29 | 0.0075 | -0.17 | 0.655 | 0.142222 | 0.981441 | 0.737959 | 1 | 0.999631 | 0.612726 |
| Ripk2_P58801_S364s_1 | 0.48 | -0.19 | -0.02 | -0.15 | 0.22 | 0.21 | -0.36 | 0.21 | 0.36 | -0.25 | -0.02 | -0.47 | 0.45 | 0.28 | 0.58 | 0.17 | 0.04 | -0.125 | 0.34 | 0.263231 | 0.888749 | 0.684105 | 1 | 0.999631 | 0.693034 |
| Rltpr_S0DHL8_S1377s_1 | 0.14 | 0.59 | 0.58 | 0.16 | 0.71 | 0.88 | -1.1 | -1.32 | 0.59 | 0.34 | -0.09 | -0.06 | 0.84 | 1.38 | 0.64 | 1.59 | -0.575 | -0.1725 | 0.745 | 0.044357 | 0.30716 | 0.557446 | 1 | 0.999631 | 0.475443 |
| Rltpr_S0DHL8_S1382s_1 | 0.11 | 1.33 | 2.47 | 0.06 | -1.09 | 1.59 | 0.24 | 0.21 | 0.42 | 1.67 | 0.24 | 0.78 | 1.98 | 0.37 | 0.61 | 1.31 | -0.755 | -0.215 | 0.075 | 0.902979 | 0.296434 | 0.719823 | 1 | 0.999631 | 0.968183 |
| Rltpr_S0DHL8_S1382s_2 | 0.58 | 1.21 | 1.95 | 1.21 | 0.7 | 2.15 | 1.12 | 1.64 | 1.19 | 1.57 | 0.38 | 0.13 | -0.71 | 0.88 | 1.09 | 1.2 | 0.165 | -0.42 | -0.6225 | 0.234663 | 0.69405 | 0.347149 | 1 | 0.999631 | 0.683301 |
| Rltpr_S0DHL8_S1385s_2 | 0.58 | 1.21 | 1.95 | 1.21 | 0.7 | 2.15 | 1.12 | 1.64 | 1.19 | 1.57 | 0.38 | 0.13 | -0.71 | 0.88 | 1.09 | 1.2 | 0.165 | -0.42 | -0.6225 | 0.234663 | 0.69405 | 0.347149 | 1 | 0.999631 | 0.683301 |
| Rltpr_S0DHL8_S955s_1 | 0.19 | 0.88 | -0.79 | -0.5 | 0.15 | 0.13 | -0.79 | -0.44 | -0.51 | 0.28 | -0.56 | -0.22 | 0.87 | 0.43 | 0.03 | 0.06 | -0.1825 | -0.1975 | 0.4025 | 0.360407 | 0.672598 | 0.641959 | 1 | 0.999631 | 0.741182 |
| Rltpr_S0DHL8_T1287t_1 | 0.3 | 0.32 | -0.74 | 1.05 | 1.44 | 0.61 | 0.65 | 2.21 | 0.3 | -0.5 | 1.4 | 0.68 | 0.24 | 0.48 | 1.44 | 1.59 | 0.995 | 0.2375 | 0.705 | 0.167634 | 0.069037 | 0.643192 | 1 | 0.999631 | 0.637399 |
| Rmdn2_Q8BSE0_S51s_1 | -0.09 | -0.02 | -0.76 | -1.49 | -0.65 | -1.27 | -1.17 | -0.19 | -1.87 | -0.75 | -0.33 | -0.96 | 0.62 | -0.34 | 0.24 | -0.43 | -0.23 | -0.3875 | 0.6125 | 0.177291 | 0.587152 | 0.408601 | 1 | 0.999631 | 0.642116 |
| Rmdn3_Q3UJU9_S46s_1 | 0.9 | 0.61 | 0.21 | -1.04 | 1.44 | 0.52 | 0.15 | -0.86 | 1.69 | 0.18 | 0.07 | -0.76 | 1.02 | 0.6 | -0.42 | 0.34 | 0.1425 | 0.125 | 0.215 | 0.669192 | 0.806379 | 0.836459 | 1 | 0.999631 | 0.883599 |
| Rnf113a1_Q8R3P8_S6s_1 | 0.64 | 2.07 | 1.15 | 0.92 | -0.07 | 0.48 | 0.92 | -1.14 | -0.55 | -0.78 | 0.42 | -0.28 | 0.74 | 0.65 | -0.24 | 1.41 | -1.1475 | -1.4925 | -0.555 | 0.240897 | 0.044506 | 0.004664 | 1 | 0.999631 | 0.684338 |
| Rnf123_Q05CH9_S675s_1 | 0.54 | -0.82 | -0.69 | 1.1 | 1.21 | 0.32 | 1.29 | 1.03 | 0.85 | -0.54 | 1.19 | 1.25 | -0.2 | 0.26 | 0.33 | 1.05 | 0.93 | 0.655 | 0.3275 | 0.524994 | 0.082024 | 0.272865 | 1 | 0.999631 | 0.822727 |
| Rnf139_Q7TMV1_S627s_1 | 2.46 | -3.37 | -0.96 | -1.12 | -3.13 | -1.7 | 3.33 | -4.84 | -2.91 | 2.68 | -2.29 | -2.87 | -2.75 | -3.33 | -3.57 | -3.49 | -0.8375 | -0.6 | -2.5375 | 0.032366 | 0.643594 | 0.697068 | 1 | 0.999631 | 0.442395 |
| Rnf14_Q9JI90_S349s_1 | -0.16 | 0.87 | 1.33 | 1.35 | -0.11 | 0.39 | 0.77 | 1.56 | 1.01 | 0.55 | 1.36 | 1.5 | 1.04 | 1.92 | 1.96 | 1.48 | -0.195 | 0.2575 | 0.7525 | 0.101079 | 0.682235 | 0.540909 | 1 | 0.999631 | 0.564448 |
| Rnf157_F7D0T8_S239s_1 | -0.23 | -1.09 | -0.61 | -0.51 | -0.71 | -0.18 | -0.72 | 0 | -0.57 | -0.33 | -0.48 | -0.9 | -1.19 | -0.32 | -0.01 | 0.1 | 0.2075 | 0.04 | 0.255 | 0.506476 | 0.509856 | 0.892473 | 1 | 0.999631 | 0.815174 |
| Rnf157_F7D0T8_S281s_1 | 0.79 | -0.35 | 1.05 | -0.94 | -1.55 | 0.73 | 0.14 | -1.31 | -0.29 | 0.01 | -1.03 | -0.52 | 0.07 | -0.45 | -1.44 | -0.67 | -0.635 | -0.595 | -0.76 | 0.174395 | 0.340274 | 0.246669 | 1 | 0.999631 | 0.642116 |
| Rnf169_E9Q7F2_S279s_1 | 1.66 | 2.17 | 5.44 | 1.99 | 1.38 | 2.29 | 3.04 | 0.78 | 1.45 | 1.85 | 4.26 | 5.15 | -0.75 | 0.1 | 4.61 | 0.23 | -0.9425 | 0.3625 | -1.7675 | 0.182777 | 0.29767 | 0.739015 | 1 | 0.999631 | 0.643861 |
| Rnf169_E9Q7F2_S390s_1 | 0.58 | 2.42 | 1.08 | 1.76 | 1.6 | 1.25 | 0.8 | 1.3 | 1.86 | 1.42 | 1.54 | 1.84 | 1.69 | 2.5 | 2.79 | 1.94 | -0.2225 | 0.205 | 0.77 | 0.122377 | 0.604645 | 0.6271 | 1 | 0.999631 | 0.590804 |
| Rnf214_Q8BFU3_S48s_1 | -0.56 | -0.27 | -0.27 | -0.31 | -0.01 | -0.38 | -0.51 | -0.09 | -0.58 | -0.75 | -0.12 | -0.21 | 0.52 | 0.52 | 0.24 | 0.27 | 0.105 | -0.0625 | 0.74 | 0.018718 | 0.679472 | 0.818527 | 1 | 0.999631 | 0.391024 |
| Rnf214_Q8BFU3_S500s_1 | -0.15 | -0.23 | 0.55 | -0.54 | -0.16 | 1.16 | 0.11 | 0.12 | 0.3 | -1.18 | -0.21 | 0.66 | 1.57 | 0.92 | 0.44 | -0.15 | 0.4 | -0.015 | 0.7875 | 0.095475 | 0.310636 | 0.973483 | 1 | 0.999631 | 0.558128 |
| Rnf217_D3YYI7_S27s_1 | 0.04 | -1.61 | -0.68 | -1.7 | -0.23 | -0.71 | 0.24 | -1.03 | -0.09 | -1.95 | -0.52 | -0.1 | -1.2 | -2.74 | -0.53 | -2.73 | 0.555 | 0.3225 | -0.8125 | 0.214104 | 0.258368 | 0.566289 | 1 | 0.999631 | 0.667209 |
| Rnf6_Q9DBU5_S183s_1 | 0.68 | -1.38 | -0.45 | -0.27 | -1.1 | -0.72 | -0.6 | -0.82 | -0.43 | 0.64 | -0.56 | -0.83 | 0.16 | -1.36 | -0.6 | -1.01 | -0.455 | 0.06 | -0.3475 | 0.49936 | 0.302326 | 0.905025 | 1 | 0.999631 | 0.813279 |
| Rnf6_Q9DBU5_S301s_1 | 0.86 | -0.86 | -1.12 | -0.38 | -1.5 | 0.11 | -0.45 | 0.1 | -0.93 | 0.45 | -0.64 | -1.9 | -0.32 | -0.84 | -1.1 | -0.92 | -0.06 | -0.38 | -0.42 | 0.374838 | 0.91066 | 0.528115 | 1 | 0.999631 | 0.74795 |
| Rnmt_D3Z4K1_S100s_1 | 1.3 | 2.28 | 0.95 | 1.79 | 1.76 | 2.09 | 1.64 | 1.37 | 2.22 | 1.94 | 1.93 | 1.65 | 0.3 | 2.29 | 1.19 | 2.21 | 0.135 | 0.355 | -0.0825 | 0.874483 | 0.703473 | 0.325081 | 1 | 0.999631 | 0.960237 |
| Rnmt_D3Z4K1_S11s_1 | 1.09 | 1.71 | 1.4 | 0.95 | 1.1 | 1.74 | 0.25 | 0.85 | 1.21 | 0.84 | 0.88 | 0.81 | 0.42 | 1.05 | 1.17 | 1.31 | -0.3025 | -0.3525 | -0.3 | 0.375946 | 0.422166 | 0.234499 | 1 | 0.999631 | 0.748307 |
| Rnmt_D3Z4K1_S11s_2 | 0.53 | 1.65 | 1.55 | 0.96 | 1.08 | 1.32 | 0.62 | 1.34 | 1.16 | 1.13 | 1.07 | 1.27 | 1.08 | 1.65 | 1.71 | 1.09 | -0.0825 | -0.015 | 0.21 | 0.565496 | 0.810724 | 0.96296 | 1 | 0.999631 | 0.84139 |
| Rnmt_D3Z4K1_S15s_1 | 0.66 | 1.43 | 2.5 | 1.78 | 0.83 | 1.16 | 0.83 | 1.79 | 1.18 | 1.6 | 1.85 | 2.15 | 1.17 | 1.16 | 2.15 | 1.49 | -0.44 | 0.1025 | -0.1 | 0.823225 | 0.324425 | 0.812709 | 1 | 0.999631 | 0.938864 |
| Rnmt_D3Z4K1_S15s_2 | 0.53 | 1.65 | 1.55 | 0.96 | 1.08 | 1.32 | 0.62 | 1.34 | 1.16 | 1.13 | 1.07 | 1.27 | 1.08 | 1.65 | 1.71 | 1.09 | -0.0825 | -0.015 | 0.21 | 0.565496 | 0.810724 | 0.96296 | 1 | 0.999631 | 0.84139 |
| Robo1_G5E843_S1403s_1 | -0.02 | -0.03 | 0.39 | -0.65 | 0.09 | 0.32 | 0.41 | 0.4 | 0.31 | 0.21 | -1.51 | 0.68 | 0.33 | 1.1 | 0.27 | 0.69 | 0.3825 | -7.9E-17 | 0.675 | 0.075407 | 0.214084 | 1 | 1 | 1 | 0.541044 |
| Robo1_G5E843_S901s_1 | -0.18 | -0.32 | -1.4 | -0.77 | 0.32 | -0.53 | -1.08 | -0.49 | 0.32 | -0.54 | -0.83 | 0.02 | -0.12 | 0.08 | -0.59 | -0.16 | 0.2225 | 0.41 | 0.47 | 0.210299 | 0.582655 | 0.312188 | 1 | 0.999631 | 0.66441 |
| Robo2_Q19AB2_S1197s_1 | -0.19 | -0.2 | -1.44 | -0.64 | -0.33 | -0.88 | -1.25 | -0.68 | -0.45 | -0.08 | -0.46 | -0.98 | 1.04 | -0.15 | -1.03 | -1.15 | -0.1675 | 0.125 | 0.295 | 0.591308 | 0.652018 | 0.738099 | 1 | 0.999631 | 0.852343 |
| Rock2_F8VPK5_S1052s_1 | 2.13 | 0.89 | 1.43 | 0.91 | 0.42 | 1.33 | 1.79 | -1.04 | 0.68 | 2.44 | -0.17 | -0.84 | 0.14 | -0.12 | -1.01 | -1.02 | -0.715 | -0.8125 | -1.8425 | 0.001527 | 0.264928 | 0.253055 | 1 | 0.999631 | 0.351308 |
| Rock2_F8VPK5_S439s_1 | 1.07 | 0.02 | 0.27 | 0.77 | -0.03 | 0.58 | 0.87 | -0.36 | 0.31 | 1.35 | -0.09 | -0.8 | -0.56 | -0.78 | -0.24 | -0.26 | -0.2675 | -0.34 | -0.9925 | 0.013816 | 0.487849 | 0.489657 | 1 | 0.999631 | 0.385092 |
| Rogdi_Q3TDK6_S227s_1 | -0.82 | -0.64 | -0.15 | -0.71 | -0.41 | -0.74 | 0.46 | 0.12 | -0.21 | -0.87 | 0.13 | 0.64 | 0.16 | -0.35 | -0.05 | -0.16 | 0.4375 | 0.5025 | 0.48 | 0.127466 | 0.218721 | 0.200643 | 1 | 0.999631 | 0.59788 |
| Ror1_Q9Z139_S436s_1 | 1 | -0.74 | 0.77 | 0.44 | -1.98 | -1.79 | -0.96 | -1.95 | -0.74 | 0.65 | -0.39 | -0.4 | 0.1 | -1.1 | 0.3 | -0.63 | -2.0375 | -0.5875 | -0.7 | 0.173083 | 0.001046 | 0.23291 | 1 | 0.999631 | 0.642116 |
| Rpap3_Q9D706_S429s_1 | -0.55 | -1.4 | -1.65 | -1.28 | -0.44 | -2.08 | -0.77 | -1.02 | -1.2 | -1.55 | -1.3 | -0.34 | -1.53 | -1.63 | -1.28 | -3.17 | 0.1425 | 0.1225 | -0.6825 | 0.174262 | 0.73584 | 0.745905 | 1 | 0.999631 | 0.642116 |
| Rph3a_P47708_S245s_1 | -0.58 | -4.52 | 1.95 | 0.39 | -2.77 | -3.74 | 3.57 | 2.56 | -3.68 | -2.28 | 2.61 | 3.84 | -4.39 | -4.08 | 1.45 | 0.73 | 0.595 | 0.8125 | -0.8825 | 0.612663 | 0.760086 | 0.676478 | 1 | 0.999631 | 0.860335 |
| Rph3a_P47708_S271s_1 | 3.18 | 1.17 | 1.58 | 2.15 | 0.81 | 1.88 | 2.48 | 1.25 | 0.88 | 1.62 | 1.41 | 0.28 | -1.36 | 0.39 | 0.8 | 0.22 | -0.415 | -0.9725 | -2.0075 | 0.006372 | 0.438146 | 0.076956 | 1 | 0.999631 | 0.351545 |
| Rph3a_P47708_S679s_1 | -0.64 | -1.17 | 0.01 | -0.91 | -0.63 | -1.74 | 0.9 | -0.85 | -0.19 | -1.16 | 0.43 | 1.26 | -0.37 | -1.33 | -0.28 | -0.45 | 0.0975 | 0.7625 | 0.07 | 0.855637 | 0.860244 | 0.174331 | 1 | 0.999631 | 0.951817 |
| Rph3a_P47708_S680s_1 | -0.26 | -0.15 | 1.02 | -0.95 | 0.15 | -0.65 | 1.9 | -0.73 | 0.88 | -0.36 | 1.44 | 3.16 | 1.69 | -0.82 | -0.2 | -0.27 | 0.2525 | 1.365 | 0.185 | 0.765381 | 0.700996 | 0.090369 | 1 | 0.999631 | 0.916326 |
| Rph3a_P47708_T188t_1 | 1.1 | 0.75 | -0.7 | 1.13 | 0.81 | 0.85 | 0.34 | 0.47 | 0.84 | 0.26 | 1.06 | -0.71 | -1.17 | 0.55 | -0.07 | 0.83 | 0.0475 | -0.2075 | -0.535 | 0.359588 | 0.913689 | 0.703803 | 1 | 0.999631 | 0.741182 |
| Rpl13_P47963_S139s_1 | 0.65 | 0.28 | 0.54 | 1.29 | -0.17 | 0.61 | 0.77 | 0.47 | 0.18 | 1.65 | 0.61 | 0.55 | -0.11 | 0.48 | 0.57 | 0.41 | -0.27 | 0.0575 | -0.3525 | 0.304907 | 0.428751 | 0.884474 | 1 | 0.999631 | 0.716239 |
| Rpl15_Q9CZM2_T197t_1 | 0.84 | 1.16 | 1.65 | 1.36 | 0.12 | 1.48 | 1.55 | 1.6 | -0.14 | 0.64 | 2.32 | -0.78 | -0.5 | 2.39 | 0.14 | 0.95 | -0.065 | -0.7425 | -0.5075 | 0.399563 | 0.870797 | 0.252594 | 1 | 0.999631 | 0.762046 |
| Rpl17_Q6ZWZ7_S141s_1 | -0.72 | -0.11 | 0.57 | 1.15 | -0.03 | -0.45 | 0.02 | 0.68 | -0.27 | -0.58 | 0.96 | 1.12 | -0.13 | -0.01 | 0.05 | 0.79 | -0.1675 | 0.085 | -0.0475 | 0.916574 | 0.711792 | 0.876676 | 1 | 0.999631 | 0.973495 |
| Rpl24_Q8BP67_T83t_1 | 0.62 | 2.62 | 1.93 | 1.76 | 1.7 | 1.31 | 1.1 | 1.72 | 1.62 | 1.26 | 1.68 | 2.44 | 0.97 | 2.44 | 1.52 | 2.34 | -0.275 | 0.0175 | 0.085 | 0.868785 | 0.529375 | 0.970131 | 1 | 0.999631 | 0.957311 |
| Rpl6_P47911_S21s_1 | 2.15 | 0.88 | 1.64 | 1.94 | 0.69 | 0.18 | 1.83 | 0.69 | 1.01 | 1.75 | 0.9 | 1.02 | 0.34 | -0.39 | 1.05 | 0.46 | -0.805 | -0.4825 | -1.2875 | 0.011139 | 0.089558 | 0.210301 | 1 | 0.999631 | 0.385092 |
| Rprd2_Q6NXI6_T332t_1 | 0.59 | 1 | 1.97 | 0.8 | -0.09 | 1.52 | 0.8 | 1.56 | 1.29 | 1.93 | 0.74 | 1.03 | 0.87 | 1.03 | 1.82 | 1.61 | -0.1425 | 0.1575 | 0.2425 | 0.551004 | 0.762155 | 0.69924 | 1 | 0.999631 | 0.836275 |
| Rprd2_Q6NXI6_T698t_1 | 0.4 | 0.67 | 0.79 | 0.64 | 0.61 | 0.34 | 0.08 | 0.78 | 0.52 | 0.51 | 0.45 | 1.02 | 1.05 | 1.16 | 0.66 | 1.21 | -0.1725 | -3.9E-17 | 0.395 | 0.182571 | 0.524864 | 1 | 1 | 1 | 0.643861 |
| Rps17_P63276_S113s_1 | -1.16 | 0.02 | 1.05 | -1.88 | 0.78 | -0.57 | 0.45 | -1.56 | 0.47 | -2.05 | 0.91 | 1.68 | 1.37 | -0.95 | 0.09 | -1.05 | 0.2675 | 0.745 | 0.3575 | 0.636778 | 0.716829 | 0.4159 | 1 | 0.999631 | 0.869841 |
| Rps2_P25444_S264s_1 | 0.33 | 1.34 | 0.56 | 0.23 | 0.33 | 0.72 | 0.19 | 0.87 | 1.05 | 1.25 | 0.87 | 0.96 | 1.28 | 1.68 | 1.04 | 1.75 | -0.0875 | 0.4175 | 0.8225 | 0.040329 | 0.794596 | 0.21491 | 1 | 0.999631 | 0.472911 |
| Rps26_P62855_T96t_1 | 0.73 | 3.66 | 4.77 | 1.47 | 2.34 | 3.92 | 3.76 | 3.2 | 3.26 | 0.35 | 3.79 | 3.51 | 2.31 | 1.38 | 3.96 | 2.13 | 0.6475 | 0.07 | -0.2125 | 0.819878 | 0.464187 | 0.947406 | 1 | 0.999631 | 0.938303 |
| Rps27_Q6ZWU9_S11s_1 | -0.51 | 4.13 | 3.22 | 1.06 | 3.95 | 2.59 | 2.07 | 2.49 | 4.23 | -0.04 | 3.53 | 4.21 | 5.1 | 2.11 | 3.23 | 1.68 | 0.8 | 1.0075 | 1.055 | 0.350182 | 0.417847 | 0.428645 | 1 | 0.999631 | 0.737144 |
| Rps28_G3UYV7_S10s_1 | 1.78 | 0.93 | 1.56 | 1.57 | 1.05 | 1.55 | 2.22 | 0.76 | 0.99 | 1.51 | 0.46 | 0.8 | -0.53 | 0.24 | 0.88 | -0.37 | -0.065 | -0.52 | -1.405 | 0.005038 | 0.864729 | 0.144448 | 1 | 0.999631 | 0.351308 |
| Rps3_P62908_T221t_1 | -0.96 | 2.04 | 1.96 | 0.87 | 1.51 | 1.53 | 0.73 | 1.73 | 1.54 | -0.23 | 2.21 | 2.39 | 2.16 | 1.9 | 1.99 | 2.47 | 0.3975 | 0.5 | 1.1525 | 0.095721 | 0.547056 | 0.539402 | 1 | 0.999631 | 0.558128 |
| Rps6ka2_Q9WUT3_S218s_1 | 0.33 | 0.37 | -2.4 | -0.94 | 0.37 | -1.57 | -1.9 | -3.15 | -0.82 | 1.49 | -1.42 | -1.74 | 0.63 | 0.66 | 0.77 | -0.94 | -0.9025 | 0.0375 | 0.94 | 0.191584 | 0.303502 | 0.965008 | 1 | 0.999631 | 0.648674 |
| Rps6ka2_Q9WUT3_S360s_1 | -0.41 | -0.32 | -0.05 | -0.42 | -0.32 | -0.62 | -0.52 | -0.72 | -0.94 | -0.69 | -0.88 | -0.73 | 0.6 | -0.91 | 0.07 | -0.9 | -0.245 | -0.51 | 0.015 | 0.970443 | 0.334616 | 0.065421 | 1 | 0.999631 | 0.989597 |
| Rps6ka2_Q9WUT3_S377s_1 | 1.18 | -0.48 | -0.61 | 0.88 | 1.65 | 1.16 | 0.57 | 1.4 | 0.69 | -0.4 | 0 | 0.44 | -0.76 | 1.33 | 0.7 | 0.36 | 0.9525 | -0.06 | 0.165 | 0.777228 | 0.074292 | 0.903261 | 1 | 0.999631 | 0.921382 |
| Rps6ka3_B1AXN9_S199s_1 | -0.58 | 0.5 | -0.37 | -0.46 | -0.12 | 0.08 | -0.88 | -0.8 | 0.27 | -0.05 | -0.52 | -0.97 | 0.72 | 0.77 | -0.05 | -0.59 | -0.2025 | -0.09 | 0.44 | 0.310439 | 0.582339 | 0.815633 | 1 | 0.999631 | 0.71924 |
| Rps6ka3_B1AXN9_S341s_1 | -1.46 | -1.01 | -1.7 | -2.35 | -1.27 | -1.29 | -1.93 | -2.87 | -1.06 | -1.3 | -1.46 | -1.31 | -0.05 | -0.97 | -1.67 | -0.96 | -0.21 | 0.3475 | 0.7175 | 0.125577 | 0.643924 | 0.317681 | 1 | 0.999631 | 0.596745 |
| Rps6ka3_B1AXN9_S387s_1 | -0.36 | -0.23 | -0.37 | 0.16 | 0.05 | 0.25 | -0.37 | -0.26 | 0.62 | -0.18 | 0.43 | 0.59 | 0.85 | 0.55 | 0.3 | 0.39 | 0.1175 | 0.565 | 0.7225 | 0.030286 | 0.670718 | 0.084455 | 1 | 0.999631 | 0.43753 |
| Rps6ka3_B1AXN9_S687s_1 | -0.77 | 0.73 | 0.1 | -0.67 | -0.14 | -0.12 | 0.03 | -1.24 | 0.98 | -0.2 | -0.13 | -0.44 | -0.17 | 1.1 | -0.85 | 0.72 | -0.215 | 0.205 | 0.3525 | 0.510672 | 0.630805 | 0.657881 | 1 | 0.999631 | 0.817683 |
| Rps6kb1_Q8BSK8_S424s_1 | 1.21 | 1.66 | -0.31 | 1.08 | 1.8 | 1.46 | 1.01 | 0.71 | 2.41 | 1.69 | 1.12 | 0.93 | 0.66 | 1.57 | 0.95 | 1.09 | 0.335 | 0.6275 | 0.1575 | 0.732739 | 0.478958 | 0.236846 | 1 | 0.999631 | 0.907277 |
| Rps6kc1_E9QMX4_S448s_1 | -0.37 | 0.34 | 1.01 | -0.3 | 0.15 | 0.63 | -0.86 | 0.68 | 0.56 | -0.01 | -0.05 | 0.49 | 1.27 | 1.66 | 0.15 | 1 | -0.02 | 0.0775 | 0.85 | 0.084291 | 0.965385 | 0.839153 | 1 | 0.999631 | 0.549732 |
| Rps6kc1_E9QMX4_S558s_1 | 0.07 | -0.16 | -0.69 | -0.1 | 1.32 | -0.44 | -0.18 | -0.17 | -1.27 | 0 | -0.29 | -0.05 | -0.03 | 0.09 | -0.32 | -0.09 | 0.3525 | -0.1825 | 0.1325 | 0.656189 | 0.416867 | 0.621579 | 1 | 0.999631 | 0.879056 |
| Rps6kc1_E9QMX4_S779s_1 | -0.88 | -0.97 | -0.43 | -0.43 | -0.43 | -0.51 | -0.49 | -0.34 | -0.89 | -0.12 | -0.1 | -0.2 | -0.66 | -0.19 | -0.22 | 0.12 | 0.235 | 0.35 | 0.44 | 0.177065 | 0.37238 | 0.271118 | 1 | 0.999631 | 0.642116 |
| Rps8_P62242_T130t_1 | -0.13 | 0.02 | 0.01 | -0.38 | 0.06 | -0.42 | -0.3 | 0.21 | 0.59 | -0.36 | 0.18 | 0.61 | 0.67 | 1.1 | 1.06 | 0.47 | 0.0075 | 0.375 | 0.945 | 0.008634 | 0.977723 | 0.247238 | 1 | 0.999631 | 0.365244 |
| Rptor_Q8K4Q0_S863s_1 | -1.21 | -0.17 | -0.3 | -1.11 | -0.33 | -0.97 | -0.75 | -0.96 | -0.41 | -1.69 | -0.32 | 0.04 | -0.22 | -1.03 | -0.4 | -1.14 | -0.055 | 0.1025 | 0 | 1 | 0.872197 | 0.821329 | 1 | 0.999631 | 1 |
| Rptor_Q8K4Q0_T706t_1 | 1.88 | -2.38 | 0.54 | 0.93 | 0.98 | 1.94 | 0.14 | 1.85 | 2 | 2.29 | 1.35 | 0.34 | -0.8 | 2.06 | 0.43 | 1.26 | 0.985 | 1.2525 | 0.495 | 0.603505 | 0.277135 | 0.178605 | 1 | 0.999631 | 0.857824 |
| Rrad_Q6PGA2_S38s_1 | 2.46 | 1.28 | 1.03 | 1.11 | 1.33 | 1.2 | 0.9 | 1.1 | 1.55 | 1.92 | 0.88 | 1.61 | 0.11 | 1.41 | 1.83 | 0.28 | -0.3375 | 0.02 | -0.5625 | 0.286378 | 0.36793 | 0.96088 | 1 | 0.999631 | 0.705972 |
| Rragb_G3UYP1_S4s_1 | 1.18 | 0.82 | 0.55 | 0.47 | -0.23 | 0.06 | -0.66 | 0.08 | 0.31 | 0.24 | -1.4 | 0.2 | -0.48 | -0.3 | -0.37 | -0.83 | -0.9425 | -0.9175 | -1.25 | 0.001947 | 0.009985 | 0.059849 | 1 | 0.999631 | 0.351308 |
| Rragc_Q99K70_S15s_1 | 0.1 | 0.23 | 0.2 | 1.33 | 0.67 | 0.19 | 0.89 | 1.58 | 0.7 | -0.08 | 1.4 | 0.67 | -0.41 | 0.39 | 0.56 | 1.71 | 0.3675 | 0.2075 | 0.0975 | 0.845874 | 0.379614 | 0.62458 | 1 | 0.999631 | 0.948565 |
| Rragd_B1AWT4_T270t_1 | 2.37 | 0.53 | 0.37 | 1.52 | 1.08 | 1.09 | 1.72 | 1.33 | 0.84 | 1.57 | 1.18 | -0.9 | -1.29 | 0.35 | 0.32 | 0.14 | 0.1075 | -0.525 | -1.3175 | 0.038938 | 0.818277 | 0.424385 | 1 | 0.999631 | 0.463185 |
| Rras2_P62071_S186s_1 | -1.34 | -0.8 | -1.91 | -1.82 | -0.97 | -1.4 | -1.59 | -1.74 | -0.9 | -1.25 | -1.36 | -1.01 | 0 | -0.97 | -1.59 | -1.99 | 0.0425 | 0.3375 | 0.33 | 0.502435 | 0.900502 | 0.318828 | 1 | 0.999631 | 0.814092 |
| Rrbp1_Q99PL5_S1162s_1 | -0.81 | -0.3 | 0.11 | -0.65 | -0.61 | -0.1 | -1.28 | -1.17 | -0.95 | -0.49 | -1.6 | -0.01 | 1.25 | -0.18 | -1.66 | -0.41 | -0.3775 | -0.35 | 0.1625 | 0.779287 | 0.312131 | 0.397417 | 1 | 0.999631 | 0.921416 |
| Rrbp1_Q99PL5_S786s_1 | 0.22 | 0.73 | 2.58 | -0.18 | -0.64 | 1.39 | 3.53 | 0.8 | -0.09 | 0.38 | 0.07 | 1.23 | 1.76 | -0.79 | 1.38 | -0.54 | 0.4325 | -0.44 | -0.385 | 0.623777 | 0.63846 | 0.479911 | 1 | 0.999631 | 0.863497 |
| Rrp1_P56183_S383s_1 | 1.29 | 1.12 | -0.93 | -1.07 | 1.65 | 1.73 | -0.6 | -1.82 | 1.58 | 1.86 | -2.01 | -1.13 | 1.65 | 0.66 | -1.09 | -1.98 | 0.1375 | -0.0275 | -0.2925 | 0.744994 | 0.882939 | 0.978069 | 1 | 0.999631 | 0.910583 |
| Rrp36_Q3UFY0_T64t_1 | 0.57 | 0.6 | 0.54 | 0.13 | 0.86 | 0.75 | -0.25 | 0.45 | 0.11 | 1.11 | -0.04 | 0.22 | 0.79 | 0.83 | 0.66 | 0.66 | -0.0075 | -0.11 | 0.275 | 0.325269 | 0.981325 | 0.741175 | 1 | 0.999631 | 0.72463 |
| Rsf1_E9PWW9_S1312s_1 | -2.14 | -1.69 | 0.25 | -0.68 | -1.85 | -2.11 | -0.96 | -1.6 | -1.43 | -1.32 | -1.28 | -2.04 | 0.52 | -0.66 | -0.73 | -0.28 | -0.565 | -0.4525 | 0.7775 | 0.187058 | 0.310836 | 0.399624 | 1 | 0.999631 | 0.646474 |
| Rsf1_E9PWW9_T1308t_1 | -0.51 | -1.1 | -1.15 | -0.68 | -0.84 | -1.58 | -0.68 | -0.99 | -0.83 | -0.41 | -1.1 | -1.07 | 0.55 | 0.02 | -2.48 | -1.18 | -0.1625 | 0.0075 | 0.0875 | 0.888762 | 0.600517 | 0.980046 | 1 | 0.999631 | 0.963505 |
| Rsph3b_Q9DA80_T100t_1 | 0.76 | 0.51 | 1.48 | 0.33 | -0.65 | 0.27 | 0.09 | 0.67 | -0.32 | 0.47 | -0.49 | 0.4 | -0.43 | 0.24 | 1.6 | 0.43 | -0.675 | -0.755 | -0.31 | 0.521675 | 0.104658 | 0.068741 | 1 | 0.999631 | 0.820623 |
| Rspry1_Q8BVR6_S50s_1 | -0.91 | 1.6 | 2.23 | -0.11 | 1.86 | 0.69 | 1.63 | 0.81 | 2.67 | 0.01 | 2.3 | 3.07 | 3.67 | 0.35 | 1.86 | 0.27 | 0.545 | 1.31 | 0.835 | 0.379562 | 0.441254 | 0.156854 | 1 | 0.999631 | 0.751933 |
| Rsrc1_Q3TGY1_S6s_1 | 1.68 | -1.21 | 0.68 | 0.32 | -1.38 | -0.51 | 2.02 | -1.1 | -0.84 | 1.72 | -0.93 | -1.94 | -1.21 | -3.02 | -0.06 | -1.1 | -0.61 | -0.865 | -1.715 | 0.043255 | 0.47984 | 0.326732 | 1 | 0.999631 | 0.475443 |
| Rsrc2_S4R2L4_S103s_1 | 2.05 | 0.97 | 1.97 | 1.51 | 0.63 | 0.91 | 2.05 | 0.4 | 0.41 | 1.99 | 1 | -0.08 | 0.59 | -0.7 | 0.61 | 0.04 | -0.6275 | -0.795 | -1.49 | 0.004558 | 0.170053 | 0.127488 | 1 | 0.999631 | 0.351308 |
| Rsrc2_S4R2L4_S104s_1 | 2 | 0.96 | 2.06 | 1.35 | 0.51 | 1.05 | 2.16 | 0.22 | 0.19 | 1.93 | 1.13 | -0.07 | 0.72 | -0.79 | 0.68 | -0.03 | -0.6075 | -0.7975 | -1.4475 | 0.008224 | 0.222842 | 0.136108 | 1 | 0.999631 | 0.365244 |
| Rsrc2_S4R2L4_S32s_1 | -0.59 | 0.67 | 2.18 | 0.11 | -0.31 | 0.43 | 0.76 | 0.95 | 0.75 | -0.38 | 1.5 | 1.88 | 1.21 | 0.24 | 0.82 | 0.92 | -0.135 | 0.345 | 0.205 | 0.721419 | 0.818999 | 0.618256 | 1 | 0.999631 | 0.905647 |
| Rtkn_D3Z4N1_S125s_1 | 1.23 | 0.33 | 0.05 | 1.35 | 0.87 | 0.59 | 0.56 | 0.88 | 0.95 | 0.6 | 0.73 | 0.19 | -0.4 | 1.08 | 1.15 | 0.49 | -0.015 | -0.1225 | -0.16 | 0.735345 | 0.966663 | 0.748979 | 1 | 0.999631 | 0.90759 |
| Rtkn_D3Z4N1_S134s_1 | -0.43 | 1.06 | 1.38 | -0.3 | -1.49 | 0.88 | 0.46 | 0.4 | 0.76 | 0.29 | 0.63 | 1.74 | 1.62 | 1.12 | -0.08 | 0.48 | -0.365 | 0.4275 | 0.3575 | 0.521342 | 0.565714 | 0.422036 | 1 | 0.999631 | 0.820259 |
| Rtkn_D3Z4N1_S56s_1 | 0.81 | 0.4 | 0.17 | 0.51 | 0.76 | 0.99 | 0.8 | 0.88 | 0.64 | 1.68 | 0.68 | 0.27 | 1.01 | 0.75 | 0.21 | 0.17 | 0.385 | 0.345 | 0.0625 | 0.847286 | 0.154406 | 0.352254 | 1 | 0.999631 | 0.948838 |
| Rtkn_Q8C6B2_S232s_1 | -0.75 | -0.27 | -1.16 | -0.01 | -0.12 | 0.16 | -0.92 | -1.07 | -0.79 | -0.72 | -0.69 | 0.16 | 0.75 | -0.2 | -0.42 | 0.24 | 0.06 | 0.0375 | 0.64 | 0.126505 | 0.88038 | 0.918866 | 1 | 0.999631 | 0.596745 |
| Rtkn_Q8C6B2_S520s_1 | 0.08 | 0.72 | 0.55 | -0.09 | 0.61 | -0.25 | 0.57 | -1.2 | -0.75 | -0.14 | -0.35 | 1.39 | -0.1 | 0.33 | 0.18 | 0.84 | -0.3825 | -0.2775 | -0.0025 | 0.994127 | 0.405518 | 0.569873 | 1 | 0.999631 | 0.997705 |
| Rtn1_Q7M6W1_S7s_1 | -0.21 | -1.17 | 0.4 | 0.73 | -0.31 | -2.42 | 0.58 | 0.32 | -2.06 | -1.36 | 0.97 | 0.51 | -0.93 | -1.61 | -0.13 | -0.43 | -0.395 | -0.4225 | -0.7125 | 0.179221 | 0.578622 | 0.572224 | 1 | 0.999631 | 0.64376 |
| Rtn1_Q8K0T0_S16s_1 | -0.2 | 0.03 | -0.48 | 0.01 | -0.26 | -0.51 | -0.68 | -0.05 | -0.31 | 0.01 | -0.74 | 0.33 | 0.86 | 0.09 | -0.86 | -0.23 | -0.215 | -0.0175 | 0.125 | 0.755539 | 0.437192 | 0.956131 | 1 | 0.999631 | 0.913663 |
| Rtn1_Q8K0T0_S352s_1 | 1.83 | 1.43 | -0.64 | 0.73 | 2.3 | 1.33 | 0.17 | 0.61 | 2.6 | 1.48 | 0.63 | 0.41 | 1.34 | 1.83 | 0.31 | -0.17 | 0.265 | 0.4425 | -0.01 | 0.987514 | 0.679812 | 0.507654 | 1 | 0.999631 | 0.995396 |
| Rtn1_Q8K0T0_S487s_1 | -0.56 | 0.4 | 0.04 | -0.4 | 0.08 | 0.09 | -0.59 | 0.04 | 0.2 | -0.43 | -0.18 | 0.26 | 0.85 | 0.07 | 0.28 | -0.13 | 0.035 | 0.0925 | 0.3975 | 0.279632 | 0.913061 | 0.777318 | 1 | 0.999631 | 0.701922 |
| Rtn1_Q8K0T0_S70s_1 | -0.56 | -0.68 | -1.01 | -0.99 | -0.54 | -0.87 | -0.92 | -1.03 | -1.09 | -0.98 | -1.18 | -0.99 | -0.12 | -0.73 | -0.79 | -0.57 | -0.03 | -0.25 | 0.2575 | 0.396193 | 0.908017 | 0.340824 | 1 | 0.999631 | 0.760566 |
| Rtn1_Q8K0T0_S92s_1 | 0.19 | -0.79 | -1.09 | -0.27 | -0.25 | -0.79 | -0.38 | -0.13 | -0.79 | -1.02 | -0.72 | -0.66 | -1.2 | -0.43 | -0.62 | -0.47 | 0.1025 | -0.3075 | -0.19 | 0.614018 | 0.768599 | 0.37561 | 1 | 0.999631 | 0.860996 |
| Rtn3_Q9ES97_S113s_1 | 0.63 | 0.37 | -0.05 | 0.61 | 0.15 | -0.1 | 0.73 | -0.1 | 0.25 | 0.35 | 0.04 | -0.25 | -0.22 | 0.33 | 0.06 | -0.57 | -0.22 | -0.2925 | -0.49 | 0.155963 | 0.481264 | 0.328196 | 1 | 0.999631 | 0.626452 |
| Rtn3_Q9ES97_S31s_1 | -0.89 | -0.7 | -0.54 | -0.98 | -1.01 | -0.64 | -1.11 | -0.65 | -0.56 | -0.3 | -0.69 | -0.42 | 0.44 | -0.36 | -0.42 | -0.24 | -0.075 | 0.285 | 0.6325 | 0.06612 | 0.773903 | 0.285777 | 1 | 0.999631 | 0.516933 |
| Rtn3_Q9ES97_S510s_1 | -0.83 | -0.31 | -0.75 | -1.23 | -0.42 | -0.85 | -0.51 | -1.56 | -0.94 | -1.91 | -1.46 | -1.02 | -0.55 | -0.56 | -0.91 | -0.84 | -0.055 | -0.5525 | 0.065 | 0.832617 | 0.875005 | 0.127535 | 1 | 0.999631 | 0.94321 |
| Rtn3_Q9ES97_S654s_1 | -1.29 | -0.36 | -0.53 | -0.91 | -0.81 | -0.79 | -0.8 | -0.73 | -0.82 | -1.09 | -0.44 | -0.27 | 0.14 | -0.53 | -0.42 | -0.36 | -0.01 | 0.1175 | 0.48 | 0.167098 | 0.97196 | 0.722623 | 1 | 0.999631 | 0.636719 |
| Rtn3_Q9ES97_S735s_1 | 0.34 | -0.32 | -1.05 | 0.48 | 0.73 | -0.06 | 0.08 | 0 | -0.08 | -0.7 | 0.19 | -0.47 | -0.94 | 0.18 | -0.53 | -0.06 | 0.325 | -0.1275 | -0.2 | 0.647872 | 0.424538 | 0.756546 | 1 | 0.999631 | 0.874615 |
| Rtn3_Q9ES97_T505t_1 | -0.77 | 0.13 | -0.25 | -0.05 | -0.4 | -0.22 | 0.05 | -0.74 | -0.47 | -0.73 | -0.31 | -0.27 | -0.23 | -0.62 | -0.67 | -0.05 | -0.0925 | -0.21 | -0.1575 | 0.629663 | 0.766101 | 0.487405 | 1 | 0.999631 | 0.866951 |
| Rtn4_Q99P72_S145s_1 | 0.22 | 0.26 | 0.13 | 0.37 | 0.37 | 0.6 | 0.39 | 0.97 | 0.34 | 0.67 | 0.38 | 0.51 | -0.44 | 0.95 | 0.57 | 0.72 | 0.3375 | 0.23 | 0.205 | 0.572572 | 0.209651 | 0.361664 | 1 | 0.999631 | 0.844402 |
| Rtn4_Q99P72_S165s_1 | -0.51 | 0.13 | 0.22 | -0.11 | -0.25 | -0.32 | -0.28 | 0.17 | -0.25 | -0.64 | -0.04 | 0.3 | 0.18 | 0.27 | 0.26 | 0.54 | -0.1025 | -0.09 | 0.38 | 0.216099 | 0.715355 | 0.776987 | 1 | 0.999631 | 0.668933 |
| Rtn4_Q99P72_S167s_1 | -1 | -0.83 | -0.93 | -0.69 | -0.96 | -1.14 | -0.92 | -0.25 | -1.15 | -0.96 | -0.78 | -0.63 | -0.93 | -0.6 | -0.55 | 0 | 0.045 | -0.0175 | 0.3425 | 0.276277 | 0.874027 | 0.946088 | 1 | 0.999631 | 0.700187 |
| Rtn4_Q99P72_S16s_1 | -0.75 | -0.29 | -0.53 | -0.81 | -0.58 | -0.66 | -0.91 | -0.58 | -0.55 | -0.49 | -0.68 | -0.3 | 0.23 | -0.19 | -0.43 | -0.27 | -0.0875 | 0.09 | 0.43 | 0.167788 | 0.731908 | 0.732872 | 1 | 0.999631 | 0.637399 |
| Rtn4_Q99P72_S223s_1 | 1.15 | -1.17 | 0.03 | 1.2 | 1.86 | 0.08 | 0.84 | 0.91 | -0.17 | -1.07 | 0.23 | -0.54 | -1.66 | 0.19 | -0.92 | -0.19 | 0.62 | -0.69 | -0.9475 | 0.15354 | 0.317826 | 0.247289 | 1 | 0.999631 | 0.625198 |
| Rtn4_Q99P72_S329s_1 | -1.29 | 1.07 | 0.6 | 0.23 | 0.32 | 0.16 | 0.99 | 0.71 | 0.55 | -0.2 | 0.51 | 1.48 | 0.93 | 0.38 | 1.52 | 0.81 | 0.3925 | 0.4325 | 0.7575 | 0.173912 | 0.447044 | 0.452758 | 1 | 0.999631 | 0.642116 |
| Rtn4_Q99P72_S344s_1 | -0.16 | -0.3 | -1.92 | -2.04 | -0.42 | -0.3 | -1.62 | -2.06 | -0.09 | 0.19 | -1.55 | -2.08 | 0.22 | -0.43 | -2.15 | -2.27 | 0.005 | 0.2225 | -0.0525 | 0.941018 | 0.993389 | 0.740708 | 1 | 0.999631 | 0.981286 |
| Rtn4_Q99P72_S690s_1 | -0.24 | -0.53 | -1.12 | -0.36 | -0.18 | -0.42 | -0.65 | -0.14 | -0.32 | -0.34 | -0.31 | -0.14 | -0.74 | -0.15 | -0.03 | 0.21 | 0.215 | 0.285 | 0.385 | 0.277604 | 0.472695 | 0.335573 | 1 | 0.999631 | 0.700874 |
| Rtn4_Q99P72_S768s_1 | -0.16 | -0.8 | -1.76 | -1.6 | -0.73 | -0.39 | -0.58 | -1.42 | -0.74 | -0.42 | -1.86 | -1.41 | -1.1 | -0.57 | -1.78 | -1.19 | 0.3 | -0.0275 | -0.08 | 0.857949 | 0.488148 | 0.953819 | 1 | 0.999631 | 0.952251 |
| Rtn4_Q99P72_S857s_1 | -1.43 | -1.38 | -1.17 | -1.2 | -1.35 | -0.58 | -0.72 | -1.47 | -0.92 | -1.45 | -0.04 | -1.02 | -0.39 | -1.23 | -0.56 | -0.36 | 0.265 | 0.4375 | 0.66 | 0.054399 | 0.382649 | 0.224134 | 1 | 0.999631 | 0.491994 |
| Rtn4_Q99P72_S953s_1 | 1.86 | 1.57 | 1.84 | 1.66 | 1.15 | 1.81 | 1.25 | 1.41 | 1.48 | 2.29 | 1.17 | 1.36 | 1.81 | 0.95 | 1.53 | 1.06 | -0.3275 | -0.1575 | -0.395 | 0.219883 | 0.231526 | 0.622377 | 1 | 0.999631 | 0.670699 |
| Rtn4_Q99P72_S961s_1 | -0.15 | 1.07 | -0.25 | -0.22 | 0.48 | 0.22 | -0.72 | 0.04 | 1.08 | 0.71 | 0.13 | -0.02 | 1.49 | 0.73 | -0.14 | 0.13 | -0.1075 | 0.3625 | 0.44 | 0.362165 | 0.793711 | 0.392251 | 1 | 0.999631 | 0.742225 |
| Rtn4_Q99P72_T430t_1 | -0.63 | -2.31 | -2.86 | -0.92 | 0.05 | -3.02 | -0.56 | -2.44 | -1.1 | -2.64 | -1.14 | -0.69 | -1.49 | -2.61 | -2.35 | -2.83 | 0.1875 | 0.2875 | -0.64 | 0.273929 | 0.813778 | 0.645798 | 1 | 0.999631 | 0.700187 |
| Rtn4_Q99P72_T488t_1 | -4.29 | -8.57 | -3.13 | -2.22 | -6.49 | -6.84 | -2.07 | -2.87 | -8.19 | -5.02 | -1.85 | -2.44 | -7.12 | -6.34 | -3.68 | -2.68 | -0.015 | 0.1775 | -0.4025 | 0.78534 | 0.992408 | 0.917221 | 1 | 0.999631 | 0.923063 |
| Rufy3_Q9D394_S27s_1 | 0.58 | 0.58 | 0.58 | 0.33 | -0.11 | 0.54 | 0.49 | 0.35 | 0.25 | 0.8 | 0.92 | 0.89 | 0.09 | 0.51 | 1.12 | 1.07 | -0.2 | 0.1975 | 0.18 | 0.586413 | 0.453114 | 0.476992 | 1 | 0.999631 | 0.849724 |
| Rufy3_Q9D394_T51t_1 | -0.93 | -0.09 | -0.29 | -0.94 | 0.11 | -0.44 | -0.68 | -0.23 | -0.43 | -0.7 | -0.29 | -0.43 | -0.02 | 0.05 | -0.1 | -0.33 | 0.2525 | 0.1 | 0.4625 | 0.16867 | 0.440667 | 0.744405 | 1 | 0.999631 | 0.637839 |
| Rufy3_Q9D394_T5t_1 | -0.59 | -0.02 | -0.1 | -0.97 | -0.4 | -0.52 | -0.96 | -0.6 | -0.27 | -0.21 | -0.36 | 0.11 | 0.86 | 0.25 | -0.24 | 0.04 | -0.2 | 0.2375 | 0.6475 | 0.102078 | 0.52204 | 0.452538 | 1 | 0.999631 | 0.564448 |
| Rundc3a_O08576_S361s_1 | -0.71 | 0.53 | -0.44 | -0.64 | 0.54 | 0.06 | 0.42 | -0.18 | 1.53 | -0.46 | 0.09 | 0.77 | 1.94 | -0.1 | -0.08 | 0.04 | 0.525 | 0.7975 | 0.765 | 0.176741 | 0.164144 | 0.130239 | 1 | 0.999631 | 0.642116 |
| Rusc1_Q8BG26_S367s_1 | -0.34 | -1.91 | -0.1 | -0.22 | -1 | -1.47 | 0.05 | -0.16 | -1.45 | -2.04 | -0.1 | -0.04 | -2.03 | -1.9 | -0.23 | 0.45 | -0.0025 | -0.265 | -0.285 | 0.671012 | 0.996132 | 0.658862 | 1 | 0.999631 | 0.884528 |
| Rusc1_Q8BG26_T355t_1 | -0.22 | 0.05 | -1.93 | -1.19 | 0.2 | -0.38 | -1.26 | -0.76 | 0.2 | -0.54 | -1.05 | -0.56 | -0.81 | -0.09 | -1.3 | -0.55 | 0.2725 | 0.335 | 0.135 | 0.786904 | 0.597368 | 0.50542 | 1 | 0.999631 | 0.92391 |
| Rusc2_Q3V1Z0_S242s_1 | 0.78 | -0.27 | -1.8 | -1.56 | -0.17 | 0.49 | -1.56 | -1.53 | -0.13 | 0 | -1.7 | -1.72 | -1.68 | -1.18 | -1.88 | -2.7 | 0.02 | -0.175 | -1.1475 | 0.086424 | 0.9771 | 0.798244 | 1 | 0.999631 | 0.55299 |
| Rusc2_Q3V1Z0_S542s_1 | -0.91 | -0.48 | -0.67 | -1.3 | -0.38 | -0.85 | -0.74 | -1.12 | -0.37 | -1.74 | -0.14 | -0.29 | 0.15 | -1.01 | -0.54 | -1.29 | 0.0675 | 0.205 | 0.1675 | 0.6695 | 0.821135 | 0.624556 | 1 | 0.999631 | 0.883626 |
| Rusc2_Q3V1Z0_S565s_1 | 0.05 | -0.48 | 0.01 | -0.63 | -0.61 | -0.48 | -0.57 | -0.53 | -0.66 | -0.32 | -0.29 | -0.36 | 0.14 | -0.32 | -0.48 | -0.26 | -0.285 | -0.145 | 0.0325 | 0.916528 | 0.302689 | 0.612265 | 1 | 0.999631 | 0.973495 |
| Rusc2_Q3V1Z0_S580s_1 | -0.68 | -0.6 | -0.41 | -1.14 | -0.75 | -1.12 | -0.76 | -1.23 | -0.66 | -0.77 | -0.92 | -1.32 | 0.33 | -0.15 | -0.72 | -0.72 | -0.2575 | -0.21 | 0.3925 | 0.280793 | 0.368016 | 0.480834 | 1 | 0.999631 | 0.701922 |
| Rusc2_Q3V1Z0_S779s_1 | -1.33 | -0.74 | -0.69 | -1 | -0.44 | -1.02 | -0.83 | -0.87 | -0.4 | 0.63 | -0.23 | 0.43 | 0.06 | -0.22 | -1.02 | 0.52 | 0.15 | 1.0475 | 0.775 | 0.068451 | 0.590494 | 0.010742 | 1 | 0.999631 | 0.523753 |
| Rusc2_Q3V1Z0_S785s_1 | -1.46 | -1.22 | -0.9 | -1.38 | -2.15 | -1.43 | -1.24 | -1.03 | -1.45 | -1.18 | -1.35 | -0.71 | 0.26 | -1.69 | -1.06 | -0.52 | -0.2225 | 0.0675 | 0.4875 | 0.279862 | 0.493183 | 0.816549 | 1 | 0.999631 | 0.701922 |
| Ryr2_E9Q401_S1495s_1 | -0.22 | -0.08 | 0.52 | -0.2 | -0.2 | -0.17 | -0.31 | -0.18 | -0.16 | -0.34 | -0.3 | -0.38 | 0.36 | -0.43 | -0.44 | -0.46 | -0.22 | -0.3 | -0.2475 | 0.466692 | 0.423185 | 0.297302 | 1 | 0.999631 | 0.795267 |
| Ryr2_E9Q401_S2362s_1 | 0.74 | 2.67 | 1.88 | -0.01 | 0.7 | 2.63 | 1.79 | 0.37 | 1.34 | 0.75 | 0.91 | 0.92 | 2.08 | 0.42 | 1.15 | 0.22 | 0.0525 | -0.34 | -0.3525 | 0.592145 | 0.939962 | 0.548227 | 1 | 0.999631 | 0.852774 |
| Ryr2_E9Q401_S2367s_1 | 0.16 | 0.73 | 0.83 | 0.82 | -0.14 | 1.09 | 0.94 | 0.07 | 0.29 | -0.21 | 0.92 | 0.34 | 0.58 | -0.01 | 0.95 | -0.87 | -0.145 | -0.3 | -0.4725 | 0.291307 | 0.6937 | 0.376631 | 1 | 0.999631 | 0.706164 |
| Ryr2_E9Q401_S2693s_1 | -0.3 | -0.36 | 0.34 | 0.18 | -0.03 | -0.01 | -0.54 | 0.06 | -0.25 | -0.59 | -0.2 | 0.37 | 0.76 | 0.01 | 0.03 | 0.4 | -0.095 | -0.1325 | 0.335 | 0.317443 | 0.745403 | 0.682109 | 1 | 0.999631 | 0.721033 |
| Ryr2_E9Q401_S2807s_1 | -0.29 | -0.31 | -0.97 | -0.88 | -0.72 | -0.9 | -0.42 | -0.82 | -1.24 | 0.07 | -1.05 | -1.74 | -0.91 | -1.19 | -1.66 | -1.06 | -0.1025 | -0.3775 | -0.5925 | 0.091027 | 0.720337 | 0.383994 | 1 | 0.999631 | 0.557085 |
| Ryr2_E9Q401_S2813s_1 | -0.87 | -0.77 | -0.34 | -0.26 | 0.01 | 0.24 | -0.1 | -0.6 | -0.67 | -1.65 | -0.97 | 0.41 | -0.82 | -0.4 | -0.65 | -0.16 | 0.4475 | -0.16 | 0.0525 | 0.864286 | 0.156699 | 0.72135 | 1 | 0.999631 | 0.955131 |
| Ryr2_E9Q401_S4549s_1 | 1.41 | 0.56 | 0.11 | 0.26 | 1.98 | 1.24 | 0.08 | 0.26 | 1.33 | 1.19 | -0.85 | -0.04 | 0.93 | -0.08 | -1.69 | -0.49 | 0.305 | -0.1775 | -0.9175 | 0.129888 | 0.543996 | 0.748126 | 1 | 0.999631 | 0.601246 |
| S1pr2_P52592_S332s_1 | 0.83 | -0.31 | 0.88 | 0.16 | -0.93 | -0.08 | 0.89 | -0.27 | -0.18 | 1.34 | -0.31 | -0.15 | -0.03 | -0.87 | 0.48 | -0.96 | -0.4875 | -0.215 | -0.735 | 0.124812 | 0.298406 | 0.647381 | 1 | 0.999631 | 0.596063 |
| S1pr5_Q91X56_S340s_1 | -0.11 | -0.73 | 0.32 | -0.08 | -1.8 | -1.1 | -0.54 | -0.18 | -1 | -0.47 | -0.37 | 0.14 | -0.45 | -0.13 | 0.2 | -0.09 | -0.755 | -0.275 | 0.0325 | 0.921282 | 0.09257 | 0.445713 | 1 | 0.999631 | 0.975032 |
| S1pr5_Q91X56_S342s_1 | 0.66 | 1.56 | 1.32 | 0.75 | -0.09 | 0.92 | 0.16 | 1.17 | 0.51 | 0.32 | -0.04 | 0.25 | 0.64 | 1.05 | 0.75 | 0.53 | -0.5325 | -0.8125 | -0.33 | 0.322143 | 0.186415 | 0.02511 | 1 | 0.999631 | 0.72427 |
| Sacs_E9QNY8_S1781s_1 | -0.01 | 0.89 | 0.03 | -0.45 | 1.88 | 1.19 | 0.21 | -0.79 | 1.82 | -1.72 | -0.16 | -0.18 | 0.62 | 1.57 | -0.3 | -0.68 | 0.5075 | -0.175 | 0.1875 | 0.728471 | 0.396072 | 0.800886 | 1 | 0.999631 | 0.905923 |
| Sacs_E9QNY8_S2513s_1 | 0.33 | 1.42 | 1.17 | 0.96 | 1.92 | 1.8 | 0.99 | 1.31 | 0.69 | 1.12 | 1.28 | 1.83 | -0.18 | 2.16 | 1.48 | 0.92 | 0.535 | 0.26 | 0.125 | 0.809388 | 0.148192 | 0.479796 | 1 | 0.999631 | 0.936582 |
| Sacs_E9QNY8_S4266s_1 | -0.49 | -1 | -0.06 | -0.48 | -0.21 | 0.04 | -0.54 | -0.44 | -0.46 | -0.19 | -0.41 | 0.07 | -0.15 | -0.55 | -0.38 | -0.26 | 0.22 | 0.26 | 0.1725 | 0.577739 | 0.465543 | 0.398413 | 1 | 0.999631 | 0.84622 |
| Safb_S4R1M2_S298s_1 | 1.97 | -1.59 | 0.16 | -0.21 | -1.95 | -0.67 | 2.23 | -2.41 | -2.65 | 1.65 | -0.92 | -2 | -1.11 | -1.87 | -2.13 | -1.63 | -0.7825 | -1.0625 | -1.7675 | 0.025042 | 0.479818 | 0.316987 | 1 | 0.999631 | 0.419285 |
| Safb_S4R1M2_S366s_1 | 0.02 | 0.81 | -0.83 | -1.05 | 0.25 | 0.38 | -1.19 | -0.74 | 0.18 | 0.58 | -1.05 | -0.89 | 0.84 | 0.79 | -0.79 | -0.71 | -0.0625 | -0.0325 | 0.295 | 0.608387 | 0.90587 | 0.95219 | 1 | 0.999631 | 0.858627 |
| Safb_S4R1M2_S626s_1 | -0.13 | 1.44 | 1.95 | 0.12 | 0.35 | 1.01 | -0.02 | 0.87 | 0.71 | 1.03 | 0.66 | 2.3 | 0.74 | 2.43 | 2.61 | 1.84 | -0.2925 | 0.33 | 1.06 | 0.101346 | 0.57556 | 0.57311 | 1 | 0.999631 | 0.564448 |
| Safb_S4R1M2_S783s_1 | 1.17 | -1.35 | -2.9 | -0.13 | 0.18 | -0.63 | -0.11 | -1.28 | -0.39 | -0.41 | -1.57 | -2.86 | -2.32 | -0.47 | -2.66 | -2.86 | 0.3425 | -0.505 | -1.275 | 0.171942 | 0.672722 | 0.581916 | 1 | 0.999631 | 0.641657 |
| Samd1_D3YXK1_S150s_1 | 0.6 | 0.55 | 1.72 | 0.59 | 0.14 | 0.11 | 0.66 | -0.45 | 0.51 | 1.92 | 1.45 | 2.88 | 0.63 | 0.14 | 1.41 | 1.08 | -0.75 | 0.825 | -0.05 | 0.903745 | 0.071632 | 0.144954 | 1 | 0.999631 | 0.968183 |
| Samd1_D3YXK1_S242s_1 | 0.24 | 1.38 | 1.3 | 1.09 | 0.28 | 1.07 | 0.66 | 0.76 | 0.78 | 1.26 | 1.13 | 1.41 | 1 | 0.82 | 0.76 | 1.45 | -0.31 | 0.1425 | 0.005 | 0.988807 | 0.375119 | 0.676132 | 1 | 0.999631 | 0.995427 |
| Samd1_D3YXK1_T107t_1 | 0.53 | 0.58 | 1.24 | 0.82 | -0.21 | 0.07 | 0.5 | 0.91 | 0.33 | 1.01 | 1 | 1.03 | 0.06 | 0.38 | 0.83 | 1.19 | -0.475 | 0.05 | -0.1775 | 0.616424 | 0.176425 | 0.87031 | 1 | 0.999631 | 0.861203 |
| Samd14_Q8K070_S108s_1 | 0.03 | -0.54 | -0.36 | -0.22 | 0.3 | -1.24 | -0.73 | 0.09 | -0.69 | 0.28 | 0.17 | 0.27 | 0.09 | -0.17 | 0.11 | 0.61 | -0.1225 | 0.28 | 0.4325 | 0.176097 | 0.752714 | 0.393325 | 1 | 0.999631 | 0.642116 |
| Samd14_Q8K070_S140s_1 | -0.48 | -0.03 | -0.56 | -1.98 | -2.09 | -1.54 | -1.47 | -0.97 | -0.08 | 0.89 | -1.54 | -1 | 1.99 | 0.33 | -1.3 | -1.43 | -0.755 | 0.33 | 0.66 | 0.413459 | 0.125905 | 0.595564 | 1 | 0.999631 | 0.768474 |
| Samd14_Q8K070_S151s_1 | 0.97 | -1.3 | -1.13 | 0.61 | 1.37 | 0.11 | 0.3 | 1.22 | 0.4 | -0.04 | 0.43 | -0.22 | -1.96 | 0.51 | -0.34 | -0.34 | 0.9625 | 0.355 | -0.32 | 0.645715 | 0.13375 | 0.53037 | 1 | 0.999631 | 0.873384 |
| Samd14_Q8K070_S165s_1 | 2.11 | -1.47 | -0.7 | 0.22 | -1.31 | -1.06 | 1.76 | -1.24 | -0.75 | 1.38 | -1.17 | -1.74 | -2.38 | -1.5 | -1.59 | -2.13 | -0.5025 | -0.61 | -1.94 | 0.019619 | 0.589881 | 0.500798 | 1 | 0.999631 | 0.391668 |
| Samd14_Q8K070_S173s_1 | -0.73 | 2.89 | 2.37 | -0.09 | 2.05 | 1.66 | 0.52 | 1.26 | 2.21 | -0.6 | 1.85 | 2.79 | 2.74 | 1.78 | 2.37 | 1.5 | 0.2625 | 0.4525 | 0.9875 | 0.243262 | 0.75181 | 0.654143 | 1 | 0.999631 | 0.685637 |
| Samd14_Q8K070_S256s_1 | 1.73 | -0.54 | 0.42 | 1.18 | 0.39 | 0.46 | 1.42 | 0.26 | 1.36 | 1.09 | -0.65 | 0.22 | -0.25 | -0.46 | 0.12 | -0.99 | -0.065 | -0.1925 | -1.0925 | 0.056479 | 0.900394 | 0.752506 | 1 | 0.999631 | 0.496238 |

| | | | | | | | | | | | | | | | | | | | | | | | | | |
|---|---|---|---|---|---|---|---|---|---|---|---|---|---|---|---|---|---|---|---|---|---|---|---|---|---|
| Samd14_Q8K070_S260s_1 | -0.36 | -0.58 | 0.43 | -0.62 | -1.99 | 0.41 | -0.13 | 0.05 | -0.43 | 1.02 | -1.2 | 0.17 | 1.54 | -0.79 | 0.81 | -0.94 | -0.1325 | 0.1725 | 0.4375 | 0.47045 | 0.807273 | 0.731594 | 1 | 0.999631 | 0.797463 |
| Samd14_Q8K070_S77s_1 | 0.13 | 0.62 | 0.28 | 0.42 | 1.29 | 1.35 | 0.5 | 1.1 | 1.91 | 1.36 | 0.46 | 0.15 | 0.67 | 2.31 | 0.62 | 0.3 | 0.6975 | 0.6075 | 0.6125 | 0.202468 | 0.035534 | 0.171802 | 1 | 0.999631 | 0.659062 |
| Samd4b_G5E8A7_S271s_1 | -0.46 | -0.04 | 0.05 | -0.75 | -0.3 | -0.15 | -0.5 | -0.14 | -0.63 | -0.7 | 0.09 | -0.68 | -0.03 | -0.13 | -0.78 | -0.38 | 0.0275 | -0.18 | -0.03 | 0.926881 | 0.922561 | 0.581114 | 1 | 0.999631 | 0.976306 |
| Samd8_D6REI9_S15s_1 | 1.32 | 0.47 | -1.3 | 1.65 | 1.66 | 0.74 | 0.99 | 1.65 | 0.93 | 0.12 | 1.22 | 0 | -0.08 | 0.6 | 0.01 | 0.02 | 0.725 | 0.0325 | -0.3975 | 0.522563 | 0.265388 | 0.960204 | 1 | 0.999631 | 0.821224 |
| Sap130_J3QNK5_S274s_1 | 0.3 | -0.69 | -1.53 | -2 | -0.28 | -0.43 | -0.31 | -0.4 | -0.57 | 0.42 | -1.8 | -1.03 | -0.06 | -0.26 | -1.24 | -0.97 | 0.625 | 0.235 | 0.3475 | 0.524795 | 0.213094 | 0.706146 | 1 | 0.999631 | 0.822727 |
| Sarm1_Q6PDS3_S548s_1 | -0.1 | -1.01 | -1.47 | -1.34 | -1.02 | -0.93 | -1.55 | -1.35 | -0.92 | -0.11 | -1.26 | 0.32 | 0 | -0.23 | -0.17 | -0.81 | -0.2325 | 0.4875 | 0.6775 | 0.10445 | 0.526329 | 0.306475 | 1 | 0.999631 | 0.568213 |
| Sarm1_Q6PDS3_S558s_1 | 0.23 | -0.72 | -0.39 | -0.7 | -0.32 | -0.14 | -0.1 | -0.73 | -0.38 | 0.47 | -0.05 | -0.06 | -1 | 0.04 | -0.09 | 0.27 | 0.0725 | 0.39 | 0.2 | 0.608148 | 0.818368 | 0.258822 | 1 | 0.999631 | 0.858527 |
| Sarm1_Q6PDS3_S572s_1 | -1.11 | -1.18 | -0.91 | -1.62 | -0.09 | -1.3 | -0.85 | -1.49 | 0.18 | -0.8 | 0.3 | 1.52 | -0.11 | -0.29 | -0.5 | -0.48 | 0.2725 | 1.505 | 0.86 | 0.013719 | 0.463076 | 0.010243 | 1 | 0.999631 | 0.385092 |
| Sash1_F8VQK5_S1020s_1 | 0.73 | 0.16 | 0.31 | 0 | 0.57 | 0.25 | 0.32 | 0.58 | 1.04 | 0.58 | 0.68 | 0.01 | -0.06 | 0.54 | 0.7 | 0.81 | 0.13 | 0.2775 | 0.1975 | 0.548902 | 0.632469 | 0.398857 | 1 | 0.999631 | 0.835034 |
| Sash1_F8VQK5_S241s_1 | 1.02 | 1.3 | 1.57 | 1.1 | -0.35 | 0.99 | 1.57 | -0.25 | 0.79 | 2.39 | -0.16 | 0.39 | -0.11 | -0.36 | 1.36 | 0.08 | -0.7575 | -0.395 | -1.005 | 0.034342 | 0.128439 | 0.459442 | 1 | 0.999631 | 0.446273 |
| Sash1_F8VQK5_S500s_1 | -0.49 | -1.12 | -0.89 | -0.29 | -0.92 | -0.31 | 0.09 | -0.25 | -0.44 | 0.01 | -0.68 | -0.28 | -0.11 | -0.31 | -0.41 | -0.05 | 0.35 | 0.35 | 0.4775 | 0.141188 | 0.297766 | 0.271063 | 1 | 0.999631 | 0.61245 |
| Sash1_F8VQK5_S723s_1 | -0.8 | 0.52 | 0.98 | 0.62 | 0.85 | 0.17 | 0.65 | 0.16 | 1.2 | -2.77 | -1.13 | 0.16 | -0.74 | 0.57 | 0.81 | 1.59 | 0.1275 | -0.965 | 0.2275 | 0.692049 | 0.762718 | 0.258665 | 1 | 0.999631 | 0.894017 |
| Sash1_F8VQK5_S805s_1 | 0.62 | 2.16 | 1.77 | 0.94 | 0.87 | 2.29 | 0.19 | 1.61 | 2.15 | 1.75 | 1.32 | 1.67 | 2.55 | 3.13 | 1.68 | 1.77 | -0.1325 | 0.35 | 0.91 | 0.082697 | 0.804118 | 0.397627 | 1 | 0.999631 | 0.548633 |
| Sash1_F8VQK5_T279t_1 | 0.07 | -0.24 | -1.39 | -0.12 | 0.56 | 0.49 | 0.33 | 0.72 | -0.47 | 0.36 | 0.78 | -0.07 | -1.27 | 0.34 | 0.47 | 0.77 | 0.945 | 0.57 | 0.4975 | 0.359306 | 0.024721 | 0.202393 | 1 | 0.999631 | 0.741182 |
| Satb1_E9PVB7_S669s_1 | 1.18 | 1.45 | 1.31 | 0.83 | 0.57 | 0.34 | 0.27 | 1.56 | 0.49 | 1.08 | 1.16 | 1.54 | 0.66 | 0.71 | 0.77 | 1.74 | -0.5075 | -0.125 | -0.2225 | 0.527332 | 0.173868 | 0.694093 | 1 | 0.999631 | 0.824554 |
| Satb2_H3BKH3_S13s_1 | 0.79 | 2.09 | 2.05 | 1.24 | 1.25 | 1.09 | 0.93 | 1.67 | 1.44 | 0.99 | 1.49 | 1.56 | 1.44 | 2.47 | 1.78 | 1.9 | -0.3075 | -0.1725 | 0.355 | 0.390267 | 0.417913 | 0.643397 | 1 | 0.999631 | 0.758901 |
| Satb2_H3BKH3_S13s_2 | 1.14 | 1.4 | 1.71 | 1.85 | 1.31 | 0.31 | 0.74 | 1.95 | 1.67 | 1.86 | 2.03 | 2.87 | 0.32 | 1.71 | 2.99 | 2.09 | -0.4475 | 0.5825 | 0.2525 | 0.641761 | 0.274424 | 0.119497 | 1 | 0.999631 | 0.871335 |
| Satb2_H3BKH3_S7s_1 | 0.86 | 2.09 | 1.86 | 1.21 | 1.09 | 1.63 | 1.27 | 1.49 | 1.56 | 1.73 | 1.52 | 1.61 | 1.16 | 1.82 | 1.79 | 1.94 | -0.135 | 0.1 | 0.1725 | 0.646594 | 0.694291 | 0.766503 | 1 | 0.999631 | 0.874181 |
| Satb2_H3BKH3_S7s_2 | 1.14 | 1.4 | 1.71 | 1.85 | 1.31 | 0.31 | 0.74 | 1.95 | 1.67 | 1.86 | 2.03 | 2.87 | 0.32 | 1.71 | 2.99 | 2.09 | -0.4475 | 0.5825 | 0.2525 | 0.641761 | 0.274424 | 0.119497 | 1 | 0.999631 | 0.871335 |
| Saysd1_Q8K190_S16s_1 | 0.52 | 0.57 | -0.4 | 1.06 | 0.83 | 0.18 | 0.28 | 0.14 | 0.22 | -0.11 | 0.65 | 0.49 | -0.35 | 0.65 | 0.55 | 0.14 | -0.08 | -0.125 | -0.19 | 0.639027 | 0.826672 | 0.738151 | 1 | 0.999631 | 0.870995 |
| Sbf1_Q6ZPE2_S1141s_1 | 0.18 | 1.38 | 0.64 | 0.41 | 0.71 | 1.11 | 0.11 | 0.56 | 1.01 | 0.6 | 0.79 | 0.42 | 0.51 | 0.89 | 0.54 | 0.69 | -0.03 | 0.0525 | 0.005 | 0.988247 | 0.93304 | 0.876131 | 1 | 0.999631 | 0.995409 |
| Sbf1_Q6ZPE2_S1228s_1 | -0.45 | 0.7 | -1.36 | -0.15 | 0.35 | -0.08 | 0.32 | 0 | -0.01 | -1.17 | 0.53 | -0.2 | -0.83 | -0.56 | -0.89 | -0.22 | 0.4625 | 0.1025 | -0.31 | 0.496506 | 0.297599 | 0.843705 | 1 | 0.999631 | 0.811248 |
| Sbf1_Q6ZPE2_S1746s_1 | -0.33 | -0.73 | -2.28 | 1.96 | -0.48 | -1.29 | 0.42 | 1.42 | -1.05 | -1.73 | 0.61 | 1.05 | 0.21 | -0.51 | 1.33 | 1.05 | 0.3625 | 0.065 | 0.865 | 0.317252 | 0.691508 | 0.945455 | 1 | 0.999631 | 0.720971 |
| Sbf1_Q6ZPE2_S559s_1 | -0.39 | 1.01 | 1.7 | 0.16 | 0 | 1.22 | 0.49 | 1.04 | 0.57 | 1.09 | 1.19 | 0.43 | 1.52 | 0.4 | 1.67 | 0.86 | 0.0675 | 0.2 | 0.4925 | 0.353316 | 0.893257 | 0.677177 | 1 | 0.999631 | 0.73949 |
| Sbf1_Q6ZPE2_T1137t_1 | -0.19 | 1.57 | 1.25 | 0.63 | -0.33 | 1.47 | 0.55 | 1.52 | 0.2 | -0.13 | 2.02 | -0.73 | 0.08 | 1.98 | 0.32 | 0.92 | -0.0125 | -0.475 | 0.01 | 0.985146 | 0.981489 | 0.463377 | 1 | 0.999631 | 0.995133 |
| Sbf1_Q6ZPE2_T1819t_1 | -0.25 | -0.82 | 0.96 | 0.06 | -0.35 | -0.05 | -0.07 | 0 | 0.1 | 0.11 | -0.04 | 1.1 | -0.05 | 0.78 | 0.43 | 0.25 | -0.105 | 0.33 | 0.365 | 0.396819 | 0.787608 | 0.4675 | 1 | 0.999631 | 0.761093 |
| Sbf2_E9Q0D4_T1077t_1 | -1.39 | 0.27 | 0.42 | 0.42 | -0.59 | 1.02 | -0.14 | 0.78 | -1.47 | -1.04 | 1.62 | -0.18 | -0.49 | 1.3 | -1.26 | -0.5 | 0.3375 | -0.1975 | -0.1675 | 0.791033 | 0.533881 | 0.784718 | 1 | 0.999631 | 0.926419 |
| Scaf1_Q5U4C3_S240s_1 | -1.12 | -0.26 | -0.63 | -1.97 | -0.27 | -0.92 | -0.88 | -1.46 | -0.81 | -1.72 | -1.59 | -0.94 | 0.19 | -0.35 | -1.63 | -1.68 | 0.1125 | -0.27 | 0.1275 | 0.818033 | 0.795276 | 0.536754 | 1 | 0.999631 | 0.938303 |
| Scaf1_Q5U4C3_S491s_1 | 1.08 | 1.13 | 1.31 | 0.9 | 0.83 | 0.68 | 0.66 | 1.25 | 1.17 | 1.37 | 1.27 | 0.63 | -0.41 | 1.04 | 1.01 | 1.21 | -0.25 | 0.005 | -0.3925 | 0.346094 | 0.352691 | 0.985802 | 1 | 0.999631 | 0.735175 |
| Scaf1_Q5U4C3_S491s_2 | -0.21 | 0.77 | 1.2 | 0.41 | 0.42 | 0.44 | 0.29 | 0.84 | 0.21 | 0.47 | 0.75 | 0.32 | 0.07 | 0.5 | 0.44 | 0.6 | -0.045 | -0.105 | -0.14 | 0.702809 | 0.897738 | 0.768412 | 1 | 0.999631 | 0.896857 |
| Scaf1_Q5U4C3_S493s_1 | -0.43 | -1.5 | -0.83 | -2.27 | -1.54 | -1.99 | -1.57 | -2.66 | -1.92 | -1.82 | -1.41 | -2.51 | -0.44 | -2.14 | -2.41 | -2.14 | -0.6825 | -0.6575 | -0.525 | 0.360192 | 0.161613 | 0.169347 | 1 | 0.999631 | 0.741182 |
| Scaf1_Q5U4C3_S493s_2 | -0.21 | 0.77 | 1.2 | 0.41 | 0.42 | 0.44 | 0.29 | 0.84 | 0.21 | 0.47 | 0.75 | 0.32 | 0.07 | 0.5 | 0.44 | 0.6 | -0.045 | -0.105 | -0.14 | 0.702809 | 0.897738 | 0.768412 | 1 | 0.999631 | 0.896857 |
| Scaf1_Q5U4C3_S577s_2 | -0.39 | -0.75 | 0.72 | -0.69 | -0.01 | -0.78 | 0.51 | -0.4 | -0.43 | 0.13 | -1.19 | -0.97 | -0.82 | -1.93 | -0.21 | -1.12 | 0.1075 | -0.3375 | -0.7425 | 0.145474 | 0.80308 | 0.454987 | 1 | 0.999631 | 0.616744 |
| Scaf1_Q5U4C3_S579s_2 | -0.39 | -0.75 | 0.72 | -0.69 | -0.01 | -0.78 | 0.51 | -0.4 | -0.43 | 0.13 | -1.19 | -0.97 | -0.82 | -1.93 | -0.21 | -1.12 | 0.1075 | -0.3375 | -0.7425 | 0.145474 | 0.80308 | 0.454987 | 1 | 0.999631 | 0.616744 |
| Scaf1_Q5U4C3_S682s_1 | -0.36 | 0.41 | 0.09 | -0.74 | -0.74 | -0.57 | -0.81 | 0.33 | -0.12 | -0.11 | -0.59 | -0.16 | 0.49 | 0.31 | 0.06 | -0.21 | -0.2975 | -0.095 | 0.3125 | 0.383186 | 0.44008 | 0.773669 | 1 | 0.999631 | 0.754425 |
| Scaf1_Q5U4C3_S912s_1 | 0.12 | 0.68 | 3.21 | -0.07 | 0.4 | 0.67 | 1.69 | 3.07 | 0.9 | 0.55 | 3.44 | 3.66 | 1.12 | 0.88 | 4.02 | 3.71 | 0.4725 | 1.1525 | 1.4475 | 0.156149 | 0.578485 | 0.250867 | 1 | 0.999631 | 0.626452 |
| Scaf1_Q5U4C3_T328t_1 | -1.58 | -2.14 | -1.47 | -1.67 | -2.31 | -2.12 | -1.83 | -2.2 | -1.99 | -2 | -1.34 | -1.86 | -1.04 | -1.73 | -1.48 | -1.54 | -0.4 | -0.0825 | 0.2675 | 0.391435 | 0.162725 | 0.780251 | 1 | 0.999631 | 0.758901 |
| Scaf1_Q5U4C3_T923t_1 | 1.26 | -0.99 | 0.37 | 0.55 | -1.28 | -0.74 | 0.6 | -0.83 | -1.08 | 0.54 | -0.4 | -1.12 | -0.69 | -0.51 | -0.03 | -1.05 | -0.86 | -0.8125 | -0.8675 | 0.104361 | 0.152481 | 0.171352 | 1 | 0.999631 | 0.568213 |
| Scaf11_E9PZM7_S857s_1 | 0.02 | -0.13 | -0.29 | 0.37 | -0.06 | -0.58 | -0.19 | -0.32 | 0.16 | 0.66 | -0.22 | 0.28 | 0.6 | 0.1 | 0.05 | -0.14 | -0.28 | 0.2275 | 0.16 | 0.605726 | 0.314838 | 0.45964 | 1 | 0.999631 | 0.85817 |
| Scaf11_E9PZM7_S859s_1 | -0.03 | -0.77 | 0.14 | -0.06 | 0.01 | -0.68 | -0.27 | -0.19 | -0.05 | -0.32 | -0.13 | -0.36 | 0.35 | -0.26 | -0.92 | 0.42 | -0.1025 | -0.035 | 0.0775 | 0.845446 | 0.738501 | 0.905664 | 1 | 0.999631 | 0.948565 |
| Scaf4_Q7TSH6_S154s_1 | 0.22 | 0.6 | 1.48 | 0.47 | 0.51 | 0.56 | 0.19 | 0.41 | 0.62 | 0.31 | 0.47 | 0.83 | 0.79 | 1.08 | 0.9 | 1.07 | -0.275 | -0.135 | 0.2675 | 0.444562 | 0.411194 | 0.692973 | 1 | 0.999631 | 0.783305 |
| Scaf8_Q6DID3_S617s_1 | -1.28 | -3.17 | -1.24 | -1.29 | -1.89 | -2.69 | -0.87 | -1.76 | -2.28 | -1.62 | -0.67 | -1.03 | -1.72 | -1.87 | -0.95 | -1.08 | -0.0575 | 0.345 | 0.34 | 0.50463 | 0.917203 | 0.533716 | 1 | 0.999631 | 0.814292 |
| Scai_B0R0E3_S2s_1 | 0.9 | 0 | -0.72 | -0.28 | 0.16 | -0.42 | -0.2 | -0.74 | 0.16 | 1.13 | -0.94 | -0.32 | 0.02 | -0.4 | -0.76 | -0.27 | -0.275 | 0.0325 | -0.3275 | 0.422926 | 0.493647 | 0.950188 | 1 | 0.999631 | 0.773293 |
| Scai_B0R0E3_S3s_1 | 0.9 | 0 | -0.72 | -0.28 | 0.16 | -0.42 | -0.2 | -0.74 | 0.16 | 1.13 | -0.94 | -0.32 | 0.02 | -0.4 | -0.76 | -0.27 | -0.275 | 0.0325 | -0.3275 | 0.422926 | 0.493647 | 0.950188 | 1 | 0.999631 | 0.773293 |
| Scamp1_Q8K021_S112s_1 | -1.52 | -0.55 | 0.24 | -0.92 | -0.81 | 0.06 | 0.24 | -0.7 | 0.18 | -1.49 | -0.13 | -0.41 | -0.54 | -0.91 | -0.45 | -1.84 | 0.385 | 0.225 | -0.2475 | 0.604366 | 0.392382 | 0.649487 | 1 | 0.999631 | 0.85817 |
| Scamp1_Q8K021_S295s_1 | -0.15 | -0.58 | -0.87 | -0.56 | -0.68 | -0.34 | -1.04 | -0.3 | -0.31 | -0.35 | -1.27 | -1.62 | -0.03 | -0.35 | -0.64 | -1.14 | -0.05 | -0.3475 | -4.9E-17 | 1 | 0.865312 | 0.374129 | 1 | 0.999631 | 1 |
| Scamp1_Q8K021_S50s_1 | 1.69 | 0.68 | 1.37 | 1.23 | 0.61 | 1.97 | 1.18 | 1.24 | 0.2 | 1.62 | 0.6 | 0.03 | -0.05 | 1.82 | 0.6 | -0.08 | 0.0075 | -0.63 | -0.67 | 0.182823 | 0.983757 | 0.155516 | 1 | 0.999631 | 0.643861 |
| Scamp1_Q8K021_T45t_1 | 1.5 | 0.4 | 0.89 | 0.58 | 0.13 | 1.26 | 0.78 | 0.69 | 0.21 | 1.37 | 0.11 | -0.51 | -0.21 | 0.41 | 0.33 | 0.09 | -0.1275 | -0.5475 | -0.6875 | 0.067693 | 0.723138 | 0.243289 | 1 | 0.999631 | 0.521762 |
| Scamp3_E9Q855_S78s_1 | -0.18 | -0.86 | -0.61 | -0.56 | -0.78 | -0.65 | -0.79 | -0.46 | -0.86 | -0.46 | -0.63 | -0.66 | -0.13 | -0.22 | -0.3 | -0.25 | -0.1175 | -0.1 | 0.3275 | 0.259748 | 0.655709 | 0.713154 | 1 | 0.999631 | 0.692624 |
| Scap_Q6GQT6_S1051s_1 | -0.17 | -1.34 | -1.32 | -1.55 | 0.06 | -2.61 | -0.51 | -0.64 | 0.63 | -0.13 | -0.86 | -1.18 | -1.36 | -0.02 | -0.54 | -1.78 | 0.17 | 0.71 | 0.17 | 0.728092 | 0.776231 | 0.16827 | 1 | 0.999631 | 0.905923 |
| Scap_Q6GQT6_S821s_1 | -0.57 | -0.01 | -1.83 | -0.61 | -0.04 | -1.08 | -0.98 | -0.92 | 0.25 | -0.35 | -1.15 | -2.87 | -0.06 | 0.19 | -1.8 | -0.42 | -7.9E-17 | -0.275 | 0.2325 | 0.67193 | 1 | 0.69283 | 1 | 0.999631 | 0.884779 |
| Scap_Q6GQT6_S850s_1 | -0.57 | 0.01 | 0.21 | 0.04 | -0.37 | -0.66 | -0.55 | 0.28 | -0.28 | 0.06 | -0.03 | 0.22 | 0.42 | 0.33 | 0.11 | 0.79 | -0.2475 | 0.07 | 0.49 | 0.140163 | 0.445247 | 0.808041 | 1 | 0.999631 | 0.611718 |
| Scap_Q6GQT6_S934s_1 | 0.3 | -0.45 | -0.41 | 0.09 | 0.04 | -0.99 | -0.92 | -0.45 | -0.52 | -0.2 | 0.64 | -1.51 | 0.01 | 0.17 | 0.5 | 0.87 | -0.4625 | -0.28 | 0.505 | 0.154615 | 0.193107 | 0.552035 | 1 | 0.999631 | 0.625677 |
| Scaper_Q8BIZ8_S184s_1 | 0.95 | 1.2 | 0.68 | 0.34 | 1.87 | 0.86 | 0.82 | -0.04 | 1.99 | 1.08 | 0.97 | 0.98 | 1.13 | 0.08 | -0.13 | 0.26 | 0.085 | 0.4625 | -0.4575 | 0.23825 | 0.841719 | 0.204483 | 1 | 0.999631 | 0.683378 |
| Sccpdh_Q8R127_S215s_1 | -0.16 | 1.27 | 0.9 | 0.34 | 0.24 | 1 | -0.03 | 0.77 | 0.98 | 0.16 | 0.21 | -0.07 | 1.1 | 0.94 | 0.89 | 0.11 | -0.0925 | -0.2675 | 0.1725 | 0.671759 | 0.816436 | 0.509111 | 1 | 0.999631 | 0.884779 |
| Sccpdh_Q8R127_S217s_1 | -0.47 | 1.17 | 2.42 | 0.41 | -0.22 | 1.74 | 0.04 | 2.06 | 0.59 | 0.7 | 1.39 | 1.36 | 0.74 | 0.46 | 2.23 | 2.41 | 0.0225 | 0.1275 | 0.5775 | 0.418517 | 0.975785 | 0.829485 | 1 | 0.999631 | 0.771707 |
| Scfd1_Q8BRF7_S300s_1 | -0.32 | -0.09 | 0.17 | -0.18 | -0.09 | -1.18 | -0.65 | 0.31 | -0.11 | -0.52 | 0.18 | 0.67 | 0.58 | -0.04 | 0.6 | 0.05 | -0.2975 | 0.16 | 0.4025 | 0.202791 | 0.421228 | 0.625575 | 1 | 0.999631 | 0.659215 |
| Scg2_Q4W8U9_S270s_1 | 0.36 | 0.63 | 0.63 | -0.02 | 0.53 | 0.67 | 0.25 | 0.63 | 1.45 | 0.95 | -0.16 | 0.9 | 2.24 | 1.59 | 1.58 | 0.41 | 0.12 | 0.385 | 1.055 | 0.029648 | 0.659841 | 0.335388 | 1 | 0.999631 | 0.43391 |
| Scg2_Q4W8U9_S492s_1 | -0.29 | -1.15 | -0.41 | -0.53 | -0.59 | -0.71 | -0.69 | -0.45 | -0.15 | -1.25 | -0.79 | -0.12 | -0.17 | -0.64 | 0.33 | 0.09 | -0.015 | 0.0175 | 0.4975 | 0.170239 | 0.957375 | 0.961607 | 1 | 0.999631 | 0.64073 |
| Scg2_Q4W8U9_S515s_1 | 0.02 | 0.43 | 0.8 | -1.29 | 2.46 | 0.95 | 2.57 | -1.48 | 4.26 | 0.52 | 1.32 | 3.45 | 5.36 | 0.84 | 1.47 | -2.18 | 1.135 | 2.3975 | 1.3825 | 0.322349 | 0.229633 | 0.0188 | 1 | 0.999631 | 0.724386 |
| Scg2_Q4W8U9_S516s_1 | -0.23 | -0.78 | 0.43 | -1.44 | 0.34 | 1.13 | 3.02 | -0.79 | 2.63 | -2.49 | 2.48 | 4.22 | 1.01 | -1.81 | -0.59 | -0.87 | 1.43 | 2.215 | -0.06 | 0.925017 | 0.092797 | 0.110394 | 1 | 0.999631 | 0.975458 |
| Scg5_P12961_S205s_1 | -0.67 | -0.17 | -0.69 | -1.29 | -0.79 | -0.47 | -1.07 | -0.78 | -0.33 | -0.14 | -0.43 | -0.81 | 0.21 | 0.29 | -0.66 | -0.08 | -0.0725 | 0.2775 | 0.645 | 0.099332 | 0.8169 | 0.402284 | 1 | 0.999631 | 0.562298 |
| Scn1a_A0A1D5RLV0_S757s_1 | 1.12 | 0.61 | 0.19 | 1.5 | 1.03 | 0.53 | 0.95 | 0.64 | 0.28 | -0.25 | 0.78 | -0.39 | -1.07 | 0.69 | -0.11 | 0.65 | -0.0675 | -0.75 | -0.815 | 0.118042 | 0.844452 | 0.08775 | 1 | 0.999631 | 0.585087 |
| Scn1a_A2APX8_S1801s_1 | -0.53 | 0.98 | 1.73 | 1.61 | -0.63 | 0.48 | 0.03 | 1.26 | 0.72 | 0.55 | 0.83 | 1.37 | 0.57 | 0.74 | 2.12 | 1.9 | -0.6625 | -0.08 | 0.385 | 0.522507 | 0.278959 | 0.876783 | 1 | 0.999631 | 0.821224 |
| Scn1a_A2APX8_S1911s_1 | 1.05 | 0.7 | -1.59 | 0.99 | 0.79 | 1.09 | 1.04 | 0.17 | 1.41 | 0.74 | 2.62 | -1.57 | -0.83 | 0.94 | 2.58 | 0.64 | 0.485 | 0.5125 | 0.545 | 0.510371 | 0.427475 | 0.587978 | 1 | 0.999631 | 0.817683 |
| Scn1a_A2APX8_S467s_1 | -0.74 | -0.38 | -1.39 | -1.12 | -1.37 | -1.05 | -0.85 | -1.62 | -1.03 | -0.72 | -2.44 | -1.76 | 0.65 | -0.84 | -2.94 | -3.13 | -0.315 | -0.58 | -0.6575 | 0.424229 | 0.343152 | 0.207343 | 1 | 0.999631 | 0.773927 |
| Scn1a_A2APX8_S467s_2 | 0.53 | 0.42 | 0.58 | 0.37 | -0.39 | 0.43 | 0.41 | 0.35 | 0.49 | 1.07 | 0.57 | -0.45 | 0.22 | 0.01 | 0.08 | 0.02 | -0.275 | -0.055 | -0.3925 | 0.154117 | 0.342149 | 0.87728 | 1 | 0.999631 | 0.625394 |
| Scn1a_A2APX8_S470s_2 | 0.58 | 0.76 | 0.52 | 0.7 | 0.1 | 0.64 | 0.5 | 0.24 | 0.73 | 1.12 | 0.54 | -0.33 | 0.85 | -0.18 | -0.2 | 0.72 | -0.27 | -0.125 | -0.3425 | 0.336258 | 0.298445 | 0.721965 | 1 | 0.999631 | 0.730499 |
| Scn1a_A2APX8_S623s_1 | 1.28 | -0.04 | -1.31 | 0.35 | 1.12 | -0.13 | 2.21 | -0.67 | 1.43 | -0.69 | 0.3 | -0.29 | -1.78 | -1.67 | -0.84 | -1.68 | 0.5625 | 0.1175 | -1.5625 | 0.015322 | 0.453377 | 0.853971 | 1 | 0.999631 | 0.388832 |
| Scn1a_A2APX8_T464t_2 | -0.38 | -2.13 | 0.83 | -1.2 | -1.88 | -2.09 | 0.23 | -0.25 | -1.5 | -0.38 | -0.54 | -0.69 | -2.6 | -1.87 | -0.53 | -1.05 | -0.2775 | -0.0575 | -0.7925 | 0.266777 | 0.712561 | 0.925259 | 1 | 0.999631 | 0.695664 |
| Scn1a_A2APX8_T465t_2 | 0.66 | 0.37 | 0.66 | -0.05 | -0.41 | 0.38 | 0.3 | 0.48 | 0.26 | 0.84 | 0.61 | -0.58 | -0.3 | 0.37 | 0.38 | -1.01 | -0.2225 | -0.1275 | -0.55 | 0.186907 | 0.485438 | 0.736018 | 1 | 0.999631 | 0.646474 |
| Scn1b_P97952_S207s_1 | -0.87 | -0.36 | -0.26 | -0.37 | -0.2 | 0 | -0.33 | 0.12 | -0.11 | -0.17 | 0.21 | 0.4 | 0.34 | 0.04 | -0.45 | 0.75 | 0.3625 | 0.5475 | 0.635 | 0.091472 | 0.195019 | 0.078714 | 1 | 0.999631 | 0.557085 |
| Scn2a1_B1AWN6_S468s_1 | -0.55 | -1.51 | -1.83 | -2.7 | -1.76 | -1.96 | -2.31 | -2.41 | -3.99 | -1.93 | -2.31 | -2.3 | 0.29 | -2.56 | -2.44 | -2.43 | -0.4625 | -0.985 | -0.1375 | 0.849689 | 0.320109 | 0.117509 | 1 | 0.999631 | 0.949525 |
| Scn2a1_B1AWN6_S468s_2 | -0.81 | 0.11 | -0.37 | 0.15 | -0.7 | -1.03 | 0.15 | 0.12 | -0.76 | -1.59 | 0.3 | -0.02 | 0.11 | -1.63 | 0.29 | -0.36 | -0.135 | -0.2875 | -0.1675 | 0.725744 | 0.726203 | 0.539469 | 1 | 0.999631 | 0.90568 |
| Scn2a1_B1AWN6_S471s_2 | -0.81 | 0.11 | -0.37 | 0.15 | -0.7 | -1.03 | 0.15 | 0.12 | -0.76 | -1.59 | 0.3 | -0.02 | 0.11 | -1.63 | 0.29 | -0.36 | -0.135 | -0.2875 | -0.1675 | 0.725744 | 0.726203 | 0.539469 | 1 | 0.999631 | 0.90568 |
| Scn2a1_B1AWN6_S526s_1 | 0.16 | -1.09 | -0.02 | -1.23 | -1.63 | -0.65 | -0.47 | -0.87 | -0.89 | 0.08 | -0.55 | -0.78 | -0.6 | -1.45 | -0.8 | -1.46 | -0.36 | 0.01 | -0.5325 | 0.233571 | 0.412432 | 0.981054 | 1 | 0.999631 | 0.683077 |
| Scn2a1_B1AWN6_S528s_1 | 0.64 | -0.72 | -0.43 | -0.79 | -1.35 | -0.48 | -0.86 | -1.43 | -0.88 | 0.39 | -1.07 | -1.5 | -0.19 | -1.07 | -1.77 | -1.88 | -0.705 | -0.44 | -0.9025 | 0.091029 | 0.104489 | 0.386673 | 1 | 0.999631 | 0.557085 |
| Scn2a1_B1AWN6_S531s_1 | 0.93 | -0.66 | -0.51 | -0.02 | -0.82 | -0.21 | 0.51 | -0.84 | -0.64 | 0.26 | -0.26 | -0.97 | -0.56 | -1.19 | -0.75 | -1.61 | -0.275 | -0.3375 | -0.9625 | 0.047965 | 0.553931 | 0.45071 | 1 | 0.999631 | 0.48343 |
| Scn2a1_B1AWN6_S540s_1 | 0.85 | 1.14 | 0.07 | 1.1 | 1.13 | 0.66 | 0.54 | 0.65 | 0.88 | 0.53 | 0.68 | 0.25 | -0.12 | 0.83 | 0.27 | 0.92 | -0.045 | -0.205 | -0.315 | 0.419242 | 0.889961 | 0.541353 | 1 | 0.999631 | 0.772152 |
| Scn2a1_B1AWN6_S568s_1 | 0.94 | -0.08 | -2.03 | 0.76 | 1.5 | -0.32 | 0.63 | 0.71 | 0.28 | -0.71 | 0.44 | -0.75 | -1.86 | 0.17 | -0.36 | 0.22 | 0.7325 | -0.0825 | -0.355 | 0.6313 | 0.301199 | 0.902147 | 1 | 0.999631 | 0.867571 |
| Scn2a1_B1AWN6_S579s_1 | 1.01 | 0.97 | -0.15 | 1.24 | 2.02 | 0.28 | 0.75 | 0.87 | 1.27 | 0.24 | 0.56 | 0.3 | 0.14 | 1.11 | 0.01 | 1.16 | 0.2125 | -0.175 | -0.1625 | 0.713181 | 0.647755 | 0.665014 | 1 | 0.999631 | 0.900989 |
| Scn2a1_B1AWN6_S623s_1 | 0.69 | -2 | -0.46 | -0.83 | -2.28 | -0.47 | -0.35 | -0.2 | -1.93 | 0.15 | -1.68 | -1.33 | -1.47 | -1.21 | -1.8 | -2.45 | -0.175 | -0.5475 | -1.0825 | 0.080821 | 0.790607 | 0.409211 | 1 | 0.999631 | 0.545579 |
| Scn2a1_B1AWN6_T1944t_1 | -0.54 | -0.63 | 2.09 | 1.09 | -0.24 | -0.76 | 0.35 | 0.86 | -0.54 | -0.91 | 0.95 | 1.23 | 1.45 | -0.2 | 1.14 | 2.14 | -0.45 | -0.32 | 0.63 | 0.394748 | 0.504444 | 0.67074 | 1 | 0.999631 | 0.759502 |
| Scn2a1_B1AWN6_T1960t_1 | -0.29 | -1.62 | -4.98 | -5.01 | -1.41 | -1.4 | -4.22 | -4.58 | -1.23 | -0.89 | -3.61 | -4.08 | -0.91 | -2.37 | -5.27 | -4.81 | 0.0725 | 0.5225 | -0.365 | 0.784144 | 0.954067 | 0.674135 | 1 | 0.999631 | 0.92266 |
| Scn2a1_B1AWN6_T1967t_1 | -0.52 | 0.4 | 0.45 | 0.16 | 0.72 | -0.08 | 0.01 | 0.52 | 0.34 | -0.28 | 0.13 | 0.76 | 0.18 | 0.55 | 0.72 | 0.94 | 0.17 | 0.115 | 0.475 | 0.183691 | 0.613628 | 0.743162 | 1 | 0.999631 | 0.644372 |
| Scn2b_Q56A07_S192s_1 | -0.11 | 1.09 | 1.26 | 0.56 | 0.38 | 0.85 | 0.19 | 0.72 | 0.66 | 1.11 | 0.65 | 1.06 | 1.44 | 1.08 | 1.01 | 1.39 | -0.165 | 0.17 | 0.53 | 0.173572 | 0.652736 | 0.642013 | 1 | 0.999631 | 0.642116 |
| Scn2b_Q56A07_T214t_1 | 2.37 | -1.68 | 0.81 | -0.34 | -0.9 | -0.34 | 0.81 | -0.62 | -1.36 | 1.25 | -0.78 | -1.34 | -1.71 | -0.95 | -0.2 | -0.47 | -0.5525 | -0.8475 | -1.1225 | 0.183459 | 0.504445 | 0.367364 | 1 | 0.999631 | 0.644372 |
| Scn3a_A2ASI5_S466s_1 | 0.25 | 0.39 | -1.2 | -0.79 | -0.68 | 0.28 | -0.63 | -0.93 | 0.19 | 0.46 | -0.46 | -0.85 | -3.07 | -0.44 | -0.12 | -1.2 | -0.1525 | 0.1725 | -0.87 | 0.221722 | 0.737548 | 0.716461 | 1 | 0.999631 | 0.672397 |
| Scn3a_A2ASI5_S466s_2 | 0.61 | 0.05 | 0.05 | -0.89 | 0.39 | -0.66 | -0.56 | 0.02 | 1.35 | 1.32 | -0.15 | -0.65 | -0.01 | 1.12 | -2.59 | -1.23 | -0.1575 | 0.5125 | -0.6325 | 0.408903 | 0.695817 | 0.365732 | 1 | 0.999631 | 0.766996 |
| Scn3a_A2ASI5_S469s_1 | 0.25 | 0.39 | -0.58 | -0.48 | -0.68 | 0.28 | -0.32 | -0.39 | 0.19 | 0.46 | -0.38 | -0.57 | -3.07 | -0.44 | 0.02 | -1.3 | -0.1725 | 0.03 | -1.0925 | 0.117344 | 0.623428 | 0.935672 | 1 | 0.999631 | 0.585087 |
| Scn3a_A2ASI5_S469s_2 | 0.61 | 0.05 | 0.05 | -0.89 | 0.39 | -0.66 | -0.56 | 0.02 | 1.35 | 1.32 | -0.15 | -0.65 | -0.01 | 1.12 | -2.59 | -1.23 | -0.1575 | 0.5125 | -0.6325 | 0.408903 | 0.695817 | 0.365732 | 1 | 0.999631 | 0.766996 |
| Scn3a_A2ASI5_S623s_1 | 1.15 | -3.05 | -1.49 | 0.06 | 0.06 | -1.29 | 0.76 | -0.76 | 0.65 | 0.26 | 0.14 | -0.41 | -3.28 | -3.18 | -1.48 | -1.04 | 0.525 | 0.9925 | -1.4125 | 0.151926 | 0.556674 | 0.246864 | 1 | 0.999631 | 0.625198 |
| Scn8a_F7D6K4_S678s_1 | -0.88 | -1.24 | -1.58 | -2.43 | -1.3 | -0.69 | -2.04 | -1.86 | -1.25 | 0.22 | -1.21 | -2.67 | 0.65 | -0.73 | -1.66 | -2.27 | 0.06 | 0.305 | 0.53 | 0.418829 | 0.891185 | 0.621569 | 1 | 0.999631 | 0.772004 |
| Scrib_Q80U72_S1263s_1 | 1.09 | 0.94 | 1.81 | 1.6 | 0.41 | 2.44 | 1.46 | 0.9 | 1.31 | 2.06 | 0.72 | 1.35 | 0.63 | 1.11 | 1.55 | 1.08 | -0.0575 | -2.4E-16 | -0.2675 | 0.441671 | 0.90086 | 1 | 1 | 1 | 0.781983 |
| Scrib_Q80U72_S1464s_1 | 0.07 | -1.17 | -0.32 | 0.33 | -1.19 | -0.9 | -1.05 | 0.17 | -0.39 | -0.73 | 0.71 | 0.32 | -0.15 | -0.71 | 0.4 | -0.29 | -0.47 | 0.25 | 0.085 | 0.837975 | 0.29931 | 0.58429 | 1 | 0.999631 | 0.946137 |
| Scrib_Q80U72_S506s_1 | -1.47 | -0.34 | -0.23 | -0.86 | -0.76 | -0.85 | -0.82 | -0.39 | -0.38 | -0.64 | -0.45 | -0.41 | 0.52 | -0.27 | -0.52 | -0.55 | 0.02 | 0.255 | 0.52 | 0.214598 | 0.952913 | 0.456322 | 1 | 0.999631 | 0.66742 |
| Scrn1_Q9CZC8_S322s_1 | 0.25 | 1.09 | 0.15 | 0.84 | 0.67 | 1.45 | 1.15 | 1.54 | 1.42 | -0.86 | -0.5 | 0.01 | 2.1 | 0.8 | 0.67 | 1.13 | 0.62 | -0.565 | 0.5925 | 0.172793 | 0.090736 | 0.289766 | 1 | 0.999631 | 0.642116 |
| Scrn1_Q9CZC8_S360s_1 | 0.46 | 1.17 | 0.45 | 0.49 | 0.35 | 0.62 | 0.35 | 0.27 | 1.73 | 1.53 | 1.31 | 1.13 | 0.74 | 0.87 | 0.42 | 0.66 | -0.245 | 0.7825 | 0.03 | 0.920979 | 0.385902 | 0.023995 | 1 | 0.999631 | 0.975032 |
| Scyl2_G5E8J9_S676s_1 | -0.08 | 0.55 | -0.03 | 0.01 | -0.28 | 0.06 | 0.02 | -0.29 | 0.69 | 0.4 | 0.46 | 0.52 | 0.89 | 0.31 | 0.42 | 0.42 | -0.235 | 0.405 | 0.3975 | 0.206132 | 0.391307 | 0.156842 | 1 | 0.999631 | 0.6605 |
| Sdc3_Q64519_S418s_1 | -0.77 | -0.02 | 0.1 | -0.43 | -0.46 | -0.03 | -0.61 | -0.88 | -0.53 | -0.51 | -0.52 | 0.11 | 0.88 | -0.47 | -0.08 | -0.08 | -0.215 | -0.0825 | 0.3425 | 0.381897 | 0.502668 | 0.794412 | 1 | 0.999631 | 0.753668 |
| Sdccag8_Q05DD9_S92s_1 | 2.38 | 0.31 | 0.53 | 0.66 | -0.41 | -0.04 | 0.97 | 0.43 | -0.38 | 1.02 | 0.56 | 0.45 | -0.41 | -1.25 | -0.21 | -0.66 | -0.7325 | -0.5575 | -1.6025 | 0.009216 | 0.1819 | 0.301116 | 1 | 0.999631 | 0.367426 |
| Sde2_Q8K1J5_S269s_1 | 0.27 | 1.62 | 0.81 | 0.98 | 0.57 | 0.81 | -0.06 | 1.36 | 0.64 | 0.98 | 1.51 | 0.76 | 0.01 | 0.91 | 2.01 | 1.54 | -0.25 | 0.0525 | 0.1975 | 0.690513 | 0.54242 | 0.886319 | 1 | 0.999631 | 0.893492 |
| Sdk1_Q3UH53_S297s_2 | -0.65 | 0.54 | 0.22 | 0.08 | -0.34 | -0.39 | 0.22 | 0.13 | -0.24 | -0.17 | 0.96 | 0.7 | 0.03 | -0.01 | 0.53 | 0.63 | -0.1425 | 0.265 | 0.2475 | 0.491078 | 0.671877 | 0.517337 | 1 | 0.999631 | 0.808323 |
| Sdk1_Q3UH53_T296t_2 | -0.65 | 0.54 | 0.22 | 0.08 | -0.34 | -0.39 | 0.22 | 0.13 | -0.24 | -0.17 | 0.96 | 0.7 | 0.03 | -0.01 | 0.53 | 0.63 | -0.1425 | 0.265 | 0.2475 | 0.491078 | 0.671877 | 0.517337 | 1 | 0.999631 | 0.808323 |
| Sdpr_Q63918_S27s_1 | -0.61 | 1.05 | -1.55 | -0.55 | 0.42 | 0.02 | -1.63 | -0.85 | -0.65 | 0.47 | -0.56 | -0.16 | 0.82 | 0.42 | -0.65 | -1.32 | -0.095 | 0.19 | 0.2325 | 0.722619 | 0.880837 | 0.731498 | 1 | 0.999631 | 0.905674 |
| Sdpr_Q63918_S293s_1 | -1.23 | -2.16 | -0.84 | -0.33 | 0.35 | -1.66 | -0.98 | -1.14 | -0.93 | -0.18 | -0.52 | 0.45 | -0.13 | -0.84 | -0.53 | -0.43 | 0.2825 | 0.845 | 0.6575 | 0.144708 | 0.59853 | 0.096474 | 1 | 0.999631 | 0.616037 |
| Sec14l2_Q99J08_T401t_1 | 0.53 | -0.23 | 0.77 | 0.34 | -0.81 | -0.15 | 0.97 | 0.11 | -1.06 | 0.97 | -0.57 | -0.32 | -2.99 | -1.67 | 0.21 | -0.74 | -0.3225 | -0.5975 | -1.65 | 0.026299 | 0.450724 | 0.220839 | 1 | 0.999631 | 0.422285 |
| Sec16a_E9QAT4_S1028s_1 | 0.11 | 0.56 | -0.12 | 0.03 | 0.32 | -0.3 | 0.23 | 0.48 | 0.51 | 0 | 0.03 | 0.25 | 0.52 | 0.87 | -0.4 | 0.39 | 0.0375 | 0.0525 | 0.2 | 0.578644 | 0.89803 | 0.852556 | 1 | 0.999631 | 0.84622 |
| Sec16a_E9QAT4_S2101s_1 | 1.72 | 1.49 | 0.41 | 1.11 | 1.44 | 2.1 | 1.05 | 1.36 | 1.81 | 1.14 | 1.14 | 0.96 | 1.72 | 2.24 | 1.08 | 1.37 | 0.305 | 0.08 | 0.42 | 0.310619 | 0.426373 | 0.828899 | 1 | 0.999631 | 0.71924 |
| Sec16a_E9QAT4_S554s_1 | -0.26 | 1 | 0.27 | 0.67 | 0.23 | 0.13 | -0.22 | 0.61 | 0.3 | 0.28 | 0.33 | 0.91 | 1.29 | 0.33 | 0.42 | 1.06 | -0.2325 | 0.035 | 0.355 | 0.37302 | 0.511328 | 0.920335 | 1 | 0.999631 | 0.747035 |
| Sec22b_O08547_S137s_1 | -0.24 | 0.33 | 0.25 | 0.54 | 0.26 | 0 | 0.01 | 1.13 | -0.06 | -0.29 | 0.96 | 0.84 | 0.32 | 0.48 | 0.88 | 1.02 | 0.13 | 0.1425 | 0.455 | 0.17448 | 0.707699 | 0.707432 | 1 | 0.999631 | 0.642116 |
| Sec22b_O08547_S175s_1 | -0.02 | -0.28 | 1.74 | -0.16 | -0.28 | 0.34 | 0.63 | 0.02 | 0.01 | 0.03 | 1.33 | 1.49 | 0.36 | 0.38 | 1.63 | 0.5 | -0.1425 | 0.395 | 0.3975 | 0.461123 | 0.769965 | 0.496085 | 1 | 0.999631 | 0.792226 |
| Sec22b_O08547_S177s_1 | 0.49 | 0.84 | 1.07 | 0.46 | 0.64 | 1.26 | 0.59 | 0.86 | 0.94 | 1.12 | 0.76 | 1.43 | 2.96 | 1.65 | 1.42 | 0.79 | 0.1225 | 0.3475 | 0.99 | 0.057422 | 0.670261 | 0.249114 | 1 | 0.999631 | 0.49679 |
| Sec23ip_G3X928_S600s_1 | -0.75 | -0.57 | -0.8 | -0.96 | -0.27 | -0.22 | -0.99 | -1.39 | 0.57 | -0.28 | -0.25 | -0.98 | 1.12 | 0.72 | -0.17 | -0.67 | 0.0525 | 0.535 | 1.02 | 0.035343 | 0.87528 | 0.160439 | 1 | 0.999631 | 0.453654 |
| Sec23ip_G3X928_S748s_1 | 0.73 | 1.85 | 1.75 | 0.96 | 0.9 | 1.59 | 1.17 | 1.21 | 1.57 | 1.52 | 2.09 | 1.78 | 1.46 | 2.04 | 1.83 | 2.1 | -0.105 | 0.4175 | 0.535 | 0.162919 | 0.761896 | 0.249103 | 1 | 0.999631 | 0.631442 |
| Sec24b_Q80ZX0_S1203s_1 | -2.28 | 0.66 | 1.93 | 0.84 | 0.02 | 0.69 | 0.62 | 0.62 | 0.98 | -0.03 | 1.61 | 1.78 | 0.71 | 1.27 | 1.62 | 1.52 | 0.2 | 0.7975 | 0.9925 | 0.235491 | 0.802363 | 0.366365 | 1 | 0.999631 | 0.683301 |
| Sec24c_G3X972_S864s_1 | 0.2 | 0.88 | -0.28 | -0.27 | 0.23 | -0.07 | -0.34 | 0.12 | 0.43 | 0.5 | 0.42 | 0.57 | 0.21 | 1.08 | 1.06 | 0.81 | -0.1475 | 0.3475 | 0.6575 | 0.106078 | 0.663245 | 0.304714 | 1 | 0.999631 | 0.568797 |
| Sec31a_S4R2A9_S502s_1 | 0.53 | 0.6 | 0.61 | 0.44 | 1.15 | 0.01 | 0.39 | 0.65 | 1.31 | 0.8 | 0.82 | 1.35 | 1.45 | 0.69 | 1.29 | 0.44 | 0.005 | 0.525 | 0.4225 | 0.210965 | 0.986827 | 0.074577 | 1 | 0.999631 | 0.665076 |
| Sec31a_S4R2A9_S877s_1 | -0.96 | -1.32 | -0.02 | -1.04 | -0.99 | -1.14 | -0.65 | -0.44 | -1.26 | -1.2 | 0.16 | -0.58 | -0.31 | -0.72 | 0.34 | -0.11 | 0.03 | 0.115 | 0.635 | 0.126339 | 0.932074 | 0.790879 | 1 | 0.999631 | 0.596745 |
| Sec61b_Q9CQS8_S13s_1 | 0.09 | -2.17 | 0.33 | 0.17 | -0.9 | 0.06 | -0.03 | -0.19 | -1.09 | 0.31 | -0.9 | -0.01 | 1.41 | 0.48 | 0.47 | 0.46 | 0.13 | -0.0275 | 1.1 | 0.084367 | 0.82138 | 0.964565 | 1 | 0.999631 | 0.549732 |
| Sec61b_Q9CQS8_S17s_1 | -2.82 | -7.38 | -2.22 | -2.51 | -3.56 | -5.66 | -2 | -2.24 | -4.53 | -2.78 | -2.5 | -2.82 | -3.17 | -3.88 | -4.09 | -3.3 | 0.3675 | 0.575 | 0.1225 | 0.907776 | 0.77129 | 0.610626 | 1 | 0.999631 | 0.96989 |
| Sec62_Q8BU14_S335s_2 | -0.56 | 0.46 | 2.67 | 2.1 | -0.81 | 0.23 | 1.15 | 2.02 | 0.63 | 0.46 | 0.8 | 1.86 | -0.2 | -0.71 | 1.02 | 1.42 | -0.52 | -0.23 | -0.785 | 0.328557 | 0.537513 | 0.748564 | 1 | 0.999631 | 0.725792 |
| Sec62_Q8BU14_S341s_1 | 1.04 | 0.88 | 0.64 | 0.59 | 0.55 | 1.2 | 0.92 | 0.48 | 1.04 | 2.2 | 0.08 | 0.43 | -0.02 | -0.57 | 1.19 | -0.52 | 0 | 0.15 | -0.7675 | 0.099059 | 1 | 0.746571 | 1 | 0.999631 | 0.562298 |
| Sec62_Q8BU14_S341s_2 | -0.86 | 0.69 | 2.16 | 1.9 | 0.03 | 1.13 | 1.36 | 1.24 | 0.54 | 1.31 | 0.86 | 1.31 | -0.53 | -1.16 | 0.86 | 0.78 | -0.0325 | 0.0325 | -0.985 | 0.207493 | 0.961351 | 0.959703 | 1 | 0.999631 | 0.6614 |
| Sec62_Q8BU14_S341s_3 | -1.05 | -1.08 | 2.02 | 0.4 | 0.71 | 0.5 | 0.54 | 1.89 | 0.88 | -0.41 | 0.42 | 1.89 | 0.85 | -1.25 | -0.11 | -0.23 | 0.8375 | 0.6225 | -0.2575 | 0.731328 | 0.256665 | 0.428706 | 1 | 0.999631 | 0.906897 |
| Sec62_Q8BU14_S345s_3 | -1.05 | -1.08 | 2.02 | 0.4 | 0.71 | 0.5 | 0.54 | 1.89 | 0.88 | -0.41 | 0.42 | 1.89 | 0.85 | -1.25 | -0.11 | -0.23 | 0.8375 | 0.6225 | -0.2575 | 0.731328 | 0.256665 | 0.428706 | 1 | 0.999631 | 0.906897 |
| Sec62_Q8BU14_T158t_1 | 1.53 | 0.31 | 0.75 | 1.12 | -0.38 | 0.72 | 1.44 | -0.4 | 0.65 | 1.58 | 0.21 | 0.25 | 0.33 | -0.18 | 0.85 | -0.27 | -0.5825 | -0.255 | -0.745 | 0.082927 | 0.251553 | 0.544121 | 1 | 0.999631 | 0.549443 |
| Sec62_Q8BU14_T343t_2 | -0.86 | 0.69 | 1.71 | 1.73 | 0.03 | 1.13 | 1.51 | 0.49 | 0.54 | 1.31 | 0.9 | 0.82 | -0.53 | -1.16 | 0.73 | 0.19 | -0.0275 | 0.075 | -1.01 | 0.148627 | 0.96473 | 0.896705 | 1 | 0.999631 | 0.62132 |
| Sec62_Q8BU14_T343t_3 | -1.05 | -1.08 | 2.02 | 0.4 | 0.71 | 0.5 | 0.54 | 1.89 | 0.88 | -0.41 | 0.42 | 1.89 | 0.85 | -1.25 | -0.11 | -0.23 | 0.8375 | 0.6225 | -0.2575 | 0.731328 | 0.256665 | 0.428706 | 1 | 0.999631 | 0.906897 |
| Secisbp2l_Q6A098_S897s_1 | 1.28 | 1.69 | -1.15 | -0.78 | 1.63 | 2.01 | -0.56 | -2.87 | 0.95 | 1.21 | 0.26 | -0.11 | 0.13 | 1.12 | -2.23 | -1.35 | -0.2075 | 0.3175 | -0.8425 | 0.356574 | 0.855753 | 0.648844 | 1 | 0.999631 | 0.740467 |
| Selplg_Q3TA56_S365s_1 | -0.46 | 0.73 | 0.3 | 0.13 | 0.44 | 0.46 | -0.44 | -0.11 | 0.72 | -0.45 | -0.14 | 0.67 | 0.75 | 1.41 | 0.48 | 0.31 | -0.0875 | 0.025 | 0.5625 | 0.162136 | 0.806087 | 0.949712 | 1 | 0.999631 | 0.631442 |
| Sema4b_Q62179_S780s_1 | 0.42 | 0.35 | 0.7 | 0.94 | -0.09 | 0.2 | 0.28 | 1.18 | -0.27 | 0.91 | 0.1 | 0.85 | 0.93 | 0.12 | -0.19 | -0.46 | -0.21 | -0.205 | -0.5025 | 0.196913 | 0.541614 | 0.567157 | 1 | 0.999631 | 0.653609 |
| Sema4c_Q64151_S761s_1 | -1.09 | -1.71 | -0.71 | -1.07 | -1.42 | -1.42 | -0.55 | 0.29 | -2.09 | -1.78 | -0.34 | 0.04 | -0.5 | -1.32 | -0.57 | 0.58 | 0.37 | 0.1025 | 0.6925 | 0.14237 | 0.414813 | 0.846103 | 1 | 0.999631 | 0.612726 |
| Sema4d_O09126_S832s_1 | -2.01 | -2.01 | -1.39 | 0.41 | -1.63 | -2.51 | -0.9 | 0.04 | -1.64 | -1.55 | 0.4 | 1.17 | -1.69 | -2.35 | -0.11 | 0.19 | 4.71E-16 | 0.845 | 0.26 | 0.725281 | 1 | 0.303751 | 1 | 0.999631 | 0.90568 |
| Sema6a_K3W4S5_S672s_1 | 1.01 | -1.43 | -1.87 | -1.25 | -1.61 | -1.02 | -1.61 | -2.45 | -1.37 | 0.24 | -2.65 | -1.66 | -1.18 | -1.7 | -1.54 | -2.56 | -0.7875 | -0.475 | -0.86 | 0.197999 | 0.233726 | 0.54403 | 1 | 0.999631 | 0.65543 |
| Sema6b_O54951_S647s_1 | -0.13 | -1.41 | -0.46 | 0.94 | -1.18 | -1.47 | -0.13 | 0.46 | -0.41 | -0.27 | -0.15 | -0.19 | -0.66 | -0.41 | -1.14 | -0.02 | -0.315 | 0.01 | -0.2925 | 0.570716 | 0.601042 | 0.983055 | 1 | 0.999631 | 0.84338 |
| Senp3_Q9EP97_S206s_1 | -0.81 | -0.17 | -0.8 | -0.39 | 0.84 | 0.17 | -0.72 | -0.82 | 0.49 | 0.83 | 0.5 | -0.46 | -1.07 | -0.27 | -2.12 | 0.15 | 0.41 | 0.8825 | -0.285 | 0.57279 | 0.340955 | 0.030821 | 1 | 0.999631 | 0.844402 |
| Senp3_Q9EP97_S226s_1 | -0.69 | 0.08 | 0.18 | -0.45 | -0.15 | -0.13 | -0.39 | 0.64 | -0.53 | -1.28 | 0.32 | -0.79 | 0.16 | -0.21 | 0.02 | 0.36 | 0.2125 | -0.35 | 0.3025 | 0.358731 | 0.537067 | 0.396839 | 1 | 0.999631 | 0.741182 |
| Senp7_Q9CX65_S12s_1 | 0.07 | 0.04 | -0.1 | 0.51 | 0.06 | -0.74 | 0.14 | 0.17 | 0.01 | -0.04 | 1.04 | -1.07 | 0.23 | -0.46 | -0.6 | 0.54 | -0.2225 | -0.145 | -0.2025 | 0.572641 | 0.478343 | 0.744595 | 1 | 0.999631 | 0.844402 |
| Senp7_Q9CX65_S214s_1 | -1.89 | -0.15 | -0.76 | -1.22 | -1.08 | -1.11 | -1.13 | -2.36 | -1.28 | 0.01 | -1.05 | -0.75 | -0.45 | -1.03 | -1.21 | 0.06 | -0.415 | 0.2375 | 0.3475 | 0.457969 | 0.379983 | 0.602418 | 1 | 0.999631 | 0.790722 |
| Sepn1_D3Z2R5_S13s_1 | 0.05 | 0.09 | 0.21 | 0.5 | 0.07 | -0.27 | 0.06 | -0.04 | 0.05 | 1.18 | 0.35 | 0.89 | 0.69 | 0.26 | 0.25 | 0.61 | -0.2575 | 0.405 | 0.24 | 0.406355 | 0.316568 | 0.236205 | 1 | 0.999631 | 0.766128 |
| Sept11_Q8C1B7_S9s_1 | -0.62 | 0.99 | -0.8 | -0.76 | 1.2 | 0.32 | -1.68 | -0.83 | 1.42 | 0.21 | -2.72 | -0.31 | 1.5 | 0.31 | -0.94 | -0.62 | 0.05 | -0.0525 | 0.36 | 0.57029 | 0.941325 | 0.95055 | 1 | 0.999631 | 0.84338 |
| Sept4_P28661_S100s_1 | -0.62 | -0.75 | 0.25 | 0.04 | -0.51 | -1 | -0.23 | 0.01 | -1.08 | -1.13 | -0.46 | -0.38 | -0.86 | -1.1 | -0.26 | -0.15 | -0.1625 | -0.4925 | -0.3225 | 0.399299 | 0.648005 | 0.18448 | 1 | 0.999631 | 0.762042 |
| Sept5_Q9Z2Q6_S327s_1 | -0.07 | 0.17 | -0.49 | -0.09 | 0.08 | -0.04 | -0.39 | 0.11 | -0.28 | -0.24 | -0.04 | 0.09 | -0.61 | 0.14 | -0.54 | 0.35 | 0.06 | 0.0025 | -0.045 | 0.895061 | 0.82448 | 0.992631 | 1 | 0.999631 | 0.965568 |
| Sept5_Q9Z2Q6_T13t_1 | 1.76 | 1.57 | 1.3 | 1.14 | 1.03 | 1.63 | 0.81 | 0.73 | 1.39 | 1.91 | 0.81 | 1 | 0.92 | 1.03 | 0.96 | 0.72 | -0.3925 | -0.165 | -0.535 | 0.083174 | 0.217791 | 0.620361 | 1 | 0.999631 | 0.549732 |
| Sept5_Q9Z2Q6_T336t_1 | 1.01 | -0.11 | -1.09 | 0.88 | 1.51 | 0.03 | 0.74 | 0.45 | 0.12 | -1.02 | -0.02 | -0.36 | -1.85 | 0 | -0.77 | -0.15 | 0.51 | -0.4925 | -0.865 | 0.16509 | 0.352695 | 0.3537 | 1 | 0.999631 | 0.633483 |
| Sept6_Q9R1T4_S411s_1 | 0.5 | 0.26 | 0.78 | 1.11 | 0.73 | 0.98 | 0.12 | 1.3 | 0.54 | -0.6 | 0.97 | 0.22 | 1.01 | 0.61 | 0.92 | 1.08 | 0.12 | -0.38 | 0.2425 | 0.43756 | 0.727165 | 0.347029 | 1 | 0.999631 | 0.781057 |
| Sept7_E9Q9F5_S334s_1 | 0.37 | 0.12 | -0.78 | -0.65 | -0.31 | -0.14 | -1.09 | -0.41 | -0.21 | 0.08 | -0.46 | -0.3 | 0.03 | 0.32 | -0.5 | -0.25 | -0.2525 | 0.0125 | 0.135 | 0.71891 | 0.500216 | 0.971186 | 1 | 0.999631 | 0.904495 |
| Sept7_E9Q9F5_S423s_1 | 1.25 | -0.6 | -0.59 | 0.66 | 0.28 | 0.2 | 0.57 | 0.56 | 0.06 | -0.09 | 0.32 | -0.59 | -0.35 | -0.57 | -0.2 | -0.69 | 0.2225 | -0.255 | -0.6325 | 0.19665 | 0.627127 | 0.598616 | 1 | 0.999631 | 0.653373 |
| Sept7_E9Q9F5_S77s_1 | -1.04 | -0.5 | -0.88 | -1.12 | -0.6 | -0.72 | -1.2 | -1.08 | -0.77 | -1.36 | -1.41 | -1.09 | -0.69 | -0.77 | -1.25 | -1.05 | -0.015 | -0.2725 | -0.055 | 0.853433 | 0.957224 | 0.356843 | 1 | 0.999631 | 0.950701 |
| Sept8_B1AQZ0_S10s_1 | 0.95 | -0.08 | 0.95 | 0.01 | 0.01 | 0.9 | 0.43 | -0.31 | 0.42 | 1.01 | -0.24 | 0.22 | 0.07 | -0.18 | 0.28 | -0.67 | -0.2 | -0.105 | -0.5825 | 0.15243 | 0.614172 | 0.792293 | 1 | 0.999631 | 0.625198 |
| Sept8_B1AQZ0_S442s_1 | 0.54 | 0.01 | 0.13 | 0.44 | 0.82 | 0.79 | 0.67 | 0.79 | 0.51 | 0.57 | 0.05 | 0.45 | 0.22 | 0.95 | 0.94 | 0.24 | 0.4875 | 0.115 | 0.3075 | 0.351681 | 0.075707 | 0.677239 | 1 | 0.999631 | 0.73814 |
| Sept8_B1AQZ0_S444s_1 | -1.16 | 0.47 | -0.26 | -0.27 | -0.36 | -0.53 | -0.43 | -0.55 | -0.28 | -0.18 | -0.19 | 0.28 | 0.83 | 0.27 | -0.46 | 0.24 | -0.1625 | 0.2125 | 0.525 | 0.24261 | 0.653434 | 0.578309 | 1 | 0.999631 | 0.685099 |
| Sept8_B1AQZ0_S446s_1 | 0.11 | 0.89 | 0.17 | 0.39 | 0.53 | 0.48 | -0.15 | 0.53 | 0.59 | 0.04 | 0.15 | 1.23 | 0.53 | 0.41 | 0.5 | 0.86 | -0.0425 | 0.1125 | 0.185 | 0.546377 | 0.888546 | 0.753552 | 1 | 0.999631 | 0.834213 |
| Sept9_Q80UG5_S30s_1 | -1.05 | -0.1 | -0.49 | -0.02 | 1.86 | -0.49 | -0.25 | -0.09 | 0.02 | 1.4 | -0.51 | -0.18 | -0.75 | 0.75 | -0.79 | 0.09 | 0.6725 | 0.5975 | 0.24 | 0.588886 | 0.233872 | 0.220086 | 1 | 0.999631 | 0.851154 |
| Sept9_Q80UG5_S330s_1 | 1.31 | -0.15 | 0.53 | 0.27 | -0.75 | 0.16 | 0.56 | -0.82 | -0.48 | 0.91 | -0.27 | -0.47 | -0.66 | -1.1 | -0.39 | -0.87 | -0.7025 | -0.5675 | -1.245 | 0.007744 | 0.138508 | 0.22255 | 1 | 0.999631 | 0.358173 |
| Serbp1_Q9CY58_S234s_1 | -0.31 | 1.22 | 4.43 | 2.34 | 0.69 | 1.19 | 2.94 | 3.43 | 1.35 | -0.59 | 4.47 | 4.83 | 0.5 | 1 | 5.01 | 3.49 | 0.1425 | 0.595 | 0.58 | 0.639409 | 0.889817 | 0.670739 | 1 | 0.999631 | 0.870995 |
| Serbp1_Q9CY58_S327s_1 | -1.32 | 1.22 | 0.54 | 0.46 | 0.86 | 0.89 | 0.27 | 0.41 | 1.41 | 0.3 | 0.34 | 0.91 | 1.27 | 1.42 | 0.73 | 1.12 | 0.3825 | 0.515 | 0.91 | 0.109734 | 0.471436 | 0.366387 | 1 | 0.999631 | 0.573624 |
| Serbp1_Q9CY58_S329s_1 | -0.5 | 1.18 | 0.61 | 0.48 | 0.49 | 0.5 | 0.2 | 0.55 | 0.89 | 0.48 | 0.28 | 0.99 | 1.11 | 1.41 | 0.71 | 1.17 | -0.0075 | 0.2175 | 0.6575 | 0.125323 | 0.983998 | 0.589246 | 1 | 0.999631 | 0.596551 |
| Serinc1_Q9QZI8_S364s_1 | 0.35 | 0.07 | 0 | 0.51 | 0.07 | 0.23 | 1.98 | -0.09 | 0.32 | 0.39 | -0.71 | 0.4 | 0.87 | 0.27 | -0.34 | 0.35 | 0.315 | -0.1325 | 0.055 | 0.871562 | 0.510005 | 0.698425 | 1 | 0.999631 | 0.959119 |
| Sertm1_Q8CD78_S106s_1 | -1.3 | -0.58 | -0.81 | -1.78 | -1.67 | -1.53 | -1.57 | -0.87 | -1.25 | -0.59 | -0.77 | -0.96 | 0.96 | -0.65 | -0.18 | -0.62 | -0.2925 | 0.225 | 0.995 | 0.051205 | 0.4142 | 0.518477 | 1 | 0.999631 | 0.489558 |
| Sertm1_Q8CD78_S107s_1 | -1.3 | -0.58 | -0.81 | -1.78 | -1.67 | -1.53 | -1.57 | -0.87 | -1.25 | -0.59 | -0.77 | -0.96 | 0.96 | -0.65 | -0.18 | -0.62 | -0.2925 | 0.225 | 0.995 | 0.051205 | 0.4142 | 0.518477 | 1 | 0.999631 | 0.489558 |
| Set_Q9EQU5_S30s_1 | 0.91 | 1.76 | 1.63 | 1.55 | 1.39 | 1.31 | 1.09 | 1.06 | 2.06 | 1.78 | 2.02 | 2.52 | 1.41 | 2.07 | 2.33 | 1.99 | -0.25 | 0.6325 | 0.4875 | 0.171017 | 0.3879 | 0.065648 | 1 | 0.999631 | 0.641312 |
| Setd1a_E9PYH6_S1159s_1 | 0.63 | 1.43 | 2.56 | 2.07 | 1.08 | 0.31 | 0.68 | 2.05 | 1.19 | 1.5 | 2.03 | 2.69 | 0.77 | 1.92 | 2.59 | 2.9 | -0.6425 | 0.18 | 0.3725 | 0.524171 | 0.235939 | 0.721272 | 1 | 0.999631 | 0.822727 |
| Setd1a_E9PYH6_S1160s_1 | 0.41 | 1.03 | 2.56 | 2.07 | 1.12 | 0 | 0.68 | 2.05 | 1.67 | 0.73 | 2.03 | 2.69 | 0.5 | 2 | 2.59 | 2.9 | -0.555 | 0.2625 | 0.48 | 0.465604 | 0.357536 | 0.654074 | 1 | 0.999631 | 0.794915 |
| Setd1a_E9PYH6_T931t_1 | 0.33 | 1.39 | 2.17 | 1.53 | 1.74 | 1.4 | 1.24 | 1.52 | 1.85 | 1.43 | 1.11 | 1.94 | 1.07 | 1.81 | 1.83 | 1.35 | 0.12 | 0.2275 | 0.16 | 0.711701 | 0.76459 | 0.597431 | 1 | 0.999631 | 0.900631 |
| Setd1b_Q8CFT2_S985s_2 | -0.97 | 1.62 | 1.42 | 1.3 | 1.07 | 0.11 | 0.56 | 2.08 | 0.94 | -0.12 | 0.9 | 1.92 | 0.82 | 1.55 | 2.58 | 1.24 | 0.1125 | 0.0675 | 0.705 | 0.287447 | 0.864625 | 0.918615 | 1 | 0.999631 | 0.705972 |
| Setd2_E9Q5F9_T1827t_1 | 0.17 | 0.32 | 0.67 | -0.93 | 0.8 | 0.21 | 2.04 | 0.77 | 1.46 | 0.09 | 1.26 | 2.28 | 1.99 | 0.32 | -0.31 | -1.14 | 0.8975 | 1.215 | 0.1575 | 0.813641 | 0.091213 | 0.043011 | 1 | 0.999631 | 0.937041 |
| Setd3_Q91WC0_S12s_1 | 0.57 | 1.81 | 1.28 | 1.91 | 1.12 | 1.33 | 1.47 | 1.67 | 1.54 | 0.93 | 2.61 | 1.52 | -0.15 | 1.68 | 1.22 | 1.85 | 0.005 | 0.2575 | -0.2425 | 0.641531 | 0.988746 | 0.574484 | 1 | 0.999631 | 0.871335 |

| | | | | | | | | | | | | | | | | | | | | | | | | |
|---|---|---|---|---|---|---|---|---|---|---|---|---|---|---|---|---|---|---|---|---|---|---|---|---|
| Setd3_Q91WC0_S21s_1 | -0.32 | 0.32 | 0.2 | -0.19 | 0.12 | 1.28 | 0.05 | -0.29 | 1.06 | 1.1 | -0.2 | 0.21 | 1.63 | 0.83 | 0.37 | -0.47 | 0.2875 | 0.54 | 0.5875 | 0.220179 | 0.462376 | 0.176335 | 1 | 0.999631 | 0.670699 |
| Setd3_Q91WC0_S571s_1 | -1.34 | 0.57 | 0.78 | 0.26 | 0.2 | -0.63 | -0.32 | 0.68 | -0.42 | 0.28 | 1.02 | 0.89 | -0.21 | 0.26 | 0.22 | 0.76 | -0.085 | 0.375 | 0.19 | 0.704126 | 0.870362 | 0.494677 | 1 | 0.999631 | 0.897149 |
| Sf1_D3YVH4_S54s_1 | -0.07 | 0.81 | 1.23 | 0.61 | -0.03 | 0.22 | 0.5 | 0.92 | -0.8 | 0.21 | 1.22 | 0.56 | 0.15 | 0.03 | 0.37 | 0.9 | -0.2425 | -0.3475 | -0.2825 | 0.455799 | 0.507736 | 0.47602 | 1 | 0.999631 | 0.790041 |
| Sf1_D3YVH4_S56s_1 | -0.07 | 0.81 | 1.23 | 0.61 | -0.03 | 0.22 | 0.5 | 0.92 | -0.8 | 0.21 | 1.22 | 0.56 | 0.15 | 0.03 | 0.37 | 0.9 | -0.2425 | -0.3475 | -0.2825 | 0.455799 | 0.507736 | 0.47602 | 1 | 0.999631 | 0.790041 |
| Sf3b1_Q99NB9_S129s_1 | 0.79 | 0.38 | 0.46 | 0.27 | 0.26 | -0.04 | 0.13 | 0.67 | 0.25 | 0.51 | 0.45 | 0.09 | 0.39 | 0.75 | 0.15 | 0.82 | -0.22 | -0.15 | 0.0525 | 0.860879 | 0.43151 | 0.574675 | 1 | 0.999631 | 0.954046 |
| Sf3b1_Q99NB9_T211t_1 | -1.99 | -0.66 | -1.14 | -2.06 | -1.37 | -1.11 | -0.68 | -1.26 | 0.75 | -2.66 | -0.65 | -1.04 | -0.12 | -1.65 | -0.92 | -2.66 | 0.3575 | 0.5625 | 0.125 | 0.830824 | 0.362395 | 0.426107 | 1 | 0.999631 | 0.942418 |
| Sf3b1_Q99NB9_T223t_1 | -1.53 | -0.94 | -0.36 | -0.51 | -2.55 | -1.56 | -0.3 | -1.15 | -2 | -0.91 | -1.19 | -1.05 | -3.09 | -1.14 | -0.38 | -1.24 | -0.555 | -0.4525 | -0.6275 | 0.294442 | 0.284405 | 0.251867 | 1 | 0.999631 | 0.708293 |
| Sf3b1_Q99NB9_T235t_1 | 0.06 | 0.24 | 0.62 | -0.44 | -0.25 | 0.61 | -0.11 | -0.88 | -0.23 | 0.37 | 0.64 | -1.02 | -0.3 | 0.73 | -0.91 | -0.15 | -0.2775 | -0.18 | -0.2775 | 0.511652 | 0.479272 | 0.675846 | 1 | 0.999631 | 0.817687 |
| Sf3b1_Q99NB9_T248t_1 | 1.05 | -1.38 | -0.18 | -0.87 | 0.61 | 0.52 | 0.06 | -0.67 | -0.58 | 0.48 | -0.19 | -1.23 | -0.3 | -0.18 | -1.79 | -1.49 | 0.475 | -0.035 | -0.595 | 0.3385 | 0.398848 | 0.952069 | 1 | 0.999631 | 0.73164 |
| Sf3b1_Q99NB9_T267t_1 | 0.33 | 1.64 | 0.61 | -0.77 | 0.66 | 0.58 | 0.1 | -1.48 | 0.44 | -0.15 | -0.21 | 0.09 | 0.99 | 0.31 | 0.65 | 0.25 | -0.4875 | -0.41 | 0.0975 | 0.845814 | 0.445982 | 0.413758 | 1 | 0.999631 | 0.948565 |
| Sf3b1_Q99NB9_T273t_1 | -0.69 | -2.75 | -1.53 | -1.78 | -3.64 | -2.27 | -0.64 | -1.7 | -3.06 | -1.82 | -1.76 | -2.11 | -2.18 | -1.95 | -2.88 | -1.38 | -0.375 | -0.5 | -0.41 | 0.422419 | 0.57955 | 0.325329 | 1 | 0.999631 | 0.773293 |
| Sf3b1_Q99NB9_T278t_1 | 1.78 | -1.15 | -0.45 | -0.4 | -1.35 | 0.36 | 0.23 | -0.17 | -1.19 | 1.01 | -0.8 | -1.02 | -1.71 | -1.47 | -0.51 | -0.32 | -0.1775 | -0.445 | -0.9475 | 0.166472 | 0.789258 | 0.541885 | 1 | 0.999631 | 0.63587 |
| Sf3b1_Q99NB9_T303t_1 | 0.08 | 1.9 | 0.34 | -0.27 | 1.43 | 1.53 | -0.31 | 0.29 | 1.78 | 0.85 | 0.66 | 1.31 | 1.99 | 0.76 | 0.47 | 0.27 | 0.2225 | 0.6375 | 0.36 | 0.530249 | 0.708597 | 0.230479 | 1 | 0.999631 | 0.826645 |
| Sf3b1_Q99NB9_T313t_1 | 0.11 | 0.45 | 0.55 | -0.13 | 0.22 | -0.46 | 0.34 | 0.56 | -0.14 | 0.15 | -0.5 | 0.1 | 0.72 | -0.6 | -0.57 | -0.29 | -0.08 | -0.3425 | -0.43 | 0.275953 | 0.802038 | 0.263868 | 1 | 0.999631 | 0.700187 |
| Sf3b1_Q99NB9_T426t_1 | 1.36 | 2.44 | 1.02 | 1.32 | 0.63 | 0.75 | 0.2 | 1.79 | 0.36 | 1.34 | 1.24 | 0.53 | 1.63 | 2.53 | 0.28 | 1.4 | -0.6925 | -0.6675 | -0.075 | 0.886469 | 0.143544 | 0.125396 | 1 | 0.999631 | 0.963505 |
| Sfn_O70456_S45s_1 | 0.09 | 0.85 | 0.65 | 0.45 | 0.07 | 0.67 | 0.25 | 0.47 | 0.16 | 0.17 | 0.17 | 0.98 | 0.6 | 0.74 | 1.52 | 1.27 | -0.145 | -0.14 | 0.5225 | 0.145219 | 0.612874 | 0.663017 | 1 | 0.999631 | 0.616182 |
| Sfr1_Q8BP27_S115s_1 | 1.84 | 0.35 | 1.88 | 1.43 | -0.29 | 0.67 | 1.78 | 0.12 | -0.05 | 1.94 | 0.55 | -0.23 | -0.04 | -0.22 | 0.53 | 0 | -0.805 | -0.8225 | -1.3075 | 0.009101 | 0.152362 | 0.165284 | 1 | 0.999631 | 0.366532 |
| Sfr1_Q8BP27_S67s_1 | 0.79 | 1.36 | 1.68 | 0.9 | 0.71 | 1.1 | 0.3 | 1.25 | 1.08 | 1.55 | 1.1 | 1.55 | 1.33 | 1.6 | 1.02 | 1.57 | -0.3425 | 0.1375 | 0.1975 | 0.547194 | 0.320031 | 0.660739 | 1 | 0.999631 | 0.834277 |
| Sfr1_Q8BP27_S67s_2 | 0.78 | 1.71 | 1.48 | 1.41 | 1.03 | 1.15 | 0.79 | 1.32 | 1.3 | 1.28 | 1.41 | 1.65 | 1.04 | 1.65 | 1.87 | 1.9 | -0.2725 | 0.065 | 0.27 | 0.439668 | 0.366746 | 0.826402 | 1 | 0.999631 | 0.781983 |
| Sfr1_Q8BP27_S71s_2 | 0.39 | 0.99 | 1.48 | 1.41 | 0.34 | 0.68 | 0.79 | 1.32 | 0.5 | 0.87 | 1.41 | 1.65 | 0.46 | 0.96 | 1.87 | 1.9 | -0.285 | 0.04 | 0.23 | 0.602153 | 0.425088 | 0.916583 | 1 | 0.999631 | 0.856988 |
| Sfswap_Q3USH5_S831s_1 | 2 | 0.76 | 1.44 | 0.9 | 0.36 | 0.84 | 1.46 | 0.46 | 1.4 | 1.92 | 0.71 | 0.28 | 0.11 | 0.5 | 0.38 | 0.03 | -0.495 | -0.1975 | -1.02 | 0.01572 | 0.220376 | 0.663028 | 1 | 0.999631 | 0.388832 |
| Sgip1_F6VMS8_S27s_1 | 0.11 | 0.21 | 2.6 | 0.67 | -0.82 | 0.26 | 3.53 | 0.05 | -0.56 | 0.65 | 1.63 | 1.09 | 0.76 | -1.07 | 1.46 | 0.45 | -0.1425 | -0.195 | -0.4975 | 0.482487 | 0.882279 | 0.770372 | 1 | 0.999631 | 0.802587 |
| Sgip1_Q8VD37_S160s_1 | 2.6 | 0.83 | 1.33 | 2.27 | 1.34 | 1.62 | 2.59 | 1.62 | 1.79 | 2.22 | 1.56 | 0.5 | -0.21 | 1.02 | 1.65 | 0.68 | 0.035 | -0.24 | -0.9725 | 0.091033 | 0.940581 | 0.643974 | 1 | 0.999631 | 0.557085 |
| Sgip1_Q8VD37_S169s_1 | -0.1 | 1.09 | -0.16 | 0.46 | 0.45 | 1.02 | -0.37 | 0.04 | 0.72 | 0.17 | 0.24 | 0.34 | 0.34 | 0.67 | 0.53 | 0.11 | -0.0375 | 0.045 | 0.09 | 0.804496 | 0.927806 | 0.898619 | 1 | 0.999631 | 0.934231 |
| Sgip1_Q8VD37_S170s_1 | 0.07 | 1.99 | 1.48 | 1.14 | 1.2 | 2.21 | 1.19 | 1.19 | 1.77 | 1.1 | 1.47 | 1.15 | 0.72 | 1.99 | 1.39 | 2.04 | 0.2775 | 0.2025 | 0.365 | 0.464193 | 0.549908 | 0.641842 | 1 | 0.999631 | 0.793831 |
| Sgip1_Q8VD37_S24s_2 | -3.36 | -1.4 | 0.79 | -1.27 | 0.03 | -1.82 | 1.51 | -1.53 | -1.84 | -4.87 | -0.13 | -0.49 | 0.08 | -2.02 | -1.94 | -1.56 | 0.8575 | -0.5225 | -0.05 | 0.952967 | 0.393968 | 0.657944 | 1 | 0.999631 | 0.984317 |
| Sgip1_Q8VD37_S264s_1 | -1.15 | -0.14 | 0.06 | -0.39 | -1.28 | -0.94 | -0.79 | -0.74 | -0.52 | -0.03 | -0.54 | 0.51 | 0.77 | -0.31 | -0.04 | 0.58 | -0.5325 | 0.26 | 0.655 | 0.120928 | 0.132679 | 0.50287 | 1 | 0.999631 | 0.588397 |
| Sgip1_Q8VD37_S265s_1 | -0.5 | 0.52 | 0.39 | -0.38 | 0.05 | -0.15 | -0.68 | -0.16 | -0.13 | -0.42 | -0.09 | 0.15 | 0.67 | 0.08 | -0.14 | 0.11 | -0.2425 | -0.13 | 0.1725 | 0.635134 | 0.480906 | 0.698818 | 1 | 0.999631 | 0.869104 |
| Sgip1_Q8VD37_S27s_1 | 0.36 | 2.3 | 2.6 | 0.67 | 0.64 | 1.95 | 3.53 | 0.05 | 1.37 | 1.63 | 1.63 | 1.09 | 2.64 | 0.48 | 1.46 | 0.45 | 0.06 | -0.0525 | -0.225 | 0.74204 | 0.94254 | 0.922461 | 1 | 0.999631 | 0.909328 |
| Sgip1_Q8VD37_S27s_1_1 | -0.38 | 1.13 | 1.04 | -0.63 | -0.3 | 0.64 | 2.23 | -1.38 | 0.17 | 0.34 | 0.27 | -0.51 | 1.32 | -0.38 | -0.05 | -1.11 | 0.0075 | -0.2225 | -0.345 | 0.582038 | 0.99234 | 0.645652 | 1 | 0.999631 | 0.847567 |
| Sgip1_Q8VD37_S300s_1 | -1.66 | -1.12 | -0.91 | -1.57 | -0.84 | -1.02 | -1.15 | -1.37 | -1.48 | -1.76 | -1.79 | -0.69 | 1.23 | -1.75 | -3.61 | -1.19 | 0.22 | -0.115 | -0.015 | 0.986322 | 0.449397 | 0.743938 | 1 | 0.999631 | 0.995135 |
| Sgip1_Q8VD37_S319s_1 | -1.12 | -1.42 | -2.34 | -1.91 | -0.58 | -0.98 | -0.2 | -3.28 | -0.96 | -1.05 | -3.51 | -1.81 | -0.5 | -0.94 | -4.52 | -2.67 | 0.4375 | -0.135 | -0.46 | 0.581954 | 0.513128 | 0.819919 | 1 | 0.999631 | 0.847567 |
| Sgip1_Q8VD37_S371s_1 | 0.02 | 3.35 | 2.35 | 0.48 | 4.97 | 2.41 | 3.24 | 1.88 | 5.17 | -0.11 | 3.99 | 5.31 | 5.34 | 1.46 | 2.06 | 1.13 | 1.575 | 2.04 | 0.9475 | 0.381319 | 0.103712 | 0.133626 | 1 | 0.999631 | 0.752924 |
| Sgip1_Q8VD37_S377s_1 | 0.31 | 0.29 | -0.53 | 1.78 | 2.03 | 1.27 | 1.61 | 1.31 | 0.57 | -2.14 | 0.74 | 0.24 | -2.94 | 0.68 | -0.02 | -0.2 | 1.0925 | -0.61 | -1.0825 | 0.201566 | 0.045146 | 0.413381 | 1 | 0.999631 | 0.658952 |
| Sgip1_Q8VD37_S398s_1 | -0.83 | 2.34 | 0.57 | -0.49 | 1.52 | 1.54 | -0.24 | 0.08 | 1.7 | -0.35 | 0.14 | 0.55 | 1.9 | 1.35 | 0.39 | 0.18 | 0.3275 | 0.1125 | 0.5575 | 0.447996 | 0.663647 | 0.879101 | 1 | 0.999631 | 0.786511 |
| Sgip1_Q8VD37_S423s_1 | -1.06 | -2.07 | -0.29 | -1 | -1.72 | -1.63 | -0.7 | -0.43 | -1.65 | -1.61 | 0.15 | -0.09 | -1.19 | -0.45 | 0.33 | -1.18 | -0.015 | 0.305 | 0.4825 | 0.341283 | 0.974394 | 0.588072 | 1 | 0.999631 | 0.733164 |
| Sgip1_Q8VD37_S424s_1 | 0.42 | 0.89 | -0.14 | 0.16 | -0.72 | -0.59 | -0.52 | -0.3 | -0.45 | -0.2 | -0.54 | -0.68 | -0.33 | -0.79 | -0.43 | 0.08 | -0.865 | -0.8 | -0.7 | 0.064871 | 0.015348 | 0.025645 | 1 | 0.999631 | 0.515898 |
| Sgip1_Q8VD37_S424s_2 | -0.27 | 0.38 | -2.19 | -2.15 | -1.2 | -0.34 | -1.9 | -1.93 | -0.92 | 0.32 | -2.23 | -2.41 | -0.37 | -0.72 | -2.7 | -1.95 | -0.285 | -0.2525 | -0.3775 | 0.615544 | 0.672558 | 0.753398 | 1 | 0.999631 | 0.861203 |
| Sgip1_Q8VD37_S429s_1 | 0.41 | -0.36 | 0.02 | -0.07 | -1.72 | -0.3 | -0.3 | -1.02 | -1.7 | -1.96 | -0.55 | -0.45 | 0.04 | -1.36 | -1.63 | -0.22 | -0.835 | -1.165 | -0.7925 | 0.098575 | 0.052851 | 0.019331 | 1 | 0.999631 | 0.562298 |
| Sgip1_Q8VD37_S437s_2 | -0.37 | -0.56 | -1.83 | -2.22 | -1.14 | -0.35 | -1.88 | -1.85 | -1.12 | 0.31 | -2.21 | -2.47 | -0.12 | -1.11 | -1.95 | -2.41 | -0.06 | -0.1275 | -0.1525 | 0.805068 | 0.911213 | 0.85471 | 1 | 0.999631 | 0.934299 |
| Sgip1_Q8VD37_S441s_1 | 0.16 | 0.2 | 0.7 | -0.19 | -0.22 | 0.45 | -0.08 | -0.41 | -0.3 | 0.5 | 0.02 | 0.29 | 0.72 | -0.01 | 0.05 | 0.61 | -0.2825 | -0.09 | 0.125 | 0.708525 | 0.377105 | 0.775614 | 1 | 0.999631 | 0.898927 |
| Sgip1_Q8VD37_S465s_1 | 0 | -0.08 | -0.51 | -1.18 | -1.86 | 0.77 | -1.79 | 0.46 | 0.31 | 1.56 | -1.85 | 0.57 | 1.06 | 1.39 | -0.61 | -0.32 | -0.1625 | 0.59 | 0.8225 | 0.144553 | 0.809352 | 0.398549 | 1 | 0.999631 | 0.616037 |
| Sgip1_Q8VD37_S484s_2 | -0.58 | -0.14 | -0.24 | -0.8 | -0.38 | -0.38 | -0.83 | -0.68 | -0.44 | -0.37 | -0.86 | -0.43 | -0.04 | 0.01 | -0.55 | -0.08 | -0.1275 | -0.085 | 0.275 | 0.373898 | 0.645326 | 0.764455 | 1 | 0.999631 | 0.747035 |
| Sgip1_Q8VD37_S490s_1 | -0.71 | -1.6 | -0.48 | -1.23 | -1.19 | -0.92 | -1.48 | -1.17 | -0.67 | -0.17 | -1.84 | -0.95 | 0.55 | -0.55 | -1.02 | -1 | -0.185 | 0.0975 | 0.5 | 0.278679 | 0.570476 | 0.821081 | 1 | 0.999631 | 0.70098 |
| Sgip1_Q8VD37_S490s_2 | 0.2 | 1.05 | 1.13 | 0.33 | 0.39 | 1.04 | -0.36 | -0.08 | 0.16 | 1 | -1.24 | 0.14 | 0.7 | 1.12 | 0.89 | 0.89 | -0.43 | -0.6625 | 0.2225 | 0.504454 | 0.291981 | 0.202924 | 1 | 0.999631 | 0.814292 |
| Sgip1_Q8VD37_S491s_1 | -0.89 | -2.06 | -2.19 | -2.93 | -1.71 | -2.01 | -2.26 | -2.6 | -2.11 | -1.89 | -3.32 | -2.66 | -0.91 | -1 | -2.99 | -2.68 | -0.1275 | -0.4775 | 0.1225 | 0.84446 | 0.775957 | 0.352678 | 1 | 0.999631 | 0.94856 |
| Sgip1_Q8VD37_S491s_2 | -0.39 | -0.13 | 0.15 | -0.31 | -0.33 | -0.04 | -0.36 | -0.16 | -0.2 | -0.36 | -0.38 | 0.02 | -0.27 | 0.2 | -0.03 | 0.54 | -0.0525 | -0.06 | 0.28 | 0.372325 | 0.836715 | 0.822035 | 1 | 0.999631 | 0.746986 |
| Sgip1_Q8VD37_S503s_1 | -0.4 | -0.32 | 0.26 | -0.29 | -0.18 | -0.46 | -0.17 | 0.96 | -0.53 | -0.06 | -0.07 | -0.92 | -0.07 | -0.05 | 0.33 | 0.76 | 0.225 | -0.2075 | 0.43 | 0.206036 | 0.545266 | 0.517995 | 1 | 0.999631 | 0.6605 |
| Sgip1_Q8VD37_S78s_1 | -1.08 | -0.57 | 0.05 | -0.63 | -0.56 | -1.15 | -0.48 | -0.27 | -1.03 | -0.94 | -0.25 | -0.24 | 0.04 | -0.73 | -0.59 | 0.21 | -0.0575 | -0.0575 | 0.29 | 0.440819 | 0.864305 | 0.870705 | 1 | 0.999631 | 0.781983 |
| Sgip1_Q8VD37_T180t_1 | 0.82 | 0.3 | 0.64 | 1.25 | 1.05 | -0.17 | 1.07 | 1.36 | 1.23 | 0.38 | 1.06 | 0.55 | -0.18 | 0.9 | 1.01 | 1.44 | 0.075 | 0.0525 | 0.04 | 0.922957 | 0.850673 | 0.87475 | 1 | 0.999631 | 0.975032 |
| Sgip1_Q8VD37_T180t_2 | -0.68 | -0.45 | -0.02 | 1.11 | -0.34 | -0.19 | 0.39 | 0.82 | -1.11 | -0.6 | 0.94 | 0.78 | -1.47 | 0.44 | 1.01 | 1.43 | 0.18 | 0.0125 | 0.3625 | 0.592563 | 0.696543 | 0.98305 | 1 | 0.999631 | 0.852841 |
| Sgip1_Q8VD37_T182t_1 | 0.8 | 0.53 | -0.41 | 0.46 | 0.83 | 0.79 | 0.13 | 0.19 | 0.67 | 0.29 | 0.17 | -0.18 | -0.75 | 0.86 | -0.14 | 0.15 | 0.14 | -0.1075 | -0.315 | 0.472101 | 0.691299 | 0.761018 | 1 | 0.999631 | 0.79887 |
| Sgip1_Q8VD37_T182t_2 | -0.68 | -0.45 | -0.02 | 1.11 | -0.34 | -0.19 | 0.39 | 0.82 | -1.11 | -0.6 | 0.94 | 0.78 | -1.47 | 0.44 | 1.01 | 1.43 | 0.18 | 0.0125 | 0.3625 | 0.592563 | 0.696543 | 0.98305 | 1 | 0.999631 | 0.852841 |
| Sgip1_Q8VD37_T247t_1 | -0.02 | -0.56 | 0.93 | 0.39 | -0.3 | 0.25 | -0.31 | -0.14 | 0.03 | 0.45 | -0.79 | 0.49 | 0.48 | 0.59 | 0.57 | 0.41 | -0.31 | -0.14 | 0.3275 | 0.379193 | 0.402812 | 0.74633 | 1 | 0.999631 | 0.751637 |
| Sgip1_Q8VD37_T258t_1 | -0.01 | -0.02 | -0.07 | -0.37 | -0.53 | -0.39 | -0.64 | -0.49 | -0.41 | 0.14 | -0.27 | -0.26 | 0.26 | -0.15 | -0.43 | -0.08 | -0.395 | -0.0825 | 0.0175 | 0.951544 | 0.124992 | 0.755526 | 1 | 0.999631 | 0.984079 |
| Sgip1_Q8VD37_T259t_1 | -0.01 | -0.02 | -0.07 | -0.37 | -0.53 | -0.39 | -0.64 | -0.49 | -0.41 | 0.14 | -0.27 | -0.26 | 0.26 | -0.15 | -0.43 | -0.08 | -0.395 | -0.0825 | 0.0175 | 0.951544 | 0.124992 | 0.755526 | 1 | 0.999631 | 0.984079 |
| Sgip1_Q8VD37_T259t_2 | -3.19 | -5.35 | -2.7 | -3.3 | -3.55 | -3.73 | -2.63 | -3.96 | -4.01 | -3.85 | -2.99 | -3.48 | -3.33 | -4.09 | -3.61 | -3.1 | 0.1675 | 0.0525 | 0.1025 | 0.858329 | 0.777809 | 0.927433 | 1 | 0.999631 | 0.952251 |
| Sgip1_Q8VD37_T25t_2 | -3.36 | -1.4 | -4.71 | -4.07 | 0.03 | -1.82 | -2.76 | -5.75 | -1.84 | -4.87 | -3.81 | -4.86 | 0.08 | -2.02 | -4.08 | -3.37 | 0.81 | -0.46 | 1.0375 | 0.310366 | 0.503992 | 0.603839 | 1 | 0.999631 | 0.71924 |
| Sgip1_Q8VD37_T263t_1 | -0.85 | -0.88 | 0.07 | -0.4 | -0.29 | -0.67 | -1.02 | -0.05 | -0.25 | -1.07 | -0.14 | 0.36 | 0.48 | -0.93 | -0.44 | -1.53 | 0.0075 | 0.24 | -0.09 | 0.848236 | 0.982523 | 0.541299 | 1 | 0.999631 | 0.949393 |
| Sgip1_Q8VD37_T263t_2 | -3.19 | -5.35 | -2.7 | -3.3 | -3.55 | -3.73 | -2.63 | -3.96 | -4.01 | -3.85 | -2.99 | -3.48 | -3.33 | -4.09 | -3.61 | -3.1 | 0.1675 | 0.0525 | 0.1025 | 0.858329 | 0.777809 | 0.927433 | 1 | 0.999631 | 0.952251 |
| Sgip1_Q8VD37_T335t_1 | -0.62 | -0.05 | -0.44 | -0.66 | 0.07 | -0.28 | -1.07 | -0.56 | -0.05 | -0.26 | -0.48 | -0.33 | 0.33 | 0.16 | -0.64 | -0.37 | -0.0175 | 0.1625 | 0.3125 | 0.361775 | 0.956848 | 0.554477 | 1 | 0.999631 | 0.742006 |
| Sgip1_Q8VD37_T409t_1 | 0.15 | 0.03 | -0.08 | 0.02 | -0.02 | -0.26 | 0.03 | -0.21 | -0.09 | -0.39 | -0.35 | -0.19 | 0.16 | -0.1 | -0.09 | -0.15 | -0.145 | -0.285 | -0.075 | 0.776051 | 0.543892 | 0.261798 | 1 | 0.999631 | 0.920998 |
| Sgip1_Q8VD37_T455t_1 | -1.67 | -1.05 | -2.12 | -1.35 | -1.94 | -1.15 | -0.81 | -0.5 | -2.23 | -1.54 | -1.91 | -0.04 | 0.18 | -1.19 | -1.04 | -1.39 | 0.4475 | 0.1175 | 0.6875 | 0.133527 | 0.270975 | 0.816641 | 1 | 0.999631 | 0.60492 |
| Sgk2_Q9QZS5_S10s_1 | -0.45 | 0.38 | 0.12 | -0.99 | 0.11 | -0.68 | -0.21 | 1.09 | 0.73 | -0.04 | -0.26 | -0.52 | -0.1 | -0.33 | -1.36 | -1.08 | 0.3125 | 0.2125 | -0.4825 | 0.281763 | 0.504693 | 0.609762 | 1 | 0.999631 | 0.702862 |
| Sgk223_Q571I4_S219s_1 | 0.73 | 0.41 | 0.17 | 1.25 | 1 | 0.59 | 0.98 | 0.75 | 0.42 | 0.93 | 1.26 | 1.33 | 1.13 | 0.8 | 0.81 | 1.1 | 0.19 | 0.345 | 0.32 | 0.339587 | 0.542747 | 0.338066 | 1 | 0.999631 | 0.732185 |
| Sgk223_Q571I4_S40s_1 | 1.44 | 1.66 | 0.63 | 1.45 | 1.42 | 1.88 | 1.4 | 1.33 | 1.84 | 1.34 | 1.74 | 1.1 | 0.61 | 1.57 | 1.22 | 1.53 | 0.2125 | 0.21 | -0.0625 | 0.863812 | 0.502241 | 0.534606 | 1 | 0.999631 | 0.955131 |
| Sgk223_Q571I4_S720s_1 | -1.43 | 0.05 | -1.27 | -0.23 | 0.03 | 0 | -0.07 | 0.29 | -0.33 | -1.34 | -0.15 | -0.41 | -0.61 | -0.52 | -0.26 | -0.27 | 0.7825 | 0.1625 | 0.305 | 0.455619 | 0.067358 | 0.717452 | 1 | 0.999631 | 0.790041 |
| Sgpp1_Q9JI99_S101s_1 | -0.44 | -0.26 | -0.29 | -0.6 | -0.35 | -0.42 | -0.7 | -0.29 | -0.31 | 0.03 | -0.34 | -0.14 | 0.97 | -0.04 | 0.27 | 0.07 | -0.0425 | 0.2075 | 0.715 | 0.047078 | 0.863279 | 0.422039 | 1 | 0.999631 | 0.480102 |
| Sgpp1_Q9JI99_S65s_1 | 1.16 | -2.03 | 0.12 | 0.43 | -1.73 | -1.42 | 0.61 | -1.06 | -1.36 | 0.01 | -0.92 | -0.38 | -1.55 | -1.86 | 0.2 | -1.03 | -0.82 | -0.5825 | -0.98 | 0.197151 | 0.293215 | 0.394468 | 1 | 0.999631 | 0.653975 |
| Sgsm1_B2RQR5_S229s_1 | 0.34 | 0.97 | 1.21 | -0.53 | 2.39 | 0.48 | 2.54 | -0.44 | 3.83 | -0.23 | 1.42 | 2.67 | 4.8 | -0.55 | 0.54 | -0.93 | 0.745 | 1.425 | 0.4675 | 0.688686 | 0.320427 | 0.112282 | 1 | 0.999631 | 0.892031 |
| Sgsm1_B2RQR5_S397s_1 | -1.13 | -1.85 | -1.27 | -1.63 | -1.31 | -1.8 | -1.74 | -0.78 | -1.56 | -1.67 | -0.87 | -0.86 | 0.21 | -0.56 | -0.3 | -0.92 | 0.0625 | 0.23 | 1.0775 | 0.010592 | 0.849566 | 0.487854 | 1 | 0.999631 | 0.383871 |
| Sgsm1_B2RQR5_S633s_1 | -1.14 | 0.41 | 0.75 | -0.47 | 0.5 | 0.9 | 0.9 | 0.03 | 0.6 | -1.65 | 0.26 | 0.67 | -0.56 | -0.14 | 0.52 | -0.4 | 0.695 | 0.0825 | -0.0325 | 0.945644 | 0.152061 | 0.895483 | 1 | 0.999631 | 0.982217 |
| Sgsm1_B2RQR5_T647t_1 | 1.23 | 1.27 | 3.2 | 2.69 | 2.63 | 2.05 | 4.29 | 3.68 | 2.19 | 1.18 | 4.28 | 3.28 | -0.4 | 0.34 | 1.9 | 0.45 | 1.065 | 0.635 | -1.525 | 0.033108 | 0.119123 | 0.400318 | 1 | 0.999631 | 0.44294 |
| Sgsm2_Q80U12_S682s_1 | -2.04 | -1.03 | 1.08 | 1.98 | -0.08 | -0.75 | -0.26 | 2.1 | -0.42 | -2.24 | 0.44 | 0.24 | -0.42 | -1.1 | 2.24 | 2.53 | 0.255 | -0.4925 | 0.815 | 0.468886 | 0.79216 | 0.610471 | 1 | 0.999631 | 0.796529 |
| Sgta_Q8BJU0_S306s_1 | 2.6 | 0.71 | 0.59 | 1.93 | 1.98 | 1.02 | 2.67 | 1.56 | 1.14 | 0.92 | 1.36 | 0.38 | -1.39 | -0.6 | 0.46 | -0.64 | 0.35 | -0.5075 | -2 | 0.005388 | 0.528308 | 0.325422 | 1 | 0.999631 | 0.351308 |
| Sgta_Q8BJU0_S306s_2 | 0.3 | 1.73 | 0.64 | -0.2 | -0.06 | 1.27 | 1.53 | 0.4 | 1.87 | 1.87 | 0.02 | -0.57 | 0.24 | -1.1 | -1.31 | -2.38 | 0.1675 | 0.18 | -1.755 | 0.015905 | 0.745133 | 0.789299 | 1 | 0.999631 | 0.388832 |
| Sgta_Q8BJU0_S308s_1 | 1.3 | 0.05 | 0.67 | 0.58 | -0.08 | -0.1 | 1.36 | 0.32 | 0.36 | 0.72 | 0.1 | -0.04 | -1.6 | -1.07 | -0.39 | -0.8 | -0.275 | -0.365 | -1.615 | 0.001957 | 0.519668 | 0.306997 | 1 | 0.999631 | 0.351308 |
| Sgta_Q8BJU0_T304t_1 | 2.67 | 0.52 | 1.1 | 1.71 | 1.44 | 1.3 | 2.78 | 1.07 | 1.53 | 1.72 | 0.85 | 0.44 | -0.73 | -0.37 | 0.63 | -1.16 | 0.1475 | -0.365 | -1.9075 | 0.006063 | 0.788851 | 0.484487 | 1 | 0.999631 | 0.351545 |
| Sgta_Q8BJU0_T304t_2 | 0.3 | 1.73 | 0.64 | -0.2 | -0.06 | 1.27 | 1.53 | 0.4 | 1.87 | 1.87 | 0.02 | -0.57 | 0.24 | -1.1 | -1.31 | -2.38 | 0.1675 | 0.18 | -1.755 | 0.015905 | 0.745133 | 0.789299 | 1 | 0.999631 | 0.388832 |
| Sgta_Q8BJU0_T81t_1 | 0.21 | 0.12 | 0.31 | -0.41 | -0.35 | -0.03 | -0.28 | -0.3 | -0.02 | 0.44 | -0.09 | 0.65 | 0.2 | -0.02 | -0.17 | 0.05 | -0.2975 | 0.1875 | -0.0425 | 0.884729 | 0.284558 | 0.547757 | 1 | 0.999631 | 0.963505 |
| Sgta_Q8BJU0_T81t_2 | 0.07 | -0.3 | -0.32 | 0.27 | -0.68 | -0.42 | 0.27 | -0.33 | -0.78 | 0.29 | -0.62 | -0.49 | -0.21 | 0.09 | -3.04 | 1.16 | -0.22 | -0.33 | -0.43 | 0.589974 | 0.477981 | 0.330479 | 1 | 0.999631 | 0.8515 |
| Sgtb_Q8VD33_S295s_1 | 1.54 | -0.02 | -0.74 | 1.23 | 0.61 | 1.02 | 1.72 | 1.13 | 1.08 | 1.01 | 0.65 | -0.18 | -0.54 | 0.06 | 0.04 | -0.46 | 0.6175 | 0.1375 | -0.7275 | 0.187269 | 0.266168 | 0.80637 | 1 | 0.999631 | 0.646474 |
| Sgtb_Q8VD33_S295s_2 | -0.47 | 0.17 | -0.49 | 0.65 | 0.51 | -0.11 | 0.74 | 0.34 | 0.9 | 0.13 | 0.49 | 0.21 | -1.25 | -1 | -0.36 | -1.03 | 0.405 | 0.4675 | -0.875 | 0.038929 | 0.27035 | 0.213018 | 1 | 0.999631 | 0.463185 |
| Sgtb_Q8VD33_S299s_1 | 0.41 | 0.12 | -0.05 | 0.11 | -0.92 | 0.32 | 0.45 | -0.34 | 0.41 | 0.33 | 0 | -0.42 | -0.89 | -0.22 | -0.58 | -0.53 | -0.27 | -0.0675 | -0.7025 | 0.032939 | 0.456029 | 0.818458 | 1 | 0.999631 | 0.44294 |
| Sgtb_Q8VD33_S304s_2 | -0.47 | 0.17 | -1.3 | 1.57 | 0.51 | -0.11 | 1.06 | 0.81 | 0.9 | 0.13 | 1.02 | 1.14 | -1.25 | -1 | 0.05 | 0.97 | 0.575 | 0.805 | -0.3 | 0.67049 | 0.345081 | 0.194675 | 1 | 0.999631 | 0.884249 |
| Sh2b1_Q91ZM2_S126s_1 | -0.02 | 0.13 | -0.58 | 0.38 | 0.84 | -0.35 | 0 | -0.35 | -0.27 | -0.4 | -0.5 | -0.21 | 0.2 | 0.34 | -0.47 | 0.17 | 0.0575 | -0.3225 | 0.0825 | 0.807568 | 0.875406 | 0.289233 | 1 | 0.999631 | 0.93582 |
| Sh2b1_Q91ZM2_S127s_1 | -0.29 | -0.21 | -0.66 | 0.17 | 0.31 | -0.43 | -0.09 | -0.21 | -0.46 | -0.64 | -0.42 | -0.22 | -0.17 | -0.1 | -0.72 | -0.03 | 0.1425 | -0.1875 | -0.0075 | 0.981151 | 0.632647 | 0.513527 | 1 | 0.999631 | 0.994608 |
| Sh2b1_Q91ZM2_S219s_1 | 0.95 | -0.38 | 0.32 | 0.95 | 0.73 | 1.21 | 1.65 | 1.2 | 1.5 | 1.02 | 0.99 | 1.17 | -0.98 | 0.98 | 1.54 | 0.29 | 0.7375 | 0.71 | -0.0025 | 0.996545 | 0.077473 | 0.077464 | 1 | 0.999631 | 0.9984 |
| Sh2b3_D3Z0T9_S102s_1 | 0.19 | 1.38 | 1.04 | 0.67 | 0.57 | 1.05 | 0.82 | 0.55 | 0.7 | 0.39 | 0.93 | 1.02 | 0.26 | 0.78 | 0.9 | 0.73 | -0.0725 | -0.06 | -0.1525 | 0.663708 | 0.823836 | 0.859153 | 1 | 0.999631 | 0.881991 |
| Sh2d3c_Q9QZS8_S189s_1 | 1.92 | -1.2 | 0.65 | -0.09 | -2 | 0.25 | 0.66 | -0.56 | -1 | 1.02 | 0 | 0.02 | -0.6 | -1.13 | 0.63 | 0 | -0.7325 | -0.31 | -0.595 | 0.387749 | 0.353349 | 0.654958 | 1 | 0.999631 | 0.757144 |
| Sh2d3c_Q9QZS8_S283s_1 | 0.2 | 0.75 | -0.12 | -0.35 | 0.84 | 0.48 | 0.01 | -0.26 | 0.82 | -0.18 | 0.57 | 1.05 | 1.21 | -0.25 | 1.01 | -1.02 | 0.1475 | 0.445 | 0.1175 | 0.828497 | 0.685968 | 0.258866 | 1 | 0.999631 | 0.941315 |
| Sh2d3c_Q9QZS8_S289s_1 | -0.81 | -0.72 | -0.76 | -0.56 | -0.54 | -0.66 | -0.87 | -0.64 | -1.06 | -0.42 | -0.84 | -0.04 | -0.46 | -0.08 | -0.54 | -0.4 | 0.035 | 0.1225 | 0.3425 | 0.224846 | 0.882731 | 0.688834 | 1 | 0.999631 | 0.67471 |
| Sh2d3c_Q9QZS8_S290s_1 | -0.81 | -0.72 | -0.76 | -0.56 | -0.54 | -0.66 | -0.87 | -0.64 | -1.06 | -0.42 | -0.84 | -0.04 | -0.46 | -0.08 | -0.54 | -0.4 | 0.035 | 0.1225 | 0.3425 | 0.224846 | 0.882731 | 0.688834 | 1 | 0.999631 | 0.67471 |
| Sh2d5_Q8JZW5_S12s_1 | 1.83 | 0.3 | -1.02 | 1.33 | 2.02 | 0.71 | 1.46 | 1.43 | 1.07 | 0.31 | 1 | 0.59 | -0.4 | 1.35 | 0.01 | 0.52 | 0.795 | 0.1325 | -0.24 | 0.715811 | 0.216761 | 0.824439 | 1 | 0.999631 | 0.902706 |
| Sh2d5_Q8JZW5_S237s_1 | 0.95 | 1.46 | -0.48 | 0.44 | 1.22 | 1.13 | -0.2 | 0.34 | 1.19 | 1.03 | 0.87 | 0.65 | 0.46 | 1.84 | 0.69 | 0.54 | 0.03 | 0.3425 | 0.29 | 0.566542 | 0.952176 | 0.432672 | 1 | 0.999631 | 0.84194 |
| Sh3bgrl2_Q8BG73_S103s_1 | 1.54 | 0.21 | 1.05 | 1.41 | 0.55 | 0.84 | 1.31 | 0.6 | 0.95 | 1.35 | 1.4 | 0.72 | -0.56 | 1.54 | 0.66 | 0.49 | -0.2275 | 0.0525 | -0.52 | 0.313488 | 0.538738 | 0.886842 | 1 | 0.999631 | 0.719861 |
| Sh3bp1_P55194_S535s_1 | 0.36 | 0.71 | 1.02 | 0.68 | 0.45 | 0.97 | 0.1 | 0.38 | 0.49 | 0.6 | -0.06 | 0.47 | 0.34 | 1.36 | 0.92 | 0.84 | -0.2175 | -0.3175 | 0.1725 | 0.599433 | 0.4661 | 0.285992 | 1 | 0.999631 | 0.856093 |
| Sh3bp4_Q921I6_S245s_1 | 0.6 | 0.78 | 0.42 | 0.34 | 1.57 | 0.48 | -0.06 | -0.14 | 1.45 | 0.73 | 0.06 | 0.67 | 0.2 | 1.58 | 0.72 | 0.94 | -0.0725 | 0.1925 | 0.325 | 0.37082 | 0.858696 | 0.578704 | 1 | 0.999631 | 0.746338 |
| Sh3bp4_Q921I6_S42s_1 | 0.31 | -0.59 | -1.12 | 0.66 | 0.93 | 0.05 | 0.63 | 1.1 | 0.09 | -0.55 | 0.94 | 0.87 | -0.67 | -0.08 | 0.6 | 0.56 | 0.8625 | 0.5225 | 0.2875 | 0.560095 | 0.081123 | 0.318559 | 1 | 0.999631 | 0.839805 |
| Sh3bp5l_Q99LH9_S30s_1 | -0.21 | 0.67 | 3.41 | 0.36 | 0.26 | 0.13 | 0.74 | 3.06 | 0.16 | 0.45 | 3.47 | 3.78 | 0.8 | 0.8 | 4.18 | 3.8 | -0.01 | 0.9075 | 1.3375 | 0.219716 | 0.991255 | 0.409251 | 1 | 0.999631 | 0.670699 |
| Sh3bp5l_Q99LH9_S361s_1 | -0.44 | 0.54 | -1.08 | -0.1 | 0.32 | 0.04 | -0.28 | 0.25 | 0.5 | 0.6 | 0.2 | 1.29 | 0.98 | 0.86 | -0.09 | 1.18 | 0.3525 | 0.9175 | 1.0025 | 0.043913 | 0.362377 | 0.048761 | 1 | 0.999631 | 0.475443 |
| Sh3bp5l_Q99LH9_S377s_1 | 0.17 | 0.16 | 0.08 | -0.05 | 0.17 | -0.23 | -0.05 | -0.09 | 0.26 | 0.68 | 0.4 | 0.33 | -0.68 | 0.22 | 0.71 | -0.23 | -0.14 | 0.3275 | -0.085 | 0.811235 | 0.562839 | 0.211412 | 1 | 0.999631 | 0.936949 |
| Sh3bp5l_Q99LH9_T13t_1 | 0.44 | 1.48 | 0.49 | -0.27 | 1.09 | 1.14 | 0.61 | -0.44 | 1.64 | 1.22 | 0.62 | 0.39 | 0.64 | 1.6 | 0.89 | 0.24 | 0.065 | 0.4325 | 0.3075 | 0.504791 | 0.893391 | 0.348263 | 1 | 0.999631 | 0.814292 |
| Sh3d19_Q91X43_S195s_1 | 1.87 | 0.85 | 0.54 | 1.14 | 2.07 | 1.93 | 1.29 | 1.29 | 1.38 | 1.23 | 0.6 | 0.9 | 0.49 | 1.51 | 0.56 | 0.11 | 0.545 | -0.0725 | -0.4325 | 0.320144 | 0.163766 | 0.842299 | 1 | 0.999631 | 0.723684 |
| Sh3gl1_Q62419_S288s_1 | 0.94 | -0.21 | 0.83 | 0.32 | 0.96 | 1.1 | 1.37 | 1.48 | 1.45 | 0.12 | 1.72 | 1.55 | 1.39 | 0.76 | 1.51 | 1.63 | 0.7575 | 0.74 | 0.8525 | 0.041064 | 0.042493 | 0.121529 | 1 | 0.999631 | 0.474979 |
| Sh3gl2_Q62420_S262s_1 | 0.22 | 1.03 | 0.35 | 0.65 | 0.61 | 0.54 | 0.25 | 0.87 | 0.53 | 0.33 | 0.72 | -1.29 | 0.03 | 0.79 | 0.35 | 0.76 | 0.005 | -0.49 | -0.08 | 0.808388 | 0.986288 | 0.317321 | 1 | 0.999631 | 0.93635 |
| Sh3gl3_J3QQ44_T243t_1 | 1.35 | 1.51 | 0.57 | 2.41 | 1.39 | 1.43 | 1.98 | 2.12 | 1.36 | 1.45 | 1.6 | -2.82 | -0.55 | 1.71 | 1.83 | 0.4 | 0.27 | -1.0625 | -0.6125 | 0.333569 | 0.522806 | 0.295119 | 1 | 0.999631 | 0.728079 |
| Sh3gl3_J3QQ44_T248t_1 | 0.84 | 2.54 | 1.81 | 1.49 | 1.12 | 2.02 | 1.33 | 1.99 | 1.94 | 2 | 2.1 | -2.51 | 0.11 | 1.74 | 1.67 | 1.35 | -0.055 | -0.7875 | -0.4525 | 0.372907 | 0.895244 | 0.448269 | 1 | 0.999631 | 0.747035 |
| Sh3glb2_Q8R3V5_S379s_1 | -1.36 | -1.16 | -0.58 | -1.26 | -1.47 | -1.15 | -1.17 | -1.18 | -1.29 | -1.1 | -0.9 | -0.9 | -0.74 | -1.06 | -1.17 | -0.46 | -0.1525 | 0.0425 | 0.2325 | 0.474172 | 0.583674 | 0.882366 | 1 | 0.999631 | 0.799827 |
| Sh3kbp1_Q8R550_S373s_1 | 0.46 | 0.91 | 0.37 | 1.7 | 1.7 | 1.28 | 1 | 1.75 | 1.36 | 0.1 | 1.11 | 1.03 | -0.22 | 1.6 | 1.58 | 0.68 | 0.5725 | 0.04 | 0.05 | 0.921099 | 0.146139 | 0.923291 | 1 | 0.999631 | 0.975032 |
| Sh3pxd2a_O89032_S377s_1 | 0.37 | 1.22 | 0.61 | 1.01 | 0.69 | 0.66 | 0.68 | 0.83 | 0.31 | 0.96 | 0.64 | 0.51 | 0.49 | 1.17 | 0.71 | 0.8 | -0.0875 | -0.1975 | -0.01 | 0.975123 | 0.753894 | 0.523443 | 1 | 0.999631 | 0.991207 |
| Sh3pxd2a_O89032_S503s_1 | -0.8 | -0.43 | 0.35 | 0.11 | -0.7 | -1.15 | -0.43 | 0.37 | -0.97 | -0.28 | -0.18 | 0.34 | -0.08 | -0.75 | 0.72 | 0.25 | -0.285 | -0.08 | 0.2275 | 0.589123 | 0.493713 | 0.837935 | 1 | 0.999631 | 0.851233 |
| Sh3pxd2a_O89032_S769s_1 | -0.21 | -0.44 | -0.28 | -0.13 | -0.62 | -0.1 | -0.69 | -0.68 | -0.21 | -0.18 | -0.68 | -0.09 | -0.05 | -0.03 | -0.56 | 0.02 | -0.2575 | -0.025 | 0.11 | 0.69822 | 0.335714 | 0.92487 | 1 | 0.999631 | 0.895731 |
| Sh3pxd2a_O89032_S950s_1 | -0.02 | 0.2 | -0.19 | 0.88 | 0.95 | 0.53 | 0.43 | 0.79 | 0.45 | -0.02 | 0.03 | 0.3 | -0.26 | 0.9 | 0.21 | 0.51 | 0.4575 | -0.0275 | 0.1225 | 0.74597 | 0.168815 | 0.931424 | 1 | 0.999631 | 0.911088 |
| Sh3pxd2a_O89032_S965s_1 | 0.53 | 0.35 | 0.29 | 1.31 | 1.27 | 1.21 | 1.23 | 0.72 | 1.63 | 0.47 | 1.03 | 0.86 | -1.15 | 0.3 | 0.74 | 0.05 | 0.4875 | 0.3775 | -0.635 | 0.19001 | 0.149086 | 0.316841 | 1 | 0.999631 | 0.648674 |
| Sh3pxd2a_O89032_S986s_1 | 0.81 | 0.24 | -0.03 | -0.15 | 1.04 | 0.63 | -0.31 | 0.43 | -0.16 | 0.26 | 0.5 | 0.01 | -0.25 | 0.87 | -0.57 | 0.48 | 0.23 | -0.065 | -0.085 | 0.836228 | 0.540267 | 0.838394 | 1 | 0.999631 | 0.945157 |
| Sh3rf1_E9QQ33_S740s_1 | -1.51 | -0.24 | -1.04 | -1.25 | -1.17 | -0.86 | -1.27 | -0.67 | -0.86 | -1.44 | -1.82 | -0.17 | -0.67 | -0.99 | -0.42 | -0.5 | 0.0175 | -0.0625 | 0.365 | 0.317284 | 0.959085 | 0.888355 | 1 | 0.999631 | 0.720971 |
| Sh3rf1_E9QQ33_S804s_1 | -0.37 | -0.03 | -0.84 | -0.31 | 0.02 | -0.38 | -0.73 | -0.82 | -0.69 | -0.59 | -0.37 | -0.38 | -0.09 | -0.07 | -0.94 | -1.19 | -0.09 | -0.12 | -0.185 | 0.623189 | 0.771788 | 0.671111 | 1 | 0.999631 | 0.862913 |
| Shank1_D3YZU4_S1283s_1 | -0.03 | 0.9 | 0.54 | 0.49 | 0.44 | -0.03 | -0.01 | 0.59 | 0.09 | 0.05 | 0.58 | 1.03 | 0.38 | -0.08 | 0.59 | 1.09 | -0.2275 | -0.0375 | 0.02 | 0.955593 | 0.463943 | 0.913038 | 1 | 0.999631 | 0.984649 |
| Shank1_D3YZU4_S1434s_1 | 0.92 | -0.59 | -0.91 | 0.41 | 0.98 | 1.12 | 0.52 | 0.6 | 0.15 | -0.3 | 0.26 | -0.9 | -1.02 | 0.21 | -0.68 | -0.8 | 0.8475 | -0.155 | -0.53 | 0.291925 | 0.078066 | 0.747804 | 1 | 0.999631 | 0.706164 |
| Shank1_D3YZU4_S1531s_1 | 1.73 | -0.84 | -3.25 | 0.74 | 2.03 | 0.09 | 0.77 | 0.33 | 0.2 | -2.16 | -0.06 | -2.14 | -3.13 | -1.07 | -1.85 | -1.15 | 1.21 | -0.635 | -1.395 | 0.189523 | 0.246616 | 0.561914 | 1 | 0.999631 | 0.648674 |
| Shank1_D3YZU4_S1899s_1 | 1.45 | -0.6 | -1.31 | -0.24 | 0.07 | -0.18 | -0.13 | -0.91 | 0.45 | -1.5 | -1.99 | -2.03 | 0.06 | -0.59 | -2.31 | -1.19 | -0.1125 | -1.0925 | -0.8325 | 0.243182 | 0.843185 | 0.159422 | 1 | 0.999631 | 0.685637 |
| Shank1_D3YZU4_S1998s_1 | 2.43 | -1.58 | -3.24 | 0.56 | 1.58 | 1.3 | 1.95 | 0.43 | 1.52 | 0.19 | -0.67 | -1.61 | -2.64 | 1.09 | -3.83 | -0.39 | 1.7725 | 0.315 | -0.985 | 0.485937 | 0.130123 | 0.793568 | 1 | 0.999631 | 0.804522 |
| Shank1_D3YZU4_S2017s_1 | 0.56 | -2.25 | 1.03 | -0.8 | -0.16 | 1.57 | -0.63 | -2.09 | -1.58 | 0.23 | -2.15 | -0.04 | -2.34 | 1.17 | -0.58 | -0.48 | 0.0375 | -0.52 | -0.1925 | 0.828447 | 0.9672 | 0.531463 | 1 | 0.999631 | 0.941315 |
| Shank1_D3YZU4_S413s_1 | -1.53 | -0.87 | -1.37 | -2.49 | -0.62 | -1.72 | -1.06 | -1.55 | -0.19 | -2.17 | -0.34 | -0.1 | 0.38 | -1.49 | -1.99 | -1.18 | 0.3275 | 0.865 | 0.495 | 0.391665 | 0.440142 | 0.142091 | 1 | 0.999631 | 0.758901 |
| Shank1_D3YZU4_S519s_1 | 3.8 | 0.08 | 0.42 | 2.1 | 1.22 | 1.53 | 2.69 | 0.47 | 0.27 | 0.65 | 0.17 | -0.84 | -3.83 | -0.87 | -0.4 | -1.27 | -0.1225 | -1.5375 | -3.1925 | 0.00835 | 0.884744 | 0.079767 | 1 | 0.999631 | 0.365244 |
| Shank1_D3YZU4_S890s_1 | 0.82 | 2.11 | 0.83 | 1.25 | 1.64 | 0.56 | 0.46 | 0.48 | 1.1 | 0.89 | 1.45 | 1.49 | 0.75 | 0.39 | 0.49 | 2.1 | -0.4675 | -0.02 | -0.32 | 0.512766 | 0.275583 | 0.956323 | 1 | 0.999631 | 0.818346 |
| Shank1_D3YZU5_S638s_1 | 0.3 | 1.16 | -0.3 | -1.1 | 0.64 | -0.3 | -0.31 | -0.67 | 0.36 | -0.65 | 0.23 | 1.52 | 1.47 | 0.66 | -1.2 | -1.1 | -0.175 | 0.35 | -0.0575 | 0.936289 | 0.734696 | 0.560343 | 1 | 0.999631 | 0.97939 |
| Shank2_D3Z5K8_S1013s_1 | 0.42 | -1.21 | 1.52 | -0.04 | -1.92 | -0.12 | 0.61 | -1.32 | -0.94 | 1.22 | -1.81 | 0.2 | -1.07 | -0.92 | 0.15 | -1.69 | -0.86 | -0.505 | -1.055 | 0.112198 | 0.244115 | 0.51489 | 1 | 0.999631 | 0.577068 |
| Shank2_D3Z5K8_S1043s_1 | -1.31 | -0.34 | -0.48 | -0.39 | 0.28 | -0.3 | 0.25 | -0.39 | -0.57 | -0.79 | 0.59 | -0.11 | -0.44 | -0.04 | -0.64 | -0.65 | 0.59 | 0.41 | 0.1875 | 0.580947 | 0.098887 | 0.313798 | 1 | 0.999631 | 0.847076 |
| Shank2_D3Z5K8_S1370s_1 | 0.73 | 0.1 | -0.57 | 0.13 | 1.84 | -0.33 | -0.22 | -0.29 | 0.99 | 0.62 | -0.44 | -0.64 | 0.05 | -0.65 | -1.3 | -0.21 | 0.1525 | 0.035 | -0.625 | 0.153742 | 0.780044 | 0.939833 | 1 | 0.999631 | 0.625198 |
| Shank2_D3Z5K8_S1603s_1 | 1.34 | -0.44 | -1.54 | 0.99 | 1.91 | 0.84 | 1.4 | 1.43 | 0.16 | -0.93 | -0.47 | -1.51 | -0.48 | -0.24 | -0.4 | -0.52 | 1.3075 | -0.775 | -0.4975 | 0.420844 | 0.060754 | 0.267055 | 1 | 0.999631 | 0.772817 |
| Shank2_D3Z5K8_S1699s_1 | 1.44 | -0.32 | -2.37 | 1.11 | 1.92 | 0.16 | 1.21 | 0.92 | 0.26 | -1.69 | 0.8 | -1.18 | -0.92 | 0.23 | -1.73 | -0.93 | 1.0875 | -0.4175 | -0.8025 | 0.3455 | 0.205116 | 0.647933 | 1 | 0.999631 | 0.734968 |
| Shank2_D3Z5K8_S1703s_1 | -1.74 | 0.43 | -0.06 | -0.93 | -1.29 | 0.04 | -1.77 | -0.69 | -0.28 | 0.22 | -0.46 | 0.02 | 0.18 | 0.12 | -0.33 | 0.3 | -0.3525 | 0.45 | 0.6425 | 0.203942 | 0.5367 | 0.361876 | 1 | 0.999631 | 0.659781 |
| Shank2_D3Z5K8_S446s_1 | 1.48 | -0.59 | -0.79 | 0.62 | 1.83 | -0.19 | 1.39 | 1.1 | 0.03 | -1.1 | 0.63 | -0.93 | -0.83 | -0.27 | -0.73 | -1.05 | 0.8525 | -0.5225 | -0.9 | 0.111437 | 0.189762 | 0.401341 | 1 | 0.999631 | 0.577068 |
| Shank2_D3Z5K8_S821s_1 | 0.6 | -1.01 | -1.67 | 0.64 | 1.36 | -0.06 | 0.71 | 0.38 | -0.02 | -1.04 | -0.37 | -1.19 | -1.94 | -0.12 | -0.83 | -0.4 | 0.9575 | -0.295 | -0.4625 | 0.473177 | 0.130408 | 0.618582 | 1 | 0.999631 | 0.799439 |
| Shank2_D3Z5K8_S951s_1 | -0.84 | -0.07 | -0.73 | -1.29 | 0.14 | -0.76 | -0.54 | -0.82 | 0 | -0.93 | 0.17 | 0.17 | -0.33 | -1.19 | -0.96 | -0.65 | 0.2375 | 0.585 | -0.05 | 0.890143 | 0.512902 | 0.151842 | 1 | 0.999631 | 0.9636 |
| Shank2_Q80Z38_S372s_1 | 0.49 | 1.68 | 0.77 | 1.31 | 1.76 | 0.62 | 0.57 | 0.72 | 1.49 | 0.45 | 1.16 | 1.24 | -0.05 | 1.26 | 0.75 | 1.77 | -0.145 | 0.0225 | -0.13 | 0.779586 | 0.714508 | 0.951937 | 1 | 0.999631 | 0.921416 |
| Shank3_Q4ACU6_S1032s_1 | -0.54 | -0.76 | 0.36 | 0.38 | -0.48 | -0.08 | -0.08 | 0.8 | -0.77 | -0.55 | -0.15 | 0.78 | -0.41 | 0.15 | 0 | 0.84 | 0.18 | -0.0325 | 0.285 | 0.495152 | 0.657897 | 0.941929 | 1 | 0.999631 | 0.810677 |
| Shank3_Q4ACU6_S1134s_1 | -0.05 | -0.78 | -1.86 | -0.15 | -0.05 | -0.68 | -0.55 | -0.34 | -1.39 | -1.05 | -1.23 | -1.7 | -1.14 | -0.69 | -1.13 | -1.2 | 0.305 | -0.6325 | -0.33 | 0.45702 | 0.483976 | 0.169086 | 1 | 0.999631 | 0.790722 |
| Shank3_Q4ACU6_S1510s_1 | 1.02 | 0.38 | -0.16 | 1.27 | 1.49 | 0.83 | 0.66 | 1.03 | 0.58 | -0.25 | 0.14 | 0.48 | -0.68 | 0.53 | -0.37 | 1 | 0.375 | -0.39 | -0.5075 | 0.312313 | 0.338267 | 0.330537 | 1 | 0.999631 | 0.719861 |
| Shank3_Q4ACU6_S1529s_1 | 2.78 | -0.33 | -2.24 | 1.61 | 2.41 | 0.24 | 2.18 | 1.34 | 0.39 | -0.92 | 0.24 | -1.13 | -2.1 | -0.23 | -0.71 | -1.46 | 1.0875 | -0.81 | -1.58 | 0.140062 | 0.308623 | 0.430682 | 1 | 0.999631 | 0.611718 |
| Shank3_Q4ACU6_S1577s_1 | 0.94 | -2.63 | -1.04 | 1.64 | 1.27 | -1.66 | 2.44 | 1.75 | -1.28 | -2.95 | 0.62 | -1.1 | -1.8 | -2.07 | -0.3 | -0.92 | 1.2225 | -0.905 | -1 | 0.284606 | 0.295414 | 0.396065 | 1 | 0.999631 | 0.705486 |
| Shank3_Q4ACU6_S351s_1 | 1.7 | 1.96 | 0.57 | 0.19 | 0.8 | 1.25 | 0.22 | 0.28 | 0.78 | 1.64 | -0.01 | -0.54 | 0.39 | 1.46 | 0.68 | -0.82 | -0.4675 | -0.6375 | -0.6775 | 0.263583 | 0.332388 | 0.291947 | 1 | 0.999631 | 0.693034 |
| Shank3_Q4ACU6_S375s_1 | 0.91 | 0.31 | -0.31 | 0.55 | 0.88 | 0.02 | 0.14 | 0.41 | 0.16 | 0.21 | 0.73 | 0.1 | 0.01 | -0.02 | -0.63 | 1.12 | -0.0025 | -0.065 | -0.245 | 0.585834 | 0.994286 | 0.848491 | 1 | 0.999631 | 0.849524 |
| Shank3_Q4ACU6_S434s_1 | -0.5 | 0.59 | 0.37 | 0.03 | -0.82 | -0.38 | -0.41 | -0.18 | -0.51 | -0.39 | -0.48 | -1.25 | 0.33 | -0.42 | -0.73 | 0.33 | -0.57 | -0.78 | -0.245 | 0.533168 | 0.100073 | 0.046967 | 1 | 0.999631 | 0.827798 |
| Shank3_Q4ACU6_S482s_1 | -0.88 | 0.66 | 0.68 | -0.09 | 0.72 | 0.24 | 1.71 | -0.22 | 1.37 | -0.75 | 1.48 | 2.29 | 2.38 | -0.17 | 0.79 | -1.25 | 0.52 | 1.005 | 0.345 | 0.647333 | 0.326841 | 0.156683 | 1 | 0.999631 | 0.874566 |
| Shank3_Q4ACU6_S685s_1 | 1.46 | 1.43 | 0.66 | 1.75 | 1.47 | 1.2 | 1.25 | 1.44 | 1.08 | 0.43 | 1.6 | 0.78 | -0.41 | 1.52 | 0.54 | 2.08 | 0.015 | -0.3525 | -0.3925 | 0.484317 | 0.960509 | 0.350649 | 1 | 0.999631 | 0.803422 |
| Shank3_Q4ACU6_S694s_1 | 0.81 | 1.16 | 0.74 | 1.15 | 1.88 | 1.1 | 0.56 | 1.08 | 1.35 | 0.68 | 1.17 | 0.88 | -0.06 | 1.66 | 0.69 | 1.83 | 0.19 | 0.055 | 0.065 | 0.885839 | 0.571634 | 0.84503 | 1 | 0.999631 | 0.963505 |
| Shank3_Q4ACU6_S801s_1 | 2.48 | 0.36 | -0.31 | 2.26 | 2.09 | 1.18 | 2.2 | 1.35 | 1.03 | -0.2 | 0.99 | 0.36 | -2.21 | 0.02 | -0.3 | 0.17 | 0.5075 | -0.6525 | -1.7775 | 0.041853 | 0.447893 | 0.343993 | 1 | 0.999631 | 0.475443 |
| Shank3_Q4ACU6_S838s_1 | -0.54 | 0.25 | -0.68 | -0.48 | 0.17 | -1.19 | -0.36 | 0.52 | 0.23 | -2.55 | -0.44 | 0.57 | 1.38 | 0.5 | -0.65 | 0.04 | 0.1475 | -0.185 | 0.68 | 0.166758 | 0.727095 | 0.778148 | 1 | 0.999631 | 0.636447 |
| Shank3_Q4ACU6_S995s_1 | -0.46 | -0.17 | -0.67 | -0.54 | 0.05 | -0.63 | -0.55 | -0.46 | -0.42 | -0.94 | -0.43 | -0.25 | 0.06 | -0.5 | -0.66 | -0.01 | 0.0625 | -0.05 | 0.1825 | 0.554417 | 0.819671 | 0.858323 | 1 | 0.999631 | 0.838079 |
| Shank3_Q4ACU6_T1234t_1 | -0.61 | -0.74 | -1.26 | -0.25 | -0.3 | -0.88 | -0.52 | -0.12 | -0.72 | -1 | -0.61 | -0.61 | -0.55 | -0.47 | -0.71 | -0.57 | 0.26 | -0.02 | 0.14 | 0.652629 | 0.419324 | 0.947254 | 1 | 0.999631 | 0.877704 |
| Shc2_Q8BMC3_S286s_1 | -0.16 | -0.02 | 0.53 | -0.16 | -0.92 | -0.64 | -0.25 | -0.3 | -0.94 | 0.41 | -0.38 | 0.89 | 0.03 | -0.28 | -0.15 | 0.12 | -0.575 | -0.0525 | -0.1175 | 0.69342 | 0.075243 | 0.904065 | 1 | 0.999631 | 0.895102 |
| Shc3_Q61120_S282s_1 | 0.51 | 0.1 | -0.3 | 0.59 | 0.74 | -0.25 | 0.26 | 0.05 | 0.22 | 0.11 | 0.27 | 0.48 | -0.03 | 0.05 | -0.4 | 0.41 | -0.025 | 0.045 | -0.2175 | 0.519732 | 0.939987 | 0.879875 | 1 | 0.999631 | 0.819491 |
| Shc3_Q61120_S354s_1 | 0.66 | 0.8 | 0.2 | 1.05 | 0.79 | 0.78 | 0.44 | 1 | 0.85 | 0.51 | 0.78 | 0.32 | 0.4 | 1.28 | 0.66 | 0.97 | 0.075 | -0.0625 | 0.15 | 0.653583 | 0.794146 | 0.833019 | 1 | 0.999631 | 0.878344 |
| Shc3_Q61120_S359s_1 | -0.82 | -0.06 | -0.48 | -0.24 | 0.06 | 0.06 | -0.44 | 0.23 | 0.04 | -0.11 | -0.72 | -1.2 | -0.16 | 0.05 | 0.57 | -0.45 | 0.3775 | -0.0975 | 0.4025 | 0.250589 | 0.213535 | 0.787923 | 1 | 0.999631 | 0.686448 |
| Shisa6_F6VQZ6_S191s_1 | -0.46 | -0.19 | 0.64 | -0.04 | -0.46 | 0.15 | -0.34 | 0.39 | -0.35 | -0.41 | -0.02 | 0.04 | 0.39 | 0.19 | 0.45 | 0.71 | -0.0525 | -0.1725 | 0.4475 | 0.19651 | 0.878108 | 0.592633 | 1 | 0.999631 | 0.653373 |
| Shisa6_F6VQZ6_S410s_1 | -1.48 | 1.15 | 0.67 | -0.8 | 2.84 | 0.01 | 0.74 | -0.36 | 3.19 | -1.92 | 0.97 | 1.49 | 3.21 | 0.05 | 0.59 | -0.46 | 0.9225 | 1.0475 | 0.9625 | 0.290324 | 0.277833 | 0.334498 | 1 | 0.999631 | 0.706164 |
| Shisa6_F6VQZ6_T190t_1 | 0.41 | 0.9 | 0.8 | 0.02 | 1.68 | 1.19 | -0.05 | 0.56 | 1.83 | 1.04 | 0.27 | 0.7 | 1.93 | 1.14 | 0.2 | 0.99 | 0.3125 | 0.4275 | 0.5325 | 0.224283 | 0.465126 | 0.298921 | 1 | 0.999631 | 0.67471 |
| Shisa6_F6VQZ6_T264t_1 | -2 | 0.81 | -0.13 | -1.43 | 0.97 | -0.12 | 0.97 | -1.03 | 1.19 | -2.63 | 0.63 | 0.23 | -0.28 | -1.67 | -1.11 | -1.89 | 0.885 | 0.5425 | -0.55 | 0.408577 | 0.228437 | 0.559571 | 1 | 0.999631 | 0.766996 |
| Shisa7_F6T9T6_S104s_1 | -4.36 | -6.64 | -2.74 | -3.38 | -5.31 | -5.81 | -1.35 | -2.57 | -6.23 | -5.07 | -2.51 | -2.08 | -5.69 | -6.09 | -2.68 | -2.48 | 0.52 | 0.3075 | 0.045 | 0.967118 | 0.658703 | 0.785656 | 1 | 0.999631 | 0.988373 |
| Shisa7_F6T9T6_S121s_1 | -0.35 | -1 | -1.15 | -1.35 | -1.25 | -1.25 | -1.13 | -1.15 | -1.79 | -0.4 | -1.49 | -1.2 | -0.17 | -0.49 | -1.17 | -1.11 | -0.2325 | -0.2575 | 0.2275 | 0.541467 | 0.430953 | 0.511094 | 1 | 0.999631 | 0.832213 |
| Shisa7_F6T9T6_S222s_1 | 1.14 | 0.13 | -0.92 | 0.99 | 1.16 | -0.66 | 0.67 | -0.04 | 0.51 | -1.02 | 0.13 | -0.04 | -1.43 | -0.21 | -0.76 | 0.29 | -0.0525 | -0.44 | -0.8625 | 0.144933 | 0.925988 | 0.420163 | 1 | 0.999631 | 0.616037 |
| Shisa7_F6T9T6_S255s_1 | -0.86 | 0.19 | -1.18 | -3.14 | -0.63 | -0.63 | -0.88 | -2.13 | -2.09 | -1.1 | -3.07 | -1.98 | -0.11 | -1.32 | -1.13 | -1.24 | 0.18 | -0.8125 | 0.2975 | 0.658547 | 0.795318 | 0.271587 | 1 | 0.999631 | 0.879726 |
| Shisa7_F6T9T6_S268s_1 | -3.64 | 1.45 | 0.96 | -1.95 | 2.98 | 0.24 | 2.52 | -1.52 | 4.11 | -1.6 | 2.88 | 3.83 | 4.55 | -0.54 | -0.3 | -1 | 1.85 | 3.1 | 1.4725 | 0.335297 | 0.194784 | 0.066141 | 1 | 0.999631 | 0.730219 |
| Shisa7_F6T9T6_S272s_1 | 0.41 | -0.83 | 0.66 | 1.08 | 0.52 | -1.97 | 0.87 | 1.36 | -0.65 | -1.47 | 0.29 | 0.52 | -0.81 | -1.41 | 0.36 | 1.61 | -0.135 | -0.6575 | -0.3925 | 0.575659 | 0.856259 | 0.262517 | 1 | 0.999631 | 0.846085 |
| Shisa7_F6T9T6_S67s_1 | 0.09 | 0.55 | -0.68 | -0.1 | 0.81 | 0.12 | -0.36 | -0.16 | 0.27 | 0.21 | -0.27 | -0.47 | -0.78 | 0.5 | -0.51 | -0.01 | 0.1375 | -0.03 | -0.165 | 0.683378 | 0.715512 | 0.931856 | 1 | 0.999631 | 0.889602 |
| Shisa7_F6T9T6_T349t_1 | -2.45 | 0.39 | -0.67 | 0.37 | 0.64 | -0.11 | -0.53 | 0.22 | 0.06 | -1.99 | -0.46 | 0.01 | 0.27 | -0.19 | -0.66 | 0.27 | 0.645 | -0.005 | 0.5125 | 0.426933 | 0.324972 | 0.994518 | 1 | 0.999631 | 0.775206 |

| | | | | | | | | | | | | | | | | | | | | | | | | | |
|---|---|---|---|---|---|---|---|---|---|---|---|---|---|---|---|---|---|---|---|---|---|---|---|---|---|
| Shisa9_E9QN38_S321s_1 | -0.29 | -0.68 | -0.22 | -1.22 | -0.41 | -0.56 | -0.45 | -0.65 | -0.86 | -0.32 | 0.22 | -0.56 | -0.48 | -0.61 | -1.63 | -0.75 | 0.085 | 0.2225 | -0.265 | 0.493112 | 0.776754 | 0.538466 | 1 | 0.999631 | 0.80955 |
| Shroom1_D3Z3D5_S293s_1 | -0.02 | -0.98 | 0.23 | 0.43 | -0.24 | -0.27 | -0.2 | 0.05 | -0.75 | -0.05 | 0.05 | 0.22 | -1.03 | -0.71 | -0.87 | -0.11 | -0.08 | -0.0475 | -0.595 | 0.156716 | 0.819298 | 0.90388 | 1 | 0.999631 | 0.626749 |
| Shroom2_A7TU71_S1047s_1 | -0.33 | 1.86 | 1.56 | 0.91 | 0.27 | 0.71 | 0.62 | 1.23 | 0.88 | 0.94 | 0.62 | 0.96 | 2.06 | 1.11 | 1.8 | 1.58 | -0.2925 | -0.15 | 0.6375 | 0.223538 | 0.5569 | 0.752172 | 1 | 0.999631 | 0.67471 |
| Shroom2_A7TU71_S1095s_1 | 0.99 | -1.63 | -0.24 | 0.05 | 2.39 | -1.04 | 5.26 | -0.89 | 6.99 | 0.09 | -0.13 | 1.93 | 1.75 | -0.74 | -0.95 | -0.23 | 1.6375 | 2.4275 | 0.165 | 0.819864 | 0.245705 | 0.126013 | 1 | 0.999631 | 0.938303 |
| Shroom2_A7TU71_S1159s_1 | 0.54 | -0.35 | -0.78 | 0.79 | 0.55 | -0.4 | 0.66 | 0.61 | -0.36 | -0.01 | 0.16 | -0.67 | -0.91 | -0.09 | -0.54 | -0.15 | 0.305 | -0.27 | -0.4725 | 0.282644 | 0.49052 | 0.522449 | 1 | 0.999631 | 0.704034 |
| Shroom2_A7TU71_S1171s_1 | -0.64 | 1.37 | 1.58 | 0.35 | 0.32 | 0.5 | 0.26 | 1.02 | 0.77 | 0.08 | 1.28 | 1.42 | 1.53 | 1.67 | 1.53 | 1.2 | -0.14 | 0.2225 | 0.8175 | 0.125422 | 0.781746 | 0.687311 | 1 | 0.999631 | 0.596551 |
| Shroom2_A7TU71_S318s_1 | -0.41 | -0.04 | -0.34 | -0.95 | -0.81 | -0.48 | -1.16 | -0.73 | 0.1 | 0.57 | -0.99 | -0.75 | 1.02 | 1 | -0.25 | -0.17 | -0.36 | 0.1675 | 0.835 | 0.067931 | 0.247102 | 0.688166 | 1 | 0.999631 | 0.522149 |
| Sik2_H3BJC9_S342s_1 | -0.08 | 0.32 | 1.24 | -0.16 | -0.06 | 0.07 | 0.21 | 0.57 | 0.05 | -0.11 | 0.7 | 0.9 | 0.94 | 0.4 | 0.53 | 0.95 | -0.1325 | 0.055 | 0.375 | 0.341075 | 0.719605 | 0.893562 | 1 | 0.999631 | 0.733032 |
| Sik2_H3BJC9_T359t_1 | 0.2 | 1.93 | 1.83 | 0.53 | 0.88 | 1.47 | -0.12 | 0.7 | 0.95 | 0.82 | 0.99 | 1.02 | 1.5 | 0.84 | 0.25 | -0.26 | -0.39 | -0.1775 | -0.54 | 0.333547 | 0.454525 | 0.687936 | 1 | 0.999631 | 0.728079 |
| Sik3_F6U6U5_S456s_1 | 0.66 | 0.66 | -0.25 | 0.72 | 1.18 | 0.25 | 0.7 | 0.84 | 0.79 | -0.08 | 0.41 | 0.44 | -0.43 | 0.66 | -0.11 | 0.55 | 0.295 | -0.0575 | -0.28 | 0.472196 | 0.392915 | 0.865511 | 1 | 0.999631 | 0.79887 |
| Sik3_F6U6U5_S684s_1 | -0.1 | -0.43 | -0.74 | -0.59 | -0.48 | -0.08 | -1.09 | -0.42 | -1.96 | -0.48 | -0.74 | -0.85 | -0.11 | -0.57 | -1.35 | -1.1 | -0.0525 | -0.5425 | -0.3175 | 0.386571 | 0.864578 | 0.173446 | 1 | 0.999631 | 0.756141 |
| Sin3a_Q60520_S941s_1 | 0.02 | 1.98 | 0.77 | 0.33 | 1.37 | 1.2 | -0.03 | 0.17 | 1.25 | 1.21 | 0.52 | 0.57 | 0.89 | 1.88 | -0.25 | 0.9 | -0.0975 | 0.1125 | 0.08 | 0.887451 | 0.850776 | 0.807234 | 1 | 0.999631 | 0.963505 |
| Sin3b_Q62141_S698s_1 | 0.17 | -0.29 | 1.93 | 0.64 | -0.47 | -0.31 | 0.35 | 1.1 | -0.01 | 0.51 | -0.87 | 1.84 | 0.14 | 0.22 | 0.91 | 2.1 | -0.445 | -0.245 | 0.23 | 0.702669 | 0.425429 | 0.713475 | 1 | 0.999631 | 0.896857 |
| Sipa1_P46062_S72s_1 | -0.06 | -1.78 | -1.41 | -1.36 | -2.08 | -3.09 | -0.61 | -1.86 | -0.55 | -1.3 | -1.5 | -0.36 | -0.36 | -3.09 | -1.52 | -1.38 | -0.7575 | 0.225 | -0.435 | 0.484641 | 0.208877 | 0.624297 | 1 | 0.999631 | 0.803422 |
| Sipa1l1_Q8C0T5_S1127s_1 | -1.17 | -1.71 | -3.04 | -2.15 | -0.54 | -0.98 | -1.64 | -2.28 | -0.83 | -1.1 | -1.26 | -2.08 | -0.68 | -1.7 | -3.11 | -1.29 | 0.6575 | 0.7 | 0.3225 | 0.589814 | 0.218665 | 0.155062 | 1 | 0.999631 | 0.851414 |
| Sipa1l1_Q8C0T5_S114s_1 | -0.18 | 0.37 | 0.3 | 0.23 | 0.23 | 0.24 | -0.31 | 0.48 | 0.01 | -0.18 | 0.07 | 0.62 | 1.22 | 0.13 | 0.37 | 0.61 | -0.02 | -0.05 | 0.4025 | 0.246224 | 0.943913 | 0.864854 | 1 | 0.999631 | 0.685751 |
| Sipa1l1_Q8C0T5_S1170s_1 | 0.76 | 0.13 | -0.43 | 2.02 | 1.5 | 0.33 | 1.37 | 1.95 | 0.41 | -0.41 | 1.15 | 0.6 | -2.25 | 0.56 | 0.36 | 0.99 | 0.6675 | -0.1825 | -0.705 | 0.381024 | 0.260291 | 0.748614 | 1 | 0.999631 | 0.752619 |
| Sipa1l1_Q8C0T5_S1184s_1 | -1.79 | -4.94 | -1.04 | 1.44 | -4.56 | -4.15 | 1.57 | 0.53 | -4.59 | -4.59 | 0.33 | -0.69 | -3.85 | -4.67 | -0.96 | -0.51 | -0.07 | -0.8025 | -0.915 | 0.520114 | 0.967802 | 0.610322 | 1 | 0.999631 | 0.819528 |
| Sipa1l1_Q8C0T5_S1249s_1 | 0.2 | 0.28 | -0.82 | 0.03 | 0.36 | 1.02 | -0.46 | 0.37 | 0.08 | 0.15 | -0.6 | 0.3 | 0.62 | 1.28 | -0.17 | 0.21 | 0.4 | 0.06 | 0.5625 | 0.196651 | 0.329454 | 0.866557 | 1 | 0.999631 | 0.653373 |
| Sipa1l1_Q8C0T5_S1372s_1 | 0.79 | 0.18 | 0.15 | 0.88 | 1.56 | 0.61 | 1.04 | 0.54 | 0.26 | -0.3 | 0.11 | -0.43 | -1.19 | 0.39 | -0.94 | 0.48 | 0.4375 | -0.59 | -0.815 | 0.106357 | 0.218002 | 0.087069 | 1 | 0.999631 | 0.568797 |
| Sipa1l1_Q8C0T5_S1384s_1 | -0.09 | -0.96 | -1.02 | -0.95 | 1.23 | -0.43 | 1.91 | -0.32 | 0.6 | -2.25 | 0.53 | -1.16 | -1.71 | -2.54 | -1.28 | -2.4 | 1.3525 | 0.185 | -1.2275 | 0.010629 | 0.036092 | 0.777346 | 1 | 0.999631 | 0.383871 |
| Sipa1l1_Q8C0T5_S148s_1 | 2.83 | 1.57 | 1.9 | 1.88 | 1.38 | 1.4 | 2.62 | 1.09 | 1.29 | 1.35 | 1.79 | 1.28 | -2.16 | -0.2 | -0.83 | 1.34 | -0.4225 | -0.6175 | -2.5075 | 0.0044 | 0.336376 | 0.09616 | 1 | 0.999631 | 0.351308 |
| Sipa1l1_Q8C0T5_S1528s_1 | -0.82 | -0.25 | -0.97 | -0.85 | 0.01 | -0.45 | -0.47 | -0.61 | -0.33 | -1.24 | -0.63 | -0.41 | -0.16 | -0.99 | -0.74 | -0.69 | 0.3425 | 0.07 | 0.0775 | 0.808986 | 0.248224 | 0.827184 | 1 | 0.999631 | 0.93635 |
| Sipa1l1_Q8C0T5_S1564s_1 | -1.57 | -0.2 | 0.05 | 0.13 | -0.22 | 0.01 | -0.16 | 0.53 | -0.46 | -1.31 | 0.74 | -0.14 | 0.08 | -0.44 | 0.55 | 0.39 | 0.4375 | 0.105 | 0.5425 | 0.246839 | 0.317719 | 0.845907 | 1 | 0.999631 | 0.685751 |
| Sipa1l1_Q8C0T5_S1582s_1 | 0.67 | 1.12 | 0.1 | 0.7 | 1.4 | 1.05 | 0.74 | 0.96 | 0.69 | 0.16 | 1.07 | -0.34 | -0.75 | 1.11 | 0.18 | 0.89 | 0.39 | -0.2525 | -0.29 | 0.534399 | 0.223999 | 0.521648 | 1 | 0.999631 | 0.828569 |
| Sipa1l1_Q8C0T5_S1600s_1 | 0.85 | 0.81 | -1.71 | 1.25 | 1.47 | 0.53 | 0.6 | 0.95 | 0.78 | -0.03 | -0.11 | -0.24 | -0.27 | 0.48 | 0.13 | 0.37 | 0.5875 | -0.2 | -0.1225 | 0.845726 | 0.366817 | 0.756704 | 1 | 0.999631 | 0.948565 |
| Sipa1l1_Q8C0T5_S1712s_1 | -1.04 | 1.99 | 2.36 | 0.04 | 1.49 | 1.06 | 1.95 | 2.48 | 1.84 | -0.84 | 3.87 | 3.43 | 2.4 | 0.47 | 2.83 | 0.73 | 0.9075 | 1.2375 | 0.77 | 0.383022 | 0.24836 | 0.295189 | 1 | 0.999631 | 0.754387 |
| Sipa1l1_Q8C0T5_S208s_1 | 0.5 | 0.01 | -0.62 | 0.04 | 0.68 | 0.17 | 0.06 | -0.11 | 0.83 | 0.2 | 0.12 | 0.05 | 0.49 | 0.03 | -0.73 | 0.08 | 0.2175 | 0.3175 | -0.015 | 0.968452 | 0.512678 | 0.359283 | 1 | 0.999631 | 0.988771 |
| Sipa1l1_Q8C0T5_S211s_1 | 1.23 | 0.14 | -0.64 | 0.13 | 1.13 | -0.04 | 0.25 | 0.05 | 0.54 | 0.06 | -0.22 | -0.96 | -1.64 | -0.82 | -0.72 | -0.42 | 0.1325 | -0.36 | -1.115 | 0.033381 | 0.769784 | 0.457498 | 1 | 0.999631 | 0.44294 |
| Sipa1l1_Q8C0T5_S288s_1 | 1.44 | 1.13 | 1.22 | 1.73 | 2.1 | 1.81 | 1.78 | 2.2 | 1.43 | 0.86 | 1.97 | 1.93 | -0.76 | 1.7 | 1.18 | 1.95 | 0.5925 | 0.1675 | -0.3625 | 0.534269 | 0.047597 | 0.623064 | 1 | 0.999631 | 0.828493 |
| Sipa1l1_Q8C0T5_S310s_1 | 0.57 | 0.98 | -0.73 | 0.32 | 3.21 | 1.1 | 1.25 | -0.21 | 2.03 | -1.31 | -0.29 | -0.94 | -0.87 | -1.25 | -0.38 | -0.97 | 1.0525 | -0.4125 | -1.1525 | 0.019451 | 0.157178 | 0.579027 | 1 | 0.999631 | 0.391668 |
| Sipa1l1_Q8C0T5_T1190t_1 | 2.33 | 1.05 | 1.95 | 1.47 | -0.21 | 1.24 | 1.98 | 0.4 | -0.44 | 1.54 | 1.61 | -0.32 | -0.49 | -0.21 | -2.1 | -0.39 | -0.8475 | -1.1025 | -2.4975 | 0.000545 | 0.125084 | 0.081295 | 1 | 0.999631 | 0.351308 |
| Sipa1l2_Q80TE4_S1406s_1 | -0.84 | -0.68 | -0.1 | -0.61 | -0.31 | -0.5 | -0.38 | -1.26 | -0.48 | -0.8 | -0.69 | 0.46 | 0.62 | -0.19 | -0.32 | -0.79 | -0.055 | 0.18 | 0.3875 | 0.313735 | 0.863414 | 0.620329 | 1 | 0.999631 | 0.719939 |
| Sipa1l2_Q80TE4_S1488s_1 | -1.49 | 0.6 | -0.55 | -1.07 | -0.22 | 0.29 | -0.52 | -0.83 | 0.58 | -0.62 | -0.51 | -0.34 | 1.03 | 0.32 | -0.78 | -1.11 | 0.3075 | 0.405 | 0.4925 | 0.426297 | 0.529036 | 0.427002 | 1 | 0.999631 | 0.774781 |
| Sipa1l2_Q80TE4_S195s_1 | 0.84 | -0.08 | -0.15 | -0.07 | -0.3 | -0.1 | -0.22 | -0.35 | 0.06 | 1.05 | 0.14 | -0.57 | -0.22 | 0.1 | 0.23 | -0.32 | -0.3775 | 0.035 | -0.1875 | 0.580808 | 0.231259 | 0.93263 | 1 | 0.999631 | 0.846995 |
| Sipa1l3_G3X9J0_S1358s_1 | -0.26 | -0.65 | -1.16 | -0.84 | -0.04 | -0.67 | -0.34 | 0.18 | -0.38 | -1.42 | -0.23 | -1.02 | -0.6 | -0.88 | -0.87 | -0.41 | 0.51 | -0.035 | 0.0375 | 0.904215 | 0.129813 | 0.923714 | 1 | 0.999631 | 0.968498 |
| Sipa1l3_G3X9J0_S94s_1 | -2.65 | -0.82 | -0.37 | -0.69 | -0.65 | -0.87 | -0.46 | -0.05 | -0.84 | 0.16 | 0.49 | 0.9 | 0.64 | -0.06 | 1.08 | -2.64 | 0.625 | 1.31 | 0.8875 | 0.308933 | 0.23666 | 0.045589 | 1 | 0.999631 | 0.718845 |
| Sirpa_P97797_S462s_1 | 0.97 | -0.1 | 1.46 | 0.4 | 0.78 | -0.43 | 0.66 | 1.31 | 0.33 | -0.73 | 1.38 | 1.1 | -0.8 | -0.93 | 0.51 | 0.58 | -0.1025 | -0.1625 | -0.8425 | 0.120069 | 0.828802 | 0.764144 | 1 | 0.999631 | 0.587703 |
| Sirpa_P97797_Y460y_1 | 0.38 | -2.62 | -0.79 | -0.54 | -1.08 | -2.52 | -0.01 | -0.89 | -1.87 | -1.26 | -0.47 | -1.66 | -0.39 | -2.18 | -0.45 | -1.64 | -0.2325 | -0.4225 | -0.2725 | 0.691635 | 0.7471 | 0.50902 | 1 | 0.999631 | 0.894017 |
| Sirt2_Q8VDQ8_S331s_1 | 0 | 1.55 | 0.68 | -0.49 | 1.67 | 0.81 | 0.67 | 0.19 | 1.6 | 0.49 | 1.34 | 2.25 | 2.92 | 1.44 | 0.22 | 0.47 | 0.4 | 0.985 | 0.8275 | 0.23733 | 0.435471 | 0.090378 | 1 | 0.999631 | 0.683301 |
| Sirt2_Q8VDQ8_S335s_1 | -2.51 | -0.14 | 0.12 | -1.8 | 0.94 | -0.67 | 0.74 | -0.65 | 3.01 | 0.22 | 0.75 | 1.56 | 3.23 | 0.22 | 0.16 | -1.63 | 1.1725 | 2.4675 | 1.5775 | 0.144721 | 0.1119 | 0.009622 | 1 | 0.999631 | 0.616037 |
| Skiv2l_Q6NZR5_S240s_2 | -0.92 | -0.63 | 0.61 | -1.55 | -0.91 | -0.75 | -0.86 | -0.12 | -0.68 | 0.02 | -0.18 | -0.23 | -0.3 | -0.59 | -0.48 | 0.12 | -0.0375 | 0.355 | 0.31 | 0.51409 | 0.935958 | 0.450513 | 1 | 0.999631 | 0.81893 |
| Skiv2l_Q6NZR5_S242s_2 | -0.92 | -0.63 | 0.61 | -1.55 | -0.91 | -0.75 | -0.86 | -0.12 | -0.68 | 0.02 | -0.18 | -0.23 | -0.3 | -0.59 | -0.48 | 0.12 | -0.0375 | 0.355 | 0.31 | 0.51409 | 0.935958 | 0.450513 | 1 | 0.999631 | 0.81893 |
| Skiv2l_Q6NZR5_S253s_1 | -0.32 | 0.16 | 0.07 | -0.43 | -0.13 | -0.48 | -0.63 | -0.32 | -0.42 | -0.16 | -0.27 | -0.05 | 0.2 | 0.15 | -0.01 | 0.25 | -0.26 | -0.095 | 0.2775 | 0.34083 | 0.349234 | 0.727603 | 1 | 0.999631 | 0.733032 |
| Skp1_Q9WTX5_T131t_1 | 0.65 | 0.99 | 1.6 | 0.15 | 0.06 | 0.74 | 0.33 | 0.74 | 0.6 | 1.32 | 0.49 | 0.97 | 0.19 | 0.95 | 0.42 | 0.56 | -0.38 | -0.0025 | -0.3175 | 0.411914 | 0.314337 | 0.99475 | 1 | 0.999631 | 0.767937 |
| Slain1_F7B673_S270s_1 | -0.9 | -1.33 | -1.99 | -1.58 | -0.97 | -1.57 | -1.04 | -2.03 | -1.35 | -3.16 | -3.22 | -2.63 | -1.95 | -2.02 | -1.85 | -1.72 | 0.0475 | -1.14 | -0.435 | 0.195065 | 0.895196 | 0.034955 | 1 | 0.999631 | 0.65277 |
| Slain2_Q8CI08_S354s_1 | 2.81 | 1.06 | 0.96 | 1.86 | 1.26 | 2.21 | 2.01 | 1.33 | 1.34 | 1.95 | 0.78 | 0.38 | -0.98 | 0.73 | 1.14 | -0.2 | 0.03 | -0.56 | -1.5 | 0.026788 | 0.948938 | 0.293081 | 1 | 0.999631 | 0.422285 |
| Slain2_Q8CI08_S87s_1 | 1.35 | -0.31 | 0.08 | 0.99 | -0.38 | -0.44 | 1.07 | -0.24 | 0.44 | 1.39 | 0.4 | 0.27 | -0.67 | -0.48 | 0.16 | -0.96 | -0.525 | 0.0975 | -1.015 | 0.045336 | 0.303558 | 0.830052 | 1 | 0.999631 | 0.476658 |
| Slbp_Q3V2L1_S7s_1 | 1.65 | 1.68 | 1.87 | 1.63 | 0.99 | 0.94 | 1.07 | 1.22 | 1.45 | 2.05 | 0.69 | 0.77 | 0.97 | 0.9 | 1.42 | 0.83 | -0.6525 | -0.4675 | -0.6775 | 0.033781 | 0.018998 | 0.212043 | 1 | 0.999631 | 0.444185 |
| Slc12a2_E9QM38_S75s_1 | 0.13 | 0.44 | 1.65 | -0.86 | -0.77 | 0.54 | -0.78 | 1.37 | -0.53 | 0.1 | -2.17 | 0.19 | 1.7 | 0.45 | 0.61 | 0.39 | -0.25 | -0.9425 | 0.4475 | 0.43022 | 0.705078 | 0.183595 | 1 | 0.999631 | 0.778001 |
| Slc12a2_E9QM38_T211t_1 | 0.08 | 2.03 | 0.49 | 1.26 | 1.28 | 1.79 | 1.12 | 1.73 | 1.48 | -0.11 | -0.76 | -0.48 | -0.53 | 1.39 | 0.58 | 0.61 | 0.515 | -0.9325 | -0.4525 | 0.414956 | 0.266108 | 0.145092 | 1 | 0.999631 | 0.7693 |
| Slc12a5_Q91V14_S1044s_1 | 0.25 | -0.34 | -1.69 | -0.61 | 0.04 | 0.45 | -0.61 | -0.89 | 0.12 | 0.06 | -0.62 | -0.76 | -0.13 | 0.2 | -0.65 | -0.63 | 0.345 | 0.2975 | 0.295 | 0.519393 | 0.478993 | 0.517693 | 1 | 0.999631 | 0.819456 |
| Slc12a5_Q91V14_S1047s_1 | 0.22 | 0.67 | 0.21 | 0.17 | -0.1 | 0.34 | -0.42 | 0.27 | 0.28 | 0.04 | -0.11 | 0.18 | 0.31 | 0.89 | 0.32 | 0.85 | -0.295 | -0.22 | 0.275 | 0.373201 | 0.317244 | 0.413774 | 1 | 0.999631 | 0.747035 |
| Slc12a5_Q91V14_S1048s_1 | -1.35 | 1.23 | 1.52 | -0.42 | -0.76 | 0.72 | -1.78 | 0.37 | -0.22 | 0.82 | 0.16 | 0.91 | 1.98 | 0.98 | 0.53 | 1.25 | -0.6075 | 0.1725 | 0.94 | 0.180311 | 0.441848 | 0.793589 | 1 | 0.999631 | 0.64376 |
| Slc12a5_Q91V14_S48s_1 | 0.31 | 1.27 | -0.89 | -1.23 | 0.5 | 0.49 | -1.4 | -0.27 | 0.76 | 0.49 | -0.4 | -0.65 | 1.2 | 0.99 | -1.65 | -1.5 | -0.035 | 0.185 | -0.105 | 0.899849 | 0.956989 | 0.760619 | 1 | 0.999631 | 0.96757 |
| Slc12a5_Q91V14_S49s_1 | 0.09 | -0.79 | -1.4 | -0.34 | 0.29 | -0.59 | -0.43 | -0.4 | -0.38 | -0.95 | -0.79 | -0.58 | -1.69 | -0.42 | -1.14 | -0.8 | 0.3275 | -0.065 | -0.4025 | 0.357211 | 0.404011 | 0.86048 | 1 | 0.999631 | 0.74093 |
| Slc12a5_Q91V14_S54s_1 | -1.52 | -0.82 | -0.22 | -0.22 | -1.45 | -1.23 | -1.04 | 0.42 | -1.92 | -0.92 | 0.11 | 0.63 | -0.58 | -2.09 | -0.72 | -0.13 | -0.13 | 0.17 | -0.185 | 0.712715 | 0.792309 | 0.773273 | 1 | 0.999631 | 0.9008 |
| Slc12a5_Q91V14_S955s_1 | 1.12 | 0.44 | -1.78 | 1.58 | 1.66 | 0.72 | 0.69 | 0.98 | 0.76 | -0.14 | 0.42 | -0.51 | -3 | 1.08 | -0.75 | 0.16 | 0.6725 | -0.2075 | -0.9675 | 0.337247 | 0.341619 | 0.769879 | 1 | 0.999631 | 0.730499 |
| Slc12a5_Q91V14_S963s_1 | -0.29 | -1.93 | -4.27 | -1.68 | 0.12 | -2.53 | -2.27 | -2.2 | -1.14 | -2.97 | -2.53 | -2.43 | -3.74 | -1.68 | -2.98 | -3.52 | 0.3225 | -0.225 | -0.9375 | 0.270214 | 0.71866 | 0.77948 | 1 | 0.999631 | 0.698438 |
| Slc12a5_Q91V14_T1029t_1 | 0.6 | 1.58 | 1.2 | 1.15 | 1.08 | 1.97 | 0.29 | 1.44 | 1.61 | 0.81 | 1.51 | 1.03 | 1.2 | 1.98 | 1.12 | 1.92 | 0.0625 | 0.1075 | 0.4225 | 0.253744 | 0.878061 | 0.745287 | 1 | 0.999631 | 0.688735 |
| Slc12a6_Q6P6P5_S50s_1 | 1.65 | 1.6 | 2.25 | 1.54 | 0.89 | 2.43 | 1.2 | 1.81 | 1.53 | 1.32 | 1.5 | 0.47 | -3 | 0.83 | 1.25 | 1.03 | -0.1775 | -0.555 | -1.7325 | 0.074214 | 0.649293 | 0.129719 | 1 | 0.999631 | 0.539897 |
| Slc14a1_Q8VHL0_S4s_1 | -0.06 | -0.86 | -0.19 | -0.47 | -1.27 | -0.64 | -0.46 | -0.52 | -0.72 | -0.33 | 0.06 | -0.13 | 0.12 | -0.54 | 0.19 | -0.27 | -0.3275 | 0.115 | 0.27 | 0.414422 | 0.306934 | 0.711933 | 1 | 0.999631 | 0.768728 |
| Slc16a1_P53986_S213s_1 | 0.87 | 0.45 | 0.52 | -0.21 | -0.26 | 1.09 | 0.76 | -0.4 | 0.34 | 1.4 | 0.61 | -0.72 | 0.54 | 0.15 | -0.15 | -1.01 | -0.11 | -3.1E-16 | -0.525 | 0.224717 | 0.796558 | 1 | 1 | 1 | 0.67471 |
| Slc16a1_P53986_S491s_1 | -0.91 | -1.59 | -1.4 | -0.06 | -1.41 | -1.77 | -1.1 | -0.34 | -1.39 | -0.86 | -1.44 | -1.1 | -0.74 | -1.63 | -0.7 | -0.37 | -0.165 | -0.2075 | 0.13 | 0.767549 | 0.710933 | 0.593865 | 1 | 0.999631 | 0.917335 |
| Slc16a1_P53986_S491s_2 | -1.27 | -1.81 | -1.18 | -1.23 | -1.55 | -1.41 | -1.11 | -2.01 | -1.2 | -1.61 | -1.09 | -0.88 | -0.23 | -0.72 | -1.64 | -1.19 | -0.1475 | 0.1775 | 0.4275 | 0.27152 | 0.625789 | 0.550165 | 1 | 0.999631 | 0.699288 |
| Slc16a8_O35308_S207s_1 | 1.8 | 1.72 | 2.26 | 0.69 | 0.4 | 1.48 | 1.07 | 1.11 | 1.23 | 0.71 | -4.71 | -0.47 | 0.28 | 0.11 | -0.01 | -0.44 | -0.6025 | -2.4275 | -1.6325 | 0.001915 | 0.158338 | 0.064909 | 1 | 0.999631 | 0.351308 |
| Slc17a7_Q3TXX4_S36s_1 | -1.18 | -1.4 | -2.7 | -2.77 | -1.27 | -0.32 | -1.96 | -2.42 | -0.56 | -0.74 | -1.9 | -1.93 | 0.92 | -0.36 | -2.75 | -2.66 | 0.52 | 0.73 | 0.8 | 0.362995 | 0.369177 | 0.18461 | 1 | 0.999631 | 0.74343 |
| Slc17a8_D3YTT3_S376s_1 | -0.07 | 0.76 | -0.57 | -0.16 | -0.27 | -0.4 | -0.59 | -0.2 | -1.61 | -0.62 | -1.35 | -1.14 | 0.46 | 0.16 | -0.51 | -0.2 | -0.355 | -1.17 | -0.0125 | 0.973994 | 0.299349 | 0.010426 | 1 | 0.999631 | 0.990749 |
| Slc1a1_P51906_S516s_1 | 0.13 | 0 | -0.64 | 0.22 | 0.88 | 0.29 | 0.5 | 0.03 | -0.04 | -0.96 | 0.52 | -0.32 | -1.16 | 0.2 | 0.48 | 0.15 | 0.4975 | -0.1275 | -0.01 | 0.981233 | 0.139062 | 0.740502 | 1 | 0.999631 | 0.994608 |
| Slc1a2_P43006_S24s_1 | 0.23 | 0.43 | 0.34 | 0.06 | 0.33 | 0.58 | 0.38 | -0.07 | 0.66 | 0.58 | 0.34 | 1.04 | 1.2 | 0.35 | 0.39 | -0.27 | 0.04 | 0.39 | 0.1525 | 0.67393 | 0.878296 | 0.174627 | 1 | 0.999631 | 0.885515 |
| Slc1a2_P43006_S25s_1 | -0.62 | 0.63 | 0.53 | 0.33 | 0.18 | 0.75 | -0.25 | 0.18 | 0.3 | 0.91 | 0.1 | 0.68 | 0.21 | 0.11 | 0.4 | 0.18 | -0.0025 | 0.28 | 0.0075 | 0.982889 | 0.994609 | 0.453783 | 1 | 0.999631 | 0.99485 |
| Slc1a2_P43006_S3s_1 | 0.27 | -0.49 | -0.47 | -0.79 | -1.13 | -0.54 | -0.52 | -1.04 | -0.66 | 0.14 | -0.6 | -1.04 | -0.18 | -0.67 | -0.93 | -0.79 | -0.4375 | -0.17 | -0.2725 | 0.433078 | 0.196768 | 0.642963 | 1 | 0.999631 | 0.779401 |
| Slc1a2_P43006_S505s_1 | -1.14 | -0.8 | -1.45 | -1.25 | -2.13 | -1.08 | -1.02 | -1.03 | -1.54 | -0.43 | -0.62 | -2.83 | -1.11 | -0.69 | -1.36 | -0.98 | -0.155 | -0.195 | 0.125 | 0.67849 | 0.649697 | 0.712848 | 1 | 0.999631 | 0.887183 |
| Slc1a2_P43006_S520s_1 | 0.72 | 1.16 | 1.28 | 1.13 | 0.59 | 1.68 | 0.69 | 1.02 | 0.44 | 0.62 | 1.1 | -0.22 | -0.34 | 0.74 | 0.05 | 0.05 | -0.0775 | -0.5875 | -0.9475 | 0.015456 | 0.809717 | 0.111402 | 1 | 0.999631 | 0.388832 |
| Slc1a2_P43006_S530s_1 | -0.94 | 0.26 | 0.04 | -0.42 | 0.49 | -0.47 | -0.38 | -0.47 | 0.58 | -0.41 | -0.23 | 0.15 | 1.55 | -0.34 | -0.07 | -0.59 | 0.0575 | 0.2875 | 0.4025 | 0.446627 | 0.876988 | 0.44627 | 1 | 0.999631 | 0.78505 |
| Slc1a2_P43006_S532s_1 | -0.67 | -0.37 | 0.05 | -0.7 | -0.37 | -0.57 | -0.6 | -0.29 | -0.75 | -0.42 | -0.98 | -0.4 | 0.3 | -0.72 | -0.36 | -0.51 | -0.035 | -0.215 | 0.1 | 0.771196 | 0.898965 | 0.479242 | 1 | 0.999631 | 0.919413 |
| Slc1a2_P43006_S542s_1 | -1.15 | -0.26 | -0.58 | -1.05 | 0.22 | 0.59 | -0.94 | -1.19 | -0.06 | -1.41 | -0.77 | -0.32 | 0.08 | -0.46 | -1 | -0.86 | 0.43 | 0.12 | 0.2 | 0.586776 | 0.362465 | 0.754119 | 1 | 0.999631 | 0.849744 |
| Slc1a2_P43006_S558s_1 | -1.9 | -1.03 | -1.04 | -1.01 | -0.7 | -2.05 | 0.23 | -0.77 | -1.08 | -3.12 | -0.49 | -0.16 | -0.94 | -1.36 | -1.06 | -0.39 | 0.4225 | 0.0325 | 0.3075 | 0.393025 | 0.395725 | 0.9589 | 1 | 0.999631 | 0.758901 |
| Slc1a2_P43006_S562s_1 | -2.59 | -3.31 | -1.92 | -2.06 | -2.07 | -3.66 | -1.93 | -2.27 | -2.84 | -4.08 | -1.35 | -1.3 | -1.25 | -3.26 | -2.04 | -2 | -0.0125 | 0.0775 | 0.3325 | 0.510566 | 0.979253 | 0.906646 | 1 | 0.999631 | 0.817683 |
| Slc1a2_P43006_T4t_1 | 0.37 | -0.41 | -0.01 | -0.68 | -1.15 | -0.55 | -0.07 | -1.19 | -0.66 | 0.16 | -0.04 | -1.09 | -0.09 | -0.58 | -0.95 | -0.45 | -0.5575 | -0.225 | -0.335 | 0.348272 | 0.15572 | 0.563844 | 1 | 0.999631 | 0.736697 |
| Slc1a2_P43006_Y522y_1 | 0.79 | 0.26 | -1.96 | -1.45 | 0.83 | 0.57 | -0.92 | -2.08 | -0.66 | -0.96 | -1.96 | -1.92 | -2.02 | -3.36 | -2.32 | -1.98 | 0.19 | -0.785 | -1.83 | 0.018321 | 0.818142 | 0.254231 | 1 | 0.999631 | 0.391024 |
| Slc1a2_P43006_Y537y_1 | -0.43 | -1.58 | -2.25 | -0.75 | -0.54 | -2.25 | -0.78 | -1.44 | -1.32 | -1.23 | -1.16 | -1.46 | -0.77 | -1.73 | -2.91 | -0.07 | 1.57E-16 | -0.04 | -0.1175 | 0.859209 | 1 | 0.924068 | 1 | 0.999631 | 0.953124 |
| Slc1a3_P56564_S4s_1 | 1.62 | 0.51 | -3.42 | -3.83 | -0.69 | 1.68 | -2.74 | -3.15 | 0.41 | 2.47 | -1.25 | -5.21 | 0.41 | 0 | -7 | -7.28 | 0.055 | 0.385 | -2.1875 | 0.31294 | 0.970671 | 0.831096 | 1 | 0.999631 | 0.719861 |
| Slc1a3_P56564_S512s_1 | -0.93 | 0.15 | 0.06 | -0.56 | -0.55 | 0.19 | -0.64 | -0.39 | -0.45 | -1.14 | -0.19 | -0.39 | 0.5 | 0.36 | -0.22 | 0 | -0.0275 | -0.2225 | 0.48 | 0.199445 | 0.937004 | 0.542943 | 1 | 0.999631 | 0.657744 |
| Slc1a4_O35874_S507s_1 | -1.11 | -0.76 | -0.88 | -1.31 | -0.58 | -1.52 | -1.39 | -1.35 | -0.72 | -0.45 | -0.94 | -0.59 | 0.34 | -0.43 | -1.05 | -1.03 | -0.195 | 0.34 | 0.4725 | 0.236089 | 0.526826 | 0.227628 | 1 | 0.999631 | 0.683301 |
| Slc1a4_O35874_T24t_1 | -0.45 | -0.92 | 0.67 | 0.22 | -1.02 | -0.8 | 0.82 | -0.52 | -0.97 | 0.54 | -0.02 | -0.37 | -0.98 | -1.19 | -0.41 | -0.36 | -0.26 | -0.085 | -0.615 | 0.166464 | 0.610428 | 0.853608 | 1 | 0.999631 | 0.63587 |
| Slc20a1_Q61609_S269s_1 | -0.21 | -0.67 | -0.45 | -1.27 | -1.16 | -0.97 | -1.45 | -1 | -2.1 | -0.74 | -0.44 | -0.97 | -0.02 | -0.13 | -0.5 | -0.85 | -0.495 | -0.4125 | 0.275 | 0.441114 | 0.132837 | 0.347051 | 1 | 0.999631 | 0.781983 |
| Slc20a2_Q80UP8_S256s_1 | 0.49 | -1.66 | -0.3 | -1.26 | -2 | -0.82 | -0.47 | -1.73 | -1.24 | 0.86 | -0.84 | -0.79 | 0.54 | -1.07 | -0.46 | -1.72 | -0.5725 | 0.18 | 0.005 | 0.993537 | 0.315672 | 0.768218 | 1 | 0.999631 | 0.997705 |
| Slc20a2_Q80UP8_S268s_1 | -0.61 | -1.81 | -1.74 | -1.37 | -1.35 | -1.68 | -1 | -1.07 | -1.43 | -1.5 | -1.31 | -1.35 | -1.09 | -1 | -1.24 | -1.59 | 0.1075 | -0.015 | 0.1525 | 0.670132 | 0.756968 | 0.963751 | 1 | 0.999631 | 0.884054 |
| Slc20a2_Q80UP8_S316s_1 | 3.07 | 1.09 | 1.09 | 1.77 | 1.39 | 1.32 | 1.72 | 1.34 | 1.05 | 0.5 | 1.23 | -0.08 | -1.25 | 1.72 | 0.5 | 1.25 | -0.3125 | -1.08 | -1.2 | 0.115069 | 0.499855 | 0.060863 | 1 | 0.999631 | 0.579336 |
| Slc23a2_Q9EPR4_S18s_1 | -1.57 | -2.89 | -1.61 | -0.91 | -2.19 | -2.92 | -1.59 | -1.76 | -1.76 | -3.87 | -1.98 | -0.79 | -1.64 | -2.19 | -2.5 | -1.7 | -0.37 | -0.355 | -0.2625 | 0.569417 | 0.449843 | 0.606123 | 1 | 0.999631 | 0.84338 |
| Slc23a2_Q9EPR4_S81s_1 | 0.31 | 0.69 | -0.91 | -1.26 | 0.38 | 0.11 | -1.12 | -0.73 | 0.06 | -0.2 | -0.66 | -0.39 | 0.85 | 0.6 | 0.02 | 0.13 | -0.0475 | -0.005 | 0.6925 | 0.17948 | 0.929875 | 0.991598 | 1 | 0.999631 | 0.64376 |
| Slc24a2_F6RT95_S43s_1 | -0.24 | 1.37 | 1.09 | 0.09 | 1.13 | 1.05 | 0.52 | 0.01 | 2.05 | -0.12 | 0.96 | 2.01 | 2.15 | -0.42 | -0.11 | -0.81 | 0.1 | 0.6475 | -0.375 | 0.586511 | 0.824413 | 0.28679 | 1 | 0.999631 | 0.849724 |
| Slc24a2_F6RT95_S65s_1 | -0.11 | -1.83 | -1.54 | -1.22 | -1.56 | -1.87 | -1.24 | -1.91 | -2.23 | -0.89 | -1.48 | -1.91 | -0.81 | -1.47 | -1.92 | -1.12 | -0.47 | -0.4525 | -0.155 | 0.728882 | 0.266188 | 0.339517 | 1 | 0.999631 | 0.905923 |
| Slc24a2_F6RT95_S68s_1 | -0.31 | -0.81 | -0.81 | -0.16 | -1.55 | -0.47 | -1.47 | 0.9 | -1.56 | 0.21 | -1.43 | -1.48 | -0.33 | -0.57 | -0.62 | -0.19 | -0.125 | -0.5425 | 0.095 | 0.752965 | 0.819429 | 0.245624 | 1 | 0.999631 | 0.913377 |
| Slc24a2_Q8BUN9_S312s_1 | 1.19 | -0.88 | 1.07 | 0.81 | -2.35 | 0.41 | 2.86 | -0.73 | -2.07 | 0.53 | -0.9 | 0.29 | -0.25 | -1.56 | 0.5 | 0.21 | -0.5 | -1.085 | -0.8225 | 0.193872 | 0.628683 | 0.138245 | 1 | 0.999631 | 0.650692 |
| Slc24a4_E9PUL6_S296s_1 | -0.84 | 0.4 | 1.61 | -0.34 | -0.27 | -0.08 | -0.48 | 0.03 | 0.18 | 0.1 | 0.97 | 2.08 | 2.33 | 0.46 | 2.03 | 0.53 | -0.4075 | 0.625 | 1.13 | 0.10635 | 0.430393 | 0.337501 | 1 | 0.999631 | 0.568797 |
| Slc25a35_Q5SWT3_T99t_1 | -0.16 | -0.22 | 0.16 | -0.54 | 0.04 | -0.39 | -0.73 | -0.77 | -0.48 | 0.25 | 0.13 | 0.9 | 0.83 | -0.55 | 0.06 | 0.51 | -0.2725 | 0.39 | 0.4025 | 0.294843 | 0.373772 | 0.285965 | 1 | 0.999631 | 0.708705 |
| Slc25a4_P48962_S149s_1 | -0.55 | -1.53 | -1.89 | -1.36 | -0.97 | -1.51 | -1.52 | -1.55 | -0.78 | -1.07 | -0.83 | -1.34 | -0.41 | -1.38 | -1.07 | -1.69 | -0.055 | 0.3275 | 0.195 | 0.637498 | 0.873984 | 0.360807 | 1 | 0.999631 | 0.870011 |
| Slc25a4_P48962_T84t_1 | -3.23 | -3.56 | -2.86 | -4.77 | -6.76 | -1.27 | -3.23 | -3.08 | -7.41 | -4.54 | -2.53 | -9.38 | -2.52 | -1.44 | -6.27 | -6.9 | 0.02 | -2.36 | -0.6775 | 0.574743 | 0.984751 | 0.103345 | 1 | 0.999631 | 0.845624 |
| Slc25a4_P48962_Y191y_1 | -0.7 | 0.92 | 0.61 | 0.33 | 0.11 | -0.04 | -0.13 | 0.62 | 1.08 | 0.42 | -0.1 | 0.76 | 0.28 | 0.89 | 0.05 | 0.66 | -0.15 | 0.25 | 0.18 | 0.665647 | 0.705384 | 0.565535 | 1 | 0.999631 | 0.88264 |
| Slc25a46_Q9CQS4_S61s_1 | -0.28 | -0.71 | -1.64 | -0.51 | -0.23 | -1.1 | -0.87 | -0.96 | -0.67 | -0.79 | -0.69 | -0.88 | -0.86 | -1.02 | -0.73 | -1.01 | -0.005 | 0.0275 | -0.12 | 0.737806 | 0.989276 | 0.936365 | 1 | 0.999631 | 0.907836 |
| Slc25a46_Q9CQS4_S94s_1 | 0.61 | 0.14 | 1.83 | 0.32 | 0.15 | 0.85 | 0.17 | 0.26 | 0.02 | 0.81 | 0.2 | 0.5 | 0.73 | 1.4 | 1.72 | 1.01 | -0.3675 | -0.3425 | 0.49 | 0.282228 | 0.38453 | 0.424529 | 1 | 0.999631 | 0.703679 |
| Slc25a46_Q9CQS4_T44t_1 | -1.27 | -2.09 | -1.67 | -2.11 | -2.41 | -2.42 | -1.59 | -2.3 | -2.82 | -1.61 | -1.69 | -2.51 | -1.82 | -1.75 | -1.85 | -0.71 | -0.395 | -0.3725 | 0.2525 | 0.508526 | 0.243186 | 0.341431 | 1 | 0.999631 | 0.816932 |
| Slc25a46_Q9CQS4_T45t_1 | -1.17 | -1.9 | -1.94 | -1.66 | -2.14 | -2.2 | -1.5 | -1.84 | -2.59 | -1.46 | -2.42 | -2 | -1.57 | -1.53 | -2.05 | -1.16 | -0.2525 | -0.45 | 0.09 | 0.784713 | 0.410531 | 0.215992 | 1 | 0.999631 | 0.922904 |
| Slc26a6_Q8CIW6_S728s_1 | -0.21 | -0.38 | 0.29 | -0.22 | -0.41 | -0.53 | -0.42 | -0.31 | -0.79 | -0.3 | 0.15 | 0.23 | -0.21 | -0.28 | -0.03 | -0.02 | -0.2875 | -0.0475 | -0.005 | 0.986031 | 0.2825 | 0.885262 | 1 | 0.999631 | 0.995133 |
| Slc29a1_E9PXM6_S273s_1 | -0.2 | -1.3 | -0.16 | -1.45 | -0.51 | -1.21 | -0.8 | -1.31 | -1.12 | -0.08 | -1.54 | -0.32 | 0.09 | -1.17 | -0.64 | -1.27 | -0.18 | 0.0125 | 0.03 | 0.947987 | 0.652771 | 0.97875 | 1 | 0.999631 | 0.983162 |
| Slc2a13_Q3UHK1_S629s_1 | -2.1 | -1.29 | -0.84 | -1.43 | -1.19 | -1.34 | -1.26 | -1.71 | -1.78 | -1.04 | -1.88 | -1.42 | 0.03 | -0.98 | -0.77 | -1.54 | 0.04 | -0.115 | 0.6 | 0.181159 | 0.902887 | 0.748346 | 1 | 0.999631 | 0.643763 |
| Slc2a13_Q3UHK1_S634s_1 | -1.19 | -1.76 | -0.74 | -2.16 | -1.21 | -1.16 | -0.49 | -1.2 | -1.01 | 0.04 | -1.08 | -0.35 | -0.53 | -0.39 | -2.19 | -0.84 | 0.4475 | 0.8625 | 0.475 | 0.349912 | 0.249788 | 0.062201 | 1 | 0.999631 | 0.736999 |
| Slc2a3_P32037_S471s_1 | 0.11 | -3.01 | -0.37 | 0.07 | -0.29 | -1.12 | 2.17 | -0.66 | -0.68 | -1.42 | 0.15 | -0.81 | -1.56 | -2.17 | -0.65 | -2.12 | 0.825 | 0.11 | -0.825 | 0.272328 | 0.372712 | 0.87859 | 1 | 0.999631 | 0.699889 |
| Slc30a1_Q60738_S501s_1 | -0.16 | 1.45 | -2.32 | -3.46 | -0.46 | -3.3 | -2.56 | -2.77 | -0.44 | 0.4 | -1.32 | -3.51 | 0.97 | -2.84 | -2.76 | -2.16 | -1.15 | -0.095 | -0.575 | 0.634184 | 0.302166 | 0.935999 | 1 | 0.999631 | 0.868569 |
| Slc30a10_Q3UVU3_S157s_1 | 0.88 | -3 | -0.15 | -1.4 | -1.73 | -0.18 | 0.17 | 0.45 | -1.5 | 0.33 | -1.64 | -1.8 | -1.13 | -2.74 | 0.22 | -2.2 | 0.595 | -0.235 | -0.545 | 0.554192 | 0.485211 | 0.782395 | 1 | 0.999631 | 0.837952 |
| Slc30a3_P97441_S10s_1 | -0.63 | -2.82 | 0.77 | -0.68 | -0.37 | -0.06 | -0.23 | 0 | 0.52 | 0.97 | -0.05 | -1.46 | -1.41 | 1.05 | -1.47 | -0.77 | 0.675 | 0.835 | 0.19 | 0.817327 | 0.322523 | 0.309851 | 1 | 0.999631 | 0.937991 |
| Slc30a3_P97441_S22s_1 | 0.97 | -0.24 | 1.22 | 1.81 | -0.83 | 1.01 | 2.38 | 2.82 | -0.17 | -0.21 | 2.2 | 1.78 | 0.41 | -0.92 | 1.66 | -0.06 | 0.405 | -0.04 | -0.6675 | 0.298772 | 0.618805 | 0.953415 | 1 | 0.999631 | 0.711958 |
| Slc30a3_P97441_S23s_1 | 0.71 | -0.61 | -2.4 | -1.44 | -0.84 | 0.5 | -0.93 | -3.21 | -0.7 | -0.49 | -2.11 | -2.03 | 0.27 | -1.22 | -2.22 | -2.94 | -0.185 | -0.3975 | -0.5925 | 0.481934 | 0.83325 | 0.573255 | 1 | 0.999631 | 0.802327 |
| Slc32a1_O35633_S95s_1 | -0.35 | -1.39 | -2.03 | -2.53 | -1.3 | -1.45 | -2.36 | -2.94 | -1.92 | -1.05 | -2.38 | -3.07 | -0.46 | -1.43 | -3.2 | -2.66 | -0.4375 | -0.53 | -0.3625 | 0.601253 | 0.440404 | 0.371751 | 1 | 0.999631 | 0.856765 |
| Slc32a1_O35633_S97s_1 | 0.37 | 0.44 | 0.29 | -0.04 | 0.24 | 0.21 | -0.44 | -0.02 | 0.25 | 0.52 | -0.3 | 0.22 | 1.31 | 0.71 | 0.36 | 0.36 | -0.2675 | -0.0925 | 0.42 | 0.216526 | 0.34377 | 0.74931 | 1 | 0.999631 | 0.669465 |
| Slc33a1_Q99J27_S42s_1 | 2.77 | 1.6 | 2.89 | 1.85 | 0.8 | 2.13 | 3.08 | 1.85 | 1.32 | 2.76 | 2.37 | 1.55 | -0.45 | 0.38 | 1.86 | 0.94 | -0.3125 | -0.2775 | -1.595 | 0.014195 | 0.557316 | 0.547859 | 1 | 0.999631 | 0.385092 |
| Slc35a5_M0QWE5_S129s_1 | 0.11 | 0.2 | 1.48 | 1.44 | 0.17 | 1 | 0.92 | 0.53 | 0.61 | 1.09 | 1.14 | 1.24 | 0 | 0.04 | 1.1 | 0.59 | -0.1525 | 0.2125 | -0.375 | 0.417967 | 0.716816 | 0.606875 | 1 | 0.999631 | 0.771539 |
| Slc35d3_Q8BGF8_S344s_1 | -0.28 | 0.02 | -1.17 | 0.17 | -0.38 | -0.49 | -0.15 | -0.55 | 0.36 | -0.08 | 0.49 | 0.64 | -0.27 | -0.15 | -1.85 | -0.52 | -0.0775 | 0.6675 | -0.3825 | 0.432743 | 0.822308 | 0.093773 | 1 | 0.999631 | 0.779247 |
| Slc35e1_Q8CD26_S363s_1 | -0.05 | 3.24 | 1.19 | 0.22 | 4.79 | 1.37 | 4.61 | 2.07 | 5.05 | -0.18 | 3.76 | 4.47 | 3.14 | -0.02 | 2.79 | 1.07 | 2.06 | 2.125 | 0.595 | 0.516066 | 0.060386 | 0.10078 | 1 | 0.999631 | 0.819449 |
| Slc35g2_D3YVE8_S4s_1 | 0.95 | -1.28 | -0.55 | -0.23 | -1.27 | -0.96 | -0.01 | -1.07 | -0.82 | 0.24 | -0.94 | -0.36 | -0.67 | -0.86 | -0.39 | -0.97 | -0.55 | -0.1925 | -0.445 | 0.356139 | 0.29283 | 0.704908 | 1 | 0.999631 | 0.74034 |
| Slc38a1_Q8K2P7_S52s_1 | -0.18 | 0.26 | 0.84 | 0.17 | -0.27 | -0.09 | 0.58 | -1.15 | 0.41 | -0.06 | 0.63 | 1.37 | 0.72 | -0.44 | 1.65 | 1.66 | -0.505 | 0.315 | 0.625 | 0.24109 | 0.239959 | 0.421627 | 1 | 0.999631 | 0.684338 |
| Slc38a7_Q8BWH0_S28s_1 | -0.64 | -0.54 | -0.53 | -0.45 | 0.48 | -0.04 | -0.54 | -0.27 | -0.27 | -1.31 | 0.03 | -0.53 | 0.61 | -0.32 | -1.06 | -0.12 | 0.4475 | 0.02 | 0.3175 | 0.413269 | 0.147524 | 0.952662 | 1 | 0.999631 | 0.768474 |
| Slc39a10_Q6P5F6_S537s_1 | 0.75 | -0.73 | -0.49 | -0.73 | 0.34 | -1.45 | -0.11 | -0.34 | 0.1 | 1.04 | -0.95 | -1.3 | 0.2 | -1.08 | -0.85 | -1.67 | -0.09 | 0.0225 | -0.55 | 0.289708 | 0.854289 | 0.969244 | 1 | 0.999631 | 0.706164 |
| Slc39a10_Q6P5F6_S593s_1 | -0.62 | 0.25 | 0.73 | -0.03 | 0.1 | 0.06 | -0.15 | 0.63 | 0.2 | -0.59 | 0.06 | 1.26 | 0.83 | 0.33 | -0.03 | 0.41 | 0.0775 | 0.15 | 0.3025 | 0.425633 | 0.826845 | 0.74611 | 1 | 0.999631 | 0.774624 |
| Slc39a10_Q6P5F6_T538t_1 | 1.17 | -1.82 | -1.11 | -0.5 | -0.49 | 0.32 | 0.39 | -1.07 | -0.7 | -0.11 | -1.35 | -1.06 | -0.35 | -1.63 | -1.71 | -1.57 | 0.3525 | -0.24 | -0.75 | 0.260172 | 0.590036 | 0.702271 | 1 | 0.999631 | 0.692624 |
| Slc39a10_Q6P5F6_T542t_1 | 1.39 | -0.38 | 0.21 | 0.85 | -0.41 | 1.93 | 1.4 | 0.3 | -0.24 | 0.67 | 0.9 | -0.93 | -1.92 | -0.54 | 0.24 | -1.06 | 0.2875 | -0.4175 | -1.3375 | 0.033768 | 0.629036 | 0.440956 | 1 | 0.999631 | 0.444185 |
| Slc39a6_D3Z7N4_S204s_1 | 0.48 | 0.19 | 0.37 | 0 | 1.01 | 0.33 | 0.48 | 0.19 | 0.87 | 0.89 | 0.51 | 0.43 | -0.05 | 0.3 | 1.08 | 0.48 | 0.2425 | 0.415 | 0.1925 | 0.564612 | 0.403296 | 0.147693 | 1 | 0.999631 | 0.84139 |
| Slc39a7_Q31125_S312s_1 | -1 | 0.23 | -0.56 | -0.75 | -0.4 | -0.15 | -0.36 | -0.1 | -0.17 | -1.08 | 0.22 | 0.54 | 0.68 | 0.01 | 0.36 | 0.15 | 0.2675 | 0.3975 | 0.82 | 0.040983 | 0.415299 | 0.373204 | 1 | 0.999631 | 0.474979 |
| Slc3a2_P10852_S2s_1 | 0.57 | -0.34 | -0.1 | -0.08 | 0.14 | -0.12 | -0.11 | -0.4 | -0.45 | -0.03 | -0.17 | -0.56 | 0.23 | 0.14 | 0.13 | -0.16 | -0.135 | -0.315 | 0.0725 | 0.814273 | 0.646722 | 0.313114 | 1 | 0.999631 | 0.937111 |
| Slc43a2_Q8CGA3_S274s_1 | 0.77 | -0.24 | -0.62 | -0.37 | -0.7 | -0.43 | -1.11 | 0.85 | 0.27 | 0.05 | -0.33 | -0.22 | 0.62 | -0.28 | -0.24 | -0.5 | -0.2325 | 0.0575 | 0.015 | 0.970851 | 0.637837 | 0.874664 | 1 | 0.999631 | 0.989618 |
| Slc45a4_F7BEQ1_S344s_1 | -0.77 | -1 | -0.33 | -1.01 | -1.23 | -0.41 | -1.17 | -0.78 | -0.82 | -0.43 | -0.99 | -0.75 | 0.9 | -0.09 | -0.19 | 0.04 | -0.12 | 0.03 | 0.9425 | 0.021549 | 0.6963 | 0.916695 | 1 | 0.999631 | 0.405125 |
| Slc45a4_F7BEQ1_S365s_1 | 1.08 | 0.56 | 0.3 | -0.09 | 0.66 | 0.47 | -0.17 | 0.42 | 0.57 | 0.42 | -0.29 | -0.25 | 0.68 | 0.62 | -0.22 | 0.99 | -0.1175 | -0.35 | 0.055 | 0.887255 | 0.72986 | 0.347037 | 1 | 0.999631 | 0.963505 |
| Slc47a1_D6RGQ3_S18s_1 | 0.03 | -2.19 | -1.83 | 0.3 | 0.06 | -2.4 | -0.67 | -0.78 | -1 | -0.34 | -1.2 | 0.12 | -0.05 | -0.72 | -0.47 | -0.28 | -0.025 | 0.3175 | 0.5425 | 0.371489 | 0.972377 | 0.619202 | 1 | 0.999631 | 0.746522 |
| Slc4a10_Q5DTL9_S1056s_1 | -0.55 | 0.13 | 0.2 | -0.03 | -0.23 | -0.08 | -0.34 | 0.02 | -0.32 | -0.24 | -0.04 | 0.09 | 0.24 | 0.07 | 0.14 | 0.75 | -0.095 | -0.065 | 0.3625 | 0.267777 | 0.729929 | 0.819701 | 1 | 0.999631 | 0.696016 |
| Slc4a10_Q5DTL9_S246s_1 | 1.99 | 0.22 | -0.54 | 0.23 | -0.33 | 0.09 | 0.63 | -0.68 | -0.19 | 0.67 | -0.93 | -0.89 | -0.92 | -1.56 | -1.03 | -1.85 | -0.5475 | -0.81 | -1.815 | 0.00701 | 0.336526 | 0.196091 | 1 | 0.999631 | 0.358173 |
| Slc4a10_Q5DTL9_S260s_1 | -0.32 | -0.07 | -1.11 | -1.1 | -0.79 | -0.04 | -1.25 | -1.12 | -1.18 | -0.08 | -0.79 | -0.43 | -0.26 | -0.76 | -1.05 | -1.16 | -0.15 | 0.03 | -0.1575 | 0.675404 | 0.701365 | 0.936949 | 1 | 0.999631 | 0.885853 |
| Slc4a10_Q5DTL9_S275s_1 | -2.38 | -3.55 | -3.82 | -2.3 | -1.39 | -4.24 | -2.26 | -2.2 | -2.39 | -3.98 | -2.96 | -2.89 | -2.19 | -2.9 | -3.28 | -3.88 | 0.49 | -0.0425 | -0.05 | 0.921002 | 0.454547 | 0.930981 | 1 | 0.999631 | 0.975032 |
| Slc4a10_Q5DTL9_S279s_1 | 1.15 | 0.75 | -0.5 | 1.68 | 2.31 | 1.19 | 1.48 | 1.7 | 1.24 | -0.14 | 1.29 | 0.17 | -1.47 | 0.63 | 0.1 | 1.09 | 0.9 | -0.13 | -0.6825 | 0.308051 | 0.091807 | 0.812755 | 1 | 0.999631 | 0.718178 |
| Slc4a10_Q5DTL9_S296s_1 | -0.68 | 3.07 | 3.12 | -0.41 | 3.23 | 2.5 | 3.77 | 1.68 | 4.83 | -0.51 | 3.56 | 4.66 | 4.02 | -0.33 | 2.33 | -1.41 | 1.52 | 1.86 | -0.1225 | 0.928562 | 0.147143 | 0.201767 | 1 | 0.999631 | 0.976781 |
| Slc4a10_Q5DTL9_S80s_2 | -0.76 | 0.9 | 2.89 | 0.53 | 0.01 | 0.83 | 1.62 | 2.73 | 0.46 | -0.52 | 2.87 | 3.42 | 0.94 | 0.62 | 3.58 | 1.26 | 0.4075 | 0.6675 | 0.71 | 0.424633 | 0.625181 | 0.526381 | 1 | 0.999631 | 0.774174 |
| Slc4a10_Q5DTL9_S88s_1 | -1.82 | -0.11 | -0.07 | -0.78 | -0.57 | -0.36 | -0.48 | -0.23 | -0.38 | -1.49 | 0 | 0.07 | 0.77 | -0.41 | -0.34 | 0.08 | 0.285 | 0.245 | 0.72 | 0.154438 | 0.496419 | 0.635133 | 1 | 0.999631 | 0.625677 |
| Slc4a10_Q5DTL9_S90s_1 | -1.26 | 0.62 | -0.37 | -0.8 | 0.01 | 0.02 | -1.22 | -0.65 | 0.3 | -0.32 | -1.91 | 0.37 | 0.9 | 0.47 | 0.14 | -0.77 | -0.0075 | 0.0625 | 0.6375 | 0.22925 | 0.987386 | 0.917611 | 1 | 0.999631 | 0.679776 |
| Slc4a2_P13808_S156s_2 | -0.53 | 1.34 | 1.3 | 0.25 | 0.01 | 0.3 | 0.2 | 0.39 | 0.32 | 0.02 | -1.3 | 1.12 | 0.72 | 0.54 | 0.33 | -0.01 | -0.365 | -0.55 | -0.195 | 0.678191 | 0.419741 | 0.380066 | 1 | 0.999631 | 0.887183 |
| Slc4a2_P13808_S158s_1 | -0.44 | 0.08 | 0.39 | -0.13 | -0.45 | 0.14 | 0.3 | -0.1 | 0.35 | -0.14 | -1.71 | -0.89 | 0.36 | -1.57 | -0.54 | -0.23 | -0.0025 | -0.5725 | -0.47 | 0.302349 | 0.993371 | 0.239163 | 1 | 0.999631 | 0.714978 |
| Slc4a2_P13808_S158s_2 | -0.53 | 1.34 | 1.3 | 0.25 | 0.01 | 0.3 | 0.2 | 0.39 | 0.32 | 0.02 | -1.3 | 1.12 | 0.72 | 0.54 | 0.33 | -0.01 | -0.365 | -0.55 | -0.195 | 0.678191 | 0.419741 | 0.380066 | 1 | 0.999631 | 0.887183 |
| Slc4a3_F8WHM3_S198s_1 | -0.27 | -0.91 | -0.81 | -0.47 | -0.22 | -0.37 | -0.75 | -0.85 | -1.05 | -1.02 | -0.77 | -0.61 | -0.48 | -0.51 | -0.13 | -0.46 | 0.0675 | -0.2475 | 0.22 | 0.456677 | 0.813832 | 0.386072 | 1 | 0.999631 | 0.790625 |
| Slc4a3_F8WHM3_S296s_1 | -0.62 | -0.98 | -0.85 | -1.06 | -0.9 | -0.89 | -1.6 | -1.16 | -0.93 | -0.52 | -0.51 | -0.44 | 0.07 | -0.53 | -0.7 | -0.31 | -0.26 | 0.2775 | 0.51 | 0.112293 | 0.358893 | 0.30905 | 1 | 0.999631 | 0.577263 |
| Slc4a4_D3Z7G8_S192s_1 | 1.26 | 0.14 | -0.86 | -0.78 | -0.49 | 0.45 | -0.88 | -0.01 | -0.32 | 0.18 | -1.15 | -0.59 | -0.74 | 0.26 | -0.57 | -0.55 | -0.1725 | -0.41 | -0.34 | 0.514005 | 0.745346 | 0.44662 | 1 | 0.999631 | 0.81893 |
| Slc4a4_D3Z7G8_S204s_1 | -0.03 | 0.49 | 0.58 | -0.09 | 0.25 | 0.34 | -0.76 | 0.37 | 0.25 | 0.51 | -0.34 | 0.91 | 1.05 | 0.78 | 0.34 | 0.58 | -0.1875 | 0.095 | 0.45 | 0.176616 | 0.595547 | 0.787534 | 1 | 0.999631 | 0.642116 |
| Slc4a4_E9Q8N8_S1029s_1 | -0.7 | -2.07 | 1.86 | 1.1 | -2.33 | -1.85 | 1.24 | 0.46 | -1.91 | -1.07 | 0.36 | -0.02 | -0.55 | -2.42 | 0.18 | 0.42 | -0.6675 | -0.7075 | -0.64 | 0.50135 | 0.534878 | 0.435417 | 1 | 0.999631 | 0.814092 |
| Slc4a4_E9Q8N8_S1060s_1 | -0.64 | 0.37 | 0.44 | -0.3 | -0.66 | -0.28 | -0.89 | 0.21 | -0.55 | 0.5 | -0.5 | 0.22 | 0.4 | 0.78 | 0 | -0.55 | -0.3725 | -0.05 | 0.19 | 0.642557 | 0.330684 | 0.897582 | 1 | 0.999631 | 0.871335 |
| Slc4a4_E9Q8N8_S239s_1 | 0.55 | 1.12 | 0.11 | -0.12 | 1.54 | 1.19 | -0.04 | -0.1 | 2.42 | 0.37 | 0.51 | 1.31 | 2.31 | 0.93 | 0.76 | -0.45 | 0.2325 | 0.7375 | 0.4725 | 0.420928 | 0.627389 | 0.172848 | 1 | 0.999631 | 0.772817 |
| Slc4a4_E9Q8N8_S245s_1 | -1.06 | -0.31 | -0.32 | -1.38 | -0.31 | -0.65 | -0.68 | -1.11 | -0.05 | -1.26 | -0.59 | 0.03 | 0.63 | -1.08 | -0.57 | -1.2 | 0.08 | 0.3 | 0.2125 | 0.660881 | 0.817553 | 0.470257 | 1 | 0.999631 | 0.880315 |
| Slc4a4_E9Q8N8_S86s_1 | 1.12 | -0.09 | -0.04 | 1.76 | 0.54 | 1.66 | 0.66 | 1.64 | 0.32 | 0.45 | -0.7 | 0.08 | -0.9 | 0.77 | -0.06 | 1.2 | 0.4375 | -0.65 | -0.435 | 0.469703 | 0.399994 | 0.210451 | 1 | 0.999631 | 0.796991 |
| Slc4a4_E9Q8N8_T1064t_1 | 0.72 | 1.05 | 0.96 | 0.73 | 1.09 | 2.05 | -0.22 | 2.27 | 1.57 | 1.67 | 0.36 | 2.12 | -1.6 | 2.12 | 1.94 | 0.45 | 0.4325 | 0.565 | -0.1375 | 0.856227 | 0.422691 | 0.178327 | 1 | 0.999631 | 0.951817 |
| Slc4a7_H6WZF5_S1072s_1 | 1.17 | 0.66 | 0.24 | 1.4 | 1.77 | 1.35 | 0.83 | 1.37 | 1.56 | 0.84 | 1.67 | 1.38 | -0.17 | 0.77 | 1.62 | 1.75 | 0.4625 | 0.495 | 0.125 | 0.800689 | 0.20899 | 0.184837 | 1 | 0.999631 | 0.931985 |
| Slc4a7_H6WZF5_S233s_1 | 0.91 | 1.33 | 0.52 | 1.86 | 3.06 | 0.7 | 1.77 | 1.41 | 1.82 | -0.28 | 1.53 | 1.27 | -0.48 | 0.6 | 0.43 | 0.74 | 0.58 | -0.07 | -0.8325 | 0.067673 | 0.28836 | 0.892241 | 1 | 0.999631 | 0.521762 |
| Slc4a7_H6WZF5_S255s_1 | -0.83 | 0.29 | -0.49 | -1.16 | -0.44 | -1.15 | -0.76 | -1.39 | 0.59 | 0.04 | -0.11 | -1.68 | 0.26 | 0.05 | -0.89 | 0.02 | -0.3875 | 0.2575 | 0.4075 | 0.34078 | 0.328198 | 0.634222 | 1 | 0.999631 | 0.733032 |
| Slc4a7_H6WZF5_S283s_1 | 1.15 | 0.75 | -0.5 | 1.68 | 2.31 | 1.19 | 1.48 | 1.7 | 1.24 | -0.14 | 1.29 | 0.17 | -1.47 | 0.63 | 0.1 | 1.09 | 0.9 | -0.13 | -0.6825 | 0.308051 | 0.091807 | 0.812755 | 1 | 0.999631 | 0.718178 |
| Slc4a7_H6WZF5_S84s_1 | 0.57 | 1.92 | 1.93 | 0.5 | 0.81 | 1.93 | 1.02 | 1.57 | 1.6 | 1.97 | 2.05 | 2.81 | 2.15 | 1.53 | 1.99 | 1.81 | 0.1025 | 0.8775 | 0.64 | 0.16 | 0.822799 | 0.082202 | 1 | 0.999631 | 0.629764 |
| Slc4a8_Q8JZR6_S194s_1 | 1.71 | -0.94 | -0.77 | 0.19 | -1.15 | 0.3 | 0.23 | -1.38 | 0.08 | 1.72 | -1.15 | -0.87 | -1.11 | -2.21 | -0.75 | -2.71 | -0.5475 | -0.1025 | -1.7425 | 0.026241 | 0.421764 | 0.895548 | 1 | 0.999631 | 0.422285 |
| Slc4a8_Q8JZR6_S207s_1 | -1.05 | 0.27 | 0.55 | -0.87 | -0.27 | -0.6 | -0.56 | 0.4 | 0.09 | -0.9 | 0.69 | 0.65 | 0.64 | -0.15 | 0.55 | 0.3 | 0.0175 | 0.4075 | 0.61 | 0.188034 | 0.968817 | 0.435817 | 1 | 0.999631 | 0.647379 |
| Slc4a8_Q8JZR6_S994s_1 | -0.25 | -0.38 | -0.62 | -0.49 | -0.61 | -0.11 | -0.64 | -0.4 | -0.59 | -0.46 | -0.3 | -0.22 | -0.65 | -0.69 | -0.29 | 0.04 | -0.005 | 0.0425 | 0.0375 | 0.899703 | 0.98442 | 0.866771 | 1 | 0.999631 | 0.967515 |
| Slc5a3_Q9JKZ2_S37s_2 | 1.6 | 1.13 | 2.06 | 1.03 | 0.28 | 1.79 | 1.3 | 0.87 | 1 | 2.39 | 0.93 | 0.63 | -1.49 | 0.6 | 0.77 | 0.86 | -0.395 | -0.2175 | -1.27 | 0.04478 | 0.338396 | 0.630649 | 1 | 0.999631 | 0.475443 |
| Slc5a3_Q9JKZ2_Y39y_2 | 1.6 | 1.13 | 2.06 | 1.03 | 0.28 | 1.79 | 1.3 | 0.87 | 1 | 2.39 | 0.93 | 0.63 | -1.49 | 0.6 | 0.77 | 0.86 | -0.395 | -0.2175 | -1.27 | 0.04478 | 0.338396 | 0.630649 | 1 | 0.999631 | 0.475443 |
| Slc5a7_Q8BGY9_S550s_1 | -0.04 | 0.3 | 12.2 | 10.66 | 0.46 | -0.74 | 6.9 | 10.96 | 0.55 | -0.33 | 12.75 | 11.63 | 0.92 | 0.25 | 12.12 | 11.83 | -1.385 | 0.37 | 0.5 | 0.896669 | 0.70122 | 0.926861 | 1 | 0.999631 | 0.966487 |
| Slc6a1_P31648_S14s_1 | -1.93 | -0.64 | -0.62 | -1.37 | -0.76 | -1.33 | -0.97 | -1.15 | -1 | -1.43 | -1.17 | -0.46 | 0.37 | -1.11 | -1.12 | -1.4 | 0.0875 | 0.125 | 0.325 | 0.513132 | 0.808928 | 0.750631 | 1 | 0.999631 | 0.818518 |

| | | | | | | | | | | | | | | | | | | | | | | | | | |
|---|---|---|---|---|---|---|---|---|---|---|---|---|---|---|---|---|---|---|---|---|---|---|---|---|---|
| Slc6a1_P31648_S591s_1 | -2.14 | -0.39 | -0.3 | -2 | -1.7 | -0.83 | -1.28 | -0.88 | -2.28 | -1.92 | -1.29 | -1.52 | -1.18 | -0.67 | -2.47 | -0.46 | 0.035 | -0.545 | 0.0125 | 0.983659 | 0.944589 | 0.30297 | 1 | 0.999631 | 0.995133 |
| Slc6a1_P31648_S592s_1 | -1.47 | -0.48 | -0.27 | -0.68 | -0.44 | -0.92 | -0.52 | -0.54 | -0.46 | -0.19 | -0.45 | -0.14 | 1.14 | -1.31 | -0.05 | 0.22 | 0.12 | 0.415 | 0.725 | 0.195323 | 0.714188 | 0.22585 | 1 | 0.999631 | 0.652992 |
| Slc6a1_P31648_T15t_1 | -1.89 | -0.42 | 0.25 | -1.47 | -0.86 | -0.73 | -0.48 | -1.35 | -0.7 | -1 | -1.07 | 0.19 | 0.61 | -1.53 | -0.32 | -1.38 | 0.0275 | 0.2375 | 0.2275 | 0.719001 | 0.955277 | 0.655778 | 1 | 0.999631 | 0.904498 |
| Slc6a11_P31650_S21s_1 | -3.07 | -7.89 | 0.63 | 0.4 | -5.68 | -5.37 | 0.21 | 0.51 | -6.11 | -4.08 | 0.04 | 0.38 | -3.7 | -6.04 | 0.56 | 0.2 | -0.1 | 0.04 | 0.2375 | 0.910907 | 0.963828 | 0.985154 | 1 | 0.999631 | 0.971293 |
| Slc6a13_P31649_T587t_1 | -0.29 | 0.37 | 0.04 | 0.74 | 0.66 | -0.41 | -0.54 | -0.44 | -0.66 | 0.15 | -0.64 | 0.53 | -0.8 | 0.39 | 0.78 | 0.68 | -0.3975 | -0.37 | 0.0475 | 0.91252 | 0.302992 | 0.351681 | 1 | 0.999631 | 0.972191 |
| Slc6a15_Q8BG16_S55s_1 | -1.67 | 3.32 | 2.63 | 0.9 | 2.82 | 3.78 | 1.29 | 1.86 | 3.62 | -0.35 | 2.17 | 2.73 | 4.21 | 3.42 | 3.19 | 1.67 | 1.1425 | 0.7475 | 1.8275 | 0.106175 | 0.297217 | 0.537089 | 1 | 0.999631 | 0.568797 |
| Slc6a15_Q8BG16_S675s_1 | 0.21 | 0.63 | -0.27 | 0.45 | 0.79 | 0.89 | 0.29 | 0.19 | 0.59 | 0.09 | 0.54 | 0.25 | -0.58 | 1.3 | 0.33 | 0.09 | 0.285 | 0.1125 | 0.03 | 0.94551 | 0.375143 | 0.710839 | 1 | 0.999631 | 0.982217 |
| Slc6a15_Q8BG16_S686s_1 | 0.15 | -0.52 | 0.26 | -0.41 | 0.86 | -0.32 | -0.38 | -0.34 | -0.16 | -1.07 | -0.35 | -0.07 | -0.17 | 0.37 | 0.47 | 0.05 | 0.085 | -0.2825 | 0.31 | 0.350503 | 0.821082 | 0.418869 | 1 | 0.999631 | 0.737333 |
| Slc6a15_Q8BG16_S701s_1 | 0.58 | 0.25 | 0.17 | -0.23 | 0.29 | 0.34 | -0.42 | -0.55 | 0.07 | 0.91 | -0.13 | 0.26 | 1.19 | 1.09 | 0.43 | 0.14 | -0.2775 | 0.085 | 0.52 | 0.166795 | 0.407119 | 0.797816 | 1 | 0.999631 | 0.636447 |
| Slc6a17_Q8BJI1_S665s_1 | -2.22 | 2.4 | 0.86 | 0.61 | 2.94 | 2.13 | 1.49 | 1.96 | 2.36 | -1.49 | 2.33 | 1.36 | 1.18 | 1.5 | 1.59 | 0.55 | 1.7175 | 0.7275 | 0.7925 | 0.366179 | 0.07399 | 0.525945 | 1 | 0.999631 | 0.745796 |
| Slc6a17_Q8BJI1_S682s_1 | 0.53 | 0.19 | -0.76 | 1.48 | 1.42 | 0.43 | 0.58 | 0.92 | 0.64 | -0.16 | 0.52 | -0.43 | -1.62 | 0.58 | -0.17 | 0.38 | 0.4775 | -0.2175 | -0.5675 | 0.367082 | 0.336435 | 0.666522 | 1 | 0.999631 | 0.745796 |
| Slc6a17_Q8BJI1_S682s_2 | 0.48 | 0.03 | 0 | 1.65 | 1.47 | -0.4 | 1.61 | 1.52 | 0.55 | -0.84 | 1.25 | 0.32 | -2.44 | -0.77 | 0.14 | 0.68 | 0.51 | -0.22 | -1.1375 | 0.126316 | 0.377336 | 0.68514 | 1 | 0.999631 | 0.596745 |
| Slc6a17_Q8BJI1_S686s_1 | 1.48 | -0.22 | -0.63 | 1 | 1.34 | -0.11 | 0.88 | 0.58 | 0.35 | 0.51 | 0.47 | -0.18 | -0.1 | 0 | 0.15 | 0.11 | 0.265 | -0.12 | -0.3675 | 0.455125 | 0.624182 | 0.809216 | 1 | 0.999631 | 0.790041 |
| Slc6a17_Q8BJI1_S686s_2 | 0.48 | 0.03 | 0 | 1.65 | 1.47 | -0.4 | 1.61 | 1.52 | 0.55 | -0.84 | 1.25 | 0.32 | -2.44 | -0.77 | 0.14 | 0.68 | 0.51 | -0.22 | -1.1375 | 0.126316 | 0.377336 | 0.68514 | 1 | 0.999631 | 0.596745 |
| Slc6a17_Q8BJI1_S701s_1 | -1.28 | -0.44 | -0.84 | -1.55 | -0.29 | -0.46 | -0.97 | -0.72 | -0.35 | -1.27 | -1.01 | -0.85 | 0.13 | -0.93 | -1.08 | -1.18 | 0.4175 | 0.1575 | 0.2625 | 0.524939 | 0.22322 | 0.654931 | 1 | 0.999631 | 0.822727 |
| Slc6a17_Q8BJI1_S715s_1 | -0.33 | -0.05 | 0.25 | 0.4 | 0.48 | -0.34 | 0.39 | 0.18 | -0.78 | -1.15 | 0 | -0.6 | 0.84 | 0.34 | -1.07 | -0.35 | 0.11 | -0.7 | -0.1275 | 0.775601 | 0.718654 | 0.061063 | 1 | 0.999631 | 0.920998 |
| Slc6a17_Q8BJI1_T722t_1 | -1.99 | -0.22 | -1.37 | -0.65 | 0.41 | -0.63 | -0.14 | -0.65 | -0.15 | -2.08 | -0.3 | -0.04 | -0.91 | -0.47 | -0.5 | -1.25 | 0.805 | 0.415 | 0.275 | 0.533719 | 0.097983 | 0.473457 | 1 | 0.999631 | 0.828143 |
| Slc6a6_O35316_T596t_1 | 0.46 | -0.23 | 0.41 | 1.53 | -0.03 | 0.07 | 1.98 | 1.98 | -0.07 | -0.33 | 1.87 | 1.12 | -1.09 | 0.2 | 0.65 | 1.03 | 0.4575 | 0.105 | -0.345 | 0.533016 | 0.458059 | 0.856104 | 1 | 0.999631 | 0.827798 |
| Slc6a7_Q6PGE7_S598s_1 | -0.88 | -3.41 | -1.33 | -1.39 | -1.48 | -4.14 | -1.08 | -0.81 | -1.78 | -2.56 | -0.64 | -1.39 | -1.67 | -1.73 | -1.11 | -0.47 | -0.125 | 0.16 | 0.5075 | 0.393587 | 0.880069 | 0.798316 | 1 | 0.999631 | 0.759138 |
| Slc6a7_Q6PGE7_S600s_1 | -1.31 | -1.86 | -2.38 | -2.13 | -0.88 | -1.55 | -1.77 | -1.74 | -1.13 | -1.74 | -2.1 | -1.69 | -0.96 | -0.95 | -1.42 | -1.95 | 0.435 | 0.255 | 0.6 | 0.130034 | 0.223727 | 0.467742 | 1 | 0.999631 | 0.601275 |
| Slc6a7_Q6PGE7_S616s_1 | 0.48 | 0.09 | -2.38 | -1.57 | 1.13 | 0.54 | -1.13 | -1.79 | 0.28 | -0.24 | -1.12 | -0.72 | -0.42 | 0.47 | -1.17 | -1.98 | 0.5325 | 0.395 | 0.07 | 0.925759 | 0.530081 | 0.556316 | 1 | 0.999631 | 0.97588 |
| Slc6a8_F6UKB4_T561t_1 | -1.18 | 3.94 | -1.61 | -1.68 | 3.45 | 0.59 | -1.53 | -2.49 | -1.07 | -0.52 | -2.32 | -0.49 | 4.25 | 1.48 | -1.77 | -1.41 | 0.1375 | -0.9675 | 0.77 | 0.640369 | 0.93162 | 0.435989 | 1 | 0.999631 | 0.871024 |
| Slc7a10_E9Q1V6_S5s_1 | -0.91 | 0.07 | -1.04 | -1.39 | -1.09 | -1.54 | -1.33 | -1.65 | -1.43 | -1.04 | -1.87 | -1.35 | 0.42 | -1.71 | -1.44 | -1.18 | -0.585 | -0.605 | -0.16 | 0.765486 | 0.128495 | 0.134832 | 1 | 0.999631 | 0.916326 |
| Slc7a10_E9Q1V6_S9s_1 | -0.65 | -1.16 | -0.03 | -0.78 | -1.72 | -0.64 | -0.74 | -0.15 | -0.76 | -0.42 | -1.81 | -0.4 | -1.53 | -1.24 | -0.92 | -1.25 | -0.1575 | -0.1925 | -0.58 | 0.10767 | 0.698904 | 0.642657 | 1 | 0.999631 | 0.569416 |
| Slc7a14_Q8BXR1_S465s_1 | -0.97 | -0.8 | -0.82 | -1.18 | 0.21 | -1.04 | -1.08 | -0.02 | -0.31 | 0.03 | 0.18 | 0.26 | 0.64 | -0.35 | 0.74 | -0.71 | 0.46 | 0.9825 | 1.0225 | 0.025744 | 0.229898 | 0.004276 | 1 | 0.999631 | 0.422114 |
| Slc7a14_Q8BXR1_S691s_1 | -0.27 | 0.06 | 1.17 | 0.04 | -1.2 | 3.42 | 0.39 | 2.18 | -2.3 | -1.01 | 1.75 | -3.76 | -1.91 | 2.99 | -0.16 | -1.24 | 0.9475 | -1.58 | -0.33 | 0.733834 | 0.315263 | 0.153856 | 1 | 0.999631 | 0.907462 |
| Slc7a2_E9QJY0_S662s_1 | -3.48 | 1.38 | 0.82 | 0.61 | 1.37 | 0.72 | 0.41 | -0.91 | 0.78 | -0.1 | 0.92 | 1.68 | 0.72 | 0.56 | 1.26 | 1.52 | 0.565 | 0.9875 | 1.1825 | 0.243007 | 0.590667 | 0.34205 | 1 | 0.999631 | 0.685637 |
| Slc7a4_Q8BLQ7_S427s_1 | -1.05 | -1.85 | -0.43 | -0.77 | -1.19 | -2.07 | -0.72 | -1.72 | -1.23 | -0.79 | -0.61 | -1.01 | -0.13 | 0.32 | -0.43 | -0.16 | -0.4 | 0.115 | 0.925 | 0.032082 | 0.352323 | 0.7518 | 1 | 0.999631 | 0.442395 |
| Slc7a4_Q8BLQ7_S626s_1 | -0.95 | -0.91 | -0.43 | -0.78 | -0.75 | -0.97 | -0.57 | -0.54 | -1.19 | -0.77 | -0.19 | -0.4 | -0.27 | -1.2 | -0.04 | -0.04 | 0.06 | 0.13 | 0.38 | 0.297915 | 0.81747 | 0.680002 | 1 | 0.999631 | 0.711242 |
| Slc7a6_Q8BGK6_S30s_1 | -1.36 | -0.54 | -1.26 | -0.55 | -1.79 | -0.72 | -1.27 | -0.39 | -1.28 | -0.89 | -4.09 | -1.89 | 0.07 | -0.7 | -1.07 | -0.7 | -0.115 | -1.11 | 0.3275 | 0.385401 | 0.767873 | 0.122339 | 1 | 0.999631 | 0.756021 |
| Slc7a6os_Q7TPE5_S256s_1 | 0.07 | 1.22 | 2.64 | 1.21 | 0.09 | 2.36 | 1.35 | -0.06 | 1.91 | 2.12 | 1.43 | 1.75 | 0 | 0.88 | 1.1 | 0.08 | -0.35 | 0.5175 | -0.77 | 0.185115 | 0.613285 | 0.32723 | 1 | 0.999631 | 0.646072 |
| Slc7a6os_Q7TPE5_S257s_1 | 0.07 | 1.22 | 2.64 | 1.21 | 0.09 | 2.36 | 1.35 | -0.06 | 1.91 | 2.12 | 1.43 | 1.75 | 0 | 0.88 | 1.1 | 0.08 | -0.35 | 0.5175 | -0.77 | 0.185115 | 0.613285 | 0.32723 | 1 | 0.999631 | 0.646072 |
| Slc7a8_Q9QXW9_S21s_1 | -0.97 | 0.73 | 0.5 | -0.52 | 1.2 | -0.05 | 1.39 | -0.29 | 1.98 | -1.28 | 1.48 | 2.35 | 3.38 | -0.54 | 0.3 | -0.18 | 0.6275 | 1.1975 | 0.805 | 0.356162 | 0.264293 | 0.160377 | 1 | 0.999631 | 0.74034 |
| Slc7a8_Q9QXW9_S21s_2 | -1.43 | 0.58 | -2 | -1.07 | -0.82 | 0.2 | 0.78 | -1.22 | 1.1 | -2.15 | 1.23 | 1.08 | 0.88 | -1.04 | 0.7 | -0.23 | 0.715 | 1.295 | 1.0575 | 0.122325 | 0.282207 | 0.157617 | 1 | 0.999631 | 0.590804 |
| Slc7a8_Q9QXW9_S527s_1 | 1.31 | 2.68 | 2.55 | 0.94 | 3.12 | 1.98 | 2.61 | 2.27 | 2.81 | 0.26 | 3.04 | 2.2 | 2.2 | 0.31 | 1.94 | 0.23 | 0.625 | 0.2075 | -0.7 | 0.272887 | 0.209669 | 0.762562 | 1 | 0.999631 | 0.700187 |
| Slc8a1_Q68FL0_S282s_1 | -0.25 | 0.69 | 0.96 | 0.54 | 0.23 | 0.87 | -0.02 | 0.58 | 0.64 | 0.95 | 0.61 | 1.28 | 1.5 | 0.93 | 1.31 | 1.34 | -0.07 | 0.385 | 0.785 | 0.043814 | 0.84278 | 0.280987 | 1 | 0.999631 | 0.475443 |
| Slc8a2_Q8K596_S266s_1 | -0.7 | -0.01 | 0.07 | -0.74 | -0.88 | 0.56 | -0.77 | 0.04 | 0.39 | 0.53 | -0.32 | -0.48 | -1.06 | -0.37 | -0.13 | -0.32 | 0.0825 | 0.375 | -0.125 | 0.723784 | 0.839358 | 0.316334 | 1 | 0.999631 | 0.905674 |
| Slc8a2_Q8K596_S394s_1 | 0.15 | 0.14 | 1.48 | 1.19 | -0.85 | 1.51 | 0.72 | 1.57 | 0.29 | 1.04 | 0.85 | 0.46 | 0.47 | -0.88 | 1.31 | 0.65 | -0.0025 | -0.08 | -0.3525 | 0.518049 | 0.996666 | 0.84188 | 1 | 0.999631 | 0.819449 |
| Slc8a2_Q8K596_S622s_1 | -0.26 | -0.22 | -0.6 | -0.13 | 0.07 | -0.41 | -0.5 | 0.12 | -0.23 | -0.31 | 0.06 | 0.41 | -0.14 | 0.04 | -0.07 | 0.67 | 0.1225 | 0.285 | 0.4275 | 0.186309 | 0.658619 | 0.329697 | 1 | 0.999631 | 0.646474 |
| Slc8a3_E9PX50_S629s_1 | 1.28 | -0.05 | -0.3 | 0.29 | 1.98 | -0.92 | -0.15 | -0.24 | 1.38 | 1.05 | -0.78 | -0.38 | -0.46 | -0.76 | 0.16 | 0.22 | -0.1375 | 0.0125 | -0.515 | 0.248184 | 0.830414 | 0.982783 | 1 | 0.999631 | 0.685751 |
| Slc9a1_Q61165_S697s_1 | -0.55 | -0.64 | -0.43 | -0.83 | -0.87 | -0.78 | -1.06 | -0.72 | -0.85 | -0.33 | -0.96 | -0.75 | -0.09 | -0.54 | -0.79 | -0.65 | -0.245 | -0.11 | 0.095 | 0.744847 | 0.329002 | 0.685633 | 1 | 0.999631 | 0.910583 |
| Slc9a1_Q61165_S790s_1 | -0.83 | 0.62 | 0.84 | 0.54 | 0.22 | 0.51 | -0.38 | 0.29 | -0.17 | -0.81 | 0.25 | 0.67 | 0.55 | 0.03 | 0.19 | 0.46 | -0.1325 | -0.3075 | 0.015 | 0.971101 | 0.753015 | 0.522251 | 1 | 0.999631 | 0.989676 |
| Slc9a3r1_P70441_S117s_1 | 0.26 | 0.37 | 0.74 | 0.77 | -0.42 | 0.23 | 0.15 | -0.45 | -0.46 | 0.66 | 0.65 | 0.86 | 0.72 | 0.55 | 0.87 | 1.33 | -0.6575 | -0.1075 | 0.3325 | 0.294643 | 0.045546 | 0.765477 | 1 | 0.999631 | 0.708556 |
| Slc9a3r1_P70441_S273s_1 | 0.13 | 0.69 | -0.17 | -0.65 | 0.12 | -0.41 | -0.8 | -0.84 | 0.29 | 0.11 | -0.35 | 0.24 | -0.78 | 0.47 | -0.78 | 0.25 | -0.4825 | 0.0725 | -0.21 | 0.634142 | 0.217418 | 0.837488 | 1 | 0.999631 | 0.868569 |
| Slc9a3r1_P70441_S275s_1 | -0.49 | -0.76 | -0.38 | -0.8 | -1.11 | -1.48 | -1.1 | -1.48 | -0.97 | -0.37 | -0.71 | -0.14 | 0.18 | -0.78 | -0.81 | 0.08 | -0.685 | 0.06 | 0.275 | 0.435092 | 0.02278 | 0.837771 | 1 | 0.999631 | 0.779781 |
| Slc9a3r1_P70441_S285s_2 | -1.31 | -1.37 | -0.8 | -0.57 | -0.94 | -2.27 | -0.82 | -0.63 | -1.16 | -0.94 | -0.71 | -1.45 | -1.26 | -1.75 | -0.29 | -0.28 | -0.1525 | -0.0525 | 0.1175 | 0.782714 | 0.716198 | 0.867519 | 1 | 0.999631 | 0.922379 |
| Slc9a3r1_P70441_S286s_1 | -0.49 | 0.01 | -0.35 | 0.1 | -0.61 | -0.07 | 0.2 | -0.88 | -0.04 | -0.37 | -0.86 | -0.33 | 0.92 | -0.05 | -1.01 | 0.78 | -0.1575 | -0.2175 | 0.3425 | 0.46381 | 0.632269 | 0.472691 | 1 | 0.999631 | 0.793747 |
| Slc9a3r1_P70441_S289s_2 | -2.02 | -0.51 | 0.27 | -0.79 | -0.52 | -0.11 | -0.9 | -1.03 | -2.53 | -1.13 | -0.46 | 0.06 | -0.03 | -0.88 | 0.19 | -0.47 | 0.1225 | -0.2525 | 0.465 | 0.369276 | 0.802328 | 0.702828 | 1 | 0.999631 | 0.746224 |
| Slc9a3r1_P70441_S297s_1 | -1.07 | 0.3 | -0.02 | -1.67 | -0.03 | 0.21 | -0.24 | -0.82 | 0.27 | 0.81 | -0.68 | 0.25 | 0.84 | -0.33 | -0.49 | -0.13 | 0.395 | 0.7775 | 0.5875 | 0.271998 | 0.419749 | 0.157395 | 1 | 0.999631 | 0.699658 |
| Slc9a3r1_P70441_S297s_2 | -1.44 | -1.23 | -0.8 | -0.57 | -0.75 | -1.7 | -0.82 | -0.63 | -0.98 | -1 | -0.71 | -1.45 | -0.25 | -1.47 | -0.29 | -0.28 | 0.035 | -0.025 | 0.4375 | 0.276821 | 0.919461 | 0.936591 | 1 | 0.999631 | 0.700187 |
| Slc9a3r1_P70441_S2s_1 | 0.23 | 0.57 | -0.53 | -0.41 | 0.43 | -0.89 | 0.53 | 1.15 | 0.18 | 0.59 | 0.09 | -0.53 | 0.65 | -1.66 | 0.27 | -0.26 | 0.34 | 0.1175 | -0.215 | 0.688345 | 0.481878 | 0.754152 | 1 | 0.999631 | 0.891814 |
| Slc9a3r2_Q9JHL1_S192s_1 | -1.49 | -0.01 | 2.02 | -0.19 | 0.19 | -0.08 | 0.81 | 1.09 | 0.29 | -0.6 | 1.44 | 2.15 | 0.27 | 0.51 | 2.5 | 1.48 | 0.42 | 0.7375 | 1.1075 | 0.174842 | 0.544446 | 0.383554 | 1 | 0.999631 | 0.642116 |
| Slc9a6_B0QZV3_S668s_1 | -0.94 | -0.71 | -0.75 | -0.61 | -0.46 | -0.53 | -0.84 | -0.51 | -0.61 | -0.46 | -1.14 | -0.62 | -0.05 | -0.09 | -0.54 | -0.34 | 0.1675 | 0.045 | 0.4975 | 0.096523 | 0.497433 | 0.868479 | 1 | 0.999631 | 0.55886 |
| Slc9a6_B0QZV3_T665t_1 | -0.59 | -0.69 | -0.53 | -0.74 | -0.5 | -0.83 | -0.65 | -0.57 | -0.88 | -0.6 | -1.55 | -0.43 | 0.01 | -0.35 | -0.32 | -0.69 | 5.89E-17 | -0.2275 | 0.3 | 0.302588 | 1 | 0.475892 | 1 | 0.999631 | 0.714978 |
| Slc9a7_Q8BLV3_S696s_1 | 0.16 | 0.66 | -0.11 | 0.14 | 0.3 | 0.5 | 0.04 | 0.25 | 0.7 | -0.18 | 0.15 | 0.35 | -0.32 | 0.51 | 0.22 | 0.23 | 0.06 | 0.0425 | -0.0525 | 0.869742 | 0.826962 | 0.891388 | 1 | 0.999631 | 0.958159 |
| Slc9b2_Q5BKR2_S62s_1 | -0.44 | -0.2 | 0.16 | -1.2 | -0.62 | -1 | -0.64 | -1.02 | -0.38 | -0.42 | -0.01 | -0.68 | 0.51 | 0.04 | 0.52 | -0.02 | -0.4 | 0.0475 | 0.6825 | 0.087306 | 0.259207 | 0.893573 | 1 | 0.999631 | 0.553616 |
| Slco2b1_D3YY31_S35s_1 | 0.52 | 1.4 | 1 | -0.18 | 0.37 | 0.62 | 0.25 | -0.21 | 1.23 | 1.48 | 0.09 | 1.11 | 2.09 | 1.53 | 0.85 | 1 | -0.4275 | 0.2925 | 0.6825 | 0.146814 | 0.288733 | 0.519269 | 1 | 0.999631 | 0.618703 |
| Slitrk5_Q810B7_S829s_1 | 0.51 | 0.11 | -0.39 | 0.19 | 0.61 | -0.19 | -0.09 | 0.18 | 0.53 | 0.05 | -0.11 | -0.15 | -0.51 | 0.43 | 0.5 | 0.4 | 0.0225 | -0.025 | 0.1 | 0.779072 | 0.942464 | 0.935749 | 1 | 0.999631 | 0.921416 |
| Slk_O54988_S189s_1 | -0.69 | 0.53 | 0.15 | 0.19 | 0.86 | -0.08 | -0.79 | -0.09 | 0.62 | 0.37 | -0.51 | 0.51 | 0.91 | 1.22 | 0.02 | -0.01 | -0.07 | 0.2025 | 0.49 | 0.259681 | 0.868137 | 0.601617 | 1 | 0.999631 | 0.692624 |
| Slmap_F8WIH0_S27s_1 | -0.4 | -0.3 | 0.12 | 0.63 | -0.52 | -0.34 | -0.27 | -0.61 | -0.7 | -0.57 | 0.27 | 0.28 | -0.47 | -0.48 | -0.43 | -0.29 | -0.4475 | -0.1925 | -0.43 | 0.200634 | 0.166132 | 0.611982 | 1 | 0.999631 | 0.658263 |
| Slmap_F8WIH0_S314s_1 | -2.65 | -0.76 | 0.01 | -0.81 | -1.76 | -0.48 | -0.72 | -0.11 | -0.99 | -2.09 | -0.64 | -0.8 | -0.74 | -1.67 | 0.07 | -0.51 | 0.285 | -0.0775 | 0.34 | 0.579727 | 0.63761 | 0.896424 | 1 | 0.999631 | 0.84667 |
| Sltm_Q8CH25_S199s_1 | 0.45 | 1.28 | 1.7 | 1.43 | 0.85 | 0.86 | 1.05 | 1.55 | 0.75 | 1.07 | 1.93 | 2.13 | 2.13 | 1.98 | 1.47 | 2.25 | -0.1375 | 0.255 | 0.7425 | 0.065155 | 0.692682 | 0.55495 | 1 | 0.999631 | 0.515898 |
| Sltm_Q8CH25_S534s_1 | -0.16 | 0.9 | 0.55 | 0.03 | 0.32 | 0.68 | -0.2 | 0.42 | 0.55 | 0.42 | 0.45 | 0.65 | 0.98 | 1.12 | 0.76 | 0.76 | -0.025 | 0.1875 | 0.575 | 0.106398 | 0.941304 | 0.552704 | 1 | 0.999631 | 0.568797 |
| Sltm_Q8CH25_S794s_1 | 0.91 | -0.03 | 0.39 | 0.84 | 0.93 | 1 | 1.9 | 0.78 | 1.14 | 0.07 | 0.53 | -0.35 | -0.51 | 0.31 | -0.69 | 0.71 | 0.625 | -0.18 | -0.5725 | 0.188139 | 0.106473 | 0.654039 | 1 | 0.999631 | 0.647379 |
| Sltm_Q8CH25_S902s_1 | 1.56 | -0.66 | 0.74 | 0.66 | -0.73 | 0.01 | 1.49 | -0.78 | -1.09 | 1.35 | -0.46 | -0.71 | -0.88 | -1.44 | -0.48 | -0.32 | -0.5775 | -0.8025 | -1.355 | 0.020658 | 0.369059 | 0.229078 | 1 | 0.999631 | 0.397234 |
| Sltm_Q8CH25_S981s_1 | -0.99 | -2.69 | -1.42 | -1.37 | -1.96 | -3.75 | -0.94 | -2.33 | -1.75 | -1.32 | -1.69 | -1.25 | -0.3 | -2.14 | -1.68 | -1.56 | -0.6275 | 0.115 | 0.1975 | 0.70119 | 0.324463 | 0.77544 | 1 | 0.999631 | 0.896835 |
| Slx4_Q6P1D7_S909s_1 | 0.98 | -0.54 | -0.76 | -0.63 | -0.38 | -0.37 | -0.15 | -0.75 | 0.43 | 0.17 | -1.39 | -0.41 | -0.21 | -1.33 | -1.29 | -1.95 | -0.175 | -0.0625 | -0.9575 | 0.088117 | 0.679517 | 0.906998 | 1 | 0.999631 | 0.553858 |
| Smad1_Q8C3Y6_T322t_1 | -0.16 | 0.28 | 2.35 | 1.29 | -0.37 | 1.01 | 1.57 | 1.02 | -0.07 | 0.61 | 2.07 | 0.4 | -0.55 | -0.06 | 1.62 | 0.34 | -0.1325 | -0.1875 | -0.6025 | 0.366291 | 0.832107 | 0.773997 | 1 | 0.999631 | 0.745796 |
| Smad2_E9PXA8_T8t_1 | 0.31 | 0.68 | 0.44 | 0.11 | 0.03 | -0.16 | -0.22 | -0.35 | 0.33 | -0.67 | -0.04 | 0.47 | 0.41 | 0.44 | 0.41 | 0.21 | -0.56 | -0.3625 | -0.0175 | 0.949379 | 0.05016 | 0.2903 | 1 | 0.999631 | 0.983795 |
| Smad3_Q8BUN5_T8t_1 | -0.11 | 0.88 | 0.51 | 0.08 | 0.3 | 0.03 | -0.23 | 0.16 | 0.07 | 0.45 | 0.19 | 0.69 | 1.29 | 1.05 | 0.26 | 0.67 | -0.275 | 0.01 | 0.4775 | 0.208781 | 0.380029 | 0.975097 | 1 | 0.999631 | 0.662575 |
| Smap_Q9R0P4_S145s_1 | -0.92 | 3.17 | 3.14 | 1.29 | 0.17 | 3.38 | 1 | 0.67 | 2.38 | 2.18 | 1.24 | 1.22 | 1.32 | 2.84 | 2.34 | 2.47 | -0.365 | 0.085 | 0.5725 | 0.520871 | 0.724454 | 0.923378 | 1 | 0.999631 | 0.82015 |
| Smap2_Q7TN29_S219s_1 | -0.53 | 0.13 | -0.04 | 0.3 | 1.31 | 0.16 | 0.38 | 0.54 | 0.38 | -0.48 | 0.5 | 0.1 | 0.17 | 0.08 | 0.38 | 0.1 | 0.6325 | 0.16 | 0.2175 | 0.472987 | 0.088802 | 0.632596 | 1 | 0.999631 | 0.79925 |
| Smap2_Q7TN29_S239s_1 | -0.31 | 0.12 | 0.1 | -0.06 | 0.23 | -0.07 | -0.45 | 0.31 | -0.15 | -0.23 | -0.19 | 0.35 | 0.02 | 0.32 | 0.19 | 0.2 | 0.0425 | -0.0175 | 0.22 | 0.425874 | 0.879388 | 0.948972 | 1 | 0.999631 | 0.77466 |
| Smarca2_Q6DIC0_S1382s_1 | 0.67 | 1.36 | 1.77 | 1.46 | 0.58 | 1.05 | 0.99 | 1.75 | 1.02 | 1.26 | 1.71 | 1.67 | 0.82 | 1.6 | 1.89 | 2.18 | -0.2225 | 0.1 | 0.3075 | 0.44736 | 0.540257 | 0.765451 | 1 | 0.999631 | 0.785754 |
| Smarca2_Q6DIC0_S166s_1 | -0.87 | 0.94 | -1.24 | -0.45 | 0.33 | 0.5 | 0.5 | 0.51 | 0.72 | -0.69 | 1.7 | 2.01 | 0.8 | 0.64 | 1.16 | 0.45 | 0.865 | 1.34 | 1.1675 | 0.034263 | 0.084455 | 0.076287 | 1 | 0.999631 | 0.446273 |
| Smarca2_Q6DIC0_S321s_1 | -0.53 | 0.61 | 0.89 | 0.3 | -0.14 | 0.88 | -0.3 | -0.04 | 0.71 | 0.77 | -1.07 | 0.11 | 0.75 | 1.61 | 0.27 | 0.1 | -0.2175 | -0.1875 | 0.365 | 0.429242 | 0.595873 | 0.708033 | 1 | 0.999631 | 0.776658 |
| Smarca4_Q3TKT4_T1390t_1 | 0.19 | 1.42 | 4.24 | 2.11 | 0.38 | 1.49 | 2.39 | 1.88 | 1.67 | 1.18 | 2.5 | 2.3 | 0.55 | 0.39 | 2.73 | 0.92 | -0.455 | -0.0775 | -0.8425 | 0.344164 | 0.585175 | 0.921926 | 1 | 0.999631 | 0.733869 |
| Smarcc1_Q3UZD0_S309s_1 | 0.45 | 1.07 | 1.35 | 1.01 | 0.44 | 0.84 | 0.86 | 0.87 | 0.91 | 1.33 | 1.06 | 1.32 | 0.56 | 0.76 | 1.15 | 1.26 | -0.2175 | 0.185 | -0.0375 | 0.908596 | 0.457721 | 0.53534 | 1 | 0.999631 | 0.969941 |
| Smarcc1_Q3UZD0_S327s_1 | 0.08 | 0.52 | 1.81 | 0.76 | -0.2 | 0.63 | 0.78 | 1.23 | 0.39 | 0.49 | 0.81 | 0.88 | 0.2 | 0.35 | 2.14 | 1.32 | -0.1825 | -0.15 | 0.21 | 0.700538 | 0.689757 | 0.707863 | 1 | 0.999631 | 0.896835 |
| Smarcc1_Q3UZD0_S327s_2 | 0.53 | 0.52 | 0.3 | 0.92 | -0.47 | 0.56 | 0.3 | 0.81 | -0.32 | 0.25 | 1.28 | 1.25 | 0.8 | 1.11 | 0.62 | 0.57 | -0.2675 | 0.0475 | 0.2075 | 0.484505 | 0.439492 | 0.909457 | 1 | 0.999631 | 0.803422 |
| Smarcc1_Q3UZD0_S329s_2 | 0.53 | 0.52 | 0.3 | 0.92 | -0.47 | 0.56 | 0.3 | 0.81 | -0.32 | 0.25 | 1.28 | 1.25 | 0.8 | 1.11 | 0.62 | 0.57 | -0.2675 | 0.0475 | 0.2075 | 0.484505 | 0.439492 | 0.909457 | 1 | 0.999631 | 0.803422 |
| Smarcc2_Q3UID0_S283s_1 | 0.27 | 1.37 | 1.07 | 1.26 | 0.63 | 0.51 | 0.6 | 0.79 | 0.82 | 0.83 | 0.86 | 1.83 | 1.15 | 1.44 | 1.15 | 1.04 | -0.36 | 0.0925 | 0.2025 | 0.54765 | 0.262955 | 0.805799 | 1 | 0.999631 | 0.834277 |
| Smarcc2_Q3UID0_S302s_1 | 0.15 | 0.94 | 0.86 | 0.24 | -0.4 | 0.21 | 0.07 | 0.45 | -0.07 | 0.81 | 0.27 | 0.94 | 0.75 | 0.07 | 0.57 | -0.11 | -0.465 | -0.06 | -0.2275 | 0.517426 | 0.168862 | 0.864266 | 1 | 0.999631 | 0.819449 |
| Smarcc2_Q3UID0_S302s_2 | 0.3 | 1.35 | 1.57 | 1.07 | 0.78 | 1.04 | 0.56 | 1.01 | 1.41 | 1.37 | 0.66 | 0.83 | 0.83 | 0.88 | 1.24 | 0.69 | -0.225 | -0.005 | -0.1625 | 0.648407 | 0.508444 | 0.989082 | 1 | 0.999631 | 0.874615 |
| Smarcc2_Q3UID0_S304s_2 | 0.42 | 1.06 | 1.67 | 1.14 | 0.71 | 0.96 | 0.56 | 1.12 | 1.62 | 1.41 | 0.69 | 0.92 | 0.69 | 0.76 | 1.3 | 0.74 | -0.235 | 0.0875 | -0.2 | 0.571334 | 0.479509 | 0.810547 | 1 | 0.999631 | 0.843666 |
| Smarcc2_Q3UID0_S306s_2 | -0.16 | 2.22 | 0.54 | 0.45 | 0.99 | 1.29 | 0.56 | -0.16 | 0.58 | 1.21 | 0.45 | -0.15 | 1.28 | 1.27 | 0.67 | 0.29 | -0.0925 | -0.24 | 0.115 | 0.828504 | 0.866422 | 0.65871 | 1 | 0.999631 | 0.941315 |
| Smc1a_Q9CU62_S956s_1 | -0.09 | -0.22 | 0.16 | 2.34 | -0.98 | -1.01 | -0.45 | 0.68 | 0.73 | -1.16 | 1.54 | 2.06 | 0.86 | 0.7 | 0.71 | 1.04 | -0.9875 | 0.245 | 0.28 | 0.620687 | 0.149002 | 0.763682 | 1 | 0.999631 | 0.861888 |
| Smc3_Q9CW03_S1067s_1 | 0.85 | 0.24 | 1.4 | 0.96 | -0.28 | 1.3 | 1.32 | 0.81 | 1.96 | 2.29 | 0.7 | 0.59 | 0.86 | -1.06 | 0.52 | -1.51 | -0.075 | 0.5225 | -1.16 | 0.068173 | 0.862998 | 0.287946 | 1 | 0.999631 | 0.522688 |
| Smchd1_Q6P5D8_S1975s_1 | -0.27 | 0.35 | 1.33 | 0.7 | -0.68 | 0.13 | 0.98 | 0.62 | 0.29 | -0.39 | 0.87 | 0.68 | -0.21 | -0.62 | 1.37 | -0.12 | -0.265 | -0.165 | -0.4225 | 0.423856 | 0.574616 | 0.704124 | 1 | 0.999631 | 0.773835 |
| Smcr8_Q3UMB5_S393s_1 | 0.56 | 0.41 | -0.34 | -1.34 | 0.76 | 0.3 | 0.07 | -1.18 | 0.62 | -1.08 | -0.51 | 0.24 | 1.33 | -0.91 | -0.34 | -0.99 | 0.165 | -0.005 | -0.05 | 0.93626 | 0.765011 | 0.992576 | 1 | 0.999631 | 0.97939 |
| Smcr8_Q3UMB5_S788s_1 | -1.29 | -0.92 | -0.4 | -0.99 | -1.14 | -0.77 | -0.74 | -0.63 | -1.53 | -0.84 | -0.04 | -0.39 | -0.43 | -0.72 | -0.02 | -0.42 | 0.08 | 0.2 | 0.5025 | 0.138382 | 0.781921 | 0.609272 | 1 | 0.999631 | 0.611124 |
| Smek1_E9Q481_S780s_1 | -0.11 | 0.77 | 0.6 | -0.13 | 0.69 | 0.78 | 0.15 | 0.31 | -0.36 | 0.98 | -0.35 | 0.54 | 0.43 | 1.16 | 1.13 | 1.01 | 0.2 | -0.08 | 0.65 | 0.086323 | 0.541344 | 0.846946 | 1 | 0.999631 | 0.55299 |
| Smg1_Q8BKX6_S3553s_1 | 0.92 | 0.57 | 0.66 | 0.45 | 1.64 | -0.26 | 0.57 | 0.02 | 0.89 | 1.03 | 0.9 | -0.09 | -0.13 | 0.18 | 0.37 | 1.42 | -0.1575 | 0.0325 | -0.19 | 0.623099 | 0.711811 | 0.921543 | 1 | 0.999631 | 0.862913 |
| Smg1_Q8BKX6_S3585s_1 | 1.29 | 1.39 | 1.66 | 1.41 | 0.32 | 1.47 | 1.54 | 1.07 | 1.41 | 1.42 | 0.7 | 0.7 | 0.84 | 1.19 | 0.66 | 0.39 | -0.3375 | -0.38 | -0.6675 | 0.044715 | 0.321621 | 0.222546 | 1 | 0.999631 | 0.475443 |
| Smg6_P61406_S331s_1 | -0.46 | 1.08 | 0.9 | 0.73 | 0.64 | 1.08 | 0.76 | 1.06 | 1.66 | 0.37 | 1.55 | 0.57 | -0.48 | 0.73 | 0.35 | 0.03 | 0.3225 | 0.475 | -0.405 | 0.364994 | 0.403972 | 0.321213 | 1 | 0.999631 | 0.745324 |
| Smg9_Q9DB90_S32s_1 | 0.27 | 0.56 | 1.03 | 1.09 | 1.3 | 0.36 | 1.11 | 1.39 | 0.7 | 0.35 | 1.53 | 1.82 | -0.24 | 1.05 | 1.78 | 2.01 | 0.3025 | 0.3625 | 0.4125 | 0.430806 | 0.383894 | 0.382166 | 1 | 0.999631 | 0.778235 |
| Smim13_E9Q942_S70s_1 | -1.32 | -0.15 | -0.77 | -0.79 | -0.39 | -1.1 | -0.94 | -0.33 | -0.73 | -0.21 | -0.25 | -1.12 | 0.23 | 0.11 | 0.34 | -0.41 | 0.0675 | 0.18 | 0.825 | 0.037038 | 0.843508 | 0.616553 | 1 | 0.999631 | 0.456881 |
| Smpd3_Q9JJY3_S173s_1 | -0.69 | -0.2 | -0.95 | -0.24 | -0.45 | -0.84 | -0.68 | 0.06 | -0.69 | -0.63 | -0.77 | -0.07 | -0.06 | -0.46 | -1.1 | -0.06 | 0.0425 | -0.02 | 0.1 | 0.779765 | 0.893376 | 0.948425 | 1 | 0.999631 | 0.921416 |
| Smpd3_Q9JJY3_S202s_1 | 1.53 | -0.25 | 0.46 | 1.08 | 0.26 | -0.01 | 1.23 | 0.89 | 0.28 | 0.11 | 1.06 | 0.11 | -0.52 | -0.17 | 0.49 | 1.22 | -0.1125 | -0.315 | -0.45 | 0.392732 | 0.807031 | 0.48243 | 1 | 0.999631 | 0.758901 |
| Smpd3_Q9JJY3_S208s_1 | 1.54 | -0.91 | -1.01 | 0.63 | 0.59 | -0.39 | 0.48 | -0.03 | 0.07 | -0.07 | -0.2 | -0.64 | -1.7 | -1.06 | -0.76 | -0.6 | 0.1 | -0.2725 | -1.0925 | 0.095408 | 0.866914 | 0.642592 | 1 | 0.999631 | 0.558128 |
| Smpd3_Q9JJY3_S289s_2 | -0.65 | 1.98 | 1.43 | 1.1 | 3.61 | 1.89 | 2.04 | 2.75 | 3.11 | -0.87 | 2.55 | 2.02 | 0.26 | 0.25 | 2.08 | -0.06 | 1.6075 | 0.7375 | -0.3325 | 0.621211 | 0.025805 | 0.42777 | 1 | 0.999631 | 0.861888 |
| Smpd3_Q9JJY3_S292s_1 | -1.33 | 1.52 | 0.66 | 0.55 | 2.17 | 1.61 | 0.88 | 2.14 | 2.41 | -1.13 | 2.22 | 1.5 | 0.19 | 1.29 | 1.63 | 0.04 | 1.35 | 0.9 | 0.4375 | 0.50329 | 0.047578 | 0.321487 | 1 | 0.999631 | 0.814092 |
| Smpd3_Q9JJY3_S292s_2 | -0.65 | 1.98 | 1.65 | 1.27 | 3.61 | 1.89 | 2.14 | 3.21 | 3.11 | -0.87 | 3.12 | 2.38 | 0.26 | 0.25 | 2.42 | 0.18 | 1.65 | 0.8725 | -0.285 | 0.689789 | 0.026927 | 0.377712 | 1 | 0.999631 | 0.892781 |
| Smtn_D3Z3Q3_S357s_1 | 0.26 | -0.39 | -1.15 | -0.21 | 3.72 | 0.61 | 0 | -1.22 | 1.26 | 4.6 | 2.14 | 4.89 | -0.44 | -0.45 | -1.47 | -0.3 | 1.15 | 3.595 | -0.2925 | 0.486453 | 0.240782 | 0.001551 | 1 | 0.999631 | 0.804724 |
| Snap23_B0R030_S110s_1 | 0.11 | 0.35 | 0.04 | -0.3 | 0.06 | -0.18 | -1.02 | -0.61 | -0.3 | 0.7 | -0.87 | -0.33 | -0.25 | 1.19 | -0.39 | 0.11 | -0.4875 | -0.25 | 0.115 | 0.776192 | 0.152284 | 0.512558 | 1 | 0.999631 | 0.921058 |
| Snap25_P60879_S115s_1 | 0.7 | -0.53 | -0.42 | -0.5 | -0.71 | 0.21 | -0.58 | -0.41 | -0.53 | -0.67 | -0.12 | -0.14 | -0.63 | -0.2 | -0.5 | -0.36 | -0.185 | -0.1775 | -0.235 | 0.518885 | 0.624411 | 0.624564 | 1 | 0.999631 | 0.819456 |
| Snap25_P60879_S130s_1 | 0.49 | 1.32 | -0.18 | -0.03 | 2.02 | 0.98 | -0.15 | 0.19 | 1.56 | 0.38 | -0.11 | -0.21 | 0.93 | 1.15 | 0.03 | 0.12 | 0.36 | 0.005 | 0.1575 | 0.722998 | 0.511613 | 0.992033 | 1 | 0.999631 | 0.905674 |
| Snap25_P60879_S25s_1 | -0.19 | 0.84 | 0.2 | 0.43 | 0.15 | 0.36 | -0.04 | 0.15 | 0.21 | 0.23 | 0.41 | 0.17 | 0.62 | 0.24 | 0.44 | 1.05 | -0.165 | -0.065 | 0.2675 | 0.439683 | 0.58081 | 0.828038 | 1 | 0.999631 | 0.781983 |
| Snap25_P60879_S28s_1 | 0.3 | 0.32 | 0.35 | 2.28 | 1.02 | 0.44 | 1.74 | 2.22 | 0.44 | -0.4 | 2.25 | 1.17 | -0.69 | 0.39 | 1.96 | 1.2 | 0.5425 | 0.0525 | -0.0975 | 0.884104 | 0.353968 | 0.937279 | 1 | 0.999631 | 0.963505 |
| Snap29_Q9ERB0_S153s_1 | -1.72 | -4.48 | 0.44 | 1.23 | -3 | -3.71 | 1.28 | -0.08 | -3.76 | -1.92 | 0.2 | 0.06 | -3.21 | -2.9 | 0.09 | -0.62 | -0.245 | -0.2225 | -0.5275 | 0.681834 | 0.868289 | 0.869157 | 1 | 0.999631 | 0.88924 |
| Snap29_Q9ERB0_S163s_1 | 1.26 | 0.78 | 1.26 | -0.24 | 0.89 | 0.23 | 0.75 | 0.63 | 0.61 | 0.59 | 0.43 | 1.06 | 1.49 | 0.24 | 0.32 | 0.75 | -0.14 | -0.0925 | -0.065 | 0.885766 | 0.720273 | 0.814452 | 1 | 0.999631 | 0.963505 |
| Snap91_E9Q9A3_S16s_1 | 0.46 | -0.54 | 0.53 | 0.41 | 0.79 | 1.41 | 0.38 | 1.8 | 1.02 | -0.2 | 1.7 | 1.43 | 0.53 | 0.95 | 2.73 | 0.77 | 0.88 | 0.7725 | 1.03 | 0.075407 | 0.052705 | 0.126899 | 1 | 0.999631 | 0.541044 |
| Snap91_E9Q9A3_S298s_1 | -0.01 | -0.3 | 0.14 | -0.54 | -0.75 | 0.42 | -0.76 | -0.69 | 0.09 | -0.3 | -0.73 | 0.06 | 1.06 | 0.54 | -0.74 | 0.46 | -0.2675 | -0.0425 | 0.5075 | 0.246642 | 0.45597 | 0.891281 | 1 | 0.999631 | 0.685751 |
| Snap91_E9Q9A3_S303s_1 | 0.01 | 0.75 | 0.29 | 0.31 | -0.24 | 0.47 | -0.01 | 0.55 | 0.29 | 0.81 | 0.13 | 0.32 | 0.36 | 0.9 | 0.66 | 0.13 | -0.1475 | 0.0475 | 0.1725 | 0.58742 | 0.629323 | 0.871647 | 1 | 0.999631 | 0.850093 |
| Snap91_E9Q9A3_S311s_1 | -0.44 | -0.36 | -0.62 | -1.06 | -0.87 | -0.6 | -1 | -0.9 | -0.6 | -0.42 | -0.8 | -0.69 | 0.31 | -0.5 | -0.62 | -0.4 | -0.2225 | -0.0075 | 0.3175 | 0.351219 | 0.418781 | 0.978373 | 1 | 0.999631 | 0.738069 |
| Snap91_E9Q9A3_S323s_1 | -1.28 | -0.19 | -0.44 | -1.36 | -0.23 | -0.7 | -0.73 | -1.04 | -0.38 | -1.28 | -0.55 | -0.02 | 0.72 | -0.61 | -0.67 | -0.51 | 0.1425 | 0.26 | 0.55 | 0.234318 | 0.694724 | 0.526501 | 1 | 0.999631 | 0.683223 |
| Snap91_E9Q9A3_S333s_1 | -1.48 | -0.29 | 0.21 | -0.72 | 0.29 | -0.23 | -0.12 | -0.28 | -0.3 | -1.45 | -0.07 | 0.92 | 1.12 | -0.6 | 0.39 | -0.83 | 0.485 | 0.345 | 0.59 | 0.289036 | 0.231984 | 0.540488 | 1 | 0.999631 | 0.706046 |
| Snap91_E9Q9A3_S587s_1 | -0.16 | 0.04 | 0.58 | 0.03 | -0.31 | 0.54 | -0.45 | -0.28 | -0.06 | -0.42 | -0.93 | 0.54 | 0.4 | 0.82 | 0.41 | 0.85 | -0.2475 | -0.34 | 0.4975 | 0.125037 | 0.449338 | 0.373252 | 1 | 0.999631 | 0.596063 |
| Snap91_E9Q9A3_S593s_1 | 0.47 | -0.29 | -1.28 | 0.06 | 0.95 | -0.15 | 0.11 | -0.07 | -0.24 | -1.2 | -0.22 | -1 | -0.72 | -0.24 | -0.67 | -0.48 | 0.47 | -0.405 | -0.2675 | 0.5176 | 0.299438 | 0.373713 | 1 | 0.999631 | 0.819449 |
| Snap91_E9Q9A3_T310t_1 | -0.45 | -0.43 | -0.52 | -0.53 | -0.9 | -0.5 | -0.59 | -0.32 | -0.72 | -0.25 | -0.15 | 0.7 | 1.01 | -0.08 | -0.08 | 0.39 | -0.095 | 0.3775 | 0.7925 | 0.035465 | 0.702863 | 0.284398 | 1 | 0.999631 | 0.453811 |
| Snap91_E9Q9A3_T310t_2 | -0.52 | -0.25 | -0.64 | -1.02 | -0.63 | -0.48 | -0.62 | -0.48 | -0.34 | -0.52 | -0.82 | -0.64 | 0.38 | -0.14 | -0.88 | -0.72 | 0.055 | 0.0275 | 0.7075 | 0.102704 | 0.835325 | 0.922522 | 1 | 0.999631 | 0.565572 |
| Snap91_E9Q9A3_T322t_1 | -1.25 | -0.23 | -0.35 | -1.28 | -0.36 | -0.43 | -0.66 | -1.11 | -0.49 | -1.39 | -0.6 | -0.14 | 0.24 | -0.63 | -0.55 | -0.88 | -0.81 | 0.1375 | 0.2175 | 0.412977 | 0.699945 | 0.618951 | 1 | 0.999631 | 0.768474 |
| Snap91_Q61548_S296s_1 | -0.49 | -0.45 | -0.36 | -0.62 | -1.5 | -0.27 | -0.64 | -1.01 | -1.11 | 0.55 | -0.07 | -1.27 | 1.11 | -0.36 | -0.54 | 0 | -0.375 | 0.005 | 0.5325 | 0.201946 | 0.255352 | 0.990799 | 1 | 0.999631 | 0.658952 |
| Snap91_Q61548_S296s_2 | -0.11 | -0.54 | 0.11 | -0.12 | 0.32 | -0.01 | 0.1 | -0.22 | -1.08 | -0.48 | -0.05 | 0.58 | -1.18 | -0.65 | 0.48 | 0.54 | 0.2125 | -0.0925 | -0.0375 | 0.933085 | 0.437833 | 0.81348 | 1 | 0.999631 | 0.978327 |
| Snap91_Q61548_S300s_1 | -0.41 | -0.53 | 0.03 | -0.35 | -0.59 | -0.52 | -0.32 | -0.23 | -0.34 | -0.4 | -0.1 | -0.12 | 0.04 | -0.24 | -0.08 | 0.25 | -0.1 | 0.075 | 0.3075 | 0.296242 | 0.698183 | 0.77611 | 1 | 0.999631 | 0.709528 |
| Snap91_Q61548_S300s_2 | -0.11 | -0.54 | 0.11 | -0.12 | 0.32 | -0.01 | 0.1 | -0.22 | -1.08 | -0.48 | -0.05 | 0.58 | -1.18 | -0.65 | 0.48 | 0.54 | 0.2125 | -0.0925 | -0.0375 | 0.933085 | 0.437833 | 0.81348 | 1 | 0.999631 | 0.978327 |
| Snapin_Q9Z266_S10s_1 | -1.13 | 0.02 | 0.82 | 1.58 | 0.22 | -0.54 | -0.61 | 1.46 | 0.12 | -0.17 | 0.86 | 0.28 | 0.12 | 0.19 | 0.71 | 0.15 | -0.19 | -0.05 | -0.03 | 0.956757 | 0.776939 | 0.929964 | 1 | 0.999631 | 0.984803 |
| Snapin_Q9Z266_S133s_1 | -0.16 | 0.62 | 0.04 | -0.04 | 0.65 | 0.77 | -0.22 | 0.12 | 0.86 | -0.06 | 0.18 | 0.19 | 0.67 | 0.9 | -0.36 | 0.34 | 0.215 | 0.1775 | 0.4525 | 0.168266 | 0.519715 | 0.583789 | 1 | 0.999631 | 0.637541 |
| Snapin_Q9Z266_T14t_1 | -0.3 | -0.04 | -0.16 | -0.81 | 0.6 | 0 | -1.57 | -0.64 | 0.71 | 0.59 | -0.13 | 0.29 | 1.03 | 0.41 | -0.28 | -0.67 | -0.075 | 0.6925 | 0.45 | 0.302479 | 0.873416 | 0.050057 | 1 | 0.999631 | 0.714978 |
| Snapin_Q9Z266_T14t_2 | 0.6 | 2.74 | 1.29 | 1.07 | 0.56 | 0.73 | 1.56 | 1.09 | 1.14 | 2.44 | 0.33 | 0.33 | 1.27 | 0.31 | 0.16 | 0.94 | -0.54 | -0.365 | -0.755 | 0.158303 | 0.26036 | 0.556935 | 1 | 0.999631 | 0.628615 |
| Snca_O55042_S42s_1 | -0.12 | -1.73 | -1.04 | -0.32 | -0.18 | 0.13 | -0.5 | 0.06 | -0.82 | -0.87 | -0.87 | -0.78 | -0.49 | -0.45 | 0.05 | -0.71 | 0.68 | -0.0325 | 0.4025 | 0.344993 | 0.112736 | 0.932933 | 1 | 0.999631 | 0.734195 |
| Snca_O55042_T81t_1 | 1.48 | 0.55 | -0.93 | 2.52 | 2.52 | 0.69 | 1.85 | 1.47 | 1.53 | -1.12 | 1.03 | -0.98 | 0.07 | 0.37 | 0.28 | 0.06 | 0.7275 | -0.79 | -0.71 | 0.295897 | 0.330241 | 0.375129 | 1 | 0.999631 | 0.709072 |
| Sncb_Q91ZZ3_S45s_1 | -0.95 | -1.85 | 0 | -1.05 | -0.23 | -0.09 | -1.12 | -1 | -1.09 | -1.09 | -1.19 | -0.31 | 1.37 | 0.11 | 0.54 | -1.34 | 0.3525 | 0.0425 | 1.1325 | 0.091179 | 0.437401 | 0.921199 | 1 | 0.999631 | 0.557085 |
| Sncb_Q91ZZ3_S63s_1 | -0.45 | 0.26 | 0.5 | -0.3 | -0.05 | -0.71 | -0.37 | -0.27 | 0.38 | 0.15 | -0.59 | -0.38 | 0.73 | -0.53 | 0.6 | 0.44 | -0.06 | -0.3175 | 0.12 | 0.780177 | 0.331939 | 0.471938 | 1 | 0.999631 | 0.921416 |
| Sncb_Q91ZZ3_S71s_1 | -0.03 | 0.43 | -0.4 | -0.36 | 0.29 | 0.64 | -0.65 | -0.18 | -0.02 | -0.87 | -0.48 | 0.06 | 0.87 | 0.44 | -0.03 | -0.79 | 0.115 | -0.2375 | 0.2125 | 0.613677 | 0.751604 | 0.486666 | 1 | 0.999631 | 0.860866 |
| Sncb_Q91ZZ3_T44t_1 | -0.95 | -1.85 | 0 | -1.05 | -0.23 | -0.09 | -1.12 | -1 | -1.09 | -1.09 | -1.19 | -0.31 | 1.37 | 0.11 | 0.54 | -1.34 | 0.3525 | 0.0425 | 1.1325 | 0.091179 | 0.437401 | 0.921199 | 1 | 0.999631 | 0.557085 |
| Snd1_Q3TJ56_S426s_1 | -0.23 | -0.05 | 0.91 | 0.15 | -0.56 | -0.26 | -0.07 | 0.01 | -0.31 | 0.02 | 0 | -0.31 | 0.56 | -0.01 | 0.67 | 0.31 | -0.415 | -0.345 | 0.1875 | 0.594697 | 0.221488 | 0.30134 | 1 | 0.999631 | 0.854138 |
| Snip1_Q8BIZ6_S144s_1 | 0.95 | -0.02 | 1.93 | 0.93 | 0.11 | 0.59 | 1.52 | -0.03 | 0.93 | 0.57 | 0.8 | 0.71 | 0.21 | -0.95 | 0.63 | -0.57 | -0.4 | -0.195 | -1.1175 | 0.050033 | 0.429509 | 0.637953 | 1 | 0.999631 | 0.488436 |
| Snip1_Q8BIZ6_S145s_1 | 1.36 | -0.34 | 1.56 | 0.84 | -0.6 | -0.21 | 1.05 | -0.41 | -0.49 | 1.09 | 0.32 | 0.09 | -1.49 | -1.51 | -0.08 | -0.59 | -0.8975 | -0.6025 | -1.7725 | 0.006521 | 0.112167 | 0.252926 | 1 | 0.999631 | 0.351545 |
| Snip1_Q8BIZ6_S18s_1 | -1.52 | 0.4 | -0.87 | -0.13 | 0.82 | 0.82 | -0.56 | 0.66 | 1.12 | -0.09 | 0.67 | 0.32 | 1.01 | 1.45 | 1.03 | 0.77 | 0.53 | 0.6 | 1.16 | 0.043298 | 0.357408 | 0.280551 | 1 | 0.999631 | 0.475443 |
| Snip1_Q8BIZ6_S33s_1 | 0.58 | 1.04 | 0.78 | 0.52 | 0.75 | 1.22 | -0.01 | 0.46 | 1.03 | 1.22 | 0.05 | 0.05 | 1.5 | 1.77 | 0.78 | 0.82 | -0.125 | -0.1675 | 0.4875 | 0.172747 | 0.703056 | 0.655536 | 1 | 0.999631 | 0.642116 |
| Snip1_Q8BIZ6_S50s_1 | -0.66 | -1.1 | -0.27 | -0.79 | -0.78 | -1.18 | -0.2 | -1.08 | -1.35 | -0.78 | 0.22 | 0.4 | -0.82 | -1.41 | 0.87 | 0.25 | -0.24 | 0.1925 | 0.2925 | 0.598156 | 0.533064 | 0.693571 | 1 | 0.999631 | 0.855276 |
| Snn_P61807_S49s_1 | 0.18 | -0.06 | 0.05 | 2.04 | -1.05 | -0.12 | 0.57 | 1.65 | -0.69 | 0.87 | 1.9 | 1.26 | 0.55 | 0.67 | 1.56 | 1.48 | -0.29 | 0.2825 | 0.5125 | 0.345487 | 0.66769 | 0.672633 | 1 | 0.999631 | 0.734968 |
| Snn_P61807_S83s_1 | -0.42 | -0.53 | -0.57 | -0.68 | -0.62 | -0.42 | -0.57 | -0.45 | -0.3 | -0.05 | -0.21 | 0.07 | -0.12 | -0.29 | -0.34 | -0.18 | 0.035 | 0.4275 | 0.3175 | 0.241733 | 0.88084 | 0.110425 | 1 | 0.999631 | 0.684831 |
| Snph_Q80U23_S200s_1 | 1.74 | -0.01 | 1.06 | 0.4 | -0.76 | -0.01 | 0.84 | -0.64 | -0.63 | 1.29 | -0.4 | -0.87 | 0.13 | -1.57 | -1.01 | -1.3 | -0.94 | -0.95 | -1.735 | 0.006425 | 0.083843 | 0.121734 | 1 | 0.999631 | 0.351545 |
| Snph_Q80U23_S200s_2 | 1.71 | 0.1 | 0.84 | 0.56 | -0.88 | 0.08 | 0.99 | -0.44 | -0.04 | 1.2 | 0.01 | -0.14 | -0.19 | -1.53 | -0.68 | -1.22 | -0.865 | -0.545 | -1.7075 | 0.003357 | 0.105105 | 0.247108 | 1 | 0.999631 | 0.351308 |
| Snph_Q80U23_S204s_1 | 2.12 | -0.49 | 0.32 | 0.15 | -1.21 | 0.11 | 1.01 | -1.22 | -0.61 | 1.46 | -0.83 | -1.32 | -0.15 | -1.59 | -1.58 | -1.21 | -0.8525 | -0.85 | -1.3075 | 0.041374 | 0.236379 | 0.264395 | 1 | 0.999631 | 0.474979 |
| Snph_Q80U23_S204s_2 | 1.71 | 0.1 | 0.84 | 0.56 | -0.88 | 0.08 | 0.99 | -0.44 | -0.04 | 1.2 | 0.01 | -0.14 | -0.19 | -1.53 | -0.68 | -1.22 | -0.865 | -0.545 | -1.7075 | 0.003357 | 0.105105 | 0.247108 | 1 | 0.999631 | 0.351308 |
| Snph_Q80U23_S20s_1 | -0.32 | -0.97 | -0.6 | -1.14 | -0.93 | -0.68 | -0.84 | -0.65 | -1.08 | -0.12 | -0.55 | -0.84 | 0.11 | -0.4 | -0.95 | -0.34 | -0.0175 | 0.11 | 0.3625 | 0.307937 | 0.949887 | 0.739083 | 1 | 0.999631 | 0.718178 |
| Snph_Q80U23_S219s_1 | -0.1 | 0.3 | -2.17 | -2.84 | -0.78 | -0.36 | -2.2 | -3.33 | -0.38 | 0.62 | -2.14 | -2.29 | 1.02 | 0.18 | -1.96 | -2.51 | -0.465 | 0.155 | 0.385 | 0.694618 | 0.603377 | 0.864188 | 1 | 0.999631 | 0.895302 |
| Snph_Q80U23_T19t_1 | -0.28 | -0.33 | -0.07 | -0.65 | -0.85 | -0.37 | -0.61 | -0.32 | -0.84 | 0.04 | -0.15 | -0.53 | 0.39 | 0.06 | -0.07 | 0.27 | -0.205 | -0.0375 | 0.495 | 0.10702 | 0.449687 | 0.901589 | 1 | 0.999631 | 0.805353 |
| Snph_Q80U23_T292t_1 | 0.36 | -0.92 | -1 | 0.33 | 0.2 | -0.71 | 0.12 | -0.5 | -0.65 | -0.74 | -0.62 | -0.99 | -1.66 | 0.16 | -0.82 | -0.53 | 0.085 | -0.4425 | -0.405 | 0.432761 | 0.843665 | 0.283311 | 1 | 0.999631 | 0.779247 |
| Snrk_Q8VDU5_S518s_1 | 0.99 | 0.12 | -0.3 | 1.14 | 1.57 | 1.26 | 0.85 | 1.25 | 0.74 | 0.12 | 0.65 | -0.19 | -0.83 | 0.97 | 1.29 | 0.88 | 0.745 | -0.1575 | 0.105 | 0.838021 | 0.078199 | 0.705563 | 1 | 0.999631 | 0.946137 |
| Snrk_Q8VDU5_S569s_1 | 1.59 | 0.62 | 1.04 | 1.14 | 0.51 | 1.37 | 1.26 | 0.57 | 1.07 | 2.07 | 0.58 | 0.25 | 0.02 | -0.11 | 0.3 | 0.2 | -0.17 | -0.105 | -0.995 | 0.008978 | 0.616546 | 0.811242 | 1 | 0.999631 | 0.365396 |
| Snrk_Q8VDU5_S596s_1 | 1.15 | 1.21 | 0.39 | 0.03 | 1.41 | 1.4 | -0.04 | -0.35 | 2.16 | 1.64 | 0.42 | -0.31 | 2.44 | 0.03 | -0.47 | -0.06 | -0.09 | 0.2825 | -0.21 | 0.746294 | 0.860371 | 0.628521 | 1 | 0.999631 | 0.911375 |
| Snrnp200_Q6P4T2_S225s_1 | -0.1 | 0.45 | -0.15 | -0.48 | -0.17 | 0 | -0.16 | -0.49 | 0.16 | 0.28 | -0.68 | -0.23 | 0.46 | 0.44 | -0.4 | -0.06 | -0.135 | -0.0475 | 0.18 | 0.604237 | 0.643375 | 0.888 | 1 | 0.999631 | 0.85817 |
| Snrnp70_Q62376_S226s_1 | 0.19 | 0.82 | 0.58 | 0.11 | 0.58 | 0.58 | -0.32 | 0.56 | 0.8 | 0.88 | 0.27 | 0.51 | 1.35 | 1.12 | 0.58 | 0.66 | -0.075 | 0.19 | 0.5025 | 0.146431 | 0.817129 | 0.526719 | 1 | 0.999631 | 0.618236 |
| Snrnp70_Q62376_S408s_1 | 0.06 | 0.06 | 0.1 | -0.06 | 0.72 | 0.91 | -0.21 | 0.03 | 0.47 | 0.11 | 0.44 | 0.97 | 0.61 | 1.17 | 0.87 | 0.76 | 0.3225 | 0.4575 | 0.8125 | 0.013119 | 0.325654 | 0.125631 | 1 | 0.999631 | 0.385092 |
| Snta1_Q61234_S183s_1 | -1.21 | -1.04 | -0.58 | -1.03 | -0.66 | -1.37 | -0.87 | -1.03 | -0.52 | -1.99 | -0.39 | -0.55 | -0.42 | -1.32 | -0.64 | -1.27 | -0.0175 | 0.1025 | 0.0525 | 0.874914 | 0.950377 | 0.80231 | 1 | 0.999631 | 0.960237 |
| Snta1_Q61234_S194s_1 | 1.26 | 0.65 | 0.49 | 1.26 | 1.11 | 1.59 | 1.05 | 1.31 | 1.57 | 1.4 | 1.13 | 1 | -0.75 | 1.19 | 0.8 | 0.99 | 0.35 | 0.36 | -0.3575 | 0.458227 | 0.259377 | 0.26016 | 1 | 0.999631 | 0.790722 |
| Snta1_Q61234_S195s_1 | 0.98 | -0.31 | 0.4 | 0.82 | -0.44 | 0.05 | 0.8 | 0.62 | 0.41 | 1.06 | 0.4 | 0.02 | -0.65 | -0.35 | 0.13 | 0.19 | -0.215 | 0 | -0.6425 | 0.118705 | 0.598348 | 1 | 1 | 1 | 0.585512 |
| Sntb1_Q99L88_S86s_1 | 0.24 | -0.54 | -0.09 | -0.13 | -1.41 | -0.29 | -0.36 | -0.34 | -1.28 | -0.74 | -0.63 | -0.59 | -1.17 | 0.34 | -0.15 | 0.53 | -0.47 | -0.68 | 0.0175 | 0.967055 | 0.195169 | 0.045041 | 1 | 0.999631 | 0.988373 |
| Sntb2_B7ZNU9_S373s_1 | 0.24 | 0.1 | -0.85 | 0.28 | 0.07 | 0.23 | -0.3 | 0.28 | -0.05 | -0.21 | -0.17 | -0.79 | -0.41 | 0.79 | 0.02 | 0.13 | 0.1275 | -0.2475 | 0.19 | 0.630641 | 0.704777 | 0.487623 | 1 | 0.999631 | 0.867509 |
| Sntb2_B7ZNU9_S75s_1 | -0.23 | -0.06 | -0.27 | -0.61 | -0.39 | -0.5 | -0.89 | -0.54 | -0.41 | -0.17 | -0.03 | 0.03 | 0.17 | 0.04 | -0.6 | -0.08 | -0.2875 | 0.1475 | 0.175 | 0.5688 | 0.286715 | 0.582906 | 1 | 0.999631 | 0.843185 |
| Sntb2_B7ZNU9_S90s_1 | 0.3 | 0.27 | 0.13 | -0.18 | -0.02 | -0.05 | -0.44 | -0.68 | 0 | 0.72 | -0.06 | -0.22 | 0.57 | 0.47 | 0.01 | 0.27 | -0.4275 | 0.09 | 0.2 | 0.492942 | 0.147054 | 0.758555 | 1 | 0.999631 | 0.80955 |
| Snurf_Q9WU12_S32s_1 | 1.42 | 1.55 | -0.46 | 1.54 | 1.81 | 1.07 | 1.33 | 1.25 | 1.65 | 1.03 | 0.3 | -0.01 | -1.25 | 0.55 | 0.08 | 0.75 | 0.3525 | -0.52 | -0.98 | 0.12983 | 0.475643 | 0.321371 | 1 | 0.999631 | 0.601246 |
| Snw1_Q9CSN1_S224s_1 | 0.47 | 0.13 | -0.19 | -1.36 | 0.09 | 0.04 | -0.33 | -1.42 | -0.17 | 0.45 | -1.02 | -1.05 | 0.13 | 0.34 | -0.71 | -0.64 | -0.1675 | -0.21 | 0.0175 | 0.970225 | 0.737105 | 0.680543 | 1 | 0.999631 | 0.989586 |
| Snw1_Q9CSN1_S224s_2 | 0.37 | -0.33 | -0.32 | -1.03 | -0.12 | -0.06 | -0.76 | -0.35 | -0.54 | 0.17 | -1.06 | -1.15 | 0.01 | -0.18 | -0.32 | -0.57 | 0.005 | -0.3175 | 0.0625 | 0.862313 | 0.98872 | 0.456588 | 1 | 0.999631 | 0.954601 |
| Snw1_Q9CSN1_S232s_2 | 0.37 | -0.33 | -0.32 | -1.03 | -0.12 | -0.06 | -0.76 | -0.35 | -0.54 | 0.17 | -1.06 | -1.15 | 0.01 | -0.18 | -0.32 | -0.57 | 0.005 | -0.3175 | 0.0625 | 0.862313 | 0.98872 | 0.456588 | 1 | 0.999631 | 0.954601 |

| | | | | | | | | | | | | | | | | | | | | | | | | | |
|---|---|---|---|---|---|---|---|---|---|---|---|---|---|---|---|---|---|---|---|---|---|---|---|---|---|
| Snx14_G3UXB6_S695s_1 | -0.22 | -0.8 | -0.07 | -0.13 | -0.8 | -0.03 | -0.57 | -0.4 | -0.32 | -0.09 | -1.18 | 0.05 | 0.45 | 0.74 | 0.16 | 0.33 | -0.145 | -0.08 | 0.725 | 0.036694 | 0.628677 | 0.82304 | 1 | 0.999631 | 0.454456 |
| Snx15_D3YVT2_S215s_1 | 0.56 | 0.63 | -0.05 | 0.69 | 1.47 | 1.25 | 0.42 | 0.58 | 1.34 | 0.39 | 0.99 | 0.63 | 0.37 | 1.63 | 0.73 | 1.29 | 0.4725 | 0.38 | 0.5475 | 0.162737 | 0.187528 | 0.25967 | 1 | 0.999631 | 0.631442 |
| Snx16_Q8BLP0_S29s_1 | 0.09 | -0.06 | -0.77 | 0.84 | 0.43 | 0.28 | 1.34 | 0.71 | 0.36 | -0.38 | 1.29 | 0.63 | -0.59 | 0.38 | 1.21 | 0.73 | 0.665 | 0.45 | 0.4075 | 0.411348 | 0.126279 | 0.344145 | 1 | 0.999631 | 0.767937 |
| Snx16_Q8BLP0_S70s_1 | -0.72 | -0.37 | -0.35 | -0.86 | -0.59 | -0.21 | -0.42 | -1.29 | -0.92 | -0.9 | 0.17 | -0.16 | -0.27 | -0.51 | -0.02 | 0.43 | -0.0525 | 0.1225 | 0.4825 | 0.155012 | 0.868377 | 0.722449 | 1 | 0.999631 | 0.6258 |
| Snx17_Q8BVL3_S409s_1 | 1.46 | 1.57 | 1.37 | 2.39 | 0.38 | 2.59 | 3.09 | 0.98 | -0.19 | 1.61 | 1.1 | 1.24 | -2.84 | 0.4 | 1.58 | 0.85 | 0.0625 | -0.7575 | -1.7 | 0.074024 | 0.919168 | 0.116104 | 1 | 0.999631 | 0.539897 |
| Snx17_Q8BVL3_S415s_1 | -0.47 | 0.83 | 1.29 | 0.13 | 0.16 | 0.85 | 0.3 | 0.5 | 0.55 | 0.73 | 0.61 | 1.11 | 1.46 | 0.2 | 1.16 | 1.33 | 0.0075 | 0.305 | 0.5925 | 0.227158 | 0.985519 | 0.467098 | 1 | 0.999631 | 0.678384 |
| Snx2_Q9CWK8_S93s_1 | -0.64 | 0.15 | -0.32 | -0.22 | 0.13 | -0.23 | -0.23 | -0.12 | -0.42 | -0.44 | -0.13 | -0.23 | -0.69 | -0.17 | -0.6 | -0.48 | 0.145 | -0.0475 | -0.2275 | 0.457734 | 0.597578 | 0.864517 | 1 | 0.999631 | 0.790722 |
| Snx2_Q9CWK8_S97s_1 | -0.65 | -0.05 | -0.38 | -0.9 | -0.63 | -0.21 | -1.16 | -0.8 | -0.47 | -0.72 | -0.98 | -0.44 | 0.81 | -0.32 | -1.31 | -0.4 | -0.205 | -0.1575 | 0.19 | 0.683217 | 0.523713 | 0.600879 | 1 | 0.999631 | 0.889602 |
| Snx2_Q9CWK8_T101t_1 | -1.45 | -1.71 | -0.04 | 0.16 | -1.3 | -0.47 | -0.82 | -0.58 | -0.88 | -0.29 | -0.01 | 0.6 | -0.54 | -0.25 | 0.73 | 0.47 | -0.0325 | 0.615 | 0.8625 | 0.127451 | 0.946458 | 0.263802 | 1 | 0.999631 | 0.59788 |
| Snx2_Q9CWK8_T104t_1 | 0.68 | 0.89 | 0.6 | 0.53 | 1.07 | -0.59 | -0.35 | -0.14 | -0.49 | -0.32 | 0 | -0.52 | 1.5 | 0.33 | 0.39 | -0.64 | -0.6775 | -1.0075 | -0.28 | 0.533653 | 0.105101 | 0.003343 | 1 | 0.999631 | 0.828143 |
| Snx25_E9Q1K0_S126s_1 | -0.32 | 0.15 | 0.04 | -0.57 | 0.07 | -0.15 | -0.21 | 0.19 | 0.02 | -0.47 | 0.67 | 0.76 | 0.94 | 0.02 | 0.28 | 0.53 | 0.15 | 0.42 | 0.6175 | 0.085449 | 0.589439 | 0.265856 | 1 | 0.999631 | 0.551124 |
| Snx27_Q3UHD6_S49s_1 | -0.63 | 0.5 | 0.92 | 0.16 | 0.27 | 0.07 | -0.36 | 0.35 | 0.3 | 0.06 | 0.21 | 0.86 | 0.79 | 0.45 | 0.7 | 0.61 | -0.155 | 0.12 | 0.4 | 0.300892 | 0.683564 | 0.758355 | 1 | 0.999631 | 0.714183 |
| Snx29_D3YZ90_S371s_1 | 1.24 | 0.67 | 0.21 | -1.73 | 1.2 | 0.52 | 3.74 | -1.33 | 2.15 | -1.13 | 0.43 | 0.71 | 0.73 | -3.56 | -0.78 | -2.72 | 0.935 | 0.4425 | -1.68 | 0.115021 | 0.385952 | 0.590399 | 1 | 0.999631 | 0.579336 |
| Snx30_Q8CE50_S67s_1 | -0.79 | -1.15 | -1.38 | 0.03 | -0.28 | -0.49 | -0.14 | 0.62 | -0.75 | -1.41 | 0.41 | 0.15 | 0.46 | -1.61 | -0.29 | -0.53 | 0.75 | 0.4225 | 0.33 | 0.517282 | 0.083992 | 0.402482 | 1 | 0.999631 | 0.819449 |
| Snx7_F8WI30_S8s_1 | -0.12 | 0.07 | -0.88 | -0.54 | -0.2 | -0.22 | -0.61 | -0.51 | -0.84 | -0.62 | -0.59 | -0.35 | 0.22 | -0.06 | -1.05 | -0.83 | -0.0175 | -0.2325 | -0.0625 | 0.87483 | 0.953435 | 0.454695 | 1 | 0.999631 | 0.960237 |
| Sod1_P08228_S99s_1 | -1.64 | 0.89 | 1.58 | 0.02 | 0.25 | 0.76 | 0.7 | 0.23 | 1.03 | -0.64 | 0.78 | 1.51 | 2.05 | 1.56 | 0.47 | 1.56 | 0.2725 | 0.4575 | 1.1975 | 0.105222 | 0.669401 | 0.539713 | 1 | 0.999631 | 0.568213 |
| Soga1_A2ACV6_S173s_1 | -0.7 | -0.94 | -1.48 | -2.57 | -1.18 | -1.59 | -1.4 | -1.78 | -1.74 | -0.24 | -1.89 | -2.32 | -0.08 | -0.79 | -1.83 | -1.91 | -0.065 | -0.125 | 0.27 | 0.631667 | 0.879241 | 0.825371 | 1 | 0.999631 | 0.867677 |
| Soga1_E1U8D0_S1015s_1 | -0.55 | 0.38 | 0.17 | -0.45 | -0.54 | -0.69 | -0.73 | 0.11 | -0.63 | -0.26 | -0.01 | 0.77 | 1.13 | -0.4 | 0.36 | 0.44 | -0.35 | 0.08 | 0.495 | 0.242284 | 0.313372 | 0.837917 | 1 | 0.999631 | 0.685099 |
| Soga3_Q6NZL0_S637s_1 | 1.11 | 1.23 | 0.98 | 0.86 | 1.58 | 1.28 | -0.09 | 0.59 | 1.2 | 0.96 | 1.29 | 1.08 | -0.12 | 1.33 | 1.12 | 1.74 | -0.205 | 0.0875 | -0.0275 | 0.947901 | 0.601778 | 0.7284 | 1 | 0.999631 | 0.983162 |
| Soga3_Q6NZL0_S639s_1 | -0.37 | 0.47 | -0.16 | -0.11 | -0.06 | 0.18 | -0.42 | -0.04 | 0.15 | 0.09 | 0.28 | 0.3 | 1.09 | 0.15 | 0.07 | 0.59 | -0.0425 | 0.2475 | 0.5175 | 0.163399 | 0.883519 | 0.389728 | 1 | 0.999631 | 0.631937 |
| Soga3_Q6NZL0_S760s_1 | 1.55 | 0.62 | 2.4 | 2.3 | -0.13 | 0.81 | 2.25 | 0.31 | 0.46 | 1.92 | 0.68 | 0.14 | -0.84 | -0.74 | 0.34 | -0.13 | -0.9075 | -0.9175 | -2.06 | 0.001605 | 0.152505 | 0.10845 | 1 | 0.999631 | 0.351308 |
| Soga3_Q6NZL0_S779s_1 | 0.48 | 0.4 | -0.04 | 0.6 | 0.34 | 0.26 | 0.24 | 0.56 | 0.65 | 0.54 | 0.59 | 0.44 | 0.13 | 0.44 | 0.44 | 0.48 | -0.01 | 0.195 | 0.0125 | 0.965194 | 0.969435 | 0.467698 | 1 | 0.999631 | 0.987856 |
| Soga3_Q6NZL0_T514t_1 | -0.15 | 0.58 | 0.21 | 0.01 | -0.1 | 0.67 | -0.14 | 0.48 | 0.1 | 0.47 | 0.07 | 0.7 | 0.91 | 0.63 | 0.09 | 0.35 | 0.065 | 0.1725 | 0.3325 | 0.312428 | 0.834986 | 0.565379 | 1 | 0.999631 | 0.719861 |
| Son_H9KV01_S1723s_1 | -0.26 | -0.4 | -0.66 | -0.57 | 0.28 | -0.1 | -0.78 | -0.41 | -0.23 | -0.51 | 0.41 | 0.17 | -0.3 | 0.72 | 0.08 | 0.38 | 0.22 | 0.4325 | 0.6925 | 0.051472 | 0.473981 | 0.171656 | 1 | 0.999631 | 0.489777 |
| Sorbs1_Q62417_S1209s_1 | 0.44 | 0.44 | 0.49 | 0.1 | 0.07 | 0.06 | 0.12 | 0.12 | 0.11 | 1.11 | 0.1 | 0.62 | 0.58 | 0.14 | 0.33 | 0.08 | -0.275 | 0.1175 | -0.085 | 0.762122 | 0.265076 | 0.712845 | 1 | 0.999631 | 0.915029 |
| Sorbs1_Q62417_S250s_1 | 1.46 | 2.64 | -0.5 | -0.62 | 3.12 | 2.26 | 0.71 | -0.53 | 4.28 | 0.69 | -0.21 | -0.1 | 2.7 | 0.72 | -0.81 | -2.38 | 0.645 | 0.42 | -0.6875 | 0.549799 | 0.513599 | 0.711794 | 1 | 0.999631 | 0.83548 |
| Sorbs1_Q62417_S254s_1 | 0.28 | 0.14 | -0.17 | -0.06 | 0.38 | 0.13 | 0.06 | 0.46 | -0.09 | 0 | 0.69 | 0.23 | -0.1 | 0.21 | -0.49 | 0.46 | 0.21 | 0.16 | -0.0275 | 0.930592 | 0.416589 | 0.582466 | 1 | 0.999631 | 0.977897 |
| Sorbs1_Q62417_S270s_1 | 0.76 | -0.67 | 0.54 | 0.17 | -0.34 | 0.3 | -0.33 | 0.58 | -0.94 | 0.41 | 0.54 | 0.66 | -0.35 | -0.48 | 0.68 | 0.55 | -0.1475 | -0.0325 | -0.1 | 0.819761 | 0.710508 | 0.944938 | 1 | 0.999631 | 0.938303 |
| Sorbs1_Q62417_S2s_1 | 1.53 | -0.67 | -0.27 | -0.37 | -0.74 | -0.46 | 0.5 | -1.22 | -1.27 | 0.6 | -1.07 | -1.22 | -1.23 | -0.72 | -1.27 | -1.04 | -0.535 | -0.795 | -1.12 | 0.044043 | 0.353881 | 0.212455 | 1 | 0.999631 | 0.475443 |
| Sorbs1_Q62417_S3s_1 | 1.53 | -0.67 | -0.27 | -0.37 | -0.74 | -0.46 | 0.5 | -1.22 | -1.27 | 0.6 | -1.07 | -1.22 | -1.23 | -0.72 | -1.27 | -1.04 | -0.535 | -0.795 | -1.12 | 0.044043 | 0.353881 | 0.212455 | 1 | 0.999631 | 0.475443 |
| Sorbs1_Q62417_S533s_1 | 0.13 | 0.5 | -0.5 | 0.55 | 0.9 | 0.89 | 0.02 | 0.81 | 0.82 | 0.37 | 0.35 | 0.57 | 1.1 | 0.94 | 0.46 | 1.05 | 0.485 | 0.3575 | 0.7175 | 0.059786 | 0.188493 | 0.284075 | 1 | 0.999631 | 0.50521 |
| Sorbs1_Q62417_S554s_1 | 0.57 | -0.33 | -0.95 | -0.26 | 0.18 | -0.09 | 0.09 | 0.37 | 0.15 | 0.06 | -0.05 | 0.16 | -0.53 | 0.14 | -0.46 | -0.89 | 0.38 | 0.3225 | -0.1925 | 0.633544 | 0.297345 | 0.371793 | 1 | 0.999631 | 0.868569 |
| Sorbs1_Q62417_S576s_1 | -0.17 | -0.35 | -0.8 | 0.01 | 0.11 | -0.3 | -0.36 | 0.44 | -0.23 | -1.32 | -0.41 | -0.95 | -1.48 | -0.01 | -0.61 | -0.37 | 0.3 | -0.4 | -0.29 | 0.462137 | 0.34601 | 0.264464 | 1 | 0.999631 | 0.792786 |
| Sorbs1_Q62417_S58s_1 | 0.41 | 0.96 | -0.78 | -0.63 | -0.07 | 0.65 | -1.12 | -0.51 | 0.26 | 0.8 | -1.35 | -1.41 | 0.64 | 1.23 | -0.93 | -0.16 | -0.2525 | -0.415 | 0.205 | 0.72373 | 0.629214 | 0.516624 | 1 | 0.999631 | 0.905674 |
| Sorbs1_Q62417_S58s_2 | 0.56 | 0.98 | 0.09 | -0.36 | 1.12 | 1.26 | -0.53 | -1.06 | 1.26 | 1.1 | 0.52 | -1.56 | 0.96 | 1.57 | 0.81 | -0.22 | -0.12 | 0.0125 | 0.4625 | 0.331503 | 0.839004 | 0.984431 | 1 | 0.999631 | 0.727302 |
| Sorbs1_Q62417_S62s_1 | -1.11 | -1.23 | -0.79 | -1.17 | -3.13 | -2.57 | -1.39 | -1.59 | -1.54 | -1.43 | -1.5 | -1.31 | 1.22 | -0.46 | -1.33 | -0.63 | -1.095 | -0.37 | 0.775 | 0.158452 | 0.024651 | 0.165102 | 1 | 0.999631 | 0.628615 |
| Sorbs1_Q62417_S635s_1 | 0.77 | 0.68 | -0.16 | 1.07 | 2.05 | 0.61 | 1.01 | 0.84 | 1.23 | 0.06 | 0.85 | -0.05 | -1.37 | 1.32 | 0.9 | 0.49 | 0.5375 | -0.0675 | -0.255 | 0.6685 | 0.212419 | 0.870153 | 1 | 0.999631 | 0.883335 |
| Sorbs1_Q62417_S638s_1 | -0.11 | -1.14 | -1.72 | -1.37 | -0.47 | -0.73 | -0.77 | -0.28 | -0.2 | -0.89 | -0.87 | -1.01 | -0.62 | -0.72 | -0.59 | -0.79 | 0.5225 | 0.3425 | 0.405 | 0.304775 | 0.189351 | 0.404355 | 1 | 0.999631 | 0.716239 |
| Sorbs1_Q62417_S666s_1 | 0.63 | -1.65 | -1.61 | -0.84 | 0.22 | 0.04 | -0.15 | -0.28 | -0.05 | -1.37 | -2.31 | -2.27 | -0.55 | 0.51 | -0.51 | -1.42 | 0.825 | -0.6325 | 0.375 | 0.538379 | 0.128861 | 0.358407 | 1 | 0.999631 | 0.830863 |
| Sorbs1_Q62417_S694s_1 | 0.96 | 0.46 | 0.28 | 1 | 0.46 | 0.47 | 0.36 | 1.38 | 0.18 | 0.47 | 0.69 | 1.25 | -0.76 | 1.19 | 1.34 | 1.77 | -0.0075 | -0.0275 | 0.21 | 0.703108 | 0.982199 | 0.934893 | 1 | 0.999631 | 0.896857 |
| Sorbs1_Q62417_S707s_1 | 0.18 | 0.18 | -0.88 | 0.07 | 0.16 | 0.23 | -0.35 | 0.26 | 0.09 | 0.1 | -0.35 | -0.58 | 0.27 | 1.05 | -0.54 | 0.19 | 0.1875 | -0.0725 | 0.355 | 0.412288 | 0.577043 | 0.83497 | 1 | 0.999631 | 0.768112 |
| Sorbs1_Q62417_S722s_1 | -0.23 | -0.13 | -0.75 | -0.48 | 0.33 | -0.52 | -0.49 | -0.66 | -0.01 | -0.35 | -0.33 | -0.82 | -0.21 | 0.28 | -0.28 | -0.3 | 0.0625 | 0.02 | 0.27 | 0.378836 | 0.842946 | 0.946184 | 1 | 0.999631 | 0.751637 |
| Sorbs1_Q62417_S791s_1 | -0.53 | -0.24 | -0.09 | -0.04 | -0.66 | -0.23 | -0.21 | -0.1 | -0.36 | 0.33 | -0.14 | 0.47 | 0.42 | 0.25 | 0.48 | 0.57 | -0.075 | 0.3 | 0.655 | 0.036072 | 0.777097 | 0.33048 | 1 | 0.999631 | 0.453963 |
| Sorbs1_Q62417_S969s_1 | 0.02 | 0.9 | -2.01 | -1.49 | -0.45 | -0.91 | -1.42 | -0.66 | -0.62 | 0.26 | -2.12 | -0.2 | 0.77 | 0.44 | -0.47 | -0.31 | -0.215 | -0.025 | 0.7525 | 0.268486 | 0.733681 | 0.973311 | 1 | 0.999631 | 0.69658 |
| Sorbs1_Q62417_S96s_1 | -0.11 | 1.35 | 0.73 | -0.18 | 0.64 | 0.05 | 0.42 | -0.13 | -0.28 | -0.8 | 0.17 | 0.4 | 0.52 | 0.23 | -0.57 | -0.26 | -0.2025 | -0.575 | -0.4675 | 0.303963 | 0.620282 | 0.215801 | 1 | 0.999631 | 0.715879 |
| Sorbs1_Q62417_T286t_1 | 0.17 | 1.06 | 0.62 | 0.36 | 0.38 | 0.84 | -0.22 | 0.7 | 0.41 | 0.24 | 0.39 | 0.16 | 0.8 | 1.09 | 0.3 | 0.79 | -0.1275 | -0.2525 | 0.1925 | 0.561067 | 0.708273 | 0.391942 | 1 | 0.999631 | 0.840285 |
| Sorbs2_B7ZWM6_S109s_1 | -0.02 | -1.67 | -0.82 | -0.73 | -1.72 | -2.66 | -0.18 | -0.5 | -1.75 | -2.09 | -0.26 | -0.19 | -1.49 | -1.82 | -0.37 | 0.06 | -0.455 | -0.2625 | -0.095 | 0.857064 | 0.454937 | 0.637535 | 1 | 0.999631 | 0.952021 |
| Sorbs2_B7ZWM6_S1111s_1 | -0.82 | 0.07 | -1.24 | -1.02 | 0.3 | -1.05 | -0.96 | -1.06 | -0.08 | -0.43 | 0.06 | 0.19 | -1.02 | -1.19 | -2.14 | -0.7 | 0.06 | 0.6875 | -0.51 | 0.253229 | 0.889097 | 0.076298 | 1 | 0.999631 | 0.688735 |
| Sorbs2_B7ZWM6_S1113s_1 | -0.16 | 0.55 | -1.18 | -0.58 | 0.25 | -0.46 | -0.51 | -0.04 | -0.17 | 0.47 | 0.64 | 0.73 | -0.12 | 0.23 | -0.39 | 0.26 | 0.1525 | 0.76 | 0.3375 | 0.421407 | 0.708941 | 0.095455 | 1 | 0.999631 | 0.772817 |
| Sorbs2_B7ZWM6_S130s_1 | -0.7 | -1.49 | -1.37 | -0.95 | -0.9 | -1.27 | -0.8 | -1.05 | -1.38 | -1.67 | -1.3 | -0.98 | -0.97 | -0.79 | -0.93 | -0.69 | 0.1225 | -0.205 | 0.2825 | 0.357803 | 0.669521 | 0.505738 | 1 | 0.999631 | 0.741099 |
| Sorbs2_B7ZWM6_S143s_1 | -0.95 | -0.89 | -0.55 | -1.22 | -0.53 | -1.12 | -1.19 | -1.1 | -1.17 | -1 | -1.46 | -0.5 | -0.66 | -0.7 | -1.44 | -0.87 | -0.0825 | -0.13 | -0.015 | 0.962001 | 0.771429 | 0.677011 | 1 | 0.999631 | 0.986594 |
| Sorbs2_B7ZWM6_S27s_1 | 1.5 | 1.4 | -1.06 | 0.35 | 2.11 | 1.95 | 0.33 | -0.31 | 1.83 | 0.34 | 0.14 | -0.29 | -0.98 | 2.53 | -0.57 | -0.24 | 0.4725 | -0.0425 | -0.3625 | 0.674739 | 0.528838 | 0.949668 | 1 | 0.999631 | 0.885515 |
| Sorbs2_B7ZWM6_S354s_1 | 1.1 | 1.72 | 0.6 | 1.4 | 3.04 | 1.74 | 0.69 | 1 | 2.02 | 2.44 | 1.34 | 2.81 | 0 | 1.68 | 1.11 | 1.38 | 0.4125 | 0.9475 | -0.1625 | 0.712934 | 0.444626 | 0.039386 | 1 | 0.999631 | 0.900854 |
| Sorbs2_B7ZWM6_S397s_1 | -1.06 | -1.96 | -0.9 | -0.93 | 0.35 | -2.85 | -0.43 | -0.69 | -1.63 | -0.35 | 1.35 | 1.35 | -0.58 | -3.1 | -1.1 | -0.71 | 0.3075 | 1.3925 | -0.16 | 0.78464 | 0.638694 | 0.065825 | 1 | 0.999631 | 0.922904 |
| Sorbs2_B7ZWM6_S40s_1 | -0.87 | -0.8 | -1.1 | -1.07 | -0.38 | -0.94 | -1.09 | -1.91 | -0.72 | -0.03 | -0.47 | -0.22 | -0.14 | -0.58 | -1.28 | -0.52 | -0.12 | 0.6 | 0.33 | 0.324493 | 0.734133 | 0.0499 | 1 | 0.999631 | 0.72463 |
| Sorbs2_B7ZWM6_S592s_1 | -1.07 | 0.61 | -1.78 | -1.35 | -1.52 | -1.11 | -1.52 | -1.63 | -0.4 | -1.77 | -1.37 | -1.4 | 0.94 | -0.92 | -1.7 | -0.77 | -0.5475 | -0.3375 | 0.285 | 0.6764 | 0.29045 | 0.546763 | 1 | 0.999631 | 0.88667 |
| Sorbs2_B7ZWM6_S645s_1 | -0.18 | -1.39 | -0.85 | -0.38 | 0.33 | -0.77 | -0.55 | 0.17 | -0.76 | -1.91 | -0.57 | -0.53 | -2.08 | -0.66 | -0.37 | -0.01 | 0.495 | -0.2425 | -0.08 | 0.873767 | 0.223098 | 0.571253 | 1 | 0.999631 | 0.959802 |
| Sorbs2_B7ZWM6_S820s_1 | 1.73 | 1.18 | -0.96 | 0.34 | 1.33 | 1.65 | 0.29 | 0.28 | 1.14 | 1.41 | 0.14 | 0.08 | -0.07 | 0.95 | -0.01 | 0.33 | 0.315 | 0.12 | -0.2725 | 0.639697 | 0.611175 | 0.845226 | 1 | 0.999631 | 0.870995 |
| Sorbs2_B7ZWM6_S82s_1 | -0.73 | -1.09 | -0.41 | -1.17 | -1.95 | -1.37 | -1.64 | -0.5 | -0.1 | -0.7 | -0.14 | -0.5 | 0.04 | -1.61 | -0.18 | -0.56 | -0.515 | 0.49 | 0.2725 | 0.520284 | 0.189922 | 0.12889 | 1 | 0.999631 | 0.819528 |
| Sorbs2_B7ZWM6_S844s_1 | -0.93 | -1.46 | -1.46 | -1.4 | -1.44 | -1.69 | -1.15 | -1.61 | -1.65 | -1.16 | -1.53 | -1.42 | -1.11 | -1.75 | -1.82 | -0.99 | -0.16 | -0.1275 | -0.105 | 0.748331 | 0.555506 | 0.64152 | 1 | 0.999631 | 0.911931 |
| Sorbs2_B7ZWM6_S99s_1 | -0.51 | -0.19 | 0.48 | 0.27 | -0.14 | 0.01 | 0.26 | 1.67 | 0.64 | -0.86 | 1.88 | 1 | 0.12 | -0.39 | -0.5 | 1.54 | 0.4375 | 0.6525 | 0.18 | 0.718539 | 0.347235 | 0.265147 | 1 | 0.999631 | 0.90414 |
| Sorbs2_D3Z080_S330s_1 | 0.16 | -0.99 | -0.3 | -1.1 | -0.61 | -0.32 | -1.25 | -1.26 | -0.82 | 1.29 | -0.59 | -0.22 | -0.9 | -1.33 | -1.4 | -1.05 | -0.3025 | 0.4725 | -0.6125 | 0.117978 | 0.443947 | 0.37837 | 1 | 0.999631 | 0.585087 |
| Sorbs2_D3Z080_S350s_1 | -0.66 | -2.13 | -0.88 | -0.63 | -0.7 | -1.08 | -1.09 | -1.31 | -1.24 | -0.53 | -1.26 | -0.42 | -0.79 | -0.73 | -0.82 | -0.83 | 0.03 | 0.2125 | 0.2825 | 0.472714 | 0.938283 | 0.617813 | 1 | 0.999631 | 0.799014 |
| Sorbs2_D3Z080_S40s_1 | 0.93 | -1.21 | -0.13 | -0.06 | -0.73 | -0.18 | 1.76 | -3.24 | -2.94 | 0.46 | -0.82 | 0.83 | -0.87 | -2.07 | -1.9 | -1.44 | -0.48 | -0.5 | -1.4525 | 0.013875 | 0.620851 | 0.55372 | 1 | 0.999631 | 0.385092 |
| Sorbs2_Q3UTJ2_S373s_1 | -0.7 | -2.13 | -0.92 | -0.65 | -0.72 | -1.14 | -1.11 | -1.35 | -1.27 | -0.52 | -1.23 | -0.41 | -0.75 | -0.73 | -0.81 | -0.87 | 0.02 | 0.2425 | 0.31 | 0.427012 | 0.958416 | 0.566819 | 1 | 0.999631 | 0.775206 |
| Sorcs1_E9PYT6_S1133s_1 | -0.95 | 0.03 | -0.87 | -0.31 | 0.22 | -1.04 | -0.98 | -0.38 | -0.6 | -0.49 | -0.91 | -0.39 | 0.24 | -0.15 | 0.07 | -0.5 | -0.02 | -0.0725 | 0.44 | 0.220449 | 0.958723 | 0.820505 | 1 | 0.999631 | 0.670752 |
| Sorcs1_E9PYT6_T1152t_1 | -0.85 | -2.36 | -0.75 | -2 | -1.85 | -1.25 | -1.29 | -2.22 | -1.26 | -0.62 | -2.58 | -0.8 | 0.65 | -1.77 | -1.43 | -1.29 | -0.1625 | 0.175 | 0.53 | 0.399053 | 0.719763 | 0.753097 | 1 | 0.999631 | 0.761767 |
| Sorcs3_Q8VI51_T175t_1 | 0.25 | -0.92 | -0.29 | -0.36 | -0.67 | -0.02 | 0.02 | -0.17 | 0.46 | 1.2 | 0.16 | -0.07 | 0.07 | 0.02 | 0 | 0.04 | 0.12 | 0.7675 | 0.3625 | 0.275283 | 0.716171 | 0.070363 | 1 | 0.999631 | 0.700187 |
| Sort1_Q6PHU5_S791s_1 | -1.55 | -0.89 | 0.32 | -1.21 | -1.86 | 1.47 | -0.93 | -0.1 | -2.3 | -0.72 | -0.3 | -0.59 | 0.27 | 0.27 | 0.35 | -0.16 | 0.4775 | -0.145 | 1.015 | 0.037724 | 0.511711 | 0.795848 | 1 | 0.999631 | 0.458331 |
| Sos1_Q62245_S1082s_1 | -0.3 | -0.22 | 1.26 | 1.03 | -0.61 | -0.74 | -0.16 | 0.83 | -1.4 | -1.3 | 0 | 1.12 | -0.11 | -1.02 | 0.46 | 1.24 | -0.6125 | -0.8375 | -0.3 | 0.60544 | 0.244518 | 0.217968 | 1 | 0.999631 | 0.85817 |
| Sos1_Q62245_S1147s_1 | 1.13 | 0.9 | 1.84 | 1.06 | 0.13 | 0.96 | 0.86 | 0.67 | 1.1 | 1.62 | 0.45 | -0.39 | 0.42 | 0.26 | 0.32 | 0.06 | -0.5775 | -0.5375 | -0.9675 | 0.01062 | 0.099408 | 0.265727 | 1 | 0.999631 | 0.383871 |
| Sos1_Q62245_S1196s_1 | -0.67 | -0.71 | -0.76 | -0.86 | -0.7 | -0.98 | -1.02 | -0.71 | -0.4 | -0.52 | -0.94 | 0.03 | -0.63 | -0.4 | -0.51 | -0.43 | -0.1025 | 0.2925 | 0.2575 | 0.333792 | 0.669631 | 0.326113 | 1 | 0.999631 | 0.728079 |
| Sos2_S4R2C4_S1003s_1 | 0.36 | -0.29 | -0.29 | 0.2 | 0 | -0.48 | 0.48 | -0.15 | 0.31 | -0.37 | 0.16 | 0.24 | -0.59 | 0.33 | -0.29 | 0.12 | -0.0325 | 0.09 | -0.1025 | 0.760098 | 0.917361 | 0.766365 | 1 | 0.999631 | 0.914614 |
| Sowaha_Q8BLS7_S242s_1 | 2.84 | 0.15 | 1.02 | 2.21 | 0.21 | -0.51 | 2.22 | -0.26 | 0.52 | 1.24 | 0.43 | -0.57 | -1.99 | -1.34 | -1.57 | -0.74 | -1.14 | -1.15 | -2.965 | 0.00059 | 0.155931 | 0.096503 | 1 | 0.999631 | 0.351308 |
| Sowaha_Q8BLS7_S258s_1 | 0.34 | 1.04 | -1.65 | -0.08 | 0.42 | 0.46 | -0.17 | -0.65 | 0.65 | 0.55 | -1.58 | -0.93 | 0.31 | 0.65 | -0.37 | -1.48 | 0.1025 | -0.24 | -0.135 | 0.83815 | 0.857809 | 0.734401 | 1 | 0.999631 | 0.946179 |
| Sox1_P53783_S325s_1 | 1.48 | 0.48 | -2.64 | -2.42 | -0.01 | 0.39 | -1.73 | -2.26 | 1.01 | 1.13 | -3.39 | -2.75 | -0.31 | 0.39 | -2.65 | -2.8 | -0.1275 | -0.225 | -0.5675 | 0.613531 | 0.902967 | 0.867844 | 1 | 0.999631 | 0.860866 |
| Sox10_Q04888_S24s_1 | 0.36 | 1.57 | 0.34 | 0.84 | 1.03 | 0.15 | 0.06 | 0.55 | 0.92 | 0.92 | 0.49 | 1.38 | 1.19 | 1.34 | 1.23 | 1.35 | -0.33 | 0.15 | 0.5 | 0.173727 | 0.392326 | 0.685515 | 1 | 0.999631 | 0.642116 |
| Sox10_Q04888_S45s_1 | 0.72 | 1.39 | 1.49 | 0.77 | 0.43 | 0.97 | 0.47 | 0.63 | 1.01 | 1.37 | 0.4 | 0.54 | 0.82 | 1.48 | 1.06 | 1.6 | -0.4675 | -0.2625 | 0.1475 | 0.664224 | 0.142586 | 0.451856 | 1 | 0.999631 | 0.882037 |
| Sox5_B2KFM4_S327s_1 | 0.52 | 0.58 | 0.7 | 0.18 | -0.32 | -0.39 | 0.24 | 0.08 | -0.43 | 0.89 | 0.61 | 0.41 | -0.53 | 0.48 | 0.89 | 0.5 | -0.5925 | -0.125 | -0.16 | 0.664852 | 0.053797 | 0.719152 | 1 | 0.999631 | 0.882387 |
| Spag9_F8WH31_S1036s_1 | -0.53 | 0.59 | 0.25 | 0.81 | 1.12 | 0.96 | 0.81 | 0.24 | 1.07 | -0.13 | -0.01 | 0.47 | -0.43 | 0.66 | 0.97 | 0.16 | 0.5025 | 0.07 | 0.06 | 0.88942 | 0.194787 | 0.864094 | 1 | 0.999631 | 0.963505 |
| Spag9_F8WH31_S1086s_1 | -1.09 | -0.24 | 0.22 | -0.23 | 0.11 | -0.64 | -0.27 | 0.38 | 0.64 | -0.25 | 0.36 | 0.86 | -0.43 | 0.52 | 0.49 | -0.04 | 0.23 | 0.7375 | 0.47 | 0.24304 | 0.539151 | 0.079493 | 1 | 0.999631 | 0.685637 |
| Spag9_F8WH31_S175s_1 | 1.81 | 2.48 | 1.7 | 0.55 | 0.69 | 1.58 | 1.23 | -0.08 | 1.2 | 2.36 | 0.02 | 0.12 | 2.38 | 0.06 | 0.64 | 0.53 | -0.78 | -0.71 | -0.7325 | 0.234025 | 0.146069 | 0.265652 | 1 | 0.999631 | 0.683077 |
| Spag9_F8WH31_S578s_1 | 2.42 | 2.45 | 1.11 | 1.73 | 1.82 | 2.3 | 1.96 | 1.61 | 1.54 | 2.19 | 1.48 | 0.29 | 0.3 | 1.54 | 1.42 | 1.15 | -0.005 | -0.5525 | -0.825 | 0.080065 | 0.989191 | 0.272292 | 1 | 0.999631 | 0.545579 |
| Sparcl1_P70663_S155s_1 | 0.57 | 0.64 | 0.58 | 0.43 | 0.37 | 0.75 | 0.43 | 0.1 | 0.82 | 1.19 | 0.86 | 0.61 | 0.7 | 1.37 | 0.79 | 0.89 | -0.1425 | 0.315 | 0.3825 | 0.199083 | 0.578409 | 0.240238 | 1 | 0.999631 | 0.657167 |
| Sparcl1_P70663_S171s_1 | 0.21 | 1.16 | 1.3 | 0.54 | 0.79 | 1.35 | 0.27 | 0.86 | 2.29 | 1.45 | 0.66 | 1.1 | 1.48 | 1.77 | 0.88 | 0.82 | 0.015 | 0.5725 | 0.435 | 0.270363 | 0.966917 | 0.20377 | 1 | 0.999631 | 0.698438 |
| Sparcl1_P70663_S406s_1 | 1.38 | 0.08 | 2.67 | 1.48 | 0.45 | 1.61 | 1.83 | -1.54 | 0.5 | 2.14 | 0.4 | 1.09 | -0.3 | -0.24 | 0.67 | 0 | -0.815 | -0.37 | -1.37 | 0.027129 | 0.33045 | 0.543741 | 1 | 0.999631 | 0.423069 |
| Sparcl1_P70663_S411s_1 | 1.36 | 0.37 | 1.51 | 0.79 | 0.3 | 1.17 | 1.42 | 0.24 | 1.22 | 1.96 | 0.58 | 0.58 | -0.32 | -0.24 | 0.63 | -0.24 | -0.225 | 0.0775 | -1.05 | 0.019025 | 0.579243 | 0.854618 | 1 | 0.999631 | 0.391024 |
| Sparcl1_P70663_S70s_1 | 0.36 | 0.23 | 2.03 | 0.82 | -0.04 | 0.75 | 0.86 | 0.24 | 0.31 | 0.71 | 0.48 | 1.27 | 2.29 | 1.37 | 1.82 | 2.05 | -0.4075 | -0.1675 | 1.0225 | 0.044312 | 0.372764 | 0.711258 | 1 | 0.999631 | 0.475443 |
| Sparcl1_P70663_S70s_2 | -0.79 | 0.96 | 1.51 | 0.34 | 0.72 | 1.26 | 0.68 | 0.94 | 1.04 | 0.87 | 0.8 | 2.05 | 0.88 | 1.36 | 1.75 | 1.29 | 0.395 | 0.685 | 0.815 | 0.128207 | 0.422039 | 0.21965 | 1 | 0.999631 | 0.598614 |
| Sparcl1_P70663_S78s_1 | -0.4 | 0.2 | 2.19 | 1.21 | 0 | 0.68 | 0.84 | 0.51 | 0.46 | 0.74 | 0.35 | 1.59 | 0.34 | 0.95 | 1.86 | 1.65 | -0.2925 | -0.015 | 0.4 | 0.514323 | 0.597048 | 0.979423 | 1 | 0.999631 | 0.819042 |
| Sparcl1_P70663_S78s_2 | -0.92 | 0.8 | 1.51 | 0.34 | 0.6 | 1.2 | 0.68 | 0.94 | 1.07 | 0.79 | 0.8 | 2.05 | 0.67 | 1.3 | 1.75 | 1.29 | 0.4225 | 0.745 | 0.82 | 0.141855 | 0.403987 | 0.195688 | 1 | 0.999631 | 0.612726 |
| Spast_Q9QYY8_S15s_1 | -0.65 | -1.64 | -1.11 | -0.82 | -0.71 | -1.95 | -0.87 | -0.15 | -2.18 | -0.81 | 0.93 | -1.2 | -3.18 | -1.89 | 0.11 | -0.16 | 0.135 | 0.24 | -0.225 | 0.751863 | 0.752546 | 0.699672 | 1 | 0.999631 | 0.912945 |
| Spast_Q9QYY8_S171s_1 | 2.19 | -0.27 | -1.75 | 0.27 | 0.12 | -0.1 | 1.15 | -0.53 | -0.38 | 0.5 | 0.61 | -0.78 | -2.08 | -1.63 | -1.29 | -1.43 | 0.05 | -0.1225 | -1.7175 | 0.038585 | 0.948718 | 0.87457 | 1 | 0.999631 | 0.461663 |
| Spast_Q9QYY8_T217t_1 | 1.66 | 0.11 | 0.46 | 1.19 | 1.94 | 1.39 | 1.21 | 1.21 | 0.82 | 0.41 | 1.44 | 0.17 | -1.58 | 1.43 | 0.97 | 1.02 | 0.5825 | -0.145 | -0.395 | 0.56816 | 0.164549 | 0.743182 | 1 | 0.999631 | 0.843185 |
| Spata18_J3QPR5_S132s_1 | 1.67 | -0.32 | 0.92 | -0.39 | -2.26 | 0.37 | 0.6 | -0.58 | -0.21 | 1.1 | -1.29 | -1.5 | -1.3 | -0.75 | -0.27 | -1.45 | -0.9375 | -0.945 | -1.4125 | 0.022937 | 0.214821 | 0.194579 | 1 | 0.999631 | 0.41165 |
| Spata2_Q8K004_S247s_1 | -0.64 | 0.27 | 0.01 | -0.32 | 0.35 | -0.36 | -0.44 | -0.34 | -0.32 | -0.45 | 0.18 | -0.08 | 0.67 | 0.52 | -0.35 | 0.35 | -0.0275 | 0.0025 | 0.4675 | 0.206583 | 0.931222 | 0.993552 | 1 | 0.999631 | 0.6605 |
| Spata2l_Q8BNN1_S318s_1 | -0.11 | 0.36 | -0.83 | 0.69 | 0.42 | -1.15 | 0.34 | 0.47 | 0.38 | 0.78 | -0.19 | -0.2 | -1.15 | -0.84 | 1.3 | -1.04 | -0.0075 | 0.165 | -0.46 | 0.455497 | 0.987613 | 0.690665 | 1 | 0.999631 | 0.790041 |
| Spata2l_Q8BNN1_S326s_1 | 0.04 | -0.21 | -4.94 | -3.14 | -0.19 | -1.32 | -2.22 | -3.38 | -0.48 | -0.54 | -2.1 | -5.43 | -0.16 | -0.39 | -5.18 | -3.62 | 0.285 | -0.075 | -0.275 | 0.849207 | 0.808765 | 0.957871 | 1 | 0.999631 | 0.949525 |
| Spats2l_E9Q106_S179s_1 | 0.14 | 0.74 | 0.39 | 0.93 | 0.65 | 0.89 | 0.98 | 1.22 | 0.73 | 0.68 | 0.98 | 0.98 | -1.06 | 0.76 | 0.98 | 0.94 | 0.385 | 0.2925 | -0.145 | 0.7719 | 0.200319 | 0.318581 | 1 | 0.999631 | 0.919413 |
| Specc1_Q5SXY1_S133s_1 | 0.85 | 1.54 | 0.21 | -0.27 | 0.54 | 2.06 | 0.12 | 0.25 | 2.24 | 2.26 | 0.24 | 0.5 | 0.23 | 1.92 | 0.76 | 0.93 | 0.16 | 0.7275 | 0.3775 | 0.46069 | 0.770385 | 0.251221 | 1 | 0.999631 | 0.791962 |
| Specc1_Q5SXY1_S355s_1 | -0.52 | 0.61 | 0.51 | 0.18 | -0.08 | -0.22 | -0.57 | 0.21 | -0.16 | -0.21 | 0.12 | 1.28 | 0.54 | 0.43 | -0.41 | 1.04 | -0.36 | 0.0625 | 0.205 | 0.621049 | 0.302036 | 0.884655 | 1 | 0.999631 | 0.861888 |
| Specc1_Q5SXY1_S55s_1 | 1.37 | 0.77 | 0.12 | 1.93 | 1.17 | 0.72 | 1.18 | 1.41 | 0.57 | 0.85 | 1.6 | 1.34 | -1.19 | 0.65 | 0.36 | 1.49 | 0.0725 | 0.0425 | -0.72 | 0.260489 | 0.860905 | 0.924101 | 1 | 0.999631 | 0.692624 |
| Specc1_Q5SXY1_S847s_1 | 0.69 | -0.23 | -1.47 | 0.71 | 0.33 | 0.09 | 0.49 | -0.41 | 0.04 | -0.46 | 0.35 | 0.23 | -0.69 | 0.21 | 0.81 | 0.35 | 0.2 | 0.115 | 0.245 | 0.662218 | 0.697478 | 0.822638 | 1 | 0.999631 | 0.881177 |
| Specc1_Q5SXY1_S9s_1 | 0.52 | 0.59 | 0.25 | 0.34 | 1.47 | 2.1 | 0.31 | 0.46 | 0.81 | 0.51 | 0.37 | 1.56 | 0.02 | 2.51 | 1.03 | 0.3 | 0.66 | 0.3875 | 0.54 | 0.320721 | 0.145104 | 0.256929 | 1 | 0.999631 | 0.723691 |
| Specc1l_Q2KN98_S171s_1 | 1.42 | 0.65 | 1.73 | 0.86 | 0.28 | 0.3 | 1.14 | 0.28 | 1.01 | 1.61 | 0.58 | 1.18 | 0.56 | 0.19 | 1.31 | 0.37 | -0.665 | -0.07 | -0.5575 | 0.168361 | 0.085292 | 0.846071 | 1 | 0.999631 | 0.637627 |
| Speg_D3Z7T0_S34s_1 | -0.67 | -0.52 | 0.49 | -0.23 | 0.67 | -1.28 | -0.08 | -1.01 | -0.17 | -1.07 | -0.51 | -0.31 | 0.06 | 0.06 | -0.45 | -0.67 | -0.1925 | -0.2825 | -0.0175 | 0.961618 | 0.693404 | 0.438523 | 1 | 0.999631 | 0.986594 |
| Speg_E9QQ25_S1177s_1 | -0.17 | 0.63 | -0.53 | -0.59 | 0.56 | 0.17 | -0.31 | -0.33 | 0.24 | 0.56 | 0.14 | -0.52 | 0.91 | -0.1 | -0.31 | -0.09 | 0.1875 | 0.27 | 0.2675 | 0.519281 | 0.615247 | 0.485537 | 1 | 0.999631 | 0.819456 |
| Speg_E9QQ25_S2020s_1 | -0.15 | 0.24 | 0.01 | -0.21 | -0.61 | -0.1 | -0.48 | 0.08 | -0.38 | 0.43 | 0.13 | 0.12 | 0.52 | 0.02 | -0.14 | 0.81 | -0.25 | 0.1025 | 0.33 | 0.319478 | 0.374959 | 0.72082 | 1 | 0.999631 | 0.722906 |
| Speg_E9QQ25_S2042s_1 | 0.21 | 0.8 | -0.04 | -0.09 | 0.35 | 0.79 | -0.34 | 0.24 | 0.96 | 0.9 | 0.4 | 0.12 | 1.16 | 1.07 | 0.1 | 0.46 | 0.04 | 0.375 | 0.4775 | 0.214582 | 0.907098 | 0.279768 | 1 | 0.999631 | 0.66742 |
| Speg_E9QQ25_S2114s_1 | 0.51 | -0.21 | -0.79 | 0.11 | -0.35 | -0.43 | -0.06 | -0.42 | -0.26 | 0.45 | -0.79 | -0.15 | -0.37 | -0.51 | -0.61 | -0.22 | -0.22 | -0.0925 | -0.3325 | 0.349382 | 0.509394 | 0.813083 | 1 | 0.999631 | 0.736792 |
| Speg_E9QQ25_S2115s_1 | -0.82 | -0.6 | -0.53 | -0.08 | -1.78 | -1.67 | -0.21 | -0.31 | -0.16 | 0.45 | -0.7 | -0.23 | 0.69 | -1.11 | -0.3 | 0.05 | -0.485 | 0.3475 | 0.34 | 0.425678 | 0.28552 | 0.310531 | 1 | 0.999631 | 0.774624 |
| Speg_E9QQ25_S2135s_1 | -0.31 | 0.02 | -0.4 | -0.46 | -0.23 | -0.04 | -0.3 | -0.57 | 0.34 | -0.17 | -0.45 | -0.14 | 0.09 | 0.25 | -0.49 | -0.68 | 0.0025 | 0.1825 | 0.08 | 0.806204 | 0.992304 | 0.527916 | 1 | 0.999631 | 0.935192 |
| Speg_E9QQ25_S2171s_1 | -0.95 | -1.58 | -0.2 | 0.17 | -0.87 | -0.95 | -0.59 | -0.2 | -1.06 | -1.56 | -0.69 | 0.51 | -0.08 | -0.75 | -0.28 | 0.05 | -0.0125 | -0.06 | 0.375 | 0.397054 | 0.976294 | 0.912536 | 1 | 0.999631 | 0.761093 |
| Speg_E9QQ25_S2327s_1 | -0.27 | -0.24 | -0.31 | 0.28 | -0.63 | -0.1 | 0.46 | 0.55 | -0.36 | -0.32 | -0.6 | 0.6 | -0.92 | -0.24 | 0.43 | 1.27 | 0.205 | -0.035 | 0.27 | 0.573594 | 0.552089 | 0.918534 | 1 | 0.999631 | 0.844597 |
| Speg_E9QQ25_S2347s_1 | 0.52 | 0.12 | 0.02 | 0.16 | 0.01 | 0.24 | 0.68 | 0.46 | 0.13 | 0.62 | 0.47 | -0.17 | 0.3 | 0.15 | -0.35 | 0.12 | 0.1425 | 0.0575 | -0.15 | 0.611201 | 0.602027 | 0.843896 | 1 | 0.999631 | 0.859461 |
| Speg_E9QQ25_S2396s_1 | 0.97 | -0.9 | -0.82 | 1.26 | 0.96 | -1.38 | -0.35 | 1.67 | 1.14 | 1.19 | 1.17 | -0.31 | -0.44 | 0.46 | 0.36 | -0.11 | 0.0975 | 0.67 | -0.06 | 0.91529 | 0.900224 | 0.29361 | 1 | 0.999631 | 0.97286 |
| Speg_E9QQ25_S2451s_1 | 0.4 | 0.24 | -0.49 | 0.84 | 0.63 | 0.79 | 0.26 | 0.25 | 0.85 | 0.3 | 0.44 | -0.34 | -0.49 | 1.34 | 0.68 | 0.6 | 0.235 | 0.065 | 0.285 | 0.54176 | 0.497288 | 0.867147 | 1 | 0.999631 | 0.832428 |
| Speg_E9QQ25_S2461s_1 | 1.4 | -0.38 | 0.33 | 1.58 | 1.01 | 0.41 | 1.38 | 0.83 | 0.35 | 0.69 | 2.09 | 1.43 | 0.03 | 0.52 | 0.75 | 1.3 | 0.175 | 0.4075 | -0.0825 | 0.870729 | 0.715223 | 0.47149 | 1 | 0.999631 | 0.958735 |
| Speg_E9QQ25_S2499s_1 | 1.42 | 0.98 | 1.14 | 1.55 | 0.74 | 1.07 | 1.45 | 1.29 | 1.67 | 1.85 | 1.67 | 1.3 | -0.03 | 0.61 | 1.22 | 1.2 | -0.135 | 0.35 | -0.5225 | 0.17681 | 0.63313 | 0.224028 | 1 | 0.999631 | 0.642116 |
| Speg_E9QQ25_S2777s_1 | 1.29 | 0.4 | -0.03 | 0.75 | 0.91 | 0.09 | 1.04 | 1.05 | -0.22 | -0.25 | 0.46 | 0.1 | -0.25 | -0.4 | -0.33 | 0.5 | 0.17 | -0.58 | -0.7225 | 0.083271 | 0.653378 | 0.130299 | 1 | 0.999631 | 0.549732 |
| Speg_E9QQ25_S2777s_2 | 0.01 | -1.02 | -0.04 | -1.38 | -1.55 | -1.6 | -0.28 | -1.85 | -0.93 | -0.12 | 0.11 | -0.95 | -0.57 | -1.34 | 0.26 | 0.38 | -0.7125 | 0.135 | 0.29 | 0.570628 | 0.155352 | 0.759297 | 1 | 0.999631 | 0.84338 |
| Speg_E9QQ25_S490s_1 | -0.23 | 0.12 | -0.07 | -0.18 | 0.1 | 0.13 | 0 | -0.16 | -0.28 | -0.42 | 0.18 | -0.49 | 0.13 | 0.29 | 0.05 | -0.06 | 0.1075 | -0.1625 | 0.1925 | 0.479767 | 0.657169 | 0.556861 | 1 | 0.999631 | 0.801333 |
| Speg_E9QQ25_S511s_1 | 0.21 | 0.59 | -0.07 | 0.45 | -0.06 | -0.21 | 0.2 | 0.46 | 0.66 | -0.09 | 0.77 | 0.46 | 0.5 | 0.34 | -0.29 | 0.77 | -0.1975 | 0.155 | 0.035 | 0.917006 | 0.492362 | 0.617643 | 1 | 0.999631 | 0.973513 |
| Speg_E9QQ25_S554s_1 | 0.82 | -0.05 | -0.52 | 0.76 | 1.09 | 0.55 | 0.7 | 0.92 | 0.19 | -0.73 | 1.27 | -0.05 | -0.36 | 1.3 | 0.78 | 0.29 | 0.5625 | -0.0825 | 0.25 | 0.597686 | 0.148407 | 0.868972 | 1 | 0.999631 | 0.855033 |
| Speg_E9QQ25_S858s_1 | 1.49 | -1.31 | -0.39 | 0.4 | -1.59 | -1.12 | 1.13 | -1.13 | -1.92 | 0.57 | -1.57 | -1.97 | -2.26 | -1.39 | -0.94 | -1.01 | -0.725 | -1.27 | -1.4475 | 0.035844 | 0.345543 | 0.114835 | 1 | 0.999631 | 0.453811 |
| Speg_E9QQ25_T2303t_1 | 0.81 | -0.93 | -1.18 | 0.35 | 1.17 | -0.55 | 0.92 | 0.96 | -0.28 | -1.38 | 0.68 | 0.14 | -1.07 | 0.07 | 0.09 | -0.08 | 0.8625 | 0.0275 | -0.01 | 0.984801 | 0.153571 | 0.963089 | 1 | 0.999631 | 0.995133 |
| Speg_E9QQ25_T2774t_1 | -0.29 | -0.76 | 0.31 | -1.63 | -2.32 | -1.22 | -0.85 | -0.49 | -1.74 | -0.31 | 0 | -0.4 | 0.66 | -2.35 | -0.22 | 0.58 | -0.6275 | -0.02 | 0.26 | 0.717778 | 0.251605 | 0.969662 | 1 | 0.999631 | 0.903626 |
| Speg_E9QQ25_T2774t_2 | 0.01 | -1.02 | -0.04 | -1.38 | -1.55 | -1.6 | -0.28 | -1.85 | -0.93 | -0.12 | 0.11 | -0.95 | -0.57 | -1.34 | 0.26 | 0.38 | -0.7125 | 0.135 | 0.29 | 0.570628 | 0.155352 | 0.759297 | 1 | 0.999631 | 0.84338 |
| Speg_E9QQ25_T2847t_1 | 0.77 | -0.22 | 0.18 | 0.49 | 0.4 | 0.82 | 0.64 | 0.81 | 0.15 | 0.29 | 0.38 | -0.05 | -0.61 | 0.23 | 0.26 | 0.3 | 0.3625 | -0.1125 | -0.26 | 0.472502 | 0.242627 | 0.712616 | 1 | 0.999631 | 0.799014 |
| Speg_E9QQ25_T453t_1 | -1.14 | 0.35 | -0.29 | -1.18 | -0.55 | -0.45 | -0.6 | -0.6 | 0.03 | -0.99 | -0.29 | 0.53 | 1.03 | -0.17 | -0.12 | -0.16 | 0.015 | 0.385 | 0.71 | 0.149257 | 0.968643 | 0.421052 | 1 | 0.999631 | 0.621356 |
| Spen_Q62504_S1267s_1 | 1.82 | 0.21 | 1.25 | 1.48 | -0.29 | 0.73 | 1.58 | 0.36 | 0.07 | 1.67 | 0.33 | 0.02 | -1.04 | -0.34 | 0.08 | -0.23 | -0.595 | -0.6675 | -1.5725 | 0.004162 | 0.244314 | 0.198953 | 1 | 0.999631 | 0.351308 |
| Spen_Q62504_S1276s_1 | 0.96 | 0.04 | 1.05 | 0.72 | -0.49 | 0.41 | 1.03 | 0.17 | -0.02 | 1.13 | 0.14 | 0.03 | 0.07 | -0.68 | 0.12 | -0.34 | -0.4125 | -0.3725 | -0.9 | 0.026513 | 0.310292 | 0.337314 | 1 | 0.999631 | 0.422285 |
| Spen_Q62504_S188s_1 | -0.03 | -3.55 | -0.99 | -0.82 | -2.48 | -3.37 | -0.03 | -1.92 | -2.86 | -1.93 | -0.93 | -0.07 | -1.35 | -3.69 | -0.14 | -2.35 | -0.6025 | -0.1 | -0.535 | 0.565127 | 0.508255 | 0.906349 | 1 | 0.999631 | 0.84139 |
| Spen_Q62504_S2128s_1 | 0.06 | -0.28 | -0.38 | -0.63 | -0.01 | -1.05 | -0.49 | -0.06 | -1.54 | -0.78 | -0.56 | -0.23 | -0.6 | -0.04 | -0.36 | 0.21 | -0.095 | -0.47 | 0.11 | 0.728265 | 0.770418 | 0.201537 | 1 | 0.999631 | 0.905923 |
| Spen_Q62504_S647s_1 | 0.73 | 0.2 | 0.2 | 0.16 | 0.66 | 0.05 | 0.26 | -0.46 | 0.16 | 0.84 | -0.12 | -0.71 | 0.24 | -0.27 | -0.38 | -0.77 | -0.195 | -0.28 | -0.6175 | 0.082225 | 0.544721 | 0.461358 | 1 | 0.999631 | 0.54792 |
| Spen_Q62504_S758s_2 | 0.48 | -0.51 | -0.12 | -0.51 | -0.38 | 0.1 | -0.4 | 0.06 | -0.53 | 0.35 | -0.14 | -0.48 | -0.58 | 0.34 | 0.73 | 0.33 | 0.01 | -0.035 | 0.37 | 0.356378 | 0.974982 | 0.920116 | 1 | 0.999631 | 0.740405 |
| Spg20_D3Z3F8_S126s_1 | 0 | 0.06 | -0.09 | 0.07 | -0.39 | -0.22 | -0.13 | 0.34 | 0.01 | 0.24 | -0.27 | 0.03 | 0.53 | 0.82 | 0.35 | 0.32 | -0.11 | -0.0075 | 0.495 | 0.093638 | 0.676376 | 0.97627 | 1 | 0.999631 | 0.558128 |
| Sphkap_E9PUC4_S1067s_1 | 1.18 | 0.03 | -0.43 | 0.11 | 1.35 | -0.18 | 0.36 | 0.21 | 1.32 | 0.43 | 0.13 | 0.39 | -3.09 | 0.55 | 0.02 | 0.22 | 0.2125 | 0.345 | -0.7975 | 0.328495 | 0.64149 | 0.428511 | 1 | 0.999631 | 0.725792 |
| Sphkap_E9PUC4_S1070s_1 | -1.82 | 1.61 | 1.25 | -0.74 | -1.34 | 0.8 | -0.27 | -0.48 | 0.6 | -0.41 | 0.88 | 1.02 | 2.17 | 1.15 | 0.89 | 0.67 | -0.3975 | 0.4475 | 1.145 | 0.160153 | 0.625175 | 0.565959 | 1 | 0.999631 | 0.629764 |
| Sphkap_E9PUC4_S1223s_1 | 1.68 | -0.59 | 0.11 | 1.22 | 0.15 | 0.45 | 1.88 | 0.31 | 0.48 | 1.09 | 0.77 | -0.16 | -1.94 | -0.96 | -0.37 | -0.66 | 0.0925 | -0.06 | -1.5875 | 0.018272 | 0.875674 | 0.911605 | 1 | 0.999631 | 0.391024 |
| Sphkap_E9PUC4_S1244s_1 | -0.26 | -0.1 | -0.22 | -0.58 | -0.13 | -0.57 | -0.5 | -0.18 | -0.18 | -0.34 | -0.3 | 0.25 | 0.14 | -0.2 | -0.41 | 0.14 | -0.055 | 0.1475 | 0.2075 | 0.479938 | 0.831603 | 0.593315 | 1 | 0.999631 | 0.801333 |
| Sphkap_E9PUC4_S1273s_1 | 0.58 | 0.57 | -0.99 | 1.57 | 1.44 | 1.54 | 1.66 | 2.28 | 1.45 | 0.12 | 1.34 | 0.67 | 0.14 | 1.11 | 1.51 | 0.94 | 1.2975 | 0.4625 | 0.4925 | 0.387225 | 0.030538 | 0.421551 | 1 | 0.999631 | 0.756827 |
| Sphkap_E9PUC4_S1369s_1 | 0.33 | -0.06 | -1.38 | 0.39 | 1.91 | 0.57 | 2.16 | 1.13 | 0.54 | -1.22 | 2.03 | 1.41 | -0.4 | -0.57 | 0.53 | 0.54 | 1.6225 | 0.87 | 0.205 | 0.67672 | 0.009831 | 0.248148 | 1 | 0.999631 | 0.886862 |
| Sphkap_E9PUC4_S1403s_1 | 0.87 | -0.81 | -4.54 | -0.55 | 0.94 | -0.14 | -0.2 | -1.18 | 0.01 | -0.84 | -1.27 | -3.17 | -2.83 | -0.66 | -2.5 | -2.44 | 1.1125 | -0.06 | -0.85 | 0.433695 | 0.30724 | 0.958211 | 1 | 0.999631 | 0.779567 |
| Sphkap_E9PUC4_S841s_1 | 1.34 | -0.94 | -0.77 | 0.4 | -0.43 | -0.7 | 1.04 | -0.28 | 0.09 | 1.51 | -0.07 | -0.95 | -0.56 | -2.43 | -2.02 | -0.74 | -0.1 | 0.1375 | -1.445 | 0.043902 | 0.866975 | 0.835588 | 1 | 0.999631 | 0.475443 |
| Spire1_D3YTL8_S229s_1 | -0.03 | 0.47 | -0.13 | -0.22 | 0.66 | 0.28 | -0.16 | 0.23 | -0.21 | -0.64 | -0.35 | -0.7 | -0.17 | 0.61 | -0.06 | 0.76 | 0.23 | -0.4975 | 0.2625 | 0.450962 | 0.442995 | 0.105684 | 1 | 0.999631 | 0.787797 |
| Spire1_D3YTL8_S275s_1 | 0.73 | -0.32 | 0.33 | -0.05 | -0.56 | 0.02 | 0.52 | 0.53 | -0.13 | 0.28 | -0.79 | -0.3 | -1.03 | -0.68 | -0.06 | 0.4 | -0.045 | -0.4075 | -0.515 | 0.228261 | 0.902019 | 0.267252 | 1 | 0.999631 | 0.679038 |
| Spire1_D3YTL8_S565s_1 | -2.07 | -1.36 | -1.84 | -0.71 | -0.83 | -3.35 | -2.09 | -1.43 | -2.52 | -3.33 | -2.18 | -2.13 | -0.34 | -0.97 | -1.34 | -2.47 | -0.43 | -1.045 | 0.215 | 0.675706 | 0.453198 | 0.028995 | 1 | 0.999631 | 0.885987 |
| Spock2_F6SLR4_S72s_1 | -0.57 | 0.86 | 0.14 | -0.37 | 0.27 | 0.8 | 0.94 | 1.27 | 0.43 | -0.94 | 1.32 | 2.37 | 1.48 | 1.47 | 2.86 | 1.82 | 0.805 | 0.78 | 1.8925 | 0.001897 | 0.062202 | 0.272926 | 1 | 0.999631 | 0.351308 |
| Sppl2b_Q3TD49_S534s_1 | 0.96 | 1.85 | 1.59 | 1.14 | 1 | 1.57 | 0.33 | 1.1 | 2.5 | 1.51 | 1.12 | 1.51 | 1.62 | 1.31 | 0.14 | 1.12 | -0.385 | 0.275 | -0.3375 | 0.409901 | 0.291851 | 0.475839 | 1 | 0.999631 | 0.767719 |
| Spred1_Q924S8_S318s_1 | 1.34 | -0.13 | -0.47 | 0.97 | 1.71 | 0.82 | 0.67 | 0.47 | 0.02 | -0.19 | 0.77 | 0.3 | -0.89 | 0.29 | 0.33 | 1.29 | 0.49 | -0.2025 | -0.1725 | 0.763488 | 0.324443 | 0.664691 | 1 | 0.999631 | 0.915649 |
| Spred2_Q5SRF8_S195s_1 | 0.94 | -1.1 | -2.97 | -2.83 | -1.17 | -1.79 | -2.41 | -2.89 | -1.4 | 0.19 | -3.38 | -3.21 | -1.09 | -1.44 | -2.9 | -3.33 | -0.575 | -0.46 | -0.7 | 0.451944 | 0.507855 | 0.668624 | 1 | 0.999631 | 0.788148 |
| Spred3_Q6P6N5_S286s_1 | 1.34 | -0.08 | 0.86 | 0.66 | -0.14 | 0.43 | 0.78 | -0.29 | 0.73 | 1.66 | 0.47 | 0.59 | 0.52 | -0.32 | 0.05 | -0.52 | -0.5 | 0.1675 | -0.7625 | 0.079541 | 0.223157 | 0.683321 | 1 | 0.999631 | 0.545579 |
| Spry1_Q9QXV9_S133s_1 | 0.53 | -1.54 | -0.65 | -0.2 | -0.1 | -0.88 | -0.74 | -0.62 | -0.12 | 0.6 | 0.78 | -0.65 | -0.68 | -0.22 | -0.79 | -0.53 | -0.12 | 0.6175 | -0.09 | 0.84133 | 0.789795 | 0.247003 | 1 | 0.999631 | 0.947261 |
| Spry2_Q9QXV8_S117s_1 | 0.65 | -1.14 | -1.32 | -0.09 | 0.36 | -0.23 | -0.04 | -0.36 | -0.47 | -0.48 | -0.31 | -0.47 | -0.82 | -1.04 | -0.42 | -0.3 | 0.4075 | 0.0425 | -0.17 | 0.723814 | 0.392331 | 0.9254 | 1 | 0.999631 | 0.905674 |
| Sptan1_P16546_S1029s_1 | -0.03 | 0.03 | -0.92 | -0.37 | 0.1 | -0.08 | -0.53 | -0.14 | 0.15 | 0.18 | -0.9 | -0.27 | 0.2 | 0.28 | -0.47 | -0.81 | 0.16 | 0.1125 | 0.1225 | 0.747092 | 0.608332 | 0.757683 | 1 | 0.999631 | 0.911931 |
| Sptan1_P16546_S1031s_1 | -0.11 | -0.54 | -1.61 | -0.28 | 0.19 | -1.3 | -0.75 | -0.3 | -0.29 | -1.37 | -0.9 | -0.65 | -1.17 | 0.03 | -0.44 | -1.12 | 0.095 | -0.1675 | -0.04 | 0.928255 | 0.832146 | 0.685985 | 1 | 0.999631 | 0.976721 |
| Sptan1_P16546_S1190s_1 | -2.12 | 1.08 | -2.82 | 1.15 | 0.48 | -0.36 | -0.97 | -0.6 | -0.74 | -1.95 | 0.68 | 0.11 | -0.09 | 2.17 | 0.97 | 2.4 | 0.315 | 0.2025 | 2.04 | 0.068764 | 0.738149 | 0.843706 | 1 | 0.999631 | 0.523794 |
| Sptan1_P16546_S1217s_1 | -0.19 | 0.29 | 0.56 | 0.25 | -0.12 | -0.05 | 0.01 | 0.21 | -0.27 | 0.07 | 0.53 | 1.03 | 0.54 | 0.43 | 0.92 | 1.13 | -0.215 | 0.1125 | 0.5275 | 0.1174 | 0.428444 | 0.75326 | 1 | 0.999631 | 0.585087 |
| Sptan1_P16546_S328s_1 | -1.78 | -0.22 | -0.01 | -0.81 | -0.97 | -0.72 | -0.74 | -0.86 | -0.9 | -1.94 | -0.92 | -0.58 | -0.07 | -1.1 | -1.06 | -0.14 | -0.1175 | -0.38 | 0.1125 | 0.812746 | 0.771169 | 0.432405 | 1 | 0.999631 | 0.937041 |
| Sptan1_P16546_S330s_1 | -1.89 | -0.57 | -0.41 | 0.13 | -0.72 | -1.56 | -1.04 | -0.56 | -0.91 | -0.92 | -1.3 | -1.1 | 0.05 | -1.28 | -1.11 | -1.57 | -0.285 | -0.3725 | -0.2925 | 0.580327 | 0.541263 | 0.401585 | 1 | 0.999631 | 0.846774 |
| Sptan1_P16546_S809s_1 | 0.66 | 0.52 | 0.12 | 0.68 | 0.49 | 0.61 | 0.04 | 1.15 | 0.54 | 0.72 | 0.72 | 0.57 | 1.01 | 0.68 | 0.24 | 0.39 | 0.0775 | 0.1425 | 0.085 | 0.783546 | 0.80556 | 0.588914 | 1 | 0.999631 | 0.922628 |
| Sptb_E9Q397_S1226s_1 | -0.48 | -0.29 | 0.38 | -0.13 | 0.01 | 0.07 | -0.87 | -0.96 | -0.27 | -0.07 | -0.51 | -0.5 | 0.13 | -0.41 | 0.07 | -0.65 | -0.3075 | -0.2075 | -0.085 | 0.799386 | 0.399031 | 0.486287 | 1 | 0.999631 | 0.931051 |
| Sptb_E9Q397_S2114s_1 | -3.76 | -1.03 | -2.17 | -0.77 | -0.12 | -1.92 | -1.15 | -1.52 | -0.84 | -0.66 | -2.14 | -2.16 | -0.1 | -1.48 | -2.61 | -1.14 | 0.755 | 0.4825 | 0.6 | 0.431888 | 0.291305 | 0.499113 | 1 | 0.999631 | 0.778676 |
| Sptb_E9Q397_T2113t_1 | 0.54 | -1.6 | -2.29 | -0.29 | 0.27 | -1.96 | -1.13 | -2.41 | -0.57 | -0.12 | -1.95 | -1.79 | -0.64 | -1.25 | -1.84 | -1.28 | -0.3975 | -0.1975 | -0.3425 | 0.582418 | 0.60384 | 0.776701 | 1 | 0.999631 | 0.847651 |
| Sptb_Q3UGX2_S2300s_1 | 0.25 | 0.75 | 0.64 | 0.05 | 0.19 | 0.41 | -0.27 | 0.38 | 0.23 | 0.68 | 0.03 | 0.4 | 0.61 | 0.41 | 0.4 | 0.17 | -0.245 | -0.0875 | -0.025 | 0.93288 | 0.414261 | 0.767234 | 1 | 0.999631 | 0.978327 |
| Sptb_Q3UGX2_T2113t_1 | 1.26 | -0.95 | 0.45 | -0.27 | -1.41 | -0.16 | 0.65 | -1.31 | -0.96 | 0.8 | -0.81 | -1.07 | -0.61 | -1.55 | -0.98 | -1.92 | -0.68 | -0.6325 | -1.3875 | 0.023154 | 0.284893 | 0.299507 | 1 | 0.999631 | 0.414424 |
| Sptbn1_Q62261_S1662s_1 | 0.6 | -0.91 | -0.16 | -1.44 | -1.93 | -0.95 | -0.75 | -1.53 | -0.82 | -0.94 | -2.21 | -2.17 | -1.54 | -0.17 | -1.83 | -2.2 | -0.8125 | -1.0575 | -0.9575 | 0.121209 | 0.122204 | 0.076089 | 1 | 0.999631 | 0.589339 |
| Sptbn1_Q62261_S2102s_1 | -0.28 | -1.64 | -0.57 | -0.75 | -1.21 | -1.58 | -0.8 | -0.69 | -1.3 | -0.37 | -1.01 | -0.99 | -1.07 | -1.31 | -1 | -0.97 | -0.26 | -0.1075 | -0.2775 | 0.442748 | 0.491939 | 0.775579 | 1 | 0.999631 | 0.782367 |

| | | | | | | | | | | | | | | | | | | | | | | | | |
|---|---|---|---|---|---|---|---|---|---|---|---|---|---|---|---|---|---|---|---|---|---|---|---|---|
| Sptbn1_Q62261_S2127s_1 | -1.84 | -1.97 | -0.33 | -0.54 | -2.32 | -1.79 | -0.79 | -0.25 | -2.56 | -2.24 | -0.12 | -0.43 | -1.68 | -2.51 | -0.32 | 0 | -0.1175 | -0.1675 | 0.0425 | 0.947949 | 0.838731 | 0.803865 | 1 | 0.999631 | 0.983162 |
| Sptbn1_Q62261_S2127s_2 | 0.34 | -0.26 | -0.96 | -0.86 | -0.16 | -0.17 | -0.79 | -0.72 | -0.25 | 0.46 | -0.99 | -1.22 | -0.15 | -0.06 | -0.65 | -1.24 | -0.025 | -0.065 | -0.09 | 0.830058 | 0.945574 | 0.889841 | 1 | 0.999631 | 0.942193 |
| Sptbn1_Q62261_S2137s_1 | -0.55 | -1.18 | -0.99 | -1.39 | -1.32 | -1.18 | -1.22 | -1.49 | -1.3 | -0.63 | -1.07 | -1.23 | -0.42 | -1.2 | -1.13 | -1.03 | -0.275 | -0.03 | 0.0825 | 0.801718 | 0.3329 | 0.92168 | 1 | 0.999631 | 0.932527 |
| Sptbn1_Q62261_S2137s_2 | 0.34 | -0.26 | -1.5 | -1.4 | -0.16 | -0.17 | -1.13 | -1.35 | -0.25 | 0.46 | -1.54 | -1.87 | -0.15 | -0.06 | -1.2 | -1.75 | 0.0025 | -0.095 | -0.085 | 0.879862 | 0.996084 | 0.881445 | 1 | 0.999631 | 0.962995 |
| Sptbn1_Q62261_S2164s_1 | -0.77 | 1.38 | 2.04 | -0.6 | -0.8 | 0.17 | -1.06 | 0.42 | 0.3 | 1 | 0.35 | 1.7 | 3.85 | 0.89 | 1.27 | 1.81 | -0.83 | 0.325 | 1.4425 | 0.109349 | 0.253906 | 0.641178 | 1 | 0.999631 | 0.572573 |
| Sptbn1_Q62261_S2164s_2 | 0.68 | 2.11 | 1.79 | 1.39 | 1.52 | 1.76 | 0.49 | 1.71 | 1.71 | 1.63 | 1.48 | 1.88 | 2.42 | 2 | 1.86 | 1.99 | -0.1225 | 0.1825 | 0.575 | 0.145855 | 0.772552 | 0.609904 | 1 | 0.999631 | 0.616968 |
| Sptbn1_Q62261_S2168s_1 | -1.38 | 0.5 | 1.97 | -1.23 | -1.59 | -0.53 | -1.79 | -0.68 | -0.4 | 0.55 | 0.06 | 1.05 | 3.15 | 0.3 | 0.64 | 1.11 | -1.1125 | 0.35 | 1.335 | 0.151888 | 0.161181 | 0.643621 | 1 | 0.999631 | 0.625198 |
| Sptbn1_Q62261_S2168s_2 | 0.68 | 2.11 | 1.79 | 1.39 | 1.52 | 1.76 | 0.49 | 1.71 | 1.71 | 1.63 | 1.48 | 1.88 | 2.42 | 2 | 1.86 | 1.99 | -0.1225 | 0.1825 | 0.575 | 0.145855 | 0.772552 | 0.609904 | 1 | 0.999631 | 0.616968 |
| Sptbn1_Q62261_S2315s_1 | 0.44 | 0.26 | 1.25 | 0.11 | 0.56 | -0.1 | 2.19 | 0.29 | 1.62 | -0.19 | 0.24 | 1.14 | 1.03 | -1.91 | -0.04 | -1.22 | 0.22 | 0.1875 | -1.05 | 0.119324 | 0.675349 | 0.690241 | 1 | 0.999631 | 0.586867 |
| Sptbn1_Q62261_S2315s_2 | 2.44 | 2.79 | 4.82 | 3.23 | 3.87 | 1.65 | 5.13 | 3 | 5.16 | 1.93 | 4.45 | 5.81 | 2.56 | 0.59 | 3.29 | 0.67 | 0.0925 | 1.0175 | -1.5425 | 0.063811 | 0.906511 | 0.259608 | 1 | 0.999631 | 0.513532 |
| Sptbn1_Q62261_S2318s_1 | 2.19 | 4.64 | 4.03 | 0.97 | 3.65 | 3.56 | 3.41 | 1.41 | 5.77 | 3.43 | 3.12 | 4.74 | 6.2 | 1.41 | 3.1 | 1.01 | 0.05 | 1.3075 | -0.0275 | 0.982089 | 0.953923 | 0.17129 | 1 | 0.999631 | 0.994846 |
| Sptbn1_Q62261_S2318s_2 | 2.31 | 2.94 | 3.49 | 1.78 | 3.87 | 1.37 | 4.24 | 1.6 | 5.23 | 2.03 | 2.97 | 4.32 | 2.95 | 0.46 | 1.91 | -0.39 | 0.14 | 1.0075 | -1.3975 | 0.08054 | 0.84888 | 0.179011 | 1 | 0.999631 | 0.545579 |
| Sptbn1_Q62261_S2339s_1 | -0.68 | -1.56 | -0.86 | -0.6 | -0.56 | -1.46 | -0.92 | -0.71 | -1.35 | -1.48 | -0.55 | -0.7 | -1 | -0.8 | -0.97 | -0.31 | 0.0125 | -0.095 | 0.155 | 0.647911 | 0.970065 | 0.789113 | 1 | 0.999631 | 0.874615 |
| Sptbn1_Q62261_S2340s_1 | -0.59 | -0.67 | -0.73 | -0.47 | -0.34 | -0.8 | -0.64 | -0.44 | -0.67 | -0.96 | -0.55 | -0.48 | -0.7 | -0.59 | -0.69 | -0.35 | 0.06 | -0.05 | 0.0325 | 0.902987 | 0.807427 | 0.844695 | 1 | 0.999631 | 0.968183 |
| Sptbn1_Q62261_S2357s_1 | 0.64 | 0.86 | -0.87 | 2.6 | 3 | 1.72 | 2.11 | 2.14 | 1.92 | -0.24 | 2.26 | 1.64 | -1.59 | 1.27 | 1.18 | 1.26 | 1.435 | 0.5875 | -0.2775 | 0.749289 | 0.056459 | 0.466051 | 1 | 0.999631 | 0.912137 |
| Sptbn1_Q62261_T2186t_1 | 0.11 | 2.17 | 0.54 | 0.83 | 0.74 | 2.18 | 0.25 | 2.09 | 0.42 | -0.07 | 1.66 | -0.43 | -0.63 | 1.72 | -0.29 | 0.93 | 0.4025 | -0.5175 | -0.48 | 0.455418 | 0.503704 | 0.385032 | 1 | 0.999631 | 0.790041 |
| Sptbn1_Q62261_T2194t_1 | -0.2 | 0.37 | -0.34 | 0.05 | 0.08 | -0.03 | -0.6 | -0.05 | -0.08 | -0.17 | -0.1 | -0.07 | 0.31 | 0.49 | 0.03 | 0.41 | -0.12 | -0.075 | 0.34 | 0.267252 | 0.680175 | 0.780729 | 1 | 0.999631 | 0.695893 |
| Sptbn1_Q62261_T2316t_2 | 2.24 | 4.45 | 2.98 | -0.46 | 2.87 | 2.49 | 4.71 | 0.94 | 4.63 | 2.94 | 3.12 | 4.94 | 4.38 | 2.58 | 0.09 | -1.45 | 0.45 | 1.605 | -0.9025 | 0.521022 | 0.684354 | 0.13053 | 1 | 0.999631 | 0.820181 |
| Sptbn1_Q62261_T2319t_1 | 2 | 2.96 | 4.53 | 0.98 | 3.29 | 2.53 | 3.74 | 2.12 | 4.95 | 1.91 | 3.64 | 5.04 | 4.34 | 0.15 | 3.28 | 0.39 | 0.3025 | 1.2675 | -0.5775 | 0.600862 | 0.682963 | 0.186922 | 1 | 0.999631 | 0.856689 |
| Sptbn1_Q62261_T2319t_2 | 2.01 | 2.1 | 2.98 | -0.46 | 2.33 | 0.98 | 4.71 | 0.94 | 3.06 | 1.62 | 3.12 | 4.94 | 3.2 | 0.98 | 0.09 | -1.45 | 0.5825 | 1.5275 | -0.9525 | 0.370555 | 0.560477 | 0.10591 | 1 | 0.999631 | 0.746338 |
| Sptbn2_Q68FG2_S1060s_1 | 0.28 | -2 | -0.47 | -1.24 | -2.58 | -1.95 | -0.13 | -1.62 | -1.56 | -0.2 | -0.88 | -1.07 | -1.22 | -2.52 | -0.95 | -0.15 | -0.7125 | -0.07 | -0.3525 | 0.577574 | 0.28133 | 0.894751 | 1 | 0.999631 | 0.84622 |
| Sptbn2_Q68FG2_S2199s_1 | 0.61 | -0.6 | -1.53 | -0.17 | 0.12 | 0.15 | -0.04 | 0.29 | -0.75 | -0.13 | -1.04 | -1.01 | -2.43 | -0.2 | -0.26 | -0.15 | 0.5525 | -0.31 | -0.3375 | 0.601503 | 0.227783 | 0.518899 | 1 | 0.999631 | 0.856765 |
| Sptbn2_Q68FG2_S2200s_1 | 0.93 | -1.08 | -1.67 | 1.5 | 0.3 | 0.58 | 1.07 | 1.06 | -0.65 | 0.03 | -0.37 | -1.48 | -3.31 | -0.53 | -3.84 | -0.57 | 0.8325 | -0.5375 | -1.9825 | 0.070489 | 0.251016 | 0.470906 | 1 | 0.999631 | 0.530612 |
| Sptbn2_Q68FG2_S2254s_1 | 0.06 | 1.38 | 0.58 | 0.56 | 0.94 | 1.57 | 0.42 | 1.11 | 0.48 | 0.55 | 0.83 | 0.56 | 1.17 | 0.92 | 0.65 | 1.25 | 0.365 | -0.04 | 0.3525 | 0.335457 | 0.344799 | 0.904488 | 1 | 0.999631 | 0.730235 |
| Sptbn2_Q68FG2_T2354t_1 | 0.46 | 0 | -0.32 | -0.09 | 0.38 | 1.01 | -0.16 | 0.31 | 0.16 | -0.61 | 0.03 | -0.4 | -0.4 | 0.95 | -1.43 | 0.23 | 0.3725 | -0.2175 | -0.175 | 0.730021 | 0.276886 | 0.489454 | 1 | 0.999631 | 0.906325 |
| Sptbn4_Q91ZE6_S1445s_1 | -0.72 | 0.3 | 0.59 | -0.14 | 0.14 | -0.19 | -0.29 | 0.19 | 0.38 | -0.19 | 0.25 | 0.74 | 0.46 | -0.05 | 1.13 | 0.07 | -0.045 | 0.2875 | 0.395 | 0.345926 | 0.895471 | 0.445551 | 1 | 0.999631 | 0.73511 |
| Sptbn4_Q91ZE6_S1577s_1 | 0.69 | 0.21 | 0.23 | 0.46 | -0.01 | 0.73 | 0.39 | 0.45 | 0.46 | 1.16 | -0.36 | -0.28 | 0.38 | 0.26 | 0.48 | 0.49 | -0.0075 | -0.1525 | 0.005 | 0.98543 | 0.978276 | 0.697317 | 1 | 0.999631 | 0.995133 |
| Sptbn4_Q91ZE6_S1863s_1 | -0.23 | -1.27 | -0.2 | 0.06 | -1.99 | -0.78 | 0.3 | -0.69 | -0.42 | -0.25 | -0.36 | -0.53 | -1.39 | -1.19 | -0.13 | -1.29 | -0.38 | 0.02 | -0.59 | 0.186702 | 0.467112 | 0.953493 | 1 | 0.999631 | 0.646474 |
| Sptbn4_Q91ZE6_S2156s_1 | 1.2 | -0.67 | -0.94 | 0.95 | 0.92 | -0.56 | 0.76 | 0.09 | 0.37 | -0.8 | 0.2 | -0.81 | -1.05 | -0.46 | -1.08 | -0.41 | 0.1675 | -0.395 | -0.885 | 0.12523 | 0.774813 | 0.501158 | 1 | 0.999631 | 0.596415 |
| Sptbn4_Q91ZE6_S2245s_1 | 0.24 | -1.33 | -1.5 | -0.91 | -1.33 | -1.07 | -0.99 | -1.9 | -1.19 | -0.01 | -1.15 | -1.53 | -2.03 | -1.66 | -2.68 | -1.46 | -0.4475 | -0.095 | -1.0825 | 0.040076 | 0.315333 | 0.846164 | 1 | 0.999631 | 0.472324 |
| Sptbn4_Q91ZE6_S2263s_1 | 0.07 | -0.12 | -0.68 | -0.08 | -0.24 | -0.27 | -0.84 | -0.95 | -0.59 | 0.2 | -1.3 | -0.74 | -0.97 | -0.49 | -1.89 | -0.24 | -0.3725 | -0.405 | -0.695 | 0.118197 | 0.242178 | 0.295318 | 1 | 0.999631 | 0.585087 |
| Sptbn4_Q91ZE6_S2296s_1 | -0.5 | -4.59 | -0.47 | -0.1 | -3.99 | -3.77 | 0.17 | -1.72 | -4.28 | -2.25 | -1.22 | -1.47 | -3.29 | -5.13 | -1.37 | -1.12 | -0.9125 | -0.89 | -1.3125 | 0.288349 | 0.464833 | 0.422787 | 1 | 0.999631 | 0.706035 |
| Sptbn4_Q91ZE6_S2297s_1 | -4.35 | -12.2 | 0.77 | 1.04 | -8.39 | -8.6 | 1.08 | -0.47 | -12.57 | -6.57 | -0.24 | -0.61 | -8.18 | -9.68 | -0.87 | -1.08 | -0.41 | -1.3125 | -1.2675 | 0.693988 | 0.903298 | 0.714514 | 1 | 0.999631 | 0.895131 |
| Sptbn4_Q91ZE6_S2391s_1 | 0.02 | 0.39 | 0.42 | -0.19 | 0.43 | 0.05 | -0.27 | 0.73 | -0.13 | -0.59 | 0.46 | 1.02 | 0.83 | 0.25 | 0.78 | 0.23 | 0.075 | 0.03 | 0.3625 | 0.261917 | 0.812064 | 0.939415 | 1 | 0.999631 | 0.692758 |
| Sqstm1_D3YZJ1_T272t_1 | -0.1 | -0.16 | -1.06 | -0.66 | -0.36 | -0.21 | -0.6 | -0.78 | -0.66 | 0.07 | -0.64 | -0.76 | 0.01 | 0.22 | -0.65 | -0.41 | 0.0075 | -0.0025 | 0.2875 | 0.424962 | 0.980839 | 0.994146 | 1 | 0.999631 | 0.774293 |
| Src_P05480_S17s_1 | -0.36 | -0.14 | -0.53 | -0.86 | -0.68 | -0.34 | -0.87 | -0.81 | -0.69 | -0.06 | -0.91 | -0.87 | 0.19 | 0.1 | -0.87 | -0.07 | -0.2025 | -0.16 | 0.31 | 0.38021 | 0.471105 | 0.611968 | 1 | 0.999631 | 0.752331 |
| Src_P05480_S74s_1 | -2.65 | -0.81 | -0.96 | -2.19 | -0.37 | -0.91 | -0.78 | -1.25 | -0.58 | -3.47 | -0.36 | -0.6 | -0.5 | -1.51 | -1.31 | -2.15 | 0.825 | 0.4 | 0.285 | 0.595143 | 0.102366 | 0.604075 | 1 | 0.999631 | 0.8545 |
| Srcap_E9Q9V7_S559s_1 | -0.6 | 0.24 | 0.04 | -0.24 | 0.23 | -0.27 | -1.6 | -0.49 | 0.56 | -0.55 | -0.13 | -0.13 | 0.61 | 0.33 | 0.12 | 0.84 | -0.3925 | 0.0775 | 0.615 | 0.080138 | 0.363594 | 0.819455 | 1 | 0.999631 | 0.545579 |
| Srcin1_B1AQX9_S1021s_1 | -0.85 | -0.38 | -0.07 | -0.53 | -1.32 | -0.85 | -0.67 | -0.54 | -0.92 | -0.5 | -0.33 | -0.68 | 0.3 | -0.74 | -1.09 | -0.31 | -0.3875 | -0.15 | -0.0025 | 0.99472 | 0.21504 | 0.608407 | 1 | 0.999631 | 0.997705 |
| Srcin1_B1AQX9_S1037s_1 | 1.11 | -0.89 | -1.27 | -0.18 | 0.87 | -0.12 | 0.41 | 0.07 | 0.03 | -0.29 | -0.58 | -1.02 | -0.84 | -0.07 | -0.93 | -0.68 | 0.615 | -0.1575 | -0.3225 | 0.543693 | 0.258949 | 0.767382 | 1 | 0.999631 | 0.832881 |
| Srcin1_B1AQX9_S1077s_1 | -0.65 | -0.27 | -0.58 | -0.38 | -0.49 | -0.82 | -0.7 | -0.59 | -0.73 | -0.55 | -0.82 | -0.26 | -0.02 | -0.22 | -0.41 | -0.21 | -0.18 | -0.12 | 0.255 | 0.35965 | 0.46818 | 0.654319 | 1 | 0.999631 | 0.741182 |
| Srcin1_B1AQX9_S1094s_1 | 0.56 | 2.08 | 1.79 | 1.24 | 0.92 | 1.86 | 0.7 | 1.07 | 1.53 | 1.41 | 1.41 | 1.15 | 1.45 | 1.59 | 1.43 | 2.02 | -0.28 | -0.0425 | 0.205 | 0.602705 | 0.506528 | 0.908792 | 1 | 0.999631 | 0.857278 |
| Srcin1_B1AQX9_S1101s_1 | 0.98 | 1.39 | 3.36 | 1.12 | 0.84 | 2.28 | 1.21 | 0.29 | 1.56 | 2.02 | 0.58 | 1.3 | 1.46 | 1.85 | 1.78 | 1.86 | -0.5575 | -0.3475 | 0.025 | 0.962338 | 0.382783 | 0.55245 | 1 | 0.999631 | 0.986642 |
| Srcin1_B1AQX9_S1152s_1 | 0.52 | 0.25 | 0.49 | -1.25 | -0.42 | 0.19 | 0.69 | -1.48 | -0.09 | -0.76 | -1.16 | -1.04 | -0.62 | -1.54 | -1.05 | -2.61 | -0.2575 | -0.765 | -1.4575 | 0.024256 | 0.654807 | 0.127535 | 1 | 0.999631 | 0.415644 |
| Srcin1_B1AQX9_S1189s_1 | 0.68 | 2.32 | 2.96 | 2.14 | 1.37 | 1.7 | 1.92 | 2.34 | 2.26 | 1.97 | 3.01 | 2.47 | 3.07 | 1.49 | 2.37 | 2.41 | -0.1925 | 0.4025 | 0.31 | 0.57069 | 0.696663 | 0.430333 | 1 | 0.999631 | 0.84338 |
| Srcin1_B1AQX9_S12s_1 | -0.2 | 0.58 | 0.69 | -0.08 | -0.2 | 0.43 | -0.48 | 0.12 | 0.11 | 0.43 | -0.06 | 0.43 | 0.92 | 0.53 | 0.16 | 1.17 | -0.28 | -0.02 | 0.4475 | 0.23671 | 0.414539 | 0.949825 | 1 | 0.999631 | 0.683301 |
| Srcin1_B1AQX9_S13s_1 | -0.09 | -0.07 | 0.85 | -0.13 | -1.31 | -0.31 | -0.32 | 0.03 | -0.78 | 0.25 | 0.43 | 0.58 | 0.85 | 0.06 | 0.26 | 1.18 | -0.6175 | -0.02 | 0.4475 | 0.261118 | 0.132963 | 0.96013 | 1 | 0.999631 | 0.69271 |
| Srcin1_B1AQX9_S199s_1 | 0.51 | 0.34 | 0.38 | 0.47 | 1.17 | 0.6 | 0.33 | 0.85 | 0.4 | -0.18 | 0.47 | 0.2 | -0.11 | 0.64 | 0.52 | 0.43 | 0.3125 | -0.2025 | -0.055 | 0.849648 | 0.268199 | 0.45425 | 1 | 0.999631 | 0.949525 |
| Srcin1_B1AQX9_S213s_1 | -1.15 | -0.56 | -0.8 | -1.21 | -0.96 | -0.75 | -1.08 | -0.76 | -0.87 | -0.71 | -0.73 | -0.47 | -0.4 | -0.66 | -0.32 | -0.48 | 0.0425 | 0.235 | 0.465 | 0.132064 | 0.873879 | 0.403755 | 1 | 0.999631 | 0.603669 |
| Srcin1_B1AQX9_S259s_1 | -0.55 | -0.16 | -0.42 | -0.51 | -0.74 | -0.21 | -0.74 | -0.45 | 0.11 | -0.06 | -0.12 | 0.3 | 1.08 | 0.29 | -0.3 | 0.28 | -0.125 | 0.4675 | 0.7475 | 0.055869 | 0.632895 | 0.095071 | 1 | 0.999631 | 0.495701 |
| Srcin1_B1AQX9_S332s_1 | -1.7 | -1.71 | 0.26 | 0.39 | -2.1 | -2.31 | -0.8 | 0.45 | -2.79 | -2.61 | 1.37 | -0.52 | -1.72 | -1.37 | 0.71 | 1.08 | -0.5 | -0.4475 | 0.365 | 0.651304 | 0.516347 | 0.652306 | 1 | 0.999631 | 0.876458 |
| Srcin1_B1AQX9_S340s_1 | -1.4 | -0.82 | -0.36 | -1.26 | -0.08 | -1.24 | -0.27 | -0.84 | -0.31 | -1.67 | -0.65 | -0.11 | 0.38 | -1.05 | -1.35 | -0.07 | 0.3525 | 0.275 | 0.4375 | 0.355759 | 0.354978 | 0.518965 | 1 | 0.999631 | 0.74034 |
| Srcin1_B1AQX9_S373s_1 | -0.48 | -0.41 | 0.29 | -0.18 | -0.76 | -0.7 | -0.08 | -0.25 | -0.87 | -0.56 | -0.65 | 0.05 | 0.1 | -1.1 | 0.07 | -0.2 | -0.2525 | -0.3125 | -0.0875 | 0.814117 | 0.41238 | 0.342969 | 1 | 0.999631 | 0.937041 |
| Srcin1_B1AQX9_S377s_1 | -1.27 | -0.91 | 2.52 | 1.06 | -0.86 | -0.58 | 0.97 | 1.79 | -0.97 | -1 | 2.02 | 2.41 | -0.35 | -1.26 | 1.95 | 1.63 | -0.02 | 0.265 | 0.1425 | 0.887299 | 0.982997 | 0.810103 | 1 | 0.999631 | 0.963505 |
| Srcin1_B1AQX9_S396s_1 | 0.23 | -0.08 | 0.74 | 0.47 | 0.59 | -0.94 | 0.26 | 1.1 | 0.1 | 0.31 | 0.37 | 0.37 | -0.67 | -0.19 | 0.19 | -0.15 | -0.0875 | -0.0525 | -0.545 | 0.118566 | 0.846216 | 0.852039 | 1 | 0.999631 | 0.585512 |
| Srcin1_B1AQX9_S398s_1 | -0.29 | 0.06 | 0.95 | -0.22 | 1.03 | 0.07 | -0.39 | 1.29 | -0.19 | -0.38 | 1.16 | 0.46 | 0.1 | 0.96 | -0.39 | 1.02 | 0.375 | 0.1375 | 0.2975 | 0.509969 | 0.42927 | 0.757372 | 1 | 0.999631 | 0.817324 |
| Srcin1_B1AQX9_S39s_1 | -1.03 | -4.86 | -2.33 | -1.85 | -3.18 | -4.48 | -0.51 | -3.75 | -4.22 | -2.59 | -1.44 | -2.51 | -2.27 | -4.68 | -2.62 | -3.68 | -0.4625 | -0.1725 | -0.795 | 0.364163 | 0.65414 | 0.843865 | 1 | 0.999631 | 0.744334 |
| Srcin1_B1AQX9_S45s_1 | 0.34 | -0.8 | -0.7 | -0.39 | 0.56 | -0.17 | -0.01 | -0.32 | 0.25 | -0.45 | -0.22 | -0.08 | 0.86 | -0.39 | -0.61 | 0.11 | 0.4025 | 0.2625 | 0.38 | 0.38224 | 0.267487 | 0.448821 | 1 | 0.999631 | 0.753668 |
| Srcin1_B1AQX9_S468s_1 | 0.35 | 0.75 | 0.38 | -0.59 | 0.39 | -0.22 | 0.06 | 0.05 | 0.35 | -0.19 | -0.03 | -0.17 | 0.96 | 0.12 | 0.86 | 0.24 | -0.1525 | -0.2325 | 0.3225 | 0.414111 | 0.659368 | 0.512227 | 1 | 0.999631 | 0.768655 |
| Srcin1_B1AQX9_S506s_1 | 0.55 | -1.07 | -0.87 | 0.06 | -0.08 | -0.55 | 0.03 | -0.2 | -0.86 | -0.3 | -0.47 | -1.08 | -1.28 | -0.8 | -1.13 | -0.88 | 0.1325 | -0.345 | -0.69 | 0.120566 | 0.744421 | 0.424582 | 1 | 0.999631 | 0.588344 |
| Srcin1_B1AQX9_S507s_1 | 0.8 | -1.13 | -0.67 | 0.19 | 0.12 | -0.09 | 0.16 | -0.15 | -0.5 | -0.1 | -0.28 | -0.96 | -1.22 | -0.64 | -0.9 | -0.63 | 0.2125 | -0.2575 | -0.645 | 0.175145 | 0.623538 | 0.57719 | 1 | 0.999631 | 0.642116 |
| Srcin1_B1AQX9_S526s_1 | -0.7 | -0.39 | -1.14 | -1.6 | -0.24 | -1.04 | -0.19 | -1.08 | -1.01 | -1.62 | -0.68 | -1.12 | -0.17 | -0.96 | -1.47 | -1.31 | 0.32 | -0.15 | -0.02 | 0.961081 | 0.40296 | 0.678728 | 1 | 0.999631 | 0.986594 |
| Srcin1_B1AQX9_S529s_1 | -1.16 | 0.65 | 1.05 | -0.48 | 0.28 | 0.55 | -0.08 | 0.19 | 0.29 | -0.64 | 0.2 | 1 | 0.81 | 0.19 | 0.93 | 0.17 | 0.22 | 0.1975 | 0.51 | 0.336567 | 0.657233 | 0.725948 | 1 | 0.999631 | 0.730499 |
| Srcin1_B1AQX9_S53s_1 | -1.07 | -1.34 | 0.08 | -0.66 | -0.06 | -0.96 | -0.21 | 0.57 | -0.91 | -1.07 | -0.4 | 0.13 | 0.22 | -0.34 | 0.49 | 0.52 | 0.5825 | 0.185 | 0.97 | 0.03165 | 0.198539 | 0.658406 | 1 | 0.999631 | 0.442395 |
| Srcin1_B1AQX9_S541s_1 | -1.83 | -5.78 | -2.3 | -1.52 | -2.92 | -5.35 | -0.94 | -1.75 | -4.88 | -4.15 | -1.1 | -1.33 | -1.07 | -5.55 | -2.39 | -2.82 | 0.1175 | -0.0075 | -0.1 | 0.931026 | 0.920365 | 0.994936 | 1 | 0.999631 | 0.977912 |
| Srcin1_B1AQX9_S546s_1 | -0.44 | -0.64 | -0.65 | -0.68 | -0.49 | -0.32 | -0.98 | -0.63 | -0.26 | -0.22 | -0.6 | -0.6 | -0.03 | -0.24 | -0.44 | -0.65 | -0.0025 | 0.1825 | 0.2625 | 0.359842 | 0.99227 | 0.479319 | 1 | 0.999631 | 0.741182 |
| Srcin1_B1AQX9_S555s_1 | -0.26 | 0.17 | 0.47 | -0.1 | 0.12 | -0.01 | -0.64 | -0.04 | 0.01 | -0.28 | -0.57 | 0.18 | 0.33 | 0.55 | 0.61 | 0.35 | -0.2125 | -0.235 | 0.39 | 0.201733 | 0.481583 | 0.445754 | 1 | 0.999631 | 0.658952 |
| Srcin1_B1AQX9_S625s_1 | 0.61 | -0.44 | 1.48 | 0.8 | -0.67 | 0.48 | 0.74 | 0.72 | -0.47 | 0.89 | -0.36 | 0.55 | -0.17 | -0.55 | 1.3 | 1.51 | -0.295 | -0.46 | -0.09 | 0.879823 | 0.550544 | 0.363808 | 1 | 0.999631 | 0.962995 |
| Srcin1_B1AQX9_S64s_1 | -1.1 | 0.89 | 0.26 | -0.42 | 2.48 | -0.33 | 2.06 | 0.42 | 2.9 | -1.17 | 1.96 | 2.46 | 2.17 | -0.24 | 0.35 | 0.07 | 1.25 | 1.63 | 0.68 | 0.291007 | 0.099526 | 0.090769 | 1 | 0.999631 | 0.706164 |
| Srcin1_B1AQX9_S654s_1 | -0.73 | -0.21 | 0.25 | -0.13 | -0.46 | -0.28 | -0.63 | -0.49 | -0.52 | -0.13 | 0.17 | -0.44 | -0.01 | -0.09 | -0.15 | -0.09 | -0.26 | -0.025 | 0.12 | 0.694371 | 0.376766 | 0.937204 | 1 | 0.999631 | 0.895302 |
| Srcin1_B1AQX9_S667s_1 | -1.18 | -1.15 | -0.46 | -0.47 | -1.14 | -0.82 | -0.24 | -0.35 | -1.3 | -1.22 | -0.61 | -0.52 | -1.88 | -0.89 | -1.04 | -0.1 | 0.1775 | -0.0975 | -0.1625 | 0.704361 | 0.595045 | 0.771269 | 1 | 0.999631 | 0.897336 |
| Srcin1_B1AQX9_S670s_1 | -0.5 | -1.25 | -0.24 | -0.86 | -0.43 | -0.06 | -0.66 | -0.08 | 0.04 | -0.08 | -0.27 | -0.53 | 0.06 | -0.27 | -0.02 | -0.63 | 0.405 | 0.5025 | 0.4975 | 0.16199 | 0.218019 | 0.13541 | 1 | 0.999631 | 0.631442 |
| Srcin1_B1AQX9_S878s_1 | -0.83 | -0.04 | -0.07 | -0.71 | -0.28 | -0.2 | -0.68 | -0.49 | -0.32 | -0.56 | -0.08 | -0.15 | 0.53 | -0.01 | -0.18 | -0.13 | -9.8E-17 | 0.135 | 0.465 | 0.185336 | 1 | 0.659889 | 1 | 0.999631 | 0.646212 |
| Srcin1_B1AQX9_S900s_1 | -1.43 | -1.1 | -1.34 | -2.04 | -1.14 | -1.34 | -1.41 | -2.08 | -1.53 | -1.13 | -1.44 | -1.56 | -0.3 | -1.57 | -2.07 | -1.96 | -0.015 | 0.0625 | 0.0025 | 0.995574 | 0.963521 | 0.834816 | 1 | 0.999631 | 0.997705 |
| Srcin1_B1AQX9_S941s_1 | 2.09 | -0.09 | -1.47 | 2.64 | 2.14 | 0.36 | 2.19 | 1.41 | 1.04 | -0.92 | 1.72 | 0.51 | -1.67 | 0.2 | 0.05 | 0.11 | 0.7325 | -0.205 | -1.12 | 0.236625 | 0.427287 | 0.830972 | 1 | 0.999631 | 0.683301 |
| Srcin1_B1AQX9_T629t_1 | 1.29 | -0.2 | 1.56 | 0.84 | -0.7 | -0.31 | 1.13 | 0.5 | -0.74 | 0.68 | 0.45 | 0.14 | -0.73 | -1.42 | 0.12 | -0.05 | -0.7175 | -0.74 | -1.3925 | 0.01824 | 0.191523 | 0.145234 | 1 | 0.999631 | 0.391024 |
| Srcin1_B1AQX9_T642t_1 | -0.38 | -1.13 | -0.47 | -0.01 | -0.89 | -1.09 | -0.4 | -0.39 | -0.48 | -0.58 | -0.43 | 0.28 | -0.69 | -0.49 | -0.73 | 0.03 | -0.195 | 0.195 | 0.0275 | 0.937194 | 0.560855 | 0.576615 | 1 | 0.999631 | 0.979415 |
| Srcin1_B1AQX9_T658t_1 | -0.99 | -0.82 | -0.4 | -0.71 | -0.88 | -1.19 | -1.1 | -1.05 | -0.93 | -1.11 | -1.08 | -1.14 | -0.47 | -1.05 | -0.8 | -0.93 | -0.325 | -0.335 | -0.0825 | 0.77819 | 0.220588 | 0.216621 | 1 | 0.999631 | 0.921416 |
| Srcin1_B1AQX9_T658t_2 | -2.51 | -0.37 | -0.63 | -0.39 | 0.39 | -2.33 | -1.49 | -0.06 | 0.25 | -0.7 | -0.27 | -0.43 | 1.42 | -0.4 | -3.29 | -0.48 | 0.1025 | 0.6875 | 0.2875 | 0.761193 | 0.886414 | 0.205942 | 1 | 0.999631 | 0.91472 |
| Srcin1_B1AQX9_T671t_2 | -1.14 | -1.21 | 1.07 | -0.29 | -0.55 | -1.49 | 0.37 | 2.2 | -1.51 | -1.29 | -0.35 | -0.69 | -1.37 | -0.92 | -2 | -0.16 | 0.525 | -0.5675 | -0.72 | 0.246716 | 0.52959 | 0.316235 | 1 | 0.999631 | 0.685751 |
| Srcin1_B1AQX9_Y430y_1 | -0.57 | 1.27 | -0.76 | -0.18 | 0.07 | 0.42 | -0.34 | -0.61 | 0.13 | -1.54 | -0.32 | -0.67 | 2.05 | 1.1 | 0.15 | -0.5 | -0.055 | -0.54 | 0.76 | 0.25795 | 0.909523 | 0.329917 | 1 | 0.999631 | 0.692624 |
| Srcin1_Q9QWI6_S46s_1 | -0.53 | 0.32 | -1.14 | -1.3 | -0.28 | 0.13 | -1.56 | -1.43 | -0.04 | 0.06 | -1.14 | -1.31 | 0.79 | 0.28 | -1.07 | -0.68 | -0.1225 | 0.055 | 0.4925 | 0.36346 | 0.813165 | 0.910543 | 1 | 0.999631 | 0.743747 |
| Srcin1_Q9QWI6_S47s_1 | -1.4 | -1.71 | -0.04 | -0.72 | -1.51 | -2.24 | -0.87 | -0.7 | -1.66 | -1.61 | -0.25 | -0.29 | -0.29 | -1.33 | -0.38 | 0.31 | -0.3625 | 0.015 | 0.545 | 0.278226 | 0.459692 | 0.976567 | 1 | 0.999631 | 0.700874 |
| Srgap1_D3YZW1_S835s_1 | 0.32 | 0.12 | 1.23 | 0.44 | -0.14 | 0.48 | 1.5 | 0.39 | 0.35 | -0.23 | 0.52 | 0.83 | -0.21 | -0.94 | 0.84 | -1.11 | 0.03 | -0.16 | -0.8825 | 0.095174 | 0.942633 | 0.659909 | 1 | 0.999631 | 0.558128 |
| Srgap1_D3YZW1_S906s_1 | 1.52 | -0.2 | 1.23 | 0.88 | 0.02 | 0.35 | 1.27 | 0.76 | 0.76 | 1.36 | 0.37 | 0.55 | -0.95 | 0.03 | 1.2 | -0.03 | -0.2575 | -0.0975 | -0.795 | 0.164273 | 0.568581 | 0.82146 | 1 | 0.999631 | 0.632578 |
| Srgap1_D3YZW1_S917s_1 | 0.92 | -0.86 | -1.55 | 0.19 | 0.89 | 1.04 | 0.16 | -0.12 | 0.39 | 0.8 | -0.37 | -0.61 | -1.2 | 0.5 | -0.3 | -0.16 | 0.8175 | 0.3775 | 0.035 | 0.95296 | 0.168798 | 0.522927 | 1 | 0.999631 | 0.984317 |
| Srgap2_Q91Z67_S206s_1 | 1.26 | -0.58 | 0.52 | 0.44 | -0.72 | -0.53 | 0.72 | -0.13 | -0.68 | 1.04 | 0.17 | -0.04 | -1.2 | -0.87 | 0.42 | -0.42 | -0.575 | -0.2875 | -0.9275 | 0.085467 | 0.241018 | 0.563907 | 1 | 0.999631 | 0.551124 |
| Srgap2_Q91Z67_S427s_1 | -1.6 | -1.91 | -0.79 | -1.23 | -1.75 | -1.72 | -1.06 | -0.99 | -1.8 | -1.39 | -0.66 | 0.02 | -1.38 | -1.61 | -0.2 | 0.15 | 0.0025 | 0.425 | 0.6225 | 0.215223 | 0.994259 | 0.364015 | 1 | 0.999631 | 0.668192 |
| Srgap2_Q91Z67_S500s_1 | -0.82 | 0.46 | 2.18 | -0.59 | -0.53 | -0.79 | 0.7 | -1.33 | -0.52 | 1.72 | -0.65 | -3.39 | -0.19 | -1.04 | -0.11 | -0.96 | -0.795 | -1.0175 | -0.8825 | 0.196313 | 0.280569 | 0.355386 | 1 | 0.999631 | 0.653373 |
| Srgap2_Q91Z67_S860s_1 | 0.12 | 0.04 | -0.98 | -0.69 | 0.48 | 0.48 | -0.35 | -0.48 | 0.12 | -0.01 | -0.05 | -0.61 | 0.28 | 0.59 | -0.37 | -0.65 | 0.41 | 0.24 | 0.34 | 0.417212 | 0.302713 | 0.501846 | 1 | 0.999631 | 0.771432 |
| Srgap2_Q91Z67_S983s_1 | 0.05 | -0.24 | 0.25 | -0.26 | -0.01 | 0.33 | -0.14 | 0.42 | -0.15 | 0.62 | -0.16 | 0.52 | 0.35 | 0.67 | 0.46 | 1.15 | 0.2 | 0.2575 | 0.7075 | 0.042376 | 0.468166 | 0.415547 | 1 | 0.999631 | 0.475443 |
| Srgap2_Q91Z67_S994s_1 | -0.42 | -0.34 | -0.71 | -0.07 | -0.56 | -0.23 | -0.04 | -0.46 | -0.32 | 0.1 | -0.62 | -0.54 | -1.23 | 0.01 | -0.41 | 0.25 | 0.0625 | 0.04 | 0.04 | 0.917062 | 0.816571 | 0.891064 | 1 | 0.999631 | 0.973513 |
| Srgap2_Q91Z67_T982t_1 | 0.05 | -0.24 | 0.25 | -0.26 | -0.01 | 0.33 | -0.14 | 0.42 | -0.15 | 0.62 | -0.16 | 0.52 | 0.35 | 0.67 | 0.46 | 1.15 | 0.2 | 0.2575 | 0.7075 | 0.042376 | 0.468166 | 0.415547 | 1 | 0.999631 | 0.475443 |
| Srgap3_E9QN14_S1054s_1 | 0.18 | 0.45 | -0.46 | -0.16 | 0.6 | -0.27 | 0.52 | 0.62 | 0.07 | -1.47 | 0.66 | -0.15 | -1.2 | 0.16 | 0.48 | 0.56 | 0.365 | -0.225 | -0.0025 | 0.995588 | 0.286619 | 0.636677 | 1 | 0.999631 | 0.997705 |
| Srgap3_E9QN14_S413s_1 | 0.93 | 0.06 | -0.16 | 1.94 | 1.23 | 1.25 | 1.01 | 1.38 | 0.5 | -0.41 | 1.34 | -0.16 | -0.83 | -0.3 | 0.17 | 0.76 | 0.525 | -0.375 | -0.7425 | 0.194541 | 0.273449 | 0.514205 | 1 | 0.999631 | 0.65187 |
| Srgap3_E9QN14_S813s_1 | -0.96 | -0.17 | -0.44 | -0.91 | -0.31 | -0.72 | -0.81 | -0.67 | -0.8 | -0.95 | -0.92 | -0.4 | 0.03 | -0.13 | -0.56 | -0.52 | -0.0075 | -0.1475 | 0.325 | 0.324801 | 0.979401 | 0.627985 | 1 | 0.999631 | 0.72463 |
| Srgap3_E9QN14_S834s_1 | 1.63 | -0.72 | -1.19 | 0.44 | 2.47 | 0.56 | 0.8 | 0.61 | 0.58 | -1.2 | 0.03 | -0.64 | -1.7 | 0.07 | -0.27 | -0.46 | 1.07 | -0.3475 | -0.63 | 0.352465 | 0.144944 | 0.603391 | 1 | 0.999631 | 0.739179 |
| Srgap3_E9QN14_S844s_1 | 0.75 | -0.09 | 0.14 | 1.07 | 0.13 | 0 | 2.36 | 0.21 | 0.01 | -0.42 | 0.8 | 0.24 | -0.09 | -1.92 | -0.43 | -1.17 | 0.2075 | -0.31 | -1.37 | 0.015382 | 0.716032 | 0.431676 | 1 | 0.999631 | 0.388832 |
| Srgap3_E9QN14_S844s_2 | 0.97 | 2.28 | 2.04 | 0.78 | 2.78 | 2.08 | 3.16 | 1.68 | 3.28 | 0.13 | 3.26 | 2.2 | 2.12 | -0.11 | 0.77 | -0.46 | 0.9075 | 0.7 | -0.9375 | 0.154402 | 0.082089 | 0.35266 | 1 | 0.999631 | 0.625677 |
| Srgap3_E9QN14_S850s_1 | 1.29 | -0.24 | 0.67 | 1.16 | -1.03 | -0.1 | 0.97 | 0.01 | -0.94 | 0.75 | 0.55 | 0.08 | -0.9 | -1.54 | 0.41 | -0.14 | -0.7575 | -0.61 | -1.2625 | 0.033424 | 0.154806 | 0.231325 | 1 | 0.999631 | 0.44294 |
| Srgap3_E9QN14_S850s_2 | 0.97 | 2.28 | 2.04 | 0.78 | 2.78 | 2.08 | 3.16 | 1.68 | 3.28 | 0.13 | 3.26 | 2.2 | 2.12 | -0.11 | 0.77 | -0.46 | 0.9075 | 0.7 | -0.9375 | 0.154402 | 0.082089 | 0.35266 | 1 | 0.999631 | 0.625677 |
| Srgap3_E9QN14_S884s_1 | 1.19 | -0.23 | -0.64 | -0.11 | 0.03 | -0.09 | 0 | 0.09 | -0.1 | 0.63 | -0.66 | -0.75 | -0.74 | -0.35 | 0.46 | -0.73 | -0.045 | -0.2725 | -0.3925 | 0.416158 | 0.910661 | 0.577723 | 1 | 0.999631 | 0.770412 |
| Srgap3_E9QN14_S895s_1 | -0.46 | -0.59 | -1.59 | -0.82 | -0.32 | -0.36 | -0.88 | -1 | -0.69 | -0.88 | -1.03 | -1.12 | -0.88 | -0.57 | -1.8 | -0.82 | 0.225 | -0.065 | -0.1525 | 0.701116 | 0.515026 | 0.84172 | 1 | 0.999631 | 0.896835 |
| Srgap3_E9QN14_S924s_1 | 2.48 | 2.3 | 2.89 | 1.7 | 1.34 | 3.03 | 5.19 | 1.21 | 1.21 | 2.96 | 3.75 | 4.29 | 1.83 | 1.39 | 2.58 | 0.19 | 0.35 | 0.71 | -0.845 | 0.131867 | 0.677137 | 0.286449 | 1 | 0.999631 | 0.603456 |
| Srgap3_E9QN14_S987s_1 | 1.52 | -0.6 | 0.07 | 1.51 | 1.75 | 0.52 | 1.71 | 1.66 | 0.36 | -1.54 | 0.57 | 0.72 | -0.78 | 0.57 | 0.91 | 1.32 | 0.785 | -0.5975 | -0.12 | 0.849261 | 0.181202 | 0.383191 | 1 | 0.999631 | 0.949525 |
| Srp68_Q8BMA6_S12s_1 | 0.66 | -0.62 | 0.84 | 1.71 | -0.61 | 0.12 | 0.54 | 0.49 | -0.18 | 0.26 | -0.25 | 0.74 | -2.16 | -0.24 | 0.09 | 0.59 | -0.5125 | -0.505 | -1.0775 | 0.139213 | 0.329627 | 0.329366 | 1 | 0.999631 | 0.611718 |
| Srp68_Q8BMA6_S26s_1 | 0.14 | 0.18 | 0.46 | 0.02 | -0.52 | 0.75 | 0.7 | -0.29 | -0.07 | 1 | 0.93 | -0.2 | 0.59 | -0.46 | -0.83 | 0.92 | -0.04 | 0.215 | -0.145 | 0.738606 | 0.912485 | 0.557787 | 1 | 0.999631 | 0.90804 |
| Srp72_E9Q740_S560s_1 | 0.28 | 1.9 | 1.96 | 0.32 | 0.18 | 1.1 | 0.16 | 0.88 | 1.08 | 1.74 | 0.94 | 1.13 | 1.81 | 1.74 | 1.27 | 1.68 | -0.535 | 0.1075 | 0.51 | 0.29689 | 0.296268 | 0.823668 | 1 | 0.999631 | 0.710119 |
| Srpk2_O54781_S487s_1 | 0.65 | 0.18 | -0.19 | -0.68 | 0.82 | -0.15 | -0.59 | 0.12 | 0.6 | 0.45 | -0.11 | -0.4 | 1.05 | 0.4 | -0.24 | 0.61 | 0.06 | 0.145 | 0.465 | 0.271062 | 0.88328 | 0.708048 | 1 | 0.999631 | 0.698674 |
| Srpk2_O54781_S490s_1 | -0.12 | 1.1 | 1.38 | 0.42 | 0.43 | 0.51 | -0.38 | 0.45 | 0.75 | 0.69 | 0.47 | 0.85 | 0.84 | 0.89 | 0.83 | 1.13 | -0.4425 | -0.005 | 0.2275 | 0.554158 | 0.286745 | 0.989308 | 1 | 0.999631 | 0.837952 |
| Srpk2_O54781_T331t_1 | -0.41 | 0.87 | 1.58 | 0.74 | 0.24 | 1.21 | 0.59 | 0.49 | 0.75 | 0.51 | 1.14 | 1.36 | 1.37 | 0.46 | 1.61 | 1.62 | -0.0625 | 0.245 | 0.57 | 0.252876 | 0.888582 | 0.586215 | 1 | 0.999631 | 0.688432 |
| Srpk2_O54781_T332t_1 | -0.32 | 0.57 | 0.61 | -0.02 | 0.42 | 0.07 | 0.43 | 0.23 | 0.14 | 0.43 | -0.24 | -0.03 | 0.36 | 0.37 | -0.63 | 0.95 | 0.0775 | -0.135 | 0.0525 | 0.89935 | 0.798761 | 0.678675 | 1 | 0.999631 | 0.967236 |
| Srrm1_E9PUK6_S220s_1 | -0.25 | -0.02 | 0.18 | -0.43 | -0.25 | -0.26 | -0.61 | -0.05 | -0.28 | 0.2 | -0.07 | -0.04 | 0.75 | 0.37 | 0.03 | 0.42 | -0.1625 | 0.0825 | 0.5225 | 0.107746 | 0.550457 | 0.762389 | 1 | 0.999631 | 0.569416 |
| Srrm1_E9PUK6_S387s_2 | -0.07 | -0.32 | 0.29 | -0.39 | -0.42 | -0.97 | -0.17 | 0.07 | -0.32 | 0.48 | -0.25 | -0.6 | -0.13 | -0.29 | -0.09 | -0.19 | -0.25 | -0.05 | -0.0525 | 0.85437 | 0.441063 | 0.879368 | 1 | 0.999631 | 0.951263 |
| Srrm1_E9PUK6_S391s_2 | -0.07 | -0.32 | 0.02 | -0.61 | -0.42 | -0.97 | -0.38 | -0.03 | -0.32 | 0.48 | -0.11 | -0.42 | -0.13 | -0.29 | 0.39 | -0.34 | -0.205 | 0.1525 | 0.1525 | 0.626216 | 0.502165 | 0.627068 | 1 | 0.999631 | 0.86465 |
| Srrm1_E9PUK6_S412s_1 | 3.14 | 3.68 | 3.22 | -0.39 | 3.14 | 3.01 | 4.63 | -0.59 | 4.4 | 3.82 | 2.74 | 3.78 | 4.32 | -0.41 | 1.29 | -0.4 | 0.135 | 1.2725 | -1.2125 | 0.337603 | 0.913235 | 0.168224 | 1 | 0.999631 | 0.730629 |
| Srrm1_E9PUK6_S429s_1 | -0.59 | -2.21 | -0.89 | -1.02 | -1.33 | -1.8 | -0.79 | -1.13 | -1.26 | -0.23 | -1.09 | -1 | -0.47 | -1.29 | -0.9 | -1.3 | -0.085 | 0.2825 | 0.1875 | 0.656771 | 0.836812 | 0.510452 | 1 | 0.999631 | 0.879056 |
| Srrm1_E9PUK6_S448s_1 | 1.46 | -0.65 | 0.95 | 0.33 | -0.81 | -0.22 | 0.9 | -0.09 | -0.73 | 0.94 | -0.23 | -0.74 | -0.61 | -1.1 | 0.3 | -0.05 | -0.5775 | -0.7125 | -0.8875 | 0.1117 | 0.293109 | 0.220065 | 1 | 0.999631 | 0.577068 |
| Srrm1_E9PUK6_S461s_1 | -6.25 | -20.04 | 1.29 | 0.71 | -7.48 | -9.07 | 0.18 | 0.18 | -9.59 | -3.71 | 0.61 | 1.2 | -6.3 | -8.68 | 0.97 | 1.5 | 2.025 | 3.2 | 2.945 | 0.528874 | 0.66453 | 0.498412 | 1 | 0.999631 | 0.825257 |
| Srrm1_E9PUK6_S550s_1 | 0.13 | 0.3 | 0.64 | -0.01 | 0.07 | 0.21 | 0.12 | 0.22 | 0.38 | 0.65 | 0.75 | 0.68 | 0.77 | 0.96 | 0.88 | 0.88 | -0.11 | 0.35 | 0.6075 | 0.052113 | 0.668764 | 0.216193 | 1 | 0.999631 | 0.489777 |
| Srrm1_E9PUK6_S561s_2 | -0.18 | 0.1 | 1.26 | 0.32 | -0.1 | 0 | 0.58 | 0.1 | 0.15 | 0.61 | -0.05 | -0.22 | 0.26 | -0.28 | 0.36 | 0.17 | -0.23 | -0.2525 | -0.2475 | 0.518772 | 0.534833 | 0.512903 | 1 | 0.999631 | 0.819456 |
| Srrm1_E9PUK6_S563s_2 | -0.18 | 0.1 | 1.26 | 0.32 | -0.1 | 0 | 0.58 | 0.1 | 0.15 | 0.61 | -0.05 | -0.22 | 0.26 | -0.28 | 0.36 | 0.17 | -0.23 | -0.2525 | -0.2475 | 0.518772 | 0.534833 | 0.512903 | 1 | 0.999631 | 0.819456 |
| Srrm1_E9PUK6_S572s_2 | -0.03 | 0 | 0.13 | -0.18 | -0.47 | -0.37 | -0.09 | 0.22 | -0.18 | -0.12 | 0.33 | 0.83 | -0.39 | -0.49 | 0.34 | -0.23 | -0.1575 | 0.235 | -0.1725 | 0.569606 | 0.556593 | 0.45572 | 1 | 0.999631 | 0.84338 |
| Srrm1_E9PUK6_S574s_2 | -0.03 | 0 | 0.13 | -0.18 | -0.47 | -0.37 | -0.09 | 0.22 | -0.18 | -0.12 | 0.33 | 0.83 | -0.39 | -0.49 | 0.34 | -0.23 | -0.1575 | 0.235 | -0.1725 | 0.569606 | 0.556593 | 0.45572 | 1 | 0.999631 | 0.84338 |
| Srrm1_E9PUK6_S591s_1 | -0.96 | -3.24 | -1.09 | -2.2 | -3.76 | -3.24 | -1.81 | -3.81 | -3.84 | -1.31 | -2.28 | -3.33 | -0.93 | -3.47 | -3.04 | -2.54 | -1.2825 | -0.8175 | -0.6225 | 0.373958 | 0.06739 | 0.254993 | 1 | 0.999631 | 0.747035 |
| Srrm1_E9PUK6_S591s_2 | 0.72 | 0.26 | 1.02 | 0.26 | 0.15 | 0.2 | 0.45 | 0.67 | 0.22 | 1.52 | 0.09 | 0.1 | 0.55 | 0.1 | 0.51 | 0.43 | -0.1975 | -0.0825 | -0.1675 | 0.589787 | 0.503776 | 0.838384 | 1 | 0.999631 | 0.851414 |
| Srrm1_E9PUK6_S602s_2 | 0.46 | 0.17 | 0.37 | -0.28 | -0.09 | -0.15 | 0.27 | -0.03 | 0.19 | 0.88 | -0.07 | -0.37 | 0.53 | 0.49 | 0.34 | 0.37 | -0.18 | -0.0225 | 0.2525 | 0.393907 | 0.518238 | 0.948948 | 1 | 0.999631 | 0.759303 |
| Srrm1_E9PUK6_S616s_1 | 0.11 | -0.03 | 0.35 | -0.13 | -0.28 | -0.35 | -0.41 | -0.07 | -0.14 | 0.39 | 0.16 | -0.09 | 0.52 | 0.6 | 0.35 | 0.49 | -0.3525 | 0.005 | 0.415 | 0.14924 | 0.179817 | 0.985233 | 1 | 0.999631 | 0.621356 |
| Srrm1_E9PUK6_S624s_1 | 0.05 | 0.06 | 0.58 | -0.21 | -0.26 | -0.25 | -0.07 | -0.17 | -0.32 | 0.22 | -0.56 | -0.28 | 0.84 | 1.51 | -0.42 | 0.16 | -0.3075 | -0.355 | 0.4025 | 0.379893 | 0.266357 | 0.261248 | 1 | 0.999631 | 0.752193 |
| Srrm1_E9PUK6_S635s_1 | -0.02 | 0.54 | 1.08 | 0.02 | 0.35 | 0.67 | 0.09 | 0.68 | 0.46 | 0.37 | 0.7 | 1.08 | 1.07 | 0.61 | 1.12 | 0.9 | 0.0425 | 0.2475 | 0.52 | 0.153973 | 0.898658 | 0.479672 | 1 | 0.999631 | 0.625198 |
| Srrm1_E9PUK6_S635s_2 | 0.21 | -0.01 | -0.03 | 0.21 | -0.07 | -0.81 | -0.34 | 0.61 | -0.29 | 0.76 | -0.29 | 0.36 | 1.22 | 0.82 | 0.6 | 0.1 | -0.2475 | 0.04 | 0.59 | 0.091964 | 0.472165 | 0.901644 | 1 | 0.999631 | 0.557085 |
| Srrm1_E9PUK6_S655s_2 | 0.82 | 0.3 | 0.6 | 0.54 | 0.01 | -0.19 | 0.93 | 0.51 | -0.08 | 1.49 | -0.55 | -0.51 | 0.16 | -0.49 | 0.1 | -0.03 | -0.25 | -0.4775 | -0.63 | 0.0545 | 0.443788 | 0.326495 | 1 | 0.999631 | 0.491994 |
| Srrm1_E9PUK6_S657s_2 | 0.82 | 0.3 | 0.6 | 0.54 | 0.01 | -0.19 | 0.93 | 0.51 | -0.08 | 1.49 | -0.55 | -0.51 | 0.16 | -0.49 | 0.1 | -0.03 | -0.25 | -0.4775 | -0.63 | 0.0545 | 0.443788 | 0.326495 | 1 | 0.999631 | 0.491994 |
| Srrm1_E9PUK6_S664s_1 | 2.26 | 0.52 | 1.56 | 1.92 | -0.38 | 1 | 1.84 | 1.71 | 0.95 | 2.4 | 1.19 | 0.27 | 0.4 | 0.35 | 1.3 | 1.03 | -0.5225 | -0.3625 | -0.795 | 0.098313 | 0.374791 | 0.508233 | 1 | 0.999631 | 0.561867 |
| Srrm1_E9PUK6_S672s_1 | 1.18 | 0.02 | 0.95 | 0.57 | -0.15 | 0.12 | 1.3 | 0.01 | 0.31 | 1.32 | 0.85 | 0.72 | -0.06 | -0.26 | 0.74 | 0.49 | -0.36 | 0.12 | -0.4525 | 0.251346 | 0.39523 | 0.739955 | 1 | 0.999631 | 0.687138 |
| Srrm1_E9PUK6_S714s_1 | 0.2 | -0.09 | -0.08 | -0.64 | -0.3 | -0.45 | -0.41 | -0.56 | -0.32 | 0.24 | -0.23 | -0.41 | 0.46 | 0.1 | -0.36 | -0.17 | -0.2775 | -0.0275 | 0.16 | 0.626533 | 0.322479 | 0.927434 | 1 | 0.999631 | 0.864779 |
| Srrm1_E9PUK6_S723s_1 | 2.95 | 1.36 | 1.97 | 1.23 | 0.54 | 0.67 | 2.04 | 0.37 | 1.77 | 2.3 | 0.14 | -0.27 | 1.06 | -0.8 | -0.48 | -0.05 | -0.9725 | -0.8925 | -1.945 | 0.004194 | 0.08188 | 0.197159 | 1 | 0.999631 | 0.351308 |
| Srrm1_E9PUK6_S725s_1 | 1.33 | -2.83 | -0.4 | -1.59 | -3.21 | -1.87 | 0.67 | -3.24 | -5.9 | 0.37 | -2.32 | -3.19 | -0.85 | -2.89 | -2.25 | -2.56 | -1.04 | -1.8875 | -1.265 | 0.163475 | 0.352968 | 0.179981 | 1 | 0.999631 | 0.631937 |
| Srrm1_E9PUK6_S731s_1 | -0.18 | -0.33 | 1.86 | 1.15 | 0.15 | -1.44 | 1.67 | 0.59 | -0.28 | -0.03 | 0.69 | 0.07 | 0.54 | 0.01 | 0.35 | 0.72 | -0.3825 | -0.5125 | -0.22 | 0.673729 | 0.60504 | 0.346217 | 1 | 0.999631 | 0.885515 |
| Srrm1_E9PUK6_S756s_1 | -0.6 | -2.03 | 0.33 | -0.53 | -1.78 | -2.22 | -0.39 | 0.32 | -2.56 | -0.92 | 0.06 | -0.12 | -1.56 | -1.86 | -0.05 | 0.17 | -0.31 | -0.1775 | -0.1175 | 0.85434 | 0.650893 | 0.796856 | 1 | 0.999631 | 0.951263 |
| Srrm1_E9PUK6_S756s_2 | 0.31 | -0.55 | -0.12 | -0.11 | -0.44 | -0.67 | -0.24 | -0.44 | -0.42 | 0.55 | -0.44 | -0.93 | -0.12 | -0.57 | -0.72 | -0.57 | -0.33 | -0.1925 | -0.3775 | 0.242631 | 0.254059 | 0.613805 | 1 | 0.999631 | 0.685099 |
| Srrm1_E9PUK6_S758s_2 | 0.31 | -0.55 | -0.12 | -0.11 | -0.44 | -0.67 | -0.24 | -0.44 | -0.42 | 0.55 | -0.44 | -0.93 | -0.12 | -0.57 | -0.72 | -0.57 | -0.33 | -0.1925 | -0.3775 | 0.242631 | 0.254059 | 0.613805 | 1 | 0.999631 | 0.685099 |
| Srrm1_E9PUK6_S774s_1 | -0.66 | -2.94 | -0.1 | -0.79 | -2.61 | -3.03 | 0.03 | -1.3 | -2.41 | -1.79 | -0.64 | -0.05 | -0.85 | -1.9 | -0.27 | -0.6 | -0.605 | -0.1 | 0.2175 | 0.736186 | 0.46317 | 0.890837 | 1 | 0.999631 | 0.90759 |
| Srrm1_E9PUK6_S787s_1 | 0.25 | 0 | 0.11 | -0.25 | 0.19 | 0.18 | -0.77 | -0.41 | -1.07 | -0.02 | -0.13 | -0.51 | 0.17 | -0.09 | -0.92 | -0.72 | -0.23 | -0.46 | -0.4175 | 0.239692 | 0.467328 | 0.172385 | 1 | 0.999631 | 0.683719 |
| Srrm1_E9PUK6_T593t_2 | 0.72 | 0.26 | 1.02 | 0.26 | 0.15 | 0.2 | 0.45 | 0.67 | 0.22 | 1.52 | 0.09 | 0.1 | 0.55 | 0.1 | 0.51 | 0.43 | -0.1975 | -0.0825 | -0.1675 | 0.589787 | 0.503776 | 0.838384 | 1 | 0.999631 | 0.851414 |
| Srrm1_E9PUK6_T633t_2 | 0.21 | -0.01 | -0.03 | 0.21 | -0.07 | -0.81 | -0.34 | 0.61 | -0.29 | 0.76 | -0.29 | 0.36 | 1.22 | 0.82 | 0.6 | 0.1 | -0.2475 | 0.04 | 0.59 | 0.091964 | 0.472165 | 0.901644 | 1 | 0.999631 | 0.557085 |
| Srrm2_Q8BTI8_S1068s_1 | -0.37 | -0.37 | -0.3 | -0.53 | -0.35 | -0.92 | -0.99 | -0.56 | -0.46 | -0.35 | -0.41 | -0.58 | 0.3 | -0.07 | -0.28 | -0.08 | -0.3125 | -0.0575 | 0.36 | 0.211235 | 0.250166 | 0.812088 | 1 | 0.999631 | 0.665474 |
| Srrm2_Q8BTI8_S1097s_1 | 1.67 | -0.31 | 0.32 | 0.67 | -0.63 | 0.16 | 0.82 | -0.49 | 0.19 | 0.95 | -0.19 | -0.17 | -0.15 | -0.07 | 0.29 | -1.45 | -0.6225 | -0.3925 | -0.9325 | 0.102296 | 0.230639 | 0.416489 | 1 | 0.999631 | 0.564448 |
| Srrm2_Q8BTI8_S1151s_1 | -0.95 | 0.9 | 0.66 | 0.1 | 0.1 | 0.36 | -0.74 | -0.08 | 0.84 | -0.22 | -0.08 | -0.18 | 1.42 | 1.8 | 0.25 | 0.48 | -0.2675 | -0.0875 | 0.81 | 0.144605 | 0.560807 | 0.851421 | 1 | 0.999631 | 0.616037 |
| Srrm2_Q8BTI8_S1179s_1 | 1.44 | -0.3 | -0.23 | -0.62 | -0.8 | 0.16 | 0.47 | -1.11 | -0.44 | 0.75 | -0.57 | -1.06 | -0.18 | -1.12 | -1.4 | -2.37 | -0.3925 | -0.4025 | -1.34 | 0.043931 | 0.48049 | 0.475661 | 1 | 0.999631 | 0.475443 |
| Srrm2_Q8BTI8_S1216s_1 | -0.18 | -0.91 | -0.73 | -0.44 | -0.71 | -0.6 | -1.09 | -0.61 | -0.14 | -0.72 | -0.76 | -0.25 | 0.47 | -0.01 | -0.66 | -0.47 | -0.1875 | 0.0975 | 0.3975 | 0.276739 | 0.507449 | 0.746786 | 1 | 0.999631 | 0.700187 |
| Srrm2_Q8BTI8_S1269s_1 | -0.41 | 0.47 | -0.07 | -0.86 | 0.25 | 0.06 | -0.8 | -0.49 | 0.38 | 0.18 | -0.43 | 0 | 0.98 | 0.48 | -0.03 | -0.21 | -0.0275 | 0.25 | 0.5225 | 0.219395 | 0.942709 | 0.494846 | 1 | 0.999631 | 0.670174 |
| Srrm2_Q8BTI8_S1278s_1 | 0.43 | 0.51 | 0.96 | 0.28 | 0 | 0.31 | 0.13 | 0.49 | 0.46 | 0.93 | 0.81 | 0.42 | 0.6 | 0.61 | 0.79 | 1.12 | -0.3125 | 0.11 | 0.235 | 0.438208 | 0.266557 | 0.700534 | 1 | 0.999631 | 0.78194 |
| Srrm2_Q8BTI8_S1305s_1 | -0.3 | 0.5 | 0.53 | -0.03 | 0.15 | 0.17 | -0.47 | 0.23 | 0.02 | 0.41 | 0.34 | 0.66 | 0.97 | 0.89 | 0.46 | 0.87 | -0.155 | 0.1825 | 0.6225 | 0.074356 | 0.623776 | 0.559885 | 1 | 0.999631 | 0.539897 |
| Srrm2_Q8BTI8_S1359s_1 | 0.18 | 0.1 | 1.36 | -0.09 | 0.01 | 0.05 | 0.21 | 1.82 | -0.49 | -0.33 | 1.46 | -0.47 | 0.13 | 0.26 | 0.78 | 0.27 | 0.135 | -0.345 | -0.0275 | 0.943675 | 0.791001 | 0.525384 | 1 | 0.999631 | 0.981629 |
| Srrm2_Q8BTI8_S1360s_1 | -0.22 | 0.07 | 0.24 | -0.22 | -0.26 | -0.47 | -0.58 | 0 | -0.46 | 0.19 | 0.16 | 0.11 | 0.5 | 0.33 | 0.1 | 0.44 | -0.295 | 0.0325 | 0.375 | 0.201002 | 0.28474 | 0.908876 | 1 | 0.999631 | 0.65842 |
| Srrm2_Q8BTI8_S1378s_1 | 0.21 | 0.07 | 0.78 | 0.27 | 0.21 | 0.41 | 0.2 | -0.11 | 0.02 | 0.81 | 0.08 | -0.19 | -0.4 | 0.04 | 0.46 | 0.27 | -0.155 | -0.1525 | -0.24 | 0.46313 | 0.57621 | 0.639668 | 1 | 0.999631 | 0.793564 |
| Srrm2_Q8BTI8_S1380s_1 | -0.03 | 0.03 | 0.01 | -0.46 | -0.2 | 0.16 | -0.28 | -0.28 | -0.27 | -0.12 | -0.2 | -0.54 | 0.22 | -0.15 | -1.27 | -0.53 | -0.0375 | -0.17 | -0.32 | 0.405027 | 0.885679 | 0.526049 | 1 | 0.999631 | 0.765844 |
| Srrm2_Q8BTI8_S1397s_1 | 0.19 | 0.41 | 1.13 | 0.59 | 0.56 | 0.09 | 0.67 | 0.13 | -0.13 | -0.04 | 0.15 | 0.11 | 0.09 | -0.36 | 0.48 | 0.09 | -0.2175 | -0.5575 | -0.505 | 0.153553 | 0.484259 | 0.082154 | 1 | 0.999631 | 0.625198 |
| Srrm2_Q8BTI8_S1400s_1 | 0.84 | -0.29 | -0.6 | -0.66 | -0.73 | 0.04 | -0.23 | -1.35 | 0.07 | 1.06 | 0.36 | -1.3 | 0.02 | 0.32 | -1.45 | -1.18 | -0.39 | 0.225 | -0.395 | 0.459515 | 0.394189 | 0.688466 | 1 | 0.999631 | 0.79152 |
| Srrm2_Q8BTI8_S1419s_1 | 0.36 | 0.99 | 1.18 | -0.78 | 1.87 | 0.71 | 1.46 | 0.46 | 3.3 | 0.05 | 1.18 | 1.7 | 3.64 | -0.18 | 1.32 | -1.11 | 0.6875 | 1.12 | 0.48 | 0.621216 | 0.200508 | 0.142185 | 1 | 0.999631 | 0.861888 |
| Srrm2_Q8BTI8_S1453s_3 | -0.63 | -1.1 | 0.32 | -0.31 | -1.6 | -2.73 | 0.43 | -1.03 | -0.21 | -1.09 | 0.36 | -0.28 | -1.71 | -2.38 | -2.43 | 0.13 | -0.8025 | 0.125 | -1.1675 | 0.078809 | 0.232334 | 0.76829 | 1 | 0.999631 | 0.545479 |
| Srrm2_Q8BTI8_S1455s_3 | -0.63 | -1.1 | 0.32 | -0.31 | -1.6 | -2.73 | 0.43 | -1.03 | -0.21 | -1.09 | 0.36 | -0.28 | -1.71 | -2.38 | -2.43 | 0.13 | -0.8025 | 0.125 | -1.1675 | 0.078809 | 0.232334 | 0.76829 | 1 | 0.999631 | 0.545479 |
| Srrm2_Q8BTI8_S1557s_1 | 2.21 | -0.27 | 0.87 | 0.63 | -0.58 | 0.28 | 1.18 | -0.38 | -1.02 | 1.24 | -0.41 | -0.31 | -0.76 | -0.26 | 0.21 | -0.7 | -0.735 | -0.985 | -1.2375 | 0.038267 | 0.23008 | 0.144651 | 1 | 0.999631 | 0.459475 |
| Srrm2_Q8BTI8_S1647s_2 | -0.39 | -0.19 | -2.71 | -1.57 | -0.68 | -1.01 | -2.02 | -2.37 | -0.4 | -0.25 | -0.9 | -2.53 | -0.13 | -0.29 | -2.28 | -3.11 | -0.305 | 0.195 | -0.2375 | 0.771929 | 0.632945 | 0.780179 | 1 | 0.999631 | 0.919413 |
| Srrm2_Q8BTI8_S1649s_1 | 0.42 | 0 | 0.61 | 0.15 | -0.41 | 0.1 | 0.04 | -0.2 | 0.09 | 0.78 | 0.12 | 0.12 | 0.23 | 0.22 | 0.02 | 0.2 | -0.4125 | -0.0175 | -0.1275 | 0.651202 | 0.151316 | 0.952833 | 1 | 0.999631 | 0.876438 |
| Srrm2_Q8BTI8_S1650s_1 | 0.42 | 0 | 0.42 | -0.08 | -0.41 | 0.1 | 0.12 | -0.42 | 0.09 | 0.78 | -0.29 | 0.43 | 0.23 | 0.22 | -0.8 | 0.28 | -0.3425 | 0.0625 | -0.2075 | 0.557215 | 0.242302 | 0.846542 | 1 | 0.999631 | 0.838305 |
| Srrm2_Q8BTI8_S1825s_1 | 2.32 | -1.5 | -0.02 | -0.84 | -1.82 | -0.42 | 3.34 | -3.58 | -4.11 | 2.04 | -1.94 | -2.03 | -0.39 | -1.82 | -1.55 | -2.53 | -0.61 | -1.5 | -1.5625 | 0.081232 | 0.6719 | 0.267937 | 1 | 0.999631 | 0.54635 |
| Srrm2_Q8BTI8_S2075s_1 | -0.71 | -1.03 | 0.58 | -0.49 | -0.54 | -0.24 | -0.38 | -0.62 | 0.1 | -0.37 | -0.84 | 0.03 | 0.17 | 0.47 | 0.45 | -0.19 | -0.0325 | 0.1425 | 0.6375 | 0.137742 | 0.930891 | 0.732462 | 1 | 0.999631 | 0.610682 |

| | | | | | | | | | | | | | | | | | | | | | | | | | |
|---|---|---|---|---|---|---|---|---|---|---|---|---|---|---|---|---|---|---|---|---|---|---|---|---|---|
| Srrm2_Q8BTI8_S2084s_1 | 0.14 | 1.05 | 0.45 | -0.18 | 0.28 | 0.06 | -0.32 | -0.48 | 0.72 | 0.86 | 0.88 | 0.68 | 1.75 | 0.18 | -0.48 | 1.43 | -0.48 | 0.42 | 0.355 | 0.520271 | 0.18688 | 0.214349 | 1 | 0.999631 | 0.819528 |
| Srrm2_Q8BTI8_S2224s_1 | 0.16 | -0.22 | 0.29 | 0.25 | 0.31 | 0.36 | -0.44 | 0.55 | 0.3 | 0.13 | -0.16 | 0.36 | -1.15 | -1.33 | -1.05 | 0.66 | 0.075 | 0.0375 | -0.8375 | 0.098083 | 0.806331 | 0.890354 | 1 | 0.999631 | 0.561078 |
| Srrm2_Q8BTI8_S2351s_1 | 1 | 0.29 | 0.65 | -0.16 | -0.15 | -0.08 | -0.08 | -0.03 | -0.13 | 0.79 | 0.43 | -0.12 | 0.58 | 0.39 | -0.55 | 0.65 | -0.53 | -0.2025 | -0.1775 | 0.657219 | 0.109094 | 0.582144 | 1 | 0.999631 | 0.879214 |
| Srrm2_Q8BTI8_S2360s_2 | -0.16 | 0.89 | 0.74 | 0.51 | 0.77 | 0.59 | 0.42 | 0.75 | 0.97 | 0.04 | 0.73 | 0.85 | 0.46 | 0.78 | 0.68 | 0.72 | 0.1375 | 0.1525 | 0.165 | 0.61153 | 0.653633 | 0.665 | 1 | 0.999631 | 0.859739 |
| Srrm2_Q8BTI8_S2381s_1 | 1.57 | -1.79 | 0.49 | -0.03 | -1.24 | -1.72 | 0.53 | -0.56 | -0.77 | 1.66 | -0.64 | -0.82 | -2.13 | -1.15 | -0.91 | -1.51 | -0.8075 | -0.2025 | -1.485 | 0.04763 | 0.296572 | 0.802775 | 1 | 0.999631 | 0.481835 |
| Srrm2_Q8BTI8_S2404s_1 | -0.1 | -0.28 | -0.29 | -0.22 | -0.36 | -0.67 | -0.54 | -0.21 | -0.35 | -0.21 | -0.13 | -0.12 | -0.46 | 0 | -0.24 | 0.07 | -0.2225 | 0.02 | 0.065 | 0.813356 | 0.371719 | 0.933994 | 1 | 0.999631 | 0.937041 |
| Srrm2_Q8BTI8_S2535s_1 | -0.14 | 0.27 | 0.08 | -0.14 | -0.27 | -0.39 | -0.59 | 0.05 | -0.06 | 0.14 | -0.08 | -0.04 | 0.41 | 0.49 | -0.07 | 0.48 | -0.3175 | -0.0275 | 0.31 | 0.296971 | 0.248681 | 0.913192 | 1 | 0.999631 | 0.710149 |
| Srrm2_Q8BTI8_S2642s_1 | -0.36 | -0.21 | 0.26 | -0.11 | -0.65 | -0.44 | -0.49 | -0.52 | -0.2 | 0.03 | 0.02 | 0.12 | 0.26 | -0.44 | -0.11 | 0.37 | -0.42 | 0.0975 | 0.125 | 0.692738 | 0.122631 | 0.714264 | 1 | 0.999631 | 0.894787 |
| Srrm2_Q8BTI8_S2646s_1 | 0.61 | 1.53 | 0.21 | 0.64 | 0.69 | 0.82 | 0.51 | 0.97 | 1.73 | 1.09 | -0.05 | 0.6 | 1.32 | 1 | 1.26 | 1.06 | -1.4E-16 | 0.095 | 0.4125 | 0.252814 | 1 | 0.835432 | 1 | 0.999631 | 0.688432 |
| Srrm2_Q8BTI8_S2648s_1 | -0.55 | -0.49 | -0.15 | -0.76 | -0.77 | -0.93 | -1.08 | -0.87 | -0.21 | -0.57 | -0.35 | -0.34 | 0.97 | -0.16 | -0.45 | -0.44 | -0.425 | 0.12 | 0.4675 | 0.247899 | 0.120111 | 0.652123 | 1 | 0.999631 | 0.685751 |
| Srrm2_Q8BTI8_S2656s_1 | 1.02 | 0.32 | 0.65 | 0.11 | 0.01 | 0.14 | 0.31 | 0.05 | 0.05 | 1.19 | 0.2 | 0.14 | 0.52 | -0.14 | 0.86 | 0.06 | -0.3975 | -0.13 | -0.2 | 0.575763 | 0.185603 | 0.721611 | 1 | 0.999631 | 0.846085 |
| Srrm2_Q8BTI8_S2656s_2 | 1.4 | 0.04 | 0.71 | 0.61 | 0.01 | 0.15 | 0.93 | 0.06 | -0.17 | 0.85 | 0.3 | -0.58 | -0.53 | -0.67 | -0.25 | -0.45 | -0.4025 | -0.59 | -1.165 | 0.007153 | 0.292841 | 0.181421 | 1 | 0.999631 | 0.358173 |
| Srrm2_Q8BTI8_S2660s_2 | 1.4 | 0.04 | 0.71 | 0.61 | 0.01 | 0.15 | 0.93 | 0.06 | -0.17 | 0.85 | 0.3 | -0.58 | -0.53 | -0.67 | -0.25 | -0.45 | -0.4025 | -0.59 | -1.165 | 0.007153 | 0.292841 | 0.181421 | 1 | 0.999631 | 0.358173 |
| Srrm2_Q8BTI8_S2681s_1 | 2.22 | -3.92 | -2.63 | -2.3 | -4.87 | -1.43 | 2.22 | -5.5 | -4.87 | 2.05 | -4.34 | -3.14 | -2.68 | -4.65 | -4.69 | -4.36 | -0.7375 | -0.9175 | -2.4375 | 0.065967 | 0.695895 | 0.604044 | 1 | 0.999631 | 0.516933 |
| Srrm2_Q8BTI8_S333s_1 | 1.86 | 0.62 | 0.73 | 0.69 | -0.08 | 1.16 | 1.07 | -0.73 | 0.49 | 1.64 | -0.2 | 0.4 | 0.67 | -0.4 | 0.28 | -0.54 | -0.62 | -0.3925 | -0.9725 | 0.041486 | 0.241881 | 0.411492 | 1 | 0.999631 | 0.474979 |
| Srrm2_Q8BTI8_S349s_1 | 0.17 | 0.21 | 0.44 | -0.09 | -0.39 | -0.26 | 0.04 | -0.31 | 0.49 | 0.37 | 0.28 | 0.64 | 0.64 | 0.23 | 0.02 | 0.82 | -0.4125 | 0.2625 | 0.245 | 0.434904 | 0.130697 | 0.325671 | 1 | 0.999631 | 0.779717 |
| Srrm2_Q8BTI8_S349s_2 | -0.41 | -0.26 | 0.75 | 0.2 | -0.01 | -0.48 | -0.26 | 0.39 | -0.13 | -0.11 | 0.32 | 0.15 | 0.49 | 0.18 | 0.05 | 0.34 | -0.16 | -0.0125 | 0.195 | 0.571528 | 0.649401 | 0.97003 | 1 | 0.999631 | 0.843675 |
| Srrm2_Q8BTI8_S351s_1 | 0.61 | -0.64 | 0.09 | -0.02 | -0.53 | -0.32 | -0.07 | -0.34 | 0.31 | 0.84 | 0.46 | 0.22 | 0.55 | 0.7 | 0.24 | 0.85 | -0.325 | 0.4475 | 0.575 | 0.121577 | 0.324245 | 0.205672 | 1 | 0.999631 | 0.590097 |
| Srrm2_Q8BTI8_S351s_2 | -0.41 | -0.26 | 0.75 | 0.2 | -0.01 | -0.48 | -0.26 | 0.39 | -0.13 | -0.11 | 0.32 | 0.15 | 0.49 | 0.18 | 0.05 | 0.34 | -0.16 | -0.0125 | 0.195 | 0.571528 | 0.649401 | 0.97003 | 1 | 0.999631 | 0.843675 |
| Srrm2_Q8BTI8_S396s_1 | -0.64 | -0.11 | -0.11 | -1.03 | 0.12 | -0.23 | -2.06 | -1.1 | -0.26 | 0.12 | -1.75 | -2.37 | -0.8 | 0.39 | -1.67 | -0.59 | -0.345 | -0.5925 | -0.195 | 0.678857 | 0.497353 | 0.321438 | 1 | 0.999631 | 0.887183 |
| Srrm2_Q8BTI8_S454s_1 | 0.74 | -0.9 | 0.72 | 0.39 | -1.3 | -1.25 | 0.65 | -1.08 | -1.41 | 0.22 | -0.61 | -1.3 | -0.73 | -1.52 | 0.35 | -0.65 | -0.9825 | -1.0125 | -0.875 | 0.114481 | 0.102247 | 0.071211 | 1 | 0.999631 | 0.579001 |
| Srrm2_Q8BTI8_S778s_1 | -0.73 | 0.55 | -0.81 | -2.7 | -0.49 | 0 | -0.28 | -0.54 | 0.71 | -0.88 | -0.71 | -0.79 | 0.65 | -1.63 | -0.67 | -2.16 | 0.595 | 0.505 | -0.03 | 0.969836 | 0.343803 | 0.468088 | 1 | 0.999631 | 0.989372 |
| Srrm2_Q8BTI8_S829s_2 | 0.25 | -0.12 | -1.54 | -1.39 | 0.22 | -0.72 | -1.66 | -0.51 | -0.03 | 0.35 | -0.64 | -0.4 | 0.13 | 0.21 | -0.87 | -0.24 | 0.0325 | 0.52 | 0.5075 | 0.316486 | 0.952427 | 0.292857 | 1 | 0.999631 | 0.720734 |
| Srrm2_Q8BTI8_S829s_3 | -0.2 | 0.29 | 0.79 | -0.12 | -0.16 | 0.09 | 0.11 | 0.09 | -0.43 | -0.02 | 0.04 | 0.49 | 0.15 | 0.61 | 0.64 | 0.89 | -0.1575 | -0.17 | 0.3825 | 0.275918 | 0.601451 | 0.61954 | 1 | 0.999631 | 0.700187 |
| Srrm2_Q8BTI8_S870s_1 | 1.45 | -3.06 | 0.41 | -0.79 | -1.57 | 0.02 | 2.74 | -2.72 | -5.44 | 1.93 | -1.76 | -2.34 | -2.28 | -2.89 | -1.71 | -3.8 | 0.115 | -1.405 | -2.1725 | 0.037069 | 0.929525 | 0.368031 | 1 | 0.999631 | 0.456881 |
| Srrm2_Q8BTI8_S928s_1 | 0.87 | 1.02 | 0.28 | -0.34 | -0.24 | 0.12 | -0.07 | 0.2 | 0.98 | 1.17 | -0.19 | 0.02 | 0.76 | 0.82 | 0.3 | 0.37 | -0.455 | 0.0375 | 0.105 | 0.78043 | 0.217163 | 0.933675 | 1 | 0.999631 | 0.921416 |
| Srrm2_Q8BTI8_T1390t_1 | -2.41 | -3.6 | -0.67 | -1.25 | -2.07 | -2.98 | -0.94 | -0.79 | -2.65 | -2.14 | -1.39 | -0.5 | -1.41 | -3.31 | -0.86 | -0.92 | 0.2875 | 0.3125 | 0.3575 | 0.640019 | 0.694645 | 0.661864 | 1 | 0.999631 | 0.870995 |
| Srrm2_Q8BTI8_T1428t_1 | 0.25 | 1.49 | 1.51 | 0.81 | 0.75 | 1.08 | 1.21 | 0.57 | 1.04 | 0.52 | -0.01 | 0.41 | 0.26 | 0.3 | -0.25 | -0.52 | -0.1125 | -0.525 | -1.0675 | 0.01993 | 0.755167 | 0.198383 | 1 | 0.999631 | 0.391668 |
| Srrm2_Q8BTI8_T1448t_1 | 0.27 | 0.54 | 0.64 | 0.67 | 0.06 | -0.02 | 0.05 | 0.17 | 0.35 | 0.67 | 0.62 | 0.34 | 0.8 | 0.7 | 0.25 | 0.72 | -0.465 | -0.035 | 0.0875 | 0.758361 | 0.077392 | 0.891746 | 1 | 0.999631 | 0.914449 |
| Srrm2_Q8BTI8_T1812t_1 | 0 | 0.71 | 1.17 | 0.51 | -0.11 | 0.55 | 0.42 | 0.83 | 0.49 | 0.86 | 0.47 | 0.93 | 1.2 | 0.78 | 1.57 | 1.66 | -0.175 | 0.09 | 0.705 | 0.075516 | 0.613984 | 0.78276 | 1 | 0.999631 | 0.541044 |
| Srrm2_Q8BTI8_T1880t_1 | -1.76 | -2.33 | -0.67 | -2.11 | -2.79 | -2.58 | -1.93 | -2.3 | -2.61 | -1.59 | -0.8 | -2.71 | -0.05 | -3.13 | -3.47 | -1.82 | -0.6825 | -0.21 | -0.4 | 0.598123 | 0.122194 | 0.699772 | 1 | 0.999631 | 0.855276 |
| Srrm2_Q8BTI8_T2056t_1 | 0.1 | -0.06 | 0.51 | 0.15 | -0.44 | -0.3 | -0.29 | -0.07 | -0.12 | 0.54 | 0.29 | 0.13 | 0.63 | 0.36 | 0.18 | 0.79 | -0.45 | 0.035 | 0.315 | 0.299426 | 0.102561 | 0.900611 | 1 | 0.999631 | 0.712852 |
| Srrm2_Q8BTI8_T2350t_1 | 0.17 | 0.13 | 0.6 | 0.13 | -0.06 | 0.27 | -0.24 | 0.2 | 0.12 | 0.58 | 0.15 | 0.42 | 0.73 | 0.71 | 0.23 | 0.73 | -0.215 | 0.06 | 0.3425 | 0.253484 | 0.423783 | 0.82408 | 1 | 0.999631 | 0.688735 |
| Srrm2_Q8BTI8_T2362t_1 | 0.94 | 0.19 | 0.94 | 0.4 | 0.3 | 0.53 | 0.48 | 0.21 | 0.35 | 1.09 | 0.62 | 0.45 | 0.18 | 0.12 | 0.39 | 0.16 | -0.2375 | 0.01 | -0.405 | 0.201224 | 0.410651 | 0.974666 | 1 | 0.999631 | 0.658939 |
| Srrm2_Q8BTI8_T2362t_2 | 0.24 | -0.53 | 0.64 | 0.43 | -0.3 | -0.13 | 0.44 | 0.55 | 0.28 | 0.16 | 0.44 | 0.54 | 0.05 | -0.84 | 0.73 | 0.53 | -0.055 | 0.16 | -0.0775 | 0.859399 | 0.877144 | 0.624318 | 1 | 0.999631 | 0.953128 |
| Srrm2_Q8BTI8_T823t_1 | 0.83 | 0.28 | -0.2 | -0.48 | 0.06 | -0.07 | -0.91 | -0.03 | 0.37 | 0.58 | -0.43 | 0.68 | 0.68 | 0.74 | 0.55 | -0.05 | -0.345 | 0.1925 | 0.3725 | 0.3361 | 0.373665 | 0.629117 | 1 | 0.999631 | 0.730466 |
| Srrm2_Q8BTI8_T823t_2 | -0.24 | -0.08 | 0.77 | -0.19 | -0.34 | -0.5 | -0.53 | -0.19 | -0.21 | -0.1 | 0.18 | -0.41 | 0.23 | 0.06 | -0.17 | 0.69 | -0.455 | -0.2 | 0.1375 | 0.698372 | 0.161659 | 0.54107 | 1 | 0.999631 | 0.895731 |
| Srrm2_Q8BTI8_T823t_3 | -0.2 | 0.29 | 0.79 | -0.12 | -0.16 | 0.09 | 0.11 | 0.09 | -0.43 | -0.02 | 0.04 | 0.49 | 0.15 | 0.61 | 0.64 | 0.89 | -0.1575 | -0.17 | 0.3825 | 0.275918 | 0.601451 | 0.61954 | 1 | 0.999631 | 0.700187 |
| Srrm2_Q8BTI8_T841t_2 | -1.38 | -0.03 | 1.48 | 0.26 | -1.46 | -0.19 | -0.15 | -0.05 | -0.5 | -0.96 | 0.52 | -0.41 | 0.38 | -0.17 | 0.12 | 1.05 | -0.545 | -0.42 | 0.2625 | 0.656767 | 0.380891 | 0.49251 | 1 | 0.999631 | 0.879056 |
| Srrm2_Q8BTI8_T841t_3 | -0.2 | 0.29 | 0.79 | -0.12 | -0.16 | 0.09 | 0.11 | 0.09 | -0.43 | -0.02 | 0.04 | 0.49 | 0.15 | 0.61 | 0.64 | 0.89 | -0.1575 | -0.17 | 0.3825 | 0.275918 | 0.601451 | 0.61954 | 1 | 0.999631 | 0.700187 |
| Srrm2_Q8BTI8_T877t_2 | 2.37 | 0.27 | 0.47 | 0.42 | -0.7 | -0.02 | 0.62 | -0.32 | 0.26 | 1.58 | -0.86 | -1 | -0.83 | -0.83 | -0.98 | -1.97 | -0.9875 | -0.8875 | -2.035 | 0.003304 | 0.08629 | 0.220709 | 1 | 0.999631 | 0.351308 |
| Srrm2_Q8BTI8_T955t_1 | -1.21 | 0.27 | 0.45 | -0.14 | -0.4 | -0.26 | -0.52 | -0.07 | -0.02 | 0.11 | -0.19 | 0.2 | 0.71 | -0.54 | 0.42 | 0.53 | -0.155 | 0.1825 | 0.4375 | 0.352846 | 0.693656 | 0.646445 | 1 | 0.999631 | 0.73949 |
| Srrm2_Q8BTI8_T973t_1 | -0.62 | -0.24 | -0.1 | -0.24 | -0.15 | -0.36 | -0.31 | -0.2 | -0.26 | -0.36 | -0.38 | -0.62 | 0.36 | -0.05 | -0.18 | -0.01 | 0.045 | -0.105 | 0.33 | 0.265516 | 0.855731 | 0.688007 | 1 | 0.999631 | 0.694352 |
| Srrm3_Q80WV7_S562s_1 | 1.12 | 0.65 | 0.11 | 0.38 | -0.13 | -0.26 | -0.24 | 0.09 | -0.14 | 0.36 | 0.38 | 0.56 | 0.62 | 0.66 | 0.35 | 0.38 | -0.7 | -0.275 | -0.0625 | 0.84355 | 0.037543 | 0.401203 | 1 | 0.999631 | 0.94821 |
| Srrm3_Q80WV7_T352t_1 | 0.51 | 0.72 | 0.33 | 0.53 | 0.12 | 0.13 | 0.13 | 0.41 | -0.33 | 0.49 | 0.5 | 0.78 | 0.39 | 0.83 | 0.51 | 0.99 | -0.325 | -0.1625 | 0.1575 | 0.58489 | 0.201094 | 0.60901 | 1 | 0.999631 | 0.849277 |
| Srrm4_Q8BKA3_S271s_1 | 0.09 | 0.3 | 0.88 | 0.59 | 0.16 | 0.04 | 0.53 | -0.46 | 0.17 | 0.56 | 0.3 | 0.75 | 1.18 | 0.35 | 0.6 | 1.11 | -0.3975 | -0.02 | 0.345 | 0.315456 | 0.22917 | 0.946078 | 1 | 0.999631 | 0.72069 |
| Srrm4_Q8BKA3_S285s_1 | 0.85 | -0.15 | 0.84 | 0.06 | 0.35 | 0.39 | 0.66 | -0.46 | 1.35 | 0.05 | 0.28 | -0.42 | -0.07 | 0.07 | 0.82 | 0.06 | -0.165 | -0.085 | -0.18 | 0.630158 | 0.659339 | 0.849583 | 1 | 0.999631 | 0.867378 |
| Srrm4_Q8BKA3_S496s_1 | 0.17 | -0.14 | 0.32 | 0.22 | -0.28 | -0.48 | 0.19 | -0.3 | 0.02 | 0.41 | 0.68 | 0.09 | -0.18 | -0.12 | 0.63 | 0.66 | -0.36 | 0.1575 | 0.105 | 0.748649 | 0.200554 | 0.576356 | 1 | 0.999631 | 0.911931 |
| Srrt_Q99MR6_S4s_1 | 1.81 | -0.41 | 0.47 | 0.15 | -2.25 | -0.27 | 0.51 | -0.27 | -0.79 | 0.76 | -1.14 | -1.09 | 0.29 | -1.89 | -1.16 | -1.68 | -1.075 | -1.07 | -1.615 | 0.02413 | 0.133587 | 0.096678 | 1 | 0.999631 | 0.415644 |
| Srrt_Q99MR6_S67s_1 | 0.75 | 1.49 | 1.64 | 1.27 | 0.87 | 1.06 | 0.53 | 1.32 | 1.16 | 1.48 | 1.52 | 1.41 | 1.17 | 1.89 | 1.77 | 1.66 | -0.3425 | 0.105 | 0.335 | 0.318974 | 0.285733 | 0.721172 | 1 | 0.999631 | 0.722245 |
| Srrt_Q99MR6_T543t_1 | 0.11 | 0.93 | 0.53 | 0.57 | 0.94 | 0.39 | 0 | 0.21 | 0.82 | 0.27 | 0.92 | 0.53 | 0.85 | 1.11 | 0.3 | 1 | -0.15 | 0.1 | 0.28 | 0.39763 | 0.635052 | 0.73925 | 1 | 0.999631 | 0.761181 |
| Srsf1_H7BX95_S249s_1 | -0.47 | -2.28 | 1.28 | -0.41 | 0.93 | -0.58 | 2.32 | -1.28 | 2.98 | -0.77 | 1.45 | 1.99 | 3.25 | -1.89 | -0.26 | -0.45 | 0.8175 | 1.8825 | 0.6325 | 0.573269 | 0.391786 | 0.067698 | 1 | 0.999631 | 0.844549 |
| Srsf10_Q3TFP0_S133s_1 | -1.25 | -0.13 | 0.55 | 0.21 | 1.06 | -0.25 | -0.68 | 0.01 | 3.85 | -1.35 | 0.05 | -0.97 | 3.03 | -0.27 | 0.48 | 0.19 | 0.19 | 0.55 | 1.0125 | 0.190413 | 0.70678 | 0.611734 | 1 | 0.999631 | 0.648674 |
| Srsf11_E9Q6E5_S462s_1 | -1.43 | 2.71 | 2.95 | 1.2 | 2.26 | 1.53 | 2.97 | 1.85 | 4.18 | -0.5 | 3.06 | 4.11 | 3.27 | -0.15 | 3.33 | 1.71 | 0.795 | 1.355 | 0.6825 | 0.540263 | 0.392567 | 0.302257 | 1 | 0.999631 | 0.831295 |
| Srsf11_E9Q6E5_S477s_1 | 1.45 | 3.18 | 2.09 | 1.67 | 1.4 | 2.11 | 1.67 | 1.27 | 2.75 | 2.71 | 2.12 | 2.08 | 2 | 1.89 | 2.01 | 1.73 | -0.485 | 0.3175 | -0.19 | 0.644071 | 0.267066 | 0.463027 | 1 | 0.999631 | 0.872436 |
| Srsf11_E9Q6E5_S510s_1 | -0.88 | 1.3 | 1.32 | 0.28 | 1.37 | 0.77 | 1.26 | 0.77 | 2.17 | 0.43 | 1.35 | 2.15 | 1.27 | 0.21 | 1.51 | -0.2 | 0.5375 | 1.02 | 0.1925 | 0.75062 | 0.305308 | 0.11659 | 1 | 0.999631 | 0.912627 |
| Srsf2_Q62093_S191s_1 | 1.27 | -0.31 | -0.52 | -0.27 | -0.72 | -0.13 | 0.44 | -1.58 | -0.86 | 1.15 | -1.29 | -2.62 | -0.88 | -0.59 | -2.14 | -1.14 | -0.54 | -0.9475 | -1.23 | 0.033618 | 0.337318 | 0.241274 | 1 | 0.999631 | 0.443263 |
| Srsf2_Q62093_S212s_1 | 1.04 | 0.47 | 0.43 | 0.08 | 0.9 | 0.96 | 0.65 | -0.56 | -0.35 | 1.34 | -0.64 | -2.01 | 0.06 | -0.63 | -0.88 | -0.44 | -0.0175 | -0.92 | -0.9775 | 0.016222 | 0.965767 | 0.176034 | 1 | 0.999631 | 0.388832 |
| Srsf2_Q62093_S220s_1 | 2.1 | 1.53 | 1.69 | 1.01 | -0.16 | 1.76 | 1.46 | 0.47 | 1.32 | 2.85 | 0.53 | 0.85 | 0.29 | 0.81 | 1.22 | 1.07 | -0.7 | -0.195 | -0.735 | 0.061964 | 0.162245 | 0.711393 | 1 | 0.999631 | 0.509355 |
| Srsf2_Q62093_S221s_1 | 2.1 | 1.53 | 1.69 | 1.01 | -0.16 | 1.76 | 1.46 | 0.47 | 1.32 | 2.85 | 0.53 | 0.85 | 0.29 | 0.81 | 1.22 | 1.07 | -0.7 | -0.195 | -0.735 | 0.061964 | 0.162245 | 0.711393 | 1 | 0.999631 | 0.509355 |
| Srsf2_Q62093_S26s_1 | 0.53 | 0.29 | 0.78 | 0.75 | 0.3 | 0.37 | 0.78 | 0.2 | 0.47 | 1.1 | 0.52 | 0.65 | 0.52 | 0.42 | 0.72 | 0.48 | -0.175 | 0.0975 | -0.0525 | 0.849139 | 0.51676 | 0.728492 | 1 | 0.999631 | 0.949525 |
| Srsf2_Q62093_T25t_1 | 0.49 | 0.22 | -0.68 | -0.51 | 0.18 | 0.24 | -0.47 | -1.45 | 0.37 | 0.99 | -0.96 | -0.16 | 0.37 | 0.33 | -0.99 | -0.76 | -0.255 | 0.18 | -0.1425 | 0.753267 | 0.584643 | 0.708639 | 1 | 0.999631 | 0.913377 |
| Srsf3_P84104_S108s_1 | 2.87 | -0.1 | 1.42 | 1.24 | -0.43 | -0.19 | 1.17 | 0.18 | -0.06 | 2.54 | -0.08 | -0.35 | -0.37 | -0.07 | 0.49 | -0.53 | -1.175 | -0.845 | -1.4775 | 0.02972 | 0.086978 | 0.305174 | 1 | 0.999631 | 0.433947 |
| Srsf4_Q542V3_S455s_1 | 0.23 | 0.37 | 0.61 | 0.38 | -0.41 | 0.52 | 0.16 | -0.83 | 0.06 | 1.16 | 0.18 | -0.42 | 0.2 | -0.08 | 0.1 | 0.25 | -0.5375 | -0.1525 | -0.28 | 0.311313 | 0.14084 | 0.680495 | 1 | 0.999631 | 0.71924 |
| Srsf7_Q8BL97_S181s_1 | -3.56 | -3.31 | -0.57 | -0.86 | -4.41 | -3.9 | -0.7 | -0.88 | -4 | -2.15 | -0.5 | -0.96 | -1.04 | -3.08 | -0.35 | -0.58 | -0.3975 | 0.1725 | 0.8125 | 0.360699 | 0.712825 | 0.857523 | 1 | 0.999631 | 0.741241 |
| Srsf7_Q8BL97_S233s_1 | -1.48 | -6.32 | -7.28 | -7.85 | -5.81 | -4.65 | -8.06 | -12.1 | -7.7 | -3.4 | -4.73 | -9.56 | -7.39 | -5.75 | -7.09 | -8.5 | -1.9225 | -0.615 | -1.45 | 0.284957 | 0.312988 | 0.71968 | 1 | 0.999631 | 0.705838 |
| Srsf9_F6RTE1_S141s_1 | 2.05 | 0.44 | 1.08 | 0.94 | -0.62 | 0.62 | 1.03 | -0.15 | 0.37 | 1.98 | -0.11 | -0.02 | -0.4 | -0.6 | 0.52 | -0.93 | -0.9075 | -0.5725 | -1.48 | 0.008219 | 0.081468 | 0.310716 | 1 | 0.999631 | 0.365244 |
| Ssbp2_Q9CYZ8_S291s_1 | 0.6 | 1.69 | 1.11 | 0.61 | 1.03 | 0.96 | 0.33 | 0.65 | 1.43 | 1.19 | 0.4 | 0.73 | 1.47 | 1.67 | 0.42 | 0.86 | -0.26 | -0.065 | 0.1025 | 0.800945 | 0.450784 | 0.861468 | 1 | 0.999631 | 0.931985 |
| Ssbp2_Q9CYZ8_T303t_1 | 0.76 | 0.33 | 0.73 | 0.72 | -0.21 | -0.25 | 0 | 0.02 | 0.34 | 0.3 | 0.29 | -0.41 | 1.39 | 0.11 | -0.08 | 1.15 | -0.745 | -0.505 | 0.0075 | 0.985166 | 0.012765 | 0.108184 | 1 | 0.999631 | 0.995133 |
| Ssbp3_B1AS37_S327s_1 | 0.18 | 0.94 | 0.33 | -0.02 | 0.9 | 0.95 | -0.09 | 1.19 | 1.13 | 0.36 | 1.29 | 0.88 | 1.21 | 1.31 | 0.78 | 0.37 | 0.38 | 0.5575 | 0.56 | 0.136606 | 0.317234 | 0.123029 | 1 | 0.999631 | 0.609985 |
| Ssbp4_Q3U4B1_S320s_1 | -0.01 | 1.27 | -1.77 | -0.17 | 0.76 | 0.5 | -0.47 | 0.37 | 1.36 | 0.31 | 0.37 | 0.77 | 0.53 | 1.09 | -0.71 | -0.05 | 0.46 | 0.8725 | 0.385 | 0.561093 | 0.457933 | 0.173562 | 1 | 0.999631 | 0.840285 |
| Ssbp4_Q3U4B1_T333t_1 | 0.61 | 1.23 | 0.08 | 0.04 | 1.43 | -0.36 | -0.07 | -0.92 | -1.01 | -0.78 | 0.11 | -1.17 | 1.06 | -0.25 | 0.1 | -0.06 | -0.47 | -1.2025 | -0.2775 | 0.511371 | 0.385932 | 0.014116 | 1 | 0.999631 | 0.817687 |
| Ssfa2_Q922B9_S667s_1 | -0.22 | -0.98 | -0.78 | 0.04 | 0.07 | 0.11 | -0.28 | -1.2 | 0.02 | -0.5 | -0.98 | 0.49 | 0.88 | 0 | 0.02 | -1.12 | 0.16 | 0.2425 | 0.43 | 0.366686 | 0.685425 | 0.554569 | 1 | 0.999631 | 0.745796 |
| Ssh2_Q5SW75_S1216s_1 | 0.94 | -0.73 | -1.46 | -0.07 | -0.04 | 0.4 | -0.26 | -1.45 | 0.07 | 0.24 | -1.55 | -1.02 | -2.05 | 1.19 | -3.62 | 0.52 | -0.0075 | -0.235 | -0.66 | 0.53349 | 0.989766 | 0.69955 | 1 | 0.999631 | 0.828143 |
| Ssh2_Q5SW75_S17s_1 | -0.22 | 0.71 | -0.33 | -0.56 | -0.19 | 0.05 | -0.24 | -0.52 | -0.95 | 1.15 | 0.24 | 0.87 | 0.27 | 1.32 | 0.63 | -0.06 | -0.125 | 0.4275 | 0.64 | 0.150544 | 0.712855 | 0.412767 | 1 | 0.999631 | 0.623175 |
| Ssh2_Q5SW75_S25s_1 | -0.36 | 0.97 | 1.08 | 0.83 | -0.3 | -0.42 | -0.3 | 0.72 | -0.66 | -0.22 | 0.23 | -0.89 | 1.27 | -0.33 | 0.04 | 1.49 | -0.705 | -1.015 | -0.0125 | 0.981055 | 0.119674 | 0.03436 | 1 | 0.999631 | 0.994608 |
| Ssh2_Q5SW75_S25s_2 | -0.97 | 0.85 | 0.51 | 1.05 | 0.14 | -1.46 | -0.75 | 0.29 | 0.78 | 0.54 | 0.62 | 2.35 | -1.07 | -1.91 | -0.02 | 1.13 | -0.805 | 0.7125 | -0.8275 | 0.259933 | 0.173575 | 0.234481 | 1 | 0.999631 | 0.692624 |
| Ssh2_Q5SW75_S461s_1 | 1.53 | 3.33 | 2.25 | 1.56 | 2.89 | 3.08 | 1.46 | 2.01 | 3.28 | 2.39 | 1.87 | 3.26 | 3.91 | 3.15 | 2.79 | 2.26 | 0.1925 | 0.5325 | 0.86 | 0.120288 | 0.715069 | 0.313795 | 1 | 0.999631 | 0.587933 |
| Ssh2_Q5SW75_S9s_1 | 0.32 | -1.06 | -0.71 | -0.6 | -0.19 | -0.17 | -0.36 | -0.81 | -0.29 | 0.19 | -0.51 | 0.25 | -0.23 | -0.06 | 0.06 | -0.31 | 0.13 | 0.4225 | 0.3775 | 0.30494 | 0.715634 | 0.274014 | 1 | 0.999631 | 0.716239 |
| Ssh2_Q5SW75_T1422t_1 | 0.29 | 0.99 | -0.23 | -0.37 | 0.39 | -0.43 | -0.47 | 0.01 | 0.32 | 0.36 | -0.32 | -0.17 | 0.58 | 0.88 | -0.43 | 0.24 | -0.295 | -0.1225 | 0.1475 | 0.730427 | 0.446811 | 0.745545 | 1 | 0.999631 | 0.906608 |
| Ssh3_Q8K330_S643s_1 | 1.44 | -0.64 | 0.35 | 1.06 | -0.31 | 0.48 | 0.48 | 0.48 | 0.71 | 1.94 | 0.27 | 0.33 | -0.71 | -0.47 | 1.14 | -0.29 | -0.27 | 0.26 | -0.635 | 0.280892 | 0.571883 | 0.641133 | 1 | 0.999631 | 0.701922 |
| Ssh3_Q8K330_S9s_1 | -0.16 | -0.07 | 0.18 | -0.65 | -0.91 | -1.21 | -0.28 | -1.46 | -0.84 | 0.87 | -0.31 | -0.11 | 0.64 | 0.61 | -0.53 | 0.17 | -0.79 | 0.0775 | 0.3975 | 0.294464 | 0.041195 | 0.848908 | 1 | 0.999631 | 0.708293 |
| Ssr3_Q9DCF9_S105s_1 | 2.46 | 1.51 | 1.65 | 1.29 | 1.02 | 2.84 | 2.11 | 1.09 | 2.35 | 2.99 | 1.78 | 0.93 | 1.33 | -0.57 | 1.34 | 1 | 0.0375 | 0.285 | -0.9525 | 0.080747 | 0.93758 | 0.560027 | 1 | 0.999631 | 0.545579 |
| Sst_P60041_S92s_1 | -0.8 | -0.73 | -0.95 | -1.05 | -0.14 | -0.88 | -0.84 | -2.26 | -0.47 | -0.3 | -1.81 | -0.38 | -0.27 | -1.1 | -0.84 | -1.51 | -0.1475 | 0.1425 | -0.0475 | 0.88793 | 0.737052 | 0.712229 | 1 | 0.999631 | 0.963505 |
| Sstr3_P30935_S245s_1 | -0.82 | 0.64 | 0.56 | 0.27 | -0.37 | 0.76 | 0.27 | 0.37 | -0.17 | 0.38 | 0.31 | 0.03 | -0.52 | 0.57 | -0.66 | 1.26 | 0.095 | -0.025 | 7.85E-17 | 1 | 0.817031 | 0.947732 | 1 | 0.999631 | 1 |
| Stac2_Q8R1B0_S221s_1 | 1.14 | -0.2 | 0.05 | 0.19 | 0.73 | -0.65 | 0.54 | -0.91 | -0.12 | -0.46 | -0.56 | -1.28 | 1.28 | 0.2 | -0.74 | -0.37 | -0.3675 | -0.9 | -0.2025 | 0.689419 | 0.451249 | 0.043838 | 1 | 0.999631 | 0.892416 |
| Stac2_Q8R1B0_S228s_1 | -0.74 | 1.36 | 1.16 | -0.43 | 0.31 | 1.05 | -0.07 | 0.12 | 1.42 | -0.15 | 0.92 | 0.86 | 1.38 | 0.69 | 0.47 | -0.45 | 0.015 | 0.425 | 0.185 | 0.757928 | 0.97797 | 0.468857 | 1 | 0.999631 | 0.914449 |
| Stac2_Q8R1B0_S274s_1 | 2.55 | 1.49 | 2.02 | 0.85 | 0.75 | 1.63 | 1.29 | 1.89 | 1.02 | 1.74 | 1.16 | 1.12 | 0.92 | 0.93 | 1.36 | 0.92 | -0.3375 | -0.4675 | -0.695 | 0.108517 | 0.440902 | 0.26798 | 1 | 0.999631 | 0.57047 |
| Stam_Q3UGN9_S156s_1 | 1 | 1.54 | 0.92 | 1.77 | 1 | 1.05 | 1.3 | 1.15 | 1.61 | 1.06 | 1.9 | 1.22 | -0.12 | 1.47 | 1.64 | 1.72 | -0.1825 | 0.14 | -0.13 | 0.783094 | 0.532453 | 0.674364 | 1 | 0.999631 | 0.922613 |
| Stambpl1_Q76N33_S76s_1 | -0.37 | 0.19 | 0.02 | -0.35 | -0.35 | -0.12 | -0.28 | -0.36 | -0.15 | 0.06 | -0.14 | 0.07 | 0.34 | 0.47 | -0.04 | 0.06 | -0.15 | 0.0875 | 0.335 | 0.272176 | 0.564101 | 0.743328 | 1 | 0.999631 | 0.699819 |
| Stard10_G3UY87_S276s_1 | 0.15 | -0.16 | 0.5 | 0.26 | -0.18 | -0.07 | -0.11 | 0.18 | -0.17 | 0.59 | -0.21 | 0.19 | -0.4 | 0.22 | 0.22 | -0.04 | -0.2325 | -0.0875 | -0.1875 | 0.539934 | 0.38359 | 0.773842 | 1 | 0.999631 | 0.831295 |
| Stard13_F8WIY7_S572s_1 | 0.45 | -0.54 | 1.13 | 1.18 | 0.79 | 1.23 | 0.57 | 1.13 | 1.06 | 1.24 | 0.86 | 1.12 | -0.22 | 0.75 | 0.82 | 1.49 | 0.375 | 0.515 | 0.155 | 0.760779 | 0.385947 | 0.233187 | 1 | 0.999631 | 0.914614 |
| Stard7_F7BBA2_S392s_1 | 1.57 | 0.26 | 0.86 | 0.36 | 0.2 | 0.44 | 0.85 | 0.53 | 0.58 | 1.7 | 0.48 | -0.04 | 0.1 | -0.35 | -0.14 | 0.11 | -0.2575 | -0.0825 | -0.8325 | 0.042724 | 0.474187 | 0.857753 | 1 | 0.999631 | 0.475443 |
| Stau2_E2QRQ3_S447s_1 | 0.24 | 0.19 | 0.41 | -0.03 | -0.13 | 0.44 | 1.19 | 1.04 | -0.06 | -0.19 | 0.66 | 0.71 | 0.99 | 0.89 | 1.35 | 1.31 | 0.4325 | 0.0775 | 0.9325 | 0.007308 | 0.231005 | 0.806133 | 1 | 0.999631 | 0.358173 |
| Steap2_Q8BWB6_S482s_1 | -0.97 | -0.36 | -0.46 | -0.73 | -0.25 | -0.34 | -0.55 | -0.59 | -0.01 | -0.44 | -1.13 | 0.07 | 0.76 | -0.48 | -0.85 | 0.08 | 0.1975 | 0.2525 | 0.5075 | 0.22415 | 0.458224 | 0.474041 | 1 | 0.999631 | 0.67471 |
| Stim1_P70302_S575s_1 | 0.63 | -0.07 | -0.76 | 2.1 | 1 | 0.11 | 1.18 | 1.72 | 0.37 | -0.6 | 1.49 | 0.47 | -0.91 | -0.07 | 0.22 | 0.9 | 0.5275 | -0.0425 | -0.44 | 0.499997 | 0.408118 | 0.949244 | 1 | 0.999631 | 0.81332 |
| Stim1_P70302_S660s_1 | 0.88 | 0.26 | 1.29 | 0.88 | 0.28 | 0.32 | 0.14 | 0.89 | 0.38 | -0.19 | 0.25 | 0.4 | -0.45 | -0.6 | 0.3 | 0.87 | -0.42 | -0.6175 | -0.7975 | 0.080059 | 0.207433 | 0.074845 | 1 | 0.999631 | 0.545579 |
| Stim1_P70302_S660s_2 | 0.25 | 0.74 | -0.29 | -0.28 | 0.82 | 0.02 | -0.07 | -0.51 | 0.25 | 0.26 | -0.39 | 0.41 | 0.3 | 0.52 | -0.6 | -0.84 | -0.04 | 0.0275 | -0.26 | 0.544541 | 0.916746 | 0.936566 | 1 | 0.999631 | 0.833306 |
| Stim1_P70302_S667s_2 | 0.25 | 0.74 | -0.79 | -0.94 | 0.82 | 0.02 | -0.16 | -0.4 | 0.25 | 0.26 | -0.16 | 0.53 | 0.3 | 0.52 | -1 | -1.61 | 0.255 | 0.405 | -0.2625 | 0.661446 | 0.585418 | 0.357582 | 1 | 0.999631 | 0.880723 |
| Stim1_P70302_S668s_1 | 0.14 | 0.77 | 0.54 | 0.25 | 0.03 | 0.55 | 0.28 | -0.47 | 1.5 | 1 | 0.85 | 0.27 | 1.2 | 1.06 | -0.02 | 0.23 | -0.3275 | 0.48 | 0.1925 | 0.60903 | 0.309134 | 0.177332 | 1 | 0.999631 | 0.858851 |
| Stim2_P83093_S523s_1 | 1.23 | -0.29 | -1.37 | 1.25 | 1.34 | 0.44 | 0.74 | 0.89 | 0.54 | -0.86 | 0.78 | -0.95 | -3.04 | 0.61 | -0.84 | 0.1 | 0.6475 | -0.3275 | -0.9975 | 0.276553 | 0.295413 | 0.640359 | 1 | 0.999631 | 0.700187 |
| Stim2_P83093_S609s_1 | -0.76 | 0.04 | 0.14 | -0.61 | -0.72 | -0.21 | -1.21 | -0.41 | -0.4 | -0.08 | -0.71 | -0.33 | 0.92 | 0.18 | -0.01 | 0.23 | -0.34 | -0.0825 | 0.6275 | 0.102774 | 0.338151 | 0.797049 | 1 | 0.999631 | 0.565572 |
| Stim2_P83093_S631s_1 | -1 | 2.45 | 2.39 | 1.04 | 2.13 | 2.18 | 1.28 | 1.83 | 2.26 | -0.93 | 2.17 | 2.72 | 1.61 | 1.41 | 2.96 | 1.99 | 0.635 | 0.335 | 0.7725 | 0.328026 | 0.396715 | 0.739354 | 1 | 0.999631 | 0.72534 |
| Stim2_P83093_S640s_1 | 0.58 | 2.03 | 2.06 | 1.28 | 0.14 | 1.96 | 0.79 | 1.24 | 1.07 | 1.61 | 1.53 | 1.74 | 1.28 | 1.86 | 1.81 | 2.75 | -0.455 | 3.93E-17 | 0.4375 | 0.353064 | 0.364332 | 1 | 1 | 1 | 0.73949 |
| Stim2_P83093_S650s_1 | -0.36 | 0.44 | 0.74 | 0.23 | -0.43 | 0.58 | -0.02 | 0.43 | -0.06 | -0.3 | -0.05 | 0.02 | 0.64 | 0.83 | 0.49 | 0.68 | -0.1225 | -0.36 | 0.3975 | 0.236544 | 0.729584 | 0.262637 | 1 | 0.999631 | 0.683301 |
| Stip1_Q60864_S481s_1 | 0.53 | 0.49 | 0.91 | 0 | 0.38 | 0.6 | 2.34 | -0.75 | 2.1 | 0.56 | 1.5 | 2.54 | 2.14 | 0.09 | -1.15 | -0.86 | 0.16 | 1.1925 | -0.4275 | 0.535155 | 0.790236 | 0.024773 | 1 | 0.999631 | 0.829113 |
| Stk10_O55098_T950t_1 | 0.18 | -0.07 | -0.06 | -0.11 | 0.41 | -0.27 | -0.43 | -0.11 | 0.09 | 0.36 | -0.48 | 0.71 | -0.32 | 0.55 | 0.33 | 0.58 | -0.085 | 0.185 | 0.3 | 0.347223 | 0.75985 | 0.566009 | 1 | 0.999631 | 0.736373 |
| Stk11ip_Q3TAA7_S388s_1 | 0.12 | 0.41 | 1.65 | 0.31 | 0.27 | -0.31 | 0.62 | -0.1 | -0.15 | 0.61 | 0.85 | -0.8 | -0.3 | 0.65 | -0.3 | -0.12 | -0.5025 | -0.495 | -0.64 | 0.155379 | 0.234194 | 0.324409 | 1 | 0.999631 | 0.626444 |
| Stk24_Q99KH8_S4s_1 | -1.13 | -0.1 | -0.43 | -0.6 | -0.08 | -0.24 | -0.82 | -0.89 | -0.21 | -0.52 | -0.63 | -0.12 | -0.03 | 0.26 | -0.6 | -0.5 | 0.0575 | 0.195 | 0.3475 | 0.33479 | 0.863585 | 0.53691 | 1 | 0.999631 | 0.72979 |
| Stk26_Q99JT2_S4s_1 | -0.76 | -0.49 | -0.31 | -0.5 | -0.91 | -0.48 | -0.55 | -0.2 | -0.48 | -0.05 | -1.36 | -0.16 | 0.61 | 0.06 | 0.36 | -0.04 | -0.02 | 0.0025 | 0.7625 | 0.024097 | 0.940444 | 0.994298 | 1 | 0.999631 | 0.415644 |
| Stk32c_Q8QZV4_S10s_2 | 1.47 | 0.59 | -0.01 | 1.38 | 1.08 | 0.83 | 1.45 | 1.24 | 0.68 | 0.53 | 1.14 | 0.31 | -0.42 | 0.97 | -0.32 | 0.88 | 0.2925 | -0.1925 | -0.58 | 0.256741 | 0.454338 | 0.634707 | 1 | 0.999631 | 0.691567 |
| Stk32c_Q8QZV4_S18s_1 | -0.25 | 0.19 | 0.34 | -0.31 | -0.51 | -0.23 | -0.35 | -0.07 | -0.36 | 0.16 | 0.08 | 0.11 | 0.35 | -0.19 | 0.17 | 0.18 | -0.2825 | 0.005 | 0.135 | 0.655574 | 0.315801 | 0.98616 | 1 | 0.999631 | 0.879056 |
| Stk32c_Q8QZV4_S18s_2 | 1.47 | 0.59 | -0.01 | 1.38 | 1.08 | 0.83 | 1.45 | 1.24 | 0.68 | 0.53 | 1.14 | 0.31 | -0.42 | 0.97 | -0.32 | 0.88 | 0.2925 | -0.1925 | -0.58 | 0.256741 | 0.454338 | 0.634707 | 1 | 0.999631 | 0.691567 |
| Stk32c_Q8QZV4_S32s_1 | 0.52 | 0.39 | -1.43 | -0.59 | 0.02 | -0.41 | -1.02 | -0.29 | -0.4 | -0.27 | -1.54 | -1.12 | 0.25 | -0.1 | -0.77 | -0.56 | -0.1475 | -0.555 | -0.0175 | 0.971617 | 0.758997 | 0.296245 | 1 | 0.999631 | 0.989788 |
| Stk32c_Q8QZV4_S69s_1 | 1.12 | 0.05 | 0.06 | 1.72 | 1.39 | 0.87 | 1.46 | 1.44 | 0.46 | -0.34 | 1.1 | 0.21 | -1.69 | 0.36 | 0.23 | 0.27 | 0.5525 | -0.38 | -0.945 | 0.132736 | 0.217104 | 0.442564 | 1 | 0.999631 | 0.604583 |
| Stk38_Q91VJ4_S281s_1 | -0.69 | 0.98 | 0.57 | 0.13 | -0.46 | 1.35 | -0.24 | 1.61 | -0.71 | -0.8 | 1.29 | -1.33 | 0.24 | 1.92 | -0.43 | -0.52 | 0.3175 | -0.635 | 0.055 | 0.927851 | 0.588575 | 0.315464 | 1 | 0.999631 | 0.976452 |
| Stk39_Q9Z1W9_S382s_1 | 1.69 | 1.11 | -0.66 | 0.39 | 0.63 | 1.76 | 0.14 | 0.35 | 1.11 | 1.52 | -0.51 | -0.67 | -0.23 | 0.85 | -0.38 | -0.4 | 0.0875 | -0.27 | -0.6725 | 0.237619 | 0.877371 | 0.689667 | 1 | 0.999631 | 0.683301 |
| Stk39_Q9Z1W9_S383s_1 | 1.64 | 1.28 | -0.66 | 0.39 | 0.79 | 1.59 | 0.14 | 0.35 | 1.29 | 1.47 | -0.51 | -0.67 | -0.19 | 0.92 | -0.38 | -0.4 | 0.055 | -0.2675 | -0.675 | 0.244026 | 0.920936 | 0.697276 | 1 | 0.999631 | 0.685637 |
| Stk39_Q9Z1W9_T366t_1 | 0.6 | 0.23 | 0 | 0.21 | -0.08 | 0.38 | -0.36 | 0.07 | 0.31 | 0.91 | 0.02 | 0.08 | -0.41 | 0.41 | 0.03 | 0.08 | -0.2575 | 0.07 | -0.2325 | 0.455531 | 0.368271 | 0.820218 | 1 | 0.999631 | 0.790041 |
| Stmn1_P54227_S16s_1 | -0.28 | 0.84 | -0.32 | 0.04 | 0.68 | 0.19 | -0.3 | -0.19 | 0.66 | 0.15 | 0.16 | 0.58 | 0.75 | 0.61 | -0.08 | 0.19 | 0.025 | 0.3175 | 0.2975 | 0.431064 | 0.945794 | 0.366968 | 1 | 0.999631 | 0.778271 |
| Stmn1_P54227_S25s_1 | -0.43 | 0.92 | -0.8 | 0.09 | 0.91 | -0.24 | 0.32 | 0.36 | 0.35 | -0.78 | 0.72 | 0.01 | 0.81 | -0.28 | -0.05 | 0.7 | 0.3925 | 0.13 | 0.35 | 0.450264 | 0.373898 | 0.78391 | 1 | 0.999631 | 0.787533 |
| Stmn1_P54227_S38s_1 | -0.3 | -0.02 | -0.14 | -0.4 | -0.1 | -0.22 | -0.51 | -0.31 | -0.08 | 0.23 | -0.03 | 0.02 | 0.67 | 0.24 | -0.03 | 0.35 | -0.07 | 0.25 | 0.5225 | 0.094771 | 0.777311 | 0.333037 | 1 | 0.999631 | 0.558128 |
| Stmn1_P54227_S3s_1 | -0.15 | -0.85 | -0.46 | -1.43 | -0.86 | -1.41 | -0.22 | -0.71 | -1.51 | -0.65 | -0.57 | -0.39 | -1.34 | -1.22 | -1.13 | -0.19 | -0.0775 | -0.0575 | -0.2475 | 0.543687 | 0.839141 | 0.882565 | 1 | 0.999631 | 0.832881 |
| Stmn1_P54227_S46s_1 | -2.51 | 2.61 | 2.14 | -0.1 | 2.81 | 1.62 | 1.39 | 1.19 | 2.93 | -1.34 | 2.83 | 3.15 | 3.86 | 1.13 | 2.19 | 0.86 | 1.2175 | 1.3575 | 1.475 | 0.219226 | 0.265218 | 0.331095 | 1 | 0.999631 | 0.669859 |
| Stmn1_P54227_S4s_1 | -0.15 | -0.85 | -0.46 | -1.43 | -0.86 | -1.41 | -0.22 | -0.71 | -1.51 | -0.65 | -0.57 | -0.39 | -1.34 | -1.22 | -1.13 | -0.19 | -0.0775 | -0.0575 | -0.2475 | 0.543687 | 0.839141 | 0.882565 | 1 | 0.999631 | 0.832881 |
| Stmn1_P54227_S63s_1 | 0.37 | -0.8 | -1.54 | 0.13 | 0.26 | -0.76 | 0.21 | -0.28 | 0.01 | -0.52 | -0.08 | -0.92 | -0.1 | -1.12 | -0.51 | -0.68 | 0.3175 | 0.0825 | -0.1425 | 0.764787 | 0.509267 | 0.86127 | 1 | 0.999631 | 0.916147 |
| Stmn2_P55821_S62s_1 | -1.4 | -0.72 | -0.77 | -1.11 | -0.95 | -1.25 | -1.2 | -0.78 | -1.03 | -1.56 | -0.86 | -0.49 | 0.28 | -0.89 | -0.84 | -0.77 | -0.045 | 0.015 | 0.445 | 0.242529 | 0.871127 | 0.963436 | 1 | 0.999631 | 0.685099 |
| Stmn2_P55821_S97s_1 | 0.8 | 1.02 | 0.34 | 0.76 | 1.45 | 1.16 | 0.8 | 1.32 | 1.02 | 0.34 | 1.23 | 0.83 | -0.14 | 1.22 | 1.02 | 0.5 | 0.4525 | 0.125 | -0.08 | 0.83116 | 0.131073 | 0.685257 | 1 | 0.999631 | 0.942552 |
| Stmn3_O70166_S50s_1 | -1.17 | -0.37 | -0.11 | -1.23 | -1.3 | -0.31 | -1.39 | -1.3 | -0.9 | -0.37 | -0.69 | -0.6 | 1.13 | -0.6 | -0.35 | -0.3 | -0.355 | 0.08 | 0.69 | 0.166451 | 0.373163 | 0.816856 | 1 | 0.999631 | 0.63587 |
| Stmn3_O70166_S53s_1 | -1.54 | -0.34 | -0.28 | -1.27 | -1.18 | -0.38 | -0.99 | -0.68 | -0.14 | -0.89 | -0.67 | -0.15 | 2.28 | -0.73 | -0.42 | -0.44 | 0.05 | 0.395 | 1.03 | 0.15785 | 0.895206 | 0.324105 | 1 | 0.999631 | 0.628615 |
| Stmn3_O70166_S81s_1 | -1.67 | 0.62 | 0.82 | -0.35 | 0.43 | -0.41 | 0.25 | 0.59 | 0.21 | -1.48 | 0.8 | 1.31 | 1.15 | -0.6 | 1.2 | 0.33 | 0.36 | 0.355 | 0.665 | 0.311221 | 0.523402 | 0.63157 | 1 | 0.999631 | 0.71924 |
| Stmn4_Q05DI3_S90s_1 | -0.5 | 0.28 | -0.6 | -1.05 | -0.54 | -0.38 | -0.91 | -0.39 | -0.21 | 0.38 | -0.34 | -0.54 | 0.74 | 0.02 | -0.5 | -1 | -0.0875 | 0.29 | 0.2825 | 0.54177 | 0.796393 | 0.438726 | 1 | 0.999631 | 0.832428 |
| Stom_P54116_S7s_1 | 2.65 | -0.27 | 0.41 | 1 | 0.17 | 0.64 | 1.91 | -0.74 | -0.73 | 1.89 | -1.04 | -1.93 | -0.76 | -1.8 | -0.57 | -1.62 | -0.4525 | -1.4 | -2.135 | 0.006461 | 0.541201 | 0.142627 | 1 | 0.999631 | 0.351545 |
| Stox2_Q499E5_S110s_1 | 0.62 | -0.23 | -0.71 | -1.17 | 0.34 | 0.15 | -0.01 | -0.83 | -0.63 | -0.13 | -0.24 | -1.76 | 0.14 | -0.05 | -1.76 | -0.01 | 0.285 | -0.3175 | -0.0475 | 0.930976 | 0.527796 | 0.53382 | 1 | 0.999631 | 0.977912 |
| Strada_Q3UUJ4_S276s_1 | -0.82 | -0.39 | 0.65 | -0.02 | -0.28 | -0.56 | -0.71 | -0.12 | -0.49 | -0.66 | -0.03 | 0.67 | -0.07 | -0.29 | 0.39 | 0.01 | -0.2725 | 0.0175 | 0.155 | 0.683747 | 0.457854 | 0.967496 | 1 | 0.999631 | 0.889602 |
| Strip1_Q8C079_S335s_1 | -0.7 | -0.68 | 0.05 | -0.54 | -0.47 | -0.62 | -0.85 | -0.83 | -0.71 | -0.53 | -0.59 | -0.33 | 0.21 | -0.51 | -0.64 | -0.14 | -0.225 | -0.0725 | 0.1975 | 0.555806 | 0.429114 | 0.799287 | 1 | 0.999631 | 0.838305 |
| Strip2_E9QN41_S328s_1 | -0.38 | -0.21 | -0.16 | -0.73 | -0.1 | -1.2 | -0.73 | 0.35 | -0.33 | -0.16 | 0.08 | 0.25 | 0.26 | -0.22 | 0.04 | 0.17 | -0.05 | 0.33 | 0.4325 | 0.156237 | 0.895014 | 0.255184 | 1 | 0.999631 | 0.626452 |
| Strip2_E9QN41_S364s_1 | -0.39 | 0.51 | 0.45 | 0.21 | 0.64 | -0.09 | -0.04 | 0.31 | 0.08 | 0.28 | 0.58 | 0.75 | 0.55 | 0.39 | 0.25 | 1.11 | 0.01 | 0.2275 | 0.38 | 0.282141 | 0.974809 | 0.478439 | 1 | 0.999631 | 0.703634 |
| Strn_F6Z700_S227s_1 | -0.78 | 0.33 | -0.48 | -0.08 | 1.39 | -0.02 | 0.07 | 0.88 | 0.41 | -1.03 | 0.41 | 0.47 | -0.41 | 0.82 | 0.29 | 0.13 | 0.8325 | 0.3175 | 0.46 | 0.247995 | 0.067954 | 0.471802 | 1 | 0.999631 | 0.685751 |
| Strn_F6Z700_S81s_1 | 1.15 | 1 | 1.15 | 1.38 | 1.75 | 1.66 | 1.14 | 2.24 | 1.68 | 0.84 | 1.72 | 1.86 | -0.59 | 1.89 | 0.92 | 1.75 | 0.5275 | 0.355 | -0.1775 | 0.741954 | 0.104411 | 0.270316 | 1 | 0.999631 | 0.909328 |
| Strn3_B2RQS1_S229s_1 | -0.84 | 0.02 | 0.52 | 0.65 | -0.94 | -0.99 | -0.05 | -0.07 | -0.39 | -1.08 | 0.47 | 0.4 | 0.34 | 0.31 | 1.16 | 1.03 | -0.6 | -0.2375 | 0.6225 | 0.15993 | 0.177369 | 0.621888 | 1 | 0.999631 | 0.629764 |
| Strn4_P58404_S206s_1 | -0.43 | -0.3 | 0.43 | 1.09 | 0.46 | -0.45 | 0.88 | 0.46 | -0.12 | -0.59 | 1.27 | 0.46 | -0.13 | -0.06 | 0.37 | 0.9 | 0.14 | 0.0575 | 0.0725 | 0.866794 | 0.750253 | 0.909294 | 1 | 0.999631 | 0.956456 |
| Strn4_P58404_S223s_1 | -0.1 | 0.58 | 0.78 | 0.01 | -0.21 | 0.1 | -0.09 | -0.37 | 0.39 | -0.06 | 0.19 | 0.57 | 1.56 | 0.93 | 0.06 | 0.48 | -0.46 | -0.045 | 0.44 | 0.293844 | 0.148272 | 0.886781 | 1 | 0.999631 | 0.708293 |
| Stt3b_Q3TDQ1_S495s_1 | -0.14 | 0.28 | -0.41 | -0.1 | 0.65 | 0.32 | -0.1 | -0.21 | 0.69 | 1.1 | -1.03 | -0.11 | 0.61 | 0.13 | -0.33 | -0.68 | 0.2575 | 0.255 | 0.025 | 0.944952 | 0.405477 | 0.592971 | 1 | 0.999631 | 0.982141 |
| Stub1_Q9WUD1_S20s_1 | -2.35 | -4.09 | -4.4 | -4.17 | -3.87 | -3.89 | -3.58 | -4.33 | -3.33 | -3.61 | -3.28 | -5.62 | -3.83 | -3.71 | -4.39 | -4.42 | -0.165 | -0.2075 | -0.335 | 0.498659 | 0.72775 | 0.752401 | 1 | 0.999631 | 0.812694 |
| Stub1_Q9WUD1_S274s_1 | -0.8 | -0.06 | -0.37 | -0.64 | -0.23 | -0.81 | -0.73 | -0.15 | -0.26 | -0.45 | -0.5 | 0.56 | 0.43 | 0.57 | -0.05 | 0.99 | -0.0125 | 0.305 | 0.9525 | 0.016699 | 0.966576 | 0.380331 | 1 | 0.999631 | 0.391024 |
| Stx12_Q9ER00_S142s_1 | -0.39 | -1.88 | -1.92 | -1.44 | -1.32 | -2.1 | -0.97 | -1.4 | -1.1 | -0.14 | -1.28 | -1.21 | -0.99 | -2.53 | -0.52 | -1.74 | -0.04 | 0.475 | -0.0375 | 0.943747 | 0.924516 | 0.295525 | 1 | 0.999631 | 0.981629 |
| Stx17_Q9D0I4_S288s_1 | -0.94 | -0.12 | -0.99 | -1.32 | 0.28 | 0.41 | -0.96 | -0.49 | 0.02 | 0.3 | -0.37 | -0.56 | 0.54 | 0.74 | -1.15 | -0.66 | 0.6525 | 0.69 | 0.71 | 0.178909 | 0.137421 | 0.076816 | 1 | 0.999631 | 0.643093 |
| Stx1a_O35526_S14s_1 | -2.33 | -1.15 | -0.64 | -1.02 | -1.39 | -1.48 | -1.26 | -0.81 | -1.41 | -1.43 | -0.72 | -0.58 | -0.54 | -1.22 | -0.54 | -0.85 | 0.05 | 0.25 | 0.4975 | 0.247961 | 0.900165 | 0.562762 | 1 | 0.999631 | 0.685751 |
| Stx1a_O35526_S59s_1 | -0.75 | 0.16 | -0.14 | -1.26 | 0.91 | 0.53 | 0.97 | -0.65 | 2.77 | -0.11 | -0.13 | 0.9 | 2.89 | 0.11 | -0.1 | -0.85 | 0.9375 | 1.355 | 1.01 | 0.208821 | 0.069773 | 0.06798 | 1 | 0.999631 | 0.662575 |
| Stx1a_O35526_S64s_1 | -0.96 | 0.86 | 1.11 | -0.41 | 1.56 | 0.16 | 1.15 | 0.32 | 2.65 | -0.59 | 2.18 | 2.66 | 3.38 | 0.01 | 0.99 | 0.28 | 0.6475 | 1.575 | 1.015 | 0.22192 | 0.256521 | 0.076683 | 1 | 0.999631 | 0.672466 |
| Stx1a_O35526_T10t_1 | -1.32 | -0.78 | 0.21 | -0.46 | -1.84 | -1.96 | -1.08 | -0.49 | -1.56 | -0.95 | -0.7 | -0.1 | 0.02 | -0.98 | -0.39 | -0.38 | -0.755 | -0.24 | 0.155 | 0.701376 | 0.120407 | 0.585408 | 1 | 0.999631 | 0.896835 |
| Stx1b_P61264_S14s_1 | -2.57 | -0.72 | -0.58 | -0.77 | -0.84 | -1.08 | -1.21 | -0.76 | -0.74 | -1.58 | -0.71 | -0.52 | 0.35 | -0.75 | -0.35 | -0.86 | 0.1875 | 0.2725 | 0.7575 | 0.164967 | 0.686432 | 0.588781 | 1 | 0.999631 | 0.633483 |
| Stx4_P70452_S117s_1 | -1.42 | -2.15 | -3.15 | -1.77 | -1.4 | -2.1 | -2.63 | -2.26 | -1.66 | -2.19 | -1.91 | -2.34 | -1.23 | -1.7 | -1.5 | -2.58 | 0.025 | 0.0975 | 0.37 | 0.434839 | 0.954782 | 0.812368 | 1 | 0.999631 | 0.779717 |
| Stx4_P70452_S15s_1 | -1.19 | 0.14 | 0.41 | -0.38 | -0.17 | -0.22 | -0.67 | -0.56 | -0.2 | -0.19 | -0.41 | -0.06 | 0.57 | -0.06 | 0.02 | -0.32 | -0.15 | 0.04 | 0.3075 | 0.464934 | 0.697497 | 0.916371 | 1 | 0.999631 | 0.794369 |
| Stx6_Q9JKK1_S2s_1 | 0.18 | -0.12 | -0.6 | 0.23 | 0.04 | -0.31 | -0.32 | 0.11 | -0.49 | -0.72 | -0.05 | -0.03 | -0.85 | -0.04 | -0.54 | 0.06 | -0.0425 | -0.245 | -0.265 | 0.451315 | 0.884185 | 0.446946 | 1 | 0.999631 | 0.788148 |
| Stx7_Q8BH40_S129s_1 | 0.26 | 0.39 | -0.13 | -1.12 | 0.36 | 0.33 | 0.29 | -0.24 | -0.99 | 1.07 | 0.41 | 1.2 | -1.25 | -1.42 | 2.11 | -0.42 | 0.335 | 0.5725 | -0.095 | 0.902159 | 0.390802 | 0.320381 | 1 | 0.999631 | 0.968183 |
| Stx7_Q8BH40_S75s_1 | -0.7 | -0.34 | -0.13 | -1.35 | -0.22 | -0.17 | -1.1 | -0.19 | -0.97 | -0.86 | 0.42 | -1.62 | 0.41 | -0.22 | -0.25 | -0.02 | 0.21 | -0.1275 | 0.61 | 0.112918 | 0.572903 | 0.791871 | 1 | 0.999631 | 0.577408 |
| Stx7_Q8BH40_T41t_1 | 0.72 | 1.89 | 0.22 | -0.41 | 1.29 | 1.4 | -0.58 | -0.07 | 1.71 | 0.84 | 0.07 | 0.17 | 1.67 | 1.33 | 0.02 | 0.36 | -0.095 | 0.0925 | 0.24 | 0.677305 | 0.878729 | 0.87051 | 1 | 0.999631 | 0.887183 |
| Stx7_Q8BH40_T79t_1 | -0.38 | 0.55 | 0.1 | -0.12 | 0.14 | -0.03 | -0.6 | 0.24 | 0.32 | 0.14 | 0.38 | 0.54 | 0.63 | 0.49 | 0.42 | 0.58 | -0.1 | 0.3075 | 0.4925 | 0.128621 | 0.755162 | 0.310976 | 1 | 0.999631 | 0.599259 |
| Stxbp1_O08599_S507s_1 | -0.59 | 0.23 | 0.22 | 0.43 | 0.13 | -0.26 | 0.17 | -0.13 | -0.01 | -0.71 | 1.11 | 0.55 | -0.91 | -0.34 | 0.17 | 0.63 | -0.095 | 0.1625 | -0.185 | 0.6584 | 0.757038 | 0.71505 | 1 | 0.999631 | 0.879726 |
| Stxbp1_O08599_S509s_1 | -1.03 | -0.97 | -1.35 | -0.05 | -1.28 | -1.05 | 1.08 | -0.23 | -1.28 | -1.85 | 0.05 | -1.05 | -0.78 | -1.1 | -1.07 | -1.75 | 0.48 | -0.1825 | -0.325 | 0.403377 | 0.394186 | 0.698775 | 1 | 0.999631 | 0.764348 |
| Stxbp1_O08599_S516s_1 | -1.28 | -1.24 | -1.18 | 0.05 | 1.65 | 0.49 | 1.59 | 1.67 | -0.03 | -3.44 | 1.27 | 0.08 | -0.65 | -0.39 | 0.74 | -2.05 | 2.2625 | 0.3825 | 0.325 | 0.590272 | 0.000383 | 0.67975 | 1 | 0.999631 | 0.851598 |
| Stxbp1_O08599_S593s_1 | -1.07 | -2.63 | -2.86 | -3.01 | -2.3 | -2.24 | -1.56 | -3.96 | -2.59 | -1.47 | -3.03 | -3.83 | -2.19 | -2.87 | -3.78 | -4.36 | -0.1225 | -0.3375 | -0.9075 | 0.152637 | 0.841224 | 0.580125 | 1 | 0.999631 | 0.625198 |
| Stxbp1_O08599_S594s_1 | -0.35 | -0.16 | 0.18 | -0.31 | -0.19 | -0.43 | -0.44 | -0.31 | -0.45 | -0.55 | -0.53 | -0.25 | 0.24 | 0.02 | -0.03 | 0.26 | -0.1825 | -0.285 | 0.2825 | 0.324881 | 0.474891 | 0.291326 | 1 | 0.999631 | 0.72463 |
| Stxbp1_O08599_S594s_1_1 | -2.08 | -4.77 | -5.61 | -6.25 | -4.32 | -4.1 | -4.16 | -8.16 | -4.57 | -2.61 | -4.61 | -7.36 | -3.81 | -4.52 | -6.91 | -7.59 | -0.5075 | -0.11 | -1.03 | 0.360081 | 0.661853 | 0.92367 | 1 | 0.999631 | 0.741182 |
| Stxbp1_O08599_Y473y_1 | -0.18 | -1.78 | -2.16 | 0.41 | -0.15 | -1.96 | 0.41 | -0.57 | -0.86 | -2.05 | -0.16 | -2.04 | -3.48 | -2.31 | -1.19 | -1.62 | 0.36 | -0.35 | -1.2225 | 0.107087 | 0.612564 | 0.614584 | 1 | 0.999631 | 0.568797 |
| Stxbp4_B1ATH0_S11s_1 | 1.49 | 1.38 | 2.21 | 1.53 | 0.75 | 1.57 | 1.35 | 1.62 | 1.26 | 1.81 | 1.33 | 0.95 | 0.2 | 0.55 | 1.08 | 1.38 | -0.33 | -0.315 | -0.85 | 0.040794 | 0.318294 | 0.336428 | 1 | 0.999631 | 0.474979 |
| Stxbp5_D3Z079_S1023s_1 | -0.94 | 0.62 | 0.41 | -0.24 | -0.42 | 0.08 | -0.41 | -0.43 | -0.1 | 0.22 | -0.19 | 0.49 | 0.6 | 0.07 | -0.39 | 0.83 | -0.2575 | 0.1425 | 0.315 | 0.486018 | 0.508443 | 0.721197 | 1 | 0.999631 | 0.804525 |

| | | | | | | | | | | | | | | | | | | | | | | | | | |
|---|---|---|---|---|---|---|---|---|---|---|---|---|---|---|---|---|---|---|---|---|---|---|---|---|---|
| Stxbp5_D3Z079_S693s_1 | -0.49 | 0.22 | -1.02 | -0.11 | 0.49 | -0.29 | -0.23 | -0.37 | -0.34 | -0.82 | -0.85 | 0.12 | -0.5 | 0.08 | 0.25 | -0.49 | 0.25 | -0.1225 | 0.185 | 0.620438 | 0.489881 | 0.745055 | 1 | 0.999631 | 0.861888 |
| Stxbp5_D3Z079_S750s_1 | -0.39 | -0.23 | -3.92 | -5.07 | -1.01 | -0.51 | -3.76 | -5.13 | -0.48 | 0.07 | -3.12 | -5.82 | -0.38 | -0.41 | -6.13 | -5.44 | -0.2 | 0.065 | -0.6875 | 0.680824 | 0.886733 | 0.96661 | 1 | 0.999631 | 0.888714 |
| Stxbp5_D3Z079_S811s_1 | -0.98 | -0.85 | -0.59 | -0.9 | -0.4 | -1.21 | -0.86 | -0.61 | -1.07 | -0.76 | -1.39 | -2.06 | 0.89 | -0.9 | -0.63 | -0.6 | 0.06 | -0.49 | 0.52 | 0.239208 | 0.828908 | 0.169406 | 1 | 0.999631 | 0.683719 |
| Stxbp5_D3Z079_S867s_1 | -1.28 | -1.67 | -0.47 | -1.36 | -1.05 | -1.57 | -0.9 | -0.42 | -1.71 | -1.35 | -0.33 | -0.33 | -0.74 | -0.89 | -0.11 | -0.4 | 0.21 | 0.265 | 0.66 | 0.090766 | 0.571566 | 0.545731 | 1 | 0.999631 | 0.557085 |
| Stxbp5_D3Z079_T749t_1 | 0.41 | 0.85 | 0.87 | 0.06 | -0.11 | 0.5 | -0.42 | -0.07 | 0.56 | 0.97 | 0.37 | 0.96 | 0.41 | 0.71 | 0.58 | 0.91 | -0.5725 | 0.1675 | 0.105 | 0.737326 | 0.098868 | 0.593379 | 1 | 0.999631 | 0.907787 |
| Stxbp5l_Q5DQR4_S1120s_1 | -0.88 | -0.42 | -1.13 | -0.85 | -0.84 | -0.78 | -0.77 | -1.64 | -1.56 | -1.06 | -1.27 | -1.06 | -1.32 | -1.32 | -1.03 | -0.3 | -0.1875 | -0.4175 | -0.1725 | 0.618342 | 0.551193 | 0.162907 | 1 | 0.999631 | 0.861888 |
| Stxbp5l_Q5DQR4_S588s_1 | -0.56 | 0.89 | -0.52 | 1.66 | 1.13 | 0.21 | 1.2 | 0.87 | 0.29 | -0.58 | 0.81 | 1.02 | -0.62 | 0.14 | 1.14 | 1.85 | 0.485 | 0.0175 | 0.26 | 0.70561 | 0.382961 | 0.976538 | 1 | 0.999631 | 0.89781 |
| Stxbp5l_Q5DQR4_S597s_1 | -2.28 | -0.09 | -0.29 | -1 | -1.27 | -0.51 | -1.05 | -0.66 | -0.64 | -2.09 | -0.52 | -0.73 | 0.2 | -0.97 | -0.25 | -0.85 | 0.0425 | -0.08 | 0.4475 | 0.408088 | 0.931355 | 0.887885 | 1 | 0.999631 | 0.766996 |
| Stxbp5l_Q5DQR4_S598s_1 | -1.45 | -0.32 | -0.99 | -2.36 | -0.51 | -0.95 | -1.99 | -1.8 | -0.5 | -0.84 | -1.61 | -1.1 | -0.05 | -1.3 | -1.28 | -1.71 | -0.0325 | 0.2675 | 0.195 | 0.711846 | 0.949635 | 0.573446 | 1 | 0.999631 | 0.900703 |
| Stxbp5l_Q5DQR4_S792s_1 | 0.45 | 0.24 | -0.78 | 0.02 | 0.91 | -0.16 | -0.44 | -0.41 | -0.21 | -0.47 | -0.26 | -0.09 | 0.06 | 0.68 | -0.87 | -0.64 | -0.0075 | -0.24 | -0.175 | 0.694632 | 0.985551 | 0.475309 | 1 | 0.999631 | 0.895302 |
| Stxbp5l_Q5DQR4_S819s_2 | 0.49 | 2.13 | 1.96 | 1.51 | 1.69 | 2.25 | 0.65 | 2.14 | 1.9 | 1.03 | 0.06 | 1.58 | 2.01 | 2.17 | 1.94 | 2.03 | 0.16 | -0.38 | 0.515 | 0.214185 | 0.743925 | 0.466296 | 1 | 0.999631 | 0.667209 |
| Stxbp5l_Q5DQR4_S822s_1 | -1.42 | -0.01 | 2.54 | 0.36 | -1.79 | -0.96 | -0.57 | 0.9 | -0.47 | -0.36 | 1.13 | 2.23 | 0.91 | -0.25 | 2.23 | 2.35 | -0.9725 | 0.265 | 0.9425 | 0.300476 | 0.275824 | 0.770349 | 1 | 0.999631 | 0.71369 |
| Stxbp5l_Q5DQR4_S822s_2 | 0.49 | 2.13 | 1.96 | 1.51 | 1.69 | 2.25 | 0.65 | 2.14 | 1.9 | 1.03 | 0.06 | 1.58 | 2.01 | 2.17 | 1.94 | 2.03 | 0.16 | -0.38 | 0.515 | 0.214185 | 0.743925 | 0.466296 | 1 | 0.999631 | 0.667209 |
| Sucla2_Q9Z2I9_S279s_1 | -0.5 | 0.09 | -0.39 | -0.75 | -0.19 | -0.73 | -0.99 | -1.11 | -0.54 | -0.45 | -0.55 | -0.42 | 0 | -0.45 | -0.25 | -0.5 | -0.3675 | -0.1025 | 0.0875 | 0.775943 | 0.265544 | 0.713491 | 1 | 0.999631 | 0.920998 |
| Suds3_Q8BR65_S234s_2 | -0.4 | 0.18 | 0.87 | -0.21 | -0.86 | -1.51 | -0.51 | 0.14 | -0.26 | 0.33 | -1.03 | -1 | -1.4 | -0.07 | 0.24 | 0.39 | -0.795 | -0.6 | -0.32 | 0.510533 | 0.092144 | 0.184582 | 1 | 0.999631 | 0.817683 |
| Suds3_Q8BR65_S236s_2 | -0.4 | 0.18 | 0.87 | -0.21 | -0.86 | -1.51 | -0.51 | 0.14 | -0.26 | 0.33 | -1.03 | -1 | -1.4 | -0.07 | 0.24 | 0.39 | -0.795 | -0.6 | -0.32 | 0.510533 | 0.092144 | 0.184582 | 1 | 0.999631 | 0.817683 |
| Sugp2_Q8CH09_S206s_1 | 0.4 | 0.76 | 0.33 | 1.09 | 1.19 | 0.62 | 0.47 | 1.15 | 0.93 | 0.3 | 1.03 | 0.99 | 0.03 | 0.84 | 0.8 | 1.31 | 0.2125 | 0.1675 | 0.1 | 0.784125 | 0.497639 | 0.593988 | 1 | 0.999631 | 0.92266 |
| Sugp2_Q8CH09_S733s_1 | -2.18 | -0.77 | 0.47 | 0 | -0.24 | -1.05 | 0.03 | 0.02 | -1.1 | -1.1 | 0.46 | 0.92 | -0.68 | -1.46 | 0.36 | 0.81 | 0.31 | 0.415 | 0.3775 | 0.585701 | 0.593014 | 0.555057 | 1 | 0.999631 | 0.849524 |
| Sugp2_Q8CH09_S838s_1 | -0.22 | 0.4 | -0.49 | -0.81 | 0.48 | -0.15 | -0.56 | -0.35 | 0.15 | 0.1 | -0.14 | -1 | 0.46 | 0.03 | -0.08 | -0.26 | 0.135 | 0.0575 | 0.3175 | 0.379071 | 0.71092 | 0.88224 | 1 | 0.999631 | 0.751637 |
| Sulf2_Q8CFG0_S843s_1 | -2.06 | 1.33 | 1.3 | 0.4 | 1.49 | 1.07 | 0.89 | 2.13 | 1.54 | -0.63 | 2.18 | 2.19 | 1.51 | 0.83 | 1.73 | 0.85 | 1.1525 | 1.0775 | 0.9875 | 0.197362 | 0.144153 | 0.25094 | 1 | 0.999631 | 0.654248 |
| Sumo1_P63166_S2s_1 | 0.69 | 1.55 | 1.45 | 0.24 | 0.39 | 1.28 | 0.46 | 1.19 | 1.46 | 1.14 | 0.89 | 1.1 | 1.4 | 1.34 | 0.9 | 1.36 | -0.1525 | 0.165 | 0.2675 | 0.48077 | 0.701929 | 0.652855 | 1 | 0.999631 | 0.801848 |
| Susd5_G3UW60_S237s_1 | 0.33 | -1.59 | -2.47 | -1.63 | -1.15 | -0.8 | -0.66 | -1.3 | -0.65 | 0.67 | -1.3 | -0.87 | -2.04 | -1.01 | -2.09 | -2.23 | 0.3625 | 0.8025 | -0.5025 | 0.409274 | 0.521188 | 0.238933 | 1 | 0.999631 | 0.766996 |
| Sv2a_Q9JIS5_S127s_1 | -0.47 | -1.12 | -0.77 | -0.79 | -1.66 | -1.42 | -0.91 | -1.1 | -1.62 | -0.52 | -0.74 | -1.07 | -0.7 | -1.27 | -1.25 | -0.79 | -0.485 | -0.2 | -0.215 | 0.483288 | 0.116021 | 0.545454 | 1 | 0.999631 | 0.803286 |
| Svil_Q8K4L2_S666s_1 | -0.88 | 0.59 | 1.3 | -0.27 | 1.89 | 0.81 | 0.99 | 0.48 | 1.87 | -0.41 | 1.76 | 2.15 | 0.87 | -0.17 | 0.79 | 0.2 | 0.8575 | 1.1575 | 0.2375 | 0.644391 | 0.126582 | 0.113261 | 1 | 0.999631 | 0.872436 |
| Svil_Q8K4L2_S857s_1 | -0.23 | -0.34 | -0.39 | -0.13 | 0.04 | -0.58 | -0.26 | -0.28 | -0.26 | 0.36 | 0 | 0.44 | -0.94 | -0.34 | -0.01 | 0.11 | 0.0025 | 0.4075 | -0.0225 | 0.944394 | 0.992136 | 0.162163 | 1 | 0.999631 | 0.982024 |
| Svop_Q8BFT9_S25s_1 | -1.52 | -1.53 | -0.24 | -1.15 | -1.09 | -1.53 | -0.77 | -0.66 | -1.25 | -0.97 | -0.78 | 0.22 | -0.11 | -1.21 | 0.4 | 0.24 | 0.0975 | 0.415 | 0.94 | 0.066551 | 0.795614 | 0.354089 | 1 | 0.999631 | 0.518767 |
| Svop_Q8BFT9_S31s_1 | -1.28 | -0.19 | -0.16 | -1.08 | -0.34 | -0.23 | -0.56 | -0.3 | -0.35 | -0.93 | -0.16 | 0.32 | 0.54 | -0.04 | 0.06 | -0.27 | 0.32 | 0.3975 | 0.75 | 0.070105 | 0.355196 | 0.335599 | 1 | 0.999631 | 0.529516 |
| Svop_Q8BFT9_S31s_2 | -2.18 | -0.7 | -0.67 | -0.17 | -0.63 | -1.51 | -1.75 | 0.01 | -1.46 | -2.4 | -1.97 | 0.09 | -0.5 | -1.32 | -0.33 | -0.3 | -0.04 | -0.505 | 0.3175 | 0.514173 | 0.941559 | 0.429801 | 1 | 0.999631 | 0.81893 |
| Svop_Q8BFT9_T534t_1 | 0.9 | -0.33 | -1.04 | 0.03 | -0.29 | 0.57 | 0.07 | -0.24 | -0.57 | 1.15 | -0.62 | -1.46 | -0.4 | -0.43 | -0.84 | -2.1 | 0.1375 | -0.265 | -0.8325 | 0.141408 | 0.754009 | 0.668853 | 1 | 0.999631 | 0.61245 |
| Syap1_Q9D5V6_T262t_1 | 0 | 0.25 | 0.33 | -0.04 | 0.06 | -0.05 | -0.52 | 0.1 | 0.1 | 0.37 | 0.07 | 0.4 | 0.37 | 0.66 | 0.18 | 0.43 | -0.2375 | 0.1 | 0.275 | 0.333358 | 0.382588 | 0.698517 | 1 | 0.999631 | 0.728079 |
| Sybu_Q8BHS8_S197s_1 | 1.13 | -0.38 | 0.58 | -0.04 | -0.64 | -0.05 | 0.1 | -0.56 | -0.04 | 1.68 | -0.52 | -0.27 | -0.4 | -0.32 | -0.72 | -0.71 | -0.61 | -0.11 | -0.86 | 0.04582 | 0.142008 | 0.843056 | 1 | 0.999631 | 0.477815 |
| Sybu_Q8BHS8_S201s_2 | 1.11 | -0.07 | 0.66 | 0.37 | -0.16 | 0.1 | 0.69 | -0.67 | 0.03 | 1 | -0.24 | -0.61 | -0.73 | -0.82 | -0.12 | -0.79 | -0.5275 | -0.4725 | -1.1325 | 0.008812 | 0.193591 | 0.282595 | 1 | 0.999631 | 0.365244 |
| Sybu_Q8BHS8_S244s_1 | 1.11 | 0.69 | 1.08 | 0.71 | 0.35 | 0.95 | 1.02 | 0.85 | 1.22 | 1.49 | 0.72 | 0.7 | 0.77 | 1.15 | 1.21 | 0.44 | -0.105 | 0.135 | -0.005 | 0.987059 | 0.704245 | 0.655234 | 1 | 0.999631 | 0.995165 |
| Sybu_Q8BHS8_S548s_1 | -0.87 | -1.29 | -0.59 | -0.47 | -0.15 | -1.33 | -0.59 | -0.6 | -0.61 | -0.69 | -0.58 | -0.35 | -0.31 | -0.65 | -1.29 | 0.13 | 0.1375 | 0.2475 | 0.275 | 0.479919 | 0.687036 | 0.396695 | 1 | 0.999631 | 0.801333 |
| Sympk_Q80X82_S1260s_1 | 0.28 | 1.85 | 1.46 | -0.04 | 2.12 | 1.51 | 0.4 | 0.77 | 1.92 | 1.4 | 1.52 | 1.89 | 2.42 | 1.71 | 1.57 | 1.47 | 0.3125 | 0.795 | 0.905 | 0.087081 | 0.570909 | 0.109387 | 1 | 0.999631 | 0.553514 |
| Sympk_Q80X82_S1276s_1 | 2.29 | -1.22 | 0.21 | 0.08 | -1.11 | 0.07 | 2.76 | -2.23 | -0.97 | 2.9 | -0.99 | -1.18 | -3.15 | -1.26 | -0.84 | -2.34 | -0.4675 | -0.4 | -2.2375 | 0.015938 | 0.674188 | 0.705578 | 1 | 0.999631 | 0.388832 |
| Sympk_Q80X82_S494s_1 | -0.35 | 0.45 | 0.4 | 0.26 | 0.25 | 0.1 | -0.43 | 0.23 | 0.2 | 0.06 | -0.47 | -0.12 | -0.15 | 0.4 | -0.13 | 1.03 | -0.1525 | -0.2725 | 0.0975 | 0.794596 | 0.617866 | 0.382853 | 1 | 0.999631 | 0.928784 |
| Syn1_O88935_S11s_1 | 0.73 | 0.46 | -0.3 | 0.08 | 0.72 | 0.52 | -0.26 | -0.05 | 0.59 | 0.42 | -0.16 | 0.25 | 0.48 | 0.94 | -0.09 | 0.58 | -0.01 | 0.0325 | 0.235 | 0.518263 | 0.977263 | 0.921302 | 1 | 0.999631 | 0.819449 |
| Syn1_O88935_S330s_1 | -0.34 | 0.2 | -1.03 | -0.7 | 0.06 | -0.69 | -0.34 | -0.68 | -1.01 | -0.43 | -0.33 | -0.6 | -0.1 | 0.53 | -0.19 | -0.43 | 0.055 | -0.125 | 0.42 | 0.276858 | 0.874379 | 0.718064 | 1 | 0.999631 | 0.700187 |
| Syn1_O88935_S332s_1 | 0.03 | -0.17 | 0.44 | -0.06 | 0.01 | -0.34 | 0.46 | 0.61 | -0.52 | -0.04 | 0.43 | 0.99 | -0.14 | 0.23 | 1.48 | 0.71 | 0.125 | 0.155 | 0.51 | 0.220228 | 0.687569 | 0.680181 | 1 | 0.999631 | 0.670699 |
| Syn1_O88935_S341s_1 | -0.39 | -0.28 | 0.03 | -0.88 | -0.58 | -0.78 | -0.68 | -0.34 | -0.52 | -0.69 | -0.47 | -0.69 | 0.01 | -0.49 | 0 | -0.31 | -0.215 | -0.2125 | 0.1825 | 0.564869 | 0.459367 | 0.465305 | 1 | 0.999631 | 0.84139 |
| Syn1_O88935_S39s_1 | 0.92 | -0.65 | 0.23 | 1.21 | 0.84 | 0.68 | 0.66 | 0.75 | 0.09 | 0.11 | -0.64 | -0.03 | -0.96 | 1.24 | 0.59 | 0.35 | 0.305 | -0.545 | -0.1225 | 0.830294 | 0.467812 | 0.238005 | 1 | 0.999631 | 0.942193 |
| Syn1_O88935_S427s_1 | 0.44 | -2.83 | -1.15 | -0.68 | -2.49 | -3.64 | 0.06 | -2.24 | -4.36 | -1.91 | -2.19 | -2.72 | -3.76 | -3.01 | -2.17 | -3.8 | -1.0225 | -1.74 | -2.13 | 0.011124 | 0.268504 | 0.044823 | 1 | 0.999631 | 0.385092 |
| Syn1_O88935_S437s_1 | 0.56 | 0.5 | -1.07 | -1.19 | 0.61 | 0.87 | -0.62 | -2.5 | 0.1 | 0.6 | -0.43 | -1.25 | 0.56 | 0.87 | -1.47 | -0.79 | -0.11 | 0.055 | 0.0925 | 0.88793 | 0.889497 | 0.923314 | 1 | 0.999631 | 0.963505 |
| Syn1_O88935_S449s_1 | 2.43 | 0.32 | -0.15 | 1.13 | 2.51 | 0.93 | 0.64 | 0.46 | 0.8 | -0.42 | 0.1 | -0.9 | 0.14 | 0.36 | -0.5 | -0.14 | 0.2025 | -1.0375 | -0.9675 | 0.104367 | 0.757485 | 0.114217 | 1 | 0.999631 | 0.568213 |
| Syn1_O88935_S510s_1 | 1.13 | -0.99 | -1.17 | 0.32 | 1.29 | -0.57 | 0.5 | 0.37 | -0.11 | -1.05 | 0.34 | -0.8 | -1.22 | -0.61 | -0.87 | -0.38 | 0.575 | -0.2275 | -0.5925 | 0.287179 | 0.351756 | 0.696214 | 1 | 0.999631 | 0.705972 |
| Syn1_O88935_S520s_1 | -0.72 | -1.45 | -0.8 | -1.86 | -0.92 | -1.49 | -0.72 | -0.2 | -1.13 | -1.75 | -0.08 | -0.97 | -0.44 | -1.13 | 0.29 | -1.02 | 0.375 | 0.225 | 0.6325 | 0.165266 | 0.347898 | 0.608192 | 1 | 0.999631 | 0.633686 |
| Syn1_O88935_S551s_1 | -0.59 | -1.88 | -2.62 | -1.42 | -0.33 | -1.3 | -1.27 | -1.68 | -1.13 | -1.55 | -1.8 | -1.76 | -1 | -1.67 | -2.22 | -2.36 | 0.4825 | 0.0675 | -0.185 | 0.713481 | 0.332422 | 0.87943 | 1 | 0.999631 | 0.900989 |
| Syn1_O88935_S551s_2 | -0.59 | -0.98 | 0.56 | 0.77 | 1.08 | 0.22 | 0.85 | 0.64 | 0.03 | -1.2 | 0.11 | 0.3 | -2 | 0.03 | 0.97 | -0.07 | 0.7575 | -0.13 | -0.2075 | 0.758708 | 0.117088 | 0.80103 | 1 | 0.999631 | 0.914449 |
| Syn1_O88935_S553s_1 | -0.69 | -2.61 | -1 | -0.85 | -0.67 | -0.79 | -0.69 | -1.16 | -1.27 | -1.44 | -1.73 | -0.92 | -1.48 | -1.02 | -0.53 | -1.59 | 0.46 | -0.0525 | 0.1325 | 0.786703 | 0.315661 | 0.909621 | 1 | 0.999631 | 0.92378 |
| Syn1_O88935_S553s_2 | -0.59 | -0.98 | 0.56 | 0.77 | 1.08 | 0.22 | 0.85 | 0.64 | 0.03 | -1.2 | 0.11 | 0.3 | -2 | 0.03 | 0.97 | -0.07 | 0.7575 | -0.13 | -0.2075 | 0.758708 | 0.117088 | 0.80103 | 1 | 0.999631 | 0.914449 |
| Syn1_O88935_S55s_1 | -1.43 | 0.03 | -0.85 | -0.16 | -0.16 | -1.4 | 0.38 | 0.52 | -0.59 | -0.23 | -0.98 | 0.35 | -0.01 | -0.55 | -0.49 | -0.95 | 0.4375 | 0.24 | 0.1025 | 0.801115 | 0.40265 | 0.584106 | 1 | 0.999631 | 0.932038 |
| Syn1_O88935_S605s_1 | 3.27 | 0.72 | 1.21 | 1.89 | 1.5 | 2.15 | 2.32 | 0.92 | 0.21 | 1.15 | 0.84 | -1.27 | -2.27 | -0.22 | -0.74 | -0.48 | -0.05 | -1.54 | -2.7 | 0.001879 | 0.931398 | 0.046888 | 1 | 0.999631 | 0.351308 |
| Syn1_O88935_S624s_1 | 0.19 | 0.54 | -0.78 | 0.46 | 0.23 | 0.79 | 0.9 | 0.91 | -0.6 | 1.21 | 0.49 | -1.4 | -0.15 | 0.5 | 0.29 | 0.26 | 0.605 | -0.1775 | 0.1225 | 0.743338 | 0.121906 | 0.765907 | 1 | 0.999631 | 0.90972 |
| Syn1_O88935_S62s_1 | 1.32 | -0.98 | -0.82 | 0.54 | 2.32 | -0.79 | 0.6 | 0.28 | 0.23 | -1.78 | 0.04 | -0.4 | -2.88 | -0.18 | -0.02 | 0.05 | 0.5875 | -0.4925 | -0.7725 | 0.337472 | 0.438823 | 0.451522 | 1 | 0.999631 | 0.730499 |
| Syn1_O88935_S642s_1 | 1.85 | -0.52 | -0.28 | 2.52 | 1.83 | 0.44 | 1.57 | 1.15 | 0.23 | -0.81 | 0.21 | -0.34 | -1.29 | 0.42 | -1.06 | 0.23 | 0.355 | -1.07 | -1.3175 | 0.110616 | 0.624909 | 0.155788 | 1 | 0.999631 | 0.575791 |
| Syn1_O88935_S642s_1_1 | 1.79 | -0.6 | 0.13 | 2.67 | 1.82 | 0.42 | 1.47 | 0.74 | 0.27 | -0.39 | -0.34 | -0.08 | -1.13 | 0.58 | -0.12 | -0.01 | 0.115 | -1.1325 | -1.1675 | 0.133713 | 0.872931 | 0.121845 | 1 | 0.999631 | 0.60492 |
| Syn1_O88935_S664s_1 | 1.39 | -0.56 | 0.69 | 0.18 | -1.27 | -1.48 | 0.55 | -0.07 | -1.46 | 0.4 | -0.71 | -0.6 | -0.78 | -0.85 | 0.14 | -0.16 | -0.9925 | -1.0175 | -0.8375 | 0.097853 | 0.111036 | 0.077579 | 1 | 0.999631 | 0.560722 |
| Syn1_O88935_S666s_1 | 2.45 | -0.47 | -0.06 | 0.85 | 0.94 | 0.98 | 2 | 0.23 | -1.12 | -0.05 | 0.15 | -1.52 | -1.81 | -0.6 | -1.26 | -1.32 | 0.345 | -1.3275 | -1.94 | 0.010722 | 0.604606 | 0.076379 | 1 | 0.999631 | 0.383871 |
| Syn1_O88935_S666s_1_1 | 2.34 | 2.53 | -0.76 | 1.32 | 3.15 | 1.83 | 0.88 | 1.09 | 2.42 | 1.35 | 0.62 | -0.18 | 2.15 | 1.64 | 0.47 | 0.44 | 0.38 | -0.305 | -0.1825 | 0.810491 | 0.635288 | 0.710414 | 1 | 0.999631 | 0.936878 |
| Syn1_O88935_S67s_1 | -0.6 | -0.26 | -0.72 | -0.18 | -0.02 | -0.59 | -0.35 | -0.11 | -0.6 | -1.19 | -0.59 | -0.35 | -0.54 | -0.24 | -0.24 | -0.07 | 0.1725 | -0.2425 | 0.1675 | 0.563123 | 0.53083 | 0.425108 | 1 | 0.999631 | 0.841038 |
| Syn1_O88935_S682s_1 | -1.03 | -0.82 | -1.8 | -0.95 | -0.12 | -1.22 | -0.94 | -1.27 | -1.07 | -1.61 | -0.79 | -1.12 | -0.58 | -1.11 | -0.73 | -1.15 | 0.2625 | 0.0025 | 0.2575 | 0.446526 | 0.479458 | 0.993965 | 1 | 0.999631 | 0.785008 |
| Syn1_O88935_S684s_1 | -1 | -1 | 0.95 | 1.24 | 0.04 | -0.87 | 0.48 | 1.99 | -0.65 | -1.75 | 2.15 | 2.03 | -0.7 | -0.86 | 2.13 | 0.57 | 0.3625 | 0.3975 | 0.2375 | 0.767802 | 0.630092 | 0.690116 | 1 | 0.999631 | 0.917335 |
| Syn1_O88935_S9s_1 | 0.45 | 0.37 | -0.07 | 0.22 | 0.63 | 0.26 | -0.07 | 0.2 | 0.39 | 0.22 | 0.11 | 0.26 | 0.4 | 0.62 | -0.01 | 0.5 | 0.0125 | 0.0025 | 0.135 | 0.647055 | 0.963451 | 0.992263 | 1 | 0.999631 | 0.874422 |
| Syn1_O88935_T339t_1 | -1.1 | -0.25 | -0.53 | -0.59 | -0.42 | -0.62 | -0.98 | -0.57 | -0.6 | -0.87 | -0.85 | -0.51 | 0.46 | -0.25 | -0.58 | -0.74 | -0.03 | -0.09 | 0.34 | 0.363152 | 0.916802 | 0.754673 | 1 | 0.999631 | 0.743604 |
| Syn1_O88935_T448t_1 | 0.37 | 0.16 | 1.3 | 1.89 | 0.48 | -0.5 | 1.94 | 1.67 | -0.1 | -0.17 | 1.75 | 0.05 | 0.35 | 0.07 | 1.18 | 0.9 | -0.0325 | -0.5475 | -0.305 | 0.51927 | 0.958371 | 0.343539 | 1 | 0.999631 | 0.819456 |
| Syn2_Q64332_S108s_1 | -2.39 | -3.24 | -3.54 | -2.51 | -2.82 | -3.27 | -2.52 | -2.72 | -3.49 | -3.43 | -3.44 | -2.78 | -3.22 | -3.67 | -2.15 | -2.6 | 0.0875 | -0.365 | 0.01 | 0.981845 | 0.802924 | 0.323535 | 1 | 0.999631 | 0.994846 |
| Syn2_Q64332_S111s_1 | -1.61 | -1.79 | 1.29 | 1.05 | -1.49 | -1.77 | 0.2 | 0.34 | -2.01 | -1.84 | 0.38 | 0.69 | -1.56 | -2.02 | 1.28 | 0.76 | -0.415 | -0.43 | -0.12 | 0.904435 | 0.633583 | 0.652023 | 1 | 0.999631 | 0.96853 |
| Syn2_Q64332_S426s_1 | -1.27 | -2.84 | 0.42 | -0.2 | -2.66 | -2.41 | -0.12 | 0.33 | -2.82 | -1.84 | -0.14 | -0.06 | -2.42 | -2.25 | 0.25 | 0.63 | -0.2425 | -0.2425 | 0.025 | 0.978427 | 0.789584 | 0.777439 | 1 | 0.999631 | 0.99279 |
| Syn2_Q64332_S426s_2 | 0.16 | 0.47 | 0.59 | 0.23 | 0.58 | 0.48 | -0.43 | 0.74 | 0.18 | 0.33 | -0.1 | 0.29 | 0.42 | 1.27 | 0.74 | 0.63 | -0.02 | -0.1875 | 0.4025 | 0.20793 | 0.951012 | 0.480483 | 1 | 0.999631 | 0.661649 |
| Syn2_Q64332_S480s_1 | -0.15 | 0.98 | 0.61 | 0.81 | 0.24 | 0.54 | 0.28 | 0.61 | 0.03 | 0.04 | 0.14 | 1.43 | 0.4 | 0.79 | 0.5 | 1.15 | -0.145 | -0.1525 | 0.1475 | 0.678637 | 0.648426 | 0.719875 | 1 | 0.999631 | 0.887183 |
| Syn2_Q64332_S483s_1 | -0.48 | 0 | -0.94 | -0.22 | 0.69 | 1.21 | 0.02 | -0.36 | -1.15 | -0.09 | -0.13 | 0.05 | 1.53 | 0.31 | -0.29 | -1.14 | 0.8 | 0.08 | 0.5125 | 0.365575 | 0.072602 | 0.828816 | 1 | 0.999631 | 0.745725 |
| Syn2_Q64332_S53s_1 | -0.86 | -5.07 | -1.33 | -0.56 | -0.99 | -2.44 | -0.75 | -0.24 | -1.5 | -2.68 | -0.25 | -0.41 | -2.13 | -1.51 | 0.08 | 0.11 | 0.85 | 0.745 | 1.0925 | 0.298187 | 0.400272 | 0.474652 | 1 | 0.999631 | 0.711311 |
| Syn2_Q64332_S546s_1 | 1.81 | 0.05 | -0.81 | 0.69 | 0.35 | 1.14 | 0.12 | 0.48 | -1.03 | 1.44 | -0.36 | -0.59 | -2.8 | 2.12 | 0.02 | -0.26 | 0.0875 | -0.57 | -0.665 | 0.503953 | 0.87257 | 0.417655 | 1 | 0.999631 | 0.814292 |
| Syn2_Q64332_S552s_1 | -0.02 | 0.36 | 0.01 | 0.68 | 0.87 | 0.84 | 0.38 | 0.95 | 0.59 | -0.95 | 0.51 | 1.03 | 0.19 | 0.75 | -0.02 | 0.98 | 0.5025 | 0.0375 | 0.2175 | 0.534208 | 0.103116 | 0.93378 | 1 | 0.999631 | 0.828493 |
| Syn2_Q64332_T422t_1 | 0.25 | 1 | 1.22 | 0.34 | -0.04 | 1 | -0.03 | 0.82 | 0.67 | 1.16 | 0.34 | 0.91 | 1.04 | 1.33 | 1.09 | 1.28 | -0.265 | 0.0675 | 0.4825 | 0.162403 | 0.490285 | 0.843392 | 1 | 0.999631 | 0.631442 |
| Syn2_Q64332_T422t_2 | 0.14 | 0.5 | 0.59 | 0.23 | 0.61 | 0.5 | -0.43 | 0.74 | 0.16 | 0.33 | -0.1 | 0.29 | 0.43 | 1.24 | 0.74 | 0.63 | -0.01 | -0.195 | 0.395 | 0.213205 | 0.975731 | 0.466511 | 1 | 0.999631 | 0.666787 |
| Syn2_Q64332_T449t_1 | 1.26 | 1.64 | 2.09 | 0.13 | 1.18 | 2.55 | 0.68 | 1.32 | 2.14 | 2.33 | 1.11 | 0.86 | 2.46 | 2.93 | 1.88 | 1.55 | 0.1525 | 0.33 | 0.925 | 0.087424 | 0.775191 | 0.531763 | 1 | 0.999631 | 0.553616 |
| Syn2_Q64332_T449t_1_1 | 0.02 | 1.15 | 1.79 | 0.47 | 0.52 | 1.07 | 0.4 | 0.84 | 0.68 | 1.1 | 0.55 | -0.61 | 1.18 | 1.45 | 0.97 | 1.68 | -0.15 | -0.4275 | 0.4625 | 0.293911 | 0.718426 | 0.406041 | 1 | 0.999631 | 0.708293 |
| Syn3_Q8JZP2_S404s_1 | 0.41 | -0.37 | -0.8 | 0.53 | 1.91 | -0.09 | 1.03 | 0.89 | 0.76 | -1.05 | -0.13 | -0.86 | -1.23 | 0.26 | -0.3 | -1.24 | 0.9925 | -0.2625 | -0.57 | 0.24764 | 0.066132 | 0.598117 | 1 | 0.999631 | 0.685751 |
| Syn3_Q8JZP2_S418s_1 | 1.26 | -1.66 | 0.11 | 1.08 | 1.13 | -1.86 | 1.28 | 1.29 | 0.54 | -2.02 | 1.5 | 0.36 | -2 | -1.7 | 1.59 | -0.98 | 0.2625 | -0.1025 | -0.97 | 0.299967 | 0.767462 | 0.906608 | 1 | 0.999631 | 0.713477 |
| Syn3_Q8JZP2_S446s_1 | 2.94 | -0.77 | -1.51 | 1.12 | 2.49 | -0.55 | 2.44 | 0.71 | 0.99 | -1.36 | 0.1 | -1.56 | -2.04 | -1.51 | -1.74 | -2.22 | 0.8275 | -0.9025 | -2.3225 | 0.022533 | 0.443947 | 0.381515 | 1 | 0.999631 | 0.407815 |
| Syn3_Q8JZP2_S461s_1 | -0.14 | -0.28 | 0.03 | -0.32 | -0.6 | -0.6 | -0.79 | -0.18 | -0.66 | -0.2 | -0.84 | -0.39 | 0 | -0.49 | -0.54 | -0.12 | -0.365 | -0.345 | -0.11 | 0.699946 | 0.180024 | 0.222609 | 1 | 0.999631 | 0.896612 |
| Syn3_Q8JZP2_S461s_2 | -0.56 | -2.2 | -0.75 | 0.31 | -0.94 | -1.34 | -0.36 | 0.03 | -1.42 | -0.99 | -0.31 | -0.56 | -0.61 | -2.63 | -0.22 | -0.26 | 0.1475 | -0.02 | -0.13 | 0.850232 | 0.789751 | 0.97021 | 1 | 0.999631 | 0.949525 |
| Syn3_Q8JZP2_S469s_1 | 0.43 | -0.55 | 0.55 | -0.25 | -0.1 | 0.1 | -0.25 | -0.12 | -0.38 | -0.29 | -0.75 | -0.62 | -0.2 | -0.81 | -0.49 | -0.22 | -0.1375 | -0.555 | -0.475 | 0.20089 | 0.67048 | 0.121899 | 1 | 0.999631 | 0.65842 |
| Syn3_Q8JZP2_S469s_2 | -0.12 | -1.74 | -0.01 | 0.81 | -0.34 | -0.81 | 0.06 | 0.21 | -0.93 | -0.94 | 0.54 | -0.15 | -0.38 | -0.91 | 0.3 | -0.36 | 0.045 | -0.105 | -0.0725 | 0.894696 | 0.93299 | 0.857911 | 1 | 0.999631 | 0.965377 |
| Syn3_Q8JZP2_S480s_1 | 1.24 | -1.19 | -1.49 | 0.54 | 1.41 | -0.97 | 0.68 | 0.35 | -0.28 | -0.46 | 0.13 | -0.51 | -1.16 | -0.24 | -0.57 | -0.19 | 0.5925 | -0.055 | -0.315 | 0.619529 | 0.424971 | 0.928696 | 1 | 0.999631 | 0.861888 |
| Syn3_Q8JZP2_S480s_2 | 0.08 | -0.54 | -0.41 | 0.62 | 0.71 | 0.03 | 0.25 | 0.47 | 0.31 | -0.19 | 0.21 | -0.19 | -1.51 | 0.33 | 0.14 | -0.28 | 0.4275 | 0.0975 | -0.2675 | 0.578272 | 0.225003 | 0.774724 | 1 | 0.999631 | 0.84622 |
| Syn3_Q8JZP2_S483s_1 | 0.79 | 0.21 | 0.29 | 1.03 | 0.48 | 0.09 | 0.2 | 0.99 | 0.3 | 0.54 | 0.19 | 0.79 | -0.53 | -0.99 | 1.13 | 0.38 | -0.14 | -0.125 | -0.5825 | 0.254399 | 0.668751 | 0.685765 | 1 | 0.999631 | 0.689457 |
| Syn3_Q8JZP2_S483s_2 | 0.04 | -0.62 | -0.41 | 0.62 | 0.71 | -0.02 | 0.25 | 0.47 | 0.33 | -0.32 | 0.21 | -0.19 | -1.59 | 0.3 | 0.14 | -0.28 | 0.445 | 0.1 | -0.265 | 0.591755 | 0.21897 | 0.777277 | 1 | 0.999631 | 0.852627 |
| Syn3_Q8JZP2_S494s_1 | -1.98 | -6.01 | -0.97 | -1.51 | -2.11 | -3.71 | 2.4 | -2.35 | -2.66 | -2.47 | -2.1 | -1.1 | -2.95 | -3.72 | -1.26 | -3.14 | 1.175 | 0.535 | -0.15 | 0.889153 | 0.437851 | 0.607647 | 1 | 0.999631 | 0.963505 |
| Syn3_Q8JZP2_S519s_1 | 2.76 | -0.05 | 0.17 | 1.42 | 0.2 | 0.58 | 1.85 | 0.21 | -0.22 | 0.36 | -0.4 | -0.47 | -1.37 | -0.73 | 0.28 | -0.89 | -0.365 | -1.2575 | -1.7525 | 0.022365 | 0.590141 | 0.063859 | 1 | 0.999631 | 0.405775 |
| Syn3_Q8JZP2_S540s_1 | 0.96 | 0.26 | 0.26 | 0.19 | 0.02 | 0.34 | 0.04 | 0.19 | -0.52 | 0.74 | -0.96 | -1.67 | -1.02 | -0.98 | -0.41 | -0.06 | -0.27 | -1.02 | -1.035 | 0.013634 | 0.344932 | 0.068794 | 1 | 0.999631 | 0.385092 |
| Syn3_Q8JZP2_S544s_2 | -0.21 | -1.15 | 0.23 | 0.64 | -1.9 | -1 | 0.05 | -0.08 | -1.69 | -1.9 | -0.32 | 0.05 | 0.25 | -1.54 | -0.48 | 1.24 | -0.61 | -0.8425 | -0.01 | 0.987372 | 0.280105 | 0.1628 | 1 | 0.999631 | 0.995351 |
| Syn3_Q8JZP2_S9s_1 | -0.26 | -0.39 | -0.34 | -0.14 | 0.01 | -0.51 | -0.4 | -0.22 | -0.24 | -0.38 | -0.03 | 0 | 0.29 | -0.07 | 0.14 | 0.05 | 0.0025 | 0.12 | 0.385 | 0.168843 | 0.991982 | 0.634033 | 1 | 0.999631 | 0.637839 |
| Syncrip_G3UZ48_S151s_1 | -1.62 | 1.27 | -0.65 | 0.23 | 0.44 | 1.43 | -0.03 | 0.11 | 1.43 | -0.83 | 1.37 | 0.86 | 1.4 | 0.46 | 0.19 | 0.46 | 0.68 | 0.9 | 0.82 | 0.199178 | 0.293493 | 0.230715 | 1 | 0.999631 | 0.657167 |
| Syndig1l_Q3USQ7_S15s_1 | 3.16 | -0.35 | -0.07 | 2 | 1.72 | 0.46 | 2.87 | 2.25 | 0.61 | 0.25 | 0.3 | 0.33 | -1.46 | -0.2 | 0.8 | -0.84 | 0.64 | -0.8125 | -1.61 | 0.079652 | 0.461107 | 0.291993 | 1 | 0.999631 | 0.545579 |
| Syne1_Q6ZWR6_S1360s_1 | 0.03 | -0.26 | -0.52 | -0.54 | 0.7 | -1.17 | -0.72 | -0.57 | -1.51 | -0.48 | -2.25 | -1.47 | -3.14 | -1.28 | -3.44 | -0.16 | -0.1175 | -1.105 | -1.6825 | 0.034875 | 0.779351 | 0.01915 | 1 | 0.999631 | 0.44933 |
| Syne1_Q6ZWR6_S1369s_1 | -0.9 | -0.38 | 0.33 | 0.15 | -0.26 | -0.22 | -0.78 | -0.59 | 0.2 | -0.94 | -0.46 | -0.73 | -0.35 | 0.68 | 0.02 | 1.26 | -0.2625 | -0.2825 | 0.6025 | 0.20139 | 0.450277 | 0.474561 | 1 | 0.999631 | 0.658952 |
| Syne1_Q6ZWR6_S6239s_1 | 1.87 | 0.82 | 0.37 | 1.43 | -0.45 | 0.66 | 1.58 | -0.13 | -0.99 | 0.96 | 0.25 | -0.04 | 0.05 | -0.71 | -0.26 | -0.46 | -0.7075 | -1.0775 | -1.4675 | 0.003805 | 0.195391 | 0.052268 | 1 | 0.999631 | 0.351308 |
| Syne1_Q6ZWR6_S8252s_1 | -0.71 | 0.19 | -0.22 | -0.33 | -0.04 | 0.25 | -1.04 | -0.84 | -0.47 | -1.12 | -1.54 | -0.56 | 1.21 | 0.97 | -0.92 | -0.32 | -0.15 | -0.655 | 0.5025 | 0.34143 | 0.691352 | 0.086538 | 1 | 0.999631 | 0.73317 |
| Syne1_Q6ZWR6_S8306s_1 | -0.38 | 1.33 | 1.48 | 1.26 | 0.79 | 0.98 | 0.94 | 1.38 | 1.07 | 0.85 | 1.48 | 2.13 | 1.43 | 1.11 | 2.08 | 1.78 | 0.1 | 0.46 | 0.6775 | 0.174281 | 0.821002 | 0.363214 | 1 | 0.999631 | 0.642116 |
| Syne2_E9QP46_S6349s_1 | 0.42 | 2.24 | 2.2 | 1.72 | 1.43 | 1.13 | 0.85 | 1.39 | 1.7 | 1.33 | 1.59 | 2.14 | 1.74 | 1.49 | 1.75 | 1.91 | -0.445 | 0.045 | 0.0775 | 0.859625 | 0.320355 | 0.92012 | 1 | 0.999631 | 0.953272 |
| Syngap1_F6SEU4_S1027s_1 | -1.32 | -1.66 | -1.46 | -1.21 | -1.83 | -1.69 | -1.65 | -1.04 | -1.9 | -1.87 | -1.72 | -1.1 | -0.91 | -1.64 | -1.63 | -1.49 | -0.14 | -0.235 | -0.005 | 0.986769 | 0.620235 | 0.430808 | 1 | 0.999631 | 0.995165 |
| Syngap1_F6SEU4_S1033s_1 | -0.77 | -0.85 | -0.43 | -0.74 | -0.63 | -1.16 | -1.05 | -0.95 | -1.09 | -0.89 | -1.35 | -1.18 | 0.04 | -0.94 | -0.84 | -0.26 | -0.25 | -0.43 | 0.1975 | 0.550386 | 0.342392 | 0.122291 | 1 | 0.999631 | 0.836038 |
| Syngap1_F6SEU4_S1053s_1 | 0.11 | 0.42 | 0.42 | 0.14 | -0.05 | 0.23 | -0.26 | 0.21 | -0.11 | 0.16 | -0.63 | -0.05 | 0.63 | 0.55 | 0.55 | 0.39 | -0.24 | -0.43 | 0.2575 | 0.346117 | 0.359488 | 0.151522 | 1 | 0.999631 | 0.735175 |
| Syngap1_F6SEU4_S1070s_1 | 1.99 | 1.9 | 0.94 | 1.9 | 2.88 | 0.29 | 1.4 | 0.06 | 1.29 | 1.27 | 2.04 | 3.01 | 0.09 | -0.59 | 1.3 | 3.35 | -0.525 | 0.22 | -0.645 | 0.419708 | 0.406642 | 0.638081 | 1 | 0.999631 | 0.772233 |
| Syngap1_F6SEU4_S1091s_1 | 1.28 | 0.02 | 0.48 | 1.22 | 1.13 | 0.68 | 0.81 | 0.56 | 1.58 | -0.31 | -0.29 | 0.23 | -0.79 | 0.13 | 0.7 | 0.32 | 0.045 | -0.4475 | -0.66 | 0.157567 | 0.898939 | 0.388293 | 1 | 0.999631 | 0.628426 |
| Syngap1_F6SEU4_S1111s_1 | 0.23 | -0.12 | -0.93 | -0.61 | 0.04 | -0.69 | -0.4 | -0.97 | 0 | -0.44 | -0.19 | -0.68 | 0.4 | -0.81 | -1.42 | 0.52 | -0.1475 | 0.03 | 0.03 | 0.953142 | 0.682835 | 0.929919 | 1 | 0.999631 | 0.984317 |
| Syngap1_F6SEU4_S1115s_1 | -0.23 | 0.09 | 0.81 | 0.12 | -0.04 | 0.17 | -0.12 | 0.19 | -0.33 | 0.22 | 0.23 | 0.34 | 0.29 | -0.03 | 1.05 | 1.21 | -0.1475 | -0.0825 | 0.4325 | 0.289745 | 0.622017 | 0.798692 | 1 | 0.999631 | 0.706164 |
| Syngap1_F6SEU4_S1118s_1 | 0.1 | 1.38 | 1.11 | 0.31 | -0.14 | 0.71 | 0.52 | 1.08 | 0.4 | 0.73 | 0.25 | 0.72 | 0.66 | 1.09 | 0.4 | 2 | -0.1825 | -0.2 | 0.3125 | 0.503009 | 0.65245 | 0.583996 | 1 | 0.999631 | 0.814092 |
| Syngap1_F6SEU4_S1135s_1 | 0.78 | 0.2 | -0.22 | 0.71 | 0.74 | -0.37 | 1.27 | 0.51 | 0.12 | 0.31 | 1.44 | 0.75 | 0.23 | -0.89 | 0.24 | 0.69 | 0.17 | 0.2875 | -0.3 | 0.483385 | 0.681969 | 0.469362 | 1 | 0.999631 | 0.803286 |
| Syngap1_F6SEU4_S1162s_1 | -0.21 | -1.63 | -2.14 | -1.62 | -1.31 | -1.86 | -0.8 | -1.61 | -1.31 | -1.92 | -2.39 | -1.28 | -1.71 | -1.91 | -2.47 | -1.2 | 0.005 | -0.325 | -0.4225 | 0.386257 | 0.991234 | 0.498989 | 1 | 0.999631 | 0.756141 |
| Syngap1_F6SEU4_S117s_1 | 0.4 | -0.09 | -0.22 | 0.36 | 0.76 | 0.1 | -0.16 | 0.43 | -0.37 | -0.89 | -0.03 | 0.42 | -1.66 | -0.1 | -0.76 | 0.01 | 0.17 | -0.33 | -0.74 | 0.10622 | 0.5858 | 0.362901 | 1 | 0.999631 | 0.568797 |
| Syngap1_F6SEU4_S1201s_1 | 0.34 | 1.2 | 1.97 | 1.51 | 0.51 | 0.52 | 0.63 | 1.86 | 1.09 | 0.83 | 1.78 | 2.69 | 0.96 | 1.65 | 2.45 | 2.44 | -0.375 | 0.3425 | 0.62 | 0.217219 | 0.419925 | 0.506208 | 1 | 0.999631 | 0.669523 |
| Syngap1_F6SEU4_S165s_1 | 1.76 | -1.47 | -1.23 | 1.51 | -0.05 | -2.13 | 0.78 | -1.09 | -0.39 | 0.12 | 0.13 | -0.61 | -2.13 | -2.71 | -0.45 | -1.27 | -0.765 | -0.33 | -1.7825 | 0.06148 | 0.416783 | 0.672862 | 1 | 0.999631 | 0.509355 |
| Syngap1_F6SEU4_S534s_1 | -0.15 | 1.41 | 0.39 | -0.75 | 0.8 | 0.98 | -0.19 | 0.31 | 0.82 | -0.42 | 0.05 | 0.37 | 1.47 | 0.45 | -0.01 | 0.03 | 0.25 | -0.02 | 0.26 | 0.630227 | 0.616759 | 0.968099 | 1 | 0.999631 | 0.867378 |
| Syngap1_F6SEU4_S535s_1 | -0.68 | 0.26 | 0.32 | -0.47 | 0.42 | 0.39 | -0.44 | -0.1 | 0.49 | 0.55 | -0.55 | -0.5 | 1.02 | 0.91 | -0.17 | 0.59 | 0.21 | 0.14 | 0.73 | 0.089367 | 0.556909 | 0.730044 | 1 | 0.999631 | 0.555695 |
| Syngap1_F6SEU4_S752s_1 | -1.85 | -1.28 | -1.72 | -0.23 | 0.28 | -1.59 | -0.41 | -0.69 | -0.96 | -2.06 | -0.92 | -0.52 | -1.82 | -1.67 | -1.17 | -0.81 | 0.6675 | 0.155 | -0.0975 | 0.824227 | 0.202908 | 0.745058 | 1 | 0.999631 | 0.939586 |
| Syngap1_F6SEU4_S766s_1 | -0.61 | 0.06 | -0.65 | 1.08 | 1.2 | -0.23 | 0.78 | 1.54 | 0.04 | -1.54 | 0.78 | 0.64 | -0.8 | -0.16 | 0.02 | 0.78 | 0.8525 | 0.01 | -0.01 | 0.983949 | 0.124517 | 0.986845 | 1 | 0.999631 | 0.995133 |
| Syngap1_F6SEU4_S779s_1 | 1.45 | -0.57 | -1.28 | 0.83 | 0.95 | -0.07 | 0.56 | 0.42 | -0.14 | -1.1 | -0.12 | -0.82 | -2.19 | -0.31 | -1.14 | 0.12 | 0.3575 | -0.6525 | -0.9875 | 0.18756 | 0.553307 | 0.300475 | 1 | 0.999631 | 0.646474 |
| Syngap1_F6SEU4_S780s_1 | 0.58 | -0.66 | -1.1 | 0.7 | 1.04 | 0 | 0.71 | 0.53 | -0.07 | -0.61 | 0.12 | -0.56 | -2.23 | -0.52 | -1.04 | 0.21 | 0.69 | -0.16 | -0.775 | 0.228307 | 0.168245 | 0.733042 | 1 | 0.999631 | 0.679038 |
| Syngap1_F6SEU4_S824s_1 | 0.67 | -1.14 | -1.32 | -0.78 | 0.32 | -0.6 | -0.39 | -0.99 | -1 | -1.85 | -1.09 | -1.08 | -2.39 | -1.41 | -1.64 | -0.79 | 0.2275 | -0.6125 | -0.915 | 0.106913 | 0.648448 | 0.217514 | 1 | 0.999631 | 0.568797 |
| Syngap1_F6SEU4_S87s_1 | 1.56 | 1.62 | 1.09 | 2.1 | 1.89 | 2.09 | 1.42 | 2.21 | 1.26 | 0.48 | 1.6 | 0.45 | -0.73 | 1.85 | 0.67 | 1.5 | 0.31 | -0.645 | -0.77 | 0.192769 | 0.343027 | 0.112225 | 1 | 0.999631 | 0.649499 |
| Syngap1_F6SEU4_S892s_1 | 0.26 | -1.3 | 0.3 | 0.11 | 0.03 | -0.67 | -0.19 | 0.42 | -1.23 | 0.98 | 0.46 | 0.44 | 0.2 | -0.33 | 0.58 | 0.25 | 0.055 | 0.32 | 0.3325 | 0.449967 | 0.899392 | 0.575068 | 1 | 0.999631 | 0.787269 |
| Syngap1_F6SEU4_S895s_1 | -3.16 | -0.77 | 0.56 | -0.33 | -1.5 | -1.47 | 0.65 | 0.68 | -0.87 | -2.39 | 0.48 | 1.21 | 0.55 | -2.9 | -0.42 | -0.72 | 0.515 | 0.5325 | 0.0525 | 0.954781 | 0.559435 | 0.586354 | 1 | 0.999631 | 0.984441 |
| Syngap1_F6SEU4_S985s_1 | -1.16 | -3.48 | -4.37 | -2.99 | -1.48 | -2.83 | -1.38 | -3.26 | -3.03 | -2.4 | -3.24 | -3.96 | -4.01 | -2.38 | -3.46 | -2.98 | 0.7625 | -0.1575 | -0.2075 | 0.759812 | 0.309187 | 0.814466 | 1 | 0.999631 | 0.914614 |
| Syngap1_F6SEU4_Y100y_1 | -0.43 | -0.96 | 0.95 | -0.24 | -0.51 | -1.14 | 2.74 | -0.26 | 0.03 | -0.89 | 0.43 | 2.27 | 0.38 | -2.47 | -0.89 | -1.22 | 0.3775 | 0.63 | -0.88 | 0.190022 | 0.650952 | 0.373496 | 1 | 0.999631 | 0.648674 |
| Syngap1_F6SEU4_Y34y_1 | 0.77 | 0.25 | -0.47 | 0.08 | 0.22 | 0 | 0.09 | 0.29 | -0.17 | 0.73 | -0.49 | -0.68 | -0.3 | 0.34 | -0.56 | 0.34 | -0.0075 | -0.31 | -0.2025 | 0.596215 | 0.980996 | 0.45753 | 1 | 0.999631 | 0.854508 |
| Syngap1_F6SEU4_Y39y_1 | -0.12 | -0.32 | -1.8 | -1.63 | -0.46 | -0.96 | -1.2 | -0.69 | -0.84 | -1.16 | -1.58 | -1.01 | -0.29 | -0.81 | -1.48 | -0.83 | 0.14 | -0.18 | 0.115 | 0.811981 | 0.755361 | 0.69211 | 1 | 0.999631 | 0.936949 |
| Syngap1_J3QQ18_S1298s_1 | 0.28 | 0.63 | -0.49 | 0.31 | 0.82 | 0.53 | 0.48 | 0.35 | -0.27 | -0.11 | -0.71 | 0.3 | -0.08 | 0.66 | -0.64 | -0.05 | 0.3625 | -0.38 | -0.21 | 0.59126 | 0.261667 | 0.297177 | 1 | 0.999631 | 0.852343 |
| Syngr3_Q8R191_S5s_1 | 1.1 | -1.44 | 0.4 | 0.07 | -0.84 | -1.04 | 1.2 | -1.01 | -0.13 | 0.42 | -0.95 | 0.56 | -0.93 | -0.76 | 0.62 | -0.03 | -0.455 | -0.0575 | -0.3075 | 0.603986 | 0.506769 | 0.921303 | 1 | 0.999631 | 0.858163 |
| Syngr3_Q8R191_T133t_1 | -0.42 | -0.07 | -0.01 | -0.74 | -0.44 | 0.46 | -0.66 | -1.14 | -0.98 | 0.13 | -0.53 | -1.45 | -0.23 | -0.15 | -2.24 | -0.49 | -0.135 | -0.3975 | -0.4675 | 0.360081 | 0.727324 | 0.324569 | 1 | 0.999631 | 0.741182 |
| Synj1_F6XIX0_S44s_1 | 0.62 | -0.9 | -0.85 | 0.92 | 1.39 | 0.26 | 0.32 | 1.06 | -0.5 | -1.09 | 0.07 | -0.32 | -1.15 | -0.32 | 0.14 | -0.21 | 0.81 | -0.4075 | -0.3325 | 0.527618 | 0.140234 | 0.429432 | 1 | 0.999631 | 0.824554 |
| Synj1_F7BQW7_S1064s_1 | 0.75 | 0.37 | -0.48 | 0.76 | 1.36 | 0.4 | 0.59 | 0.58 | 0.43 | -0.31 | 0.49 | -0.34 | -1.4 | 0.59 | 0.23 | 0.22 | 0.3825 | -0.2825 | -0.44 | 0.39469 | 0.322846 | 0.472335 | 1 | 0.999631 | 0.759502 |
| Synj1_F7BQW7_S1068s_1 | 0.02 | -0.18 | -0.55 | 0.21 | 0.52 | -0.09 | -0.25 | 0.06 | 0.4 | -0.43 | -0.04 | -0.09 | -2.16 | 0.27 | -0.05 | -0.11 | 0.185 | 0.085 | -0.3875 | 0.4796 | 0.538038 | 0.781622 | 1 | 0.999631 | 0.801333 |
| Synj1_F7BQW7_S1099s_1 | -0.37 | -0.32 | -0.11 | -0.25 | -0.78 | -0.62 | -0.55 | -0.68 | -0.65 | -0.06 | -0.64 | -0.24 | -0.02 | -0.13 | -0.27 | -0.12 | -0.395 | -0.135 | 0.1275 | 0.628079 | 0.116243 | 0.618347 | 1 | 0.999631 | 0.865998 |
| Synj1_F7BQW7_S1162s_1 | 0.78 | -0.26 | -2.59 | 0.58 | 3.45 | 0.47 | 2.24 | 0.93 | 1.24 | -3.04 | 1.85 | -0.63 | -3.7 | -0.56 | -0.73 | -1.9 | 2.145 | 0.2275 | -1.35 | 0.161028 | 0.034548 | 0.843552 | 1 | 0.999631 | 0.631211 |
| Synj1_F7BQW7_S1175s_1 | 0.11 | -2.34 | -2.86 | -1.19 | -1.21 | -1.8 | -0.59 | -2.48 | -2.1 | -1.73 | -3.26 | -4.36 | -2.75 | -2.4 | -4.39 | -2.64 | 0.05 | -1.2925 | -1.475 | 0.060224 | 0.941867 | 0.12322 | 1 | 0.999631 | 0.505962 |
| Synj1_F7BQW7_S1190s_1 | 0.51 | 0.24 | 0.44 | 0.55 | 1.33 | 0.03 | 0.12 | 0.5 | 0.59 | 0.22 | 0.35 | 0.55 | -0.79 | 0.78 | 0.74 | 0.99 | 0.06 | -0.0075 | -0.005 | 0.990634 | 0.859783 | 0.976267 | 1 | 0.999631 | 0.996087 |
| Synj1_F7BQW7_S11s_1 | 0.86 | -0.33 | -1 | 0.37 | -0.25 | -1.07 | 1.3 | -2.43 | -3.03 | -0.74 | -0.08 | 0.65 | -0.29 | -0.54 | 0.3 | 0.09 | -0.5875 | -0.775 | -0.085 | 0.849483 | 0.452836 | 0.336515 | 1 | 0.999631 | 0.949525 |
| Synj1_F7BQW7_S1308s_1 | -1.1 | -0.35 | -2.45 | -0.96 | -0.03 | 0.1 | -0.97 | -2.61 | -0.38 | -0.17 | -3.42 | -0.31 | 0.28 | -0.62 | -0.23 | -0.18 | 0.3375 | 0.145 | 1.0275 | 0.05028 | 0.621695 | 0.854644 | 1 | 0.999631 | 0.488571 |
| Synj1_F7BQW7_T1063t_1 | 0.33 | 0.59 | -0.39 | 0.92 | 1.2 | 1.08 | 0.88 | 0.51 | 0.46 | 0.13 | 0.27 | -0.79 | -0.39 | 0.82 | 0.42 | 0.72 | 0.555 | -0.345 | 0.03 | 0.941587 | 0.134326 | 0.402064 | 1 | 0.999631 | 0.9813 |
| Synm_Q70IV5_S1075s_1 | 0.45 | 0.18 | -1.19 | -0.63 | 1.15 | 1.04 | -1.35 | -0.16 | 0.57 | 1.22 | -0.15 | 1.11 | -0.22 | 0.76 | -0.91 | -1.28 | 0.4675 | 0.985 | -0.115 | 0.833073 | 0.461644 | 0.060272 | 1 | 0.999631 | 0.943363 |
| Synm_Q70IV5_S172s_1 | 0.44 | -0.51 | -0.64 | 0.24 | 0.63 | 0.05 | -0.22 | -0.24 | 0.16 | 1.57 | 0.45 | 1.29 | 0.39 | 0.38 | -0.36 | 0.03 | 0.1725 | 0.985 | 0.2275 | 0.539263 | 0.633905 | 0.042417 | 1 | 0.999631 | 0.830899 |
| Synpo_Q8CC35_S19s_1 | -1.16 | -0.52 | -0.87 | -0.96 | -0.9 | -1.09 | -0.96 | -1.26 | -1.23 | -1.67 | -1.03 | -1 | -0.42 | -1.2 | -1.13 | -0.57 | -0.175 | -0.355 | 0.0475 | 0.882146 | 0.50659 | 0.239602 | 1 | 0.999631 | 0.963505 |
| Synpo_Q8CC35_S328s_1 | -0.43 | 0.01 | -0.82 | -1.1 | -0.3 | 0 | -1.05 | -0.96 | -0.55 | -0.19 | -1.39 | -1.06 | 0.37 | 0.07 | -0.81 | -0.84 | 0.0075 | -0.2125 | 0.2825 | 0.495977 | 0.9838 | 0.580022 | 1 | 0.999631 | 0.811036 |
| Synpo_Q8CC35_S331s_1 | -1.33 | -2.86 | -2.35 | -1.31 | -2 | -2.81 | -1.05 | -1.54 | -3.16 | -3.08 | -1.45 | -1.03 | -0.82 | -2.19 | -3.16 | -1.27 | 0.1125 | -0.2175 | 0.1025 | 0.862268 | 0.823037 | 0.722077 | 1 | 0.999631 | 0.954601 |
| Synpo_Q8CC35_S433s_1 | -0.77 | -0.65 | -0.3 | -0.57 | -0.53 | -0.03 | -0.71 | -0.12 | -0.53 | -0.23 | -0.49 | -0.7 | 0.14 | 0.01 | -0.18 | 0.23 | 0.225 | 0.085 | 0.6225 | 0.045037 | 0.423388 | 0.746092 | 1 | 0.999631 | 0.475465 |
| Synpo_Q8CC35_S501s_1 | -0.48 | 0.22 | -0.03 | -0.15 | 0.24 | 0.14 | -0.3 | -0.23 | 0.05 | -0.4 | -0.05 | -0.13 | 0.42 | 0.5 | 0.1 | 0.16 | 0.0725 | -0.0225 | 0.405 | 0.186435 | 0.795515 | 0.935055 | 1 | 0.999631 | 0.646474 |
| Synpo_Q8CC35_S566s_1 | -0.09 | -1.38 | -0.45 | -0.36 | -0.3 | 0.35 | -0.54 | -0.73 | -0.52 | -0.69 | -2.08 | -0.97 | -1.5 | 0.69 | 0.19 | -1.19 | 0.265 | -0.495 | 0.1175 | 0.832609 | 0.490979 | 0.280008 | 1 | 0.999631 | 0.94321 |
| Synpo_Q8CC35_S594s_1 | -0.42 | -1.82 | -2 | -1.28 | -0.45 | -1.86 | -1.2 | -1.3 | -2.06 | -2.11 | -2.02 | -1.14 | -1.68 | -1.69 | -0.81 | -1.37 | 0.1775 | -0.4525 | -0.0075 | 0.985824 | 0.690647 | 0.301593 | 1 | 0.999631 | 0.995133 |
| Synpo_Q8CC35_S615s_1 | 1.15 | -0.92 | -1.1 | 1 | 1.44 | 0.53 | 0.9 | 0.99 | -0.47 | -0.89 | 0.22 | -0.6 | -1.66 | 0.1 | -0.24 | 0.01 | 0.9325 | -0.4675 | -0.48 | 0.469456 | 0.129489 | 0.437239 | 1 | 0.999631 | 0.796837 |
| Synpo_Q8CC35_S643s_1 | -1.12 | -0.59 | -1.43 | -0.59 | -0.83 | -1.44 | -1.26 | -0.94 | -1.54 | -1.64 | -0.76 | -1.49 | -0.52 | -1.22 | -1.82 | 0.1 | -0.185 | -0.425 | 0.0675 | 0.88338 | 0.551583 | 0.226385 | 1 | 0.999631 | 0.963505 |
| Synpo_Q8CC35_S654s_1 | 0.05 | -0.18 | -2 | -0.49 | 0.81 | -0.68 | -0.32 | -0.48 | -1.23 | -1.5 | -1.06 | -0.7 | -1.33 | -0.68 | -1.02 | -0.82 | 0.4875 | -0.4675 | -0.3075 | 0.519366 | 0.366446 | 0.336176 | 1 | 0.999631 | 0.819456 |
| Synpo_Q8CC35_S819s_1 | 0.24 | -0.73 | -0.91 | 0.38 | 3.87 | 0.44 | -0.35 | -0.79 | 1.91 | 4.13 | 3.2 | 4.45 | -1.55 | 1.12 | 0.51 | 0.56 | 1.0475 | 3.6775 | 0.415 | 0.501978 | 0.287983 | 0.000138 | 1 | 0.472282 | 0.814092 |
| Synpo_Q8CC35_S892s_1 | 0.46 | 0.8 | -0.04 | 0.68 | 1.57 | 1.11 | -0.22 | -0.4 | 1.44 | 2.42 | 1.16 | 2.41 | 0.5 | -0.05 | 0.36 | 0.8 | 0.04 | 1.3825 | -0.0725 | 0.826203 | 0.93499 | 0.005441 | 1 | 0.999631 | 0.940686 |
| Synpo_Q8CC35_T308t_1 | 0.68 | -0.31 | -0.44 | -0.22 | -0.28 | 0.18 | -1.19 | -0.83 | -1.04 | 0.43 | -1.26 | -1.17 | -0.91 | 0.59 | -0.35 | 0.37 | -0.4575 | -0.6875 | -0.0025 | 0.995399 | 0.268643 | 0.159011 | 1 | 0.999631 | 0.997705 |
| Synpo_Q8CC35_T503t_1 | -0.57 | -0.99 | 0.69 | -0.26 | -0.67 | -0.9 | 0.17 | 1.01 | -1.07 | -0.7 | -0.22 | 0.97 | -1.11 | -0.48 | -1.07 | 1.85 | 0.185 | 0.0275 | 0.08 | 0.908514 | 0.723786 | 0.958731 | 1 | 0.999631 | 0.969941 |
| Synpo_Q8CC35_T530t_1 | -0.11 | -0.26 | -0.01 | -0.41 | -0.49 | -0.37 | -0.83 | 0.61 | -1.31 | 0.19 | -1.73 | -0.13 | -0.72 | -0.21 | 0.56 | 0.69 | -0.0725 | -0.5475 | 0.2775 | 0.471468 | 0.836113 | 0.24886 | 1 | 0.999631 | 0.798132 |
| Synrg_V9GXD9_S743s_1 | -0.51 | 0.75 | 1.16 | 1 | 0.85 | 0.27 | 0.03 | 0.72 | 0.6 | 0.1 | 0.64 | 1.23 | 0.89 | 0.76 | 1.27 | 1.13 | -0.1325 | 0.0425 | 0.4125 | 0.330895 | 0.753243 | 0.922947 | 1 | 0.999631 | 0.727302 |
| Synrg_V9GXD9_S803s_1 | -0.39 | 0.63 | -0.52 | 0.24 | 1.11 | 0.74 | 0.1 | 1.17 | 0.6 | -0.03 | 1.06 | 0.81 | 0.42 | 0.61 | 0.64 | 0.35 | 0.79 | 0.62 | 0.515 | 0.155381 | 0.060378 | 0.126777 | 1 | 0.999631 | 0.626444 |
| Synrg_V9GXD9_S920s_1 | -0.24 | -1.36 | -4.19 | -2.86 | -2.14 | -2.32 | -1.39 | -3.67 | -1.67 | -1.11 | -3.23 | -3.83 | -1.33 | -2.03 | -3.61 | -3.1 | -0.2175 | -0.2975 | -0.355 | 0.683916 | 0.799502 | 0.750018 | 1 | 0.999631 | 0.889602 |
| Synrg_V9GXD9_S989s_1 | 0.2 | 1.25 | 1.19 | 1.49 | 1.01 | 0.96 | 0.99 | 0.98 | 1.29 | 0.73 | 1.67 | 1.45 | -0.39 | 1.31 | 2.16 | 1.7 | -0.0475 | 0.2525 | 0.1625 | 0.777769 | 0.88479 | 0.5047 | 1 | 0.999631 | 0.921382 |
| Syp_Q62277_Y81y_1 | -0.54 | 2 | 1.59 | -0.25 | 2.86 | 0.95 | 1.78 | 0.77 | 3.49 | -1.38 | 1.73 | 2.75 | 4.86 | 0.15 | 0.57 | -0.21 | 0.89 | 0.9475 | 0.6425 | 0.576502 | 0.226652 | 0.387606 | 1 | 0.999631 | 0.846204 |
| Syt1_P46096_S264s_1 | -4.3 | -5.14 | 3.03 | 2.13 | -2.62 | -4.32 | 1.86 | 2 | -3.01 | -6.19 | 2.52 | 2.81 | -1.89 | -3.9 | 2.11 | 1.85 | 0.3 | 0.1025 | 0.6125 | 0.775967 | 0.893319 | 0.968225 | 1 | 0.999631 | 0.920998 |
| Syt1_P46096_T128t_1 | -0.75 | 4.86 | 4.27 | 1.5 | 3.61 | 3.86 | 2.62 | 2.83 | 4.18 | 0.41 | 4.34 | 4.81 | 4.54 | 2.73 | 3.74 | 2.4 | 0.76 | 0.965 | 0.8825 | 0.459429 | 0.509942 | 0.497646 | 1 | 0.999631 | 0.79152 |
| Syt1_P46096_T201t_1 | -0.83 | -0.93 | -1.06 | -0.98 | -1.12 | -0.97 | -1.1 | -1.04 | -1.07 | -0.81 | -1.1 | -0.8 | -0.39 | -0.74 | -0.69 | -1.02 | -0.1075 | 0.005 | 0.24 | 0.397342 | 0.643404 | 0.983874 | 1 | 0.999631 | 0.761181 |
| Syt1_P46096_T285t_1 | -0.15 | -0.28 | -0.55 | -0.66 | -0.75 | -0.44 | -0.52 | -0.96 | -0.42 | 0.09 | -1.14 | -0.27 | -0.48 | -0.83 | -1.01 | -0.95 | -0.2575 | -0.025 | -0.4075 | 0.179685 | 0.343316 | 0.940226 | 1 | 0.999631 | 0.64376 |

| | | | | | | | | | | | | | | | | | | | | | | | | | |
|---|---|---|---|---|---|---|---|---|---|---|---|---|---|---|---|---|---|---|---|---|---|---|---|---|---|
| Syt1_P46096_T334t_1 | -1.33 | -0.72 | -0.76 | -1.16 | -0.53 | -0.82 | -1.17 | -0.85 | -0.49 | -0.8 | -1.66 | -0.58 | -0.96 | -0.88 | -0.83 | 0.23 | 0.15 | 0.11 | 0.3825 | 0.308485 | 0.595139 | 0.751954 | 1 | 0.999631 | 0.718534 |
| Syt10_Q9R0N4_S169s_1 | 2.28 | -2.45 | -0.67 | 0.18 | -0.24 | -1.51 | 0.72 | -0.74 | 0.01 | 1.16 | -2.53 | 0 | -0.59 | -2.13 | 0.7 | -1.07 | -0.2775 | -0.175 | -0.6075 | 0.539071 | 0.767901 | 0.871038 | 1 | 0.999631 | 0.830899 |
| Syt10_Q9R0N4_S218s_1 | 1.82 | -1.27 | 1.02 | 0.47 | -0.54 | 0.15 | 2.07 | 0.18 | -0.69 | 1.18 | -0.71 | -0.59 | 0.14 | -0.7 | 0.12 | -0.99 | -0.045 | -0.7125 | -0.8675 | 0.197863 | 0.952516 | 0.329913 | 1 | 0.999631 | 0.655274 |
| Syt11_Q9R0N3_S113s_1 | -0.55 | -1.23 | -1.08 | -0.93 | -0.86 | -1.15 | -0.85 | -0.91 | -1.01 | -0.92 | -0.99 | -0.76 | -0.24 | -1.04 | -0.76 | -0.64 | 0.005 | 0.0275 | 0.2775 | 0.384132 | 0.984825 | 0.91844 | 1 | 0.999631 | 0.754721 |
| Syt11_Q9R0N3_S133s_1 | -1.3 | -0.41 | -0.41 | -0.64 | -0.24 | -0.66 | -0.84 | -0.55 | -0.01 | -1.7 | -0.84 | -0.2 | 0.48 | 0 | -0.24 | -0.08 | 0.1175 | 0.0025 | 0.73 | 0.049481 | 0.700653 | 0.995394 | 1 | 0.999631 | 0.487521 |
| Syt11_Q9R0N3_S416s_1 | 2.45 | 1.5 | 2.73 | 1.7 | 0.33 | 1.97 | 2.44 | 1.72 | 1.65 | 2.91 | 1.3 | 1.13 | 1.38 | 0.77 | 2 | 1.23 | -0.48 | -0.3475 | -0.75 | 0.090442 | 0.353956 | 0.474258 | 1 | 0.999631 | 0.557085 |
| Syt11_Q9R0N3_T139t_1 | 1.52 | -0.78 | -0.82 | -0.2 | -0.52 | -0.89 | -0.29 | 0 | -0.64 | 0.27 | -0.68 | -0.54 | -0.26 | 0.54 | 0.18 | 0.61 | -0.355 | -0.3275 | 0.3375 | 0.539394 | 0.512007 | 0.554787 | 1 | 0.999631 | 0.830899 |
| Syt14_B7ZP31_S195s_2 | 0.86 | -1.78 | -1.05 | 0.15 | -0.87 | -1.4 | 0.65 | 0.05 | -2.07 | -0.94 | -1.07 | -0.72 | -2.1 | -2.7 | -1.01 | -0.52 | 0.0625 | -0.745 | -1.1275 | 0.125097 | 0.92524 | 0.235795 | 1 | 0.999631 | 0.596063 |
| Syt14_B7ZP31_S197s_2 | 0.86 | -1.78 | -1.05 | 0.15 | -0.87 | -1.4 | 0.65 | 0.05 | -2.07 | -0.94 | -1.07 | -0.72 | -2.1 | -2.7 | -1.01 | -0.52 | 0.0625 | -0.745 | -1.1275 | 0.125097 | 0.92524 | 0.235795 | 1 | 0.999631 | 0.596063 |
| Syt17_Q920M7_S100s_1 | 0.8 | 0.91 | -0.36 | 0.69 | 1.09 | 1.42 | 0.53 | 1.21 | 1.45 | 0.65 | 0.7 | 0.85 | 0.46 | 1.19 | 0.77 | 0.83 | 0.5525 | 0.4025 | 0.3025 | 0.423351 | 0.157277 | 0.294988 | 1 | 0.999631 | 0.77354 |
| Syt17_Q920M7_S114s_1 | 0.63 | -0.78 | -1.01 | -0.32 | 1.23 | 0.84 | -0.53 | 0.04 | 0.5 | 0 | -1.22 | -1.06 | -0.78 | 0.36 | -0.63 | -0.37 | 0.765 | -0.075 | 0.015 | 0.973025 | 0.151549 | 0.8849 | 1 | 0.999631 | 0.990551 |
| Syt17_Q920M7_S115s_1 | 0.63 | -0.78 | -1.01 | -0.32 | 1.23 | 0.84 | -0.53 | 0.04 | 0.5 | 0 | -1.22 | -1.06 | -0.78 | 0.36 | -0.63 | -0.37 | 0.765 | -0.075 | 0.015 | 0.973025 | 0.151549 | 0.8849 | 1 | 0.999631 | 0.990551 |
| Syt17_Q920M7_S84s_1 | 1.93 | -3.94 | -3.41 | -2.73 | -2.45 | -1.98 | 1.22 | -5.1 | -6.45 | 1.2 | -3.68 | -3.1 | -1.83 | -3.16 | -3.56 | -3.32 | -0.04 | -0.97 | -0.93 | 0.440137 | 0.97981 | 0.58432 | 1 | 0.999631 | 0.781983 |
| Syt3_G3X9Y1_S290s_1 | 1.8 | 0.24 | 0.74 | 0.22 | -0.95 | 0.35 | 0.78 | -0.93 | -0.44 | 1.41 | -0.79 | -0.65 | -0.37 | -1.24 | -0.16 | -0.67 | -0.9375 | -0.8675 | -1.36 | 0.010923 | 0.103779 | 0.158549 | 1 | 0.999631 | 0.385092 |
| Syt3_G3X9Y1_S586s_1 | -0.56 | -2.13 | -2.93 | -4.27 | -2 | -3.78 | -2.45 | -3.53 | -3.22 | -2.11 | -1.34 | -5.16 | -2.91 | -1.49 | -4.44 | -2.64 | -0.4675 | -0.485 | -0.3975 | 0.642862 | 0.54998 | 0.623322 | 1 | 0.999631 | 0.871518 |
| Syt4_P40749_S135s_1 | 0.73 | -0.19 | -1 | -0.89 | 0.17 | 0.09 | -0.82 | -0.92 | -0.54 | 0.9 | -0.84 | -0.24 | 0.94 | 0.64 | -0.29 | -0.35 | -0.0325 | 0.1575 | 0.5725 | 0.263872 | 0.944801 | 0.761371 | 1 | 0.999631 | 0.693392 |
| Syt5_Q9R0N5_S95s_1 | 0.08 | 0.36 | 0.9 | 0.35 | 0.72 | 1.01 | 0.48 | 0.39 | 0.81 | 0.6 | -0.15 | -0.05 | 0.41 | 0.42 | 0.28 | 1.04 | 0.2275 | -0.12 | 0.115 | 0.722347 | 0.442301 | 0.724122 | 1 | 0.999631 | 0.905674 |
| Syt7_Q9R0N7_S107s_1 | 0.52 | 0.84 | 0.76 | 0.18 | 0.45 | 0.88 | 0.15 | 0.68 | 0.15 | 0.66 | 0.7 | -0.14 | 0.35 | 0.68 | 0.48 | 0.63 | -0.035 | -0.2325 | -0.04 | 0.889665 | 0.903466 | 0.467293 | 1 | 0.999631 | 0.963505 |
| Syt7_Q9R0N7_S129s_1 | -1.28 | -0.71 | -1.21 | -1.02 | -0.89 | -1.26 | -1.14 | -0.92 | -0.98 | -0.82 | -1.32 | -2.25 | -0.25 | -0.45 | -0.72 | -1.44 | 0.0025 | -0.2875 | 0.34 | 0.340877 | 0.992329 | 0.446012 | 1 | 0.999631 | 0.733032 |
| Syt7_Q9R0N7_S172s_1 | -1.18 | -1.7 | -0.5 | 0.5 | -1.03 | -1.37 | -0.29 | 0.28 | -2.13 | 0.07 | -0.55 | 0.25 | 0.19 | -0.99 | 0.27 | 0.11 | 0.1175 | 0.13 | 0.615 | 0.260269 | 0.831639 | 0.841022 | 1 | 0.999631 | 0.692624 |
| Syt7_Q9R0N7_S52s_1 | -0.38 | -0.62 | 1.81 | 0.1 | -1.11 | -0.77 | -0.23 | 1.2 | -0.77 | -0.53 | 1.43 | -0.61 | 0.18 | -0.91 | 1.76 | 1.78 | -0.455 | -0.3475 | 0.475 | 0.530928 | 0.49978 | 0.609285 | 1 | 0.999631 | 0.826822 |
| Syt7_Q9R0N7_S61s_1 | -1.29 | -3.21 | -1.59 | -1.61 | -2.08 | -2.52 | -1.48 | -1.85 | -2.94 | -2.27 | -1.63 | -1.49 | -1.32 | -2.4 | -1.39 | -1.8 | -0.0575 | -0.1575 | 0.1975 | 0.684591 | 0.9015 | 0.760138 | 1 | 0.999631 | 0.889888 |
| Syt7_Q9R0N7_S61s_2 | -3.54 | -3.22 | -2.71 | -1.85 | -2.41 | -2.92 | -2.08 | -2.07 | -2.36 | -2.26 | -2.14 | -1.93 | -1.24 | -2.28 | -1.45 | -1.82 | 0.46 | 0.6575 | 1.1325 | 0.025429 | 0.284994 | 0.120653 | 1 | 0.999631 | 0.422114 |
| Syt7_Q9R0N7_T253t_1 | -0.23 | 0.66 | -0.39 | 0.2 | 0.63 | 0.61 | 0.14 | 0.15 | 0.63 | 0.15 | -0.08 | -0.24 | -0.11 | 0.56 | -0.01 | 0.24 | 0.3225 | 0.055 | 0.11 | 0.748455 | 0.327185 | 0.873239 | 1 | 0.999631 | 0.911931 |
| Syt7_Q9R0N7_T262t_1 | 0.72 | -0.12 | -0.76 | -1.12 | -0.85 | 0.6 | -0.69 | -0.96 | -0.56 | -0.03 | -1.29 | -1.12 | -0.13 | 0.4 | -0.6 | -1.45 | -0.155 | -0.43 | -0.125 | 0.813449 | 0.760188 | 0.377 | 1 | 0.999631 | 0.937041 |
| Syt7_Q9R0N7_T58t_1 | -0.72 | -1.36 | -0.71 | -1.22 | -1.49 | -0.87 | -1 | -1 | -1.06 | 0.08 | -0.32 | -0.79 | -0.38 | -0.58 | -0.57 | -0.05 | -0.0875 | 0.48 | 0.6075 | 0.071059 | 0.762728 | 0.186346 | 1 | 0.999631 | 0.531307 |
| Syt7_Q9R0N7_T58t_2 | -3.54 | -3.22 | -2.71 | -1.85 | -2.41 | -2.92 | -2.08 | -2.07 | -2.36 | -2.26 | -2.14 | -1.93 | -1.24 | -2.28 | -1.45 | -1.82 | 0.46 | 0.6575 | 1.1325 | 0.025429 | 0.284994 | 0.120653 | 1 | 0.999631 | 0.422114 |
| Syt9_Q9R0N9_S207s_1 | 0.08 | -0.29 | 1.16 | 0.66 | -0.92 | -0.43 | 1.12 | -0.35 | 0.16 | 1.12 | -0.29 | -0.24 | 1.3 | -1.55 | -0.35 | -0.89 | -0.5475 | -0.215 | -0.775 | 0.231386 | 0.295877 | 0.633711 | 1 | 0.999631 | 0.681565 |
| Syt9_Q9R0N9_T159t_1 | 0.93 | -1.2 | 0.32 | 0.18 | -1.9 | -0.12 | 0.12 | 0 | -2.22 | 0.47 | -0.27 | -2.22 | -3 | -0.37 | -1.08 | -0.53 | -0.5325 | -1.1175 | -1.3025 | 0.076582 | 0.380005 | 0.148333 | 1 | 0.999631 | 0.542546 |
| Sytl5_Q80T23_S347s_1 | 0.98 | 0.96 | -0.35 | 0.5 | 0.9 | 0.58 | 0.47 | 0.53 | 0.64 | -0.52 | -0.15 | 0.49 | 0.83 | 0.96 | 0.45 | -0.74 | 0.0975 | -0.4075 | -0.1475 | 0.759925 | 0.782793 | 0.341817 | 1 | 0.999631 | 0.914614 |
| Syvn1_E9QQ99_S557s_1 | -0.11 | 0.77 | -1.21 | -0.58 | -1.71 | 0.1 | -0.58 | 0 | 0.37 | 0.91 | -0.56 | -0.58 | 0.14 | 1.2 | 0.24 | 0.57 | -0.265 | 0.3175 | 0.82 | 0.105839 | 0.626815 | 0.546067 | 1 | 0.999631 | 0.568797 |
| Szrd1_Q6NXN1_S18s_1 | -0.56 | -0.07 | 0.5 | -0.05 | 0.03 | -1.13 | -0.32 | -0.17 | -0.29 | -0.48 | 0.45 | 0.59 | 0.78 | -0.27 | 0.76 | -0.02 | -0.3525 | 0.1125 | 0.3575 | 0.359158 | 0.33808 | 0.761646 | 1 | 0.999631 | 0.741182 |
| Szrd1_Q6NXN1_S20s_1 | -1.02 | -1.29 | 0.03 | -0.28 | -1.09 | -1.67 | -0.62 | -0.53 | -1.33 | -1.61 | -0.13 | 0.14 | -0.3 | -0.88 | 0.51 | -0.3 | -0.3375 | 0.2925 | 0.3975 | 0.365353 | 0.414779 | 0.854299 | 1 | 0.999631 | 0.745574 |
| Szt2_A2A9C3_T2432t_1 | -0.81 | 0.29 | -0.21 | -0.73 | 0.21 | -1.16 | -0.71 | -1.51 | -0.45 | -0.59 | 0.09 | -0.26 | -0.16 | -0.44 | -0.18 | -0.48 | -0.4275 | 0.0625 | 0.05 | 0.882015 | 0.343019 | 0.854315 | 1 | 0.999631 | 0.963505 |
| Tab3_Q571K4_S103s_1 | -0.09 | -0.38 | -0.7 | -0.51 | -0.48 | 0.68 | -0.53 | -1 | -0.51 | -0.26 | -1.23 | -0.8 | 0.24 | 0.27 | 0.02 | -0.29 | 0.0875 | -0.28 | 0.48 | 0.127292 | 0.822055 | 0.378518 | 1 | 0.999631 | 0.59788 |
| Tacc1_F8VQ95_S119s_1 | -1.09 | -0.92 | -0.58 | -0.7 | -0.79 | -0.85 | -0.55 | -1.48 | -0.96 | 0.11 | -0.88 | -0.93 | -0.09 | -0.22 | -0.37 | -0.77 | -0.095 | 0.1575 | 0.46 | 0.144026 | 0.74833 | 0.638044 | 1 | 0.999631 | 0.616037 |
| Tacc1_F8VQ95_S403s_1 | -0.86 | 0.45 | 0.75 | 0.22 | -0.18 | -0.35 | 0.14 | 0.13 | -0.27 | -0.97 | 0.55 | 1.24 | 0.79 | -0.56 | 0.63 | -0.03 | -0.205 | -0.0025 | 0.0675 | 0.88389 | 0.595008 | 0.996379 | 1 | 0.999631 | 0.963505 |
| Tacc1_F8VQ95_S45s_1 | -1.34 | 0.24 | 0.7 | -0.1 | -0.48 | 0.19 | -1.27 | 0.52 | -0.38 | 0.14 | -1.31 | 1.26 | -0.48 | -0.03 | -0.76 | 0.36 | -0.135 | 0.0525 | -0.1025 | 0.832846 | 0.803634 | 0.933064 | 1 | 0.999631 | 0.94321 |
| Tacc2_E9QL08_S213s_1 | 0.24 | 0.5 | 0.61 | 0.16 | 0.26 | 0.22 | -0.09 | 0.85 | 0.18 | 0.63 | 0.63 | 0.93 | 0.47 | 0.82 | 0.81 | 1.2 | -0.0675 | 0.215 | 0.4475 | 0.152438 | 0.817645 | 0.45313 | 1 | 0.999631 | 0.625198 |
| Tacc2_E9QL08_S240s_1 | -0.68 | 0.33 | 0.33 | -0.47 | -0.14 | -1.45 | -0.29 | -2.72 | -0.7 | -0.05 | -1.04 | -0.89 | 0.15 | -0.04 | -1.37 | 0.83 | -1.0275 | -0.5475 | 0.015 | 0.976346 | 0.108294 | 0.162322 | 1 | 0.999631 | 0.991938 |
| Tacc2_E9QL08_S305s_1 | -0.54 | -0.5 | -0.22 | -0.66 | -0.7 | -0.78 | -0.47 | -0.94 | -0.46 | -0.18 | -0.6 | -0.31 | -0.21 | -0.62 | -0.48 | -0.86 | -0.2425 | 0.0925 | -0.0625 | 0.828207 | 0.348259 | 0.72142 | 1 | 0.999631 | 0.941315 |
| Tacc2_E9QL08_S396s_1 | -1.6 | 0.41 | 1.31 | -1.7 | -2.2 | -0.61 | -2.01 | -1.35 | -0.56 | 0.48 | -0.43 | 0.12 | 2.2 | 0 | 0.04 | 0.42 | -1.1475 | 0.2975 | 1.06 | 0.202486 | 0.140544 | 0.672131 | 1 | 0.999631 | 0.659062 |
| Tacc2_E9QL08_S600s_1 | -1.23 | 0.16 | -0.72 | -0.83 | -0.59 | -0.43 | -0.69 | -0.5 | -0.59 | -0.97 | 0.4 | 0.36 | 0.58 | -0.56 | -0.84 | -0.63 | 0.1025 | 0.455 | 0.2925 | 0.509194 | 0.760693 | 0.320054 | 1 | 0.999631 | 0.816932 |
| Tada3_Q8R0L9_S279s_1 | 2.39 | 0.02 | 1.65 | 1.28 | -1.21 | -0.21 | 1.78 | 0.23 | 0.03 | 1.52 | 0.43 | 0.96 | -0.62 | -1.24 | 0.73 | -0.4 | -1.1875 | -0.6 | -1.7175 | 0.014542 | 0.116117 | 0.289517 | 1 | 0.999631 | 0.385392 |
| Taf10_Q8K0H5_S44s_1 | -0.04 | 0.8 | 0.14 | -0.17 | -0.57 | -0.31 | -0.53 | 0.05 | -0.13 | 0.16 | 0.11 | -0.02 | 0.42 | 0.33 | 0.34 | 0.01 | -0.5225 | -0.1525 | 0.0925 | 0.771871 | 0.118465 | 0.614426 | 1 | 0.999631 | 0.919413 |
| Taf15_Q8BQ46_S373s_1 | 1.02 | 2.38 | 2.41 | 0.33 | -0.07 | 3.01 | 0.36 | 1.02 | 1.63 | 3.36 | 0.85 | -0.63 | 0.84 | 0.79 | 1.81 | 0.84 | -0.455 | -0.2325 | -0.465 | 0.395049 | 0.548244 | 0.785587 | 1 | 0.999631 | 0.759939 |
| Taf3_Q5HZG4_T56t_1 | -0.23 | -0.89 | 0.6 | -0.11 | -0.72 | -0.89 | -0.56 | -0.08 | -0.13 | -0.51 | -0.09 | -0.06 | 0.93 | 0.44 | -0.87 | 0.17 | -0.405 | -0.04 | 0.325 | 0.499256 | 0.288971 | 0.910871 | 1 | 0.999631 | 0.813279 |
| Taf6_Q62311_S653s_1 | -0.59 | 0.13 | -0.71 | 0.77 | -0.24 | -0.95 | -0.93 | 0.62 | -0.04 | -0.69 | 0.63 | 0.81 | 1.21 | 0.14 | 0.69 | 0.82 | -0.275 | 0.2775 | 0.815 | 0.077118 | 0.569457 | 0.557863 | 1 | 0.999631 | 0.542683 |
| Tagln3_Q9R1Q8_S163s_1 | 0.01 | 0.34 | -0.6 | 0.11 | -0.06 | 0.24 | -0.5 | -0.12 | -0.78 | -0.25 | -0.27 | -0.05 | -0.41 | -0.16 | -0.41 | 0.04 | -0.075 | -0.3025 | -0.2 | 0.530844 | 0.807919 | 0.35072 | 1 | 0.999631 | 0.826822 |
| Taldo1_Q93092_S227s_1 | -1.58 | 1.75 | 1.65 | 1.2 | 0.85 | 1.29 | 0.42 | 0.94 | 0.74 | 0.11 | 0.97 | 1.31 | 0.92 | 1.14 | 1.32 | 1.04 | 0.12 | 0.0275 | 0.35 | 0.620161 | 0.866329 | 0.970025 | 1 | 0.999631 | 0.861888 |
| Tanc1_E9QAF9_S211s_1 | -0.18 | 0.25 | -0.22 | -0.35 | 0.52 | 0 | -0.46 | -0.25 | -0.12 | -0.63 | -0.74 | -0.41 | -1.46 | 0.22 | 0.15 | 0.35 | 0.0775 | -0.35 | -0.06 | 0.893123 | 0.800399 | 0.234521 | 1 | 0.999631 | 0.964596 |
| Tanc1_E9QAF9_S438s_1 | -1.23 | -0.09 | -0.64 | -1.29 | 0.19 | 0.5 | -0.77 | -0.43 | -0.66 | 0.44 | -0.09 | 0.29 | -0.04 | 0.81 | -0.15 | 0.06 | 0.685 | 0.8075 | 1.0575 | 0.017723 | 0.115867 | 0.062095 | 1 | 0.999631 | 0.391024 |
| Tanc1_E9QAF9_S455s_1 | -0.73 | -0.02 | -0.44 | -0.21 | -0.1 | -0.83 | -0.44 | -1.1 | -0.53 | -0.83 | -0.09 | -0.85 | -0.03 | -0.12 | -0.41 | 0.1 | -0.2675 | -0.225 | 0.235 | 0.43646 | 0.408135 | 0.468024 | 1 | 0.999631 | 0.780748 |
| Tanc2_A2A690_S220s_1 | 0.84 | -0.44 | -0.77 | 0.82 | 0.44 | -0.02 | 0.63 | 0.81 | 0.09 | -0.55 | 0.31 | 0.25 | -0.76 | 0.08 | 1.05 | -0.41 | 0.3525 | -0.0875 | -0.1225 | 0.820048 | 0.433922 | 0.847053 | 1 | 0.999631 | 0.938303 |
| Tanc2_A2A690_S238s_1 | -0.29 | -0.58 | -0.15 | 1.29 | 0.59 | 0.88 | 0.49 | 1.19 | 0.48 | -1.07 | 0.61 | -0.4 | -0.66 | 0.23 | 0.56 | 0.65 | 0.72 | -0.1625 | 0.1275 | 0.796364 | 0.12307 | 0.76197 | 1 | 0.999631 | 0.929895 |
| Tanc2_A2A690_S254s_1 | -0.4 | 0.22 | 0.2 | 0.66 | 0.82 | 0.63 | -0.35 | 1.43 | 0.05 | -0.58 | 0.42 | 0.02 | -0.73 | 0.99 | -0.22 | 0.82 | 0.4625 | -0.1925 | 0.045 | 0.9222 | 0.289715 | 0.578309 | 1 | 0.999631 | 0.975032 |
| Tanc2_A2A690_S268s_1 | 0.01 | -0.37 | 0.46 | 0.15 | -0.45 | 0.87 | 0.51 | 0.03 | -0.67 | 0.26 | 0.01 | -0.41 | -0.88 | 0.99 | -0.21 | -0.42 | 0.1775 | -0.265 | -0.1925 | 0.662092 | 0.623199 | 0.423965 | 1 | 0.999631 | 0.881128 |
| Tanc2_A2A690_S400s_1 | -0.07 | -1.63 | -1.21 | -1.46 | -1.98 | -1.89 | -0.73 | -1.91 | -1.7 | -0.42 | -1.1 | -1.64 | -1.23 | -1.66 | -1.63 | -1.66 | -0.535 | -0.1225 | -0.4525 | 0.267488 | 0.249479 | 0.786273 | 1 | 0.999631 | 0.695932 |
| Tanc2_A2A690_T434t_1 | -0.56 | -0.43 | -0.53 | -0.54 | -1.16 | -0.89 | -0.74 | -0.16 | -1.12 | -0.47 | -0.89 | -0.92 | 0.38 | -0.45 | -0.69 | -0.21 | -0.2225 | -0.335 | 0.2725 | 0.399571 | 0.445976 | 0.220741 | 1 | 0.999631 | 0.762046 |
| Tanc2_E9Q2K2_S21s_1 | 0.28 | 0.1 | -0.25 | 0.83 | 0.77 | 0.36 | 1.96 | -0.03 | 0.32 | -0.96 | 0.35 | -0.7 | -1.43 | -2.02 | -1.81 | -1.57 | 0.525 | -0.4875 | -1.9475 | 0.000167 | 0.274872 | 0.2567 | 1 | 0.999631 | 0.351308 |
| Tanc2_E9Q2K2_S22s_1 | 0.63 | -2.6 | -1.36 | 0.01 | -1.71 | -4.08 | 0.52 | -1.08 | -2.02 | -1.37 | -1.1 | -2.63 | -1.99 | -3.98 | -1.39 | -2.24 | -0.7575 | -0.95 | -1.57 | 0.07188 | 0.4675 | 0.201911 | 1 | 0.999631 | 0.532005 |
| Tanc2_F6VP19_S1124s_1 | 1.38 | 0.72 | -0.28 | 0.09 | 0.32 | 1.67 | 0.46 | 0.11 | 1.57 | 1.69 | 0.06 | -0.11 | 0.47 | 0.88 | 0.17 | 0.14 | 0.1625 | 0.325 | -0.0625 | 0.881076 | 0.735381 | 0.562992 | 1 | 0.999631 | 0.963364 |
| Tanc2_F6VP19_S589s_1 | -1.93 | -1.61 | -1.37 | -2.61 | -1.95 | -2.25 | -1.56 | -2.79 | -1.41 | -3.15 | -1.93 | -1.14 | -0.88 | -3.29 | -2.42 | -3.09 | -0.2575 | -0.0275 | -0.54 | 0.349038 | 0.508419 | 0.95566 | 1 | 0.999631 | 0.736748 |
| Tanc2_F6VP19_S623s_1 | 0.62 | -0.13 | -0.36 | -0.8 | -0.22 | -0.22 | 1.23 | -1.44 | -0.1 | -1.56 | -0.11 | -0.66 | -0.76 | -2.17 | -1.31 | -2.04 | 0.005 | -0.44 | -1.4025 | 0.009723 | 0.992959 | 0.337124 | 1 | 0.999631 | 0.374514 |
| Tanc2_F6VP19_S664s_1 | -0.2 | 0.44 | 0.12 | -0.37 | -0.22 | 0.27 | -0.71 | 0.2 | -0.22 | 0.5 | 0.27 | 0.04 | 1.36 | 0.75 | -0.05 | 0.5 | -0.1125 | 0.15 | 0.6425 | 0.114558 | 0.73365 | 0.626901 | 1 | 0.999631 | 0.579001 |
| Tanc2_F6VP19_S677s_1 | -1.49 | -0.65 | -0.38 | -1.47 | -1.72 | -0.86 | -1.84 | -0.98 | -1.3 | -0.21 | -0.78 | -0.52 | 0.55 | -0.51 | -0.47 | -0.05 | -0.3525 | 0.295 | 0.8775 | 0.049611 | 0.374537 | 0.451067 | 1 | 0.999631 | 0.487521 |
| Tanc2_F6VP19_S677s_2 | 0.34 | 0.17 | -2.19 | -0.54 | 1.4 | 0.2 | -0.05 | -0.78 | -0.1 | -0.29 | 0.26 | -0.43 | -0.86 | 0.16 | 0 | -1.13 | 0.7475 | 0.415 | 0.0975 | 0.870932 | 0.270057 | 0.459125 | 1 | 0.999631 | 0.958735 |
| Tanc2_F6VP19_S692s_1 | 0.25 | -0.17 | -0.14 | -0.25 | -0.25 | 0.3 | -0.44 | -0.12 | -0.29 | 0.23 | -0.25 | 0.17 | -0.19 | 0.38 | -0.65 | 0.28 | -0.05 | 0.0425 | 0.0325 | 0.922218 | 0.85678 | 0.878078 | 1 | 0.999631 | 0.975032 |
| Tanc2_F6VP19_S971s_1 | -0.63 | -0.67 | -0.22 | -0.19 | -0.07 | -0.64 | -0.55 | -0.09 | 0.59 | -0.61 | 0.54 | 0.05 | -0.02 | -0.56 | 0.28 | 0.22 | 0.09 | 0.57 | 0.4075 | 0.217843 | 0.747968 | 0.125699 | 1 | 0.999631 | 0.669859 |
| Tanc2_F6VP19_S974s_1 | -2.14 | 1.75 | -0.24 | -0.67 | 0.85 | 1.28 | 0.53 | 1.24 | 1.01 | -1.24 | 1.42 | 2.06 | 2.14 | 1.34 | 1.97 | 1.1 | 1.3 | 1.1375 | 1.9625 | 0.02263 | 0.09771 | 0.241387 | 1 | 0.999631 | 0.408305 |
| Tanc2_F6VP19_T1123t_1 | 0.46 | -0.38 | -0.19 | 0.08 | 0.03 | 0.58 | 0.56 | -0.18 | 0.52 | 0.46 | 0.23 | -0.07 | 0.32 | 0.15 | 0.37 | 0.24 | 0.255 | 0.2925 | -0.2775 | 0.361899 | 0.426778 | 0.343704 | 1 | 0.999631 | 0.742006 |
| Taok1_Q5F2E8_S421s_1 | -0.14 | 0.03 | -0.01 | -0.45 | -0.29 | -0.17 | -0.42 | -0.31 | -0.2 | 0.26 | 0.16 | 0.02 | 0.49 | 0.24 | -0.24 | 0.12 | -0.155 | 0.2025 | 0.295 | 0.332724 | 0.534473 | 0.451964 | 1 | 0.999631 | 0.728079 |
| Taok1_Q5F2E8_S9s_1 | 0.03 | 1.21 | 0.38 | 0.77 | 0.93 | 1.52 | 0.1 | 0.9 | 1.08 | 0.45 | 0.89 | 1.35 | 1.37 | 1.26 | 0.95 | 1.07 | 0.265 | 0.345 | 0.565 | 0.118463 | 0.505976 | 0.341679 | 1 | 0.999631 | 0.585447 |
| Taok2_Q6ZQ29_S1018s_1 | 0.61 | -1.15 | -0.9 | -0.62 | 1.82 | -0.22 | -0.04 | -0.31 | 0.5 | -0.24 | -0.35 | 0.31 | -0.36 | 0.14 | -0.66 | -0.73 | 0.8275 | 0.57 | 0.1125 | 0.798722 | 0.176242 | 0.213195 | 1 | 0.999631 | 0.93091 |
| Taok2_Q6ZQ29_S493s_1 | 0.89 | 1.19 | 1.34 | 1.24 | 0.86 | 1.83 | 1.29 | 1.69 | 1.54 | 2.33 | 1.75 | 1.65 | -0.27 | 1.9 | 2.24 | 1.54 | 0.2525 | 0.6525 | 0.1875 | 0.725605 | 0.410315 | 0.044756 | 1 | 0.999631 | 0.90568 |
| Taok3_Q8BYC6_S324s_1 | -1.15 | 0.43 | 0.65 | 0.35 | 0.06 | -1.31 | 0.14 | 1.21 | -0.08 | 0.34 | 1.17 | 0.99 | 0.41 | -1.13 | 0.22 | 0.21 | -0.045 | 0.535 | -0.1425 | 0.782494 | 0.939882 | 0.283209 | 1 | 0.999631 | 0.922325 |
| Tardbp_H3BJV1_S35s_1 | 0.56 | 0.78 | -0.46 | 0.64 | 0.27 | 0.64 | -0.29 | -0.5 | 0.5 | 1.29 | -0.04 | -1.53 | 1.03 | 1.13 | -0.67 | 0.26 | -0.35 | -0.325 | 0.0575 | 0.906036 | 0.382735 | 0.590982 | 1 | 0.999631 | 0.969331 |
| Tbc1d1_Q60949_S660s_1 | 0.14 | 1.38 | 0.44 | 0.97 | 0.43 | -0.01 | 0.11 | 0.83 | 0.87 | 0.22 | 1.11 | 1.51 | 0.64 | 0.8 | 1.38 | 0.84 | -0.3925 | 0.195 | 0.1825 | 0.619532 | 0.28861 | 0.62705 | 1 | 0.999631 | 0.861888 |
| Tbc1d10a_Q5SPX8_S441s_1 | 0.43 | -3.72 | 0.46 | 2.18 | 0.17 | -1.21 | 1.72 | 0.6 | -0.23 | -1.09 | 0.33 | 1.56 | -1.04 | -1.02 | 0.99 | 0.79 | 0.4825 | 0.305 | 0.0925 | 0.936546 | 0.686018 | 0.795457 | 1 | 0.999631 | 0.97939 |
| Tbc1d10a_Q5SPX8_S52s_1 | 0.59 | 0.95 | 0.03 | 0.41 | 0.34 | 0.7 | 0.07 | 0.18 | 0.62 | 1.04 | 0.54 | 0.69 | 0.22 | 0.66 | 0.68 | 0.86 | -0.1725 | 0.2275 | 0.11 | 0.730939 | 0.567694 | 0.453009 | 1 | 0.999631 | 0.906897 |
| Tbc1d10b_Q8BHL3_S22s_1 | 0.64 | 3.16 | 2.81 | 0.14 | 3.71 | 1.3 | 2.38 | 0.97 | 5.14 | 0.03 | 3.7 | 4.28 | 5.56 | 1.2 | 1.77 | 0.15 | 0.4025 | 1.6 | 0.4825 | 0.68553 | 0.638033 | 0.190722 | 1 | 0.999631 | 0.890608 |
| Tbc1d10b_Q8BHL3_S644s_1 | 2.17 | 0.33 | -1.49 | 1.05 | 2.57 | 0.85 | 1.51 | 0.44 | 0.89 | -0.71 | -0.74 | -0.56 | 0.22 | 0.95 | -1.35 | -1.21 | 0.8275 | -0.795 | -0.8625 | 0.30983 | 0.306249 | 0.310087 | 1 | 0.999631 | 0.71896 |
| Tbc1d10b_Q8BHL3_S647s_1 | 1.88 | -0.75 | -1.84 | 1.63 | 2.48 | 0.81 | 1.33 | 1.18 | 0.85 | -0.53 | 0.78 | -0.09 | -2.28 | 0.55 | -0.39 | -0.32 | 1.22 | 0.0225 | -0.84 | 0.378746 | 0.173751 | 0.978794 | 1 | 0.999631 | 0.751637 |
| Tbc1d10b_Q8BHL3_S664s_1 | 1.3 | -0.45 | -1.1 | 0.4 | 1.38 | -0.27 | 0.72 | 0.41 | 0.32 | -0.8 | 0.01 | -0.3 | -0.99 | 0.01 | -0.69 | -0.59 | 0.5225 | -0.23 | -0.6025 | 0.27027 | 0.369003 | 0.668357 | 1 | 0.999631 | 0.698438 |
| Tbc1d10b_Q8BHL3_S673s_1 | 0.12 | -0.21 | -0.85 | 0.11 | -0.11 | -1.05 | -0.45 | -0.23 | -0.16 | -0.19 | -0.2 | -0.76 | -0.04 | 0.17 | -1.12 | -0.29 | -0.2525 | -0.12 | -0.1125 | 0.774389 | 0.46738 | 0.712954 | 1 | 0.999631 | 0.920998 |
| Tbc1d10b_Q8BHL3_S693s_1 | -0.94 | -0.69 | -1.95 | -2.11 | -0.65 | -1.23 | -0.5 | -1.01 | -0.54 | -2.01 | -0.98 | -0.73 | -1.24 | -1.18 | -1.41 | -2.47 | 0.575 | 0.3575 | -0.1525 | 0.74135 | 0.171141 | 0.452107 | 1 | 0.999631 | 0.909067 |
| Tbc1d10b_Q8BHL3_T136t_1 | 1.25 | 0.54 | -1.23 | -0.04 | 1.5 | 0.57 | -0.16 | 0.19 | 0.68 | 0.05 | -0.04 | -0.45 | -1.45 | 0.96 | -0.7 | -0.47 | 0.395 | -0.07 | -0.545 | 0.411906 | 0.49912 | 0.895916 | 1 | 0.999631 | 0.767937 |
| Tbc1d13_Q8R3D1_S184s_1 | 1.67 | 0.43 | 1.11 | 0.94 | 0.34 | 0.65 | 1.46 | 0.53 | 0.43 | 1.31 | 0.99 | 0.67 | -0.89 | 0.62 | 0.68 | 0.66 | -0.2925 | -0.1875 | -0.77 | 0.117093 | 0.439795 | 0.600305 | 1 | 0.999631 | 0.585087 |
| Tbc1d14_E9Q2V7_S296s_1 | 1.26 | -0.82 | 1.87 | 1.26 | 0.17 | 0.7 | 0.91 | 0.82 | 0.74 | 1.35 | -0.23 | 0.29 | 0.17 | 0.86 | 1.19 | 0.57 | -0.2425 | -0.355 | -0.195 | 0.735881 | 0.66603 | 0.566966 | 1 | 0.999631 | 0.90759 |
| Tbc1d14_E9Q2V7_S613s_1 | 0.53 | 0.06 | -0.19 | -0.94 | -1.01 | -1.26 | -0.45 | -1.83 | 0.1 | -0.13 | -0.96 | -0.42 | 0.38 | -3.1 | -1.46 | -1.15 | -1.0025 | -0.2175 | -1.1975 | 0.107074 | 0.035166 | 0.586454 | 1 | 0.999631 | 0.568797 |
| Tbc1d15_Q9CXF4_S205s_1 | -0.44 | 0.09 | 0.72 | 0.94 | -0.42 | -0.52 | 0.61 | 1.13 | -0.33 | -0.34 | 1.77 | 1.12 | -0.61 | 0.47 | 0.35 | 0.24 | -0.1275 | 0.2275 | -0.215 | 0.606024 | 0.791553 | 0.689462 | 1 | 0.999631 | 0.85817 |
| Tbc1d15_Q9CXF4_S655s_1 | 1.47 | 2.65 | 3.55 | 0.09 | 1.93 | 1.19 | 1.91 | 1.34 | 0.81 | 0.12 | 3.46 | 4.4 | 2.88 | -2.01 | 2.78 | 0.22 | -0.3475 | 0.2575 | -0.9725 | 0.416351 | 0.614728 | 0.813965 | 1 | 0.999631 | 0.77063 |
| Tbc1d15_Q9CXF4_S662s_1 | 0.28 | 0.09 | -1.11 | 0.93 | 0.11 | 0.05 | 1 | 1.2 | 0.01 | -0.22 | 0.95 | 0.21 | -0.29 | -1.49 | 0.32 | 0.77 | 0.5425 | 0.19 | -0.22 | 0.711465 | 0.283453 | 0.691185 | 1 | 0.999631 | 0.900631 |
| Tbc1d22a_Q8R5A6_S166s_1 | -0.75 | -1.77 | -1.63 | -1.73 | -2.32 | -1.79 | -0.96 | -2.43 | -2.35 | -1.42 | -2.36 | -2.26 | -2.04 | -2.02 | -3.08 | -1.46 | -0.405 | -0.6275 | -0.68 | 0.134002 | 0.33862 | 0.108793 | 1 | 0.999631 | 0.60492 |
| Tbc1d22b_Q80VE5_S141s_1 | 0.52 | 0.96 | 1.43 | 1.41 | 0.23 | 1.88 | 1.55 | 1.84 | 1.56 | 0.65 | 1.79 | 0.93 | 0.01 | 0.72 | 1.57 | -0.1 | 0.295 | 0.1525 | -0.53 | 0.249542 | 0.502531 | 0.681518 | 1 | 0.999631 | 0.686359 |
| Tbc1d22b_Q80VE5_S58s_1 | 0.92 | 1.37 | 0.53 | 1.61 | 1.51 | 1.6 | 1.16 | 1.46 | 1.68 | 0.9 | 1.53 | 1.31 | 0.12 | 2.19 | 1.47 | 1.44 | 0.325 | 0.2475 | 0.1975 | 0.68166 | 0.311326 | 0.472322 | 1 | 0.999631 | 0.88924 |
| Tbc1d22b_Q80VE5_T117t_1 | 0.57 | 1.36 | 0.52 | 0.81 | 0.87 | 1.05 | 0.67 | 1.14 | 1.27 | 0.69 | 1.36 | 0.52 | -0.51 | 1.87 | 0.57 | 0.94 | 0.1175 | 0.145 | -0.0975 | 0.846748 | 0.686682 | 0.665268 | 1 | 0.999631 | 0.948838 |
| Tbc1d23_Q8K0F1_S468s_1 | 1.14 | 1.6 | 1.32 | 1.56 | 0.64 | 1.16 | 0.9 | 0.85 | 1 | 1.27 | 1.1 | 0.42 | 1.23 | 1.3 | 0.94 | 0.85 | -0.5175 | -0.4575 | -0.325 | 0.268046 | 0.069423 | 0.144575 | 1 | 0.999631 | 0.696016 |
| Tbc1d23_Q8K0F1_S469s_2 | 0.23 | 0.49 | 1.7 | 1.19 | 0.45 | 0.64 | 0.77 | 0.84 | 1.55 | 0.87 | 1.2 | 2.33 | 0.15 | 0.7 | 1.49 | 2.4 | -0.2275 | 0.585 | 0.2825 | 0.610604 | 0.538066 | 0.213354 | 1 | 0.999631 | 0.859267 |
| Tbc1d25_S4R2S6_S378s_1 | 0.03 | 0.41 | 1.03 | 0.65 | 0.42 | 0.37 | 0.1 | 0.9 | 0.22 | 0.54 | 1.6 | 0.94 | 0.6 | 0.89 | 0.7 | 1.26 | -0.0825 | 0.295 | 0.3325 | 0.325873 | 0.795261 | 0.44961 | 1 | 0.999631 | 0.724678 |
| Tbc1d30_Q69ZT9_S624s_1 | 2.24 | -0.02 | 0.18 | 0.31 | -0.47 | -0.23 | 0.27 | -0.26 | 0.26 | 2.48 | -0.31 | -0.74 | -0.24 | 0.74 | -1.05 | -1.24 | -0.85 | -0.255 | -1.125 | 0.096631 | 0.121058 | 0.744577 | 1 | 0.999631 | 0.55886 |
| Tbc1d5_Q80XQ2_S546s_1 | 1.64 | 1.33 | 0.26 | 0.66 | -0.17 | 0.94 | 0.02 | 0.52 | 1.04 | 0.86 | 0.78 | -0.35 | 0.24 | 1.17 | 0.72 | 0.94 | -0.645 | -0.39 | -0.205 | 0.60791 | 0.134557 | 0.385435 | 1 | 0.999631 | 0.858527 |
| Tbc1d5_Q80XQ2_S565s_1 | -0.4 | -0.37 | -0.15 | -0.07 | -0.44 | -0.35 | -0.61 | -0.04 | -0.87 | 0 | -0.3 | -0.2 | -0.27 | 0.08 | -0.06 | -0.04 | -0.1125 | -0.095 | 0.195 | 0.476813 | 0.661674 | 0.743613 | 1 | 0.999631 | 0.799947 |
| Tbc1d5_Q80XQ2_S578s_1 | -1.76 | 0.42 | -0.59 | -1.92 | -0.37 | -0.38 | -1.24 | -1.64 | -0.07 | -1.08 | -0.83 | -1.04 | 0.63 | -0.67 | -1.37 | -1.79 | 0.055 | 0.2075 | 0.1625 | 0.810959 | 0.923923 | 0.707872 | 1 | 0.999631 | 0.936949 |
| Tbc1d7_Q9D0K0_S124s_1 | -2 | -0.04 | 0.07 | -0.91 | -1.2 | 0.06 | -1.64 | -1.89 | -1.09 | -2.18 | -2.1 | -1.12 | -0.11 | -1.16 | -0.29 | -1.13 | -0.4475 | -0.9025 | 0.0475 | 0.927475 | 0.452314 | 0.112833 | 1 | 0.999631 | 0.976306 |
| Tbc1d9_Q3UYK3_S238s_1 | -0.23 | -0.77 | -0.61 | -0.65 | -0.65 | -0.75 | -0.76 | -0.28 | -0.35 | 0.15 | -0.32 | -0.78 | 1.01 | -0.58 | 0.29 | 0.27 | -0.045 | 0.24 | 0.8125 | 0.054988 | 0.864242 | 0.431075 | 1 | 0.999631 | 0.492066 |
| Tbc1d9b_Q5SVR0_S464s_1 | -0.91 | -0.3 | -0.28 | -1.05 | 0.08 | -0.43 | -0.75 | -0.67 | -0.1 | -0.94 | -0.97 | -0.76 | 0.53 | -0.32 | -0.33 | -0.41 | 0.1925 | -0.0575 | 0.5025 | 0.177074 | 0.553183 | 0.86363 | 1 | 0.999631 | 0.642116 |
| Tbc1d9b_Q5SVR0_T1082t_1 | 0.51 | 1.71 | 1.61 | -0.7 | 2.8 | 1.85 | 1.8 | 2.46 | 3.54 | 0.23 | 1.77 | 2.29 | 2.89 | 2.33 | 1.98 | 0.68 | 1.445 | 1.175 | 1.1875 | 0.095106 | 0.026422 | 0.15584 | 1 | 0.999631 | 0.558128 |
| Tbcb_Q9D1E6_S110s_1 | -0.2 | -0.18 | -0.06 | -0.3 | -0.03 | 0.01 | -0.35 | 0.13 | -0.18 | 0.03 | -0.11 | 0.62 | -0.27 | 0.46 | 0.12 | 0.56 | 0.125 | 0.1475 | 0.4025 | 0.199997 | 0.611986 | 0.342783 | 1 | 0.999631 | 0.658081 |
| Tbcb_Q9D1E6_S173s_1 | 1.06 | 1.56 | 1.3 | 1.08 | 1 | 1.09 | 0.93 | 1.3 | 1.54 | 1.74 | 0.91 | 0.91 | 0.99 | 0.69 | 1.06 | 0.54 | -0.17 | 0.025 | -0.43 | 0.160304 | 0.509591 | 0.9359 | 1 | 0.999631 | 0.629764 |
| Tbcel_D3YVP4_S18s_1 | 0.21 | 0.82 | 0.87 | -0.04 | 0.26 | 0.26 | -0.42 | 0.77 | 0.73 | 0.56 | 0.61 | 0.79 | 1.36 | 1.16 | 0.97 | 0.77 | -0.2475 | 0.2075 | 0.6 | 0.094282 | 0.494153 | 0.50041 | 1 | 0.999631 | 0.558128 |
| Tbcel_D3YVP4_S41s_1 | -0.3 | -0.24 | -0.26 | 0.15 | 0.24 | 0.23 | 0.19 | 0.51 | 0.46 | -0.11 | 0.43 | 0.44 | 0.34 | -0.26 | -0.35 | 0.47 | 0.455 | 0.4675 | 0.2125 | 0.508779 | 0.093187 | 0.114998 | 1 | 0.999631 | 0.816932 |
| Tbck_E9Q1W7_S890s_1 | -0.53 | -0.22 | -0.28 | -0.21 | -0.34 | 0.04 | -0.63 | -0.17 | -0.23 | -0.21 | -0.63 | 0.43 | -0.15 | 0.16 | -0.17 | 0.57 | 0.035 | 0.15 | 0.4125 | 0.186821 | 0.89381 | 0.624038 | 1 | 0.999631 | 0.646474 |
| Tbr1_Q64336_S549s_1 | 1.04 | 1.32 | 0.59 | 0.49 | 1.28 | 0.98 | 0.4 | 0.51 | 1.07 | 1.51 | 1.02 | 0.46 | 0.77 | 0.25 | 0.21 | 0.51 | -0.0675 | 0.155 | -0.425 | 0.202361 | 0.836295 | 0.647621 | 1 | 0.999631 | 0.659062 |
| Tbr1_Q64336_S640s_1 | 0.13 | 1.26 | 1.17 | 0.33 | 0.63 | 0.41 | 0.26 | 1.57 | 0.66 | 0.4 | 1.08 | 1.86 | 1.03 | 0.92 | 1.69 | 2.19 | -0.005 | 0.2775 | 0.735 | 0.10794 | 0.990287 | 0.522605 | 1 | 0.999631 | 0.569416 |
| Tbr1_Q64336_S84s_1 | -0.27 | 0.46 | 0.9 | 0.32 | 0.24 | 0.01 | 0.44 | 0.74 | 0.23 | 0.18 | 0.92 | 0.95 | 0.53 | 0.54 | -0.01 | 0.44 | 0.005 | 0.2175 | 0.0225 | 0.947171 | 0.987862 | 0.545364 | 1 | 0.999631 | 0.982895 |
| Tcaf3_Q6QR59_Y458y_2 | -0.69 | -2.44 | -2.63 | -1.28 | -1.12 | -2.2 | -1.88 | -1.32 | -2.1 | -0.96 | -1.48 | -0.28 | -1.4 | -1.44 | -1.72 | -1.11 | 0.13 | 0.555 | 0.3425 | 0.473311 | 0.793271 | 0.333164 | 1 | 0.999631 | 0.799534 |
| Tcea1_E9PYD5_S111s_1 | 0.79 | 1.36 | 1.54 | 1.12 | 0.7 | 0.89 | 0.79 | 1.08 | 1.28 | 1.35 | 1.39 | 1.76 | 1.23 | 1.54 | 1.65 | 1.79 | -0.3375 | 0.2425 | 0.35 | 0.264612 | 0.233041 | 0.404705 | 1 | 0.999631 | 0.6935 |
| Tceal3_Q8R0A5_S120s_1 | 2.37 | 1.48 | 1.97 | 0.11 | -0.57 | 2.46 | 4.01 | -3.36 | -0.49 | 2.66 | -2.66 | -0.39 | 0.04 | -0.36 | -0.21 | -2.18 | -0.8475 | -1.7025 | -2.1425 | 0.006323 | 0.56234 | 0.124309 | 1 | 0.999631 | 0.351545 |
| Tceal3_Q8R0A5_S124s_1 | 0.8 | 2.07 | 2.33 | 1.47 | 0.33 | 2.83 | 1.6 | 0.45 | 1.51 | 2.22 | 1.05 | 1.34 | 0.32 | 0.57 | 0.83 | 1.48 | -0.365 | -0.1375 | -0.8675 | 0.067104 | 0.552919 | 0.746074 | 1 | 0.999631 | 0.520345 |
| Tceal5_Q8CCT4_S23s_1 | 1.94 | 0.95 | 2.45 | 2.1 | -0.26 | 1.14 | 3.3 | 0.77 | -0.54 | 1.46 | 1.51 | 1.26 | -0.57 | -0.53 | 1.83 | 0.52 | -0.6225 | -0.9375 | -1.5475 | 0.024638 | 0.39591 | 0.110249 | 1 | 0.999631 | 0.417291 |
| Tceal5_Q8CCT4_T24t_1 | 1.95 | 0.8 | 2.45 | 2.1 | -0.3 | 1.07 | 3.3 | 0.77 | -0.73 | 1.54 | 1.51 | 1.26 | -0.65 | -0.7 | 1.83 | 0.52 | -0.615 | -0.93 | -1.575 | 0.029193 | 0.412113 | 0.142089 | 1 | 0.999631 | 0.430938 |
| Tceb3_Q8C877_S125s_1 | 1.62 | 0.24 | 1.17 | 0.3 | -0.05 | 0.51 | 1.35 | 0.49 | 0.51 | 1.7 | 0.39 | 0.03 | 0.03 | -0.53 | 0.39 | -0.23 | -0.2575 | -0.175 | -0.9175 | 0.045659 | 0.557102 | 0.714212 | 1 | 0.999631 | 0.477252 |
| Tceb3_Q8C877_S195s_1 | 0.22 | 1.47 | 1.4 | 1.66 | 1.15 | 1.91 | 0.57 | 1.04 | 1.6 | 1.12 | 1.7 | 1.82 | 0.77 | 1.87 | 1.84 | 1.5 | -0.02 | 0.3725 | 0.3075 | 0.47611 | 0.962299 | 0.342173 | 1 | 0.999631 | 0.799882 |
| Tcf12_Q61286_T313t_1 | -1.32 | 0.53 | -0.25 | -0.24 | -0.29 | -0.02 | -0.98 | -0.12 | -0.11 | 0.08 | -0.67 | -0.36 | 0.23 | -0.41 | -0.46 | -0.8 | -0.0325 | 0.055 | -0.04 | 0.927539 | 0.939636 | 0.895207 | 1 | 0.999631 | 0.976306 |
| Tcf20_Q9EPQ8_S261s_1 | -0.52 | 0.13 | -0.15 | -0.61 | -0.76 | 0.23 | -0.1 | -0.51 | -0.69 | -0.43 | -0.28 | -0.58 | 0.15 | -0.19 | -1.11 | 0.16 | 0.0025 | -0.2075 | 0.04 | 0.916218 | 0.99383 | 0.472429 | 1 | 0.999631 | 0.973495 |
| Tcf20_Q9EPQ8_S587s_1 | 0.63 | -0.28 | -0.61 | -1.41 | 0.21 | 0.33 | -0.28 | -0.44 | 0.32 | 1.02 | -0.71 | -1.4 | 0.77 | 0.08 | -0.83 | -0.49 | 0.3725 | 0.225 | 0.3 | 0.56647 | 0.413193 | 0.716976 | 1 | 0.999631 | 0.84194 |
| Tcf20_Q9EPQ8_S588s_1 | -0.22 | -1.06 | -0.67 | -1.24 | -1.3 | -2.01 | -0.61 | -0.97 | -1.05 | -0.51 | -0.28 | -0.24 | -1.05 | -1.49 | -1.67 | -0.79 | -0.4525 | -0.2775 | -0.4525 | 0.222236 | 0.285321 | 0.421632 | 1 | 0.999631 | 0.672826 |
| Tcf20_Q9EPQ8_S603s_1 | 0.24 | 0.01 | -0.04 | -0.05 | -0.42 | -0.07 | -0.18 | -0.08 | 0.09 | 0.69 | -0.41 | -0.47 | 0.97 | 0.18 | -0.34 | -0.25 | -0.2275 | -0.065 | 0.1 | 0.780263 | 0.359281 | 0.843924 | 1 | 0.999631 | 0.921416 |
| Tcf20_Q9EPQ8_S669s_1 | -0.51 | -0.92 | -0.06 | -0.68 | -1.27 | -1.33 | -0.6 | -0.71 | -0.66 | 0.11 | -0.67 | -0.62 | 0.4 | -0.49 | -0.32 | -0.39 | -0.435 | 0.0825 | 0.3425 | 0.32467 | 0.187134 | 0.797674 | 1 | 0.999631 | 0.72463 |
| Tcf20_Q9EPQ8_T1699t_1 | -0.42 | -1.17 | -0.21 | -0.48 | -1.13 | -0.94 | -0.75 | -0.45 | -1.32 | -0.44 | -0.17 | -0.52 | -0.65 | -0.37 | -0.37 | -0.17 | -0.2475 | -0.0425 | 0.18 | 0.573164 | 0.431596 | 0.905422 | 1 | 0.999631 | 0.844516 |
| Tcf4_Q60722_S212s_1 | -0.13 | 0.41 | 0.08 | -1 | -0.25 | 0.09 | -0.26 | -0.52 | 0.31 | 0.09 | -0.53 | -0.97 | 0.85 | 0.25 | 0.18 | -0.32 | -0.075 | -0.115 | 0.4 | 0.336603 | 0.832051 | 0.78554 | 1 | 0.999631 | 0.730499 |
| Tcf4_Q60722_S555s_1 | -0.5 | 3.4 | 3.99 | 1.61 | 1.71 | 3.6 | 2.18 | 1.7 | 2.74 | 1.08 | 3.01 | 2.58 | 1.95 | 1.44 | 2.18 | 2.27 | 0.1725 | 0.2275 | -0.165 | 0.852402 | 0.856454 | 0.811343 | 1 | 0.999631 | 0.950222 |
| Tcof1_H3BL37_S1252s_1 | 1.64 | 0.67 | 0.38 | 0.25 | -0.37 | 0.58 | 0.38 | -0.81 | 0.5 | 1.5 | 0.04 | -0.57 | -0.37 | 0 | -0.13 | -0.92 | -0.79 | -0.3675 | -1.09 | 0.019728 | 0.097326 | 0.47558 | 1 | 0.999631 | 0.391668 |
| Tcof1_H3BL37_S593s_1 | -0.26 | 0.66 | 1.37 | 1.38 | 0.48 | -0.47 | 0.34 | 0.46 | 0.31 | 1.12 | 0.82 | 1.2 | 1.46 | 1.16 | 1.11 | 0.77 | -0.585 | 0.075 | 0.3375 | 0.433654 | 0.20221 | 0.862924 | 1 | 0.999631 | 0.779567 |
| Tcof1_H3BL37_S855s_1 | 2.97 | 1.22 | 2.53 | 2.4 | 0.26 | 1.73 | 2.03 | 1.33 | 1.21 | 2.38 | 1.25 | 1.07 | 1.55 | 0.75 | 1.86 | 0.82 | -0.9425 | -0.8025 | -1.035 | 0.044527 | 0.085988 | 0.110342 | 1 | 0.999631 | 0.475443 |
| Tcof1_H3BL37_S879s_1 | 0.71 | 1.72 | 1.22 | 1.04 | 0.25 | 1.12 | 0.32 | 0.88 | 1.31 | 1.27 | 1.08 | 1 | 1.24 | 1.61 | 1.12 | 1.35 | -0.53 | -0.0075 | 0.1575 | 0.623941 | 0.139345 | 0.980036 | 1 | 0.999631 | 0.863524 |
| Tcp11l1_Q8BTG3_S44s_1 | 1.58 | -1.26 | 0.37 | 0.13 | -1.27 | -0.32 | 2.77 | -1.18 | -0.98 | 1.03 | -0.17 | -0.59 | 1.09 | -1.42 | -0.64 | -1.16 | -0.205 | -0.3825 | -0.7375 | 0.316846 | 0.830806 | 0.561712 | 1 | 0.999631 | 0.720778 |
| Tcp11l1_Q8BTG3_S55s_1 | 1.38 | 0.8 | 0.83 | 1.12 | 0.49 | 0.84 | 1.01 | 0.69 | 1.44 | 1.76 | 0.18 | 0.34 | 0.11 | -0.22 | 0.17 | -0.36 | -0.275 | -0.1025 | -1.1075 | 0.003722 | 0.320203 | 0.807373 | 1 | 0.999631 | 0.351308 |
| Tcp11l2_Q8K1H7_S16s_1 | 1.39 | -2.49 | 1.71 | 1.47 | -0.03 | 0.24 | 1.8 | 0.07 | 0.62 | 1.73 | -0.74 | 0.99 | -1.52 | -2.47 | 0.17 | -1.41 | 1.96E-16 | -0.3 | -2.0525 | 0.066995 | 1 | 0.8942 | 1 | 0.999631 | 0.520345 |
| Tdrd3_Q91W18_S349s_1 | 0.8 | 1.02 | 0.33 | 0.67 | 0.93 | 0.51 | 1.36 | 0.33 | 0.33 | 0.7 | 0.7 | 0.49 | 0.61 | 0.24 | 1.33 | 1.22 | 1.61 | 0.0775 | -0.08 | 0.395 | 0.303022 | 0.808623 | 0.764972 | 1 | 0.999631 | 0.714978 |
| Tecpr1_Q80VP0_S272s_1 | 0.12 | -0.05 | -0.48 | -0.01 | 1.36 | 0.85 | -0.69 | -0.54 | 0.61 | 0.11 | -0.56 | 0.71 | 0.25 | 1.23 | -0.07 | 0.16 | 0.35 | 0.3225 | 0.4975 | 0.19126 | 0.483983 | 0.373776 | 1 | 0.999631 | 0.648674 |
| Tecpr1_Q80VP0_S303s_1 | -3.32 | -5.28 | -2.86 | -4.05 | -2.89 | -3.76 | -2.85 | -3.83 | -4.53 | -2.81 | -2.93 | -4.75 | -3.84 | -3.59 | -3.16 | -3.92 | 0.545 | 0.1225 | 0.25 | 0.634791 | 0.329561 | 0.852918 | 1 | 0.999631 | 0.868886 |
| Tenm1_Q9WTS4_S145s_1 | -0.3 | -0.98 | -0.1 | -0.18 | -0.97 | -0.6 | -0.38 | -0.36 | -0.87 | 0.67 | -0.83 | 0.22 | 0.13 | 0.77 | 1.11 | 0.56 | -0.1875 | 0.1875 | 1.0325 | 0.012956 | 0.54262 | 0.666078 | 1 | 0.999631 | 0.385092 |
| Tenm1_Q9WTS4_S59s_1 | 1.94 | 0.03 | -0.06 | 2.22 | 1.25 | 0.93 | 3.05 | 0.94 | 1.41 | 0.16 | 1.78 | 0.61 | -2.95 | -2.17 | -0.08 | -0.66 | 0.51 | -0.0425 | -2.4975 | 0.009314 | 0.472945 | 0.947076 | 1 | 0.999631 | 0.367426 |
| Tenm2_Q9WTS5_S41s_1 | 0.88 | 0.26 | -0.71 | 0.32 | 0.78 | 0.4 | 0.23 | 0.64 | 0.63 | -0.35 | -0.16 | 0.03 | -1.44 | 0.26 | -0.22 | 0.59 | 0.325 | -0.15 | -0.39 | 0.462151 | 0.390025 | 0.7108 | 1 | 0.999631 | 0.792786 |
| Tenm2_Q9WTS5_S42s_1 | -0.32 | -0.57 | -0.82 | -0.06 | -0.49 | -1.98 | 0.33 | -0.39 | -0.43 | -1.28 | 0 | -0.19 | -0.83 | -0.79 | -0.88 | -0.01 | -0.19 | -0.0325 | -0.185 | 0.582386 | 0.695517 | 0.927953 | 1 | 0.999631 | 0.847651 |
| Tenm4_F6UP10_S21s_1 | 0.41 | -0.43 | 0.55 | -0.92 | -1.37 | -0.22 | 1.09 | -0.99 | -0.44 | 0.46 | -0.57 | -0.92 | -0.26 | -2.04 | -0.79 | -2.43 | -0.275 | -0.27 | -1.2825 | 0.045471 | 0.64049 | 0.5508 | 1 | 0.999631 | 0.477252 |
| Tenm4_Q3UHK6_T2737t_2 | -1.31 | -1.86 | -1.56 | -1.53 | -2.6 | -1.33 | -0.63 | -0.72 | -2.99 | 0.16 | -0.92 | -0.44 | -1.23 | -0.78 | -0.42 | -0.13 | 0.245 | 0.5175 | 0.925 | 0.018357 | 0.590263 | 0.417737 | 1 | 0.999631 | 0.391024 |
| Tenm4_Q3UHK6_T2738t_2 | -1.31 | -1.86 | -1.56 | -1.53 | -2.6 | -1.33 | -0.63 | -0.72 | -2.99 | 0.16 | -0.92 | -0.44 | -1.23 | -0.78 | -0.42 | -0.13 | 0.245 | 0.5175 | 0.925 | 0.018357 | 0.590263 | 0.417737 | 1 | 0.999631 | 0.391024 |
| Tenm4_Q3UHK6_Y2735y_2 | -1.31 | -1.86 | -1.56 | -1.53 | -2.6 | -1.33 | -0.63 | -0.72 | -2.99 | 0.16 | -0.92 | -0.44 | -1.23 | -0.78 | -0.42 | -0.13 | 0.245 | 0.5175 | 0.925 | 0.018357 | 0.590263 | 0.417737 | 1 | 0.999631 | 0.391024 |
| Terf2_D3YZ08_S398s_1 | 1.63 | 1.52 | 1.26 | 1.06 | 0.45 | 0.94 | 0.37 | 0.62 | 1.4 | 1.59 | 1 | 1.83 | 1.45 | 1.26 | 1.75 | 1.3 | -0.7725 | 0.0875 | 0.0725 | 0.802727 | 0.016511 | 0.768485 | 1 | 0.999631 | 0.932746 |
| Terf2ip_Q91VL8_S43s_1 | -1.2 | 1.4 | 0.75 | -0.29 | -1.15 | -0.43 | -0.67 | -0.17 | -0.08 | 0.11 | -0.19 | 0.43 | 2.12 | 0.5 | 0.21 | 1.03 | -0.77 | -0.0975 | 0.8 | 0.229642 | 0.189088 | 0.858366 | 1 | 0.999631 | 0.680148 |
| Tex2_B1ATR2_S165s_1 | -2.01 | -0.13 | -0.37 | -1.91 | -0.47 | -0.16 | -0.41 | -1.91 | 0.43 | -2.32 | 0.33 | -0.29 | -1.16 | -0.91 | -1.03 | -1.67 | 0.3675 | 0.6425 | -0.0875 | 0.861391 | 0.529095 | 0.381431 | 1 | 0.999631 | 0.954096 |
| Tex2_B1ATR2_S221s_1 | 2.14 | 1.13 | -0.08 | 1.99 | 1.25 | 1.75 | 2.06 | 0.99 | 2.27 | 1.1 | 1.62 | 0.92 | -1.5 | 0.25 | -0.19 | 0.4 | 0.2175 | 0.1825 | -1.555 | 0.026578 | 0.678641 | 0.740342 | 1 | 0.999631 | 0.422285 |
| Tex2_B1ATR2_S265s_1 | 0.56 | 0.95 | -1.06 | 0.03 | 0.56 | 0.08 | -0.06 | 0.52 | 0.25 | -0.62 | -0.01 | 0.43 | -1.05 | 0.88 | -0.57 | -1.02 | 0.155 | -0.1075 | -0.56 | 0.344613 | 0.730079 | 0.820946 | 1 | 0.999631 | 0.73415 |
| Tex2_B1ATR2_S265s_2 | 0.64 | 0.01 | -1.2 | 0.95 | 0.09 | 0.08 | 0.86 | 1.62 | -0.61 | -0.6 | 1.54 | 0.33 | -2.29 | -0.46 | 0.1 | 0.76 | 0.5625 | 0.065 | -0.5725 | 0.431282 | 0.321052 | 0.91767 | 1 | 0.999631 | 0.778271 |
| Tex2_B1ATR2_S269s_1 | -1.59 | -0.43 | -0.88 | -0.66 | -1.05 | -0.42 | -1.44 | -0.68 | -0.61 | -0.37 | -1 | -0.47 | -0.71 | -0.47 | -0.41 | -0.29 | -0.0075 | 0.2775 | 0.42 | 0.228669 | 0.983329 | 0.416176 | 1 | 0.999631 | 0.679038 |
| Tex2_B1ATR2_S269s_2 | 0.64 | 0.01 | -1.2 | 0.95 | 0.09 | 0.08 | 0.86 | 1.62 | -0.61 | -0.6 | 1.54 | 0.33 | -2.29 | -0.46 | 0.1 | 0.76 | 0.5625 | 0.065 | -0.5725 | 0.431282 | 0.321052 | 0.91767 | 1 | 0.999631 | 0.778271 |
| Tex2_B1ATR2_S294s_1 | 1.18 | -0.71 | -1.44 | 1.61 | 2.71 | -0.87 | 1.08 | 0.27 | 0.6 | -0.88 | 0.15 | -0.06 | -0.9 | -0.16 | -0.29 | -0.99 | 0.6375 | -0.2075 | -0.745 | 0.288131 | 0.488809 | 0.769846 | 1 | 0.999631 | 0.706035 |
| Tex2_B1ATR2_S791s_1 | 0.96 | 0.42 | 2.3 | 0.6 | 0.43 | 0.86 | 1.08 | 0.85 | 1.46 | 1.85 | 0.5 | 0.26 | 0.45 | -0.14 | 0.61 | 0.15 | -0.265 | -0.0525 | -0.8025 | 0.101228 | 0.54702 | 0.921365 | 1 | 0.999631 | 0.564448 |
| Tex2_B1ATR2_S795s_1 | 0.07 | -1.23 | -0.01 | -1.36 | -0.94 | -0.47 | -1.45 | -1.14 | -0.65 | -0.62 | -0.97 | 0.33 | 0.33 | -0.72 | -0.73 | -1.2 | -0.3675 | 0.155 | 0.0525 | 0.913833 | 0.399831 | 0.737846 | 1 | 0.999631 | 0.972551 |
| Tex2_B1ATR2_S803s_1 | -1.55 | -1.79 | -1.17 | -2.18 | -2.21 | -2.07 | -1.81 | -1.9 | -2.1 | -0.95 | -1.86 | -1.02 | 0.09 | -1.56 | -1.98 | -1.71 | -0.325 | 0.19 | 0.3825 | 0.445883 | 0.28777 | 0.62094 | 1 | 0.999631 | 0.784402 |
| Tex264_E9Q137_S200s_1 | 0.53 | -1.02 | 0.83 | 0.82 | 0 | 0.02 | 0.84 | 0.94 | 0.2 | 0.13 | 0.54 | 1.04 | 0.61 | 0.3 | 0.62 | 0.68 | 0.16 | 0.1875 | 0.2625 | 0.562569 | 0.740912 | 0.692004 | 1 | 0.999631 | 0.841038 |
| Tex264_E9Q137_S205s_1 | -0.25 | -0.24 | -0.6 | 0 | -0.26 | -0.88 | -0.07 | -1.3 | 0.2 | -0.51 | -0.46 | 0.26 | 0.2 | -0.21 | -0.88 | -0.47 | -0.355 | 0.145 | -0.0675 | 0.838518 | 0.31379 | 0.640981 | 1 | 0.999631 | 0.946365 |
| Tfeb_Q3UKG7_S180s_1 | 1.21 | -0.66 | 0.72 | 0.33 | -0.7 | 0.39 | 0.37 | 0.05 | 0.94 | 1.25 | -1.24 | 0.96 | 0.33 | 1.03 | 1.23 | 0.68 | -0.3725 | 0.0775 | 0.4175 | 0.358838 | 0.420364 | 0.902386 | 1 | 0.999631 | 0.741182 |
| Tgfb1i1_E9Q1D5_S155s_1 | 0.45 | 0.2 | 0.24 | 0.28 | 0.75 | 0.02 | -0.05 | -0.23 | -0.15 | 1.31 | 0.2 | 1.19 | -0.43 | 0.07 | 0.3 | -0.02 | -0.17 | 0.345 | -0.3125 | 0.290712 | 0.564187 | 0.380773 | 1 | 0.999631 | 0.706164 |
| Tgfbrap1_Q3UR70_S853s_1 | 2.23 | 5.35 | 2.05 | 0.68 | 7.3 | 2.5 | 3.24 | 0.94 | 8.38 | 1.52 | 2.07 | 2.8 | 6.76 | 2.99 | 0.38 | -1.25 | 0.9175 | 1.115 | -0.3575 | 0.831469 | 0.523776 | 0.487427 | 1 | 0.999631 | 0.942694 |
| Tgfbrap1_Q3UR70_S853s_2 | 1.69 | 1.85 | 0.97 | 0.14 | 0.89 | 1.24 | 1.81 | -0.37 | 1.78 | 1.43 | 1.42 | 1.49 | 1.56 | -0.54 | 1.02 | 1.1 | -0.27 | 0.3675 | -0.3775 | 0.503575 | 0.628156 | 0.378661 | 1 | 0.999631 | 0.814092 |

| | | | | | | | | | | | | | | | | | | | | | | | | | |
|---|---|---|---|---|---|---|---|---|---|---|---|---|---|---|---|---|---|---|---|---|---|---|---|---|---|
| Tgfbrap1_Q3UR70_S855s_2 | 1.69 | 1.85 | 0.97 | 0.14 | 0.89 | 1.24 | 1.81 | -0.37 | 1.78 | 1.43 | 1.42 | 1.49 | 1.56 | -0.54 | 1.02 | 1.1 | -0.27 | 0.3675 | -0.3775 | 0.503575 | 0.628156 | 0.378661 | 1 | 0.999631 | 0.814092 |
| Th_P24529_S19s_1 | 2.3 | -1.44 | -0.13 | 1.72 | -0.95 | -0.59 | 2.25 | 0.13 | -1.32 | 0.18 | -2.02 | 0.76 | -0.67 | -1.06 | 1.2 | -1.38 | -0.4025 | -1.2125 | -1.09 | 0.239786 | 0.677337 | 0.213063 | 1 | 0.999631 | 0.683719 |
| Th_P24529_S31s_1 | 1.18 | -0.77 | -0.92 | -0.01 | -0.29 | -1.3 | -0.34 | 0.27 | -0.4 | 0.29 | -0.84 | -1.39 | -0.29 | -1.08 | -0.61 | -1.84 | -0.285 | -0.455 | -0.825 | 0.154008 | 0.597073 | 0.418836 | 1 | 0.999631 | 0.625198 |
| Th_P24529_S40s_1 | 0.31 | -0.35 | 0.06 | 0.91 | -0.3 | -0.32 | 0.51 | 1.82 | -0.37 | -0.02 | -0.16 | 0.88 | 0.15 | -0.48 | 1.05 | -0.37 | 0.195 | -0.15 | -0.145 | 0.742632 | 0.711632 | 0.705514 | 1 | 0.999631 | 0.90972 |
| Thada_A8C756_S78s_1 | 0.07 | -1.01 | -1.04 | -3.09 | -0.96 | -1.16 | -1.63 | -1.57 | -0.72 | 0.13 | -1.09 | -0.26 | 0.39 | -0.59 | -1.69 | -0.85 | -0.0625 | 0.7825 | 0.5825 | 0.412483 | 0.918615 | 0.240883 | 1 | 0.999631 | 0.768114 |
| Thoc1_Q8R3N6_S560s_1 | 1.23 | 0.09 | -2.57 | -2.04 | -0.67 | 0.06 | -1.19 | -3.86 | -0.03 | 1.22 | -3.27 | -4.01 | -0.34 | -1.35 | -3.09 | -3.86 | -0.5925 | -0.7 | -1.3375 | 0.211622 | 0.579331 | 0.597588 | 1 | 0.999631 | 0.665474 |
| Thoc2_B1AZI6_S1417s_1 | 0.07 | 0.32 | 0.31 | 0.12 | -0.45 | 0.23 | 0.24 | -0.03 | 0.18 | 0.78 | -0.1 | 0.12 | 0.2 | 0.47 | 0.32 | 0.55 | -0.2075 | 0.04 | 0.18 | 0.505631 | 0.446642 | 0.889345 | 1 | 0.999631 | 0.814836 |
| Thra_P63058_S12s_1 | 0.36 | 0.72 | 1.88 | 0.89 | 0.23 | 0.59 | 2.03 | 1.4 | 1.73 | 1.11 | 1.12 | 1.55 | 0.11 | -1.41 | 1.03 | -1.05 | 0.1 | 0.415 | -1.2925 | 0.05005 | 0.838141 | 0.292296 | 1 | 0.999631 | 0.488436 |
| Thrap3_Q569Z6_S190s_1 | -0.26 | 0.47 | 2.19 | 1.46 | 0.19 | 1.71 | 1.53 | 0.56 | 0.46 | 0.12 | 0.92 | 0.92 | -1.1 | -0.18 | 1.06 | 0.7 | 0.0325 | -0.36 | -0.845 | 0.21302 | 0.95617 | 0.505948 | 1 | 0.999631 | 0.666787 |
| Thrap3_Q569Z6_S238s_2 | -0.51 | 0.27 | -0.08 | -0.45 | -0.81 | -0.61 | -1 | -1.05 | -0.53 | -0.19 | 0.49 | -1.32 | -0.46 | -0.14 | -1.09 | -0.23 | -0.675 | -0.195 | -0.2875 | 0.410675 | 0.03682 | 0.644756 | 1 | 0.999631 | 0.767937 |
| Thrap3_Q569Z6_S243s_1 | -0.21 | -0.01 | -0.45 | -1.22 | -0.73 | -0.72 | -0.58 | -0.74 | -0.41 | 0.14 | -1.17 | -1 | 0.26 | -0.25 | -1.19 | -0.45 | -0.22 | -0.1375 | 0.065 | 0.875851 | 0.493728 | 0.736587 | 1 | 0.999631 | 0.960853 |
| Thrap3_Q569Z6_S243s_2 | -0.51 | 0.27 | -0.08 | -0.45 | -0.81 | -0.61 | -1 | -1.05 | -0.53 | -0.19 | 0.49 | -1.32 | -0.46 | -0.14 | -1.09 | -0.23 | -0.675 | -0.195 | -0.2875 | 0.410675 | 0.03682 | 0.644756 | 1 | 0.999631 | 0.767937 |
| Thrap3_Q569Z6_S248s_1 | 1.06 | 0.15 | 0.86 | 0.55 | -0.59 | 0.03 | 0.76 | -0.19 | 0.28 | 1.28 | 0.66 | 0.29 | -0.34 | -0.22 | 0.27 | 0.06 | -0.6525 | -0.0275 | -0.7125 | 0.048151 | 0.099647 | 0.936948 | 1 | 0.999631 | 0.48343 |
| Thrap3_Q569Z6_S264s_1 | -0.98 | -1.42 | -0.56 | -1.37 | -1.38 | -2.67 | -0.81 | -0.95 | -1.63 | -1.27 | -0.84 | -0.87 | 0.06 | -1.75 | -0.46 | -0.86 | -0.37 | -0.07 | 0.33 | 0.456341 | 0.4213 | 0.830929 | 1 | 0.999631 | 0.790326 |
| Thrap3_Q569Z6_S320s_1 | 0.15 | 0.21 | 1.08 | 0.68 | -0.35 | -0.12 | 0.19 | 0.44 | -0.1 | 0.59 | 0.57 | 0.75 | 0.3 | -0.05 | 0.65 | 0.57 | -0.49 | -0.0775 | -0.1625 | 0.631716 | 0.153771 | 0.817899 | 1 | 0.999631 | 0.867677 |
| Thrap3_Q569Z6_S532s_1 | 3.05 | 0.2 | 1.45 | 0.73 | -0.69 | 0.09 | 1.71 | -0.51 | 1.24 | 3.73 | -0.08 | -0.64 | -0.3 | -0.28 | 0.05 | -0.56 | -1.2075 | -0.295 | -1.63 | 0.017425 | 0.121752 | 0.76745 | 1 | 0.999631 | 0.391024 |
| Thrap3_Q569Z6_S55s_1 | 1.14 | -1.04 | 0.06 | -0.62 | -1.84 | -1.95 | -0.12 | -1.8 | -2.2 | 0.7 | -1.22 | -1.96 | -1.39 | -2.32 | -1.44 | -1.9 | -1.3125 | -1.055 | -1.6475 | 0.007773 | 0.046515 | 0.165852 | 1 | 0.999631 | 0.358173 |
| Thrap3_Q569Z6_S572s_1 | -1.82 | 1.61 | 1.08 | 0.61 | 0.94 | 1.17 | 0.21 | 0.83 | 1.1 | -0.19 | 1.12 | 1.13 | 0.82 | 1.24 | 0.63 | 1.16 | 0.4175 | 0.42 | 0.5925 | 0.397038 | 0.551934 | 0.568994 | 1 | 0.999631 | 0.761093 |
| Thrap3_Q569Z6_S669s_1 | -0.46 | -3.07 | 1.15 | 0.4 | -3.06 | -2.62 | 0.82 | -0.69 | -3.5 | -1.42 | 0.28 | -0.32 | -3.19 | -5.01 | -0.33 | -0.96 | -0.8925 | -0.745 | -1.8775 | 0.139754 | 0.425419 | 0.490509 | 1 | 0.999631 | 0.611718 |
| Thrap3_Q569Z6_S679s_1 | -0.04 | 0.18 | 0.39 | -0.15 | -0.3 | -0.44 | -0.68 | -0.45 | -0.14 | 0.41 | 0.04 | 0.25 | 0.44 | 0.21 | -0.31 | 0.33 | -0.5625 | 0.045 | 0.0725 | 0.812114 | 0.049384 | 0.869649 | 1 | 0.999631 | 0.936949 |
| Thrap3_Q569Z6_S695s_1 | 0.22 | -2.43 | 1.01 | -0.08 | 3.06 | 0.26 | 0.86 | -0.49 | 3.72 | 1.15 | 2.29 | 2.12 | 4.02 | -0.13 | 1.6 | 0.99 | 1.2425 | 2.64 | 1.94 | 0.071852 | 0.197629 | 0.007858 | 1 | 0.999631 | 0.532005 |
| Thrap3_Q569Z6_S860s_1 | -0.91 | 0.46 | -0.33 | -0.1 | -0.06 | 0.14 | -0.33 | -0.02 | 0.03 | -0.06 | 0.42 | 0.4 | -0.21 | 0.29 | 0.31 | 0.31 | 0.1525 | 0.4175 | 0.395 | 0.288319 | 0.652434 | 0.249627 | 1 | 0.999631 | 0.706035 |
| Thrap3_Q569Z6_S935s_1 | -1.81 | -1.89 | -0.05 | -0.27 | -1.5 | -2.56 | -0.7 | -0.36 | -2.1 | -1.76 | 0.04 | 0.43 | -1.78 | -1.21 | -0.79 | -0.22 | -0.275 | 0.1575 | 0.005 | 0.992737 | 0.659939 | 0.825008 | 1 | 0.999631 | 0.997601 |
| Tiam1_G3UWG2_S1407s_1 | 0.05 | 0.5 | -0.1 | 0.51 | 0.64 | 0.27 | 0.29 | 1.6 | 0.09 | -0.49 | 0.74 | 1.17 | 0.37 | 0.72 | 0.92 | 1.28 | 0.46 | 0.1375 | 0.5825 | 0.097054 | 0.230363 | 0.735541 | 1 | 0.999631 | 0.55886 |
| Tiam1_G3UWG2_S1519s_1 | -0.86 | -1 | -2.32 | -0.89 | -0.28 | -1.34 | -1.26 | -0.89 | -1.13 | -1.54 | -2.87 | -1.58 | -0.82 | -0.39 | -1.89 | -2.31 | 0.325 | -0.5125 | -0.085 | 0.873687 | 0.448424 | 0.311491 | 1 | 0.999631 | 0.959802 |
| Tiam1_G3UWG2_S358s_1 | -0.39 | -0.64 | 0.01 | -0.57 | -0.37 | -0.9 | -0.26 | -0.34 | -0.31 | -0.3 | -0.19 | 0.18 | 0.14 | 0.07 | -0.08 | -0.2 | -0.07 | 0.2425 | 0.38 | 0.20651 | 0.805351 | 0.397991 | 1 | 0.999631 | 0.6605 |
| Tiam1_G3UWG2_T1405t_1 | -0.17 | 0.41 | -0.34 | 0.33 | 0.73 | 0.44 | 0.13 | 0.07 | 0.25 | -0.15 | 0.71 | -0.12 | -0.12 | 0.68 | 0.06 | 0.3 | 0.285 | 0.115 | 0.1725 | 0.601746 | 0.355696 | 0.725879 | 1 | 0.999631 | 0.856765 |
| Tiam2_Q6ZPF3_S20s_1 | 1.35 | 0.63 | 2.37 | -1.3 | 3.66 | 1.76 | 3.85 | 0.6 | 5.86 | 1.9 | 3.46 | 3.77 | 5.56 | 1.8 | 1.04 | 0.28 | 1.705 | 2.985 | 1.4075 | 0.253691 | 0.097417 | 0.011544 | 1 | 0.999631 | 0.688735 |
| Tiam2_Q6ZPF3_S440s_1 | -1 | -0.35 | -2.36 | -0.92 | -0.55 | -0.43 | -0.52 | -1.13 | -0.36 | -1.04 | -3.15 | -0.47 | 0.03 | -0.4 | -0.77 | -1.32 | 0.5 | -0.0975 | 0.5425 | 0.286905 | 0.274431 | 0.887956 | 1 | 0.999631 | 0.705972 |
| Tiam2_Q6ZPF3_S600s_1 | -0.33 | 0.22 | 0.2 | 0.32 | 0.4 | 0.07 | -0.04 | 0.03 | 0.13 | -0.59 | 0.76 | 0.9 | -0.53 | 0.76 | 0.8 | 1.22 | 0.0125 | 0.1975 | 0.46 | 0.287991 | 0.962949 | 0.613652 | 1 | 0.999631 | 0.706002 |
| Tjp1_B9EHJ3_S1534s_1 | -0.75 | 0.45 | -0.68 | -0.4 | 0 | 0.65 | -0.38 | -0.57 | 0.76 | -0.27 | -0.49 | -0.47 | 0.63 | 0.4 | -0.32 | -0.32 | 0.27 | 0.2275 | 0.4425 | 0.279046 | 0.497128 | 0.583684 | 1 | 0.999631 | 0.701431 |
| Tjp1_B9EHJ3_S175s_1 | 0.56 | 0.46 | 0.22 | 0.06 | 0.62 | 0.16 | -0.42 | 0.08 | 0.88 | 0.76 | -0.56 | -0.28 | 0.37 | 0.19 | 0.28 | -0.11 | -0.215 | -0.125 | -0.1425 | 0.617874 | 0.483017 | 0.752289 | 1 | 0.999631 | 0.861861 |
| Tjp1_B9EHJ3_S212s_1 | 1.29 | -0.73 | -0.03 | 0.99 | 1.29 | 0.36 | 1.31 | 1.17 | 0.63 | 0.02 | 1.12 | -0.03 | -1.33 | 0.32 | 1.12 | 0.99 | 0.6525 | 0.055 | -0.105 | 0.872557 | 0.202177 | 0.913978 | 1 | 0.999631 | 0.959468 |
| Tjp1_B9EHJ3_S353s_1 | -0.45 | -1.47 | 0.7 | 1.2 | -1.74 | -1.96 | 0.35 | 0.92 | -1.9 | -1.43 | -0.15 | 0.98 | -1.35 | -1.27 | 1 | 1.06 | -0.6025 | -0.62 | -0.135 | 0.863955 | 0.469421 | 0.433135 | 1 | 0.999631 | 0.955131 |
| Tjp1_B9EHJ3_S617s_1 | -0.25 | 0.21 | 0.12 | 0.06 | 0.48 | -0.26 | 0.07 | -0.21 | 0.3 | 0.02 | -0.18 | 0.14 | 0.19 | -0.3 | -0.62 | -0.03 | -0.015 | 0.035 | -0.225 | 0.464484 | 0.95711 | 0.894112 | 1 | 0.999631 | 0.794196 |
| Tjp1_B9EHJ3_S622s_1 | -0.24 | 0.37 | 0.06 | 0 | 0.6 | -0.13 | -0.24 | -0.14 | 0.41 | 0.72 | 0.12 | -0.06 | 0.13 | 0.08 | 0.09 | 0.31 | -0.025 | 0.25 | 0.105 | 0.706271 | 0.932839 | 0.404301 | 1 | 0.999631 | 0.898066 |
| Tjp1_B9EHJ3_S899s_1 | 0.69 | -1.77 | -0.8 | -1.34 | -0.76 | -0.65 | -0.11 | -0.53 | -1.43 | -0.22 | 0.09 | -0.53 | -0.81 | 0.03 | -0.84 | 0.44 | 0.2925 | 0.2825 | 0.51 | 0.383882 | 0.574113 | 0.626424 | 1 | 0.999631 | 0.754721 |
| Tjp1_B9EHJ3_S912s_1 | 0.18 | 1.11 | 1.26 | 0.69 | 0.23 | 0.93 | 0.21 | 0.69 | 0.57 | 1.15 | 0.81 | 1.08 | 0.65 | 1.2 | 1.07 | 1.44 | -0.295 | 0.0925 | 0.28 | 0.431771 | 0.391132 | 0.778993 | 1 | 0.999631 | 0.778604 |
| Tjp1_B9EHJ3_T354t_1 | -0.28 | -1.46 | -2.81 | -2.87 | -1.31 | -1.28 | -2.2 | -2.79 | -1.16 | -0.98 | -2.28 | -3.06 | -1.82 | -0.75 | -3.31 | -2.93 | -0.04 | -0.015 | -0.3475 | 0.642138 | 0.950316 | 0.982931 | 1 | 0.999631 | 0.871335 |
| Tjp1_P39447_S912s_1 | 0.52 | 1.91 | 1.85 | 1.19 | 0.69 | 1.71 | 0.61 | 1.23 | 1.19 | 1.96 | 1.26 | 1.67 | 0.91 | 1.98 | 1.56 | 2.04 | -0.3075 | 0.1525 | 0.255 | 0.554108 | 0.463758 | 0.696863 | 1 | 0.999631 | 0.837952 |
| Tjp2_Q9Z0U1_S107s_1 | -0.44 | -0.3 | 0.05 | -0.67 | -0.39 | -0.82 | -0.7 | 0.03 | -0.52 | -0.41 | -0.44 | 0.09 | -0.41 | -0.57 | 0.03 | 0.38 | -0.13 | 0.02 | 0.1975 | 0.557341 | 0.669216 | 0.94506 | 1 | 0.999631 | 0.838305 |
| Tjp2_Q9Z0U1_S1136s_1 | -0.45 | 0.51 | 0.34 | -0.19 | -0.28 | 0.19 | -0.47 | 0.14 | 0.46 | 0.44 | 0.07 | 0.66 | 1.18 | 0.66 | 0.48 | 0.58 | -0.1575 | 0.355 | 0.6725 | 0.070748 | 0.627228 | 0.279224 | 1 | 0.999631 | 0.530826 |
| Tjp2_Q9Z0U1_S213s_1 | -2.24 | -6.36 | -7.38 | -6.17 | -4.12 | -4.53 | -6.08 | -6.44 | -6.01 | -3.73 | -4.22 | -6.11 | -4.57 | -5.11 | -5.91 | -8.13 | 0.245 | 0.52 | -0.3925 | 0.736903 | 0.821413 | 0.639465 | 1 | 0.999631 | 0.907709 |
| Tjp2_Q9Z0U1_S393s_1 | 0.44 | 0.54 | 1.08 | 0.52 | 0.25 | 0.54 | 0.01 | 0.37 | 0.69 | 1.28 | 0.33 | 0.92 | 0.93 | 0.63 | 1.06 | 1.22 | -0.3525 | 0.16 | 0.315 | 0.306562 | 0.216395 | 0.61139 | 1 | 0.999631 | 0.717667 |
| Tjp2_Q9Z0U1_S395s_1 | -0.19 | -2.63 | -0.52 | -1.15 | -0.28 | -0.41 | -1.05 | -0.77 | -0.6 | 0.21 | -0.59 | -0.98 | 1.09 | -0.18 | -0.58 | -0.05 | 0.495 | 0.6325 | 1.1925 | 0.067564 | 0.358241 | 0.268287 | 1 | 0.999631 | 0.521762 |
| Tjp2_Q9Z0U1_S404s_1 | 0.46 | -0.34 | 0.67 | 0.51 | -1.02 | 0.07 | 0.4 | 0.05 | -0.11 | 0.48 | -0.05 | -0.11 | -0.81 | -1.03 | 0.13 | -1.13 | -0.45 | -0.2725 | -1.035 | 0.023564 | 0.266587 | 0.409418 | 1 | 0.999631 | 0.415644 |
| Tjp2_Q9Z0U1_S421s_1 | 0.44 | -7.35 | -3.15 | -2.27 | -2.13 | -6.68 | -1.75 | -4.09 | -8.31 | -0.2 | -2.96 | -4.43 | -2.52 | -3.29 | -4.32 | -6.43 | -0.58 | -0.8925 | -1.0575 | 0.494696 | 0.72852 | 0.653241 | 1 | 0.999631 | 0.810404 |
| Tjp2_Q9Z0U1_S684s_1 | -0.74 | 0.25 | 0.02 | -0.41 | 0.48 | -0.76 | -0.32 | -0.58 | 0.2 | -0.77 | -0.07 | 0.39 | -0.67 | -0.11 | 0.05 | -0.29 | -0.075 | 0.1575 | -0.035 | 0.917294 | 0.839451 | 0.668548 | 1 | 0.999631 | 0.973536 |
| Tjp2_Q9Z0U1_S988s_1 | -0.7 | -0.09 | 0.63 | 0.91 | 0.5 | -0.31 | 0.89 | 1.09 | 0.04 | -0.5 | 0.81 | 0.9 | -0.33 | -0.33 | 0.89 | 1.16 | 0.355 | 0.125 | 0.16 | 0.754304 | 0.445854 | 0.7924 | 1 | 0.999631 | 0.913377 |
| Tkt_P40142_T287t_1 | 0.21 | -0.04 | -0.15 | -1.98 | 0.61 | 0.64 | -0.51 | -0.77 | 0.12 | 1.06 | -0.46 | -0.61 | 2.67 | 0.34 | -0.96 | -0.76 | 0.4825 | 0.5175 | 0.8125 | 0.347002 | 0.405061 | 0.380193 | 1 | 0.999631 | 0.736174 |
| Tle1_Q5SQA7_S31s_1 | -0.22 | 0.62 | 0.47 | 0.32 | 1.2 | 1.38 | 0.53 | 0.59 | 1.34 | 0.53 | 0.69 | 0.49 | 0.85 | 1.24 | 0.71 | 0.91 | 0.6275 | 0.465 | 0.63 | 0.064667 | 0.078539 | 0.174814 | 1 | 0.999631 | 0.515625 |
| Tle3_E0CY76_S202s_1 | -0.15 | 0.83 | 1.92 | 0.93 | 1.11 | 0.36 | 0.97 | 0.9 | 1.32 | 0.63 | 1.7 | 1.89 | 1.48 | 1.28 | 1.34 | 1.39 | -0.0475 | 0.5025 | 0.49 | 0.272956 | 0.914232 | 0.313726 | 1 | 0.999631 | 0.700187 |
| Tle3_E0CY76_S262s_1 | 0.47 | 0.46 | 0.57 | 0.16 | -0.41 | 0.11 | 0.4 | 0.74 | -0.19 | 0.61 | 0.46 | 0.59 | 1.01 | 0.12 | 0.44 | 0.52 | -0.205 | -0.0475 | 0.1075 | 0.724805 | 0.516534 | 0.870892 | 1 | 0.999631 | 0.905674 |
| Tle3_E0CY76_S285s_1 | -0.68 | 0.92 | 0.8 | -0.15 | 0.49 | 0.49 | -0.06 | 0.46 | 0.33 | -0.24 | 0.08 | 0.57 | 0.55 | -0.14 | 0.61 | 0.47 | 0.1225 | -0.0375 | 0.15 | 0.727881 | 0.764508 | 0.929414 | 1 | 0.999631 | 0.905923 |
| Tle4_Q62441_S292s_1 | -0.45 | 1.36 | 0.97 | 0.49 | 1.22 | -0.07 | 0.29 | 1.04 | 0.82 | -0.1 | 1.36 | 1.4 | 1.02 | 1.21 | 1.07 | 1.31 | 0.0275 | 0.2775 | 0.56 | 0.195685 | 0.953506 | 0.580113 | 1 | 0.999631 | 0.653056 |
| Tln1_P26039_S2040s_1 | 0.5 | 0.22 | 0.41 | 0.41 | 2.21 | 0.07 | 0.36 | -0.24 | 0.19 | 1.84 | 2.25 | 3.17 | 0.39 | 1.44 | 0.23 | 1.37 | 0.215 | 1.4775 | 0.4725 | 0.218124 | 0.677702 | 0.026646 | 1 | 0.999631 | 0.669859 |
| Tln2_E9PUM4_S473s_1 | -0.16 | -0.48 | -0.33 | -0.68 | 0.02 | -0.42 | -0.31 | 0.85 | 0.08 | -0.46 | -0.47 | 0.27 | 0.39 | 0.62 | -0.28 | -0.71 | 0.4475 | 0.2675 | 0.4175 | 0.273146 | 0.210655 | 0.378835 | 1 | 0.999631 | 0.700187 |
| Tln2_E9PUM4_T1843t_1 | -0.42 | -0.47 | -0.66 | -0.5 | -0.59 | -0.33 | -0.88 | -0.78 | -0.37 | -0.08 | -0.64 | -0.27 | -0.29 | -0.3 | -0.59 | -0.29 | -0.1325 | 0.1725 | 0.145 | 0.586667 | 0.600161 | 0.508451 | 1 | 0.999631 | 0.849724 |
| Tmcc1_F8WJ98_S131s_1 | 2.45 | -0.21 | 0.62 | 1.92 | 1.19 | 1.97 | 2.29 | 1.44 | 0.97 | 0.59 | 1.3 | -0.2 | -1.62 | 0.06 | 0.05 | 0.88 | 0.5275 | -0.53 | -1.3525 | 0.080662 | 0.383893 | 0.403107 | 1 | 0.999631 | 0.545579 |
| Tmcc1_F8WJ98_S419s_1 | -0.18 | 0.4 | -0.64 | -0.26 | 0.43 | -0.64 | -0.49 | -0.47 | -0.56 | -0.01 | -0.47 | -0.2 | 0.03 | 0.21 | -0.64 | -0.4 | -0.1225 | -0.14 | -0.03 | 0.931341 | 0.729044 | 0.656879 | 1 | 0.999631 | 0.978006 |
| Tmcc1_F8WJ98_S450s_1 | -0.89 | -0.63 | -1.34 | -2.09 | 0.29 | 0.34 | -0.21 | 0.54 | -0.29 | -0.55 | 0.66 | 0.46 | 1.9 | 0.08 | 0.85 | -1.19 | 1.4775 | 1.3075 | 1.6475 | 0.027909 | 0.002748 | 0.012276 | 1 | 0.999631 | 0.423977 |
| Tmcc1_F8WJ98_S451s_1 | -0.93 | -1.44 | -0.14 | -1.22 | -1.05 | -0.96 | -0.43 | -0.16 | -1.68 | -1.4 | -0.08 | -0.03 | -1.17 | 0.2 | -0.5 | -0.7 | 0.2825 | 0.135 | 0.39 | 0.360631 | 0.454945 | 0.784437 | 1 | 0.999631 | 0.741241 |
| Tmcc2_Q80W04_S119s_1 | -1.14 | -3.11 | -0.93 | -1.42 | -3.66 | -2.45 | -2.14 | -1.44 | -1.58 | -0.53 | -1.4 | -0.66 | 0.68 | -0.87 | -0.8 | -0.86 | -0.7725 | 0.6075 | 1.1875 | 0.062189 | 0.226159 | 0.265084 | 1 | 0.999631 | 0.509361 |
| Tmcc2_Q80W04_S126s_1 | 0.05 | -0.36 | 0.9 | 0.79 | -0.09 | 1.47 | 1.07 | 0.78 | 1.22 | 2.3 | -0.14 | 0.05 | -1.3 | -0.05 | 0.46 | -1.88 | 0.4625 | 0.5125 | -1.0375 | 0.093728 | 0.304118 | 0.392594 | 1 | 0.999631 | 0.558128 |
| Tmcc2_Q80W04_S189s_1 | 0.68 | 0.14 | 0.05 | 1.21 | 1.05 | 1.15 | 0.76 | 1.76 | 0.94 | 0.49 | 0.91 | 0.67 | -0.74 | 1.01 | 1.18 | 1.05 | 0.66 | 0.2325 | 0.105 | 0.835453 | 0.094049 | 0.495361 | 1 | 0.999631 | 0.944699 |
| Tmcc2_Q80W04_S435s_1 | 0.26 | 1.54 | -0.26 | 0.54 | 1.43 | 1.2 | 0.21 | 0.67 | 0.95 | 0.77 | 0.42 | 0.49 | 0.46 | 1.47 | 0.3 | 0.44 | 0.3575 | 0.1375 | 0.1475 | 0.748629 | 0.435438 | 0.736106 | 1 | 0.999631 | 0.911931 |
| Tmcc2_Q80W04_S75s_1 | -0.77 | -1.28 | -1.45 | 0.3 | 0.47 | -0.54 | 0.37 | 0.07 | -0.69 | -1.32 | 0.09 | -0.6 | -2.46 | 0.39 | -1.06 | 0.21 | 0.8925 | 0.17 | 0.07 | 0.918497 | 0.067445 | 0.719203 | 1 | 0.999631 | 0.974207 |
| Tmcc3_Q8R310_S216s_1 | 0.26 | 1.54 | 0.16 | 0.99 | 1.43 | 1.2 | 0.51 | 1.14 | 0.95 | 0.77 | 0.86 | 1.02 | 0.46 | 1.47 | 0.67 | 0.94 | 0.3325 | 0.1625 | 0.1475 | 0.719955 | 0.402179 | 0.654987 | 1 | 0.999631 | 0.904808 |
| Tmed8_Q3UHI4_S21s_1 | 2.34 | 0.85 | 1.22 | 1.41 | 0.25 | 1.16 | 2.41 | 0.31 | 0.53 | 1.71 | 0.75 | 0.19 | -0.52 | 0.03 | 0.46 | -0.28 | -0.4225 | -0.66 | -1.5325 | 0.003344 | 0.447134 | 0.163078 | 1 | 0.999631 | 0.351308 |
| Tmeff2_Q9QYM9_T358t_1 | 0.24 | -2.22 | -0.41 | -0.66 | -2.84 | 1 | 1.07 | -0.36 | -2.42 | -0.54 | -0.91 | -2.94 | -3.6 | -1.82 | -1.36 | -2.74 | 0.48 | -0.94 | -1.6175 | 0.029562 | 0.600436 | 0.197177 | 1 | 0.999631 | 0.433885 |
| Tmem100_Q9CQG9_S121s_1 | -0.11 | 0.35 | -0.23 | -0.27 | -0.33 | 0.57 | -0.59 | 0.04 | 0.25 | 0.22 | -0.15 | 0.12 | 0.63 | 0.46 | 0.14 | 0.27 | -0.0125 | 0.175 | 0.44 | 0.156251 | 0.969787 | 0.527326 | 1 | 0.999631 | 0.626452 |
| Tmem100_Q9CQG9_S15s_1 | -0.09 | -0.82 | -1.6 | -1.3 | -0.37 | -0.8 | -1.47 | -1.19 | -0.1 | 0.24 | -1.37 | -0.77 | -0.46 | -0.96 | -1.08 | -1.14 | -0.005 | 0.4525 | 0.0425 | 0.913726 | 0.990212 | 0.347702 | 1 | 0.999631 | 0.972551 |
| Tmem11_Q8BK08_S16s_1 | 0.73 | -0.72 | 0.06 | -0.02 | -0.78 | -0.48 | 0.46 | -0.79 | -0.33 | 0.59 | -0.27 | -0.65 | -0.27 | -0.75 | -0.73 | -1.05 | -0.41 | -0.1775 | -0.7125 | 0.08212 | 0.336513 | 0.664137 | 1 | 0.999631 | 0.54792 |
| Tmem121_Q80XA0_S310s_1 | 0.49 | 0.44 | -1.93 | -1.21 | -0.08 | -1.47 | -1.63 | -1.98 | -0.81 | -0.13 | -1.24 | -2.43 | -0.34 | -0.46 | 0.37 | -1.81 | -0.7375 | -0.6 | -0.0075 | 0.991133 | 0.276917 | 0.394827 | 1 | 0.999631 | 0.996392 |
| Tmem127_G3UXI2_S17s_1 | -0.17 | -2.42 | -0.51 | -0.74 | -1.41 | -0.5 | -0.95 | -0.41 | -0.75 | -0.55 | -0.22 | -0.16 | -0.2 | -0.62 | -0.38 | -0.22 | 0.1425 | 0.54 | 0.605 | 0.236251 | 0.781533 | 0.29015 | 1 | 0.999631 | 0.683301 |
| Tmem132a_Q922P8_S883s_1 | -1.34 | -0.82 | -0.02 | -0.63 | -0.65 | -0.65 | -0.91 | 0.43 | -0.71 | -0.74 | 0.25 | 0.6 | 0.5 | -0.49 | 0.31 | 0.67 | 0.2575 | 0.5525 | 0.95 | 0.036005 | 0.52994 | 0.221987 | 1 | 0.999631 | 0.453963 |
| Tmem132a_Q922P8_S938s_1 | -1.83 | -0.23 | -0.24 | 0.12 | -0.57 | -0.24 | -0.72 | 0.54 | -0.68 | -0.55 | -0.77 | 0.44 | 1.32 | -0.14 | -0.58 | 1.38 | 0.2975 | 0.155 | 1.04 | 0.110019 | 0.547031 | 0.754057 | 1 | 0.999631 | 0.573646 |
| Tmem132c_E9QP73_S563s_1 | -0.66 | -0.42 | 0.8 | 0.49 | -0.9 | -0.11 | 0.37 | 1.07 | 0.28 | 0.35 | -0.18 | -0.33 | -1.13 | -1.57 | 0.01 | -1.8 | 0.055 | -0.0225 | -1.175 | 0.040744 | 0.913605 | 0.955243 | 1 | 0.999631 | 0.474979 |
| Tmem132d_Q76HP3_S1035s_1 | 1.28 | -0.82 | -1.05 | -1.05 | -0.26 | 0.6 | -0.76 | -1.8 | -0.28 | 0.45 | -1.4 | -1.49 | 0.32 | -0.65 | -2.06 | -1.66 | -0.145 | -0.27 | -0.6025 | 0.392842 | 0.829239 | 0.682907 | 1 | 0.999631 | 0.758901 |
| Tmem132d_Q76HP3_S554s_1 | -0.78 | -1.23 | -0.12 | 0.08 | -1.92 | 0.03 | 0.53 | -1.51 | -1.64 | 0.2 | -1.33 | -0.82 | -1.58 | -3.57 | -1.29 | -2.64 | -0.205 | -0.385 | -1.7575 | 0.009834 | 0.732984 | 0.432737 | 1 | 0.999631 | 0.375381 |
| Tmem135_Q8C8G3_S198s_1 | -0.9 | 0.39 | -0.96 | -2 | -1.07 | 0.52 | -0.72 | -0.29 | -0.61 | -0.39 | -0.5 | -0.97 | 0.66 | 0.88 | 1.24 | 0.26 | 0.4775 | 0.25 | 1.6275 | 0.008768 | 0.393636 | 0.607799 | 1 | 0.999631 | 0.365244 |
| Tmem145_E9PUE6_S511s_1 | -1.5 | -2.04 | -1 | -1.26 | -1.99 | -2.34 | -1.13 | -1.24 | -2.14 | -2.4 | -0.83 | -1.45 | -1.24 | -2.22 | -1.11 | -1.01 | -0.225 | -0.255 | 0.055 | 0.887187 | 0.557753 | 0.548256 | 1 | 0.999631 | 0.963505 |
| Tmem151a_Q6GQT5_S323s_1 | -0.41 | -0.11 | -0.57 | -1.47 | -0.26 | -0.15 | -0.51 | -1.08 | -0.27 | -0.61 | -2.05 | 0.51 | 0.25 | -0.94 | -1.27 | -0.46 | 0.14 | 0.035 | 0.035 | 0.936996 | 0.709615 | 0.95038 | 1 | 0.999631 | 0.97939 |
| Tmem151b_Q68FE7_S486s_1 | 0.5 | -0.38 | 0.28 | -0.2 | -0.37 | 1.26 | 0.23 | 0.34 | 0.41 | 0.67 | 0.27 | 1.11 | -1.08 | 1.45 | 0.3 | 0.56 | 0.315 | 0.565 | 0.2575 | 0.628431 | 0.437247 | 0.110569 | 1 | 0.999631 | 0.865998 |
| Tmem151b_Q68FE7_S488s_1 | 0.16 | 0.04 | 0.04 | -0.18 | -0.57 | 0.25 | -0.23 | -0.34 | 1.03 | 1.15 | -0.26 | -0.08 | 1.11 | 0.13 | -0.52 | -0.4 | -0.2375 | 0.445 | 0.065 | 0.871352 | 0.395867 | 0.270626 | 1 | 0.999631 | 0.959003 |
| Tmem163_Q8C996_S11s_1 | -0.57 | -0.87 | -0.87 | -1.11 | -1.07 | -1.09 | -1.16 | -1.09 | -0.64 | -0.27 | -1.01 | -0.7 | 0.21 | -0.77 | -1.09 | -0.66 | -0.2475 | 0.2 | 0.2775 | 0.439888 | 0.324357 | 0.484489 | 1 | 0.999631 | 0.781983 |
| Tmem163_Q8C996_S45s_1 | -0.13 | -0.32 | 0.72 | -0.69 | -0.51 | -0.29 | -0.06 | -0.93 | 0.19 | -0.93 | -0.2 | 0.4 | 0.44 | -1.21 | -0.1 | -0.82 | -0.3425 | -0.03 | -0.3175 | 0.500714 | 0.365444 | 0.94316 | 1 | 0.999631 | 0.813968 |
| Tmem163_Q8C996_S56s_1 | -1.12 | -0.81 | -0.46 | -0.82 | -0.84 | -0.87 | -1.06 | -0.65 | -0.85 | -0.77 | -0.66 | -0.33 | 0.01 | -0.48 | -0.74 | -0.25 | -0.0525 | 0.15 | 0.4375 | 0.17561 | 0.841078 | 0.591383 | 1 | 0.999631 | 0.642116 |
| Tmem163_Q8C996_S7s_1 | 0.5 | -0.8 | 0.75 | 0.1 | -0.86 | 0.34 | 0.48 | -0.38 | -1.41 | 0.26 | -1.79 | -0.57 | -0.87 | -0.4 | -0.08 | -0.47 | -0.2425 | -1.015 | -0.5925 | 0.16154 | 0.59108 | 0.081077 | 1 | 0.999631 | 0.631442 |
| Tmem169_D3YUR6_S15s_1 | -0.72 | 0.18 | 0.43 | -0.22 | 0.43 | 0.07 | -0.04 | 0.3 | 0.87 | -0.65 | 0.3 | 0.24 | 1.41 | 0.31 | -2.06 | -0.24 | 0.2725 | 0.2725 | -0.0625 | 0.926998 | 0.404355 | 0.51112 | 1 | 0.999631 | 0.976306 |
| Tmem169_D3YUR6_S20s_1 | 1.12 | 1.32 | 0.66 | 1.25 | 0.57 | 0.28 | 0.44 | 0.18 | 0.25 | 0.48 | -0.32 | -0.87 | 0.58 | -0.56 | 1.75 | 0.11 | -0.72 | -1.2025 | -0.6175 | 0.226703 | 0.021446 | 0.008074 | 1 | 0.999631 | 0.677921 |
| Tmem169_D3YUR6_S20s_2 | 0.47 | 1.08 | 1.45 | 0.71 | -0.35 | 0.9 | 1.02 | 1.91 | 0.55 | 1.59 | 2.26 | 1.35 | 1.01 | 0.05 | 0.49 | 1.73 | -0.0575 | 0.51 | -0.1075 | 0.802334 | 0.905395 | 0.240427 | 1 | 0.999631 | 0.932677 |
| Tmem169_D3YUR6_S69s_1 | -0.54 | 2.21 | 1.29 | 1.54 | 1.46 | 2.35 | 0.51 | 1.26 | 2.15 | 1.07 | 0.16 | 1.06 | 1.87 | 1.15 | 0.63 | -0.47 | 0.27 | -0.015 | -0.33 | 0.630705 | 0.668317 | 0.981401 | 1 | 0.999631 | 0.867509 |
| Tmem169_D3YUR6_T62t_1 | -0.45 | 2.24 | 2.58 | 0.8 | 0.85 | 2.48 | 1.24 | 0.18 | 2.35 | 1.97 | 0.99 | 1.78 | 1.35 | 0.08 | 1.13 | 1.37 | -0.105 | 0.48 | -0.31 | 0.649278 | 0.887962 | 0.482559 | 1 | 0.999631 | 0.874768 |
| Tmem184b_Q8BG09_S382s_1 | 1.67 | 1.12 | 1.72 | 0.62 | -1.1 | 0.75 | 1.69 | 0.5 | -0.89 | 2.03 | 0.92 | 0.58 | 0.48 | -0.41 | -0.01 | 0.07 | -0.8225 | -0.6225 | -1.25 | 0.006029 | 0.176666 | 0.311784 | 1 | 0.999631 | 0.351545 |
| Tmem19_Q91W52_S294s_1 | -1.2 | -0.58 | 0.07 | -0.64 | -0.31 | -0.84 | -0.74 | -0.29 | -0.24 | -0.4 | 0.13 | 0.04 | 0.36 | -0.54 | -0.97 | 0.04 | 0.0425 | 0.47 | 0.31 | 0.458356 | 0.89897 | 0.183375 | 1 | 0.999631 | 0.790722 |
| Tmem200b_D3Z0P8_S177s_1 | 1.34 | -0.14 | -1.79 | 0.44 | 1.46 | -0.6 | 0.43 | -0.82 | -0.8 | -0.46 | -0.99 | -1.22 | -1.16 | -1.31 | -2.01 | -1 | 0.155 | -0.83 | -1.3325 | 0.056141 | 0.834193 | 0.198332 | 1 | 0.999631 | 0.495761 |
| Tmem200b_D3Z0P8_S5s_1 | 1.37 | -1.55 | -0.44 | -0.79 | -1.38 | -1.05 | 0.44 | -1.88 | -2.24 | 0.78 | -1.03 | -1.62 | -1.71 | -2.15 | -2.53 | -2.36 | -0.615 | -0.675 | -1.835 | 0.010403 | 0.391582 | 0.401898 | 1 | 0.999631 | 0.383871 |
| Tmem200c_J3QK46_S487s_1 | 0.14 | -0.77 | -0.52 | 0.53 | 0.59 | -0.87 | 0.67 | 0.02 | -0.27 | -1.28 | 1.18 | 0.52 | -0.64 | -0.94 | -0.68 | -0.41 | 0.2575 | 0.1925 | -0.5125 | 0.180763 | 0.569156 | 0.732166 | 1 | 0.999631 | 0.64376 |
| Tmem201_A2A8U2_S454s_1 | 0.15 | 0.44 | -0.1 | 0.48 | 1.03 | 0.63 | 0.07 | 0.06 | 0.62 | -0.28 | 0.86 | 0.28 | 0.16 | 0.39 | -0.46 | 0.68 | 0.205 | 0.1275 | -0.05 | 0.88359 | 0.526096 | 0.702381 | 1 | 0.999631 | 0.963505 |
| Tmem229a_B9EJI9_S4s_1 | -0.46 | -1.54 | 0.78 | -0.62 | -0.49 | -0.9 | -0.29 | 0.71 | -0.97 | -0.79 | 0.32 | 0.46 | 0.33 | -1.1 | 0.94 | -0.18 | 0.2175 | 0.215 | 0.4575 | 0.44421 | 0.68846 | 0.700724 | 1 | 0.999631 | 0.783224 |
| Tmem245_D3YWD3_S12s_1 | 0.38 | -0.25 | 0.22 | -0.03 | -0.52 | -0.21 | -0.17 | 0.1 | -0.19 | 0.35 | 0.58 | -0.26 | -0.44 | -0.01 | -0.27 | 0.24 | -0.28 | 0.04 | -0.2 | 0.515579 | 0.321616 | 0.89822 | 1 | 0.999631 | 0.819449 |
| Tmem245_D3YWD3_S329s_1 | 0.21 | -0.58 | 0.48 | -0.35 | -0.05 | -1.28 | -0.18 | -0.5 | -0.46 | 1.45 | 0.13 | 0.39 | -1.11 | 0.05 | -0.69 | 0.72 | -0.4425 | 0.4375 | -0.1975 | 0.672398 | 0.26304 | 0.350183 | 1 | 0.999631 | 0.885053 |
| Tmem25_Q9DCF1_S297s_1 | 0.22 | 0.27 | 1.56 | 0.55 | -0.36 | -0.97 | -0.18 | -0.31 | -0.82 | -1.96 | -0.15 | -0.99 | 0.45 | -2.87 | 1.08 | 0.11 | -1.105 | -1.63 | -0.9575 | 0.254553 | 0.013903 | 0.006044 | 1 | 0.999631 | 0.689457 |
| Tmem261_Q9CQ00_S36s_1 | 0.24 | 0.36 | 1.12 | 0.55 | 0.36 | 0.33 | 0.65 | 0.5 | 0.09 | 0.73 | 0.86 | 1.7 | 0.53 | 0.62 | 1.68 | 1.67 | -0.1075 | 0.2775 | 0.5575 | 0.182842 | 0.70675 | 0.490824 | 1 | 0.999631 | 0.643861 |
| Tmem38a_Q3TMP8_S289s_1 | 0.63 | 0.94 | 1.65 | -0.77 | -1.7 | 1.9 | -0.35 | -0.33 | 0.73 | 2.94 | 0.17 | 0.71 | 2.52 | 1.33 | 0.17 | -0.17 | -0.7325 | 0.525 | 0.35 | 0.620948 | 0.364174 | 0.465461 | 1 | 0.999631 | 0.861888 |
| Tmem40_E9Q6I9_S104s_1 | -1.85 | -0.94 | -1.21 | -0.64 | -0.99 | -1.14 | -1.1 | -2.36 | -0.63 | -2.38 | -1.01 | -0.3 | -0.15 | -0.8 | -1.38 | -0.89 | -0.2375 | 0.08 | 0.355 | 0.373864 | 0.56759 | 0.872404 | 1 | 0.999631 | 0.747035 |
| Tmem47_Q9JJG6_S6s_1 | -0.83 | -1.66 | -0.52 | -2 | -1.37 | -1.14 | -0.32 | -1.33 | -1.18 | -1.08 | -0.7 | -0.14 | 0.17 | -2.29 | -0.84 | -2.1 | 0.2125 | 0.4775 | -0.0125 | 0.983655 | 0.615283 | 0.273779 | 1 | 0.999631 | 0.995133 |
| Tmem51_Q99LG1_S114s_1 | -2.53 | -1.88 | -1.35 | -0.87 | -0.63 | -1.19 | -1.41 | -2.3 | -0.15 | -1.23 | -1.29 | -1.34 | 1.09 | -1.05 | -1.18 | -2.11 | 0.275 | 0.655 | 0.845 | 0.233068 | 0.564668 | 0.166987 | 1 | 0.999631 | 0.683035 |
| Tmem51_Q99LG1_S151s_1 | 0.24 | 1.33 | 0.62 | 0.95 | 0.26 | 1.34 | -0.36 | -1.32 | 1.21 | 1.05 | -1.89 | 0.5 | 0.31 | 1.06 | -1.18 | -0.71 | -0.805 | -0.5675 | -0.915 | 0.104986 | 0.168812 | 0.409861 | 1 | 0.999631 | 0.568213 |
| Tmem51_Q99LG1_S153s_1 | -0.2 | -0.14 | 0.18 | -0.97 | 0.29 | 0.17 | -0.82 | -0.89 | 0.25 | -0.51 | -0.5 | -0.06 | 1.08 | 0.54 | -0.6 | 0.31 | -0.03 | 0.0775 | 0.615 | 0.17757 | 0.940402 | 0.823308 | 1 | 0.999631 | 0.642116 |
| Tmem51_Q99LG1_T230t_1 | -0.87 | -0.85 | -0.98 | -0.97 | -0.94 | -0.29 | -0.91 | -1.44 | -0.47 | -0.89 | -2.2 | -1.25 | -0.41 | -1.08 | -0.49 | -0.92 | 0.0225 | -0.285 | 0.1925 | 0.509484 | 0.940272 | 0.4693 | 1 | 0.999631 | 0.816932 |
| Tmem55a_Q9CZX7_S47s_1 | -0.29 | -0.17 | -0.52 | 0.05 | -0.36 | -0.13 | -0.11 | -0.14 | -0.75 | -0.43 | -0.16 | 0.28 | -1.92 | 0.86 | 0.58 | 0.1 | 0.0475 | -0.0325 | 0.1375 | 0.814545 | 0.850136 | 0.917326 | 1 | 0.999631 | 0.937111 |
| Tmem55a_Q9CZX7_T22t_1 | -1.55 | -1.61 | -1.17 | -1.75 | -2.46 | -1.36 | -1.4 | -1.59 | -2.12 | -0.99 | -1.38 | -2.27 | -1.22 | -1.44 | -0.62 | -1.13 | -0.1825 | -0.17 | 0.4175 | 0.196635 | 0.581206 | 0.639397 | 1 | 0.999631 | 0.653373 |
| Tmem55b_E0CYE6_S169s_1 | -1.22 | 0.2 | -0.39 | -1.15 | 0.05 | -0.02 | -0.66 | -0.79 | -0.18 | -1.08 | -0.46 | -0.37 | 0.57 | 0.11 | -0.44 | -0.41 | 0.285 | 0.1175 | 0.5975 | 0.181046 | 0.485444 | 0.769993 | 1 | 0.999631 | 0.64376 |
| Tmem63c_D6RI74_S94s_2 | -1.01 | 0.67 | 0.02 | -0.02 | 0.35 | 0.67 | 0.04 | 0.43 | 0.49 | 0.97 | 0.76 | 0.35 | 1.77 | 0.85 | 0.43 | 0.68 | 0.4575 | 0.7275 | 1.0175 | 0.044849 | 0.249319 | 0.087084 | 1 | 0.999631 | 0.475443 |
| Tmem74b_A2AU34_S198s_1 | -0.16 | -0.23 | -0.02 | 0.05 | -0.21 | -0.01 | -0.57 | 0.28 | 0.16 | 0.08 | -0.1 | 0.47 | 0.34 | 0.4 | 0.77 | 0.97 | -0.0375 | 0.2425 | 0.71 | 0.030835 | 0.891738 | 0.362877 | 1 | 0.999631 | 0.439946 |
| Tmem74b_A2AU34_S3s_1 | -0.88 | -0.84 | -0.72 | -0.8 | -1.05 | -0.07 | -1.48 | -1.07 | -0.95 | -0.23 | -0.94 | -0.83 | 0.41 | -0.17 | -0.35 | -0.65 | -0.1075 | 0.0725 | 0.62 | 0.072259 | 0.750797 | 0.793308 | 1 | 0.999631 | 0.532251 |
| Tmem74b_A2AU34_S62s_1 | -0.99 | -0.14 | -1.3 | -1.5 | -0.37 | -0.31 | -1.29 | -0.72 | -0.72 | -0.39 | -0.45 | -0.7 | 0.49 | -0.31 | -0.02 | -0.76 | 0.31 | 0.4175 | 0.8325 | 0.067858 | 0.429324 | 0.251162 | 1 | 0.999631 | 0.522149 |
| Tmem88_Q9D0N8_S145s_1 | -0.23 | -0.86 | -2 | -1.58 | -0.37 | 0.27 | -0.74 | 0.08 | -0.25 | 0.15 | -0.2 | -1.37 | 0.09 | 0.14 | -0.26 | 0.18 | 0.9775 | 0.75 | 1.205 | 0.015331 | 0.048061 | 0.148605 | 1 | 0.999631 | 0.388832 |
| Tmem88_Q9D0N8_S148s_1 | -0.22 | -1.19 | -0.99 | -0.17 | 0.04 | 1.42 | -0.16 | 0.75 | 0.21 | -0.31 | 1.06 | -0.27 | 0.14 | 0.77 | 0.57 | 1.31 | 1.155 | 0.815 | 1.34 | 0.005865 | 0.023048 | 0.075132 | 1 | 0.999631 | 0.351545 |
| Tmpo_Q61033_S155s_1 | 2.04 | -1.1 | -0.43 | -0.39 | -1.66 | 0.11 | 1.33 | -1.55 | -1.87 | 2.35 | -1.6 | -2.56 | -1.97 | -2.29 | -2.01 | -3.1 | -0.4725 | -0.95 | -2.3725 | 0.004716 | 0.586878 | 0.405731 | 1 | 0.999631 | 0.351308 |
| Tmpo_Q61033_T159t_1 | -0.24 | 0.47 | -0.01 | -0.5 | -0.32 | 0.11 | -0.77 | -0.58 | 0.34 | 0.64 | -0.59 | -0.01 | 0.73 | 0.09 | -0.62 | -0.42 | -0.32 | 0.165 | 0.015 | 0.969548 | 0.337425 | 0.653034 | 1 | 0.999631 | 0.989176 |
| Tmtc1_D6RG80_S168s_1 | -0.03 | -0.72 | -0.61 | -3.13 | -1.86 | -2.07 | -0.68 | -1.92 | -0.21 | -0.98 | -1.15 | -1.49 | -1.95 | -1.7 | -3.62 | -3.82 | -0.51 | 0.165 | -1.65 | 0.054274 | 0.455545 | 0.803233 | 1 | 0.999631 | 0.491994 |
| Tmub1_D3Z5X7_S71s_1 | 0.25 | 0.43 | 0.32 | 0.26 | -0.08 | 0.2 | 0.2 | 0.05 | 0.3 | 0.22 | 0.42 | 0.36 | 0.28 | 1.28 | 0.55 | 1.06 | -0.2225 | 0.01 | 0.4775 | 0.156026 | 0.355558 | 0.966725 | 1 | 0.999631 | 0.626452 |
| Tmx1_Q8VBT0_S245s_1 | -0.57 | -0.07 | 0.44 | 0.6 | -0.58 | -0.4 | -0.36 | 0.03 | -0.46 | 0.41 | -0.13 | 0.55 | 0.4 | 0.01 | 0.28 | 0.7 | -0.4275 | -0.0075 | 0.2475 | 0.490918 | 0.219636 | 0.984151 | 1 | 0.999631 | 0.808319 |
| Tmx2_D3Z2J6_S205s_1 | -0.39 | 0.12 | -0.59 | -1.04 | -0.44 | -0.2 | -0.67 | -1.35 | -0.59 | -0.28 | -0.7 | -0.38 | 0.3 | 0.16 | -1.11 | 0.35 | -0.19 | -0.0125 | 0.4 | 0.363658 | 0.605956 | 0.96873 | 1 | 0.999631 | 0.743747 |
| Tmx4_Q8C0L0_S304s_1 | 1.62 | 0.46 | 1.5 | 0.69 | -0.76 | 0.5 | 1.65 | -0.7 | -0.56 | 0.74 | 0.21 | -0.53 | 0.15 | -0.6 | 0.13 | -0.19 | -0.895 | -1.1025 | -1.195 | 0.009338 | 0.147767 | 0.026201 | 1 | 0.999631 | 0.367426 |
| Tnc_Q80YX0_S72s_1 | -0.03 | 0.48 | 0.35 | -0.14 | 0.49 | 0.7 | -0.07 | 0.38 | 0.32 | 0.54 | 0.49 | 0.34 | 0.91 | 1.03 | 0.54 | 0.82 | 0.21 | 0.2575 | 0.66 | 0.045719 | 0.476515 | 0.350817 | 1 | 0.999631 | 0.477252 |
| Tnfrsf19_Q9JLL3_S273s_1 | 0.74 | 0.11 | -0.09 | 0.44 | 0.23 | 0.18 | 0.47 | 0.38 | 0.33 | 0.53 | -0.02 | -0.12 | -0.42 | -0.1 | 0.29 | 0.75 | 0.015 | -0.12 | -0.17 | 0.639466 | 0.95695 | 0.697166 | 1 | 0.999631 | 0.870995 |
| Tnfrsf21_Q9EPU5_S565s_1 | 0.55 | 0.34 | 1.3 | 0.68 | 0.24 | 1.32 | 0.42 | 1.29 | 0.98 | 1.51 | 0.73 | 0.92 | 0.53 | 0.05 | 1.15 | -0.02 | 0.1 | 0.3175 | -0.29 | 0.449573 | 0.787026 | 0.338003 | 1 | 0.999631 | 0.787118 |
| Tnik_E9PUL9_S519s_1 | 1.05 | -0.17 | -1.51 | 1.62 | 1.37 | 0.59 | 2.5 | 1.38 | 0.21 | -2 | 1.28 | -0.78 | -4.4 | -0.51 | -1.18 | -0.87 | 1.2125 | -0.57 | -1.9875 | 0.063974 | 0.110644 | 0.512404 | 1 | 0.999631 | 0.513532 |
| Tnik_E9PUL9_S581s_1 | 1.85 | 0.65 | -0.74 | 1.02 | 0.82 | 1.01 | 1.13 | 0.29 | 0.33 | 0.2 | 0.13 | -1.62 | -1.22 | 0.92 | -1.38 | -0.46 | 0.1175 | -0.935 | -1.23 | 0.091574 | 0.824131 | 0.167285 | 1 | 0.999631 | 0.557085 |
| Tnik_E9PUL9_S611s_1 | -0.83 | -0.59 | 0.28 | -0.06 | -0.51 | -0.77 | -0.34 | -0.3 | -0.56 | -0.04 | -0.02 | 0.81 | -0.01 | -0.44 | 0.32 | 0.84 | -0.18 | 0.3475 | 0.4775 | 0.245157 | 0.577763 | 0.38768 | 1 | 0.999631 | 0.685751 |
| Tnik_E9PUL9_S649s_1 | 1.13 | 0.93 | 0.79 | 1.06 | 0.99 | 1.08 | 0.86 | 1.08 | 0.52 | 0.15 | 1.09 | 0.37 | 0.17 | 0.9 | 0.22 | 0.48 | 0.025 | -0.445 | -0.535 | 0.094171 | 0.91637 | 0.155023 | 1 | 0.999631 | 0.558128 |
| Tnik_E9PUL9_S651s_1 | 1.61 | -0.41 | -2.04 | 0.88 | 2.06 | 0.41 | 0.87 | 1.07 | 0.63 | -0.57 | 0.62 | -0.57 | -2.06 | 0.24 | -0.45 | -0.37 | 1.0925 | 0.0175 | -0.67 | 0.420399 | 0.175823 | 0.981812 | 1 | 0.999631 | 0.77267 |
| Tnik_E9PUL9_S659s_1 | 0.28 | 0.28 | -0.19 | -0.07 | -0.09 | 0.29 | -0.08 | -0.08 | 0.73 | 0.74 | 0.43 | 0.7 | 0.3 | 0.87 | 0.49 | 0.81 | -0.065 | 0.575 | 0.5425 | 0.089502 | 0.80216 | 0.050669 | 1 | 0.999631 | 0.555926 |
| Tnik_E9PUL9_S678s_1 | 2.19 | -0.19 | -1.29 | 1.82 | 1.37 | 1.01 | 1.97 | 1.21 | 0.62 | 0.48 | 1.16 | 0.04 | -2.69 | -0.38 | -0.44 | -0.41 | 0.7575 | -0.0575 | -1.6125 | 0.087626 | 0.325033 | 0.939463 | 1 | 0.999631 | 0.553787 |
| Tnik_E9PUL9_S740s_1 | -0.74 | -0.64 | -0.72 | -1.06 | -0.95 | -1.18 | -0.87 | -0.69 | -0.86 | -0.56 | -0.68 | -0.21 | -1.39 | 0.13 | -1.02 | -0.34 | -0.1325 | 0.2125 | 0.135 | 0.726867 | 0.603213 | 0.442706 | 1 | 0.999631 | 0.905774 |
| Tnik_E9PUL9_S925s_1 | -0.48 | -0.76 | -0.39 | -0.31 | -0.52 | -0.58 | -0.42 | -0.94 | -0.77 | -0.62 | -0.15 | -0.16 | -0.99 | -0.66 | -0.7 | -0.06 | -0.13 | 0.06 | -0.1175 | 0.706579 | 0.616565 | 0.831465 | 1 | 0.999631 | 0.898087 |
| Tnik_E9PUL9_T552t_1 | 0.17 | -0.11 | -0.24 | 0.54 | 1.03 | 0.68 | 0.07 | 0.48 | 0.37 | -0.01 | 0.17 | 0.29 | -0.73 | 0.88 | 0.4 | 0.58 | 0.475 | 0.115 | 0.1925 | 0.640746 | 0.155565 | 0.686676 | 1 | 0.999631 | 0.871305 |
| Tnk2_D3Z3U5_S757s_1 | 1.03 | 0.25 | 0.5 | 1.14 | 1.09 | 0.51 | 0.79 | -0.69 | 0.06 | 0.77 | -0.98 | 0.36 | -0.14 | 0.31 | -0.52 | -0.88 | -0.305 | -0.6775 | -1.0375 | 0.01809 | 0.488213 | 0.138854 | 1 | 0.999631 | 0.391024 |
| Tnk2_D3Z3U5_S785s_1 | 1.71 | -0.31 | 0.08 | 0.98 | 1.56 | 1.9 | 1.28 | 0.85 | 0.8 | 0.53 | 0.91 | -0.55 | -0.74 | 0.87 | 1.5 | 1.24 | 0.7825 | -0.1925 | 0.1025 | 0.867483 | 0.127087 | 0.715878 | 1 | 0.999631 | 0.9567 |
| Tnks1bp1_P58871_S1063s_1 | -0.29 | 0.18 | 1.17 | 0.51 | -0.1 | -0.12 | 0.38 | 0.28 | 0.15 | -0.28 | 0.66 | 1.35 | 0.44 | 0.32 | 1.47 | 1.11 | -0.2825 | 0.0775 | 0.4425 | 0.309331 | 0.43724 | 0.86478 | 1 | 0.999631 | 0.718845 |
| Tnks1bp1_P58871_S1131s_1 | 1.55 | 0.22 | -0.81 | 0.85 | 1.66 | 1.16 | 1.06 | 1.38 | 0.83 | 0.2 | 1.17 | -0.12 | -0.96 | 0.96 | 0.32 | 0.65 | 0.8625 | 0.0675 | -0.21 | 0.725807 | 0.102595 | 0.900473 | 1 | 0.999631 | 0.90568 |
| Tnks1bp1_P58871_S1290s_1 | 1.34 | -0.29 | -0.63 | 0.45 | 0.98 | 0.44 | 0.44 | 0.05 | 0.34 | 0.56 | -0.07 | -0.48 | -0.67 | 0.05 | 0.06 | -0.2 | 0.26 | -0.13 | -0.4075 | 0.387926 | 0.573473 | 0.784719 | 1 | 0.999631 | 0.757344 |
| Tnks1bp1_P58871_S1375s_1 | -0.48 | -0.33 | -0.21 | -0.05 | -0.18 | -0.28 | -0.33 | -0.21 | -0.75 | -0.38 | 0.04 | -0.18 | -0.11 | -0.14 | -0.4 | 0.11 | 0.0175 | -0.05 | 0.1325 | 0.63602 | 0.941795 | 0.860142 | 1 | 0.999631 | 0.869397 |
| Tnks1bp1_P58871_S1437s_1 | -0.55 | -0.57 | 0.15 | -0.62 | -0.66 | -0.95 | -0.44 | -0.42 | -0.89 | -0.76 | -0.65 | 0.2 | 0.33 | -1.1 | -0.14 | 0 | -0.22 | -0.1275 | 0.17 | 0.663147 | 0.45696 | 0.714689 | 1 | 0.999631 | 0.88184 |
| Tnks1bp1_P58871_S1611s_1 | -0.28 | 0.09 | 0.85 | 0.25 | -0.24 | 0.46 | 0.04 | -0.02 | -0.01 | 0.39 | 0.42 | 0.48 | 0.92 | 0.05 | 0.38 | 0.41 | -0.1675 | 0.0925 | 0.2125 | 0.54957 | 0.606546 | 0.772895 | 1 | 0.999631 | 0.835379 |
| Tnks1bp1_P58871_S1657s_1 | 1 | -0.49 | 0.85 | 1.05 | 1.09 | -0.42 | 1.4 | 0.69 | 0.45 | -0.64 | 1.3 | 0.95 | -1.84 | -1.04 | 0.36 | -0.86 | 0.0875 | -0.0875 | -1.4475 | 0.022563 | 0.862579 | 0.867436 | 1 | 0.999631 | 0.407815 |
| Tnks1bp1_P58871_S220s_1 | -0.13 | -0.6 | -3.14 | -1.42 | -0.22 | -0.49 | -0.9 | -0.86 | -0.03 | -0.45 | -1.06 | -1.71 | -0.45 | -1.17 | -1.84 | -3.03 | 0.705 | 0.51 | -0.3 | 0.691882 | 0.267137 | 0.457786 | 1 | 0.999631 | 0.894017 |
| Tnks1bp1_P58871_S429s_1 | 0.6 | -0.5 | -0.2 | 0.76 | 1.3 | -0.49 | 0.55 | 0.23 | 0.38 | -0.47 | 0.26 | 0.49 | -0.31 | 0.22 | -0.42 | 0.35 | 0.2325 | -7.9E-17 | -0.205 | 0.601741 | 0.616175 | 1 | 1 | 1 | 0.856765 |
| Tnks1bp1_P58871_S539s_1 | -0.47 | 0.06 | 0.54 | 0.42 | 0.05 | 0.22 | 0.08 | 1.08 | -0.08 | -0.78 | 0.47 | 0.88 | 0.01 | 0.44 | 0.55 | 1.04 | 0.22 | -0.015 | 0.3725 | 0.314186 | 0.542589 | 0.971881 | 1 | 0.999631 | 0.72001 |
| Tnks1bp1_P58871_S539s_2 | -0.55 | -0.09 | 0.32 | 0.21 | -0.43 | -0.03 | 0.08 | -0.22 | -0.64 | -1 | 0.01 | -0.38 | -0.6 | -0.74 | -0.51 | 0.91 | -0.1225 | -0.475 | -0.2075 | 0.636198 | 0.67741 | 0.179908 | 1 | 0.999631 | 0.869397 |
| Tnks1bp1_P58871_S568s_1 | 1.3 | -0.42 | -1.02 | 0.68 | 1.84 | -0.82 | 0.82 | 0.22 | -0.02 | -1.02 | -0.34 | -0.22 | -0.91 | -0.63 | -1.16 | -0.49 | 0.38 | -0.535 | -0.9325 | 0.095302 | 0.578695 | 0.32608 | 1 | 0.999631 | 0.558128 |
| Tnks1bp1_P58871_S602s_1 | -1.53 | -0.94 | -0.39 | -0.82 | -1.11 | -0.92 | -0.93 | -0.67 | -1.11 | -1.07 | -0.42 | -0.3 | 0.02 | -0.51 | -0.6 | 0.08 | 0.0125 | 0.195 | 0.6675 | 0.080757 | 0.96764 | 0.584376 | 1 | 0.999631 | 0.545579 |
| Tnks1bp1_P58871_S763s_1 | 1.37 | -1.48 | -1.46 | 0.4 | 1.19 | -1.56 | 1.04 | 0.1 | -0.45 | -2.38 | 0.17 | -0.56 | -0.83 | -1.41 | -0.93 | -1.22 | 0.485 | -0.5125 | -0.805 | 0.231907 | 0.560836 | 0.519898 | 1 | 0.999631 | 0.682192 |
| Tnks1bp1_P58871_S866s_1 | 0.03 | 1.37 | 1.05 | 0.6 | 0.84 | 1.05 | 0.7 | 1.14 | 1.21 | 0.12 | 1.89 | 1.19 | 0.51 | 1.27 | 1.35 | 1.63 | 0.17 | 0.34 | 0.4275 | 0.300139 | 0.620896 | 0.46264 | 1 | 0.999631 | 0.713499 |
| Tnks1bp1_P58871_S887s_1 | 1.78 | -0.62 | -0.41 | 0.71 | 2.04 | 0.77 | 0.97 | 0.43 | 1.75 | -0.24 | 0.46 | -0.48 | 0.29 | 1.09 | 0.34 | 0.23 | 0.6875 | 0.0075 | 0.1225 | 0.823655 | 0.262967 | 0.99105 | 1 | 0.999631 | 0.939038 |
| Tnks1bp1_P58871_S911s_1 | 2.13 | 1.35 | -1.37 | 0.5 | 1.04 | 2.62 | 1.44 | -0.71 | 1.09 | 3.15 | -0.2 | -2.01 | -0.28 | -0.35 | -1.7 | -1.95 | 0.445 | -0.145 | -1.7225 | 0.044625 | 0.616929 | 0.898173 | 1 | 0.999631 | 0.475443 |
| Tnks1bp1_P58871_S912s_1 | 2.39 | 0.74 | 2.37 | 2.08 | 1.91 | 2.08 | 2.08 | 0.6 | 1.72 | 2.26 | 1.56 | 0.93 | -2.31 | -0.08 | 0.8 | 0.91 | -0.2275 | -0.2775 | -2.065 | 0.018169 | 0.64936 | 0.553232 | 1 | 0.999631 | 0.391024 |
| Tnks1bp1_P58871_S927s_1 | 0.8 | 0.42 | -0.55 | 1.4 | 1.13 | 0.15 | 0.7 | 1.11 | -0.01 | -0.62 | 0.64 | -0.94 | -1.59 | -0.35 | 1.48 | -0.21 | 0.255 | -0.75 | -0.685 | 0.320406 | 0.576213 | 0.162913 | 1 | 0.999631 | 0.723684 |
| Tnks1bp1_P58871_T131t_1 | 1.62 | 1.86 | 0.47 | 1.01 | 2.33 | 2.97 | 1.12 | 1.39 | 1.41 | 1.8 | 0.89 | 0.31 | 0.14 | 1.33 | 0.24 | 1.6 | 0.7125 | -0.1375 | -0.4125 | 0.394268 | 0.174094 | 0.757048 | 1 | 0.999631 | 0.759303 |
| Tnks1bp1_P58871_T533t_1 | -0.78 | -0.44 | -1.88 | -1.33 | -1.47 | -0.46 | -1.38 | -1.72 | -0.24 | -0.21 | -0.9 | -1.19 | 0.43 | -1.33 | -0.58 | -1.06 | -0.15 | 0.4725 | 0.4725 | 0.341143 | 0.719501 | 0.263862 | 1 | 0.999631 | 0.733032 |
| Tnks1bp1_P58871_T533t_2 | -0.55 | -0.09 | 0.32 | 0.21 | -0.43 | -0.03 | 0.08 | -0.22 | -0.64 | -1 | 0.01 | -0.38 | -0.6 | -0.74 | -0.51 | 0.91 | -0.1225 | -0.475 | -0.2075 | 0.636198 | 0.67741 | 0.179908 | 1 | 0.999631 | 0.869397 |
| Tnpo1_J3QMX2_S24s_1 | 0.46 | -1.06 | -0.86 | -1.32 | -1.3 | -1.56 | -0.96 | -1.12 | -1.28 | 0.19 | -0.42 | -0.01 | -0.45 | -1.14 | -0.81 | -0.51 | -0.54 | 0.315 | -0.0325 | 0.940196 | 0.212988 | 0.524499 | 1 | 0.999631 | 0.981146 |
| Tnrc18_F6Z7L1_T1852t_1 | 1.34 | 1.17 | 1.43 | 1.02 | 1.09 | 0.98 | 0.94 | 1.37 | 0.66 | 1.58 | 1.54 | 1.42 | 0.6 | 1.57 | 1.47 | 1.61 | -0.145 | 0.06 | 0.0725 | 0.826181 | 0.567577 | 0.844424 | 1 | 0.999631 | 0.940686 |
| Tnrc6b_Q8BKI2_S1313s_1 | 1.53 | 3.6 | 3.36 | 1.58 | 4.89 | 3.31 | 4.34 | 2.63 | 6.56 | 1.26 | 5 | 5.67 | 4.4 | 2.19 | 3.57 | 1.44 | 1.275 | 2.105 | 0.3825 | 0.617878 | 0.081996 | 0.082699 | 1 | 0.999631 | 0.861861 |
| Tnrc6b_Q8BKI2_S1793s_1 | -0.11 | 0.33 | 0.53 | 0.01 | -0.01 | -0.33 | -0.06 | 0.07 | 0.2 | -0.7 | 0.29 | -0.18 | -0.66 | 0.48 | -0.65 | 0.32 | -0.2725 | -0.2875 | -0.3175 | 0.408938 | 0.320265 | 0.385935 | 1 | 0.999631 | 0.766996 |

| | | | | | | | | | | | | | | | | | | | | | | | | | |
|---|---|---|---|---|---|---|---|---|---|---|---|---|---|---|---|---|---|---|---|---|---|---|---|---|---|
| Tnrc6c_B1ATC3_T986t_1 | 0.57 | 0.72 | 1.23 | 1.45 | 0.25 | 0.71 | 0.68 | 0.62 | 0.68 | 0.81 | 1.4 | 1.28 | 1.3 | 1.14 | 1.01 | 1.4 | -0.4275 | 0.05 | 0.22 | 0.489415 | 0.173955 | 0.878344 | 1 | 0.999631 | 0.807405 |
| Tns1_E9Q0S6_S1054s_1 | 0.17 | 0.21 | -0.2 | -0.21 | 0.76 | 0.06 | -0.01 | -0.85 | 0.02 | 1.11 | 0.02 | -0.65 | 0.25 | -0.59 | -0.02 | -0.28 | -0.0025 | 0.1325 | -0.1525 | 0.623178 | 0.994578 | 0.738565 | 1 | 0.999631 | 0.862913 |
| Tns1_E9Q0S6_S1062s_1 | -0.2 | 0.18 | 0.11 | 0.72 | -0.13 | 0.41 | -0.06 | -0.07 | -0.28 | 0.12 | 0.93 | 1 | 0.36 | 0.48 | 1.27 | 0.97 | -0.165 | 0.24 | 0.5675 | 0.124943 | 0.579535 | 0.538085 | 1 | 0.999631 | 0.596063 |
| Tns1_E9Q0S6_S1067s_1 | 0.38 | -0.6 | 0.69 | 0.3 | -0.93 | -0.23 | -0.02 | 0.31 | -1.2 | -0.79 | 0.28 | 0.33 | -0.65 | -0.04 | 1.06 | 0.17 | -0.41 | -0.5375 | -0.0575 | 0.898049 | 0.307613 | 0.260915 | 1 | 0.999631 | 0.966853 |
| Tns1_E9Q0S6_S1118s_1 | -0.27 | 0.43 | -0.79 | -1.62 | 0.41 | -0.23 | -0.8 | -0.28 | 0.35 | 0.18 | -1.37 | -2.24 | 0.92 | -0.82 | -0.01 | -0.75 | 0.3375 | -0.2075 | 0.3975 | 0.476633 | 0.482117 | 0.760239 | 1 | 0.999631 | 0.799947 |
| Tns1_E9Q0S6_S1126s_1 | 0.21 | 0.43 | -0.39 | -0.53 | 0.17 | -0.29 | -0.6 | -0.58 | -0.34 | 0.14 | -0.86 | -0.08 | 0.69 | -0.05 | -0.85 | -0.29 | -0.255 | -0.215 | -0.055 | 0.893744 | 0.450333 | 0.546659 | 1 | 0.999631 | 0.965164 |
| Tns1_E9Q0S6_S1346s_1 | 0.28 | 0.27 | 0.56 | 0.63 | 1.2 | 0.11 | -0.16 | -0.12 | 0.47 | 0.89 | 0.76 | 1.66 | 0.02 | 0.26 | 0.39 | 0.75 | -0.1775 | 0.51 | -0.08 | 0.78545 | 0.622139 | 0.141535 | 1 | 0.999631 | 0.923063 |
| Tns1_E9Q0S6_S1363s_1 | 0.63 | -0.06 | 0.02 | 0.16 | 0.36 | -0.16 | 0.27 | 0.23 | -0.7 | 0.5 | 0.81 | 0.73 | -0.75 | -0.29 | -0.11 | 0.18 | -0.0125 | 0.1475 | -0.43 | 0.207096 | 0.963973 | 0.711275 | 1 | 0.999631 | 0.660822 |
| Tns1_E9Q0S6_S1397s_1 | -0.01 | -1.66 | -2.09 | -1.42 | -0.53 | -0.81 | -1.67 | -1.13 | -1.11 | -0.54 | -1.13 | -0.32 | -1.04 | -1.68 | -2.38 | -2.45 | 0.26 | 0.52 | -0.5925 | 0.276307 | 0.59406 | 0.289858 | 1 | 0.999631 | 0.700187 |
| Tns1_E9Q0S6_S1468s_1 | -5.56 | -5.47 | -3.77 | -4.37 | -2.68 | -4.59 | -0.84 | -3.56 | -1.87 | -5.23 | -1.96 | -1.91 | -3.9 | -6.6 | -2.05 | -4.1 | 1.875 | 2.05 | 0.63 | 0.484292 | 0.037498 | 0.030072 | 1 | 0.999631 | 0.803422 |
| Tns1_E9Q0S6_S154s_1 | -0.2 | -0.51 | 0.04 | -1.26 | -0.55 | -0.97 | -0.52 | -0.99 | -0.33 | -1.41 | -1.41 | -1.87 | 0.33 | -0.2 | 0.39 | -1.91 | -0.275 | -0.7725 | 0.135 | 0.810251 | 0.431261 | 0.098159 | 1 | 0.999631 | 0.936878 |
| Tns1_E9Q0S6_S792s_1 | -0.39 | -0.44 | -0.02 | -0.61 | 0.55 | -0.74 | 0.14 | -0.37 | -0.31 | -0.38 | -0.32 | 0.45 | -0.58 | 0.01 | -0.11 | 0.13 | 0.26 | 0.225 | 0.2275 | 0.458138 | 0.455019 | 0.467135 | 1 | 0.999631 | 0.790722 |
| Tns1_E9Q0S6_S847s_1 | -0.83 | -0.33 | 0.97 | 0.72 | 0.09 | 0.01 | -0.28 | -0.11 | -0.32 | -0.35 | 0.22 | 0.62 | -0.65 | -0.44 | 0.61 | 0.79 | -0.205 | -0.09 | -0.055 | 0.916969 | 0.633351 | 0.848138 | 1 | 0.999631 | 0.973513 |
| Tns2_Q8CGB6_S1064s_1 | 0.48 | 0.3 | 0.35 | -0.38 | 0.83 | 0.46 | 0.55 | -1.01 | 2.5 | 0.18 | 0.86 | 1.84 | 2.96 | 0.55 | -0.76 | -0.29 | 0.02 | 1.1575 | 0.4275 | 0.570062 | 0.963962 | 0.046489 | 1 | 0.999631 | 0.84338 |
| Tns2_Q8CGB6_S802s_1 | 1.08 | -0.75 | 0.13 | 0.18 | -0.05 | -0.53 | -0.05 | -0.84 | -0.37 | 0.98 | 0.25 | 0.32 | -0.35 | -0.65 | 0 | -0.11 | -0.5275 | 0.135 | -0.4375 | 0.306807 | 0.22599 | 0.766656 | 1 | 0.999631 | 0.717667 |
| Tns2_Q8CGB6_S908s_1 | 1.26 | 0.2 | 0.62 | 0.44 | 0.94 | 0.66 | 0 | -0.57 | 0.87 | 0.71 | 0.15 | 1.12 | 0.7 | 1.63 | 0.6 | 0.3 | -0.3725 | 0.0825 | 0.1775 | 0.653735 | 0.372529 | 0.812306 | 1 | 0.999631 | 0.878344 |
| Tns2_Q8CGB6_Y460y_1 | -0.15 | 0.76 | 0.44 | 0.1 | 0.22 | 0.45 | -0.53 | -0.19 | 0.23 | 0.76 | -0.02 | 0.64 | 0.98 | 0.13 | 0.53 | 0.56 | -0.3 | 0.115 | 0.2625 | 0.439311 | 0.37872 | 0.723928 | 1 | 0.999631 | 0.781983 |
| Tns3_Q5SSZ5_S934s_1 | 1.03 | -0.07 | 0.71 | 0.57 | -0.34 | -0.14 | 0.33 | -0.25 | -0.02 | 0.97 | -0.19 | -0.48 | -0.79 | -0.96 | -0.36 | -0.71 | -0.66 | -0.49 | -1.265 | 0.003375 | 0.063666 | 0.241126 | 1 | 0.999631 | 0.351308 |
| Tollip_Q9QZ06_T25t_1 | -1.42 | -0.16 | -0.26 | -0.46 | -0.11 | -1.65 | -0.97 | 0.03 | -1.17 | -1.06 | -0.04 | 0.3 | 0.28 | -0.42 | -0.32 | 0.39 | -0.1 | 0.0825 | 0.5575 | 0.171126 | 0.830136 | 0.856323 | 1 | 0.999631 | 0.641448 |
| Tom1_Q3UDC3_S160s_1 | -0.66 | 0.12 | -0.14 | -0.09 | 0.3 | -0.07 | -0.36 | -0.12 | -0.15 | -0.11 | -0.22 | -0.36 | 0.27 | 0.21 | 0.04 | 0.1 | 0.13 | -0.0175 | 0.3475 | 0.25029 | 0.653406 | 0.949466 | 1 | 0.999631 | 0.686417 |
| Tom1_Q3UDC3_S176s_1 | 0.77 | 0.12 | -0.49 | 0.44 | 0.65 | 0.64 | 0.54 | 0.02 | 0.26 | 0.42 | 0.19 | -0.19 | 0.23 | 0.48 | 0.07 | -0.04 | 0.2525 | -0.04 | -0.025 | 0.942876 | 0.466158 | 0.906878 | 1 | 0.999631 | 0.981629 |
| Tom1_Q3UDC3_S180s_1 | 3.12 | -0.75 | -0.21 | 1.42 | -0.37 | 0.88 | 2.22 | 0.17 | -0.03 | 1.45 | 0.14 | -1.22 | -2.38 | -0.97 | -0.28 | -0.84 | -0.17 | -0.81 | -2.0125 | 0.03693 | 0.849893 | 0.376432 | 1 | 0.999631 | 0.456822 |
| Tom1_Q3UDC3_S183s_1 | 0.82 | -0.68 | 0.03 | -1.28 | -1.62 | -1.29 | -0.7 | -1.77 | -0.91 | 1.19 | -1.99 | -1.31 | -1.67 | -1.99 | -1.17 | -2.18 | -1.0675 | -0.4775 | -1.475 | 0.012034 | 0.049201 | 0.51686 | 1 | 0.999631 | 0.385092 |
| Tom1_Q3UDC3_S400s_1 | 2.45 | -0.68 | 0.24 | 1.8 | 1.25 | 1.64 | 2.11 | 1.5 | 0.82 | 1.25 | 1.76 | 0.33 | 0.1 | 0.08 | 1.03 | -0.02 | 0.6725 | 0.0875 | -0.655 | 0.343161 | 0.319876 | 0.899462 | 1 | 0.999631 | 0.73348 |
| Tom1_Q3UDC3_S486s_1 | 0.11 | 0.17 | -0.8 | 0.28 | 0.31 | -0.07 | -0.01 | 0.39 | 0.22 | 0.08 | 0.2 | -0.06 | -0.48 | 0.12 | 0 | 0.37 | 0.215 | 0.17 | 0.0625 | 0.861349 | 0.508409 | 0.595965 | 1 | 0.999631 | 0.954096 |
| Tom1_Q3UDC3_S496s_1 | 0.08 | 0.4 | 0.56 | -0.18 | 0.66 | 0.54 | -0.33 | 0.02 | 0.7 | 1.08 | 0.28 | 1.17 | 0.72 | 0.98 | 0.94 | 0.75 | 0.0075 | 0.5925 | 0.6325 | 0.052613 | 0.981688 | 0.089595 | 1 | 0.999631 | 0.489777 |
| Tom1l1_Q8BZR6_S93s_1 | -1.31 | 0.31 | 0.52 | -0.31 | 0.3 | 0.18 | -0.19 | -0.08 | -0.19 | -1.38 | -0.91 | 0.38 | 0.18 | -0.36 | -0.45 | -0.56 | 0.25 | -0.3275 | -0.1 | 0.821861 | 0.556434 | 0.539587 | 1 | 0.999631 | 0.938464 |
| Tom1l2_Q5SXA5_S110s_1 | -0.26 | 0.01 | 0.07 | -0.25 | -0.23 | -0.04 | -0.45 | -0.11 | -0.24 | -0.05 | -0.16 | 0.17 | 0.04 | 0.34 | 0.03 | 0.26 | -0.1 | 0.0375 | 0.275 | 0.323879 | 0.688406 | 0.883729 | 1 | 0.999631 | 0.724485 |
| Tom1l2_Q5SXA5_S344s_1 | 2.42 | -0.52 | -1.51 | 0.62 | 1.09 | 1.18 | 1.01 | -0.47 | 1.17 | 0.72 | -0.35 | -1.39 | -1.55 | 0.07 | -1.1 | -1.69 | 0.45 | -0.215 | -1.32 | 0.128037 | 0.582094 | 0.808668 | 1 | 0.999631 | 0.598425 |
| Tom1l2_Q5SXA5_S429s_1 | -0.53 | -0.43 | -0.24 | -0.42 | -0.53 | -0.59 | -0.65 | -0.69 | -0.63 | -0.63 | -0.4 | -0.24 | 0.09 | -0.43 | -0.43 | -0.03 | -0.21 | -0.07 | 0.205 | 0.47277 | 0.37795 | 0.782238 | 1 | 0.999631 | 0.799014 |
| Tomm20_Q9DCC8_S138s_1 | -0.01 | 1.05 | 1.12 | 0.12 | 0.26 | 1.2 | 1.38 | 2.04 | 0.33 | 0.62 | 2.58 | 2.82 | 1.31 | -0.57 | 1.34 | 0.85 | 0.65 | 1.0175 | 0.1625 | 0.75139 | 0.176143 | 0.137329 | 1 | 0.999631 | 0.912912 |
| Tomm22_Q9CPQ3_S15s_1 | -0.53 | -0.32 | -0.13 | -0.94 | -0.74 | -0.82 | -1.03 | -0.68 | -0.79 | -0.77 | -0.72 | -0.51 | 0.26 | -0.45 | -0.36 | -0.13 | -0.3375 | -0.2175 | 0.31 | 0.343328 | 0.239219 | 0.446375 | 1 | 0.999631 | 0.73348 |
| Tomm34_Q9CYG7_S8s_1 | -0.91 | -0.05 | 0.54 | 0.13 | 0.42 | 0.27 | -0.11 | 0.61 | 0.14 | -0.22 | 0.63 | 1.27 | 0.01 | -0.76 | 1.02 | 0.92 | 0.37 | 0.5275 | 0.37 | 0.463445 | 0.321755 | 0.24812 | 1 | 0.999631 | 0.793615 |
| Tomm70a_Q9CZW5_S105s_2 | -0.3 | -0.69 | -0.82 | -1.17 | -1.61 | -0.7 | -1.03 | -0.99 | -1.21 | -0.25 | -2.86 | -2.73 | -1.12 | -0.08 | -1.14 | -0.62 | -0.3375 | -1.0175 | 0.005 | 0.988879 | 0.297374 | 0.112723 | 1 | 0.999631 | 0.995427 |
| Tomm70a_Q9CZW5_S108s_1 | -1.08 | -6.44 | -4.3 | -2.49 | -4.31 | -3.54 | -2.16 | -3.62 | -5 | -3.51 | -4.13 | -3.1 | -4.34 | -4.26 | -2.62 | -3.52 | 0.17 | -0.3575 | -0.1075 | 0.918069 | 0.87334 | 0.736447 | 1 | 0.999631 | 0.973954 |
| Tomm70a_Q9CZW5_S94s_1 | -0.64 | -1.87 | -2.2 | -1.49 | -2.13 | -2.11 | -1.55 | -2.01 | -2.04 | -1.16 | -2.59 | -2.26 | -1.16 | -1.72 | -1.9 | -1.56 | -0.4 | -0.4625 | -0.035 | 0.929753 | 0.303627 | 0.314649 | 1 | 0.999631 | 0.97754 |
| Tomm70a_Q9CZW5_S94s_2 | -0.3 | -0.69 | -0.9 | -1.1 | -1.61 | -0.7 | -0.98 | -1.12 | -1.21 | -0.25 | -2.53 | -1.61 | -1.12 | -0.08 | -0.88 | -0.85 | -0.355 | -0.6525 | 0.015 | 0.96526 | 0.269593 | 0.196906 | 1 | 0.999631 | 0.987856 |
| Top1_Q04750_S114s_1 | 0.3 | 0.7 | 0.36 | 0.42 | 0.27 | 0.06 | 0.1 | 0.58 | 0.31 | 0.69 | 0.66 | 0.56 | 0.84 | 0.78 | 0.63 | 0.88 | -0.1925 | 0.11 | 0.3375 | 0.226639 | 0.460761 | 0.66929 | 1 | 0.999631 | 0.677921 |
| Top2b_Q64511_S1537s_1 | -0.02 | -0.25 | 0.05 | -1 | -1.37 | -0.37 | -0.44 | -0.96 | -0.32 | 0.18 | -0.19 | -1.27 | -0.06 | -0.44 | -0.61 | -0.89 | -0.48 | -0.095 | -0.195 | 0.582604 | 0.203072 | 0.813542 | 1 | 0.999631 | 0.847801 |
| Top2b_Q64511_S1539s_1 | -0.07 | -2.25 | -1.02 | -0.95 | -2.34 | -2.41 | -0.96 | -1.34 | -1.73 | -0.85 | -1.12 | -1.08 | -1.72 | -3.1 | -1.39 | -1.78 | -0.69 | -0.1225 | -0.925 | 0.114587 | 0.214956 | 0.794442 | 1 | 0.999631 | 0.579001 |
| Topors_Q80Z37_S1025s_1 | 0.47 | 1.06 | 1.69 | 1.31 | 0.84 | 0.65 | 0.71 | 1.15 | 1.04 | 0.95 | 1.28 | 1.29 | 1.42 | 1.19 | 1.63 | 1.31 | -0.295 | 0.0075 | 0.255 | 0.458227 | 0.373574 | 0.981602 | 1 | 0.999631 | 0.790722 |
| Tor1aip1_Q921T2_S140s_1 | -0.06 | -0.53 | -0.49 | -0.2 | -0.18 | 0.3 | -0.4 | -0.32 | -0.09 | -0.2 | -0.59 | -0.37 | -0.73 | 0.12 | 0.51 | -0.39 | 0.17 | 0.0075 | 0.1975 | 0.577843 | 0.544109 | 0.977743 | 1 | 0.999631 | 0.84622 |
| Tor1aip1_Q921T2_S151s_1 | 0.79 | 1.12 | 2.57 | 1.31 | 0.32 | 1.37 | 0.85 | 0.69 | 0.92 | 2.21 | 0.71 | 0.97 | 1.04 | 0.58 | 1.13 | 1.55 | -0.64 | -0.245 | -0.3725 | 0.406482 | 0.164825 | 0.62115 | 1 | 0.999631 | 0.766226 |
| Tor1aip1_Q921T2_S253s_1 | 0.67 | -0.34 | 0.8 | 0.68 | -0.13 | 0 | 1.02 | -0.06 | -0.02 | 0.22 | 1.36 | -0.48 | -0.33 | -1.21 | 0.01 | -0.43 | -0.245 | -0.1825 | -0.9425 | 0.03623 | 0.533508 | 0.692806 | 1 | 0.999631 | 0.453963 |
| Tox2_E9Q0B9_S199s_1 | 0.27 | 0.53 | 1.25 | -0.07 | -0.38 | 0.85 | 0.27 | 0.87 | 0.09 | 1 | -0.01 | 0.78 | 0.17 | 0.18 | 0.8 | 0.99 | -0.0925 | -0.03 | 0.04 | 0.917093 | 0.820762 | 0.938929 | 1 | 0.999631 | 0.973513 |
| Tox2_E9Q0B9_S506s_1 | 1.66 | 1 | 0.61 | -0.12 | 0.8 | 1.38 | 0.38 | 0.8 | 1.19 | 2.06 | -0.58 | 0.13 | 0.38 | 1.07 | 0.84 | -0.56 | 0.0525 | -0.0875 | -0.355 | 0.480915 | 0.900621 | 0.888522 | 1 | 0.999631 | 0.801848 |
| Tp53bp1_P70399_S1622s_1 | -0.38 | 1.39 | 0.17 | -0.38 | -0.57 | 1.03 | -0.87 | 0.14 | 0.94 | 0.95 | 0.63 | 0.98 | 2.06 | 0.87 | 0.93 | 0.33 | -0.2675 | 0.675 | 0.8475 | 0.128697 | 0.626184 | 0.137523 | 1 | 0.999631 | 0.599259 |
| Tp53bp1_P70399_S1669s_1 | 0.85 | 1.55 | 1.64 | 0.74 | 0.53 | 1.07 | 0.73 | 1.1 | 1.38 | 1.66 | 1.4 | 1.67 | 1.21 | 1.85 | 1.61 | 1.95 | -0.3375 | 0.3325 | 0.46 | 0.203316 | 0.30453 | 0.301044 | 1 | 0.999631 | 0.659781 |
| Tp53bp1_P70399_S332s_1 | -0.23 | 0.72 | 1.5 | 0.17 | -0.08 | 0.79 | 0.23 | 0.5 | 0.73 | 0.32 | 0.33 | 1.23 | 1.42 | 0.82 | 0.55 | 1.12 | -0.18 | 0.1125 | 0.4375 | 0.320139 | 0.666472 | 0.794196 | 1 | 0.999631 | 0.723684 |
| Tp53bp1_P70399_S381s_1 | -0.48 | 0.21 | -0.04 | -0.49 | -0.15 | -0.34 | -0.47 | 0.08 | -0.18 | -0.35 | -0.67 | -0.1 | 0.93 | -0.14 | 0.03 | -0.12 | -0.02 | -0.125 | 0.375 | 0.309052 | 0.944128 | 0.672914 | 1 | 0.999631 | 0.718845 |
| Tp53bp1_P70399_S429s_1 | -0.76 | -1.3 | 0.12 | -0.22 | -0.48 | -0.86 | -0.62 | -1.09 | -0.5 | -0.98 | -0.98 | 0.12 | 0.23 | -0.29 | -1.01 | -0.22 | -0.2225 | -0.045 | 0.2175 | 0.605192 | 0.542683 | 0.913146 | 1 | 0.999631 | 0.85817 |
| Tp53bp1_P70399_S533s_1 | -0.05 | 1.06 | 1.17 | 0.51 | 0.59 | 0.62 | 0.58 | 0.81 | 1.09 | 0.96 | 0.98 | 1.46 | 0.98 | 1.06 | 1.26 | 1.12 | -0.0225 | 0.45 | 0.4325 | 0.230632 | 0.945331 | 0.213049 | 1 | 0.999631 | 0.681463 |
| Tp53bp1_P70399_S552s_1 | -0.52 | 0.67 | 0.77 | 0.53 | 0.17 | 0.37 | 0.45 | 0.41 | 0.43 | 0.41 | 0.84 | 1.27 | 0.81 | 1.1 | 1.45 | 1.55 | -0.0125 | 0.375 | 0.865 | 0.042807 | 0.970639 | 0.338802 | 1 | 0.999631 | 0.475443 |
| Tp53bp1_P70399_T323t_1 | 1.95 | 1.03 | 2.42 | 0.91 | 0.14 | 1.71 | 2.22 | -0.26 | 1.27 | 2.51 | -0.13 | 0.83 | -0.03 | -0.31 | -0.6 | -0.18 | -0.625 | -0.4575 | -1.8575 | 0.000997 | 0.333301 | 0.452441 | 1 | 0.999631 | 0.351308 |
| Tpbgl_Q8C013_S375s_1 | 1.13 | -0.38 | -0.72 | 0.27 | 0.13 | -0.05 | 0.18 | 0.01 | 0.9 | 1.01 | 0.51 | 0.11 | -0.18 | 0.83 | 0.21 | -0.12 | -0.0075 | 0.5575 | 0.11 | 0.81195 | 0.985409 | 0.23216 | 1 | 0.999631 | 0.936949 |
| Tpd52_F8WHQ1_S155s_1 | -1.28 | -0.58 | -0.9 | -1.06 | -0.03 | -0.91 | -0.71 | -1.5 | -1.24 | -2.37 | -0.96 | -0.5 | -0.72 | -0.97 | -1.04 | -1.14 | 0.1675 | -0.3125 | -0.0125 | 0.965702 | 0.643926 | 0.469034 | 1 | 0.999631 | 0.988111 |
| Tpd52_F8WHQ1_S186s_1 | 0.33 | 0.43 | -0.27 | 0.4 | 0.43 | 0.52 | 0.03 | 0.45 | -0.09 | 0.52 | 0.19 | 0.35 | -0.24 | 0.45 | 0.4 | 0.77 | 0.135 | 0.02 | 0.1225 | 0.716716 | 0.631993 | 0.94557 | 1 | 0.999631 | 0.903179 |
| Tpd52_F8WHQ1_S198s_1 | -8.33 | -10.96 | -4.29 | -4.1 | -9.4 | -7.3 | -3.62 | -5.17 | -10.93 | -5.67 | -3.56 | -4.73 | -9.39 | -6.81 | -3.5 | -3.96 | 0.5475 | 0.6975 | 1.005 | 0.579659 | 0.756606 | 0.723714 | 1 | 0.999631 | 0.84667 |
| Tpd52l1_B2RUC1_S142s_1 | 0.51 | -0.58 | 2.06 | -0.36 | -0.81 | 0.09 | 2.05 | -0.15 | 1.63 | -0.19 | 1.94 | 2.37 | 1.65 | -1.87 | 0.7 | -0.44 | -0.1125 | 1.03 | -0.3975 | 0.63718 | 0.881323 | 0.179714 | 1 | 0.999631 | 0.870011 |
| Tpd52l2_F6VQ81_S140s_1 | -0.33 | -0.59 | -0.73 | 0.04 | 0.02 | 0.13 | 0.59 | -0.4 | -0.39 | -0.14 | -0.45 | -0.25 | -1.88 | 0.11 | -0.56 | 0.43 | 0.4875 | 0.095 | -0.0725 | 0.88736 | 0.146167 | 0.735123 | 1 | 0.999631 | 0.963505 |
| Tpd52l2_Q8BKP1_S12s_1 | -0.26 | 0.18 | -0.99 | 0.03 | 0.19 | -0.15 | -0.38 | -0.26 | -0.17 | -0.04 | 0.05 | 0.01 | -0.12 | 0.08 | 0.28 | 0.07 | 0.11 | 0.2225 | 0.3375 | 0.331508 | 0.738903 | 0.495307 | 1 | 0.999631 | 0.727302 |
| Tpd52l2_Q8BKP1_S148s_1 | 0.92 | 2.38 | 2.01 | 0.03 | 3.43 | 0.74 | 2.78 | 2.06 | 4.87 | 0.99 | 4.13 | 5.09 | 4.12 | -0.16 | 1.94 | -0.62 | 0.9175 | 2.435 | -0.015 | 0.988406 | 0.207774 | 0.026236 | 1 | 0.999631 | 0.995409 |
| Tpd52l2_Q8BKP1_S186s_1 | -0.17 | 0.38 | -0.3 | -0.57 | -0.21 | -0.39 | -0.63 | 0.01 | -0.15 | -0.55 | -0.06 | 0.21 | 0.58 | 0.32 | -0.42 | -0.1 | -0.14 | 0.0275 | 0.26 | 0.467895 | 0.645315 | 0.93074 | 1 | 0.999631 | 0.795565 |
| Tpgs1_Q99MS8_S279s_1 | 0.29 | 0.25 | 1.08 | 0.07 | -0.25 | -0.06 | -0.02 | 0.07 | -0.22 | 0.47 | 0.45 | 0.17 | 0.48 | 0.61 | 0.31 | 1.08 | -0.4875 | -0.205 | 0.1975 | 0.566967 | 0.126848 | 0.537496 | 1 | 0.999631 | 0.842178 |
| Tph2_Q8CGV2_S19s_1 | -0.19 | -0.64 | -0.76 | -0.34 | -0.66 | -0.42 | -0.65 | -0.74 | -0.36 | 0.08 | -0.7 | -0.27 | 0.22 | -0.15 | -0.54 | -0.11 | -0.135 | 0.17 | 0.3375 | 0.282646 | 0.602553 | 0.563478 | 1 | 0.999631 | 0.704034 |
| Tpi1_P17751_S245s_1 | 1.4 | 1.35 | 1.62 | 0.97 | 0.78 | 1.11 | 2.09 | 0.84 | 0.85 | 1.43 | 1.34 | 1.26 | 0.15 | 0.94 | 0.97 | 0.28 | -0.13 | -0.115 | -0.75 | 0.042739 | 0.716865 | 0.684123 | 1 | 0.999631 | 0.475443 |
| Tpi1_P17751_S262s_1 | -0.08 | 1.51 | 0.31 | 0.09 | 0.89 | 0.46 | -0.38 | 0.03 | 0.72 | 0.19 | 0.33 | 0.99 | 1.5 | 0.88 | -0.47 | 0.36 | -0.2075 | 0.1 | 0.11 | 0.832694 | 0.639065 | 0.80788 | 1 | 0.999631 | 0.94321 |
| Tpi1_P17751_T228t_1 | 0.13 | 0.03 | -0.41 | 0.26 | -0.75 | 0.19 | 0.33 | 0.33 | 0.02 | 0.24 | 0.27 | 0.28 | -0.25 | 0.2 | 0.14 | 0.32 | 0.0225 | 0.2 | 0.1 | 0.73821 | 0.946555 | 0.464062 | 1 | 0.999631 | 0.908007 |
| Tpm1_G5E8R0_S6s_1 | 0.2 | -0.27 | 1.09 | 0.44 | -1.68 | -0.88 | -0.27 | -0.37 | -1.1 | 1.12 | 0.09 | 0.58 | -1.34 | -0.78 | 0.78 | 0.44 | -1.165 | -0.1925 | -0.59 | 0.287879 | 0.01957 | 0.711205 | 1 | 0.999631 | 0.705972 |
| Tpm3_D3Z6I8_S51s_1 | 0.07 | 1.01 | 0.97 | 0.84 | 0.47 | 1.05 | 0.17 | 0.66 | 0.85 | 1.03 | 1.02 | 1.04 | 0.34 | 0.91 | 1.36 | 1.01 | -0.135 | 0.2625 | 0.1825 | 0.611852 | 0.682941 | 0.392393 | 1 | 0.999631 | 0.859904 |
| Tpm3_P21107_S207s_1 | 0.35 | -0.04 | -0.01 | -0.2 | 0.28 | -0.2 | -0.11 | -0.74 | -0.04 | 0.25 | 0.05 | -0.97 | -0.59 | -0.16 | -0.34 | -0.08 | -0.2175 | -0.2025 | -0.3175 | 0.283571 | 0.477062 | 0.555354 | 1 | 0.999631 | 0.704942 |
| Tppp_Q7TQD2_S151s_1 | 0.14 | 1.51 | 1.75 | 0.72 | -0.13 | 1.49 | -0.15 | 1.1 | 1.98 | 1.9 | 1.24 | 1.4 | 1.93 | 1.65 | 1.78 | 1.85 | -0.4525 | 0.6 | 0.7725 | 0.076562 | 0.394297 | 0.172638 | 1 | 0.999631 | 0.542546 |
| Tppp_Q7TQD2_S159s_1 | 2.09 | 1.46 | 0.65 | 1.92 | 2.23 | 3.03 | 1.49 | 1.48 | 2.64 | 1.88 | 1.4 | 0.42 | -1.69 | 2.47 | 1.07 | 0.78 | 0.5275 | 0.055 | -0.8725 | 0.292687 | 0.275315 | 0.9171 | 1 | 0.999631 | 0.706671 |
| Tppp_Q7TQD2_S182s_1 | 3.01 | -0.43 | -0.51 | 1.3 | 2.08 | 2.45 | 1.5 | 2.07 | 1.19 | 0.61 | 0.62 | -0.06 | -1.31 | 0.78 | 1.42 | -0.66 | 1.1825 | -0.2525 | -0.785 | 0.392992 | 0.140266 | 0.743169 | 1 | 0.999631 | 0.758901 |
| Tppp_Q7TQD2_S19s_1 | 0.5 | -4.17 | -2.35 | -1.41 | -2.8 | -3.12 | 0 | -3.12 | -4.5 | -1.64 | -2.63 | -4.2 | -6.85 | -3.67 | -2.97 | -4.64 | -0.4025 | -1.385 | -2.675 | 0.032542 | 0.704307 | 0.195885 | 1 | 0.999631 | 0.442395 |
| Tppp_Q7TQD2_S19s_2 | 1.88 | 1.73 | 1.42 | 1.84 | 2.28 | 2.01 | 1.71 | 2.38 | 2.23 | 1.53 | 1.51 | 1.49 | -0.52 | 2.16 | 1.56 | 1.46 | 0.3775 | -0.0275 | -0.5525 | 0.328257 | 0.187298 | 0.924944 | 1 | 0.999631 | 0.725441 |
| Tppp_Q7TQD2_S31s_1 | -0.73 | -0.27 | -0.34 | -0.42 | -0.43 | -0.6 | -0.8 | -0.71 | -0.7 | -0.52 | -0.56 | -0.28 | 0.01 | -0.32 | -0.57 | -0.11 | -0.195 | -0.075 | 0.1925 | 0.507499 | 0.442816 | 0.773312 | 1 | 0.999631 | 0.81618 |
| Tppp_Q7TQD2_T15t_1 | 0.11 | -3.75 | -3.64 | -3.74 | -2.39 | -3.24 | -2.64 | -4.32 | -2.58 | -0.58 | -2.82 | -3.68 | -1.49 | -3.58 | -3.23 | -4.31 | -0.3925 | 0.34 | -0.3975 | 0.681838 | 0.666317 | 0.734597 | 1 | 0.999631 | 0.88924 |
| Tppp_Q7TQD2_T15t_2 | 1.88 | 1.73 | 1.42 | 1.84 | 2.28 | 2.01 | 1.71 | 2.38 | 2.23 | 1.53 | 1.51 | 1.49 | -0.52 | 2.16 | 1.56 | 1.46 | 0.3775 | -0.0275 | -0.5525 | 0.328257 | 0.187298 | 0.924944 | 1 | 0.999631 | 0.725441 |
| Tpr_Q7M739_S2031s_1 | -0.04 | -1.03 | -0.42 | 0.45 | -1.08 | -0.79 | -0.22 | -0.44 | -0.54 | -1.45 | -1.62 | -0.19 | 1.15 | 0.08 | 0.19 | 0.59 | -0.3725 | -0.69 | 0.7625 | 0.088708 | 0.33853 | 0.153718 | 1 | 0.999631 | 0.553858 |
| Tpr_Q7M739_S2067s_1 | 0.86 | 1.91 | 1.39 | 0.94 | 1.1 | 0.53 | 0.59 | 1.42 | 0.57 | 1.11 | 1.95 | 1.18 | 1.23 | 0.75 | 1.43 | 1.58 | -0.365 | -0.0725 | -0.0275 | 0.938238 | 0.312003 | 0.85256 | 1 | 0.999631 | 0.979905 |
| Tpr_Q7M739_S379s_1 | 0.06 | 0.06 | -0.3 | -0.42 | -0.06 | -0.42 | -0.52 | -0.99 | 0.07 | -0.22 | -0.8 | -0.34 | -0.09 | 0.18 | -0.93 | -0.32 | -0.3475 | -0.1725 | -0.14 | 0.677916 | 0.256346 | 0.565642 | 1 | 0.999631 | 0.887183 |
| Tprgl_B1AX95_S10s_1 | -1.06 | -0.46 | -0.89 | -1.61 | -0.57 | -0.87 | -2.26 | -0.69 | -0.87 | -1.12 | -0.64 | -0.48 | -0.07 | -0.64 | -0.47 | -0.58 | -0.0925 | 0.2275 | 0.565 | 0.118156 | 0.835377 | 0.494557 | 1 | 0.999631 | 0.585087 |
| Tprgl_B1AX95_S14s_1 | -1.03 | 0.39 | -0.61 | -0.48 | 0.58 | 0.54 | -0.06 | 0.01 | 0.51 | -1.02 | -0.29 | -0.07 | 0.18 | 1.01 | -0.97 | -0.2 | 0.7 | 0.215 | 0.4375 | 0.381932 | 0.079225 | 0.62153 | 1 | 0.999631 | 0.753668 |
| Tprgl_B1AX95_T34t_1 | -0.72 | -0.06 | 1.41 | -0.12 | 0.17 | 1.01 | 0.64 | 0.38 | 0.4 | -0.45 | -0.3 | 0.53 | 0.01 | 0.37 | 0.86 | -0.04 | 0.4225 | -0.0825 | 0.1725 | 0.721627 | 0.375071 | 0.866595 | 1 | 0.999631 | 0.905674 |
| Tprn_A2AI08_S457s_1 | -0.91 | -1.4 | 0.72 | 0.06 | -1.2 | -0.79 | -0.35 | 0.09 | -1.45 | -0.71 | 0.34 | 1.03 | -0.39 | -0.67 | -0.66 | 0.34 | -0.18 | 0.185 | 0.0375 | 0.941024 | 0.726803 | 0.777495 | 1 | 0.999631 | 0.981286 |
| Tra2a_E9QP00_S14s_1 | -0.42 | -0.92 | -0.29 | 0.36 | -1.29 | -2.05 | 0.01 | -0.49 | -1.39 | -0.21 | -0.52 | -1.77 | -1.18 | -0.92 | 0.57 | -0.54 | -0.6375 | -0.655 | -0.2 | 0.66582 | 0.215758 | 0.163028 | 1 | 0.999631 | 0.88264 |
| Tra2a_E9QP00_S14s_2 | 0.3 | -0.34 | 0.03 | -0.93 | 0.02 | -0.17 | 0.91 | 0.43 | 0.48 | 0.1 | -0.21 | 0.47 | 1.34 | 0.14 | -0.27 | -0.63 | 0.5325 | 0.445 | 0.38 | 0.442647 | 0.177056 | 0.225032 | 1 | 0.999631 | 0.782367 |
| Tra2a_E9QP00_S20s_1 | 2.69 | 1.22 | 2.07 | 2.15 | 0.46 | 1.49 | 2.45 | 1.38 | 1.74 | 2.4 | 1.36 | 1.05 | 1.52 | 0.35 | 1.1 | 0.78 | -0.5875 | -0.395 | -1.095 | 0.022016 | 0.240251 | 0.361397 | 1 | 0.999631 | 0.405544 |
| Tra2a_E9QP00_S236s_1 | 1.79 | 2.05 | 0.98 | 2.51 | 0.18 | 0.17 | 1.82 | 1.06 | 0.46 | 0.99 | 1.89 | 1.26 | 0.45 | -1.03 | 0.03 | 1.44 | -1.025 | -0.6825 | -1.61 | 0.015482 | 0.056594 | 0.140654 | 1 | 0.999631 | 0.388832 |
| Tra2a_E9QP00_S260s_1 | -1.03 | -1.92 | -1.28 | -1.51 | -2.29 | -1.74 | -0.93 | -1.79 | -2.29 | -0.75 | -2.05 | -2.58 | -0.23 | -1.88 | -2.01 | -2.03 | -0.2525 | -0.4825 | -0.1025 | 0.826529 | 0.49079 | 0.289187 | 1 | 0.999631 | 0.940834 |
| Tra2a_E9QP00_S260s_2 | -0.36 | 0.47 | 1.1 | 0.23 | 0.06 | 1 | 0.18 | 0.73 | 0.64 | 0.82 | 0.59 | 0.76 | 0.96 | 0.59 | 1.29 | 0.96 | 0.1325 | 0.3425 | 0.59 | 0.138306 | 0.732184 | 0.336784 | 1 | 0.999631 | 0.611124 |
| Tra2a_E9QP00_S262s_2 | -0.58 | 0.47 | 1.1 | 0.23 | -0.07 | 1 | 0.18 | 0.73 | 0.53 | 0.59 | 0.59 | 0.76 | 0.87 | 0.49 | 1.29 | 0.96 | 0.155 | 0.3125 | 0.5975 | 0.162965 | 0.713491 | 0.413932 | 1 | 0.999631 | 0.631442 |
| Tra2a_E9QP00_S2s_1 | 0.78 | 0.13 | -0.15 | 0 | 0.15 | -0.17 | 0.13 | 0.28 | -0.08 | 0.75 | -0.19 | 0.2 | 0.31 | 0 | 0.16 | 0.37 | -0.0925 | -0.02 | 0.02 | 0.948905 | 0.753511 | 0.952966 | 1 | 0.999631 | 0.983587 |
| Tra2a_E9QP00_S2s_2 | 0.3 | -0.34 | 0.03 | -0.93 | 0.02 | -0.17 | 0.91 | 0.43 | 0.48 | 0.1 | -0.21 | 0.47 | 1.34 | 0.14 | -0.27 | -0.63 | 0.5325 | 0.445 | 0.38 | 0.442647 | 0.177056 | 0.225032 | 1 | 0.999631 | 0.782367 |
| Tra2a_E9QP00_T200t_1 | -0.27 | 2.37 | 1.64 | -0.22 | 3.22 | 0.82 | 3.51 | 0.35 | 4.91 | -0.94 | 2.93 | 4.28 | 5.4 | -0.59 | 0.23 | 0.07 | 1.095 | 1.915 | 0.3975 | 0.759609 | 0.24572 | 0.151459 | 1 | 0.999631 | 0.914614 |
| Tra2b_P62996_S29s_2 | 1.46 | -0.39 | 0.01 | 0.95 | 0 | -1.93 | 1.32 | -2.03 | -0.4 | 0.65 | -0.85 | -0.04 | -0.08 | -3.01 | -2.64 | -1.46 | -1.1675 | -0.6675 | -2.305 | 0.007528 | 0.166054 | 0.204013 | 1 | 0.999631 | 0.358173 |
| Tra2b_P62996_S2s_1 | 1.83 | 0.21 | 0.29 | 0.14 | -0.7 | -0.51 | 1.05 | -0.12 | -0.01 | 0.84 | -0.17 | 0.19 | 0.22 | -1.51 | 0.02 | -0.04 | -0.6875 | -0.405 | -0.945 | 0.100736 | 0.209189 | 0.38089 | 1 | 0.999631 | 0.564448 |
| Tra2b_P62996_S99s_2 | -0.61 | -0.18 | -0.13 | -0.23 | -0.83 | -1.16 | 1.96 | -0.87 | 0.35 | -0.91 | -0.37 | 0.23 | -0.52 | -2.48 | -1.25 | -3.28 | 0.0625 | 0.1125 | -1.595 | 0.018668 | 0.92446 | 0.748535 | 1 | 0.999631 | 0.391024 |
| Tra2b_P62996_T201t_1 | -0.37 | 1.09 | 1.44 | 0.78 | 0.88 | 0.81 | 0.12 | 0.99 | 0.84 | 0.99 | 0.81 | 1.12 | 1.32 | 1.45 | 1.14 | 1.38 | -0.035 | 0.205 | 0.5875 | 0.177675 | 0.935158 | 0.617266 | 1 | 0.999631 | 0.642116 |
| Tra2b_P62996_T33t_2 | 1.46 | -0.39 | 0.01 | 0.95 | 0 | -1.93 | 1.32 | -2.03 | -0.4 | 0.65 | -0.85 | -0.04 | -0.08 | -3.01 | -2.64 | -1.46 | -1.1675 | -0.6675 | -2.305 | 0.007528 | 0.166054 | 0.204013 | 1 | 0.999631 | 0.358173 |
| Traf3_Q3UHJ1_S243s_1 | -1.19 | -0.49 | 2.75 | 1.79 | -0.49 | -0.39 | 0.53 | 2.54 | -0.23 | -1.04 | 3.3 | 3.75 | 0.49 | -0.26 | 3.08 | 2.96 | -0.1675 | 0.73 | 0.8525 | 0.434838 | 0.867282 | 0.579329 | 1 | 0.999631 | 0.779717 |
| Traf3_Q3UHJ1_S330s_1 | 0.09 | 0.18 | 0 | -0.97 | -0.44 | 0.47 | -0.3 | -0.64 | -0.33 | 0.22 | -0.67 | -0.41 | 0.83 | 0.19 | 0.27 | 0.16 | -0.0525 | -0.1225 | 0.5375 | 0.158127 | 0.889061 | 0.734157 | 1 | 0.999631 | 0.628615 |
| Trafd1_Q3UDK1_S450s_1 | 0.91 | -0.16 | -0.52 | 0.46 | 1.11 | 0.75 | 0.71 | 0.29 | 0.68 | 0.32 | 0.09 | 0.43 | -1.27 | 0.83 | -0.26 | -0.44 | 0.5425 | 0.2075 | -0.4575 | 0.379966 | 0.171104 | 0.577212 | 1 | 0.999631 | 0.752193 |
| Tram1_Q91V04_S365s_1 | -0.69 | -2.83 | -1.12 | -1.71 | -1.86 | -1.91 | -1.11 | -2.06 | -2.03 | -1.17 | -1.28 | -0.9 | -0.57 | -1.38 | -1.23 | -1.08 | -0.1475 | 0.2425 | 0.5225 | 0.292617 | 0.760385 | 0.627731 | 1 | 0.999631 | 0.706671 |
| Trappc10_Q3TLI0_S707s_1 | -0.97 | -0.58 | -1.06 | -0.72 | -0.27 | -0.7 | -0.99 | -0.74 | -0.47 | -0.98 | -0.65 | -0.62 | 0.25 | -0.13 | -0.81 | -0.62 | 0.1575 | 0.1525 | 0.505 | 0.153614 | 0.568554 | 0.572226 | 1 | 0.999631 | 0.625198 |
| Trappc12_Q8C0C3_S10s_1 | -1.26 | -0.22 | 0.12 | -0.74 | -0.94 | -0.93 | -0.79 | -0.46 | -0.41 | -0.64 | -0.47 | -0.4 | -0.05 | -0.87 | 0.14 | -0.36 | -0.255 | 0.045 | 0.24 | 0.551113 | 0.473691 | 0.897008 | 1 | 0.999631 | 0.836275 |
| Trappc12_Q8C0C3_S140s_1 | -0.13 | 2.83 | 2.82 | 0.6 | 2.2 | 2.31 | 1.46 | 1.43 | 2.74 | 0.17 | 2.78 | 2.81 | 3.08 | 1.89 | 2.45 | 1.15 | 0.32 | 0.595 | 0.6125 | 0.427716 | 0.651626 | 0.501221 | 1 | 0.999631 | 0.775391 |
| Trappc12_Q8C0C3_S309s_1 | -0.24 | -0.87 | 0.1 | -0.02 | -0.53 | -0.61 | 0.06 | 0.14 | -0.49 | -0.23 | 0.15 | 0.06 | -0.62 | -0.63 | 0.08 | -0.18 | 0.0225 | 0.13 | -0.08 | 0.814917 | 0.945822 | 0.685727 | 1 | 0.999631 | 0.937235 |
| Trappc12_Q8C0C3_T143t_1 | -1.22 | 0.9 | 1.82 | 0.24 | 0.75 | 1.17 | 0.95 | 1.44 | 1.2 | -0.72 | 1.94 | 1.83 | 1.96 | 0.88 | 1.69 | 1.57 | 0.6425 | 0.6275 | 1.09 | 0.10133 | 0.294637 | 0.429901 | 1 | 0.999631 | 0.564448 |
| Trappc4_Q9ES56_S219s_1 | -0.35 | -0.63 | -1.24 | -0.22 | 0.77 | -1 | -0.7 | -0.23 | 0.37 | -1.02 | 0.45 | 0.31 | 1.71 | 0.02 | -1.09 | -0.96 | 0.32 | 0.6375 | 0.53 | 0.401534 | 0.470945 | 0.153138 | 1 | 0.999631 | 0.763221 |
| Trappc8_E9PY51_S657s_1 | -1.6 | -1.01 | -0.07 | -1.06 | -0.64 | -1.23 | -0.99 | -0.5 | -0.43 | -1.53 | -1.13 | -0.43 | -1.44 | -0.65 | -0.7 | -0.43 | 0.095 | 0.055 | 0.13 | 0.749762 | 0.800002 | 0.896103 | 1 | 0.999631 | 0.912137 |
| Trappc9_Q3U0M1_S646s_1 | 0.93 | 0.7 | 1.06 | 0.68 | 0.07 | 1.88 | 0.67 | 1.82 | 0.59 | 0.84 | 0.58 | 0.45 | -0.58 | 1.14 | 0.52 | 0.12 | 0.2675 | -0.2275 | -0.5425 | 0.193256 | 0.548387 | 0.384574 | 1 | 0.999631 | 0.649965 |
| Trex1_Q91XB0_S78s_1 | -0.78 | 0.61 | 0.43 | 0.18 | 0.38 | 0.71 | -0.49 | 0.14 | 0.95 | 1.12 | 1.07 | 0.6 | 1.14 | 1.42 | 0.87 | 1.6 | 0.075 | 0.825 | 1.1475 | 0.012549 | 0.852389 | 0.04372 | 1 | 0.999631 | 0.385092 |
| Trhde_Q8K093_S11s_1 | 0.15 | -3.16 | -3.41 | -3.08 | -1.79 | -2.33 | -1.85 | -3.47 | -2.08 | -0.78 | -2.1 | -4.28 | -2.31 | -2.5 | -3.29 | -4.08 | 0.015 | 0.065 | -0.67 | 0.419143 | 0.985231 | 0.946178 | 1 | 0.999631 | 0.772152 |
| Trim2_Q9ESN6_S424s_1 | -1.71 | -0.1 | 1.17 | -0.54 | 0.42 | 0.6 | 0.26 | 0.34 | 1.54 | -0.94 | 0.72 | 1.23 | 0.49 | -0.28 | 1 | 0.48 | 0.7 | 0.9325 | 0.7175 | 0.244961 | 0.222629 | 0.215512 | 1 | 0.999631 | 0.685751 |
| Trim2_Q9ESN6_S424s_2 | -2.01 | -0.2 | 0.05 | -1.83 | -0.43 | -0.75 | -0.43 | -0.1 | -0.13 | -1.66 | 0.3 | 0.17 | 0.36 | -0.52 | -0.12 | -0.11 | 0.57 | 0.6675 | 0.9 | 0.114605 | 0.283492 | 0.306379 | 1 | 0.999631 | 0.579001 |
| Trim2_Q9ESN6_S428s_1 | -1.46 | 0.06 | 0.31 | -0.32 | -0.54 | -0.77 | -0.01 | -0.2 | -0.22 | -1.04 | 0.08 | 0.53 | 0.28 | -0.1 | 0.13 | 0.21 | -0.0275 | 0.19 | 0.4825 | 0.262119 | 0.947965 | 0.698587 | 1 | 0.999631 | 0.692889 |
| Trim2_Q9ESN6_S440s_1 | 2.16 | 1.52 | 1.77 | 1.35 | -1.53 | 1.95 | 3.56 | 0.21 | 0.44 | 1.83 | -0.59 | 0.69 | 0.96 | 1.2 | 2.07 | 0.84 | -0.6525 | -1.1075 | -0.4325 | 0.260294 | 0.501425 | 0.048595 | 1 | 0.999631 | 0.692624 |
| Trim2_Q9ESN6_S456s_1 | 2.58 | 0.72 | 0.52 | 1.99 | 2.07 | 2.14 | 2.37 | 2.23 | 1.15 | 0.4 | 2.11 | 0.11 | -1.12 | 1.57 | 0.84 | 0.39 | 0.75 | -0.51 | -1.0325 | 0.14835 | 0.139381 | 0.410081 | 1 | 0.999631 | 0.62132 |
| Trim24_E9Q1U8_S675s_1 | 0.54 | 0.18 | 1.1 | 0.36 | 0.03 | 0.9 | 0.01 | 0.3 | 0.42 | 0.93 | 0.27 | 0.34 | 0.35 | 1.01 | 0.11 | 0.89 | -0.235 | -0.055 | 0.045 | 0.897698 | 0.481404 | 0.860828 | 1 | 0.999631 | 0.966758 |
| Trim24_E9Q1U8_S742s_1 | 0.7 | 1.27 | 1.52 | 1.23 | 0.3 | 0.49 | 1 | 1.24 | 1.25 | 1.66 | 1.33 | 1.75 | 1.16 | 1.4 | 1.06 | 1.66 | -0.4225 | 0.3175 | 0.14 | 0.654222 | 0.211914 | 0.294654 | 1 | 0.999631 | 0.878537 |
| Trim28_Q62318_S23s_1 | -0.29 | 0.37 | 0.45 | 0.09 | 0.15 | -0.09 | -0.35 | 0.1 | -0.02 | -0.1 | 0.25 | 0.47 | 0.67 | 0.41 | 0.41 | 0.79 | -0.2025 | -0.005 | 0.415 | 0.186659 | 0.478181 | 0.986421 | 1 | 0.999631 | 0.646474 |
| Trim28_Q62318_S473s_1 | -0.28 | 1.49 | 1.43 | 0.33 | 0.37 | 0.69 | 0.48 | 0.71 | 0.79 | 0.18 | 1.17 | 1.38 | 1.15 | 1.14 | 0.98 | 1.98 | -0.18 | 0.1375 | 0.57 | 0.248393 | 0.677971 | 0.777012 | 1 | 0.999631 | 0.685751 |
| Trim28_Q62318_S51s_1 | 0.2 | 0.89 | 0.76 | 0.42 | 0.23 | 0.51 | 0.02 | 0.57 | 0.45 | 0.55 | 0.52 | 0.56 | 0.93 | 0.94 | 1.15 | 0.52 | -0.235 | -0.0475 | 0.3175 | 0.31156 | 0.413467 | 0.860303 | 1 | 0.999631 | 0.71924 |
| Trim28_Q62318_S594s_1 | -0.05 | 0.75 | 0.47 | 0.1 | 0.47 | 0.52 | -0.38 | 0.2 | 0.65 | 0.14 | 0.43 | 0.04 | 0.61 | 0.51 | 0.58 | 0.73 | -0.115 | -0.0025 | 0.29 | 0.340485 | 0.721334 | 0.993397 | 1 | 0.999631 | 0.733032 |
| Trim28_Q62318_S697s_1 | 1.6 | 0.77 | 1.48 | 1.55 | -0.1 | 0.3 | 1.07 | 0.63 | 0.83 | 1.4 | 0.82 | 0.68 | 0.28 | 0.18 | 0.93 | 0.54 | -0.875 | -0.4175 | -0.8675 | 0.02362 | 0.028632 | 0.205817 | 1 | 0.999631 | 0.415644 |
| Trim3_Q9R1R2_S437s_1 | -0.13 | -1.4 | 1.36 | 1.13 | -0.4 | -0.85 | 1.37 | 1.68 | -0.74 | -0.07 | 1.65 | 1.18 | -1.1 | -1.6 | 1.34 | 1.32 | 0.21 | 0.265 | -0.25 | 0.774336 | 0.789088 | 0.722746 | 1 | 0.999631 | 0.920998 |
| Trim3_Q9R1R2_S529s_1 | 1.04 | 0.76 | -0.11 | 0 | 0.75 | -0.05 | -0.19 | 0.31 | 1.94 | 2.56 | 0.97 | 0.97 | 1.49 | 1.86 | 0.88 | 0.55 | -0.2175 | 1.1875 | 0.7725 | 0.090886 | 0.560447 | 0.027896 | 1 | 0.999631 | 0.557085 |
| Trim3_Q9R1R2_S7s_1 | -0.93 | 0.56 | 0.48 | 0.28 | -0.2 | 0.37 | 0.26 | 0.52 | 0.58 | 0.48 | 0.43 | 0.8 | 1.07 | 0.45 | 0.71 | 1.04 | 0.14 | 0.475 | 0.72 | 0.096848 | 0.720087 | 0.22976 | 1 | 0.999631 | 0.55886 |
| Trim32_Q8CH72_S330s_1 | 0.11 | -0.42 | -0.52 | -1.83 | -0.48 | -0.4 | -1.03 | -1.02 | -0.01 | -0.97 | -0.54 | 0.02 | 0.92 | -0.76 | -0.64 | -0.63 | -0.0675 | 0.29 | 0.3875 | 0.477168 | 0.876789 | 0.534301 | 1 | 0.999631 | 0.800086 |
| Trim46_D3YXA6_S604s_1 | -1.35 | -1.65 | -1.68 | -1.34 | -0.7 | -1.29 | -1.28 | -1.75 | -1.29 | -2.01 | -0.86 | -1.59 | -0.61 | -1.31 | -1.13 | -1.52 | 0.25 | 0.0675 | 0.3625 | 0.258535 | 0.410993 | 0.832802 | 1 | 0.999631 | 0.692624 |
| Trim56_Q80VI1_S454s_1 | -0.27 | 0.33 | -0.56 | -0.61 | 0.05 | 0.44 | -0.79 | -0.3 | 0.18 | 0.29 | -0.48 | -0.34 | 1.05 | 0.77 | 0.36 | 0.44 | 0.1275 | 0.19 | 0.9325 | 0.018364 | 0.725268 | 0.57464 | 1 | 0.999631 | 0.391024 |
| Trim66_E9PZP2_S767s_1 | -0.35 | -0.34 | -0.29 | -0.22 | -0.52 | 0.09 | -0.47 | -1.18 | -0.53 | 0.42 | -0.86 | 0.13 | 0.94 | -0.07 | -0.02 | -0.19 | -0.22 | 0.09 | 0.465 | 0.184181 | 0.489023 | 0.791852 | 1 | 0.999631 | 0.644602 |
| Trim67_Q505D9_S275s_1 | 1.37 | -0.5 | -0.22 | 0.68 | -0.28 | 0.37 | 0.47 | 0.16 | -0.02 | 1.7 | 0.02 | -0.06 | -0.78 | -0.61 | -0.37 | -0.21 | -0.1525 | 0.0775 | -0.825 | 0.088856 | 0.732479 | 0.889907 | 1 | 0.999631 | 0.553932 |
| Trim67_Q505D9_T38t_1 | 0.15 | 1 | -0.16 | -0.79 | -0.19 | 0.02 | -1.11 | -1.91 | -1.3 | 1.14 | -3.92 | 0.19 | -0.08 | 1.43 | 0.2 | -0.91 | -0.8475 | -1.0225 | 0.11 | 0.845564 | 0.137483 | 0.322487 | 1 | 0.999631 | 0.948565 |
| Trim9_E9Q524_S46s_1 | -0.17 | -0.49 | -0.76 | -0.22 | -0.29 | -0.3 | -0.73 | -0.28 | -0.44 | -0.1 | -0.48 | -0.54 | -0.6 | 0.04 | -0.67 | -0.23 | 0.01 | 0.02 | 0.045 | 0.884326 | 0.970305 | 0.941684 | 1 | 0.999631 | 0.963505 |
| Trim9_E9Q524_T41t_1 | -0.69 | -1.65 | -2.26 | -1.36 | -0.67 | -1.32 | -1.37 | -1.69 | -1.28 | -1.18 | -1.39 | -1.32 | -1.32 | -1.21 | -1.96 | -1.67 | 0.2275 | 0.1975 | -0.05 | 0.899279 | 0.568881 | 0.587574 | 1 | 0.999631 | 0.967236 |
| Trim9_E9Q524_T443t_1 | -1.15 | -0.24 | 0.44 | -0.04 | -1.42 | 0.14 | -0.27 | -0.84 | -0.2 | 0.55 | -0.19 | -0.04 | 0.56 | -0.52 | 0.55 | 0.05 | -0.35 | 0.2775 | 0.4075 | 0.35341 | 0.451173 | 0.485779 | 1 | 0.999631 | 0.73949 |
| Trio_Q0KL02_S1724s_1 | 1.62 | 0.41 | 0.3 | 0.97 | 0.65 | 0.53 | 0.46 | 0.44 | 0.45 | 0.31 | 0.71 | 0.69 | -0.6 | 0.49 | 1.4 | 0.95 | -0.305 | -0.285 | -0.265 | 0.600132 | 0.381353 | 0.429347 | 1 | 0.999631 | 0.856618 |
| Trio_Q0KL02_S1818s_1 | -1.42 | -0.08 | -1.46 | -1.24 | -1.36 | -0.76 | -0.6 | -1.28 | -0.29 | -1.6 | -1.44 | -0.98 | 0.81 | 0.4 | -0.47 | -0.81 | 0.05 | -0.0275 | 1.0325 | 0.054538 | 0.897229 | 0.949679 | 1 | 0.999631 | 0.491994 |
| Trio_Q0KL02_S1845s_1 | 0.24 | -0.2 | -0.08 | -0.23 | 0.99 | 0.43 | -0.01 | 0.59 | -0.05 | -0.25 | 0.57 | 0.59 | 0.37 | 0.42 | 0.85 | 0.31 | 0.5675 | 0.2825 | 0.555 | 0.077144 | 0.081264 | 0.37089 | 1 | 0.999631 | 0.542683 |
| Trio_Q0KL02_S2282s_1 | -1.11 | -0.51 | -0.47 | -0.71 | -0.49 | -0.94 | -0.89 | -0.69 | -0.59 | -0.91 | -0.89 | -0.21 | -0.55 | -0.43 | -0.73 | -0.42 | -0.0525 | 0.05 | 0.1675 | 0.563254 | 0.846189 | 0.86674 | 1 | 0.999631 | 0.841038 |
| Trio_Q0KL02_S2458s_1 | 0.72 | -0.79 | -1.69 | -0.36 | 1.48 | -0.74 | 0.18 | -0.41 | -0.53 | -1.81 | -0.92 | -1.56 | -0.92 | -0.98 | -0.91 | -1.14 | 0.6575 | -0.675 | -0.4575 | 0.358085 | 0.309222 | 0.230069 | 1 | 0.999631 | 0.741182 |
| Trio_Q0KL02_T2560t_1 | 0.19 | 0.2 | -0.09 | -0.15 | 0.6 | 0.13 | -0.53 | -0.34 | 0.68 | 0.45 | -0.23 | -0.16 | 0.83 | 0.26 | -0.24 | 0.34 | -0.0725 | 0.1475 | 0.26 | 0.425053 | 0.819974 | 0.636344 | 1 | 0.999631 | 0.774293 |
| Triobp_Q99KW3_S1101s_1 | 0.96 | 0.2 | -0.52 | -0.07 | -0.04 | -0.41 | 0.08 | -0.89 | -0.34 | 0.41 | -1.08 | -0.41 | -0.06 | -0.13 | -0.48 | -0.49 | -0.4575 | -0.4975 | -0.4325 | 0.261474 | 0.256512 | 0.267318 | 1 | 0.999631 | 0.69271 |
| Trip12_G5E870_S1049s_1 | -2.26 | -4.3 | -6.43 | -4.68 | -3.02 | -4.64 | -4.98 | -5 | -4.04 | -2.95 | -3.68 | -4.98 | -2.49 | -4.74 | -4.84 | -5.43 | 0.0075 | 0.505 | 0.0425 | 0.963218 | 0.992943 | 0.548828 | 1 | 0.999631 | 0.987149 |
| Trip12_G5E870_S239s_1 | 1.28 | -1.6 | 0.74 | 0.08 | -1.82 | -0.04 | 1.05 | -2.19 | -2.98 | 0.71 | -1.57 | -2.69 | -1.09 | -1.82 | -1.01 | -1.8 | -0.875 | -1.7575 | -1.555 | 0.025842 | 0.320131 | 0.077614 | 1 | 0.999631 | 0.422248 |
| Trip12_G5E870_S312s_1 | -0.28 | 0.54 | 0.41 | 0.1 | -0.04 | 0.08 | -0.21 | 0.72 | 0.14 | 0.03 | 0.18 | 0.36 | 0.38 | 0.19 | 0.56 | 0.61 | -0.055 | -0.015 | 0.2425 | 0.434602 | 0.86381 | 0.958097 | 1 | 0.999631 | 0.779717 |
| Trip12_G5E870_S77s_1 | 1.89 | -0.21 | 1.1 | 0.6 | -0.75 | 0.15 | 0.83 | -0.27 | -0.21 | 1.59 | 0.16 | -0.17 | -0.53 | -1.02 | -0.47 | -0.76 | -0.855 | -0.5025 | -1.54 | 0.006598 | 0.12171 | 0.382924 | 1 | 0.999631 | 0.351545 |
| Trmt10a_Q8C1Z8_S22s_1 | 1.63 | 2.16 | 2.1 | 1.48 | 0.92 | 2.25 | 1.4 | 1.05 | 2.52 | 3.12 | 1.31 | 1.33 | 0.75 | 1.68 | 1.51 | 0.8 | -0.4375 | 0.2275 | -0.6575 | 0.084486 | 0.248407 | 0.627408 | 1 | 0.999631 | 0.549801 |
| Trmt10a_Q8C1Z8_S324s_1 | 0.69 | 2.66 | 1.68 | 0.87 | 1.73 | 1.86 | 0.35 | 0.57 | 1.13 | 2.21 | 0.35 | 1.72 | 2.98 | 2.34 | 1.68 | 1.57 | -0.3475 | -0.1225 | 0.6675 | 0.222109 | 0.529169 | 0.825922 | 1 | 0.999631 | 0.672638 |
| Trpc5_Q9QX29_S770s_1 | 0.18 | 0.86 | 0.18 | -0.23 | -0.43 | 0.94 | -0.23 | -0.16 | -0.95 | 1.1 | 0.09 | -0.28 | 0.12 | 0.02 | 0.57 | -0.15 | -0.2175 | -0.2575 | -0.1075 | 0.751766 | 0.580901 | 0.585299 | 1 | 0.999631 | 0.912945 |
| Trpc6_Q3UZG1_S736s_1 | -0.84 | -0.51 | -1.23 | 0.27 | -0.68 | -0.78 | -0.6 | -1.74 | -1.3 | -0.56 | -1.1 | -0.58 | -1 | -0.6 | -0.9 | -0.19 | -0.3725 | -0.3075 | -0.095 | 0.809784 | 0.377991 | 0.43517 | 1 | 0.999631 | 0.936643 |
| Trpm3_Q5F4S6_S1292s_1 | 0.2 | 0.16 | -0.09 | 0.37 | 0.34 | -0.08 | -0.19 | 0.55 | 0.46 | 0.59 | 0.14 | -0.06 | 0.23 | 0.09 | 0.85 | 0.63 | -0.005 | 0.1225 | 0.29 | 0.349227 | 0.985736 | 0.6598 | 1 | 0.999631 | 0.736792 |
| Trpm3_Q5F4S6_S1629s_1 | -0.69 | 0.6 | -0.61 | -0.55 | 0.9 | 0.11 | -0.5 | 0.32 | 0.25 | -0.39 | -1.09 | -0.4 | 0.31 | 0.71 | -0.36 | -0.04 | 0.52 | -0.095 | 0.4675 | 0.263936 | 0.231789 | 0.819265 | 1 | 0.999631 | 0.693392 |
| Trpm3_Q5F4S6_T102t_1 | 1.06 | 0.72 | -0.71 | -0.46 | 1.3 | 1.75 | -0.2 | 0.76 | 0.79 | 1.15 | -0.85 | -1.29 | -0.13 | 1.79 | -1.08 | -0.73 | 0.75 | -0.2025 | -0.19 | 0.782579 | 0.196578 | 0.760896 | 1 | 0.999631 | 0.922325 |
| Trpm4_Q7TN37_S527s_1 | 0.16 | -0.04 | 0.82 | 0.23 | 0.19 | -0.01 | -0.22 | -0.42 | 0.07 | -0.14 | -0.14 | 0.53 | 0.42 | 0.29 | 0.82 | -0.03 | -0.4075 | -0.2125 | 0.0825 | 0.802493 | 0.188089 | 0.498939 | 1 | 0.999631 | 0.932677 |
| Trpm7_Q923J1_T555t_1 | -0.3 | -0.32 | -0.42 | -1 | -1.53 | 0 | -0.38 | -0.63 | -0.17 | -0.39 | -0.37 | 0 | 0.23 | 0.34 | -0.31 | -0.44 | -0.125 | 0.2775 | 0.465 | 0.179278 | 0.742046 | 0.339229 | 1 | 0.999631 | 0.64376 |
| Trps1_Q80V18_S584s_1 | 0.24 | 1.27 | 2.14 | 0.35 | 0.51 | 0.51 | 0.56 | 0.47 | 0.83 | 0.76 | 1.56 | 2.03 | 1.23 | 0.78 | 1.85 | 1.67 | -0.4875 | 0.295 | 0.3825 | 0.440211 | 0.278376 | 0.565477 | 1 | 0.999631 | 0.781983 |
| Trpv2_Q9WTR1_S15s_1 | -0.77 | -0.42 | -0.25 | -0.45 | -0.61 | -0.67 | -0.64 | -0.46 | -0.94 | -0.48 | -0.21 | -0.38 | -0.12 | -0.83 | -0.48 | -0.07 | -0.1225 | -0.03 | 0.0975 | 0.750574 | 0.621056 | 0.91565 | 1 | 0.999631 | 0.912627 |
| Trpv2_Q9WTR1_S47s_1 | -1.07 | -0.27 | -0.27 | -0.83 | -0.59 | -0.15 | -0.88 | -0.76 | -0.45 | -1.21 | -0.71 | -0.7 | 0.13 | 0.14 | -0.51 | -0.43 | 0.015 | -0.1575 | 0.4425 | 0.20787 | 0.961595 | 0.622939 | 1 | 0.999631 | 0.661649 |
| Trpv2_Q9WTR1_T14t_1 | -1.8 | -0.09 | -0.13 | 0.12 | 0.07 | -0.78 | 0.35 | 0.27 | -0.69 | 0.03 | 0.27 | 0.86 | 0.83 | 0.85 | 0.65 | 0.91 | 0.4525 | 0.5925 | 1.285 | 0.015741 | 0.363268 | 0.267663 | 1 | 0.999631 | 0.388832 |
| Trrap_F7CGG2_S1350s_1 | 1.65 | 0.61 | 2.01 | 1.19 | 0.2 | 1.63 | 1.08 | 0.53 | 1.37 | 1.75 | 0.09 | 0.88 | -0.08 | 1.18 | 1.47 | 0.84 | -0.505 | -0.3425 | -0.5125 | 0.271657 | 0.256151 | 0.460587 | 1 | 0.999631 | 0.699288 |
| Tsc1_Q9EP53_S1068s_1 | -0.06 | 1.59 | 0.39 | 0.04 | 1.2 | 0.6 | -0.16 | 0.12 | 0.64 | -0.26 | -0.04 | 0.61 | 1.27 | 0.97 | 0.49 | -0.01 | -0.05 | -0.2525 | 0.19 | 0.683515 | 0.913774 | 0.567976 | 1 | 0.999631 | 0.889602 |
| Tsc1_Q9EP53_S1133s_1 | -1.43 | -1.49 | -1.54 | -1.3 | -1.58 | -1.51 | -1.13 | -1.53 | -2.02 | -1.42 | -1.77 | -1.28 | -1.05 | -1.59 | -1.36 | -1.61 | 0.0025 | -0.1825 | 0.0375 | 0.893063 | 0.991868 | 0.514517 | 1 | 0.999631 | 0.964596 |
| Tsc1_Q9EP53_S321s_1 | -0.45 | -0.16 | -0.69 | -0.68 | -0.02 | 0.36 | -0.88 | -0.67 | 0.23 | -0.34 | -0.56 | -0.69 | 0.28 | 0.39 | -0.73 | -0.16 | 0.1925 | 0.155 | 0.44 | 0.220792 | 0.579839 | 0.616857 | 1 | 0.999631 | 0.671424 |
| Tsc1_Q9EP53_S511s_1 | 1.02 | 0.61 | 1.2 | 1.41 | 1.05 | 1.29 | 2.31 | 1.41 | 1.57 | 0.68 | 1.24 | 0.49 | 0.63 | 0.05 | 0.78 | 0.69 | 0.455 | -0.065 | -0.5225 | 0.126524 | 0.215703 | 0.850282 | 1 | 0.999631 | 0.596745 |
| Tsc1_Q9EP53_S580s_1 | 0.04 | -1.47 | -0.43 | 0.72 | 0.23 | 0.23 | 1.07 | 0.41 | -0.25 | -0.87 | 0.4 | 1.12 | -1.46 | -0.31 | 0.64 | 1.12 | 0.77 | 0.385 | 0.2825 | 0.668044 | 0.130434 | 0.509247 | 1 | 0.999631 | 0.883098 |

| | | | | | | | | | | | | | | | | | | | | | | | | | |
|---|---|---|---|---|---|---|---|---|---|---|---|---|---|---|---|---|---|---|---|---|---|---|---|---|---|
| Tsc2_Q3UHB2_S1132s_1 | 0.24 | 0.63 | -0.48 | 0.98 | 0.23 | 0.41 | 0.6 | 0.74 | -0.3 | 0.74 | 0.31 | 1.17 | -2.4 | 0.59 | 0.31 | -0.13 | 0.1525 | 0.1375 | -0.75 | 0.277813 | 0.670938 | 0.754499 | 1 | 0.999631 | 0.700874 |
| Tsc2_Q3UHB2_S1254s_1 | -0.48 | 0.54 | 0.49 | -0.18 | -0.03 | 0.42 | 0.28 | 0.89 | 0.6 | 0.39 | -0.99 | 1.4 | 0.94 | 0.76 | 1.34 | 0.82 | 0.2975 | 0.2575 | 0.8725 | 0.027813 | 0.400645 | 0.624145 | 1 | 0.999631 | 0.423977 |
| Tsc2_Q3UHB2_S1366s_1 | 0.36 | -0.41 | -0.57 | 1.66 | 1.36 | -1.39 | 2.19 | 0.71 | 0.23 | -0.75 | 0.37 | -0.09 | -0.52 | 0.03 | -0.52 | -0.68 | 0.4575 | -0.32 | -0.6825 | 0.198805 | 0.571644 | 0.549565 | 1 | 0.999631 | 0.656905 |
| Tsc2_Q3UHB2_S1413s_1 | -0.19 | 0.01 | -0.13 | -0.04 | 0.03 | -0.81 | -0.71 | -0.08 | 0.2 | -0.21 | 0.03 | 0.58 | 0.67 | 0.21 | 0.15 | 0.33 | -0.305 | 0.2375 | 0.4275 | 0.141802 | 0.307648 | 0.397613 | 1 | 0.999631 | 0.612726 |
| Tsc2_Q3UHB2_S664s_1 | -0.49 | -0.43 | -0.08 | -0.9 | -0.42 | -0.69 | -0.51 | -0.54 | -0.46 | -0.6 | -0.28 | -0.33 | 0.4 | -0.1 | -0.46 | -0.3 | -0.065 | 0.0575 | 0.36 | 0.286607 | 0.809761 | 0.837335 | 1 | 0.999631 | 0.705972 |
| Tsc2_Q3UHB2_S981s_1 | 1.23 | 1.09 | -0.23 | 1.03 | 1.72 | 1.41 | 1.03 | 1.13 | 1.46 | 0.88 | 1.45 | 1.09 | -0.39 | 1.54 | 0.65 | 0.74 | 0.5425 | 0.44 | -0.145 | 0.771869 | 0.180121 | 0.271603 | 1 | 0.999631 | 0.919413 |
| Tsc2_Q3UHB2_T1271t_1 | -0.84 | 0.42 | -0.95 | -0.6 | -0.26 | -0.13 | -1.14 | -0.93 | -0.46 | -0.6 | -1.07 | -1.32 | 0.13 | -0.06 | -1.31 | -0.47 | -0.1225 | -0.37 | 0.065 | 0.88444 | 0.76114 | 0.353432 | 1 | 0.999631 | 0.963505 |
| Tsc22d2_E9Q7M2_S587s_1 | 0.51 | 1.41 | 0.5 | 0.71 | 0.03 | 1.72 | 0.39 | 1 | 0.97 | 0.78 | 0.59 | 0.97 | 0.73 | 1.48 | 1.28 | 1.58 | 0.0025 | 0.045 | 0.485 | 0.182786 | 0.995278 | 0.882679 | 1 | 0.999631 | 0.643861 |
| Tsc22d4_D3YZZ4_S38s_1 | 0.03 | 0.58 | 0.57 | -0.27 | -0.1 | 1.25 | -0.49 | 2.25 | 0.9 | 1.15 | 0.16 | 0.74 | 1.66 | 1.15 | 1.42 | 0.82 | 0.5 | 0.51 | 1.035 | 0.011765 | 0.413127 | 0.158495 | 1 | 0.999631 | 0.385092 |
| Tsc22d4_Q9EQN3_S62s_1 | 0.18 | -0.22 | 0.62 | -0.61 | -0.56 | -0.07 | -0.17 | -0.62 | -0.03 | 0.11 | -0.29 | -0.34 | 0.26 | -0.39 | 0.24 | -0.65 | -0.3475 | -0.13 | -0.1275 | 0.740077 | 0.314312 | 0.698346 | 1 | 0.999631 | 0.908664 |
| Tsnax_Q9QZE7_T165t_1 | 1.19 | -2.36 | -0.53 | 0.34 | -1.11 | -1.83 | 0.12 | -0.95 | -1.35 | -0.18 | -1.08 | -1.26 | -1.19 | -2.69 | -1.21 | -2.6 | -0.6025 | -0.6275 | -1.5825 | 0.060029 | 0.432389 | 0.390146 | 1 | 0.999631 | 0.505461 |
| Tspan13_Q9D8C2_S143s_1 | 0.13 | 0.84 | 0.54 | 0.37 | 0.6 | 0.64 | 0.14 | -0.18 | 0.4 | 0.65 | 0.37 | 1.04 | 0.64 | 1.13 | 1.31 | 0.58 | -0.17 | 0.145 | 0.445 | 0.183823 | 0.580652 | 0.625866 | 1 | 0.999631 | 0.644372 |
| Tspyl1_O88852_S51s_1 | 0.19 | 0.56 | 0.13 | 0.17 | 0.1 | 0.08 | -0.11 | 0.25 | 0.16 | 0.52 | -0.12 | 0.24 | 0.64 | 0.56 | 0.04 | 0.41 | -0.1825 | -0.0625 | 0.15 | 0.605568 | 0.468831 | 0.81859 | 1 | 0.999631 | 0.85817 |
| Tspyl2_Q7TQI8_S504s_1 | 0.35 | 0.3 | 0.04 | 0.34 | 0.1 | -0.11 | -0.14 | -0.53 | 1.15 | 0.41 | -0.2 | -1.21 | 1.05 | 0.34 | 0.03 | -0.76 | -0.4275 | -0.22 | -0.0925 | 0.819052 | 0.122346 | 0.65058 | 1 | 0.999631 | 0.938303 |
| Tspyl4_Q8VD63_S116s_1 | 0.39 | 3.17 | 2.68 | 0.9 | 1.7 | 2.38 | 2.03 | 1.29 | 2.89 | 0.91 | 2.62 | 2.11 | 1.87 | 2.48 | 2.13 | 1.65 | 0.065 | 0.3475 | 0.2475 | 0.695069 | 0.918814 | 0.628524 | 1 | 0.999631 | 0.895345 |
| Tspyl4_Q8VD63_S397s_1 | 2.51 | -0.04 | 1.37 | 2.02 | 0.96 | 1.36 | 2.42 | 1.32 | 1.11 | 1.86 | 0.92 | 0.01 | -2.36 | 0.31 | 1.02 | -0.19 | 0.05 | -0.49 | -1.77 | 0.04764 | 0.931096 | 0.429767 | 1 | 0.999631 | 0.481835 |
| Tspyl4_Q8VD63_S87s_1 | 0.21 | 1.04 | 1.14 | 0.89 | 0.69 | 0.45 | 0.64 | 0.35 | 1.24 | 1.12 | 0.68 | 0.95 | 0.76 | 1.36 | 0.73 | 0.72 | -0.2875 | 0.1775 | 0.0725 | 0.827638 | 0.339196 | 0.570132 | 1 | 0.999631 | 0.941202 |
| Tssc4_D3Z6S3_T124t_1 | 1.09 | 1.28 | 1.28 | 0.81 | 0.94 | 1.18 | 0.73 | 1.12 | 1.05 | 1.52 | 1.08 | 1.51 | 1.34 | 1.64 | 1.29 | 1.36 | -0.1225 | 0.175 | 0.2925 | 0.305985 | 0.637061 | 0.528604 | 1 | 0.999631 | 0.716954 |
| Ttbk1_Q6PCN3_S1222s_1 | 1.48 | 0.57 | -0.48 | 1.19 | 1.99 | 1.3 | 1.39 | 1.35 | 1.33 | 0.89 | 1.36 | 0.58 | -1.78 | 0.88 | 0.61 | 0.87 | 0.8175 | 0.35 | -0.545 | 0.437312 | 0.092984 | 0.453596 | 1 | 0.999631 | 0.781057 |
| Ttbk1_Q6PCN3_S433s_1 | 0.96 | 0.05 | -0.68 | 0.78 | 1.06 | 0.17 | 0.34 | 0.84 | 0.27 | 0.1 | 0.54 | -0.2 | -1.25 | 0.37 | 0.05 | 0.55 | 0.325 | -0.1 | -0.3475 | 0.510621 | 0.449882 | 0.808581 | 1 | 0.999631 | 0.817683 |
| Ttbk1_Q6PCN3_S441s_1 | 0.27 | 0.72 | 0.19 | -0.56 | -2.19 | 0.18 | -1.15 | -0.45 | 0.48 | 0.88 | -0.11 | 0.23 | -0.76 | 0.11 | -0.49 | 0.38 | -1.0575 | 0.215 | -0.345 | 0.396 | 0.070103 | 0.56133 | 1 | 0.999631 | 0.760506 |
| Ttbk1_Q6PCN3_S441s_2 | 1.14 | -0.22 | -1.11 | 1.13 | 1.01 | -0.02 | 1.13 | 1.49 | 0.22 | 0.63 | 1.11 | 0.21 | -2.07 | 0.42 | 0.34 | 0.66 | 0.6675 | 0.3075 | -0.3975 | 0.594808 | 0.266791 | 0.577461 | 1 | 0.999631 | 0.854138 |
| Ttbk1_Q6PCN3_S456s_1 | 0.61 | -0.74 | 0.22 | -0.9 | -2.05 | -0.81 | -0.79 | -1.35 | -1.13 | 1.26 | -1.24 | -1.32 | -1.24 | -1.88 | -1.3 | -2.2 | -1.0475 | -0.405 | -1.4525 | 0.007423 | 0.041621 | 0.537763 | 1 | 0.999631 | 0.358173 |
| Ttbk1_Q6PCN3_S484s_1 | 0.34 | 0.82 | -0.06 | 0.29 | 0.62 | 0.65 | -0.13 | 0.36 | 0.55 | 0.74 | 0.17 | 0.84 | 0.1 | 0.41 | 0.94 | 0.71 | 0.0275 | 0.2275 | 0.1925 | 0.563509 | 0.929369 | 0.462685 | 1 | 0.999631 | 0.841193 |
| Ttbk1_Q6PCN3_S638s_1 | -0.26 | -0.33 | -0.08 | -1.27 | -1.03 | -2.25 | -0.34 | -1.12 | -1.74 | -1.16 | 0.04 | -0.76 | -2.7 | -1.3 | -0.67 | 0.24 | -0.7 | -0.42 | -0.6225 | 0.321412 | 0.15033 | 0.36228 | 1 | 0.999631 | 0.72408 |
| Ttbk1_Q6PCN3_S655s_1 | 2.46 | -0.1 | 0.39 | 1.5 | 0.31 | 0.55 | 2.03 | 0.58 | 0.43 | 1.5 | 0.4 | -0.56 | -2.06 | -0.69 | 0.09 | -0.52 | -0.195 | -0.62 | -1.8575 | 0.016678 | 0.755107 | 0.345642 | 1 | 0.999631 | 0.391024 |
| Ttbk1_Q6PCN3_S827s_1 | -0.87 | 0.41 | -2.01 | -1.56 | -0.47 | 0.05 | -1.31 | -1 | -0.06 | -0.08 | -0.43 | -0.81 | 0.29 | 0.62 | -0.6 | -1.84 | 0.325 | 0.6625 | 0.625 | 0.36781 | 0.561989 | 0.223781 | 1 | 0.999631 | 0.745968 |
| Ttbk1_Q6PCN3_T325t_1 | -0.95 | -0.35 | 0.05 | -0.48 | -0.51 | -0.35 | -0.66 | -0.82 | -0.61 | 0.24 | 0.62 | -0.38 | -0.41 | -0.47 | -0.59 | -0.27 | -0.1525 | 0.4 | -0.0025 | 0.993566 | 0.608675 | 0.300057 | 1 | 0.999631 | 0.997705 |
| Ttbk1_Q6PCN3_T444t_2 | 1.14 | -0.22 | -1.11 | 1.13 | 1.01 | -0.02 | 1.13 | 1.49 | 0.22 | 0.63 | 1.11 | 0.21 | -2.07 | 0.42 | 0.34 | 0.66 | 0.6675 | 0.3075 | -0.3975 | 0.594808 | 0.266791 | 0.577461 | 1 | 0.999631 | 0.854138 |
| Ttbk2_Q3UVR3_S1181s_1 | 1.35 | 0.21 | 0.26 | 0 | 0.2 | 1.91 | 0.68 | 1.02 | 0.6 | 1.57 | 0.66 | 0.25 | -0.64 | 0.78 | 0.57 | 0.31 | 0.4975 | 0.315 | -0.2 | 0.650979 | 0.288761 | 0.458207 | 1 | 0.999631 | 0.876297 |
| Ttc14_E9PXW3_S570s_1 | 1.04 | -0.79 | 0.18 | 0.35 | -1.01 | -0.5 | 0.77 | -0.24 | -0.1 | 0.45 | 0.39 | 0.68 | -1.31 | -0.88 | -0.03 | -0.08 | -0.44 | 0.16 | -0.77 | 0.130558 | 0.387374 | 0.701767 | 1 | 0.999631 | 0.602177 |
| Ttc26_E0CXQ7_S14s_1 | 0.34 | -1 | 1.48 | 0.4 | -0.4 | 0.05 | 0.31 | 0.73 | 0.4 | 0.8 | 0.26 | 0.83 | 0.59 | 0.46 | 1.16 | 0.79 | -0.1325 | 0.2675 | 0.445 | 0.388487 | 0.799142 | 0.59558 | 1 | 0.999631 | 0.757772 |
| Ttc3_G5E8T2_S1903s_1 | -0.8 | 0.34 | 0.73 | 0.33 | 0.47 | 0.02 | 0.34 | 0.77 | 0.5 | -0.18 | 0.99 | 1.58 | 1.19 | 0.78 | -0.19 | 0.22 | 0.25 | 0.5725 | 0.35 | 0.442254 | 0.514171 | 0.249568 | 1 | 0.999631 | 0.782337 |
| Ttc7b_E9Q6P5_S158s_1 | 0.5 | -0.17 | -1.49 | -0.62 | -0.94 | -0.62 | -0.74 | 0.58 | -0.47 | -0.72 | -0.02 | 0.3 | -0.24 | 0.64 | -0.04 | -0.71 | 0.015 | 0.2175 | 0.3575 | 0.468368 | 0.976293 | 0.63973 | 1 | 0.999631 | 0.795782 |
| Ttc7b_E9Q6P5_S160s_1 | -0.3 | -0.72 | 0.19 | -0.65 | -0.96 | -0.84 | -0.2 | -0.94 | -1.1 | -0.64 | 0.5 | -0.47 | -0.9 | -0.16 | -0.64 | -0.62 | -0.365 | -0.0575 | -0.21 | 0.530487 | 0.273155 | 0.887211 | 1 | 0.999631 | 0.826702 |
| Ttc7b_E9Q6P5_S202s_1 | -2.23 | -2.76 | -4.01 | -3.17 | -1.66 | -3.67 | -3.32 | -3.44 | -2.98 | -3.3 | -2.85 | -3.07 | -0.35 | -2.8 | -3.36 | -3.69 | 0.02 | -0.0075 | 0.4925 | 0.511829 | 0.97074 | 0.985019 | 1 | 0.999631 | 0.817687 |
| Ttc7b_E9Q6P5_S629s_1 | -0.71 | -0.28 | -0.49 | -0.85 | -0.86 | -0.4 | -1.03 | -0.67 | -0.64 | -0.43 | -0.61 | -0.6 | 0.71 | 0.01 | -0.87 | -0.42 | -0.1575 | 0.0125 | 0.44 | 0.274115 | 0.567332 | 0.961621 | 1 | 0.999631 | 0.700187 |
| Ttc7b_E9Q6P5_S630s_1 | -0.42 | -0.34 | -0.8 | -0.26 | -0.57 | -0.14 | -0.68 | -0.01 | -0.85 | -0.29 | -0.79 | -0.26 | -0.13 | 0.09 | -0.5 | -0.12 | 0.105 | -0.0925 | 0.29 | 0.330588 | 0.710054 | 0.747888 | 1 | 0.999631 | 0.727302 |
| Ttc7b_E9Q6P5_S678s_1 | -1.75 | -1.77 | -1.84 | -2.89 | -1.68 | -1.9 | -2.48 | -2.99 | -2.14 | -2.16 | -2.05 | -1.68 | -0.03 | -1.84 | -1.59 | -1.82 | -0.2 | 0.055 | 0.7425 | 0.155955 | 0.624581 | 0.872328 | 1 | 0.999631 | 0.626452 |
| Ttc9_Q3V038_S15s_1 | -1.48 | 0.04 | 1.04 | 0.01 | -0.14 | -0.05 | 0.01 | 0.72 | 0.04 | -0.63 | 0.82 | 1.2 | 0.31 | 0.36 | 0.36 | 0.17 | 0.2325 | 0.455 | 0.3975 | 0.433534 | 0.654291 | 0.456269 | 1 | 0.999631 | 0.779567 |
| Ttc9b_Q9D6E4_S226s_1 | 1.06 | 0.4 | 0.8 | 0.82 | -0.55 | 0.02 | 1.66 | -0.12 | -0.44 | 0.32 | 0.46 | -0.21 | -0.13 | -0.38 | -0.13 | -0.45 | -0.5175 | -0.7375 | -1.0425 | 0.00428 | 0.293776 | 0.03966 | 1 | 0.999631 | 0.351308 |
| Ttc9b_Q9D6E4_S27s_1 | -0.22 | 0.35 | -0.01 | -0.42 | -0.08 | -0.08 | -0.08 | -0.91 | -0.03 | -0.41 | 0.09 | -0.33 | -0.12 | -0.18 | 0.55 | 0.61 | -0.2125 | -0.095 | 0.29 | 0.397607 | 0.506094 | 0.743472 | 1 | 0.999631 | 0.761181 |
| Tti1_Q91V83_S783s_1 | 0.51 | 0.05 | 0.47 | 1.27 | -0.27 | -0.26 | 0.64 | 0.02 | -0.24 | 0.71 | 0.44 | -0.83 | -0.19 | -1 | -0.58 | -0.92 | -0.5425 | -0.555 | -1.2475 | 0.005935 | 0.152062 | 0.216142 | 1 | 0.999631 | 0.351545 |
| Ttll12_F2Z423_S11s_1 | -0.26 | -1.91 | 0.4 | 0.44 | -1.75 | -1.54 | 0.07 | 0.63 | -2.08 | -0.37 | 0.08 | 0.42 | -1.72 | -1.83 | 0.75 | 0.42 | -0.315 | -0.155 | -0.2625 | 0.733455 | 0.659408 | 0.823863 | 1 | 0.999631 | 0.907462 |
| Ttll7_E9Q6C6_S410s_1 | 0.25 | 0.67 | 0.3 | -0.13 | -0.01 | -0.13 | -0.24 | -0.43 | 0.59 | 1.38 | 1.41 | -0.14 | 0.99 | 0.14 | 1.14 | 1.52 | -0.475 | 0.5375 | 0.675 | 0.095461 | 0.107842 | 0.211394 | 1 | 0.999631 | 0.558128 |
| Ttll9_D6RG93_S81s_2 | -1.48 | -2.45 | 1.56 | 2.6 | -1.47 | -2.95 | -0.66 | 3.27 | -1.37 | -3.4 | 1.5 | 0.96 | -2.15 | -3.18 | 1.94 | 2.87 | -0.51 | -0.635 | -0.1875 | 0.906949 | 0.737661 | 0.652824 | 1 | 0.999631 | 0.969801 |
| Ttll9_D6RG93_Y87y_2 | -1.48 | -2.45 | 1.56 | 2.6 | -1.47 | -2.95 | -0.66 | 3.27 | -1.37 | -3.4 | 1.5 | 0.96 | -2.15 | -3.18 | 1.94 | 2.87 | -0.51 | -0.635 | -0.1875 | 0.906949 | 0.737661 | 0.652824 | 1 | 0.999631 | 0.969801 |
| Ttyh1_Q9D3A9_S449s_1 | -1.48 | -1.18 | -0.69 | -0.94 | -1.1 | -0.66 | -1.09 | -1.17 | -0.93 | -1.13 | -1.32 | -0.31 | -0.15 | -0.68 | -0.62 | -0.55 | 0.0675 | 0.15 | 0.5725 | 0.08512 | 0.811711 | 0.650428 | 1 | 0.999631 | 0.550778 |
| Ttyh2_Q3TH73_S442s_1 | 0.07 | 0.09 | 1.46 | 0.28 | 0.55 | -0.31 | -0.22 | 0.11 | 0.3 | 0.58 | -0.05 | 0.29 | 0.34 | -0.33 | 0.75 | 0.38 | -0.4425 | -0.195 | -0.19 | 0.649602 | 0.275494 | 0.609063 | 1 | 0.999631 | 0.875089 |
| Ttyh3_Q6P5F7_S496s_1 | -1.35 | -0.82 | -0.56 | -1.75 | -0.94 | -1.43 | -1.09 | -1.53 | -0.66 | -1.28 | -0.87 | -0.24 | 0.63 | -1.4 | -1 | -0.98 | -0.1275 | 0.3575 | 0.4325 | 0.396657 | 0.706488 | 0.346873 | 1 | 0.999631 | 0.761029 |
| Ttyh3_Q6P5F7_S519s_1 | -0.87 | -1.49 | -1.32 | -1.57 | -1.74 | -1.88 | -1.24 | -1.96 | -2.17 | -0.73 | -2.19 | -1.64 | -0.75 | -1.84 | -1 | -1.34 | -0.3925 | -0.37 | 0.08 | 0.816516 | 0.201739 | 0.356975 | 1 | 0.999631 | 0.937777 |
| Ttyh3_Q6P5F7_S522s_1 | 0.05 | -0.8 | -1.78 | -1.77 | -1.89 | -0.53 | -1.18 | -1.84 | -0.36 | 0.49 | -2.49 | -1.42 | -0.59 | -0.7 | -1.91 | -0.96 | -0.285 | 0.13 | 0.035 | 0.944959 | 0.577945 | 0.851963 | 1 | 0.999631 | 0.982141 |
| Ttyh3_Q6P5F7_T427t_1 | -0.62 | -1.24 | -1.23 | 0.7 | -1.33 | -0.73 | -0.28 | 0.79 | -1.37 | -0.1 | -0.24 | -0.21 | -1.07 | -1.3 | -0.97 | -0.82 | 0.21 | 0.1175 | -0.4425 | 0.348868 | 0.718331 | 0.819061 | 1 | 0.999631 | 0.736697 |
| Tuba4a_P68368_S158s_1 | -0.85 | -1.31 | 0.28 | -0.31 | -1.24 | -1.31 | -0.31 | -0.51 | -1.09 | -0.73 | 0.12 | 0.35 | -0.25 | -0.65 | 0.26 | 0.54 | -0.295 | 0.21 | 0.5225 | 0.250215 | 0.489914 | 0.655623 | 1 | 0.999631 | 0.686417 |
| Tuba4a_P68368_S277s_1 | -0.08 | -0.29 | -0.5 | -0.93 | 0.7 | -0.39 | 0.06 | -0.59 | -1.24 | -2.23 | -0.78 | 0.31 | 1.37 | -0.17 | -1.58 | -1.52 | 0.395 | -0.535 | -0.025 | 0.969103 | 0.289914 | 0.319696 | 1 | 0.999631 | 0.98892 |
| Tuba4a_P68368_S340s_1 | 0.23 | 1.9 | 0.6 | 0.35 | 1.09 | 1.7 | 0.09 | 0.73 | 1.49 | 1.34 | 0.6 | 0.59 | 1.85 | 2.14 | 0.81 | 0.88 | 0.1325 | 0.235 | 0.65 | 0.206823 | 0.784256 | 0.600456 | 1 | 0.999631 | 0.660763 |
| Tuba4a_P68368_T223t_1 | -0.51 | 0.37 | -0.01 | -0.54 | 0.14 | -0.34 | 0.2 | -0.85 | 0.07 | -0.69 | 0.25 | 0.85 | 0.8 | -0.65 | -0.22 | -0.93 | -0.04 | 0.2925 | -0.0775 | 0.860376 | 0.910073 | 0.468972 | 1 | 0.999631 | 0.953592 |
| Tuba4a_P68368_Y103y_1 | 3.46 | -3.7 | -1.18 | -2.27 | -1.69 | -2.26 | 1.49 | -3.22 | -1.92 | 2.91 | -2.26 | -1.25 | -3.01 | -3.85 | -2.85 | -2.32 | -0.4975 | 0.2925 | -2.085 | 0.14111 | 0.75197 | 0.860242 | 1 | 0.999631 | 0.61245 |
| Tuba4a_P68368_Y357y_1 | 0.18 | 0.02 | 0.03 | -0.09 | -0.15 | 0.26 | -1 | -0.46 | 0.76 | 0.37 | 0.86 | -0.13 | 0.39 | 0.65 | -1.45 | 0.96 | -0.3725 | 0.43 | 0.1025 | 0.84261 | 0.260774 | 0.179416 | 1 | 0.999631 | 0.948041 |
| Tubb3_Q9ERD7_S48s_1 | 0.01 | 0.14 | 0.73 | -0.3 | -0.48 | 1.07 | 0.33 | 0.78 | -2.05 | 0.12 | 0.19 | -1.64 | 0.09 | 0.9 | -1.62 | -0.81 | 0.28 | -0.99 | -0.505 | 0.366724 | 0.493345 | 0.108518 | 1 | 0.999631 | 0.745796 |
| Tubb4b_P68372_S338s_1 | -0.4 | 0.72 | 2.22 | 0.69 | 0.04 | 0.67 | 0.26 | 0.88 | 0.37 | -1.27 | 1.32 | 1.15 | 0.02 | 0.33 | 1.22 | 0.27 | -0.345 | -0.415 | -0.3475 | 0.535644 | 0.518451 | 0.562712 | 1 | 0.999631 | 0.829362 |
| Tubb4b_P68372_T219t_1 | -0.99 | -0.53 | -0.46 | -0.6 | -0.87 | -0.37 | -0.63 | -0.51 | -1.21 | -0.89 | -0.69 | -0.69 | -0.49 | 0.16 | -1.04 | 0.11 | 0.05 | -0.225 | 0.33 | 0.367428 | 0.84855 | 0.420899 | 1 | 0.999631 | 0.745967 |
| Tubgcp3_P58854_Y542y_1 | -0.2 | 0.47 | -0.82 | -0.47 | 0.09 | 0.29 | -0.56 | -0.65 | 0.54 | 0.7 | 0 | -0.17 | 0.52 | 0.51 | -0.49 | 0.29 | 0.0475 | 0.5225 | 0.4625 | 0.254551 | 0.899233 | 0.180881 | 1 | 0.999631 | 0.689457 |
| Tubgcp5_Q8BKN5_S182s_1 | 0.46 | 0.96 | 1.42 | 0.61 | 0.03 | 0.99 | 0.56 | 1.02 | 1.29 | 0.6 | 0.93 | 1.45 | 1.25 | 1.15 | 1.79 | 1.72 | -0.2125 | 0.205 | 0.615 | 0.091568 | 0.54353 | 0.544187 | 1 | 0.999631 | 0.557085 |
| Tusc1_Q673H1_S146s_1 | 2.15 | 0.07 | 0.58 | 0.63 | 0.53 | 0.67 | 1.68 | 0.04 | 1.1 | 1.82 | 0.33 | 0.33 | -0.1 | -0.89 | -0.06 | -0.42 | -0.1275 | 0.0375 | -1.225 | 0.026001 | 0.808063 | 0.944107 | 1 | 0.999631 | 0.422285 |
| Tusc5_Q8C838_S74s_1 | -0.05 | -0.64 | 0.2 | -0.93 | -0.88 | 0.08 | -0.79 | -1.02 | 0.07 | 1.35 | -0.53 | -0.28 | 2.01 | 0.1 | -0.86 | 0.18 | -0.2975 | 0.5075 | 0.7125 | 0.249526 | 0.43604 | 0.299351 | 1 | 0.999631 | 0.686359 |
| Twf1_Q91YR1_T349t_1 | 1.5 | -0.51 | -0.02 | 0.72 | -0.75 | -0.06 | 0.85 | 0.74 | -0.21 | 0.54 | 0.63 | -1.46 | -1.06 | -1.19 | -0.04 | -1.01 | -0.2275 | -0.5475 | -1.2475 | 0.028041 | 0.670929 | 0.368896 | 1 | 0.999631 | 0.423977 |
| Twf2_Q9Z0P5_S165s_1 | 1.13 | -0.75 | -1.2 | 0.4 | 0.46 | -0.05 | 0.25 | -0.19 | 0.26 | -1.14 | 0.22 | -0.15 | 0.38 | -0.08 | -0.47 | -0.66 | 0.2225 | -0.0975 | -0.1025 | 0.84988 | 0.666373 | 0.864911 | 1 | 0.999631 | 0.949525 |
| Txlna_Q6PAM1_S515s_1 | -0.42 | -1.14 | 1.07 | 0.82 | -0.99 | -2.02 | 0.34 | 0.1 | 0.13 | -0.63 | 0.59 | 1.1 | -0.44 | -0.77 | 0.55 | 1.21 | -0.725 | 0.215 | 0.055 | 0.929858 | 0.293991 | 0.713605 | 1 | 0.999631 | 0.97754 |
| Txlna_Q6PAM1_S515s_2 | 0.34 | 0.54 | 0.92 | -0.43 | 0.15 | -0.5 | -0.28 | 0.01 | 0.22 | 0.47 | -0.14 | 0.14 | 0.39 | 0.42 | -0.11 | 0.59 | -0.4975 | -0.17 | -0.02 | 0.956484 | 0.176355 | 0.62915 | 1 | 0.999631 | 0.984803 |
| Txnrd3_Q99MD6_S16s_1 | 0.46 | 1.2 | 1.29 | 0.78 | 0.09 | 0.43 | 0.77 | 0.67 | 0.77 | 0.79 | 0.85 | 1.25 | 0.18 | 0.99 | 1.29 | 1.03 | -0.4425 | -0.0175 | -0.06 | 0.867171 | 0.168698 | 0.953428 | 1 | 0.999631 | 0.956562 |
| Txnrd3_Q99MD6_S50s_1 | 1.93 | 0.74 | -2.94 | -1.53 | 1.34 | 1.36 | -1.36 | -2.99 | 1.96 | 1.65 | -1.26 | -1.72 | 0.23 | 1.27 | -2.32 | -2.72 | 0.0375 | 0.6075 | -0.435 | 0.726006 | 0.977046 | 0.627664 | 1 | 0.999631 | 0.90568 |
| Txnrd3_Q99MD6_T15t_1 | 0.46 | 1.2 | 1.29 | 0.78 | 0.09 | 0.43 | 0.77 | 0.67 | 0.77 | 0.79 | 0.85 | 1.25 | 0.18 | 0.99 | 1.29 | 1.03 | -0.4425 | -0.0175 | -0.06 | 0.867171 | 0.168698 | 0.953428 | 1 | 0.999631 | 0.956562 |
| Tyro3_P55144_S804s_1 | 0.13 | -0.88 | -0.16 | -0.22 | -0.96 | -1.31 | -0.29 | -1.33 | -0.88 | 0.02 | -0.49 | -0.33 | -0.61 | -0.92 | -0.84 | -0.45 | -0.69 | -0.1375 | -0.4225 | 0.208016 | 0.073869 | 0.681255 | 1 | 0.999631 | 0.661649 |
| Tyro3_P55144_S855s_1 | -0.65 | -0.11 | -0.21 | -0.8 | -0.09 | -0.71 | -0.65 | -0.17 | -0.45 | -0.28 | -0.13 | -0.18 | 0.33 | -0.15 | -0.5 | -0.44 | 0.0375 | 0.1825 | 0.2525 | 0.448642 | 0.899492 | 0.518364 | 1 | 0.999631 | 0.786796 |
| U2af1_Q9D883_S61s_1 | -0.33 | 0.52 | 1.22 | 0.7 | 0.34 | 0.09 | 0.29 | 0.03 | 1.18 | 0.27 | 0.77 | 0.48 | 0.71 | -0.23 | -1.21 | 0.44 | -0.34 | 0.1475 | -0.6 | 0.255799 | 0.351805 | 0.708432 | 1 | 0.999631 | 0.690775 |
| U2surp_Q6NV83_S202s_1 | -0.85 | 2.7 | 2.3 | 1.46 | -0.22 | 1.51 | -0.3 | 1.32 | 1.93 | 1.62 | 2.1 | 2.86 | 0.31 | 2.4 | 1.94 | 3.23 | -0.825 | 0.725 | 0.5675 | 0.517798 | 0.322769 | 0.339753 | 1 | 0.999631 | 0.819449 |
| U2surp_Q6NV83_S67s_1 | -0.58 | 0.63 | 1.73 | -0.44 | -1.19 | 0.73 | -0.28 | -0.83 | -0.29 | 0.65 | -1.2 | -0.21 | 1 | 0.35 | 0.34 | 0.77 | -0.7275 | -0.5975 | 0.28 | 0.598873 | 0.253253 | 0.331491 | 1 | 0.999631 | 0.855702 |
| U2surp_Q6NV83_S810s_1 | 0.6 | 1.35 | 2.8 | 1.2 | -0.64 | 2.01 | 0.16 | 0.28 | -0.07 | -0.11 | -0.27 | 0.68 | -0.23 | 1.33 | 2.46 | 1.46 | -1.035 | -1.43 | -0.2325 | 0.721654 | 0.135311 | 0.01474 | 1 | 0.999631 | 0.905674 |
| Uba1_Q02053_S13s_1 | 2.02 | 2.63 | 3.15 | 2.15 | 1.27 | 2.85 | 2.56 | 1.91 | 2.59 | 2.91 | 1.96 | 1.62 | 1.41 | 1.89 | 2.3 | 2.15 | -0.34 | -0.2175 | -0.55 | 0.156862 | 0.435258 | 0.59072 | 1 | 0.999631 | 0.626839 |
| Uba1_Q02053_S810s_1 | -0.54 | 0.58 | 0.32 | -0.4 | 0.88 | -0.22 | 0.53 | 0.64 | 0.7 | -0.36 | 0.98 | 1.28 | 1.06 | 0.16 | 0.64 | 0.69 | 0.4675 | 0.66 | 0.6475 | 0.105245 | 0.23308 | 0.159542 | 1 | 0.999631 | 0.568213 |
| Uba1_Q02053_S816s_1 | -1.52 | -1.72 | 0.91 | -0.32 | -0.43 | -1.17 | 0.17 | -1.37 | -0.6 | -0.65 | -0.68 | 0.36 | -0.31 | -0.79 | -1.01 | -0.23 | -0.0375 | 0.27 | 0.0775 | 0.894375 | 0.952569 | 0.654215 | 1 | 0.999631 | 0.965376 |
| Uba1_Q02053_S835s_1 | 0.37 | 1.47 | 0.09 | -0.18 | 0.25 | 1.52 | -0.52 | -0.57 | 0.46 | 1.51 | -0.48 | -0.05 | 0.44 | 1.1 | -0.56 | -0.16 | -0.2675 | -0.0775 | -0.2325 | 0.638855 | 0.632428 | 0.882723 | 1 | 0.999631 | 0.870943 |
| Uba1_Q02053_S876s_1 | -2.21 | -0.57 | -0.4 | -1.3 | -0.71 | -0.63 | -0.53 | -2.16 | -0.5 | -1.25 | -0.34 | -0.08 | -0.19 | -0.38 | -0.86 | -1.41 | 0.1125 | 0.5775 | 0.41 | 0.402213 | 0.829983 | 0.235509 | 1 | 0.999631 | 0.763561 |
| Uba3_Q3TL72_S363s_1 | -0.58 | 0.55 | 0.19 | -0.2 | 0.28 | 0.2 | -0.88 | -0.55 | -0.23 | -0.17 | -0.46 | -0.82 | -0.12 | -0.01 | -0.59 | -0.74 | -0.2275 | -0.41 | -0.355 | 0.329938 | 0.558642 | 0.238511 | 1 | 0.999631 | 0.726756 |
| Ubac1_F6Z1E7_S18s_1 | 0.13 | 0.43 | 0.63 | 0.27 | 0.19 | 0.74 | 0.25 | 0.54 | 0.67 | 0.97 | 0.76 | 1.49 | 0.72 | 1.43 | 0.98 | 0.8 | 0.065 | 0.6075 | 0.6175 | 0.061574 | 0.806743 | 0.062443 | 1 | 0.999631 | 0.509355 |
| Ubap1_Q8BH48_S146s_1 | -0.39 | -0.22 | -0.62 | -0.05 | -0.62 | -0.26 | -0.81 | 0.17 | -0.34 | 0.65 | 0.33 | -0.65 | 0.8 | 0.5 | -0.27 | 0.27 | -0.06 | 0.3175 | 0.645 | 0.074465 | 0.844627 | 0.384179 | 1 | 0.999631 | 0.540038 |
| Ubap2l_Q80X50_S467s_1 | 0.25 | 0.98 | 1.42 | 0.73 | 0.47 | 0.56 | 0.65 | 0.82 | 0.28 | 0.21 | 1.56 | 2.07 | 0.72 | 1.03 | 1.59 | 1.05 | -0.22 | 0.185 | 0.2525 | 0.484246 | 0.484256 | 0.711567 | 1 | 0.999631 | 0.803422 |
| Ubap2l_Q80X50_S609s_1 | 0.65 | 1.02 | 0.29 | 0.49 | 0.91 | 0.84 | 0.23 | 0.72 | 1.48 | 1.24 | 0.93 | 1.01 | 1.36 | 1.73 | 1.1 | 0.97 | 0.0625 | 0.5525 | 0.6775 | 0.053692 | 0.829278 | 0.078809 | 1 | 0.999631 | 0.4909 |
| Ubap2l_Q80X50_S992s_1 | -0.1 | -0.37 | -0.95 | -0.84 | 1.92 | 0.07 | 2.26 | -0.56 | 1.57 | -2.93 | 0.2 | -0.18 | -1.11 | -1.99 | -1.18 | -2.82 | 1.4875 | 0.23 | -1.21 | 0.0209 | 0.040884 | 0.785068 | 1 | 0.999631 | 0.399638 |
| Ube2e2_H3BJB5_S11s_1 | 1.08 | -0.08 | 2.52 | 1.33 | -2.27 | -1.01 | 1.84 | 0.88 | -0.87 | 0.66 | 0.8 | 0.32 | -1.43 | -2.88 | 0.65 | 0.75 | -1.3525 | -0.985 | -1.94 | 0.049857 | 0.165415 | 0.123608 | 1 | 0.999631 | 0.488436 |
| Ube2e3_B2FDH7_S8s_1 | 1.67 | 0.88 | 1.81 | 1.19 | 1.08 | 0.67 | 1.19 | 0.87 | 0.98 | 2.29 | 0.37 | 1.19 | 0.38 | -0.49 | 1.18 | -0.74 | -0.435 | -0.18 | -1.305 | 0.019416 | 0.174551 | 0.688308 | 1 | 0.999631 | 0.391668 |
| Ube2j1_F6SPK0_S210s_1 | 0.16 | 0.36 | 0.02 | 0.16 | 0.08 | 0.64 | 0 | 0.44 | -0.13 | 0.31 | 0.41 | -0.13 | -0.05 | 0.98 | -0.03 | 0.24 | 0.115 | -0.06 | 0.11 | 0.737372 | 0.665653 | 0.82407 | 1 | 0.999631 | 0.907787 |
| Ube2o_Q6ZPJ3_S110s_1 | -0.59 | -1.09 | -1.3 | -1.21 | -1.19 | -0.84 | -0.69 | -1.43 | -0.48 | -0.31 | -1.68 | -0.79 | 0.09 | -0.38 | -0.74 | -1.34 | 0.01 | 0.2325 | 0.455 | 0.246362 | 0.973079 | 0.534876 | 1 | 0.999631 | 0.685751 |
| Ube2o_Q6ZPJ3_S1196s_1 | -0.8 | -0.06 | 0.31 | -0.17 | -0.08 | -0.07 | -0.65 | 0.41 | -0.07 | -0.24 | -0.16 | 0.48 | 0.23 | 0.12 | 0.32 | 1.33 | 0.0825 | 0.1825 | 0.68 | 0.106754 | 0.812422 | 0.586341 | 1 | 0.999631 | 0.568797 |
| Ube2o_Q6ZPJ3_S317s_1 | -2.14 | -0.52 | -1.21 | -0.96 | -0.66 | -0.95 | -1.08 | 0.21 | -0.66 | -1.76 | -2.31 | -0.47 | -1.53 | -1.09 | -0.6 | -0.76 | 0.5875 | -0.0925 | 0.2125 | 0.610902 | 0.200216 | 0.859607 | 1 | 0.999631 | 0.859461 |
| Ube2o_Q6ZPJ3_S376s_1 | 1.56 | 0.65 | 2.7 | 0.83 | -0.16 | 0.32 | 2.78 | 1.36 | 0.67 | 0.21 | 1.96 | 1.79 | -0.49 | -1.27 | 2.49 | 0.5 | -0.36 | -0.2775 | -1.1275 | 0.187481 | 0.612959 | 0.633489 | 1 | 0.999631 | 0.646474 |
| Ube2o_Q6ZPJ3_S510s_1 | 1.26 | 2.18 | 0.43 | 2.13 | 2.33 | 1.49 | 1.32 | 1.8 | 2.33 | 1.32 | 2.04 | 1.78 | 0.45 | 2.51 | 1.6 | 1.95 | 0.235 | 0.3675 | 0.1275 | 0.819092 | 0.60652 | 0.428025 | 1 | 0.999631 | 0.938303 |
| Ube2o_Q6ZPJ3_S833s_1 | -0.21 | -0.43 | -0.3 | -0.76 | -0.68 | -0.02 | -1.06 | -0.49 | -0.02 | -0.17 | -0.59 | 0.08 | 0.48 | 0.06 | -0.05 | 0.18 | -0.1375 | 0.25 | 0.5925 | 0.062713 | 0.654627 | 0.387842 | 1 | 0.999631 | 0.512833 |
| Ube2o_Q6ZPJ3_S836s_1 | -0.54 | -0.57 | -0.48 | -0.4 | -0.21 | -0.43 | -0.79 | -0.58 | -0.7 | -0.48 | -0.47 | -0.43 | -0.14 | -0.21 | -0.61 | -0.18 | -0.005 | -0.0225 | 0.2125 | 0.440617 | 0.984008 | 0.925943 | 1 | 0.999631 | 0.781983 |
| Ube2o_Q6ZPJ3_S893s_1 | 0.14 | 0.93 | 1.19 | 0.31 | 0.42 | 0.77 | 0.13 | 0.39 | 1.1 | 0.9 | 0.17 | 0.94 | 1.53 | 0.73 | 0.51 | 1 | -0.215 | 0.135 | 0.3 | 0.429226 | 0.514188 | 0.707214 | 1 | 0.999631 | 0.776658 |
| Ube2ql1_E9QMD2_S66s_1 | 0.03 | 0.14 | 1.62 | 1.47 | -1.18 | 1.86 | 1.89 | 1.68 | 0.43 | -0.12 | 0.87 | -0.17 | -2.9 | -0.14 | 0.92 | 0.25 | 0.2475 | -0.5625 | -1.2825 | 0.140334 | 0.743607 | 0.251641 | 1 | 0.999631 | 0.611934 |
| Ube2v1_Q9CZY3_S104s_1 | -0.63 | -0.36 | 0.3 | -0.33 | 0.01 | -0.13 | -0.28 | -0.36 | -0.28 | -0.22 | 0.02 | 0.38 | 0.53 | -0.34 | 0.4 | 0.66 | 0.065 | 0.23 | 0.5675 | 0.131591 | 0.821176 | 0.46845 | 1 | 0.999631 | 0.603456 |
| Ube3a_O08759_S8s_1 | 0.17 | 0.59 | 0.24 | -0.31 | 0.01 | -0.26 | -0.25 | -0.29 | 0.13 | 0.68 | 0.09 | 0.23 | 1.2 | 0.54 | 0.31 | 0.78 | -0.37 | 0.11 | 0.535 | 0.133742 | 0.206167 | 0.717481 | 1 | 0.999631 | 0.60492 |
| Ube3b_Q9ES34_S421s_1 | -0.42 | 1.18 | 0.42 | 0.07 | -1.99 | 0.06 | -0.45 | 0.36 | -1.06 | -0.92 | 0.47 | 0.47 | 0.18 | -0.89 | -0.02 | 0.2 | -0.8175 | -0.5725 | -0.445 | 0.315653 | 0.171958 | 0.277515 | 1 | 0.999631 | 0.720734 |
| Ube4b_G3UZD6_S101s_1 | -1.11 | -1.25 | -0.5 | -1.05 | -0.78 | -0.85 | 0.12 | 0.09 | -0.28 | -1.69 | 0.41 | 0.54 | -1.7 | 0.06 | -0.9 | -0.24 | 0.6225 | 0.7225 | 0.2825 | 0.517157 | 0.095987 | 0.176798 | 1 | 0.999631 | 0.819449 |
| Ube4b_G3UZD6_S105s_1 | -0.68 | 0.29 | -0.26 | 0.74 | 0.39 | 0.29 | 0.86 | -0.15 | -0.19 | -0.02 | 0.63 | -0.65 | -1.99 | -0.19 | -0.13 | 0.35 | 0.325 | -0.08 | -0.5125 | 0.368376 | 0.407264 | 0.846935 | 1 | 0.999631 | 0.745968 |
| Ube4b_G3UZD6_S31s_1 | -0.18 | 0.58 | 0.34 | -0.74 | 0.96 | 0.89 | -0.29 | -0.22 | 0.94 | 0.42 | 0.06 | 0.66 | 1.75 | 1.72 | -0.3 | 0.26 | 0.335 | 0.52 | 0.8575 | 0.144946 | 0.451907 | 0.18538 | 1 | 0.999631 | 0.616037 |
| Ubn1_Q4G0F8_S493s_1 | -0.33 | -1.15 | 0.3 | -0.6 | -0.41 | 0.49 | 0.49 | -0.44 | -0.11 | 0.53 | -0.46 | -1.37 | -1.09 | -0.03 | -0.52 | -1.2 | 0.4775 | 0.0925 | -0.265 | 0.531488 | 0.25412 | 0.8467 | 1 | 0.999631 | 0.827175 |
| Ubn2_Q80WC1_S777s_2 | -0.56 | -1.15 | -0.48 | -1.12 | -1.2 | -1.19 | -1.67 | -0.64 | -1.69 | -0.67 | -0.67 | -1.38 | -0.56 | -1.25 | -0.67 | -0.35 | -0.3475 | -0.275 | 0.12 | 0.719654 | 0.295915 | 0.441548 | 1 | 0.999631 | 0.904782 |
| Ubn2_Q80WC1_S780s_2 | -0.56 | -1.15 | -0.48 | -1.12 | -1.2 | -1.19 | -1.67 | -0.64 | -1.69 | -0.67 | -0.67 | -1.38 | -0.56 | -1.25 | -0.67 | -0.35 | -0.3475 | -0.275 | 0.12 | 0.719654 | 0.295915 | 0.441548 | 1 | 0.999631 | 0.904782 |
| Ubqln2_Q9QZM0_S25s_1 | 0.11 | 0.34 | -0.17 | -0.28 | -0.19 | -0.06 | -0.45 | -0.16 | -0.16 | 0.48 | -0.48 | -0.11 | 0.6 | 0.05 | 0.07 | -0.06 | -0.215 | -0.0675 | 0.165 | 0.590291 | 0.422581 | 0.828229 | 1 | 0.999631 | 0.851598 |
| Ubqln4_Q99NB8_S5s_1 | 1.14 | -1.94 | -1.65 | -0.49 | -0.75 | -0.78 | 0.75 | -0.86 | -0.27 | -1 | -0.94 | -2.1 | -1.95 | -1.77 | -1.21 | -2.56 | 0.325 | -0.3425 | -1.1375 | 0.11453 | 0.647152 | 0.630126 | 1 | 0.999631 | 0.579001 |
| Ubr4_A2AN08_S178s_1 | -0.01 | 0.89 | -0.22 | -0.25 | 0.27 | 0.43 | -0.57 | -0.09 | 0.49 | 1 | 0.77 | 0.56 | 1.42 | 0.18 | -0.19 | 0.69 | -0.0925 | 0.6025 | 0.4225 | 0.350772 | 0.80148 | 0.099728 | 1 | 0.999631 | 0.737443 |
| Ubr4_A2AN08_S181s_1 | -0.45 | -0.26 | -0.15 | -0.52 | -1.2 | -0.71 | -0.7 | -0.41 | -0.59 | -1.04 | -0.05 | -0.17 | -0.32 | -0.76 | -0.07 | -0.1 | -0.41 | -0.1175 | 0.0325 | 0.912012 | 0.15666 | 0.704492 | 1 | 0.999631 | 0.971964 |
| Ubr4_A2AN08_S457s_1 | 1.92 | 1.22 | 1.86 | 1.32 | 0.01 | 1.38 | 2.14 | -0.01 | 0.41 | 1.87 | 1.45 | 1.26 | 0.7 | 0.53 | 1.06 | 1 | -0.7 | -0.3325 | -0.7575 | 0.033058 | 0.199924 | 0.389925 | 1 | 0.999631 | 0.44294 |
| Ubr4_F6SSP6_S1439s_1 | -3.24 | 4.74 | 5.5 | 2.26 | 4.38 | 3.67 | 2.67 | 4.88 | 4.74 | 0.76 | 6.36 | 6.96 | 4.94 | 2.84 | 5.82 | 4.07 | 1.585 | 2.39 | 2.1025 | 0.243667 | 0.367624 | 0.259903 | 1 | 0.999631 | 0.685637 |
| Ubr4_F6SSP6_S1442s_1 | 0.15 | 0.55 | -0.88 | -0.01 | 0.24 | 1.04 | -0.34 | 0.68 | 0.96 | 1.2 | -0.16 | -0.01 | 0.32 | -0.41 | 0.02 | -2.24 | 0.4525 | 0.545 | -0.53 | 0.381044 | 0.295645 | 0.241693 | 1 | 0.999631 | 0.752619 |
| Ubr4_F6SSP6_S1464s_1 | 0.93 | 0.48 | 0.85 | 1.06 | 0.36 | 0.03 | 0.35 | 0.95 | 0.69 | 0.15 | 0.16 | 0.64 | -0.62 | 0.57 | 0.78 | 0.52 | -0.4075 | -0.42 | -0.5175 | 0.191144 | 0.189277 | 0.163177 | 1 | 0.999631 | 0.648674 |
| Ubr4_F6SSP6_S1570s_1 | -0.99 | -1.66 | -2.09 | -1.63 | -1.46 | -1.43 | -1.48 | -1.81 | -1.97 | -1.14 | -1.24 | -2.51 | -2.87 | -1.86 | -1.85 | -0.92 | 0.0475 | -0.1225 | -0.2825 | 0.538558 | 0.875665 | 0.762625 | 1 | 0.999631 | 0.830863 |
| Ubr4_F6SSP6_T1444t_1 | -0.23 | -0.24 | 0.39 | 0.62 | -0.14 | -0.03 | 0.78 | 0.03 | 0.21 | -0.1 | 0.01 | 0.29 | -0.5 | -0.88 | 0.48 | -0.8 | 0.025 | -0.0325 | -0.56 | 0.186415 | 0.941142 | 0.915568 | 1 | 0.999631 | 0.646474 |
| Ubr4_F6SSP6_T1569t_1 | -0.99 | -1.66 | -2.09 | -1.63 | -1.46 | -1.43 | -1.48 | -1.81 | -1.97 | -1.14 | -1.24 | -2.51 | -2.87 | -1.86 | -1.85 | -0.92 | 0.0475 | -0.1225 | -0.2825 | 0.538558 | 0.875665 | 0.762625 | 1 | 0.999631 | 0.830863 |
| Ubr5_E9Q2H1_S1543s_1 | 1.09 | 2.18 | 1.6 | 1.4 | 0.94 | 1.81 | 1.4 | 1.59 | 1.13 | 0.86 | 2.08 | 0.43 | 0.64 | 0.94 | 0.99 | 1.3 | -0.1325 | -0.4425 | -0.6 | 0.097848 | 0.692473 | 0.307595 | 1 | 0.999631 | 0.560722 |
| Ubxn1_Q922Y1_T65t_1 | -0.2 | 1.35 | 1.46 | 0.36 | 0.41 | 1.19 | 0.06 | 1.27 | 0.55 | 1.45 | 0.94 | 1.39 | 0.91 | 0.17 | 1.52 | 0.66 | -0.01 | 0.34 | 0.0725 | 0.878947 | 0.983134 | 0.452083 | 1 | 0.999631 | 0.962598 |
| Ubxn2b_Q0KL01_S216s_1 | 1.61 | 0.63 | 0.62 | 2.14 | 0.98 | 1.03 | 2.04 | 0.99 | 1.89 | 1.25 | 1.47 | 1.43 | -1.88 | 0.61 | 1.2 | 0.39 | 0.01 | 0.26 | -1.17 | 0.113111 | 0.982024 | 0.529069 | 1 | 0.999631 | 0.577408 |
| Ubxn4_Q8VCH8_T487t_1 | 0.21 | 1.02 | 0.19 | 1.01 | 1.53 | 2.16 | 0.43 | 1.37 | 0.85 | -0.45 | 0.07 | -1.19 | -2.15 | 1.73 | 0.33 | -0.76 | 0.765 | -0.7875 | -0.82 | 0.290017 | 0.095269 | 0.120537 | 1 | 0.999631 | 0.706164 |
| Ubxn6_Q99PL6_S36s_1 | 1.22 | 0.2 | 0.9 | 1.74 | 1.57 | 1.47 | 1.51 | 1.63 | 1.29 | 0.58 | 1.74 | 1.24 | -0.68 | 1.8 | 1.7 | 1.69 | 0.53 | 0.1975 | 0.1125 | 0.855804 | 0.155508 | 0.631345 | 1 | 0.999631 | 0.951817 |
| Ubxn6_Q99PL6_S75s_1 | 0.52 | -0.47 | -0.05 | 0.64 | 0.27 | 0.08 | 0.07 | -0.03 | 0.15 | 0.04 | -0.21 | 0.25 | -1.49 | -0.12 | 0.35 | -0.03 | -0.0625 | -0.1025 | -0.4825 | 0.31673 | 0.843453 | 0.756122 | 1 | 0.999631 | 0.720734 |
| Ubxn6_Q99PL6_S78s_1 | 1.73 | -0.98 | -2.52 | 0.83 | 0.97 | 1.91 | 0.8 | 0.25 | 0.95 | 0.4 | 0.58 | -1.02 | -2.37 | 0.61 | -0.23 | -1.18 | 1.2175 | 0.4625 | -0.5575 | 0.572206 | 0.185641 | 0.611526 | 1 | 0.999631 | 0.844319 |
| Uck2_Q99PM9_S254s_1 | 3.24 | 0.29 | 1.04 | 1.16 | 0.5 | 1.3 | 3 | -0.33 | -0.14 | 1.57 | 0.71 | 0.22 | -0.66 | -1.71 | 0.21 | -1.04 | -0.315 | -0.8425 | -2.2325 | 0.007154 | 0.70457 | 0.220132 | 1 | 0.999631 | 0.358173 |
| Ufd1l_P70362_S209s_1 | -0.39 | 0.84 | 2.31 | 1.44 | 0.7 | 1.19 | 0.3 | 0.84 | 0.53 | 0.34 | 0.08 | 1.75 | 0.68 | 1.01 | 1.79 | 0.9 | -0.2925 | -0.375 | 0.045 | 0.936875 | 0.595845 | 0.545472 | 1 | 0.999631 | 0.97939 |
| Ufd1l_P70362_S247s_1 | 0.45 | 0.15 | -0.37 | 0.96 | 1.32 | 0.56 | 0.84 | 1.55 | 0.82 | -0.44 | 0.9 | 0.28 | -1.23 | 0.51 | 1.39 | 0.73 | 0.77 | 0.0925 | 0.0525 | 0.927148 | 0.062955 | 0.825577 | 1 | 0.999631 | 0.976306 |
| Ufd1l_P70362_S299s_1 | 0.15 | 0.61 | 0.24 | 1.62 | 1.18 | 0.59 | 0.89 | 1.57 | 0.54 | 0.19 | 2.94 | 2.34 | -0.04 | 0.86 | 1.48 | 1.58 | 0.4025 | 0.8475 | 0.315 | 0.521081 | 0.327821 | 0.226995 | 1 | 0.999631 | 0.820181 |
| Uhrf1bp1_B2KF50_S929s_1 | -1.75 | 0.54 | 1.06 | -0.75 | 0.61 | -0.1 | 0.1 | -0.33 | 1.24 | -1.64 | 0.9 | 1.83 | 3.39 | -0.23 | 0.12 | 0.32 | 0.295 | 0.8075 | 1.125 | 0.23246 | 0.625778 | 0.362818 | 1 | 0.999631 | 0.682449 |
| Uhrf1bp1l_A2RSJ4_S1080s_1 | 2.22 | 0.93 | 0.24 | 2.34 | 1.32 | 1.98 | 1.72 | 1.46 | 1.74 | 1.02 | 1.12 | 0.24 | 1.65 | 0.11 | 0.51 | -0.19 | 0.1875 | -0.4025 | -0.9125 | 0.147466 | 0.707207 | 0.471788 | 1 | 0.999631 | 0.620049 |
| Uhrf1bp1l_A2RSJ4_S1336s_1 | -0.27 | -1.29 | -1.27 | -0.83 | -0.5 | -0.65 | -0.8 | -0.44 | -0.26 | -0.45 | 0.06 | -0.33 | 0.11 | -0.63 | -0.56 | -0.72 | 0.3175 | 0.67 | 0.465 | 0.212661 | 0.317709 | 0.060464 | 1 | 0.999631 | 0.66658 |
| Uhrf1bp1l_A2RSJ4_S774s_1 | -0.37 | 0.37 | -1.02 | -0.5 | 0.17 | -0.46 | -0.65 | -0.48 | -0.08 | -0.84 | -0.18 | 0.42 | -0.15 | 0.06 | -0.66 | -0.23 | 0.025 | 0.21 | 0.135 | 0.714727 | 0.94483 | 0.601387 | 1 | 0.999631 | 0.901895 |
| Uhrf1bp1l_A2RSJ4_S879s_1 | -2.02 | 0.08 | 0.09 | -2.02 | -2.35 | 0.69 | -1.37 | -0.98 | -1.11 | -1.75 | -0.32 | -7.16 | -0.12 | 1.47 | -1.66 | -1.92 | -0.035 | -1.6175 | 0.41 | 0.634748 | 0.963674 | 0.271055 | 1 | 0.999631 | 0.868886 |
| Uhrf1bp1l_A2RSJ4_S890s_1 | -0.98 | 0.8 | 0.19 | -1.76 | -0.01 | -0.23 | -0.48 | 0.11 | -0.19 | -0.39 | -0.6 | -1.02 | 0.73 | 0.62 | -0.73 | -0.38 | 0.285 | -0.1125 | 0.4975 | 0.427399 | 0.601893 | 0.839802 | 1 | 0.999631 | 0.77521 |
| Uhrf1bp1l_A2RSJ4_S914s_1 | 2.92 | -2.13 | 0.79 | 0.56 | -0.17 | 0.35 | 1.49 | -0.04 | -0.2 | 1.16 | -0.14 | -0.6 | -1.59 | -0.99 | -0.8 | -2.1 | -0.1275 | -0.48 | -1.905 | 0.062548 | 0.893227 | 0.6176 | 1 | 0.999631 | 0.51189 |
| Uhrf1bp1l_A2RSJ4_S934s_1 | 0.6 | 1.85 | 2.05 | 1.6 | 1.01 | 1.49 | 1.11 | 1.73 | 1.63 | 1.33 | 2.03 | 2.47 | 0.69 | 1.67 | 1.62 | 2.77 | -0.19 | 0.34 | 0.1625 | 0.749535 | 0.616951 | 0.418392 | 1 | 0.999631 | 0.912137 |
| Uhrf1bp1l_A2RSJ4_S952s_1 | 1.08 | -0.11 | 0.49 | 0.6 | -0.34 | 0.49 | 0.44 | -0.02 | -0.01 | 0.76 | 0.31 | 0.05 | -0.63 | -0.62 | 0 | -0.69 | -0.3725 | -0.2375 | -1 | 0.015998 | 0.296461 | 0.495209 | 1 | 0.999631 | 0.388832 |
| Uhrf1bp1l_A2RSJ4_S988s_1 | 1.51 | -0.56 | -0.79 | -0.15 | -1.51 | 0.04 | 0.49 | -1.48 | -0.06 | 1.25 | -1.54 | -0.8 | -1.43 | -1.82 | -1.18 | -3.32 | -0.6175 | -0.29 | -1.94 | 0.011647 | 0.356289 | 0.68114 | 1 | 0.999631 | 0.385092 |
| Uhrf1bp1l_Q8C318_S326s_1 | -1.35 | 0.45 | 0.95 | -1.37 | -1.76 | -0.43 | -1.69 | -1.19 | 0.13 | 0.89 | -0.28 | 0.71 | 2.47 | -0.12 | -0.43 | 0.56 | -0.9375 | 0.6925 | 0.95 | 0.238877 | 0.148062 | 0.266399 | 1 | 0.999631 | 0.683719 |
| Uhrf1bp1l_Q8C318_S326s_2 | -0.56 | -1.04 | -1.06 | -0.55 | -0.98 | -1.02 | -0.8 | -0.53 | 0.04 | -0.07 | -0.71 | -0.18 | -0.83 | -0.75 | -1.23 | -0.35 | -0.03 | 0.5725 | 0.0125 | 0.968503 | 0.912042 | 0.078941 | 1 | 0.999631 | 0.988771 |
| Uhrf1bp1l_Q8C318_S330s_1 | 3.05 | 1.38 | 1.78 | 2.93 | 1.3 | 2.05 | 2.88 | 2.24 | 2.49 | 2.56 | 0.99 | 1.56 | -2.17 | 0.84 | 1.65 | 1.41 | -0.1675 | -0.385 | -1.8525 | 0.050182 | 0.736142 | 0.471251 | 1 | 0.999631 | 0.488453 |
| Uhrf1bp1l_Q8C318_S330s_2 | 0.88 | -0.11 | -0.07 | 0.86 | -0.27 | 0.51 | 0.87 | -0.1 | 0.69 | 0.93 | 0 | -0.06 | -1.35 | -0.13 | -0.67 | -0.1 | -0.1375 | -1.2E-16 | -0.9525 | 0.042925 | 0.726291 | 1 | 1 | 1 | 0.475443 |
| Uhrf1bp1l_Q8C318_S335s_1 | -0.5 | 1.52 | 2.78 | 0.36 | -2.6 | 0.44 | -0.09 | -0.3 | 0.05 | 2.55 | 0.87 | 1.33 | 0.62 | -0.61 | 1.03 | 0.45 | -1.6775 | 0.16 | -0.6675 | 0.353803 | 0.073076 | 0.836987 | 1 | 0.999631 | 0.739673 |
| Uhrf1bp1l_Q8C318_T205t_1 | 1.1 | 1.37 | -0.94 | 0.47 | 1.88 | 1.19 | 0.28 | 0.93 | 0.69 | 0.68 | 0.14 | 0.9 | -0.49 | 1.21 | -0.64 | 0.98 | 0.57 | 0.1025 | -0.235 | 0.712634 | 0.322388 | 0.840787 | 1 | 0.999631 | 0.9008 |
| Uhrf2_Q7TMI3_S668s_1 | 2.99 | 1.56 | 2.39 | 1.8 | 0.28 | 1.03 | 1.34 | 1.17 | 1.84 | 2.5 | 0.02 | 0.42 | 2.96 | 0.53 | 0.66 | 0.05 | -1.23 | -0.99 | -1.135 | 0.105256 | 0.011418 | 0.127035 | 1 | 0.999631 | 0.568213 |
| Ulk1_Q6PB82_S450s_1 | -0.8 | -0.56 | -0.59 | -0.89 | -0.66 | -0.64 | -0.24 | -0.26 | -0.67 | -1.25 | -0.4 | -0.25 | 0.19 | -0.71 | -0.42 | 0.03 | 0.26 | 0.0675 | 0.4825 | 0.14568 | 0.320241 | 0.82536 | 1 | 0.999631 | 0.616968 |
| Ulk1_Q6PB82_S628s_1 | -0.38 | 0.09 | -0.04 | 0.55 | 0.86 | -0.07 | 0.14 | 0.15 | 0.17 | -0.08 | 0.73 | 0.59 | 0.22 | 0.26 | 0.45 | 0.66 | 0.215 | 0.2975 | 0.3425 | 0.285216 | 0.512696 | 0.370048 | 1 | 0.999631 | 0.705851 |
| Ulk1_Q6PB82_S643s_1 | 0.27 | -0.13 | -0.13 | 0.36 | 0.42 | 0.57 | -0.5 | 0.62 | 1.01 | -0.04 | 0.84 | 0.76 | 1.2 | 0.73 | 0.2 | 0.91 | 0.185 | 0.55 | 0.6675 | 0.062984 | 0.581091 | 0.114179 | 1 | 0.999631 | 0.512978 |
| Ulk1_Q6PB82_S763s_1 | -0.99 | -0.04 | -0.01 | 0.11 | -0.17 | -0.34 | -0.62 | 0.39 | -0.27 | -0.29 | -0.58 | 0.14 | -0.29 | 0.67 | -0.28 | -0.62 | 0.0475 | -0.0175 | 0.1025 | 0.798369 | 0.894163 | 0.95892 | 1 | 0.999631 | 0.930748 |
| Ulk2_Q9QY01_S322s_1 | -0.99 | -0.11 | -0.96 | -0.72 | -1.56 | -1.67 | -0.8 | -1.29 | -0.5 | -0.66 | -1.82 | -0.04 | 0.03 | -1.58 | -0.8 | -0.12 | -0.635 | -0.06 | 0.0775 | 0.856833 | 0.075461 | 0.888874 | 1 | 0.999631 | 0.951933 |
| Ulk2_Q9QY01_S430s_1 | -1.98 | -2.02 | -3.09 | -1.79 | -0.5 | -0.75 | -1.76 | -1.65 | -0.86 | -2.04 | -0.63 | -2.36 | 0.04 | -1.24 | -0.73 | -2.31 | 1.055 | 0.7475 | 1.16 | 0.052789 | 0.03111 | 0.153925 | 1 | 0.999631 | 0.489777 |
| Ulk2_Q9QY01_S600s_1 | -0.03 | -0.02 | 0.53 | 0.34 | 0.18 | -0.48 | -0.3 | 0.16 | -0.12 | 0.17 | 0.59 | 0.17 | -0.09 | -0.41 | -0.28 | 0.81 | -0.315 | -0.0025 | -0.1975 | 0.584944 | 0.290525 | 0.993085 | 1 | 0.999631 | 0.849277 |
| Ulk2_Q9QY01_S772s_1 | -1.31 | -0.19 | -0.58 | -1.32 | -0.37 | -0.96 | -0.35 | -1.54 | -0.59 | -0.37 | -1.46 | -0.49 | 0.34 | -0.74 | 0.13 | -0.33 | 0.045 | 0.1225 | 0.7 | 0.101468 | 0.910741 | 0.754477 | 1 | 0.999631 | 0.564448 |
| Ulk2_Q9QY01_S781s_1 | -0.98 | 0.61 | 0.26 | 0.22 | 0.44 | 0.35 | -0.82 | -0.03 | 0.23 | -0.08 | -0.1 | -0.61 | 2.07 | 1.25 | 0.77 | 0.57 | -0.0425 | -0.1675 | 1.1375 | 0.034145 | 0.922789 | 0.676964 | 1 | 0.999631 | 0.446273 |
| Ulk3_Q3U3Q1_S464s_1 | 1.37 | 1.17 | 0.84 | 1.34 | 1.73 | 1.48 | 1.52 | 2.3 | 1.53 | 0.88 | 1.21 | 2.18 | 2.13 | 1.42 | 1.65 | 2.48 | 0.5775 | 0.27 | 0.74 | 0.048768 | 0.073128 | 0.441293 | 1 | 0.999631 | 0.485306 |
| Unc119_B1AQD9_S37s_1 | -1.92 | -2.81 | -1.02 | 0.44 | -2.13 | -3.16 | -0.9 | 1.25 | -1.77 | -3.72 | -2.93 | 2.08 | -0.59 | -1.95 | 0.37 | -1.45 | 0.0925 | -0.2575 | 0.4225 | 0.578706 | 0.926975 | 0.836649 | 1 | 0.999631 | 0.84622 |
| Unc13a_Q4KUS2_S241s_1 | -1.01 | -1.25 | -0.19 | 0.56 | -1.48 | -0.69 | -0.16 | -0.65 | 0.06 | -0.43 | -0.53 | -0.94 | 0.02 | -0.05 | 0.16 | 0.06 | -0.2725 | 0.0125 | 0.52 | 0.239368 | 0.566269 | 0.977884 | 1 | 0.999631 | 0.683719 |
| Unc13a_Q4KUS2_S244s_1 | -1.08 | -0.78 | -1.37 | -1.58 | -1.42 | -0.55 | -1.81 | -1.85 | -1.01 | -0.96 | -2.05 | -1.98 | 0.01 | -0.68 | -1.79 | -1.03 | -0.205 | -0.2975 | 0.33 | 0.443811 | 0.580498 | 0.43138 | 1 | 0.999631 | 0.782802 |
| Unc13a_Q4KUS2_S249s_1 | -0.83 | -1.17 | -0.53 | -2.86 | -0.37 | -0.54 | -1.56 | -2.05 | -1.12 | -0.39 | -2.21 | -1.42 | 1.58 | -0.15 | -1.85 | -1.9 | 0.2175 | 0.0625 | 0.7675 | 0.375034 | 0.71599 | 0.915262 | 1 | 0.999631 | 0.748001 |
| Unc13a_Q4KUS2_S255s_1 | -1.14 | -1.36 | -0.39 | -2.39 | -2.9 | -1.13 | -2.35 | -2.08 | -2.16 | -0.38 | -1.47 | -1.8 | -0.19 | -1.54 | -1 | -1.61 | -0.795 | -0.1325 | 0.235 | 0.642431 | 0.147021 | 0.801872 | 1 | 0.999631 | 0.871335 |
| Unc13a_Q4KUS2_S255s_2 | 0.33 | -1.1 | -1.56 | -0.65 | -0.9 | -1.36 | -0.32 | -1.6 | -1.33 | 0.38 | -1.22 | -1.55 | -1.32 | -1.51 | -1.24 | -1.79 | -0.3 | -0.185 | -0.72 | 0.118218 | 0.527241 | 0.738946 | 1 | 0.999631 | 0.585087 |
| Unc13a_Q4KUS2_S259s_1 | -0.79 | -2.46 | -2 | -1.91 | -2.31 | -1.83 | -1.17 | -1.34 | -2.5 | -0.68 | -1.18 | -1.96 | -1.17 | -2.91 | -1.35 | -1.61 | 0.1275 | 0.21 | 0.03 | 0.952631 | 0.767127 | 0.680443 | 1 | 0.999631 | 0.984317 |
| Unc13a_Q4KUS2_S385s_1 | 1.2 | 1.6 | 1.02 | 1.9 | 1.5 | 1.61 | 0.84 | 0.95 | 1.9 | 0.74 | 1.01 | 1.3 | 0.78 | 1.44 | 1.16 | 1.02 | -0.205 | -0.1925 | -0.33 | 0.319112 | 0.529714 | 0.589851 | 1 | 0.999631 | 0.722397 |
| Unc13a_Q4KUS2_S440s_1 | 0.36 | -0.44 | -0.4 | -0.53 | -0.86 | -0.06 | -0.26 | -0.64 | -0.48 | 0.2 | -0.62 | -0.84 | -0.14 | -0.74 | -0.83 | -1.34 | -0.2025 | -0.1825 | -0.51 | 0.184884 | 0.532719 | 0.600562 | 1 | 0.999631 | 0.646072 |

| | | | | | | | | | | | | | | | | | | | | | | | | |
|---|---|---|---|---|---|---|---|---|---|---|---|---|---|---|---|---|---|---|---|---|---|---|---|---|
| Unc13a_Q4KUS2_T416t_1 | 0.99 | -0.67 | 1.01 | -0.03 | -1.93 | 0.42 | 1.74 | -1.9 | -2.87 | -0.29 | -0.73 | -0.09 | -0.36 | -1.75 | 0.4 | -0.95 | -0.7425 | -1.32 | -0.99 | 0.104876 | 0.399075 | 0.076545 | 1 | 0.999631 | 0.568213 |
| Unc13b_E9Q263_S1250s_1 | 2.04 | -0.85 | 0.62 | 0.49 | -1.51 | -0.39 | 1.27 | -0.81 | -0.63 | 1.83 | -0.61 | -0.74 | -0.33 | -1.12 | -0.8 | -1.06 | -0.935 | -0.6125 | -1.4025 | 0.031416 | 0.222842 | 0.426952 | 1 | 0.999631 | 0.441911 |
| Unc13c_Q8K0T7_T511t_1 | -0.08 | -1.67 | -0.35 | -0.36 | -0.99 | -1.27 | -0.41 | -0.91 | -0.66 | -0.43 | -1.12 | -0.69 | -0.69 | -1.76 | -0.53 | -1.79 | -0.28 | -0.11 | -0.5775 | 0.245167 | 0.492661 | 0.782812 | 1 | 0.999631 | 0.685751 |
| Unc5a_F7CVI0_S143s_1 | 1.58 | -0.66 | 0.31 | 0.72 | -0.92 | 0.83 | 1.77 | -0.11 | -0.73 | 0.72 | -0.13 | -1.23 | -0.28 | -0.41 | -0.08 | -0.85 | -0.095 | -0.83 | -0.8925 | 0.086794 | 0.886093 | 0.171651 | 1 | 0.999631 | 0.553514 |
| Unc5a_F7CVI0_S155s_1 | 0.15 | -0.66 | -1.01 | -0.99 | 0.44 | -0.32 | -0.59 | -0.68 | -0.37 | -1.52 | -0.49 | -1.54 | -0.67 | -0.86 | -1.9 | -0.13 | 0.34 | -0.3525 | -0.2625 | 0.567711 | 0.385404 | 0.411548 | 1 | 0.999631 | 0.843037 |
| Unc5c_E9PVI4_S447s_1 | -0.19 | -0.27 | -0.28 | -0.17 | -0.01 | 0.06 | -0.39 | 0.05 | -0.24 | 0.12 | -0.49 | 0.22 | 0 | 0.37 | -0.06 | 0.22 | 0.155 | 0.13 | 0.36 | 0.199206 | 0.529297 | 0.635509 | 1 | 0.999631 | 0.657167 |
| Unc5c_E9PVI4_S638s_1 | -0.76 | 1.07 | 0.48 | -0.31 | -0.13 | 1.21 | -0.75 | 0.12 | 0.07 | 0.83 | 0.04 | -0.26 | 1.2 | 1.29 | -0.47 | -0.19 | -0.0075 | 0.05 | 0.3375 | 0.554863 | 0.988766 | 0.912892 | 1 | 0.999631 | 0.838109 |
| Unc79_E5CYJ9_S1759s_1 | -0.55 | -0.2 | -1.54 | -0.64 | 0.25 | 0.37 | -0.29 | -0.69 | -0.1 | -1.1 | -0.24 | -0.8 | 0.6 | 0.47 | -0.54 | -1.22 | 0.6425 | 0.1725 | 0.56 | 0.275566 | 0.121502 | 0.658101 | 1 | 0.999631 | 0.700187 |
| Unc79_E5CYJ9_S689s_1 | 0.35 | -0.42 | -0.96 | -0.31 | -0.19 | -0.81 | -0.82 | -0.08 | 0.06 | -0.01 | -0.62 | -0.41 | -0.82 | -0.17 | -0.39 | -0.96 | -0.14 | 0.09 | -0.25 | 0.503379 | 0.697026 | 0.79842 | 1 | 0.999631 | 0.814092 |
| Unc79_G8JL63_S1078s_1 | 1.36 | 0.44 | 0.56 | 1.06 | 0.94 | 0.57 | 0.58 | 0.11 | 0.09 | 1.39 | 0.33 | -0.1 | 0.44 | -0.48 | 0.09 | 1.21 | -0.305 | -0.4275 | -0.54 | 0.222654 | 0.354212 | 0.30662 | 1 | 0.999631 | 0.673692 |
| Unc80_Q8BLN6_S1259s_1 | 2.35 | 0.58 | 0.42 | 0.89 | 0.51 | 2.07 | 0.98 | -0.14 | 1.07 | 1.53 | -0.22 | -0.64 | 0.74 | 1.19 | -0.26 | -0.16 | -0.205 | -0.625 | -0.6825 | 0.217438 | 0.725472 | 0.32308 | 1 | 0.999631 | 0.669718 |
| Unc80_Q8BLN6_S166s_1 | 1.3 | -1.29 | -0.84 | -0.42 | -3.07 | -2.85 | 1.78 | -1.63 | -4.62 | 0.52 | -1.74 | -0.81 | -2.72 | -2.88 | -2.92 | -0.85 | -1.13 | -1.35 | -2.03 | 0.012294 | 0.307653 | 0.221223 | 1 | 0.999631 | 0.385092 |
| Unc80_Q8BLN6_S246s_1 | -0.39 | -2.31 | -0.99 | -3.43 | -2.36 | -0.35 | -0.83 | -1.35 | -1.71 | 0.22 | -0.65 | -0.18 | -0.01 | -3.98 | -0.23 | 0.33 | 0.5575 | 1.2 | 0.8075 | 0.443817 | 0.4407 | 0.115677 | 1 | 0.999631 | 0.782802 |
| Unc80_Q8BLN6_S246s_2 | -0.8 | -0.8 | -0.6 | -1.19 | -1.04 | -0.98 | -0.95 | -1.2 | -1.05 | -0.56 | -1.91 | -1.5 | 0.09 | -1.17 | -0.75 | -0.37 | -0.195 | -0.4075 | 0.2975 | 0.40633 | 0.447033 | 0.265286 | 1 | 0.999631 | 0.766128 |
| Unc80_Q8BLN6_S257s_1 | -0.13 | -1.16 | 0.14 | -0.95 | -0.72 | -0.7 | -0.75 | -0.88 | -1.1 | -0.54 | -0.05 | -0.33 | 0.58 | -1.33 | 0.05 | -0.04 | -0.2375 | 0.02 | 0.34 | 0.495881 | 0.499358 | 0.959883 | 1 | 0.999631 | 0.811036 |
| Unc80_Q8BLN6_S257s_2 | -0.8 | -0.8 | -0.6 | -1.19 | -1.04 | -0.98 | -0.95 | -1.2 | -1.05 | -0.56 | -1.91 | -1.5 | 0.09 | -1.17 | -0.75 | -0.37 | -0.195 | -0.4075 | 0.2975 | 0.40633 | 0.447033 | 0.265286 | 1 | 0.999631 | 0.766128 |
| Unc80_Q8BLN6_S295s_1 | -1.1 | -0.78 | -0.54 | -0.3 | -1.02 | -1.1 | -0.75 | -0.81 | -1.1 | -0.29 | -0.72 | -0.79 | -1 | -1.16 | -0.06 | -0.27 | -0.24 | -0.045 | 0.0575 | 0.87491 | 0.393833 | 0.883831 | 1 | 0.999631 | 0.960237 |
| Unc80_Q8BLN6_S3110s_1 | -0.28 | -0.47 | 0.01 | 0.37 | -0.62 | -1.18 | -0.09 | -0.66 | -0.62 | -0.22 | -1.44 | -0.55 | -0.21 | -0.49 | 0.13 | -0.39 | -0.545 | -0.615 | -0.1475 | 0.642238 | 0.122892 | 0.107279 | 1 | 0.999631 | 0.871335 |
| Unc80_Q8BLN6_S316s_1 | -0.21 | 0.1 | -0.34 | 0.78 | -0.57 | -0.51 | -0.21 | 0.83 | -0.05 | -0.04 | 1.03 | 0.34 | 0.01 | 0.54 | -0.16 | 1.62 | -0.1975 | 0.2375 | 0.42 | 0.376698 | 0.631904 | 0.534994 | 1 | 0.999631 | 0.748858 |
| Unc80_Q8BLN6_S3203s_1 | -1.06 | -0.54 | -0.74 | 0.08 | -0.73 | -0.61 | -0.37 | 0.34 | 0.12 | -0.63 | -1.54 | -0.33 | 0.44 | -0.8 | 0.11 | -0.83 | 0.2225 | -0.03 | 0.295 | 0.484645 | 0.542656 | 0.943812 | 1 | 0.999631 | 0.803422 |
| Unc80_Q8BLN6_S415s_1 | -0.73 | -1.31 | -0.58 | -1.11 | -0.57 | -1.16 | -0.66 | -0.01 | -0.05 | -1.2 | -0.18 | -0.27 | -1.47 | -1.18 | -1.75 | -0.96 | 0.3325 | 0.5075 | -0.4075 | 0.224395 | 0.327413 | 0.170133 | 1 | 0.999631 | 0.67471 |
| Unc80_Q8BLN6_S526s_1 | 0.21 | -0.19 | -0.16 | -0.19 | -0.12 | -0.61 | -0.06 | -0.56 | -0.59 | -0.3 | -0.19 | -0.98 | -0.96 | -0.52 | -0.8 | -0.4 | -0.255 | -0.4325 | -0.5875 | 0.063096 | 0.353449 | 0.157647 | 1 | 0.999631 | 0.512978 |
| Unc80_Q8BLN6_T1014t_1 | 0.24 | -1.91 | 0.18 | 0.33 | -0.2 | -0.01 | 0.17 | 0.09 | 0.14 | 0.19 | 0.06 | -0.75 | -0.73 | 0.51 | 0.66 | -0.06 | 0.3025 | 0.2 | 0.385 | 0.508764 | 0.556714 | 0.713541 | 1 | 0.999631 | 0.816932 |
| Unk_Q8BL48_S240s_1 | -0.5 | 0.56 | 1.46 | 0.55 | 0.18 | 0.58 | 0.21 | 0.74 | 0.69 | 0.82 | 1.03 | 1.26 | 1.24 | 0.92 | 1.49 | 0.98 | -0.09 | 0.4325 | 0.64 | 0.159148 | 0.830306 | 0.319227 | 1 | 0.999631 | 0.629514 |
| Unk_Q8BL48_S371s_1 | -0.35 | 1.02 | 1.71 | 0.16 | 0.3 | 0.35 | 0.76 | 0.86 | 1.38 | 0.93 | 1.64 | 1.67 | -0.22 | 1.42 | 2.36 | 0.44 | -0.0675 | 0.77 | 0.365 | 0.57822 | 0.883025 | 0.126635 | 1 | 0.999631 | 0.84622 |
| Unkl_Q5FWH2_S15s_1 | -0.74 | 1.45 | -0.28 | 0.37 | 0.3 | 0.34 | -0.34 | 0.21 | 0.72 | 0.41 | 0.44 | 0.38 | 1.42 | 0.75 | 0.72 | 0.54 | -0.0725 | 0.2875 | 0.6575 | 0.202669 | 0.878779 | 0.541536 | 1 | 0.999631 | 0.659215 |
| Uqcrc2_Q9DB77_S169s_1 | -0.24 | -0.57 | 0.45 | -0.27 | -1.23 | -0.64 | 0.04 | -0.55 | -0.4 | -0.36 | -0.77 | -0.37 | -0.31 | -0.62 | -0.97 | 0.13 | -0.4375 | -0.3175 | -0.285 | 0.441661 | 0.243606 | 0.31512 | 1 | 0.999631 | 0.781983 |
| Urb1_E9PU96_S17s_1 | -0.17 | -0.65 | -0.33 | -0.52 | -0.73 | -1.02 | -0.41 | -1.04 | -1.82 | -1.1 | -0.16 | 0.19 | 0.12 | -2.48 | -1.27 | 0.07 | -0.3825 | -0.305 | -0.4725 | 0.421797 | 0.181369 | 0.509282 | 1 | 0.999631 | 0.772942 |
| Uri1_Q3TLD5_S341s_1 | -0.44 | 0.03 | 0.73 | 0.71 | -0.2 | -0.24 | 0.95 | 0.52 | -0.46 | -0.51 | -0.4 | -0.25 | 0.54 | -0.91 | 0.35 | -0.38 | 1.18E-16 | -0.6625 | -0.3575 | 0.426222 | 1 | 0.073437 | 1 | 0.999631 | 0.774781 |
| Usp10_P52479_S209s_1 | -1.16 | -0.78 | -0.15 | -0.24 | 0.05 | -0.85 | -0.31 | -0.35 | -0.67 | -0.74 | -0.32 | 0.27 | -0.19 | -0.05 | 0.35 | -0.66 | 0.2175 | 0.2175 | 0.445 | 0.238623 | 0.523963 | 0.552524 | 1 | 0.999631 | 0.683403 |
| Usp11_Q99K46_S906s_1 | -1.76 | -0.63 | -1.6 | -0.28 | -0.55 | -0.75 | -0.99 | -1.41 | -1.28 | -0.85 | 0.16 | 0.6 | 0.78 | -0.13 | 0.01 | -0.2 | 0.1425 | 0.725 | 1.1825 | 0.019945 | 0.727394 | 0.192478 | 1 | 0.999631 | 0.391668 |
| Usp14_E9PYI8_S108s_1 | -0.8 | -0.02 | -0.01 | -0.63 | 0.19 | -0.7 | -0.51 | -0.49 | 0.36 | 0.52 | -0.36 | -0.1 | 1.35 | 0.06 | -0.37 | 0.29 | -0.0125 | 0.47 | 0.6975 | 0.127376 | 0.969442 | 0.184279 | 1 | 0.999631 | 0.59788 |
| Usp14_E9PYI8_S358s_1 | -1.13 | -1.82 | -1.28 | -1.11 | -1.38 | -1.62 | -0.99 | -1.2 | -1.34 | -1.23 | -1.38 | -0.92 | -1.07 | -2.12 | -1.38 | -1.7 | 0.0375 | 0.1175 | -0.2325 | 0.502316 | 0.896142 | 0.682568 | 1 | 0.999631 | 0.814092 |
| Usp14_E9PYI8_T200t_1 | 0.14 | 1.21 | 1.76 | 1.04 | 0.81 | 0.97 | 0.87 | 2.4 | 1.02 | 0.81 | 1.2 | 0.02 | -0.84 | 0.81 | 1.35 | 0.99 | 0.225 | -0.275 | -0.46 | 0.411267 | 0.641718 | 0.523166 | 1 | 0.999631 | 0.767937 |
| Usp15_Q8R5H1_S229s_1 | 0.54 | 0.55 | -0.55 | 0.61 | 1.16 | 0.68 | 0.64 | 0.89 | 0.58 | 0.35 | 0.09 | 0.02 | -1.05 | 0.73 | 0.31 | 0.53 | 0.555 | -0.0275 | -0.1575 | 0.741917 | 0.126431 | 0.93697 | 1 | 0.999631 | 0.909328 |
| Usp15_Q8R5H1_S621s_1 | 0.64 | 0.14 | -0.12 | -0.04 | 0.05 | 0.12 | -0.29 | 0.54 | -0.13 | 0.09 | -0.04 | -0.38 | 0.24 | 0.3 | -0.13 | 0.13 | -0.05 | -0.27 | -0.02 | 0.946907 | 0.868624 | 0.357885 | 1 | 0.999631 | 0.982839 |
| Usp15_Q8R5H1_T226t_1 | 1.57 | -0.62 | -0.53 | 1.52 | 1.58 | 1.03 | 1.55 | 0.98 | 0.67 | 0.63 | 1.05 | -0.3 | -1.4 | 0.4 | 0.87 | 0.09 | 0.8 | 0.0275 | -0.495 | 0.482752 | 0.186777 | 0.964226 | 1 | 0.999631 | 0.802885 |
| Usp15_Q8R5H1_T622t_1 | 0.44 | -0.01 | 0.47 | -0.07 | -0.15 | 0.18 | -0.41 | 0.98 | -0.16 | 0.07 | 0.66 | -0.09 | 0.31 | 0.18 | -0.21 | 0.27 | -0.0575 | -0.0875 | -0.07 | 0.813642 | 0.872626 | 0.775315 | 1 | 0.999631 | 0.937041 |
| Usp20_Q8C6M1_S112s_1 | -0.55 | -0.47 | -0.2 | -0.49 | -0.44 | -0.3 | -0.66 | -0.2 | -0.57 | -0.09 | -0.41 | -0.12 | 0.57 | -0.09 | -0.37 | -0.31 | 0.0275 | 0.13 | 0.3775 | 0.249681 | 0.912056 | 0.622043 | 1 | 0.999631 | 0.686359 |
| Usp20_Q8C6M1_S369s_1 | 0.62 | -0.65 | -0.37 | -0.22 | -0.55 | -0.39 | -0.16 | -0.21 | -0.51 | 0.4 | -0.41 | -0.79 | -0.7 | -0.68 | -0.46 | -0.57 | -0.1725 | -0.1725 | -0.4475 | 0.210594 | 0.602877 | 0.660453 | 1 | 0.999631 | 0.664726 |
| Usp20_Q8C6M1_S374s_2 | -0.3 | -0.01 | -1.03 | 0.25 | -0.59 | -0.42 | -0.11 | 0.55 | -0.86 | -0.08 | 0.7 | 0.49 | 0.13 | 0.21 | -1.76 | 0.2 | 0.13 | 0.335 | -0.0325 | 0.950706 | 0.736064 | 0.453643 | 1 | 0.999631 | 0.984079 |
| Usp20_Q8C6M1_S410s_1 | -0.06 | -0.63 | -0.01 | -0.74 | -0.69 | -0.16 | -0.45 | -0.5 | -0.83 | 0.27 | -0.22 | -0.18 | -0.18 | -0.44 | -0.41 | -0.41 | -0.09 | 0.12 | -3.9E-17 | 1 | 0.756919 | 0.724946 | 1 | 0.999631 | 1 |
| Usp20_Q8C6M1_T378t_1 | -1.16 | -1.37 | 0.83 | -0.21 | -1.11 | -0.36 | 0.28 | 0.05 | -1.26 | -0.8 | 0.64 | -0.28 | -0.54 | -1.15 | -0.58 | -0.15 | 0.1925 | 0.0525 | -0.1275 | 0.804881 | 0.72318 | 0.929247 | 1 | 0.999631 | 0.934231 |
| Usp20_Q8C6M1_T378t_2 | -0.3 | -0.01 | -1.03 | 0.25 | -0.59 | -0.42 | -0.11 | 0.55 | -0.86 | -0.08 | 0.7 | 0.49 | 0.13 | 0.21 | -1.76 | 0.2 | 0.13 | 0.335 | -0.0325 | 0.950706 | 0.736064 | 0.453643 | 1 | 0.999631 | 0.984079 |
| Usp22_Q5DU02_T147t_1 | 0.02 | 1.69 | 0.91 | 0.86 | 0.37 | 1.18 | 0.41 | 1.06 | 0.22 | 0.31 | 0.51 | 1.35 | 0.61 | 1.43 | 1.01 | 1.21 | -0.115 | -0.2725 | 0.195 | 0.63183 | 0.776357 | 0.528653 | 1 | 0.999631 | 0.867717 |
| Usp24_E9PV45_S1139s_1 | 1.32 | 0.09 | 0.95 | 0.57 | -0.02 | 0.29 | 1.3 | 0.01 | 0.47 | 1.43 | 0.85 | 0.72 | -0.08 | -0.16 | 0.74 | 0.49 | -0.3375 | 0.135 | -0.485 | 0.219932 | 0.416091 | 0.711402 | 1 | 0.999631 | 0.670699 |
| Usp24_E9PV45_S1297s_1 | 2.12 | 0.72 | -0.66 | 1.75 | 2.22 | 1 | 1.79 | 1.67 | 1.64 | 0.54 | 1.31 | 0.3 | -1.33 | 0.94 | 0.26 | 0.77 | 0.6875 | -0.035 | -0.8225 | 0.266553 | 0.272937 | 0.955701 | 1 | 0.999631 | 0.695664 |
| Usp24_E9PV45_S1923s_1 | 0.69 | 0.22 | 1.56 | 2.18 | -0.83 | -0.37 | 2.18 | 0.64 | -0.03 | 1.47 | 0.72 | 1.26 | -1 | -4.18 | -0.25 | -0.12 | -0.7575 | -0.3075 | -2.55 | 0.016895 | 0.297596 | 0.556517 | 1 | 0.999631 | 0.391024 |
| Usp24_E9PV45_S1941s_1 | -0.18 | 0.89 | 1.07 | -0.67 | 0.89 | 0.7 | 3.04 | -0.6 | 1.71 | -0.98 | 0.9 | 1.26 | 0.68 | -1.63 | -0.16 | -3 | 0.73 | 0.445 | -1.305 | 0.125683 | 0.346985 | 0.499969 | 1 | 0.999631 | 0.596745 |
| Usp24_E9PV45_S2045s_1 | -1.44 | 0.02 | 0.08 | -0.92 | -0.31 | -0.46 | -0.37 | -0.26 | -0.06 | -1.15 | -0.21 | 0.07 | 0.79 | -0.07 | -0.04 | -0.3 | 0.215 | 0.2275 | 0.66 | 0.158964 | 0.57917 | 0.617638 | 1 | 0.999631 | 0.629092 |
| Usp24_E9PV45_S2559s_1 | 0.3 | 0.6 | 0.72 | 0.47 | 0.4 | 0.51 | 0.15 | 1.07 | 0.53 | 0.56 | 0.44 | 0.97 | 0.44 | 1.34 | 1.49 | 1.25 | 0.01 | 0.1025 | 0.6075 | 0.087913 | 0.972265 | 0.700301 | 1 | 0.999631 | 0.553787 |
| Usp25_P57080_T61t_1 | -0.77 | -0.52 | -0.48 | -0.24 | -0.15 | -1.13 | -0.86 | -0.47 | -0.59 | -0.67 | -1.02 | 0.27 | 0.13 | -0.07 | -2.04 | -0.41 | -0.15 | 7.85E-17 | -0.095 | 0.84567 | 0.621966 | 1 | 1 | 1 | 0.948565 |
| Usp31_E9Q6Y8_S1103s_1 | -0.13 | -0.09 | 0.27 | 0.38 | -0.08 | 0.13 | 0.41 | -0.06 | -0.54 | -0.03 | 0.41 | 0.42 | -0.72 | -0.91 | 0.58 | -0.39 | -0.0075 | -0.0425 | -0.4675 | 0.24459 | 0.977574 | 0.894384 | 1 | 0.999631 | 0.685751 |
| Usp31_E9Q6Y8_S1127s_1 | 1.62 | 0.9 | 0.92 | 1.15 | 0.89 | 1.63 | 0.94 | 1.33 | 1.55 | 1.74 | 0.75 | 0.66 | 1.32 | 1.14 | 1.21 | 0.52 | 0.05 | 0.0275 | -0.1 | 0.758418 | 0.868795 | 0.938525 | 1 | 0.999631 | 0.914449 |
| Usp31_E9Q6Y8_S49s_1 | -0.33 | 0.35 | 0.35 | 0.01 | 0.28 | 1.11 | 0.1 | 0.11 | 0.35 | 0.06 | 0.38 | 0.36 | -0.2 | 0.07 | 0.34 | 0.27 | 0.305 | 0.1925 | 0.025 | 0.934399 | 0.367587 | 0.49504 | 1 | 0.999631 | 0.978794 |
| Usp31_E9Q6Y8_S793s_1 | 1 | 0.05 | 0.17 | 1.47 | 0.76 | 1.74 | 0.77 | 1.26 | 1.39 | 1.52 | 1.16 | 0.98 | -3.35 | 1.33 | 1.86 | 0.79 | 0.46 | 0.59 | -0.515 | 0.627066 | 0.279586 | 0.14582 | 1 | 0.999631 | 0.86524 |
| Usp32_F8VPZ3_S1372s_1 | -0.58 | -0.34 | 0.24 | -0.54 | -0.97 | -0.73 | -0.66 | -0.27 | -0.55 | -0.98 | -0.39 | 0.27 | 0.61 | -0.21 | 0.07 | 0.31 | -0.3525 | -0.1075 | 0.5 | 0.153171 | 0.258416 | 0.76331 | 1 | 0.999631 | 0.625198 |
| Usp32_F8VPZ3_S1394s_1 | 1.54 | 0.14 | 0.16 | -0.31 | -1 | 0.49 | 1.74 | -0.81 | 0.18 | 2.87 | -0.63 | -1.59 | -0.09 | -0.35 | -0.7 | -1.75 | -0.2775 | -0.175 | -1.105 | 0.053594 | 0.6799 | 0.846338 | 1 | 0.999631 | 0.4909 |
| Usp32_F8VPZ3_T1326t_1 | -1.02 | -0.57 | -0.88 | 0.17 | 0.36 | -0.42 | 0.28 | -0.75 | -0.46 | -1.8 | -0.1 | 1.18 | -0.35 | -0.08 | 0.66 | -0.43 | 0.4425 | 0.28 | 0.525 | 0.200967 | 0.270977 | 0.646391 | 1 | 0.999631 | 0.65842 |
| Usp35_M0QWN7_S929s_1 | 1.84 | 1.54 | 0.77 | -0.12 | 0.89 | 2.6 | 0.52 | 0.63 | 0.78 | 0.17 | -0.9 | 0.35 | 2.24 | 0.52 | -0.48 | 0.2 | 0.1525 | -0.9075 | -0.3875 | 0.555381 | 0.797106 | 0.111057 | 1 | 0.999631 | 0.838109 |
| Usp35_M0QWN7_T738t_1 | 0.89 | -0.12 | 0.98 | 0.45 | -0.59 | 0.22 | 0.87 | -0.07 | 0.04 | 1.18 | 0.18 | -0.16 | 0.63 | -0.27 | 0.61 | 0.18 | -0.4425 | -0.24 | -0.2625 | 0.485388 | 0.283589 | 0.553945 | 1 | 0.999631 | 0.804287 |
| Usp36_B0QZF8_S264s_1 | 1.09 | -0.21 | -0.48 | 0.49 | -0.5 | -0.3 | 0.25 | -0.01 | -0.69 | 0.3 | -1.26 | -0.61 | -0.56 | 0.04 | -0.64 | -1.03 | -0.3625 | -0.7875 | -0.77 | 0.095793 | 0.37106 | 0.11509 | 1 | 0.999631 | 0.558128 |
| Usp39_Q3TIX9_S81s_1 | 1.22 | 0.3 | 1.44 | 0.71 | -0.47 | 0.43 | 1.28 | -0.26 | -0.09 | 1.41 | -0.14 | -0.5 | -0.61 | -0.98 | -0.56 | -1.08 | -0.6725 | -0.7475 | -1.725 | 0.000563 | 0.161438 | 0.140702 | 1 | 0.999631 | 0.351308 |
| Usp4_P35123_S655s_1 | -0.96 | 1.9 | 1.49 | 1.01 | 1.2 | 0.85 | 0.17 | 0.81 | 1.47 | 1.08 | 0.75 | 1.26 | 1.66 | 2.06 | 1.05 | 1.61 | -0.1025 | 0.28 | 0.735 | 0.24307 | 0.865137 | 0.639456 | 1 | 0.999631 | 0.685637 |
| Usp4_P35123_S680s_1 | -0.08 | 1.18 | 1.17 | 0.79 | 0.46 | 1.04 | 0.49 | 1.37 | 1.01 | 1.37 | 1.04 | 1.43 | 2.29 | 1.79 | 1.06 | 1.2 | 0.075 | 0.4475 | 0.82 | 0.075778 | 0.844405 | 0.22407 | 1 | 0.999631 | 0.541044 |
| Usp42_B2RQC2_S936s_1 | 1.89 | 0.29 | 0.72 | 0.68 | -0.77 | -0.48 | 0.83 | 0.55 | -0.77 | 0.85 | 0.49 | 0.33 | -0.54 | -0.51 | -0.03 | -1.07 | -0.8625 | -0.67 | -1.4325 | 0.00625 | 0.103342 | 0.179322 | 1 | 0.999631 | 0.351545 |
| Usp43_Q8BUM9_S967s_1 | -0.88 | -1.51 | -0.85 | -0.81 | -1.32 | -1.4 | -1.18 | -0.57 | -1.12 | -1.33 | -0.31 | 0.1 | -0.73 | -1.46 | -0.9 | -0.32 | -0.105 | 0.3475 | 0.16 | 0.647247 | 0.733759 | 0.385205 | 1 | 0.999631 | 0.874565 |
| Usp43_Q8BUM9_S970s_1 | 0.07 | 0.34 | -0.31 | -0.9 | -0.3 | -0.73 | -0.69 | -0.55 | -0.33 | -0.19 | -0.51 | -0.5 | 1.11 | 0.39 | -0.2 | -0.33 | -0.3675 | -0.1825 | 0.4425 | 0.318952 | 0.27878 | 0.584297 | 1 | 0.999631 | 0.722245 |
| Usp47_Q8BY87_S812s_1 | -1.05 | -0.43 | -0.68 | 0.06 | -0.35 | -0.31 | -1.43 | -0.84 | 0.12 | 0.08 | -1.03 | -1.19 | 1.21 | 0 | -1.11 | 0.92 | -0.2075 | 0.02 | 0.78 | 0.168096 | 0.577285 | 0.962278 | 1 | 0.999631 | 0.637399 |
| Usp47_Q8BY87_S829s_1 | 2.09 | -0.44 | -0.9 | 0.3 | -0.54 | 0.25 | 1.06 | -1.59 | 0.45 | 0.94 | -0.64 | -2.04 | -0.82 | -2.08 | -0.77 | -2.3 | -0.4675 | -0.585 | -1.755 | 0.026826 | 0.542777 | 0.4798 | 1 | 0.999631 | 0.422285 |
| Usp5_Q3U4W8_S728s_1 | 1.2 | 0.04 | 0.48 | 0.18 | -0.89 | 0.37 | 0.84 | -1.18 | -0.1 | 0.87 | -0.07 | -0.68 | -0.02 | -1.12 | -0.7 | -1.27 | -0.69 | -0.47 | -1.2525 | 0.010459 | 0.199815 | 0.275647 | 1 | 0.999631 | 0.383871 |
| Usp5_Q3U4W8_S756s_1 | 0.2 | -0.27 | 0.4 | 0.31 | -0.31 | -0.59 | -0.22 | 0.2 | -0.46 | -0.38 | 0.23 | 0.58 | 0.13 | -0.06 | 0.15 | 1.36 | -0.39 | -0.1675 | 0.235 | 0.549098 | 0.201695 | 0.621859 | 1 | 0.999631 | 0.835034 |
| Usp5_Q3U4W8_S760s_1 | -0.55 | -0.43 | -0.1 | -0.97 | 0.15 | -0.31 | -0.88 | -0.71 | -0.19 | 0.05 | -0.51 | -0.25 | 0.59 | 0.34 | -0.83 | -0.5 | 0.075 | 0.2875 | 0.4125 | 0.320259 | 0.821229 | 0.34158 | 1 | 0.999631 | 0.723684 |
| Usp5_Q3U4W8_S762s_1 | -1.7 | -0.96 | -0.12 | -0.93 | -1.34 | -1.7 | -0.87 | -0.64 | -0.81 | -0.04 | -0.88 | -0.36 | -0.33 | -0.38 | -0.38 | -0.41 | -0.21 | 0.405 | 0.5525 | 0.155522 | 0.60517 | 0.317448 | 1 | 0.999631 | 0.626444 |
| Usp5_Q3U4W8_T623t_1 | -0.53 | -0.17 | -0.34 | -0.37 | -0.22 | -0.44 | -0.67 | -0.29 | -0.2 | -0.34 | -0.53 | -0.31 | -0.26 | 0.33 | -0.17 | 0.04 | -0.0525 | 0.0075 | 0.3375 | 0.249496 | 0.832579 | 0.975984 | 1 | 0.999631 | 0.686359 |
| Usp53_P15975_S490s_1 | 0.81 | 0.98 | 0.32 | 0.04 | 0.47 | 0.45 | -0.03 | -0.43 | -0.06 | 0.78 | 0.35 | 0.03 | 0.36 | 0 | -1.2 | 0.5 | -0.4225 | -0.2625 | -0.6225 | 0.182689 | 0.233382 | 0.44251 | 1 | 0.999631 | 0.643861 |
| Usp54_F6ZXV1_S74s_1 | -1.06 | -0.22 | -0.9 | -0.81 | -1.25 | -0.51 | -0.71 | -1.23 | -0.52 | -0.88 | -0.79 | 0.14 | -0.05 | 0.06 | -1.37 | -0.68 | -0.1775 | 0.235 | 0.2375 | 0.557158 | 0.574824 | 0.494951 | 1 | 0.999631 | 0.838305 |
| Usp6nl_Q80XC3_S405s_1 | 1.39 | -1.78 | 0.08 | -0.62 | -4.31 | -0.38 | 2.59 | -2.73 | -2.1 | 1.41 | -1.09 | -1.96 | -2.82 | -2.28 | -1.6 | -3.2 | -0.975 | -0.7025 | -2.2425 | 0.006973 | 0.489912 | 0.447958 | 1 | 0.999631 | 0.358173 |
| Usp6nl_Q80XC3_S544s_1 | 0.1 | 0.15 | -1.16 | -0.66 | 0.01 | 0.28 | -0.61 | -0.4 | -0.6 | -0.2 | -2.75 | -0.44 | -0.48 | -1.26 | 0.23 | -0.38 | 0.2125 | -0.605 | -0.08 | 0.856432 | 0.583976 | 0.332707 | 1 | 0.999631 | 0.951817 |
| Usp8_Q80U87_S603s_1 | 2.08 | 0.24 | -0.76 | 1.55 | 0.14 | 1.23 | 1.7 | 0.71 | 0.41 | 2.25 | -0.34 | -2.04 | -2.07 | -1.05 | -0.45 | -2.18 | 0.1675 | -0.7075 | -2.215 | 0.008266 | 0.796029 | 0.463057 | 1 | 0.999631 | 0.365244 |
| Usp8_Q80U87_S680s_1 | -0.22 | -0.06 | -0.17 | -0.4 | 0.3 | -0.15 | -0.64 | -0.03 | -0.21 | -0.36 | 0.24 | 0.53 | 0.62 | 0.12 | 1.69 | 0 | 0.0825 | 0.2625 | 0.82 | 0.068885 | 0.772041 | 0.385644 | 1 | 0.999631 | 0.523934 |
| Usp9x_P70398_S2443s_1 | -1.05 | -0.45 | -0.24 | -0.52 | 0.14 | -0.5 | -0.95 | -0.7 | -0.46 | -1.06 | -0.63 | -0.36 | 0.71 | 0.33 | -0.49 | -0.1 | 0.0625 | -0.0625 | 0.6775 | 0.084621 | 0.850145 | 0.837214 | 1 | 0.999631 | 0.549982 |
| Utrn_E9Q6R7_S933s_1 | 0.02 | 0.43 | -0.21 | 0.43 | 0.32 | 0.02 | -0.49 | -0.28 | -0.42 | 0.18 | -0.53 | 0.47 | -0.17 | 0 | -0.9 | 0.75 | -0.275 | -0.2425 | -0.2475 | 0.538898 | 0.370552 | 0.476133 | 1 | 0.999631 | 0.830899 |
| Uvrag_Q8K245_T517t_1 | -0.53 | -0.18 | -0.64 | -0.78 | 0.64 | -0.47 | -0.31 | -0.07 | 0.23 | -0.49 | -0.71 | -0.13 | -0.08 | 0.16 | -0.14 | -0.37 | 0.48 | 0.2575 | 0.425 | 0.163949 | 0.159926 | 0.415235 | 1 | 0.999631 | 0.632208 |
| Vamp2_B0QZN5_S75s_1 | -0.82 | -1.16 | -0.67 | -1.07 | -0.49 | -0.13 | -0.73 | 0.08 | -1.46 | -0.49 | -0.01 | -0.8 | -0.54 | 0.14 | -0.41 | -0.45 | 0.6125 | 0.24 | 0.615 | 0.062173 | 0.055607 | 0.507923 | 1 | 0.999631 | 0.509361 |
| Vamp2_B0QZN5_S80s_1 | -0.39 | 0.28 | -0.21 | -0.87 | 0.27 | 0.37 | -1.31 | -0.06 | 0.71 | 0.18 | -0.05 | -1.05 | 1.16 | 0.2 | -1.15 | -0.43 | 0.115 | 0.245 | 0.2425 | 0.64017 | 0.794688 | 0.576888 | 1 | 0.999631 | 0.870995 |
| Vapa_Q9WV55_S166s_1 | -0.69 | -1.44 | -0.69 | -0.03 | -0.16 | -0.71 | -0.57 | 0.22 | -0.46 | -0.96 | -0.68 | -1.3 | -1.13 | -0.67 | -0.63 | -0.82 | 0.4075 | -0.1375 | -0.1 | 0.781294 | 0.290391 | 0.710146 | 1 | 0.999631 | 0.921661 |
| Vapa_Q9WV55_S3s_1 | 0.76 | 0.26 | 1.31 | 1.33 | -0.14 | 0.25 | 0.26 | 0.77 | 0.06 | 0.87 | 0.3 | 0.56 | -0.27 | -0.11 | 1.05 | 1.13 | -0.63 | -0.4675 | -0.465 | 0.314451 | 0.094283 | 0.200882 | 1 | 0.999631 | 0.720156 |
| Vapb_Q8BH80_S144s_1 | 0.33 | 0.72 | 0.12 | 0.64 | 0.54 | 0.31 | 0.29 | 0.5 | 0.78 | -0.09 | 0.45 | 0.57 | 0.8 | 0.9 | 0.25 | 0.65 | -0.0425 | -0.025 | 0.1975 | 0.51823 | 0.869836 | 0.934483 | 1 | 0.999631 | 0.819449 |
| Vapb_Q8BH80_S206s_1 | 0.71 | -0.73 | -0.41 | -0.27 | -0.08 | -0.24 | -0.4 | 0.27 | 0.34 | -0.08 | 0.36 | 0.21 | -0.29 | 0.44 | -0.9 | -0.03 | 0.0625 | 0.3825 | -0.02 | 0.962635 | 0.86352 | 0.302263 | 1 | 0.999631 | 0.986848 |
| Vash1_Q8C1W1_S323s_1 | 0.76 | 0.23 | -4.86 | -5.33 | -0.44 | 0.5 | -3.38 | -5.9 | 0.76 | 1.83 | -3.93 | -5.9 | 1.1 | 0.25 | -5.25 | -7.08 | -0.005 | 0.49 | -0.445 | 0.836477 | 0.997829 | 0.813746 | 1 | 0.999631 | 0.945333 |
| Vash1_Q8C1W1_S334s_1 | 1.63 | 1.3 | 2.12 | 1.42 | 0.83 | 1.55 | 2.3 | 1.25 | 1.28 | 1.91 | 1.02 | 1.36 | 0.17 | 0.88 | 1.11 | 0.29 | -0.135 | -0.225 | -1.005 | 0.015216 | 0.719517 | 0.487583 | 1 | 0.999631 | 0.388832 |
| Vasp_P70460_S317s_1 | 0.04 | 0.5 | 1.4 | 0.85 | 0 | 0.1 | -0.03 | 1.1 | 0.6 | 0.55 | 1.03 | 0.91 | 0.1 | 0.59 | 0.81 | 1.31 | -0.405 | 0.075 | 0.005 | 0.990106 | 0.323481 | 0.830139 | 1 | 0.999631 | 0.995892 |
| Vat1l_Q80TB8_S31s_1 | 2.98 | -1.92 | -0.24 | -0.41 | -0.48 | 0.46 | 1.36 | -0.61 | 0.37 | 2.42 | -1.79 | -1.47 | -0.75 | -1.12 | 0.48 | -1.11 | 0.08 | -0.22 | -0.7275 | 0.447318 | 0.934364 | 0.855911 | 1 | 0.999631 | 0.785754 |
| Vat1l_Q80TB8_S394s_1 | -0.9 | -1.55 | 1.08 | 0.88 | -0.99 | -1.98 | -1 | -1.01 | -0.58 | -0.46 | -0.36 | 1.51 | 0.24 | 0.1 | 0.14 | -1.55 | -1.1225 | 0.15 | -0.145 | 0.834202 | 0.097492 | 0.836365 | 1 | 0.999631 | 0.943909 |
| Vcam1_Q3UPN1_S166s_1 | -0.52 | -1.41 | -0.26 | -0.5 | -1.41 | -1.36 | -0.21 | -1.77 | -1 | -0.47 | -0.3 | -0.62 | -0.34 | -1 | 0.1 | 0.24 | -0.515 | 0.075 | 0.4225 | 0.30531 | 0.237317 | 0.82524 | 1 | 0.999631 | 0.716239 |
| Vcl_Q64727_S290s_1 | 0.47 | 0.26 | 1.07 | 0.74 | 1.24 | 1.03 | 0.32 | 0.02 | -0.28 | 2.03 | 1.94 | 2.96 | -1.33 | 1.46 | 1.04 | 1.03 | 0.0175 | 1.0275 | -0.085 | 0.887306 | 0.961281 | 0.132582 | 1 | 0.999631 | 0.963505 |
| Vcl_Q64727_S346s_1 | -0.23 | -0.41 | 0.39 | 0.65 | 1.64 | 0.42 | -0.26 | -0.37 | 0.39 | 1.11 | 1.29 | 2.73 | -0.09 | -0.51 | 0.73 | 0.13 | 0.2575 | 1.28 | -0.035 | 0.928486 | 0.60478 | 0.031454 | 1 | 0.999631 | 0.976781 |
| Vcp_Q01853_S702s_1 | -0.34 | 1.05 | 0.34 | -0.16 | -0.67 | 0.24 | -0.6 | -0.09 | 0.17 | 0.5 | 0.42 | 0.57 | 0.74 | 0.53 | 0.1 | 0.2 | -0.5025 | 0.1925 | 0.17 | 0.656546 | 0.215569 | 0.593171 | 1 | 0.999631 | 0.879056 |
| Vcp_Q01853_S748s_1 | 0.4 | -0.06 | -0.1 | 0.05 | -0.66 | 0.01 | 0.29 | -0.6 | 0.03 | 0.78 | -0.08 | -0.24 | 0.39 | -0.27 | -0.69 | -0.47 | -0.3125 | 0.05 | -0.3325 | 0.328687 | 0.329995 | 0.874208 | 1 | 0.999631 | 0.725922 |
| Vcpip1_Q8CDG3_S1007s_1 | 0.24 | 0.09 | -0.45 | 0.06 | -0.09 | 0.01 | -0.51 | -0.56 | 0.57 | 0.17 | -0.36 | -0.74 | 0.66 | 0.49 | -0.17 | -0.97 | -0.2725 | -0.075 | 0.0175 | 0.966372 | 0.350711 | 0.834602 | 1 | 0.999631 | 0.988351 |
| Vcpip1_Q8CDG3_S1196s_1 | -1.08 | -0.01 | 0.04 | 0.17 | -0.09 | 0.34 | -0.3 | 0.17 | -0.03 | 0.29 | -0.25 | 0.46 | -0.08 | -0.07 | -0.06 | 0.41 | 0.25 | 0.3375 | 0.27 | 0.461862 | 0.48179 | 0.362438 | 1 | 0.999631 | 0.792786 |
| Vcpip1_Q8CDG3_S746s_1 | 0.71 | 0.71 | 0.62 | 0.62 | 0.67 | 1.01 | 0.42 | 0.8 | 0.69 | 1.21 | 0.95 | 1.13 | -0.31 | 1.53 | 0.94 | 0.82 | 0.06 | 0.33 | 0.08 | 0.843867 | 0.809764 | 0.214895 | 1 | 0.999631 | 0.948343 |
| Vcpip1_Q8CDG3_T760t_1 | -0.16 | 0.56 | 1.5 | 1.77 | 1.58 | 2.55 | 1.86 | 2.12 | 1.72 | 0.63 | 2.06 | 1.44 | -0.66 | 1.38 | 1.59 | 1.71 | 1.11 | 0.545 | 0.0875 | 0.891486 | 0.036821 | 0.298259 | 1 | 0.999631 | 0.964253 |
| Vdac1_Q60932_S104s_1 | -1.02 | -1.04 | -0.13 | -0.42 | -1.05 | -1.5 | -0.62 | -0.28 | -1.13 | -1.06 | -0.28 | 0.29 | -0.31 | -0.84 | -0.26 | 0.39 | -0.21 | 0.1075 | 0.3975 | 0.305317 | 0.571027 | 0.794994 | 1 | 0.999631 | 0.716239 |
| Vdac1_Q60932_S215s_1 | -0.61 | -0.9 | -0.17 | -0.98 | -0.99 | -0.79 | -1.02 | -0.78 | -0.65 | -0.58 | -0.7 | -0.77 | -1.25 | -0.46 | -0.27 | -0.15 | -0.23 | -0.01 | 0.1325 | 0.712658 | 0.416056 | 0.971706 | 1 | 0.999631 | 0.9008 |
| Vdac1_Q60932_S240s_1 | 0.35 | 2.66 | 0.41 | 0.02 | 1.4 | 1.28 | -0.52 | -0.22 | 0.96 | 1.69 | -0.2 | -0.02 | 1.26 | 1.57 | 0.06 | 0.3 | -0.375 | -0.2525 | -0.0625 | 0.921829 | 0.592055 | 0.707747 | 1 | 0.999631 | 0.975032 |
| Vdac1_Q60932_S57s_1 | 0.47 | -1.15 | -0.57 | -1.77 | -0.27 | -1.2 | -0.71 | -1.52 | -0.81 | -0.46 | -2.12 | -0.65 | -0.71 | -1.06 | -0.86 | -1.92 | -0.17 | -0.255 | -0.3825 | 0.466809 | 0.740555 | 0.650789 | 1 | 0.999631 | 0.795267 |
| Vdac1_Q60932_T107t_1 | -1 | -1.61 | 0.8 | -0.22 | -0.69 | -2.03 | 0.19 | -0.05 | -1.53 | -0.73 | 0.29 | -0.55 | -0.88 | -1.02 | -0.66 | 0.88 | -0.1375 | -0.1225 | 0.0875 | 0.887237 | 0.830935 | 0.834474 | 1 | 0.999631 | 0.963505 |
| Vdac1_Q60932_T250t_1 | -1.11 | -1.24 | -0.6 | -1.89 | -2.07 | -1.02 | -1.11 | -1.6 | -1.95 | -1.7 | -0.44 | -1.73 | -0.71 | -1.32 | -2.26 | -2.11 | -0.24 | -0.245 | -0.39 | 0.393407 | 0.527101 | 0.573862 | 1 | 0.999631 | 0.759134 |
| Vdac2_G3UX26_S104s_1 | -1.56 | -1.63 | -1.02 | -1.32 | -1.68 | -2.27 | -1.58 | -1.94 | -1.7 | -1.53 | -0.82 | -0.41 | -0.93 | -2.14 | -0.59 | -0.91 | -0.485 | 0.2675 | 0.24 | 0.547493 | 0.112486 | 0.468362 | 1 | 0.999631 | 0.834277 |
| Vdac2_G3UX26_S240s_1 | -0.72 | -0.33 | -0.51 | -0.74 | -0.98 | -1 | -0.74 | -0.25 | -1.39 | -1.27 | -1.5 | -1.33 | -0.73 | -0.49 | -0.51 | -0.48 | -0.1675 | -0.7975 | 0.0225 | 0.933714 | 0.553985 | 0.009692 | 1 | 0.999631 | 0.978577 |
| Vdac2_G3UX26_T250t_1 | -1.33 | -1.55 | -1.12 | -1.68 | -1.75 | -0.9 | -1.42 | -1.63 | -2.2 | -1.77 | -0.77 | -3.95 | -1.28 | -1.61 | -3.34 | -2.78 | -0.005 | -0.7525 | -0.8325 | 0.109986 | 0.986391 | 0.238171 | 1 | 0.999631 | 0.573646 |
| Vdac2_G3UX26_T57t_1 | -0.41 | 0.2 | 0.96 | 0.26 | -0.35 | -0.56 | 0.33 | 0.29 | 0.05 | 0.49 | 0.95 | 0.99 | -0.61 | -0.97 | 0.29 | -0.2 | -0.325 | 0.3675 | -0.625 | 0.149546 | 0.395922 | 0.34498 | 1 | 0.999631 | 0.621721 |
| Vgf_Q0VGU4_S423s_1 | 1.21 | -1.7 | -0.47 | -1.11 | -1.54 | -0.72 | 0.14 | -1.83 | -0.95 | 0.18 | -1.4 | -1.22 | -1.11 | -1.93 | -1.53 | -2.1 | -0.47 | -0.33 | -1.15 | 0.08175 | 0.496319 | 0.613426 | 1 | 0.999631 | 0.54792 |
| Vill_G5E8H0_S232s_1 | -0.83 | -0.52 | -0.76 | -0.46 | -0.19 | -0.16 | -1.05 | 0 | -0.17 | -0.29 | -0.16 | -0.42 | -0.36 | -0.14 | -0.51 | 0.6 | 0.2925 | 0.3825 | 0.54 | 0.128768 | 0.35578 | 0.152033 | 1 | 0.999631 | 0.599316 |
| Vim_P20152_S10s_1 | 2.3 | -0.69 | 0.95 | 1.93 | 1.59 | -0.23 | 2.56 | 0.01 | 0.28 | 0.67 | 0.21 | 0.8 | -0.84 | -0.67 | -0.6 | -0.96 | -0.14 | -0.6325 | -1.89 | 0.010769 | 0.864427 | 0.320608 | 1 | 0.999631 | 0.383871 |
| Vim_P20152_S39s_1 | 2.29 | 1.48 | 0.82 | 2.2 | 1.85 | 1.7 | 2.19 | 1.06 | 2.17 | 2.08 | 1.38 | 1.59 | -1.23 | 1.44 | 1.53 | 1.85 | 0.0025 | 0.1075 | -0.8 | 0.271052 | 0.995193 | 0.790364 | 1 | 0.999631 | 0.698674 |
| Vim_P20152_S73s_1 | 0.84 | 0.5 | 2.15 | 1.11 | 1.27 | 2.12 | 0.93 | 1.32 | 0.93 | 1.75 | 1.58 | 2.46 | 3.57 | 1.01 | 2.43 | 1.72 | 0.26 | 0.53 | 1.0325 | 0.10706 | 0.547901 | 0.267681 | 1 | 0.999631 | 0.568797 |
| Vipas39_Q8BGQ1_S119s_1 | 0.73 | -0.64 | -0.13 | 0.63 | 0.54 | -0.12 | 0.2 | -0.49 | 0.24 | -0.35 | 0.13 | 0.05 | -0.83 | -0.31 | -0.45 | 0.02 | -0.115 | -0.13 | -0.54 | 0.193566 | 0.77293 | 0.729323 | 1 | 0.999631 | 0.650305 |
| Vipas39_Q8BGQ1_S77s_1 | 1.67 | -0.63 | 2.35 | -0.41 | -0.84 | 0.23 | 0.74 | -0.66 | -1.06 | 0.98 | -0.71 | -2.16 | -2.19 | -1.33 | -0.65 | -2.32 | -0.8775 | -1.4825 | -2.3675 | 0.00904 | 0.249313 | 0.110663 | 1 | 0.999631 | 0.365535 |
| Vps13c_Q8BX70_S1891s_1 | -1 | -0.57 | -0.62 | -0.59 | -0.72 | -0.4 | -1.23 | -0.31 | -0.34 | -0.59 | -0.33 | -0.77 | -0.47 | -0.26 | -1.4 | -0.95 | 0.03 | 0.1875 | -0.075 | 0.825455 | 0.919526 | 0.485077 | 1 | 0.999631 | 0.940357 |
| Vps13c_Q8BX70_S2442s_1 | 2.66 | 0.66 | -0.36 | -0.36 | 0.3 | 1.24 | -0.1 | -0.5 | 1.46 | 1.9 | -0.25 | 0.32 | 0.54 | 0.41 | -0.12 | -0.73 | -0.415 | 0.2075 | -0.625 | 0.371116 | 0.56221 | 0.786545 | 1 | 0.999631 | 0.746338 |
| Vps13c_Q8BX70_S736s_1 | -0.64 | 0.06 | 0.6 | -0.82 | -0.25 | -0.25 | -0.45 | -0.44 | -0.08 | -0.54 | -0.02 | 0.64 | 0.16 | 0.11 | 0.55 | -0.14 | -0.1475 | 0.2 | 0.37 | 0.351962 | 0.680962 | 0.630811 | 1 | 0.999631 | 0.738579 |
| Vps13c_Q8BX70_S841s_1 | 1.96 | -1.02 | -0.01 | 0.24 | -1.56 | -0.53 | 0.62 | -0.76 | -1.42 | 1.05 | -1.19 | -1.4 | -1.27 | -1.27 | -0.59 | -1.5 | -0.85 | -1.0325 | -1.45 | 0.032473 | 0.230077 | 0.195469 | 1 | 0.999631 | 0.442395 |
| Vps13c_Q8BX70_S841s_2 | 0.76 | -0.51 | 0.36 | -0.14 | -0.89 | -0.75 | 0.55 | -1.11 | -0.75 | 0.46 | -1.18 | -1.32 | -0.77 | -1.47 | -1.6 | -2.05 | -0.6675 | -0.815 | -1.59 | 0.002761 | 0.161566 | 0.110257 | 1 | 0.999631 | 0.351308 |
| Vps13d_B1ART2_S1043s_1 | -0.79 | -0.03 | -0.68 | -2.17 | 0.5 | -0.09 | -1.08 | -1.67 | -0.05 | -0.16 | -1.53 | 0.05 | 0.38 | 0.25 | -0.49 | 0.13 | 0.3325 | 0.495 | 0.985 | 0.061724 | 0.581866 | 0.372211 | 1 | 0.999631 | 0.509355 |
| Vps13d_B1ART2_S1725s_1 | -0.44 | 0.33 | 0.02 | -0.28 | 0.13 | -0.18 | -0.61 | -0.11 | -0.06 | -1.08 | -0.09 | 0.65 | 0.88 | -1.19 | 0.29 | 0.2 | -0.1 | -0.0525 | 0.1375 | 0.767 | 0.736084 | 0.896671 | 1 | 0.999631 | 0.917066 |
| Vps13d_B1ART2_S2080s_1 | 0.07 | 1.18 | 0.41 | -0.7 | 0.27 | 0.93 | -0.28 | -0.21 | 0.47 | 1.15 | -0.51 | -0.07 | 0.71 | 1.02 | -0.16 | 0.95 | -0.0625 | 0.02 | 0.39 | 0.411968 | 0.891894 | 0.968174 | 1 | 0.999631 | 0.767937 |
| Vps13d_B1ART2_S2435s_1 | 0.14 | -0.2 | -0.95 | -0.56 | -0.25 | 1.25 | 0.13 | -0.86 | 0.03 | 0.32 | -0.85 | -0.34 | 0.41 | 0.4 | -0.42 | 0.39 | 0.46 | 0.1825 | 0.5875 | 0.127153 | 0.34643 | 0.625223 | 1 | 0.999631 | 0.59788 |
| Vps13d_B1ART2_S2862s_1 | 0.74 | -0.43 | 0.63 | 1 | 0.24 | -0.64 | 0.27 | 0.66 | -0.56 | 0.21 | 1.14 | 1.18 | 0.42 | 0.39 | 0 | 1.11 | -0.3525 | 0.0075 | -0.005 | 0.990246 | 0.405213 | 0.987928 | 1 | 0.999631 | 0.995892 |
| Vps13d_B1ART2_S2865s_1 | -1.06 | -0.31 | -1.26 | -1.1 | -0.66 | -1.42 | -1.24 | -1.41 | -0.32 | -0.16 | -0.53 | -1.11 | 0.01 | -0.64 | -1.1 | 0.13 | -0.25 | 0.4025 | 0.5325 | 0.191417 | 0.44651 | 0.25571 | 1 | 0.999631 | 0.648674 |
| Vps13d_B1ART2_S2920s_1 | -0.69 | -1.23 | -0.77 | -1.26 | -0.56 | -0.05 | -1.22 | -1.25 | -0.71 | -0.78 | -1.5 | -1 | -0.02 | -0.88 | -0.04 | -1.42 | 0.2175 | -0.01 | 0.3975 | 0.330725 | 0.541206 | 0.973815 | 1 | 0.999631 | 0.727302 |
| Vps13d_B1ART2_S3011s_1 | -1.93 | -0.61 | 0.56 | -1.4 | -0.21 | -0.97 | 0.29 | -1.22 | -0.08 | -2.38 | -0.99 | -0.47 | -0.34 | -1.99 | -0.76 | -1.68 | 0.3175 | -0.135 | -0.3475 | 0.56878 | 0.589169 | 0.838543 | 1 | 0.999631 | 0.843185 |
| Vps26a_P40336_S315s_1 | -0.5 | -0.56 | -0.09 | -0.3 | -0.38 | 0.11 | -0.55 | 0.04 | 0.48 | -0.5 | -0.17 | 0.57 | -0.02 | 0.84 | 0.14 | 0.12 | 0.1675 | 0.4575 | 0.6325 | 0.065814 | 0.549154 | 0.188173 | 1 | 0.999631 | 0.516933 |
| Vps26b_Q8C0E2_S302s_1 | 1.31 | -0.36 | -0.46 | -0.4 | 0.08 | -0.2 | 0.16 | -0.3 | -0.2 | 0.66 | -0.45 | -0.94 | -0.18 | 1 | -0.38 | -0.69 | -0.0875 | -0.255 | -0.085 | 0.873046 | 0.840206 | 0.621411 | 1 | 0.999631 | 0.959648 |
| Vps26b_Q8C0E2_S304s_1 | 0.41 | -0.14 | -0.12 | 0.29 | -0.04 | 0.17 | 0.21 | 0.14 | -0.08 | 0.44 | -0.02 | -0.24 | -0.67 | 0.14 | -0.11 | 0.12 | 0.01 | -0.085 | -0.24 | 0.458995 | 0.969124 | 0.769375 | 1 | 0.999631 | 0.79152 |
| Vps26b_Q8C0E2_S330s_1 | -0.05 | -1.18 | -0.36 | 0.06 | -0.69 | -1.07 | -0.18 | -0.28 | -1.02 | 0.03 | 0.14 | 0.05 | -1.56 | -0.95 | -0.09 | -0.9 | -0.1725 | 0.1825 | -0.4925 | 0.261348 | 0.640008 | 0.652412 | 1 | 0.999631 | 0.69271 |
| Vps26b_Q8C0E2_S334s_1 | 1.27 | -0.01 | -0.79 | 1.07 | 1.38 | 0.12 | 0.84 | 0.38 | 0.61 | 0.05 | 1.17 | -0.26 | -1.62 | -0.79 | 0.29 | -0.63 | 0.295 | 0.0075 | -1.0725 | 0.085079 | 0.571457 | 0.988844 | 1 | 0.999631 | 0.550778 |
| Vps26b_Q8C0E2_T325t_1 | 3.03 | -0.42 | -2.59 | 2.55 | 3.17 | 1.2 | 2.75 | 2.29 | 1.85 | -0.23 | 1.39 | 0.05 | -4.66 | 1.04 | 0.4 | -0.25 | 1.71 | 0.1225 | -1.51 | 0.341512 | 0.171237 | 0.919303 | 1 | 0.999631 | 0.73317 |
| Vps35_Q9EQH3_S783s_1 | -0.55 | -0.24 | -0.14 | 0.03 | -0.37 | 0.09 | -0.49 | 0.19 | -0.7 | -0.54 | 0.17 | 0.22 | -0.39 | 0 | 0.22 | 0.44 | 0.08 | 0.0125 | 0.2925 | 0.355702 | 0.778699 | 0.969188 | 1 | 0.999631 | 0.74034 |
| Vps35_Q9EQH3_S7s_1 | 1.07 | -1.8 | 0.51 | -0.05 | -1.39 | -0.68 | 0.55 | -1.63 | -1.02 | 0.09 | -2.03 | -1 | -0.73 | -2.19 | -0.34 | -1.71 | -0.72 | -0.9225 | -1.175 | 0.105101 | 0.317105 | 0.194367 | 1 | 0.999631 | 0.568213 |
| Vps45_P97390_S441s_1 | -0.25 | -1.22 | -0.43 | -0.61 | -0.91 | -1.88 | -0.45 | -0.59 | -0.94 | -0.54 | -0.85 | -0.58 | -0.73 | -1.14 | -0.09 | 0.35 | -0.33 | -0.1 | 0.225 | 0.586626 | 0.409417 | 0.743379 | 1 | 0.999631 | 0.849724 |
| Vps4a_Q8VEJ9_S97s_1 | 1.73 | 1.13 | 1.38 | 1.25 | -0.03 | 1.28 | 1.94 | -0.46 | 0.31 | 2.21 | 0.37 | 0.16 | 0.27 | 0.36 | 0.88 | -0.69 | -0.69 | -0.61 | -1.1675 | 0.011826 | 0.213581 | 0.224354 | 1 | 0.999631 | 0.385092 |
| Vps4b_P46467_S102s_1 | 0.99 | -0.35 | 0.83 | 1.25 | -0.43 | 0.24 | 1 | 0.05 | 0.19 | 1.56 | 0.08 | 0.16 | -0.36 | -0.65 | 0.16 | -0.96 | -0.465 | -0.1825 | -1.1325 | 0.024246 | 0.312669 | 0.705269 | 1 | 0.999631 | 0.415644 |
| Vps51_Q3UVL4_S640s_1 | 2.47 | -3.23 | -1.61 | -1.77 | -3.29 | -1.86 | 0.3 | -2.36 | -6.21 | 2.48 | -1.46 | -0.61 | -3.89 | -4.6 | -0.24 | -1.8 | -0.7675 | -0.415 | -1.5975 | 0.246837 | 0.536352 | 0.821804 | 1 | 0.999631 | 0.685751 |
| Vps51_Q3UVL4_S652s_1 | 1.56 | 0.84 | -2.34 | 2.26 | 2.53 | 1.22 | 1.56 | 1.65 | 1.66 | 0.05 | 1.35 | 0 | -3.98 | 0.47 | -0.22 | 0.22 | 1.16 | 0.185 | -1.4575 | 0.253557 | 0.222389 | 0.846516 | 1 | 0.999631 | 0.688735 |
| Vsnl1_P62761_T146t_1 | -0.47 | -0.63 | -0.56 | -0.91 | -0.75 | -0.57 | -0.81 | -1.1 | -1.39 | -0.06 | -0.92 | -0.98 | 0.13 | -0.46 | -0.62 | -1.07 | -0.165 | -0.195 | 0.1375 | 0.682631 | 0.524089 | 0.570005 | 1 | 0.999631 | 0.889594 |
| Vti1b_F6UHS3_S80s_1 | 0.44 | -0.29 | -1.56 | 0.79 | 0.91 | 0.4 | 0.56 | 0.86 | -0.15 | -1.06 | 0.39 | -0.86 | -1.2 | 0.46 | -0.64 | -0.42 | 0.8375 | -0.265 | -0.295 | 0.609643 | 0.118874 | 0.642667 | 1 | 0.999631 | 0.858923 |
| Vwa5b1_A9Z1V5_S664s_1 | 0.65 | -1.91 | -0.52 | -0.01 | -1.46 | -0.59 | -0.44 | 0.27 | -1.62 | -2.21 | -1.54 | -1.39 | -1.08 | -2.33 | -0.75 | -1.57 | -0.1075 | -1.2425 | -0.985 | 0.118211 | 0.855006 | 0.040182 | 1 | 0.999631 | 0.585087 |
| Vwa5b1_A9Z1V5_S804s_1 | -0.48 | -0.92 | 0.02 | -0.35 | 0.02 | -0.68 | -1.13 | 0.38 | -0.01 | -0.06 | -0.92 | -0.97 | -0.05 | 0.31 | 0.1 | -0.42 | 0.08 | -0.0575 | 0.4175 | 0.218802 | 0.840354 | 0.873107 | 1 | 0.999631 | 0.669859 |
| Vwa5b2_E0CZ01_S1017s_1 | -0.71 | -0.17 | -0.99 | -1.66 | -1.1 | -1.33 | -1.01 | -1.95 | -0.05 | -2.15 | -1.31 | -1.07 | -0.06 | -1 | -1.66 | -1.58 | -0.465 | -0.2625 | -0.1925 | 0.684166 | 0.246349 | 0.604694 | 1 | 0.999631 | 0.889649 |
| Vwa5b2_E0CZ01_S759s_1 | 2.07 | 0.28 | -0.82 | 0.03 | 0.55 | 0.1 | 0.33 | -0.28 | 0.87 | 1.19 | -0.86 | -1.57 | -1.8 | -0.42 | -1.33 | -0.72 | -0.215 | -0.4825 | -1.4575 | 0.037432 | 0.709689 | 0.546629 | 1 | 0.999631 | 0.458129 |
| Wasf1_Q8R5H6_S310s_1 | 0.86 | -0.1 | 0.69 | 1.61 | 0.71 | 1.08 | 1.89 | 0.18 | 0.7 | 0.62 | 0.14 | -0.43 | -0.65 | 1.55 | 0.56 | 0.32 | 0.2 | -0.5075 | -0.32 | 0.554043 | 0.676134 | 0.259918 | 1 | 0.999631 | 0.837952 |
| Wasf1_Q8R5H6_S321s_1 | -0.05 | -1.24 | 0.2 | 1.05 | -0.16 | -0.66 | 0.48 | 1.21 | -0.7 | -1.14 | 0.56 | 0 | -1.44 | -0.53 | 0.14 | 0.22 | 0.2275 | -0.31 | -0.3925 | 0.491526 | 0.689709 | 0.581448 | 1 | 0.999631 | 0.80893 |
| Wasf1_Q8R5H6_T323t_1 | -0.2 | -0.87 | -0.79 | -0.22 | -0.05 | -0.45 | -0.38 | -0.34 | -0.65 | 0.04 | -1 | -0.67 | -0.73 | 0.48 | -0.17 | 0.25 | 0.215 | -0.05 | 0.4775 | 0.211862 | 0.45083 | 0.880674 | 1 | 0.999631 | 0.665852 |
| Wasf3_Q8VHI6_S229s_1 | -0.68 | -1.12 | -0.93 | 0 | -0.64 | -0.95 | 0.41 | -1.37 | -0.79 | -0.94 | -2.18 | -2.28 | -0.35 | -0.53 | -1.89 | -0.86 | 0.045 | -0.865 | -0.225 | 0.60358 | 0.918463 | 0.081934 | 1 | 0.999631 | 0.857824 |
| Wbp11_Q923D5_S237s_1 | 0.67 | 1.38 | 1.24 | 0.45 | 0.11 | -0.16 | 0.67 | -0.1 | -0.31 | 0.92 | 0.87 | 1.11 | 1.3 | 1.06 | 0.75 | 1.38 | -0.805 | -0.2875 | 0.1875 | 0.577803 | 0.03397 | 0.482084 | 1 | 0.999631 | 0.84622 |
| Wbp4_Q61048_S97s_1 | -0.18 | 1.03 | 1.15 | 0.43 | 0.75 | 0.12 | -0.33 | -0.07 | 0.87 | -0.49 | 0.45 | -0.06 | 0.37 | 1.22 | -0.52 | 0.53 | -0.49 | -0.415 | -0.2075 | 0.656098 | 0.229361 | 0.343112 | 1 | 0.999631 | 0.879056 |
| Wdfy3_G3UYW1_S1942s_1 | -0.27 | -0.2 | 0.61 | 0.78 | -0.69 | 0.14 | -0.05 | -0.07 | -0.08 | 0.1 | 0.48 | -0.99 | -0.54 | 0.13 | -0.53 | -0.82 | -0.3975 | -0.3525 | -0.67 | 0.099293 | 0.276385 | 0.407638 | 1 | 0.999631 | 0.562298 |
| Wdfy3_G3UYW1_S2277s_1 | -1.13 | 0.67 | 0.14 | -0.1 | 2.03 | 0.59 | 1.41 | -0.47 | 1.72 | -1.7 | 1.05 | 0.56 | -0.85 | -0.72 | 0.31 | -0.97 | 0.995 | 0.5125 | -0.4525 | 0.346351 | 0.120091 | 0.492152 | 1 | 0.999631 | 0.735175 |
| Wdfy3_G3UYW1_S2434s_1 | 1.17 | 1.39 | 0.79 | 1.61 | 1.02 | 0.66 | 0.92 | 0.85 | 0.98 | 1.12 | 0.84 | 1.14 | -0.83 | 0.89 | 0.15 | 0.75 | -0.3775 | -0.22 | -1 | 0.041334 | 0.192739 | 0.443523 | 1 | 0.999631 | 0.474979 |
| Wdr11_Q8K1X1_S405s_1 | -0.78 | 0.31 | 0.29 | -0.66 | -0.49 | 0.19 | -0.25 | -0.86 | 0.3 | -1.37 | 0.2 | 1.5 | 0.54 | -0.24 | -0.08 | -0.14 | -0.1425 | 0.3675 | 0.23 | 0.549057 | 0.709492 | 0.544171 | 1 | 0.999631 | 0.835034 |
| Wdr13_Q91V09_S70s_1 | 0.3 | 1.07 | 1 | 0.38 | -0.07 | 0.5 | -0.15 | 0.67 | 0.54 | 1.15 | 0.52 | 0.41 | -0.09 | 1.13 | 0.58 | 0.61 | -0.45 | -0.0325 | -0.13 | 0.723569 | 0.192708 | 0.919324 | 1 | 0.999631 | 0.905674 |
| Wdr13_Q91V09_S79s_1 | 1.44 | 1.21 | -0.16 | 1.62 | 2.02 | 1.4 | 1.08 | 1.41 | 1.43 | 0.51 | 1.71 | 0.75 | -3.05 | 1.58 | 1.09 | 1 | 0.45 | 0.0725 | -0.8725 | 0.384926 | 0.318248 | 0.878616 | 1 | 0.999631 | 0.75538 |
| Wdr17_H3BLI8_S309s_1 | -0.3 | -0.79 | 0.18 | 0 | -0.83 | -0.72 | -0.51 | -0.35 | -0.68 | -0.21 | 0.08 | -0.09 | 0.33 | -0.26 | -0.35 | 0.1 | -0.375 | 0.0025 | 0.1825 | 0.588135 | 0.230714 | 0.993839 | 1 | 0.999631 | 0.850634 |
| Wdr20_Q3UWE6_S357s_1 | 0.16 | 0.04 | -0.39 | 0.23 | -0.22 | -0.06 | 1.28 | -0.46 | -0.51 | -0.04 | -0.16 | -0.18 | -0.31 | -0.17 | -0.19 | -0.23 | 0.125 | -0.2325 | -0.235 | 0.409439 | 0.762605 | 0.406885 | 1 | 0.999631 | 0.766996 |
| Wdr20_Q3UWE6_S434s_1 | 0.08 | -0.59 | -0.93 | -0.25 | -0.58 | -0.89 | -0.06 | -0.58 | -0.3 | -0.27 | -0.51 | 0.38 | -0.65 | -0.01 | 0.16 | 0.92 | -0.105 | 0.2475 | 0.5275 | 0.216112 | 0.745827 | 0.469854 | 1 | 0.999631 | 0.668933 |
| Wdr20_Q3UWE6_S465s_1 | 0.95 | 1.14 | 0.18 | 0.88 | 1.63 | 1.72 | 0.47 | 1.23 | 0.93 | 0.32 | 0.23 | 0.7 | -0.15 | 1.61 | 1.02 | 0.66 | 0.475 | -0.2425 | -0.0025 | 0.995372 | 0.220557 | 0.459967 | 1 | 0.999631 | 0.997705 |

| | | | | | | | | | | | | | | | | | | | | | | | | | |
|---|---|---|---|---|---|---|---|---|---|---|---|---|---|---|---|---|---|---|---|---|---|---|---|---|---|
| Wdr25_E9Q349_S60s_1 | 0.82 | 1.49 | 0.04 | 1.06 | 1.29 | 0.56 | 0.62 | 0.61 | 1.23 | 1.47 | 0.41 | 1.54 | -1.84 | 1.22 | -0.31 | 0.02 | -0.0825 | 0.31 | -1.08 | 0.111415 | 0.823307 | 0.454764 | 1 | 0.999631 | 0.577068 |
| Wdr26_E0CYH4_S101s_1 | 0.37 | 0.19 | 1.2 | -0.03 | 0.92 | 0.75 | 0.63 | 0.85 | 1.02 | 1.1 | 0.69 | 1.05 | 1.05 | 0.35 | 0.87 | 0.67 | 0.355 | 0.5325 | 0.3025 | 0.407494 | 0.286165 | 0.134944 | 1 | 0.999631 | 0.766996 |
| Wdr33_Q8K4P0_S1204s_1 | 0.81 | 2.38 | 3.04 | 0.15 | 2.55 | 1.78 | 1.95 | 1.72 | 3.4 | 1.33 | 2.62 | 3.12 | 4.01 | 1.79 | 2.03 | 1.46 | 0.405 | 1.0225 | 0.7275 | 0.357872 | 0.522346 | 0.178581 | 1 | 0.999631 | 0.741099 |
| Wdr37_Q8CBE3_S30s_1 | 1.12 | -2.58 | -0.41 | -0.17 | -2.91 | 0.64 | 1.98 | -2.81 | -2.18 | 1.5 | -1.3 | -0.8 | -2.26 | -1.32 | 0.37 | -0.43 | -0.265 | -0.185 | -0.4 | 0.631216 | 0.830712 | 0.845786 | 1 | 0.999631 | 0.867571 |
| Wdr37_Q8CBE3_S31s_1 | 1.3 | -1.48 | 0.37 | 0.42 | -2.18 | 0.26 | 3.3 | -2.52 | -2.77 | 1.7 | -1.05 | -0.57 | -1.3 | -2.45 | -0.26 | -1.8 | -0.4375 | -0.825 | -1.605 | 0.034367 | 0.727079 | 0.392488 | 1 | 0.999631 | 0.446273 |
| Wdr4_E9Q156_S440s_1 | 0.3 | 1.23 | 2.8 | 1.36 | 0.86 | -0.01 | 1.22 | 2.4 | 0.53 | 0.27 | 1.31 | 2.79 | 1.65 | 1.31 | 1.7 | 3.04 | -0.305 | -0.1975 | 0.5025 | 0.401584 | 0.636803 | 0.773241 | 1 | 0.999631 | 0.763221 |
| Wdr44_Q6NVE8_S405s_1 | 0.31 | 1.21 | 1.35 | 0.74 | 0.79 | 0.99 | 0.7 | 1.06 | 1.28 | 1.27 | 0.99 | 1.19 | 1 | 0.63 | 1.53 | 0.36 | -0.0175 | 0.28 | -0.0225 | 0.952979 | 0.954651 | 0.379741 | 1 | 0.999631 | 0.984317 |
| Wdr44_Q6NVE8_S50s_1 | -0.3 | 1.79 | 1.54 | -0.25 | 0.47 | -0.06 | 0.66 | -0.25 | 0.91 | -0.48 | 0.91 | 1.62 | 1.23 | -0.01 | 0.09 | 0.52 | -0.49 | 0.045 | -0.2375 | 0.68239 | 0.384869 | 0.944171 | 1 | 0.999631 | 0.889581 |
| Wdr44_Q6NVE8_S563s_1 | -0.1 | 0.63 | 0.61 | 0.43 | 0.27 | 0.07 | -0.02 | 0.07 | -0.42 | 0.72 | 1.04 | 0.39 | 1.23 | 0.75 | 0.58 | 0.32 | -0.295 | 0.04 | 0.3275 | 0.333468 | 0.292299 | 0.915984 | 1 | 0.999631 | 0.728079 |
| Wdr44_Q6NVE8_S576s_1 | -1.01 | 1.33 | 2.96 | 1.36 | 0.29 | 1.21 | 1.73 | 2.22 | 0.48 | 0.78 | 1.19 | 1.55 | 1.15 | 0.83 | 2.05 | 2.9 | 0.2025 | -0.16 | 0.5725 | 0.489791 | 0.80025 | 0.831346 | 1 | 0.999631 | 0.807612 |
| Wdr47_Q8CGF6_S292s_1 | 2.33 | 0.57 | 0.7 | 0.9 | 1.53 | 0.86 | 0.79 | 2.03 | 1.28 | 0.2 | 2.4 | 0.87 | 0.05 | 1.2 | 1.6 | 2 | 0.1775 | 0.0625 | 0.0875 | 0.872619 | 0.710664 | 0.912145 | 1 | 0.999631 | 0.959468 |
| Wdr47_Q8CGF6_S297s_1 | 1.01 | 0.08 | -0.54 | 0.35 | 1.09 | 0.46 | 0.62 | 0.37 | 0.36 | 0.07 | 0.64 | 0.29 | -1.04 | 0.3 | 0.44 | 0.53 | 0.41 | 0.115 | -0.1675 | 0.725885 | 0.289427 | 0.756251 | 1 | 0.999631 | 0.90568 |
| Wdr47_Q8CGF6_S304s_1 | 0.56 | 0.02 | -1.05 | 0.08 | 0.3 | -0.28 | -0.58 | -0.34 | 0.08 | 0.18 | -1.08 | -0.72 | -0.93 | -0.19 | -1.27 | -0.69 | -0.1275 | -0.2875 | -0.6725 | 0.134071 | 0.746986 | 0.526752 | 1 | 0.999631 | 0.60492 |
| Wdr47_Q8CGF6_S359s_1 | 0.15 | 0.66 | 0.39 | -0.22 | 0.34 | 0.29 | 0.18 | -0.32 | 0.42 | 0.77 | 0.86 | 1.53 | 1.2 | 0.57 | -0.26 | 0.47 | -0.1225 | 0.65 | 0.25 | 0.521132 | 0.686023 | 0.081957 | 1 | 0.999631 | 0.820181 |
| Wdr47_Q8CGF6_S374s_1 | -0.35 | -0.03 | -1.04 | -0.59 | -0.07 | 0.06 | -0.86 | -0.66 | -0.55 | 0.24 | -0.54 | -0.57 | 0.68 | 0.09 | 0 | 0.35 | 0.12 | 0.1475 | 0.7825 | 0.037903 | 0.726561 | 0.66387 | 1 | 0.999631 | 0.459387 |
| Wdr47_Q8CGF6_T285t_1 | 0.14 | 0.3 | 0.33 | 0.92 | 1.55 | 1.18 | 0.39 | 0.97 | 0.96 | -0.15 | 0.45 | 0.98 | -1.13 | 0.9 | 1.17 | 1.16 | 0.6 | 0.1375 | 0.1025 | 0.850159 | 0.098084 | 0.698426 | 1 | 0.999631 | 0.949525 |
| Wdr59_Q8C0M0_S603s_1 | -0.94 | 0.9 | 0.84 | -2.27 | 0.79 | 0.14 | 1.01 | -1.28 | 1.7 | -1.7 | 0.96 | 1.5 | 0.14 | -1.31 | 0.16 | -3.24 | 0.5325 | 0.9825 | -0.695 | 0.4712 | 0.513065 | 0.315142 | 1 | 0.999631 | 0.797981 |
| Wdr7_Q920I9_S1151s_1 | -0.75 | -0.02 | -1.02 | -1.02 | -0.7 | -0.85 | -1.19 | -0.51 | -0.19 | -1.18 | -1.34 | -0.31 | 0.02 | -0.23 | -1.24 | -0.88 | -0.11 | -0.0525 | 0.12 | 0.763912 | 0.733851 | 0.893718 | 1 | 0.999631 | 0.915728 |
| Wdr7_Q920I9_S1455s_1 | -0.83 | -0.54 | -0.51 | -0.99 | -0.83 | -0.61 | -1.12 | -0.68 | -0.85 | -0.59 | -1.02 | -0.88 | 0.02 | -0.2 | -0.69 | -0.27 | -0.0925 | -0.1175 | 0.4325 | 0.168093 | 0.725823 | 0.658703 | 1 | 0.999631 | 0.637399 |
| Wdr7_Q920I9_S716s_1 | -0.14 | 0.13 | 0.78 | -0.73 | -0.89 | 0.58 | -0.56 | -0.5 | 0.54 | 0.24 | -0.66 | 0.15 | 0.28 | 1.52 | 1.21 | -0.25 | -0.3525 | 0.0575 | 0.68 | 0.190623 | 0.427445 | 0.88892 | 1 | 0.999631 | 0.648674 |
| Wdr7_Q920I9_S772s_1 | -0.66 | -5.76 | -4.58 | -2.61 | -1.74 | -4.45 | -4.27 | -5.38 | -3.01 | -1.64 | -4.84 | -3.21 | -4.36 | -4.96 | -3.93 | -6.15 | -0.5575 | 0.2275 | -1.4475 | 0.18559 | 0.63458 | 0.838439 | 1 | 0.999631 | 0.646224 |
| Wdr7_Q920I9_S772s_2 | -0.7 | -4.18 | -3.28 | -2.11 | -2.73 | -3.63 | -2.1 | -3.24 | -2.48 | -2.41 | -2.17 | -4.38 | -2.28 | -4.52 | -3.93 | -5.82 | -0.3575 | -0.2925 | -1.57 | 0.107467 | 0.624026 | 0.71506 | 1 | 0.999631 | 0.569416 |
| Wdr7_Q920I9_S776s_1 | -0.11 | -0.66 | 0.12 | -0.29 | -0.99 | -0.45 | 0 | -1.16 | -1 | 0.11 | -0.69 | -0.87 | -0.96 | -1.63 | -0.88 | -1.75 | -0.415 | -0.3775 | -1.07 | 0.010011 | 0.245715 | 0.285857 | 1 | 0.999631 | 0.377907 |
| Wdr7_Q920I9_S776s_2 | -0.7 | -4.18 | -3.28 | -2.11 | -2.73 | -3.63 | -2.1 | -3.24 | -2.48 | -2.41 | -2.17 | -4.38 | -2.28 | -4.52 | -3.93 | -5.82 | -0.3575 | -0.2925 | -1.57 | 0.107467 | 0.624026 | 0.71506 | 1 | 0.999631 | 0.569416 |
| Wdr7_Q920I9_S935s_1 | -0.23 | -0.28 | 0.96 | -1.21 | -0.12 | 0.04 | -0.24 | -0.99 | -0.26 | 0.35 | -1.54 | -0.14 | 0.44 | 0.43 | -0.21 | -2.32 | -0.1375 | -0.2075 | -0.225 | 0.748498 | 0.77253 | 0.709388 | 1 | 0.999631 | 0.911931 |
| Wdr7_Q920I9_T1195t_1 | -0.02 | 0.21 | -0.56 | -0.82 | -1.02 | -0.41 | -0.81 | -0.31 | -0.88 | -1.27 | 0.05 | -2.05 | 0.5 | 0.68 | -1.34 | -1.72 | -0.34 | -0.74 | -0.1725 | 0.775046 | 0.317939 | 0.145372 | 1 | 0.999631 | 0.920998 |
| Wdr7_Q920I9_T926t_1 | 0.66 | 0.5 | 0.76 | 0.96 | 0.58 | 0.93 | 0.7 | 0.91 | 0.57 | 0.77 | 1.26 | 0.96 | -0.88 | 0.96 | 0.58 | 1.64 | 0.06 | 0.17 | -0.145 | 0.777899 | 0.810283 | 0.543021 | 1 | 0.999631 | 0.921382 |
| Wdr77_D6RIL4_T5t_1 | -1.09 | 0.93 | 0.63 | -0.07 | 0.48 | 0.29 | -0.01 | 0.63 | 0.73 | -0.58 | 0.58 | 1.04 | 1.53 | 0.9 | 0.79 | 0.78 | 0.2475 | 0.3425 | 0.9 | 0.080724 | 0.58735 | 0.52522 | 1 | 0.999631 | 0.545579 |
| Wdr81_K4DI77_S1570s_1 | -0.59 | 0.6 | -0.53 | -0.43 | -0.32 | -0.81 | -0.55 | -0.4 | -0.69 | -1.99 | -0.34 | -0.31 | -0.46 | -0.71 | -0.55 | -0.41 | -0.2825 | -0.595 | -0.295 | 0.405028 | 0.411495 | 0.224156 | 1 | 0.999631 | 0.765844 |
| Wdr91_S4R1X1_S257s_1 | -0.38 | -2.29 | -0.51 | -0.5 | -1 | -0.79 | -0.13 | -0.51 | -0.61 | -0.93 | -1.73 | -1.75 | -2.54 | -0.06 | -0.36 | -0.43 | 0.3125 | -0.335 | 0.0725 | 0.911808 | 0.512169 | 0.516046 | 1 | 0.999631 | 0.971848 |
| Wdtc1_Q80ZK9_S227s_1 | 0.7 | -1.52 | 1.47 | -1.1 | -0.97 | 0.86 | -0.44 | 1.27 | 0.01 | 0.43 | 0.8 | 0.75 | 0.93 | 0.68 | 1.02 | -0.08 | 0.2925 | 0.61 | 0.75 | 0.281692 | 0.70779 | 0.367205 | 1 | 0.999631 | 0.702855 |
| Wfs1_P56695_S158s_1 | -0.46 | -0.2 | -0.08 | 0.28 | 0.07 | -0.1 | -0.11 | -0.35 | 0.24 | 0.1 | 0.67 | 0.6 | 1.1 | 0.45 | 0.72 | 0.42 | -0.0075 | 0.5175 | 0.7875 | 0.028101 | 0.977771 | 0.100554 | 1 | 0.999631 | 0.423977 |
| Wfs1_P56695_S236s_1 | 1.91 | 1.04 | 1.48 | 1.14 | 0.79 | 1.15 | 0.9 | -0.02 | 1.3 | 1.56 | 0.45 | 1.06 | 0.56 | 0.27 | 0.02 | -0.13 | -0.6875 | -0.3 | -1.2125 | 0.003764 | 0.073529 | 0.398434 | 1 | 0.999631 | 0.351308 |
| Wfs1_P56695_S32s_1 | -0.99 | -0.99 | 1.13 | 0.89 | -0.68 | -1.59 | 0.37 | 0.9 | -1.2 | -1.16 | 0.91 | 1.35 | -0.3 | -0.8 | 1.34 | 1.43 | -0.26 | -0.035 | 0.4075 | 0.572917 | 0.714443 | 0.964232 | 1 | 0.999631 | 0.844402 |
| Wfs1_P56695_S3s_1 | -0.65 | -1.28 | -0.75 | -0.42 | -1.3 | -0.94 | -1.12 | -0.88 | -1.18 | -0.8 | -1.06 | -0.89 | -0.75 | -0.52 | -1.24 | -0.38 | -0.285 | -0.2075 | 0.0525 | 0.874769 | 0.32737 | 0.478074 | 1 | 0.999631 | 0.960237 |
| Wfs1_P56695_S3s_2 | 0.04 | -0.96 | -0.64 | -1.2 | -0.62 | -0.43 | -0.78 | -0.69 | -1.51 | -0.17 | -0.88 | -0.7 | 0.38 | -0.1 | 0.32 | -1.25 | 0.06 | -0.125 | 0.5275 | 0.266602 | 0.853208 | 0.754328 | 1 | 0.999631 | 0.695664 |
| Wfs1_P56695_S9s_2 | 0.04 | -0.96 | -0.64 | -1.2 | -0.62 | -0.43 | -0.78 | -0.69 | -1.51 | -0.17 | -0.88 | -0.7 | 0.38 | -0.1 | 0.32 | -1.25 | 0.06 | -0.125 | 0.5275 | 0.266602 | 0.853208 | 0.754328 | 1 | 0.999631 | 0.695664 |
| Wfs1_P56695_T30t_1 | -1.63 | -2.12 | -2.14 | -1.76 | -1.19 | -2.84 | -2.38 | -2.99 | -2.64 | -2.84 | -1.62 | -1.71 | -0.29 | -2.31 | -1.05 | -2.61 | -0.4375 | -0.29 | 0.3475 | 0.513364 | 0.314185 | 0.43789 | 1 | 0.999631 | 0.818661 |
| Wfs1_P56695_T5t_1 | -0.65 | -1.28 | -0.75 | -0.42 | -1.3 | -0.94 | -1.12 | -0.88 | -1.18 | -0.8 | -1.06 | -0.89 | -0.75 | -0.52 | -1.24 | -0.38 | -0.285 | -0.2075 | 0.0525 | 0.874769 | 0.32737 | 0.478074 | 1 | 0.999631 | 0.960237 |
| Wfs1_P56695_T5t_2 | 0.04 | -0.96 | -0.64 | -1.2 | -0.62 | -0.43 | -0.78 | -0.69 | -1.51 | -0.17 | -0.88 | -0.7 | 0.38 | -0.1 | 0.32 | -1.25 | 0.06 | -0.125 | 0.5275 | 0.266602 | 0.853208 | 0.754328 | 1 | 0.999631 | 0.695664 |
| Wipf2_Q6PEV3_S155s_1 | 0.27 | -0.79 | 0.53 | 0.62 | -0.05 | -1.19 | 0.08 | 0.86 | -0.53 | -0.78 | -0.24 | 0.35 | 0.21 | -0.09 | -0.09 | -0.21 | -0.2325 | -0.4575 | -0.2025 | 0.59257 | 0.643341 | 0.282883 | 1 | 0.999631 | 0.852841 |
| Wipf2_Q6PEV3_S175s_1 | 1.14 | 2.76 | 1.96 | 0.57 | 2.73 | 1.62 | 2.68 | 0.41 | 4.63 | 1.2 | 2.59 | 3.51 | 3.51 | 0.18 | 0.8 | -0.72 | 0.2525 | 1.375 | -0.665 | 0.460352 | 0.698209 | 0.097879 | 1 | 0.999631 | 0.791962 |
| Wipf3_D3Z5R4_S161s_1 | 1.65 | -3.36 | -1.13 | 0.37 | 2.65 | 0.65 | 1.85 | 0.47 | -0.06 | -3.94 | 0.29 | -0.68 | -1.03 | -1.02 | -0.87 | -1.09 | 2.0225 | -0.48 | -0.385 | 0.678836 | 0.072825 | 0.6992 | 1 | 0.999631 | 0.887183 |
| Wipf3_D3Z5R4_S211s_1 | -0.21 | 0.47 | -0.43 | -0.71 | 0.08 | -0.06 | -0.91 | -0.67 | -0.37 | -0.19 | -0.95 | -0.38 | 0.53 | -0.05 | -0.47 | -0.29 | -0.17 | -0.2525 | 0.15 | 0.689008 | 0.644552 | 0.469272 | 1 | 0.999631 | 0.892221 |
| Wipf3_D3Z5R4_T106t_1 | 0 | 0.02 | 0.54 | -0.07 | 0.02 | 0.25 | -0.16 | 0.3 | -0.44 | -0.38 | 0.1 | 0.15 | 0.71 | -0.15 | 0.67 | 0.78 | -0.02 | -0.265 | 0.38 | 0.268469 | 0.940785 | 0.375773 | 1 | 0.999631 | 0.69658 |
| Wipf3_D3Z5R4_T240t_1 | -0.07 | 0.21 | 0.3 | 0.75 | 0.82 | 0.6 | 0.51 | 0.96 | -0.52 | -0.11 | 0.03 | -0.52 | -0.98 | 0.5 | 0.62 | 0.11 | 0.425 | -0.5775 | -0.235 | 0.576049 | 0.152436 | 0.077661 | 1 | 0.999631 | 0.846204 |
| Wipf3_D3Z5R4_T338t_1 | -0.49 | 0.63 | -0.49 | -0.68 | -0.08 | 0.63 | -1.16 | -0.41 | -0.3 | 0.49 | -0.84 | -0.95 | 0.54 | 0.97 | 0.77 | -1 | 0.0025 | -0.1425 | 0.5775 | 0.275368 | 0.995643 | 0.746497 | 1 | 0.999631 | 0.700187 |
| Wipf3_D3Z5R4_T357t_1 | 0.5 | 0.61 | -0.39 | 0.7 | 1.08 | 0.52 | 0.02 | 1.01 | 0.31 | 0.06 | 0.04 | 0.01 | -0.64 | 0.43 | 0.37 | 0.34 | 0.3025 | -0.25 | -0.23 | 0.558151 | 0.422393 | 0.442568 | 1 | 0.999631 | 0.838323 |
| Wipi2_Q80W47_S395s_1 | 0.17 | -0.17 | 0.17 | 0.26 | -0.19 | -0.09 | -0.25 | 0.33 | -0.04 | 0.15 | -0.03 | 0.72 | 0.19 | 0.8 | 0.13 | 0.31 | -0.1575 | 0.0925 | 0.25 | 0.403581 | 0.552855 | 0.749637 | 1 | 0.999631 | 0.764562 |
| Wnk1_P83741_S1037s_1 | -0.71 | 0.26 | 0.16 | -0.86 | 0.76 | -0.18 | 0.31 | 0.16 | 0.15 | -0.52 | 0.37 | 0.52 | 0.11 | 0.39 | -0.24 | 0.82 | 0.55 | 0.4175 | 0.5575 | 0.178498 | 0.158571 | 0.29656 | 1 | 0.999631 | 0.642903 |
| Wnk1_P83741_S165s_1 | 0.59 | 1.17 | 0.87 | 0.51 | 0.82 | 0.83 | 0.63 | 1.03 | 0.92 | -0.08 | 0.01 | 0.89 | 0.92 | 0.88 | 1.89 | 0.95 | 0.0425 | -0.35 | 0.375 | 0.293386 | 0.873428 | 0.329984 | 1 | 0.999631 | 0.707696 |
| Wnk1_P83741_S168s_1 | 0.99 | 1.28 | -0.04 | 2.65 | 3 | 1.53 | 1.21 | 1.88 | 1.1 | 0.16 | 2.19 | 0.55 | -0.69 | 1.67 | 1.33 | 0.63 | 0.685 | -0.22 | -0.485 | 0.4805 | 0.278008 | 0.7312 | 1 | 0.999631 | 0.801848 |
| Wnk1_P83741_S2222s_1 | -0.09 | 0.71 | 0.4 | -0.16 | 0.11 | 0.15 | -0.29 | 0.33 | 0.11 | 0.47 | 0.2 | 0.58 | 0.78 | 0.61 | 0.44 | 0.72 | -0.14 | 0.125 | 0.4225 | 0.195851 | 0.647119 | 0.683811 | 1 | 0.999631 | 0.653145 |
| Wnk1_P83741_S2273s_1 | -0.52 | -1.63 | -1.87 | -1.13 | -2.65 | -1.91 | -1.01 | -2.67 | -2.45 | -0.41 | -2.09 | -1.61 | -1.97 | -1.49 | -3.13 | -1.82 | -0.7725 | -0.3525 | -0.815 | 0.100759 | 0.124844 | 0.491936 | 1 | 0.999631 | 0.564448 |
| Wnk1_P83741_S2276s_1 | -0.19 | -0.87 | -0.61 | -0.65 | -1.86 | -1.25 | -0.65 | -0.71 | -2.18 | -0.33 | -1.51 | -0.97 | -1.29 | -0.47 | -1.03 | -0.65 | -0.5375 | -0.6675 | -0.28 | 0.388508 | 0.144865 | 0.137299 | 1 | 0.999631 | 0.757772 |
| Wnk1_P83741_S2616s_1 | 0.57 | 0.18 | 0.51 | 0.8 | 0.64 | 0.69 | 0.51 | 0.97 | 0.23 | -0.04 | 0.51 | 0.39 | -0.59 | 0.44 | 0.8 | 0.64 | 0.1875 | -0.2425 | -0.1925 | 0.611566 | 0.480861 | 0.390015 | 1 | 0.999631 | 0.859739 |
| Wnk1_P83741_T71t_1 | 2.01 | 0.04 | 0.95 | 1.92 | -0.41 | 1.13 | 1.52 | 0.92 | 0.79 | 1.6 | 0.12 | 0.22 | -0.79 | -0.45 | 0.99 | -0.55 | -0.44 | -0.5475 | -1.43 | 0.02843 | 0.447048 | 0.320324 | 1 | 0.999631 | 0.427072 |
| Wnk2_E9Q6Q2_S125s_1 | -0.48 | -0.22 | -1.72 | -1.58 | -1.22 | -0.75 | -1.36 | 0.59 | -0.76 | -0.44 | -1.88 | -1.23 | 0.32 | -0.67 | -0.45 | -0.34 | 0.315 | -0.0775 | 0.715 | 0.127639 | 0.562369 | 0.870192 | 1 | 0.999631 | 0.598244 |
| Wnk2_E9Q6Q2_S1456s_1 | 0.32 | -0.98 | -0.72 | 0.22 | -0.09 | -0.51 | 0.04 | 0.46 | -0.51 | -1.22 | -0.42 | -0.56 | -0.82 | -0.56 | -0.88 | 0.2 | 0.265 | -0.3875 | -0.225 | 0.597608 | 0.504258 | 0.335596 | 1 | 0.999631 | 0.855033 |
| Wnk2_E9Q6Q2_S1660s_1 | 0.32 | 0.14 | -0.41 | 0.4 | 0.82 | 0.65 | 0.23 | -0.26 | 0.38 | -0.21 | -0.03 | 0.37 | 0.17 | 0.28 | -0.52 | 0.87 | 0.2475 | 0.015 | 0.0875 | 0.816499 | 0.470829 | 0.960848 | 1 | 0.999631 | 0.937777 |
| Wnk2_E9Q6Q2_S1687s_1 | 0.79 | 0.43 | -1.16 | 0.84 | 0.57 | 0.09 | 0.68 | 0.61 | 0.66 | 0.24 | 0.82 | 0.01 | -0.49 | -0.02 | -0.17 | 0.64 | 0.2625 | 0.2075 | -0.235 | 0.64249 | 0.577407 | 0.66985 | 1 | 0.999631 | 0.871335 |
| Wnk2_E9Q6Q2_S988s_1 | -0.89 | 1.53 | 1.41 | -0.19 | 2.5 | 0.95 | 1.35 | 1.5 | 2.13 | -1.13 | 1.76 | 2.07 | 2.37 | -1 | 1.87 | 0.41 | 1.11 | 0.7425 | 0.4475 | 0.596569 | 0.095922 | 0.397275 | 1 | 0.999631 | 0.854508 |
| Wnk2_E9Q6Q2_S990s_1 | -0.61 | -0.68 | -0.09 | -0.38 | 0.62 | 0.98 | -0.43 | -0.53 | 0.56 | -0.6 | 0.15 | -0.94 | 1.4 | -0.58 | -0.93 | -0.65 | 0.6 | 0.2325 | 0.25 | 0.633337 | 0.159715 | 0.551111 | 1 | 0.999631 | 0.868569 |
| Wnk4_F6Z709_S18s_1 | 1.76 | 0.69 | 1.4 | 1.81 | 0.94 | 0.45 | 1.3 | 0.09 | 1.28 | 2.56 | 0.33 | 1.38 | -0.64 | -0.27 | 0.56 | -0.73 | -0.72 | -0.0275 | -1.685 | 0.00205 | 0.084868 | 0.955956 | 1 | 0.999631 | 0.351308 |
| Wrnip1_Q91XU0_S153s_1 | 1.32 | 1.65 | 1.38 | 1.29 | 1.11 | 1.19 | 0.89 | 1.04 | 1.39 | 1.69 | 1.04 | 0.81 | 0.6 | 1.49 | 0.79 | 1.17 | -0.3525 | -0.1775 | -0.3975 | 0.218024 | 0.167514 | 0.548701 | 1 | 0.999631 | 0.669859 |
| Wrnip1_Q91XU0_S465s_1 | 1.91 | 0.75 | 1.74 | 1.78 | 1.04 | 1.1 | 2.02 | 0.88 | 1.48 | 2.29 | 1.69 | 1.87 | 0.49 | 0.45 | 1.46 | 0.8 | -0.285 | 0.2875 | -0.745 | 0.078006 | 0.463657 | 0.425502 | 1 | 0.999631 | 0.543121 |
| Wrnip1_Q91XU0_S65s_1 | 1.56 | 1.61 | 1.98 | 1.59 | 1.03 | 1.56 | 1.31 | 1.45 | 1.51 | 2.21 | 1.25 | 0.75 | 0.41 | 1.26 | 1.46 | 1.11 | -0.3475 | -0.255 | -0.625 | 0.07913 | 0.199812 | 0.479287 | 1 | 0.999631 | 0.545579 |
| Wrnip1_Q91XU0_S75s_1 | 0.91 | 2.57 | 1.63 | 1.89 | 1.13 | 1.55 | 1.02 | 1.56 | 2.25 | 2.28 | 2.03 | 2.3 | 2.08 | 1.28 | 1.71 | 2.44 | -0.435 | 0.465 | 0.1275 | 0.767932 | 0.272108 | 0.23264 | 1 | 0.999631 | 0.917335 |
| Wtap_E0CYH0_S297s_1 | -2.49 | -0.69 | -0.12 | -0.66 | -1.15 | -1.76 | -0.54 | -0.62 | -0.61 | -2.76 | -1.08 | -0.85 | -0.6 | -3.04 | 0.7 | -0.1 | -0.0275 | -0.335 | 0.23 | 0.782234 | 0.959497 | 0.603519 | 1 | 0.999631 | 0.922325 |
| Wtip_Q7TQJ8_S29s_1 | 2.16 | 0.15 | 1.01 | 1.93 | 2.58 | 0.47 | 1.56 | -0.3 | 1.13 | 3.33 | 0.79 | 1.83 | -2.05 | 0.09 | 0.48 | -0.03 | -0.235 | 0.4575 | -1.69 | 0.025748 | 0.734424 | 0.489647 | 1 | 0.999631 | 0.422114 |
| Wwc1_Q5SXA9_S891s_1 | -0.27 | -1.28 | -1.2 | -0.55 | 0 | 0.14 | -0.18 | -0.71 | -0.12 | -0.38 | -0.36 | -0.01 | -0.1 | 0.03 | -0.03 | -0.54 | 0.6375 | 0.6075 | 0.665 | 0.075609 | 0.087516 | 0.083487 | 1 | 0.999631 | 0.541044 |
| Wwc2_Q6NXJ0_T999t_1 | -0.42 | 2.08 | 1.95 | 1.07 | -1.13 | 2.61 | 0.93 | 2.28 | -1.23 | -0.09 | 2.8 | -3.41 | -1.42 | 2.82 | -0.3 | 0.66 | 0.0025 | -1.6525 | -0.73 | 0.434559 | 0.997753 | 0.193795 | 1 | 0.999631 | 0.779717 |
| Wwp2_Q9DBH0_S211s_1 | 1.95 | -1.02 | 1.19 | 0.62 | -0.19 | 0.62 | 1.38 | -0.07 | -0.29 | 1.53 | -0.25 | -0.7 | -1.59 | -0.53 | 0.46 | -0.91 | -0.25 | -0.6125 | -1.3275 | 0.074267 | 0.701078 | 0.399508 | 1 | 0.999631 | 0.539897 |
| Xkr4_Q5GH67_S207s_1 | -0.41 | -0.35 | -0.66 | -1.12 | -0.35 | -0.79 | -1.05 | -0.89 | -0.76 | -1.11 | -1.22 | -1.09 | -0.73 | -0.37 | -3.85 | -0.47 | -0.135 | -0.41 | -0.72 | 0.348348 | 0.650498 | 0.177686 | 1 | 0.999631 | 0.736697 |
| Xkr4_Q5GH67_S520s_1 | -0.63 | 0.17 | -0.64 | -0.85 | -0.31 | -0.49 | -0.64 | -0.68 | -0.46 | -0.44 | -0.4 | -0.48 | 0.33 | -0.09 | -0.77 | -0.25 | -0.0425 | 0.0425 | 0.2925 | 0.431437 | 0.887964 | 0.887689 | 1 | 0.999631 | 0.778288 |
| Xkr4_Q5GH67_S522s_1 | -0.5 | 0.58 | -0.55 | -0.42 | 0.04 | -0.17 | -0.57 | 0.08 | 0.07 | -0.28 | -0.16 | -0.02 | 0.15 | 0.24 | -0.42 | -0.12 | 0.0675 | 0.125 | 0.185 | 0.607972 | 0.843359 | 0.706736 | 1 | 0.999631 | 0.858527 |
| Xkr4_Q5GH67_S87s_1 | 0.23 | 0.67 | 0.98 | 0.74 | 0.43 | 0.8 | 0.17 | 0.93 | 1.17 | 1.19 | 0.97 | 0.71 | 1.73 | 1.45 | 1.42 | 1.42 | -0.0725 | 0.355 | 0.85 | 0.014361 | 0.808073 | 0.230824 | 1 | 0.999631 | 0.385092 |
| Xkr4_Q5GH67_T208t_1 | -0.42 | -1.32 | -0.39 | -0.93 | -1.37 | -0.97 | -1.24 | -0.51 | -1.3 | -0.43 | -1.19 | -0.36 | -0.65 | -0.62 | -0.54 | -0.18 | -0.2575 | -0.055 | 0.2675 | 0.419459 | 0.445941 | 0.879739 | 1 | 0.999631 | 0.772152 |
| Xkr4_Q5GH67_T579t_1 | 0.81 | -0.44 | 0.88 | 1.11 | 0.4 | 0.76 | 0.38 | 1.08 | 0.07 | 1.1 | -0.72 | 0.38 | -0.45 | 0.94 | 1.19 | 0.92 | 0.065 | -0.3825 | 0.06 | 0.902762 | 0.868909 | 0.442709 | 1 | 0.999631 | 0.968183 |
| Xkr6_E9Q6C8_T106t_1 | 1.58 | 1.18 | 0.56 | -1.68 | 1.3 | 0.26 | 2.04 | -1.36 | 3.19 | 1.5 | 0.35 | 1.8 | 3.17 | -0.71 | -0.41 | -0.62 | 0.15 | 1.3 | -0.0525 | 0.958711 | 0.867097 | 0.136063 | 1 | 0.999631 | 0.98569 |
| Xkr6_E9Q6C8_T539t_1 | 0.81 | -1.35 | -0.41 | 0 | -1.31 | -0.57 | -0.39 | -1.05 | -0.81 | 0.66 | -0.92 | -0.85 | -0.71 | -1.17 | -0.58 | -0.93 | -0.5925 | -0.2425 | -0.61 | 0.20615 | 0.229191 | 0.658708 | 1 | 0.999631 | 0.6605 |
| Xkr6_E9Q6C8_T569t_1 | 1.43 | -0.67 | 1.04 | 0.37 | -0.73 | 0.82 | -0.01 | 0.51 | 0.27 | 1.73 | -0.52 | 0.64 | 2.26 | 1.83 | 0.88 | 0.53 | -0.395 | -0.0125 | 0.8325 | 0.16304 | 0.461449 | 0.983298 | 1 | 0.999631 | 0.631442 |
| Xkr7_Q5GH64_S154s_1 | -0.28 | 0 | -0.29 | -0.67 | -0.82 | -0.01 | -0.37 | -0.67 | -0.97 | 0.79 | -0.29 | -1.38 | 0.12 | -0.39 | -0.55 | -0.86 | -0.1575 | -0.1525 | -0.11 | 0.735689 | 0.594712 | 0.748855 | 1 | 0.999631 | 0.90759 |
| Xkr7_Q5GH64_T496t_1 | -0.53 | -0.08 | -0.36 | -0.57 | -0.48 | -0.47 | -0.77 | -0.61 | -0.87 | -0.38 | -0.98 | -0.37 | 0.26 | -0.46 | -0.55 | -0.4 | -0.1975 | -0.265 | 0.0975 | 0.753591 | 0.438918 | 0.364868 | 1 | 0.999631 | 0.913377 |
| Xpo4_Q9ESJ0_S521s_1 | 0.09 | 2 | 1.32 | -0.94 | 1.07 | 0.85 | 0.78 | 1.56 | 1.74 | 0.4 | 1.95 | 2.45 | 1.77 | 1.3 | 1.19 | 0.15 | 0.4475 | 1.0175 | 0.485 | 0.468162 | 0.46896 | 0.168703 | 1 | 0.999631 | 0.795565 |
| Xpr1_D3Z2B4_S600s_1 | 0.74 | 1.42 | 1.64 | 1.82 | 1.44 | 1.8 | 1.33 | 2.29 | 1.52 | 0.83 | 1.56 | 2 | -0.12 | 1.78 | 2.05 | 1.97 | 0.31 | 0.0725 | 0.015 | 0.977529 | 0.385422 | 0.843506 | 1 | 0.999631 | 0.992174 |
| Xrcc1_Q60596_S484s_1 | 0.07 | 3.38 | 2.3 | 0.38 | 2.07 | 1.91 | 1.18 | 1.55 | 2.49 | -0.51 | 2.28 | 2.74 | 2.25 | 1.32 | 1.86 | 1.73 | 0.145 | 0.2175 | 0.2575 | 0.720348 | 0.839938 | 0.818729 | 1 | 0.999631 | 0.904858 |
| Xrn1_F8VQ87_T1041t_1 | 0.32 | 1.86 | -0.31 | 1.27 | 0.67 | 2 | 1.48 | 2.42 | 1.94 | 0.59 | 1.68 | -0.21 | 0.03 | 1.2 | -0.07 | 0.15 | 0.8575 | 0.215 | -0.4575 | 0.398642 | 0.14979 | 0.732604 | 1 | 0.999631 | 0.761726 |
| Xrn2_Q9DBR1_S475s_1 | 0.14 | 1.15 | 0.23 | 0.38 | 0.59 | 0.39 | -0.3 | 0.15 | 0.45 | 0.31 | -0.07 | 0.36 | 0.69 | 1.03 | 0.39 | 0.29 | -0.2675 | -0.2125 | 0.125 | 0.718096 | 0.43471 | 0.509299 | 1 | 0.999631 | 0.903915 |
| Xrn2_Q9DBR1_S499s_1 | 0.06 | 0.87 | 1.14 | 0.77 | 1.11 | -0.26 | 0.52 | 0.82 | 0.5 | 0.47 | 1.04 | 1.26 | 0.8 | 0.95 | 2.41 | 1.42 | -0.1625 | 0.1075 | 0.685 | 0.139364 | 0.67441 | 0.756037 | 1 | 0.999631 | 0.611718 |
| Yaf2_Q99LW6_S173s_1 | 2.14 | 0.9 | 0.82 | 1.2 | 0.77 | 1.7 | 1.37 | 1.15 | 1.35 | 1.89 | 0.23 | 0.55 | -0.71 | 1.17 | 0.69 | 0.1 | -0.0175 | -0.26 | -0.9525 | 0.074928 | 0.962837 | 0.581731 | 1 | 0.999631 | 0.540556 |
| Yap1_G3UYA6_S94s_1 | -0.06 | -0.19 | -0.05 | -0.07 | -0.23 | -0.41 | -0.29 | -0.1 | -0.13 | 0.41 | -0.46 | 0.2 | 0.45 | 0.27 | 0.38 | -0.01 | -0.165 | 0.0975 | 0.365 | 0.194816 | 0.485795 | 0.732911 | 1 | 0.999631 | 0.652362 |
| Yars_Q91WQ3_S6s_1 | 0.58 | -0.29 | 0.64 | 0.56 | -0.39 | -0.35 | -0.37 | -0.88 | -0.78 | 0.45 | -0.48 | 0.09 | -0.04 | -0.32 | -0.81 | -0.16 | -0.87 | -0.5525 | -0.705 | 0.062076 | 0.017877 | 0.166504 | 1 | 0.999631 | 0.509355 |
| Ybx1_P62960_S163s_1 | 0.11 | 2.26 | 1.48 | 0.84 | 0.71 | 1.01 | 0.88 | 0.63 | 1.36 | 0.33 | 0.44 | 0.62 | 1.15 | 0.72 | 0.56 | -0.78 | -0.365 | -0.485 | -0.76 | 0.203151 | 0.425514 | 0.334845 | 1 | 0.999631 | 0.659781 |
| Ybx1_P62960_S172s_1 | 0.86 | 1.69 | 1.39 | 0.77 | 0.96 | 2.12 | 1.41 | 0.87 | 2.51 | 2.28 | 0.01 | 1.29 | 1.5 | 0.79 | 0.36 | -0.13 | 0.1625 | 0.345 | -0.5475 | 0.213578 | 0.666593 | 0.543694 | 1 | 0.999631 | 0.667146 |
| Ybx1_P62960_S174s_1 | 1.07 | 2.02 | 2.68 | 2.8 | 1.57 | 1.39 | 2.32 | 1.51 | 2.28 | 2.19 | 1.79 | 2.13 | 0.63 | 0.37 | 2.21 | 1.4 | -0.445 | -0.045 | -0.99 | 0.089743 | 0.322828 | 0.913789 | 1 | 0.999631 | 0.556851 |
| Ybx1_P62960_S312s_1 | -0.24 | -1.32 | -0.39 | -0.58 | -0.63 | -1.8 | -0.67 | -0.36 | -0.85 | 0.86 | -0.21 | -0.59 | 0.06 | 0.82 | -0.38 | 0.12 | -0.2325 | 0.435 | 0.7875 | 0.061252 | 0.566655 | 0.336397 | 1 | 0.999631 | 0.509355 |
| Ybx3_Q9JKB3_S259s_1 | 0.63 | 1.82 | 1.73 | 2.18 | 1.2 | 1.19 | 1.32 | 0.34 | 1.64 | 2.06 | 1.35 | 1.11 | 1.16 | 1.63 | 1.27 | 2.2 | -0.5775 | -0.05 | -0.025 | 0.952628 | 0.176719 | 0.901209 | 1 | 0.999631 | 0.984317 |
| Yeats2_Q3TUF7_S467s_1 | -1.07 | 1.22 | 1.07 | 0.05 | 0.33 | 0.12 | 0.08 | -0.37 | 0.73 | -0.82 | -0.34 | 0.66 | 0.82 | 0.35 | 0.29 | -0.15 | -0.2775 | -0.26 | 0.01 | 0.984987 | 0.591265 | 0.664023 | 1 | 0.999631 | 0.995133 |
| Yes1_Q04736_S26s_1 | -1.08 | -2.24 | -2.9 | -2.88 | -1.77 | -2.93 | -2.11 | -2.7 | -2.54 | -1.91 | -3.44 | -3.3 | -1.97 | -2.76 | -3.96 | -3.22 | -0.1025 | -0.5225 | -0.7025 | 0.223821 | 0.83004 | 0.329314 | 1 | 0.999631 | 0.67471 |
| Ythdc1_D6RG95_S321s_1 | 0.17 | 0.87 | 0.52 | 0.51 | 0.34 | 0.26 | -0.54 | 0.52 | 0.53 | 0.54 | 0.44 | 0.38 | 1.22 | 1.4 | 0.77 | 0.95 | -0.3725 | -0.045 | 0.5675 | 0.084756 | 0.263681 | 0.865282 | 1 | 0.999631 | 0.550457 |
| Ythdc2_B2RR83_S1104s_1 | 0.57 | 1.29 | 0.43 | -0.13 | 0.42 | 0.11 | -0.54 | 0.42 | 0.83 | 0.49 | 0.12 | 0.53 | 0.58 | 0.98 | 0.54 | 0.02 | -0.4375 | -0.0475 | -0.01 | 0.97928 | 0.26869 | 0.894926 | 1 | 0.999631 | 0.993455 |
| Ythdc2_B2RR83_S1105s_1 | 1.43 | 1.28 | 0.45 | 0.47 | 0.08 | 0.63 | -0.38 | 0.19 | 0.58 | 1.4 | 0.49 | 0.34 | 1.15 | -0.41 | -0.08 | 0.14 | -0.7775 | -0.205 | -0.7075 | 0.125692 | 0.051839 | 0.588931 | 1 | 0.999631 | 0.596745 |
| Ythdc2_B2RR83_S1284s_1 | -0.89 | 0.14 | 0.27 | 0.01 | -0.64 | -0.37 | -0.08 | -2.05 | -0.17 | -0.49 | 0.56 | 0.04 | -1.06 | 0.29 | -0.52 | 0.16 | -0.6675 | 0.1025 | -0.165 | 0.696808 | 0.188163 | 0.782312 | 1 | 0.999631 | 0.895731 |
| Ywhae_P62259_S65s_1 | 0.42 | 0.95 | -1.96 | -2.53 | 0.31 | 0.64 | -2.14 | -2.24 | 0.71 | 0.65 | -1.74 | -2.82 | 1.84 | 1.09 | -2.21 | -2.09 | -0.0775 | -0.02 | 0.4375 | 0.705544 | 0.937869 | 0.984947 | 1 | 0.999631 | 0.89781 |
| Ywhaz_P63101_S28s_1 | 0.76 | 0.3 | 0.2 | 0.24 | -0.84 | -0.34 | 1.26 | 0.25 | -0.4 | 0.94 | 0.66 | 0.92 | 0.2 | 0.43 | 0.4 | 0.87 | -0.2925 | 0.155 | 0.1 | 0.738688 | 0.522882 | 0.67641 | 1 | 0.999631 | 0.90804 |
| Ywhaz_P63101_S37s_1 | -0.08 | -1.3 | 0.06 | -1.17 | -1.07 | -1.4 | -0.11 | -0.43 | -0.89 | -0.56 | -0.84 | -0.25 | -0.94 | -0.95 | 0.43 | -0.18 | -0.13 | -0.0125 | 0.2125 | 0.656296 | 0.771462 | 0.974941 | 1 | 0.999631 | 0.879056 |
| Ywhaz_P63101_S64s_1 | 0.32 | 0.56 | 0.34 | -0.11 | -0.36 | 0.5 | -0.24 | -0.2 | -0.04 | 1 | -0.14 | 0.33 | 0.37 | 0.57 | 0.17 | 0.42 | -0.3525 | 0.01 | 0.105 | 0.715374 | 0.259577 | 0.976485 | 1 | 0.999631 | 0.902377 |
| Zbed3_Q9D0L1_S168s_1 | 1.81 | -0.48 | -0.71 | 2.48 | 1.97 | 1.81 | 2.01 | 1.33 | 1.86 | 0.71 | 1.03 | 0.06 | -3.56 | -0.16 | 0.02 | -0.56 | 1.005 | 0.14 | -1.84 | 0.088668 | 0.185839 | 0.8576 | 1 | 0.999631 | 0.553858 |
| Zbtb11_G5E8B9_S473s_1 | 0.58 | 2.27 | 3.56 | 0.93 | 0.68 | 2.59 | 2.74 | 0.68 | 1.11 | 2.02 | 3.09 | 3.99 | 1.26 | -0.25 | 2.55 | 1.34 | -0.1625 | 0.7175 | -0.61 | 0.440417 | 0.83484 | 0.386486 | 1 | 0.999631 | 0.781983 |
| Zbtb11_G5E8B9_S493s_1 | -0.4 | 0.63 | 1.65 | 0.21 | 0.57 | 0.91 | 0.72 | -0.58 | 1.52 | -1.16 | 0.92 | 1.08 | 0.92 | 0.06 | 1.89 | -0.05 | -0.1175 | 0.0675 | 0.1825 | 0.751036 | 0.817827 | 0.918579 | 1 | 0.999631 | 0.912766 |
| Zbtb16_Q3UQ17_S307s_1 | -1.1 | 1.98 | -0.32 | -1.22 | 0.18 | -1.54 | -0.98 | -1.64 | -0.16 | -0.32 | -1.12 | -0.64 | 1.91 | -0.09 | -0.85 | -0.75 | -0.83 | -0.395 | 0.22 | 0.795859 | 0.283082 | 0.569562 | 1 | 0.999631 | 0.929517 |
| Zbtb18_H7BX69_S223s_1 | 0.6 | -0.15 | 1.32 | 0.91 | 1.71 | 1.6 | -0.36 | 0.97 | -0.1 | 1.14 | 2.06 | 1.47 | 1.35 | 0.34 | 2.28 | 1.49 | 0.31 | 0.4725 | 0.695 | 0.175992 | 0.559195 | 0.372636 | 1 | 0.999631 | 0.642116 |
| Zbtb20_Q8K0L9_S359s_1 | -0.8 | 0.91 | 1.03 | 0.59 | -0.26 | -0.18 | 0.17 | 0.15 | 0.21 | 0.6 | -0.35 | 1.03 | 1.37 | 0.49 | 0.42 | 0.97 | -0.4625 | -0.06 | 0.38 | 0.424358 | 0.295057 | 0.902449 | 1 | 0.999631 | 0.773927 |
| Zbtb20_Q8K0L9_T232t_1 | 0.1 | 0.14 | 0.98 | 0.38 | -0.74 | -0.25 | 0.39 | 0.68 | -0.42 | -0.21 | -0.1 | 0.71 | -1.01 | -0.19 | 0.79 | 1 | -0.38 | -0.405 | -0.2525 | 0.608186 | 0.339736 | 0.271311 | 1 | 0.999631 | 0.858527 |
| Zbtb22_Q9Z0G7_S602s_1 | -0.02 | 1.26 | 0.98 | -0.11 | 1.02 | 0.07 | 1.07 | 0.84 | 1.61 | 0.75 | 0.98 | 1.24 | 1.08 | 1.48 | 1.38 | 1.01 | 0.2225 | 0.6175 | 0.71 | 0.095415 | 0.594894 | 0.150016 | 1 | 0.999631 | 0.558128 |
| Zbtb44_Q8R0A2_S161s_1 | -0.65 | 1.54 | 2.13 | 1.09 | 1.44 | 1.41 | 0.91 | 2.16 | 1.78 | -0.09 | 2.16 | 2.33 | 1.92 | 2.06 | 1.35 | 1.88 | 0.4525 | 0.5175 | 0.775 | 0.19471 | 0.450257 | 0.481215 | 1 | 0.999631 | 0.652222 |
| Zbtb45_Q52KG4_S435s_1 | -0.01 | -0.21 | -0.22 | 0.65 | 0.15 | -0.34 | -0.23 | 0.54 | 0.79 | 0.97 | -0.21 | 1.29 | 2.32 | 0.73 | 0.72 | 0.31 | -0.0225 | 0.6575 | 0.9675 | 0.065594 | 0.945361 | 0.122311 | 1 | 0.999631 | 0.516933 |
| Zbtb7a_D3Z4C3_S331s_1 | 2.17 | 0.04 | 1.26 | 0.65 | -0.67 | 1.24 | 2.92 | -1.37 | 0.38 | 2.91 | -0.34 | -0.51 | -0.2 | -1.78 | -0.34 | -1.92 | -0.5 | -0.42 | -2.09 | 0.005127 | 0.591439 | 0.602254 | 1 | 0.999631 | 0.351308 |
| Zc2hc1a_Q8BJH1_S222s_1 | 0.89 | 0.2 | 0.21 | 1.03 | 0.96 | 1.5 | 1.13 | 1.32 | 0.45 | 0.71 | 0.87 | 0.73 | -1.1 | 1.31 | 1.08 | 0.89 | 0.645 | 0.1075 | -0.0375 | 0.94602 | 0.058357 | 0.726793 | 1 | 0.999631 | 0.982217 |
| Zc2hc1a_Q8BJH1_S251s_1 | 0.44 | -0.15 | -0.13 | -0.07 | 0.22 | 1.16 | 0.13 | 0.93 | 0.31 | -0.41 | 0.04 | 0.8 | 0.69 | 0.91 | 1.39 | 0.23 | 0.5875 | 0.1625 | 0.7825 | 0.042243 | 0.101679 | 0.631566 | 1 | 0.999631 | 0.475443 |
| Zc2hc1a_Q8BJH1_T242t_1 | -0.39 | 1.06 | 1.32 | 0.67 | 0.7 | 1.17 | 0.31 | 1.24 | 0.82 | 0.9 | 1.03 | 1.29 | 0.6 | 1.5 | 1.62 | 1.56 | 0.19 | 0.345 | 0.655 | 0.165187 | 0.658222 | 0.399674 | 1 | 0.999631 | 0.633618 |
| Zc2hc1a_Q8BJH1_T243t_1 | -0.31 | 1.21 | 1.32 | 0.67 | 0.89 | 1.2 | 0.31 | 1.24 | 0.91 | 0.98 | 1.03 | 1.29 | 0.91 | 1.95 | 1.62 | 1.56 | 0.1875 | 0.33 | 0.7875 | 0.095522 | 0.659836 | 0.413119 | 1 | 0.999631 | 0.558128 |
| Zc3h12a_Q5D1E7_S394s_1 | 0.85 | -0.08 | 1.09 | 0.98 | -0.08 | 0.73 | 0.79 | 0.51 | 0.76 | 1.58 | 0.97 | 1.08 | -0.62 | 0.22 | 0.85 | 0.27 | -0.2225 | 0.3875 | -0.53 | 0.224051 | 0.538514 | 0.290721 | 1 | 0.999631 | 0.67471 |
| Zc3h12b_G3X9I7_S702s_1 | 1.15 | 0.93 | 1.13 | 0.18 | 0.23 | 1.18 | 0.73 | -0.1 | 0.44 | 0.75 | 0.48 | -0.11 | 0.72 | 0.7 | -0.22 | 0.43 | -0.3375 | -0.4575 | -0.44 | 0.243851 | 0.380736 | 0.196553 | 1 | 0.999631 | 0.685637 |
| Zc3h12c_E9Q1I3_S511s_1 | 0.35 | -0.14 | 0.65 | 0.45 | -0.23 | -0.25 | 0.31 | 0.2 | -0.1 | 0.29 | 0.45 | -0.42 | -0.29 | -0.82 | 0.33 | -0.25 | -0.32 | -0.2725 | -0.585 | 0.116549 | 0.288015 | 0.401694 | 1 | 0.999631 | 0.583887 |
| Zc3h13_E9Q784_S110s_1 | 0.85 | 0.9 | 2 | 1.21 | 0.29 | 1.27 | 1.29 | 1.08 | 1.07 | 1.85 | 1.26 | 0.61 | -0.05 | 0.89 | 1.6 | 1.4 | -0.2575 | -0.0425 | -0.28 | 0.539609 | 0.494641 | 0.91266 | 1 | 0.999631 | 0.83098 |
| Zc3h13_E9Q784_S1273s_1 | 1.05 | -0.43 | 0.61 | 0.59 | -0.53 | -0.71 | 0.52 | -0.03 | -0.71 | 1.35 | -0.98 | -0.84 | -1.4 | -1.32 | -0.64 | -2.85 | -0.6425 | -0.75 | -2.0075 | 0.003364 | 0.145876 | 0.216748 | 1 | 0.999631 | 0.351308 |
| Zc3h13_E9Q784_S1499s_1 | 0.72 | 0.34 | 2.68 | 2.32 | 0.71 | 0.61 | 2.07 | 0.38 | 0.84 | 0.84 | 2.08 | 1.07 | 0.89 | 0.88 | 1.84 | 1.34 | -0.5725 | -0.3075 | -0.2775 | 0.629942 | 0.36889 | 0.606676 | 1 | 0.999631 | 0.86722 |
| Zc3h13_E9Q784_S198s_1 | 0.13 | 0.21 | -0.03 | -0.24 | 0.13 | 0.3 | 0.13 | -0.62 | -0.23 | 0.5 | -0.04 | 0.89 | 0.54 | 0.93 | 1.28 | -0.69 | -0.0325 | 0.2625 | 0.4975 | 0.276784 | 0.912326 | 0.43124 | 1 | 0.999631 | 0.700187 |
| Zc3h13_E9Q784_S242s_1 | -0.4 | -0.45 | -0.27 | -0.48 | -0.53 | -1.05 | -0.64 | -0.3 | -0.7 | -0.31 | -0.86 | -0.12 | -0.19 | -0.85 | 0.1 | 0.02 | -0.23 | -0.0975 | 0.17 | 0.588717 | 0.392797 | 0.725933 | 1 | 0.999631 | 0.851127 |
| Zc3h13_E9Q784_S316s_3 | -0.92 | -0.43 | 0.06 | 0.37 | 0.06 | 0.97 | 0.64 | -0.01 | -0.2 | 1.4 | 0.57 | 0.47 | 0.74 | 0.63 | -0.89 | 0.55 | 0.645 | 0.79 | 0.4875 | 0.311493 | 0.115232 | 0.092342 | 1 | 0.999631 | 0.71924 |
| Zc3h13_E9Q784_S317s_3 | -0.92 | -0.43 | 0.06 | 0.37 | 0.06 | 0.97 | 0.64 | -0.01 | -0.2 | 1.4 | 0.57 | 0.47 | 0.74 | 0.63 | -0.89 | 0.55 | 0.645 | 0.79 | 0.4875 | 0.311493 | 0.115232 | 0.092342 | 1 | 0.999631 | 0.71924 |
| Zc3h13_E9Q784_S324s_3 | -0.92 | -0.43 | 0.06 | 0.37 | 0.06 | 0.97 | 0.64 | -0.01 | -0.2 | 1.4 | 0.57 | 0.47 | 0.74 | 0.63 | -0.89 | 0.55 | 0.645 | 0.79 | 0.4875 | 0.311493 | 0.115232 | 0.092342 | 1 | 0.999631 | 0.71924 |
| Zc3h13_E9Q784_S703s_1 | -0.86 | 0.35 | -0.39 | -0.63 | 0.25 | 0 | 0.29 | -0.41 | 1.03 | 0 | 0.23 | 0.61 | 0.99 | 0.46 | -0.07 | -0.21 | 0.415 | 0.85 | 0.675 | 0.117288 | 0.242791 | 0.043171 | 1 | 0.999631 | 0.585087 |
| Zc3h13_E9Q784_S704s_1 | -0.45 | 0.61 | 0.98 | 0.81 | 0.59 | 0.48 | 0.69 | 0.59 | 1.11 | 0.61 | 0.88 | 1.04 | 1.01 | 1.06 | 1.83 | 1.53 | 0.1 | 0.4225 | 0.87 | 0.051437 | 0.776864 | 0.268199 | 1 | 0.999631 | 0.489777 |
| Zc3h13_E9Q784_S77s_1 | 0.34 | 0.46 | 0.98 | 0.73 | 0.2 | 0.65 | 0.31 | 0.74 | 0.48 | 0.89 | 0.8 | 0.48 | 0.64 | 1.15 | 0.84 | 0.75 | -0.1525 | 0.035 | 0.2175 | 0.465877 | 0.585477 | 0.899814 | 1 | 0.999631 | 0.795249 |
| Zc3h13_E9Q784_S907s_1 | 1.62 | 0.64 | 1.72 | 0.9 | 0.1 | 0.97 | 1.83 | 0.47 | -0.48 | 1.45 | 0.12 | 0.04 | 0.21 | -0.73 | -0.17 | -0.87 | -0.3775 | -0.9375 | -1.61 | 0.002101 | 0.405531 | 0.071529 | 1 | 0.999631 | 0.351308 |
| Zc3h13_E9Q784_T1109t_1 | 0.65 | 1.27 | 0.75 | 0.74 | 0.8 | 0.81 | 0.67 | 0.34 | 0.59 | 1.01 | 0.8 | 0.94 | 1.12 | 1.63 | 1.02 | 1.08 | -0.1975 | -0.0175 | 0.36 | 0.250782 | 0.47151 | 0.948992 | 1 | 0.999631 | 0.686697 |
| Zc3h13_E9Q784_T363t_1 | -0.12 | 0.37 | 1.47 | 0.98 | 0.14 | 0.69 | 0.47 | 1.22 | -0.14 | 0.56 | 1.19 | 1.57 | 0.51 | 0.96 | 1.95 | 2.13 | -0.045 | 0.12 | 0.7125 | 0.17585 | 0.913306 | 0.806014 | 1 | 0.999631 | 0.642116 |
| Zc3h14_Q8BJ05_S309s_1 | -1.85 | 0.77 | 1.05 | 1.21 | 0.58 | 0.58 | 0.42 | 0.87 | 0.7 | 0.22 | 1.46 | 0.39 | 1.38 | 0.23 | 0.67 | 0.86 | 0.3175 | 0.3975 | 0.49 | 0.474937 | 0.626789 | 0.566881 | 1 | 0.999631 | 0.799827 |
| Zc3h14_Q8BJ05_S515s_1 | -0.36 | 0.58 | 0.57 | -0.41 | 0.21 | -0.16 | -0.44 | -0.16 | 0.22 | -0.11 | -0.51 | -0.16 | 0.86 | 1.13 | -0.27 | -0.29 | -0.2325 | -0.235 | 0.2625 | 0.570587 | 0.501554 | 0.51007 | 1 | 0.999631 | 0.84338 |
| Zc3h18_H3BIW0_S507s_1 | 1.23 | -0.22 | 1.11 | 0.84 | 0.02 | 0.37 | 0.47 | 0.09 | 0.24 | 1.51 | -0.25 | -0.22 | -0.22 | 0.4 | 0.25 | -0.14 | -0.5025 | -0.42 | -0.6675 | 0.11251 | 0.192601 | 0.410534 | 1 | 0.999631 | 0.577306 |
| Zc3h18_H3BIW0_S64s_1 | 1.85 | 1.51 | 2.57 | 1.77 | 0.9 | 1.94 | 2.1 | 0.57 | 1.7 | 2.65 | 0.78 | 0.96 | -0.83 | -0.73 | 0.9 | -0.34 | -0.5475 | -0.4025 | -2.175 | 0.000753 | 0.22466 | 0.39804 | 1 | 0.999631 | 0.351308 |
| Zc3h18_H3BIW0_S64s_2 | 1.19 | 1.02 | 1.38 | 0.72 | 0.11 | 0.66 | 0.76 | 0.69 | 1.25 | 0.87 | 1.25 | 1.56 | 0.6 | 0.6 | 1.45 | -0.51 | -0.5225 | 0.155 | -0.5425 | 0.228451 | 0.08976 | 0.59261 | 1 | 0.999631 | 0.679038 |
| Zc3h18_H3BIW0_S71s_2 | 0.51 | 0.8 | 1.38 | 0.72 | 0.15 | 0.93 | 0.76 | 0.69 | 1.21 | 0.5 | 1.25 | 1.56 | 0.25 | 0.22 | 1.45 | -0.51 | -0.22 | 0.2775 | -0.5 | 0.278663 | 0.480867 | 0.420157 | 1 | 0.999631 | 0.70098 |
| Zc3h18_H3BIW0_S75s_1 | -1.47 | 2.24 | 1.48 | -0.17 | 0.68 | 1.59 | 1.53 | -0.01 | 1.42 | -1.52 | 1.28 | 2.04 | 1.22 | 0.17 | 0.82 | -0.06 | 0.4275 | 0.285 | 0.0175 | 0.981913 | 0.595478 | 0.774388 | 1 | 0.999631 | 0.994846 |
| Zc3h18_H3BIW0_S80s_1 | -0.66 | 0.39 | 0.7 | -1.3 | -1.28 | -0.85 | -0.49 | -0.8 | -0.35 | -1.31 | -0.51 | -0.18 | -0.03 | -1.01 | -1.2 | -0.81 | -0.6375 | -0.37 | -0.545 | 0.295567 | 0.194975 | 0.465591 | 1 | 0.999631 | 0.708945 |
| Zc3h18_H3BIW0_S840s_1 | 1.41 | 1.45 | 1.73 | 0.98 | 0.53 | 1.77 | 1.86 | 0.18 | 0.67 | 1.1 | 0.24 | 0.47 | 0.49 | 0.11 | 1.13 | -0.41 | -0.3075 | -0.7725 | -1.0625 | 0.01988 | 0.492341 | 0.02958 | 1 | 0.999631 | 0.391668 |
| Zc3h18_H3BIW0_S849s_1 | 1.15 | 0.13 | 0.71 | 1.01 | 0.02 | 0.36 | 0.84 | 1.09 | 0.32 | 1.58 | 0.89 | -0.03 | 0.37 | 0.16 | 0.07 | 0.65 | -0.1725 | -0.06 | -0.4375 | 0.207601 | 0.629665 | 0.886717 | 1 | 0.999631 | 0.6614 |
| Zc3h18_H3BIW0_S890s_1 | 0.57 | -0.15 | 0.94 | 0.45 | -0.16 | 0 | 0.45 | 0.3 | 0.04 | 0.92 | 0.47 | 0.18 | -0.33 | 0.11 | 0.27 | 0.4 | -0.305 | -0.05 | -0.34 | 0.331152 | 0.34666 | 0.883845 | 1 | 0.999631 | 0.727302 |
| Zc3h3_Q8CHP0_S212s_1 | 0.94 | -0.43 | 1.25 | -0.05 | -1.69 | -0.43 | 1.64 | -0.54 | -0.28 | -0.15 | -0.43 | -0.16 | -1.89 | -2.29 | -1.7 | -1.36 | -0.6825 | -0.6825 | -2.2375 | 0.000524 | 0.345997 | 0.121523 | 1 | 0.999631 | 0.351308 |
| Zc3h4_E9Q8K8_S1221s_1 | 0.5 | 2 | 0.76 | 0.13 | 1.03 | 0.89 | 0.47 | 0.51 | 1.62 | 1.11 | 0.74 | 0.57 | 1.9 | 2.04 | 0.97 | 0.32 | -0.1225 | 0.1625 | 0.46 | 0.400823 | 0.772477 | 0.722379 | 1 | 0.999631 | 0.763209 |
| Zc3h4_E9Q8K8_S1227s_1 | -0.11 | 0.88 | 0.97 | 0.99 | 0.92 | 0.66 | -0.04 | 1.39 | 1.06 | 0.47 | 1.06 | 1.64 | 1.65 | 1.17 | 1.81 | 1.33 | 0.05 | 0.375 | 0.8075 | 0.043456 | 0.901099 | 0.335614 | 1 | 0.999631 | 0.475443 |
| Zc3h4_E9Q8K8_S868s_1 | 0.78 | 1 | -0.63 | -0.9 | 0.11 | 0.35 | -0.48 | -0.52 | 1.13 | 2.05 | -2.28 | -1.96 | 0.94 | 0.66 | -1.68 | -0.3 | -0.1975 | -0.3275 | -0.1575 | 0.816936 | 0.692112 | 0.750384 | 1 | 0.999631 | 0.937777 |
| Zc3h6_Q8BYK8_S1150s_1 | 0.81 | 1.35 | 2.25 | 1.16 | 0.56 | 1.68 | 0.68 | 1.73 | 1.21 | 1.47 | 1.68 | 1.47 | 0.86 | 1.83 | 1.02 | 1.58 | -0.23 | 0.065 | -0.07 | 0.862649 | 0.59613 | 0.855451 | 1 | 0.999631 | 0.954766 |
| Zc3h7b_F8VPP8_S372s_1 | 0.37 | -0.36 | -0.19 | 0.73 | -0.24 | -0.69 | 0.43 | 0.19 | -0.35 | 0.48 | -0.17 | 1.02 | -0.38 | 0.08 | 0.03 | 1.05 | -0.215 | 0.1075 | 0.0575 | 0.888892 | 0.565214 | 0.793427 | 1 | 0.999631 | 0.963505 |
| Zc3hc1_H3BKM2_S301s_1 | 1.26 | 2.26 | 2.26 | 1.86 | 1.65 | 1.97 | 1.68 | 1.16 | 2.33 | 2.23 | 1.97 | 2.01 | 1.67 | 1.87 | 1.78 | 2.29 | -0.295 | 0.225 | -0.0075 | 0.982319 | 0.382568 | 0.481107 | 1 | 0.999631 | 0.994846 |
| Zc3hc1_H3BKM2_S351s_1 | 0.08 | 0.34 | 0.01 | 0.34 | 0.01 | -0.69 | -0.36 | 0.23 | 0.41 | -0.42 | -0.01 | 0.29 | 0.26 | -0.28 | -0.5 | 0.18 | -0.395 | -0.125 | -0.2775 | 0.370891 | 0.196001 | 0.667953 | 1 | 0.999631 | 0.746338 |
| Zc3hc1_H3BKM2_S352s_1 | 0.14 | 0.57 | 0.01 | 0.34 | 0.27 | -0.32 | -0.36 | 0.23 | 0.47 | -0.03 | -0.01 | 0.29 | 0.39 | 0.42 | -0.5 | 0.18 | -0.31 | -0.085 | -0.1425 | 0.662583 | 0.292888 | 0.757927 | 1 | 0.999631 | 0.881311 |
| Zc3hc1_H3BKM2_S367s_1 | 0.22 | 1.35 | 2.28 | 0.09 | 0.56 | 1.29 | 1.6 | 0.42 | 1.83 | 0.75 | 0.86 | 2.27 | 0.36 | 0.44 | 1.09 | -0.5 | -0.0175 | 0.4425 | -0.6375 | 0.274539 | 0.974232 | 0.454294 | 1 | 0.999631 | 0.700187 |
| Zcchc6_E9PUA2_S705s_1 | 0.82 | 0.85 | 0.68 | 0.36 | 0.91 | -0.41 | 0.63 | 0.39 | 0.74 | 1.28 | 0.2 | -0.38 | 0.62 | 0.29 | 0.42 | -0.29 | -0.2975 | -0.2175 | -0.4175 | 0.204255 | 0.391612 | 0.580566 | 1 | 0.999631 | 0.659781 |
| Zdbf2_Q5SS00_S144s_1 | 1.42 | -0.97 | 0.31 | 2.06 | 1.15 | 0.53 | 1.49 | 2.13 | 0.8 | 0.01 | 0.53 | 1.86 | -1.18 | 0.81 | 1.1 | 1.32 | 0.62 | 0.095 | -0.1925 | 0.802528 | 0.360853 | 0.890107 | 1 | 0.999631 | 0.932677 |
| Zdhhc14_Q8BQQ1_S351s_1 | 1.44 | 0.23 | -0.05 | 0.51 | 0.83 | 0.17 | 0.21 | 0.64 | 0.31 | 1.02 | 0.24 | -0.24 | -0.77 | 0.41 | -0.64 | 0.73 | -0.07 | -0.2 | -0.6 | 0.230466 | 0.852658 | 0.634838 | 1 | 0.999631 | 0.681463 |
| Zdhhc14_Q8BQQ1_T19t_1 | -0.9 | -1.12 | -0.19 | -0.21 | -1.37 | -0.98 | -0.52 | -1.12 | -1.41 | -0.69 | 0.13 | 0.01 | -0.27 | -1.26 | -0.63 | -0.21 | -0.3925 | 0.115 | 0.0125 | 0.973535 | 0.259105 | 0.788553 | 1 | 0.999631 | 0.990676 |
| Zdhhc18_Q5Y5T2_S34s_1 | 0.25 | -0.77 | -1.5 | -0.94 | 0.02 | -1.18 | -0.39 | -0.37 | -0.53 | -0.72 | -0.24 | -0.94 | -1.68 | 0.03 | -0.4 | -1.19 | 0.26 | 0.1325 | -0.07 | 0.88987 | 0.552597 | 0.743708 | 1 | 0.999631 | 0.9636 |
| Zdhhc18_Q5Y5T2_S358s_1 | -0.41 | 0.14 | -0.38 | -1.17 | -0.19 | -0.26 | -1.35 | -1.11 | -0.15 | 0.17 | -0.94 | -0.53 | 0.26 | 0.19 | -0.58 | -0.11 | -0.2725 | 0.0925 | 0.395 | 0.302428 | 0.503606 | 0.808855 | 1 | 0.999631 | 0.714978 |
| Zdhhc23_B7ZNY9_S242s_1 | -0.44 | -0.33 | 1.16 | 1.05 | -0.59 | -0.24 | 0.8 | 1.13 | -0.32 | 0.11 | 1.58 | 2 | -0.92 | 0.41 | 1.67 | 1.83 | -0.085 | 0.4825 | 0.3875 | 0.575444 | 0.876459 | 0.456332 | 1 | 0.999631 | 0.846085 |
| Zdhhc5_Q8VDZ4_S372s_1 | -0.1 | -2.06 | -1.04 | -0.55 | -1.22 | -1.01 | -0.27 | -1.22 | -2.12 | -0.76 | -1.68 | -1.51 | -2.28 | -1.26 | -0.91 | -1.62 | 0.0075 | -0.58 | -0.58 | 0.256274 | 0.986932 | 0.249803 | 1 | 0.999631 | 0.691343 |

| | | | | | | | | | | | | | | | | | | | | | | | | | |
|---|---|---|---|---|---|---|---|---|---|---|---|---|---|---|---|---|---|---|---|---|---|---|---|---|---|
| Zdhhc5_Q8VDZ4_S380s_1 | -0.24 | -0.25 | -0.04 | -0.54 | -0.06 | -0.24 | -0.6 | 0.18 | -0.36 | -0.27 | -0.37 | 0.52 | 0.04 | 0.05 | 0.11 | -0.15 | 0.0875 | 0.1475 | 0.28 | 0.316113 | 0.753139 | 0.633462 | 1 | 0.999631 | 0.720734 |
| Zdhhc5_Q8VDZ4_S398s_1 | 0.11 | 1.22 | 0.92 | 0.6 | 0.5 | 1.36 | 0.04 | 0.74 | 0.68 | 0.82 | 0.81 | 0.99 | 0.85 | 1.38 | 0.86 | 1.5 | -0.0525 | 0.1125 | 0.435 | 0.231932 | 0.889436 | 0.719042 | 1 | 0.999631 | 0.682192 |
| Zdhhc5_Q8VDZ4_S409s_1 | -1.5 | -2.9 | -3.17 | -2.23 | -1.3 | -3.43 | -2.61 | -2.67 | -2.3 | -2.45 | -2.26 | -2.39 | -1.38 | -2.74 | -3.27 | -3.42 | -0.0525 | 0.1 | -0.2525 | 0.649813 | 0.921658 | 0.798806 | 1 | 0.999631 | 0.875258 |
| Zdhhc5_Q8VDZ4_S554s_1 | 0.84 | 1.54 | 1.89 | 0.81 | 0.64 | 1.79 | 0.74 | 1.36 | 1.17 | 2.09 | 1.17 | 1.07 | 0.95 | 2.06 | 1.49 | 1.87 | -0.1375 | 0.105 | 0.3225 | 0.418117 | 0.72464 | 0.782841 | 1 | 0.999631 | 0.771665 |
| Zdhhc5_Q8VDZ4_S621s_1 | -0.51 | 0.17 | -0.98 | -0.54 | 0.36 | -0.91 | -0.58 | -0.71 | 0.03 | -0.15 | -0.54 | -0.38 | -0.03 | -0.29 | -0.62 | -0.05 | 0.005 | 0.205 | 0.2175 | 0.526386 | 0.989527 | 0.532032 | 1 | 0.999631 | 0.824261 |
| Zdhhc5_Q8VDZ4_S694s_1 | -0.74 | -1.07 | -0.41 | -0.62 | -0.93 | -1.26 | -0.82 | -0.59 | -1.1 | -1.18 | -0.56 | -0.27 | -0.46 | -0.95 | -0.54 | -0.22 | -0.19 | -0.0675 | 0.1675 | 0.585703 | 0.500741 | 0.832347 | 1 | 0.999631 | 0.849524 |
| Zdhhc8_Q5Y5T5_S335s_1 | -1.02 | -1.31 | -0.94 | -0.78 | -0.92 | -0.83 | -1.02 | -0.83 | -0.95 | -1.15 | -0.8 | -0.73 | -0.19 | -0.95 | -1.09 | -0.17 | 0.1125 | 0.105 | 0.4125 | 0.238109 | 0.650259 | 0.691953 | 1 | 0.999631 | 0.683301 |
| Zdhhc8_Q5Y5T5_S672s_1 | -0.74 | -0.61 | -0.33 | -0.54 | -1.03 | -0.57 | -1.11 | -0.43 | -0.81 | -0.28 | -0.31 | -0.3 | 0.2 | -0.29 | -0.31 | 0.36 | -0.23 | 0.13 | 0.545 | 0.09216 | 0.411723 | 0.629419 | 1 | 0.999631 | 0.557085 |
| Zdhhc8_Q5Y5T5_S679s_1 | 2.13 | -2.03 | -0.44 | 1.89 | -0.2 | 0.38 | 2.08 | 0.49 | -1.07 | -0.6 | -0.14 | -1.18 | -1.79 | -2.07 | -0.55 | -0.88 | 0.3 | -1.135 | -1.71 | 0.084325 | 0.753611 | 0.221045 | 1 | 0.999631 | 0.549732 |
| Zfand2b_E0CY18_S163s_1 | 1.51 | 0.55 | -0.64 | 0.96 | 2.09 | 0.84 | 1.57 | 1.69 | 0.79 | -1.73 | 0.52 | 0 | -1.09 | 1.01 | -0.04 | 0.1 | 0.9525 | -0.7 | -0.6 | 0.31168 | 0.078456 | 0.298763 | 1 | 0.999631 | 0.71924 |
| Zfand2b_E0CY18_S173s_1 | 0.99 | -0.04 | -0.39 | 1.07 | 0.27 | 0.32 | 0.97 | 0.95 | 0.85 | 0.84 | 1.19 | 0.49 | 0.07 | 0.13 | 0.79 | 0.47 | 0.22 | 0.435 | -0.0425 | 0.918893 | 0.596818 | 0.296695 | 1 | 0.999631 | 0.974324 |
| Zfand2b_E0CY18_T185t_1 | 0.3 | -0.55 | -0.03 | -0.82 | 0.45 | 0.29 | -1 | -0.13 | -0.91 | -0.56 | -0.13 | 0.76 | -0.3 | 0.72 | -0.21 | 0.21 | 0.1775 | 0.065 | 0.38 | 0.329738 | 0.667096 | 0.881611 | 1 | 0.999631 | 0.726674 |
| Zfand5_O88878_S128s_1 | -1.4 | 0.31 | -0.4 | -0.45 | -0.92 | -0.27 | -0.28 | -0.65 | -0.35 | -0.32 | -1.02 | -0.67 | 0.3 | 0.15 | -1.26 | -1.42 | -0.045 | -0.105 | -0.0725 | 0.892622 | 0.90862 | 0.793026 | 1 | 0.999631 | 0.964461 |
| Zfand5_O88878_S48s_1 | 0.33 | 0.48 | -0.13 | -0.66 | -0.09 | 0.32 | -0.36 | -0.31 | 0.25 | 0.78 | -0.49 | 0.06 | 0.52 | 0.75 | -0.6 | 0.08 | -0.115 | 0.145 | 0.1825 | 0.658423 | 0.733508 | 0.708135 | 1 | 0.999631 | 0.879726 |
| Zfand5_O88878_S48s_2 | 1.19 | -0.19 | 0.98 | 0.57 | -0.22 | -1.43 | 0.51 | 0.53 | -0.96 | 0.57 | -0.65 | 1.11 | 1.73 | 0.32 | -0.92 | 1.39 | -0.79 | -0.62 | -0.0075 | 0.99017 | 0.147956 | 0.266321 | 1 | 0.999631 | 0.995892 |
| Zfand5_O88878_S58s_2 | 0.71 | 0.05 | 0.98 | 0.57 | 0.21 | -0.98 | 0.51 | 0.53 | -0.81 | 1.31 | -0.65 | 1.11 | 1.3 | -0.69 | -0.92 | 1.39 | -0.51 | -0.3375 | -0.3075 | 0.608063 | 0.229992 | 0.544477 | 1 | 0.999631 | 0.858527 |
| Zfml_E9QML5_S1571s_1 | -0.09 | -0.5 | 0.67 | 0.1 | 0.43 | -1.07 | -0.15 | -0.67 | 0.08 | -0.67 | 0.46 | -0.6 | 0.34 | 0.28 | 0.31 | 0.72 | -0.41 | -0.2275 | 0.3675 | 0.286502 | 0.327013 | 0.558214 | 1 | 0.999631 | 0.705972 |
| Zfml_E9QML5_S606s_1 | 2.03 | 2.1 | 2.54 | 1.56 | 0.41 | 2.69 | 1.2 | 1.4 | 1.79 | 2.51 | 1.43 | 0.91 | 1.19 | 1.69 | 1.35 | 1.88 | -0.6325 | -0.3975 | -0.53 | 0.131133 | 0.212515 | 0.336047 | 1 | 0.999631 | 0.602916 |
| Zfml_E9QML5_S881s_1 | -0.07 | 0.07 | 2.79 | 2.23 | -0.29 | 0.04 | 1.25 | 2.85 | -0.68 | 0.26 | 1.66 | 2.1 | 1.12 | 1.68 | 3.86 | 3.89 | -0.2925 | -0.42 | 1.3825 | 0.143385 | 0.741692 | 0.623354 | 1 | 0.999631 | 0.614578 |
| Zfp219_Q4VA98_S643s_1 | 1.4 | 1.42 | 1.68 | 0.85 | 1.76 | 1 | 0.6 | 0.91 | 1.85 | 1.99 | 0.1 | 1.29 | 1.12 | 0.97 | 1.81 | 1.18 | -0.27 | -0.03 | -0.0675 | 0.837604 | 0.432362 | 0.947251 | 1 | 0.999631 | 0.946036 |
| Zfp362_E9QA68_S145s_1 | -0.19 | 0.33 | 0.55 | 0.35 | 0.37 | -0.17 | -0.17 | 0.69 | 0.18 | -0.46 | 0.38 | 0.93 | -0.37 | 0.06 | 1.39 | 1 | -0.08 | -0.0025 | 0.26 | 0.558308 | 0.800172 | 0.994462 | 1 | 0.999631 | 0.838323 |
| Zfp37_Q6P3Z4_S586s_1 | 0.55 | 0.65 | 1.68 | -0.29 | 1.44 | 0.57 | 4.21 | 0.48 | 3.19 | 0.81 | 2.58 | 3.71 | 2.24 | -1.2 | 1.5 | -2.75 | 1.0275 | 1.925 | -0.7 | 0.509797 | 0.237747 | 0.016126 | 1 | 0.999631 | 0.817176 |
| Zfp385b_G5E8Y8_S185s_1 | -0.35 | -0.05 | 0.25 | 0.34 | -0.75 | -0.62 | 0.14 | -1.58 | 0.09 | 0.27 | 0.01 | 1 | -1.14 | 0.53 | 1.35 | -0.12 | -0.75 | 0.295 | 0.1075 | 0.836012 | 0.080823 | 0.379157 | 1 | 0.999631 | 0.945122 |
| Zfp407_G3UVV3_S221s_1 | 1.8 | -1.97 | 1.1 | 1.49 | -1.77 | -0.61 | 1.25 | -0.45 | -1.68 | 1.32 | -0.52 | -0.14 | -2.92 | -1.6 | -1 | -0.59 | -1 | -0.86 | -2.1325 | 0.032215 | 0.294346 | 0.364364 | 1 | 0.999631 | 0.442395 |
| Zfp532_S4R1H5_S293s_1 | -3.55 | -9.83 | -4.1 | -3.41 | -6.95 | -6.24 | -2.7 | -4.14 | -20.82 | -3.99 | -3.21 | -5.06 | -6.67 | -6.53 | -5.81 | -4.23 | 0.215 | -3.0475 | -0.5875 | 0.671846 | 0.889336 | 0.426456 | 1 | 0.999631 | 0.884779 |
| Zfp532_S4R1H5_S297s_1 | -0.24 | -0.14 | 0.08 | -0.01 | -0.49 | -0.68 | -0.2 | 0.4 | -0.43 | 0.24 | 0.26 | 0.35 | 0.46 | -0.16 | 0.14 | 0.43 | -0.165 | 0.1825 | 0.295 | 0.316565 | 0.591413 | 0.526216 | 1 | 0.999631 | 0.720734 |
| Zfp532_S4R1H5_S306s_1 | 0.96 | -0.02 | 0.78 | -0.49 | -1.33 | 0.29 | 1.4 | -1.68 | -1.69 | 0.61 | -0.7 | -1 | -0.38 | -1.68 | -0.46 | -0.79 | -0.6375 | -1.0025 | -1.135 | 0.028576 | 0.375257 | 0.092103 | 1 | 0.999631 | 0.427571 |
| Zfp532_S4R1H5_S333s_1 | 0.16 | -0.05 | -0.37 | -0.01 | -1.06 | -0.38 | -0.21 | -0.09 | -0.5 | 0.25 | 0.38 | -0.24 | 0.62 | -0.26 | -0.42 | 0.63 | -0.3675 | 0.04 | 0.21 | 0.557637 | 0.243994 | 0.895816 | 1 | 0.999631 | 0.838305 |
| Zfp532_S4R1H5_S349s_1 | 0.63 | 1.72 | 0.93 | 0.4 | 0.97 | 1.47 | -0.07 | 0.7 | 0.57 | 0.57 | 0.28 | 0.32 | 2.28 | 1.32 | 0.15 | 0.49 | -0.1525 | -0.485 | 0.14 | 0.78956 | 0.720657 | 0.178681 | 1 | 0.999631 | 0.925593 |
| Zfp644_Q3UMJ0_S376s_1 | 0.89 | 0.28 | 2.13 | 1.4 | 0.35 | 0.66 | 1.38 | 0.44 | 0.26 | 1.93 | 0.95 | 0.47 | 0.2 | -0.25 | 0.95 | -0.2 | -0.4675 | -0.2725 | -1 | 0.055741 | 0.305773 | 0.595691 | 1 | 0.999631 | 0.495418 |
| Zfp646_Q6NV66_S601s_1 | 0.85 | -1.39 | 0.15 | 0.52 | -0.27 | -0.42 | 0.85 | -0.21 | -0.32 | 0.36 | -0.34 | -0.85 | -0.41 | -1.34 | -0.99 | -1.16 | -0.045 | -0.32 | -1.0075 | 0.071374 | 0.932406 | 0.542393 | 1 | 0.999631 | 0.531405 |
| Zfp651_E9PZ11_T406t_1 | 1.44 | 1.71 | 1.7 | 1.53 | 1.21 | 1.25 | 1.32 | 1.67 | 1.6 | 2.24 | 1.6 | 1.29 | 0.66 | 0.87 | 1.16 | 1.66 | -0.2325 | 0.0875 | -0.5075 | 0.130795 | 0.360328 | 0.765764 | 1 | 0.999631 | 0.602177 |
| Zfp704_Q9ERQ3_S521s_1 | -1.38 | 0.92 | 0.96 | -2.12 | 1.87 | -0.44 | 2.13 | -0.25 | 2.16 | -1.32 | 1.44 | 1.47 | 2.5 | -1.06 | 0.15 | -2.65 | 1.2325 | 1.3425 | 0.14 | 0.902165 | 0.193523 | 0.18241 | 1 | 0.999631 | 0.968183 |
| Zfp740_D3Z606_S19s_1 | 0.32 | 0.76 | 0.68 | 0.52 | -0.07 | -0.07 | -0.09 | 0.33 | 0.64 | 0.77 | 0.48 | 0.66 | 0.67 | 1.09 | 0.32 | 0.69 | -0.545 | 0.0675 | 0.1225 | 0.68027 | 0.054618 | 0.79012 | 1 | 0.999631 | 0.888282 |
| Zfp777_G5E8L5_S615s_1 | 1.36 | -0.15 | 0.81 | 0.42 | 0.21 | -0.24 | 0.64 | 0.36 | -0.24 | 1.28 | -0.24 | -0.43 | -0.31 | -0.22 | -0.63 | -0.99 | -0.3675 | -0.5175 | -1.1475 | 0.014129 | 0.346442 | 0.302845 | 1 | 0.999631 | 0.385092 |
| Zfpm1_O35615_S681s_1 | 0.36 | 0.9 | 2.16 | 1.79 | 0.06 | 1.11 | 1.41 | 1.87 | 0.43 | 0.27 | 0.6 | 0.94 | 0.69 | 0.35 | 1.73 | 1.04 | -0.19 | -0.7425 | -0.35 | 0.479221 | 0.716518 | 0.111432 | 1 | 0.999631 | 0.801052 |
| Zfpm2_Q8CCH7_S1057s_1 | -0.1 | -0.04 | 2.21 | 0.91 | 0.58 | 0.97 | 0.58 | 1.62 | 0.34 | 0.53 | 1.46 | 1.71 | 1.35 | 1.99 | 1.67 | 1.59 | 0.1925 | 0.265 | 0.905 | 0.108838 | 0.724871 | 0.651423 | 1 | 0.999631 | 0.571276 |
| Zfr_O88532_S1054s_1 | -0.23 | 1 | 1.29 | 0.7 | 0.61 | 1.06 | 0.37 | 0.94 | 0.55 | 1.04 | 0.56 | 0.7 | 1.23 | 0.66 | 0.81 | 1.29 | 0.055 | 0.0225 | 0.3075 | 0.440862 | 0.884586 | 0.952006 | 1 | 0.999631 | 0.781983 |
| Zfr2_E9Q5M4_S130s_1 | 0.58 | 1.76 | 1.95 | 1.4 | 1.28 | 1.43 | 0.74 | 2.15 | -1.84 | -1.32 | -2.16 | -2.02 | -1.93 | -2.15 | -2.04 | -1.61 | -0.0225 | -3.2575 | -3.355 | 4.67E-06 | 0.956932 | 9.05E-06 | 1 | 0.09263 | 0.047748 |
| Zfyve1_Q810J8_S81s_1 | 0.19 | -0.98 | -0.01 | 0.15 | -1.55 | -0.93 | 0.7 | 0.22 | -0.73 | 1.05 | -0.64 | -0.22 | -1.07 | -2.3 | 0 | -0.83 | -0.2275 | 0.0275 | -0.8875 | 0.112099 | 0.674407 | 0.953884 | 1 | 0.999631 | 0.577068 |
| Zfyve19_Q9DAZ9_S219s_1 | 0.35 | 0.63 | 1.82 | 0.26 | 0.18 | 0.47 | 0.2 | 1.07 | 0.63 | 0.46 | 0.72 | 1.02 | 0.21 | 0.83 | 0.78 | 1.35 | -0.285 | -0.0575 | 0.0275 | 0.949562 | 0.496705 | 0.883977 | 1 | 0.999631 | 0.983798 |
| Zfyve19_Q9DAZ9_S94s_1 | 2.74 | -0.62 | -1.26 | 1.46 | 2.19 | 2.26 | 1.74 | 1.32 | -1.41 | 0.38 | -0.39 | -1.49 | 0.63 | 0.78 | -0.42 | -1.27 | 1.2975 | -1.3075 | -0.65 | 0.475392 | 0.14101 | 0.166205 | 1 | 0.999631 | 0.799882 |
| Zfyve19_Q9DAZ9_T377t_1 | 1.43 | 2.38 | 1.89 | -0.92 | 1.31 | 2.35 | 2.84 | -0.12 | 1.52 | 2.84 | 1.67 | 1.63 | 1.37 | 2.35 | 0.05 | -2.13 | 0.4 | 0.72 | -0.785 | 0.455096 | 0.640774 | 0.32153 | 1 | 0.999631 | 0.790041 |
| Zfyve26_Q5DU37_S605s_2 | -1.47 | 0.19 | 0.73 | 0.19 | 0.09 | -0.69 | 0.05 | 0.24 | -1.09 | -1.18 | 1.26 | 0.19 | -0.7 | 0.2 | 0.26 | 1.05 | 0.0125 | -0.115 | 0.2925 | 0.599998 | 0.979651 | 0.864166 | 1 | 0.999631 | 0.856618 |
| Zfyve28_Q6ZPK7_S686s_1 | -0.51 | -0.28 | 0.72 | 0.03 | -0.43 | -0.47 | -0.06 | 0.51 | -0.5 | -0.01 | 0.16 | 0.25 | 0.27 | -0.33 | 0.36 | 0.57 | -0.1025 | -0.015 | 0.2275 | 0.544189 | 0.781754 | 0.96606 | 1 | 0.999631 | 0.833226 |
| Zkscan1_Q8BGS3_S13s_1 | 0.48 | 0.51 | 0.87 | 0.45 | 1.45 | 0.69 | 0.28 | 1.29 | 1.18 | 1.6 | 1.12 | 0.86 | 1.72 | 0.05 | 1.71 | 1.08 | 0.35 | 0.6125 | 0.5625 | 0.199736 | 0.302819 | 0.050472 | 1 | 0.999631 | 0.657804 |
| Zmym2_Q9CU65_S837s_1 | 1.42 | -0.34 | 0.17 | 0.15 | 1.01 | -0.75 | 0.55 | 0.44 | 0.03 | -0.22 | 0.02 | -0.6 | -0.62 | -0.39 | 0.15 | 0.06 | -0.0375 | -0.5425 | -0.55 | 0.217544 | 0.939939 | 0.20738 | 1 | 0.999631 | 0.669718 |
| Zmynd8_Q3UH28_S407s_2 | -1.19 | -1.17 | 0.03 | -0.21 | -1.44 | -2.11 | -0.41 | -1.03 | -1.82 | -0.95 | -0.17 | -0.17 | -1.11 | -1.52 | -0.38 | -0.1 | -0.6125 | -0.1425 | -0.1425 | 0.753752 | 0.203066 | 0.768958 | 1 | 0.999631 | 0.913377 |
| Zmynd8_Q3UH28_S419s_2 | -1.19 | -1.17 | 0.03 | -0.21 | -1.44 | -2.11 | -0.41 | -1.03 | -1.82 | -0.95 | -0.17 | -0.17 | -1.11 | -1.52 | -0.38 | -0.1 | -0.6125 | -0.1425 | -0.1425 | 0.753752 | 0.203066 | 0.768958 | 1 | 0.999631 | 0.913377 |
| Zmynd8_Q3UH28_S465s_1 | 1.5 | 1.74 | 1.67 | 1.52 | 0.22 | 1.14 | 1.35 | 0.42 | 2.07 | 1.9 | 1.1 | 1.61 | 2.8 | 1.42 | 0.11 | 0.55 | -0.825 | 0.0625 | -0.3875 | 0.489268 | 0.027755 | 0.834069 | 1 | 0.999631 | 0.807292 |
| Zmynd8_Q3UH28_S524s_1 | -0.54 | 0.17 | -1.54 | -1.02 | -0.04 | -0.24 | -1.14 | -1.13 | -0.18 | -0.06 | -0.63 | -0.52 | -0.16 | 0.23 | -0.9 | -0.69 | 0.095 | 0.385 | 0.3525 | 0.436369 | 0.8327 | 0.348074 | 1 | 0.999631 | 0.780748 |
| Zmynd8_Q3UH28_S629s_1 | -0.59 | -2.22 | -0.12 | -0.86 | -1.09 | -0.29 | -0.46 | -0.56 | -1.55 | -0.04 | -0.09 | 0.06 | -0.99 | -1.67 | 0.11 | 0.15 | 0.3475 | 0.5425 | 0.3475 | 0.553213 | 0.459146 | 0.33535 | 1 | 0.999631 | 0.837597 |
| Zmynd8_Q3UH28_S734s_1 | -0.85 | -1.2 | -0.47 | -0.96 | -1.12 | -1.51 | -1.25 | -0.61 | -0.91 | -0.72 | -0.91 | -0.51 | 0.2 | -0.34 | -1.09 | -0.47 | -0.2525 | 0.1075 | 0.445 | 0.231247 | 0.413531 | 0.700477 | 1 | 0.999631 | 0.681463 |
| Znf148_Q61624_S665s_1 | 0.54 | 1.77 | 1.73 | 1.13 | 0.62 | 1.17 | 0.63 | 1.18 | 1.19 | 1.19 | 1.13 | 0.46 | 0.39 | 1.1 | 1.16 | 1.08 | -0.3925 | -0.3 | -0.36 | 0.353857 | 0.286436 | 0.424243 | 1 | 0.999631 | 0.739673 |
| Znf148_Q61624_S784s_1 | -1.36 | -0.95 | -1.5 | -2.63 | -1.5 | -0.92 | -1.46 | -2.18 | -0.35 | -0.74 | -1.87 | -3.35 | -0.56 | -0.3 | -1.49 | -2 | 0.095 | 0.0325 | 0.5225 | 0.318527 | 0.826482 | 0.961941 | 1 | 0.999631 | 0.722158 |
| Znf318_Q99PP2_S1591s_1 | 0.25 | 1.94 | 1.21 | 0.94 | 0.62 | 1.47 | 0.99 | 1.32 | 2.53 | 1.54 | 1.89 | 1.93 | -0.19 | 0.54 | 1.05 | 1.41 | 0.015 | 0.8875 | -0.3825 | 0.432036 | 0.970134 | 0.053868 | 1 | 0.999631 | 0.778808 |
| Znf318_Q99PP2_S2140s_1 | -0.33 | 0.65 | 1.1 | 0.89 | 0.17 | -0.38 | -0.02 | 0.52 | 0.1 | -0.32 | 0.4 | 1.51 | 0.29 | 0.65 | 0.95 | 1.22 | -0.505 | -0.155 | 0.2 | 0.6183 | 0.205295 | 0.748429 | 1 | 0.999631 | 0.861888 |
| Znf318_Q99PP2_S2206s_1 | -0.22 | 2.16 | 1.87 | 1.75 | 1.66 | 1.42 | 1.36 | 1.26 | 1.59 | 0.78 | 1.84 | 1.83 | 2 | 2.01 | 2.17 | 2.21 | 0.035 | 0.12 | 0.7075 | 0.192225 | 0.945518 | 0.828484 | 1 | 0.999631 | 0.648844 |
| Znf318_Q99PP2_S41s_1 | 0.68 | -0.16 | 1.5 | 0.72 | 0.28 | -0.19 | 0.39 | 1.45 | 0.57 | -0.03 | 0.73 | 0.98 | 1.35 | 0.51 | 0.85 | 1.18 | -0.2025 | -0.1225 | 0.2875 | 0.485561 | 0.663788 | 0.765173 | 1 | 0.999631 | 0.804287 |
| Znf318_Q99PP2_S79s_2 | 0.94 | 0.91 | 1.31 | 1.1 | 0.9 | 0.86 | 1.1 | 0.47 | 1.24 | 1.75 | 0.75 | 0.59 | 0.49 | -0.11 | 0.17 | 0.78 | -0.2325 | 0.0175 | -0.7325 | 0.036541 | 0.385592 | 0.957664 | 1 | 0.999631 | 0.453963 |
| Znf318_Q99PP2_S81s_2 | 0.94 | 0.91 | 1.31 | 1.1 | 0.9 | 0.86 | 1.1 | 0.47 | 1.24 | 1.75 | 0.75 | 0.59 | 0.49 | -0.11 | 0.17 | 0.78 | -0.2325 | 0.0175 | -0.7325 | 0.036541 | 0.385592 | 0.957664 | 1 | 0.999631 | 0.453963 |
| Znf365_Q8BG89_S146s_1 | -0.88 | -1.34 | -0.82 | -0.82 | -0.74 | -1.05 | -0.89 | -0.87 | -0.88 | -0.97 | -0.61 | -0.75 | 0.03 | -1.01 | -1.2 | -0.09 | 0.0775 | 0.1625 | 0.3975 | 0.305063 | 0.761144 | 0.544268 | 1 | 0.999631 | 0.716239 |
| Znf365_Q8BG89_S329s_1 | 0.32 | 1.19 | 0.71 | -0.09 | 2.27 | 1.59 | 2.05 | 1.02 | 2.05 | -0.46 | 1.14 | 0.62 | 0.4 | -0.44 | 0.67 | -1.01 | 1.2 | 0.305 | -0.6275 | 0.19785 | 0.01211 | 0.580817 | 1 | 0.999631 | 0.655274 |
| Znf365_Q8BG89_T206t_1 | -1.32 | -0.38 | -0.87 | -0.99 | -0.48 | -0.72 | -0.92 | -0.93 | -1.12 | -1.66 | -0.09 | -1.75 | -0.86 | -0.24 | -1.64 | -1.34 | 0.1275 | -0.265 | -0.13 | 0.740479 | 0.663596 | 0.540078 | 1 | 0.999631 | 0.90883 |
| Znf385a_Q8VD12_S133s_1 | -0.33 | -0.58 | 1.32 | 0.62 | -0.21 | -0.24 | 0.52 | 1.03 | 0.27 | -0.17 | 1.1 | 2.3 | -0.08 | 0.09 | 1.75 | 1.99 | 0.0175 | 0.6175 | 0.68 | 0.294407 | 0.972131 | 0.339926 | 1 | 0.999631 | 0.708293 |
| Znf385a_Q8VD12_S140s_1 | 0.09 | 0.73 | 0.6 | -0.12 | 0.09 | 0 | -0.4 | 1.16 | -0.09 | 0.2 | -0.19 | 1.12 | 1.12 | 0.41 | 1.18 | 0.99 | -0.1125 | -0.065 | 0.6 | 0.098866 | 0.776706 | 0.864851 | 1 | 0.999631 | 0.562298 |
| Znf451_Q8C0P7_S1032s_1 | 0.98 | 2.23 | 2.41 | 0.71 | 0.87 | 2.75 | 1.17 | 1.57 | 2.25 | 2.77 | 1.51 | 1.04 | 0.91 | 2.35 | 2.21 | 2.56 | 0.0075 | 0.31 | 0.425 | 0.434024 | 0.989066 | 0.567589 | 1 | 0.999631 | 0.779717 |
| Znf592_Q8BHZ4_S332s_1 | 0.37 | 0.24 | 1.32 | 0.22 | -0.31 | -0.62 | 0.32 | -0.04 | -0.01 | 0.5 | 0.34 | 0.29 | 0.85 | 0.24 | -0.31 | 0.43 | -0.7 | -0.2575 | -0.235 | 0.548345 | 0.073465 | 0.447337 | 1 | 0.999631 | 0.834633 |
| Znf592_Q8BHZ4_S346s_1 | 0 | -0.58 | -0.5 | -0.71 | -1 | -0.75 | -0.98 | -0.58 | -0.79 | 1.23 | -0.68 | -1.06 | -0.63 | -0.02 | -1.06 | -1.08 | -0.38 | 0.1225 | -0.25 | 0.481055 | 0.186026 | 0.812062 | 1 | 0.999631 | 0.801848 |
| Znf609_Q8BZ47_S413s_1 | 1.34 | 1.8 | -0.43 | 0.69 | 2.24 | 0.63 | 0.8 | 0.3 | 0.97 | -0.16 | 0.88 | -0.94 | 2.05 | -1.02 | -0.33 | 0.9 | 0.1425 | -0.6625 | -0.45 | 0.543551 | 0.808051 | 0.288305 | 1 | 0.999631 | 0.832881 |
| Znf609_Q8BZ47_S491s_1 | -0.13 | 1.59 | 1.94 | 0.07 | 0.37 | 1.07 | 0.63 | 0.56 | 0.62 | 1.57 | 0.64 | 0.82 | 1.01 | 1.08 | 0.32 | 0.53 | -0.21 | 0.045 | -0.1325 | 0.800722 | 0.681021 | 0.9326 | 1 | 0.999631 | 0.931985 |
| Znf622_Q91VY9_S145s_1 | 2.47 | 0.02 | 2.47 | 1.81 | -0.27 | 0.9 | 2.66 | 0.66 | -0.83 | 1.42 | 0.76 | 0.74 | -0.62 | -0.91 | 1.96 | 1.19 | -0.705 | -1.17 | -1.2875 | 0.127196 | 0.3529 | 0.10667 | 1 | 0.999631 | 0.59788 |
| Znf687_Q9D2D7_S228s_1 | -0.64 | -2.43 | -1.85 | -1.72 | -2.02 | -1.19 | -1 | -1.68 | -1.57 | -1 | -1.66 | -1.59 | -1.52 | -1.45 | -2.65 | -2.88 | 0.1875 | 0.205 | -0.465 | 0.366727 | 0.664993 | 0.620032 | 1 | 0.999631 | 0.745796 |
| Znf750_Q8BH05_S373s_1 | 1.19 | 1.02 | 1.29 | 1.01 | 0.84 | 1.48 | 0.41 | 1.56 | 1.2 | 1.49 | 0.98 | 0.63 | 0.84 | 1.37 | 1 | 0.82 | -0.055 | -0.0525 | -0.12 | 0.67005 | 0.865945 | 0.853926 | 1 | 0.999631 | 0.884054 |
| Znf830_Q8R1N0_S216s_1 | -0.99 | 0.06 | 2.06 | 1.26 | -0.25 | 0.03 | -0.43 | 1.93 | 1.11 | -0.16 | 2.38 | 3.26 | 0.44 | 0.59 | 3.06 | 2.95 | -0.2775 | 1.05 | 1.1625 | 0.193072 | 0.71502 | 0.247674 | 1 | 0.999631 | 0.649713 |
| Znrf2_Q71FD5_S18s_1 | 0.21 | 1.11 | -0.23 | -0.69 | 0.34 | -0.23 | -0.39 | -0.48 | 0.5 | 0.55 | -1.47 | -1.14 | 0.4 | 1.07 | 0.54 | 0.21 | -0.29 | -0.49 | 0.455 | 0.306855 | 0.496212 | 0.421143 | 1 | 0.999631 | 0.717667 |
| Znrf2_Q71FD5_S75s_1 | 0.28 | 0.37 | 0.17 | -0.53 | 0.05 | 0.32 | 0.07 | -0.22 | 0.64 | 0.58 | 0.32 | 0.62 | 0.82 | 0.24 | 0.92 | 0.44 | -0.0175 | 0.4675 | 0.5325 | 0.132256 | 0.953093 | 0.139599 | 1 | 0.999631 | 0.604009 |
| Zranb2_E9PUD0_S106s_1 | -0.02 | 1.26 | 0.39 | 1.78 | 0.99 | 1.18 | 0.65 | 1.57 | 0.88 | 0.34 | 1.75 | 0.71 | -0.44 | 1.49 | -0.06 | 2.22 | 0.245 | 0.0675 | -0.05 | 0.940512 | 0.580162 | 0.889136 | 1 | 0.999631 | 0.981146 |
| Zranb2_E9PUD0_S116s_1 | -0.96 | 1.59 | 0.73 | 1.11 | 0.74 | 0.27 | -0.22 | 0.94 | 0.72 | 0.34 | 0.38 | 0.67 | 1.77 | 1.53 | 0.79 | 1.42 | -0.185 | -0.09 | 0.76 | 0.188915 | 0.741538 | 0.864363 | 1 | 0.999631 | 0.648465 |
| Zranb2_E9PUD0_S151s_1 | -0.53 | 0.52 | 1.18 | 0.45 | 0.32 | 0.33 | -0.11 | 0.62 | 0.28 | 0.26 | 0.52 | 0.69 | 0.71 | 0.79 | 0.38 | 1.14 | -0.115 | 0.0325 | 0.35 | 0.397979 | 0.7692 | 0.932881 | 1 | 0.999631 | 0.761708 |
| Zranb2_E9PUD0_S83s_1 | -0.01 | 1.3 | 1.29 | 0.99 | 0.74 | 0.82 | 0.2 | 1.1 | 1.05 | 0.78 | 1.01 | 1.68 | 0.36 | 1.59 | 1.4 | 1.63 | -0.1775 | 0.2375 | 0.3525 | 0.426562 | 0.639624 | 0.54072 | 1 | 0.999631 | 0.775085 |
| Zrsr1_Q64707_S418s_1 | 1.82 | -0.6 | 1.72 | 0.93 | -1.01 | 0.28 | 1.34 | 0.05 | -0.25 | 1.3 | -0.45 | -0.33 | -1.09 | -1.5 | -0.86 | -0.63 | -0.8025 | -0.9 | -1.9875 | 0.004494 | 0.24102 | 0.174555 | 1 | 0.999631 | 0.351308 |
| Zswim8_Q3UHH1_S1060s_1 | -0.79 | -0.24 | -1.37 | -0.72 | -0.23 | -0.47 | -0.44 | -0.84 | -0.67 | -0.53 | -0.29 | -0.59 | -0.37 | -0.55 | 0.27 | -0.38 | 0.285 | 0.26 | 0.5225 | 0.158536 | 0.376478 | 0.412949 | 1 | 0.999631 | 0.628615 |
| Zswim8_Q3UHH1_S48s_1 | 1.79 | 0.38 | 2.1 | 1.49 | 0.69 | 1.36 | 1.5 | 1.31 | 2.37 | 2.38 | 1.3 | 0.89 | 0.25 | -0.78 | 1.04 | 0.76 | -0.225 | 0.295 | -1.1225 | 0.05238 | 0.589452 | 0.562026 | 1 | 0.999631 | 0.489777 |
| Zswim8_Q3UHH1_S53s_1 | 0.74 | -0.88 | -2.14 | -3.18 | -0.91 | -1.38 | -1.34 | -4.28 | -0.88 | 0.33 | -2.54 | -3.59 | -0.98 | -1.53 | -3.72 | -5.16 | -0.6125 | -0.305 | -1.4825 | 0.196256 | 0.538811 | 0.770546 | 1 | 0.999631 | 0.653373 |
| Zswim8_Q3UHH1_S564s_1 | 1.74 | 0.23 | 0.84 | 1.47 | 0.15 | 0.41 | 1.74 | 0.42 | 0.75 | 1.76 | 1.05 | 0.55 | -1.29 | -0.1 | 0.86 | 0.25 | -0.39 | -0.0425 | -1.14 | 0.053657 | 0.414979 | 0.921009 | 1 | 0.999631 | 0.4909 |
| Zwint_Q9CQU5_S216s_1 | 0.68 | 0.2 | -1.19 | 1 | 2.66 | 1.49 | 1.27 | 1.6 | 1.07 | -0.15 | 0.7 | -0.36 | -0.87 | 1.14 | 0.22 | -0.35 | 1.5825 | 0.1425 | -0.1375 | 0.81627 | 0.013282 | 0.795043 | 1 | 0.999631 | 0.937777 |
| Zyx_Q7TQE2_S241s_1 | 0.87 | 0.33 | -1.47 | -0.12 | 0.93 | 0.55 | 0.09 | -0.02 | 0.12 | 0.36 | 0.01 | -0.76 | -1.1 | 0.71 | -0.61 | -1.23 | 0.485 | 0.03 | -0.46 | 0.455626 | 0.352862 | 0.954084 | 1 | 0.999631 | 0.790041 |
| Zyx_Q7TQE2_S305s_1 | 0.1 | -0.33 | -1.33 | -0.37 | 0.1 | -0.43 | -0.31 | -0.21 | -0.47 | -1.18 | -0.23 | -0.55 | -1.05 | -0.01 | -0.83 | -0.76 | 0.27 | -0.125 | -0.18 | 0.654991 | 0.449054 | 0.745192 | 1 | 0.999631 | 0.878992 |
| Zzef1_E9Q5W5_S1475s_1 | -0.82 | -0.05 | -1.1 | 0.58 | -0.26 | -0.79 | -0.42 | 0.4 | -0.53 | -0.37 | -0.08 | -0.77 | 0.57 | 0.51 | 2.07 | 0.14 | 0.08 | -0.09 | 1.17 | 0.050562 | 0.856339 | 0.827769 | 1 | 0.999631 | 0.489558 |
| Zzef1_E9Q5W5_S1488s_1 | 0.46 | -0.44 | -0.3 | 0.15 | 0.07 | 0.25 | 0.49 | 0.27 | 0.36 | 0.19 | 0.97 | 0.06 | -0.66 | -0.14 | 0.42 | 0.75 | 0.3025 | 0.4275 | 0.125 | 0.754252 | 0.315594 | 0.223296 | 1 | 0.999631 | 0.913377 |
| Zzef1_E9Q5W5_S2407s_1 | 0.25 | 0.58 | -0.11 | -0.87 | -0.12 | 0.07 | -0.95 | -0.21 | 0.42 | 0.33 | -0.64 | -0.08 | 1.51 | 0.31 | 0.13 | 0.06 | -0.265 | 0.045 | 0.54 | 0.254165 | 0.503474 | 0.911351 | 1 | 0.999631 | 0.689137 |
| Zzef1_E9Q5W5_T2066t_1 | -0.61 | 0.08 | -1.33 | -1.18 | -1.1 | -0.53 | -1.22 | -1.09 | -1.48 | -0.78 | -0.6 | -0.05 | -1.28 | 0.02 | 0.14 | -0.23 | -0.225 | 0.0325 | 0.4225 | 0.361613 | 0.549709 | 0.940156 | 1 | 0.999631 | 0.741948 |

TMT all phosphosites synaptoneurosomes

| phosphosite SYP | | normalized protein expression SYP | | | | | | | | | | | | | | | | LogFC over WT SYP | | | p-value / WT SYP | | | adjusted p-value / WT SYP | | |
|---|---|---|---|---|---|---|---|---|---|---|---|---|---|---|---|---|---|---|---|---|---|---|---|---|---|---|
| id | name | SYP_WT | SYP_WT | SYP_WT | SYP_WT | SYP_MNK1 | SYP_MNK1 | SYP_MNK1 | SYP_MNK1 | SYP_MNK2 | SYP_MNK2 | SYP_MNK2 | SYP_MNK2 | SYP_MNKD | SYP_MNKD | SYP_MNKD | SYP_MNKD | logFC.SYP_ | logFC.SYP_ | logFC.SYP_ | P.Value.SY | P.Value.SY | P.Value.SY | adj.P.Val.S | adj.P.Val.S | adj.P.Val.SYP_MNKDKO.over.SYP_WT |
| 0610010F05Rik_M0QWF3_S116s_1 | 0610010F0 | -0.36 | -0.42 | 0.61 | -0.52 | -0.39 | 0.61 | 0.27 | -0.26 | -0.36 | -1.38 | -0.24 | -0.33 | 0.94 | -0.46 | -0.08 | 0.38 | 0.23 | -0.405 | 0.3675 | 0.37472 | 0.51879 | 0.302157 | 0.989223 | 0.997988 | 0.994666 |
| 0610011F06Rik_G5E8X1_S149s_1 | 0610011F0 | -0.2 | 0.12 | 0.12 | -0.12 | 0.34 | -0.28 | 0.29 | 0.11 | 0.01 | 0.14 | -0.23 | 0.06 | -0.21 | 0.41 | 0.23 | 0.35 | 0.135 | 0.015 | 0.215 | 0.418981 | 0.567598 | 0.946235 | 0.989223 | 0.997988 | 0.994666 |
| 1110002E22Rik_E0CYV9_S1338s_1 | 1110002E2 | -0.6 | -0.14 | 0.08 | -0.23 | 0.41 | -1.03 | -1.01 | -0.26 | 0.11 | -0.55 | 0.16 | -0.42 | 1.52 | 0.31 | -0.15 | -0.36 | -0.25 | 0.0475 | 0.5525 | 0.22515 | 0.501965 | 0.865743 | 0.989223 | 0.997988 | 0.994666 |
| 1110037F02Rik_E9PZY8_S1628s_1 | 1110037F0 | 0.44 | -0.85 | -0.2 | -0.52 | -0.03 | -0.81 | 0.37 | -0.08 | 0.03 | -0.44 | -1.29 | -1 | -0.05 | 0.64 | 0.6 | -0.27 | 0.145 | -0.3925 | 0.5125 | 0.193965 | 0.692307 | 0.339645 | 0.994893 | 0.997988 | 0.994666 |
| 1700021K19Rik_D3Z7B1_S137s_1 | 1700021K1 | 0.07 | -0.27 | 1.03 | 0.18 | 0.05 | -0.32 | 0.7 | 0.43 | -0.2 | -0.11 | 1.12 | 0.84 | -0.15 | -0.14 | 0.63 | 0.6 | -0.0375 | 0.16 | -0.0175 | 0.962425 | 0.915812 | 0.706236 | 0.997335 | 0.997988 | 0.999507 |
| 1700112E06Rik_E0CYG4_T63t_1 | 1700112E0 | 0.46 | 0.85 | 1.1 | 0.93 | 0.57 | 0.23 | 0.99 | 0.65 | 0.54 | 0.49 | 0.46 | 0.61 | 0.57 | 0.3 | 0.53 | 0.6 | -0.225 | -0.31 | -0.335 | 0.211967 | 0.392141 | 0.204695 | 0.989223 | 0.997988 | 0.994666 |
| 2010300C02Rik_E9Q3M9_S1094s_1 | 2010300C0 | -2.15 | -2.53 | -2.42 | -4.07 | -2.16 | -2.8 | -1.75 | -2.62 | -3.8 | -2.01 | -0.4 | -2.35 | -1.96 | -1.1 | -2.79 | -2.66 | 0.46 | 0.6525 | 0.665 | 0.237705 | 0.329761 | 0.386904 | 0.989223 | 0.997988 | 0.994666 |
| 2010300C02Rik_E9Q3M9_S1123s_1 | 2010300C0 | -1.8 | -0.6 | -0.61 | 0.5 | -1.73 | -3.63 | 0.4 | 0.1 | -3.27 | -0.82 | 0.53 | -0.08 | -2.16 | -1.47 | -0.35 | -0.79 | -0.5875 | -0.2825 | -0.565 | 0.331598 | 0.521566 | 0.740536 | 0.989223 | 0.997988 | 0.994666 |
| 2010300C02Rik_E9Q3M9_S113s_1 | 2010300C0 | -0.74 | -1.25 | -1.64 | -2.53 | -1.21 | -0.64 | -1.35 | -1.89 | -1.93 | -0.94 | 0.32 | -1.92 | -3.18 | -0.45 | -2.12 | -1.91 | 0.2675 | 0.4225 | -0.375 | 0.546405 | 0.539031 | 0.486229 | 0.989223 | 0.997988 | 0.994666 |
| 2010300C02Rik_E9Q3M9_S162s_1 | 2010300C0 | -1.71 | -0.88 | -0.9 | -0.54 | -0.79 | -1.95 | -0.23 | -0.52 | -1.37 | -1.13 | -1.14 | -0.71 | -1 | -0.55 | -0.81 | -0.43 | 0.135 | -0.08 | 0.31 | 0.350965 | 0.752226 | 0.799868 | 0.995762 | 0.997988 | 0.994666 |
| 2010300C02Rik_E9Q3M9_S189s_1 | 2010300C0 | -1.58 | -1.51 | -0.67 | -0.34 | -1.27 | -1.41 | -0.26 | -0.56 | -0.8 | -1.43 | -0.75 | -0.75 | -0.37 | -0.61 | -0.65 | -0.84 | 0.15 | 0.0925 | 0.4075 | 0.26386 | 0.707589 | 0.798054 | 0.995437 | 0.997988 | 0.994666 |
| 2010300C02Rik_E9Q3M9_S325s_1 | 2010300C0 | 2.81 | 0.94 | 1.1 | 1.1 | 0.64 | 3.41 | 1.87 | -0.06 | 1.43 | 2.72 | 0.04 | 0.94 | 0.77 | 1.02 | 0.83 | 1.43 | -0.0225 | -0.205 | -0.475 | 0.3092 | 0.976694 | 0.752047 | 0.999187 | 0.997988 | 0.994666 |
| 2010300C02Rik_E9Q3M9_S465s_1 | 2010300C0 | 2.06 | 1.44 | 4.3 | 5.02 | 0.09 | 3.15 | 3.66 | 4.83 | 1.56 | 1.78 | 4.45 | 4.19 | 1.23 | -0.53 | 3.83 | 4.02 | -0.2725 | -0.21 | -1.0675 | 0.384995 | 0.811806 | 0.836678 | 0.995762 | 0.997988 | 0.994666 |
| 2010300C02Rik_E9Q3M9_S47s_1 | 2010300C0 | -0.42 | -0.09 | -0.19 | 0.17 | 0.49 | 0.17 | 1.58 | 0.15 | -0.39 | 0.01 | -0.03 | 0.21 | -0.39 | 0.34 | -0.29 | 0.23 | 0.73 | 0.0825 | 0.105 | 0.717781 | 0.065372 | 0.741813 | 0.989223 | 0.997988 | 0.996966 |
| 2010300C02Rik_E9Q3M9_S507s_1 | 2010300C0 | 2.93 | 2.77 | 2.63 | 2.24 | 2.42 | 2.5 | 2.2 | 2.51 | 0.71 | 2.89 | 1.32 | 2.55 | 0.24 | 0.44 | 2.59 | 2.64 | -0.235 | -0.775 | -1.165 | 0.082879 | 0.329874 | 0.147667 | 0.989223 | 0.997988 | 0.994666 |
| 2010300C02Rik_E9Q3M9_S570s_1 | 2010300C0 | 2.7 | 2.67 | 2.89 | 2.68 | 2.46 | 2.31 | 2.29 | 2.69 | 1.14 | 2.77 | 2.17 | 2.45 | 0.69 | 0.98 | 2.8 | 2.59 | -0.2975 | -0.6025 | -0.97 | 0.083467 | 0.178047 | 0.125715 | 0.989223 | 0.997988 | 0.994666 |
| 2010300C02Rik_E9Q3M9_S572s_1 | 2010300C0 | 2.44 | 2.31 | 2.74 | 2.15 | 1.99 | 2.11 | 1.46 | 2.3 | 1.21 | 2.38 | 1.59 | 2.54 | 0.71 | 0.81 | 2.96 | 2.54 | -0.445 | -0.48 | -0.655 | 0.252315 | 0.118136 | 0.198206 | 0.989223 | 0.997988 | 0.994666 |
| 2010300C02Rik_E9Q3M9_S612s_1 | 2010300C0 | 4.02 | 3.65 | 2.91 | 1.61 | 3.21 | 3.64 | 0.81 | 0.18 | 1.27 | 3.85 | 1.28 | 2.87 | 0.47 | 0.75 | -1.57 | 2.69 | -1.0875 | -0.73 | -2.4625 | 0.019941 | 0.235238 | 0.338981 | 0.989223 | 0.997988 | 0.994666 |
| 2010300C02Rik_E9Q3M9_S712s_1 | 2010300C0 | 0.29 | 0.1 | -0.78 | -0.79 | 0.44 | 0.04 | -0.27 | -0.75 | 0.27 | 0.26 | -0.33 | -0.49 | -0.45 | 0.74 | -0.64 | -0.68 | 0.16 | 0.2225 | 0.0375 | 0.931138 | 0.669797 | 0.539861 | 0.992876 | 0.997988 | 0.999507 |
| 2010300C02Rik_E9Q3M9_S730s_1 | 2010300C0 | -1.91 | -0.07 | 0.61 | 0.56 | 0.24 | -1.45 | 1.01 | 0.18 | -1.72 | -0.33 | 0.9 | 0.16 | -0.75 | -1.21 | 0.91 | 1.19 | 0.1975 | -0.045 | 0.2375 | 0.750475 | 0.775724 | 0.950551 | 0.995762 | 0.997988 | 0.9971 |
| 2010300C02Rik_E9Q3M9_S898s_1 | 2010300C0 | -2.44 | -2.68 | -2.82 | -5.35 | -1.42 | -2.28 | -2.73 | -3.02 | -3.96 | -2.8 | -0.3 | -2.61 | -1.97 | -0.77 | -3.32 | -2.64 | 0.96 | 0.905 | 1.1475 | 0.160603 | 0.178357 | 0.332715 | 0.989223 | 0.997988 | 0.994666 |
| 2010300C02Rik_E9Q3M9_S918s_1 | 2010300C0 | 0.33 | -1.17 | -0.83 | -0.54 | -0.44 | 0.65 | -0.48 | -0.37 | -0.54 | -0.62 | 0.84 | -0.65 | -0.1 | 0.31 | -0.36 | -0.32 | 0.3925 | 0.31 | 0.435 | 0.261948 | 0.342781 | 0.507966 | 0.989223 | 0.997988 | 0.994666 |
| 2010300C02Rik_E9Q3M9_S92s_1 | 2010300C0 | -0.92 | -0.71 | -0.81 | -0.7 | -0.15 | -0.42 | -0.19 | -0.44 | -0.99 | -0.71 | -1.56 | -1.11 | -0.9 | -0.25 | -1.08 | -0.87 | 0.485 | -0.3075 | 0.01 | 0.970774 | 0.037541 | 0.246487 | 0.989223 | 0.997988 | 0.999507 |
| 2010300C02Rik_E9Q3M9_S95s_1 | 2010300C0 | 0.09 | -0.37 | -0.15 | 0.21 | 0.46 | 0.1 | 0.7 | 0.72 | -0.53 | -0.25 | -0.04 | 0.03 | -1.06 | 0.27 | -0.47 | 0.01 | 0.55 | -0.1425 | -0.2575 | 0.468067 | 0.050244 | 0.576001 | 0.989223 | 0.997988 | 0.994666 |
| 2010300C02Rik_E9Q3M9_S961s_1 | 2010300C0 | 0.02 | 0.22 | 0.31 | 0.24 | 0.28 | 0.03 | 0.29 | 0.05 | 0.02 | -0.02 | 0.12 | 0.24 | 0.26 | 0.56 | 0.02 | -0.09 | -0.035 | -0.1075 | -0.01 | 0.96914 | 0.864685 | 0.615481 | 0.996493 | 0.997988 | 0.999507 |
| 2010300C02Rik_E9Q3M9_T660t_1 | 2010300C0 | 0 | 0.54 | 0.55 | 0.72 | 0.2 | -0.27 | 0.86 | 0.77 | 0 | 0.4 | 0.54 | 0.51 | -0.14 | 0.43 | 0.48 | 0.62 | -0.0625 | -0.09 | -0.105 | 0.722307 | 0.845717 | 0.733245 | 0.996349 | 0.997988 | 0.9971 |
| 2010300C02Rik_E9Q3M9_T959t_1 | 2010300C0 | -0.62 | 0.15 | 0.37 | 0.58 | -0.53 | -0.83 | 0.85 | 0.7 | -0.27 | 0.15 | 0.43 | 0.4 | 0.64 | 0.5 | 0.33 | 0.24 | -0.0725 | 0.0575 | 0.3075 | 0.352109 | 0.875793 | 0.86226 | 0.996493 | 0.997988 | 0.994666 |
| 2010300C02Rik_F7B0R9_S599s_1 | 2010300C0 | -1.98 | -1.69 | -2.46 | -3.4 | -1.98 | -2.19 | -1.41 | -1.76 | -2.14 | -1.77 | 0.72 | -1.61 | -1.62 | -1.46 | -2.15 | -2.66 | 0.5475 | 1.1825 | 0.41 | 0.373654 | 0.188382 | 0.107666 | 0.989223 | 0.997988 | 0.994666 |
| 2310035C23Rik_G3X9J4_S141s_1 | 2310035C2 | 0.21 | 0.35 | 0.05 | 0.83 | 0.09 | 0.2 | 0.69 | 0.38 | 0.32 | 0.29 | 0.59 | 0.76 | -0.1 | 0.16 | 0.65 | 0.89 | -0.02 | 0.13 | 0.04 | 0.902102 | 0.938965 | 0.625623 | 0.999187 | 0.997988 | 0.999507 |
| 2310035C23Rik_G3X9J4_S180s_1 | 2310035C2 | 0.37 | 0.67 | -0.2 | 0.15 | 0.5 | 0.27 | 0.16 | 0.37 | 0.27 | 0.2 | -0.58 | 0.05 | -0.18 | 0.81 | -1.27 | -0.15 | 0.0775 | -0.2625 | -0.445 | 0.336 | 0.759014 | 0.397856 | 0.995762 | 0.997988 | 0.994666 |
| 2310035C23Rik_G3X9J4_S182s_1 | 2310035C2 | -0.3 | -0.32 | -0.12 | -0.36 | 0.32 | -0.06 | 0.03 | -0.04 | -0.69 | -0.31 | -0.24 | -0.38 | -0.84 | 0.35 | -1.03 | -0.45 | 0.3375 | -0.13 | -0.2175 | 0.531123 | 0.130037 | 0.561507 | 0.989223 | 0.997988 | 0.994666 |
| 2310035C23Rik_G3X9J4_S193s_1 | 2310035C2 | -1 | -1.04 | -0.43 | -2.45 | -2.26 | -0.01 | -1.84 | -1.16 | 1.15 | -1.2 | -0.76 | -0.55 | 0.47 | -3.08 | -0.44 | -1.16 | -0.0875 | 0.89 | 0.1775 | 0.81752 | 0.881114 | 0.170517 | 0.996493 | 0.997988 | 0.999507 |
| 2310035C23Rik_G3X9J4_S243s_1 | 2310035C2 | -0.01 | 0.37 | -0.19 | -0.48 | 0.49 | -0.32 | -0.01 | -0.41 | -0.34 | 0.23 | -0.66 | -0.4 | -0.02 | 0.31 | -0.6 | -0.39 | 0.015 | -0.215 | -0.0975 | 0.759243 | 0.959608 | 0.477962 | 0.999187 | 0.997988 | 0.9971 |
| 2310035C23Rik_G3X9J4_S244s_1 | 2310035C2 | -0.01 | 0.37 | -0.43 | -0.62 | 0.49 | -0.32 | -0.19 | -0.61 | -0.34 | 0.23 | -0.84 | -0.43 | -0.02 | 0.31 | -0.7 | -0.45 | 0.015 | -0.1725 | -0.0425 | 0.902396 | 0.963833 | 0.608605 | 0.999187 | 0.997988 | 0.999507 |
| 2310035C23Rik_G3X9J4_S422s_1 | 2310035C2 | 1.46 | 0.24 | 1.51 | 1.98 | -0.65 | 2.31 | 0.17 | 1.76 | 1.5 | 1.13 | 0.95 | 2.44 | 0.39 | -1.88 | 1.1 | 1.13 | -0.4 | 0.2075 | -1.1125 | 0.144022 | 0.565157 | 0.663042 | 0.989223 | 0.997988 | 0.994666 |
| 2310035C23Rik_G3X9J4_S56s_1 | 2310035C2 | 0.42 | 0.84 | 0.04 | 0.42 | 0.54 | 0.38 | 0.83 | 0.55 | 0.44 | 0.47 | 0 | 0.01 | -0.3 | 0.31 | 0.4 | 0.35 | 0.145 | -0.2 | -0.24 | 0.425952 | 0.561187 | 0.463252 | 0.989223 | 0.997988 | 0.994666 |
| 2310035C23Rik_G3X9J4_T143t_1 | 2310035C2 | 0.1 | 0.23 | 0.32 | 0.17 | 0.24 | 0.18 | 0.6 | 0.74 | 0.05 | 0.32 | -0.2 | 0.44 | -0.22 | 0.72 | 0.25 | 0.49 | 0.235 | -0.0525 | 0.105 | 0.7109 | 0.310821 | 0.825391 | 0.989223 | 0.997988 | 0.996966 |
| 4833439L19Rik_D3Z1F7_S164s_1 | 4833439L1 | -0.63 | 0.6 | 0.67 | 0.92 | 0.01 | -1.04 | -0.69 | 0.73 | 0.1 | -0.42 | 1.85 | 0.57 | 1.13 | 0.67 | 0.76 | 0.45 | -0.6375 | 0.135 | 0.3625 | 0.360144 | 0.207514 | 0.807198 | 0.989223 | 0.997988 | 0.994666 |
| 4931406P16Rik_Q8C5X1_S662s_1 | 4931406P1 | -1.09 | -1.29 | -0.88 | -0.87 | -0.53 | -1.16 | -0.83 | -0.76 | -0.64 | -1.1 | -0.26 | -0.2 | 0.2 | -0.27 | -0.42 | -0.44 | 0.2125 | 0.4825 | 0.8 | 0.013683 | 0.37575 | 0.11228 | 0.989223 | 0.997988 | 0.994666 |
| 5031439G07Rik_B1APX2_S30s_1 | 5031439G0 | -0.17 | 0.03 | -0.35 | 0.42 | 0.2 | -0.56 | -0.41 | 0.39 | 0.72 | 0.07 | 1.46 | 0.42 | 0.35 | 0.1 | -1.03 | -0.31 | -0.0775 | 0.685 | -0.205 | 0.578236 | 0.79985 | 0.07813 | 0.995762 | 0.997988 | 0.995259 |
| 5031439G07Rik_B1APX2_S311s_2 | 5031439G0 | -0.95 | -0.49 | -0.8 | -0.18 | 0.12 | -0.36 | -0.6 | -0.82 | -0.28 | 0.24 | -1.52 | -0.41 | -0.37 | -0.96 | -0.09 | -0.2 | 0.19 | 0.1125 | 0.2 | 0.525373 | 0.521115 | 0.780961 | 0.989223 | 0.997988 | 0.994666 |
| 5031439G07Rik_B1APX2_S322s_1 | 5031439G0 | 0.18 | 0.62 | 0.1 | -0.52 | 0.46 | -0.17 | 1.56 | -0.36 | 0.59 | 0.31 | 1.22 | 1.79 | 0.7 | 1.29 | 0.93 | 0.6 | 0.2775 | 0.8825 | 0.785 | 0.034747 | 0.54879 | 0.051787 | 0.989223 | 0.997988 | 0.994666 |
| 5031439G07Rik_B1APX2_S322s_2 | 5031439G0 | -0.2 | -0.38 | 0 | 0.43 | 0.22 | -0.28 | 0.05 | -0.19 | 0.13 | 0.65 | -0.66 | -0.3 | -0.02 | -0.59 | 0.53 | 0.51 | -0.0125 | -0.0075 | 0.145 | 0.678232 | 0.961374 | 0.982793 | 0.999187 | 0.997988 | 0.996159 |
| 5031439G07Rik_B1APX2_S347s_1 | 5031439G0 | -1.66 | -1.55 | -2.71 | -2.38 | -0.42 | -1.41 | -2.38 | -2.65 | -1.03 | -1.49 | -1.31 | -3.33 | -1.27 | -0.45 | -2.25 | -2.21 | 0.36 | 0.285 | 0.53 | 0.292958 | 0.499743 | 0.606431 | 0.989223 | 0.997988 | 0.994666 |
| 5031439G07Rik_B1APX2_S405s_1 | 5031439G0 | 0.15 | 0.11 | -0.05 | -0.03 | 0.76 | -0.61 | 0.25 | -0.16 | -0.6 | -0.54 | -1.02 | 0.13 | -1.59 | 0.22 | 0.03 | -0.24 | 0.015 | -0.5525 | -0.44 | 0.302062 | 0.961849 | 0.08172 | 0.999187 | 0.997988 | 0.994666 |
| 5031439G07Rik_B1APX2_T336t_1 | 5031439G0 | 0.53 | 0.19 | 0.53 | 0.68 | 0.24 | 0.26 | 0.32 | 0.72 | 0.32 | 0.4 | 0.35 | 0.73 | 0.19 | 0.33 | 0.4 | 0.39 | -0.0975 | -0.0325 | -0.155 | 0.526849 | 0.672188 | 0.889172 | 0.992876 | 0.997988 | 0.994666 |
| 5730455P16Rik_J3KMQ6_S19s_1 | 5730455P1 | -2.13 | -0.95 | -0.92 | -1.65 | -2.44 | -1.56 | -1.8 | -1.98 | -0.73 | -1.4 | 0.1 | -1.09 | -0.69 | -0.94 | -1.04 | -1.58 | -0.5325 | 0.6325 | 0.35 | 0.353689 | 0.151486 | 0.162026 | 0.989223 | 0.997988 | 0.994666 |
| 6330403A02Rik_J3QP43_S27s_1 | 6330403A0 | 0.13 | 0.34 | 0.46 | 0.53 | 0.66 | 0.37 | 0.96 | 0.68 | -0.08 | -0.05 | -0.31 | 0.53 | 0.34 | 0.8 | 0.6 | 0.32 | 0.3025 | -0.3425 | 0.15 | 0.557419 | 0.204373 | 0.213896 | 0.989223 | 0.997988 | 0.995259 |
| 6330403A02Rik_J3QP43_S28s_1 | 6330403A0 | 0.89 | 1.03 | 0.49 | 0.75 | 0.64 | 0.56 | 0.71 | 0.48 | 0.22 | 0.66 | 0.22 | 0.38 | 0.74 | 1.02 | 0.39 | 0.02 | -0.1925 | -0.42 | -0.2475 | 0.423705 | 0.385172 | 0.106358 | 0.989223 | 0.997988 | 0.994666 |
| 6330403A02Rik_J3QP43_S29s_1 | 6330403A0 | 0.74 | 0.89 | 0.95 | 1.33 | 0.85 | 0.89 | 1.04 | 0.32 | 0.24 | 0.32 | -1.09 | 0.67 | 0.86 | 0.69 | 1.31 | -0.03 | -0.2025 | -0.9425 | -0.27 | 0.435661 | 0.433955 | 0.040656 | 0.989223 | 0.997988 | 0.994666 |
| 6430548M08Rik_Q3TA40_S208s_1 | 6430548M | 1.1 | 0.14 | 1.22 | 1.46 | 1.17 | 1.93 | 1.64 | 1.73 | 0.63 | 0.96 | 2.29 | 2.3 | -0.35 | 1.56 | 0.69 | 1.69 | 0.6375 | 0.565 | -0.0825 | 0.873626 | 0.084943 | 0.271602 | 0.989223 | 0.997988 | 0.999507 |
| 6430548M08Rik_Q3TA40_S275s_1 | 6430548M | 1.1 | 1.1 | 1.02 | 1.05 | 0.76 | 0.73 | 0.58 | 0.85 | 0.61 | 1.16 | 1.07 | 0.85 | 0.07 | 1.16 | 0.83 | 0.85 | -0.3375 | -0.145 | -0.34 | 0.268606 | 0.110227 | 0.526667 | 0.989223 | 0.997988 | 0.994666 |
| 6430548M08Rik_Q3TA40_S72s_1 | 6430548M | -1.06 | -0.58 | -0.75 | -0.2 | -0.62 | -0.94 | -0.25 | -0.51 | -1.27 | -0.6 | 0.86 | -0.5 | -1.07 | -0.05 | -1.21 | -0.29 | 0.0675 | 0.27 | -0.0075 | 0.983424 | 0.803377 | 0.561381 | 0.995762 | 0.997988 | 1 |
| 8030462N17Rik_Q0VAW6_S145s_1 | 8030462N1 | -0.51 | -0.91 | -0.41 | -0.38 | -0.92 | -0.51 | -0.94 | -0.72 | 0.75 | -0.22 | 0.29 | -0.86 | 0.3 | -0.85 | -0.66 | -0.85 | -0.22 | 0.5425 | 0.0375 | 0.911573 | 0.354515 | 0.170222 | 0.989223 | 0.997988 | 0.999507 |
| 8030462N17Rik_Q0VAW6_S66s_1 | 8030462N1 | -0.64 | -0.66 | 0.03 | -0.22 | -1.12 | -1.2 | -0.44 | 0.24 | -1.25 | -0.86 | 0.46 | 0.14 | -0.62 | -0.33 | 0.17 | -0.47 | -0.2575 | -0.005 | 0.06 | 0.84196 | 0.492633 | 0.990651 | 0.989223 | 0.997988 | 0.999507 |
| A0A0G2JDV2_S170s_1 | A0A0G2JD\ | -0.21 | 0.07 | -0.4 | 0.17 | -0.36 | -0.02 | -0.85 | 0.54 | 0.64 | 0.12 | -1.53 | -0.24 | 1.25 | 0.42 | 0.31 | -0.51 | -0.08 | -0.16 | 0.46 | 0.258715 | 0.808709 | 0.728712 | 0.995762 | 0.997988 | 0.994666 |
| A0A0G2JFH6_S27s_1 | A0A0G2JFH | -2.48 | 1.08 | 1.88 | 0.91 | 1.03 | -2.31 | 3.18 | 1.85 | 1.3 | 0.63 | 0.4 | 1.88 | 1.99 | 2.14 | 2.84 | 3.03 | 0.59 | 0.705 | 2.1525 | 0.032783 | 0.65206 | 0.444526 | 0.990358 | 0.997988 | 0.994666 |
| A0A0G2JFH6_S28s_1 | A0A0G2JFH | -1.19 | 0.39 | 1.74 | 1.67 | -0.13 | -0.23 | 0.19 | 0.03 | 1.7 | -0.83 | 2.8 | 1.56 | 1.63 | 0.99 | 0.52 | -0.41 | -0.6875 | 0.655 | 0.03 | 0.966869 | 0.282657 | 0.478833 | 0.989223 | 0.997988 | 0.999507 |
| A0A0G2JGJ4_S77s_1 | A0A0G2JGJ | 0.14 | 0.75 | -0.01 | -1.62 | 0.24 | 0.66 | -0.18 | -1.51 | 1.11 | 0.6 | -1.99 | -1.15 | 1.46 | -0.02 | -0.53 | -0.44 | -0.0125 | -0.1725 | 0.3025 | 0.628444 | 0.983784 | 0.827673 | 0.999187 | 0.997988 | 0.995259 |
| A0A0J9YUM2_S765s_1 | A0A0J9YUN | 0.71 | 0.1 | 0.39 | 0.73 | 0.28 | 1.54 | -0.08 | 1.19 | 0.77 | -0.41 | 1.09 | 0.2 | -0.3 | 0.3 | -0.35 | -0.14 | 0.25 | -0.07 | -0.605 | 0.056569 | 0.52994 | 0.850037 | 0.989223 | 0.997988 | 0.994666 |
| A0A0J9YUM2_S766s_1 | A0A0J9YUN | 0.64 | 0.24 | 0.49 | 0.94 | 0.52 | 1.38 | -0.05 | 1.67 | 0.68 | 0.22 | 1.2 | 0.51 | -0.83 | 0.4 | -0.09 | 0.11 | 0.3025 | 0.075 | -0.68 | 0.068169 | 0.460494 | 0.799414 | 0.989223 | 0.997988 | 0.994666 |
| A0A0J9YVH8_S1262s_1 | A0A0J9YVH | 0.22 | 1.37 | 1.52 | 0.56 | 0.29 | 0.33 | 0.65 | 1.06 | -0.03 | 0.73 | 2.03 | 0.76 | 1.03 | 1.02 | -0.41 | 0.7 | -0.335 | -0.045 | -0.3325 | 0.467019 | 0.362767 | 0.928087 | 0.989223 | 0.997988 | 0.994666 |
| A0A0N4SUT6_T239t_1 | A0A0N4SU | -0.81 | 0.69 | -0.07 | 0.53 | -0.07 | 0.74 | -0.01 | -1.07 | 0.33 | -0.82 | 1.25 | 0.31 | 0.86 | 1.08 | 0.29 | -0.69 | -0.1875 | 0.1825 | 0.3 | 0.5479 | 0.689231 | 0.721092 | 0.994672 | 0.997988 | 0.994666 |
| A0A0N4SW73_S1188s_1 | A0A0N4SW | -5.27 | -4.25 | -3.91 | -3.85 | -4.03 | -4.59 | -3.39 | -3.89 | -4.2 | -3.99 | -2.22 | -3.93 | -2.5 | -2.87 | -3.46 | -4.05 | 0.345 | 0.735 | 1.1 | 0.035852 | 0.393806 | 0.185862 | 0.989223 | 0.997988 | 0.994666 |
| A0A0N4SW93_S866s_1 | A0A0N4SW | 0.2 | -0.76 | -0.49 | -0.39 | -0.04 | 0.62 | 0.22 | 0.3 | -0.98 | -1.38 | -0.09 | -0.36 | -1.79 | 0.23 | -0.63 | -0.58 | 0.635 | -0.3425 | -0.3325 | 0.466437 | 0.044614 | 0.360558 | 0.989223 | 0.997988 | 0.994666 |
| A0A0R4J006_S143s_1 | A0A0R4J00 | 1.1 | 1 | 0.42 | 0.81 | 1.46 | -0.18 | 0.76 | 0.59 | 0.34 | 0.83 | -0.23 | 0.71 | 0.63 | 0.33 | 0.4 | 0.98 | -0.175 | -0.42 | -0.2475 | 0.393503 | 0.633558 | 0.202495 | 0.989223 | 0.997988 | 0.994666 |
| A0A0R4J0J1_S543s_1 | A0A0R4J0J | 1.24 | 0.33 | 0.32 | 0.79 | 0.59 | 1 | 0.96 | 0.65 | 0.77 | 0.94 | 0.09 | 0.8 | 0.08 | 0.62 | 0.67 | 0.74 | 0.13 | -0.02 | -0.1425 | 0.653897 | 0.643547 | 0.950066 | 0.990258 | 0.997988 | 0.995259 |
| A0A0R4J0K0_T176t_1 | A0A0R4J0K | 0.11 | -0.14 | -0.36 | 0.38 | -0.36 | 0.39 | 0.62 | 0.16 | -0.7 | 0.49 | 0.57 | 0.32 | -0.14 | 0.43 | -0.07 | -0.13 | 0.205 | 0.1725 | 0.025 | 0.9299 | 0.488865 | 0.624775 | 0.989223 | 0.997988 | 0.999507 |
| A0A0R4J0M1_S162s_1 | A0A0R4J0N | 0.59 | 0.01 | 0.46 | 0.71 | 0.68 | 0.53 | 0.97 | 0.88 | -0.32 | 0.4 | -0.02 | 0.31 | -0.67 | 0.65 | 0.05 | 0.38 | 0.3225 | -0.35 | -0.34 | 0.346729 | 0.207158 | 0.227761 | 0.989223 | 0.997988 | 0.994666 |
| A0A0R4J1E3_S613s_1 | A0A0R4J1E | 0.49 | 0.39 | 0.29 | 0.45 | 0.33 | 0.17 | 0.59 | 0.27 | -0.13 | 0.39 | -0.26 | 0.47 | -0.78 | 0.72 | 0 | 0.26 | -0.065 | -0.2875 | -0.355 | 0.321434 | 0.754188 | 0.279914 | 0.995762 | 0.997988 | 0.994666 |
| A0A0R4J216_S30s_1 | A0A0R4J21 | 0.06 | 0.6 | 0.86 | 0.72 | 0.57 | 0.76 | -0.49 | 0.67 | 1.44 | 0.48 | 0.98 | 1.26 | 1.03 | 0.26 | 0.41 | 0.52 | -0.1825 | 0.48 | -0.005 | 0.986772 | 0.598901 | 0.144208 | 0.989223 | 0.997988 | 1 |
| A0A0R4J216_T23t_1 | A0A0R4J21 | 0.92 | 0.59 | 0.54 | 0.2 | -0.22 | 0.5 | 0.59 | 0.79 | 1.02 | 0.04 | 1.35 | -1.59 | 1.04 | 0.66 | 0.42 | 0.83 | -0.1475 | -0.3575 | 0.175 | 0.533001 | 0.617183 | 0.566041 | 0.989223 | 0.997988 | 0.994666 |
| A0A0U1RPY2_S57s_1 | A0A0U1RP\ | -2.01 | -1.54 | -1.95 | -4.68 | -2.07 | -1.84 | -2.13 | -2.08 | 2.37 | -1.51 | -2.05 | -0.82 | 3.05 | -1.72 | 2.56 | -1.8 | 0.515 | 2.0425 | 3.0675 | 0.038095 | 0.430151 | 0.08418 | 0.989223 | 0.997988 | 0.994666 |
| A0A140LHX7_S67s_1 | A0A140LH) | -0.53 | 0.28 | 0.18 | 0.2 | 0.19 | -0.52 | 0.41 | -0.17 | -0.41 | -0.15 | 0.12 | 0.33 | -0.42 | 0.34 | -0.02 | -0.09 | -0.055 | -0.06 | -0.08 | 0.792326 | 0.855419 | 0.837297 | 0.996349 | 0.997988 | 0.999507 |
| A0A140LHX7_S68s_1 | A0A140LH) | -0.12 | 0.73 | 0.24 | -0.29 | 0.32 | -0.15 | 0.15 | -0.39 | 0.38 | 0.31 | -0.14 | -0.08 | 0.58 | 0.96 | -0.39 | -0.15 | -0.1575 | -0.0225 | 0.11 | 0.78096 | 0.601992 | 0.940319 | 0.989223 | 0.997988 | 0.998226 |
| A0A140LI67_S906s_1 | A0A140LI6 | -0.01 | -0.13 | -0.96 | -0.3 | -2.84 | -0.94 | -3.13 | -2.5 | 1.99 | -0.7 | 0.26 | -1.46 | 1.68 | -0.24 | 0.23 | -0.33 | -2.0025 | 0.3725 | 0.685 | 0.181597 | 0.002363 | 0.596366 | 0.989223 | 0.997988 | 0.994666 |
| A0A140LIW3_S1289s_1 | A0A140LIW | -0.41 | -1.42 | 0.59 | 0.05 | -0.21 | -0.43 | -0.26 | 0.55 | 2.05 | 0.18 | 0.21 | 0.61 | 2.94 | -1.14 | -0.51 | 0.2 | 0.21 | 1.06 | 0.67 | 0.449908 | 0.640875 | 0.087295 | 0.989223 | 0.997988 | 0.994666 |
| A0A140LIW3_S1290s_1 | A0A140LIW | -0.41 | -1.42 | 0.26 | -0.05 | -0.21 | -0.43 | -0.6 | 0.4 | 2.05 | 0.18 | -0.55 | 0.58 | 2.94 | -1.14 | -1.08 | -0.33 | 0.195 | 0.97 | 0.5025 | 0.581114 | 0.633853 | 0.133279 | 0.989223 | 0.997988 | 0.995259 |
| A0A140LIW3_S1350s_1 | A0A140LIW | 1.41 | -0.38 | -0.68 | 0.26 | 1.04 | 1.36 | 0.54 | 0.69 | 0.12 | 1.38 | 0.55 | 0.28 | 1.09 | 2.1 | -0.12 | 0.29 | 0.755 | 0.43 | 0.6875 | 0.278289 | 0.127803 | 0.40671 | 0.989223 | 0.997988 | 0.994666 |
| A0A140LIW3_S1450s_1 | A0A140LIW | 0.28 | 0.48 | 0.57 | 0.75 | 0.8 | 0.22 | 0.85 | 0.74 | 0.24 | 0.57 | 0.45 | 0.71 | 0.3 | 0.57 | 0.53 | 0.75 | 0.1325 | -0.0275 | 0.0175 | 0.94405 | 0.584727 | 0.905941 | 0.989223 | 0.997988 | 0.999507 |
| A0A140LJD2_S105s_1 | A0A140LJD | 0.38 | 0.69 | 0.03 | -0.03 | 0.63 | 0.25 | 0.46 | 0.13 | 0.16 | 0.44 | 0.32 | 0.28 | 0.27 | 0.68 | -0.01 | 0.08 | 0.1 | 0.0325 | -0.0125 | 0.965991 | 0.695107 | 0.896934 | 0.994893 | 0.997988 | 0.999507 |
| A0A1B0GQW2_S50s_1 | A0A1B0GQ | -0.62 | -0.29 | -0.16 | -0.28 | -0.56 | -0.47 | -0.71 | -0.47 | -0.27 | -0.31 | -0.81 | -0.22 | 1.28 | 0.27 | -0.4 | -0.91 | -0.215 | -0.065 | 0.3975 | 0.402629 | 0.324734 | 0.792527 | 0.989223 | 0.997988 | 0.994666 |
| A0A1B0GR85_S1132s_1 | A0A1B0GR | 0.78 | 0.41 | 1.24 | 0.82 | 1.11 | 0.62 | 0.35 | 0.91 | 0.46 | 0.72 | 1.11 | 1.2 | 0.5 | 0.88 | 0.32 | 0.81 | -0.065 | 0.06 | -0.185 | 0.523602 | 0.813964 | 0.835213 | 0.995762 | 0.997988 | 0.994666 |
| A0A1B0GR85_S1194s_1 | A0A1B0GR | -1.09 | 0.54 | -0.78 | 0.02 | 0.16 | -0.03 | 0.08 | -0.35 | 0.49 | 0.18 | -0.44 | -0.16 | 0.59 | 0.41 | -0.19 | -0.47 | 0.2925 | 0.345 | 0.4125 | 0.359 | 0.448162 | 0.415835 | 0.989223 | 0.997988 | 0.994666 |
| A0A1B0GR85_S1194s_2 | A0A1B0GR | 0.5 | -1.41 | -0.35 | 0.2 | -1.58 | 0.49 | -0.04 | -0.51 | -0.2 | 0.2 | -0.24 | 0.16 | -0.14 | -0.33 | -0.84 | -0.73 | -0.145 | 0.245 | -0.245 | 0.582295 | 0.792242 | 0.568313 | 0.995762 | 0.997988 | 0.995259 |
| A0A1B0GR85_S1202s_2 | A0A1B0GR | 0.5 | -1.41 | -0.35 | 0.2 | -1.58 | 0.49 | -0.04 | -0.51 | -0.2 | 0.2 | -0.24 | 0.16 | -0.14 | -0.33 | -0.84 | -0.73 | -0.145 | 0.245 | -0.245 | 0.582295 | 0.792242 | 0.568313 | 0.995762 | 0.997988 | 0.995259 |
| A0A1B0GR85_S1331s_1 | A0A1B0GR | 0.43 | 0.58 | 0.84 | 0.33 | 0.63 | 0.05 | 0.66 | 0.33 | 0.58 | 0.52 | 0.84 | 0.5 | 0.6 | 0.84 | 0.11 | 0.61 | -0.1275 | 0.065 | -0.005 | 0.98542 | 0.603165 | 0.779003 | 0.989223 | 0.997988 | 1 |
| A0A1B0GR85_S1454s_1 | A0A1B0GR | -0.79 | -0.49 | -0.29 | -0.12 | -0.34 | -0.91 | -1.08 | -0.68 | -0.85 | -0.58 | -0.75 | -0.31 | -0.14 | -0.31 | -0.38 | -0.15 | -0.33 | -0.2 | 0.1775 | 0.497285 | 0.22832 | 0.442267 | 0.989223 | 0.997988 | 0.994666 |
| A0A1B0GS09_S77s_1 | A0A1B0GS( | -0.08 | 0.03 | 0.03 | 0.36 | 0.29 | -0.1 | 0.15 | 0.01 | -0.45 | 0.53 | 0.49 | 0.16 | -0.12 | 0.69 | -0.08 | -0.1 | 0.0025 | 0.0975 | 0.0125 | 0.965511 | 0.990834 | 0.737902 | 0.999187 | 0.997988 | 0.999507 |
| A0A1D5RLB2_S38s_1 | A0A1D5RLI | 0.66 | 0.65 | 0.53 | 0.52 | -0.67 | 0.89 | -0.16 | 0.35 | 0.95 | 0.35 | 0.27 | -0.09 | 0.57 | -0.58 | 0.6 | -0.07 | -0.4875 | -0.22 | -0.46 | 0.182712 | 0.177251 | 0.430981 | 0.989223 | 0.997988 | 0.994666 |
| A0A1D5RLL0_S106s_1 | A0A1D5RLI | 1.22 | 1.06 | 0.33 | 0.21 | 0.17 | 0.62 | 0.3 | 0.48 | 0.55 | 0.94 | 0.62 | 0.33 | 0.64 | 0.34 | 0.31 | 0.72 | -0.3125 | -0.095 | -0.2025 | 0.532948 | 0.309423 | 0.763639 | 0.989223 | 0.997988 | 0.994666 |
| A0A1W2P772_S197s_1 | A0A1W2P7 | 0.57 | 0.22 | 0.83 | 0.33 | 0.21 | 0.08 | 0.69 | 0.7 | -0.16 | 0.29 | -0.39 | 0.72 | 0.13 | 0.22 | 0.56 | 0.7 | -0.0675 | -0.3725 | -0.085 | 0.757661 | 0.795184 | 0.2531 | 0.995762 | 0.997988 | 0.9971 |
| A0A1W2P772_S207s_1 | A0A1W2P7 | -0.9 | -0.97 | -0.06 | -0.26 | 0.1 | -0.21 | -0.14 | 0.44 | -0.8 | -1.2 | -0.02 | -0.18 | -0.34 | -0.25 | -0.88 | -0.35 | 0.595 | -0.0025 | 0.0925 | 0.771462 | 0.069705 | 0.994543 | 0.989223 | 0.997988 | 0.9971 |
| A0A1W2P772_S210s_1 | A0A1W2P7 | -0.15 | -0.1 | -1.34 | -0.11 | -0.39 | -0.27 | -0.43 | -0.53 | 0.33 | -0.66 | 0.32 | -1.46 | -0.48 | -1.51 | -0.78 | -0.5 | 0.02 | 0.0575 | -0.3925 | 0.335227 | 0.950613 | 0.908051 | 0.999187 | 0.997988 | 0.994666 |
| A0A1W2P832_S760s_1 | A0A1W2P8 | -0.62 | -0.95 | -0.87 | -1.05 | -0.72 | -0.83 | -0.06 | -0.43 | -1.1 | -0.27 | -0.26 | -0.9 | -1.55 | -0.37 | -0.66 | -0.46 | 0.3625 | 0.24 | 0.1125 | 0.73215 | 0.171707 | 0.407304 | 0.989223 | 0.997988 | 0.9971 |
| A0A1W2P832_S776s_1 | A0A1W2P8 | -3.09 | -4.04 | -3.44 | -4.16 | -2.87 | -5.32 | -4.89 | -3.24 | -5.06 | -2.69 | -1.57 | -4.48 | -3.51 | -2.6 | -4.08 | -2.19 | -0.3975 | 0.2325 | 0.5875 | 0.236637 | 0.502929 | 0.758777 | 0.989223 | 0.997988 | 0.994666 |
| A0A1Y7VKX4_S609s_1 | A0A1Y7VK) | 0.48 | -0.23 | -0.55 | -0.67 | -0.05 | 0.53 | -1.12 | -1.43 | 0.79 | -0.04 | -0.29 | -0.37 | 1.11 | -0.14 | -0.74 | -0.26 | -0.275 | 0.265 | 0.235 | 0.609317 | 0.573122 | 0.486852 | 0.989223 | 0.997988 | 0.995259 |
| A0A1Y7VLQ2_S103s_2 | A0A1Y7VL( | 1.3 | 0.67 | 0.59 | -1.16 | 0.17 | -0.03 | -0.38 | -1.38 | -0.15 | 0.66 | -0.29 | -1.55 | -0.01 | 1.42 | -0.68 | 0.4 | -0.755 | -0.6825 | -0.0675 | 0.913717 | 0.204282 | 0.298368 | 0.989223 | 0.997988 | 0.999507 |
| A0A1Y7VLQ2_S95s_2 | A0A1Y7VL( | 1.3 | 0.67 | 0.59 | -1.16 | 0.17 | -0.03 | -0.38 | -1.38 | -0.15 | 0.66 | -0.29 | -1.55 | -0.01 | 1.42 | -0.68 | 0.4 | -0.755 | -0.6825 | -0.0675 | 0.913717 | 0.204282 | 0.298368 | 0.989223 | 0.997988 | 0.999507 |
| A0A1Y7VNP1_S418s_2 | A0A1Y7VN | -0.57 | -0.4 | -1.48 | -2.73 | -0.66 | -1.08 | -1.13 | -1.34 | -0.77 | -0.31 | -0.92 | -0.82 | -0.76 | -0.08 | -1.53 | -1.44 | 0.2425 | 0.59 | 0.3425 | 0.558343 | 0.633303 | 0.270119 | 0.989223 | 0.997988 | 0.995259 |
| A0A1Y7VNP1_S421s_2 | A0A1Y7VN | -1.08 | -1.37 | -1.48 | -2.73 | -1.11 | -1.54 | -1.13 | -1.34 | -1.21 | -1.23 | -0.92 | -0.82 | -1.4 | -0.55 | -1.53 | -1.44 | 0.385 | 0.62 | 0.435 | 0.32143 | 0.315203 | 0.131012 | 0.989223 | 0.997988 | 0.994666 |
| A0A1Y7VNZ6_S556s_1 | A0A1Y7VN. | -2.46 | -0.83 | -2.17 | -1.39 | -0.12 | -1.9 | -2.07 | -1.56 | -0.47 | -0.68 | -0.99 | -1.48 | 0.84 | -0.21 | -0.4 | -2.04 | 0.3 | 0.8075 | 1.26 | 0.071973 | 0.571553 | 0.081659 | 0.989223 | 0.997988 | 0.994666 |
| A0A286YCB8_S126s_1 | A0A286YCE | -0.56 | -0.39 | -0.54 | 0.18 | 0.26 | -0.42 | -0.11 | -0.31 | 0.07 | -0.98 | -0.22 | -0.2 | -0.59 | 0.05 | -0.18 | 0.17 | 0.1825 | -0.005 | 0.19 | 0.532701 | 0.505364 | 0.987355 | 0.989223 | 0.997988 | 0.994666 |
| A0A286YCB8_S127s_1 | A0A286YCE | 1.38 | 3.37 | 0.32 | -0.5 | 2.48 | 2.02 | -0.16 | -0.88 | 2.55 | 2.57 | -0.3 | -0.23 | 2.29 | 3.62 | -0.37 | -0.03 | -0.2775 | 0.005 | 0.235 | 0.8307 | 0.784044 | 0.996122 | 0.995762 | 0.998367 | 0.999507 |
| A0A286YCB8_S131s_1 | A0A286YCE | 0.66 | 1.48 | 0.42 | -0.1 | 1.07 | 0.61 | 0.98 | 0.72 | 0.52 | 0.86 | 0.29 | -0.13 | 0.24 | 1.2 | -0.21 | 0.4 | 0.23 | -0.23 | -0.2075 | 0.634824 | 0.514616 | 0.559105 | 0.989223 | 0.997988 | 0.995259 |
| A0A286YCX6_S971s_1 | A0A286YC) | -0.12 | 0 | 0.58 | -0.12 | 0.06 | 0.07 | 0.48 | 0.03 | 0.3 | 0.8 | 0.24 | -0.34 | 0.16 | 0 | 0.23 | 0.01 | 0.075 | 0.165 | 0.015 | 0.955275 | 0.767309 | 0.605824 | 0.995762 | 0.997988 | 0.999507 |
| A0A286YCX6_T968t_1 | A0A286YC) | -0.14 | 0.69 | 0.6 | 0.41 | 0.29 | -0.9 | 0.47 | 1.31 | -0.19 | -0.44 | -2.09 | 0.73 | -0.22 | 0.94 | 0.26 | 1.39 | -0.0975 | -0.8875 | 0.2025 | 0.620358 | 0.831474 | 0.143882 | 0.995762 | 0.997988 | 0.995259 |
| A0A286YD83_S49s_1 | A0A286YD8 | 1.36 | 1.34 | 2.03 | 1.04 | 1.41 | 0.98 | 1.71 | 1.94 | 0.62 | 1.37 | 1.15 | 2.07 | 0.7 | 1.31 | 1.38 | 0.69 | 0.0675 | -0.14 | -0.4225 | 0.214402 | 0.829127 | 0.708166 | 0.995762 | 0.997988 | 0.994666 |
| A0A286YDE4_S719s_1 | A0A286YDI | -0.28 | -0.9 | -0.65 | -1.54 | -0.63 | 0.87 | -1.12 | -0.74 | 0.08 | -1.45 | -2.09 | -0.59 | -0.32 | -1.04 | -1.15 | -1.15 | 0.4375 | -0.17 | -0.0725 | 0.8395 | 0.366632 | 0.740091 | 0.989223 | 0.997988 | 0.999507 |
| A0A286YDL2_S718s_1 | A0A286YDI | 0.62 | 0.48 | -0.06 | 0.76 | 0.54 | 0.54 | -0.17 | 0.49 | 0.44 | -0.09 | -0.44 | 0.8 | -0.77 | 0.82 | 0.16 | 0.29 | -0.1 | -0.2725 | -0.325 | 0.411001 | 0.725011 | 0.438729 | 0.995437 | 0.997988 | 0.994666 |
| A0A286YDL2_S741s_1 | A0A286YDI | -0.28 | -0.06 | 0.14 | 0.2 | 0.74 | 0.71 | 0.39 | 1.08 | -0.77 | -0.82 | 1.49 | 0.35 | -0.13 | 0.84 | -0.33 | 0.24 | 0.73 | 0.0625 | 0.155 | 0.634257 | 0.012734 | 0.904511 | 0.989223 | 0.997988 | 0.995259 |
| A0A286YDL2_S741s_2 | A0A286YDI | 0 | -0.71 | 0.25 | 0.23 | -0.1 | 0.75 | 0.29 | 0.45 | -0.36 | -1.2 | 0.97 | 0.36 | -1.57 | -0.18 | -1.21 | -0.5 | 0.405 | 1.57E-16 | -0.8075 | 0.06644 | 0.208034 | 1 | 0.989223 | 1 | 0.994666 |
| A0A286YDL2_S745s_1 | A0A286YDI | 0.38 | 0.39 | 0.61 | 0.7 | 0.8 | 0.69 | 0.25 | 0.36 | 0.45 | -0.22 | 0.81 | 0.87 | -0.5 | 0.77 | -0.01 | 0 | 0.005 | -0.0425 | -0.455 | 0.17895 | 0.982601 | 0.887852 | 0.999187 | 0.997988 | 0.994666 |
| A0A286YDL2_S745s_2 | A0A286YDI | 0 | -0.71 | 0.52 | 1.05 | -0.1 | 0.75 | 0.52 | 0.86 | -0.36 | -1.2 | 1.06 | 0.84 | -1.57 | -0.18 | 0.12 | 0.17 | 0.2925 | -0.13 | -0.58 | 0.281189 | 0.484153 | 0.827267 | 0.989223 | 0.997988 | 0.994666 |
| A0A286YDL2_S750s_1 | A0A286YDI | 1.9 | 0.1 | 1.02 | 2.37 | 0.73 | 0.86 | 1.43 | 2.28 | 1.48 | 1.94 | 1.55 | 1.21 | 0.35 | 0.45 | 1.76 | 1.71 | -0.0225 | 0.1975 | -0.28 | 0.631552 | 0.967532 | 0.690327 | 0.999187 | 0.997988 | 0.995259 |
| A0A2C9F2C1_S114s_1 | A0A2C9F2C | -0.34 | -0.1 | 0.16 | 0.12 | 0.3 | 0.01 | 0.25 | -0.15 | -0.52 | -0.28 | 0.06 | 0.08 | -1.28 | 0.3 | -1.2 | 0.11 | 0.1425 | -0.125 | -0.4775 | 0.282511 | 0.540927 | 0.626203 | 0.989223 | 0.997988 | 0.994666 |
| A0A2I3BPI6_T9t_1 | A0A2I3BPI6 | 0.42 | -0.53 | 0.41 | 0.11 | -1.57 | -0.28 | -0.09 | 0.23 | 0.48 | -0.25 | -0.68 | 0.08 | 1.01 | -1.11 | 0.14 | 0.19 | -0.53 | -0.195 | -0.045 | 0.923868 | 0.235295 | 0.578789 | 0.989223 | 0.997988 | 0.999507 |
| A0A2I3BQ39_S187s_1 | A0A2I3BQ3 | -0.75 | -1.11 | -1.79 | -1.93 | -0.18 | -0.95 | -1.21 | -2.22 | -1.2 | -1.26 | 0.79 | -1.25 | -1.55 | -0.57 | -1.94 | -0.43 | 0.255 | 0.665 | 0.2725 | 0.54925 | 0.589228 | 0.236511 | 0.989223 | 0.997988 | 0.994666 |
| A0A2I3BQP6_S426s_1 | A0A2I3BQF | 0.08 | 0.17 | -0.42 | -0.26 | 0.13 | 0.13 | -0.09 | -0.2 | 0.03 | 0.27 | 0.15 | -0.26 | 0.22 | 0.42 | -0.43 | -0.16 | 0.1 | 0.155 | 0.12 | 0.688747 | 0.669974 | 0.543415 | 0.992876 | 0.997988 | 0.996159 |
| A0A2I3BRN2_S374s_1 | A0A2I3BRN | -0.54 | -1.12 | -1.19 | -1.09 | -1.15 | 0.01 | -0.16 | -0.84 | 2.48 | -0.72 | -3.61 | -0.56 | 3.09 | -1.61 | 2.23 | -0.59 | 0.45 | 0.3825 | 1.765 | 0.098354 | 0.189008 | 0.729158 | 0.989223 | 0.997988 | 0.994666 |
| A0A2R8VHB6_S125s_1 | A0A2R8VH | -1.42 | -0.8 | -0.2 | -0.41 | -0.31 | -1 | -0.26 | -0.38 | -1.25 | -1.7 | 0.21 | -0.29 | -0.84 | -0.29 | -0.32 | -0.62 | 0.22 | -0.05 | 0.19 | 0.574876 | 0.509282 | 0.917822 | 0.989223 | 0.997988 | 0.995259 |
| A0A2R8VHB6_S130s_1 | A0A2R8VH | -1.24 | 0.24 | -1.27 | -1.61 | 0.42 | 1.71 | -1.38 | -1.24 | 1.85 | 0.84 | -0.8 | -1.27 | 0.38 | -0.54 | -1.87 | -1.01 | 0.8475 | 1.125 | 0.21 | 0.715308 | 0.268969 | 0.156903 | 0.989223 | 0.997988 | 0.996966 |
| A0A2R8VHB6_S148s_1 | A0A2R8VH | -1.84 | -2.27 | -1.27 | -2.17 | -2.45 | -1.12 | -1.17 | -1.15 | 1.55 | -2.15 | -1.51 | -0.79 | 1.48 | -2.32 | 0.95 | -1.66 | 0.415 | 1.1625 | 1.5 | 0.103476 | 0.296764 | 0.147341 | 0.989223 | 0.997988 | 0.994666 |
| A0A2R8VHB6_S148s_2 | A0A2R8VH | 0.17 | 1.26 | 0.81 | 0.5 | -0.68 | 0.39 | 0.36 | 0.35 | 0.72 | 0.22 | 0.61 | 0.9 | -1.26 | 0.25 | 0.43 | 0.77 | -0.58 | -0.0725 | -0.6375 | 0.207174 | 0.124057 | 0.814071 | 0.989223 | 0.997988 | 0.994666 |
| A0A2R8W6P0_S335s_1 | A0A2R8W6 | 0.64 | 0.78 | 0.91 | 0.77 | 0.59 | 0.87 | 1.18 | 0.88 | 0.19 | 0.47 | 0.5 | 0.95 | 0.01 | 0.94 | 0.72 | 0.91 | 0.105 | -0.2475 | -0.13 | 0.657604 | 0.635025 | 0.327488 | 0.989223 | 0.997988 | 0.995259 |
| A0A3B2WBC6_S10s_1 | A0A3B2WE | -0.55 | -0.25 | 0.49 | 0.41 | -0.48 | -0.7 | 0.37 | 0.11 | -0.23 | 0.18 | 0.07 | -0.07 | -0.46 | -0.16 | 0.08 | 0.28 | -0.2 | -0.0375 | -0.09 | 0.789457 | 0.576917 | 0.902149 | 0.989223 | 0.997988 | 0.999507 |
| A0A3B2WCD2_S1010s_1 | A0A3B2WC | 0.34 | -0.54 | -0.64 | -0.29 | 0.46 | 0.32 | -0.22 | -0.35 | 1.54 | -0.25 | -3.19 | -0.56 | 1.8 | 0.29 | -0.52 | -0.46 | 0.335 | -0.3325 | 0.56 | 0.315438 | 0.301114 | 0.708736 | 0.989223 | 0.997988 | 0.994666 |
| A0A3B2WCD2_S1019s_1 | A0A3B2WC | 0.34 | 0.24 | 0.3 | 0.09 | 1.09 | -0.02 | 0.15 | -0.04 | -0.38 | 0.06 | 0.1 | 0.02 | -0.22 | 0.49 | 0.27 | -0.03 | 0.0525 | -0.2925 | -0.115 | 0.66342 | 0.860632 | 0.217057 | 0.996493 | 0.997988 | 0.996159 |
| A0A3B2WCD2_S1217s_1 | A0A3B2WC | -0.62 | -0.82 | -0.24 | -0.82 | -1.23 | -0.01 | -1.01 | -1.03 | 2.42 | -0.65 | -1.68 | -0.08 | 3.25 | -1.22 | 1.29 | -0.39 | -0.195 | 0.6275 | 1.3575 | 0.145388 | 0.547442 | 0.434165 | 0.989223 | 0.997988 | 0.994666 |
| A0A3B2WCD2_S1301s_1 | A0A3B2WC | 1.96 | 1.45 | 1.92 | 0.65 | 1.15 | 2.01 | 0.23 | 1.08 | 0.86 | 1.26 | 0.9 | 1.75 | 1.29 | 0.86 | 0.48 | 2.19 | -0.3775 | -0.3025 | -0.29 | 0.533842 | 0.404882 | 0.427437 | 0.989223 | 0.997988 | 0.994666 |
| A0A3B2WCD2_S1379s_1 | A0A3B2WC | 0.06 | 0.06 | -0.83 | -0.13 | -0.15 | 0.67 | 0.2 | 0.33 | -0.31 | -0.05 | 0.01 | -0.06 | -0.68 | 0.3 | -0.84 | 0.26 | 0.4725 | 0.1075 | -0.03 | 0.937463 | 0.136461 | 0.698748 | 0.989223 | 0.997988 | 0.999507 |
| A0A3B2WCD2_S877s_1 | A0A3B2WC | 1.41 | 2 | 2.21 | 1.44 | 1.92 | 2.05 | 1.88 | 1.86 | 0.71 | 1.87 | 0.56 | 1.8 | 0.44 | 0.64 | 1.84 | 2 | 0.1625 | -0.53 | -0.535 | 0.242918 | 0.53143 | 0.206574 | 0.989223 | 0.997988 | 0.994666 |
| A0A3B2WCD2_S898s_1 | A0A3B2WC | -0.21 | 0.77 | 0.94 | 0.74 | 0.52 | -0.16 | 1.46 | 1.21 | 0.58 | 0.3 | 1.27 | 0.65 | 0.29 | 1.02 | 1.07 | 1.59 | 0.1975 | 0.14 | 0.4325 | 0.278481 | 0.643414 | 0.688326 | 0.990258 | 0.997988 | 0.994666 |
| A0A3B2WCD2_S951s_1 | A0A3B2WC | 1.06 | 0.62 | 1.02 | 1.24 | 1.48 | 0.84 | 0.97 | 1.21 | 0.43 | 0.89 | 1.58 | 1.34 | 0.18 | 1.53 | 1.51 | 0.94 | 0.14 | 0.075 | 0.055 | 0.880343 | 0.577821 | 0.812824 | 0.989223 | 0.997988 | 0.999507 |
| A0A3Q4EBK4_S301s_1 | A0A3Q4EB | 0.86 | 1.25 | -0.62 | -0.32 | 0.84 | 0.03 | 0.94 | -0.5 | 0.7 | 0.78 | -0.95 | 1.37 | -0.37 | 0.71 | -0.48 | 1.21 | 0.035 | 0.1825 | -0.025 | 0.964635 | 0.945957 | 0.766698 | 0.999187 | 0.997988 | 0.999507 |
| A0A494B917_S42s_1 | A0A494B9 | 0.33 | 0.53 | 0.17 | 1.36 | 0.67 | 0.72 | 0.78 | 0.6 | 0.6 | 1.25 | 1.65 | 1.07 | -0.47 | 0.76 | 0.26 | 0.83 | 0.095 | 0.545 | -0.2525 | 0.5367 | 0.747743 | 0.150596 | 0.995762 | 0.997988 | 0.994666 |
| A0A494BAF6_S191s_1 | A0A494BAI | 0.45 | -0.22 | 0.04 | -0.58 | -0.65 | 0.42 | -0.68 | -0.81 | -0.22 | -0.57 | -0.88 | -0.36 | 0.66 | -0.19 | -0.52 | -0.9 | -0.3525 | -0.43 | -0.16 | 0.692013 | 0.337191 | 0.174965 | 0.989223 | 0.997988 | 0.996159 |
| A0A498WGM0_S167s_1 | A0A498WG | 0.29 | 0.24 | -0.47 | -0.38 | -0.49 | 1.13 | -0.86 | 0.3 | 1.53 | 0.59 | -0.17 | -0.13 | 1.93 | -0.25 | -0.12 | 0.13 | 0.1 | 0.535 | 0.5025 | 0.34062 | 0.825309 | 0.236786 | 0.995762 | 0.997988 | 0.994666 |
| A0A571BDA2_S895s_1 | A0A571BD. | 0.03 | 0.95 | 0.39 | 0.33 | 0.37 | 0.38 | 0.87 | 0.97 | -0.13 | 0.69 | 0.48 | -0.05 | 0.03 | 1.08 | 0.81 | 0.38 | 0.2225 | -0.1775 | 0.15 | 0.65843 | 0.439528 | 0.571476 | 0.989223 | 0.997988 | 0.995259 |
| A0A571BDA2_S971s_1 | A0A571BD. | 0.41 | 0.85 | 0.97 | 0.57 | 1.08 | 0.54 | 0.86 | 0.84 | 0.45 | 0.71 | 0.56 | 0.94 | 1.06 | 0.95 | 1.02 | 0.89 | 0.13 | -0.035 | 0.28 | 0.277323 | 0.586735 | 0.886574 | 0.989223 | 0.997988 | 0.994666 |
| A0A571BEK8_S23s_1 | A0A571BEI | -1.36 | -2.62 | -1.56 | -3.16 | -1.62 | -0.82 | -1.52 | -3.17 | 0.13 | -0.48 | -1.42 | -2.09 | 0.79 | -1.64 | -2.08 | -1.3 | 0.3925 | 1.21 | 1.1175 | 0.129313 | 0.509204 | 0.068552 | 0.989223 | 0.997988 | 0.994666 |
| A0A571BGD8_S21s_1 | A0A571BGI | -0.87 | -0.51 | 0.08 | -0.29 | -0.88 | -0.92 | 0.11 | 0.18 | -0.95 | -0.72 | 0.22 | 0.32 | -0.22 | -1.05 | -0.03 | 0.12 | 0.02 | 0.115 | 0.1025 | 0.773357 | 0.955656 | 0.764238 | 0.999187 | 0.997988 | 0.99798 |
| A0A5F8MPK1_S6s_1 | A0A5F8MP | -0.68 | -0.87 | -1.16 | -1.25 | 0.02 | -0.29 | -0.33 | -0.77 | -0.39 | -0.69 | -0.64 | -0.32 | 0.36 | 0.11 | -0.45 | -0.49 | 0.6475 | 0.48 | 0.8725 | 0.016591 | 0.030233 | 0.073071 | 0.989223 | 0.997988 | 0.994666 |
| A0A5F8MQ17_S313s_1 | A0A5F8MC | -1.13 | 0.23 | -0.31 | -0.93 | 0.19 | -0.93 | -0.17 | -1.16 | -0.71 | -0.69 | -0.73 | -0.51 | -0.49 | 0.16 | -0.45 | -0.29 | 0.0175 | -0.125 | 0.2675 | 0.470271 | 0.966738 | 0.710039 | 0.999187 | 0.997988 | 0.994666 |
| A0A5F8MQ82_S377s_1 | A0A5F8MC | -0.73 | 0.2 | 0.31 | 0.83 | 1.08 | -0.22 | 0.6 | 0.22 | -0.09 | 0.5 | -0.37 | 0.05 | 0.05 | 0.29 | 0.3 | 0.75 | 0.2675 | -0.13 | 0.195 | 0.604549 | 0.516792 | 0.731039 | 0.989223 | 0.997988 | 0.995259 |
| A0A5H1ZRL1_S6s_1 | A0A5H1ZRI | 0.68 | 0.82 | 0.39 | 1.16 | 0 | 1.04 | 0.3 | 1.81 | 1.03 | 0.87 | 1 | -0.29 | 0.91 | 1.38 | 0.34 | 0.72 | 0.025 | -0.11 | 0.075 | 0.813538 | 0.951872 | 0.763043 | 0.999187 | 0.997988 | 0.999507 |
| A0A6I8MWZ7_S543s_1 | A0A6I8MW | -0.35 | -0.2 | -0.11 | -0.29 | -0.18 | -0.11 | -0.28 | -0.46 | 0.03 | 0.04 | 0.65 | 0.32 | 0.74 | -0.02 | -0.23 | -0.2 | -0.02 | 0.4975 | 0.31 | 0.310833 | 0.92214 | 0.062965 | 0.997335 | 0.997988 | 0.994666 |
| A0A7N9VR94_S286s_1 | A0A7N9VR | 2.54 | 1.79 | 1.48 | 0.84 | 1.77 | 2.32 | 0.99 | 1.37 | 2.86 | 2.26 | 0.42 | 1.69 | 2.5 | 0.44 | 2.04 | 1.52 | -0.05 | 0.145 | -0.0375 | 0.943422 | 0.907031 | 0.802358 | 0.997038 | 0.997988 | 0.999507 |
| A0A7N9VR94_S3383s_1 | A0A7N9VR | 0.19 | -0.25 | -0.28 | 0.12 | 0.53 | -0.07 | 0.58 | -0.12 | -0.2 | 0.13 | -0.11 | -0.16 | 0.52 | 0.6 | 0.41 | 0.17 | 0.285 | -0.03 | 0.48 | 0.087464 | 0.302349 | 0.898104 | 0.989223 | 0.997988 | 0.994666 |
| A0A7N9VR94_S55s_1 | A0A7N9VR | 0.59 | 0.02 | 0.64 | -0.18 | 1.16 | 0.28 | 0.35 | 0.27 | 0.28 | 0.67 | -0.52 | 0.47 | 0.69 | 0.89 | 0.05 | 0.2 | 0.2475 | -0.0425 | 0.19 | 0.565705 | 0.439423 | 0.902767 | 0.989223 | 0.997988 | 0.995259 |
| A230065H16Rik_G3UW55_S11s_1 | A230065H1 | 0.05 | -0.4 | 0.34 | 0.38 | -0.18 | 0.26 | 0.18 | 0.51 | 0.31 | -0.15 | 1.86 | -0.08 | 1.05 | 0.84 | 0.48 | 0.78 | 0.1 | 0.3925 | 0.695 | 0.0345 | 0.713057 | 0.419945 | 0.995437 | 0.997988 | 0.994666 |
| A2AG58-2_A2AG58_S205s_1 | A2AG58-2_ | 0.43 | 0.42 | -0.36 | 0.04 | 0.29 | -0.03 | 0.09 | 0.14 | 0.13 | 0.16 | 0.3 | 0.31 | 1.01 | 0.07 | 0.36 | -0.2 | -0.01 | 0.0925 | 0.1775 | 0.61461 | 0.968468 | 0.7225 | 0.999187 | 0.997988 | 0.995259 |
| A830010M20Rik_Q3TEG0_T427t_1 | A830010M | 0.11 | 0.78 | 0.89 | 0.12 | 0.4 | -0.18 | -0.39 | -0.1 | 0.35 | 0.55 | 0.55 | 0.36 | 1.06 | 0.35 | 0.25 | 0.63 | -0.5425 | -0.0225 | 0.0975 | 0.762816 | 0.091345 | 0.934248 | 0.989223 | 0.997988 | 0.9971 |
| Aagab_Q8R2R3_S211s_1 | Aagab_Q8I | -1.3 | -0.46 | -0.64 | 0.14 | -1 | -1.35 | -0.08 | -0.87 | -0.62 | -0.91 | 0.36 | 0.2 | 0.19 | -0.04 | -0.17 | -0.21 | -0.26 | 0.3225 | 0.5075 | 0.162734 | 0.507771 | 0.449735 | 0.989223 | 0.997988 | 0.994666 |
| Aak1_Q3UHJ0_S18s_1 | Aak1_Q3UI | 0.42 | -0.02 | 0.73 | 0.68 | 0.26 | 0.58 | 0.48 | 0.6 | 0.71 | 0.39 | 0.26 | 0.55 | 0.16 | 0.34 | 0.71 | 0.5 | 0.0275 | 0.025 | -0.025 | 0.929574 | 0.911429 | 0.92392 | 0.997335 | 0.997988 | 0.999507 |
| Aak1_Q3UHJ0_S21s_1 | Aak1_Q3UI | 0.31 | 0.29 | 0.08 | 0.42 | 0.87 | 0.32 | 0.52 | 0.45 | -0.05 | 0.66 | -0.38 | 0.35 | 0.4 | 0.07 | 0.24 | 0.43 | 0.265 | -0.13 | 0.01 | 0.96681 | 0.250754 | 0.651179 | 0.989223 | 0.997988 | 0.999507 |
| Aak1_Q3UHJ0_S621s_2 | Aak1_Q3UI | -0.5 | -0.52 | 0.27 | 0.05 | -0.27 | -0.79 | 0.21 | 0.16 | -0.68 | -0.95 | 0.32 | -0.3 | -1.32 | -0.05 | 0 | 0.15 | 0.0025 | -0.2275 | -0.13 | 0.746158 | 0.993766 | 0.524667 | 0.999278 | 0.997988 | 0.9971 |
| Aak1_Q3UHJ0_S622s_1 | Aak1_Q3UI | -0.07 | 0.58 | 0.67 | 1.28 | 1.24 | 0 | 0.03 | -0.3 | -0.04 | 0.42 | 0.79 | 1.09 | 0.49 | 0.72 | 0.79 | 0.36 | -0.3725 | -0.05 | -0.025 | 0.940125 | 0.382243 | 0.893768 | 0.989223 | 0.997988 | 0.999507 |
| Aak1_Q3UHJ0_S622s_2 | Aak1_Q3UI | 0.32 | 0.22 | 0.16 | 0.01 | 0.45 | -0.01 | 0.24 | -0.14 | 0.09 | 0.19 | 0.15 | -0.1 | -0.14 | 0.54 | 0.13 | 0.23 | -0.0425 | -0.095 | 0.0125 | 0.961285 | 0.850966 | 0.6594 | 0.996349 | 0.997988 | 0.999507 |
| Aak1_Q3UHJ0_S635s_1 | Aak1_Q3UI | -0.54 | -1.09 | -1.86 | -0.83 | 0.26 | -0.15 | 0.23 | -0.26 | -1.66 | -1.31 | -0.33 | -1.27 | -1.5 | -0.32 | -1.56 | -0.85 | 1.1 | -0.0625 | 0.0225 | 0.956226 | 0.007068 | 0.875592 | 0.989223 | 0.997988 | 0.999507 |
| Aak1_Q3UHJ0_S650s_1 | Aak1_Q3UI | -0.68 | -1.8 | -2.26 | -1.62 | 0.66 | -0.15 | 0.02 | -1.11 | -1.59 | -0.99 | 1.04 | -1.83 | -1.3 | -0.2 | -2.32 | -1.46 | 1.445 | 0.7475 | 0.27 | 0.602165 | 0.010471 | 0.276651 | 0.989223 | 0.997988 | 0.995259 |
| Aak1_Q3UHJ0_S668s_1 | Aak1_Q3UI | -1.34 | -0.56 | -1.53 | -1.97 | -1.43 | -0.69 | -1.96 | -1.79 | -0.03 | -1.2 | -1.8 | -1.9 | 0.2 | -2.18 | -1.67 | -1.55 | -0.1175 | 0.1175 | 0.05 | 0.927766 | 0.765741 | 0.811317 | 0.995762 | 0.997988 | 0.999507 |
| Aak1_Q3UHJ0_S668s_2 | Aak1_Q3UI | -1.1 | -1.18 | -1.26 | -2.88 | -0.85 | -0.33 | -1.89 | -1.03 | 0.59 | -1.98 | -0.38 | -1.62 | -0.12 | -1.79 | -1.5 | -1.99 | 0.58 | 0.7575 | 0.255 | 0.647554 | 0.256317 | 0.268282 | 0.989223 | 0.997988 | 0.995259 |
| Aak1_Q3UHJ0_S676s_1 | Aak1_Q3UI | -0.3 | -0.03 | -0.43 | -0.85 | -0.22 | 1.1 | -0.61 | -0.37 | 1.48 | 0.22 | -1.87 | -0.57 | 1.67 | -1.25 | -0.83 | -0.6 | 0.3775 | 0.2175 | 0.15 | 0.807541 | 0.360349 | 0.740787 | 0.989223 | 0.997988 | 0.999507 |
| Aak1_Q3UHJ0_S676s_2 | Aak1_Q3UI | -0.44 | -0.94 | -0.91 | -1.45 | -0.73 | 0.59 | -1.16 | -0.59 | 1.09 | -1.29 | -0.56 | -0.95 | -0.26 | -1.11 | -0.91 | -1.18 | 0.4625 | 0.5075 | 0.07 | 0.833724 | 0.274382 | 0.349193 | 0.989223 | 0.997988 | 0.999507 |
| Aak1_Q3UHJ0_S680s_1 | Aak1_Q3UI | 0.72 | 0.69 | 0.48 | 0.45 | 0.52 | 0.33 | 0.79 | 0.5 | 0.02 | 0.48 | 0.22 | 0.36 | -0.48 | 0.77 | -0.05 | 0.44 | -0.05 | -0.315 | -0.415 | 0.223478 | 0.815343 | 0.184657 | 0.995762 | 0.997988 | 0.994666 |
| Aak1_Q3UHJ0_S729s_1 | Aak1_Q3UI | 0.14 | -0.23 | -0.22 | 0.25 | 0.22 | 0.03 | 0.56 | 0.06 | -0.71 | -0.19 | -0.33 | -0.01 | -1.23 | 0.22 | -0.38 | -0.2 | 0.2325 | -0.295 | -0.3825 | 0.297199 | 0.34361 | 0.273005 | 0.989223 | 0.997988 | 0.994666 |
| Aak1_Q3UHJ0_S795s_1 | Aak1_Q3UI | -2.72 | -2.52 | -2.12 | -2.71 | -1.56 | -2.1 | -1.24 | -2.98 | -3.51 | -3.02 | 0.37 | -2.15 | -3.12 | -1.42 | -2.27 | -2.64 | 0.5475 | 0.44 | 0.155 | 0.695316 | 0.185135 | 0.576816 | 0.989223 | 0.997988 | 0.996159 |
| Aak1_Q3UHJ0_S848s_1 | Aak1_Q3UI | 0.61 | -0.31 | 1.31 | 0.79 | 0.59 | -0.17 | 0.21 | 1.09 | -0.7 | 0.57 | 1.9 | 0.67 | -0.08 | 0.35 | 1.17 | 1.17 | -0.17 | 0.01 | 0.0525 | 0.906603 | 0.681069 | 0.986209 | 0.993613 | 0.997988 | 0.999507 |
| Aak1_Q3UHJ0_S936s_1 | Aak1_Q3UI | 0.94 | 0.66 | 0.53 | 0.45 | 0.46 | 1.15 | 0.95 | 0.74 | -0.05 | 1.18 | -0.73 | 0.9 | -0.03 | 0.04 | 0.8 | 0.9 | 0.18 | -0.32 | -0.2175 | 0.498114 | 0.467704 | 0.471727 | 0.989223 | 0.997988 | 0.994666 |
| Aak1_Q3UHJ0_T604t_1 | Aak1_Q3UI | -0.07 | 0.92 | 0.42 | 0.14 | 0.67 | 0.1 | -0.04 | -0.16 | 0.36 | 0.38 | 0.51 | -0.22 | 0.16 | 0.79 | 0.15 | -0.01 | -0.21 | -0.095 | -0.08 | 0.804621 | 0.495808 | 0.756218 | 0.989223 | 0.997988 | 0.999507 |
| Aak1_Q3UHJ0_T618t_1 | Aak1_Q3UI | 0.65 | 0.83 | 1.14 | 1.12 | 0.33 | 0.32 | 0.59 | 0.49 | 0.42 | 0.99 | 0.97 | 0.86 | 1.09 | 0.87 | 0.85 | 0.83 | -0.5025 | -0.125 | -0.025 | 0.91997 | 0.044415 | 0.62161 | 0.989223 | 0.997988 | 0.999507 |
| Aak1_Q3UHJ0_T618t_2 | Aak1_Q3UI | 0.15 | 0.07 | 0.19 | 0.02 | 0.3 | -0.17 | 0.23 | -0.04 | -0.06 | -0.03 | 0.2 | -0.16 | -0.36 | 0.42 | 0.09 | 0.2 | -0.0275 | -0.12 | -0.02 | 0.939544 | 0.897581 | 0.575805 | 0.996493 | 0.997988 | 0.999507 |
| Aak1_Q3UHJ0_T672t_2 | Aak1_Q3UI | 0.1 | -0.78 | 0.05 | -0.29 | -0.51 | 1.25 | -0.44 | -0.05 | 1.57 | -0.94 | -0.58 | -0.22 | -0.27 | -0.71 | -0.24 | -0.37 | 0.2925 | 0.1875 | -0.1675 | 0.573349 | 0.502095 | 0.733631 | 0.989223 | 0.997988 | 0.995259 |
| Aak1_Q3UHJ0_T679t_1 | Aak1_Q3UI | 0.72 | 0.69 | 0.48 | 0.45 | 0.52 | 0.33 | 0.79 | 0.5 | 0.02 | 0.48 | 0.22 | 0.36 | -0.48 | 0.77 | -0.05 | 0.44 | -0.05 | -0.315 | -0.415 | 0.223478 | 0.815343 | 0.184657 | 0.995762 | 0.997988 | 0.994666 |
| Aatk_B1AZF9_S662s_1 | Aatk_B1AZ | 0.13 | 0.79 | 1.28 | 0.26 | 0.41 | -0.19 | 0.13 | 0.4 | -2.15 | 0.68 | 0.28 | 0.43 | -1.1 | -1.29 | 0.27 | 0.37 | -0.4275 | -0.805 | -1.0525 | 0.053932 | 0.197514 | 0.23189 | 0.989223 | 0.997988 | 0.994666 |

| | | | | | | | | | | | | | | | | | | | | | | | | | | |
|---|---|---|---|---|---|---|---|---|---|---|---|---|---|---|---|---|---|---|---|---|---|---|---|---|---|---|
| Aatk_B1AZF9_S670s_1 | Aatk_B1AZ | -0.08 | -2.25 | -0.52 | 0.04 | 0.22 | -0.26 | 1 | -0.55 | -0.88 | -0.32 | -0.84 | -1 | -1.04 | 0.26 | 0.34 | 0.63 | 0.805 | -0.0575 | 0.75 | 0.224763 | 0.17773 | 0.911142 | 0.989223 | 0.997988 | 0.994666 |
| Aatk_B1AZF9_S987s_1 | Aatk_B1AZ | -0.4 | -0.7 | -0.42 | -0.4 | -0.01 | -0.57 | 0.07 | -0.58 | -0.42 | -0.42 | -0.01 | -0.69 | -0.58 | -0.05 | 0.17 | -0.03 | 0.2075 | 0.095 | 0.3575 | 0.202554 | 0.412312 | 0.695254 | 0.989223 | 0.997988 | 0.994666 |
| Abca2_P41234_S2381s_1 | Abca2_P41 | -0.87 | -0.08 | 0.56 | -0.41 | -0.88 | -1.04 | 0.42 | -0.1 | -1.72 | 0.22 | -0.48 | 0.44 | -0.86 | -2.92 | 0.25 | 0.23 | -0.2 | -0.185 | -0.625 | 0.393051 | 0.643456 | 0.728966 | 0.990258 | 0.997988 | 0.994666 |
| Abcb1a_P21447_S667s_1 | Abcb1a_P2 | -1.12 | -1.87 | -2.25 | -2.46 | -2.7 | -0.95 | -3.28 | -2.52 | -1.78 | -1.22 | -1.42 | -3.1 | -0.35 | -3.13 | -2.25 | -2.81 | -0.4375 | 0.045 | -0.21 | 0.73731 | 0.414453 | 0.926406 | 0.989223 | 0.997988 | 0.9971 |
| Abcc4_E9Q236_T561t_1 | Abcc4_E9C | 0.01 | 0.93 | 0.86 | 1.11 | 0.35 | 0.34 | 0.71 | 0.55 | 0.95 | 1.14 | 0.36 | 0.95 | 0.53 | 1.12 | 0.64 | 0.89 | -0.24 | 0.1225 | 0.0675 | 0.834723 | 0.417076 | 0.706817 | 0.989223 | 0.997988 | 0.999507 |
| Abcd2_Q61285_S58s_1 | Abcd2_Q6 | 0.27 | 0.63 | 0.93 | 0.68 | 0.2 | 0.5 | -0.24 | 0.39 | 0.78 | 1.07 | 1.12 | 0.99 | 1.3 | -0.42 | 0.26 | 1.19 | -0.415 | 0.3625 | -0.045 | 0.915793 | 0.136307 | 0.155782 | 0.989223 | 0.997988 | 0.999507 |
| Abcd3_P55096_S424s_1 | Abcd3_P55 | 0.72 | 0.43 | 0.15 | 0.47 | 0.76 | 0.37 | 0.87 | 0.41 | 0.19 | 0.54 | -0.23 | 0.91 | 0.03 | 0.52 | 0.43 | -0.06 | 0.16 | -0.09 | -0.2125 | 0.441793 | 0.506875 | 0.769164 | 0.989223 | 0.997988 | 0.994666 |
| Abcf1_Q6P542_S194s_1 | Abcf1_Q6P | -2.32 | -0.72 | -0.29 | 0.17 | -1.43 | -1.66 | -0.29 | 0.08 | -1.01 | -1.63 | -0.02 | -0.09 | -1.71 | -0.99 | 0.14 | -0.84 | -0.035 | 0.1025 | -0.06 | 0.920753 | 0.954499 | 0.866205 | 0.999187 | 0.997988 | 0.999507 |
| Abhd17b_Q7M759_S282s_1 | Abhd17b_( | 0.11 | 0.85 | 1.21 | 0.63 | 0.53 | -0.14 | 0.63 | 0.24 | -0.05 | 0.45 | 1.67 | 1.54 | 0.41 | 0.12 | 1.16 | 0.19 | -0.385 | 0.2025 | -0.23 | 0.524316 | 0.230925 | 0.660081 | 0.989223 | 0.997988 | 0.994666 |
| Abi1_B7ZCU4_S183s_1 | Abi1_B7ZC | 0.44 | 0.58 | 0.55 | 0.5 | 0.33 | 0.33 | 0.27 | 0.19 | 0.44 | 0.84 | 0.47 | 0.28 | 0.72 | 0.64 | 0.21 | 0.33 | -0.2375 | -0.01 | -0.0425 | 0.86297 | 0.237639 | 0.964615 | 0.989223 | 0.997988 | 0.999507 |
| Abi1_B7ZCU4_S222s_1 | Abi1_B7ZC | -1.09 | -2.17 | -2.86 | -4.45 | -1.59 | -1.37 | -2.16 | -2.7 | -1 | -2.11 | -0.98 | -2.3 | 0.56 | -2.77 | -2.29 | -2.19 | 0.6875 | 1.045 | 0.97 | 0.298239 | 0.322938 | 0.165094 | 0.989223 | 0.997988 | 0.994666 |
| Abi1_B7ZCU4_S225s_1 | Abi1_B7ZC | 0.03 | -0.17 | -0.67 | -0.23 | -1.11 | 0.24 | -0.4 | -0.74 | 1.39 | 0.18 | -0.09 | -0.63 | 0.56 | -0.25 | -0.37 | -0.69 | -0.2425 | 0.4725 | 0.0725 | 0.831037 | 0.471552 | 0.296306 | 0.989223 | 0.997988 | 0.999507 |
| Abi1_B7ZCU4_S231s_1 | Abi1_B7ZC | -2.83 | -4.41 | -1.47 | -2.09 | -1.73 | -1.98 | -1.32 | -1.47 | -3.25 | -3.76 | 0.47 | -1.49 | -1.26 | -0.99 | -1.42 | -1.81 | 1.075 | 0.6925 | 1.33 | 0.048805 | 0.091005 | 0.500134 | 0.989223 | 0.997988 | 0.994666 |
| Abi2_A0A087WRS3_S191s_1 | Abi2_A0A0 | -1.17 | -1.29 | -2.38 | -1.63 | -2.69 | -1.64 | -2.48 | -1.58 | 0.09 | -1.85 | -3.29 | -1.76 | 1.43 | -3.46 | -2.54 | -2.45 | -0.48 | -0.085 | -0.1375 | 0.887882 | 0.230424 | 0.899509 | 0.989223 | 0.997988 | 0.999507 |
| Abi2_A0A087WRS3_S195s_1 | Abi2_A0A0 | -1.91 | -2.78 | -2.37 | -2.18 | -2.81 | -1.18 | -3.95 | -2.1 | 0.25 | -1.93 | -4.23 | -1.92 | 0.69 | -2.96 | -2.99 | -2.94 | -0.2 | 0.3525 | 0.26 | 0.751947 | 0.718372 | 0.672747 | 0.995437 | 0.997988 | 0.9971 |
| Abi2_Q6AXH6_S183s_1 | Abi2_Q6AX | 0.46 | 0.65 | 0.51 | 0.68 | 0.3 | 0.17 | 0.43 | 0.32 | 0.21 | 0.62 | 0.62 | 0.39 | 0.51 | 0.8 | 0.47 | 0.39 | -0.27 | -0.115 | -0.0325 | 0.891758 | 0.197864 | 0.606885 | 0.989223 | 0.997988 | 0.999507 |
| Abi2_Q6AXH6_S216s_1 | Abi2_Q6AX | 0.62 | -0.79 | -0.77 | -0.49 | -0.93 | 0.22 | -0.77 | 0.18 | -0.74 | -1.01 | 1.63 | -1.12 | -1.33 | 0.01 | -1.4 | -1.49 | 0.0325 | 0.0475 | -0.695 | 0.161697 | 0.939264 | 0.942842 | 0.999187 | 0.997988 | 0.994666 |
| Abi2_Q6AXH6_S227s_1 | Abi2_Q6AX | 0.39 | -0.21 | -0.13 | -0.5 | 0.26 | 0.14 | -0.46 | -0.11 | 0.08 | -0.11 | -0.42 | -0.53 | 0.18 | 0.25 | -0.83 | -0.62 | 0.07 | -0.1325 | -0.1425 | 0.691385 | 0.802846 | 0.640855 | 0.995762 | 0.997988 | 0.996159 |
| Abl1_P00520_S709s_1 | Abl1_P005 | -2.67 | -1.56 | -3.3 | -3.58 | -1.64 | -3 | -3.95 | -3.23 | 0.37 | -1.4 | -0.48 | -2.68 | -1.16 | -1.23 | -2.79 | -3.43 | -0.1775 | 1.73 | 0.625 | 0.350403 | 0.763955 | 0.03525 | 0.995762 | 0.997988 | 0.994666 |
| Abl1_P00520_S713s_1 | Abl1_P005 | -3.73 | -2.75 | -3.61 | -6.8 | -2.93 | -4.37 | -4.82 | -5.08 | -0.2 | -1.51 | -2.67 | -4.79 | -0.03 | -2.76 | -2.76 | -2.78 | -0.0775 | 1.93 | 2.14 | 0.050906 | 0.929474 | 0.121334 | 0.997641 | 0.997988 | 0.994666 |
| Abl1_P00520_S903s_1 | Abl1_P005 | -0.49 | -0.62 | 0.26 | 0.64 | -1.32 | -0.93 | 0.19 | 0.52 | -0.02 | 0.65 | 1.05 | 1.07 | -0.72 | -0.51 | 0.53 | 0.75 | -0.3325 | 0.74 | 0.065 | 0.88752 | 0.502855 | 0.08679 | 0.989223 | 0.997988 | 0.999507 |
| Abl1_P00520_Y179y_1 | Abl1_P005 | -0.33 | 0.26 | 0.27 | 0.37 | 0.35 | -0.45 | -0.1 | 0.29 | -0.21 | -0.01 | 0.47 | -0.32 | 0.8 | 0.08 | 0.54 | 0.33 | -0.12 | -0.16 | 0.295 | 0.322984 | 0.670447 | 0.577679 | 0.992876 | 0.997988 | 0.994666 |
| Abl2_F8VQH0_S632s_1 | Abl2_F8VQ | -1.51 | -0.59 | -1.64 | -0.91 | -0.99 | -1.4 | -0.73 | -0.68 | -1.77 | -1.38 | -0.64 | -0.5 | -1.23 | -1.35 | -0.57 | -0.41 | 0.2125 | 0.09 | 0.2725 | 0.461132 | 0.505215 | 0.818277 | 0.989223 | 0.997988 | 0.994666 |
| Abl2_F8VQH0_S785s_1 | Abl2_F8VQ | -0.45 | 0.05 | 0.37 | -0.7 | 0.41 | 0.01 | -0.17 | 0.26 | -0.25 | -0.22 | 0.4 | -0.02 | -0.02 | 1.25 | -0.05 | -0.01 | 0.31 | 0.16 | 0.475 | 0.258143 | 0.313686 | 0.614315 | 0.989223 | 0.997988 | 0.994666 |
| Abl2_F8VQH0_S819s_1 | Abl2_F8VQ | -0.7 | 0.03 | -0.5 | -1.91 | -0.22 | 0.1 | -0.22 | 0.02 | 1.49 | 0.01 | 1.22 | 0.62 | 2.19 | 0.26 | 0.25 | -0.14 | 0.69 | 1.605 | 1.41 | 0.040927 | 0.111401 | 0.009428 | 0.989223 | 0.997988 | 0.994666 |
| Abl2_F8VQH0_S822s_1 | Abl2_F8VQ | 1.66 | -0.27 | 1.29 | 1.35 | 0.74 | 1.53 | 0.6 | 1.75 | 1.43 | 0.71 | 1.35 | 1.63 | 0.83 | 1.51 | 0.88 | 0.99 | 0.1475 | 0.2725 | 0.045 | 0.92007 | 0.758497 | 0.555073 | 0.995762 | 0.997988 | 0.999507 |
| Abl2_F8VQH0_S936s_1 | Abl2_F8VQ | 0.4 | -0.84 | -0.51 | -0.62 | 0.58 | 0.5 | -0.41 | -0.35 | -0.55 | -0.82 | -0.8 | 0.05 | -1.62 | 0.34 | -1.36 | 0 | 0.4725 | -0.1375 | -0.2675 | 0.611186 | 0.226576 | 0.698734 | 0.989223 | 0.997988 | 0.995259 |
| Abl2_F8VQH0_Y439y_1 | Abl2_F8VQ | 0.82 | 0.49 | 0.9 | 0.66 | 1.1 | 0.46 | 0.57 | 0.77 | -0.28 | 0.69 | 0.14 | 0.15 | 0.45 | -0.23 | 0.32 | 0.39 | 0.0075 | -0.5425 | -0.485 | 0.097457 | 0.974654 | 0.073227 | 0.999187 | 0.997988 | 0.994666 |
| Ablim1_E9Q9C4_S304s_1 | Ablim1_E9 | -0.55 | -0.65 | -1.95 | -2.44 | -1.17 | -0.49 | -1.65 | -2.25 | -0.98 | -0.57 | 1.05 | -1.31 | 0.57 | -0.34 | -1.59 | -2.87 | 0.0075 | 0.945 | 0.34 | 0.665594 | 0.988985 | 0.16455 | 0.999187 | 0.997988 | 0.996159 |
| Ablim1_E9Q9C4_S319s_1 | Ablim1_E9 | -0.28 | -0.87 | -1.41 | -1.08 | -0.1 | -0.12 | -0.21 | -1.2 | -0.64 | -0.29 | -1.86 | -1.01 | -0.31 | -0.26 | -1.48 | -1.83 | 0.5025 | -0.04 | -0.06 | 0.894671 | 0.180602 | 0.921757 | 0.989223 | 0.997988 | 0.999507 |
| Ablim1_E9Q9C4_S319s_2 | Ablim1_E9 | 1.2 | 1.59 | -0.24 | 1.05 | 1.52 | 0.34 | 0.58 | 1.16 | 1.04 | 0.86 | -0.04 | 1.33 | -1.02 | 1.1 | 0.09 | 0.35 | -3.1E-16 | -0.1025 | -0.77 | 0.18287 | 1 | 0.827836 | 1 | 0.997988 | 0.994666 |
| Ablim1_E9Q9C4_S378s_1 | Ablim1_E9 | 0.68 | 0.33 | 0.96 | 0.41 | -0.25 | 1.1 | -0.32 | 0.26 | 0.52 | 0.9 | -0.09 | 0.58 | 0.83 | -1.85 | 0.07 | 0.62 | -0.3975 | -0.1175 | -0.6775 | 0.257354 | 0.281349 | 0.690669 | 0.989223 | 0.997988 | 0.994666 |
| Ablim1_E9Q9C4_S383s_1 | Ablim1_E9 | 0.99 | 0.84 | 0.58 | 0.32 | 1.31 | 0.76 | 0.87 | 0.62 | 1 | 1.04 | 0.27 | 0.64 | 0.46 | 1.61 | 0.49 | 0.2 | 0.2075 | 0.055 | 0.0075 | 0.983702 | 0.431679 | 0.845714 | 0.989223 | 0.997988 | 1 |
| Ablim1_E9Q9C4_S387s_1 | Ablim1_E9 | 1.11 | 0.8 | 1.34 | 0.51 | 1.06 | 0.91 | 0.39 | 1.37 | 1.38 | 1.16 | 1.14 | 1.65 | -0.6 | 1.08 | 1.71 | 0.93 | -0.0075 | 0.3925 | -0.16 | 0.747116 | 0.979941 | 0.174123 | 0.999187 | 0.997988 | 0.9971 |
| Ablim1_E9Q9C4_S404s_1 | Ablim1_E9 | -0.58 | -0.39 | -0.42 | -0.22 | -0.08 | -0.03 | -0.35 | -0.08 | -0.82 | -0.6 | 0.27 | 0.09 | -0.43 | -0.4 | -0.39 | -0.28 | 0.2675 | 0.1375 | 0.0275 | 0.905752 | 0.219637 | 0.65736 | 0.989223 | 0.997988 | 0.999507 |
| Ablim1_E9Q9C4_S407s_1 | Ablim1_E9 | 0.46 | -0.61 | 0.44 | 0.12 | 0.29 | 0.49 | 0.22 | 1.05 | 0.35 | 0.05 | -0.01 | 0.49 | -0.29 | 0.07 | -0.31 | 0.38 | 0.41 | 0.1175 | -0.14 | 0.678598 | 0.227079 | 0.706307 | 0.989223 | 0.997988 | 0.996159 |
| Ablim1_E9Q9C4_S410s_1 | Ablim1_E9 | 0.57 | 0.46 | 0.7 | 0.08 | 0.88 | 0.89 | 0.38 | 0.47 | 0.89 | 0.84 | 0.95 | 0.53 | -0.28 | 0.35 | 0.8 | 0.31 | 0.2025 | 0.35 | -0.1575 | 0.615556 | 0.422268 | 0.174843 | 0.989223 | 0.997988 | 0.995259 |
| Ablim1_E9Q9C4_S425s_1 | Ablim1_E9 | -0.99 | -1.04 | -1 | -0.39 | -0.37 | -0.6 | -0.3 | -0.21 | 0.08 | -1.17 | -0.31 | -0.45 | 0.35 | -0.22 | -0.15 | -0.93 | 0.485 | 0.3925 | 0.6175 | 0.095342 | 0.068466 | 0.251152 | 0.989223 | 0.997988 | 0.994666 |
| Ablim1_E9Q9C4_S442s_1 | Ablim1_E9 | -2.26 | -1.12 | -0.16 | -0.86 | 0.29 | -1.25 | -0.64 | -1.23 | -2.03 | -2.03 | 0.08 | -0.27 | -1.12 | -0.44 | -1.32 | -1.09 | 0.3925 | 0.0375 | 0.1075 | 0.815577 | 0.454476 | 0.953756 | 0.989223 | 0.997988 | 0.999507 |
| Ablim1_E9Q9C4_S491s_1 | Ablim1_E9 | 0.16 | 0.26 | -0.19 | 0.52 | 0.44 | 0.25 | 0.3 | -0.08 | 0.2 | 0.37 | -0.21 | 0.24 | -1.21 | -0.27 | 0.14 | -0.09 | 0.04 | -0.0375 | -0.545 | 0.150848 | 0.870176 | 0.885125 | 0.996493 | 0.997988 | 0.994666 |
| Ablim1_E9Q9C4_S492s_1 | Ablim1_E9 | 0.34 | 0.4 | 0.01 | 0.68 | 0.55 | 0.54 | 0.59 | 0.32 | 0.17 | 0.36 | -0.89 | 0.67 | -0.77 | 0.65 | 0.21 | 0.35 | 0.1425 | -0.28 | -0.2475 | 0.498327 | 0.536253 | 0.459253 | 0.989223 | 0.997988 | 0.994666 |
| Ablim1_E9Q9C4_S544s_1 | Ablim1_E9 | 0.53 | -0.76 | -0.69 | -0.18 | 1.2 | 0.21 | -0.31 | 0.02 | -0.93 | -0.06 | -0.29 | 0.12 | -0.16 | 0.1 | -1.11 | -0.19 | 0.555 | -0.015 | -0.065 | 0.872001 | 0.206103 | 0.968488 | 0.989223 | 0.997988 | 0.999507 |
| Ablim1_E9Q9C4_S559s_1 | Ablim1_E9 | -0.1 | -0.21 | -0.14 | -0.01 | 0.63 | 0.02 | 0.01 | 0.17 | -0.45 | -0.24 | -0.05 | -0.26 | -0.63 | 0.59 | -0.33 | 0.24 | 0.3225 | -0.135 | 0.0825 | 0.798814 | 0.180562 | 0.53384 | 0.989223 | 0.997988 | 0.999507 |
| Ablim1_E9Q9C4_S573s_1 | Ablim1_E9 | 0.15 | 0.42 | 0.37 | 1.1 | 0.84 | 0.46 | 1.2 | 0.9 | -0.65 | 0.81 | 0.95 | 0.95 | -0.71 | 0.65 | 0.29 | 0.8 | 0.34 | 0.005 | -0.2525 | 0.534626 | 0.25414 | 0.990691 | 0.989223 | 0.997988 | 0.994666 |
| Ablim1_E9Q9C4_S610s_1 | Ablim1_E9 | -1.05 | -1.54 | -1.76 | -1.03 | -0.45 | -1.66 | -0.73 | -1.18 | -1.72 | -0.78 | -0.28 | -1.59 | -1.53 | -1.07 | -1.25 | -1.36 | 0.34 | 0.2525 | 0.0425 | 0.880225 | 0.318562 | 0.520357 | 0.989223 | 0.997988 | 0.999507 |
| Ablim1_E9Q9C4_T303t_1 | Ablim1_E9 | -0.78 | 0.13 | -0.05 | -0.69 | -0.16 | -1.13 | -0.3 | -0.41 | -1.38 | -1.1 | -0.44 | -0.59 | -0.79 | -0.47 | -0.43 | -0.13 | -0.1525 | -0.53 | -0.1075 | 0.733623 | 0.641624 | 0.141068 | 0.989223 | 0.997988 | 0.9971 |
| Ablim2_E9Q0W6_S313s_1 | Ablim2_E9 | 0.64 | 0.97 | 0.65 | 0.86 | 0.38 | 0.15 | 1.12 | 0.62 | -0.38 | 0.9 | -0.98 | 0.64 | 0.58 | -0.41 | 0.89 | 0.93 | -0.2125 | -0.735 | -0.2825 | 0.430001 | 0.434193 | 0.116616 | 0.989223 | 0.997988 | 0.994666 |
| Ablim2_E9Q0W6_S366s_1 | Ablim2_E9 | -2.44 | -1.61 | -0.76 | -3.08 | -2.53 | -1.69 | -2.83 | -2.46 | -0.43 | -2.12 | -1.43 | -1.72 | -0.76 | -3.98 | -1.77 | -1.82 | -0.405 | 0.5475 | -0.11 | 0.88339 | 0.435684 | 0.350348 | 0.989223 | 0.997988 | 0.999507 |
| Ablim2_E9Q0W6_S402s_1 | Ablim2_E9 | 0.34 | -0.36 | 0.8 | 0.8 | -1.34 | 0.81 | 0.37 | 0.12 | 2.15 | -0.43 | -1.98 | 0.68 | 1.79 | -1.37 | 0.6 | 0.65 | -0.405 | -0.29 | 0.0225 | 0.972042 | 0.422464 | 0.723023 | 0.989223 | 0.997988 | 0.999507 |
| Ablim2_E9Q0W6_S457s_1 | Ablim2_E9 | -0.97 | -0.91 | -0.26 | -0.85 | -0.88 | -1.77 | -0.3 | -2.06 | -0.75 | -0.9 | 0.3 | -1.37 | -0.15 | -0.53 | -0.18 | -0.6 | -0.505 | 0.0675 | 0.3825 | 0.193376 | 0.24316 | 0.862472 | 0.989223 | 0.997988 | 0.994666 |
| Ablim2_E9Q0W6_S486s_1 | Ablim2_E9 | 1.44 | -0.6 | 0.47 | 1.13 | 1.22 | 2.31 | 0.88 | 1.45 | 0.49 | 0.28 | 0.54 | 0.21 | -0.36 | 0.83 | 0.6 | 0.77 | 0.855 | -0.23 | -0.15 | 0.764908 | 0.112269 | 0.60597 | 0.989223 | 0.997988 | 0.9971 |
| Ablim2_E9Q0W6_S487s_1 | Ablim2_E9 | 1.44 | -0.6 | 0.44 | 0.23 | 1.22 | 2.31 | 0.28 | 0.9 | 0.49 | 0.28 | 0.95 | 0.57 | -0.36 | 0.83 | -0.05 | 0.16 | 0.8 | 0.195 | -0.2325 | 0.623066 | 0.162586 | 0.652057 | 0.989223 | 0.997988 | 0.995259 |
| Ablim2_E9Q0W6_S511s_1 | Ablim2_E9 | 0.06 | 0.46 | 0.37 | 0.46 | 0.23 | -0.09 | 0.63 | 0.48 | -0.22 | 0.27 | 0.34 | 0.54 | -0.99 | 0.14 | 0.46 | 0.39 | -0.025 | -0.105 | -0.3375 | 0.372485 | 0.918649 | 0.683372 | 0.997335 | 0.997988 | 0.994666 |
| Ablim2_E9Q0W6_T397t_1 | Ablim2_E9 | -0.51 | -1.98 | -0.56 | -0.35 | -1.43 | -1.26 | 0.08 | -0.27 | -0.64 | -1.92 | -0.15 | -0.42 | -1.03 | -1.14 | -0.44 | -0.63 | 0.13 | 0.0675 | 0.04 | 0.923034 | 0.789861 | 0.894966 | 0.995762 | 0.997988 | 0.999507 |
| Ablim3_Q69ZX8_S282s_1 | Ablim3_Q6 | 2.74 | 1.88 | 2.02 | 1.04 | 1.31 | 2.61 | 3.66 | 2.08 | 0.71 | 2.51 | 0.76 | 2.43 | 0.74 | 0.72 | 2.63 | 3.15 | 0.495 | -0.3175 | -0.11 | 0.865905 | 0.375645 | 0.576627 | 0.989223 | 0.997988 | 0.999507 |
| Ablim3_Q69ZX8_S337s_1 | Ablim3_Q6 | 0.89 | 0.73 | 1.37 | 1.14 | 0.92 | 0.86 | 1.14 | 1.28 | 1.09 | 0.96 | 1.92 | 1.44 | 0.64 | 1.08 | 1.19 | 1.23 | 0.0175 | 0.32 | 0.0025 | 0.992784 | 0.941392 | 0.29767 | 0.999187 | 0.997988 | 1 |
| Ablim3_Q69ZX8_S372s_1 | Ablim3_Q6 | -1.01 | -0.81 | -0.62 | 0.05 | 0.18 | -0.48 | -0.54 | -0.74 | -0.96 | -0.97 | -0.75 | 0.04 | 0.25 | 0.35 | -0.05 | -0.08 | 0.2025 | -0.0625 | 0.715 | 0.04129 | 0.53064 | 0.857457 | 0.989223 | 0.997988 | 0.994666 |
| Ablim3_Q69ZX8_S373s_1 | Ablim3_Q6 | 0.79 | 0.72 | 0.2 | 0.85 | 0.66 | 1.54 | 0.73 | 1.13 | 0.57 | 1.02 | 1.18 | 1.03 | 0.73 | 1.29 | 1.07 | 0.62 | 0.375 | 0.31 | 0.2875 | 0.329126 | 0.209012 | 0.256595 | 0.989223 | 0.997988 | 0.994666 |
| Ablim3_Q69ZX8_S408s_1 | Ablim3_Q6 | 0.86 | 0.69 | 0.1 | -0.43 | 0.69 | 1.3 | 0.3 | 0.6 | 0.27 | 0.79 | 0.16 | -0.06 | 0.37 | 0.94 | 0.1 | -0.19 | 0.4175 | -0.015 | 0 | 1 | 0.263106 | 0.966504 | 0.989223 | 0.997988 | 1 |
| Ablim3_Q69ZX8_S502s_1 | Ablim3_Q6 | 1.05 | 0.05 | 0.66 | 0.79 | 0.47 | 0.49 | 0.61 | 0.73 | 0.56 | 0.61 | 0.88 | 0.93 | -1.32 | 1.26 | 0.58 | 0.66 | -0.0625 | 0.1075 | -0.3425 | 0.540177 | 0.814055 | 0.703984 | 0.995762 | 0.997988 | 0.994666 |
| Ablim3_Q69ZX8_T542t_1 | Ablim3_Q6 | 1.17 | 1.02 | 0.23 | -0.3 | 0.95 | 1.11 | 0.68 | 0.18 | 0.67 | 1.11 | -0.71 | 0.46 | 0.2 | -0.1 | -0.37 | 0.27 | 0.2 | -0.1475 | -0.53 | 0.192685 | 0.609183 | 0.764194 | 0.989223 | 0.997988 | 0.994666 |
| Abr_Q5SSL4_S53s_1 | Abr_Q5SSL | 0.01 | 0.71 | 0.23 | 0.3 | 0.87 | 0.65 | 0.46 | 0.34 | -0.13 | -0.44 | 0.04 | 0.25 | -0.17 | 0.71 | 0.1 | 0.16 | 0.2675 | -0.3825 | -0.1125 | 0.706866 | 0.292928 | 0.175645 | 0.989223 | 0.997988 | 0.996966 |
| Abr_Q5SSL4_S72s_1 | Abr_Q5SSL | 0.23 | 0.09 | -0.13 | 0.69 | 0.13 | 0.45 | 1.08 | 0.42 | -0.53 | 0.28 | -1.08 | 0.26 | -0.63 | -0.68 | -0.27 | 0.18 | 0.3 | -0.4875 | -0.57 | 0.095461 | 0.31979 | 0.217938 | 0.989223 | 0.997988 | 0.994666 |
| Abr_Q5SSL4_T294t_1 | Abr_Q5SSL | 2.57 | 0.94 | 1.48 | 2.04 | 0.4 | 2.09 | 0.33 | 1.62 | 2.25 | 2.95 | 1.08 | 1.28 | 2.08 | 0.83 | 1.3 | 1.22 | -0.6475 | 0.1325 | -0.4 | 0.366129 | 0.227882 | 0.800279 | 0.989223 | 0.997988 | 0.994666 |
| Acap2_D4AFX6_S386s_1 | Acap2_D4A | -0.27 | -1.1 | -0.17 | -0.4 | -1.13 | 0.55 | -0.13 | -0.23 | -0.3 | 0.36 | -0.39 | -0.2 | 0.57 | 0.22 | -0.77 | -0.61 | 0.25 | 0.3525 | 0.3375 | 0.401367 | 0.527193 | 0.266775 | 0.989223 | 0.997988 | 0.994666 |
| Acap2_D4AFX6_S388s_2 | Acap2_D4A | -0.43 | -0.5 | 0.89 | 0.75 | -0.91 | 0.06 | -0.61 | 0.3 | 0.4 | -0.58 | 0.99 | 0.15 | -1.21 | -0.36 | 0.13 | 0.39 | -0.4675 | 0.0625 | -0.44 | 0.377468 | 0.30187 | 0.894351 | 0.989223 | 0.997988 | 0.994666 |
| Acap2_D4AFX6_S391s_1 | Acap2_D4A | 0.71 | 0.5 | 0.87 | 0.85 | -0.51 | 0.64 | -0.11 | 0.73 | 0.63 | 0.7 | 0.89 | 0.43 | 1.28 | 0.13 | 0.08 | 0.73 | -0.545 | -0.07 | -0.1775 | 0.598173 | 0.11794 | 0.75906 | 0.989223 | 0.997988 | 0.995259 |
| Acap2_D4AFX6_S391s_2 | Acap2_D4A | 0.04 | -0.24 | 0.89 | 0.75 | -0.68 | 0.12 | -0.61 | 0.3 | 1.26 | 0.32 | 0.99 | 0.15 | -2.03 | 0.21 | 0.13 | 0.39 | -0.5775 | 0.32 | -0.685 | 0.256004 | 0.139497 | 0.413415 | 0.989223 | 0.997988 | 0.994666 |
| Acap2_D4AFX6_S528s_1 | Acap2_D4A | -0.02 | 0.17 | 0.97 | 1.02 | 0.51 | -0.45 | 1 | 1.33 | -0.61 | -0.02 | 0.51 | 1.03 | -0.41 | 0.14 | 0.98 | 0.93 | 0.0625 | -0.3075 | -0.125 | 0.76916 | 0.887582 | 0.481573 | 0.996493 | 0.997988 | 0.9971 |
| Acap2_D4AFX6_S559s_1 | Acap2_D4A | -0.33 | -1.81 | -1.06 | -0.05 | -1.51 | 0.27 | -0.21 | -0.51 | -2.74 | -2.35 | 0.89 | 0.53 | -2.73 | -3.02 | -0.56 | -1.04 | 0.3225 | -0.105 | -1.025 | 0.142187 | 0.523248 | 0.907934 | 0.989223 | 0.997988 | 0.994666 |
| Acap2_D4AFX6_S562s_1 | Acap2_D4A | 0.46 | 0.04 | 0.53 | 1.07 | -0.5 | 0.78 | 0.12 | 1.08 | 1.07 | -0.43 | 0 | 1.4 | 1.23 | -0.22 | 0.11 | 1.23 | -0.155 | -0.015 | 0.0625 | 0.883802 | 0.696762 | 0.974021 | 0.994893 | 0.997988 | 0.999507 |
| Acap2_D4AFX6_S568s_1 | Acap2_D4A | 0.47 | -1.33 | -0.45 | -0.89 | -1.42 | 1.14 | -0.95 | 0.49 | -1.54 | -2.02 | -0.88 | 0.04 | -2.33 | -3.77 | 0.14 | 0.3 | 0.365 | -0.55 | -0.865 | 0.362529 | 0.568028 | 0.32798 | 0.989223 | 0.997988 | 0.994666 |
| Acap2_D4AFX6_S580s_1 | Acap2_D4A | -0.49 | -0.7 | 1.99 | 1.69 | -0.25 | -1.68 | 1.07 | -0.03 | -1.05 | -0.6 | 3.22 | 1.21 | -1.54 | 0.36 | -0.52 | 1.38 | -0.845 | 0.0725 | -0.7025 | 0.406275 | 0.300554 | 0.945383 | 0.989223 | 0.997988 | 0.994666 |
| Acap3_Q6NXL5_S383s_1 | Acap3_Q6N | -0.48 | -0.13 | -0.28 | -0.14 | -0.06 | -0.54 | -0.06 | 0.18 | -0.34 | -0.06 | 0.77 | 0.11 | 0.5 | 0.81 | -0.02 | 0.13 | 0.1375 | 0.3775 | 0.6125 | 0.051942 | 0.567668 | 0.218371 | 0.989223 | 0.997988 | 0.994666 |
| Acap3_Q6NXL5_S383s_2 | Acap3_Q6N | -0.15 | 0.74 | 0.48 | 0.43 | 0.77 | -0.07 | -0.16 | -0.01 | -0.35 | 0.99 | 1.21 | 0.26 | -1.43 | 0.87 | 0.47 | 0.72 | -0.2425 | 0.1525 | -0.2175 | 0.682571 | 0.436593 | 0.704016 | 0.989223 | 0.997988 | 0.996159 |
| Acap3_Q6NXL5_S388s_1 | Acap3_Q6N | -0.1 | 0.54 | 0.26 | -0.58 | 0.9 | -0.18 | -0.15 | -0.07 | -0.38 | 0.48 | 0.65 | -0.36 | 0.81 | 0.68 | -0.72 | 0.04 | 0.095 | 0.0675 | 0.1725 | 0.684673 | 0.78914 | 0.85579 | 0.995762 | 0.997988 | 0.996159 |
| Acin1_F6RJ39_S414s_1 | Acin1_F6RJ | -2.41 | -0.79 | 0 | -1.25 | -1.42 | -1.27 | -1.42 | -1.27 | -1.42 | 0.12 | 0.16 | -0.5 | 0.27 | 0.41 | -0.69 | 0.26 | -0.2325 | 0.7025 | 1.175 | 0.047804 | 0.621905 | 0.241355 | 0.989223 | 0.997988 | 0.994666 |
| Acin1_F6RJ39_S425s_1 | Acin1_F6RJ | 0.4 | 0.21 | 0.02 | 0.57 | 0.17 | 0.35 | 0.44 | 0.68 | -0.14 | 0.38 | 0.08 | 0.31 | 0.43 | 0.31 | 0.8 | 0.59 | 0.11 | -0.1425 | 0.2325 | 0.381073 | 0.637703 | 0.56615 | 0.989223 | 0.997988 | 0.994666 |
| Acin1_F6RJ39_S437s_1 | Acin1_F6RJ | -1.6 | -3.27 | -2.58 | -6.02 | -3.82 | -1.96 | -2.44 | -2.65 | 2.54 | -3.43 | -2.9 | -1.67 | 3.63 | -4.18 | 2.5 | -4.02 | 0.65 | 2.0025 | 2.85 | 0.168129 | 0.472722 | 0.18827 | 0.989223 | 0.997988 | 0.994666 |
| Acin1_F6RJ39_S541s_1 | Acin1_F6RJ | 1.16 | 0.83 | 1.13 | -0.01 | 0.68 | 0.63 | -0.26 | 0.34 | 0.69 | 1.1 | 0.65 | 0.15 | 1.6 | 1.23 | 0.9 | 0.21 | -0.43 | -0.13 | 0.2075 | 0.611671 | 0.238281 | 0.711573 | 0.989223 | 0.997988 | 0.995259 |
| Acin1_F6RJ39_S601s_1 | Acin1_F6RJ | -0.64 | 0.68 | 0.1 | -0.81 | -0.8 | -0.54 | -0.39 | -0.6 | -0.89 | -1.19 | -0.49 | -0.27 | 0.74 | 0.25 | 0.74 | -0.85 | -0.415 | -0.5425 | 0.3875 | 0.432531 | 0.260114 | 0.196421 | 0.989223 | 0.997988 | 0.994666 |
| Acin1_F6RJ39_S603s_1 | Acin1_F6RJ | 1.17 | 0.52 | 0.56 | 0.2 | 0.58 | 0.76 | 0.5 | 0.25 | 0.77 | 0.25 | 0.47 | 0.22 | 1.53 | 0.84 | 1.52 | 0.03 | -0.09 | -0.185 | 0.3675 | 0.381692 | 0.739887 | 0.523258 | 0.995762 | 0.997988 | 0.994666 |
| Acin1_F6RJ39_S675s_1 | Acin1_F6RJ | 2.36 | 0.98 | 1.05 | 0.73 | -0.33 | 1.83 | -0.64 | 0.13 | 1.69 | 0.79 | -0.69 | 0.3 | 2.48 | -0.91 | 0.55 | -0.18 | -1.0325 | -0.7575 | -0.795 | 0.289667 | 0.106908 | 0.204988 | 0.989223 | 0.997988 | 0.994666 |
| Acin1_F6RJ39_S784s_1 | Acin1_F6RJ | 0.17 | 0.02 | -0.53 | -0.61 | -1.44 | -0.29 | -1.41 | -0.76 | 0.08 | -1.1 | -1.04 | -1.34 | 0.87 | -1.62 | -0.38 | -1.12 | -0.7375 | -0.6125 | -0.325 | 0.547998 | 0.054963 | 0.130332 | 0.989223 | 0.997988 | 0.994666 |
| Acin1_F6RJ39_S832s_1 | Acin1_F6RJ | -2.09 | -2.21 | -1.1 | -2.67 | -2.9 | -2.21 | -2.1 | -1.95 | -2.51 | -2.71 | -0.25 | -1.87 | -0.97 | -3.06 | -1.92 | -2.31 | -0.2725 | 0.1825 | -0.0475 | 0.92622 | 0.481492 | 0.759327 | 0.989223 | 0.997988 | 0.999507 |
| Acin1_F6RJ39_S923s_1 | Acin1_F6RJ | -1.05 | -1.45 | -0.55 | -0.43 | 0.16 | -0.51 | 0.18 | -0.85 | -1.33 | -1.84 | 0.44 | -0.69 | -0.78 | 0.33 | -0.22 | -0.88 | 0.615 | 0.015 | 0.4825 | 0.224538 | 0.103987 | 0.976562 | 0.989223 | 0.997988 | 0.994666 |
| Acin1_F6RJ39_S937s_1 | Acin1_F6RJ | 0.4 | 0.1 | -0.21 | -0.17 | -0.62 | -0.23 | -0.43 | 0.14 | -0.22 | -0.64 | -0.18 | -0.48 | 0.63 | 0.13 | 0.07 | -0.81 | -0.315 | -0.41 | -0.025 | 0.944143 | 0.248491 | 0.129216 | 0.989223 | 0.997988 | 0.999507 |
| Acin1_Q9JIX8_S858s_1 | Acin1_Q9JI | -2.45 | -2.76 | -1.38 | -3.09 | -3.56 | -2.59 | -2.78 | -2.5 | -3.02 | -3.35 | -0.6 | -2.18 | -1.09 | -4.03 | -2.31 | -2.86 | -0.4375 | 0.1325 | -0.1525 | 0.813243 | 0.311644 | 0.83877 | 0.989223 | 0.997988 | 0.999507 |
| Acly_Q3TS02_S455s_1 | Acly_Q3TS( | 0.09 | -0.9 | 0.14 | -0.1 | -1.22 | 0.08 | -0.98 | -0.03 | -0.06 | -0.48 | 0.8 | 0.44 | 0.12 | -0.59 | -0.32 | -0.22 | -0.345 | 0.3675 | -0.06 | 0.854015 | 0.391438 | 0.341299 | 0.989223 | 0.997988 | 0.999507 |
| Acly_Q3TS02_S481s_1 | Acly_Q3TS( | -1.33 | -1.4 | -1.37 | -1.11 | -1.11 | -1.47 | -1.12 | -0.91 | -1.31 | -0.97 | -0.88 | -1.36 | -0.63 | -0.75 | -1.34 | -1.31 | 0.15 | 0.1725 | 0.295 | 0.302163 | 0.501547 | 0.463655 | 0.989223 | 0.997988 | 0.994666 |
| Aco2_Q99KI0_S559s_1 | Aco2_Q99I | 0.79 | 1.08 | 0.39 | 0.22 | 0.56 | 0.33 | 0.1 | 0.33 | 0.88 | 1.04 | 1.01 | 0.44 | 1.31 | 1.17 | 0.18 | -0.23 | -0.29 | 0.2225 | -0.0125 | 0.976295 | 0.285847 | 0.444767 | 0.989223 | 0.997988 | 1 |
| Acot1_O55137_S416s_1 | Acot1_O55 | -1.01 | -2.21 | -1.65 | -1.12 | -1.29 | -1.71 | -0.72 | -1.39 | -1.61 | -0.52 | -0.37 | -1.61 | -0.87 | -0.73 | -1.76 | -0.72 | 0.22 | 0.47 | 0.4775 | 0.233466 | 0.530897 | 0.285821 | 0.989223 | 0.997988 | 0.994666 |
| Acot1_O55137_S56s_1 | Acot1_O55 | -0.97 | -0.21 | 0.42 | 0.07 | 0.18 | -0.88 | -0.35 | -0.34 | -0.29 | -0.35 | 0.25 | 0.48 | -0.26 | 0.81 | 0.76 | 1.02 | -0.175 | 0.195 | 0.755 | 0.092831 | 0.631915 | 0.598471 | 0.989223 | 0.997988 | 0.994666 |
| Acot11_F6SS52_S14s_1 | Acot11_F6 | -1.26 | -1.44 | -1.3 | -1.16 | -0.76 | -1.23 | -1.01 | -1.24 | -1.52 | -1.37 | -0.73 | -0.84 | -0.8 | -0.22 | -1.52 | -1.04 | 0.23 | 0.175 | 0.395 | 0.239151 | 0.305377 | 0.514094 | 0.989223 | 0.997988 | 0.994666 |
| Acot11_F6SS52_S20s_1 | Acot11_F6 | 1.56 | 1.92 | 1.65 | 1.45 | 1.22 | 1.57 | 1.4 | 0.92 | 1.25 | 1.26 | 1.39 | 0.01 | 0.09 | 1.59 | 0.07 | 1.23 | -0.3675 | -0.6675 | -0.9 | 0.049306 | 0.147358 | 0.085017 | 0.989223 | 0.997988 | 0.994666 |
| Acot7_Q91V12_S285s_1 | Acot7_Q91 | -1.95 | -1.35 | -2.95 | -3.25 | 0.1 | -1.03 | -2.52 | -3.53 | -2.24 | -1.66 | -2.64 | -2.87 | -0.85 | -0.27 | -3.75 | -2.83 | 0.63 | 0.0225 | 0.45 | 0.585859 | 0.438081 | 0.963067 | 0.989223 | 0.997988 | 0.995259 |
| Acot7_Q91V12_S2s_1 | Acot7_Q91 | 0.28 | -0.1 | 0.39 | 0.82 | 0.59 | 0.33 | 1.08 | 1.06 | -0.32 | -0.65 | -0.08 | 0.86 | -0.5 | 0.44 | 0.44 | 0.31 | 0.4175 | -0.395 | -0.175 | 0.602721 | 0.176291 | 0.313099 | 0.989223 | 0.997988 | 0.995259 |
| Acsbg1_Q99PU5_S55s_1 | Acsbg1_Q9 | 1.46 | 1.15 | 0.12 | -0.58 | 0.66 | -1.46 | 0.41 | -0.08 | 0.66 | 0.87 | 0.73 | 0.16 | 0.43 | 1.33 | -0.42 | -0.38 | -0.655 | 0.0675 | -0.2975 | 0.607234 | 0.28805 | 0.88603 | 0.989223 | 0.997988 | 0.995259 |
| Acsbg1_Q99PU5_S70s_1 | Acsbg1_Q9 | 0.62 | 0.95 | 0.09 | 0.24 | 0.99 | 1.14 | 1.13 | 0.78 | 0.96 | 1.1 | -0.05 | 0.36 | 0.18 | 1.39 | 0.06 | 0.86 | 0.535 | 0.1175 | 0.1475 | 0.699476 | 0.063673 | 0.735616 | 0.989223 | 0.997988 | 0.996159 |
| Acsl6_Q5F2C5_S364s_1 | Acsl6_Q5F | -0.14 | 0.27 | 0.32 | 0.86 | 1.4 | -0.29 | 0.52 | 0.17 | 0.77 | 0.24 | 0.25 | 0.26 | -0.04 | 0.82 | 0.34 | 0.34 | 0.1225 | 0.0525 | 0.0375 | 0.906282 | 0.758178 | 0.855816 | 0.995762 | 0.997988 | 0.999507 |
| Acss2_Q9QXG4_S265s_1 | Acss2_Q9C | 0.41 | 0.78 | 0.08 | 0 | 0.41 | 0.7 | 0.57 | 0.14 | 0.71 | 0.75 | 0.44 | 0.21 | 0.27 | 0.58 | 0.37 | 0.36 | 0.1375 | 0.21 | 0.0775 | 0.777509 | 0.602763 | 0.450682 | 0.989223 | 0.997988 | 0.99798 |
| Acss2_Q9QXG4_S267s_1 | Acss2_Q9C | -0.47 | -0.18 | -0.23 | -0.31 | -0.11 | -0.26 | 0.14 | -0.32 | -0.17 | 0.22 | -0.32 | 0 | -0.19 | 0.08 | -0.28 | -0.54 | 0.16 | 0.23 | 0.065 | 0.797273 | 0.463421 | 0.330295 | 0.989223 | 0.997988 | 0.999507 |
| Acss2_Q9QXG4_S30s_1 | Acss2_Q9C | -0.44 | -0.19 | -0.4 | -0.67 | 0.15 | -0.74 | -0.05 | -0.62 | -1.03 | -0.23 | -0.9 | -0.26 | -0.6 | 0.12 | -0.42 | -0.66 | 0.11 | -0.18 | 0.035 | 0.900693 | 0.691149 | 0.527776 | 0.994893 | 0.997988 | 0.999507 |
| Actb_P60710_S233s_1 | Actb_P607 | -0.21 | -0.2 | 0.03 | -0.42 | -0.14 | 0.14 | -0.18 | -0.13 | -0.06 | 0.09 | -0.25 | -0.91 | -0.48 | -0.3 | -0.12 | -0.2 | 0.1225 | -0.0825 | -0.075 | 0.759277 | 0.569984 | 0.773607 | 0.989223 | 0.997988 | 0.9971 |
| Actb_P60710_S239s_1 | Actb_P607 | -0.07 | -0.03 | -0.66 | 0.05 | 1.32 | -0.38 | 0.56 | 0.24 | -0.49 | 1.3 | -0.1 | -0.21 | -0.3 | 1.11 | 0.72 | -0.25 | 0.6125 | 0.3025 | 0.4975 | 0.228996 | 0.132171 | 0.480334 | 0.989223 | 0.997988 | 0.994666 |
| Actb_P60710_S52s_1 | Actb_P607 | -1.36 | 2.14 | -2.25 | -2.49 | 0.78 | -0.55 | -1.72 | -0.99 | -2.46 | 0.64 | -3.92 | -2.45 | 0.18 | -1.65 | -1.33 | -1 | 0.37 | -1.0575 | 0.04 | 0.967914 | 0.720326 | 0.410786 | 0.995437 | 0.997988 | 0.999507 |
| Actb_P60710_T203t_1 | Actb_P607 | 0.27 | 0.2 | 0.34 | 0.23 | 0.06 | -0.39 | -0.24 | -0.2 | -0.53 | -0.21 | -0.33 | -0.31 | -0.27 | -0.72 | -0.25 | -0.1 | -0.4525 | -0.605 | -0.595 | 0.037372 | 0.051074 | 0.016272 | 0.989223 | 0.997988 | 0.994666 |
| Actb_P60710_Y198y_1 | Actb_P607 | -0.26 | -0.46 | 0.4 | 0.35 | -0.23 | -0.55 | -0.33 | -0.42 | 0.21 | -0.4 | 0.87 | -0.01 | 1.89 | 1.51 | -0.29 | 0.1 | -0.39 | 0.16 | 0.795 | 0.159974 | 0.170696 | 0.654751 | 0.989223 | 0.997988 | 0.994666 |
| Actr3_Q99JY9_S418s_1 | Actr3_Q99 | 7.38 | 4 | 5.91 | 2.33 | 4.76 | 7.5 | 6.66 | 5.2 | -0.51 | 6.62 | 0.65 | 5.99 | -2.35 | 0.91 | 5.72 | 6.65 | 1.125 | -1.7175 | -2.1725 | 0.300351 | 0.319144 | 0.36541 | 0.989223 | 0.997988 | 0.994666 |
| Acvr2a_P27038_S184s_1 | Acvr2a_P2 | -0.68 | 0.22 | -1.46 | -0.39 | -0.03 | -1.17 | -0.3 | -1.16 | -0.68 | -0.97 | -0.41 | -1.16 | -1.19 | -0.25 | 0.23 | -0.5 | -0.0875 | -0.2275 | 0.15 | 0.737404 | 0.838951 | 0.560754 | 0.995762 | 0.997988 | 0.9971 |
| Acy3_Q91XE4_S7s_1 | Acy3_Q91X | 1.5 | 0.44 | 0.62 | -0.65 | 0.1 | 1.25 | -2.88 | -0.61 | 3.18 | 0.88 | 0.71 | -0.93 | 2.76 | -1.02 | 1.75 | 0.32 | -1.0125 | 0.4825 | 0.475 | 0.568311 | 0.251682 | 0.572836 | 0.989223 | 0.997988 | 0.995259 |
| Adam22_Q9R1V6_S772s_1 | Adam22_Q | 0.28 | 0.39 | 0.76 | 0.47 | 0.76 | 0.3 | 0.89 | 0.63 | -0.22 | 0.51 | -0.62 | 1.03 | 0.34 | -0.93 | 1.04 | 0.76 | 0.17 | -0.3 | -0.1725 | 0.695227 | 0.474119 | 0.443901 | 0.989223 | 0.997988 | 0.996159 |
| Adam22_Q9R1V6_S774s_1 | Adam22_Q | 0.29 | 0.5 | -0.73 | -1.03 | 0.67 | 0.42 | -0.58 | -0.32 | -0.46 | 0.3 | -1.2 | -0.93 | 0.26 | -0.6 | -0.34 | -0.36 | 0.29 | -0.33 | -0.0175 | 0.966521 | 0.521748 | 0.492696 | 0.989223 | 0.997988 | 0.999507 |
| Adam22_Q9R1V6_S796s_1 | Adam22_Q | 0.34 | 0.31 | 0.84 | 1.14 | 0.49 | 0.55 | 0.75 | 0.87 | 0.72 | 0.5 | 0.28 | 0.85 | 0.95 | 0.34 | 0.74 | 0.52 | 0.0075 | -0.07 | -0.02 | 0.947062 | 0.97743 | 0.806462 | 0.999187 | 0.997988 | 0.999507 |
| Adam22_Q9R1V6_S813s_1 | Adam22_Q | -0.09 | 0.38 | 0.45 | 0.53 | 0.47 | 0.38 | 0.62 | 0.31 | 0.32 | 0.98 | 0.8 | 0.55 | 0.98 | 0.49 | 0.54 | 0.25 | 0.1275 | 0.345 | 0.2475 | 0.39061 | 0.582302 | 0.212044 | 0.989223 | 0.997988 | 0.994666 |
| Adam22_Q9R1V6_S819s_1 | Adam22_Q | 0.18 | 0.76 | 0.22 | 0.83 | 0.67 | -0.17 | 1.08 | 0.25 | -0.17 | 0.58 | -0.37 | 0.41 | -0.44 | 0.95 | 0.19 | -0.62 | -0.04 | -0.385 | -0.4775 | 0.251797 | 0.903477 | 0.242649 | 0.996493 | 0.997988 | 0.994666 |
| Adam22_Q9R1V6_S867s_1 | Adam22_Q | -0.48 | 0.86 | 0.04 | 0.62 | -0.05 | -0.72 | -0.33 | -0.07 | 0.89 | 0.63 | 1.51 | -0.18 | 1.12 | 0.4 | 1.04 | 0.69 | -0.5525 | 0.4525 | 0.5525 | 0.154552 | 0.132785 | 0.323688 | 0.989223 | 0.997988 | 0.994666 |
| Adam22_Q9R1V6_S867s_2 | Adam22_Q | 0.33 | 0.83 | 1.3 | 0.84 | -0.28 | 0.04 | 0.64 | 0.94 | 1.22 | 0.76 | 2.01 | 0.48 | 0.26 | 0.09 | 1.57 | 0.82 | -0.49 | 0.2925 | -0.14 | 0.724475 | 0.178346 | 0.45989 | 0.989223 | 0.997988 | 0.9971 |
| Adam22_Q9R1V6_S871s_2 | Adam22_Q | 0.94 | 1.46 | 1.3 | 0.84 | 0.6 | 0.65 | 0.64 | 0.94 | 1.56 | 1.1 | 2.01 | 0.48 | 0.88 | 0.78 | 1.57 | 0.82 | -0.4275 | 0.1525 | -0.1225 | 0.683622 | 0.095132 | 0.67924 | 0.989223 | 0.997988 | 0.996159 |
| Adam22_Q9R1V6_S874s_1 | Adam22_Q | 0.05 | 0.01 | 0.52 | 0.64 | -0.16 | 0.24 | -0.03 | 0.07 | 0.55 | -0.74 | 0.12 | -0.29 | 1.35 | -0.45 | 0.08 | 0.48 | -0.275 | -0.395 | 0.06 | 0.884536 | 0.275653 | 0.262772 | 0.989223 | 0.997988 | 0.999507 |
| Adcy2_Q3V1Q3_S1089s_1 | Adcy2_Q3V | 0.95 | 0.28 | 0.57 | 0.63 | 0.8 | 0.52 | 0.13 | 0.39 | 0.64 | 0.62 | 0.52 | 0.32 | 0.06 | -0.07 | 0.49 | 0.29 | -0.1475 | -0.0825 | -0.415 | 0.150631 | 0.560906 | 0.732049 | 0.989223 | 0.997988 | 0.994666 |
| Adcy2_Q3V1Q3_S476s_1 | Adcy2_Q3V | 0.03 | 0.29 | 0.6 | 0.69 | 0.06 | 0.08 | 0.59 | 0.61 | -0.06 | 0.37 | 0.21 | 0.64 | -0.45 | 0.15 | 0.24 | 0.25 | -0.0675 | -0.1125 | -0.355 | 0.24346 | 0.79717 | 0.677477 | 0.995762 | 0.997988 | 0.994666 |
| Adcy5_P84309_S137s_1 | Adcy5_P84 | 1.11 | 0.91 | 0.04 | 0.18 | 0.61 | 0.22 | 0.53 | -0.03 | 1.08 | 0.91 | 0.1 | 0.34 | 1.04 | 0.71 | 0.54 | -0.52 | -0.2275 | 0.0475 | -0.1175 | 0.782518 | 0.480826 | 0.895646 | 0.989223 | 0.997988 | 0.998274 |
| Adcy5_P84309_S156s_1 | Adcy5_P84 | 0.26 | 0.64 | 0.89 | 1.15 | 0.44 | 0.53 | 0.9 | 1.03 | 0.09 | 0.56 | 0.38 | 0.36 | -0.3 | 0.38 | 0.99 | 0.72 | -0.01 | -0.3875 | -0.2875 | 0.433884 | 0.971028 | 0.175907 | 0.999187 | 0.997988 | 0.994666 |
| Adcy5_P84309_S49s_1 | Adcy5_P84 | 1.79 | -0.92 | -0.8 | 0.11 | 0.1 | 1.1 | -1.02 | -1.08 | 2.04 | 0.93 | 0.25 | -2.07 | 1.5 | -0.06 | -0.82 | 0.07 | -0.27 | 0.2425 | 0.1275 | 0.856817 | 0.706598 | 0.798003 | 0.995437 | 0.997988 | 0.999507 |
| Adcy5_P84309_S8s_1 | Adcy5_P84 | 0.15 | 0.82 | 0.61 | 0.91 | 0.24 | 0.36 | 1.29 | 1.26 | 0.54 | 0.69 | 1.25 | 0.98 | 0.49 | 0.6 | 1.13 | 1.02 | 0.165 | 0.2425 | 0.1875 | 0.530126 | 0.626429 | 0.399375 | 0.989223 | 0.997988 | 0.994666 |
| Adcy5_P84309_T62t_1 | Adcy5_P84 | -0.82 | -2.56 | -0.78 | -0.47 | -2.11 | -0.15 | -2.78 | -2.03 | 0.8 | -1.16 | -0.39 | 0.77 | 0.87 | -2.28 | -1.6 | -3.22 | -0.61 | 1.1625 | -0.4 | 0.648477 | 0.36221 | 0.084719 | 0.989223 | 0.997988 | 0.995259 |
| Adcy5_P84309_T665t_1 | Adcy5_P84 | 1.48 | 0.71 | -1.25 | -1.14 | 1.09 | 1.4 | -0.84 | -1.24 | 1 | 0.92 | -0.1 | -0.91 | -1.01 | 1.21 | -0.89 | -1.13 | 0.1525 | 0.2775 | -0.405 | 0.605233 | 0.854349 | 0.706503 | 0.996349 | 0.997988 | 0.995259 |
| Adcy6_F8VQ52_S53s_1 | Adcy6_F8V | -0.15 | 0.62 | 0.66 | 0.36 | 0.66 | 0.47 | 0.24 | 0.08 | 0.67 | 0.13 | 0.44 | 0.47 | 0.8 | 0.52 | 0.21 | 0.37 | -0.01 | 0.055 | 0.1025 | 0.726476 | 0.9703 | 0.840582 | 0.999187 | 0.997988 | 0.9971 |
| Adcy6_F8VQ52_S576s_1 | Adcy6_F8V | 0.46 | 0.62 | 0.56 | 0.47 | 0.87 | 0.51 | 0.52 | 0.28 | 0.31 | 0.4 | 0.28 | 0.65 | -0.05 | 1.03 | 0.07 | 0.4 | 0.0175 | -0.1175 | -0.165 | 0.589684 | 0.935923 | 0.587344 | 0.99868 | 0.997988 | 0.995259 |
| Adcy8_P97490_S67s_1 | Adcy8_P97 | 0.54 | -0.24 | 0.12 | 0.83 | 0.61 | 0.43 | 0.69 | 0.84 | 0.46 | 0.85 | -0.6 | 0.4 | -0.25 | 1.16 | -0.31 | 0.43 | 0.33 | -0.035 | -0.055 | 0.894934 | 0.261962 | 0.928294 | 0.989223 | 0.997988 | 0.999507 |
| Adcy9_P51830_S1273s_1 | Adcy9_P51 | -0.16 | 0.36 | 0.17 | -0.02 | 0.39 | -0.07 | 0.49 | 0.16 | -0.09 | -0.13 | 0.47 | -0.06 | -0.43 | 0.07 | -0.31 | 0.13 | 0.155 | -0.04 | -0.2225 | 0.416277 | 0.517851 | 0.874469 | 0.989223 | 0.997988 | 0.994666 |
| Adcy9_P51830_S354s_1 | Adcy9_P51 | -0.25 | -0.57 | 1.66 | 1.31 | -0.77 | 0.15 | 0.54 | 0.93 | 1.87 | -0.01 | 0.47 | 0.21 | 2.11 | -0.61 | 0.69 | 0.85 | -0.325 | 0.0975 | 0.2225 | 0.752033 | 0.587517 | 0.878042 | 0.989223 | 0.997988 | 0.9971 |
| Adcy9_P51830_S39s_1 | Adcy9_P51 | -0.03 | -2.48 | -1.05 | 0.12 | -1.61 | 0.28 | -1.73 | -0.4 | -0.76 | -1.48 | -2.53 | -1.5 | 0.3 | -2.05 | -1.43 | -0.7 | -0.005 | -0.7075 | -0.11 | 0.875278 | 0.99414 | 0.284073 | 0.999278 | 0.997988 | 0.999507 |
| Adcy9_P51830_S49s_1 | Adcy9_P51 | -0.44 | 0.02 | 0.41 | 0.95 | 0.34 | 0.06 | 0.85 | 1.23 | -0.69 | -0.52 | 0.53 | -0.05 | -0.82 | 0.46 | 0.21 | 1.24 | 0.385 | -0.4175 | 0.0375 | 0.939114 | 0.328702 | 0.311222 | 0.989223 | 0.997988 | 0.999507 |
| Adcy9_P51830_S610s_1 | Adcy9_P51 | 0.44 | 0.84 | 1.29 | 1.09 | 0.82 | 0.11 | 1.38 | 1.1 | 0.44 | 0.92 | 1.28 | 1.03 | 0.48 | 0.76 | 0.47 | 0.69 | -0.0625 | 0.0025 | -0.315 | 0.273389 | 0.852495 | 0.993252 | 0.996349 | 0.997988 | 0.994666 |
| Adcy9_P51830_S613s_1 | Adcy9_P51 | 0.67 | 0.38 | 1.15 | 1.01 | 0.72 | 0.34 | 1.14 | 0.71 | 0.48 | 0.67 | 0.55 | 0.84 | -0.1 | 0.62 | 0.38 | 0.53 | -0.075 | -0.1675 | -0.445 | 0.159624 | 0.786316 | 0.521147 | 0.995762 | 0.997988 | 0.994666 |
| Adcy9_P51830_S678s_1 | Adcy9_P51 | 1.97 | 1.82 | 1.94 | 0.83 | 1.54 | 1.81 | 1.72 | 1.64 | -0.64 | 2.12 | -0.91 | 1.66 | 0.15 | -0.32 | 1.22 | 1.9 | 0.0375 | -1.0825 | -0.9025 | 0.116294 | 0.901019 | 0.167272 | 0.996493 | 0.997988 | 0.994666 |
| Adcy9_P51830_T1275t_1 | Adcy9_P51 | -1.02 | 0.19 | 0.22 | 0.33 | -1.48 | -1.1 | 0.11 | 0.59 | -0.4 | -0.97 | 1.15 | 0.16 | -0.58 | -0.25 | -0.29 | 0.12 | -0.4 | 0.055 | -0.18 | 0.627916 | 0.457567 | 0.915219 | 0.989223 | 0.997988 | 0.995259 |
| Adcyap1r1_E9Q3E8_S438s_1 | Adcyap1r1_ | 0.66 | 0.18 | 1.3 | 0.63 | 2.21 | 0.64 | 1.19 | 0.62 | 1.55 | 0.4 | 1.26 | 1.29 | 0.58 | 0.56 | 0.51 | 1.02 | 0.4725 | 0.4325 | -0.025 | 0.936052 | 0.273452 | 0.241242 | 0.989223 | 0.997988 | 0.999507 |
| Add1_F8WGR0_S12s_1 | Add1_F8W | 0.42 | 0.2 | 0.9 | 0.68 | 0.15 | 0.12 | 0.33 | 0.85 | 0.3 | 0.13 | 0.96 | 0.75 | 0.64 | 0.42 | 0.36 | -0.02 | -0.1875 | -0.015 | -0.2 | 0.483734 | 0.492828 | 0.958807 | 0.989223 | 0.997988 | 0.994666 |
| Add1_F8WGR0_S353s_2 | Add1_F8W | 1.45 | 1.24 | 0.83 | 0.81 | 0.36 | 0.57 | 0.11 | 0.12 | 1.39 | 1.06 | 0.66 | 0.43 | 0.76 | 0.59 | 0.59 | 0.85 | -0.7925 | -0.1975 | -0.385 | 0.168904 | 0.009803 | 0.519069 | 0.989223 | 0.997988 | 0.994666 |
| Add1_F8WGR0_S355s_1 | Add1_F8W | 0.83 | 0.8 | 0.52 | 0.95 | 0.41 | 0.2 | 0.34 | 0.62 | 0.98 | 0.54 | 0.84 | 0.48 | 1.33 | 0.55 | 0.79 | 0.54 | -0.3825 | -0.065 | 0.0275 | 0.922356 | 0.103773 | 0.78483 | 0.989223 | 0.997988 | 0.999507 |
| Add1_F8WGR0_S355s_2 | Add1_F8W | 1.03 | 1.15 | 0.93 | 0.79 | 0.41 | 0.76 | 0.02 | 0.16 | 0.97 | 0.87 | 0.75 | 0.44 | -0.06 | 0.09 | 0.59 | 1.01 | -0.6375 | -0.2175 | -0.5675 | 0.092471 | 0.024393 | 0.3608 | 0.989223 | 0.997988 | 0.994666 |
| Add1_F8WGR0_S366s_1 | Add1_F8W | 0.89 | 0.45 | 0.27 | 0.49 | 0.52 | 0.2 | 0.51 | 0.1 | -0.1 | 0.4 | 0.07 | 0.22 | -0.15 | 0.58 | -0.06 | 0.09 | -0.1925 | -0.3775 | -0.41 | 0.171287 | 0.425398 | 0.150839 | 0.989223 | 0.997988 | 0.994666 |
| Add1_F8WGR0_S457s_2 | Add1_F8W | 0.76 | 0.68 | 0.07 | -0.7 | -2.02 | 0.43 | -1.13 | 0.22 | 2.75 | -1.13 | 1.44 | 0.03 | 1.17 | -0.6 | 1.01 | -0.9 | -0.8275 | 0.57 | -0.0325 | 0.955357 | 0.189771 | 0.487771 | 0.989223 | 0.997988 | 0.999507 |
| Add1_F8WGR0_S464s_1 | Add1_F8W | 0.88 | -0.47 | -0.45 | 0.38 | -0.95 | 1.18 | -0.74 | 0.72 | 2.62 | 0.67 | -1.1 | -0.6 | 2.65 | -0.61 | 1.1 | -0.75 | -0.0325 | 0.3125 | 0.5125 | 0.510502 | 0.953817 | 0.696138 | 0.999187 | 0.997988 | 0.994666 |
| Add1_F8WGR0_S465s_1 | Add1_F8W | 0.97 | -0.76 | 0.66 | 0.6 | -1.11 | 1.19 | -0.26 | 0.38 | 2.82 | 0.77 | 2.09 | 1.16 | 2.8 | -0.5 | 1.19 | -0.17 | -0.3175 | 1.3425 | 0.4625 | 0.541476 | 0.573691 | 0.035942 | 0.989223 | 0.997988 | 0.994666 |
| Add1_F8WGR0_S465s_2 | Add1_F8W | 0.76 | 0.68 | 0.07 | -0.7 | -2.02 | 0.43 | -1.13 | 0.22 | 2.75 | -1.13 | 1.44 | 0.03 | 1.17 | -0.6 | 1.01 | -0.9 | -0.8275 | 0.57 | -0.0325 | 0.955357 | 0.189771 | 0.487771 | 0.989223 | 0.997988 | 0.999507 |
| Add1_F8WGR0_S617s_1 | Add1_F8W | 0.53 | 0.05 | 0.89 | 1.05 | -0.7 | 0.47 | -0.21 | 0.71 | 0.49 | 0.39 | 0.59 | 0.81 | 0.53 | -0.77 | 0.52 | 0.66 | -0.5625 | -0.06 | -0.395 | 0.343873 | 0.163583 | 0.83426 | 0.989223 | 0.997988 | 0.994666 |
| Add1_F8WGR0_S631s_1 | Add1_F8W | 0.69 | 1.78 | 0.21 | 0.88 | 1.19 | 0.33 | 1.05 | 0.59 | 1.5 | 1.38 | 0.79 | 1 | 0.98 | 1.52 | 0.81 | 1.32 | -0.1 | 0.2775 | 0.2675 | 0.487888 | 0.790807 | 0.464941 | 0.995762 | 0.997988 | 0.994666 |
| Add1_F8WGR0_T358t_1 | Add1_F8W | 0.58 | 0.55 | 1.32 | 1.27 | 0.17 | 0.59 | 0.52 | 0.93 | 0.23 | 0.43 | 1.64 | 0.76 | 0.88 | -0.27 | 1.35 | 1.21 | -0.3775 | -0.165 | -0.1375 | 0.744904 | 0.21569 | 0.66592 | 0.989223 | 0.997988 | 0.9971 |
| Add1_F8WGR0_T358t_2 | Add1_F8W | 1.05 | 1.15 | 0.93 | 0.79 | 0.43 | 0.77 | 0.02 | 0.16 | 0.97 | 0.9 | 0.75 | 0.44 | -0.03 | 0.11 | 0.59 | 1.01 | -0.635 | -0.215 | -0.56 | 0.092898 | 0.025601 | 0.368649 | 0.989223 | 0.997988 | 0.994666 |
| Add1_F8WGR0_T641t_1 | Add1_F8W | 0.81 | 1.06 | 0.82 | 0.64 | 0.58 | 0.37 | 0.66 | 0.48 | 0.74 | 1.11 | 0.63 | 0.76 | 1 | 1.15 | 0.78 | 0.74 | -0.31 | -0.0225 | 0.085 | 0.732002 | 0.16278 | 0.922346 | 0.989223 | 0.997988 | 0.9971 |
| Add1_F8WGR0_T641t_2 | Add1_F8W | 0.23 | -0.22 | -0.04 | 0.05 | 0.76 | -0.59 | 0.61 | 0.3 | 0 | 0.17 | -0.03 | 0.39 | -0.09 | 0.79 | 0.15 | 0.43 | 0.265 | 0.1275 | 0.315 | 0.28483 | 0.427 | 0.583539 | 0.989223 | 0.997988 | 0.994666 |
| Add1_F8WGR0_T645t_1 | Add1_F8W | -0.51 | -0.76 | -0.42 | 0.18 | -0.03 | -0.07 | -0.13 | 0.27 | -0.65 | 0.16 | -0.76 | -0.15 | 0.08 | -0.06 | 0.49 | 0.23 | 0.3875 | 0.0275 | 0.5625 | 0.082428 | 0.16593 | 0.93185 | 0.989223 | 0.997988 | 0.994666 |
| Add1_F8WGR0_T645t_2 | Add1_F8W | 0.23 | -0.22 | -0.12 | -0.04 | 0.76 | -0.59 | 0.82 | 0.43 | 0 | 0.17 | -0.4 | 0.6 | -0.09 | 0.79 | 0.15 | 0.38 | 0.3925 | 0.13 | 0.345 | 0.243991 | 0.271601 | 0.644317 | 0.989223 | 0.997988 | 0.994666 |
| Add1_F8WHZ9_S631s_1 | Add1_F8W | 0.69 | 1.78 | 0.21 | 0.88 | 1.19 | 0.33 | 1.05 | 0.59 | 1.5 | 1.38 | 0.79 | 1 | 0.98 | 1.52 | 0.81 | 1.32 | -0.1 | 0.2775 | 0.2675 | 0.487888 | 0.790807 | 0.464941 | 0.995762 | 0.997988 | 0.994666 |
| Add1_F8WHZ9_T641t_1 | Add1_F8W | 0.5 | 1.07 | 0.45 | 0.29 | 0.54 | 0.11 | 0.36 | 0.11 | 0.5 | 0.85 | 0.19 | 0.51 | 0.92 | 1.12 | 0.47 | 0.46 | -0.2975 | -0.065 | 0.165 | 0.584318 | 0.258503 | 0.812126 | 0.989223 | 0.997988 | 0.995259 |
| Add1_F8WHZ9_T641t_2 | Add1_F8W | 0.23 | -0.22 | -0.04 | 0.05 | 0.76 | -0.59 | 0.61 | 0.3 | 0 | 0.17 | -0.03 | 0.39 | -0.09 | 0.79 | 0.15 | 0.43 | 0.265 | 0.1275 | 0.315 | 0.28483 | 0.427 | 0.583539 | 0.989223 | 0.997988 | 0.994666 |
| Add1_F8WHZ9_T645t_1 | Add1_F8W | -0.51 | -0.76 | -0.42 | 0.18 | -0.03 | -0.07 | -0.13 | 0.27 | -0.65 | 0.16 | -0.76 | -0.15 | 0.08 | -0.06 | 0.49 | 0.23 | 0.3875 | 0.0275 | 0.5625 | 0.082428 | 0.16593 | 0.93185 | 0.989223 | 0.997988 | 0.994666 |
| Add1_F8WHZ9_T645t_2 | Add1_F8W | 0.23 | -0.22 | -0.12 | -0.04 | 0.76 | -0.59 | 0.82 | 0.43 | 0 | 0.17 | -0.4 | 0.6 | -0.09 | 0.79 | 0.15 | 0.38 | 0.3925 | 0.13 | 0.345 | 0.243991 | 0.271601 | 0.644317 | 0.989223 | 0.997988 | 0.994666 |
| Add1_Q9QYC0_S600s_1 | Add1_Q9Q | 0.69 | 1.78 | 0.21 | 0.88 | 1.19 | 0.33 | 1.05 | 0.59 | 1.5 | 1.38 | 0.79 | 1 | 0.98 | 1.52 | 0.81 | 1.32 | -0.1 | 0.2775 | 0.2675 | 0.487888 | 0.790807 | 0.464941 | 0.995762 | 0.997988 | 0.994666 |
| Add1_Q9QYC0_S628s_1 | Add1_Q9Q | -2.69 | -0.81 | -2.11 | -2.8 | -0.95 | -2.64 | -2.63 | -2.84 | -2.23 | -2.44 | -0.36 | -2.73 | -2.29 | -0.89 | -3.02 | -3.59 | -0.1625 | 0.1625 | -0.345 | 0.60714 | 0.776208 | 0.79996 | 0.995762 | 0.997988 | 0.995259 |
| Add1_Q9QYC0_T610t_1 | Add1_Q9Q | 1.06 | 1.29 | 1.64 | 1.63 | 0.75 | 0.53 | 1.47 | 1.47 | 0.95 | 1.31 | 1.7 | 1.7 | 1.04 | 1.28 | 1.6 | 1.36 | -0.35 | 0.01 | -0.085 | 0.753589 | 0.26671 | 0.971503 | 0.989223 | 0.997988 | 0.9971 |
| Add1_Q9QYC0_T610t_2 | Add1_Q9Q | 0.23 | -0.22 | 0.29 | 0.56 | 0.76 | -0.59 | 0.86 | 0.61 | 0 | 0.17 | 0.12 | 0.69 | -0.09 | 0.79 | 0.49 | 0.5 | 0.195 | 0.03 | 0.2075 | 0.500035 | 0.599755 | 0.913512 | 0.989223 | 0.997988 | 0.994666 |
| Add1_Q9QYC0_T614t_1 | Add1_Q9Q | -0.51 | -0.76 | -0.42 | 0.18 | -0.03 | -0.07 | -0.13 | 0.27 | -0.65 | 0.16 | -0.76 | -0.15 | 0.08 | -0.06 | 0.49 | 0.23 | 0.3875 | 0.0275 | 0.5625 | 0.082428 | 0.16593 | 0.93185 | 0.989223 | 0.997988 | 0.994666 |
| Add1_Q9QYC0_T614t_2 | Add1_Q9Q | 0.23 | -0.22 | 0.36 | 0.66 | 0.76 | -0.59 | 0.97 | 0.71 | 0 | 0.17 | 0.07 | 0.8 | -0.09 | 0.79 | 0.56 | 0.51 | 0.205 | 0.0025 | 0.185 | 0.558884 | 0.599686 | 0.993322 | 0.989223 | 0.997988 | 0.995259 |
| Add2_Q9QYB8_S11s_1 | Add2_Q9Q | 0.72 | 0.42 | 0.51 | 0.59 | 0.28 | 0.65 | 1.12 | 0.9 | 0.38 | 1.1 | 0.67 | 0.58 | -0.45 | 0.46 | 0.69 | 0.98 | 0.1775 | 0.1225 | -0.14 | 0.687673 | 0.481694 | 0.618918 | 0.989223 | 0.997988 | 0.996159 |
| Add2_Q9QYB8_S528s_1 | Add2_Q9Q | 0.98 | 0.05 | 0.38 | 1.18 | 0.29 | 0.7 | 0.87 | 1.25 | 1.97 | 0.98 | 1.17 | 0.63 | 1.56 | 1.02 | 0.77 | 0.53 | 0.13 | 0.54 | 0.3225 | 0.38671 | 0.700197 | 0.186759 | 0.994893 | 0.997988 | 0.994666 |
| Add2_Q9QYB8_S530s_1 | Add2_Q9Q | 1.6 | 0.31 | 0.39 | 1.6 | 1.1 | 1.14 | 0.88 | 1.68 | 0.51 | 0.64 | 1.6 | 0.89 | -0.02 | 1.16 | 1.17 | 0.74 | 0.225 | -0.065 | -0.2125 | 0.63549 | 0.564895 | 0.878437 | 0.989223 | 0.997988 | 0.995259 |
| Add2_Q9QYB8_S532s_1 | Add2_Q9Q | 0.03 | 0.6 | 0.54 | 0.22 | -0.1 | 1.02 | 0.77 | 0.78 | 0.21 | 0.38 | 0.75 | 0.62 | -0.13 | 0.99 | -0.1 | 0.37 | 0.27 | 0.1425 | -0.065 | 0.845596 | 0.383732 | 0.576632 | 0.989223 | 0.997988 | 0.999507 |
| Add2_Q9QYB8_S539s_1 | Add2_Q9Q | 0.21 | -0.24 | -0.76 | -0.88 | 0.46 | -0.1 | 0.26 | -0.81 | 0.73 | 0.36 | -0.49 | -0.33 | 0.58 | 0.3 | -0.7 | -0.74 | 0.37 | 0.485 | 0.2775 | 0.514396 | 0.330612 | 0.227625 | 0.989223 | 0.997988 | 0.994666 |
| Add2_Q9QYB8_S594s_1 | Add2_Q9Q | 0.09 | 1.06 | 0.29 | 0.01 | 0.54 | -0.62 | -0.06 | 0.11 | 0.38 | 0.22 | 1.3 | 0.81 | 0.77 | 1.07 | 0.49 | -0.08 | -0.37 | 0.315 | 0.2 | 0.58656 | 0.297167 | 0.384797 | 0.989223 | 0.997988 | 0.995259 |
| Add2_Q9QYB8_S598s_1 | Add2_Q9Q | -1.12 | -2.32 | -2.27 | -1.43 | 0.27 | -0.63 | -1.28 | -1.9 | -0.68 | -2.01 | -3.19 | -1.98 | 0.23 | -0.35 | -0.62 | -2.04 | 0.9 | -0.18 | 1.09 | 0.064098 | 0.10096 | 0.74445 | 0.989223 | 0.997988 | 0.994666 |
| Add2_Q9QYB8_S598s_2 | Add2_Q9Q | -2.47 | -1.23 | 0.46 | 0.28 | -1.15 | -3.1 | -0.34 | 0.03 | -2.29 | -2.22 | 0.35 | 0.12 | -1.9 | -1.86 | 0.38 | 0.35 | -0.4 | -0.27 | -0.0175 | 0.98323 | 0.641552 | 0.760836 | 0.989223 | 0.997988 | 1 |
| Add2_Q9QYB8_S598s_3 | Add2_Q9Q | -2.07 | -1 | -2.12 | -1.84 | -1.24 | -1.75 | -1.66 | -1.89 | -2.03 | -1.53 | -0.86 | -1.43 | -1.52 | -1.14 | -1.42 | -1.35 | 0.1225 | 0.295 | 0.4 | 0.230074 | 0.69658 | 0.425964 | 0.994893 | 0.997988 | 0.994666 |
| Add2_Q9QYB8_S602s_1 | Add2_Q9Q | 0.33 | 0.98 | 1.23 | 0.96 | 0.09 | -0.25 | 0.36 | 0.89 | 0.49 | 0.75 | 1.99 | 1.11 | 1.85 | 0.5 | 1.09 | 1.14 | -0.6025 | 0.21 | 0.27 | 0.460919 | 0.086487 | 0.586241 | 0.989223 | 0.997988 | 0.994666 |
| Add2_Q9QYB8_S602s_2 | Add2_Q9Q | 0.82 | 0.93 | 1.27 | 1.11 | -0.07 | 0.24 | 0.31 | 0.72 | 0.89 | 1.19 | 1.06 | 1.01 | 0.51 | -0.08 | 1.18 | 0.95 | -0.7325 | 0.005 | -0.3925 | 0.256986 | 0.014319 | 0.982138 | 0.989223 | 0.997988 | 0.994666 |

| | | | | | | | | | | | | | | | | | | | | | | | | | | |
|---|---|---|---|---|---|---|---|---|---|---|---|---|---|---|---|---|---|---|---|---|---|---|---|---|---|---|
| Add2_Q9QYB8_S602s_3 | Add2_Q9Q | -2.07 | -1 | -2.12 | -1.84 | -1.24 | -1.75 | -1.66 | -1.89 | -2.03 | -1.53 | -0.86 | -1.43 | -1.52 | -1.14 | -1.42 | -1.35 | 0.1225 | 0.295 | 0.4 | 0.230074 | 0.69658 | 0.425964 | 0.994893 | 0.997988 | 0.994666 |
| Add2_Q9QYB8_S606s_2 | Add2_Q9Q | 0.96 | 1.06 | 0.98 | 0.79 | 0.02 | 0.4 | 0.02 | 0.46 | 1.03 | 1.34 | 0.75 | 0.66 | 0.65 | -0.02 | 0.87 | 0.68 | -0.7225 | -0.0025 | -0.4025 | 0.174078 | 0.008183 | 0.991823 | 0.989223 | 0.997988 | 0.994666 |
| Add2_Q9QYB8_S606s_3 | Add2_Q9Q | -2.07 | -1 | -2.12 | -1.84 | -1.24 | -1.75 | -1.66 | -1.89 | -2.03 | -1.53 | -0.86 | -1.43 | -1.52 | -1.14 | -1.42 | -1.35 | 0.1225 | 0.295 | 0.4 | 0.230074 | 0.69658 | 0.425964 | 0.994893 | 0.997988 | 0.994666 |
| Add2_Q9QYB8_S60s_1 | Add2_Q9Q | -1.31 | -0.53 | -0.23 | -0.14 | -0.11 | -1.21 | 0.7 | -0.28 | -1.63 | -1.39 | -1.1 | -0.17 | -0.66 | -0.71 | -0.27 | -0.22 | 0.3275 | -0.52 | 0.0875 | 0.793496 | 0.467104 | 0.225483 | 0.989223 | 0.997988 | 0.999507 |
| Add2_Q9QYB8_S614s_2 | Add2_Q9Q | 2.37 | 0.33 | 0.42 | 0.4 | 0.33 | 2.41 | -0.5 | 0.04 | 2.37 | 0.98 | -1.97 | 0.17 | 1.05 | -0.68 | -0.42 | 0 | -0.31 | -0.4925 | -0.8925 | 0.145377 | 0.664451 | 0.594034 | 0.991343 | 0.997988 | 0.994666 |
| Add2_Q9QYB8_S618s_1 | Add2_Q9Q | 1.4 | 1.42 | 1.76 | 1.16 | 0.25 | 1.03 | -0.2 | 0.17 | 3.46 | 1.77 | 0.01 | 1.26 | 3.89 | -0.27 | 1.26 | 1.19 | -1.1225 | 0.19 | 0.0825 | 0.91512 | 0.004599 | 0.773028 | 0.989223 | 0.997988 | 0.999507 |
| Add2_Q9QYB8_S618s_2 | Add2_Q9Q | 2.37 | 0.33 | 0.42 | 0.4 | 0.33 | 2.41 | -0.5 | 0.04 | 2.37 | 0.98 | -1.97 | 0.17 | 1.05 | -0.68 | -0.42 | 0 | -0.31 | -0.4925 | -0.8925 | 0.145377 | 0.664451 | 0.594034 | 0.991343 | 0.997988 | 0.994666 |
| Add2_Q9QYB8_S623s_1 | Add2_Q9Q | 2.62 | 1.87 | 1.96 | 2.11 | 0.32 | 1.99 | 0.59 | 1.4 | 2.42 | 2.4 | 2.07 | 1.5 | 2.16 | 0.92 | 1.62 | 1.6 | -1.065 | -0.0425 | -0.565 | 0.121408 | 0.023365 | 0.890083 | 0.989223 | 0.997988 | 0.994666 |
| Add2_Q9QYB8_S62s_1 | Add2_Q9Q | -0.81 | -0.34 | 0.21 | -0.24 | 0.1 | -0.8 | 0.47 | -0.61 | -0.74 | -0.72 | -1.3 | -0.58 | -0.14 | -0.63 | -0.42 | -0.13 | 0.085 | -0.54 | -0.035 | 0.907565 | 0.814357 | 0.099495 | 0.995762 | 0.997988 | 0.999507 |
| Add2_Q9QYB8_S692s_1 | Add2_Q9Q | 0.01 | 0.57 | 0.58 | 0.51 | 0.1 | -0.24 | 0.12 | 0.1 | -0.19 | 0.35 | 0.55 | 0.25 | 0.12 | 0.21 | 0.27 | 0.26 | -0.3975 | -0.1775 | -0.2025 | 0.429844 | 0.114124 | 0.512577 | 0.989223 | 0.997988 | 0.994666 |
| Add2_Q9QYB8_S692s_2 | Add2_Q9Q | 2.79 | 2.33 | -1.29 | -0.91 | 1.64 | 2.49 | -1.16 | -1.19 | 1.86 | 1.86 | 0.08 | -0.98 | 1.06 | 2.23 | -1.45 | -1.18 | -0.285 | -0.025 | -0.565 | 0.638872 | 0.816331 | 0.982169 | 0.995762 | 0.997988 | 0.995259 |
| Add2_Q9QYB8_S696s_1 | Add2_Q9Q | -1.33 | -1.07 | -1.1 | -0.89 | -1.14 | -1.91 | -0.38 | -1.05 | -1.45 | -0.69 | -0.67 | -0.54 | -0.54 | -1.13 | -0.75 | -0.91 | -0.0225 | 0.26 | 0.265 | 0.31727 | 0.946306 | 0.362099 | 0.999187 | 0.997988 | 0.994666 |
| Add2_Q9QYB8_S700s_1 | Add2_Q9Q | 2.2 | 1.87 | 1.28 | 1.25 | 1.27 | 1.76 | 1.26 | 1.25 | 1.48 | 1.98 | 0.86 | 1.17 | 1.68 | 1.72 | 1.65 | 1.45 | -0.265 | -0.2775 | -0.025 | 0.933707 | 0.375665 | 0.434854 | 0.989223 | 0.997988 | 0.999507 |
| Add2_Q9QYB8_T561t_1 | Add2_Q9Q | 0.23 | 0.76 | 0.84 | 0.48 | 0.31 | 0.27 | 1.02 | 0.66 | 0.23 | 0.7 | 0.27 | 0.75 | 0.16 | 0.64 | 0.36 | 1.04 | -0.0125 | -0.09 | -0.0275 | 0.926562 | 0.962705 | 0.731024 | 0.999187 | 0.997988 | 0.999507 |
| Add3_Q9QYB5_S42s_1 | Add3_Q9Q | 0.3 | 0.4 | 0.56 | 0.27 | 0.49 | 0.46 | 0.1 | 0.27 | 0.13 | 0.31 | 0.53 | 0.71 | 0.02 | 0.31 | 0.36 | 0.47 | -0.0525 | 0.0375 | -0.0925 | 0.70423 | 0.804229 | 0.872767 | 0.995762 | 0.997988 | 0.996159 |
| Add3_Q9QYB5_S461s_1 | Add3_Q9Q | -0.63 | -0.91 | -0.46 | -1.68 | -1.12 | 0.86 | -0.64 | 0.16 | 2.57 | -1.26 | -1.21 | -0.22 | 2.99 | -1.95 | 2.5 | -0.62 | 0.735 | 0.89 | 1.65 | 0.146205 | 0.146113 | 0.302457 | 0.989223 | 0.997988 | 0.994666 |
| Add3_Q9QYB5_S641s_1 | Add3_Q9Q | 0.28 | 0.47 | 1.34 | 0.98 | 0.25 | 0.13 | 0.51 | 1 | 0.4 | 0.61 | 0.86 | 1.03 | -0.24 | 0.46 | 0.99 | 1.08 | -0.295 | -0.0425 | -0.195 | 0.624426 | 0.371372 | 0.891452 | 0.989223 | 0.997988 | 0.995259 |
| Add3_Q9QYB5_S645s_1 | Add3_Q9Q | -0.5 | -0.28 | -0.24 | -0.48 | -0.5 | -0.68 | -0.47 | -0.49 | -0.72 | 0.18 | -0.14 | -0.4 | -0.17 | -0.5 | -0.6 | -0.76 | -0.16 | 0.105 | -0.1325 | 0.602 | 0.435793 | 0.692965 | 0.989223 | 0.997988 | 0.995259 |
| Add3_Q9QYB5_S649s_1 | Add3_Q9Q | 1.4 | 1.48 | 1.86 | 1.6 | 1.24 | 1.22 | 1.48 | 1.56 | 1.39 | 1.5 | 1.56 | 1.51 | 1.14 | 1.59 | 1.59 | 1.57 | -0.21 | -0.095 | -0.1125 | 0.661359 | 0.351062 | 0.666863 | 0.989223 | 0.997988 | 0.995671 |
| Add3_Q9QYB5_S64s_1 | Add3_Q9Q | -0.55 | 0.23 | -0.53 | -2.59 | 1.15 | -0.57 | 0.18 | -1.75 | -0.08 | -0.28 | -1.76 | -1.73 | -0.21 | 0.27 | -2.21 | -1.41 | 0.6125 | -0.1025 | -0.03 | 0.967449 | 0.425218 | 0.880675 | 0.989223 | 0.997988 | 0.999507 |
| Add3_Q9QYB5_T12t_1 | Add3_Q9Q | 0.17 | 0.49 | 0.35 | 0.36 | 0.61 | 0.2 | 0.55 | 0.4 | 0.15 | 0.53 | 0.21 | 0.29 | 0.58 | 0.67 | 0.25 | -0.04 | 0.0975 | -0.0475 | 0.0225 | 0.932782 | 0.647104 | 0.828758 | 0.99033 | 0.997988 | 0.999507 |
| Adgra1_Q8C4G9_S537s_1 | Adgra1_Q8 | -1.2 | -0.32 | 0.52 | -0.2 | 0.11 | -1.08 | 0.1 | 0.12 | -0.72 | -0.26 | 1.54 | -2.12 | -0.98 | -0.79 | -1.26 | 0.52 | 0.1125 | -0.09 | -0.3275 | 0.517551 | 0.795885 | 0.904126 | 0.995762 | 0.997988 | 0.994666 |
| Adgra1_Q8C4G9_T239t_1 | Adgra1_Q8 | 4.05 | 2.75 | 3.25 | 1.03 | 3.65 | 4.83 | 3.72 | 2.28 | 0.24 | 4.16 | 1.12 | 3.48 | 0.12 | 0.18 | 3.07 | 3.68 | 0.85 | -0.52 | -1.0075 | 0.322217 | 0.258919 | 0.605341 | 0.989223 | 0.997988 | 0.994666 |
| Adgrb1_E9Q4H5_S336s_1 | Adgrb1_E9 | -0.57 | -0.78 | -0.52 | -0.33 | 0.84 | -0.66 | -0.15 | 0 | -0.58 | -0.64 | -1.78 | -0.74 | -0.09 | 0.84 | -0.44 | -0.2 | 0.5575 | -0.385 | 0.5775 | 0.10925 | 0.120041 | 0.254525 | 0.989223 | 0.997988 | 0.994666 |
| Adgrb1_E9Q4H5_S349s_1 | Adgrb1_E9 | 0.45 | 0.62 | 0.9 | 0.24 | 1.73 | 1.5 | 0.94 | 1.54 | -2 | 1.49 | 1.03 | 1.49 | 0.63 | -0.8 | 1.97 | 0.5 | 0.875 | -0.05 | 0.0225 | 0.966839 | 0.008068 | 0.947749 | 0.989223 | 0.997988 | 0.999507 |
| Adgrb1_E9Q4H5_S364s_1 | Adgrb1_E9 | -2.11 | -1.28 | -0.59 | -1.43 | -0.11 | -1.57 | -0.39 | -1.8 | -2.78 | -2.16 | -1.76 | -1.29 | -1.21 | -0.66 | -1.75 | -0.78 | 0.385 | -0.645 | 0.2525 | 0.535083 | 0.433139 | 0.159616 | 0.989223 | 0.997988 | 0.994666 |
| Adgrb1_E9Q4H5_S527s_1 | Adgrb1_E9 | 0.29 | 0.96 | 0.53 | 0.18 | 1.04 | -0.47 | 0.55 | 0.43 | -1.38 | 0.22 | -2.32 | 0.26 | -0.67 | -0.96 | -0.46 | 0.25 | -0.1025 | -1.295 | -0.95 | 0.019448 | 0.775006 | 0.055149 | 0.995762 | 0.997988 | 0.994666 |
| Adgrb1_E9Q4H5_T308t_1 | Adgrb1_E9 | -0.57 | 0.16 | 0.1 | 0 | -0.42 | 0.15 | 0.39 | 0.6 | -0.26 | 0.42 | 0.67 | 0.02 | -0.82 | 0.47 | -0.09 | -0.18 | 0.2575 | 0.29 | -0.0775 | 0.822482 | 0.401314 | 0.351748 | 0.989223 | 0.997988 | 0.999507 |
| Adgrb2_B1ASC0_S1131s_1 | Adgrb2_B1 | -0.24 | -1.02 | 0.85 | 0.05 | -2.85 | -0.69 | -0.29 | -0.03 | 1.07 | -0.3 | 0.62 | -1.01 | 1.38 | -2.02 | 1.14 | 0.98 | -0.875 | 0.185 | 0.46 | 0.562119 | 0.208501 | 0.741444 | 0.989223 | 0.997988 | 0.995259 |
| Adgrb2_B1ASC0_S1415s_1 | Adgrb2_B1 | 0.58 | 0.59 | 0.92 | 0.48 | 1.02 | 0.51 | 0.81 | 0.66 | 0.58 | 0.59 | 0.89 | 0.84 | 0.82 | 1.11 | 0.85 | 0.24 | 0.1075 | 0.0825 | 0.1125 | 0.691826 | 0.635498 | 0.717113 | 0.989223 | 0.997988 | 0.996159 |
| Adgrl1_H7BX15_S1128s_1 | Adgrl1_H7I | -0.15 | -0.28 | -0.03 | -0.11 | -0.37 | -0.12 | -0.46 | -0.55 | -0.27 | -0.34 | -0.31 | -0.46 | 0.59 | -0.22 | -0.28 | -0.17 | -0.2325 | -0.2025 | 0.1225 | 0.668608 | 0.283165 | 0.342248 | 0.989223 | 0.997988 | 0.996159 |
| Adgrl1_H7BX15_S1146s_1 | Adgrl1_H7I | -1.39 | -3.45 | -1.97 | -1.43 | -2.1 | -1.11 | -1.76 | -1.11 | -2.61 | -2.31 | -0.76 | -0.91 | -1.66 | -3.54 | -1.03 | -1.35 | 0.54 | 0.4125 | 0.165 | 0.804687 | 0.292337 | 0.509181 | 0.989223 | 0.997988 | 0.999507 |
| Adgrl1_H7BX15_S1171s_1 | Adgrl1_H7I | 0.05 | -0.59 | 0.09 | 0.1 | 0.67 | 0.44 | 0.66 | 0.41 | 0.56 | 0.4 | -1.48 | 0.05 | -0.86 | 0.82 | -0.28 | 0.63 | 0.6325 | -0.03 | 0.165 | 0.698776 | 0.025823 | 0.949276 | 0.989223 | 0.997988 | 0.996159 |
| Adgrl1_H7BX15_S1213s_1 | Adgrl1_H7I | -0.36 | 0.38 | 1.45 | 1.46 | 0.29 | -0.49 | 1.68 | 2.13 | -0.5 | -0.41 | 1.05 | 1.75 | -1.38 | 0.19 | 0.89 | 0.86 | 0.17 | -0.26 | -0.5925 | 0.357078 | 0.798623 | 0.688172 | 0.995762 | 0.997988 | 0.994666 |
| Adgrl1_H7BX15_S1219s_1 | Adgrl1_H7I | 1.56 | 1.37 | 0.33 | 1.27 | 1.06 | 1.78 | 1.16 | 0.95 | 1.67 | 1.74 | 0.64 | 1.09 | 0.04 | 0.71 | -0.19 | -0.09 | 0.105 | 0.1525 | -1.015 | 0.018639 | 0.756586 | 0.690413 | 0.995762 | 0.997988 | 0.994666 |
| Adgrl1_H7BX15_S1222s_1 | Adgrl1_H7I | 0.45 | -0.13 | 0.46 | 0.59 | 0.17 | 0.41 | 1.03 | 0.56 | -0.23 | -0.15 | -0.26 | 1.11 | -0.82 | 0.6 | -0.17 | -0.44 | 0.2 | -0.225 | -0.55 | 0.154203 | 0.479333 | 0.552087 | 0.989223 | 0.997988 | 0.994666 |
| Adgrl1_H7BX15_S1374s_1 | Adgrl1_H7I | -1.1 | -0.21 | -0.41 | -0.84 | -0.49 | -0.78 | -0.63 | -1.39 | -1.45 | -0.91 | -1.13 | -0.08 | -0.29 | -0.28 | -0.79 | -1.96 | -0.1825 | -0.2525 | -0.19 | 0.664536 | 0.553313 | 0.495055 | 0.989223 | 0.997988 | 0.996159 |
| Adgrl1_H7BX15_S1384s_1 | Adgrl1_H7I | 0.24 | -0.83 | -0.9 | -0.85 | -0.27 | 0.93 | -0.4 | -0.5 | -1.08 | 0.08 | 0.6 | -0.2 | -1.25 | 0.06 | -1.51 | -1.13 | 0.525 | 0.435 | -0.3725 | 0.402719 | 0.222963 | 0.329625 | 0.989223 | 0.997988 | 0.994666 |
| Adgrl1_H7BX15_S1444s_1 | Adgrl1_H7I | 0.03 | -0.09 | -0.03 | 0.66 | 0.41 | 0.06 | 0.71 | 0.57 | 0.3 | 0.48 | -0.12 | 0.43 | -0.32 | 0.39 | -0.01 | 0.29 | 0.295 | 0.13 | -0.055 | 0.85402 | 0.285803 | 0.63915 | 0.989223 | 0.997988 | 0.999507 |
| Adgrl1_H7BX15_T1172t_1 | Adgrl1_H7I | 1.65 | 0.84 | -0.05 | -0.04 | 1.61 | 1.67 | 0.29 | 0.14 | 0.5 | 1.13 | 0.44 | -0.04 | -0.15 | 0.68 | 0.04 | 0.48 | 0.3275 | -0.0925 | -0.3375 | 0.451318 | 0.538531 | 0.838638 | 0.989223 | 0.997988 | 0.994666 |
| Adgrl2_Q0P6A0_S145s_1 | Adgrl2_Q0I | 0.2 | 0.34 | -0.08 | -0.23 | 0.34 | 0.07 | 0.07 | -0.37 | -0.09 | -0.02 | -0.58 | 0 | -0.28 | 0.38 | -0.41 | -0.39 | -0.03 | -0.23 | -0.2325 | 0.436653 | 0.905019 | 0.381993 | 0.996493 | 0.997988 | 0.994666 |
| Adgrl2_Q0P6A0_S175s_1 | Adgrl2_Q0I | 0.96 | 1.18 | -0.08 | -0.45 | 1.82 | 1.06 | 0.01 | -0.07 | 0.06 | 0.8 | -0.6 | -0.16 | -0.74 | 0.97 | -0.39 | 0.07 | 0.3025 | -0.3775 | -0.425 | 0.413288 | 0.580837 | 0.429207 | 0.989223 | 0.997988 | 0.994666 |
| Adgrl2_Q0P6A0_S180s_1 | Adgrl2_Q0I | 0.41 | 0.76 | 1 | 0.85 | 0.46 | 0.65 | 0.44 | 0.79 | 0.25 | 0.57 | 0.97 | 1.25 | -0.43 | 0.45 | 0.5 | 0.61 | -0.17 | 0.005 | -0.4725 | 0.16173 | 0.462574 | 0.986476 | 0.989223 | 0.997988 | 0.994666 |
| Adgrl3_F6WMY5_S558s_1 | Adgrl3_F6\ | 2.21 | 0.43 | -0.07 | 0.55 | -0.39 | 2.53 | -0.26 | 0.83 | 2.31 | 0.43 | -0.56 | 0.04 | 1.89 | -0.6 | 0.52 | 0.49 | -0.1025 | -0.225 | -0.205 | 0.750124 | 0.88913 | 0.753123 | 0.996493 | 0.997988 | 0.9971 |
| Adgrl3_F6WMY5_S576s_1 | Adgrl3_F6\ | -1.06 | -2.22 | -2 | -3.14 | -1.95 | -0.71 | -1.94 | -1.83 | -1.26 | -1.76 | -1.79 | -1.73 | 0.15 | -5.14 | -1.66 | -2.16 | 0.4975 | 0.47 | -0.0975 | 0.924087 | 0.315541 | 0.293248 | 0.989223 | 0.997988 | 0.999507 |
| Adgrl3_F6WMY5_S590s_1 | Adgrl3_F6\ | 0.93 | 0.43 | 0.01 | 0.57 | 0.6 | 0.71 | 0.4 | 0.33 | 0.39 | 1 | -0.78 | 0.72 | -0.93 | 0.49 | -0.25 | 1 | 0.025 | -0.1525 | -0.4075 | 0.376673 | 0.923108 | 0.720366 | 0.997335 | 0.997988 | 0.994666 |
| Adgrl3_F6WMY5_S776s_1 | Adgrl3_F6\ | 1.07 | 0.25 | 0.37 | -0.35 | 0.67 | 1.75 | -0.38 | -0.27 | -1.11 | -0.07 | -0.76 | -0.62 | 0.66 | 0.3 | -0.6 | -0.6 | 0.1075 | -0.975 | -0.395 | 0.367519 | 0.837155 | 0.024986 | 0.995762 | 0.997988 | 0.994666 |
| Adgrl3_F6WMY5_S821s_1 | Adgrl3_F6\ | -2.28 | -2.29 | -1.66 | -1.87 | -0.76 | -2.87 | -1.32 | -2.39 | -3.06 | -2.63 | -0.32 | -1.75 | -1.03 | -1.44 | -1.53 | -0.9 | 0.19 | 0.085 | 0.8 | 0.019333 | 0.687279 | 0.882505 | 0.994672 | 0.997988 | 0.994666 |
| Adgrl3_F6WMY5_S858s_1 | Adgrl3_F6\ | 0.52 | 0.36 | 1.52 | -0.09 | 0.76 | 1.42 | 0.02 | 0.81 | 1.36 | 0.83 | 0.57 | 1.14 | 1.47 | -2.4 | 2.86 | 0.54 | 0.175 | 0.3975 | 0.04 | 0.968451 | 0.679229 | 0.315434 | 0.993464 | 0.997988 | 0.999507 |
| Adh6a_E9Q4Y1_T3t_3 | Adh6a_E9( | 0.04 | 1.16 | 0.54 | 0.06 | 0.91 | -0.33 | 0.6 | 0.18 | -0.9 | 0.08 | -0.61 | 0.51 | -0.17 | -0.27 | 0.29 | 0.46 | -0.11 | -0.68 | -0.3725 | 0.299372 | 0.766965 | 0.123214 | 0.995762 | 0.997988 | 0.994666 |
| Adh6a_E9Q4Y1_T7t_3 | Adh6a_E9( | 0.04 | 1.16 | 0.54 | 0.06 | 0.91 | -0.33 | 0.6 | 0.18 | -0.9 | 0.08 | -0.61 | 0.51 | -0.17 | -0.27 | 0.29 | 0.46 | -0.11 | -0.68 | -0.3725 | 0.299372 | 0.766965 | 0.123214 | 0.995762 | 0.997988 | 0.994666 |
| Adh6a_E9Q4Y1_T9t_3 | Adh6a_E9( | 0.04 | 1.16 | 0.54 | 0.06 | 0.91 | -0.33 | 0.6 | 0.18 | -0.9 | 0.08 | -0.61 | 0.51 | -0.17 | -0.27 | 0.29 | 0.46 | -0.11 | -0.68 | -0.3725 | 0.299372 | 0.766965 | 0.123214 | 0.995762 | 0.997988 | 0.994666 |
| Adra1d_A2ANQ2_S444s_1 | Adra1d_A2 | -0.52 | -1.02 | 0.04 | -0.18 | -1.18 | 0.84 | -1 | -0.05 | 1.18 | -0.75 | -0.52 | -0.54 | 0.88 | -1.16 | -0.66 | -0.43 | 0.0725 | 0.2625 | 0.0775 | 0.869697 | 0.880044 | 0.586765 | 0.996493 | 0.997988 | 0.999507 |
| Adra1d_P97714_S281s_1 | Adra1d_P9 | -0.5 | -0.54 | -1.36 | -0.56 | -0.88 | -0.45 | -0.77 | -0.57 | -1.17 | -0.41 | 0.49 | -1.53 | -1.15 | 0.08 | -0.5 | -0.25 | 0.0725 | 0.085 | 0.285 | 0.434182 | 0.788902 | 0.856425 | 0.995762 | 0.997988 | 0.994666 |
| Adra2a_Q01338_S255s_1 | Adra2a_Q0 | 0.38 | 0.41 | 0.09 | 0.66 | 0.44 | 0.68 | 0.98 | 0.33 | 0.76 | 0.67 | 0.6 | 0.2 | 1.25 | 0.58 | 1.04 | 0.19 | 0.2225 | 0.1725 | 0.38 | 0.246097 | 0.375893 | 0.491453 | 0.989223 | 0.997988 | 0.994666 |
| Adra2a_Q01338_S339s_1 | Adra2a_Q0 | -0.1 | 0.42 | -0.39 | -1.37 | -0.43 | -0.21 | -0.25 | -1 | 0.2 | 0.3 | -0.51 | -0.95 | 0.55 | 0.16 | -1.35 | -0.88 | -0.1125 | 0.12 | -0.02 | 0.970417 | 0.779571 | 0.794085 | 0.995762 | 0.997988 | 0.999507 |
| Adra2a_Q01338_S346s_1 | Adra2a_Q0 | 0.01 | 0.56 | 0.45 | 0.4 | 0.77 | 0.56 | 0.55 | 1.12 | 1.11 | 0.72 | 0.73 | 0.9 | 0.83 | 1.11 | 1.06 | 0.34 | 0.395 | 0.51 | 0.48 | 0.118244 | 0.128226 | 0.055895 | 0.989223 | 0.997988 | 0.994666 |
| Adra2a_Q01338_S360s_1 | Adra2a_Q0 | 0.64 | 0.07 | -0.94 | -0.06 | 0.23 | 1.35 | -1.11 | -0.21 | 1.22 | 0.57 | -0.41 | -0.45 | 1.47 | -0.74 | -0.41 | -0.15 | 0.1375 | 0.305 | 0.115 | 0.833505 | 0.802142 | 0.539119 | 0.995762 | 0.997988 | 0.999507 |
| Adra2a_Q01338_S362s_1 | Adra2a_Q0 | 1.33 | 0.3 | 2.09 | 2.98 | 0.86 | 1.51 | 1.27 | 1.43 | 1.25 | 0.28 | 3.72 | 2.93 | 1.8 | 0.26 | 2.1 | 2.98 | -0.4075 | 0.37 | 0.11 | 0.877941 | 0.451547 | 0.668339 | 0.989223 | 0.997988 | 0.999507 |
| Adra2c_Q01337_S257s_1 | Adra2c_Q0 | 1.44 | 0.81 | 1.17 | 1.52 | 0.82 | 1.02 | 0.85 | 1.24 | 1.21 | 1.47 | 1.65 | 0.78 | 0.54 | 0.53 | 1.29 | 1 | -0.2525 | 0.0425 | -0.395 | 0.213983 | 0.319696 | 0.884177 | 0.989223 | 0.997988 | 0.994666 |
| Adra2c_Q01337_S287s_1 | Adra2c_Q0 | 0.81 | 0.49 | 1.41 | 2.51 | -0.05 | 0.86 | 1.09 | 2.39 | -0.19 | 0.14 | 1.55 | 1.13 | -1.12 | 0.29 | 1.4 | 0.89 | -0.2325 | -0.6475 | -0.94 | 0.16312 | 0.699602 | 0.264311 | 0.994893 | 0.997988 | 0.994666 |
| Adrb1_P34971_S274s_1 | Adrb1_P34 | -0.19 | 0.6 | 1.13 | 0.65 | 0.82 | -0.31 | 0.83 | 0.6 | -0.24 | 0.4 | 0.13 | 1.27 | 0.25 | 0.28 | 1.05 | 0.81 | -0.0625 | -0.1575 | 0.05 | 0.889969 | 0.868134 | 0.705073 | 0.996493 | 0.997988 | 0.999507 |
| Adrbk1_Q7TS64_S628s_1 | Adrbk1_Q7 | -0.48 | -0.84 | -0.71 | -0.25 | -0.08 | -0.38 | -0.16 | -0.28 | -0.86 | -0.84 | -0.63 | -0.66 | -1.2 | -0.02 | -0.58 | -0.5 | 0.345 | -0.1775 | -0.005 | 0.987578 | 0.150761 | 0.456268 | 0.989223 | 0.997988 | 1 |
| AF529169_Q8K3V7_S118s_1 | AF529169_ | 0.5 | -0.19 | 0.09 | 0.65 | -0.01 | 0.82 | 0.03 | 0.47 | 0.48 | 0.14 | 1.16 | -0.5 | 0.33 | -0.14 | 0.61 | 0.62 | 0.065 | 0.0575 | 0.0925 | 0.768275 | 0.828323 | 0.884141 | 0.995762 | 0.997988 | 0.9971 |
| Afap1_Q80YS6_S549s_1 | Afap1_Q80 | -1.58 | -0.78 | -1.13 | -0.85 | -0.84 | 0.23 | -1.85 | -1.4 | 0.29 | -0.32 | -0.74 | -0.97 | 0.73 | -0.28 | -1.42 | -1.1 | 0.12 | 0.65 | 0.5675 | 0.262527 | 0.790794 | 0.087509 | 0.995762 | 0.997988 | 0.994666 |
| Afap1_Q80YS6_S669s_1 | Afap1_Q80 | -0.49 | -0.61 | 0.16 | 0.34 | 0.27 | -0.16 | 0.53 | -0.01 | -0.17 | -0.17 | 0.17 | 0.36 | 0.34 | 0.43 | 0.11 | -0.3 | 0.3075 | 0.1975 | 0.295 | 0.37932 | 0.324178 | 0.523058 | 0.989223 | 0.997988 | 0.994666 |
| Afap1l1_Q8BZI0_S329s_1 | Afap1l1_Q | 0.38 | -0.53 | -0.57 | -0.19 | 0.24 | -0.15 | -0.57 | -0.23 | -0.22 | 0.3 | 1.93 | -1.91 | -0.34 | -0.3 | -0.92 | -0.38 | 0.05 | 0.2525 | -0.2575 | 0.421613 | 0.867651 | 0.732626 | 0.996493 | 0.997988 | 0.994666 |
| Afap1l1_Q8BZI0_S747s_1 | Afap1l1_Q | -3.06 | -1.66 | -2.18 | -2.71 | -1.02 | -2.87 | -1.9 | -2.83 | -3.04 | -2.65 | -1.88 | -2.27 | -0.69 | -1.08 | -2.72 | -2.62 | 0.2475 | -0.0575 | 0.625 | 0.279846 | 0.617462 | 0.88404 | 0.989223 | 0.997988 | 0.994666 |
| Afap1l2_Q5DTU0_S484s_1 | Afap1l2_Q | -0.33 | -0.6 | -0.47 | -0.22 | 0.08 | -0.84 | -0.28 | -0.55 | -0.59 | -0.28 | -0.57 | -0.07 | -0.52 | -0.08 | -0.65 | -0.47 | 0.0075 | 0.0275 | -0.025 | 0.921823 | 0.977061 | 0.907742 | 0.999187 | 0.997988 | 0.999507 |
| Aff4_Q9ESC8_S1059s_1 | Aff4_Q9ES | -0.23 | 0.28 | 0.36 | 1.11 | 0.52 | 0.13 | 0.76 | 0.17 | -0.06 | 0.2 | 0.04 | 0.45 | -0.4 | 0.46 | 0.01 | 0.54 | 0.015 | -0.2225 | -0.2275 | 0.543874 | 0.96329 | 0.498658 | 0.999187 | 0.997988 | 0.994666 |
| Aff4_Q9ESC8_S544s_1 | Aff4_Q9ES | -0.73 | -1.21 | -1.82 | -5.09 | -2.35 | -0.38 | -1.99 | -2.7 | 1.59 | -1.01 | -1.85 | -2.17 | 2.22 | -2.38 | -0.2 | -1.72 | 0.3575 | 1.3525 | 1.6925 | 0.188998 | 0.710839 | 0.255729 | 0.995437 | 0.997988 | 0.994666 |
| Aff4_Q9ESC8_S614s_1 | Aff4_Q9ES | -0.55 | -2.3 | -1.16 | -0.82 | -2.88 | -0.92 | -2.05 | -1.65 | 0.82 | -1.83 | -3.07 | -1.53 | 1.54 | -2.54 | -0.93 | -0.23 | -0.6675 | -0.195 | 0.6675 | 0.423373 | 0.213254 | 0.808142 | 0.989223 | 0.997988 | 0.994666 |
| Aff4_Q9ESC8_S615s_1 | Aff4_Q9ES | -0.55 | -2.3 | -1.16 | -0.82 | -2.88 | -0.92 | -2.05 | -1.65 | 0.82 | -1.83 | -3.07 | -1.53 | 1.54 | -2.54 | -0.93 | -0.23 | -0.6675 | -0.195 | 0.6675 | 0.423373 | 0.213254 | 0.808142 | 0.989223 | 0.997988 | 0.994666 |
| Aftph_H3BKM5_S240s_1 | Aftph_H3B | 0.39 | 0.6 | 0.99 | 0.95 | 0.96 | 0.95 | 1.48 | 0.36 | 0.28 | 1.05 | 1.23 | 0.77 | -1.4 | 0.81 | 1.06 | 0.88 | 0.205 | 0.1 | -0.395 | 0.482077 | 0.495125 | 0.734981 | 0.989223 | 0.997988 | 0.994666 |
| Aftph_H3BKM5_S525s_1 | Aftph_H3B | 6.92 | 3.63 | 4 | 0.48 | 4.73 | 7.2 | 4.83 | 3.53 | 0.95 | 6.02 | 0.43 | 3.78 | 1.32 | 0.52 | 4.15 | 4.94 | 1.315 | -0.9625 | -1.025 | 0.487952 | 0.328293 | 0.555106 | 0.989223 | 0.997988 | 0.994666 |
| Agap1_E9Q3K3_S124s_1 | Agap1_E9C | 0.69 | 1.22 | -0.79 | 0.27 | 0.6 | 1.03 | 0.52 | 0.53 | 1.43 | 0.63 | 0.46 | 0.54 | 0.29 | 0.36 | -0.19 | 0.6 | 0.3225 | 0.4175 | -0.0825 | 0.853277 | 0.4503 | 0.376114 | 0.989223 | 0.997988 | 0.999507 |
| Agap1_E9Q3K3_S152s_1 | Agap1_E9C | 0.3 | -0.78 | -0.96 | -1.38 | -0.88 | 0.34 | -0.7 | -0.92 | 0.51 | -0.88 | -0.03 | -0.74 | 0.51 | -1.4 | -1.23 | -1.62 | 0.165 | 0.42 | -0.23 | 0.681992 | 0.706849 | 0.374186 | 0.995437 | 0.997988 | 0.996159 |
| Agap1_E9Q3K3_S292s_1 | Agap1_E9C | 1 | -0.23 | 0.06 | 0.88 | -0.25 | 0.52 | 0.07 | 0.61 | 1.33 | 0.75 | 0.82 | 0.19 | 0.44 | 1.58 | 0.6 | 0.41 | -0.19 | 0.345 | 0.33 | 0.431299 | 0.601389 | 0.380803 | 0.989223 | 0.997988 | 0.994666 |
| Agap1_E9Q3K3_S295s_1 | Agap1_E9C | -0.66 | 0.51 | -0.68 | -0.4 | 0.16 | -0.14 | -0.61 | -0.77 | 0.32 | 0.23 | 0.01 | -0.34 | 1.32 | 0.44 | 0.23 | -0.65 | -0.0325 | 0.3625 | 0.6425 | 0.195208 | 0.926589 | 0.297879 | 0.997335 | 0.997988 | 0.994666 |
| Agap1_E9Q3K3_S295s_2 | Agap1_E9C | 0.18 | -0.93 | -0.79 | -0.03 | -0.59 | -0.25 | -0.7 | -0.56 | 0.62 | -0.47 | -0.6 | -0.18 | 1.46 | 0.27 | -0.22 | -0.78 | -0.1325 | 0.235 | 0.575 | 0.281782 | 0.670908 | 0.54973 | 0.992876 | 0.997988 | 0.994666 |
| Agap1_E9Q3K3_S57s_1 | Agap1_E9C | -0.13 | -1.68 | -1.33 | -1.2 | -1.26 | 0.52 | -1.54 | -0.94 | 1.45 | -1.56 | -2.05 | -0.99 | 0.39 | -1.72 | -1.54 | -1.43 | 0.28 | 0.2975 | 0.01 | 0.985501 | 0.590893 | 0.695628 | 0.989223 | 0.997988 | 1 |
| Agap1_E9Q3K3_S591s_1 | Agap1_E9C | -0.79 | -0.95 | -0.11 | -0.95 | -1.07 | -0.09 | -0.98 | -0.66 | 0.51 | -0.71 | -0.92 | -0.9 | 1.4 | -1.18 | 0.56 | -1.17 | 7.85E-17 | 0.195 | 0.6025 | 0.339526 | 1 | 0.623848 | 1 | 0.997988 | 0.994666 |
| Agap1_E9Q3K3_S674s_1 | Agap1_E9C | 0.95 | 0.4 | -0.92 | -0.51 | 0.17 | 0.39 | -0.48 | -0.66 | 0.12 | 0.68 | 0.87 | -0.98 | -1.07 | 0.47 | -0.25 | -0.79 | -0.125 | 0.1925 | -0.39 | 0.452881 | 0.785935 | 0.727475 | 0.995762 | 0.997988 | 0.994666 |
| Agap1_E9Q3K3_S720s_1 | Agap1_E9C | -1.15 | -1.28 | -1.36 | -1.44 | -0.92 | -1.88 | -0.91 | -1.04 | -1.34 | -0.8 | -0.14 | -0.82 | 0.4 | -0.02 | -0.43 | -0.93 | 0.12 | 0.5325 | 1.0625 | 0.009349 | 0.667344 | 0.097775 | 0.99272 | 0.997988 | 0.994666 |
| Agap1_E9Q3K3_S92s_1 | Agap1_E9C | -1.6 | -1.56 | -1.08 | -0.94 | -0.82 | -2.22 | -0.56 | -0.37 | -1.39 | -1.41 | 1.11 | -1.06 | -0.52 | -0.8 | -0.54 | -1.04 | 0.3025 | 0.6075 | 0.57 | 0.068313 | 0.484531 | 0.307842 | 0.989223 | 0.997988 | 0.994666 |
| Agap1_E9Q3K3_T1035t_1 | Agap1_E9C | -1.33 | -2.74 | -2.46 | -3.11 | -2.38 | -1.89 | -0.67 | -1.58 | -3.31 | -2.08 | 0.79 | -1.94 | -2.6 | -2.06 | -3.12 | -2 | 0.78 | 0.775 | -0.035 | 0.938208 | 0.132527 | 0.368983 | 0.989223 | 0.997988 | 0.999507 |
| Agap1_E9Q3K3_T224t_1 | Agap1_E9C | -0.6 | -1.21 | -1.09 | -0.59 | 0.16 | -0.88 | -0.41 | -0.64 | -1.12 | -1.02 | 0.19 | -0.44 | -0.18 | -0.48 | -0.47 | -0.56 | 0.43 | 0.275 | 0.45 | 0.121078 | 0.174536 | 0.447582 | 0.989223 | 0.997988 | 0.994666 |
| Agap2_Q3UHD9_S1148s_1 | Agap2_Q3I | -0.14 | -0.47 | -0.88 | -1.57 | 0.12 | -0.01 | -0.39 | -1.67 | -0.77 | -0.2 | -1.37 | -1.58 | -0.96 | -0.03 | -0.95 | -0.29 | 0.2775 | -0.215 | 0.2075 | 0.602792 | 0.561347 | 0.617116 | 0.989223 | 0.997988 | 0.995259 |
| Agap2_Q3UHD9_S1171s_1 | Agap2_Q3I | 0.03 | -0.98 | -0.16 | -0.77 | -0.88 | 0.61 | -1.53 | -0.48 | 0.32 | -0.86 | -1.05 | 0 | 0.64 | -0.86 | -1.22 | -0.99 | -0.1 | 0.0725 | -0.1375 | 0.7692 | 0.832122 | 0.857949 | 0.995762 | 0.997988 | 0.9971 |
| Agap2_Q3UHD9_S1171s_2 | Agap2_Q3I | -0.91 | -3.12 | -1.39 | -1 | -2.82 | -0.76 | -1.97 | -0.5 | -1.72 | -2.7 | -1.01 | -0.78 | -3.56 | -2.06 | -2.41 | -2.16 | 0.0925 | 0.0525 | -0.9425 | 0.124133 | 0.888793 | 0.931838 | 0.996493 | 0.997988 | 0.994666 |
| Agap2_Q3UHD9_S1175s_2 | Agap2_Q3I | -1.09 | -2.98 | -1.39 | -1 | -2.75 | -0.8 | -1.97 | -0.5 | -1.9 | -3.08 | -1.01 | -0.78 | -4.75 | -2.2 | -2.41 | -2.16 | 0.11 | -0.0775 | -1.265 | 0.094886 | 0.859509 | 0.902758 | 0.996349 | 0.997988 | 0.994666 |
| Agap2_Q3UHD9_S149s_1 | Agap2_Q3I | -1.46 | 1.08 | 0.33 | -0.5 | 0.43 | -0.6 | 0.28 | -0.43 | -0.1 | 0.83 | 0.28 | 0.38 | 0.84 | 1.4 | 0.26 | 0.34 | 0.0575 | 0.485 | 0.8475 | 0.154244 | 0.915985 | 0.378474 | 0.997335 | 0.997988 | 0.994666 |
| Agap2_Q3UHD9_S168s_1 | Agap2_Q3I | 0.17 | -1.96 | -0.88 | -2.1 | -1.14 | 0.62 | -1.32 | -0.39 | -0.13 | -0.87 | 0.74 | -0.7 | -0.09 | -0.7 | -0.02 | -0.7 | 0.635 | 0.9525 | 0.815 | 0.145203 | 0.315749 | 0.132076 | 0.989223 | 0.997988 | 0.994666 |
| Agap2_Q3UHD9_S211s_1 | Agap2_Q3I | -0.85 | -0.1 | -0.01 | -0.21 | 0.37 | -0.69 | 0.38 | -0.32 | -0.74 | -0.52 | -0.5 | 0.18 | -0.87 | 0.39 | -0.34 | -0.12 | 0.2275 | -0.1025 | 0.0575 | 0.870199 | 0.501902 | 0.741283 | 0.989223 | 0.997988 | 0.999507 |
| Agap2_Q3UHD9_S26s_1 | Agap2_Q3I | 0.27 | 0.97 | 0.03 | -0.21 | 0.71 | 0.19 | 0.72 | 0.33 | 0.83 | 0.31 | 0.17 | 0.3 | -0.17 | 0.88 | 0.41 | -0.48 | 0.2225 | 0.1375 | -0.105 | 0.793848 | 0.476644 | 0.670515 | 0.989223 | 0.997988 | 0.999507 |
| Agap2_Q3UHD9_S33s_1 | Agap2_Q3I | 0.68 | -0.07 | 0.04 | 0.43 | 0.15 | 0.59 | 0.79 | 0.72 | -0.24 | 0.33 | 1 | 0.43 | -0.42 | 0.18 | 0.2 | 0.32 | 0.2925 | 0.11 | -0.2 | 0.511446 | 0.291579 | 0.740504 | 0.989223 | 0.997988 | 0.994666 |
| Agap2_Q3UHD9_S349s_1 | Agap2_Q3I | -0.49 | 0.57 | -1.1 | -0.09 | 0.12 | -1.31 | -0.89 | -0.92 | -0.34 | -0.13 | -1.06 | -0.73 | 0.18 | 0.56 | 0.5 | -0.97 | -0.4725 | -0.2875 | 0.345 | 0.476499 | 0.295143 | 0.482313 | 0.989223 | 0.997988 | 0.994666 |
| Agap2_Q3UHD9_S368s_1 | Agap2_Q3I | 0.37 | 0.31 | -0.01 | 0.17 | -0.29 | -0.13 | 0.23 | 0.63 | -0.13 | -0.67 | 1.07 | -0.44 | -1.19 | -0.36 | -0.42 | 0.72 | -0.1 | -0.2525 | -0.5225 | 0.218 | 0.708018 | 0.527532 | 0.995437 | 0.997988 | 0.994666 |
| Agap2_Q3UHD9_S424s_1 | Agap2_Q3I | -0.19 | 0.33 | 0.03 | 0.01 | 0.69 | 0.48 | 0.65 | -0.24 | -0.26 | 0.4 | -0.39 | 0.93 | 0.09 | 0.61 | 0.81 | 0.26 | 0.35 | 0.125 | 0.3975 | 0.173846 | 0.228258 | 0.716596 | 0.989223 | 0.997988 | 0.994666 |
| Agap2_Q3UHD9_S632s_1 | Agap2_Q3I | 0.32 | -1.24 | -0.39 | -1.28 | 1.25 | -0.8 | -0.32 | 0.37 | -1.66 | -1.74 | 0.36 | -0.9 | 0.45 | -0.54 | -1.28 | -0.7 | 0.7725 | -0.3375 | 0.13 | 0.793846 | 0.17014 | 0.557933 | 0.989223 | 0.997988 | 0.999507 |
| Agap2_Q3UHD9_S642s_1 | Agap2_Q3I | -3.7 | -5.49 | -2.53 | -4.17 | -3.15 | -3.92 | -2.55 | -3.49 | -7.14 | -5.93 | 1.77 | -2.9 | -3.41 | -1.36 | -5.21 | -3.17 | 0.695 | 0.4225 | 0.685 | 0.441344 | 0.26762 | 0.815368 | 0.989223 | 0.997988 | 0.994666 |
| Agap2_Q3UHD9_S650s_1 | Agap2_Q3I | -3.07 | -2.39 | -3.33 | -9.01 | -2.37 | -1.76 | -1.36 | -1.8 | 1.04 | -2.91 | -4.98 | -2.06 | 1.67 | -2.48 | -0.02 | -4.29 | 2.6275 | 2.2225 | 3.17 | 0.092385 | 0.072549 | 0.218118 | 0.989223 | 0.997988 | 0.994666 |
| Agap2_Q3UHD9_S921s_1 | Agap2_Q3I | 0.28 | 0.19 | 0.59 | 1 | 0.51 | 0.08 | 0.92 | 1.07 | 0.05 | 0.05 | 0.42 | 0.48 | -0.22 | 0.46 | 0.67 | 0.57 | 0.13 | -0.265 | -0.145 | 0.651739 | 0.674594 | 0.344537 | 0.992876 | 0.997988 | 0.995259 |
| Agap2_Q3UHD9_S99s_1 | Agap2_Q3I | 1.98 | 0.11 | 0.56 | 1 | -0.49 | 0.75 | 1.02 | 1.34 | -0.49 | 0.49 | -1.27 | 1.09 | -1.9 | -0.52 | -0.38 | 0.26 | -0.2575 | -0.9575 | -1.5475 | 0.019444 | 0.619732 | 0.137511 | 0.989223 | 0.997988 | 0.994666 |
| Agap2_Q3UHD9_T350t_1 | Agap2_Q3I | 0.2 | 0.22 | 0.67 | 0.94 | 0.17 | 0.23 | 1.12 | 0.86 | 0.46 | 1.3 | 0.24 | 0.57 | -0.13 | -0.13 | 1.13 | 0.8 | 0.0875 | 0.135 | -0.09 | 0.814637 | 0.780512 | 0.675178 | 0.995762 | 0.997988 | 0.999507 |
| Agap2_Q3UHD9_T629t_1 | Agap2_Q3I | 0.15 | 0.61 | -1.97 | 0.44 | 0.36 | -0.64 | 0.17 | 0.81 | -0.29 | 0.83 | -0.08 | -0.12 | 0.02 | 0.39 | 0.29 | -0.44 | 0.3675 | 0.2775 | 0.2575 | 0.656926 | 0.544847 | 0.643767 | 0.989223 | 0.997988 | 0.995259 |
| Agap2_Q3UHD9_T803t_1 | Agap2_Q3I | 0.62 | 1.05 | -0.2 | -0.44 | 1.14 | 0.34 | 0.17 | -0.29 | 0.12 | 1.01 | 0.28 | 0.04 | 0.63 | 2.53 | 0.01 | -0.26 | 0.0825 | 0.105 | 0.47 | 0.475207 | 0.8486 | 0.797009 | 0.996349 | 0.997988 | 0.994666 |
| Agap3_Q8VHH5_S395s_1 | Agap3_Q8\ | -1.91 | -1.53 | -0.91 | -1.76 | -1.7 | -0.75 | -1.4 | -1.12 | -0.87 | -1.39 | -0.28 | -0.28 | -0.7 | -1.9 | -2.04 | -1.72 | 0.285 | 0.8225 | -0.0625 | 0.871717 | 0.377413 | 0.043605 | 0.989223 | 0.997988 | 0.999507 |
| Agap3_Q8VHH5_S478s_1 | Agap3_Q8\ | -1.85 | -3.11 | -2.62 | -2.74 | -1.34 | -1.21 | -1.18 | -0.8 | -2.14 | -2.44 | 1 | -0.95 | -2.62 | -1.43 | -1.86 | -3.05 | 1.4475 | 1.4475 | 0.34 | 0.448659 | 0.000986 | 0.076253 | 0.989223 | 0.997988 | 0.994666 |
| Agap3_Q8VHH5_S573s_1 | Agap3_Q8\ | -0.5 | -0.26 | -0.67 | -1.04 | -0.83 | -1.09 | -0.42 | -1.56 | 0.23 | 0.06 | -0.56 | -0.94 | 0.46 | -0.54 | -0.54 | -0.83 | -0.3575 | 0.315 | 0.255 | 0.477882 | 0.265206 | 0.363379 | 0.989223 | 0.997988 | 0.994666 |
| Agfg1_Q8K2K6_T177t_2 | Agfg1_Q8K | -1.09 | -2.4 | -1.34 | -2.94 | -2.61 | -0.01 | -2.2 | -1.59 | -1.13 | -2.56 | 2.77 | -0.97 | -1.27 | -0.47 | -1.82 | -3.23 | 0.34 | 1.47 | 0.245 | 0.709473 | 0.597214 | 0.193564 | 0.989223 | 0.997988 | 0.996966 |
| Agps_Q8C0I1_S52s_1 | Agps_Q8C( | 1.22 | 1 | 1.44 | 1.35 | 0.82 | 0.89 | 1.77 | 1.41 | 0.88 | 1.5 | 1.91 | 1.5 | 0.77 | 1.15 | 2.14 | 1.87 | -0.03 | 0.195 | 0.23 | 0.523744 | 0.914688 | 0.496075 | 0.997335 | 0.997988 | 0.994666 |
| Agps_Q8C0I1_S57s_1 | Agps_Q8C( | 0.38 | 0.37 | 0.45 | 0.54 | 0.13 | 0.26 | 0.39 | 0.8 | 0.66 | 0.85 | 1.53 | 0.76 | 0.42 | 0.83 | 0.98 | 1.31 | -0.04 | 0.515 | 0.45 | 0.124872 | 0.860709 | 0.076766 | 0.996493 | 0.997988 | 0.994666 |
| Ahctf1_Q8CJF7_S1218s_1 | Ahctf1_Q8 | 0.73 | 0.08 | 0.83 | 1.62 | 0.25 | -0.06 | 1.33 | 1.67 | -0.05 | 0.16 | 0.96 | 1.42 | 0.79 | 0.69 | 0.97 | 1.38 | -0.0175 | -0.1925 | 0.1425 | 0.701238 | 0.970994 | 0.669642 | 0.999187 | 0.997988 | 0.996159 |
| Ahctf1_Q8CJF7_S1381s_1 | Ahctf1_Q8 | -0.71 | -0.39 | 1.98 | 1.99 | -0.81 | -0.99 | 1.08 | 1.81 | -1.21 | -1.13 | 0.95 | 1.62 | 0.14 | -0.99 | 2.33 | 2.15 | -0.445 | -0.66 | 0.19 | 0.841604 | 0.614862 | 0.476165 | 0.989223 | 0.997988 | 0.999507 |
| Ahctf1_Q8CJF7_S1507s_1 | Ahctf1_Q8 | 1.18 | 0.92 | 0.83 | 0.38 | 0.51 | 0.75 | 0.54 | 0.59 | -0.18 | 0.31 | -0.52 | 0.58 | 0.63 | 0.4 | 0.41 | 0.16 | -0.23 | -0.78 | -0.4275 | 0.144651 | 0.350358 | 0.03613 | 0.989223 | 0.997988 | 0.994666 |
| Ahcyl2_F8WGT1_S109s_1 | Ahcyl2_F8\ | 0.46 | 1.06 | 0.81 | 0.61 | 0.72 | 0.51 | 1.39 | 0.91 | 0.97 | 0.97 | 0.76 | 1.04 | 0.04 | 0.9 | 1.08 | 1.46 | 0.1475 | 0.2 | 0.135 | 0.704369 | 0.588261 | 0.403532 | 0.989223 | 0.997988 | 0.996159 |
| Ahdc1_Q6PAL7_S1060s_1 | Ahdc1_Q6I | 0.54 | 0.17 | 0.37 | -0.3 | 0.12 | 0.05 | -0.56 | 0.2 | -0.14 | 0.83 | -0.34 | -0.23 | 1.2 | 0.95 | -0.32 | 0.4 | -0.2425 | -0.165 | 0.3625 | 0.366663 | 0.403172 | 0.632977 | 0.989223 | 0.997988 | 0.994666 |
| Ahdc1_Q6PAL7_S1180s_1 | Ahdc1_Q6I | 0.36 | 0.74 | 1 | 0.94 | 0.5 | 0.04 | 1.1 | 0.41 | 0.1 | 0.53 | 0.56 | 1.31 | 0.03 | 0.69 | 0.74 | 0.81 | -0.2475 | -0.135 | -0.1925 | 0.520138 | 0.40572 | 0.673805 | 0.989223 | 0.997988 | 0.994666 |
| Ahdc1_Q6PAL7_S1467s_1 | Ahdc1_Q6I | 0.54 | -0.04 | 0.18 | -0.1 | -1.79 | 0.51 | -1.4 | -0.56 | 0.68 | -0.04 | -0.91 | 0.18 | 1.5 | -0.86 | -0.46 | -0.57 | -0.955 | -0.1675 | -0.2425 | 0.64439 | 0.074995 | 0.652462 | 0.989223 | 0.997988 | 0.995259 |
| Ahdc1_Q6PAL7_S892s_1 | Ahdc1_Q6I | 0.01 | 1.03 | 0.88 | 0.9 | 0.6 | 0.94 | 0.93 | 0.62 | 0.86 | 0.75 | 0.37 | 1.37 | 0.78 | 0.84 | 0.94 | 0.53 | 0.0675 | 0.1325 | 0.0675 | 0.825732 | 0.812959 | 0.692925 | 0.995762 | 0.997988 | 0.999507 |
| Ahdc1_Q6PAL7_S99s_1 | Ahdc1_Q6I | -0.9 | -0.38 | 0.2 | 0.35 | -0.55 | -0.38 | 0.71 | 1.16 | 0.25 | -1.15 | 2.02 | 0.61 | 0.68 | 0.8 | 0.57 | 0.71 | 0.4175 | 0.615 | 0.8725 | 0.02404 | 0.382796 | 0.353706 | 0.989223 | 0.997988 | 0.994666 |
| Ahi1_E9QP54_S26s_1 | Ahi1_E9QP | -2.23 | -1.46 | -2.18 | -4.41 | -2.84 | -1.15 | -2.36 | -3.38 | 0.28 | -2.54 | 0.29 | -3.74 | 0.23 | -3.64 | -1.29 | -2.73 | 0.1375 | 1.1425 | 0.7125 | 0.449529 | 0.844604 | 0.296335 | 0.996349 | 0.997988 | 0.994666 |
| Ahi1_E9QP54_S974s_1 | Ahi1_E9QP | -0.32 | 0.36 | -1.35 | -0.33 | -0.47 | -0.87 | 0.09 | -0.35 | -0.29 | -0.59 | 1.26 | -0.96 | 0.62 | -0.26 | 0.92 | 0.31 | 0.01 | 0.265 | 0.8075 | 0.084867 | 0.979581 | 0.636143 | 0.999187 | 0.997988 | 0.994666 |
| Ahi1_E9QP54_T122t_1 | Ahi1_E9QP | -0.74 | 0.11 | -0.44 | -0.14 | -1.25 | -0.74 | 0.14 | -0.21 | 0.29 | -0.46 | 0.63 | -0.69 | 0.89 | -0.5 | 1.29 | 0.49 | -0.2125 | 0.245 | 0.845 | 0.070392 | 0.554007 | 0.511923 | 0.989223 | 0.997988 | 0.994666 |
| Ahi1_E9QP54_T5t_1 | Ahi1_E9QP | -1.62 | -1.52 | -4.21 | -4.09 | -1.43 | -2.73 | -4.04 | -3.49 | -0.12 | -1.16 | -2.37 | -3.54 | 1.02 | -2.29 | -3.27 | -3.64 | -0.0625 | 1.0625 | 0.815 | 0.474142 | 0.93893 | 0.271026 | 0.999187 | 0.997988 | 0.994666 |
| Ahnak_E9Q616_S136s_1 | Ahnak_E9C | 0.26 | -1.06 | -1.03 | -1.13 | -1.06 | -0.27 | -0.36 | -0.71 | -1 | -0.68 | -0.54 | -1.05 | 0.1 | -0.06 | 0.61 | -1.03 | 0.14 | -0.0775 | 0.645 | 0.18519 | 0.708267 | 0.832177 | 0.995437 | 0.997988 | 0.994666 |
| Ahnak_E9Q616_S5504s_1 | Ahnak_E9C | 0.74 | 0.73 | 0.34 | 0.51 | 0.79 | 0.72 | 0.11 | 0.29 | 0.51 | 0.75 | -0.09 | -0.81 | -0.28 | 0.43 | -0.79 | 0.47 | -0.1025 | -0.49 | -0.6225 | 0.099434 | 0.681485 | 0.20627 | 0.993613 | 0.997988 | 0.994666 |
| Ahsa2_F6S1R2_S175s_1 | Ahsa2_F6S | -1.23 | -1.92 | -1.94 | -1.96 | -1.57 | -0.57 | -0.07 | -0.46 | -1.67 | -1.1 | 1.23 | -1.11 | -0.76 | -0.65 | -3.43 | -2.18 | 1.095 | 1.1 | 0.0075 | 0.990327 | 0.012848 | 0.098477 | 0.989223 | 0.997988 | 1 |
| Ahsg_P29699_S138s_1 | Ahsg_P296 | -1.4 | -0.87 | -3.44 | -3.37 | -2.44 | -1.33 | -4.22 | -3.66 | 0.24 | -1.1 | -1.95 | -3.54 | 0.21 | 0.44 | -4.33 | -4.4 | -0.6425 | 0.6825 | 0.25 | 0.847506 | 0.434782 | 0.463105 | 0.989223 | 0.997988 | 0.999507 |
| Aida_Q8C4Q6_S161s_1 | Aida_Q8C4 | -0.36 | 0.95 | 0.04 | -0.84 | 0.66 | 0.14 | 0.08 | 0.26 | 0.01 | 0.21 | -0.33 | -0.04 | 0.21 | 0.54 | -0.85 | 0.18 | 0.3375 | 0.015 | 0.0725 | 0.876479 | 0.395415 | 0.969659 | 0.989223 | 0.997988 | 0.999507 |
| Aimp1_Q3UZG4_S147s_1 | Aimp1_Q3I | -1.19 | -3.83 | -1.98 | -1.48 | -2.41 | -2.05 | -2.06 | -2.27 | -2.71 | -2.36 | -2.29 | -1.88 | -1.06 | -1.14 | -1.62 | -1.89 | -0.0775 | -0.19 | 0.6925 | 0.246482 | 0.886341 | 0.739008 | 0.996493 | 0.997988 | 0.994666 |
| Aimp2_Q8R010_S36s_1 | Aimp2_Q8I | -0.66 | 0.62 | -0.5 | -0.27 | 0.47 | -0.15 | -0.46 | 0.02 | -0.26 | 0.03 | -0.06 | -0.28 | -0.58 | -0.37 | -0.48 | -0.04 | 0.1725 | 0.06 | -0.165 | 0.632102 | 0.622397 | 0.852691 | 0.989223 | 0.997988 | 0.995259 |
| Akap1_O08715_S101s_1 | Akap1_O0I | -0.47 | 0.22 | -1.08 | -1.04 | 0.53 | 0.52 | -0.5 | -0.4 | -0.54 | -0.5 | 0.06 | -0.49 | -0.45 | 0.32 | -0.6 | -0.85 | 0.63 | 0.225 | 0.1975 | 0.624752 | 0.139478 | 0.526673 | 0.989223 | 0.997988 | 0.995259 |
| Akap1_O08715_S428s_1 | Akap1_O0I | -0.01 | -1.69 | -1.39 | -1.36 | 0.61 | 1.08 | -1.72 | -0.98 | -0.1 | -0.43 | -1.53 | -1.05 | -0.28 | -0.31 | -1.17 | -1.51 | 0.86 | 0.335 | 0.295 | 0.532714 | 0.219704 | 0.481563 | 0.989223 | 0.997988 | 0.994666 |
| Akap1_O08715_S55s_1 | Akap1_O0I | 0.15 | -0.43 | -0.53 | 0.23 | 0.17 | 0.42 | 0.36 | 0.16 | -0.31 | -0.1 | 0.2 | -0.57 | -0.65 | 0.15 | -0.35 | -0.29 | 0.4225 | -0.05 | -0.14 | 0.652968 | 0.125527 | 0.865861 | 0.989223 | 0.997988 | 0.995259 |
| Akap1_O08715_T487t_1 | Akap1_O0I | -0.39 | 0.3 | 0.01 | 0.35 | 0.03 | -0.45 | 0.41 | 0.2 | -0.14 | 0.2 | 0.24 | 0.6 | 0.34 | 0.67 | 0.21 | 0.22 | -0.02 | 0.1575 | 0.2925 | 0.309933 | 0.943731 | 0.576314 | 0.999187 | 0.997988 | 0.994666 |
| Akap10_O88845_S187s_1 | Akap10_O | 0.7 | 0.6 | -0.33 | -1.18 | 1.25 | 0.71 | -0.1 | 0.11 | 0.61 | 0.58 | -0.12 | -0.1 | 0.26 | 0.16 | -0.87 | -0.58 | 0.545 | 0.295 | -0.205 | 0.679603 | 0.284994 | 0.53005 | 0.989223 | 0.997988 | 0.996159 |
| Akap10_O88845_S281s_1 | Akap10_O | -0.72 | -2.1 | -2.46 | -3.64 | -1.76 | 0.37 | -4.07 | -2.35 | -0.32 | -1.33 | -3.25 | -2.24 | -0.24 | -2.98 | -2.89 | -2.98 | 0.2775 | 0.445 | -0.0425 | 0.957603 | 0.770413 | 0.570105 | 0.995762 | 0.997988 | 0.999507 |
| Akap11_E9Q777_S910s_1 | Akap11_E9 | 0.63 | 0.48 | -0.26 | 0.64 | 0.37 | 1.03 | -0.7 | 0.03 | 0.46 | 0.98 | 0.14 | -0.14 | 0.08 | 0.5 | -0.18 | -0.55 | -0.19 | -0.0125 | -0.41 | 0.24859 | 0.63846 | 0.970738 | 0.989223 | 0.997988 | 0.994666 |
| Akap11_E9Q777_S912s_1 | Akap11_E9 | 0.63 | 0.48 | -0.26 | 0.64 | 0.37 | 1.03 | -0.7 | 0.03 | 0.46 | 0.98 | 0.14 | -0.14 | 0.08 | 0.5 | -0.18 | -0.55 | -0.19 | -0.0125 | -0.41 | 0.24859 | 0.63846 | 0.970738 | 0.989223 | 0.997988 | 0.994666 |
| Akap12_Q9WTQ5_S1292s_1 | Akap12_Q9 | 1.14 | 1.46 | 0.5 | 0.9 | 0.71 | 0.02 | 1.56 | 0.49 | 0.6 | 1.3 | 0.67 | 0.61 | 0.64 | 0.65 | 2.16 | 1.06 | -0.305 | -0.205 | 0.1275 | 0.757644 | 0.422811 | 0.502746 | 0.989223 | 0.997988 | 0.9971 |
| Akap12_Q9WTQ5_S1645s_1 | Akap12_Q9 | 1.03 | 1.15 | 0.05 | -0.47 | 0.66 | 0.5 | -0.37 | -0.62 | 1.01 | 1.25 | -0.63 | -0.19 | 1.2 | 0.89 | -0.23 | -0.8 | -0.3975 | -0.08 | -0.175 | 0.756513 | 0.403341 | 0.885507 | 0.989223 | 0.997988 | 0.9971 |
| Akap12_Q9WTQ5_S234s_1 | Akap12_Q9 | 0.2 | 0.5 | -0.75 | -0.05 | 0.67 | 0.31 | -0.16 | -0.27 | 0.01 | 0.31 | 0.15 | -0.19 | -0.3 | 0.01 | -0.38 | -0.48 | 0.1625 | 0.095 | -0.2625 | 0.432512 | 0.641605 | 0.765024 | 0.989223 | 0.997988 | 0.994666 |
| Akap12_Q9WTQ5_S270s_1 | Akap12_Q9 | 1.68 | 2.18 | 0.11 | -0.73 | 1.39 | 0.27 | 0.63 | 0.05 | 1.4 | 1.74 | -1.01 | -0.25 | 1.03 | 1.99 | 0.74 | 0.17 | -0.225 | -0.34 | 0.1725 | 0.80411 | 0.731476 | 0.686778 | 0.995762 | 0.997988 | 0.999507 |
| Akap12_Q9WTQ5_S350s_1 | Akap12_Q9 | -1.26 | -0.37 | -0.48 | -1.55 | -0.36 | -1.16 | -0.99 | -1.29 | -0.95 | -0.91 | -0.38 | -0.71 | -0.16 | -0.49 | -0.29 | -0.61 | -0.035 | 0.1775 | 0.5275 | 0.146543 | 0.921531 | 0.602498 | 0.997335 | 0.997988 | 0.994666 |
| Akap12_Q9WTQ5_S467s_1 | Akap12_Q9 | 0.63 | 0.79 | -0.07 | -0.03 | 0.68 | 0.61 | 0.45 | 0.29 | 0.45 | 0.86 | -0.41 | -0.22 | 0.44 | 0.78 | 0.14 | 0.11 | 0.1775 | -0.16 | 0.0375 | 0.906469 | 0.52543 | 0.671056 | 0.989223 | 0.997988 | 0.999507 |
| Akap12_Q9WTQ5_S507s_1 | Akap12_Q9 | -3.12 | -3.67 | -2.92 | -2.27 | -2.37 | -3.11 | -1.01 | -2.21 | -3.84 | -2.35 | -2.83 | -3.03 | -3.57 | -3.43 | -3.21 | -1.78 | 0.82 | -0.0175 | -0.0025 | 0.995849 | 0.114143 | 0.966457 | 0.989223 | 0.997988 | 1 |
| Akap12_Q9WTQ5_S543s_1 | Akap12_Q9 | -2.47 | -0.79 | -3.07 | -3.55 | -0.87 | -1.96 | -3.16 | -4.27 | -2.13 | -1.63 | -1.13 | -3.48 | -0.81 | -1.26 | -3.78 | -4.1 | -0.095 | 0.3775 | -0.0175 | 0.984621 | 0.908714 | 0.596789 | 0.997335 | 0.997988 | 1 |
| Akap12_Q9WTQ5_S584s_1 | Akap12_Q9 | 0.29 | 0.45 | 0.84 | 0.63 | 0.49 | 0.15 | -0.01 | 0.46 | 0.43 | 0.83 | 0.91 | 0.75 | 0.29 | 1.12 | 0.64 | 0.5 | -0.28 | 0.1775 | 0.085 | 0.766287 | 0.256076 | 0.469259 | 0.989223 | 0.997988 | 0.9971 |
| Akap12_Q9WTQ5_S598s_1 | Akap12_Q9 | -2.95 | -4.33 | -2.57 | -3.4 | -1.48 | -3.2 | -1.19 | -2.7 | -4.13 | -3.09 | -1.27 | -2.29 | -3.64 | -1.75 | -2.68 | -1.62 | 1.17 | 0.6175 | 0.89 | 0.135403 | 0.05788 | 0.35398 | 0.989223 | 0.997988 | 0.994666 |
| Akap12_Q9WTQ5_S613s_1 | Akap12_Q9 | 0.45 | 0.76 | 1.24 | 0.16 | 0.44 | 0.47 | 0.02 | 0.54 | 1.09 | 1.12 | 0.32 | 0.55 | 0.6 | 0.89 | 0.98 | 0.74 | -0.285 | 0.1175 | 0.15 | 0.623897 | 0.337741 | 0.722165 | 0.989223 | 0.997988 | 0.995259 |
| Akap12_Q9WTQ5_S682s_1 | Akap12_Q9 | -0.51 | -1.02 | -1.13 | -1.41 | -2.93 | -0.8 | -2.39 | -0.84 | 0.69 | -1.06 | 0.83 | 0.18 | 0.97 | -1.56 | 1.2 | -1.36 | -0.7225 | 1.1775 | 0.83 | 0.242664 | 0.188885 | 0.026039 | 0.989223 | 0.997988 | 0.994666 |
| Akap12_Q9WTQ5_S684s_1 | Akap12_Q9 | -1.06 | -0.71 | -0.41 | -1.93 | -2.1 | -0.5 | -2.35 | -1.16 | 1.73 | -0.45 | -1.03 | -1.17 | 2.34 | -2.1 | -0.06 | -0.41 | -0.5 | 0.7975 | 0.97 | 0.272945 | 0.326055 | 0.256872 | 0.989223 | 0.997988 | 0.994666 |
| Akap12_Q9WTQ5_S786s_1 | Akap12_Q9 | -4.26 | -3.72 | -4.25 | -8.52 | -3.35 | -4.39 | -3.43 | -3.83 | -5.59 | -3.57 | -1.29 | -5.45 | -3.59 | -2.85 | -3.16 | -3.9 | 1.4375 | 1.2125 | 1.8125 | 0.093832 | 0.169253 | 0.36862 | 0.989223 | 0.997988 | 0.994666 |
| Akap12_Q9WTQ5_S885s_1 | Akap12_Q9 | -0.33 | -0.5 | -0.37 | -0.2 | -0.09 | 0.65 | -0.66 | -0.25 | 1.99 | -0.19 | -0.83 | -0.67 | 1.58 | -1.21 | 0.1 | 0.09 | 0.2625 | 0.425 | 0.49 | 0.370153 | 0.396449 | 0.48524 | 0.989223 | 0.997988 | 0.994666 |
| Akap5_H3BIV5_S149s_1 | Akap5_H3I | 1.57 | -0.1 | 0.69 | 1.21 | 1.76 | 0.51 | 1.33 | 0.94 | -0.06 | -0.52 | -0.32 | 0.54 | -0.7 | 0.85 | 0.75 | 0.32 | 0.2925 | -0.9325 | -0.5375 | 0.282205 | 0.497765 | 0.049651 | 0.989223 | 0.997988 | 0.994666 |
| Akap5_H3BIV5_S178s_1 | Akap5_H3I | 0.53 | 0.36 | -0.05 | 0.49 | 0.48 | 0.34 | 0.77 | 0.48 | -0.09 | 0.7 | 0.45 | 0.33 | -0.35 | 0.39 | -0.15 | 0.33 | 0.185 | 0.015 | -0.2775 | 0.353939 | 0.434821 | 0.955669 | 0.989223 | 0.997988 | 0.994666 |
| Akap5_H3BIV5_S249s_1 | Akap5_H3I | 0.58 | -1.71 | -0.89 | -0.19 | 0.41 | 0.69 | 0.33 | 0.45 | -0.76 | -1.12 | 0.15 | -0.8 | -1.04 | 0.23 | -0.82 | -1.41 | 1.0225 | -0.08 | -0.2075 | 0.710203 | 0.048551 | 0.878382 | 0.989223 | 0.997988 | 0.996966 |
| Akap5_H3BIV5_S258s_1 | Akap5_H3I | 0.37 | 0.2 | 0.06 | 0.4 | 0.3 | -0.15 | 0.36 | 0.04 | -0.3 | 0.27 | -0.22 | -0.11 | -0.51 | 0.23 | 0.04 | 0.23 | -0.12 | -0.3475 | -0.26 | 0.356326 | 0.595415 | 0.166861 | 0.989223 | 0.997988 | 0.994666 |
| Akap5_H3BIV5_S269s_1 | Akap5_H3I | 1.81 | -1.16 | 0.14 | 0.6 | 0.2 | 1.05 | 0.33 | 1.1 | 0.09 | 0.39 | 0.86 | 0.72 | -0.66 | 0.04 | 0.35 | 0.54 | 0.3225 | 0.1675 | -0.28 | 0.646982 | 0.586864 | 0.775198 | 0.989223 | 0.997988 | 0.995259 |
| Akap5_H3BIV5_S277s_1 | Akap5_H3I | 0.37 | 0.56 | -0.08 | 0.46 | -0.21 | -0.38 | -0.36 | -0.05 | 0.71 | 0.52 | -0.27 | -0.28 | 0.2 | 0.54 | -0.6 | -0.51 | -0.5775 | -0.1575 | -0.42 | 0.240486 | 0.031395 | 0.628213 | 0.989223 | 0.997988 | 0.994666 |
| Akap5_H3BIV5_S29s_1 | Akap5_H3I | 1.86 | 0.69 | 1.45 | 2.48 | 0.47 | 1.75 | 0.64 | 2.14 | 1.67 | 1.1 | 0.86 | 1.18 | 1.72 | 0.2 | 0.45 | 1.07 | -0.37 | -0.4175 | -0.76 | 0.140368 | 0.473075 | 0.319771 | 0.989223 | 0.997988 | 0.994666 |
| Akap5_H3BIV5_S75s_1 | Akap5_H3I | 5.57 | 2.3 | 4.8 | 1.4 | 1.76 | 6.03 | 5.42 | 4.67 | 2.88 | 2.67 | -0.34 | 5.69 | 1.99 | -0.16 | 6.08 | 5.68 | 0.9525 | -0.7925 | -0.12 | 0.937993 | 0.427452 | 0.57012 | 0.989223 | 0.997988 | 0.999507 |
| Akap5_H3BIV5_T227t_1 | Akap5_H3I | 1 | 0.33 | 0.38 | 0.94 | 0.52 | 0.12 | 0.91 | 0.33 | 0.17 | 0.61 | 0.4 | 0.47 | -0.78 | 0.82 | 0.4 | 0.47 | -0.1925 | -0.25 | -0.435 | 0.290347 | 0.497952 | 0.355353 | 0.989223 | 0.997988 | 0.994666 |
| Akap6_E9Q9K8_S1070s_1 | Akap6_E9C | 1.37 | 1.25 | 1.14 | 1 | 1.14 | 0.89 | 1.24 | 0.98 | 0.56 | 1.07 | 1.1 | 1.21 | -0.72 | 1.02 | 0.83 | 1.22 | -0.1275 | -0.205 | -0.6025 | 0.193464 | 0.550245 | 0.41164 | 0.989223 | 0.997988 | 0.994666 |
| Akap6_E9Q9K8_S423s_1 | Akap6_E9C | 1.4 | 0.64 | -2.27 | -1.65 | 0.63 | 0.85 | -1.06 | -0.82 | 0.26 | 1.52 | -0.38 | -0.88 | -1.83 | 0.95 | -0.76 | -1.45 | 0.37 | 0.6 | -0.3025 | 0.748135 | 0.674724 | 0.513869 | 0.992876 | 0.997988 | 0.9971 |
| Akap6_E9Q9K8_S480s_1 | Akap6_E9C | -0.68 | -2.08 | -1.39 | -1.4 | -0.63 | -1.25 | -1.24 | -1.15 | -1.55 | -1.43 | -0.08 | -1.09 | -1.47 | -0.65 | -1.86 | -1.57 | 0.32 | 0.35 | 1.57E-16 | 1 | 0.346511 | 0.422452 | 0.989223 | 0.997988 | 1 |
| Akap6_E9Q9K8_S533s_1 | Akap6_E9C | -0.51 | -0.32 | -0.92 | -0.55 | 0.27 | -0.09 | -0.36 | 0.01 | -0.37 | -1.01 | -0.08 | 0.03 | -0.09 | -0.1 | -1.65 | -0.61 | 0.5325 | 0.2175 | -0.0375 | 0.924437 | 0.051072 | 0.478931 | 0.989223 | 0.997988 | 0.999507 |
| Akap6_E9Q9K8_T2234t_1 | Akap6_E9C | -0.49 | -0.38 | -1.02 | 0.55 | 0.43 | -0.53 | -0.54 | -0.28 | -0.51 | 0.06 | 0.09 | -0.32 | 0.71 | 0.76 | 0.54 | -0.02 | 0.105 | 0.165 | 0.8325 | 0.053431 | 0.786245 | 0.654563 | 0.995762 | 0.997988 | 0.994666 |
| Akap9_E9QQ10_T361t_1 | Akap9_E9C | 0.53 | 1.1 | 0.95 | 1 | 1.08 | 0.79 | 0.54 | 1.33 | 0.94 | 1.24 | -0.54 | 0.95 | 0.43 | 0.75 | 0.34 | 1.4 | 0.04 | -0.2475 | -0.165 | 0.607414 | 0.878222 | 0.553981 | 0.996493 | 0.997988 | 0.995259 |
| Akt1s1_E9QKI5_S118s_1 | Akt1s1_E9( | 1.23 | 1.19 | 0.92 | 1.21 | 0.48 | 0.82 | 1.24 | 1.09 | 1.75 | 1.53 | 1.11 | 0.95 | 1.05 | 0.62 | 1.23 | 1.25 | -0.23 | 0.1975 | -0.1 | 0.703183 | 0.357518 | 0.459158 | 0.989223 | 0.997988 | 0.996159 |
| Akt1s1_E9QKI5_S143s_1 | Akt1s1_E9( | 0.23 | 1.16 | 0.02 | 0.22 | -0.73 | 0.45 | -0.84 | 0.03 | 0.81 | 0.81 | 1.15 | -0.03 | -0.31 | 0.31 | 0.08 | 0.18 | -0.68 | 0.2775 | -0.3425 | 0.31297 | 0.105822 | 0.457152 | 0.989223 | 0.997988 | 0.994666 |
| Akt1s1_E9QKI5_S229s_1 | Akt1s1_E9( | 0.78 | 1.5 | 0.71 | 0.7 | 1.1 | 0.57 | 0.24 | 0.45 | 1.52 | 1.68 | 0.83 | 0.96 | 0.93 | 0.54 | 0.72 | 1.65 | -0.3325 | 0.325 | 0.0375 | 0.913137 | 0.274748 | 0.316982 | 0.989223 | 0.997988 | 0.999507 |
| Akt1s1_E9QKI5_S230s_1 | Akt1s1_E9( | 0.78 | 1.5 | 0.71 | 0.7 | 1.1 | 0.57 | 0.24 | 0.45 | 1.52 | 1.68 | 0.83 | 0.96 | 0.93 | 0.54 | 0.72 | 1.65 | -0.3325 | 0.325 | 0.0375 | 0.913137 | 0.274748 | 0.316982 | 0.989223 | 0.997988 | 0.999507 |
| Akt1s1_E9QKI5_S239s_1 | Akt1s1_E9( | -0.34 | 0.55 | 0.52 | 0.58 | 0.96 | 0.21 | 0.32 | 0.45 | 0.22 | 0.59 | -0.63 | 0.98 | -0.46 | -0.18 | 1.24 | -0.32 | 0.1575 | -0.0375 | -0.2575 | 0.564376 | 0.603953 | 0.926249 | 0.989223 | 0.997988 | 0.995259 |
| Akt3_Q9WUA6_S34s_1 | Akt3_Q9W | 0 | 0.01 | -0.21 | 0.15 | -0.34 | -1.15 | -0.4 | -1.23 | -0.39 | -0.26 | -0.17 | -0.39 | 0.19 | 0.19 | -0.14 | -0.21 | -0.7675 | -0.29 | 0.02 | 0.935598 | 0.02122 | 0.19867 | 0.989223 | 0.997988 | 0.999507 |
| Akt3_Q9WUA6_T447t_1 | Akt3_Q9W | -1.12 | -0.87 | -0.94 | -1.28 | -0.71 | -1.13 | 0.11 | -0.96 | -1.38 | -0.56 | -1.84 | -1.28 | -1.16 | -1.43 | -0.25 | -0.8 | 0.38 | -0.2125 | 0.1425 | 0.656826 | 0.238485 | 0.502604 | 0.989223 | 0.997988 | 0.995259 |
| Alad_P10518_S215s_1 | Alad_P105 | 0.78 | 0.69 | 0 | -0.05 | 0.57 | 0.58 | -0.59 | -0.02 | 1.21 | 0.89 | 0.89 | 0.16 | 1.41 | 0.98 | 0.36 | -0.25 | -0.22 | 0.4325 | 0.27 | 0.526844 | 0.540407 | 0.217654 | 0.989223 | 0.997988 | 0.994666 |

| | | | | | | | | | | | | | | | | | | | | | | | | | | |
|---|---|---|---|---|---|---|---|---|---|---|---|---|---|---|---|---|---|---|---|---|---|---|---|---|---|---|
| Aldh1a1_P24549_S2s_1 | Aldh1a1_P | 0.62 | 0.41 | 0.91 | 1.01 | 0.67 | 1.14 | 0.82 | 0.69 | 0.92 | 1.25 | 0.19 | 0.46 | 0.76 | 1.31 | 0.46 | 1.01 | 0.0925 | -0.0325 | 0.1475 | 0.617429 | 0.701027 | 0.915916 | 0.99521 | 0.997988 | 0.995259 |
| Aldh1a1_P24549_S3s_1 | Aldh1a1_P | 0.62 | 0.41 | 0.91 | 1.01 | 0.67 | 1.14 | 0.82 | 0.69 | 0.92 | 1.25 | 0.19 | 0.46 | 0.76 | 1.31 | 0.46 | 1.01 | 0.0925 | -0.0325 | 0.1475 | 0.617429 | 0.701027 | 0.915916 | 0.99521 | 0.997988 | 0.995259 |
| Aldh1l1_Q8R0Y6_T544t_1 | Aldh1l1_Q | -0.21 | 0.92 | 0.93 | 0.56 | 0.6 | -0.08 | 0.81 | 0.37 | -0.22 | 0.75 | 0.96 | 0.76 | 0.27 | 0.3 | 0.74 | 0.6 | -0.125 | 0.0125 | -0.0725 | 0.826948 | 0.710945 | 0.973785 | 0.995437 | 0.997988 | 0.999507 |
| Aldoa_P05064_S39s_1 | Aldoa_P05 | 0.2 | 0.46 | -0.41 | -0.23 | 0.43 | -0.03 | -0.19 | -0.64 | -0.1 | 0.41 | -0.84 | -0.75 | 0.19 | 0.71 | -0.55 | -0.17 | -0.1125 | -0.325 | 0.04 | 0.911212 | 0.721139 | 0.383294 | 0.995437 | 0.997988 | 0.999507 |
| Aldoa_P05064_S46s_1 | Aldoa_P05 | 0.13 | 0.29 | 0.37 | 0.56 | 0.43 | -0.15 | 0 | 0.48 | -0.18 | -0.02 | 0.62 | -0.14 | 0.06 | -0.11 | -0.16 | -0.02 | -0.1475 | -0.2675 | -0.395 | 0.122394 | 0.547628 | 0.330522 | 0.989223 | 0.997988 | 0.994666 |
| Aldoc_P05063_S36s_1 | Aldoc_P05 | -0.44 | 0.74 | -1.79 | 0.57 | 0.69 | -0.28 | 0.07 | 0.48 | -0.05 | 0.43 | -0.82 | -0.42 | -0.16 | 0.88 | -1.88 | 0.08 | 0.47 | 0.015 | -0.04 | 0.95637 | 0.409753 | 0.979609 | 0.989223 | 0.997988 | 0.999507 |
| Aldoc_P05063_S39s_1 | Aldoc_P05 | -0.1 | 0.62 | 0.01 | -0.01 | 0.4 | 0.33 | -0.15 | 0.02 | 0.07 | -0.19 | -0.81 | 0.06 | -0.42 | 1.04 | -0.31 | 0.36 | 0.02 | -0.3475 | 0.0375 | 0.922919 | 0.938509 | 0.26811 | 0.999187 | 0.997988 | 0.999507 |
| Aldoc_P05063_S45s_1 | Aldoc_P05 | 0.57 | 0.51 | 0.23 | 0.04 | 0.68 | 0.14 | -0.12 | 0.08 | 0.37 | 0.44 | -0.71 | 0.11 | 0.52 | 0.59 | 0.01 | 0.15 | -0.1425 | -0.285 | -0.02 | 0.941519 | 0.586723 | 0.384268 | 0.989223 | 0.997988 | 0.999507 |
| Alkbh3_Q8K1E6_S28s_1 | Alkbh3_Q8 | -2.4 | -4.43 | -1.14 | -1.27 | -3.4 | -2.31 | -1.19 | -1.35 | -2.86 | -3.11 | -2.01 | -1.38 | -2.01 | -4.52 | -0.87 | -0.96 | 0.2475 | -0.03 | 0.22 | 0.824861 | 0.757578 | 0.968836 | 0.995762 | 0.997988 | 0.999507 |
| Alkbh5_Q3TSG4_S362s_1 | Alkbh5_Q3 | -1.86 | -1.96 | -3 | -2.6 | -2.03 | -2.56 | -2.27 | -1.56 | -2.35 | -1.87 | -1.07 | -2.37 | -1.69 | -1.11 | -1.76 | -2.34 | 0.25 | 0.44 | 0.63 | 0.125054 | 0.477049 | 0.291783 | 0.989223 | 0.997988 | 0.994666 |
| Alkbh5_Q3TSG4_S385s_1 | Alkbh5_Q3 | -1.14 | -1.53 | -1.67 | -1.29 | -2.55 | -1.52 | -1.5 | -2.38 | -1.04 | -1.1 | -1.14 | -2.15 | -0.38 | -0.54 | -1.34 | -1.69 | -0.58 | 0.05 | 0.42 | 0.261478 | 0.093612 | 0.875659 | 0.989223 | 0.997988 | 0.994666 |
| Alkbh5_Q3TSG4_S65s_1 | Alkbh5_Q3 | -3.95 | -3.1 | -4.25 | -8.32 | -3.9 | -3.7 | -3.74 | -5.79 | -3.53 | -2.17 | -0.88 | -3.62 | -1.33 | -3.12 | -5.26 | -6.51 | 0.6225 | 2.355 | 0.85 | 0.549003 | 0.572499 | 0.069512 | 0.989223 | 0.997988 | 0.994666 |
| Als2_Q920R0_S1016s_1 | Als2_Q920 | 0.87 | 0.37 | 0.48 | 0.03 | 1.1 | 0.49 | 0.36 | 0.49 | 0.55 | 0.52 | -0.2 | 0.56 | -0.6 | 0.68 | -0.14 | -0.27 | 0.1725 | -0.08 | -0.52 | 0.162461 | 0.538256 | 0.787143 | 0.989223 | 0.997988 | 0.994666 |
| Als2_Q920R0_S1329s_1 | Als2_Q920 | 4.19 | 3.13 | 4.38 | 0.9 | 3.2 | 4.28 | 4.71 | 3.72 | 0.32 | 4.34 | 0.34 | 4.58 | 0.55 | 0.85 | 4.86 | 5.17 | 0.8275 | -0.755 | -0.2925 | 0.819031 | 0.290216 | 0.552369 | 0.989223 | 0.997988 | 0.999507 |
| Als2_Q920R0_S1458s_1 | Als2_Q920 | -0.41 | 0.37 | -0.15 | -0.4 | -1.34 | 0.26 | -0.93 | -0.17 | 1.2 | 0.14 | -1.04 | -0.26 | -0.82 | -0.24 | -0.78 | -0.19 | -0.3975 | 0.1575 | -0.36 | 0.257079 | 0.324032 | 0.741275 | 0.989223 | 0.997988 | 0.994666 |
| Als2_Q920R0_S460s_1 | Als2_Q920 | -1.53 | -0.67 | -0.95 | -1.83 | -0.2 | -1.54 | -1.16 | -1.62 | -1.51 | -1.37 | -0.81 | -1.3 | -0.62 | -0.63 | -1.14 | -1 | 0.115 | -0.0025 | 0.3975 | 0.252337 | 0.77562 | 0.993946 | 0.995762 | 0.997988 | 0.994666 |
| Als2_Q920R0_S477s_1 | Als2_Q920 | -0.08 | 0.82 | 1.28 | 1.44 | 0.99 | -0.32 | 1.13 | 0.9 | 0.1 | 0.59 | 0.59 | 1.29 | -0.4 | 1.56 | 0.47 | 1.85 | -0.19 | -0.2225 | 0.005 | 0.992998 | 0.672413 | 0.593251 | 0.992876 | 0.997988 | 1 |
| Als2_Q920R0_S486s_1 | Als2_Q920 | 0.39 | 0.71 | 0.58 | 0.23 | 1.21 | 0.41 | 1.1 | 0.54 | 0.89 | 1.37 | -0.35 | 0.49 | 0.43 | 1.04 | 0.69 | 0.04 | 0.3375 | 0.1225 | 0.0725 | 0.808212 | 0.228494 | 0.749305 | 0.989223 | 0.997988 | 0.999507 |
| Als2_Q920R0_T504t_1 | Als2_Q920 | 0.08 | 0.63 | 0.67 | 0.79 | 1 | 0.09 | 1.36 | 0.78 | 0.17 | 0.22 | 0.17 | 1 | -0.21 | 1.08 | 0.72 | 1.11 | 0.265 | -0.1525 | 0.1325 | 0.719022 | 0.421367 | 0.611526 | 0.989223 | 0.997988 | 0.9971 |
| Amer2_Q8CCJ4_S223s_1 | Amer2_Q8 | -3.1 | -2.34 | -1.58 | -2.62 | -2.35 | -2.93 | -2.35 | -2.89 | -3.07 | -2.33 | -1.87 | -1.28 | -2.75 | -2.32 | -1.81 | -3.25 | -0.22 | 0.2725 | -0.1225 | 0.778456 | 0.541214 | 0.566084 | 0.989223 | 0.997988 | 0.99798 |
| Amer2_Q8CCJ4_S227s_1 | Amer2_Q8 | -1.73 | -0.94 | -0.91 | -1.21 | -1.35 | -1.89 | -0.64 | -0.62 | -1.51 | -1.74 | -0.42 | -0.2 | -1.45 | -1.07 | -0.61 | -0.08 | 0.0725 | 0.23 | 0.395 | 0.301677 | 0.83999 | 0.587412 | 0.995762 | 0.997988 | 0.994666 |
| Amer2_Q8CCJ4_S244s_1 | Amer2_Q8 | -0.24 | -0.79 | -1.32 | -0.92 | -1.63 | -0.05 | -1.41 | -1.71 | 1.96 | -0.28 | -0.41 | -1.3 | 1.84 | -1.48 | -1.24 | -1.1 | -0.3825 | 0.81 | 0.3225 | 0.657953 | 0.377906 | 0.239853 | 0.989223 | 0.997988 | 0.995259 |
| Amer2_Q8CCJ4_S269s_1 | Amer2_Q8 | 3.32 | 0.64 | 3.08 | -0.99 | 2.92 | 3.9 | 3.21 | 1.79 | 1.05 | 3.6 | -2.28 | 2.23 | 1.63 | 0.11 | 2.7 | 3.21 | 1.4425 | -0.3625 | 0.4 | 0.710327 | 0.160695 | 0.798696 | 0.989223 | 0.997988 | 0.996966 |
| Amer2_Q8CCJ4_S280s_1 | Amer2_Q8 | 0.65 | 0.72 | 0.68 | 0.82 | 0.55 | 0.48 | 0.37 | 0.61 | 0.58 | 0.61 | 0.37 | 0.3 | 0.62 | 0.69 | 0.67 | 0.64 | -0.215 | -0.2525 | -0.0625 | 0.781217 | 0.290074 | 0.25347 | 0.989223 | 0.997988 | 0.998226 |
| Amer2_Q8CCJ4_S71s_1 | Amer2_Q8 | -0.84 | -0.82 | 0.17 | 0.19 | 0.01 | -0.42 | 0.68 | 0.61 | -0.32 | -0.7 | 0.39 | 0.56 | -0.81 | 0.17 | 0.14 | 0.48 | 0.545 | 0.3075 | 0.32 | 0.439609 | 0.176197 | 0.460744 | 0.989223 | 0.997988 | 0.994666 |
| Amer2_Q8CCJ4_S98s_1 | Amer2_Q8 | 0.35 | 0.4 | -0.11 | -0.12 | 0.77 | 0.06 | 0.26 | 0.32 | -0.07 | -0.14 | -0.79 | -0.49 | -1.13 | -0.2 | -0.74 | -0.05 | 0.2225 | -0.5025 | -0.66 | 0.068914 | 0.396955 | 0.093176 | 0.989223 | 0.997988 | 0.994666 |
| Amer2_Q8CCJ4_T100t_1 | Amer2_Q8 | 0.35 | 0.4 | -0.11 | -0.12 | 0.77 | 0.06 | 0.26 | 0.32 | -0.07 | -0.14 | -0.79 | -0.49 | -1.13 | -0.2 | -0.74 | -0.05 | 0.2225 | -0.5025 | -0.66 | 0.068914 | 0.396955 | 0.093176 | 0.989223 | 0.997988 | 0.994666 |
| Amer3_Q6NS69_S17s_1 | Amer3_Q6 | -0.64 | -1.02 | -0.65 | -1.4 | -0.45 | -0.96 | -1.34 | -1.4 | -0.41 | -0.65 | 0.26 | -1.26 | -0.37 | -1.03 | -1.6 | -1.3 | -0.11 | 0.4125 | -0.1475 | 0.673789 | 0.719495 | 0.281916 | 0.995437 | 0.997988 | 0.996159 |
| Amot_Q8VHG2_S1083s_1 | Amot_Q8V | 0.41 | -0.99 | 0.01 | -0.35 | 0.15 | 0.19 | 0.62 | 0.05 | -0.24 | 0.53 | -0.49 | -0.15 | -0.51 | 0.1 | -0.23 | 0.08 | 0.4825 | 0.1425 | 0.09 | 0.801021 | 0.169507 | 0.704647 | 0.989223 | 0.997988 | 0.999507 |
| Amot_Q8VHG2_S149s_1 | Amot_Q8V | -0.72 | -0.89 | -0.66 | -0.17 | -1.24 | -0.5 | -1.45 | -0.46 | 1.15 | -1.04 | -0.51 | -0.08 | 1.37 | -1.05 | 0.29 | -0.29 | -0.3025 | 0.49 | 0.69 | 0.192241 | 0.348016 | 0.310784 | 0.989223 | 0.997988 | 0.994666 |
| Amot_Q8VHG2_S693s_1 | Amot_Q8V | 0.03 | -0.26 | 0.11 | 1.07 | -0.12 | -0.25 | 1.14 | 0.72 | -0.01 | -0.05 | 0.06 | 1.07 | -0.56 | -0.71 | 0.39 | 0.46 | 0.135 | 0.03 | -0.3425 | 0.423573 | 0.747993 | 0.939218 | 0.995762 | 0.997988 | 0.994666 |
| Amotl1_Q9D4H4_S645s_1 | Amotl1_Q9 | -0.5 | -0.16 | 0.27 | -0.09 | -0.88 | 0.22 | -0.69 | -1.23 | 2.21 | 0.04 | -1.01 | -0.26 | 2.61 | -1.64 | 0.31 | 0.27 | -0.525 | 0.365 | 0.5075 | 0.52099 | 0.156997 | 0.574418 | 0.989223 | 0.997988 | 0.994666 |
| Amotl2_B8JK81_S50s_1 | Amotl2_B8 | -1.33 | 0.15 | 0.04 | -0.06 | 0.05 | -0.79 | -0.16 | -0.29 | 0.74 | -0.2 | 0.17 | -0.35 | 0.68 | 0.1 | 0.52 | 0.41 | 0.0025 | 0.39 | 0.7275 | 0.081612 | 0.994748 | 0.359748 | 0.99937 | 0.997988 | 0.994666 |
| Amotl2_B8JK81_S51s_1 | Amotl2_B8 | -1.33 | 0.15 | 0.04 | -0.06 | 0.05 | -0.79 | -0.16 | -0.29 | 0.74 | -0.2 | 0.17 | -0.35 | 0.68 | 0.1 | 0.52 | 0.41 | 0.0025 | 0.39 | 0.7275 | 0.081612 | 0.994748 | 0.359748 | 0.99937 | 0.997988 | 0.994666 |
| Ampd2_A2AE27_S3s_1 | Ampd2_A2 | -1.22 | 0.07 | -0.52 | 0.85 | -0.52 | -0.87 | 1.5 | 0.46 | -0.21 | 0.72 | 0.7 | 1.14 | -1.12 | -2.39 | 0.27 | -0.91 | 0.3475 | 0.7925 | -0.8325 | 0.209836 | 0.575806 | 0.134501 | 0.989223 | 0.997988 | 0.994666 |
| Ampd2_Q9DBT5_S12s_1 | Ampd2_Q9 | -0.81 | -0.94 | -0.04 | 0.48 | -0.39 | 0.28 | -0.09 | 0.07 | -0.81 | -0.56 | 0.4 | -0.29 | -0.91 | -0.73 | 0.85 | 0.03 | 0.295 | 0.0125 | 0.1375 | 0.781725 | 0.422241 | 0.976036 | 0.989223 | 0.997988 | 0.998226 |
| Ampd2_Q9DBT5_S19s_1 | Ampd2_Q9 | -1.03 | -1.06 | -0.8 | -1.61 | -1.12 | -1.4 | -1.26 | -1.35 | -1.61 | -1.35 | 0.85 | -1.1 | -1.15 | -0.44 | -0.94 | -1.62 | -0.1575 | 0.3225 | 0.0875 | 0.794981 | 0.523487 | 0.55725 | 0.989223 | 0.997988 | 0.999507 |
| Ampd2_Q9DBT5_S53s_1 | Ampd2_Q9 | -0.74 | -2.5 | -0.98 | -2.99 | -0.5 | -0.63 | -1.91 | -1.52 | -1.97 | -2.28 | 1.26 | -1.31 | -3.14 | 0.47 | -2.62 | -1.82 | 0.6625 | 0.7275 | 0.025 | 0.976612 | 0.274164 | 0.411479 | 0.989223 | 0.997988 | 1 |
| Ampd2_Q9DBT5_S87s_1 | Ampd2_Q9 | -1.23 | -0.79 | -1.25 | -1.3 | -0.71 | -1.61 | -1.19 | -1.44 | -1.78 | -1.36 | -0.92 | -1.35 | -0.85 | -0.69 | -1.22 | -1.37 | -0.095 | -0.21 | 0.11 | 0.693221 | 0.726291 | 0.444697 | 0.995437 | 0.997988 | 0.996159 |
| Ampd3_O08739_S85s_1 | Ampd3_O0 | -1.04 | 0.33 | -1.24 | -1.25 | -0.05 | -0.4 | -0.45 | -2.24 | -1.52 | -0.18 | -0.8 | -1.71 | -0.3 | -0.3 | -1.75 | -2.36 | 0.015 | -0.2525 | -0.3775 | 0.527116 | 0.978643 | 0.60794 | 0.999187 | 0.997988 | 0.994666 |
| Amph_Q7TQF7_S244s_1 | Amph_Q7T | -0.17 | 0.39 | -1.03 | -0.37 | 0.73 | -0.44 | 0.24 | -0.36 | -1.01 | 0.17 | -0.32 | -0.07 | -0.4 | 0.54 | -1.43 | -0.39 | 0.3375 | -0.0125 | -0.125 | 0.793292 | 0.395828 | 0.974359 | 0.989223 | 0.997988 | 0.999507 |
| Amph_Q7TQF7_S250s_1 | Amph_Q7T | -0.27 | 0.38 | 0.49 | 0.81 | 0.29 | 0.16 | 1.03 | 0.51 | -0.15 | 0.03 | 0.69 | 0.7 | -0.57 | 0.27 | 0.28 | 0.62 | 0.145 | -0.035 | -0.2025 | 0.580148 | 0.646232 | 0.917349 | 0.99033 | 0.997988 | 0.995259 |
| Amph_Q7TQF7_S252s_1 | Amph_Q7T | 0.46 | 0.5 | 0.49 | 0.53 | 0.61 | 0.23 | 0.99 | 0.53 | -0.12 | 0.32 | 0.43 | 0.64 | -0.31 | 0.57 | 0.48 | 0.53 | 0.095 | -0.1775 | -0.1775 | 0.536985 | 0.682372 | 0.471448 | 0.994114 | 0.997988 | 0.994666 |
| Amph_Q7TQF7_S262s_1 | Amph_Q7T | 0.63 | 1.81 | 0.31 | 0.14 | 0.87 | 0.48 | 0.38 | -0.44 | 1.12 | 1.58 | -0.03 | 0.03 | 1.43 | 1.58 | 0.38 | -0.02 | -0.4 | -0.0475 | 0.12 | 0.814606 | 0.372327 | 0.926468 | 0.989223 | 0.997988 | 0.999507 |
| Amph_Q7TQF7_S262s_2 | Amph_Q7T | 0.2 | 0.25 | 0.34 | 0.7 | 0.56 | 0.33 | 0.93 | 0.77 | -0.02 | 0.72 | 0.07 | 0.64 | -0.36 | 0.56 | 0.39 | 0.65 | 0.275 | -0.02 | -0.0625 | 0.840289 | 0.266621 | 0.942173 | 0.989223 | 0.997988 | 0.999507 |
| Amph_Q7TQF7_S272s_1 | Amph_Q7T | 0.38 | -0.57 | 0.45 | 0.65 | 0.32 | -0.41 | 1 | 0.64 | -1.68 | -0.07 | -0.25 | 0.68 | -0.27 | 0.19 | 0.7 | 0.3 | 0.16 | -0.5575 | 0.0025 | 0.994474 | 0.684077 | 0.304089 | 0.994379 | 0.997988 | 1 |
| Amph_Q7TQF7_S272s_2 | Amph_Q7T | 0.3 | 0.33 | 0.38 | 0.73 | 0.71 | 0.52 | 0.98 | 0.75 | -0.03 | 0.81 | 0.07 | 0.64 | -0.32 | 0.6 | 0.42 | 0.72 | 0.305 | -0.0625 | -0.08 | 0.795728 | 0.191395 | 0.824121 | 0.989223 | 0.997988 | 0.999507 |
| Amph_Q7TQF7_S500s_1 | Amph_Q7T | 0.37 | 1.04 | 0.59 | 0.51 | 1.01 | 0.25 | 0.45 | 0.24 | 0.17 | 0.67 | 0.76 | 0.39 | 0.78 | 1.1 | 0.49 | 0.49 | -0.14 | -0.13 | 0.0875 | 0.756628 | 0.609604 | 0.62225 | 0.989223 | 0.997988 | 0.9971 |
| Amph_Q7TQF7_T260t_1 | Amph_Q7T | 1.24 | 1.45 | 0.86 | 0.06 | 1.29 | 0.3 | 0.13 | 0.49 | 0.05 | 1.22 | -1.21 | -0.16 | 1.3 | 1.37 | 0.32 | -0.52 | -0.35 | -0.9275 | -0.285 | 0.580088 | 0.377332 | 0.114027 | 0.989223 | 0.997988 | 0.995259 |
| Anapc5_Q3TWF7_T217t_1 | Anapc5_Q | -0.21 | 0.19 | 0.55 | 0.54 | 0.64 | 0.6 | -0.04 | 0.24 | 0.9 | 0.61 | -0.61 | 0.48 | 1.12 | 0.9 | 0.86 | 0.08 | 0.0925 | 0.0775 | 0.4725 | 0.1757 | 0.740128 | 0.838515 | 0.995762 | 0.997988 | 0.994666 |
| Ank1_Q02357_S1804s_1 | Ank1_Q02 | 0.49 | -0.22 | 0.37 | 0.4 | -0.43 | 0.62 | -0.77 | -0.34 | 1.86 | -0.06 | -1.21 | -0.73 | 2.05 | -1.01 | 0.06 | -0.16 | -0.49 | -0.295 | -0.025 | 0.967167 | 0.176962 | 0.642288 | 0.989223 | 0.997988 | 0.999507 |
| Ank1_Q02357_S897s_1 | Ank1_Q02 | 0.65 | -0.48 | -1.15 | 0.04 | 0.1 | -0.19 | 0.33 | 0.34 | -0.07 | -0.08 | -0.13 | 0.59 | -1 | 0.28 | -0.33 | -0.61 | 0.38 | 0.3125 | -0.18 | 0.692853 | 0.342219 | 0.456217 | 0.989223 | 0.997988 | 0.996159 |
| Ank1_Q02357_T1421t_1 | Ank1_Q02 | 1.5 | 0.1 | 0.84 | 1.28 | 0.29 | 1.32 | -0.05 | 1.28 | 1.21 | 0.32 | 0.73 | 0.48 | 0.61 | 1.08 | -0.81 | -0.77 | -0.22 | -0.245 | -0.9025 | 0.116087 | 0.61714 | 0.515176 | 0.989223 | 0.997988 | 0.994666 |
| Ank1_Q02357_T907t_1 | Ank1_Q02 | 1.08 | 0.59 | 0.97 | 1.61 | 1.23 | 1.44 | 1.12 | 1.61 | 0.76 | 0.45 | 0.6 | 1.25 | 0.52 | 1.29 | -0.21 | 0.71 | 0.2875 | -0.2975 | -0.485 | 0.228965 | 0.311315 | 0.346764 | 0.989223 | 0.997988 | 0.994666 |
| Ank2_Q8C8R3_S1454s_1 | Ank2_Q8C | -0.99 | 0.32 | -0.21 | 0.17 | 0.6 | 0.15 | 0.38 | 0.14 | -0.41 | 0.33 | 0.12 | -0.04 | -0.51 | 0.39 | 0.29 | 0.21 | 0.495 | 0.1775 | 0.2725 | 0.473685 | 0.151953 | 0.612716 | 0.989223 | 0.997988 | 0.994666 |
| Ank2_Q8C8R3_S1456s_1 | Ank2_Q8C | -0.37 | 0.23 | 0.08 | -0.36 | 0.05 | -0.5 | -0.03 | -0.39 | -0.2 | -0.35 | 0.44 | 0 | -1.13 | 0.27 | 0.27 | 0.21 | -0.1125 | 0.0775 | 0.01 | 0.979467 | 0.661694 | 0.782954 | 0.990434 | 0.997988 | 1 |
| Ank2_Q8C8R3_S1727s_1 | Ank2_Q8C | 0.63 | 0.45 | 1.5 | 0.83 | -0.66 | 1.29 | 0.3 | 0.53 | 2.26 | 0.2 | 1.25 | 0.92 | 1.94 | -0.72 | 1.5 | 1.05 | -0.4875 | 0.305 | 0.09 | 0.87576 | 0.279262 | 0.515988 | 0.989223 | 0.997988 | 0.999507 |
| Ank2_Q8C8R3_S1727s_2 | Ank2_Q8C | 0.33 | 1.01 | 1.01 | 0.06 | -1.46 | 1.46 | -0.54 | -0.22 | 2.35 | 1.42 | 1.11 | 0.7 | 0.17 | -0.49 | 1.11 | 1.49 | -0.7925 | 0.7925 | -0.0325 | 0.946616 | 0.198987 | 0.084159 | 0.989223 | 0.997988 | 0.999507 |
| Ank2_Q8C8R3_S1728s_1 | Ank2_Q8C | 0.07 | 0.75 | -1.89 | -0.26 | -0.75 | 0.73 | -2.6 | -1.53 | 1.68 | 0.19 | 1.13 | -1.09 | 1.73 | -0.08 | -0.32 | -1.93 | -0.705 | 0.81 | 0.1825 | 0.824841 | 0.379445 | 0.290543 | 0.989223 | 0.997988 | 0.999507 |
| Ank2_Q8C8R3_S1728s_2 | Ank2_Q8C | 0.33 | 1.01 | 1.01 | 0.06 | -1.46 | 1.46 | -0.54 | -0.22 | 2.35 | 1.42 | 1.11 | 0.7 | 0.17 | -0.49 | 1.11 | 1.49 | -0.7925 | 0.7925 | -0.0325 | 0.946616 | 0.198987 | 0.084159 | 0.989223 | 0.997988 | 0.999507 |
| Ank2_Q8C8R3_S1731s_1 | Ank2_Q8C | -1.64 | -1.71 | -1.2 | -2.32 | -2.56 | -0.1 | -1.46 | -1.46 | 0.18 | -1.36 | -0.2 | -0.52 | -0.94 | -2.21 | -2.19 | -1.48 | 0.3225 | 1.2425 | 0.0125 | 0.974588 | 0.52864 | 0.011399 | 0.989223 | 0.997988 | 0.999507 |
| Ank2_Q8C8R3_S1750s_1 | Ank2_Q8C | 0.95 | -0.81 | -0.7 | 0.63 | 0.63 | 1.52 | 1.02 | 1.14 | -0.36 | -0.27 | 0.4 | -0.03 | -1.7 | -0.06 | -0.72 | -0.31 | 1.06 | -0.0825 | -0.715 | 0.204761 | 0.040244 | 0.858086 | 0.989223 | 0.997988 | 0.994666 |
| Ank2_Q8C8R3_S1805s_1 | Ank2_Q8C | 3.68 | 2.42 | 4.74 | 3.35 | 2.72 | 4.3 | 3.79 | 3.99 | -0.74 | 3.12 | -1.1 | 4.07 | 0.6 | -1.13 | 4.92 | 5.15 | 0.1525 | -2.21 | -1.1625 | 0.418855 | 0.775598 | 0.099079 | 0.995762 | 0.997988 | 0.994666 |
| Ank2_Q8C8R3_S1821s_1 | Ank2_Q8C | 6.86 | 1.56 | 4.22 | 0.44 | 4.75 | 7.23 | 5.42 | 4.66 | -0.13 | 4.01 | 0.72 | 5.96 | -5.32 | -1.83 | 5.31 | 6.85 | 2.245 | -0.63 | -2.0175 | 0.469921 | 0.11767 | 0.721056 | 0.989223 | 0.997988 | 0.994666 |
| Ank2_Q8C8R3_S1835s_1 | Ank2_Q8C | 7.8 | 4.12 | 4.91 | 1.6 | 4.76 | 8.79 | 4.81 | 4.35 | -0.96 | 6.49 | 0.62 | 4.93 | -1.83 | -0.11 | 5.3 | 5.73 | 1.07 | -1.8375 | -2.335 | 0.25184 | 0.455815 | 0.34383 | 0.989223 | 0.997988 | 0.994666 |
| Ank2_Q8C8R3_S1844s_1 | Ank2_Q8C | 0.46 | 0.54 | 0.83 | 0.96 | 1.08 | 0.38 | 0.84 | 1.06 | -0.09 | 0.56 | 0.93 | 1.02 | 0.09 | 0.63 | 0.75 | 0.78 | 0.1425 | -0.0925 | -0.135 | 0.630367 | 0.578509 | 0.767031 | 0.989223 | 0.997988 | 0.995259 |
| Ank2_Q8C8R3_S1847s_1 | Ank2_Q8C | -0.1 | 0.06 | 0.84 | 0.85 | 0.65 | 0.12 | 0.76 | 0.85 | -0.37 | -0.01 | 0.86 | 0.68 | 0.29 | 0.72 | 0.6 | 0.59 | 0.1825 | -0.1225 | 0.1375 | 0.666083 | 0.567698 | 0.751263 | 0.989223 | 0.997988 | 0.996159 |
| Ank2_Q8C8R3_S1856s_1 | Ank2_Q8C | 1.79 | 0.53 | 0.12 | 0.69 | 1.17 | 1.92 | 0.71 | 1.01 | 0.63 | 1.34 | 1.03 | 0.62 | -0.1 | 0.69 | 0.95 | 0.44 | 0.42 | 0.1225 | -0.2875 | 0.498774 | 0.328395 | 0.757269 | 0.989223 | 0.997988 | 0.994666 |
| Ank2_Q8C8R3_S1870s_1 | Ank2_Q8C | 8.6 | 4.2 | 5.9 | 0.87 | 4.52 | 9.32 | 6.77 | 5.9 | 0.84 | 7.62 | 2.23 | 6.9 | 0.77 | -2.05 | 9.07 | 6.98 | 1.735 | -0.495 | -1.2 | 0.647515 | 0.302312 | 0.80751 | 0.989223 | 0.997988 | 0.995259 |
| Ank2_Q8C8R3_S1879s_1 | Ank2_Q8C | 2.8 | 2.46 | 2.76 | 2.54 | 2.66 | 2.97 | 2.11 | 3.06 | 1.44 | 3.02 | 3.07 | 3.12 | 1.23 | 1.08 | 3.23 | 2.9 | 0.06 | 0.0225 | -0.53 | 0.326932 | 0.825068 | 0.956252 | 0.995762 | 0.997988 | 0.994666 |
| Ank2_Q8C8R3_S1893s_1 | Ank2_Q8C | 7.66 | 2 | 1.83 | -1.25 | 3.37 | 8.37 | 3.02 | 1.61 | 1.47 | 6.47 | -0.68 | 2.93 | -0.07 | 0.47 | 4.1 | 2.95 | 1.5325 | -0.0125 | -0.6975 | 0.699659 | 0.455798 | 0.995192 | 0.989223 | 0.998021 | 0.996159 |
| Ank2_Q8C8R3_S1953s_1 | Ank2_Q8C | 5.41 | 2.19 | 2.59 | 0.86 | 3.47 | 5.64 | 4.01 | 3.13 | 1.07 | 4.52 | 0.91 | 3.28 | 0.03 | 1.21 | 3.39 | 3.37 | 1.3 | -0.3175 | -0.7625 | 0.493453 | 0.19667 | 0.781067 | 0.989223 | 0.997988 | 0.994666 |
| Ank2_Q8C8R3_S1964s_1 | Ank2_Q8C | 0.48 | -0.13 | 0.11 | 0.43 | 0.58 | 0.57 | 0.44 | 0.7 | 0.1 | 0.19 | 0.69 | -0.07 | 0.24 | 0.44 | 0 | 0.43 | 0.35 | 0.005 | 0.055 | 0.837219 | 0.150724 | 0.985393 | 0.989223 | 0.997988 | 0.999507 |
| Ank2_Q8C8R3_S2118s_1 | Ank2_Q8C | -0.44 | 0.29 | -0.53 | -0.54 | 0.79 | 0.64 | 0 | -0.57 | 0.06 | 0.45 | -1.26 | -0.58 | 0.16 | -0.8 | -0.47 | -0.6 | 0.52 | -0.0275 | -0.1225 | 0.710772 | 0.178928 | 0.947375 | 0.989223 | 0.997988 | 0.996966 |
| Ank2_Q8C8R3_S2163s_1 | Ank2_Q8C | -0.61 | -0.71 | -0.8 | -1.23 | -0.1 | 0.37 | -0.38 | -0.51 | -0.83 | -0.89 | 0.75 | -1.19 | -1.2 | 0.28 | -1.88 | -0.99 | 0.6825 | 0.2975 | -0.11 | 0.809792 | 0.031751 | 0.50674 | 0.989223 | 0.997988 | 0.999507 |
| Ank2_Q8C8R3_S2204s_1 | Ank2_Q8C | 0.26 | 0.15 | -0.1 | 0.04 | 0.8 | 0.29 | -0.19 | -0.57 | -0.36 | 0.49 | -0.17 | 0.13 | -0.03 | 0.87 | -0.64 | -0.45 | -0.005 | -0.065 | -0.15 | 0.683318 | 0.987561 | 0.805671 | 0.999187 | 0.997988 | 0.996159 |
| Ank2_Q8C8R3_S2266s_1 | Ank2_Q8C | -0.06 | 1.07 | 0.29 | -0.43 | 0.96 | 0.21 | 0.14 | -0.51 | 0.63 | 0.33 | 0.14 | -0.48 | 0.61 | 0.87 | 0.89 | 0.14 | -0.0175 | -0.0625 | 0.41 | 0.295471 | 0.966529 | 0.874432 | 0.999187 | 0.997988 | 0.994666 |
| Ank2_Q8C8R3_S2392s_1 | Ank2_Q8C | 0.48 | 0.61 | 0.38 | 0.52 | 1.03 | 0.25 | 0.88 | 0.41 | 0.38 | 0.83 | 0.21 | 0.31 | -0.08 | 1.03 | 0.56 | 0.38 | 0.145 | -0.065 | -0.025 | 0.933011 | 0.564266 | 0.782711 | 0.989223 | 0.997988 | 0.999507 |
| Ank2_Q8C8R3_S2450s_1 | Ank2_Q8C | -0.34 | 0.84 | 1.96 | 1.85 | 0.23 | -0.27 | 1.78 | 1.24 | -0.21 | 0.69 | 1.57 | 1.84 | 0 | 0.26 | 1.92 | 1.92 | -0.3325 | -0.105 | -0.0525 | 0.937547 | 0.601231 | 0.870492 | 0.989223 | 0.997988 | 0.999507 |
| Ank2_Q8C8R3_S2455s_2 | Ank2_Q8C | 0.44 | 1.56 | 0.3 | 0.11 | 0.46 | -0.12 | 0.23 | 0.13 | 0.49 | 0.4 | 1.16 | 0.63 | -1.2 | 1.1 | 0.43 | 0.18 | -0.4275 | 0.0675 | -0.475 | 0.390614 | 0.240555 | 0.856852 | 0.989223 | 0.997988 | 0.994666 |
| Ank2_Q8C8R3_S2458s_2 | Ank2_Q8C | 0.44 | 1.56 | 0.3 | 0.11 | 0.46 | -0.12 | 0.23 | 0.13 | 0.49 | 0.4 | 1.16 | 0.63 | -1.2 | 1.1 | 0.43 | 0.18 | -0.4275 | 0.0675 | -0.475 | 0.390614 | 0.240555 | 0.856852 | 0.989223 | 0.997988 | 0.994666 |
| Ank2_Q8C8R3_S2524s_1 | Ank2_Q8C | 0.83 | 1.32 | 1.36 | 1.17 | 1.1 | 0.79 | 1.33 | 1.24 | 0.78 | 1.09 | 1.09 | 0.94 | 0.49 | 1.13 | 1.16 | 1.08 | -0.055 | -0.195 | -0.205 | 0.468871 | 0.816633 | 0.413226 | 0.995762 | 0.997988 | 0.994666 |
| Ank2_Q8C8R3_S2739s_1 | Ank2_Q8C | 2.51 | 2.39 | 0.82 | 1.55 | 3.03 | 2.14 | 0.47 | 2.9 | 2.1 | 1.97 | 3.06 | 1.4 | 1.38 | 2.85 | 1.4 | 1.84 | 0.3175 | 0.315 | 0.05 | 0.91972 | 0.616954 | 0.528789 | 0.989223 | 0.997988 | 0.999507 |
| Ank2_Q8C8R3_S2746s_1 | Ank2_Q8C | 2.05 | 2.39 | 2.84 | 2.49 | 1.72 | 2.51 | 2.41 | 2.29 | 1.65 | 2.15 | 2.78 | 2.64 | 1.5 | 1.9 | 2.5 | 2.75 | -0.21 | -0.1375 | -0.28 | 0.437484 | 0.456188 | 0.677081 | 0.989223 | 0.997988 | 0.994666 |
| Ank2_Q8C8R3_S2769s_1 | Ank2_Q8C | -0.13 | -0.12 | -0.18 | 0.03 | 0.78 | -1.19 | 0.49 | -0.9 | -0.48 | -0.05 | 0.19 | 0.38 | -0.9 | -1.27 | 0.09 | -0.58 | -0.105 | 0.11 | -0.565 | 0.114054 | 0.819445 | 0.67178 | 0.995762 | 0.997988 | 0.994666 |
| Ank2_Q8C8R3_S2783s_1 | Ank2_Q8C | -1.09 | -0.44 | -1.53 | -0.76 | -0.38 | -0.98 | -0.29 | -1.1 | -0.71 | -0.32 | -1.27 | -0.66 | -0.04 | 0.09 | -0.44 | -0.65 | 0.2675 | 0.215 | 0.695 | 0.058982 | 0.417603 | 0.519079 | 0.989223 | 0.997988 | 0.994666 |
| Ank2_Q8C8R3_S27s_1 | Ank2_Q8C | -0.94 | -0.19 | -0.17 | -0.4 | 0.39 | -0.91 | 0 | -0.26 | -1.14 | -1.33 | -0.84 | -0.64 | -0.53 | -0.65 | -0.11 | 0.34 | 0.23 | -0.5625 | 0.1875 | 0.572171 | 0.497835 | 0.073643 | 0.989223 | 0.997988 | 0.995259 |
| Ank2_Q8C8R3_S2852s_1 | Ank2_Q8C | 0.87 | 0.8 | 0.46 | 0.41 | 1.01 | 0.36 | 0.32 | 0.27 | 0.78 | 0.68 | 0.48 | 0.28 | 0.03 | 0.97 | -0.23 | 0.02 | -0.145 | -0.08 | -0.4375 | 0.20602 | 0.57926 | 0.742488 | 0.989223 | 0.997988 | 0.994666 |
| Ank2_Q8C8R3_S2852s_2 | Ank2_Q8C | -1.35 | -0.85 | -1.72 | -2.17 | -0.36 | -1.46 | -1.86 | -1.79 | -0.56 | -0.82 | -1.26 | -2.32 | -1.6 | -0.65 | -1.66 | -2.21 | 0.155 | 0.2825 | -0.0075 | 0.985898 | 0.714563 | 0.54192 | 0.995437 | 0.997988 | 1 |
| Ank2_Q8C8R3_S2855s_1 | Ank2_Q8C | -2.07 | -0.65 | -0.6 | -0.85 | -0.46 | -2.43 | -0.84 | -1.01 | -1.85 | -1.28 | -0.42 | -0.99 | -0.31 | -0.92 | -0.5 | -0.55 | -0.1425 | -0.0925 | 0.4725 | 0.236491 | 0.77837 | 0.834214 | 0.995762 | 0.997988 | 0.994666 |
| Ank2_Q8C8R3_S29s_1 | Ank2_Q8C | 0.05 | 0.5 | 0.22 | 0.07 | 0.71 | 0.08 | 0.24 | 0.19 | 0.01 | 0.2 | -0.21 | 0.29 | 0.17 | 0.33 | -0.1 | 0.26 | 0.095 | -0.1375 | -0.045 | 0.858144 | 0.692677 | 0.567766 | 0.994893 | 0.997988 | 0.999507 |
| Ank2_Q8C8R3_S3060s_1 | Ank2_Q8C | 0.01 | -0.22 | 0.75 | 1 | 0.33 | 0.38 | 0.92 | 1.36 | -0.26 | 0.13 | 0.59 | 1.09 | -1.05 | -0.43 | 0.25 | 0.18 | 0.3625 | 0.0025 | -0.6475 | 0.146986 | 0.343929 | 0.995098 | 0.989223 | 0.998021 | 0.994666 |
| Ank2_Q8C8R3_S30s_1 | Ank2_Q8C | 0 | 0.49 | 0.61 | 0.52 | 0.8 | 0.56 | 0.38 | 0.61 | -1.05 | -0.08 | -0.9 | 0.44 | 0.38 | -0.78 | 0.46 | 0.39 | 0.1825 | -0.8025 | -0.2925 | 0.418926 | 0.442533 | 0.059648 | 0.989223 | 0.997988 | 0.994666 |
| Ank2_Q8C8R3_S3218s_1 | Ank2_Q8C | 1.25 | 0.25 | 0.41 | 0.7 | 0.35 | 1.4 | 0.31 | 0.64 | 0.71 | 0.61 | 1.17 | 0.53 | 0.46 | 0.77 | 0.65 | 0.93 | 0.0225 | 0.1025 | 0.05 | 0.868245 | 0.947301 | 0.734263 | 0.999187 | 0.997988 | 0.999507 |
| Ank2_Q8C8R3_S3254s_1 | Ank2_Q8C | 0 | 0.53 | 0.42 | 0.14 | 0.7 | 0.07 | -0.14 | 0.27 | -0.35 | 0.36 | 1.06 | -0.54 | 0.28 | 0.8 | 1.24 | 0.48 | -0.0475 | -0.14 | 0.4275 | 0.179757 | 0.856972 | 0.718157 | 0.996349 | 0.997988 | 0.994666 |
| Ank2_Q8C8R3_S3264s_1 | Ank2_Q8C | -1.53 | -1.07 | -1.88 | -1.55 | -1.13 | -1.66 | -0.89 | -1.55 | -1.45 | -1.54 | -0.65 | -1.81 | -1.6 | -0.43 | -1.71 | -1.6 | 0.2 | 0.145 | 0.1725 | 0.640028 | 0.482121 | 0.658156 | 0.989223 | 0.997988 | 0.995259 |
| Ank2_Q8C8R3_S3343s_1 | Ank2_Q8C | 0.12 | -0.08 | -0.6 | 0.39 | 0.27 | 0.15 | 0.65 | 0.39 | -0.64 | -0.23 | -1.19 | -0.41 | -0.57 | 0.61 | -0.29 | -0.33 | 0.4075 | -0.575 | -0.1025 | 0.775295 | 0.161056 | 0.101288 | 0.989223 | 0.997988 | 0.99798 |
| Ank2_Q8C8R3_S3371s_1 | Ank2_Q8C | 0.75 | 0.81 | 0.52 | 0.86 | -0.05 | 1.15 | -0.58 | 0.56 | 1.01 | 1.36 | 0.69 | 0.49 | 1.35 | -0.36 | 0.44 | 0.81 | -0.465 | 0.1525 | -0.175 | 0.648023 | 0.233966 | 0.570048 | 0.989223 | 0.997988 | 0.995259 |
| Ank2_Q8C8R3_S3390s_1 | Ank2_Q8C | 0.43 | 0.82 | 1.07 | 0.65 | 1.17 | 0.38 | 0.43 | 0.72 | 0.18 | 0.61 | 1.32 | 0.88 | 0.2 | 1.08 | 0.72 | 0.64 | -0.0675 | 0.005 | -0.0825 | 0.778419 | 0.802048 | 0.987087 | 0.995762 | 0.997988 | 0.99798 |
| Ank2_Q8C8R3_S3407s_1 | Ank2_Q8C | 0.45 | 0.72 | 0.43 | 0.5 | 0.67 | 0.35 | 0.13 | 0.62 | 0.28 | 0.76 | 0.68 | 0.67 | -0.07 | 1.04 | 0.34 | 0.48 | -0.0825 | 0.0725 | -0.0775 | 0.796448 | 0.71378 | 0.750362 | 0.995437 | 0.997988 | 0.999507 |
| Ank2_Q8C8R3_S3439s_1 | Ank2_Q8C | 0.59 | 0.47 | 0.74 | 0.96 | -0.65 | 1.46 | -0.31 | 0.14 | 2.13 | 0.61 | -2.3 | 0.56 | 2.23 | -2.65 | 0.18 | 0.52 | -0.53 | -0.44 | -0.62 | 0.492265 | 0.251254 | 0.59842 | 0.989223 | 0.997988 | 0.994666 |
| Ank2_Q8C8R3_S34s_1 | Ank2_Q8C | -0.25 | 0.27 | 0.3 | -0.15 | 0.83 | -0.07 | 0.19 | -0.14 | -1.07 | -0.26 | -0.99 | -0.1 | -0.11 | -0.14 | -0.07 | -0.01 | 0.16 | -0.6475 | -0.125 | 0.624001 | 0.586117 | 0.065598 | 0.989223 | 0.997988 | 0.995259 |
| Ank2_Q8C8R3_S34s_2 | Ank2_Q8C | -0.95 | -0.32 | -0.13 | -0.58 | 0.79 | -1.31 | -0.41 | -0.52 | -2 | -1.21 | -1.07 | -0.42 | -3.09 | -0.15 | -0.52 | -0.46 | 0.1325 | -0.68 | -0.56 | 0.390414 | 0.763546 | 0.096659 | 0.995762 | 0.997988 | 0.994666 |
| Ank2_Q8C8R3_S3714s_1 | Ank2_Q8C | 0.26 | 0.55 | 0.13 | 0.41 | 0.55 | 0.08 | 0.58 | 0.45 | -0.15 | 0.32 | 0.26 | 0.45 | 0.26 | 0.18 | 0.42 | 0.54 | 0.0775 | -0.1175 | 0.0125 | 0.959235 | 0.732655 | 0.628856 | 0.995762 | 0.997988 | 0.999507 |
| Ank2_Q8C8R3_S3720s_1 | Ank2_Q8C | 0.5 | 0.22 | 0.62 | 0.83 | 0.83 | 0.28 | 0.92 | 0.67 | 0.03 | 0.39 | 0.21 | 0.58 | -0.37 | 0.68 | 0.17 | 0.46 | 0.1325 | -0.24 | -0.3075 | 0.338016 | 0.596457 | 0.34821 | 0.989223 | 0.997988 | 0.994666 |
| Ank2_Q8C8R3_S3792s_1 | Ank2_Q8C | -0.13 | 0.4 | 0.07 | -0.16 | 0.29 | -0.29 | -0.12 | -0.43 | -0.23 | 0.13 | -0.26 | 0.19 | -0.45 | 0.08 | -0.16 | -0.38 | -0.1825 | -0.0875 | -0.2725 | 0.32158 | 0.480425 | 0.726766 | 0.989223 | 0.997988 | 0.994666 |
| Ank2_Q8C8R3_S3808s_1 | Ank2_Q8C | 0.9 | 1.06 | -2.61 | -6.84 | 0.36 | 0.5 | -1.1 | -0.88 | 2.29 | 1.13 | -0.5 | -0.79 | 2.4 | 0.74 | 3.68 | -3.13 | 1.5925 | 2.405 | 2.795 | 0.190482 | 0.340894 | 0.189306 | 0.989223 | 0.997988 | 0.994666 |
| Ank2_Q8C8R3_S3814s_1 | Ank2_Q8C | 0.23 | 0.53 | 0.08 | 0.12 | 0.68 | 0.21 | 0.48 | 0.15 | -0.24 | 0.3 | -0.75 | 0.06 | -0.41 | 0.75 | -0.41 | 0.16 | 0.14 | -0.3975 | -0.2175 | 0.519733 | 0.550877 | 0.195872 | 0.989223 | 0.997988 | 0.994666 |
| Ank2_Q8C8R3_S3878s_1 | Ank2_Q8C | -1.87 | -1.51 | -0.91 | -1.21 | -1.41 | -1.27 | -0.94 | -1.08 | -1.06 | -1.23 | -0.55 | -0.74 | -0.41 | -2.11 | -1.32 | -1.15 | 0.2 | 0.48 | 0.1275 | 0.755113 | 0.467693 | 0.132166 | 0.989223 | 0.997988 | 0.9971 |
| Ank2_Q8C8R3_S3879s_1 | Ank2_Q8C | -4.06 | -3.48 | -3.23 | -6.54 | -3.12 | -3.8 | -2.75 | -5.03 | -3.02 | -3.83 | -1.64 | -3.16 | -1.87 | -4.05 | -3.04 | -3.22 | 0.6525 | 1.415 | 1.2825 | 0.125198 | 0.419841 | 0.100433 | 0.989223 | 0.997988 | 0.994666 |
| Ank2_Q8C8R3_S3921s_1 | Ank2_Q8C | 1.23 | -0.87 | 0.2 | 0.14 | -1.28 | 1.68 | -0.76 | 0.68 | 3.52 | 0.32 | 1.89 | 0.26 | 1.55 | -0.32 | 2.17 | -0.73 | -0.095 | 1.3225 | 0.4925 | 0.507612 | 0.892751 | 0.120244 | 0.996493 | 0.997988 | 0.994666 |
| Ank2_Q8C8R3_S3s_2 | Ank2_Q8C | 0.56 | 0.57 | -1.34 | -1.08 | 0.18 | 0.8 | -0.67 | -1.11 | 1.09 | 0.96 | -1.76 | -0.69 | 1.26 | -0.08 | -0.14 | -1.26 | 0.1225 | 0.2225 | 0.2675 | 0.68495 | 0.837916 | 0.772341 | 0.995762 | 0.997988 | 0.996159 |
| Ank2_Q8C8R3_S44s_1 | Ank2_Q8C | 0.18 | 0.44 | 0.56 | 0.54 | 0.7 | 0.18 | 0.34 | 0.35 | 0.01 | 0.29 | -0.01 | 0.58 | 0.2 | 0.26 | 0.43 | 0.62 | -0.0375 | -0.2125 | -0.0525 | 0.831946 | 0.866638 | 0.394693 | 0.996493 | 0.997988 | 0.999507 |
| Ank2_Q8C8R3_S49s_1 | Ank2_Q8C | -0.25 | 0.27 | 0.3 | -0.15 | 0.83 | -0.07 | 0.19 | -0.14 | -1.07 | -0.26 | -0.99 | -0.1 | -0.11 | -0.14 | -0.07 | -0.01 | 0.16 | -0.6475 | -0.125 | 0.624001 | 0.586117 | 0.065598 | 0.989223 | 0.997988 | 0.995259 |
| Ank2_Q8C8R3_S49s_2 | Ank2_Q8C | -0.95 | -0.32 | -0.13 | -0.58 | 0.79 | -1.31 | -0.41 | -0.52 | -2 | -1.21 | -1.07 | -0.42 | -3.09 | -0.15 | -0.52 | -0.46 | 0.1325 | -0.68 | -0.56 | 0.390414 | 0.763546 | 0.096659 | 0.995762 | 0.997988 | 0.994666 |
| Ank2_Q8C8R3_S588s_1 | Ank2_Q8C | -0.99 | 0.32 | 0.01 | 0.57 | 0.6 | 0.15 | 0.45 | 0.31 | -0.41 | 0.33 | 0.52 | 0.17 | -0.51 | 0.39 | 0.39 | 0.25 | 0.4 | 0.175 | 0.1525 | 0.709339 | 0.276958 | 0.659847 | 0.989223 | 0.997988 | 0.996966 |
| Ank2_Q8C8R3_S590s_1 | Ank2_Q8C | -0.08 | 0.3 | 0.31 | 0.07 | 0.06 | -0.4 | 0.08 | -0.04 | -0.06 | -0.17 | 0.69 | 0.09 | -1.09 | 0.32 | 0.27 | 0.31 | -0.225 | -0.0125 | -0.1975 | 0.60279 | 0.332615 | 0.963223 | 0.989223 | 0.997988 | 0.995259 |
| Ank2_Q8C8R3_S6s_2 | Ank2_Q8C | 0.56 | 0.57 | -1.34 | -1.08 | 0.18 | 0.8 | -0.67 | -1.11 | 1.09 | 0.96 | -1.76 | -0.69 | 1.26 | -0.08 | -0.14 | -1.26 | 0.1225 | 0.2225 | 0.2675 | 0.68495 | 0.837916 | 0.772341 | 0.995762 | 0.997988 | 0.996159 |
| Ank2_Q8C8R3_S7s_1 | Ank2_Q8C | -3.83 | -5.17 | -4.52 | -8.07 | -2.01 | -5.08 | -4.05 | -4.24 | -7 | -5.57 | 0.27 | -4.69 | -3.06 | -2.04 | -4.48 | -4 | 1.5525 | 1.15 | 2.0025 | 0.056577 | 0.139252 | 0.478421 | 0.989223 | 0.997988 | 0.994666 |
| Ank2_Q8C8R3_S892s_1 | Ank2_Q8C | -0.92 | -1.46 | -1.29 | -1.81 | -0.46 | 0.41 | -1.12 | -0.64 | -1.55 | -1.71 | 0.98 | -0.99 | -0.88 | -0.65 | -1.49 | -1.23 | 0.9175 | 0.5525 | 0.3075 | 0.339613 | 0.029879 | 0.362935 | 0.989223 | 0.997988 | 0.994666 |
| Ank2_Q8C8R3_S939s_1 | Ank2_Q8C | 0.56 | -2.31 | -1.6 | -1.31 | -1.68 | 1.1 | -1.4 | -0.29 | 1.32 | -1.79 | -0.42 | -0.27 | -0.31 | -2.33 | -1.45 | -1.38 | 0.5975 | 0.875 | -0.2025 | 0.761045 | 0.445404 | 0.284129 | 0.989223 | 0.997988 | 0.9971 |
| Ank2_Q8C8R3_T2123t_1 | Ank2_Q8C | 1.09 | 0.32 | 0.12 | 1.04 | 0.49 | 0.68 | 0.62 | 1.17 | 0.15 | 0.69 | 0.97 | 0.83 | -0.38 | 0.74 | 0.69 | 0.52 | 0.0975 | 0.0175 | -0.25 | 0.512036 | 0.752697 | 0.95767 | 0.995762 | 0.997988 | 0.994666 |
| Ank2_Q8C8R3_T2260t_1 | Ank2_Q8C | 0.41 | 0.71 | 1.07 | 0.98 | 0.4 | 0.2 | 0.61 | 1 | 0.55 | 0.57 | 1.35 | 0.92 | 0.6 | 0.79 | 1.12 | 0.81 | -0.24 | 0.055 | 0.0375 | 0.889872 | 0.38314 | 0.847939 | 0.989223 | 0.997988 | 0.999507 |
| Ank2_Q8C8R3_T2361t_1 | Ank2_Q8C | 0.83 | 0.53 | -0.08 | 0.49 | 0.56 | 0.86 | 0.39 | 0.6 | 0.27 | 1.04 | -0.81 | 0.21 | -0.67 | 0.12 | -0.25 | -0.45 | 0.16 | -0.265 | -0.755 | 0.033083 | 0.544565 | 0.529439 | 0.989223 | 0.997988 | 0.994666 |
| Ank2_Q8C8R3_T2570t_1 | Ank2_Q8C | 0.16 | 0.55 | -0.11 | -0.26 | 0.82 | -0.42 | 0.24 | -0.27 | -0.18 | 0.41 | -0.14 | -0.17 | -0.51 | 0.78 | -0.02 | -0.22 | 0.0075 | -0.105 | -0.0775 | 0.827546 | 0.98232 | 0.708603 | 0.999187 | 0.997988 | 0.999507 |
| Ank2_Q8C8R3_T2805t_1 | Ank2_Q8C | -3.3 | -3.16 | -1.07 | -1.51 | -2.98 | -2.94 | -1.64 | -1.14 | -0.95 | -3.09 | 0.74 | -1.07 | -1.6 | -0.93 | -0.99 | -1.24 | 0.085 | 1.1675 | 1.07 | 0.074811 | 0.896034 | 0.197201 | 0.996493 | 0.997988 | 0.994666 |
| Ank2_Q8C8R3_T3063t_1 | Ank2_Q8C | 0.31 | -0.42 | -1.24 | -1.01 | -2.08 | 0.99 | -1.4 | -2.17 | -1.08 | -0.44 | -2.25 | -1.28 | -0.56 | -4.75 | -2.45 | -1.28 | -0.575 | -0.6725 | -1.67 | 0.077438 | 0.431137 | 0.187214 | 0.989223 | 0.997988 | 0.994666 |
| Ank2_Q8C8R3_T3078t_1 | Ank2_Q8C | 0.46 | 0.41 | 0.27 | 0.57 | 0.35 | 0.35 | 0.58 | 0.53 | 0.43 | 0.57 | -0.02 | 0.52 | -0.61 | 0.5 | -0.08 | 0.31 | 0.025 | -0.0525 | -0.3975 | 0.217359 | 0.901901 | 0.824734 | 0.996493 | 0.997988 | 0.994666 |
| Ank2_Q8C8R3_T3746t_1 | Ank2_Q8C | 1.44 | 0.8 | 0.44 | 0.29 | 1.25 | 1.43 | 0.94 | 0.62 | -0.64 | 1.37 | -1 | 0.78 | -0.74 | -0.45 | 0.46 | 0.44 | 0.3175 | -0.615 | -0.815 | 0.068394 | 0.34057 | 0.29779 | 0.989223 | 0.997988 | 0.994666 |
| Ank2_Q8C8R3_T3767t_1 | Ank2_Q8C | -1.08 | -0.25 | -0.65 | -0.52 | -0.21 | -1.06 | -0.22 | -0.96 | -0.81 | -0.62 | -0.37 | -0.57 | -1.11 | -0.13 | -0.58 | -0.55 | 0.0125 | 0.0325 | 0.0325 | 0.917885 | 0.967641 | 0.900776 | 0.999187 | 0.997988 | 0.999507 |
| Ank2_Q8C8R3_T3773t_1 | Ank2_Q8C | 0.76 | 1.55 | 0.83 | 0.66 | 0.86 | 0.23 | 0.6 | 0.42 | 0.59 | 0.91 | 0.67 | 0.59 | 0.56 | 1.23 | 0.3 | 0.35 | -0.4225 | -0.26 | -0.34 | 0.322239 | 0.154523 | 0.353225 | 0.989223 | 0.997988 | 0.994666 |
| Ank2_Q8C8R3_T3784t_1 | Ank2_Q8C | 0.09 | 0.2 | 0.51 | 0.71 | 0.02 | -0.34 | 0.78 | 0.25 | -0.1 | 0.28 | 0.53 | 0.6 | -0.17 | 0.05 | 0.24 | -0.02 | -0.2 | -0.05 | -0.3525 | 0.200637 | 0.509062 | 0.853449 | 0.989223 | 0.997988 | 0.994666 |
| Ank2_Q8C8R3_T3813t_1 | Ank2_Q8C | 0.35 | 0.39 | 0.06 | 0.43 | 0.58 | 0.16 | 0.57 | 0.38 | 0.05 | 0.38 | -0.73 | 0.34 | -0.28 | 0.56 | -0.29 | 0.37 | 0.115 | -0.2975 | -0.2175 | 0.472178 | 0.601738 | 0.344722 | 0.989223 | 0.997988 | 0.994666 |
| Ank2_Q8C8R3_T5t_1 | Ank2_Q8C | -1.46 | -1.58 | 0.22 | -0.78 | -2.74 | -1.5 | -0.55 | -0.26 | -1.91 | -1.01 | 0.04 | -0.65 | -0.34 | -0.31 | 0.27 | 0.59 | -0.3625 | 0.0175 | 0.9525 | 0.060895 | 0.561909 | 0.974037 | 0.989223 | 0.997988 | 0.994666 |
| Ank2_Q8C8R3_T5t_2 | Ank2_Q8C | 0.56 | 0.57 | -1.34 | -1.08 | 0.18 | 0.8 | -0.67 | -1.11 | 1.09 | 0.96 | -1.76 | -0.69 | 1.26 | -0.08 | -0.14 | -1.26 | 0.1225 | 0.2225 | 0.2675 | 0.68495 | 0.837916 | 0.772341 | 0.995762 | 0.997988 | 0.996159 |
| Ank3_A0A087WNU5_S1186s_1 | Ank3_A0A | 0.64 | 0.98 | -1.03 | -1.45 | -0.12 | 0.37 | -1.35 | -0.87 | 0.45 | 0.99 | -1.41 | -1.04 | 0.17 | 0.51 | -1.72 | -1.51 | -0.2775 | -0.0375 | -0.4225 | 0.570117 | 0.663246 | 0.959979 | 0.990434 | 0.997988 | 0.995259 |
| Ank3_A0A087WNU5_S1316s_1 | Ank3_A0A | 0.76 | 0.97 | 0.74 | 0.73 | 0.81 | 0.52 | 0.54 | 0.57 | -0.19 | 0.75 | 0.34 | 0.57 | 0.07 | -0.43 | 0.61 | 0.77 | -0.19 | -0.4325 | -0.545 | 0.117632 | 0.361851 | 0.134894 | 0.989223 | 0.997988 | 0.994666 |
| Ank3_A0A087WNU5_S1338s_1 | Ank3_A0A | 0.58 | 0.35 | 1.31 | 0.76 | -0.84 | 0.71 | 0.01 | 0.46 | 1.69 | 0.75 | 1.05 | 1.2 | 1.1 | -0.81 | 0.84 | 0.72 | -0.665 | 0.4225 | -0.2875 | 0.537412 | 0.110601 | 0.201791 | 0.989223 | 0.997988 | 0.994666 |
| Ank3_A0A087WNU5_S1378s_1 | Ank3_A0A | 2.17 | 0.93 | 1.01 | 2.77 | 0.32 | 1.13 | 0.92 | 2.24 | 2.11 | 2.29 | 3.12 | 1.19 | 0.99 | 1.45 | 1.51 | 1.4 | -0.5675 | 0.4575 | -0.3825 | 0.406921 | 0.312418 | 0.418676 | 0.989223 | 0.997988 | 0.994666 |
| Ank3_A0A087WNU5_S1380s_1 | Ank3_A0A | 1.3 | 0.17 | 1.24 | 1.19 | 0.24 | 1.76 | 0.88 | 1.33 | 2.37 | 1.08 | 1.09 | 2.05 | 2.51 | 0.12 | 2.49 | 0.74 | 0.0775 | 0.6725 | 0.49 | 0.428511 | 0.848098 | 0.134015 | 0.996349 | 0.997988 | 0.994666 |
| Ank3_A0A087WNU5_S1390s_1 | Ank3_A0A | -0.75 | -0.01 | -0.48 | -0.8 | 0.48 | 0.22 | -0.37 | -0.66 | 0.58 | -0.26 | -0.12 | -0.16 | 0.47 | 0.19 | -0.88 | -1.45 | 0.4275 | 0.52 | 0.0925 | 0.843126 | 0.214473 | 0.111497 | 0.989223 | 0.997988 | 0.999507 |
| Ank3_A0A087WNU5_S2266s_1 | Ank3_A0A | -1.68 | -1.17 | -2.11 | -2.62 | -0.89 | -1.14 | -3.41 | -2.25 | -1.8 | -2.71 | 0.56 | -1.75 | -0.26 | -0.79 | -1.86 | -2.48 | -0.0275 | 0.47 | 0.5475 | 0.3305 | 0.962583 | 0.500591 | 0.999187 | 0.997988 | 0.994666 |
| Ank3_A0A087WNU5_S2266s_2 | Ank3_A0A | -1.73 | -2.47 | -1.74 | -2.58 | -4.05 | -2 | -4.64 | -1.95 | -3.11 | -2.49 | -0.54 | -2.06 | -2.09 | -2.34 | -3.6 | -2.92 | -1.03 | 0.08 | -0.6075 | 0.161355 | 0.137344 | 0.884319 | 0.989223 | 0.997988 | 0.994666 |
| Ank3_A0A087WNU5_S2704s_1 | Ank3_A0A | 0.09 | 0.48 | 0.64 | 0.6 | 0.13 | 0.62 | 0.12 | 0.28 | 0.6 | -0.19 | 1.5 | 0.79 | 0.09 | -0.41 | 0.12 | 0.52 | -0.165 | 0.2225 | -0.3725 | 0.225008 | 0.494391 | 0.55702 | 0.989223 | 0.997988 | 0.994666 |
| Ank3_A0A087WNU5_T2270t_1 | Ank3_A0A | 0.78 | -0.35 | 0.87 | 0.52 | -0.38 | 0.52 | -0.14 | 0.15 | 0.4 | 0.54 | 1.37 | 0.74 | -0.12 | -0.2 | -0.03 | -0.08 | -0.4175 | 0.3075 | -0.5625 | 0.108813 | 0.243327 | 0.405282 | 0.989223 | 0.997988 | 0.994666 |
| Ank3_A0A087WNU5_T2270t_2 | Ank3_A0A | -1.73 | -2.47 | -1.74 | -2.58 | -4.05 | -2 | -4.64 | -1.95 | -3.11 | -2.49 | -0.54 | -2.06 | -2.09 | -2.34 | -3.6 | -2.92 | -1.03 | 0.08 | -0.6075 | 0.161355 | 0.137344 | 0.884319 | 0.989223 | 0.997988 | 0.994666 |
| Ank3_G5E8K5_S1648s_1 | Ank3_G5E | 0.93 | -0.24 | -0.46 | 0.9 | -0.24 | 0.88 | 0.73 | 1.12 | 0.16 | -0.51 | 0.74 | 0.26 | -0.37 | -0.03 | -0.24 | -0.06 | 0.34 | -0.12 | -0.4575 | 0.256155 | 0.451056 | 0.783231 | 0.989223 | 0.997988 | 0.994666 |
| Ank3_G5E8K5_S1656s_1 | Ank3_G5E | -0.08 | 1.29 | 0.54 | 0.83 | 1.12 | 3.29 | -0.3 | 0.27 | 2.98 | 1.42 | -1.51 | 0.79 | 0.62 | 1.41 | 0.06 | 0.4 | 0.45 | 0.275 | -0.0225 | 0.956031 | 0.545052 | 0.751372 | 0.989223 | 0.997988 | 0.999507 |
| Ank3_G5E8K5_S27s_1 | Ank3_G5E | -0.29 | -0.36 | -0.93 | -0.32 | -0.08 | -2.08 | -1.12 | -2.16 | -1.77 | -1.33 | -0.48 | -0.85 | -1.25 | -1.4 | -1.12 | -0.68 | -0.885 | -0.6325 | -0.6375 | 0.04913 | 0.086661 | 0.088744 | 0.989223 | 0.997988 | 0.994666 |
| Ank3_G5E8K5_S507s_1 | Ank3_G5E | 0.5 | 0.51 | 0.19 | 0.54 | 1.21 | 1.04 | 0.66 | 0.5 | 0.42 | 0.95 | 0.35 | 0.96 | 0.37 | 1.03 | 0.57 | 0.63 | 0.4175 | 0.235 | 0.215 | 0.417688 | 0.113495 | 0.368083 | 0.989223 | 0.997988 | 0.994666 |
| Ank3_G5E8K5_S606s_1 | Ank3_G5E | 1.16 | 0.93 | 0.2 | 0.52 | 1.13 | 0.37 | 0.88 | -0.05 | 0.39 | 0.55 | -1.53 | 0.21 | -0.23 | 0.62 | -0.15 | 0.47 | -0.12 | -0.7975 | -0.525 | 0.145648 | 0.727696 | 0.134987 | 0.995569 | 0.997988 | 0.994666 |
| Ank3_G5E8K5_S919s_1 | Ank3_G5E | 2.23 | 1.24 | 0.7 | 0.32 | 1.9 | 1.5 | 0.28 | 1.19 | -0.17 | 1.46 | -1.61 | 0.46 | -0.97 | -0.88 | -0.04 | 1.35 | 0.095 | -1.0875 | -1.2575 | 0.065851 | 0.847768 | 0.140923 | 0.996349 | 0.997988 | 0.994666 |
| Ank3_G5E8K5_S921s_1 | Ank3_G5E | 1.49 | -0.13 | 0.84 | -1.03 | 1.94 | 2.11 | 1.75 | 1.63 | -1.42 | 0.29 | -0.72 | 1.92 | -0.45 | -0.84 | 1.46 | 0.97 | 1.565 | -0.275 | -0.0075 | 0.991425 | 0.012217 | 0.735734 | 0.989223 | 0.997988 | 1 |
| Ank3_G5E8K5_T1201t_1 | Ank3_G5E | -0.38 | 0.49 | 0.17 | -0.31 | 0.21 | -0.4 | -0.4 | -0.51 | -0.51 | -0.39 | 0.2 | 0.06 | 0.43 | 0.08 | 0.02 | -0.26 | -0.2675 | -0.1525 | 0.075 | 0.80689 | 0.370668 | 0.619279 | 0.989223 | 0.997988 | 0.999507 |
| Ank3_S4R162_S33s_1 | Ank3_S4R1 | 0.71 | 0.49 | 1.23 | -0.14 | 0.32 | 2.24 | 0.71 | 0.23 | 1.27 | 0.23 | -0.21 | 0.82 | 1.46 | -2.01 | 1.14 | 1.2 | 0.3025 | -0.045 | -0.125 | 0.871097 | 0.549367 | 0.91505 | 0.989223 | 0.997988 | 0.999507 |
| Ank3_S4R162_S73s_1 | Ank3_S4R1 | 0.21 | -0.3 | 0.55 | 1.44 | -1.62 | 0.29 | -0.03 | 0.91 | 1.93 | -0.21 | 1.23 | 0.44 | 1.24 | -1.84 | 0.27 | -0.27 | -0.5875 | 0.3725 | -0.625 | 0.359923 | 0.32806 | 0.503005 | 0.989223 | 0.997988 | 0.994666 |
| Ank3_S4R162_S87s_1 | Ank3_S4R1 | -0.67 | -0.83 | -0.52 | -0.8 | -0.05 | -1.01 | 0.04 | -0.28 | -1.33 | -0.81 | 0.02 | -0.07 | -1.04 | -0.43 | -0.64 | -0.37 | 0.38 | 0.1575 | 0.085 | 0.747315 | 0.200121 | 0.650214 | 0.989223 | 0.997988 | 0.9971 |
| Ank3_S4R162_T32t_1 | Ank3_S4R1 | 2.19 | 1.03 | 0.96 | 0.89 | 0.45 | 2.08 | 0.28 | 1.12 | 0.6 | 1.43 | -0.73 | 1.33 | 1.17 | -1.25 | 1.45 | 1.68 | -0.285 | -0.61 | -0.505 | 0.458243 | 0.551091 | 0.279165 | 0.989223 | 0.997988 | 0.994666 |
| Ank3_S4R1S2_S1411s_1 | Ank3_S4R1 | -0.67 | -0.83 | -0.52 | -0.8 | -0.05 | -1.01 | 0.04 | -0.28 | -1.33 | -0.81 | 0.02 | -0.07 | -1.04 | -0.43 | -0.64 | -0.37 | 0.38 | 0.1575 | 0.085 | 0.747315 | 0.200121 | 0.650214 | 0.989223 | 0.997988 | 0.9971 |
| Ank3_S4R1S2_S1421s_1 | Ank3_S4R1 | 1.01 | -0.24 | 0.36 | 0.8 | 0.59 | 0.34 | 0.83 | 1.11 | -0.23 | 0.07 | -0.14 | 0.37 | -1.06 | -0.61 | 0.27 | 0.52 | 0.235 | -0.465 | -0.7025 | 0.142822 | 0.484169 | 0.182809 | 0.989223 | 0.997988 | 0.994666 |
| Ank3_S4R2K9_S1645s_1 | Ank3_S4R2 | 0.49 | 1.13 | 0.68 | 0.9 | 1.05 | 0.18 | 0.75 | 0.8 | 0.33 | 1.14 | 1.19 | 0.75 | 0.93 | 1.13 | 1.27 | 0.79 | -0.105 | 0.0525 | 0.23 | 0.397749 | 0.699755 | 0.855868 | 0.994893 | 0.997988 | 0.994666 |
| Ank3_S4R2K9_S1679s_1 | Ank3_S4R2 | 1.78 | 0.58 | 0.73 | 1.01 | 0.82 | 1.71 | 0.06 | 0.76 | 1.47 | 0.7 | -0.87 | 0.68 | 0.31 | 0.48 | 0.46 | 0.14 | -0.1875 | -0.53 | -0.6775 | 0.059803 | 0.650066 | 0.324073 | 0.990358 | 0.997988 | 0.994666 |
| Ank3_S4R2K9_S1853s_1 | Ank3_S4R2 | 2.95 | -0.91 | -0.36 | 0.97 | 1.46 | 2.13 | 1.15 | 1.64 | 0.33 | -0.12 | 0.62 | 0.45 | -2.29 | 0.15 | -1.13 | -0.93 | 0.9325 | -0.3425 | -1.7125 | 0.074418 | 0.24606 | 0.662408 | 0.989223 | 0.997988 | 0.994666 |
| Ank3_S4R2K9_S2113s_1 | Ank3_S4R2 | -1.74 | -2.88 | -2.25 | -1.87 | -1.54 | -2.69 | -1.26 | -1.71 | -2.42 | -2.5 | -0.89 | -1.88 | -2.04 | -1.53 | -2.71 | -2.21 | 0.385 | 0.2625 | 0.0625 | 0.866515 | 0.336353 | 0.551614 | 0.989223 | 0.997988 | 0.999507 |
| Ank3_S4R2K9_S2215s_1 | Ank3_S4R2 | -0.05 | -0.98 | 0.29 | -1.24 | -1.95 | 0.35 | -0.51 | -0.4 | 3.1 | -0.58 | 0.19 | 0.08 | 1.94 | -1.59 | 1.34 | -1.44 | -0.1325 | 1.1925 | 0.5575 | 0.525513 | 0.808405 | 0.160918 | 0.995762 | 0.997988 | 0.994666 |
| Ank3_S4R2K9_S2246s_1 | Ank3_S4R2 | 0.28 | 0.68 | -0.15 | 0.62 | 0.63 | 0.11 | 0.17 | 0.21 | 0.86 | 0.19 | 1 | 0.18 | 0.76 | 1.05 | 0.69 | 0.41 | -0.0775 | 0.2 | 0.37 | 0.230701 | 0.772811 | 0.534611 | 0.995762 | 0.997988 | 0.994666 |
| Ank3_S4R2K9_S2330s_1 | Ank3_S4R2 | 0.79 | -1.28 | -0.59 | -0.62 | -0.96 | 1.27 | -1.06 | -0.68 | 1.91 | -0.66 | -0.76 | -0.6 | 1.49 | -1.18 | -0.31 | -0.66 | 0.0675 | 0.3975 | 0.26 | 0.69192 | 0.913731 | 0.574914 | 0.997335 | 0.997988 | 0.996159 |
| Ankhd1_F7DE17_S168s_1 | Ankhd1_F7 | -1.65 | -1.44 | -0.68 | -2.57 | -0.4 | -1.7 | -0.74 | -0.72 | -1.93 | -1.52 | -0.79 | -1.36 | -0.85 | -0.86 | -0.33 | -1.14 | 0.695 | 0.185 | 0.79 | 0.086385 | 0.145771 | 0.675587 | 0.989223 | 0.997988 | 0.994666 |
| Ankhd1_F7DE17_T29t_1 | Ankhd1_F7 | -0.06 | -0.6 | -0.38 | -0.51 | -0.96 | -0.62 | -1.01 | -0.7 | -0.53 | -0.26 | -0.48 | -0.63 | -0.03 | -0.66 | -0.63 | -0.47 | -0.435 | -0.0875 | -0.06 | 0.825955 | 0.082407 | 0.709156 | 0.989223 | 0.997988 | 0.999507 |
| Ankib1_Q6ZPS6_S738s_1 | Ankib1_Q6 | 0.67 | 0.72 | 0.84 | 0.9 | 0.74 | 0.34 | 0.77 | 0.93 | 0.7 | 0.79 | 1.22 | 0.8 | 0.93 | 1.13 | 0.9 | 0.6 | -0.0875 | 0.095 | 0.1075 | 0.661948 | 0.69394 | 0.678057 | 0.994893 | 0.997988 | 0.995804 |
| Ankrd11_E9Q4F7_S1676s_1 | Ankrd11_E | 0 | -1.63 | -0.77 | -2.05 | -0.78 | -0.61 | 0.22 | -1.45 | -1.58 | -0.19 | -0.33 | -1.07 | 0.2 | -0.51 | 0.66 | 0.29 | 0.4575 | 0.32 | 1.2725 | 0.026286 | 0.389233 | 0.545388 | 0.989223 | 0.997988 | 0.994666 |
| Ankrd12_G5E893_T1349t_1 | Ankrd12_G | -1.64 | -0.31 | -0.46 | 0.43 | -0.62 | -0.85 | 0.35 | -1.07 | 0.41 | -0.05 | 2 | -2.47 | 1.02 | 0.22 | 0.2 | 0.11 | -0.0525 | 0.4675 | 0.8825 | 0.081225 | 0.914281 | 0.607555 | 0.997335 | 0.997988 | 0.994666 |
| Ankrd13c_Q3UX43_S411s_1 | Ankrd13c_ | 0.64 | 0.99 | 0.17 | 0.08 | 0.86 | 0.51 | 0.86 | 0.52 | 0.1 | 0.81 | -0.19 | 0.24 | -0.1 | 0.95 | 0.44 | 0.26 | 0.2175 | -0.23 | -0.0825 | 0.808464 | 0.435415 | 0.485561 | 0.989223 | 0.997988 | 0.999507 |
| Ankrd13d_Q6PD24_T556t_1 | Ankrd13d_ | 0.78 | 0.67 | -0.53 | 0.17 | 0.36 | 0.45 | 0.36 | 0.02 | 0.75 | 0.99 | -0.55 | 0 | 0.8 | 0.74 | 0.23 | -0.08 | 0.025 | 0.025 | 0.15 | 0.694394 | 0.938742 | 0.955217 | 0.999187 | 0.997988 | 0.996159 |
| Ankrd17_E9QKG6_S1804s_1 | Ankrd17_E | -0.33 | -0.04 | 0.13 | 0.52 | 0.06 | -0.12 | 0.29 | 0.29 | -0.47 | 0.27 | -0.03 | -0.21 | 0.28 | -0.43 | 0.61 | 0.76 | 0.06 | -0.18 | 0.235 | 0.506014 | 0.814643 | 0.531661 | 0.995762 | 0.997988 | 0.994666 |
| Ankrd17_E9QKG6_S2150s_1 | Ankrd17_E | -1.12 | -0.28 | -0.57 | 0.32 | -0.41 | -1.34 | 0.06 | 0.44 | -0.92 | 0.09 | -0.39 | -0.6 | -0.58 | -0.98 | 0.13 | 0.45 | 0.1 | -0.0425 | 0.1675 | 0.701601 | 0.825892 | 0.909395 | 0.995762 | 0.997988 | 0.996159 |
| Ankrd27_Q3UMR0_S634s_1 | Ankrd27_C | 0.23 | 0.25 | 0.44 | 0.44 | -0.33 | 0.72 | 0.02 | 0.02 | 0.87 | 0.65 | 0.61 | 0.57 | 0.8 | 0.67 | -0.1 | -1.15 | -0.2325 | 0.335 | -0.285 | 0.52377 | 0.398611 | 0.14211 | 0.989223 | 0.997988 | 0.994666 |
| Ankrd34a_B2RW11_S149s_1 | Ankrd34a_ | 0.51 | 0.4 | 0.16 | -0.02 | 0.64 | 0.4 | 0.33 | -0.2 | 0.46 | 0.57 | -0.28 | 0.03 | -0.29 | 0.41 | -0.12 | -0.02 | 0.03 | -0.0675 | -0.2675 | 0.342334 | 0.908616 | 0.81024 | 0.997335 | 0.997988 | 0.994666 |
| Ankrd34a_B2RW11_S183s_1 | Ankrd34a_ | 0.16 | 0.4 | 0.55 | 0.98 | 0.06 | 0.44 | 0.76 | 0.34 | 0.23 | 0.61 | 0.26 | 0.66 | -0.24 | 0.37 | 0.45 | 0.85 | -0.1225 | -0.0825 | -0.165 | 0.615828 | 0.649611 | 0.757753 | 0.990358 | 0.997988 | 0.995259 |

| | | | | | | | | | | | | | | | | | | | | | | | | | | |
|---|---|---|---|---|---|---|---|---|---|---|---|---|---|---|---|---|---|---|---|---|---|---|---|---|---|---|
| Ankrd34a_B2RW11_S225s_1 | Ankrd34a_ | 0.58 | -0.31 | -0.19 | 0.41 | 0.2 | 0.48 | 0 | 0.39 | 0.04 | -0.32 | 0.43 | 0.39 | -1.5 | 0.11 | -0.63 | -0.39 | 0.145 | 0.0125 | -0.725 | 0.099451 | 0.606757 | 0.968124 | 0.989223 | 0.997988 | 0.994666 |
| Ankrd34a_B2RW11_S246s_1 | Ankrd34a_ | -0.2 | 0.18 | 0.1 | 0 | -0.3 | -0.37 | 0.07 | -0.23 | -0.25 | -0.31 | -0.01 | -0.03 | -0.68 | -0.38 | -0.09 | -0.08 | -0.2275 | -0.17 | -0.3275 | 0.23055 | 0.310089 | 0.450481 | 0.989223 | 0.997988 | 0.994666 |
| Ankrd34a_B2RW11_S391s_1 | Ankrd34a_ | 0.21 | -1.24 | 0.4 | 0.38 | 0.42 | 0.63 | 0.7 | 0.66 | -0.33 | 0.39 | 0.49 | 0.46 | -0.18 | 0.82 | 0.32 | 1.01 | 0.665 | 0.315 | 0.555 | 0.246821 | 0.11198 | 0.469529 | 0.989223 | 0.997988 | 0.994666 |
| Ankrd34a_B2RW11_S408s_1 | Ankrd34a_ | 0.18 | 0.67 | 0.19 | -0.06 | 0.58 | 0.44 | -0.31 | -0.28 | 0.3 | 0.61 | -0.11 | 0.12 | 0.81 | 0.54 | -0.59 | -0.44 | -0.1375 | -0.015 | -0.165 | 0.674634 | 0.651464 | 0.956048 | 0.990358 | 0.997988 | 0.996159 |
| Ankrd34a_B2RW11_S460s_1 | Ankrd34a_ | -0.54 | -0.8 | -1.03 | -0.41 | -0.25 | -0.39 | -0.08 | -0.18 | -0.29 | -0.32 | 0.02 | -0.47 | -1.28 | 0.03 | -0.58 | -0.65 | 0.47 | 0.43 | 0.075 | 0.824403 | 0.064184 | 0.109703 | 0.989223 | 0.997988 | 0.999507 |
| Ankrd34a_B2RW11_S472s_1 | Ankrd34a_ | 4.45 | 1.83 | 3.35 | -1.48 | 3.04 | 5.83 | 3.09 | 2.64 | -2.77 | 3.92 | -0.79 | 2.82 | 0.8 | -3.72 | 3.06 | 3.41 | 1.6125 | -1.2425 | -1.15 | 0.524213 | 0.22546 | 0.486442 | 0.989223 | 0.997988 | 0.994666 |
| Ankrd34a_B2RW11_T315t_1 | Ankrd34a_ | -0.61 | -0.22 | -0.88 | -0.99 | 0.2 | -0.23 | -0.24 | -0.71 | -0.48 | -0.17 | -0.36 | -0.23 | -0.32 | 0.47 | -0.12 | -0.32 | 0.43 | 0.365 | 0.6025 | 0.075713 | 0.155805 | 0.176013 | 0.989223 | 0.997988 | 0.994666 |
| Ankrd34b_Q3UUF8_S293s_1 | Ankrd34b_ | 4.35 | 2.44 | 0.7 | -0.09 | 1.93 | 4.78 | 1.54 | 0.49 | 1.22 | 3.64 | -0.12 | 0.22 | 1.26 | 0.94 | 0.23 | 1.25 | 0.335 | -0.61 | -0.93 | 0.310585 | 0.772925 | 0.596541 | 0.995762 | 0.997988 | 0.994666 |
| Ankrd34c_Q8BLB8_S231s_1 | Ankrd34c_ | 0.33 | 0.25 | -0.72 | 0.18 | 0.07 | -0.74 | 0.06 | -1.13 | 0.03 | 1.08 | -0.39 | -1.39 | 0.68 | 0.05 | 0.31 | 0.09 | -0.445 | -0.1775 | 0.2725 | 0.413951 | 0.257469 | 0.73776 | 0.989223 | 0.997988 | 0.994666 |
| Ankrd34c_Q8BLB8_T347t_1 | Ankrd34c_ | -0.39 | -0.71 | -1.62 | -1.51 | -0.54 | -1.09 | -2.37 | -1.34 | -2.13 | -0.63 | -0.48 | -1.77 | -0.69 | -0.29 | -1.23 | -2.1 | -0.2775 | -0.195 | -0.02 | 0.96628 | 0.545451 | 0.68727 | 0.989223 | 0.997988 | 0.999507 |
| Ankrd50_F7BE84_S1101s_1 | Ankrd50_F | 0.38 | -0.24 | 0.22 | -0.11 | -0.83 | -0.12 | 0.12 | -0.27 | 0.37 | -0.1 | -0.29 | -0.7 | -0.04 | -0.2 | 1.68 | -0.77 | -0.3375 | -0.2425 | 0.105 | 0.838175 | 0.249692 | 0.430987 | 0.989223 | 0.997988 | 0.999507 |
| Ankrd50_F7BE84_S1279s_1 | Ankrd50_F | -0.39 | -1 | -1.59 | -2.34 | -1.1 | -0.59 | -1.27 | -1.51 | 2.44 | -1.64 | -2.32 | -1 | 3.41 | -2.25 | 2.02 | -2.38 | 0.2125 | 0.7 | 1.53 | 0.264977 | 0.625827 | 0.4914 | 0.989223 | 0.997988 | 0.994666 |
| Ankrd63_A2ARS0_S193s_1 | Ankrd63_A | -0.18 | 0.45 | 0.41 | 0.5 | -0.57 | -1.32 | 0.37 | -0.45 | 0.53 | 0.7 | 0.8 | -0.44 | 1.34 | 0.65 | 1.21 | 0.41 | -0.7875 | 0.1025 | 0.6075 | 0.08315 | 0.058876 | 0.766274 | 0.989223 | 0.997988 | 0.994666 |
| Ankrd63_A2ARS0_S193s_2 | Ankrd63_A | 0.88 | -1.54 | -0.92 | 0.51 | -0.4 | 0.13 | 0.45 | 0.6 | 0.05 | -0.44 | 0.97 | -0.56 | -0.91 | -0.2 | -0.29 | -0.57 | 0.4625 | 0.2725 | -0.225 | 0.684943 | 0.414182 | 0.658678 | 0.989223 | 0.997988 | 0.996159 |
| Ankrd63_A2ARS0_S203s_1 | Ankrd63_A | 0.14 | -2.02 | -1.19 | 0.24 | -1.16 | -0.4 | -0.01 | 0.34 | -0.45 | -1.18 | -0.33 | -0.99 | -0.35 | -1.13 | -1.04 | -1.15 | 0.4 | -0.03 | -0.21 | 0.69718 | 0.48843 | 0.955785 | 0.989223 | 0.997988 | 0.996159 |
| Ankrd63_A2ARS0_S203s_2 | Ankrd63_A | 1.02 | -1.44 | -1.02 | 0.51 | -0.34 | 0.22 | 0.46 | 0.61 | 0.18 | -0.38 | 1 | -0.57 | -0.88 | -0.08 | -0.29 | -0.46 | 0.47 | 0.29 | -0.195 | 0.731523 | 0.41413 | 0.645727 | 0.989223 | 0.997988 | 0.9971 |
| Ankrd63_A2ARS0_S268s_1 | Ankrd63_A | -0.38 | -0.93 | -0.72 | -0.5 | -0.44 | -1.19 | -0.37 | -0.84 | -0.9 | -0.27 | -0.76 | -0.51 | -0.71 | -0.22 | -0.98 | -0.15 | -0.0775 | 0.0225 | 0.1175 | 0.693322 | 0.773573 | 0.929666 | 0.995762 | 0.997988 | 0.996159 |
| Ankrd63_A2ARS0_S293s_1 | Ankrd63_A | 3.01 | -1.23 | -0.38 | 1.54 | 0.18 | 2.21 | 1.2 | 2.58 | 1.26 | -0.15 | 0.14 | 0.22 | -1.54 | -0.72 | -0.26 | -0.3 | 0.8075 | -0.3675 | -1.44 | 0.125706 | 0.404557 | 0.680851 | 0.989223 | 0.997988 | 0.994666 |
| Ankrd63_A2ARS0_S341s_1 | Ankrd63_A | -1.9 | -2.54 | -3 | -1.07 | -1.45 | -2.33 | -1.23 | -1.29 | -2.26 | -2.74 | 0.29 | -1.62 | -2.99 | -1.38 | -1.23 | -1.99 | 0.5525 | 0.545 | 0.23 | 0.670538 | 0.245118 | 0.448938 | 0.989223 | 0.997988 | 0.996159 |
| Ankrd63_A2ARS0_T342t_1 | Ankrd63_A | -1.9 | -2.54 | -3 | -1.07 | -1.45 | -2.33 | -1.23 | -1.29 | -2.26 | -2.74 | 0.29 | -1.62 | -2.99 | -1.38 | -1.23 | -1.99 | 0.5525 | 0.545 | 0.23 | 0.670538 | 0.245118 | 0.448938 | 0.989223 | 0.997988 | 0.996159 |
| Anks1_Q3UHP6_S663s_1 | Anks1_Q3U | 0.99 | 0.26 | 0.03 | 0.19 | 0.99 | 1.22 | -0.06 | 0.18 | 0.31 | 1.22 | 0.33 | -0.7 | 0.58 | 0.48 | -0.79 | -0.16 | 0.215 | -0.0775 | -0.34 | 0.395914 | 0.56462 | 0.858126 | 0.989223 | 0.997988 | 0.994666 |
| Anks1_Q3UHP6_S903s_1 | Anks1_Q3U | 1.42 | 0.64 | 2.14 | 0.37 | 1.5 | 1.88 | 2.63 | 2.25 | 0.88 | 1.73 | -1 | 2.53 | 1.18 | -1.55 | 2.83 | 3.35 | 0.9225 | -0.1075 | 0.31 | 0.761568 | 0.059545 | 0.888478 | 0.989223 | 0.997988 | 0.9971 |
| Anks1b_S4R286_S1222s_1 | Anks1b_S4 | -1.83 | -2.31 | -1.71 | -0.95 | -0.32 | -2.37 | -0.18 | -0.54 | -1.83 | -1.67 | 0.18 | -0.12 | -0.19 | -0.99 | -0.07 | -0.65 | 0.8475 | 0.84 | 1.225 | 0.00809 | 0.134743 | 0.149959 | 0.989223 | 0.997988 | 0.994666 |
| Anks1b_S4R286_S364s_1 | Anks1b_S4 | -0.94 | -0.85 | 1.5 | -0.06 | -0.05 | -1.24 | 0.22 | 0.85 | -1.02 | -0.87 | 1.73 | 1.2 | -0.7 | 0.04 | 0.47 | -0.25 | 0.0325 | 0.3475 | -0.0225 | 0.968347 | 0.959159 | 0.66781 | 0.999187 | 0.997988 | 0.999507 |
| Anks1b_S4R286_S510s_1 | Anks1b_S4 | 0.2 | 0.08 | 0.53 | 0.95 | 0.73 | -0.1 | 0.67 | 0.44 | 0.22 | 0.8 | 0.5 | 0.43 | -0.11 | -0.28 | 0.46 | 0.62 | -0.005 | 0.0475 | -0.2675 | 0.428498 | 0.986573 | 0.865385 | 0.999187 | 0.997988 | 0.994666 |
| Anks1b_S4R286_S589s_1 | Anks1b_S4 | -0.51 | 1.25 | -1.24 | -3.22 | 0.92 | 1.21 | -1.44 | -1.85 | 0.17 | 1.04 | -2.04 | -1.58 | -2.39 | -0.33 | -1.87 | -1.12 | 0.64 | 0.3275 | -0.4975 | 0.583606 | 0.545961 | 0.752446 | 0.989223 | 0.997988 | 0.995259 |
| Anks1b_S4R286_T968t_1 | Anks1b_S4 | -1.21 | -0.84 | -0.59 | -0.42 | -0.72 | -1.74 | 0.19 | -0.67 | -0.57 | -0.46 | -0.95 | -0.44 | 0.17 | -0.44 | -0.03 | -0.35 | 0.03 | 0.16 | 0.6025 | 0.062184 | 0.941748 | 0.554774 | 0.999187 | 0.997988 | 0.994666 |
| Anln_Q8K298_S637s_1 | Anln_Q8K2 | -1.11 | -0.44 | -0.06 | -0.84 | -0.86 | -1.53 | -0.38 | -1.2 | -1.63 | -1.82 | 0.28 | 0.88 | -1.05 | -1.19 | -0.17 | -0.13 | -0.38 | 0.04 | -0.0225 | 0.952627 | 0.282492 | 0.95087 | 0.989223 | 0.997988 | 0.999507 |
| Anln_Q8K298_S656s_1 | Anln_Q8K2 | -0.41 | -0.23 | -0.37 | -1.44 | -0.29 | -0.78 | -2.78 | -1.53 | -0.01 | -0.49 | -0.31 | -1.32 | -0.3 | -1.79 | -1.33 | -1.31 | -0.7325 | 0.08 | -0.57 | 0.195253 | 0.204512 | 0.839377 | 0.989223 | 0.997988 | 0.994666 |
| Ano8_Q6PB70_S637s_1 | Ano8_Q6P | 0.02 | 0.41 | 0.33 | 0.14 | 0.62 | -0.37 | 0.71 | 0.48 | -0.69 | -0.09 | -0.31 | -0.04 | -0.22 | 0.82 | 0.22 | 0.45 | 0.135 | -0.5075 | 0.0925 | 0.75668 | 0.645774 | 0.066176 | 0.99033 | 0.997988 | 0.9971 |
| Antxr1_Q9CZ52_S561s_1 | Antxr1_Q9 | 0.21 | 0.61 | 0.6 | 0.94 | 0.65 | 0.71 | 0.71 | 0.85 | 0.03 | 0.15 | 0.45 | 0.88 | 0.73 | 0.59 | 0.75 | 0.49 | 0.14 | -0.2125 | 0.05 | 0.847909 | 0.54718 | 0.467044 | 0.989223 | 0.997988 | 0.999507 |
| Anxa7_Q07076_S311s_1 | Anxa7_Q07 | -0.42 | 0.26 | 0.99 | 0.65 | -0.27 | -0.5 | 0.34 | 0.57 | 0.28 | -0.05 | 0.86 | 0.82 | 0.97 | 0.33 | 0.75 | 0.81 | -0.335 | 0.1075 | 0.345 | 0.347125 | 0.391926 | 0.776895 | 0.989223 | 0.997988 | 0.994666 |
| Ap1g1_Q8CBB7_S235s_1 | Ap1g1_Q8 | -0.52 | -0.13 | -0.12 | -0.18 | -0.15 | -0.57 | 0.35 | -0.17 | -0.37 | -0.25 | -0.12 | -0.06 | -0.88 | 0.21 | -0.38 | -0.07 | 0.1025 | 0.0375 | -0.0425 | 0.890126 | 0.69456 | 0.866882 | 0.994893 | 0.997988 | 0.999507 |
| Ap2a1_P17426_S517s_1 | Ap2a1_P17 | 0.02 | -1.68 | -0.22 | 0.63 | -0.45 | -0.95 | 0.93 | 0.49 | -0.1 | -0.75 | -0.58 | 1.39 | -0.38 | 0.11 | -0.56 | 0.58 | 0.3175 | 0.3025 | 0.25 | 0.63167 | 0.589494 | 0.632594 | 0.989223 | 0.997988 | 0.995259 |
| Ap2a1_P17426_S518s_1 | Ap2a1_P17 | 0.02 | -1.68 | -0.22 | 0.63 | -0.45 | -0.95 | 0.93 | 0.49 | -0.1 | -0.75 | -0.58 | 1.39 | -0.38 | 0.11 | -0.56 | 0.58 | 0.3175 | 0.3025 | 0.25 | 0.63167 | 0.589494 | 0.632594 | 0.989223 | 0.997988 | 0.995259 |
| Ap2a1_P17426_S611s_1 | Ap2a1_P17 | -1.33 | -1.7 | -1.26 | -1.46 | -0.58 | -1.25 | -0.53 | -0.9 | -1.81 | -1.69 | -1.72 | -1.17 | -0.75 | -0.7 | -1.17 | -0.79 | 0.6225 | -0.16 | 0.585 | 0.041835 | 0.030635 | 0.526065 | 0.989223 | 0.997988 | 0.994666 |
| Ap2a1_P17426_S626s_1 | Ap2a1_P17 | 0.2 | 0.26 | 0.3 | 0.33 | 0.63 | -0.13 | -0.02 | 0.04 | -0.23 | -0.18 | 0.44 | 0.65 | 0.42 | 0.58 | 0.38 | 0.15 | -0.1425 | -0.1025 | 0.11 | 0.642469 | 0.554758 | 0.711831 | 0.989223 | 0.997988 | 0.995259 |
| Ap2a1_P17426_S631s_1 | Ap2a1_P17 | -0.12 | -0.03 | -0.51 | -0.91 | 1.17 | 0.36 | -0.3 | -0.43 | -0.51 | -0.92 | 0.35 | 0.32 | 0.13 | 1.41 | -0.35 | 0.13 | 0.5925 | 0.2025 | 0.7225 | 0.114197 | 0.163916 | 0.595334 | 0.989223 | 0.997988 | 0.994666 |
| Ap2a1_P17426_S655s_1 | Ap2a1_P17 | 0.47 | 1.21 | -1.34 | 1.87 | 0.59 | 0.74 | 0.92 | 2.3 | 0.56 | 1.01 | 3.13 | 2.24 | 0.36 | 0.81 | 1.82 | 0.87 | 0.585 | 1.1825 | 0.4125 | 0.547915 | 0.413542 | 0.167997 | 0.989223 | 0.997988 | 0.994666 |
| Ap2m1_P84091_S45s_1 | Ap2m1_P8 | 0.59 | 0.93 | 1.16 | 0.81 | 0.55 | 0.56 | 0.57 | 0.51 | 0.48 | 0.48 | 0.78 | 0.47 | 0.73 | 0.64 | 0.14 | 0.26 | -0.325 | -0.32 | -0.43 | 0.138756 | 0.153837 | 0.193102 | 0.989223 | 0.997988 | 0.994666 |
| Ap2m1_P84091_T156t_1 | Ap2m1_P8 | 0.07 | 0.55 | 0.53 | -0.04 | 0.87 | 0.1 | 0.98 | 0.13 | -0.14 | 0.12 | -0.36 | 0.9 | 0.16 | 1.09 | -0.01 | 0.11 | 0.2425 | -0.1475 | 0.06 | 0.857677 | 0.432098 | 0.662332 | 0.989223 | 0.997988 | 0.999507 |
| Ap3b2_Q9JME5_S272s_1 | Ap3b2_Q9 | -0.38 | -0.1 | 0.22 | -0.11 | 0.06 | -0.43 | -0.11 | 0.29 | -0.66 | -0.16 | -0.22 | 0.04 | -0.68 | -0.26 | -0.09 | -0.28 | 0.045 | -0.1575 | -0.235 | 0.38867 | 0.857718 | 0.547108 | 0.996349 | 0.997988 | 0.994666 |
| Ap3b2_Q9JME5_S8s_1 | Ap3b2_Q9 | -0.17 | -0.29 | -2.63 | -1.64 | -0.24 | -0.76 | -1.61 | -1.86 | -0.09 | -0.15 | 0.95 | -1.82 | 0.74 | 0.46 | -2.26 | -1.83 | 0.065 | 0.905 | 0.46 | 0.59172 | 0.916501 | 0.238646 | 0.997335 | 0.997988 | 0.995259 |
| Ap3d1_O54774_S760s_1 | Ap3d1_O5 | 3.32 | 4.41 | 4.4 | 2.34 | 4.1 | 2.94 | 4.03 | 4.04 | 1.66 | 4.4 | 1.17 | 4.46 | 1.32 | 1.57 | 4.38 | 4.54 | 0.16 | -0.695 | -0.665 | 0.457037 | 0.758371 | 0.44448 | 0.995762 | 0.997988 | 0.994666 |
| Ap3d1_O54774_S825s_1 | Ap3d1_O5 | 1.21 | 0.72 | 0.79 | 1.14 | 1.06 | 1.47 | 0.7 | 1.12 | 1.71 | 1.18 | 0.54 | 0.95 | 1.06 | 0.34 | 0.81 | 1 | 0.1225 | 0.13 | -0.1625 | 0.566879 | 0.631259 | 0.674561 | 0.989223 | 0.997988 | 0.995259 |
| Ap4e1_Q80V94_S675s_1 | Ap4e1_Q8 | -0.37 | -0.26 | -0.28 | 0.02 | 0.21 | 0.29 | 0.18 | 0 | -0.04 | 0.13 | -1.41 | -0.33 | -0.27 | 0.16 | 0.1 | 0.81 | 0.3925 | -0.19 | 0.4225 | 0.182274 | 0.085324 | 0.605714 | 0.989223 | 0.997988 | 0.994666 |
| Apba1_B2RUJ5_S246s_1 | Apba1_B2R | -2.2 | -1.36 | -2.44 | -4.68 | -3.15 | -1.75 | -4.33 | -3.78 | -0.49 | -2.26 | -1.99 | -2.93 | 0.72 | -1.83 | -1.68 | -3.05 | -0.5825 | 0.7525 | 1.21 | 0.212434 | 0.466251 | 0.349445 | 0.989223 | 0.997988 | 0.994666 |
| Apba1_B2RUJ5_S289s_1 | Apba1_B2R | 0.77 | 0.94 | 1.27 | 0.97 | 0.04 | 0.42 | 0.05 | 0.33 | 0.84 | 1.18 | 1.23 | 1.25 | 1.47 | 0.26 | 1.17 | 0.51 | -0.7775 | 0.1375 | -0.135 | 0.689869 | 0.006141 | 0.560153 | 0.989223 | 0.997988 | 0.996159 |
| Apba1_B2RUJ5_S317s_1 | Apba1_B2R | -0.17 | -0.03 | -0.29 | -0.11 | -0.37 | -0.32 | 0.15 | 0.15 | 0.5 | 0.52 | 0.06 | 0.32 | 0.9 | 0.78 | 0.13 | 0.11 | 0.0525 | 0.5 | 0.63 | 0.050959 | 0.819111 | 0.048739 | 0.995762 | 0.997988 | 0.994666 |
| Apba1_B2RUJ5_S317s_2 | Apba1_B2R | -0.03 | 0.38 | 0.38 | 0.01 | 0.49 | 0.21 | 0.25 | 0.41 | 0.17 | 0.63 | 0.88 | 0.47 | -0.58 | 0.73 | -0.05 | 0.14 | 0.155 | 0.3525 | -0.125 | 0.707927 | 0.485417 | 0.191329 | 0.989223 | 0.997988 | 0.996966 |
| Apba1_B2RUJ5_S372s_1 | Apba1_B2R | -1.71 | -2.48 | -1.42 | -1.01 | -0.95 | -1.6 | -0.51 | -0.75 | -1.09 | 0.01 | -1.03 | -1.08 | -1.81 | -0.05 | -1.37 | -1.43 | 0.7025 | 0.8575 | 0.49 | 0.314937 | 0.089697 | 0.059351 | 0.989223 | 0.997988 | 0.994666 |
| Apba1_B2RUJ5_S391s_1 | Apba1_B2R | 0.05 | 0.38 | 0.05 | -0.24 | -0.48 | 0.32 | -0.75 | -0.78 | 0.9 | 0.13 | -0.59 | -0.1 | 1.28 | -0.55 | -0.15 | -0.08 | -0.4825 | 0.025 | 0.065 | 0.876297 | 0.140666 | 0.943103 | 0.989223 | 0.997988 | 0.999507 |
| Apba1_B2RUJ5_S573s_1 | Apba1_B2R | -0.62 | -0.39 | 0.74 | 0.52 | -1.42 | 0.05 | -0.48 | 0.32 | 0.09 | -0.38 | 0.74 | 0.96 | 0.33 | -1.22 | 0.38 | 0.73 | -0.445 | 0.29 | -0.0075 | 0.98836 | 0.356543 | 0.51374 | 0.989223 | 0.997988 | 1 |
| Apba1_B2RUJ5_T309t_1 | Apba1_B2R | -1.03 | -0.19 | -0.58 | -1.01 | -0.51 | -1.22 | 0.51 | -0.53 | -0.87 | -0.66 | -0.29 | -0.08 | -0.36 | -0.13 | -1.17 | -0.1 | 0.265 | 0.2275 | 0.2625 | 0.460171 | 0.506891 | 0.462187 | 0.989223 | 0.997988 | 0.994666 |
| Apba1_B2RUJ5_T309t_2 | Apba1_B2R | -0.03 | 0.38 | 0.38 | 0.01 | 0.49 | 0.21 | 0.25 | 0.41 | 0.17 | 0.63 | 0.88 | 0.47 | -0.58 | 0.73 | -0.05 | 0.14 | 0.155 | 0.3525 | -0.125 | 0.707927 | 0.485417 | 0.191329 | 0.989223 | 0.997988 | 0.996966 |
| Apba1_B2RUJ5_T333t_1 | Apba1_B2R | 0.31 | 0.82 | -0.49 | -0.17 | -0.03 | 0.12 | -0.55 | -0.67 | 0.04 | 0.73 | -0.38 | -0.36 | 0.05 | 0.42 | -0.37 | -0.16 | -0.4 | -0.11 | -0.1325 | 0.712008 | 0.267538 | 0.777009 | 0.989223 | 0.997988 | 0.996966 |
| Apba1_B2RUJ5_T375t_1 | Apba1_B2R | -4.78 | -2.52 | -0.93 | -0.28 | -1.86 | -4.33 | -0.69 | -0.39 | -2.55 | -2.31 | -0.57 | -0.52 | -3.11 | -1.72 | -0.04 | -0.53 | 0.31 | 0.64 | 0.7775 | 0.467017 | 0.788869 | 0.529578 | 0.995762 | 0.997988 | 0.994666 |
| Apba2_P98084_S479s_1 | Apba2_P98 | -0.56 | 0.04 | -0.09 | -0.98 | 0.63 | -0.9 | -0.34 | 0.01 | -0.35 | -0.27 | -0.5 | -0.74 | 0.6 | 0.09 | -0.16 | -0.06 | 0.2475 | -0.0675 | 0.515 | 0.142285 | 0.525216 | 0.819975 | 0.989223 | 0.997988 | 0.994666 |
| Apba2_P98084_S481s_1 | Apba2_P98 | 0.74 | -0.83 | 0.27 | 1.07 | 0.05 | 0.6 | 0.37 | 0.61 | -0.01 | 0.01 | 1.03 | 0.48 | -1.07 | -0.06 | 0.45 | 0.54 | 0.095 | 0.065 | -0.3475 | 0.509366 | 0.818758 | 0.88783 | 0.995762 | 0.997988 | 0.994666 |
| Apba2_P98084_T307t_1 | Apba2_P98 | -0.64 | -1.71 | 0.1 | -0.56 | -2.08 | 0.57 | -0.79 | -0.14 | 0.87 | -1.67 | -1.29 | 0.21 | 0.51 | -2.33 | -0.12 | -1.08 | 0.0925 | 0.2325 | -0.0525 | 0.935785 | 0.878306 | 0.719448 | 0.996493 | 0.997988 | 0.999507 |
| Apbb1ip_Q8R5A3_S507s_1 | Apbb1ip_Q | -0.69 | -1.39 | -1.19 | -0.56 | -1.96 | -0.27 | -1.75 | -1.78 | -0.8 | -2.24 | -1.92 | -0.63 | 0.85 | -1.22 | -0.36 | -1.79 | -0.4825 | -0.44 | 0.3275 | 0.562706 | 0.265631 | 0.324913 | 0.989223 | 0.997988 | 0.995259 |
| Apbb2_F6V8Y6_S413s_1 | Apbb2_F6V | 0.66 | 1.83 | 1.43 | 0.86 | 1.37 | 1.52 | 0.53 | 1.1 | 1.04 | 1.15 | 0.52 | 1.26 | 1.2 | 0.81 | 1.02 | 0.93 | -0.065 | -0.2025 | -0.205 | 0.531613 | 0.85177 | 0.549871 | 0.996349 | 0.997988 | 0.994666 |
| Apbb3_Q8R1C9_S111s_1 | Apbb3_Q8 | 0.47 | 0.24 | 0.97 | 0.01 | 0.16 | 0.66 | -0.47 | 0.04 | 0.43 | -0.05 | -0.5 | 0.67 | 0.95 | -0.14 | 0.4 | 0.05 | -0.325 | -0.285 | -0.1075 | 0.757076 | 0.327803 | 0.421193 | 0.989223 | 0.997988 | 0.9971 |
| Apc_B2RUG9_S1030s_1 | Apc_B2RU | -1.22 | -0.73 | -1.71 | -2.43 | -0.43 | -1.75 | -1.85 | -1.89 | -1.75 | -1.5 | -0.27 | -1.08 | -0.64 | -0.2 | -2.02 | -1.77 | 0.0425 | 0.3725 | 0.365 | 0.497104 | 0.92759 | 0.431216 | 0.997416 | 0.997988 | 0.994666 |
| Apc_B2RUG9_S1040s_1 | Apc_B2RU | 2.77 | 0.53 | -0.01 | -2.55 | 0.76 | 2.1 | -0.41 | -1.2 | 2.33 | 2.81 | -1.31 | 0 | 1.85 | 0.42 | 0.44 | -1.28 | 0.1275 | 0.7725 | 0.1725 | 0.875061 | 0.909826 | 0.550606 | 0.997335 | 0.997988 | 0.999507 |
| Apc_B2RUG9_S1370s_1 | Apc_B2RU | 1.51 | 1.22 | 0.63 | 0.49 | 0.57 | 0.59 | 1.14 | 0.57 | 1.12 | 1.71 | 0.98 | 0.82 | 0.61 | 0.88 | 0.51 | 0.3 | -0.245 | 0.195 | -0.3875 | 0.242098 | 0.426621 | 0.561611 | 0.989223 | 0.997988 | 0.994666 |
| Apc_B2RUG9_S1435s_1 | Apc_B2RU | 1.11 | 1.09 | 0.93 | 1.4 | 0.97 | 1.09 | 0.61 | 1.35 | 0.22 | 1.36 | 0.61 | 0.62 | 0.27 | 0.74 | 0.46 | 1.19 | -0.1275 | -0.43 | -0.4675 | 0.132959 | 0.603879 | 0.17276 | 0.989223 | 0.997988 | 0.994666 |
| Apc_B2RUG9_S1772s_1 | Apc_B2RU | 0.63 | 1 | 1.98 | 1.19 | 0.62 | 0.49 | 1.37 | 1.36 | 0.82 | 1.08 | 2.17 | 1.84 | 1.22 | 1.07 | 2.48 | 1.33 | -0.24 | 0.2775 | 0.325 | 0.453164 | 0.516366 | 0.511642 | 0.989223 | 0.997988 | 0.994666 |
| Apc_B2RUG9_S2260s_1 | Apc_B2RU | 0.25 | 0.2 | 0.43 | 0.35 | 0.14 | 0.36 | -0.14 | 0.46 | 0.34 | 0.39 | 0.9 | 0.61 | 0.59 | 0.12 | 0.43 | 0.34 | -0.1025 | 0.2525 | 0.0625 | 0.79562 | 0.649532 | 0.292411 | 0.990358 | 0.997988 | 0.999507 |
| Apc_B2RUG9_S2283s_1 | Apc_B2RU | -1.48 | -1.23 | -0.99 | -1.15 | -1.15 | -1.31 | -1.32 | -1.55 | -1.45 | -1.27 | -1.23 | -1.25 | -1.05 | -0.88 | -1.26 | -1.18 | -0.12 | -0.0875 | 0.12 | 0.631092 | 0.587299 | 0.695065 | 0.989223 | 0.997988 | 0.995259 |
| Apc_B2RUG9_S2290s_1 | Apc_B2RU | -2.59 | -2.58 | -1.64 | -0.82 | -3.33 | -1.39 | -1.78 | -1.83 | -1.84 | -4.14 | -0.46 | -0.78 | -0.35 | -1.39 | -1.02 | -1.62 | -0.175 | 0.1025 | 0.8125 | 0.118719 | 0.74881 | 0.901954 | 0.995762 | 0.997988 | 0.994666 |
| Apc_B2RUG9_S2449s_1 | Apc_B2RU | -0.53 | -0.57 | 0.58 | 0.59 | 0.32 | -0.67 | 0.52 | 0.93 | -1 | -0.69 | 0.81 | 0.62 | -0.84 | 0.4 | 0.27 | 0.3 | 0.2575 | -0.0825 | 0.015 | 0.972324 | 0.564162 | 0.874997 | 0.989223 | 0.997988 | 0.999507 |
| Apc_B2RUG9_S2674s_1 | Apc_B2RU | 0.15 | 0.08 | 0.87 | 0.95 | -0.16 | 0.29 | 0.88 | 0.66 | -0.12 | 0.5 | 0.23 | 0.59 | -0.13 | 0.7 | 0.74 | 0.24 | -0.095 | -0.2125 | -0.125 | 0.716404 | 0.775769 | 0.502565 | 0.995762 | 0.997988 | 0.996966 |
| Apc_B2RUG9_S2710s_1 | Apc_B2RU | 0 | 0.4 | 0.44 | 0.06 | -0.19 | 0.02 | 0.43 | 0.4 | 0.13 | 0.45 | 0.25 | 0.4 | 0.17 | -0.3 | 0.38 | 0.01 | -0.06 | 0.0825 | -0.16 | 0.557945 | 0.808616 | 0.721594 | 0.995762 | 0.997988 | 0.995259 |
| Apc_B2RUG9_S2771s_1 | Apc_B2RU | 8.2 | 4.44 | 5.3 | 1.77 | 4.79 | 9.47 | 5.02 | 4.51 | -0.4 | 7.55 | 0.66 | 5.37 | -0.2 | -1.94 | 5.32 | 6.24 | 1.02 | -1.6325 | -2.5725 | 0.231457 | 0.506689 | 0.425151 | 0.989223 | 0.997988 | 0.994666 |
| Apc_B2RUG9_S2829s_1 | Apc_B2RU | -1.42 | -0.67 | -0.89 | -0.67 | -2.21 | -0.61 | -0.89 | -1.06 | 1.63 | -0.94 | 0.77 | -0.77 | 2.36 | -2.05 | 1.08 | -1.38 | -0.28 | 1.085 | 0.915 | 0.331998 | 0.471568 | 0.093016 | 0.989223 | 0.997988 | 0.994666 |
| Apc_B2RUG9_S742s_1 | Apc_B2RU | 0.46 | 1.21 | 0.1 | 0.13 | 1.05 | 1.39 | 0.39 | 0.9 | 0.82 | 0 | 1.31 | -0.12 | 0.94 | 0.75 | 0.54 | 0.93 | 0.4575 | 0.0275 | 0.315 | 0.339577 | 0.197859 | 0.947628 | 0.989223 | 0.997988 | 0.994666 |
| Apc_B2RUG9_S838s_1 | Apc_B2RU | -0.09 | -0.59 | -0.39 | -1.37 | -1.41 | 1.09 | -1.26 | 0.28 | 1.88 | -0.35 | -1.9 | -0.65 | 1.52 | -0.75 | 0.19 | -1.11 | 0.285 | 0.355 | 0.5725 | 0.346065 | 0.634244 | 0.63669 | 0.989223 | 0.997988 | 0.994666 |
| Apc_B2RUG9_S983s_1 | Apc_B2RU | -2.11 | -2.49 | -0.8 | -0.9 | -1.25 | -2.69 | -1.09 | -0.41 | -1.93 | -1.96 | 1.71 | -0.5 | -2.44 | -1.49 | -1.12 | -0.45 | 0.215 | 0.905 | 0.2 | 0.717418 | 0.709864 | 0.30575 | 0.995437 | 0.997988 | 0.996966 |
| Apc_B2RUG9_T1437t_1 | Apc_B2RU | -0.16 | 0.85 | -0.48 | -0.51 | 0.2 | -0.58 | -0.45 | -0.9 | 0.1 | 0.07 | 0.24 | -0.81 | 0.27 | 0.76 | 0.14 | -0.79 | -0.3575 | -0.025 | 0.17 | 0.702612 | 0.362245 | 0.949736 | 0.989223 | 0.997988 | 0.996159 |
| Apc2_D3YTR0_S1230s_1 | Apc2_D3YT | 0.42 | 0.08 | 0.53 | 0.22 | 0.18 | -0.45 | 0.94 | 0.36 | -0.12 | 0.23 | 0.2 | 0.3 | -0.41 | 0.61 | 0.14 | 0.73 | -0.055 | -0.16 | -0.045 | 0.889274 | 0.862973 | 0.49582 | 0.996493 | 0.997988 | 0.999507 |
| Apc2_D3YTR0_S1834s_1 | Apc2_D3YT | 0.2 | 0.19 | 0.77 | 0.36 | -0.84 | 0.5 | -0.56 | 0.03 | 2.7 | 0.38 | -0.49 | 0.42 | 2.62 | -0.95 | 0.3 | 0.46 | -0.5975 | 0.3725 | 0.2275 | 0.737274 | 0.101674 | 0.560147 | 0.989223 | 0.997988 | 0.9971 |
| Apc2_D3YTR0_S1880s_1 | Apc2_D3YT | -0.83 | -0.31 | -1.2 | -1.55 | -0.01 | -0.45 | 0.04 | -0.98 | 0.03 | 0.84 | -0.89 | -0.59 | 0.9 | 0 | -0.19 | -0.78 | 0.6225 | 0.82 | 0.955 | 0.049217 | 0.105694 | 0.093803 | 0.989223 | 0.997988 | 0.994666 |
| Apc2_D3YTR0_S1890s_1 | Apc2_D3YT | -0.28 | 0.29 | 0.35 | 0.23 | 0.17 | -0.14 | 0.71 | 0.32 | -0.11 | -0.08 | -0.25 | 0.44 | -0.36 | 0.22 | 0.03 | 0.21 | 0.1175 | -0.1475 | -0.1225 | 0.661194 | 0.665177 | 0.586099 | 0.991354 | 0.997988 | 0.995671 |
| Apc2_D3YTR0_S2010s_1 | Apc2_D3YT | -0.8 | 0.41 | 0.88 | 0.74 | 0.68 | -0.22 | -0.26 | 0.37 | -0.31 | 0.17 | 2.33 | 1.07 | 1.06 | 0.8 | 1.28 | 1.1 | -0.165 | 0.5075 | 0.7525 | 0.085756 | 0.697231 | 0.430889 | 0.994893 | 0.997988 | 0.994666 |
| Apc2_D3YTR0_S2128s_1 | Apc2_D3YT | -0.85 | -0.61 | 0.49 | -1.04 | -0.41 | -0.57 | 0.06 | -0.19 | 0.37 | -0.55 | 0.65 | 0.97 | 0.98 | -0.25 | 0.23 | -0.5 | 0.225 | 0.8625 | 0.6175 | 0.198724 | 0.541694 | 0.084038 | 0.989223 | 0.997988 | 0.994666 |
| Apc2_D3YTR0_S2172s_1 | Apc2_D3YT | 1.06 | 2.01 | 0.58 | 0.08 | 1.98 | -0.13 | 0.16 | -0.18 | 1.86 | 1.59 | 1.19 | 0.18 | 2.2 | 2.41 | 0.4 | 0.29 | -0.475 | 0.2725 | 0.3925 | 0.540399 | 0.428214 | 0.600594 | 0.989223 | 0.997988 | 0.994666 |
| Apc2_D3YTR0_S2205s_1 | Apc2_D3YT | 0.82 | 1.06 | 1.03 | -0.09 | 0.8 | 0.49 | 0.27 | 0.14 | -0.48 | 0.59 | -1.04 | 0.38 | 1.23 | -1.19 | 0.56 | 0.38 | -0.28 | -0.8425 | -0.46 | 0.402541 | 0.392673 | 0.086783 | 0.989223 | 0.997988 | 0.994666 |
| Apc2_D3YTR0_T128t_1 | Apc2_D3YT | -0.62 | -0.28 | -1.15 | -1.08 | -0.54 | -0.44 | -1.45 | -0.99 | -0.43 | -0.04 | -0.41 | -0.55 | 0.16 | -0.68 | -0.87 | -1.09 | -0.0725 | 0.425 | 0.1625 | 0.656864 | 0.822281 | 0.156934 | 0.995762 | 0.997988 | 0.995259 |
| Api5_O35841_S464s_1 | Api5_O358 | -2.32 | -2.58 | -1.78 | -4.94 | -2.56 | -2.42 | -2.97 | -3.05 | -2.08 | -2.27 | -3.75 | -3.6 | -0.52 | -2.8 | -4.15 | -2.91 | 0.155 | -0.02 | 0.31 | 0.73157 | 0.807516 | 0.978391 | 0.995762 | 0.997988 | 0.9971 |
| Aplf_Q9D842_S23s_1 | Aplf_Q9D8 | -1.84 | -1.11 | -1.65 | -2.34 | -1.12 | -1.93 | -1.07 | -1.75 | -1.7 | -1.15 | -0.35 | -2.1 | -1.84 | -0.99 | -0.55 | -0.87 | 0.2675 | 0.41 | 0.6725 | 0.107503 | 0.440264 | 0.364437 | 0.989223 | 0.997988 | 0.994666 |
| Aplp2_Q60709_T724t_1 | Aplp2_Q60 | -0.28 | 0 | 0.05 | 0.39 | 0.12 | -0.15 | 0.58 | 0.46 | -0.55 | 0.23 | 0.88 | 0.34 | -0.89 | -0.15 | -0.01 | 0.15 | 0.2125 | 0.185 | -0.265 | 0.412286 | 0.426647 | 0.593086 | 0.989223 | 0.997988 | 0.994666 |
| Apoe_P08226_T140t_1 | Apoe_P082 | -0.24 | 0.23 | -1.18 | -0.23 | 0.23 | 0.03 | -0.74 | -1.69 | 0.63 | 0.18 | 0.21 | 0.06 | 0.62 | 0.66 | 1.1 | -0.62 | -0.1875 | 0.625 | 0.795 | 0.108456 | 0.700176 | 0.092732 | 0.994893 | 0.997988 | 0.994666 |
| App_Q3TWF3_S441s_1 | App_Q3TW | -0.26 | 0.32 | -0.3 | -1.35 | 0.84 | -0.52 | 0.3 | -0.8 | -0.1 | -0.54 | -0.82 | -0.89 | 0.04 | 0.67 | -0.59 | -1.71 | 0.3525 | -0.19 | -1.6E-16 | 1 | 0.463576 | 0.629529 | 0.989223 | 0.997988 | 1 |
| App_Q3TWF3_T725t_1 | App_Q3TW | 0.54 | 0.74 | 0.09 | -0.02 | 0.69 | 0.36 | 0.55 | 0.51 | 0.68 | 0.68 | -0.37 | 0.11 | 0.33 | 0.97 | 0.49 | 0.34 | 0.19 | -0.0625 | 0.195 | 0.51769 | 0.454209 | 0.851526 | 0.989223 | 0.997988 | 0.994666 |
| Appl1_Q8K3H0_S401s_1 | Appl1_Q8K | -0.36 | -0.75 | -0.73 | -1.08 | -0.28 | -0.4 | -0.29 | -0.81 | -0.69 | -0.1 | -0.46 | -1.16 | -1.27 | -1.44 | -1.31 | -0.78 | 0.285 | 0.1275 | -0.47 | 0.121156 | 0.270697 | 0.675988 | 0.989223 | 0.997988 | 0.994666 |
| Aqp4_P55088_S254s_1 | Aqp4_P55C | 0.53 | 1.26 | 1.16 | 0.67 | 1.37 | 0.56 | 0.41 | 0.55 | 0.7 | 0.83 | 1.05 | 0.72 | 0.61 | 1.21 | 1.24 | 0.81 | -0.1825 | -0.08 | 0.0625 | 0.834444 | 0.553201 | 0.760181 | 0.989223 | 0.997988 | 0.999507 |
| Aqp4_P55088_S263s_1 | Aqp4_P55C | -1.58 | -1.14 | -1.04 | -1.08 | -1 | -1.86 | -0.53 | -0.89 | -1.75 | -1.22 | -0.73 | -0.97 | -1.28 | -0.96 | -1.04 | -1.03 | 0.14 | 0.0425 | 0.1325 | 0.602982 | 0.665252 | 0.885165 | 0.991354 | 0.997988 | 0.995259 |
| Araf_P04627_S255s_1 | Araf_P046 | -0.32 | -0.65 | -1.59 | -1.4 | -0.52 | -0.8 | -1.4 | -0.62 | -0.07 | -0.61 | 0.65 | -0.36 | 0.3 | -0.43 | 0.06 | -0.53 | 0.155 | 0.8925 | 0.84 | 0.047911 | 0.66731 | 0.049993 | 0.99272 | 0.997988 | 0.994666 |
| Araf_P04627_S580s_1 | Araf_P046 | -0.57 | -0.41 | -1.26 | -0.22 | -0.82 | -0.84 | -1.29 | -0.93 | -0.95 | -0.98 | -0.37 | -1.09 | 0.36 | -0.25 | -1.15 | -0.89 | -0.355 | -0.2325 | 0.1325 | 0.746972 | 0.23169 | 0.46274 | 0.989223 | 0.997988 | 0.9971 |
| Arcn1_Q5XJY5_S493s_1 | Arcn1_Q5X | -0.12 | 0.72 | 0.65 | 0.67 | 0.19 | 0.04 | 0.02 | 0.37 | 0.4 | 0.27 | 0.95 | 0.55 | 1.16 | 0.68 | 1.01 | 1.35 | -0.325 | 0.0625 | 0.57 | 0.085938 | 0.235872 | 0.830931 | 0.989223 | 0.997988 | 0.994666 |
| Arel1_E9PZX5_S337s_1 | Arel1_E9PZ | -0.5 | 0.07 | -0.21 | 0.27 | -0.44 | -1.1 | 0.19 | 0.25 | -0.39 | -0.65 | 0.77 | -0.21 | -0.29 | 0.37 | 0.45 | 0.52 | -0.1825 | -0.0275 | 0.355 | 0.264847 | 0.611795 | 0.939562 | 0.989223 | 0.997988 | 0.994666 |
| Arf6_P62331_S38s_1 | Arf6_P623 | -0.15 | -1.75 | -0.1 | 0.22 | -0.09 | -0.07 | 0.42 | 0.12 | -0.84 | 0.12 | -1.44 | -0.22 | -0.95 | -0.27 | -0.89 | 0.07 | 0.54 | -0.15 | -0.065 | 0.892985 | 0.230766 | 0.774954 | 0.989223 | 0.997988 | 0.999507 |
| Arfgap1_V9GWV1_S150s_1 | Arfgap1_V9 | -0.38 | -0.44 | -0.38 | -0.03 | -0.03 | 0.57 | -0.12 | 0.42 | 1.18 | -0.32 | 0.44 | 0.86 | 1.8 | 0.04 | 0.77 | 0.26 | 0.5175 | 0.8475 | 1.025 | 0.029305 | 0.062006 | 0.036173 | 0.989223 | 0.997988 | 0.994666 |
| Arfgap1_V9GWV1_S159s_1 | Arfgap1_V9 | 0.58 | -0.61 | 0.98 | 0.2 | -0.22 | 0.88 | -0.08 | 0.68 | 1.42 | 0.23 | 0.34 | 0.93 | 0.85 | 0.45 | -0.39 | 0.52 | 0.0275 | 0.4425 | 0.07 | 0.869409 | 0.947058 | 0.314423 | 0.999187 | 0.997988 | 0.999507 |
| Arfgap1_V9GWV1_S256s_1 | Arfgap1_V9 | -0.02 | 0.4 | 0.04 | 0.6 | 0.96 | 0.39 | 0.7 | 0.04 | -0.76 | -0.89 | 0.07 | 0.9 | -0.91 | -0.11 | 0.21 | 0.58 | 0.2675 | -0.425 | -0.3125 | 0.407208 | 0.354852 | 0.335498 | 0.989223 | 0.997988 | 0.994666 |
| Arfgap1_V9GWV1_S330s_1 | Arfgap1_V9 | 1.16 | 0.83 | -1.08 | 0.48 | 0.76 | 1.17 | 0.67 | 0.26 | 0.46 | 1.1 | -0.18 | 0.18 | 0.46 | 0.84 | 0.15 | 0.27 | 0.3675 | 0.0425 | 0.0825 | 0.866782 | 0.457801 | 0.935645 | 0.989223 | 0.997988 | 0.999507 |
| Arfgap1_V9GWV1_S330s_2 | Arfgap1_V9 | 0.13 | -0.3 | -0.19 | 0.48 | 0.4 | -0.43 | 0.18 | 0.13 | 0.16 | -0.65 | 0.41 | -0.42 | -1.19 | 0.96 | -0.26 | 0.27 | 0.04 | -0.155 | -0.085 | 0.855804 | 0.887603 | 0.638459 | 0.996493 | 0.997988 | 0.999507 |
| Arfgap1_V9GWV1_S335s_2 | Arfgap1_V9 | 0.13 | -0.3 | -0.21 | 0.22 | 0.4 | -0.43 | 0.46 | 0.21 | 0.16 | -0.65 | 0.18 | -0.56 | -1.19 | 0.96 | -0.28 | 0.04 | 0.2 | -0.1775 | -0.0775 | 0.863039 | 0.477204 | 0.555634 | 0.989223 | 0.997988 | 0.999507 |
| Arfgap1_V9GWV1_S347s_1 | Arfgap1_V9 | 0.76 | 0.27 | 0.92 | 1.25 | 0.66 | 0.46 | 1.14 | 1.13 | 0.15 | 0.58 | 0.52 | 0.77 | -0.47 | 0.8 | 0.68 | 0.7 | 0.0475 | -0.295 | -0.3725 | 0.338063 | 0.871621 | 0.320204 | 0.996493 | 0.997988 | 0.994666 |
| Arfgap1_V9GWV1_T135t_1 | Arfgap1_V9 | 0.2 | 0.52 | 0.16 | 0.59 | 1.15 | 0 | 0.79 | 0.49 | -0.25 | 0.35 | -0.2 | 0.48 | -0.31 | 0.61 | 0.29 | 0.35 | 0.24 | -0.2725 | -0.1325 | 0.65098 | 0.423369 | 0.331719 | 0.989223 | 0.997988 | 0.995259 |
| Arfgap3_Q9D8S3_S403s_1 | Arfgap3_Q | 0.22 | -1.7 | -0.79 | 0.52 | -2.39 | 1.42 | -0.59 | 0.62 | 2.08 | -1.19 | -0.06 | 0.74 | 1.49 | -1.56 | 0.49 | -0.49 | 0.2025 | 0.83 | 0.42 | 0.571305 | 0.810844 | 0.293662 | 0.995762 | 0.997988 | 0.995259 |
| Arfgef1_G3X9K3_S1076s_1 | Arfgef1_G3 | -0.73 | -1.24 | 0.73 | 1.76 | -1.04 | 0.2 | 0.39 | 1.72 | 0.23 | -0.73 | 3.1 | 0.95 | 0.65 | -1.01 | 1.11 | 1.68 | 0.1875 | 0.7575 | 0.4775 | 0.550782 | 0.809483 | 0.429501 | 0.995762 | 0.997988 | 0.994666 |
| Arfgef1_G3X9K3_S1550s_1 | Arfgef1_G3 | -0.29 | 0.59 | -0.04 | -0.55 | 0.5 | 0.02 | 0.46 | -0.18 | -1.16 | 0.16 | -2.07 | 0.45 | -1.03 | -1.01 | -0.25 | -0.55 | 0.2725 | -0.5825 | -0.6375 | 0.088128 | 0.394544 | 0.332753 | 0.989223 | 0.997988 | 0.994666 |
| Arfgef1_G3X9K3_S1563s_1 | Arfgef1_G3 | -1.56 | -2.46 | -1.74 | -2.66 | -1.6 | -1.92 | -1.43 | -2.48 | -3.17 | -1.68 | -2 | -2.13 | -2.47 | -1.14 | -1.8 | -1.69 | 0.2475 | -0.14 | 0.33 | 0.409432 | 0.490744 | 0.735253 | 0.989223 | 0.997988 | 0.994666 |
| Arfgef1_G3X9K3_S1566s_1 | Arfgef1_G3 | 0.79 | 0.09 | 0.34 | 0.16 | 0.31 | 0.7 | 0.28 | 0.97 | -0.18 | 0.15 | 0.87 | 0.96 | -0.32 | -0.13 | 0.01 | 0.44 | 0.22 | 0.105 | -0.345 | 0.256164 | 0.424138 | 0.757304 | 0.989223 | 0.997988 | 0.994666 |
| Arfgef1_G3X9K3_S234s_1 | Arfgef1_G3 | -0.96 | -1.7 | 0.78 | 1.01 | -1.47 | -0.66 | 1.26 | -0.85 | -0.76 | -1.08 | -2.29 | 1.86 | 1.15 | -1.61 | 1.13 | 0.37 | -0.2125 | -0.35 | 0.4775 | 0.562514 | 0.784309 | 0.718613 | 0.995762 | 0.997988 | 0.995259 |
| Arfgef1_G3X9K3_S52s_1 | Arfgef1_G3 | -0.91 | -0.79 | -0.42 | -0.68 | -1.52 | -0.67 | -1.43 | -1.28 | -0.15 | -0.78 | -1.08 | -0.55 | 0.48 | -1.7 | -0.7 | -0.77 | -0.525 | 0.06 | 0.0275 | 0.950932 | 0.071531 | 0.828151 | 0.989223 | 0.997988 | 0.999507 |
| Arfgef1_G3X9K3_S661s_1 | Arfgef1_G3 | 0.19 | 0.48 | 0.35 | 0.38 | 0.45 | 0.28 | 0.85 | 0.54 | 0.17 | 0.25 | 0.16 | 1.07 | -0.11 | 0.9 | -0.11 | 0.44 | 0.18 | 0.0625 | -0.07 | 0.819898 | 0.422736 | 0.82337 | 0.989223 | 0.997988 | 0.999507 |
| Arfgef2_A2A5R2_S621s_1 | Arfgef2_A2 | -0.48 | 0.18 | 0.16 | 0.46 | 0.62 | -0.16 | 0.49 | 0.54 | -0.11 | 0.47 | -0.39 | 0.14 | -0.18 | 1.05 | -0.14 | 0.75 | 0.2925 | -0.0525 | 0.29 | 0.456633 | 0.334507 | 0.863883 | 0.989223 | 0.997988 | 0.994666 |
| Arfgef2_F6YCJ0_S129s_1 | Arfgef2_F6 | 0.31 | 1.04 | 1.9 | 1.78 | -0.01 | 0.18 | 1.27 | 1.95 | -1 | 0.74 | 1.48 | 1.77 | -1.16 | -0.52 | 0.69 | 1.61 | -0.41 | -0.51 | -1.1025 | 0.115167 | 0.452071 | 0.444428 | 0.989223 | 0.997988 | 0.994666 |
| Arfgef2_F6YCJ0_S138s_1 | Arfgef2_F6 | 0.89 | 0.91 | -0.19 | 0.6 | 0.25 | 0.57 | 0.41 | 0.26 | 0.89 | 1.14 | -0.36 | 0.19 | 0.73 | 0.88 | 0.34 | 0.23 | -0.18 | -0.0875 | -0.0075 | 0.982244 | 0.54614 | 0.834494 | 0.989223 | 0.997988 | 1 |
| Arfgef2_F6YCJ0_S187s_1 | Arfgef2_F6 | -1.31 | -0.9 | -0.51 | -1.92 | -2.17 | 0.05 | -1.13 | -0.85 | 1.56 | -1.01 | -2.65 | -0.28 | 1.99 | -2.33 | 0.33 | -1.18 | 0.135 | 0.565 | 0.8625 | 0.331468 | 0.788014 | 0.497982 | 0.995762 | 0.997988 | 0.994666 |
| Arfgef2_F6YCJ0_T155t_1 | Arfgef2_F6 | -2.11 | -1.51 | -0.3 | -0.88 | -2.1 | -1.87 | -0.38 | -1.18 | -0.88 | -1.92 | -1.85 | -1.07 | -0.14 | -2.21 | -0.32 | -0.48 | -0.1825 | -0.23 | 0.4125 | 0.475168 | 0.717878 | 0.614997 | 0.995437 | 0.997988 | 0.994666 |
| Arfgef3_Q3UGY8_S1736s_1 | Arfgef3_Q3 | -1.07 | -0.61 | 0.24 | -1.12 | -1.59 | -0.4 | -1.79 | -0.86 | 0.63 | -1.61 | -0.23 | 0.09 | 1.34 | -2.56 | -0.1 | -0.48 | -0.52 | 0.36 | 0.19 | 0.804354 | 0.241492 | 0.504636 | 0.989223 | 0.997988 | 0.999507 |
| Arfgef3_Q3UGY8_S1957s_1 | Arfgef3_Q3 | -0.27 | -0.2 | 0.24 | -0.06 | -1.11 | -0.11 | -0.53 | -0.08 | 1.82 | -0.34 | 0.4 | 0.42 | 1.25 | -1.86 | 0.49 | 0.4 | -0.385 | 0.6475 | 0.1425 | 0.817807 | 0.210566 | 0.172326 | 0.989223 | 0.997988 | 0.999507 |
| Arfgef3_Q3UGY8_S2054s_1 | Arfgef3_Q3 | -0.95 | -1.02 | -0.49 | -0.34 | -0.31 | -0.97 | 0.01 | -0.73 | -1.02 | -0.62 | -0.25 | -0.19 | -1.1 | -0.46 | -1.11 | -0.37 | 0.2 | 0.18 | -0.06 | 0.848072 | 0.509803 | 0.547819 | 0.989223 | 0.997988 | 0.999507 |
| Arfgef3_Q3UGY8_S2088s_1 | Arfgef3_Q3 | -0.07 | 0.35 | -0.01 | -0.22 | 0.09 | -0.2 | -0.13 | -0.34 | -0.77 | -0.13 | -0.13 | -0.42 | 0.11 | -0.87 | -0.93 | 0.14 | -0.1575 | -0.375 | -0.4 | 0.269739 | 0.495359 | 0.173249 | 0.989223 | 0.997988 | 0.994666 |
| Arfgef3_Q3UGY8_S471s_1 | Arfgef3_Q3 | 0.53 | 0.49 | 1.45 | 1.31 | 0.59 | 0.75 | 1.17 | 1.24 | -0.2 | 0.06 | 0.75 | 1.18 | -1.25 | 0.44 | 0.75 | 1.07 | -0.0075 | -0.4975 | -0.6925 | 0.216757 | 0.981008 | 0.235405 | 0.999187 | 0.997988 | 0.994666 |
| Arglu1_Q3UL36_S75s_1 | Arglu1_Q3 | 0.48 | -1.16 | -0.58 | -0.62 | -2.16 | -0.19 | -0.71 | -0.13 | 1 | -0.93 | -1.99 | -1.24 | 1.86 | -1.08 | 0.97 | -0.8 | -0.3275 | -0.32 | 0.7075 | 0.326913 | 0.540919 | 0.627935 | 0.989223 | 0.997988 | 0.994666 |
| Arhgap1_Q5FWK3_S223s_1 | Arhgap1_Q | 2.4 | 0.75 | -0.29 | -0.89 | -0.13 | 2.38 | -1.02 | -0.59 | 3.96 | 2.21 | -1.12 | 0.33 | 3.46 | -0.05 | 0.73 | 0.53 | -0.3325 | 0.8525 | 0.675 | 0.473895 | 0.715671 | 0.468959 | 0.995437 | 0.997988 | 0.994666 |
| Arhgap1_Q5FWK3_S27s_1 | Arhgap1_Q | 0.62 | -0.82 | 0.02 | 0.04 | 0.44 | 0.5 | -0.19 | 0.42 | -0.84 | -0.9 | -0.12 | 0.22 | -1.48 | 0.3 | -0.65 | -0.3 | 0.3275 | -0.375 | -0.4975 | 0.294626 | 0.351235 | 0.361943 | 0.989223 | 0.997988 | 0.994666 |
| Arhgap1_Q5FWK3_S51s_1 | Arhgap1_Q | -0.03 | 0.32 | 0.08 | -0.17 | 0.32 | -0.63 | 0.2 | -0.07 | 0.19 | 0.24 | 0.79 | 0.16 | 0.56 | 0.4 | 0.35 | -0.04 | -0.095 | 0.295 | 0.2675 | 0.320703 | 0.730043 | 0.262315 | 0.995762 | 0.997988 | 0.994666 |
| Arhgap10_Q6Y5D8_S488s_1 | Arhgap10_ | 0.32 | 0.14 | 0.41 | 0.32 | 0.67 | 0.8 | 1.2 | 0.41 | 0.56 | 1.03 | 0.56 | 0.69 | -0.23 | 0.77 | 0.32 | 0.46 | 0.4725 | 0.4125 | 0.0325 | 0.910036 | 0.073473 | 0.094821 | 0.989223 | 0.997988 | 0.999507 |
| Arhgap12_B2RUJ8_S199s_1 | Arhgap12_ | 0.78 | 0.49 | 0.93 | 0.5 | 0.6 | 1.31 | -1.12 | 0.46 | 1.87 | 1.32 | -0.41 | 0.17 | 0.87 | 0.55 | -0.9 | -2.16 | -0.3625 | 0.0625 | -1.085 | 0.115301 | 0.459648 | 0.900668 | 0.989223 | 0.997988 | 0.994666 |
| Arhgap12_B2RUJ8_S213s_1 | Arhgap12_ | -0.12 | -0.74 | 0.33 | 0.09 | -0.09 | -0.14 | 0.69 | -0.6 | -0.64 | 0.21 | -0.17 | 0.48 | -0.33 | 0.26 | 0.45 | -0.1 | 0.075 | 0.08 | 0.18 | 0.588213 | 0.831874 | 0.819213 | 0.995762 | 0.997988 | 0.995259 |
| Arhgap17_E9QAJ9_S669s_1 | Arhgap17_ | 0.77 | 1.07 | 0.19 | -0.04 | 0.33 | 0.09 | -0.14 | -0.44 | -0.15 | 0.63 | 0.7 | 0.14 | 0.85 | -0.65 | 0.03 | 0.14 | -0.5375 | -0.1675 | -0.405 | 0.32972 | 0.117544 | 0.628806 | 0.989223 | 0.997988 | 0.994666 |
| Arhgap17_E9QAJ9_S671s_1 | Arhgap17_ | 0.52 | 0.48 | 0.26 | 0.02 | 0 | -0.44 | -0.08 | -0.51 | -0.01 | 0.24 | 0.94 | 0.29 | 0.7 | -0.72 | 0.21 | 0.45 | -0.5775 | 0.045 | -0.16 | 0.655529 | 0.034593 | 0.87355 | 0.989223 | 0.997988 | 0.995259 |
| Arhgap20_Q6IFT4_S46s_1 | Arhgap20_ | -1.85 | -2 | -1.93 | -2.61 | -0.77 | -1.45 | -1.43 | -1.91 | -1.56 | -1.49 | -1.07 | -1.82 | -1.18 | -0.78 | -1.37 | -2.25 | 0.7075 | 0.6125 | 0.7025 | 0.08531 | 0.044202 | 0.053821 | 0.989223 | 0.997988 | 0.994666 |
| Arhgap21_B7ZCJ0_S1090s_2 | Arhgap21_ | 2.82 | -0.03 | 1.46 | 1.52 | 0.72 | 3.66 | 0.42 | 1.17 | 3.01 | 1.65 | -0.54 | 1.97 | -0.67 | -0.34 | 0.18 | 1.44 | 0.05 | 0.08 | -1.29 | 0.079092 | 0.951379 | 0.924298 | 0.999187 | 0.997988 | 0.994666 |
| Arhgap21_B7ZCJ0_S1093s_2 | Arhgap21_ | 2.82 | -0.03 | 1.46 | 1.52 | 0.72 | 3.66 | 0.42 | 1.17 | 3.01 | 1.65 | -0.54 | 1.97 | -0.67 | -0.34 | 0.18 | 1.44 | 0.05 | 0.08 | -1.29 | 0.079092 | 0.951379 | 0.924298 | 0.999187 | 0.997988 | 0.994666 |
| Arhgap21_B7ZCJ0_S1502s_1 | Arhgap21_ | -0.75 | -0.98 | -0.82 | -0.76 | -0.9 | -0.45 | -1.23 | -0.74 | -0.73 | -0.76 | 0.08 | -0.5 | 0.05 | -0.92 | -0.3 | -0.18 | -0.0025 | 0.35 | 0.49 | 0.112444 | 0.991619 | 0.207248 | 0.999187 | 0.997988 | 0.994666 |
| Arhgap21_B7ZCJ0_S1729s_1 | Arhgap21_ | -2.1 | -2.27 | -1.35 | -2.31 | -1.5 | -3.18 | -1.22 | -2.25 | -2.43 | -1.97 | -1.19 | -2.36 | -1.4 | -0.91 | -1.65 | -1.35 | -0.03 | 0.02 | 0.68 | 0.056417 | 0.947763 | 0.956834 | 0.999187 | 0.997988 | 0.994666 |
| Arhgap21_B7ZCJ0_S42s_1 | Arhgap21_ | 0.55 | 0.76 | 0.28 | 0.61 | 0.97 | 0.52 | 0.39 | 0.04 | -0.15 | 0.32 | 0.16 | 0.63 | -0.83 | 0.89 | 0.22 | 0.09 | -0.07 | -0.31 | -0.4575 | 0.249266 | 0.790706 | 0.24912 | 0.995762 | 0.997988 | 0.994666 |
| Arhgap21_B7ZCJ0_S436s_1 | Arhgap21_ | 1.12 | 0.01 | 0.65 | 1.32 | 0.53 | 1.23 | 0.79 | 1.77 | 1.21 | 1.43 | 1.21 | 0.88 | 0.78 | 0.93 | 0.86 | 0.75 | 0.305 | 0.4075 | 0.055 | 0.868873 | 0.438042 | 0.242508 | 0.989223 | 0.997988 | 0.999507 |
| Arhgap21_B7ZCJ0_S470s_1 | Arhgap21_ | 0.3 | 0.6 | 0.88 | 0.45 | -0.52 | -0.16 | 0.87 | -0.21 | 0.27 | 0.82 | -0.23 | -0.27 | -0.08 | 0.72 | 0.45 | -0.23 | -0.5625 | -0.41 | -0.3425 | 0.284121 | 0.118432 | 0.214609 | 0.989223 | 0.997988 | 0.994666 |
| Arhgap21_B7ZCJ0_S478s_1 | Arhgap21_ | 0.26 | 0.72 | 0.64 | 0.51 | -0.03 | 0.27 | 0.46 | 0.22 | 0.21 | 0.44 | -0.38 | 0.43 | 0.39 | 0.16 | -0.11 | 0.41 | -0.3025 | -0.3575 | -0.32 | 0.235782 | 0.198939 | 0.212095 | 0.989223 | 0.997988 | 0.994666 |
| Arhgap21_B7ZCJ0_S687s_1 | Arhgap21_ | 0.28 | 0.63 | -0.1 | 0.61 | 0.81 | 0.86 | 1.3 | 0.5 | 1.01 | 0.72 | 0.91 | 0.69 | 0.24 | 0.71 | 0.62 | 0.48 | 0.5125 | 0.4775 | 0.1575 | 0.57562 | 0.088742 | 0.089063 | 0.989223 | 0.997988 | 0.995259 |
| Arhgap21_B7ZCJ0_T62t_1 | Arhgap21_ | 0.56 | 0.72 | 1.13 | 0.82 | 1.21 | 0.28 | 1.63 | 0.62 | -0.43 | 0.48 | -0.01 | -0.14 | 0 | -0.5 | -0.14 | 0.16 | 0.1275 | -0.8325 | -0.9275 | 0.006675 | 0.702782 | 0.013643 | 0.995437 | 0.997988 | 0.994666 |
| Arhgap21_B7ZCJ0_T686t_1 | Arhgap21_ | 0.28 | 0.63 | -0.1 | 0.61 | 0.81 | 0.86 | 1.3 | 0.5 | 1.01 | 0.72 | 0.91 | 0.69 | 0.24 | 0.71 | 0.62 | 0.48 | 0.5125 | 0.4775 | 0.1575 | 0.57562 | 0.088742 | 0.089063 | 0.989223 | 0.997988 | 0.995259 |
| Arhgap22_Q8BL80_S22s_1 | Arhgap22_ | 0.4 | -0.6 | -0.34 | -0.8 | -0.12 | -0.14 | -1.28 | -0.33 | -1.27 | -0.76 | -0.55 | -1.92 | -0.21 | 0.08 | -0.74 | -0.48 | -0.1325 | -0.79 | -0.0025 | 0.994253 | 0.723403 | 0.074022 | 0.995437 | 0.997988 | 1 |
| Arhgap22_Q8BL80_S387s_1 | Arhgap22_ | 1.15 | 0.49 | 0.64 | 0.52 | 0.93 | -0.06 | -0.03 | -0.26 | 0.58 | 0.5 | 1.18 | 0.04 | 1.85 | -0.07 | 0.08 | -0.38 | -0.555 | -0.125 | -0.33 | 0.512104 | 0.109923 | 0.690455 | 0.989223 | 0.997988 | 0.994666 |
| Arhgap23_F8VQ11_S1182s_1 | Arhgap23_ | 0.24 | -0.41 | 0.17 | -0.28 | -0.19 | -0.24 | -0.43 | -0.58 | 0.45 | 0.42 | 0.17 | -0.44 | 0.6 | 0.76 | -0.5 | 0.63 | -0.29 | 0.22 | 0.4425 | 0.237256 | 0.255159 | 0.471556 | 0.989223 | 0.997988 | 0.994666 |
| Arhgap23_F8VQ11_S145s_1 | Arhgap23_ | 0.43 | 0.43 | 0.24 | -0.21 | 0.54 | 0.7 | 0.06 | 0.23 | 0.34 | 0.42 | 0.15 | 0.12 | 0.31 | 0.4 | 0.12 | 0.13 | 0.16 | 0.035 | 0.0175 | 0.946845 | 0.541097 | 0.886909 | 0.989223 | 0.997988 | 0.999507 |
| Arhgap23_F8VQ11_S155s_1 | Arhgap23_ | 0.51 | 0.61 | 0.21 | 0.24 | 0.76 | 0.36 | 0.31 | 0.28 | 0.22 | 0.48 | -0.13 | 0.1 | -0.06 | 0.88 | -0.05 | -0.08 | 0.035 | -0.225 | -0.22 | 0.483799 | 0.877765 | 0.365894 | 0.996493 | 0.997988 | 0.994666 |
| Arhgap23_F8VQ11_S166s_1 | Arhgap23_ | 0.81 | 0.28 | 0.44 | 0.67 | 1.05 | 0.69 | 0.91 | 0.18 | 0.81 | 0.83 | 0.33 | 0.12 | -0.02 | 0.81 | 0.4 | 0.65 | 0.1575 | -0.0275 | -0.09 | 0.754684 | 0.560207 | 0.919044 | 0.989223 | 0.997988 | 0.9971 |
| Arhgap23_F8VQ11_S215s_1 | Arhgap23_ | -0.06 | -0.49 | 0.4 | -0.39 | -0.29 | -0.26 | -0.48 | -0.22 | -0.22 | -0.14 | 0.88 | 0.02 | -0.49 | 0.06 | -0.93 | -0.66 | -0.1775 | 0.27 | -0.37 | 0.279194 | 0.498176 | 0.437905 | 0.989223 | 0.997988 | 0.994666 |
| Arhgap23_F8VQ11_S344s_1 | Arhgap23_ | 0.26 | 0.3 | 1.55 | 0.5 | 0.29 | -0.34 | 1.02 | 1.25 | 0.11 | -0.57 | 0.96 | 0.52 | 0.23 | 0.63 | 0.23 | -0.62 | -0.0975 | -0.3975 | -0.535 | 0.208376 | 0.825825 | 0.369499 | 0.995762 | 0.997988 | 0.994666 |
| Arhgap23_F8VQ11_S388s_1 | Arhgap23_ | 6.17 | 2 | 3.55 | 1.88 | 2.68 | 6.92 | 4.3 | 3.51 | 0.38 | 4.39 | 2.13 | 3.47 | 0.08 | 0.39 | 4.07 | 4.65 | 0.9525 | -0.8075 | -1.1025 | 0.420129 | 0.422813 | 0.493543 | 0.989223 | 0.997988 | 0.994666 |
| Arhgap23_F8VQ11_S400s_1 | Arhgap23_ | 0.24 | -0.1 | -0.54 | -1.15 | 1.38 | -0.82 | 0.02 | -0.36 | -1.77 | -1.36 | -0.74 | -1 | -1.98 | 0.46 | -0.28 | -0.51 | 0.4425 | -0.83 | -0.19 | 0.730115 | 0.397121 | 0.051929 | 0.989223 | 0.997988 | 0.9971 |
| Arhgap23_F8VQ11_S413s_1 | Arhgap23_ | -0.97 | -0.26 | -0.6 | -0.85 | -0.18 | -0.13 | -0.73 | -0.96 | -0.78 | -0.12 | -0.93 | -0.6 | -0.39 | -0.32 | -0.35 | -1.04 | 0.17 | 0.0625 | 0.145 | 0.62727 | 0.559704 | 0.826175 | 0.989223 | 0.997988 | 0.995259 |
| Arhgap23_F8VQ11_T127t_1 | Arhgap23_ | -0.73 | -0.97 | -0.25 | -0.65 | -0.17 | -0.97 | -0.01 | -0.03 | -1.06 | -0.45 | -1.55 | -0.65 | -1 | -1.32 | -0.36 | -0.83 | 0.355 | -0.2775 | -0.2275 | 0.464604 | 0.251769 | 0.390763 | 0.989223 | 0.997988 | 0.994666 |
| Arhgap23_F8VQ11_T897t_1 | Arhgap23_ | -1.06 | -0.43 | -1.19 | -1.46 | -0.33 | -1 | -0.72 | -1.38 | -0.73 | -0.81 | -1.24 | -1.33 | -0.64 | -0.49 | -1.54 | -1.44 | 0.1775 | 0.0075 | 0.0075 | 0.983711 | 0.586791 | 0.980234 | 0.989223 | 0.997988 | 1 |
| Arhgap24_D3Z5T4_T241t_1 | Arhgap24_ | 0.03 | 0.45 | 0.38 | 0.57 | -0.49 | 0.67 | -0.88 | 0.15 | 1.62 | 0.04 | -0.31 | 0.12 | 2.49 | -0.8 | 0.01 | 0.94 | -0.495 | 0.01 | 0.3025 | 0.641479 | 0.191243 | 0.981476 | 0.989223 | 0.997988 | 0.995259 |
| Arhgap25_Q8BYW1_S274s_1 | Arhgap25_ | 0.61 | 1.4 | 1.35 | 0.91 | 1.09 | 0.82 | 1.02 | 1.12 | 0.26 | 1.04 | 0.59 | 1.36 | 0.65 | 0.59 | 0.75 | 0.48 | -0.055 | -0.255 | -0.45 | 0.129485 | 0.828441 | 0.448448 | 0.995762 | 0.997988 | 0.994666 |
| Arhgap26_F7D661_S18s_1 | Arhgap26_ | -0.68 | 0.09 | -1.26 | -0.38 | -1.4 | -1.82 | -2.22 | -0.66 | -0.57 | -1.37 | -1.9 | 0.01 | -0.29 | -1.07 | 0.46 | -1.09 | -0.9675 | -0.4 | 0.06 | 0.89438 | 0.040757 | 0.415844 | 0.989223 | 0.997988 | 0.999507 |
| Arhgap31_A6X8Z5_S1163s_1 | Arhgap31_ | 0.07 | -0.75 | -0.77 | -0.25 | -0.19 | 0.03 | -0.77 | 0.1 | -0.31 | 0.5 | 0.86 | -0.69 | 0.69 | 0.07 | -0.74 | 0.02 | 0.2175 | 0.515 | 0.435 | 0.262774 | 0.482723 | 0.22596 | 0.989223 | 0.997988 | 0.994666 |
| Arhgap31_A6X8Z5_S1242s_1 | Arhgap31_ | 1.61 | 2.33 | 1.67 | 0.43 | 0.26 | 1.19 | -0.23 | -0.31 | -0.16 | 0.64 | 1 | 0.77 | -0.14 | 0.58 | 0.79 | 1.31 | -1.2825 | -0.9475 | -0.875 | 0.091298 | 0.023545 | 0.060169 | 0.989223 | 0.997988 | 0.994666 |
| Arhgap31_A6X8Z5_S765s_1 | Arhgap31_ | 0.73 | 1.01 | 0.09 | -0.26 | 0.66 | -0.53 | 0.51 | 0.46 | 0.65 | 0.66 | 0.11 | 0.6 | 0.02 | 0.29 | 0.51 | -0.47 | -0.1175 | 0.1125 | -0.305 | 0.425588 | 0.761446 | 0.740804 | 0.995762 | 0.997988 | 0.994666 |
| Arhgap31_A6X8Z5_S850s_1 | Arhgap31_ | -0.42 | 0.41 | -0.38 | -0.25 | -0.02 | -0.19 | 0.5 | 0.64 | 0.4 | -0.01 | 0.63 | 0.53 | 0.22 | 0.16 | 0.3 | 0.44 | 0.3925 | 0.5475 | 0.44 | 0.141182 | 0.21393 | 0.081924 | 0.989223 | 0.997988 | 0.994666 |
| Arhgap32_Q811P8_S1405s_1 | Arhgap32_ | -0.23 | -0.18 | -0.93 | -0.48 | -0.1 | -0.1 | -0.13 | -0.65 | -0.83 | -0.35 | -0.3 | -1.04 | -0.5 | 0.06 | -1.01 | -0.65 | 0.21 | -0.175 | -0.07 | 0.829548 | 0.434828 | 0.554311 | 0.989223 | 0.997988 | 0.999507 |
| Arhgap32_Q811P8_S1588s_1 | Arhgap32_ | -0.85 | -0.32 | -0.25 | -1.48 | -0.3 | -1.11 | -0.51 | -1.7 | -0.55 | -1.11 | -0.29 | -0.74 | -0.36 | -1.44 | -3.01 | -1.85 | -0.18 | 0.0525 | -0.94 | 0.123482 | 0.659877 | 0.880442 | 0.990434 | 0.997988 | 0.994666 |
| Arhgap32_Q811P8_S1823s_1 | Arhgap32_ | 1.04 | 0.05 | 1.16 | 1.38 | -0.42 | 1.53 | -0.1 | 0.65 | 2.33 | 0.35 | 0.91 | 1.54 | 1.5 | -0.58 | 0.33 | 0.34 | -0.4925 | 0.375 | -0.51 | 0.313888 | 0.321945 | 0.450738 | 0.989223 | 0.997988 | 0.994666 |
| Arhgap32_Q811P8_S1876s_1 | Arhgap32_ | -0.05 | -0.21 | 0.91 | -2.76 | -0.44 | 0.69 | 0.76 | 0.18 | -0.5 | 1.01 | -0.69 | 0.53 | 1 | -1.13 | 0.26 | 0.37 | 0.825 | 0.615 | 0.6525 | 0.422138 | 0.275674 | 0.44401 | 0.989223 | 0.997988 | 0.994666 |
| Arhgap32_Q811P8_S47s_1 | Arhgap32_ | 0.59 | 0.45 | 0.42 | 0.76 | 0.56 | -0.4 | 0.1 | 0.73 | 0.17 | 0.46 | 0.32 | 0.78 | -0.94 | 0.54 | 0 | 0.32 | -0.3075 | -0.1225 | -0.575 | 0.134868 | 0.308991 | 0.609808 | 0.989223 | 0.997988 | 0.994666 |
| Arhgap32_Q811P8_S613s_1 | Arhgap32_ | 0.55 | 0.1 | 0.28 | 0.07 | -0.24 | 0.55 | 0.09 | 0.49 | -0.86 | -0.05 | 1.36 | 0.3 | 0.09 | 0.32 | -0.23 | -0.18 | -0.0275 | -0.0625 | -0.25 | 0.355322 | 0.916534 | 0.890533 | 0.997335 | 0.997988 | 0.994666 |
| Arhgap32_Q811P8_S655s_1 | Arhgap32_ | 0.42 | -1.82 | -0.28 | 0.25 | 0.39 | 0.88 | 0.57 | 0.66 | -0.54 | -0.96 | 0.25 | 0.26 | -2.39 | 0.42 | -0.57 | -0.43 | 0.9825 | 0.11 | -0.385 | 0.584405 | 0.064794 | 0.841351 | 0.989223 | 0.997988 | 0.995259 |
| Arhgap32_Q811P8_S682s_1 | Arhgap32_ | 0.55 | 0.5 | -2.22 | -2.52 | -0.02 | 0.92 | -3.66 | -2.91 | 0.45 | 0.26 | -0.72 | -1.58 | 0.39 | -1.07 | -2.87 | -2.67 | -0.495 | 0.525 | -0.6325 | 0.526295 | 0.679488 | 0.543163 | 0.993464 | 0.997988 | 0.994666 |
| Arhgap32_Q811P8_S856s_1 | Arhgap32_ | -0.71 | -0.14 | 1.59 | 1.25 | 0.03 | -0.52 | 1.03 | 1.38 | -0.66 | -0.31 | 1.69 | 1.28 | 0.18 | 0.71 | 0.9 | 0.47 | -0.0175 | 0.0025 | 0.0675 | 0.899089 | 0.977685 | 0.997221 | 0.999187 | 0.999272 | 0.999507 |

| | | | | | | | | | | | | | | | | | | | | | | | | | | |
|---|---|---|---|---|---|---|---|---|---|---|---|---|---|---|---|---|---|---|---|---|---|---|---|---|---|---|
| Arhgap32_Q811P8_S892s_1 | Arhgap32_ | 0.35 | -2.24 | -2.22 | -1.58 | -0.64 | 0.77 | -0.63 | -0.27 | -2.1 | -1.6 | 0.43 | -1.45 | -2.88 | -0.46 | -3.04 | -2.77 | 1.23 | 0.2425 | -0.865 | 0.2754 | 0.071802 | 0.743756 | 0.989223 | 0.997988 | 0.994666 |
| Arhgap32_Q811P8_S935s_1 | Arhgap32_ | 0.13 | -0.04 | -0.57 | -0.2 | 0.65 | 0.36 | 0.35 | -0.16 | -0.08 | 0.46 | -0.88 | -0.3 | -0.28 | 0.29 | -0.77 | -0.32 | 0.47 | -0.03 | -0.1 | 0.751062 | 0.105319 | 0.928875 | 0.989223 | 0.997988 | 0.9971 |
| Arhgap32_Q811P8_S952s_1 | Arhgap32_ | 1.24 | 0.65 | 1.33 | 1.5 | 0.95 | 1.22 | 1.24 | 1.69 | 0.84 | 0.95 | 1.21 | 1.46 | 0.38 | 1.03 | 0.89 | 1.06 | 0.095 | -0.065 | -0.34 | 0.277272 | 0.732728 | 0.817545 | 0.995762 | 0.997988 | 0.994666 |
| Arhgap33_Q80YF9_S1144s_1 | Arhgap33_ | 2.08 | 1.52 | 2.64 | 0.83 | 0.84 | 2.32 | 2.1 | 2.12 | 0.72 | 2.15 | 2.5 | 2.36 | 0.67 | 0.9 | 3.18 | 2.1 | 0.0775 | 0.165 | -0.055 | 0.930636 | 0.870489 | 0.754457 | 0.996493 | 0.997988 | 0.999507 |
| Arhgap33_Q80YF9_S594s_1 | Arhgap33_ | 0.64 | -0.32 | 0.46 | 0.47 | -0.21 | 0.95 | -0.27 | 0.43 | 0.86 | 0.27 | -0.35 | 0.52 | 1.08 | -0.38 | -0.47 | 0.99 | -0.0875 | 0.0125 | -0.0075 | 0.986937 | 0.807452 | 0.971444 | 0.995762 | 0.997988 | 1 |
| Arhgap33_Q80YF9_S660s_1 | Arhgap33_ | 1.07 | 0.91 | 0.1 | 0.03 | 0.92 | 0.67 | 0.09 | 0.11 | 0.9 | 1.03 | -0.06 | 0.21 | 0.45 | 1.26 | 0.13 | -0.04 | -0.08 | -0.0075 | -0.0775 | 0.846697 | 0.816307 | 0.984284 | 0.995762 | 0.997988 | 0.999507 |
| Arhgap33_Q80YF9_S887s_1 | Arhgap33_ | 0.07 | 0.12 | 0.15 | 0.29 | 0.92 | -0.28 | 0.77 | 0.23 | 0.11 | 0.37 | 0.61 | 0.69 | 0.01 | 0.9 | -0.18 | 0.69 | 0.2525 | 0.2875 | 0.1975 | 0.535268 | 0.411189 | 0.236159 | 0.989223 | 0.997988 | 0.994666 |
| Arhgap33_Q80YF9_S949s_1 | Arhgap33_ | 1.06 | -0.7 | 0.58 | 1.28 | -0.14 | 1.37 | 0.77 | 1.72 | 0.35 | 0.17 | 1.42 | 1.28 | 0.57 | 0.49 | 0.23 | 0.51 | 0.375 | 0.25 | -0.105 | 0.811678 | 0.496428 | 0.627114 | 0.989223 | 0.997988 | 0.999507 |
| Arhgap33_Q80YF9_S949s_2 | Arhgap33_ | 0.99 | 0.55 | 0.58 | 0.48 | 0.35 | 0.37 | 0.17 | -0.17 | 0.41 | 0.58 | 0.96 | 0.08 | 0.52 | 1.78 | 0.94 | -0.2 | -0.47 | -0.1425 | 0.11 | 0.796262 | 0.072634 | 0.604081 | 0.989223 | 0.997988 | 0.999507 |
| Arhgap33_Q80YF9_S998s_1 | Arhgap33_ | 0.96 | -0.66 | 0.68 | 0.67 | -1.25 | 1.01 | -0.04 | 0.92 | -0.14 | -0.86 | -1.12 | 1.44 | 0.01 | -3.94 | -1.47 | -0.09 | -0.2525 | -0.5825 | -1.785 | 0.063986 | 0.662083 | 0.360275 | 0.990434 | 0.997988 | 0.994666 |
| Arhgap33_Q80YF9_T943t_1 | Arhgap33_ | 0.54 | 0.81 | 0.67 | 0.97 | 0.93 | 0.79 | 1.16 | 1.06 | 0.58 | 1.56 | 1.22 | 1.32 | 0.82 | 1.45 | 1.29 | 0.94 | 0.2375 | 0.4225 | 0.3775 | 0.178869 | 0.286692 | 0.156145 | 0.989223 | 0.997988 | 0.994666 |
| Arhgap33_Q80YF9_T943t_2 | Arhgap33_ | 0.99 | 0.55 | 0.58 | 0.48 | 0.35 | 0.37 | 0.17 | -0.17 | 0.41 | 0.58 | 0.96 | 0.08 | 0.52 | 1.78 | 0.94 | -0.2 | -0.47 | -0.1425 | 0.11 | 0.796262 | 0.072634 | 0.604081 | 0.989223 | 0.997988 | 0.999507 |
| Arhgap35_Q91YM2_S1134s_1 | Arhgap35_ | 0.4 | 0.17 | -0.05 | -0.16 | 0.07 | -0.63 | -0.12 | 0.13 | -0.26 | 0.14 | -0.91 | 0.4 | 0.2 | -1.76 | -0.37 | -0.2 | -0.2275 | -0.2475 | -0.6225 | 0.176487 | 0.39315 | 0.465074 | 0.989223 | 0.997988 | 0.994666 |
| Arhgap35_Q91YM2_S1134s_2 | Arhgap35_ | -0.15 | -1.84 | 1.58 | 1.09 | -0.69 | -0.6 | 0.44 | 0.89 | -0.45 | -0.39 | 2.36 | -0.79 | -0.98 | -2.41 | 1.49 | 0.95 | -0.16 | 0.0125 | -0.4075 | 0.6917 | 0.831353 | 0.989324 | 0.995762 | 0.997988 | 0.996159 |
| Arhgap35_Q91YM2_S1142s_1 | Arhgap35_ | -0.54 | -0.01 | -0.99 | -0.38 | -0.44 | 0.02 | -0.31 | 0.12 | 0.25 | 0.16 | -1.49 | -0.48 | 0.04 | -0.23 | -0.03 | -0.26 | 0.3275 | 0.09 | 0.36 | 0.229903 | 0.258123 | 0.836489 | 0.989223 | 0.997988 | 0.994666 |
| Arhgap35_Q91YM2_S1142s_2 | Arhgap35_ | -0.15 | -1.84 | 1.58 | 1.09 | -0.69 | -0.6 | 0.44 | 0.89 | -0.45 | -0.39 | 2.36 | -0.79 | -0.98 | -2.41 | 1.49 | 0.95 | -0.16 | 0.0125 | -0.4075 | 0.6917 | 0.831353 | 0.989324 | 0.995762 | 0.997988 | 0.996159 |
| Arhgap35_Q91YM2_S975s_1 | Arhgap35_ | 0.7 | 0.52 | 0.8 | 0.54 | 0.44 | 0.27 | 0.99 | 0.61 | 0.47 | 0.42 | 0.52 | 0.98 | -0.59 | 0.96 | 0.5 | 0.95 | -0.0625 | -0.0425 | -0.185 | 0.63187 | 0.791872 | 0.8568 | 0.995762 | 0.997988 | 0.995259 |
| Arhgap39_H7BX46_S117s_1 | Arhgap39_ | -1.61 | -2.12 | -0.59 | -3.02 | -2.64 | -0.74 | -1.99 | -1.46 | 2.65 | -0.91 | -1.38 | -0.22 | 3.65 | -2.88 | 2.33 | -1.86 | 0.1275 | 1.87 | 2.145 | 0.157385 | 0.826409 | 0.066892 | 0.995762 | 0.997988 | 0.994666 |
| Arhgap39_H7BX46_S125s_1 | Arhgap39_ | -1.6 | -2.22 | -2.19 | -6.45 | -2.37 | -2.03 | -1.4 | -1.48 | 3.71 | -1.9 | -1.9 | -1.01 | 4.81 | -3.02 | 3.66 | -3.11 | 1.295 | 2.84 | 3.7 | 0.096463 | 0.211485 | 0.089425 | 0.989223 | 0.997988 | 0.994666 |
| Arhgap39_H7BX46_S171s_1 | Arhgap39_ | 0.35 | 0.44 | 0.46 | 0.36 | 0.43 | 0.34 | 0.67 | 0.39 | 0.21 | 0.63 | 0.37 | 0.17 | 0.41 | 0.81 | 0.34 | 0.39 | 0.055 | -0.0575 | 0.085 | 0.725542 | 0.784144 | 0.794174 | 0.995762 | 0.997988 | 0.9971 |
| Arhgap39_H7BX46_S297s_1 | Arhgap39_ | -0.5 | -0.12 | -0.23 | -1.1 | -0.87 | -0.06 | -1.54 | -1.85 | 1.1 | -0.56 | -1.13 | -0.96 | 2.13 | -1.93 | -1.79 | -1.38 | -0.5925 | 0.1 | -0.255 | 0.769337 | 0.188662 | 0.847271 | 0.989223 | 0.997988 | 0.9971 |
| Arhgap39_H7BX46_S364s_1 | Arhgap39_ | 0.34 | -0.42 | -0.41 | 0.06 | -0.06 | 0.59 | 0.03 | -0.11 | 0.17 | -0.58 | 0.24 | 0.49 | -0.06 | 0.07 | -0.26 | -0.07 | 0.22 | 0.1875 | 0.0275 | 0.92125 | 0.442468 | 0.566015 | 0.989223 | 0.997988 | 0.999507 |
| Arhgap39_H7BX46_S382s_1 | Arhgap39_ | -0.43 | 0.37 | 0.14 | -0.75 | -0.36 | -0.39 | -0.25 | -0.03 | -0.29 | -0.16 | 0.69 | -0.75 | -0.61 | 0.38 | -0.91 | -0.82 | -0.09 | 0.04 | -0.3225 | 0.427418 | 0.761601 | 0.919194 | 0.995762 | 0.997988 | 0.994666 |
| Arhgap39_H7BX46_S599s_1 | Arhgap39_ | 0.89 | 0.81 | 0.88 | 0.94 | 0.5 | 0.54 | 0.66 | 1 | 0.45 | 0.51 | 1.14 | 0.97 | 0.05 | 0.88 | 0.33 | 0.59 | -0.205 | -0.1125 | -0.4175 | 0.144955 | 0.348928 | 0.652412 | 0.989223 | 0.997988 | 0.994666 |
| Arhgap39_H7BX46_S639s_1 | Arhgap39_ | -0.6 | -0.28 | 0.71 | 0.82 | 0.11 | -0.43 | 0.49 | 1.42 | 0.62 | 0.07 | -0.49 | 0.76 | -0.84 | 0.81 | 0 | 0.76 | 0.235 | 0.0775 | 0.02 | 0.967909 | 0.630152 | 0.860271 | 0.989223 | 0.997988 | 0.999507 |
| Arhgap39_H7BX46_S685s_1 | Arhgap39_ | -0.51 | -0.6 | -0.17 | 0.31 | -0.06 | -0.8 | 0.59 | 0.35 | -0.56 | -0.3 | -0.09 | 0.76 | -0.82 | 0.57 | -0.08 | 0.09 | 0.2625 | 0.195 | 0.1825 | 0.626582 | 0.477391 | 0.593867 | 0.989223 | 0.997988 | 0.995259 |
| Arhgap39_H7BX46_T532t_1 | Arhgap39_ | -1.16 | -2.3 | -1.95 | -1.58 | -0.98 | -1.19 | -0.66 | -0.53 | -2.94 | -2.14 | 1.63 | -1.18 | -1.35 | -0.79 | -0.67 | -1.49 | 0.9075 | 0.59 | 0.6725 | 0.078614 | 0.014368 | 0.521379 | 0.989223 | 0.997988 | 0.994666 |
| Arhgap44_Q5SSM3_S598s_1 | Arhgap44_ | 1.73 | -0.58 | -0.16 | -0.81 | -0.16 | 1.85 | -0.16 | -0.62 | 1.21 | 0.77 | -1 | 0.05 | 0.44 | -0.08 | -0.25 | -0.56 | 0.1825 | 0.2125 | -0.1575 | 0.781125 | 0.795582 | 0.755255 | 0.995762 | 0.997988 | 0.998226 |
| Arhgap44_Q5SSM3_S606s_1 | Arhgap44_ | -0.01 | 0.59 | 0.21 | 0.33 | 0.7 | -0.27 | 0.51 | 0.39 | -0.46 | 1.1 | 0.09 | 0.46 | -0.72 | 0.18 | 0.31 | 0.33 | 0.0525 | 0.0175 | -0.255 | 0.439685 | 0.851587 | 0.96136 | 0.996349 | 0.997988 | 0.994666 |
| Arhgap44_Q5SSM3_S636s_1 | Arhgap44_ | -0.13 | -0.08 | -0.27 | -0.21 | 0.08 | -0.52 | 0.26 | 0.36 | 0.17 | -0.28 | 0.4 | -0.04 | -0.71 | 0.24 | -0.24 | 0.25 | 0.2175 | 0.235 | 0.0575 | 0.846356 | 0.402365 | 0.337125 | 0.989223 | 0.997988 | 0.999507 |
| Arhgap44_Q5SSM3_S805s_1 | Arhgap44_ | -0.99 | -1.27 | -1.15 | -2.11 | -1.07 | -0.86 | -1.38 | -1.3 | -2.61 | -1.63 | -0.19 | -1.35 | -1.93 | -1.07 | -1.35 | -1.41 | 0.2275 | -0.065 | -0.06 | 0.860708 | 0.456096 | 0.900619 | 0.989223 | 0.997988 | 0.999507 |
| Arhgap5_E9PYT0_S1219s_1 | Arhgap5_E | 0.84 | 0.26 | 1.32 | 0.61 | 0.81 | 0.25 | 1.01 | 1.06 | -0.13 | 0.39 | 0.78 | 1.01 | -1.06 | 0.25 | 0.75 | 1.75 | 0.025 | -0.245 | -0.335 | 0.563383 | 0.93553 | 0.489565 | 0.99868 | 0.997988 | 0.995259 |
| Arhgap5_E9PYT0_S968s_1 | Arhgap5_E | -0.43 | 0.43 | 0.84 | -0.08 | 0.43 | -0.95 | 1.49 | 1.02 | 0.17 | 0.38 | 0.26 | -0.73 | 0.13 | -0.53 | 0.8 | 0.1 | 0.3075 | -0.17 | -0.065 | 0.870017 | 0.573467 | 0.657296 | 0.989223 | 0.997988 | 0.999507 |
| Arhgap5_E9PYT0_T1218t_1 | Arhgap5_E | 0.84 | 0.26 | 0.58 | -0.08 | 0.81 | 0.25 | 0.5 | 0.57 | -0.13 | 0.39 | 0.2 | 0.01 | -1.06 | 0.25 | -0.14 | 0.83 | 0.1325 | -0.2825 | -0.43 | 0.337351 | 0.627465 | 0.32751 | 0.989223 | 0.997988 | 0.994666 |
| Arhgdia_Q99PT1_S34s_1 | Arhgdia_Q | -2.38 | -1.73 | -0.01 | -0.05 | -2.69 | -3.14 | 0.38 | 0.1 | -3.26 | -2.4 | -0.6 | -0.05 | -4.1 | -2.26 | -0.48 | -0.29 | -0.295 | -0.535 | -0.74 | 0.437904 | 0.756427 | 0.536171 | 0.995762 | 0.997988 | 0.994666 |
| Arhgdia_Q99PT1_Y133y_1 | Arhgdia_Q | -0.99 | 0.59 | 0.15 | 0.36 | -0.7 | -0.41 | -1.43 | 0.44 | 0.77 | -0.73 | 0.51 | -0.1 | 1.16 | -0.21 | 0.1 | 0.01 | -0.5525 | 0.085 | 0.2375 | 0.60188 | 0.268242 | 0.854509 | 0.989223 | 0.997988 | 0.995259 |
| Arhgef1_E9PUF7_S961s_1 | Arhgef1_E | -0.28 | -0.04 | 0.01 | 0.68 | 0 | -0.28 | 1.22 | 0.49 | -0.27 | -0.34 | 0.54 | 1.62 | -0.65 | -0.52 | -0.3 | 0.48 | 0.265 | 0.295 | -0.34 | 0.348869 | 0.49014 | 0.539849 | 0.989223 | 0.997988 | 0.994666 |
| Arhgef1_E9PUF7_T429t_1 | Arhgef1_E | -0.02 | -0.56 | -0.01 | -0.35 | -0.96 | -0.22 | -0.61 | -0.03 | 0.39 | -0.39 | 0.92 | 0.09 | 0.28 | -0.53 | -0.46 | 0.18 | -0.22 | 0.4875 | 0.1025 | 0.738585 | 0.442311 | 0.163574 | 0.989223 | 0.997988 | 0.9971 |
| Arhgef10_Q8C033_S7s_1 | Arhgef10_ | 0.11 | 0.01 | -0.34 | -0.45 | -0.01 | -0.24 | -0.61 | -0.74 | 0.15 | 0.12 | 0.29 | -0.38 | 1.17 | 0.27 | -0.3 | 0.29 | -0.2325 | 0.2125 | 0.525 | 0.165594 | 0.385923 | 0.427428 | 0.989223 | 0.997988 | 0.994666 |
| Arhgef10_Q8C033_S852s_1 | Arhgef10_ | 1.02 | 1.33 | 0.35 | -0.31 | 1.6 | 0.71 | 0.3 | -0.18 | 1.63 | 1.07 | 0.54 | 0.34 | 0.15 | 1.06 | 0.42 | 0.63 | 0.01 | 0.2975 | -0.0325 | 0.937307 | 0.983496 | 0.512447 | 0.999187 | 0.997988 | 0.999507 |
| Arhgef11_Q68FM7_S1125s_1 | Arhgef11_ | 4.47 | 2.38 | 3.82 | 1.62 | 2.67 | 4.66 | 4.28 | 3.06 | 1.15 | 3.97 | 1.39 | 3.86 | -0.29 | 0.13 | 3.85 | 4.56 | 0.595 | -0.48 | -1.01 | 0.414985 | 0.411674 | 0.59341 | 0.989223 | 0.997988 | 0.994666 |
| Arhgef11_Q68FM7_S1353s_1 | Arhgef11_ | -3.49 | -2.86 | -2.78 | -3.07 | -1.74 | -2.47 | -1.81 | -2.72 | -3.73 | -3.7 | -1.23 | -3.22 | -1.85 | 0.14 | -2.54 | -2 | 0.865 | 0.08 | 1.4875 | 0.023612 | 0.01858 | 0.887452 | 0.989223 | 0.997988 | 0.994666 |
| Arhgef11_Q68FM7_S1488s_1 | Arhgef11_ | -0.22 | 0.07 | -0.18 | 0.04 | 0.22 | -0.28 | 0.6 | 0.06 | -0.35 | 0.37 | -0.33 | 0.08 | -0.34 | -0.04 | -0.1 | 0.3 | 0.2225 | 0.015 | 0.0275 | 0.914532 | 0.387763 | 0.953466 | 0.989223 | 0.997988 | 0.999507 |
| Arhgef11_Q68FM7_S253s_1 | Arhgef11_ | 2.16 | 1.08 | 0.34 | -0.53 | 0.47 | 2.49 | 0.32 | 0.55 | -0.46 | 0.35 | 0.32 | 0.17 | -0.93 | -0.59 | -0.22 | -0.21 | 0.195 | -0.6675 | -1.25 | 0.045024 | 0.773955 | 0.248899 | 0.995762 | 0.997988 | 0.994666 |
| Arhgef11_Q68FM7_S295s_1 | Arhgef11_ | 0.49 | 0.43 | 0.73 | 0.23 | 0.36 | 0 | 0.69 | 0.25 | -0.67 | 0.63 | 0.78 | 0.04 | -0.67 | 0.2 | -0.11 | 0.57 | -0.145 | -0.275 | -0.4725 | 0.169838 | 0.551055 | 0.449625 | 0.989223 | 0.997988 | 0.994666 |
| Arhgef11_Q68FM7_S311s_1 | Arhgef11_ | -1.36 | -1.46 | -1.11 | -0.58 | -0.39 | -0.81 | -0.78 | -0.76 | -0.63 | -1.95 | -1.06 | -0.69 | -1.44 | -0.32 | -1.24 | -0.9 | 0.4425 | 0.045 | 0.1525 | 0.661123 | 0.120721 | 0.903201 | 0.989223 | 0.997988 | 0.995671 |
| Arhgef11_Q68FM7_S585s_1 | Arhgef11_ | -3.12 | -3.68 | -2.07 | -1.31 | -1.23 | -2.87 | -1.57 | -2.81 | -5.64 | -3.04 | 0.21 | -1.94 | -2.13 | -2.05 | -3.06 | -2.1 | 0.425 | -0.0575 | 0.21 | 0.699118 | 0.488162 | 0.960605 | 0.989223 | 0.997988 | 0.996159 |
| Arhgef11_Q68FM7_S663s_1 | Arhgef11_ | 0.52 | -0.08 | 0.83 | 0.65 | 0.67 | -0.08 | 0.71 | 0.72 | -0.55 | -0.02 | 0.38 | 0.65 | -0.53 | 1.16 | 0.42 | 0.67 | 0.025 | -0.365 | -0.05 | 0.902622 | 0.933841 | 0.304958 | 0.998562 | 0.997988 | 0.999507 |
| Arhgef11_Q68FM7_S665s_1 | Arhgef11_ | -0.38 | 0.11 | -0.15 | -0.04 | 0.69 | -0.33 | 0.03 | 0.01 | -0.46 | -0.42 | -0.32 | -0.12 | -0.6 | 0.44 | -0.13 | -0.24 | 0.215 | -0.215 | -0.0175 | 0.953479 | 0.442826 | 0.357666 | 0.989223 | 0.997988 | 0.999507 |
| Arhgef12_F8VQN6_S1177s_1 | Arhgef12_ | 0.94 | 1.04 | -0.04 | 0.26 | 1.03 | 1.75 | -0.31 | -0.9 | 1.03 | 1.39 | -1.87 | -0.14 | 0.28 | 1.7 | -0.84 | -0.62 | -0.1575 | -0.4475 | -0.42 | 0.476108 | 0.790298 | 0.528875 | 0.995762 | 0.997988 | 0.994666 |
| Arhgef12_F8VQN6_S1328s_1 | Arhgef12_ | -0.22 | -0.31 | 0.44 | -0.2 | -0.18 | -0.44 | -0.59 | 0.6 | -0.16 | 0.09 | 1.42 | -0.15 | 0.41 | 0.11 | -0.15 | -0.68 | -0.08 | 0.3725 | -0.005 | 0.98789 | 0.807114 | 0.374532 | 0.995762 | 0.997988 | 1 |
| Arhgef12_F8VQN6_S190s_1 | Arhgef12_ | 0.39 | 0.51 | 1.33 | 0.93 | 0.95 | 0.63 | 0.69 | 0.89 | 0.21 | 0.58 | 0.56 | 0.9 | -0.17 | 0.67 | 0.19 | 0.56 | -1.8E-16 | -0.2275 | -0.4775 | 0.169362 | 1 | 0.451547 | 1 | 0.997988 | 0.994666 |
| Arhgef12_F8VQN6_S342s_1 | Arhgef12_ | -0.12 | 1.26 | 1.39 | 1.19 | 0.47 | 0.69 | 1.44 | 1.03 | 0.07 | 0.5 | 0.82 | 1.62 | 0.39 | 0.76 | 0.91 | 1.04 | -0.0225 | -0.1775 | -0.155 | 0.69217 | 0.954687 | 0.700901 | 0.999187 | 0.997988 | 0.996159 |
| Arhgef12_F8VQN6_T737t_1 | Arhgef12_ | 0.15 | -1.84 | -0.33 | -1.52 | -1.24 | -0.34 | -0.47 | -0.94 | -1.11 | -0.02 | -0.73 | -0.25 | -0.67 | -1.41 | -0.63 | -0.69 | 0.1375 | 0.3575 | 0.035 | 0.942461 | 0.773969 | 0.482194 | 0.995762 | 0.997988 | 0.999507 |
| Arhgef17_Q80U35_S142s_1 | Arhgef17_ | 0.57 | 0.94 | -0.47 | -0.67 | 0.22 | 0.57 | -0.54 | -1.59 | 0.66 | 0.59 | -0.86 | -0.1 | 0.29 | 0.5 | -0.77 | -0.24 | -0.4275 | -0.02 | -0.1475 | 0.752344 | 0.450583 | 0.967945 | 0.989223 | 0.997988 | 0.9971 |
| Arhgef17_Q80U35_S152s_1 | Arhgef17_ | -0.11 | 0.81 | 1.25 | -0.02 | 0.48 | -0.15 | 0.45 | 0.06 | 0.91 | 0.28 | -1.11 | 0.92 | 0.93 | 0.52 | 0.36 | 0.71 | -0.2725 | -0.2325 | 0.1475 | 0.691833 | 0.457311 | 0.667672 | 0.989223 | 0.997988 | 0.996159 |
| Arhgef17_Q80U35_S2005s_1 | Arhgef17_ | -1.23 | -0.75 | -2.86 | -1.66 | -0.49 | -1.99 | -1.72 | -2.29 | -1.3 | 0.03 | -1.14 | -3.29 | -1.16 | -1.75 | -1.78 | -1.39 | 0.0025 | 0.2 | 0.105 | 0.819518 | 0.996323 | 0.787036 | 0.99973 | 0.997988 | 0.999507 |
| Arhgef18_Q6P9R4_S577s_1 | Arhgef18_ | -0.49 | 0.07 | -0.24 | 0.37 | 0.03 | -0.07 | -0.03 | -0.09 | 0.11 | 0.06 | -0.25 | -0.08 | -0.56 | -0.25 | -0.39 | 0.08 | 0.0325 | 0.0325 | -0.2075 | 0.488505 | 0.895476 | 0.902485 | 0.996493 | 0.997988 | 0.994666 |
| Arhgef2_H3BJU7_S617s_1 | Arhgef2_H | -0.28 | 0 | 0.09 | -0.32 | 0.14 | -0.49 | 0.07 | -0.12 | -0.43 | -0.64 | -0.27 | -0.24 | -0.95 | 0.43 | -0.39 | -0.49 | 0.0275 | -0.2675 | -0.2225 | 0.517617 | 0.908597 | 0.26597 | 0.997335 | 0.997988 | 0.994666 |
| Arhgef2_H3BJU7_S856s_1 | Arhgef2_H | -0.59 | -0.66 | -0.55 | -0.2 | -0.23 | -0.54 | -0.52 | -1.15 | -1.08 | -0.56 | 0.27 | -0.49 | -1.51 | -0.22 | -0.71 | -0.23 | -0.11 | 0.035 | -0.1675 | 0.634172 | 0.679001 | 0.91384 | 0.993464 | 0.997988 | 0.995259 |
| Arhgef2_H3BJU7_S902s_1 | Arhgef2_H | -0.04 | 0.96 | 0.13 | -0.22 | 0.46 | 0.26 | -0.98 | -0.31 | 0.51 | 0.44 | 0.35 | 0.16 | 0.54 | 0.41 | 0.24 | -0.13 | -0.35 | 0.1575 | 0.0575 | 0.864157 | 0.391038 | 0.61062 | 0.989223 | 0.997988 | 0.999507 |
| Arhgef2_H3BJU7_S911s_1 | Arhgef2_H | -1.36 | 1.01 | -0.56 | -1.89 | 0.97 | -2 | -1.19 | -2.45 | -1.56 | -0.41 | -2.78 | -1.43 | 0.5 | -0.12 | -0.83 | -0.19 | -0.4675 | -0.845 | 0.54 | 0.396781 | 0.589331 | 0.257182 | 0.989223 | 0.997988 | 0.994666 |
| Arhgef2_H3BJU7_S926s_1 | Arhgef2_H | -1.84 | -0.71 | -0.77 | -1.65 | -0.75 | -2.38 | -1.44 | -1.91 | -1.33 | -1.25 | -1.37 | -1.25 | -0.09 | -0.67 | -1.18 | -1.17 | -0.3775 | -0.0575 | 0.465 | 0.259994 | 0.389294 | 0.858443 | 0.989223 | 0.997988 | 0.994666 |
| Arhgef2_H3BJU7_S926s_2 | Arhgef2_H | -3.08 | -4.59 | -1.38 | -1.77 | -2.38 | -3.28 | -1.1 | -1.35 | -3.54 | -3.27 | 0.66 | -1.19 | -3.23 | -1.1 | -1.95 | -1.81 | 0.6775 | 0.87 | 0.6825 | 0.376733 | 0.390327 | 0.427335 | 0.989223 | 0.997988 | 0.994666 |
| Arhgef2_H3BJU7_S930s_2 | Arhgef2_H | -3.08 | -4.59 | -1.38 | -1.77 | -2.38 | -3.28 | -1.1 | -1.35 | -3.54 | -3.27 | 0.66 | -1.19 | -3.23 | -1.1 | -1.95 | -1.81 | 0.6775 | 0.87 | 0.6825 | 0.376733 | 0.390327 | 0.427335 | 0.989223 | 0.997988 | 0.994666 |
| Arhgef2_H3BJU7_T766t_1 | Arhgef2_H | 1.07 | 0.63 | -0.19 | -0.37 | 0.57 | 0.37 | 0.14 | -0.14 | 0.51 | 1.01 | 0.68 | 0.26 | 0.47 | 1.14 | -0.31 | 0.01 | -0.05 | 0.33 | 0.0425 | 0.924831 | 0.891933 | 0.394588 | 0.996493 | 0.997988 | 0.999507 |
| Arhgef25_G5E825_T603t_1 | Arhgef25_ | 0.96 | 1.17 | -0.92 | 0.88 | 0.43 | 0.57 | 0.12 | 0.96 | 1.21 | 0.76 | -1.02 | 0.16 | 0.34 | 0.33 | -0.15 | -0.05 | -0.0025 | -0.245 | -0.405 | 0.407055 | 0.995805 | 0.695663 | 0.999616 | 0.997988 | 0.994666 |
| Arhgef26_D3YYY8_S123s_1 | Arhgef26_ | 0.9 | 0.96 | 0.05 | 0.57 | 0.12 | -0.28 | 1.33 | 0.38 | 0.82 | 0.58 | 0.74 | 1.61 | 0.84 | 0.4 | 0.64 | 1.25 | -0.2325 | 0.3175 | 0.1625 | 0.615737 | 0.553404 | 0.352742 | 0.989223 | 0.997988 | 0.995259 |
| Arhgef26_D3YYY8_S199s_1 | Arhgef26_ | 0.44 | 0.65 | 1.11 | -0.07 | 0.17 | 0.31 | 1.34 | -0.21 | -0.05 | 0.08 | -2.5 | -0.32 | -1.07 | 0.36 | 0.29 | 0.55 | -0.13 | -1.23 | -0.5 | 0.269564 | 0.743808 | 0.065211 | 0.995762 | 0.997988 | 0.994666 |
| Arhgef26_D3YYY8_S202s_1 | Arhgef26_ | 0.44 | 0.65 | 1.11 | -0.07 | 0.17 | 0.31 | 1.34 | -0.21 | -0.05 | 0.08 | -2.5 | -0.32 | -1.07 | 0.36 | 0.29 | 0.55 | -0.13 | -1.23 | -0.5 | 0.269564 | 0.743808 | 0.065211 | 0.995762 | 0.997988 | 0.994666 |
| Arhgef26_D3YYY8_S221s_1 | Arhgef26_ | -0.14 | -0.19 | -0.4 | 0.52 | 0.36 | 0.01 | 0.44 | 0.18 | -0.04 | 0.33 | -0.25 | -0.91 | -0.06 | 0.27 | 0.32 | 0.07 | 0.3 | -0.165 | 0.2025 | 0.488407 | 0.275688 | 0.634567 | 0.989223 | 0.997988 | 0.994666 |
| Arhgef28_G5E8P2_S1198s_1 | Arhgef28_ | -0.77 | -0.53 | 0.34 | -1.59 | -1.84 | 0.21 | -1.01 | -0.81 | 2.33 | -1.5 | 0.36 | 0.3 | 1.96 | -1.83 | 1.2 | -0.59 | -0.225 | 1.01 | 0.8225 | 0.336145 | 0.670349 | 0.218832 | 0.992876 | 0.997988 | 0.994666 |
| Arhgef4_E9PYK8_S1277s_1 | Arhgef4_E | -0.33 | -0.31 | -0.46 | -0.15 | -0.23 | -0.78 | -0.51 | 0.01 | 1.08 | 0.46 | -0.77 | 0.2 | 0.04 | 0.56 | -0.28 | 0.13 | -0.065 | 0.555 | 0.425 | 0.141751 | 0.790357 | 0.178815 | 0.995762 | 0.997988 | 0.994666 |
| Arhgef40_S4R189_S1089s_1 | Arhgef40_ | -0.62 | -0.22 | -0.37 | -0.67 | -0.08 | -0.04 | -0.61 | -0.94 | 1.28 | -0.46 | -1.57 | -0.21 | 1.67 | -1.37 | -0.2 | -0.56 | 0.0525 | 0.23 | 0.355 | 0.554971 | 0.850524 | 0.678272 | 0.996349 | 0.997988 | 0.99525 |
| Arhgef40_S4R189_S1494s_1 | Arhgef40_ | 0.54 | -0.15 | 0.19 | 0.35 | 0.36 | 0.34 | 0.95 | 0.27 | -0.17 | -0.08 | -0.35 | 0.77 | -0.21 | 0.23 | 1.54 | 0.6 | 0.2475 | -0.19 | 0.3075 | 0.454182 | 0.35739 | 0.554625 | 0.989223 | 0.997988 | 0.994666 |
| Arhgef6_F6WMJ3_S248s_1 | Arhgef6_F | 0.69 | 0.48 | 1.11 | 1.07 | 0.83 | 0.68 | 0.68 | 0.82 | 0.62 | 0.8 | 0.74 | 0.86 | 0.49 | 0.82 | 0.97 | 0.82 | -0.085 | -0.0825 | -0.0625 | 0.817443 | 0.714415 | 0.73455 | 0.995437 | 0.997988 | 0.999507 |
| Arhgef6_F6WMJ3_S511s_1 | Arhgef6_F | 0.52 | 0.68 | 0.52 | 0.25 | 0.5 | 0.61 | 0.68 | 0.46 | -0.21 | 0.47 | 0.33 | 0.25 | 0.03 | 0.78 | 0.44 | 0.39 | 0.07 | -0.2825 | -0.0825 | 0.758668 | 0.737267 | 0.272772 | 0.995762 | 0.997988 | 0.9971 |
| Arhgef6_F6WMJ3_S663s_1 | Arhgef6_F | 1.49 | 0.68 | 0.28 | -0.29 | 0.78 | 1.41 | 0.4 | 0.5 | 0.32 | 0.63 | 0.54 | 0.68 | -0.34 | 0.77 | 0.18 | -0.14 | 0.2325 | 0.0025 | -0.4225 | 0.346618 | 0.57962 | 0.994813 | 0.989223 | 0.998021 | 0.994666 |
| Arhgef6_F6WMJ3_S703s_1 | Arhgef6_F | 0.1 | 0.28 | -0.11 | -0.13 | 0.6 | -0.25 | -0.08 | -0.1 | -0.47 | -0.16 | -0.77 | -0.15 | -0.07 | 0.25 | -0.01 | 0.17 | 0.0075 | -0.4225 | 0.05 | 0.838383 | 0.977073 | 0.118018 | 0.999187 | 0.997988 | 0.999507 |
| Arhgef7_Q9ES28_S103s_1 | Arhgef7_Q | 0.05 | 0.08 | 0.08 | 0.07 | 0.38 | -0.14 | 0.46 | 0.33 | -0.32 | 0.03 | -0.22 | 0.13 | -0.5 | -0.22 | -0.28 | 0.14 | 0.1875 | -0.165 | -0.285 | 0.269126 | 0.405801 | 0.458717 | 0.989223 | 0.997988 | 0.994666 |
| Arhgef7_Q9ES28_S176s_1 | Arhgef7_Q | -1.29 | -0.74 | -0.73 | -1.63 | -0.05 | -0.7 | -0.91 | -0.53 | -1.04 | -0.37 | -0.54 | -0.53 | 0.13 | 0.19 | -1.08 | -0.63 | 0.55 | 0.4775 | 0.75 | 0.079038 | 0.099098 | 0.139555 | 0.989223 | 0.997988 | 0.994666 |
| Arhgef7_Q9ES28_S445s_1 | Arhgef7_Q | 0.04 | 0.34 | 0.56 | 0.41 | 0.23 | -0.1 | 0.44 | 0.34 | -0.04 | 0.16 | 0.27 | 0.53 | 0.23 | 0.4 | 0.07 | 0.14 | -0.11 | -0.1075 | -0.1275 | 0.609594 | 0.638424 | 0.659928 | 0.989223 | 0.997988 | 0.995259 |
| Arhgef7_Q9ES28_S621s_1 | Arhgef7_Q | 0.47 | -0.09 | 0.63 | 0.66 | 0.28 | 0.27 | 0.21 | 0.35 | 0.25 | 0.35 | 0.31 | 0.39 | -0.39 | 0.61 | -0.09 | -0.01 | -0.14 | -0.0925 | -0.3875 | 0.244526 | 0.564843 | 0.713209 | 0.989223 | 0.997988 | 0.994666 |
| Arhgef7_Q9ES28_S665s_1 | Arhgef7_Q | -0.14 | 0.07 | -0.35 | 0.22 | 0.1 | -0.46 | 0.58 | 0.39 | -0.56 | 0.16 | 0.38 | -0.02 | -0.45 | 0.84 | 0.09 | 0.16 | 0.2025 | 0.04 | 0.21 | 0.531573 | 0.489467 | 0.888219 | 0.989223 | 0.997988 | 0.994666 |
| Arhgef7_Q9ES28_S719s_1 | Arhgef7_Q | 0.94 | 0.4 | 0.53 | 0.34 | -1.02 | 1.43 | -0.74 | -0.1 | 2.36 | 0.69 | -0.2 | 0.39 | 1.5 | -0.61 | -0.44 | -0.8 | -0.66 | 0.2575 | -0.64 | 0.23502 | 0.21964 | 0.627374 | 0.989223 | 0.997988 | 0.994666 |
| Arhgef7_Q9ES28_S80s_1 | Arhgef7_Q | 1.44 | 0.9 | 0.07 | -0.18 | 0.92 | 1.27 | -0.47 | -0.57 | 1.15 | 0.53 | 0.1 | -0.13 | 1.06 | 1.21 | -0.24 | -0.4 | -0.27 | -0.145 | -0.15 | 0.776281 | 0.62234 | 0.747926 | 0.989223 | 0.997988 | 0.99798 |
| Arhgef7_Q9ES28_S83s_1 | Arhgef7_Q | 0.2 | -0.16 | -1.8 | -0.93 | 0.01 | 1.02 | -0.72 | -2.08 | 0.11 | -0.04 | -1.73 | -2 | 0.38 | -0.28 | -0.16 | -0.09 | 0.23 | -0.2425 | 0.635 | 0.18352 | 0.741455 | 0.707304 | 0.995762 | 0.997988 | 0.994666 |
| Arid1a_E9QAQ7_S1760s_1 | Arid1a_E9 | -0.61 | -1.15 | -0.74 | -1.17 | -1.24 | -1.33 | -0.64 | -0.89 | -1.64 | -1.3 | -1.12 | -0.75 | -2.12 | -0.82 | -1.29 | -1.26 | -0.1075 | -0.285 | -0.455 | 0.203406 | 0.68258 | 0.328172 | 0.994175 | 0.997988 | 0.994666 |
| Arid1a_E9QAQ7_S365s_1 | Arid1a_E9 | -2.29 | -1.86 | -0.51 | -2.13 | -3.11 | -2.97 | -2.13 | -1.53 | 1.75 | -1.96 | 0 | -2.75 | 2.17 | -1.23 | -0.4 | -0.52 | -0.7375 | 0.9575 | 1.7025 | 0.04538 | 0.166277 | 0.333807 | 0.989223 | 0.997988 | 0.994666 |
| Arid1a_E9QAQ7_S770s_1 | Arid1a_E9 | -0.83 | 0.09 | -0.4 | 0.42 | -0.28 | -0.23 | -0.82 | 0.47 | -0.09 | -0.09 | 0.43 | -0.16 | -0.52 | 0.43 | 0.36 | 0.95 | -0.035 | 0.2025 | 0.485 | 0.256644 | 0.925415 | 0.544402 | 0.997335 | 0.997988 | 0.994666 |
| Arid1b_E9Q6R4_S1549s_1 | Arid1b_E9 | -0.28 | -1.25 | -0.42 | -0.23 | -0.35 | -0.82 | -0.18 | 0.01 | -0.89 | -0.65 | 0.05 | -0.28 | -0.8 | -0.38 | -0.93 | -0.41 | 0.21 | 0.1025 | -0.085 | 0.791938 | 0.509479 | 0.761248 | 0.989223 | 0.997988 | 0.999507 |
| Arid1b_E9Q6R4_S1755s_1 | Arid1b_E9 | -0.81 | -0.97 | 0.03 | -0.52 | -1.59 | -0.31 | -0.55 | -0.83 | -0.91 | -0.98 | -0.14 | -1.18 | -0.28 | -0.3 | -0.48 | -0.81 | -0.2525 | -0.235 | 0.1 | 0.745622 | 0.482406 | 0.492287 | 0.989223 | 0.997988 | 0.9971 |
| Arid1b_E9Q6R4_S1756s_1 | Arid1b_E9 | -0.81 | -0.97 | 0.03 | -0.52 | -1.59 | -0.31 | -0.55 | -0.83 | -0.91 | -0.98 | -0.14 | -1.18 | -0.28 | -0.3 | -0.48 | -0.81 | -0.2525 | -0.235 | 0.1 | 0.745622 | 0.482406 | 0.492287 | 0.989223 | 0.997988 | 0.9971 |
| Arid4a_F8VPQ2_S867s_1 | Arid4a_F8 | 0.09 | 0.43 | -0.28 | -0.14 | -0.07 | -0.1 | 0.56 | -0.55 | -0.08 | 0.36 | -0.03 | -0.19 | 0.09 | 0.6 | 0.38 | -0.24 | -0.065 | -0.01 | 0.1825 | 0.547288 | 0.828696 | 0.969429 | 0.995762 | 0.997988 | 0.994666 |
| Arid5b_Q8BM75_S789s_1 | Arid5b_Q8 | 0.7 | 0.95 | 0.76 | 0.69 | 0.16 | 0.63 | 0.91 | 0.52 | 0.72 | 0.67 | 1.1 | 0.51 | 0.51 | 0.75 | 0.86 | 1.18 | -0.22 | -0.025 | 0.05 | 0.845435 | 0.362347 | 0.914336 | 0.989223 | 0.997988 | 0.999507 |
| Arl13b_Q640N2_S328s_1 | Arl13b_Q6 | 0.49 | 0.22 | 0.88 | 0.34 | 0.63 | 0.28 | 0.09 | 0.24 | 0.94 | 0.38 | 0.71 | 0.39 | 0.62 | 0.85 | 0.26 | 0.17 | -0.1725 | 0.1225 | -0.0075 | 0.979081 | 0.486687 | 0.64194 | 0.989223 | 0.997988 | 1 |
| Arl2_Q9D0J4_S45s_1 | Arl2_Q9D0 | -0.22 | 0.12 | -0.29 | 0.33 | 0.68 | -0.38 | 0.64 | 0.19 | 0.32 | 0.15 | 0.42 | 0.21 | 0.21 | 0.41 | 0.43 | 0.09 | 0.2975 | 0.29 | 0.3 | 0.271156 | 0.34581 | 0.248115 | 0.989223 | 0.997988 | 0.994666 |
| Arl6ip6_Q8BH07_T19t_1 | Arl6ip6_Q8 | -0.97 | -0.1 | -0.61 | -1.27 | -0.28 | -1.68 | -1.55 | -1.38 | -1.95 | -1.42 | -0.74 | -1.99 | 0.29 | 0.04 | -1.05 | -1.39 | -0.485 | -0.7875 | 0.21 | 0.652038 | 0.236793 | 0.067263 | 0.989223 | 0.997988 | 0.995259 |
| Armcx4_E9PWM3_S34s_1 | Armcx4_E9 | -0.05 | -0.88 | -0.27 | 0.59 | -1.7 | 0.39 | -1.18 | -1.8 | 1.27 | -0.65 | -1.78 | -2 | 1.66 | -1.38 | 0.58 | 0.55 | -0.92 | -0.6375 | 0.505 | 0.434438 | 0.111126 | 0.389929 | 0.989223 | 0.997988 | 0.994666 |
| Arpc1a_Q9R0Q6_S236s_1 | Arpc1a_Q9 | 0.72 | 0.16 | 0.57 | 0.4 | 0.86 | 0.32 | 0.45 | 0.31 | 0.18 | 0.17 | 0.48 | 0.31 | -0.15 | 0.66 | 0.56 | 0.82 | 0.0225 | -0.1775 | 0.01 | 0.973738 | 0.925539 | 0.453936 | 0.997335 | 0.997988 | 0.999507 |
| Arpc5_Q9CPW4_S77s_1 | Arpc5_Q9C | -0.21 | 0.68 | -0.15 | -0.01 | 0.29 | 0.07 | 0.32 | -0.06 | 0.51 | 0.02 | -0.15 | 0.24 | 0.9 | 0.49 | -0.07 | 0.14 | 0.0775 | 0.0775 | 0.2875 | 0.399003 | 0.772729 | 0.791813 | 0.995762 | 0.997988 | 0.994666 |
| Arpc5l_Q9D898_S64s_1 | Arpc5l_Q9 | 0.5 | 0.01 | 1.52 | -1.03 | 0.1 | 1.18 | 1.78 | 0.62 | 1.47 | -0.19 | -3.07 | 1.41 | 1.67 | -0.45 | 1.25 | 1.73 | 0.67 | -0.345 | 0.8 | 0.245991 | 0.262478 | 0.742336 | 0.989223 | 0.997988 | 0.994666 |
| Arpp19_P56212_S2s_1 | Arpp19_P5 | -0.36 | -0.19 | 0.02 | -0.57 | 0.26 | -0.08 | -0.13 | -0.03 | -0.75 | 0.16 | 0.26 | 0.63 | -1.02 | 0.02 | 0.09 | -0.16 | 0.28 | 0.35 | 0.0075 | 0.981729 | 0.240479 | 0.316245 | 0.989223 | 0.997988 | 1 |
| Arpp19_P56212_S7s_1 | Arpp19_P5 | 1.4 | 1.33 | 2.12 | 2.21 | 0.92 | 1.17 | 1.61 | 1.85 | 1.39 | 1.97 | 1.34 | 1.78 | 1.39 | 0.74 | 1.48 | 1.88 | -0.3775 | -0.145 | -0.3925 | 0.287618 | 0.263274 | 0.643149 | 0.989223 | 0.997988 | 0.994666 |
| Arpp19_P56212_T22t_1 | Arpp19_P5 | 2.24 | 2.32 | 1.89 | 2.33 | 1.39 | 1.96 | 1.17 | 1.74 | 1.89 | 1.84 | 1.2 | 2.28 | 1.91 | 0.61 | 2.1 | 1.95 | -0.63 | -0.3925 | -0.5525 | 0.166918 | 0.032532 | 0.199586 | 0.989223 | 0.997988 | 0.994666 |
| Arpp19_P56212_T6t_1 | Arpp19_P5 | 2.24 | 2.32 | 1.89 | 2.33 | 1.39 | 1.96 | 1.17 | 1.74 | 1.89 | 1.84 | 1.2 | 2.28 | 1.91 | 0.61 | 2.1 | 1.95 | -0.63 | -0.3925 | -0.5525 | 0.166918 | 0.032532 | 0.199586 | 0.989223 | 0.997988 | 0.994666 |
| Arpp21_E0CYU3_S32s_1 | Arpp21_E0 | -2.87 | -2.95 | -0.24 | -0.28 | -2.54 | -3.28 | 0.1 | 0.03 | -3.33 | -2.59 | 0.11 | -0.19 | -2.98 | -2.51 | 0.01 | -0.04 | 0.1625 | 0.085 | 0.205 | 0.83127 | 0.871275 | 0.933172 | 0.996493 | 0.997988 | 0.999507 |
| Arpp21_Q9DCB4_S346s_1 | Arpp21_Q | -1.71 | -1.3 | -1.16 | -2.05 | -2.16 | -0.91 | -2.47 | -2.05 | -0.63 | -1.37 | -0.75 | -2.18 | 0.49 | -1.99 | -1.35 | -1.81 | -0.3425 | 0.3225 | 0.39 | 0.489663 | 0.384429 | 0.4337 | 0.989223 | 0.997988 | 0.994666 |
| Arpp21_Q9DCB4_S362s_1 | Arpp21_Q | -0.05 | 0.45 | -1.26 | -1.03 | -0.15 | -0.85 | -0.49 | -1.31 | 0.29 | 0.53 | -0.7 | -0.69 | 1.44 | -1.66 | -0.43 | -0.45 | -0.2275 | 0.33 | 0.1975 | 0.771581 | 0.611176 | 0.505164 | 0.989223 | 0.997988 | 0.9971 |
| Arpp21_Q9DCB4_S381s_1 | Arpp21_Q | -2.57 | -0.4 | -1.7 | -2.01 | -0.57 | -2.03 | -1.5 | -1.55 | -2.35 | -1.54 | -1.04 | -1.61 | -0.79 | -0.55 | -1.52 | -1.75 | 0.2575 | 0.035 | 0.5175 | 0.324008 | 0.612475 | 0.944344 | 0.989223 | 0.997988 | 0.994666 |
| Arpp21_Q9DCB4_S408s_1 | Arpp21_Q | -0.1 | -0.37 | 0.13 | -1.81 | -1.31 | -0.09 | -1.94 | -1.65 | 0.51 | -0.67 | -0.4 | -2.12 | 0.87 | -1.52 | -0.78 | -1.33 | -0.71 | -0.1325 | -0.1525 | 0.809209 | 0.209602 | 0.834904 | 0.989223 | 0.997988 | 0.999507 |
| Arpp21_Q9DCB4_T87t_1 | Arpp21_Q | -1.23 | -2.49 | -1.26 | -2 | -2.67 | -1.8 | -2.13 | -1.19 | -1.03 | -1.94 | -0.3 | -0.71 | -0.99 | -1.9 | -1.31 | -1.82 | -0.2025 | 0.75 | 0.24 | 0.538616 | 0.627152 | 0.119588 | 0.989223 | 0.997988 | 0.994666 |
| Arrb1_J3QNU6_S413s_1 | Arrb1_J3Q | 1.47 | 1.2 | 0.97 | 0.85 | 0.83 | 1.25 | 0.44 | 0.92 | 1.22 | 1.5 | 1.69 | 1.08 | 0.36 | 1.3 | 0.71 | 0.42 | -0.2625 | 0.25 | -0.425 | 0.191197 | 0.330804 | 0.348703 | 0.989223 | 0.997988 | 0.994666 |
| Arvcf_P98203_S273s_1 | Arvcf_P98 | 0.19 | 0.46 | 0.62 | 0.53 | 0.78 | 0.09 | 0.37 | 0.18 | 0.33 | 0.45 | 0.36 | 0.32 | 0.55 | 0.94 | 0.82 | 0.48 | -0.095 | -0.085 | 0.2475 | 0.340403 | 0.696098 | 0.695568 | 0.994893 | 0.997988 | 0.994666 |
| Arvcf_P98203_S281s_1 | Arvcf_P98 | 1.05 | 0.77 | 0.8 | 1.34 | 0.42 | 1.83 | 0.18 | 0.87 | 3.04 | 0.98 | 0.4 | 0.89 | 2.79 | 0.31 | 1.42 | 0.94 | -0.165 | 0.3375 | 0.375 | 0.469429 | 0.664646 | 0.547221 | 0.991343 | 0.997988 | 0.994666 |
| Arvcf_P98203_S283s_1 | Arvcf_P98 | -0.43 | -0.22 | 0.7 | 0.69 | -0.82 | 0.6 | -0.33 | 0.33 | 1.26 | -0.26 | -0.56 | 0.57 | 1.69 | -1.36 | 0.42 | 0.11 | -0.24 | 0.0675 | 0.03 | 0.961902 | 0.567846 | 0.888565 | 0.989223 | 0.997988 | 0.999507 |
| Arvcf_P98203_S873s_1 | Arvcf_P98 | -0.71 | -1.08 | -0.28 | 0.04 | -0.43 | -0.52 | -0.3 | -0.13 | -0.26 | -1.26 | 0.5 | 0.04 | 0.27 | -0.3 | -0.17 | -0.32 | 0.1625 | 0.2625 | 0.3775 | 0.2628 | 0.577783 | 0.548627 | 0.989223 | 0.997988 | 0.994666 |
| Arvcf_P98203_T573t_1 | Arvcf_P98 | 0.06 | 1.93 | 2.07 | 1.17 | 1.18 | 1.23 | 2.38 | 2.34 | -0.71 | 1.4 | 0.63 | 2.03 | -1.1 | -0.5 | 1.66 | 2.81 | 0.475 | -0.47 | -0.59 | 0.515735 | 0.370272 | 0.493283 | 0.989223 | 0.997988 | 0.994666 |
| Asap1_E9QMJ1_S1010s_1 | Asap1_E9C | 6.64 | 0.72 | 2.98 | -0.86 | 2.01 | 6.87 | 5.62 | 4 | 0.11 | 3.09 | 0.93 | 3.62 | -0.96 | -1.01 | 3.19 | 3.7 | 2.255 | -0.4325 | -1.14 | 0.524824 | 0.19498 | 0.786847 | 0.989223 | 0.997988 | 0.994666 |
| Asap1_E9QMJ1_S1024s_1 | Asap1_E9C | -0.01 | 0.52 | 0.29 | 0.06 | 0.29 | -0.19 | 0.56 | 0.18 | 0.37 | 0.1 | -0.26 | -0.28 | -0.84 | 0.66 | -0.4 | 0.39 | -0.005 | -0.2325 | -0.2625 | 0.496946 | 0.984143 | 0.384858 | 0.999187 | 0.997988 | 0.994666 |
| Asap1_E9QMJ1_S1111s_1 | Asap1_E9C | -1.93 | -0.6 | -1.65 | -1.5 | -1.17 | -2.03 | -0.79 | -1.37 | -0.58 | -0.47 | -1.38 | -2.36 | 0.04 | -0.83 | -1.45 | -0.52 | 0.08 | 0.2225 | 0.73 | 0.108631 | 0.832736 | 0.653979 | 0.995762 | 0.997988 | 0.994666 |
| Asap1_E9QMJ1_S505s_1 | Asap1_E9C | 0.49 | 0.76 | 0.46 | 0.56 | 0.65 | 0.6 | 0.78 | 0.44 | 0.34 | 0.77 | 0.03 | 0.52 | 0.02 | 0.69 | 0.3 | 0.52 | 0.05 | -0.1525 | -0.185 | 0.482655 | 0.808773 | 0.541583 | 0.995762 | 0.997988 | 0.994666 |
| Asap1_E9QMJ1_S819s_1 | Asap1_E9C | -0.14 | -0.38 | 0.1 | 0.53 | 0.34 | -0.68 | 0.18 | 0.19 | -0.2 | -0.2 | -0.11 | 0.2 | -0.18 | 0.55 | -0.41 | -0.12 | -0.02 | -0.105 | -0.0675 | 0.836285 | 0.949827 | 0.701143 | 0.999187 | 0.997988 | 0.999507 |
| Asap1_E9QMJ1_S823s_1 | Asap1_E9C | 0.42 | -0.95 | -1.17 | 0.61 | -0.51 | -0.25 | 0.23 | 0.25 | -1.94 | -1.33 | 1.52 | -0.15 | -1.76 | 0.26 | -1.19 | -0.67 | 0.2025 | -0.2025 | -0.5675 | 0.337941 | 0.661563 | 0.798922 | 0.990434 | 0.997988 | 0.994666 |
| Asap2_D3YX85_S712s_1 | Asap2_D3Y | -0.12 | -1.74 | -0.55 | -0.66 | -0.37 | 0.06 | -0.83 | -0.28 | 0.14 | -1.43 | 0.09 | -0.45 | 0.14 | -0.89 | -0.83 | -0.67 | 0.4125 | 0.355 | 0.205 | 0.626119 | 0.29418 | 0.463318 | 0.989223 | 0.997988 | 0.995259 |
| Asb3_D6RDR7_S35s_1 | Asb3_D6RI | -1.47 | -1.92 | -0.88 | -1.09 | -1.6 | -1.54 | -1.6 | -0.85 | -1.4 | -1.14 | -0.06 | -1.73 | -0.98 | -1.41 | -0.79 | -0.66 | -0.0575 | 0.2575 | 0.38 | 0.260164 | 0.853602 | 0.54332 | 0.996349 | 0.997988 | 0.994666 |
| Ash1l_Q99MY8_S1678s_1 | Ash1l_Q99 | 0.21 | -0.64 | 0.47 | 0.43 | -1.6 | 0.25 | -0.63 | -0.31 | 2.26 | 0.53 | 0.03 | -0.31 | 2.64 | -1.82 | 0.98 | -0.3 | -0.69 | 0.51 | 0.2575 | 0.765636 | 0.140045 | 0.387327 | 0.989223 | 0.997988 | 0.9971 |
| Asic1_Q6NXK8_S497s_1 | Asic1_Q6N | 5.42 | 0.98 | 1.88 | 1.7 | 1.26 | 6.56 | 2.24 | 2.88 | 1.23 | 3.16 | 2.22 | 2.22 | 1.6 | 1.04 | 2.52 | 3.83 | 0.74 | -0.2875 | -0.2475 | 0.807358 | 0.575145 | 0.761645 | 0.989223 | 0.997988 | 0.999507 |
| Aspscr1_Q8VBT9_S116s_1 | Aspscr1_Q | 0.28 | 0.45 | 0.41 | 0.55 | 0.28 | 0.55 | 0.84 | 0.84 | 0.79 | 0.77 | 0.17 | 0.71 | 0.83 | 0.36 | 0.37 | 0.35 | 0.205 | 0.1875 | 0.055 | 0.824501 | 0.374685 | 0.444997 | 0.989223 | 0.997988 | 0.999507 |
| Aspscr1_Q8VBT9_S202s_1 | Aspscr1_Q | -3.97 | -2.62 | -0.66 | -0.49 | -1.83 | -3.13 | 0.04 | -0.18 | -3.02 | -1.89 | 0.37 | -0.73 | -1.35 | -0.97 | -0.31 | -0.43 | 0.66 | 0.6175 | 1.17 | 0.15213 | 0.500708 | 0.532796 | 0.989223 | 0.997988 | 0.994666 |
| Astn1_Q61137_S192s_1 | Astn1_Q61 | 0.47 | 0.28 | -0.53 | 0.13 | 0.38 | 0.53 | 0.55 | 0.2 | -0.19 | 0.52 | -2.21 | -0.12 | -1.07 | 0.03 | -0.23 | 0.09 | 0.3275 | -0.5875 | -0.3825 | 0.310592 | 0.247673 | 0.32523 | 0.989223 | 0.997988 | 0.994666 |
| Astn1_Q61137_S274s_1 | Astn1_Q61 | 0.21 | 0.1 | 0.33 | 0.4 | -0.06 | 0.14 | 0.3 | 0.35 | -0.26 | 0.51 | 0.26 | 0.44 | -0.07 | 0.6 | -0.43 | -0.05 | -0.0775 | -0.0225 | -0.2475 | 0.405932 | 0.715866 | 0.92987 | 0.995437 | 0.997988 | 0.994666 |
| Astn1_Q61137_S893s_1 | Astn1_Q61 | -1.74 | -0.35 | 0.64 | -0.84 | -1.1 | 0.4 | -1.28 | -1.24 | 1.86 | -0.05 | 0.39 | -0.14 | 1.58 | -0.96 | -0.01 | -0.46 | -0.2325 | 1.0875 | 0.61 | 0.370764 | 0.686909 | 0.105666 | 0.994672 | 0.997988 | 0.994666 |
| Astn1_Q61137_S898s_1 | Astn1_Q61 | -0.62 | -0.72 | 0.52 | 0.43 | -2 | 0.97 | -0.62 | -0.05 | 2.14 | 0.05 | 1.25 | 0.83 | 1.14 | -1.67 | 0.11 | 0.29 | -0.3275 | 1.165 | 0.065 | 0.915845 | 0.603177 | 0.045479 | 0.989223 | 0.997988 | 0.999507 |
| Atad1_Q9D5T0_S322s_1 | Atad1_Q9D | -0.33 | -0.21 | -1.72 | -0.65 | -0.19 | -0.66 | -0.52 | -0.43 | 0.01 | -0.03 | -0.28 | -0.3 | -0.91 | 0.68 | -0.72 | 0.45 | 0.2775 | 0.5775 | 0.6025 | 0.247213 | 0.444388 | 0.137616 | 0.989223 | 0.997988 | 0.994666 |
| Atat1_Q8K341_S249s_1 | Atat1_Q8K | 0.01 | -0.8 | 0.1 | -0.53 | -0.68 | 0.61 | -0.71 | -0.17 | 0.94 | -0.66 | -0.44 | 0.07 | 1.26 | -0.5 | -0.05 | 0.43 | 0.0675 | 0.2825 | 0.59 | 0.190978 | 0.855512 | 0.49667 | 0.996349 | 0.997988 | 0.994666 |
| Atat1_Q8K341_S253s_1 | Atat1_Q8K | -1.1 | -1.96 | -0.67 | -1.48 | -2.15 | -0.82 | -0.99 | -0.8 | -1.43 | -1.28 | -0.71 | -0.63 | -0.64 | -4.57 | -0.96 | -0.42 | 0.1125 | 0.29 | -0.345 | 0.699928 | 0.781924 | 0.421548 | 0.995762 | 0.997988 | 0.996159 |
| Atat1_Q8K341_T238t_1 | Atat1_Q8K | -0.64 | -0.58 | -1.21 | -1.1 | -0.59 | -0.83 | -1.04 | -0.53 | 0.58 | -0.16 | -0.45 | -0.32 | 1.37 | 0.09 | -0.96 | -0.68 | 0.135 | 0.795 | 0.8375 | 0.127398 | 0.595665 | 0.028839 | 0.989223 | 0.997988 | 0.994666 |
| Atcay_Q8BHE3_S370s_1 | Atcay_Q8B | 0.1 | -0.85 | 1.29 | 1.88 | -1.22 | 0.09 | 1.21 | 1.43 | 0.5 | -0.26 | 1.76 | 1.59 | 0.66 | -0.94 | 1.32 | 1.37 | -0.2275 | 0.2925 | -0.0025 | 0.997248 | 0.763609 | 0.676533 | 0.995762 | 0.997988 | 1 |
| Atcay_Q8BHE3_S46s_1 | Atcay_Q8B | 0.06 | 0.38 | -0.18 | 0.17 | 0.63 | 0.25 | 0.24 | -0.07 | 0.25 | 0.57 | -0.31 | 0.11 | 0.36 | 0.65 | -0.03 | 0.02 | 0.155 | 0.0475 | 0.1425 | 0.610219 | 0.531937 | 0.86183 | 0.989223 | 0.997988 | 0.995259 |
| Atf2_P16951_S149s_1 | Atf2_P169 | -0.53 | -1.55 | -0.7 | -0.63 | -0.68 | -1.77 | -0.36 | -2.48 | -1.3 | -0.03 | -1.82 | -2.55 | -1.12 | -1.44 | -1.03 | -0.38 | -0.47 | -0.5725 | -0.14 | 0.692121 | 0.357146 | 0.304156 | 0.989223 | 0.997988 | 0.996159 |
| Atf2_P16951_S150s_1 | Atf2_P169 | -0.53 | -1.55 | -0.7 | -0.63 | -0.68 | -1.77 | -0.36 | -2.48 | -1.3 | -0.03 | -1.82 | -2.55 | -1.12 | -1.44 | -1.03 | -0.38 | -0.47 | -0.5725 | -0.14 | 0.692121 | 0.357146 | 0.304156 | 0.989223 | 0.997988 | 0.996159 |
| Atf2_P16951_S212s_2 | Atf2_P169 | -2.77 | -1.95 | -3.12 | -3.28 | -3.06 | -3.59 | -2 | -2.72 | -2.51 | -2.2 | -3.82 | -4.87 | -1.4 | -1.59 | -1.56 | -3.83 | -0.0625 | -0.57 | 0.685 | 0.266868 | 0.88238 | 0.372169 | 0.996493 | 0.997988 | 0.994666 |
| Atf2_P16951_S94s_1 | Atf2_P169 | -1.51 | -1.67 | -1.15 | -0.96 | -0.66 | -1.86 | -0.38 | -1.12 | -1.48 | -1.06 | -0.53 | -0.86 | -1.35 | -0.77 | -1.09 | -0.67 | 0.3175 | 0.34 | 0.3525 | 0.246046 | 0.387966 | 0.270493 | 0.989223 | 0.997988 | 0.994666 |
| Atf2_P16951_T53t_1 | Atf2_P169 | 0.21 | -0.14 | -0.63 | -1.62 | -1.56 | -1.21 | -0.63 | -0.2 | -0.37 | 0.31 | 0.35 | -1.13 | -0.19 | 0.12 | -1.29 | 0.24 | -0.355 | 0.335 | 0.265 | 0.59878 | 0.451153 | 0.506501 | 0.989223 | 0.997988 | 0.995259 |
| Atg16l1_Q8C0J2_S303s_1 | Atg16l1_Q | -0.85 | -0.97 | -0.94 | -0.67 | -0.42 | -1 | -0.2 | -0.73 | -1.59 | -0.54 | 0.02 | -0.75 | -0.99 | -0.13 | -0.87 | -0.88 | 0.27 | 0.1425 | 0.14 | 0.62386 | 0.29002 | 0.688781 | 0.989223 | 0.997988 | 0.995259 |
| Atg2a_Q6P4T0_S1243s_1 | Atg2a_Q6P | 0.2 | 0.28 | 1.64 | 1.27 | -0.95 | 0.09 | 1.55 | 1.44 | 0.3 | 0.92 | 0.88 | -0.03 | 0.1 | 1.25 | 0.05 | 1.41 | -0.315 | -0.33 | -0.145 | 0.766137 | 0.613493 | 0.438661 | 0.989223 | 0.997988 | 0.9971 |
| Atg2b_Q80XK6_S1571s_1 | Atg2b_Q8C | -1.4 | -0.47 | -0.5 | -0.51 | -0.02 | -1.35 | -0.67 | 0.01 | -1.08 | -2.06 | 0.56 | -0.63 | -1.17 | -0.58 | -0.38 | -0.2 | 0.2125 | -0.0825 | 0.1375 | 0.689746 | 0.582356 | 0.879926 | 0.989223 | 0.997988 | 0.996159 |
| Atg2b_Q80XK6_S1581s_1 | Atg2b_Q8C | -2.63 | -0.99 | -0.76 | -0.97 | -0.85 | -3.17 | -0.8 | -1.98 | -1.37 | -2.15 | -2.98 | -0.6 | -0.14 | -1.23 | -3.83 | -0.11 | -0.3625 | -0.4375 | 0.01 | 0.990683 | 0.568797 | 0.481568 | 0.989223 | 0.997988 | 1 |
| Atg2b_Q80XK6_S1741s_1 | Atg2b_Q8C | -0.25 | -0.55 | -0.85 | -0.61 | 0.03 | 0.25 | -0.09 | -0.3 | -0.39 | -0.59 | 0.04 | -0.88 | -1.03 | -0.36 | -0.37 | -0.94 | 0.5375 | 0.11 | -0.11 | 0.704094 | 0.044681 | 0.69603 | 0.989223 | 0.997988 | 0.996159 |
| Atg2b_Q80XK6_S239s_1 | Atg2b_Q8C | -0.03 | 0.45 | -1.04 | -1.29 | 0.64 | 0.04 | -0.34 | -0.7 | 0.29 | 0.47 | -1.42 | -0.82 | -0.32 | 0.58 | -0.68 | -1.78 | 0.3875 | 0.1075 | -0.0725 | 0.901371 | 0.414148 | 0.848946 | 0.989223 | 0.997988 | 0.999507 |
| Atg2b_Q80XK6_S240s_1 | Atg2b_Q8C | -0.03 | 0.45 | -1.04 | -1.29 | 0.64 | 0.04 | -0.34 | -0.7 | 0.29 | 0.47 | -1.42 | -0.82 | -0.32 | 0.58 | -0.68 | -1.78 | 0.3875 | 0.1075 | -0.0725 | 0.901371 | 0.414148 | 0.848946 | 0.989223 | 0.997988 | 0.999507 |
| Atg2b_Q80XK6_S255s_1 | Atg2b_Q8C | 0.26 | -0.41 | -0.09 | -0.02 | 0.24 | 0.31 | 0.42 | 0.2 | -0.78 | -0.49 | -0.23 | 0.3 | -1.33 | 0.37 | 0.26 | 0.04 | 0.3575 | -0.235 | -0.1 | 0.810705 | 0.1381 | 0.447862 | 0.989223 | 0.997988 | 0.999507 |
| Atg2b_Q80XK6_S379s_1 | Atg2b_Q8C | -0.82 | -1.03 | -0.19 | -0.62 | -0.66 | -0.62 | -0.11 | -0.14 | -0.88 | -1.48 | 0.72 | 0.05 | -1.14 | -0.53 | -0.41 | -0.33 | 0.2825 | 0.2675 | 0.0625 | 0.840478 | 0.31508 | 0.588688 | 0.989223 | 0.997988 | 0.999507 |
| Atg2b_Q80XK6_S885s_1 | Atg2b_Q8C | -1.51 | -1.13 | -3.11 | -3.28 | 0.04 | -2.06 | -3.25 | -4.09 | -1.08 | -1.23 | -0.92 | -2.9 | -1.2 | -0.24 | -3.01 | -2.33 | -0.0825 | 0.725 | 0.5625 | 0.449137 | 0.927825 | 0.281327 | 0.997447 | 0.997988 | 0.994666 |
| Atg9a_Q68FE2_S735s_1 | Atg9a_Q68 | 0.29 | -0.03 | 0.91 | 0.89 | -1.64 | 0.51 | -0.64 | 0.24 | 1.47 | 0.13 | 1.23 | 0.56 | 0.99 | -1.09 | 0.95 | 0.79 | -0.8975 | 0.3325 | -0.105 | 0.839518 | 0.094901 | 0.399685 | 0.989223 | 0.997988 | 0.999507 |
| Atg9a_Q68FE2_S735s_2 | Atg9a_Q68 | 0.82 | 1.26 | 1.61 | 1.45 | -1.18 | 1.16 | 0.38 | 0.3 | 1.61 | 0.95 | 2.16 | 0.94 | 0.7 | -0.38 | 1.36 | 1.65 | -1.12 | 0.13 | -0.4525 | 0.342444 | 0.039548 | 0.714087 | 0.989223 | 0.997988 | 0.994666 |
| Atg9a_Q68FE2_S738s_2 | Atg9a_Q68 | 0.66 | 1.26 | 1.53 | 1.54 | -1.38 | 1.14 | 0.56 | -0.04 | 1.63 | 1.04 | 2.39 | 0.94 | 0.41 | -0.2 | 1.16 | 1.53 | -1.1775 | 0.2525 | -0.5225 | 0.245015 | 0.04703 | 0.524457 | 0.989223 | 0.997988 | 0.994666 |
| Atg9a_Q68FE2_S741s_2 | Atg9a_Q68 | 1.18 | 1.26 | 1.81 | 1.19 | -0.74 | 1.21 | -0.16 | 1.03 | 1.57 | 0.72 | 1.51 | 0.91 | 1.3 | -0.9 | 1.84 | 1.93 | -1.025 | -0.1825 | -0.3175 | 0.609454 | 0.047692 | 0.547679 | 0.989223 | 0.997988 | 0.995259 |
| Atg9a_Q68FE2_S761s_1 | Atg9a_Q68 | -0.42 | -0.81 | -0.96 | -0.93 | -0.3 | -0.44 | -1.01 | -0.64 | -1.08 | -0.61 | 0.36 | -1.17 | -0.32 | 0.43 | -0.79 | -0.52 | 0.1825 | 0.155 | 0.48 | 0.169534 | 0.476382 | 0.682344 | 0.989223 | 0.997988 | 0.994666 |
| Atl1_Q8BH66_S10s_1 | Atl1_Q8BH | -0.42 | -1.26 | -1.09 | -1.04 | 0.2 | -0.65 | -0.65 | -0.76 | -2.09 | -0.89 | -1.2 | -0.92 | -2.89 | -0.81 | -1.31 | -1.38 | 0.4875 | -0.3225 | -0.645 | 0.189418 | 0.139276 | 0.369059 | 0.989223 | 0.997988 | 0.994666 |
| Atl1_Q8BH66_S15s_1 | Atl1_Q8BH | 0.94 | 1.28 | -0.96 | -1.22 | 1.79 | 1.58 | -0.4 | -0.93 | -0.82 | 0.82 | -1.25 | -0.67 | -0.49 | 0.66 | -0.59 | -0.57 | 0.5 | -0.49 | -0.2575 | 0.68905 | 0.546269 | 0.493487 | 0.989223 | 0.997988 | 0.996159 |
| Atl1_Q8BH66_S18s_1 | Atl1_Q8BH | 1.73 | -0.01 | 0.66 | -0.06 | 1.66 | 1.86 | 0.53 | 0.56 | -1.75 | 0.57 | -0.48 | 0 | -2.38 | -0.64 | 0.35 | 0.88 | 0.5725 | -0.995 | -1.0275 | 0.186075 | 0.269724 | 0.119716 | 0.989223 | 0.997988 | 0.994666 |
| Atn1_O35126_S646s_1 | Atn1_O351 | -3.04 | -0.84 | -0.83 | -0.47 | -1.52 | -1.72 | -0.42 | -1.34 | -2.64 | -1.58 | -0.32 | -0.81 | -0.75 | -1.31 | -0.77 | -1.18 | 0.045 | -0.0425 | 0.2925 | 0.603268 | 0.938804 | 0.9515 | 0.999187 | 0.997988 | 0.995259 |
| Atn1_O35126_T654t_1 | Atn1_O351 | -2.22 | -0.88 | -2.89 | -3.3 | -2.27 | -3.71 | -4.4 | -3.04 | -2.29 | -2.05 | -1.07 | -3.2 | -0.18 | -1.58 | -2.31 | -2.88 | -1.0325 | 0.17 | 0.585 | 0.415066 | 0.122566 | 0.786352 | 0.989223 | 0.997988 | 0.994666 |
| Atp11c_F6Q8D3_S1105s_1 | Atp11c_F6 | -0.76 | 0 | 0.56 | -0.17 | -1.51 | -0.74 | -1.05 | -0.15 | 0.57 | -0.39 | -0.05 | -0.18 | 1.49 | -1.47 | -0.48 | -0.21 | -0.77 | 0.08 | -0.075 | 0.902418 | 0.069347 | 0.821716 | 0.989223 | 0.997988 | 0.999507 |
| Atp13a1_Q9EPE9_S22s_1 | Atp13a1_Q | 0.63 | 0.12 | 1.05 | 0.43 | -0.03 | 0.8 | 0.67 | 0.68 | 2.26 | 0.52 | -1.15 | 0.85 | 2.7 | -1.81 | 2.96 | 1.72 | -0.0275 | 0.0625 | 0.835 | 0.400784 | 0.926273 | 0.924397 | 0.997335 | 0.997988 | 0.994666 |
| Atp13a1_Q9EPE9_S896s_1 | Atp13a1_Q | 1.29 | 0.8 | 0.88 | 0.14 | 1.03 | 0.79 | 1.03 | 0.68 | 1.46 | 1.82 | 0.52 | 1.14 | 1.1 | 1.97 | 1.69 | 1.73 | 0.105 | 0.4575 | 0.845 | 0.030229 | 0.714453 | 0.237949 | 0.995437 | 0.997988 | 0.994666 |
| Atp1a1_Q8VDN2_S16s_1 | Atp1a1_Q8 | 0.61 | 0.5 | -0.28 | -0.54 | 1.18 | 0.45 | -0.56 | -0.78 | -0.55 | -0.11 | 0.06 | -0.74 | -0.45 | 0.76 | -0.6 | -0.08 | -3E-16 | -0.4075 | -0.165 | 0.693039 | 1 | 0.267339 | 1 | 0.997988 | 0.996159 |
| Atp1a1_Q8VDN2_S207s_1 | Atp1a1_Q8 | 1.34 | 1.43 | 1.05 | 1.21 | 1.1 | 1.25 | 0.55 | 1.26 | 0.54 | 1.12 | 1 | 0.91 | 1.42 | 0.92 | 0.91 | 0.93 | -0.2175 | -0.365 | -0.2125 | 0.414737 | 0.387515 | 0.149419 | 0.989223 | 0.997988 | 0.994666 |
| Atp1a1_Q8VDN2_S228s_1 | Atp1a1_Q8 | 0.41 | 0.77 | 1.09 | 0.67 | 1.07 | 0.2 | 0.86 | 0.55 | -0.04 | 0.43 | -0.02 | 0.49 | -0.41 | 0.85 | 0.4 | 0.69 | -0.065 | -0.52 | -0.3525 | 0.321442 | 0.813289 | 0.073793 | 0.995762 | 0.997988 | 0.994666 |
| Atp1a1_Q8VDN2_S40s_1 | Atp1a1_Q8 | 1.23 | 1.32 | 2.21 | 1.51 | 0.62 | 1.25 | 0.35 | 0.6 | 1.65 | 0.58 | 0.98 | 1.17 | 1.38 | -0.05 | 1 | 0.97 | -0.8625 | -0.4725 | -0.7425 | 0.081581 | 0.01948 | 0.182433 | 0.989223 | 0.997988 | 0.994666 |
| Atp1a1_Q8VDN2_T485t_1 | Atp1a1_Q8 | 0.63 | 1.62 | 1.01 | 0.37 | 0.97 | 0.91 | 0.37 | 1.05 | -0.01 | 0.19 | 0.09 | 0.27 | 0.37 | 0.84 | 0.77 | 1.26 | -0.0825 | -0.7725 | -0.0975 | 0.783543 | 0.799685 | 0.031958 | 0.995762 | 0.997988 | 0.998491 |
| Atp1a1_Q8VDN2_Y684y_1 | Atp1a1_Q8 | 4.36 | 2.41 | 3.71 | 1.18 | 2.49 | 3.99 | 3.69 | 3.24 | -0.02 | 3.86 | 0.68 | 3.56 | -0.76 | 0.02 | 3.1 | 3.52 | 0.4375 | -0.895 | -1.445 | 0.215789 | 0.528604 | 0.411876 | 0.989223 | 0.997988 | 0.994666 |
| Atp1a2_Q6PIE5_S10s_1 | Atp1a2_Q6 | -0.04 | 1.22 | 1.24 | 0.94 | 1.19 | 1.08 | 0.48 | 0.81 | 0.49 | 0.74 | 1.87 | 0.68 | 1.45 | 1.5 | 0.93 | 1.19 | 0.05 | 0.105 | 0.4275 | 0.247523 | 0.88457 | 0.804695 | 0.996493 | 0.997988 | 0.994666 |
| Atp1a2_Q6PIE5_S439s_1 | Atp1a2_Q6 | 1.29 | 1.55 | 2.14 | 1.72 | 1.78 | 1.39 | 0.77 | 1.65 | 0.88 | 1.11 | 1.5 | 1.92 | 1.42 | 1.76 | 1.54 | 1.7 | -0.2775 | -0.3225 | -0.07 | 0.800176 | 0.377305 | 0.326256 | 0.989223 | 0.997988 | 0.999507 |
| Atp1a2_Q6PIE5_S464s_1 | Atp1a2_Q6 | -1.13 | -1.58 | -2.17 | -1.79 | -0.63 | -1.4 | -2.01 | -2.29 | -2.13 | -1.06 | -1.53 | -2.12 | -1.57 | -1.47 | -2.66 | -2.72 | 0.085 | -0.0425 | -0.4375 | 0.29645 | 0.835131 | 0.903812 | 0.995762 | 0.997988 | 0.994666 |
| Atp1a2_Q6PIE5_S496s_1 | Atp1a2_Q6 | 0.26 | 0.64 | 0.42 | 0.38 | 0.56 | 0.15 | 0.68 | 0.3 | 0.04 | 0.44 | 0.13 | 0.6 | -0.03 | 0.74 | 0.34 | 0.66 | -0.0025 | -0.1225 | 0.0025 | 0.992747 | 0.991134 | 0.610947 | 0.999187 | 0.997988 | 1 |
| Atp1a2_Q6PIE5_S58s_1 | Atp1a2_Q6 | 0.76 | 0.76 | 1.14 | 0.37 | 0.64 | 0.05 | -0.49 | 0.58 | -0.84 | 0.39 | -0.24 | 0.5 | 0.13 | -0.14 | 0.4 | 0.31 | -0.5625 | -0.805 | -0.5825 | 0.059441 | 0.105629 | 0.047865 | 0.989223 | 0.997988 | 0.994666 |

| | | | | | | | | | | | | | | | | | | | | | | | | | | |
|---|---|---|---|---|---|---|---|---|---|---|---|---|---|---|---|---|---|---|---|---|---|---|---|---|---|---|
| Atp1a3_Q6PIC6_S206s_1 | Atp1a3_Q6 | -0.05 | 1.32 | 0.43 | 0.62 | 1.18 | 0.66 | 0.61 | 0.84 | 0.9 | 0.85 | 1.58 | 0 | 1.17 | 0.37 | 0.8 | -0.1 | 0.2425 | 0.2525 | -0.02 | 0.960171 | 0.461899 | 0.552997 | 0.989223 | 0.997988 | 0.999507 |
| Atp1a3_Q6PIC6_S207s_1 | Atp1a3_Q6 | 1.57 | 0.79 | 1.04 | 0.86 | 1.26 | 0.65 | 0.32 | 0.35 | 0.45 | 1.09 | 0.92 | 0.41 | 1.11 | 1.83 | 0.53 | 0.59 | -0.42 | -0.3475 | -0.05 | 0.892243 | 0.18824 | 0.249646 | 0.989223 | 0.997988 | 0.999507 |
| Atp1a3_Q6PIC6_S212s_1 | Atp1a3_Q6 | 1.82 | 1.76 | 1.38 | 0.33 | 1.89 | 1.35 | 0.47 | 0.04 | 0.95 | 1.52 | 0.85 | 0.47 | 1.08 | 1.77 | 0.03 | 0.44 | -0.385 | -0.375 | -0.4925 | 0.326789 | 0.44626 | 0.364851 | 0.989223 | 0.997988 | 0.994666 |
| Atp1a3_Q6PIC6_S218s_1 | Atp1a3_Q6 | 0.6 | 0.81 | 0.94 | 1.05 | 0.68 | 0.52 | 0.77 | 0.65 | 0.68 | 0.72 | 1.23 | 0.41 | -0.03 | 0.44 | 1.18 | 0.79 | -0.195 | -0.09 | -0.255 | 0.435422 | 0.366059 | 0.731423 | 0.989223 | 0.997988 | 0.994666 |
| Atp1a3_Q6PIC6_S37s_1 | Atp1a3_Q6 | -1.01 | -0.95 | -1.22 | -1.61 | -1.08 | -0.83 | -1.61 | -1.4 | -1.42 | -1.35 | -0.29 | -1.58 | -0.47 | -0.66 | -1.38 | -1.29 | -0.0325 | 0.0375 | 0.2475 | 0.445397 | 0.904152 | 0.913793 | 0.996493 | 0.997988 | 0.994666 |
| Atp1a3_Q6PIC6_S442s_1 | Atp1a3_Q6 | 0.68 | 1.1 | 1.49 | 0.93 | 1.28 | 0.53 | 0.68 | 1.08 | 0.64 | 1.25 | 1.25 | 0.76 | 0.54 | 0.79 | 0.46 | 1.82 | -0.1575 | -0.075 | -0.1475 | 0.694786 | 0.576112 | 0.791516 | 0.989223 | 0.997988 | 0.996159 |
| Atp1a3_Q6PIC6_S474s_1 | Atp1a3_Q6 | 0.47 | 0.9 | 0.2 | -0.55 | 0.94 | 0.18 | 0.26 | -0.92 | -0.72 | 0.32 | -1.03 | -0.23 | 0.25 | 0.91 | 0.19 | -0.53 | -0.14 | -0.67 | -0.05 | 0.905508 | 0.759798 | 0.13358 | 0.995762 | 0.997988 | 0.999507 |
| Atp1a3_Q6PIC6_S56s_1 | Atp1a3_Q6 | -1.13 | -1.21 | -0.75 | -0.54 | -1.55 | -1 | -0.88 | -0.68 | -1.76 | -1.16 | -1.26 | -1.8 | -1.42 | -1.26 | -0.78 | -0.8 | -0.12 | -0.5875 | -0.1575 | 0.594007 | 0.669866 | 0.060646 | 0.992876 | 0.997988 | 0.995259 |
| Atp1a3_Q6PIC6_S643s_1 | Atp1a3_Q6 | 0.09 | 0.4 | 0.14 | -0.26 | 0.98 | -0.4 | 0.42 | -0.1 | -0.37 | -0.68 | -1.25 | -0.33 | 0.28 | 1 | -0.25 | 0.06 | 0.1325 | -0.75 | 0.18 | 0.594916 | 0.696833 | 0.028672 | 0.994893 | 0.997988 | 0.995259 |
| Atp1a3_Q6PIC6_S712s_1 | Atp1a3_Q6 | 0.5 | 0.86 | -0.11 | -0.06 | 0.95 | -0.33 | 0.77 | -0.01 | 0.07 | 0.35 | -0.4 | 0.16 | -0.19 | 0.66 | 0.07 | 0.06 | 0.0475 | -0.2525 | -0.1475 | 0.659925 | 0.899693 | 0.430002 | 0.996493 | 0.997988 | 0.995259 |
| Atp1a3_Q6PIC6_T368t_1 | Atp1a3_Q6 | -2.9 | -1.1 | -1.39 | -2.63 | -1.38 | -2.47 | -2.11 | -2.58 | -1.52 | -2.82 | -2.78 | -2.51 | 0.56 | -2.37 | -2.1 | -1.66 | -0.13 | -0.4025 | 0.6125 | 0.401792 | 0.787359 | 0.435342 | 0.995762 | 0.997988 | 0.994666 |
| Atp1a3_Q6PIC6_T370t_1 | Atp1a3_Q6 | -0.06 | -0.22 | 0.64 | 1.22 | -0.73 | 0.56 | -0.23 | 0.95 | 0.98 | -0.03 | 0.73 | 0.83 | 1.56 | -1 | 1.43 | 1.02 | -0.2575 | 0.2325 | 0.3575 | 0.56722 | 0.58482 | 0.563954 | 0.989223 | 0.997988 | 0.995259 |
| Atp1a3_Q6PIC6_T71t_1 | Atp1a3_Q6 | 0.63 | 0.8 | 1.37 | 1.74 | 1.06 | 0.33 | 1.03 | 0.84 | -1.09 | 0.22 | -1.8 | 1.72 | -0.46 | 0 | 1.87 | 1.65 | -0.32 | -1.3725 | -0.37 | 0.531055 | 0.332181 | 0.08764 | 0.989223 | 0.997988 | 0.994666 |
| Atp2a2_O55143_S553s_1 | Atp2a2_O5 | 0.66 | 0.47 | 0.48 | 1.12 | 0.99 | 0.21 | -0.57 | 0.07 | 0.19 | 0.56 | 1.46 | 0.27 | -0.35 | 1.57 | 0.72 | 1.1 | -0.5075 | -0.0625 | 0.0775 | 0.857192 | 0.175612 | 0.856567 | 0.989223 | 0.997988 | 0.999507 |
| Atp2a2_O55143_S661s_1 | Atp2a2_O5 | 1.29 | 0.87 | 1.18 | 0.75 | 1.18 | 1.37 | 1.58 | 1.3 | -0.41 | 1.34 | 0.02 | 1.18 | 0.36 | -0.24 | 2.32 | 1.79 | 0.335 | -0.49 | 0.035 | 0.950501 | 0.166608 | 0.277453 | 0.989223 | 0.997988 | 0.999507 |
| Atp2a2_O55143_S663s_1 | Atp2a2_O5 | 1.48 | 1 | 2.16 | 0.18 | 1.64 | 1.15 | 1.82 | 1.77 | -1.75 | 1.17 | -2.69 | 1.23 | -0.19 | -1.51 | 1.98 | 2.27 | 0.39 | -1.715 | -0.5675 | 0.519817 | 0.365416 | 0.10128 | 0.989223 | 0.997988 | 0.994666 |
| Atp2b1_G5E829_S1139s_1 | Atp2b1_G5 | 0.77 | 0.2 | 0.46 | 0.74 | -0.06 | 0.69 | 0.91 | 0.77 | 0.57 | 0.46 | 0.87 | 0.6 | -0.54 | 0.53 | 0.19 | 0.42 | 0.035 | 0.0825 | -0.3925 | 0.241057 | 0.902886 | 0.734072 | 0.996493 | 0.997988 | 0.994666 |
| Atp2b1_G5E829_S1140s_1 | Atp2b1_G5 | 0.77 | 0.2 | 0.46 | 0.74 | -0.06 | 0.69 | 0.91 | 0.77 | 0.57 | 0.46 | 0.87 | 0.6 | -0.54 | 0.53 | 0.19 | 0.42 | 0.035 | 0.0825 | -0.3925 | 0.241057 | 0.902886 | 0.734072 | 0.996493 | 0.997988 | 0.994666 |
| Atp2b1_G5E829_S1177s_1 | Atp2b1_G5 | 0.21 | 0.87 | 1.17 | 1.06 | -0.07 | 0.12 | -0.53 | 0.17 | 1.88 | 0.91 | -0.16 | 0.68 | 2.27 | -0.39 | 0.46 | 0.63 | -0.905 | -3.1E-16 | -0.085 | 0.87755 | 0.01175 | 1 | 0.989223 | 1 | 0.999507 |
| Atp2b1_G5E829_S1178s_1 | Atp2b1_G5 | 0.5 | 0.92 | 1.2 | 1.06 | -0.12 | 0.58 | 0.19 | 0 | 2.34 | 1.15 | -0.52 | 0.81 | 2.49 | -0.43 | 0.81 | 0.73 | -0.7575 | 0.025 | -0.02 | 0.972 | 0.015885 | 0.964436 | 0.989223 | 0.997988 | 0.999507 |
| Atp2b1_G5E829_S1178s_2 | Atp2b1_G5 | 0.7 | -1.27 | 0.55 | 0.82 | 0.15 | 0.99 | 0.14 | 0.95 | 1.73 | 0.12 | 0.14 | 0.52 | -0.51 | -1 | -0.01 | 0.22 | 0.3575 | 0.4275 | -0.525 | 0.331649 | 0.482004 | 0.462641 | 0.989223 | 0.997988 | 0.994666 |
| Atp2b1_G5E829_S1182s_2 | Atp2b1_G5 | 0.7 | -1.27 | 0.55 | 0.82 | 0.15 | 0.99 | 0.14 | 0.95 | 1.73 | 0.12 | 0.14 | 0.52 | -0.51 | -1 | -0.01 | 0.22 | 0.3575 | 0.4275 | -0.525 | 0.331649 | 0.482004 | 0.462641 | 0.989223 | 0.997988 | 0.994666 |
| Atp2b1_G5E829_S17s_1 | Atp2b1_G5 | 1.79 | 0.72 | 1.27 | 1.93 | 0.95 | 1.82 | 1.39 | 1.97 | 1.38 | 1.46 | 1.6 | 1.63 | 1.26 | 1.06 | 0.94 | 0.8 | 0.105 | 0.09 | -0.4125 | 0.233207 | 0.769441 | 0.774526 | 0.995762 | 0.997988 | 0.994666 |
| Atp2b1_G5E829_S338s_1 | Atp2b1_G5 | 0.5 | 0.47 | 1.72 | 0.52 | 0.63 | 0.38 | 1.28 | 0.74 | -0.26 | 0.49 | 0.23 | 1.2 | -0.17 | -0.26 | 1.05 | 1.23 | -0.045 | -0.3875 | -0.34 | 0.481952 | 0.900126 | 0.370161 | 0.996493 | 0.997988 | 0.994666 |
| Atp2b2_Q3UHH0_S1100s_1 | Atp2b2_Q3 | 1.3 | 1.47 | 1.09 | 1.37 | 1.05 | 1.17 | 0.49 | 1.15 | 0.51 | 0.74 | 0.56 | 1.08 | 0.19 | 1.38 | 1.04 | 0.94 | -0.3425 | -0.585 | -0.42 | 0.202777 | 0.180496 | 0.033492 | 0.989223 | 0.997988 | 0.994666 |
| Atp2b2_Q3UHH0_S1101s_1 | Atp2b2_Q3 | 1.3 | 1.47 | 1.31 | 1.16 | 1.05 | 1.17 | 1.03 | 0.93 | 0.51 | 0.74 | 0.84 | 1.24 | 0.19 | 1.38 | 0.86 | 0.85 | -0.265 | -0.4775 | -0.49 | 0.135854 | 0.208327 | 0.076848 | 0.989223 | 0.997988 | 0.994666 |
| Atp2b2_Q3UHH0_S1111s_1 | Atp2b2_Q3 | 0.1 | 0.16 | 0.04 | -0.2 | 0.18 | 0.22 | 0.41 | -0.19 | -0.77 | 0.1 | -0.95 | 0.3 | -1.28 | 0.38 | -0.3 | -0.23 | 0.13 | -0.355 | -0.3825 | 0.316978 | 0.570166 | 0.311724 | 0.989223 | 0.997988 | 0.994666 |
| Atp2b2_Q3UHH0_S1114s_1 | Atp2b2_Q3 | 1.31 | 1.47 | 1.46 | 0.96 | 1.8 | 1.36 | 1.23 | 1.26 | -0.11 | 0.77 | 0.21 | 1.42 | -0.83 | 0.76 | 0.62 | 1.42 | 0.1125 | -0.7275 | -0.8075 | 0.11118 | 0.643449 | 0.072282 | 0.990258 | 0.997988 | 0.994666 |
| Atp2b2_Q3UHH0_S1151s_1 | Atp2b2_Q3 | -0.66 | 1.13 | 0.26 | 0.42 | 0.11 | 0.78 | -0.13 | -0.51 | 0.55 | 0.92 | 0.61 | 0.4 | 1.74 | -0.3 | -0.23 | 0.14 | -0.225 | 0.3325 | 0.05 | 0.928435 | 0.604336 | 0.399092 | 0.989223 | 0.997988 | 0.999507 |
| Atp2b2_Q3UHH0_S1152s_1 | Atp2b2_Q3 | 0.59 | 0.74 | 0.85 | 0.98 | 0.3 | 0.44 | 0.5 | 0.57 | 0.8 | 0.24 | 0.57 | 0.97 | 1.15 | 0.12 | 0.84 | 0.47 | -0.3375 | -0.145 | -0.145 | 0.630663 | 0.128857 | 0.568007 | 0.989223 | 0.997988 | 0.995259 |
| Atp2b2_Q3UHH0_S1156s_1 | Atp2b2_Q3 | 2.11 | 0.36 | 0.15 | -0.35 | 0.34 | 3.31 | -0.22 | 0.05 | 2.58 | 1.98 | -0.33 | 0.18 | 0.77 | -0.11 | 0.03 | 0.38 | 0.3025 | 0.535 | -0.3 | 0.575905 | 0.72374 | 0.50196 | 0.995437 | 0.997988 | 0.995259 |
| Atp2b2_Q3UHH0_S1162s_1 | Atp2b2_Q3 | 0.99 | -0.05 | 0.85 | 0.9 | 0.61 | 1.75 | 0.75 | 1.37 | -0.92 | 0.66 | 0.85 | 0.93 | 0.23 | 1.46 | -0.38 | 0.41 | 0.4475 | -0.2925 | -0.2425 | 0.58705 | 0.232364 | 0.542436 | 0.989223 | 0.997988 | 0.995259 |
| Atp2b2_Q3UHH0_S1175s_1 | Atp2b2_Q3 | 0.84 | 0.78 | 0.35 | 0.48 | 0.32 | 0.68 | 0.41 | 0.03 | -0.12 | 0.5 | -0.21 | 0.39 | -0.09 | 0.19 | 0 | 0.5 | -0.2525 | -0.4725 | -0.4625 | 0.108814 | 0.311025 | 0.107825 | 0.989223 | 0.997988 | 0.994666 |
| Atp2b2_Q3UHH0_S18s_1 | Atp2b2_Q3 | 0.99 | -0.03 | -1.13 | -0.45 | -0.27 | 1.43 | 0.23 | 0.13 | -0.48 | 0.42 | 0.37 | -0.71 | -1.44 | -0.58 | -1.02 | -0.4 | 0.535 | 0.055 | -0.705 | 0.164772 | 0.32064 | 0.912206 | 0.989223 | 0.997988 | 0.994666 |
| Atp2b2_Q3UHH0_S19s_1 | Atp2b2_Q3 | 1.89 | -0.41 | -0.36 | 0.03 | 0.57 | 2.79 | 0.4 | 0.88 | -0.68 | 0.25 | 0.63 | -0.21 | -0.4 | -0.21 | -0.46 | 0.12 | 0.8725 | -0.29 | -0.525 | 0.328686 | 0.222236 | 0.610648 | 0.989223 | 0.997988 | 0.994666 |
| Atp2b2_Q3UHH0_T1113t_1 | Atp2b2_Q3 | 1.2 | 1.32 | 1.37 | 1.04 | 1.59 | 1.38 | 1.1 | 1.15 | 0 | 0.66 | 0.39 | 1.37 | -0.47 | 0.65 | 0.73 | 1.32 | 0.0725 | -0.6275 | -0.675 | 0.109749 | 0.743284 | 0.07906 | 0.995762 | 0.997988 | 0.994666 |
| Atp2b2_S4R1C4_S1107s_1 | Atp2b2_S4 | 1.7 | 1.77 | 1.58 | 1.27 | 1.62 | 1.62 | 1.23 | 1.38 | -0.07 | 1.17 | -0.87 | 0.98 | 0.69 | 0.81 | 1.39 | 1.66 | -0.1175 | -1.2775 | -0.4425 | 0.176755 | 0.605846 | 0.021128 | 0.989223 | 0.997988 | 0.994666 |
| Atp2b2_S4R1C4_S1116s_2 | Atp2b2_S4 | 0.32 | 1.43 | 1.23 | 0.48 | 0.65 | 0.52 | 0.51 | -0.33 | 0.74 | 0.92 | 0.31 | 1.28 | -0.86 | 0.77 | 0.42 | 0.93 | -0.5275 | -0.0525 | -0.55 | 0.259763 | 0.160391 | 0.882003 | 0.989223 | 0.997988 | 0.994666 |
| Atp2b2_S4R1C4_S1117s_2 | Atp2b2_S4 | 0.22 | 1.04 | -0.79 | -2.53 | 0.62 | 0.32 | 0.02 | -0.08 | 0.46 | 0.81 | -0.62 | -0.55 | -0.75 | 0.65 | -1.42 | -0.96 | 0.735 | 0.54 | -0.105 | 0.89369 | 0.30342 | 0.483845 | 0.989223 | 0.997988 | 0.999507 |
| Atp2b2_S4R1C4_S1130s_1 | Atp2b2_S4 | -0.85 | 1.19 | 1.18 | 0.77 | 0.57 | -0.59 | 0.87 | 0.92 | 0.66 | 1.16 | 0.25 | 0.65 | 1.07 | 0.5 | 1.44 | 1.86 | -0.13 | 0.1075 | 0.645 | 0.239647 | 0.810654 | 0.826485 | 0.995762 | 0.997988 | 0.994666 |
| Atp2b2_S4R1C4_S1130s_2 | Atp2b2_S4 | 0.03 | 0.2 | -0.79 | -2.53 | 0.55 | -0.05 | 0.02 | -0.08 | -0.08 | 0.6 | -0.62 | -0.55 | -0.57 | 0.43 | -1.42 | -0.96 | 0.8825 | 0.61 | 0.1425 | 0.830207 | 0.150052 | 0.341671 | 0.989223 | 0.997988 | 0.999507 |
| Atp2b2_S4R1C4_S1135s_1 | Atp2b2_S4 | -0.55 | 0.3 | -1.11 | 0.07 | 0.69 | -0.19 | -1.55 | -0.13 | 0.66 | 0.35 | -1.51 | -2.84 | 0.26 | 0.61 | 1.28 | 0.12 | 0.0275 | -0.5125 | 0.89 | 0.05352 | 0.95723 | 0.520336 | 0.999187 | 0.997988 | 0.994666 |
| Atp2b2_S4R1C4_S1142s_1 | Atp2b2_S4 | 0 | 0.7 | 0.94 | 0.8 | 0.98 | 0.8 | 0.89 | 0.39 | 0.76 | 0.92 | 0.8 | 0.29 | 0.71 | 0.76 | 1.13 | 0.54 | 0.155 | 0.0825 | 0.175 | 0.566557 | 0.584822 | 0.779202 | 0.989223 | 0.997988 | 0.995259 |
| Atp2b3_A2ALL9_S1124s_1 | Atp2b3_A2 | 0.99 | 0.75 | 0.5 | 0.51 | 1.04 | 0.97 | 0.58 | 0.09 | 0.65 | 1.08 | 0.27 | 0.47 | 1.01 | 0.97 | 0.88 | 0.61 | -0.0175 | -0.07 | 0.18 | 0.48532 | 0.95047 | 0.79419 | 0.999187 | 0.997988 | 0.994666 |
| Atp2b3_A2ALL9_S1126s_1 | Atp2b3_A2 | 0.54 | 0.55 | 2.15 | 1.7 | 0.91 | 0.51 | 1.34 | 1.46 | 0.27 | 0.52 | 2.33 | 1.87 | 0.62 | 0.53 | 1.56 | 1.66 | -0.18 | 0.0125 | -0.1425 | 0.768584 | 0.680552 | 0.983218 | 0.993464 | 0.997988 | 0.9971 |
| Atp2b3_Q8C048_S315s_1 | Atp2b3_Q8 | 2.2 | 0.41 | 0.89 | -0.13 | -0.64 | 2.47 | 0.04 | 0.33 | 0.47 | 0.85 | 0.05 | 0.46 | 2.02 | -3.48 | 0.84 | 0.17 | -0.2925 | -0.385 | -0.955 | 0.400808 | 0.691524 | 0.444086 | 0.994893 | 0.997988 | 0.994666 |
| Atp2b3_Q8C048_S316s_1 | Atp2b3_Q8 | 2.06 | 0.26 | 1.3 | 0.21 | -0.52 | 2.93 | 0.38 | 1.14 | 2 | 0.54 | -0.34 | 0.59 | 2.46 | -2.04 | 0.58 | 1.48 | 0.025 | -0.26 | -0.3375 | 0.717368 | 0.973445 | 0.6673 | 0.999187 | 0.997988 | 0.996966 |
| Atp2b3_Q8C048_S336s_1 | Atp2b3_Q8 | -0.13 | 0.11 | 0.73 | -0.12 | -0.43 | 0.46 | -0.72 | -1.04 | 0.6 | -0.32 | -1.15 | 0.51 | 0.46 | -1.02 | -0.05 | -0.67 | -0.58 | -0.2375 | -0.4675 | 0.253669 | 0.145128 | 0.593444 | 0.989223 | 0.997988 | 0.994666 |
| Atp2b3_Q8C048_S578s_1 | Atp2b3_Q8 | -2.65 | -2.1 | -0.95 | -2.5 | -1.2 | -2.27 | -1.01 | -2.31 | -2.83 | -2.67 | -1.99 | -1.27 | -1.01 | -1.17 | -2.23 | -1.41 | 0.3525 | -0.14 | 0.595 | 0.212725 | 0.466436 | 0.7775 | 0.989223 | 0.997988 | 0.994666 |
| Atp2b3_Q8C048_S8s_1 | Atp2b3_Q8 | 2.33 | 1.63 | 0.64 | 0.85 | 1.75 | 2.36 | 1.05 | 0.88 | 1.06 | 2.23 | 0.81 | 0.8 | 0.2 | 1.28 | 0.86 | 1.25 | 0.1475 | -0.1375 | -0.465 | 0.314524 | 0.756992 | 0.778062 | 0.995762 | 0.997988 | 0.994666 |
| Atp2b3_Q8C048_T1079t_1 | Atp2b3_Q8 | -0.2 | 0.61 | 0.88 | 1.49 | 0.93 | 0.24 | 1.01 | 1.24 | 0.44 | 0.8 | 1.28 | 1.05 | -0.26 | 1.15 | 0.77 | 1.12 | 0.16 | 0.1975 | 2.36E-16 | 1 | 0.687715 | 0.618163 | 0.994672 | 0.997988 | 1 |
| Atp2b4_E9Q828_S1065s_1 | Atp2b4_E9 | 1.67 | 1.29 | 1.01 | 1.1 | 0.43 | 2.66 | 0 | 0.31 | 3 | 1.85 | 0.58 | 0.82 | 2.89 | 0.4 | 0.52 | 0.7 | -0.4175 | 0.295 | -0.14 | 0.803313 | 0.466494 | 0.582697 | 0.989223 | 0.997988 | 0.999507 |
| Atp2b4_E9Q828_S329s_1 | Atp2b4_E9 | 0.63 | 0.97 | 1.04 | 0.38 | 1.04 | 0.72 | 1.48 | 0.88 | -0.45 | 1.08 | -0.41 | 1.23 | -0.26 | -0.22 | 1.2 | 1.25 | 0.275 | -0.3925 | -0.2625 | 0.555308 | 0.318738 | 0.405756 | 0.989223 | 0.997988 | 0.995259 |
| Atp2c1_Q3UZR5_S944s_1 | Atp2c1_Q3 | -0.27 | 0.64 | 0.32 | -0.46 | 0.4 | -2.31 | 0.3 | -0.46 | -0.26 | 0.87 | 1.13 | -2.15 | -0.01 | 0.87 | -0.34 | 0.13 | -0.575 | -0.16 | 0.105 | 0.781233 | 0.354044 | 0.821882 | 0.989223 | 0.997988 | 0.998226 |
| Atp6v1a_P50516_S384s_1 | Atp6v1a_P | -0.27 | -0.06 | 0.13 | -0.08 | -0.01 | -0.56 | 0.02 | -0.33 | -0.01 | -0.38 | -0.84 | -0.11 | -0.1 | 0.4 | 0.08 | -0.07 | -0.15 | -0.265 | 0.1475 | 0.561785 | 0.524225 | 0.330925 | 0.989223 | 0.997988 | 0.995259 |
| Atp6v1a_P50516_S568s_1 | Atp6v1a_P | 0.68 | 0.59 | 0.05 | 0.34 | 0.8 | -0.5 | -0.19 | 0.73 | -0.24 | -0.03 | -0.56 | -0.02 | -0.64 | 0.94 | 0.09 | 0.55 | -0.205 | -0.6275 | -0.18 | 0.639301 | 0.568033 | 0.034458 | 0.989223 | 0.997988 | 0.995259 |
| Atp6v1b2_P62814_S498s_1 | Atp6v1b2_ | 0.33 | 0.22 | 0.51 | 0.26 | 0.65 | 0.13 | 0.67 | -0.04 | -0.41 | -0.06 | -0.48 | 0.13 | 0.14 | 0.41 | -0.01 | 0.31 | 0.0225 | -0.535 | -0.1175 | 0.628959 | 0.928044 | 0.049618 | 0.997447 | 0.997988 | 0.995259 |
| Atp6v1e1_P50518_T195t_1 | Atp6v1e1_ | -0.34 | 0.32 | 0.8 | 0.51 | -0.36 | -0.11 | -0.36 | -1.32 | 0.08 | -0.76 | 0.55 | -0.71 | 1.12 | 0.11 | 0.4 | 0.66 | -0.86 | -0.5325 | 0.25 | 0.483497 | 0.036319 | 0.202833 | 0.989223 | 0.997988 | 0.994666 |
| Atp6v1g2_Q9WTT4_S65s_1 | Atp6v1g2_ | 0.63 | 0.63 | 1.69 | 0.29 | 0.31 | -0.14 | 1.05 | 0.9 | 0.5 | 0.54 | -0.28 | -0.1 | -0.1 | 0.27 | 0.27 | 0.49 | -0.28 | -0.645 | -0.5775 | 0.128725 | 0.484686 | 0.1116 | 0.989223 | 0.997988 | 0.994666 |
| Atp6v1h_Q8BVE3_S483s_1 | Atp6v1h_Q | 0.84 | 0.59 | 1.7 | 2.26 | -1.13 | 0.29 | 1.45 | 2.3 | 1.8 | 1.44 | 2.13 | 2.18 | 1.82 | 0.42 | 1.82 | 1.13 | -0.62 | 0.54 | -0.05 | 0.918074 | 0.408643 | 0.214622 | 0.989223 | 0.997988 | 0.999507 |
| Atp6v1h_Q8BVE3_S58s_1 | Atp6v1h_Q | 0.82 | 0.65 | 0.39 | 0.44 | 0.81 | 1.36 | 0.55 | 0.09 | 0.84 | 0.95 | 1.09 | 0.33 | 0.74 | 0.61 | 0.75 | 0.78 | 0.1275 | 0.2275 | 0.145 | 0.548871 | 0.67711 | 0.39106 | 0.99315 | 0.997988 | 0.994666 |
| Atp6v1h_Q8BVE3_S59s_1 | Atp6v1h_Q | 0.82 | 0.65 | 0.52 | 0.57 | 0.81 | 1.36 | 0.62 | 0.14 | 0.84 | 0.95 | 1.29 | 0.32 | 0.74 | 0.61 | 0.95 | 0.81 | 0.0925 | 0.21 | 0.1375 | 0.564733 | 0.750632 | 0.444372 | 0.995762 | 0.997988 | 0.995259 |
| Atp7a_Q64430_S356s_1 | Atp7a_Q64 | 0.43 | 1.3 | 0.41 | 0.59 | 0.52 | -0.31 | 0.36 | 0.49 | -0.48 | 0.05 | -1.25 | -1.14 | -0.12 | 0.82 | -0.11 | 0.56 | -0.4175 | -1.3875 | -0.395 | 0.273353 | 0.195671 | 0.004551 | 0.989223 | 0.997988 | 0.994666 |
| Atp8a1_P70704_S1109s_1 | Atp8a1_P7 | -0.11 | 0.16 | -0.07 | 0.94 | 0.38 | -0.43 | 2.23 | 0.94 | -0.54 | -0.16 | 0.34 | 0.22 | -0.43 | 0.06 | 1.21 | 1.04 | 0.55 | -0.265 | 0.24 | 0.596468 | 0.331442 | 0.438839 | 0.989223 | 0.997988 | 0.995259 |
| Atp8a1_P70704_S1111s_1 | Atp8a1_P7 | 0.11 | 0.36 | -0.23 | 0.36 | 0.29 | 0.25 | 0.71 | 0.58 | 0.05 | 0.28 | -0.39 | 0.57 | -0.62 | 0.34 | 0.06 | 0.34 | 0.3075 | -0.0225 | -0.12 | 0.705121 | 0.224135 | 0.938299 | 0.989223 | 0.997988 | 0.996159 |
| Atp8a1_P70704_S29s_1 | Atp8a1_P7 | 0.78 | 0.9 | 2.89 | 2.95 | 0.19 | 0.71 | 2.14 | 3.23 | 0.36 | 1.24 | 2.88 | 3.47 | 0.24 | 0.03 | 3.03 | 3.28 | -0.3125 | 0.1075 | -0.235 | 0.800132 | 0.697057 | 0.89744 | 0.994893 | 0.997988 | 0.999507 |
| Atp8a1_P70704_T28t_1 | Atp8a1_P7 | -1.72 | -0.96 | 0.08 | 1.92 | -1.64 | -1.79 | 0.16 | 1.96 | -1.91 | -1.33 | 1.32 | 1.43 | -2.04 | -1.86 | 0.25 | 0.43 | -0.1575 | 0.0475 | -0.635 | 0.487243 | 0.87748 | 0.963419 | 0.996493 | 0.997988 | 0.994666 |
| Atp8a2_P98200_S421s_1 | Atp8a2_P9 | 0.38 | -0.44 | 0.46 | 0.18 | -0.75 | 0.66 | -0.15 | -0.1 | 0.28 | 0.11 | 0.82 | 0.18 | 0.24 | -0.58 | 0.49 | 0.42 | -0.23 | 0.2025 | -0.0025 | 0.994292 | 0.52062 | 0.50446 | 0.989223 | 0.997988 | 1 |
| Atp8a2_P98200_S422s_1 | Atp8a2_P9 | 0.67 | 0.61 | 0.46 | 0.18 | -0.6 | 0.81 | -0.15 | -0.1 | 1.13 | 0.98 | 0.82 | 0.18 | 1.11 | -0.3 | 0.49 | 0.42 | -0.49 | 0.2975 | -0.05 | 0.883897 | 0.157654 | 0.310542 | 0.989223 | 0.997988 | 0.999507 |
| Atp8a2_P98200_S435s_1 | Atp8a2_P9 | -0.12 | -0.44 | -0.38 | -1.01 | -0.2 | -0.12 | -0.53 | -0.55 | 0.62 | 0.2 | -1.24 | -0.77 | 0.34 | -0.21 | -0.51 | -0.38 | 0.1375 | 0.19 | 0.2975 | 0.357065 | 0.605134 | 0.674154 | 0.989223 | 0.997988 | 0.994666 |
| Atp8a2_P98200_S6s_1 | Atp8a2_P9 | 0.78 | 1.08 | 0.29 | 0.81 | 0.78 | 0.49 | 0.76 | 0.57 | 0.73 | 0.75 | 0.54 | 0.87 | 0.48 | 0.93 | 0.96 | -0.05 | -0.09 | -0.0175 | -0.16 | 0.629564 | 0.711355 | 0.944491 | 0.995437 | 0.997988 | 0.995259 |
| Atp8a2_P98200_T1108t_1 | Atp8a2_P9 | -0.75 | 0.7 | -0.23 | -0.03 | 0.04 | 0.28 | 0.03 | -0.01 | 0.07 | 0.4 | -0.23 | -0.92 | -1.21 | 0.13 | -0.37 | -0.61 | 0.1625 | -0.0925 | -0.4375 | 0.302275 | 0.615566 | 0.820217 | 0.989223 | 0.997988 | 0.994666 |
| Atrx_Q61687_S1290s_1 | Atrx_Q616 | -0.37 | -0.15 | 0.09 | 0.4 | -0.28 | -0.13 | -0.49 | -0.6 | -0.44 | 0.32 | 0.74 | -0.07 | 0.1 | 0.04 | 0.55 | 0.22 | -0.3675 | 0.145 | 0.235 | 0.409333 | 0.167099 | 0.660027 | 0.989223 | 0.997988 | 0.994666 |
| Atrx_Q61687_S315s_1 | Atrx_Q616 | 2.95 | 1.48 | 1.09 | 0.23 | 1.95 | 2.64 | 1.7 | 1.07 | -0.55 | 3.06 | -0.24 | 1.13 | 0.78 | -1.25 | 2.13 | 2.06 | 0.4025 | -0.5875 | -0.5075 | 0.556452 | 0.497853 | 0.512781 | 0.989223 | 0.997988 | 0.995259 |
| Atrx_Q61687_S590s_1 | Atrx_Q616 | -0.09 | 0.7 | 0.1 | -0.52 | 0.37 | -0.58 | 0.82 | 0.2 | -0.15 | 0.26 | 0.29 | -0.05 | -0.35 | 0.75 | 0.9 | 0.25 | 0.155 | 0.04 | 0.34 | 0.393119 | 0.682922 | 0.897353 | 0.994272 | 0.997988 | 0.994666 |
| Atrx_Q61687_S665s_1 | Atrx_Q616 | 1.87 | 1.4 | 0.65 | 0.45 | 1.34 | 1.26 | 0.1 | -0.23 | 0.89 | 1.48 | -0.68 | 0.64 | 1.77 | 1.12 | 0.28 | 0.56 | -0.475 | -0.51 | -0.16 | 0.724571 | 0.334198 | 0.344728 | 0.989223 | 0.997988 | 0.9971 |
| Atrx_Q61687_S801s_1 | Atrx_Q616 | 1.81 | 1.21 | 0.29 | -0.12 | 1.28 | 1.07 | -0.49 | -0.06 | 1.55 | 0.61 | -2.18 | 0.02 | 1.39 | 0.94 | 0.18 | -0.01 | -0.3475 | -0.7975 | -0.1725 | 0.737443 | 0.534437 | 0.336581 | 0.989223 | 0.997988 | 0.9971 |
| Atrx_Q61687_S912s_1 | Atrx_Q616 | 1.15 | 0.5 | 0.69 | 0.16 | -1 | 1.86 | -0.67 | -0.17 | 2.74 | 1.15 | -0.07 | 0.61 | 2.35 | -0.73 | 0.4 | -0.08 | -0.62 | 0.4825 | -0.14 | 0.82412 | 0.317288 | 0.416409 | 0.989223 | 0.997988 | 0.999507 |
| Atxn1_J3QPR1_S213s_1 | Atxn1_J3Q | -0.62 | -0.18 | -0.06 | 0.22 | -0.54 | -1.94 | 0.86 | 1.35 | -0.47 | -1.11 | 0.31 | -1.42 | 0.16 | -0.02 | -0.78 | -0.94 | 0.0925 | -0.5125 | -0.235 | 0.51013 | 0.890745 | 0.235422 | 0.996493 | 0.997988 | 0.994666 |
| Atxn1_J3QPR1_S759s_1 | Atxn1_J3Q | -1.32 | -3.31 | -0.94 | -2.32 | -2.39 | -2.06 | -1.05 | -1.47 | -3.24 | -1.35 | -0.27 | -0.97 | -1.38 | -3 | -0.83 | -0.17 | 0.23 | 0.515 | 0.6275 | 0.392396 | 0.677925 | 0.494345 | 0.993395 | 0.997988 | 0.994666 |
| Atxn1_J3QPR1_S81s_1 | Atxn1_J3Q | 0.15 | 0.27 | 0.53 | 0.44 | 0.24 | 0.14 | 0.7 | 0.08 | -0.13 | 0.69 | 1.25 | 0.63 | -0.66 | 0.4 | 0.21 | 0.58 | -0.0575 | 0.2625 | -0.215 | 0.518523 | 0.806692 | 0.42549 | 0.995762 | 0.997988 | 0.994666 |
| Atxn1l_P0C7T6_T328t_1 | Atxn1l_P0C | -1.41 | -0.47 | -1.11 | -0.86 | -0.74 | -0.65 | -1.26 | -1.15 | -0.6 | -0.87 | -1.12 | -1.51 | 1.21 | -0.66 | -1.82 | -1.68 | 0.0125 | -0.0625 | 0.225 | 0.731836 | 0.964763 | 0.840788 | 0.999187 | 0.997988 | 0.9971 |
| Atxn2_F6U2C2_S190s_1 | Atxn2_F6U | -3.13 | -1.08 | 0.05 | -0.23 | -3.01 | -2.02 | -0.81 | 0.08 | -0.37 | -1.96 | 1.41 | -0.25 | -0.29 | -1.86 | -0.72 | -1.62 | -0.3425 | 0.805 | -0.025 | 0.972314 | 0.691406 | 0.373498 | 0.994893 | 0.997988 | 0.999507 |
| Atxn2_F6U2C2_S276s_1 | Atxn2_F6U | -1.99 | -2.09 | -2.16 | -2.47 | -1.93 | -2.37 | -2.12 | -1.56 | -1.58 | -1.56 | -1.52 | -1.5 | -0.22 | -1.8 | -1.91 | -1.97 | 0.1825 | 0.6375 | 0.7025 | 0.125782 | 0.477911 | 0.015751 | 0.989223 | 0.997988 | 0.994666 |
| Atxn2_F6U2C2_S356s_1 | Atxn2_F6U | -0.01 | -0.19 | -1.02 | 0.3 | 0.42 | -0.61 | 0.73 | 0.72 | -0.51 | 0 | 0.37 | -0.6 | -0.83 | 0.06 | 0.17 | 1.41 | 0.545 | 0.045 | 0.4325 | 0.404483 | 0.200962 | 0.903169 | 0.989223 | 0.997988 | 0.994666 |
| Atxn2_F6U2C2_S388s_1 | Atxn2_F6U | 0.43 | 0.25 | -0.06 | -0.87 | 0.7 | 0.12 | -0.24 | 0.41 | 0.08 | 0.08 | 1.05 | 0.41 | -1.68 | 0.2 | 0.51 | 0.15 | 0.31 | 0.4675 | -0.1425 | 0.790428 | 0.388743 | 0.231212 | 0.989223 | 0.997988 | 0.999507 |
| Atxn2_F6U2C2_S403s_1 | Atxn2_F6U | -0.98 | -1.31 | -1.15 | -1.08 | -1 | -1.11 | -0.26 | -0.78 | -1.27 | -1.58 | 1.32 | -0.8 | -0.89 | 0.07 | -0.68 | -1 | 0.3425 | 0.5475 | 0.505 | 0.124999 | 0.199538 | 0.374562 | 0.989223 | 0.997988 | 0.994666 |
| Atxn2_F6U2C2_S444s_1 | Atxn2_F6U | -0.49 | 0.07 | -1.13 | -0.39 | -0.18 | 0.33 | -1.82 | -0.75 | 0.83 | 0.12 | -0.14 | -1.52 | 1.14 | -0.71 | -0.93 | 0 | -0.12 | 0.3075 | 0.36 | 0.476904 | 0.803688 | 0.554834 | 0.995762 | 0.997988 | 0.994666 |
| Atxn2_F6U2C2_S519s_1 | Atxn2_F6U | -0.4 | -0.67 | 0.13 | -0.71 | -0.56 | -0.12 | 0.1 | -0.14 | -0.49 | -0.57 | -0.39 | 0.19 | -0.3 | -0.44 | 0.38 | -0.44 | 0.2325 | 0.0975 | 0.2125 | 0.513903 | 0.408567 | 0.744989 | 0.989223 | 0.997988 | 0.994666 |
| Atxn2_F6U2C2_S551s_1 | Atxn2_F6U | -0.87 | -0.99 | 0.38 | 0.09 | 0.15 | -0.06 | -0.27 | -0.45 | -0.2 | 0.17 | 0.03 | 0.23 | 0.43 | 0.33 | -0.78 | -0.33 | 0.19 | 0.405 | 0.26 | 0.556307 | 0.603402 | 0.28389 | 0.989223 | 0.997988 | 0.995259 |
| Atxn2_F6U2C2_T240t_1 | Atxn2_F6U | -6.57 | -5.39 | -4.16 | -11 | -1.95 | -6.34 | -3.72 | -5.73 | -3.25 | -5.47 | -0.9 | -4.41 | -3.6 | -2.85 | -4.9 | -4.73 | 2.345 | 3.2725 | 2.76 | 0.064971 | 0.150963 | 0.060009 | 0.989223 | 0.997988 | 0.994666 |
| Atxn2_F6U2C2_T326t_1 | Atxn2_F6U | -0.99 | -0.31 | -0.92 | -1.09 | -0.16 | -0.51 | -0.39 | -0.63 | -1.13 | -1.32 | -1.38 | -1.28 | -1.58 | -1.1 | -0.43 | -1.92 | 0.405 | -0.45 | -0.43 | 0.276794 | 0.137197 | 0.103413 | 0.989223 | 0.997988 | 0.994666 |
| Atxn2_F6U2C2_T401t_1 | Atxn2_F6U | -0.98 | -1.31 | -1.15 | -1.08 | -1 | -1.11 | -0.26 | -0.78 | -1.27 | -1.58 | 1.32 | -0.8 | -0.89 | 0.07 | -0.68 | -1 | 0.3425 | 0.5475 | 0.505 | 0.124999 | 0.199538 | 0.374562 | 0.989223 | 0.997988 | 0.994666 |
| Atxn2_F6U2C2_T431t_1 | Atxn2_F6U | -0.72 | -1.26 | -0.6 | -0.86 | -0.94 | -1.46 | -0.34 | -0.62 | -1.13 | -0.65 | -0.72 | -0.77 | -0.29 | -1.05 | -0.48 | -0.09 | 0.02 | 0.0425 | 0.3825 | 0.232776 | 0.947316 | 0.866091 | 0.999187 | 0.997988 | 0.994666 |
| Atxn2_F6U2C2_T521t_1 | Atxn2_F6U | -1.26 | -1.63 | 0.18 | -0.46 | -1 | -1.62 | 0.21 | -0.23 | -2.01 | -0.98 | -1.34 | -0.44 | -0.75 | -1.98 | -0.38 | 0.4 | 0.1325 | -0.4 | 0.115 | 0.844678 | 0.799965 | 0.426023 | 0.995762 | 0.997988 | 0.999507 |
| Atxn2l_Q7TQH0_S109s_1 | Atxn2l_Q7 | -0.44 | 0.43 | -0.61 | -0.53 | 0.43 | -0.39 | 0.05 | -0.38 | -0.18 | -0.19 | -0.37 | -0.65 | -0.08 | -0.3 | -0.35 | -0.43 | 0.215 | -0.06 | -0.0025 | 0.993524 | 0.512262 | 0.843166 | 0.989223 | 0.997988 | 1 |
| Atxn2l_Q7TQH0_S304s_1 | Atxn2l_Q7 | -2.1 | -1.7 | -3.63 | -3.31 | -0.58 | -1.33 | -3.07 | -2.89 | 2.28 | -1.62 | -0.7 | -3.61 | 2.49 | 0.01 | -2.13 | -1.91 | 0.7175 | 1.7725 | 2.3 | 0.045274 | 0.302688 | 0.14882 | 0.989223 | 0.997988 | 0.994666 |
| Atxn2l_Q7TQH0_S337s_1 | Atxn2l_Q7 | -3.33 | -3.92 | -2.86 | -4.4 | -2.5 | -3.52 | -2.56 | -2.9 | -4.59 | -2.82 | -1.67 | -2.36 | -2.28 | -2.89 | -0.8 | -2.93 | 0.7575 | 0.7675 | 1.4025 | 0.029234 | 0.080678 | 0.251985 | 0.989223 | 0.997988 | 0.994666 |
| Atxn2l_Q7TQH0_S388s_1 | Atxn2l_Q7 | -2.7 | -4.18 | -2.21 | -2.69 | -1.35 | -2.22 | -2.24 | -2.19 | -1.4 | -1.73 | 0.13 | -1.81 | -0.41 | -2.13 | -1.55 | -1.55 | 0.945 | 1.7425 | 1.535 | 0.014762 | 0.058566 | 0.012949 | 0.989223 | 0.997988 | 0.994666 |
| Atxn2l_Q7TQH0_S591s_1 | Atxn2l_Q7 | -1.06 | -0.28 | -0.48 | -0.5 | -0.04 | -1.13 | 0.13 | -0.62 | -1.28 | -0.42 | -0.32 | -0.43 | -0.85 | -0.71 | -0.2 | -0.19 | 0.165 | -0.0325 | 0.0925 | 0.758688 | 0.628613 | 0.917104 | 0.989223 | 0.997988 | 0.9971 |
| Atxn2l_Q7TQH0_S631s_1 | Atxn2l_Q7 | -2.41 | -1.43 | -1.39 | -1.47 | -1.05 | -1.89 | -0.7 | -2.06 | -1.94 | -1.27 | -1.5 | -1.08 | -1.49 | -1.03 | -0.88 | -1.13 | 0.25 | 0.2275 | 0.5425 | 0.118364 | 0.531108 | 0.497912 | 0.989223 | 0.997988 | 0.994666 |
| Atxn2l_Q7TQH0_S681s_1 | Atxn2l_Q7 | -1.91 | -1.84 | -1.84 | -2.34 | -1.53 | -2.01 | -1.38 | -2.28 | -1.75 | -1.37 | -1.27 | -1.17 | -0.32 | -1.94 | -1.4 | -1.08 | 0.1825 | 0.5925 | 0.7975 | 0.057303 | 0.516553 | 0.038264 | 0.989223 | 0.997988 | 0.994666 |
| Atxn7l2_B7ZCC4_S514s_1 | Atxn7l2_B7 | -0.88 | -0.93 | -1 | -0.98 | -0.17 | -0.48 | -0.72 | -0.77 | -0.67 | -0.48 | -0.93 | -1.08 | -0.23 | 0.15 | 0.56 | 0.28 | 0.4125 | 0.1575 | 1.1375 | 0.00163 | 0.089821 | 0.501708 | 0.989223 | 0.997988 | 0.994666 |
| Aurkaip1_Q9DCJ7_S51s_1 | Aurkaip1_Q | -3.59 | -2.2 | -1.97 | -2.84 | -0.99 | -1.5 | -2.42 | -2.37 | -3.44 | -1.6 | -1.14 | -3.35 | -1.45 | -0.33 | -1.92 | -2.03 | 0.83 | 0.2675 | 1.2175 | 0.034607 | 0.100974 | 0.673836 | 0.989223 | 0.997988 | 0.994666 |
| Avl9_Q80U56_S368s_1 | Avl9_Q80U | 0.43 | 0.47 | 0.88 | 0.86 | 0.76 | 0.71 | 0.74 | 0.78 | 0.49 | 0.57 | 1.28 | 0.06 | 0.66 | 0.78 | 0.79 | 1.36 | 0.0875 | -0.06 | 0.2375 | 0.402894 | 0.686237 | 0.848032 | 0.994672 | 0.997988 | 0.994666 |
| Axin1_F6XWC2_S445s_1 | Axin1_F6X | -1.43 | -1.83 | -1.31 | -2.35 | -3.14 | -1.24 | -1.7 | -1.19 | 1.02 | -0.75 | -2.81 | -0.66 | 1.2 | -1.27 | 0.81 | -0.78 | -0.0875 | 0.93 | 1.72 | 0.015176 | 0.853515 | 0.226275 | 0.996349 | 0.997988 | 0.994666 |
| Axin2_O88566_S454s_1 | Axin2_O88 | -0.23 | -0.33 | -1.77 | -2 | -0.45 | -0.04 | -0.94 | -1.66 | 0.12 | -0.38 | 0.42 | -1.46 | 0.91 | -0.08 | -2.61 | -1.9 | 0.31 | 0.7575 | 0.1625 | 0.843571 | 0.561154 | 0.208365 | 0.989223 | 0.997988 | 0.999507 |
| B630019K06Rik_Q7TNS5_S41s_1 | B630019K0 | 2.68 | 1.02 | 1.43 | 1.55 | 0.12 | 2.14 | 0.64 | 1.09 | 1.97 | 2.35 | 0.37 | 1.25 | 2.16 | 0.55 | 1.36 | 1.78 | -0.6725 | -0.185 | -0.2075 | 0.663507 | 0.207497 | 0.725298 | 0.989223 | 0.997988 | 0.996159 |
| B630019K06Rik_Q7TNS5_S42s_1 | B630019K0 | 0.36 | -0.19 | -0.56 | 0.49 | -0.36 | 0.34 | -1.08 | 0.41 | 0.56 | 0.53 | 0.1 | 0.83 | 1.6 | -0.59 | 1.46 | 0.99 | -0.1975 | 0.48 | 0.84 | 0.133111 | 0.629965 | 0.155245 | 0.989223 | 0.997988 | 0.994666 |
| B630019K06Rik_Q7TNS5_S97s_1 | B630019K0 | 1.45 | 0.27 | 1.6 | 1.82 | 1.07 | 0.85 | 2.12 | 1.72 | 0.66 | 0.45 | 2.56 | 1.99 | -0.92 | 0.47 | 1.84 | 1.97 | 0.155 | 0.13 | -0.445 | 0.519347 | 0.718305 | 0.819694 | 0.995437 | 0.997988 | 0.994666 |
| Babam1_Q3UI43_S48s_1 | Babam1_Q | 0.58 | 0.17 | -0.13 | -0.05 | 0.98 | 1.06 | -0.35 | 0.16 | 0.3 | 0.53 | 1.13 | 0.23 | -0.11 | 0.35 | -0.58 | -0.3 | 0.32 | 0.405 | -0.3025 | 0.338261 | 0.395588 | 0.197792 | 0.989223 | 0.997988 | 0.994666 |
| Bad_Q3U9H3_S128s_1 | Bad_Q3U9 | 0.75 | 0.16 | 0.67 | 0.5 | -0.18 | 0.53 | -0.48 | -0.13 | 1 | 0.74 | 0.6 | 0.48 | 1.68 | -0.36 | 0.28 | 0.44 | -0.585 | 0.185 | -0.01 | 0.981766 | 0.063033 | 0.463443 | 0.989223 | 0.997988 | 1 |
| Bad_Q3U9H3_S94s_1 | Bad_Q3U9 | -0.15 | 0.22 | -0.52 | -0.41 | 0.33 | -0.79 | 0.26 | -0.53 | -0.94 | -0.08 | -0.64 | -0.31 | -0.67 | 0.52 | -0.42 | -0.29 | 0.0325 | -0.2775 | 0 | 1 | 0.922473 | 0.356212 | 0.997335 | 0.997988 | 1 |
| Bag2_Q91YN9_S20s_1 | Bag2_Q91Y | -0.95 | -1.7 | -2.09 | -6.03 | -0.76 | -1.31 | -1.83 | -1.31 | 0.34 | -2.58 | -5.15 | -1.3 | 1.86 | -2.17 | 0.95 | -1.35 | 1.39 | 0.52 | 2.515 | 0.073591 | 0.187341 | 0.714793 | 0.989223 | 0.997988 | 0.994666 |
| Bag3_Q9JLV1_S179s_1 | Bag3_Q9JL | 0.28 | 0.87 | -1.74 | -0.04 | -0.04 | -0.76 | -0.51 | -0.84 | 1.11 | -0.05 | -0.56 | -0.61 | -0.66 | -0.87 | -0.12 | -0.65 | -0.38 | 0.13 | -0.4175 | 0.447962 | 0.482542 | 0.835839 | 0.989223 | 0.997988 | 0.994666 |
| Bag3_Q9JLV1_S297s_1 | Bag3_Q9JL | 0.34 | -0.92 | -2.44 | -1.82 | -0.41 | 1.29 | -1.79 | -1.55 | -1.11 | 0.57 | -0.07 | -0.98 | -0.87 | -1.4 | -1.63 | -0.79 | 0.595 | 0.8125 | 0.0375 | 0.948755 | 0.468492 | 0.235007 | 0.989223 | 0.997988 | 0.999507 |
| Bag3_Q9JLV1_S390s_1 | Bag3_Q9JL | 0.34 | 0.4 | -0.07 | 0.07 | 0.15 | 0.13 | 0.32 | -0.01 | -0.14 | 0.55 | -0.24 | 0.16 | -0.32 | 0.07 | -0.18 | 0.76 | -0.0375 | -0.1025 | -0.1025 | 0.745425 | 0.864004 | 0.703972 | 0.996493 | 0.997988 | 0.9971 |
| Bag4_Q8CI61_S175s_1 | Bag4_Q8CI | -0.72 | -0.28 | -0.56 | -1.1 | -0.36 | -0.52 | -1.55 | -1.94 | -0.09 | -0.09 | -1.75 | -1.35 | 0.61 | -0.36 | -1.25 | -0.99 | -0.4275 | -0.155 | 0.1675 | 0.703807 | 0.305877 | 0.728737 | 0.989223 | 0.997988 | 0.996159 |
| Bag4_Q8CI61_S175s_2 | Bag4_Q8CI | -2.68 | -2.3 | -2.35 | -2.12 | -2.48 | -3.39 | -1.35 | -1.65 | -1.77 | -5.12 | 1.01 | -2.04 | -2.66 | -2.02 | -1.56 | -1.22 | 0.145 | 0.3825 | 0.4975 | 0.187065 | 0.744059 | 0.730341 | 0.995762 | 0.997988 | 0.994666 |
| Bag4_Q8CI61_S179s_2 | Bag4_Q8CI | -2.68 | -2.3 | -2.35 | -2.12 | -2.48 | -3.39 | -1.35 | -1.65 | -1.77 | -5.12 | 1.01 | -2.04 | -2.66 | -2.02 | -1.56 | -1.22 | 0.145 | 0.3825 | 0.4975 | 0.187065 | 0.744059 | 0.730341 | 0.995762 | 0.997988 | 0.994666 |
| Bag6_Q3UF95_S1085s_1 | Bag6_Q3U | -0.2 | 0.01 | 0.17 | 0.04 | -0.53 | 0.08 | 0.05 | -0.08 | -0.07 | 0 | -0.22 | 0.24 | -0.01 | 0.48 | -0.15 | -1.09 | -0.125 | -0.0175 | -0.1975 | 0.587403 | 0.594195 | 0.938172 | 0.989223 | 0.997988 | 0.995259 |
| Bag6_Q3UF95_S968s_2 | Bag6_Q3U | 2.23 | 0.63 | 1.84 | 2.53 | -0.17 | 1.63 | 1.85 | 2.24 | 2.03 | 2.51 | 2.24 | 1.35 | 1.06 | 1.85 | 1.27 | 1.48 | -0.42 | 0.225 | -0.3925 | 0.383432 | 0.493513 | 0.629633 | 0.989223 | 0.997988 | 0.994666 |
| Bag6_Q3UF95_S977s_1 | Bag6_Q3U | -0.86 | -0.69 | -0.45 | -0.78 | -0.56 | -0.71 | -0.8 | -0.94 | -0.13 | -0.64 | -0.22 | -0.53 | 0.29 | 0.25 | -0.86 | -0.87 | -0.0575 | 0.315 | 0.3975 | 0.288906 | 0.78887 | 0.207193 | 0.995762 | 0.997988 | 0.994666 |
| Bag6_Q3UF95_S977s_2 | Bag6_Q3U | 2.23 | 0.63 | 1.84 | 2.53 | -0.17 | 1.63 | 1.85 | 2.24 | 2.03 | 2.51 | 2.24 | 1.35 | 1.06 | 1.85 | 1.27 | 1.48 | -0.42 | 0.225 | -0.3925 | 0.383432 | 0.493513 | 0.629633 | 0.989223 | 0.997988 | 0.994666 |
| Bag6_Q3UF95_T967t_1 | Bag6_Q3U | -2.76 | -1.6 | -0.37 | -0.69 | -2.05 | -1.31 | -1.14 | -1.03 | 2.2 | -0.82 | -2.2 | -0.72 | 1.81 | -0.48 | -0.67 | -0.49 | -0.0275 | 0.97 | 1.3975 | 0.074657 | 0.958548 | 0.319669 | 0.999187 | 0.997988 | 0.994666 |
| Bai3_Q80ZF8_S1220s_1 | Bai3_Q80Z | -0.26 | 0.21 | 0.88 | -0.11 | 0.07 | 0.05 | -0.19 | -0.15 | 0.17 | -0.14 | 1.06 | 0.49 | -0.59 | 0.27 | 0.28 | -0.26 | -0.235 | 0.215 | -0.255 | 0.481108 | 0.427858 | 0.563261 | 0.989223 | 0.997988 | 0.994666 |
| Bai3_Q80ZF8_S1411s_1 | Bai3_Q80Z | 0.18 | 0.65 | 1.7 | 1.4 | 0.34 | 0.16 | 1.53 | 0.73 | 0.18 | 0.34 | 0.74 | 1.97 | 0.55 | -0.54 | 1.18 | 0.41 | -0.2925 | -0.175 | -0.5825 | 0.239538 | 0.505954 | 0.727911 | 0.989223 | 0.997988 | 0.994666 |
| Bai3_Q80ZF8_S751s_1 | Bai3_Q80Z | 0.1 | 0.96 | 0.21 | 0.45 | 0.85 | -0.04 | 0.97 | 0.1 | 0.04 | 0.34 | 0.43 | 0.79 | 0 | 0.9 | 0.67 | 0.98 | 0.04 | -0.03 | 0.2075 | 0.537162 | 0.903563 | 0.91774 | 0.996493 | 0.997988 | 0.994666 |
| Bai3_Q80ZF8_T1215t_1 | Bai3_Q80Z | 0.31 | -0.29 | -1.11 | -1.35 | -0.25 | 0.04 | -1.32 | -0.99 | -0.75 | -0.27 | 0.03 | -0.77 | -0.09 | 0.16 | -0.81 | -1.27 | -0.02 | 0.17 | 0.1075 | 0.823105 | 0.965339 | 0.686587 | 0.999187 | 0.997988 | 0.999507 |
| Baiap2_Q8BKX1_S324s_1 | Baiap2_Q8 | 0.04 | 0.66 | 0.43 | 0.48 | 1.21 | 0.7 | 0.14 | 0.24 | 0.5 | 0.74 | 1.07 | 0.1 | 1.22 | 1.24 | 0.39 | 0.01 | 0.17 | 0.2 | 0.3125 | 0.393886 | 0.575944 | 0.494004 | 0.989223 | 0.997988 | 0.994666 |
| Baiap2_Q8BKX1_S326s_1 | Baiap2_Q8 | 0.55 | 0.85 | 1.05 | 0.75 | 0.89 | 0.31 | 0.59 | 0.64 | 0.53 | 0.81 | 1.12 | 0.82 | 1.2 | 1.04 | 1.05 | 0.84 | -0.1925 | 0.02 | 0.2325 | 0.358043 | 0.414808 | 0.934093 | 0.989223 | 0.997988 | 0.994666 |
| Baiap2_Q8BKX1_S367s_1 | Baiap2_Q8 | -2.53 | -3.72 | -3.38 | -5.25 | -1.73 | -2.8 | -2.65 | -3.3 | -4.15 | -3.06 | -0.21 | -3.07 | -2.24 | -0.93 | -3.4 | -3.19 | 1.1 | 1.0975 | 1.28 | 0.098589 | 0.086694 | 0.243113 | 0.989223 | 0.997988 | 0.994666 |
| Baiap2_Q8BKX1_S396s_1 | Baiap2_Q8 | -3.65 | -2.21 | -2.12 | -2.09 | -1.18 | -3.31 | -1.34 | -1.74 | -2.89 | -2.83 | -2.9 | -2.12 | -2.19 | -1.56 | -2.12 | -2.61 | 0.625 | -0.1675 | 0.3975 | 0.366689 | 0.277702 | 0.688055 | 0.989223 | 0.997988 | 0.994666 |
| Baiap2_Q8BKX1_S515s_1 | Baiap2_Q8 | 1.05 | 0.76 | 0.89 | 0.87 | 0.68 | 1.28 | 0.86 | 1.26 | 0.15 | 0.51 | -0.17 | 1.3 | -1.76 | 0.36 | -0.1 | 0.93 | 0.1275 | -0.445 | -1.035 | 0.080937 | 0.586999 | 0.213315 | 0.989223 | 0.997988 | 0.994666 |
| Baiap2_Q8BKX1_T341t_1 | Baiap2_Q8 | -1.32 | -1.73 | -1.18 | -1.09 | -1.32 | -1.4 | 0.49 | -0.14 | -3.61 | -1.53 | 0.22 | -0.9 | -2.36 | -1.56 | -0.64 | -0.23 | 0.7375 | -0.125 | 0.1325 | 0.781142 | 0.127166 | 0.864684 | 0.989223 | 0.997988 | 0.998226 |
| Baiap2_Q8BKX1_T521t_1 | Baiap2_Q8 | 0.49 | 0.19 | 0.26 | 0.37 | 0.27 | 0.21 | 0.65 | 0.73 | 0.36 | 0.21 | 0.02 | 0.36 | -0.21 | 0.68 | 0.3 | 0.43 | 0.1375 | -0.09 | -0.0275 | 0.921256 | 0.547569 | 0.681963 | 0.989223 | 0.997988 | 0.999507 |
| Baiap2l2_Q80Y61_S459s_1 | Baiap2l2_C | -0.09 | -0.22 | 0.58 | 1.31 | -0.05 | -0.83 | 0.42 | 0.7 | 1.22 | 1.87 | 0.19 | 0.31 | 0.79 | 1.12 | 0.69 | 1.04 | -0.335 | 0.5025 | 0.515 | 0.197752 | 0.466218 | 0.328465 | 0.989223 | 0.997988 | 0.994666 |
| Baiap3_S4R1K2_S215s_1 | Baiap3_S4 | -0.84 | -0.86 | -1.97 | -2.56 | -0.72 | -0.56 | -2.15 | -2.5 | -0.61 | -0.64 | -1.54 | -2.07 | 0.54 | -1.02 | -1.35 | -1.07 | 0.075 | 0.3425 | 0.8325 | 0.159404 | 0.897771 | 0.515087 | 0.996493 | 0.997988 | 0.994666 |
| Basp1_Q91XV3_S128s_1 | Basp1_Q91 | -0.03 | 0.98 | 0.68 | 0.25 | 0.42 | 0.76 | -0.05 | 0.36 | 1.12 | 0.58 | 0.86 | 0.53 | 1.04 | 1.29 | 0.69 | 0.65 | -0.0975 | 0.3025 | 0.4475 | 0.182811 | 0.747572 | 0.329032 | 0.995762 | 0.997988 | 0.994666 |
| Basp1_Q91XV3_S128s_2 | Basp1_Q91 | 0.13 | 0.46 | 0.46 | 0.75 | 0.35 | 0.31 | 0.62 | 0.36 | 0.11 | 0.65 | 0.96 | 0.62 | -0.42 | 0.87 | 0.84 | 0.72 | -0.04 | 0.135 | 0.0525 | 0.883773 | 0.859252 | 0.623333 | 0.996349 | 0.997988 | 0.999507 |
| Basp1_Q91XV3_S131s_1 | Basp1_Q91 | 0.44 | 0.92 | 1.76 | 1.5 | 0.72 | 0.47 | 0.98 | 1.4 | 0.32 | 0.77 | 1.9 | 2.26 | 0.87 | 1.02 | 1.29 | 1.6 | -0.2625 | 0.1575 | 0.04 | 0.911714 | 0.467692 | 0.758639 | 0.989223 | 0.997988 | 0.999507 |
| Basp1_Q91XV3_S131s_2 | Basp1_Q91 | 0.13 | 0.46 | 0.59 | 0.62 | 0.35 | 0.31 | 0.57 | 0.23 | 0.11 | 0.65 | 0.97 | 0.81 | -0.42 | 0.87 | 0.48 | 0.77 | -0.085 | 0.185 | -0.025 | 0.942386 | 0.700802 | 0.504547 | 0.995169 | 0.997988 | 0.999507 |
| Basp1_Q91XV3_S167s_1 | Basp1_Q91 | 0.54 | 0.93 | 0.03 | 0.57 | 0.78 | 0.62 | 0.36 | 0.61 | 0.31 | 0.36 | 0.99 | 0.59 | 0.75 | 0.87 | -0.4 | 0.4 | 0.075 | 0.045 | -0.1125 | 0.757531 | 0.770146 | 0.875786 | 0.995762 | 0.997988 | 0.9971 |
| Basp1_Q91XV3_S169s_2 | Basp1_Q91 | 0.77 | 1.26 | 0.98 | 0.93 | 0.63 | 1.39 | 0.5 | 0.77 | 0.75 | 0.66 | 1.12 | 0.88 | -0.48 | 0.7 | 0.69 | 0.81 | -0.1625 | -0.1325 | -0.555 | 0.138095 | 0.545932 | 0.575113 | 0.989223 | 0.997988 | 0.994666 |
| Basp1_Q91XV3_S191s_1 | Basp1_Q91 | 0 | 0.28 | 0.16 | 0.64 | -0.24 | -0.12 | 0.46 | 0.84 | -0.36 | -1.01 | 0.57 | 0.46 | 0.02 | -0.56 | 0.52 | 0.16 | -0.035 | -0.355 | -0.235 | 0.460916 | 0.909422 | 0.379624 | 0.997335 | 0.997988 | 0.994666 |
| Basp1_Q91XV3_S218s_1 | Basp1_Q91 | 0.91 | 0.68 | 0.8 | 1.16 | 0.67 | 0.68 | 0.97 | 1.32 | 0.63 | 0.86 | 0.4 | 0.92 | 0.75 | 1.26 | 0.73 | 1.33 | 0.0225 | -0.185 | 0.13 | 0.637562 | 0.926826 | 0.449787 | 0.997335 | 0.997988 | 0.995259 |
| Basp1_Q91XV3_S40s_1 | Basp1_Q91 | -1.43 | 0.33 | -1.6 | -1.79 | -0.53 | -0.73 | -1.3 | -2.05 | -0.97 | -0.25 | -0.87 | -1.43 | 0.08 | -0.12 | -0.8 | -1.76 | -0.03 | 0.2425 | 0.4725 | 0.429696 | 0.95568 | 0.637937 | 0.999187 | 0.997988 | 0.994666 |
| Basp1_Q91XV3_S40s_2 | Basp1_Q91 | -1.18 | -0.8 | 1.41 | 1.23 | -0.95 | -1.44 | 1.26 | 1.08 | -1.75 | -1.03 | 0.42 | 0.99 | -1.74 | -1.41 | 0.72 | 1.52 | -0.1775 | -0.5075 | -0.3925 | 0.668579 | 0.832704 | 0.541752 | 0.995762 | 0.997988 | 0.996159 |
| Basp1_Q91XV3_S92s_1 | Basp1_Q91 | -0.53 | -0.05 | -1.07 | -0.53 | -0.26 | -0.61 | -0.4 | -0.61 | -0.84 | -0.2 | -0.39 | -0.41 | -1.51 | -0.3 | -0.63 | -0.61 | 0.075 | 0.085 | -0.2175 | 0.54828 | 0.780305 | 0.771573 | 0.995762 | 0.997988 | 0.994666 |
| Basp1_Q91XV3_T132t_1 | Basp1_Q91 | 0.16 | 0.72 | 0.39 | 0.83 | 0.32 | -0.58 | 0.25 | 0.59 | 0.19 | 0.03 | 1.38 | 1.43 | 0.96 | 0.26 | 1.1 | 0.56 | -0.38 | 0.2325 | 0.195 | 0.525482 | 0.243825 | 0.566896 | 0.989223 | 0.997988 | 0.994666 |
| Basp1_Q91XV3_T132t_2 | Basp1_Q91 | 1.04 | 0.03 | 0.38 | 0.83 | 0.81 | -0.36 | 0.68 | 0.39 | 0.46 | 1.25 | 1.07 | 0.61 | 0 | 1.3 | 0.94 | 0.8 | -0.19 | 0.2775 | 0.19 | 0.61359 | 0.588836 | 0.398344 | 0.989223 | 0.997988 | 0.995259 |
| Basp1_Q91XV3_T183t_1 | Basp1_Q91 | -1.15 | -0.74 | 1.06 | -1.07 | -1.15 | -1.02 | -0.37 | -0.09 | -1.54 | -1.01 | 0.13 | 0.29 | -0.09 | -0.63 | 0.72 | -0.4 | -0.1825 | -0.0575 | 0.375 | 0.502809 | 0.729212 | 0.926381 | 0.995762 | 0.997988 | 0.994666 |
| Basp1_Q91XV3_T31t_1 | Basp1_Q91 | 0.47 | 0.53 | 1.1 | 0.76 | -0.7 | 1.31 | -0.44 | 0.46 | 2.27 | 0.8 | 0.45 | 0.84 | 2.23 | -1.61 | 0.59 | 0.53 | -0.5575 | 0.375 | -0.28 | 0.696001 | 0.232904 | 0.381689 | 0.989223 | 0.997988 | 0.996159 |
| Basp1_Q91XV3_T31t_2 | Basp1_Q91 | 0.03 | -0.02 | 1.24 | 1.19 | -0.95 | 0.69 | 0.47 | 1.04 | 2.15 | 0.48 | 1.39 | 1.27 | 0.6 | -0.63 | 1.13 | 0.68 | -0.2975 | 0.7125 | -0.165 | 0.736293 | 0.56385 | 0.151875 | 0.989223 | 0.997988 | 0.9971 |
| Basp1_Q91XV3_T36t_1 | Basp1_Q91 | 0.81 | 0.52 | 0.68 | 0.76 | -0.37 | 1.4 | 0.18 | 0.51 | 2.22 | 1.07 | 0.01 | 0.68 | 2.13 | -0.27 | 0.75 | 0.49 | -0.2625 | 0.3025 | 0.0825 | 0.864031 | 0.484562 | 0.5068 | 0.989223 | 0.997988 | 0.999507 |
| Basp1_Q91XV3_T36t_2 | Basp1_Q91 | -0.36 | -0.28 | 1.3 | 1.2 | -0.95 | 0.07 | 0.79 | 1.06 | 1.25 | 0.02 | 1.07 | 1.17 | 0.01 | -0.89 | 0.99 | 1.01 | -0.2225 | 0.4125 | -0.185 | 0.753569 | 0.699197 | 0.423024 | 0.994893 | 0.997988 | 0.9971 |
| Baz1b_Q9Z277_S1464s_1 | Baz1b_Q9Z | 2.11 | 1.95 | 0.8 | 1.48 | 1.19 | 1.69 | 0.02 | 1.48 | 1.32 | 2.02 | 0.88 | 1.59 | 0.35 | 1.04 | 1.27 | 0.82 | -0.49 | -0.1325 | -0.715 | 0.07966 | 0.286321 | 0.7296 | 0.989223 | 0.997988 | 0.994666 |
| Baz1b_Q9Z277_S1464s_2 | Baz1b_Q9Z | 0.62 | 0.4 | 0.72 | 0.61 | -0.63 | 0.06 | 0.88 | 0.61 | -0.35 | 0.05 | 0.91 | 0.32 | -0.66 | -0.52 | 0.59 | 0.95 | -0.3575 | -0.355 | -0.4975 | 0.242328 | 0.313707 | 0.26671 | 0.989223 | 0.997988 | 0.994666 |
| Baz1b_Q9Z277_S1468s_2 | Baz1b_Q9Z | 0.62 | 0.4 | 0.72 | 0.61 | -0.63 | 0.06 | 0.88 | 0.61 | -0.35 | 0.05 | 0.91 | 0.32 | -0.66 | -0.52 | 0.59 | 0.95 | -0.3575 | -0.355 | -0.4975 | 0.242328 | 0.313707 | 0.26671 | 0.989223 | 0.997988 | 0.994666 |
| BC023829_Q3TYW3_S305s_1 | BC023829_ | 0.12 | 0.76 | 0.39 | 0.56 | 1.17 | 0.34 | 0.41 | 0.48 | 0 | 0.23 | 0.65 | 0.5 | 0.89 | 0.98 | 0.45 | 0.59 | 0.1425 | -0.1125 | 0.27 | 0.332436 | 0.606125 | 0.669371 | 0.989223 | 0.997988 | 0.994666 |
| Bcam_Q9R069_S615s_1 | Bcam_Q9R | 0.99 | -0.06 | -2.01 | -1.03 | -0.42 | 0.02 | -2.22 | -1.28 | 0.37 | -0.11 | -1.5 | -2.41 | 0.9 | 0.5 | -1.86 | -1.75 | -0.4475 | -0.385 | -0.025 | 0.976623 | 0.535507 | 0.635834 | 0.989223 | 0.997988 | 1 |
| Bcan_Q61361_S29s_1 | Bcan_Q613 | -8.25 | -8.08 | -5.83 | -11.61 | -6.07 | -8.42 | -3.01 | -4.2 | -4.31 | -7.32 | -6.71 | -4.06 | -3.17 | -4.73 | -3.7 | -5.88 | 3.0175 | 2.8425 | 4.0725 | 0.00561 | 0.059386 | 0.048123 | 0.989223 | 0.997988 | 0.994666 |
| Bcan_Q61361_S587s_1 | Bcan_Q613 | -0.77 | 0.12 | 0.28 | 0.04 | -0.06 | -0.53 | -0.4 | 0.47 | -0.66 | -0.31 | 1.24 | 0.05 | -0.65 | 0.12 | 0.42 | 0.06 | -0.0475 | 0.1625 | 0.07 | 0.843255 | 0.886477 | 0.722133 | 0.996493 | 0.997988 | 0.999507 |
| Bcar1_Q61140_S572s_1 | Bcar1_Q61 | -0.46 | 0.48 | 0.14 | 0.1 | 0.27 | -0.11 | 0.04 | -1.1 | 0.27 | 0.04 | -1.39 | -0.66 | -0.44 | 0.87 | -0.99 | -0.49 | -0.29 | -0.5 | -0.3275 | 0.458742 | 0.426429 | 0.24746 | 0.989223 | 0.997988 | 0.994666 |
| Bcar1_Q61140_S643s_1 | Bcar1_Q61 | -0.08 | -0.24 | 0.04 | 0.06 | -0.6 | -0.14 | -0.04 | -0.57 | -0.44 | 0.06 | -0.63 | 0.18 | -0.13 | 0.53 | -0.33 | -1.31 | -0.2825 | -0.1525 | -0.255 | 0.523435 | 0.243259 | 0.571422 | 0.989223 | 0.997988 | 0.994666 |
| Bcar3_Q9QZK2_S164s_1 | Bcar3_Q9C | 0.94 | 0.55 | -0.1 | -0.04 | 0.82 | 0.59 | 0.3 | -0.2 | 0.03 | 0.89 | 0.19 | 0.18 | 0.36 | -0.01 | 0.07 | -0.2 | 0.04 | -0.015 | -0.2825 | 0.389095 | 0.905921 | 0.96437 | 0.996513 | 0.997988 | 0.994666 |
| Bcar3_Q9QZK2_S245s_1 | Bcar3_Q9C | 0.18 | 0.48 | 1.19 | 0.73 | 0.18 | -0.41 | 0.72 | 0.84 | -0.05 | 0.28 | 0.45 | 0.81 | 0.01 | 0.29 | 0.16 | 0.5 | -0.3125 | -0.2725 | -0.405 | 0.197546 | 0.390814 | 0.392399 | 0.989223 | 0.997988 | 0.994666 |
| Bcar3_Q9QZK2_S250s_1 | Bcar3_Q9C | 0.15 | -0.15 | -0.17 | 0.18 | -0.17 | -0.07 | 0.08 | 0.23 | -0.67 | 0.26 | 0.7 | -0.71 | -0.43 | -0.27 | -0.02 | 0.13 | 0.015 | -0.1075 | -0.15 | 0.565287 | 0.945161 | 0.771691 | 0.999187 | 0.997988 | 0.995259 |
| Bcas1_Q80YN3_S316s_1 | Bcas1_Q80 | -0.04 | 0.35 | 0.73 | 0.2 | -0.83 | 0.08 | -0.74 | -0.38 | 2.07 | -0.44 | -0.71 | 0.48 | 1.61 | -0.31 | 0.25 | 0.5 | -0.7775 | 0.04 | 0.2025 | 0.638956 | 0.022878 | 0.946274 | 0.989223 | 0.997988 | 0.995259 |
| Bcas1_Q80YN3_S388s_1 | Bcas1_Q80 | 0.41 | 0.19 | -0.68 | 0.47 | 0.49 | 0.81 | 0.22 | -0.32 | 0.94 | 0.8 | 0.94 | -0.06 | 0.93 | 0.17 | 0.36 | 0.44 | 0.2025 | 0.5575 | 0.3775 | 0.288625 | 0.573934 | 0.154109 | 0.989223 | 0.997988 | 0.994666 |
| Bcas1_Q80YN3_S390s_1 | Bcas1_Q80 | 0.41 | 0.19 | -0.68 | 0.47 | 0.49 | 0.81 | 0.22 | -0.32 | 0.94 | 0.8 | 0.94 | -0.06 | 0.93 | 0.17 | 0.36 | 0.44 | 0.2025 | 0.5575 | 0.3775 | 0.288625 | 0.573934 | 0.154109 | 0.989223 | 0.997988 | 0.994666 |
| Bcas1_Q80YN3_S414s_1 | Bcas1_Q80 | 0.8 | 0.98 | 1.46 | 1.18 | 1.35 | -0.08 | 1.16 | 1 | -0.04 | 0.75 | 1.69 | 1.15 | 0.14 | 0.68 | 1.08 | 0.98 | -0.2475 | -0.2175 | -0.385 | 0.232024 | 0.487568 | 0.581364 | 0.989223 | 0.997988 | 0.994666 |
| Bcas1_Q80YN3_S425s_1 | Bcas1_Q80 | 1.16 | 1.29 | 0.74 | 0.51 | 1.16 | 0.96 | 1.02 | 0.92 | 0.81 | 1.56 | 0.77 | 0.99 | 0.51 | 1.62 | 0.18 | 1.13 | 0.09 | 0.1075 | -0.065 | 0.865063 | 0.717053 | 0.718946 | 0.995437 | 0.997988 | 0.999507 |
| Bcas1_Q80YN3_S443s_1 | Bcas1_Q80 | 1.39 | 0.87 | -0.13 | 0.61 | 1.52 | 0.77 | -0.48 | 0.57 | 1.4 | -0.2 | 0.27 | -0.8 | -0.84 | 1.61 | -0.09 | 0.69 | -0.09 | -0.5175 | -0.3425 | 0.548525 | 0.851197 | 0.337895 | 0.996349 | 0.997988 | 0.994666 |
| Bcas1_Q80YN3_S497s_1 | Bcas1_Q80 | 1.11 | 1.29 | 1.38 | 0.8 | 0.44 | 1.05 | 0.4 | 1.54 | -1.38 | 0.76 | -1.5 | 1.33 | 0.53 | -3.05 | 1.08 | 1.06 | -0.2875 | -1.3425 | -1.24 | 0.180629 | 0.374746 | 0.070133 | 0.989223 | 0.997988 | 0.994666 |
| Bcas1_Q80YN3_S525s_1 | Bcas1_Q80 | 0.69 | 1.43 | 0.59 | 0.77 | 0.78 | 0.41 | 0.73 | 0.72 | 0.54 | 0.54 | 0.33 | 0.5 | 0.89 | 0.8 | 0.96 | 0.71 | -0.21 | -0.3925 | -0.03 | 0.914021 | 0.424751 | 0.158043 | 0.989223 | 0.997988 | 0.999507 |
| Bcas1_Q80YN3_S525s_2 | Bcas1_Q80 | -1.18 | -0.79 | -0.07 | -0.21 | -0.85 | -0.97 | -0.58 | -0.93 | -1.86 | -1.24 | 0.05 | -0.57 | -1.24 | -0.61 | -0.55 | -0.52 | -0.27 | -0.3425 | -0.1675 | 0.627615 | 0.375951 | 0.468468 | 0.989223 | 0.997988 | 0.995259 |
| Bcas1_Q80YN3_S601s_1 | Bcas1_Q80 | -1.35 | -1.27 | -0.25 | 0.15 | -0.6 | -1.74 | 0.53 | 0.38 | -1.99 | -1.98 | 0.24 | 0.36 | -1.9 | -1 | 0.33 | 0.61 | 0.3225 | -0.1625 | 0.19 | 0.763374 | 0.579954 | 0.812619 | 0.989223 | 0.997988 | 0.9971 |
| Bcas1_Q80YN3_S615s_1 | Bcas1_Q80 | -0.11 | 0.19 | 0.5 | 0.66 | 0.38 | -0.42 | 0.6 | 0.2 | -0.4 | 0.14 | 0.02 | 0.49 | -0.18 | 0.35 | 0.55 | 0.12 | -0.12 | -0.2475 | -0.1 | 0.73623 | 0.691736 | 0.408796 | 0.994893 | 0.997988 | 0.9971 |

| | | | | | | | | | | | | | | | | | | | | | | | | | | |
|---|---|---|---|---|---|---|---|---|---|---|---|---|---|---|---|---|---|---|---|---|---|---|---|---|---|---|
| Bcas1_Q80YN3_T379t_1 | Bcas1_Q80 | 1.29 | 1.29 | 0.93 | 1.4 | 1.83 | 0.61 | 1.29 | 1.07 | 1.19 | 1.16 | 1.85 | 1.3 | 1.19 | 1.53 | 1.69 | 1.48 | -0.0275 | 0.1475 | 0.245 | 0.347314 | 0.926568 | 0.571617 | 0.997335 | 0.997988 | 0.994666 |
| Bcas1_Q80YN3_T523t_1 | Bcas1_Q80 | 0.73 | 1.47 | 1.06 | 1.03 | 1.52 | -0.2 | 0.7 | 0.96 | 0.18 | 1.07 | 1.34 | 1.11 | 0.13 | 0.97 | 1.04 | 0.95 | -0.3275 | -0.1475 | -0.3 | 0.351926 | 0.3986 | 0.650862 | 0.989223 | 0.997988 | 0.994666 |
| Bcas1_Q80YN3_T523t_2 | Bcas1_Q80 | -1.18 | -0.79 | -0.07 | -0.21 | -0.85 | -0.97 | -0.58 | -0.93 | -1.86 | -1.24 | 0.05 | -0.57 | -1.24 | -0.61 | -0.55 | -0.52 | -0.27 | -0.3425 | -0.1675 | 0.627615 | 0.375951 | 0.468468 | 0.989223 | 0.997988 | 0.995259 |
| Bcas1_Q80YN3_T591t_1 | Bcas1_Q80 | 0.86 | 1.26 | 0.84 | 0.76 | 1.13 | 0.78 | 0.99 | 0.81 | 0.96 | 1.29 | 0.69 | 1.23 | 0.75 | 1.38 | 1.44 | 1.28 | -0.0025 | 0.1125 | 0.2825 | 0.319472 | 0.991036 | 0.656003 | 0.999187 | 0.997988 | 0.994666 |
| Bcas3_Q0VF62_S503s_1 | Bcas3_Q0V | 1.01 | 0.41 | 0.95 | 0.86 | 0.38 | 0.05 | 0.79 | 0.91 | 0.56 | 0.67 | 1.18 | 1.47 | -0.13 | 0.88 | 1.03 | 1.03 | -0.275 | 0.1625 | -0.105 | 0.760689 | 0.33378 | 0.585865 | 0.989223 | 0.997988 | 0.9971 |
| Bcas3_Q0VF62_S518s_1 | Bcas3_Q0V | -0.47 | 0.5 | 0.04 | -0.49 | 0.58 | -0.15 | 0.03 | -0.4 | 0.76 | 0.28 | -0.25 | 0.03 | 0.88 | 0.9 | -0.45 | -0.22 | 0.12 | 0.31 | 0.3825 | 0.37816 | 0.713977 | 0.371809 | 0.995437 | 0.997988 | 0.994666 |
| Bcas3_Q0VF62_S550s_1 | Bcas3_Q0V | 0.39 | 0.77 | 0.9 | 0.94 | 0.77 | 0 | 0.83 | 0.47 | 0.72 | 0.45 | 0.87 | 0.88 | 0.49 | 0.58 | 1.04 | 0.52 | -0.2325 | -0.02 | -0.0925 | 0.731713 | 0.398241 | 0.934265 | 0.989223 | 0.997988 | 0.9971 |
| Bcas3_Q0VF62_S854s_1 | Bcas3_Q0V | 0.26 | 0.1 | 0.52 | 0.29 | 0.25 | -0.17 | 0.42 | 0.63 | -0.73 | 0.03 | 0.46 | 0.27 | -0.02 | -0.32 | 0.16 | 0.32 | -0.01 | -0.285 | -0.2575 | 0.336681 | 0.967872 | 0.368492 | 0.999187 | 0.997988 | 0.994666 |
| Bcas3_Q0VF62_S862s_1 | Bcas3_Q0V | 0.22 | 0.66 | 0.47 | 0.1 | 1.42 | 1.34 | 0.76 | 0.38 | 0.92 | 0.53 | 0.81 | 0.78 | 1.38 | 0.56 | 0.83 | -0.4 | 0.6125 | 0.3975 | 0.23 | 0.56817 | 0.066102 | 0.120501 | 0.989223 | 0.997988 | 0.995259 |
| Bcas3_Q0VF62_S871s_1 | Bcas3_Q0V | 0.41 | 0.49 | 0.9 | 1.07 | 0.72 | 0.42 | 0.09 | 1.85 | 0.25 | 0.28 | 0.92 | 1.02 | 0.21 | 0.4 | 0.54 | 0.88 | 0.0525 | -0.1 | -0.21 | 0.469007 | 0.895159 | 0.738795 | 0.996493 | 0.997988 | 0.994666 |
| Bckdha_Q3U3J1_S338s_1 | Bckdha_Q3 | 5.47 | -0.82 | -2.15 | -2.76 | -1.17 | 5.39 | 0.59 | -2.06 | -0.3 | -0.25 | -4.35 | -2.11 | 4.04 | -2.38 | -1.56 | -1.51 | 0.7525 | -1.6875 | -0.2875 | 0.888321 | 0.726384 | 0.372306 | 0.995437 | 0.997988 | 0.999507 |
| Bckdha_Q3U3J1_S348s_1 | Bckdha_Q3 | 0.02 | 0.67 | 0.58 | 0.12 | 0.67 | -0.02 | 0.63 | 0.21 | -0.39 | 0.24 | -0.13 | 0.28 | -0.74 | 0.42 | 0.64 | 0.51 | 0.025 | -0.3475 | -0.14 | 0.710009 | 0.927055 | 0.233666 | 0.997335 | 0.997988 | 0.996966 |
| Bckdha_Q3U3J1_T339t_1 | Bckdha_Q3 | 0.58 | 1.51 | 0.94 | 0.8 | 0.44 | 0.57 | 0.22 | 0.56 | 1.23 | 0.75 | 1.22 | 0.54 | 1.3 | 0.29 | 0.74 | 1.31 | -0.51 | -0.0225 | -0.0475 | 0.891177 | 0.076576 | 0.940511 | 0.989223 | 0.997988 | 0.999507 |
| Bckdk_D3Z7R0_S31s_1 | Bckdk_D3Z | -0.69 | -0.61 | -0.32 | -0.21 | -0.68 | -0.73 | -0.54 | -0.34 | 0.18 | -0.47 | -0.23 | -0.05 | 0.41 | -0.24 | -0.11 | -0.24 | -0.115 | 0.315 | 0.4125 | 0.157775 | 0.611756 | 0.231427 | 0.989223 | 0.997988 | 0.994666 |
| Bcl11a_Q5STS9_S205s_1 | Bcl11a_Q5 | 0.12 | 0.01 | -2.35 | -1.7 | 0.43 | -0.01 | -0.14 | -1.14 | -0.76 | -0.26 | -1.08 | -0.7 | -0.33 | 0.31 | -0.82 | -0.62 | 0.765 | 0.28 | 0.615 | 0.324093 | 0.239571 | 0.63615 | 0.989223 | 0.997988 | 0.994666 |
| Bcl11a_Q5STS9_S328s_1 | Bcl11a_Q5 | -0.65 | 0.06 | -0.32 | -0.77 | 0.2 | -0.16 | -0.13 | 0.28 | -0.31 | -0.11 | 0.45 | 1.27 | -0.64 | 0.04 | -0.24 | -0.46 | 0.4675 | 0.745 | 0.095 | 0.751215 | 0.101872 | 0.087508 | 0.989223 | 0.997988 | 0.9971 |
| Bcl11a_Q5STS9_S431s_1 | Bcl11a_Q5 | -1.33 | -0.85 | -1.02 | -1.39 | -0.78 | -1.88 | -0.82 | -1.68 | -1.23 | -1.16 | -1.63 | -1.67 | 1.15 | -1.45 | -0.66 | -1.09 | -0.1425 | -0.275 | 0.635 | 0.264001 | 0.663073 | 0.29616 | 0.990434 | 0.997988 | 0.994666 |
| Bcl11a_Q5STS9_S432s_1 | Bcl11a_Q5 | -1.81 | -1.51 | -0.83 | -2.46 | -1.18 | -1.9 | -1.71 | -1.85 | -0.85 | -1.26 | -1.83 | -1.81 | 1.4 | -1.6 | -0.34 | -0.68 | -0.0075 | 0.215 | 1.3475 | 0.060222 | 0.983841 | 0.600687 | 0.999187 | 0.997988 | 0.994666 |
| Bcl11a_Q5STS9_S608s_1 | Bcl11a_Q5 | 0.37 | -0.22 | -1.26 | -0.42 | 0.22 | -0.41 | -0.24 | -0.81 | 0.11 | 0.45 | 0.11 | -1.14 | 0.75 | 0.22 | 0.73 | -0.03 | 0.0725 | 0.265 | 0.8 | 0.067983 | 0.851508 | 0.571663 | 0.996349 | 0.997988 | 0.994666 |
| Bcl11a_Q5STS9_S718s_1 | Bcl11a_Q5 | -1.73 | -2.13 | -0.9 | -1.52 | -0.55 | -2.72 | -0.92 | -1.68 | -0.89 | -2.29 | -1.35 | -1.07 | 0.19 | -1.49 | -0.78 | -1.79 | 0.1025 | 0.17 | 0.6025 | 0.23377 | 0.837078 | 0.672855 | 0.995762 | 0.997988 | 0.994666 |
| Bcl11a_Q5STS9_S86s_1 | Bcl11a_Q5 | -0.88 | -0.24 | -0.43 | -0.29 | -0.61 | -1.03 | 0.37 | -0.31 | -0.49 | -1.32 | -0.87 | -0.23 | 0.32 | -0.05 | 0.31 | 0.27 | 0.065 | -0.2675 | 0.6725 | 0.028248 | 0.846764 | 0.400443 | 0.996349 | 0.997988 | 0.994666 |
| Bcl11b_Q99PV8_S309s_1 | Bcl11b_Q9 | 0.14 | 0.5 | 0.85 | 1.47 | 0.04 | -0.23 | 0.93 | 0.99 | -0.13 | 0.75 | 1.02 | 0.95 | 1.05 | 0.2 | 1.36 | 1.4 | -0.3075 | -0.0925 | 0.2625 | 0.519234 | 0.452416 | 0.81228 | 0.989223 | 0.997988 | 0.994666 |
| Bcl11b_Q99PV8_S690s_1 | Bcl11b_Q9 | -0.8 | -0.89 | -1.5 | -1.87 | -0.19 | -1.45 | -0.61 | -1.39 | -0.33 | -0.72 | -0.77 | -1.48 | 0.81 | -0.08 | -0.18 | -0.51 | 0.355 | 0.44 | 1.275 | 0.008537 | 0.370238 | 0.242023 | 0.989223 | 0.997988 | 0.994666 |
| Bcl11b_Q99PV8_T344t_1 | Bcl11b_Q9 | 0.02 | 0.74 | 0.16 | 0.56 | 0.48 | 0.26 | 0.68 | 0.5 | -0.07 | 0.34 | -0.29 | 0.2 | -0.17 | 0.28 | 0.57 | 0.08 | 0.11 | -0.325 | -0.18 | 0.545775 | 0.658481 | 0.254745 | 0.990434 | 0.997988 | 0.994666 |
| Bcl11b_Q99PV8_T672t_1 | Bcl11b_Q9 | 0.92 | 0.78 | 0.46 | 0.36 | 1.2 | 0.28 | 1.24 | 0.11 | 0.35 | 1.23 | 0.23 | 0.2 | 0.86 | 1.2 | 1.05 | 0.85 | 0.0775 | -0.1275 | 0.36 | 0.185666 | 0.816794 | 0.683358 | 0.995762 | 0.997988 | 0.994666 |
| Bcl2l13_P59017_S35s_1 | Bcl2l13_P5 | 0.09 | 0.35 | -0.32 | -0.27 | 0.61 | -0.26 | 0.31 | -0.16 | 0.11 | 0.28 | -0.66 | 0.08 | -0.31 | -0.13 | -0.19 | -0.14 | 0.1625 | -0.01 | -0.155 | 0.557154 | 0.576825 | 0.973468 | 0.989223 | 0.997988 | 0.995259 |
| Bcl2l13_P59017_S377s_1 | Bcl2l13_P5 | 1.53 | 0.73 | 1.36 | 1.35 | 0.82 | 1.74 | 1.1 | 1.54 | 0.42 | 0.47 | 0.77 | 0.73 | 0.7 | 1.34 | 0.7 | 0.58 | 0.0575 | -0.645 | -0.4125 | 0.196717 | 0.847127 | 0.03283 | 0.996349 | 0.997988 | 0.994666 |
| Bcl2l13_P59017_S387s_1 | Bcl2l13_P5 | 5.61 | 2.05 | 2.97 | 2.76 | 1.77 | 6.68 | 2.53 | 2.44 | 3.7 | 2.37 | 0.34 | 2.89 | 3.1 | 1.71 | 2.19 | 2.49 | 0.0075 | -1.0225 | -0.975 | 0.208484 | 0.994901 | 0.291259 | 0.999395 | 0.997988 | 0.994666 |
| Bcl7a_Q9CXE2_S207s_1 | Bcl7a_Q9C | -0.53 | -0.81 | -1.63 | -1.46 | -0.75 | -1.06 | -1.6 | -3.06 | 0.14 | -0.36 | -0.05 | -0.77 | 0.45 | 0.43 | -2.13 | -1.94 | -0.51 | 0.8475 | 0.31 | 0.651659 | 0.341998 | 0.033555 | 0.989223 | 0.997988 | 0.995259 |
| Bcl9_Q9D219_S291s_1 | Bcl9_Q9D2 | 0.97 | 0.25 | 0.31 | 0.71 | -0.29 | 0.66 | 0.9 | -0.61 | 0.17 | 0.46 | -0.68 | 0.25 | 0.64 | 1.08 | -0.1 | 0.49 | -0.395 | -0.51 | -0.0325 | 0.922772 | 0.324463 | 0.145558 | 0.989223 | 0.997988 | 0.999507 |
| Bcl9_Q9D219_S686s_1 | Bcl9_Q9D2 | -3.61 | -3.42 | -2.94 | -3.78 | -2.27 | -4.15 | -3.49 | -2.83 | -4.03 | -2.64 | -2.24 | -3.47 | -1.71 | -2.02 | -2.89 | -3.12 | 0.2525 | 0.3425 | 1.0025 | 0.02792 | 0.553711 | 0.434427 | 0.989223 | 0.997988 | 0.994666 |
| Bcl9l_Q67FY2_S1014s_1 | Bcl9l_Q67F | -0.48 | -0.86 | -1.54 | -1.44 | 0.76 | -0.66 | -2.28 | -1.91 | -0.44 | -0.42 | -0.32 | -0.73 | 0.48 | 0.28 | -0.92 | -1.19 | 0.0575 | 0.6025 | 0.7425 | 0.138653 | 0.929374 | 0.07205 | 0.997639 | 0.997988 | 0.994666 |
| Bcl9l_Q67FY2_S116s_1 | Bcl9l_Q67F | -2.65 | -2.91 | -1.94 | -2.12 | -2.82 | -3.88 | -2.76 | -2.14 | -2.85 | -2.1 | -0.66 | -2.27 | -1.26 | -2.2 | -2.71 | -2.71 | -0.495 | 0.435 | 0.185 | 0.654619 | 0.243105 | 0.38433 | 0.989223 | 0.997988 | 0.995259 |
| Bclaf1_F8WI22_S177s_1 | Bclaf1_F8V | 1.1 | 0.82 | 0.75 | 0.74 | 0.61 | 0.92 | 0.76 | 0.95 | 0.27 | 0.61 | 0.39 | 0.9 | 0.94 | 0.31 | 1.34 | 0.92 | -0.0425 | -0.31 | 0.025 | 0.932272 | 0.841835 | 0.223372 | 0.996064 | 0.997988 | 0.999507 |
| Bclaf1_F8WI22_S221s_1 | Bclaf1_F8V | -1.16 | -2.07 | -0.28 | -0.79 | -0.26 | -0.57 | 0.47 | 0.32 | -1.37 | -0.08 | -0.49 | -0.2 | -0.32 | 0.57 | -0.02 | -0.47 | 1.065 | 0.54 | 1.015 | 0.040167 | 0.030284 | 0.256097 | 0.989223 | 0.997988 | 0.994666 |
| Bclaf1_F8WI22_S261s_2 | Bclaf1_F8V | -0.48 | -0.3 | -0.64 | -0.18 | -0.87 | -0.36 | -0.56 | 0.18 | -1.11 | -1.17 | -0.17 | -0.79 | -2.03 | -0.15 | -0.11 | -1.15 | -0.0025 | -0.41 | -0.46 | 0.325684 | 0.992847 | 0.185393 | 0.999187 | 0.997988 | 0.994666 |
| Bclaf1_F8WI22_S261s_3 | Bclaf1_F8V | 1.62 | 1.39 | 2.03 | 1.19 | 0.81 | 1.8 | 2.04 | 3.38 | 0.86 | 0.93 | 0.53 | 2.9 | -1.15 | 0.36 | 3.97 | 2.6 | 0.45 | -0.2525 | -0.1125 | 0.910829 | 0.388271 | 0.635514 | 0.989223 | 0.997988 | 0.999507 |
| Bclaf1_F8WI22_S263s_3 | Bclaf1_F8V | 0.44 | 1.79 | 2.44 | 1.3 | -0.57 | 2 | 1.18 | 2.97 | 1.06 | 0.88 | 2.18 | 2.31 | 1.59 | 1.3 | 3.47 | 2.54 | -0.0975 | 0.115 | 0.7325 | 0.235598 | 0.897128 | 0.826405 | 0.996493 | 0.997988 | 0.994666 |
| Bclaf1_F8WI22_S267s_1 | Bclaf1_F8V | 1.16 | 0.09 | -0.24 | 0.69 | -0.93 | 0.8 | -0.49 | 1.28 | 0.97 | 0.31 | 0.84 | 0.06 | 0.62 | -0.1 | 0.91 | -0.92 | -0.26 | 0.12 | -0.2975 | 0.546617 | 0.638029 | 0.75412 | 0.989223 | 0.997988 | 0.994666 |
| Bclaf1_F8WI22_S267s_2 | Bclaf1_F8V | 1.62 | 1.09 | -0.22 | 0.14 | 0 | 1.35 | -0.06 | 0.68 | 0.04 | 0.45 | 0.72 | -0.01 | -2.24 | 0.13 | 0.34 | -0.44 | -0.165 | -0.3575 | -1.21 | 0.089053 | 0.7388 | 0.427487 | 0.995762 | 0.997988 | 0.994666 |
| Bclaf1_F8WI22_S267s_3 | Bclaf1_F8V | 1.62 | 1.39 | 2.6 | 1.34 | 0.81 | 1.8 | 0.7 | 2.77 | 0.86 | 0.93 | 2.73 | 2.03 | -1.15 | 0.36 | 3.23 | 2.52 | -0.2175 | -0.1 | -0.4975 | 0.589389 | 0.675016 | 0.843494 | 0.992876 | 0.997988 | 0.995259 |
| Bclaf1_F8WI22_S284s_2 | Bclaf1_F8V | 0.61 | 0.44 | -0.05 | 0.1 | 0.34 | 0.82 | 0.02 | 0.34 | 1.12 | 0.44 | 0.11 | -0.19 | -0.2 | 1.11 | 0.26 | -0.22 | 0.105 | 0.095 | -0.0375 | 0.918626 | 0.695951 | 0.779883 | 0.994893 | 0.997988 | 0.999507 |
| Bclaf1_F8WI22_S289s_1 | Bclaf1_F8V | -0.36 | 0.84 | 0.03 | -0.41 | 0.11 | -0.25 | 0.12 | 0.08 | 0.38 | -1.44 | 0.17 | -0.22 | 1.29 | 0.36 | 0.48 | -0.15 | -0.01 | -0.3025 | 0.47 | 0.274466 | 0.974885 | 0.528548 | 0.999187 | 0.997988 | 0.994666 |
| Bclaf1_F8WI22_S289s_2 | Bclaf1_F8V | 0.9 | 0.92 | -0.05 | 0.1 | 0.68 | 0.67 | 0.02 | 0.34 | 0.77 | 0.99 | 0.11 | -0.19 | -0.33 | 1.18 | 0.26 | -0.22 | -0.04 | -0.0475 | -0.245 | 0.568397 | 0.899675 | 0.900811 | 0.996493 | 0.997988 | 0.995259 |
| Bclaf1_F8WI22_S395s_1 | Bclaf1_F8V | 3 | 1.51 | 3.46 | 0.54 | 0.91 | 2.99 | 4 | 3.71 | 0.87 | 2.26 | -0.78 | 3.43 | 0.75 | 0.88 | 5.02 | 4.12 | 0.775 | -0.6825 | 0.565 | 0.615205 | 0.369704 | 0.499472 | 0.989223 | 0.997988 | 0.995259 |
| Bclaf1_F8WI22_S494s_1 | Bclaf1_F8V | 2.41 | 1.15 | 0.51 | 0.22 | 0.35 | 1.92 | -0.32 | 0.51 | 1.62 | 1.58 | -0.21 | 0.07 | 2.16 | -0.52 | 0.77 | 0.16 | -0.4575 | -0.3075 | -0.43 | 0.527608 | 0.456289 | 0.625684 | 0.989223 | 0.997988 | 0.994666 |
| Bclaf1_F8WI22_S510s_1 | Bclaf1_F8V | 0.45 | 0.43 | 0.04 | -0.29 | 0.43 | -0.03 | 0.29 | -0.01 | -0.56 | -0.32 | -0.84 | -0.29 | -0.46 | 0.41 | 0.51 | -0.19 | 0.0125 | -0.66 | -0.09 | 0.787039 | 0.961532 | 0.035827 | 0.999187 | 0.997988 | 0.999507 |
| Bclaf1_F8WI22_S646s_1 | Bclaf1_F8V | 1.97 | 0.56 | 0.97 | 0.96 | -0.37 | 1.9 | -0.37 | 1.16 | 2.05 | 0.59 | -0.83 | 0.62 | 2.05 | -0.09 | 1.33 | 0.34 | -0.535 | -0.5075 | -0.2075 | 0.696911 | 0.365728 | 0.410421 | 0.989223 | 0.997988 | 0.996159 |
| Bclaf1_F8WI22_S656s_1 | Bclaf1_F8V | 0.39 | 0.24 | 0.05 | -0.39 | 0.13 | -0.06 | 0.22 | 0.32 | -0.62 | -0.77 | -0.49 | -0.37 | 0.35 | 0.63 | 0.37 | -0.31 | 0.08 | -0.635 | 0.1875 | 0.553846 | 0.746391 | 0.032857 | 0.995762 | 0.997988 | 0.994666 |
| Bclaf1_F8WI22_S759s_1 | Bclaf1_F8V | 0.33 | 0.14 | 0.82 | 0.49 | -1.61 | 0.92 | -0.54 | -0.19 | 0.08 | -0.8 | 0.39 | -0.86 | 1.97 | -5.3 | 1.02 | 0.4 | -0.8 | -0.7425 | -0.9225 | 0.519126 | 0.13195 | 0.062721 | 0.989223 | 0.997988 | 0.994666 |
| Bclaf1_F8WI22_T256t_2 | Bclaf1_F8V | 3.44 | 2.5 | 2.27 | 2.09 | 1.01 | 2.98 | 2.42 | 3.34 | 1.27 | 1.96 | 5.21 | 3.48 | -2.63 | 0.54 | 2.88 | 2.93 | -0.1375 | 0.405 | -1.645 | 0.179927 | 0.798443 | 0.625418 | 0.995762 | 0.997988 | 0.994666 |
| Bclaf1_F8WI22_T256t_3 | Bclaf1_F8V | 2.58 | 1.71 | 2.44 | 1.3 | 1.6 | 2.39 | 1.18 | 2.97 | 0.81 | 1.73 | 2.18 | 2.31 | -3.81 | 0.49 | 3.47 | 2.54 | 0.0275 | -0.25 | -1.335 | 0.35733 | 0.95288 | 0.573561 | 0.999187 | 0.997988 | 0.994666 |
| Bclaf1_F8WI22_T305t_1 | Bclaf1_F8V | 0.62 | 0.34 | 0.02 | 0.47 | 0.42 | 0.2 | -0.76 | 0.44 | -0.19 | -0.14 | -1.73 | -0.71 | 1.28 | 0.83 | -1.2 | -0.4 | -0.2875 | -1.055 | -0.235 | 0.665203 | 0.385093 | 0.022457 | 0.989223 | 0.997988 | 0.996159 |
| Bclaf1_F8WI22_T339t_1 | Bclaf1_F8V | 0.14 | 0.88 | -0.14 | -0.04 | 0.08 | 0.62 | -1.33 | 0.11 | 1.16 | 0.4 | 0.38 | 0.99 | 1.47 | 0.13 | 0.76 | -0.4 | -0.34 | 0.5225 | 0.28 | 0.539627 | 0.454288 | 0.138862 | 0.989223 | 0.997988 | 0.994666 |
| Bcor_Q8CGN4_S1391s_1 | Bcor_Q8CG | -2.01 | -1.71 | -2.26 | -2.95 | -4.69 | -2.09 | -2.24 | -2.97 | -0.21 | -2.44 | -1.96 | -2 | 1.77 | -3.1 | -0.63 | -1.97 | -0.765 | 0.58 | 1.25 | 0.206086 | 0.211436 | 0.28369 | 0.989223 | 0.997988 | 0.994666 |
| Bcr_Q6PAJ1_S356s_1 | Bcr_Q6PAJ | -0.27 | -1.45 | -0.81 | -0.7 | 0.27 | -0.3 | -0.19 | -0.16 | -0.9 | -0.92 | 0 | -0.69 | -1.03 | 0.06 | -0.46 | -0.69 | 0.7125 | 0.18 | 0.2775 | 0.447397 | 0.036974 | 0.603064 | 0.989223 | 0.997988 | 0.994666 |
| Bcr_Q6PAJ1_S384s_1 | Bcr_Q6PAJ | 0.11 | 1.16 | -0.22 | -0.3 | 0.24 | -0.44 | 0.65 | -0.62 | -0.1 | 1.01 | -0.47 | -0.45 | 1.34 | 0.47 | 0.63 | 0.42 | -0.23 | -0.19 | 0.5275 | 0.211316 | 0.590404 | 0.682538 | 0.989223 | 0.997988 | 0.994666 |
| Bcr_Q6PAJ1_S697s_1 | Bcr_Q6PAJ | -0.67 | -1.6 | -1.16 | -1.23 | -1.25 | -0.56 | -1.33 | -1.35 | -0.41 | -1.06 | -2.15 | -1.32 | 0.47 | -2.04 | -1.24 | -1.27 | 0.0425 | -0.07 | 0.145 | 0.783315 | 0.885632 | 0.862412 | 0.996493 | 0.997988 | 0.998491 |
| Begain_F8WIG2_S16s_1 | Begain_F8\ | -1.46 | -1.64 | -1.15 | -1.8 | -1.83 | -1.05 | -1.35 | -1.22 | -0.59 | -2.11 | -0.18 | -0.86 | 0.15 | -1.33 | -1.12 | -1.21 | 0.15 | 0.5775 | 0.635 | 0.124262 | 0.573309 | 0.199107 | 0.989223 | 0.997988 | 0.994666 |
| Begain_F8WIG2_S265s_1 | Begain_F8\ | 0.61 | 0.82 | 0.56 | 1.02 | 0.59 | 0.59 | 0.73 | 0.7 | 0.37 | 0.56 | 0.18 | 0.79 | -0.15 | 0.83 | -0.17 | -0.16 | -0.1 | -0.2775 | -0.665 | 0.059995 | 0.638244 | 0.276816 | 0.989223 | 0.997988 | 0.994666 |
| Begain_F8WIG2_S460s_1 | Begain_F8\ | 1.19 | -0.93 | -1.17 | 0.07 | -3.3 | 1.35 | -0.4 | 0.25 | 1.14 | -1.01 | 0.75 | -0.76 | -0.23 | -1.79 | -1.5 | -1.48 | -0.315 | 0.24 | -1.04 | 0.101973 | 0.747768 | 0.727405 | 0.995762 | 0.997988 | 0.994666 |
| Begain_F8WIG2_S482s_1 | Begain_F8\ | -0.19 | 0.22 | 0.74 | 0.63 | -0.36 | 0.08 | 0.49 | 0.5 | -1.48 | -0.42 | -0.59 | 0.68 | 0.17 | -3.02 | 0.52 | 0.32 | -0.1725 | -0.8025 | -0.8525 | 0.284002 | 0.583957 | 0.113211 | 0.989223 | 0.997988 | 0.994666 |
| Begain_F8WIG2_S492s_1 | Begain_F8\ | 1.14 | 0.8 | 0.71 | 0.9 | 0.93 | 1.01 | 0.48 | 0.83 | 0.77 | 0.86 | 0.73 | 0.91 | 0.38 | 0.99 | 0.12 | 0.05 | -0.075 | -0.07 | -0.5025 | 0.115734 | 0.742109 | 0.748453 | 0.995762 | 0.997988 | 0.994666 |
| Begain_F8WIG2_S529s_1 | Begain_F8\ | 0.48 | -0.58 | -0.06 | 0.58 | 0.05 | 0.43 | 0.3 | 0.65 | -0.17 | -0.16 | 0.7 | 0.62 | -0.68 | -0.21 | -0.61 | -1.13 | 0.2525 | 0.1425 | -0.7625 | 0.054662 | 0.429025 | 0.699361 | 0.989223 | 0.997988 | 0.994666 |
| Begain_F8WIG2_S579s_1 | Begain_F8\ | -0.31 | 0.55 | 0.33 | 0.59 | 0.17 | -0.23 | 0 | -0.49 | 0.84 | 0.28 | 1.64 | 0.13 | 0.92 | 0.46 | 0 | 0.23 | -0.4275 | 0.4325 | 0.1125 | 0.732475 | 0.157753 | 0.293569 | 0.989223 | 0.997988 | 0.9971 |
| Begain_F8WIG2_S589s_1 | Begain_F8\ | -0.75 | 0.02 | -0.31 | -0.95 | 0.3 | -0.88 | -0.19 | -0.99 | -0.89 | -0.99 | -0.62 | -0.45 | -0.47 | 0.14 | -1.3 | -1.37 | 0.0575 | -0.24 | -0.2525 | 0.551283 | 0.876076 | 0.422752 | 0.996493 | 0.997988 | 0.994666 |
| Begain_F8WIG2_Y495y_1 | Begain_F8\ | 1.17 | -0.1 | 1.21 | 1.52 | 0.34 | 1.23 | 0.93 | 1.22 | 0.77 | 0.81 | 1.34 | 1.19 | -0.85 | 0.53 | 0.35 | 0.77 | -0.02 | 0.0775 | -0.75 | 0.146126 | 0.960004 | 0.84134 | 0.999187 | 0.997988 | 0.994666 |
| Begain_F8WIG2_Y614y_1 | Begain_F8\ | 1.29 | 0.14 | -0.32 | 0.72 | -2.56 | -0.14 | 1.14 | 0.67 | 1.15 | 0 | 1.28 | 0.65 | 0.57 | 1.78 | 0.25 | 0.47 | -0.68 | 0.3125 | 0.31 | 0.515394 | 0.394409 | 0.483599 | 0.989223 | 0.997988 | 0.994666 |
| Bet1_O35623_S50s_1 | Bet1_O356 | -0.2 | -1.3 | -1.79 | 0.26 | -0.24 | -0.27 | 0.57 | 0.17 | -0.94 | -0.9 | -2.6 | -2.01 | -2.56 | 0.29 | -0.88 | -0.82 | 0.815 | -0.855 | -0.235 | 0.72964 | 0.112137 | 0.165885 | 0.989223 | 0.997988 | 0.9971 |
| Bicd1_G3UXK5_S546s_1 | Bicd1_G3U | 0.22 | -0.24 | 0.02 | 0.97 | 0.04 | 0.18 | 0.6 | 1.15 | 0.09 | 0.17 | 1.14 | 0.9 | 0.37 | 0.42 | 0.66 | 0.71 | 0.25 | 0.3325 | 0.2975 | 0.364735 | 0.491842 | 0.38609 | 0.989223 | 0.997988 | 0.994666 |
| Bicd2_D3Z390_S504s_1 | Bicd2_D3Z | 0.18 | 0.32 | 0.31 | 0.13 | 0.43 | 0.07 | 0.65 | 0.24 | 0.16 | 0.23 | 0.24 | 0.2 | 0.15 | 0.95 | 0.33 | 0.16 | 0.1125 | -0.0275 | 0.1625 | 0.560118 | 0.612061 | 0.892884 | 0.989223 | 0.997988 | 0.995259 |
| Bin1_Q6P1B9_S245s_1 | Bin1_Q6P1 | 1.68 | 1.15 | 1.12 | -0.05 | 1.72 | 1.99 | 1.52 | 1.3 | -0.49 | 1.47 | -2.65 | 1.26 | -0.47 | -1.23 | 1.85 | 1.69 | 0.6575 | -1.0775 | -0.515 | 0.502367 | 0.111775 | 0.253644 | 0.989223 | 0.997988 | 0.994666 |
| Bin1_Q6P1B9_S265s_1 | Bin1_Q6P1 | 0.61 | 0.93 | 0.74 | 0.74 | 0.38 | 0.08 | 0.85 | 0.7 | 0.07 | 0.62 | 0.45 | 0.76 | 0.43 | 0.78 | 0.75 | 0.51 | -0.2525 | -0.28 | -0.1375 | 0.570826 | 0.316455 | 0.268477 | 0.989223 | 0.997988 | 0.995259 |
| Bin1_Q6P1B9_S265s_2 | Bin1_Q6P1 | 0.86 | 1.15 | 0.56 | 0.34 | 0.65 | 0.62 | 0.91 | 0.51 | 0.55 | 0.79 | 0.32 | 0.77 | 0.09 | 0.86 | 0.71 | 0.81 | -0.055 | -0.12 | -0.11 | 0.721052 | 0.826957 | 0.656058 | 0.995762 | 0.997988 | 0.9971 |
| Bin1_Q6P1B9_S267s_1 | Bin1_Q6P1 | 0.37 | 0.67 | 1.18 | 0.91 | 0.09 | -0.09 | 1.07 | 0.74 | -0.12 | 0.5 | 0.67 | 0.96 | 0.5 | 0.2 | 1.07 | 0.77 | -0.33 | -0.28 | -0.1475 | 0.634683 | 0.333075 | 0.387124 | 0.989223 | 0.997988 | 0.995259 |
| Bin1_Q6P1B9_S267s_2 | Bin1_Q6P1 | 0.89 | 1.5 | 0.7 | 0.89 | 0.67 | 0.79 | 1.27 | 0.88 | 0.99 | 1.04 | 0.4 | 1.25 | 0.56 | 1.62 | 0.99 | 1.05 | -0.0925 | -0.075 | 0.06 | 0.852991 | 0.726484 | 0.799272 | 0.995437 | 0.997988 | 0.999507 |
| Bin1_Q6P1B9_S273s_1 | Bin1_Q6P1 | -0.26 | 0.7 | 0.21 | 0.57 | -0.16 | -0.5 | 0 | -0.13 | -0.54 | 0.21 | -0.23 | -0.02 | -0.2 | 0.57 | -0.28 | 0.26 | -0.5025 | -0.45 | -0.2175 | 0.518457 | 0.095505 | 0.163741 | 0.989223 | 0.997988 | 0.994666 |
| Bin1_Q6P1B9_S273s_2 | Bin1_Q6P1 | 0.92 | 0.84 | 0.36 | -0.61 | 0.63 | 0.43 | 0.35 | -0.07 | 0.2 | 0.63 | 0.2 | 0.03 | -0.25 | 0.22 | 0.3 | 0.47 | -0.0425 | -0.1125 | -0.1925 | 0.626524 | 0.909515 | 0.766899 | 0.997335 | 0.997988 | 0.995259 |
| Bin1_Q6P1B9_S293s_1 | Bin1_Q6P1 | 5.05 | 0.48 | 1.37 | 1.26 | 0.37 | 4.9 | 4.85 | 1.42 | 0.32 | 3.47 | 1.15 | 1.78 | -0.3 | 0.11 | 1.55 | 2.82 | 0.845 | -0.36 | -0.995 | 0.36841 | 0.530345 | 0.738304 | 0.989223 | 0.997988 | 0.994666 |
| Bin2_S4R2J8_S393s_1 | Bin2_S4R2 | 0.08 | -1.3 | -1.49 | -0.91 | -1.41 | -0.37 | 1.03 | -1.05 | -1.05 | -0.7 | 0.09 | -1.07 | -0.3 | -1.06 | -0.67 | -0.47 | 0.455 | 0.2225 | 0.28 | 0.484107 | 0.437899 | 0.607924 | 0.989223 | 0.997988 | 0.994666 |
| Bin2_S4R2J8_S415s_1 | Bin2_S4R2 | -0.73 | 0.29 | -1.59 | -2.51 | -0.46 | -0.23 | 0.05 | -0.06 | 0.5 | -0.19 | -1.17 | -1.44 | 0.48 | -0.74 | -1.27 | -0.53 | 0.96 | 0.56 | 0.62 | 0.341401 | 0.106316 | 0.416415 | 0.989223 | 0.997988 | 0.994666 |
| Bin2_S4R2J8_S415s_2 | Bin2_S4R2 | 0.37 | -0.55 | 0.49 | 0.26 | -0.11 | -0.11 | -0.39 | -1.12 | 1.14 | 0.03 | 0.49 | -0.6 | 0.33 | 1.59 | 0.18 | -1.08 | -0.575 | 0.1225 | 0.1125 | 0.838164 | 0.117144 | 0.773951 | 0.989223 | 0.997988 | 0.999507 |
| Bin2_S4R2J8_S428s_1 | Bin2_S4R2 | -0.13 | -0.09 | -0.62 | -0.54 | -0.62 | 0.05 | -1.64 | -1.51 | -0.08 | -0.12 | 0.86 | -0.7 | -0.44 | 0.24 | -0.7 | -1.16 | -0.585 | 0.335 | -0.17 | 0.630927 | 0.170249 | 0.365808 | 0.989223 | 0.997988 | 0.995259 |
| Bin2_S4R2J8_S448s_1 | Bin2_S4R2 | -1.11 | -1.17 | -1.7 | -0.53 | -2.05 | -0.24 | -2.87 | -0.84 | -0.7 | -1.97 | -0.28 | -0.59 | -0.63 | -2.86 | -1.65 | -1.19 | -0.3725 | 0.2425 | -0.455 | 0.375294 | 0.520842 | 0.576667 | 0.989223 | 0.997988 | 0.994666 |
| Bin2_S4R2J8_T392t_1 | Bin2_S4R2 | -0.74 | -0.93 | -1.56 | -0.87 | -1.39 | -2.12 | -1.42 | -0.51 | -0.89 | -1.4 | 0.47 | -1.33 | -0.18 | -1.24 | -0.33 | -0.68 | -0.335 | 0.2375 | 0.4175 | 0.23289 | 0.377283 | 0.602431 | 0.989223 | 0.997988 | 0.994666 |
| Birc6_S4R1L5_S2216s_1 | Birc6_S4R1 | -1.32 | -1.03 | -0.37 | -1.62 | -2.36 | -0.45 | -1.01 | -1.62 | 1.7 | -0.85 | -2.41 | -0.6 | 1.45 | -1.69 | 0.51 | -1.28 | -0.275 | 0.545 | 0.8325 | 0.255293 | 0.550311 | 0.497893 | 0.989223 | 0.997988 | 0.994666 |
| Birc6_S4R1L5_S552s_2 | Birc6_S4R1 | -1.5 | -1.39 | -1.34 | -0.58 | -0.73 | -1.54 | -0.58 | -1.54 | -0.62 | -1.02 | -0.16 | -0.9 | -0.65 | 0.1 | -1.25 | -1.03 | 0.105 | 0.5275 | 0.495 | 0.211964 | 0.757077 | 0.120387 | 0.995762 | 0.997988 | 0.994666 |
| Birc6_S4R1L5_S556s_1 | Birc6_S4R1 | 0.48 | -0.03 | -0.08 | -0.5 | 0.17 | 0.42 | -0.37 | 0.34 | 0.66 | 0.07 | 0.63 | 0.37 | 0.92 | 0.67 | -0.08 | -0.36 | 0.1725 | 0.465 | 0.32 | 0.407515 | 0.56259 | 0.133257 | 0.989223 | 0.997988 | 0.994666 |
| Birc6_S4R1L5_S556s_2 | Birc6_S4R1 | -1.5 | -1.39 | -1.34 | -0.58 | -0.73 | -1.54 | -0.58 | -1.54 | -0.62 | -1.02 | -0.16 | -0.9 | -0.65 | 0.1 | -1.25 | -1.03 | 0.105 | 0.5275 | 0.495 | 0.211964 | 0.757077 | 0.120387 | 0.995762 | 0.997988 | 0.994666 |
| Bles03_Q8VD62_S59s_1 | Bles03_Q8' | -1.76 | -0.07 | 0.37 | -0.07 | -1.3 | -2.19 | -0.18 | -0.88 | -3.54 | -0.84 | 0.26 | -0.68 | -2.28 | -1.08 | 0.92 | 0.6 | -0.755 | -0.8175 | -0.0775 | 0.921042 | 0.204714 | 0.343022 | 0.989223 | 0.997988 | 0.999507 |
| Bloc1s4_Q8VED2_S11s_1 | Bloc1s4_Q8 | 0.36 | -0.99 | 0.52 | -0.04 | -0.19 | -0.4 | -0.08 | -0.31 | 0.99 | 0.78 | -0.42 | -0.23 | 1 | 0.69 | 0.07 | 1.08 | -0.2075 | 0.3175 | 0.7475 | 0.094391 | 0.55523 | 0.502494 | 0.989223 | 0.997988 | 0.994666 |
| Bmp2k_Q91Z96_S1087s_1 | Bmp2k_Q9 | -2.01 | -0.49 | -1.19 | -1.2 | -0.84 | -1.39 | -1.24 | -0.78 | -0.46 | -0.94 | -2.32 | -0.86 | 0.11 | -1.29 | -0.15 | -1.51 | 0.16 | 0.0775 | 0.5125 | 0.306044 | 0.647159 | 0.872724 | 0.99033 | 0.997988 | 0.994666 |
| Bmp2k_Q91Z96_T644t_1 | Bmp2k_Q9 | -0.63 | 0.43 | 1.42 | 0.87 | -0.09 | 0.18 | -0.2 | 0.52 | 0.82 | 0.1 | 0.43 | 1.47 | 1.09 | 1.17 | 0.43 | 0.71 | -0.42 | 0.1825 | 0.3275 | 0.476485 | 0.347286 | 0.713134 | 0.989223 | 0.997988 | 0.994666 |
| Bmpr2_O35607_S515s_1 | Bmpr2_O3 | 0.56 | 0.58 | 0.66 | 0.69 | 0.6 | 0.93 | 0.84 | 0.73 | 0.79 | 1.47 | 1.4 | 0.87 | 0.59 | 1.39 | 0.55 | 0.6 | 0.1525 | 0.51 | 0.16 | 0.573395 | 0.453735 | 0.06796 | 0.989223 | 0.997988 | 0.995259 |
| Bmpr2_O35607_S586s_1 | Bmpr2_O3 | -0.41 | -0.87 | 0.14 | 0.47 | -0.41 | 0.83 | -0.1 | 1.25 | -1.05 | -0.56 | 1.82 | -0.27 | -0.46 | -0.38 | -0.45 | 0.53 | 0.56 | 0.1525 | -0.0225 | 0.954148 | 0.237955 | 0.810531 | 0.989223 | 0.997988 | 0.999507 |
| Bmpr2_O35607_S680s_1 | Bmpr2_O3 | 0.58 | -0.85 | -0.09 | 0.42 | 0.5 | 0.34 | 0.58 | 0.32 | 0.17 | 0.45 | 1.73 | 1.1 | 0.33 | 1.69 | 0.41 | -1.08 | 0.42 | 0.8475 | 0.3225 | 0.590632 | 0.230441 | 0.089091 | 0.989223 | 0.997988 | 0.995259 |
| Bmpr2_O35607_S862s_1 | Bmpr2_O3 | -0.22 | 0.74 | -0.42 | 0.11 | 0.6 | 0.49 | 0.25 | -0.29 | 0.2 | 1.1 | -0.57 | -0.13 | 0.3 | 0.85 | -0.31 | -0.42 | 0.21 | 0.0975 | 0.0525 | 0.894657 | 0.530995 | 0.818702 | 0.989223 | 0.997988 | 0.999507 |
| Bmpr2_O35607_S863s_1 | Bmpr2_O3 | 0.09 | 0.51 | 0.18 | 0.34 | 0.48 | 0.28 | 0.49 | 0.43 | 0.32 | 0.95 | 0.17 | 0.33 | 0.21 | 0.91 | -0.14 | 0.14 | 0.14 | 0.1625 | -1.6E-16 | 1 | 0.507774 | 0.538403 | 0.989223 | 0.997988 | 1 |
| Bmpr2_O35607_T517t_1 | Bmpr2_O3 | 0.47 | 0.67 | 0.04 | 0.73 | 0.5 | 0.33 | 1.04 | 0.66 | 0.6 | 0.7 | 0.36 | 0.55 | 0.69 | 0.92 | 0.99 | 1.01 | 0.155 | 0.075 | 0.425 | 0.134389 | 0.560577 | 0.76294 | 0.989223 | 0.997988 | 0.994666 |
| Bnip3_O55003_S79s_1 | Bnip3_O55 | 0.06 | 0.05 | -0.42 | -0.4 | 0.18 | -0.52 | -0.55 | -0.99 | 0.11 | 0.15 | -0.14 | -0.47 | -0.15 | 0.31 | -0.48 | -0.64 | -0.2925 | 0.09 | -0.0625 | 0.838216 | 0.343274 | 0.731117 | 0.989223 | 0.997988 | 0.999507 |
| Bnip3_O55003_S88s_1 | Bnip3_O55 | 0.72 | 0.96 | 0.71 | 0.54 | 0.73 | 0.39 | 0.35 | 0.45 | 0.25 | 0.77 | 0.36 | 0.68 | -1.97 | 0.66 | 0.52 | 0.45 | -0.2525 | -0.2175 | -0.8175 | 0.183271 | 0.258771 | 0.372826 | 0.989223 | 0.997988 | 0.994666 |
| Bnip3_O55003_T66t_1 | Bnip3_O55 | 0.6 | -0.37 | -0.13 | -0.08 | -1.15 | -0.12 | -2.06 | -0.19 | 1.96 | -0.68 | -0.66 | 0.48 | 2.55 | -0.98 | 0.28 | -0.15 | -0.885 | 0.27 | 0.42 | 0.552572 | 0.083304 | 0.655465 | 0.989223 | 0.997988 | 0.994666 |
| Bnip3l_Q91Z78_S165s_1 | Bnip3l_Q9 | -0.87 | -0.57 | -0.76 | -0.74 | 0.76 | -1.22 | -0.33 | -0.6 | -1.82 | -0.69 | -0.34 | -0.85 | -0.15 | 0.45 | -0.3 | -0.53 | 0.3875 | -0.19 | 0.6025 | 0.060542 | 0.346924 | 0.581435 | 0.989223 | 0.997988 | 0.994666 |
| Bnip3l_Q91Z78_S167s_1 | Bnip3l_Q9 | -5.41 | -4.61 | -3.36 | -4.2 | -3.69 | -7.7 | -3.7 | -3.57 | -11.22 | -4.41 | -2.85 | -3.97 | -3.38 | -3.89 | -3.84 | -4.18 | -0.27 | -1.2175 | 0.5725 | 0.219973 | 0.776903 | 0.478905 | 0.995762 | 0.997988 | 0.994666 |
| Bod1l_E9Q6J5_S1989s_1 | Bod1l_E9Q | 0.41 | 0.26 | -0.01 | 0.84 | 1.18 | -0.41 | 0.78 | 0.37 | 0.85 | 0.82 | 1.19 | 1.4 | 0.13 | 1.43 | 0.47 | 0.51 | 0.105 | 0.69 | 0.26 | 0.472369 | 0.78017 | 0.032124 | 0.995762 | 0.997988 | 0.994666 |
| Bod1l_E9Q6J5_S2898s_1 | Bod1l_E9Q | -1.87 | -1.6 | -0.1 | -2.46 | -2.33 | -1.49 | -1.65 | -1.65 | -0.71 | -1.36 | 0.87 | -1.34 | 0.55 | -1.54 | -1.11 | -0.92 | -0.2725 | 0.8725 | 0.7525 | 0.240385 | 0.583014 | 0.206806 | 0.989223 | 0.997988 | 0.994666 |
| Bod1l_E9Q6J5_T2890t_1 | Bod1l_E9Q | -1.43 | -3.41 | -4.97 | -12.84 | -2.79 | -5.4 | -5.35 | -5.41 | -1.89 | -2 | -4.87 | -5.52 | 0.3 | -0.44 | -6.04 | -4.9 | 0.925 | 2.0925 | 2.8925 | 0.268465 | 0.675647 | 0.378472 | 0.992876 | 0.997988 | 0.994666 |
| Bola1_Q9D8S9_S81s_1 | Bola1_Q9D | -1.39 | -0.88 | -1.4 | -1.72 | -0.45 | -1.54 | -1.91 | -2.49 | -1.76 | -0.6 | -0.88 | -2.19 | -1.46 | -0.64 | -1.43 | -1.01 | -0.25 | -0.01 | 0.2125 | 0.502213 | 0.569656 | 0.980323 | 0.989223 | 0.997988 | 0.994666 |
| Bop1_P97452_S559s_1 | Bop1_P974 | 1.09 | 1.14 | 0.53 | 1.05 | -0.11 | 1.03 | 0.78 | 0.86 | 1.36 | 1.4 | 2.16 | 1.94 | 1.62 | 1.1 | 1.32 | 1.11 | -0.3125 | 0.7625 | 0.335 | 0.238099 | 0.328509 | 0.024857 | 0.989223 | 0.997988 | 0.994666 |
| Bptf_E9Q6A7_S228s_1 | Bptf_E9Q6 | 0.07 | -0.48 | -0.27 | -0.47 | -0.24 | -0.59 | -0.06 | -0.36 | -0.64 | -0.61 | -0.12 | -0.53 | -0.3 | -0.15 | -0.08 | -0.31 | -0.025 | -0.1875 | 0.0775 | 0.758773 | 0.916169 | 0.461414 | 0.997335 | 0.997988 | 0.9971 |
| Bptf_E9Q6A7_S2353s_1 | Bptf_E9Q6 | -0.12 | -0.42 | -0.58 | -0.58 | -0.3 | -0.8 | -0.55 | -0.76 | -1.14 | -0.22 | 0.44 | -0.88 | 0.09 | -0.13 | -0.2 | -0.27 | -0.1775 | -0.025 | 0.2975 | 0.250525 | 0.450681 | 0.946956 | 0.989223 | 0.997988 | 0.994666 |
| Braf_P28028_S348s_1 | Braf_P280 | -1.69 | -1.11 | -2.75 | -2.82 | -0.92 | -2.1 | -1.96 | -3.08 | -2.35 | -1.51 | -2.58 | -2.79 | -1.56 | -0.73 | -3.07 | -2.69 | 0.0775 | -0.215 | 0.08 | 0.896665 | 0.887732 | 0.654228 | 0.996493 | 0.997988 | 0.999507 |
| Braf_P28028_S714s_1 | Braf_P280 | -0.68 | -0.43 | -0.08 | -0.16 | -0.02 | -1.09 | -0.32 | -0.2 | -0.64 | -0.04 | 0.03 | -0.38 | -0.17 | -0.01 | -0.17 | -0.23 | -0.07 | 0.08 | 0.1925 | 0.45463 | 0.814423 | 0.765292 | 0.995762 | 0.997988 | 0.994666 |
| Braf_P28028_S735s_1 | Braf_P280 | 0.36 | 0.25 | -0.26 | 0.14 | 0.13 | -0.1 | 0.34 | 0.19 | -0.24 | 0.04 | 0.13 | -0.08 | -0.47 | 0.07 | -0.18 | 0.4 | 0.0175 | -0.16 | -0.1675 | 0.572898 | 0.940392 | 0.511921 | 0.999187 | 0.997988 | 0.995259 |
| Braf_P28028_T384t_1 | Braf_P280 | -0.36 | 0.35 | 0.68 | 0.31 | -0.06 | -0.24 | 0.8 | 0.39 | 0.03 | 0.01 | 1.02 | 0.21 | -0.12 | -0.54 | 0.77 | 0.75 | -0.0225 | 0.0725 | -0.03 | 0.940036 | 0.945584 | 0.832193 | 0.999187 | 0.997988 | 0.999507 |
| Braf_P28028_T790t_1 | Braf_P280 | -1.88 | -1.6 | -1.98 | -1.68 | -0.97 | -1.43 | -1.17 | -2.4 | -1.54 | -2.41 | -0.8 | -2.21 | -1.3 | -0.71 | -2.38 | -2.08 | 0.2925 | 0.045 | 0.1675 | 0.673999 | 0.394297 | 0.905505 | 0.989223 | 0.997988 | 0.996159 |
| Brap_Q99MP8_S86s_1 | Brap_Q99M | -1.65 | -1.42 | -1.13 | -1.36 | -1.3 | -1.38 | -1.42 | -0.82 | -1.94 | -1.01 | -0.67 | -0.47 | -0.43 | -2.18 | -1.84 | -1.7 | 0.16 | 0.3675 | -0.1475 | 0.714937 | 0.510666 | 0.316189 | 0.989223 | 0.997988 | 0.996966 |
| Brap_Q99MP8_S86s_2 | Brap_Q99M | -0.23 | -1.67 | -0.94 | -0.57 | -1.39 | -0.5 | -0.81 | 0.05 | -0.65 | -0.83 | 0.45 | -0.18 | -1.5 | -0.82 | -1.67 | -0.79 | 0.19 | 0.55 | -0.3425 | 0.393351 | 0.645828 | 0.205735 | 0.99033 | 0.997988 | 0.994666 |
| Brd1_G5E8P1_T885t_1 | Brd1_G5E8 | 5.66 | 2.35 | 5.42 | 0.45 | 3.08 | 6.27 | 5.24 | 3.62 | 1.14 | 5.48 | -0.27 | 5.03 | 1.36 | -1.46 | 5.99 | 5.98 | 1.0825 | -0.625 | -0.5025 | 0.791781 | 0.396423 | 0.706722 | 0.989223 | 0.997988 | 0.999507 |
| Brd3_Q8K2F0_S262s_1 | Brd3_Q8K2 | -0.01 | -0.1 | -0.05 | 0.13 | -0.14 | -0.12 | -0.08 | -0.09 | 0.03 | -0.02 | -0.02 | -0.34 | -0.25 | -0.44 | 0.28 | -0.13 | -0.1 | -0.08 | -0.1275 | 0.626114 | 0.609026 | 0.712948 | 0.989223 | 0.997988 | 0.995259 |
| Brd4_Q3UH70_S603s_1 | Brd4_Q3UH | -0.77 | -0.33 | -0.84 | -0.75 | -0.53 | -1.39 | -0.61 | -0.75 | -0.72 | -1.03 | -0.22 | -0.85 | -0.86 | 0.12 | 0.15 | -0.79 | -0.1475 | -0.0325 | 0.3275 | 0.344236 | 0.587197 | 0.903581 | 0.989223 | 0.997988 | 0.994666 |
| Brd8_D3YZC7_S547s_1 | Brd8_D3YZ | -1.8 | -2.65 | -3.35 | -6.09 | -4.7 | -2.94 | -4.07 | -3.82 | -0.95 | -3.42 | -1.95 | -5.63 | -1.33 | -3.86 | -2.09 | -3.84 | -0.41 | 0.485 | 0.6925 | 0.486081 | 0.638374 | 0.689048 | 0.989223 | 0.997988 | 0.994666 |
| Brd8_D3YZC7_T264t_1 | Brd8_D3YZ | -0.88 | -1.31 | -0.52 | -1.3 | -0.82 | -0.6 | -1.75 | -1.57 | -1.78 | -1.51 | -2.59 | -4.6 | -0.41 | -1.75 | -0.66 | -1.82 | -0.1825 | -1.6175 | -0.1575 | 0.703463 | 0.597537 | 0.033129 | 0.989223 | 0.997988 | 0.996159 |
| Brd9_Q3UQU0_S565s_1 | Brd9_Q3U( | 0.52 | 1.1 | -0.39 | -0.39 | -0.81 | 0.38 | -1.29 | -1.11 | 1.16 | -0.08 | -1.38 | -0.68 | 2.2 | -1.12 | 0.03 | 0.12 | -0.9175 | -0.455 | 0.0975 | 0.889108 | 0.083053 | 0.453258 | 0.989223 | 0.997988 | 0.999507 |
| Brf1_G3X8S2_S358s_1 | Brf1_G3X8 | -3.12 | -2.74 | -1.45 | -6.2 | -2.74 | -3.34 | -2.4 | -3.33 | -1.41 | -2.17 | -0.77 | -2.63 | 0.74 | -4.71 | -2.48 | -1.58 | 0.425 | 1.6325 | 1.37 | 0.307322 | 0.637495 | 0.115796 | 0.989223 | 0.997988 | 0.994666 |
| Brf1_G3X8S2_S552s_1 | Brf1_G3X8 | -2.28 | -1.71 | -1.47 | -1.83 | -2.07 | -2.75 | -2.9 | -2.17 | -2.03 | -1.46 | -1.04 | -1.4 | -0.01 | -2.61 | -1.72 | -1.41 | -0.65 | 0.34 | 0.385 | 0.472538 | 0.049813 | 0.27945 | 0.989223 | 0.997988 | 0.994666 |
| Brsk1_Q5RJI5_S384s_1 | Brsk1_Q5R | -1.1 | -2.58 | -1.34 | -0.82 | -0.82 | -1.28 | -0.87 | -1.05 | -1.13 | -2.02 | 0.34 | -1.05 | -0.55 | -1.15 | -1.03 | -1.37 | 0.455 | 0.495 | 0.435 | 0.317223 | 0.260278 | 0.39844 | 0.989223 | 0.997988 | 0.994666 |
| Brsk1_Q5RJI5_S399s_1 | Brsk1_Q5R | 0.06 | 0.36 | 0.28 | 0.53 | 0.34 | 0.16 | 0.72 | 0.52 | 0.32 | 0.38 | 0.59 | 0.36 | 0.08 | 0.43 | -0.09 | 0.49 | 0.1275 | 0.105 | -0.08 | 0.762737 | 0.58251 | 0.639188 | 0.989223 | 0.997988 | 0.9971 |
| Brsk1_Q5RJI5_S427s_1 | Brsk1_Q5R | -0.58 | -0.7 | -0.64 | -0.74 | 0.2 | -0.5 | -0.89 | -0.95 | -0.15 | -0.78 | -0.74 | -0.69 | 1.35 | 0.28 | -0.1 | -0.63 | 0.13 | 0.075 | 0.89 | 0.057054 | 0.660338 | 0.753687 | 0.990434 | 0.997988 | 0.994666 |
| Brsk1_Q5RJI5_S432s_1 | Brsk1_Q5R | -0.69 | -0.26 | -1.98 | -1.61 | 0.11 | -0.99 | -0.94 | -0.77 | 0.66 | 0.76 | -0.82 | -1.73 | -0.12 | 0.2 | -0.91 | -1.93 | 0.4875 | 0.8525 | 0.445 | 0.442755 | 0.288077 | 0.214658 | 0.989223 | 0.997988 | 0.994666 |
| Brsk1_Q5RJI5_S450s_1 | Brsk1_Q5R | 0.44 | 0.36 | 0.04 | 0.17 | 0.99 | 0.6 | 0.8 | 0.56 | 0.09 | 0.23 | 0.03 | 0.41 | -0.53 | 0.78 | -0.16 | 0.42 | 0.485 | -0.0625 | -0.125 | 0.714744 | 0.050375 | 0.78266 | 0.989223 | 0.997988 | 0.996966 |
| Brsk1_Q5RJI5_S499s_1 | Brsk1_Q5R | -1.08 | -0.77 | -3.28 | -3.03 | -0.31 | -2.3 | -1.69 | -2.2 | -1.31 | -1.08 | -1.79 | -2.55 | -0.54 | -0.13 | -1.61 | -2.29 | 0.415 | 0.3575 | 0.8975 | 0.235757 | 0.557075 | 0.589895 | 0.989223 | 0.997988 | 0.994666 |
| Brsk1_Q5RJI5_S508s_1 | Brsk1_Q5R | -0.49 | -1.88 | -1.84 | -1.22 | -0.61 | -0.98 | -0.49 | -1.29 | -1.8 | -1.17 | -0.98 | -0.95 | -1.21 | -0.5 | -1.25 | -0.93 | 0.515 | 0.1325 | 0.385 | 0.328049 | 0.184742 | 0.730925 | 0.989223 | 0.997988 | 0.994666 |
| Brsk1_Q5RJI5_S519s_1 | Brsk1_Q5R | -0.92 | -1.32 | -1.96 | -1.75 | -0.3 | 0.04 | -0.58 | -1.28 | -0.31 | -0.34 | -1.79 | -1.83 | -0.08 | -0.41 | -1.35 | -0.76 | 0.9575 | 0.42 | 0.8375 | 0.047107 | 0.023659 | 0.377778 | 0.989223 | 0.997988 | 0.994666 |
| Brsk1_Q5RJI5_S519s_2 | Brsk1_Q5R | 0.86 | 1.23 | 0.87 | 0.08 | 1.51 | 0.86 | 0.73 | 1.07 | 0.46 | 0.99 | 0.57 | 1.22 | 0.04 | 0.29 | -0.29 | 0.5 | 0.2825 | 0.05 | -0.625 | 0.086999 | 0.378997 | 0.878327 | 0.989223 | 0.997988 | 0.994666 |
| Brsk1_Q5RJI5_S527s_1 | Brsk1_Q5R | 0.42 | 0.04 | 0.84 | 0.67 | 0.69 | 0.18 | 1.05 | 0.73 | 0.28 | 0.88 | 0.26 | 0.76 | 0.55 | 1.35 | 0.28 | 1.07 | 0.17 | 0.0525 | 0.32 | 0.352993 | 0.552868 | 0.853871 | 0.989223 | 0.997988 | 0.994666 |
| Brsk1_Q5RJI5_S751s_1 | Brsk1_Q5R | -1.74 | -1.13 | -0.8 | -1.02 | -0.3 | -0.47 | -0.51 | -0.51 | -2 | -1.8 | 0.09 | -0.54 | -0.6 | -0.97 | -0.84 | -0.56 | 0.725 | 0.11 | 0.43 | 0.163821 | 0.018002 | 0.828544 | 0.989223 | 0.997988 | 0.994666 |
| Brsk1_Q5RJI5_T535t_1 | Brsk1_Q5R | 0.03 | 0.01 | 0.99 | 1.07 | 0.19 | 0.51 | 0.9 | 0.81 | -0.01 | 0.33 | 0.97 | 0.68 | -0.21 | 0.85 | 0.36 | 0.84 | 0.0775 | -0.0325 | -0.065 | 0.868938 | 0.819603 | 0.929817 | 0.995762 | 0.997988 | 0.999507 |
| Brsk1_Q5RJI5_T739t_1 | Brsk1_Q5R | -1.7 | -1.32 | -1.07 | -1.87 | -2.41 | -0.75 | -1.89 | -2.38 | 0.29 | -3.05 | -2.7 | -1.04 | 0.78 | -2.24 | -1.32 | -1.81 | -0.3675 | -0.135 | 0.3425 | 0.588734 | 0.379633 | 0.850412 | 0.989223 | 0.997988 | 0.995259 |
| Brsk2_G3UWV4_S368s_1 | Brsk2_G3U | -1.88 | -2.24 | -1.71 | -1.95 | -1.08 | -1.99 | -1.49 | -1.64 | -1.51 | -1.92 | -0.48 | -1.58 | -0.94 | -1.55 | -2.06 | -1.35 | 0.395 | 0.5725 | 0.47 | 0.154554 | 0.158898 | 0.124614 | 0.989223 | 0.997988 | 0.994666 |
| Brsk2_G3UWV4_S383s_1 | Brsk2_G3U | 0.17 | 0.2 | 0.11 | -0.75 | 0.17 | -0.06 | 0.38 | -0.48 | -0.5 | 0.27 | -0.03 | 0.15 | -0.64 | 0.69 | -0.61 | -0.04 | 0.07 | 0.04 | -0.0825 | 0.834682 | 0.82251 | 0.89904 | 0.995762 | 0.997988 | 0.999507 |
| Brsk2_G3UWV4_S394s_1 | Brsk2_G3U | -0.6 | -0.59 | 0.3 | 1.04 | -1.43 | -0.18 | 0.56 | 0.74 | -0.8 | -0.02 | -0.51 | 0.85 | 0.18 | 0.02 | 0.42 | 0.95 | -0.115 | -0.1575 | 0.355 | 0.424573 | 0.839685 | 0.754829 | 0.995762 | 0.997988 | 0.994666 |
| Brsk2_G3UWV4_S417s_1 | Brsk2_G3U | -1.87 | -1.4 | -1.52 | -1.49 | -0.87 | -1.48 | -0.8 | -1.54 | -1.91 | -1.71 | -1.72 | -1.49 | -1.76 | -1.81 | -1.38 | -1.87 | 0.3975 | -0.1375 | -0.135 | 0.600994 | 0.158907 | 0.554797 | 0.989223 | 0.997988 | 0.995259 |
| Brsk2_G3UWV4_S458s_1 | Brsk2_G3U | -1.29 | -2.03 | -1.13 | -1.44 | -2.55 | -1.1 | -1.04 | -1.4 | -1.04 | -2.22 | -0.83 | -1.1 | -0.85 | -1.66 | -1.82 | -1.74 | -0.05 | 0.175 | -0.045 | 0.893446 | 0.897925 | 0.642865 | 0.996493 | 0.997988 | 0.999507 |
| Brsk2_G3UWV4_S462s_1 | Brsk2_G3U | -0.6 | -2 | -0.58 | -0.97 | -0.86 | -0.72 | -0.51 | -0.06 | -1.59 | -1.36 | -0.17 | -1.43 | -2 | -0.42 | -0.89 | -1.68 | 0.5 | -0.1 | -0.21 | 0.657735 | 0.198664 | 0.823781 | 0.989223 | 0.997988 | 0.995259 |
| Brsk2_G3UWV4_S478s_1 | Brsk2_G3U | 0.78 | 0.71 | 1.26 | 1.28 | 0.9 | 0.31 | 1.13 | 1.54 | 0.28 | 0.46 | 0.74 | 1.39 | -0.01 | 0.23 | 0.59 | 0.63 | -0.0375 | -0.29 | -0.6475 | 0.045772 | 0.905302 | 0.37211 | 0.996493 | 0.997988 | 0.994666 |
| Brsk2_G3UWV4_S512s_1 | Brsk2_G3U | -2.59 | -3.65 | -3.09 | -3.79 | -1.06 | -3.29 | -1.68 | -2.88 | -4.2 | -3.05 | -2.43 | -2.81 | -2.65 | -2 | -2.11 | -1.77 | 1.0525 | 0.1575 | 1.1475 | 0.009387 | 0.072821 | 0.728187 | 0.989223 | 0.997988 | 0.994666 |
| Brsk2_G3UWV4_S536s_1 | Brsk2_G3U | 0.35 | 0.62 | 0.46 | 0.22 | 0.74 | 0.59 | 0.62 | 0.47 | 0.74 | 0.54 | 0.09 | 0.55 | 0.32 | 1.15 | 0.11 | 0.61 | 0.1925 | 0.0675 | 0.135 | 0.655685 | 0.365394 | 0.781769 | 0.989223 | 0.997988 | 0.995259 |
| Brsk2_G3UWV4_T454t_1 | Brsk2_G3U | -3.07 | -2.32 | -1.4 | -3.16 | -1.72 | -2.18 | -3.29 | -1.81 | -3.93 | -3.36 | -0.81 | -2.21 | -0.43 | -2.01 | -2.41 | -1.46 | 0.2375 | -0.09 | 0.91 | 0.122786 | 0.636273 | 0.900567 | 0.989223 | 0.997988 | 0.994666 |
| Brsk2_G3UWV4_T466t_1 | Brsk2_G3U | -0.97 | -0.21 | -0.82 | -2.93 | -0.71 | -0.62 | -1.19 | -1.42 | -1.81 | -1.03 | -0.3 | -1.97 | -1.18 | -0.74 | -1.47 | -1.48 | 0.2475 | -0.045 | 0.015 | 0.97884 | 0.659256 | 0.943908 | 0.990434 | 0.997988 | 1 |
| Brsk2_G3UWV4_T470t_1 | Brsk2_G3U | -2.39 | -2.6 | -2.54 | -3.55 | -1.89 | -1.95 | -1.93 | -2.51 | -1.82 | -2.34 | -2.08 | -2.02 | -1.16 | -1.81 | -3.79 | -2.24 | 0.7 | 0.705 | 0.52 | 0.371949 | 0.050017 | 0.048209 | 0.989223 | 0.997988 | 0.994666 |
| Brsk2_G3UWV4_T486t_1 | Brsk2_G3U | 0.06 | 0.03 | 0.36 | 0.58 | 0.48 | 0.36 | 1.11 | 0.38 | -0.06 | 0.29 | 0.32 | 0.57 | -0.24 | 0.85 | 0.36 | 0.48 | 0.325 | 0.0225 | 0.105 | 0.738058 | 0.240119 | 0.929566 | 0.989223 | 0.997988 | 0.9971 |
| Brwd1_Q921C3_S1751s_1 | Brwd1_Q9 | 0.65 | 0.53 | 0.86 | 0.78 | -0.53 | 1.4 | -0.53 | 0.21 | 1.17 | 0.5 | 0.77 | 1.07 | 0.72 | -0.09 | 0.53 | 1.05 | -0.5675 | 0.1725 | -0.1525 | 0.621271 | 0.212089 | 0.490049 | 0.989223 | 0.997988 | 0.995259 |
| Brwd1_Q921C3_S1830s_1 | Brwd1_Q9 | 0.13 | 0.29 | 0.48 | 1.23 | 0.79 | -0.61 | 1.11 | 0.83 | 0.1 | 0 | -0.51 | 0.58 | 0.17 | 0.55 | 0.21 | 0.79 | -0.0025 | -0.49 | -0.1025 | 0.754537 | 0.995366 | 0.181267 | 0.999616 | 0.997988 | 0.9971 |
| Brwd1_Q921C3_S1857s_1 | Brwd1_Q9 | 0.83 | 0.84 | 1.46 | 1.78 | -0.15 | 0.63 | 0.84 | 1.5 | 0.67 | 0.85 | 2.62 | 1.54 | 1.1 | 0.3 | 1.86 | 0.99 | -0.5225 | 0.1925 | -0.165 | 0.683003 | 0.210428 | 0.687059 | 0.989223 | 0.997988 | 0.996159 |
| Brwd1_Q921C3_S2035s_1 | Brwd1_Q9 | -1.65 | -0.52 | 0.58 | 0.25 | 0.1 | -0.68 | 0.09 | 0.98 | -0.66 | -0.49 | 0.34 | 0.83 | -0.44 | -2.25 | 1.41 | 1.82 | 0.4575 | 0.34 | 0.47 | 0.613601 | 0.408787 | 0.548542 | 0.989223 | 0.997988 | 0.995259 |
| Bscl2_Q9Z2E9_S280s_1 | Bscl2_Q9Z | 1.84 | 1.67 | 3.63 | 1.33 | 2.03 | 3.12 | 2.82 | 3.23 | 0.69 | 2.41 | 0.35 | 3.69 | 2.75 | -3.68 | 6.63 | 4.47 | 0.6825 | -0.3325 | 0.425 | 0.827382 | 0.21715 | 0.690437 | 0.989223 | 0.997988 | 0.999507 |
| Bsg_K3W4Q8_S201s_1 | Bsg_K3W4 | -3.93 | -3.51 | -1.97 | -3.44 | -3.35 | -4 | -1.46 | -2.73 | -1.93 | -3.42 | -3.26 | -1.68 | 0.62 | -3.38 | -0.27 | -2.29 | 0.3275 | 0.64 | 1.8825 | 0.056945 | 0.59723 | 0.279131 | 0.989223 | 0.997988 | 0.994666 |
| Bsg_K3W4Q8_T203t_1 | Bsg_K3W4 | 0.72 | 0.24 | 0.93 | -0.05 | -0.74 | 1.61 | 0.31 | 0.37 | 4.88 | 1.08 | 0.43 | 0.52 | 4.61 | -1.23 | 0.6 | 0.76 | -0.0725 | 1.2675 | 0.725 | 0.507976 | 0.881746 | 0.207728 | 0.996493 | 0.997988 | 0.994666 |
| Bsn_O88737_S105s_1 | Bsn_O8873 | 0.71 | 0.53 | 0.79 | 0.34 | 0.64 | 0.65 | -0.06 | 0.22 | 0.85 | 1.19 | -0.46 | 0.57 | 1.32 | -0.37 | 0.63 | 0.82 | -0.23 | -0.055 | 0.0075 | 0.984339 | 0.37527 | 0.883511 | 0.989223 | 0.997988 | 1 |
| Bsn_O88737_S1100s_1 | Bsn_O8873 | 0.85 | 1.41 | 0.8 | 0.7 | 1.13 | 0.39 | 0.81 | 0.35 | 1.18 | 1.56 | 0.05 | 0.38 | 1.21 | 1.28 | 0.94 | 0.73 | -0.27 | -0.1475 | 0.1 | 0.722778 | 0.347668 | 0.70329 | 0.989223 | 0.997988 | 0.9971 |
| Bsn_O88737_S1108s_1 | Bsn_O8873 | 0.4 | 1.54 | 1.08 | 0.91 | 0.31 | 0.45 | 1.08 | 0.42 | 1.23 | 1.53 | 1.06 | 1.23 | 0.69 | 1.38 | 0.81 | 1.02 | -0.4175 | 0.28 | -0.0075 | 0.981537 | 0.201559 | 0.3569 | 0.989223 | 0.997988 | 1 |
| Bsn_O88737_S1114s_1 | Bsn_O8873 | 0.18 | 0.96 | 1.19 | 0.93 | 0.25 | 0.45 | 0.81 | 0.35 | 0.56 | 1.02 | 1.4 | 1.39 | 0.21 | 1.18 | 0.97 | 0.87 | -0.35 | 0.2775 | -0.0075 | 0.982341 | 0.237377 | 0.400443 | 0.989223 | 0.997988 | 1 |
| Bsn_O88737_S1137s_1 | Bsn_O8873 | -1.09 | -1.01 | -0.14 | 0.51 | -0.61 | -1.19 | 0.62 | 0.14 | -1.11 | -0.55 | -0.16 | -0.1 | -1.43 | -0.77 | 0.14 | 0.19 | 0.1725 | -0.0475 | -0.035 | 0.945463 | 0.733418 | 0.912656 | 0.995762 | 0.997988 | 0.999507 |
| Bsn_O88737_S1173s_1 | Bsn_O8873 | 1.41 | 0.35 | 0.27 | 0.78 | 1.17 | 1.46 | 1.31 | 1.23 | 0.31 | 0.75 | -0.14 | 0.54 | -0.63 | 0.54 | 0.32 | 0.61 | 0.59 | -0.3375 | -0.4925 | 0.232775 | 0.070388 | 0.337236 | 0.989223 | 0.997988 | 0.994666 |
| Bsn_O88737_S121s_1 | Bsn_O8873 | -0.36 | -0.72 | 0.12 | 0 | -1.01 | 0.09 | -0.51 | -0.35 | 0.67 | -0.43 | -1.26 | -0.25 | 1.07 | -1.11 | -0.35 | 0.11 | -0.205 | -0.0775 | 0.17 | 0.720265 | 0.518139 | 0.856257 | 0.989223 | 0.997988 | 0.9971 |
| Bsn_O88737_S1236s_1 | Bsn_O8873 | -0.08 | -0.06 | -0.43 | -0.37 | 0.18 | 0.71 | -0.6 | -0.37 | -1.12 | -0.01 | -1.41 | -0.25 | -1.52 | -0.95 | -1 | -0.01 | 0.215 | -0.4625 | -0.635 | 0.099558 | 0.509776 | 0.221064 | 0.989223 | 0.997988 | 0.994666 |
| Bsn_O88737_S1369s_1 | Bsn_O8873 | 1.87 | 1.03 | 0.76 | 1.08 | 0.98 | 2.48 | 2.06 | 2.5 | -0.07 | 1.42 | 1.06 | 1.54 | 0.1 | -0.69 | 1.9 | 1.53 | 0.82 | -0.1975 | -0.475 | 0.433415 | 0.070093 | 0.64551 | 0.989223 | 0.997988 | 0.994666 |
| Bsn_O88737_S1370s_1 | Bsn_O8873 | 2.48 | 1.23 | 0.6 | 0.8 | 2.96 | 2.88 | 1.44 | 1.57 | -1.28 | 2.17 | -0.39 | 1.4 | -0.62 | -1.03 | 1.67 | 1.31 | 0.935 | -0.8025 | -0.945 | 0.205809 | 0.105277 | 0.33087 | 0.989223 | 0.997988 | 0.994666 |
| Bsn_O88737_S138s_2 | Bsn_O8873 | -0.33 | -1.09 | -1.15 | -1.29 | -1.02 | 1.2 | -1.49 | -0.19 | -0.04 | -1.36 | 0.32 | -0.85 | -1.15 | -1.28 | -2.48 | -1.39 | 0.59 | 0.4825 | -0.61 | 0.139001 | 0.311099 | 0.275905 | 0.989223 | 0.997988 | 0.994666 |
| Bsn_O88737_S1407s_1 | Bsn_O8873 | 0.26 | -0.72 | -0.79 | -0.27 | 0.64 | 0.56 | 0.24 | 0.25 | -0.85 | -0.55 | -0.06 | -0.12 | -1.1 | -0.34 | -0.24 | -0.09 | 0.8025 | -0.015 | -0.0625 | 0.861038 | 0.02011 | 0.963742 | 0.989223 | 0.997988 | 0.999507 |
| Bsn_O88737_S142s_1 | Bsn_O8873 | 0.17 | -1.32 | 1.19 | 1.52 | -0.38 | 0.75 | -0.17 | 0.38 | -0.23 | -0.73 | 0.84 | 0.58 | -0.82 | -0.96 | 1.16 | 0.44 | -0.245 | -0.275 | -0.435 | 0.553882 | 0.690939 | 0.680454 | 0.994893 | 0.997988 | 0.994666 |
| Bsn_O88737_S142s_2 | Bsn_O8873 | -0.33 | -1.09 | -1.15 | -1.29 | -1.02 | 1.2 | -1.49 | -0.19 | -0.04 | -1.36 | 0.32 | -0.85 | -1.15 | -1.28 | -2.48 | -1.39 | 0.59 | 0.4825 | -0.61 | 0.139001 | 0.311099 | 0.275905 | 0.989223 | 0.997988 | 0.994666 |
| Bsn_O88737_S1460s_1 | Bsn_O8873 | 1.34 | 0.13 | 0.04 | 0.8 | 0.88 | 1.11 | 0.16 | 0.53 | 0.24 | 0.58 | 0.08 | -0.01 | -0.11 | 1.05 | -0.2 | 0.35 | 0.0925 | -0.355 | -0.305 | 0.471675 | 0.800401 | 0.322174 | 0.995762 | 0.997988 | 0.994666 |
| Bsn_O88737_S1482s_1 | Bsn_O8873 | 0.76 | 1.27 | 0.88 | 0.78 | 1.02 | 0.56 | 0.88 | 0.7 | 0.5 | 1.26 | 0.39 | 0.81 | 0.78 | 0.88 | 0.54 | 0.86 | -0.1325 | -0.1825 | -0.1575 | 0.537039 | 0.568912 | 0.519226 | 0.989223 | 0.997988 | 0.994666 |
| Bsn_O88737_S151s_1 | Bsn_O8873 | -0.4 | 0.02 | -0.45 | -0.48 | -0.05 | -0.39 | 0.49 | 0.09 | -0.28 | -0.05 | 1.73 | -0.28 | -0.53 | 0.77 | 0.19 | 0.06 | 0.3625 | 0.6075 | 0.45 | 0.196141 | 0.19039 | 0.222225 | 0.989223 | 0.997988 | 0.994666 |
| Bsn_O88737_S1650s_1 | Bsn_O8873 | -0.68 | -0.73 | -0.67 | -0.83 | 0.27 | -0.18 | -0.23 | -0.29 | -0.87 | -0.57 | 1.39 | -0.03 | 0.73 | -0.06 | -0.35 | -0.54 | 0.62 | 0.7075 | 0.6725 | 0.062854 | 0.017824 | 0.162753 | 0.989223 | 0.997988 | 0.994666 |
| Bsn_O88737_S1990s_1 | Bsn_O8873 | 0.1 | 0.77 | 0.54 | 0.19 | 0.48 | -0.03 | 0.64 | 0.48 | -0.15 | 0.49 | -0.11 | 0.66 | -0.06 | 0.32 | 0.84 | 0.71 | -0.0075 | -0.1775 | 0.0525 | 0.865562 | 0.977088 | 0.556302 | 0.999187 | 0.997988 | 0.999507 |
| Bsn_O88737_S2046s_1 | Bsn_O8873 | -0.29 | -0.09 | -0.28 | -0.15 | 0.58 | 0.04 | 0.21 | -0.12 | -0.75 | -0.63 | -0.41 | -0.12 | -0.87 | 0.43 | -0.47 | -0.06 | 0.38 | -0.275 | -0.04 | 0.902384 | 0.125957 | 0.262235 | 0.989223 | 0.997988 | 0.999507 |

| | | | | | | | | | | | | | | | | | | | | | | | | | | |
|---|---|---|---|---|---|---|---|---|---|---|---|---|---|---|---|---|---|---|---|---|---|---|---|---|---|---|
| Bsn_O88737_S241s_1 | Bsn_O8873 | -1.05 | -3.19 | -1.99 | -1.41 | -0.5 | -0.11 | -0.45 | -0.19 | -1.65 | -1.17 | 0.2 | -0.7 | -0.49 | -0.71 | -2.58 | -1.91 | 1.5975 | 1.08 | 0.4875 | 0.439318 | 0.005279 | 0.082259 | 0.989223 | 0.997988 | 0.994666 |
| Bsn_O88737_S245s_1 | Bsn_O8873 | -1.11 | -2.17 | -1.07 | -1.5 | -0.3 | -0.36 | -0.25 | 0.13 | -1.16 | -1.68 | -0.77 | -0.39 | -1.96 | -1.57 | -0.47 | -0.87 | 1.2675 | 0.4625 | 0.245 | 0.563353 | 0.001979 | 0.242831 | 0.989223 | 0.997988 | 0.995259 |
| Bsn_O88737_S2578s_1 | Bsn_O8873 | -0.35 | 0.05 | 0.59 | -0.24 | -0.59 | 0.22 | -0.4 | 0.66 | -0.67 | -0.29 | 0.69 | 0.69 | -1.37 | -0.95 | -1.17 | -0.02 | -0.04 | 0.0925 | -0.89 | 0.039532 | 0.9104 | 0.81819 | 0.997335 | 0.997988 | 0.994666 |
| Bsn_O88737_S2659s_1 | Bsn_O8873 | 2.53 | 1.55 | -0.16 | -1.02 | 0.01 | 2.68 | 0.67 | 0.15 | 1.85 | 2.22 | -0.61 | -0.09 | 1.36 | 0.2 | 0.73 | 1.48 | 0.1525 | 0.1175 | 0.2175 | 0.775258 | 0.86263 | 0.900885 | 0.996493 | 0.997988 | 0.99798 |
| Bsn_O88737_S2805s_1 | Bsn_O8873 | -0.47 | -0.42 | -0.66 | -0.77 | 0.26 | 0.59 | -0.39 | -0.17 | -1.5 | -0.05 | -0.3 | -0.04 | -0.57 | -0.03 | -1.01 | -0.33 | 0.6525 | 0.1075 | 0.095 | 0.744631 | 0.036765 | 0.769574 | 0.989223 | 0.997988 | 0.9971 |
| Bsn_O88737_S2808s_1 | Bsn_O8873 | 1.77 | 1.35 | -0.41 | 0.05 | 1.8 | 1.19 | 0.87 | 0.44 | 0.21 | 1.2 | -0.05 | -0.01 | 1.15 | 0.49 | -0.21 | 0.99 | 0.385 | -0.3525 | -0.085 | 0.878514 | 0.479259 | 0.527385 | 0.989223 | 0.997988 | 0.999507 |
| Bsn_O88737_S2811s_1 | Bsn_O8873 | -0.25 | -1.46 | -0.94 | -0.51 | -0.34 | 0.28 | 0.41 | -0.17 | -0.14 | -1.18 | -0.67 | -0.26 | -0.79 | -0.12 | -0.67 | -0.47 | 0.835 | 0.2275 | 0.2775 | 0.421114 | 0.02884 | 0.536457 | 0.989223 | 0.997988 | 0.994666 |
| Bsn_O88737_S2822s_1 | Bsn_O8873 | 0.2 | 0.14 | -0.18 | -0.16 | 0.22 | 0.7 | -0.08 | 0.5 | -1.92 | -0.4 | 0.19 | 0.38 | -1.08 | -1.11 | -1.09 | -0.61 | 0.335 | -0.4375 | -0.9725 | 0.003845 | 0.204475 | 0.391517 | 0.989223 | 0.997988 | 0.994666 |
| Bsn_O88737_S284s_1 | Bsn_O8873 | -0.44 | -0.49 | 0.3 | 0.31 | 0.82 | 0.73 | 0.84 | 0.1 | 0.06 | -0.28 | -0.3 | 0.87 | -0.31 | -0.32 | 0.39 | 1.05 | 0.7025 | 0.1675 | 0.2825 | 0.487849 | 0.042413 | 0.646356 | 0.989223 | 0.997988 | 0.994666 |
| Bsn_O88737_S2858s_1 | Bsn_O8873 | 0.41 | 0.45 | 0.26 | 0.59 | 0.77 | 0.36 | 1.22 | 0.55 | 0.41 | 0.85 | 0.52 | 0.31 | 0.27 | 1.11 | 0.53 | 1.22 | 0.2975 | 0.095 | 0.355 | 0.254038 | 0.255811 | 0.682287 | 0.989223 | 0.997988 | 0.994666 |
| Bsn_O88737_S2860s_1 | Bsn_O8873 | 0.36 | 1.31 | 1.42 | 0.8 | 0.97 | 0.33 | 1.15 | 1.04 | 0.26 | 0.79 | 0.75 | 1.03 | 0.75 | 1.46 | 1.38 | 0.89 | -0.1 | -0.265 | 0.1475 | 0.664107 | 0.755784 | 0.418757 | 0.995762 | 0.997988 | 0.996159 |
| Bsn_O88737_S2866s_1 | Bsn_O8873 | -0.7 | -1.28 | -1.14 | -1.86 | -1.02 | -0.76 | -1.12 | -1.13 | -1.34 | -0.79 | -1.02 | -1.51 | -1.12 | -0.81 | -1.44 | -1.22 | 0.2375 | 0.08 | 0.0975 | 0.761097 | 0.415805 | 0.801778 | 0.989223 | 0.997988 | 0.9971 |
| Bsn_O88737_S2908s_1 | Bsn_O8873 | 0.19 | 0.53 | 0.36 | 0.25 | 0.59 | 0.03 | 0.55 | 0.24 | -0.13 | 0.44 | -0.36 | 0.14 | 0.12 | 0.27 | 0.04 | 0.35 | 0.02 | -0.31 | -0.1375 | 0.567563 | 0.930355 | 0.246479 | 0.997844 | 0.997988 | 0.995259 |
| Bsn_O88737_S301s_1 | Bsn_O8873 | 0.3 | -0.18 | -0.41 | -0.41 | 0.41 | 0.57 | 0.26 | 0.05 | -0.7 | -0.14 | 0.05 | -0.31 | -0.6 | 0.42 | -0.84 | -0.6 | 0.4975 | -0.1 | -0.23 | 0.521055 | 0.075086 | 0.72338 | 0.989223 | 0.997988 | 0.994666 |
| Bsn_O88737_S3022s_1 | Bsn_O8873 | 0.35 | 0.67 | 0.8 | 1.01 | 0.67 | 0.64 | 0.84 | 0.94 | -0.14 | 0.74 | 0.54 | 0.7 | -0.98 | 0.51 | 0.41 | 0.63 | 0.065 | -0.2475 | -0.565 | 0.1858 | 0.778179 | 0.405201 | 0.995762 | 0.997988 | 0.994666 |
| Bsn_O88737_S3158s_1 | Bsn_O8873 | -0.14 | 0.08 | -0.29 | -0.46 | 0.13 | 0.2 | 0.09 | 0.02 | -0.33 | 0.17 | 0.19 | 0.04 | -0.77 | 0.1 | 0.06 | 0.19 | 0.3125 | 0.22 | 0.0975 | 0.751388 | 0.169226 | 0.381023 | 0.989223 | 0.997988 | 0.9971 |
| Bsn_O88737_S3218s_1 | Bsn_O8873 | -0.28 | -0.1 | -0.27 | -0.39 | -0.33 | -0.24 | 0.58 | 0.08 | -1.23 | -0.03 | -0.74 | -0.22 | -0.78 | -0.74 | 0.01 | -0.33 | 0.2825 | -0.295 | -0.2 | 0.476718 | 0.295949 | 0.35461 | 0.989223 | 0.997988 | 0.994666 |
| Bsn_O88737_S3258s_1 | Bsn_O8873 | 0.25 | -0.21 | 1.23 | 1.12 | -0.66 | 0.57 | 0.34 | 0.69 | 1.7 | -0.2 | 0.32 | 0.24 | 1.5 | -1.51 | 1.05 | 1.1 | -0.3625 | -0.0825 | -0.0625 | 0.927885 | 0.414581 | 0.870345 | 0.989223 | 0.997988 | 0.999507 |
| Bsn_O88737_S3301s_1 | Bsn_O8873 | 0.71 | 0.75 | 0.8 | 0.57 | -0.78 | 0.58 | -0.4 | 0.35 | 0.55 | 0.57 | 1.38 | 0.9 | -0.81 | 0.3 | -0.01 | -0.11 | -0.77 | 0.1425 | -0.865 | 0.016058 | 0.040849 | 0.591658 | 0.989223 | 0.997988 | 0.994666 |
| Bsn_O88737_S3382s_1 | Bsn_O8873 | -1.61 | -0.81 | -0.97 | -1.78 | 0.01 | -1.48 | -0.94 | -1.71 | -2.1 | -1.66 | -1.26 | -1.28 | -1.46 | -0.52 | -1.47 | -1.23 | 0.2625 | -0.2825 | 0.1225 | 0.729879 | 0.541601 | 0.404584 | 0.989223 | 0.997988 | 0.9971 |
| Bsn_O88737_S3460s_1 | Bsn_O8873 | -0.26 | -1.54 | -0.88 | -1.14 | -1.04 | -0.23 | -1.21 | -0.59 | -2.01 | -1.13 | 0.63 | -0.63 | -1.1 | -1.94 | -0.95 | -0.49 | 0.1875 | 0.17 | -0.165 | 0.686885 | 0.595162 | 0.764264 | 0.989223 | 0.997988 | 0.996159 |
| Bsn_O88737_S3466s_1 | Bsn_O8873 | -0.86 | -0.98 | -0.9 | -1.99 | -2.06 | 0.12 | -1.98 | -1.45 | -0.01 | -0.76 | 0.66 | -1.83 | -0.31 | -1.41 | -2.1 | -1.38 | -0.16 | 0.6975 | -0.1175 | 0.792701 | 0.759649 | 0.228443 | 0.995762 | 0.997988 | 0.999507 |
| Bsn_O88737_S3512s_1 | Bsn_O8873 | 0.99 | 1.4 | 1.46 | 1.42 | 0.56 | 0.58 | 0.88 | 1.3 | 0.79 | 1.19 | 1.28 | 1.21 | 0.38 | 1.08 | 1.03 | 1.07 | -0.4875 | -0.2 | -0.4275 | 0.151492 | 0.082528 | 0.414754 | 0.989223 | 0.997988 | 0.994666 |
| Bsn_O88737_S3512s_2 | Bsn_O8873 | 0.56 | 1.01 | 2.04 | 1.52 | 0.93 | 0.55 | 1.28 | 1.8 | 0.76 | 0.51 | 1.44 | 1.51 | -0.48 | 1.2 | 1.4 | 1.83 | -0.1425 | -0.2275 | -0.295 | 0.597169 | 0.722259 | 0.574529 | 0.995437 | 0.997988 | 0.995259 |
| Bsn_O88737_S3516s_1 | Bsn_O8873 | 1.67 | 1.6 | 1.68 | 1.83 | 0.92 | 0.93 | 1.41 | 1.44 | 2.21 | 2.21 | 2.52 | 2.13 | 0.72 | 0.73 | 1.7 | 1.72 | -0.52 | 0.5725 | -0.4775 | 0.170634 | 0.044473 | 0.024579 | 0.989223 | 0.997988 | 0.994666 |
| Bsn_O88737_S3516s_2 | Bsn_O8873 | 1.32 | 1.33 | 2.01 | 1.63 | 1.83 | 1.25 | 1.42 | 1.68 | 0.95 | 1.19 | 0.75 | 1.97 | 0.29 | 1.28 | 1.96 | 1.89 | -0.0275 | -0.3575 | -0.2175 | 0.605637 | 0.915185 | 0.301155 | 0.997335 | 0.997988 | 0.995259 |
| Bsn_O88737_S3519s_1 | Bsn_O8873 | 0.59 | 0.75 | 1.36 | 1.26 | 1.46 | 1.15 | 1.8 | 1.59 | 0.16 | 0.57 | -0.15 | 1.25 | 0.63 | 0.62 | 1.52 | 1.85 | 0.51 | -0.5325 | 0.165 | 0.666065 | 0.08696 | 0.16962 | 0.989223 | 0.997988 | 0.996159 |
| Bsn_O88737_S3519s_2 | Bsn_O8873 | 0.72 | 0.93 | 1.14 | 1.45 | 1.13 | 0.77 | 0.79 | 1.49 | 0.67 | 0.64 | 1.04 | 1.4 | -0.5 | 1.13 | 1.34 | 1.6 | -0.015 | -0.1225 | -0.1675 | 0.726382 | 0.955915 | 0.669176 | 0.999187 | 0.997988 | 0.9971 |
| Bsn_O88737_S3545s_1 | Bsn_O8873 | 0.26 | 0.18 | 0.33 | 0.48 | -0.16 | 0 | 0.16 | 0.15 | -0.49 | 0.24 | 0.01 | -0.13 | -1.67 | 0.15 | 0.02 | 0.16 | -0.275 | -0.405 | -0.6475 | 0.165691 | 0.20468 | 0.124459 | 0.989223 | 0.997988 | 0.994666 |
| Bsn_O88737_S3730s_1 | Bsn_O8873 | -3.35 | -2.61 | -2.45 | -2.57 | -2.64 | -2.48 | -2.43 | -2.77 | -0.6 | -3.62 | -2.32 | -1.74 | 0.81 | -1.28 | -1.92 | -1.47 | 0.165 | 0.675 | 1.78 | 0.012509 | 0.537109 | 0.28109 | 0.989223 | 0.997988 | 0.994666 |
| Bsn_O88737_S435s_1 | Bsn_O8873 | -1.99 | -2.97 | -2.84 | -9.39 | -3.33 | -1.45 | -3.44 | -2.72 | 1.08 | -3.14 | -3.74 | -1.67 | 2.64 | -2.36 | 1.63 | -3.16 | 1.5625 | 2.43 | 3.985 | 0.060822 | 0.316733 | 0.190187 | 0.989223 | 0.997988 | 0.994666 |
| Bsn_O88737_S572s_1 | Bsn_O8873 | -2.13 | -3.79 | -3.51 | -11.59 | -2.83 | -3.22 | -1.95 | -3.02 | 1.51 | -2.84 | -3.46 | -1.96 | 2.79 | -3.19 | 2.58 | -3.67 | 2.5 | 3.5675 | 4.8825 | 0.063423 | 0.197003 | 0.115331 | 0.989223 | 0.997988 | 0.994666 |
| Bsn_O88737_S817s_1 | Bsn_O8873 | 0.52 | 1.33 | 1.59 | 0.47 | 1.01 | 1.17 | -1.3 | 0.74 | 0.6 | 2.02 | 0.79 | -1.33 | 1.49 | -0.18 | -1 | -0.32 | -0.5725 | -0.4575 | -0.98 | 0.103058 | 0.333446 | 0.504469 | 0.989223 | 0.997988 | 0.994666 |
| Bsn_O88737_S90s_1 | Bsn_O8873 | -1.25 | -3.74 | -2.66 | -4.82 | -1.82 | 0.26 | -1.34 | -0.82 | 0.18 | -2.44 | -5.81 | -1.21 | -0.17 | -2.21 | -0.22 | -2.22 | 2.1875 | 0.7975 | 1.9125 | 0.045144 | 0.017629 | 0.545195 | 0.989223 | 0.997988 | 0.994666 |
| Bsn_O88737_T2595t_1 | Bsn_O8873 | -0.03 | 0.74 | 4.85 | 4.18 | -0.07 | 0.5 | 2.35 | 4.02 | -0.65 | 0.25 | 6.75 | 5.19 | 0.77 | -0.36 | 3.78 | 3.73 | -0.735 | 0.45 | -0.455 | 0.742851 | 0.579446 | 0.81336 | 0.989223 | 0.997988 | 0.9971 |
| Bsn_O88737_T3254t_1 | Bsn_O8873 | 3.29 | 2.02 | 3.19 | 2.3 | 1.16 | 3.98 | 1.55 | 2.54 | 2.13 | 2.34 | 0.71 | 2.91 | 1.41 | -2.4 | 2.33 | 2.37 | -0.3925 | -0.6775 | -1.7725 | 0.108297 | 0.535515 | 0.21974 | 0.989223 | 0.997988 | 0.994666 |
| Bsn_O88737_T3896t_1 | Bsn_O8873 | 0.16 | 0.42 | 0.18 | 0.42 | 0.77 | -0.29 | 0 | 0.68 | -0.13 | 0.4 | 0.72 | 0.44 | -0.89 | 0.32 | -0.57 | 0.7 | -0.005 | 0.0625 | -0.405 | 0.312982 | 0.98649 | 0.809303 | 0.999187 | 0.997988 | 0.994666 |
| Bsn_O88737_T55t_1 | Bsn_O8873 | -0.99 | 0.61 | 0.08 | 0.36 | 0.79 | -0.33 | 0.64 | -0.24 | 0.48 | 0.25 | 0.31 | -0.17 | 0.05 | 0.71 | 0.3 | -0.92 | 0.2 | 0.2025 | 0.02 | 0.966275 | 0.644264 | 0.598592 | 0.99033 | 0.997988 | 0.999507 |
| Btbd10_E9Q6L8_S44s_1 | Btbd10_E9 | -0.15 | -0.22 | 0.47 | 0.13 | -0.66 | 0.16 | -0.89 | -0.61 | 1.48 | 0.03 | -0.85 | -0.31 | 1.83 | -0.82 | -0.01 | -0.13 | -0.5575 | 0.03 | 0.16 | 0.769578 | 0.089404 | 0.951437 | 0.989223 | 0.997988 | 0.9971 |
| Btbd10_E9Q6L8_S71s_1 | Btbd10_E9 | 0.22 | -0.36 | 0.36 | 0.54 | -0.49 | 0.54 | -0.14 | -0.7 | 1.49 | 0.3 | -0.21 | 1 | 2.19 | -0.63 | 0.07 | 0.45 | -0.3875 | 0.455 | 0.33 | 0.571978 | 0.271546 | 0.289721 | 0.989223 | 0.997988 | 0.995259 |
| Btbd9_Q8C726_T578t_1 | Btbd9_Q8C | -1.36 | 0.34 | -0.4 | -1.74 | -0.43 | 0.38 | -0.38 | 0.33 | 0.29 | -0.08 | -1.45 | -0.83 | -0.59 | -0.77 | -1.73 | 0.32 | 0.765 | 0.2725 | 0.0975 | 0.866258 | 0.13538 | 0.630752 | 0.989223 | 0.997988 | 0.999507 |
| Bzw1_Q9CQC6_S411s_1 | Bzw1_Q9C | -0.1 | -0.26 | 0.16 | -0.64 | -2.47 | 0.42 | -3.28 | -0.87 | 1.37 | -0.36 | 0.59 | -0.58 | -0.73 | -1.56 | -0.25 | -0.65 | -1.34 | 0.465 | -0.5875 | 0.119531 | 0.095692 | 0.32591 | 0.989223 | 0.997988 | 0.994666 |
| Bzw1_Q9CQC6_S413s_1 | Bzw1_Q9C | -0.61 | -0.53 | 1.51 | 0.75 | -2.3 | 1.03 | -0.46 | 0.1 | 0.73 | 0.75 | 0.77 | 0.33 | 0.73 | -1.21 | 0.79 | 0.2 | -0.6875 | 0.365 | -0.1525 | 0.808554 | 0.377963 | 0.468246 | 0.989223 | 0.997988 | 0.999507 |
| C0HKD9_S94s_1 | C0HKD9_S | -1.68 | -2.24 | -2.7 | -3.28 | -2.2 | -3.34 | -3.05 | -2.36 | -4.35 | -2.22 | -0.98 | -3.28 | -2.75 | -1.96 | -2.44 | -2.88 | -1.3225 | -1.8625 | -0.755 | 0.935542 | 0.531306 | 0.746509 | 0.989223 | 0.997988 | 0.999507 |
| C2cd2_E9Q3C1_S436s_1 | C2cd2_E9C | -0.24 | 0.03 | 0.12 | -0.04 | 0.67 | -0.3 | 0.27 | -0.4 | -0.14 | -0.38 | 0.31 | -0.4 | 0.42 | 0.83 | 0.36 | 0.79 | -0.2625 | -0.2325 | -0.0325 | 0.030432 | 0.751673 | 0.638036 | 0.995762 | 0.997988 | 0.994666 |
| C2cd2l_Q80X80_S374s_1 | C2cd2l_Q8 | 0.7 | 0.44 | 0.03 | 0.61 | 0.48 | 0.53 | 1.24 | 0.65 | -0.64 | 0.83 | -0.88 | 0.43 | 0.02 | -0.76 | 1.04 | 0.96 | 0.0925 | -0.12 | 0.6325 | 0.769045 | 0.31584 | 0.251518 | 0.989223 | 0.997988 | 0.9971 |
| C2cd2l_Q80X80_S396s_1 | C2cd2l_Q8 | -0.67 | -0.38 | -0.65 | 0.04 | -0.34 | -0.26 | 0.35 | -0.28 | -0.22 | 0.15 | -0.19 | -0.15 | 0.21 | 0.55 | -0.02 | 0 | 0.28 | -0.51 | -0.13 | 0.058909 | 0.313436 | 0.241241 | 0.989223 | 0.997988 | 0.994666 |
| C2cd2l_Q80X80_S411s_1 | C2cd2l_Q8 | -0.29 | -0.08 | -0.23 | 0.02 | 0.09 | -0.63 | 0.6 | -0.13 | -0.26 | -0.11 | -0.67 | -0.23 | 0.01 | 0.36 | 0.27 | 0.21 | 0.2825 | 0.3125 | 0.6 | 0.157428 | 0.664916 | 0.466778 | 0.991343 | 0.997988 | 0.994666 |
| C2cd2l_Q80X80_S468s_1 | C2cd2l_Q8 | 1.91 | 1.25 | 1.51 | 2.8 | 1.35 | 2.11 | 1.88 | 2.8 | 1.83 | 1.27 | 2.17 | 1.47 | 2.07 | 1.45 | 2.38 | 2.12 | 0.1275 | -0.1725 | 0.3575 | 0.731099 | 0.696666 | 0.644221 | 0.994893 | 0.997988 | 0.9971 |
| C2cd2l_Q80X80_S48s_2 | C2cd2l_Q8 | 0.79 | 1.61 | 1.57 | 1.64 | 0.79 | 0.51 | 0.52 | 1.07 | 0.53 | 1.44 | 1.33 | 1.48 | 0.67 | 1.07 | 1.08 | 1.61 | 0.1675 | -0.1825 | 0.1375 | 0.37511 | 0.034095 | 0.532011 | 0.989223 | 0.997988 | 0.994666 |
| C2cd2l_Q80X80_S59s_1 | C2cd2l_Q8 | 1.11 | 1.43 | 0.98 | 0.82 | 0.27 | 1.26 | 0.45 | 1.16 | 0.62 | 1.01 | 0.94 | -0.38 | 1.95 | 1.19 | 0.41 | 0.04 | -0.68 | -0.2075 | -0.295 | 0.668892 | 0.335612 | 0.157917 | 0.989223 | 0.997988 | 0.996159 |
| C2cd2l_Q80X80_S613s_1 | C2cd2l_Q8 | -0.33 | 0.44 | 0.89 | 0.6 | 0.05 | -0.58 | 0.71 | 0.14 | -0.62 | 0.16 | 0.72 | 0.18 | -0.63 | 0.51 | 1.35 | 1.32 | -0.3 | -0.5375 | -0.1875 | 0.638898 | 0.392735 | 0.455883 | 0.989223 | 0.997988 | 0.995259 |
| C2cd2l_Q80X80_S619s_1 | C2cd2l_Q8 | 0.68 | 1.28 | 1.91 | 0.33 | 0.83 | 0.15 | 1.01 | 0.03 | 0.59 | 1.18 | -0.36 | 1.12 | 0.23 | 1.02 | 1.12 | 0.6 | -0.32 | -0.29 | 0.2375 | 0.456686 | 0.201126 | 0.388419 | 0.989223 | 0.997988 | 0.994666 |
| C2cd2l_Q80X80_S623s_1 | C2cd2l_Q8 | 0.32 | 1.27 | -0.02 | 0.13 | 1.02 | 0.51 | 0.74 | -0.69 | 0.41 | 1.12 | 0.72 | 0.99 | -0.02 | 1.31 | 0.34 | 0.82 | -0.545 | -0.4175 | -0.3075 | 0.650272 | 0.946214 | 0.283567 | 0.999187 | 0.997988 | 0.995259 |
| C2cd2l_Q80X80_S660s_1 | C2cd2l_Q8 | -0.08 | 0.38 | -0.35 | 0.26 | 0.34 | -0.12 | 0.12 | -0.02 | 0.12 | 0.43 | -0.17 | 0.69 | -1.35 | 0.3 | -0.19 | 0.22 | -0.03 | 0.385 | 0.1875 | 0.465674 | 0.912407 | 0.469912 | 0.997335 | 0.997988 | 0.994666 |
| C2cd2l_Q80X80_S662s_1 | C2cd2l_Q8 | 0.68 | 0.7 | -0.25 | 0.3 | 0.69 | 0.47 | 0.11 | 0.11 | 0.68 | 0.48 | 0.46 | 0.14 | -1.8 | 0.84 | -0.57 | -0.76 | 0.0275 | 0.215 | -0.3075 | 0.113216 | 0.965924 | 0.778296 | 0.999187 | 0.997988 | 0.994666 |
| C2cd2l_Q80X80_T388t_1 | C2cd2l_Q8 | -1.69 | -1.46 | -0.77 | -2.39 | -0.28 | -1.8 | -1.42 | -1.67 | -2.34 | -0.81 | -1.96 | -1.28 | -2.95 | 0.1 | -0.48 | -1.57 | -0.0125 | 0.0825 | -0.93 | 0.603257 | 0.530727 | 0.965144 | 0.989223 | 0.997988 | 0.995259 |
| C2cd2l_Q80X80_T417t_1 | C2cd2l_Q8 | 1.57 | 1.31 | 1.45 | 1.7 | 1.05 | 0.87 | 1.81 | 1.75 | 1.58 | 2.06 | 2.45 | 1.29 | 1.77 | 1.75 | 2.36 | 1.91 | 0.285 | -0.02 | 0.3525 | 0.11842 | 0.632858 | 0.286431 | 0.989223 | 0.997988 | 0.994666 |
| C2cd2l_Q80X80_T417t_2 | C2cd2l_Q8 | 1.88 | 0.21 | -0.03 | 0.68 | 1.05 | 1.6 | 0.84 | -0.07 | 1.18 | 1.04 | 1.23 | 0.1 | -0.5 | 1.38 | 0.4 | 0.32 | -0.1375 | 0.3375 | 0.44 | 0.596304 | 0.735689 | 0.671972 | 0.995762 | 0.997988 | 0.995259 |
| C2cd2l_Q80X80_T428t_1 | C2cd2l_Q8 | 1.29 | -0.67 | -1.65 | -1.62 | 0.22 | 1.17 | -0.96 | -1.08 | 0.45 | 0.19 | -1.1 | -1.3 | -0.38 | 0.71 | -0.88 | -1.51 | 0.17 | 0.2025 | -0.285 | 0.842356 | 0.517195 | 0.762938 | 0.989223 | 0.997988 | 0.999507 |
| C2cd2l_Q80X80_T428t_2 | C2cd2l_Q8 | 1.88 | 0.21 | -0.46 | -0.06 | 1.05 | 1.6 | -0.25 | 0.24 | 1.18 | 1.04 | 1.88 | -0.86 | -0.5 | 1.38 | 0.28 | -0.51 | 0.5 | 0.2225 | 0.1475 | 0.710917 | 0.652244 | 0.556471 | 0.990358 | 0.997988 | 0.996966 |
| C2cd4cC2CD4 family_Q5HZI2_S153s_1 | C2cd4cC2C | 0.01 | -0.88 | 0.47 | -0.04 | -0.54 | -0.48 | 2.03 | 1 | -1.28 | -1.22 | 1.31 | 0.86 | -1.11 | -1.17 | -0.71 | 0.75 | 0.2675 | 0.4175 | -0.23 | 0.379078 | 0.327808 | 0.96707 | 0.989223 | 0.997988 | 0.994666 |
| C2cd4cC2CD4 family_Q5HZI2_S175s_1 | C2cd4cC2C | 0.26 | 0.08 | -1.01 | -0.86 | -1.07 | 0.18 | -0.51 | -0.51 | -0.42 | -0.44 | -0.51 | -1.01 | -1.4 | -0.47 | -1.18 | -1.35 | 0.6125 | 0.0275 | -0.45 | 0.095403 | 0.810911 | 0.561115 | 0.995762 | 0.997988 | 0.994666 |
| C2cd4cC2CD4 family_Q5HZI2_S203s_1 | C2cd4cC2C | 0.67 | -1 | -0.12 | -0.08 | -0.07 | 0.71 | 0.3 | 0.65 | -0.56 | -0.49 | 2.59 | 0.45 | -1.12 | 0.14 | -0.22 | 0.19 | -0.095 | -0.2125 | -0.7175 | 0.788023 | 0.183134 | 0.396928 | 0.989223 | 0.997988 | 0.999507 |
| C2cd4cC2CD4 family_Q5HZI2_S205s_1 | C2cd4cC2C | 1.18 | -0.02 | 0.26 | 0.18 | 0.1 | 1.37 | -0.09 | 0.28 | 1.46 | -0.31 | 2.24 | 0.15 | 0.21 | 1.07 | -0.32 | -0.07 | 0.53 | 0.63 | -0.12 | 0.664088 | 0.970394 | 0.421189 | 0.999187 | 0.997988 | 0.996159 |
| C2cd4cC2CD4 family_Q5HZI2_S259s_1 | C2cd4cC2C | 2.21 | 1.06 | 0.21 | 0.11 | 0.52 | 3.49 | 2.13 | 0.51 | 1.58 | 2.31 | 1.09 | 0.52 | 1.71 | -0.39 | 2.26 | 2.96 | 0.015 | 0.485 | -0.1775 | 0.351012 | 0.327788 | 0.410858 | 0.989223 | 0.997988 | 0.994666 |
| C2cd4cC2CD4 family_Q5HZI2_T282t_1 | C2cd4cC2C | -0.48 | -1.59 | -1.48 | -0.43 | -0.83 | -0.22 | -1.3 | -0.6 | -0.31 | -0.89 | 1.17 | -0.13 | 0.09 | 0.3 | -0.92 | -0.99 | 0.765 | 0.4775 | 0.7375 | 0.190844 | 0.500547 | 0.084109 | 0.989223 | 0.997988 | 0.994666 |
| C2cd5_E9Q026_S295s_1 | C2cd5_E9C | 0.55 | 0.16 | 0.78 | 1.15 | 0.36 | 1.67 | 1.07 | 0.9 | 0 | -0.6 | 0.4 | 0.99 | -0.98 | 0.4 | 0.77 | 1.05 | 0.2575 | 0.955 | 0.615 | 0.467714 | 0.336679 | 0.25814 | 0.989223 | 0.997988 | 0.994666 |
| C2cd5_Q7TPS5_S284s_1 | C2cd5_Q7T | 6.33 | 2.6 | 2.29 | 1.57 | 2.97 | 6.38 | 3.55 | 1.03 | 1.96 | 3.05 | 2.01 | 1.4 | 1.06 | 2.72 | 1.14 | 1.73 | 0.34 | -0.4625 | -0.35 | 0.144328 | 0.828883 | 0.285987 | 0.995762 | 0.997988 | 0.994666 |
| C2cd5_Q7TPS5_T330t_1 | C2cd5_Q7T | -2.27 | -1.83 | -0.51 | -1.03 | -1.46 | -1.65 | -4.14 | -1.04 | -2.12 | -2.34 | 0.69 | -1.47 | -0.62 | -1.47 | -2.42 | -1.91 | 0.285 | -1.0925 | -1.535 | 0.706099 | 0.361242 | 0.889001 | 0.989223 | 0.997988 | 0.996331 |
| C2cd5_Q7TPS5_T809t_1 | C2cd5_Q7T | -0.28 | 0.15 | -0.64 | 0.01 | 0.02 | 0.12 | 0.12 | -0.2 | -0.13 | 0.21 | -0.11 | -0.45 | -0.51 | 0.1 | -0.58 | -0.12 | -0.6625 | 0.1 | -0.195 | 0.770576 | 0.417843 | 0.799526 | 0.989223 | 0.997988 | 0.9971 |
| C530008M17Rik_E9Q5L4_S1076s_1 | C530008M | -0.29 | 0.5 | 0.21 | 0.43 | 0.45 | 0.48 | 0.76 | 0.59 | -0.02 | 0.3 | -0.61 | 0.45 | -0.44 | 0.28 | 0.52 | 0.49 | 0.205 | 0.07 | -0.0875 | 1 | 0.174277 | 0.575004 | 0.989223 | 0.997988 | 1 |
| C530008M17Rik_E9Q5L4_S1100s_1 | C530008M | -0.22 | 0.42 | 0.17 | -0.31 | 0.31 | 0.16 | 0.24 | -0.38 | -0.3 | -0.18 | -0.19 | -0.3 | 0.04 | 0.08 | -0.02 | -0.16 | 0.3575 | -0.1825 | 2.16E-16 | 0.910976 | 0.804527 | 0.316118 | 0.995762 | 0.997988 | 0.999507 |
| C530008M17Rik_E9Q5L4_S1259s_1 | C530008M | -0.35 | 0.05 | -0.9 | -0.74 | 0.51 | 0.2 | 0.43 | -0.23 | -0.04 | -0.08 | -0.52 | 0.11 | -1.56 | 0.5 | -0.1 | 0.54 | 0.0675 | -0.2575 | -0.03 | 0.517872 | 0.035541 | 0.250144 | 0.989223 | 0.997988 | 0.994666 |
| C530008M17Rik_E9Q5L4_S571s_1 | C530008M | 0.1 | -0.43 | 1.17 | 1.08 | -0.55 | 0.5 | -0.36 | 0.03 | 1.41 | -0.27 | 0.12 | 1.04 | 1.53 | -1.65 | 0.6 | 0.75 | 0.7125 | 0.3525 | 0.33 | 0.80627 | 0.200108 | 0.853882 | 0.989223 | 0.997988 | 0.999507 |
| C530008M17Rik_E9Q5L4_S730s_1 | C530008M | 0.62 | 1.12 | -1.55 | -0.35 | 1.81 | 1.03 | 1.25 | -0.79 | 0.76 | 1.71 | -0.27 | -0.82 | 0.85 | 0.42 | 0.47 | -0.36 | -0.575 | 0.095 | -0.1725 | 0.516802 | 0.245928 | 0.60083 | 0.989223 | 0.997988 | 0.994666 |
| C530008M17Rik_E9Q5L4_S867s_1 | C530008M | -0.18 | 0.63 | 0.79 | -0.9 | 0.96 | -0.25 | 0.17 | 0.4 | -0.75 | -0.24 | -0.37 | -0.13 | -0.47 | 0.46 | 0.05 | 0.49 | 0.865 | 0.385 | 0.385 | 0.91425 | 0.593871 | 0.279701 | 0.989223 | 0.997988 | 0.999507 |
| C530008M17Rik_E9Q5L4_T1033t_1 | C530008M | -0.22 | 0.3 | -0.06 | -0.06 | -0.02 | -0.04 | 0.59 | -0.05 | 0.18 | 0.31 | -0.12 | -0.08 | 0.61 | 0.3 | -0.14 | -0.28 | 0.235 | -0.4575 | 0.0475 | 0.656217 | 0.604257 | 0.729658 | 0.989223 | 0.997988 | 0.995259 |
| C78339_Q3URQ4_T70t_1 | C78339_Q3 | 0.42 | -0.51 | -0.37 | -1.25 | 0.13 | 0.22 | -1.03 | -1.36 | -0.57 | 0.12 | -0.14 | -0.39 | 0.8 | 2.76 | -0.89 | -0.59 | 0.13 | 0.0825 | 0.1325 | 0.251615 | 0.86512 | 0.631737 | 0.996493 | 0.997988 | 0.994666 |
| Cables1_Q3U148_T376t_1 | Cables1_Q3 | -0.23 | -0.17 | -0.17 | 1.34 | 0.38 | 0.13 | 0.29 | 1.73 | 0.06 | 0.56 | 0.96 | 1.32 | -0.2 | 1.13 | 1.36 | 1.04 | -0.0825 | 0.1825 | 0.9475 | 0.214462 | 0.379017 | 0.256109 | 0.989223 | 0.997988 | 0.994666 |
| Cables2_Q8K3M5_S103s_1 | Cables2_Q | 0.54 | 0.48 | 0.48 | 0.31 | 0.92 | 0.74 | 0.22 | 0.9 | 0.29 | 0.19 | -0.16 | 0.6 | -0.19 | 0.68 | 0.07 | 0.64 | 0.44 | 0.5325 | 0.64 | 0.60136 | 0.321945 | 0.373949 | 0.989223 | 0.997988 | 0.995259 |
| Cabp1_D3Z1M4_S101s_1 | Cabp1_D3Z | -0.16 | -0.04 | -0.69 | -0.48 | 0.03 | -0.16 | -0.38 | -1.04 | -0.32 | 0.01 | -0.29 | -0.19 | -0.32 | -0.01 | -0.62 | -0.51 | 0.2425 | -0.2225 | -0.1525 | 0.935668 | 0.880966 | 0.55841 | 0.996493 | 0.997988 | 0.999507 |
| Cachd1_Z4YK18_S1169s_1 | Cachd1_Z4 | 1.33 | 1.57 | 1.56 | 1.54 | 0.59 | 1.29 | 1.45 | 0.65 | 1.37 | 1.81 | 2.03 | 1.91 | 0.72 | 0.74 | 1.76 | 1.55 | -0.045 | 0.145 | -0.0225 | 0.352004 | 0.085972 | 0.261103 | 0.989223 | 0.997988 | 0.994666 |
| Cacna1a_E9Q1R5_S1899s_1 | Cacna1a_E | 0.83 | 1.01 | 0.91 | 0.96 | 0.76 | 0.51 | 1.06 | 0.91 | 0.34 | 0.95 | 0.68 | 0.84 | 0.1 | 0.82 | 0.59 | 0.72 | -0.505 | 0.28 | -0.3075 | 0.180179 | 0.589523 | 0.345614 | 0.989223 | 0.997988 | 0.994666 |
| Cacna1a_E9Q1R5_S1934s_1 | Cacna1a_E | -0.22 | -0.89 | -1.09 | -0.63 | 0.26 | 0.24 | -0.24 | -0.61 | -1.24 | -0.65 | -0.68 | -0.67 | -1.11 | 0.15 | -1.1 | -0.72 | -0.1175 | -0.225 | -0.37 | 0.973039 | 0.065344 | 0.719735 | 0.989223 | 0.997988 | 0.999507 |
| Cacna1a_E9Q1R5_S1951s_1 | Cacna1a_E | 0.18 | 0.57 | 0.17 | 0.08 | 1.05 | 0.59 | 0.41 | 0.37 | 0.13 | 0.59 | -0.01 | 0.41 | 0.02 | 0.87 | 0.3 | 0.05 | 0.62 | -0.1025 | 0.0125 | 0.837485 | 0.175624 | 0.904167 | 0.989223 | 0.997988 | 0.999507 |
| Cacna1a_E9Q1R5_S1969s_1 | Cacna1a_E | 0.12 | -0.42 | -0.09 | 0.21 | -0.12 | 0.1 | 0.78 | 0.02 | 0.68 | -0.06 | 0.71 | 0.07 | 0.18 | 0.55 | 0.74 | 0.21 | 0.355 | 0.03 | 0.06 | 0.117869 | 0.400971 | 0.195665 | 0.989223 | 0.997988 | 0.994666 |
| Cacna1a_E9Q1R5_S1981s_1 | Cacna1a_E | -0.46 | -0.57 | -1.13 | -1.09 | -0.53 | -0.63 | -0.76 | -0.84 | -0.24 | -0.53 | -1.41 | -1.25 | 0.39 | 0.31 | -1.22 | -1.01 | 0.24 | 0.395 | 0.465 | 0.349054 | 0.621195 | 0.89678 | 0.989223 | 0.997988 | 0.994666 |
| Cacna1a_E9Q1R5_S2173s_1 | Cacna1a_E | -0.31 | 0.7 | 0.74 | 1.13 | -0.36 | 1.31 | 1.46 | 0.98 | 1.83 | 0.39 | 0.93 | 1.85 | 1.98 | 1.08 | 1.3 | 0.98 | 0.1225 | -0.045 | 0.43 | 0.073796 | 0.55727 | 0.15566 | 0.989223 | 0.997988 | 0.994666 |
| Cacna1a_E9Q1R5_S2205s_1 | Cacna1a_E | -0.57 | -0.69 | -0.21 | -0.42 | -0.13 | -0.24 | 0.43 | -0.35 | -1.39 | -0.56 | -0.43 | -0.41 | -0.76 | 0.18 | 0 | -0.05 | 0.2825 | 0.685 | 0.77 | 0.301825 | 0.141049 | 0.455315 | 0.989223 | 0.997988 | 0.994666 |
| Cacna1a_E9Q1R5_S2226s_1 | Cacna1a_E | 0.08 | 0.48 | 0.33 | 0.39 | 1.03 | -0.14 | 0.43 | 0.03 | -0.03 | 0.22 | 0.45 | 0.38 | 0.61 | 0.81 | 0.38 | 0.43 | 0.4 | -0.225 | 0.315 | 0.34986 | 0.953268 | 0.779323 | 0.999187 | 0.997988 | 0.994666 |
| Cacna1a_E9Q1R5_S2256s_1 | Cacna1a_E | 0.19 | 0.36 | 0.75 | 0.34 | 0.25 | -0.11 | 0.43 | 0.44 | -0.25 | -0.16 | 0.35 | 0.54 | -0.05 | 0.09 | 0.49 | 0.43 | 0.0175 | -0.065 | 0.2375 | 0.53008 | 0.517315 | 0.313072 | 0.989223 | 0.997988 | 0.994666 |
| Cacna1a_E9Q1R5_S705s_1 | Cacna1a_E | 1.11 | 1.47 | 0.95 | 0.91 | 1.06 | 1.14 | 0.96 | 0.87 | 1.1 | 1.28 | 0.42 | 0.96 | 0.88 | 1.46 | 0.56 | 0.87 | -0.1575 | -0.29 | -0.17 | 0.571485 | 0.64813 | 0.544869 | 0.99033 | 0.997988 | 0.995259 |
| Cacna1a_E9Q1R5_S745s_1 | Cacna1a_E | 0.28 | 0.52 | -0.17 | -1.1 | 1.05 | 0.17 | -0.76 | -0.96 | -1.61 | -0.18 | -1.04 | -0.84 | 0 | -0.44 | -1.26 | -1.08 | -0.1025 | -0.17 | -0.1675 | 0.218439 | 0.988714 | 0.099857 | 0.999187 | 0.997988 | 0.994666 |
| Cacna1a_E9Q1R5_S820s_1 | Cacna1a_E | 0.35 | 0.55 | 0.38 | 0.29 | 1.01 | -0.14 | 0.42 | 0.27 | 0.26 | 0.38 | 0.25 | 0.18 | 0.31 | 0.77 | 0.11 | 0.1 | -0.0075 | -0.8 | -0.5775 | 0.790282 | 0.992877 | 0.552473 | 0.999187 | 0.997988 | 0.999507 |
| Cacna1a_E9Q1R5_T1888t_1 | Cacna1a_E | 0.43 | 0.5 | 0.11 | 0.45 | 1.08 | 0.22 | 0.59 | 0.2 | -0.33 | 0.1 | -0.31 | 0.64 | -0.44 | 0.21 | 0.4 | 0.39 | -0.0025 | -0.125 | -0.07 | 0.427873 | 0.578915 | 0.24923 | 0.989223 | 0.997988 | 0.994666 |
| Cacna1a_E9Q1R5_T986t_1 | Cacna1a_E | 0.73 | 1.2 | -0.25 | 0.25 | 0.78 | 0.74 | 0.31 | 0.01 | 0.08 | 0.27 | -0.2 | 0.06 | 0.74 | -0.23 | -0.23 | -0.03 | 0.15 | -0.3475 | -0.2325 | 0.303615 | 0.950141 | 0.231733 | 0.999187 | 0.997988 | 0.994666 |
| Cacna1b_O55017_S1907s_1 | Cacna1b_C | 0.25 | 0.37 | -0.13 | -0.82 | 0.48 | 0.04 | 1.28 | 0.04 | -0.34 | 0.15 | 0.09 | 0.25 | 0.05 | 0.81 | 0.72 | 0.93 | -0.0225 | -0.43 | -0.42 | 0.072106 | 0.181926 | 0.712538 | 0.989223 | 0.997988 | 0.994666 |
| Cacna1b_O55017_S1929s_1 | Cacna1b_C | 0.04 | -0.37 | -0.35 | -0.03 | 1.03 | -0.33 | 0.23 | 0.09 | -0.98 | -0.34 | -0.55 | 0.14 | -0.78 | 0.05 | -0.5 | -0.61 | 0.5425 | 0.12 | 0.71 | 0.332234 | 0.19666 | 0.400413 | 0.989223 | 0.997988 | 0.994666 |
| Cacna1b_O55017_S1951s_1 | Cacna1b_C | 1.12 | 0.8 | 1.08 | 0.28 | 1.14 | 1.16 | 1.36 | 1.23 | 0.04 | 1.18 | -0.01 | 1.05 | -0.08 | 0.61 | 0.63 | 1.24 | 0.4325 | -0.255 | -0.2825 | 0.542694 | 0.13726 | 0.505276 | 0.989223 | 0.997988 | 0.994666 |
| Cacna1b_O55017_S2007s_1 | Cacna1b_C | 1.26 | -0.71 | 0.19 | 1.12 | 0.49 | 1.56 | 0.89 | 1.25 | 0.37 | 0.19 | 0.86 | 0.67 | -0.24 | 0.29 | 0.01 | 0.11 | 0.4025 | -0.255 | -0.22 | 0.36605 | 0.238364 | 0.900712 | 0.989223 | 0.997988 | 0.994666 |
| Cacna1b_O55017_S2062s_1 | Cacna1b_C | -0.09 | -0.33 | -0.59 | -0.14 | 0.37 | 0.16 | 0.32 | -0.14 | -0.38 | 0.62 | -0.54 | -0.19 | -0.24 | 0.07 | -0.22 | -0.06 | 0.5825 | 0.0575 | -0.4225 | 0.489043 | 0.070794 | 0.601535 | 0.989223 | 0.997988 | 0.994666 |
| Cacna1b_O55017_S2221s_1 | Cacna1b_C | -1.78 | -1.58 | -0.96 | -1.07 | -0.4 | -1.04 | -0.33 | -0.73 | -2.43 | -1.96 | -0.29 | -0.99 | -1.2 | -0.47 | -0.77 | -0.85 | 0.465 | 0.165 | 0.175 | 0.110961 | 0.029742 | 0.886576 | 0.989223 | 0.997988 | 0.994666 |
| Cacna1b_O55017_S2244s_1 | Cacna1b_C | 0.1 | -0.32 | 0.2 | 0.22 | 0.26 | -0.04 | 0.36 | 0.37 | -0.14 | -0.14 | -0.13 | 0.32 | -0.49 | 0.1 | 0 | 0.25 | 0.7225 | -0.07 | 0.525 | 0.762745 | 0.426876 | 0.7699 | 0.989223 | 0.997988 | 0.9971 |
| Cacna1b_O55017_S424s_1 | Cacna1b_C | -0.64 | -0.55 | -1.46 | -0.74 | -0.32 | -0.11 | -0.98 | -0.82 | -1.5 | -0.65 | -2.01 | -0.46 | -0.8 | -1.92 | -1.12 | -0.88 | 0.1875 | -0.0725 | -0.085 | 0.362029 | 0.362379 | 0.463059 | 0.989223 | 0.997988 | 0.994666 |
| Cacna1b_O55017_S446s_1 | Cacna1b_C | 1.5 | 0.68 | -1.23 | -0.02 | 0.83 | 2 | 0.12 | -0.54 | -0.44 | 0.85 | -1.2 | -0.28 | -0.68 | 0.11 | -0.43 | 0.28 | 0.29 | -0.3075 | -0.3325 | 0.476165 | 0.599424 | 0.449614 | 0.989223 | 0.997988 | 0.994666 |
| Cacna1b_O55017_S745s_1 | Cacna1b_C | 0.63 | 0.92 | 1.22 | 1.29 | 0.67 | 0.64 | 1.78 | 0.55 | 0.69 | 0.98 | 1.79 | 1.46 | 0.27 | 1.02 | 1.5 | 1.87 | 0.37 | -0.5 | -0.4125 | 0.700344 | 0.755044 | 0.504443 | 0.995762 | 0.997988 | 0.996159 |
| Cacna1b_O55017_S770s_1 | Cacna1b_C | 1.02 | 0.64 | 0.91 | 0.51 | 0.39 | 1.29 | 0.33 | 0.62 | 0.96 | 0.88 | 0.7 | 0.75 | 1.44 | -0.16 | 1.06 | 0.97 | -0.105 | 0.215 | 0.15 | 0.879533 | 0.692435 | 0.819255 | 0.994893 | 0.997988 | 0.999507 |
| Cacna1b_O55017_S773s_2 | Cacna1b_C | 0.52 | 0.85 | 0.65 | -0.12 | -0.05 | 1.06 | -0.11 | 0.33 | 0.98 | -0.18 | -1.35 | 0.05 | 0.81 | 0.03 | 0.35 | 0.82 | -0.1125 | 0.0525 | 0.0575 | 0.932953 | 0.629474 | 0.243476 | 0.989223 | 0.997988 | 0.999507 |
| Cacna1b_O55017_S783s_1 | Cacna1b_C | -0.02 | 0.46 | -0.65 | -0.52 | 0.34 | 0.21 | -0.33 | -0.56 | -0.54 | 0.26 | -0.42 | -0.83 | -0.43 | -0.34 | -1.15 | -0.61 | -0.1675 | -0.6 | 0.0275 | 0.212101 | 0.773877 | 0.577532 | 0.995762 | 0.997988 | 0.994666 |
| Cacna1b_O55017_S875s_1 | Cacna1b_C | 4.28 | 0.2 | 1.79 | 2.41 | 0.14 | 3.65 | 1.59 | 3.38 | 1.73 | 2.4 | 1.81 | 2.17 | 0.74 | 0.5 | 1.95 | 2.6 | 0.0975 | -0.2 | -0.45 | 0.410071 | 0.984305 | 0.852795 | 0.999187 | 0.997988 | 0.994666 |
| Cacna1b_O55017_S915s_1 | Cacna1b_C | 1.52 | 1.11 | 0.83 | 1.44 | 0.26 | 1.33 | 0.43 | 0.9 | 2.25 | 1.72 | 1.77 | 1.08 | 1.6 | 0.47 | 1.27 | 0.73 | 0.02 | -0.1425 | -0.7225 | 0.543004 | 0.133998 | 0.155737 | 0.989223 | 0.997988 | 0.994666 |
| Cacna1b_O55017_T1935t_1 | Cacna1b_C | -1.93 | -2.61 | -1.44 | -2.28 | -0.94 | -0.65 | -1.44 | -0.78 | -1.88 | -3.03 | 0.66 | -1.12 | -2.2 | -0.63 | -1.85 | -1.91 | -0.495 | 0.48 | -0.2075 | 0.33866 | 0.006035 | 0.336526 | 0.989223 | 0.997988 | 0.994666 |
| Cacna1c_F7C376_S1825s_1 | Cacna1c_F | -0.13 | -0.81 | 0.96 | 0.63 | -1.44 | 0.15 | -0.19 | 0.36 | 1.45 | -0.16 | -0.81 | 1.05 | 2.22 | -2.43 | 0.76 | 0.31 | 1.1125 | 0.7225 | 0.4175 | 0.954331 | 0.400124 | 0.715626 | 0.989223 | 0.997988 | 0.999507 |
| Cacna1c_F7C376_S2208s_1 | Cacna1c_F | 0.92 | -0.67 | -0.65 | 0.78 | 0.19 | 1.42 | 0.39 | 0.79 | 0.04 | -0.11 | 1.53 | 0.16 | -0.64 | 0.49 | -0.68 | -1.19 | -0.4425 | 0.22 | 0.0525 | 0.270842 | 0.224804 | 0.567852 | 0.989223 | 0.997988 | 0.994666 |
| Cacna1c_F7C376_S840s_1 | Cacna1c_F | 0.83 | 0.85 | 0.8 | 0.99 | 0.77 | 0.66 | 0.57 | 0.8 | 0.63 | 0.83 | 0.81 | 0.86 | 0.31 | 0.95 | 0.89 | 0.78 | 0.6025 | 0.31 | -0.6 | 0.600904 | 0.406064 | 0.683606 | 0.989223 | 0.997988 | 0.995259 |
| Cacna1c_F7C376_S863s_1 | Cacna1c_F | -2.09 | -0.41 | -0.63 | -0.55 | -1.61 | -0.43 | -0.32 | -0.9 | -0.2 | -0.62 | -0.59 | -0.43 | -1.27 | -0.79 | -0.12 | -0.47 | -0.1675 | -0.085 | -0.135 | 0.570531 | 0.81821 | 0.269375 | 0.995762 | 0.997988 | 0.995259 |
| Cacna1c_F7C376_S863s_2 | Cacna1c_F | -2.01 | -0.6 | -0.22 | -0.09 | -0.03 | -0.44 | -0.35 | -0.55 | -0.2 | -0.79 | 0.68 | 0.35 | -0.92 | -0.13 | 0 | 0.42 | 0.105 | 0.46 | 0.2575 | 0.264359 | 0.376986 | 0.172061 | 0.989223 | 0.997988 | 0.994666 |
| Cacna1c_F7C376_T867t_2 | Cacna1c_F | -2.01 | -0.6 | -0.22 | -0.09 | -0.03 | -0.44 | -0.35 | -0.55 | -0.2 | -0.79 | 0.68 | 0.35 | -0.92 | -0.13 | 0 | 0.42 | 0.3875 | 0.74 | 0.5725 | 0.264359 | 0.376986 | 0.172061 | 0.989223 | 0.997988 | 0.994666 |
| Cacna1d_Q99246_S1684s_1 | Cacna1d_C | -1.07 | -2.04 | -1.86 | -0.68 | -1.25 | -0.68 | -1.27 | -2.59 | -0.87 | -2.02 | -1.21 | -2.64 | -0.75 | -0.97 | -1.45 | -1.38 | 0.3875 | 0.74 | 0.5725 | 0.474159 | 0.941557 | 0.57814 | 0.999187 | 0.997988 | 0.994666 |
| Cacna1e_Q61290_S1051s_1 | Cacna1e_C | 0.66 | 0.57 | 0.43 | 0.68 | 0.72 | 0.54 | 0.94 | 0.66 | 0.44 | 0.58 | 0.14 | 0.72 | 0.12 | 0.82 | 0.38 | 0.65 | -0.035 | -0.2725 | 0.275 | 0.724546 | 0.535767 | 0.621635 | 0.989223 | 0.997988 | 0.9971 |
| Cacna1e_Q61290_S1099s_1 | Cacna1e_C | -0.49 | 1.34 | 0.81 | 0.32 | 0.97 | 0.63 | 0.31 | 0.1 | 1.09 | 0.23 | 0.72 | 0.45 | 0.93 | 0.98 | 0.57 | 0.47 | 0.13 | -0.115 | -0.0925 | 0.558905 | 0.985486 | 0.762678 | 0.999187 | 0.997988 | 0.995259 |
| Cacna1e_Q61290_S15s_1 | Cacna1e_C | -2.25 | -1.63 | -0.02 | -10.42 | -3.78 | -1.17 | -0.94 | -2.67 | 0.8 | -1.56 | -2.06 | -0.41 | 2.23 | -2.13 | 0.48 | -2.54 | 0.0075 | 0.1275 | 0.2425 | 0.183902 | 0.491377 | 0.208061 | 0.989223 | 0.997988 | 0.994666 |
| Cacna1e_Q61290_S2054s_1 | Cacna1e_C | 6.75 | 2.44 | 6.19 | 3.26 | 4.15 | 7.35 | 7.15 | 6.72 | 0.29 | 5.03 | 1.75 | 7.1 | -1.72 | 0.46 | 6.03 | 6.85 | 1.44 | 2.7725 | 3.09 | 0.391988 | 0.156368 | 0.500111 | 0.989223 | 0.997988 | 0.994666 |
| Cacna1e_Q61290_S2073s_1 | Cacna1e_C | 1.19 | 0.33 | 0.47 | 1.14 | 1.03 | 0.72 | 0.75 | 1.01 | 0.69 | 0.61 | 0.8 | 1.07 | -1.06 | 0.91 | -0.05 | -0.03 | 1.6825 | -1.1175 | -1.755 | 0.086278 | 0.731401 | 0.972452 | 0.995762 | 0.997988 | 0.994666 |
| Cacna1e_Q61290_S2097s_1 | Cacna1e_C | 0.66 | 0.32 | 1.87 | 0.78 | 1.24 | 0.09 | 0.85 | 2.31 | 0.61 | 0.26 | 0.03 | 0.33 | -0.01 | 0.62 | 0.48 | 2.61 | 0.095 | 0.01 | -0.84 | 0.977028 | 0.680499 | 0.126638 | 0.993464 | 0.997988 | 1 |
| Cacna1e_Q61290_S2099s_1 | Cacna1e_C | 0.86 | 0.92 | 0.35 | 0.28 | 0.76 | 1.61 | 0.21 | 0.46 | 1.09 | 1.47 | 0.23 | 0.3 | 0.36 | 0.61 | -0.1 | 0.12 | 0.215 | -0.6 | 0.0175 | 0.244487 | 0.654106 | 0.637341 | 0.990434 | 0.997988 | 0.994666 |
| Cacna1e_Q61290_S20s_1 | Cacna1e_C | -1.08 | -1.44 | -0.46 | -2.42 | -2.64 | -0.5 | -1.12 | -1.43 | 1.3 | -0.56 | -2.29 | -0.13 | 2.39 | -1.63 | 1.84 | -0.72 | 0.1575 | 0.17 | -0.355 | 0.073789 | 0.895033 | 0.239276 | 0.996493 | 0.997988 | 0.994666 |
| Cacna1e_Q61290_S737s_1 | Cacna1e_C | 0.03 | 0.54 | 0.3 | 0.18 | 0.6 | 0.05 | 0.43 | 0.23 | 0.08 | 0.41 | -0.09 | 0.2 | 0.25 | 0.63 | 0.21 | 0.39 | -0.0725 | 0.93 | 1.82 | 0.672376 | 0.780621 | 0.638216 | 0.995762 | 0.997988 | 0.996159 |
| Cacna1e_Q61290_S740s_1 | Cacna1e_C | 0.63 | 0.56 | 1.24 | 0.63 | 0.79 | 0.74 | 0.83 | 1.66 | 1.14 | 0.87 | 0.56 | 0.71 | 0.03 | 1.05 | 1.04 | 1.08 | 0.065 | -0.1125 | 0.1075 | 0.917573 | 0.427274 | 0.835311 | 0.989223 | 0.997988 | 0.999507 |
| Cacna1e_Q61290_S793s_1 | Cacna1e_C | 1.71 | 1.12 | 2.22 | 0.94 | 1.63 | 1.96 | -0.24 | 1.63 | 0.06 | 1.88 | 1.21 | 1.83 | 0.15 | -0.07 | 0.68 | 0.98 | 0.24 | 0.055 | 0.035 | 0.020586 | 0.633744 | 0.606311 | 0.989223 | 0.997988 | 0.994666 |
| Cacna1e_Q61290_S794s_1 | Cacna1e_C | 2.99 | 2.14 | 3.42 | 0.58 | 2.79 | 3.09 | 2.75 | 2.33 | 0.66 | 2.8 | 0.74 | 3.05 | 0.97 | 0.12 | 3.07 | 3.2 | -0.2525 | -0.2525 | -1.0625 | 0.613342 | 0.437845 | 0.561758 | 0.989223 | 0.997988 | 0.995259 |
| Cacna1e_Q61290_S826s_1 | Cacna1e_C | -0.46 | -0.04 | -0.49 | -0.31 | -0.3 | -0.25 | 0.42 | -0.45 | -0.49 | -0.3 | -0.44 | -0.64 | -0.7 | -0.33 | -0.13 | -0.5 | 0.4575 | -0.47 | -0.4425 | 0.72956 | 0.50167 | 0.533705 | 0.989223 | 0.997988 | 0.9971 |
| Cacna1e_Q61290_S852s_1 | Cacna1e_C | 1.4 | -1.8 | -0.05 | 0.74 | -1.2 | 2.33 | -0.23 | 0.98 | 0.59 | -0.55 | 0.22 | 0.12 | -1.08 | -1.08 | -1.19 | -0.28 | 0.18 | -0.1425 | -0.09 | 0.156565 | 0.658331 | 0.972868 | 0.990434 | 0.997988 | 0.994666 |
| Cacna1e_Q61290_S856s_1 | Cacna1e_C | 2.3 | 0.86 | 1.72 | 2.15 | 1.92 | 2.43 | 2.08 | 2.53 | 1.36 | 1.46 | 1.48 | 2.32 | 0.7 | 1.47 | 1.54 | 1.64 | 0.3975 | 0.0225 | -0.98 | 0.304169 | 0.195073 | 0.79437 | 0.989223 | 0.997988 | 0.994666 |
| Cacna1e_Q61290_S873s_1 | Cacna1e_C | 1.07 | -0.74 | -0.65 | 0.15 | 0.83 | 1.26 | 0.7 | 0.8 | 0.31 | -0.02 | 0.76 | 0.34 | -0.88 | 0.27 | -0.61 | -0.47 | 0.4825 | -0.1025 | -0.42 | 0.426205 | 0.046171 | 0.382748 | 0.989223 | 0.997988 | 0.994666 |
| Cacna1e_Q61290_S948s_1 | Cacna1e_C | 2.09 | 0.3 | 0.19 | 0.3 | 0.28 | 2.18 | 0.15 | 0.26 | 1.09 | 0.97 | 0.72 | 0.48 | -1.23 | -0.2 | 0.12 | 0.03 | 0.94 | 0.39 | -0.38 | 0.069505 | 0.996663 | 0.835507 | 0.999864 | 0.997988 | 0.994666 |
| Cacna1e_Q61290_S988s_1 | Cacna1e_C | -1.36 | -1.64 | -2.17 | -3.23 | -0.83 | -2 | -1.46 | -2.02 | -2.72 | -2.04 | -0.15 | -1.61 | -1.25 | -0.96 | -2.71 | -1.58 | -0.0025 | 0.095 | -1.04 | 0.377371 | 0.276129 | 0.457458 | 0.989223 | 0.997988 | 0.994666 |
| Cacna1e_Q61290_T2225t_1 | Cacna1e_C | -1.08 | -1.89 | -1.84 | -2.68 | -1.66 | -1.83 | -0.97 | -1.36 | -2.31 | -1.57 | 1.41 | -1.98 | -1.58 | -0.76 | -1.51 | -2.27 | 0.5225 | 0.47 | 0.475 | 0.444281 | 0.277338 | 0.363301 | 0.989223 | 0.997988 | 0.994666 |
| Cacna1e_Q61290_T29t_1 | Cacna1e_C | 0.13 | -0.16 | -0.2 | 0.35 | 0.52 | 0.3 | 0.12 | 0.32 | 0.26 | 0.27 | 0.44 | 0.12 | -0.64 | 0.84 | 0.29 | 0.15 | 0.4175 | 0.76 | 0.3425 | 0.717042 | 0.233195 | 0.31735 | 0.989223 | 0.997988 | 0.996966 |
| Cacna1e_Q61290_T441t_1 | Cacna1e_C | 1.95 | 1.45 | 2.51 | 2.39 | 1.92 | 1.57 | 2.66 | 2.82 | 0.95 | 1.36 | 1.33 | 2.54 | -0.13 | 0.4 | 2.54 | 2.8 | 0.285 | 0.2425 | 0.13 | 0.347343 | 0.657002 | 0.220837 | 0.990434 | 0.997988 | 0.994666 |
| Cacna1g_Q5SUG2_S1063s_1 | Cacna1g_C | -0.45 | -0.54 | 0.68 | 0.41 | -1.67 | -0.12 | 0.51 | 0.47 | -1.43 | -0.44 | -2.25 | 0.31 | 0.84 | -2.35 | 1.56 | 0.39 | 0.1675 | -0.53 | -0.6725 | 0.915488 | 0.674104 | 0.121806 | 0.992876 | 0.997988 | 0.999507 |
| Cacna1g_Q5SUG2_S2068s_1 | Cacna1g_C | 0.87 | 0.27 | -0.69 | 0.47 | 0.37 | 1.65 | 0.52 | 0.62 | 0.89 | 1.2 | 0.23 | 0.83 | 1.55 | -0.92 | 0.82 | 1.27 | -0.2275 | -0.9775 | 0.085 | 0.45325 | 0.202515 | 0.175823 | 0.989223 | 0.997988 | 0.994666 |
| Cacna1g_Q5SUG2_S467s_1 | Cacna1g_C | 0.9 | 0.4 | 0.48 | 0.31 | 0.91 | 0.91 | 0.79 | 0.05 | 0.65 | 0.58 | -0.28 | 0.24 | 0.61 | 1 | 0.32 | 0.32 | 0.56 | 0.5575 | 0.45 | 0.88752 | 0.6136 | 0.44977 | 0.989223 | 0.997988 | 0.999507 |
| Cacna1g_Q5SUG2_S518s_1 | Cacna1g_C | 1.49 | -0.15 | 0.7 | 1.06 | 0.17 | 1.81 | 0.79 | 1.21 | 0.87 | 0.41 | 0.14 | 0.67 | 0.09 | 0.22 | 0.36 | 0.64 | 0.1425 | -0.225 | 0.04 | 0.258681 | 0.632105 | 0.516927 | 0.989223 | 0.997988 | 0.994666 |
| Cacna1h_E9Q6P9_S1203s_1 | Cacna1h_E | -0.45 | -1.21 | 0.49 | 0.07 | -1.22 | -0.48 | 0.62 | -0.14 | 0.45 | -1.24 | -0.39 | 0 | 0.68 | -1.68 | 1.26 | 0.61 | 0.22 | -0.2525 | -0.4475 | 0.468786 | 0.950834 | 0.967094 | 0.999187 | 0.997988 | 0.994666 |
| Cacna1h_E9Q6P9_S2195s_1 | Cacna1h_E | 1.16 | 0.01 | -0.41 | 0.58 | 0.54 | 0.3 | 0.52 | 0.65 | 0.65 | 1.23 | 0.86 | -0.03 | 0.93 | 1.02 | -0.03 | 0.25 | -0.03 | -0.02 | 0.4925 | 0.626893 | 0.635271 | 0.426312 | 0.989223 | 0.997988 | 0.995259 |
| Cacna1h_E9Q6P9_T1172t_1 | Cacna1h_E | -1.31 | -0.61 | -0.91 | -0.2 | -0.58 | -1.23 | 0.04 | 0.03 | -0.16 | -1.03 | 0.49 | 0.32 | 0.06 | 0.37 | -0.91 | -0.01 | 0.1675 | 0.3425 | 0.2075 | 0.118043 | 0.399534 | 0.13088 | 0.989223 | 0.997988 | 0.994666 |
| Cacna1h_E9Q6P9_T751t_1 | Cacna1h_E | 0.43 | 0.58 | 0.78 | 0.7 | 1.61 | 0.54 | 1.01 | 0.22 | 0.87 | 0.48 | 1.08 | 0.59 | 0.63 | 1.26 | 0.98 | 1.1 | 0.3225 | 0.6625 | 0.635 | 0.173476 | 0.49828 | 0.584585 | 0.989223 | 0.997988 | 0.994666 |
| Cacna1i_E9Q7P2_S1017s_1 | Cacna1i_E9 | 0.26 | 0.91 | 0.41 | 0.14 | 0.58 | 0.89 | 0.12 | 0.09 | 0.06 | 0.29 | 0.02 | 0.34 | -0.5 | 0.53 | -0.35 | 0.28 | 0.2225 | 0.1325 | 0.37 | 0.211705 | 0.972255 | 0.338101 | 0.999187 | 0.997988 | 0.994666 |
| Cacna1i_E9Q7P2_S1744s_1 | Cacna1i_E9 | 0.63 | -0.44 | 0.59 | 0.22 | 0.04 | 0.45 | 0.39 | 0.09 | 0.54 | -0.36 | 0.19 | 0.34 | 0.74 | -0.05 | -0.26 | -0.52 | -0.01 | -0.2525 | -0.44 | 0.48163 | 0.979739 | 0.829124 | 0.999187 | 0.997988 | 0.994666 |
| Cacna1i_E9Q7P2_S1745s_1 | Cacna1i_E9 | -1.91 | 0.84 | 0.99 | 0.71 | 1.53 | 0.4 | 0.69 | 0.94 | 1.12 | 1.26 | 1.29 | 0.77 | 1.7 | 1.35 | 0.3 | 0.3 | -0.0075 | -0.0725 | -0.2725 | 0.294703 | 0.275921 | 0.160325 | 0.989223 | 0.997988 | 0.994666 |
| Cacna1i_E9Q7P2_S2037s_1 | Cacna1i_E9 | 2.16 | -0.93 | -0.32 | 0.28 | 0.46 | 1.86 | 0.12 | 0.93 | 0.14 | -0.54 | 1.52 | 0.04 | -0.22 | 0.27 | -0.51 | -0.29 | 0.7325 | 0.9525 | 0.755 | 0.444769 | 0.429839 | 0.991659 | 0.989223 | 0.997988 | 0.994666 |
| Cacna1i_E9Q7P2_S2157s_1 | Cacna1i_E9 | -1.56 | -1.11 | -0.25 | -0.47 | -0.32 | -0.2 | -0.14 | 0.12 | -1.44 | -0.8 | 1.36 | -0.03 | -0.77 | 1.14 | -1.15 | -1.73 | 0.545 | -0.0075 | -0.485 | 0.725802 | 0.051307 | 0.327067 | 0.989223 | 0.997988 | 0.9971 |
| Cacna1i_E9Q7P2_S895s_1 | Cacna1i_E9 | 0.97 | -1.09 | -0.46 | 0.63 | -0.21 | 1.31 | 0.02 | 0.91 | -0.8 | -0.83 | 1.25 | -0.11 | -1.65 | 0.56 | -0.55 | 0.07 | 0.7125 | 0.62 | 0.22 | 0.514268 | 0.371737 | 0.828265 | 0.989223 | 0.997988 | 0.994666 |
| Cacna1i_E9Q7P2_T469t_1 | Cacna1i_E9 | -0.3 | -0.59 | -0.79 | -1.64 | -0.39 | -0.54 | -1.22 | -1.43 | -0.06 | -0.47 | -1.39 | -1.68 | 0.1 | -0.75 | -1.03 | -1.01 | 0.495 | -0.135 | -0.405 | 0.693831 | 0.862808 | 0.878391 | 0.996493 | 0.997988 | 0.996159 |
| Cacnb1_A2A543_S464s_1 | Cacnb1_A2 | 0.19 | 0.57 | -0.29 | 0.29 | -0.58 | -0.15 | -0.09 | -0.06 | 0.37 | 0.39 | 0.76 | 0.01 | -0.2 | 0.83 | -0.06 | -0.22 | -0.065 | -0.07 | 0.1575 | 0.764697 | 0.143674 | 0.503903 | 0.989223 | 0.997988 | 0.9971 |
| Cacnb1_Q8R3Z5_S186s_1 | Cacnb1_Q8 | -1.26 | 0.74 | 0.64 | 0.34 | 0.6 | -0.5 | 1.57 | 1.49 | 0.44 | -0.23 | 0.9 | 0.47 | 1.83 | 0.3 | -0.78 | 0.79 | -0.41 | 0.1925 | -0.1025 | 0.520146 | 0.277178 | 0.572798 | 0.989223 | 0.997988 | 0.994666 |
| Cacnb1_Q8R3Z5_S436s_1 | Cacnb1_Q8 | -0.23 | 0.44 | -0.36 | -1.53 | -0.5 | 0.31 | -0.81 | -2.08 | 0.62 | 0.24 | -0.77 | -0.24 | 0.07 | -0.84 | -2.72 | -1.47 | 0.675 | 0.28 | 0.42 | 0.223938 | 0.54857 | 0.435417 | 0.989223 | 0.997988 | 0.994666 |
| Cacnb1_Q8R3Z5_S53s_2 | Cacnb1_Q8 | -1.44 | -0.51 | -1.98 | -0.82 | 0.38 | -0.01 | -0.26 | -0.04 | -1.02 | -1.17 | -0.59 | -1.42 | -2.81 | -0.05 | -1.81 | -1.52 | -0.35 | 0.3825 | -0.82 | 0.552489 | 0.006872 | 0.715658 | 0.989223 | 0.997988 | 0.994666 |
| Cacnb1_Q8R3Z5_S68s_1 | Cacnb1_Q8 | -0.64 | 0.18 | -0.87 | -0.19 | -0.07 | -1.08 | -0.69 | -0.07 | -0.04 | 0.59 | 0.22 | 0.62 | 0.82 | -0.74 | 1.92 | 0.24 | 1.205 | 0.1375 | -0.36 | 0.117208 | 0.77793 | 0.042047 | 0.995762 | 0.997988 | 0.994666 |
| Cacnb1_Q8R3Z5_T205t_1 | Cacnb1_Q8 | 0.31 | 0.85 | 0.97 | 0.61 | 0.2 | 0.03 | 0.01 | 0.03 | 0.37 | 0.93 | -0.2 | 0.21 | 1.22 | -0.62 | 0.08 | 0.89 | -0.0975 | 0.7275 | 0.94 | 0.504118 | 0.021742 | 0.266177 | 0.989223 | 0.997988 | 0.994666 |
| Cacnb1_Q8R3Z5_T217t_1 | Cacnb1_Q8 | -0.25 | 0.54 | 1.01 | 0.89 | 0.24 | -1.02 | 1.35 | 1.33 | -0.07 | -0.31 | 0.78 | 1.11 | -0.95 | 0.44 | 0.42 | 1 | -0.6175 | -0.3575 | -0.2925 | 0.510258 | 0.898119 | 0.693512 | 0.996493 | 0.997988 | 0.994666 |
| Cacnb1_Q8R3Z5_T499t_1 | Cacnb1_Q8 | -0.84 | -0.21 | 0.1 | -0.52 | -0.22 | -1.24 | -2.2 | -1.88 | -1.65 | -0.04 | -2.94 | -2.22 | 0.57 | 0.51 | -1.5 | 0.52 | -0.0725 | -0.17 | -0.32 | 0.452354 | 0.04527 | 0.046338 | 0.989223 | 0.997988 | 0.994666 |
| Cacnb1_Q8R3Z5_T540t_1 | Cacnb1_Q8 | 0.36 | 0.93 | 0.76 | 0.94 | 0.26 | 0.33 | 0.92 | 0.71 | 0.32 | 0.66 | 0.01 | 0.8 | -0.08 | 0.8 | 0.83 | 0.99 | -1.0175 | -1.345 | 0.3925 | 0.728222 | 0.462229 | 0.29439 | 0.989223 | 0.997988 | 0.9971 |
| Cacnb2_Q8CC27_S485s_1 | Cacnb2_Q8 | 0.58 | 0.35 | 0.8 | 1.4 | 0.77 | 0.66 | 1.16 | 1.29 | 0.08 | 0.51 | 1.16 | 0.5 | -0.37 | 0.64 | 0.82 | 1.14 | -0.1925 | -0.3 | -0.1125 | 0.578751 | 0.53275 | 0.519772 | 0.989223 | 0.997988 | 0.995259 |
| Cacnb2_Q8CC27_S508s_1 | Cacnb2_Q8 | 1.92 | 1.34 | -1.71 | 0.23 | 1.52 | 1.63 | -0.43 | -0.99 | 0.39 | 1.37 | -0.08 | -0.66 | 0 | 2.07 | -0.36 | -0.13 | 0.1875 | -0.22 | -0.225 | 0.953538 | 0.988973 | 0.814532 | 0.999187 | 0.997988 | 0.999507 |
| Cacnb2_Q8CC27_S510s_1 | Cacnb2_Q8 | 1.92 | 1.34 | 1.29 | 1.48 | 1.52 | 1.63 | 1.63 | 2.1 | 0.39 | 1.37 | 0.8 | 1.72 | 0 | 2.07 | 1.28 | 1.35 | -0.0125 | -0.19 | -0.05 | 0.460253 | 0.404354 | 0.226073 | 0.989223 | 0.997988 | 0.994666 |
| Cacnb2_Q8CC27_S552s_1 | Cacnb2_Q8 | 1.13 | 0.25 | 0.97 | 1.28 | 0.35 | 1.68 | 0.24 | 1.43 | 0.58 | 0.6 | 1.26 | 0.68 | -0.61 | -0.15 | 1.27 | 1.76 | 0.2125 | -0.4375 | -0.3325 | 0.54871 | 0.966152 | 0.684352 | 0.999187 | 0.997988 | 0.994666 |
| Cacnb2_Q8CC27_S570s_1 | Cacnb2_Q8 | 1.2 | -0.08 | 0.56 | 0.31 | -0.9 | 1.11 | -0.07 | -0.16 | 0.92 | 0.2 | 0.29 | 1.21 | -0.34 | -0.48 | 0.61 | 0.58 | 0.0175 | -0.1275 | -0.34 | 0.327162 | 0.290565 | 0.672474 | 0.989223 | 0.997988 | 0.994666 |
| Cacnb2_Q8CC27_S69s_1 | Cacnb2_Q8 | -0.8 | -0.59 | -0.44 | -1.82 | -0.9 | -0.98 | -0.61 | -0.86 | -0.18 | -0.66 | -0.81 | -0.78 | 0.09 | -0.73 | -1.15 | -0.65 | -0.5025 | 0.1575 | -0.405 | 0.464654 | 0.820957 | 0.402354 | 0.995762 | 0.997988 | 0.994666 |
| Cacnb2_Q8CC27_S83s_1 | Cacnb2_Q8 | -1.12 | 2.15 | 0.06 | -0.32 | -0.52 | -1.59 | 0.21 | -0.19 | 0.2 | 0.26 | -0.15 | 0.06 | 0.43 | -0.18 | 0.43 | -0.66 | 0.075 | 0.305 | 0.3025 | 0.779968 | 0.322392 | 0.875718 | 0.989223 | 0.997988 | 0.998226 |
| Cacnb2_Q8CC27_T549t_1 | Cacnb2_Q8 | -0.55 | -0.78 | 1.29 | 2.24 | -0.28 | -1.15 | 1.03 | 0.49 | -1.13 | -0.71 | 3.58 | 2.19 | 0.21 | 0.31 | 1.66 | 0.06 | -0.715 | -0.1 | -0.1875 | 0.989037 | 0.49455 | 0.716155 | 0.989223 | 0.997988 | 1 |
| Cacnb3_D3Z3Z3_S151s_1 | Cacnb3_D3 | 0.26 | 0.83 | 0.67 | 0.52 | 0.98 | 0.23 | 0.32 | 0.42 | 0.33 | 0.66 | 0.89 | 0.8 | 0.97 | 0.92 | 0.47 | 0.26 | -0.5275 | 0.4325 | 0.01 | 0.765903 | 0.749805 | 0.688757 | 0.995762 | 0.997988 | 0.9971 |
| Cacnb3_D3Z3Z3_S31s_1 | Cacnb3_D3 | 0.75 | 1.66 | 1.72 | 1.02 | 1.63 | 1.35 | 0.88 | 1.35 | 0.56 | 0.63 | 1.94 | 1.73 | -0.71 | 1.66 | 1.22 | 1.24 | -0.0825 | 0.1 | 0.085 | 0.429492 | 0.960931 | 0.86394 | 0.999187 | 0.997988 | 0.994666 |
| Cacnb3_D3Z3Z3_S392s_1 | Cacnb3_D3 | 1.2 | 1.31 | 0.99 | 0.96 | 1.46 | 1.09 | 0.89 | 0.94 | 0.84 | 0.92 | 1.32 | 0.77 | 0.74 | 1.28 | 1.12 | 0.86 | 0.015 | -0.0725 | -0.435 | 0.653834 | 0.930561 | 0.525003 | 0.997844 | 0.997988 | 0.995259 |
| Cacnb3_D3Z3Z3_S402s_1 | Cacnb3_D3 | 1.07 | 0.93 | 1 | 1.37 | 1.04 | 1.02 | 1.08 | 1.1 | 1.08 | 1.22 | 1.45 | 0.95 | -0.77 | 1 | 1.17 | 0.91 | -0.02 | -0.1525 | -0.115 | 0.269565 | 0.875098 | 0.726276 | 0.996493 | 0.997988 | 0.994666 |
| Cacnb3_D3Z3Z3_S406s_1 | Cacnb3_D3 | 0.19 | 0.53 | 2.66 | 1.88 | 1.09 | -0.96 | 2.11 | 1.98 | -0.34 | 0.32 | 2.82 | 2.56 | -0.45 | 0.39 | 2.43 | 2.61 | -0.0325 | 0.0825 | -0.515 | 0.933372 | 0.745454 | 0.977093 | 0.995762 | 0.997988 | 0.999507 |
| Cacnb4_Q8R0S4_S16s_1 | Cacnb4_Q8 | -0.1 | 0.76 | 0.82 | 1.19 | 0.7 | 0.19 | 0.61 | 0.91 | 0.54 | 0.05 | 1.56 | 1.27 | 1.27 | 0.78 | 0.8 | 0.71 | -0.26 | 0.025 | -0.07 | 0.515224 | 0.841343 | 0.662628 | 0.995932 | 0.997988 | 0.994666 |

| | | | | | | | | | | | | | | | | | | | | | | | | | | |
|---|---|---|---|---|---|---|---|---|---|---|---|---|---|---|---|---|---|---|---|---|---|---|---|---|---|---|
| Cacnb4_Q8R0S4_S38s_1 | Cacnb4_Q8 | 0.59 | 0.09 | 0.7 | 0.63 | 1.46 | 0.83 | 1.07 | 1.09 | -0.1 | -0.12 | -0.32 | 0.5 | -0.84 | 1.12 | -0.2 | 0.64 | -0.065 | 0.1875 | 0.2225 | 0.476331 | 0.03271 | 0.09157 | 0.989223 | 0.997988 | 0.994666 |
| Cacnb4_Q8R0S4_S423s_1 | Cacnb4_Q8 | 0.26 | 0.41 | 0.42 | 0.23 | 0.76 | 0.45 | 0.97 | 0.93 | -1.82 | 0.32 | -0.32 | 0.51 | -0.73 | -0.15 | -0.35 | 0.37 | 0.61 | -0.5125 | -0.3225 | 0.092403 | 0.064217 | 0.209722 | 0.989223 | 0.997988 | 0.994666 |
| Cacnb4_Q8R0S4_S443s_2 | Cacnb4_Q8 | 1.44 | 1.17 | 1.83 | 1.49 | 0.29 | 1.77 | -0.58 | 1.33 | 1.91 | 0.69 | 2.38 | 1.63 | -1.08 | 0 | 0.97 | 1.32 | 0.4475 | -0.6575 | -0.545 | 0.044976 | 0.143125 | 0.65954 | 0.989223 | 0.997988 | 0.994666 |
| Cacnb4_Q8R0S4_S447s_1 | Cacnb4_Q8 | 1.66 | 2.99 | 3 | 2.15 | 1.71 | 2.22 | 1.16 | 2.05 | 3.12 | 2.11 | 2.68 | 2.13 | 1.92 | 1.45 | 2.49 | 2.34 | -0.78 | 0.17 | -1.18 | 0.33923 | 0.114856 | 0.882316 | 0.989223 | 0.997988 | 0.994666 |
| Cacnb4_Q8R0S4_S447s_2 | Cacnb4_Q8 | 1.68 | 1.24 | 1.83 | 1.49 | 0.72 | 1.87 | -0.58 | 1.33 | 2.14 | 0.98 | 2.38 | 1.63 | -1.83 | 0.81 | 0.97 | 1.32 | -0.665 | 0.06 | -0.4 | 0.085794 | 0.168104 | 0.530012 | 0.989223 | 0.997988 | 0.994666 |
| Cacnb4_Q8R0S4_S507s_1 | Cacnb4_Q8 | -0.67 | -1.01 | -0.09 | -0.07 | -0.87 | 0 | -0.34 | -0.92 | 2.15 | -0.02 | -0.18 | -0.72 | 2.11 | -3.31 | 1.56 | 0.09 | -0.725 | 0.2225 | -1.2425 | 0.59708 | 0.82589 | 0.231858 | 0.995762 | 0.997988 | 0.995259 |
| Cacnb4_Q8R0S4_S50s_1 | Cacnb4_Q8 | -0.8 | -0.59 | -0.44 | -1.82 | -0.9 | -0.98 | -0.61 | -0.86 | -0.18 | -0.66 | -0.81 | -0.78 | 0.09 | -0.73 | -1.15 | -0.65 | -0.0725 | 0.7675 | 0.5725 | 0.464654 | 0.820957 | 0.402354 | 0.995762 | 0.997988 | 0.994666 |
| Cacnb4_Q8R0S4_S64s_1 | Cacnb4_Q8 | -1.12 | 2.15 | 1.68 | 0.7 | -0.52 | -1.59 | 0.83 | 0.29 | 0.2 | 0.26 | 0.63 | 1.45 | 0.43 | -0.18 | 1.16 | 0.54 | 0.075 | 0.305 | 0.3025 | 0.600974 | 0.182486 | 0.757142 | 0.989223 | 0.997988 | 0.995259 |
| Cacng2_O88602_S249s_1 | Cacng2_O8 | -0.94 | -0.29 | 0.14 | -1.51 | 0.03 | -0.85 | -0.23 | -1.58 | 2.6 | -0.31 | -2.65 | -1.01 | 2.94 | -1.34 | 1.12 | -1.27 | -1.1 | -0.2175 | -0.365 | 0.303625 | 0.98731 | 0.763153 | 0.999187 | 0.997988 | 0.994666 |
| Cacng2_O88602_S253s_1 | Cacng2_O8 | 3.43 | 3.7 | 2.59 | 1.79 | 3.36 | 3.87 | 1.74 | 1.85 | 1.19 | 2.91 | 0.49 | 2.12 | 1.26 | 0.62 | 1.97 | 2.33 | -0.0075 | 0.3075 | 1.0125 | 0.031226 | 0.779196 | 0.080403 | 0.995762 | 0.997988 | 0.994666 |
| Cacng2_O88602_Y288y_1 | Cacng2_O8 | 0.68 | 0.25 | 0.9 | 0.05 | -0.84 | 0.41 | -0.67 | 0.24 | -0.52 | -0.55 | 0.05 | 0.19 | 0.25 | -1.19 | -1.03 | -0.67 | -0.1725 | -1.2 | -1.3325 | 0.015214 | 0.087073 | 0.050873 | 0.989223 | 0.997988 | 0.994666 |
| Cacng3_Q9JJV5_S248s_1 | Cacng3_Q9 | 3.6 | 3.74 | 4.09 | 3.21 | 3.78 | 3.95 | 2.95 | 3.74 | 0.6 | 3.04 | 1.58 | 3.99 | -0.11 | 0.58 | 3.03 | 3.27 | -0.685 | -0.6775 | -1.13 | 0.028348 | 0.858368 | 0.07814 | 0.996349 | 0.997988 | 0.994666 |
| Cacng3_Q9JJV5_S302s_1 | Cacng3_Q9 | 0.03 | -1.31 | -0.97 | -1.74 | 0.63 | 0.14 | -1.4 | -0.06 | -1.86 | -1.5 | 1.86 | 1.31 | -0.9 | 0.49 | -1.32 | -2.31 | -0.055 | -1.3575 | -1.9675 | 0.984098 | 0.139496 | 0.310264 | 0.989223 | 0.997988 | 1 |
| Cacng4_Q9JJV4_S228s_1 | Cacng4_Q9 | 0.9 | 0.48 | 0.54 | 0.91 | -0.56 | 1.03 | 0.12 | 0.01 | 1.31 | 0.83 | -1.24 | -0.03 | 1.93 | -2.1 | 0.8 | 0.08 | 0.825 | 0.95 | -0.0125 | 0.491388 | 0.136436 | 0.368453 | 0.989223 | 0.997988 | 0.994666 |
| Cacng4_Q9JJV4_S253s_1 | Cacng4_Q9 | 0.6 | -0.21 | 0.74 | 1.02 | -1.21 | -0.02 | 0.8 | 0.7 | 3.36 | 0.17 | -1.26 | 0.27 | 3.72 | -0.76 | 3.16 | 0.36 | -0.5575 | -0.49 | -0.53 | 0.280905 | 0.349836 | 0.91315 | 0.989223 | 0.997988 | 0.994666 |
| Cacng4_Q9JJV4_S259s_1 | Cacng4_Q9 | 2.73 | 2.69 | 2.97 | 2.52 | 2.54 | 2.36 | 2.55 | 2.86 | 0.7 | 2.57 | 2.37 | 2.58 | 0.44 | 0.46 | 3.14 | 2.82 | -0.47 | 0.0975 | 1.0825 | 0.152064 | 0.506128 | 0.157178 | 0.989223 | 0.997988 | 0.994666 |
| Cacng8_S4R2F1_S251s_1 | Cacng8_S4 | 0.82 | 0.97 | 0.51 | 0.65 | 1.58 | 0.98 | 1.06 | 0.7 | 0.58 | 1.18 | 0.76 | 0.5 | 0.99 | 0.46 | 1.05 | 0.66 | -0.15 | -0.6725 | -1.0125 | 0.843173 | 0.206935 | 0.94492 | 0.989223 | 0.997988 | 0.999507 |
| Cacng8_S4R2F1_S254s_1 | Cacng8_S4 | 0.95 | 0.9 | 1.23 | 0.97 | 1.33 | 0.76 | 1.33 | 1.05 | 0.72 | 0.96 | 0.51 | 1.08 | 0.71 | 0.65 | 1.13 | 1.26 | 0.3425 | 0.0175 | 0.0525 | 0.776575 | 0.650006 | 0.417986 | 0.990358 | 0.997988 | 0.99798 |
| Cacng8_S4R2F1_S257s_1 | Cacng8_S4 | 0.49 | -0.41 | 0.34 | 0.41 | -0.15 | 0.35 | 0.36 | 0.24 | -0.45 | 0.26 | 1.38 | 0.47 | -0.85 | 0.17 | 0.47 | 0.3 | 0.105 | -0.195 | -0.075 | 0.627169 | 0.978421 | 0.62472 | 0.999187 | 0.997988 | 0.995259 |
| Cacng8_S4R2F1_S290s_1 | Cacng8_S4 | 1.53 | 1.42 | 1.4 | 1.41 | 1.66 | 1.59 | 1.53 | 1.92 | 0.58 | 1.49 | 0.73 | 1.47 | -0.09 | 0.61 | 1.08 | 1.46 | -0.0075 | 0.2075 | -0.185 | 0.085954 | 0.267143 | 0.217296 | 0.989223 | 0.997988 | 0.994666 |
| Cacng8_S4R2F1_S357s_1 | Cacng8_S4 | -0.25 | -0.7 | 0.76 | 0.75 | 0.36 | 0.35 | 0.97 | 0.99 | -0.99 | -0.4 | 0.37 | 0.82 | -1.54 | 0.14 | 0.09 | 0.61 | 0.235 | -0.3725 | -0.675 | 0.571429 | 0.202593 | 0.710584 | 0.989223 | 0.997988 | 0.995259 |
| Cacng8_S4R2F1_T415t_1 | Cacng8_S4 | -2.23 | -1.39 | -1.84 | -0.57 | -1.05 | -1.23 | -1.96 | -2.42 | -0.23 | -2.35 | -1.55 | -0.63 | 0.51 | -0.66 | -0.84 | -1.71 | 0.5275 | -0.19 | -0.315 | 0.146823 | 0.725715 | 0.567636 | 0.995437 | 0.997988 | 0.994666 |
| Cactin_Q9CS00_S148s_1 | Cactin_Q9 | 0.96 | 1.09 | 1.18 | 0.73 | 1.49 | 1.12 | 0.92 | 1.4 | 0.36 | 0.57 | 1.26 | 0.31 | 0.74 | 0.66 | -0.13 | 0.71 | -0.1575 | 0.3175 | 0.8325 | 0.119135 | 0.314508 | 0.223872 | 0.989223 | 0.997988 | 0.994666 |
| Cactin_Q9CS00_S96s_1 | Cactin_Q9 | -3.97 | -3.02 | -2.25 | -2.92 | -5.41 | -3.11 | -4 | -3.57 | -1.49 | -2.96 | -1.27 | -2.63 | -1.26 | -2.45 | -2.28 | -2.74 | 0.2425 | -0.365 | -0.495 | 0.088941 | 0.100241 | 0.089391 | 0.989223 | 0.997988 | 0.994666 |
| Cad_E9QAI5_T1770t_1 | Cad_E9QAI | 1.04 | 0.07 | -0.29 | 0.53 | 0.77 | 0.58 | 0.77 | 0.26 | 0.41 | 1 | -0.42 | 0.15 | -0.64 | 0.67 | -0.28 | 0.15 | -0.9825 | 0.9525 | 0.8575 | 0.38339 | 0.435745 | 0.897393 | 0.989223 | 0.997988 | 0.994666 |
| Cadm2_G3UZM4_S423s_1 | Cadm2_G3 | 1.99 | 1.67 | 0.19 | 0.68 | 1.89 | 1.92 | 0.51 | -0.29 | 1.73 | 1.79 | -0.78 | -1.19 | -0.16 | 2.23 | -1.31 | -0.53 | 0.2575 | -0.0525 | -0.3625 | 0.18579 | 0.838376 | 0.365836 | 0.995762 | 0.997988 | 0.994666 |
| Cadm4_Q8R464_S361s_1 | Cadm4_Q8 | 0.41 | 0.84 | 0.65 | 0.42 | 0.6 | 0.35 | 0.89 | 0.46 | 0.22 | 0.49 | 0.84 | 0.64 | 0.12 | 0.84 | 0.97 | 0.74 | -0.125 | -0.745 | -1.075 | 0.760727 | 0.982679 | 0.894638 | 0.999187 | 0.997988 | 0.9971 |
| Cadps_Q80TJ1_S375s_1 | Cadps_Q80 | 0.46 | 0.71 | 0.92 | 0.66 | 0.69 | 1.4 | 2.08 | 1.3 | 0.1 | 0.77 | 0.22 | 1.49 | -0.12 | -0.41 | 1.69 | 1.93 | -0.005 | -0.0325 | 0.0875 | 0.880109 | 0.054661 | 0.902619 | 0.989223 | 0.997988 | 0.999507 |
| Cadps_Q80TJ1_S486s_1 | Cadps_Q80 | -1.83 | -2.03 | -0.16 | -1.44 | -1.34 | -2.34 | -1.57 | -0.2 | -1.58 | -1.65 | 0.19 | -1.41 | -0.48 | -1.88 | -0.71 | -2.02 | 0.68 | -0.0425 | 0.085 | 0.863146 | 0.996377 | 0.653873 | 0.99973 | 0.997988 | 0.999507 |
| Cadps_Q80TJ1_S89s_1 | Cadps_Q80 | 0.77 | -0.76 | -1.27 | 0.34 | -0.21 | 0.53 | 0.01 | 0.99 | -0.75 | -0.55 | 2.06 | 0.12 | -1.29 | -0.16 | -1.37 | -1.57 | 0.0025 | 0.2525 | 0.0925 | 0.127975 | 0.278184 | 0.534704 | 0.989223 | 0.997988 | 0.994666 |
| Cadps_Q80TJ1_S96s_1 | Cadps_Q80 | 0.13 | 0.73 | -2.37 | -2.14 | 0.68 | -0.03 | -1.54 | -2.53 | -0.44 | 0.32 | -2.4 | -2.16 | -0.08 | 1.22 | -1.77 | -2.55 | 0.56 | 0.45 | -0.8675 | 0.906904 | 0.950525 | 0.777611 | 0.999187 | 0.997988 | 0.999507 |
| Cadps2_S4R1C6_S161s_1 | Cadps2_S4 | 0.39 | 0.88 | 1.06 | 1.34 | 0.75 | 0.28 | 0.99 | 1.03 | 0.28 | 0.41 | 0.86 | 0.37 | -0.17 | 0.55 | 0.64 | 0.67 | 0.0575 | -0.2575 | 0.1175 | 0.152574 | 0.598753 | 0.150747 | 0.989223 | 0.997988 | 0.994666 |
| Cadps2_S4R1C6_S39s_1 | Cadps2_S4 | -0.84 | -1.05 | -0.73 | -0.34 | -0.97 | -0.95 | -0.37 | -0.57 | -1.39 | -1.07 | -1.32 | -0.3 | -1.51 | 0.04 | -0.92 | -1.37 | -0.155 | -0.4375 | -0.495 | 0.61118 | 0.923108 | 0.391541 | 0.997335 | 0.997988 | 0.995259 |
| Cadps2_S4R1C6_T811t_1 | Cadps2_S4 | 0.96 | 0.97 | 2.09 | 1.72 | 0.85 | 1.1 | 0.82 | 1.84 | 0.89 | 0.83 | 1.5 | 2.16 | 1.3 | 0.11 | 1.48 | 2.06 | 0.025 | -0.28 | -0.2 | 0.679333 | 0.446246 | 0.827568 | 0.989223 | 0.997988 | 0.996159 |
| Calcoco1_E9Q7U2_S563s_1 | Calcoco1_E | 0.13 | 0.13 | 0.63 | 1.02 | 0.05 | 0.07 | 0.87 | 0.61 | -0.55 | 0.33 | 0.09 | 0.73 | -0.26 | 0.35 | 0.35 | 0.72 | -0.2825 | -0.09 | -0.1975 | 0.578459 | 0.805259 | 0.370254 | 0.995762 | 0.997988 | 0.995259 |
| Calml3_P0DP28_S82s_1 | Calml3_P0I | -0.49 | 0.12 | 0.09 | -0.66 | -0.45 | 0.07 | -1.25 | -0.36 | -1.42 | -0.24 | 0.35 | -0.36 | -0.78 | -1.38 | -1.08 | -0.06 | -0.0775 | -0.3275 | -0.1875 | 0.132917 | 0.452984 | 0.659778 | 0.989223 | 0.997988 | 0.994666 |
| Camk1_Q91YS8_S324s_1 | Camk1_Q9 | 0.25 | 0.65 | -0.17 | 0.52 | 0.68 | -0.18 | 0.39 | -0.81 | -0.29 | -0.03 | -1.95 | 0.68 | -0.24 | -0.04 | 0.57 | 0.15 | -0.2625 | -0.1825 | -0.59 | 0.514029 | 0.437334 | 0.212809 | 0.989223 | 0.997988 | 0.994666 |
| Camk1_Q91YS8_T177t_1 | Camk1_Q9 | -0.09 | 0.53 | 0.83 | 0.51 | 0.38 | 0.14 | 1.65 | 1.63 | 0.77 | 1.09 | 0.07 | 1.34 | 0.18 | 0.02 | 1.57 | 1.82 | -0.2925 | -0.71 | -0.2025 | 0.357848 | 0.249842 | 0.303917 | 0.989223 | 0.997988 | 0.994666 |
| Camk1_Q91YS8_T339t_1 | Camk1_Q9 | -0.23 | -0.52 | -0.34 | -0.41 | 0.72 | -1.36 | 0.4 | -1.08 | -0.83 | -0.52 | -0.3 | -0.22 | -0.87 | -0.09 | -0.5 | -0.25 | 0.505 | 0.3725 | 0.4525 | 0.845551 | 0.925788 | 0.697823 | 0.997335 | 0.997988 | 0.999507 |
| Camk1d_Q8BW96_S319s_1 | Camk1d_Q | 1.41 | 0.72 | 0.66 | 0.5 | 0.44 | 0.32 | 0.66 | 0.69 | 0.7 | 0.48 | -0.08 | 0.47 | -0.08 | 0.66 | 0.22 | 0.37 | 0.045 | -0.0925 | -0.0525 | 0.111211 | 0.282288 | 0.175706 | 0.989223 | 0.997988 | 0.994666 |
| Camk1d_Q8BW96_S349s_1 | Camk1d_Q | 0.58 | 0.14 | 0.36 | 0.59 | -1.18 | 0.74 | -0.63 | 0.09 | 0.91 | -0.07 | 0.18 | -0.26 | 1.12 | -1.11 | -0.02 | 0.01 | -0.295 | -0.43 | -0.53 | 0.368716 | 0.133004 | 0.471311 | 0.989223 | 0.997988 | 0.994666 |
| Camk1d_Q8BW96_T172t_1 | Camk1d_Q | 0.33 | 1.04 | 0.1 | -0.66 | 1.36 | 0.35 | 0.07 | -0.07 | 0.07 | 0.43 | -1.19 | 0.12 | 0.1 | 1.08 | -0.65 | -0.21 | -0.6625 | -0.2275 | -0.4175 | 0.800277 | 0.616163 | 0.475188 | 0.989223 | 0.997988 | 0.999507 |
| Camk2a_F8WIS9_S272s_1 | Camk2a_F8 | 0.81 | 2.27 | 2.38 | -0.27 | 2.02 | -0.71 | 2.54 | 0.57 | -1.07 | 3.2 | 0.14 | 1.88 | 0.31 | -2.02 | 2.66 | 2.19 | 0.225 | -0.345 | -0.1225 | 0.635174 | 0.819857 | 0.795331 | 0.995762 | 0.997988 | 0.995259 |
| Camk2a_F8WIS9_S275s_1 | Camk2a_F8 | -1.23 | -1.69 | -2.51 | -3.39 | -2.2 | -1.48 | -2.79 | -2.84 | 0.19 | -1.46 | -2.45 | -3.52 | 1.29 | -2 | -2.15 | -2.78 | -0.1925 | -0.26 | -0.5125 | 0.388881 | 0.814102 | 0.632311 | 0.995762 | 0.997988 | 0.994666 |
| Camk2a_F8WIS9_S275s_2 | Camk2a_F8 | -2.38 | -1.78 | -2.34 | -1.8 | -2.36 | -2.59 | -4.25 | -2.37 | 0 | -1.37 | -2.22 | -3.1 | 0.5 | -2.02 | -1.97 | -2.02 | -0.1225 | 0.395 | 0.795 | 0.256531 | 0.096259 | 0.521536 | 0.989223 | 0.997988 | 0.994666 |
| Camk2a_F8WIS9_S279s_1 | Camk2a_F8 | -2.31 | -0.32 | -1.2 | -2.8 | -1.73 | -1.98 | -1.92 | -2.63 | 0.24 | -0.61 | -0.47 | -1.42 | 1.99 | -0.65 | -1.09 | -1.55 | -0.8175 | 0.4025 | 0.6975 | 0.142484 | 0.453978 | 0.094442 | 0.989223 | 0.997988 | 0.994666 |
| Camk2a_F8WIS9_S279s_2 | Camk2a_F8 | -2.38 | -1.78 | -2.34 | -1.8 | -2.36 | -2.59 | -4.25 | -2.37 | 0 | -1.37 | -2.22 | -3.1 | 0.5 | -2.02 | -1.97 | -2.02 | -0.4075 | 1.0925 | 1.3325 | 0.256531 | 0.096259 | 0.521536 | 0.989223 | 0.997988 | 0.994666 |
| Camk2a_F8WIS9_S481s_1 | Camk2a_F8 | -0.55 | 0.33 | 0.16 | -0.16 | -0.16 | -1.03 | -0.41 | 0.37 | 0.27 | 0.06 | 0.61 | -0.04 | 1.11 | 0.58 | 0.42 | -0.05 | -0.8175 | 0.4025 | 0.6975 | 0.120751 | 0.478051 | 0.343482 | 0.989223 | 0.997988 | 0.994666 |
| Camk2a_F8WIS9_S485s_1 | Camk2a_F8 | -0.01 | 0.16 | 0.48 | 0.44 | 0.09 | 0.03 | 1.1 | 0.54 | 0.28 | -0.07 | 0.91 | 1.9 | 0.54 | 0.2 | 1.74 | 1.03 | -0.2525 | 0.28 | 0.57 | 0.126543 | 0.567315 | 0.278902 | 0.989223 | 0.997988 | 0.994666 |
| Camk2a_F8WIS9_T253t_1 | Camk2a_F8 | -1.86 | -0.85 | -1.1 | -1 | -1.18 | -3.12 | -1.03 | -1.75 | -2.23 | -0.78 | -1.29 | -1.05 | -1.02 | -1.69 | -0.77 | -1.16 | 0.1725 | 0.4875 | 0.61 | 0.899098 | 0.259512 | 0.728998 | 0.989223 | 0.997988 | 0.999507 |
| Camk2a_F8WIS9_T286t_1 | Camk2a_F8 | -3.46 | -2.17 | -2.88 | -4.49 | -2.38 | -3.49 | -3.17 | -4.03 | -3.92 | -2.27 | -1.23 | -3.55 | -1.14 | -1.79 | -3.18 | -3.3 | -0.5675 | -0.135 | 0.0425 | 0.19024 | 0.974247 | 0.480448 | 0.999187 | 0.997988 | 0.994666 |
| Camk2a_P11798_S330s_1 | Camk2a_P1 | 0.68 | 1.25 | 1.79 | 1.79 | 1.12 | 0.91 | 1.08 | 1.86 | 0.55 | 1.05 | 1.89 | 1.54 | 0.27 | 1.33 | 0.22 | 0.98 | -0.0175 | 0.5075 | 0.8975 | 0.108233 | 0.695251 | 0.76106 | 0.994893 | 0.997988 | 0.994666 |
| Camk2a_P11798_S331s_1 | Camk2a_P1 | 1.94 | 1.32 | -0.43 | -0.92 | 0.17 | 1.24 | -0.92 | -1.2 | 1.71 | 2.09 | -0.17 | -0.72 | 0.99 | 0.27 | -0.69 | -0.63 | -0.135 | -0.12 | -0.6775 | 0.492659 | 0.405972 | 0.772708 | 0.989223 | 0.997988 | 0.994666 |
| Camk2a_P11798_S333s_2 | Camk2a_P1 | 0.55 | 0.94 | 1.59 | 1.82 | -0.62 | 0.44 | 0.79 | 1.86 | 0.79 | 0.9 | 2.11 | 1.27 | -0.63 | 0.13 | 0.82 | 1.2 | -0.655 | 0.25 | -0.4925 | 0.101774 | 0.272183 | 0.917747 | 0.989223 | 0.997988 | 0.994666 |
| Camk2a_P11798_T334t_1 | Camk2a_P1 | -0.32 | 1.71 | 0.99 | 0.42 | -0.58 | 0.16 | 0.18 | 0.44 | 0.14 | 0.77 | 1.2 | 1.1 | 0.76 | 0.09 | 0.61 | 0.51 | -0.6075 | 0.0425 | -0.845 | 0.641725 | 0.172765 | 0.827206 | 0.989223 | 0.997988 | 0.995259 |
| Camk2a_P11798_T334t_2 | Camk2a_P1 | 1.14 | 1.16 | 0.89 | 0.39 | -0.48 | 1.08 | -0.33 | 0.14 | 1.05 | 1.5 | 1.49 | 0.92 | -0.28 | 0.41 | 0.98 | 0.79 | -0.65 | 0.1025 | -0.2075 | 0.257129 | 0.063165 | 0.242129 | 0.989223 | 0.997988 | 0.994666 |
| Camk2a_P11798_T336t_1 | Camk2a_P1 | -0.01 | 1.39 | 1.22 | 0.23 | 0.38 | -0.54 | -0.31 | -0.07 | -0.02 | 0.56 | 0.53 | 0.7 | 0.78 | 0.31 | 0.84 | 0.71 | -0.7925 | 0.345 | -0.42 | 0.901388 | 0.053394 | 0.498503 | 0.989223 | 0.997988 | 0.999507 |
| Camk2a_P11798_T336t_2 | Camk2a_P1 | 0.6 | 1.16 | 0.92 | 0.79 | -0.59 | 0.62 | -0.03 | 0.68 | 0.69 | 1.03 | 1.57 | 0.98 | -0.3 | 0.13 | 0.93 | 0.8 | -0.8425 | -0.265 | -0.0475 | 0.188122 | 0.059177 | 0.471661 | 0.989223 | 0.997988 | 0.994666 |
| Camk2a_P11798_T337t_1 | Camk2a_P1 | 1.88 | 1.86 | 1.85 | 0.76 | 1.25 | 1.67 | -0.1 | 0.72 | 1.26 | 1.97 | 1.75 | 1.46 | 0.17 | 0.99 | 1.19 | 0.81 | -0.6975 | 0.2 | -0.4775 | 0.055112 | 0.136596 | 0.946872 | 0.989223 | 0.997988 | 0.994666 |
| Camk2a_P11798_T337t_2 | Camk2a_P1 | 0.27 | 0.94 | 0.93 | 0.4 | -0.84 | 0.17 | -0.28 | 0.13 | 0.52 | 0.67 | 1.43 | 0.86 | -0.66 | -0.28 | 0.96 | 0.79 | -0.7025 | 0.0225 | -0.7975 | 0.327163 | 0.0219 | 0.445813 | 0.989223 | 0.997988 | 0.994666 |
| Camk2b_P28652_S331s_1 | Camk2b_P2 | -0.52 | 0.08 | -0.41 | -0.18 | -0.29 | -0.37 | 0.12 | -0.4 | -0.24 | -0.15 | -0.29 | -0.27 | -0.11 | 0.42 | -0.71 | -0.64 | -0.84 | 0.235 | -0.4325 | 0.994019 | 0.926051 | 0.931117 | 0.997335 | 0.997988 | 1 |
| Camk2b_P28652_S363s_1 | Camk2b_P2 | -1.36 | 0.16 | -1.58 | -0.73 | -0.23 | -0.39 | -0.68 | -1.03 | -0.16 | -0.35 | -0.6 | -1.25 | 1.27 | 0.19 | -1.44 | -0.37 | 0.0225 | 0.02 | -0.0025 | 0.224625 | 0.476817 | 0.519852 | 0.989223 | 0.997988 | 0.994666 |
| Camk2b_P28652_S367s_1 | Camk2b_P2 | 0.69 | 0.71 | 1.42 | 0.67 | 0.27 | 0.78 | 0.24 | 0.62 | 0.54 | 0.78 | 0.33 | 0.85 | 1.47 | -1.05 | 0.91 | 1.14 | 0.295 | 0.2875 | 0.79 | 0.645878 | 0.165029 | 0.376831 | 0.989223 | 0.997988 | 0.995259 |
| Camk2b_P28652_S371s_1 | Camk2b_P2 | -1.19 | -0.43 | 0.37 | -0.01 | 0.07 | -1.23 | 0.39 | 0.93 | -1.07 | -0.61 | 0.56 | 1.27 | -0.24 | 0.32 | 0.62 | 0.82 | -0.395 | -0.2475 | -0.255 | 0.114639 | 0.498165 | 0.547757 | 0.989223 | 0.997988 | 0.994666 |
| Camk2b_P28652_T287t_1 | Camk2b_P2 | -1.33 | -1.02 | -0.93 | -1.25 | -1.16 | -1.24 | -0.94 | -1.34 | -1.72 | -0.97 | 0.07 | -1.37 | -0.32 | -0.47 | -0.99 | -1.14 | 0.355 | 0.3525 | 0.695 | 0.186322 | 0.863153 | 0.734573 | 0.996493 | 0.997988 | 0.994666 |
| Camk2b_P28652_T382t_1 | Camk2b_P2 | -1.51 | 0.06 | -0.43 | -0.62 | -0.56 | -2.44 | -0.11 | -0.97 | -0.75 | -1.22 | -0.92 | -1.09 | -0.39 | -0.46 | -1.08 | -1.02 | -0.0375 | 0.135 | 0.4025 | 0.77152 | 0.474599 | 0.313917 | 0.989223 | 0.997988 | 0.9971 |
| Camk2b_P28652_T400t_1 | Camk2b_P2 | -0.44 | 1.04 | 1.42 | 1.02 | -1.09 | -0.67 | -0.1 | 0.51 | -0.88 | 0.22 | 0.57 | 0.68 | 0.85 | -0.68 | 1.08 | 1 | -0.395 | -0.37 | -0.1125 | 0.717009 | 0.048492 | 0.248643 | 0.989223 | 0.997988 | 0.996966 |
| Camk2b_P28652_T400t_2 | Camk2b_P2 | 1.01 | 1.18 | 0.66 | 0.99 | -0.48 | 1.05 | -0.53 | -0.37 | 1.43 | 1.85 | -0.11 | 0.41 | -0.07 | 0.65 | 0.25 | 0.69 | -1.0975 | -0.6125 | -0.1975 | 0.065524 | 0.021217 | 0.884415 | 0.989223 | 0.997988 | 0.994666 |
| Camk2b_P28652_T401t_2 | Camk2b_P2 | 0.65 | 0.79 | 1.23 | 1.01 | -0.66 | 0.3 | 0.2 | 0.32 | 1.04 | 1.32 | 1.47 | 0.87 | -0.47 | 0.03 | 1.12 | 1.03 | -1.0425 | -0.065 | -0.58 | 0.24842 | 0.013422 | 0.331843 | 0.989223 | 0.997988 | 0.994666 |
| Camk2b_Q5SVI1_S427s_1 | Camk2b_Q | 0.67 | 0.74 | 0.55 | 0.25 | 0.31 | 0.21 | 1.01 | 0.34 | -0.25 | 1.06 | -0.1 | 0.56 | -0.36 | 0.03 | -0.22 | 0.91 | -0.88 | 0.255 | -0.4925 | 0.195439 | 0.745189 | 0.49704 | 0.995762 | 0.997988 | 0.994666 |
| Camk2b_Q5SVJ0_T400t_1 | Camk2b_Q | -0.44 | 1.04 | 1.42 | 1.02 | -1.09 | -0.67 | -0.1 | 0.51 | -0.88 | 0.22 | 0.57 | 0.68 | 0.85 | -0.68 | 1.08 | 1 | -0.085 | -0.235 | -0.4625 | 0.717009 | 0.048492 | 0.248643 | 0.989223 | 0.997988 | 0.996966 |
| Camk2b_Q5SVJ0_T400t_2 | Camk2b_Q | 1.01 | 1.18 | 0.66 | 0.99 | -0.48 | 1.05 | -0.53 | -0.37 | 1.43 | 1.85 | -0.11 | 0.41 | -0.07 | 0.65 | 0.25 | 0.69 | -1.0975 | -0.6125 | -0.1975 | 0.065524 | 0.021217 | 0.884415 | 0.989223 | 0.997988 | 0.994666 |
| Camk2b_Q5SVJ0_T401t_2 | Camk2b_Q | 0.65 | 0.79 | 1.23 | 1.01 | -0.66 | 0.3 | 0.2 | 0.32 | 1.04 | 1.32 | 1.47 | 0.87 | -0.47 | 0.03 | 1.12 | 1.03 | -1.0425 | -0.065 | -0.58 | 0.24842 | 0.013422 | 0.331843 | 0.989223 | 0.997988 | 0.994666 |
| Camk2d_E9Q1T1_S235s_1 | Camk2d_E9 | 0.27 | 0.45 | 0.76 | 0.77 | 0.65 | 0.26 | 0.94 | 0.58 | 0.18 | 0.28 | 0.77 | 0.82 | -0.21 | 0.75 | 0.62 | 0.61 | -0.88 | 0.255 | -0.4925 | 0.697388 | 0.854831 | 0.85116 | 0.996349 | 0.997988 | 0.996159 |
| Camk2d_E9Q1T1_S276s_1 | Camk2d_E9 | -1 | 0.54 | -0.54 | -1.51 | -0.78 | -0.86 | -0.99 | -1.41 | 0.24 | 0.11 | -0.77 | -0.78 | 1.66 | -0.12 | 0 | -0.66 | 0.045 | -0.05 | -0.12 | 0.184165 | 0.386495 | 0.507712 | 0.989223 | 0.997988 | 0.994666 |
| Camk2d_E9Q1T1_S276s_2 | Camk2d_E9 | -3.08 | 0.3 | -1.36 | -2.68 | -1.77 | -3.11 | -4.02 | -2.4 | 0.21 | -0.05 | -3.67 | -2.34 | 1.46 | -1.05 | -2.82 | -2.78 | -0.3825 | 0.3275 | 0.8475 | 0.710702 | 0.178742 | 0.8193 | 0.989223 | 0.997988 | 0.996966 |
| Camk2d_E9Q1T1_S280s_1 | Camk2d_E9 | -3.04 | 0.84 | -0.04 | -1.92 | -1.04 | -3.33 | -1.53 | -1.88 | -0.43 | -0.35 | -0.83 | -1.23 | 2.12 | 0.59 | -0.24 | -0.3 | -1.12 | 0.2425 | 0.4075 | 0.109202 | 0.316411 | 0.683625 | 0.989223 | 0.997988 | 0.994666 |
| Camk2d_E9Q1T1_S280s_2 | Camk2d_E9 | -3.08 | 0.3 | -1.36 | -2.68 | -1.77 | -3.11 | -4.02 | -2.4 | 0.21 | -0.05 | -3.67 | -2.34 | 1.46 | -1.05 | -2.82 | -2.78 | -0.905 | 0.33 | 1.5825 | 0.710702 | 0.178742 | 0.8193 | 0.989223 | 0.997988 | 0.996966 |
| Camk2d_E9Q1T1_S338s_1 | Camk2d_E9 | 0.55 | 0.69 | 1.39 | 0.31 | 0.95 | 0.88 | 0.58 | 0.49 | 0.85 | 0.67 | 0.62 | 0.63 | 1.15 | 0.78 | 0.75 | 0.6 | -1.12 | 0.2425 | 0.4075 | 0.785731 | 0.972331 | 0.881797 | 0.999187 | 0.997988 | 0.999507 |
| Camk2d_E9Q1T1_S524s_1 | Camk2d_E9 | 0.17 | 0.64 | -0.18 | 0.09 | 0.21 | 0.26 | 1.1 | 0.04 | 0.15 | 0.78 | 0.53 | 0.41 | 0.06 | 0.61 | 0.56 | 0.74 | -0.01 | -0.0425 | 0.085 | 0.301447 | 0.480617 | 0.30577 | 0.989223 | 0.997988 | 0.994666 |
| Camk2d_E9Q1T1_S528s_1 | Camk2d_E9 | -0.71 | -0.79 | -1.24 | 1.41 | -0.75 | -0.91 | 0.41 | 0.37 | -0.87 | 0.02 | 0.14 | 0.47 | -0.77 | -1.13 | 0.79 | 0.66 | 0.2225 | 0.2875 | 0.3125 | 0.749906 | 0.854931 | 0.652727 | 0.996349 | 0.997988 | 0.9971 |
| Camk2d_E9Q1T1_T287t_1 | Camk2d_E9 | -3.46 | -2.17 | -2.89 | -4.55 | -2.38 | -3.49 | -3.18 | -4.07 | -3.92 | -2.27 | -1.23 | -3.57 | -1.14 | -1.79 | -3.19 | -3.32 | 0.1125 | 0.2725 | 0.22 | 0.19136 | 0.981975 | 0.474686 | 0.999187 | 0.997988 | 0.994666 |
| Camk2g_Q923T9_S334s_1 | Camk2g_Q | -2.26 | -1.83 | -1.27 | -2.89 | -1.83 | -2.39 | -1.52 | -2.11 | -2.48 | -2.94 | -1.31 | -1.46 | -0.31 | -1.04 | -2.5 | -2.33 | -0.0125 | 0.52 | 0.9075 | 0.379424 | 0.792929 | 0.975717 | 0.995762 | 0.997988 | 0.994666 |
| Camk2g_Q923T9_S349s_1 | Camk2g_Q | -0.98 | -0.35 | -0.69 | -0.83 | -0.83 | -1.34 | 0.03 | -0.64 | -0.39 | -0.77 | -0.65 | -0.78 | -1.05 | -0.75 | -1.57 | -0.45 | 0.1 | 0.015 | 0.5175 | 0.456813 | 0.957119 | 0.789802 | 0.999187 | 0.997988 | 0.994666 |
| Camk2g_Q923T9_T287t_1 | Camk2g_Q | -4.33 | -2.07 | -3.16 | -4.2 | -3.67 | -3.91 | -3.83 | -4.6 | -5.03 | -3.33 | -1.73 | -3.94 | -1.7 | -2 | -3.14 | -3.43 | 0.0175 | 0.065 | -0.2425 | 0.178164 | 0.290591 | 0.9306 | 0.989223 | 0.997988 | 0.994666 |
| Camk4_P08414_S337s_1 | Camk4_P08 | -0.89 | -0.91 | -0.69 | -1.11 | -0.68 | -0.94 | -0.61 | -0.41 | -0.19 | -0.52 | -1.53 | -0.21 | 0.45 | 0.67 | -0.81 | -0.31 | -0.5625 | -0.0675 | 0.8725 | 0.03452 | 0.296636 | 0.413027 | 0.989223 | 0.997988 | 0.994666 |
| Camk4_P08414_T196t_1 | Camk4_P08 | -0.47 | -0.2 | -1.13 | -0.36 | -0.24 | 0.15 | -0.42 | -0.88 | -0.28 | 0.17 | -0.99 | -1.27 | -0.18 | 0.16 | -0.37 | -0.08 | 0.24 | 0.2875 | 0.9 | 0.176673 | 0.543176 | 0.892416 | 0.989223 | 0.997988 | 0.994666 |
| Camkk1_Q8VBY2_S458s_1 | Camkk1_Q | -0.26 | -0.16 | -0.33 | -0.45 | -0.1 | -0.59 | -0.31 | -0.46 | -1.16 | -0.5 | -1.46 | -0.59 | -1.12 | -0.99 | -0.79 | -1.02 | 0.1925 | -0.0525 | 0.4225 | 0.015795 | 0.763241 | 0.051206 | 0.995762 | 0.997988 | 0.994666 |
| Camkk1_Q8VBY2_S475s_1 | Camkk1_Q | -0.12 | -0.16 | -0.29 | -0.81 | 0.58 | -0.53 | -0.08 | -0.86 | -1.33 | -0.58 | -0.36 | -1.24 | -0.86 | 0.43 | -0.43 | -0.52 | -0.065 | -0.6275 | -0.68 | 1 | 0.728092 | 0.120176 | 0.995762 | 0.997988 | 1 |
| Camkk1_Q8VBY2_S492s_1 | Camkk1_Q | 0.06 | 0.86 | -0.07 | 0.08 | 1.07 | 0.91 | 0.3 | -0.14 | -0.69 | 0.22 | -0.26 | -0.45 | -1.47 | -1.4 | 0.08 | 0.1 | 0.1225 | -0.5325 | 1.57E-16 | 0.079203 | 0.399688 | 0.122058 | 0.989223 | 0.997988 | 0.994666 |
| Camkk1_Q8VBY2_S82s_1 | Camkk1_Q | -0.26 | -2.54 | 0.29 | 0.55 | -0.29 | -0.43 | 0.52 | -0.25 | -2.13 | -1.28 | -1.39 | 0.92 | -2.49 | -0.8 | -0.12 | 0.41 | 0.3025 | -0.5275 | -0.905 | 0.755128 | 0.566294 | 0.578445 | 0.989223 | 0.997988 | 0.9971 |
| Camkk1_Q8VBY2_T108t_1 | Camkk1_Q | -3.08 | -3.21 | -1.95 | -3.92 | -2.76 | -3.43 | -2.93 | -2.8 | -1.95 | -3.38 | -0.33 | -2.15 | -1.22 | -1.24 | -2.71 | -3.16 | 0.3775 | -0.48 | -0.26 | 0.1301 | 0.88498 | 0.133567 | 0.996493 | 0.997988 | 0.994666 |
| Camkk2_Q8C078_S129s_1 | Camkk2_Q | 0.36 | -0.14 | 0.88 | 0.86 | 0.62 | 0.42 | 0.55 | 1.06 | 1.17 | 0.09 | 0.39 | 1.52 | -0.25 | 0.05 | -0.15 | 1.37 | 0.06 | 1.0875 | 0.9575 | 0.595342 | 0.571595 | 0.464013 | 0.989223 | 0.997988 | 0.995259 |
| Camkk2_Q8C078_S133s_1 | Camkk2_Q | 1.01 | 0.08 | -0.11 | 0.14 | 0.09 | 0.26 | 0.89 | 0.45 | -0.13 | 0.06 | -0.29 | 0.38 | -0.25 | 0.59 | -0.35 | 0.04 | 0.1725 | 0.3025 | -0.235 | 0.448937 | 0.656036 | 0.396886 | 0.990434 | 0.997988 | 0.994666 |
| Camkk2_Q8C078_S495s_1 | Camkk2_Q | -0.04 | 0.08 | 0.09 | -0.27 | 0.49 | -0.11 | 0.47 | 0.11 | -0.65 | 0.05 | -0.77 | -0.09 | -0.34 | -0.48 | -0.25 | 0.08 | 0.1425 | -0.275 | -0.2725 | 0.411527 | 0.262537 | 0.248253 | 0.989223 | 0.997988 | 0.994666 |
| Camkk2_Q8C078_S511s_1 | Camkk2_Q | -0.03 | -0.65 | 0.02 | -0.11 | 0.6 | -0.54 | 1.05 | 0.47 | -0.56 | -0.79 | 0.13 | 0.42 | -0.64 | 0.49 | 0.69 | -0.14 | 0.275 | -0.33 | -0.2125 | 0.429274 | 0.13308 | 0.982516 | 0.989223 | 0.997988 | 0.994666 |
| Camkk2_Q8C078_S522s_1 | Camkk2_Q | -1.78 | -2.52 | -0.67 | -3.29 | -2.22 | -0.55 | -1.8 | -1.54 | 1.54 | -1.53 | -1.5 | -0.66 | 2.18 | -1.24 | 0.76 | -1.83 | 0.5875 | -0.0075 | 0.2925 | 0.053873 | 0.374688 | 0.08668 | 0.989223 | 0.997988 | 0.994666 |
| Camkk2_Q8C078_S572s_1 | Camkk2_Q | -1.15 | -0.98 | -1.52 | -0.81 | -0.85 | -1.89 | -0.42 | -1.67 | -1.52 | -0.76 | -0.91 | -1.43 | -1.16 | -0.2 | -1.3 | -1.62 | 0.5375 | 1.5275 | 2.0325 | 0.901544 | 0.803258 | 0.88956 | 0.995762 | 0.997988 | 0.999507 |
| Camkk2_Q8C078_S99s_1 | Camkk2_Q | -0.54 | -0.74 | -0.83 | -0.92 | -0.24 | -0.32 | -0.15 | -0.37 | -0.72 | -0.65 | 0.19 | -0.92 | -0.94 | 0.11 | -0.92 | -0.76 | -0.0925 | -0.04 | 0.045 | 0.679744 | 0.03749 | 0.443854 | 0.989223 | 0.997988 | 0.996159 |
| Camkv_Q3UHL1_S387s_1 | Camkv_Q3 | 0.94 | 0.73 | 1.09 | 0.46 | -0.03 | 1.25 | 0.02 | 0.62 | 0.62 | 0.44 | 0.19 | 1.13 | -0.01 | -1.61 | 0.51 | 0.82 | 0.4875 | 0.2325 | 0.13 | 0.118118 | 0.326891 | 0.472349 | 0.989223 | 0.997988 | 0.994666 |
| Camkv_Q3UHL1_S392s_1 | Camkv_Q3 | -0.04 | -0.17 | 0.32 | 0.09 | -1 | 0.5 | -0.87 | -0.23 | 0.62 | -0.1 | -0.13 | 0.36 | 0.14 | -1.25 | -0.11 | -0.29 | -0.34 | -0.21 | -0.8775 | 0.241617 | 0.228807 | 0.612194 | 0.989223 | 0.997988 | 0.994666 |
| Camkv_Q3UHL1_S436s_1 | Camkv_Q3 | -0.77 | -1.52 | -0.63 | -2.23 | -1.35 | 0.04 | -1.19 | -1.27 | 1.04 | -1.44 | -0.93 | -0.64 | 1.12 | -1.53 | -0.31 | -1.15 | -0.45 | 0.1375 | -0.4275 | 0.213071 | 0.460595 | 0.205334 | 0.989223 | 0.997988 | 0.994666 |
| Camkv_Q3UHL1_S436s_2 | Camkv_Q3 | -1.87 | -0.71 | -1.44 | -3.05 | -2.19 | -0.5 | -2.11 | -1.58 | 0.59 | -1.1 | -0.78 | -0.58 | -1.09 | -1.58 | -0.92 | -1.61 | 0.345 | 0.795 | 0.82 | 0.355102 | 0.759972 | 0.043545 | 0.995762 | 0.997988 | 0.994666 |
| Camkv_Q3UHL1_S506s_1 | Camkv_Q3 | -0.03 | -0.86 | -0.61 | 0.49 | -1.07 | 0.67 | 0.39 | -0.23 | 0.54 | -0.74 | 0.81 | 0.22 | 0.74 | -0.12 | 0.66 | 0.8 | 0.1725 | 1.3 | 0.4675 | 0.068878 | 0.674582 | 0.309861 | 0.992876 | 0.997988 | 0.994666 |
| Camkv_Q3UHL1_T430t_1 | Camkv_Q3 | 0.74 | 0.54 | -0.03 | 0.52 | 0.75 | 0.79 | 1.02 | 0.73 | 0.19 | 0.73 | 0.04 | 0.23 | -0.67 | 0.84 | -0.11 | -0.2 | 0.1925 | 0.46 | 0.7725 | 0.22226 | 0.140727 | 0.602981 | 0.989223 | 0.997988 | 0.994666 |
| Camkv_Q3UHL1_T438t_1 | Camkv_Q3 | 0.42 | -0.85 | -0.35 | -0.71 | -0.42 | 0.86 | -0.68 | -0.69 | 1.93 | -0.49 | -2.47 | -0.28 | 1.62 | -1.09 | 0.43 | -0.52 | 0.38 | -0.145 | -0.4775 | 0.429604 | 0.74983 | 0.957235 | 0.995762 | 0.997988 | 0.994666 |
| Camkv_Q3UHL1_T438t_2 | Camkv_Q3 | -0.71 | -0.13 | 0.91 | 0.99 | -1.14 | 0.13 | 1.26 | 1.08 | 1.15 | -0.31 | 0.56 | 1.34 | -0.49 | -0.69 | 0.95 | 1.19 | 0.14 | 0.045 | 0.4825 | 0.965768 | 0.912336 | 0.427151 | 0.997335 | 0.997988 | 0.999507 |
| Camkv_Q3UHL1_T441t_1 | Camkv_Q3 | -0.26 | 0.21 | 0.72 | -0.22 | -0.77 | 0.13 | -0.35 | -0.44 | 2.71 | -0.06 | -1.04 | 0.54 | 2.63 | -0.9 | 0.44 | 0.1 | 0.0675 | 0.42 | -0.025 | 0.518419 | 0.156876 | 0.57006 | 0.989223 | 0.997988 | 0.994666 |
| Camkv_Q3UHL1_T441t_2 | Camkv_Q3 | -0.93 | 0.05 | -1.26 | -2.88 | -1.88 | 0.12 | -1.96 | -1.55 | 2.12 | -0.86 | -0.73 | -0.48 | 0 | -1.12 | -0.82 | -1.4 | -0.47 | 0.425 | 0.455 | 0.501956 | 0.927607 | 0.152273 | 0.997416 | 0.997988 | 0.994666 |
| Camkv_Q3UHL1_T446t_1 | Camkv_Q3 | 0.49 | 0.47 | 0.33 | 0.71 | 0.5 | 0.45 | 0.79 | 0.56 | 0.39 | 0.58 | 0.3 | 0.72 | -0.04 | 0.92 | 0.25 | 0.3 | -0.0625 | 1.2675 | 0.42 | 0.622727 | 0.722241 | 0.991158 | 0.995437 | 0.997988 | 0.995259 |
| Camkv_Q3UHL1_T446t_2 | Camkv_Q3 | 1.17 | 0.13 | 0.99 | 1.25 | 0.32 | 0.72 | 1.48 | 1.36 | 0.93 | 0.82 | 0.67 | 1.47 | -0.76 | 0.18 | 1.05 | 1.29 | 0.075 | -0.0025 | -0.1425 | 0.386112 | 0.818424 | 0.793285 | 0.995762 | 0.997988 | 0.994666 |
| Camkv_Q3UHL1_T470t_1 | Camkv_Q3 | 0.44 | 0.49 | 0.47 | 0.76 | 0.75 | 0.58 | 1.02 | 1.05 | 0.23 | 0.43 | 0.37 | 0.45 | -0.44 | 0.4 | 0.42 | 0.57 | 0.085 | 0.0875 | -0.445 | 0.328418 | 0.182807 | 0.434751 | 0.989223 | 0.997988 | 0.994666 |
| Camsap1_A0A0A0MQE5_S1069s_1 | Camsap1_/ | 0.39 | 0.14 | 0.38 | 0.21 | 0.37 | 0.42 | 0.6 | 0.55 | -0.78 | 0.2 | -0.48 | -0.13 | 0.08 | 0.02 | -0.52 | 0.32 | 0.31 | -0.17 | -0.3025 | 0.280155 | 0.323053 | 0.060634 | 0.989223 | 0.997988 | 0.994666 |
| Camsap1_A0A0A0MQE5_S1133s_1 | Camsap1_/ | 0.13 | -0.25 | -0.43 | -1.7 | 0.44 | 0.44 | 0.25 | -0.77 | -0.71 | 0.27 | -0.09 | -0.27 | -0.25 | 0.31 | -0.17 | -0.25 | 0.205 | -0.5775 | -0.305 | 0.275027 | 0.176796 | 0.41203 | 0.989223 | 0.997988 | 0.994666 |
| Camsap1_A0A0A0MQE5_S1407s_1 | Camsap1_/ | 1.48 | -0.65 | -0.11 | -0.75 | 0.93 | 1.61 | 0.42 | -0.51 | -1.3 | 1.01 | -0.63 | -0.62 | -1.44 | -0.69 | -0.86 | -0.06 | 0.6525 | 0.3625 | 0.4725 | 0.188811 | 0.322197 | 0.563176 | 0.989223 | 0.997988 | 0.994666 |
| Camsap1_A0A0A0MQE5_S550s_1 | Camsap1_/ | -0.07 | 0.62 | 0.59 | 0.08 | 0.69 | -1.12 | 0.28 | -0.35 | 0.03 | 0.62 | 0.75 | 0.29 | 0.88 | 0.36 | -0.04 | 0.6 | 0.62 | -0.3775 | -0.755 | 0.645927 | 0.31119 | 0.683367 | 0.989223 | 0.997988 | 0.995259 |
| Camsap1_A0A0A0MQE5_S550s_2 | Camsap1_/ | 1.12 | 0.38 | 0.61 | 0.31 | -0.48 | 0.31 | 0.77 | 1.64 | 0.59 | 0.85 | 0.45 | 1.05 | 0.04 | -0.2 | 0.67 | -0.24 | -0.43 | 0.1175 | 0.145 | 0.122013 | 0.91998 | 0.643189 | 0.997335 | 0.997988 | 0.994666 |
| Camsap1_A0A0A0MQE5_S553s_1 | Camsap1_/ | 0.18 | -0.51 | -0.32 | 0.03 | -0.99 | -0.09 | -1.27 | 0.12 | -0.08 | -0.41 | 1.3 | 0.13 | 0.58 | -0.87 | 0.65 | -1.86 | -0.045 | 0.13 | -0.5375 | 0.703343 | 0.290206 | 0.344057 | 0.989223 | 0.997988 | 0.996159 |
| Camsap1_A0A0A0MQE5_S553s_2 | Camsap1_/ | 1.12 | 0.38 | 0.61 | 0.31 | -0.48 | 0.31 | 0.77 | 1.64 | 0.59 | 0.85 | 0.45 | 1.05 | 0.04 | -0.2 | 0.67 | -0.24 | -0.4025 | 0.39 | -0.22 | 0.122013 | 0.91998 | 0.643189 | 0.997335 | 0.997988 | 0.994666 |
| Camsap1_A0A0A0MQE5_S626s_1 | Camsap1_/ | -1.78 | -1.84 | -1.09 | -0.96 | -0.84 | -0.65 | -0.63 | -0.91 | -0.59 | -0.82 | 0.75 | -0.16 | 0.46 | -0.08 | -0.84 | -1.1 | -0.045 | 0.13 | -0.5375 | 0.033663 | 0.036731 | 0.014665 | 0.989223 | 0.997988 | 0.994666 |
| Camsap1_A0A0A0MQE5_S718s_1 | Camsap1_/ | -0.05 | -0.15 | -0.34 | 0.35 | 0.08 | -0.47 | 0.89 | 0.14 | -0.19 | -0.37 | -0.35 | 0.18 | -1.2 | -1.42 | 0.13 | 0.43 | 0.66 | 1.2125 | 1.0275 | 0.331775 | 0.52981 | 0.605867 | 0.989223 | 0.997988 | 0.994666 |
| Camsap1_A0A0A0MQE5_T559t_1 | Camsap1_/ | -0.04 | -1.33 | -0.48 | 0.23 | 0.18 | 0.4 | 0.73 | 0.89 | -0.81 | -0.99 | 0.53 | 0.23 | -1.08 | -0.03 | 0.46 | 0.28 | 0.2075 | -0.135 | -0.4675 | 0.508325 | 0.026913 | 0.763796 | 0.989223 | 0.997988 | 0.994666 |
| Camsap1_V9GXD7_S104s_1 | Camsap1_\ | -1.56 | -1.54 | -1.4 | 0.02 | -1.29 | -1.79 | -0.4 | -0.43 | -1.86 | -1.59 | 0.34 | -0.44 | -0.67 | -0.98 | -0.73 | -1.31 | 0.955 | 0.145 | 0.3125 | 0.632489 | 0.763671 | 0.692972 | 0.995762 | 0.997988 | 0.995259 |
| Camsap2_H7BX08_S1009s_1 | Camsap2_I | -1.54 | -1.1 | -1.17 | -1.09 | -0.49 | -1.31 | -0.25 | -0.86 | -1.43 | -0.97 | -0.37 | -0.96 | -0.56 | -0.35 | -1.03 | -1.29 | 0.1425 | 0.2325 | 0.1975 | 0.186878 | 0.107928 | 0.324201 | 0.989223 | 0.997988 | 0.994666 |
| Camsap2_H7BX08_S1302s_1 | Camsap2_I | -0.13 | -0.42 | -0.7 | 0.18 | 0.22 | -0.35 | 0.72 | -0.51 | -0.5 | -0.2 | 0.4 | -0.48 | -0.77 | 0.37 | 0.31 | 0.25 | 0.4975 | 0.2925 | 0.4175 | 0.398427 | 0.411555 | 0.817726 | 0.989223 | 0.997988 | 0.994666 |
| Camsap2_H7BX08_S1308s_1 | Camsap2_I | -0.13 | -0.28 | 0.05 | -0.19 | 0.09 | -0.42 | 0.28 | -0.34 | -0.42 | 0.07 | -0.41 | -0.12 | -0.76 | 0.19 | -0.72 | -0.48 | 0.2875 | 0.0725 | 0.3075 | 0.315868 | 0.869804 | 0.723126 | 0.996493 | 0.997988 | 0.994666 |
| Camsap2_H7BX08_S408s_1 | Camsap2_I | -1.45 | -1.98 | -1.8 | -0.71 | -0.22 | -2.06 | -0.57 | -1.13 | -1.73 | -2.1 | -1.08 | -1.3 | -2.81 | -0.74 | -1.16 | -0.77 | 0.04 | -0.0825 | -0.305 | 0.827404 | 0.29816 | 0.854931 | 0.989223 | 0.997988 | 0.999507 |
| Camsap2_H7BX08_S589s_1 | Camsap2_I | 0.53 | 1.39 | 0.93 | 0.65 | 0.88 | 0.75 | 1 | 0.27 | 0.5 | 1.13 | 1.38 | 0.83 | 0.29 | -0.24 | 0.68 | 0.82 | 0.49 | -0.0675 | 0.115 | 0.174422 | 0.598983 | 0.781353 | 0.989223 | 0.997988 | 0.994666 |
| Camsap2_H7BX08_S590s_1 | Camsap2_I | 0.43 | 0.46 | 0.54 | 0.39 | 0.35 | 0.06 | 0.24 | 0.27 | 0.23 | 0.5 | 0.62 | 0.31 | 0.45 | 0.32 | 0.18 | 0.53 | -0.15 | 0.085 | -0.4875 | 0.71602 | 0.272636 | 0.852714 | 0.989223 | 0.997988 | 0.996966 |
| Camsap2_H7BX08_S674s_1 | Camsap2_I | 0.16 | -0.24 | 0.01 | -0.51 | -0.08 | 0.1 | -1.5 | -0.36 | 1.8 | -0.07 | 0.52 | -0.45 | 2.09 | -0.94 | -0.01 | -1.51 | -0.225 | -0.04 | -0.085 | 0.941601 | 0.415166 | 0.240595 | 0.989223 | 0.997988 | 0.999507 |
| Camsap2_H7BX08_S853s_1 | Camsap2_I | -0.48 | -1.44 | -1.6 | -1.86 | -0.47 | -0.45 | -0.99 | -1.04 | -1.07 | -1.18 | 0.37 | -1.39 | -1.49 | 0.08 | -2.35 | -2.3 | -0.315 | 0.595 | 0.0525 | 0.772757 | 0.104054 | 0.285078 | 0.989223 | 0.997988 | 0.997893 |
| Camsap2_H7BX08_S922s_1 | Camsap2_I | 0.33 | 0.44 | -0.39 | -0.13 | 0.44 | 0.4 | 0.89 | 0.03 | -0.51 | 0.36 | -1.79 | 0.28 | -0.37 | 0.21 | -0.16 | 0.19 | 0.6075 | 0.5275 | -0.17 | 0.752841 | 0.216013 | 0.355104 | 0.989223 | 0.997988 | 0.9971 |
| Camsap3_Q80VC9_S1095s_1 | Camsap3_( | -1.08 | -1.18 | -0.87 | -0.9 | -0.54 | -1.06 | -0.58 | -0.89 | -0.72 | -0.98 | 0.44 | -0.35 | -0.41 | -0.15 | -1.25 | -0.61 | 0.3775 | -0.4775 | -0.095 | 0.206722 | 0.302213 | 0.10005 | 0.989223 | 0.997988 | 0.994666 |
| Camsap3_Q80VC9_S345s_1 | Camsap3_( | 0.34 | 0.36 | 1.99 | 1.55 | 0.92 | 0.59 | 1.61 | 1.87 | -0.35 | 0.5 | -0.35 | 1.53 | -0.68 | -1.08 | 1.58 | 1.25 | 0.24 | 0.605 | 0.4025 | 0.278731 | 0.694224 | 0.219417 | 0.994893 | 0.997988 | 0.994666 |
| Camsap3_Q80VC9_S346s_1 | Camsap3_( | -0.06 | 1.05 | 1.07 | 1.83 | 0.24 | 1.65 | 2.1 | 1.51 | -0.06 | 1.52 | 0.5 | 2.14 | -2.42 | 0.66 | 0.52 | 1.45 | 0.1875 | -0.7275 | -0.7925 | 0.279989 | 0.436911 | 0.92774 | 0.989223 | 0.997988 | 0.994666 |
| Camsap3_Q80VC9_S359s_1 | Camsap3_( | 0.35 | 0.03 | 0.95 | 1.08 | 0.21 | 0.72 | 1.48 | 0.82 | -0.46 | 0.8 | 0.56 | 1.28 | 0.5 | 0.65 | 0.72 | 1.5 | 0.4025 | 0.0525 | -0.92 | 0.509349 | 0.571155 | 0.893814 | 0.989223 | 0.997988 | 0.994666 |
| Camsap3_Q80VC9_S380s_1 | Camsap3_( | 1.38 | -1.19 | 2.78 | 1.47 | 1.18 | 0.99 | 2.68 | 1.98 | -1.2 | 1.17 | 0.22 | 1.14 | 0.48 | -1.45 | 1.08 | 1.9 | 0.205 | -0.0575 | 0.24 | 0.529235 | 0.462609 | 0.391246 | 0.989223 | 0.997988 | 0.994666 |
| Camsap3_Q80VC9_S443s_1 | Camsap3_( | -1.13 | -0.94 | 0 | -0.85 | -0.17 | -0.17 | -0.21 | -0.21 | -0.78 | -0.8 | -0.49 | -0.4 | -1.04 | -0.01 | -0.99 | -1.07 | 0.5975 | -0.7775 | -0.6075 | 0.899276 | 0.082373 | 0.7141 | 0.989223 | 0.997988 | 0.999507 |
| Camsap3_Q80VC9_S560s_1 | Camsap3_( | 0.38 | 0.3 | 0.06 | 0.42 | 0.22 | 0.21 | 0.85 | 0.21 | 0.22 | 0.46 | -0.01 | 0.36 | 0.06 | 0.75 | 0.77 | 0.33 | 0.54 | 0.1125 | -0.0475 | 0.499421 | 0.73384 | 0.886968 | 0.995762 | 0.997988 | 0.994666 |
| Camsap3_Q80VC9_S598s_1 | Camsap3_( | -0.01 | 0.73 | -0.11 | 0.16 | 0.82 | 0.54 | 0.21 | 0.28 | 0.31 | 0.49 | -0.01 | 0.32 | -0.22 | 1.02 | 0.26 | 0.23 | 0.0825 | -0.0325 | 0.1875 | 0.710242 | 0.336134 | 0.755017 | 0.989223 | 0.997988 | 0.996966 |
| Camsap3_Q80VC9_S779s_1 | Camsap3_( | -0.3 | -0.52 | 0.06 | 0.21 | 0.16 | -0.42 | 0.2 | 0.21 | 0.17 | 0.09 | 0.78 | -0.09 | 1.1 | 0.41 | 0.08 | 0.34 | 0.27 | 0.085 | 0.13 | 0.078268 | 0.519956 | 0.223236 | 0.989223 | 0.997988 | 0.994666 |
| Camsap3_Q80VC9_S785s_1 | Camsap3_( | -0.53 | 0.35 | -1.02 | -0.34 | -0.05 | -1.07 | 0.05 | -0.56 | 0.11 | -0.27 | 1.9 | 0.34 | 1.34 | 0.25 | -1.28 | -0.44 | 0.175 | 0.375 | 0.62 | 0.542264 | 0.952232 | 0.107717 | 0.999187 | 0.997988 | 0.994666 |
| Camsap3_Q80VC9_S824s_1 | Camsap3_( | -0.19 | -0.37 | -0.38 | -0.99 | 0.48 | 0.03 | 0.38 | -0.25 | -0.59 | 0.51 | 0.16 | -0.24 | 0.42 | 0.01 | -1.24 | -1.23 | -0.0225 | 0.905 | 0.3525 | 0.951134 | 0.04195 | 0.193637 | 0.989223 | 0.997988 | 0.999507 |
| Camsap3_Q80VC9_T842t_1 | Camsap3_( | -1.91 | -1.49 | -0.85 | -0.71 | -1.13 | -1.73 | -0.36 | -0.87 | -1.54 | -1.4 | -0.84 | -0.84 | -1.43 | -0.82 | -1.13 | -0.49 | 0.6425 | 0.4425 | -0.0275 | 0.464431 | 0.578186 | 0.808689 | 0.989223 | 0.997988 | 0.994666 |
| Camta1_A2A891_S1228s_1 | Camta1_A2 | -1.46 | -1.53 | -1.37 | -0.58 | -1.58 | -2.18 | -0.82 | -1.23 | -2.3 | -0.94 | -0.29 | -1 | -0.33 | -0.15 | -0.5 | 0.24 | 0.2175 | 0.085 | 0.2725 | 0.008219 | 0.550442 | 0.822268 | 0.989223 | 0.997988 | 0.994666 |
| Cand1_Q6ZQ38_S1219s_1 | Cand1_Q6 | 0.23 | 0.56 | 2.52 | 1.89 | 0.64 | 0.49 | 1.67 | 2.46 | 0.12 | 1.55 | 2.86 | 2.47 | 1.02 | 0.53 | 0.75 | 2.18 | -0.2175 | 0.1025 | 1.05 | 0.763824 | 0.981107 | 0.543176 | 0.999187 | 0.997988 | 0.9971 |
| Cand1_Q6ZQ38_S1220s_1 | Cand1_Q6 | 0.91 | 0.44 | 0.24 | 0.57 | 0.52 | -0.63 | -0.38 | -0.19 | 0.13 | 0.59 | 0.22 | 0.13 | 0.3 | -0.45 | -1.08 | -0.19 | 0.015 | 0.45 | -0.18 | 0.027461 | 0.040952 | 0.294803 | 0.989223 | 0.997988 | 0.994666 |
| Cand1_Q6ZQ38_S1226s_1 | Cand1_Q6 | 1.85 | 0.07 | 1.07 | 0.74 | 1.29 | 1.65 | 0.32 | 1.12 | 0.75 | 0.23 | -1.66 | 1.03 | -1.41 | 0.71 | -0.46 | 0.85 | -0.71 | -0.2725 | -0.895 | 0.113947 | 0.710739 | 0.211585 | 0.995437 | 0.997988 | 0.994666 |
| Cand1_Q6ZQ38_S1230s_1 | Cand1_Q6 | 0.72 | 0.93 | 1.31 | 0.74 | 0.36 | 1.26 | 0.7 | 0.98 | 0.71 | 0.84 | 0.66 | 0.73 | 0.61 | 0.76 | 0.85 | 0.89 | 0.1625 | -0.845 | -1.01 | 0.56765 | 0.717055 | 0.425318 | 0.995437 | 0.997988 | 0.995259 |
| Cand1_Q6ZQ38_S558s_1 | Cand1_Q6 | 0.57 | 0.67 | 0.31 | 0.28 | 0.47 | -0.38 | 0.43 | 0.76 | 0.39 | 0.6 | 1.07 | 0.54 | 0.82 | 0.53 | 0.52 | 0.77 | -0.1 | -0.19 | -0.1475 | 0.418463 | 0.639455 | 0.449238 | 0.989223 | 0.997988 | 0.994666 |
| Cand1_Q6ZQ38_T1229t_1 | Cand1_Q6 | 0.72 | 0.93 | 1.31 | 0.74 | 0.36 | 1.26 | 0.7 | 0.98 | 0.71 | 0.84 | 0.66 | 0.73 | 0.61 | 0.76 | 0.85 | 0.89 | -0.1375 | 0.1925 | 0.2025 | 0.56765 | 0.717055 | 0.425318 | 0.995437 | 0.997988 | 0.995259 |
| Canx_P35564_S563s_1 | Canx_P355 | -1.78 | -1.11 | -0.44 | -0.59 | -1.91 | -1.88 | -0.42 | -0.63 | -1.46 | -1.41 | 0.09 | -0.76 | -1.45 | -1.46 | 0.48 | 0.21 | -0.1 | -0.19 | -0.1475 | 0.45344 | 0.622458 | 0.83408 | 0.989223 | 0.997988 | 0.994666 |
| Canx_P35564_S582s_1 | Canx_P355 | 1.06 | 0.55 | 0.71 | 0.8 | 0.93 | 0.74 | 0.95 | 1.18 | 1.02 | 1.07 | 1.37 | 0.66 | 0.51 | 1.29 | 1.52 | 1.45 | -0.23 | 0.095 | 0.425 | 0.204257 | 0.453146 | 0.335992 | 0.989223 | 0.997988 | 0.994666 |
| Cap1_P40124_S307s_1 | Cap1_P401 | 1.72 | 0.72 | -0.19 | -1.16 | -0.1 | 2.5 | -1.26 | -0.38 | 1.7 | 1.05 | -2.14 | -0.3 | 1.99 | -0.16 | -0.66 | -1.37 | 0.17 | 0.25 | 0.4125 | 0.700911 | 0.925572 | 0.833738 | 0.997335 | 0.997988 | 0.996159 |
| Cap1_P40124_S309s_1 | Cap1_P401 | -0.8 | -0.67 | -1.07 | -2.68 | -1 | -0.21 | -1.05 | -2.16 | -0.81 | -0.51 | -1.93 | -1.48 | 0.03 | -1.7 | -1.28 | -1.21 | -0.0825 | -0.195 | -0.3225 | 0.633321 | 0.719163 | 0.816558 | 0.995437 | 0.997988 | 0.995259 |
| Cap1_P40124_S34s_1 | Cap1_P401 | -0.75 | -0.95 | -1.16 | -0.82 | -0.92 | -0.95 | -0.82 | -0.85 | -0.58 | -0.45 | -0.57 | -1.03 | -0.28 | -0.37 | -0.84 | -1.11 | 0.2 | 0.1225 | 0.265 | 0.359288 | 0.864831 | 0.291259 | 0.996493 | 0.997988 | 0.994666 |
| Cap2_D3YTR7_S145s_1 | Cap2_D3Y | -3.05 | -2.19 | -2.14 | -6.34 | -3.06 | -1.56 | -4.43 | -3.51 | -0.45 | -2.62 | -3.36 | -2.79 | 0.46 | -2.75 | -2.86 | -4.08 | 0.035 | 0.2625 | 0.27 | 0.360647 | 0.772589 | 0.295197 | 0.995762 | 0.997988 | 0.994666 |
| Cap2_D3YTR7_S149s_1 | Cap2_D3Y | -2.15 | -0.68 | -0.15 | 0.31 | -0.44 | -2.33 | 0.04 | 0.05 | -2.23 | -1.6 | -0.25 | 0.34 | -1.31 | -0.76 | -0.15 | 0.4 | 0.29 | 1.125 | 1.1225 | 0.721335 | 0.997091 | 0.710282 | 0.999926 | 0.997988 | 0.9971 |
| Cap2_D3YTR7_S199s_1 | Cap2_D3Y | 7.56 | 3.25 | 4.54 | 1.23 | 3.49 | 7.61 | 3.62 | 3.22 | -0.28 | 6.59 | 0.16 | 3.71 | -0.49 | -0.18 | 4.22 | 4.62 | -0.0025 | -0.2675 | 0.2125 | 0.219506 | 0.814277 | 0.389948 | 0.995762 | 0.997988 | 0.994666 |
| Cap2_D3YTR7_T198t_1 | Cap2_D3Y | 5.74 | 3.72 | 5.35 | 2.3 | 3.78 | 5.9 | 4.48 | 4.41 | 0.57 | 5.08 | 1.68 | 4.74 | 0.63 | -0.29 | 4.89 | 5.05 | 0.34 | -1.6 | -2.1025 | 0.235801 | 0.647658 | 0.309348 | 0.99033 | 0.997988 | 0.994666 |
| Capn5_O08688_T44t_1 | Capn5_O0 | -0.38 | 0.6 | 0.58 | -0.31 | 0.39 | 0.62 | -0.36 | -0.28 | 0.19 | -0.08 | -0.31 | 0.55 | 1.1 | -0.16 | 0.2 | 0.24 | 0.365 | -1.26 | -1.7075 | 0.572156 | 0.933785 | 0.91917 | 0.998562 | 0.997988 | 0.995259 |
| Capza2_P47754_T63t_1 | Capza2_P4 | 0.31 | -0.36 | 0.03 | -0.51 | 0.45 | -0.05 | 0.47 | -0.12 | -1.18 | -0.28 | -0.82 | -0.54 | -1.21 | 0.04 | -0.12 | -0.24 | -0.03 | -0.035 | 0.2225 | 0.494925 | 0.270119 | 0.086515 | 0.989223 | 0.997988 | 0.994666 |
| Capzb_P47757_S263s_1 | Capzb_P47 | -2.13 | -2.81 | -2.89 | -3.31 | -1.19 | -2.9 | -1.61 | -2.24 | -3.28 | -2.5 | -0.55 | -2.37 | -2.11 | -0.98 | -2.29 | -1.87 | 0.32 | -0.5725 | -0.25 | 0.030726 | 0.083956 | 0.306013 | 0.989223 | 0.997988 | 0.994666 |
| Carhsp1_Q9CR86_S31s_1 | Carhsp1_Q | 1.03 | 1.35 | 0.34 | -0.11 | 1.15 | 1.15 | -0.01 | -0.04 | -0.07 | 1.12 | 0.58 | 0.01 | -0.19 | 0.3 | 0.34 | 0.22 | 0.8 | 0.61 | 0.9725 | 0.211814 | 0.839279 | 0.568918 | 0.995762 | 0.997988 | 0.994666 |
| Carhsp1_Q9CR86_S42s_1 | Carhsp1_Q | -1.12 | -0.9 | -1.29 | -0.84 | -0.03 | -1.42 | -0.56 | -0.28 | -1.96 | -1.17 | -0.34 | -0.43 | -1.15 | -0.28 | -0.92 | -1.09 | -0.09 | -0.2425 | -0.485 | 0.54743 | 0.181405 | 0.873145 | 0.989223 | 0.997988 | 0.994666 |
| Carhsp1_Q9CR86_S53s_1 | Carhsp1_Q | -2.59 | -2 | -2.38 | -2.93 | -0.8 | -3.47 | -1.93 | -2.91 | -3.81 | -3.09 | -1.9 | -3.09 | -2.47 | -1.69 | -2.24 | -2.05 | 0.465 | 0.0625 | 0.1775 | 0.259102 | 0.723747 | 0.264601 | 0.995437 | 0.997988 | 0.994666 |
| Cars_Q9ER72_S19s_1 | Cars_Q9ER | 1.52 | 1.36 | 0.57 | 0.61 | 0.69 | 0.98 | 0.55 | 1.07 | 0.63 | 1.74 | 1.08 | 0.75 | 0.43 | 1.37 | 0.64 | 0.95 | 0.1975 | -0.4975 | 0.3625 | 0.63478 | 0.526135 | 0.922912 | 0.989223 | 0.997988 | 0.995259 |
| Cars_Q9ER72_S307s_1 | Cars_Q9ER | -0.11 | -0.03 | -1.21 | -0.2 | 0.14 | -0.13 | -0.37 | 0.29 | -0.44 | -0.6 | -0.48 | 0.31 | -0.34 | -0.18 | 0.12 | -0.19 | -0.1925 | 0.035 | -0.1675 | 0.476507 | 0.271827 | 0.812317 | 0.989223 | 0.997988 | 0.994666 |
| Cartpt_P56388_S48s_1 | Cartpt_P56 | -0.13 | 0.32 | -0.41 | -0.41 | 0.31 | -0.75 | 0.6 | -0.05 | -0.66 | 0.12 | -0.12 | 0.08 | -0.5 | -1.09 | 0.81 | 0.58 | 0.37 | 0.085 | 0.24 | 0.816877 | 0.592664 | 0.96589 | 0.989223 | 0.997988 | 0.999507 |
| Casc3_Q8K3W3_S262s_1 | Casc3_Q8K | 0.03 | 0.1 | 0.91 | 0.51 | 0.48 | -1.28 | 0.51 | 0.71 | -0.12 | 0.66 | -0.11 | 0.01 | -1.08 | -0.67 | 0.73 | 1.2 | 0.185 | 0.0125 | 0.1075 | 0.531615 | 0.551494 | 0.38127 | 0.989223 | 0.997988 | 0.994666 |
| Casc3_Q8K3W3_S44s_1 | Casc3_Q8K | -0.4 | 0.91 | 0.63 | 0.64 | 0.24 | 0.44 | 0.52 | 0.41 | 0.31 | 0.22 | 0.93 | 0.5 | 0.3 | 0.36 | 1.46 | 1.26 | -0.2825 | -0.2775 | -0.3425 | 0.349137 | 0.891949 | 0.896589 | 0.996493 | 0.997988 | 0.994666 |
| Casc4_E9PZJ6_S231s_1 | Casc4_E9P | 0.16 | -0.99 | -2.78 | -0.8 | -0.74 | -1.3 | -0.48 | -0.21 | -1.19 | -0.34 | -1.24 | -1.28 | 0.2 | -0.54 | -1.38 | -0.2 | -0.0425 | 0.045 | 0.4 | 0.338303 | 0.480918 | 0.880426 | 0.989223 | 0.997988 | 0.994666 |
| Casc4_E9PZJ6_S232s_1 | Casc4_E9P | 0.16 | -0.99 | -2.78 | -0.8 | -0.74 | -1.3 | -0.48 | -0.21 | -1.19 | -0.34 | -1.24 | -1.28 | 0.2 | -0.54 | -1.38 | -0.2 | 0.42 | 0.09 | 0.6225 | 0.338303 | 0.480918 | 0.880426 | 0.989223 | 0.997988 | 0.994666 |
| Cask_O70589_S571s_1 | Cask_O705 | 0.64 | 0.57 | 0.61 | 0.81 | 0.28 | 0.38 | 0.48 | 0.72 | 0.17 | 0.54 | 0.65 | 0.67 | 0.39 | -0.01 | 0.71 | 0.59 | 0.42 | 0.09 | 0.6225 | 0.376935 | 0.370688 | 0.515166 | 0.989223 | 0.997988 | 0.994666 |
| Cask_O70589_S582s_1 | Cask_O705 | -0.08 | 0.03 | 0.18 | -0.74 | -0.68 | 0.46 | -0.12 | -0.36 | 2.26 | 0.17 | -1.16 | 0.71 | 2.51 | -2.65 | 1.46 | 0.16 | -0.1925 | -0.15 | -0.2375 | 0.599376 | 0.945037 | 0.34432 | 0.999187 | 0.997988 | 0.995259 |
| Cask_O70589_S583s_1 | Cask_O705 | -1.59 | -0.79 | 0.33 | -0.49 | -0.48 | -1.28 | 0.7 | -0.06 | -0.43 | 0.17 | -1.27 | 0.6 | -0.23 | 0 | 0.05 | 0.33 | -0.0225 | 0.6475 | 0.5225 | 0.129192 | 0.501308 | 0.453895 | 0.989223 | 0.997988 | 0.994666 |
| Cask_O70589_T182t_1 | Cask_O705 | 1.1 | 0.75 | 1.41 | 1.18 | 1.28 | 1.36 | 1.13 | 1.29 | 0.8 | 1.52 | 1.29 | 1.47 | 1.02 | 1.76 | 0.75 | 2.13 | 0.355 | 0.4025 | 0.6725 | 0.416531 | 0.49744 | 0.559119 | 0.989223 | 0.997988 | 0.994666 |
| Caskin1_Q6P9K8_S1012s_1 | Caskin1_Q | 0.51 | 0.38 | 0.35 | 0.47 | 0.34 | 0.66 | 0.24 | 0.63 | 0.43 | 0.47 | 0.85 | 0.45 | 0.35 | 1.11 | 0.49 | 0.5 | 0.155 | 0.16 | 0.305 | 0.492587 | 0.849902 | 0.580527 | 0.996349 | 0.997988 | 0.994666 |
| Caskin1_Q6P9K8_S1188s_1 | Caskin1_Q | 0.25 | 0.2 | -0.48 | -0.96 | 0.62 | 0 | 0.07 | -0.99 | -0.09 | 0.4 | -0.58 | -0.63 | -0.03 | 0.31 | -0.78 | -0.83 | 0.04 | 0.1225 | 0.185 | 0.834867 | 0.682836 | 0.952885 | 0.994272 | 0.997988 | 0.999507 |

| | | | | | | | | | | | | | | | | | | | | | | | | | | |
|---|---|---|---|---|---|---|---|---|---|---|---|---|---|---|---|---|---|---|---|---|---|---|---|---|---|---|
| Caskin1_Q6P9K8_S1233s_1 | Caskin1_Q( | 1.76 | 1.69 | 0.43 | 1.19 | 1.31 | 2.21 | 1.76 | 1.78 | 0.92 | 2.16 | 1.64 | 1.41 | 0.62 | 2.24 | 1.45 | 1.61 | 0.1725 | 0.0225 | -0.085 | 0.633445 | 0.185682 | 0.511238 | 0.989223 | 0.997988 | 0.995259 |
| Caskin1_Q6P9K8_S1288s_1 | Caskin1_Q( | -0.4 | -0.14 | 0.24 | 0.39 | -0.14 | 0.07 | 1.13 | 1.04 | -0.21 | 0.51 | 0.68 | 1.65 | 0.31 | -0.25 | 1.22 | 0.33 | 0.4975 | 0.265 | 0.2125 | 0.320562 | 0.193958 | 0.151644 | 0.989223 | 0.997988 | 0.994666 |
| Caskin1_Q6P9K8_S1330s_1 | Caskin1_Q( | 4.16 | 3.41 | 0.61 | 1.15 | 3.4 | 2.06 | 0.65 | 0.89 | 3.53 | 1.99 | 0.68 | 0.3 | 3.47 | 3.76 | 0.67 | 0.68 | 0.5025 | 0.635 | 0.38 | 0.858291 | 0.533004 | 0.483714 | 0.989223 | 0.997988 | 0.999507 |
| Caskin1_Q6P9K8_S358s_1 | Caskin1_Q( | -0.17 | 0.44 | 0.46 | 0.44 | 0.44 | -0.17 | 0.81 | 0.35 | 0 | 0.25 | 0.35 | 0.42 | -0.26 | 0.56 | -0.16 | 0.54 | -0.5825 | -0.7075 | -0.1875 | 0.700872 | 0.820527 | 0.881818 | 0.995762 | 0.997988 | 0.996159 |
| Caskin1_Q6P9K8_S390s_1 | Caskin1_Q( | -1.31 | -1.33 | -0.91 | -0.78 | -0.4 | -1.28 | -0.33 | -0.18 | -1.2 | -1.05 | 1.99 | 0.54 | -0.14 | 0.47 | -0.18 | 0.06 | 0.065 | -0.0375 | -0.1225 | 0.002524 | 0.106221 | 0.121148 | 0.989223 | 0.997988 | 0.994666 |
| Caskin1_Q6P9K8_S423s_1 | Caskin1_Q( | -0.85 | -1.01 | -1.6 | -0.68 | -0.56 | -1.14 | -0.98 | -0.85 | -0.89 | -1.22 | 0.68 | -0.37 | -1.4 | -0.22 | -1.22 | -0.74 | 0.535 | 1.1525 | 1.135 | 0.69647 | 0.581145 | 0.210075 | 0.989223 | 0.997988 | 0.996159 |
| Caskin1_Q6P9K8_S432s_1 | Caskin1_Q( | 0.63 | 0.34 | -0.4 | 0.27 | 1.16 | 0.51 | 0 | 0.2 | 0.62 | 0.79 | -0.09 | 0.15 | 0.6 | 0.39 | 0.14 | 0.31 | 0.1525 | 0.585 | 0.14 | 0.618736 | 0.456194 | 0.630017 | 0.989223 | 0.997988 | 0.995259 |
| Caskin1_Q6P9K8_S579s_1 | Caskin1_Q( | 0.78 | 0.89 | 0.66 | 1.1 | 0.99 | 0.74 | 0.73 | 0.37 | 0.85 | 0.78 | 0.8 | 0.86 | -0.09 | 0.82 | 0.59 | 0.57 | 0.2575 | 0.1575 | 0.15 | 0.202502 | 0.522111 | 0.871153 | 0.989223 | 0.997988 | 0.994666 |
| Caskin1_Q6P9K8_S720s_1 | Caskin1_Q( | -0.21 | -1 | -0.76 | -0.09 | 0.09 | -0.33 | -0.37 | 0.04 | -0.02 | 0.08 | -0.23 | -0.11 | 0.59 | 0.77 | -0.17 | -0.23 | -0.15 | -0.035 | -0.385 | 0.060076 | 0.208797 | 0.136762 | 0.989223 | 0.997988 | 0.994666 |
| Caskin1_Q6P9K8_S750s_1 | Caskin1_Q( | 0.48 | -0.02 | -0.42 | 0.38 | 0.13 | 0.43 | 0.54 | 0.08 | -0.1 | 0.22 | 1.17 | -0.12 | -0.78 | 0.34 | -0.76 | -0.31 | 0.3725 | 0.445 | 0.755 | 0.200032 | 0.493644 | 0.61732 | 0.989223 | 0.997988 | 0.994666 |
| Caskin1_Q6P9K8_S757s_1 | Caskin1_Q( | -0.99 | -2.23 | -1.52 | -0.47 | -1.24 | -0.16 | -0.66 | -1.17 | -1.11 | -1.3 | -0.41 | -1.15 | -0.6 | -0.21 | -1.03 | -0.83 | 0.19 | 0.1875 | -0.4825 | 0.150617 | 0.264429 | 0.464602 | 0.989223 | 0.997988 | 0.994666 |
| Caskin1_Q6P9K8_S820s_1 | Caskin1_Q( | -3.02 | -1.92 | -2.29 | -5.15 | -2.04 | -3.41 | -1.92 | -1.66 | -0.93 | -2.43 | 1.04 | -2.39 | -0.64 | -2.71 | -4.66 | -2.65 | 0.495 | 0.31 | 0.635 | 0.653767 | 0.263865 | 0.072379 | 0.989223 | 0.997988 | 0.995259 |
| Caskin1_Q6P9K8_S822s_1 | Caskin1_Q( | -2.93 | -2.01 | -2.38 | -3.49 | -1.46 | -1.5 | -2.3 | -2.33 | -0.09 | -2.55 | -0.62 | -1.48 | -0.28 | -1.52 | -1.2 | -2.71 | 0.8375 | 1.9175 | 0.43 | 0.042047 | 0.06348 | 0.025262 | 0.989223 | 0.997988 | 0.994666 |
| Caskin1_Q6P9K8_S918s_1 | Caskin1_Q( | -1.46 | -1.19 | -1.29 | -0.91 | -0.26 | -1.15 | -0.53 | -0.89 | -0.97 | -0.63 | 0.14 | -0.83 | 0.02 | 0.88 | -0.96 | -0.65 | 0.805 | 1.5175 | 1.275 | 0.032683 | 0.086412 | 0.062323 | 0.989223 | 0.997988 | 0.994666 |
| Caskin1_Q6P9K8_S965s_1 | Caskin1_Q( | 0.69 | -0.96 | -0.55 | -0.45 | -1.11 | 1.56 | -0.75 | -0.53 | 1.55 | -0.57 | -2.53 | -0.3 | 0.3 | -0.95 | -1.11 | -1.32 | 0.505 | 0.64 | 1.035 | 0.361446 | 0.859459 | 0.857714 | 0.996349 | 0.997988 | 0.994666 |
| Caskin1_Q6P9K8_T1197t_1 | Caskin1_Q( | 1.68 | 1.85 | 2.62 | 0.89 | 2.09 | 1.7 | 2.33 | 1.83 | -0.7 | 1.69 | -0.66 | 2.63 | -0.34 | -0.41 | 2.8 | 2.81 | 0.11 | -0.145 | -0.4525 | 0.532643 | 0.54733 | 0.230877 | 0.989223 | 0.997988 | 0.994666 |
| Caskin1_Q6P9K8_T670t_1 | Caskin1_Q( | 0.04 | 0.07 | -0.14 | -0.4 | 0.46 | 0.33 | -0.19 | 0.06 | -0.25 | 0.36 | 0.27 | 0.33 | -0.37 | 0.17 | -0.04 | 0.12 | 0.2275 | -1.02 | -0.545 | 0.767142 | 0.278685 | 0.275913 | 0.989223 | 0.997988 | 0.9971 |
| Caskin1_Q6P9K8_T708t_1 | Caskin1_Q( | -0.38 | -0.43 | 0.02 | 0.12 | -0.15 | -0.66 | 0.31 | 0.15 | -0.75 | -0.51 | -0.19 | 0.21 | -0.65 | -0.14 | -0.06 | 0.13 | 0.2725 | 0.285 | 0.0775 | 0.965352 | 0.780442 | 0.629233 | 0.995762 | 0.997988 | 0.999507 |
| Caskin1_Q6P9K8_T745t_1 | Caskin1_Q( | -0.04 | -1.97 | 0.5 | 1.21 | -0.27 | 0.06 | 0.81 | 0.59 | -0.37 | -1.39 | 0.64 | 0.28 | -0.58 | -0.59 | 0.47 | 0.17 | 0.08 | -0.1425 | -0.0125 | 0.930491 | 0.56603 | 0.853588 | 0.989223 | 0.997988 | 0.999507 |
| Caskin1_Q6P9K8_T759t_1 | Caskin1_Q( | 0.33 | -0.74 | -0.86 | -1.4 | 0.82 | -1.2 | 0.19 | 0.2 | -1.66 | -0.09 | 2.94 | -2.18 | -0.97 | 0.58 | -1.25 | -0.39 | 0.3725 | -0.135 | -0.0575 | 0.75452 | 0.210774 | 0.693867 | 0.989223 | 0.997988 | 0.9971 |
| Caskin1_Q6P9K8_T996t_1 | Caskin1_Q( | -3.94 | -4.54 | -1.65 | -3.11 | -1.8 | -4.61 | -1.42 | -2.83 | -5.3 | -3.2 | 0.75 | -2.06 | -3.39 | -1.87 | -2.3 | -2.6 | 0.67 | 0.42 | 0.16 | 0.244962 | 0.443425 | 0.492997 | 0.989223 | 0.997988 | 0.994666 |
| Caskin1_Q6P9K8_Y457y_1 | Caskin1_Q( | 0.13 | 0.19 | -1.88 | -0.24 | -0.26 | 0.25 | 0.04 | -0.22 | 0.38 | -0.49 | 0.64 | 0.02 | 0.26 | 0.56 | -1.17 | -0.58 | 0.645 | 0.8575 | 0.77 | 0.706346 | 0.39606 | 0.267479 | 0.989223 | 0.997988 | 0.996541 |
| Caskin2_Q8VHK1_S694s_1 | Caskin2_Q8 | 0.87 | 1.04 | 0.91 | 0.65 | 0.13 | 0.83 | 0.27 | 0.19 | 0.74 | 1.11 | 1.24 | 0.79 | 1.25 | 0.22 | 0.75 | 0.65 | 0.4025 | 0.5875 | 0.2175 | 0.611321 | 0.058309 | 0.665424 | 0.989223 | 0.997988 | 0.995259 |
| Casp3_P70677_S26s_1 | Casp3_P70 | -0.05 | -0.22 | -0.88 | -0.18 | 0.34 | -0.01 | 0.51 | 0.06 | -0.18 | 0.22 | -0.21 | -0.03 | -0.19 | 0.2 | -0.03 | -0.05 | -0.5125 | 0.1025 | -0.15 | 0.277878 | 0.060154 | 0.309198 | 0.989223 | 0.997988 | 0.994666 |
| Cast_Q921U7_S117s_1 | Cast_Q921 | 1.13 | 1.77 | 1.18 | 1.09 | 1.48 | 1.11 | 1.36 | 1.57 | 0.62 | 1.2 | 0.54 | 1.13 | 0.84 | 1.68 | 1.29 | 1.29 | 0.5575 | 0.2825 | 0.315 | 0.953109 | 0.724466 | 0.16241 | 0.995437 | 0.997988 | 0.999507 |
| Cbfa2t2_O70374_S235s_1 | Cbfa2t2_O | -0.42 | -0.65 | -0.33 | -0.82 | -0.58 | -0.31 | -0.77 | -0.72 | -0.3 | 0.21 | -0.08 | -1.81 | 0.45 | 0.03 | -0.08 | -0.21 | 0.0875 | -0.42 | -0.0175 | 0.04731 | 0.861086 | 0.893229 | 0.996493 | 0.997988 | 0.994666 |
| Cbfa2t2_O70374_S380s_1 | Cbfa2t2_O | 2.42 | -0.31 | 0.55 | -5.14 | 0.08 | 1.54 | 0.27 | 0.84 | -0.03 | 2.17 | -0.88 | 0.03 | 0.69 | -1.02 | 1.22 | 0.05 | -0.04 | 0.06 | 0.6025 | 0.557557 | 0.366423 | 0.538472 | 0.989223 | 0.997988 | 0.995259 |
| Cbfa2t2_O70374_S522s_1 | Cbfa2t2_O | -3.03 | -2.79 | -3.08 | -2.21 | -3.06 | -3.16 | -2.95 | -2.83 | -1.43 | -3.76 | -1.96 | -3.14 | -0.65 | -2.99 | -1.86 | -2.3 | 1.3025 | 0.9425 | 0.855 | 0.124181 | 0.402415 | 0.700225 | 0.989223 | 0.997988 | 0.994666 |
| Cbl_P22682_S617s_1 | Cbl_P2268 | -0.28 | 0.16 | 0.64 | 0.4 | 0.33 | -0.72 | 0.8 | 0.55 | -1.05 | -0.32 | 1.5 | -0.32 | -0.59 | -0.13 | 0.22 | 0.37 | -0.2225 | 0.205 | 0.8275 | 0.435097 | 0.978917 | 0.60826 | 0.999187 | 0.997988 | 0.994666 |
| Cbl_P22682_S861s_1 | Cbl_P2268 | 0.74 | 1.39 | 0.37 | 1.05 | 1.14 | 0.91 | 1.1 | 0.89 | 0.4 | 1.16 | 0.05 | -0.14 | 0.31 | 0.28 | 1.14 | 0.6 | 0.01 | -0.2775 | -0.2625 | 0.371966 | 0.651072 | 0.181513 | 0.990358 | 0.997988 | 0.994666 |
| Cbx3_D3Z313_S93s_1 | Cbx3_D3Z3 | 2.52 | 0.8 | -2.1 | -1.15 | -0.77 | 2.56 | -2.33 | -0.61 | 3.15 | 1.74 | -0.69 | -1.71 | 2.63 | -0.46 | -0.08 | -2.45 | 0.1225 | -0.52 | -0.305 | 0.93222 | 0.807119 | 0.649755 | 0.995762 | 0.997988 | 0.999507 |
| Cbx3_D3Z313_S95s_1 | Cbx3_D3Z3 | 0.78 | -0.44 | 0.16 | 0.21 | -1.88 | 0.54 | -1.29 | -0.35 | 1.03 | 0.29 | 2.52 | 0.6 | 1.12 | -1 | 1.03 | -0.09 | -0.305 | 0.605 | -0.1075 | 0.867702 | 0.108994 | 0.098154 | 0.989223 | 0.997988 | 0.999507 |
| Cbx5_Q61686_S110s_1 | Cbx5_Q616 | -0.8 | 0.31 | 0.7 | -0.19 | -1.11 | -0.56 | -0.96 | -0.82 | 1.65 | 0.16 | -0.38 | -0.35 | 2.09 | 1.22 | 0.16 | 0.07 | -0.9225 | 0.9325 | 0.0875 | 0.127241 | 0.030716 | 0.623347 | 0.989223 | 0.997988 | 0.994666 |
| Cbx5_Q61686_S92s_1 | Cbx5_Q616 | 0 | 0.15 | 0.23 | 0.17 | 0.48 | -0.33 | 0.13 | 0.04 | 0.01 | -0.26 | -0.04 | 0.15 | -0.44 | 0.52 | 0.22 | -0.43 | -0.8675 | 0.265 | 0.88 | 0.578849 | 0.810338 | 0.434223 | 0.995762 | 0.997988 | 0.995259 |
| Cby1_Q9D1C2_S20s_1 | Cby1_Q9D | 0.04 | 0.21 | -0.52 | 0.02 | -0.13 | 0.44 | -0.52 | 0.08 | 0.14 | 0.82 | -0.47 | -0.27 | 0.23 | 0.14 | -0.74 | -0.57 | -0.0575 | -0.1725 | -0.17 | 0.606011 | 0.916767 | 0.733321 | 0.997335 | 0.997988 | 0.995259 |
| Cby1_Q9D1C2_S9s_1 | Cby1_Q9D | -0.87 | -0.78 | -0.28 | -0.6 | -0.21 | -0.27 | -0.22 | -0.5 | -0.69 | 0.09 | -1.25 | -0.38 | -0.89 | -0.44 | -0.94 | -1.16 | 0.03 | 0.1175 | -0.1725 | 0.427849 | 0.165207 | 0.821786 | 0.989223 | 0.997988 | 0.994666 |
| Cc2d1a_E9PX94_S267s_1 | Cc2d1a_E9 | 0.34 | 0.44 | 0.32 | 0.91 | 0.44 | 0.11 | 0.52 | 0.83 | 0.34 | 0.34 | 0.07 | 0.72 | -0.1 | 0.46 | 0.61 | 0.8 | 0.3325 | 0.075 | -0.225 | 0.841354 | 0.914142 | 0.605088 | 0.997335 | 0.997988 | 0.999507 |
| Cc2d1a_E9PX94_S390s_1 | Cc2d1a_E9 | -0.43 | -0.38 | 0.02 | -0.07 | -0.69 | -0.8 | -0.48 | -0.39 | -1.59 | 0.14 | 0.5 | -0.25 | -1.76 | -1.81 | -0.13 | -0.24 | -0.0275 | -0.135 | -0.06 | 0.119583 | 0.122158 | 0.850771 | 0.989223 | 0.997988 | 0.994666 |
| Ccar2_Q8VDP4_S298s_1 | Ccar2_Q8V | -0.34 | 0.21 | 0 | 0.05 | 0.44 | -1.22 | -0.05 | -1.28 | 0.36 | 0.06 | 0.49 | -0.68 | 0.91 | -0.18 | 0.35 | -0.1 | -0.375 | -0.085 | -0.77 | 0.419105 | 0.247029 | 0.806924 | 0.989223 | 0.997988 | 0.994666 |
| Ccar2_Q8VDP4_S612s_1 | Ccar2_Q8V | -0.09 | -0.16 | -3.7 | -13.35 | 0.51 | 0.14 | -3.92 | -4.09 | -0.2 | 0.25 | -2.41 | -3.63 | -0.66 | -0.26 | -4.22 | -3.68 | -0.5075 | 0.0775 | 0.265 | 0.455733 | 0.395471 | 0.330576 | 0.989223 | 0.997988 | 0.994666 |
| Ccar2_Q8VDP4_S625s_1 | Ccar2_Q8V | -1.91 | -2.41 | -3.64 | -3.66 | -6.1 | -2.11 | -3.58 | -2.91 | -0.47 | -2.12 | -3.55 | -2.06 | 0.32 | -3.39 | -2.67 | -4.39 | 2.485 | 2.8275 | 2.12 | 0.700388 | 0.372597 | 0.2377 | 0.989223 | 0.997988 | 0.996159 |
| Ccdc115_Q8VE99_T90t_1 | Ccdc115_C | 0.4 | 0.73 | 1.63 | 0.79 | 0.94 | 0.3 | 0.98 | 2.44 | 0.88 | 0.67 | 2.74 | -0.08 | 0.85 | 0.67 | 2.1 | 2.63 | -0.77 | 0.855 | 0.3725 | 0.20825 | 0.568291 | 0.783249 | 0.989223 | 0.997988 | 0.994666 |
| Ccdc120_A2AEV7_S253s_1 | Ccdc120_A | 0.71 | 0.48 | 0.11 | -1.13 | -0.26 | 0.24 | -2.02 | -1.29 | 0.98 | 0.52 | -2.17 | -0.45 | 2.56 | -1.5 | -1.5 | -1.6 | 0.2775 | 0.165 | 0.675 | 0.568994 | 0.158774 | 0.65885 | 0.989223 | 0.997988 | 0.995259 |
| Ccdc120_A2AEV7_S283s_1 | Ccdc120_A | 0.15 | -0.12 | -0.01 | 0.04 | 0.7 | 0.07 | -0.65 | -0.25 | 0.79 | 0.41 | 1.02 | 0.05 | 0.68 | -0.42 | -0.89 | 0.04 | -0.875 | -0.3225 | -0.5525 | 0.655812 | 0.878208 | 0.070053 | 0.996493 | 0.997988 | 0.995259 |
| Ccdc132_Q8CI71_S15s_1 | Ccdc132_C | 0.45 | 0.69 | 0.04 | -0.21 | 0.38 | 0.21 | 0.09 | -0.16 | 0.29 | 0.51 | -0.16 | -0.15 | -0.07 | 0.71 | -0.23 | -0.31 | -0.0475 | 0.5525 | -0.1625 | 0.530047 | 0.680737 | 0.690982 | 0.993508 | 0.997988 | 0.994666 |
| Ccdc132_Q8CI71_S494s_1 | Ccdc132_C | 1.09 | 0.45 | -0.8 | -1.16 | 0.84 | 0.7 | -0.64 | -0.88 | 1.66 | 1.1 | -1.75 | -0.75 | 1.67 | 0.3 | -0.34 | -0.63 | -0.1125 | -0.12 | -0.2175 | 0.594156 | 0.858091 | 0.841164 | 0.996349 | 0.997988 | 0.995259 |
| Ccdc132_Q8CI71_S498s_1 | Ccdc132_C | 1.54 | 1.86 | 0.92 | 0.59 | 0.56 | 1.8 | -0.54 | -0.8 | 3.37 | 1.52 | -1.34 | 1.06 | 2.78 | 0.03 | 0.03 | 0.72 | 0.11 | 0.17 | 0.355 | 0.602322 | 0.12507 | 0.933308 | 0.989223 | 0.997988 | 0.995259 |
| Ccdc136_D3Z165_T102t_1 | Ccdc136_D | -0.06 | 0.71 | -0.6 | -0.74 | 1.49 | 0.28 | 0.33 | -0.98 | -0.53 | -0.3 | -1.74 | -0.87 | -0.6 | 1.32 | -0.55 | 0.21 | -0.9725 | -0.075 | -0.3375 | 0.609949 | 0.414944 | 0.14431 | 0.989223 | 0.997988 | 0.995259 |
| Ccdc177_Q3UHB8_S209s_1 | Ccdc177_C | 0.21 | 0.32 | 0.56 | 0.67 | -0.36 | 0.31 | -0.15 | 0.3 | 0.66 | -0.37 | -0.47 | 0.21 | 0.81 | -0.77 | -0.63 | 0.72 | 0.4525 | -0.6875 | 0.2675 | 0.356672 | 0.125502 | 0.193201 | 0.989223 | 0.997988 | 0.994666 |
| Ccdc177_Q3UHB8_S209s_2 | Ccdc177_C | 0.75 | 0.35 | 0.73 | 0.87 | -0.12 | 0.35 | -0.29 | 0.2 | 1.1 | 0.5 | 0.21 | 0.45 | 0.56 | -0.46 | 0.73 | 0.81 | -0.415 | -0.4325 | -0.4075 | 0.452154 | 0.025189 | 0.690415 | 0.989223 | 0.997988 | 0.994666 |
| Ccdc177_Q3UHB8_S211s_1 | Ccdc177_C | 0.85 | 1.14 | 1.17 | 0.73 | 0.47 | 0.63 | 0.07 | 0.43 | 0.73 | 0.33 | 0.02 | 0.36 | 1.54 | -0.07 | 0.6 | 0.62 | -0.64 | -0.11 | -0.265 | 0.423893 | 0.03243 | 0.035386 | 0.989223 | 0.997988 | 0.994666 |
| Ccdc177_Q3UHB8_S211s_2 | Ccdc177_C | 0.88 | 1 | 0.83 | 0.98 | -0.13 | 1.19 | -0.29 | 0.49 | 1.23 | 0.85 | -0.06 | 0.67 | 0.79 | -0.25 | 0.48 | 0.83 | -0.5725 | -0.6125 | -0.3 | 0.159496 | 0.102779 | 0.426288 | 0.989223 | 0.997988 | 0.994666 |
| Ccdc177_Q3UHB8_S273s_1 | Ccdc177_C | 0.13 | 0.13 | 0.03 | 0.34 | 0.77 | 0.69 | -0.04 | 0.84 | 0.84 | 0.31 | 0.41 | 0.44 | 0.6 | 1.14 | 0.49 | 0.53 | -0.6075 | -0.25 | -0.46 | 0.066563 | 0.144094 | 0.161337 | 0.989223 | 0.997988 | 0.994666 |
| Ccdc177_Q3UHB8_S281s_1 | Ccdc177_C | -1.86 | -0.27 | 0.16 | 0.14 | -0.74 | -0.86 | 0.06 | -0.26 | -0.78 | -0.31 | -0.23 | -0.03 | 0.07 | -0.49 | 0.63 | -0.36 | 0.4075 | 0.3425 | 0.5325 | 0.417973 | 0.987588 | 0.802021 | 0.999187 | 0.997988 | 0.994666 |
| Ccdc177_Q3UHB8_S303s_1 | Ccdc177_C | -0.39 | -0.24 | -0.71 | -0.59 | -0.4 | -0.87 | -0.44 | -0.69 | -0.8 | -0.6 | -1.39 | -0.9 | -1.12 | -0.11 | -0.76 | -0.91 | 0.0075 | 0.12 | 0.42 | 0.429605 | 0.609953 | 0.11926 | 0.989223 | 0.997988 | 0.994666 |
| Ccdc177_Q3UHB8_S310s_1 | Ccdc177_C | 1.19 | 0.32 | 0.54 | 1.41 | 0.69 | 0.44 | 0.93 | 1.66 | 0.44 | 0.4 | 1.24 | 1.1 | 0.12 | 0.08 | 1.45 | 0.78 | -0.1175 | -0.44 | -0.2425 | 0.538966 | 0.858804 | 0.842924 | 0.996349 | 0.997988 | 0.994666 |
| Ccdc177_Q3UHB8_S423s_1 | Ccdc177_C | 1.38 | -0.13 | 0.23 | 0.36 | -1.54 | 1.47 | -1.08 | -0.31 | 3.15 | 1.48 | -0.47 | 0.78 | 3.35 | -1.79 | 2.22 | 0.04 | 0.065 | -0.07 | -0.2575 | 0.633856 | 0.226563 | 0.308527 | 0.989223 | 0.997988 | 0.995259 |
| Ccdc177_Q3UHB8_S90s_1 | Ccdc177_C | -0.45 | 0.11 | -0.15 | 0.16 | 0.43 | -0.28 | -0.02 | 0.15 | -0.77 | -0.24 | -0.42 | -0.41 | -0.73 | 0.02 | -0.09 | -0.44 | -0.825 | 0.775 | 0.495 | 0.440698 | 0.555637 | 0.15857 | 0.989223 | 0.997988 | 0.994666 |
| Ccdc177_Q3UHB8_T208t_1 | Ccdc177_C | 0.82 | 1.77 | 0.25 | 0.02 | 0.53 | 2.42 | -0.37 | -0.13 | 2.07 | 1.74 | -0.88 | -0.3 | 1.15 | 0.75 | -0.2 | -0.54 | 0.1525 | -0.3775 | -0.2275 | 0.422676 | 0.876143 | 0.938419 | 0.996493 | 0.997988 | 0.994666 |
| Ccdc177_Q3UHB8_T208t_2 | Ccdc177_C | 0.91 | 1.12 | 0.87 | 1.02 | -0.13 | 1.34 | -0.29 | 0.59 | 1.26 | 0.92 | -0.17 | 0.75 | 0.83 | -0.2 | 0.39 | 0.84 | -0.1025 | -0.0575 | -0.425 | 0.11915 | 0.13011 | 0.394612 | 0.989223 | 0.997988 | 0.994666 |
| Ccdc25_Q78PG9_S204s_1 | Ccdc25_Q7 | -0.59 | -0.03 | -0.26 | -1.01 | -0.65 | -0.5 | -1.39 | -0.01 | -0.8 | -0.48 | -4.2 | -0.54 | 0.09 | 0.51 | -1.2 | -0.12 | -0.6025 | -0.29 | -0.515 | 0.492132 | 0.644759 | 0.230955 | 0.99033 | 0.997988 | 0.994666 |
| Ccdc6_D3YZP9_S233s_1 | Ccdc6_D3Y | -0.67 | -0.47 | -0.24 | 0.03 | -0.3 | -0.73 | -0.62 | -0.17 | -0.46 | -0.51 | -0.22 | -0.66 | -0.53 | -0.14 | -0.5 | -0.1 | -0.165 | -1.0325 | 0.2925 | 0.941709 | 0.645257 | 0.620207 | 0.99033 | 0.997988 | 0.999507 |
| Ccdc6_D3YZP9_S237s_1 | Ccdc6_D3Y | 0.78 | -0.04 | 0.35 | 0.71 | 0.81 | 0.96 | 0.6 | 0.96 | 0.58 | 0.49 | 0.3 | 0.25 | 0.12 | 0.88 | 0.51 | 0.55 | -0.1175 | -0.125 | 0.02 | 0.830391 | 0.161277 | 0.865508 | 0.989223 | 0.997988 | 0.999507 |
| Ccdc6_D3YZP9_S316s_1 | Ccdc6_D3Y | -1.4 | -1.03 | -0.66 | -0.62 | -1 | -1.25 | -0.08 | -0.54 | -0.83 | -1.68 | 1.08 | -2.34 | -0.61 | 0.04 | -1.23 | -1.52 | 0.3825 | -0.045 | 0.065 | 0.806955 | 0.526167 | 0.982628 | 0.989223 | 0.997988 | 0.999507 |
| Ccdc6_D3YZP9_S345s_1 | Ccdc6_D3Y | -0.46 | -0.64 | -0.97 | 0.01 | -0.41 | -0.43 | -0.44 | -0.22 | -1.27 | -0.44 | -0.92 | 0.06 | -1.48 | -0.3 | -0.93 | -0.45 | 0.21 | -0.015 | 0.0975 | 0.452453 | 0.593069 | 0.727051 | 0.989223 | 0.997988 | 0.994666 |
| Ccdc6_D3YZP9_S360s_1 | Ccdc6_D3Y | -0.48 | -1.19 | -1.72 | -1.9 | -0.56 | -0.57 | -1.2 | -1.8 | -0.56 | -0.76 | -1.64 | -2.92 | -0.92 | 0.19 | -2.29 | -2.28 | 0.14 | -0.1275 | -0.275 | 0.996762 | 0.489928 | 0.797838 | 0.989223 | 0.997988 | 1 |
| Ccdc6_D3YZP9_S45s_1 | Ccdc6_D3Y | -2.82 | -1.43 | -1.37 | -2.01 | -1.07 | -2 | -1.31 | -2.43 | -2.59 | -0.97 | -0.33 | -2.77 | -1.43 | -0.45 | -1.87 | -1.97 | 0.29 | -0.1475 | -0.0025 | 0.319228 | 0.637687 | 0.70046 | 0.989223 | 0.997988 | 0.994666 |
| Ccdc85a_E0CX32_S216s_1 | Ccdc85a_E | 0.18 | -0.67 | 0.62 | -0.08 | -1.57 | 0.32 | -1.41 | -1.13 | 0.87 | 0 | -0.01 | 0.13 | -0.35 | -2.43 | 0.45 | -0.06 | 0.205 | 0.2425 | 0.4775 | 0.340994 | 0.066095 | 0.513234 | 0.989223 | 0.997988 | 0.994666 |
| Ccdc85a_E0CX32_S240s_1 | Ccdc85a_E | -1.72 | -1.36 | -1.09 | -2.29 | -2.17 | -1.47 | -1.63 | -1.69 | 2.04 | -1.82 | -2.12 | -0.73 | 2.53 | -2.3 | 1.94 | -1.75 | -0.96 | 0.235 | -0.61 | 0.142034 | 0.693825 | 0.28884 | 0.994893 | 0.997988 | 0.994666 |
| Ccdc85a_E0CX32_S250s_1 | Ccdc85a_E | -0.02 | 0.8 | -4.32 | -4.03 | 0.55 | -0.03 | -4.61 | -7.89 | 1.28 | 0.48 | -3.23 | -4.78 | 1.15 | 1.32 | -5.86 | -5.43 | -0.125 | 0.9575 | 1.72 | 0.878383 | 0.593003 | 0.847572 | 0.989223 | 0.997988 | 0.999507 |
| Ccdc85a_E0CX32_S273s_1 | Ccdc85a_E | -2.4 | -2.57 | -1.24 | -12.75 | -2.06 | -2.29 | -1.81 | -2.24 | -1.09 | -2.15 | -2.2 | -1.92 | 0.2 | -1.88 | -0.39 | -1.08 | -1.1025 | 0.33 | -0.3125 | 0.114334 | 0.266374 | 0.23581 | 0.989223 | 0.997988 | 0.994666 |
| Ccdc85a_E0CX32_S284s_1 | Ccdc85a_E | -1.45 | -1.58 | -0.47 | -1.95 | -3.4 | -0.79 | -1.75 | -1.18 | 2.96 | -1.48 | -1.07 | 0.19 | 2.9 | -1.33 | 2.67 | -1 | 2.64 | 2.9 | 3.9525 | 0.058682 | 0.4835 | 0.129902 | 0.989223 | 0.997988 | 0.994666 |
| Ccdc85a_E0CX32_S308s_1 | Ccdc85a_E | 5.56 | 2.52 | 3.88 | 1.24 | 3.09 | 5.84 | 4.13 | 3.35 | 0.95 | 5.06 | 0.89 | 3.76 | 1.23 | 1.18 | 4.11 | 3.97 | -0.4175 | 1.5125 | 2.1725 | 0.532002 | 0.414481 | 0.606015 | 0.989223 | 0.997988 | 0.994666 |
| Ccdc88a_Q5SNZ0_S1332s_1 | Ccdc88a_C | 0.54 | 0.97 | 1.25 | 0.48 | 0.74 | -0.1 | 0.54 | 0.69 | 0.02 | 0.9 | 0.43 | 0.97 | 0.12 | 0.25 | 0.99 | 0.98 | 0.8025 | -0.635 | -0.6775 | 0.506131 | 0.261696 | 0.475511 | 0.989223 | 0.997988 | 0.994666 |
| Ccdc88a_Q5SNZ0_S1362s_1 | Ccdc88a_C | -2.32 | -2.75 | -1.65 | -3.62 | -0.48 | -2.99 | -1.84 | -2.52 | -3.63 | -2.88 | -0.04 | -1.69 | -1.06 | -0.88 | -2.45 | -2 | -0.3425 | -0.23 | -0.225 | 0.084052 | 0.318257 | 0.511898 | 0.989223 | 0.997988 | 0.994666 |
| Ccdc88a_Q5SNZ0_S1505s_1 | Ccdc88a_C | 0.53 | 0.52 | 0.3 | -0.57 | -0.25 | 1.24 | -0.81 | -0.31 | 1.03 | -0.03 | -0.82 | 0.12 | 0.39 | -0.54 | -0.43 | -0.75 | 0.6275 | 0.525 | 0.9875 | 0.184298 | 0.63366 | 0.787268 | 0.989223 | 0.997988 | 0.994666 |
| Ccdc88a_Q5SNZ0_S1594s_1 | Ccdc88a_C | -0.01 | 0.4 | 0.34 | 0.17 | 0.32 | -0.46 | 0.11 | -0.11 | 0.11 | -0.11 | 0.27 | 0.3 | -0.57 | 0 | -0.04 | 0.75 | -0.2275 | -0.12 | -0.5275 | 0.566593 | 0.309804 | 0.719989 | 0.989223 | 0.997988 | 0.995259 |
| Ccdc88a_Q5SNZ0_S1726s_1 | Ccdc88a_C | 0.2 | 0.41 | -2.59 | -0.44 | 0.88 | -0.02 | 0.32 | -0.15 | 0.93 | -0.15 | -1.09 | -0.64 | -0.31 | 0.75 | -0.22 | -0.44 | -0.26 | -0.0825 | -0.19 | 0.416301 | 0.200811 | 0.616749 | 0.989223 | 0.997988 | 0.994666 |
| Ccdc88c_Q6VGS5_S1563s_1 | Ccdc88c_C | 0.07 | -0.17 | 0.13 | -0.06 | -0.75 | -0.32 | 0.57 | 0.86 | 0.05 | 0.81 | 0.41 | -0.43 | 1.12 | -1.33 | 0.99 | 1 | 0.8625 | 0.3675 | 0.55 | 0.420894 | 0.794935 | 0.485578 | 0.995762 | 0.997988 | 0.994666 |
| Ccdc88c_Q6VGS5_S1675s_1 | Ccdc88c_C | -2.77 | -2.41 | -1.7 | -1.44 | -3.81 | -1.73 | -2.63 | -2.16 | -0.61 | -2.51 | -2.06 | -3.11 | 0.51 | -3.71 | -1.23 | -1.73 | 0.0975 | 0.2175 | 0.4525 | 0.511325 | 0.327143 | 0.989451 | 0.989223 | 0.997988 | 0.994666 |
| Ccdc88c_Q6VGS5_S227s_1 | Ccdc88c_C | -0.18 | 0.21 | 0 | 0.24 | -0.02 | 0.1 | 0.11 | -0.95 | -0.18 | -0.35 | -0.3 | -0.55 | -0.46 | -0.53 | -0.4 | 0.09 | -0.5025 | 0.0075 | 0.54 | 0.16253 | 0.398114 | 0.094972 | 0.989223 | 0.997988 | 0.994666 |
| Ccdc92_Q8VDN4_S194s_1 | Ccdc92_Q8 | -1.24 | -1.39 | -0.95 | -0.93 | -0.59 | -1.48 | -0.58 | -0.85 | -2.49 | -1.18 | -1.18 | -1.12 | -1.48 | -0.07 | -0.66 | -0.83 | -0.2575 | -0.4125 | -0.3925 | 0.300501 | 0.372842 | 0.327313 | 0.989223 | 0.997988 | 0.994666 |
| Ccdc92_Q8VDN4_S261s_1 | Ccdc92_Q8 | -1.63 | -1.23 | -2 | -5.96 | -0.89 | -2.73 | -3.15 | -2.7 | 1.46 | -3.81 | -3.32 | -2.75 | 1.71 | -2.7 | -1.68 | -2.21 | 0.2525 | -0.365 | 0.3675 | 0.262743 | 0.746415 | 0.67512 | 0.995762 | 0.997988 | 0.994666 |
| Ccdc97_Q9DBT3_S210s_1 | Ccdc97_Q9 | -0.43 | -0.42 | -0.12 | 0.66 | 0.02 | -0.13 | -0.72 | 0.52 | 0.19 | 0.29 | -0.69 | -0.2 | -1.66 | -0.4 | 0.14 | -0.24 | 0.3375 | 0.6 | 1.485 | 0.321239 | 1 | 0.943566 | 1 | 0.997988 | 0.994666 |
| Cckbr_P56481_S257s_1 | Cckbr_P56 | 0.46 | 0.5 | 0.51 | -0.01 | -0.89 | 1.29 | -0.25 | -1.33 | 2.04 | 0.55 | -0.67 | -0.49 | 2.08 | 0.3 | -0.2 | -0.13 | -3.9E-17 | -0.025 | -0.4625 | 0.774953 | 0.234276 | 0.98973 | 0.989223 | 0.997988 | 0.99798 |
| Cckbr_P56481_S443s_1 | Cckbr_P56 | -1.25 | -0.9 | -0.08 | 0.01 | 0.18 | 0.22 | 0.52 | 0.64 | -0.72 | -0.01 | -0.43 | 0.44 | -0.04 | 0.76 | 0.02 | 0.12 | -0.66 | -0.0075 | 0.1475 | 0.065048 | 0.018209 | 0.357868 | 0.989223 | 0.997988 | 0.994666 |
| Cckbr_P56481_T433t_1 | Cckbr_P56 | 0.12 | 0.7 | 0.55 | 0.69 | 0.4 | 0.73 | 0.86 | 0.45 | 0.1 | 0.43 | 0.19 | 0.76 | 0.2 | 0.93 | 0.3 | 0.47 | 0.945 | 0.375 | 0.77 | 0.888327 | 0.693751 | 0.585617 | 0.994893 | 0.997988 | 0.999507 |
| Ccm2_Q8K2Y9_S164s_1 | Ccm2_Q8K | -0.73 | 0.29 | 0.63 | 0.79 | -0.53 | -1.16 | 0.73 | 0.59 | 0.01 | -0.26 | 0.39 | -0.22 | -1.03 | -2.07 | 1.31 | 1.29 | 0.095 | -0.145 | -0.04 | 0.648714 | 0.519122 | 0.488744 | 0.989223 | 0.997988 | 0.995259 |
| Ccm2_Q8K2Y9_S393s_1 | Ccm2_Q8K | -0.53 | -1.62 | -1.4 | -1.24 | -1.07 | -1.26 | -0.7 | -1.33 | -2.04 | -1.11 | -1.04 | -1.36 | -1.61 | -1.07 | -1.58 | -1.25 | -0.3375 | -0.265 | -0.37 | 0.574838 | 0.720462 | 0.585389 | 0.995437 | 0.997988 | 0.995259 |
| Ccm2_Q8K2Y9_S417s_1 | Ccm2_Q8K | -1.1 | -2.09 | -2.29 | -4 | -2.58 | -1.02 | -2.02 | -2.93 | 3 | -1.58 | -4.7 | -2.28 | 2.91 | -2.36 | 1.51 | -2.84 | 0.1075 | -0.19 | -0.18 | 0.126864 | 0.721271 | 0.518382 | 0.995437 | 0.997988 | 0.994666 |
| Ccm2_Q8K2Y9_S425s_1 | Ccm2_Q8K | -0.3 | -0.11 | 0.13 | 0.33 | -0.08 | 0.13 | 0.05 | -0.26 | -0.08 | -0.89 | 0.1 | -0.34 | -0.8 | 0.35 | -0.09 | -0.61 | 0.2325 | 0.98 | 2.175 | 0.378774 | 0.822235 | 0.304081 | 0.995762 | 0.997988 | 0.994666 |
| Ccnk_Q3U3M5_S329s_1 | Ccnk_Q3U | -0.69 | -1.22 | -0.55 | -0.41 | -1.59 | -2.08 | -0.12 | -0.38 | -0.87 | -1.25 | -0.29 | -0.98 | -0.27 | -0.92 | 0.36 | -0.61 | -0.0525 | -0.315 | -0.3 | 0.3261 | 0.492098 | 0.671666 | 0.989223 | 0.997988 | 0.994666 |
| Ccnk_Q3U3M5_S341s_1 | Ccnk_Q3U | 0.57 | -0.52 | -0.57 | -1.04 | -0.41 | 0.26 | -0.35 | -0.87 | -0.17 | 0.35 | 0.68 | -0.6 | 0.79 | 0.76 | -0.64 | -0.9 | -0.325 | -0.13 | 0.3575 | 0.463066 | 0.904755 | 0.305919 | 0.996493 | 0.997988 | 0.994666 |
| Ccnl1_Q52KE7_S358s_1 | Ccnl1_Q52 | -2.08 | -2.68 | -2.26 | -1.94 | -1.6 | -3.53 | -1.1 | -1.29 | -3.23 | -2.87 | -0.24 | -2.27 | -1.72 | -1.1 | -1.89 | -1.69 | 0.0475 | 0.455 | 0.3925 | 0.054879 | 0.501381 | 0.888771 | 0.989223 | 0.997988 | 0.994666 |
| Ccnl1_Q52KE7_S380s_1 | Ccnl1_Q52 | -0.75 | -1.36 | -1.43 | -2.04 | -1.47 | -1.07 | -1.1 | -1 | -1.25 | -2.08 | -0.47 | -2.59 | 0.6 | -1.46 | 0.36 | -0.32 | 0.36 | 0.0875 | 0.64 | 0.037839 | 0.449276 | 0.688803 | 0.989223 | 0.997988 | 0.994666 |
| Ccnl2_Q9JJA7_S366s_1 | Ccnl2_Q9JJ | -0.38 | -0.55 | -0.72 | -0.8 | -0.61 | -0.88 | -0.38 | -0.67 | -1.01 | -0.41 | -0.98 | -1.67 | 0.51 | 0.56 | -0.12 | -0.78 | 0.235 | -0.2025 | 1.19 | 0.090657 | 0.919494 | 0.210918 | 0.997335 | 0.997988 | 0.994666 |
| Ccnt2_Q7TQK0_S549s_1 | Ccnt2_Q7T | 4.42 | 1.67 | 0.67 | 0.18 | 0.75 | 3.54 | 0.49 | 0.17 | 1.4 | 2.53 | -1.13 | 0.67 | 2.04 | 0.15 | 1.04 | 1.57 | -0.0225 | -0.405 | 0.655 | 0.556767 | 0.639739 | 0.427548 | 0.989223 | 0.997988 | 0.995259 |
| Ccnt2_Q7TQK0_S596s_1 | Ccnt2_Q7T | -0.23 | 0.03 | 0.68 | 0.48 | 0.63 | -0.64 | 0.39 | 0.54 | -0.17 | 0.13 | 1.78 | -0.17 | -0.42 | 0.42 | 1.04 | 0.83 | -0.4975 | -0.8675 | -0.535 | 0.566041 | 0.977735 | 0.753495 | 0.999187 | 0.997988 | 0.995259 |
| Ccnt2_Q7TQK0_S657s_1 | Ccnt2_Q7T | -1.59 | -3.17 | -2.25 | -3.89 | -3.39 | -2.13 | -3.26 | -2.38 | 0.75 | -2.21 | -1.79 | -2.97 | 1.27 | -2.96 | -1.14 | -1.21 | -0.01 | 0.1525 | 0.2275 | 0.076762 | 0.904016 | 0.189958 | 0.996493 | 0.997988 | 0.994666 |
| Ccny_Q8BGU5_S25s_1 | Ccny_Q8B( | 1 | -0.56 | 0.5 | 0.98 | 0.24 | 1.28 | 0.87 | 1.31 | -0.5 | -0.92 | -0.14 | 1.1 | -0.95 | 0.25 | 0.92 | 0.74 | -0.065 | 1.17 | 1.715 | 0.647458 | 0.304144 | 0.278366 | 0.989223 | 0.997988 | 0.995259 |
| Ccny_Q8BGU5_S324s_1 | Ccny_Q8B( | 0.55 | 0.94 | 0.55 | 0.86 | 0.27 | 0.66 | 1.43 | 0.31 | 0.54 | 0.71 | 0.99 | 0.37 | 0.67 | 1.2 | 0.74 | 0.13 | 0.445 | -0.595 | -0.24 | 0.894537 | 0.851955 | 0.768557 | 0.996349 | 0.997988 | 0.999507 |
| Ccny_Q8BGU5_S326s_1 | Ccny_Q8B( | 0.41 | 0.68 | 0.76 | 0.7 | 0.9 | 0.16 | 0.94 | 0.61 | 0.27 | 0.59 | 0.4 | 0.5 | 0.26 | 0.85 | 0.53 | 0.55 | -0.0575 | -0.0725 | -0.04 | 0.722848 | 0.952406 | 0.377284 | 0.999187 | 0.997988 | 0.9971 |
| Ccny_Q8BGU5_S33s_1 | Ccny_Q8B( | -0.23 | 0.09 | 0.31 | 0.65 | -0.62 | -0.53 | -0.07 | 0.52 | -0.45 | 0.14 | 0.05 | 0.38 | -1.13 | -0.51 | -0.31 | -0.53 | 0.015 | -0.1975 | -0.09 | 0.023147 | 0.267046 | 0.558288 | 0.989223 | 0.997988 | 0.994666 |
| Ccny_Q8BGU5_S45s_1 | Ccny_Q8B( | -0.93 | -0.32 | -0.73 | -2 | -0.75 | -0.65 | -0.92 | -1.35 | 0.28 | -0.66 | -0.31 | -1.04 | 0.72 | -0.4 | -0.18 | -2.06 | -0.38 | -0.175 | -0.825 | 0.414313 | 0.838757 | 0.222066 | 0.995762 | 0.997988 | 0.994666 |
| Ccny_Q8BGU5_S83s_1 | Ccny_Q8B( | -1.8 | -1.83 | -2.38 | -1.98 | -1.62 | -2.34 | -1.51 | -1.27 | -2.77 | -1.37 | -0.83 | -2.27 | -1.81 | -1.6 | -2.45 | -2.66 | 0.0775 | 0.5625 | 0.515 | 0.687483 | 0.302551 | 0.672356 | 0.989223 | 0.997988 | 0.996159 |
| Ccny_Q8BGU5_S99s_1 | Ccny_Q8B( | -0.57 | 0.08 | -0.69 | -0.76 | -0.01 | -0.66 | 0.16 | -0.48 | -0.82 | -0.82 | -1.33 | -1.08 | -0.77 | -0.4 | -0.47 | -0.45 | 0.3125 | 0.1875 | -0.1325 | 0.894989 | 0.433512 | 0.085326 | 0.989223 | 0.997988 | 0.999507 |
| Ccny_Q8BGU5_T37t_1 | Ccny_Q8B( | -2.05 | -1.64 | -1.55 | -2.82 | -1.55 | -2.59 | -2.2 | -2.52 | -2.24 | -3.01 | -2.08 | -1.87 | -1.34 | -2.15 | -2.35 | -1.74 | 0.2375 | -0.5275 | -0.0375 | 0.753243 | 0.590839 | 0.465279 | 0.989223 | 0.997988 | 0.9971 |
| Ccnyl1_E9Q226_S274s_1 | Ccnyl1_E9( | 0.32 | 0.74 | 0.61 | 0.32 | 0.92 | 0.11 | 0.95 | 0.39 | 0.51 | 0.72 | 0.74 | 0.52 | 0.14 | 0.98 | 0.24 | 0.44 | -0.2 | -0.285 | 0.12 | 0.868559 | 0.727846 | 0.582987 | 0.995569 | 0.997988 | 0.999507 |
| Ccnyl1_E9Q226_S276s_1 | Ccnyl1_E9( | 0.22 | 0.5 | 0.49 | 0.59 | 0.71 | 0.17 | 0.93 | 0.33 | 0.43 | 0.67 | 0.4 | 0.34 | 0.04 | 0.79 | 0.15 | 0.56 | 0.095 | 0.125 | -0.0475 | 0.813555 | 0.733338 | 0.963729 | 0.995762 | 0.997988 | 0.999507 |
| Ccnyl1_E9Q226_S52s_1 | Ccnyl1_E9( | -0.36 | 0.13 | -0.2 | 0.25 | 0.56 | -0.07 | 0.23 | -0.3 | 0.25 | -0.09 | 0.25 | -0.52 | -0.47 | 0.05 | -0.32 | 0.05 | 0.085 | 0.01 | -0.065 | 0.645763 | 0.586846 | 0.950419 | 0.989223 | 0.997988 | 0.995259 |
| Ccp110_Q7TSH4_S176s_1 | Ccp110_Q | -1 | -1.47 | -0.45 | -1.53 | -2.26 | -1.17 | -1.8 | -1.48 | 1.5 | -0.94 | -1.04 | -1.67 | 1.73 | -2.2 | 0.29 | -0.99 | 0.15 | 0.0175 | -0.1275 | 0.307442 | 0.127512 | 0.401563 | 0.989223 | 0.997988 | 0.994666 |
| Ccs_Q9WU84_S267s_1 | Ccs_Q9WU | -0.35 | -0.24 | -2.8 | -0.85 | 0.63 | -0.27 | -2.33 | -0.3 | -0.38 | -0.38 | 0.09 | 0.56 | 0.46 | 0.75 | -0.8 | -0.55 | -0.565 | 0.575 | 0.82 | 0.132891 | 0.520066 | 0.103906 | 0.989223 | 0.997988 | 0.994666 |
| Ccsap_Q8QZT2_S129s_1 | Ccsap_Q8C | -1.4 | -1.51 | -1.99 | -2.5 | -1.54 | -1.57 | -2.97 | -2.36 | -1.08 | -1.17 | -0.9 | -1 | 0.5 | -1.1 | -1.98 | -2.42 | 0.4925 | 1.0325 | 1.025 | 0.35102 | 0.528067 | 0.021423 | 0.989223 | 0.997988 | 0.994666 |
| Ccsap_Q8QZT2_S222s_1 | Ccsap_Q8C | 2.67 | -1 | 2.19 | -8.48 | 0.48 | 2.6 | 0.59 | 0.92 | -1.69 | 1.53 | -1.22 | 1.09 | 0.43 | -0.78 | 0.53 | 1.99 | -0.26 | 0.8125 | 0.6 | 0.457292 | 0.31668 | 0.645886 | 0.989223 | 0.997988 | 0.994666 |
| Ccser1_Q8C0C4_S375s_1 | Ccser1_Q8 | -0.05 | -1.2 | 1.15 | 0.79 | -0.37 | 0 | 0.62 | 0.58 | -0.31 | -0.55 | 0.57 | 1.22 | -0.64 | -0.51 | -0.02 | 0.14 | 2.3025 | 1.0825 | 1.6975 | 0.416999 | 0.946541 | 0.921084 | 0.999187 | 0.997988 | 0.994666 |
| Ccser2_Q3UHI0_S222s_1 | Ccser2_Q3 | -0.75 | -0.62 | 0.47 | -0.39 | 0.06 | -0.47 | 0.39 | -1.13 | -0.62 | -0.44 | 0.15 | -0.72 | -0.41 | 0.46 | 0.34 | 0.01 | 0.035 | 0.06 | -0.4325 | 0.259146 | 0.932105 | 0.809037 | 0.998475 | 0.997988 | 0.994666 |
| Cct2_P80314_S3s_1 | Cct2_P803 | -1.25 | -0.97 | -2.41 | -1.55 | -0.43 | -1.7 | -1.46 | -1.21 | -0.46 | -1.18 | -1.13 | -1.16 | 0.22 | -0.74 | -1.43 | -0.91 | 0.035 | -0.085 | 0.4225 | 0.091595 | 0.398047 | 0.149784 | 0.989223 | 0.997988 | 0.994666 |
| Ccz1_Q8C1Y8_S264s_1 | Ccz1_Q8C1 | 6.63 | 4.65 | 3.71 | 1.27 | 4.88 | 7.49 | 4.16 | 2.89 | 0.35 | 6.11 | -0.18 | 4.05 | 0.01 | 2.4 | 3.81 | 4.47 | 0.345 | 0.5625 | 0.83 | 0.292866 | 0.537366 | 0.374941 | 0.989223 | 0.997988 | 0.994666 |
| Cd200_Q80VX2_S270s_1 | Cd200_Q8( | 1.04 | -0.75 | -0.11 | 0.4 | -0.43 | 1.56 | 0.65 | 0.62 | 1.91 | -0.3 | 0.36 | 0.08 | 2.12 | -0.33 | 1.31 | 0.63 | 0.79 | -1.4825 | -1.3925 | 0.202578 | 0.381424 | 0.522726 | 0.989223 | 0.997988 | 0.994666 |
| Cd248_Q91V98_S754s_1 | Cd248_Q9 | -1.88 | -1.51 | -1.08 | -1.86 | -0.67 | -1.82 | -1.89 | -1.55 | -1.67 | -1.9 | -0.74 | -1.16 | -0.26 | -0.52 | -0.81 | -0.78 | 0.455 | 0.3675 | 0.7875 | 0.007178 | 0.77075 | 0.532194 | 0.995762 | 0.997988 | 0.994666 |
| Cd2ap_Q9JLQ0_S404s_1 | Cd2ap_Q9J | -0.62 | -0.62 | -0.98 | -1.49 | -1.38 | -1.7 | -0.4 | -0.94 | -1.34 | -0.65 | 1 | -0.69 | 0.52 | -0.8 | -1.3 | -0.86 | 0.1 | 0.215 | 0.99 | 0.472836 | 0.6152 | 0.330496 | 0.989223 | 0.997988 | 0.994666 |
| Cd2ap_Q9JLQ0_S510s_1 | Cd2ap_Q9J | -0.12 | 0.42 | -0.56 | 0.14 | 0.07 | -0.14 | -0.11 | 0.31 | -0.02 | 0.88 | 0.98 | 0.4 | 0.42 | -0.24 | 0.44 | -0.67 | -0.1775 | 0.5075 | 0.3175 | 0.961609 | 0.818947 | 0.103294 | 0.995762 | 0.997988 | 0.999507 |
| Cd2ap_Q9JLQ0_S580s_1 | Cd2ap_Q9J | 0.37 | 1.1 | -1.33 | -0.96 | 0.42 | 0.03 | -1.37 | -1.95 | -0.88 | 0.71 | -0.33 | -0.42 | 0.41 | -0.51 | -1.15 | -0.81 | 0.0625 | 0.59 | 0.0175 | 0.608702 | 0.473182 | 0.966941 | 0.989223 | 0.997988 | 0.995259 |
| Cd34_Q64314_S343s_1 | Cd34_Q643 | 0.29 | 0.12 | 0.16 | 0.04 | -0.42 | -0.65 | 0.46 | -0.23 | 0.26 | -0.58 | -1.46 | 0.77 | -0.39 | 0.67 | -0.02 | 0.11 | -0.5125 | -0.025 | -0.31 | 0.837704 | 0.216489 | 0.397985 | 0.989223 | 0.997988 | 0.999507 |
| Cd44_Q80X37_S544s_1 | Cd44_Q80 | -1.75 | -1 | -0.08 | -1.47 | -0.87 | -2.2 | -0.39 | -1.19 | -1.87 | -1.32 | -0.84 | -0.94 | -1.21 | -0.93 | -0.71 | -0.57 | -0.3625 | -0.405 | -0.06 | 0.584057 | 0.85752 | 0.693924 | 0.996349 | 0.997988 | 0.995259 |
| Cdc23_Q8BGZ4_S460s_1 | Cdc23_Q8 | 0.73 | 0.37 | 0.98 | 0.56 | 0.7 | 0.25 | 0.57 | 0.69 | 0.44 | 0.67 | 0.38 | 1.15 | -0.58 | 0.63 | 0.7 | 0.8 | -0.0875 | -0.1675 | 0.22 | 0.467238 | 0.650459 | 1 | 0.990358 | 1 | 0.994666 |
| Cdc23_Q8BGZ4_S470s_1 | Cdc23_Q8 | -0.77 | -1.25 | -0.04 | -0.06 | -0.67 | -1.34 | 0.37 | -0.1 | -0.67 | -0.8 | -0.91 | 0.53 | -0.99 | -0.8 | -0.44 | 0.34 | -0.1075 | -4.9E-17 | -0.2725 | 0.889958 | 0.830055 | 0.875996 | 0.995762 | 0.997988 | 0.999507 |
| Cdc23_Q8BGZ4_T478t_1 | Cdc23_Q8 | 1.03 | 1.45 | 0.54 | 0.71 | 1.23 | 1.15 | 0.68 | 0.64 | 0.4 | 1.08 | 0.46 | 0.57 | 0.98 | 1.25 | -0.18 | 0.07 | 0.095 | 0.0675 | 0.0575 | 0.332614 | 0.979012 | 0.3161 | 0.999187 | 0.997988 | 0.994666 |
| Cdc27_Q8R568_S371s_1 | Cdc27_Q8 | -0.67 | 0.11 | 0.6 | -0.3 | 0.87 | 0.14 | 0.72 | 0.4 | -0.85 | 0.86 | 0.78 | 0.46 | -0.19 | 0.18 | 0.74 | 0.67 | -0.0075 | -0.305 | -0.4025 | 0.278042 | 0.094864 | 0.421752 | 0.989223 | 0.997988 | 0.994666 |
| Cdc42bpa_F6Q5A5_S234s_1 | Cdc42bpa_ | -0.38 | 0.19 | -0.91 | -1.49 | -0.64 | -0.22 | -0.96 | -2.4 | 0.03 | -0.44 | -1.17 | -1.3 | -0.52 | -1.19 | -0.61 | -0.81 | 0.5975 | 0.3775 | 0.415 | 0.73436 | 0.45626 | 0.87402 | 0.989223 | 0.997988 | 0.9971 |
| Cdc42bpa_F6Q5A5_S945s_1 | Cdc42bpa_ | -1.15 | -1.24 | -1 | -1.41 | -0.4 | -0.97 | -0.77 | -0.99 | -1.17 | -1.19 | -0.25 | -0.99 | -0.2 | -0.44 | -1.06 | -0.66 | -0.4075 | -0.0725 | -0.135 | 0.051946 | 0.099033 | 0.308172 | 0.989223 | 0.997988 | 0.994666 |
| Cdc42bpa_F6Q5A5_S967s_1 | Cdc42bpa_ | 0.6 | 0.39 | -0.91 | 0.05 | 0.33 | 0.61 | -0.16 | -0.22 | 0.01 | 0.35 | 0.2 | -0.41 | -0.41 | -0.06 | 0.01 | 0.27 | 0.4175 | 0.3 | 0.61 | 0.83251 | 0.777472 | 0.989421 | 0.995762 | 0.997988 | 0.999507 |
| Cdc42bpb_Q7TT50_S1692s_1 | Cdc42bpb_ | 0.82 | -0.69 | 0.36 | 0.91 | -3.39 | -1.36 | 0.2 | 0.6 | 0.6 | 0.18 | 0.77 | 1.35 | 2.09 | -2.77 | 1.35 | 0.39 | 0.1075 | 0.005 | -0.08 | 0.931195 | 0.140641 | 0.391808 | 0.989223 | 0.997988 | 0.999507 |
| Cdc42ep1_Q91W92_S121s_1 | Cdc42ep1_ | -1.11 | -0.95 | -0.45 | -0.82 | -0.26 | -1.43 | -0.22 | -0.82 | -1.11 | -0.89 | -1.1 | -0.82 | -1.04 | -1.01 | -1.26 | -0.48 | -1.3375 | 0.375 | -0.085 | 0.690891 | 0.649386 | 0.545931 | 0.990343 | 0.997988 | 0.996159 |
| Cdc42ep2_Q8JZX9_S137s_1 | Cdc42ep2_ | 1.05 | 1.47 | 1.55 | 1.31 | 1.53 | 0.01 | 1.3 | 1.73 | 0.34 | 0.77 | 0.97 | 1.93 | -0.89 | 0.71 | 1.05 | 1.2 | 0.15 | -0.1475 | -0.115 | 0.105634 | 0.606346 | 0.35844 | 0.989223 | 0.997988 | 0.994666 |
| Cdc42ep4_Q9JM96_S116s_1 | Cdc42ep4_ | 0.3 | -1.11 | -1.01 | -2.42 | 0.58 | 0.39 | -1.54 | -1.62 | -0.57 | -0.36 | -0.92 | -2.43 | -1.06 | -0.04 | -3.08 | -3.08 | -0.2025 | -0.3425 | -0.8275 | 0.373151 | 0.481388 | 0.987873 | 0.989223 | 0.997988 | 0.994666 |
| Cdc42ep4_Q9JM96_S140s_1 | Cdc42ep4_ | 0.01 | 0.8 | 0.41 | 0.39 | -0.05 | -0.12 | -0.01 | 0.38 | 0.27 | 0.61 | -0.06 | 0.31 | 1.15 | 0.26 | 0.43 | 0.26 | 0.5125 | -0.01 | -0.755 | 0.699541 | 0.183917 | 0.658457 | 0.989223 | 0.997988 | 0.996159 |
| Cdc42ep4_Q9JM96_S64s_1 | Cdc42ep4_ | 0.25 | 0.53 | 0.86 | 0.32 | 0.07 | 0.29 | 0.4 | -0.05 | -0.16 | 1.13 | 0.35 | 1.14 | -0.49 | 0.13 | -0.23 | 0.07 | -0.3525 | -0.12 | 0.1225 | 0.047801 | 0.21005 | 0.727867 | 0.989223 | 0.997988 | 0.994666 |
| Cdc5l_Q6A068_T385t_1 | Cdc5l_Q6A | 0 | -0.01 | -0.61 | 0.3 | -0.79 | -0.64 | 0.82 | 0.13 | -0.23 | -0.48 | 0.03 | 0.52 | 0.57 | 0.41 | -0.14 | 0.21 | -0.3125 | 0.125 | -0.62 | 0.275393 | 0.920667 | 0.899541 | 0.997335 | 0.997988 | 0.994666 |
| Cdh10_P70408_T701t_1 | Cdh10_P7( | 0.15 | 0.36 | -0.13 | -0.26 | 0.68 | 0.16 | 0.03 | -0.18 | -0.01 | -0.01 | -0.04 | -0.03 | 0.58 | 0.56 | -0.33 | -0.09 | -0.04 | 0.04 | 0.3425 | 0.639011 | 0.602113 | 0.821746 | 0.989223 | 0.997988 | 0.995259 |
| Cdh12_Q5RJH3_S649s_1 | Cdh12_Q5 | -0.38 | 0.24 | 0.39 | -0.53 | 0.4 | -0.62 | -0.34 | -0.78 | 0.12 | -0.56 | 0.45 | 0.47 | -0.06 | -0.46 | -0.05 | 0.1 | 0.1425 | -0.0525 | 0.15 | 0.878265 | 0.45411 | 0.587099 | 0.989223 | 0.997988 | 0.999507 |
| Cdh22_Q9WTP5_S705s_1 | Cdh22_Q9 | 0.56 | 0.66 | 0.38 | 0.25 | 0.21 | 0.87 | -0.05 | 0.46 | 0.43 | 0.81 | 0.57 | 0.14 | 1.23 | 0.68 | 0.04 | 0.46 | -0.265 | 0.19 | -0.0475 | 0.658129 | 0.732906 | 0.918989 | 0.995762 | 0.997988 | 0.995259 |
| Cdip1_F6X7Q6_T17t_1 | Cdip1_F6X | 0.59 | -1.58 | -0.98 | -0.35 | 0.53 | 0.75 | 0 | 0.48 | -0.88 | -1.06 | 0.35 | -0.3 | -1.77 | -0.2 | -1.33 | -1.15 | -0.09 | 0.025 | 0.14 | 0.32966 | 0.047493 | 0.837858 | 0.989223 | 0.997988 | 0.994666 |
| Cdk10_Q3UMM4_T196t_1 | Cdk10_Q3 | 0.1 | -0.17 | -0.87 | -0.25 | -0.01 | -0.72 | 0.14 | -0.38 | -0.73 | 0.08 | -0.51 | -0.28 | 0.33 | 0 | 0.36 | -0.37 | 1.02 | 0.1075 | -0.5325 | 0.250957 | 0.856457 | 0.837693 | 0.996349 | 0.997988 | 0.994666 |
| Cdk11b_P24788_S47s_1 | Cdk11b_P2 | -2.07 | -0.65 | -1.19 | -3.64 | -1.5 | -1.06 | -1.65 | -1.54 | -2.55 | -2.02 | -0.67 | -3.11 | -1.59 | 0.31 | -0.77 | -1.1 | 0.055 | -0.0625 | 0.3775 | 0.134691 | 0.45724 | 0.789991 | 0.989223 | 0.997988 | 0.994666 |
| Cdk11b_P24788_S578s_1 | Cdk11b_P2 | -0.1 | -0.18 | -0.45 | -0.6 | -0.34 | -0.85 | -0.27 | -0.44 | -0.68 | -0.41 | -0.11 | -0.86 | 0.03 | 0.04 | 0.39 | -0.25 | 0.45 | -0.2 | 1.1 | 0.17223 | 0.556186 | 0.494985 | 0.989223 | 0.997988 | 0.994666 |
| Cdk11b_P24788_S65s_1 | Cdk11b_P2 | 0.73 | -0.43 | 0.96 | 0.83 | -0.5 | 0.16 | 1.39 | 1.67 | 0.51 | 0.29 | -0.35 | -0.91 | 1.13 | 0.74 | 1.98 | 0.42 | -0.1425 | -0.1825 | 0.385 | 0.246594 | 0.773517 | 0.171203 | 0.995762 | 0.997988 | 0.994666 |
| Cdk11b_P24788_S72s_1 | Cdk11b_P2 | -3.96 | -5.26 | -4.17 | -3.58 | -3.02 | -4.04 | -2.99 | -3.32 | -2.85 | -3.61 | -3.52 | -2.37 | -1.78 | -2.85 | -1.36 | -2.54 | 0.1575 | -0.6375 | 0.545 | 0.001423 | 0.052836 | 0.027317 | 0.989223 | 0.997988 | 0.994666 |
| Cdk12_Q14AX6_S1079s_1 | Cdk12_Q1 | -0.86 | -2.01 | -2.13 | -1.81 | -2.02 | -2.29 | -1.89 | -1.62 | -1.63 | -1.53 | -1.17 | -2.19 | -1.63 | -1.52 | -1.4 | -1.9 | 0.9 | 1.155 | 2.11 | 0.792585 | 0.45202 | 0.843263 | 0.989223 | 0.997988 | 0.999507 |
| Cdk13_Q69ZA1_S326s_1 | Cdk13_Q6 | 1.14 | -0.16 | 0.61 | -4.84 | 0.46 | 0.76 | -1.74 | -0.25 | 0.54 | 0.03 | -1.54 | 0.17 | 1.5 | -1.68 | -0.53 | -0.35 | -0.2525 | 0.0725 | 0.09 | 0.676826 | 0.627271 | 0.6303 | 0.989223 | 0.997988 | 0.996159 |
| Cdk13_Q69ZA1_T1087t_1 | Cdk13_Q6 | -0.16 | -1.32 | -0.3 | 0.43 | -0.81 | -0.92 | -1.49 | -2.04 | 0.81 | -0.8 | -1.34 | -1.62 | 1.1 | -0.52 | -0.78 | -1.04 | 0.62 | 0.6125 | 0.5475 | 0.960645 | 0.046277 | 0.509511 | 0.989223 | 0.997988 | 0.999507 |
| Cdk13_Q69ZA1_T588t_1 | Cdk13_Q6 | -2.14 | -2.65 | -1.1 | -1.39 | -2.58 | -3.35 | -1.06 | -0.91 | -2.97 | -2.29 | -0.21 | -1.63 | -0.67 | -2.63 | -0.87 | -1.75 | -0.9775 | -0.4 | 0.0275 | 0.52785 | 0.80143 | 0.942639 | 0.995762 | 0.997988 | 0.994666 |
| Cdk13_Q69ZA1_T871t_1 | Cdk13_Q6 | 0.15 | -0.8 | -1.03 | -0.87 | -0.5 | -0.52 | -0.88 | -1.01 | -1.29 | -0.6 | -2.44 | -1.1 | 0.16 | -1.81 | -1.11 | -0.59 | -0.155 | 0.045 | 0.34 | 0.674266 | 0.774582 | 0.137883 | 0.995762 | 0.997988 | 0.996159 |
| Cdk14_D3YV23_S32s_1 | Cdk14_D3Y | -0.91 | -0.7 | -1.38 | -2.67 | 0.05 | -1.02 | -0.61 | -1.63 | -1.38 | -0.43 | -1.45 | -1.58 | -1.48 | -0.61 | -1.78 | -1.49 | -0.09 | -0.72 | -0.2 | 0.8771 | 0.252966 | 0.674923 | 0.989223 | 0.997988 | 0.999507 |
| Cdk16_Q04735_S103s_1 | Cdk16_Q0 | -0.08 | -1.22 | -1.37 | -0.88 | -0.29 | 0.15 | -0.25 | -0.6 | -1.49 | -1.13 | 1.13 | -0.46 | -1.89 | -0.24 | -1.28 | -0.84 | 0.6125 | 0.205 | 0.075 | 0.693458 | 0.081182 | 0.505982 | 0.989223 | 0.997988 | 0.996159 |
| Cdk16_Q04735_S43s_1 | Cdk16_Q0 | -0.86 | 0.45 | -0.35 | -0.46 | 1.59 | 0.21 | 0.11 | -0.12 | 2.2 | 1.91 | -0.9 | -0.59 | 2.07 | 1.27 | 0.14 | 0.13 | 0.64 | 0.4 | -0.175 | 0.038992 | 0.115957 | 0.231555 | 0.989223 | 0.997988 | 0.994666 |
| Cdk16_Q04735_S60s_1 | Cdk16_Q0 | 0.11 | 0.33 | -0.52 | -0.47 | 0.52 | 0.13 | -0.06 | -0.11 | 0.16 | 0.61 | 0.5 | -0.33 | -0.26 | 0.91 | -0.74 | -0.53 | 0.7525 | 0.96 | 1.2075 | 0.966914 | 0.380131 | 0.269729 | 0.989223 | 0.997988 | 0.999507 |
| Cdk16_Q04735_S75s_1 | Cdk16_Q0 | -0.69 | -1.36 | -0.61 | -1.06 | -0.23 | -1.12 | -0.03 | -0.4 | -0.9 | -1.12 | 0.48 | -0.34 | -1.13 | 0.02 | -0.79 | -0.72 | 0.2575 | 0.3725 | -0.0175 | 0.421235 | 0.14497 | 0.263 | 0.989223 | 0.997988 | 0.994666 |
| Cdk16_Q04735_S84s_1 | Cdk16_Q0 | -1.24 | -0.9 | -1.46 | -1.98 | -0.05 | -1.38 | -0.84 | -1.41 | -0.95 | -0.48 | -1.03 | -1.28 | -0.26 | -0.37 | -1.61 | -1.14 | 0.485 | 0.46 | 0.275 | 0.190808 | 0.232002 | 0.166556 | 0.989223 | 0.997988 | 0.994666 |
| Cdk17_Q8K0D0_S38s_1 | Cdk17_Q8 | 1.19 | 1.91 | 0.75 | 0.09 | 1.7 | 1.56 | 0.63 | 0.37 | 0.47 | 1.49 | 1.39 | 0.47 | -1.37 | 0.09 | -0.23 | 0.76 | 0.475 | 0.46 | 0.55 | 0.054373 | 0.864547 | 0.947407 | 0.996493 | 0.997988 | 0.994666 |
| Cdk17_Q8K0D0_S53s_1 | Cdk17_Q8 | 0.12 | -0.42 | -1.12 | -0.86 | 0.16 | 0.5 | -0.72 | -0.64 | -0.55 | -0.29 | 0.52 | -0.72 | -0.46 | 0.07 | -1.12 | -0.97 | 0.08 | -0.03 | -1.1725 | 0.898585 | 0.32664 | 0.432406 | 0.989223 | 0.997988 | 0.999507 |
| Cdk17_Q8K0D0_S81s_1 | Cdk17_Q8 | -0.29 | 0.19 | 0.12 | 1.09 | 0.61 | -0.23 | 0.6 | -0.23 | 0.11 | 0.59 | -0.3 | 0.82 | -0.1 | 1.1 | 0.38 | 0.37 | 0.395 | 0.31 | -0.05 | 0.684186 | 0.80877 | 0.943124 | 0.995762 | 0.997988 | 0.996159 |
| Cdk17_Q8K0D0_S96s_1 | Cdk17_Q8 | -0.52 | -1.07 | -1.11 | -0.77 | -0.04 | -0.58 | -0.39 | -0.39 | -0.5 | -0.65 | -0.03 | -0.47 | -0.65 | -0.16 | -0.64 | -0.15 | -0.09 | 0.0275 | 0.16 | 0.117932 | 0.055601 | 0.105394 | 0.989223 | 0.997988 | 0.994666 |
| Cdk18_Q04899_S109s_1 | Cdk18_Q0 | -1.05 | -0.93 | 0.38 | 0.03 | -0.87 | -1.05 | 0.52 | 0.33 | -1.15 | -1.1 | 0.38 | -0.21 | -0.94 | -0.02 | 0 | 1 | 0.5175 | 0.455 | 0.4675 | 0.430333 | 0.7998 | 0.792517 | 0.995762 | 0.997988 | 0.994666 |
| Cdk18_Q04899_S66s_1 | Cdk18_Q0 | -1.32 | -1.56 | -1.57 | -1.46 | -0.24 | -1.75 | 0.01 | -0.55 | -1.96 | -1.55 | 0.02 | -1.34 | -2.07 | -0.09 | -0.88 | -0.55 | 0.125 | -0.1275 | 0.4025 | 0.193133 | 0.050087 | 0.528084 | 0.989223 | 0.997988 | 0.994666 |
| Cdk5r1_P61809_T138t_1 | Cdk5r1_P6 | -2.5 | -4.27 | -0.69 | -0.98 | -2.53 | -2.5 | -0.99 | -1.23 | -3.1 | -2.86 | 0.1 | -0.97 | -2.53 | -2.45 | -1.4 | -1.05 | 0.845 | 0.27 | 0.58 | 0.751955 | 0.71151 | 0.686499 | 0.995437 | 0.997988 | 0.9971 |
| Cdkl5_Q3UTQ8_S306s_1 | Cdkl5_Q3U | 0.85 | -0.41 | 0.62 | 0.83 | 1 | 1.21 | 0.66 | 1.14 | -0.93 | 0.19 | 0.12 | 1.2 | -0.71 | -0.99 | 0.58 | 0.66 | 0.2975 | 0.4025 | 0.2525 | 0.253399 | 0.135428 | 0.515263 | 0.989223 | 0.997988 | 0.994666 |
| Cdkl5_Q3UTQ8_S377s_1 | Cdkl5_Q3U | 0 | -0.15 | -0.08 | 0.63 | 0.22 | 0.3 | -0.36 | 1.78 | -0.49 | -0.25 | 0.37 | 1.12 | -1.12 | 0.13 | 0.4 | 0.36 | 0.53 | -0.3275 | -0.5875 | 0.698742 | 0.408148 | 0.82677 | 0.989223 | 0.997988 | 0.996159 |
| Cdkl5_Q3UTQ8_S407s_1 | Cdkl5_Q3U | 1.24 | 0.22 | 0.48 | 0.95 | 1.57 | 1.46 | 1.51 | 1.55 | 0.12 | 0.1 | 0.56 | 1.19 | -1.35 | 0.59 | 0.35 | 0.01 | 0.385 | 0.0875 | -0.1575 | 0.106266 | 0.014402 | 0.523445 | 0.989223 | 0.997988 | 0.994666 |
| Cdkl5_Q3UTQ8_S497s_1 | Cdkl5_Q3U | -2.67 | -2.71 | -2.72 | -2.78 | -2.45 | -3.16 | -1.7 | -4.49 | -3.66 | -3.12 | 1.52 | -1.19 | -1.3 | -1.96 | -1.61 | -1.3 | 0.8 | -0.23 | -0.8225 | 0.001201 | 0.670514 | 0.299123 | 0.992876 | 0.997988 | 0.994666 |
| Cdkl5_Q3UTQ8_S681s_1 | Cdkl5_Q3U | -3.76 | -2.67 | -3.56 | -1.97 | -2.65 | -2.8 | -4.61 | -3.31 | -2.55 | -3.56 | -0.22 | -4.35 | -0.63 | -1.08 | -3.32 | -2.89 | -0.23 | 1.1075 | 1.1775 | 0.171394 | 0.525935 | 0.715894 | 0.989223 | 0.997988 | 0.994666 |
| Cdkl5_Q3UTQ8_S691s_1 | Cdkl5_Q3U | 1.06 | 0.5 | -0.01 | 0.46 | 0.69 | 1.26 | 0.13 | 0.62 | 0.48 | 0.18 | 0.3 | 0.69 | -0.14 | 1.12 | 0.26 | 0.35 | -0.3525 | 0.32 | 1.01 | 0.773796 | 0.602853 | 0.757182 | 0.989223 | 0.997988 | 0.99798 |
| Cdkl5_Q3UTQ8_S761s_1 | Cdkl5_Q3U | -0.86 | -0.84 | 0.06 | -0.22 | -0.1 | -0.57 | 0.39 | -0.01 | -1.55 | -0.93 | -0.98 | -0.46 | -2.03 | -0.25 | -0.66 | -0.45 | 0.1725 | -0.09 | -0.105 | 0.405615 | 0.236383 | 0.154528 | 0.989223 | 0.997988 | 0.994666 |
| Cdkl5_Q3UTQ8_Y171y_1 | Cdkl5_Q3U | 1.59 | 0.5 | -0.88 | 1.39 | 0.14 | 1.94 | 0.43 | 0.3 | 2.01 | 1.84 | 1.63 | 0.14 | -1.53 | 1.12 | -0.01 | -1.06 | 0.3925 | -0.515 | -0.3825 | 0.182526 | 0.932825 | 0.258544 | 0.998562 | 0.997988 | 0.994666 |
| Cdkn1b_P46414_S10s_1 | Cdkn1b_P4 | -1.21 | -1.46 | -1.91 | -3.62 | -1.34 | -1.37 | -2.33 | -2.25 | -3.12 | -0.84 | -2.48 | -1.67 | -1.12 | -3.31 | -2.33 | -2.19 | 0.0525 | 0.755 | -1.02 | 0.768845 | 0.679438 | 0.972999 | 0.993464 | 0.997988 | 0.9971 |
| Cdkn1b_P46414_S178s_1 | Cdkn1b_P4 | 0.13 | 0.09 | -2.15 | 0.33 | -0.06 | -1.76 | -0.78 | -0.81 | -1.21 | -0.75 | 0.05 | -0.37 | -0.74 | -0.35 | 0.4 | 0.23 | 0.2275 | 0.0225 | -0.1875 | 0.629851 | 0.463849 | 0.774074 | 0.989223 | 0.997988 | 0.995259 |
| Cdkn2aip_Q8BI72_S169s_1 | Cdkn2aip_( | -2.49 | -5.23 | -2.47 | -8.61 | -4.08 | -3.68 | -5.11 | -4.42 | -0.47 | -3.53 | -2.05 | -3.14 | 0.41 | -4.85 | -2.64 | -3.44 | -0.4525 | -0.17 | 0.285 | 0.208426 | 0.767812 | 0.113924 | 0.995762 | 0.997988 | 0.994666 |
| Cds1_P98191_S56s_1 | Cds1_P981 | 0.73 | 0.69 | 0.39 | -0.79 | 0.43 | 0.37 | -0.98 | -0.84 | 0.57 | 0.51 | 0.37 | -0.22 | 0.45 | 0.77 | -0.73 | 0.47 | 0.3775 | 2.4025 | 2.07 | 0.974422 | 0.303203 | 0.894986 | 0.989223 | 0.997988 | 0.999507 |
| Cds2_Q6PBC0_S20s_1 | Cds2_Q6PE | 0.73 | 0.36 | -0.03 | -0.28 | -0.42 | 0.92 | -1.21 | -0.8 | 1.17 | 0.99 | 0.09 | -0.66 | 0.12 | 0.19 | -0.3 | 0.16 | -0.51 | 0.0525 | -0.015 | 0.620879 | 0.245307 | 0.660422 | 0.989223 | 0.997988 | 0.995259 |

| | | | | | | | | | | | | | | | | | | | | | | | | | | |
|---|---|---|---|---|---|---|---|---|---|---|---|---|---|---|---|---|---|---|---|---|---|---|---|---|---|---|
| Cds2_Q6PBC0_S32s_1 | Cds2_Q6PE | 0.37 | 0.08 | 0 | -0.52 | -0.81 | 0.42 | -1.05 | -0.72 | 1.39 | 0.34 | -0.16 | -0.52 | 1.65 | -0.43 | -0.11 | -0.23 | -0.5725 | 0.2025 | -0.1525 | 0.629959 | 0.181182 | 0.529107 | 0.989223 | 0.997988 | 0.995259 |
| Cds2_Q6PBC0_S32s_2 | Cds2_Q6PE | -2.63 | -1.32 | -0.39 | -1.17 | -2.3 | -1.09 | -1.5 | -0.81 | -1.79 | -2.79 | -0.51 | -1.09 | -1.02 | -2.32 | -0.93 | -1.16 | -0.5225 | 0.28 | 0.2375 | 0.969814 | 0.926664 | 0.786264 | 0.997335 | 0.997988 | 0.999507 |
| Cds2_Q6PBC0_S36s_2 | Cds2_Q6PE | -2.63 | -1.32 | -0.93 | -1.54 | -2.3 | -1.09 | -2.21 | -1.03 | -1.79 | -2.79 | -0.18 | -1.81 | -1.02 | -2.32 | -0.95 | -1.4 | -0.0475 | -0.1675 | 0.02 | 0.695338 | 0.910222 | 0.949898 | 0.997335 | 0.997988 | 0.996159 |
| Cdv3_Q4VAA2_S51s_1 | Cdv3_Q4V | 0.8 | -0.63 | -0.38 | 0.11 | -0.35 | 1.13 | 0.11 | 0.03 | -0.43 | 0.21 | -0.19 | 0.53 | -0.29 | -1.59 | 0.59 | 0.1 | -0.0525 | -0.0375 | 0.1825 | 0.607202 | 0.550486 | 0.88583 | 0.989223 | 0.997988 | 0.995259 |
| Cdv3_Q4VAA2_S58s_1 | Cdv3_Q4V | 1.74 | -0.74 | 0.01 | 0.49 | -0.57 | 2.45 | 0.02 | -0.39 | 1.57 | 0.2 | -0.71 | -0.72 | 0.52 | -0.99 | -0.02 | -2.09 | 0.255 | 0.055 | -0.2725 | 0.165067 | 0.997392 | 0.670388 | 1 | 0.997988 | 0.994666 |
| Cecr6_Q99MX7_S521s_1 | Cecr6_Q99 | -0.15 | 0.43 | 0.35 | -0.07 | 1.08 | -0.05 | 0.61 | 0.14 | -0.25 | -0.32 | -0.08 | 0.57 | 0.09 | 1.03 | 0.02 | -0.2 | 0.0025 | -0.29 | -1.02 | 0.781909 | 0.339658 | 0.588527 | 0.989223 | 0.997988 | 0.998226 |
| Cecr6_Q99MX7_S527s_1 | Cecr6_Q99 | 0.49 | 0.84 | 0.4 | 0 | 0.84 | 0.32 | 0.57 | 0.4 | 0.57 | 1.09 | 0.38 | 0.16 | 0.02 | 0.49 | 0.56 | 0.67 | 0.305 | -0.16 | 0.095 | 0.993162 | 0.699301 | 0.697924 | 0.994893 | 0.997988 | 1 |
| Cecr6_Q99MX7_S548s_1 | Cecr6_Q99 | 0.82 | 0.86 | 0.7 | 0.49 | 1.03 | 0.94 | 0.94 | 1.12 | -0.08 | 0.8 | 0.37 | 0.93 | -0.01 | 0.11 | 0.81 | 1.11 | 0.1 | 0.1175 | 0.0025 | 0.519957 | 0.180009 | 0.469165 | 0.989223 | 0.997988 | 0.994666 |
| Celf4_Q7TSY6_S164s_1 | Celf4_Q7TS | -1.28 | -0.08 | -0.43 | -2.14 | -1.41 | -1.94 | -2.33 | -2.23 | 0.82 | -1.93 | -1.8 | -2.43 | 1.81 | -1.74 | -1 | -1.7 | 0.29 | -0.2125 | -0.2125 | 0.70069 | 0.056948 | 0.650234 | 0.989223 | 0.997988 | 0.996159 |
| Celsr2_F7BHW1_S637s_1 | Celsr2_F7B | 0.78 | 0.96 | 0.76 | 1.33 | 0.35 | 1.09 | 1.05 | 1.48 | 0.75 | 0.59 | 1.11 | 1.73 | 0.27 | 0.21 | 0.54 | 0.07 | -0.995 | -0.3525 | 0.325 | 0.025035 | 0.905899 | 0.782297 | 0.996513 | 0.997988 | 0.994666 |
| Celsr2_F7BHW1_S808s_1 | Celsr2_F7B | 0.51 | 0.14 | -0.62 | 0.14 | 0.47 | 0.15 | -0.23 | -0.4 | -0.29 | 0.3 | 0.87 | -0.99 | -0.79 | -0.51 | -1.34 | -0.97 | 0.035 | 0.0875 | -0.685 | 0.017285 | 0.888593 | 0.875593 | 0.996493 | 0.997988 | 0.994666 |
| Cend1_Q9JKC6_S109s_1 | Cend1_Q9J | -1.07 | -0.17 | -0.24 | -0.34 | -1.98 | 0.24 | -1.52 | -1.39 | 1.4 | -1.09 | -0.17 | -0.36 | 0.91 | -2.32 | -0.2 | -0.91 | -0.045 | -0.07 | -0.945 | 0.784744 | 0.168278 | 0.454392 | 0.989223 | 0.997988 | 0.99912 |
| Cend1_Q9JKC6_S10s_1 | Cend1_Q9J | -0.21 | -0.02 | -0.19 | -0.26 | -0.34 | -1.09 | -0.73 | -0.71 | 1.54 | -0.3 | -2.35 | -0.28 | 2.2 | -0.77 | 0.46 | -0.37 | -0.7075 | 0.4 | -0.175 | 0.372253 | 0.039318 | 0.804577 | 0.989223 | 0.997988 | 0.994666 |
| Cend1_Q9JKC6_S90s_1 | Cend1_Q9J | -0.67 | -0.48 | 0.04 | 0.2 | -0.37 | -0.81 | 0.49 | 0.24 | -0.75 | -0.41 | -0.22 | 0.18 | -0.52 | 0.11 | 0.08 | 0.03 | -0.5475 | -0.1775 | 0.55 | 0.624385 | 0.74894 | 0.818142 | 0.995762 | 0.997988 | 0.995259 |
| Cenpa_D6RJ71_S15s_1 | Cenpa_D6F | 0.87 | 0.96 | 1.82 | 1.25 | -0.01 | 0.09 | 0.51 | 0.39 | 1.08 | 0.84 | -0.53 | 0.92 | 1.68 | -0.54 | 1.09 | 1.8 | 0.115 | -0.0725 | 0.1525 | 0.687784 | 0.006031 | 0.149059 | 0.989223 | 0.997988 | 0.996159 |
| Cenpv_Q9CXS4_S18s_1 | Cenpv_Q9C | -0.74 | -0.3 | 0.58 | -0.05 | -1.14 | -0.62 | -1 | 0.07 | 0.07 | -0.68 | -0.82 | 0.42 | 0.78 | -1.09 | -0.04 | 0.12 | -0.98 | -0.6475 | -0.2175 | 0.878923 | 0.172445 | 0.756453 | 0.989223 | 0.997988 | 0.999507 |
| Cenpv_Q9CXS4_S6s_1 | Cenpv_Q9C | -3.7 | -3.89 | -3.94 | -11.45 | -2.82 | -3.71 | -3.46 | -4.45 | -2.9 | -3.48 | -2.12 | -3.94 | -1.37 | -3.15 | -4.03 | -3.01 | -0.545 | -0.125 | 0.07 | 0.11789 | 0.216512 | 0.144901 | 0.989223 | 0.997988 | 0.994666 |
| Cep128_F7CIB0_S536s_1 | Cep128_F7 | -2.37 | -3.12 | -0.43 | 0.03 | -2.93 | -1.9 | 0.08 | -0.26 | -1.31 | -2.39 | -1.67 | 0.01 | -0.02 | -3.05 | 0.09 | -0.13 | 2.135 | 2.635 | 2.855 | 0.464436 | 0.806807 | 0.87007 | 0.995762 | 0.997988 | 0.994666 |
| Cep131_B1AXI9_S47s_1 | Cep131_B1 | -1.83 | -1.22 | -1.94 | -1.32 | -1.28 | -1.53 | -1.16 | -1.36 | -1.04 | -1.83 | -0.75 | -1.61 | -0.33 | -0.8 | -1.38 | -0.99 | 0.22 | 0.1325 | 0.695 | 0.054199 | 0.342662 | 0.423841 | 0.989223 | 0.997988 | 0.994666 |
| Cep131_B1AXI9_S78s_1 | Cep131_B1 | -0.41 | -0.5 | 0.59 | 0.18 | -0.23 | -1.04 | 0.33 | -0.41 | -0.13 | -0.6 | 1.03 | 0.66 | 0.18 | 0.41 | -0.05 | 0.4 | 0.245 | 0.27 | 0.7025 | 0.413573 | 0.427656 | 0.533475 | 0.989223 | 0.997988 | 0.994666 |
| Cep170_H7BX26_S1048s_1 | Cep170_H7 | -0.41 | 0.37 | -1 | -0.54 | -0.07 | -0.53 | -0.66 | -0.26 | 0.33 | -0.63 | -0.46 | -0.79 | -0.08 | 0.74 | -0.78 | -1.45 | -0.3025 | 0.275 | 0.27 | 0.996128 | 0.963278 | 0.984302 | 0.999187 | 0.997988 | 1 |
| Cep170_H7BX26_S1155s_1 | Cep170_H7 | -3.14 | -2.99 | -2.3 | -4.54 | -3.13 | -3.47 | -3.45 | -3.94 | -2.97 | -3.35 | -0.43 | -2.58 | -0.91 | -3.41 | -3.99 | -3.71 | 0.015 | 0.0075 | 0.0025 | 0.753352 | 0.58472 | 0.228864 | 0.989223 | 0.997988 | 0.9971 |
| Cep170_H7BX26_S1157s_1 | Cep170_H7 | -3.77 | -0.89 | -1.67 | -8.64 | -2.4 | -2.29 | -3.65 | -3.03 | -2.06 | -1.67 | -0.18 | -2.81 | -0.45 | -1.51 | -2.78 | -3.1 | -0.255 | 0.91 | 0.2375 | 0.2765 | 0.556202 | 0.216828 | 0.989223 | 0.997988 | 0.994666 |
| Cep170_H7BX26_S1215s_1 | Cep170_H7 | -2.71 | -2.48 | -1.16 | -1.34 | -3.61 | -2.26 | -1.92 | -0.92 | 2.25 | -2.49 | -0.28 | -1.16 | 2.5 | -1.46 | -0.61 | -1.9 | 0.9 | 2.0625 | 1.7825 | 0.120842 | 0.676042 | 0.139297 | 0.992974 | 0.997988 | 0.994666 |
| Cep170_H7BX26_S1260s_1 | Cep170_H7 | -0.8 | -0.3 | -0.68 | 0.08 | -0.03 | -0.94 | -0.1 | -0.55 | -0.54 | -0.22 | 0.65 | 0.21 | -0.15 | 0.38 | 0.7 | -0.09 | -0.255 | 1.5025 | 1.555 | 0.076807 | 0.948585 | 0.21272 | 0.999187 | 0.997988 | 0.994666 |
| Cep170_H7BX26_S1296s_1 | Cep170_H7 | -2.21 | -2.3 | -1.67 | -2.92 | 0.02 | -0.98 | -2.59 | -0.75 | -1.71 | -1.99 | -0.16 | -1.23 | -2.36 | -0.54 | -0.58 | -1.82 | 0.02 | 0.45 | 0.635 | 0.080244 | 0.0509 | 0.051877 | 0.989223 | 0.997988 | 0.994666 |
| Cep170_H7BX26_S1511s_1 | Cep170_H7 | -0.25 | -0.84 | -0.63 | -0.26 | -0.69 | -1.1 | 0.1 | -0.09 | -0.45 | -0.3 | 0.5 | -0.21 | 0.25 | 0.02 | 0.06 | -0.08 | 1.2 | 1.0025 | 0.95 | 0.05526 | 0.877443 | 0.221482 | 0.996493 | 0.997988 | 0.994666 |
| Cep170_H7BX26_S1512s_1 | Cep170_H7 | -0.25 | -0.84 | -0.63 | -0.26 | -0.69 | -1.1 | 0.1 | -0.09 | -0.45 | -0.3 | 0.5 | -0.21 | 0.25 | 0.02 | 0.06 | -0.08 | 0.05 | 0.38 | 0.5575 | 0.05526 | 0.877443 | 0.221482 | 0.996493 | 0.997988 | 0.994666 |
| Cep170_H7BX26_S1512s_2 | Cep170_H7 | 0.13 | 0.25 | -0.53 | 0.08 | 0.4 | -0.4 | 0.53 | 0.78 | 0.72 | 0.52 | 0.52 | -0.07 | -1.28 | 0.51 | 0.17 | -0.29 | 0.05 | 0.38 | 0.5575 | 0.630201 | 0.299872 | 0.153492 | 0.989223 | 0.997988 | 0.995259 |
| Cep170_H7BX26_S1519s_1 | Cep170_H7 | -1.82 | -2.1 | 1.04 | 0.7 | -2.43 | -1.58 | 0.63 | 0.8 | -1.2 | -1.08 | 1.94 | 1.2 | -0.43 | -0.49 | 0.67 | 0.67 | 0.345 | 0.44 | -0.205 | 0.414952 | 0.919877 | 0.459022 | 0.997335 | 0.997988 | 0.994666 |
| Cep170_H7BX26_S1519s_2 | Cep170_H7 | 0.13 | 0.25 | -0.53 | 0.08 | 0.4 | -0.4 | 0.53 | 0.78 | 0.72 | 0.52 | 0.52 | -0.07 | -1.28 | 0.51 | 0.17 | -0.29 | -0.1 | 0.76 | 0.65 | 0.630201 | 0.299872 | 0.153492 | 0.989223 | 0.997988 | 0.995259 |
| Cep170_H7BX26_S378s_1 | Cep170_H7 | 3.37 | 1.22 | 2.21 | 1.61 | 0.91 | 3.89 | 2.29 | 1.95 | 3.62 | 3.01 | 1.33 | 2.24 | 2.42 | 0.32 | 3.37 | 2.56 | 0.345 | 0.44 | -0.205 | 0.927566 | 0.818193 | 0.478332 | 0.995762 | 0.997988 | 0.999507 |
| Cep170_H7BX26_S443s_1 | Cep170_H7 | -2.13 | -1.28 | -1.43 | -1.59 | -0.86 | -1.68 | -0.63 | -1.56 | -1.87 | -1.27 | -1.11 | -1.67 | -1.61 | -0.8 | -1.65 | -1.48 | 0.1575 | 0.4475 | 0.065 | 0.491557 | 0.216313 | 0.668706 | 0.989223 | 0.997988 | 0.994666 |
| Cep170_H7BX26_S829s_1 | Cep170_H7 | -2.44 | -1.97 | 0.18 | 0.42 | -1.76 | -2.76 | 0.01 | 0.05 | -2.74 | -1.75 | -0.5 | -0.2 | -1.65 | -1.39 | -0.22 | -0.37 | 0.425 | 0.1275 | 0.2225 | 0.950512 | 0.852714 | 0.680588 | 0.996349 | 0.997988 | 0.999507 |
| Cep170_H7BX26_S914s_1 | Cep170_H7 | -1.98 | -0.54 | -1.56 | -4 | -1.95 | -1.08 | -1.73 | -2.22 | -0.66 | -1.55 | 0.72 | -2.04 | -1.43 | -0.86 | -1.76 | -2.06 | -0.1625 | -0.345 | 0.045 | 0.481379 | 0.685639 | 0.198807 | 0.994624 | 0.997988 | 0.994666 |
| Cep170_H7BX26_S914s_2 | Cep170_H7 | -1.81 | -1.9 | -1.39 | -2.87 | -1.97 | -1.6 | -1.5 | -2.42 | -2.34 | -2.5 | -0.11 | -0.06 | -2.26 | -0.94 | -1.95 | -2.36 | 0.275 | 1.1375 | 0.4925 | 0.794623 | 0.746478 | 0.288064 | 0.995762 | 0.997988 | 0.999507 |
| Cep170_H7BX26_S950s_1 | Cep170_H7 | -3.37 | -3.43 | -1.9 | -3.41 | -2.21 | -3.65 | -2.75 | -2.42 | -5.3 | -2.83 | 0.43 | -2.06 | -3.12 | -2.8 | -1.74 | -2.76 | 0.12 | 0.74 | 0.115 | 0.373071 | 0.55928 | 0.592706 | 0.989223 | 0.997988 | 0.994666 |
| Cep170_H7BX26_S985s_1 | Cep170_H7 | -1.3 | -1.34 | -0.47 | -1.5 | -0.84 | -0.14 | -1.59 | -1.17 | -1.75 | -1.25 | -3.12 | -1.14 | -0.85 | -3.74 | -1.52 | -1.63 | 0.27 | 0.5875 | 0.4225 | 0.218275 | 0.566553 | 0.193333 | 0.989223 | 0.997988 | 0.994666 |
| Cep170_H7BX26_T394t_1 | Cep170_H7 | -3.2 | -1.88 | -1.93 | -1.1 | -0.34 | -1.82 | -1.01 | -1.38 | -1.47 | -1.84 | -0.87 | -1.49 | -1.65 | -0.77 | -1.19 | -1.25 | 0.2175 | -0.6625 | -0.7825 | 0.10042 | 0.096125 | 0.205586 | 0.989223 | 0.997988 | 0.994666 |
| Cep170_H7BX26_T912t_1 | Cep170_H7 | -0.4 | -0.39 | -0.44 | -0.7 | -0.62 | -0.9 | -1.08 | -1.18 | -0.5 | -0.99 | -0.42 | -0.35 | -0.5 | -0.69 | -0.65 | -0.76 | 0.89 | 0.61 | 0.8125 | 0.482448 | 0.063424 | 0.7355 | 0.989223 | 0.997988 | 0.994666 |
| Cep170_H7BX26_T912t_2 | Cep170_H7 | -1.81 | -1.9 | -1.39 | -2.87 | -1.97 | -1.6 | -1.5 | -2.42 | -2.34 | -2.5 | -0.11 | -0.06 | -2.26 | -0.94 | -1.95 | -2.36 | -0.4625 | -0.0825 | -0.1675 | 0.794623 | 0.746478 | 0.288064 | 0.995762 | 0.997988 | 0.999507 |
| Cep170b_Q80U49_S1032s_1 | Cep170b_C | -0.12 | -0.15 | -0.2 | -0.21 | 0.08 | -0.58 | 0.27 | -0.28 | -0.23 | -0.02 | -0.31 | -0.05 | -0.83 | 0.43 | -0.21 | -0.34 | 0.12 | 0.74 | 0.115 | 0.829124 | 0.864136 | 0.933314 | 0.996493 | 0.997988 | 0.999507 |
| Cep170b_Q80U49_S1101s_1 | Cep170b_C | -3 | -1.94 | -1.26 | -3.23 | -1.57 | -2.94 | -2.17 | -2.72 | -2.91 | -2.48 | -0.3 | -1.98 | -0.66 | -1.47 | -2.06 | -1.96 | 0.0425 | 0.0175 | -0.0675 | 0.142797 | 0.988169 | 0.513597 | 0.999187 | 0.997988 | 0.994666 |
| Cep170b_Q80U49_S1122s_1 | Cep170b_C | -0.26 | -0.06 | -1.26 | -1.84 | -0.9 | -1.19 | -1.48 | -1.54 | -0.67 | -0.2 | 1.12 | -1.11 | -0.46 | -0.14 | -2.63 | -2.59 | 0.0075 | 0.44 | 0.82 | 0.405194 | 0.330246 | 0.292163 | 0.989223 | 0.997988 | 0.994666 |
| Cep170b_Q80U49_S1166s_1 | Cep170b_C | -1.22 | 0.33 | 1.15 | 0.68 | 0.01 | -1.35 | -0.42 | 0.12 | 0 | -0.81 | 1.59 | 0.56 | 1.38 | 1 | 0.99 | 0.13 | -0.4225 | 0.64 | -0.6 | 0.251768 | 0.261003 | 0.878232 | 0.989223 | 0.997988 | 0.994666 |
| Cep170b_Q80U49_S1347s_1 | Cep170b_C | -0.75 | -0.58 | 0.01 | 0.29 | -0.29 | -1.23 | 0.61 | 0.02 | -0.24 | -0.12 | -0.11 | 0.34 | 0.11 | 0.04 | 0.22 | 0.47 | -0.645 | 0.1 | 0.64 | 0.159714 | 0.935061 | 0.474446 | 0.99868 | 0.997988 | 0.994666 |
| Cep170b_Q80U49_S1533s_1 | Cep170b_C | 0.41 | 0.64 | -0.1 | 0.1 | 0.32 | 0.53 | 1.03 | 0.65 | 0.47 | -0.02 | 1.18 | 1.16 | -0.54 | 0.65 | 0.75 | 0.41 | 0.035 | 0.225 | 0.4675 | 0.878941 | 0.18668 | 0.233033 | 0.989223 | 0.997988 | 0.999507 |
| Cep170b_Q80U49_S371s_1 | Cep170b_C | -1.7 | -0.46 | -1.5 | -0.89 | -0.07 | -0.71 | -0.41 | -0.63 | -2.38 | -0.76 | -0.95 | -0.54 | -0.14 | -0.6 | -0.47 | -0.75 | 0.37 | 0.435 | 0.055 | 0.085705 | 0.061893 | 0.966612 | 0.989223 | 0.997988 | 0.994666 |
| Cep170b_Q80U49_S478s_1 | Cep170b_C | -1.02 | -1.26 | -1.22 | -1.93 | -0.27 | -1.74 | -0.98 | -0.9 | -0.86 | -0.31 | -0.59 | -1.13 | -0.87 | -0.18 | -1.29 | -0.28 | 0.6825 | -0.02 | 0.6475 | 0.072262 | 0.298596 | 0.060369 | 0.989223 | 0.997988 | 0.994666 |
| Cep170b_Q80U49_S534s_1 | Cep170b_C | 0.3 | -0.28 | 0.45 | 1.1 | 0.07 | 0.23 | 0.99 | 1.11 | 0.11 | -0.09 | 1.9 | 1.01 | -0.12 | -0.2 | 1.21 | 0.85 | 0.385 | 0.635 | 0.7025 | 0.923352 | 0.586219 | 0.506316 | 0.989223 | 0.997988 | 0.999507 |
| Cep170b_Q80U49_S819s_1 | Cep170b_C | 0.73 | 0.76 | 0.67 | 0.84 | -0.34 | 0.98 | 0.68 | 0.92 | 1.26 | 0.37 | -0.17 | 0.64 | 0.86 | -0.28 | 0.44 | 1.12 | 0.2075 | 0.34 | 0.0425 | 0.53382 | 0.559412 | 0.495399 | 0.989223 | 0.997988 | 0.994666 |
| Cep170b_Q80U49_S840s_1 | Cep170b_C | 1.12 | -0.32 | -0.89 | 0.08 | -0.01 | 1.7 | -0.25 | -0.35 | 1.19 | 0.25 | 0.23 | -1.15 | 0.36 | 0.58 | -0.98 | -1.71 | -0.19 | -0.225 | -0.215 | 0.493533 | 0.634965 | 0.822153 | 0.989223 | 0.997988 | 0.994666 |
| Cep170b_Q80U49_S843s_1 | Cep170b_C | 0.74 | -0.43 | -0.99 | -0.3 | 0.02 | 1.29 | -0.43 | -0.44 | 0.69 | 0.13 | 0.73 | -0.84 | 0.04 | 0.29 | -1.19 | -1.65 | 0.275 | 0.1325 | -0.435 | 0.491714 | 0.482835 | 0.395048 | 0.989223 | 0.997988 | 0.994666 |
| Cep170b_Q80U49_S965s_1 | Cep170b_C | 0.17 | 0.53 | 0.71 | 0.29 | 0.14 | -0.59 | -0.25 | 0.1 | -0.13 | 1.3 | 0.34 | 0.47 | 0.78 | 0.53 | -0.11 | 0.02 | 0.355 | 0.4225 | -0.3825 | 0.692338 | 0.048522 | 0.837293 | 0.989223 | 0.997988 | 0.996159 |
| Cep170b_Q80U49_S981s_1 | Cep170b_C | 0.57 | 0.95 | -0.22 | -0.08 | 0.47 | 0.53 | 0.41 | 0.23 | 0.2 | 0.54 | 0.09 | -0.16 | 0.38 | 0.79 | 0.22 | -0.05 | -0.575 | 0.07 | -0.12 | 0.932471 | 0.731241 | 0.681709 | 0.995762 | 0.997988 | 0.999507 |
| Cep250_E9Q5A8_S2322s_1 | Cep250_E9 | 0.97 | 0.06 | 0 | 0.19 | -0.01 | 0.69 | -0.53 | -0.09 | 1.83 | 0.86 | 0.6 | 0.08 | 1.77 | -0.97 | 1.03 | 0.35 | 0.105 | -0.1375 | 0.03 | 0.677116 | 0.407145 | 0.222841 | 0.989223 | 0.997988 | 0.996159 |
| Cep41_H9KUZ8_S286s_1 | Cep41_H9K | -4.84 | -4.45 | -3.42 | -4.37 | -4.43 | -4.28 | -3.1 | -4.98 | -3.61 | -4.27 | -2.68 | -4.32 | -2.38 | -2.55 | -3.52 | -3.58 | -0.29 | 0.5375 | 0.24 | 0.014799 | 0.875303 | 0.255577 | 0.996493 | 0.997988 | 0.994666 |
| Cep68_Q3TRP8_S419s_1 | Cep68_Q3T | 0.52 | -0.41 | -0.13 | 0.13 | 0.6 | 0.48 | -0.2 | 0 | 0.05 | 0.52 | 1.28 | 0.05 | 1.35 | 0.86 | 0.31 | -0.05 | 0.0725 | 0.55 | 1.2625 | 0.14536 | 0.525083 | 0.236204 | 0.989223 | 0.997988 | 0.994666 |
| Cep97_Q9CZ62_S449s_1 | Cep97_Q9C | -0.49 | -0.6 | 0.79 | 0.35 | -0.54 | 0.36 | 0.17 | 0.2 | 0.2 | -0.08 | 1.22 | 0.06 | -0.13 | 0.7 | 0.87 | 1.3 | 0.1925 | 0.4475 | 0.59 | 0.15084 | 0.926715 | 0.444356 | 0.997335 | 0.997988 | 0.994666 |
| Cep97_Q9CZ62_S725s_1 | Cep97_Q9C | 5.68 | 3.97 | 4.76 | 1.51 | 2.59 | 6.19 | 3.65 | 3.11 | 2.35 | 5.07 | -0.5 | 4.28 | 1.96 | 0.28 | 4 | 4.6 | 0.035 | 0.3375 | 0.6725 | 0.286387 | 0.926831 | 0.388064 | 0.997335 | 0.997988 | 0.994666 |
| Cetn2_B1AUQ7_S11s_1 | Cetn2_B1A | -1.61 | -0.64 | -1.78 | -1.71 | -0.47 | -1.86 | -1.21 | -2.01 | -1.28 | -1.47 | -1.04 | -1.79 | -0.62 | -0.56 | -0.84 | -1.34 | -0.095 | -1.18 | -1.27 | 0.114742 | 0.909778 | 0.904454 | 0.997335 | 0.997988 | 0.994666 |
| Cfl1_P18760_S24s_1 | Cfl1_P1876 | -0.83 | -1.64 | -0.07 | -0.92 | -1.2 | -0.72 | -0.83 | -0.7 | -0.23 | -1 | -1.03 | -0.77 | 1.23 | -1.13 | -0.91 | -1.1 | 0.0475 | 0.04 | 0.595 | 0.521904 | 0.994218 | 0.775397 | 0.999278 | 0.997988 | 0.994666 |
| Cfl1_P18760_S3s_1 | Cfl1_P1876 | -1.39 | -2.82 | -2.17 | -2.1 | -1.51 | -1.94 | -1.01 | -1.51 | -2.81 | -2.87 | -0.27 | -1.94 | -3.16 | -1.54 | -2.94 | -2.38 | 0.0025 | 0.1075 | 0.3875 | 0.403349 | 0.099112 | 0.810472 | 0.989223 | 0.997988 | 0.994666 |
| Cfl1_P18760_T25t_1 | Cfl1_P1876 | 0.25 | 0.26 | 0.32 | -0.27 | -0.73 | 0.67 | -0.87 | -0.52 | 2.07 | 0.48 | -0.59 | -0.09 | 1.86 | -0.69 | -0.16 | -0.22 | 0.6275 | 0.1475 | -0.385 | 0.915431 | 0.197764 | 0.555297 | 0.989223 | 0.997988 | 0.999507 |
| Cfl2_P45591_S3s_1 | Cfl2_P4559 | -1.26 | -3.19 | -2.11 | -1.87 | -1.3 | -1.52 | -1.39 | -1.48 | -2.43 | -2.72 | -0.54 | -1.89 | -3.62 | -1.43 | -2.77 | -2.04 | -0.5025 | 0.3275 | 0.0575 | 0.53569 | 0.106428 | 0.714406 | 0.989223 | 0.997988 | 0.994666 |
| Cgn_D3YUW7_S208s_1 | Cgn_D3YU | -1.85 | -1.91 | -1.19 | -1.15 | -2.81 | -1.01 | -1.54 | -1.63 | 0 | -1.88 | -1.02 | -1.16 | -0.05 | -2.82 | -0.6 | -1.68 | 0.685 | 0.2125 | -0.3575 | 0.68904 | 0.591828 | 0.251577 | 0.989223 | 0.997988 | 0.996159 |
| Cgnl1_D3Z3H4_S203s_1 | Cgnl1_D3Z | 0.13 | 1.01 | 1.12 | 1.13 | 0.18 | 0.06 | 0.62 | 1.05 | 0.4 | 0.6 | 0.31 | 0.7 | 0.6 | 0.47 | 1.45 | 1.59 | -0.2225 | 0.51 | 0.2375 | 0.643375 | 0.288284 | 0.264161 | 0.989223 | 0.997988 | 0.995259 |
| Cgnl1_D3Z3H4_S257s_1 | Cgnl1_D3Z | -0.29 | 0.28 | 0.43 | 0.57 | -1.27 | -0.67 | 0.8 | -0.01 | -0.68 | -0.11 | -0.19 | 0.84 | 0.33 | -0.33 | 0.76 | -0.14 | -0.37 | -0.345 | 0.18 | 0.787552 | 0.253541 | 0.458902 | 0.989223 | 0.997988 | 0.999507 |
| Champ1_Q8K327_S110s_1 | Champ1_Q | 2.77 | -0.08 | 0.4 | 0.35 | -0.57 | 1.55 | 0.51 | 0.21 | 1.68 | 2.05 | 0.01 | 0.37 | 1.58 | 0.42 | 1.23 | 1.14 | -0.535 | -0.2825 | -0.0925 | 0.711137 | 0.53219 | 0.818865 | 0.989223 | 0.997988 | 0.996966 |
| Champ1_Q8K327_S204s_1 | Champ1_Q | 0.69 | -0.03 | -0.65 | -0.08 | -0.47 | -0.37 | -0.92 | -0.6 | -0.36 | -0.15 | -1.21 | -1.48 | 1.08 | -0.35 | 0.05 | -0.03 | -0.435 | 0.1675 | 0.2325 | 0.623091 | 0.095632 | 0.086246 | 0.989223 | 0.997988 | 0.995259 |
| Champ1_Q8K327_S265s_1 | Champ1_Q | 0.59 | -1.36 | 0.07 | -0.35 | -2.37 | 0.11 | -1.55 | -0.58 | 1.11 | -0.96 | -1.59 | -0.77 | 1.72 | -1.58 | 0.24 | -0.81 | -0.5725 | -0.7825 | 0.205 | 0.832899 | 0.192763 | 0.656014 | 0.989223 | 0.997988 | 0.999507 |
| Champ1_Q8K327_S405s_1 | Champ1_Q | 0.85 | 0.45 | 0.39 | -0.94 | 0.65 | 0.68 | 0.05 | -0.19 | 0.47 | -0.75 | -0.8 | 0.45 | 0.05 | 0.03 | 0.37 | 0.6 | -0.835 | -0.29 | 0.155 | 0.856138 | 0.794968 | 0.49392 | 0.995762 | 0.997988 | 0.999507 |
| Champ1_Q8K327_S416s_1 | Champ1_Q | 0.1 | -0.13 | 0.22 | -0.18 | -0.84 | -0.08 | 0.44 | 0.25 | -0.58 | -0.14 | 0.04 | -0.06 | -1.07 | 0.27 | 0.46 | 0.23 | 0.11 | -0.345 | 0.075 | 0.936885 | 0.848834 | 0.452486 | 0.996349 | 0.997988 | 0.999507 |
| Champ1_Q8K327_S425s_1 | Champ1_Q | -0.22 | 0.28 | 0.88 | 0.57 | -0.49 | -0.86 | 0.29 | 0.35 | 0.05 | -0.23 | -0.07 | 0.41 | 0.45 | 0.07 | 1.57 | 1.29 | -0.06 | -0.1875 | -0.03 | 0.282101 | 0.159491 | 0.287066 | 0.989223 | 0.997988 | 0.994666 |
| Champ1_Q8K327_S532s_1 | Champ1_Q | -0.15 | -0.28 | 0.62 | -0.3 | -0.66 | -0.95 | -0.47 | -0.27 | -0.75 | -0.83 | 0.48 | -0.44 | 0.01 | -0.13 | -0.71 | -0.38 | -0.555 | -0.3375 | 0.4675 | 0.396964 | 0.079567 | 0.354938 | 0.989223 | 0.997988 | 0.994666 |
| Chat_Q8BQV2_S366s_1 | Chat_Q8BC | 0.09 | 0.1 | -0.52 | -0.24 | 0.45 | 0.12 | -0.58 | -0.22 | 0.31 | 0.35 | -0.83 | -0.28 | 0.04 | 0.51 | -0.53 | -0.31 | -0.56 | -0.3575 | -0.275 | 0.826467 | 0.773064 | 0.929067 | 0.995762 | 0.997988 | 0.999507 |
| Chchd2_Q9D1L0_S45s_1 | Chchd2_Q9 | 0.28 | 0.57 | 0.11 | 0.12 | 0.25 | 0.17 | 0.38 | 0.17 | 0.63 | 0.46 | 0.11 | -0.22 | 0.39 | -0.02 | -0.05 | 0.35 | 0.085 | 0.03 | 0.07 | 0.694043 | 0.897726 | 0.927101 | 0.996493 | 0.997988 | 0.996159 |
| Chchd2_Q9D1L0_T13t_1 | Chchd2_Q9 | -0.03 | 0.25 | -0.25 | -0.22 | -0.42 | -0.05 | -1.11 | -0.86 | 0.54 | 0.06 | -0.64 | -0.25 | 1.27 | -0.5 | -0.11 | 0.09 | -0.0275 | -0.025 | -0.1025 | 0.539342 | 0.08515 | 0.974172 | 0.989223 | 0.997988 | 0.994666 |
| Chchd3_Q9D9P1_S112s_1 | Chchd3_Q9 | -0.89 | 0.02 | 0.66 | 1.13 | -2.95 | 1.02 | -0.59 | 0.62 | 0.62 | 0.94 | 0.74 | 0.39 | 0.76 | -1.19 | 1.15 | 0.07 | -0.5475 | -0.01 | 0.25 | 0.957712 | 0.42356 | 0.326261 | 0.989223 | 0.997988 | 0.999507 |
| Chchd6_E9Q4M4_S164s_1 | Chchd6_E9 | 0.62 | 0.61 | 0.24 | 0.25 | -0.29 | 0.63 | -1.2 | -0.25 | 0.81 | 0.24 | -0.46 | -0.21 | 1.52 | -0.35 | -0.26 | -0.21 | -0.705 | 0.4425 | -0.0325 | 0.573786 | 0.087866 | 0.317342 | 0.989223 | 0.997988 | 0.995259 |
| Chchd6_E9Q4M4_S172s_1 | Chchd6_E9 | -0.14 | 0.78 | -0.8 | 0.9 | 0.06 | -0.24 | -0.08 | 0.4 | 1.07 | 0.25 | 0.73 | 0.4 | 1.31 | 0.04 | 0.77 | -0.66 | -0.7075 | -0.335 | -0.255 | 0.742159 | 0.712962 | 0.333582 | 0.995437 | 0.997988 | 0.9971 |
| Chd1_P40201_S1690s_1 | Chd1_P402 | -0.14 | 0.66 | -1.29 | -1.11 | -1.13 | -1.93 | -0.45 | -1.68 | -0.43 | 0.14 | -0.68 | 0.03 | 0.83 | 0.12 | 0.96 | -0.75 | -0.15 | 0.4275 | 0.18 | 0.192699 | 0.130518 | 0.619175 | 0.989223 | 0.997988 | 0.994666 |
| Chd2_E9PZM4_S1365s_1 | Chd2_E9PZ | 0.47 | 0.48 | 2.13 | 0.67 | 0.14 | 0.58 | 0.74 | 1.32 | -0.7 | -0.08 | 0.24 | 1.16 | 0.59 | -0.91 | 1.4 | 1.39 | -0.8275 | 0.235 | 0.76 | 0.604643 | 0.584514 | 0.157732 | 0.989223 | 0.997988 | 0.995259 |
| Chd3_B1AR17_S142s_1 | Chd3_B1AF | 0.85 | -0.32 | 0.49 | -0.27 | -1.25 | 0.66 | -0.68 | -0.31 | 1.14 | 0.75 | 0.72 | 0.21 | 1.17 | -2.16 | 0.36 | 0.45 | -0.2425 | -0.7825 | -0.32 | 0.740436 | 0.224947 | 0.172135 | 0.989223 | 0.997988 | 0.9971 |
| Chd3_B1AR17_S1651s_1 | Chd3_B1AF | 2.53 | 0.71 | 0.55 | -0.69 | 0.74 | 1.78 | 0.07 | 0.55 | 0.81 | 0.83 | -1.2 | 1.2 | 0.35 | -0.21 | -0.52 | 0.98 | -0.5825 | 0.5175 | -0.2325 | 0.359979 | 0.988045 | 0.636628 | 0.999187 | 0.997988 | 0.994666 |
| Chd4_E9QAS4_S1329s_1 | Chd4_E9Q | -4.47 | -6.27 | -4.19 | -7.76 | -5.69 | -4.59 | -6.6 | -4.27 | -10.98 | -4.97 | -1.61 | -4.06 | -2.84 | -7.35 | -3.6 | -3.96 | 0.01 | -0.365 | -0.625 | 0.288763 | 0.656562 | 0.88663 | 0.990434 | 0.997988 | 0.994666 |
| Chd4_E9QAS4_S1515s_1 | Chd4_E9Q | -0.04 | -0.92 | -0.08 | -0.05 | -0.85 | -0.11 | -0.47 | -0.51 | 0.16 | -0.15 | -0.3 | -0.57 | 0.86 | -0.77 | -0.12 | -0.1 | 0.385 | 0.2675 | 1.235 | 0.55616 | 0.473743 | 0.849219 | 0.989223 | 0.997988 | 0.995259 |
| Chd5_E9PYL1_S1633s_1 | Chd5_E9PY | -1.03 | -1.7 | -1.85 | -2.22 | -1.65 | -1.48 | -1.35 | -2.97 | -1.77 | -1.73 | -1.34 | -2.56 | -1.74 | -1.89 | -2.92 | -2.74 | -0.2125 | 0.0575 | 0.24 | 0.140115 | 0.703274 | 0.683163 | 0.995437 | 0.997988 | 0.994666 |
| Chd6_A3KFM7_S2688s_1 | Chd6_A3KF | -1.06 | -1.38 | -4.01 | -1.36 | -0.53 | -1.93 | -1.5 | -2.72 | -0.56 | -0.25 | -2.77 | -3.69 | -0.2 | -0.32 | -2.19 | -2.93 | -0.1625 | -0.15 | -0.6225 | 0.528819 | 0.698574 | 0.888167 | 0.994893 | 0.997988 | 0.994666 |
| Chd8_F7AL76_S659s_1 | Chd8_F7AL | -0.82 | -0.45 | -1.75 | -1.16 | -0.05 | -0.57 | -1.7 | -1.16 | -0.8 | -0.51 | -0.47 | -1.26 | -0.85 | -0.19 | -0.89 | -1.26 | 0.2825 | 0.135 | 0.5425 | 0.511953 | 0.68323 | 0.420845 | 0.994272 | 0.997988 | 0.994666 |
| Chd9_Q8BYH8_S2078s_1 | Chd9_Q8B | 1.4 | 0.98 | 0.59 | 0.89 | -0.23 | 1.14 | -0.23 | 0.78 | 1.44 | 1.03 | -0.58 | 0.69 | 1.26 | -0.28 | 0.93 | 0.87 | 0.175 | 0.285 | 0.2475 | 0.491385 | 0.139087 | 0.482876 | 0.989223 | 0.997988 | 0.994666 |
| Chd9_Q8BYH8_S2734s_1 | Chd9_Q8B | 0.92 | 0.08 | 0 | -0.8 | -2.45 | 1.04 | -3 | -2.49 | 2.4 | -0.19 | -0.16 | -0.27 | 2.31 | -0.79 | 0.22 | -0.69 | -0.6 | -0.32 | -0.27 | 0.766526 | 0.064394 | 0.559574 | 0.989223 | 0.997988 | 0.9971 |
| Chga_P26339_S282s_1 | Chga_P263 | -0.05 | 0.38 | 0.19 | -0.2 | -0.24 | -0.46 | -0.71 | -0.51 | 1.32 | 0.31 | -0.4 | 0.8 | 1.51 | -1.31 | 0.48 | 0.4 | -1.775 | 0.395 | 0.2125 | 0.732013 | 0.035551 | 0.288203 | 0.989223 | 0.997988 | 0.9971 |
| Chga_P26339_S350s_1 | Chga_P263 | -2.35 | -1.79 | -1.26 | -1.14 | -2.11 | -2.62 | -0.91 | -0.84 | -2.19 | -2.26 | -0.5 | -0.99 | -1.28 | -1.56 | -0.16 | -0.77 | -0.56 | 0.4275 | 0.19 | 0.120201 | 0.97513 | 0.760153 | 0.999187 | 0.997988 | 0.994666 |
| Chga_P26339_S414s_1 | Chga_P263 | -1.55 | -1.65 | -1.13 | -3.82 | -3.15 | -1.26 | -3.09 | -1.89 | -0.23 | -1.2 | -1.41 | -1.11 | 0.14 | -2.32 | -0.52 | -1.67 | 0.015 | 0.15 | 0.6925 | 0.216205 | 0.647362 | 0.108021 | 0.99033 | 0.997988 | 0.994666 |
| Chgb_P16014_S100s_1 | Chgb_P160 | 0.41 | 0.48 | 0.47 | 0.6 | 0.89 | 0.76 | 0.47 | 0.72 | 0.52 | 1.1 | 0.57 | 0.59 | 0.55 | 0.51 | 1.03 | 0.9 | -0.31 | 1.05 | 0.945 | 0.316164 | 0.300687 | 0.390771 | 0.989223 | 0.997988 | 0.994666 |
| Chgb_P16014_S129s_1 | Chgb_P160 | -0.55 | -0.72 | -0.41 | -0.41 | -1.42 | -0.42 | -0.86 | -0.48 | 0.72 | -0.49 | -0.68 | -0.22 | 1.15 | -1.51 | 0.02 | 0.07 | 0.22 | 0.205 | 0.2575 | 0.389496 | 0.340228 | 0.310223 | 0.989223 | 0.997988 | 0.994666 |
| Chgb_P16014_S220s_1 | Chgb_P160 | 0.88 | 1.04 | 0.82 | 0.37 | 1.39 | 0.56 | 0.44 | 0.73 | 0.49 | 1.26 | 0.88 | 0.76 | 0.21 | 1.53 | 0.99 | 0.83 | -0.2725 | 0.355 | 0.455 | 0.742513 | 0.993054 | 0.797082 | 0.999187 | 0.997988 | 0.9971 |
| Chgb_P16014_S234s_2 | Chgb_P160 | -2.62 | -1.5 | 0.39 | -0.21 | -3.24 | -1.84 | -0.88 | -0.62 | 0.8 | -2.5 | 1.25 | 0.31 | 0.49 | -2.59 | 1 | 0.51 | 0.0025 | 0.07 | 0.1125 | 0.377818 | 0.409144 | 0.333097 | 0.989223 | 0.997988 | 0.994666 |
| Chgb_P16014_S237s_1 | Chgb_P160 | -1.03 | -0.78 | -0.25 | -1.89 | -2.14 | -0.59 | -1.34 | -1.96 | 1.47 | -0.33 | -0.51 | -0.07 | 2.08 | -2.43 | 0.11 | -0.8 | -0.66 | 0.95 | 0.8375 | 0.415945 | 0.271287 | 0.05719 | 0.989223 | 0.997988 | 0.994666 |
| Chgb_P16014_S237s_2 | Chgb_P160 | -2.69 | -1.49 | 0.19 | -0.38 | -3.02 | -1.65 | -1.08 | -0.76 | 0.88 | -2.45 | 1.38 | 0.12 | 0.48 | -3.6 | 0.84 | 0.35 | -0.52 | 1.1275 | 0.7275 | 0.568882 | 0.459476 | 0.270642 | 0.989223 | 0.997988 | 0.995259 |
| Chgb_P16014_S300s_1 | Chgb_P160 | -1.74 | -1.37 | -0.13 | 0.3 | -1.25 | -1.54 | 0.8 | 0.97 | -0.85 | -1.05 | 0.5 | 0.43 | -0.78 | -0.27 | 1.26 | 0.64 | -0.535 | 1.075 | 0.61 | 0.143738 | 0.511634 | 0.409779 | 0.989223 | 0.997988 | 0.994666 |
| Chgb_P16014_S301s_1 | Chgb_P160 | 0.57 | 0.24 | -0.56 | -0.66 | 0.49 | 0.11 | -0.4 | -0.11 | 0.64 | 0.77 | 0 | -0.75 | 0.31 | 0.65 | 0.47 | -0.17 | 0.48 | 0.4925 | 0.9475 | 0.274575 | 0.725502 | 0.551303 | 0.995437 | 0.997988 | 0.994666 |
| Chgb_P16014_S308s_1 | Chgb_P160 | 0.02 | -0.78 | -1.44 | -1.5 | -0.91 | -0.4 | -0.89 | -0.92 | 0.39 | 0.53 | -0.15 | -0.96 | 0.17 | 0.18 | 0 | -0.48 | 0.125 | 0.2675 | 0.4175 | 0.045584 | 0.696833 | 0.086728 | 0.994893 | 0.997988 | 0.994666 |
| Chgb_P16014_S308s_2 | Chgb_P160 | -1.13 | -0.62 | -2.54 | -0.95 | 0.24 | -0.18 | -0.94 | -1.15 | 0.53 | -0.42 | -1.31 | -1.51 | 0.09 | 0.16 | -0.1 | -1.6 | 0.145 | 0.8775 | 0.8925 | 0.110496 | 0.127004 | 0.291601 | 0.989223 | 0.997988 | 0.994666 |
| Chgb_P16014_S342s_1 | Chgb_P160 | -1.28 | -1.38 | -1.01 | -1 | -1.79 | -1.05 | -1.35 | -1.14 | -0.66 | -0.72 | -0.06 | -0.82 | -1.44 | -1.6 | -0.56 | -0.62 | 0.8025 | 0.6325 | 0.9475 | 0.733044 | 0.512047 | 0.042836 | 0.989223 | 0.997988 | 0.9971 |
| Chgb_P16014_S365s_1 | Chgb_P160 | 0.63 | 0.09 | 0 | 0.08 | -1.13 | 1.53 | -0.34 | 0 | 3.19 | 0.86 | -1.07 | 0.21 | 3.17 | -0.88 | 2.15 | 0.25 | -0.165 | 0.6025 | 0.1125 | 0.251155 | 0.724796 | 0.465896 | 0.995437 | 0.997988 | 0.994666 |
| Chgb_P16014_S378s_1 | Chgb_P160 | -1.38 | -2.14 | -1.41 | -3.94 | -3.24 | -0.81 | -2.09 | -1.62 | 0.74 | -1.08 | -1.65 | -1.34 | 0.44 | -2.93 | -0.23 | -1.46 | -0.185 | 0.5975 | 0.9725 | 0.184836 | 0.690191 | 0.082035 | 0.994672 | 0.997988 | 0.994666 |
| Chgb_P16014_S390s_1 | Chgb_P160 | 5.36 | 1.3 | 2.05 | 0.36 | 3.16 | 5.61 | 2.94 | 2.23 | 0.29 | 5.84 | 0.06 | 1.95 | 2.15 | 1.07 | 2.76 | 3.25 | 0.2775 | 1.385 | 1.1725 | 0.968976 | 0.29679 | 0.877091 | 0.989223 | 0.997988 | 0.999507 |
| Chgb_P16014_S408s_1 | Chgb_P160 | 3.5 | 0.1 | -1.16 | -2.53 | -0.31 | 3.93 | -1.69 | -1.21 | -0.33 | 3.17 | -2.06 | -1.19 | 1.5 | -1.46 | 0.25 | -0.95 | 1.2175 | -0.2325 | 0.04 | 0.909252 | 0.896457 | 0.957628 | 0.996493 | 0.997988 | 0.999507 |
| Chgb_P16014_S428s_1 | Chgb_P160 | -1.33 | -1.01 | -2.42 | -1.97 | -0.07 | -1.2 | -1.49 | -1.56 | -0.65 | -0.49 | -0.57 | -1.58 | 0.29 | 0.06 | -1.06 | -2.4 | 0.2025 | -0.08 | -0.1425 | 0.172591 | 0.192851 | 0.056989 | 0.989223 | 0.997988 | 0.994666 |
| Chgb_P16014_S99s_1 | Chgb_P160 | 0.86 | 0.99 | 0.6 | 0.64 | 0.85 | 0.8 | 0.8 | 0.81 | 0.72 | 1.22 | 0.61 | 1.03 | 0.82 | 0.87 | 1.36 | 0.97 | 0.6025 | 0.86 | 0.905 | 0.376548 | 0.835813 | 0.62119 | 0.995762 | 0.997988 | 0.994666 |
| Chic1_Q8CBW7_S80s_1 | Chic1_Q8C | 0.24 | 0.4 | -0.33 | -0.94 | 0.15 | -0.25 | -2.49 | -0.83 | 1.23 | 0.12 | -0.87 | 0.07 | 1.25 | 1.15 | -3.16 | -0.23 | 0.0425 | 0.1225 | 0.2325 | 0.923376 | 0.253326 | 0.555984 | 0.989223 | 0.997988 | 0.999507 |
| Chm_Q3UR39_S622s_1 | Chm_Q3UF | -0.17 | 0.47 | -0.87 | -0.5 | -0.15 | -0.38 | -0.64 | -0.95 | 0.64 | 0.26 | -0.38 | -0.54 | 0.21 | 0.99 | -1 | -1.06 | -0.6975 | 0.295 | -0.09 | 0.92157 | 0.446111 | 0.511796 | 0.989223 | 0.997988 | 0.999507 |
| Chmp7_Q8R1T1_S232s_1 | Chmp7_Q8 | -0.53 | -0.44 | 0.22 | 0.22 | -0.02 | -0.24 | 0.24 | 0.32 | -0.56 | -0.24 | 0.19 | 0.38 | -0.46 | 0.26 | 0.17 | 0.23 | -0.2625 | 0.2625 | 0.0525 | 0.567319 | 0.462239 | 0.816103 | 0.989223 | 0.997988 | 0.995259 |
| Chrm1_P12657_S244s_1 | Chrm1_P12 | -2.56 | -0.06 | 1.68 | 1.75 | -0.81 | -1.71 | -0.24 | 1.32 | -0.72 | -1.18 | 0.12 | 1.53 | 0.55 | -1.48 | 0.93 | -0.65 | 0.2075 | 0.075 | 0.1825 | 0.716826 | 0.588668 | 0.798018 | 0.989223 | 0.997988 | 0.996966 |
| Chrm1_P12657_S251s_1 | Chrm1_P12 | -2.71 | 0.22 | -1.04 | -1.01 | -1.33 | -2.02 | -0.26 | -2.29 | -0.09 | -2.11 | -0.4 | -3.53 | 0.05 | -0.99 | -0.11 | 0.42 | -0.5625 | -0.265 | -0.365 | 0.134707 | 0.612209 | 0.656216 | 0.989223 | 0.997988 | 0.994666 |
| Chrm1_P12657_S254s_1 | Chrm1_P12 | -0.82 | -0.18 | -0.94 | -1.71 | -0.72 | -1.03 | -1.03 | -1.49 | -0.21 | -1.05 | -0.33 | -0.85 | 0.33 | -0.56 | -0.83 | -0.46 | -0.34 | -0.3975 | 0.9775 | 0.20944 | 0.661799 | 0.432052 | 0.990434 | 0.997988 | 0.994666 |
| Chrm1_P12657_S451s_1 | Chrm1_P12 | -0.61 | -2.01 | -0.16 | -0.06 | -0.68 | 0.31 | -0.52 | -0.79 | 1.33 | -0.69 | -3.61 | 0.05 | 1.61 | -1.94 | -0.15 | -0.66 | -0.155 | 0.3025 | 0.5325 | 0.581156 | 0.545718 | 0.984111 | 0.989223 | 0.997988 | 0.995259 |
| Chrm2_Q9ERZ4_S232s_1 | Chrm2_Q9 | 1.31 | 0.65 | 0.07 | -0.25 | 0.12 | 0.22 | 0.14 | 0.43 | 0.75 | 1.47 | 1.15 | 1.09 | -0.03 | 0.8 | 0.47 | 0.47 | 0.29 | -0.02 | 0.425 | 0.964448 | 0.540594 | 0.103103 | 0.989223 | 0.997988 | 0.999507 |
| Chrm3_Q9ERZ3_S288s_1 | Chrm3_Q9 | 0.41 | 0.83 | -0.57 | 0.71 | 0.54 | 0.05 | -0.19 | -0.2 | -0.13 | 1.08 | 0.48 | -0.01 | 0.75 | 0.57 | -0.01 | -0.11 | -0.2175 | 0.67 | -0.0175 | 0.908496 | 0.421016 | 0.980688 | 0.989223 | 0.997988 | 0.999507 |
| Chrm3_Q9ERZ3_S384s_1 | Chrm3_Q9 | 0.02 | 0.84 | 0.75 | 0.16 | 0.63 | 0.08 | 0.51 | -0.15 | -0.1 | 0.62 | -0.4 | 0.51 | 0.05 | 1.02 | 0.47 | 0.04 | -0.295 | 0.01 | -0.045 | 0.889928 | 0.563195 | 0.410659 | 0.989223 | 0.997988 | 0.999507 |
| Chrm3_Q9ERZ3_S472s_1 | Chrm3_Q9 | -1.27 | -0.72 | -0.1 | -0.96 | -1.32 | -0.12 | -0.76 | -1.49 | 0.37 | -0.1 | -4.57 | -0.8 | 1.64 | -3.06 | -0.57 | -1.11 | -0.175 | -0.285 | -0.0475 | 0.988594 | 0.679394 | 0.616184 | 0.993464 | 0.997988 | 1 |
| Chrm3_Q9ERZ3_S577s_1 | Chrm3_Q9 | -0.47 | -1 | -0.35 | -0.6 | -0.57 | -0.55 | -0.39 | -0.51 | -1.16 | -0.46 | -0.31 | -1.69 | -1.01 | -0.66 | -0.1 | 0.08 | -0.16 | -0.5125 | -0.0125 | 0.582865 | 0.660179 | 0.415519 | 0.990434 | 0.997988 | 0.995259 |
| Chrm4_P32211_S262s_1 | Chrm4_P32 | 1.1 | 0.05 | 0.8 | 0.74 | 0.3 | 0.77 | 0.46 | 1.05 | 0.54 | 0.68 | 2.41 | 1.03 | 0.38 | 0.58 | 0.31 | 0.01 | 0.1 | -0.3 | 0.1825 | 0.268347 | 0.927239 | 0.300978 | 0.997335 | 0.997988 | 0.994666 |
| Chrm4_P32211_T320t_1 | Chrm4_P32 | 1 | 0.91 | 0.57 | 0.96 | 1.97 | 1.53 | 0.92 | 1.26 | 0.43 | 1.41 | 0.58 | 1.59 | -0.57 | 1.55 | 0.63 | 1.14 | -0.0275 | 0.4925 | -0.3525 | 0.7059 | 0.069209 | 0.668396 | 0.989223 | 0.997988 | 0.996324 |
| Chrna4_O70174_S468s_1 | Chrna4_O7 | 0.26 | 0.07 | -0.03 | 0.3 | 0.31 | 0.23 | 0.46 | 0.04 | 0.12 | 0.19 | -0.62 | 0.16 | 0.01 | 0.28 | 0.42 | 0.55 | 0.56 | 0.1425 | -0.1725 | 0.516659 | 0.60917 | 0.4917 | 0.989223 | 0.997988 | 0.994666 |
| Chrna4_O70174_S491s_1 | Chrna4_O7 | -0.87 | -0.8 | 0.51 | 0.25 | -0.17 | -0.66 | 0.22 | 0.65 | -1.22 | -0.56 | -0.09 | -0.45 | -1.1 | -0.7 | -0.17 | -0.59 | 0.11 | -0.1875 | 0.165 | 0.324181 | 0.5811 | 0.408565 | 0.989223 | 0.997988 | 0.994666 |
| Chrna4_O70174_S543s_1 | Chrna4_O7 | -0.23 | 0.12 | 0.62 | 0.24 | 1.09 | 0.09 | 0.9 | 0.35 | 0.11 | 0.11 | -0.72 | 0.55 | -0.54 | 0.3 | 0.02 | 0.18 | 0.2375 | -0.3525 | -0.4125 | 0.528121 | 0.198081 | 0.607647 | 0.989223 | 0.997988 | 0.994666 |
| Chrnb2_Q9ERK7_S395s_1 | Chrnb2_Q9 | -0.06 | 0.27 | 0.75 | 0.34 | 0.34 | -0.58 | 0.72 | 0.91 | -0.14 | 0.14 | 0.17 | 0.32 | 0.08 | -0.4 | 0.4 | 0.76 | 0.42 | -0.175 | -0.1975 | 0.732218 | 0.950985 | 0.442202 | 0.999187 | 0.997988 | 0.9971 |
| Chst15_Q91XQ5_S185s_1 | Chst15_Q9 | 0.51 | 0.78 | 0.68 | 0.56 | -0.91 | 0.95 | -0.2 | -0.39 | 2.8 | 0.53 | -1.7 | 0.89 | 3.01 | -0.82 | 0.55 | 0.1 | 0.0225 | -0.2025 | -0.115 | 0.915437 | 0.070685 | 0.997577 | 0.989223 | 0.999531 | 0.999507 |
| Cic_Q924A2_S1343s_1 | Cic_Q924A | -1.12 | -0.88 | -0.99 | -1.04 | -1.77 | -0.81 | -3.04 | -1.38 | 0.54 | 0.02 | -0.19 | -1.73 | 1.2 | -0.96 | -1.59 | -1.48 | -0.77 | -0.0025 | 0.0775 | 0.617036 | 0.120889 | 0.177461 | 0.989223 | 0.997988 | 0.995259 |
| Cic_Q924A2_S1809s_1 | Cic_Q924A | 0.14 | 0.34 | 0.05 | 0.35 | 0.52 | -0.53 | 0.49 | 0.17 | 0.28 | 0.83 | 0.4 | 0.56 | -0.17 | 1.08 | -0.32 | -0.55 | -0.7425 | 0.6675 | 0.3 | 0.588659 | 0.841225 | 0.222648 | 0.995932 | 0.997988 | 0.995259 |
| Cic_Q924A2_S2148s_1 | Cic_Q924A | -2.4 | -1.98 | -2.78 | -2.47 | -2.25 | -3.23 | -2.45 | -2.97 | -2.11 | -2.11 | -1.45 | -2.93 | -0.97 | -1.44 | -2.91 | -2.48 | -0.0575 | 0.2975 | -0.21 | 0.33514 | 0.30929 | 0.477664 | 0.989223 | 0.997988 | 0.994666 |
| Cic_Q924A2_S2195s_1 | Cic_Q924A | -0.19 | -0.12 | 0.34 | 0.55 | -0.96 | -0.54 | -0.16 | 0.48 | 0.17 | 0.12 | 0.45 | 0.52 | 0.15 | 0.03 | 0 | 0.81 | -0.3175 | 0.2575 | 0.4575 | 0.743978 | 0.233862 | 0.527741 | 0.989223 | 0.997988 | 0.9971 |
| Cisd3_Z4YKM2_S71s_1 | Cisd3_Z4YK | 0.79 | 0.44 | 0.5 | 0.3 | 1.09 | 0.51 | 0.28 | 0.81 | 0.59 | 0.89 | 0.12 | 0.42 | 0.49 | 1.56 | 0.33 | 0.74 | -0.44 | 0.17 | 0.1025 | 0.420377 | 0.52425 | 0.992277 | 0.989223 | 0.997988 | 0.994666 |
| Cit_E9QL53_S1348s_1 | Cit_E9QL5 | -0.24 | 0.2 | -0.21 | -0.13 | 0.45 | -0.06 | 0.16 | 0.12 | 0.58 | 0.15 | -0.47 | -0.49 | -0.06 | 0.19 | -0.39 | 0.15 | 0.165 | -0.0025 | 0.2725 | 0.797698 | 0.262811 | 0.904145 | 0.989223 | 0.997988 | 0.999507 |
| Cit_E9QL53_S1458s_1 | Cit_E9QL5 | 1.35 | 0.96 | 0.18 | 0.8 | 0.66 | 1.86 | 0.34 | 0.13 | 1.59 | 0.89 | 0.93 | 0.68 | 1.21 | 0.5 | -0.09 | 0.24 | 0.2625 | 0.0375 | 0.0675 | 0.361002 | 0.861799 | 0.554403 | 0.996493 | 0.997988 | 0.994666 |
| Cit_E9QL53_S1608s_1 | Cit_E9QL5 | 0.36 | 0.36 | 0.44 | 0.54 | 0.89 | -0.02 | 0.59 | 0.42 | -0.69 | -0.71 | -0.83 | -0.02 | -0.25 | 1.08 | 0.6 | 0.95 | -0.075 | 0.2 | -0.3575 | 0.617825 | 0.857648 | 0.003607 | 0.996349 | 0.997988 | 0.995259 |
| Cit_E9QL53_S1883s_1 | Cit_E9QL5 | 0.62 | 0.65 | 1.06 | 0.47 | 0.56 | 0.9 | 0.39 | 0.14 | 0.94 | 0.4 | -0.19 | 0.72 | 1.09 | 0.34 | 0.23 | 0.37 | 0.045 | -0.9875 | 0.17 | 0.521761 | 0.436287 | 0.460573 | 0.989223 | 0.997988 | 0.994666 |
| Cit_E9QL53_S1966s_1 | Cit_E9QL5 | 0.4 | -0.36 | -0.47 | -0.28 | 0.21 | 0.24 | -0.38 | -0.37 | 0.78 | 0.31 | 0.08 | -0.34 | 0.55 | -0.46 | -0.93 | -0.66 | -0.2025 | -0.2325 | -0.1925 | 0.614079 | 0.725638 | 0.260788 | 0.995437 | 0.997988 | 0.995259 |
| Cit_E9QL53_S479s_1 | Cit_E9QL5 | -0.19 | -0.06 | -0.43 | -0.43 | 0.71 | -0.73 | 0.24 | 0.26 | -0.38 | -0.14 | -0.65 | -0.34 | -0.56 | 0.27 | -1.08 | -0.69 | 0.1025 | 0.385 | -0.1975 | 0.485688 | 0.244336 | 0.669052 | 0.989223 | 0.997988 | 0.994666 |
| Cit_E9QL53_S698s_1 | Cit_E9QL5 | 7.49 | 3.43 | 5.78 | -1.4 | 3.9 | 8.12 | 5.16 | 4.74 | 0.41 | 6.72 | -4.63 | 5.84 | 1.39 | -4.04 | 7.75 | 5.86 | 0.3975 | -0.1 | -0.2375 | 0.696776 | 0.375274 | 0.543595 | 0.989223 | 0.997988 | 0.996159 |
| Ciz1_F7AT58_S192s_1 | Ciz1_F7AT | 1.26 | -0.34 | -0.42 | 0.76 | 1.53 | -1.32 | 0.61 | -0.8 | 0.37 | -0.87 | 1.15 | -0.36 | 0.02 | 1.64 | -0.95 | 1.14 | 1.655 | -1.74 | -1.085 | 0.818933 | 0.651157 | 0.665864 | 0.990358 | 0.997988 | 0.999507 |
| Ckap4_Q8BMK4_S367s_1 | Ckap4_Q8E | 0.49 | 0.39 | -0.15 | -0.09 | 0.77 | 0.5 | 0.36 | 0.07 | 0.55 | 0.99 | 0.39 | 0.08 | 0.26 | 0.97 | 0.64 | 0.79 | -0.31 | -0.2425 | 0.1475 | 0.108015 | 0.329997 | 0.261158 | 0.989223 | 0.997988 | 0.994666 |
| Ckap4_Q8BMK4_S68s_1 | Ckap4_Q8E | -0.93 | -1.71 | -1.31 | -1.95 | -2.01 | -0.23 | -0.97 | -1.9 | -0.15 | -0.23 | -0.21 | -1.62 | 1.37 | -2.86 | -1.48 | -1.2 | 0.265 | 0.3425 | 0.505 | 0.593244 | 0.659281 | 0.048895 | 0.990434 | 0.997988 | 0.995259 |
| Ckap5_Z4YL78_S816s_1 | Ckap5_Z4Y | -0.03 | 0.03 | 0.54 | 0.5 | -0.18 | -0.1 | 0.33 | 0.38 | 0.3 | 0.35 | 0.44 | -0.15 | 0.84 | 0.27 | 1.01 | 0.82 | 0.1975 | 0.9225 | 0.4325 | 0.126298 | 0.559173 | 0.92437 | 0.989223 | 0.997988 | 0.994666 |
| Ckb_Q04447_S199s_1 | Ckb_Q044 | -0.27 | 0.18 | -0.09 | -0.26 | 0.02 | -0.25 | -0.06 | -0.25 | -0.11 | -0.1 | -0.65 | -0.2 | -0.33 | 0.14 | -0.67 | -0.29 | -0.1525 | -0.025 | 0.475 | 0.526696 | 0.908211 | 0.533252 | 0.997335 | 0.997988 | 0.994666 |
| Ckb_Q04447_T322t_1 | Ckb_Q044 | -0.7 | 0.39 | 0.19 | 0.34 | -0.11 | -0.9 | 0.37 | -0.21 | -0.84 | -0.31 | 0.59 | 0.01 | -0.51 | -0.02 | 0.09 | 0.18 | -0.025 | -0.155 | -0.1775 | 0.72156 | 0.467499 | 0.626488 | 0.989223 | 0.997988 | 0.9971 |
| Ckb_Q04447_T35t_1 | Ckb_Q044 | -0.68 | -0.38 | -0.08 | -0.9 | 0.18 | -0.73 | 0.03 | -0.67 | -0.62 | -0.51 | -0.88 | -1.03 | -0.49 | 0.34 | -0.58 | -0.43 | -0.2675 | -0.1925 | -0.12 | 0.501036 | 0.502469 | 0.369012 | 0.989223 | 0.997988 | 0.994666 |
| Ckmt1_P30275_S319s_1 | Ckmt1_P30 | 0.07 | 0.16 | -0.5 | -0.24 | 0.07 | 0.09 | -0.15 | -0.58 | -0.32 | -0.45 | -0.39 | -0.46 | -0.89 | -0.15 | -0.31 | -0.59 | 0.2125 | -0.25 | 0.22 | 0.237653 | 0.954554 | 0.264641 | 0.999187 | 0.997988 | 0.994666 |
| Clasp1_E9Q6L0_S676s_1 | Clasp1_E9 | 0.52 | -2.22 | -0.31 | -0.09 | -1.76 | 0.57 | -0.63 | -0.2 | 1.62 | -0.15 | -0.76 | -0.06 | 0.79 | -1.6 | -1.24 | -0.43 | -0.015 | -0.2775 | -0.3575 | 0.893413 | 0.97641 | 0.341905 | 0.999187 | 0.997988 | 0.999507 |
| Clasp1_E9QKH0_S1111s_2 | Clasp1_E9 | -0.77 | -0.02 | -0.11 | 0.1 | -0.43 | -0.71 | -0.05 | 0.26 | 0.2 | 0.35 | 0.3 | -0.49 | -0.43 | -0.12 | -0.2 | -1.26 | 0.02 | 0.6875 | -0.095 | 0.401662 | 0.916073 | 0.361675 | 0.997335 | 0.997988 | 0.994666 |
| Clasp1_E9QNY9_S1118s_1 | Clasp1_E9 | -0.78 | -0.23 | -0.67 | -1.13 | -0.29 | -0.86 | -1.1 | -1.37 | -1.29 | -0.59 | -2.48 | -0.66 | -0.19 | -2.47 | -0.5 | -0.69 | -0.0325 | 0.29 | -0.3025 | 0.614449 | 0.522747 | 0.244925 | 0.989223 | 0.997988 | 0.995259 |
| Clasp1_E9QNY9_S1184s_1 | Clasp1_E9 | 1.02 | 0.88 | -2.67 | -1.78 | 0.48 | 1.56 | -0.82 | -1.18 | 1.31 | 0.82 | -0.76 | -0.98 | -0.8 | 0.68 | -0.81 | -1.86 | -0.2025 | -0.5525 | -0.26 | 0.948812 | 0.511448 | 0.454741 | 0.989223 | 0.997988 | 0.999507 |
| Clasp1_E9QNY9_S1211s_1 | Clasp1_E9 | 0.65 | 0.05 | 0.54 | 0.14 | -0.59 | 0.98 | -0.76 | -0.19 | 1.48 | 0.57 | 0.16 | 0.77 | 1.02 | -2.42 | 0.49 | 0.52 | 0.6475 | 0.735 | -0.06 | 0.5384 | 0.246195 | 0.250921 | 0.989223 | 0.997988 | 0.994666 |
| Clasp1_E9QNY9_S572s_1 | Clasp1_E9 | -0.33 | -0.12 | -0.07 | -1.1 | -0.03 | -0.3 | 0.08 | -0.13 | 2.97 | -0.25 | -1.51 | -0.05 | 3.36 | -0.78 | 1.83 | -0.6 | -0.485 | 0.4 | -0.4425 | 0.155251 | 0.29178 | 0.43277 | 0.989223 | 0.997988 | 0.994666 |
| Clasp1_E9QNY9_S594s_1 | Clasp1_E9 | 0.66 | -0.33 | -0.09 | -0.19 | 0.14 | 0.42 | 0.25 | 0.35 | -0.56 | 0.22 | 1.03 | 0.65 | 0.4 | 0.78 | 0.26 | 0.07 | 0.31 | 0.695 | 1.3575 | 0.266487 | 0.319802 | 0.432168 | 0.989223 | 0.997988 | 0.994666 |
| Clasp1_E9QNY9_S598s_1 | Clasp1_E9 | -0.99 | -1.26 | -1.09 | -0.86 | -0.65 | -0.79 | -0.8 | -0.81 | -1.17 | -0.75 | -0.06 | -0.63 | -0.11 | -0.38 | -0.8 | -0.67 | 0.2775 | 0.3225 | 0.365 | 0.059618 | 0.182672 | 0.192291 | 0.989223 | 0.997988 | 0.994666 |
| Clasp1_E9QNY9_S600s_1 | Clasp1_E9 | -0.94 | -1.09 | -0.82 | -0.42 | -0.48 | -0.66 | -0.36 | -0.17 | -0.75 | -0.66 | 0.48 | -0.11 | -0.09 | -0.25 | -0.49 | -0.94 | 0.2875 | 0.3975 | 0.56 | 0.227065 | 0.122235 | 0.125783 | 0.989223 | 0.997988 | 0.994666 |
| Clasp1_E9QNY9_S620s_1 | Clasp1_E9 | 0.17 | 0.85 | 1.02 | 1.21 | 1.23 | 0.97 | 1.21 | 0.98 | 0.14 | 0.84 | -0.53 | 0.95 | 0.27 | -0.14 | 0.19 | 1.47 | 0.4 | 0.5575 | 0.375 | 0.392365 | 0.315419 | 0.273208 | 0.989223 | 0.997988 | 0.994666 |
| Clasp1_E9QNY9_S646s_1 | Clasp1_E9 | -0.82 | -0.74 | -1.21 | -0.69 | -1.69 | 0.19 | -1.49 | -0.94 | 0.18 | -0.48 | -1.31 | -0.33 | 0.5 | -0.98 | -0.64 | -0.96 | 0.285 | -0.4625 | -0.365 | 0.377436 | 0.778202 | 0.290316 | 0.995762 | 0.997988 | 0.994666 |
| Clasp1_E9QNY9_S646s_2 | Clasp1_E9 | -1.14 | -0.76 | -1.07 | -0.54 | -1.72 | -1.3 | -0.96 | -0.45 | 0.69 | -2.04 | -0.76 | -0.4 | -0.82 | 0.01 | -0.72 | -0.7 | -0.1175 | 0.38 | 0.345 | 0.299196 | 0.476168 | 0.644583 | 0.989223 | 0.997988 | 0.994666 |
| Clasp1_E9QNY9_S647s_2 | Clasp1_E9 | -1.14 | -0.76 | -1.19 | -1.53 | -1.72 | -1.3 | -1.21 | -0.58 | 0.69 | -2.04 | -0.66 | -1.05 | -0.82 | 0.01 | -1.37 | -1.52 | -0.23 | 0.25 | 0.32 | 0.559632 | 0.876278 | 0.480303 | 0.996493 | 0.997988 | 0.995259 |
| Clasp1_E9QNY9_S685s_1 | Clasp1_E9 | -0.1 | -1.4 | -0.65 | -1.12 | -1.28 | 0.37 | -0.96 | -0.94 | 1.7 | -1.01 | -1.47 | -0.03 | 0.95 | -1.46 | -0.59 | -0.35 | -0.0475 | 0.39 | 0.23 | 0.404129 | 0.792276 | 0.379699 | 0.995762 | 0.997988 | 0.994666 |
| Clasp1_E9QNY9_S779s_1 | Clasp1_E9 | -0.59 | -1.48 | -2.03 | -1.55 | -0.27 | -1.22 | -2.34 | -2.17 | 0.18 | -0.07 | -0.18 | -1.31 | 0.01 | -0.24 | -1.42 | -2.18 | 0.115 | 0.615 | 0.455 | 0.41733 | 0.865068 | 0.033169 | 0.996493 | 0.997988 | 0.994666 |
| Clasp2_F7DCH5_S1294s_1 | Clasp2_F7D | 0.05 | 0.67 | 0.95 | 0.39 | 1.1 | 0.67 | 0.11 | 0.53 | 0.24 | 0.81 | 2.33 | 0.51 | 1 | 0.91 | 0.61 | 0.55 | -0.0875 | 1.0675 | 0.455 | 0.394879 | 0.773383 | 0.352552 | 0.995762 | 0.997988 | 0.994666 |
| Clasp2_F7DCH5_S1307s_1 | Clasp2_F7D | -0.73 | 0.4 | 1.01 | 0.65 | -1.27 | 0.43 | 1.03 | 1.37 | 0.5 | -0.05 | 0.47 | 1.32 | 0.7 | -0.79 | 1.45 | 1 | 0.0875 | 0.4575 | 0.2525 | 0.650546 | 0.926116 | 0.617182 | 0.997335 | 0.997988 | 0.995259 |
| Clasp2_F7DCH5_S14s_1 | Clasp2_F7D | -0.06 | 0.15 | -0.55 | -0.76 | -0.21 | 0.06 | 0.08 | -0.04 | -0.73 | 0.29 | -1.54 | -0.39 | -0.81 | -1.18 | -0.41 | -0.27 | 0.0575 | 0.2275 | 0.2575 | 0.292609 | 0.311157 | 0.503984 | 0.989223 | 0.997988 | 0.994666 |
| Clasp2_F7DCH5_S370s_1 | Clasp2_F7D | -0.12 | 0.25 | 0.62 | 0.03 | 0.27 | 0.58 | 0.6 | 0.01 | -0.58 | -0.09 | 0.12 | 0.38 | -0.36 | 0.63 | -0.7 | 0.6 | 0.2775 | -0.2875 | -0.3625 | 0.694106 | 0.520324 | 0.435379 | 0.989223 | 0.997988 | 0.996159 |
| Clasp2_F7DCH5_S374s_1 | Clasp2_F7D | -1.36 | -1.09 | -0.77 | -0.72 | -0.76 | -1.28 | -0.65 | -0.76 | -1.28 | -0.82 | -0.07 | -0.5 | -0.3 | -0.61 | -0.87 | -0.71 | 0.17 | -0.2375 | -0.1525 | 0.208122 | 0.635692 | 0.339335 | 0.989223 | 0.997988 | 0.994666 |
| Clasp2_F7DCH5_S376s_1 | Clasp2_F7D | -0.7 | -1.01 | -0.61 | -0.66 | -0.56 | -1.1 | -0.49 | -0.72 | -0.98 | -0.86 | -0.06 | -0.51 | -0.15 | -0.17 | -0.66 | -0.52 | 0.1225 | 0.3175 | 0.3625 | 0.174904 | 0.906536 | 0.610921 | 0.996975 | 0.997988 | 0.994666 |
| Clasp2_F7DCH5_S413s_1 | Clasp2_F7D | -0.02 | -0.09 | 0.14 | -1.45 | 0 | -0.63 | 0.12 | -0.44 | -1.61 | -0.25 | -0.04 | -0.71 | -1.17 | 0.55 | 0.04 | -0.17 | 0.0275 | 0.1425 | 0.37 | 0.733022 | 0.766279 | 0.540373 | 0.995762 | 0.997988 | 0.9971 |
| Clasp2_F7DCH5_S426s_1 | Clasp2_F7D | -3.34 | -2.12 | -1.39 | -4.41 | -2.73 | -2.17 | -1.74 | -2.84 | -0.65 | -3.04 | -0.65 | -1.33 | 0.26 | -2.38 | -1.85 | -1.61 | 0.1175 | -0.2975 | 0.1675 | 0.093918 | 0.48915 | 0.100066 | 0.989223 | 0.997988 | 0.994666 |
| Clasp2_F7DCH5_S461s_1 | Clasp2_F7D | -2.26 | -2.09 | -1.93 | -2.62 | -1.53 | -2.08 | -1.79 | -2.15 | -1.6 | -1.38 | -1.01 | -1.79 | 0.02 | -2.01 | -1.78 | -1.77 | 0.445 | 1.3975 | 1.42 | 0.101903 | 0.209454 | 0.018357 | 0.989223 | 0.997988 | 0.994666 |
| Clasp2_F7DCH5_S465s_1 | Clasp2_F7D | -1.08 | -1.58 | -0.64 | -0.65 | -1.5 | -0.67 | -0.91 | -1.08 | 0.42 | -1.19 | -0.31 | -0.13 | 0.89 | -1.62 | -1.05 | -0.77 | 0.3375 | 0.78 | 0.84 | 0.522959 | 0.863324 | 0.11264 | 0.996493 | 0.997988 | 0.994666 |
| Clasp2_F7DCH5_S531s_1 | Clasp2_F7D | -2.62 | -2.66 | -3.47 | -4.81 | -1.67 | -1.46 | -2.6 | -4.47 | -0.52 | -2.68 | -2.93 | -3.33 | -0.09 | -1.83 | -2.79 | -2.98 | -0.0525 | 0.685 | 0.35 | 0.073156 | 0.279744 | 0.183214 | 0.989223 | 0.997988 | 0.994666 |
| Clasp2_F7DCH5_S535s_1 | Clasp2_F7D | -0.02 | -0.37 | -0.17 | -0.11 | -1.04 | 0.61 | -0.78 | 0.02 | 1.43 | -0.36 | 0.33 | 0.3 | 0.45 | -0.09 | -0.94 | -0.99 | 0.84 | 1.025 | 1.4675 | 0.551437 | 0.730626 | 0.146071 | 0.995762 | 0.997988 | 0.994666 |
| Clasp2_F7DCH5_S591s_1 | Clasp2_F7D | -3.58 | -4.04 | -2.3 | -2.5 | -1.9 | -4.88 | -1.9 | -2.01 | -6.02 | -4.41 | -0.62 | -2.31 | -2.97 | -1.53 | -2.34 | -2.15 | -0.13 | 0.5925 | -0.225 | 0.104914 | 0.564412 | 0.831451 | 0.989223 | 0.997988 | 0.994666 |

| | | | | | | | | | | | | | | | | | | | | | | | | | | |
|---|---|---|---|---|---|---|---|---|---|---|---|---|---|---|---|---|---|---|---|---|---|---|---|---|---|---|
| Clasp2_F7DCH5_S635s_1 | Clasp2_F7D | -0.32 | -0.44 | -0.68 | -0.31 | -0.65 | -0.51 | -0.05 | 0.03 | -0.13 | 0.16 | 0.13 | 0.02 | -0.35 | 0.13 | -0.33 | -0.39 | 0.4325 | -0.235 | 0.8575 | 0.435991 | 0.568101 | 0.051644 | 0.989223 | 0.997988 | 0.994666 |
| Clasp2_F7DCH5_S641s_1 | Clasp2_F7D | -2.09 | -2.07 | -0.53 | -1.4 | -1.25 | -1.43 | -0.13 | -1.9 | -0.6 | -2.74 | 0.6 | -2.32 | -0.73 | -0.07 | -1.45 | -0.69 | 0.1425 | 0.4825 | 0.2025 | 0.106204 | 0.481917 | 0.738018 | 0.989223 | 0.997988 | 0.994666 |
| Clasp2_F7DCH5_S905s_1 | Clasp2_F7D | -0.51 | -0.97 | -0.67 | -0.44 | -0.54 | -0.76 | -0.49 | -0.8 | -0.25 | -0.55 | 0.07 | 0.38 | -0.01 | -1.13 | -0.22 | 0.32 | 0.345 | 0.2575 | 0.7875 | 0.294246 | 1 | 0.07297 | 1 | 0.997988 | 0.994666 |
| Clasp2_F7DCH5_S968s_1 | Clasp2_F7D | 0.67 | -0.23 | 0.07 | 0.46 | 0.31 | 0.95 | -0.02 | 0.4 | 0.13 | 0.01 | 1.26 | 0.65 | -0.64 | 0.56 | -0.4 | -0.69 | 9.81E-17 | 0.56 | 0.3875 | 0.173538 | 0.586817 | 0.460642 | 0.989223 | 0.997988 | 0.994666 |
| Clasp2_F7DCH5_T564t_1 | Clasp2_F7D | -0.26 | -1.01 | -0.16 | 0.67 | -0.91 | -0.46 | 0.98 | -0.54 | -0.7 | -0.48 | 0.61 | 0.12 | -1.11 | -0.17 | -0.24 | 0.63 | 0.1675 | 0.27 | -0.535 | 0.945301 | 0.931545 | 0.860008 | 0.998293 | 0.997988 | 0.999507 |
| Clasp2_Q8BRT1_S8s_1 | Clasp2_Q8 | -0.06 | 0.15 | 0.85 | 0.85 | -0.21 | 0.06 | 1.13 | 1.44 | -0.73 | 0.29 | 0.14 | 1.04 | -0.81 | -1.18 | 0.66 | 1.08 | -0.0425 | 0.0775 | -0.0325 | 0.368511 | 0.719827 | 0.539547 | 0.995437 | 0.997988 | 0.994666 |
| Clasrp_Q8CFC7_S294s_1 | Clasrp_Q8 | 1.74 | 0.9 | 0.44 | -0.01 | 0.24 | 1.56 | 0.07 | 0.13 | 1.03 | 0.42 | 0.03 | 0.27 | 1.09 | 1.08 | 0.04 | -0.2 | 0.1575 | -0.2625 | -0.51 | 0.585427 | 0.577484 | 0.441173 | 0.989223 | 0.997988 | 0.995259 |
| Clasrp_Q8CFC7_S541s_1 | Clasrp_Q8 | 1.78 | 1.15 | 0.66 | 0.53 | 0.53 | 0.62 | 0.02 | 0.91 | 0.9 | 0.31 | 0.11 | -0.05 | 1.08 | 0.9 | 0.51 | -0.12 | -0.2675 | -0.33 | -0.265 | 0.285572 | 0.161188 | 0.074785 | 0.989223 | 0.997988 | 0.994666 |
| Clcn6_O35454_S684s_1 | Clcn6_O35 | -0.62 | -0.11 | 0.57 | 0.27 | 0.42 | -0.7 | 0.21 | 0.4 | -0.68 | -1 | -0.01 | 0.24 | -0.47 | 0.3 | 0.38 | 0.62 | -0.51 | -0.7125 | -0.4375 | 0.627705 | 0.88028 | 0.328003 | 0.996493 | 0.997988 | 0.995259 |
| Cldn10_Q9Z0S6_S147s_1 | Cldn10_Q9 | -0.02 | 0.09 | 1.07 | -0.64 | -0.86 | -0.24 | 0.9 | -0.07 | 0.73 | -0.1 | -1.49 | -0.37 | 0.79 | 0.61 | 0.36 | 0.12 | 0.055 | -0.39 | 0.18 | 0.389219 | 0.683495 | 0.430138 | 0.994272 | 0.997988 | 0.994666 |
| Cldn11_Q60771_S194s_1 | Cldn11_Q6 | 0.06 | 0.1 | -0.33 | 0.37 | 0.15 | -0.49 | -1.42 | -0.77 | -0.51 | -0.26 | -1.12 | -0.5 | 0.17 | -0.9 | -0.55 | -0.02 | -0.1925 | -0.4325 | 0.345 | 0.268471 | 0.080346 | 0.043961 | 0.989223 | 0.997988 | 0.994666 |
| Cldn11_Q60771_S196s_1 | Cldn11_Q6 | -1.05 | -1.04 | 0.16 | -1.16 | -0.54 | -1.72 | -1.91 | -0.84 | -1.09 | -0.95 | -2.98 | -0.72 | 0.24 | -1.52 | -0.06 | -0.55 | -0.6825 | -0.6475 | -0.375 | 0.532034 | 0.281039 | 0.256077 | 0.989223 | 0.997988 | 0.994666 |
| Cldn11_Q60771_S197s_1 | Cldn11_Q6 | -1.28 | -0.35 | 0.77 | 1.24 | -0.45 | -1.71 | -0.67 | -0.14 | -1.25 | -0.98 | 5.78 | -0.12 | 0.08 | -1.62 | 0.18 | 0.45 | -0.48 | -0.6625 | 0.3 | 0.628792 | 0.17985 | 0.620369 | 0.989223 | 0.997988 | 0.995259 |
| Cldn11_Q60771_S198s_1 | Cldn11_Q6 | -0.28 | -0.12 | -0.4 | -0.36 | -0.18 | -0.9 | -0.62 | -1.03 | -0.23 | -0.38 | 0.38 | -0.67 | 0.68 | -0.23 | -0.28 | -0.39 | -0.8375 | 0.7625 | -0.3225 | 0.454597 | 0.144653 | 0.817113 | 0.989223 | 0.997988 | 0.994666 |
| Cldn12_Q9ET43_S228s_1 | Cldn12_Q9 | -1.05 | -1.33 | -2.82 | -0.39 | -0.73 | -0.83 | -1.51 | 0.2 | 0.06 | -1.29 | -2.9 | -0.76 | -1.95 | -1.08 | -2.14 | -2.43 | -0.3925 | 0.065 | 0.235 | 0.3695 | 0.243931 | 0.809831 | 0.989223 | 0.997988 | 0.994666 |
| Cldn5_O54942_T207t_1 | Cldn5_O54 | -1.34 | -0.59 | 0.4 | 0 | -1.03 | -0.88 | -0.81 | -0.28 | 1.17 | -0.47 | -0.22 | 0.98 | 2.01 | -1.63 | 0.42 | 0.42 | 0.68 | 0.175 | -0.5025 | 0.366395 | 0.366018 | 0.17762 | 0.989223 | 0.997988 | 0.994666 |
| Clec16a_Z4YLG3_S406s_1 | Clec16a_Z4 | 1.69 | 0.08 | 0.57 | 0.94 | 1.05 | 1.6 | 0.17 | 0.79 | 0.68 | 0.52 | 0.82 | 0.67 | -3.39 | -0.62 | 0.46 | 0.55 | -0.3675 | 0.7475 | 0.6875 | 0.093683 | 0.846586 | 0.681063 | 0.996349 | 0.997988 | 0.994666 |
| Clec2l_P0C7M9_S29s_1 | Clec2l_P0C | 0.36 | 0.55 | 0.05 | -0.1 | -0.94 | 0.77 | -0.99 | -0.66 | 2.33 | 0.31 | -1.18 | 0 | 2.11 | -0.3 | 0.02 | -0.54 | 0.0825 | -0.1475 | -1.57 | 0.851442 | 0.134066 | 0.823794 | 0.989223 | 0.997988 | 0.999507 |
| Clec2l_P0C7M9_S47s_1 | Clec2l_P0C | 0.31 | 0.77 | 1.53 | 1.55 | 1.12 | 0.15 | 1.16 | 0.97 | 0.36 | 0.31 | 1.69 | 1.63 | 0.14 | 0.95 | 1.45 | 0.93 | -0.67 | 0.15 | 0.1075 | 0.674767 | 0.616579 | 0.927334 | 0.989223 | 0.997988 | 0.996159 |
| Clec2l_P0C7M9_S52s_1 | Clec2l_P0C | 0.87 | 0.79 | 0.99 | 0.63 | 0.71 | 0.33 | 0.5 | 0.69 | 0.17 | 0.6 | -0.36 | 0.52 | -0.4 | -0.73 | 0.56 | 0.57 | -0.19 | -0.0425 | -0.1725 | 0.045932 | 0.237918 | 0.061425 | 0.989223 | 0.997988 | 0.994666 |
| Clint1_Q5SUH6_S227s_1 | Clint1_Q5S | 0.39 | 1 | 1.24 | -4.29 | -0.25 | 1.13 | -0.65 | -0.41 | 1.66 | 0.59 | -1.03 | 0.61 | -0.37 | -1.33 | 1.46 | -0.04 | -0.2625 | -0.5875 | -0.82 | 0.779592 | 0.753012 | 0.488453 | 0.995762 | 0.997988 | 0.998226 |
| Clint1_Q5SUH6_S299s_1 | Clint1_Q5S | -2.22 | -1.07 | -3.88 | -3.31 | -2.35 | -2.61 | -3.67 | -4.16 | -0.86 | -1.32 | -3.16 | -3.13 | -0.19 | -1.52 | -3.34 | -3.75 | 0.37 | 0.8725 | 0.345 | 0.644404 | 0.397016 | 0.520582 | 0.989223 | 0.997988 | 0.995259 |
| Clint1_Q5SUH6_S457s_1 | Clint1_Q5S | 0.83 | 0.63 | -0.78 | 0.1 | 1.55 | 0.39 | 0.75 | -0.22 | 0.09 | 0.2 | -0.4 | 0.05 | -0.92 | 0.66 | -0.65 | -0.06 | -0.5775 | 0.5025 | 0.42 | 0.372714 | 0.385532 | 0.588926 | 0.989223 | 0.997988 | 0.994666 |
| Clint1_Q5SUH6_T308t_1 | Clint1_Q5S | -4.47 | -4.81 | -1.67 | -2.44 | -4.19 | -5.59 | -2.17 | -3.08 | -3.8 | -3.63 | -2.64 | -2.29 | -1.33 | -4.92 | -1.92 | -2 | 0.4225 | -0.21 | -0.4375 | 0.41519 | 0.658137 | 0.735921 | 0.990434 | 0.997988 | 0.994666 |
| Clint1_Q5SUH7_S457s_1 | Clint1_Q5S | 0.3 | -0.43 | 0.11 | -0.37 | -0.62 | 0.7 | -0.49 | 0.06 | 0.31 | -0.47 | 0.31 | -0.03 | 0.25 | -1.53 | -0.08 | 0.17 | -0.41 | 0.2575 | 0.805 | 0.653661 | 0.977299 | 0.670312 | 0.999187 | 0.997988 | 0.995259 |
| Clip1_D3Z3M7_S1116s_1 | Clip1_D3Z3 | -1.49 | 0.01 | -0.93 | -1.08 | -0.31 | -0.69 | -0.65 | -1.23 | -1.58 | -0.71 | -1.99 | -1.16 | -1.25 | -0.05 | -1.46 | -1.06 | 0.01 | 0.1275 | -0.2 | 0.850359 | 0.677994 | 0.256497 | 0.993395 | 0.997988 | 0.999507 |
| Clip1_D3Z3M7_S194s_1 | Clip1_D3Z3 | -0.41 | -0.19 | -0.19 | -0.77 | 0.06 | -0.25 | -0.71 | -0.47 | -1.03 | -0.33 | -0.41 | -0.62 | -0.32 | -0.4 | -1.22 | -0.77 | 0.1525 | -0.4875 | -0.0825 | 0.357252 | 0.855864 | 0.446394 | 0.996349 | 0.997988 | 0.994666 |
| Clip1_D3Z3M7_S347s_1 | Clip1_D3Z3 | -0.75 | -1.2 | -0.77 | -0.92 | -0.29 | -0.35 | 0.22 | -0.43 | -1.23 | -0.96 | -0.03 | -0.87 | -1.01 | -0.05 | -0.84 | -1.03 | 0.0475 | -0.2075 | -0.2875 | 0.568638 | 0.01624 | 0.661084 | 0.989223 | 0.997988 | 0.995259 |
| Clip2_Q9Z0H8_S196s_1 | Clip2_Q9Z | 0.49 | 0.34 | 0.08 | 0.42 | 0.39 | 0.52 | -0.06 | 0.13 | -0.24 | 0 | 0.43 | -0.39 | -0.59 | 0.24 | 0.26 | 0.21 | 0.6975 | 0.1375 | 0.1775 | 0.315311 | 0.706499 | 0.170496 | 0.995437 | 0.997988 | 0.994666 |
| Clip2_Q9Z0H8_S203s_1 | Clip2_Q9Z | 0.55 | 1.38 | 1.55 | 1.31 | 0.24 | 1.24 | 0.5 | 1.2 | 1.41 | 1.09 | 2.11 | 1.19 | 1.15 | 0.72 | 1.14 | 1.32 | -0.0875 | -0.3825 | -0.3025 | 0.711452 | 0.255443 | 0.463851 | 0.989223 | 0.997988 | 0.996966 |
| Clip2_Q9Z0H8_S208s_1 | Clip2_Q9Z | 0.56 | 0.45 | 0.23 | 0.64 | -0.11 | 0.68 | 0.59 | 0.36 | 0.13 | 0.76 | 1 | 0.63 | 0.75 | -0.01 | 0.89 | 0.68 | -0.4025 | 0.2525 | -0.115 | 0.711131 | 0.722329 | 0.550911 | 0.995437 | 0.997988 | 0.996966 |
| Clip2_Q9Z0H8_S212s_1 | Clip2_Q9Z | -0.05 | -0.64 | -0.81 | 0.12 | -0.7 | 0.27 | -0.95 | -0.13 | -0.51 | -0.67 | 0.83 | -0.39 | -0.44 | -0.59 | -0.85 | -0.26 | -0.09 | 0.16 | 0.1075 | 0.544865 | 0.927132 | 0.69454 | 0.997335 | 0.997988 | 0.994666 |
| Clip2_Q9Z0H8_S39s_1 | Clip2_Q9Z | -0.86 | -0.12 | -0.4 | 0.19 | -1.05 | -1.13 | -0.33 | -0.4 | -0.67 | 0.48 | 0.15 | 0.71 | -0.47 | -0.56 | 0.32 | 0.11 | -0.0325 | 0.16 | -0.19 | 0.669105 | 0.20151 | 0.240167 | 0.989223 | 0.997988 | 0.996159 |
| Clip2_Q9Z0H8_S43s_1 | Clip2_Q9Z | -1.42 | -1.32 | -1.71 | -1.76 | -1.41 | -0.73 | -0.92 | -1.03 | -0.03 | -0.9 | -4.05 | -1.65 | 0.69 | -1.03 | -0.68 | -1.34 | -0.43 | 0.465 | 0.1475 | 0.054972 | 0.051328 | 0.892632 | 0.989223 | 0.997988 | 0.994666 |
| Clip2_Q9Z0H8_S50s_1 | Clip2_Q9Z | 1.06 | 0.44 | 0.68 | 0.34 | 1.56 | 1.35 | 0.05 | 0.49 | 0.77 | 1.17 | 0.45 | 0.95 | 0.85 | 1.13 | 0.77 | 0.15 | 0.53 | -0.105 | 0.9625 | 0.762086 | 0.545775 | 0.463752 | 0.989223 | 0.997988 | 0.9971 |
| Clip2_Q9Z0H8_S924s_1 | Clip2_Q9Z | 1.61 | 1.09 | 0.12 | -0.64 | 0.68 | 3.34 | 0.51 | -0.47 | 0.63 | 2.05 | 0.03 | 0.17 | 1.08 | 0.85 | 0.44 | 0.3 | 0.2325 | 0.205 | 0.095 | 0.807757 | 0.576366 | 0.778618 | 0.989223 | 0.997988 | 0.999507 |
| Clip2_Q9Z0H8_S974s_1 | Clip2_Q9Z | -2.54 | -2.09 | -1.99 | -2.4 | -1.38 | -2.84 | -0.8 | -2.03 | -3.31 | -2.54 | -1.13 | -1.86 | -2.5 | -0.71 | -2.44 | -1.67 | 0.47 | 0.175 | 0.1225 | 0.337867 | 0.27011 | 0.922438 | 0.989223 | 0.997988 | 0.994666 |
| Clip3_B9EHT4_S399s_1 | Clip3_B9EH | 0.71 | 0.86 | 0.95 | 1.17 | 0.76 | 0.18 | 1.1 | 1.32 | 0.71 | 1.25 | 1.13 | 1.1 | 0.78 | 1.33 | 1.25 | 1.28 | 0.4925 | 0.045 | 0.425 | 0.371893 | 0.779367 | 0.602644 | 0.995762 | 0.997988 | 0.994666 |
| Clip4_Q8CI96_S580s_1 | Clip4_Q8CI | -1.97 | -1.92 | -2.42 | -2.37 | -2.02 | -2.32 | -2.21 | -2.27 | -1.66 | -2.51 | -1.1 | -1.93 | -1.97 | -2.68 | -1.57 | -1.84 | -0.0825 | 0.125 | 0.2375 | 0.628771 | 0.876946 | 0.293092 | 0.996493 | 0.997988 | 0.995259 |
| Clip4_Q8CI96_S608s_1 | Clip4_Q8CI | -1.85 | -3.03 | -1.47 | -1.6 | -1.31 | -2.54 | 0.39 | -0.39 | -2.01 | -1.95 | -2.44 | -0.93 | -1.26 | -0.87 | -1.38 | -1.23 | -0.035 | 0.37 | 0.155 | 0.061133 | 0.135732 | 0.73612 | 0.989223 | 0.997988 | 0.994666 |
| Clk2_D3Z7L4_S17s_1 | Clk2_D3Z7 | -1.62 | -2.69 | -2.8 | -2.13 | -3.33 | -1.94 | -2.33 | -1.69 | -0.25 | -2.76 | -0.91 | -2.63 | 0.47 | -3.27 | 0.97 | -1.97 | 1.025 | 0.155 | 0.8025 | 0.159428 | 0.976648 | 0.296033 | 0.999187 | 0.997988 | 0.994666 |
| Clk2_D3Z7L4_S50s_1 | Clk2_D3Z7 | -0.87 | -1.33 | -1.12 | -1.82 | -2.99 | -1.5 | -1.85 | -1.46 | -1.45 | -1.47 | -0.71 | -1.62 | -0.24 | -2.28 | -0.74 | -1.68 | -0.0125 | 0.6725 | 1.36 | 0.916807 | 0.118423 | 0.930908 | 0.989223 | 0.997988 | 0.999507 |
| Clk3_O35492_S157s_1 | Clk3_O354 | -0.69 | -0.17 | -0.35 | -0.45 | -0.29 | -0.91 | 0.49 | 0.15 | -0.42 | -0.1 | -0.16 | -0.61 | -0.14 | 0.24 | 0.77 | 0.1 | -0.665 | -0.0275 | 0.05 | 0.044861 | 0.414896 | 0.704035 | 0.989223 | 0.997988 | 0.994666 |
| Clmn_Q8C5W0_S358s_1 | Clmn_Q8C | -0.1 | -0.05 | 0.17 | 0.49 | 0.15 | -0.42 | 0.89 | 0.71 | -0.45 | -0.4 | 0.45 | 0.29 | -0.25 | 0.56 | 0.39 | 0.61 | 0.275 | 0.0925 | 0.6575 | 0.510626 | 0.543269 | 0.613619 | 0.989223 | 0.997988 | 0.994666 |
| Clmn_Q8C5W0_S419s_1 | Clmn_Q8C | -1.44 | -0.81 | -0.48 | -2.3 | -0.12 | -2.16 | -1.49 | -1.8 | -2.32 | -2.22 | -0.17 | -1.44 | -0.57 | -0.24 | -0.58 | -1.6 | 0.205 | -0.155 | 0.2 | 0.299303 | 0.803598 | 0.634809 | 0.995762 | 0.997988 | 0.994666 |
| Cln3_D3YVA8_S14s_1 | Cln3_D3YV | -0.22 | 0.63 | 0.19 | -0.19 | -0.09 | 0.31 | -0.01 | -0.37 | 0.4 | 0.06 | 0.89 | 0.03 | 0.31 | 0.77 | -0.13 | -0.93 | -0.135 | -0.28 | 0.51 | 0.813849 | 0.612524 | 0.447812 | 0.989223 | 0.997988 | 0.999507 |
| Clns1a_Q923F1_S3s_1 | Clns1a_Q9 | -0.85 | -0.27 | -0.47 | -0.62 | -0.14 | -0.66 | -0.33 | -0.53 | -0.92 | -0.47 | -0.42 | -0.76 | -0.35 | -0.09 | -0.75 | -0.25 | -0.1425 | 0.2425 | -0.0975 | 0.485492 | 0.564066 | 0.717235 | 0.989223 | 0.997988 | 0.994666 |
| Clstn3_D3Z601_S841s_1 | Clstn3_D3Z | -0.98 | -0.92 | -0.64 | -1.23 | -0.61 | -0.44 | -0.8 | -1.5 | -0.48 | -0.26 | -0.63 | -0.49 | 0.68 | -0.75 | -0.41 | -0.73 | 0.1375 | -0.09 | 0.1925 | 0.113311 | 0.719534 | 0.066704 | 0.995437 | 0.997988 | 0.994666 |
| Clstn3_D3Z601_S914s_1 | Clstn3_D3Z | 0.41 | 0.41 | 0.07 | 0.18 | 0.95 | -0.11 | 0.56 | 0.27 | 1.12 | 0.35 | 0.75 | 0.73 | 0.24 | 1.14 | 0.76 | 0.52 | 0.105 | 0.4775 | 0.64 | 0.183477 | 0.591571 | 0.088225 | 0.989223 | 0.997988 | 0.994666 |
| Clta_B1AWD8_S105s_1 | Clta_B1AW | -0.63 | 0.11 | 0.63 | 0.12 | 0.26 | -0.73 | -0.02 | -0.48 | -1.66 | -0.81 | -1.15 | -0.06 | -0.42 | -0.06 | -0.17 | 0.11 | 0.15 | 0.47 | 0.3975 | 0.558283 | 0.395306 | 0.040141 | 0.989223 | 0.997988 | 0.995259 |
| Cltc_Q68FD5_S576s_1 | Cltc_Q68F | -0.14 | 0.2 | 0.12 | 0.29 | 0.33 | -0.06 | -0.05 | 0.37 | 0.11 | -0.11 | -0.49 | -0.02 | 0.72 | 0.05 | -0.14 | -0.41 | -0.3 | -0.9775 | -0.1925 | 0.840912 | 0.895062 | 0.324925 | 0.996493 | 0.997988 | 0.999507 |
| Cltc_Q68FD5_S67s_1 | Cltc_Q68F | 0.52 | 0.46 | 0.5 | 0.13 | 0.93 | 0.16 | 0.56 | 0.1 | 0.28 | 0.77 | 0.53 | 0.11 | 0.37 | 1.14 | 0.28 | 0.1 | 0.03 | -0.245 | -0.0625 | 0.818501 | 0.893604 | 0.935658 | 0.996493 | 0.997988 | 0.999507 |
| Cltc_Q68FD5_T235t_1 | Cltc_Q68F | 1.17 | 1.05 | 2.63 | 1.37 | 0.82 | 2.02 | 2.58 | 2.17 | -0.53 | 1.02 | -0.45 | 2.28 | -0.09 | 0.32 | 3.42 | 2.97 | 0.035 | 0.02 | 0.07 | 0.906465 | 0.484954 | 0.180105 | 0.989223 | 0.997988 | 0.999507 |
| Cltc_Q68FD5_T302t_1 | Cltc_Q68F | 0.62 | 2.21 | 0.82 | 0.66 | 2.4 | 1.43 | 0.77 | 0.11 | 0.54 | 1.33 | -0.51 | 0.29 | 1.01 | 1.85 | 0.68 | 0.67 | 0.3425 | -0.975 | 0.1 | 0.956241 | 0.857789 | 0.209069 | 0.996349 | 0.997988 | 0.999507 |
| Cluh_Z4YLI8_S1287s_1 | Cluh_Z4YL | -0.59 | -0.13 | 0.02 | 0.05 | -0.29 | -0.29 | -0.18 | -0.11 | 0.47 | -0.08 | 0.33 | 0.2 | 0.59 | -0.17 | -0.02 | -0.03 | 0.1 | -0.665 | -0.025 | 0.392757 | 0.811048 | 0.151749 | 0.995762 | 0.997988 | 0.994666 |
| Clvs1_Q9D4C9_S34s_1 | Clvs1_Q9D | 8.16 | 4.48 | 5.62 | 1.21 | 5.3 | 9.11 | 6.41 | 5.17 | -3.87 | 7.37 | -0.07 | 6.01 | -0.04 | 0.12 | 7.3 | 8.11 | -0.055 | 0.3925 | 0.255 | 0.660903 | 0.281065 | 0.348346 | 0.989223 | 0.997988 | 0.995671 |
| Clvs1_Q9D4C9_S41s_1 | Clvs1_Q9D | -1.14 | -1.84 | -1.54 | -0.79 | -2.15 | -0.35 | -1.06 | -1.44 | 0.39 | -0.93 | -1.65 | -1.53 | 0.51 | -2.38 | -1.93 | -1.25 | 1.63 | -2.5075 | -0.995 | 0.915629 | 0.85298 | 0.427377 | 0.996349 | 0.997988 | 0.999507 |
| Clvs1_Q9D4C9_S50s_1 | Clvs1_Q9D | -1.28 | -0.92 | -1.62 | -2.22 | -0.65 | -0.99 | -0.99 | -2.26 | -1.17 | -0.93 | -0.23 | -1.81 | -1 | -0.48 | -2.44 | -1.42 | 0.0775 | 0.3975 | 0.065 | 0.714796 | 0.504694 | 0.274462 | 0.989223 | 0.997988 | 0.996966 |
| Clvs2_Q8BG92_S302s_1 | Clvs2_Q8B | -3.25 | -2.46 | -1.83 | -2.24 | -2.54 | -4.3 | -2.03 | -1.84 | -2.56 | -2.42 | -1.27 | -1.89 | -2.66 | -1.84 | -1.75 | -2.27 | 0.2875 | 0.475 | 0.175 | 0.415838 | 0.685086 | 0.333191 | 0.994624 | 0.997988 | 0.994666 |
| Cmc2_F8WHQ0_S47s_1 | Cmc2_F8W | 1.11 | 0.39 | 0.2 | 0.51 | 1.25 | 0.52 | -0.15 | 0.41 | 0.92 | 0.77 | 0.44 | 1.02 | 1.06 | 1.03 | 0.38 | 0.79 | -0.2325 | 0.41 | 0.315 | 0.402551 | 0.897744 | 0.415795 | 0.996493 | 0.997988 | 0.994666 |
| Cmip_Q9D486_S349s_1 | Cmip_Q9D | 3.84 | 3.79 | 4.08 | 2.34 | 2.77 | 4.21 | 2.39 | 3.3 | 0.08 | 3.75 | -1.14 | 3.66 | 0.87 | -1.11 | 3.44 | 3.55 | -0.045 | 0.235 | 0.2625 | 0.102563 | 0.504308 | 0.120389 | 0.989223 | 0.997988 | 0.994666 |
| Cmpk1_Q9DBP5_S147s_1 | Cmpk1_Q9 | -1.59 | -1.12 | -0.6 | -0.45 | -3.11 | -1.1 | -1.31 | -0.22 | -0.08 | -1.12 | -1.58 | -0.2 | 0.11 | -2.49 | -0.58 | -1.22 | -0.345 | -1.925 | -1.825 | 0.85219 | 0.411075 | 0.654348 | 0.989223 | 0.997988 | 0.999507 |
| Cmtm4_Q8CJ61_S194s_1 | Cmtm4_Q8 | 2.36 | 1.43 | 1.91 | 2.15 | 2.02 | 2.2 | 1.73 | 2.16 | 1.84 | 1.87 | 1.38 | 2.19 | 1.19 | 2.08 | 2.14 | 1.93 | -0.495 | 0.195 | -0.105 | 0.704477 | 0.809346 | 0.636581 | 0.995762 | 0.997988 | 0.996159 |
| Cmtm4_Q8CJ61_S199s_1 | Cmtm4_Q8 | 1.15 | 1.52 | 1.15 | 0.56 | 1.84 | 1.55 | 0.98 | 0.59 | 0.12 | 0.86 | 1.39 | 0.75 | 0.15 | 1.39 | 1.06 | 0.69 | 0.065 | -0.1425 | -0.1275 | 0.453265 | 0.677408 | 0.3727 | 0.99315 | 0.997988 | 0.994666 |
| Cmtm6_Q9CZ69_S8s_1 | Cmtm6_Q9 | -0.17 | -0.05 | 0.09 | -0.04 | 0.2 | -0.27 | -1.62 | 0.1 | -0.48 | -0.69 | -0.21 | -1 | 0.32 | 0.28 | -0.51 | -0.09 | 0.145 | -0.315 | -0.2725 | 0.879077 | 0.391202 | 0.050077 | 0.989223 | 0.997988 | 0.999507 |
| Cmtm7_Q9ESD6_S2s_1 | Cmtm7_Q9 | -0.24 | 0.48 | 0.66 | -0.3 | -2.37 | -1.14 | 0.3 | -0.86 | 2.4 | 0.22 | 1.99 | 0.41 | 1.46 | -0.56 | 0.45 | -0.16 | -0.355 | -0.5525 | 0.0425 | 0.759463 | 0.054854 | 0.073195 | 0.989223 | 0.997988 | 0.9971 |
| Cnbp_P53996_T165t_1 | Cnbp_P539 | -3.67 | -2.87 | -3.32 | -6.08 | -2.03 | -3.02 | -4.76 | -4.31 | -2.43 | -6.01 | -3.47 | -1.2 | -2.61 | -4.01 | -2.01 | -0.35 | -1.1675 | 1.105 | 0.1475 | 0.081962 | 0.585357 | 0.524274 | 0.989223 | 0.997988 | 0.994666 |
| Cnksr2_Q80YA9_S12s_1 | Cnksr2_Q8 | -0.48 | 0.01 | -0.26 | -0.4 | 0.5 | -1.12 | 0.91 | -0.17 | -1.22 | 0.34 | -0.82 | 0.25 | -0.48 | 0.93 | 0.53 | -0.19 | 0.455 | 0.7075 | 1.74 | 0.207801 | 0.474151 | 0.841771 | 0.989223 | 0.997988 | 0.994666 |
| Cnksr2_Q80YA9_S325s_1 | Cnksr2_Q8 | 3.3 | 0.41 | 0.42 | 1.98 | 0.89 | 3.27 | 1.18 | 2.62 | 2.21 | 1.75 | 1.11 | 1.66 | 0.55 | 1.58 | 1.11 | 0.66 | 0.3125 | -0.08 | 0.48 | 0.412787 | 0.560416 | 0.815393 | 0.989223 | 0.997988 | 0.994666 |
| Cnksr2_Q80YA9_S325s_2 | Cnksr2_Q8 | 1.49 | 1.05 | -0.22 | -0.97 | 0.76 | 1.48 | -0.04 | -0.08 | 0.69 | 1.55 | 0.02 | -0.17 | 0.59 | 0.8 | 0.02 | -0.43 | 0.4625 | 0.155 | -0.5525 | 0.873216 | 0.750921 | 0.768198 | 0.995762 | 0.997988 | 0.999507 |
| Cnksr2_Q80YA9_S390s_1 | Cnksr2_Q8 | -0.27 | 0.14 | 0.41 | 0.04 | 0.29 | -0.33 | 0.64 | 0.05 | -0.14 | 0.08 | 0.32 | 0.4 | -0.06 | 0.53 | 0.07 | 0.15 | 0.1925 | 0.185 | -0.0925 | 0.735985 | 0.770399 | 0.740337 | 0.995762 | 0.997988 | 0.9971 |
| Cnksr2_Q80YA9_S430s_1 | Cnksr2_Q8 | -0.82 | -1.26 | -0.26 | -1.16 | -2.61 | -0.34 | -1.39 | -0.35 | -1.27 | -1.65 | 1.87 | -0.14 | 0.55 | -1.86 | -0.96 | -0.74 | 0.0825 | 0.085 | 0.0925 | 0.81145 | 0.577965 | 0.441872 | 0.989223 | 0.997988 | 0.999507 |
| Cnksr2_Q80YA9_S467s_1 | Cnksr2_Q8 | -0.67 | -1.16 | -1.63 | -1.68 | -0.86 | -0.53 | -1.06 | -0.82 | 0.16 | -1.92 | 2.77 | 0.03 | -0.09 | -0.34 | -0.63 | -1.59 | -0.2975 | 0.5775 | 0.1225 | 0.150871 | 0.131951 | 0.105621 | 0.989223 | 0.997988 | 0.994666 |
| Cnksr2_Q80YA9_S503s_1 | Cnksr2_Q8 | -1.52 | -0.8 | -0.29 | -0.67 | -0.88 | -1.01 | 0.61 | -0.83 | -0.51 | -0.47 | -0.22 | 0.32 | -0.67 | -0.53 | -0.09 | -0.29 | 0.4675 | 1.545 | 0.6225 | 0.21685 | 0.504762 | 0.104023 | 0.989223 | 0.997988 | 0.994666 |
| Cnksr2_Q80YA9_S505s_1 | Cnksr2_Q8 | -0.34 | -0.55 | -0.82 | -0.79 | -0.41 | -0.45 | -0.22 | -0.7 | -0.57 | -0.25 | -1.12 | -0.68 | -0.06 | -0.17 | -0.92 | -0.93 | 0.2925 | 0.6 | 0.425 | 0.737643 | 0.437024 | 0.911605 | 0.989223 | 0.997988 | 0.9971 |
| Cnksr2_Q80YA9_S702s_1 | Cnksr2_Q8 | 0.37 | 0.54 | -0.05 | -0.09 | 0.24 | 0.31 | 0.63 | -0.1 | 0.35 | 0.75 | 0.2 | 0.32 | 0.07 | 0.61 | 0.62 | 0.32 | 0.18 | -0.03 | 0.105 | 0.456627 | 0.768597 | 0.424794 | 0.995762 | 0.997988 | 0.994666 |
| Cnksr2_Q80YA9_S756s_1 | Cnksr2_Q8 | -0.84 | -0.93 | -0.62 | -0.24 | -0.81 | -0.78 | -0.24 | -0.01 | -1.51 | -0.95 | -0.02 | -0.25 | -1.15 | -0.46 | -0.19 | -0.48 | 0.0775 | 0.2125 | 0.2125 | 0.777941 | 0.493389 | 0.94717 | 0.989223 | 0.997988 | 0.99798 |
| Cnksr2_Q80YA9_S906s_1 | Cnksr2_Q8 | 0.09 | 0.3 | 0.4 | 0.14 | 0.36 | -0.22 | 0.48 | 0.05 | -0.02 | 0.05 | 0.21 | 0.27 | 0.04 | 0.5 | 0.07 | 0.43 | 0.1975 | -0.025 | 0.0875 | 0.912831 | 0.786186 | 0.630286 | 0.995762 | 0.997988 | 0.999507 |
| Cnksr2_Q80YA9_T339t_2 | Cnksr2_Q8 | 1.24 | 0.97 | -0.22 | -0.97 | 0.61 | 1.21 | -0.04 | -0.08 | 0.51 | 1.33 | 0.02 | -0.17 | 0.5 | 0.72 | 0.02 | -0.43 | -0.065 | -0.105 | 0.0275 | 0.922092 | 0.754904 | 0.768503 | 0.995762 | 0.997988 | 0.999507 |
| Cnksr2_Q80YA9_T809t_1 | Cnksr2_Q8 | -3.14 | -4.98 | -2.87 | -4.7 | -1.72 | -2.52 | -3.45 | -2.99 | -5.53 | -3.65 | -1.06 | -2.14 | -2.23 | -1.75 | -3.01 | -2.53 | 0.17 | 0.1675 | -0.0525 | 0.018757 | 0.055004 | 0.407428 | 0.989223 | 0.997988 | 0.994666 |
| Cnnm1_Q0GA42_S850s_1 | Cnnm1_Q0 | 0.49 | -0.26 | -0.49 | -0.11 | 0.34 | 0.61 | 0.23 | 0.74 | -0.68 | -0.83 | 0.26 | 0.02 | -2.24 | 0.2 | -0.68 | -0.76 | 1.2525 | 0.8275 | 1.5425 | 0.154605 | 0.063058 | 0.545632 | 0.989223 | 0.997988 | 0.994666 |
| Cnnm1_Q0GA42_T824t_1 | Cnnm1_Q0 | -1.06 | -0.85 | -1.05 | -1.83 | -0.69 | -0.92 | -2.07 | -1.98 | 0.46 | -0.46 | -3.04 | -0.88 | 0.25 | -0.95 | -0.53 | -0.95 | 0.5725 | -0.215 | -0.7775 | 0.106442 | 0.589745 | 0.755387 | 0.989223 | 0.997988 | 0.994666 |
| Cnnm3_Q32NY4_S667s_1 | Cnnm3_Q3 | 1.41 | 1.57 | 1.07 | 0.96 | 1.1 | 1.14 | 1.85 | 1.09 | 1.3 | 1.56 | 1.58 | 0.96 | 1.12 | 0.31 | 1.56 | 1.31 | -0.2175 | 0.2175 | 0.6525 | 0.604934 | 0.876582 | 0.714117 | 0.996493 | 0.997988 | 0.995259 |
| Cnnm4_Q69ZF7_S663s_1 | Cnnm4_Q6 | 2.16 | -0.3 | -0.15 | -0.19 | 1.28 | 2.24 | -0.35 | 0.37 | -0.64 | 1.35 | 0.78 | 0.05 | -1.64 | 0.39 | -0.83 | -0.11 | 0.0425 | 0.0975 | -0.1775 | 0.186128 | 0.488369 | 0.994004 | 0.989223 | 0.997988 | 0.994666 |
| Cnnm4_Q69ZF7_S766s_1 | Cnnm4_Q6 | 0.88 | 0.69 | 0.39 | 0.44 | 0.09 | 0.17 | -0.59 | 0.68 | 0.66 | -0.55 | 0.96 | -0.45 | 0.8 | 0.76 | -0.5 | 0.25 | 0.505 | 0.005 | -0.9275 | 0.446656 | 0.119408 | 0.280807 | 0.989223 | 0.997988 | 0.994666 |
| Cnot10_Q8BH15_S520s_1 | Cnot10_Q8 | -1.27 | -1.06 | 0.05 | -0.93 | -1.49 | -2.55 | -1.38 | -1.78 | -0.8 | -1.06 | -0.73 | -1.7 | -0.35 | -0.76 | -1.97 | 0.11 | -0.5125 | -0.445 | -0.2725 | 0.905113 | 0.025757 | 0.475998 | 0.989223 | 0.997988 | 0.999507 |
| Cnot2_E9Q8D5_S165s_1 | Cnot2_E9Q | -0.52 | -0.49 | -0.73 | -1.05 | -0.61 | -0.54 | -0.51 | -0.59 | -0.38 | -0.49 | -0.86 | -0.78 | 0.29 | -1.97 | -0.65 | -0.16 | -0.9975 | -0.27 | 0.06 | 0.876345 | 0.542284 | 0.778667 | 0.989223 | 0.997988 | 0.999507 |
| Cnot3_F6YBV1_S256s_1 | Cnot3_F6Y | -1.33 | -1.87 | -0.54 | -1.75 | -2.15 | -0.45 | -1.71 | -1.59 | 0.02 | -1.15 | -0.34 | -0.54 | 0.79 | -2.65 | -1.05 | -1.87 | 0.135 | 0.07 | 0.075 | 0.803141 | 0.816579 | 0.048107 | 0.995762 | 0.997988 | 0.999507 |
| Cnp_P16330_S22s_1 | Cnp_P1633 | -1.29 | -2.73 | -2.52 | -13.23 | -2.9 | -1.63 | -2.87 | -3.11 | 2.57 | -2.52 | -4.11 | -1.6 | 3.56 | -4.38 | 3.1 | -3.98 | -0.1025 | 0.87 | 0.1775 | 0.156543 | 0.344429 | 0.214572 | 0.989223 | 0.997988 | 0.994666 |
| Cnp_P16330_S382s_1 | Cnp_P1633 | 0.53 | 1.34 | 1.2 | 0.61 | 1.47 | 0.3 | 1.03 | 0.25 | 0.17 | 0.01 | -0.47 | 0.75 | -0.63 | 0.86 | 0.94 | 1.25 | 2.315 | 3.5275 | 4.5175 | 0.493145 | 0.661831 | 0.039868 | 0.990434 | 0.997988 | 0.994666 |
| Cnp_P16330_S408s_1 | Cnp_P1633 | 0.98 | 1.47 | 1.36 | 1.11 | 1.42 | 0.54 | 0.65 | 0.36 | 0.14 | 0.17 | 0.89 | 0.35 | 0.65 | 0.75 | 1.34 | 0.65 | -0.1575 | -0.805 | -0.315 | 0.192531 | 0.117478 | 0.010642 | 0.989223 | 0.997988 | 0.994666 |
| Cnp_P16330_S4s_1 | Cnp_P1633 | -0.15 | 0.31 | -3.6 | -2.51 | -1.18 | -0.87 | -2.83 | -2.66 | 1.93 | -0.29 | -2.21 | -2.56 | 2.3 | 0.98 | -1.88 | -2.19 | -0.4875 | -0.8425 | -0.3825 | 0.313957 | 0.667689 | 0.567647 | 0.992842 | 0.997988 | 0.994666 |
| Cnr1_P47746_S317s_1 | Cnr1_P477 | -1.56 | -1.4 | -1.88 | -2.32 | -0.79 | -1.66 | -1.56 | -2.24 | -1.65 | -1.37 | -1.25 | -1.91 | -1.13 | -0.46 | -1.88 | -1.98 | -0.3975 | 0.705 | 1.29 | 0.312823 | 0.530742 | 0.413679 | 0.989223 | 0.997988 | 0.994666 |
| Cnr1_P47746_S442s_1 | Cnr1_P477 | 7.78 | 3.7 | 5.87 | 0.92 | 5.1 | 8.57 | 6.26 | 4.85 | 1.31 | 7.12 | -1.79 | 5.94 | 2.68 | -0.74 | 7.21 | 7.08 | 0.2275 | 0.245 | 0.4275 | 0.804932 | 0.280252 | 0.523955 | 0.989223 | 0.997988 | 0.999507 |
| Cnst_Q8CBC4_S279s_1 | Cnst_Q8CB | 0.15 | -0.08 | 0.93 | 0.7 | -0.18 | -0.96 | 0.55 | 0.59 | 0.11 | 0 | 1.69 | 0.73 | 0.34 | 0.78 | 1.37 | 0.99 | 1.6275 | -1.4225 | -0.51 | 0.221818 | 0.318556 | 0.638113 | 0.989223 | 0.997988 | 0.994666 |
| Cntnap1_O54991_S1384s_1 | Cntnap1_O | 0.4 | 1.01 | 1.5 | 0.77 | 1.02 | 0.59 | 1.55 | 1.13 | 0.63 | 0.94 | -0.52 | 1.12 | -0.98 | -0.11 | 1.16 | 1.32 | -0.425 | 0.2075 | 0.445 | 0.311592 | 0.634235 | 0.384437 | 0.989223 | 0.997988 | 0.994666 |
| Coasy_Q9DBL7_S177s_1 | Coasy_Q9D | -1.92 | -2.26 | -1.27 | -2.3 | -2.69 | -0.86 | -2.64 | -2.08 | -0.33 | -1.76 | -3.39 | -1.18 | 0.49 | -2.68 | -2.24 | -2.66 | 0.1525 | -0.3775 | -0.5725 | 0.816823 | 0.77528 | 0.665446 | 0.995762 | 0.997988 | 0.999507 |
| Cobl_G3UWY3_S1121s_1 | Cobl_G3UV | 0.38 | -2.29 | -0.9 | 0.31 | 0.12 | 0.55 | -0.08 | 0.26 | -1.13 | -1.82 | 1.01 | -0.2 | -2.05 | -0.57 | -1.42 | -1.33 | -0.13 | 0.2725 | 0.165 | 0.27278 | 0.169525 | 0.908585 | 0.989223 | 0.997988 | 0.994666 |
| Cobl_G3UWY3_S669s_1 | Cobl_G3UV | -0.31 | -0.69 | 0.01 | 0.11 | 0.27 | -0.49 | 0.22 | 0.05 | -0.38 | -0.62 | 0.49 | 0.16 | 0.1 | 0.2 | 0.16 | 0.29 | 0.8375 | 0.09 | -0.7175 | 0.157151 | 0.421648 | 0.691896 | 0.989223 | 0.997988 | 0.994666 |
| Cobl_G3UWY3_S819s_1 | Cobl_G3UV | 0.37 | 0.25 | -0.36 | 0.25 | 0.53 | 0.24 | 0.62 | 0.42 | 0.07 | 0.44 | -0.12 | 0 | 0.52 | 0.92 | -0.21 | 0.21 | 0.2325 | 0.1325 | 0.4075 | 0.488304 | 0.205688 | 0.910129 | 0.989223 | 0.997988 | 0.994666 |
| Cobl_Q5NBX1_S269s_1 | Cobl_Q5NE | -1.02 | -0.55 | -0.56 | -0.43 | -0.97 | -0.39 | -0.31 | -1.24 | -0.26 | -0.42 | 0.49 | 0.18 | -0.08 | -1.22 | -0.74 | -0.15 | 0.325 | -0.03 | 0.2325 | 0.783394 | 0.763418 | 0.05107 | 0.995762 | 0.997988 | 0.998491 |
| Cobll1_B1AZ15_S234s_1 | Cobll1_B1A | -0.64 | -1.52 | -0.66 | -1.53 | -1.13 | -1.44 | -0.56 | -1.53 | -1.31 | -1 | -1.24 | -1.49 | -0.64 | 0.09 | -2.24 | -0.95 | -0.0875 | 0.6375 | 0.0925 | 0.767456 | 0.81998 | 0.57815 | 0.995762 | 0.997988 | 0.9971 |
| Cobll1_B1AZ15_S281s_1 | Cobll1_B1A | 3.81 | 2.25 | 3.32 | -2.26 | 4.42 | 4.05 | 3.28 | 2.99 | -1.86 | 3.69 | -0.48 | 2.91 | 1.49 | -2.22 | 3.62 | 3.87 | -0.0775 | -0.1725 | 0.1525 | 0.957537 | 0.144286 | 0.670658 | 0.989223 | 0.997988 | 0.999507 |
| Cog3_E9QL65_S212s_1 | Cog3_E9QL | -0.32 | -0.38 | -0.22 | -0.08 | -0.33 | -0.91 | 0.36 | 0.06 | -0.14 | 0.07 | 0.01 | 0.15 | -0.18 | -0.2 | -0.01 | 0.48 | 1.905 | -0.715 | -0.09 | 0.317995 | 0.882387 | 0.223396 | 0.996493 | 0.997988 | 0.994666 |
| Coil_E9Q284_S149s_1 | Coil_E9Q2 | 2.06 | 1.73 | 2.32 | 2.01 | 0.47 | 1.55 | 0.74 | 2.53 | 0.9 | 1.65 | 1.64 | 2.24 | 1.38 | 0.32 | 2.19 | 2.36 | 0.045 | 0.2725 | 0.2725 | 0.327413 | 0.138923 | 0.215966 | 0.989223 | 0.997988 | 0.994666 |
| Coil_E9Q284_S174s_1 | Coil_E9Q2 | 1.32 | 1.2 | 1.7 | 1.5 | -0.68 | 1.2 | 0.36 | 1.08 | 0.93 | 0.59 | 1.55 | 1.34 | 0.75 | 0.05 | 1.19 | 1.14 | -0.7075 | -0.4225 | -0.4675 | 0.07271 | 0.048159 | 0.271372 | 0.989223 | 0.997988 | 0.994666 |
| Col15a1_O35206_S125s_1 | Col15a1_O | 1.49 | -1.01 | 0.66 | 0.55 | -0.42 | 1.47 | -0.39 | 0.75 | -0.03 | -0.36 | 0.23 | 0.59 | -0.08 | -0.41 | -0.04 | 0.4 | -0.94 | -0.3275 | -0.6475 | 0.386494 | 0.910118 | 0.550384 | 0.997335 | 0.997988 | 0.994666 |
| Col4a3bp_Q9EQG9_S123s_1 | Col4a3bp_ | 0.89 | 1.41 | 0.32 | 0.15 | 1.06 | 0.85 | 0.64 | 0.51 | 0.42 | 2.04 | -0.49 | 0.53 | 1.37 | -1.56 | 1.09 | 0.72 | -0.07 | -0.315 | -0.455 | 0.662474 | 0.823058 | 0.902831 | 0.995762 | 0.997988 | 0.995804 |
| Col4a3bp_Q9EQG9_S123s_2 | Col4a3bp_ | -0.43 | -0.37 | -0.53 | -1.54 | -0.36 | -1.51 | -0.55 | -0.31 | -2.18 | -0.34 | -2.15 | 0.3 | -3.71 | -2.13 | -0.93 | 0.4 | 0.0725 | -0.0675 | -0.2875 | 0.295002 | 0.927209 | 0.554983 | 0.997335 | 0.997988 | 0.994666 |
| Col4a3bp_Q9EQG9_S126s_1 | Col4a3bp_ | -0.34 | -0.36 | 0.45 | -0.2 | -0.06 | -0.25 | -0.01 | 0.62 | -1.86 | -0.29 | -1.08 | -0.12 | -0.5 | -3.36 | 0.37 | 0.25 | 0.035 | -0.375 | -0.875 | 0.386766 | 0.530161 | 0.118877 | 0.989223 | 0.997988 | 0.994666 |
| Col4a3bp_Q9EQG9_S126s_2 | Col4a3bp_ | -0.43 | -0.37 | -0.53 | -1.54 | -0.36 | -1.51 | -0.55 | -0.31 | -2.18 | -0.34 | -2.15 | 0.3 | -3.71 | -2.13 | -0.93 | 0.4 | 0.1875 | -0.725 | -0.6975 | 0.295002 | 0.927209 | 0.554983 | 0.997335 | 0.997988 | 0.994666 |
| Comt_O88587_S218s_1 | Comt_O88 | -0.15 | 0.28 | -1.64 | -2.34 | 0.07 | -0.45 | -1.65 | -1.71 | -0.1 | 0.35 | -1.31 | -1.77 | 0.32 | 0.25 | -1.74 | -1.66 | 0.035 | -0.375 | -0.875 | 0.733788 | 0.967305 | 0.72161 | 0.999187 | 0.997988 | 0.9971 |
| Copa_F8WHL2_S173s_1 | Copa_F8W | -0.5 | 1.33 | 0.59 | 0.31 | 1.22 | 0.28 | 0.59 | 0 | -0.26 | 0.83 | 0.39 | 0.18 | 0.48 | 0.47 | 0.65 | 0.34 | 0.0275 | 0.255 | 0.255 | 0.89363 | 0.835493 | 0.731761 | 0.995762 | 0.997988 | 0.999507 |
| Copb2_O55029_S860s_1 | Copb2_O5 | -3.55 | -3.25 | -2.92 | -2.79 | -3.47 | -2.1 | -2.06 | -3.3 | -3.22 | -3.1 | -3.7 | -2.85 | -1.25 | -2.12 | -2.72 | -3.24 | 0.09 | -0.1475 | 0.0525 | 0.10179 | 0.335643 | 0.757912 | 0.989223 | 0.997988 | 0.994666 |
| Copg1_Q9QZE5_T584t_1 | Copg1_Q9 | 0.09 | -0.18 | -0.11 | 0.09 | 0.46 | -0.26 | 0.48 | -0.01 | -0.21 | -0.02 | -0.47 | 0.24 | 0.02 | 0.87 | -0.52 | 0.62 | 0.395 | -0.09 | 0.795 | 0.439171 | 0.444443 | 0.721773 | 0.989223 | 0.997988 | 0.994666 |
| Cops3_O88543_S410s_1 | Cops3_O8 | 0.4 | -0.01 | 0.81 | 0.91 | 0.36 | 0.33 | 0.31 | 0.54 | 0.12 | 1.07 | 0.5 | 0.55 | 0.67 | 1.86 | 0.34 | 0.33 | 0.195 | -0.0875 | 0.275 | 0.519202 | 0.591493 | 0.918386 | 0.989223 | 0.997988 | 0.994666 |
| Cops3_O88543_S423s_1 | Cops3_O8 | 0.27 | -0.25 | 0.38 | 0.37 | -0.85 | -0.13 | 0.02 | 0.74 | 0.51 | 0.3 | 0.05 | -0.1 | 0.7 | -0.85 | -0.23 | 0.76 | -0.1425 | 0.0325 | 0.2725 | 0.814773 | 0.494633 | 0.992444 | 0.989223 | 0.997988 | 0.999507 |
| Cops5_O35864_S284s_1 | Cops5_O3 | 0.15 | 0.61 | -0.81 | -0.22 | 0.59 | -0.17 | -0.65 | 0.03 | 0.3 | -0.62 | -0.77 | -0.95 | -1.11 | 0.26 | 0.04 | -1.41 | -0.2475 | -0.0025 | -0.0975 | 0.329878 | 0.963643 | 0.291031 | 0.999187 | 0.997988 | 0.994666 |
| Coro1a_O89053_S426s_1 | Coro1a_O8 | 1.68 | 0.12 | 0.02 | 0.8 | 0.9 | 1.67 | 0.54 | 1.25 | 0.63 | 0.04 | -0.04 | 0.5 | 0.33 | 1.45 | 0.15 | -0.26 | 0.0175 | -0.4425 | -0.4875 | 0.637099 | 0.323101 | 0.37641 | 0.989223 | 0.997988 | 0.995259 |
| Coro1a_O89053_T418t_1 | Coro1a_O8 | -1.21 | -2.21 | -2.35 | -1.31 | -1.9 | -1.53 | -2.09 | -1.72 | -1.35 | -1.89 | 0.43 | -1.42 | -0.5 | -1.22 | -2.67 | -2.58 | 0.435 | -0.3725 | -0.2375 | 0.960872 | 0.903515 | 0.214245 | 0.996493 | 0.997988 | 0.999507 |
| Coro1a_O89053_T424t_1 | Coro1a_O8 | -0.6 | -2.19 | -0.73 | -1.14 | -0.99 | -1.06 | -0.84 | -0.01 | -0.46 | -0.5 | 0.28 | -0.11 | -1.08 | -1.22 | -0.68 | -0.3 | -0.04 | 0.7125 | 0.0275 | 0.415735 | 0.302815 | 0.035969 | 0.989223 | 0.997988 | 0.994666 |
| Coro7_Q9D2V7_S459s_1 | Coro7_Q9D | 0.33 | -0.11 | -0.42 | -0.34 | 0.2 | 0.18 | 0.39 | -0.39 | -1.19 | -0.05 | -0.59 | 0.06 | -0.92 | -0.15 | 0.24 | 0.06 | 0.44 | 0.9675 | 0.345 | 0.865973 | 0.414012 | 0.38824 | 0.989223 | 0.997988 | 0.999507 |
| Cotl1_Q9CQI6_S141s_1 | Cotl1_Q9C | 0.55 | 0.33 | 0.45 | 1.1 | -1.02 | 0.55 | -0.22 | 0.23 | 1.3 | 0.54 | 0.07 | 0.61 | 1.5 | -1.39 | 0.51 | -0.23 | 0.23 | -0.3075 | -0.0575 | 0.389753 | 0.078398 | 0.945555 | 0.989223 | 0.997988 | 0.994666 |
| Cox6a1_Q9DCW5_S28s_1 | Cox6a1_Q9 | -0.53 | -0.64 | -1.11 | -1.01 | -2.29 | -1.13 | -1.55 | -1.57 | 0.14 | -1.27 | -0.82 | -0.79 | 1.21 | -2.81 | -1.71 | -1.59 | -0.7225 | 0.0225 | -0.51 | 0.603417 | 0.022091 | 0.691563 | 0.989223 | 0.997988 | 0.995259 |
| Cpeb2_E9Q5X2_S514s_1 | Cpeb2_E9C | 0.03 | 0.08 | 0.65 | 0.16 | -0.42 | 0.18 | 0.52 | -0.15 | 0.31 | -0.07 | 0.23 | -0.02 | -0.43 | 0.48 | -0.32 | 0.2 | -0.8125 | 0.1375 | -0.4025 | 0.434586 | 0.490616 | 0.636903 | 0.989223 | 0.997988 | 0.994666 |
| Cpeb2_E9Q5X2_S517s_1 | Cpeb2_E9C | 0.97 | -0.23 | 0.04 | 1.1 | -0.09 | 0.08 | 0.8 | 1.17 | 0.53 | 1.04 | 0.53 | 0.19 | 0.47 | 0.81 | 0.71 | 0.75 | -0.1975 | -0.1175 | -0.2475 | 0.557561 | 0.962212 | 0.787437 | 0.999187 | 0.997988 | 0.995259 |
| Cpeb3_D3Z1R6_S197s_1 | Cpeb3_D3Z | -0.51 | -0.1 | -0.33 | -0.34 | -0.27 | -0.42 | -0.59 | -0.4 | 0.59 | -0.31 | 0.18 | -0.8 | 1.3 | 0.23 | -0.56 | -0.23 | 0.02 | 0.1025 | 0.215 | 0.241928 | 0.635398 | 0.488635 | 0.989223 | 0.997988 | 0.994666 |
| Cpeb3_D3Z1R6_S291s_1 | Cpeb3_D3Z | -1.15 | -1.66 | -0.49 | -0.65 | -1.06 | -0.55 | -0.13 | -0.45 | -1.12 | -0.92 | -0.09 | -0.91 | -0.83 | -0.65 | -0.87 | -0.67 | -0.1 | 0.235 | 0.505 | 0.471971 | 0.210394 | 0.5329 | 0.989223 | 0.997988 | 0.994666 |
| Cpeb3_D3Z1R6_S413s_1 | Cpeb3_D3Z | -0.7 | -0.42 | -1.38 | -1.12 | -0.87 | -1.08 | -0.97 | -1.02 | -0.63 | -0.23 | -0.9 | -0.98 | -1.89 | -0.29 | -1.09 | -0.47 | 0.44 | 0.2275 | 0.2325 | 0.942864 | 0.762664 | 0.481156 | 0.995762 | 0.997988 | 0.999507 |
| Cpeb4_Q5SU48_S97s_1 | Cpeb4_Q5 | -0.27 | 0.04 | -0.27 | 0.07 | 0.14 | 0 | 0.44 | 0.63 | -0.79 | -0.06 | -0.09 | 0.22 | -0.59 | -0.89 | 0.29 | 0.42 | -0.08 | 0.22 | -0.03 | 0.81387 | 0.110403 | 0.798554 | 0.989223 | 0.997988 | 0.999507 |
| Cpeb4_Q5SU48_S99s_1 | Cpeb4_Q5 | 0.09 | 0.33 | 0.44 | 0.12 | 0.45 | 0.46 | 0.68 | 0.57 | -0.09 | 0.8 | 0.55 | 0.25 | -0.81 | 0.15 | 1 | 0.99 | 0.41 | -0.0725 | -0.085 | 0.8395 | 0.177359 | 0.624519 | 0.989223 | 0.997988 | 0.999507 |
| Cplx1_P63040_S93s_1 | Cplx1_P63 | -0.11 | -0.11 | -0.57 | -0.41 | 0.19 | -0.44 | -0.27 | -0.13 | -0.68 | -0.19 | -0.75 | -0.39 | -0.5 | 0.45 | -0.72 | -0.19 | 0.295 | 0.1325 | 0.0875 | 0.853041 | 0.571087 | 0.424803 | 0.989223 | 0.997988 | 0.999507 |
| Cplx2_P84086_T95t_1 | Cplx2_P84 | -1.47 | -1.25 | -0.13 | -0.03 | -1.79 | -1 | -0.04 | -0.08 | -1.37 | -1.72 | 0.12 | -0.04 | -2.18 | -0.18 | -0.35 | -0.04 | 0.1375 | -0.2025 | 0.06 | 0.954839 | 0.988283 | 0.952967 | 0.999187 | 0.997988 | 0.999507 |
| Cplx3_Q8R1B5_T113t_1 | Cplx3_Q8R | -1.41 | -0.41 | -1.98 | -2.57 | -0.86 | -1.8 | -2.48 | -3.2 | -0.75 | -1.28 | -0.66 | -2.66 | -1.21 | -0.89 | -1.98 | -2.24 | -0.0075 | -0.0325 | 0.0325 | 0.980911 | 0.424555 | 0.670777 | 0.989223 | 0.997988 | 1 |
| Cpne1_B7ZCP4_T469t_1 | Cpne1_B7Z | 0.81 | -0.19 | -0.09 | -0.99 | 0.1 | 0.07 | -1.08 | -0.67 | 0.74 | 0.06 | -1.31 | -0.69 | 1.54 | 0.56 | 0.04 | -0.89 | -0.4925 | 0.255 | 0.0125 | 0.46465 | 0.528894 | 0.731435 | 0.989223 | 0.997988 | 0.994666 |
| Cpne5_Q8JZW4_T138t_1 | Cpne5_Q8J | -2.06 | -1.02 | -1.59 | -0.96 | -0.3 | -2.81 | -0.46 | -1.21 | -2.48 | -1.27 | -1.99 | -1.1 | -1.3 | 0.19 | -0.8 | -0.81 | -0.28 | -0.185 | 0.4275 | 0.100878 | 0.708601 | 0.465346 | 0.995437 | 0.997988 | 0.994666 |
| Cpne6_Q3UYN2_S330s_1 | Cpne6_Q3 | -0.79 | -1.06 | -1.3 | -0.72 | -1.49 | -0.59 | -0.91 | -0.85 | -0.54 | -0.37 | 0.59 | -1.25 | -0.12 | -0.92 | -0.54 | -0.59 | 0.2125 | -0.3025 | 0.7275 | 0.15941 | 0.978013 | 0.173758 | 0.999187 | 0.997988 | 0.994666 |
| Cpne6_Q3UYN2_S553s_1 | Cpne6_Q3 | 0.94 | 0.07 | 0.78 | 0.05 | 0.87 | 1.38 | 0.2 | 0.24 | 1.53 | 0.31 | 0.85 | 1.25 | 1.31 | 0.55 | 0.21 | 0.65 | 0.0075 | 0.575 | 0.425 | 0.539543 | 0.561478 | 0.172197 | 0.989223 | 0.997988 | 0.994666 |
| Cpne6_Q3UYN2_S553s_2 | Cpne6_Q3 | 0.29 | 0.32 | 0.32 | 0.29 | -1.26 | -0.02 | -0.05 | 0.03 | -0.34 | 0.43 | -0.06 | -0.15 | -1.5 | -0.39 | 0.32 | 0.19 | 0.2125 | 0.525 | 0.22 | 0.141998 | 0.077468 | 0.193134 | 0.989223 | 0.997988 | 0.994666 |
| Cpne6_Q3UYN2_S555s_1 | Cpne6_Q3 | 0.18 | 0.64 | 0.5 | 0.52 | 0.04 | 0.22 | 0.25 | 0.05 | 0.42 | 0.77 | 0.47 | 0.4 | 0.61 | 0.73 | 0.47 | -0.21 | -0.63 | -0.335 | -0.65 | 0.839557 | 0.154927 | 0.810427 | 0.989223 | 0.997988 | 0.999507 |
| Cpne6_Q3UYN2_S555s_2 | Cpne6_Q3 | -0.35 | -0.1 | 0.31 | 0.07 | 0.88 | 0.21 | 0.02 | 0.13 | -0.31 | 0.09 | -0.01 | -0.07 | -0.97 | 0.71 | 0.4 | 0.33 | -0.32 | 0.055 | -0.06 | 0.737852 | 0.254312 | 0.813761 | 0.989223 | 0.997988 | 0.9971 |
| Cpne6_Q3UYN2_S555s_3 | Cpne6_Q3 | -1.27 | -0.96 | 0.54 | -0.1 | -1.24 | -0.62 | 0.49 | 0.21 | -0.14 | -1 | 1.84 | 0.58 | 0.39 | -0.88 | 0.01 | 0.62 | 0.3275 | -0.0575 | 0.135 | 0.34608 | 0.762163 | 0.263183 | 0.995762 | 0.997988 | 0.994666 |
| Cpne6_Q3UYN2_S68s_1 | Cpne6_Q3 | -0.11 | -0.25 | -1.07 | -0.39 | -0.41 | 0.47 | -0.7 | -0.53 | -0.34 | 0.21 | 0.4 | -0.89 | 0.25 | -0.94 | -0.43 | -0.06 | 0.1575 | 0.7675 | 0.4825 | 0.656875 | 0.636649 | 0.424575 | 0.989223 | 0.997988 | 0.995259 |
| Cpne6_Q3UYN2_T547t_1 | Cpne6_Q3 | -0.6 | -0.44 | -0.13 | -0.3 | -0.31 | -0.43 | -0.19 | -0.42 | -0.7 | -0.4 | -0.28 | -0.11 | -0.65 | -0.03 | -0.14 | -0.55 | 0.1625 | 0.3 | 0.16 | 0.926198 | 0.887806 | 0.983516 | 0.996493 | 0.997988 | 0.999507 |
| Cpne6_Q3UYN2_T547t_2 | Cpne6_Q3 | -0.26 | -0.03 | 0.32 | -0.03 | 0.92 | 0.15 | -0.07 | 0 | -0.46 | 0.18 | -0.02 | -0.13 | -0.98 | 0.69 | 0.51 | 0.3 | 0.03 | -0.005 | 0.025 | 0.747158 | 0.395768 | 0.671487 | 0.989223 | 0.997988 | 0.9971 |
| Cpne6_Q3UYN2_T547t_3 | Cpne6_Q3 | -1.27 | -0.96 | 0.54 | -0.1 | -1.24 | -0.62 | 0.49 | 0.21 | -0.14 | -1 | 1.84 | 0.58 | 0.39 | -0.88 | 0.01 | 0.62 | 0.25 | -0.1075 | 0.13 | 0.34608 | 0.762163 | 0.263183 | 0.995762 | 0.997988 | 0.994666 |
| Cpne6_Q3UYN2_T551t_2 | Cpne6_Q3 | -0.51 | -0.32 | 1.23 | 1.18 | -1.19 | 0.41 | 1.37 | 1.17 | 0.62 | -0.35 | 1.41 | 1.67 | -1.34 | 0.05 | 1.5 | 1.86 | 0.1575 | 0.7675 | 0.4825 | 0.873676 | 0.945735 | 0.466675 | 0.999187 | 0.997988 | 0.999507 |
| Cpne6_Q3UYN2_T551t_3 | Cpne6_Q3 | -1.27 | -0.96 | 0.54 | -0.1 | -1.24 | -0.62 | 0.49 | 0.21 | -0.14 | -1 | 1.84 | 0.58 | 0.39 | -0.88 | 0.01 | 0.62 | 0.045 | 0.4425 | 0.1225 | 0.34608 | 0.762163 | 0.263183 | 0.995762 | 0.997988 | 0.994666 |
| Cpox_P36552_S99s_1 | Cpox_P365 | 2.41 | 1.06 | 0.75 | 2.43 | 1.06 | 1.92 | 1.36 | 2.07 | 2.83 | 2.91 | 2.79 | 1.71 | 1.29 | 1.65 | 2.04 | 1.56 | 0.1575 | 0.7675 | 0.4825 | 0.951706 | 0.897339 | 0.096641 | 0.996493 | 0.997988 | 0.999507 |
| Cpsf7_Q8BTV2_S48s_1 | Cpsf7_Q8B | -2.2 | -1.68 | -1.73 | -1.58 | -2.11 | -1.69 | -1.43 | -2.75 | -1.78 | -1.83 | -0.69 | -1.01 | -0.46 | -1.38 | -1.56 | -1.53 | -0.06 | 0.8975 | -0.0275 | 0.115647 | 0.552928 | 0.185473 | 0.989223 | 0.997988 | 0.994666 |
| Cpsf7_Q8BTV2_T194t_1 | Cpsf7_Q8B | -1.01 | -0.6 | -0.99 | -0.96 | -1.59 | -1.84 | -1.11 | -1.08 | -1.8 | -1.15 | -0.3 | -0.36 | -1.09 | -1.49 | -0.41 | -0.37 | -0.1975 | 0.47 | 0.565 | 0.87964 | 0.071808 | 0.973404 | 0.989223 | 0.997988 | 0.999507 |
| Cracr2b_E9Q2F1_S224s_1 | Cracr2b_E9 | -0.48 | -0.16 | -0.24 | -1.48 | -0.6 | -1.45 | -1.27 | -2.02 | -0.23 | -0.19 | -1.6 | -0.32 | 0.74 | -1.61 | -0.89 | -1.04 | -0.515 | -0.0125 | 0.05 | 0.840508 | 0.091149 | 0.990942 | 0.989223 | 0.997988 | 0.999507 |
| Crebbp_F8VPR5_S2350s_1 | Crebbp_F8 | -2.33 | -3.59 | -3.14 | -1.07 | -2.54 | -2.19 | -1.55 | -2.76 | -2.69 | -2.3 | -1.19 | -1.47 | -0.95 | -2.8 | -1.7 | -0.76 | -0.745 | 0.005 | -0.11 | 0.155891 | 0.624393 | 0.315747 | 0.989223 | 0.997988 | 0.994666 |
| Creg2_Q8BGC9_S109s_2 | Creg2_Q8B | -0.2 | -0.94 | -0.62 | -0.48 | -2.08 | 0.17 | -1.46 | -0.19 | 1.06 | -0.37 | -0.25 | -0.73 | 0.94 | -2.45 | 0.1 | -1.01 | 0.2725 | 0.62 | 0.98 | 0.946654 | 0.517356 | 0.256866 | 0.989223 | 0.997988 | 0.999507 |
| Creg2_Q8BGC9_S111s_2 | Creg2_Q8B | -0.2 | -0.94 | -0.62 | -0.48 | -2.08 | 0.17 | -1.46 | -0.19 | 1.06 | -0.37 | -0.25 | -0.73 | 0.94 | -2.45 | 0.1 | -1.01 | -0.33 | 0.4875 | -0.045 | 0.946654 | 0.517356 | 0.256866 | 0.989223 | 0.997988 | 0.999507 |
| Crhr1_P35347_S386s_1 | Crhr1_P35 | 1.64 | 1.45 | 1.65 | 1.31 | 2.05 | 1.28 | 1.4 | 1.82 | -0.11 | 1.38 | -0.43 | 2.11 | 0.11 | -0.88 | 1.08 | 1.44 | -0.33 | 0.4875 | -0.045 | 0.054344 | 0.622524 | 0.191445 | 0.989223 | 0.997988 | 0.994666 |
| Crhr1_P35347_S396s_1 | Crhr1_P35 | -0.31 | -0.27 | -1.25 | -1.32 | -0.99 | -0.53 | -0.32 | -1.87 | 0.02 | -0.37 | -0.74 | -0.59 | 0.16 | 0.01 | -0.29 | -0.94 | 0.125 | -0.775 | -1.075 | 0.199879 | 0.742105 | 0.306204 | 0.995762 | 0.997988 | 0.994666 |
| Crhr1_P35347_S400s_1 | Crhr1_P35 | 1.25 | 1.14 | 0.83 | 0.77 | 1.24 | 0.92 | 0.35 | 0.35 | 1.35 | 1.37 | 0.48 | 0.9 | 0.93 | 1.3 | 0.19 | 0.45 | -0.14 | 0.3675 | 0.5225 | 0.392287 | 0.332051 | 0.923914 | 0.989223 | 0.997988 | 0.994666 |
| Crhr1_P35347_S405s_1 | Crhr1_P35 | -0.31 | 0.02 | -0.65 | -1.44 | 0.25 | -1.03 | -0.36 | -0.53 | -0.71 | -0.96 | -1 | -0.88 | -0.14 | 0.53 | -0.17 | -0.38 | -0.2825 | 0.0275 | -0.28 | 0.170991 | 0.654943 | 0.398936 | 0.990434 | 0.997988 | 0.994666 |
| Crip2_Q9DCT8_S115s_1 | Crip2_Q9D | -0.82 | -0.98 | -0.77 | -0.82 | -0.81 | -1.27 | -0.41 | -0.46 | -1.24 | -0.62 | 0.18 | -0.6 | -1.56 | -0.12 | -0.31 | -0.63 | 0.1775 | -0.2925 | 0.555 | 0.587071 | 0.668707 | 0.399095 | 0.992876 | 0.997988 | 0.995259 |
| Crk_Q8JZR2_S40s_1 | Crk_Q8JZR | 1.66 | 1.13 | 0.75 | 0.51 | 3.09 | 3.14 | 0.42 | 0.83 | 0.86 | 3.08 | 0.29 | 0.81 | 1.5 | 1.42 | -0.07 | 0.49 | 0.11 | 0.2775 | 0.1925 | 0.690399 | 0.225159 | 0.686822 | 0.989223 | 0.997988 | 0.996159 |
| Crkl_P47941_S107s_1 | Crkl_P4794 | 0.31 | 0.21 | 0.96 | 0.64 | 0.38 | 0.13 | 0.76 | 0.76 | -0.08 | 0.23 | 0.54 | 0.96 | -0.1 | 0.38 | 0.14 | 0.97 | 0.8575 | 0.2475 | -0.1775 | 0.58106 | 0.933846 | 0.707922 | 0.998562 | 0.997988 | 0.995259 |
| Crmp1_P97427_S495s_1 | Crmp1_P9 | -0.48 | -0.23 | 0.76 | -0.19 | 0.27 | -0.42 | 0.68 | 0.29 | -1.21 | -0.15 | -0.28 | 0.1 | -1.23 | 0.01 | -0.32 | 0.16 | -0.0225 | -0.1175 | -0.1825 | 0.461431 | 0.505113 | 0.385603 | 0.989223 | 0.997988 | 0.994666 |
| Crmp1_P97427_S540s_1 | Crmp1_P9 | -0.33 | -0.3 | -1.31 | -0.35 | -0.03 | -0.79 | -0.86 | -0.97 | -1.01 | -0.33 | 0.94 | -0.44 | 0.2 | -1.44 | -1.33 | -0.9 | 0.24 | -0.35 | -0.31 | 0.507068 | 0.78813 | 0.437836 | 0.995762 | 0.997988 | 0.994666 |
| Crmp1_P97427_S542s_1 | Crmp1_P9 | 0.31 | 1.1 | -0.28 | -0.52 | 0.61 | 0.35 | -0.3 | 0.1 | 0.71 | 1.04 | 1.47 | -0.26 | 0.38 | 0.77 | -0.26 | -0.25 | -0.09 | 0.3625 | -0.295 | 0.98619 | 0.924335 | 0.247164 | 0.997335 | 0.997988 | 1 |
| Crmp1_P97427_S570s_1 | Crmp1_P9 | -0.93 | -2.05 | -1.77 | -1.81 | -1.75 | -1.69 | -1.68 | -2.1 | -1.32 | -1.22 | 0.3 | -1.15 | -1.87 | -0.65 | -2.27 | -1.81 | 0.0375 | 0.5875 | 0.0075 | 0.981126 | 0.576071 | 0.098524 | 0.989223 | 0.997988 | 1 |
| Crmp1_P97427_S8s_1 | Crmp1_P9 | -3.5 | -5.17 | -2.62 | -1.79 | -3.88 | -4.68 | -1.67 | -1.9 | -5.21 | -4.29 | -2.21 | -2.1 | -3.35 | -3.16 | -2.67 | -3.12 | -0.165 | 0.7925 | -0.01 | 0.769234 | 0.791759 | 0.845542 | 0.995762 | 0.997988 | 0.9971 |
| Crmp1_P97427_T509t_1 | Crmp1_P9 | -1.55 | -1.29 | -1.09 | -0.83 | -0.55 | -1.59 | -0.22 | -0.88 | -1.74 | -1.25 | -0.65 | -0.79 | -1.18 | -0.82 | -0.99 | -0.9 | 0.2375 | -0.1825 | 0.195 | 0.42034 | 0.276537 | 0.796396 | 0.989223 | 0.997988 | 0.994666 |
| Crmp1_Q6P1J1_T27t_1 | Crmp1_Q6 | -2.04 | -2.11 | -1.63 | -1.64 | -4.6 | -2.9 | -2.05 | -1.65 | -1 | -2.98 | -1.81 | -1.55 | -0.96 | -3.84 | -1.15 | -1.49 | 0.38 | 0.0825 | 0.2175 | 0.993547 | 0.13786 | 0.962511 | 0.989223 | 0.997988 | 1 |
| Crocc_Q8CJ40_S1043s_1 | Crocc_Q8C | -0.15 | -0.16 | 0.29 | -0.71 | -0.24 | -1.04 | 0.24 | -0.42 | 0.69 | -0.22 | -1.36 | -0.71 | 1.26 | -0.43 | -0.29 | -0.42 | -0.945 | 0.02 | -0.005 | 0.636987 | 0.594712 | 0.636872 | 0.989223 | 0.997988 | 0.995259 |
| Crocc_Q8CJ40_S1289s_1 | Crocc_Q8C | -1.1 | -1.78 | -1.39 | -2.87 | -2.73 | -1.08 | -1.19 | -2.22 | 1.77 | -1.53 | -2.71 | -0.83 | 2.99 | -2.8 | 1.33 | -0.91 | -0.1825 | -0.2175 | 0.2125 | 0.116592 | 0.968661 | 0.305596 | 0.999187 | 0.997988 | 0.994666 |
| Crocc_Q8CJ40_S1304s_1 | Crocc_Q8C | -0.02 | 0.42 | -0.63 | 0.43 | 0.47 | 0.32 | 0.54 | 0.32 | -0.57 | 0.75 | 0.39 | -0.42 | 0.02 | 0.49 | 0.03 | 0.38 | -0.02 | 0.96 | 1.9375 | 0.580252 | 0.226753 | 0.975119 | 0.989223 | 0.997988 | 0.995259 |
| Crocc_Q8CJ40_S1319s_1 | Crocc_Q8C | 0.31 | 0.82 | 0.64 | 1.1 | 0.12 | 0.08 | 1.2 | 0.35 | 0.57 | 0.51 | -0.08 | 1.04 | 0.45 | 0.73 | 0.08 | 1.21 | 0.3625 | -0.0125 | 0.18 | 0.762801 | 0.395238 | 0.514768 | 0.989223 | 0.997988 | 0.9971 |
| Crocc_Q8CJ40_S1404s_1 | Crocc_Q8C | 0.53 | 0.87 | 0.68 | -0.41 | 0.51 | 0.62 | 0.46 | 0.21 | 0.71 | 0.34 | 0.06 | 0.27 | 0.27 | 0.89 | 0.8 | 0.05 | -0.28 | -0.2075 | -0.1 | 0.818285 | 0.917675 | 0.829454 | 0.997335 | 0.997988 | 0.999507 |
| Crocc_Q8CJ40_S337s_2 | Crocc_Q8C | 0.06 | 0 | -0.42 | -0.03 | -0.14 | -1.23 | -0.04 | -1.03 | 0.26 | 0.11 | -0.4 | 0.03 | 0.23 | -0.29 | 0.74 | 0.48 | 0.0325 | -0.0725 | 0.085 | 0.221952 | 0.147143 | 0.699751 | 0.989223 | 0.997988 | 0.994666 |
| Crocc_Q8CJ40_S337s_3 | Crocc_Q8C | 0.25 | 0 | -0.24 | -0.37 | -0.42 | -0.38 | -0.13 | -0.76 | -0.23 | 0.02 | 0.29 | -0.4 | -2.17 | -0.78 | 0.51 | 0.77 | -0.5125 | 0.0975 | 0.3875 | 0.602892 | 0.200168 | 0.969911 | 0.989223 | 0.997988 | 0.995259 |
| Crocc_Q8CJ40_T330t_2 | Crocc_Q8C | 0.06 | 0 | -0.42 | -0.03 | -0.14 | -1.23 | -0.04 | -1.03 | 0.26 | 0.11 | -0.4 | 0.03 | 0.23 | -0.29 | 0.74 | 0.48 | -0.3325 | 0.01 | -0.3275 | 0.221952 | 0.147143 | 0.699751 | 0.989223 | 0.997988 | 0.994666 |
| Crocc_Q8CJ40_T330t_3 | Crocc_Q8C | 0.25 | 0 | -0.24 | -0.37 | -0.42 | -0.38 | -0.13 | -0.76 | -0.23 | 0.02 | 0.29 | -0.4 | -2.17 | -0.78 | 0.51 | 0.77 | -0.5125 | 0.0975 | 0.3875 | 0.602892 | 0.200168 | 0.969911 | 0.989223 | 0.997988 | 0.995259 |
| Cry2_Q9R194_S557s_1 | Cry2_Q9R1 | -1.17 | -0.21 | -2.22 | -4.19 | -1.01 | -0.95 | -5.03 | -3.72 | -1.07 | -0.45 | -2.56 | -2.91 | -0.45 | -0.3 | -2.5 | -2.77 | -0.3325 | 0.01 | -0.3275 | 0.639451 | 0.527207 | 0.827587 | 0.989223 | 0.997988 | 0.995259 |

| | | | | | | | | | | | | | | | | | | | | | | | | | | |
|---|---|---|---|---|---|---|---|---|---|---|---|---|---|---|---|---|---|---|---|---|---|---|---|---|---|---|
| Cs_Q9CZU6_S226s_1 | Cs_Q9CZU | 0.74 | 1 | 0.24 | 0.08 | 0.63 | 0.27 | 0.05 | 0.17 | 0.64 | 0.64 | 1.03 | 0.49 | 0.67 | 0.93 | 0.17 | 0.27 | -0.73 | 0.2 | 0.4425 | 0.987644 | 0.414235 | 0.527007 | 0.989223 | 0.997988 | 1 |
| Csdc2_Q91YQ3_S48s_1 | Csdc2_Q91 | -1.62 | -0.86 | -1.38 | -1.79 | -1.33 | -1.26 | -1.32 | -1.07 | -1.55 | -1.66 | 0.21 | -1.94 | -1.15 | 0.04 | -1.22 | -1.61 | -0.235 | 0.185 | -0.005 | 0.313137 | 0.524382 | 0.722396 | 0.989223 | 0.997988 | 0.994666 |
| Csde1_Q91W50_S123s_1 | Csde1_Q91 | 0.12 | 0.41 | -0.54 | 0.14 | 0.87 | 0.43 | -0.13 | -0.13 | -0.05 | 0.24 | 0.01 | -0.1 | 0.48 | 0.32 | 0.2 | -0.14 | 0.1675 | 0.1775 | 0.4275 | 0.548653 | 0.492392 | 0.978007 | 0.989223 | 0.997988 | 0.994666 |
| Csnk1a1_E9Q2U6_T349t_1 | Csnk1a1_E | -0.59 | -0.42 | 1.55 | 1.87 | -1.58 | 0.18 | 0.58 | 1.32 | -0.06 | 0.23 | -0.01 | 1 | 0.01 | -0.84 | 0.43 | 1.03 | 0.2275 | -0.0075 | 0.1825 | 0.515892 | 0.544059 | 0.620845 | 0.989223 | 0.997988 | 0.994666 |
| Csnk1d_Q9DC28_S331s_1 | Csnk1d_Q9 | -1.76 | -2.74 | -1.67 | -1.75 | -1.36 | -1.91 | -1.71 | -1.99 | -1.38 | -2.46 | 1.26 | -0.32 | -0.51 | -1.35 | -1.59 | -1.52 | -0.4775 | -0.3125 | -0.445 | 0.073252 | 0.450679 | 0.119225 | 0.989223 | 0.997988 | 0.994666 |
| Csnk1d_Q9DC28_T347t_1 | Csnk1d_Q9 | -0.34 | 0.36 | 0.25 | 0.28 | 0.21 | -0.33 | 0.44 | -0.43 | 0.21 | -0.04 | 0.92 | 0.4 | 0.17 | 0.35 | 0.05 | 1.05 | 0.2375 | 1.255 | 0.7375 | 0.414545 | 0.576338 | 0.440263 | 0.989223 | 0.997988 | 0.994666 |
| Csnk1e_Q9JMK2_S363s_1 | Csnk1e_Q9 | 0.1 | -1.21 | 0.01 | -0.86 | -0.54 | 0.42 | -0.84 | -0.72 | 1.73 | 0.66 | -1.73 | -0.08 | 1.31 | -1.94 | -0.2 | -0.27 | -0.165 | 0.235 | 0.2675 | 0.746787 | 0.865097 | 0.385851 | 0.996493 | 0.997988 | 0.9971 |
| Csnk1e_Q9JMK2_S389s_1 | Csnk1e_Q9 | -0.3 | 0.37 | 0.57 | 0.13 | -0.68 | 0.09 | 0.55 | 0.69 | -0.49 | 0.4 | 0.71 | 0.68 | 0.06 | 0.2 | 0.5 | 0.52 | 0.07 | 0.635 | 0.215 | 0.660648 | 0.93343 | 0.707508 | 0.998562 | 0.997988 | 0.995671 |
| Csnk1g1_Q8BTH8_S361s_1 | Csnk1g1_Q | 0.51 | -0.78 | 0.59 | 0.97 | -0.35 | 1.39 | -0.06 | 0.78 | 0.98 | 0.21 | 1.97 | 0.88 | 0.91 | 1.01 | 0.09 | 0.16 | -0.03 | 0.1325 | 0.1275 | 0.620763 | 0.815621 | 0.188678 | 0.995762 | 0.997988 | 0.995259 |
| Csnk1g1_Q8BTH8_T214t_1 | Csnk1g1_Q | -0.72 | -0.37 | -0.41 | -0.66 | -0.27 | -0.6 | -0.9 | -0.87 | -0.36 | -1.03 | -2.58 | -0.71 | 0.71 | -0.83 | -0.95 | -0.5 | 0.1175 | 0.6875 | 0.22 | 0.711248 | 0.616486 | 0.205049 | 0.989223 | 0.997988 | 0.996966 |
| Csnk2b_G3UXG7_S158s_1 | Csnk2b_G3 | 1.14 | 0.94 | 1.03 | 1.08 | 0.94 | 1 | 1.16 | 1.57 | 1 | 1.18 | 0.72 | 1.02 | 0.77 | 0.93 | 0.98 | 1.02 | -0.12 | -0.63 | 0.1475 | 0.596575 | 0.59963 | 0.758262 | 0.989223 | 0.997988 | 0.995259 |
| Cspg5_Q71M36_S467s_1 | Cspg5_Q71 | -0.05 | 0.91 | 0.35 | 0.63 | 0.76 | 0.03 | 0.96 | 0.86 | -0.25 | 0.37 | 0.29 | 0.83 | 0.14 | 0.06 | 0.59 | 0.56 | 0.12 | -0.0675 | -0.1225 | 0.688941 | 0.54214 | 0.648418 | 0.989223 | 0.997988 | 0.996159 |
| Cspg5_Q71M36_S467s_1_1 | Cspg5_Q71 | 0.21 | 0.68 | 0.27 | 0.11 | 0.64 | 0.53 | 0.72 | 0.49 | 0.1 | 0.93 | -0.23 | 0.31 | 0.31 | 0.46 | 0.19 | 0.13 | 0.1925 | -0.15 | -0.1225 | 0.859187 | 0.229412 | 0.896979 | 0.989223 | 0.997988 | 0.999507 |
| Cspg5_Q71M36_S475s_1 | Cspg5_Q71 | -0.27 | -0.03 | 0.28 | 0.28 | 0.26 | -0.23 | 0.61 | 0.43 | -0.09 | 0.14 | -0.13 | 0.2 | 0.47 | 0.14 | 0.29 | 0.15 | 0.2775 | -0.04 | -0.045 | 0.45035 | 0.456941 | 0.884409 | 0.989223 | 0.997988 | 0.994666 |
| Cspg5_Q71M36_S475s_1_1 | Cspg5_Q71 | -0.35 | -0.08 | 0.81 | 0.79 | 0.72 | -0.08 | 0.93 | 0.45 | -0.39 | 0.41 | 0.76 | 1.31 | -0.13 | 0.53 | 0.61 | 0.91 | 0.2025 | -0.035 | 0.1975 | 0.627029 | 0.564609 | 0.609816 | 0.989223 | 0.997988 | 0.995259 |
| Cspg5_Q71M36_S541s_1 | Cspg5_Q71 | 0.21 | 0.3 | 0.33 | 0.75 | 0.62 | 1.16 | 0.98 | 0.51 | 0.51 | 0.89 | 0.51 | 0.96 | -0.34 | 0.56 | 0.13 | 0.79 | 0.2125 | 0.23 | 0.1875 | 0.727896 | 0.118611 | 0.21705 | 0.989223 | 0.997988 | 0.9971 |
| Cspg5_Q71M36_S543s_1 | Cspg5_Q71 | 0.32 | 0.31 | 0.16 | 0.71 | 0.84 | 0.26 | 0.78 | 0.49 | 0.03 | 0.62 | 0.05 | 0.79 | -0.9 | 0.45 | 0.13 | 0.55 | 0.42 | 0.32 | -0.1125 | 0.401512 | 0.379595 | 0.992859 | 0.989223 | 0.997988 | 0.994666 |
| Cspp1_B2RX88_S806s_1 | Cspp1_B2R | 0.13 | 0.82 | -1.56 | -1 | -0.48 | 0.12 | -1.01 | -1.35 | -0.01 | 0.23 | -0.26 | -1.3 | 0.56 | -0.86 | -1.16 | -0.5 | 0.2175 | -0.0025 | -0.3175 | 0.883791 | 0.625022 | 0.90786 | 0.989223 | 0.997988 | 0.999507 |
| Csrp1_P97315_S192s_1 | Csrp1_P97 | 0.71 | 0.94 | 0.46 | 0.46 | 0.81 | 0.46 | 0.64 | 0.52 | 0.05 | 0.68 | 0.17 | 0.52 | -0.4 | 0.77 | 0.14 | 0.4 | -0.2775 | 0.0675 | -0.0875 | 0.214192 | 0.875091 | 0.278824 | 0.996493 | 0.997988 | 0.994666 |
| Csrp1_P97315_S81s_1 | Csrp1_P97 | -1.65 | -0.54 | -2.73 | -4.5 | -0.52 | -0.8 | -2.31 | -2.63 | -1.01 | -1.23 | -1.07 | -2.41 | -1.51 | -0.4 | -2.22 | -2.53 | -0.035 | -0.2875 | -0.415 | 0.425154 | 0.37271 | 0.269855 | 0.989223 | 0.997988 | 0.994666 |
| Cstf2_F6ZKC7_S145s_1 | Cstf2_F6ZK | 0.88 | 0.55 | -2.04 | -2.11 | 0.38 | -0.03 | -1.35 | -1.61 | 0.92 | 0.84 | -1.89 | -1.86 | 0.93 | 0.3 | -1 | -1.83 | 0.79 | 0.925 | 0.69 | 0.754648 | 0.973336 | 0.855325 | 0.999187 | 0.997988 | 0.9971 |
| Cstf3_Q99LI7_S691s_1 | Cstf3_Q99 | -1.96 | -2.75 | -3.07 | -3.45 | -3.56 | -1.51 | -3.3 | -3.43 | -0.8 | -2.64 | -1.23 | -2.44 | -0.41 | -2.58 | -1.15 | -3.17 | 0.0275 | 0.1825 | 0.28 | 0.150733 | 0.78635 | 0.071399 | 0.995762 | 0.997988 | 0.994666 |
| Ctage5_H3BK44_S496s_1 | Ctage5_H3 | -0.52 | 0.35 | -0.99 | -0.71 | 0.22 | -0.53 | 0.22 | -0.46 | -0.27 | -0.2 | -0.47 | -1.03 | -0.55 | 0.59 | -0.06 | -0.02 | -0.1425 | 1.03 | 0.98 | 0.252244 | 0.364544 | 0.944194 | 0.989223 | 0.997988 | 0.994666 |
| Ctage5_H3BK44_S543s_1 | Ctage5_H3 | -1.05 | -2.27 | -1.22 | -1.07 | 0.25 | -0.01 | 0.41 | 0.7 | -1.2 | -1.11 | 1.31 | -0.4 | -1.67 | -0.28 | -0.94 | 0.68 | 0.33 | -0.025 | 0.4575 | 0.138927 | 0.000498 | 0.104139 | 0.989223 | 0.997988 | 0.994666 |
| Ctage5_H3BK44_S593s_1 | Ctage5_H3 | -4.38 | -1.47 | -1.76 | -1.42 | -1.78 | -2.8 | 0.26 | -0.81 | -4.66 | -1.46 | -0.37 | -0.07 | -2.63 | -0.45 | -0.17 | -0.36 | 1.74 | 1.0525 | 0.85 | 0.119435 | 0.264539 | 0.582867 | 0.989223 | 0.997988 | 0.994666 |
| Ctage5_H3BK44_S636s_1 | Ctage5_H3 | 0.36 | 0.74 | 0.2 | 0.76 | 0.88 | 0.17 | 0.32 | 0.39 | -0.09 | 0.32 | 0.33 | 0.15 | -0.89 | 0.04 | 0.69 | 0.94 | 0.975 | 0.6175 | 1.355 | 0.460755 | 0.771736 | 0.195439 | 0.995762 | 0.997988 | 0.994666 |
| Ctdp1_Q7TSG2_S502s_1 | Ctdp1_Q7T | -0.97 | -0.62 | -1.43 | -0.61 | -0.37 | -0.46 | -0.5 | -0.61 | -0.11 | -0.33 | -0.76 | -0.63 | -0.07 | 0.33 | -1.66 | -0.81 | -0.075 | -0.3375 | -0.32 | 0.44992 | 0.120781 | 0.144211 | 0.989223 | 0.997988 | 0.994666 |
| Ctdspl2_Q8BG15_S9s_1 | Ctdspl2_Q8 | 0.94 | 0.04 | 0.06 | -0.22 | 0.06 | 0.69 | -0.05 | -0.13 | 0.48 | 0.24 | 0.11 | 0.3 | 1.46 | 0.65 | -0.14 | -0.32 | 0.4225 | 0.45 | 0.355 | 0.655733 | 0.848155 | 0.798073 | 0.996349 | 0.997988 | 0.995259 |
| Ctif_Q6PEE2_S18s_1 | Ctif_Q6PEE | -1.63 | -1.28 | -1.32 | -2.23 | -0.67 | -2.03 | -1.54 | -1.71 | -1.61 | -1.51 | -1.07 | -1.58 | -1.73 | -0.82 | -1.54 | -1.7 | -0.0625 | 0.0775 | 0.2075 | 0.625254 | 0.725256 | 0.562608 | 0.995437 | 0.997988 | 0.995259 |
| Ctnna2_E0CXB9_S275s_1 | Ctnna2_E0 | 0.41 | 0.84 | 1.16 | 0.36 | 0.6 | 0.2 | 0.88 | 0.75 | 0.45 | 0.73 | 0.5 | 0.67 | -0.21 | 0.65 | 0.18 | 0.47 | 0.1275 | 0.1725 | 0.1675 | 0.205339 | 0.759774 | 0.691902 | 0.995762 | 0.997988 | 0.994666 |
| Ctnna2_E0CXB9_S667s_1 | Ctnna2_E0 | 1.78 | 2.62 | 1.38 | 1.93 | 1.62 | 1.58 | 1.63 | 1.62 | 2.08 | 2.26 | 2.8 | 1.56 | 1.23 | 2.06 | 1.16 | 1.35 | -0.085 | -0.105 | -0.42 | 0.202448 | 0.291468 | 0.509976 | 0.989223 | 0.997988 | 0.994666 |
| Ctnna2_E0CXB9_S667s_2 | Ctnna2_E0 | 0.65 | 0.86 | 0.97 | 1.33 | -0.28 | 0.37 | 1.06 | 1.51 | 0.41 | 0.54 | 1.33 | 1.81 | -2.48 | 0.73 | -1.15 | 1.7 | -0.315 | 0.2475 | -0.4775 | 0.15719 | 0.47979 | 0.849672 | 0.989223 | 0.997988 | 0.994666 |
| Ctnna2_E0CXB9_S914s_1 | Ctnna2_E0 | 0.35 | 0.23 | -0.19 | -0.07 | 0.59 | 0.29 | 0.5 | 0.08 | -0.36 | 0.17 | -0.48 | -0.04 | -0.7 | 0.26 | -0.29 | 0.37 | -0.2875 | 0.07 | -1.2525 | 0.602559 | 0.248562 | 0.336255 | 0.989223 | 0.997988 | 0.995259 |
| Ctnna2_E0CXB9_S952s_1 | Ctnna2_E0 | 4.63 | 3.08 | 5.02 | 2.57 | 3.41 | 4.99 | 5.42 | 4.62 | 0.26 | 4.57 | 0.84 | 5.41 | -0.42 | 0.61 | 4.84 | 5.68 | 0.285 | -0.2575 | -0.17 | 0.420608 | 0.245788 | 0.408365 | 0.989223 | 0.997988 | 0.994666 |
| Ctnna2_E0CXB9_T274t_1 | Ctnna2_E0 | -0.05 | 0.8 | 1.31 | 0.89 | 0.42 | 0.14 | 1.34 | 0.72 | 0.8 | 0.94 | 0.18 | 0.87 | -0.1 | 0.6 | 0.41 | 0.7 | 0.785 | -1.055 | -1.1475 | 0.364033 | 0.826486 | 0.908953 | 0.995762 | 0.997988 | 0.994666 |
| Ctnna2_E0CXB9_T666t_2 | Ctnna2_E0 | 0.53 | 0.5 | 0.41 | 1.47 | -2.29 | 0.15 | 0.78 | 1.4 | -0.11 | 0.4 | 0.79 | 1.67 | -3.57 | -0.22 | -6.13 | 1.42 | -0.0825 | -0.04 | -0.335 | 0.077087 | 0.346003 | 0.926988 | 0.989223 | 0.997988 | 0.994666 |
| Ctnna2_E0CXB9_T670t_1 | Ctnna2_E0 | 0.18 | 0.77 | 1.53 | 0.83 | 1.73 | 0.4 | 1.42 | 0.73 | 0.34 | 0.72 | -0.17 | 1.67 | -0.92 | 0.59 | 1 | 1.5 | -0.7175 | -0.04 | -2.8525 | 0.605196 | 0.545453 | 0.68299 | 0.989223 | 0.997988 | 0.995259 |
| Ctnna2_E0CXB9_T670t_2 | Ctnna2_E0 | 0.85 | 1.36 | 1.11 | 0.78 | 1.39 | 0.58 | 1.18 | 1.53 | 0.89 | 0.78 | 1.7 | 1.69 | -1.4 | 1.88 | 1.48 | 1.93 | 0.2425 | -0.1875 | -0.285 | 0.941871 | 0.609865 | 0.452324 | 0.989223 | 0.997988 | 0.999507 |
| Ctnnb1_E9Q6A9_S191s_1 | Ctnnb1_E9 | 1.2 | 0.15 | 0.23 | 0.23 | 0.95 | 0.88 | 0.4 | 0.41 | 0.63 | 0.56 | -0.42 | 0.15 | -0.38 | 0.48 | -0.01 | -0.21 | 0.145 | 0.24 | -0.0525 | 0.183521 | 0.508259 | 0.53987 | 0.989223 | 0.997988 | 0.994666 |
| Ctnnb1_Q02248_S552s_1 | Ctnnb1_Q0 | -0.05 | 0.38 | 0.45 | 0.27 | 0.78 | -0.12 | 1.3 | 0.2 | -0.76 | -0.17 | -0.6 | 1.1 | -0.17 | 0.23 | 0.07 | 1.21 | 0.2075 | -0.2225 | -0.4825 | 0.836684 | 0.422315 | 0.39443 | 0.989223 | 0.997988 | 0.999507 |
| Ctnnb1_Q02248_S675s_1 | Ctnnb1_Q0 | 0.64 | 0.88 | 0.55 | -0.43 | 1.1 | 0.65 | 1.32 | 0.24 | 0.17 | 0.41 | -1 | -0.49 | -0.19 | 0.7 | -0.64 | 0.02 | 0.2775 | -0.37 | 0.0725 | 0.295817 | 0.275779 | 0.15552 | 0.989223 | 0.997988 | 0.994666 |
| Ctnnb1_Q02248_T551t_1 | Ctnnb1_Q0 | 0.19 | 0.42 | 1.28 | 1.29 | 0.72 | 0.08 | 2.59 | 1.18 | -0.32 | 0.06 | 0.7 | 2.63 | -0.07 | 0.11 | 1.21 | 2.85 | 0.4175 | -0.6375 | -0.4375 | 0.727137 | 0.529634 | 0.966205 | 0.989223 | 0.997988 | 0.9971 |
| Ctnnbip1_Q9JJN6_S10s_1 | Ctnnbip1_ | 0.5 | 0.52 | 0.64 | 0.62 | 0.89 | 0.24 | 0.62 | 0.28 | 0.04 | 0.78 | -0.22 | 0.53 | 0.25 | 1.06 | 0.07 | 0.77 | 0.3475 | -0.0275 | 0.23 | 0.912514 | 0.787287 | 0.320395 | 0.995762 | 0.997988 | 0.999507 |
| Ctnnbl1_Q9CWL8_S545s_1 | Ctnnbl1_Q | -0.74 | -1.5 | -1.44 | -1.5 | -1.28 | -1.44 | -1.01 | -1.5 | -1.34 | -0.95 | -1.97 | -1.69 | -0.88 | -0.53 | -1.33 | -1.7 | -0.0625 | -0.2875 | -0.0325 | 0.597016 | 0.962011 | 0.549256 | 0.999187 | 0.997988 | 0.995259 |
| Ctnnd1_P30999_S269s_1 | Ctnnd1_P3 | -0.87 | 0.6 | -0.56 | -0.4 | 0.69 | 0.82 | -0.35 | 0.15 | 1.09 | 0.21 | 1.61 | -0.48 | 1.25 | 1.41 | -0.42 | -0.64 | -0.0125 | -0.1925 | 0.185 | 0.238036 | 0.137256 | 0.107695 | 0.989223 | 0.997988 | 0.994666 |
| Ctnnd1_P30999_S312s_1 | Ctnnd1_P3 | -1.02 | -1.21 | -0.43 | -0.01 | -1.17 | -0.54 | -0.56 | -0.87 | 2.14 | -0.75 | -1.07 | 0.01 | 2.14 | -1.31 | 0.14 | -0.03 | 0.635 | 0.915 | 0.7075 | 0.208535 | 0.718578 | 0.297784 | 0.995437 | 0.997988 | 0.994666 |
| Ctnnd1_P30999_S346s_1 | Ctnnd1_P3 | -0.39 | -0.11 | 0.15 | 0.07 | 0.86 | -0.74 | 0.9 | 0.35 | -0.87 | -0.65 | 0.01 | 0.49 | -0.55 | 0.16 | -0.17 | 0.41 | -0.1175 | 0.75 | 0.9025 | 0.913911 | 0.302926 | 0.598853 | 0.989223 | 0.997988 | 0.999507 |
| Ctnnd1_P30999_S349s_1 | Ctnnd1_P3 | 0 | -0.45 | -0.59 | -0.38 | 0.27 | -0.18 | 0.05 | -0.58 | -0.4 | -0.13 | -0.5 | -0.16 | -0.67 | 0.11 | -0.7 | -0.46 | 0.4125 | -0.185 | 0.0325 | 0.798191 | 0.368119 | 0.810905 | 0.989223 | 0.997988 | 0.999507 |
| Ctnnd1_P30999_S47s_1 | Ctnnd1_P3 | -0.82 | -0.68 | 0.43 | 0.47 | -0.43 | -0.78 | 1.05 | 0.85 | -0.7 | -0.09 | 1.27 | 1.19 | -0.12 | 0.24 | 0.87 | 1.05 | 0.245 | 0.0575 | -0.075 | 0.153032 | 0.540907 | 0.319258 | 0.989223 | 0.997988 | 0.994666 |
| Ctnnd1_P30999_T310t_1 | Ctnnd1_P3 | 0.39 | -0.14 | 0.58 | 0.5 | -0.52 | 0.95 | -0.37 | -0.23 | 2.25 | 0.33 | -0.06 | 0.76 | 2.56 | -0.85 | 0.61 | 0.82 | 0.3225 | 0.5675 | 0.66 | 0.490362 | 0.322318 | 0.342218 | 0.989223 | 0.997988 | 0.994666 |
| Ctnnd2_B7ZNF6_S1054s_1 | Ctnnd2_B7 | -1.56 | -1.77 | -1.27 | -1.86 | -0.04 | -1.6 | -1.01 | -1.29 | -2.22 | -1.36 | -0.3 | -1.71 | -0.72 | -0.78 | -1.34 | -0.94 | -0.375 | 0.4875 | 0.4525 | 0.033588 | 0.105757 | 0.605885 | 0.989223 | 0.997988 | 0.994666 |
| Ctnnd2_B7ZNF6_S1056s_1 | Ctnnd2_B7 | 0.63 | 1.08 | 0.74 | 0.94 | 0.92 | 0.76 | 0.63 | 0.88 | 0.65 | 0.78 | 1.18 | 0.76 | 0.29 | 0.34 | 0.15 | 0.53 | 0.63 | 0.2175 | 0.67 | 0.058659 | 0.816015 | 0.983317 | 0.995762 | 0.997988 | 0.994666 |
| Ctnnd2_B7ZNF6_S1061s_1 | Ctnnd2_B7 | 2.1 | 1.02 | 1 | 1.46 | 0.64 | 1.48 | 1.22 | 1.36 | 2.19 | 2.6 | 1.45 | 1.54 | 0.91 | 0.66 | 1.17 | 1.34 | -0.05 | -0.005 | -0.52 | 0.280088 | 0.508555 | 0.171888 | 0.989223 | 0.997988 | 0.994666 |
| Ctnnd2_B7ZNF6_S1068s_1 | Ctnnd2_B7 | 0.05 | -0.17 | 0.73 | 0.2 | -1.03 | 0.24 | -0.77 | -0.42 | 1.65 | -0.29 | -1.18 | 0.3 | 1.39 | -2.76 | -1.4 | 0.49 | -0.22 | 0.55 | -0.375 | 0.367926 | 0.065017 | 0.885806 | 0.989223 | 0.997988 | 0.994666 |
| Ctnnd2_B7ZNF6_S1068s_2 | Ctnnd2_B7 | -0.64 | -0.4 | -0.57 | -1.23 | -1.18 | 0.04 | -1.82 | -1.66 | 1.43 | -1.21 | -1.17 | -0.63 | -0.68 | -2.8 | -1.83 | -1.18 | -0.6975 | -0.0825 | -0.7725 | 0.078429 | 0.316822 | 0.598926 | 0.989223 | 0.997988 | 0.994666 |
| Ctnnd2_B7ZNF6_S1072s_1 | Ctnnd2_B7 | 0.7 | 0.43 | 1.17 | 1.12 | 0.43 | 1.42 | 0.31 | 0.61 | 2.71 | 1.02 | -0.67 | 0.72 | 2.26 | -2.08 | -0.54 | 0.36 | -0.445 | 0.315 | -0.9125 | 0.308244 | 0.614654 | 0.88986 | 0.989223 | 0.997988 | 0.994666 |
| Ctnnd2_B7ZNF6_S1072s_2 | Ctnnd2_B7 | 0.36 | 0.39 | -0.57 | -1.23 | -0.6 | 0.98 | -1.82 | -1.66 | 2.22 | 0.05 | -1.17 | -0.63 | -0.03 | -1.9 | -1.83 | -1.18 | -0.1625 | 0.09 | -0.855 | 0.098724 | 0.449986 | 0.616565 | 0.989223 | 0.997988 | 0.994666 |
| Ctnnd2_B7ZNF6_S1123s_1 | Ctnnd2_B7 | 0.46 | 1.12 | 0.75 | 0.62 | 1.05 | 0.15 | 0.71 | 0.61 | 0.42 | 0.49 | 0.28 | 0.68 | -1.42 | 0.18 | 0.47 | 1.05 | -0.5125 | 0.38 | -0.9725 | 0.214523 | 0.694897 | 0.285058 | 0.994893 | 0.997988 | 0.994666 |
| Ctnnd2_B7ZNF6_S1216s_1 | Ctnnd2_B7 | 0.04 | 0.56 | 0.26 | 0.41 | 0.81 | -0.47 | 0.72 | 1.03 | -0.27 | 1.09 | 0.64 | 0.51 | -0.3 | 0.36 | -0.63 | 0.25 | -0.1075 | -0.27 | -0.6675 | 0.221758 | 0.567098 | 0.597297 | 0.989223 | 0.997988 | 0.994666 |
| Ctnnd2_B7ZNF6_S200s_1 | Ctnnd2_B7 | 0.7 | -0.05 | -0.4 | -0.57 | 0.36 | 0.45 | 0.54 | 0.09 | -0.31 | 0.07 | 2.28 | -0.56 | 0.32 | -0.41 | -0.86 | 0.07 | 0.205 | 0.175 | -0.3975 | 0.722772 | 0.184798 | 0.490398 | 0.989223 | 0.997988 | 0.9971 |
| Ctnnd2_B7ZNF6_S263s_1 | Ctnnd2_B7 | 1.01 | 1.14 | 0.23 | 0.45 | 0.5 | 1.09 | 0.47 | 0.33 | 1.06 | 1.64 | 0.92 | 0.6 | 1.39 | 1.25 | 0.14 | 0.38 | 0.44 | 0.45 | -0.14 | 0.832799 | 0.714903 | 0.309758 | 0.995437 | 0.997988 | 0.999507 |
| Ctnnd2_B7ZNF6_S263s_2 | Ctnnd2_B7 | 0.35 | 0.3 | 0.32 | 0.29 | 0.68 | 0.23 | 0.58 | 0.42 | 0.33 | 0.52 | 0.4 | 0.59 | -0.48 | 0.09 | -0.33 | 0.1 | -0.11 | 0.3475 | 0.0825 | 0.090615 | 0.440204 | 0.487842 | 0.989223 | 0.997988 | 0.994666 |
| Ctnnd2_B7ZNF6_S272s_1 | Ctnnd2_B7 | 0.36 | 1.33 | 1.41 | 1.15 | 0.1 | 0.58 | 0.45 | 0.82 | 1.04 | 0.75 | 1.89 | 1.41 | 0.9 | 0.19 | 0.32 | 0.97 | 0.1625 | 0.145 | -0.47 | 0.1964 | 0.084909 | 0.55995 | 0.989223 | 0.997988 | 0.994666 |
| Ctnnd2_B7ZNF6_S272s_2 | Ctnnd2_B7 | 0.35 | 0.3 | 0.32 | 0.29 | 0.68 | 0.23 | 0.58 | 0.42 | 0.33 | 0.52 | 0.4 | 0.59 | -0.48 | 0.09 | -0.33 | 0.1 | -0.575 | 0.21 | -0.4675 | 0.090615 | 0.440204 | 0.487842 | 0.989223 | 0.997988 | 0.994666 |
| Ctnnd2_B7ZNF6_S281s_1 | Ctnnd2_B7 | 1.47 | 0.03 | 0.06 | 0.56 | 1.21 | 1.45 | 0.58 | 0.83 | 0.41 | 0.67 | 0.9 | 0.77 | -0.05 | 0.58 | -0.04 | 0.23 | 0.1625 | 0.145 | -0.47 | 0.370115 | 0.218783 | 0.665344 | 0.989223 | 0.997988 | 0.994666 |
| Ctnnd2_B7ZNF6_S311s_1 | Ctnnd2_B7 | 0.54 | 0.82 | 0.55 | 0.38 | 0.72 | 0.16 | 0.68 | 0.33 | 0.16 | 1.04 | -1.31 | 0.47 | 0.02 | -0.99 | -0.22 | 0.3 | 0.4875 | 0.1575 | -0.35 | 0.036028 | 0.671207 | 0.331138 | 0.992876 | 0.997988 | 0.994666 |
| Ctnnd2_B7ZNF6_S323s_1 | Ctnnd2_B7 | 0.06 | 0.48 | 0.1 | -0.13 | 0.69 | 0.24 | 1.26 | 0.13 | -0.63 | 0.03 | -0.4 | 0.81 | -0.7 | -0.73 | -0.25 | 0.34 | -0.1 | -0.4825 | -0.795 | 0.177556 | 0.164764 | 0.624382 | 0.989223 | 0.997988 | 0.994666 |
| Ctnnd2_B7ZNF6_S32s_1 | Ctnnd2_B7 | 0.56 | 0.81 | 0.57 | 0.84 | 0.68 | 0.86 | 1.04 | 0.72 | 0.28 | 0.73 | -0.02 | 0.55 | -0.57 | 0.44 | -0.25 | 0.51 | 0.4525 | -0.175 | -0.4625 | 0.063046 | 0.542336 | 0.239239 | 0.989223 | 0.997988 | 0.994666 |
| Ctnnd2_B7ZNF6_S394s_1 | Ctnnd2_B7 | -1.12 | -1.04 | -1.19 | -0.92 | -1.11 | -0.24 | -0.86 | -0.48 | -0.81 | -0.89 | -0.81 | -0.36 | -0.31 | -0.17 | -1.97 | -0.72 | 0.13 | -0.31 | -0.6625 | 0.511785 | 0.145759 | 0.153454 | 0.989223 | 0.997988 | 0.994666 |
| Ctnnd2_B7ZNF6_S411s_1 | Ctnnd2_B7 | 0.67 | 0.7 | 0.53 | 0.67 | 0.94 | 0.67 | 0.96 | 0.79 | 0.67 | 0.78 | 0.37 | 0.77 | -0.63 | 0.33 | -0.61 | 0.43 | 0.395 | 0.35 | 0.275 | 0.042562 | 0.337337 | 0.981718 | 0.989223 | 0.997988 | 0.994666 |
| Ctnnd2_B7ZNF6_S434s_1 | Ctnnd2_B7 | 0.04 | -0.17 | 0.63 | 0.99 | 0.27 | -0.32 | 1.06 | 0.78 | -0.66 | 0.03 | 1.23 | 0.88 | -0.53 | 0.18 | 0.38 | 1.18 | 0.1975 | 0.005 | -0.7625 | 0.871947 | 0.847922 | 0.995816 | 0.996349 | 0.998158 | 0.999507 |
| Ctnnd2_B7ZNF6_S436s_1 | Ctnnd2_B7 | -0.34 | -0.22 | 0.79 | 0.96 | -0.04 | 0.45 | 1 | 1.18 | -0.33 | -0.18 | 0.92 | 0.8 | -0.41 | -0.45 | 0.14 | 0.63 | 0.075 | -0.0025 | -0.07 | 0.453862 | 0.407795 | 0.991116 | 0.989223 | 0.997988 | 0.994666 |
| Ctnnd2_B7ZNF6_S457s_1 | Ctnnd2_B7 | 0.48 | 0.26 | 0.73 | 0.82 | 0.85 | 0.77 | 0.85 | 0.78 | 0.32 | 0.61 | 0.36 | 0.78 | 0 | 0.18 | -0.23 | 0.37 | 0.35 | 0.005 | -0.32 | 0.092704 | 0.287153 | 0.822998 | 0.989223 | 0.997988 | 0.994666 |
| Ctnnd2_B7ZNF6_S470s_1 | Ctnnd2_B7 | 0.58 | 0 | -0.09 | 0.63 | 0.51 | 1.04 | 0.64 | 1.32 | 0.06 | 0.01 | 0.54 | 0.63 | 0.09 | 0.12 | 0.02 | 0.68 | 0.24 | -0.055 | -0.4925 | 0.862095 | 0.065653 | 0.91806 | 0.989223 | 0.997988 | 0.999507 |
| Ctnnd2_B7ZNF6_S486s_1 | Ctnnd2_B7 | 0.2 | 0.34 | 0.46 | 0.14 | 0.03 | 0.08 | 0.4 | 0.44 | 0.31 | 0.13 | -0.38 | -0.08 | -0.21 | 0.73 | -0.45 | 0.48 | 0.5975 | 0.03 | -0.0525 | 0.656046 | 0.827679 | 0.254466 | 0.995762 | 0.997988 | 0.995259 |
| Ctnnd2_B7ZNF6_S528s_1 | Ctnnd2_B7 | 2.7 | 0.95 | 1.37 | 1.94 | 0.16 | 2.17 | 1.26 | 1.83 | 2.57 | 2.91 | 3.5 | 1.32 | 0.59 | 0.79 | 0.67 | 1.05 | -0.0475 | -0.29 | -0.1475 | 0.03474 | 0.472163 | 0.155411 | 0.989223 | 0.997988 | 0.994666 |
| Ctnnd2_B7ZNF6_S530s_1 | Ctnnd2_B7 | 0.87 | 1.15 | 1.58 | 1.25 | 0.8 | 1.2 | 1.05 | 1 | 1.37 | 1.1 | 1.05 | 1.51 | 0.57 | 0.66 | 0.63 | 1.34 | -0.385 | 0.835 | -0.965 | 0.18659 | 0.407351 | 0.859467 | 0.989223 | 0.997988 | 0.994666 |
| Ctnnd2_B7ZNF6_S533s_1 | Ctnnd2_B7 | 0.95 | 1.87 | 1.71 | 1.17 | 1.5 | 1.04 | 1.25 | 1.31 | 0.94 | 1.67 | 0.64 | 1.63 | 1.32 | 0.9 | 1.16 | 1.9 | -0.2 | 0.045 | -0.4125 | 0.757567 | 0.589648 | 0.56299 | 0.989223 | 0.997988 | 0.9971 |
| Ctnnd2_B7ZNF6_S7s_1 | Ctnnd2_B7 | 0.68 | 0.3 | 1.47 | 1.05 | 0.51 | 1.14 | 1.07 | 0.89 | -0.07 | 0.67 | 1.09 | 1.63 | 0.01 | 0.24 | 1.21 | 1.55 | -0.15 | -0.205 | -0.105 | 0.780777 | 0.9288 | 0.915947 | 0.997639 | 0.997988 | 0.998226 |
| Ctnnd2_B7ZNF6_T1052t_1 | Ctnnd2_B7 | 0.39 | 0.69 | 1.1 | 0.94 | 0.76 | -0.09 | 1.06 | 0.98 | 0.65 | 0.32 | 1.6 | 1.13 | 0.43 | 0.47 | 1.13 | 1.03 | 0.0275 | -0.045 | -0.1225 | 0.960187 | 0.749489 | 0.670935 | 0.995762 | 0.997988 | 0.999507 |
| Ctnnd2_B7ZNF6_T468t_1 | Ctnnd2_B7 | 1.54 | 1.53 | 0.49 | 0.69 | 1.09 | 2.45 | 0.48 | 1.15 | 1.42 | 1.03 | 0.85 | 0.78 | 0.82 | 1 | 0.31 | 0.5 | -0.1025 | 0.145 | -0.015 | 0.261408 | 0.621225 | 0.898561 | 0.989223 | 0.997988 | 0.994666 |
| Ctps1_P70698_S562s_1 | Ctps1_P70 | 1.98 | 0.48 | 0.9 | 2.34 | -0.15 | 2.17 | 1.32 | 1.86 | 1.56 | 2.01 | 2.34 | 1.16 | 1.57 | 0.14 | 2.29 | 1.45 | 0.23 | -0.0425 | -0.405 | 0.913561 | 0.836135 | 0.484154 | 0.995762 | 0.997988 | 0.999507 |
| Ctr9_Q62018_S925s_1 | Ctr9_Q620 | -1.48 | -2.25 | -1.73 | -1.16 | -2.45 | -1.68 | -1.94 | -1.57 | -1.47 | -2.32 | -0.5 | -1.34 | -1.23 | -1.69 | -0.71 | -1.38 | -0.125 | 0.3425 | -0.0625 | 0.257004 | 0.43064 | 0.565673 | 0.989223 | 0.997988 | 0.994666 |
| Cttn_Q921L6_S380s_1 | Cttn_Q921 | 0.41 | 0.56 | 0.61 | 0.77 | 0.67 | 0.4 | 1.16 | 0.85 | -0.31 | 0.79 | 0.01 | 0.62 | -0.25 | 0.78 | 0.54 | 0.55 | -0.255 | 0.2475 | 0.4025 | 0.545806 | 0.455407 | 0.323341 | 0.989223 | 0.997988 | 0.994666 |
| Cttn_Q921L6_S380s_2 | Cttn_Q921 | 0.44 | 0.43 | 0.38 | 0.43 | 0.98 | 0.38 | 0.81 | 0.5 | -0.07 | 0.21 | -0.22 | 0.51 | -1 | 1.01 | -0.08 | -0.16 | 0.1825 | -0.31 | -0.1825 | 0.265283 | 0.282375 | 0.219567 | 0.989223 | 0.997988 | 0.994666 |
| Cttn_Q921L6_S381s_1 | Cttn_Q921 | 0.28 | 0.41 | 0.8 | 0.93 | 0.68 | 0.05 | 1.27 | 1.01 | -0.03 | 0.55 | 0.15 | 0.85 | -0.29 | 0.53 | 0.69 | 0.65 | 0.2475 | -0.3125 | -0.4775 | 0.520603 | 0.646769 | 0.451764 | 0.99033 | 0.997988 | 0.994666 |
| Cttn_Q921L6_T286t_1 | Cttn_Q921 | -1.28 | 0.12 | -1.01 | -1.41 | -0.33 | -0.36 | -0.42 | -0.44 | -0.41 | -0.53 | 1.28 | 0.33 | -0.02 | -0.34 | -0.79 | 0.02 | 0.1475 | -0.225 | -0.21 | 0.151189 | 0.171439 | 0.06183 | 0.989223 | 0.997988 | 0.994666 |
| Cttn_Q921L6_T364t_1 | Cttn_Q921 | 0.5 | 2.14 | 1.53 | 1.12 | 0.88 | 0.39 | 1.02 | 0.87 | 0.6 | 1.26 | 1 | 1.31 | 1.33 | 1.48 | 1.11 | 1.05 | 0.5075 | 1.0625 | 0.6125 | 0.831591 | 0.170121 | 0.47274 | 0.989223 | 0.997988 | 0.999507 |
| Cttn_Q921L6_T364t_2 | Cttn_Q921 | 0.45 | 0.43 | 0.36 | 0.36 | 0.96 | 0.39 | 0.73 | 0.54 | -0.08 | 0.23 | -0.19 | 0.56 | -0.98 | 0.99 | -0.13 | -0.16 | -0.5325 | -0.28 | -0.08 | 0.266694 | 0.257463 | 0.291406 | 0.989223 | 0.997988 | 0.994666 |
| Cttnbp2_B9EJA2_S1529s_1 | Cttnbp2_B | -0.34 | -0.5 | 0.16 | 0.36 | 0.02 | 0.51 | -0.06 | -0.61 | 2.33 | 0.02 | -0.76 | -0.01 | 2.18 | -0.96 | 0.52 | 0.14 | 0.255 | -0.27 | -0.47 | 0.384692 | 0.888392 | 0.462773 | 0.996493 | 0.997988 | 0.994666 |
| Cttnbp2_B9EJA2_S367s_1 | Cttnbp2_B | -0.38 | -0.21 | 0.85 | 0.49 | 0.35 | -0.49 | 0.48 | 0.57 | 0.24 | 0.36 | -0.28 | 0.97 | 0.59 | 0.11 | 0.96 | 1 | 0.045 | 0.475 | 0.55 | 0.221755 | 0.914398 | 0.72903 | 0.997335 | 0.997988 | 0.994666 |
| Cttnbp2_B9EJA2_S465s_1 | Cttnbp2_B | -0.36 | -0.73 | -0.77 | 0.32 | 0.18 | -0.38 | 0.25 | -0.59 | -0.11 | 0.08 | 2.07 | 1.31 | 0.5 | -0.09 | 0.61 | 0.83 | 0.04 | 0.135 | 0.4775 | 0.034489 | 0.46202 | 0.04469 | 0.989223 | 0.997988 | 0.994666 |
| Cttnbp2_B9EJA2_S472s_1 | Cttnbp2_B | 0.82 | -0.12 | 0.4 | 0.26 | 0.35 | 1.62 | -0.15 | 0.31 | 1.47 | 0.3 | -1.99 | 0.29 | 0.54 | -0.84 | 0.23 | 0.32 | 0.25 | 1.2225 | 0.8475 | 0.470834 | 0.638882 | 0.636306 | 0.989223 | 0.997988 | 0.994666 |
| Cttnbp2_B9EJA2_T15t_1 | Cttnbp2_B | -2.27 | -1.15 | -1.16 | -0.26 | -1.42 | -1.15 | -1.28 | -1.05 | -0.74 | -1.86 | -0.43 | -1.36 | -0.36 | -0.27 | -0.86 | -1.53 | 0.1925 | -0.3225 | -0.2775 | 0.355395 | 0.970242 | 0.819183 | 0.999187 | 0.997988 | 0.994666 |
| Cttnbp2nl_Q99LJ0_S477s_1 | Cttnbp2nl_ | 0.63 | 0.42 | 0.96 | 0.75 | 0.9 | 0.77 | 0.93 | 1.06 | 0.53 | 0.85 | 2.62 | 0.65 | -0.1 | 0.61 | 1.67 | 0.21 | -0.015 | 0.1125 | 0.455 | 0.819892 | 0.315765 | 0.336129 | 0.989223 | 0.997988 | 0.999507 |
| Cttnbp2nl_Q99LJ0_S481s_1 | Cttnbp2nl_ | 0.4 | 0.65 | 0.29 | 0.24 | 0.35 | 0.3 | 0.53 | 0.37 | 0.07 | 0.9 | 0.1 | 0.2 | -0.18 | 0.03 | 0.7 | 0.56 | 0.225 | 0.4725 | -0.0925 | 0.691058 | 0.971308 | 0.776695 | 0.999187 | 0.997988 | 0.996159 |
| Ctu1_Q99J10_S378s_1 | Ctu1_Q99J | -0.26 | 0.7 | 1.14 | 0.96 | 0.01 | -0.47 | 0.66 | 0.38 | 0.7 | -0.29 | 2.4 | 1.77 | 0.28 | 1.26 | 1.09 | 0.32 | -0.0075 | -0.0775 | -0.1175 | 0.801114 | 0.2233 | 0.414155 | 0.989223 | 0.997988 | 0.999507 |
| Cul3_F6UY44_S80s_1 | Cul3_F6UY | -1.1 | -1.17 | -0.72 | -1.13 | -0.12 | -1.66 | -0.33 | -1.27 | -0.92 | -0.78 | -0.92 | -1.88 | -0.41 | -0.12 | -0.56 | -0.7 | -0.49 | 0.51 | 0.1025 | 0.047506 | 0.624804 | 0.758971 | 0.989223 | 0.997988 | 0.994666 |
| Cul4b_J3QJX0_S126s_1 | Cul4b_J3Q | 1.17 | 1.36 | 0.3 | 0.52 | 0.22 | 0.2 | -0.25 | 0.26 | 1.16 | 0.53 | -0.01 | 0.49 | 1.2 | 1.13 | 0.32 | 0.36 | 0.185 | -0.095 | 0.5825 | 0.817842 | 0.035544 | 0.421984 | 0.989223 | 0.997988 | 0.999507 |
| Cul4b_J3QJX0_T32t_1 | Cul4b_J3Q | 0.32 | 0.65 | -0.83 | 0 | -0.69 | -0.34 | -0.71 | -0.6 | 0.6 | 0.98 | 0.08 | -1.33 | 0.4 | 0.12 | 0.09 | 0.14 | -0.73 | -0.295 | -0.085 | 0.66753 | 0.09073 | 0.931478 | 0.989223 | 0.997988 | 0.996159 |
| Cul9_E9QP09_S1983s_1 | Cul9_E9QP | -1.4 | -0.63 | 0.03 | 0.49 | -0.05 | -0.22 | 0.92 | -0.01 | 0.15 | 0.25 | -0.24 | 0.47 | 0.01 | 0.09 | -0.35 | -0.07 | -0.62 | 0.0475 | 0.1525 | 0.485054 | 0.253066 | 0.229838 | 0.989223 | 0.997988 | 0.994666 |
| Cul9_E9QP09_S2440s_1 | Cul9_E9QP | -0.11 | 0.22 | 0.33 | 0.15 | 0.34 | -0.36 | 0.45 | 0.05 | -0.29 | 0.14 | -0.05 | 0.14 | 0.21 | 0.59 | 0.26 | 0.48 | 0.5375 | 0.535 | 0.2975 | 0.349794 | 0.914348 | 0.49027 | 0.997335 | 0.997988 | 0.994666 |
| Cul9_E9QP09_S584s_1 | Cul9_E9QP | 0.22 | 0.58 | 0.15 | 0.01 | 0.4 | -0.11 | -0.06 | -0.08 | 0.41 | 0.37 | 0.52 | 0.04 | 0.48 | 0.76 | 0.19 | 0.26 | -0.0275 | -0.1625 | 0.2375 | 0.500689 | 0.40506 | 0.695887 | 0.989223 | 0.997988 | 0.994666 |
| Cul9_E9QP09_S592s_1 | Cul9_E9QP | 3.17 | 1.09 | 3.16 | -0.05 | 0.23 | 3.87 | 2.07 | 2.68 | 0.85 | 2.58 | 0.41 | 3.06 | 1.54 | -2.92 | 3.24 | 3.82 | -0.2025 | 0.095 | 0.1825 | 0.775335 | 0.699182 | 0.897226 | 0.994893 | 0.997988 | 0.99798 |
| Cwc22_B1AYU7_S39s_1 | Cwc22_B1 | -0.24 | -0.47 | 0.43 | 0.43 | -2.55 | 0.09 | -1.23 | -0.4 | 1.4 | -0.56 | -0.1 | 0.41 | 2.06 | -1.51 | 0.85 | -0.57 | 0.37 | -0.1175 | -0.4225 | 0.815737 | 0.08405 | 0.589426 | 0.989223 | 0.997988 | 0.999507 |
| Cwc22_B1AYU7_S61s_1 | Cwc22_B1 | -1.83 | -0.88 | -1.16 | -1.41 | -1.84 | -1.87 | -1.69 | -1.41 | -1.16 | -1.45 | -0.54 | -1.55 | -0.48 | -0.59 | -1.03 | -1.24 | -1.06 | 0.25 | 0.17 | 0.151775 | 0.178719 | 0.66065 | 0.989223 | 0.997988 | 0.994666 |
| Cwc22_B1AYU7_S824s_1 | Cwc22_B1 | 0.49 | 0.21 | -0.19 | -0.95 | -2.19 | 0.27 | -2.05 | -0.93 | 1.1 | 0.08 | -0.44 | -1.06 | 1.29 | -0.3 | -1.28 | -1.59 | -0.3825 | 0.145 | 0.485 | 0.582306 | 0.081961 | 0.953791 | 0.989223 | 0.997988 | 0.995259 |
| Cx3cl1_O35188_S262s_1 | Cx3cl1_O3 | -0.36 | -0.27 | -0.05 | 0.51 | 0.16 | -0.54 | 0.74 | 0.06 | -0.34 | -0.38 | 0.43 | 0.04 | -1.19 | 0.53 | 0.59 | 0.57 | -1.115 | 0.03 | -0.36 | 0.718422 | 0.661254 | 0.948104 | 0.990434 | 0.997988 | 0.9971 |
| Cx3cl1_O35188_S384s_1 | Cx3cl1_O3 | -1.19 | 0.69 | -1 | -0.19 | -0.23 | -0.33 | 0.35 | -0.31 | -0.04 | 0.43 | -0.04 | -0.47 | -0.02 | 0.48 | -0.22 | -0.12 | 0.1475 | -0.02 | 0.1675 | 0.323786 | 0.504149 | 0.393009 | 0.989223 | 0.997988 | 0.994666 |
| Cxadr_P97792_S332s_1 | Cxadr_P97 | 0.94 | 0.52 | 0.66 | 0.73 | 0.9 | 1.01 | 0.91 | 1.42 | 1.52 | 1.28 | 1.74 | 1.46 | 1.14 | 1.84 | 0.86 | 1.2 | 0.2925 | 0.3925 | 0.4525 | 0.086239 | 0.151658 | 0.006629 | 0.989223 | 0.997988 | 0.994666 |
| Cyb561_Q60720_S246s_2 | Cyb561_Q6 | -0.2 | -0.45 | 0.64 | 0.68 | -0.08 | 0.38 | 0.3 | 0.52 | 0.33 | 0.31 | 0.84 | 0.36 | -0.05 | 0.09 | 0.98 | 1.06 | 0.3475 | 0.7875 | 0.5475 | 0.401141 | 0.731912 | 0.395297 | 0.995762 | 0.997988 | 0.994666 |
| Cyb561_Q60720_S248s_2 | Cyb561_Q6 | -0.2 | -0.45 | 0.64 | 0.68 | -0.08 | 0.38 | 0.3 | 0.52 | 0.33 | 0.31 | 0.84 | 0.36 | -0.05 | 0.09 | 0.98 | 1.06 | 0.1125 | 0.2925 | 0.3525 | 0.401141 | 0.731912 | 0.395297 | 0.995762 | 0.997988 | 0.994666 |
| Cyb5b_Q9CQX2_S33s_1 | Cyb5b_Q9 | 0.81 | 1.15 | 0.84 | 1 | 0.51 | 0.79 | 1.24 | 1.23 | 0.58 | 1.13 | 0.58 | 1.31 | 0.37 | 0.84 | 0.77 | 1.16 | 0.1125 | 0.2925 | 0.3525 | 0.54421 | 0.976133 | 0.850793 | 0.999187 | 0.997988 | 0.994666 |
| Cyb5b_Q9CQX2_T3t_1 | Cyb5b_Q9 | 0.96 | 0.22 | 1.2 | 1.51 | -0.25 | 0.96 | 1.05 | 1.76 | 1.8 | 0.95 | 1.36 | 0.73 | 1.56 | 0.18 | 0.87 | 0.82 | -0.0075 | -0.05 | -0.165 | 0.774153 | 0.841955 | 0.525832 | 0.996092 | 0.997988 | 0.99798 |
| Cyfip1_Q7TMB8_S1233s_1 | Cyfip1_Q7 | 1.11 | 1.85 | 0.74 | 0.82 | 0.33 | 0.78 | 0 | 0.53 | 1.61 | 1.26 | 0.61 | 0.6 | 0.98 | 0.4 | 0.26 | 0.53 | -0.0925 | 0.2375 | -0.115 | 0.105003 | 0.044925 | 0.763656 | 0.989223 | 0.997988 | 0.994666 |
| Cyfip1_Q7TMB8_S872s_1 | Cyfip1_Q7 | -0.43 | 0.22 | 0.63 | 0.05 | 0.52 | -0.51 | 0.74 | 0.01 | 0.28 | -0.05 | -0.38 | 0.27 | -0.48 | -0.04 | -0.07 | 0.64 | -0.72 | -0.11 | -0.5875 | 0.764214 | 0.838217 | 0.77515 | 0.995762 | 0.997988 | 0.9971 |
| Cyfip1_Q7TMB8_T1234t_1 | Cyfip1_Q7 | 0.82 | 1.21 | 0.82 | 0.92 | 0.73 | 0.6 | 0.14 | 0.69 | 0.54 | 0.65 | 0.35 | 0.57 | 0.24 | 0.31 | 0.29 | 0.62 | 0.0725 | -0.0875 | -0.105 | 0.039384 | 0.111851 | 0.087188 | 0.989223 | 0.997988 | 0.994666 |
| Cyld_Z4YJJ5_S344s_1 | Cyld_Z4YJJ | 1.03 | -0.15 | 0.93 | 0.2 | -0.44 | 1.39 | -0.21 | 0.66 | 2.52 | 0.65 | 0.51 | 0.08 | 2.94 | -1.26 | 0.55 | -0.95 | -0.4025 | -0.415 | -0.5775 | 0.835255 | 0.746237 | 0.445963 | 0.995762 | 0.997988 | 0.999507 |
| Cyld_Z4YJJ5_S397s_1 | Cyld_Z4YJJ | 0.08 | 0.54 | 0.48 | 0.4 | 0.28 | -0.48 | 0.66 | 0.51 | 0.08 | 0.28 | 0.45 | 0.67 | 0.25 | 0.53 | 0.57 | 0.54 | -0.1525 | 0.4375 | -0.1825 | 0.693644 | 0.65867 | 0.983591 | 0.990434 | 0.997988 | 0.996159 |
| Cyld_Z4YJJ5_S398s_1 | Cyld_Z4YJJ | -1.07 | -1.08 | 0.07 | 0.28 | -0.7 | -1.53 | 0.43 | -0.41 | -1.18 | -0.61 | -0.28 | 0.3 | -0.87 | -0.37 | 0.15 | -0.32 | -0.1325 | -0.005 | 0.0975 | 0.815734 | 0.837105 | 0.986909 | 0.995762 | 0.997988 | 0.999507 |
| Cys1_Q8R4T1_S116s_1 | Cys1_Q8R4 | 0.78 | 1.21 | 0.71 | 0.5 | 0 | 0.69 | 0.15 | 0.04 | 1.62 | 1.02 | 0.85 | 0.22 | 0.93 | 0.62 | -1.88 | 0.25 | -0.1025 | 0.0075 | 0.0975 | 0.191048 | 0.050718 | 0.710822 | 0.989223 | 0.997988 | 0.994666 |
| Cys1_Q8R4T1_T55t_1 | Cys1_Q8R4 | -1.31 | -1.83 | -0.11 | 0.43 | -0.99 | -0.13 | 0 | 0.66 | -0.13 | -1.76 | -0.94 | 0.46 | 0.14 | -0.61 | 0.09 | -0.05 | -0.58 | 0.1275 | -0.82 | 0.264415 | 0.307683 | 0.861865 | 0.989223 | 0.997988 | 0.994666 |
| D17Wsu92e_B2KF52_S215s_1 | D17Wsu92 | -0.99 | -1.17 | -0.67 | -1.12 | -0.2 | -0.87 | -0.12 | -1.04 | -0.75 | -0.79 | -1.12 | -1.58 | -0.94 | 0.19 | -1.88 | -0.81 | 0.59 | 0.1125 | 0.5975 | 0.768285 | 0.1604 | 0.79341 | 0.989223 | 0.997988 | 0.9971 |
| D17Wsu92e_B2KF52_S265s_1 | D17Wsu92 | 0.57 | 0.56 | -0.08 | 0.22 | 0.99 | 0.14 | 0.31 | 0.22 | -0.17 | 1.39 | -1.03 | -0.27 | 0.31 | -0.15 | -0.09 | 0.22 | 0.43 | -0.0725 | 0.1275 | 0.383708 | 0.731302 | 0.504999 | 0.995762 | 0.997988 | 0.994666 |
| D2Wsu81e_Q3UHX9_S338s_1 | D2Wsu81e | -0.23 | -0.12 | 0.08 | -0.15 | -0.5 | -0.17 | -0.42 | -0.44 | -0.16 | -0.16 | -0.28 | -0.02 | 0.3 | 0.33 | -0.4 | -1.21 | 0.0975 | -0.3375 | -0.245 | 0.715456 | 0.200681 | 0.814084 | 0.989223 | 0.997988 | 0.996966 |
| D430041D05Rik_F6ZGR6_S1044s_1 | D430041D | 0.15 | 1.06 | 1.33 | 1.1 | 1 | -0.05 | 0.7 | 0.79 | 0.78 | 1.01 | -1.5 | -0.05 | 0.75 | 0.34 | 1.03 | -0.02 | -0.2775 | -0.05 | -0.14 | 0.310068 | 0.399549 | 0.163869 | 0.989223 | 0.997988 | 0.994666 |
| D430041D05Rik_F6ZGR6_S1070s_1 | D430041D | -1.14 | -1.27 | -1.04 | -0.91 | -0.98 | -1.71 | 0 | -1.11 | -1.63 | -1.43 | -1.04 | -0.55 | -1.74 | -0.91 | -1.88 | -0.94 | -0.3 | -0.85 | -0.385 | 0.39173 | 0.698729 | 0.804947 | 0.994893 | 0.997988 | 0.994666 |
| D430041D05Rik_F6ZGR6_S1088s_1 | D430041D | 0.32 | 0.56 | 0.57 | 0.41 | 0.82 | 0.51 | 0.72 | 0.32 | 0.65 | 0.6 | 0.11 | 0.56 | 0.37 | 0.54 | 0.1 | 0.58 | 0.14 | -0.0725 | -0.2775 | 0.783983 | 0.561394 | 0.948515 | 0.989223 | 0.997988 | 0.998711 |
| D430041D05Rik_F6ZGR6_S1104s_1 | D430041D | -0.97 | -1.34 | -1.05 | -1.57 | -0.77 | -1.06 | -0.82 | -0.99 | -1.63 | -1.62 | -0.14 | -0.71 | -1.5 | -0.38 | -1.56 | -1.77 | 0.1275 | 0.015 | -0.0675 | 0.847685 | 0.18264 | 0.598394 | 0.989223 | 0.997988 | 0.999507 |
| D430041D05Rik_F6ZGR6_S1108s_1 | D430041D | 0.77 | -0.06 | -0.73 | -1.34 | -1.65 | 1.17 | -0.39 | -0.48 | 2.05 | -1.18 | -2.07 | -0.07 | 2.06 | 0.5 | 0.1 | -1.11 | 0.3225 | 0.2075 | -0.07 | 0.319768 | 0.996936 | 0.979708 | 0.999868 | 0.997988 | 0.994666 |
| D430041D05Rik_F6ZGR6_S1226s_1 | D430041D | 1.68 | -0.42 | 0.97 | 2.3 | -0.82 | 0.58 | 0.35 | 2.2 | 1.81 | 2.11 | 1.85 | 1.62 | 1.43 | -1.95 | 0.52 | -0.45 | 0.0025 | 0.0225 | 0.7275 | 0.152789 | 0.465121 | 0.214674 | 0.989223 | 0.997988 | 0.994666 |
| D430041D05Rik_F6ZGR6_S726s_1 | D430041D | -0.31 | -0.75 | 0.85 | 0.54 | -0.7 | 1.25 | -0.66 | 0.1 | 1.65 | -1.33 | -0.99 | -0.06 | 1.19 | -0.3 | 0.34 | 0.07 | -0.555 | 0.715 | -1.245 | 0.607312 | 0.8733 | 0.701533 | 0.996493 | 0.997988 | 0.995259 |
| D430041D05Rik_F6ZGR6_S762s_1 | D430041D | 2.74 | 3 | 1.37 | 1.44 | 2.23 | 2.57 | 0.3 | 0.74 | 3.13 | 2.51 | 0.82 | 1.14 | 2.77 | 0.93 | 1.09 | 0.32 | -0.085 | -0.265 | 0.2425 | 0.184197 | 0.292357 | 0.70867 | 0.989223 | 0.997988 | 0.994666 |
| D430041D05Rik_F6ZGR6_S785s_1 | D430041D | 0.57 | 0.71 | 0.6 | 0.42 | 0.69 | 0.4 | 1.09 | 0.56 | 0.23 | 0.83 | 0.42 | 0.59 | -0.18 | 0.69 | 0.24 | 0.68 | -0.6775 | -0.2375 | -0.86 | 0.45637 | 0.637593 | 0.805618 | 0.989223 | 0.997988 | 0.994666 |
| D430041D05Rik_F6ZGR6_S927s_1 | D430041D | -0.3 | 0.21 | 0.08 | 0.54 | -0.23 | -0.71 | 0.58 | 0.05 | -1.31 | -0.35 | -0.26 | -0.5 | -0.87 | 0.06 | -0.44 | -0.37 | 0.11 | -0.0575 | -0.2175 | 0.109331 | 0.529808 | 0.044646 | 0.989223 | 0.997988 | 0.994666 |
| D430041D05Rik_F6ZGR6_S993s_2 | D430041D | 0.39 | 1.38 | 0.95 | -0.16 | 0.29 | -0.15 | 1.93 | 1.77 | -0.53 | 0.74 | -0.56 | 0.67 | -0.14 | 0.83 | 1.34 | 1.59 | -0.21 | -0.7375 | -0.5375 | 0.587374 | 0.570739 | 0.251784 | 0.989223 | 0.997988 | 0.995259 |
| D430041D05Rik_F6ZGR6_S994s_1 | D430041D | 0.77 | 0.99 | 1.6 | 1.15 | 2.09 | 0.78 | 1.19 | 0.85 | 0.41 | 1 | 0.35 | 0.36 | 0.19 | 0.33 | 2.02 | 1.32 | 0.32 | -0.56 | 0.265 | 0.720657 | 0.775725 | 0.059981 | 0.995762 | 0.997988 | 0.9971 |
| D430041D05Rik_F6ZGR6_S994s_2 | D430041D | 0.39 | 1.38 | 2.14 | 1.76 | 0.29 | -0.15 | 1.58 | 1.79 | -0.53 | 0.74 | 3.21 | 1.43 | -0.14 | 0.83 | 2.5 | 2.57 | 0.1 | -0.5975 | -0.1625 | 0.973706 | 0.337872 | 0.791496 | 0.989223 | 0.997988 | 0.999507 |
| D430042O09Rik_F6XQI1_T1128t_1 | D430042O | -3.21 | -2.11 | -2.59 | -9.85 | -3.3 | -1.83 | -3.95 | -4.05 | -0.96 | -2.32 | -2.01 | -2.89 | 0.18 | -4.25 | -1.8 | -2.74 | -0.54 | -0.205 | 0.0225 | 0.210932 | 0.479534 | 0.164508 | 0.989223 | 0.997988 | 0.994666 |
| D630045J12Rik_D3YTS3_S1197s_1 | D630045J1 | -0.58 | -1.68 | -0.81 | -0.72 | -0.05 | -0.95 | -0.09 | -0.91 | -2.18 | -1.6 | -1.35 | -0.95 | -1.44 | -0.29 | -1.52 | -0.76 | 1.1575 | 2.395 | 2.2875 | 0.8887 | 0.224356 | 0.144078 | 0.989223 | 0.997988 | 0.999507 |
| D630045J12Rik_D3YTS3_S1303s_1 | D630045J1 | 0.49 | 1.24 | 0.32 | 0.81 | 0.06 | -0.37 | 0.45 | 0.79 | 0.25 | 1.23 | 0.54 | 0.82 | 0.08 | -0.11 | -0.62 | 0.13 | 0.4475 | -0.5725 | -0.055 | 0.022427 | 0.168726 | 0.987516 | 0.989223 | 0.997988 | 0.994666 |
| D630045J12Rik_D3YTS3_S1429s_1 | D630045J1 | -0.27 | -0.68 | -0.2 | -0.48 | 0.04 | -0.37 | 0.29 | -0.85 | -0.56 | -0.44 | -1.88 | -1.22 | -0.92 | -0.3 | -1.88 | -0.52 | -0.4825 | -0.005 | -0.845 | 0.211831 | 0.538678 | 0.113497 | 0.989223 | 0.997988 | 0.994666 |
| D630045J12Rik_D3YTS3_S1587s_1 | D630045J1 | 0.85 | 0.62 | 0.36 | 0.44 | 1.08 | 0.49 | 0.46 | 0.45 | 0.29 | 1.11 | 0.77 | 0.51 | 0.52 | 0.77 | 0.39 | 0.82 | 0.185 | -0.6175 | -0.4975 | 0.821992 | 0.831904 | 0.702641 | 0.995762 | 0.997988 | 0.999507 |
| D630045J12Rik_D3YTS3_S1799s_1 | D630045J1 | 0.55 | 0.38 | 0.35 | -0.04 | 0.47 | 1.19 | 0.17 | 0.45 | 1.4 | 0.82 | 1.15 | 0.1 | 0.81 | 0.55 | 0.62 | 0.22 | 0.0525 | 0.1025 | 0.0575 | 0.379063 | 0.371287 | 0.118996 | 0.989223 | 0.997988 | 0.994666 |
| Dab2ip_F6RK07_S1062s_1 | Dab2ip_F6 | 0.83 | 0.21 | -0.8 | -0.8 | 1.17 | 1.54 | -0.39 | -0.67 | 0.15 | 0.35 | 0.02 | -0.15 | 0.63 | 1.19 | -0.55 | -0.27 | 0.26 | 0.5575 | 0.24 | 0.468539 | 0.375471 | 0.573692 | 0.989223 | 0.997988 | 0.994666 |
| Dab2ip_F6RK07_S641s_1 | Dab2ip_F6 | 0.64 | -0.94 | -1.01 | -1.04 | 0.47 | -0.02 | 0.32 | 0.35 | -0.15 | -0.19 | 0.51 | -0.21 | -0.63 | 0.46 | -1.01 | -0.36 | 0.5525 | 0.2325 | 0.39 | 0.680228 | 0.055553 | 0.203813 | 0.989223 | 0.997988 | 0.996159 |
| Dab2ip_F6RK07_S657s_1 | Dab2ip_F6 | 0.1 | 0.3 | 0.04 | 0.35 | 0.44 | -0.04 | 0.56 | 0.37 | 0.13 | 0.33 | 0.19 | 0.26 | -0.43 | 0.47 | -0.05 | 0.2 | 0.8675 | 0.5775 | 0.2025 | 0.597479 | 0.557486 | 0.888018 | 0.989223 | 0.997988 | 0.995259 |
| Dab2ip_F6RK07_S789s_1 | Dab2ip_F6 | -1.75 | -2.09 | -0.25 | -0.77 | -1.1 | -2.19 | -0.56 | -0.07 | -2.65 | -1.8 | 0.09 | -0.2 | -1.5 | -0.85 | -0.79 | -0.65 | 0.135 | 0.03 | -0.15 | 0.558386 | 0.676961 | 0.914951 | 0.99315 | 0.997988 | 0.995259 |
| Dab2ip_F6RK07_S872s_1 | Dab2ip_F6 | 0.11 | 0.33 | 0.65 | 0.55 | 0.5 | -0.28 | 0.26 | 0.3 | -0.03 | 0.12 | 0.78 | 0.71 | 0.3 | 0.75 | 0.19 | 0.18 | 0.235 | 0.075 | 0.2675 | 0.838282 | 0.411741 | 0.958074 | 0.989223 | 0.997988 | 0.999507 |
| Dab2ip_F6RK07_S889s_1 | Dab2ip_F6 | 1.25 | 0.32 | 1.45 | 0.82 | 0.27 | 0.92 | 0.87 | 1.07 | 0.39 | 0.69 | 0.94 | 0.65 | 0.02 | 0.42 | 1.32 | 0.95 | -0.215 | -0.015 | -0.055 | 0.476165 | 0.582908 | 0.356269 | 0.989223 | 0.997988 | 0.994666 |
| Dact1_D3Z5V0_S492s_1 | Dact1_D3Z | 0.19 | 0.29 | 0.93 | 0.86 | 1.5 | 0.25 | -0.02 | 0.75 | -0.31 | -0.73 | -1.23 | 0.55 | 0.68 | -0.39 | 0.63 | 0.74 | -0.1775 | -0.2925 | -0.2825 | 0.670163 | 0.889087 | 0.036606 | 0.996493 | 0.997988 | 0.996159 |
| Dact3_Q0PHV7_S10s_1 | Dact3_Q0P | 0.9 | 1.88 | 0.77 | 0.78 | 2.79 | 2.05 | 1.36 | 0.83 | 1.27 | 2.07 | -0.95 | 0.94 | 1.67 | -0.03 | 0.74 | 0.66 | 0.0525 | -0.9975 | -0.1525 | 0.463442 | 0.170746 | 0.693298 | 0.989223 | 0.997988 | 0.994666 |
| Dact3_Q0PHV7_S165s_1 | Dact3_Q0P | 0.02 | 0.53 | -0.57 | -0.65 | 0 | -0.04 | -0.83 | -0.09 | -0.36 | -0.14 | 0.68 | -0.64 | -0.1 | 0.37 | -0.84 | -1.73 | 0.675 | -0.25 | -0.3225 | 0.427969 | 0.833187 | 0.894303 | 0.995762 | 0.997988 | 0.994666 |
| Dact3_Q0PHV7_S217s_1 | Dact3_Q0P | 0.91 | -2.62 | -0.8 | -1.17 | 0.42 | -0.26 | -0.76 | -0.91 | -0.45 | 0.8 | -1.02 | -1.01 | 0.46 | 0.79 | -1.73 | -0.44 | -0.0725 | 0.0525 | -0.4075 | 0.403889 | 0.441137 | 0.51296 | 0.989223 | 0.997988 | 0.994666 |
| Dact3_Q0PHV7_S237s_1 | Dact3_Q0P | -0.54 | -0.67 | -1.77 | -1.72 | -0.24 | -0.5 | -0.64 | -1.45 | -0.42 | -0.6 | -1.15 | -2.35 | -0.62 | -0.59 | -1.47 | -1.3 | 0.5425 | 0.5 | 0.69 | 0.658646 | 0.264246 | 0.929934 | 0.989223 | 0.997988 | 0.995259 |
| Dact3_Q0PHV7_S268s_1 | Dact3_Q0P | -0.18 | -0.79 | 0.45 | -0.26 | -0.65 | 1.08 | -0.4 | -0.14 | 2.58 | 0.15 | -0.39 | 1.09 | 2.79 | -0.79 | 1.1 | -0.39 | 0.4675 | 0.045 | 0.18 | 0.265717 | 0.700136 | 0.123928 | 0.994893 | 0.997988 | 0.994666 |
| Dact3_Q0PHV7_S290s_1 | Dact3_Q0P | -0.78 | -0.55 | -0.32 | -0.33 | -0.91 | -0.1 | -0.36 | -0.53 | -0.44 | -0.32 | -0.32 | 0.39 | 0.29 | 0.81 | -0.58 | -1.91 | 0.1675 | 1.0525 | 0.8725 | 0.791879 | 0.937377 | 0.257689 | 0.999187 | 0.997988 | 0.999507 |
| Dact3_Q0PHV7_S370s_1 | Dact3_Q0P | -1 | -0.01 | 0.65 | 0.14 | -1.79 | -1.14 | -1.61 | -0.44 | -0.27 | -0.79 | 0.34 | 0.14 | 1.17 | -0.85 | -0.16 | 0.05 | 0.02 | 0.3225 | 0.1475 | 0.833397 | 0.019902 | 0.829564 | 0.989223 | 0.997988 | 0.999507 |
| Dact3_Q0PHV7_S370s_2 | Dact3_Q0P | -3.04 | -2.15 | -2.23 | -3.53 | -2.6 | -3.36 | -4.8 | -3.24 | -3.14 | -2.67 | -1.93 | -2.97 | -1.98 | -3 | -3.69 | -3.62 | -1.19 | -0.09 | 0.1075 | 0.500213 | 0.165871 | 0.885847 | 0.989223 | 0.997988 | 0.994666 |
| Dact3_Q0PHV7_S374s_1 | Dact3_Q0P | 1.76 | 1.64 | 2.02 | 1.75 | 1.02 | 1.95 | 1.04 | 1.47 | 2.24 | 1.88 | 1.31 | 1.77 | 2.27 | 1.02 | 1.78 | 1.87 | -0.7625 | 0.06 | -0.335 | 0.857554 | 0.146793 | 0.977649 | 0.989223 | 0.997988 | 0.999507 |
| Dact3_Q0PHV7_S374s_2 | Dact3_Q0P | -3.04 | -2.15 | -2.23 | -3.53 | -2.6 | -3.36 | -4.8 | -3.24 | -3.14 | -2.67 | -1.93 | -2.97 | -1.98 | -3 | -3.69 | -3.62 | -0.4225 | 0.0075 | -0.0575 | 0.500213 | 0.165871 | 0.885847 | 0.989223 | 0.997988 | 0.994666 |
| Dact3_Q0PHV7_S409s_1 | Dact3_Q0P | -2.28 | -2.22 | -0.79 | -0.43 | -1.49 | -2.21 | -0.45 | -0.19 | -3.06 | -1.75 | 0.76 | -0.01 | -1.53 | -1.27 | 0.13 | -0.37 | -0.7625 | 0.06 | -0.335 | 0.255578 | 0.568223 | 0.63714 | 0.989223 | 0.997988 | 0.994666 |
| Dact3_Q0PHV7_S6s_1 | Dact3_Q0P | -1.43 | -1.14 | -0.88 | -1.7 | -0.82 | -2.15 | -0.56 | -1.43 | -1.81 | -1.94 | -0.18 | -1.15 | -1.35 | -0.66 | -1.09 | -0.97 | 0.345 | 0.415 | 0.67 | 0.369538 | 0.90177 | 0.96739 | 0.996493 | 0.997988 | 0.994666 |
| Dag1_Q62165_T788t_1 | Dag1_Q62 | 1.47 | -0.19 | -1.21 | 0.02 | 1.02 | 0.46 | 0.46 | 0.34 | 0.32 | 0.39 | -0.91 | -0.73 | 0.23 | 1.24 | -0.27 | 0.05 | 0.0475 | 0.0175 | 0.27 | 0.623591 | 0.308341 | 0.669834 | 0.989223 | 0.997988 | 0.995259 |
| Dagla_Q6WQJ1_S1032s_1 | Dagla_Q6W | -2.17 | -2.45 | -2.61 | -2.52 | -1.95 | -2.48 | -2.19 | -2.57 | -1.05 | -1.97 | -3.99 | -2.89 | -0.16 | -1.55 | -3.03 | -3.02 | 0.5475 | -0.255 | 0.29 | 0.436954 | 0.559669 | 0.948833 | 0.989223 | 0.997988 | 0.994666 |
| Dagla_Q6WQJ1_S744s_1 | Dagla_Q6W | 1.55 | -1.12 | 0.01 | 0.96 | 0.48 | 1.57 | 0.94 | 1.42 | 0 | -0.55 | -0.39 | 0.42 | -1.58 | 0.79 | -1.16 | -0.5 | 0.14 | -0.0375 | 0.4975 | 0.19 | 0.207832 | 0.412015 | 0.989223 | 0.997988 | 0.994666 |
| Dagla_Q6WQJ1_S954s_1 | Dagla_Q6W | 0.3 | -0.46 | 0.41 | 0 | 0.48 | 0.2 | 0.33 | -0.06 | -0.61 | -0.95 | -0.13 | 0.11 | -1.33 | 0.43 | -1.17 | -0.98 | 0.7525 | -0.48 | -0.9625 | 0.08587 | 0.519687 | 0.189752 | 0.989223 | 0.997988 | 0.994666 |
| Dap_Q91XC8_S2s_1 | Dap_Q91X | 0 | 0.3 | -0.37 | -1.21 | -0.71 | 0.3 | -0.92 | -0.96 | 1.04 | -0.06 | -0.77 | -0.59 | 1.16 | -1.62 | -0.34 | -0.53 | 0.175 | -0.4575 | -0.825 | 0.983389 | 0.55041 | 0.649751 | 0.989223 | 0.997988 | 1 |
| Dap_Q91XC8_S3s_1 | Dap_Q91X | 0 | 0.3 | -0.37 | -1.21 | -0.71 | 0.3 | -0.92 | -0.96 | 1.04 | -0.06 | -0.77 | -0.59 | 1.16 | -1.62 | -0.34 | -0.53 | -0.2525 | 0.225 | -0.0125 | 0.983389 | 0.55041 | 0.649751 | 0.989223 | 0.997988 | 1 |
| Dapk1_Q80YE7_S326s_1 | Dapk1_Q8 | -0.73 | 0.12 | -0.63 | -3.07 | -0.15 | -0.42 | -2.37 | -1.6 | -0.25 | -0.29 | -3.11 | -1.59 | 1.72 | -0.37 | -0.48 | -0.78 | -0.2525 | 0.225 | -0.0125 | 0.187538 | 0.939402 | 0.786937 | 0.999187 | 0.997988 | 0.994666 |
| Dapk2_Q8VDF3_S349s_1 | Dapk2_Q8 | 0.97 | 1.23 | 0.79 | 1.19 | -0.22 | 1.48 | -0.2 | 1.12 | 1.04 | 0.27 | 1.09 | 0.4 | -0.25 | 0.24 | 1.42 | -0.02 | -0.0575 | -0.2325 | 1.1 | 0.102204 | 0.260936 | 0.244553 | 0.989223 | 0.997988 | 0.994666 |
| Dars_Q922B2_S249s_1 | Dars_Q922 | -0.73 | -0.63 | -0.88 | 0.23 | -0.36 | -1.84 | 0.24 | -0.07 | -2.26 | -0.18 | -1.23 | -1.33 | -0.87 | -0.57 | 0.18 | -0.28 | -0.5 | -0.345 | -0.6975 | 0.744539 | 0.991731 | 0.137467 | 0.999187 | 0.997988 | 0.9971 |
| Daxx_O35613_S515s_1 | Daxx_O356 | -1.72 | 0.79 | 0.02 | -0.11 | -0.17 | -1.06 | -0.37 | -1.43 | 1.12 | 0.12 | 2.09 | 0.38 | 1.88 | 1.31 | 0.07 | -0.03 | -0.005 | -0.7475 | 0.1175 | 0.121517 | 0.368521 | 0.085526 | 0.989223 | 0.997988 | 0.994666 |
| Dbi_M0QWU8_S2s_1 | Dbi_M0QW | 0.17 | 0.95 | 1.05 | 2.18 | 0.21 | 0.62 | 1.13 | -1.1 | 0.45 | 1.32 | 1.64 | 1.97 | 0.91 | -0.62 | 0.85 | 0.69 | -0.5025 | 1.1825 | 1.0625 | 0.241123 | 0.148996 | 0.606845 | 0.989223 | 0.997988 | 0.994666 |
| Dbi_M0QWU8_Y29y_1 | Dbi_M0QW | -0.47 | 0.05 | -0.12 | 0.38 | 0.15 | 0.1 | 0.09 | 0.07 | -0.06 | 0.31 | -0.05 | -0.02 | -0.68 | 0.41 | -0.13 | 0.31 | -0.8725 | 0.2575 | -0.63 | 0.958953 | 0.55906 | 0.746452 | 0.989223 | 0.997988 | 0.999507 |
| Dbn1_F7CPL2_S78s_1 | Dbn1_F7C | 1.15 | 0.24 | 0.25 | -0.1 | 1.12 | 1.04 | 0.29 | 0.23 | 0.42 | 0.89 | 0.76 | -0.08 | 0.4 | 1.15 | 0.07 | -0.22 | 0.1425 | 0.085 | 0.0175 | 0.930823 | 0.43294 | 0.753152 | 0.989223 | 0.997988 | 0.999507 |
| Dbn1_F7CPL2_S79s_1 | Dbn1_F7C | -0.27 | -0.13 | 0.01 | -0.13 | 0.65 | -0.13 | 0.3 | 0.19 | -0.29 | -0.29 | 0.37 | -0.2 | -0.11 | 0.32 | -0.07 | -0.26 | 0.285 | 0.1125 | -0.035 | 0.690476 | 0.13284 | 0.911237 | 0.989223 | 0.997988 | 0.996159 |
| Dbn1_Q9QXS6_S241s_1 | Dbn1_Q9Q | -0.3 | 0.3 | 0.81 | -0.15 | -0.5 | 0.62 | -0.69 | -0.39 | -1.25 | -0.58 | -1.72 | -0.35 | 0.58 | -0.67 | -0.52 | -0.39 | 0.3825 | 0.0275 | 0.1 | 0.300417 | 0.298164 | 0.017278 | 0.989223 | 0.997988 | 0.994666 |
| Dbn1_Q9QXS6_S344s_1 | Dbn1_Q9Q | 0.31 | 0.01 | 0.57 | 0.81 | -0.8 | 1.1 | -0.51 | 0.11 | 0.98 | 0.35 | -1.2 | 0.64 | 1.67 | -2.09 | 0.32 | 0.61 | -0.405 | -1.14 | -0.415 | 0.682373 | 0.308586 | 0.633271 | 0.989223 | 0.997988 | 0.996159 |

| | | | | | | | | | | | | | | | | | | | | | | | | | | |
|---|---|---|---|---|---|---|---|---|---|---|---|---|---|---|---|---|---|---|---|---|---|---|---|---|---|---|
| Dbn1_Q9QXS6_S385s_1 | Dbn1_Q9Q | -0.18 | 0.66 | 0.14 | 0.3 | 0.74 | 0.35 | 0.67 | 0.27 | -0.24 | -0.34 | -0.11 | 0.08 | -0.62 | 0.02 | 0.2 | 0.3 | -0.45 | -0.2325 | -0.2975 | 0.431712 | 0.299556 | 0.167741 | 0.989223 | 0.997988 | 0.994666 |
| Dbn1_Q9QXS6_S387s_1 | Dbn1_Q9Q | 0.73 | 1.02 | 0.49 | 0.39 | 0.95 | 0.64 | 0.84 | 0.41 | -0.03 | 0.82 | -0.37 | 0.19 | -1.17 | 0.57 | 0.06 | 0.54 | 0.2775 | -0.3825 | -0.255 | 0.147566 | 0.830338 | 0.137122 | 0.995762 | 0.997988 | 0.994666 |
| Dbn1_Q9QXS6_S658s_1 | Dbn1_Q9Q | 0.33 | 0.56 | 0.27 | 0.46 | 0.55 | 0.27 | 0.5 | 0.34 | 0.05 | 0.58 | -0.01 | 0.3 | -0.54 | 0.65 | 0.01 | 0.25 | 0.0525 | -0.505 | -0.6575 | 0.32783 | 0.961033 | 0.467837 | 0.999187 | 0.997988 | 0.994666 |
| Dbn1_Q9QXS6_T394t_1 | Dbn1_Q9Q | -0.18 | -0.11 | -0.24 | 0.01 | -0.24 | -1.38 | 0.4 | 0.3 | 0.15 | -0.08 | -0.35 | -0.21 | 1.65 | 1.13 | -0.45 | 0.26 | 0.01 | -0.175 | -0.3125 | 0.112364 | 0.801236 | 0.973304 | 0.995762 | 0.997988 | 0.994666 |
| Dbndd1_H3BK17_S6s_1 | Dbndd1_H | 1.41 | 1.42 | -0.48 | -0.2 | 0.93 | 1.08 | -0.04 | 0.33 | 0.74 | 0.5 | -0.28 | -0.95 | 0.17 | 1.52 | -0.59 | -0.84 | -0.1 | 0.0075 | 0.7775 | 0.481645 | 0.942623 | 0.373241 | 0.999187 | 0.997988 | 0.994666 |
| Dbnl_Q62418_S141s_1 | Dbnl_Q624 | 0.06 | 0.13 | 0.92 | 1.23 | 0.5 | 0.45 | 1.22 | 1.28 | -0.2 | -0.49 | 1.91 | 1.57 | 0.51 | 0.92 | 1.91 | 1.66 | 0.0375 | -0.535 | -0.4725 | 0.146213 | 0.452584 | 0.855064 | 0.989223 | 0.997988 | 0.994666 |
| Dbnl_Q62418_S235s_1 | Dbnl_Q624 | -0.46 | -1.27 | -0.1 | -0.53 | -1.12 | 0.23 | -2.18 | -0.16 | 0.73 | -0.49 | -0.83 | -0.02 | 0.99 | -2.61 | -0.68 | 0.06 | 0.2775 | 0.1125 | 0.665 | 0.966546 | 0.686231 | 0.303105 | 0.994672 | 0.997988 | 0.999507 |
| Dbnl_Q62418_S277s_1 | Dbnl_Q624 | -2.24 | -2.07 | -2.09 | -2.05 | -1.23 | -1.43 | -1.41 | -1.35 | -2.35 | -2.13 | -0.11 | -1.58 | -1.39 | -0.79 | -1.34 | -1.81 | -0.2175 | 0.4375 | 0.03 | 0.020762 | 0.003338 | 0.253826 | 0.989223 | 0.997988 | 0.994666 |
| Dbnl_Q62418_T299t_1 | Dbnl_Q624 | -2.54 | -2.34 | -2.4 | -4.94 | -0.94 | -2.34 | -2.29 | -2.61 | -2.03 | -3 | -0.64 | -2.58 | -2.04 | -1.08 | -2.85 | -1.76 | 0.7575 | 0.57 | 0.78 | 0.112837 | 0.144808 | 0.196725 | 0.989223 | 0.997988 | 0.994666 |
| Dbp_Q60925_S82s_1 | Dbp_Q609 | 0.51 | 0.64 | 0.33 | 0.22 | 0.45 | 0.3 | 0.85 | 0.65 | -0.33 | 0.24 | -0.25 | 0.22 | -0.98 | 0.78 | 0.81 | 1.27 | 1.01 | 0.9925 | 1.1225 | 0.92554 | 0.551435 | 0.096403 | 0.989223 | 0.997988 | 0.999507 |
| Dbr1_Q923B1_S505s_1 | Dbr1_Q92 | 1.83 | -1.45 | 1 | 0.18 | 1.5 | 0.21 | 0.62 | -0.38 | 1.77 | 0.2 | -0.33 | 0.68 | 1.48 | -2.83 | 1.33 | 0.56 | 0.1375 | -0.455 | 0.045 | 0.81095 | 0.89038 | 0.798894 | 0.996493 | 0.997988 | 0.999507 |
| Dbr1_Q923B1_S520s_1 | Dbr1_Q92 | -2.19 | -1.73 | -1.62 | -1.36 | -2.61 | -1.99 | -3.52 | -1.13 | -1.62 | -1.86 | -0.35 | -1.85 | -0.29 | -2.41 | -1.68 | -2.62 | 0.0975 | 0.19 | -0.255 | 0.961597 | 0.248873 | 0.45151 | 0.989223 | 0.997988 | 0.999507 |
| Dcaf10_A2AKB9_S100s_1 | Dcaf10_A2 | -1.02 | -1.54 | -1.09 | -1.44 | -1.76 | -1.21 | -1.85 | -2 | -0.07 | -0.92 | -1.11 | -2.17 | 0.31 | -0.85 | -0.57 | -0.63 | -0.5875 | 0.305 | -0.025 | 0.027706 | 0.124102 | 0.640426 | 0.989223 | 0.997988 | 0.994666 |
| Dcaf6_Q9DC22_S584s_1 | Dcaf6_Q9D | -0.76 | 0.37 | -0.49 | 0.01 | 0.68 | -0.58 | -0.02 | -0.16 | -0.16 | 0.2 | -0.85 | -0.56 | 0.37 | 0.71 | -1.42 | 0.39 | -0.4325 | 0.205 | 0.8375 | 0.655494 | 0.587736 | 0.725407 | 0.989223 | 0.997988 | 0.995259 |
| Dcaf8_Q8N7N5_S100s_1 | Dcaf8_Q8N | 0.12 | -0.75 | -0.17 | -0.64 | -2.57 | 0.78 | -1.38 | -0.54 | 0.87 | 0.18 | -0.87 | -0.01 | 0.07 | -4.03 | 0.12 | -0.2 | 0.1975 | -0.125 | 0.23 | 0.477044 | 0.393222 | 0.33631 | 0.989223 | 0.997988 | 0.994666 |
| Dcbld2_Q91ZV3_S582s_1 | Dcbld2_Q9 | 1.45 | 1.04 | 0.83 | 0.97 | 1.08 | 1.86 | 0.78 | 0.96 | 0.45 | 0.89 | 1.31 | 0.68 | -0.28 | 0.71 | 0.63 | 0.17 | -0.5675 | 0.4025 | -0.65 | 0.034312 | 0.744332 | 0.398915 | 0.995762 | 0.997988 | 0.994666 |
| Dcbld2_Q91ZV3_T738t_1 | Dcbld2_Q9 | 1.57 | 2.28 | 1.74 | 2 | 1.58 | 1.92 | 1.92 | 1.6 | 1.21 | 1.22 | 1.17 | 1.54 | 0.32 | -0.25 | 1.11 | 2.26 | 0.0975 | -0.24 | -0.765 | 0.074371 | 0.562686 | 0.034211 | 0.989223 | 0.997988 | 0.994666 |
| Dcc_P70211_T1135t_1 | Dcc_P7021 | -2.34 | -3.88 | -2.07 | -1.18 | -1.55 | -2.01 | -1.26 | -1.03 | -1.56 | -2.88 | -0.15 | -1.32 | -1.93 | -0.81 | -2.55 | -1.54 | -0.1425 | -0.6125 | -1.0375 | 0.294379 | 0.121036 | 0.232668 | 0.989223 | 0.997988 | 0.994666 |
| Dclk1_Q1EDH1_S25s_1 | Dclk1_Q1E | -0.81 | -0.37 | -0.1 | 0.22 | 0.13 | -0.52 | -0.22 | 0 | -0.16 | -0.57 | 0.25 | 0.1 | 0.04 | 0.14 | 0.19 | 0.24 | 0.905 | 0.89 | 0.66 | 0.173969 | 0.699305 | 0.591532 | 0.994893 | 0.997988 | 0.994666 |
| Dclk1_Q1EDH1_S30s_1 | Dclk1_Q1E | 0.56 | -0.56 | 0.05 | -0.11 | 0.85 | 0.31 | -0.15 | 0.18 | 0.59 | 0.59 | 0.44 | 0.21 | 0.62 | 1.28 | 0.13 | 0.14 | 0.1125 | 0.17 | 0.4175 | 0.159753 | 0.346963 | 0.131304 | 0.989223 | 0.997988 | 0.994666 |
| Dclk1_Q1EDH1_S33s_1 | Dclk1_Q1E | -0.85 | -0.69 | -1.47 | -1.65 | 0.48 | -0.31 | -1.09 | -1.4 | 0 | -0.09 | -0.4 | -0.51 | 0.46 | 1.46 | -1.14 | -0.89 | 0.3125 | 0.4725 | 0.5575 | 0.08113 | 0.214081 | 0.012886 | 0.989223 | 0.997988 | 0.994666 |
| Dclk1_Q1EDH1_S40s_1 | Dclk1_Q1E | -4.8 | -4.07 | -2.83 | -1.98 | -3.2 | -4.59 | -2.02 | -2.1 | -2.77 | -3.42 | -1.56 | -2.61 | -0.69 | -1.88 | -2.01 | -2.25 | 0.585 | 0.915 | 1.1375 | 0.024097 | 0.565107 | 0.234097 | 0.989223 | 0.997988 | 0.994666 |
| Dclk1_Q1EDH1_S414s_1 | Dclk1_Q1E | 0.3 | -0.13 | 0.42 | -0.29 | -0.33 | 0.43 | -0.49 | 0.16 | 0.21 | -0.01 | 0.16 | 0.09 | 0.64 | 0.51 | 0.13 | 0.06 | 0.4425 | 0.83 | 1.7125 | 0.38072 | 0.658247 | 0.880982 | 0.990434 | 0.997988 | 0.994666 |
| Dclk1_Q1EDH1_S45s_1 | Dclk1_Q1E | -2.84 | -2.22 | -1.73 | -1.47 | -2.08 | -2.48 | -1.2 | -1.33 | -2.19 | -2.01 | -1.15 | -1.49 | -0.59 | -1.72 | -1.53 | -1.18 | -0.1325 | 0.0375 | 0.26 | 0.065998 | 0.481522 | 0.369756 | 0.989223 | 0.997988 | 0.994666 |
| Dclk1_Q1EDH1_S45s_2 | Dclk1_Q1E | -0.4 | -0.49 | -1.25 | -1.69 | -0.35 | -0.51 | -1.02 | -1.11 | -1.03 | -0.06 | -0.15 | -0.48 | -0.77 | -1.28 | -1.86 | -0.95 | 0.2925 | 0.355 | 0.81 | 0.522906 | 0.56325 | 0.191078 | 0.989223 | 0.997988 | 0.994666 |
| Dclk1_Q1EDH1_S46s_1 | Dclk1_Q1E | -3.11 | -2.91 | -1.83 | -1.96 | -1.95 | -2.71 | -1.53 | -2.7 | -2.35 | -3.47 | -1.55 | -2.34 | -0.77 | -1.65 | -2.37 | -1.75 | 0.21 | 0.5275 | -0.2575 | 0.095148 | 0.583186 | 0.958816 | 0.989223 | 0.997988 | 0.994666 |
| Dclk1_Q1EDH1_S46s_2 | Dclk1_Q1E | -1.27 | -1.42 | -1.8 | -1.63 | -1.41 | -1.46 | -1.28 | -1.64 | -0.9 | -1.07 | -1.15 | -0.81 | -0.54 | -1.19 | -1.61 | -1.69 | 0.23 | 0.025 | 0.8175 | 0.416625 | 0.711586 | 0.039707 | 0.995437 | 0.997988 | 0.994666 |
| Dclk1_Q1EDH1_S48s_2 | Dclk1_Q1E | -1.17 | -1.26 | -1.59 | -1.65 | -1.54 | -1.51 | -1.18 | -1.45 | -0.8 | -1.29 | -0.77 | -0.69 | -0.49 | -1.18 | -1.69 | -1.41 | 0.0825 | 0.5475 | 0.2725 | 0.495765 | 0.991139 | 0.060526 | 0.999187 | 0.997988 | 0.994666 |
| Dclk1_Q9JLM8_S307s_1 | Dclk1_Q9JI | -0.3 | 0.03 | 0.1 | 0.1 | -0.34 | -0.36 | 0.03 | 0.01 | 0.21 | 0.24 | 0.4 | -0.03 | 0.04 | 0.08 | 0.1 | 0.21 | -0.0025 | 0.53 | 0.225 | 0.602574 | 0.516578 | 0.343955 | 0.989223 | 0.997988 | 0.995259 |
| Dclk1_Q9JLM8_S307s_2 | Dclk1_Q9JI | -2 | -1.71 | -2.71 | -0.6 | -2.06 | -1.75 | -1.1 | -1.74 | -1.34 | -1.68 | -0.61 | -1.59 | -1.01 | -0.12 | -1.45 | -0.95 | -0.1475 | 0.2225 | 0.125 | 0.102128 | 0.837479 | 0.356946 | 0.995762 | 0.997988 | 0.994666 |
| Dclk1_Q9JLM8_S323s_2 | Dclk1_Q9JI | -2 | -1.71 | -2.71 | -0.6 | -2.06 | -1.75 | -1.1 | -1.74 | -1.34 | -1.68 | -0.61 | -1.59 | -1.01 | -0.12 | -1.45 | -0.95 | 0.0925 | 0.45 | 0.8725 | 0.102128 | 0.837479 | 0.356946 | 0.995762 | 0.997988 | 0.994666 |
| Dclk1_Q9JLM8_S32s_1 | Dclk1_Q9JI | -2.17 | -2.24 | -3.75 | -2.69 | -2.01 | -2.24 | -2.24 | -2.97 | -2.6 | -1.32 | -2.47 | -2.68 | -1.54 | -1.65 | -2.77 | -2.32 | 0.0925 | 0.45 | 0.8725 | 0.178438 | 0.398261 | 0.349109 | 0.989223 | 0.997988 | 0.994666 |
| Dclk1_Q9JLM8_S363s_1 | Dclk1_Q9JI | 0.04 | -0.47 | -0.03 | 1.07 | -0.76 | -0.3 | 0.37 | 0.18 | 0.24 | 0.24 | 0.94 | -0.32 | -0.02 | 0.12 | 0.86 | 0.27 | 0.3475 | 0.445 | 0.6425 | 0.691991 | 0.487354 | 0.76554 | 0.989223 | 0.997988 | 0.996159 |
| Dclk1_Q9JLM8_S364s_1 | Dclk1_Q9JI | 0.76 | 0.11 | -0.64 | -0.09 | 0.09 | 0.42 | -0.24 | -0.13 | 0.48 | 0.87 | 0.06 | -0.56 | -0.18 | 0.02 | -0.62 | -0.51 | -0.28 | 0.1225 | 0.155 | 0.3245 | 1 | 0.669572 | 1 | 0.997988 | 0.994666 |
| Dclk1_Q9JLM8_S392s_1 | Dclk1_Q9JI | -0.02 | 0.1 | 0.12 | 0.33 | 0.09 | -0.32 | 0.61 | 0.44 | 0.1 | 0.21 | 0.88 | 0.5 | -0.54 | 0.82 | -0.18 | 0 | -3.9E-17 | 0.1775 | -0.3575 | 0.748821 | 0.784126 | 0.276657 | 0.995762 | 0.997988 | 0.9971 |
| Dclk1_Q9JLM8_S736s_1 | Dclk1_Q9JI | -0.1 | 0.31 | 0.43 | 0.34 | 0.32 | -0.42 | 0.55 | 0.53 | 0.12 | -0.12 | 0.24 | 0.14 | -0.73 | 0.07 | 0.17 | 0.56 | 0.0725 | 0.29 | -0.1075 | 0.501103 | 1 | 0.524897 | 1 | 0.997988 | 0.994666 |
| Dclk1_Q9JLM8_S742s_1 | Dclk1_Q9JI | 0.57 | 0.54 | 0.47 | 0.52 | 0.71 | 0.42 | 0.92 | 0.58 | 0.65 | 0.92 | 0.64 | 0.83 | 0.18 | 1.26 | 0.71 | 0.25 | 1.57E-16 | -0.15 | -0.2275 | 0.806982 | 0.532562 | 0.277747 | 0.989223 | 0.997988 | 0.999507 |
| Dclk1_Q9JLM8_S77s_1 | Dclk1_Q9JI | -0.47 | 0.15 | -0.57 | -0.82 | -0.16 | -0.44 | -0.08 | -0.8 | -0.87 | -0.22 | -0.76 | -0.22 | -0.19 | -0.03 | -0.66 | -0.57 | 0.1325 | 0.235 | 0.075 | 0.833758 | 0.843782 | 0.768767 | 0.996349 | 0.997988 | 0.999507 |
| Dclk2_Q6PGN3_S343s_1 | Dclk2_Q6P | -1.97 | -1 | -1.45 | -3.16 | -1.11 | -2.16 | -1.81 | -2.16 | -1.93 | -0.72 | -1.6 | -1.89 | -0.4 | -2.74 | -1.95 | -1.72 | 0.0575 | -0.09 | 0.065 | 0.753781 | 0.861075 | 0.486785 | 0.996493 | 0.997988 | 0.9971 |
| Dclk2_Q6PGN3_S346s_1 | Dclk2_Q6P | -1.15 | -0.6 | -1.33 | -1.11 | -0.48 | -0.65 | -0.52 | -0.51 | -0.39 | -0.47 | -1.5 | -0.43 | 1.23 | -1.54 | -0.92 | -0.64 | 0.085 | 0.36 | 0.1925 | 0.32085 | 0.052138 | 0.308522 | 0.989223 | 0.997988 | 0.994666 |
| Dcp1a_Q91YD3_S542s_1 | Dcp1a_Q9 | 0.32 | 0.56 | -0.27 | 0.24 | -0.01 | 0.35 | -0.1 | 0.09 | 0.96 | 0.2 | -0.64 | -0.3 | 0.01 | 0.93 | 0.04 | -0.16 | 0.5075 | 0.35 | 0.58 | 0.982276 | 0.610778 | 0.687339 | 0.989223 | 0.997988 | 1 |
| Dcps_Q9DAR7_T21t_1 | Dcps_Q9D | -2.05 | -1.28 | -3.01 | -2.75 | -0.88 | -2.25 | -2.04 | -3.87 | -1.08 | -1.74 | 0.15 | -2.61 | -0.19 | -2.64 | -3.35 | -2.89 | -0.13 | -0.1575 | -0.0075 | 0.994453 | 0.98454 | 0.159078 | 0.999187 | 0.997988 | 1 |
| Dctn1_E9Q586_S2s_1 | Dctn1_E9Q | -1.34 | -3.13 | -1.13 | -2.29 | -3.73 | -0.4 | -1.42 | -1.22 | -0.08 | -2.27 | -0.34 | -0.53 | -0.55 | -1.62 | -1.82 | -1.13 | 0.0125 | 0.9525 | 0.005 | 0.195726 | 0.708246 | 0.085477 | 0.995437 | 0.997988 | 0.994666 |
| Dctn4_Q8CBY8_S196s_1 | Dctn4_Q8C | -1.36 | -1.74 | -1.25 | -1.39 | -0.91 | -1.42 | -0.72 | -0.56 | -2.19 | -0.73 | 0.58 | -1.21 | -1.26 | -0.91 | -1.08 | -0.98 | 0.28 | 1.1675 | 0.6925 | 0.15192 | 0.066514 | 0.327728 | 0.989223 | 0.997988 | 0.994666 |
| Dcx_O88809_S306s_1 | Dcx_O888 | -0.62 | -1.54 | -1.07 | -1.8 | -1.67 | -1.44 | -1.45 | -0.75 | -0.95 | 0.76 | -1.23 | -1.66 | -0.25 | -0.57 | -1.63 | -1.51 | 0.5325 | 0.5475 | 0.3775 | 0.534538 | 0.835068 | 0.384439 | 0.995762 | 0.997988 | 0.994666 |
| Dcx_O88809_S332s_1 | Dcx_O888 | 0.67 | 0.11 | 0.07 | 0.22 | 0.55 | -0.17 | 0.45 | 0.16 | 0.94 | 1.91 | 0.47 | 0.73 | 1.03 | 0.52 | 0.36 | 0.45 | -0.07 | 0.4875 | 0.2675 | 0.268251 | 0.938878 | 0.060965 | 0.999187 | 0.997988 | 0.994666 |
| Dcx_O88809_S332s_2 | Dcx_O888 | 0.25 | -0.35 | -1.89 | -1.43 | 0.67 | 0.86 | -0.78 | -2 | 0.39 | 1.43 | -1.13 | -0.79 | -1.63 | 0.81 | -1.31 | -1.75 | -0.02 | 0.745 | 0.3225 | 0.868324 | 0.464211 | 0.247072 | 0.989223 | 0.997988 | 0.999507 |
| Dcx_O88809_S339s_1 | Dcx_O888 | -0.27 | -0.08 | -0.43 | -0.03 | -0.22 | -1.57 | -0.08 | 0.01 | -0.42 | 0.11 | 0.77 | -0.39 | -0.27 | 0.2 | -0.48 | 0.14 | 0.5425 | 0.83 | -0.115 | 0.714818 | 0.489861 | 0.499617 | 0.989223 | 0.997988 | 0.996966 |
| Dda1_D3Z6F8_S30s_1 | Dda1_D3Z | -2.12 | -2.44 | -2.28 | -3.06 | -1.93 | -2.76 | -2.52 | -2.57 | -2.14 | -1.77 | -0.73 | -2.08 | -0.78 | -2.15 | -1.72 | -1.88 | -0.2625 | 0.22 | 0.1 | 0.047715 | 0.919842 | 0.065401 | 0.997335 | 0.997988 | 0.994666 |
| Ddah1_Q9CWS0_S33s_1 | Ddah1_Q9 | 0.65 | 0.58 | 0.76 | 0.63 | 0.82 | 0.04 | -0.57 | 0.63 | 0.85 | -0.16 | 0.91 | 0.43 | -0.2 | 1.09 | 0.21 | 0.19 | 0.03 | 0.795 | 0.8425 | 0.31528 | 0.215154 | 0.617169 | 0.989223 | 0.997988 | 0.994666 |
| Ddhd1_F8WHP5_S707s_1 | Ddhd1_F8 | -0.24 | 0.19 | 0.43 | 0.65 | 0.35 | 0 | 0.69 | -0.35 | -0.06 | -0.31 | -0.67 | 0.67 | -1.05 | -0.16 | -0.72 | 0.25 | -0.425 | -0.1475 | -0.3325 | 0.090471 | 0.785816 | 0.337404 | 0.995762 | 0.997988 | 0.994666 |
| Ddhd1_F8WHP5_T710t_1 | Ddhd1_F8 | -0.72 | -0.3 | 0.26 | 0.35 | -0.16 | -0.66 | 1.03 | 0.58 | -0.36 | -0.09 | -0.58 | 0.69 | -0.32 | -0.21 | 0.16 | 0.59 | -0.085 | -0.35 | -0.6775 | 0.656991 | 0.488908 | 0.963102 | 0.989223 | 0.997988 | 0.995259 |
| Ddhd2_Q80Y98_S447s_1 | Ddhd2_Q8 | 0.32 | 1.08 | -1.09 | -0.79 | 0.81 | -0.47 | -0.34 | -0.5 | -0.99 | -0.66 | -0.77 | -0.54 | -0.1 | 1.08 | 0.07 | -0.27 | 0.3 | 0.0175 | 0.1575 | 0.566212 | 0.992557 | 0.221085 | 0.999187 | 0.997988 | 0.995259 |
| Ddi2_A2ADY9_S121s_1 | Ddi2_A2A | -0.49 | 0.28 | 0.38 | 0.36 | 0.32 | -0.19 | 0.55 | 0.61 | -0.1 | 0 | 0.21 | 0.65 | -0.11 | 0.29 | -0.01 | 0.74 | -0.005 | -0.62 | 0.315 | 0.771218 | 0.53211 | 0.850038 | 0.989223 | 0.997988 | 0.9971 |
| Ddi2_A2ADY9_S128s_1 | Ddi2_A2A | 0.86 | 0.24 | 0.93 | 0.78 | 1.07 | 0.83 | 1.86 | 1 | 0 | 0.7 | -0.59 | 1.32 | -0.12 | 0.34 | 0.91 | 1 | 0.19 | 0.0575 | 0.095 | 0.619889 | 0.130813 | 0.432543 | 0.989223 | 0.997988 | 0.995259 |
| Ddn_Q80TS7_S138s_1 | Ddn_Q80T | 0.6 | 0.69 | 0.68 | 0.68 | 0.9 | 0.89 | 0.69 | 0.99 | 0.65 | 0.73 | 1.09 | 1.23 | 0.5 | 0.78 | 0.19 | 0.39 | 0.4875 | -0.345 | -0.17 | 0.429996 | 0.313449 | 0.278035 | 0.989223 | 0.997988 | 0.994666 |
| Ddn_Q80TS7_S354s_1 | Ddn_Q80T | 0.56 | 0.67 | 0.9 | 0.34 | 0.2 | 0.63 | 0.5 | 0.68 | 0.23 | 0.94 | 0.02 | 0.95 | 0.06 | 0.53 | 0.01 | 0.12 | 0.205 | 0.2625 | -0.1975 | 0.121219 | 0.622277 | 0.786619 | 0.989223 | 0.997988 | 0.994666 |
| Ddn_Q80TS7_S388s_1 | Ddn_Q80T | 0.09 | 0.41 | -1.22 | -0.07 | 0.07 | 1.01 | 1.25 | 0.81 | -1.29 | 0.53 | 1.06 | -0.24 | -0.89 | -0.21 | -0.58 | 0.48 | -0.115 | -0.0825 | -0.4375 | 0.819669 | 0.038342 | 0.711575 | 0.989223 | 0.997988 | 0.999507 |
| Ddn_Q80TS7_S524s_1 | Ddn_Q80T | 0.25 | -0.58 | -0.84 | -2.61 | -0.85 | 0.74 | -1.5 | -1.86 | 0.08 | -0.73 | -0.12 | -1.29 | -0.24 | -1.1 | -2.42 | -1.26 | 0.9825 | 0.2125 | -0.1025 | 0.647293 | 0.915556 | 0.49251 | 0.997335 | 0.997988 | 0.995259 |
| Ddn_Q80TS7_S77s_1 | Ddn_Q80T | 0.51 | 0.68 | 0.83 | 0.76 | 0.62 | 0.8 | 0.67 | 0.93 | 0.68 | 0.76 | 1.01 | 0.94 | 0.51 | 0.78 | 0.36 | 0.2 | 0.0775 | 0.43 | -0.31 | 0.36763 | 0.771831 | 0.490924 | 0.995762 | 0.997988 | 0.994666 |
| Ddx17_Q3U741_S522s_1 | Ddx17_Q3 | -1.06 | -2.74 | -1.4 | -2.31 | -3.16 | -1.23 | -1.95 | -1.95 | 0.02 | -3.37 | -1.93 | -2.19 | 0.81 | -1.5 | -1.96 | -1.73 | 0.06 | 0.1525 | -0.2325 | 0.263522 | 0.703397 | 0.988953 | 0.995437 | 0.997988 | 0.994666 |
| Ddx20_Q9JJY4_S472s_1 | Ddx20_Q9 | -1.84 | -1.2 | -0.34 | -0.96 | -1.39 | -0.68 | -1.01 | -1.56 | -0.71 | -0.68 | -1.34 | -1.8 | 0.06 | -0.67 | -0.72 | -1.87 | -0.195 | 0.01 | 0.7825 | 0.558009 | 0.836828 | 0.906897 | 0.995762 | 0.997988 | 0.995259 |
| Ddx20_Q9JJY4_S496s_1 | Ddx20_Q9 | -1.97 | -2.5 | -2.02 | -1.23 | -1.99 | -1.12 | -1.76 | -2.23 | 0.36 | -1.91 | -1.52 | -1.18 | -0.23 | -2.94 | -1.54 | -0.77 | -0.075 | -0.0475 | 0.285 | 0.354031 | 0.66354 | 0.124399 | 0.990661 | 0.997988 | 0.994666 |
| Ddx21_Q9JIK5_S243s_1 | Ddx21_Q9 | 0.86 | 0.64 | 0.72 | 0.64 | -1.26 | 1.15 | -0.16 | 0.66 | 1.85 | 0.21 | -0.37 | 0.87 | 0.56 | -2.07 | 0.68 | 0.68 | 0.155 | 0.8675 | 0.56 | 0.242072 | 0.224984 | 0.870158 | 0.989223 | 0.997988 | 0.994666 |
| Ddx23_D3Z0M9_S108s_1 | Ddx23_D3 | 0.44 | -0.77 | 0.37 | 0.41 | -1.48 | 0.03 | -0.38 | 0.15 | -0.03 | -1.58 | -0.58 | 0.26 | 1.27 | -1.34 | 0.79 | 0.46 | -0.6175 | -0.075 | -0.7525 | 0.756471 | 0.248676 | 0.231538 | 0.989223 | 0.997988 | 0.9971 |
| Ddx24_F8WJA0_S126s_1 | Ddx24_F8 | 0.01 | -0.02 | 0.43 | 0.31 | -0.87 | -0.12 | 0.28 | -0.27 | -0.03 | 0.2 | 0.87 | 0.15 | -0.77 | -0.06 | 0.67 | 0.26 | -0.5325 | -0.595 | 0.1825 | 0.656551 | 0.166499 | 0.681173 | 0.989223 | 0.997988 | 0.995259 |
| Ddx27_H3BLJ6_S133s_1 | Ddx27_H3 | -0.36 | -0.61 | -2.57 | -3.7 | -1.11 | -0.35 | -2.23 | -5.05 | -0.38 | -0.32 | -4.2 | -3.58 | 0.92 | -1.54 | -4.22 | -2.45 | -0.4275 | 0.115 | -0.1575 | 0.991363 | 0.739191 | 0.786988 | 0.995762 | 0.997988 | 1 |
| Ddx3x_Q62167_S612s_1 | Ddx3x_Q6 | -1.02 | -3.71 | -2.48 | -1.76 | -1.17 | -0.73 | -1.24 | -1.51 | -6.31 | -1.69 | -5.06 | -1.91 | -3 | -7.75 | -1.7 | -0.36 | -0.375 | -0.31 | -0.0125 | 0.517557 | 0.069825 | 0.206178 | 0.989223 | 0.997988 | 0.994666 |
| Ddx51_Q6P9R1_S77s_1 | Ddx51_Q6 | 1.45 | 0.48 | 1 | 1.09 | 0.93 | 0.76 | 0.15 | 1.02 | 0.66 | 0.52 | 1.04 | 1.34 | 0.91 | 0.5 | 1.19 | 0.56 | 1.08 | -1.5 | -0.96 | 0.494393 | 0.350659 | 0.709975 | 0.989223 | 0.997988 | 0.994666 |
| Ddx54_Q8K4L0_S774s_1 | Ddx54_Q8 | 0.42 | 0.28 | 1.02 | 0.47 | -1.14 | 0.33 | -0.6 | 0.29 | 0.54 | -0.11 | 0.76 | 0.73 | 0 | -1.78 | 0.61 | 0.73 | -0.29 | -0.115 | -0.215 | 0.255014 | 0.053777 | 0.821796 | 0.989223 | 0.997988 | 0.994666 |
| Ddx59_Q9DBN9_S150s_1 | Ddx59_Q9 | -2.9 | -1.52 | -3.15 | -4.37 | -2.19 | -1.72 | -2.71 | -2.96 | -2.27 | -1.69 | -2.22 | -2.55 | -0.84 | -3.03 | -3.26 | -1.78 | -0.8275 | -0.0675 | -0.6575 | 0.310297 | 0.322894 | 0.17874 | 0.989223 | 0.997988 | 0.994666 |
| Dek_E9Q8Y1_S33s_1 | Dek_E9Q8 | -5.23 | -4.24 | -3.03 | -3.77 | -4.99 | -4.95 | -4.18 | -3.19 | -5.27 | -4.75 | -1.64 | -3.38 | -3.92 | -3.36 | -2.77 | -4.63 | 0.59 | 0.8025 | 0.7575 | 0.484703 | 0.64597 | 0.712223 | 0.99033 | 0.997988 | 0.994666 |
| Dek_E9Q8Y1_S55s_1 | Dek_E9Q8 | -2.57 | -2.48 | -4.13 | -3.39 | -2.06 | -3.21 | -3.3 | -2.63 | -2.79 | -3.12 | -0.31 | -2.49 | -1.98 | -3.22 | -3.01 | -1.75 | -0.26 | 0.3075 | 0.3975 | 0.215496 | 0.454646 | 0.175795 | 0.989223 | 0.997988 | 0.994666 |
| Dennd1a_A2ALU3_S86s_1 | Dennd1a_ | -1.94 | -0.91 | -0.93 | -1.83 | -0.16 | -1.86 | -0.79 | -1.6 | -1.37 | -1.42 | -1.35 | -1.01 | -1.14 | -0.49 | -1.06 | -1.3 | 0.3425 | 0.965 | 0.6525 | 0.272981 | 0.507665 | 0.722238 | 0.989223 | 0.997988 | 0.994666 |
| Dennd1a_Q8K382_S473s_1 | Dennd1a_ | 0.2 | 0.08 | 0.3 | -0.37 | 0.37 | 0.18 | 0.13 | 0.19 | 0.08 | -0.06 | 0.08 | -0.06 | 0.94 | 0.45 | -0.09 | 0.55 | 0.3 | 0.115 | 0.405 | 0.209051 | 0.481319 | 0.858963 | 0.989223 | 0.997988 | 0.994666 |
| Dennd1a_Q8K382_T707t_1 | Dennd1a_ | 0.22 | 0.08 | -1.29 | -0.93 | 0.09 | 0.24 | -0.29 | -0.24 | -0.3 | -0.03 | -0.25 | -1.33 | -0.28 | -0.02 | -1.04 | -1.28 | 0.165 | -0.0425 | 0.41 | 0.705686 | 0.278424 | 0.995595 | 0.989223 | 0.998132 | 0.996159 |
| Dennd4a_E9Q8V6_S1053s_1 | Dennd4a_ | -0.23 | 0.1 | -0.41 | -1.05 | 0 | -0.64 | -1.03 | -0.99 | 0.21 | -0.35 | -0.49 | -1.07 | 0.73 | 0.17 | -1.6 | -0.75 | 0.43 | 0.0025 | -0.175 | 0.947249 | 0.444309 | 0.940109 | 0.989223 | 0.997988 | 0.999507 |
| Dennd4a_E9Q8V6_S1251s_1 | Dennd4a_ | -0.78 | -0.8 | -0.93 | -0.99 | -0.33 | -0.77 | -1.54 | -1.28 | -0.48 | -0.9 | -0.58 | -0.61 | 0.35 | 0.09 | -1.13 | -1.2 | -0.2675 | -0.0275 | 0.035 | 0.338228 | 0.726242 | 0.302705 | 0.995437 | 0.997988 | 0.994666 |
| Dennd4a_E9Q8V6_S1252s_1 | Dennd4a_ | -0.65 | -0.47 | -0.27 | -0.93 | -0.38 | -0.75 | -0.87 | -0.83 | -0.1 | -0.95 | -0.54 | -0.48 | 0.52 | -0.04 | -0.75 | -0.46 | -0.105 | 0.2325 | 0.4025 | 0.263741 | 0.601671 | 0.821587 | 0.989223 | 0.997988 | 0.994666 |
| Dennd4a_E9Q8V6_S1512s_1 | Dennd4a_ | -1.78 | -1.48 | -1.17 | -0.78 | -1.59 | -1.01 | -1.06 | -0.83 | -0.39 | -0.67 | -1.68 | -1.25 | 0.23 | -0.59 | -0.79 | -1.05 | -0.1275 | 0.0625 | 0.3975 | 0.065929 | 0.547117 | 0.416856 | 0.989223 | 0.997988 | 0.994666 |
| Dennd4a_E9Q8V6_S732s_1 | Dennd4a_ | -0.22 | 0.64 | 0.94 | -0.15 | 0.97 | 0.18 | 1.28 | 0.78 | -0.09 | 0.77 | 3.39 | 1.19 | 0.19 | 1.41 | 1.18 | 0.86 | 0.18 | 0.305 | 0.7525 | 0.151691 | 0.193662 | 0.180714 | 0.989223 | 0.997988 | 0.994666 |
| Dennd4b_Q3U1Y4_S736s_1 | Dennd4b_ | 0.76 | 0.29 | 0.34 | 0.67 | 0.19 | 1.73 | -0.38 | 0.2 | 2.08 | 0.37 | -0.56 | 0.02 | 2.19 | -0.45 | 0.45 | 0.12 | 0.5 | 1.0125 | 0.6075 | 0.907795 | 0.85542 | 0.944459 | 0.996349 | 0.997988 | 0.999507 |
| Dennd4b_Q3U1Y4_S930s_1 | Dennd4b_ | -0.05 | -0.32 | 0.03 | -0.15 | 0.12 | -0.36 | 0.56 | -0.07 | 0.08 | -0.18 | -0.98 | 0.18 | 0.03 | -0.34 | -0.19 | -0.1 | -0.08 | -0.0375 | 0.0625 | 0.90883 | 0.479249 | 0.740211 | 0.989223 | 0.997988 | 0.999507 |
| Dennd4b_Q3U1Y4_S951s_1 | Dennd4b_ | -0.86 | -1.03 | -1.44 | -1.82 | -0.8 | -0.67 | -1.36 | -1.77 | -0.88 | -1.97 | -1.16 | -1.55 | -0.19 | -1.16 | -1.36 | -1.61 | 0.185 | -0.1025 | -0.0275 | 0.596446 | 0.687007 | 0.763497 | 0.994672 | 0.997988 | 0.995259 |
| Dennd4b_Q3U1Y4_S961s_1 | Dennd4b_ | -0.18 | -0.54 | -0.04 | -0.33 | -0.99 | -0.22 | -0.87 | -0.48 | 0.2 | -0.51 | -1.82 | -1.13 | 0.87 | -1.26 | -0.26 | -0.17 | 0.1375 | -0.1025 | 0.2075 | 0.877886 | 0.176765 | 0.229523 | 0.989223 | 0.997988 | 0.999507 |
| Dennd4c_E9Q449_S1323s_1 | Dennd4c_ | -0.19 | -0.86 | -0.92 | -0.54 | -0.33 | 0.4 | -0.37 | -0.52 | 0.08 | -0.53 | -1.12 | -0.93 | 0.33 | -1.17 | -0.25 | -0.06 | -0.3675 | -0.5425 | 0.0675 | 0.376973 | 0.173119 | 0.994052 | 0.989223 | 0.997988 | 0.994666 |
| Dennd5a_Q6PAL8_S193s_1 | Dennd5a_ | -0.1 | -0.06 | 0.25 | 0.26 | 0.53 | 0.02 | 0.27 | 0.06 | -0.2 | 0.27 | 0.28 | 0.19 | -0.4 | 0.35 | 0.05 | 0.44 | 0.4225 | 0.0025 | 0.34 | 0.937198 | 0.56591 | 0.841407 | 0.989223 | 0.997988 | 0.999507 |
| Dennd5b_A2RSQ0_S178s_1 | Dennd5b_ | -0.38 | -0.06 | 0.45 | 0.16 | 0.3 | -0.19 | -0.15 | -0.32 | -0.39 | -0.08 | -0.21 | -0.34 | -0.07 | 0.31 | -0.19 | 0.04 | 0.1325 | 0.0475 | 0.0225 | 0.94338 | 0.620729 | 0.26508 | 0.989223 | 0.997988 | 0.999507 |
| Dennd6a_Q8BH65_S16s_1 | Dennd6a_ | -0.43 | -0.19 | 0.26 | -1.23 | -0.63 | -0.71 | -0.81 | -0.82 | -0.45 | -1.03 | 1.16 | -0.76 | 0.03 | -0.83 | -1.63 | -1.2 | -0.1325 | -0.2975 | -0.02 | 0.280166 | 0.306088 | 0.813714 | 0.989223 | 0.997988 | 0.994666 |
| Denr_E9Q0G1_S73s_1 | Denr_E9Q | -0.53 | -0.52 | -1.16 | 1.22 | -0.33 | -1.18 | -0.92 | 1.35 | -0.99 | -1.29 | -0.27 | 1.1 | -0.3 | -0.47 | 1.2 | -1.26 | -0.345 | 0.1275 | -0.51 | 0.951164 | 0.973426 | 0.863411 | 0.999187 | 0.997988 | 0.999507 |
| Denr_E9Q0G1_S9s_1 | Denr_E9Q | -0.49 | 0.42 | 0.39 | -0.9 | -0.67 | -0.42 | 0.85 | 1.07 | 0.06 | 0.25 | 1.43 | 0.2 | 0.77 | 0.86 | -0.11 | 1.22 | -0.0225 | -0.115 | 0.04 | 0.077721 | 0.488771 | 0.177399 | 0.989223 | 0.997988 | 0.994666 |
| Depdc5_P61460_S1496s_1 | Depdc5_P | -1.01 | -1 | -0.95 | -0.51 | -1.25 | -0.85 | -0.82 | -0.28 | -0.62 | -1.02 | -0.27 | -0.49 | -0.34 | -0.74 | -0.83 | -0.76 | 0.3525 | 0.63 | 0.83 | 0.451567 | 0.804748 | 0.321942 | 0.995762 | 0.997988 | 0.994666 |
| Dfnb31_Q80VW5_S246s_1 | Dfnb31_Q | 2.38 | 0.16 | 0.95 | 1.85 | 0.33 | 1.29 | 0.76 | 0.99 | 1.07 | 2.42 | 0.3 | 0.9 | 1.01 | 0.24 | 0.82 | 1.47 | 0.0675 | 0.2675 | 0.2 | 0.394987 | 0.326416 | 0.789247 | 0.989223 | 0.997988 | 0.994666 |
| Dfnb31_Q80VW5_S685s_1 | Dfnb31_Q | -0.48 | 0.38 | 0.01 | 0.03 | 0.74 | -0.46 | 0.12 | -0.37 | 0.38 | -0.74 | 0.13 | 0.2 | 0.78 | 0.3 | 0.33 | 0.95 | -0.4925 | -0.1625 | -0.45 | 0.068712 | 0.946453 | 0.981862 | 0.999187 | 0.997988 | 0.994666 |
| Dgkb_Q6NS52_S418s_1 | Dgkb_Q6N | -0.92 | -1.19 | -1.13 | -0.76 | -0.57 | -1.04 | 0.27 | -0.36 | -1.49 | -1.17 | -1.19 | -0.81 | -1.42 | -0.68 | -0.5 | -0.46 | 0.0225 | 0.0075 | 0.605 | 0.44582 | 0.088231 | 0.510537 | 0.989223 | 0.997988 | 0.994666 |
| Dgkb_Q6NS52_S94s_1 | Dgkb_Q6N | 1.29 | -0.07 | -0.68 | 0.78 | 0.91 | 1.14 | 0.26 | 0.95 | 0.08 | 0.01 | 0.53 | -0.14 | -1.04 | 0.53 | -0.05 | -0.04 | 0.575 | -0.165 | 0.235 | 0.361308 | 0.293826 | 0.638985 | 0.989223 | 0.997988 | 0.994666 |
| Dgkb_Q6NS52_S95s_1 | Dgkb_Q6N | 1.15 | -1.95 | -0.74 | 0.19 | 0.6 | 1.24 | 0.57 | 0.96 | -0.48 | -1.03 | 0.59 | -0.02 | -1.88 | 0.29 | -0.17 | -0.15 | 0.485 | -0.21 | -0.48 | 0.847476 | 0.077257 | 0.879195 | 0.989223 | 0.997988 | 0.999507 |
| Dgkb_Q6NS52_T117t_1 | Dgkb_Q6N | 0.65 | -0.93 | -0.89 | -0.16 | 0.37 | 0.27 | 0.49 | 0.52 | -0.45 | -0.68 | -0.04 | -0.7 | -0.13 | -0.01 | 0.23 | -0.23 | 1.18 | 0.1025 | -0.14 | 0.457311 | 0.069146 | 0.736442 | 0.989223 | 0.997988 | 0.994666 |
| Dgkd_E9PUQ8_S44s_1 | Dgkd_E9P | -0.89 | -3.93 | -2.96 | -2.78 | -3.25 | -1.22 | -2.83 | -1.78 | -3.6 | -4.83 | 0.54 | -3.08 | -2.84 | -1.67 | -3.64 | -3.64 | 0.745 | -0.135 | 0.2975 | 0.663872 | 0.597487 | 0.92935 | 0.989223 | 0.997988 | 0.996159 |
| Dgkg_F2Z409_S98s_1 | Dgkg_F2Z4 | -0.17 | 0.27 | -0.64 | 0.49 | -0.73 | 0.2 | -0.23 | -0.11 | 0.57 | 0.31 | -0.69 | 0.25 | 0.6 | 0.7 | 0.1 | -0.69 | 0.37 | -0.1025 | -0.3075 | 0.642886 | 0.53566 | 0.745997 | 0.989223 | 0.997988 | 0.995259 |
| Dgkg_F2Z409_T87t_1 | Dgkg_F2Z4 | -0.76 | -1.39 | -0.51 | -1.09 | -1.95 | -0.1 | -1.38 | -1.14 | 1.32 | -0.91 | -1.11 | -0.61 | 1.12 | -2.23 | -0.08 | -0.86 | -0.205 | 0.1225 | 0.19 | 0.52199 | 0.62092 | 0.282261 | 0.989223 | 0.997988 | 0.994666 |
| Dgkh_D3YXJ0_S1067s_1 | Dgkh_D3Y | 0.1 | 0.53 | -0.39 | -0.01 | -0.33 | -0.52 | 0.2 | 0.18 | 0.16 | 0.23 | 0.06 | 0.64 | -0.06 | 0.28 | 0.09 | -0.12 | -0.205 | 0.61 | 0.425 | 0.971877 | 0.552197 | 0.449808 | 0.989223 | 0.997988 | 0.999507 |
| Dgkh_D3YXJ0_S677s_1 | Dgkh_D3Y | 0.15 | 0.16 | -0.31 | -0.32 | 0.22 | -0.06 | -0.48 | -0.19 | 0.21 | 0.35 | -0.6 | 0.02 | 0.09 | 0.53 | -1.03 | -0.29 | -0.175 | 0.215 | -0.01 | 0.800115 | 0.851173 | 0.798237 | 0.996349 | 0.997988 | 0.999507 |
| Dgki_D3YWQ0_S705s_1 | Dgki_D3YW | -0.05 | -0.03 | 0.08 | -0.23 | 0.1 | -0.35 | 0.31 | 0.03 | -0.32 | -0.28 | 0.19 | -0.24 | -0.53 | 0.15 | -0.06 | 0.11 | -0.0475 | 0.075 | -0.095 | 0.924484 | 0.727226 | 0.65095 | 0.995569 | 0.997988 | 0.999507 |
| Dgki_D3YWQ0_S906s_1 | Dgki_D3YW | -0.58 | -0.11 | -0.47 | -0.09 | -0.47 | -0.62 | 0.2 | -0.16 | -0.36 | -0.25 | -0.96 | -0.34 | -0.21 | -0.23 | -0.55 | -0.37 | 0.08 | -0.105 | -0.025 | 0.913968 | 0.85064 | 0.539731 | 0.996349 | 0.997988 | 0.999507 |
| Dgki_D3YWQ0_T704t_1 | Dgki_D3YW | 0.12 | 0.26 | 0.58 | -0.52 | 0.25 | 0.18 | -0.21 | -1.64 | 0.09 | 0.43 | -0.19 | -0.2 | -0.03 | 0.53 | -1.21 | -1.46 | 0.05 | -0.165 | -0.0275 | 0.211826 | 0.327548 | 0.80226 | 0.989223 | 0.997988 | 0.994666 |
| Dgkq_D6RI30_S17s_1 | Dgkq_D6RI | 0.32 | -0.06 | -0.62 | -0.59 | 0.61 | -0.12 | 0.07 | -0.22 | 0.23 | 0.08 | 0.28 | -0.51 | -0.42 | 0.69 | -0.19 | -0.53 | -0.465 | -0.0775 | -0.6525 | 0.739322 | 0.313993 | 0.428502 | 0.989223 | 0.997988 | 0.9971 |
| Dgkq_D6RI30_S22s_1 | Dgkq_D6RI | 0.23 | 0.35 | -0.08 | -0.27 | 0.17 | 0.3 | 0.64 | -0.08 | 0.19 | 0.28 | -0.76 | 0.09 | -0.2 | 1 | -0.39 | -0.05 | 0.3225 | 0.2575 | 0.125 | 0.928899 | 0.44466 | 0.730949 | 0.989223 | 0.997988 | 0.999507 |
| Dgkq_D6RI30_S39s_1 | Dgkq_D6RI | 0.57 | 0.22 | 0.33 | 0.04 | -0.4 | 1.53 | -0.27 | 0.27 | 2.7 | 0.18 | -1.4 | 0.72 | 1.94 | -0.38 | -0.01 | 0.57 | 0.2 | -0.1075 | 0.0325 | 0.629167 | 0.986034 | 0.734337 | 0.999187 | 0.997988 | 0.995259 |
| Dgkz_Q80UP3_S706s_1 | Dgkz_Q80U | -2.3 | -3.27 | -2.15 | -2.85 | -2.52 | -2.21 | -1.58 | -1.75 | -2.05 | -2.68 | -0.21 | -1.87 | -1.18 | -2.08 | -1.83 | -2.04 | -0.0075 | 0.26 | 0.24 | 0.035394 | 0.0913 | 0.110713 | 0.989223 | 0.997988 | 0.994666 |
| Dhrs13_B0QZT2_S291s_1 | Dhrs13_B0 | -1.02 | -1.11 | -0.05 | -0.3 | -1.37 | -0.78 | -0.42 | -0.55 | -0.39 | -0.56 | -0.25 | -1.29 | 0.78 | -0.99 | -0.35 | -0.36 | 0.6275 | 0.94 | 0.86 | 0.388279 | 0.641509 | 0.994471 | 0.989223 | 0.997988 | 0.994666 |
| Dhrs13_B0QZT2_S307s_1 | Dhrs13_B0 | -0.24 | -0.25 | 0.31 | 0.42 | 0.1 | 0.41 | -0.28 | 0.29 | 0.03 | 0.18 | 0.36 | -0.27 | 0.2 | 0.34 | -0.4 | 0.03 | -0.16 | -0.0025 | 0.39 | 0.953508 | 0.797928 | 0.956647 | 0.995762 | 0.997988 | 0.999507 |
| Dhx29_Q8BN72_S198s_1 | Dhx29_Q8 | -0.82 | 0.04 | -0.5 | -0.34 | 0.04 | -0.54 | 0.29 | -0.17 | -0.77 | -0.52 | -1.1 | -0.54 | -1.17 | -0.21 | -0.41 | -0.25 | 0.07 | 0.015 | -0.0175 | 0.749891 | 0.290682 | 0.256013 | 0.989223 | 0.997988 | 0.9971 |
| Dhx30_Q99PU8_S226s_1 | Dhx30_Q9 | -0.87 | -1.6 | -0.74 | -0.48 | -0.54 | -1.6 | -0.31 | -0.41 | -1.56 | -0.67 | -0.49 | -0.69 | -1.78 | 0.51 | -0.28 | -0.53 | 0.31 | -0.3275 | -0.105 | 0.431233 | 0.583652 | 0.843083 | 0.989223 | 0.997988 | 0.994666 |
| Dhx32_Q8BZS9_S12s_1 | Dhx32_Q8 | -2.08 | -0.8 | -0.5 | 0.37 | -1.06 | -1.25 | -0.4 | -0.03 | -0.58 | 0.03 | 0.53 | -0.53 | 0.3 | 0.08 | 0.22 | 0.47 | 0.2075 | 0.07 | 0.4025 | 0.060674 | 0.898286 | 0.265429 | 0.996493 | 0.997988 | 0.994666 |
| Dhx9_O70133_S136s_1 | Dhx9_O70 | 0.01 | 0.43 | 0.22 | 0.76 | -0.13 | 0.04 | 0.36 | 1.11 | -0.45 | 0.02 | 0.75 | 0.71 | -0.67 | 0.14 | 0.96 | 0.67 | 0.0675 | 0.615 | 1.02 | 0.841064 | 0.975698 | 0.779188 | 0.999187 | 0.997988 | 0.999507 |
| Dhx9_O70133_S137s_1 | Dhx9_O70 | 0.72 | 0.57 | 0.37 | 0.53 | 0.45 | 0.31 | 0.46 | 0.89 | 0.02 | 0.6 | 0.36 | 0.57 | -0.17 | 0.45 | 0.88 | 0.39 | -0.01 | -0.0975 | -0.08 | 0.588639 | 0.929241 | 0.507547 | 0.997639 | 0.997988 | 0.995259 |
| Diap1_E9PV41_S22s_1 | Diap1_E9P | 0.62 | 0.61 | 0.94 | 0.73 | 1.04 | 0.3 | 0.2 | 0.63 | -0.17 | 0.68 | 1.61 | 1.15 | 0.43 | 0.72 | 1.23 | 0.96 | -0.02 | -0.16 | -0.16 | 0.687677 | 0.48248 | 0.812102 | 0.989223 | 0.997988 | 0.996159 |
| Dido1_Q8C9B9_S1726s_1 | Dido1_Q8 | -0.46 | -0.96 | 0.04 | 0.18 | -0.84 | 0.18 | -0.3 | -0.58 | -1.4 | -0.8 | 0.43 | 0.25 | -0.06 | -1.56 | -0.06 | -0.31 | -0.1825 | 0.0925 | 0.11 | 0.651894 | 0.804768 | 0.86771 | 0.995762 | 0.997988 | 0.995259 |
| Dip2b_Q3UH60_S50s_1 | Dip2b_Q3 | 0.9 | 0.17 | -0.55 | -0.22 | 0.16 | 0.73 | 0.92 | 1.07 | -0.11 | -0.51 | 0.16 | -0.11 | -0.99 | 0.77 | -1.76 | -0.56 | -0.085 | -0.08 | -0.1975 | 0.229767 | 0.103671 | 0.543932 | 0.989223 | 0.997988 | 0.994666 |
| Dip2b_Q3UH60_S782s_1 | Dip2b_Q3 | -0.79 | 0.22 | -0.44 | -0.23 | -1.13 | 0.14 | 0.2 | -0.05 | 0.56 | 0.09 | 0.16 | -1.1 | -0.12 | -1.31 | -0.52 | 0.78 | 0.645 | -0.2175 | -0.71 | 0.969987 | 0.787423 | 0.565826 | 0.995762 | 0.997988 | 0.999507 |
| Dip2b_Q3UH60_T70t_1 | Dip2b_Q3 | 0.11 | -0.86 | -0.57 | -0.29 | -0.38 | 0.06 | 0.49 | -0.82 | 0.03 | 0.1 | 1.48 | -0.13 | 0.18 | -0.69 | -0.53 | 0.14 | 0.1 | 0.2375 | 0.0175 | 0.60466 | 0.499884 | 0.091237 | 0.989223 | 0.997988 | 0.995259 |
| Dip2c_B7ZMZ7_T123t_1 | Dip2c_B7Z | -0.92 | -3.02 | -3.92 | -2.22 | -2.44 | -1 | -3.42 | -0.77 | -1.62 | -2.69 | 0.8 | -1.74 | -1.56 | -1.54 | -3.29 | -2.41 | 0.24 | 0.7725 | 0.1775 | 0.640784 | 0.440284 | 0.188004 | 0.989223 | 0.997988 | 0.995259 |
| Diras2_Q5PR73_S126s_1 | Diras2_Q5 | -0.2 | -0.6 | 0.6 | 0.57 | -0.9 | 0.18 | 0.98 | 0.65 | 0.35 | 0.41 | 0.29 | 0.54 | 1.71 | -0.88 | 0.98 | 0.13 | 0.6125 | 1.2075 | 0.32 | 0.503566 | 0.773634 | 0.362138 | 0.995762 | 0.997988 | 0.994666 |
| Diras2_Q5PR73_S175s_1 | Diras2_Q5 | 1.22 | 0.77 | -1.07 | -1.77 | 1.71 | 1.06 | -0.14 | -1.26 | 0.42 | 0.53 | -2.28 | -1.45 | -0.07 | 1.39 | -1.25 | -0.87 | 0.135 | 0.305 | 0.3925 | 0.987765 | 0.517389 | 0.58969 | 0.989223 | 0.997988 | 1 |
| Diras2_Q5PR73_S35s_1 | Diras2_Q5 | 0.56 | 0.87 | 0.92 | 0.82 | 1.39 | 0.88 | 1.12 | 0.69 | 0.49 | 0.62 | -0.26 | 0.96 | -0.03 | 1.25 | 0.45 | 0.91 | 0.555 | -0.4825 | 0.0125 | 0.656983 | 0.351821 | 0.282813 | 0.989223 | 0.997988 | 0.995259 |
| Dis3l2_Q8CI75_S864s_1 | Dis3l2_Q8 | -0.62 | -0.42 | 0.87 | 0.74 | -0.87 | -0.49 | 0.3 | 0.78 | 0.97 | -0.22 | 0.43 | 0.78 | 0.55 | -0.75 | 0.97 | 0.91 | 0.2275 | -0.34 | -0.1475 | 0.597 | 0.667912 | 0.446438 | 0.992842 | 0.997988 | 0.995259 |
| Disp2_Q8CIP5_S1119s_1 | Disp2_Q8C | 0.45 | 0.32 | 0.77 | 0.81 | 0.34 | -0.07 | 0.62 | 1.03 | 0.64 | 0.51 | 1.53 | 1.29 | 0.56 | 0.52 | 0.69 | 0.89 | -0.2125 | 0.3475 | 0.2775 | 0.76087 | 0.712368 | 0.21183 | 0.995437 | 0.997988 | 0.9971 |
| Disp2_Q8CIP5_S1121s_1 | Disp2_Q8C | 0.23 | -0.02 | 0.11 | 0.23 | 0.12 | 0.62 | 0.19 | 0.74 | -0.19 | -0.3 | 0.87 | 0.41 | -0.19 | -0.07 | 0.18 | 0.16 | -0.1075 | 0.405 | 0.0775 | 0.62677 | 0.251994 | 0.847718 | 0.989223 | 0.997988 | 0.995259 |
| Disp2_Q8CIP5_S1317s_1 | Disp2_Q8C | 0.15 | 0.97 | 0.42 | 0.05 | 0.2 | 0.8 | 0.27 | 0.73 | 0.68 | 0.92 | 0.63 | 0.76 | -0.32 | 0.51 | -0.27 | 0.23 | 0.28 | 0.06 | -0.1175 | 0.28981 | 0.723101 | 0.219691 | 0.995437 | 0.997988 | 0.994666 |
| Disp2_Q8CIP5_S1345s_1 | Disp2_Q8C | 0.2 | -0.37 | -0.74 | 0.02 | -0.35 | 0.19 | -0.87 | -0.32 | 0.7 | 0.66 | 0.33 | -0.01 | 0.5 | 0.39 | 0.11 | 0.18 | 0.1025 | 0.35 | -0.36 | 0.103971 | 0.717787 | 0.058845 | 0.995437 | 0.997988 | 0.994666 |
| Disp2_Q8CIP5_S6s_1 | Disp2_Q8C | -0.49 | -0.5 | 0.39 | 0.17 | -1.04 | -0.15 | -0.62 | -0.34 | 2.06 | -0.35 | 0.75 | -0.4 | 2.17 | -1.49 | 0.41 | 0.32 | -0.115 | 0.6425 | 0.5175 | 0.515393 | 0.194997 | 0.293887 | 0.989223 | 0.997988 | 0.994666 |
| Dixdc1_Q80Y83_S13s_1 | Dixdc1_Q8 | -1.82 | -1.4 | -2.83 | -1.74 | -1.49 | -1.78 | -2.92 | -2.48 | -1.19 | -0.87 | -1.76 | -2.68 | -0.62 | -1.24 | -2.22 | -1.79 | -0.43 | 0.6225 | 0.46 | 0.301921 | 0.60667 | 0.504608 | 0.989223 | 0.997988 | 0.994666 |
| Dixdc1_Q80Y83_S335s_1 | Dixdc1_Q8 | 0.75 | -0.9 | -0.31 | -0.29 | -1.37 | 0.88 | -0.78 | 0.18 | 0.67 | -0.48 | -0.18 | 0.02 | 0.96 | -0.93 | -0.52 | -0.44 | -0.22 | 0.3225 | 0.48 | 0.929004 | 0.876658 | 0.639446 | 0.996493 | 0.997988 | 0.999507 |
| Dixdc1_Q80Y83_S50s_1 | Dixdc1_Q8 | 0.07 | 0.84 | 0.57 | 0.8 | -1.03 | 0.99 | -0.09 | -0.19 | 2.87 | 1.07 | -0.01 | 0.6 | 2.28 | -1 | 0.66 | 0.47 | -0.085 | 0.195 | -0.045 | 0.958752 | 0.151379 | 0.354056 | 0.989223 | 0.997988 | 0.999507 |
| Dkc1_Q9ESX5_S508s_1 | Dkc1_Q9E | 0.71 | 0.51 | 0.33 | 0.34 | -0.48 | 0.71 | -0.35 | 0.25 | 0.69 | 0.28 | 0.01 | 0.19 | 0.8 | -0.24 | 0.53 | 0.76 | -0.65 | 0.5625 | 0.0325 | 0.974324 | 0.17824 | 0.472841 | 0.989223 | 0.997988 | 0.999507 |
| Dkk3_Q9QUN9_S295s_1 | Dkk3_Q9Q | -1.37 | -0.99 | -0.6 | -0.16 | -1 | -1.26 | -0.45 | -0.49 | -0.89 | -0.52 | -0.6 | -0.4 | -0.3 | -0.45 | -0.62 | -0.68 | -0.44 | -0.18 | -0.01 | 0.412848 | 0.952318 | 0.573469 | 0.999187 | 0.997988 | 0.994666 |
| Dlc1_Q9R0Z9_S584s_1 | Dlc1_Q9R0 | -0.72 | -0.4 | -0.88 | -0.84 | -0.53 | -0.39 | -0.79 | -1.1 | 0.17 | -0.06 | 0.37 | -0.13 | 0.48 | -0.58 | -1.15 | -0.39 | -0.02 | 0.1775 | 0.2675 | 0.42951 | 0.975897 | 0.008298 | 0.999187 | 0.997988 | 0.994666 |
| Dlg1_D3Z3B8_S149s_1 | Dlg1_D3Z3 | -0.97 | -0.97 | -0.43 | -0.15 | -0.13 | -0.27 | -0.09 | -0.27 | -0.4 | -0.82 | -0.58 | -0.51 | 0.09 | 0.54 | -0.66 | -0.31 | 0.0075 | 0.7975 | 0.3 | 0.150132 | 0.11508 | 0.84955 | 0.989223 | 0.997988 | 0.994666 |
| Dlg1_D3Z3B8_S40s_1 | Dlg1_D3Z3 | 0.99 | 0.91 | 1.09 | 0.97 | 1.04 | 1.33 | 0.91 | 1.15 | 0.7 | 1.23 | 1.25 | 1.04 | 0.3 | 0.91 | 1.08 | 1.09 | 0.44 | 0.0525 | 0.545 | 0.599427 | 0.571899 | 0.777759 | 0.989223 | 0.997988 | 0.995259 |
| Dlg1_D3Z3B8_S41s_1 | Dlg1_D3Z3 | -0.27 | 0.07 | 0.47 | -0.09 | 0.56 | 0.29 | 0.44 | 0.19 | -0.41 | -0.4 | 0.2 | -0.09 | 0.1 | 0.31 | -0.42 | -0.88 | 0.1175 | 0.065 | -0.145 | 0.444036 | 0.199849 | 0.42676 | 0.989223 | 0.997988 | 0.994666 |
| Dlg1_D3Z3B8_S487s_1 | Dlg1_D3Z3 | 0.22 | 0.78 | -0.23 | -1.19 | 0.68 | 0.31 | 2.14 | -0.7 | 0.63 | 0.07 | -1.58 | -1.12 | 0.28 | 0.02 | -0.41 | -0.11 | 0.325 | -0.22 | -0.2675 | 0.90819 | 0.281945 | 0.518433 | 0.989223 | 0.997988 | 0.999507 |
| Dlg1_D3Z3B8_S492s_1 | Dlg1_D3Z3 | 0.3 | -0.21 | -0.27 | 0.1 | 0.38 | 0.44 | -0.12 | 0.51 | -0.36 | 0.15 | -1.04 | 0.27 | -0.31 | -1.03 | 0.4 | 0.03 | 0.7125 | -0.395 | 0.05 | 0.565725 | 0.220041 | 0.518787 | 0.989223 | 0.997988 | 0.995259 |
| Dlg1_E9Q9H0_S102s_1 | Dlg1_E9Q9 | -0.07 | 0.38 | 0.19 | 0.55 | 0.33 | -0.24 | 0.83 | -0.01 | 0.24 | -0.04 | -0.2 | 0.38 | -0.74 | 0.45 | -0.1 | 0.41 | 0.3225 | -0.225 | -0.2075 | 0.458811 | 0.905526 | 0.51838 | 0.996513 | 0.997988 | 0.994666 |
| Dlg1_E9Q9H0_S122s_1 | Dlg1_E9Q9 | 0.63 | 0.75 | 0.63 | 0.47 | 1.11 | 0.63 | 0.96 | 0.77 | -0.23 | 0.41 | 0.5 | 1.1 | -0.2 | -0.46 | 0.41 | 0.62 | -0.035 | -0.1675 | -0.2575 | 0.116396 | 0.266218 | 0.577485 | 0.989223 | 0.997988 | 0.994666 |
| Dlg2_E9Q2L2_S28s_1 | Dlg2_E9Q2 | 1.18 | 1.35 | 1.38 | 0.89 | 1.34 | 1.27 | 1.09 | 1.05 | -0.34 | 1.02 | -0.75 | 1.28 | 0.22 | -0.97 | 1.11 | 1.3 | 0.2475 | -0.175 | -0.5275 | 0.141873 | 0.954618 | 0.090447 | 0.999187 | 0.997988 | 0.994666 |
| Dlg2_E9Q2L2_S318s_1 | Dlg2_E9Q2 | -1.58 | -0.78 | -0.84 | -1.36 | -0.17 | -1.39 | 0.07 | -0.94 | -2.12 | -1.75 | -0.94 | -1.07 | -0.87 | -0.44 | -1.91 | -0.67 | -0.0125 | -0.8975 | -0.785 | 0.668921 | 0.185063 | 0.364766 | 0.989223 | 0.997988 | 0.996159 |
| Dlg2_E9Q2L2_S323s_1 | Dlg2_E9Q2 | -1.04 | -0.37 | 0.2 | -0.45 | 0.19 | -0.73 | 0.07 | -0.74 | -0.79 | -0.74 | -0.84 | 0.01 | -0.05 | 0.59 | -1.22 | -1.34 | 0.5325 | -0.33 | 0.1675 | 0.857677 | 0.752718 | 0.611759 | 0.995762 | 0.997988 | 0.999507 |
| Dlg2_E9Q2L2_S328s_1 | Dlg2_E9Q2 | 0.83 | 0.59 | 0.89 | 1.3 | 1.09 | 1.25 | 1.38 | 1.38 | 1 | 1.09 | 0.92 | 1.3 | 0.74 | 1.05 | 0.7 | 0.46 | 0.1125 | -0.175 | -0.09 | 0.551593 | 0.136112 | 0.4843 | 0.989223 | 0.997988 | 0.994666 |
| Dlg2_E9Q2L2_S360s_1 | Dlg2_E9Q2 | 2.72 | 1.51 | 1.65 | -0.17 | 1.71 | 2.89 | 1.59 | 1.48 | -0.53 | 2.79 | -0.95 | 1.67 | 0.95 | -1.49 | 2.04 | 1.82 | 0.3725 | 0.175 | -0.165 | 0.503049 | 0.428566 | 0.478501 | 0.989223 | 0.997988 | 0.994666 |
| Dlg2_E9Q2L2_S365s_1 | Dlg2_E9Q2 | 6.51 | 2.79 | 4.95 | 2.01 | 2.87 | 7.17 | 5.48 | 4.4 | 1.24 | 5.13 | 2.71 | 4.82 | -0.4 | 1.01 | 5.03 | 5.5 | 0.49 | -0.6825 | -0.5975 | 0.412689 | 0.444282 | 0.627228 | 0.989223 | 0.997988 | 0.994666 |
| Dlg2_E9Q2L2_S406s_1 | Dlg2_E9Q2 | -0.69 | -0.85 | -1.33 | -0.5 | 0.66 | 0.62 | -0.36 | 0.53 | -0.16 | -0.7 | 0.36 | -0.19 | -0.33 | 0.91 | -1.38 | -1.32 | 0.915 | -0.59 | -1.28 | 0.557773 | 0.003643 | 0.055919 | 0.989223 | 0.997988 | 0.995259 |
| Dlg2_E9Q2L2_S414s_1 | Dlg2_E9Q2 | 0.41 | 0.12 | 0.51 | 0.39 | 0.63 | 0.23 | 0.27 | 0.55 | -0.35 | -0.2 | 0.17 | 0.3 | -0.43 | 0.43 | -0.06 | 0.21 | 1.205 | 0.67 | 0.3125 | 0.271097 | 0.77558 | 0.154822 | 0.995762 | 0.997988 | 0.994666 |
| Dlg2_E9Q2L2_S45s_1 | Dlg2_E9Q2 | -0.23 | 0.36 | 0.24 | -0.51 | 0.82 | -0.58 | -0.47 | -0.92 | -1.33 | -0.49 | -1.83 | -0.7 | -0.25 | 0.47 | -0.9 | -0.28 | 0.0625 | -0.3775 | -0.32 | 0.579701 | 0.544149 | 0.018373 | 0.989223 | 0.997988 | 0.995259 |
| Dlg2_E9Q2L2_S528s_1 | Dlg2_E9Q2 | -1.15 | -0.1 | 0.37 | 0.43 | -0.82 | -0.19 | 0.18 | 0.56 | -0.26 | -0.15 | -0.86 | 0.3 | 0.92 | -1.04 | 0.1 | 0.3 | -0.2525 | -1.0525 | -0.205 | 0.723886 | 0.918541 | 0.760699 | 0.997335 | 0.997988 | 0.9971 |
| Dlg2_E9Q2L2_S530s_1 | Dlg2_E9Q2 | -0.68 | -0.62 | -0.78 | -0.72 | -0.34 | -0.56 | -1.06 | -0.66 | -0.63 | -0.41 | -1.44 | -0.11 | 0.21 | -0.8 | -1.1 | -1.18 | 0.045 | -0.13 | 0.1825 | 0.960196 | 0.844846 | 0.868543 | 0.996349 | 0.997988 | 0.999507 |
| Dlg2_E9Q2L2_S656s_1 | Dlg2_E9Q2 | -3.55 | -3.22 | -2.4 | -6.61 | -2.15 | -3.31 | -4.36 | -2.54 | -3.12 | -3.04 | -1.24 | -2.98 | -1.56 | -5.16 | -2.7 | -3.36 | 0.045 | 0.0525 | -0.0175 | 0.474409 | 0.356191 | 0.162187 | 0.989223 | 0.997988 | 0.994666 |
| Dlg2_E9Q2L2_S65s_1 | Dlg2_E9Q2 | 0.31 | 0.58 | 0.04 | 0.18 | 1.04 | 0.5 | 0.7 | 0.22 | 0.04 | 0.83 | -0.93 | 0.07 | -0.21 | 0.27 | -0.23 | 0.17 | 0.855 | 1.35 | 0.75 | 0.310624 | 0.211288 | 0.478024 | 0.989223 | 0.997988 | 0.994666 |
| Dlg2_E9Q2L2_T317t_1 | Dlg2_E9Q2 | -1.79 | -1.88 | -0.34 | -1.16 | -0.49 | -1.5 | -0.32 | -1.27 | -1.98 | -1.84 | -0.81 | -1.02 | -0.9 | -0.35 | -1.22 | -1.15 | 0.3375 | -0.275 | -0.2775 | 0.355567 | 0.368582 | 0.78724 | 0.989223 | 0.997988 | 0.994666 |
| Dlg3_Q52KF7_S139s_1 | Dlg3_Q52K | -0.42 | -0.08 | 0.42 | 0.49 | 0.17 | -0.13 | 0.25 | 0.2 | -0.52 | -0.35 | -0.35 | 0.28 | -0.26 | 0.41 | -0.14 | 0.31 | 0.3975 | -0.12 | 0.3875 | 0.943758 | 0.941451 | 0.294643 | 0.999187 | 0.997988 | 0.999507 |
| Dlg3_Q52KF7_S368s_1 | Dlg3_Q52K | 1.06 | -0.22 | 0.1 | 0.84 | 0.72 | 1.51 | 1.1 | 1.4 | 0.66 | 0.24 | 0.43 | 0.76 | -0.03 | 0.5 | 0.03 | 0.24 | 0.02 | -0.3375 | -0.0225 | 0.468222 | 0.059659 | 0.82114 | 0.989223 | 0.997988 | 0.994666 |
| Dlg3_Q52KF7_S491s_1 | Dlg3_Q52K | 0.76 | 0.34 | -0.98 | -0.7 | -0.85 | 0.3 | -1.23 | -0.57 | 1.04 | 0.16 | -0.36 | -1.1 | 1.43 | -0.5 | -1.29 | -1.76 | 0.7375 | 0.0775 | -0.26 | 0.599078 | 0.372568 | 0.887297 | 0.989223 | 0.997988 | 0.995259 |
| Dlg3_Q52KF7_S493s_1 | Dlg3_Q52K | 0.27 | 0.35 | 0.89 | 0.21 | -0.25 | 0.57 | -0.19 | 0.28 | 1.03 | 0.27 | -0.4 | 0.59 | 1.27 | -0.04 | 0.29 | 0.5 | -0.4425 | 0.08 | -0.385 | 0.830019 | 0.265754 | 0.870664 | 0.989223 | 0.997988 | 0.999507 |
| Dlg3_Q52KF7_S495s_1 | Dlg3_Q52K | 0.02 | -0.44 | -0.86 | -0.63 | -0.36 | 0.19 | -0.7 | -0.65 | 0.3 | -0.17 | -1.66 | 0.18 | 0.29 | -0.76 | -0.9 | -0.66 | -0.3275 | -0.0575 | 0.075 | 0.932709 | 0.746415 | 0.764596 | 0.995762 | 0.997988 | 0.999507 |
| Dlg3_Q52KF7_T365t_1 | Dlg3_Q52K | 1.2 | 0.37 | 0.04 | 0.87 | 0.88 | 1.47 | 1.17 | 1.61 | 0.65 | 0.5 | 0.71 | 0.78 | -0.24 | 0.99 | 0.17 | 0.42 | 0.0975 | 0.14 | -0.03 | 0.45957 | 0.062176 | 0.894809 | 0.989223 | 0.997988 | 0.994666 |
| Dlg4_Q62108_S139s_1 | Dlg4_Q621 | 0.75 | 0.7 | 1.25 | 0.72 | 0.72 | 0.38 | 0.29 | 0.56 | 0.26 | -0.02 | 0.58 | 0.82 | 1.02 | 0.72 | 0.47 | 0.84 | 0.6625 | 0.04 | -0.285 | 0.729703 | 0.141039 | 0.135757 | 0.989223 | 0.997988 | 0.9971 |
| Dlg4_Q62108_S292s_1 | Dlg4_Q621 | 1.42 | 1.11 | 1.32 | 1.21 | 0.83 | 1.49 | 0.19 | 0.85 | 1.47 | 0.84 | 0.5 | 0.8 | 1.57 | 0.64 | 0.51 | 0.53 | -0.3675 | -0.445 | -0.0925 | 0.172596 | 0.177737 | 0.204053 | 0.989223 | 0.997988 | 0.994666 |
| Dlg4_Q62108_S412s_1 | Dlg4_Q621 | 1.01 | 1.01 | 1.49 | 1.34 | 0.59 | 0.85 | 1.39 | 1.16 | 1.09 | 1.3 | 1.24 | 1.25 | 0.97 | 0.82 | 0.73 | 0.93 | -0.425 | -0.3625 | -0.4525 | 0.181829 | 0.418915 | 0.973777 | 0.989223 | 0.997988 | 0.994666 |
| Dlg4_Q62108_S415s_1 | Dlg4_Q621 | 0.75 | 1.17 | 1.21 | 1.03 | 0.97 | 0.64 | 1.17 | 0.96 | 0.67 | 1.12 | 0.74 | 1.15 | 0.66 | 0.9 | 0.85 | 0.96 | -0.215 | 0.0075 | -0.35 | 0.428569 | 0.647394 | 0.624709 | 0.99033 | 0.997988 | 0.994666 |
| Dlg4_Q62108_S419s_1 | Dlg4_Q621 | -0.23 | -0.11 | -0.12 | -0.42 | 0.36 | -0.27 | -0.02 | -0.12 | -0.78 | -0.37 | -0.3 | 0.09 | -0.69 | 0.18 | -0.87 | -0.66 | -0.105 | -0.12 | -0.1975 | 0.351867 | 0.375543 | 0.645102 | 0.989223 | 0.997988 | 0.994666 |
| Dlg4_Q62108_S507s_1 | Dlg4_Q621 | 0.35 | 0.08 | 2.03 | 1.83 | 0.12 | 0.39 | 1.39 | 2 | -0.32 | -0.39 | 2.21 | 2.29 | -0.31 | -0.5 | 1.66 | 1.47 | 0.2075 | -0.12 | -0.29 | 0.475865 | 0.869695 | 0.876716 | 0.996493 | 0.997988 | 0.994666 |
| Dlg4_Q62108_S510s_1 | Dlg4_Q621 | -0.84 | -0.38 | -0.04 | -0.39 | -0.6 | -0.54 | -0.82 | -0.66 | -1.06 | -1.17 | -0.53 | 0.3 | -0.85 | -1.31 | -0.68 | -0.87 | -0.0975 | -0.125 | -0.4925 | 0.096575 | 0.325945 | 0.595279 | 0.989223 | 0.997988 | 0.994666 |
| Dlg4_Q62108_S603s_1 | Dlg4_Q621 | -0.87 | -0.61 | -1.08 | -0.57 | 0.27 | -0.5 | -0.23 | -1.03 | -0.68 | 0.03 | -1.82 | -0.95 | -0.65 | -0.61 | -0.58 | -0.34 | -0.2425 | -0.2025 | -0.515 | 0.355793 | 0.212024 | 0.855656 | 0.989223 | 0.997988 | 0.994666 |
| Dlg4_Q62108_S70s_1 | Dlg4_Q621 | -0.43 | 0.12 | 0.4 | 0.16 | 0.84 | -0.31 | 0.93 | 0.77 | -0.1 | -0.39 | 0.24 | 0.55 | -0.28 | 0.04 | 0.02 | -0.12 | 0.41 | -0.0725 | 0.2375 | 0.59352 | 0.173454 | 0.967338 | 0.989223 | 0.997988 | 0.995259 |
| Dlg5_E9Q9I2_S109s_1 | Dlg5_E9Q9 | 0.51 | 0.43 | 1.02 | 0.24 | 0.34 | 0.33 | 0.85 | 0.68 | 0.03 | 0.86 | 1.25 | 0.49 | 0.05 | 0.28 | 1.07 | 1.22 | 0.495 | 0.0125 | -0.1475 | 0.769969 | 1 | 0.746899 | 1 | 0.997988 | 0.9971 |
| Dlg5_E9Q9I2_S1209s_1 | Dlg5_E9Q9 | -2.99 | -1.56 | -1.31 | -2.05 | -1.06 | -2.71 | -0.58 | -1.68 | -1.98 | -1.63 | -0.82 | -1.46 | -1.24 | -1.37 | -0.53 | -1.14 | 1.96E-17 | 0.1075 | 0.105 | 0.051435 | 0.390194 | 0.259802 | 0.989223 | 0.997988 | 0.994666 |
| Dlg5_E9Q9I2_S818s_1 | Dlg5_E9Q9 | -0.91 | 0.02 | 0.23 | 0.82 | 0.76 | -1.52 | 0.54 | 0.12 | -0.81 | -0.13 | -0.67 | 0.52 | -0.83 | 0.4 | 0.12 | 1.39 | 0.47 | 0.505 | 0.9075 | 0.671622 | 0.90847 | 0.494449 | 0.997335 | 0.997988 | 0.996159 |
| Dlgap1_Q9D415_S169s_1 | Dlgap1_Q9 | 0.46 | 0.82 | 1.11 | 0.99 | 0.58 | 0.61 | 0.6 | 1.13 | 0.54 | 0.64 | 1.38 | 0.82 | 1.2 | 0.55 | 1.02 | 0.81 | -0.065 | -0.3125 | 0.23 | 0.857562 | 0.648164 | 1 | 0.99033 | 1 | 0.999507 |
| Dlgap1_Q9D415_S389s_1 | Dlgap1_Q9 | -0.84 | 0.63 | 0.35 | -0.24 | 0.46 | -0.27 | 0.23 | -0.09 | -0.72 | 0.16 | -0.22 | -0.01 | 0.63 | 0.73 | -0.4 | -0.01 | -0.115 | -5.9E-17 | 0.05 | 0.537015 | 0.76705 | 0.653128 | 0.995762 | 0.997988 | 0.994666 |

| | | | | | | | | | | | | | | | | | | | | | | | | | | |
|---|---|---|---|---|---|---|---|---|---|---|---|---|---|---|---|---|---|---|---|---|---|---|---|---|---|---|
| Dlgap1_Q9D415_S403s_1 | Dlgap1_Q9 | -1.03 | -0.94 | -0.87 | -1.09 | -0.85 | -0.28 | -0.56 | -1.23 | 1.73 | -0.98 | -2.79 | -0.41 | 2.49 | -2.38 | 0.37 | -0.9 | 0.1075 | -0.1725 | 0.2625 | 0.344143 | 0.341935 | 0.65758 | 0.989223 | 0.997988 | 0.994666 |
| Dlgap1_Q9D415_S418s_1 | Dlgap1_Q9 | -0.92 | -0.27 | -0.55 | -1.57 | 0.02 | -1.33 | -0.43 | -0.95 | -0.63 | -0.36 | 0.84 | -0.93 | 0.44 | 0.08 | -1.57 | -1.45 | 0.2525 | 0.37 | 0.8775 | 0.711477 | 0.694956 | 0.243726 | 0.994893 | 0.997988 | 0.996966 |
| Dlgap1_Q9D415_S421s_1 | Dlgap1_Q9 | 0.16 | 0.39 | 0.7 | 0.5 | 1.11 | 0.59 | 0.33 | 0.64 | -0.08 | -0.11 | 0.53 | 0.42 | 0.24 | 0.53 | 0.33 | -0.3 | 0.155 | 0.5575 | 0.2025 | 0.41244 | 0.373064 | 0.360132 | 0.989223 | 0.997988 | 0.994666 |
| Dlgap1_Q9D415_S437s_1 | Dlgap1_Q9 | -0.02 | 1.03 | 0.36 | 0.28 | 1.66 | 0.52 | 0.92 | 0.28 | -0.09 | 0.34 | -0.74 | 0.49 | 0.27 | 0.54 | 0.04 | 0.23 | 0.23 | -0.2475 | -0.2375 | 0.640238 | 0.259178 | 0.273992 | 0.989223 | 0.997988 | 0.995259 |
| Dlgap1_Q9D415_S509s_1 | Dlgap1_Q9 | 0.58 | 0.1 | 0.13 | 0.64 | 0.28 | 1.1 | -0.04 | 0.65 | 1.06 | 0.32 | 0.86 | 0.51 | 0.25 | 0.93 | -0.12 | -0.37 | 0.4325 | -0.4125 | -0.1425 | 0.588033 | 0.660423 | 0.255866 | 0.990434 | 0.997988 | 0.995259 |
| Dlgap1_Q9D415_S516s_1 | Dlgap1_Q9 | -1.71 | -0.69 | -0.65 | -0.53 | -0.92 | -1.19 | -1.22 | -0.24 | -0.84 | -1.5 | 0.02 | -0.66 | -0.57 | -1.12 | -1.55 | -1.45 | 0.135 | 0.325 | -0.19 | 0.461311 | 0.99439 | 0.715213 | 0.999278 | 0.997988 | 0.994666 |
| Dlgap1_Q9D415_S52s_1 | Dlgap1_Q9 | -0.63 | 0.84 | -1.35 | -1.25 | -0.15 | 0.4 | 0.43 | 0.02 | 0.31 | 0.04 | -0.65 | -2.05 | -0.37 | 1.06 | -1.22 | -0.17 | 0.0025 | 0.15 | -0.2775 | 0.504899 | 0.13507 | 0.987935 | 0.989223 | 0.997988 | 0.994666 |
| Dlgap1_Q9D415_S937s_1 | Dlgap1_Q9 | -1.41 | -1.1 | 0.31 | -0.15 | -1.31 | 0.22 | -0.46 | -0.1 | 0.14 | -1.16 | -0.94 | -0.1 | 0.85 | -1.45 | -0.32 | -0.42 | 0.7725 | 0.01 | 0.4225 | 0.658988 | 0.716004 | 0.880901 | 0.995437 | 0.997988 | 0.995259 |
| Dlgap1_Q9D415_S963s_1 | Dlgap1_Q9 | 0.74 | 0.21 | -0.26 | 0.08 | 0.53 | -0.58 | 0.29 | 0.2 | -0.64 | 0.47 | -1.02 | 0.06 | -1.14 | 0.12 | 0.25 | 0.2 | 0.175 | 0.0725 | 0.2525 | 0.410772 | 0.801899 | 0.247514 | 0.995762 | 0.997988 | 0.994666 |
| Dlgap1_Q9D415_T436t_1 | Dlgap1_Q9 | 0.05 | 0.1 | 0.01 | -1.06 | 0.97 | 0.17 | 0.91 | 0.33 | 0.44 | 0.5 | -1.14 | -0.31 | 0.36 | 0.41 | 0.33 | -0.09 | -0.0825 | -0.475 | -0.335 | 0.180451 | 0.038546 | 0.830669 | 0.989223 | 0.997988 | 0.994666 |
| Dlgap2_Q0VF59_S1041s_1 | Dlgap2_Q0 | -1.04 | -2.66 | -1.63 | -3.26 | -1.96 | 0.52 | -0.86 | -0.01 | 2.07 | -1.79 | -3.08 | -0.36 | 2.22 | -2.08 | 0.92 | -3.17 | 0.82 | 0.0975 | 0.4775 | 0.188799 | 0.035782 | 0.223737 | 0.989223 | 0.997988 | 0.994666 |
| Dlgap2_Q0VF59_S1048s_1 | Dlgap2_Q0 | -0.09 | 0.24 | -0.03 | -0.11 | 0.6 | -0.13 | 0.45 | 0.02 | -0.37 | -0.02 | -0.33 | -0.29 | -0.54 | 0.35 | -0.56 | -0.08 | 1.57 | 1.3575 | 1.62 | 0.483584 | 0.361417 | 0.26841 | 0.989223 | 0.997988 | 0.994666 |
| Dlgap2_Q0VF59_S249s_1 | Dlgap2_Q0 | 6.3 | 1.59 | 2.77 | -1.32 | 1.25 | 7.36 | 3.9 | 3.75 | 1.56 | 3.58 | -0.37 | 3.34 | 1.32 | -0.51 | 3.66 | 3.69 | 0.2325 | -0.255 | -0.21 | 0.854371 | 0.328365 | 0.846726 | 0.989223 | 0.997988 | 0.999507 |
| Dlgap2_Q0VF59_S390s_1 | Dlgap2_Q0 | -4.83 | -8.1 | -2.23 | -5.57 | -3.51 | -5.63 | -3.57 | -4.41 | -7.86 | -4.54 | 0.57 | -4.33 | -3.64 | -1.51 | -4.53 | -3.83 | 1.73 | -0.3075 | -0.295 | 0.152087 | 0.430226 | 0.53916 | 0.989223 | 0.997988 | 0.994666 |
| Dlgap2_Q0VF59_S454s_2 | Dlgap2_Q0 | -0.18 | 1.23 | 0.58 | 0.91 | -1.02 | 0.73 | -0.13 | 0.14 | 1.6 | 0.44 | 1.6 | 0.16 | 0.4 | 0.26 | 1.11 | 0.39 | 0.9025 | 1.1425 | 1.805 | 0.800518 | 0.136785 | 0.502539 | 0.989223 | 0.997988 | 0.999507 |
| Dlgap2_Q0VF59_S457s_1 | Dlgap2_Q0 | 0.63 | 0.61 | 0.86 | 0.53 | -0.62 | 0.72 | -0.52 | -0.16 | 1.47 | 0.39 | 0.41 | -0.6 | 2.44 | -0.88 | 0.27 | 0.33 | -0.705 | 0.315 | -0.095 | 0.852088 | 0.032118 | 0.571017 | 0.989223 | 0.997988 | 0.999507 |
| Dlgap2_Q0VF59_S461s_1 | Dlgap2_Q0 | -3.43 | -2.31 | -2.72 | -4.44 | -1.75 | -2.74 | -2.25 | -3.03 | -3.14 | -2.48 | -1.84 | -3.08 | -2.01 | -1.66 | -2.39 | -3.15 | -0.8025 | -0.24 | -0.1175 | 0.106228 | 0.141668 | 0.274253 | 0.989223 | 0.997988 | 0.994666 |
| Dlgap2_Q0VF59_S668s_1 | Dlgap2_Q0 | 0.81 | 0.1 | -0.1 | 0.25 | 0.16 | 0.83 | 0.16 | 0.45 | 0.51 | 0.24 | 0.36 | 0.02 | -0.24 | 0.76 | -0.35 | -0.05 | 0.7825 | 0.59 | 0.9225 | 0.505528 | 0.637353 | 0.949324 | 0.989223 | 0.997988 | 0.994666 |
| Dlgap2_Q0VF59_S746s_1 | Dlgap2_Q0 | 1.11 | 0.46 | 0.87 | 0.64 | -0.55 | 1.14 | -0.71 | 0.02 | 1.47 | 0.66 | 0.57 | 0.4 | 0.91 | -1.07 | 0.15 | 0.18 | 0.135 | 0.0175 | -0.235 | 0.115337 | 0.08345 | 0.987136 | 0.989223 | 0.997988 | 0.994666 |
| Dlgap2_Q0VF59_S777s_1 | Dlgap2_Q0 | -0.65 | 0 | -0.59 | -0.93 | 0.39 | -0.75 | -0.02 | -0.8 | -0.53 | -0.43 | -1.61 | -0.71 | -0.99 | 0.3 | -1.03 | -0.9 | -0.795 | 0.005 | -0.7275 | 0.771423 | 0.486703 | 0.434697 | 0.989223 | 0.997988 | 0.9971 |
| Dlgap2_Q0VF59_S807s_1 | Dlgap2_Q0 | -1.08 | -0.62 | -0.65 | -1.69 | 2.64 | -0.93 | -0.47 | 0.56 | -1.68 | -1.53 | 0.23 | -1.46 | -1.51 | 0.24 | -2.72 | -0.02 | 0.2475 | -0.2775 | -0.1125 | 0.990937 | 0.07054 | 0.83711 | 0.989223 | 0.997988 | 1 |
| Dlgap2_Q0VF59_S812s_1 | Dlgap2_Q0 | 3.11 | 1.22 | 1.12 | 1.02 | 1.16 | 3.35 | 2.49 | 1.22 | 0.28 | 2.31 | 0.1 | 1.33 | 0.91 | -0.1 | 1.2 | 1.61 | 1.46 | -0.1 | 0.0075 | 0.23008 | 0.503455 | 0.357956 | 0.989223 | 0.997988 | 0.994666 |
| Dlgap2_Q0VF59_S984s_1 | Dlgap2_Q0 | 0.65 | 0.53 | 0.49 | 0.29 | 0.27 | 0.23 | -0.03 | 0.45 | 0.17 | 0.6 | 0.62 | 0.42 | 0.56 | 0.3 | 0.39 | 0.28 | 0.4375 | -0.6125 | -0.7125 | 0.652031 | 0.247837 | 0.8694 | 0.989223 | 0.997988 | 0.995259 |
| Dlgap2_Q0VF59_T460t_1 | Dlgap2_Q0 | -5.52 | -5.87 | -3.29 | -5.01 | -3.96 | -6.07 | -3.77 | -3.74 | -6.98 | -4.89 | -1.87 | -4.28 | -4.85 | -3.21 | -4.24 | -4.09 | -0.26 | -0.0375 | -0.1075 | 0.194988 | 0.454515 | 0.693234 | 0.989223 | 0.997988 | 0.994666 |
| Dlgap2_Q0VF59_T744t_1 | Dlgap2_Q0 | 0.48 | 0.57 | 0.38 | 0.58 | -0.28 | 0.95 | 0.52 | 0.66 | 2.05 | 1.03 | 0.44 | 0.63 | 2.52 | -0.29 | 0.85 | 1.38 | 0.5375 | 0.4175 | 0.825 | 0.27648 | 0.891999 | 0.172128 | 0.996493 | 0.997988 | 0.994666 |
| Dlgap2_Q0VF59_T816t_1 | Dlgap2_Q0 | 3 | 0.7 | 2.67 | 1.2 | 1.52 | 3.67 | 2.91 | 2.71 | 0.28 | 2.47 | 1.19 | 3.1 | 0.84 | 0.36 | 2.35 | 2.51 | -0.04 | 0.535 | 0.6125 | 0.589313 | 0.221752 | 0.860932 | 0.989223 | 0.997988 | 0.995259 |
| Dlgap3_B1AS06_S185s_1 | Dlgap3_B1 | 4.65 | 0.37 | 5.85 | 2.24 | 1.43 | 5.23 | 7.51 | 6.91 | -0.3 | 2.15 | 0.94 | 6.13 | -0.06 | -1.24 | 6.21 | 6.84 | 0.81 | -0.1325 | -0.3775 | 0.869407 | 0.224647 | 0.522138 | 0.989223 | 0.997988 | 0.999507 |
| Dlgap3_B1AS06_S407s_2 | Dlgap3_B1 | -0.29 | 0.7 | -0.37 | -0.01 | -1.38 | 0.76 | -0.57 | -0.83 | 1.63 | -0.28 | 0.56 | -0.44 | 0.96 | -1.29 | 0.04 | -0.23 | 1.9925 | -1.0475 | -0.34 | 0.781746 | 0.297035 | 0.479428 | 0.989223 | 0.997988 | 0.998226 |
| Dlgap3_B1AS06_S410s_1 | Dlgap3_B1 | -1.49 | -0.83 | 0.54 | -0.08 | -0.71 | -1.28 | -0.25 | -0.39 | -0.51 | -1.24 | 0.5 | 0.41 | 2.24 | -0.04 | 0.63 | -0.06 | -0.5125 | 0.36 | -0.1375 | 0.092615 | 0.678362 | 0.649842 | 0.993464 | 0.997988 | 0.994666 |
| Dlgap3_B1AS06_S410s_2 | Dlgap3_B1 | -0.29 | 0.7 | -0.37 | -0.01 | -1.38 | 0.76 | -0.57 | -0.83 | 1.63 | -0.28 | 0.56 | -0.44 | 0.96 | -1.29 | 0.04 | -0.23 | -0.1925 | 0.255 | 1.1575 | 0.781746 | 0.297035 | 0.479428 | 0.989223 | 0.997988 | 0.998226 |
| Dlgap3_B1AS06_S429s_1 | Dlgap3_B1 | -0.81 | -0.72 | -0.16 | -1.29 | -1.16 | 0.67 | -0.84 | -0.2 | 1.3 | -1.11 | -1.61 | -1.46 | 2.02 | -0.87 | -0.22 | -1.34 | -0.5125 | 0.36 | -0.1375 | 0.367881 | 0.416121 | 0.969439 | 0.989223 | 0.997988 | 0.994666 |
| Dlgap3_B1AS06_S559s_1 | Dlgap3_B1 | -0.12 | 0.28 | 0.02 | -0.06 | 0.29 | 0.15 | 0.44 | 0.16 | 0.09 | -0.01 | -0.58 | -0.06 | -0.08 | 0.14 | -0.46 | -0.56 | 0.3625 | 0.025 | 0.6425 | 0.333811 | 0.291995 | 0.501 | 0.989223 | 0.997988 | 0.994666 |
| Dlgap3_B1AS06_S58s_1 | Dlgap3_B1 | 0.71 | 0.66 | 0.75 | 0.66 | 0.86 | 0.4 | 0.84 | 0.87 | 0.08 | 0.72 | 0.15 | 0.43 | -0.01 | 0.88 | 0.56 | 0.34 | 0.23 | -0.17 | -0.27 | 0.366715 | 0.823896 | 0.163235 | 0.995762 | 0.997988 | 0.994666 |
| Dlgap3_B1AS06_S712s_1 | Dlgap3_B1 | 6.25 | 1.91 | 4.52 | 0.4 | 2.26 | 7.34 | 4.3 | 3.74 | 2.58 | 4.01 | -0.25 | 3.78 | 2.72 | 0.85 | 4.07 | 5.14 | 0.0475 | -0.35 | -0.2525 | 0.956557 | 0.438244 | 0.606652 | 0.989223 | 0.997988 | 0.999507 |
| Dlgap3_B1AS06_S753s_1 | Dlgap3_B1 | 0.26 | 0.73 | 1 | 0.56 | 0.93 | 0.1 | 1.05 | 0.85 | -0.12 | 0.57 | 0.49 | 0.78 | -0.33 | 0.41 | 0.68 | 0.9 | 1.14 | -0.74 | -0.075 | 0.521307 | 0.746451 | 0.483356 | 0.995762 | 0.997988 | 0.994666 |
| Dlgap3_B1AS06_S771s_1 | Dlgap3_B1 | -0.5 | -0.94 | -1.89 | -1.81 | -0.89 | -0.79 | -1.58 | 0.09 | -1.26 | -2.16 | 0.55 | -1.01 | -1.19 | -0.16 | -3.12 | -2.9 | 0.095 | -0.2075 | -0.2225 | 0.434674 | 0.289606 | 0.603522 | 0.989223 | 0.997988 | 0.994666 |
| Dlgap3_B1AS06_S779s_1 | Dlgap3_B1 | 0.43 | -0.14 | 0.31 | -1.36 | -0.04 | 0.54 | -2.47 | -0.52 | -0.3 | -1.08 | -0.48 | -1.41 | -0.34 | -0.53 | 0.15 | -0.67 | 0.4925 | 0.315 | -0.5575 | 0.719969 | 0.529558 | 0.197334 | 0.989223 | 0.997988 | 0.9971 |
| Dlgap3_B1AS06_S783s_1 | Dlgap3_B1 | 1.29 | 0.5 | 0.84 | 1.13 | 0.91 | 1.23 | 0.25 | 0.97 | 1.8 | 1.04 | 0.34 | 0.75 | 1.3 | -0.51 | 0.47 | 0.81 | -0.4325 | -0.6275 | -0.1575 | 0.326667 | 0.73729 | 0.906939 | 0.995762 | 0.997988 | 0.994666 |
| Dlgap3_B1AS06_S919s_1 | Dlgap3_B1 | 0.82 | 0.34 | 0.8 | 0.6 | 0.46 | 0.68 | 0.14 | 0.83 | 0.3 | 0.08 | 0.94 | 0.29 | 0.19 | 0.43 | 0.09 | 0.14 | -0.1 | 0.0425 | -0.4225 | 0.112079 | 0.649848 | 0.394861 | 0.990358 | 0.997988 | 0.994666 |
| Dlgap3_B1AS06_S922s_1 | Dlgap3_B1 | -0.21 | 0.9 | 0.68 | -0.01 | 0.21 | -0.45 | -0.15 | -0.27 | -1.37 | -0.15 | 0.06 | 0.16 | 0.24 | -0.23 | 0.45 | 0.72 | -0.1125 | -0.2375 | -0.4275 | 0.900212 | 0.136069 | 0.148381 | 0.989223 | 0.997988 | 0.999507 |
| Dlgap4_B1AZP2_S678s_1 | Dlgap4_B1 | -1.66 | -0.47 | -0.42 | -1.28 | -1.86 | -0.33 | -1.71 | -1.47 | 0.17 | -0.1 | -0.49 | -0.65 | 2.36 | 0.03 | -1.09 | -1.72 | -0.505 | -0.665 | -0.045 | 0.321016 | 0.386747 | 0.086357 | 0.989223 | 0.997988 | 0.994666 |
| Dlgap4_B1AZP2_S681s_1 | Dlgap4_B1 | 0.29 | 0.11 | 0.85 | 0.9 | -0.09 | 1.21 | -0.06 | 0.7 | 0.41 | 0.02 | 0.67 | 0.66 | 0.84 | -0.4 | 0.52 | 0.33 | -0.385 | 0.69 | 0.8525 | 0.549831 | 0.791131 | 0.740327 | 0.995762 | 0.997988 | 0.994666 |
| Dlgap4_B7ZNS2_S156s_1 | Dlgap4_B7 | 1.5 | 0.42 | 0.33 | 0.82 | 0.47 | 1.7 | -0.05 | 0.34 | 1.26 | 0.77 | 0.33 | 0.48 | 1.28 | 0.26 | 0.08 | -0.06 | -0.0975 | -0.0975 | -0.215 | 0.365418 | 0.726803 | 0.869897 | 0.995569 | 0.997988 | 0.994666 |
| Dlgap4_B7ZNS2_S35s_1 | Dlgap4_B7 | 0.89 | -0.28 | 0.15 | 0.49 | 0.3 | 1.13 | 0.45 | 0.51 | 0.01 | 0.1 | -0.51 | 0.53 | -1.05 | 1.03 | -1.09 | -1.53 | -0.1525 | -0.0575 | -0.3775 | 0.116104 | 0.386869 | 0.426011 | 0.989223 | 0.997988 | 0.994666 |
| Dlgap4_B7ZNS2_S377s_1 | Dlgap4_B7 | -1.41 | -0.5 | -1.17 | -1.86 | -0.86 | -0.62 | -1.15 | -1.46 | 0.36 | -0.39 | -1.57 | -0.63 | 0.71 | -0.92 | -1.5 | -0.9 | 0.285 | -0.28 | -0.9725 | 0.277927 | 0.53869 | 0.170476 | 0.989223 | 0.997988 | 0.994666 |
| Dlgap4_B7ZNS2_S405s_1 | Dlgap4_B7 | -0.55 | 0.12 | 0.53 | -0.56 | -0.3 | -0.16 | -1.13 | -0.04 | 0.07 | -0.23 | 1.16 | 0 | 1.19 | -0.17 | -0.46 | -0.73 | 0.2125 | 0.6775 | 0.5825 | 0.880088 | 0.426698 | 0.378847 | 0.989223 | 0.997988 | 0.999507 |
| Dlgap4_B7ZNS2_S405s_2 | Dlgap4_B7 | -0.3 | -1.33 | 0.21 | -0.35 | -1.25 | -0.14 | -1.63 | -0.78 | -1.22 | -0.95 | -0.77 | 0 | 0.22 | -0.57 | -1.66 | -1.54 | -0.2925 | 0.365 | 0.0725 | 0.396367 | 0.254987 | 0.481255 | 0.989223 | 0.997988 | 0.994666 |
| Dlgap4_B7ZNS2_S410s_1 | Dlgap4_B7 | 0.96 | -0.58 | 0.81 | 0.73 | -0.5 | 0.82 | -0.53 | -0.04 | 0.11 | -0.19 | 0.41 | 0.91 | 1.57 | 0.56 | 0.22 | 0.03 | -0.5075 | -0.2925 | -0.445 | 0.80853 | 0.241967 | 0.685762 | 0.989223 | 0.997988 | 0.999507 |
| Dlgap4_B7ZNS2_S410s_2 | Dlgap4_B7 | -0.3 | -1.33 | 0.21 | -0.35 | -1.25 | -0.14 | -1.63 | -0.78 | -1.22 | -0.95 | -0.77 | 0 | 0.22 | -0.57 | -1.66 | -1.54 | -0.5425 | -0.17 | 0.115 | 0.396367 | 0.254987 | 0.481255 | 0.989223 | 0.997988 | 0.994666 |
| Dlgap4_B7ZNS2_S415s_1 | Dlgap4_B7 | 1.19 | 0.64 | 1.2 | 1.39 | 1.35 | 1.13 | 1.62 | 1.5 | 0.81 | 1.14 | 1.08 | 1.32 | -0.09 | 0.95 | 0.91 | 0.98 | -0.5075 | -0.2925 | -0.445 | 0.239246 | 0.255644 | 0.946217 | 0.989223 | 0.997988 | 0.994666 |
| Dlgap4_B7ZNS2_S517s_1 | Dlgap4_B7 | 0.63 | 0.82 | 0.79 | 0.96 | 0.4 | 0.71 | 1.4 | 0.77 | 0.37 | 0.68 | 1.08 | 1.31 | 0.76 | 0.53 | 0.58 | 0.94 | 0.295 | -0.0175 | -0.4175 | 0.68884 | 0.939913 | 0.82719 | 0.999187 | 0.997988 | 0.996159 |
| Dlgap4_B7ZNS2_S649s_1 | Dlgap4_B7 | -2.44 | -0.6 | -2.14 | -1.28 | 0.52 | -0.72 | -0.37 | -0.71 | -0.67 | -0.25 | 0.27 | -1.94 | -0.07 | 0 | -1.63 | -1.5 | 0.02 | 0.06 | -0.0975 | 0.170213 | 0.020244 | 0.121554 | 0.989223 | 0.997988 | 0.994666 |
| Dlgap4_B7ZNS2_S715s_1 | Dlgap4_B7 | 0.14 | 0.19 | -0.45 | -0.34 | 0.29 | 0.39 | 0.24 | -0.27 | 0.22 | -0.17 | -0.56 | -0.06 | 0.01 | 0.95 | -0.28 | -0.63 | 1.295 | 0.9675 | 0.815 | 0.74288 | 0.310235 | 0.921957 | 0.989223 | 0.997988 | 0.9971 |
| Dlgap4_B7ZNS2_S956s_1 | Dlgap4_B7 | -0.19 | 0.37 | -0.21 | -0.18 | 0.77 | 0.09 | 0.66 | -0.33 | 0.02 | 0.23 | -0.63 | -0.02 | -0.71 | 0.66 | 0 | 0.14 | 0.2775 | -0.0275 | 0.1275 | 0.829003 | 0.2778 | 0.866467 | 0.989223 | 0.997988 | 0.999507 |
| Dlgap4_B7ZNS2_T543t_1 | Dlgap4_B7 | 0.35 | 0.28 | 0.28 | 0.28 | 0.85 | 0.38 | 0.36 | 0.41 | 0.79 | 0.13 | 0.51 | 0.22 | 0.7 | 0.53 | 0.2 | 0.08 | 0.35 | -0.0475 | 0.075 | 0.752829 | 0.356029 | 0.630996 | 0.989223 | 0.997988 | 0.9971 |
| Dlgap4_I1E4X0_S228s_1 | Dlgap4_I1E | -0.46 | -0.01 | 0.55 | 0.8 | 0.16 | -0.15 | 0.92 | 0.96 | -0.91 | -1.15 | 0.67 | 0.33 | -0.76 | 0.86 | 0.23 | 0.15 | 0.2025 | 0.115 | 0.08 | 0.816853 | 0.517033 | 0.345094 | 0.989223 | 0.997988 | 0.999507 |
| Dmap1_Q9JI44_S455s_1 | Dmap1_Q9 | -1.41 | -1.16 | -2.94 | -4.52 | -2.77 | -1.14 | -3.42 | -2.88 | -0.38 | -1.67 | -3.52 | -2.61 | 0.23 | -2.81 | -1.31 | -2.83 | 0.2525 | -0.485 | -0.1 | 0.384721 | 0.955306 | 0.613921 | 0.999187 | 0.997988 | 0.994666 |
| Dmd_P11531_S3606s_1 | Dmd_P115 | 0.76 | 0.57 | 0.35 | 0.46 | -0.05 | 0.75 | -0.52 | 0.36 | 0.15 | 0.8 | -0.18 | 0.17 | 0.75 | -0.29 | 0.59 | 0.63 | -0.045 | 0.4625 | 0.8275 | 0.710819 | 0.2134 | 0.293817 | 0.989223 | 0.997988 | 0.996966 |
| Dmtn_Q9WV69_S11s_1 | Dmtn_Q9V | 0.47 | -0.28 | -0.58 | 0.14 | 0.03 | 0.07 | -0.03 | 0.49 | 0.14 | -0.49 | 0.23 | -0.03 | -0.19 | 0.46 | 0.11 | -0.17 | -0.4 | -0.3 | -0.115 | 0.722032 | 0.490307 | 0.936364 | 0.989223 | 0.997988 | 0.9971 |
| Dmtn_Q9WV69_S152s_1 | Dmtn_Q9V | -2.01 | -2.17 | -2.48 | -3.92 | -2.18 | -1.04 | -2.83 | -3.04 | 1.49 | -2.07 | -1.36 | -1.14 | 1.54 | -1.29 | -2.71 | -2.14 | 0.2025 | 0.025 | 0.115 | 0.125317 | 0.514 | 0.039808 | 0.989223 | 0.997988 | 0.994666 |
| Dmtn_Q9WV69_S156s_1 | Dmtn_Q9V | 0.13 | 0.15 | -0.1 | -0.18 | -0.95 | 0.46 | -0.2 | -0.15 | 1.48 | 0.18 | -0.47 | 0.53 | 1.35 | -0.9 | 0.8 | -0.1 | 0.3725 | 1.875 | 1.495 | 0.553219 | 0.512128 | 0.308811 | 0.989223 | 0.997988 | 0.994666 |
| Dmtn_Q9WV69_S21s_1 | Dmtn_Q9V | 1.53 | 1.94 | 1.46 | 1.41 | 1.62 | 1.87 | 1.81 | 0.49 | 1.03 | 1.43 | 0.93 | 1.58 | 0.58 | 1.44 | 1.74 | 1.54 | -0.21 | 0.43 | 0.2875 | 0.433711 | 0.693981 | 0.212256 | 0.994893 | 0.997988 | 0.994666 |
| Dmtn_Q9WV69_S226s_1 | Dmtn_Q9V | 0.95 | 0.88 | 1.09 | 0.66 | -0.27 | 0.59 | 0.31 | 0.64 | 0.76 | 0.81 | 0.88 | 0.86 | -1.08 | 0.72 | 0.88 | 0.81 | -0.1375 | -0.3425 | -0.26 | 0.243545 | 0.055297 | 0.754291 | 0.989223 | 0.997988 | 0.994666 |
| Dmtn_Q9WV69_S26s_1 | Dmtn_Q9V | 0.61 | 0.16 | 0.93 | 0.81 | -0.11 | 0.35 | 0.9 | 0.34 | 0.02 | 0.08 | 0.23 | 0.09 | 0.36 | 0.4 | 0.19 | 0.55 | -0.5775 | -0.0675 | -0.5625 | 0.363637 | 0.391971 | 0.060965 | 0.989223 | 0.997988 | 0.994666 |
| Dmtn_Q9WV69_S279s_1 | Dmtn_Q9V | 0.18 | 0.29 | -0.76 | -0.8 | 0.34 | 0.26 | -0.53 | -1.2 | -0.29 | 0.05 | -1.26 | -1.14 | 0.17 | 0.34 | -0.24 | -0.67 | -0.2575 | -0.5225 | -0.2525 | 0.654815 | 0.981812 | 0.372785 | 0.999187 | 0.997988 | 0.995259 |
| Dmtn_Q9WV69_S287s_1 | Dmtn_Q9V | -1.03 | -1.13 | -0.84 | -0.84 | -0.31 | -1.12 | -1 | -0.95 | -0.56 | -0.81 | -0.81 | -1.54 | 0.86 | -0.96 | -0.72 | -0.43 | -0.01 | -0.3875 | 0.1725 | 0.141566 | 0.64902 | 0.913754 | 0.99033 | 0.997988 | 0.994666 |
| Dmtn_Q9WV69_S289s_1 | Dmtn_Q9V | 0.29 | 0.43 | 0.78 | 0.25 | 0.6 | 0.19 | 0.17 | -0.11 | 0.46 | 0.94 | 1.15 | 0.62 | 0.67 | 0.35 | 0.02 | 0.45 | 0.115 | 0.03 | 0.6475 | 0.809724 | 0.374891 | 0.196961 | 0.989223 | 0.997988 | 0.999507 |
| Dmtn_Q9WV69_S303s_1 | Dmtn_Q9V | -0.45 | -0.68 | -1.73 | -0.69 | -0.41 | -0.24 | -0.94 | -0.65 | -1.21 | -1.07 | -0.85 | -1.27 | -1.49 | -0.35 | -1.39 | -0.89 | -0.225 | 0.355 | -0.065 | 0.719769 | 0.338297 | 0.520465 | 0.989223 | 0.997988 | 0.9971 |
| Dmtn_Q9WV69_S307s_1 | Dmtn_Q9V | -0.73 | -0.73 | 0.1 | 0.24 | -0.2 | -0.33 | 0.64 | 0.13 | -1.37 | -0.44 | -0.92 | -0.1 | -1.46 | -0.32 | -0.18 | 0.15 | 0.3275 | -0.2125 | -0.1425 | 0.690162 | 0.335611 | 0.282473 | 0.989223 | 0.997988 | 0.996159 |
| Dmtn_Q9WV69_S307s_1_1 | Dmtn_Q9V | 0.16 | 0.79 | 1.02 | 0.92 | 0.54 | 0.49 | 1.08 | 1.06 | 0.16 | 0.82 | -0.1 | 0.74 | -0.24 | 0.02 | 0.7 | 1.06 | 0.34 | -0.4275 | -0.1725 | 0.377945 | 0.805827 | 0.33885 | 0.995762 | 0.997988 | 0.994666 |
| Dmtn_Q9WV69_S315s_1 | Dmtn_Q9V | 1.4 | 1.1 | 1.21 | 0.91 | 1.06 | 1.25 | 1.13 | 1.39 | 1.06 | 1.69 | 0.56 | 1.28 | 0.3 | 0.51 | 1.2 | 1.11 | 0.07 | -0.3175 | -0.3375 | 0.235043 | 0.808968 | 0.979873 | 0.995762 | 0.997988 | 0.994666 |
| Dmtn_Q9WV69_S333s_1 | Dmtn_Q9V | -0.93 | -1.18 | -1 | -1.12 | -0.21 | -0.58 | -0.12 | -0.79 | -1.6 | -1.38 | -0.58 | -1.1 | -1.27 | -0.64 | -0.85 | -0.64 | 0.0525 | -0.0075 | -0.375 | 0.432072 | 0.022454 | 0.701395 | 0.989223 | 0.997988 | 0.994666 |
| Dmtn_Q9WV69_S87s_1 | Dmtn_Q9V | 0.28 | -0.82 | -0.36 | -0.29 | -0.66 | 0.81 | -0.53 | -0.21 | -0.42 | -0.42 | -0.87 | -0.6 | -0.36 | -1.21 | -0.55 | -0.43 | 0.6325 | -0.1075 | 0.2075 | 0.324755 | 0.70121 | 0.351487 | 0.99521 | 0.997988 | 0.994666 |
| Dmtn_Q9WV69_S92s_1 | Dmtn_Q9V | 0.07 | 0.68 | 0.61 | 0.5 | 0.65 | -0.78 | 0.69 | 0.21 | 0.01 | 0.66 | -0.01 | 0.56 | 0.19 | 0.99 | 0.56 | 0.47 | 0.15 | -0.28 | -0.34 | 0.760802 | 0.461766 | 0.567474 | 0.989223 | 0.997988 | 0.9971 |
| Dmtn_Q9WV69_S92s_2 | Dmtn_Q9V | 0.51 | 0.11 | 0.28 | 0.89 | 0.71 | 0.43 | 1.04 | 0.28 | -0.18 | 0.25 | 1.13 | 0.56 | -1.17 | 0.77 | 0.18 | -0.44 | -0.2725 | -0.16 | 0.0875 | 0.186278 | 0.547855 | 0.982488 | 0.989223 | 0.997988 | 0.994666 |
| Dmtn_Q9WV69_S96s_2 | Dmtn_Q9V | 0.43 | 0.01 | 0.27 | -0.04 | 0.71 | 0.41 | 1.23 | 0.92 | -0.17 | 0.33 | 0.74 | 0.51 | -0.99 | 0.73 | 0.04 | 0.01 | 0.1675 | -0.0075 | -0.6125 | 0.570172 | 0.02872 | 0.508929 | 0.989223 | 0.997988 | 0.995259 |
| Dmwd_E9Q6J8_S111s_1 | Dmwd_E9C | 0.87 | 1.13 | 2.55 | 2.3 | 0.44 | 0.72 | 1.61 | 2.55 | 0.85 | 0.73 | 2.18 | 2.19 | 0.1 | 0.93 | 1.39 | 1.5 | 0.65 | 0.185 | -0.22 | 0.164349 | 0.507607 | 0.678125 | 0.989223 | 0.997988 | 0.994666 |
| Dmwd_E9Q6J8_S487s_1 | Dmwd_E9C | -0.73 | -0.67 | -0.94 | 0.04 | -0.16 | -1.25 | -0.09 | -0.21 | -0.61 | -0.69 | -0.22 | -0.8 | -0.01 | 0.01 | 0.02 | -0.39 | -0.3825 | -0.225 | -0.7325 | 0.130253 | 0.674833 | 0.986315 | 0.992876 | 0.997988 | 0.994666 |
| Dmwd_E9Q6J8_S537s_1 | Dmwd_E9C | 5.07 | 1.51 | 3.48 | -2.13 | 2.27 | 5.11 | 4.78 | 3.38 | -2.86 | 3.99 | -0.17 | 3.82 | -0.09 | -1.8 | 4.76 | 4.74 | 0.1475 | -0.005 | 0.4825 | 0.967326 | 0.20842 | 0.691292 | 0.989223 | 0.997988 | 0.999507 |
| Dmwd_E9Q6J8_S634s_1 | Dmwd_E9C | 0.63 | 0.75 | 0.95 | 0.39 | 0.88 | 0.27 | 0.95 | 0.99 | 0.36 | 0.35 | 1.28 | 0.45 | 0.64 | 0.89 | 0.87 | 0.95 | 1.9025 | -0.7875 | -0.08 | 0.532804 | 0.719643 | 0.812757 | 0.995437 | 0.997988 | 0.994666 |
| Dmwd_E9Q6J8_T453t_1 | Dmwd_E9C | -0.69 | -0.37 | -0.19 | -0.58 | -0.12 | 0.08 | 0.59 | 0.16 | -0.22 | -0.54 | 0.29 | 0.52 | 0.41 | 0.52 | 0.64 | 0.33 | 0.0925 | -0.07 | 0.1575 | 0.003845 | 0.026535 | 0.145743 | 0.989223 | 0.997988 | 0.994666 |
| Dmwd_E9Q6J8_T457t_1 | Dmwd_E9C | 0.91 | 0.19 | -0.4 | 0.32 | 0.78 | 0.61 | 0.02 | 0.13 | 0.42 | 0.87 | 0.38 | 0.21 | 0.67 | 0.73 | 0.31 | -0.68 | 0.635 | 0.47 | 0.9325 | 0.995253 | 0.698168 | 0.517689 | 0.994893 | 0.997988 | 1 |
| Dmwd_E9Q6J8_T461t_1 | Dmwd_E9C | -0.22 | 0.73 | 0.75 | 0.34 | 0.48 | 0.55 | 0.26 | 0.04 | 0.67 | 0.45 | 1.04 | -0.13 | 0.62 | 1.42 | 0.56 | 0.81 | 0.13 | 0.215 | 0.0025 | 0.200948 | 0.814029 | 0.758993 | 0.995762 | 0.997988 | 0.994666 |
| Dmwd_Q08274_S659s_1 | Dmwd_Q08 | 0.14 | 0.41 | 0.33 | 0.46 | 0.63 | 0.45 | 0.9 | 0.53 | 0.16 | 0.42 | 0.97 | 0.7 | 0.13 | 0.73 | 0.35 | 0.9 | -0.0675 | 0.1075 | 0.4525 | 0.487471 | 0.195273 | 0.386751 | 0.989223 | 0.997988 | 0.994666 |
| Dmxl1_Q6PNC0_S1255s_1 | Dmxl1_Q6I | 0.41 | 0.78 | 1.4 | 0.27 | 0.55 | 0.44 | 0.77 | 0.52 | 0.85 | -0.04 | 0.1 | 0.42 | 0.77 | 1.11 | 0.66 | 0.54 | 0.2925 | 0.2275 | 0.1925 | 0.865568 | 0.620949 | 0.277812 | 0.989223 | 0.997988 | 0.999507 |
| Dmxl1_Q6PNC0_S1286s_1 | Dmxl1_Q6I | -0.85 | -1.01 | -1.34 | -0.49 | -0.63 | -1.03 | -0.88 | -0.9 | -1.25 | -1.27 | -1.03 | -0.69 | -0.88 | -0.91 | -0.56 | -0.71 | -0.145 | -0.3825 | 0.055 | 0.571905 | 0.803686 | 0.620954 | 0.995762 | 0.997988 | 0.995259 |
| Dmxl1_Q6PNC0_S1900s_1 | Dmxl1_Q6I | -0.28 | 0.72 | 0.8 | 0.43 | 1.06 | -0.73 | 0.23 | 0.79 | 0.03 | -0.19 | 0.03 | 0.13 | 0.15 | -0.19 | 0.48 | 0.49 | 0.0625 | -0.1375 | 0.1575 | 0.581834 | 0.854986 | 0.18205 | 0.996349 | 0.997988 | 0.995259 |
| Dmxl1_Q6PNC0_S206s_1 | Dmxl1_Q6I | -2.02 | -1.67 | -2.89 | -4.51 | -1.77 | -1.6 | -4.37 | -4.17 | -1.41 | -1.4 | -3.28 | -2.95 | -0.4 | -1.31 | -3.16 | -4.61 | -0.08 | -0.4175 | -0.185 | 0.685055 | 0.810298 | 0.485532 | 0.995762 | 0.997988 | 0.996159 |
| Dmxl1_Q6PNC0_S2372s_1 | Dmxl1_Q6I | 1.41 | -0.32 | 0.62 | 0.57 | 0.85 | 1.2 | 0.51 | 0.68 | -0.1 | 0.21 | 0.43 | 0.82 | 0.06 | 0.66 | 0.03 | 0.01 | -0.205 | 0.5125 | 0.4025 | 0.348531 | 0.527287 | 0.569194 | 0.989223 | 0.997988 | 0.994666 |
| Dmxl1_Q6PNC0_S916s_1 | Dmxl1_Q6I | 0.25 | 0.54 | 0.02 | 0.27 | 0.64 | 0.23 | 0.81 | -0.1 | 0.02 | 0.19 | -0.31 | -0.02 | 0.12 | 0.71 | -0.12 | -0.31 | 0.24 | -0.23 | -0.38 | 0.579215 | 0.647486 | 0.225932 | 0.99033 | 0.997988 | 0.995259 |
| Dmxl1_Q6PNC0_S922s_1 | Dmxl1_Q6I | 1.77 | 0.69 | 0.3 | 0.7 | 0.96 | 1.22 | 0.81 | 1.25 | 0.45 | 0.31 | 0.25 | 0.93 | -0.93 | 1.51 | 0.5 | 0.46 | 0.125 | -0.3 | -0.17 | 0.391832 | 0.568888 | 0.308973 | 0.989223 | 0.997988 | 0.994666 |
| Dmxl1_Q6PNC0_T661t_1 | Dmxl1_Q6I | 0.42 | 0.48 | 0.56 | 0.18 | -0.28 | 0.74 | -0.4 | -0.35 | 1.36 | -0.12 | -1.42 | 0.31 | 1.82 | -0.09 | 0.54 | 0.39 | 0.195 | -0.38 | -0.48 | 0.543849 | 0.139062 | 0.490492 | 0.989223 | 0.997988 | 0.994666 |
| Dmxl2_Q8BPN8_S1141s_1 | Dmxl2_Q8I | 0.07 | 0.75 | 0.6 | 0.32 | 0.59 | 0.08 | 0.51 | 0.12 | 0.05 | 0.37 | -0.17 | 0.02 | -0.33 | 0.68 | -0.07 | 0.47 | -0.4825 | -0.3775 | 0.255 | 0.451378 | 0.666048 | 0.175823 | 0.991946 | 0.997988 | 0.994666 |
| Dmxl2_Q8BPN8_S1144s_1 | Dmxl2_Q8I | -0.44 | 0.31 | 0.62 | 0.16 | 0.26 | 0.31 | 0.55 | 0.42 | -0.32 | -0.01 | 0.68 | 0.18 | 0 | 0.15 | 0.56 | 0.51 | -0.11 | -0.3675 | -0.2475 | 0.650668 | 0.422338 | 0.927509 | 0.989223 | 0.997988 | 0.995259 |
| Dmxl2_Q8BPN8_S1152s_1 | Dmxl2_Q8I | -1.13 | 0.22 | 0.07 | -1.06 | 0.36 | -0.97 | -0.5 | -1.09 | -1.35 | -1.28 | -1.09 | -0.63 | -0.5 | -0.08 | -0.61 | -0.42 | 0.2225 | -0.03 | 0.1425 | 0.852002 | 0.868881 | 0.145319 | 0.996493 | 0.997988 | 0.999507 |
| Dmxl2_Q8BPN8_S1288s_1 | Dmxl2_Q8I | 0.58 | 1 | 0.51 | 0.33 | 0.68 | 0.43 | 0.6 | 0.09 | 0.25 | 0.5 | 0.19 | 0.16 | -0.05 | 0.44 | -0.08 | 0.1 | -0.075 | -0.6125 | 0.0725 | 0.089735 | 0.539172 | 0.196929 | 0.989223 | 0.997988 | 0.994666 |
| Dmxl2_Q8BPN8_S1399s_1 | Dmxl2_Q8I | -1.89 | -1.38 | -2.54 | -4.3 | -0.64 | -1.59 | -2.95 | -2.92 | -2.29 | -1.23 | -0.37 | -2.83 | -0.86 | -1.4 | -2.61 | -2.79 | -0.155 | -0.33 | -0.5025 | 0.395429 | 0.504795 | 0.277797 | 0.989223 | 0.997988 | 0.994666 |
| Dmxl2_Q8BPN8_S1403s_1 | Dmxl2_Q8I | 1.03 | 0.93 | 0.67 | 0.5 | 1.04 | 0.86 | 0.56 | 0.39 | -0.1 | 0.72 | 0.07 | 0.68 | 0.14 | 0.63 | -0.03 | 0.5 | 0.5025 | 0.8475 | 0.6125 | 0.113133 | 0.778108 | 0.15167 | 0.995762 | 0.997988 | 0.994666 |
| Dmxl2_Q8BPN8_S1856s_1 | Dmxl2_Q8I | 0.55 | 0.46 | 0.9 | 0.48 | 0.71 | 0.14 | 1.03 | 0.92 | 0.12 | 0.35 | 0.7 | 1.08 | 0.04 | 1.06 | 0.39 | 0.43 | -0.07 | -0.44 | -0.4725 | 0.694781 | 0.702242 | 0.901549 | 0.995437 | 0.997988 | 0.996159 |
| Dmxl2_Q8BPN8_S1925s_1 | Dmxl2_Q8I | -0.93 | -0.93 | 1.38 | 0.94 | -1.19 | -1.07 | 1.08 | 0.99 | -1.37 | -1.78 | 1.53 | 1.42 | -0.41 | -1.79 | 1.15 | 1.23 | 0.1025 | -0.035 | -0.1175 | 0.932746 | 0.831944 | 0.862174 | 0.995762 | 0.997988 | 0.999507 |
| Dmxl2_Q8BPN8_S2010s_1 | Dmxl2_Q8I | 0.08 | -0.13 | 1.05 | 0.25 | -0.8 | -0.34 | -1.18 | -0.18 | 2.73 | -0.46 | 1.1 | 0.29 | 2.48 | -1.61 | 0.77 | -0.23 | -0.1625 | -0.165 | -0.07 | 0.959787 | 0.021681 | 0.37419 | 0.989223 | 0.997988 | 0.999507 |
| Dmxl2_Q8BPN8_S221s_1 | Dmxl2_Q8I | -0.24 | 0.25 | -0.68 | -0.61 | -0.15 | -0.27 | 0.27 | -0.09 | -0.34 | 0.09 | -2.59 | -0.97 | -0.75 | -0.33 | -0.62 | 0.27 | -0.9375 | 0.6025 | 0.04 | 0.913203 | 0.362991 | 0.288241 | 0.989223 | 0.997988 | 0.999507 |
| Dmxl2_Q8BPN8_S2395s_1 | Dmxl2_Q8I | -0.07 | 0.27 | -0.18 | 0.08 | 0.27 | 0.32 | 0.92 | 0.38 | 0.01 | 0.48 | 0.26 | 0.76 | -0.07 | 0.52 | 0.56 | 0.91 | 0.26 | -0.6325 | -0.0375 | 0.144248 | 0.090494 | 0.191593 | 0.989223 | 0.997988 | 0.994666 |
| Dmxl2_Q8BPN8_S2626s_1 | Dmxl2_Q8I | 0.6 | 0.76 | 0.68 | 0.38 | 0.37 | 0.58 | 0.63 | 0.51 | 0.33 | 0.63 | -0.26 | 0.78 | 0.19 | 0.52 | 0.47 | 0.35 | 0.4475 | 0.3525 | 0.455 | 0.366598 | 0.691461 | 0.425612 | 0.994893 | 0.997988 | 0.994666 |
| Dmxl2_Q8BPN8_S2697s_1 | Dmxl2_Q8I | -0.14 | 0.24 | 0.58 | 0.03 | -0.74 | -0.16 | -0.35 | 0.49 | -0.57 | -0.21 | 0.98 | 0.4 | -0.55 | -0.89 | 0.45 | -0.11 | -0.0825 | -0.235 | -0.2225 | 0.222918 | 0.262798 | 0.94218 | 0.989223 | 0.997988 | 0.994666 |
| Dmxl2_Q8BPN8_S2698s_1 | Dmxl2_Q8I | -0.29 | 0.15 | 0.51 | -0.16 | -0.73 | -0.16 | -0.59 | -0.01 | -0.44 | -0.08 | 1.08 | 0.06 | -0.03 | -0.86 | -0.3 | -0.38 | -0.3675 | -0.0275 | -0.4525 | 0.168426 | 0.155474 | 0.785848 | 0.989223 | 0.997988 | 0.994666 |
| Dmxl2_Q8BPN8_S423s_1 | Dmxl2_Q8I | 0.42 | 0.48 | 0.66 | 0.8 | -1.03 | 0.38 | -0.66 | 0.24 | 0.39 | 0.32 | 1.1 | 0.41 | -0.48 | -1.11 | 0.5 | 0.33 | -0.425 | 0.1025 | -0.445 | 0.071877 | 0.034776 | 0.894751 | 0.989223 | 0.997988 | 0.994666 |
| Dmxl2_Q8BPN8_S423s_2 | Dmxl2_Q8I | 0.7 | 1.34 | 1.1 | 0.25 | 0.34 | 0.65 | -0.23 | 0.46 | 0.66 | 0.58 | 0.54 | 0.42 | -0.5 | 0.4 | 0.89 | 1.04 | -0.8575 | -0.035 | -0.78 | 0.364346 | 0.115197 | 0.31864 | 0.989223 | 0.997988 | 0.994666 |
| Dmxl2_Q8BPN8_S434s_1 | Dmxl2_Q8I | -0.24 | 0.69 | 0.44 | -0.25 | -0.34 | 0.34 | -0.18 | 0.08 | 0.56 | 0.14 | 0.28 | 0.52 | 0.75 | -0.96 | 0.81 | 0.1 | -0.5425 | -0.2975 | -0.39 | 0.973958 | 0.547252 | 0.478778 | 0.989223 | 0.997988 | 0.999507 |
| Dmxl2_Q8BPN8_S434s_2 | Dmxl2_Q8I | 0.7 | 1.34 | 1.1 | 0.25 | 0.34 | 0.65 | -0.23 | 0.46 | 0.66 | 0.58 | 0.54 | 0.42 | -0.5 | 0.4 | 0.89 | 1.04 | -0.185 | 0.215 | 0.015 | 0.364346 | 0.115197 | 0.31864 | 0.989223 | 0.997988 | 0.994666 |
| Dmxl2_Q8BPN8_S451s_1 | Dmxl2_Q8I | 1.39 | 1.23 | 1.01 | 0.64 | 1.29 | 1.29 | 1.02 | 0.87 | 0.95 | 0.91 | 0.55 | 0.68 | -0.3 | 1.38 | 0.6 | 0.68 | -0.5425 | -0.2975 | -0.39 | 0.240842 | 0.841431 | 0.268408 | 0.995932 | 0.997988 | 0.994666 |
| Dmxl2_Q8BPN8_S462s_1 | Dmxl2_Q8I | -0.15 | -0.54 | -0.05 | -0.52 | -2.53 | 0.54 | -1.37 | -0.68 | 1.71 | -0.76 | 0.45 | -0.94 | 1.29 | -2.04 | -0.07 | -0.62 | 0.05 | -0.295 | -0.4775 | 0.943377 | 0.254166 | 0.463582 | 0.989223 | 0.997988 | 0.999507 |
| Dmxl2_Q8BPN8_S463s_1 | Dmxl2_Q8I | -0.92 | -0.77 | -0.44 | -0.43 | -1.71 | -0.05 | -1 | -1.35 | 1.45 | -1.33 | 0.93 | -0.52 | 1.26 | -2.22 | 0.4 | -0.9 | -0.695 | 0.43 | -0.045 | 0.691558 | 0.310783 | 0.217768 | 0.989223 | 0.997988 | 0.996159 |
| Dmxl2_Q8BPN8_S481s_1 | Dmxl2_Q8I | 1.6 | 1.34 | 1.31 | 0.8 | 1.84 | 1.8 | 2.12 | 0.87 | -0.92 | 1.54 | -1.19 | 2.52 | -0.54 | -0.67 | 1.58 | 1.14 | -0.3875 | 0.7725 | 0.275 | 0.135068 | 0.249235 | 0.36133 | 0.989223 | 0.997988 | 0.994666 |
| Dnaja1_P63037_S112s_1 | Dnaja1_P6 | 0.44 | 0.44 | 2.23 | 0.46 | 0.87 | 0.25 | 0.62 | -0.22 | 2.18 | 0.51 | 1.45 | 1.17 | 0.02 | -0.12 | 0.18 | 0.96 | 0.395 | -0.775 | -0.885 | 0.211479 | 0.28932 | 0.417667 | 0.989223 | 0.997988 | 0.994666 |
| Dnaja1_P63037_S335s_1 | Dnaja1_P6 | 1.36 | 1.45 | 1.06 | 0.61 | 1.81 | 1.14 | 0.79 | 0.49 | 1.18 | 1.61 | 1.14 | 0.74 | 0.68 | 2.09 | 0.84 | 0.88 | -0.5125 | 0.435 | -0.6325 | 0.994854 | 0.856314 | 0.873966 | 0.996349 | 0.997988 | 1 |
| Dnajb1_Q9QYJ3_S16s_1 | Dnajb1_Q9 | 1.08 | 1.07 | 0.66 | 0.49 | 0.98 | 1.14 | -0.53 | -0.29 | 1.56 | 1.11 | -0.01 | 0.4 | 1.98 | 0.97 | 0.58 | 0.21 | -0.0625 | 0.0475 | 0.0025 | 0.789269 | 0.260646 | 0.875967 | 0.989223 | 0.997988 | 0.999507 |
| Dnajb2_Q9QYI5_S311s_1 | Dnajb2_Q9 | -0.34 | 0.07 | 0.95 | 1.18 | -0.8 | -0.26 | 1.01 | 0.73 | 0.43 | -0.19 | 2.23 | 1.92 | 1.21 | -0.99 | 1.88 | 1.54 | -0.5 | -0.06 | 0.11 | 0.509439 | 0.564346 | 0.324631 | 0.989223 | 0.997988 | 0.994666 |
| Dnajb2_Q9QYI5_S78s_1 | Dnajb2_Q9 | 0.55 | 0.88 | 1.57 | 1.11 | -0.1 | 0 | 0.95 | 1.08 | 0.48 | 0.16 | 1.91 | 0.91 | 0.96 | -0.02 | 1.68 | 0.89 | -0.295 | 0.6325 | 0.445 | 0.715917 | 0.16485 | 0.703971 | 0.989223 | 0.997988 | 0.996966 |
| Dnajb4_Q9D832_S148s_1 | Dnajb4_Q9 | -2 | -1.39 | -2.01 | -3.5 | -1.1 | -0.69 | -1.62 | -2.56 | -0.12 | -1.92 | -1.83 | -1.72 | 0.45 | -2.44 | -1.76 | -1.25 | -0.545 | -0.1625 | -0.15 | 0.176992 | 0.201102 | 0.171754 | 0.989223 | 0.997988 | 0.994666 |
| Dnajc2_Q3TRX6_S47s_1 | Dnajc2_Q3 | -1.27 | -1.03 | -0.54 | -1.8 | -0.67 | -1.54 | 0.35 | -1.04 | -1.58 | -2.15 | 0.31 | -1.2 | -0.53 | -0.6 | -1.8 | -0.81 | 0.7325 | 0.8275 | 0.975 | 0.577235 | 0.343442 | 0.992675 | 0.989223 | 0.997988 | 0.995259 |
| Dnajc5_P60904_S10s_1 | Dnajc5_P6I | -0.59 | -0.16 | -0.3 | -0.43 | -0.03 | -0.79 | 0.03 | -0.52 | -0.48 | -0.42 | -1.49 | -0.48 | -0.55 | -0.09 | -0.47 | -0.37 | 0.435 | 0.005 | 0.225 | 1 | 0.872056 | 0.277028 | 0.996493 | 0.997988 | 1 |
| Dnajc5_P60904_S12s_1 | Dnajc5_P6I | -1.28 | -0.31 | 0.25 | 0.28 | -0.39 | -1.17 | 0.66 | 0.42 | -0.88 | -0.68 | -0.49 | 0.22 | -0.42 | -0.49 | 0.06 | 0 | 0.0425 | -0.3475 | 1.18E-16 | 0.895046 | 0.774653 | 0.652787 | 0.995762 | 0.997988 | 0.999507 |
| Dnajc5_P60904_S15s_1 | Dnajc5_P6I | -1.7 | -0.43 | -0.99 | -0.82 | -0.45 | -1.36 | 0.1 | -1.65 | -0.72 | -1.8 | -1.09 | -0.92 | -0.56 | -0.29 | -0.84 | -0.45 | 0.145 | -0.1925 | 0.0525 | 0.195002 | 0.748691 | 0.686907 | 0.995762 | 0.997988 | 0.994666 |
| Dnajc5_P60904_T11t_1 | Dnajc5_P6I | -0.73 | 0.08 | -0.24 | -0.25 | -0.64 | -0.22 | 0.15 | -0.3 | -0.15 | -0.74 | -0.83 | -0.41 | -0.51 | -0.21 | -0.54 | -0.35 | 0.145 | -0.1475 | 0.45 | 0.666002 | 0.905189 | 0.387416 | 0.996493 | 0.997988 | 0.996159 |
| Dnajc6_B1B0B7_S38s_1 | Dnajc6_B1I | -0.04 | 1.06 | 0.29 | 0.83 | 0.14 | 0.6 | 0.84 | 0.72 | -0.02 | 0.57 | 0.3 | 0.6 | -0.05 | 0.67 | 0.42 | 0.44 | 0.0325 | -0.2475 | -0.1175 | 0.621732 | 0.897961 | 0.591315 | 0.996493 | 0.997988 | 0.995259 |
| Dnajc6_B1B0B7_S39s_1 | Dnajc6_B1I | 0.08 | 0.51 | 0.41 | 0.41 | 0.61 | 0.11 | 0.7 | 0.46 | 0.29 | 0.58 | 0.25 | 0.45 | -0.04 | 0.87 | 0.31 | 0.33 | 0.04 | -0.1725 | -0.165 | 0.957884 | 0.615938 | 0.858891 | 0.989223 | 0.997988 | 0.999507 |
| Dnajc6_B1B0B7_T403t_1 | Dnajc6_B1I | 0.35 | 0.62 | 0.42 | -0.04 | 1.16 | 0.58 | 1.03 | 0.45 | 0.13 | 0.52 | -0.11 | -0.23 | 0.23 | -0.57 | 0.64 | -0.21 | 0.1175 | 0.04 | 0.015 | 0.358848 | 0.103742 | 0.352118 | 0.989223 | 0.997988 | 0.994666 |
| Dnajc6_Q80TZ3_S591s_2 | Dnajc6_Q8 | -0.73 | -2.73 | -2.16 | -2.22 | -1.17 | -0.75 | -1.42 | -0.91 | -2.89 | -2.91 | 1.58 | -1.79 | -3.26 | -0.6 | -1.53 | -2.72 | 0.4675 | -0.26 | -0.315 | 0.918778 | 0.060137 | 0.652056 | 0.989223 | 0.997988 | 0.999507 |
| Dnajc6_Q80TZ3_S595s_2 | Dnajc6_Q8 | -0.73 | -2.73 | -2.16 | -2.22 | -1.17 | -0.75 | -1.42 | -0.91 | -2.89 | -2.91 | 1.58 | -1.79 | -3.26 | -0.6 | -1.53 | -2.72 | 0.8975 | 0.4575 | -0.0675 | 0.918778 | 0.060137 | 0.652056 | 0.989223 | 0.997988 | 0.999507 |
| Dnajc6_Q80TZ3_S734s_1 | Dnajc6_Q8 | -2.19 | -2.22 | -1.28 | -0.73 | -1.28 | -1.22 | -0.37 | -0.98 | -2.79 | -2.93 | -0.25 | -0.68 | -2.49 | -0.83 | -1.34 | -1.21 | 0.8975 | 0.4575 | -0.0675 | 0.777556 | 0.135359 | 0.93529 | 0.989223 | 0.997988 | 0.99798 |
| Dnajc6_Q80TZ3_S739s_1 | Dnajc6_Q8 | -0.04 | 0.28 | 0.23 | 0.63 | 0.46 | 0.35 | 0.1 | 0.74 | 0.21 | 0.64 | 0.83 | -0.02 | -0.67 | 0.96 | -0.46 | 0.03 | 0.6425 | -0.0575 | 0.1375 | 0.441332 | 0.583587 | 0.626666 | 0.989223 | 0.997988 | 0.994666 |
| Dnajc6_Q80TZ3_S769s_1 | Dnajc6_Q8 | -2.22 | -1.46 | -2.1 | -2.26 | -1.61 | -1.28 | -1.59 | -1.79 | -1.4 | -1.55 | -1.96 | -1.81 | -0.52 | -1.58 | -2.88 | -2.5 | 0.1375 | 0.14 | -0.31 | 0.789037 | 0.116567 | 0.253658 | 0.989223 | 0.997988 | 0.999507 |
| Dnajc6_Q80TZ3_S770s_1 | Dnajc6_Q8 | -0.19 | -1.47 | -1.89 | -1.47 | -1.67 | -0.5 | -0.77 | -1.31 | -0.59 | 0.03 | -1.5 | -1.62 | -0.68 | -0.27 | -1.24 | -1.21 | 0.4425 | 0.33 | 0.14 | 0.358175 | 0.654485 | 0.512372 | 0.990434 | 0.997988 | 0.994666 |
| Dnajc6_Q80TZ3_T590t_1 | Dnajc6_Q8 | -0.66 | -1.8 | 1.02 | 1.05 | 0.04 | -0.46 | 0.42 | 1.16 | -1.69 | -2.12 | 2.04 | 0.96 | -1.9 | -0.23 | 1.13 | 0.49 | 0.1925 | 0.335 | 0.405 | 0.97143 | 0.57381 | 0.92245 | 0.989223 | 0.997988 | 0.999507 |
| Dnajc8_E9PZT4_S35s_1 | Dnajc8_E9I | -4.41 | -4.12 | -3.32 | -3.17 | -4.23 | -5.47 | -5.14 | -5.06 | -5.49 | -4.14 | -4 | -3.34 | -2.31 | -5.36 | -3.41 | -3.38 | 0.3875 | -0.105 | -0.03 | 0.826003 | 0.010565 | 0.350898 | 0.989223 | 0.997988 | 0.999507 |
| Dnal1_D3YYU7_S56s_1 | Dnal1_D3Y | -1.04 | -1.39 | -1.19 | -1.33 | -1.2 | -0.83 | -0.02 | -0.87 | -1.93 | -1.46 | -0.62 | -0.89 | -0.99 | -0.42 | -1.21 | -1.01 | -1.22 | -0.4875 | 0.14 | 0.244495 | 0.107323 | 0.969604 | 0.989223 | 0.997988 | 0.994666 |
| Dnm1_P39053_S512s_1 | Dnm1_P39 | 1.25 | 0.81 | 1.08 | 0.54 | 1.27 | 0.7 | 1.02 | 0.49 | 0.2 | 0.37 | 0.01 | 0.72 | -0.18 | 1 | 0.35 | 0.52 | 0.5075 | 0.0125 | 0.33 | 0.155584 | 0.854506 | 0.052679 | 0.996349 | 0.997988 | 0.994666 |
| Dnm1_P39053_S61s_1 | Dnm1_P39 | 0.45 | 0.29 | 1.49 | 0.57 | -0.22 | 0.08 | 0.13 | 0.54 | 0.16 | -0.52 | 0 | 0.57 | 1.13 | 0.3 | 0.51 | 0.51 | -0.05 | -0.595 | -0.4975 | 0.804073 | 0.105708 | 0.100161 | 0.989223 | 0.997988 | 0.999507 |
| Dnm1_P39053_S644s_1 | Dnm1_P39 | -0.95 | -0.33 | 0.04 | -0.3 | 1.14 | 0.06 | 0.13 | 0.02 | 0.84 | -0.17 | -0.14 | 0.28 | -0.1 | 0.24 | -0.62 | 0.65 | -0.5675 | -0.6475 | -0.0875 | 0.255767 | 0.058973 | 0.105978 | 0.989223 | 0.997988 | 0.994666 |
| Dnm1_P39053_S774s_1 | Dnm1_P39 | -1.13 | -1.79 | -1.64 | -2.15 | -2.68 | -0.38 | -2.04 | -1.57 | -0.45 | -2.18 | 1.03 | -1.55 | -0.06 | -1.44 | -2.38 | -2.2 | 0.7225 | 0.5875 | 0.4275 | 0.766886 | 0.983618 | 0.203129 | 0.999187 | 0.997988 | 0.9971 |
| Dnm1_P39053_S774s_2 | Dnm1_P39 | -2.5 | -3.66 | -2.48 | -4.55 | -4.18 | -2 | -2.18 | -1.98 | -0.85 | -2.9 | 0.99 | -2.28 | -0.41 | -2.09 | -2.75 | -2.78 | 0.01 | 0.89 | 0.1575 | 0.080134 | 0.288044 | 0.043436 | 0.989223 | 0.997988 | 0.994666 |
| Dnm1_P39053_S777s_1 | Dnm1_P39 | -1.24 | -1.49 | -0.01 | -0.02 | -2.01 | -1.56 | -0.92 | -1.04 | -0.07 | -1.42 | -1.22 | -0.54 | 0.72 | -1.77 | -0.32 | 0.09 | 0.7125 | 2.0375 | 1.29 | 0.542205 | 0.138868 | 0.797226 | 0.989223 | 0.997988 | 0.994666 |
| Dnm1_P39053_S778s_1 | Dnm1_P39 | -1.34 | -0.88 | -2.54 | -2.71 | -1.6 | -0.49 | -2.89 | -2.63 | -0.06 | -1.66 | -1.59 | -1.77 | 0.71 | -1.91 | -2.67 | -1.5 | -0.6925 | -0.1225 | 0.37 | 0.493712 | 0.955618 | 0.300607 | 0.999187 | 0.997988 | 0.994666 |
| Dnm1_P39053_S778s_2 | Dnm1_P39 | -2.54 | -4.12 | -2.24 | -4.12 | -4.79 | -1.98 | -1.71 | -1.71 | -1.22 | -3.34 | 1.15 | -1.82 | -0.77 | -2.15 | -2.76 | -2.53 | -0.035 | 0.5975 | 0.525 | 0.074217 | 0.380837 | 0.062717 | 0.989223 | 0.997988 | 0.994666 |
| Dnm1_P39053_S795s_1 | Dnm1_P39 | -1.74 | -2.63 | -1.85 | -2.55 | -0.35 | -1.43 | -1 | -1.32 | -1.87 | -2.41 | 0.88 | 0.24 | -1.58 | 0.16 | -1.82 | -1.18 | 0.7075 | 1.9475 | 1.2025 | 0.044074 | 0.006662 | 0.086208 | 0.989223 | 0.997988 | 0.994666 |
| Dnm1_P39053_S822s_1 | Dnm1_P39 | 0.49 | 0.54 | 0.16 | 0.02 | 0.72 | 0.71 | 0.19 | 0.05 | 0.43 | 0.8 | 1.6 | 0.86 | -0.2 | 0.93 | -0.02 | 0.33 | 1.1675 | 1.4025 | 1.0875 | 0.895605 | 0.6628 | 0.070668 | 0.990434 | 0.997988 | 0.999507 |
| Dnm1_P39053_S847s_1 | Dnm1_P39 | -1.59 | -1.52 | -1.81 | -2.47 | -0.14 | -1.17 | -1.65 | -2.2 | -1.82 | -1.69 | -1.56 | -2.02 | -1.12 | -0.14 | -2 | -1.68 | 0.115 | 0.62 | -0.0425 | 0.19473 | 0.239297 | 0.793118 | 0.989223 | 0.997988 | 0.994666 |
| Dnm1_P39053_S851s_1 | Dnm1_P39 | -0.79 | -2.48 | -1.92 | -0.71 | -0.48 | -0.39 | -0.07 | -0.89 | -1.16 | -1.55 | -0.47 | -0.57 | -2.92 | -0.81 | -1.88 | -1.24 | 0.5575 | 0.075 | 0.6125 | 0.683185 | 0.040916 | 0.277898 | 0.989223 | 0.997988 | 0.996159 |
| Dnm1_P39053_S853s_1 | Dnm1_P39 | -2.39 | -3.29 | -1.83 | -0.58 | -1.17 | -1.92 | -0.12 | -0.49 | -3.13 | -2.84 | 0.37 | 0 | -2.88 | -0.96 | -0.89 | -0.55 | 1.0175 | 0.5375 | -0.2375 | 0.324683 | 0.100885 | 0.519798 | 0.989223 | 0.997988 | 0.994666 |
| Dnm1_P39053_S857s_1 | Dnm1_P39 | -2.12 | -2.22 | -2.05 | -1.37 | -1.56 | -1.86 | -0.79 | -1.22 | -2.65 | -1.93 | 0.8 | -1 | -1.9 | -0.76 | -1.93 | -1.68 | 1.0975 | 0.6225 | 0.7025 | 0.31541 | 0.091289 | 0.299527 | 0.989223 | 0.997988 | 0.994666 |
| Dnm1_P39053_T404t_1 | Dnm1_P39 | 0.76 | 0.85 | 0.3 | 1.23 | 0.73 | 0.64 | -0.09 | -0.42 | 0.57 | 0.62 | -2.61 | 0.44 | 0.34 | 1.03 | -0.11 | 1.4 | 0.5825 | 0.745 | 0.3725 | 0.7628 | 0.122584 | 0.181549 | 0.989223 | 0.997988 | 0.9971 |
| Dnm1_P39053_T776t_2 | Dnm1_P39 | -2.37 | -2.34 | -3.21 | -6.42 | -2.6 | -2.1 | -3.42 | -2.68 | 0.28 | -1.63 | 0.79 | -3.35 | 0.65 | -1.85 | -2.81 | -3.78 | -0.57 | -1.03 | -0.12 | 0.18479 | 0.324294 | 0.051457 | 0.989223 | 0.997988 | 0.994666 |
| Dnm1l_Q8K1M6_S532s_1 | Dnm1l_Q8I | -0.48 | -1.39 | -1.62 | -1.08 | -0.15 | -1.22 | -0.18 | -0.85 | -1.71 | -0.86 | -0.23 | -0.35 | -0.92 | -0.08 | -0.76 | -0.83 | 0.885 | 2.6075 | 1.6375 | 0.174453 | 0.154732 | 0.39811 | 0.989223 | 0.997988 | 0.994666 |
| Dnm1l_Q8K1M6_S535s_1 | Dnm1l_Q8I | 1.51 | -0.3 | -1.2 | -1.45 | 1.35 | 1.71 | -1 | -1.16 | 1.03 | 0.58 | -0.23 | -0.79 | -0.73 | 1.37 | -1.67 | -1.56 | 0.5425 | 0.355 | 0.495 | 0.737141 | 0.511002 | 0.479042 | 0.989223 | 0.997988 | 0.9971 |
| Dnm1l_Q8K1M6_S622s_1 | Dnm1l_Q8I | -0.4 | -1.41 | -0.9 | -0.51 | 0.01 | -0.29 | 0.21 | -0.2 | -0.76 | -0.59 | -0.34 | -0.5 | -1.14 | 0.59 | -0.73 | -0.35 | 0.585 | 0.5075 | -0.2875 | 0.365837 | 0.026703 | 0.38509 | 0.989223 | 0.997988 | 0.994666 |
| Dnm3_E9QLL2_S763s_1 | Dnm3_E9C | 0.82 | 1.28 | 1 | 0.91 | 0.58 | 0.42 | 0.29 | 0.64 | 0.89 | 0.94 | 0.82 | 0.61 | 1.5 | 0.66 | 0.96 | 0.67 | 0.7375 | 0.2575 | 0.3975 | 0.849587 | 0.03651 | 0.415724 | 0.989223 | 0.997988 | 0.999507 |
| Dnm3_E9QLL2_S763s_2 | Dnm3_E9C | 0.02 | -1.11 | -0.33 | -0.32 | -0.88 | 0.55 | -0.52 | -0.29 | -0.03 | -1.05 | -0.35 | -0.62 | -0.55 | -1.38 | -0.99 | -0.63 | -0.52 | -0.1875 | -0.055 | 0.20492 | 0.692666 | 0.820557 | 0.994893 | 0.997988 | 0.994666 |
| Dnm3_E9QLL2_S767s_1 | Dnm3_E9C | 0.08 | -0.14 | 1.11 | 0.63 | 0.18 | 0.6 | 0.44 | 0.35 | 0.73 | 0 | 0.24 | 0.78 | 0.99 | -0.17 | 0.34 | 0.9 | 0.15 | -0.0775 | -0.4525 | 0.81116 | 0.929884 | 0.960487 | 0.997796 | 0.997988 | 0.999507 |
| Dnm3_E9QLL2_S767s_2 | Dnm3_E9C | 0.4 | -0.84 | 0.21 | 0.4 | -0.72 | 1.05 | -0.1 | 0.23 | 0.32 | -0.68 | -0.21 | -0.05 | -0.32 | -1.27 | -0.61 | -0.1 | -0.0275 | 0.0175 | 0.095 | 0.14551 | 0.870247 | 0.596609 | 0.996493 | 0.997988 | 0.994666 |
| Dnm3_E9QLL2_T769t_2 | Dnm3_E9C | -0.85 | -1.69 | -1.3 | -1.82 | -1.19 | -0.64 | -1.27 | -1.19 | -0.83 | -1.87 | -0.55 | -1.65 | -1.07 | -1.6 | -1.6 | -1.56 | 0.0725 | -0.1975 | -0.6175 | 0.890491 | 0.25768 | 0.625658 | 0.989223 | 0.997988 | 0.999507 |
| Dnmt3a_G3UXY2_S28s_1 | Dnmt3a_G | 1.4 | 0.97 | 1.29 | 1.36 | -0.28 | 0.96 | 0.27 | 0.73 | 1.48 | 1.82 | 1.53 | 1.08 | 1.88 | 0.33 | 1.47 | 1.22 | 0.3425 | 0.19 | -0.0425 | 0.934367 | 0.022012 | 0.389533 | 0.989223 | 0.997988 | 0.999507 |
| Dnmt3a_G3UXY2_T46t_1 | Dnmt3a_G | 0.79 | 0.02 | -0.1 | -0.85 | 0 | 0.07 | -0.75 | -0.63 | 0.34 | 0.46 | 0.43 | -0.5 | 1.72 | 1.18 | 0.28 | -0.53 | -0.835 | 0.2225 | -0.03 | 0.227873 | 0.455119 | 0.592241 | 0.989223 | 0.997988 | 0.994666 |
| Dnpep_Q8BPW9_T201t_1 | Dnpep_Q8I | -3.48 | -2.43 | -2.91 | -4.27 | -2.01 | -3.39 | -2.94 | -3.3 | -3.3 | -2.49 | -1.24 | -3.23 | -1.4 | -2.14 | -3.03 | -3.19 | -0.2925 | 0.2175 | 0.6975 | 0.144349 | 0.446907 | 0.236037 | 0.989223 | 0.997988 | 0.994666 |
| Doc2a_H3BJT3_S156s_1 | Doc2a_H3E | 1.17 | -0.2 | -0.14 | 0.65 | 0.45 | 0.88 | 0.86 | 1.03 | -0.08 | -0.04 | -0.09 | 0.03 | -0.78 | 0.21 | 0.8 | 0.41 | 0.3625 | 0.7075 | 0.8325 | 0.647178 | 0.237041 | 0.250893 | 0.989223 | 0.997988 | 0.995259 |
| Dock1_Q8BUR4_S1858s_1 | Dock1_Q8E | 0.24 | 0.94 | -0.26 | -0.19 | 0.44 | 0.71 | 0.36 | 0.29 | 0.66 | 0.32 | -0.87 | -1.78 | 0.32 | 0.82 | -0.52 | -0.43 | 0.435 | -0.415 | -0.21 | 0.7484 | 0.397835 | 0.311649 | 0.989223 | 0.997988 | 0.9971 |
| Dock11_A2AF67_S1095s_1 | Dock11_A2 | 0.53 | -0.45 | 2.42 | 2.53 | -1.26 | 1.16 | 1.62 | 2.64 | 1.19 | -0.06 | 3.59 | 2.65 | 1.19 | -0.4 | 2.82 | 2.3 | 0.2675 | -0.6 | -0.135 | 0.8058 | 0.820104 | 0.546638 | 0.995762 | 0.997988 | 0.999507 |
| Dock3_Q8CIQ7_S1655s_1 | Dock3_Q8C | -0.42 | -0.21 | -0.13 | 0.15 | -0.11 | -0.47 | -0.13 | 0.08 | -0.12 | 0.15 | 0.01 | 0.13 | 0.16 | -0.09 | -0.52 | -0.3 | -0.2175 | 0.585 | 0.22 | 0.897824 | 0.982993 | 0.406449 | 0.999187 | 0.997988 | 0.999507 |
| Dock3_Q8CIQ7_S1763s_1 | Dock3_Q8C | 2.03 | 0.64 | 0.18 | -0.22 | 1.35 | 1.75 | 0.15 | 0.11 | -0.61 | 1.67 | 0.59 | 0.48 | 0.6 | 0.85 | 0.25 | 0.08 | -0.005 | 0.195 | -0.035 | 0.667917 | 0.752807 | 0.839588 | 0.995762 | 0.997988 | 0.996159 |
| Dock3_Q8CIQ7_S1765s_1 | Dock3_Q8C | -0.85 | -0.23 | 1.45 | 2.63 | 1.13 | -1.02 | 1.33 | 2.33 | -0.08 | -0.54 | 2.81 | 1.5 | 0.39 | 0.81 | 0.88 | 1.62 | 0.1825 | -0.125 | -0.2125 | 0.813381 | 0.834374 | 0.859431 | 0.995762 | 0.997988 | 0.999507 |
| Dock3_Q8CIQ7_T200t_1 | Dock3_Q8C | -0.55 | -1.07 | 0.21 | 0.06 | -0.4 | -0.75 | 0.13 | 0.23 | -0.12 | -1.41 | 0.16 | 0.51 | 0.11 | -0.18 | 0.53 | 0.12 | 0.1925 | 0.1725 | 0.175 | 0.19623 | 0.70505 | 0.800625 | 0.995437 | 0.997988 | 0.994666 |
| Dock4_P59764_S1608s_1 | Dock4_P59 | 0.73 | 0.71 | 0.84 | 0.82 | 1.09 | 1.13 | 0.9 | 0.95 | 0.8 | 0.58 | 0.21 | 0.94 | -0.07 | 0.41 | -0.08 | 0.78 | 0.14 | 0.1225 | 0.4825 | 0.09553 | 0.236858 | 0.562169 | 0.989223 | 0.997988 | 0.994666 |
| Dock4_P59764_S1629s_1 | Dock4_P59 | 0.31 | 0.44 | 0.47 | 0.22 | 0.44 | 0.03 | 0.22 | 0.12 | 0.55 | 0.48 | 0.93 | 0.56 | 0.68 | 0.56 | 0.12 | 0.24 | 0.2425 | -0.1425 | -0.515 | 0.874479 | 0.458633 | 0.246091 | 0.989223 | 0.997988 | 0.999507 |
| Dock4_P59764_S1709s_1 | Dock4_P59 | -0.41 | -1.06 | -0.54 | -0.88 | 0.2 | -0.38 | -0.25 | -0.2 | -0.68 | -0.38 | -0.13 | -0.28 | -0.81 | 0.09 | -0.48 | -0.29 | -0.1575 | 0.27 | 0.04 | 0.262202 | 0.04609 | 0.19096 | 0.989223 | 0.997988 | 0.994666 |
| Dock4_P59764_S1726s_1 | Dock4_P59 | 0.92 | -2.44 | -1.6 | -1.42 | -1.05 | 0.91 | -1.12 | -0.41 | -1.67 | -0.74 | -1.32 | -1.27 | -1.2 | -2.82 | -2.16 | -1.92 | 0.565 | 0.355 | 0.35 | 0.229084 | 0.353421 | 0.86456 | 0.989223 | 0.997988 | 0.994666 |
| Dock4_P59764_T1869t_1 | Dock4_P59 | -0.91 | -1.36 | -0.74 | -0.77 | -0.59 | -1.28 | -0.47 | -0.9 | -1.32 | -1.02 | -0.32 | -1.02 | -1.15 | -0.17 | -0.9 | -0.6 | 0.7175 | -0.115 | -0.89 | 0.445329 | 0.621607 | 0.932885 | 0.989223 | 0.997988 | 0.994666 |
| Dock5_B2RY04_S1832s_1 | Dock5_B2F | 0.56 | 0.78 | 1.68 | 0.53 | 1.28 | 0.52 | 0.64 | 0.61 | -0.25 | 0.61 | 0.42 | 0.64 | 0.31 | 0.2 | 0 | 1.81 | 0.135 | 0.025 | 0.24 | 0.521633 | 0.706752 | 0.157451 | 0.995437 | 0.997988 | 0.994666 |
| Dock6_Q8VDR9_S178s_1 | Dock6_Q8\ | -0.15 | -0.42 | 1.07 | 0.47 | -0.71 | 0.72 | 0.62 | 1.03 | 0.19 | 0.67 | 1.61 | 0.9 | -1.94 | 0.35 | 1.72 | 0.54 | -0.125 | -0.5325 | -0.3075 | 0.919412 | 0.715107 | 0.188411 | 0.995437 | 0.997988 | 0.999507 |
| Dock7_Q8R1A4_S1382s_1 | Dock7_Q8F | -0.79 | -0.67 | -0.4 | -1.25 | 0.15 | -0.97 | -0.48 | -0.76 | -1.25 | -0.85 | -0.07 | -0.58 | -0.68 | 0.01 | -0.69 | -0.64 | 0.1725 | 0.6 | -0.075 | 0.372438 | 0.416596 | 0.784848 | 0.989223 | 0.997988 | 0.994666 |

| | | | | | | | | | | | | | | | | | | | | | | | | | | |
|---|---|---|---|---|---|---|---|---|---|---|---|---|---|---|---|---|---|---|---|---|---|---|---|---|---|---|
| Dock7_Q8R1A4_S1428s_1 | Dock7_Q8F | -0.41 | 0.57 | 0.38 | -0.78 | 0.77 | 0.9 | -0.91 | -0.31 | 1.2 | 0.7 | 0.18 | 0.1 | 0.78 | 0.88 | -1.04 | -0.31 | 0.2625 | 0.09 | 0.2775 | 0.7933 | 0.728689 | 0.160467 | 0.995762 | 0.997988 | 0.999507 |
| Dock7_Q8R1A4_S180s_1 | Dock7_Q8F | 0.15 | -1.05 | -0.29 | -0.26 | -0.09 | -0.7 | -0.44 | -0.22 | 0.04 | 0.84 | 0.49 | 0.03 | 0.25 | 1.05 | -0.22 | -0.53 | 0.1725 | 0.605 | 0.1375 | 0.254667 | 1 | 0.059318 | 1 | 0.997988 | 0.994666 |
| Dock7_Q8R1A4_S182s_1 | Dock7_Q8F | -0.69 | -1.12 | -0.92 | -0.85 | -0.81 | -0.35 | -1.03 | -0.73 | -0.53 | -1.02 | 0.44 | -0.69 | -0.59 | -0.56 | -0.8 | -1.04 | 1.18E-16 | 0.7125 | 0.5 | 0.562877 | 0.490732 | 0.216945 | 0.989223 | 0.997988 | 0.995259 |
| Dock7_Q8R1A4_S30s_1 | Dock7_Q8F | 0.88 | 0.1 | -0.06 | 0.69 | 1.47 | 0.63 | 0.15 | 0.1 | 1.21 | 0.95 | 0.28 | 0.39 | -0.33 | 0.2 | -0.62 | 0.37 | 0.165 | 0.445 | 0.1475 | 0.179728 | 0.629046 | 0.378132 | 0.989223 | 0.997988 | 0.994666 |
| Dock7_Q8R1A4_S864s_1 | Dock7_Q8F | -0.59 | -1.41 | -0.84 | -0.75 | -0.11 | -0.41 | 0.01 | -0.9 | -0.37 | -0.16 | -1.41 | -0.57 | -0.1 | -0.24 | -1.21 | -0.89 | 0.185 | 0.305 | -0.4975 | 0.418623 | 0.091464 | 0.441363 | 0.989223 | 0.997988 | 0.994666 |
| Dock7_Q8R1A4_S900s_1 | Dock7_Q8F | -2.8 | -1.23 | -0.19 | -0.59 | -2.15 | -3.88 | 0 | -0.11 | 0.94 | -2.29 | 0.08 | -0.83 | 0.74 | 3.11 | -0.99 | 0.87 | 0.545 | 0.27 | 0.2875 | 0.036349 | 0.725172 | 0.407475 | 0.995437 | 0.997988 | 0.994666 |
| Dock7_Q8R1A4_S929s_1 | Dock7_Q8F | -0.22 | 0.08 | -0.11 | -0.38 | 0.26 | 0.72 | 0.24 | 0.44 | 0.84 | 0.95 | 0.62 | 0.23 | 0.72 | 1.32 | -0.52 | -0.65 | -0.3325 | 0.6775 | 2.135 | 0.433305 | 0.028945 | 0.00989 | 0.989223 | 0.997988 | 0.994666 |
| Dock7_Q8R1A4_T214t_1 | Dock7_Q8F | -0.95 | -0.56 | -0.59 | -0.45 | -1.02 | -0.74 | 0 | -0.2 | 0.39 | -1.57 | 1.73 | -0.62 | 0.92 | -0.63 | -2.23 | -0.64 | 0.5725 | 0.8175 | 0.375 | 0.989947 | 0.613108 | 0.352086 | 0.989223 | 0.997988 | 1 |
| Dock9_E9QMR2_S1191s_1 | Dock9_E9C | 0.37 | -0.25 | 1.33 | 1.18 | -0.17 | 0.27 | 1.45 | 0.57 | -0.03 | 0.42 | 0.74 | 0.79 | 0.12 | 0.32 | 0.28 | 1.24 | 0.1475 | 0.62 | -0.0075 | 0.703725 | 0.784926 | 0.66573 | 0.995762 | 0.997988 | 0.996159 |
| Dock9_E9QMR2_S924s_1 | Dock9_E9C | 0.85 | 1.01 | 1.44 | 1.25 | 1.54 | 0.4 | 1.11 | 1.17 | 0.91 | 0.81 | 0.79 | 1.37 | 0.71 | 1.44 | 1.49 | 1.54 | -0.1275 | -0.1775 | -0.1675 | 0.5999 | 0.781709 | 0.519314 | 0.995762 | 0.997988 | 0.995259 |
| Dopey2_Q3UHQ6_S601s_1 | Dopey2_Q3 | -0.66 | -1.57 | -0.34 | -0.18 | -1.12 | -1.27 | -0.36 | -0.56 | -2.61 | -0.83 | -0.22 | 0.71 | -0.27 | -1.34 | 0.29 | -0.16 | -0.0825 | -0.1675 | 0.1575 | 0.487754 | 0.7083 | 0.942285 | 0.995437 | 0.997988 | 0.994666 |
| Dos_E9PW07_S136s_1 | Dos_E9PW | -0.06 | -1.12 | -0.02 | -0.09 | -0.5 | -0.04 | 1.04 | 0.72 | -1.68 | -1.45 | 0.37 | 0.16 | -1.43 | -0.21 | -0.67 | 0.6 | -0.14 | -0.05 | 0.3175 | 0.82666 | 0.157781 | 0.556237 | 0.989223 | 0.997988 | 0.999507 |
| Dos_E9PW07_S282s_1 | Dos_E9PW | -0.1 | -0.11 | 0.28 | 0.41 | 0.35 | -0.56 | 0.46 | 0.48 | -0.25 | 0.15 | -0.14 | 0.65 | -0.79 | -0.31 | 0.25 | 0.38 | 0.6275 | -0.3275 | -0.105 | 0.487166 | 0.837406 | 0.951438 | 0.995762 | 0.997988 | 0.994666 |
| Dos_E9PW07_S320s_1 | Dos_E9PW | -0.86 | -1.3 | -1.7 | -2.25 | -1.05 | 0 | -1.63 | -2.32 | 0.59 | -1.21 | -4.47 | -2.03 | 0.86 | -1.44 | -0.78 | -0.97 | 0.0625 | -0.0175 | -0.2375 | 0.105932 | 0.5974 | 0.793717 | 0.989223 | 0.997988 | 0.994666 |
| Dos_E9PW07_S485s_1 | Dos_E9PW | -0.41 | -0.46 | 0.18 | 0.27 | -0.17 | -0.63 | 0.71 | 0.62 | -0.67 | -0.12 | 0.28 | 0.22 | -0.92 | -0.04 | 0.48 | 0.74 | 0.2775 | -0.2525 | 0.945 | 0.6826 | 0.524881 | 0.91891 | 0.989223 | 0.997988 | 0.996159 |
| Dos_E9PW07_S527s_1 | Dos_E9PW | 0.76 | 1.02 | 0.58 | 1.48 | 0.81 | 0.75 | 1.71 | 1.39 | 1.18 | 1.59 | 1.01 | 1.07 | 1.38 | 1.32 | 1.69 | 1.52 | 0.2375 | 0.0325 | 0.17 | 0.095173 | 0.524664 | 0.384642 | 0.989223 | 0.997988 | 0.994666 |
| Dos_E9PW07_S544s_1 | Dos_E9PW | -1.46 | -1.69 | -0.94 | -1.17 | -1.14 | -1.36 | -2.04 | -1.03 | -1.92 | -1.43 | -0.14 | -2.04 | -1.53 | -1.02 | -1.86 | -1.16 | 0.205 | 0.2525 | 0.5175 | 0.800808 | 0.79819 | 0.880107 | 0.995762 | 0.997988 | 0.999507 |
| Dos_E9PW07_S617s_1 | Dos_E9PW | -0.36 | -1.25 | -0.97 | -0.62 | -0.98 | -0.04 | -1.2 | -0.69 | -0.72 | -0.47 | 0.2 | -1.18 | -0.1 | -0.73 | -0.45 | -0.8 | -0.0775 | -0.0675 | -0.0775 | 0.373498 | 0.825805 | 0.480226 | 0.995762 | 0.997988 | 0.994666 |
| Dos_E9PW07_T600t_1 | Dos_E9PW | -0.47 | 0.69 | 0.42 | 0.6 | 1.42 | -0.2 | 0.92 | 0.8 | -0.27 | 0.45 | 1.44 | 0.34 | 0.89 | 0.72 | 1.73 | 1.76 | 0.0725 | 0.2575 | 0.28 | 0.032208 | 0.315738 | 0.677386 | 0.989223 | 0.997988 | 0.994666 |
| Dos_E9PW07_T698t_1 | Dos_E9PW | -0.92 | 0.47 | 0.3 | -0.3 | -0.52 | -1.41 | 0.13 | -0.4 | -1.24 | -0.18 | 0.52 | 0.51 | -0.26 | 1.22 | -0.31 | 0.28 | 0.425 | 0.18 | 0.965 | 0.459823 | 0.317495 | 0.975656 | 0.989223 | 0.997988 | 0.994666 |
| Dpf1_D3YYG6_T3t_1 | Dpf1_D3YY | -0.12 | 1.41 | -0.11 | -0.45 | -0.58 | -1.73 | 0.46 | -0.68 | 0.26 | 1.33 | -0.52 | 0.56 | -0.72 | -0.04 | -2.41 | 0.01 | -0.4375 | 0.015 | 0.345 | 0.150193 | 0.163549 | 0.671186 | 0.989223 | 0.997988 | 0.994666 |
| Dpf2_F8WIP7_T204t_1 | Dpf2_F8WI | -3.26 | -5.07 | -4.76 | -3.48 | -3.34 | -4.99 | -9.39 | -4.22 | -4.39 | -3.68 | -1.48 | -3.12 | -2.91 | -3.1 | -3.43 | -4.79 | -0.815 | 0.225 | -0.9725 | 0.31957 | 0.286829 | 0.182584 | 0.989223 | 0.997988 | 0.994666 |
| Dpp10_E9QN98_S28s_1 | Dpp10_E9C | -0.06 | 0.86 | -0.79 | 0.09 | 0.85 | 0.26 | 0.79 | 0.1 | 0.32 | 0.15 | 1.72 | -0.05 | 0.63 | 0.73 | 0.76 | 0.07 | -1.3425 | 0.975 | 0.585 | 0.198284 | 0.229019 | 0.317886 | 0.989223 | 0.997988 | 0.994666 |
| Dpp10_E9QN98_T11t_1 | Dpp10_E9C | 0.17 | -0.31 | 0.46 | 0.17 | -0.78 | 0.57 | -1.02 | -0.38 | 2.13 | 0.08 | -1.05 | 0.2 | 2.13 | -0.95 | -0.94 | 0.08 | 0.475 | 0.51 | 0.5225 | 0.949185 | 0.185768 | 0.726236 | 0.989223 | 0.997988 | 0.999507 |
| Dpp6_Q80VM5_S13s_1 | Dpp6_Q80 | -0.06 | 0.21 | 2.44 | 0.82 | -0.83 | 0.22 | 5.25 | 2.48 | 0.39 | 0.38 | 2.13 | 2.52 | 0.2 | -0.31 | 2.86 | 3.73 | -0.525 | 0.2175 | -0.0425 | 0.44552 | 0.465897 | 0.489119 | 0.989223 | 0.997988 | 0.994666 |
| Dpp8_Q80YA7_S568s_1 | Dpp8_Q80 | -3.16 | -2.81 | -4.48 | -2.51 | -1.61 | -2.89 | -4.52 | -2.51 | -1.75 | -2.13 | -1.27 | -2.43 | -0.16 | -2.7 | -1.13 | -2.13 | 0.9275 | 0.5025 | 0.7675 | 0.022954 | 0.591877 | 0.018319 | 0.989223 | 0.997988 | 0.994666 |
| Dpy19l1_A6X919_S22s_1 | Dpy19l1_A | 1.03 | 0.84 | -0.02 | 0.36 | 0.91 | 0.45 | 0.62 | 0.3 | 1.14 | 0.56 | 0.27 | -0.09 | -0.04 | 1.28 | 0.6 | 0.8 | 0.3575 | 1.345 | 1.71 | 0.776454 | 0.952986 | 0.820081 | 0.999187 | 0.997988 | 0.99798 |
| Dpy19l1_A6X919_S23s_1 | Dpy19l1_A | -0.29 | -0.33 | -1.08 | 0.91 | 0.26 | -0.64 | -0.55 | 0.46 | 0.17 | -0.18 | 1.9 | 1.74 | -0.23 | 0.55 | 0.66 | 1.03 | 0.0175 | -0.0825 | 0.1075 | 0.160293 | 0.862548 | 0.098819 | 0.996493 | 0.997988 | 0.994666 |
| Dpy19l1_A6X919_S46s_1 | Dpy19l1_A | -1.16 | -1.18 | -1.5 | 0 | -0.49 | 0.92 | -0.19 | -0.45 | 1.94 | -0.71 | -1.89 | 0.15 | 2.29 | -0.74 | 0.44 | 1.12 | 0.08 | 1.105 | 0.7 | 0.022048 | 0.062601 | 0.300692 | 0.989223 | 0.997988 | 0.994666 |
| Dpysl2_O08553_S517s_2 | Dpysl2_O0 | -2.64 | -2.01 | -1.96 | -1.59 | -2.11 | -2.58 | -2.12 | -2.08 | -1.52 | -1.87 | -1.58 | -1.88 | -0.59 | -2.14 | -1.69 | -1.89 | 0.9075 | 0.8325 | 1.7375 | 0.264436 | 0.545726 | 0.255556 | 0.989223 | 0.997988 | 0.994666 |
| Dpysl2_O08553_S518s_1 | Dpysl2_O0 | -0.76 | -0.74 | -0.36 | -0.67 | -0.83 | -0.95 | -1.28 | -1.07 | 0.19 | -0.66 | -2.03 | -1.28 | 1.07 | -1.08 | -0.48 | -0.8 | -0.1725 | 0.3375 | 0.4725 | 0.514359 | 0.094922 | 0.501973 | 0.989223 | 0.997988 | 0.994666 |
| Dpysl2_O08553_S518s_2 | Dpysl2_O0 | -4.14 | -4.23 | -1.25 | -2.95 | -3.08 | -4.65 | -1.97 | -1.71 | -3.69 | -4.52 | -1.21 | -1.87 | -2.12 | -3.62 | -1.42 | -1.48 | -0.4 | -0.3125 | 0.31 | 0.219357 | 0.730677 | 0.728622 | 0.995762 | 0.997988 | 0.994666 |
| Dpysl2_O08553_S540s_1 | Dpysl2_O0 | 0.66 | -0.17 | 1.03 | 0.51 | 0.24 | 0.26 | 2.36 | 0.37 | -0.14 | 0.5 | 0.74 | 1.24 | -0.11 | 0.6 | 0.92 | 1.54 | 0.29 | 0.32 | 0.9825 | 0.589174 | 0.570135 | 0.840029 | 0.989223 | 0.997988 | 0.995259 |
| Dpysl2_O08553_S542s_1 | Dpysl2_O0 | -0.63 | -0.52 | -0.51 | -0.67 | -0.2 | 0.03 | 0.7 | -0.61 | -0.24 | -0.5 | 1.37 | 0.68 | -0.34 | 0.21 | -0.59 | 0.13 | 0.3 | 0.0775 | 0.23 | 0.140318 | 0.086964 | 0.055103 | 0.989223 | 0.997988 | 0.994666 |
| Dpysl2_O08553_S570s_1 | Dpysl2_O0 | -0.95 | -2.57 | -1.29 | -1.76 | -0.82 | -0.54 | -1 | -0.8 | -1.55 | -1.21 | 0.19 | -0.71 | -2.42 | 0.18 | -1.81 | -1.87 | 0.5625 | 0.91 | 0.435 | 0.790006 | 0.038497 | 0.118238 | 0.989223 | 0.997988 | 0.999507 |
| Dpysl2_O08553_S62s_1 | Dpysl2_O0 | -1.19 | 0.03 | -0.6 | -2.08 | 0.51 | -0.55 | -2.47 | -1.78 | 0.38 | -1.11 | -1.08 | -1.33 | 1.56 | -4.86 | -0.76 | -1.46 | 0.8525 | 0.8225 | 0.1625 | 0.729165 | 0.872915 | 0.751761 | 0.996493 | 0.997988 | 0.9971 |
| Dpysl2_O08553_T509t_1 | Dpysl2_O0 | -0.96 | -0.59 | -0.74 | -0.58 | -0.39 | -0.92 | -0.15 | -0.58 | -1.24 | -0.61 | -0.71 | -0.68 | -0.84 | -0.47 | -0.62 | -0.73 | -0.1125 | 0.175 | -0.42 | 0.82956 | 0.407349 | 0.709314 | 0.989223 | 0.997988 | 0.999507 |
| Dpysl2_O08553_T512t_1 | Dpysl2_O0 | -0.84 | -0.32 | -1.27 | -5.11 | -0.68 | -0.04 | -3.88 | -3.49 | 0.64 | -0.44 | -3.44 | -3.33 | 0.9 | -1.35 | -1.86 | -2.85 | 0.2075 | -0.0925 | 0.0525 | 0.612635 | 0.912766 | 0.853601 | 0.997335 | 0.997988 | 0.995259 |
| Dpysl2_O08553_T514t_1 | Dpysl2_O0 | -0.96 | -0.89 | -0.56 | -1.46 | -0.68 | -0.52 | -1.82 | -1.22 | 0.52 | -1.2 | -2.44 | -2.02 | 0.88 | -1.99 | -1.58 | -0.05 | -0.1375 | 0.2425 | 0.595 | 0.654739 | 0.791842 | 0.611496 | 0.995762 | 0.997988 | 0.995259 |
| Dpysl2_O08553_T514t_2 | Dpysl2_O0 | -3.04 | -2.58 | -1.85 | -3.51 | -2.37 | -3.19 | -2.19 | -2.22 | -2.21 | -2.36 | -1.77 | -2.2 | -1.24 | -2.56 | -1.78 | -1.95 | -0.0925 | -0.3175 | 0.2825 | 0.0737 | 0.538975 | 0.134123 | 0.989223 | 0.997988 | 0.994666 |
| Dpysl3_E9PWE8_S101s_1 | Dpysl3_E9F | -1.88 | -1.46 | -1.1 | -1.28 | -1.11 | -1.36 | -0.95 | -1.4 | -1.54 | -1.15 | -0.11 | -1.29 | -0.95 | -0.22 | -1.22 | -1.04 | 0.2525 | 0.61 | 0.8625 | 0.100907 | 0.38358 | 0.281874 | 0.989223 | 0.997988 | 0.994666 |
| Dpysl3_E9PWE8_S585s_1 | Dpysl3_E9F | -1.31 | -1.69 | -1.71 | -2 | -0.98 | 0.03 | -0.27 | -2.05 | -1.3 | -0.83 | -0.26 | -1.73 | 0.16 | -1.19 | -3.12 | -2.2 | 0.225 | 0.4075 | 0.5725 | 0.889432 | 0.082034 | 0.097002 | 0.989223 | 0.997988 | 0.999507 |
| Dpysl3_E9PWE8_S631s_2 | Dpysl3_E9F | -0.1 | -1.06 | -0.56 | -0.4 | -0.53 | -0.36 | 0.22 | -0.1 | 0.45 | 0.29 | 0.48 | -0.23 | 0.63 | 0.09 | 0.09 | -0.21 | 0.86 | 0.6475 | 0.09 | 0.054399 | 0.260831 | 0.026334 | 0.989223 | 0.997988 | 0.994666 |
| Dpysl3_E9PWE8_S631s_3 | Dpysl3_E9F | -2.96 | -1.75 | -4.27 | -2.08 | -2.35 | -2.75 | -1.68 | -3.5 | -1.59 | -1.93 | -1.65 | -2.62 | -2.48 | -2.01 | -2.89 | -2.04 | 0.3375 | 0.7775 | 0.68 | 0.466753 | 0.74844 | 0.171057 | 0.995762 | 0.997988 | 0.994666 |
| Dpysl3_E9PWE8_S635s_1 | Dpysl3_E9F | -0.02 | 0.34 | 0.11 | 0.12 | 0.33 | -0.24 | 0.14 | 0.14 | 0.65 | 0.52 | 1.21 | 0.28 | 1.14 | 0.77 | 0.51 | 0.09 | 0.195 | 0.8175 | 0.41 | 0.123956 | 0.840584 | 0.076007 | 0.995762 | 0.997988 | 0.994666 |
| Dpysl3_E9PWE8_S652s_1 | Dpysl3_E9F | 2.5 | 0.54 | 0.33 | 0.49 | 0.91 | 2.86 | 1.16 | 0.74 | 0.3 | 2 | 0.11 | 1.09 | -0.16 | 0.18 | 0.7 | 0.14 | -0.045 | 0.5275 | 0.49 | 0.166015 | 0.479144 | 0.88316 | 0.989223 | 0.997988 | 0.994666 |
| Dpysl3_E9PWE8_S654s_1 | Dpysl3_E9F | -1.65 | -1.01 | 0.38 | -0.05 | 0.19 | -0.8 | 1.47 | 0.68 | -0.08 | -1.06 | 0.47 | 0.78 | -0.35 | 0.49 | 0.87 | 1.53 | 0.4525 | -0.09 | -0.75 | 0.051902 | 0.127202 | 0.295913 | 0.989223 | 0.997988 | 0.994666 |
| Dpysl3_E9PWE8_T621t_1 | Dpysl3_E9F | -0.95 | -0.01 | -0.51 | -0.11 | 0.01 | -0.62 | 0.32 | -0.06 | -0.5 | -0.46 | -0.78 | -0.07 | 0.07 | 0.12 | -0.06 | -0.01 | 0.9675 | 0.61 | 1.2175 | 0.164148 | 0.333888 | 0.847622 | 0.989223 | 0.997988 | 0.994666 |
| Dpysl3_E9PWE8_T622t_1 | Dpysl3_E9F | -0.9 | -0.27 | -0.57 | -0.6 | 0.04 | -0.81 | 0.02 | -0.45 | -0.99 | -0.16 | -0.91 | -0.49 | -0.49 | 0.09 | -0.39 | -0.56 | 0.3075 | -0.0575 | 0.425 | 0.381402 | 0.319659 | 0.852486 | 0.989223 | 0.997988 | 0.994666 |
| Dpysl3_E9PWE8_T627t_1 | Dpysl3_E9F | -2.09 | -2.76 | -2.62 | -6.35 | -3.45 | -1.85 | -2.41 | -2.27 | 1.26 | -2.04 | -3.42 | -1.57 | 1.89 | -3.34 | -0.55 | -3.06 | 0.285 | -0.0525 | 0.2475 | 0.129224 | 0.299774 | 0.123689 | 0.989223 | 0.997988 | 0.994666 |
| Dpysl3_E9PWE8_T627t_2 | Dpysl3_E9F | -5.13 | -7.76 | -1.7 | 0.92 | -6.85 | -4.73 | -0.87 | 0.98 | -2.92 | -4.56 | 1.45 | 0.59 | -2.59 | -5.49 | -0.84 | -1.39 | 0.96 | 2.0125 | 2.19 | 0.652295 | 0.8048 | 0.333399 | 0.995762 | 0.997988 | 0.995259 |
| Dpysl3_E9PWE8_T627t_3 | Dpysl3_E9F | -2.96 | -1.75 | -4.27 | -2.08 | -2.35 | -2.75 | -1.68 | -3.5 | -1.59 | -1.93 | -1.65 | -2.62 | -2.48 | -2.01 | -2.89 | -2.04 | 0.55 | 2.0575 | 0.84 | 0.466753 | 0.74844 | 0.171057 | 0.995762 | 0.997988 | 0.994666 |
| Dpysl3_E9PWE8_T632t_1 | Dpysl3_E9F | -2.48 | -2.94 | -3.19 | -3.73 | -2.37 | -2.88 | -2.77 | -3.01 | -0.46 | -2.7 | -2.94 | -2.95 | 0.68 | -2.16 | -1.7 | -2.35 | 0.195 | 0.8175 | 0.41 | 0.02877 | 0.309698 | 0.194462 | 0.989223 | 0.997988 | 0.994666 |
| Dpysl3_E9PWE8_T632t_2 | Dpysl3_E9F | -2.6 | -3.93 | -1.32 | -0.76 | -3.22 | -2.67 | -0.19 | -0.49 | -1.42 | -2.07 | 0.29 | -0.63 | -1.04 | -2.53 | -0.24 | -0.88 | 0.3275 | 0.8225 | 1.7025 | 0.217554 | 0.575334 | 0.150257 | 0.989223 | 0.997988 | 0.994666 |
| Dpysl3_E9PWE8_T637t_1 | Dpysl3_E9F | 0.46 | 0.45 | -0.02 | -0.22 | 0.12 | 0.06 | -0.31 | 0 | 0.23 | 0.64 | 0.58 | 0.02 | 1.04 | 0.73 | 0.34 | 0.03 | 0.51 | 1.195 | 0.98 | 0.273495 | 0.434384 | 0.47855 | 0.989223 | 0.997988 | 0.994666 |
| Dpysl4_Q3TMU8_S570s_1 | Dpysl4_Q3 | -0.56 | -1.31 | -1.21 | -1.21 | -1.02 | -0.58 | -1.27 | -1.34 | -0.95 | -1.27 | -0.68 | -0.39 | -1.36 | -0.2 | -2.03 | -1.03 | -0.2 | 0.2 | 0.3675 | 0.843241 | 0.942929 | 0.408549 | 0.999187 | 0.997988 | 0.999507 |
| Dpysl4_Q3TMU8_S572s_1 | Dpysl4_Q3 | -0.42 | 0.37 | 0.57 | 0.39 | 0.15 | -0.34 | 0.29 | 0.53 | 0.31 | 0.43 | 0.54 | 0.27 | 0.13 | 0.8 | 0.5 | 0 | 0.02 | 0.25 | -0.0825 | 0.69292 | 0.820035 | 0.571426 | 0.995762 | 0.997988 | 0.996159 |
| Dpysl5_Q9EQF6_T509t_1 | Dpysl5_Q9 | -2.05 | -1.69 | -1.55 | -1.05 | -1.79 | -1.91 | -1.14 | -1.2 | -1.43 | -1.7 | -0.53 | -0.9 | -1.07 | -1.58 | -1.97 | -1.73 | -0.07 | 0.16 | 0.13 | 0.993862 | 0.807405 | 0.224067 | 0.995762 | 0.997988 | 1 |
| Dpysl5_Q9EQF6_T509t_2 | Dpysl5_Q9 | -2.51 | -2.72 | -2.85 | -2.44 | -2.22 | -3.35 | -2.44 | -2.62 | -3.6 | -2.05 | -0.62 | -2.3 | -2.81 | -2.12 | -2.75 | -3.1 | 0.075 | 0.445 | -0.0025 | 0.824783 | 0.924822 | 0.402065 | 0.997335 | 0.997988 | 0.999507 |
| Dpysl5_Q9EQF6_T514t_1 | Dpysl5_Q9 | -1.54 | -1.57 | -1.9 | -1.37 | -1.64 | -1.45 | -1.03 | -1.66 | -2 | -1.23 | -0.77 | -1.5 | -1.47 | -1.61 | -2.04 | -2.08 | -0.0275 | 0.4875 | -0.065 | 0.460436 | 0.54388 | 0.487695 | 0.989223 | 0.997988 | 0.994666 |
| Dpysl5_Q9EQF6_T514t_2 | Dpysl5_Q9 | -2.51 | -2.72 | -2.85 | -2.44 | -2.22 | -3.35 | -2.44 | -2.62 | -3.6 | -2.05 | -0.62 | -2.3 | -2.81 | -2.12 | -2.75 | -3.1 | 0.15 | 0.22 | -0.205 | 0.824783 | 0.924822 | 0.402065 | 0.997335 | 0.997988 | 0.999507 |
| Drd1_Q61616_S441s_1 | Drd1_Q616 | -0.34 | -1.56 | -0.25 | 0.4 | -1.43 | -0.21 | -0.77 | 0.76 | 0.08 | -1.03 | -0.07 | -0.87 | 0.51 | -0.8 | 0.74 | 0.08 | -0.0275 | 0.4875 | -0.065 | 0.273017 | 0.964197 | 0.940779 | 0.999187 | 0.997988 | 0.994666 |
| Drd1_Q61616_T446t_1 | Drd1_Q616 | -0.05 | -0.42 | -0.44 | -0.42 | -1.28 | -1.19 | -0.36 | -0.54 | 0.69 | 0.23 | -0.03 | -0.58 | 1.5 | 0.05 | 0.06 | -0.63 | 0.025 | -0.035 | 0.57 | 0.216651 | 0.096915 | 0.212165 | 0.989223 | 0.997988 | 0.994666 |
| Drd5_Q8BLD9_S385s_1 | Drd5_Q8BL | 0.18 | 0.13 | 0.59 | 0.61 | 0.43 | -0.14 | 0.52 | -0.47 | -0.18 | 0.94 | 0 | 1.39 | -0.58 | 0.23 | 1.32 | 0.95 | -0.51 | 0.41 | 0.5775 | 0.812859 | 0.33533 | 0.687293 | 0.989223 | 0.997988 | 0.999507 |
| Drosha_Q5HZJ0_T272t_1 | Drosha_Q5 | -3.76 | -3.02 | -3.52 | -4.67 | -2.29 | -3.53 | -3.45 | -3.57 | -2.93 | -4.04 | -4.11 | -3.15 | -1.33 | -1.43 | -2.15 | -2.43 | -0.2925 | 0.16 | 0.1025 | 0.001458 | 0.24018 | 0.677828 | 0.989223 | 0.997988 | 0.994666 |
| Dst_S4R1P5_S106s_1 | Dst_S4R1P | -0.49 | -0.65 | 0.02 | -0.04 | -0.13 | -0.44 | 0.87 | -0.05 | -0.21 | -0.07 | 0 | 0.22 | -0.63 | 0 | 0.68 | 0.05 | 0.5325 | 0.185 | 1.9075 | 0.373918 | 0.307385 | 0.299897 | 0.989223 | 0.997988 | 0.994666 |
| Dst_S4R1P5_S140s_1 | Dst_S4R1P | 0.33 | 0.53 | 0.46 | 0.34 | -0.87 | 0.54 | -0.37 | -0.22 | 2.52 | 0.75 | -0.13 | 0.15 | 2.69 | -0.69 | 0.92 | 0.26 | 0.3525 | 0.275 | 0.315 | 0.559575 | 0.063774 | 0.467777 | 0.989223 | 0.997988 | 0.995259 |
| Dst_S4R1P5_S4072s_1 | Dst_S4R1P | 1.41 | 0.42 | 0.61 | 0.93 | 0.71 | 1.06 | 1.06 | 1.15 | -0.05 | 0.17 | -0.07 | 0.91 | 0.08 | 0.73 | 0.94 | 1.37 | -0.645 | 0.4075 | 0.38 | 0.864581 | 0.583333 | 0.099988 | 0.989223 | 0.997988 | 0.999507 |
| Dst_S4R1P5_S5775s_1 | Dst_S4R1P | -0.25 | -0.06 | 0.43 | -0.14 | 0.3 | 0.43 | 0.93 | 0.33 | -0.06 | -0.59 | -0.4 | 0.18 | -0.27 | 0.09 | -0.42 | 0.28 | 0.1525 | -0.6025 | -0.0625 | 0.79589 | 0.077733 | 0.45506 | 0.989223 | 0.997988 | 0.999507 |
| Dst_S4R1P5_S77s_1 | Dst_S4R1P | 0.6 | 0.72 | 1.26 | 1.28 | 0.8 | 1.09 | 1.8 | 1.68 | 1.43 | 1.84 | 1.87 | 1.33 | 1.13 | 1.46 | 2.53 | 1.61 | 0.5025 | -0.2125 | -0.075 | 0.076609 | 0.248429 | 0.040015 | 0.989223 | 0.997988 | 0.994666 |
| Dstn_Q9R0P5_S3s_1 | Dstn_Q9R0 | 0.7 | 1.21 | 1.31 | 0.63 | 0.75 | 0.82 | 0.91 | 0.87 | 0.52 | 0.75 | 2.12 | 1.01 | 0.51 | 0.61 | 1.22 | 3.12 | 0.3775 | 0.6525 | 0.7175 | 0.492218 | 0.606849 | 0.727865 | 0.989223 | 0.997988 | 0.994666 |
| Dtd1_Q9DD18_S197s_1 | Dtd1_Q9DI | 1.44 | 1.67 | 0.87 | 1.12 | 0.99 | 1.23 | -0.11 | 0.93 | 1.26 | 1.09 | 1.33 | 0.42 | 0.46 | 1.39 | 0.58 | 0.56 | -0.125 | 0.1375 | 0.4025 | 0.127942 | 0.162704 | 0.425024 | 0.989223 | 0.997988 | 0.994666 |
| Dtd1_Q9DD18_S205s_1 | Dtd1_Q9DI | -1.91 | -1.58 | -1.07 | -1.15 | -1.75 | -3.1 | -1.44 | -1.63 | -2.61 | -1.68 | -1.13 | -1.63 | -2.93 | -2.31 | -1.63 | -1.49 | -0.515 | -0.25 | -0.5275 | 0.118416 | 0.19719 | 0.379379 | 0.989223 | 0.997988 | 0.994666 |
| Dtna_Q8CFR5_S500s_1 | Dtna_Q8CF | -1.13 | -2 | -0.99 | -5.32 | -2.4 | -0.11 | -2.03 | -1.58 | 3.18 | -1.37 | -1.4 | -0.53 | 3.56 | -2.93 | 2.48 | -2.17 | -0.5525 | -0.335 | -0.6625 | 0.139189 | 0.405054 | 0.099871 | 0.989223 | 0.997988 | 0.994666 |
| Dtna_Q9D2N4_S306s_1 | Dtna_Q9D2 | -4.27 | -6.09 | -3.4 | -4.3 | -4.28 | -4.77 | -4.11 | -3.19 | -3.67 | -4.22 | -2.52 | -3.66 | -1.45 | -5.66 | -2.13 | -3.82 | 0.83 | 2.33 | 2.595 | 0.212429 | 0.472527 | 0.128637 | 0.989223 | 0.997988 | 0.994666 |
| Dtna_Q9D2N4_S308s_1 | Dtna_Q9D2 | -3.89 | -4.83 | -4.91 | -6.8 | -4.31 | -3.79 | -5.36 | -5.84 | 0.44 | -1.52 | -1.98 | -4.28 | 1.83 | -2.7 | -2.46 | -5.61 | 0.4275 | 0.9975 | 1.25 | 0.067171 | 0.679225 | 0.009381 | 0.993464 | 0.997988 | 0.994666 |
| Dtna_Q9D2N4_T501t_1 | Dtna_Q9D2 | 1.36 | 0.63 | 0.05 | 0.49 | 1.45 | 0.99 | 0.26 | 0.81 | 1.21 | 1.1 | 1.91 | -0.1 | 0.97 | 1.71 | 0.83 | 0.2 | 0.2825 | 3.2725 | 2.8725 | 0.4816 | 0.505747 | 0.410911 | 0.989223 | 0.997988 | 0.994666 |
| Dtnb_G3UXK0_S364s_1 | Dtnb_G3U) | -0.67 | -0.75 | -0.7 | -1.04 | -0.06 | -0.8 | -0.72 | -1.07 | -0.88 | -0.75 | -0.14 | -0.62 | -0.63 | -0.04 | -0.92 | -0.84 | 0.245 | 0.3975 | 0.295 | 0.530131 | 0.640687 | 0.455549 | 0.989223 | 0.997988 | 0.994666 |
| Dtnb_G3UXK0_S501s_2 | Dtnb_G3U) | -1.1 | -0.63 | -0.59 | 0.12 | -1.49 | 0.08 | -0.98 | -1.25 | 1.38 | -0.78 | -1.8 | -0.01 | 2.03 | -1.43 | -0.12 | -0.05 | 0.1275 | 0.1925 | 0.1825 | 0.3461 | 0.388742 | 0.704265 | 0.989223 | 0.997988 | 0.994666 |
| Dtnb_G3UXK0_T504t_2 | Dtnb_G3U) | -1.18 | -1.13 | 0.45 | -0.3 | -1.13 | 0.3 | -1.07 | -1.05 | 0.46 | -1.35 | -1.42 | -0.42 | 0.73 | -1.42 | 0.12 | 0.56 | -0.36 | 0.2475 | 0.6575 | 0.360386 | 0.681411 | 0.794369 | 0.993613 | 0.997988 | 0.994666 |
| Dtnb_G3UXK0_T507t_2 | Dtnb_G3U) | -0.75 | -0.21 | -0.75 | 0 | -2.66 | 0.16 | -0.94 | -1.3 | 1.26 | 0.1 | -1.92 | 0.16 | 2.1 | -0.45 | -0.09 | -0.11 | -0.1975 | -0.1425 | 0.5375 | 0.185854 | 0.192422 | 0.604885 | 0.989223 | 0.997988 | 0.994666 |
| Dtnb_G3UZJ3_S494s_1 | Dtnb_G3U. | 0.85 | 0.41 | -0.13 | 0.31 | 1.21 | 2.33 | -0.03 | -1.33 | 3.12 | 0.97 | -1.64 | -0.57 | 3.75 | -0.88 | -0.12 | -0.01 | -0.7575 | 0.3275 | 0.79 | 0.726239 | 0.7965 | 0.905966 | 0.995762 | 0.997988 | 0.9971 |
| Dtx4_Q6PDK8_S273s_1 | Dtx4_Q6PD | 0.31 | -0.43 | 0.59 | -0.19 | 0.68 | 0.02 | -0.18 | 0.06 | 1.33 | 0.32 | 0.78 | 0.45 | 1.38 | 0.26 | 0.53 | 0.5 | 0.185 | 0.11 | 0.325 | 0.12336 | 0.811668 | 0.083457 | 0.995762 | 0.997988 | 0.994666 |
| Dtx4_Q6PDK8_T72t_1 | Dtx4_Q6PD | -0.78 | -1.32 | -0.74 | -1.26 | -0.36 | -0.66 | -0.16 | -0.75 | -0.42 | -1.02 | -0.49 | -0.62 | 0.6 | 0.34 | -0.54 | -0.01 | 0.075 | 0.65 | 0.5975 | 0.006937 | 0.057988 | 0.168612 | 0.989223 | 0.997988 | 0.994666 |
| Dync1h1_Q9JHU4_S1228s_1 | Dync1h1_C | 0.2 | 1.36 | 0.09 | -0.03 | 0.83 | -0.22 | 0.1 | 0.5 | -0.35 | -0.13 | -0.48 | -0.05 | 0.12 | 0.35 | -0.55 | 0.31 | 0.5425 | 0.3875 | 1.1225 | 0.386614 | 0.790004 | 0.087584 | 0.995762 | 0.997988 | 0.994666 |
| Dync1h1_Q9JHU4_S4366s_1 | Dync1h1_C | 0.4 | 0.51 | 0.56 | 0.37 | 0.55 | 0.41 | 0.48 | 0.51 | -0.03 | 0.31 | 0.15 | 0.56 | 0.05 | 0.17 | 0.38 | 0.48 | -0.1025 | -0.6575 | -0.3475 | 0.436405 | 0.887625 | 0.367128 | 0.996493 | 0.997988 | 0.994666 |
| Dync1i1_O88485_S142s_1 | Dync1i1_O | -0.29 | 0.12 | 0.14 | 0.22 | -0.03 | -0.29 | 0.46 | 0.06 | 0.1 | 0.33 | -0.08 | 0.13 | -0.61 | 0.75 | 0.06 | 0.01 | 0.0275 | -0.2125 | -0.19 | 0.988167 | 0.992013 | 0.757145 | 0.999187 | 0.997988 | 1 |
| Dync1i1_O88485_S50s_1 | Dync1i1_O | -0.48 | -0.26 | -0.58 | -1.23 | -1.11 | -0.93 | -0.23 | -0.22 | 0.23 | -0.45 | 0.49 | -0.58 | -0.4 | 0.48 | 0.1 | -0.27 | 0.0025 | 0.0725 | 0.005 | 0.087028 | 0.963133 | 0.134107 | 0.999187 | 0.997988 | 0.994666 |
| Dync1i1_O88485_S80s_1 | Dync1i1_O | -0.49 | -0.23 | -0.24 | -0.15 | -0.03 | -0.54 | 0.06 | -0.06 | -0.65 | -0.26 | -0.57 | -0.37 | -0.33 | 0.11 | -0.47 | -0.23 | 0.015 | 0.56 | 0.615 | 0.851219 | 0.560115 | 0.416041 | 0.989223 | 0.997988 | 0.999507 |
| Dync1i2_Q3TPJ8_S81s_1 | Dync1i2_Q | -0.32 | 0.19 | -0.13 | -0.04 | 0.6 | 0.09 | 0.28 | 0.11 | 0.02 | 0.09 | -0.35 | 0.04 | -0.19 | 0.4 | -0.2 | -0.33 | 0.135 | -0.185 | 0.0475 | 0.98555 | 0.16042 | 0.915571 | 0.989223 | 0.997988 | 1 |
| Dync1i2_Q3TPJ8_S95s_2 | Dync1i2_Q | -1.01 | 0.14 | 0.26 | 0.41 | -0.35 | -0.56 | -0.21 | 0.73 | -0.36 | 0.3 | 1.81 | 0 | 0.78 | -0.71 | 0.45 | 0.45 | 0.345 | 0.025 | -0.005 | 0.519319 | 0.90819 | 0.374932 | 0.997335 | 0.997988 | 0.994666 |
| Dync1i2_Q3TPJ8_S98s_1 | Dync1i2_Q | -0.17 | -1.05 | -0.2 | -0.38 | 0.03 | -0.37 | -0.09 | -0.14 | 1.18 | 0.52 | 1.1 | -0.09 | 0.68 | 0.43 | 1.22 | -0.48 | -0.0475 | 0.4875 | 0.2925 | 0.048798 | 0.265475 | 0.012355 | 0.989223 | 0.997988 | 0.994666 |
| Dync1i2_Q3TPJ8_S98s_2 | Dync1i2_Q | -1.01 | 0.14 | 0.08 | 0.12 | -0.35 | -0.56 | -0.47 | 0.39 | -0.36 | 0.3 | 1.34 | -0.53 | 0.78 | -0.71 | 0.34 | 0.58 | 0.3075 | 1.1275 | 0.9125 | 0.346333 | 0.821841 | 0.468325 | 0.995762 | 0.997988 | 0.994666 |
| Dync1i2_Q3TPJ8_T174t_1 | Dync1i2_Q | -0.89 | -1.53 | -0.28 | -0.6 | 0.16 | 0.05 | -0.01 | 0.52 | -1.54 | -0.73 | 0.81 | -0.99 | -1.3 | 0.03 | -0.07 | 0.27 | -0.08 | 0.355 | 0.415 | 0.219152 | 0.008958 | 0.689879 | 0.989223 | 0.997988 | 0.994666 |
| Dync1i2_Q3TPJ8_T89t_1 | Dync1i2_Q | -0.2 | -0.68 | 0.68 | 0.15 | 0.14 | 0.11 | -0.4 | 0.65 | 0.66 | -0.51 | 0.82 | 0.89 | 0.65 | 0.5 | 0.39 | -0.3 | 1.005 | 0.2125 | 0.5575 | 0.397603 | 0.701316 | 0.278906 | 0.99521 | 0.997988 | 0.994666 |
| Dync1li1_Q8R1Q8_S207s_1 | Dync1li1_C | 0.48 | 0.44 | 0.85 | 0.97 | 0.39 | 0.46 | 1.22 | 1.19 | 0.24 | 0.61 | 0.85 | 0.97 | -0.94 | 0.51 | 0.84 | 0.84 | 0.1375 | 0.4775 | 0.3225 | 0.403343 | 0.656784 | 0.94794 | 0.990434 | 0.997988 | 0.994666 |
| Dync1li1_Q8R1Q8_S405s_1 | Dync1li1_C | -3.2 | -2.75 | -3.17 | -4.05 | -2.21 | -3.15 | -3.44 | -3.43 | -0.68 | -2.15 | -4.86 | -3.62 | -0.27 | -3.9 | -2.44 | -2.08 | 0.13 | -0.0175 | -0.3725 | 0.139235 | 0.547409 | 0.583575 | 0.989223 | 0.997988 | 0.994666 |
| Dync1li1_Q8R1Q8_S421s_1 | Dync1li1_C | -0.91 | -1.27 | -0.74 | 0.21 | -0.41 | -0.79 | 0.28 | -0.26 | -1.43 | -1.28 | 0.26 | -1.42 | -1.3 | -0.3 | 0.41 | 0.19 | 0.235 | 0.465 | 1.12 | 0.377587 | 0.324666 | 0.557397 | 0.989223 | 0.997988 | 0.994666 |
| Dync1li1_Q8R1Q8_S516s_1 | Dync1li1_C | -0.29 | 0.4 | 0.98 | 0.83 | 0.08 | -0.32 | 0.45 | 0.82 | 0.11 | 0.16 | 1.17 | 0.73 | 0.86 | 0.52 | 0.5 | 0.14 | 0.3825 | -0.29 | 0.4275 | 0.943034 | 0.551266 | 0.870435 | 0.989223 | 0.997988 | 0.999507 |
| Dync1li1_Q8R1Q8_T512t_1 | Dync1li1_C | -0.74 | 0.01 | 0.06 | 0.21 | 0.16 | -0.48 | 0.58 | -0.2 | -0.1 | -0.21 | 0 | -0.53 | 0.69 | 0.05 | -0.11 | 0.13 | -0.2225 | 0.0625 | 0.025 | 0.354272 | 0.69042 | 0.742201 | 0.994672 | 0.997988 | 0.994666 |
| Dync1li2_Q6PDL0_S194s_1 | Dync1li2_C | 0.24 | 0.33 | 0.13 | 0.02 | 0.47 | 0.08 | 0.44 | 0.2 | -0.24 | 0.5 | -0.41 | 0.08 | -0.09 | -0.19 | 0.24 | 0.31 | 0.13 | -0.095 | 0.305 | 0.656258 | 0.584392 | 0.470819 | 0.989223 | 0.997988 | 0.995259 |
| Dync1li2_Q6PDL0_S407s_1 | Dync1li2_C | -1.7 | -0.68 | -0.41 | -0.73 | -1.1 | -1.13 | -0.7 | -1.16 | -0.69 | -0.5 | 0.54 | -0.23 | -0.3 | -0.97 | 0.27 | 0.02 | 0.1175 | -0.1975 | -0.1125 | 0.134577 | 0.655603 | 0.117892 | 0.990434 | 0.997988 | 0.994666 |
| Dync1li2_Q6PDL0_S487s_1 | Dync1li2_C | 0.16 | -0.15 | 0.03 | 0.01 | 0.36 | 0.37 | -0.57 | -0.08 | 0.25 | 0.43 | -0.06 | 0.17 | 0.56 | 0.36 | -0.34 | -0.43 | -0.1425 | 0.66 | 0.635 | 0.93569 | 0.978016 | 0.419065 | 0.999187 | 0.997988 | 0.999507 |
| Dync1li2_Q6PDL0_T376t_1 | Dync1li2_C | -3.21 | -3.16 | -2.73 | -4.36 | -2.21 | -3.13 | -3.96 | -3.73 | -0.74 | -2.79 | -3.57 | -2.65 | -0.04 | -3.58 | -2.99 | -2.36 | 0.0075 | 0.185 | 0.025 | 0.160612 | 0.823817 | 0.166918 | 0.995762 | 0.997988 | 0.994666 |
| Dyrk1a_F6U6X3_S491s_1 | Dyrk1a_F6 | -0.57 | -0.84 | -0.63 | 0.06 | -1.55 | -0.73 | 0.14 | -1.58 | 1.27 | -0.17 | -2.99 | 0.3 | 1.96 | -2.2 | 1.13 | 0.46 | 0.1075 | 0.9275 | 1.1225 | 0.319078 | 0.321672 | 0.906733 | 0.989223 | 0.997988 | 0.994666 |
| Dyrk1a_F6U6X3_S720s_1 | Dyrk1a_F6 | 0.8 | 1.18 | -0.64 | 0.57 | 1.28 | 1.46 | 0.26 | 0.53 | 1.32 | 0.95 | -0.38 | 0.1 | -0.05 | 0.79 | 0.42 | 0.37 | -0.435 | 0.0975 | 0.8325 | 0.823411 | 0.383707 | 0.969117 | 0.989223 | 0.997988 | 0.999507 |
| Dyrk1a_F6U6X3_Y283y_1 | Dyrk1a_F6 | 0.3 | 0.62 | 0.07 | 0.06 | 0.41 | 0.1 | -0.03 | -0.1 | -0.18 | 0.29 | -0.25 | 0 | 0.28 | -0.41 | -0.01 | -0.16 | 0.405 | 0.02 | -0.095 | 0.241137 | 0.489733 | 0.255498 | 0.989223 | 0.997988 | 0.994666 |
| Dyrk4_Q8BI55_Y344y_1 | Dyrk4_Q8B | 0.5 | 0.31 | -1.15 | 0.18 | 0.56 | 0.87 | -0.03 | 0.66 | 0.35 | 0.18 | 0.61 | 0.49 | -0.06 | 0.38 | 0.25 | 0.11 | -0.1675 | -0.2975 | -0.3375 | 0.598283 | 0.192301 | 0.267 | 0.989223 | 0.997988 | 0.995259 |
| Dzank1_E9Q4Z6_S769s_1 | Dzank1_E9 | -0.03 | 0.15 | 0.65 | 0.44 | 0.56 | 0.28 | 0.57 | 0.43 | -0.24 | 0.33 | -0.04 | 0.3 | 0.58 | 0.06 | 0.17 | 0.5 | 0.555 | 0.4475 | 0.21 | 0.928123 | 0.508542 | 0.427236 | 0.989223 | 0.997988 | 0.999507 |
| Dzip3_F6Z1P8_S98s_1 | Dzip3_F6Z1 | -1.65 | -1.22 | -1.96 | -0.54 | -1.79 | -2.05 | -1.16 | -1.06 | -2.03 | -1.4 | 0.04 | -0.47 | -0.18 | -0.63 | -0.3 | -1.22 | 0.1575 | -0.215 | 0.025 | 0.079073 | 0.652377 | 0.474483 | 0.990358 | 0.997988 | 0.994666 |
| E030010N08Rik_F6XLV1_T56t_1 | E030010NC | 0.2 | 1.21 | 1.14 | 0.71 | 0.82 | 0.04 | 0.72 | 0.24 | 0.64 | -0.1 | 0.85 | 0.39 | 1.16 | -0.31 | 1.16 | 1.34 | -0.1725 | 0.3775 | 0.76 | 0.95924 | 0.271029 | 0.282979 | 0.989223 | 0.997988 | 0.999507 |
| E9Q0B3_S233s_1 | E9Q0B3_S2 | 0.93 | -0.98 | 1.04 | 0.61 | -0.51 | 1.38 | -0.24 | 0.94 | 1.77 | -0.78 | 2.16 | 1.25 | 0.78 | -0.57 | 0.16 | 0.16 | -0.36 | -0.37 | 0.0225 | 0.603728 | 0.989779 | 0.346135 | 0.999187 | 0.997988 | 0.995259 |
| Ebag9_Q9D0V7_S36s_1 | Ebag9_Q9D | 0.04 | 0.23 | -0.29 | -0.5 | 0.41 | 0.2 | 0.48 | -0.16 | -0.24 | 0.35 | 0.21 | -0.31 | -0.25 | 0.61 | 0.07 | -0.06 | -0.0075 | 0.7 | -0.2675 | 0.471471 | 0.191543 | 0.639974 | 0.989223 | 0.997988 | 0.994666 |
| Ece2_E9Q1S5_S27s_1 | Ece2_E9Q1 | 0.83 | 0.86 | 0.08 | 0.1 | 0.35 | 0.39 | 0.16 | -0.03 | 0.33 | 1.08 | -0.15 | -0.37 | 0.09 | 0.31 | 0.72 | 0.19 | 0.3625 | 0.1325 | 0.2225 | 0.6546 | 0.376596 | 0.533494 | 0.989223 | 0.997988 | 0.995259 |
| Edc3_Q8K2D3_S131s_1 | Edc3_Q8K2 | 0.21 | 0.46 | -0.06 | -0.13 | 0.15 | 0.31 | 0.05 | 0.42 | 0.43 | 0.81 | 0.07 | 0.11 | 0.76 | -1.1 | 0.33 | 0.05 | -0.25 | -0.245 | -0.14 | 0.793084 | 0.629336 | 0.400912 | 0.989223 | 0.997988 | 0.999507 |
| Edc4_G5E896_S6s_1 | Edc4_G5E8 | -0.39 | -0.15 | 0.06 | 0.19 | -0.12 | 0.14 | 0.41 | 0.02 | 0.01 | 0.84 | -0.28 | -0.04 | -0.44 | 0.4 | -0.43 | 0.11 | 0.1125 | 0.235 | -0.11 | 0.953871 | 0.443432 | 0.512905 | 0.989223 | 0.997988 | 0.999507 |
| Edc4_G5E896_S727s_1 | Edc4_G5E8 | 1.51 | 0.84 | 0.14 | 1.27 | 1.35 | -0.13 | -0.91 | -0.04 | 0.01 | -0.09 | 0.23 | -0.19 | 0.9 | 0.9 | 0.78 | 0.04 | 0.185 | 0.205 | -0.0175 | 0.460091 | 0.112404 | 0.017895 | 0.989223 | 0.997988 | 0.994666 |
| Eef1a1_P10126_T176t_1 | Eef1a1_P1( | 0.03 | 0.02 | 0.11 | 0.58 | -0.24 | -0.36 | 0.36 | -0.14 | -0.08 | 0.5 | 0.35 | -0.07 | -0.34 | 0.33 | 0.49 | 0.06 | -0.8725 | -0.95 | -0.285 | 0.864241 | 0.292568 | 0.969615 | 0.989223 | 0.997988 | 0.999507 |
| Eef1b2_O70251_S82s_1 | Eef1b2_O7 | -0.19 | -0.31 | 0.17 | -0.54 | 0.47 | -0.19 | -0.22 | -0.14 | -1.59 | -0.32 | -0.73 | -0.52 | -0.71 | 0.39 | -0.52 | -0.82 | -0.28 | -0.01 | -0.05 | 0.570533 | 0.464518 | 0.115788 | 0.989223 | 0.997988 | 0.995259 |
| Eef1b2_O70251_S83s_1 | Eef1b2_O7 | -3.17 | -2.75 | -1.54 | -1.43 | -3.58 | -4.63 | -0.99 | -0.89 | -3.14 | -2 | -0.72 | -0.9 | -2.58 | -1.57 | -1.83 | -1.06 | 0.1975 | -0.5725 | -0.1975 | 0.372604 | 0.739023 | 0.417638 | 0.995762 | 0.997988 | 0.994666 |
| Eef1d_D3YZT9_S152s_1 | Eef1d_D3Y | -2.17 | -2.12 | -1.26 | -1.57 | -1.71 | -2.52 | -1.39 | -1.78 | -1.95 | -1.78 | -0.34 | -1.21 | -0.56 | -1.4 | -1.02 | -1.67 | -0.3 | 0.5325 | 0.4625 | 0.106652 | 0.833893 | 0.286286 | 0.995762 | 0.997988 | 0.994666 |
| Eef1d_D3YZT9_S71s_1 | Eef1d_D3Y | 0.58 | 0.15 | 0.27 | 0.5 | 0.78 | 0.25 | 0.72 | 0.07 | 0.25 | 0.57 | 0.43 | 0.55 | -0.21 | 0.84 | -0.07 | 0.44 | -0.07 | 0.46 | 0.6175 | 0.690738 | 0.753659 | 0.740852 | 0.995762 | 0.997988 | 0.996159 |
| Eef1d_D3YZT9_T166t_1 | Eef1d_D3Y | -5.69 | -3.73 | -4.26 | -5.93 | -3.93 | -4.31 | -7.31 | -3.96 | -5.25 | -4.58 | -2.43 | -5.49 | -2.31 | -4.78 | -4.56 | -4.12 | 0.08 | 0.075 | -0.125 | 0.189522 | 0.976521 | 0.557949 | 0.999187 | 0.997988 | 0.994666 |
| Eef2_P58252_T57t_1 | Eef2_P582 | -2.3 | -3.06 | -1.36 | -0.6 | -1.95 | -2.65 | -1.01 | 0.22 | 0.61 | -0.91 | -0.84 | -0.99 | -1.14 | -2.1 | -1.3 | -0.62 | 0.025 | 0.465 | 0.96 | 0.355007 | 0.509319 | 0.055613 | 0.989223 | 0.997988 | 0.994666 |
| Eef2k_F6ZEU2_S42s_1 | Eef2k_F6ZE | 5.33 | 1.97 | 3.41 | 0.89 | 1.43 | 5.5 | 3.01 | 2.92 | 1.05 | 5.41 | 0.24 | 4.06 | 0.41 | -2.79 | 4.79 | 4.88 | 0.4825 | 1.2975 | 0.54 | 0.550914 | 0.775246 | 0.877286 | 0.995762 | 0.997988 | 0.994666 |
| Eefsec_Q9JHW4_T532t_1 | Eefsec_Q9J | -2.04 | -2 | -2.18 | -2.62 | -1.04 | -2.79 | -1.54 | -2.13 | -2.52 | -1.55 | -1.05 | -1.73 | -1.42 | -1.33 | -1.37 | -1.05 | 0.315 | -0.21 | -1.0775 | 0.005806 | 0.400668 | 0.180314 | 0.989223 | 0.997988 | 0.994666 |
| Eepd1_Q3TGW2_S106s_2 | Eepd1_Q3T | -1.33 | -1.15 | -0.09 | 0.25 | -1.18 | -1.02 | -0.86 | 0.3 | 0.97 | -1.52 | 0.25 | -0.08 | 0.54 | -0.6 | -0.22 | -0.48 | 0.335 | 0.4975 | 0.9175 | 0.398295 | 0.817119 | 0.425627 | 0.995762 | 0.997988 | 0.994666 |
| Eepd1_Q3TGW2_S110s_1 | Eepd1_Q3T | -0.55 | 0.41 | 0.51 | -0.18 | -0.5 | -0.85 | -0.39 | -0.49 | 0.28 | -0.27 | -1.36 | -0.75 | 1.16 | 0.43 | 0.06 | -0.47 | -0.11 | 0.485 | 0.39 | 0.561118 | 0.065478 | 0.196063 | 0.989223 | 0.997988 | 0.995259 |
| Eepd1_Q3TGW2_S16s_1 | Eepd1_Q3T | 1.85 | 2.71 | 6.43 | 4.87 | 2.28 | 2.82 | 4.53 | 5.5 | 0.64 | 2.61 | 2.34 | 6.31 | 1.24 | -0.96 | 5.09 | 5.6 | -0.605 | -0.5725 | 0.2475 | 0.455993 | 0.868499 | 0.480599 | 0.996493 | 0.997988 | 0.994666 |
| Eepd1_Q3TGW2_S25s_1 | Eepd1_Q3T | 0.38 | -0.99 | -2.04 | -0.77 | 0.19 | 1.23 | -1.03 | -0.94 | -0.55 | -0.85 | -0.67 | -1.73 | -0.84 | -0.86 | -2.12 | -1.36 | -0.1825 | -0.99 | -1.2225 | 0.423071 | 0.283344 | 0.856652 | 0.989223 | 0.997988 | 0.994666 |
| Efhd2_Q8C845_S74s_1 | Efhd2_Q8C | -0.85 | -1.74 | -1.51 | -0.82 | -1.48 | -0.8 | -1.22 | -1.18 | -1.34 | -1.06 | -0.67 | -0.93 | -1.38 | -1.1 | -1.52 | -1.51 | 0.7175 | -0.095 | -0.44 | 0.634045 | 0.840283 | 0.460877 | 0.995762 | 0.997988 | 0.995259 |
| Efhd2_Q8C845_S76s_1 | Efhd2_Q8C | -3.15 | -3.64 | -0.92 | -0.44 | -2.1 | -3.61 | -0.72 | -1.14 | -3.29 | -2.59 | -0.29 | -0.75 | -3.59 | -1.91 | -1.21 | -0.84 | 0.06 | 0.23 | -0.1475 | 0.864519 | 0.870299 | 0.746476 | 0.996493 | 0.997988 | 0.999507 |
| Efnb1_P52795_S280s_1 | Efnb1_P52 | 0.1 | 0.17 | 0.46 | 0.97 | 0.23 | -0.25 | 0.79 | 0.55 | -0.49 | -0.38 | 0.03 | 0.12 | 0.01 | 0.22 | 0.24 | 0.46 | 0.145 | 0.3075 | 0.15 | 0.509789 | 0.764055 | 0.062986 | 0.995762 | 0.997988 | 0.994666 |
| Efnb1_P52795_S286s_1 | Efnb1_P52 | 0.82 | 0.63 | 0.86 | 0.78 | 0.88 | 0.39 | 1.18 | 1.17 | 0.01 | 0.46 | 0.38 | 0.82 | -0.47 | 0.69 | 0.85 | 0.81 | -0.095 | -0.605 | -0.1925 | 0.396217 | 0.59774 | 0.177709 | 0.989223 | 0.997988 | 0.994666 |
| Efnb1_P52795_S337s_1 | Efnb1_P52 | 0.28 | 0.09 | -0.74 | -0.31 | 0.56 | 0.63 | 0.53 | 0.56 | 0.14 | -0.3 | 1 | 0.5 | -0.36 | 0.03 | -1.21 | 0.1 | 0.1325 | -0.355 | -0.3025 | 0.626452 | 0.020123 | 0.190298 | 0.989223 | 0.997988 | 0.995259 |
| Efnb3_O35393_S268s_1 | Efnb3_O35 | 7.05 | 2.53 | 5.32 | 1.87 | 4.6 | 7.22 | 5.81 | 4.09 | 0.04 | 5.68 | -0.46 | 3.78 | -0.22 | -0.13 | 5.19 | 6.15 | 0.74 | 0.505 | -0.19 | 0.426483 | 0.317027 | 0.26441 | 0.989223 | 0.997988 | 0.994666 |
| Efnb3_O35393_S274s_1 | Efnb3_O35 | 0.63 | 0.41 | 0.83 | 0.69 | 0.42 | 0.96 | 0.96 | 1.14 | 0.73 | 0.9 | 1.25 | 1.05 | 0.47 | 0.85 | 0.82 | 0.94 | 1.2375 | -1.9325 | -1.445 | 0.605294 | 0.354071 | 0.166454 | 0.989223 | 0.997988 | 0.995259 |
| Efnb3_O35393_S332s_1 | Efnb3_O35 | 0.33 | 0.76 | -0.03 | -0.29 | 0.74 | 0.35 | 0.42 | 0.12 | 0.3 | 0.58 | -0.79 | -0.44 | 0.49 | 0.57 | 0.36 | 0.57 | 0.23 | 0.3425 | 0.13 | 0.318221 | 0.466514 | 0.480822 | 0.989223 | 0.997988 | 0.994666 |
| Efr3a_Q8BG67_S220s_1 | Efr3a_Q8B | 0.15 | 0.49 | 0.9 | 1.11 | 0.03 | -0.44 | 1.59 | -0.17 | -1.32 | 0.53 | 2.2 | -0.54 | -1.73 | 0.2 | 1.47 | 0.18 | 0.215 | -0.28 | 0.305 | 0.327739 | 0.390169 | 0.536672 | 0.989223 | 0.997988 | 0.994666 |
| Efr3b_Q6ZQ18_S214s_1 | Efr3b_Q6Z | 0.21 | 0.81 | 0.6 | 0.72 | 0.58 | 0.31 | 0.41 | 0.63 | 0.36 | 0.52 | 1.04 | 0.57 | 0.59 | 1.21 | 0.51 | 0.43 | -0.41 | -0.445 | -0.6325 | 0.731413 | 0.655488 | 0.886075 | 0.990434 | 0.997988 | 0.9971 |
| Efr3b_Q6ZQ18_S216s_1 | Efr3b_Q6Z | 0.14 | 1.07 | 0.49 | 0.78 | 0.85 | 0.4 | 0.61 | 0.85 | 0.51 | 0.68 | 0.97 | 1.05 | 0.74 | 1.32 | 1.04 | 0.75 | -0.1025 | 0.0375 | 0.1 | 0.273388 | 0.831006 | 0.525601 | 0.995762 | 0.997988 | 0.994666 |
| Eftud1_Q8C0D5_S474s_1 | Eftud1_Q8 | -0.16 | 0.42 | 0.52 | 0.1 | -0.01 | 0.57 | 0.55 | 0.58 | 0.26 | 0.19 | -0.42 | 0.62 | -0.33 | 0.58 | -0.18 | 0.04 | 0.0575 | 0.1825 | 0.3425 | 0.5362 | 0.444569 | 0.849702 | 0.989223 | 0.997988 | 0.994666 |
| Eftud1_Q8C0D5_S59s_1 | Eftud1_Q8 | 0.62 | 0.39 | 1.02 | 0.56 | 0.08 | 0.9 | -0.41 | 0.86 | 0.6 | 0.32 | -0.34 | 1.2 | 0.19 | -0.28 | 0.38 | 0.68 | 0.2025 | -0.0575 | -0.1925 | 0.200565 | 0.413213 | 0.575466 | 0.989223 | 0.997988 | 0.994666 |
| Ehbp1_Q69ZW3_S1022s_1 | Ehbp1_Q6 | -1.35 | -1.31 | -1.38 | -1.32 | -1.04 | -0.89 | -0.86 | -0.9 | -2.45 | -0.89 | 0.22 | -1.23 | -0.97 | 0.34 | -1.06 | -1.92 | -0.29 | -0.2025 | -0.405 | 0.346514 | 0.053067 | 0.626063 | 0.989223 | 0.997988 | 0.994666 |
| Ehbp1_Q69ZW3_S1038s_1 | Ehbp1_Q6 | -0.06 | -0.16 | -0.07 | 0.08 | -0.38 | -0.33 | -0.61 | -0.22 | 1.03 | -0.27 | -0.22 | -0.14 | 2.01 | -0.41 | 0.07 | -0.47 | 0.4175 | 0.2525 | 0.4375 | 0.521018 | 0.131233 | 0.652056 | 0.989223 | 0.997988 | 0.994666 |
| Ehbp1_Q69ZW3_S335s_1 | Ehbp1_Q6 | -0.66 | -0.59 | -0.16 | -2.24 | -0.67 | -0.4 | -0.69 | -1.4 | 0.12 | -0.29 | -1.58 | -0.46 | 0.66 | -3.16 | 0.65 | -0.26 | -0.3325 | 0.1525 | 0.3525 | 0.665741 | 0.793347 | 0.511545 | 0.995762 | 0.997988 | 0.996159 |
| Ehbp1_Q69ZW3_S588s_1 | Ehbp1_Q6 | 0.53 | -0.31 | -0.55 | 0.44 | 0.12 | 1.17 | 0.33 | 0.54 | -0.09 | 0.45 | -0.61 | -0.44 | 0.04 | 0.95 | 0.33 | 0.24 | 0.1225 | 0.36 | 0.385 | 0.32551 | 0.169791 | 0.587815 | 0.989223 | 0.997988 | 0.994666 |
| Ehbp1_Q69ZW3_S764s_1 | Ehbp1_Q6 | -0.11 | -1.5 | -1.24 | -1.51 | -0.39 | 0.01 | -1.07 | -1.08 | -0.33 | -0.37 | -1.52 | -1.04 | -1.21 | 0.09 | -1.38 | -1.13 | 0.5125 | -0.2 | 0.3625 | 0.691416 | 0.2791 | 0.524852 | 0.989223 | 0.997988 | 0.996159 |
| Ehbp1_Q69ZW3_S765s_1 | Ehbp1_Q6 | 0.15 | -0.37 | -0.24 | 0.07 | 0.59 | 0.46 | 0.03 | 0.19 | 0.11 | -0.08 | -0.75 | 0.11 | -0.24 | 0.8 | -0.21 | -0.36 | 0.4575 | 0.275 | 0.1825 | 0.776935 | 0.111008 | 0.84768 | 0.989223 | 0.997988 | 0.99798 |
| Ehbp1l1_G5E8Y6_S502s_1 | Ehbp1l1_G | 0.21 | 0.6 | -0.01 | 0.23 | 0.08 | 0.04 | -0.17 | -0.07 | 0.47 | 0.13 | -0.13 | 0.07 | 1.25 | 1.16 | -0.06 | -0.11 | 0.415 | -0.055 | 0.095 | 0.456709 | 0.216664 | 0.628084 | 0.989223 | 0.997988 | 0.994666 |
| Ehbp1l1_G5E8Y6_S518s_1 | Ehbp1l1_G | 0.63 | -1.27 | 0.29 | -1.93 | -0.34 | 2.05 | 1.5 | 1 | -1.67 | 0.75 | -1.72 | 0.42 | 1.37 | -3.43 | 1.44 | 1.33 | -0.2875 | -0.1225 | 0.3025 | 0.526504 | 0.042195 | 0.985125 | 0.989223 | 0.997988 | 0.994666 |
| Ehd1_Q9WVK4_S456s_1 | Ehd1_Q9W | 0.13 | 0.17 | -0.13 | 0.33 | 0.51 | 0.13 | 0.47 | 0.41 | -0.25 | 0.37 | 0.03 | 0 | -0.34 | 0.6 | 0.06 | 0.26 | 1.6225 | 0.015 | 0.7475 | 0.944732 | 0.259911 | 0.718457 | 0.989223 | 0.997988 | 0.999507 |
| Ehd3_Q9QXY6_S456s_1 | Ehd3_Q9Q | 0.47 | 0.89 | 0.51 | 0.72 | 0.97 | 0.37 | 1.07 | 0.57 | 0.3 | 0.71 | 0.16 | 0.57 | -1.29 | 0.71 | 0.58 | 0.7 | 0.255 | -0.0875 | 0.02 | 0.335211 | 0.696739 | 0.38976 | 0.994893 | 0.997988 | 0.994666 |
| Ehd3_Q9QXY6_S479s_1 | Ehd3_Q9Q | -1.09 | 0.52 | -0.06 | -0.26 | 0.21 | -1.04 | -0.7 | -0.92 | -1.23 | -0.59 | -0.96 | -0.64 | -0.43 | 0.29 | -0.31 | -0.49 | 0.0975 | -0.2125 | -0.4725 | 0.974353 | 0.360045 | 0.115527 | 0.989223 | 0.997988 | 0.999507 |
| Ei24_Q61070_S330s_1 | Ei24_Q610 | -1.58 | -1.83 | -0.2 | -0.78 | -1.39 | -1.97 | 0.28 | -1.09 | -1.36 | -1.47 | -1.53 | -0.63 | 0.45 | 0.39 | 0.08 | -0.97 | -0.39 | -0.6325 | -0.0125 | 0.044306 | 0.919977 | 0.721424 | 0.997335 | 0.997988 | 0.994666 |
| Eif2b4_D3Z780_S63s_1 | Eif2b4_D3Z | -1.04 | -1.72 | -0.72 | -0.26 | -0.82 | -0.79 | -0.04 | -0.18 | -0.24 | -0.39 | 1.53 | 0.67 | -0.02 | 0.15 | 1.01 | 0.17 | 0.055 | -0.15 | 1.085 | 0.009359 | 0.210722 | 0.025838 | 0.989223 | 0.997988 | 0.994666 |
| Eif2b5_Q8CHW4_S540s_1 | Eif2b5_Q8C | -0.04 | 0.03 | 0.24 | -0.04 | -0.23 | -0.57 | -0.33 | -0.4 | -0.39 | 0.02 | 0.08 | -0.08 | -0.18 | -0.11 | -0.58 | -0.3 | 0.4775 | 1.3275 | 1.2625 | 0.187643 | 0.060978 | 0.540215 | 0.989223 | 0.997988 | 0.994666 |
| Eif2s2_Q99L45_S2s_1 | Eif2s2_Q99 | -0.3 | -0.08 | -0.54 | -0.5 | 0.09 | -0.31 | -0.11 | -0.48 | -0.31 | -0.09 | -0.56 | -0.68 | -0.37 | 0.08 | -0.57 | -0.38 | -0.43 | -0.14 | -0.34 | 0.865776 | 0.519056 | 0.823715 | 0.989223 | 0.997988 | 0.999507 |
| Eif3a_P23116_S1159s_1 | Eif3a_P231 | -4.22 | -5.51 | -3.44 | -2.31 | -5.09 | -3.44 | -2.22 | -1.82 | -2.96 | -5.17 | -0.15 | -1.2 | -0.46 | -5.01 | -0.41 | -2.16 | 0.1525 | -0.055 | 0.045 | 0.116949 | 0.410301 | 0.206982 | 0.989223 | 0.997988 | 0.994666 |
| Eif3a_P23116_S1300s_1 | Eif3a_P231 | -2.97 | -3.51 | -3.7 | -4.65 | -1.6 | -3.62 | -2.87 | -3.99 | -3.39 | -3.57 | -1.36 | -2.82 | -2.2 | -1.59 | -2.62 | -2.87 | 0.7275 | 1.5 | 1.86 | 0.010217 | 0.245898 | 0.128282 | 0.989223 | 0.997988 | 0.994666 |
| Eif3a_P23116_T574t_1 | Eif3a_P231 | -3.52 | -2.06 | -2.49 | -3.09 | -4.11 | -4.11 | -3.36 | -3.48 | -4.76 | -3.51 | -3.37 | -3.43 | -0.94 | -3.4 | -2.69 | -2.71 | 0.6875 | 0.9225 | 1.3875 | 0.535802 | 0.025162 | 0.051926 | 0.989223 | 0.997988 | 0.994666 |
| Eif3a_P23116_T982t_1 | Eif3a_P231 | -1.72 | -1.24 | -1.92 | -1.73 | -1.04 | -1.16 | -1.98 | -2.69 | -0.67 | -1.06 | -1.04 | -1.96 | 0.34 | -0.98 | -1.88 | -2.07 | -0.975 | -0.9775 | 0.355 | 0.3532 | 0.869921 | 0.181418 | 0.996493 | 0.997988 | 0.994666 |
| Eif3b_Q8JZQ9_S111s_1 | Eif3b_Q8JZ | -2.01 | -1.35 | -1.21 | -0.75 | -1.83 | -1.1 | -0.45 | -0.31 | -1.43 | -1.39 | -0.87 | 0.12 | -0.12 | -0.97 | 0.14 | 0.13 | -0.065 | 0.47 | 0.505 | 0.014612 | 0.337875 | 0.325702 | 0.989223 | 0.997988 | 0.994666 |
| Eif3b_Q8JZQ9_S120s_1 | Eif3b_Q8JZ | -1.27 | -0.87 | -0.61 | -0.48 | -1.25 | 0.02 | -0.33 | -0.29 | 1.28 | -0.8 | 0 | -0.63 | 1.41 | 1.35 | -0.22 | -0.39 | 0.4075 | 0.4375 | 1.125 | 0.020557 | 0.314959 | 0.132795 | 0.989223 | 0.997988 | 0.994666 |
| Eif3b_Q8JZQ9_S120s_2 | Eif3b_Q8JZ | -1.55 | -1.24 | -1.39 | -1.05 | -1.3 | -1.51 | -1.07 | -1.32 | -0.77 | -1.08 | -0.55 | -1.2 | -0.74 | -0.8 | -0.46 | -0.93 | 0.345 | 0.77 | 1.345 | 0.044968 | 0.973154 | 0.133825 | 0.999187 | 0.997988 | 0.994666 |
| Eif3b_Q8JZQ9_S123s_2 | Eif3b_Q8JZ | -1.55 | -1.24 | -1.39 | -1.05 | -1.3 | -1.51 | -1.07 | -1.32 | -0.77 | -1.08 | -0.55 | -1.2 | -0.74 | -0.8 | -0.46 | -0.93 | 0.0075 | 0.4075 | 0.575 | 0.044968 | 0.973154 | 0.133825 | 0.999187 | 0.997988 | 0.994666 |
| Eif3b_Q8JZQ9_S75s_1 | Eif3b_Q8JZ | -1.08 | -0.56 | -1.14 | -0.82 | -0.04 | -1.53 | -0.15 | -0.75 | 0.16 | -0.13 | -0.17 | -0.78 | 0.42 | -0.22 | 0.01 | -0.53 | 0.0075 | 0.4075 | 0.575 | 0.020734 | 0.444723 | 0.040089 | 0.989223 | 0.997988 | 0.994666 |
| Eif3b_Q8JZQ9_S75s_2 | Eif3b_Q8JZ | -0.86 | -0.77 | 0.15 | 0.35 | -0.77 | -1.05 | -0.12 | -0.11 | -0.33 | -0.42 | 0.28 | -0.3 | -0.13 | 0.16 | -0.02 | 0.5 | 0.2825 | 0.67 | 0.82 | 0.275237 | 0.54992 | 0.803084 | 0.989223 | 0.997988 | 0.994666 |
| Eif3b_Q8JZQ9_S79s_2 | Eif3b_Q8JZ | -0.86 | -0.77 | 0.15 | 0.35 | -0.77 | -1.05 | -0.12 | -0.11 | -0.33 | -0.42 | 0.28 | -0.3 | -0.13 | 0.16 | -0.02 | 0.5 | -0.23 | 0.09 | 0.41 | 0.275237 | 0.54992 | 0.803084 | 0.989223 | 0.997988 | 0.994666 |
| Eif3b_Q8JZQ9_S88s_1 | Eif3b_Q8JZ | -1.11 | -0.12 | -0.21 | -0.89 | -0.87 | -1.34 | -0.49 | 0.12 | -1.01 | -2.36 | -0.37 | -1.15 | -0.03 | -0.96 | 0.03 | -0.87 | -0.23 | 0.09 | 0.41 | 0.741247 | 0.870696 | 0.188767 | 0.996493 | 0.997988 | 0.9971 |
| Eif3b_Q8JZQ9_S90s_1 | Eif3b_Q8JZ | -0.21 | -0.25 | -0.39 | -0.54 | -0.68 | -0.7 | -0.91 | -1.12 | -0.84 | -0.52 | -0.73 | -0.42 | -0.39 | -1.16 | -0.75 | -0.33 | -0.0625 | -0.64 | 0.125 | 0.287347 | 0.041093 | 0.233832 | 0.989223 | 0.997988 | 0.994666 |
| Eif3c_Q8R1B4_S166s_1 | Eif3c_Q8R1 | -4.55 | -4.57 | -2.78 | -3.56 | -5.01 | -4.07 | -2.67 | -3.61 | -3.42 | -4.15 | -1.56 | -3.26 | -2.06 | -3.53 | -3.29 | -2.85 | -0.505 | -0.28 | -0.31 | 0.091965 | 0.965797 | 0.245923 | 0.999187 | 0.997988 | 0.994666 |
| Eif3c_Q8R1B4_S39s_1 | Eif3c_Q8R1 | -0.82 | -0.79 | -0.65 | -0.62 | -0.65 | -0.92 | -0.09 | -0.81 | -0.84 | -0.53 | -0.75 | -0.76 | -0.92 | -0.24 | -0.24 | -0.32 | 0.025 | 0.7675 | 0.9325 | 0.290031 | 0.681542 | 1 | 0.993613 | 1 | 0.994666 |
| Eif3e_P60229_S203s_1 | Eif3e_P602 | -0.7 | 2 | -0.14 | 0.86 | 2.05 | 0.42 | 0.27 | 0.51 | 0.8 | 0.11 | 0.26 | 0.21 | 0.71 | 0.97 | 0.07 | 1.47 | 0.1025 | 7.85E-17 | 0.29 | 0.621068 | 0.634614 | 0.777809 | 0.989223 | 0.997988 | 0.995259 |
| Eif3f_Q9DCH4_S262s_1 | Eif3f_Q9DC | -2.81 | -3.07 | -4.77 | -5.6 | -2.88 | -1.76 | -4.33 | -2.92 | -2.61 | -2.31 | -1.73 | -4.45 | -1.24 | -2.42 | -2.99 | -2.63 | 0.3075 | -0.16 | 0.3 | 0.029337 | 0.168801 | 0.132174 | 0.989223 | 0.997988 | 0.994666 |
| Eif3g_Q9Z1D1_S42s_1 | Eif3g_Q9Z1 | -1.06 | -1.67 | -0.82 | -1.14 | -0.61 | -1.48 | -0.2 | -0.79 | -1.18 | -1.16 | -0.26 | -0.71 | -0.18 | -0.68 | -0.72 | -0.68 | 1.09 | 1.2875 | 1.7425 | 0.059732 | 0.243029 | 0.28841 | 0.989223 | 0.997988 | 0.994666 |
| Eif3j1_Q3UGC7_T112t_1 | Eif3j1_Q3U | -0.23 | 0.25 | 0.56 | 0.08 | 0.27 | -0.13 | 1 | 0.67 | -0.26 | -0.1 | 0.27 | 0.65 | -0.19 | 0.06 | 0.75 | 0.88 | 0.4025 | 0.345 | 0.6075 | 0.543619 | 0.369677 | 0.933687 | 0.989223 | 0.997988 | 0.994666 |
| Eif4a2_Q8BTU6_S2s_1 | Eif4a2_Q8E | -2.14 | -0.85 | -0.68 | -0.47 | -3.71 | -1.39 | -1.37 | 0.46 | 1.45 | -1.98 | 0.38 | 0.31 | 1.39 | -1.2 | 0.97 | -1.27 | 0.2875 | -0.025 | 0.21 | 0.179542 | 0.569399 | 0.166097 | 0.989223 | 0.997988 | 0.994666 |
| Eif4a3_Q91VC3_S12s_1 | Eif4a3_Q91 | 0.15 | 1.71 | 0.95 | 0.65 | 1.1 | 2.15 | -1.08 | 0.15 | 2.26 | 0.94 | 0.73 | -0.16 | 2.01 | 2.3 | -0.16 | 0.76 | -0.4675 | 1.075 | 1.0075 | 0.549564 | 0.673365 | 0.888447 | 0.992876 | 0.997988 | 0.994666 |

| | | | | | | | | | | | | | | | | | | | | | | | | | | |
|---|---|---|---|---|---|---|---|---|---|---|---|---|---|---|---|---|---|---|---|---|---|---|---|---|---|---|
| Eif4b_Q8BGD9_S207s_1 | Eif4b_Q8B | -1.58 | -1.44 | -2.47 | -2.84 | -0.8 | -1.18 | -1.51 | -2.08 | -0.99 | -1.24 | -0.06 | -2.09 | -1.24 | -0.44 | -2.39 | -2.49 | -0.285 | 0.0775 | 0.3625 | 0.431244 | 0.121282 | 0.07694 | 0.989223 | 0.997988 | 0.994666 |
| Eif4b_Q8BGD9_S263s_1 | Eif4b_Q8B | 1.59 | 0.43 | 0.63 | 0.87 | 0.62 | 2.85 | 0.13 | 0.51 | 3.22 | 2.08 | 1.47 | 2.23 | 1.41 | 0.05 | 0.95 | 1.24 | 0.69 | 0.9875 | 0.4425 | 0.935321 | 0.804786 | 0.010256 | 0.995762 | 0.997988 | 0.999507 |
| Eif4b_Q8BGD9_S406s_1 | Eif4b_Q8B | 6.83 | -1.48 | 2.32 | -1.59 | -0.34 | 6.87 | 6.61 | -1.01 | 0.36 | 5.81 | -0.65 | 2.65 | 2.7 | -0.29 | 3.84 | 4.48 | 0.1475 | 1.37 | 0.0325 | 0.549903 | 0.547849 | 0.806465 | 0.989223 | 0.997988 | 0.994666 |
| Eif4b_Q8BGD9_S409s_1 | Eif4b_Q8B | -3.65 | -4.79 | -2.36 | -7.13 | -4.68 | -2.29 | -3.87 | -3.43 | -1.1 | -4.84 | -2.68 | -3.05 | -0.83 | -4.42 | -1.34 | -3.73 | 1.5125 | 0.5225 | 1.1625 | 0.12633 | 0.36037 | 0.184468 | 0.989223 | 0.997988 | 0.994666 |
| Eif4b_Q8BGD9_S422s_2 | Eif4b_Q8B | -0.65 | -1.47 | -0.74 | -1.5 | -1.91 | 0.12 | -1.91 | -0.19 | 1.3 | -0.5 | -0.77 | -2.04 | 0.7 | -0.86 | -0.79 | -1.35 | 0.915 | 1.565 | 1.9025 | 0.295693 | 0.82652 | 0.382583 | 0.995762 | 0.997988 | 0.994666 |
| Eif4b_Q8BGD9_S424s_1 | Eif4b_Q8B | -0.66 | 0.01 | 0.02 | 0.87 | -1.2 | -0.42 | -0.68 | -0.35 | 2.16 | -1.07 | -1.23 | 0.55 | 2.65 | -1.24 | 0.34 | -0.66 | 0.1175 | 0.5875 | 0.515 | 0.792132 | 0.072242 | 0.955666 | 0.989223 | 0.997988 | 0.999507 |
| Eif4b_Q8BGD9_S445s_1 | Eif4b_Q8B | -2.28 | -2.11 | -2.12 | -2.8 | -1.77 | -2.74 | -1.99 | -2.48 | -2.66 | -2.01 | -1.06 | -2.36 | -1.82 | -1.65 | -2.35 | -2.47 | -0.7225 | 0.0425 | 0.2125 | 0.420313 | 0.783342 | 0.437073 | 0.995762 | 0.997988 | 0.994666 |
| Eif4b_Q8BGD9_S495s_2 | Eif4b_Q8B | 0.72 | -0.69 | 0.32 | 0.08 | -2.75 | 0.98 | -2.24 | -0.59 | 1.43 | -0.65 | -0.74 | 0.14 | 1.34 | -0.05 | -1.09 | -0.17 | 0.0825 | 0.305 | 0.255 | 0.853449 | 0.13395 | 0.908109 | 0.989223 | 0.997988 | 0.999507 |
| Eif4b_Q8BGD9_S497s_1 | Eif4b_Q8B | 0.09 | -0.28 | -0.69 | -1.06 | -0.91 | 0.38 | -1.69 | -1.49 | 0.88 | 0.44 | -2.01 | -1.57 | 0.55 | -1.32 | -1.89 | -2.94 | -1.2575 | -0.0625 | -0.1 | 0.207123 | 0.373951 | 0.907234 | 0.989223 | 0.997988 | 0.994666 |
| Eif4b_Q8BGD9_S497s_2 | Eif4b_Q8B | -1.12 | -0.6 | 0.74 | 0.01 | -1.62 | -0.5 | -4.02 | 0.02 | 0.08 | -0.29 | -0.11 | 0.42 | -0.4 | -1.44 | -1.96 | 0.17 | -0.4425 | -0.08 | -0.915 | 0.266342 | 0.156431 | 0.528648 | 0.989223 | 0.997988 | 0.994666 |
| Eif4b_Q8BGD9_S498s_2 | Eif4b_Q8B | -0.51 | 0.07 | -0.15 | -0.27 | -1.32 | 0.18 | -1.29 | -0.8 | 1.76 | -0.06 | -1.6 | -0.22 | 0.15 | -1.71 | -0.49 | -0.39 | -1.2875 | 0.2675 | -0.665 | 0.350132 | 0.130787 | 0.771836 | 0.989223 | 0.997988 | 0.994666 |
| Eif4b_Q8BGD9_S504s_1 | Eif4b_Q8B | 0.25 | 0.96 | 1.23 | 1.14 | -0.83 | 0.89 | 0.3 | 0.28 | 2.14 | 1.11 | -0.14 | 0.88 | 2.76 | -1.99 | 0.52 | 1.07 | -0.5925 | 0.185 | -0.395 | 0.730648 | 0.094956 | 0.834174 | 0.989223 | 0.997988 | 0.9971 |
| Eif4b_Q8BGD9_S504s_2 | Eif4b_Q8B | -0.42 | -0.03 | -0.19 | -0.24 | -0.93 | 0.28 | -1.22 | -0.88 | 1.43 | -0.32 | -1.65 | -0.25 | 0.53 | -1.56 | -0.44 | -0.42 | -0.735 | 0.1025 | -0.305 | 0.560334 | 0.19685 | 0.969262 | 0.989223 | 0.997988 | 0.995259 |
| Eif4b_Q8BGD9_S93s_1 | Eif4b_Q8B | -2.84 | -1.54 | -1.9 | -2.36 | -0.65 | -2.92 | -1.13 | -2.58 | -2.56 | -2.1 | -1.23 | -2.39 | -1.78 | -0.26 | -2.6 | -2.21 | -0.4675 | 0.0225 | -0.2525 | 0.418626 | 0.545988 | 0.824085 | 0.989223 | 0.997988 | 0.994666 |
| Eif4b_Q8BGD9_T420t_2 | Eif4b_Q8B | -0.15 | -0.94 | -0.53 | -1.26 | -1.51 | 0.63 | -1.75 | 0.03 | 2.02 | 0.24 | -0.61 | -1.96 | 1.28 | -0.21 | -0.64 | -1.36 | 0.34 | 0.09 | 0.4475 | 0.394388 | 0.901585 | 0.41676 | 0.996493 | 0.997988 | 0.994666 |
| Eif4b_Q8BGD9_Y211y_1 | Eif4b_Q8B | -1.22 | -1.88 | 0.01 | 0.14 | -1.58 | -1.3 | -0.67 | 0.36 | -1.17 | -2.68 | 1.45 | -0.82 | -0.67 | -0.67 | 0.4 | -1.33 | 0.07 | 0.6425 | 0.4875 | 0.761668 | 0.918107 | 0.938339 | 0.997335 | 0.997988 | 0.9971 |
| Eif4e2_M0QWD1_S2s_1 | Eif4e2_M0 | 1.48 | 1.06 | 0.54 | 0.9 | 0.03 | 1.56 | -0.51 | 1.26 | 1.43 | 0.97 | 1.39 | 0.97 | 1.92 | 1.13 | 0.82 | 0.32 | -0.06 | -0.0675 | 0.17 | 0.894462 | 0.409442 | 0.495905 | 0.989223 | 0.997988 | 0.999507 |
| Eif4enif1_Q9EST3_S539s_1 | Eif4enif1_ | -1.43 | -0.22 | -0.5 | -1.29 | -0.98 | -2.06 | -1.2 | -1.78 | -1.58 | -0.52 | 0.63 | 0.01 | -0.32 | -0.28 | -1.12 | -1.35 | -0.41 | 0.195 | 0.0525 | 0.820224 | 0.114098 | 0.351819 | 0.989223 | 0.997988 | 0.999507 |
| Eif4enif1_Q9EST3_S562s_1 | Eif4enif1_ | 0.36 | -0.56 | -1.33 | -0.66 | -0.07 | 0.89 | -1.84 | -0.91 | 1.06 | -0.5 | -1.28 | -2.35 | 0.37 | 0.74 | -1.49 | -0.27 | -0.645 | 0.495 | 0.0925 | 0.493484 | 0.914554 | 0.759537 | 0.997335 | 0.997988 | 0.994666 |
| Eif4enif1_Q9EST3_S77s_1 | Eif4enif1_ | -1.11 | -0.67 | -0.01 | -0.48 | -0.8 | -1.53 | -0.77 | -0.71 | -1.23 | -0.9 | 0.56 | -0.16 | -0.28 | -0.42 | -0.53 | -0.3 | 0.065 | -0.22 | 0.385 | 0.538952 | 0.241348 | 0.761464 | 0.989223 | 0.997988 | 0.994666 |
| Eif4enif1_Q9EST3_S925s_1 | Eif4enif1_ | -2.25 | -1.59 | -2.06 | -2.46 | -1.28 | -2.66 | -2.44 | -2.24 | -1.87 | -1.58 | -0.89 | -1.77 | 0 | -0.52 | -1.97 | -2.25 | -0.385 | 0.135 | 0.185 | 0.117916 | 0.855104 | 0.103477 | 0.996349 | 0.997988 | 0.994666 |
| Eif4g1_E9PVC6_S1149s_1 | Eif4g1_E9P | -0.84 | -1.86 | 0.42 | 0.62 | -1.08 | -0.9 | -0.16 | 0.82 | -2.16 | -1.14 | 1 | 0.45 | -1.35 | -2.74 | 2.21 | -0.16 | -0.065 | 0.5625 | 0.905 | 0.927242 | 0.894652 | 0.954176 | 0.996493 | 0.997988 | 0.999507 |
| Eif4g1_E9PVC6_S1164s_1 | Eif4g1_E9P | 0.64 | -0.46 | -1.14 | -1.71 | -0.08 | 0.77 | -1.75 | -1.56 | 0.31 | 0.29 | -1.15 | -1.25 | 0.09 | -0.41 | -1.13 | -1.51 | 0.085 | -0.0475 | -0.095 | 0.898769 | 0.985629 | 0.72165 | 0.999187 | 0.997988 | 0.999507 |
| Eif4g1_E9PVC6_S1184s_1 | Eif4g1_E9P | -0.32 | -0.77 | -0.2 | 0.15 | -0.75 | -1.11 | -0.58 | -0.69 | -1.36 | -0.45 | -0.32 | -0.49 | -0.52 | -0.45 | -0.5 | -0.57 | 0.0125 | 0.2175 | -0.0725 | 0.421467 | 0.087159 | 0.277744 | 0.989223 | 0.997988 | 0.994666 |
| Eif4g1_E9PVC6_S1550s_1 | Eif4g1_E9P | -1.82 | -2.3 | -2.27 | -3.32 | -2.89 | -1.81 | -1.5 | -2.3 | 2.17 | -1.91 | -2.02 | -2.2 | 1.11 | -2.12 | 1.11 | -2.57 | -0.4975 | -0.37 | -0.225 | 0.073732 | 0.474877 | 0.16419 | 0.989223 | 0.997988 | 0.994666 |
| Eif4g1_E9PVC6_T1166t_1 | Eif4g1_E9P | -1.47 | -2.35 | -1.76 | -2.79 | -1.86 | -2.14 | -3.01 | -2.77 | -2.83 | -2.04 | -1.21 | -2.03 | -1.91 | -2.65 | -2.22 | -3.01 | 0.3025 | 1.4375 | 1.81 | 0.375494 | 0.373688 | 0.880231 | 0.989223 | 0.997988 | 0.994666 |
| Eif4g1_E9PVC6_T165t_1 | Eif4g1_E9P | 0.02 | 0.49 | -2.07 | -0.25 | -0.47 | -1.76 | 0.3 | -0.96 | -0.45 | 0.65 | -1.05 | -0.94 | 0.23 | -1.39 | -0.11 | 0.6 | -0.3525 | 0.065 | -0.355 | 0.657695 | 0.669454 | 0.99358 | 0.992876 | 0.997988 | 0.995259 |
| Eif4g1_E9PVC6_T167t_1 | Eif4g1_E9P | -1.1 | -0.72 | -0.43 | 0.51 | -0.83 | -1.22 | 1.06 | 0.63 | -0.2 | -0.41 | 1.44 | 0.53 | -0.29 | 0.43 | 0.92 | 0.92 | -0.27 | 0.005 | 0.285 | 0.057391 | 0.558092 | 0.152278 | 0.989223 | 0.997988 | 0.994666 |
| Eif4g2_F7CBP1_T469t_1 | Eif4g2_F7C | -0.82 | -0.26 | -0.73 | -0.87 | -0.18 | -0.98 | -0.63 | -1.08 | -1.3 | -0.62 | -1.03 | -1.24 | -0.75 | -0.68 | -0.44 | -0.52 | 0.345 | 0.775 | 0.93 | 0.779549 | 0.866085 | 0.181885 | 0.996493 | 0.997988 | 0.998226 |
| Eif4g3_Z4YKC4_S237s_1 | Eif4g3_Z4Y | 0.28 | 0.64 | 1.31 | 0.64 | 0.87 | -0.03 | 1.26 | 0.79 | 0.07 | 0.47 | 1 | 0.92 | -0.6 | 0.7 | 0.51 | 1.52 | -0.0475 | -0.3775 | 0.0725 | 0.693722 | 0.988544 | 0.756059 | 0.999187 | 0.997988 | 0.996159 |
| Eif4g3_Z4YKC4_S274s_1 | Eif4g3_Z4Y | -1.58 | -1.28 | -1.17 | -1.29 | -1.22 | -2.03 | -1.24 | -1.63 | -1.62 | -1.06 | -1.07 | -1.44 | -0.9 | -0.73 | -1.86 | -1.71 | 0.005 | -0.1025 | -0.185 | 0.928628 | 0.448815 | 0.894261 | 0.989223 | 0.997988 | 0.999507 |
| Eif4g3_Z4YKC4_S288s_1 | Eif4g3_Z4Y | -0.56 | -0.66 | -0.54 | -0.87 | -0.32 | -0.79 | -0.51 | -0.9 | -0.66 | -0.37 | -0.39 | -0.29 | -1.15 | -0.32 | -1.32 | -0.79 | -0.2 | 0.0325 | 0.03 | 0.432291 | 0.904285 | 0.312708 | 0.996493 | 0.997988 | 0.994666 |
| Eif4g3_Z4YKC4_S479s_1 | Eif4g3_Z4Y | -1.81 | -0.85 | -1.22 | -1.2 | -2.36 | -1.72 | -1.3 | -2.19 | -1.19 | -1.49 | -0.82 | -1.13 | -0.79 | -0.98 | -1.35 | -0.55 | 0.0275 | 0.23 | -0.2375 | 0.276528 | 0.080003 | 0.69732 | 0.989223 | 0.997988 | 0.994666 |
| Eif4g3_Z4YKC4_T175t_1 | Eif4g3_Z4Y | -3.69 | -2.72 | -2.92 | -4.88 | -2.09 | -4.67 | -3.55 | -4.41 | -4.94 | -2.73 | -1.3 | -2.85 | -1.95 | -2.57 | -3.5 | -3.02 | -0.6225 | 0.1125 | 0.3525 | 0.170883 | 0.849577 | 0.461776 | 0.996349 | 0.997988 | 0.994666 |
| Eif5b_Q05D44_S137s_2 | Eif5b_Q05 | -0.04 | 0.46 | 1.1 | 0.08 | 0.02 | -0.98 | 0.46 | 0.61 | -0.43 | 0 | 0.89 | 0.18 | -1.22 | 0.42 | 0.41 | 0.65 | -0.1275 | 0.5975 | 0.7925 | 0.490904 | 0.383973 | 0.532395 | 0.989223 | 0.997988 | 0.994666 |
| Eif5b_Q05D44_S139s_2 | Eif5b_Q05 | -0.04 | 0.46 | 1.1 | 0.08 | 0.02 | -0.98 | 0.46 | 0.61 | -0.43 | 0 | 0.89 | 0.18 | -1.22 | 0.42 | 0.41 | 0.65 | -0.3725 | -0.24 | -0.335 | 0.490904 | 0.383973 | 0.532395 | 0.989223 | 0.997988 | 0.994666 |
| Eif5b_Q05D44_S165s_1 | Eif5b_Q05 | 3.12 | 1.5 | 0.26 | -0.66 | 0.67 | 3.41 | 0.15 | 0.05 | 1.66 | 2.65 | -1.31 | 0.38 | 1.03 | -1.77 | 3.02 | 0.36 | -0.3725 | -0.24 | -0.335 | 0.722899 | 0.987839 | 0.840492 | 0.999187 | 0.997988 | 0.9971 |
| Elac2_B1ATP7_S191s_1 | Elac2_B1A | -4.92 | -6.93 | -4.35 | -10.13 | -7.48 | -5.77 | -4.12 | -3.91 | -2 | -4.29 | -3.72 | -4.23 | -0.15 | -5.33 | -2.65 | -4.63 | 0.015 | -0.21 | -0.395 | 0.045359 | 0.353662 | 0.034207 | 0.989223 | 0.997988 | 0.994666 |
| Elavl2_B1AXZ0_S119s_1 | Elavl2_B1A | -2.02 | -1.55 | -1.58 | -2.25 | -3.09 | -1.55 | -1.95 | -2.97 | -1.03 | -2.25 | -3.35 | -2.62 | 0.39 | -2.65 | -1.72 | -2.55 | 1.2625 | 3.0225 | 3.3925 | 0.74027 | 0.198644 | 0.354105 | 0.989223 | 0.997988 | 0.9971 |
| Elavl2_B1AXZ0_S221s_1 | Elavl2_B1A | 0.91 | 0.32 | 0.37 | -0.42 | 0.77 | 0.23 | 0.79 | -0.17 | 0.5 | 0.93 | -0.15 | 0.46 | 0.12 | 0.05 | 0.89 | -0.06 | -0.54 | -0.4625 | 0.2175 | 0.902866 | 0.758604 | 0.700486 | 0.995762 | 0.997988 | 0.999507 |
| Elavl4_Q61701_S36s_1 | Elavl4_Q61 | 0.01 | 0.47 | -0.33 | -0.35 | 0.22 | -0.05 | -0.41 | -0.28 | 0.11 | 0.08 | -0.24 | -0.5 | 0.52 | 0.27 | 0.17 | -0.28 | 0.11 | 0.14 | -0.045 | 0.482724 | 0.771797 | 0.761093 | 0.995762 | 0.997988 | 0.994666 |
| Elfn1_Q8C8T7_S773s_1 | Elfn1_Q8C | 0.52 | 0.77 | -0.37 | 0.3 | 1.54 | -0.14 | 1.16 | 0.31 | 0.11 | 0.22 | 0.46 | 0.35 | -1.15 | 0.55 | 0.79 | 0.69 | -0.08 | -0.0875 | 0.22 | 0.863268 | 0.349647 | 0.946313 | 0.989223 | 0.997988 | 0.999507 |
| Elfn1_Q8C8T7_T517t_1 | Elfn1_Q8C | -0.38 | -0.81 | -1.19 | 0.51 | -1.59 | 0.17 | 0.08 | -2.22 | 0.07 | -0.57 | -4.19 | -0.87 | -0.85 | -0.41 | 0.82 | 1.91 | 0.4125 | -0.02 | -0.085 | 0.220565 | 0.504703 | 0.322514 | 0.989223 | 0.997988 | 0.994666 |
| Elfn2_Q68FM6_S451s_1 | Elfn2_Q68 | 0.51 | -0.08 | -0.84 | -0.2 | 0.87 | 0.68 | 0.09 | -0.04 | -0.01 | -0.35 | -0.38 | -1.03 | -1.31 | -0.05 | -1.38 | -1.13 | -0.4225 | -0.9225 | 0.835 | 0.074898 | 0.143911 | 0.429701 | 0.989223 | 0.997988 | 0.994666 |
| Elfn2_Q68FM6_S471s_1 | Elfn2_Q68 | 0.18 | -1.12 | -0.07 | 0.61 | 1.11 | 0.93 | 1.05 | 0.71 | -0.93 | -0.86 | 0.09 | 0.37 | -1.61 | 0.47 | 0.17 | 0.13 | 0.5525 | -0.29 | -0.815 | 0.842692 | 0.017787 | 0.623644 | 0.989223 | 0.997988 | 0.999507 |
| Elfn2_Q68FM6_S639s_1 | Elfn2_Q68 | -1.55 | -3.57 | -1.62 | -1.94 | -2.73 | -0.97 | -2.08 | -0.46 | -3.21 | -2.83 | 1.09 | -0.89 | -2.64 | -1.49 | -1.1 | -1.79 | 1.05 | -0.2325 | -0.11 | 0.446572 | 0.341102 | 0.471038 | 0.989223 | 0.997988 | 0.994666 |
| Elfn2_Q68FM6_S647s_1 | Elfn2_Q68 | -0.92 | -1.27 | -2.17 | -0.6 | 0.01 | -0.64 | -1.31 | -1.07 | -0.72 | -0.92 | -2.31 | -1.36 | -1.92 | -0.29 | -1.58 | -1.94 | 0.61 | 0.71 | 0.415 | 0.695803 | 0.265723 | 0.85149 | 0.989223 | 0.997988 | 0.996159 |
| Elfn2_Q68FM6_S671s_1 | Elfn2_Q68 | -0.06 | -0.34 | 0.2 | 0.44 | -0.28 | -0.07 | 0.36 | 0.18 | -0.07 | -0.5 | 1 | 0.42 | -0.34 | 0.28 | -0.22 | -0.25 | 0.4875 | -0.0875 | -0.1925 | 0.510186 | 0.962297 | 0.682494 | 0.999187 | 0.997988 | 0.994666 |
| Elfn2_Q68FM6_S750s_1 | Elfn2_Q68 | 0.48 | 0.2 | -0.05 | 0.04 | 0.4 | 0.27 | 0.19 | 0.5 | 0.3 | -0.05 | 1.12 | 0.69 | 0.52 | 0.37 | 0.48 | 0.05 | -0.0125 | 0.1525 | -0.1925 | 0.475421 | 0.44273 | 0.280664 | 0.989223 | 0.997988 | 0.994666 |
| Elfn2_Q68FM6_T493t_1 | Elfn2_Q68 | -0.2 | -1 | -0.86 | 0.07 | 0.1 | -0.08 | 0.24 | -0.43 | -0.6 | -1.11 | -1.55 | -0.6 | -0.95 | -0.01 | -0.87 | -0.05 | 0.1725 | 0.3475 | 0.1875 | 0.941964 | 0.169482 | 0.211906 | 0.989223 | 0.997988 | 0.999507 |
| Elfn2_Q68FM6_T655t_1 | Elfn2_Q68 | 0.89 | 0.98 | 0.42 | 0.19 | 0.98 | 0.53 | 0.36 | -0.16 | 0.53 | 0.73 | -0.24 | 0.04 | -0.52 | 0.63 | -0.3 | 0.1 | 0.455 | -0.4675 | 0.0275 | 0.089249 | 0.548373 | 0.284271 | 0.989223 | 0.997988 | 0.994666 |
| Elmo1_Q8BPU7_S344s_1 | Elmo1_Q8 | 0.26 | 0.79 | 0.22 | 0.38 | 0.53 | 0.35 | 0.43 | 0.14 | 0.13 | 0.36 | 0.32 | 0.19 | 0.18 | 0.16 | 0.36 | 0.48 | -0.1925 | -0.355 | -0.6425 | 0.64821 | 0.827932 | 0.492444 | 0.995762 | 0.997988 | 0.995259 |
| Elmsan1_E9Q2I4_S456s_1 | Elmsan1_E | -1.28 | 0.13 | -0.13 | -0.08 | -0.47 | -0.63 | -0.32 | 0.47 | -0.67 | 0.38 | 0.19 | -0.24 | 0.32 | 0.31 | -1.27 | -1.6 | -0.05 | -0.1625 | -0.1175 | 0.693217 | 0.792367 | 0.522874 | 0.995762 | 0.997988 | 0.996159 |
| Emb_P21995_S312s_1 | Emb_P219 | 0.01 | 0.51 | 0.94 | 0.94 | 0.69 | -0.27 | 1.27 | 1.22 | 0.58 | 0.45 | 2.67 | 0.9 | 0.4 | 1.37 | 1.77 | 1.8 | 0.1025 | 0.255 | -0.22 | 0.092063 | 0.75237 | 0.30876 | 0.995762 | 0.997988 | 0.994666 |
| Emc4_Q9CZX9_S36s_1 | Emc4_Q9C | -0.15 | 0.37 | 0.22 | 0.49 | 0.46 | -0.29 | 0.76 | 0.34 | -0.76 | 0.04 | -0.17 | 0.15 | -0.63 | 0.47 | 1.05 | 1.25 | 0.1275 | 0.55 | 0.735 | 0.494046 | 0.770316 | 0.174546 | 0.995762 | 0.997988 | 0.994666 |
| Eml1_D3Z4J9_S115s_1 | Eml1_D3Z4 | -3.47 | -4.6 | -4.02 | -9.94 | -4.16 | -4.2 | -2.56 | -2.87 | 3.52 | -3.6 | -1.69 | -2.73 | 4.04 | -3.24 | 2.77 | -4.77 | 0.085 | -0.4175 | 0.3025 | 0.041783 | 0.147016 | 0.043187 | 0.989223 | 0.997988 | 0.994666 |
| Eml1_D3Z4J9_S154s_1 | Eml1_D3Z4 | -0.25 | -0.16 | 0.59 | 0.55 | 0.12 | 0.01 | 0.81 | -0.02 | -0.11 | 0.7 | 0.21 | 0.36 | 0.37 | 0 | 0.07 | 0.15 | 2.06 | 4.3825 | 5.2075 | 0.907057 | 0.880093 | 0.732017 | 0.996493 | 0.997988 | 0.999507 |
| Eml3_Q8VC03_S157s_1 | Eml3_Q8V | -0.82 | -1.79 | -1.28 | -2.2 | -2.26 | -0.66 | -1.52 | -1.92 | 0.96 | -1.65 | -1.83 | -1.63 | 1.31 | -3.44 | -1.32 | -1.45 | 0.0475 | 0.1075 | -0.035 | 0.739666 | 0.875625 | 0.471276 | 0.996493 | 0.997988 | 0.9971 |
| Eml3_Q8VC03_S177s_1 | Eml3_Q8V | -1.02 | -0.47 | -1.43 | -2.16 | 0.09 | -1.03 | -1.12 | -2.35 | -1.15 | -0.9 | -0.46 | -2.27 | -0.24 | 0.13 | -1.77 | -1.76 | -0.0675 | 0.485 | 0.2975 | 0.528701 | 0.762409 | 0.879521 | 0.995762 | 0.997988 | 0.994666 |
| Eml4_F8WJ93_S146s_1 | Eml4_F8W | -0.02 | -0.23 | -0.29 | 0.61 | -0.28 | 0.24 | 0.57 | 0.95 | -0.69 | 0.07 | 0.75 | 0.81 | 0.48 | 0.12 | 0.08 | 0.32 | 0.1675 | 0.075 | 0.36 | 0.435076 | 0.312428 | 0.592666 | 0.989223 | 0.997988 | 0.994666 |
| Eml4_F8WJ93_S908s_1 | Eml4_F8W | 0.41 | 0.61 | 0.68 | 0.83 | 0.37 | 0.29 | 1.05 | 0.73 | 0.4 | 0.76 | 0.46 | 0.47 | 0.29 | 0.91 | 0.83 | 0.48 | 0.3525 | 0.2175 | 0.2325 | 0.984913 | 0.928621 | 0.625895 | 0.99758 | 0.997988 | 1 |
| Enah_J3QNM3_S129s_1 | Enah_J3Q | 0.06 | 0.99 | 1.04 | 1.84 | 0.38 | 0.2 | 1.6 | 1.8 | 0.4 | 0.21 | 2.14 | 0.71 | -0.47 | 0.33 | 0.99 | 1.44 | -0.0225 | -0.11 | -0.005 | 0.436605 | 0.980131 | 0.824708 | 0.999187 | 0.997988 | 0.994666 |
| Enah_J3QNM3_S240s_1 | Enah_J3Q | 0.25 | 0.36 | 0.3 | 0.89 | 0.55 | 0.14 | 0.65 | 0.1 | -0.06 | -0.29 | 0.96 | 0.96 | -0.57 | 0.57 | -0.15 | 0 | 0.0125 | -0.1175 | -0.41 | 0.155742 | 0.72605 | 0.876604 | 0.995437 | 0.997988 | 0.994666 |
| Enah_J3QNM3_S339s_1 | Enah_J3Q | 1.02 | 0.18 | 1.05 | 1.19 | 1.08 | 1.09 | 0.84 | 1.34 | 0.79 | 0.87 | 1.55 | 1.1 | 0.95 | 1.15 | 0.69 | 0.77 | -0.09 | -0.0575 | -0.4875 | 0.922198 | 0.431992 | 0.497971 | 0.989223 | 0.997988 | 0.999507 |
| Enah_J3QNM3_S339s_2 | Enah_J3Q | -0.89 | -0.55 | -0.19 | 0.17 | -0.43 | 0.02 | -0.01 | 0.33 | -0.18 | -0.85 | 0.88 | -1.42 | -0.86 | -0.21 | -0.85 | 0.34 | 0.2275 | 0.2175 | 0.03 | 0.937369 | 0.271695 | 0.957003 | 0.989223 | 0.997988 | 0.999507 |
| Enah_J3QNM3_S340s_1 | Enah_J3Q | -0.57 | -0.46 | -1.29 | -0.19 | -0.28 | -0.35 | -0.86 | -0.5 | -0.59 | -0.7 | 0.21 | -0.41 | -0.02 | 0.43 | -1 | -0.64 | 0.3425 | -0.0275 | -0.03 | 0.433371 | 0.661164 | 0.451648 | 0.990434 | 0.997988 | 0.994666 |
| Enah_J3QNM3_S340s_2 | Enah_J3Q | -0.89 | -0.55 | -0.19 | 0.17 | -0.43 | 0.02 | -0.01 | 0.33 | -0.18 | -0.85 | 0.88 | -1.42 | -0.86 | -0.21 | -0.85 | 0.34 | 0.13 | 0.255 | 0.32 | 0.937369 | 0.271695 | 0.957003 | 0.989223 | 0.997988 | 0.999507 |
| Enah_J3QNM3_S727s_1 | Enah_J3Q | -0.86 | -0.41 | -0.39 | -1.11 | 0.2 | -0.39 | -0.36 | -0.88 | -0.06 | -0.41 | 0.57 | -0.54 | 0.28 | 0.09 | -0.11 | -0.76 | 0.3425 | -0.0275 | -0.03 | 0.109666 | 0.287407 | 0.103117 | 0.989223 | 0.997988 | 0.994666 |
| Enah_J3QNM3_T708t_1 | Enah_J3Q | 0.41 | 0.54 | 0.29 | 0.14 | -0.45 | -0.63 | -0.75 | -0.65 | 0.86 | 0.71 | 1.15 | -0.08 | 0.41 | -0.05 | 0.23 | 0.34 | 0.335 | 0.5825 | 0.5675 | 0.65266 | 0.001084 | 0.324167 | 0.989223 | 0.997988 | 0.995259 |
| Eno1_P17182_S263s_1 | Eno1_P171 | -0.81 | -0.45 | -0.92 | -0.98 | -0.37 | -0.93 | -0.66 | -0.55 | 0.15 | -0.57 | -1.72 | -0.79 | 0.55 | -0.55 | -0.57 | -0.43 | -0.965 | 0.315 | -0.1125 | 0.129698 | 0.496651 | 0.885853 | 0.989223 | 0.997988 | 0.994666 |
| Eno2_P17183_S79s_1 | Eno2_P171 | -2.43 | -1.43 | -1.28 | -0.38 | -1.54 | -2.71 | -1.12 | -1.23 | -0.79 | -1.74 | -1.5 | -1.38 | -1.42 | -0.72 | -0.16 | -0.03 | 0.1625 | 0.0575 | 0.54 | 0.134107 | 0.597756 | 0.951074 | 0.989223 | 0.997988 | 0.994666 |
| Enox1_Q8BHR2_S403s_1 | Enox1_Q8 | -3.31 | -4.19 | -4.3 | -2.68 | -3.32 | -3.67 | -2.43 | -2.64 | -3.32 | -3.27 | -1.87 | -2.81 | -3.12 | -2.6 | -3.02 | -3.67 | -0.27 | 0.0275 | 0.7975 | 0.252442 | 0.19955 | 0.123045 | 0.989223 | 0.997988 | 0.994666 |
| Ensa_P60840_S43s_1 | Ensa_P608 | -4.96 | -4.86 | -1.5 | -1.57 | -3.21 | -6.04 | -0.45 | -1.24 | -6.26 | -4.61 | -0.7 | -0.85 | -3.67 | -2.23 | -1.1 | -1.38 | 0.605 | 0.8025 | 0.5175 | 0.270086 | 0.719961 | 0.936673 | 0.995437 | 0.997988 | 0.994666 |
| Ep300_B2RWS6_S500s_1 | Ep300_B2 | -1.39 | 0.08 | -0.28 | -2.17 | -2.72 | -0.55 | -1.69 | -1.89 | 1.38 | -0.36 | -0.03 | -2.03 | 1.13 | -1.86 | -1.62 | -0.77 | 0.4875 | 0.1175 | 1.1275 | 0.83248 | 0.224704 | 0.392339 | 0.989223 | 0.997988 | 0.999507 |
| Ep400_Q8CHI8_S721s_1 | Ep400_Q8 | -2.44 | -2.33 | -1.54 | -0.9 | -3.55 | -2.58 | -2.22 | -2.58 | -2.27 | -2.84 | -1.85 | -1.38 | -0.22 | -2.67 | -0.6 | -1.12 | -0.7725 | 0.68 | 0.16 | 0.288767 | 0.055262 | 0.540666 | 0.989223 | 0.997988 | 0.994666 |
| Ep400_Q8CHI8_S728s_1 | Ep400_Q8 | -1.95 | -2.44 | -2.39 | -2.56 | -2.81 | -2.27 | -1.8 | -2.64 | -1.97 | -1.75 | -0.91 | -2.13 | -0.22 | -3.23 | -1.08 | -1.39 | -0.93 | -0.2825 | 0.65 | 0.171928 | 0.876794 | 0.074562 | 0.996493 | 0.997988 | 0.994666 |
| Ep400_Q8CHI8_S923s_1 | Ep400_Q8 | 1.1 | -0.68 | -1.75 | -1.62 | 0.65 | 1.08 | -0.61 | -1.6 | 0.09 | 0.42 | -2.85 | -1.23 | -0.64 | 0.23 | -0.36 | 0.01 | -0.045 | 0.645 | 0.855 | 0.388742 | 0.438828 | 0.860334 | 0.989223 | 0.997988 | 0.994666 |
| Ep400_Q8CHI8_T922t_1 | Ep400_Q8 | 1.1 | -0.68 | -1.75 | -1.62 | 0.65 | 1.08 | -0.61 | -1.6 | 0.09 | 0.42 | -2.85 | -1.23 | -0.64 | 0.23 | -0.36 | 0.01 | 0.6175 | -0.155 | 0.5475 | 0.388742 | 0.438828 | 0.860334 | 0.989223 | 0.997988 | 0.994666 |
| Epb4.1_P48193_S182s_1 | Epb4.1_P4 | -1.26 | -1.7 | -0.71 | -1.76 | -0.76 | -1.77 | -1.44 | -2.12 | -1.32 | -0.83 | -1.35 | -2.66 | -0.79 | -0.61 | -1.47 | -1.76 | 0.6175 | -0.155 | 0.5475 | 0.601688 | 0.658748 | 0.682892 | 0.990434 | 0.997988 | 0.995259 |
| Epb4.1_P48193_S543s_1 | Epb4.1_P4 | 1.23 | 0.7 | 0.91 | 1.47 | 0.53 | 0.74 | 0.51 | 1.41 | 1.01 | 0.63 | 1.12 | 0.89 | 0.38 | -0.14 | 0.11 | 0.56 | -0.165 | -0.1825 | 0.2 | 0.015911 | 0.35752 | 0.535651 | 0.989223 | 0.997988 | 0.994666 |
| Epb4.1_P48193_S556s_1 | Epb4.1_P4 | -0.05 | -0.48 | -0.62 | 0.01 | 0.19 | 0.22 | 0.49 | 0.2 | -0.94 | 0.26 | -0.34 | -1.07 | -0.92 | -0.43 | -0.71 | -0.02 | -0.28 | -0.165 | -0.85 | 0.450065 | 0.039175 | 0.510076 | 0.989223 | 0.997988 | 0.994666 |
| Epb4.1_P48193_S86s_1 | Epb4.1_P4 | -2.8 | -3.43 | -3.54 | -2.99 | -2.09 | -2.81 | -1.67 | -2.49 | -2.73 | -2.07 | -0.33 | -2.51 | -2.53 | -2.38 | -1.69 | -2.95 | 0.56 | -0.2375 | -0.235 | 0.041499 | 0.015536 | 0.036594 | 0.989223 | 0.997988 | 0.994666 |
| Epb4.1_P48193_S87s_1 | Epb4.1_P4 | -1.19 | -3.08 | -2.45 | -2.81 | -0.71 | -1.52 | -0.28 | -1.27 | -3.49 | -1.92 | -1.88 | -1.45 | -2.88 | -3.22 | -2.11 | -1.78 | 0.925 | 1.28 | 0.8025 | 0.819756 | 0.011556 | 0.727576 | 0.989223 | 0.997988 | 0.999507 |
| Epb4.1_P48193_S97s_1 | Epb4.1_P4 | 1.64 | 1.17 | 0.91 | 1 | 1.34 | 1.84 | 0.33 | 1.19 | 1.64 | 1.38 | 0.8 | 0.97 | -0.37 | 1.63 | 0.46 | 0.7 | 1.4375 | 0.1975 | -0.115 | 0.20735 | 0.988723 | 0.952483 | 0.999187 | 0.997988 | 0.994666 |
| Epb4.1_P48193_T62t_1 | Epb4.1_P4 | 1.41 | -0.48 | -0.86 | -0.06 | -0.08 | 0.91 | -0.15 | -0.32 | 0.67 | 0 | -0.4 | -0.15 | 0.41 | -0.42 | -0.35 | -0.03 | -0.005 | 0.0175 | -0.575 | 0.842188 | 0.86599 | 0.957173 | 0.996493 | 0.997988 | 0.999507 |
| Epb4.1l1_A2AUK5_S441s_1 | Epb4.1l1_A | -0.99 | 1.67 | 0.01 | 0.32 | 1.4 | 1.84 | 0.4 | 0.48 | 0.75 | 0.84 | -0.51 | 0.51 | 0.5 | 1.3 | -0.16 | -1.08 | 0.0875 | 0.0275 | -0.1 | 0.866669 | 0.205011 | 0.802809 | 0.989223 | 0.997988 | 0.999507 |
| Epb4.1l1_A2AUK5_S441s_2 | Epb4.1l1_A | -0.65 | -0.12 | -1.41 | -1.05 | -0.22 | -0.59 | -0.77 | -0.34 | -0.53 | -0.35 | -0.52 | -1.58 | -1.71 | -0.32 | -2.02 | -1.85 | 0.7775 | 0.145 | -0.1125 | 0.171634 | 0.317851 | 0.873969 | 0.989223 | 0.997988 | 0.994666 |
| Epb4.1l1_E9PV14_S1077s_1 | Epb4.1l1_E | 0.03 | -1.14 | -0.44 | -1.13 | -0.24 | 0.46 | -1.21 | -0.61 | 0.32 | -0.55 | -0.5 | -0.39 | 0.01 | -1.02 | -0.61 | -0.43 | 0.3275 | 0.0625 | -0.6675 | 0.674993 | 0.531099 | 0.295856 | 0.989223 | 0.997988 | 0.996159 |
| Epb4.1l1_E9PV14_S1228s_1 | Epb4.1l1_E | 0.78 | -0.22 | -0.2 | 0.31 | 0.21 | 0.23 | 0.39 | 0.54 | -0.07 | 0.25 | -0.17 | -0.2 | -0.52 | 0.39 | 0.07 | 0.03 | 0.27 | 0.39 | 0.1575 | 0.608735 | 0.541731 | 0.478932 | 0.989223 | 0.997988 | 0.995259 |
| Epb4.1l1_E9PV14_S1255s_1 | Epb4.1l1_E | 0.86 | -0.75 | 0.23 | 0.56 | -1.76 | 1.8 | 0.04 | 0.21 | 2.36 | -0.13 | -0.01 | 0.28 | 0.26 | -1.68 | -0.14 | -0.19 | 0.175 | -0.215 | -0.175 | 0.220365 | 0.830265 | 0.524599 | 0.995762 | 0.997988 | 0.994666 |
| Epb4.1l1_E9PV14_S1325s_1 | Epb4.1l1_E | 3 | 2.2 | 1.89 | 1.49 | 2.33 | 2.98 | 2.08 | 2 | -0.03 | 2.16 | 0.76 | 1.95 | -1.15 | 0.03 | 1.62 | 1.75 | -0.1525 | 0.4 | -0.6625 | 0.041506 | 0.597093 | 0.121707 | 0.989223 | 0.997988 | 0.994666 |
| Epb4.1l1_E9PV14_S430s_1 | Epb4.1l1_E | -1.76 | -0.66 | 0.05 | 0.11 | -0.22 | -1.01 | -0.11 | 0.84 | -1.98 | -1.04 | 0.81 | 0.51 | -0.94 | -0.44 | -0.39 | -0.28 | 0.2025 | -0.935 | -1.5825 | 0.906546 | 0.410721 | 0.843798 | 0.989223 | 0.997988 | 0.999507 |
| Epb4.1l1_E9PV14_S440s_1 | Epb4.1l1_E | -0.75 | -0.63 | 0.51 | 0.81 | -0.78 | -0.33 | 0.95 | 0.85 | -1.26 | -1.01 | -0.24 | 1.12 | -0.84 | -0.88 | 0.57 | -0.85 | 0.44 | 0.14 | 0.0525 | 0.347053 | 0.724509 | 0.587743 | 0.995437 | 0.997988 | 0.994666 |
| Epb4.1l1_E9PV14_S713s_1 | Epb4.1l1_E | -0.67 | -1.2 | 1.52 | 0.9 | -0.01 | -0.4 | 1.33 | 1.23 | -0.87 | -0.25 | 1.7 | -0.38 | -0.62 | -0.22 | 1.06 | 1.2 | 0.1875 | -0.3325 | -0.485 | 0.757541 | 0.562753 | 0.908977 | 0.989223 | 0.997988 | 0.9971 |
| Epb4.1l1_E9PV14_T1217t_1 | Epb4.1l1_E | 0.06 | -1.38 | -0.62 | -0.71 | -0.37 | 0.43 | -0.27 | -0.38 | -0.63 | -0.47 | -0.15 | -0.7 | -0.42 | -1.05 | -0.54 | -0.83 | 0.4 | -0.0875 | 0.2175 | 0.893388 | 0.168436 | 0.608347 | 0.989223 | 0.997988 | 0.999507 |
| Epb4.1l1_E9PV14_T1257t_1 | Epb4.1l1_E | -1.45 | 0.15 | 0.12 | 0.64 | -0.51 | 1.56 | -1.18 | -0.62 | 2.68 | 0.52 | 0.18 | 0.7 | 1.2 | 0.57 | 0.09 | -0.37 | 0.515 | 0.175 | -0.0475 | 0.348506 | 0.937169 | 0.105142 | 0.999187 | 0.997988 | 0.994666 |
| Epb4.1l1_Q9Z2H5_S378s_1 | Epb4.1l1_C | 0.06 | 0.31 | 0.4 | 0.25 | 0.51 | 0 | 0.44 | 0.13 | 0.37 | 0.14 | 0.71 | 0.4 | 0.53 | 0.71 | 0.51 | 0.39 | -0.0525 | 1.155 | 0.5075 | 0.25514 | 0.946573 | 0.522638 | 0.999187 | 0.997988 | 0.994666 |
| Epb4.1l1_Q9Z2H5_S430s_1 | Epb4.1l1_C | -0.12 | 0.23 | -0.03 | -0.68 | 0.16 | -0.68 | -0.63 | -0.88 | -0.18 | -0.04 | -0.04 | -0.63 | -0.23 | 0.09 | -0.47 | -0.65 | 0.015 | 0.15 | 0.28 | 0.592943 | 0.273723 | 0.799627 | 0.989223 | 0.997988 | 0.995259 |
| Epb4.1l1_Q9Z2H5_S461s_1 | Epb4.1l1_C | -1.77 | -1.84 | -1.52 | -2.9 | -2.51 | -1.01 | -1.37 | -1.44 | 1.49 | -1.95 | -1.72 | -0.91 | 1.43 | -1.95 | 0.91 | -2.07 | -0.3575 | -0.0725 | -0.165 | 0.088971 | 0.326914 | 0.128462 | 0.989223 | 0.997988 | 0.994666 |
| Epb4.1l1_Q9Z2H5_S461s_2 | Epb4.1l1_C | 0.14 | 1.17 | 0.95 | 0.58 | -0.7 | 0.43 | 0.41 | 0.4 | 0.64 | 0.18 | 0.59 | 0.93 | -1.34 | 0.27 | 0.63 | 0.99 | 0.425 | 1.235 | 1.5875 | 0.291074 | 0.132773 | 0.686404 | 0.989223 | 0.997988 | 0.994666 |
| Epb4.1l1_Q9Z2H5_S466s_1 | Epb4.1l1_C | 0.01 | 0.53 | 0.53 | -0.12 | -1.18 | 0.11 | -0.47 | -0.43 | 1.22 | 0.24 | 0.45 | 0.18 | 0.92 | -0.57 | 0.16 | 0 | -0.575 | -0.125 | -0.5725 | 0.767138 | 0.047471 | 0.386976 | 0.989223 | 0.997988 | 0.9971 |
| Epb4.1l1_Q9Z2H5_S466s_2 | Epb4.1l1_C | 0.01 | 1.03 | 0.32 | 0.22 | -0.6 | 0.26 | 0.08 | 0 | 0.49 | -0.01 | 0.6 | 0.49 | -1.41 | 0.08 | 0.18 | 0.1 | -0.73 | 0.285 | -0.11 | 0.154848 | 0.160518 | 0.993339 | 0.989223 | 0.997988 | 0.994666 |
| Epb4.1l1_Q9Z2H5_S510s_1 | Epb4.1l1_C | 2.03 | -0.82 | 1.77 | -1.02 | 1.73 | 3.06 | 3.99 | 3.42 | -4.26 | 0.31 | 0.68 | 2.61 | -2.03 | -1.25 | 2.18 | 3.04 | -0.46 | -0.0025 | -0.6575 | 0.996896 | 0.011277 | 0.654779 | 0.989223 | 0.997988 | 1 |
| Epb4.1l1_Q9Z2H5_S544s_1 | Epb4.1l1_C | 0.83 | -0.13 | -0.5 | -0.19 | 0.63 | 0.95 | 0.04 | -0.19 | -0.05 | 0.28 | -0.18 | -0.45 | -0.05 | 0.54 | -0.61 | -0.78 | 2.56 | -0.655 | -0.005 | 0.586567 | 0.361771 | 0.765703 | 0.989223 | 0.997988 | 0.995259 |
| Epb4.1l1_Q9Z2H5_S544s_2 | Epb4.1l1_C | -0.17 | -0.13 | -0.27 | -0.19 | 0.18 | -0.2 | -0.05 | 0.02 | -0.54 | -0.41 | -0.08 | -0.79 | -0.28 | 0.76 | -0.55 | -1.15 | 0.355 | -0.1025 | -0.2275 | 0.77649 | 0.390065 | 0.28206 | 0.989223 | 0.997988 | 0.99798 |
| Epb4.1l1_Q9Z2H5_S546s_1 | Epb4.1l1_C | 0.19 | -0.74 | -0.03 | 0.41 | 0.4 | 0.03 | 0.54 | 0.55 | -0.09 | -0.27 | 0.17 | 0.1 | -0.47 | 0.16 | 0.01 | -0.15 | 0.1775 | -0.265 | -0.115 | 0.830378 | 0.183117 | 0.94767 | 0.989223 | 0.997988 | 0.999507 |
| Epb4.1l1_Q9Z2H5_S546s_2 | Epb4.1l1_C | -0.17 | -0.13 | -0.27 | -0.19 | 0.18 | -0.2 | -0.05 | 0.02 | -0.54 | -0.41 | -0.08 | -0.79 | -0.28 | 0.76 | -0.55 | -1.15 | 0.4225 | 0.02 | -0.07 | 0.77649 | 0.390065 | 0.28206 | 0.989223 | 0.997988 | 0.99798 |
| Epb4.1l1_Q9Z2H5_S639s_1 | Epb4.1l1_C | -2.09 | -3.35 | -3.17 | -3.37 | -1.63 | -2.26 | -1.87 | -2.5 | -3.78 | -4.03 | -1.26 | -2.59 | -4.47 | -1.32 | -3.19 | -3.24 | 0.1775 | -0.265 | -0.115 | 0.926162 | 0.026489 | 0.90054 | 0.989223 | 0.997988 | 0.999507 |
| Epb4.1l1_Q9Z2H5_S666s_1 | Epb4.1l1_C | -0.21 | 0.16 | 0.13 | 0.26 | 0.2 | -0.25 | 0.08 | 0.14 | -0.13 | -0.19 | 0.2 | 0.2 | -0.36 | 0.47 | 0.39 | -0.07 | 0.93 | 0.08 | -0.06 | 0.938163 | 0.850034 | 0.783196 | 0.996349 | 0.997988 | 0.999507 |
| Epb4.1l1_Q9Z2H5_S671s_1 | Epb4.1l1_C | -0.38 | 0.1 | -0.04 | -0.11 | -0.25 | -0.51 | -0.04 | -0.26 | -0.52 | 0.44 | 0.62 | -0.2 | -0.14 | 0.15 | 0.01 | -0.14 | -0.0425 | -0.065 | 0.0225 | 0.752313 | 0.485728 | 0.547319 | 0.989223 | 0.997988 | 0.9971 |
| Epb4.1l1_Q9Z2H5_S677s_1 | Epb4.1l1_C | -0.31 | 0.04 | 0.54 | 0.64 | -0.06 | -0.42 | 0.38 | 0.63 | -0.36 | -0.62 | 1.12 | 0.45 | -0.73 | 0.17 | 0.47 | 0.48 | -0.1575 | 0.1925 | 0.0775 | 0.731272 | 0.774748 | 0.855589 | 0.995762 | 0.997988 | 0.9971 |
| Epb4.1l1_Q9Z2H5_S684s_1 | Epb4.1l1_C | 1.18 | -0.14 | -0.53 | -0.19 | 0.01 | 0.66 | 1.1 | -0.31 | -0.32 | 0.14 | -0.45 | -0.41 | -0.61 | -0.46 | -1.23 | -0.99 | -0.095 | -0.08 | -0.13 | 0.052759 | 0.538418 | 0.403386 | 0.989223 | 0.997988 | 0.994666 |
| Epb4.1l1_Q9Z2H5_S69s_1 | Epb4.1l1_C | 0.66 | -0.19 | 0.47 | -4.95 | 0.73 | 0.51 | 0.02 | 0.3 | -3.58 | 0.06 | 0.66 | 0.48 | -1.47 | -0.66 | -0.29 | -0.05 | 0.285 | -0.34 | -0.9025 | 0.744549 | 0.244878 | 0.779533 | 0.989223 | 0.997988 | 0.9971 |
| Epb4.1l1_Q9Z2H5_S75s_1 | Epb4.1l1_C | -2.19 | -1.59 | -1.66 | -2.46 | -0.63 | -2.66 | -2.03 | -2.02 | -3.05 | -2.43 | 0.03 | -1.24 | -2.3 | -1.48 | -1.52 | -1.98 | 1.3925 | 0.4075 | 0.385 | 0.639454 | 0.754407 | 0.641955 | 0.995762 | 0.997988 | 0.995259 |
| Epb4.1l1_Q9Z2H5_S782s_1 | Epb4.1l1_C | 1.52 | -2.12 | -1.69 | 0.13 | 0.61 | 1.77 | 0.67 | 1.22 | -0.89 | -1.2 | -0.37 | -0.53 | -3.29 | -0.03 | -1.87 | -1.33 | 0.14 | 0.3025 | 0.155 | 0.264677 | 0.061926 | 0.788305 | 0.989223 | 0.997988 | 0.994666 |
| Epb4.1l1_Q9Z2H5_S868s_1 | Epb4.1l1_C | 0.28 | -0.04 | 0.55 | 0.53 | -1.1 | 0.96 | 0.08 | 0.2 | 2.21 | 0.34 | 0.12 | 0.41 | 1.63 | -0.86 | 0.4 | 0.3 | 1.6075 | -0.2075 | -1.09 | 0.940013 | 0.491937 | 0.368712 | 0.989223 | 0.997988 | 0.999507 |
| Epb4.1l1_Q9Z2H5_T30t_1 | Epb4.1l1_C | -2.72 | -2.19 | -3.64 | -1.96 | -3.1 | -2.1 | -1.43 | -1.95 | -1.12 | -2.23 | -2.26 | -2.65 | -0.55 | -0.65 | -2.44 | -2.48 | -0.295 | 0.44 | 0.0375 | 0.091781 | 0.321295 | 0.253497 | 0.989223 | 0.997988 | 0.994666 |
| Epb4.1l1_Q9Z2H5_T475t_1 | Epb4.1l1_C | 0.85 | 0.29 | -0.03 | 0.23 | 0.35 | 0.71 | 0.11 | 0.34 | 0.47 | 0.23 | 0.24 | -0.16 | 0.03 | 0.66 | 0.08 | -0.05 | 0.4825 | 0.5625 | 1.0975 | 0.61288 | 0.873518 | 0.618063 | 0.996493 | 0.997988 | 0.995259 |
| Epb4.1l1_Q9Z2H5_T475t_2 | Epb4.1l1_C | -0.94 | -0.38 | -0.33 | -0.13 | -0.22 | -0.42 | -0.23 | -0.39 | -1.02 | -1.52 | 0.61 | 0.07 | -2.33 | -0.87 | -0.26 | -0.91 | 0.0425 | -0.14 | -0.155 | 0.179173 | 0.596234 | 0.967401 | 0.989223 | 0.997988 | 0.994666 |
| Epb4.1l1_Q9Z2H5_T489t_1 | Epb4.1l1_C | 0.41 | -0.88 | -0.14 | 0.42 | 0.54 | 0.46 | 0.4 | 0.6 | -0.46 | -0.79 | 0.07 | 0.04 | -1.11 | 0.13 | -0.32 | -0.21 | 0.13 | -0.02 | -0.6475 | 0.425916 | 0.115699 | 0.532226 | 0.989223 | 0.997988 | 0.994666 |
| Epb4.1l1_Q9Z2H5_T685t_1 | Epb4.1l1_C | 0.43 | -0.46 | -0.37 | 0.18 | -0.16 | 0.68 | 0.04 | 0.4 | -0.02 | -0.25 | 0.01 | -0.12 | -0.82 | -0.07 | -0.94 | -0.65 | 0.5475 | -0.2375 | -0.33 | 0.112054 | 0.34746 | 0.884964 | 0.989223 | 0.997988 | 0.994666 |
| Epb4.1l2_Q80UE4_S116s_1 | Epb4.1l2_C | -1.78 | -1.46 | -1.51 | -2.25 | -0.91 | -1.84 | -1.82 | -2.29 | -1.94 | -1.36 | -0.62 | -1.81 | -1.36 | -1.07 | -1.93 | -2 | 0.295 | -0.04 | -0.565 | 0.629675 | 0.919201 | 0.388391 | 0.997335 | 0.997988 | 0.995259 |
| Epb4.1l2_Q80UE4_S379s_1 | Epb4.1l2_C | 0.06 | 0.73 | -0.89 | -0.29 | 0.12 | -0.26 | -0.26 | -1.23 | -0.13 | 0.78 | 0.68 | -0.36 | 0.62 | 0.56 | -0.21 | -1.35 | 0.035 | 0.3175 | 0.16 | 0.996246 | 0.468797 | 0.438332 | 0.989223 | 0.997988 | 1 |
| Epb4.1l2_Q80UE4_S38s_1 | Epb4.1l2_C | 2.15 | 2.2 | 0.09 | 0.52 | 1.31 | 1.59 | 0.69 | 0.55 | 2.5 | 2.54 | 0.23 | 0.32 | 0.89 | 2.24 | 0.1 | 0.44 | -0.31 | 0.34 | 0.0025 | 0.622211 | 0.706921 | 0.835874 | 0.995437 | 0.997988 | 0.995259 |
| Epb4.1l2_Q80UE4_S492s_1 | Epb4.1l2_C | 0.41 | 1.3 | 0.52 | 0.13 | 1.09 | 0.76 | -0.3 | -0.01 | 0.13 | 0.58 | -0.15 | -0.29 | 0.49 | 1 | 0.76 | 0.38 | -0.205 | 0.1575 | -0.3225 | 0.837308 | 0.606644 | 0.145873 | 0.989223 | 0.997988 | 0.999507 |
| Epb4.1l2_Q80UE4_S543s_1 | Epb4.1l2_C | 0.48 | 0.7 | 0.04 | 0.04 | 0.75 | 0.32 | 0.66 | 0.41 | -0.05 | 0.18 | -0.31 | 0.03 | -0.62 | 0.56 | -0.1 | 0.05 | -0.205 | -0.5225 | 0.0675 | 0.317545 | 0.38941 | 0.197778 | 0.989223 | 0.997988 | 0.994666 |
| Epb4.1l2_Q80UE4_S628s_1 | Epb4.1l2_C | -0.74 | -0.18 | -0.26 | 0.07 | -0.67 | -0.83 | -0.56 | -0.44 | 0.27 | -0.36 | 0.74 | -0.42 | 1.08 | -0.27 | 0.05 | -0.67 | 0.22 | -0.3525 | -0.3425 | 0.43833 | 0.182636 | 0.340992 | 0.989223 | 0.997988 | 0.994666 |
| Epb4.1l2_Q80UE4_S86s_1 | Epb4.1l2_C | -1.31 | -1.22 | -1.7 | -2.13 | -0.26 | -1.62 | -0.72 | -1.74 | -2.24 | -1.12 | -3 | -1.2 | -0.75 | -0.91 | -1.57 | -0.98 | -0.3475 | 0.335 | 0.325 | 0.119339 | 0.224259 | 0.529286 | 0.989223 | 0.997988 | 0.994666 |
| Epb4.1l3_D0VYV6_S451s_1 | Epb4.1l3_D | 0.54 | 0.34 | 0.37 | 0.47 | 0.33 | 0.52 | 0.59 | 0.18 | -0.02 | 0.25 | -0.03 | 0.22 | 0.04 | 0.12 | 0.16 | 0.41 | 0.505 | -0.3 | 0.5375 | 0.307558 | 0.904562 | 0.152791 | 0.996493 | 0.997988 | 0.994666 |
| Epb4.1l4b_B1AXV1_S114s_1 | Epb4.1l4b_ | 0.22 | 0.51 | 1.56 | 0.32 | 0.52 | 0.45 | 0.9 | 1.02 | 0.34 | 0.45 | 0.69 | 1.38 | 0.63 | 0.81 | 0.63 | 0.49 | -0.025 | -0.325 | -0.2475 | 0.971228 | 0.838525 | 0.872066 | 0.995762 | 0.997988 | 0.999507 |
| Epb41l3_Q9WV92_S428s_1 | Epb41l3_Q | -1.19 | -1.51 | -1.36 | -1.69 | -0.68 | -1.36 | -0.56 | -1.75 | -1.34 | -1.34 | -0.3 | -1.33 | -0.37 | -0.8 | -1.19 | -1.52 | 0.07 | 0.0625 | -0.0125 | 0.165483 | 0.286482 | 0.267424 | 0.989223 | 0.997988 | 0.994666 |
| Epb41l3_Q9WV92_S451s_1 | Epb41l3_Q | -0.05 | 0.16 | -0.34 | -0.03 | 0.03 | -0.29 | -0.07 | -0.22 | -0.28 | -0.12 | -0.54 | -0.11 | -0.31 | -0.01 | -0.28 | -0.15 | 0.35 | 0.36 | 0.4675 | 0.620115 | 0.7391 | 0.409691 | 0.995762 | 0.997988 | 0.995259 |
| Epb41l3_Q9WV92_S456s_1 | Epb41l3_Q | 0.01 | -0.98 | -0.63 | -0.02 | 0.43 | 0.22 | 0.26 | 0.21 | -1.06 | -1.25 | -0.76 | -0.25 | -1.42 | -0.11 | -0.65 | -0.24 | -0.0725 | -0.1975 | -0.1225 | 0.611824 | 0.03407 | 0.236056 | 0.989223 | 0.997988 | 0.995259 |
| Epb41l3_Q9WV92_S470s_1 | Epb41l3_Q | 0.32 | 0.06 | 0.16 | -0.17 | 0.39 | 0.01 | 0.3 | 0.05 | -1.06 | -0.66 | 0.08 | 0.15 | -1.19 | -0.15 | -0.58 | 0.14 | 0.685 | -0.425 | -0.2 | 0.139812 | 0.670894 | 0.186141 | 0.992876 | 0.997988 | 0.994666 |
| Epb41l3_Q9WV92_S486s_1 | Epb41l3_Q | 1.21 | 0.2 | -0.35 | -0.52 | 1.36 | 1.26 | 0.51 | -0.06 | 0.46 | 0.04 | -0.31 | -0.31 | -0.73 | 0.73 | -0.72 | -0.38 | 0.095 | -0.465 | -0.5375 | 0.41505 | 0.204231 | 0.6965 | 0.989223 | 0.997988 | 0.994666 |
| Epb41l3_Q9WV92_S601s_1 | Epb41l3_Q | 1.59 | -1.48 | -0.63 | 0.61 | 0.58 | 1.22 | 0.16 | 1.07 | -0.71 | -0.94 | 0.93 | -0.18 | -2.44 | 0.01 | -0.78 | -0.61 | 0.6325 | -0.165 | -0.41 | 0.217844 | 0.266457 | 0.729858 | 0.989223 | 0.997988 | 0.994666 |
| Epb41l3_Q9WV92_S63s_1 | Epb41l3_Q | 1.96 | -0.46 | -0.66 | 0.07 | 0.85 | 1.57 | 0.33 | 0.13 | 0.32 | -0.12 | -0.98 | -0.63 | -1.48 | 0.04 | -1.48 | -0.98 | 0.735 | -0.2475 | -0.9775 | 0.082089 | 0.424843 | 0.351044 | 0.989223 | 0.997988 | 0.994666 |
| Epb41l3_Q9WV92_S655s_1 | Epb41l3_Q | 1.17 | 1.89 | 1.14 | 0.95 | 0.91 | 0.24 | 0.98 | 0.32 | -0.32 | 0.82 | 0.6 | 0.35 | -0.24 | 0.77 | 0.5 | 0.23 | 0.4925 | -0.58 | -1.2025 | 0.015656 | 0.049045 | 0.021859 | 0.989223 | 0.997988 | 0.994666 |
| Epb41l3_Q9WV92_S804s_1 | Epb41l3_Q | 1.2 | -0.39 | -0.42 | 0.44 | 0.45 | 0.69 | 0.43 | 0.58 | 0.11 | -0.05 | 0.12 | 0.06 | -0.97 | -0.12 | -0.33 | 0.08 | -0.675 | -0.925 | -0.9725 | 0.236177 | 0.396563 | 0.705673 | 0.989223 | 0.997988 | 0.994666 |
| Epb41l3_Q9WV92_S96s_1 | Epb41l3_Q | -0.03 | -1.43 | -0.68 | -0.28 | 0.38 | 0.07 | 0.4 | 0.12 | -1.29 | -1.54 | -1.25 | -0.23 | -2.79 | -0.29 | -0.84 | -0.31 | 0.33 | -0.1475 | -0.5425 | 0.462492 | 0.02664 | 0.271566 | 0.989223 | 0.997988 | 0.994666 |
| Epb41l3_Q9WV92_T409t_1 | Epb41l3_Q | -2.19 | -2.8 | -1.57 | -2.04 | -1.66 | -3.12 | -2.29 | -2.61 | -3.47 | -3.42 | -1.03 | -3.02 | -2.22 | -2.25 | -2.77 | -1.57 | 0.8475 | -0.4725 | -0.4525 | 0.88774 | 0.48999 | 0.327125 | 0.989223 | 0.997988 | 0.999507 |
| Epb41l3_Q9WV92_T493t_1 | Epb41l3_Q | -0.31 | -0.51 | 0.38 | 0.75 | 0.19 | -0.58 | 0.84 | 0.75 | -0.65 | -0.4 | -0.03 | 0.57 | -0.87 | 0.29 | 0.3 | 0.37 | -0.27 | -0.585 | -0.0525 | 0.895278 | 0.597102 | 0.606489 | 0.989223 | 0.997988 | 0.999507 |
| Epb41l3_Q9WV92_T495t_1 | Epb41l3_Q | 1.83 | 0.91 | -0.12 | 0.04 | 1.15 | 1.54 | 0.29 | 0.01 | 0.79 | 1.09 | -0.45 | -0.18 | 0.33 | 1.32 | -0.39 | -0.39 | 0.2225 | -0.205 | -0.055 | 0.431271 | 0.874656 | 0.520106 | 0.996493 | 0.997988 | 0.994666 |
| Epb41l3_Q9WV92_T518t_1 | Epb41l3_Q | 0.11 | -0.96 | -0.49 | -0.21 | -0.04 | -0.16 | -0.07 | -0.07 | -1.03 | -0.91 | -0.34 | -0.54 | -2.27 | -0.51 | -0.65 | -0.35 | 0.0825 | -0.3525 | -0.4475 | 0.261367 | 0.279327 | 0.321705 | 0.989223 | 0.997988 | 0.994666 |
| Epb41l3_Q9WV92_T64t_1 | Epb41l3_Q | 1.09 | -1.06 | -0.51 | -0.23 | 0.22 | 0.69 | -0.08 | -0.18 | -0.46 | -0.7 | -0.55 | -0.23 | -1.72 | -0.29 | -1.12 | -0.57 | 0.3025 | -0.3175 | -0.5575 | 0.173753 | 0.467771 | 0.499401 | 0.989223 | 0.997988 | 0.994666 |
| Epb41l3_Q9WV92_T687t_1 | Epb41l3_Q | 1.53 | 1.24 | -0.59 | 1.65 | 1.34 | 1.22 | 2.06 | 1.41 | 0.42 | 0.96 | 0.72 | 1.06 | -1.67 | 1.05 | 1.27 | 1.39 | 0.34 | -0.3075 | -0.7475 | 0.576026 | 0.294339 | 0.742505 | 0.989223 | 0.997988 | 0.995259 |
| Epb41l3_Q9WV92_T725t_1 | Epb41l3_Q | 2.24 | 1.54 | 2.08 | 2 | 1.54 | 1.83 | 1.4 | 1.98 | 0.4 | 1.54 | 1.64 | 2.15 | 0.74 | 0.16 | 2.13 | 2.11 | 0.55 | -0.1675 | -0.4475 | 0.189311 | 0.289994 | 0.20201 | 0.989223 | 0.997988 | 0.994666 |
| Epb41l3_Q9WV92_T923t_1 | Epb41l3_Q | 0.45 | 0.7 | 1.03 | 0.85 | 0.05 | 0.35 | 0.06 | 0.53 | 0.8 | 0.67 | 1.21 | 0.71 | 0.88 | 0.33 | 0.84 | 0.52 | -0.2775 | -0.5325 | -0.68 | 0.670622 | 0.054687 | 0.719234 | 0.989223 | 0.997988 | 0.996159 |
| Epb41l4a_P52963_S402s_1 | Epb41l4a_ | 0.98 | -0.36 | -0.31 | -1.03 | -0.9 | 1.1 | -0.61 | -0.38 | 2.87 | 0.92 | -1.52 | -1.05 | 2.11 | -1.65 | -0.18 | -0.82 | -0.51 | 0.09 | -0.115 | 0.955263 | 0.974719 | 0.616898 | 0.999187 | 0.997988 | 0.999507 |
| Epb41l4a_P52963_S541s_1 | Epb41l4a_ | 1.04 | 1.37 | 1.3 | 0.99 | 0.97 | 0.23 | 0.34 | 0.62 | 1.82 | 1.76 | 1.82 | 0.77 | 2.93 | 1.16 | 2.67 | 1.16 | -0.0175 | 0.485 | 0.045 | 0.110543 | 0.027484 | 0.252948 | 0.989223 | 0.997988 | 0.994666 |
| Epb41l4a_P52963_S543s_1 | Epb41l4a_ | 1.04 | 1.37 | 1.3 | 0.99 | 0.97 | 0.23 | 0.34 | 0.62 | 1.82 | 1.76 | 1.82 | 0.77 | 2.93 | 1.16 | 2.67 | 1.16 | -0.635 | 0.3675 | 0.805 | 0.110543 | 0.027484 | 0.252948 | 0.989223 | 0.997988 | 0.994666 |
| Epha5_Q6PFV6_Y649y_1 | Epha5_Q6 | 1.34 | 1.23 | 1.22 | 1.41 | 0.95 | 1.57 | 0.98 | 1.45 | 0.5 | 1.1 | 1.66 | 1.52 | 0.68 | 0.98 | 0.84 | 0.96 | -0.635 | 0.3675 | 0.805 | 0.08702 | 0.790939 | 0.729211 | 0.995762 | 0.997988 | 0.994666 |
| Epha7_Q61772_S946s_1 | Epha7_Q61 | 0.59 | 0.72 | 1.06 | 1.15 | 0.68 | 0.29 | 1.2 | 1.07 | 0.34 | 0.66 | 0.05 | 1.45 | -0.53 | 0.45 | 1.11 | 1.15 | -0.0625 | -0.105 | -0.435 | 0.427921 | 0.803271 | 0.469187 | 0.995762 | 0.997988 | 0.994666 |
| Epn1_Q80VP1_S418s_1 | Epn1_Q80 | 0.69 | 0.56 | -0.26 | 0.58 | 1.04 | 0.51 | 0.64 | 0.55 | 0.18 | 0.03 | -0.6 | -0.03 | -0.47 | 0.18 | -0.49 | 0.19 | -0.07 | -0.255 | -0.335 | 0.128447 | 0.317227 | 0.135631 | 0.989223 | 0.997988 | 0.994666 |
| Epn1_Q80VP1_S419s_1 | Epn1_Q80 | 0.25 | -0.26 | -0.55 | -0.27 | 0.36 | 0.58 | 0.37 | 0.01 | -0.07 | -0.23 | -0.52 | -0.38 | -1.22 | 0.44 | -0.68 | -0.56 | 0.2925 | -0.4975 | -0.54 | 0.456505 | 0.059441 | 0.722198 | 0.989223 | 0.997988 | 0.994666 |
| Epn1_Q80VP1_S434s_1 | Epn1_Q80 | 0.42 | 0.63 | 0.91 | 1.17 | 0.86 | 0.4 | 0.76 | 0.43 | 0.46 | 0.31 | 1.46 | 1.05 | -0.38 | 0.44 | -0.04 | 0.48 | 0.5375 | -0.0925 | -0.2975 | 0.059864 | 0.508555 | 0.910764 | 0.989223 | 0.997988 | 0.994666 |
| Epn1_Q80VP1_S453s_1 | Epn1_Q80 | 0.51 | 0.61 | 0.49 | 0.48 | 0.52 | 0.37 | 0.66 | 0.5 | 0.4 | 0.54 | 0.02 | 0.35 | -0.25 | 0.65 | 0.28 | 0.55 | -0.17 | 0.0375 | -0.6575 | 0.451481 | 0.959573 | 0.390858 | 0.999187 | 0.997988 | 0.994666 |
| Epn1_Q80VP1_T463t_1 | Epn1_Q80 | 0.3 | 0.45 | 0.35 | 0.27 | 0.74 | 0.34 | 0.62 | 0.07 | -0.05 | 0.34 | 0.02 | 0.07 | -0.55 | 0.59 | 0.06 | 0.47 | -0.01 | -0.195 | -0.215 | 0.52675 | 0.665389 | 0.267953 | 0.991397 | 0.997988 | 0.994666 |
| Epn1_Q80VP1_T469t_1 | Epn1_Q80 | 0.37 | 0.2 | 0.43 | 1.16 | 0.56 | 0.11 | 1.21 | 0.89 | -0.23 | 0.14 | 0.24 | 0.71 | -0.89 | 0.44 | 0.76 | 0.41 | 0.1 | -0.2475 | -0.2 | 0.401615 | 0.643767 | 0.320284 | 0.99033 | 0.997988 | 0.994666 |
| Epn1_Q80VP1_T493t_1 | Epn1_Q80 | -2.23 | -2.81 | -2.06 | -1.28 | -2.19 | -2.83 | -1.34 | -2.04 | -3.5 | -1.96 | -0.79 | -1.81 | -0.38 | -1.18 | -1.88 | -1.85 | 0.1525 | -0.325 | -0.36 | 0.117434 | 0.99044 | 0.891981 | 0.999187 | 0.997988 | 0.994666 |
| Epn2_Q5NCM5_S173s_1 | Epn2_Q5N | -0.09 | -0.33 | -0.68 | -0.33 | -0.26 | 0.15 | 0.01 | -0.51 | -0.34 | -0.59 | -0.73 | -0.39 | -0.07 | -0.07 | -0.36 | -0.53 | -0.005 | 0.08 | 0.7725 | 0.704586 | 0.417811 | 0.520302 | 0.989223 | 0.997988 | 0.996159 |
| Epn2_Q5NCM5_S192s_2 | Epn2_Q5N | 0.26 | -0.97 | -0.6 | -0.42 | -0.32 | 0.24 | -0.29 | -0.38 | 0.38 | 0.13 | -0.56 | -0.36 | -2.52 | -0.26 | -0.64 | -0.71 | 0.205 | -0.155 | 0.1 | 0.274019 | 0.441462 | 0.359756 | 0.989223 | 0.997988 | 0.994666 |
| Epn2_Q5NCM5_S195s_1 | Epn2_Q5N | -1.36 | -0.47 | -0.19 | 0.08 | -0.67 | -0.34 | -0.86 | -0.33 | -1.66 | -1.2 | 0.12 | -1.74 | 0.1 | -0.73 | -0.4 | 0.22 | 0.245 | 0.33 | -0.6 | 0.477855 | 0.849829 | 0.225078 | 0.996349 | 0.997988 | 0.994666 |
| Epn2_Q5NCM5_S199s_1 | Epn2_Q5N | -1.36 | -1.22 | -1.9 | -1.17 | -1.3 | -1.46 | -0.8 | -1.73 | -0.63 | -0.85 | -1.73 | -1.77 | -0.13 | -0.79 | -0.91 | -0.95 | -0.065 | -0.635 | 0.2825 | 0.040832 | 0.755408 | 0.637523 | 0.995762 | 0.997988 | 0.994666 |
| Epn2_Q5NCM5_S199s_2 | Epn2_Q5N | -0.26 | -1.06 | -0.6 | -0.42 | -0.4 | -0.16 | -0.29 | -0.38 | -0.1 | -0.15 | -0.56 | -0.36 | -2.25 | -0.49 | -0.64 | -0.71 | 0.09 | 0.1675 | 0.7175 | 0.331299 | 0.271307 | 0.285823 | 0.989223 | 0.997988 | 0.994666 |
| Epn2_Q5NCM5_S478s_1 | Epn2_Q5N | -0.75 | -1.23 | -1.04 | -0.88 | -1.01 | -0.88 | -1.29 | -0.64 | -0.97 | -1.79 | 0.37 | -1.07 | 0.41 | -0.61 | -1.67 | -1.46 | 0.2775 | 0.2925 | -0.4375 | 0.760458 | 0.932962 | 0.80523 | 0.998562 | 0.997988 | 0.9971 |
| Epn2_Q5NCM5_S482s_1 | Epn2_Q5N | 0.54 | 0.55 | 0.65 | 1.03 | 0.56 | 0.2 | 0.91 | 0.97 | 0.48 | 0.6 | 0.64 | 0.55 | 0.1 | 0.98 | 0.55 | 0.57 | 0.02 | 0.11 | 0.1425 | 0.620717 | 0.900732 | 0.581324 | 0.996493 | 0.997988 | 0.995259 |
| Epn2_Q5NCM5_T504t_1 | Epn2_Q5N | -2.26 | -2.38 | -1.94 | -1.73 | -1.75 | -2.65 | -1.04 | -2.16 | -2.39 | -2.05 | -0.55 | -1.92 | -2.36 | -1.49 | -2.01 | -1.74 | -0.0325 | -0.125 | -0.1425 | 0.557522 | 0.630746 | 0.415843 | 0.989223 | 0.997988 | 0.995259 |
| Epn2_Q5NCM6_S192s_1 | Epn2_Q5N | -0.78 | -0.89 | -0.16 | 0.06 | -1.09 | -0.19 | -0.77 | -0.2 | 0.35 | 0.23 | 0.5 | -0.18 | 0.9 | -0.38 | 0.28 | -0.11 | 0.1775 | 0.35 | 0.1775 | 0.128692 | 0.717764 | 0.054228 | 0.995437 | 0.997988 | 0.994666 |
| Epn2_Q5NCM6_S192s_2 | Epn2_Q5N | -0.46 | -0.25 | -0.89 | -1.07 | -1.16 | -0.83 | -1.9 | -1.2 | 0.8 | -0.49 | -1.13 | -1.12 | 1.13 | -0.32 | -0.58 | -0.9 | -0.12 | 0.6675 | 0.615 | 0.300562 | 0.07771 | 0.698031 | 0.989223 | 0.997988 | 0.994666 |
| Epn2_Q5NCM6_S195s_1 | Epn2_Q5N | -0.75 | -0.48 | -1.98 | -1.73 | -0.12 | -0.81 | -1.21 | -1.79 | -0.39 | -0.84 | -1.78 | -2.39 | 0.96 | -0.37 | -1.05 | -1.28 | -0.605 | 0.1825 | 0.5 | 0.184632 | 0.593531 | 0.831442 | 0.989223 | 0.997988 | 0.994666 |
| Epn2_Q5NCM6_S195s_2 | Epn2_Q5N | -0.46 | -0.34 | -0.89 | -1.07 | -1 | -0.78 | -1.9 | -1.2 | 0.65 | -0.62 | -1.13 | -1.12 | 0.92 | -0.24 | -0.58 | -0.9 | 0.2525 | -0.115 | 0.8 | 0.26958 | 0.11777 | 0.758827 | 0.989223 | 0.997988 | 0.994666 |
| Epn3_Q91W69_S257s_1 | Epn3_Q91 | -1.21 | 0 | 0.85 | 0.66 | -0.21 | -0.72 | -0.25 | 0.01 | 0.58 | -0.42 | 2.25 | -1.66 | 0.68 | 1.79 | 1.31 | 0.35 | -0.53 | 0.135 | 0.49 | 0.09555 | 0.429055 | 0.89389 | 0.989223 | 0.997988 | 0.994666 |
| Eprs_Q8CGC7_S944s_1 | Eprs_Q8CG | -2.79 | -3.19 | -1.31 | -2.29 | -1.99 | -2.73 | -1.03 | -1.16 | -4.07 | -2.29 | -1.71 | -2.41 | -1.64 | -1.6 | -1.57 | -2.19 | -0.3675 | 0.1125 | 0.9575 | 0.155228 | 0.216928 | 0.706441 | 0.989223 | 0.997988 | 0.994666 |
| Eps15_F6W2Q5_S324s_1 | Eps15_F6W | -1.08 | 0.29 | 0.32 | 0.65 | -0.55 | -3.14 | 0.71 | 0.72 | 0.15 | 0.73 | -0.37 | 0.45 | -0.25 | 0.85 | 0.64 | 0.43 | 0.6675 | -0.225 | 0.645 | 0.408571 | 0.483344 | 0.656658 | 0.989223 | 0.997988 | 0.994666 |
| Eps15l1_Q60902_S229s_1 | Eps15l1_Q | 1.05 | 0.64 | 0.81 | 1.2 | 0.2 | 0.7 | 0.86 | 1.04 | 0.46 | 0.79 | 0.56 | 0.33 | 0.13 | 0.87 | 0.16 | 0.35 | -0.61 | 0.195 | 0.3725 | 0.079879 | 0.40493 | 0.131792 | 0.989223 | 0.997988 | 0.994666 |
| Eps15l1_Q60902_S353s_1 | Eps15l1_Q | 0.72 | 0.76 | 0.56 | 0.05 | 1.01 | 0.43 | 0.91 | -0.33 | 0.32 | 0.73 | 0.56 | -0.08 | 0.28 | 0.99 | -0.15 | -0.42 | -0.225 | -0.39 | -0.5475 | 0.357494 | 0.959981 | 0.62778 | 0.999187 | 0.997988 | 0.994666 |
| Eps15l1_Q60902_T364t_1 | Eps15l1_Q | -0.99 | -0.29 | -0.01 | -0.53 | -0.77 | -0.29 | 0.21 | -0.32 | -0.52 | -0.44 | -0.59 | -0.29 | -0.67 | -0.17 | -0.54 | -0.33 | -0.0175 | -0.14 | -0.3475 | 0.926452 | 0.601102 | 0.985396 | 0.989223 | 0.997988 | 0.999507 |
| Eps8_Q08509_S624s_1 | Eps8_Q085 | -0.37 | 0.44 | -0.55 | -0.36 | 0.52 | 0.6 | 0.62 | 0.15 | -0.77 | 0.28 | -0.45 | 0.24 | -0.11 | -0.28 | -0.1 | -0.41 | 0.1625 | -0.005 | 0.0275 | 0.959673 | 0.034444 | 0.92133 | 0.989223 | 0.997988 | 0.999507 |

| | | | | | | | | | | | | | | | | | | | | | | | | | | |
|---|---|---|---|---|---|---|---|---|---|---|---|---|---|---|---|---|---|---|---|---|---|---|---|---|---|---|
| Eps8l1_E9Q4X5_S300s_1 | Eps8l1_E9( | 1.69 | 0.4 | 1.72 | 2.36 | 1.3 | 0.42 | 1.72 | 1.3 | -0.12 | 0.18 | 1.94 | 1.51 | -0.34 | 0.7 | 1.53 | 1.63 | 0.6825 | 0.035 | -0.015 | 0.259693 | 0.44394 | 0.280021 | 0.989223 | 0.997988 | 0.994666 |
| Eps8l1_E9Q4X5_S685s_1 | Eps8l1_E9( | 0.84 | -0.48 | -0.16 | 1.64 | 1.05 | -0.22 | 0.5 | -0.14 | -0.68 | -0.1 | 0.33 | -0.41 | 0.77 | 0.98 | -0.75 | 0.26 | -0.3575 | -0.665 | -0.6625 | 0.799186 | 0.754006 | 0.198514 | 0.995762 | 0.997988 | 0.999507 |
| Eps8l2_Q99K30_S242s_1 | Eps8l2_Q9! | -1.87 | -1.59 | -1.45 | -1.89 | -3.28 | -0.88 | -3.31 | -2.73 | -0.54 | -1.82 | -1.57 | -2.85 | 0.49 | -3.81 | -1.38 | -2.15 | -0.1625 | -0.675 | -0.145 | 0.98743 | 0.133134 | 0.991405 | 0.989223 | 0.997988 | 1 |
| Erbb2ip_Q80TH2_S1013s_1 | Erbb2ip_Q8 | 0.85 | -0.28 | -0.92 | -0.38 | -1.86 | 1.06 | -1.88 | -0.81 | 1.95 | -0.03 | -0.38 | -0.61 | 0.89 | -1.87 | -1.37 | -1.32 | -0.85 | 0.005 | -0.0125 | 0.273933 | 0.332045 | 0.515887 | 0.989223 | 0.997988 | 0.994666 |
| Erbb2ip_Q80TH2_S1125s_1 | Erbb2ip_Q8 | -2.26 | -0.54 | -2.06 | -1.77 | -1.22 | -2.37 | -2.41 | -1.53 | -2.54 | -2.8 | -0.7 | -2.4 | -1.3 | -1.61 | -2.58 | -2.02 | -0.69 | 0.415 | -0.735 | 0.633904 | 0.623441 | 0.432661 | 0.989223 | 0.997988 | 0.995259 |
| Erbb2ip_Q80TH2_S1209s_1 | Erbb2ip_Q8 | 6.89 | 3.41 | 3.66 | 0.75 | 1.47 | 7.27 | 2.18 | 2.23 | 3.9 | 5.12 | 0.66 | 3.13 | 1.74 | 2.36 | 3.65 | 3.74 | -0.225 | -0.4525 | -0.22 | 0.495879 | 0.804092 | 0.730089 | 0.995762 | 0.997988 | 0.994666 |
| Erbb2ip_Q80TH2_S849s_1 | Erbb2ip_Q8 | 1.37 | 0.43 | 0.24 | -0.03 | 0.14 | 0.87 | -0.24 | -0.1 | -0.16 | 0.73 | 0.49 | 0.09 | -0.28 | 0.17 | -0.07 | -0.07 | -0.39 | -0.475 | -0.805 | 0.129814 | 0.390496 | 0.565843 | 0.989223 | 0.997988 | 0.994666 |
| Erbb2ip_Q80TH2_S854s_1 | Erbb2ip_Q8 | 0.69 | 0.53 | 0.71 | 0.83 | -0.02 | 0.23 | 0.33 | 0.41 | 0.45 | 0.63 | 0.72 | 0.23 | 0.39 | 0.05 | 0.71 | 0.53 | -0.335 | -0.215 | -0.565 | 0.30769 | 0.055562 | 0.428834 | 0.989223 | 0.997988 | 0.994666 |
| Erbb4_Q61527_S997s_1 | Erbb4_Q61 | 0.18 | 0.51 | 0.76 | 0.6 | 0.53 | -0.09 | 0.13 | -0.28 | 0 | 0.29 | 0.88 | 0.43 | 0.31 | 0.27 | -0.06 | 0.18 | -0.4525 | -0.1825 | -0.27 | 0.206243 | 0.117133 | 0.683926 | 0.989223 | 0.997988 | 0.994666 |
| Erbb4_Q61527_T699t_1 | Erbb4_Q61 | 0.41 | 0.47 | -0.35 | 0.34 | 0.82 | -0.24 | 0.27 | -0.08 | 0.19 | 0.65 | -1.64 | -0.05 | -0.14 | 0.48 | -0.52 | 0.17 | -0.44 | -0.1125 | -0.3375 | 0.508449 | 0.937326 | 0.400969 | 0.999187 | 0.997988 | 0.994666 |
| Erc1_V9GWT6_S17s_1 | Erc1_V9GV | 0.31 | -0.19 | -0.85 | -1.09 | -1.27 | 0.24 | -2.48 | -1.4 | 1.89 | -0.24 | -1.04 | -0.55 | 1.83 | -1.91 | -0.9 | -0.65 | -0.025 | -0.43 | -0.22 | 0.950047 | 0.202533 | 0.478577 | 0.989223 | 0.997988 | 0.999507 |
| Erc1_V9GWT6_S21s_1 | Erc1_V9GV | -1.56 | -0.42 | -1.06 | -1.7 | -0.69 | -1.07 | -2.24 | -0.27 | -0.47 | -0.29 | 1.52 | -2.18 | 0.34 | -1.12 | -1.63 | -0.96 | -0.7725 | 0.47 | 0.0475 | 0.486335 | 0.804569 | 0.271679 | 0.995762 | 0.997988 | 0.994666 |
| Erc1_V9GWT6_S37s_1 | Erc1_V9GV | 0.14 | 0.32 | 0.35 | -0.01 | 0.04 | 0.89 | -0.47 | -0.14 | 2.03 | 0.35 | 0.97 | 0.18 | 2.6 | -0.21 | 0.47 | -0.44 | 0.1175 | 0.83 | 0.3425 | 0.526091 | 0.706505 | 0.128994 | 0.995437 | 0.997988 | 0.994666 |
| Erc1_V9GWT6_S415s_1 | Erc1_V9GV | -1.77 | -2.64 | -1.72 | -3.39 | -2.8 | -1.55 | -2.51 | -1.8 | -0.97 | -1.71 | -1.18 | -1.41 | -0.2 | -2.86 | -1.78 | -1.51 | -0.12 | 0.6825 | 0.405 | 0.21763 | 0.641761 | 0.029497 | 0.989223 | 0.997988 | 0.994666 |
| Erc1_V9GWT6_S416s_1 | Erc1_V9GV | -1.77 | -2.64 | -1.72 | -3.39 | -2.8 | -1.55 | -2.51 | -1.8 | -0.97 | -1.71 | -1.18 | -1.41 | -0.2 | -2.86 | -1.78 | -1.51 | 0.215 | 1.0625 | 0.7925 | 0.21763 | 0.641761 | 0.029497 | 0.989223 | 0.997988 | 0.994666 |
| Erc1_V9GWT6_S702s_1 | Erc1_V9GV | 0.16 | -0.21 | -0.13 | 0.43 | -0.26 | 0.71 | -0.02 | 0 | 1.86 | 0.33 | -0.31 | 0.38 | 1.72 | -0.59 | 0.43 | 0.17 | 0.215 | 1.0625 | 0.7925 | 0.448115 | 0.875385 | 0.292078 | 0.996493 | 0.997988 | 0.994666 |
| Erc1_V9GWT6_S937s_1 | Erc1_V9GV | 0.04 | 0.04 | 0.09 | 0.49 | 0.27 | -0.25 | 0.42 | 0.64 | 0.33 | 0.38 | 0.17 | 0.41 | -1.2 | 0.93 | -0.12 | -0.33 | 0.045 | 0.5025 | 0.37 | 0.442178 | 0.692345 | 0.490304 | 0.994893 | 0.997988 | 0.994666 |
| Erc2_Q3UHT7_S14s_1 | Erc2_Q3UH | -0.33 | -0.28 | -0.03 | 0.16 | 0.49 | -0.57 | -0.07 | 0.29 | 0 | 0.14 | 0.54 | 0.92 | 0.12 | 0.47 | 0.32 | -0.15 | 0.105 | 0.1575 | -0.345 | 0.263478 | 0.595026 | 0.094459 | 0.989223 | 0.997988 | 0.994666 |
| Erc2_Q3UHT7_S329s_1 | Erc2_Q3UH | 1.49 | 0.93 | 1.63 | 1.17 | 0.51 | 2.2 | -0.02 | 0.76 | 2.26 | 0.81 | 0.72 | 1.13 | 1.32 | 0.51 | 0.91 | 0.73 | 0.155 | 0.52 | 0.31 | 0.163934 | 0.352571 | 0.847162 | 0.989223 | 0.997988 | 0.994666 |
| Erc2_Q3UHT7_S966s_1 | Erc2_Q3UH | 1.23 | 1.15 | 0.6 | 0.88 | 0.68 | 0.74 | 0.5 | 0.8 | 0.73 | 1.41 | 0.21 | 0.93 | 0.08 | 1.12 | 0.6 | 0.64 | -0.4425 | -0.075 | -0.4375 | 0.269151 | 0.236488 | 0.649181 | 0.989223 | 0.997988 | 0.994666 |
| Ercc5_E9QM61_S341s_1 | Ercc5_E9QI | -3.85 | -3.73 | -3.61 | -4.67 | -2.56 | -4.28 | -4.39 | -5.08 | -4.96 | -2.78 | -2.13 | -4.19 | -1.96 | -2.74 | -3.87 | -4.36 | -0.285 | -0.145 | -0.355 | 0.204196 | 0.832836 | 0.47981 | 0.995762 | 0.997988 | 0.994666 |
| Ercc5_E9QM61_S384s_1 | Ercc5_E9QI | -2.35 | -1 | -2.06 | -2.64 | -2.52 | -2.78 | -2.9 | -2.79 | -1.93 | -2.2 | -0.41 | -3.21 | -1.01 | -0.93 | -2.84 | -2.55 | -0.1125 | 0.45 | 0.7325 | 0.751613 | 0.06767 | 0.903796 | 0.989223 | 0.997988 | 0.9971 |
| Erg_E9PY05_S222s_1 | Erg_E9PY0! | -1.61 | -2.3 | -1.44 | -1.7 | -1.87 | -0.95 | -0.79 | -1.89 | -0.64 | -1.54 | -3.53 | -1.79 | 1.09 | -3.28 | -1.58 | -1.8 | -0.735 | 0.075 | 0.18 | 0.652887 | 0.284345 | 0.846695 | 0.989223 | 0.997988 | 0.995259 |
| Erg_E9PY05_S88s_1 | Erg_E9PY0! | -2.86 | -1.84 | -3.24 | -1.68 | -2.87 | -1.17 | -4.34 | -3.57 | -0.57 | -1.42 | -1.69 | -2.78 | -0.06 | -2.41 | -1.73 | -2.48 | 0.3875 | -0.1125 | 0.37 | 0.252304 | 0.404844 | 0.17535 | 0.989223 | 0.997988 | 0.994666 |
| Ermn_Q5EBJ4_S211s_1 | Ermn_Q5EI | 2.83 | 3 | 5.95 | 2.5 | 2.26 | 2.64 | 6.18 | 4.77 | 2.4 | 2.56 | 1.03 | 5.56 | 1.68 | 2.5 | 6.26 | 7.05 | -0.5825 | 0.79 | 0.735 | 0.553592 | 0.710476 | 0.53774 | 0.995437 | 0.997988 | 0.994666 |
| Ermn_Q5EBJ4_S223s_1 | Ermn_Q5EI | 0.03 | 0.58 | 0.53 | 0.41 | 0.54 | -0.14 | 0.67 | 0.49 | -0.37 | 0.22 | 0.18 | 0.27 | -0.15 | 0.05 | 0.36 | 0.61 | 0.3925 | -0.6825 | 0.8025 | 0.552756 | 0.992465 | 0.248845 | 0.999187 | 0.997988 | 0.994666 |
| Ermn_Q5EBJ4_S227s_1 | Ermn_Q5EI | 0.76 | 1.1 | 0.82 | 0.77 | 0.77 | 0.23 | 0.6 | 0.46 | 0.37 | 0.98 | 0.52 | 0.58 | 0.69 | 0.71 | 0.89 | 0.9 | 0.0025 | -0.3125 | -0.17 | 0.784259 | 0.143616 | 0.310427 | 0.989223 | 0.997988 | 0.998711 |
| Ermn_Q5EBJ4_S230s_1 | Ermn_Q5EI | 0.35 | 0.53 | 0.53 | 0.41 | 0.09 | 0.2 | 0.84 | 0.01 | -0.66 | 0.16 | -0.3 | -0.21 | -0.36 | 0.77 | 0.46 | 1.14 | -0.3475 | -0.25 | -0.065 | 0.892477 | 0.502866 | 0.017799 | 0.989223 | 0.997988 | 0.999507 |
| Ermn_Q5EBJ4_S9s_1 | Ermn_Q5EI | -1.33 | -0.1 | -0.62 | 0.16 | -0.91 | -1.01 | -1.12 | -0.2 | 0.13 | -0.26 | 0.06 | 0.02 | 0.23 | -0.37 | -0.47 | -0.7 | -0.17 | -0.7075 | 0.0475 | 0.713078 | 0.384477 | 0.214502 | 0.989223 | 0.997988 | 0.996966 |
| Ermn_Q5EBJ4_T234t_1 | Ermn_Q5EI | 0.75 | 0.92 | 0.67 | 0.68 | 1.29 | 0.94 | 0.99 | 0.61 | 0.33 | 1.07 | -0.16 | 0.57 | 0.15 | 0.82 | 0.78 | 0.44 | -0.3375 | 0.46 | 0.145 | 0.438794 | 0.385069 | 0.328799 | 0.989223 | 0.997988 | 0.994666 |
| Ermn_Q5EBJ4_T257t_1 | Ermn_Q5EI | -2.11 | -2.02 | -1.36 | -1.61 | -1.32 | -2.97 | -1.28 | -2.56 | -2.83 | -2.46 | -1.78 | -2.58 | -1.44 | -1.69 | -1.41 | -1.9 | 0.2025 | -0.3025 | -0.2075 | 0.565384 | 0.559976 | 0.068904 | 0.989223 | 0.997988 | 0.995259 |
| Esyt2_Q3TZZ7_S667s_1 | Esyt2_Q3T. | -0.08 | -0.71 | -0.45 | -0.06 | -0.33 | -0.06 | 0.23 | 0.13 | -0.85 | -0.65 | -1.36 | -0.08 | -0.79 | -1.07 | -0.32 | -0.22 | -0.2575 | -0.6375 | 0.165 | 0.383521 | 0.229533 | 0.235423 | 0.989223 | 0.997988 | 0.994666 |
| Etl4_E9QAU4_S1100s_1 | Etl4_E9QAI | -0.45 | 0.49 | 0.21 | -0.46 | 0.9 | 0.27 | 0.13 | -0.61 | 0.02 | 0.11 | 0.16 | 0.04 | 0.45 | 1.05 | -0.1 | -0.05 | 0.3175 | -0.41 | -0.275 | 0.31396 | 0.559293 | 0.641128 | 0.989223 | 0.997988 | 0.994666 |
| Etl4_E9QAU4_S1123s_1 | Etl4_E9QAI | 0.88 | 0.24 | 0.22 | 0.72 | 0.87 | 1.9 | 0.56 | 0.98 | -0.3 | -0.05 | 1.25 | 1.28 | -0.11 | 0.11 | 0.48 | 0.56 | 0.225 | 0.135 | 0.39 | 0.397341 | 0.123091 | 0.945253 | 0.989223 | 0.997988 | 0.994666 |
| Etl4_E9QAU4_S1138s_1 | Etl4_E9QAI | -0.84 | 0.49 | -0.36 | -1.95 | 0.13 | 0.31 | -1.38 | -0.44 | 0.24 | -0.06 | 1.69 | -0.78 | 0.82 | 1.32 | -0.97 | -0.77 | 0.5625 | 0.03 | -0.255 | 0.280751 | 0.57849 | 0.178445 | 0.989223 | 0.997988 | 0.994666 |
| Etl4_E9QAU4_S1164s_1 | Etl4_E9QAI | 0.41 | 0.63 | 1.07 | 0.65 | 0.7 | -0.24 | 1.07 | 0.63 | -0.01 | 0.86 | 0.73 | 0.44 | -0.69 | 0.4 | 0.47 | 0.8 | 0.32 | 0.9375 | 0.765 | 0.248589 | 0.6442 | 0.521447 | 0.99033 | 0.997988 | 0.994666 |
| Etl4_E9QAU4_S1698s_1 | Etl4_E9QAI | 2.41 | 1.83 | 0.76 | 0.82 | 1.48 | 1.69 | 0.41 | 0.75 | 0.93 | 2.01 | -0.07 | 1.39 | 1.16 | 0.39 | 0.8 | 0.66 | -0.15 | -0.185 | -0.445 | 0.125977 | 0.432224 | 0.485205 | 0.989223 | 0.997988 | 0.994666 |
| Etl4_E9QAU4_S1813s_1 | Etl4_E9QAI | 2.29 | 0.3 | 1.6 | 1.3 | 0.02 | 2.84 | -0.21 | 1.13 | 2.17 | 0.93 | 0.88 | 1.56 | 0.92 | 0.35 | 0.05 | 0.16 | -0.3725 | -0.39 | -0.7025 | 0.04621 | 0.552389 | 0.979443 | 0.989223 | 0.997988 | 0.994666 |
| Etl4_E9QAU4_S1847s_1 | Etl4_E9QAI | 0.36 | -0.57 | -0.42 | 0.11 | 0.66 | 1.08 | 0.73 | 0.73 | -0.42 | -0.3 | 1.3 | 0.44 | -1.01 | 0.09 | -0.62 | -0.12 | -0.4275 | 0.0125 | -1.0025 | 0.432717 | 0.00727 | 0.390694 | 0.989223 | 0.997988 | 0.994666 |
| Etl4_E9QAU4_S1859s_1 | Etl4_E9QAI | 0.56 | 0.45 | 0.11 | 0.47 | 1.07 | 0.86 | 0.94 | 0.62 | 0.27 | 0.73 | 0.23 | 0.06 | -0.35 | 0.9 | 0.22 | 0.32 | 0.93 | 0.385 | -0.285 | 0.698043 | 0.055276 | 0.764074 | 0.989223 | 0.997988 | 0.996159 |
| Etl4_E9QAU4_S1894s_1 | Etl4_E9QAI | 0.6 | 0.59 | 0.56 | 0.7 | 1.34 | 1 | 1.21 | 0.96 | -0.13 | 0.54 | 0.28 | 0.79 | -0.42 | 0.35 | 0.4 | 0.83 | 0.475 | -0.075 | -0.125 | 0.317493 | 0.029962 | 0.367624 | 0.989223 | 0.997988 | 0.994666 |
| Etl4_E9QAU4_S1919s_1 | Etl4_E9QAI | -0.23 | 0.3 | 0.31 | 0.3 | 0.83 | -0.24 | 0.45 | 0.18 | 0 | 0.52 | 0.43 | 0.43 | 0.61 | 0.84 | 0.37 | 0.31 | 0.515 | -0.2425 | -0.3225 | 0.199635 | 0.64473 | 0.491846 | 0.99033 | 0.997988 | 0.994666 |
| Etl4_E9QAU4_S1950s_1 | Etl4_E9QAI | 0.61 | 0.72 | -0.46 | -0.66 | 0.59 | 1.04 | -0.44 | -1.14 | 0.67 | 0.7 | -0.55 | -0.41 | 0.8 | 1.02 | -1.02 | -0.8 | 0.135 | 0.175 | 0.3625 | 0.928419 | 0.941985 | 0.914922 | 0.999187 | 0.997988 | 0.999507 |
| Etl4_E9QAU4_S364s_1 | Etl4_E9QAI | 0.14 | -0.33 | 0.56 | 0.66 | 0.29 | -0.13 | 0.58 | 0.79 | 0.11 | 0.12 | 0.89 | 1 | -0.61 | 1.04 | 0.78 | 0.33 | -0.04 | 0.05 | -0.0525 | 0.764101 | 0.694601 | 0.440115 | 0.994893 | 0.997988 | 0.9971 |
| Etl4_E9QAU4_S410s_1 | Etl4_E9QAI | 0.11 | -0.55 | -0.69 | -1.13 | -0.24 | -0.52 | -0.06 | -0.92 | -0.27 | 0.27 | -0.46 | -1.12 | -0.46 | 0.89 | -0.72 | -1.51 | 0.125 | 0.2725 | 0.1275 | 0.826708 | 0.693497 | 0.661888 | 0.994893 | 0.997988 | 0.999507 |
| Etl4_E9QAU4_S412s_1 | Etl4_E9QAI | 0.59 | -0.13 | 0.12 | 0.33 | 0.92 | 0.52 | 0.82 | 0.59 | 0.04 | -0.11 | 0.11 | 0.25 | -0.99 | 0.42 | -0.49 | -0.54 | 0.13 | 0.17 | 0.115 | 0.105344 | 0.070211 | 0.535723 | 0.989223 | 0.997988 | 0.994666 |
| Etl4_E9QAU4_T521t_1 | Etl4_E9QAI | -0.62 | -0.34 | 0.77 | 0.51 | -0.91 | 0.31 | -1.05 | -0.83 | 1.67 | -0.42 | -0.96 | -0.61 | 1.95 | -1.45 | -0.31 | -0.23 | 0.485 | -0.155 | -0.6275 | 0.897837 | 0.129486 | 0.796906 | 0.989223 | 0.997988 | 0.999507 |
| Etl4_E9QAU4_T918t_1 | Etl4_E9QAI | 7.73 | 4.29 | 5.42 | 2.76 | 3.92 | 7.95 | 6.52 | 5.24 | 2.72 | 6.24 | 3.45 | 5.96 | 2.5 | 1.09 | 6.25 | 6.82 | -0.7 | -0.16 | -0.09 | 0.559422 | 0.469115 | 0.704478 | 0.989223 | 0.997988 | 0.995259 |
| Etv6_E9Q8J8_S126s_1 | Etv6_E9Q8 | -2.56 | -3.29 | -2.36 | -3.57 | -2.86 | -3.85 | -3.07 | -3.63 | -4.79 | -2.27 | -1.25 | -3.12 | -0.16 | -1.81 | -2.57 | -2.96 | 0.8575 | -0.4575 | -0.885 | 0.11027 | 0.282132 | 0.903484 | 0.989223 | 0.997988 | 0.994666 |
| Eva1a_D3Z511_S118s_1 | Eva1a_D3Z | 1.11 | -0.25 | 0.69 | 1.26 | 0.27 | 0.95 | 0.55 | 1.23 | 0.12 | 0.15 | 0.99 | 1.34 | -1.58 | 0.93 | -1.06 | -0.43 | -0.4075 | 0.0875 | 1.07 | 0.058265 | 0.902625 | 0.90535 | 0.996493 | 0.997988 | 0.994666 |
| Evc2_Q8K1G2_S331s_1 | Evc2_Q8K1 | 0.75 | 0.81 | 0.57 | 0.53 | 0.66 | -1.1 | -0.47 | 0.16 | 0.77 | -0.02 | 0.45 | 0.04 | 0.6 | 0.23 | -0.23 | -0.35 | 0.0475 | -0.0525 | -1.2375 | 0.064799 | 0.046427 | 0.197753 | 0.989223 | 0.997988 | 0.994666 |
| Evi5l_H3BJ67_S579s_1 | Evi5l_H3BJ | 0.5 | -0.16 | -0.44 | -0.58 | -0.04 | 0.2 | -0.92 | -0.43 | -0.04 | -0.32 | -1.2 | -0.22 | -0.07 | -0.07 | -0.79 | -0.6 | -0.8525 | -0.355 | -0.6025 | 0.534654 | 0.713507 | 0.455851 | 0.995437 | 0.997988 | 0.994666 |
| Evi5l_H3BJ67_S638s_1 | Evi5l_H3BJ | 0.55 | 0.38 | 0.16 | 0.12 | -0.1 | 0.57 | 0.4 | 0.82 | -0.07 | 1.14 | -0.07 | 0.74 | 0.53 | 0.98 | 0.31 | 1.21 | -0.1275 | -0.275 | -0.2125 | 0.146419 | 0.651833 | 0.697082 | 0.990358 | 0.997988 | 0.994666 |
| Evi5l_H3BJX1_S394s_1 | Evi5l_H3BJ | 0.45 | 0.59 | -0.61 | -0.41 | -0.2 | 0 | 0.28 | 0.41 | 0.52 | 0.9 | 1.49 | 0.78 | 0.61 | 0.48 | 1.03 | 0.54 | 0.12 | 0.1325 | 0.455 | 0.087496 | 0.728919 | 0.032798 | 0.995762 | 0.997988 | 0.994666 |
| Evl_E9PVP4_S117s_1 | Evl_E9PVP | 0.42 | -0.94 | -0.55 | 0.68 | -0.1 | 0.15 | 0 | 0.2 | 0.41 | 1.26 | 0.59 | -1.3 | 0.32 | 0.45 | 0.65 | 0.85 | 0.1175 | 0.9175 | 0.66 | 0.12706 | 0.676694 | 0.583451 | 0.99315 | 0.997988 | 0.994666 |
| Evl_E9PVP4_S137s_1 | Evl_E9PVP | 2.52 | 0.73 | 1.64 | 2.49 | -0.03 | 2.86 | 0.46 | 2.18 | 3.85 | 2.07 | 1.93 | 1.55 | 2.52 | 0.53 | 1.95 | 1.36 | 0.16 | 0.3375 | 0.665 | 0.647839 | 0.506927 | 0.415299 | 0.989223 | 0.997988 | 0.995259 |
| Evl_E9PVP4_S231s_1 | Evl_E9PVP | 1.05 | 0.06 | -1.29 | -0.5 | -0.13 | 0.92 | -1.52 | -1.23 | 0.41 | 0.34 | -0.5 | -0.53 | 0.25 | 0.2 | -0.79 | -0.23 | -0.4775 | 0.505 | -0.255 | 0.957348 | 0.629096 | 0.847296 | 0.989223 | 0.997988 | 0.999507 |
| Evl_E9PVP4_S291s_1 | Evl_E9PVP | -0.07 | -0.09 | 0.11 | -0.3 | -0.41 | 0.16 | -0.39 | -0.5 | -0.28 | -0.59 | 0.29 | -1.04 | 0.06 | -0.26 | 0.49 | -0.53 | -0.32 | 0.1 | 0.0275 | 0.9264 | 0.416751 | 0.335126 | 0.989223 | 0.997988 | 0.999507 |
| Evl_E9PVP4_S316s_1 | Evl_E9PVP | 0.93 | 1 | 0.98 | 1.01 | 1 | 0.8 | 1.33 | 1 | 0.99 | 0.84 | 1.35 | 1.2 | 0.71 | 1.33 | 1.04 | 1.16 | -0.1975 | -0.3175 | 0.0275 | 0.748151 | 0.804 | 0.607591 | 0.995762 | 0.997988 | 0.9971 |
| Exoc1_Q5PPR2_S448s_1 | Exoc1_Q5P | 0.41 | -1.56 | -1.4 | -1.42 | -0.05 | 0.92 | -0.74 | -1.5 | -0.25 | -0.99 | 1.86 | -1.18 | -1.11 | -1 | -1.5 | -1.22 | 0.0525 | 0.115 | 0.08 | 0.644806 | 0.309175 | 0.274574 | 0.989223 | 0.997988 | 0.995259 |
| Exoc1_Q5PPR2_S455s_1 | Exoc1_Q5P | 0.96 | 0.21 | -0.58 | 0.37 | 0.67 | 0.83 | -0.35 | 0.65 | 0.26 | 0.37 | 1.56 | -0.08 | -0.15 | 0.92 | 0.1 | 0.41 | 0.65 | 0.8525 | -0.215 | 0.841142 | 0.602496 | 0.534613 | 0.989223 | 0.997988 | 0.999507 |
| Exoc1_Q5PPR2_S458s_1 | Exoc1_Q5P | 1.18 | -0.73 | -0.55 | -0.19 | 0.09 | 1.24 | -0.38 | -0.11 | 0.08 | 0.91 | 0.15 | 0.3 | -0.26 | 0.36 | -0.37 | -0.06 | 0.21 | 0.2875 | 0.08 | 0.98222 | 0.583801 | 0.352448 | 0.989223 | 0.997988 | 1 |
| Exoc1_Q5PPR2_S469s_1 | Exoc1_Q5P | -1.02 | -2.37 | -0.18 | -0.5 | -0.99 | -0.38 | 0.49 | 0.27 | -1.63 | -1.83 | 2.09 | 0.91 | -0.38 | -0.54 | -0.27 | 0.22 | 0.2825 | 0.4325 | -0.01 | 0.135464 | 0.129796 | 0.356123 | 0.989223 | 0.997988 | 0.994666 |
| Exoc1_Q6P1Y9_S470s_1 | Exoc1_Q6P | -0.78 | -1.48 | -1.24 | -1.36 | -0.84 | -0.5 | -1.08 | -1.14 | 0.08 | -1.03 | -1.44 | -0.68 | -0.28 | -1.66 | -0.87 | -0.5 | 0.865 | 0.9025 | 0.775 | 0.300386 | 0.23078 | 0.241055 | 0.989223 | 0.997988 | 0.994666 |
| Exoc2_Q9D4H1_S432s_1 | Exoc2_Q9C | -0.36 | -1.41 | -0.73 | -1.34 | -0.93 | 0.72 | -1.16 | -0.33 | 0.77 | -1.12 | -0.69 | -0.44 | 1.13 | -1.02 | -1.03 | -0.74 | 0.325 | 0.4475 | 0.3875 | 0.323921 | 0.258472 | 0.217999 | 0.989223 | 0.997988 | 0.994666 |
| Exoc7_O35250_S283s_1 | Exoc7_O35 | -2.95 | -3.39 | -1.44 | -2.59 | -2.58 | -3.05 | -1.29 | -1.26 | -4.18 | -2.75 | 1.35 | -0.98 | -2.02 | -1.25 | -1.65 | -1.37 | 0.535 | 0.59 | 0.545 | 0.041837 | 0.338356 | 0.401016 | 0.989223 | 0.997988 | 0.994666 |
| Exoc8_Q6PGF7_S28s_1 | Exoc8_Q6P | -3.31 | -3.89 | 1.04 | 2.34 | -2.99 | -2.49 | 1.43 | 2.07 | -3.01 | -3.31 | 1.09 | 1.63 | -1.4 | -3.31 | 0.76 | 1.43 | 0.5475 | 0.9525 | 1.02 | 0.841154 | 0.791116 | 0.975195 | 0.995762 | 0.997988 | 0.999507 |
| Exosc5_Q9CRA8_S23s_1 | Exosc5_Q9 | -1.1 | -1.35 | -1.76 | -2.99 | -0.51 | -1.1 | -1.15 | -2.12 | -1.16 | -1.17 | -0.46 | -2.19 | -0.5 | -0.87 | -1.07 | -1.21 | 0.46 | 0.055 | 0.325 | 0.067334 | 0.256393 | 0.29717 | 0.989223 | 0.997988 | 0.994666 |
| Exph5_Q0VAV2_S1071s_1 | Exph5_Q0\ | 1.23 | 1.41 | 1.87 | 1.05 | 1.15 | 1.17 | 0.8 | 0.83 | 1.12 | 1.62 | 0.82 | 1.87 | 0.32 | 0.77 | 1.42 | 1.38 | 0.58 | 0.555 | 0.8875 | 0.248308 | 0.139241 | 0.919842 | 0.989223 | 0.997988 | 0.994666 |
| Exph5_Q0VAV2_S1117s_1 | Exph5_Q0\ | 1.26 | 1.66 | 1.61 | 1.4 | 2.29 | 2.29 | 1.34 | 1.03 | 0.85 | 2.04 | 2.23 | 2.64 | 1.09 | -0.54 | 2.17 | 1.64 | -0.4025 | -0.0325 | -0.4175 | 0.481547 | 0.464086 | 0.264324 | 0.989223 | 0.997988 | 0.994666 |
| Exph5_Q0VAV2_S331s_1 | Exph5_Q0\ | -0.68 | 0.43 | 0.55 | -0.39 | 0.73 | -0.26 | -1.25 | -0.19 | 0.62 | 0.54 | 1 | 0.01 | 0.83 | 1.21 | -0.13 | 0.09 | 0.255 | 0.4575 | -0.3925 | 0.243267 | 0.64038 | 0.153787 | 0.989223 | 0.997988 | 0.994666 |
| Exph5_Q0VAV2_S915s_1 | Exph5_Q0\ | -0.35 | 0.57 | 1.35 | 0.5 | 0.63 | -0.31 | 0.26 | -0.1 | -0.12 | -0.49 | 2.94 | 0.3 | 1.08 | 0.59 | 0.56 | 0.46 | -0.22 | 0.565 | 0.5225 | 0.689394 | 0.323342 | 0.85441 | 0.989223 | 0.997988 | 0.996159 |
| Eya1_Q8C9D0_S280s_1 | Eya1_Q8C! | 2.45 | 2.58 | 1.87 | 0.79 | 3.12 | 2.49 | 1.57 | 1.48 | -0.06 | 2.35 | 0.44 | 1.76 | 0.54 | -0.24 | 1.43 | 1.51 | -0.3975 | 0.14 | 0.155 | 0.063455 | 0.639949 | 0.225897 | 0.989223 | 0.997988 | 0.994666 |
| Eya4_Q8BY78_S338s_1 | Eya4_Q8BY | 0.07 | 0.72 | 0.59 | 0.4 | 0.39 | 0.02 | 0.64 | 0.54 | 0.55 | 0.14 | -0.02 | 0.12 | 0.65 | 0.38 | 0.01 | 0.87 | 0.2425 | -0.8 | -1.1125 | 0.912753 | 0.850243 | 0.346966 | 0.996349 | 0.997988 | 0.999507 |
| F11r_O88792_S285s_1 | F11r_O887 | 0.3 | -0.03 | -0.55 | -0.48 | -0.35 | 0.09 | -0.81 | -0.69 | 0.19 | 0.25 | -0.21 | -0.3 | -0.4 | 0.35 | -0.82 | -0.54 | -0.0475 | -0.2475 | 0.0325 | 0.64415 | 0.421859 | 0.554675 | 0.989223 | 0.997988 | 0.995259 |
| Fadd_Q61160_T204t_1 | Fadd_Q611 | 0.02 | -0.14 | 0.29 | 0.2 | -0.86 | 1 | -0.25 | -0.8 | 1.53 | 0.67 | 0.81 | 0.75 | 0.55 | -0.15 | -0.41 | -0.6 | -0.25 | 0.1725 | -0.1625 | 0.448296 | 0.452951 | 0.011967 | 0.989223 | 0.997988 | 0.994666 |
| Faf1_P54731_T268t_1 | Faf1_P547: | -0.49 | -0.6 | 0.04 | 0.39 | -0.26 | 0.42 | -1.1 | -0.87 | 1.91 | -0.3 | -3.21 | -0.52 | 2.26 | -1.78 | -1.04 | -0.36 | -0.32 | 0.8475 | -0.245 | 0.935414 | 0.475258 | 0.701739 | 0.989223 | 0.997988 | 0.999507 |
| Fahd2_Q3TC72_S129s_1 | Fahd2_Q3T | 0.03 | 0.17 | 0.25 | 0.01 | 0.23 | -1.48 | 0.26 | -0.14 | 0.14 | 0.17 | 0.23 | 0.5 | 0.3 | 0.17 | 0.26 | 0.29 | -0.2875 | -0.365 | -0.065 | 0.544523 | 0.331202 | 0.510029 | 0.989223 | 0.997988 | 0.994666 |
| Fam102a_Q78T81_S244s_1 | Fam102a_( | 0.25 | -0.81 | -1.16 | 0.28 | -0.62 | 0.65 | -0.54 | -0.27 | -0.04 | -1.17 | -2.2 | -0.72 | -0.05 | -1.09 | -1.48 | -0.53 | -0.3975 | 0.145 | 0.14 | 0.369803 | 0.708884 | 0.234758 | 0.995437 | 0.997988 | 0.994666 |
| Fam104a_A2A6P4_S62s_2 | Fam104a_/ | -2.96 | -2.29 | -1.4 | -2.94 | -1.32 | -3.76 | -2.68 | -2.9 | -1.43 | -1.46 | 0.63 | -3.27 | -1.07 | -3.54 | -3.04 | -5.35 | 0.165 | -0.6725 | -0.4275 | 0.323189 | 0.636375 | 0.216823 | 0.989223 | 0.997988 | 0.994666 |
| Fam107a_Q78TU8_T138t_1 | Fam107a_( | 1.03 | 0.18 | -0.33 | 0.37 | 0.75 | 0.81 | 0.77 | -0.11 | 0.27 | 0.72 | 0.54 | 0.44 | -0.62 | 0.67 | 0 | 0.39 | -0.2675 | 1.015 | -0.8525 | 0.616724 | 0.502789 | 0.581345 | 0.989223 | 0.997988 | 0.995259 |
| Fam110b_Q8C739_S172s_1 | Fam110b_( | -0.1 | -0.81 | -0.67 | -0.44 | -0.94 | -0.29 | -0.34 | -0.22 | -0.69 | -0.1 | 0.15 | -1.05 | 0.12 | -0.29 | 0.07 | -0.25 | 0.2425 | 0.18 | -0.2025 | 0.150318 | 0.831189 | 0.806162 | 0.995762 | 0.997988 | 0.994666 |
| Fam110b_Q8C739_S234s_1 | Fam110b_( | 0.92 | 0.78 | -1.04 | -1.75 | -0.06 | 1.06 | -0.89 | -0.9 | 0.19 | 1.08 | 0.5 | -1.44 | 0.61 | 0.79 | -0.73 | -1.21 | 0.0575 | 0.0825 | 0.4175 | 0.851901 | 0.916185 | 0.645202 | 0.997335 | 0.997988 | 0.999507 |
| Fam110b_Q8C739_S95s_1 | Fam110b_( | 0.03 | 0.37 | -0.98 | -0.93 | 0.29 | -0.39 | -0.75 | -1.18 | -0.58 | 0.13 | -0.43 | -0.91 | 0.12 | 0.49 | -0.91 | -0.53 | 0.075 | 0.355 | 0.1375 | 0.707253 | 0.765115 | 0.861461 | 0.995762 | 0.997988 | 0.996966 |
| Fam114a1_Q9D281_T197t_1 | Fam114a1_ | 0.37 | -0.02 | 0.11 | 0.03 | 0.21 | -0.53 | 0 | -0.3 | -0.27 | 0.35 | 0 | -0.07 | 1.07 | 0.67 | 0.13 | 0.18 | -0.13 | -0.07 | 0.17 | 0.213913 | 0.274559 | 0.619795 | 0.989223 | 0.997988 | 0.994666 |
| Fam114a2_Q8VE88_S112s_1 | Fam114a2_ | 0.29 | 0.15 | 0.11 | 0.75 | 0.47 | 0.27 | 0.83 | 0.17 | 0.01 | 0.02 | 0.26 | 0.39 | -0.35 | 0.17 | 0.57 | 0.52 | -0.2775 | -0.12 | 0.39 | 0.754509 | 0.670823 | 0.538648 | 0.992876 | 0.997988 | 0.9971 |
| Fam117b_Q3U3E2_S10s_1 | Fam117b_( | 0.02 | 0.34 | 0.28 | 0.71 | 0.57 | -0.26 | 0.66 | 1.25 | -0.15 | 1.02 | 0.41 | 1.64 | 0.03 | -0.1 | 1.28 | 1.8 | 0.11 | -0.155 | -0.0975 | 0.387176 | 0.533432 | 0.347442 | 0.989223 | 0.997988 | 0.994666 |
| Fam117b_Q3U3E2_S176s_1 | Fam117b_( | -0.14 | 0.2 | 0.07 | -0.04 | -1.02 | 0.94 | -1.2 | -0.86 | 2.13 | -0.27 | -0.74 | -0.13 | 2.11 | -2.05 | -0.38 | -0.39 | 0.2175 | 0.3925 | 0.415 | 0.793487 | 0.249539 | 0.704802 | 0.989223 | 0.997988 | 0.999507 |
| Fam117b_Q3U3E2_S386s_1 | Fam117b_( | -0.9 | 0.11 | 0.84 | 0.19 | -0.84 | -0.36 | -0.29 | 0.59 | -0.78 | -0.27 | 0.96 | 0.56 | 0.12 | -1.12 | -0.22 | 1.21 | -0.5575 | 0.225 | -0.2 | 0.910366 | 0.520512 | 0.908674 | 0.989223 | 0.997988 | 0.999507 |
| Fam117b_Q3U3E2_S444s_1 | Fam117b_( | -2.14 | -2.29 | 1.93 | 1.78 | -1.06 | -2.79 | 1.75 | 2.36 | -1.63 | -2.26 | 2.33 | 1.52 | -0.81 | -1.45 | 1.78 | 2.11 | -0.285 | 0.0575 | -0.0625 | 0.646981 | 0.865264 | 0.90523 | 0.996493 | 0.997988 | 0.995259 |
| Fam117b_Q3U3E2_T451t_1 | Fam117b_( | 0.25 | 0.12 | -1.1 | -0.71 | 0.69 | -0.1 | 0.1 | -0.26 | -0.26 | -0.01 | -0.82 | -1.2 | -0.23 | 0.1 | -0.57 | -0.62 | 0.245 | 0.17 | 0.5875 | 0.937174 | 0.23522 | 0.611269 | 0.989223 | 0.997988 | 0.999507 |
| FAM120A_Q6A0A9_S382s_1 | FAM120A_ | 0.48 | -0.55 | 0.05 | -0.46 | -0.16 | 0.43 | 1.21 | 0.6 | -0.71 | 0.2 | 0.53 | 0.18 | -0.06 | 0.17 | -0.59 | -0.95 | 0.4675 | -0.2125 | 0.03 | 0.524567 | 0.105919 | 0.644906 | 0.989223 | 0.997988 | 0.994666 |
| Fam122a_Q9DB52_S144s_1 | Fam122a_( | -0.68 | -1.17 | -4.48 | -3.06 | -0.61 | -1.21 | -4.14 | -4.97 | -1.06 | -0.77 | -1.61 | -3.84 | -1.43 | -0.79 | -5.73 | -4.37 | 0.64 | 0.17 | -0.2375 | 0.56631 | 0.747112 | 0.596279 | 0.995762 | 0.997988 | 0.995259 |
| Fam122a_Q9DB52_S267s_1 | Fam122a_( | -0.32 | 0.02 | -2.79 | -1.2 | 0.27 | -0.77 | -0.86 | -1.92 | -0.82 | -0.19 | -0.56 | -1.2 | -1.3 | -0.38 | -1.5 | -1.14 | -0.385 | 0.5275 | -0.7325 | 0.990225 | 0.711614 | 0.535261 | 0.995437 | 0.997988 | 1 |
| Fam126a_Q6P9N1_S433s_1 | Fam126a_( | 1.12 | 0.47 | -0.09 | 0.39 | 1.1 | 1.12 | 1.07 | 0.91 | 0.38 | 0.55 | -0.01 | 0.07 | 0.11 | 1.35 | 0.15 | 0.32 | 0.2525 | 0.38 | -0.0075 | 0.979699 | 0.067588 | 0.47932 | 0.989223 | 0.997988 | 1 |
| Fam126a_Q6P9N1_S453s_1 | Fam126a_( | 0.13 | -0.17 | -0.45 | -0.04 | 0.11 | 0.42 | 0.8 | 0.75 | -0.09 | -0.28 | -0.28 | -0.2 | -0.82 | -0.15 | -0.21 | -0.12 | 0.5775 | -0.225 | 0.01 | 0.500735 | 0.027361 | 0.72746 | 0.989223 | 0.997988 | 0.994666 |
| Fam126b_Q8C729_S321s_1 | Fam126b_( | -3.23 | -3.8 | -2.65 | -4.02 | -1.58 | -3.36 | -2.79 | -3.03 | -4.62 | -3.49 | -1.07 | -3.1 | -2.49 | -1.72 | -3.22 | -2.68 | 0.6525 | -0.08 | -0.1925 | 0.06119 | 0.135247 | 0.624736 | 0.989223 | 0.997988 | 0.994666 |
| Fam126b_Q8C729_S346s_1 | Fam126b_( | 0.66 | 0.22 | 0.67 | 0.92 | 1.27 | 0.11 | 1.33 | 0.97 | 0.06 | 0.56 | -0.2 | 1.05 | -0.99 | -0.14 | 0.72 | 1.19 | 0.735 | 0.355 | 0.8975 | 0.389257 | 0.367398 | 0.462258 | 0.989223 | 0.997988 | 0.994666 |
| Fam126b_Q8C729_S575s_1 | Fam126b_( | 0.08 | 0.41 | 0.37 | 0.36 | 0.73 | -0.07 | 0.75 | 0.19 | -0.41 | 0.58 | 1.13 | 0.26 | 0.25 | 0.34 | 0.79 | -0.05 | 0.3025 | -0.25 | -0.4225 | 0.91997 | 0.71933 | 0.806707 | 0.995437 | 0.997988 | 0.999507 |
| Fam126b_Q8C729_S577s_1 | Fam126b_( | -0.11 | 0.32 | 0.37 | 0.36 | 0.44 | 0.25 | 0.75 | 0.19 | 0.56 | 0.32 | 1.13 | 0.26 | 0.31 | 0.34 | 0.79 | -0.05 | 0.095 | 0.085 | 0.0275 | 0.691913 | 0.475087 | 0.254603 | 0.989223 | 0.997988 | 0.996159 |
| Fam126b_Q8C729_T306t_1 | Fam126b_( | 0.62 | -0.39 | -0.03 | 0.54 | 0.4 | 0.8 | 0.48 | 0.71 | -0.39 | 0.22 | 0.37 | 0.41 | -0.63 | 0.3 | -0.21 | 0.12 | 0.1725 | 0.3325 | 0.1125 | 0.412063 | 0.175858 | 0.921122 | 0.989223 | 0.997988 | 0.994666 |
| Fam126b_Q8C729_T30t_1 | Fam126b_( | 0.61 | 0.38 | 0.21 | -0.33 | 2.03 | 1.64 | 0.81 | 0.98 | -0.16 | 1.51 | -1.94 | -0.6 | 1.7 | 1.63 | -0.82 | 1.1 | 0.4125 | -0.0325 | -0.29 | 0.249835 | 0.008438 | 0.449157 | 0.989223 | 0.997988 | 0.994666 |
| Fam129a_Q3UW53_S595s_1 | Fam129a_( | 0.19 | -0.28 | -1.01 | -1.8 | -1.15 | 0.03 | -1.6 | -1.16 | -0.62 | 0.1 | -6.5 | -0.82 | 0.58 | -4.49 | -1.52 | -0.4 | 1.1475 | -0.515 | 0.685 | 0.481645 | 0.632756 | 0.384081 | 0.989223 | 0.997988 | 0.994666 |
| Fam131b_E9Q8P8_S130s_1 | Fam131b_I | 0.7 | -0.7 | -0.49 | 0.69 | 0.55 | 1.11 | 0.5 | 1.11 | 0.41 | -0.82 | 1.4 | 0.53 | -0.05 | 0.89 | -0.5 | 0.15 | -0.245 | -1.235 | -0.7325 | 0.874331 | 0.078024 | 0.550965 | 0.989223 | 0.997988 | 0.999507 |
| Fam131b_E9Q8P8_S145s_1 | Fam131b_I | 2.6 | 1.76 | 1.36 | 1.04 | 1.14 | 2.13 | 0.63 | 1.1 | 1.21 | 2.62 | 0.86 | 1.17 | -2.08 | 0.49 | 1.49 | 1.05 | 0.7675 | 0.33 | 0.0725 | 0.084163 | 0.324973 | 0.647448 | 0.989223 | 0.997988 | 0.994666 |
| Fam131b_E9Q8P8_S325s_1 | Fam131b_I | 0.63 | 1.46 | 1.71 | 1.26 | 1.72 | 0.66 | 1.17 | 1.03 | -0.24 | 0.64 | 1.01 | 1.19 | -0.85 | 0.71 | 1.54 | 1.43 | -0.44 | -0.225 | -1.4525 | 0.326662 | 0.716363 | 0.142896 | 0.995437 | 0.997988 | 0.994666 |
| Fam131b_E9Q8P8_S350s_1 | Fam131b_I | 0.74 | 0.68 | 0.35 | 0.58 | 0.56 | 0.79 | 0.65 | 0.58 | 0.94 | 1.09 | -0.1 | 0.72 | 0.09 | 1.29 | 0.12 | 0.53 | -0.12 | -0.615 | -0.5575 | 0.809838 | 0.781898 | 0.810125 | 0.995762 | 0.997988 | 0.999507 |
| Fam131b_E9Q8P8_S75s_1 | Fam131b_I | 0.71 | 1.09 | 0.8 | 0.54 | 1.08 | 0.67 | 0.76 | 0.64 | 0.2 | 0.75 | 0.18 | 0.62 | 0.29 | 0.7 | 0.51 | 0.66 | 0.0575 | 0.075 | -0.08 | 0.348523 | 0.991291 | 0.195985 | 0.999187 | 0.997988 | 0.994666 |
| Fam131c_A2ADB2_S35s_1 | Fam131c_/ | 1.35 | 1.28 | 1.01 | 1.09 | 1.79 | 1.22 | 1.72 | 1.31 | 0.57 | 1.73 | 1.17 | 1.36 | 0.16 | 1.27 | 1.72 | 1.76 | 0.0025 | -0.3475 | -0.245 | 0.908387 | 0.184903 | 0.932741 | 0.989223 | 0.997988 | 0.999507 |
| Fam133b_Q9CVI2_S82s_1 | Fam133b_( | 0.4 | 0.81 | -0.29 | 0.44 | -0.13 | 0.23 | -0.7 | -0.24 | 0.36 | 0.41 | -0.08 | 0.49 | -0.85 | 0.65 | -1.11 | -0.04 | 0.3275 | 0.025 | 0.045 | 0.155504 | 0.106998 | 0.881257 | 0.989223 | 0.997988 | 0.994666 |
| Fam134c_Q9CQV4_S26s_1 | Fam134c_( | -0.42 | -0.69 | -0.59 | -0.69 | 0.48 | -0.29 | 0.05 | -0.64 | -0.59 | -0.63 | -1.49 | -0.65 | -0.38 | -0.03 | -0.65 | -0.51 | -0.55 | -0.045 | -0.6775 | 0.428915 | 0.103277 | 0.394252 | 0.989223 | 0.997988 | 0.994666 |
| Fam13c_G3X9S1_S48s_1 | Fam13c_G: | -0.71 | 0.22 | -0.14 | 0.23 | -0.3 | -0.9 | -0.78 | -0.31 | -0.05 | -0.28 | 0.5 | -1.09 | 0.36 | 0.33 | -1.13 | -0.57 | 0.4975 | -0.2425 | 0.205 | 0.719025 | 0.136134 | 0.743229 | 0.989223 | 0.997988 | 0.9971 |
| Fam149b1_Q6NSV7_S59s_1 | Fam149b1_ | -0.98 | -0.78 | 0.47 | 0.19 | 0.04 | -0.89 | -0.04 | 0.17 | -0.76 | -0.66 | 0.89 | -0.02 | -0.52 | -1.31 | 1.05 | 0.31 | -0.4725 | -0.13 | -0.1525 | 0.783567 | 0.817086 | 0.780042 | 0.995762 | 0.997988 | 0.998491 |
| Fam160a2_F8WHJ1_S482s_1 | Fam160a2_ | -0.67 | -0.67 | 2.13 | 0.96 | -0.89 | -1.59 | 0.9 | 1.35 | -0.77 | -0.63 | 0.24 | 1.26 | -1.53 | -0.49 | 0.63 | 2.31 | 0.095 | 0.1375 | 0.1575 | 0.82414 | 0.565331 | 0.581872 | 0.989223 | 0.997988 | 0.999507 |
| Fam160a2_F8WHJ1_S529s_1 | Fam160a2_ | 0.6 | -0.72 | 0.69 | 1.45 | -0.62 | 1.01 | -0.04 | 0.73 | 0.35 | 0.67 | 1.08 | 1.02 | 0.11 | -0.72 | 0.47 | 0.75 | -0.495 | -0.4125 | -0.2075 | 0.501426 | 0.658895 | 0.554508 | 0.990434 | 0.997988 | 0.994666 |
| Fam160a2_F8WHJ1_S789s_1 | Fam160a2_ | -1.57 | -1.52 | -2.42 | -6.67 | -1.75 | -1.67 | -1.42 | -1.87 | -0.64 | -2.65 | 0.69 | -2.44 | -0.26 | -1.59 | -2.03 | -2.04 | -0.235 | 0.275 | -0.3525 | 0.185422 | 0.217759 | 0.185617 | 0.989223 | 0.997988 | 0.994666 |
| Fam160a2_F8WHJ1_S834s_1 | Fam160a2_ | 0.02 | 0.66 | 0.07 | 0.51 | 0.17 | -0.17 | -0.4 | 0.26 | -0.57 | 0.33 | -0.84 | -0.35 | -0.67 | 0.14 | -0.56 | 0.54 | 1.3675 | 1.785 | 1.565 | 0.223616 | 0.208625 | 0.062303 | 0.989223 | 0.997988 | 0.994666 |
| Fam160b1_Q8CDM8_S552s_1 | Fam160b1_ | -0.84 | -1.46 | -0.11 | -3 | -1.56 | -1.45 | -1.3 | -1.96 | 2.04 | -1.39 | -2.92 | -0.72 | 2.58 | -2.42 | 1.27 | -1.59 | -0.35 | -0.6725 | -0.4525 | 0.270367 | 0.705687 | 0.572116 | 0.995437 | 0.997988 | 0.994666 |
| Fam160b1_Q8CDM8_T551t_1 | Fam160b1_ | -0.84 | -1.46 | -0.11 | -3 | -1.56 | -1.45 | -1.3 | -1.96 | 2.04 | -1.39 | -2.92 | -0.72 | 2.58 | -2.42 | 1.27 | -1.59 | -0.215 | 0.605 | 1.3125 | 0.270367 | 0.705687 | 0.572116 | 0.995437 | 0.997988 | 0.994666 |
| Fam163b_Q8BUM6_S79s_1 | Fam163b_( | 0.62 | -0.6 | -2.13 | -0.04 | 0.73 | -0.72 | 0 | 0.61 | -0.82 | -0.22 | 0.75 | -0.18 | -1.59 | 0.59 | -0.01 | -0.27 | -0.215 | 0.605 | 1.3125 | 0.745999 | 0.267543 | 0.498444 | 0.989223 | 0.997988 | 0.9971 |
| Fam169a_Q5XG69_S359s_1 | Fam169a_( | -0.95 | -2.06 | -2.25 | -2.61 | -1.68 | -1.37 | -3.21 | -2.54 | -1.5 | -1.82 | -1.89 | -1.95 | 0.19 | -2.56 | -2.28 | -2.55 | 0.6925 | 0.42 | 0.2175 | 0.805293 | 0.645965 | 0.639629 | 0.99033 | 0.997988 | 0.999507 |
| Fam169a_Q5XG69_S512s_1 | Fam169a_( | -1.88 | -2.62 | -0.71 | -0.53 | -1.52 | -3.26 | -0.99 | -0.68 | -8.58 | -1.12 | -0.22 | -0.98 | -2.12 | -2.43 | -0.6 | -0.96 | -0.2325 | 0.1775 | 0.1675 | 0.878529 | 0.791883 | 0.470775 | 0.995762 | 0.997988 | 0.999507 |
| Fam169a_Q5XG69_T358t_1 | Fam169a_( | -2.55 | -5.67 | -2.3 | -4.14 | -3.45 | -2.72 | -2.27 | -2.58 | -3.23 | -3.79 | -2.51 | -2.74 | -1.09 | -5.18 | -4.4 | -3.28 | -0.1775 | -1.29 | -0.0925 | 0.863114 | 0.227627 | 0.432081 | 0.989223 | 0.997988 | 0.999507 |
| Fam171a1_A2ATL1_S346s_1 | Fam171a1_ | 1.88 | 1.18 | 1.03 | 2.33 | 1.47 | 2.99 | 1.47 | 1.96 | 1.88 | 0.84 | 0.66 | 1.25 | 0.9 | 1.72 | 1.2 | 1.68 | 0.91 | 0.5975 | 0.1775 | 0.547501 | 0.413748 | 0.284346 | 0.989223 | 0.997988 | 0.994666 |
| Fam171a1_A2ATL1_S400s_1 | Fam171a1_ | -0.81 | 0.04 | -0.35 | 0.17 | 0.07 | -0.13 | 0.33 | -0.2 | -0.43 | -0.31 | 0.74 | 0.44 | -0.62 | -0.02 | 0.53 | 0.6 | 0.3675 | -0.4475 | -0.23 | 0.350577 | 0.379731 | 0.357888 | 0.989223 | 0.997988 | 0.994666 |
| Fam171a1_A2ATL1_S699s_1 | Fam171a1_ | -0.61 | 0.82 | 0.77 | 1.06 | 0.54 | -0.33 | 1.08 | 1.3 | -0.24 | 0.06 | -0.19 | 1.29 | 0.16 | 0.7 | 0.96 | 0.78 | 0.255 | 0.3475 | 0.36 | 0.737046 | 0.776328 | 0.573263 | 0.995762 | 0.997988 | 0.9971 |
| Fam171a1_A2ATL1_S724s_1 | Fam171a1_ | 1.34 | 0.48 | 1.43 | 0.45 | 1.45 | 1.44 | 2.07 | 1.85 | -0.56 | 1.28 | 0.03 | 1.77 | -0.49 | 0.22 | 1.33 | 1.93 | 0.1375 | -0.28 | 0.14 | 0.751387 | 0.035108 | 0.599057 | 0.989223 | 0.997988 | 0.9971 |
| Fam171a1_A2ATL1_T323t_1 | Fam171a1_ | -1.06 | -1.08 | -0.33 | -0.82 | -0.56 | -0.32 | 0.58 | -0.75 | -0.92 | -0.59 | 1.08 | 0.69 | -1.04 | -1.32 | 0.04 | 0.75 | 0.7775 | -0.295 | -0.1775 | 0.387156 | 0.130489 | 0.095108 | 0.989223 | 0.997988 | 0.994666 |
| Fam171a2_A2A699_S261s_1 | Fam171a2_ | 0.43 | 0.06 | -0.54 | -0.02 | 0.69 | 0.36 | 0.27 | 0.18 | -0.14 | 0.23 | 0.36 | -0.07 | -0.97 | 0.35 | -0.57 | -0.41 | 0.56 | 0.8875 | 0.43 | 0.307593 | 0.169942 | 0.691726 | 0.989223 | 0.997988 | 0.994666 |
| Fam171a2_A2A699_S593s_1 | Fam171a2_ | 0.63 | -0.52 | -0.09 | -0.41 | -0.05 | 1.34 | -0.68 | -0.46 | 1.27 | 0.09 | -1.57 | -0.11 | 0.95 | -0.22 | -0.43 | -0.29 | 0.3925 | 0.1125 | -0.3825 | 0.808492 | 0.779485 | 0.976143 | 0.995762 | 0.997988 | 0.999507 |
| Fam171a2_A2A699_T378t_1 | Fam171a2_ | 0.63 | 0.41 | 1.13 | -0.02 | 0.22 | 0.6 | 0.71 | -0.59 | -0.63 | 0.73 | -0.33 | 0.95 | 0.37 | -0.02 | 0.43 | -0.66 | 0.135 | 0.0175 | 0.1 | 0.189229 | 0.425431 | 0.426378 | 0.989223 | 0.997988 | 0.994666 |
| Fam171b_Q14CH0_S410s_1 | Fam171b_( | 0.18 | -0.49 | -0.28 | -0.96 | 1.36 | 0.4 | -0.25 | 1.19 | 0.04 | 0.61 | 0.94 | 0.82 | 0.29 | -0.27 | 0.49 | 1.29 | -0.3025 | -0.3575 | -0.5075 | 0.062461 | 0.028725 | 0.014126 | 0.989223 | 0.997988 | 0.994666 |
| Fam171b_Q14CH0_S432s_1 | Fam171b_( | 0.47 | 0.61 | 0.37 | 0.03 | -0.39 | 0.6 | -0.37 | -0.36 | 0.92 | 0.15 | -1.42 | 0.16 | 1.2 | -1.62 | 0.13 | 0.06 | 1.0625 | 0.99 | 0.8375 | 0.44564 | 0.11906 | 0.395429 | 0.989223 | 0.997988 | 0.994666 |
| Fam171b_Q14CH0_S497s_1 | Fam171b_( | 3.39 | 1.23 | 0.53 | 0.1 | 2.33 | 2.88 | 1.6 | 0.83 | 0 | 2.52 | -0.52 | 1 | 0.39 | -1.3 | 1.08 | 1.36 | -0.5 | -0.4175 | -0.4275 | 0.279227 | 0.433925 | 0.527173 | 0.989223 | 0.997988 | 0.994666 |
| Fam171b_Q14CH0_S698s_1 | Fam171b_( | -0.34 | -0.29 | -0.56 | 0.16 | -0.31 | 0.22 | -0.45 | -0.27 | 0.01 | -0.2 | -0.15 | 0.11 | -0.59 | -1.11 | -0.62 | -0.63 | 0.5975 | -0.5625 | -0.93 | 0.108404 | 0.832237 | 0.424711 | 0.995762 | 0.997988 | 0.994666 |
| Fam171b_Q14CH0_S788s_1 | Fam171b_( | 1.26 | 0.81 | 1 | 0.96 | 1.06 | 1.18 | 1.08 | 0.93 | 0.95 | 1.75 | 0.7 | 1.3 | 0.73 | 1.08 | 1.06 | 1.1 | 0.055 | 0.2 | -0.48 | 0.951578 | 0.793328 | 0.568297 | 0.995762 | 0.997988 | 0.999507 |
| Fam171b_Q14CH0_T787t_1 | Fam171b_( | 0.64 | 1.59 | 0.9 | 1.61 | 1.3 | 0.84 | 0.76 | 0.66 | 0.54 | 1.32 | 0.98 | 0.82 | 0.55 | 1.49 | 1.6 | 0.72 | 0.055 | 0.1675 | -0.015 | 0.801706 | 0.346056 | 0.411559 | 0.989223 | 0.997988 | 0.999507 |
| Fam177a1_Q8BR63_S151s_1 | Fam177a1_ | 0.93 | 1.13 | 0.52 | 0.56 | 1.25 | 0.21 | 0.21 | -0.14 | 0.74 | 0.29 | 0.8 | 0.14 | -0.24 | 1.12 | 1.22 | 1.11 | -0.295 | -0.27 | -0.095 | 0.964112 | 0.255375 | 0.302966 | 0.989223 | 0.997988 | 0.999507 |
| Fam184a_F6RSY8_S671s_1 | Fam184a_I | 0.01 | 0.76 | -0.18 | 0.03 | 0.98 | 0.22 | -0.37 | -0.4 | 1.06 | 0.16 | -0.91 | -0.52 | -0.22 | 0.1 | -0.37 | 0.12 | -0.4025 | -0.2925 | 0.0175 | 0.419033 | 0.899583 | 0.652734 | 0.996493 | 0.997988 | 0.994666 |
| Fam189a1_Q6A044_S455s_1 | Fam189a1_ | -0.39 | -1.39 | -0.64 | -0.88 | 0.12 | -0.1 | 0.01 | -0.05 | -1.15 | -0.84 | 0.25 | -0.72 | -0.25 | -0.46 | 0.45 | -0.05 | -0.0475 | -0.2075 | -0.2475 | 0.044894 | 0.011125 | 0.579366 | 0.989223 | 0.997988 | 0.994666 |
| Fam189a2_E9QP68_S586s_1 | Fam189a2_ | 0.22 | 0.57 | 0.84 | 0.21 | 1.46 | -0.69 | 1.02 | 0.08 | 0.3 | 0.57 | -0.23 | 0.53 | 0.02 | 0.88 | 0.45 | 1.12 | 0.82 | 0.21 | 0.7475 | 0.633447 | 0.987199 | 0.562025 | 0.999187 | 0.997988 | 0.995259 |
| Fam195b_Q3UGS4_S21s_1 | Fam195b_( | -0.6 | -0.54 | -0.94 | -0.97 | -0.59 | -0.96 | -1.35 | -1.23 | -1.57 | -1.23 | -1.39 | -0.89 | -1.44 | -1.12 | -1.27 | -1.61 | 0.0075 | -0.1675 | 0.1575 | 0.040934 | 0.30458 | 0.071028 | 0.989223 | 0.997988 | 0.994666 |
| Fam196a_E9PZW7_S177s_1 | Fam196a_E | 0.13 | -0.8 | -1.12 | -0.09 | -0.71 | -0.23 | 0.87 | 0.15 | 0.04 | -0.34 | 0.3 | -0.43 | -0.61 | -0.31 | 0.18 | -0.01 | -0.27 | -0.5075 | -0.5975 | 0.444945 | 0.262488 | 0.319417 | 0.989223 | 0.997988 | 0.994666 |
| Fam207a_P58468_S38s_1 | Fam207a_I | 0 | 0.5 | -0.29 | -0.34 | 0.77 | 0.74 | -0.21 | -0.06 | 0.18 | 1.14 | -0.54 | 0.28 | 0.1 | 0.28 | -1.33 | -0.7 | 0.49 | 0.3625 | 0.2825 | 0.374699 | 0.316398 | 0.457343 | 0.989223 | 0.997988 | 0.994666 |
| Fam208a_Q69ZR9_S1108s_1 | Fam208a_( | -0.77 | 0.45 | -0.56 | -0.14 | 0.84 | -0.48 | 0.37 | -0.03 | -0.7 | -0.12 | -0.53 | -0.15 | -0.41 | 0.4 | -0.59 | 0.29 | 0.3425 | 0.2975 | -0.38 | 0.642705 | 0.274747 | 0.716378 | 0.989223 | 0.997988 | 0.995259 |
| Fam208a_Q69ZR9_S339s_1 | Fam208a_( | -0.14 | -0.96 | 0.06 | -1.12 | -1.22 | 0.39 | -0.5 | -0.38 | -0.11 | -0.55 | -0.02 | -1.18 | 0.9 | -1.29 | 0.16 | -0.3 | 0.43 | -0.12 | 0.1775 | 0.4338 | 0.788599 | 0.849268 | 0.995762 | 0.997988 | 0.994666 |
| Fam208a_Q69ZR9_S962s_1 | Fam208a_( | 2.14 | 1.2 | 1.8 | -0.08 | 1.38 | 2.04 | 1.45 | 1.3 | 0.65 | 2.02 | 0.28 | 0.82 | 1.31 | 0.41 | 2.23 | 2.09 | 0.1125 | 0.075 | 0.4075 | 0.678724 | 0.564533 | 0.574144 | 0.989223 | 0.997988 | 0.996159 |
| Fam21_Q6PGL7_S747s_1 | Fam21_Q6 | 0.62 | 1.04 | 0.68 | 0.34 | 1.11 | 0.25 | 1.11 | 0.75 | 0.61 | 1.03 | 0.08 | 0.13 | 0.27 | 0.96 | 0.12 | 0.39 | 0.2775 | -0.3225 | 0.245 | 0.436494 | 0.635535 | 0.499793 | 0.989223 | 0.997988 | 0.994666 |
| Fam21_Q6PGL7_S981s_1 | Fam21_Q6 | 0.64 | 1.04 | -0.04 | 0.38 | 1.13 | 0.56 | 0.27 | -0.14 | 0.32 | 1.01 | 1.11 | 0.01 | -0.08 | 0.56 | 0.85 | 0.84 | 0.135 | -0.2075 | -0.235 | 0.91375 | 0.886892 | 0.766406 | 0.996493 | 0.997988 | 0.999507 |
| Fam212b_Q80VY2_S262s_1 | Fam212b_( | 1 | 0.71 | 3.32 | 0.51 | 1.26 | 1.1 | 3.25 | 2.96 | 0.48 | 1.05 | 1.7 | 3.02 | 1.1 | 1.62 | 3.99 | 4.22 | -0.05 | 0.1075 | 0.0375 | 0.16004 | 0.328468 | 0.815995 | 0.989223 | 0.997988 | 0.994666 |
| Fam217b_A2AJW5_S106s_1 | Fam217b_/ | -0.13 | 1.21 | 0.66 | 0.18 | -0.4 | -0.22 | 0.33 | 0.25 | -0.15 | 0.61 | 0.51 | -0.82 | 0.95 | 1.18 | 1.68 | 1.2 | 0.7575 | 0.1775 | 1.3475 | 0.053074 | 0.179488 | 0.318475 | 0.989223 | 0.997988 | 0.994666 |
| Fam219a_Q9D772_S44s_1 | Fam219a_( | -1.1 | -1.37 | -1.11 | -0.58 | -0.75 | -1.5 | -0.73 | -1.11 | -1.32 | -1.11 | 0.46 | -1.03 | -0.71 | -0.9 | -0.79 | -1.4 | -0.49 | -0.4425 | 0.7725 | 0.759437 | 0.950302 | 0.504331 | 0.999187 | 0.997988 | 0.9971 |
| Fam222a_Q6PGH4_S313s_1 | Fam222a_( | -2.17 | -1.61 | -1.64 | -3.95 | -2.11 | -1.39 | -3.41 | -3.22 | -0.16 | -1.42 | -1.48 | -2.92 | 0.16 | -2.1 | -3.3 | -3.61 | 0.0175 | 0.29 | 0.09 | 0.884011 | 0.769229 | 0.251591 | 0.995762 | 0.997988 | 0.999507 |
| Fam49b_Q921M7_S117s_1 | Fam49b_Q | -0.14 | 0.72 | 0.84 | 0.08 | -0.26 | -0.56 | -0.82 | 0.14 | 0.23 | 0.55 | -3.06 | 0.11 | 0.82 | -1.79 | -1.49 | -1.6 | -0.19 | 0.8475 | 0.13 | 0.041879 | 0.043195 | 0.260421 | 0.989223 | 0.997988 | 0.994666 |
| Fam53b_Q3U0R8_S168s_1 | Fam53b_Q | -1.32 | -0.43 | -0.39 | -0.08 | 0.32 | -1.73 | -0.3 | -0.69 | -1.61 | -1.18 | -0.46 | -1 | -0.48 | -0.61 | -0.72 | -0.76 | -0.75 | -0.9175 | -1.39 | 0.785336 | 0.923517 | 0.189152 | 0.997335 | 0.997988 | 0.999507 |
| Fam53b_Q3U0R8_S180s_1 | Fam53b_Q | -0.24 | -1.38 | -1.87 | -0.93 | -0.6 | -1.92 | -1.51 | -2.28 | 0.19 | -1.86 | -3.21 | -2.59 | 0.74 | -1.17 | -2.37 | -0.85 | -0.045 | -0.5075 | -0.0875 | 0.769655 | 0.323156 | 0.313261 | 0.989223 | 0.997988 | 0.9971 |
| Fam53c_Q8BXQ8_S122s_1 | Fam53c_Q! | -1.47 | -1.73 | -1.53 | -1.59 | -0.77 | -2.51 | -1.2 | -2.32 | -1.88 | -1.25 | -0.47 | -1.06 | -1.42 | -1.02 | -2.29 | -1.91 | -0.4725 | -0.7625 | 0.1925 | 0.806667 | 0.769365 | 0.219911 | 0.995762 | 0.997988 | 0.999507 |
| Fam53c_Q8BXQ8_S162s_1 | Fam53c_Q! | -1.18 | -0.03 | 0.3 | -0.46 | -0.68 | -0.64 | -0.3 | -0.62 | -0.92 | -0.58 | -0.82 | 0.04 | 0.4 | -1.12 | -0.11 | -0.59 | -0.12 | 0.415 | -0.08 | 0.977585 | 0.524773 | 0.560555 | 0.989223 | 0.997988 | 1 |
| Fam53c_Q8BXQ8_S273s_1 | Fam53c_Q! | -1.31 | -1.56 | -2.79 | -3.06 | -1.33 | -0.74 | -3.03 | -2.37 | -0.86 | -1.48 | 0.11 | -2.66 | -0.21 | -1.18 | -2.36 | -2.18 | -0.2175 | -0.2275 | -0.0125 | 0.265 | 0.606423 | 0.170033 | 0.989223 | 0.997988 | 0.994666 |
| Fam57b_Q7TNV1_S215s_1 | Fam57b_Q | 0.22 | 0.62 | 0.53 | 0.33 | 0.15 | 0.02 | 0.06 | 0.37 | 0.06 | 0.25 | 0.43 | 0.52 | 0.92 | 0.17 | 0.38 | 0.45 | 0.3125 | 0.9575 | 0.6975 | 0.838943 | 0.220563 | 0.63691 | 0.989223 | 0.997988 | 0.999507 |
| Fam57b_Q7TNV1_S215s_2 | Fam57b_Q | 0.23 | 0.49 | -0.14 | 0.23 | -0.7 | 0.61 | -0.83 | -0.55 | 1.36 | 0.23 | -0.26 | -0.45 | 1.03 | -0.97 | -0.27 | 0.1 | -0.275 | -0.11 | 0.055 | 0.596011 | 0.133624 | 0.966581 | 0.989223 | 0.997988 | 0.995259 |
| Fam57b_Q7TNV1_S219s_2 | Fam57b_Q | 0.23 | 0.49 | -0.14 | 0.23 | -0.7 | 0.61 | -0.83 | -0.55 | 1.36 | 0.23 | -0.26 | -0.45 | 1.03 | -0.97 | -0.27 | 0.1 | -0.57 | 0.0175 | -0.23 | 0.596011 | 0.133624 | 0.966581 | 0.989223 | 0.997988 | 0.995259 |
| Fam63a_Q76LS9_S103s_1 | Fam63a_Q | 0.55 | 0.28 | 0.49 | 0.92 | -0.05 | 0.62 | 0.84 | 0.77 | 0.1 | 0.72 | 0.29 | 0.65 | 0.12 | 0.73 | 0.47 | 0.36 | -0.57 | 0.0175 | -0.23 | 0.607922 | 0.957164 | 0.649565 | 0.999187 | 0.997988 | 0.995259 |
| Fam63b_Q6PDI6_S21s_1 | Fam63b_Q | 0.2 | 0.75 | 0.5 | 0.06 | 0.3 | -0.04 | -0.34 | -0.56 | -0.04 | 0.15 | -0.08 | 0.09 | 0.85 | 0.18 | 0.12 | 0.3 | -0.015 | -0.12 | -0.14 | 0.959223 | 0.078911 | 0.17805 | 0.989223 | 0.997988 | 0.999507 |
| Fam63b_Q6PDI6_S82s_1 | Fam63b_Q | 0.34 | 0.81 | -0.2 | -0.36 | 0.88 | 0.38 | -0.27 | -1.59 | 0.62 | 0.51 | 2.05 | 0.27 | 0.97 | 1.34 | -0.31 | -0.74 | -0.5375 | -0.3475 | -0.015 | 0.751626 | 0.585351 | 0.146983 | 0.989223 | 0.997988 | 0.9971 |
| Fam65a_Q68FE6_S347s_1 | Fam65a_Q | -0.31 | 0.36 | 0.3 | 0.11 | -0.2 | -0.8 | 0.31 | 0.18 | -0.59 | 0.09 | -0.21 | 0.27 | -0.42 | -0.11 | -0.21 | -0.16 | -0.2975 | 0.715 | 0.1675 | 0.215876 | 0.442437 | 0.442452 | 0.989223 | 0.997988 | 0.994666 |
| Fam65a_Q68FE6_S755s_2 | Fam65a_Q | 0.35 | -0.19 | 0.33 | 0.87 | 0.89 | -0.88 | 0.06 | 0.92 | -0.52 | -0.47 | 0.14 | -0.09 | -0.91 | -0.97 | 0.59 | -0.21 | -0.2425 | -0.225 | -0.34 | 0.115589 | 0.835977 | 0.085099 | 0.995762 | 0.997988 | 0.994666 |
| Fam65a_Q68FE6_S875s_1 | Fam65a_Q | 1.37 | 2.62 | 4.68 | 2.85 | 3.08 | 1.26 | 2.64 | 4.33 | 1.39 | 1.77 | 1.7 | 4.82 | 1.4 | 0.81 | 4.44 | 4.48 | -0.0925 | -0.575 | -0.715 | 0.924882 | 0.948446 | 0.624471 | 0.999187 | 0.997988 | 0.999507 |
| Fam65b_Q80U16_S441s_1 | Fam65b_Q | 1.93 | 2.32 | 1.4 | 1.22 | 1.21 | 1.87 | 0.63 | 1.45 | 0.71 | 1.94 | 1.8 | 1.51 | 1.02 | -0.16 | 1.76 | 1.91 | -0.0525 | -0.46 | -0.0975 | 0.263979 | 0.252031 | 0.550712 | 0.989223 | 0.997988 | 0.994666 |
| Fam73b_Q8BK03_S276s_1 | Fam73b_Q | 0.36 | 0.73 | -0.42 | 0.04 | 0.29 | 0.7 | -0.3 | -0.4 | 0.09 | 0.52 | -0.17 | -0.1 | -0.24 | 0.16 | -1.35 | 0.17 | -0.4275 | -0.2275 | -0.585 | 0.267456 | 0.768099 | 0.772029 | 0.995762 | 0.997988 | 0.994666 |
| Fam76b_D6RG49_S193s_1 | Fam76b_D | -1.51 | -1.76 | -1.77 | -1.38 | -0.75 | -0.46 | -2.08 | -2.09 | -0.66 | -0.46 | 0.09 | -3.45 | -0.5 | -0.66 | -0.08 | -2.96 | -0.105 | -0.0925 | -0.4925 | 0.365005 | 0.539629 | 0.505077 | 0.989223 | 0.997988 | 0.994666 |
| Fam83h_Q148V8_S1041s_1 | Fam83h_Q | 0.67 | 0.34 | 1.16 | -0.26 | -1.57 | 0.92 | -1.92 | -0.42 | 1.99 | 0.84 | 0.03 | 0.6 | 1.85 | 0.19 | -0.84 | 0.68 | 0.26 | 0.485 | 0.555 | 0.989687 | 0.076683 | 0.429727 | 0.989223 | 0.997988 | 1 |
| Fam83h_Q148V8_S948s_1 | Fam83h_Q | 0.34 | 0.27 | 0.68 | 0.11 | -0.12 | 0.6 | -0.26 | 0.73 | 0.45 | 0.65 | 0.55 | 0.59 | 0.34 | -0.21 | 0.46 | 0.57 | -1.225 | 0.3875 | -0.0075 | 0.833136 | 0.709629 | 0.367678 | 0.995437 | 0.997988 | 0.999507 |
| Fam84a_Q9D650_S67s_1 | Fam84a_Q | 2.36 | 1.72 | 0.28 | 0.65 | 2.51 | 2.43 | 0.93 | 0.77 | -0.47 | 2.43 | 1.72 | 0.27 | 0.3 | -0.62 | 0.79 | 1.06 | -0.1125 | 0.21 | -0.06 | 0.144012 | 0.502028 | 0.718835 | 0.989223 | 0.997988 | 0.994666 |
| Fam91a1_Q3UVG3_S671s_1 | Fam91a1_( | -0.76 | -0.56 | 0.11 | -0.36 | -1.51 | -0.28 | -0.98 | -0.53 | 0.16 | -0.58 | -0.1 | -0.05 | 1.15 | -1.53 | -0.81 | -0.42 | 0.4075 | -0.265 | -0.87 | 0.98553 | 0.218424 | 0.397082 | 0.989223 | 0.997988 | 1 |
| Farp1_F8VPU2_S20s_1 | Farp1_F8V | 0.53 | 0.11 | 0.83 | 0.31 | 0.34 | 0.66 | 0.94 | 1.13 | 0.04 | 0.04 | -0.35 | 0.74 | -0.2 | 0.17 | 0.2 | 0.12 | -0.4325 | 0.25 | -0.01 | 0.188349 | 0.253721 | 0.304474 | 0.989223 | 0.997988 | 0.994666 |
| Farp1_F8VPU2_S373s_1 | Farp1_F8V | -0.46 | -0.39 | -0.02 | -0.62 | 0.29 | -0.31 | 0.33 | -0.2 | -0.67 | -0.47 | -0.26 | 0.28 | -0.21 | 0.3 | 0.06 | 0.32 | 0.3225 | -0.3275 | -0.3725 | 0.092913 | 0.145879 | 0.748225 | 0.989223 | 0.997988 | 0.994666 |
| Farp1_F8VPU2_S389s_1 | Farp1_F8V | 0.3 | -0.01 | 0.5 | 0.74 | 0.14 | -0.17 | 0.64 | 0.79 | -0.35 | 0.14 | 0.32 | 0.83 | -0.15 | 0.47 | 0.33 | 0.77 | 0.4 | 0.0925 | 0.49 | 0.928321 | 0.913234 | 0.64656 | 0.997335 | 0.997988 | 0.999507 |
| Farp1_F8VPU2_S403s_1 | Farp1_F8V | 1.26 | 0.59 | 0.57 | 0.85 | 0.53 | 0.68 | 0.81 | 1.12 | 0.32 | 0.6 | 0.13 | 0.9 | -0.27 | 0.41 | 0.4 | 0.58 | -0.0325 | -0.1475 | -0.0275 | 0.10337 | 0.898925 | 0.260697 | 0.996493 | 0.997988 | 0.994666 |

| | | | | | | | | | | | | | | | | | | | | | | | | | | |
|---|---|---|---|---|---|---|---|---|---|---|---|---|---|---|---|---|---|---|---|---|---|---|---|---|---|---|
| Farp1_F8VPU2_S427s_1 | Farp1_F8V | 0.55 | -0.07 | -0.58 | 0.28 | 0.79 | 0.29 | 0.42 | -0.02 | 0.08 | -0.09 | -0.08 | -0.18 | -0.18 | 0.19 | -0.4 | -0.09 | -0.0325 | -0.33 | -0.5375 | 0.60869 | 0.314347 | 0.702219 | 0.989223 | 0.997988 | 0.995259 |
| Farp1_F8VPU2_S875s_1 | Farp1_F8V | 0.58 | 0.18 | 0.69 | 1.65 | 0.78 | 0.01 | 1.56 | 1.54 | 0.04 | -0.13 | 0.34 | 1.42 | -0.04 | 0.23 | 0.91 | 1.33 | 0.325 | -0.1125 | -0.165 | 0.700914 | 0.662786 | 0.434747 | 0.990434 | 0.997988 | 0.996159 |
| Farp1_F8VPU2_S892s_1 | Farp1_F8V | -0.5 | -0.37 | -0.65 | -0.45 | -0.3 | -0.35 | -0.33 | -0.51 | -0.62 | -0.45 | 0.4 | -0.03 | -0.19 | 0.34 | -0.32 | -0.15 | 0.1975 | -0.3575 | -0.1675 | 0.135079 | 0.551759 | 0.281043 | 0.989223 | 0.997988 | 0.994666 |
| Farp1_F8VPU2_T24t_1 | Farp1_F8V | 0.67 | -0.49 | -0.2 | 0.26 | 0.21 | 0.52 | 0.78 | 0.57 | -0.29 | -0.44 | -0.17 | 0.41 | -0.75 | 0.38 | -0.93 | -0.28 | 0.12 | 0.3175 | 0.4125 | 0.267806 | 0.153884 | 0.591004 | 0.989223 | 0.997988 | 0.994666 |
| Fasn_P19096_S1577s_1 | Fasn_P190 | -0.91 | 0.02 | 0.48 | -0.89 | -0.5 | -0.97 | 0.25 | -0.9 | -0.69 | -0.67 | 0.51 | 0.24 | -0.57 | -0.19 | -0.27 | -0.27 | 0.46 | -0.1825 | -0.455 | 1 | 0.628648 | 0.700444 | 0.989223 | 0.997988 | 1 |
| Fat1_F2Z4A3_T153t_1 | Fat1_F2Z4 | -0.06 | 0.38 | -0.69 | -0.24 | 0.32 | -0.65 | 0.44 | -0.31 | -0.47 | -0.6 | -0.37 | -0.64 | -0.85 | 0.03 | -1.51 | -0.01 | -0.205 | 0.1725 | 1.77E-16 | 0.324084 | 0.76667 | 0.212045 | 0.995762 | 0.997988 | 0.994666 |
| Fbxl17_Q9QZN1_S42s_1 | Fbxl17_Q9 | 0.79 | 0.88 | 1.22 | 1.13 | -0.36 | 1.02 | 0.54 | 0.63 | 1.45 | 0.57 | -0.26 | 1.01 | 2.2 | -0.52 | 0.62 | 0.6 | 0.1025 | -0.3675 | -0.4325 | 0.600168 | 0.115002 | 0.422318 | 0.989223 | 0.997988 | 0.995259 |
| Fbxl20_Q9CZV8_T403t_1 | Fbxl20_Q9 | 2.66 | 2.38 | 2.68 | 1.63 | 2.54 | 2.35 | 2.4 | 2.68 | 0.25 | 2.49 | 0.08 | 3.06 | 0.4 | 0.09 | 2.94 | 2.82 | -0.5475 | -0.3125 | -0.28 | 0.294342 | 0.593036 | 0.248057 | 0.989223 | 0.997988 | 0.994666 |
| Fbxl4_V9GWU9_S45s_1 | Fbxl4_V9G | 0.23 | 0.19 | 1.68 | 1.77 | 0.45 | -0.36 | 1.55 | 0.89 | -0.24 | 0.32 | 0.73 | 0.03 | -0.11 | 0.68 | 1.18 | 0.8 | 0.155 | -0.8675 | -0.775 | 0.505386 | 0.537567 | 0.128058 | 0.989223 | 0.997988 | 0.994666 |
| Fbxl4_V9GWU9_T44t_1 | Fbxl4_V9G | 0.23 | 0.19 | 1.68 | 1.77 | 0.45 | -0.36 | 1.55 | 0.89 | -0.24 | 0.32 | 0.73 | 0.03 | -0.11 | 0.68 | 1.18 | 0.8 | -0.335 | -0.7575 | -0.33 | 0.505386 | 0.537567 | 0.128058 | 0.989223 | 0.997988 | 0.994666 |
| Fbxo10_F6Q7Q7_S165s_1 | Fbxo10_F6 | -0.02 | -0.5 | 0.16 | -0.62 | -1.63 | -0.11 | -0.94 | -1.05 | 1.77 | 0.03 | 0.16 | -0.47 | 1.38 | -1.45 | 0.38 | -0.23 | -0.335 | -0.7575 | -0.33 | 0.644409 | 0.083024 | 0.229054 | 0.989223 | 0.997988 | 0.995259 |
| Fbxo30_Q8BJL1_S573s_2 | Fbxo30_Q8 | -1.14 | -0.18 | -0.06 | -0.69 | 0.02 | -1.3 | 0.24 | -0.71 | -0.87 | -0.46 | -0.96 | -0.4 | 0.13 | 0.09 | -0.42 | -0.13 | -0.6875 | 0.6175 | 0.265 | 0.200223 | 0.845744 | 0.62669 | 0.996349 | 0.997988 | 0.994666 |
| Fbxo30_Q8BJL1_T574t_2 | Fbxo30_Q8 | -1.14 | -0.18 | -0.06 | -0.69 | 0.02 | -1.3 | 0.24 | -0.71 | -0.87 | -0.46 | -0.96 | -0.4 | 0.13 | 0.09 | -0.42 | -0.13 | 0.08 | -0.155 | 0.435 | 0.200223 | 0.845744 | 0.62669 | 0.996349 | 0.997988 | 0.994666 |
| Fbxo31_Q3TQF0_S33s_2 | Fbxo31_Q3 | 0.02 | -0.11 | -0.86 | -0.21 | -1.13 | -0.3 | -0.32 | -0.06 | -0.12 | 0.45 | 1.04 | 0.85 | -0.68 | 0.98 | -0.11 | -0.91 | 0.08 | -0.155 | 0.435 | 0.807792 | 0.613461 | 0.03237 | 0.989223 | 0.997988 | 0.999507 |
| Fbxo31_Q3TQF0_T37t_1 | Fbxo31_Q3 | 1.31 | 0.58 | -0.15 | 0.38 | 0.14 | 0.53 | -0.56 | -0.59 | 1.41 | 1.43 | -0.06 | -0.47 | 0.6 | 0.89 | 0.29 | 1.1 | -0.1625 | 0.845 | 0.11 | 0.609741 | 0.124476 | 0.929604 | 0.989223 | 0.997988 | 0.995259 |
| Fbxo31_Q3TQF0_T37t_2 | Fbxo31_Q3 | 0.02 | -0.11 | -0.86 | -0.21 | -1.13 | -0.3 | -0.32 | -0.06 | -0.12 | 0.45 | 1.04 | 0.85 | -0.68 | 0.98 | -0.11 | -0.91 | -0.65 | 0.0475 | 0.19 | 0.807792 | 0.613461 | 0.03237 | 0.989223 | 0.997988 | 0.999507 |
| Fbxo4_Q8CHQ0_S11s_1 | Fbxo4_Q8C | 0.53 | 0.54 | 0.66 | 0.7 | 0.27 | 0.45 | 0.17 | 0.52 | 0.43 | 0.53 | 0.4 | 0.46 | -0.09 | 0.87 | 0.25 | 0.31 | -0.1625 | 0.845 | 0.11 | 0.344309 | 0.231389 | 0.462526 | 0.989223 | 0.997988 | 0.994666 |
| Fbxo41_Q6NS60_S363s_1 | Fbxo41_Q6 | 0.96 | 0.34 | 0.59 | 0.8 | 0.46 | 1.35 | -1.52 | 0.3 | 1.93 | 0.72 | -1.33 | 0.99 | 1.51 | -1.08 | -0.43 | 0.13 | -0.255 | -0.1525 | -0.2725 | 0.245307 | 0.357082 | 0.881341 | 0.989223 | 0.997988 | 0.994666 |
| Fbxo41_Q6NS60_S476s_1 | Fbxo41_Q6 | 4.32 | 3.05 | 2.86 | 0.58 | 3.6 | 4.77 | 2.57 | 2.73 | 0.7 | 4.74 | 1.12 | 3.15 | 2.69 | -0.4 | 3.99 | 3.93 | -0.525 | -0.095 | -0.64 | 0.893138 | 0.387342 | 0.797928 | 0.989223 | 0.997988 | 0.999507 |
| Fbxo41_Q6NS60_S476s_2 | Fbxo41_Q6 | 2.4 | 2.39 | 2.19 | 1 | 2.75 | 2.13 | 1.95 | 2.23 | 0.3 | 3.07 | 0.53 | 2.29 | -0.38 | 0.34 | 3 | 3 | 0.715 | -0.275 | -0.15 | 0.547986 | 0.472902 | 0.516106 | 0.989223 | 0.997988 | 0.994666 |
| Fbxo41_Q6NS60_S838s_1 | Fbxo41_Q6 | -1.13 | -1.22 | -1.32 | -2.02 | -0.89 | -0.79 | -1.11 | -1.7 | 1.04 | -1 | -0.98 | -0.71 | 1.47 | -1.97 | -0.67 | -0.32 | 0.27 | -0.4475 | -0.505 | 0.138562 | 0.342581 | 0.068046 | 0.989223 | 0.997988 | 0.994666 |
| Fbxo41_Q6NS60_T477t_1 | Fbxo41_Q6 | 2.82 | 2.68 | 2.87 | 1.18 | 2.95 | 2.87 | 2.37 | 2.33 | 1.56 | 3.58 | 1.07 | 3.39 | 3.76 | 0.78 | 3.63 | 3.27 | 0.3 | 1.01 | 1.05 | 0.51742 | 0.5626 | 0.98534 | 0.989223 | 0.997988 | 0.994666 |
| Fbxo41_Q6NS60_T477t_2 | Fbxo41_Q6 | 2.4 | 2.39 | 2.19 | 1 | 2.75 | 2.13 | 1.95 | 2.23 | 0.3 | 3.07 | 0.53 | 2.29 | -0.38 | 0.34 | 3 | 3 | 0.2425 | 0.0125 | 0.4725 | 0.547986 | 0.472902 | 0.516106 | 0.989223 | 0.997988 | 0.994666 |
| Fbxo42_Q6PDJ6_S365s_1 | Fbxo42_Q6 | -0.44 | -0.9 | -1.13 | -1.24 | -0.01 | -0.47 | -0.7 | -0.79 | -0.88 | -0.43 | -0.03 | -0.8 | -0.69 | -0.09 | -1.04 | -1.22 | 0.27 | -0.4475 | -0.505 | 0.625018 | 0.148235 | 0.214543 | 0.989223 | 0.997988 | 0.995259 |
| Fbxo42_Q6PDJ6_S373s_1 | Fbxo42_Q6 | 1.65 | 0.79 | 0.99 | 1.72 | 0.61 | 1.5 | 1.4 | 1.67 | 1.22 | 1.25 | 1.49 | 1.57 | -0.15 | 0.67 | 0.91 | 0.35 | 0.435 | 0.3925 | 0.1675 | 0.037288 | 0.982289 | 0.746144 | 0.999187 | 0.997988 | 0.994666 |
| Fbxo42_Q6PDJ6_S429s_1 | Fbxo42_Q6 | 0.08 | -0.79 | -0.89 | -0.99 | -0.87 | 0.67 | -1.44 | -0.45 | 0.33 | -0.49 | -0.02 | -1.04 | -0.41 | -0.91 | -1.41 | -1.24 | 0.0075 | 0.095 | -0.8425 | 0.345508 | 0.791212 | 0.387123 | 0.995762 | 0.997988 | 0.994666 |
| Fbxo42_Q6PDJ6_S488s_1 | Fbxo42_Q6 | -0.5 | -1.39 | -2.11 | -1.04 | -1.58 | 0.36 | -2.27 | -1.02 | 0.33 | -0.56 | -1.99 | -2.09 | 0.19 | -0.65 | -1.22 | -1.48 | 0.125 | 0.3425 | -0.345 | 0.338832 | 0.820779 | 0.767627 | 0.995762 | 0.997988 | 0.994666 |
| Fbxo42_Q6PDJ6_S578s_1 | Fbxo42_Q6 | -0.53 | 0.43 | 0 | -0.11 | 0.64 | -0.36 | 0.29 | 0.49 | 0.06 | 0.36 | 1.09 | 0.78 | 0.01 | 0.55 | 0.25 | 0.17 | 0.1325 | 0.1825 | 0.47 | 0.324617 | 0.324713 | 0.081562 | 0.989223 | 0.997988 | 0.994666 |
| Fbxo9_Q8BK06_S22s_1 | Fbxo9_Q8E | -0.02 | 0.01 | 0.15 | -0.41 | -1.28 | 0.83 | -1.66 | -0.47 | 1.62 | -0.22 | -0.34 | 0.17 | 1.41 | -2.28 | -0.67 | -0.54 | 0.3175 | 0.625 | 0.2975 | 0.514191 | 0.277808 | 0.413023 | 0.989223 | 0.997988 | 0.994666 |
| Fcgr1_P26151_S347s_1 | Fcgr1_P261 | 0.33 | -0.03 | -0.88 | -1.31 | 0.22 | -1.03 | -1.72 | -0.91 | -1.05 | -0.93 | 0.18 | -0.36 | -0.25 | 0.35 | -1.5 | -1.31 | -0.5775 | 0.375 | -0.4525 | 0.704409 | 0.449517 | 0.881554 | 0.989223 | 0.997988 | 0.996159 |
| Fcho1_Q8K285_S498s_1 | Fcho1_Q8K | -2.55 | -1.32 | -1.53 | -1.84 | -2.29 | -2.04 | -1.44 | -2.13 | -1.24 | -2.02 | -1.33 | -1.74 | -0.69 | -1.47 | -1.23 | -1.75 | -0.3875 | -0.0675 | -0.205 | 0.178716 | 0.62437 | 0.512382 | 0.989223 | 0.997988 | 0.994666 |
| Fcho2_Q3UQN2_S394s_1 | Fcho2_Q3U | 7.37 | 2.74 | 4.75 | 0.61 | 3.85 | 7.23 | 5.96 | 3.58 | 0.76 | 5.6 | 1.02 | 4.87 | -2.03 | -1.21 | 4.49 | 5.24 | -0.165 | 0.2275 | 0.525 | 0.28416 | 0.382729 | 0.631988 | 0.989223 | 0.997988 | 0.994666 |
| Fcho2_Q3UQN2_S403s_1 | Fcho2_Q3U | 0.38 | -0.1 | 0.61 | 1.38 | -0.67 | 0.92 | 0.29 | 0.59 | 1.09 | 0.34 | 0.3 | 1.18 | 0.85 | -1.18 | 0.29 | -0.03 | 1.2875 | -0.805 | -2.245 | 0.258879 | 0.516432 | 0.682496 | 0.989223 | 0.997988 | 0.994666 |
| Fcho2_Q3UQN2_S487s_1 | Fcho2_Q3U | -0.08 | 0.48 | -0.1 | 0.39 | 0.79 | 0.79 | 0.55 | -0.03 | 0 | 0.88 | 0.12 | -0.31 | 0.07 | 0.89 | 0.23 | 0.36 | -0.285 | 0.16 | -0.585 | 0.47617 | 0.229857 | 1 | 0.989223 | 1 | 0.994666 |
| Fcho2_Q3UQN2_T682t_1 | Fcho2_Q3U | -1.68 | -1.3 | -1.9 | -2.08 | -1.45 | -0.58 | -2.07 | -1.9 | -1.4 | -0.81 | -1.01 | -2.15 | -0.09 | -0.98 | -1.07 | -1.18 | 0.3525 | 0 | 0.215 | 0.021854 | 0.519483 | 0.27841 | 0.989223 | 0.997988 | 0.994666 |
| Fchsd2_Q3USJ8_S630s_1 | Fchsd2_Q3 | -0.66 | -0.92 | -0.78 | -1.04 | -0.48 | -1.34 | -1.49 | -1.3 | -0.75 | -1.65 | -0.62 | -1.4 | -1.02 | -0.15 | -2.3 | -1.04 | 0.24 | 0.3975 | 0.91 | 0.533393 | 0.293516 | 0.406856 | 0.989223 | 0.997988 | 0.994666 |
| Fdx1l_Q9CPW2_S51s_1 | Fdx1l_Q9C | 0.6 | 0.58 | 0.31 | 0.35 | 0.09 | 0.54 | -0.37 | -0.01 | -0.19 | 0.6 | 0.03 | 0.23 | 0.21 | 0.39 | 0.22 | 0.61 | -0.3025 | -0.255 | -0.2775 | 0.676154 | 0.14331 | 0.268286 | 0.989223 | 0.997988 | 0.996159 |
| Fer_Q3TZJ5_S411s_1 | Fer_Q3TZJ | 0.29 | 0.59 | 0.78 | 0.75 | 0.35 | 0.03 | 0.42 | 0.58 | -0.11 | 0.65 | 1.13 | 0.52 | 0.21 | 0.54 | 0.89 | 0.13 | -0.3975 | -0.2925 | -0.1025 | 0.574434 | 0.285265 | 0.860081 | 0.989223 | 0.997988 | 0.995259 |
| Fez1_Q8K0X8_S298s_1 | Fez1_Q8K0 | -2.71 | -2.38 | -1.59 | -2.28 | -0.56 | -2.44 | -1.03 | -2.48 | -2.96 | -1.94 | -0.6 | -1.87 | -1.21 | -0.65 | -1.13 | -1.2 | -0.2575 | -0.055 | -0.16 | 0.003982 | 0.238162 | 0.43933 | 0.989223 | 0.997988 | 0.994666 |
| Fez1_Q8K0X8_S316s_1 | Fez1_Q8K0 | -0.47 | -0.39 | 0.18 | 0.27 | 0.45 | -0.47 | 1.04 | 0.21 | -0.3 | -0.09 | -0.19 | 0.89 | -0.36 | -0.09 | 0.59 | 0.64 | 0.6125 | 0.3975 | 1.1925 | 0.399002 | 0.275349 | 0.609237 | 0.989223 | 0.997988 | 0.994666 |
| Fez2_Q3TN06_S200s_1 | Fez2_Q3TN | -0.08 | -0.29 | -0.17 | -0.26 | -0.77 | -0.72 | -0.82 | -0.25 | -0.26 | -0.25 | 0.19 | -0.31 | 0.28 | -0.37 | 0.49 | 0.33 | 0.41 | 0.18 | 0.2975 | 0.188914 | 0.07336 | 0.851532 | 0.989223 | 0.997988 | 0.994666 |
| Fgd6_Q69ZL1_S1167s_1 | Fgd6_Q692 | 1.45 | 0.36 | 1.15 | 0.39 | -0.51 | 2.16 | -0.05 | 0.41 | 1.77 | 0.62 | 0.35 | 0.24 | 1.15 | -0.51 | 0.8 | 0.4 | -0.44 | 0.0425 | 0.3825 | 0.401706 | 0.566142 | 0.830754 | 0.989223 | 0.997988 | 0.994666 |
| Fgd6_Q69ZL1_S583s_1 | Fgd6_Q692 | -1.41 | -0.95 | -1.08 | -1.56 | -0.82 | -0.86 | -1.42 | -1.94 | -1.87 | -1.1 | -0.3 | -1.92 | -0.91 | -0.68 | -1.91 | -1.15 | -0.335 | -0.0925 | -0.3775 | 0.796203 | 0.97478 | 0.906261 | 0.999187 | 0.997988 | 0.999507 |
| Fgf12_P61329_S212s_1 | Fgf12_P61 | -0.68 | 0.32 | -0.12 | -0.37 | 0.47 | -0.83 | -0.05 | -0.65 | -1.45 | -0.57 | -1.2 | -0.35 | -0.69 | 0.27 | -0.11 | 0.04 | -0.01 | -0.0475 | 0.0875 | 0.787404 | 0.884505 | 0.07761 | 0.996493 | 0.997988 | 0.999507 |
| Fgf12_P61329_S226s_1 | Fgf12_P61 | 0.74 | 0.65 | 0.53 | 1.08 | 0.39 | 0.81 | -0.24 | -0.94 | 1.39 | 0.94 | -0.75 | 0.51 | 1.9 | -0.51 | -0.29 | -0.35 | -0.0525 | -0.68 | 0.09 | 0.314511 | 0.079949 | 0.620266 | 0.989223 | 0.997988 | 0.994666 |
| Fgf12_P61329_S227s_1 | Fgf12_P61 | -1.16 | -0.41 | -0.35 | 0.04 | -0.44 | -0.49 | -0.2 | -0.37 | 0.33 | -0.38 | -0.73 | 0.25 | 1.17 | -1.07 | -0.45 | -0.26 | -0.745 | -0.2275 | -0.5625 | 0.534154 | 0.743035 | 0.368627 | 0.995762 | 0.997988 | 0.994666 |
| Fgf12_P61329_T229t_1 | Fgf12_P61 | -0.16 | -0.4 | -0.04 | -0.54 | 0.18 | -0.44 | -0.16 | -0.59 | 1.24 | 0.61 | -0.81 | 0.27 | 1.57 | -0.43 | -0.38 | -0.29 | 0.095 | 0.3375 | 0.3175 | 0.407677 | 0.898916 | 0.179378 | 0.996493 | 0.997988 | 0.994666 |
| Fhod3_Q76LL6_S345s_1 | Fhod3_Q76 | -1.16 | -1.83 | -2 | -1.79 | -0.76 | -1.32 | -1.52 | -1.66 | -1.95 | -0.93 | -0.66 | -2.67 | -1.05 | -1.28 | -1.85 | -1.51 | 0.0325 | 0.6125 | 0.4025 | 0.385118 | 0.220091 | 0.764654 | 0.989223 | 0.997988 | 0.994666 |
| Fkbp15_Q6P9Q6_S1091s_1 | Fkbp15_Q6 | -0.67 | 0.46 | -1.44 | -1.36 | 0.4 | -0.33 | -0.72 | -0.89 | -0.88 | -0.79 | -0.98 | -0.37 | -0.54 | -0.31 | -1.78 | -1.41 | 0.38 | 0.1425 | 0.2725 | 0.625928 | 0.454399 | 0.9955 | 0.989223 | 0.998132 | 0.995259 |
| Fkbp1a_P26883_S9s_1 | Fkbp1a_P2 | -0.05 | 0.01 | 0.28 | 0.58 | 0.49 | -0.29 | 0.47 | 0.56 | -0.29 | 0.57 | 0.68 | 0.35 | 0.33 | 0.62 | 0.31 | 0.29 | 0.3675 | -0.0025 | -0.2575 | 0.491345 | 0.716632 | 0.684223 | 0.995437 | 0.997988 | 0.994666 |
| Fkbp8_O35465_S286s_1 | Fkbp8_O35 | 1.28 | 0.09 | -0.37 | 1.52 | 0.21 | 1.01 | 0.56 | 1.59 | 0.25 | 1.18 | 2.04 | 0.24 | -0.96 | 0.29 | 1.19 | 0.51 | 0.1025 | 0.1225 | 0.1825 | 0.529909 | 0.672143 | 0.609494 | 0.992876 | 0.997988 | 0.994666 |
| Flad1_Q8R123_S468s_1 | Flad1_Q8R | -1.58 | -2.07 | -1.8 | -2.9 | -2.15 | -1.84 | -2.6 | -1.65 | -1.21 | -1.5 | -0.77 | -2.53 | -0.34 | -2.43 | -1.62 | -2.01 | 0.2125 | 0.2975 | -0.3725 | 0.347057 | 0.938286 | 0.218375 | 0.999187 | 0.997988 | 0.994666 |
| Flcn_Q8QZS3_S571s_1 | Flcn_Q8QZ | -0.94 | -1.28 | -1.29 | -3.88 | -1.8 | -1.17 | -2.56 | -2.38 | 1.19 | -0.94 | -1.86 | -2.09 | 2.17 | -1.76 | -0.68 | -1.92 | 0.0275 | 0.585 | 0.4875 | 0.220441 | 0.845038 | 0.318587 | 0.996349 | 0.997988 | 0.994666 |
| Flcn_Q8QZS3_S62s_1 | Flcn_Q8QZ | 3.64 | 2.12 | 2.51 | 0.28 | 1.64 | 4 | 2 | 2.37 | 0.26 | 3.67 | 0.11 | 2.76 | 1.31 | -0.45 | 3.27 | 3.46 | -0.13 | 0.9225 | 1.3 | 0.811504 | 0.63439 | 0.66375 | 0.989223 | 0.997988 | 0.999507 |
| Flcn_Q8QZS3_S62s_2 | Flcn_Q8QZ | 1.68 | 0.28 | 2.15 | -0.31 | 0.72 | 2.4 | 1.58 | 0.99 | 0.76 | 2.38 | -0.36 | 1.37 | 1.68 | -0.97 | 1.63 | 2.12 | 0.365 | -0.4375 | -0.24 | 0.837418 | 0.448339 | 0.904721 | 0.989223 | 0.997988 | 0.999507 |
| Flcn_Q8QZS3_S73s_2 | Flcn_Q8QZ | 1.68 | 0.28 | 2.15 | -0.31 | 0.72 | 2.4 | 1.58 | 0.99 | 0.76 | 2.38 | -0.36 | 1.37 | 1.68 | -0.97 | 1.63 | 2.12 | 0.4725 | 0.0875 | 0.165 | 0.837418 | 0.448339 | 0.904721 | 0.989223 | 0.997988 | 0.999507 |
| Flcn_Q8QZS3_T313t_1 | Flcn_Q8QZ | -2.59 | -3.65 | -2.89 | -3.94 | -2.22 | -3.51 | -2.46 | -2.6 | -3.17 | -3.5 | -0.14 | -2.19 | -0.94 | -2.57 | -1.77 | -2.69 | 0.4725 | 0.0875 | 0.165 | 0.02548 | 0.182651 | 0.189866 | 0.989223 | 0.997988 | 0.994666 |
| Flii_Q9JJ28_S436s_1 | Flii_Q9JJ28 | -2.4 | -2.81 | -1.85 | -2.81 | -2.54 | -1.55 | -1.69 | -2.21 | -0.29 | -2.65 | -3.68 | -2.06 | 0.39 | -1.67 | -0.92 | -1.43 | 0.57 | 1.0175 | 1.275 | 0.009611 | 0.180222 | 0.660897 | 0.989223 | 0.997988 | 0.994666 |
| Flna_B7FAV1_S1435s_1 | Flna_B7FA | 0.09 | -0.7 | -0.54 | -0.13 | -1.02 | -1 | -0.64 | -0.65 | -0.4 | -0.92 | 0.18 | -0.14 | -0.54 | -0.46 | 0.11 | -0.14 | 0.47 | 0.2975 | 1.56 | 0.834124 | 0.075435 | 1 | 0.989223 | 1 | 0.999507 |
| Flna_B7FAV1_S2088s_1 | Flna_B7FA | -2.35 | -2.27 | -2.71 | -2.7 | -1.47 | -2.29 | -2.6 | -2.77 | -4.88 | -2.47 | -3.46 | -3.1 | -1.53 | -2.75 | -2.88 | -2.54 | -0.5075 | 7.85E-17 | 0.0625 | 0.81623 | 0.492657 | 0.07526 | 0.989223 | 0.997988 | 0.999507 |
| Flrt3_Q8BGT1_S644s_1 | Flrt3_Q8BG | -0.06 | 1.05 | 0.06 | 0.32 | 0.69 | 0.84 | 0.82 | 1.52 | 0.41 | 1.14 | -1.06 | 1.49 | 0.41 | -1.72 | 0.27 | 0.59 | 0.225 | -0.97 | 0.0825 | 0.417294 | 0.078984 | 0.789064 | 0.989223 | 0.997988 | 0.994666 |
| Flt3_Q3UEW6_S988s_1 | Flt3_Q3UE | 1.02 | 1.48 | 1.03 | 1.61 | 0.11 | 0.75 | 0.72 | 0.77 | 1.2 | 1.5 | 2.01 | 0.6 | 1.35 | 1.05 | 0.23 | 1.05 | 0.625 | 0.1525 | -0.455 | 0.281198 | 0.024552 | 0.902757 | 0.989223 | 0.997988 | 0.994666 |
| Fmn1_E9Q7P6_S168s_1 | Fmn1_E9Q | -0.08 | 0.59 | 0.22 | -0.26 | 0.1 | -0.25 | -0.03 | -0.01 | 0.76 | 0.51 | 1.18 | 0.56 | 0.73 | 0.23 | 0.42 | 0.2 | -0.6975 | 0.0425 | -0.365 | 0.352113 | 0.520271 | 0.050055 | 0.989223 | 0.997988 | 0.994666 |
| Fmn1_E9Q7P6_S588s_1 | Fmn1_E9Q | 0.25 | -1 | 0.35 | 0.14 | 0.27 | 0.29 | 0.73 | 0.51 | -0.24 | -0.65 | 0.93 | 0.5 | -0.38 | 0.42 | -0.13 | 0.31 | -0.165 | 0.635 | 0.2775 | 0.753204 | 0.152982 | 0.66242 | 0.989223 | 0.997988 | 0.9971 |
| Fmn2_Q9JL04_S16s_1 | Fmn2_Q9JI | 2.93 | 2.34 | 2.27 | 1.26 | 1.16 | 3.23 | 0.73 | 2.14 | 1.21 | 2.08 | 0.91 | 2.4 | 1.02 | -0.61 | 1.48 | 1.24 | 0.515 | 0.2 | 0.12 | 0.025646 | 0.517042 | 0.260825 | 0.989223 | 0.997988 | 0.994666 |
| Fmn2_Q9JL04_S373s_1 | Fmn2_Q9JI | 0.35 | -0.83 | 1.45 | 0.3 | -0.74 | 0.74 | 0.93 | 1.11 | 0.72 | -0.81 | 2.09 | 0.53 | -0.86 | -1.07 | -0.02 | -0.33 | -0.385 | -0.55 | -1.4175 | 0.099396 | 0.734593 | 0.645789 | 0.995762 | 0.997988 | 0.994666 |
| Fmn2_Q9JL04_S405s_1 | Fmn2_Q9JI | -0.46 | -0.05 | -0.6 | -0.19 | -0.93 | 0.08 | -0.28 | -0.04 | 1.45 | -0.75 | -0.31 | 0.08 | 1.36 | 0.19 | -0.09 | -0.56 | 0.1925 | 0.315 | -0.8875 | 0.215294 | 0.910276 | 0.357331 | 0.997335 | 0.997988 | 0.994666 |
| Fmn2_Q9JL04_S409s_1 | Fmn2_Q9JI | 0.11 | 0.04 | -1.47 | -1.17 | -0.54 | -0.05 | -1.36 | -0.79 | 0.77 | 0.14 | -0.77 | 0 | 1.5 | 0.33 | 0.15 | -0.1 | 0.0325 | 0.4425 | 0.55 | 0.054896 | 0.891097 | 0.20006 | 0.996493 | 0.997988 | 0.994666 |
| Fmn2_Q9JL04_S409s_2 | Fmn2_Q9JI | -0.32 | -0.33 | -0.59 | -0.89 | -0.46 | -0.67 | -1.06 | -0.66 | 0.39 | -0.23 | -1.17 | -0.65 | 0.38 | -0.12 | -0.97 | -0.44 | -0.0625 | 0.6575 | 1.0925 | 0.48448 | 0.469023 | 0.748592 | 0.989223 | 0.997988 | 0.994666 |
| Fmn2_Q9JL04_S413s_2 | Fmn2_Q9JI | -0.04 | -0.37 | -0.59 | -0.89 | -0.3 | -0.55 | -1.06 | -0.66 | 0.55 | -0.06 | -1.17 | -0.65 | 0.19 | -0.3 | -0.97 | -0.44 | -0.18 | 0.1175 | 0.245 | 0.783383 | 0.543312 | 0.731882 | 0.989223 | 0.997988 | 0.998491 |
| Fmn2_Q9JL04_S493s_1 | Fmn2_Q9JI | 0.51 | 0.86 | 0.11 | 0.39 | 0.88 | 0.75 | 0.45 | 0.33 | 0.49 | 0.97 | 0.41 | 0.46 | 0.42 | 1 | 0.62 | 0.34 | -0.17 | 0.14 | 0.0925 | 0.657826 | 0.598439 | 0.665779 | 0.989223 | 0.997988 | 0.995259 |
| Fmn2_Q9JL04_S614s_1 | Fmn2_Q9JI | -0.46 | 0.09 | -0.05 | 0.08 | 0.16 | -0.37 | 0.16 | 0.06 | -0.45 | -0.06 | 0.45 | 0.12 | -0.2 | 0.37 | 0.12 | 0.06 | 0.135 | 0.115 | 0.1275 | 0.522055 | 0.719746 | 0.721315 | 0.995437 | 0.997988 | 0.994666 |
| Fmn2_Q9JL04_S629s_1 | Fmn2_Q9JI | 0.08 | 0.32 | 0 | -0.05 | 0.69 | -0.06 | -0.05 | -0.13 | 0.85 | 0.52 | 2.27 | 0.07 | 0.58 | 0.97 | 0.05 | -0.57 | 0.0875 | 0.1 | 0.1725 | 0.643922 | 0.923198 | 0.09517 | 0.997335 | 0.997988 | 0.995259 |
| Fmn2_Q9JL04_S69s_1 | Fmn2_Q9JI | -0.87 | -0.85 | -2.19 | -2.05 | -0.8 | -1.63 | -0.86 | -1.6 | -1.55 | -3.37 | -0.07 | -1.62 | -1.13 | 0.29 | -2.2 | -1.23 | 0.025 | 0.84 | 0.17 | 0.471185 | 0.519255 | 0.814586 | 0.989223 | 0.997988 | 0.994666 |
| Fmn2_Q9JL04_S720s_2 | Fmn2_Q9JI | -0.45 | -0.41 | -0.3 | -0.58 | 0.08 | 0.91 | 0.01 | -0.35 | -1.31 | -0.38 | 0.81 | -0.33 | -2.67 | -0.5 | -0.57 | -0.81 | 0.2675 | -0.1625 | 0.4225 | 0.176289 | 0.069532 | 0.756727 | 0.989223 | 0.997988 | 0.994666 |
| Fmn2_Q9JL04_S724s_1 | Fmn2_Q9JI | 1.17 | 1.12 | 1.09 | 0.95 | 0.82 | 0.72 | 0.31 | 0.64 | 1.35 | 1.58 | 1.54 | 1.34 | 1.15 | 0.93 | 1.17 | 0.52 | 0.5975 | 0.1325 | -0.7025 | 0.591972 | 0.055273 | 0.105288 | 0.989223 | 0.997988 | 0.995259 |
| Fmn2_Q9JL04_S724s_2 | Fmn2_Q9JI | -0.45 | -0.41 | -0.3 | -0.58 | 0.08 | 0.91 | 0.01 | -0.35 | -1.31 | -0.38 | 0.81 | -0.33 | -2.67 | -0.5 | -0.57 | -0.81 | -0.46 | 0.37 | -0.14 | 0.176289 | 0.069532 | 0.756727 | 0.989223 | 0.997988 | 0.994666 |
| Fmnl1_Q9JL26_S995s_1 | Fmnl1_Q9J | 1.6 | 1.67 | 2.35 | 2.67 | 1.38 | 1.43 | 1.57 | 2.38 | 1.26 | 1.83 | 2.47 | 2.31 | 1.26 | 1.93 | 2.38 | 2.21 | 0.5975 | 0.1325 | -0.7025 | 0.735216 | 0.292441 | 0.783235 | 0.989223 | 0.997988 | 0.9971 |
| Fmnl1_Q9JL26_T483t_1 | Fmnl1_Q9J | -0.44 | 0.59 | 0.25 | 0.77 | 0.73 | -0.02 | 0.73 | 0.61 | 0.21 | 0.74 | -0.25 | 0.56 | -0.14 | 0.36 | 0.33 | 0.83 | -0.3825 | -0.105 | -0.1275 | 0.883397 | 0.511864 | 0.949747 | 0.989223 | 0.997988 | 0.999507 |
| Fmnl2_E9PXE6_S183s_1 | Fmnl2_E9P | 0.54 | 0.51 | 0.2 | 0.46 | 0.56 | 0.51 | 1.04 | 0.87 | 0.42 | 0.9 | 0.8 | 0.83 | -0.03 | 0.44 | 0.66 | 1.04 | 0.22 | 0.0225 | 0.0525 | 0.738438 | 0.184329 | 0.199739 | 0.989223 | 0.997988 | 0.9971 |
| Fmnl2_E9PXE6_S981s_1 | Fmnl2_E9P | 1.01 | 1.11 | 0.12 | 1.74 | 1.52 | 0.7 | 1.61 | 1.5 | 0.72 | 1.47 | 1.23 | 1.62 | -0.31 | 1.1 | 0.68 | 1.2 | 0.3175 | 0.31 | 0.1 | 0.485175 | 0.390115 | 0.500868 | 0.989223 | 0.997988 | 0.994666 |
| Fnbp1_Q80TY0_S217s_1 | Fnbp1_Q80 | 0.59 | 0.59 | 1.09 | 0.64 | 0.36 | 0.45 | 0.25 | 0.43 | 0.42 | 0.55 | 0.54 | 0.38 | 0.97 | 0.32 | 0.8 | 0.7 | 0.3375 | 0.265 | -0.3275 | 0.911774 | 0.12942 | 0.280564 | 0.989223 | 0.997988 | 0.999507 |
| Fnbp1_Q80TY0_S220s_1 | Fnbp1_Q80 | 1.56 | 0.48 | 0.74 | 0.8 | 0.37 | 0.83 | 0.07 | 0.52 | 1.23 | 0.12 | 0.25 | 0.94 | 1.15 | 0.08 | 0.46 | 0.8 | -0.355 | -0.255 | -0.03 | 0.448823 | 0.164021 | 0.481795 | 0.989223 | 0.997988 | 0.994666 |
| Fnbp1_Q80TY0_S402s_1 | Fnbp1_Q80 | -2.1 | -1.18 | -1.08 | -1.1 | -0.97 | -3.15 | -1.47 | -1.73 | -3.38 | -0.56 | -2.36 | -1.53 | -2.93 | -2.83 | -1.02 | -2.18 | -0.4475 | -0.26 | -0.2725 | 0.094376 | 0.350966 | 0.33274 | 0.989223 | 0.997988 | 0.994666 |
| Fnbp1_Q80TY0_S422s_1 | Fnbp1_Q80 | -3.06 | -1.12 | -1.96 | -2.73 | -2.34 | -3.45 | -1.84 | -2.78 | -2.51 | -2.26 | -1.8 | -2.61 | -1.38 | -2 | -2.54 | -2.88 | -0.465 | -0.5925 | -0.875 | 0.972633 | 0.452282 | 0.86347 | 0.989223 | 0.997988 | 0.999507 |
| Fnbp1_Q80TY0_S451s_1 | Fnbp1_Q80 | 0.5 | 1.06 | 0.86 | 0.74 | -0.28 | -0.07 | 0.08 | 0.61 | 0.6 | 0.82 | 1.43 | 0.36 | 0.91 | 0.16 | 1.4 | 0.92 | -0.385 | -0.0775 | 0.0175 | 0.859779 | 0.023996 | 0.966514 | 0.989223 | 0.997988 | 0.999507 |
| Fnbp1_Q80TY0_S451s_2 | Fnbp1_Q80 | -0.07 | 0.46 | -0.75 | -0.58 | -0.93 | -0.7 | -1.28 | -0.92 | 0.55 | -0.4 | -0.6 | -1.18 | -1.09 | -0.94 | -0.77 | -0.72 | -0.705 | 0.0125 | 0.0575 | 0.07401 | 0.043036 | 0.694394 | 0.989223 | 0.997988 | 0.994666 |
| Fnbp1_Q80TY0_S455s_2 | Fnbp1_Q80 | -0.07 | 0.46 | -0.75 | -0.58 | -0.93 | -0.7 | -1.28 | -0.92 | 0.55 | -0.4 | -0.6 | -1.18 | -1.09 | -0.94 | -0.77 | -0.72 | -0.7225 | -0.1725 | -0.645 | 0.07401 | 0.043036 | 0.694394 | 0.989223 | 0.997988 | 0.994666 |
| Fnbp1l_E9PUK3_S153s_1 | Fnbp1l_E9I | -0.09 | -0.57 | 0.57 | -0.51 | -1.42 | 0.59 | -1.12 | -0.56 | 0.56 | 0.51 | -2.45 | 0.43 | 1.74 | -3.66 | 0.71 | -0.26 | -0.7225 | -0.1725 | -0.645 | 0.834776 | 0.329199 | 0.90129 | 0.989223 | 0.997988 | 0.999507 |
| Fnbp1l_E9PUK3_S295s_1 | Fnbp1l_E9I | -0.01 | -0.28 | -0.23 | -0.24 | 0.62 | 0.06 | 0.11 | 0.04 | -0.12 | 0.14 | -0.06 | -0.02 | -0.28 | 0.47 | -0.14 | -0.05 | -0.4775 | -0.0875 | -0.2175 | 0.482195 | 0.107144 | 0.417701 | 0.989223 | 0.997988 | 0.994666 |
| Fnbp1l_E9PUK3_S430s_1 | Fnbp1l_E9I | 1.89 | -0.11 | 2.08 | -0.97 | -0.24 | 2.14 | 2.4 | 1.52 | 0.73 | 1.53 | 1.25 | 1.99 | 1 | -0.56 | 1.95 | 1.81 | 0.3975 | 0.175 | 0.19 | 0.695259 | 0.390146 | 0.373501 | 0.989223 | 0.997988 | 0.996159 |
| Fnbp1l_E9PUK3_S443s_1 | Fnbp1l_E9I | -0.01 | 0.41 | 1.79 | 1.01 | -0.11 | 0.12 | 2.19 | 2 | -0.07 | 0.6 | 1.17 | 2.17 | -0.75 | 0.03 | 2.11 | 2.28 | 0.7325 | 0.6525 | 0.3275 | 0.876272 | 0.697421 | 0.768046 | 0.994893 | 0.997988 | 0.999507 |
| Fnbp4_Q6ZQ03_S487s_1 | Fnbp4_Q6Z | -0.36 | -0.13 | -2.57 | -3.15 | -0.88 | -1.11 | -2.98 | -3.16 | -1.71 | -0.64 | -1.22 | -3.08 | -0.7 | 0.13 | -2.73 | -2.93 | 0.25 | 0.1675 | 0.1175 | 0.995748 | 0.574634 | 0.893877 | 0.989223 | 0.997988 | 1 |
| Fndc3a_E0CXY0_S202s_1 | Fndc3a_E0 | -0.54 | 0.85 | -0.1 | -0.51 | -0.04 | -0.67 | -0.52 | -1.07 | 0.25 | 0.43 | -0.69 | -0.49 | 1.36 | -0.29 | 0.42 | 0.15 | -0.48 | -0.11 | -0.005 | 0.306776 | 0.208102 | 0.904453 | 0.989223 | 0.997988 | 0.994666 |
| Fndc3a_E0CXY0_S213s_1 | Fndc3a_E0 | -1.5 | -1.23 | -0.49 | -0.6 | -1.14 | -0.71 | -0.54 | 0.38 | -1.31 | -0.87 | -1.98 | -1.38 | 0.2 | -0.13 | -1.45 | 0.17 | -0.5 | -0.05 | 0.485 | 0.168624 | 0.263287 | 0.238494 | 0.989223 | 0.997988 | 0.994666 |
| Fndc4_Q3TR08_S209s_1 | Fndc4_Q3T | 0.05 | 0.36 | 0.05 | -0.68 | 0.24 | 0.7 | -0.91 | -0.38 | 1.52 | 0.02 | -0.3 | 0.26 | 1.74 | -0.26 | -0.1 | -0.38 | 0.4525 | -0.43 | 0.6525 | 0.556296 | 0.935166 | 0.342204 | 0.99868 | 0.997988 | 0.995259 |
| Fnip1_F6RJ64_S237s_1 | Fnip1_F6R | 0.39 | -0.55 | 1.19 | 1.66 | -0.96 | 0.31 | -0.14 | -1.74 | 0.77 | 0.47 | 1.86 | 0.52 | -1.37 | -0.12 | 0.24 | 0.37 | -0.0325 | 0.43 | 0.305 | 0.145513 | 0.050526 | 0.669322 | 0.989223 | 0.997988 | 0.994666 |
| Foxk1_P42128_S199s_1 | Foxk1_P42 | -1.57 | -0.9 | -1.66 | -1.16 | -0.94 | -1.66 | -0.74 | -1.91 | -1.23 | -1.51 | -1.27 | -1.51 | -1.17 | -0.48 | -1.85 | -1.52 | -1.305 | 0.2325 | -0.8925 | 0.853362 | 0.976431 | 0.82489 | 0.999187 | 0.997988 | 0.999507 |
| Foxk1_P42128_S431s_1 | Foxk1_P42 | -0.26 | -1.26 | -2.5 | -2.19 | -1.04 | -0.96 | -2.51 | -2.62 | -1.28 | -0.52 | -1.34 | -3.32 | -0.74 | -0.78 | -3.22 | -2.88 | 0.01 | -0.0575 | 0.0675 | 0.636731 | 0.705082 | 0.929347 | 0.995437 | 0.997988 | 0.995259 |
| Foxk1_P42128_S87s_1 | Foxk1_P42 | -2.37 | -1.36 | -1.64 | -1.87 | -1.57 | -1.86 | -1.42 | -1.69 | -0.8 | -0.97 | -0.24 | -1.57 | 0.1 | -1.02 | -1.63 | -1.83 | -0.23 | -0.0625 | -0.3525 | 0.149267 | 0.526486 | 0.028701 | 0.989223 | 0.997988 | 0.994666 |
| Foxk2_Q3UCQ1_S389s_1 | Foxk2_Q3U | -1.62 | -1.02 | -3.27 | -2.24 | -0.51 | -1.18 | -2.08 | -2.3 | -0.55 | -0.3 | -1.93 | -3.52 | -0.65 | 0.2 | -1.71 | -1.21 | 0.175 | 0.915 | 0.715 | 0.063245 | 0.372859 | 0.561438 | 0.989223 | 0.997988 | 0.994666 |
| Fra10ac1_Q8BP78_S243s_1 | Fra10ac1_( | -3.36 | -3.72 | -2.37 | -4.13 | -4.97 | -3.43 | -4 | -3.52 | -1.46 | -2.78 | -1.97 | -3.31 | -0.97 | -5.25 | -2.06 | -3.2 | 0.52 | 0.4625 | 1.195 | 0.549203 | 0.238567 | 0.077585 | 0.989223 | 0.997988 | 0.994666 |
| Frmd3_Q8BHD4_S63s_1 | Frmd3_Q8I | 0.7 | 0.13 | -0.23 | -0.47 | 1.79 | -1.33 | 0.32 | -0.04 | -0.18 | 0.41 | -0.19 | -0.6 | -0.34 | 1.61 | 0.3 | 0.29 | -0.585 | 1.015 | 0.525 | 0.363725 | 0.804463 | 0.620162 | 0.995762 | 0.997988 | 0.994666 |
| Frmd4a_Q3TB04_S372s_1 | Frmd4a_Q | -1.39 | -0.72 | -1.37 | -1.78 | -1.32 | -0.03 | -2.18 | -1.51 | 0.67 | -1.55 | -0.65 | -0.98 | 0.21 | -0.59 | -0.62 | -0.85 | 0.1525 | -0.1725 | 0.4325 | 0.033469 | 0.905693 | 0.184475 | 0.996513 | 0.997988 | 0.994666 |
| Frmd4a_Q3TB04_S385s_1 | Frmd4a_Q | 1.33 | 1.14 | 0.77 | 0.73 | 1.3 | 1.23 | 0.86 | 0.92 | 0.57 | 1.34 | 0.14 | 1.02 | 0.78 | 0.35 | 0.89 | 0.34 | 0.055 | 0.6875 | 0.8525 | 0.175174 | 0.72836 | 0.495746 | 0.995762 | 0.997988 | 0.994666 |
| Frmd4a_Q3TB04_S402s_1 | Frmd4a_Q | -0.74 | -0.97 | -1.3 | -0.56 | -2.22 | 0.3 | -1.1 | -1.24 | 1.54 | -0.13 | -2.37 | -1.02 | 1.51 | -1.05 | -0.84 | -1.07 | 0.085 | -0.225 | -0.4025 | 0.380564 | 0.729542 | 0.598434 | 0.995762 | 0.997988 | 0.994666 |
| Frmd4a_Q3TB04_S48s_1 | Frmd4a_Q | -0.34 | -0.62 | 0.33 | 0.49 | -0.78 | -0.33 | 0.65 | 0.8 | 0.33 | 0.32 | 0.17 | 0.48 | 0.45 | -0.96 | 1.01 | 0.46 | -0.1725 | 0.3975 | 0.53 | 0.567069 | 0.783994 | 0.260321 | 0.995762 | 0.997988 | 0.995259 |
| Frmd4a_Q3TB04_S491s_1 | Frmd4a_Q | -0.38 | -0.14 | -0.75 | -0.61 | -0.93 | -0.39 | -0.37 | -0.04 | -0.66 | -0.11 | -0.09 | -0.31 | 0.02 | -0.67 | -0.36 | -0.61 | 0.12 | 0.36 | 0.275 | 0.817997 | 0.889483 | 0.495342 | 0.996493 | 0.997988 | 0.999507 |
| Frmd4a_Q3TB04_S514s_1 | Frmd4a_Q | -0.75 | -0.82 | -0.55 | -0.51 | -0.29 | -1.06 | -0.48 | -0.44 | -0.41 | -1.32 | -1.27 | -0.85 | -0.42 | -0.63 | -0.4 | 0.27 | 0.0375 | 0.1775 | 0.065 | 0.221733 | 0.714909 | 0.288879 | 0.995437 | 0.997988 | 0.994666 |
| Frmd4a_Q3TB04_S65s_1 | Frmd4a_Q | 0.49 | -0.39 | 0.47 | 0.22 | 0.63 | -0.08 | 1.12 | 0.66 | -0.3 | 0.68 | 0.51 | 0.87 | -0.26 | 0.07 | -0.03 | 0.68 | 0.09 | -0.305 | 0.3625 | 0.80209 | 0.262694 | 0.489709 | 0.989223 | 0.997988 | 0.999507 |
| Frmd5_F6XM46_S270s_1 | Frmd5_F6X | -0.07 | -1.48 | -0.77 | 1 | 0.02 | -0.7 | -0.45 | -0.3 | 0.74 | -0.39 | 0.93 | -0.99 | 1.38 | -0.23 | 0.8 | -0.82 | 0.385 | 0.2425 | -0.0825 | 0.360217 | 0.956363 | 0.531855 | 0.999187 | 0.997988 | 0.994666 |
| Frmd8_Q3UFK8_S21s_1 | Frmd8_Q3I | 1.45 | 0.83 | 1.31 | 1.09 | 0.15 | 1.34 | 0.47 | 0.81 | 2.2 | 1.26 | 0.7 | 1.23 | 2.35 | 0.1 | 1.65 | 0.84 | -0.0275 | 0.4025 | 0.6125 | 0.892935 | 0.145978 | 0.618384 | 0.989223 | 0.997988 | 0.999507 |
| Frmd8_Q3UFK8_S24s_1 | Frmd8_Q3I | 0.61 | 0.77 | 0.88 | 0.09 | 0.63 | -0.15 | -0.42 | -0.24 | 0.66 | -0.04 | 0.25 | 0.46 | 1.32 | -0.99 | -0.01 | 0.47 | -0.4775 | 0.1775 | 0.065 | 0.432036 | 0.065485 | 0.375369 | 0.989223 | 0.997988 | 0.994666 |
| Frs2_Q8C180_S428s_1 | Frs2_Q8C1 | -0.27 | -1.37 | -0.24 | -0.28 | -1.1 | 0.13 | -0.09 | -0.32 | -0.67 | -1.19 | 0.94 | -0.3 | -0.55 | -0.53 | 0.11 | -0.4 | -0.6325 | -0.255 | -0.39 | 0.573531 | 0.607573 | 0.640288 | 0.989223 | 0.997988 | 0.995259 |
| Fry_E9Q8I9_S1936s_1 | Fry_E9Q8I | 0.43 | -0.28 | -0.37 | -1.56 | 0.22 | 0.56 | 0.82 | 0.33 | -2.8 | -0.75 | -1.78 | 0.12 | -2.2 | -1.77 | -0.52 | -0.03 | 0.195 | 0.235 | 0.1975 | 0.270213 | 0.046653 | 0.229221 | 0.989223 | 0.997988 | 0.994666 |
| Fry_E9Q8I9_S1940s_1 | Fry_E9Q8I | -0.38 | -0.32 | 0.13 | 0.28 | -0.99 | -0.55 | -0.27 | -0.48 | 0.56 | -0.09 | -1.24 | -0.01 | 1.03 | -1.14 | -0.32 | -0.12 | 0.9275 | -0.8575 | -0.685 | 0.887691 | 0.086399 | 0.763601 | 0.989223 | 0.997988 | 0.999507 |
| Fry_E9Q8I9_S1955s_1 | Fry_E9Q8I | -0.83 | -0.65 | 0.61 | 0.54 | -1.38 | -1.15 | 0.21 | 0.35 | -1.35 | -0.5 | 0.52 | 0.57 | -0.8 | -0.64 | 0.79 | 0.99 | -0.5 | -0.1225 | -0.065 | 0.764339 | 0.450485 | 0.845756 | 0.989223 | 0.997988 | 0.9971 |
| Fry_E9Q8I9_S2459s_1 | Fry_E9Q8I | -1.21 | -1.01 | -0.62 | -0.19 | -0.93 | -1.4 | -0.24 | -0.48 | -1.18 | -1.29 | 0.21 | -0.5 | -0.14 | -1.07 | -0.21 | -0.01 | -0.41 | -0.1075 | 0.1675 | 0.278149 | 0.988431 | 0.868789 | 0.999187 | 0.997988 | 0.994666 |
| Fry_E9Q8I9_S2515s_1 | Fry_E9Q8I | -3.58 | -2.67 | -4.93 | -7.66 | -3.04 | -5.22 | -4.2 | -6.8 | -5.6 | -2.77 | -1.37 | -5.15 | -1.68 | -2.1 | -7.34 | -3.45 | -0.005 | 0.0675 | 0.4 | 0.467784 | 0.927857 | 0.452683 | 0.997447 | 0.997988 | 0.994666 |
| Fry_E9Q8I9_S2815s_1 | Fry_E9Q8I | -0.71 | -0.07 | 0.37 | -0.13 | -0.56 | -0.27 | -0.96 | -0.32 | 0.55 | -0.04 | -0.55 | 0.03 | 1.13 | -0.36 | -0.37 | 0.1 | -0.105 | 0.9875 | 1.0675 | 0.536503 | 0.208198 | 0.695265 | 0.989223 | 0.997988 | 0.994666 |
| Fry_E9Q8I9_S3019s_1 | Fry_E9Q8I | 0.69 | 0.48 | 0.21 | -1.89 | 0.98 | 0.13 | 0.21 | -0.16 | -1.43 | 0.31 | -1.07 | -1.29 | -0.62 | -0.68 | -1.31 | 0.81 | -0.3925 | 0.1325 | 0.26 | 0.632273 | 0.476186 | 0.271864 | 0.989223 | 0.997988 | 0.995259 |
| Fry_E9Q8I9_T1965t_1 | Fry_E9Q8I | 0.16 | 1.22 | 0.71 | 0.28 | 0.38 | 0.83 | 0.47 | -1.31 | -0.22 | 0.32 | 2.46 | 1.29 | 0.49 | 0.15 | 1.16 | 1.75 | 0.4175 | -0.7425 | -0.3225 | 0.493916 | 0.322418 | 0.531261 | 0.989223 | 0.997988 | 0.994666 |
| Fryl_F8VQ05_T1960t_1 | Fryl_F8VQ | -0.92 | -0.67 | -1.27 | -2.06 | -0.03 | -1.28 | -1.37 | -1.47 | -1.3 | -1.07 | -0.95 | -1.65 | -1.3 | -0.15 | -1.97 | -1.38 | -0.5 | 0.37 | 0.295 | 0.948833 | 0.654982 | 0.971785 | 0.990434 | 0.997988 | 0.999507 |
| Fscn1_Q61553_T403t_1 | Fscn1_Q61 | -0.01 | 0.64 | 0.27 | 0.07 | -0.41 | 0.2 | -0.31 | -0.1 | 0.38 | -0.39 | 0.29 | 0.27 | 0.77 | 0.95 | 0.18 | 0.23 | 0.1925 | -0.0125 | 0.03 | 0.348292 | 0.140172 | 0.709019 | 0.989223 | 0.997988 | 0.994666 |
| Fsd1_Q7TPM6_S300s_1 | Fsd1_Q7TP | -3.12 | -3.91 | -4.32 | -4.29 | -3.1 | -3.04 | -4.93 | -3.67 | -3.15 | -2.73 | -1.59 | -3.02 | -1.64 | -2.99 | -4.63 | -3.98 | -0.3975 | -0.105 | 0.29 | 0.360329 | 0.640966 | 0.01514 | 0.989223 | 0.997988 | 0.994666 |
| Fsd1_Q7TPM6_S469s_1 | Fsd1_Q7TP | -0.42 | -0.22 | 0.7 | 0.21 | -1.3 | -1.06 | -0.44 | 0.78 | 0.91 | -0.85 | 1.17 | 0.62 | 1.25 | -0.27 | 0.13 | -0.65 | 0.225 | 1.2875 | 0.6 | 0.918237 | 0.256174 | 0.426494 | 0.989223 | 0.997988 | 0.999507 |
| Fsd1_Q7TPM6_S477s_1 | Fsd1_Q7TP | 1.1 | 0.82 | 0.97 | 1.24 | 0.32 | 0.47 | -0.19 | 0.48 | 2.47 | 0.86 | 1.2 | 0.33 | 2.82 | 0.97 | 1.08 | 0.74 | -0.5725 | 0.395 | 0.0475 | 0.436796 | 0.010607 | 0.684291 | 0.989223 | 0.997988 | 0.994666 |
| Fsd1_Q7TPM6_S477s_2 | Fsd1_Q7TP | 0.72 | 0.77 | 0.73 | -0.26 | -0.26 | 1.18 | -1.4 | -0.28 | 2.12 | 1.47 | -0.17 | -0.64 | 1.04 | 0.28 | -0.79 | -0.08 | -0.7625 | 0.1825 | 0.37 | 0.405515 | 0.220491 | 0.748695 | 0.989223 | 0.997988 | 0.994666 |
| Fsd1l_E0CXZ1_S314s_1 | Fsd1l_E0C | -2.9 | -1.63 | -2.05 | -4.02 | -2.06 | -2.37 | -3.58 | -2.69 | -0.01 | -1.83 | -4.71 | -2.28 | 1.64 | -2.87 | -0.92 | -2.86 | -0.68 | 0.205 | -0.3775 | 0.198402 | 0.964353 | 0.651297 | 0.999187 | 0.997988 | 0.994666 |
| Fsd1l_E0CXZ1_S319s_1 | Fsd1l_E0C | -0.52 | 0.16 | -1.05 | -1.09 | 0.52 | -0.88 | -0.47 | -0.81 | -1.05 | -0.95 | 0.16 | -0.57 | 0.45 | 1.04 | -0.8 | -0.91 | -0.025 | 0.4425 | 1.3975 | 0.292229 | 0.606497 | 0.95496 | 0.989223 | 0.997988 | 0.994666 |
| Fth1_P09528_S5s_1 | Fth1_P095 | -0.01 | 0.3 | -2.43 | 1.83 | -0.43 | 0.22 | -2.43 | -1.86 | 2.23 | 0.65 | -0.75 | -1 | 2.17 | 0.19 | 1.94 | -1.74 | 0.215 | 0.0225 | 0.57 | 0.517813 | 0.277358 | 0.724136 | 0.989223 | 0.997988 | 0.994666 |
| Ftsj3_Q9DBE9_S576s_1 | Ftsj3_Q9DI | 1.27 | 1.05 | 0.76 | -0.41 | 0.18 | 0.38 | -0.7 | -0.93 | 0.79 | -0.1 | -0.16 | -0.32 | 1.34 | 0.48 | 0.28 | -0.42 | -1.0475 | 0.36 | 0.7175 | 0.618528 | 0.066149 | 0.182088 | 0.989223 | 0.997988 | 0.995259 |
| Fubp1_Q3TUE1_S628s_1 | Fubp1_Q3T | -1.15 | -1.34 | 0.96 | 1.19 | -3.42 | -1.48 | 0.43 | 1.22 | -1.51 | -0.77 | -0.15 | 1 | -1.28 | -1.44 | 0.84 | 0.93 | -0.935 | -0.615 | -0.2475 | 0.853098 | 0.500639 | 0.7232 | 0.989223 | 0.997988 | 0.999507 |
| Fxr1_Q61584_S409s_1 | Fxr1_Q615 | -0.92 | -0.07 | 0.07 | -0.6 | -0.29 | -0.3 | 0.68 | -0.01 | 0.29 | 0.04 | 0.2 | -0.16 | -0.4 | 1.01 | -1.53 | 0.81 | -0.7275 | -0.2725 | -0.1525 | 0.547958 | 0.249692 | 0.133501 | 0.989223 | 0.997988 | 0.994666 |
| Fxr2_Q6P5B5_S477s_1 | Fxr2_Q6P5 | -0.09 | 0.55 | 0.5 | 0.88 | -0.81 | -0.03 | 0.32 | 0.85 | 1.53 | 0.3 | 1.88 | 0.07 | 2.21 | 0.14 | 1.37 | 1.44 | 0.4 | 0.4725 | 0.3525 | 0.095588 | 0.345302 | 0.314613 | 0.989223 | 0.997988 | 0.994666 |
| Fxr2_Q6P5B5_S603s_1 | Fxr2_Q6P5 | 0.57 | 0.31 | 0.88 | 1.13 | 0.33 | -0.18 | 1.2 | 0.92 | 0.42 | 1.08 | 0.75 | 0.56 | -0.09 | 1.04 | 1.22 | 1.51 | -0.3775 | 0.485 | 0.83 | 0.62351 | 0.664775 | 0.94308 | 0.991343 | 0.997988 | 0.995259 |
| Fxyd7_P59648_S60s_1 | Fxyd7_P59 | -0.58 | 0.29 | -2.38 | -4.68 | 0 | -0.12 | -1.85 | -3.07 | 4.12 | -0.28 | -3.31 | -1.38 | 4.06 | -0.47 | 1.89 | -2.31 | -0.155 | -0.02 | 0.1975 | 0.108882 | 0.614428 | 0.344234 | 0.989223 | 0.997988 | 0.994666 |
| Fxyd7_P59648_S72s_1 | Fxyd7_P59 | 0.37 | 1 | 0.84 | 0.63 | 0.62 | 0.45 | 1.14 | 0.5 | 0.06 | 0.64 | -0.3 | 0.92 | -0.13 | 0.76 | 0.04 | 0.62 | 0.5775 | 1.625 | 2.63 | 0.23084 | 0.900033 | 0.26906 | 0.996493 | 0.997988 | 0.994666 |
| Fxyd7_P59648_S73s_1 | Fxyd7_P59 | 0.3 | 1.05 | 0.89 | 0.84 | 1.64 | 0.55 | 0.52 | 0.59 | 0.06 | 0.21 | 0.19 | 0.9 | 0.2 | 1.13 | 1.11 | 0.86 | -0.0325 | -0.38 | -0.3875 | 0.863248 | 0.866891 | 0.166668 | 0.996493 | 0.997988 | 0.999507 |
| Fxyd7_P59648_T62t_1 | Fxyd7_P59 | -1.62 | -1.58 | -1.6 | -2.02 | -0.94 | -1.4 | -1.84 | -1.7 | 1.83 | -0.43 | -2.41 | -1.23 | 2.8 | -1.59 | 1.74 | -2.07 | 0.055 | -0.43 | 0.055 | 0.092874 | 0.391276 | 0.177972 | 0.989223 | 0.997988 | 0.994666 |
| Fyn_D3YZ57_S21s_1 | Fyn_D3YZ5 | 4.27 | 3.41 | 3.92 | 3.06 | 2.76 | 4.12 | 2.63 | 3.64 | 1.75 | 3.56 | 2.87 | 3.89 | 1.35 | 0.66 | 3.56 | 3.77 | 0.235 | 1.145 | 1.925 | 0.096155 | 0.382456 | 0.22479 | 0.989223 | 0.997988 | 0.994666 |
| Fyn_D3YZ57_Y185y_1 | Fyn_D3YZ5 | 0.98 | 1.05 | 1.68 | 1.29 | 1.15 | 1 | 0.96 | 1.12 | 1.01 | 1.09 | 1.16 | 1.2 | 0.64 | 1.1 | 1.38 | 1.12 | -0.3775 | -0.6475 | -1.33 | 0.51706 | 0.420936 | 0.583287 | 0.989223 | 0.997988 | 0.994666 |
| Fyttd1_Q91Z49_S23s_1 | Fyttd1_Q9 | -1.39 | -1.75 | -2.54 | -3.54 | -2.62 | -1.35 | -1.99 | -1.94 | -1.03 | -2.01 | -2.25 | -1 | 0.61 | -2.2 | -0.67 | -2.03 | -0.1925 | -0.135 | -0.19 | 0.113968 | 0.512206 | 0.195044 | 0.989223 | 0.997988 | 0.994666 |
| Fzd3_Q61086_S588s_1 | Fzd3_Q610 | -0.31 | -1.43 | -1.12 | -0.71 | -0.44 | -0.11 | -0.87 | -0.29 | -0.52 | -1.15 | -1.42 | -0.2 | -0.43 | -0.86 | -0.54 | -0.96 | 0.33 | 0.7325 | 1.2325 | 0.547716 | 0.159287 | 0.852661 | 0.989223 | 0.997988 | 0.994666 |
| Fzd3_Q61086_S665s_1 | Fzd3_Q610 | 0.6 | 0.44 | -0.43 | -0.03 | 0.03 | 0.98 | 1.12 | 0.32 | 0.55 | 0.64 | -0.05 | 0.64 | 0.62 | -0.45 | 0.85 | 0.58 | 0.465 | 0.07 | 0.195 | 0.512653 | 0.204954 | 0.356449 | 0.989223 | 0.997988 | 0.994666 |
| Fzd3_Q61086_T542t_1 | Fzd3_Q610 | 1.34 | -0.34 | 0.56 | 1.44 | 0.83 | 1.39 | 0.7 | 0.79 | -0.93 | -0.22 | 0.31 | 0.89 | -2.15 | -0.95 | -0.01 | 0.45 | 0.4675 | 0.3 | 0.255 | 0.049327 | 0.673533 | 0.185778 | 0.992876 | 0.997988 | 0.994666 |
| G3bp1_P97855_T268t_1 | G3bp1_P97 | -0.62 | -0.39 | -0.74 | -1.18 | 0.2 | -0.73 | 0.08 | -1 | -0.77 | -0.53 | -0.97 | -0.29 | -1.09 | 0.39 | -0.7 | -0.54 | 0.1775 | -0.7375 | -1.415 | 0.512129 | 0.29742 | 0.738206 | 0.989223 | 0.997988 | 0.994666 |
| G3bp2_P97379_S225s_1 | G3bp2_P97 | -1 | -0.45 | 0.36 | 0.2 | -0.2 | -0.78 | 0.41 | 0.39 | -1.37 | -0.63 | 0.21 | 0.41 | -0.63 | -0.4 | 0.13 | 0.51 | 0.37 | 0.0925 | 0.2475 | 0.759871 | 0.662159 | 0.801681 | 0.990434 | 0.997988 | 0.9971 |
| G3bp2_P97379_T227t_1 | G3bp2_P97 | -0.5 | -0.47 | -0.83 | -0.7 | -0.4 | -0.57 | -0.23 | -0.76 | -0.73 | -0.57 | -0.77 | -0.8 | -0.48 | -0.07 | -0.51 | -0.5 | 0.1775 | -0.1225 | 0.125 | 0.359493 | 0.550875 | 0.671549 | 0.989223 | 0.997988 | 0.994666 |
| Gab2_Q3ZB57_S211s_1 | Gab2_Q3ZI | 0.1 | -1.42 | -0.24 | -0.66 | -0.56 | 0.18 | -0.77 | -1.02 | -0.57 | -0.72 | -0.03 | -1.09 | 0.61 | -1.3 | -0.54 | -0.21 | 0.135 | -0.0925 | 0.235 | 0.690882 | 0.975107 | 0.903985 | 0.999187 | 0.997988 | 0.996159 |
| Gab2_Q3ZB57_S402s_1 | Gab2_Q3ZI | 0.19 | -0.76 | -1.21 | -1.12 | -0.06 | -0.22 | -0.67 | -0.91 | -0.13 | 0.19 | 0.27 | -1.78 | -0.01 | 0.02 | -0.81 | -1.2 | 0.0125 | -0.0475 | 0.195 | 0.606366 | 0.488219 | 0.504995 | 0.989223 | 0.997988 | 0.995259 |
| Gab2_Q3ZB57_T388t_1 | Gab2_Q3ZI | -2.41 | -2.32 | -1.79 | -1.21 | -0.73 | -3.4 | -0.72 | -1.54 | -3.02 | -2.7 | -1.43 | -1.84 | -1.3 | -1.56 | -0.98 | -1.15 | 0.26 | 0.3625 | 0.225 | 0.067567 | 0.588488 | 0.48544 | 0.989223 | 0.997988 | 0.994666 |
| Gabarapl2_P60521_S10s_1 | Gabarapl2_ | 0.11 | 0.44 | 0.22 | -0.97 | -0.6 | 0.77 | 0.32 | 0.05 | 1.54 | 0.37 | -1.99 | 0.99 | 1.71 | -0.33 | 1.35 | 0.7 | 0.335 | -0.315 | 0.685 | 0.103015 | 0.651065 | 0.712758 | 0.990358 | 0.997988 | 0.994666 |
| Gabbr1_Q9WV18_S877s_1 | Gabbr1_Q9 | -0.24 | -0.13 | 1.23 | -0.65 | -1.91 | 1.38 | -0.4 | 0.17 | 2.53 | 0.07 | -0.13 | 1.32 | 3.34 | -1.24 | 2.64 | -0.91 | 0.185 | 0.2775 | 0.9075 | 0.413512 | 0.730147 | 0.20365 | 0.995762 | 0.997988 | 0.994666 |
| Gabbr1_Q9WV18_S909s_1 | Gabbr1_Q9 | 0.24 | 0.72 | 0.34 | -0.61 | 0.54 | -0.27 | 0.35 | -0.29 | -0.05 | 0.08 | 0.28 | 0.57 | 0.87 | 0.79 | -0.34 | -0.12 | -0.2425 | 0.895 | 0.905 | 0.76041 | 0.79898 | 0.886828 | 0.995762 | 0.997988 | 0.9971 |
| Gabbr1_Q9WV18_S949s_1 | Gabbr1_Q9 | 0.44 | -1.07 | -0.39 | 0.19 | 0.3 | 1.07 | 0.08 | 0.71 | -0.14 | -0.5 | 0.8 | 0.13 | -1.5 | 0.19 | -0.95 | -0.48 | -0.09 | 0.0475 | 0.1275 | 0.324905 | 0.079947 | 0.514128 | 0.989223 | 0.997988 | 0.994666 |
| Gabbr1_Q9WV18_T872t_1 | Gabbr1_Q9 | 0.75 | 0.78 | -0.09 | 0.07 | 1.15 | 0.83 | 0.74 | 0.79 | 0.14 | 0.3 | 0.95 | 0.6 | -0.04 | 0.89 | 0.97 | 0.76 | 0.7475 | 0.28 | -0.4775 | 0.455258 | 0.099784 | 0.706918 | 0.989223 | 0.997988 | 0.994666 |
| Gabbr1_Q9WV18_T929t_1 | Gabbr1_Q9 | 3.06 | 1.72 | 2.45 | 1.47 | 2.2 | 3.01 | 3.21 | 2.26 | 0.37 | 2.82 | 0.5 | 2.51 | 0.6 | 0.28 | 2.46 | 3 | 0.5 | 0.12 | 0.2675 | 0.398653 | 0.257135 | 0.363399 | 0.989223 | 0.997988 | 0.994666 |
| Gabbr2_Q80T41_S770s_1 | Gabbr2_Q8 | 4.08 | 1 | 2.18 | 3.04 | 1.13 | 3.52 | 1.82 | 3.28 | 1.93 | 1.93 | 2.59 | 2.22 | 1.37 | 1.29 | 1.54 | 1.75 | 0.495 | -0.625 | -0.59 | 0.09683 | 0.856914 | 0.511797 | 0.996349 | 0.997988 | 0.994666 |
| Gabbr2_Q80T41_S775s_1 | Gabbr2_Q8 | 0.18 | 0.62 | 1.23 | 1.38 | -0.3 | 0.65 | 0.74 | 1.37 | 1.24 | 1.07 | 1.16 | 1.38 | 1.4 | 0.58 | 0.83 | 0.73 | -0.1375 | -0.4075 | -1.0875 | 0.927404 | 0.575078 | 0.272492 | 0.989223 | 0.997988 | 0.999507 |
| Gabbr2_Q80T41_S778s_2 | Gabbr2_Q8 | 0.6 | -0.8 | 1.53 | 0.89 | 0.92 | 0.53 | 0.58 | 0.6 | 0.5 | -0.01 | 3.13 | 1.56 | 1.05 | 1.2 | 1.21 | 0.11 | -0.2375 | 0.36 | 0.0325 | 0.522764 | 0.825219 | 0.344149 | 0.995762 | 0.997988 | 0.994666 |
| Gabbr2_Q80T41_S783s_1 | Gabbr2_Q8 | 1.3 | 1.12 | 0.08 | 0.58 | 1.25 | 1.41 | 0.29 | 0.42 | 2.02 | 1.15 | -0.46 | 0.65 | 1.7 | 1.06 | -0.44 | -0.48 | 0.1025 | 0.74 | 0.3375 | 0.586146 | 0.850846 | 0.897653 | 0.996349 | 0.997988 | 0.995259 |
| Gabbr2_Q80T41_S792s_1 | Gabbr2_Q8 | -0.57 | 0.25 | 0.76 | -0.58 | 0.38 | 0.54 | -0.15 | 0.46 | 0.38 | 0.04 | -0.85 | 0.48 | 0.69 | 0.64 | -0.34 | 0.35 | 0.0725 | 0.07 | -0.31 | 0.37559 | 0.355226 | 0.91275 | 0.989223 | 0.997988 | 0.994666 |
| Gabbr2_Q80T41_S883s_1 | Gabbr2_Q8 | 1.31 | 1.33 | 1.3 | 1.24 | 1.3 | 1.28 | 1.57 | 1.28 | 0.5 | 1.41 | 0.54 | 1.25 | 0 | 0.2 | 1.24 | 1.31 | 0.3425 | 0.0475 | 0.37 | 0.120989 | 0.754236 | 0.215587 | 0.995762 | 0.997988 | 0.994666 |
| Gabbr2_Q80T41_T762t_1 | Gabbr2_Q8 | -0.41 | -2.9 | -1.25 | -0.32 | -1.09 | -0.59 | -0.54 | -0.68 | -2.11 | -2.62 | -0.45 | -1.32 | -2.48 | -1.14 | -1.79 | -1.46 | 0.0625 | -0.37 | -0.6075 | 0.419122 | 0.38007 | 0.560422 | 0.989223 | 0.997988 | 0.994666 |
| Gabra2_P26048_T393t_1 | Gabra2_P2 | -3.4 | -2.77 | -2.09 | -3.51 | -0.87 | 0.75 | -0.69 | -1.3 | -0.6 | -2.59 | -1.9 | 3.43 | 0.21 | -2.01 | -0.54 | -0.31 | 0.495 | -0.405 | -0.4975 | 0.001891 | 0.000872 | 0.062333 | 0.989223 | 0.997988 | 0.994666 |
| Gabra3_B1AVY2_S474s_1 | Gabra3_B1 | 0.69 | 0.59 | 0.62 | 0.83 | 0.49 | -0.14 | 0.37 | 0.57 | 0.14 | 0.3 | 0.96 | 0.56 | 0.25 | 0.76 | 0.32 | 0.17 | 2.415 | 2.5275 | 2.28 | 0.242108 | 0.152368 | 0.459503 | 0.989223 | 0.997988 | 0.994666 |
| Gabra3_B1AVY2_T468t_1 | Gabra3_B1 | -0.26 | 0.47 | 0.41 | 1.17 | -0.07 | -1.09 | 0.45 | -0.01 | -0.82 | -0.04 | -0.36 | 0.28 | -0.33 | -0.5 | 0.34 | 1.31 | -0.36 | -0.1925 | -0.3075 | 0.616529 | 0.155266 | 0.098414 | 0.989223 | 0.997988 | 0.995259 |
| Gabra4_Q3USX9_T449t_1 | Gabra4_Q3 | 0.75 | 1.3 | 0.15 | 1.02 | 0.55 | 1.15 | 0.46 | -0.18 | 0.93 | 0.03 | 1.65 | 1.05 | -0.44 | -1.43 | 0.76 | 0.85 | -0.6275 | -0.6825 | -0.2425 | 0.139017 | 0.403078 | 0.788421 | 0.989223 | 0.997988 | 0.994666 |
| Gabrd_P22933_S390s_1 | Gabrd_P22 | 0.93 | -0.14 | -0.02 | 0.05 | 0.89 | 0.81 | 0.4 | 0.08 | 0.01 | 0.59 | -0.59 | 0.2 | -0.13 | 0.95 | -0.42 | -0.48 | -0.31 | 0.11 | -0.87 | 0.589331 | 0.306513 | 0.672326 | 0.989223 | 0.997988 | 0.995259 |
| Gabrg2_P22723_S393s_1 | Gabrg2_P2 | -0.41 | -0.61 | -0.15 | 0.11 | -0.29 | -0.38 | 0.31 | -0.03 | -0.84 | -0.21 | -0.32 | 0.14 | -0.48 | 0.08 | 0.1 | 0.2 | 0.34 | -0.1525 | -0.225 | 0.415646 | 0.532334 | 0.886255 | 0.989223 | 0.997988 | 0.994666 |
| Gabrg2_P22723_T395t_1 | Gabrg2_P2 | 1.27 | 0.9 | -0.43 | 0.45 | 0.61 | 1.07 | 1.23 | 0.56 | 1.04 | 0.41 | 0.38 | 0.75 | 1.05 | 0.89 | 0.42 | 0.35 | 0.1675 | -0.0425 | 0.24 | 0.750581 | 0.420454 | 0.806092 | 0.989223 | 0.997988 | 0.9971 |
| Gad1_P48318_S527s_1 | Gad1_P48 | 0.68 | 0.8 | -0.31 | -0.39 | 0.05 | 0.43 | -0.85 | -0.8 | 1.24 | 0.76 | 0.48 | -0.4 | 1.48 | 0.86 | 0.1 | 0.45 | 0.32 | 0.0975 | 0.13 | 0.23736 | 0.267417 | 0.476527 | 0.989223 | 0.997988 | 0.994666 |
| Gad1_P48318_S55s_1 | Gad1_P48 | 1.43 | 0.69 | -0.28 | -0.03 | 1.31 | 1.53 | -0.52 | -0.38 | 1.71 | 0.82 | -2.97 | -0.69 | 1.03 | 0.42 | 0.39 | 0.61 | -0.4875 | 0.325 | 0.5275 | 0.699785 | 0.956429 | 0.454459 | 0.999187 | 0.997988 | 0.996159 |
| Gad1_P48318_S77s_1 | Gad1_P48 | 1.35 | 0.01 | -0.27 | 0.75 | 0.38 | 1.71 | 0.59 | 0.76 | -0.04 | -0.27 | 0.7 | 0.22 | -0.98 | 1.13 | 0.17 | 0.61 | 0.0325 | -0.735 | 0.16 | 0.674402 | 0.374984 | 0.46504 | 0.989223 | 0.997988 | 0.996159 |
| Gak_F6SLK1_S112s_1 | Gak_F6SLK | -2.97 | -1.48 | -2.6 | -4.61 | -1.54 | -2.6 | -4.23 | -3.07 | -2.69 | -1.93 | -0.05 | -2.31 | -2.4 | -1.51 | -2.07 | -3.9 | 0.4 | -0.3075 | -0.2275 | 0.548338 | 0.941469 | 0.158852 | 0.999187 | 0.997988 | 0.994666 |
| Gap43_P06837_S122s_1 | Gap43_P06 | 0.45 | 0.04 | 0.63 | 0.61 | -1.06 | 1.2 | -0.24 | 0.44 | 2.25 | 0.35 | 0.7 | 0.77 | 2.09 | -1.33 | 0.78 | -0.08 | 0.055 | 1.17 | 0.445 | 0.91848 | 0.46163 | 0.19684 | 0.989223 | 0.997988 | 0.999507 |
| Gap43_P06837_S128s_1 | Gap43_P06 | 1.65 | 0.69 | 0.42 | 0 | -1.26 | 1.85 | 0.03 | -0.06 | 3.51 | 1.06 | 0.84 | 0.94 | 3.26 | -0.19 | 0.9 | -0.42 | -0.3475 | 0.585 | -0.0675 | 0.80625 | 0.4055 | 0.198212 | 0.989223 | 0.997988 | 0.999507 |
| Gap43_P06837_S133s_1 | Gap43_P06 | -0.29 | -0.65 | 0.09 | -0.36 | -1.4 | -0.14 | -0.65 | -0.77 | 1.86 | -0.17 | -0.47 | -0.18 | 2.79 | -1.46 | 1.56 | -0.57 | -0.55 | 0.8975 | 0.1975 | 0.320628 | 0.189187 | 0.297255 | 0.989223 | 0.997988 | 0.994666 |

| | | | | | | | | | | | | | | | | | | | | | | | | | | |
|---|---|---|---|---|---|---|---|---|---|---|---|---|---|---|---|---|---|---|---|---|---|---|---|---|---|---|
| Gap43_P06837_S142s_1 | Gap43_P06 | 0.75 | -0.48 | -0.13 | 1.71 | -1.01 | -0.03 | -0.36 | 1.23 | 2.47 | 0.94 | 1.44 | 0.55 | 2.84 | -0.74 | 2.31 | 0.11 | -0.4375 | 0.5625 | 0.8825 | 0.44916 | 0.41393 | 0.157052 | 0.989223 | 0.997988 | 0.994666 |
| Gap43_P06837_S192s_1 | Gap43_P06 | -1.85 | -0.5 | -1.32 | -1.52 | -0.09 | -1.84 | -0.87 | -1.49 | -1.22 | -0.85 | -0.17 | -0.85 | -0.69 | -0.45 | -0.2 | -1.26 | -0.505 | 0.8875 | 0.6675 | 0.114349 | 0.618143 | 0.17873 | 0.989223 | 0.997988 | 0.994666 |
| Gap43_P06837_S193s_1 | Gap43_P06 | -1.54 | -0.39 | -1.27 | -1.42 | 0.09 | -1.75 | -0.86 | -1.45 | -1.11 | -0.81 | -0.08 | -0.76 | -0.41 | -0.22 | -0.18 | -1.25 | 0.225 | 0.525 | 0.6475 | 0.115397 | 0.719019 | 0.210098 | 0.995437 | 0.997988 | 0.994666 |
| Gap43_P06837_S86s_1 | Gap43_P06 | 0.6 | 1.04 | 0.76 | 0.57 | 0.17 | 0.91 | 0.56 | 0.91 | 0.97 | 1.15 | 1.1 | 0.61 | 1.23 | 0.78 | 1.5 | 0.84 | 0.1625 | 0.465 | 0.64 | 0.236306 | 0.684631 | 0.388005 | 0.994624 | 0.997988 | 0.994666 |
| Gap43_P06837_S96s_1 | Gap43_P06 | 0.85 | 1.02 | 1.06 | 0.59 | 0.74 | 0.88 | 0.22 | 0.57 | 0.94 | 1.16 | 1.44 | 1.01 | 1.41 | 0.67 | 1.01 | 0.73 | -0.105 | 0.215 | 0.345 | 0.788314 | 0.268082 | 0.297874 | 0.989223 | 0.997988 | 0.999507 |
| Gap43_P06837_S96s_2 | Gap43_P06 | 0.45 | 1.06 | 1.25 | 0.45 | -0.08 | 0.91 | -0.35 | 0.29 | 0.82 | 0.35 | 1.23 | 0.93 | 0.47 | 0.28 | 0.92 | 0.78 | -0.2775 | 0.2575 | 0.075 | 0.541821 | 0.103655 | 0.922904 | 0.989223 | 0.997988 | 0.994666 |
| Gap43_P06837_T171t_1 | Gap43_P06 | 0.8 | 0.82 | 0.41 | 0.7 | 1.15 | 1.09 | 1.4 | 0.9 | -0.96 | 1.31 | -0.1 | 1.24 | -0.05 | -1.42 | 1.07 | 1.4 | -0.61 | 0.03 | -0.19 | 0.469759 | 0.066871 | 0.556064 | 0.989223 | 0.997988 | 0.994666 |
| Gap43_P06837_T172t_1 | Gap43_P06 | 0.7 | 0.76 | 0.3 | 0.21 | 0.97 | 0.71 | 1.22 | 0.63 | -0.58 | 1.12 | -1.51 | 0.72 | -0.03 | -0.56 | 0.79 | 0.81 | 0.4525 | -0.31 | -0.4325 | 0.529801 | 0.142919 | 0.344061 | 0.989223 | 0.997988 | 0.994666 |
| Gap43_P06837_T95t_1 | Gap43_P06 | 0.82 | 1.04 | 1.36 | 1.66 | 0.7 | 0.77 | 1.3 | 1.66 | 0.74 | 1.3 | 2.11 | 1.43 | 1.15 | 0.57 | 1.74 | 1.5 | 0.39 | -0.555 | -0.24 | 0.953836 | 0.718711 | 0.620586 | 0.995437 | 0.997988 | 0.999507 |
| Gap43_P06837_T95t_2 | Gap43_P06 | 1.66 | 0.18 | -0.25 | -0.07 | 0.29 | 0.34 | 0.5 | 0.12 | 0.9 | 1.29 | -0.09 | 0.34 | 0.32 | 0.89 | -0.37 | -0.41 | -0.1125 | 0.175 | 0.02 | 0.591937 | 0.872332 | 0.647378 | 0.996493 | 0.997988 | 0.995259 |
| Gapdh_P16858_S208s_1 | Gapdh_P16 | -0.57 | -0.02 | -1.62 | -1.52 | 0.29 | -0.74 | -1.42 | -1.05 | -0.18 | 0.1 | -2.17 | -1.41 | -0.05 | 0.04 | -1.69 | -2.04 | -0.0675 | 0.23 | -0.2725 | 0.996705 | 0.680327 | 0.976757 | 0.993464 | 0.997988 | 1 |
| Gapdh_P16858_T209t_1 | Gapdh_P16 | -0.35 | -0.73 | -0.72 | -0.16 | -0.68 | -1.07 | -0.28 | -0.4 | -1.31 | -0.46 | -0.31 | -0.21 | -0.51 | -0.18 | -0.35 | -0.04 | 0.2025 | 0.0175 | -0.0025 | 0.41766 | 0.663066 | 0.795565 | 0.990434 | 0.997988 | 0.994666 |
| Gapvd1_Q6PAR5_S1041s_1 | Gapvd1_Q6 | -1.19 | -0.8 | -0.79 | -0.34 | -1.18 | -0.91 | -1.3 | -0.4 | 0.46 | -0.33 | -1.04 | -0.59 | 0.97 | 0.03 | -0.08 | -0.47 | -0.1175 | -0.0825 | 0.22 | 0.035356 | 0.571035 | 0.287427 | 0.989223 | 0.997988 | 0.994666 |
| Gapvd1_Q6PAR5_S1041s_2 | Gapvd1_Q6 | -2.49 | -1.3 | -1.46 | -2.55 | -1.36 | -1.54 | -1.81 | -0.66 | -1.21 | -1.36 | -1.99 | -0.97 | -0.87 | -0.57 | -2.36 | -1.16 | -0.1675 | 0.405 | 0.8925 | 0.169133 | 0.150318 | 0.17486 | 0.989223 | 0.997988 | 0.994666 |
| Gapvd1_Q6PAR5_S1045s_1 | Gapvd1_Q6 | -0.52 | -0.22 | -0.18 | 0.42 | -0.66 | -0.85 | 0.21 | 0.17 | -0.6 | 0.43 | -0.01 | 0.53 | -0.35 | 0.52 | 0.45 | 0.13 | 0.6075 | 0.5675 | 0.71 | 0.347212 | 0.648238 | 0.539509 | 0.99033 | 0.997988 | 0.994666 |
| Gapvd1_Q6PAR5_S1045s_2 | Gapvd1_Q6 | -2.49 | -1.3 | -1.46 | -2.55 | -1.36 | -1.54 | -1.81 | -0.66 | -1.21 | -1.36 | -1.99 | -0.97 | -0.87 | -0.57 | -2.36 | -1.16 | -0.1575 | 0.2125 | 0.3125 | 0.169133 | 0.150318 | 0.17486 | 0.989223 | 0.997988 | 0.994666 |
| Gapvd1_Q6PAR5_S881s_1 | Gapvd1_Q6 | -0.9 | -0.89 | -1.06 | -4.54 | 0.11 | -0.47 | -0.8 | -0.6 | -1.2 | -0.59 | -1.5 | -1.04 | -0.12 | -0.79 | -1.4 | -1.07 | 0.6075 | 0.5675 | 0.71 | 0.243928 | 0.10527 | 0.361938 | 0.989223 | 0.997988 | 0.994666 |
| Gapvd1_Q6PAR5_S893s_1 | Gapvd1_Q6 | 0.06 | 0.89 | 0.04 | 0.19 | 0.16 | -0.45 | -0.2 | -0.1 | 0.07 | -0.24 | 0.08 | -0.8 | -0.65 | 0.45 | 0.63 | 0.22 | 1.4075 | 0.765 | 1.0025 | 0.719749 | 0.133474 | 0.129505 | 0.989223 | 0.997988 | 0.9971 |
| Gapvd1_Q6PAR5_T741t_1 | Gapvd1_Q6 | -0.17 | -0.21 | -1.19 | 0.09 | -0.14 | -0.12 | -1.49 | -2.11 | 0.21 | 0.25 | -1.68 | -0.75 | -0.32 | -0.09 | -0.98 | -1.32 | -0.4425 | -0.5175 | -0.1325 | 0.454835 | 0.270392 | 0.808594 | 0.989223 | 0.997988 | 0.994666 |
| Garem_Q3UFT3_S493s_1 | Garem_Q3 | -0.15 | -0.18 | 0.35 | -0.1 | 0.24 | 0.52 | 0.22 | -0.08 | 0.14 | 0.28 | -0.19 | -0.47 | -0.06 | 0.14 | -0.19 | -0.27 | -0.595 | -0.1225 | -0.3075 | 0.770888 | 0.321144 | 0.881678 | 0.989223 | 0.997988 | 0.9971 |
| Garem_Q3UFT3_S698s_1 | Garem_Q3 | -0.26 | -0.33 | -0.14 | -0.12 | 1.4 | 0.11 | 0.24 | 0 | -0.03 | 0.85 | -0.26 | -0.34 | -0.13 | 0.38 | -0.31 | 0.27 | 0.245 | -0.04 | -0.075 | 0.330111 | 0.078239 | 0.399655 | 0.989223 | 0.997988 | 0.994666 |
| Garem_Q3UFT3_S721s_1 | Garem_Q3 | 0.22 | 0.87 | 0.35 | 0.07 | -0.2 | 0.76 | 0.7 | -0.09 | 0.38 | 0.55 | 0.38 | -0.11 | -0.06 | 0.17 | -0.05 | 0.29 | 0.65 | 0.2675 | 0.265 | 0.307889 | 0.791817 | 0.780556 | 0.995762 | 0.997988 | 0.994666 |
| Garem_Q3UFT3_T738t_1 | Garem_Q3 | 0.68 | 0.89 | -0.07 | 0.33 | 0.79 | 0.58 | 0.89 | 0.66 | 0.69 | 1.14 | 0.61 | 0.66 | 0.14 | 0.74 | 0.62 | 0.48 | -0.085 | -0.0775 | -0.29 | 0.90206 | 0.319627 | 0.288613 | 0.989223 | 0.997988 | 0.999507 |
| Gareml_Q6PAJ3_S509s_1 | Gareml_Q6 | 0.01 | 0.36 | -0.21 | 0.11 | 0.5 | 0.31 | -0.19 | 0.07 | 1.05 | 1.03 | -0.11 | -0.04 | 1.02 | 0.57 | -0.31 | 0.24 | 0.2725 | 0.3175 | 0.0375 | 0.368504 | 0.67401 | 0.262048 | 0.992876 | 0.997988 | 0.994666 |
| Gareml_Q6PAJ3_T507t_1 | Gareml_Q6 | -0.51 | 0.15 | 0.23 | 1.02 | 0.46 | -0.15 | 1.49 | 0.81 | -0.35 | 0.24 | 1.76 | 1.07 | 0.5 | 0.29 | 1.16 | 0.14 | 0.105 | 0.415 | 0.3125 | 0.454027 | 0.33747 | 0.391411 | 0.989223 | 0.997988 | 0.994666 |
| Garnl3_A2AUQ6_S26s_1 | Garnl3_A2A | 0.64 | 0.81 | 0 | 0.19 | 0.43 | 0.59 | 0.44 | 0.05 | 0.3 | 0.86 | -0.35 | -0.05 | 0.1 | 0.98 | -0.17 | 0.46 | 0.43 | 0.4575 | 0.3 | 0.844903 | 0.902784 | 0.523408 | 0.996493 | 0.997988 | 0.999507 |
| Garnl3_B1AW31_S26s_1 | Garnl3_B1A | -3.98 | -2.7 | -3.77 | -4.63 | -2.3 | -3.39 | -2.31 | -4.81 | -3.23 | -4.52 | -1.47 | -3.09 | -2.75 | -2.02 | -3.09 | -3.78 | -0.0325 | -0.22 | -0.0675 | 0.117637 | 0.383135 | 0.317361 | 0.989223 | 0.997988 | 0.994666 |
| Garnl3_Q3V0G7_S6s_1 | Garnl3_Q3 | 0.16 | -0.3 | -0.18 | -0.7 | -0.13 | -0.2 | -0.09 | 0.1 | 0.21 | 0.61 | 0.83 | -0.26 | 0.27 | 0.39 | 0.05 | 0.81 | 0.5675 | 0.6925 | 0.86 | 0.057629 | 0.485468 | 0.089835 | 0.989223 | 0.997988 | 0.994666 |
| Garnl3_Q3V0G7_S844s_3 | Garnl3_Q3 | -1.91 | -1.16 | -1.27 | -1.23 | -1.1 | -0.68 | -1.08 | -1.15 | -0.87 | -0.66 | -0.23 | -0.6 | -1.87 | -0.94 | -1.39 | -1.07 | 0.175 | 0.6025 | 0.635 | 0.814062 | 0.152585 | 0.015381 | 0.989223 | 0.997988 | 0.999507 |
| Garnl3_Q3V0G7_S848s_1 | Garnl3_Q3 | -0.44 | 0.28 | -0.25 | -0.6 | -0.02 | -0.15 | -0.28 | -0.06 | 0.8 | 0.81 | 0.26 | -0.35 | 1.57 | -0.11 | 0.53 | -0.51 | 0.39 | 0.8025 | 0.075 | 0.208591 | 0.62437 | 0.097459 | 0.989223 | 0.997988 | 0.994666 |
| Garnl3_Q3V0G7_S848s_2 | Garnl3_Q3 | 0.25 | 0.03 | 0.32 | 0.23 | 0.02 | 0.18 | -0.37 | -0.23 | 1.1 | 0.76 | 0.02 | 0.09 | 0.55 | -0.07 | 0.63 | 0.12 | 0.125 | 0.6325 | 0.6225 | 0.710957 | 0.188791 | 0.363208 | 0.989223 | 0.997988 | 0.996966 |
| Garnl3_Q3V0G7_S848s_3 | Garnl3_Q3 | -1.91 | -1.16 | -1.27 | -1.23 | -1.1 | -0.68 | -1.08 | -1.15 | -0.87 | -0.66 | -0.23 | -0.6 | -1.87 | -0.94 | -1.39 | -1.07 | -0.3075 | 0.285 | 0.1 | 0.814062 | 0.152585 | 0.015381 | 0.989223 | 0.997988 | 0.999507 |
| Garnl3_Q3V0G7_T851t_1 | Garnl3_Q3 | -0.17 | -0.43 | -0.43 | -0.27 | -0.46 | -0.45 | -0.84 | -0.63 | -0.05 | -0.09 | -0.03 | -0.44 | 1.18 | -0.31 | -0.14 | -0.94 | 0.39 | 0.8025 | 0.075 | 0.540527 | 0.221704 | 0.447024 | 0.989223 | 0.997988 | 0.994666 |
| Garnl3_Q3V0G7_T851t_2 | Garnl3_Q3 | 0.25 | 0.03 | 0.32 | 0.23 | 0.02 | 0.18 | -0.37 | -0.23 | 1.1 | 0.76 | 0.02 | 0.09 | 0.55 | -0.07 | 0.63 | 0.12 | -0.27 | 0.1725 | 0.2725 | 0.710957 | 0.188791 | 0.363208 | 0.989223 | 0.997988 | 0.996966 |
| Garnl3_Q3V0G7_T851t_3 | Garnl3_Q3 | -1.91 | -1.16 | -1.27 | -1.23 | -1.1 | -0.68 | -1.08 | -1.15 | -0.87 | -0.66 | -0.23 | -0.6 | -1.87 | -0.94 | -1.39 | -1.07 | -0.3075 | 0.285 | 0.1 | 0.814062 | 0.152585 | 0.015381 | 0.989223 | 0.997988 | 0.999507 |
| Garnl3_Q3V0G7_T901t_1 | Garnl3_Q3 | -0.26 | -0.13 | -0.24 | 0.27 | 0.46 | -0.73 | 0.75 | 0.84 | 0.35 | 0.26 | 1.21 | 0.06 | -0.74 | -0.44 | 0.74 | 0.83 | 0.39 | 0.8025 | 0.075 | 0.656304 | 0.279614 | 0.101006 | 0.989223 | 0.997988 | 0.995259 |
| Gars_Q9CZD3_S643s_1 | Gars_Q9CZ | -1.21 | -1.32 | -0.13 | -1.19 | -1.57 | -1.55 | -1.96 | -2.03 | -0.7 | -1.65 | -0.18 | -0.78 | 0.28 | -2.28 | -1.38 | -0.47 | 0.42 | 0.56 | 0.1875 | 1 | 0.028364 | 0.74138 | 0.989223 | 0.997988 | 1 |
| Gas2_P11862_S283s_1 | Gas2_P118 | 1.68 | 0.54 | -0.21 | -3.18 | -0.47 | 1.62 | -1.78 | -2.1 | 2.94 | 1.32 | -1.34 | -1.1 | 2.47 | -0.21 | 1.5 | -1.46 | -0.815 | 0.135 | 1.57E-16 | 0.466544 | 0.73554 | 0.561836 | 0.995762 | 0.997988 | 0.994666 |
| Gas2l1_Q8JZP9_S292s_1 | Gas2l1_Q8 | -0.86 | -0.58 | -0.36 | -0.69 | 0.05 | -1.15 | -0.59 | -0.51 | -0.93 | -0.64 | -0.17 | -0.66 | 0.01 | -0.12 | -0.09 | -0.31 | -0.39 | 0.7475 | 0.8675 | 0.068284 | 0.805685 | 0.930352 | 0.995762 | 0.997988 | 0.994666 |
| Gas2l1_Q8JZP9_S297s_1 | Gas2l1_Q8 | -2.84 | -3.67 | -2.48 | -4.07 | -2.84 | -2.57 | -2.8 | -4 | -1.26 | -2.96 | -3.35 | -2.91 | 0.38 | -2.45 | -2.75 | -2.58 | 0.0725 | 0.0225 | 0.495 | 0.080776 | 0.641786 | 0.257598 | 0.989223 | 0.997988 | 0.994666 |
| Gas2l1_Q8JZP9_S306s_1 | Gas2l1_Q8 | 0.07 | 0.03 | 0.11 | 0.48 | 0.31 | 0.06 | 0.34 | 0.22 | 0.32 | 0.25 | 0.38 | 0.42 | 0.87 | 0.81 | 0.6 | 0.24 | 0.2125 | 0.645 | 1.415 | 0.112395 | 0.780783 | 0.449201 | 0.995762 | 0.997988 | 0.994666 |
| Gas2l1_Q8JZP9_S352s_1 | Gas2l1_Q8 | -0.23 | -0.07 | -0.43 | -0.19 | 0.08 | 0.03 | -0.2 | 0.23 | -0.22 | -0.07 | -0.16 | -0.36 | 0.65 | 0.02 | 0.27 | 0.08 | 0.06 | 0.17 | 0.4575 | 0.087673 | 0.233325 | 0.898717 | 0.989223 | 0.997988 | 0.994666 |
| Gas2l1_Q8JZP9_S352s_2 | Gas2l1_Q8 | -0.04 | 0.02 | -1.53 | -0.87 | 0.14 | -0.01 | -0.48 | -1.39 | 0.84 | 0.37 | 0.61 | -1.56 | 1.28 | 1.02 | -0.43 | -0.69 | 0.265 | 0.0275 | 0.485 | 0.139919 | 0.717301 | 0.285586 | 0.995437 | 0.997988 | 0.994666 |
| Gas2l1_Q8JZP9_S482s_1 | Gas2l1_Q8 | -1.42 | -1.61 | -2.2 | -2.61 | -0.37 | -1.45 | -1.36 | -1.84 | -1.75 | -2.15 | 1.23 | -1.54 | -0.93 | -0.16 | -2.23 | -1.9 | 0.17 | 0.67 | 0.9 | 0.220829 | 0.103183 | 0.236684 | 0.989223 | 0.997988 | 0.994666 |
| Gas7_B1ATI9_S312s_1 | Gas7_B1AT | -1.4 | -4.11 | -2.51 | -4.85 | -1.99 | -2.65 | -1.7 | -1.47 | -1.86 | -2.79 | 2.39 | -2.14 | -1.1 | -0.57 | -2.6 | -2.61 | 0.705 | 0.9075 | 0.655 | 0.09529 | 0.104742 | 0.113763 | 0.989223 | 0.997988 | 0.994666 |
| Gas7_B1ATI9_S47s_1 | Gas7_B1AT | -2.3 | -1.8 | -1.62 | -1.31 | -0.99 | -1.64 | -2.15 | -1.53 | -0.48 | -1.16 | -1.12 | -1.6 | 0.48 | -2 | -0.87 | -1.49 | 1.265 | 2.1175 | 1.4975 | 0.164606 | 0.585041 | 0.07023 | 0.989223 | 0.997988 | 0.994666 |
| Gas7_B1ATI9_S53s_1 | Gas7_B1AT | -1.65 | -2.06 | -2 | -1.1 | -1.72 | -1.17 | -1.13 | -1.3 | -1.64 | -1.93 | -1.39 | -1.71 | -1.23 | -1.09 | -2.12 | -1.39 | 0.18 | 0.6675 | 0.7875 | 0.487869 | 0.217554 | 0.904302 | 0.989223 | 0.997988 | 0.994666 |
| Gas7_B1ATI9_S88s_1 | Gas7_B1AT | -0.17 | -0.83 | -0.55 | -0.22 | 0.62 | 0.89 | -0.09 | -0.02 | -0.78 | -0.18 | 0.25 | -0.63 | 0.21 | 0.1 | -0.26 | -0.14 | 0.3725 | 0.035 | 0.245 | 0.148303 | 0.02643 | 0.731363 | 0.989223 | 0.997988 | 0.994666 |
| Gas7_B1ATI9_S92s_1 | Gas7_B1AT | 0.07 | -1.31 | -0.46 | -0.81 | -0.91 | 0.94 | -0.82 | 0.21 | -0.84 | -1.22 | 0.33 | 0.31 | -0.83 | 0.26 | -0.52 | 0.11 | 0.7925 | 0.1075 | 0.42 | 0.346975 | 0.334653 | 0.565066 | 0.989223 | 0.997988 | 0.994666 |
| Gas7_B1ATI9_S98s_1 | Gas7_B1AT | 3.49 | 0.53 | 0.39 | 0.9 | 0.35 | 3.48 | 0.14 | 0.76 | 2.12 | 2.35 | 0.22 | 0.91 | 1.68 | -1.91 | 0.13 | 0.32 | 0.4825 | 0.2725 | 0.3825 | 0.184041 | 0.875237 | 0.926994 | 0.996493 | 0.997988 | 0.994666 |
| Gas7_B1ATI9_S99s_1 | Gas7_B1AT | -0.83 | -2.02 | -0.66 | -0.33 | -1.97 | -0.22 | -1.4 | -0.65 | -0.03 | -1.5 | -0.56 | -0.32 | 0.49 | -1.71 | -1.18 | -0.94 | -0.145 | 0.0725 | -1.2725 | 0.821168 | 0.839035 | 0.449471 | 0.995762 | 0.997988 | 0.999507 |
| Gatad2b_Q8VHR5_S335s_1 | Gatad2b_C | -0.97 | -0.38 | -0.69 | -0.89 | -0.21 | -0.56 | -0.53 | -0.61 | -0.51 | -0.14 | -0.92 | -0.16 | -0.68 | -0.19 | -0.94 | -0.41 | -0.1 | 0.3575 | 0.125 | 0.536076 | 0.288372 | 0.296738 | 0.989223 | 0.997988 | 0.994666 |
| Gatad2b_Q8VHR5_S339s_1 | Gatad2b_C | 0.26 | -0.41 | 0.44 | -0.22 | -0.01 | -0.33 | 0.19 | -0.16 | -0.79 | -0.2 | -0.12 | -0.24 | -1.01 | -0.37 | 0.03 | 0.2 | 0.255 | 0.3 | 0.1775 | 0.405494 | 0.725716 | 0.246278 | 0.995437 | 0.997988 | 0.994666 |
| Gatad2b_Q8VHR5_S487s_1 | Gatad2b_C | 0.1 | -0.64 | 0.29 | 0.49 | 0.13 | -0.75 | 0.45 | 0.07 | -0.59 | 0.46 | 0.37 | 0.81 | -0.63 | -0.41 | 0.45 | 0.68 | -0.095 | -0.355 | -0.305 | 0.926591 | 0.811006 | 0.605376 | 0.995762 | 0.997988 | 0.999507 |
| Gatad2b_Q8VHR5_T121t_1 | Gatad2b_C | 0.81 | 0.96 | -0.78 | -0.13 | 0.41 | -0.53 | 0.82 | -0.91 | -0.27 | 0.87 | -0.42 | -0.34 | -0.71 | 0.75 | -0.54 | -0.25 | -0.085 | 0.2025 | -0.0375 | 0.425936 | 0.611076 | 0.601016 | 0.989223 | 0.997988 | 0.994666 |
| Gba2_Q69ZF3_S893s_1 | Gba2_Q69Z | -0.01 | -0.61 | -0.22 | 0.18 | -0.89 | -0.06 | 0.15 | 0.6 | -0.75 | -0.56 | 0.64 | 0.41 | 0.08 | -0.18 | 0.88 | 0.29 | -0.2675 | -0.255 | -0.4025 | 0.205332 | 0.746559 | 0.796337 | 0.995762 | 0.997988 | 0.994666 |
| Gbf1_Q6DFZ1_S1318s_1 | Gbf1_Q6DF | -0.61 | -0.76 | -1.06 | -0.6 | -0.59 | -0.19 | -0.77 | -0.99 | -0.63 | 0.25 | -0.33 | -0.95 | -0.22 | -0.12 | -0.99 | -0.63 | 0.115 | 0.1 | 0.4325 | 0.372735 | 0.631694 | 0.286546 | 0.989223 | 0.997988 | 0.994666 |
| Gbf1_Q6DFZ1_S1786s_1 | Gbf1_Q6DF | -1.2 | -0.96 | -2.3 | -0.65 | 0.15 | -0.5 | -0.78 | -0.5 | -0.53 | -0.36 | 0.56 | -0.95 | 1 | 0.08 | -1.87 | -0.73 | 0.1225 | 0.3425 | 0.2675 | 0.183083 | 0.050186 | 0.061956 | 0.989223 | 0.997988 | 0.994666 |
| Gbf1_Q6DFZ1_S1791s_1 | Gbf1_Q6DF | -0.77 | -0.53 | -0.6 | -0.68 | 0.01 | -1.36 | -0.69 | -0.62 | -0.25 | -1.29 | -0.16 | -0.59 | 0.09 | 0.31 | -0.08 | -1.11 | 0.87 | 0.9575 | 0.8975 | 0.219868 | 0.947876 | 0.809632 | 0.999187 | 0.997988 | 0.994666 |
| Gcfc2_S4R198_S169s_1 | Gcfc2_S4R1 | -0.25 | 0.29 | -0.37 | -0.51 | -1.68 | 0.45 | -1.43 | -1.09 | 1.9 | -0.83 | -0.51 | -0.53 | -0.73 | -0.69 | -1.24 | -0.16 | -0.02 | 0.0725 | 0.4475 | 0.152262 | 0.147079 | 0.719143 | 0.989223 | 0.997988 | 0.994666 |
| Gdap1l1_Q8VE33_S139s_1 | Gdap1l1_Q | 0.52 | 0.47 | -0.01 | -0.03 | 0.56 | 0.72 | 0.21 | 0.15 | 0.79 | 0.56 | -0.85 | 0 | -0.06 | 0.42 | -0.39 | 0.16 | -0.7275 | 0.2175 | -0.495 | 0.490828 | 0.50447 | 0.77541 | 0.989223 | 0.997988 | 0.994666 |
| Gdap1l1_Q8VE33_S37s_1 | Gdap1l1_Q | -0.62 | -0.49 | -1.38 | -1.53 | -0.35 | -0.94 | -0.62 | -1.38 | -1.52 | -0.82 | -1.76 | -1 | -1.22 | 0.04 | -1.95 | -2.05 | 0.1725 | -0.1125 | -0.205 | 0.576869 | 0.601597 | 0.455161 | 0.989223 | 0.997988 | 0.995259 |
| Gdap1l1_Q8VE33_S38s_1 | Gdap1l1_Q | -0.62 | -0.49 | -1.38 | -1.53 | -0.35 | -0.94 | -0.62 | -1.38 | -1.52 | -0.82 | -1.76 | -1 | -1.22 | 0.04 | -1.95 | -2.05 | 0.1825 | -0.27 | -0.29 | 0.576869 | 0.601597 | 0.455161 | 0.989223 | 0.997988 | 0.995259 |
| Gdi1_P50396_S121s_1 | Gdi1_P503 | -1.01 | -0.14 | -0.67 | -0.88 | -0.15 | -0.53 | 0.1 | -0.35 | -0.08 | -0.78 | -1.45 | -1.51 | -0.86 | 0.35 | -0.22 | -0.75 | 0.1825 | -0.27 | -0.29 | 0.407779 | 0.131307 | 0.476716 | 0.989223 | 0.997988 | 0.994666 |
| Gemin5_E9PUU4_S1410s_2 | Gemin5_E9 | -4.12 | -4.66 | -2.67 | -3.95 | -3.32 | -5.19 | -2.98 | -3.49 | -5.94 | -4.5 | -1.29 | -2.81 | -2.93 | -3.54 | -3.08 | -2.98 | 0.4425 | -0.28 | 0.305 | 0.125244 | 0.85683 | 0.824184 | 0.996349 | 0.997988 | 0.994666 |
| Gemin5_E9PUU4_S1414s_1 | Gemin5_E9 | -0.19 | 0.28 | 0.73 | 0.53 | 0.25 | -0.54 | 0.37 | 0.22 | 0.26 | 0.24 | 0.55 | 0.24 | 0.93 | 0.14 | 1.05 | 0.74 | 0.105 | 0.215 | 0.7175 | 0.264729 | 0.403039 | 0.955754 | 0.989223 | 0.997988 | 0.994666 |
| Gemin5_Q8BX17_S1410s_2 | Gemin5_Q8 | -2.09 | -3.07 | -1.08 | -1.17 | -2.52 | -1.59 | -0.81 | -1.22 | -3.25 | -3.74 | -2.37 | -0.73 | -2.06 | -1.49 | -0.66 | -0.36 | -0.2625 | -0.015 | 0.3775 | 0.224528 | 0.557486 | 0.368317 | 0.989223 | 0.997988 | 0.994666 |
| Gemin5_Q8BX17_S1414s_2 | Gemin5_Q8 | -2.09 | -3.07 | -1.08 | -1.17 | -2.52 | -1.59 | -0.81 | -1.22 | -3.25 | -3.74 | -2.37 | -0.73 | -2.06 | -1.49 | -0.66 | -0.36 | 0.3175 | -0.67 | 0.71 | 0.224528 | 0.557486 | 0.368317 | 0.989223 | 0.997988 | 0.994666 |
| Gemin7_D3Z684_S3s_1 | Gemin7_D3 | -0.5 | 0.16 | -0.59 | -0.34 | -0.65 | -0.45 | 0.27 | -0.12 | 1.73 | 0.04 | 0.72 | -0.25 | 0.88 | 0.54 | 0.29 | -0.56 | 0.3175 | -0.67 | 0.71 | 0.126798 | 0.783819 | 0.078369 | 0.995762 | 0.997988 | 0.994666 |
| Gfap_P03995_S12s_1 | Gfap_P039 | -0.36 | 0.33 | -0.38 | -0.68 | 0.62 | -0.53 | -0.55 | -0.71 | -0.1 | 0.08 | -0.56 | -0.04 | 1.01 | -0.14 | 0.34 | 0.26 | 0.08 | 0.8775 | 0.605 | 0.092896 | 0.956718 | 0.693323 | 0.999187 | 0.997988 | 0.994666 |
| Gfap_P03995_S390s_1 | Gfap_P039 | 0.79 | -0.72 | 0.54 | 0.18 | -0.78 | 0.89 | -0.57 | 0.35 | 0.33 | -0.43 | 0.19 | -0.17 | 0.22 | -1.52 | 0.02 | 0.23 | -0.02 | 0.1175 | 0.64 | 0.373669 | 0.637043 | 0.56832 | 0.989223 | 0.997988 | 0.994666 |
| Gfap_P03995_T40t_1 | Gfap_P039 | 0.03 | -0.46 | -0.23 | -0.18 | 0.12 | 0.25 | 0.45 | -0.57 | -0.32 | 0.18 | 0.39 | 0.23 | 0.16 | 0.14 | 0.99 | 1.01 | -0.225 | -0.2175 | -0.46 | 0.030894 | 0.342033 | 0.215515 | 0.989223 | 0.997988 | 0.994666 |
| Gfpt1_P47856_S243s_1 | Gfpt1_P47 | -0.97 | -0.45 | -1.33 | -1.38 | -0.55 | -1.71 | -0.59 | -0.92 | -0.61 | -0.28 | -1.41 | -1.04 | -0.43 | -0.21 | -1.59 | -0.96 | 0.2725 | 0.33 | 0.785 | 0.547567 | 0.795679 | 0.570226 | 0.995762 | 0.997988 | 0.994666 |
| Gga3_S4R2D2_S159s_1 | Gga3_S4R2 | -0.18 | 0.08 | 0.38 | -0.12 | 0.29 | -0.49 | 0.2 | -0.06 | -0.19 | -0.03 | -0.08 | -0.12 | 0 | 0.3 | -0.53 | -0.25 | 0.09 | 0.1975 | 0.235 | 0.581856 | 0.834279 | 0.531239 | 0.995762 | 0.997988 | 0.995259 |
| Ggt7_Q99JP7_S83s_1 | Ggt7_Q99J | 0.41 | 1 | 0.39 | 0.26 | 0.46 | 0 | 0.39 | -0.04 | 0.35 | 0.66 | -0.04 | 0.15 | 0.56 | 0.75 | 0.76 | 0.56 | -0.055 | -0.145 | -0.16 | 0.595673 | 0.2466 | 0.405124 | 0.989223 | 0.997988 | 0.995259 |
| Gigyf1_Q99MR1_S137s_1 | Gigyf1_Q9 | -1.4 | -2.1 | -1.35 | -1.61 | -1.05 | -1.34 | -2.1 | -1.11 | -1.84 | -1.38 | -0.22 | -1.5 | -1.66 | -0.67 | -1.7 | -1.98 | -0.3125 | -0.235 | 0.1425 | 0.754438 | 0.49861 | 0.34488 | 0.989223 | 0.997988 | 0.9971 |
| Gigyf1_Q99MR1_S157s_1 | Gigyf1_Q9 | 0.06 | -0.35 | -0.16 | -0.09 | -0.58 | 0.26 | -0.85 | -0.32 | -0.55 | 0.03 | 0.37 | -0.43 | -0.59 | -0.09 | -0.22 | -0.51 | 0.215 | 0.38 | 0.1125 | 0.401167 | 0.413217 | 0.971506 | 0.989223 | 0.997988 | 0.994666 |
| Gigyf1_Q99MR1_S24s_1 | Gigyf1_Q9 | -1.25 | -1.32 | -0.34 | -0.55 | -0.39 | -0.96 | 0.59 | -0.48 | -1.64 | -0.8 | -0.62 | -0.18 | -0.13 | -1.27 | -0.26 | -0.29 | -0.2375 | -0.01 | -0.2175 | 0.329981 | 0.181589 | 0.888585 | 0.989223 | 0.997988 | 0.994666 |
| Gigyf1_Q99MR1_S413s_1 | Gigyf1_Q9 | -0.67 | 0.5 | -0.19 | 0.29 | -0.19 | -0.75 | 0.81 | -0.1 | -0.31 | 0.15 | 0.03 | -0.13 | 0.07 | 0.02 | 0.01 | 0.88 | 0.555 | 0.055 | 0.3775 | 0.473102 | 0.920182 | 0.879103 | 0.997335 | 0.997988 | 0.994666 |
| Gigyf1_Q99MR1_S540s_1 | Gigyf1_Q9 | -0.38 | -0.11 | 0.24 | 0.31 | 0.12 | -0.56 | 0.46 | 0.29 | 0.1 | 0.21 | 0.27 | 0 | 0.11 | 0.56 | 0.06 | 0.58 | -0.04 | -0.0475 | 0.2625 | 0.290184 | 0.834951 | 0.602888 | 0.995762 | 0.997988 | 0.994666 |
| Gigyf1_Q99MR1_S755s_1 | Gigyf1_Q9 | -1.48 | 0.08 | -1.24 | -1.23 | -0.95 | -1.18 | -0.36 | -1.3 | -1.28 | -1.13 | -1.52 | -0.74 | -0.3 | -0.21 | -0.52 | -1.29 | 0.0625 | 0.13 | 0.3125 | 0.374539 | 0.959706 | 0.611038 | 0.999187 | 0.997988 | 0.994666 |
| Gigyf1_Q99MR1_S756s_1 | Gigyf1_Q9 | -1.48 | 0.08 | -1.24 | -1.23 | -0.95 | -1.18 | -0.36 | -1.3 | -1.28 | -1.13 | -1.52 | -0.74 | -0.3 | -0.21 | -0.52 | -1.29 | 0.02 | -0.2 | 0.3875 | 0.374539 | 0.959706 | 0.611038 | 0.999187 | 0.997988 | 0.994666 |
| Gigyf2_G3UWR2_S161s_1 | Gigyf2_G3 | -1.68 | -2.27 | -3.23 | -3.03 | -1.95 | -1.75 | -2.65 | -2.99 | -2.18 | -1.69 | -1.67 | -2.8 | -1.42 | -1.41 | -3.08 | -2.98 | 0.02 | -0.2 | 0.3875 | 0.548873 | 0.61843 | 0.29564 | 0.989223 | 0.997988 | 0.994666 |
| Gigyf2_G3UWR2_S202s_1 | Gigyf2_G3 | -6.11 | -7.14 | -5.18 | -8.71 | -3.64 | -4.96 | -5.15 | -5.92 | -7.04 | -5.23 | -1.74 | -5.07 | -3.79 | -4.04 | -5.28 | -4.89 | 0.2175 | 0.4675 | 0.33 | 0.011571 | 0.036494 | 0.112785 | 0.989223 | 0.997988 | 0.994666 |
| Gigyf2_G3UWR2_S237s_1 | Gigyf2_G3 | -2.19 | -1.47 | -1.76 | -2.25 | -1.07 | -2.69 | -1.49 | -2.49 | -2.55 | -1.77 | -1.77 | -2.42 | -1.34 | -1.54 | -1.64 | -1.92 | 1.8675 | 2.015 | 2.285 | 0.303788 | 0.966061 | 0.505704 | 0.999187 | 0.997988 | 0.994666 |
| Gigyf2_G3UWR2_T25t_1 | Gigyf2_G3 | -0.42 | -0.34 | -0.83 | -2.12 | 0.22 | -0.38 | 0 | 0.04 | -1 | 0.28 | -0.68 | -0.77 | -0.25 | -0.17 | -3.29 | -0.15 | -0.0175 | -0.21 | 0.3075 | 0.961479 | 0.051707 | 0.426018 | 0.989223 | 0.997988 | 0.999507 |
| Git1_Q5F258_S361s_1 | Git1_Q5F2 | 0.6 | 0.54 | 0.81 | 0.79 | 0.68 | 0.19 | 0.87 | 0.62 | 0.35 | 0.61 | 0.31 | 0.71 | -0.01 | 0.58 | 0.57 | 0.65 | 0.8975 | 0.385 | -0.0375 | 0.378011 | 0.683394 | 0.406344 | 0.994272 | 0.997988 | 0.994666 |
| Git1_Q5F258_S362s_1 | Git1_Q5F2 | 0.47 | 0.67 | 0.79 | 0.7 | 0.68 | 0.32 | 0.8 | 0.61 | 0.4 | 0.6 | 0.33 | 0.68 | 0.11 | 0.77 | 0.48 | 0.54 | -0.095 | -0.19 | -0.2375 | 0.483293 | 0.79892 | 0.486328 | 0.995762 | 0.997988 | 0.994666 |
| Git1_Q5F258_S561s_1 | Git1_Q5F2 | -0.49 | 0.52 | -1.34 | -0.68 | 1.13 | -1.27 | -0.29 | -0.48 | 0.56 | 0.51 | -0.07 | -0.27 | 0 | 0.86 | -1.51 | 0 | -0.055 | -0.155 | -0.1825 | 0.563488 | 0.636253 | 0.136778 | 0.989223 | 0.997988 | 0.995259 |
| Git2_F7BIK4_S360s_1 | Git2_F7BIK | -0.44 | -0.08 | -0.46 | -0.22 | -0.64 | -0.67 | -1.15 | -0.91 | 0.57 | -0.07 | -0.31 | 0.01 | 1.33 | -0.83 | -0.61 | -0.87 | 0.27 | 0.68 | 0.335 | 0.913298 | 0.036963 | 0.212165 | 0.989223 | 0.997988 | 0.999507 |
| Gja1_P23242_S255s_1 | Gja1_P232 | 0.04 | -0.38 | -1.35 | -1.4 | -0.3 | 0.01 | -0.77 | -1.11 | -0.32 | 0.15 | -2.8 | -1.19 | -1.1 | -0.31 | -1.74 | -1.46 | -0.5425 | 0.35 | 0.055 | 0.416043 | 0.582425 | 0.691305 | 0.989223 | 0.997988 | 0.994666 |
| Gja1_P23242_S306s_1 | Gja1_P232 | -0.03 | 0.46 | 0.28 | -0.41 | -0.02 | -0.35 | -0.36 | 0.21 | -0.08 | 0.68 | -1.98 | -0.05 | -0.34 | -0.44 | -0.82 | 0.17 | 0.23 | -0.2675 | -0.38 | 0.203079 | 0.462244 | 0.443737 | 0.989223 | 0.997988 | 0.994666 |
| Gja1_P23242_S314s_1 | Gja1_P232 | 1.04 | -0.15 | 0.37 | 0.98 | 0.34 | 0.45 | -0.06 | 0.25 | 0.78 | 0.88 | -0.57 | 0.51 | 0.57 | 0.28 | 0.9 | 0.13 | -0.205 | -0.4325 | -0.4325 | 0.800264 | 0.334074 | 0.707546 | 0.989223 | 0.997988 | 0.999507 |
| Gja1_P23242_S328s_1 | Gja1_P232 | -0.71 | -0.06 | -0.17 | -0.34 | -0.18 | -0.99 | -0.13 | -0.65 | -0.9 | -0.18 | -1.11 | -0.42 | 0.15 | 0.02 | 0.2 | -0.05 | -0.315 | -0.16 | -0.09 | 0.144218 | 0.55765 | 0.280483 | 0.989223 | 0.997988 | 0.994666 |
| Gja1_P23242_S364s_1 | Gja1_P232 | 1.21 | 1.34 | 0.79 | 0.8 | 0.88 | 1.1 | 1.04 | 1.03 | 1.53 | 1.62 | 1.4 | 1.17 | 0.09 | 0.98 | 0.37 | 0.44 | -0.1675 | -0.3325 | 0.4 | 0.082307 | 0.921063 | 0.136998 | 0.997335 | 0.997988 | 0.994666 |
| Gja1_P23242_S365s_1 | Gja1_P232 | -0.07 | 0.32 | 2.3 | 2.27 | -0.06 | -0.05 | 2.28 | 2.7 | 0.08 | 0.42 | 3.76 | 3.06 | -0.85 | -0.17 | 2.13 | 2.42 | -0.0225 | 0.395 | -0.565 | 0.721773 | 0.988193 | 0.531672 | 0.999187 | 0.997988 | 0.9971 |
| Gja1_P23242_S373s_1 | Gja1_P232 | 0.07 | 0.76 | 1.03 | 0.24 | 0.94 | -0.23 | 0.98 | 0.31 | 0.43 | 0.88 | 0.76 | 0.45 | 0.25 | 0.38 | 0.81 | 0.48 | 0.0125 | 0.625 | -0.3225 | 0.883991 | 0.944853 | 0.721167 | 0.999187 | 0.997988 | 0.999507 |
| Gjc2_Q8BQU6_T190t_1 | Gjc2_Q8BC | -1.6 | -0.24 | 1.22 | 0.55 | -0.44 | -1.54 | -0.58 | 0.49 | 0.43 | -1.32 | 0.94 | -0.17 | 1.13 | 0.29 | 0.3 | 0.27 | -0.025 | 0.105 | -0.045 | 0.390776 | 0.45015 | 0.985749 | 0.989223 | 0.997988 | 0.994666 |
| Gjc3_Q921C1_S249s_1 | Gjc3_Q921 | 1.94 | 1.83 | 0.75 | 0.09 | 1.96 | 1.35 | -0.09 | 0.01 | 0.76 | 0.67 | 0.16 | 0.28 | 0.78 | 1.14 | 0.24 | 0.39 | -0.5 | -0.0125 | 0.515 | 0.288407 | 0.569616 | 0.153109 | 0.989223 | 0.997988 | 0.994666 |
| Glcci1_E9QKK4_S137s_1 | Glcci1_E9C | -0.49 | -1.87 | -1.77 | -1.55 | -1.63 | -0.24 | -2.22 | -1.46 | -0.68 | -0.57 | -0.71 | -1.74 | 0.15 | -1.3 | -1.57 | -1.85 | -0.345 | -0.685 | -0.515 | 0.591832 | 0.946202 | 0.248925 | 0.999187 | 0.997988 | 0.995259 |
| Glcci1_E9QKK4_S140s_1 | Glcci1_E9C | -1.21 | -2.81 | -2.24 | -2.84 | -2.2 | -1.64 | -2.72 | -3.3 | -2.11 | -1.3 | -2.79 | -2.87 | -0.91 | -2.83 | -3.21 | -2.11 | 0.0325 | 0.495 | 0.2775 | 0.986255 | 0.693523 | 0.987948 | 0.994893 | 0.997988 | 1 |
| Glcci1_E9QKK4_S215s_1 | Glcci1_E9C | -0.17 | -1.01 | -0.43 | -0.18 | 0.35 | -0.66 | 0.53 | 0.3 | -0.56 | -0.24 | 0.38 | 0.3 | -1.72 | 0.02 | 0.06 | 0.18 | -0.19 | 0.0075 | 0.01 | 0.861694 | 0.113998 | 0.219551 | 0.989223 | 0.997988 | 0.999507 |
| Glcci1_E9QKK4_S26s_1 | Glcci1_E9C | -0.62 | 0.53 | 1.3 | 0.83 | -0.1 | 0.6 | 0.39 | 1.01 | -0.4 | 0.98 | 1.65 | 1.19 | -0.37 | -2.01 | 1.29 | 1.52 | 0.5775 | 0.4175 | 0.0825 | 0.621346 | 0.936626 | 0.539476 | 0.998821 | 0.997988 | 0.995259 |
| Glcci1_E9QKK4_S336s_1 | Glcci1_E9C | -0.37 | -0.44 | -0.9 | -1.06 | 0.28 | -0.04 | -0.75 | -0.4 | -0.31 | -0.8 | -0.75 | -0.79 | 0.16 | 0.09 | -0.79 | -0.78 | -0.035 | 0.345 | -0.4025 | 0.308439 | 0.149126 | 0.910578 | 0.989223 | 0.997988 | 0.994666 |
| Glcci1_E9QKK4_S389s_1 | Glcci1_E9C | -0.01 | 0.5 | -0.23 | -0.21 | -0.01 | 0.18 | -0.3 | -0.05 | -0.17 | 0.2 | 0.21 | -0.28 | 0.3 | -0.09 | -0.25 | 0.26 | 0.465 | 0.03 | 0.3625 | 0.882559 | 0.819499 | 0.933557 | 0.995762 | 0.997988 | 0.999507 |
| Glod4_F6ZTG3_S109s_1 | Glod4_F6Z | 0.5 | 0.57 | 0.65 | 0.71 | 0.55 | 0.96 | -0.13 | 0.69 | 0.56 | 1.02 | 1.22 | 0.59 | 0.34 | 0.83 | -0.09 | -0.05 | -0.0575 | -0.0225 | 0.0425 | 0.244431 | 0.744274 | 0.343818 | 0.995762 | 0.997988 | 0.994666 |
| Glrb_P48168_T391t_1 | Glrb_P481 | -0.31 | -0.46 | 0.11 | 0.8 | 0.02 | 0.43 | 0.9 | 0.26 | 0.24 | -0.44 | -0.5 | 0.11 | -0.29 | 0.03 | -0.03 | 0.68 | -0.09 | 0.24 | -0.35 | 0.865655 | 0.298857 | 0.608133 | 0.989223 | 0.997988 | 0.999507 |
| Gltscr1l_Q8CHH5_S930s_1 | Gltscr1l_Q8 | 0.09 | 0.33 | -1.31 | -0.71 | -0.72 | -1.16 | -1.2 | -1.38 | 1.88 | -1.35 | -0.04 | -2.26 | 1.84 | -1.68 | -0.31 | 0.48 | 0.3675 | -0.1825 | 0.0625 | 0.516806 | 0.09165 | 0.960793 | 0.989223 | 0.997988 | 0.994666 |
| Gltscr2_Q8BK35_S305s_1 | Gltscr2_Q8 | -0.18 | 0.09 | 0.81 | 0.32 | 0.38 | -1.2 | 0.4 | 1.22 | -0.15 | 0.3 | 0.13 | 0.42 | -0.44 | -0.68 | 1 | 1.7 | -0.715 | -0.0425 | 0.4825 | 0.810286 | 0.904743 | 0.769261 | 0.996493 | 0.997988 | 0.999507 |
| Glud1_P26443_S450s_1 | Glud1_P26 | 0.34 | 1.17 | 2.36 | 1.47 | 0.82 | 0.96 | 1.64 | 1.69 | 0.09 | 0.86 | 0.93 | 1.85 | 0.64 | -1.04 | 1.73 | 1.76 | -0.06 | -0.085 | 0.135 | 0.427898 | 0.896962 | 0.443081 | 0.996493 | 0.997988 | 0.994666 |
| Glul_P15105_S19s_1 | Glul_P151C | -0.54 | -0.5 | -1.9 | -1.27 | -0.05 | -0.24 | -0.02 | -0.49 | -0.03 | -0.29 | -0.98 | 0.23 | 0.14 | -0.22 | -1.38 | -1.1 | -0.0575 | -0.4025 | -0.5625 | 0.390556 | 0.034642 | 0.085137 | 0.989223 | 0.997988 | 0.994666 |
| Glul_P15105_S343s_1 | Glul_P151C | -0.12 | -1.01 | 0.41 | 0.48 | -1.11 | -0.25 | 0.15 | 0.78 | -0.21 | -0.21 | 0.41 | 0.06 | -0.39 | -0.33 | -0.08 | 0.29 | 0.8525 | 0.785 | 0.4125 | 0.862001 | 0.921625 | 0.848174 | 0.997335 | 0.997988 | 0.999507 |
| Gm13889_Q3UPL5_S210s_1 | Gm13889_ | -2.7 | -3.22 | -2.53 | -2.98 | -1.34 | -2.8 | -2.4 | -2.81 | -4.29 | -2.2 | -1.61 | -3.51 | -2.28 | -2.09 | -2.77 | -3.19 | -0.0475 | 0.0725 | -0.0675 | 0.415398 | 0.184372 | 0.937914 | 0.989223 | 0.997988 | 0.994666 |
| Gm15800_E9Q2E4_S1496s_1 | Gm15800_ | 0.59 | 1.07 | 0.63 | 0.79 | 0.44 | -0.29 | 0.3 | 0.97 | 0.01 | 1.02 | 1.29 | 0.85 | 0.49 | 0.5 | 1.34 | 1.27 | 0.52 | -0.045 | 0.275 | 0.677646 | 0.193403 | 0.944585 | 0.989223 | 0.997988 | 0.996159 |
| Gm15800_E9Q2E4_S1498s_1 | Gm15800_ | 1.43 | 1.6 | 0.33 | 0.84 | 1.72 | 1.88 | 0.39 | 0.49 | 0.95 | 1.58 | 0.88 | 0.81 | 1.26 | 2.38 | 0.46 | 0.56 | -0.415 | 0.0225 | 0.13 | 0.818248 | 0.878011 | 0.988717 | 0.996493 | 0.997988 | 0.999507 |
| Gm15800_E9Q2E4_S1562s_1 | Gm15800_ | 0.13 | 0.42 | 0.06 | 0.11 | -0.13 | -0.44 | 0.74 | -0.51 | -0.28 | 0.18 | -0.91 | 0.49 | -1.26 | -0.05 | 0.05 | 0.41 | 0.07 | 0.005 | 0.115 | 0.321434 | 0.409747 | 0.368238 | 0.989223 | 0.997988 | 0.994666 |
| Gm15800_E9Q2E4_S1916s_1 | Gm15800_ | 1.01 | 0.93 | 1.2 | 0.46 | 1.04 | 1.04 | 1.68 | 1.05 | -0.59 | 1.43 | -1.49 | 0.75 | -0.44 | -0.57 | 1.38 | 0.99 | -0.265 | -0.31 | -0.3925 | 0.272641 | 0.274767 | 0.179375 | 0.989223 | 0.997988 | 0.994666 |
| Gm15800_E9Q2E4_T2503t_1 | Gm15800_ | 0.56 | 0.94 | 0.62 | 0.64 | 0.78 | 0.53 | 0.35 | 0.5 | 0.6 | 0.85 | 1.05 | 0.45 | 1.03 | 0.98 | 0.53 | 0.36 | 0.3025 | -0.875 | -0.56 | 0.897651 | 0.493135 | 0.843996 | 0.989223 | 0.997988 | 0.999507 |
| Gm20521_D3Z5F7_S177s_1 | Gm20521_ | 0.28 | 0.53 | -0.44 | -0.11 | 0.19 | 0.32 | -0.12 | -0.65 | 0.02 | 0.14 | -0.47 | -0.42 | -0.34 | 0.64 | -0.36 | -0.4 | -0.15 | 0.0475 | 0.035 | 0.616165 | 0.684712 | 0.420983 | 0.994624 | 0.997988 | 0.995259 |
| Gm27021_M0QWE2_T124t_1 | Gm27021_ | 1.2 | 1.3 | 1.08 | 0.45 | 1.27 | 1.41 | 1.9 | 0.43 | 0.73 | 1.63 | 0.73 | 0.91 | 0.54 | 0.75 | 0.55 | 0.7 | -0.13 | -0.2475 | -0.18 | 0.201895 | 0.500426 | 0.981134 | 0.989223 | 0.997988 | 0.994666 |
| Gm27029_Q3THG9_S88s_1 | Gm27029_ | -1.17 | -1.11 | -0.27 | -0.53 | -1.04 | -1.43 | 0.19 | -0.1 | -0.39 | -0.65 | -0.44 | -0.52 | -0.37 | -0.32 | 0.04 | 0.65 | 0.245 | -0.0075 | -0.3725 | 0.050852 | 0.677684 | 0.346743 | 0.993224 | 0.997988 | 0.994666 |
| Gm28036_F6YJB6_S336s_1 | Gm28036_ | -2.63 | -2.33 | -1.12 | -1.4 | -2.19 | -2.05 | -0.98 | -1.32 | -1.74 | -1.58 | -1.05 | -0.04 | -2.07 | -1.78 | -2.77 | -2.32 | 0.175 | 0.27 | 0.77 | 0.392097 | 0.59291 | 0.147865 | 0.989223 | 0.997988 | 0.994666 |
| Gm7694_J3QNH8_S221s_1 | Gm7694_J3 | 0.54 | 0.92 | 0.83 | 0.28 | -0.2 | 0.48 | -0.13 | -0.3 | 0.45 | 0.58 | 1 | 0.63 | 1.11 | 0.85 | 0.64 | 0.54 | 0.235 | 0.7675 | -0.365 | 0.607268 | 0.029509 | 0.930038 | 0.989223 | 0.997988 | 0.995259 |
| Gm9774_Q9JKV1_S211s_1 | Gm9774_C | -3.09 | -3.6 | -2.92 | -2.88 | -3.81 | -3.61 | -5.04 | -2.53 | -3.92 | -3.43 | -1.11 | -2.6 | -2.41 | -5.86 | -2.29 | -4.29 | -0.68 | 0.0225 | 0.1425 | 0.449262 | 0.227205 | 0.54556 | 0.989223 | 0.997988 | 0.994666 |
| Gm9774_Q9JKV1_S213s_1 | Gm9774_C | -0.71 | 0.37 | -0.69 | -1.63 | -0.77 | -1.48 | -1.29 | -1.35 | 0.46 | 0.03 | -0.63 | -1.35 | -0.08 | -0.22 | -0.86 | -1.44 | -0.625 | 0.3575 | -0.59 | 0.975533 | 0.201802 | 0.584185 | 0.989223 | 0.997988 | 1 |
| Gm9774_Q9JKV1_S213s_2 | Gm9774_C | -1.73 | -0.27 | -1.3 | 0.27 | -0.73 | -0.54 | -1.84 | -3.4 | -1.57 | -1.23 | 0.3 | -0.83 | -0.53 | -2.27 | -0.68 | -2.02 | -0.5575 | 0.2925 | 0.015 | 0.30815 | 0.236977 | 0.894582 | 0.989223 | 0.997988 | 0.994666 |
| Gm996_A2AJA9_S109s_1 | Gm996_A2 | 1.07 | 1.36 | 1.15 | 0.84 | 1.32 | 0.8 | 1.02 | 0.75 | -1.24 | 0.73 | -3.92 | 1.22 | -0.36 | -2.57 | 0.68 | 0.93 | -0.87 | -0.075 | -0.6175 | 0.070187 | 0.579051 | 0.091328 | 0.989223 | 0.997988 | 0.994666 |
| Gm996_A2AJA9_S286s_1 | Gm996_A2 | -0.39 | 0.43 | 0.34 | 0.06 | -0.58 | 0.15 | 0.19 | -0.48 | -0.44 | -0.08 | -0.99 | 0.65 | -0.22 | 0.4 | -0.64 | -1.01 | -0.1325 | -1.9075 | -1.435 | 0.220153 | 0.344987 | 0.414576 | 0.989223 | 0.997988 | 0.994666 |
| Gm996_A2AJA9_S435s_1 | Gm996_A2 | 0.33 | 0.2 | 0.85 | 0.3 | 0.32 | 0.12 | 0.68 | 0.24 | -1.75 | -0.23 | -1.41 | 0.57 | -1.08 | -1.68 | -0.18 | -0.08 | -0.29 | -0.325 | -0.4775 | 0.016598 | 0.747413 | 0.053968 | 0.995762 | 0.997988 | 0.994666 |
| Gm996_A2AJA9_S466s_1 | Gm996_A2 | 0.84 | 0.76 | 1.29 | 0.66 | 0.94 | 0.26 | 0.88 | 0.19 | 0.39 | 0.66 | 0.82 | 0.91 | 0.4 | 1.02 | 0.22 | -0.07 | -0.08 | -1.125 | -1.175 | 0.143839 | 0.268352 | 0.453223 | 0.989223 | 0.997988 | 0.994666 |
| Gm996_A2AJA9_S509s_1 | Gm996_A2 | 0.56 | 0.73 | 0.87 | 0.03 | 0.15 | 1.15 | -0.66 | -0.37 | 0.53 | -0.54 | -1.92 | -0.14 | 1.58 | -1.56 | -0.74 | -0.06 | -0.32 | -0.1925 | -0.495 | 0.254179 | 0.266627 | 0.063148 | 0.989223 | 0.997988 | 0.994666 |
| Gm996_A2AJA9_S560s_1 | Gm996_A2 | 1.04 | 0.72 | 1.02 | 0.26 | 0.77 | 1.11 | 0.53 | 1.33 | 1.1 | 0.89 | 0.27 | 0.65 | 0.69 | 0.53 | 0.03 | 0.97 | -0.48 | -1.065 | -0.7425 | 0.523034 | 0.545103 | 0.91259 | 0.989223 | 0.997988 | 0.994666 |
| Gm996_A2AJA9_S697s_1 | Gm996_A2 | 2.04 | 1.69 | 2.63 | 1.47 | 1.4 | 2.51 | 1.74 | 1.54 | 3.33 | 1.71 | -0.84 | 3 | 2.41 | -0.76 | 3.79 | 2.37 | 0.175 | -0.0325 | -0.205 | 0.995427 | 0.653232 | 0.856445 | 0.990434 | 0.997988 | 1 |
| Gm996_A2AJA9_S761s_1 | Gm996_A2 | 2.15 | 1.75 | 0.9 | 1.53 | 1.78 | 1.52 | 1.38 | 1.7 | 1.53 | 1.87 | 1.67 | 1.51 | 1.4 | 1.7 | 1.48 | 0.26 | -0.16 | -0.1575 | -0.005 | 0.380501 | 0.966794 | 0.839649 | 0.999187 | 0.997988 | 0.994666 |
| Gm996_A2AJA9_S821s_1 | Gm996_A2 | 1 | 1.06 | 1.26 | 1.08 | 0.8 | 0.7 | 0.97 | 0.85 | 0.62 | 0.9 | 0.82 | 1.21 | 1.04 | 1.11 | 0.79 | 0.8 | 0.0125 | 0.0625 | -0.3725 | 0.492826 | 0.200046 | 0.368609 | 0.989223 | 0.997988 | 0.994666 |
| Gmps_Q3THK7_S332s_1 | Gmps_Q3T | -0.47 | -0.2 | -0.45 | -0.55 | -0.32 | -0.63 | -0.43 | -0.54 | -0.12 | -0.06 | -0.36 | -0.65 | -0.33 | -0.64 | -0.41 | -0.47 | -0.27 | -0.2125 | -0.165 | 0.849894 | 0.7639 | 0.618724 | 0.995762 | 0.997988 | 0.999507 |
| Gnao1_P18872_S29s_1 | Gnao1_P18 | 1.54 | 0.95 | 1.88 | 1.74 | 1.09 | 1.19 | 2.01 | 1.73 | 1.1 | 1.27 | 1.29 | 1.37 | 0.95 | 1.27 | 1.59 | 1.49 | -0.0625 | 0.12 | -0.045 | 0.513027 | 0.943266 | 0.331849 | 0.999187 | 0.997988 | 0.994666 |
| Gnas_Z4YKV1_S337s_1 | Gnas_Z4YK | 0.14 | 0.17 | 0.44 | 0.02 | -0.52 | 0.38 | 0.08 | -0.27 | 1.63 | -0.1 | -1.02 | -0.22 | 2.34 | -0.97 | 0.76 | 0.65 | -0.0225 | -0.27 | -0.2025 | 0.425511 | 0.310315 | 0.81996 | 0.989223 | 0.997988 | 0.994666 |
| Gnaz_O70443_S27s_1 | Gnaz_O704 | 0.63 | 0.09 | 0.26 | 1.15 | 1.71 | 2.12 | 2.48 | 3.11 | 0.7 | 0.16 | -5.54 | 2.97 | -2.57 | -1.01 | 0.56 | 1.22 | -0.275 | -0.12 | 0.5025 | 0.226597 | 0.000741 | 0.55032 | 0.989223 | 0.997988 | 0.994666 |
| Gnb5_P62881_S63s_1 | Gnb5_P628 | -2.01 | -1.07 | -0.97 | -2.27 | -1.13 | -1.76 | -1.8 | -2.09 | -1.09 | -2.22 | -1.89 | -1.38 | -0.28 | -1.82 | -1.14 | -1.56 | 1.8225 | -0.96 | -0.9825 | 0.413688 | 0.760965 | 0.873749 | 0.995762 | 0.997988 | 0.994666 |
| Gng12_Q9DAS9_S10s_1 | Gng12_Q9 | 0.48 | 0.95 | 0.89 | 0.48 | 1.15 | 0.57 | 0.16 | 0.29 | 0.1 | 0.25 | 0.32 | 0.56 | 1.07 | 0.76 | 0.83 | 0.96 | -0.115 | -0.065 | 0.38 | 0.427265 | 0.585238 | 0.130824 | 0.989223 | 0.997988 | 0.994666 |
| Gng12_Q9DAS9_S10s_2 | Gng12_Q9 | 0.84 | 1.41 | 1.34 | 0.58 | 1.03 | 0.61 | 0.35 | 0.59 | -0.19 | 0.31 | 0.54 | 0.76 | -0.01 | -0.57 | 0.98 | 1.56 | -0.1575 | -0.3925 | 0.205 | 0.278673 | 0.178746 | 0.053037 | 0.989223 | 0.997988 | 0.994666 |
| Gng12_Q9DAS9_S7s_2 | Gng12_Q9 | 0.84 | 1.41 | 1.34 | 0.58 | 1.03 | 0.61 | 0.35 | 0.59 | -0.19 | 0.31 | 0.54 | 0.76 | -0.01 | -0.57 | 0.98 | 1.56 | -0.3975 | -0.6875 | -0.5525 | 0.278673 | 0.178746 | 0.053037 | 0.989223 | 0.997988 | 0.994666 |
| Gng12_Q9DAS9_T8t_1 | Gng12_Q9 | 0.63 | 0.87 | 1.19 | 0.51 | 0.69 | 0.18 | 0.39 | 0.51 | 0.14 | 0.11 | 0.7 | 0.75 | 1.3 | 1.01 | 1.07 | 1.14 | -0.3975 | -0.6875 | -0.5525 | 0.226019 | 0.167914 | 0.203246 | 0.989223 | 0.997988 | 0.994666 |
| Gng2_P63213_T52t_1 | Gng2_P632 | -0.01 | 0.74 | 0.53 | 0.13 | 0.89 | 0.09 | 0.87 | -0.02 | -0.41 | -0.1 | -0.94 | 0.36 | 0.46 | 0.95 | 0.09 | 0.62 | -0.3575 | -0.375 | 0.33 | 0.554746 | 0.728987 | 0.096437 | 0.995762 | 0.997988 | 0.99525 |
| Gng3_P63216_S12s_1 | Gng3_P632 | -0.57 | -0.11 | 0.41 | -0.42 | 0.88 | -0.36 | 0.72 | -0.62 | -0.69 | -0.18 | -0.29 | 0.26 | 0.36 | 0.95 | 0.12 | 0.31 | 0.11 | -0.62 | 0.1825 | 0.086984 | 0.437311 | 0.869833 | 0.989223 | 0.997988 | 0.994666 |
| Gng3_P63216_T10t_1 | Gng3_P632 | 0.16 | 0.71 | 0.34 | 0.37 | 1.29 | 0.36 | 0.39 | 0.65 | -0.51 | -0.1 | 0.24 | 0.73 | 0.38 | 1.1 | 0.76 | 1.27 | 0.3275 | -0.0525 | 0.6075 | 0.125718 | 0.334977 | 0.348741 | 0.989223 | 0.997988 | 0.994666 |
| Gng3_P63216_T5t_1 | Gng3_P632 | 0.46 | 0.84 | 0.78 | 0.7 | 1.22 | 0.53 | 1.2 | 0.61 | 0.21 | 0.68 | 0.44 | 0.96 | 0.53 | 1.68 | 0.89 | 1.02 | 0.2775 | -0.305 | 0.4825 | 0.293662 | 0.452861 | 0.630634 | 0.989223 | 0.997988 | 0.994666 |
| Gng3_P63216_T5t_2 | Gng3_P632 | -0.37 | 0.68 | -1.06 | -1.06 | 1.03 | 0.13 | -0.12 | -1.6 | -0.44 | 0.57 | -1.3 | 0.12 | 0.23 | 0.79 | 0.19 | -0.87 | 0.195 | -0.1225 | 0.335 | 0.303687 | 0.610736 | 0.723491 | 0.989223 | 0.997988 | 0.994666 |
| Gng4_P50153_S12s_1 | Gng4_P501 | -1.53 | -0.31 | 0.06 | 0.08 | -0.72 | -0.36 | -1.28 | -0.57 | -1.42 | -1 | -1.28 | -0.11 | 1.31 | -2.13 | 0.42 | 0.76 | 0.3125 | 0.19 | 0.5375 | 0.500921 | 0.458448 | 0.268642 | 0.989223 | 0.997988 | 0.994666 |
| Gng4_P50153_S12s_2 | Gng4_P501 | 0.61 | 0.4 | -0.57 | -0.32 | -0.03 | -0.54 | -0.65 | -0.8 | 0.3 | -0.27 | 0.13 | -0.3 | 0.39 | -0.9 | 0.43 | 0.32 | -0.3075 | -0.5275 | 0.515 | 0.943567 | 0.135977 | 0.848021 | 0.989223 | 0.997988 | 0.999507 |
| Gng4_P50153_S9s_1 | Gng4_P501 | 0.7 | 1.43 | 0.3 | 0.07 | 0.13 | 0.7 | -0.09 | 0.38 | 1.19 | 0.93 | 0.07 | 0.12 | 1.44 | 0.98 | 0.27 | 0.59 | -0.535 | -0.065 | 0.03 | 0.626301 | 0.332659 | 0.907033 | 0.989223 | 0.997988 | 0.995259 |
| Gng4_P50153_S9s_2 | Gng4_P501 | 0.15 | 0.49 | -0.57 | -0.32 | 0.59 | -0.24 | -0.65 | -0.8 | 0.11 | -0.02 | 0.13 | -0.3 | 0.45 | -0.42 | 0.43 | 0.32 | -0.345 | -0.0475 | 0.195 | 0.464035 | 0.580901 | 0.886439 | 0.989223 | 0.997988 | 0.994666 |
| Gnl1_P36916_S51s_1 | Gnl1_P369 | -0.94 | -0.6 | -0.69 | -1.01 | -1.07 | -1.22 | -1.12 | -0.87 | -0.14 | -0.73 | -0.96 | -0.72 | -0.17 | -0.2 | -0.96 | -1.12 | -0.2125 | 0.0425 | 0.2575 | 0.537462 | 0.246611 | 0.517993 | 0.989223 | 0.997988 | 0.994666 |
| Gnl1_P36916_S51s_2 | Gnl1_P369 | -1.49 | -1.45 | -1.86 | -1.97 | -1.27 | -1.56 | -1.02 | -1.41 | -0.56 | -1.37 | -0.8 | -0.69 | -0.14 | -1.13 | -0.39 | -0.72 | -0.26 | 0.1725 | 0.1975 | 0.005177 | 0.139772 | 0.012781 | 0.989223 | 0.997988 | 0.994666 |
| Gnl1_P36916_S68s_1 | Gnl1_P369 | -1.49 | -1.83 | -0.31 | -0.71 | -0.71 | -2.5 | -0.26 | -0.26 | -2.12 | -1.62 | 1.84 | -0.81 | -1.52 | -0.74 | -0.37 | -0.6 | 0.3775 | 0.8375 | 1.0975 | 0.518875 | 0.790252 | 0.631162 | 0.995762 | 0.997988 | 0.994666 |
| Gnl1_P36916_S89s_1 | Gnl1_P369 | -1.29 | -1.02 | 0.99 | -1.53 | -0.99 | -0.63 | -1.09 | -1 | -0.44 | -1.31 | -0.6 | -0.25 | -1.24 | -2.43 | -0.84 | -1.47 | 0.1525 | 0.4075 | 0.2775 | 0.218952 | 0.687116 | 0.913056 | 0.994672 | 0.997988 | 0.994666 |
| Gnl1_P36916_T48t_2 | Gnl1_P369 | -1.65 | -2.09 | -1.82 | -2.21 | -1.68 | -1.85 | -1.28 | -2.04 | -0.95 | -2.12 | -1.24 | -1.03 | -0.16 | -1.74 | -0.6 | -1.25 | -0.215 | 0.0625 | -0.7825 | 0.025011 | 0.381344 | 0.087052 | 0.989223 | 0.997988 | 0.994666 |
| Gnl1_P36916_T50t_2 | Gnl1_P369 | -1.49 | -1.45 | -1.16 | 0.72 | -1.27 | -1.56 | -1.05 | -1.77 | -0.56 | -1.37 | -0.49 | 0.43 | -0.14 | -1.13 | -0.61 | -0.14 | 0.23 | 0.6075 | 1.005 | 0.531243 | 0.27623 | 0.559559 | 0.989223 | 0.997988 | 0.994666 |
| Gnl3_Q8CI11_S472s_1 | Gnl3_Q8CI | 0.67 | 0.73 | 0.63 | 0.89 | 0.02 | 0.53 | 0.68 | 0.8 | 0.33 | 0.93 | 0.41 | 1.01 | 1.38 | 0.23 | 1.06 | 0.87 | -0.5675 | 0.3475 | 0.34 | 0.614755 | 0.371418 | 0.81383 | 0.989223 | 0.997988 | 0.995259 |
| Gnpat_Q05DV2_S12s_1 | Gnpat_Q05 | 0.56 | 0.2 | -0.29 | -0.03 | 1.49 | 0.76 | 1.13 | -2.47 | -0.72 | 0.35 | -1.05 | 0.39 | 0.51 | 1.63 | 1.44 | 1.1 | -0.2225 | -0.06 | 0.155 | 0.010837 | 0.88477 | 0.376482 | 0.996493 | 0.997988 | 0.994666 |
| Golga3_E9QP99_S1479s_1 | Golga3_E9 | 3.65 | 1.77 | 2.55 | -0.01 | 1.12 | 3.41 | 2.58 | 2.54 | 1.17 | 3.14 | 0.22 | 2.31 | 1.55 | 0.25 | 3.05 | 3.11 | 0.1175 | -0.3675 | 1.06 | 1 | 0.59603 | 0.75318 | 0.989223 | 0.997988 | 1 |
| Golga3_E9QP99_S285s_1 | Golga3_E9 | -0.38 | 0.38 | -0.26 | 0.56 | -0.1 | -0.07 | 0.29 | -0.39 | -0.08 | 0.02 | -1.19 | -1.63 | 0.95 | 0.79 | -0.8 | -0.08 | 0.4225 | -0.28 | 1.57E-16 | 0.758576 | 0.633361 | 0.105166 | 0.989223 | 0.997988 | 0.9971 |
| Golga3_E9QP99_S312s_1 | Golga3_E9 | -2.6 | -0.85 | -4.35 | -1.22 | -1.03 | -1.25 | -1.32 | -1.47 | -2.59 | -1.72 | -2.6 | -1.73 | -0.16 | -1.1 | -2.29 | -1.55 | -0.1425 | -0.795 | 0.14 | 0.24124 | 0.183051 | 0.898686 | 0.989223 | 0.997988 | 0.994666 |
| Golga3_E9QP99_T1391t_1 | Golga3_E9 | -1.97 | -1.4 | -0.73 | -1.12 | -1.92 | -2.33 | -1.26 | -0.74 | -2.31 | -1.6 | -0.68 | -0.94 | -0.91 | -0.65 | -1.15 | -1.08 | 0.9875 | 0.095 | 0.98 | 0.289414 | 0.540567 | 0.858623 | 0.989223 | 0.997988 | 0.994666 |
| GOLGA7B_Q9D428_S14s_1 | GOLGA7B_ | -0.69 | -0.25 | 0.89 | 0.57 | -0.47 | 0.01 | 0.18 | 0.58 | 0.23 | -0.2 | -2.36 | -0.16 | 1.08 | -0.7 | 0.66 | 0.76 | -0.2575 | -0.0775 | 0.3575 | 0.531132 | 0.892162 | 0.250702 | 0.996493 | 0.997988 | 0.994666 |
| GOLGA7B_Q9D428_S16s_1 | GOLGA7B_ | 0.09 | 0.27 | 0.49 | -0.3 | 0.77 | -0.39 | -0.9 | -0.43 | -0.03 | -0.1 | -2.1 | -0.79 | 2.04 | -1.08 | 0.29 | 0.01 | -0.055 | -0.7525 | 0.32 | 0.77074 | 0.339718 | 0.090444 | 0.989223 | 0.997988 | 0.9971 |
| Golim4_D3YVW2_S568s_1 | Golim4_D3 | -1.68 | -1.62 | -3.45 | -7.55 | -0.51 | -1.32 | -4.36 | -5.02 | -1.94 | -1.88 | -0.55 | -4.96 | -3.35 | -0.89 | -3.11 | -4.1 | -0.375 | -0.8925 | 0.1775 | 0.596804 | 0.614427 | 0.405295 | 0.989223 | 0.997988 | 0.995259 |
| Golph3_Q9CRA5_S35s_1 | Golph3_Q9 | -1.05 | -1.9 | -0.67 | -0.16 | -2.32 | -0.56 | -1.56 | -1.56 | 0.94 | -0.86 | 0.31 | -0.44 | 1.8 | -2.95 | -0.04 | -1.05 | 0.7725 | 1.2425 | 0.7125 | 0.678195 | 0.260801 | 0.093308 | 0.989223 | 0.997988 | 0.996159 |
| Gon4l_E9Q507_S1394s_1 | Gon4l_E9C | -1.78 | -1.52 | -0.83 | -0.82 | -0.96 | -1.88 | -1.33 | -1.38 | -0.27 | -1.97 | -0.68 | -1.03 | 1.26 | -1.24 | -0.44 | -1.21 | -0.555 | 0.9325 | 0.385 | 0.176736 | 0.643461 | 0.561241 | 0.990258 | 0.997988 | 0.994666 |
| Gorab_Q8BRM2_S335s_1 | Gorab_Q8E | 0.27 | 0.4 | 0.03 | 0.35 | -0.01 | 0.56 | -0.71 | -0.91 | 0.92 | 0.19 | -0.72 | -0.67 | 1.02 | -0.62 | -0.52 | 0.3 | -0.15 | 0.25 | 0.83 | 0.589408 | 0.152945 | 0.412104 | 0.989223 | 0.997988 | 0.995259 |
| Gorasp2_Q99JX3_S350s_1 | Gorasp2_Q | 0.57 | 0.63 | 0.53 | 0.61 | 0.5 | 0.37 | 1.06 | 0.83 | 0.19 | 0.71 | 0.25 | 0.57 | -0.15 | 0.57 | 0.38 | 0.62 | -0.53 | -0.3325 | -0.2175 | 0.400273 | 0.652259 | 0.501318 | 0.990358 | 0.997988 | 0.994666 |
| Gorasp2_Q99JX3_T154t_1 | Gorasp2_Q | -0.05 | -1.8 | -0.97 | -0.14 | -0.12 | 0 | -0.36 | -0.71 | -1.52 | -0.63 | -0.8 | -0.77 | -1.26 | -1.02 | -1.54 | -0.98 | 0.105 | -0.155 | -0.23 | 0.295209 | 0.30341 | 0.667913 | 0.989223 | 0.997988 | 0.994666 |
| Gorasp2_Q99JX3_T157t_1 | Gorasp2_Q | -0.72 | -1.9 | -3.42 | -1.96 | -0.97 | -2.18 | -1.75 | -1.81 | -1.81 | -1.34 | 0.15 | -2.91 | -3.09 | 0.14 | -2.85 | -1.95 | 0.4425 | -0.19 | -0.46 | 0.938526 | 0.561003 | 0.494314 | 0.989223 | 0.997988 | 0.999507 |
| Gpalpp1_Q69ZC8_S104s_1 | Gpalpp1_Q | -2.06 | -1.89 | -2.29 | -1.41 | -1.76 | -2.43 | -0.99 | -1.33 | -2.19 | -1.46 | -1.37 | -1.93 | -1.76 | -1.73 | -2.39 | -1.77 | 0.3225 | 0.5225 | 0.0625 | 1 | 0.436733 | 0.570519 | 0.989223 | 0.997988 | 1 |
| Gpatch8_A2A6A1_S1018s_1 | Gpatch8_A | 0.21 | 0.8 | -0.89 | -0.47 | -0.01 | -0.16 | -1.55 | -0.86 | -1.63 | -0.07 | -0.76 | -1.61 | 0.03 | -0.14 | 0.11 | -2.09 | 0.285 | 0.175 | 2.36E-16 | 0.467053 | 0.258347 | 0.088329 | 0.989223 | 0.997988 | 0.994666 |
| Gpatch8_A2A6A1_S1039s_1 | Gpatch8_A | 0.16 | -0.66 | -0.07 | -0.06 | -0.94 | -0.19 | -0.92 | 0.25 | 0.27 | -0.4 | 0.39 | -0.11 | 0.9 | 0.61 | -0.16 | -1.11 | -0.5575 | -0.93 | -0.435 | 0.641918 | 0.405142 | 0.513092 | 0.989223 | 0.997988 | 0.995259 |
| Gpatch8_A2A6A1_S1179s_1 | Gpatch8_A | 0.93 | -0.14 | -0.2 | 0.38 | -2.37 | 0.78 | -1.32 | -0.52 | 0.64 | -0.51 | 1.14 | 0.53 | 0.86 | -1.5 | 0.39 | -0.61 | -0.2925 | 0.195 | 0.2175 | 0.411377 | 0.108988 | 0.627546 | 0.989223 | 0.997988 | 0.994666 |
| Gpatch8_A2A6A1_S1179s_2 | Gpatch8_A | 0.24 | -0.31 | -0.26 | -0.83 | -2 | 0.31 | -1.64 | -0.58 | 1.09 | -0.96 | -0.09 | 0.02 | 0.39 | -1.93 | -0.19 | -0.41 | -1.1 | 0.2075 | -0.4575 | 0.632409 | 0.205335 | 0.50763 | 0.989223 | 0.997988 | 0.995259 |

| | | | | | | | | | | | | | | | | | | | | | | | | | | |
|---|---|---|---|---|---|---|---|---|---|---|---|---|---|---|---|---|---|---|---|---|---|---|---|---|---|---|
| Gpatch8_A2A6A1_S1185s_2 | Gpatch8_A | 0.24 | -0.31 | -0.26 | -0.83 | -2 | 0.31 | -1.64 | -0.58 | 1.09 | -0.96 | -0.09 | 0.02 | 0.39 | -1.93 | -0.19 | -0.41 | -0.6875 | 0.305 | -0.245 | 0.632409 | 0.205335 | 0.50763 | 0.989223 | 0.997988 | 0.995259 |
| Gpatch8_A2A6A1_S451s_1 | Gpatch8_A | 1.07 | 0.31 | -0.06 | -0.14 | -1.41 | 0.42 | -1.7 | -0.56 | 1.76 | -0.38 | -1.27 | -0.62 | 2.57 | -1.97 | -0.09 | -1.19 | -0.6875 | 0.305 | -0.245 | 0.608292 | 0.050595 | 0.518811 | 0.989223 | 0.997988 | 0.995259 |
| Gpatch8_A2A6A1_S648s_1 | Gpatch8_A | 0.38 | 0.44 | -0.58 | -0.36 | -1.44 | -0.7 | -2.27 | -0.61 | -0.71 | -0.86 | -0.76 | -0.54 | 0.28 | 0.05 | -0.44 | -1.13 | -1.1075 | -0.4225 | -0.465 | 0.498042 | 0.018321 | 0.045959 | 0.989223 | 0.997988 | 0.994666 |
| Gpatch8_A2A6A1_S735s_1 | Gpatch8_A | 0.41 | 0.34 | -0.14 | -0.25 | -0.76 | -0.29 | -0.61 | 0.03 | -0.3 | -0.39 | 0.32 | -0.3 | 0.82 | 0.23 | 0.16 | -0.61 | -1.225 | -0.6875 | -0.28 | 0.868134 | 0.099793 | 0.3744 | 0.989223 | 0.997988 | 0.999507 |
| Gpatch8_A2A6A1_S753s_1 | Gpatch8_A | 0.52 | 0.4 | 0.45 | 0.75 | -0.64 | -0.13 | 0.39 | 0.12 | -0.59 | -0.04 | -0.38 | 0.86 | 0.54 | -0.31 | 0.25 | 0.65 | -0.4975 | -0.2575 | 0.06 | 0.409207 | 0.051708 | 0.127087 | 0.989223 | 0.997988 | 0.994666 |
| Gpbp1_Q6NXH3_S114s_1 | Gpbp1_Q6 | -2.72 | -1.53 | -1.53 | -1.64 | -0.96 | -2.78 | -2.97 | -1.18 | 0.09 | -1.39 | -1.44 | -1.22 | -0.36 | -0.28 | -1.64 | -1.17 | -0.595 | -0.5675 | -0.2475 | 0.042523 | 0.828511 | 0.07927 | 0.995762 | 0.997988 | 0.994666 |
| Gpbp1l1_Q6NZP2_T353t_1 | Gpbp1l1_C | -0.46 | -1.99 | -0.94 | -0.62 | -1.94 | -1.2 | -0.61 | -0.25 | -1.02 | -1.72 | 0.29 | -0.78 | -0.44 | -2.44 | -0.37 | -0.58 | -0.1175 | 0.865 | 0.9925 | 0.93565 | 0.995733 | 0.701668 | 0.999616 | 0.997988 | 0.999507 |
| Gpd2_Q64521_S415s_1 | Gpd2_Q64 | 1.58 | 0.94 | 0.91 | 1.07 | -0.94 | 1.44 | -0.36 | -0.28 | 1.98 | 0.63 | -0.05 | 0.55 | 2.3 | -0.72 | 0.53 | -0.04 | 0.0025 | 0.195 | 0.045 | 0.329548 | 0.038815 | 0.438702 | 0.989223 | 0.997988 | 0.994666 |
| Gpd2_Q64521_T413t_1 | Gpd2_Q64 | -0.04 | 0.38 | 1.02 | 0.68 | -2.2 | 0.3 | -0.27 | -0.1 | 1.39 | -0.18 | -0.63 | 0.25 | 1.41 | -1.85 | 0.33 | 0.28 | -1.16 | -0.3475 | -0.6075 | 0.477347 | 0.072336 | 0.521198 | 0.989223 | 0.997988 | 0.994666 |
| Gphn_Q8BUV3_S262s_2 | Gphn_Q8B | 0.81 | 0.46 | 0.49 | 0.6 | 0.47 | 0.94 | 0.31 | 0.33 | 0.76 | 1.07 | 0.98 | -0.29 | 0.22 | -0.04 | 0.96 | 0.14 | -1.0775 | -0.3025 | -0.4675 | 0.374637 | 0.743254 | 0.906897 | 0.995762 | 0.997988 | 0.994666 |
| Gphn_Q8BUV3_S268s_2 | Gphn_Q8B | -1.01 | -0.97 | 0.06 | -0.06 | -0.75 | -0.86 | -0.21 | -0.81 | 0.08 | 0.93 | 0.64 | -1.09 | -0.69 | 0.4 | 0.44 | -1.09 | -0.0775 | 0.04 | -0.27 | 0.578757 | 0.627775 | 0.224369 | 0.989223 | 0.997988 | 0.995259 |
| Gphn_Q8BUV3_S270s_1 | Gphn_Q8B | 0.54 | 0.68 | 1.09 | 1.11 | 0.72 | 0.89 | 0.91 | 1.02 | 0.44 | 0.72 | 1.28 | 1.26 | 0.02 | 0.54 | 1 | 1.13 | -0.1625 | 0.635 | 0.26 | 0.584136 | 0.896727 | 0.812537 | 0.996493 | 0.997988 | 0.995259 |
| Gphn_Q8BUV3_S270s_2 | Gphn_Q8B | 0.17 | 0.32 | 0.82 | 1.06 | 0.01 | 0.49 | 0.68 | 1.36 | 0.65 | 1.11 | 2.56 | 0.83 | -0.53 | 0.22 | 0.89 | -0.03 | 0.03 | 0.07 | -0.1825 | 0.246769 | 0.903528 | 0.156771 | 0.996493 | 0.997988 | 0.994666 |
| Gphn_Q8BUV3_S280s_1 | Gphn_Q8B | -0.26 | -1.41 | -0.9 | -0.69 | -1.15 | 0.4 | 0.27 | -0.38 | 0.08 | -0.2 | -0.02 | -0.41 | 0.73 | 0.32 | -0.32 | -0.55 | 0.0425 | 0.695 | -0.455 | 0.049392 | 0.166009 | 0.046755 | 0.989223 | 0.997988 | 0.994666 |
| Gphn_Q8BUV3_S283s_1 | Gphn_Q8B | 0.72 | 0.17 | -0.69 | -1.67 | 0.39 | 1.01 | -0.14 | -1.22 | 0.31 | 0.87 | 0.39 | -0.92 | 0.8 | -0.18 | -0.81 | -1.33 | 0.6 | 0.6775 | 0.86 | 0.984144 | 0.550868 | 0.382412 | 0.989223 | 0.997988 | 1 |
| Gphn_Q8BUV3_S338s_1 | Gphn_Q8B | 6.05 | 0.76 | 1.05 | 1.6 | 0.94 | 7.77 | 7.46 | 0.2 | 1.08 | 2.69 | -0.35 | -1.76 | 1.57 | 0.31 | -1.13 | 2.33 | 0.3775 | 0.53 | -0.0125 | 0.22333 | 0.405594 | 0.177924 | 0.989223 | 0.997988 | 0.994666 |
| Gphn_Q8BUV3_T286t_1 | Gphn_Q8B | -0.1 | -0.47 | -0.91 | -0.54 | -0.34 | -0.07 | -0.61 | -0.44 | 0.17 | 0.15 | 0.16 | -0.37 | 0.38 | -0.6 | -0.65 | -0.61 | 1.7275 | -1.95 | -1.595 | 0.690002 | 0.584825 | 0.07454 | 0.989223 | 0.997988 | 0.996159 |
| Gpi_P06745_T109t_1 | Gpi_P0674 | 2.06 | 1.67 | 2.21 | 1.81 | 1.85 | 2.43 | 1.59 | 2.44 | 2.16 | 2.41 | 2.02 | 1.69 | 2.55 | 2.51 | 2.1 | 1.89 | 0.14 | 0.5325 | 0.135 | 0.262643 | 0.620238 | 0.612281 | 0.989223 | 0.997988 | 0.994666 |
| Gpkow_Q56A08_S348s_1 | Gpkow_Q5 | 3.7 | 0.49 | 2.08 | -0.36 | 0.72 | 3.16 | 1.92 | 1.69 | -0.75 | 3.38 | -2.59 | 2.06 | -1.48 | -2.22 | 3.01 | 2.54 | 0.14 | 0.1325 | 0.325 | 0.472184 | 0.659704 | 0.506412 | 0.990434 | 0.997988 | 0.994666 |
| Gpm6a_P35802_S256s_1 | Gpm6a_P3 | -3.28 | -1.98 | -3.74 | -3.44 | -1.32 | -3.07 | -2.99 | -4.67 | -2.3 | -2.76 | -3.81 | -3.38 | -0.74 | -1.52 | -3.48 | -3.93 | 0.395 | -0.9525 | -1.015 | 0.374019 | 0.888226 | 0.921877 | 0.996493 | 0.997988 | 0.994666 |
| Gpm6a_P35802_S267s_1 | Gpm6a_P3 | -2.41 | -1.98 | -1.61 | -1.4 | -1.48 | -2.42 | -1.34 | -1.54 | -1.51 | -2.64 | -1.83 | -1.67 | -1.16 | -0.5 | -1.36 | -1.4 | 0.0975 | 0.0475 | 0.6925 | 0.050744 | 0.648006 | 0.858459 | 0.99033 | 0.997988 | 0.994666 |
| Gpm6a_P35802_S267s_2 | Gpm6a_P3 | -0.78 | -1.17 | -0.3 | -0.47 | -0.35 | -0.99 | -1.04 | -0.71 | 0.22 | -0.4 | 0.57 | -0.61 | -1.29 | 0.36 | -0.6 | -1.26 | 0.155 | -0.0625 | 0.745 | 0.967294 | 0.744001 | 0.099707 | 0.995762 | 0.997988 | 0.999507 |
| Gpm6a_P35802_T268t_2 | Gpm6a_P3 | 0.38 | -0.71 | -0.3 | -0.47 | 0.19 | 0.1 | -1.04 | -0.71 | 1.16 | 0.71 | 0.57 | -0.61 | -0.59 | 1.21 | -0.6 | -1.26 | -0.0925 | 0.625 | -0.0175 | 0.948195 | 0.810529 | 0.116171 | 0.995762 | 0.997988 | 0.999507 |
| Gpm6b_P35803_S16s_1 | Gpm6b_P3 | 0.01 | -0.2 | -0.87 | -0.99 | -0.93 | 0.04 | -1.08 | -0.64 | 0.34 | 0.07 | -0.23 | -0.27 | 0.89 | -0.49 | -0.56 | -0.7 | -0.09 | 0.7325 | -0.035 | 0.501532 | 0.691201 | 0.145651 | 0.994893 | 0.997988 | 0.994666 |
| Gpm6b_P35803_S278s_1 | Gpm6b_P3 | 0.41 | 0.32 | 0.94 | 0.73 | 0.4 | 0.23 | 0.62 | 0.71 | -0.02 | 0.21 | 0.32 | 0.79 | -0.15 | 0.29 | 0.65 | 0.7 | -0.14 | 0.49 | 0.2975 | 0.459236 | 0.652108 | 0.333568 | 0.990358 | 0.997988 | 0.994666 |
| Gpm6b_P35803_S280s_1 | Gpm6b_P3 | 1.94 | 1.92 | 2.06 | 1.89 | 1.72 | 1.97 | 1.74 | 1.96 | 1.28 | 1.89 | 1.11 | 2.13 | 0.78 | 0.16 | 1.68 | 1.83 | -0.11 | -0.275 | -0.2275 | 0.059808 | 0.602737 | 0.247309 | 0.989223 | 0.997988 | 0.994666 |
| Gpm6b_P35803_T13t_1 | Gpm6b_P3 | -1.04 | -0.66 | -0.92 | -1.82 | -0.41 | 0.69 | -1.97 | -2.6 | 1.59 | -0.29 | -2.2 | -2.39 | 1.16 | 0.99 | -2.01 | -1.17 | -0.105 | -0.35 | -0.84 | 0.264233 | 0.95688 | 0.738487 | 0.999187 | 0.997988 | 0.994666 |
| Gpm6b_P35803_T288t_1 | Gpm6b_P3 | 2.16 | 1.79 | 2.1 | 1.91 | 1.55 | 2.22 | 1.3 | 1.88 | 1.54 | 2.01 | 1.48 | 2.07 | 0.68 | 1.5 | 1.5 | 1.56 | 0.0375 | 0.2875 | 0.8525 | 0.041201 | 0.350517 | 0.400503 | 0.989223 | 0.997988 | 0.994666 |
| Gpr101_Q80T62_S241s_1 | Gpr101_Q8 | 0.84 | 1.02 | -0.25 | 1.19 | 0.91 | 0.2 | 1.11 | 1.02 | 0.93 | 0.22 | 0.43 | 0.45 | 0.74 | 0.95 | 1.87 | 1.4 | -0.2525 | -0.215 | -0.68 | 0.210628 | 0.770874 | 0.602296 | 0.995762 | 0.997988 | 0.994666 |
| Gpr101_Q80T62_S378s_1 | Gpr101_Q8 | 0.06 | 0.1 | -0.13 | 0.27 | 0.13 | 0.09 | 0.65 | 0.19 | 0.58 | 0.4 | 1.67 | -0.44 | 1.31 | -0.14 | 1.19 | 0.93 | 0.11 | -0.1925 | 0.54 | 0.06368 | 0.416637 | 0.283249 | 0.989223 | 0.997988 | 0.994666 |
| Gpr101_Q80T62_T509t_1 | Gpr101_Q8 | 0.27 | 0.41 | -0.2 | 0.89 | -0.55 | 0.02 | 0.62 | 0.75 | 1.19 | 0.69 | 1.19 | 1 | 1.37 | -0.22 | 1.36 | 1.02 | 0.19 | 0.4775 | 0.7475 | 0.230845 | 0.720139 | 0.044236 | 0.995437 | 0.997988 | 0.994666 |
| Gpr137c_E9Q343_S351s_1 | Gpr137c_E | 0 | 0.17 | 0.76 | 0.32 | -0.25 | -0.47 | 0.03 | -0.02 | 0.54 | -0.3 | 0.87 | 0.03 | 0.3 | -0.03 | 0.7 | -0.22 | -0.1325 | 0.675 | 0.54 | 0.689803 | 0.078121 | 0.933747 | 0.989223 | 0.997988 | 0.996159 |
| Gpr153_Q8K0Z9_S229s_1 | Gpr153_Q8 | 1.01 | 1.77 | 1.09 | 0.79 | 0.98 | 1.95 | -0.16 | 0.05 | 2.59 | 1.34 | 0.43 | 0.34 | 2.61 | 1.13 | -0.01 | 0.55 | -0.49 | -0.0275 | -0.125 | 0.864428 | 0.355429 | 0.984825 | 0.989223 | 0.997988 | 0.999507 |
| Gpr155_A2AWR3_S835s_1 | Gpr155_A2 | -0.34 | -0.04 | 0.41 | 0.33 | -0.62 | 0.2 | -0.31 | -0.09 | 1.71 | -0.27 | -0.45 | 0.12 | 1.87 | -1.38 | 0.34 | 0.38 | -0.46 | 0.01 | -0.095 | 0.733714 | 0.30789 | 0.704435 | 0.989223 | 0.997988 | 0.9971 |
| Gpr155_A2AWR3_S835s_2 | Gpr155_A2 | -0.33 | 0.54 | -0.09 | -0.23 | -1.32 | 0.69 | -0.93 | -1.15 | 1.97 | 0.05 | -0.79 | -0.17 | 1.05 | -0.55 | -0.67 | 0.13 | -0.295 | 0.1875 | 0.2125 | 0.967794 | 0.185477 | 0.614602 | 0.989223 | 0.997988 | 0.999507 |
| Gpr155_A2AWR3_S839s_1 | Gpr155_A2 | 0.53 | 0.26 | 0.05 | 0.18 | -0.27 | 0.78 | -0.47 | -0.66 | 1.56 | 0.67 | -0.59 | -0.01 | 1.7 | -1 | -0.02 | 0.21 | -0.65 | 0.2925 | 0.0175 | 0.95101 | 0.249417 | 0.737847 | 0.989223 | 0.997988 | 0.999507 |
| Gpr155_A2AWR3_S839s_2 | Gpr155_A2 | -0.33 | 0.54 | -0.09 | -0.23 | -1.32 | 0.69 | -0.93 | -1.15 | 1.97 | 0.05 | -0.79 | -0.17 | 1.05 | -0.55 | -0.67 | 0.13 | -0.41 | 0.1525 | -0.0325 | 0.967794 | 0.185477 | 0.614602 | 0.989223 | 0.997988 | 0.999507 |
| Gpr155_A2AWR3_S860s_1 | Gpr155_A2 | -0.97 | -0.21 | 0.98 | 0.71 | -0.07 | -0.89 | 0.34 | -0.96 | -0.11 | -0.16 | 0.8 | -0.02 | 0.32 | 0.1 | 0.22 | 1.32 | -0.65 | 0.2925 | 0.0175 | 0.473321 | 0.311592 | 1 | 0.989223 | 1 | 0.994666 |
| Gpr158_Q8C419_S1059s_1 | Gpr158_Q8 | -0.12 | -0.56 | -1.14 | -0.36 | 0.57 | -0.01 | -0.01 | -0.81 | -0.88 | -0.61 | -1.37 | -0.74 | -2.14 | -1.17 | -0.84 | 0.27 | -0.5225 | 9.81E-17 | 0.3625 | 0.414022 | 0.20008 | 0.267653 | 0.989223 | 0.997988 | 0.994666 |
| Gpr158_Q8C419_S239s_1 | Gpr158_Q8 | 0.99 | -1.62 | -2.03 | -2.36 | -0.75 | 2.05 | -0.78 | -0.39 | 3.71 | -0.21 | -0.95 | -1 | 2.75 | -1.32 | 2.56 | -2.2 | 0.48 | -0.355 | -0.425 | 0.207864 | 0.170075 | 0.190686 | 0.989223 | 0.997988 | 0.994666 |
| Gpr158_Q8C419_S249s_1 | Gpr158_Q8 | 2.04 | 0.21 | -0.49 | 1.61 | 1.36 | 1.84 | 0.61 | 1.71 | 1.04 | 0.97 | 0.4 | 1.26 | -1.33 | 1.65 | 1.14 | 0.68 | 1.2875 | 1.6425 | 1.7025 | 0.694259 | 0.369467 | 0.895423 | 0.989223 | 0.997988 | 0.996159 |
| Gpr158_Q8C419_S692s_1 | Gpr158_Q8 | -1.26 | -1.02 | -1.8 | -0.78 | -0.63 | -1.63 | 0.16 | -0.84 | -1.37 | -1.43 | -2.34 | -0.9 | -2.13 | -0.36 | -1.52 | -0.61 | 0.5375 | 0.075 | -0.3075 | 0.894169 | 0.258109 | 0.441885 | 0.989223 | 0.997988 | 0.999507 |
| Gpr158_Q8C419_S819s_1 | Gpr158_Q8 | 2.35 | -1.67 | 0.01 | 0.47 | -2.24 | 1.9 | 0.44 | 1.53 | 0.08 | -1.16 | 2.64 | -0.27 | -1.03 | -1.87 | 0.02 | -0.09 | 0.48 | -0.295 | 0.06 | 0.230907 | 0.913028 | 0.974649 | 0.997335 | 0.997988 | 0.994666 |
| Gpr158_Q8C419_S881s_1 | Gpr158_Q8 | -0.82 | -2.24 | -1.72 | -1.82 | -1.49 | -0.35 | -1.9 | -1.09 | -0.39 | -1.96 | -0.91 | -1.17 | -0.51 | -1.38 | -1.98 | -1.38 | 0.1175 | 0.0325 | -1.0325 | 0.432541 | 0.309049 | 0.228472 | 0.989223 | 0.997988 | 0.994666 |
| Gpr158_Q8C419_S902s_1 | Gpr158_Q8 | -0.55 | -1.92 | -1.63 | -2.11 | -1.06 | -1.35 | -1.63 | -1.72 | -2.09 | -0.44 | 1.58 | -1.32 | -0.28 | -0.24 | -2.54 | -2.04 | 0.4425 | 0.5425 | 0.3375 | 0.658769 | 0.763075 | 0.223629 | 0.995762 | 0.997988 | 0.995259 |
| Gpr158_Q8C419_S944s_1 | Gpr158_Q8 | 0.35 | -0.44 | -0.28 | 0.1 | -0.01 | 0.51 | 0.42 | 0.38 | -0.26 | 0.3 | -0.21 | 0.09 | -0.36 | 0.32 | -0.17 | 0.31 | 0.1125 | 0.985 | 0.2775 | 0.762566 | 0.156793 | 0.863596 | 0.989223 | 0.997988 | 0.9971 |
| Gpr158_Q8C419_S946s_1 | Gpr158_Q8 | -1.14 | -0.51 | -0.89 | -0.32 | 0.14 | -0.08 | 0.36 | -0.11 | -1.81 | -0.14 | 1.18 | 0.09 | 0.43 | -1.04 | 0.12 | 0.63 | 0.3925 | 0.0475 | 0.0925 | 0.097198 | 0.012663 | 0.368577 | 0.989223 | 0.997988 | 0.994666 |
| Gpr158_Q8C419_S976s_1 | Gpr158_Q8 | 1.36 | 0.24 | 0.33 | 0.73 | 1.46 | 1.22 | 1.33 | 0.43 | 0.64 | 0.72 | 0.92 | 0.69 | 0.27 | 1.13 | 0.63 | 0.71 | 0.7925 | 0.545 | 0.75 | 0.953532 | 0.220706 | 0.7971 | 0.989223 | 0.997988 | 0.999507 |
| Gpr162_Q3UN16_S255s_1 | Gpr162_Q3 | 0.8 | -0.39 | 2.51 | 2.6 | 0.11 | 0.16 | 0.13 | 1.78 | 0.55 | -0.81 | 0.54 | 2.09 | 1.71 | -0.89 | 1.85 | 1.97 | 0.445 | 0.0775 | 0.02 | 0.801149 | 0.269398 | 0.355654 | 0.989223 | 0.997988 | 0.999507 |
| Gpr162_Q3UN16_S256s_1 | Gpr162_Q3 | 0.92 | 0.75 | 1.45 | 1.25 | 0.36 | 1.63 | 0.47 | 1 | 2.35 | 0.73 | 0.57 | 1.44 | 2.35 | 0.23 | 1.12 | 1.04 | -0.835 | -0.7875 | -0.22 | 0.837847 | 0.505981 | 0.673434 | 0.989223 | 0.997988 | 0.999507 |
| Gpr162_Q3UN16_S413s_1 | Gpr162_Q3 | 0.45 | -0.43 | -0.29 | 0.17 | 0.3 | 0.62 | 0.16 | 0.38 | -0.3 | -0.59 | 0.3 | -0.32 | -0.72 | 0.42 | -0.56 | -0.34 | -0.2275 | 0.18 | 0.0925 | 0.443125 | 0.169506 | 0.519575 | 0.989223 | 0.997988 | 0.994666 |
| Gpr162_Q3UN16_S534s_1 | Gpr162_Q3 | 1.59 | -2.36 | -1.32 | -0.37 | -1.38 | 2.12 | 0.18 | 1.06 | -2 | -2.27 | 0.61 | -0.6 | -5.41 | -0.65 | -2.33 | -2.27 | 0.39 | -0.2025 | -0.275 | 0.094447 | 0.269059 | 0.637346 | 0.989223 | 0.997988 | 0.994666 |
| Gpr162_Q3UN16_S542s_1 | Gpr162_Q3 | 1.34 | 0.67 | 0.71 | 1.77 | 0.73 | 1.12 | 1.37 | 1.42 | 0.34 | 0.24 | -0.48 | 1.18 | -1.1 | -0.32 | 0.59 | 1.77 | 1.11 | -0.45 | -2.05 | 0.169802 | 0.907237 | 0.083219 | 0.997038 | 0.997988 | 0.994666 |
| Gpr162_Q3UN16_S561s_1 | Gpr162_Q3 | 1.8 | 0.45 | 0.42 | 0.08 | -0.09 | 2.52 | -0.58 | 0.3 | 1.3 | 1.1 | 0.91 | 0.27 | 1.49 | -0.04 | -0.03 | 0.57 | 0.0375 | -0.8025 | -0.8875 | 0.702731 | 0.828277 | 0.630882 | 0.995762 | 0.997988 | 0.996159 |
| Gpr162_Q3UN16_S565s_1 | Gpr162_Q3 | 0.96 | 1.55 | 0.22 | 0.42 | 0.73 | 0.9 | -0.48 | -0.2 | 0.8 | 0.7 | 0.59 | 0.38 | 2.28 | 0.79 | 0.47 | 0.54 | -0.15 | 0.2075 | -0.19 | 0.639164 | 0.218601 | 0.613282 | 0.989223 | 0.997988 | 0.995259 |
| Gpr162_Q3UN16_T587t_1 | Gpr162_Q3 | -0.89 | -0.21 | -0.43 | -0.46 | -0.19 | -1.01 | -0.88 | -1.82 | -0.13 | -0.73 | -1.1 | -0.56 | -0.9 | 0.06 | -1.09 | -1.1 | -0.55 | -0.17 | 0.2325 | 0.456669 | 0.207003 | 0.650652 | 0.989223 | 0.997988 | 0.994666 |
| Gpr17_Q6NS65_S323s_1 | Gpr17_Q6 | 0.36 | 0.86 | 0.61 | 0.5 | 0.82 | 0.49 | 0.83 | 0.87 | 0.5 | 0.7 | 0.78 | 0.44 | 0.51 | 0.86 | 0.67 | 0.82 | -0.4775 | -0.1325 | -0.26 | 0.596892 | 0.451399 | 0.922018 | 0.989223 | 0.997988 | 0.995259 |
| Gpr17_Q6NS65_S332s_1 | Gpr17_Q6 | 1.03 | 0.58 | 0.86 | 0.79 | -0.58 | 1.87 | -0.02 | 0.78 | 2.28 | 1.61 | 0.99 | 0.65 | 2.76 | 0.55 | 0.99 | 0.73 | 0.17 | 0.0225 | 0.1325 | 0.378396 | 0.545345 | 0.156514 | 0.989223 | 0.997988 | 0.994666 |
| Gpr26_Q8BZA7_S329s_1 | Gpr26_Q8 | 1.49 | 1.58 | 0.6 | 0.37 | 1.78 | 2.89 | 0.68 | 0.36 | 1.57 | 2.23 | -0.41 | 0.63 | 1.36 | 0.63 | 0.39 | 0.19 | -0.3025 | 0.5675 | 0.4425 | 0.374145 | 0.480983 | 0.993314 | 0.989223 | 0.997988 | 0.994666 |
| Gpr34_Q9R1K6_S341s_1 | Gpr34_Q9 | -1.02 | -1.56 | -1 | -1.61 | -1.84 | -0.34 | -1.52 | -1.02 | 0.6 | -1.57 | -1.55 | -1.58 | 0.75 | -1.93 | -1.64 | -1.87 | 0.4175 | -0.005 | -0.3675 | 0.836627 | 0.747111 | 0.608595 | 0.995762 | 0.997988 | 0.999507 |
| Gpr37l1_Q99JG2_S471s_1 | Gpr37l1_Q | -1.01 | -0.43 | -0.67 | -0.87 | -0.64 | -0.85 | -0.59 | -1.03 | -1 | -0.5 | -0.88 | -1.09 | -0.66 | -0.67 | -1.29 | -1.11 | 0.1175 | 0.2725 | 0.125 | 0.508946 | 0.889254 | 0.63044 | 0.996493 | 0.997988 | 0.994666 |
| Gpr56_Q8K209_S681s_1 | Gpr56_Q8 | 0.54 | 0.33 | 0.71 | 1.2 | 0.75 | 0.82 | 0.81 | 1.23 | 0.61 | 0.78 | 1.52 | 1.22 | 1.17 | 1.17 | 1.48 | 1.31 | -0.0325 | -0.1225 | -0.1875 | 0.058546 | 0.437584 | 0.294271 | 0.989223 | 0.997988 | 0.994666 |
| Gpr61_Q8C010_S258s_1 | Gpr61_Q8 | -0.47 | -0.16 | -0.58 | -0.19 | -1.01 | 0.11 | -0.62 | -0.87 | 1.36 | -0.55 | -0.87 | -1.09 | 1.48 | -0.97 | 0.48 | 0.11 | 0.2075 | 0.3375 | 0.5875 | 0.222697 | 0.412637 | 0.906225 | 0.989223 | 0.997988 | 0.994666 |
| Gprc5b_D2DFA9_S337s_1 | Gprc5b_D2 | 1.25 | 0.88 | 0.51 | 0.55 | 1.2 | 0.18 | 0.49 | 0.43 | 0.83 | 0.73 | 0.72 | 0.76 | -0.34 | 1.01 | 0.38 | 0.62 | -0.2475 | 0.0625 | 0.625 | 0.302997 | 0.467799 | 0.88044 | 0.989223 | 0.997988 | 0.994666 |
| Gprin1_Q3UNH4_S166s_1 | Gprin1_Q3 | -0.05 | 0.07 | -0.1 | -0.49 | 0.45 | -0.46 | 0.28 | 0.04 | -0.21 | -0.14 | 1.1 | 0.5 | 0.02 | 0.4 | 0.89 | 0.31 | -0.2225 | -0.0375 | -0.38 | 0.082768 | 0.427839 | 0.211235 | 0.989223 | 0.997988 | 0.994666 |
| Gprin1_Q3UNH4_S182s_1 | Gprin1_Q3 | 0.51 | -0.22 | -0.13 | 0.21 | 0.74 | 0.64 | 1.12 | 0.66 | -0.18 | 0.42 | -0.03 | 0.5 | -0.78 | 0.75 | 0.08 | 0.25 | 0.22 | 0.455 | 0.5475 | 0.962888 | 0.020225 | 0.765654 | 0.989223 | 0.997988 | 0.999507 |
| Gprin1_Q3UNH4_S190s_1 | Gprin1_Q3 | 0.95 | 0.45 | 0.32 | 0.65 | 1.17 | 0.06 | 0.99 | 0.94 | -0.52 | 0.46 | 0.3 | 0.49 | -0.53 | 0.46 | 0.36 | 0.18 | 0.6975 | 0.085 | -0.0175 | 0.153888 | 0.522506 | 0.206319 | 0.989223 | 0.997988 | 0.994666 |
| Gprin1_Q3UNH4_S219s_1 | Gprin1_Q3 | 1.04 | 1.15 | 0.69 | 0.76 | 1.19 | 0.69 | 0.82 | 0.81 | 0.96 | 1.09 | 0.66 | 0.58 | 0.68 | 1.46 | 0.59 | 0.51 | 0.1975 | -0.41 | -0.475 | 0.742691 | 0.887554 | 0.720859 | 0.996493 | 0.997988 | 0.9971 |
| Gprin1_Q3UNH4_S369s_1 | Gprin1_Q3 | 0.63 | -0.12 | -0.46 | 0.45 | 0.82 | 0.77 | 0.95 | 0.52 | -0.48 | 0.05 | 0.69 | 0.78 | -0.97 | 0.62 | 0.89 | 1.13 | -0.0325 | -0.0875 | -0.1 | 0.566831 | 0.052766 | 0.729505 | 0.989223 | 0.997988 | 0.995259 |
| Gprin1_Q3UNH4_S433s_1 | Gprin1_Q3 | 0.37 | 0.25 | -0.15 | 0.61 | -1.01 | 0.58 | 0.87 | 0.12 | 1.67 | 0.77 | -0.06 | 0.47 | 1.57 | -1.06 | 0.18 | -0.32 | 0.64 | 0.135 | 0.2925 | 0.741488 | 0.757455 | 0.279334 | 0.995762 | 0.997988 | 0.9971 |
| Gprin1_Q3UNH4_S446s_1 | Gprin1_Q3 | 0.57 | -0.62 | -0.77 | -0.2 | 0.68 | 0.77 | 0.44 | 0.28 | -0.77 | -0.11 | -1.24 | -0.61 | -0.94 | 0.36 | 0.01 | -0.46 | -0.13 | 0.4425 | -0.1775 | 0.995171 | 0.035201 | 0.281429 | 0.989223 | 0.997988 | 1 |
| Gprin1_Q3UNH4_S468s_1 | Gprin1_Q3 | 1.49 | 0.21 | 1.28 | 1.44 | 0.63 | 2.44 | 0.99 | 1.86 | 1.29 | 1.2 | 1.04 | 1.81 | 0.72 | 0.77 | 0.66 | 0.98 | 0.7975 | -0.4275 | -0.0025 | 0.358897 | 0.434827 | 0.524455 | 0.989223 | 0.997988 | 0.994666 |
| Gprin1_Q3UNH4_S495s_1 | Gprin1_Q3 | 0.29 | 0.7 | 0.17 | 0.54 | 0.65 | 0.35 | 0.72 | 0.31 | 0.48 | 0.98 | 0.35 | 0.48 | 0.59 | 1.03 | 0.51 | 0.37 | 0.375 | 0.23 | -0.3225 | 0.468928 | 0.722756 | 0.565762 | 0.995437 | 0.997988 | 0.994666 |
| Gprin1_Q3UNH4_S519s_1 | Gprin1_Q3 | 0.4 | -0.34 | -0.37 | 0.45 | 0.43 | 0.23 | 1.08 | 0.79 | -0.45 | -0.27 | 0.28 | 0.28 | -0.71 | 0.55 | 0.1 | 0.05 | 0.0825 | 0.1475 | 0.2 | 0.918465 | 0.080994 | 0.815908 | 0.989223 | 0.997988 | 0.999507 |
| Gprin1_Q3UNH4_S53s_1 | Gprin1_Q3 | 2.18 | 0.94 | 0.2 | 1.07 | 0.81 | 2.29 | 0.65 | 1.2 | 1.68 | 1.8 | 1.14 | 0.36 | 0.07 | 1.73 | 0.47 | 0.56 | 0.5975 | -0.075 | -0.0375 | 0.452155 | 0.781442 | 0.765778 | 0.995762 | 0.997988 | 0.994666 |
| Gprin1_Q3UNH4_S576s_1 | Gprin1_Q3 | 0.81 | 0.38 | 0.67 | 1.21 | 1.19 | 0.92 | 1.9 | 1.39 | 0.16 | 0.44 | 0.86 | 1.12 | 0.17 | 1.04 | 0.99 | 0.7 | 0.14 | 0.1475 | -0.39 | 0.892534 | 0.073963 | 0.69316 | 0.989223 | 0.997988 | 0.999507 |
| Gprin1_Q3UNH4_S57s_1 | Gprin1_Q3 | 1.91 | 0.76 | 0.78 | 0.28 | 0.78 | 1.17 | -0.58 | 0.4 | 0.87 | 2.03 | 0.19 | 0.09 | 0.33 | 1.57 | 0.58 | 0.52 | 0.5825 | -0.1225 | -0.0425 | 0.677021 | 0.31369 | 0.794339 | 0.989223 | 0.997988 | 0.996159 |
| Gprin1_Q3UNH4_S618s_2 | Gprin1_Q3 | -0.23 | 0.15 | -0.2 | -1.08 | -1.18 | 1 | -0.87 | -0.52 | 2.04 | -0.53 | -0.84 | -0.14 | 1.11 | -0.8 | 0.55 | -0.24 | -0.49 | -0.1375 | -0.1825 | 0.312079 | 0.916808 | 0.46634 | 0.997335 | 0.997988 | 0.994666 |
| Gprin1_Q3UNH4_S620s_1 | Gprin1_Q3 | 0.71 | -1.32 | -0.3 | -0.39 | -0.11 | 1.63 | -0.76 | 0.18 | 2.17 | -0.32 | -1.4 | -0.09 | 1.5 | -0.84 | 0.49 | -0.11 | -0.0525 | 0.4725 | 0.495 | 0.335047 | 0.351189 | 0.591515 | 0.989223 | 0.997988 | 0.994666 |
| Gprin1_Q3UNH4_S620s_2 | Gprin1_Q3 | -0.31 | -0.26 | -0.68 | -1.61 | -1.37 | 0.95 | -0.94 | -0.44 | 1.95 | -0.06 | -0.41 | 0.17 | 1.11 | -1.13 | 0.46 | -0.17 | 0.56 | 0.415 | 0.585 | 0.164839 | 0.624093 | 0.071612 | 0.989223 | 0.997988 | 0.994666 |
| Gprin1_Q3UNH4_S622s_1 | Gprin1_Q3 | 0.33 | -0.91 | 0.05 | -0.15 | -0.45 | 1.22 | -0.49 | -0.19 | 1.8 | -0.01 | -1.02 | -0.49 | 1.46 | -1.12 | -0.01 | 0.01 | 0.265 | 1.1275 | 0.7825 | 0.643849 | 0.671447 | 0.695197 | 0.992876 | 0.997988 | 0.995259 |
| Gprin1_Q3UNH4_S622s_2 | Gprin1_Q3 | -0.27 | -0.06 | -0.46 | -1.37 | -1.28 | 0.97 | -0.91 | -0.47 | 1.99 | -0.27 | -0.59 | 0.04 | 1.11 | -0.97 | 0.5 | -0.2 | 0.1925 | 0.24 | 0.255 | 0.217065 | 0.821317 | 0.184525 | 0.995762 | 0.997988 | 0.994666 |
| Gprin1_Q3UNH4_S649s_1 | Gprin1_Q3 | 1.08 | 0.86 | 0.84 | 0.77 | 1.14 | 1.23 | 0.95 | 0.71 | 1.19 | 1.21 | 0.02 | 1.66 | 1.11 | 0.55 | 1.54 | 0.96 | 0.1175 | 0.8325 | 0.65 | 0.597868 | 0.588701 | 0.718251 | 0.989223 | 0.997988 | 0.995259 |
| Gprin1_Q3UNH4_S668s_1 | Gprin1_Q3 | -1.1 | -0.76 | 0.47 | 0.7 | -0.29 | -1.98 | 0.81 | 0.67 | -1.7 | -0.68 | 0.36 | 1.16 | -0.98 | -0.22 | 0.7 | 1.01 | 0.12 | 0.1325 | 0.1525 | 0.608714 | 0.971065 | 0.950884 | 0.999187 | 0.997988 | 0.995259 |
| Gprin1_Q3UNH4_S675s_1 | Gprin1_Q3 | 1.54 | -0.3 | 0.3 | 1.53 | 0.13 | 0.84 | 0.93 | 1.75 | -0.2 | 0.52 | 0.72 | 1.5 | -0.08 | 0.95 | 0.5 | 1.2 | -0.025 | -0.0425 | 0.3 | 0.805665 | 0.779268 | 0.805308 | 0.995762 | 0.997988 | 0.999507 |
| Gprin1_Q3UNH4_S691s_1 | Gprin1_Q3 | 0.66 | 0.37 | 0.51 | 0.96 | 1 | 0.63 | 0.78 | 0.87 | 0.38 | 0.69 | 0.79 | 0.45 | 0.61 | 1.12 | 0.55 | 0.42 | 0.145 | -0.1325 | -0.125 | 0.85754 | 0.399978 | 0.844489 | 0.989223 | 0.997988 | 0.999507 |
| Gprin1_Q3UNH4_S714s_1 | Gprin1_Q3 | 0.16 | 0.74 | 0.3 | 0.08 | 1.06 | -0.11 | 0.36 | -0.21 | -0.51 | -0.09 | -0.47 | 0.11 | -0.37 | 1.18 | 0.21 | 0.08 | 0.195 | -0.0475 | 0.05 | 0.904491 | 0.892668 | 0.062103 | 0.996493 | 0.997988 | 0.999507 |
| Gprin1_Q3UNH4_S764s_1 | Gprin1_Q3 | 0.82 | -0.89 | -0.26 | 0.53 | 0.98 | 1.12 | 0.69 | 1.04 | -0.17 | -0.44 | 0.26 | 0.34 | -0.21 | 0.33 | -0.36 | -0.54 | -0.045 | -0.56 | -0.045 | 0.569094 | 0.039541 | 0.900516 | 0.989223 | 0.997988 | 0.995259 |
| Gprin1_Q3UNH4_S764s_2 | Gprin1_Q3 | 0.98 | -1.27 | -0.31 | 0.13 | 1.4 | 0.95 | 0.6 | 0.77 | 0.3 | -0.44 | 0.24 | 0.09 | -1.44 | 0.27 | -0.61 | 0.12 | 0.9075 | -0.0525 | -0.245 | 0.600019 | 0.045469 | 0.728168 | 0.989223 | 0.997988 | 0.995259 |
| Gprin1_Q3UNH4_S771s_1 | Gprin1_Q3 | 0.24 | 0.98 | 0.69 | 0.36 | 0.87 | -0.15 | 0.5 | 0.28 | 0.43 | 0.84 | 0.67 | 0.39 | 1.42 | 1.48 | 0.78 | 0.46 | 1.0475 | 0.165 | -0.2975 | 0.187177 | 0.521655 | 0.954348 | 0.989223 | 0.997988 | 0.994666 |
| Gprin1_Q3UNH4_S771s_2 | Gprin1_Q3 | 0.98 | -1.27 | -0.31 | 0.13 | 1.4 | 0.95 | 0.6 | 0.77 | 0.3 | -0.44 | 0.24 | 0.09 | -1.44 | 0.27 | -0.61 | 0.12 | -0.1925 | 0.015 | 0.4675 | 0.600019 | 0.045469 | 0.728168 | 0.989223 | 0.997988 | 0.995259 |
| Gprin1_Q3UNH4_S817s_1 | Gprin1_Q3 | -0.18 | -1.09 | -1.56 | -0.41 | 0.58 | 0.02 | -0.22 | -0.81 | -0.82 | -1.16 | -0.11 | -0.44 | -1.8 | 0.31 | -0.85 | -0.92 | 1.0475 | 0.165 | -0.2975 | 0.992099 | 0.111164 | 0.650985 | 0.989223 | 0.997988 | 1 |
| Gprin1_Q3UNH4_S917s_1 | Gprin1_Q3 | 0.06 | -2.22 | -1.94 | -0.78 | -0.65 | 0.48 | -0.16 | 0.58 | -1.09 | -1.14 | 0.78 | -0.94 | -0.84 | 0.76 | -0.57 | -0.52 | 0.7025 | 0.1775 | -0.005 | 0.137557 | 0.038837 | 0.342262 | 0.989223 | 0.997988 | 0.994666 |
| Gprin1_Q3UNH4_T256t_1 | Gprin1_Q3 | 0.71 | 0.58 | 0.9 | 1.28 | 0.66 | 1.19 | 1.02 | 1.39 | 1.93 | 0.77 | 0.98 | 1.15 | 1.51 | 0.28 | 1.43 | 1.24 | 1.2825 | 0.6225 | 0.9275 | 0.48711 | 0.460673 | 0.307512 | 0.989223 | 0.997988 | 0.994666 |
| Gprin1_Q3UNH4_T795t_1 | Gprin1_Q3 | 0.33 | 0.61 | 0.41 | 0.42 | 0.76 | 0.3 | 0.87 | 0.47 | -0.08 | 0.3 | -0.2 | 0.45 | 0.15 | 0.72 | 0.26 | 0.47 | 0.1975 | 0.34 | 0.2475 | 0.865671 | 0.489472 | 0.205909 | 0.989223 | 0.997988 | 0.999507 |
| Gprin3_Q8BWS5_S327s_1 | Gprin3_Q8 | 0.7 | 0.62 | 0.68 | 0.8 | 1.13 | 0.57 | 1.03 | 0.51 | 1.03 | 1.44 | 0.25 | 0.66 | 0.92 | 1.55 | 0.57 | 0.58 | 0.1575 | -0.325 | -0.0425 | 0.495242 | 0.639665 | 0.628479 | 0.989223 | 0.997988 | 0.994666 |
| Gprin3_Q8BWS5_S572s_1 | Gprin3_Q8 | 1.51 | -0.29 | -0.23 | -1.58 | 0.84 | -0.09 | -0.24 | -0.23 | -0.21 | -0.58 | 3.05 | -1.33 | 0.29 | 1.36 | -2.36 | -0.29 | 0.11 | 0.145 | 0.205 | 0.907504 | 0.722713 | 0.710463 | 0.995437 | 0.997988 | 0.999507 |
| Gps1_G3UXW9_S275s_1 | Gps1_G3U | 0.6 | 1.21 | 0.86 | 0.12 | 0.91 | 0.92 | 0.63 | -0.21 | 0.77 | 0.63 | 0.45 | 0.15 | 0.83 | 0.98 | 0.49 | 0.16 | 0.2175 | 0.38 | -0.1025 | 0.804478 | 0.702691 | 0.519848 | 0.995437 | 0.997988 | 0.999507 |
| Gps1_G3UXW9_S4s_1 | Gps1_G3U | -0.39 | -0.12 | 0.34 | 0.46 | -1.4 | -1.5 | -0.6 | -0.42 | 0.14 | -1.4 | -0.28 | -0.72 | 0.9 | -1.51 | -0.81 | 0.39 | -0.135 | -0.1975 | -0.0825 | 0.547459 | 0.011934 | 0.125054 | 0.989223 | 0.997988 | 0.994666 |
| Gps1_G3UXW9_S509s_1 | Gps1_G3U | 0.34 | 0.58 | -0.04 | 0.79 | 0.58 | 0.44 | -0.23 | 0.66 | -0.04 | 0.42 | 0.97 | 0.49 | 0.83 | 0.91 | 0.25 | 0.32 | -1.0525 | -0.6375 | -0.33 | 0.601768 | 0.852758 | 0.890197 | 0.996349 | 0.997988 | 0.995259 |
| Gps1_G3UXW9_S518s_1 | Gps1_G3U | -1.23 | -1.55 | -1.63 | -0.47 | -1.56 | -0.25 | -0.48 | 0.32 | -1.3 | -0.95 | 0.32 | 0.02 | -0.32 | -0.47 | -0.64 | 0.29 | -0.055 | 0.0425 | 0.16 | 0.025256 | 0.126728 | 0.125047 | 0.989223 | 0.997988 | 0.994666 |
| Gpsm1_F6SX81_S293s_1 | Gpsm1_F6 | 1.11 | -0.11 | -0.39 | 0.09 | 1.32 | 1.03 | 0.71 | 1.45 | 0.67 | 0.28 | 1.67 | 0.85 | -0.73 | 0.64 | 0.18 | 0.58 | 0.7275 | 0.7425 | 0.935 | 0.986517 | 0.024747 | 0.131377 | 0.989223 | 0.997988 | 1 |
| Gpsm1_F6SX81_S315s_1 | Gpsm1_F6 | 0.48 | -0.53 | -0.96 | -0.67 | 0.85 | 0.43 | 0.06 | -0.78 | 0.53 | 0.47 | -0.67 | -1.41 | -0.27 | 0.26 | -0.47 | -0.71 | 0.9525 | 0.6925 | -0.0075 | 0.752071 | 0.221903 | 0.77609 | 0.989223 | 0.997988 | 0.9971 |
| Gpsm1_F6SX81_S367s_1 | Gpsm1_F6 | -0.32 | 0.49 | -0.04 | 0.01 | 0.2 | 0.45 | 0.66 | 0.76 | 0.12 | 0.42 | 0.54 | 0.17 | 0.27 | 0.04 | 0.03 | 0.28 | 0.56 | 0.15 | 0.1225 | 0.658416 | 0.087616 | 0.303135 | 0.989223 | 0.997988 | 0.995259 |
| Gpsm1_F6SX81_S477s_1 | Gpsm1_F6 | -0.06 | 0.5 | 0.15 | 0.55 | 0.2 | -0.22 | 0.79 | 0.23 | 0.1 | 0.27 | 0.34 | 0.75 | -0.16 | 0.31 | 0.69 | 0.35 | 0.4825 | 0.2775 | 0.12 | 0.966016 | 0.902448 | 0.761991 | 0.996493 | 0.997988 | 0.999507 |
| Gpsm3_G3UZQ1_S38s_1 | Gpsm3_G3 | 0.32 | 0.56 | -0.1 | 0.05 | -0.19 | 0.64 | -0.55 | 0.09 | 0.91 | 0.32 | -0.41 | -0.47 | 1.34 | -0.28 | -0.44 | 0.06 | -0.035 | 0.08 | 0.0125 | 0.929745 | 0.503381 | 0.744618 | 0.989223 | 0.997988 | 0.999507 |
| Gramd1a_F8WJ74_S11s_1 | Gramd1a_ | 0.07 | 0.2 | 0.51 | 0.21 | 0.33 | -0.04 | 0.44 | 0.18 | 0.53 | 0.67 | 0.56 | 0.33 | 1.59 | 0.74 | 1 | 0.86 | -0.21 | -0.12 | -0.0375 | 0.017921 | 0.928493 | 0.238118 | 0.99758 | 0.997988 | 0.994666 |
| Gramd1a_F8WJ74_S11s_2 | Gramd1a_ | 0.14 | -1.33 | -0.79 | -0.49 | -0.55 | 0.01 | -0.56 | -0.62 | -0.13 | 0.12 | -0.54 | -0.49 | -0.68 | -0.35 | -0.16 | -0.21 | -0.02 | 0.275 | 0.8 | 0.458372 | 0.589076 | 0.330065 | 0.989223 | 0.997988 | 0.994666 |
| Gramd1a_F8WJ74_S269s_1 | Gramd1a_ | 0.5 | 1.19 | 0.45 | 0.53 | 0.94 | -0.24 | 0.99 | 0.35 | 0.14 | 0.97 | -0.79 | 0.34 | -0.1 | 0.71 | 0.31 | 0.61 | 0.1875 | 0.3575 | 0.2675 | 0.365866 | 0.647716 | 0.230584 | 0.99033 | 0.997988 | 0.994666 |
| Gramd1a_F8WJ74_T279t_1 | Gramd1a_ | 0.57 | -0.59 | 0.11 | -0.04 | 0.42 | 0.16 | 0.38 | 0.3 | 0.38 | 0.01 | 0.67 | -0.1 | 0.23 | 0.67 | 0.79 | 0.76 | -0.1575 | -0.5025 | -0.285 | 0.085782 | 0.297042 | 0.488735 | 0.989223 | 0.997988 | 0.994666 |
| Gramd1b_Q80TI0_S165s_1 | Gramd1b_ | 0.16 | -0.44 | -0.02 | -0.22 | -0.03 | 0.3 | 0.12 | -0.16 | -0.29 | -0.32 | -0.24 | -0.26 | -0.53 | -0.49 | 0.27 | 0.56 | 0.3025 | 0.2275 | 0.6 | 0.807709 | 0.431075 | 0.524156 | 0.989223 | 0.997988 | 0.999507 |
| Gramd1b_Q80TI0_S174s_1 | Gramd1b_ | -0.1 | -1.21 | 0.7 | 0.71 | 0.49 | 0.19 | 0.63 | 0.78 | -0.51 | -0.29 | 1.27 | 0.38 | -0.26 | 0.47 | 0.74 | 0.26 | 0.1875 | -0.1475 | 0.0825 | 0.565324 | 0.276202 | 0.737917 | 0.989223 | 0.997988 | 0.995259 |
| Gramd1b_Q80TI0_S194s_1 | Gramd1b_ | 0.44 | 0.59 | 1.27 | 1.69 | 1.13 | 1.14 | 1.44 | 1.69 | 1.06 | 0.88 | 1.62 | 1.35 | 2.41 | 1.65 | 2.22 | 2.08 | 0.4975 | 0.1875 | 0.2775 | 0.01232 | 0.302995 | 0.51677 | 0.989223 | 0.997988 | 0.994666 |
| Gramd1b_Q80TI0_S197s_1 | Gramd1b_ | 0.47 | -0.58 | 0.77 | 0.77 | 0.65 | 0.64 | 0.92 | 0.86 | -0.19 | 0.43 | 1.11 | 1.06 | 0.29 | 0.99 | 0.9 | 0.68 | 0.3525 | 0.23 | 1.0925 | 0.351284 | 0.24127 | 0.573147 | 0.989223 | 0.997988 | 0.994666 |
| Gramd1b_Q80TI0_S418s_1 | Gramd1b_ | -0.36 | 0.33 | 0.44 | 0.52 | 0.54 | 0.15 | 1.02 | 0.87 | 0.31 | -0.01 | -0.15 | -0.52 | -0.36 | 0.95 | 0.98 | 1.4 | 0.41 | 0.245 | 0.3575 | 0.250664 | 0.194095 | 0.298575 | 0.989223 | 0.997988 | 0.994666 |
| Gramd1b_Q80TI0_S694s_1 | Gramd1b_ | 0.38 | 0.57 | 1.33 | 1.28 | 0.94 | 0.58 | 1.75 | 1.72 | 0.05 | 0.54 | 1.11 | 1.39 | 0.4 | 0.13 | 1.69 | 1.7 | 0.4125 | -0.325 | 0.51 | 0.846476 | 0.349466 | 0.761801 | 0.989223 | 0.997988 | 0.999507 |
| Gramd1b_Q80TI0_T200t_1 | Gramd1b_ | -0.94 | -0.19 | -0.41 | -0.66 | -0.48 | -0.53 | -0.93 | -1.19 | -0.75 | -0.25 | -0.52 | -0.23 | 0.35 | -0.79 | 0.04 | 0.32 | 0.3575 | -0.1175 | 0.09 | 0.147807 | 0.404873 | 0.672934 | 0.989223 | 0.997988 | 0.994666 |
| Grasp_Q9JJA9_S384s_1 | Grasp_Q9J | 1.52 | 2.06 | 1.63 | 0.91 | 1.73 | 1.38 | 1.11 | 0.95 | 0.54 | 1.15 | 0.63 | 1.6 | -0.13 | 0.9 | 0.96 | 1.04 | -0.2325 | 0.1125 | 0.53 | 0.049404 | 0.452494 | 0.146952 | 0.989223 | 0.997988 | 0.994666 |
| Grasp_Q9JJA9_S93s_1 | Grasp_Q9J | -0.4 | 0.5 | 0.28 | -0.09 | 0.58 | -0.08 | -0.02 | -0.07 | 0.11 | -0.05 | 0.7 | 0.38 | 0.49 | 0.67 | 0.37 | -0.03 | -0.2375 | -0.55 | -0.8375 | 0.335031 | 0.916695 | 0.482683 | 0.997335 | 0.997988 | 0.994666 |
| Grasp_Q9JJA9_T76t_1 | Grasp_Q9J | -2.34 | -0.76 | -1.31 | -0.91 | 0.41 | -1.61 | -2.17 | -1.96 | -1.93 | -2.8 | -0.92 | -1.48 | -1.39 | -1.67 | -1.91 | -3.05 | 0.03 | 0.2125 | 0.3025 | 0.186501 | 0.996759 | 0.378217 | 0.999864 | 0.997988 | 0.994666 |
| Grb10_B1ATS7_S50s_1 | Grb10_B1A | -2.66 | -2.86 | -2.5 | -6 | -2.52 | -2.2 | -2.31 | -2.84 | -0.21 | -3.23 | -2.04 | -2.6 | 0.11 | -2.46 | -2.37 | -1.61 | -0.0025 | -0.4525 | -0.675 | 0.055546 | 0.18508 | 0.138611 | 0.989223 | 0.997988 | 0.994666 |
| Grb10_B1ATS7_S96s_1 | Grb10_B1A | 0.06 | 0.87 | -0.75 | -0.29 | 0.43 | 0.09 | -0.1 | -0.26 | -0.1 | -0.06 | -0.43 | -0.58 | -0.18 | -0.31 | -0.91 | -0.65 | 1.0375 | 1.485 | 1.9225 | 0.2334 | 0.854487 | 0.482374 | 0.996349 | 0.997988 | 0.994666 |
| Grb14_Q9JLM9_S364s_1 | Grb14_Q9J | -0.45 | -0.42 | -0.63 | -0.87 | -0.13 | -0.18 | -0.59 | -0.53 | 0.83 | 0.44 | -0.95 | -0.1 | 2 | -0.96 | -0.34 | -0.03 | 0.0675 | -0.265 | -0.485 | 0.220409 | 0.322898 | 0.129324 | 0.989223 | 0.997988 | 0.994666 |
| Gria2_G5E8H1_Y876y_1 | Gria2_G5E | 0.99 | 0.42 | 0.58 | 0.36 | 0.45 | 1.1 | 0.7 | 0.51 | 0.55 | -0.57 | -0.86 | 0.26 | -0.66 | 1.1 | -0.83 | -0.09 | 0.235 | 0.6475 | 0.76 | 0.138532 | 0.690127 | 0.070525 | 0.994672 | 0.997988 | 0.994666 |
| Gria2_P23819_S899s_1 | Gria2_P23 | 0.62 | 0.69 | 0.52 | 0.3 | 0.6 | 0.57 | 0.39 | 0.53 | 0.48 | 0.38 | 0.61 | 0.35 | -0.18 | 0.58 | 0.66 | 0.32 | 0.1025 | -0.7425 | -0.7075 | 0.512681 | 0.961294 | 0.723536 | 0.999187 | 0.997988 | 0.994666 |
| Gria2_P23819_T912t_1 | Gria2_P23 | 0.49 | 0.5 | 1.04 | 0.89 | 0.98 | 0.47 | 1.37 | 0.79 | -0.1 | 0.21 | 0.49 | 0.71 | -0.26 | 0.96 | 0.25 | 0.7 | -0.01 | -0.0775 | -0.1875 | 0.359376 | 0.532877 | 0.170429 | 0.989223 | 0.997988 | 0.994666 |
| Gria3_B0QZW1_Y871y_1 | Gria3_B0Q | 1.58 | 0.49 | 0.75 | 0.68 | 1.07 | 1.55 | 0.36 | 0.88 | 0.64 | 0.45 | -0.14 | 0.67 | 0.25 | -0.98 | 0.06 | 0.33 | 0.1725 | -0.4025 | -0.3175 | 0.034442 | 0.795929 | 0.177988 | 0.995762 | 0.997988 | 0.994666 |
| Gria3_B0QZW1_Y881y_1 | Gria3_B0Q | 0.6 | 0.38 | 0.59 | 0.62 | 0.59 | 0.88 | 0.61 | 0.65 | 0.38 | 0.51 | 0.45 | 0.82 | 0.29 | 0.51 | 0.11 | 0.21 | 0.09 | -0.47 | -0.96 | 0.277872 | 0.511482 | 0.972984 | 0.989223 | 0.997988 | 0.994666 |
| Gria4_G5E863_T875t_1 | Gria4_G5E | 0.51 | 0.3 | -0.01 | 0.08 | 0.41 | 0.46 | 0.5 | 0.05 | 0.15 | -0.25 | 0.66 | 0.18 | 0.03 | 0.08 | 0.31 | -0.14 | 0.135 | -0.0075 | -0.2675 | 0.560466 | 0.561746 | 0.898599 | 0.989223 | 0.997988 | 0.995259 |
| Grid1_Q61627_S882s_1 | Grid1_Q61 | 0.54 | 0.29 | -0.07 | -0.16 | 0.53 | 1.07 | 0.19 | 0.52 | 0.48 | 0.12 | -0.52 | -0.01 | -0.32 | 0.89 | -0.49 | -0.06 | 0.135 | -0.035 | -0.15 | 0.69507 | 0.150786 | 0.661994 | 0.989223 | 0.997988 | 0.996159 |
| Grid2ip_Q0QWG9_S45s_1 | Grid2ip_Q0 | -0.24 | -1.16 | 0.22 | -0.13 | -2.56 | 0.26 | -0.98 | -0.77 | 2.65 | 0.27 | -0.59 | -0.13 | 2.01 | -2.62 | -0.09 | -0.67 | 0.4275 | -0.1325 | -0.145 | 0.986306 | 0.259473 | 0.231851 | 0.989223 | 0.997988 | 1 |
| Grik5_Q61626_S961s_1 | Grik5_Q61 | 0.56 | 0.83 | 1.36 | 0.82 | 0.59 | 0.6 | 0.72 | 0.76 | 0.57 | 0.74 | 0.59 | 0.92 | 0.74 | 0.95 | 0.79 | 0.6 | -0.685 | 0.8775 | -0.015 | 0.65235 | 0.358712 | 0.470573 | 0.989223 | 0.997988 | 0.995259 |
| Grik5_Q61626_Y891y_1 | Grik5_Q61 | -0.31 | 0.79 | 0.12 | 0.18 | 0.32 | -1.13 | -0.26 | 0.72 | 0.37 | -0.14 | 0.01 | -0.1 | 0.95 | -0.25 | 0.4 | 0.94 | -0.225 | -0.1875 | -0.1225 | 0.4145 | 0.519939 | 0.59184 | 0.989223 | 0.997988 | 0.994666 |
| Grin2a_P35436_S1025s_1 | Grin2a_P3 | 1.8 | 1.27 | 2.05 | 2.12 | 1.08 | 1.99 | 1.38 | 2.62 | 1.6 | 1.37 | 1.09 | 2 | 0.71 | 1.54 | 1.18 | 1.43 | -0.2825 | -0.16 | 0.315 | 0.08524 | 0.911397 | 0.350016 | 0.997335 | 0.997988 | 0.994666 |
| Grin2a_P35436_S1198s_1 | Grin2a_P3 | 3.7 | 2.56 | -1.12 | -2.48 | 1.61 | 4 | -1.46 | -1.81 | 3.45 | 3.68 | -4.01 | -0.87 | 2.8 | 0.16 | 1.34 | -2.18 | -0.0425 | -0.295 | -0.595 | 0.930528 | 0.962822 | 0.960135 | 0.999187 | 0.997988 | 0.999507 |
| Grin2a_P35436_S1204s_1 | Grin2a_P3 | 0.47 | 1.53 | 1 | 0.05 | -1.08 | 0.87 | -0.25 | -0.25 | 2.09 | 0.56 | -0.46 | 0.46 | 2.34 | -0.46 | 0.89 | 0.42 | -0.08 | -0.1025 | -0.135 | 0.954066 | 0.072433 | 0.860753 | 0.989223 | 0.997988 | 0.999507 |
| Grin2a_P35436_S1459s_1 | Grin2a_P3 | 0.48 | 0.52 | 0.98 | 0.96 | 0.4 | 1.31 | 1.09 | 0.66 | 1.57 | 0.65 | -0.02 | 0.89 | 0.79 | 0.98 | 0.19 | 0.52 | -0.94 | -0.1 | 0.035 | 0.691859 | 0.645921 | 0.91811 | 0.99033 | 0.997988 | 0.996159 |
| Grin2a_P35436_S882s_1 | Grin2a_P3 | 1.11 | 1.17 | 1.24 | 1.12 | 1.24 | 0.9 | 1.52 | 1.08 | 0.95 | 1.11 | 1.2 | 1.35 | 0.61 | 1.4 | 0.74 | 0.93 | 0.13 | 0.0375 | -0.115 | 0.379367 | 0.909863 | 0.97198 | 0.997335 | 0.997988 | 0.994666 |
| Grin2a_P35436_S890s_1 | Grin2a_P3 | 0.38 | 0.68 | 0.63 | 0.24 | 0.84 | 0.03 | 0.67 | 0.71 | 0.02 | 0.41 | -0.3 | -0.02 | 0.16 | 0.81 | -0.1 | 0.11 | 0.025 | -0.0075 | -0.24 | 0.422796 | 0.757755 | 0.097458 | 0.995762 | 0.997988 | 0.994666 |
| Grin2a_P35436_S929s_1 | Grin2a_P3 | -1.15 | -0.25 | -0.4 | -0.6 | 0.88 | -1.7 | -0.42 | -0.44 | -1.4 | -2.53 | -0.82 | -0.55 | -1.38 | 0.06 | -0.79 | -0.63 | 0.08 | -0.455 | -0.2375 | 0.820135 | 0.726404 | 0.141039 | 0.995437 | 0.997988 | 0.999507 |
| Grin2b_G3X9V4_S1116s_1 | Grin2b_G3 | -0.87 | -0.96 | -0.48 | -0.69 | -1.63 | -0.19 | -0.95 | -0.52 | 1.89 | -0.76 | -2.14 | -0.77 | 1.89 | -2.5 | 0.21 | -1.29 | 0.18 | -0.725 | -0.085 | 0.69778 | 0.829314 | 0.689807 | 0.995762 | 0.997988 | 0.996159 |
| Grin2b_G3X9V4_S1143s_1 | Grin2b_G3 | -0.21 | 0.33 | 1.16 | 1.3 | 0.74 | 0.02 | 1.42 | 1.32 | -0.44 | 0.16 | 0.7 | 1.39 | -1.11 | 0.49 | 0.86 | 0.92 | -0.0725 | 0.305 | 0.3275 | 0.52397 | 0.610888 | 0.699868 | 0.989223 | 0.997988 | 0.994666 |
| Grin2b_G3X9V4_S1166s_1 | Grin2b_G3 | -0.96 | -1.55 | -0.33 | -1.04 | -2.74 | 0.94 | -1.3 | -0.66 | -0.26 | -1.72 | 0.31 | -0.63 | 0.69 | -1.72 | -1.26 | -0.8 | 0.23 | -0.1925 | -0.355 | 0.714763 | 0.966128 | 0.411865 | 0.999187 | 0.997988 | 0.996966 |
| Grin2b_G3X9V4_S1284s_1 | Grin2b_G3 | -0.07 | -0.98 | -0.52 | -1 | -1.25 | 0.36 | -0.35 | -0.77 | 1.15 | -0.62 | -0.22 | 0.22 | 1.07 | -2.06 | -0.21 | -2.47 | 0.03 | 0.395 | 0.1975 | 0.717442 | 0.722288 | 0.096861 | 0.995437 | 0.997988 | 0.996966 |
| Grin2b_G3X9V4_S1303s_1 | Grin2b_G3 | 8.08 | 0.53 | 2.39 | -0.78 | 2.05 | 8.6 | 4.08 | 2.41 | 0.27 | 5.2 | -0.76 | 2.35 | 1.8 | 0.43 | 3.15 | 3.78 | 0.14 | 0.775 | -0.275 | 0.882166 | 0.419207 | 0.700557 | 0.989223 | 0.997988 | 0.999507 |
| Grin2b_G3X9V4_S882s_1 | Grin2b_G3 | 0.95 | 0.88 | -0.51 | -0.48 | 0.06 | 0.39 | -0.59 | -0.6 | 0.05 | 0.84 | -0.89 | -1.47 | 0.04 | 0.4 | -0.8 | -0.97 | 1.73 | -0.79 | -0.265 | 0.289894 | 0.385621 | 0.347045 | 0.989223 | 0.997988 | 0.994666 |
| Grin2b_G3X9V4_S882s_2 | Grin2b_G3 | 0.03 | 0.5 | 1.09 | 0.36 | 0.76 | 0.04 | 1.95 | 1.69 | 0.71 | 0.3 | -0.65 | -0.18 | 0.56 | 0.58 | 1.59 | 1.6 | -0.395 | -0.5775 | -0.5425 | 0.14943 | 0.200378 | 0.248648 | 0.989223 | 0.997988 | 0.994666 |
| Grin2b_G3X9V4_S886s_1 | Grin2b_G3 | -0.12 | 0.19 | 0.07 | 0.33 | 0.05 | -0.03 | 0.33 | 0.21 | 0.51 | 0.51 | 0.14 | 0.61 | 0.44 | 0.49 | 0.04 | 0.3 | 0.615 | -0.45 | 0.5875 | 0.433854 | 0.917264 | 0.186678 | 0.997335 | 0.997988 | 0.994666 |
| Grin2b_G3X9V4_S886s_2 | Grin2b_G3 | 0.03 | 0.5 | 1.09 | 0.36 | 0.76 | 0.04 | 1.95 | 1.69 | 0.71 | 0.3 | -0.65 | -0.18 | 0.56 | 0.58 | 1.59 | 1.6 | 0.0225 | 0.325 | 0.2 | 0.14943 | 0.200378 | 0.248648 | 0.989223 | 0.997988 | 0.994666 |
| Grin2b_G3X9V4_S917s_1 | Grin2b_G3 | 0.5 | 0.88 | 0.62 | 0.48 | 0.8 | 0.4 | 0.93 | 0.75 | -0.08 | 0.88 | 0.06 | 0.83 | 0.1 | 0.5 | 0.76 | 0.76 | 0.615 | -0.45 | 0.5875 | 0.739195 | 0.659556 | 0.521744 | 0.990434 | 0.997988 | 0.9971 |
| Grin2b_G3X9V4_S929s_1 | Grin2b_G3 | -2.18 | -2.02 | -2.03 | -3.12 | -1.58 | -2.2 | -2.4 | -1.67 | -2.56 | -3.06 | -0.78 | -2.46 | -1.88 | -1.61 | -3.03 | -2.51 | 0.1 | -0.1975 | -0.09 | 0.846948 | 0.282962 | 0.815388 | 0.989223 | 0.997988 | 0.999507 |
| Grin2b_G3X9V4_S929s_2 | Grin2b_G3 | -0.18 | -0.76 | -1.21 | -0.91 | -0.48 | -1.11 | -1.83 | -1.96 | -0.55 | -1.41 | -0.37 | -1.25 | -0.59 | -0.7 | -2.85 | -2.03 | 0.375 | 0.1225 | 0.08 | 0.176077 | 0.16316 | 0.711549 | 0.989223 | 0.997988 | 0.994666 |
| Grin2b_G3X9V4_S930s_1 | Grin2b_G3 | -1.39 | -0.83 | -1.09 | -0.07 | -0.66 | -1.31 | -0.28 | -1.19 | -1.6 | -0.84 | -0.44 | -0.7 | -0.04 | -0.53 | -0.39 | 0.14 | -0.58 | -0.13 | -0.7775 | 0.093776 | 0.967296 | 0.895462 | 0.999187 | 0.997988 | 0.994666 |
| Grin2b_G3X9V4_S930s_2 | Grin2b_G3 | -0.43 | -0.59 | -1.14 | -0.69 | -0.53 | -0.47 | -1.33 | -1.37 | -0.35 | -1.48 | -1.29 | -0.92 | -0.56 | -0.48 | -2.64 | -1.47 | -0.015 | -0.05 | 0.64 | 0.265483 | 0.497228 | 0.364927 | 0.989223 | 0.997988 | 0.994666 |
| Grin2b_G3X9V4_S940s_1 | Grin2b_G3 | 0.85 | -0.44 | -0.84 | -0.93 | -0.55 | 1.47 | -1.03 | -0.72 | 1.03 | 0.11 | -0.88 | -0.8 | 0.52 | -1.24 | -0.95 | -1.45 | -0.2125 | -0.2975 | -0.575 | 0.439321 | 0.831926 | 0.716063 | 0.995762 | 0.997988 | 0.994666 |
| Grin2b_G3X9V4_Y1039y_1 | Grin2b_G3 | 4.8 | 4.11 | 7.6 | 3.28 | 4.39 | 5.14 | 7.63 | 7.25 | 0.7 | 3.94 | 1.72 | 8.43 | -0.14 | 0.68 | 6.17 | 7.9 | 0.1325 | 0.205 | -0.44 | 0.496854 | 0.290763 | 0.469869 | 0.989223 | 0.997988 | 0.994666 |
| Grin2b_G3X9V4_Y1049y_1 | Grin2b_G3 | 0.38 | -0.21 | 1.79 | 1.89 | 0.99 | 0.25 | -0.18 | 0.79 | 1.1 | -0.69 | 1.96 | 0.31 | 0.23 | -0.33 | 0.14 | 0.76 | 1.155 | -1.25 | -1.295 | 0.172825 | 0.357938 | 0.674565 | 0.989223 | 0.997988 | 0.994666 |
| Grin2b_G3X9V4_Y1070y_1 | Grin2b_G3 | 0.14 | -0.49 | -0.19 | 0.42 | -0.22 | 0.09 | 0.65 | 0.76 | 0.55 | 0.18 | -0.5 | 0.55 | -0.48 | 0.36 | -0.44 | 0.16 | -0.5 | -0.2925 | -0.7625 | 0.832378 | 0.28939 | 0.510319 | 0.989223 | 0.997988 | 0.999507 |
| Grin2b_G3X9V4_Y1225y_1 | Grin2b_G3 | 3.17 | 1.86 | 3.37 | 1.62 | 2.87 | 3.15 | 2.48 | 3.36 | 0.12 | 2.98 | 1.84 | 3.3 | 0.88 | -0.08 | 2.97 | 2.71 | 0.35 | 0.225 | -0.07 | 0.262394 | 0.323248 | 0.560842 | 0.989223 | 0.997988 | 0.994666 |
| Grin2b_G3X9V4_Y932y_1 | Grin2b_G3 | 0.53 | 0.2 | 0.34 | 1.26 | 0.31 | 0.68 | 0.47 | 1.09 | 0.86 | -0.25 | 0.42 | 1.08 | 0.19 | 0.6 | 0.13 | -0.16 | 0.46 | -0.445 | -0.885 | 0.247381 | 0.858998 | 0.884986 | 0.996349 | 0.997988 | 0.994666 |
| Grin2b_G3X9V4_Y932y_2 | Grin2b_G3 | -0.81 | -0.39 | 0.1 | 0.44 | -0.66 | -0.1 | -0.27 | -0.6 | -0.33 | -1.93 | 0.06 | 0.44 | -0.58 | -0.41 | -1.09 | -0.47 | 0.055 | -0.055 | -0.3925 | 0.194508 | 0.455485 | 0.616973 | 0.989223 | 0.997988 | 0.994666 |
| Grip1_H7BX90_S380s_1 | Grip1_H7B | -0.26 | 0.28 | 1.39 | 0.78 | 1.11 | -0.04 | 0.83 | 0.82 | 0.77 | 0.58 | 1.22 | 1.04 | 0.91 | 1.11 | 0.68 | 1.95 | -0.2425 | -0.275 | -0.4725 | 0.183824 | 0.748012 | 0.365473 | 0.995762 | 0.997988 | 0.994666 |
| Grip1_H7BX90_S43s_1 | Grip1_H7B | 0.47 | 0.52 | 0.5 | 0.27 | 1.01 | 0.26 | 0.47 | -0.21 | 0.79 | 1.03 | -0.23 | 0.36 | 0.93 | 1.05 | 0.34 | 0.63 | 0.1325 | 0.355 | 0.615 | 0.2766 | 0.842673 | 0.879827 | 0.996349 | 0.997988 | 0.994666 |
| Grip1_H7BX90_S709s_1 | Grip1_H7B | 0.33 | 0.24 | 0.06 | -0.72 | -0.03 | 0.93 | -0.44 | -0.48 | 2.53 | 0.22 | -0.72 | -0.08 | 3.01 | -1.19 | 0.66 | -0.73 | -0.0575 | 0.0475 | 0.2975 | 0.593395 | 0.964394 | 0.458057 | 0.999187 | 0.997988 | 0.995259 |
| Grip1_H7BX90_S794s_1 | Grip1_H7B | 1.63 | 1.61 | 0.64 | 0.66 | 2 | 1.84 | 0.48 | 0.53 | 1.51 | 1.78 | 0.31 | 0.64 | 1.47 | 2.06 | 0.8 | 0.88 | 0.0175 | 0.51 | 0.46 | 0.682999 | 0.86661 | 0.862864 | 0.996493 | 0.997988 | 0.996159 |
| Grip1_H7BX90_S885s_1 | Grip1_H7B | 0.28 | -0.53 | -0.36 | -0.46 | -0.43 | 0.71 | -0.78 | -0.42 | 0.05 | -0.15 | 0.02 | -0.38 | 0.85 | -0.87 | -0.22 | 0.11 | 0.0775 | -0.075 | 0.1675 | 0.566848 | 0.918937 | 0.574611 | 0.997335 | 0.997988 | 0.995259 |
| Grip1_H7BX90_S928s_1 | Grip1_H7B | 0.58 | -0.06 | 0.34 | 0.04 | 0.57 | 0.49 | 0.82 | 0.55 | -0.11 | -0.25 | -0.63 | 0.01 | -0.03 | 1.09 | -0.03 | 0.19 | 0.0375 | 0.1525 | 0.235 | 0.813867 | 0.127211 | 0.101457 | 0.989223 | 0.997988 | 0.999507 |
| Grip1_H7BX90_T904t_1 | Grip1_H7B | -0.32 | 0.43 | -0.19 | -0.19 | 1.15 | -1.17 | -0.05 | -0.29 | 0.18 | -0.46 | 0.37 | -0.54 | 0.59 | 0.9 | -0.27 | 0.24 | 0.3825 | -0.47 | 0.08 | 0.221326 | 0.961768 | 0.886326 | 0.999187 | 0.997988 | 0.994666 |
| Grip2_E0CXE0_S10s_1 | Grip2_E0C | -0.1 | 0.54 | 0.57 | 0.8 | 0.81 | -0.06 | 0.37 | 0.24 | 0.06 | 0.07 | 1.41 | 1.49 | 0.02 | 0.96 | 1.48 | 0.91 | -0.0225 | -0.045 | 0.4325 | 0.313921 | 0.701878 | 0.48623 | 0.995437 | 0.997988 | 0.994666 |
| Grip2_E0CXE0_S773s_1 | Grip2_E0C | -1.38 | -0.89 | 0.23 | 0.26 | -0.18 | -1.44 | -0.15 | 0.19 | -1.31 | -1.08 | 0.4 | -0.01 | -0.72 | -0.68 | 0.03 | 0.34 | -0.1125 | 0.305 | 0.39 | 0.690479 | 0.920209 | 0.91895 | 0.997335 | 0.997988 | 0.996159 |
| Grip2_E0CXE0_S774s_1 | Grip2_E0C | 0.4 | 0.62 | 2.62 | 2.25 | 0.52 | 0.25 | 1.92 | 1.9 | -0.21 | 0.58 | 1.27 | 2.11 | 0.39 | 0.21 | 2 | 2.44 | 0.05 | -0.055 | 0.1875 | 0.765266 | 0.612158 | 0.437627 | 0.989223 | 0.997988 | 0.9971 |
| Grip2_E0CXE0_S810s_1 | Grip2_E0C | -0.01 | -0.35 | -0.24 | 0.7 | 0.46 | -1.59 | -0.38 | 0.59 | 0.14 | -0.77 | -1.53 | -1.66 | 0.21 | -0.14 | -0.07 | 0.25 | -0.325 | -0.535 | -0.2125 | 0.903588 | 0.617153 | 0.05614 | 0.989223 | 0.997988 | 0.999507 |
| Grip2_E0CXS4_S43s_1 | Grip2_E0C | 0.58 | 0.79 | 1.52 | 1.62 | 0.78 | 0.51 | 1.16 | 1.98 | 0.87 | 1.29 | 1.46 | 1.43 | 0.72 | 0.72 | 1.4 | 1.51 | -0.255 | -0.98 | 0.0375 | 0.911628 | 0.959799 | 0.675955 | 0.999187 | 0.997988 | 0.999507 |
| Grip2_E0CXS4_S996s_1 | Grip2_E0C | 0.81 | 0.63 | 0.92 | 0.8 | 0.33 | 0.17 | 0.77 | 1.18 | 0.56 | 0.71 | 0.87 | 0.75 | -1.25 | 0.12 | 0.64 | 1.24 | -0.02 | 0.135 | -0.04 | 0.250374 | 0.522793 | 0.752206 | 0.989223 | 0.997988 | 0.994666 |
| Gripap1_Q8VD04_S237s_1 | Gripap1_Q | -3.41 | -3.81 | -2.92 | -2.94 | -3.12 | -2.98 | -2.94 | -2.98 | -4.74 | -3.67 | -4.63 | -3.43 | -1.07 | -5.03 | -3.32 | -2.75 | -0.1775 | -0.0675 | -0.6025 | 0.761902 | 0.32726 | 0.055331 | 0.989223 | 0.997988 | 0.9971 |
| Gripap1_Q8VD04_S657s_1 | Gripap1_Q | -0.32 | -0.1 | -0.17 | 0.12 | 0.11 | -0.19 | 0.31 | 0.09 | -0.15 | 0.23 | -0.53 | -0.05 | -0.48 | 0.31 | -0.23 | -0.28 | 0.265 | -0.8475 | 0.2275 | 0.848676 | 0.384292 | 0.976379 | 0.989223 | 0.997988 | 0.999507 |
| Grm1_P97772_S969s_1 | Grm1_P97 | 1.13 | 0.84 | 0.64 | 1.44 | 0.81 | 0.65 | 1.11 | 1.15 | 1.02 | 1.16 | -0.06 | 0.71 | 0.32 | 0.73 | 1.09 | 1.82 | 0.1975 | -0.0075 | -0.0525 | 0.952575 | 0.752029 | 0.383652 | 0.995762 | 0.997988 | 0.999507 |
| Grm1_P97772_T988t_1 | Grm1_P97 | -0.62 | 0.75 | -0.14 | -0.4 | 0.34 | -0.28 | -0.41 | -0.17 | -0.75 | 0.31 | -0.54 | -0.06 | 0.11 | 0.22 | -0.52 | -0.91 | -0.0825 | -0.305 | -0.0225 | 0.672341 | 0.936625 | 0.684471 | 0.998821 | 0.997988 | 0.996159 |
| Grm2_Q14BI2_S843s_1 | Grm2_Q14 | -0.05 | 0.14 | 0.74 | 1.3 | 0.95 | 0.12 | 1.14 | 1.42 | 0.24 | 0.64 | 0.77 | 0.68 | 0.13 | 0.79 | 0.77 | 1.24 | -0.0275 | -0.1575 | -0.1725 | 0.612134 | 0.359026 | 0.884872 | 0.989223 | 0.997988 | 0.995259 |

| | | | | | | | | | | | | | | | | | | | | | | | | | | |
|---|---|---|---|---|---|---|---|---|---|---|---|---|---|---|---|---|---|---|---|---|---|---|---|---|---|---|
| Grm2_Q14BI2_S871s_1 | Grm2_Q14 | -0.65 | 0.7 | 0.42 | 0.45 | 0.34 | -0.1 | -0.05 | 0.03 | -0.1 | 0.39 | 0.65 | 0.24 | 0.41 | 0.75 | 0.1 | 0.27 | 0.375 | 0.05 | 0.2 | 0.670271 | 0.595572 | 0.853828 | 0.989223 | 0.997988 | 0.996159 |
| Grm3_Q9QYS2_S845s_1 | Grm3_Q9C | 0.43 | 0.35 | 0.64 | 0.72 | 0.57 | 0.24 | 0.86 | 0.78 | 0.15 | 0.86 | -0.08 | 0.43 | -0.12 | 0.49 | 0.16 | 0.68 | -0.175 | 0.065 | 0.1525 | 0.411424 | 0.741407 | 0.484485 | 0.995762 | 0.997988 | 0.994666 |
| Grm3_Q9QYS2_S878s_1 | Grm3_Q9C | 0.74 | 0.18 | 1.13 | 1.43 | 0.55 | 0.21 | 0.86 | 0.29 | 0.29 | 0.73 | 0.14 | 0.94 | -0.13 | 0.72 | 0.53 | 0.75 | 0.0775 | -0.195 | -0.2325 | 0.282769 | 0.240713 | 0.331623 | 0.989223 | 0.997988 | 0.994666 |
| Grm5_E9QMC2_S1005s_1 | Grm5_E9Q | 2.91 | 2.13 | 2.03 | 1.88 | 2.44 | 2.84 | 1.81 | 2.25 | 1.2 | 2.27 | 2.22 | 2.51 | 1.51 | 1.23 | 2.77 | 2.58 | -0.3925 | -0.345 | -0.4025 | 0.625923 | 0.765306 | 0.620318 | 0.995762 | 0.997988 | 0.995259 |
| Grm5_E9QMC2_S1141s_1 | Grm5_E9Q | 0.56 | 1.25 | 0.89 | 0.79 | 0.98 | 0.52 | 1.38 | 0.91 | 0.52 | 1.07 | 0.25 | 1.2 | 0.45 | 0.83 | 1.05 | 0.73 | 0.0975 | -0.1875 | -0.215 | 0.695925 | 0.781201 | 0.712301 | 0.995762 | 0.997988 | 0.996159 |
| Grm5_E9QMC2_S1153s_1 | Grm5_E9Q | 1.39 | 0.6 | 1.3 | 1.44 | 1.47 | 1.19 | 1.69 | 1.67 | 0.11 | 1.07 | 0.86 | 1.29 | -0.13 | 0.76 | 1.11 | 0.88 | 0.075 | -0.1125 | -0.1075 | 0.165949 | 0.250695 | 0.320171 | 0.989223 | 0.997988 | 0.994666 |
| Grm5_E9QMC2_S839s_1 | Grm5_E9Q | -1.69 | -3.5 | -2.12 | -2.02 | -1.8 | -2.01 | -0.63 | -1.23 | -3.13 | -2.69 | -0.09 | -1.13 | -3.55 | -1.48 | -1.84 | -1.54 | 0.3225 | -0.35 | -0.5275 | 0.692592 | 0.076013 | 0.438982 | 0.989223 | 0.997988 | 0.996159 |
| Grm5_E9QMC2_S860s_1 | Grm5_E9Q | -0.01 | -1.23 | -0.39 | 0.21 | 0.53 | -0.15 | 0.59 | 0.06 | -1.3 | -0.87 | 0.7 | -0.65 | -1.78 | 0.37 | -0.29 | -0.59 | 0.915 | 0.5725 | 0.23 | 0.674883 | 0.116142 | 0.728837 | 0.989223 | 0.997988 | 0.996159 |
| Grm5_E9QMC2_S901s_1 | Grm5_E9Q | -0.66 | -0.77 | -0.94 | -1.36 | -0.76 | -0.44 | -0.63 | -0.61 | -1.82 | -1.2 | 1.43 | -0.48 | -2.14 | -0.27 | -2.11 | -1.69 | 0.6125 | -0.175 | -0.2175 | 0.19252 | 0.193821 | 0.530512 | 0.989223 | 0.997988 | 0.994666 |
| Grm5_E9QMC2_T840t_1 | Grm5_E9Q | -0.38 | -2.09 | -1.52 | -0.75 | -0.79 | -0.23 | -0.43 | -0.33 | -1.22 | -1.4 | -0.37 | -0.9 | -1.76 | -0.62 | -2.17 | -1.89 | 0.3225 | 0.415 | -0.62 | 0.393951 | 0.083226 | 0.625616 | 0.989223 | 0.997988 | 0.994666 |
| Grm5_Q3UVX5_S870s_1 | Grm5_Q3U | -0.49 | -2.81 | -1.85 | -1.84 | -2.05 | 0.34 | -1.99 | -1.05 | -0.5 | -2.08 | -0.54 | -0.95 | -0.8 | -2.48 | -2.25 | -1.78 | 0.74 | 0.2125 | -0.425 | 0.886118 | 0.399376 | 0.211835 | 0.989223 | 0.997988 | 0.999507 |
| Grm7_Q68ED2_S900s_1 | Grm7_Q68 | 1.03 | 0.73 | 0.44 | 0.29 | 0.31 | 0.43 | 0.59 | 0.27 | 0.82 | 1.13 | 0.42 | 0.4 | 0.44 | 0.83 | 0.44 | 0.57 | 0.56 | 0.73 | -0.08 | 0.847502 | 0.369296 | 0.807356 | 0.989223 | 0.997988 | 0.999507 |
| Gsg1l_D3Z7H4_S274s_1 | Gsg1l_D3Z | 0.36 | 0.3 | 0.74 | 0.34 | 0.66 | 0.42 | 0.52 | 0.71 | 0.37 | 0.12 | 0.76 | 0.59 | -0.44 | 0.67 | 0.27 | 0.27 | -0.2225 | 0.07 | -0.0525 | 0.438913 | 0.513444 | 0.919844 | 0.989223 | 0.997988 | 0.994666 |
| Gsk3a_D3Z7E5_S39s_1 | Gsk3a_D3Z | 0.38 | -0.42 | -0.28 | -1.46 | 0.17 | -0.62 | -0.63 | -0.04 | -0.66 | -0.18 | -1.18 | -0.59 | -0.24 | -0.29 | -0.39 | -0.05 | 0.1425 | 0.025 | -0.2425 | 0.61109 | 0.689813 | 0.625794 | 0.994672 | 0.997988 | 0.995259 |
| Gsk3a_D3Z7E5_Y279y_1 | Gsk3a_D3Z | -0.81 | -1.33 | -1.34 | -1.69 | -1.57 | -0.58 | -1.27 | -1.39 | -1.22 | -0.61 | -0.42 | -1.96 | -0.72 | -0.99 | -1.34 | -1.58 | 0.165 | -0.2075 | 0.2025 | 0.668438 | 0.76797 | 0.544476 | 0.995762 | 0.997988 | 0.996159 |
| Gsk3b_E9QAQ5_S25s_1 | Gsk3b_E9C | -0.79 | -1.33 | -1.51 | -1.47 | -0.72 | -0.95 | -1.02 | -2.28 | -1.21 | -0.83 | -0.26 | -1.21 | -0.46 | -0.45 | -1.78 | -2.15 | 0.09 | 0.24 | 0.135 | 0.887012 | 0.931776 | 0.223486 | 0.998317 | 0.997988 | 0.999507 |
| Gsk3b_E9QAQ5_S402s_1 | Gsk3b_E9C | -0.01 | 0.34 | 0.56 | 0.45 | -0.15 | -0.05 | 0.49 | 0.59 | 0.18 | 0.1 | 0.56 | 0.68 | 0.52 | 0.42 | 0.58 | 0.56 | 0.0325 | 0.3975 | 0.065 | 0.461134 | 0.668972 | 0.860884 | 0.992876 | 0.997988 | 0.994666 |
| Gsk3b_E9QAQ5_S9s_1 | Gsk3b_E9C | -3.3 | -3.11 | -2.46 | -3.41 | -2.54 | -3.21 | -2.85 | -2.83 | -3.55 | -2.56 | 1.41 | -0.88 | -1.98 | -1.3 | -1.89 | -3.14 | -0.115 | 0.045 | 0.185 | 0.043158 | 0.463535 | 0.112915 | 0.989223 | 0.997988 | 0.994666 |
| Gsk3b_E9QAQ5_T433t_1 | Gsk3b_E9C | -0.63 | -1 | 0.57 | -0.59 | -1.62 | 0.45 | -0.99 | -0.5 | 1.93 | -0.09 | 1.72 | 0.33 | 2.8 | -2.28 | -0.07 | -0.79 | 0.2125 | 1.675 | 0.9925 | 0.737622 | 0.620117 | 0.032606 | 0.989223 | 0.997988 | 0.9971 |
| Gsn_P13020_S307s_1 | Gsn_P1302 | 0.84 | 0.64 | 0.64 | -0.17 | 0.98 | 0.57 | 0.76 | 0.8 | 0.57 | 0.5 | 0.32 | 0.66 | 0.27 | 0.7 | 0.17 | 0.66 | -0.2525 | 1.385 | 0.3275 | 0.904677 | 0.309691 | 0.929874 | 0.989223 | 0.997988 | 0.999507 |
| Gspt2_Q149F3_S195s_1 | Gspt2_Q14 | -1.63 | -1.24 | -2.6 | -3.27 | -0.04 | -1.65 | -1.71 | -2.54 | -2.53 | -0.88 | 0.73 | -1.91 | -0.88 | -0.16 | -2.51 | -1.76 | 0.29 | 0.025 | -0.0375 | 0.192114 | 0.276177 | 0.194918 | 0.989223 | 0.997988 | 0.994666 |
| Gsto1_O09131_S139s_1 | Gsto1_O09 | 0.63 | 0.55 | 0.47 | 0.25 | -0.23 | 0.79 | -0.39 | -0.2 | 0.52 | 0.58 | -0.61 | 0.12 | 0.93 | -1.12 | 0.01 | 0.12 | 0.7 | 1.0375 | 0.8575 | 0.266866 | 0.136973 | 0.322447 | 0.989223 | 0.997988 | 0.994666 |
| Gtf2f1_Q3THK3_T331t_1 | Gtf2f1_Q3 | -2.97 | -2.05 | -3.3 | -4.42 | -2.01 | -3.55 | -4.97 | -3.4 | -0.82 | -1.91 | -2.21 | -4.55 | -0.28 | -2.52 | -2.33 | -3.78 | -0.4825 | -0.3225 | -0.49 | 0.235388 | 0.666379 | 0.337556 | 0.992005 | 0.997988 | 0.994666 |
| Gtf2i_Q3UHU8_S783s_1 | Gtf2i_Q3U | -1.17 | -0.81 | -0.82 | -1.18 | -0.49 | -1.81 | -0.29 | -1.78 | -0.99 | -1.28 | -0.77 | -1.34 | -0.74 | -0.35 | -1.23 | -0.33 | -0.2975 | 0.8125 | 0.9575 | 0.281068 | 0.809232 | 0.686314 | 0.995762 | 0.997988 | 0.994666 |
| Gtf3c1_Q8K284_S1794s_1 | Gtf3c1_Q8 | 0.54 | 0.18 | 0.12 | 0.46 | 0.67 | -0.15 | 0.84 | 0.11 | 0.4 | 0.52 | 0.44 | 0.33 | -0.42 | 0.02 | 0.28 | 0.1 | -0.0975 | -0.1 | 0.3325 | 0.239752 | 0.882019 | 0.660713 | 0.996493 | 0.997988 | 0.994666 |
| Gtf3c4_Q8BMQ2_S17s_1 | Gtf3c4_Q8 | -0.46 | 0.31 | 0.42 | -0.23 | -0.06 | -0.6 | -0.86 | -0.44 | 0.76 | -0.64 | -0.42 | 0.22 | 0.94 | -0.26 | 0.58 | 0.48 | 0.0425 | 0.0975 | -0.33 | 0.25026 | 0.116778 | 0.937728 | 0.989223 | 0.997988 | 0.994666 |
| Gtpbp1_O08582_S24s_1 | Gtpbp1_O( | 1.25 | 1.82 | 0.13 | -0.16 | 1.62 | 1.34 | 0.44 | 0.65 | 0.13 | 0.9 | -0.1 | 0.01 | -0.14 | 1.59 | 0.15 | -0.28 | -0.5 | -0.03 | 0.425 | 0.466171 | 0.614543 | 0.299427 | 0.989223 | 0.997988 | 0.994666 |
| Gtpbp1_O08582_S580s_1 | Gtpbp1_O( | 0.03 | 0.06 | 0.19 | 0.2 | -0.08 | -0.41 | 0.53 | 0.35 | -0.1 | 0.25 | 0.16 | 0.19 | -0.02 | 0.46 | 0.09 | 0.12 | 0.2525 | -0.525 | -0.43 | 0.860381 | 0.93186 | 0.981318 | 0.998317 | 0.997988 | 0.999507 |
| Gtpbp1_O08582_S69s_1 | Gtpbp1_O( | -0.02 | 0.82 | 0.37 | 0.13 | 0.57 | -0.27 | 0.69 | 0.04 | 0.1 | 0.17 | -0.24 | 0.45 | 0.21 | 0.26 | 0.21 | 0.43 | -0.0225 | 0.005 | 0.0425 | 0.862596 | 0.827786 | 0.473987 | 0.995762 | 0.997988 | 0.999507 |
| Gtpbp1_O08582_S8s_1 | Gtpbp1_O( | 0.57 | 0.69 | 0.77 | 0.26 | 0.28 | 0.69 | 0.92 | 0.61 | 0.21 | 0.85 | 0.75 | 0.43 | -0.28 | 0.53 | 0.38 | 0.4 | -0.0675 | -0.205 | -0.0475 | 0.287073 | 0.826517 | 0.960891 | 0.995762 | 0.997988 | 0.994666 |
| Gucy1a2_F8VQK3_S41s_1 | Gucy1a2_F | 0.57 | 0.43 | 0.47 | 0.28 | 0.77 | 0.41 | 0.14 | 0.26 | 0.46 | 0.18 | 0.01 | 0.38 | 0.64 | 0.44 | 0.44 | 0.28 | 0.0525 | -0.0125 | -0.315 | 0.957769 | 0.851831 | 0.429749 | 0.996349 | 0.997988 | 0.999507 |
| Gys1_Q9Z1E4_S645s_1 | Gys1_Q9Z1 | 0.75 | 0.72 | 0.87 | 0.81 | 0.33 | 1.02 | 0.82 | 0.75 | 0.5 | 0.75 | 0.6 | 1 | 0.23 | 0.9 | 0.69 | 0.47 | -0.0425 | -0.18 | 0.0125 | 0.407954 | 0.800471 | 0.736419 | 0.995762 | 0.997988 | 0.994666 |
| Gys1_Q9Z1E4_S653s_2 | Gys1_Q9Z1 | 2.77 | 2.97 | 4.54 | 3.2 | 0.93 | 2.97 | 2.72 | 3.81 | 4.4 | 1.87 | 4.92 | 3.93 | -0.63 | 1.99 | 3.95 | 4.99 | -0.0575 | -0.075 | -0.215 | 0.485623 | 0.254611 | 0.562744 | 0.989223 | 0.997988 | 0.994666 |
| Gys1_Q9Z1E4_S657s_2 | Gys1_Q9Z1 | 2.77 | 2.97 | 4.54 | 3.2 | 0.93 | 2.97 | 2.72 | 3.81 | 4.4 | 1.87 | 4.92 | 3.93 | -0.63 | 1.99 | 3.95 | 4.99 | -0.7625 | 0.41 | -0.795 | 0.485623 | 0.254611 | 0.562744 | 0.989223 | 0.997988 | 0.994666 |
| Gys1_Q9Z1E4_S698s_1 | Gys1_Q9Z1 | -1.19 | -2.83 | -3.3 | -12.87 | -2.22 | -2.2 | -1.6 | -2.56 | 2.21 | -2.32 | -2.64 | -1.76 | 2.42 | -2.72 | 1.56 | -4.12 | -0.7625 | 0.41 | -0.795 | 0.125215 | 0.21961 | 0.141986 | 0.989223 | 0.997988 | 0.994666 |
| H1f0_P10922_S92s_1 | H1f0_P109 | 1.76 | 2.15 | 1.09 | 1.78 | 1.59 | 2.66 | 1.09 | 2.57 | 0.98 | 0.45 | 1.15 | 0.35 | 1.16 | 2.89 | 1.27 | 2.2 | 2.9025 | 3.92 | 4.3325 | 0.683582 | 0.505664 | 0.014019 | 0.989223 | 0.997988 | 0.996159 |
| H2afx_P27661_S140s_1 | H2afx_P27 | 0.66 | 0.86 | 0.04 | 0.19 | -0.08 | 1.28 | 0.38 | -0.56 | 2.25 | 2.07 | 0.66 | 0.02 | 2.2 | 0.5 | 0.56 | 0.55 | 0.2825 | -0.9625 | 0.185 | 0.266645 | 0.66216 | 0.154419 | 0.990434 | 0.997988 | 0.994666 |
| H2afy_Q9QZQ8_T178t_1 | H2afy_Q9C | 0.81 | 0.67 | 0.31 | 0.26 | 0.64 | 0.42 | 0.68 | 0.68 | 0.89 | 0.74 | -1.1 | -0.11 | 0.48 | 0.67 | -0.11 | 0.48 | -0.1825 | 0.8125 | 0.515 | 0.646907 | 0.685014 | 0.383986 | 0.994624 | 0.997988 | 0.995259 |
| Habp4_E9QKB2_S109s_1 | Habp4_E9( | -0.12 | 0.68 | 0.38 | -1.43 | 0.79 | 0.22 | 0.05 | -0.37 | -1.09 | 0.09 | -1.22 | 0.12 | -1.75 | -0.71 | 0.66 | 0.04 | 0.0925 | -0.4075 | -0.1325 | 0.617486 | 0.545508 | 0.469911 | 0.989223 | 0.997988 | 0.995259 |
| Habp4_E9QKB2_S75s_1 | Habp4_E9( | -4.6 | -4.14 | -4.28 | -4.98 | -1.9 | -4.84 | -2.89 | -7.46 | -5.06 | -2.91 | -2.01 | -3.64 | -1.88 | -3.64 | -3.16 | -3 | 0.295 | -0.4025 | -0.3175 | 0.003695 | 0.831591 | 0.100664 | 0.995762 | 0.997988 | 0.994666 |
| Habp4_E9QKB2_S7s_1 | Habp4_E9( | -0.02 | 0.43 | 0 | 0.03 | 0.72 | 0.1 | 0.31 | 0.3 | 0.32 | 0.68 | 0.02 | 0.12 | -0.23 | 0.41 | -0.1 | 0.16 | 0.2275 | 1.095 | 1.58 | 0.852148 | 0.310534 | 0.4949 | 0.989223 | 0.997988 | 0.999507 |
| Hacd3_Q8K2C9_S114s_1 | Hacd3_Q8I | 0.03 | 0.24 | 0.26 | 0.14 | 0.57 | -0.39 | 0.85 | 0.08 | -0.54 | 0.19 | -0.2 | 0.04 | -1.24 | 0.55 | 0.26 | 0.48 | 0.2475 | 0.175 | -0.05 | 0.714684 | 0.716549 | 0.249693 | 0.995437 | 0.997988 | 0.996966 |
| Hars2_G5E823_S66s_1 | Hars2_G5E | 1.91 | 1.77 | 1.75 | 1.37 | 0.99 | 0.92 | 2.08 | 1.98 | 0.2 | 1.36 | 2.84 | 2.01 | 0.25 | 1.06 | 1.53 | 1.96 | 0.11 | -0.295 | -0.155 | 0.223406 | 0.544179 | 0.854094 | 0.989223 | 0.997988 | 0.994666 |
| Hbbt1_E9Q223_S88s_1 | Hbbt1_E9C | -1.06 | -1.51 | -2.44 | -0.54 | -1.38 | 0.21 | -1.35 | -0.68 | -1.62 | -2.21 | -2.53 | -1.37 | -0.47 | -0.61 | -1.62 | -1.5 | -0.2075 | -0.0975 | -0.5 | 0.486523 | 0.260271 | 0.256421 | 0.989223 | 0.997988 | 0.994666 |
| Hcar1_E9PZR8_S331s_1 | Hcar1_E9P | -0.51 | -0.45 | -0.72 | 0.06 | -1.16 | -0.22 | -1.57 | -1.11 | 1.18 | -0.4 | -0.73 | -0.45 | 1.58 | -1.39 | -0.2 | -0.58 | 0.5875 | -0.545 | 0.3375 | 0.665533 | 0.095461 | 0.500311 | 0.989223 | 0.997988 | 0.996159 |
| Hcfc1_Q61191_S3s_1 | Hcfc1_Q61 | 1.12 | 0.8 | 0.76 | 0.96 | 0.29 | -0.71 | 0.32 | 1.17 | 0.85 | 1.62 | 2.05 | 0.13 | 0.54 | 0.88 | 0.84 | 1.13 | -0.61 | 0.305 | 0.2575 | 0.805948 | 0.118818 | 0.554125 | 0.989223 | 0.997988 | 0.999507 |
| Hcls1_P49710_S333s_1 | Hcls1_P49 | -0.23 | -1.08 | 0.72 | 0.83 | -0.48 | -0.57 | 1.08 | 0.62 | -0.01 | -0.05 | 0.33 | 0.84 | -0.26 | 0.86 | 0 | 0.85 | -0.6425 | 0.2525 | -0.0625 | 0.551909 | 0.851662 | 0.645456 | 0.996349 | 0.997988 | 0.994666 |
| Hcn1_O88704_S53s_1 | Hcn1_O88 | 6.22 | 1.36 | 3.6 | 1.61 | 4.24 | 6.8 | 5.06 | 5.41 | -2.04 | 3.13 | 0.33 | 5.11 | -3.07 | -1.42 | 4.99 | 4.98 | 0.1025 | 0.2175 | 0.3025 | 0.382017 | 0.066848 | 0.363941 | 0.989223 | 0.997988 | 0.994666 |
| Hcn1_O88704_S69s_1 | Hcn1_O88 | -0.92 | 0.6 | -0.95 | 0.36 | 0.13 | 0.74 | 0.36 | 0.49 | 0.14 | 0.35 | 0.95 | 0.77 | -0.07 | 0.75 | 0.28 | 1.45 | 2.18 | -1.565 | -1.8275 | 0.121163 | 0.134664 | 0.101226 | 0.989223 | 0.997988 | 0.994666 |
| Hcn1_O88704_S810s_1 | Hcn1_O88 | -0.64 | -0.65 | -1.08 | -0.44 | 0.12 | -0.8 | -0.22 | -0.22 | -1.06 | -0.81 | -0.43 | 0.66 | -1.19 | -0.42 | 0.38 | -0.25 | 0.6575 | 0.78 | 0.83 | 0.379134 | 0.146793 | 0.471505 | 0.989223 | 0.997988 | 0.994666 |
| Hcn1_O88704_S867s_1 | Hcn1_O88 | -1.36 | -1.47 | -0.53 | -1.45 | -0.59 | -0.58 | -1.2 | -0.53 | -0.88 | -0.49 | -1.25 | -1.31 | -0.34 | -0.3 | -1.7 | -0.9 | 0.4225 | 0.2925 | 0.3325 | 0.34191 | 0.136459 | 0.500657 | 0.989223 | 0.997988 | 0.994666 |
| Hcn1_O88704_S9s_1 | Hcn1_O88 | 1.7 | 2.37 | 0.82 | 0.01 | 1.47 | 2.45 | -0.42 | -0.28 | 2.42 | 1.81 | -0.86 | -0.13 | 2.52 | 0.91 | 0.03 | 0.9 | 0.4775 | 0.22 | 0.3925 | 0.837628 | 0.583115 | 0.619743 | 0.989223 | 0.997988 | 0.999507 |
| Hcn1_O88704_T39t_1 | Hcn1_O88 | 1.25 | 1.35 | 1.37 | 1.42 | 1.92 | 1.18 | 0.87 | 1.18 | 0.94 | 1.68 | 1.8 | 1.15 | 1.52 | 1.73 | 1.16 | 1.18 | -0.42 | -0.415 | -0.135 | 0.843237 | 0.823532 | 0.867105 | 0.995762 | 0.997988 | 0.999507 |
| Hcn2_O88703_S119s_1 | Hcn2_O88 | 0.56 | 0.54 | 0.53 | 0.48 | 0.69 | 0.32 | 0.34 | 0.5 | 0.77 | 0.69 | 0.7 | 0.45 | 0.57 | 0.93 | 0.73 | 0.82 | -0.06 | 0.045 | 0.05 | 0.324012 | 0.74961 | 0.553573 | 0.995762 | 0.997988 | 0.994666 |
| Hcn2_O88703_S131s_1 | Hcn2_O88 | 0.9 | 1.47 | 0.79 | 0.46 | 0.89 | 0.91 | 0.65 | 0.62 | 0.43 | 0.83 | 1.9 | 1.4 | 0.03 | 0.46 | 1.41 | 0.82 | -0.065 | 0.125 | 0.235 | 0.553884 | 0.609855 | 0.547411 | 0.989223 | 0.997988 | 0.994666 |
| Hcn2_O88703_S134s_1 | Hcn2_O88 | 1.71 | 1.21 | 0.93 | 0.81 | 1.13 | 1.21 | 0.54 | 0.97 | 0.66 | 1.04 | 0.44 | 0.31 | 0.19 | 0.79 | 0.78 | 0.85 | -0.1375 | 0.235 | -0.225 | 0.12158 | 0.481104 | 0.089062 | 0.989223 | 0.997988 | 0.994666 |
| Hcn2_O88703_S67s_1 | Hcn2_O88 | 1.38 | 0.61 | 1.05 | 1.3 | 0.42 | 1.1 | 1.17 | 1.29 | 0.94 | 1.27 | 2.34 | 0.59 | 1.08 | -0.24 | 1.85 | 1.25 | -0.2025 | -0.5525 | -0.5125 | 0.827302 | 0.757458 | 0.627987 | 0.995762 | 0.997988 | 0.999507 |
| Hcn2_O88703_S726s_1 | Hcn2_O88 | -0.89 | -1.44 | -0.28 | -0.29 | -1.13 | -0.3 | 0.13 | -0.24 | -1.45 | -1.13 | -0.84 | 0.27 | -1.8 | -0.3 | -0.91 | -1.35 | -0.09 | 0.2 | -0.1 | 0.396223 | 0.377304 | 0.889129 | 0.989223 | 0.997988 | 0.994666 |
| Hcn2_O88703_S750s_1 | Hcn2_O88 | -0.12 | 0.27 | 0.95 | 0.4 | 0.39 | -0.18 | 0.35 | 0.54 | -0.04 | 0.34 | 0.51 | 0.7 | -0.47 | 0.16 | 0.78 | 0.75 | 0.34 | -0.0625 | -0.365 | 0.854881 | 0.737222 | 0.993499 | 0.995762 | 0.997988 | 0.999507 |
| Hcn2_O88703_S771s_1 | Hcn2_O88 | 0.64 | 0.35 | 0.57 | 0.32 | 0.58 | 0.61 | 0.45 | 0.42 | 0.55 | 0.48 | -0.07 | 0.28 | 0.38 | 0.64 | 0.27 | 0.68 | -0.1 | 0.0025 | -0.07 | 0.927331 | 0.826129 | 0.514398 | 0.995762 | 0.997988 | 0.999507 |
| Hcn2_O88703_S795s_1 | Hcn2_O88 | -0.48 | 0.04 | 0.24 | 0.46 | 0.33 | 0.09 | 0.64 | 0.81 | 0.08 | -0.03 | -0.16 | 0.39 | -0.14 | 0.23 | 0.87 | 0.57 | 0.045 | -0.16 | 0.0225 | 0.354578 | 0.184681 | 0.98587 | 0.989223 | 0.997988 | 0.994666 |
| Hcn2_O88703_S818s_1 | Hcn2_O88 | -1.46 | -1.26 | -0.45 | -1.64 | -1.97 | -0.33 | -1.02 | -1.1 | 1.1 | -1.2 | -3.76 | -0.78 | 2.13 | -2.51 | 0.86 | -1.47 | 0.4025 | 0.005 | 0.3175 | 0.329714 | 0.811236 | 0.96298 | 0.995762 | 0.997988 | 0.994666 |
| Hcn2_O88703_S834s_1 | Hcn2_O88 | 0.26 | 0.27 | 0.5 | 0.57 | -0.73 | 0.73 | -0.49 | -0.1 | 1.64 | 0 | -0.83 | 0.13 | 1.81 | -0.96 | 0.08 | 0.33 | 0.0975 | 0.0425 | 0.955 | 0.874169 | 0.129363 | 0.737691 | 0.989223 | 0.997988 | 0.999507 |
| Hcn2_O88703_S840s_1 | Hcn2_O88 | 0.53 | 0.35 | 0.61 | 0.13 | 0.63 | 0.03 | 0.82 | 0.56 | 0.09 | -0.28 | -0.32 | 0.62 | 0.74 | 0.82 | 0.23 | 0.12 | -0.5475 | -0.165 | -0.085 | 0.797628 | 0.680264 | 0.212261 | 0.993464 | 0.997988 | 0.999507 |
| Hcn2_O88703_S842s_1 | Hcn2_O88 | 0.66 | 0.99 | 0.97 | 0.46 | 0.88 | 0.36 | 1.05 | 0.51 | 0.27 | 0.91 | 0.45 | 0.79 | -0.14 | 0.54 | 0.51 | 0.92 | 0.105 | -0.3775 | 0.0725 | 0.325521 | 0.785187 | 0.531958 | 0.995762 | 0.997988 | 0.994666 |
| Hcn2_O88703_S90s_1 | Hcn2_O88 | 0.72 | 0.53 | 0.57 | 0.8 | -0.14 | 0.79 | -0.03 | 0.36 | 1.09 | 0.75 | 1.23 | 0.05 | 1.19 | 0.2 | 0.43 | 0.47 | -0.07 | -0.165 | -0.3125 | 0.777481 | 0.146768 | 0.684811 | 0.989223 | 0.997988 | 0.99798 |
| Hcn2_O88703_T70t_1 | Hcn2_O88 | -0.31 | 0.77 | 0.15 | 0.02 | -1.04 | -0.11 | -0.53 | -0.24 | 1.73 | 0.12 | -0.24 | -0.09 | 1.24 | -1.2 | 0 | -0.44 | -0.41 | 0.125 | -0.0825 | 0.624487 | 0.071372 | 0.646422 | 0.989223 | 0.997988 | 0.995259 |
| Hcn2_O88703_T763t_1 | Hcn2_O88 | 0.15 | -0.11 | 0.81 | 1.08 | 0.11 | -0.38 | 1.17 | 1.08 | 0.4 | 0.05 | 0.94 | 0.85 | -0.04 | 0.49 | 1.39 | 1.25 | -0.6375 | 0.2225 | -0.2575 | 0.505364 | 0.977321 | 0.828945 | 0.999187 | 0.997988 | 0.994666 |
| Hcn3_O88705_S33s_1 | Hcn3_O88 | 0.52 | 0.52 | 0.89 | 0.59 | 0.23 | 1 | 0.24 | 0.44 | 0.53 | 2.21 | -0.64 | 0.05 | 0.73 | 0.74 | 0.63 | 0.03 | 0.0125 | 0.0775 | 0.29 | 0.723223 | 0.552697 | 0.87018 | 0.989223 | 0.997988 | 0.9971 |
| Hcn3_O88705_S633s_1 | Hcn3_O88 | 0.63 | 0.8 | 0.83 | 0.74 | 0.37 | 0.48 | 1.03 | 0.75 | 0.65 | 0.49 | 0.52 | 0.57 | -0.1 | 0.52 | 0.48 | 0.26 | -0.1525 | -0.0925 | -0.0975 | 0.096885 | 0.688383 | 0.360543 | 0.994672 | 0.997988 | 0.994666 |
| Hcn3_O88705_S648s_1 | Hcn3_O88 | 0.22 | -0.21 | 0.65 | 0.88 | 0.92 | -0.06 | 0.84 | 0.7 | 0.3 | 0.88 | 0.36 | 0.67 | 0.09 | 0.51 | 0.99 | 1.43 | -0.0925 | -0.1925 | -0.46 | 0.353043 | 0.527769 | 0.592443 | 0.989223 | 0.997988 | 0.994666 |
| Hcn3_O88705_S654s_1 | Hcn3_O88 | 0.78 | 0.57 | 0.49 | 0.71 | 1.13 | 0.79 | 0.95 | 1.1 | 0.7 | 0.5 | 1.29 | 1.02 | 0.04 | 1.14 | 0.45 | 1.13 | 0.215 | 0.1675 | 0.37 | 0.871001 | 0.113265 | 0.360208 | 0.989223 | 0.997988 | 0.999507 |
| Hcn3_O88705_S720s_1 | Hcn3_O88 | -2.96 | -2.28 | 0.27 | -0.34 | -1.01 | -1.56 | 0.1 | -0.21 | -0.53 | -1.88 | -0.25 | -0.32 | -1.23 | -0.82 | -0.66 | -0.19 | 0.355 | 0.24 | 0.0525 | 0.407231 | 0.391605 | 0.456166 | 0.989223 | 0.997988 | 0.994666 |
| Hcn3_O88705_S733s_1 | Hcn3_O88 | -1.6 | -2.57 | -2.91 | -5.24 | -2.51 | -0.07 | -2.6 | -1.5 | 3.51 | -3.4 | -0.79 | -0.42 | 3.64 | -1.72 | 3.09 | -2.48 | 0.6575 | 0.5825 | 0.6025 | 0.033164 | 0.120713 | 0.073372 | 0.989223 | 0.997988 | 0.994666 |
| Hcn3_O88705_T16t_1 | Hcn3_O88 | -1.4 | -1.6 | -1.16 | -1.92 | -1.95 | -0.3 | -1.13 | -2.07 | -0.34 | -1.97 | -3.12 | -0.76 | 0.6 | -0.41 | -1.24 | -1.36 | 1.41 | 2.805 | 3.7125 | 0.072714 | 0.708058 | 0.96306 | 0.995437 | 0.997988 | 0.994666 |
| Hcn4_B2RY58_S1026s_1 | Hcn4_B2RY | 0.3 | 0.41 | 0.6 | 0.65 | 0.28 | 0.32 | 0.37 | 0.8 | 0.84 | 0.91 | 0.7 | 0.51 | 0.83 | 0.8 | 0.89 | 0.79 | 0.1575 | -0.0275 | 0.9175 | 0.171185 | 0.83346 | 0.282678 | 0.995762 | 0.997988 | 0.994666 |
| Hcn4_B2RY58_S918s_1 | Hcn4_B2RY | 0.41 | 0.68 | 0.7 | -2.01 | 0.77 | 0.83 | 1.22 | 1.54 | -0.33 | 0.46 | 0.96 | -0.63 | 0.16 | 0.24 | 0.93 | 1.13 | -0.0475 | 0.25 | 0.3375 | 0.30453 | 0.084509 | 0.802048 | 0.989223 | 0.997988 | 0.994666 |
| Hcn4_B2RY58_S936s_1 | Hcn4_B2RY | 0.54 | 0.85 | -0.2 | 0.21 | -0.12 | 1.13 | -0.81 | 0.64 | 0.17 | 0.14 | 1.24 | 0.16 | 0.25 | 0.9 | 0.12 | 0.68 | 1.145 | 0.17 | 0.67 | 0.678655 | 0.756341 | 0.830895 | 0.995762 | 0.997988 | 0.996159 |
| Hcn4_B2RY58_T1071t_1 | Hcn4_B2RY | 0.72 | 0.14 | 0.78 | 0.81 | 0.54 | 0.42 | 0.61 | 0.57 | 0.94 | 1.13 | 1.42 | 0.18 | 0.98 | 0.04 | 0.98 | 1.28 | -0.14 | 0.0775 | 0.1375 | 0.551519 | 0.741622 | 0.369769 | 0.995762 | 0.997988 | 0.994666 |
| Hdac4_Q6NZM9_S465s_1 | Hdac4_Q6I | -0.73 | -0.4 | -0.85 | -0.98 | 0.07 | -0.67 | -0.48 | -0.99 | -0.53 | -0.59 | -1.72 | -0.99 | -0.72 | 0.19 | -1.11 | -0.91 | -0.0775 | 0.305 | 0.2075 | 0.766942 | 0.445107 | 0.51071 | 0.989223 | 0.997988 | 0.9971 |
| Hdac4_Q6NZM9_S629s_1 | Hdac4_Q6I | 0.25 | 0.61 | 0.06 | 0.07 | 0.92 | 0.11 | 0.46 | 0.38 | 0.1 | 0.47 | -0.55 | -0.21 | 0.09 | 0.68 | 0.29 | 0.56 | 0.2225 | -0.2175 | 0.1025 | 0.56538 | 0.407567 | 0.330956 | 0.989223 | 0.997988 | 0.995259 |
| Hdac4_Q6NZM9_S630s_1 | Hdac4_Q6I | -0.05 | 0.17 | -0.54 | -0.46 | 0.22 | -0.64 | 0.19 | 0.25 | 0.06 | 0.38 | 0.5 | -0.96 | -0.04 | 0.22 | -0.35 | 0.46 | 0.22 | -0.295 | 0.1575 | 0.345157 | 0.458262 | 0.572096 | 0.989223 | 0.997988 | 0.994666 |
| Hdac4_Q6NZM9_T463t_1 | Hdac4_Q6I | -1.19 | -1.03 | -0.46 | -0.43 | -0.14 | -1.23 | 0.49 | -0.46 | -1.46 | -1.18 | -2.09 | -0.83 | -1.01 | -0.09 | -1.03 | -0.67 | 0.225 | 0.215 | 0.2925 | 0.816037 | 0.277292 | 0.103828 | 0.989223 | 0.997988 | 0.999507 |
| Hdac5_Q3UJF1_S652s_1 | Hdac5_Q3l | -2.09 | -1.65 | -0.12 | -0.77 | -0.99 | -2.57 | -0.15 | -0.14 | -1.06 | -3.81 | -0.39 | -0.08 | 0.38 | -0.65 | -0.06 | 0.33 | 0.4425 | -0.6125 | 0.0775 | 0.035105 | 0.761536 | 0.834992 | 0.995762 | 0.997988 | 0.994666 |
| Hdac5_Q3UJF1_S662s_1 | Hdac5_Q3l | -0.68 | -0.14 | -1.05 | -0.53 | -0.39 | -0.03 | -0.03 | 0.16 | -0.37 | -0.64 | -0.05 | -0.93 | -0.19 | 0.7 | -0.87 | 0.02 | 0.195 | -0.1775 | 1.1575 | 0.203694 | 0.071688 | 0.736249 | 0.989223 | 0.997988 | 0.994666 |
| Hdac5_Q3UJF1_T488t_1 | Hdac5_Q3l | -1.9 | -1.44 | -2.52 | -2.31 | -1 | -2.42 | -1.41 | -2.93 | -1.41 | -1.28 | -2.09 | -2.25 | -1.57 | -0.96 | -3.05 | -2.15 | 0.5275 | 0.1025 | 0.515 | 0.819466 | 0.826697 | 0.428877 | 0.995762 | 0.997988 | 0.999507 |
| Hdac6_Q9Z2V5_S1148s_1 | Hdac6_Q9z | 0 | 0.28 | 0.06 | 0.59 | 0.43 | -0.01 | 0.54 | 0.51 | 0.21 | 0.56 | 0.94 | 0.45 | -0.02 | 0.61 | 0.4 | 0.17 | 0.1025 | 0.285 | 0.11 | 0.834255 | 0.58532 | 0.263312 | 0.989223 | 0.997988 | 0.999507 |
| Hdac6_Q9Z2V5_S43s_1 | Hdac6_Q9z | -2.1 | -1.98 | -3.33 | -3.4 | -2.07 | -2.32 | -2.02 | -2.63 | -2.42 | -1.59 | -2.19 | -2.62 | -2.04 | -1.27 | -3.09 | -2.77 | 0.135 | 0.3075 | 0.0575 | 0.440351 | 0.278048 | 0.266691 | 0.989223 | 0.997988 | 0.994666 |
| Hdac6_Q9Z2V5_S59s_1 | Hdac6_Q9z | -0.1 | -0.26 | -0.34 | -0.09 | 0.06 | -0.59 | 0.77 | -0.45 | -0.28 | -0.43 | -0.22 | -1.08 | -0.78 | 0.26 | -1.49 | -1 | 0.4425 | 0.4975 | 0.41 | 0.173527 | 0.657893 | 0.271205 | 0.990434 | 0.997988 | 0.994666 |
| Hdac6_Q9Z2V5_S909s_1 | Hdac6_Q9z | 1.46 | 1.19 | -0.75 | 0.58 | 1.73 | 1.61 | 0.23 | 0.85 | 0.58 | 1.13 | 1.16 | 0.51 | -0.2 | 1.37 | 0.09 | 0.11 | 0.145 | -0.305 | -0.555 | 0.621171 | 0.385192 | 0.649767 | 0.989223 | 0.997988 | 0.995259 |
| Hdac9_Q99N13_S218s_1 | Hdac9_Q99 | -0.63 | -0.89 | -0.85 | -1.37 | -0.65 | -0.92 | -1.45 | -1.13 | -1.17 | -1.17 | -0.7 | -1.6 | -0.32 | -0.51 | -0.88 | -0.86 | 0.485 | 0.225 | -0.2775 | 0.315365 | 0.706452 | 0.441994 | 0.995437 | 0.997988 | 0.994666 |
| Hdgf_E0CXA0_S101s_1 | Hdgf_E0CX | -6.21 | -5.76 | -5.68 | -2.26 | -7.01 | -7.14 | -5.19 | -4.86 | -3.89 | -7.51 | -3.85 | -4.15 | -1.94 | -3.47 | -7.51 | -8.47 | -0.1025 | -0.225 | 0.2925 | 0.812346 | 0.273215 | 0.909344 | 0.989223 | 0.997988 | 0.999507 |
| Hdgf_E0CXA0_S133s_1 | Hdgf_E0CX | -4.55 | -3.7 | -0.54 | 0.45 | -3.51 | -4.86 | -0.27 | 0.19 | -3.94 | -3.76 | 0.4 | -0.4 | -2.62 | -2.82 | 0.39 | 1.1 | -1.0725 | 0.1275 | -0.37 | 0.42642 | 0.985128 | 0.911534 | 0.999187 | 0.997988 | 0.994666 |
| Hdgf_E0CXA0_S174s_1 | Hdgf_E0CX | -1.45 | -1.01 | -2.87 | -1.93 | -2.82 | -1.29 | -2.31 | -2.2 | -2.09 | -0.85 | -2.39 | -4.39 | -1.02 | -0.73 | -2.12 | -1.2 | -0.0275 | 0.16 | 1.0975 | 0.268579 | 0.477702 | 0.420331 | 0.989223 | 0.997988 | 0.994666 |
| Hdgfrp2_Q3UMU9_S576s_1 | Hdgfrp2_Q | -2.22 | -1.75 | -1.93 | -0.83 | -2.54 | -2.8 | -1.49 | -3.23 | -1.78 | -1.86 | -0.99 | -1.8 | -0.91 | -2.21 | -1.3 | -2.33 | -0.34 | -0.615 | 0.5475 | 0.991068 | 0.087484 | 0.840098 | 0.989223 | 0.997988 | 1 |
| Hdgfrp2_Q3UMU9_S628s_2 | Hdgfrp2_Q | -1.24 | -2.02 | -0.49 | -0.92 | -2.24 | -2.45 | -1.58 | -2.69 | -1.52 | -1.34 | -0.95 | -1.22 | -1.45 | -2.83 | -2.46 | -1.66 | -0.8325 | 0.075 | -0.005 | 0.061099 | 0.019865 | 0.801232 | 0.989223 | 0.997988 | 0.994666 |
| Hdgfrp2_Q3UMU9_S635s_1 | Hdgfrp2_Q | -1.92 | -3.07 | -1.26 | -1.98 | -5.17 | -3.58 | -3.21 | -2.08 | -1.58 | -2.38 | -1.32 | -1 | -0.84 | -2.89 | -2.85 | -1.8 | -1.0725 | -0.09 | -0.9325 | 0.947247 | 0.04923 | 0.30188 | 0.989223 | 0.997988 | 0.999507 |
| Hdgfrp2_Q3UMU9_S635s_2 | Hdgfrp2_Q | -1.31 | -2.15 | -0.49 | -0.92 | -1.78 | -2.14 | -1.58 | -2.69 | -0.53 | -1.24 | -0.95 | -1.22 | -0.48 | -1.63 | -2.46 | -1.66 | -1.4525 | 0.4875 | -0.0375 | 0.508322 | 0.067241 | 0.555467 | 0.989223 | 0.997988 | 0.994666 |
| Hdgfrp2_Q3UMU9_S659s_1 | Hdgfrp2_Q | 0.14 | 0 | -0.29 | -1.24 | -0.97 | 0.34 | -0.71 | -0.77 | 0.98 | -0.1 | 1.74 | -0.82 | 1.13 | -0.14 | -1.09 | -0.71 | -0.83 | 0.2325 | -0.34 | 0.787153 | 0.66143 | 0.201045 | 0.990434 | 0.997988 | 0.999507 |
| Hdgfrp2_Q3UMU9_S661s_1 | Hdgfrp2_Q | -0.27 | 0.19 | -1.17 | -0.81 | -0.21 | -0.04 | -3.06 | -3.17 | 0.22 | -0.45 | 0.36 | -1.5 | 0.56 | -1.11 | -0.9 | -3.18 | -0.18 | 0.7975 | 0.145 | 0.391858 | 0.188794 | 0.725796 | 0.989223 | 0.997988 | 0.994666 |
| Hdlbp_Q8VDJ3_S944s_1 | Hdlbp_Q8\ | -1.72 | -2.07 | -1.87 | -3.98 | -2.03 | -1.64 | -2.31 | -2.26 | 0.45 | -1.75 | -2.22 | -1.9 | 0.99 | -2.42 | -0.25 | -1.21 | -1.105 | 0.1725 | -0.6425 | 0.056357 | 0.492425 | 0.169873 | 0.989223 | 0.997988 | 0.994666 |
| Heatr5b_Q8C547_S1737s_1 | Heatr5b_Q | -0.37 | -0.05 | 0.11 | -0.87 | 0.05 | -0.61 | -0.74 | -0.56 | -0.14 | 0.03 | -1.51 | -0.5 | 0.24 | -0.59 | -0.44 | -0.56 | 0.35 | 1.055 | 1.6875 | 0.897936 | 0.576732 | 0.563841 | 0.989223 | 0.997988 | 0.999507 |
| Heatr5b_Q8C547_S1739s_1 | Heatr5b_Q | -0.01 | 0.25 | -0.07 | 0.05 | 0.51 | -0.15 | -0.17 | -0.54 | -0.58 | 0.23 | 0.72 | -0.38 | 0.03 | 0.14 | 0.32 | -0.38 | -0.17 | -0.235 | -0.0425 | 0.916278 | 0.601404 | 0.861227 | 0.989223 | 0.997988 | 0.999507 |
| Hebp1_Q9R257_S111s_1 | Hebp1_Q9I | 0.83 | 1.01 | 0.98 | 0.85 | 0.39 | 0.06 | 0.38 | 0.62 | -0.25 | 0.36 | 0.46 | 0.65 | -0.23 | 0.58 | 0.45 | 0.11 | -0.1425 | -0.0575 | -0.0275 | 0.028813 | 0.027075 | 0.041261 | 0.989223 | 0.997988 | 0.994666 |
| Hectd1_F8WIE5_S357s_1 | Hectd1_F8 | -0.65 | 0.08 | -0.64 | -1.17 | -0.47 | -1.1 | -1.83 | -1.56 | -0.25 | 0.23 | -2.19 | -1.32 | 1.74 | -2.27 | -0.14 | -0.64 | -0.555 | -0.6125 | -0.69 | 0.72776 | 0.117566 | 0.607545 | 0.989223 | 0.997988 | 0.9971 |
| Hectd1_F8WIE5_S358s_1 | Hectd1_F8 | -1.62 | -2.08 | -0.77 | -2.35 | -2.21 | -1.41 | -1.69 | -1.97 | 0.23 | -1.91 | -0.33 | -0.53 | 0.83 | -1.94 | 0.25 | -1.3 | -0.645 | -0.2875 | 0.2675 | 0.103812 | 0.761644 | 0.069806 | 0.995762 | 0.997988 | 0.994666 |
| Hecw1_Q8K4P8_S1071s_1 | Hecw1_Q8 | -2.03 | -2.71 | -2.91 | -6.63 | -2.18 | -2.63 | -1.83 | -2.88 | -2.45 | -2.73 | -0.32 | -2.06 | -1.27 | -1.18 | -1.58 | -2.23 | -0.115 | 1.07 | 1.165 | 0.054126 | 0.217315 | 0.129353 | 0.989223 | 0.997988 | 0.994666 |
| Hecw1_Q8K4P8_S1199s_1 | Hecw1_Q8 | 0.52 | 0.12 | 0.9 | 0.79 | -0.04 | 1.19 | 0.87 | -0.26 | 1.28 | 1.61 | 1.37 | 1.16 | 1.82 | 1.41 | 0.9 | 0.96 | 1.19 | 1.68 | 2.005 | 0.055314 | 0.709257 | 0.014868 | 0.995437 | 0.997988 | 0.994666 |
| Hecw1_Q8K4P8_S396s_1 | Hecw1_Q8 | 0.93 | 1.01 | 1.81 | 1.75 | 1.01 | -0.28 | 1.07 | 1.66 | 0.55 | 1.15 | 2.47 | 1.97 | -0.79 | 0.91 | 1.94 | 1.92 | -0.1425 | 0.7725 | 0.69 | 0.544303 | 0.265952 | 0.73141 | 0.989223 | 0.997988 | 0.994666 |
| Hecw1_Q8K4P8_S718s_1 | Hecw1_Q8 | -1.31 | -2.1 | -0.17 | -1.3 | -1.44 | -1.28 | -1.41 | -1.35 | -2.52 | -1.93 | -0.31 | -0.7 | -1.39 | -0.72 | -1.17 | -1.38 | -0.51 | 0.16 | -0.38 | 0.896703 | 0.699308 | 0.808397 | 0.994893 | 0.997988 | 0.999507 |
| Hecw1_Q8K4P8_S786s_1 | Hecw1_Q8 | -0.33 | -0.98 | -0.65 | -1.37 | -0.34 | -0.66 | -1.89 | -1.76 | 0.05 | -0.35 | -0.2 | -1.17 | 0.02 | -0.08 | -0.27 | -0.24 | -0.15 | -0.145 | 0.055 | 0.040698 | 0.444398 | 0.264162 | 0.989223 | 0.997988 | 0.994666 |
| Hecw1_Q8K4P8_S869s_1 | Hecw1_Q8 | -1.89 | -1.87 | -1.58 | -2.75 | -0.56 | -1.75 | -1.65 | -1.4 | -1.38 | -2.58 | -1.77 | -2.02 | -0.1 | -0.13 | -1.53 | -2.97 | -0.33 | 0.415 | 0.69 | 0.220653 | 0.087629 | 0.816368 | 0.989223 | 0.997988 | 0.994666 |
| Hecw1_Q8K4P8_S937s_1 | Hecw1_Q8 | 0.08 | 0.57 | 0.21 | -0.3 | 0.7 | 0.07 | -0.07 | -0.03 | -0.21 | 0.15 | 0.46 | 0.05 | 0.78 | 0.9 | -0.02 | 0.07 | 0.6825 | 0.085 | 0.84 | 0.39308 | 0.923443 | 0.921433 | 0.997335 | 0.997988 | 0.994666 |
| Hecw2_Q6I6G8_S1046s_1 | Hecw2_Q6 | 1.49 | 0.01 | 0.93 | 0.77 | -1.01 | 3.43 | 0.61 | 0.65 | 2.16 | 1.43 | 1.7 | 0.18 | 2.16 | -1.43 | 0.61 | -0.41 | 0.0275 | -0.0275 | 0.2925 | 0.446622 | 0.887113 | 0.267262 | 0.996493 | 0.997988 | 0.994666 |
| Hecw2_Q6I6G8_S1048s_1 | Hecw2_Q6 | 6.59 | 1.12 | 2.98 | -0.91 | 0.81 | 6.64 | 4.43 | 2.41 | 0.72 | 5.33 | -3.31 | 3.24 | 2.6 | -2.35 | 4.71 | 4.8 | 0.12 | 0.5675 | -0.5675 | 0.997979 | 0.520973 | 0.656857 | 0.989223 | 0.997988 | 1 |
| Hecw2_Q6I6G8_S1173s_1 | Hecw2_Q6 | 0.24 | -2.33 | -0.31 | 0.19 | -0.54 | -0.17 | -0.7 | 0.48 | -0.32 | -1.42 | 0.25 | -0.38 | -0.31 | -1.26 | -1.39 | -0.43 | 1.1275 | -0.95 | -0.005 | 0.628973 | 0.590924 | 0.893532 | 0.989223 | 0.997988 | 0.995259 |
| Hecw2_Q6I6G8_S1173s_2 | Hecw2_Q6 | -0.06 | 0.02 | 0.52 | 0.85 | -0.25 | -0.38 | -0.55 | 0.46 | 0.7 | -0.23 | 1.65 | 0.26 | 0.73 | 0.38 | 0.16 | 0.34 | 0.32 | 0.085 | -0.295 | 0.818067 | 0.136926 | 0.554029 | 0.989223 | 0.997988 | 0.999507 |
| Hecw2_Q6I6G8_S1176s_2 | Hecw2_Q6 | 0.57 | 1.03 | 0.8 | 0.9 | 0.45 | -0.01 | -0.39 | 0.62 | 1.29 | 0.43 | 1.76 | 0.28 | 1.01 | 1.4 | 0.28 | 0.6 | -0.5125 | 0.2625 | 0.07 | 0.993638 | 0.040557 | 0.757981 | 0.989223 | 0.997988 | 1 |
| Hecw2_Q6I6G8_S1181s_2 | Hecw2_Q6 | -0.89 | -1.59 | -0.59 | 0.21 | -1.23 | -0.81 | -0.87 | -0.47 | -0.07 | -1.13 | 1.27 | -0.14 | 0.38 | -1.3 | -0.18 | 0.02 | -0.6575 | 0.115 | -0.0025 | 0.377587 | 0.740105 | 0.240427 | 0.995762 | 0.997988 | 0.994666 |
| Hecw2_Q6I6G8_S406s_1 | Hecw2_Q6 | 1.56 | 0.7 | 1.85 | 2.31 | 1.03 | 1.46 | 1.94 | 2.34 | 1.08 | 2.47 | 1.94 | 1.61 | 0.65 | 1.29 | 1.76 | 1.7 | -0.13 | 0.6975 | 0.445 | 0.54964 | 0.835079 | 0.696385 | 0.995762 | 0.997988 | 0.994666 |
| Hecw2_Q6I6G8_S48s_1 | Hecw2_Q6 | -0.62 | -1.34 | -0.08 | 0.03 | -0.32 | -0.08 | -0.17 | 0.02 | -0.31 | 0.05 | 1.1 | -0.21 | 0.35 | 0.88 | -0.18 | -0.35 | 0.0875 | 0.17 | -0.255 | 0.130171 | 0.28634 | 0.155338 | 0.989223 | 0.997988 | 0.994666 |
| Hecw2_Q6I6G8_S596s_1 | Hecw2_Q6 | 0.13 | 0.16 | 0.25 | 0.03 | -0.23 | -0.82 | 0.08 | 0.45 | -0.75 | -0.28 | 0.02 | 0.65 | -1.81 | -0.01 | 0.51 | 0.56 | 0.365 | 0.66 | 0.6775 | 0.531855 | 0.371581 | 0.479136 | 0.989223 | 0.997988 | 0.994666 |
| Hecw2_Q6I6G8_S858s_1 | Hecw2_Q6 | -0.43 | -0.04 | -0.04 | -0.07 | 0.07 | -0.23 | -0.3 | 0.09 | -0.63 | -0.55 | 0.44 | -0.01 | 0.32 | 0.64 | -0.24 | -0.34 | -0.2725 | -0.2325 | -0.33 | 0.442381 | 0.813696 | 0.889058 | 0.995762 | 0.997988 | 0.994666 |
| Hepacam_Q640R3_S280s_1 | Hepacam_( | 0.51 | 0.63 | 0.88 | 0.84 | 0.34 | 0.26 | 0.62 | 0.42 | 0.39 | 0.8 | 0.44 | 0.4 | 0.2 | 1.09 | 0.61 | 0.81 | 0.0525 | -0.0425 | 0.24 | 0.894369 | 0.175848 | 0.376729 | 0.989223 | 0.997988 | 0.999507 |
| Hepacam_Q640R3_S301s_2 | Hepacam_( | 1.24 | 1.56 | 1.51 | 1.83 | 0.75 | 1.16 | 0.77 | 1.33 | 1.23 | 2.93 | 1.92 | 1.61 | -0.6 | 1.32 | 2.01 | 2.33 | -0.305 | -0.2075 | -0.0375 | 0.658495 | 0.054175 | 0.329212 | 0.989223 | 0.997988 | 0.995259 |
| Hepacam_Q640R3_S320s_1 | Hepacam_( | 2.74 | 2.48 | 2.47 | 2.49 | 2.36 | 2.11 | 1.81 | 2.29 | 1.22 | 2.4 | 1.42 | 2.36 | 1.44 | 0.06 | 2.49 | 2.28 | -0.5325 | 0.3875 | -0.27 | 0.085651 | 0.096183 | 0.063778 | 0.989223 | 0.997988 | 0.994666 |
| Hepacam_Q640R3_S321s_1 | Hepacam_( | 0.2 | 0.24 | 0.91 | 0.39 | 0.2 | 0.14 | 0.02 | 0.56 | -1.74 | 0.44 | 0.37 | 0.48 | 0.05 | -1.45 | 0.61 | 1.1 | -0.4025 | -0.695 | -0.9775 | 0.510204 | 0.428037 | 0.315113 | 0.989223 | 0.997988 | 0.994666 |
| Hepacam_Q640R3_S345s_1 | Hepacam_( | 0.27 | 0.05 | 0.21 | 0.77 | 0.26 | -0.02 | 1.04 | 0.63 | 0.41 | 0.26 | 0.02 | 0.6 | -0.37 | 0.4 | 0.31 | 0.49 | -0.205 | -0.5475 | -0.3575 | 0.70251 | 0.615176 | 0.992387 | 0.989223 | 0.997988 | 0.996159 |
| Hepacam_Q640R3_S352s_1 | Hepacam_( | -2.11 | -2.03 | -1.74 | -1.8 | -0.28 | -2.06 | -0.42 | -1.81 | -1.83 | -1.51 | -1.06 | -1.39 | -2.18 | -0.19 | -1.42 | -1.12 | 0.1525 | -0.0025 | -0.1175 | 0.124065 | 0.103441 | 0.088704 | 0.989223 | 0.997988 | 0.994666 |
| Hepacam_Q640R3_S366s_2 | Hepacam_( | -0.11 | -1.23 | 0.1 | 0.25 | -0.71 | 0.71 | -0.73 | -0.7 | 2.72 | 0.37 | 0.32 | 0.31 | 0.67 | -1.69 | 0.82 | 0.25 | 0.7775 | 0.4725 | 0.6925 | 0.671394 | 0.809471 | 0.085623 | 0.995762 | 0.997988 | 0.996159 |
| Hepacam_Q640R3_S370s_2 | Hepacam_( | -0.11 | -1.23 | 0.1 | 0.25 | -0.71 | 0.71 | -0.73 | -0.7 | 2.72 | 0.37 | 0.32 | 0.31 | 0.67 | -1.69 | 0.82 | 0.25 | -0.11 | 1.1775 | 0.26 | 0.671394 | 0.809471 | 0.085623 | 0.995762 | 0.997988 | 0.996159 |
| Hepacam_Q640R3_T297t_1 | Hepacam_( | -0.62 | 0.45 | 0.15 | -0.31 | 0.65 | -0.5 | -0.58 | -0.51 | -0.42 | -0.16 | -0.28 | -0.51 | -0.11 | 0.12 | 0.29 | 0.39 | -0.11 | 1.1775 | 0.26 | 0.424243 | 0.683647 | 0.385828 | 0.994272 | 0.997988 | 0.994666 |
| Hepacam_Q640R3_T297t_2 | Hepacam_( | 1.24 | 1.56 | 1.51 | 1.83 | 0.75 | 1.16 | 0.77 | 1.33 | 1.23 | 2.93 | 1.92 | 1.61 | -0.6 | 1.32 | 2.01 | 2.33 | -0.1525 | -0.26 | 0.255 | 0.658495 | 0.054175 | 0.329212 | 0.989223 | 0.997988 | 0.995259 |
| Hepacam_Q640R3_T378t_1 | Hepacam_( | 8.31 | 3.38 | 3.71 | 0.32 | 3.48 | 9.23 | 3.28 | 2.02 | 1.77 | 7.14 | 2.43 | 2.9 | 1.47 | -0.63 | 4.42 | 4.26 | -0.5325 | 0.3875 | -0.27 | 0.385911 | 0.771066 | 0.835617 | 0.995762 | 0.997988 | 0.994666 |
| Herc1_E9PZP8_S133s_1 | Herc1_E9P | -1.57 | -1.71 | -0.61 | -0.74 | -1.35 | -2.52 | -0.8 | -0.86 | -0.53 | -1.12 | -1.23 | -0.85 | 0.56 | -0.71 | -0.35 | -1.11 | 0.5725 | -0.37 | -1.55 | 0.11431 | 0.623004 | 0.514504 | 0.989223 | 0.997988 | 0.994666 |
| Herc1_E9PZP8_S1406s_1 | Herc1_E9P | -0.66 | -1.67 | -1.25 | -3.43 | -1.92 | -0.54 | -1.68 | -1.38 | -0.29 | -2 | -2.4 | -1.09 | -0.13 | -2.77 | -1.33 | -1.84 | -0.225 | 0.225 | 0.755 | 0.745749 | 0.53723 | 0.656179 | 0.989223 | 0.997988 | 0.9971 |
| Herc1_E9PZP8_S1428s_1 | Herc1_E9P | 1.43 | 1.29 | 0.64 | 1.41 | 1 | 1.32 | 1.04 | 1.15 | 1.38 | 1.29 | 0.67 | 0.86 | 0.47 | 1.55 | 1.29 | 0.58 | 0.3725 | 0.3075 | 0.235 | 0.536779 | 0.798053 | 0.631058 | 0.995762 | 0.997988 | 0.994666 |
| Herc1_E9PZP8_S1491s_1 | Herc1_E9P | -0.15 | -0.37 | 0.07 | -0.24 | -0.38 | -0.31 | -0.44 | -0.2 | -0.76 | -0.45 | 0.25 | -0.12 | -0.5 | -0.13 | -0.46 | -0.05 | -0.065 | -0.1425 | -0.22 | 0.659949 | 0.452021 | 0.732581 | 0.989223 | 0.997988 | 0.995259 |
| Herc1_E9PZP8_S2718s_1 | Herc1_E9P | -1.91 | -0.95 | -1.06 | -0.91 | -1.26 | -1.66 | -1.52 | -1.79 | -1.33 | -1.14 | -0.77 | -1.99 | 0 | -0.74 | -1.56 | -0.19 | -0.16 | -0.0975 | -0.1125 | 0.186228 | 0.247965 | 0.781297 | 0.989223 | 0.997988 | 0.994666 |
| Herc1_E9PZP8_S2718s_2 | Herc1_E9P | 0.45 | -0.13 | -1.24 | -2.75 | 0.02 | -0.23 | -1.04 | -1.13 | 0.88 | 0.51 | -1.53 | 0.34 | 1.34 | 0.42 | -1.12 | -0.55 | -0.35 | -0.1 | 0.585 | 0.250002 | 0.633716 | 0.241428 | 0.989223 | 0.997988 | 0.994666 |
| Herc1_E9PZP8_S2721s_1 | Herc1_E9P | -0.65 | 0.66 | 0.15 | -0.34 | -0.16 | -0.51 | -0.71 | -0.25 | -1.51 | -0.51 | -0.09 | -0.24 | 0.39 | -0.65 | -0.18 | -0.55 | 0.3225 | 0.9675 | 0.94 | 0.600825 | 0.281441 | 0.218138 | 0.989223 | 0.997988 | 0.995259 |
| Herc1_E9PZP8_S2736s_1 | Herc1_E9P | 1.1 | 0.51 | 0.18 | 0.06 | 0 | 0.75 | -0.19 | -0.19 | 0.95 | 1.22 | 0.51 | 0.11 | 1.06 | 0.6 | 0.34 | -0.05 | -0.3625 | -0.5425 | -0.2025 | 0.943979 | 0.281592 | 0.509895 | 0.989223 | 0.997988 | 0.999507 |
| Herc1_E9PZP8_S2745s_1 | Herc1_E9P | -0.08 | 0.03 | 1.14 | -0.15 | 0.03 | 0.14 | 0.4 | -0.34 | 0.9 | 0.6 | -0.71 | -0.74 | 0.59 | 0.22 | -0.12 | 0.51 | -0.37 | 0.235 | 0.025 | 0.858952 | 0.609035 | 0.655704 | 0.989223 | 0.997988 | 0.999507 |
| Herc1_E9PZP8_S4855s_1 | Herc1_E9P | 0.28 | 0.62 | 1.09 | 1.26 | -0.24 | 0.31 | 0.09 | 1.4 | 0.25 | 0.68 | 0.94 | -0.3 | -0.23 | -0.68 | 0.63 | 0.87 | -0.1775 | -0.2225 | 0.065 | 0.140838 | 0.308537 | 0.263879 | 0.989223 | 0.997988 | 0.994666 |
| Herc1_E9PZP8_T2699t_1 | Herc1_E9P | 1.16 | 1.35 | 1.53 | 1.38 | 1.43 | 1.24 | 1.5 | 1.45 | 0.52 | 1.18 | 0.83 | 1.3 | 0.68 | 1.4 | 0.81 | 1.08 | -0.4225 | -0.42 | -0.665 | 0.19696 | 0.808018 | 0.149996 | 0.995762 | 0.997988 | 0.994666 |
| Herpud2_Q9JJC9_S90s_1 | Herpud2_C | -0.2 | -0.89 | 0.37 | 0 | -0.66 | 0.1 | 0.03 | -0.27 | 1.3 | 0.11 | -1.26 | -0.03 | 2.05 | -0.81 | 0.74 | 1.04 | 0.05 | -0.3975 | -0.3625 | 0.139587 | 0.951257 | 0.700628 | 0.999187 | 0.997988 | 0.994666 |
| Hgs_Q3UMA3_S240s_1 | Hgs_Q3UM | -0.17 | 0.39 | 0.16 | 0.11 | 0.47 | -0.16 | 0.23 | 0.03 | -0.25 | 0.19 | -0.65 | -0.02 | -0.17 | -0.01 | -0.12 | 0.02 | -0.02 | 0.21 | 0.935 | 0.440463 | 0.93395 | 0.27758 | 0.998569 | 0.997988 | 0.994666 |
| Hgsnat_Q3UDW8_S206s_1 | Hgsnat_Q3 | 0.03 | 0.25 | 0.26 | 0.2 | 0.32 | -0.26 | 0.23 | -0.18 | -0.03 | 0.19 | -0.27 | 0.24 | 0.32 | -0.34 | 0.55 | 0.48 | 0.02 | -0.305 | -0.1925 | 0.812518 | 0.49883 | 0.508913 | 0.989223 | 0.997988 | 0.999507 |
| Hid1_Q8R1F6_S592s_1 | Hid1_Q8R1 | -1.1 | -1.11 | 0.14 | -0.07 | -1.43 | -1.08 | 0.04 | 0.03 | -1.37 | -0.9 | 0.17 | 0.1 | -0.07 | -1.85 | 0.05 | 0.07 | -0.1575 | -0.1525 | 0.0675 | 0.87358 | 0.872453 | 0.941634 | 0.996493 | 0.997988 | 0.999507 |
| Hid1_Q8R1F6_S669s_1 | Hid1_Q8R1 | -2.01 | -2.19 | -0.63 | 0 | -1.09 | -2.6 | 0.11 | -0.08 | -2.76 | -1.69 | -0.67 | -0.73 | -2.51 | -0.77 | -0.56 | -0.85 | -0.075 | 0.035 | 0.085 | 0.955758 | 0.685363 | 0.699039 | 0.994624 | 0.997988 | 0.999507 |
| Hid1_Q8R1F6_T590t_1 | Hid1_Q8R1 | -2.28 | -1.95 | -0.27 | -0.58 | -2.88 | -1.51 | -0.76 | -0.88 | -2.18 | -2.12 | -0.01 | -0.72 | -0.01 | -3.7 | -0.94 | -0.91 | 0.2925 | -0.255 | 0.035 | 0.884857 | 0.702292 | 0.984916 | 0.995437 | 0.997988 | 0.999507 |
| Hint1_B0R1E3_S45s_1 | Hint1_B0R | -1.56 | -0.73 | -0.37 | -0.43 | -0.5 | -1.67 | -0.22 | -0.52 | -1.98 | -0.08 | -0.94 | 0.47 | -2.7 | -2.64 | -0.64 | -0.69 | -0.2375 | 0.0125 | -0.12 | 0.151377 | 0.911449 | 0.801361 | 0.997335 | 0.997988 | 0.994666 |
| Hip1r_Q9JKY5_S19s_1 | Hip1r_Q9J | -1.87 | -0.47 | 0 | -0.22 | -1.97 | -0.94 | -1.18 | -0.29 | -1.55 | 0.06 | 0.16 | -0.28 | 0.24 | 0.03 | -1.36 | -0.74 | 0.045 | 0.14 | -0.895 | 0.727793 | 0.373626 | 0.659517 | 0.989223 | 0.997988 | 0.9971 |
| Hipk2_Q9QZR5_Y354y_1 | Hipk2_Q9C | -1.45 | -1.28 | -1.79 | -1.74 | -1.98 | -1.39 | -1.62 | -2.31 | -1.28 | -1.42 | -0.98 | -2.03 | -0.02 | -1.25 | -1.96 | -1.97 | -0.455 | 0.2375 | 0.1825 | 0.567135 | 0.355747 | 0.64238 | 0.989223 | 0.997988 | 0.995259 |
| Hipk3_Q9ERH7_Y359y_1 | Hipk3_Q9E | -0.89 | 0.31 | -0.44 | -0.57 | 0.42 | -0.86 | -0.63 | -1.11 | -0.02 | -0.36 | -0.8 | -0.57 | 0.88 | -2.86 | -0.4 | -0.39 | -0.26 | 0.1375 | 0.265 | 0.688226 | 0.715918 | 0.902998 | 0.995437 | 0.997988 | 0.996159 |
| Hirip3_Q8BLH7_S575s_1 | Hirip3_Q8E | -0.36 | -1.52 | -1.52 | -1.59 | -2.69 | -2.55 | -3.58 | -1.92 | -2.9 | -1.96 | 0.19 | -1.78 | -0.98 | -1.56 | -2.66 | -2.83 | -0.1475 | -0.04 | -0.295 | 0.155128 | 0.007373 | 0.576564 | 0.989223 | 0.997988 | 0.994666 |
| Hirip3_Q8BLH7_S80s_1 | Hirip3_Q8E | -2.85 | -4.86 | -2.58 | -2.37 | -3.36 | -3.38 | -2.5 | -3.17 | -1.97 | -2 | -2.36 | -3.11 | -1.33 | -3.16 | -3.79 | -3.2 | -1.4375 | -0.365 | -0.76 | 0.674882 | 0.909562 | 0.188948 | 0.997335 | 0.997988 | 0.996159 |
| Hist1h1a_P43275_S2s_1 | Hist1h1a_F | 1.72 | 1.37 | 1.78 | 1.78 | 1.26 | 1.39 | 1.72 | 1.93 | 1.09 | 2.03 | 1.32 | 1.93 | 0.8 | 1.33 | 0.89 | 1.56 | 0.0625 | 0.805 | 0.295 | 0.0918 | 0.720494 | 0.811421 | 0.995437 | 0.997988 | 0.994666 |
| Hist1h1a_P43275_T4t_1 | Hist1h1a_F | 1.72 | 1.37 | 1.78 | 1.78 | 1.26 | 1.39 | 1.72 | 1.93 | 1.09 | 2.03 | 1.32 | 1.93 | 0.8 | 1.33 | 0.89 | 1.56 | -0.0875 | -0.07 | -0.5175 | 0.0918 | 0.720494 | 0.811421 | 0.995437 | 0.997988 | 0.994666 |
| Hist1h1b_P43276_S18s_1 | Hist1h1b_F | 2.27 | 1.18 | 2.27 | 2.16 | 0.69 | 1.36 | 1.65 | 1.94 | 1.44 | 1.95 | 1.59 | 2.19 | 1.28 | 1.29 | 2.04 | 2.04 | -0.0875 | -0.07 | -0.5175 | 0.408608 | 0.154829 | 0.600203 | 0.989223 | 0.997988 | 0.994666 |
| Hist1h1b_P43276_S2s_1 | Hist1h1b_F | 0.3 | 0.55 | 0.78 | 0.5 | 0.25 | -0.09 | 0.39 | 0.87 | 0.28 | 0.48 | 1.1 | 0.86 | 0.41 | 0.44 | 0.98 | 1.04 | -0.56 | -0.1775 | -0.3075 | 0.509429 | 0.510608 | 0.586099 | 0.989223 | 0.997988 | 0.994666 |
| Hist1h1b_P43276_T4t_1 | Hist1h1b_F | 0.3 | 0.55 | 0.78 | 0.5 | 0.25 | -0.09 | 0.39 | 0.87 | 0.28 | 0.48 | 1.1 | 0.86 | 0.41 | 0.44 | 0.98 | 1.04 | -0.1775 | 0.1475 | 0.185 | 0.509429 | 0.510608 | 0.586099 | 0.989223 | 0.997988 | 0.994666 |
| Hist1h1c_P15864_S36s_1 | Hist1h1c_P | 0.43 | 0.02 | -0.28 | -1.14 | 0.98 | 0.49 | -0.07 | -1.72 | 0.05 | 0.51 | -0.49 | -0.61 | 0.72 | 0.18 | -1.65 | -0.09 | -0.1775 | 0.1475 | 0.185 | 0.953764 | 0.787837 | 0.795485 | 0.995762 | 0.997988 | 0.999507 |
| Hist1h1c_P15864_T146t_1 | Hist1h1c_P | -1.9 | -2.03 | -0.49 | -2.38 | -2.78 | -0.84 | -1.91 | -1.37 | 0.43 | -1.81 | -3.77 | -0.05 | 1.7 | -2.34 | 1.27 | -1.1 | 0.1625 | 0.1075 | 0.0325 | 0.110156 | 0.962519 | 0.665777 | 0.999187 | 0.997988 | 0.994666 |
| Hist1h1d_P43277_S2s_1 | Hist1h1d_F | 1.13 | 1.18 | 1.04 | 0.97 | 0.89 | 0.86 | 0.81 | 0.82 | 0.89 | 1.04 | 0.41 | 0.62 | 0.54 | 0.98 | 0.75 | 0.63 | -0.025 | 0.4 | 1.5825 | 0.161851 | 0.244246 | 0.174305 | 0.989223 | 0.997988 | 0.994666 |
| Hist1h1d_P43277_T18t_1 | Hist1h1d_F | 1.05 | 1.4 | 0.61 | 0.55 | 1.5 | 0.69 | 1.14 | 0.26 | 1.09 | 0.66 | -0.35 | 1 | 0.83 | 1.25 | 0.6 | 1.79 | -0.235 | -0.34 | -0.355 | 0.549593 | 0.988293 | 0.442772 | 0.999187 | 0.997988 | 0.994666 |
| Hist1h1d_P43277_T4t_1 | Hist1h1d_F | 1.13 | 1.18 | 0.91 | 1.21 | 0.89 | 0.86 | 0.82 | 0.76 | 0.89 | 1.04 | 0.21 | 0.61 | 0.54 | 0.98 | 0.94 | 0.81 | -0.005 | -0.3025 | 0.215 | 0.25311 | 0.189407 | 0.132469 | 0.989223 | 0.997988 | 0.994666 |
| Hist1h1e_P43274_S2s_1 | Hist1h1e_F | 1.52 | 1.43 | 0.42 | 0.62 | 1.33 | 0.55 | 0.13 | 0.32 | 1.21 | 1.25 | -0.38 | 0.57 | 0.45 | 1.59 | 0.41 | 0.58 | -0.275 | -0.42 | -0.29 | 0.553543 | 0.285954 | 0.467051 | 0.989223 | 0.997988 | 0.994666 |
| Hist1h1e_P43274_T18t_1 | Hist1h1e_F | 1.02 | 1.27 | 0.71 | 0.52 | 0.84 | 0.57 | 0.77 | -0.03 | 0.77 | 0.99 | 0.11 | 1 | 0.75 | 1.09 | 0.91 | 1.1 | -0.415 | -0.335 | -0.24 | 0.761987 | 0.253487 | 0.596068 | 0.989223 | 0.997988 | 0.9971 |
| Hist1h1e_P43274_T4t_1 | Hist1h1e_F | 1.52 | 1.43 | 0.42 | 0.62 | 1.33 | 0.55 | 0.13 | 0.32 | 1.21 | 1.25 | -0.38 | 0.57 | 0.45 | 1.59 | 0.41 | 0.58 | -0.3425 | -0.1625 | 0.0825 | 0.553543 | 0.285954 | 0.467051 | 0.989223 | 0.997988 | 0.994666 |
| Hist1h4a_P62806_S48s_1 | Hist1h4a_F | 0.33 | 0.66 | 0.63 | 0.14 | 0.6 | 0.29 | 0.66 | 0.16 | 0.35 | 0.79 | 0.15 | 0.28 | -0.1 | 0.68 | 0.18 | 0.7 | -0.415 | -0.335 | -0.24 | 0.800507 | 0.958325 | 0.852861 | 0.999187 | 0.997988 | 0.999507 |
| Hivep2_Q3UHF7_S1077s_1 | Hivep2_Q3 | 0.02 | 0.68 | -0.06 | -0.12 | 0.5 | -0.43 | 0.26 | 0.03 | 0.25 | 0.15 | 0.46 | 0.02 | 0.38 | 0.8 | 0.03 | -0.17 | -0.0125 | -0.0475 | -0.075 | 0.691152 | 0.892966 | 0.737339 | 0.996493 | 0.997988 | 0.996159 |
| Hivep2_Q3UHF7_S169s_1 | Hivep2_Q3 | -0.91 | 0.55 | -1.09 | -1.58 | -0.97 | -0.67 | -1.56 | -1.08 | 0.19 | 0.4 | 0.9 | -2.61 | 0.54 | -0.8 | -1.45 | -2.14 | -0.04 | 0.09 | 0.13 | 0.756269 | 0.501962 | 0.561652 | 0.989223 | 0.997988 | 0.9971 |
| Hivep2_Q3UHF7_S1921s_1 | Hivep2_Q3 | -1.22 | -0.96 | -0.06 | -0.4 | -1.64 | -0.46 | -0.77 | -0.68 | 0.26 | -1.09 | -0.13 | -0.38 | 0.64 | -0.81 | 0.06 | -0.74 | -0.3125 | 0.4775 | -0.205 | 0.31184 | 0.53786 | 0.412166 | 0.989223 | 0.997988 | 0.994666 |
| Hivep2_Q3UHF7_S2085s_1 | Hivep2_Q3 | -4.75 | -0.97 | -1.49 | -9.12 | -2.63 | -4.07 | -4.66 | -6.16 | -0.97 | -1.14 | -1.54 | -2.68 | 1.8 | -1.76 | -2.38 | -2.41 | -0.2275 | 0.325 | 0.4475 | 0.136757 | 0.862669 | 0.158851 | 0.996493 | 0.997988 | 0.994666 |
| Hivep2_Q3UHF7_S2295s_1 | Hivep2_Q3 | -1.12 | -0.1 | -0.01 | -0.44 | -1.65 | -1.37 | -0.71 | -0.99 | -0.17 | -0.34 | 0.88 | -0.01 | 0.36 | -0.56 | 0.41 | -0.13 | -0.2975 | 2.5 | 2.895 | 0.246616 | 0.04397 | 0.199482 | 0.996493 | 0.997988 | 0.994666 |
| Hivep2_Q3UHF7_S314s_1 | Hivep2_Q3 | -0.43 | 0.7 | 1.19 | 0.99 | -0.22 | 0.06 | 0.91 | 1.1 | -0.04 | 0.83 | 1.27 | 0.36 | -0.1 | 0.75 | 1.32 | 0.63 | -0.7625 | 0.5075 | 0.4375 | 0.933792 | 0.740248 | 0.986511 | 0.995762 | 0.997988 | 0.999507 |
| Hivep2_Q3UHF7_S395s_1 | Hivep2_Q3 | -1.26 | -0.34 | 0.08 | -0.84 | -0.33 | -1.33 | -0.47 | -0.41 | -0.05 | 0.16 | 1.01 | -0.38 | -0.04 | 0.16 | -0.6 | -0.25 | -0.15 | -0.0075 | 0.0375 | 0.273189 | 0.902816 | 0.085281 | 0.996493 | 0.997988 | 0.994666 |
| Hivep2_Q3UHF7_S41s_1 | Hivep2_Q3 | -1.85 | -0.51 | 0.26 | -0.5 | -0.97 | -0.36 | -0.86 | -0.11 | -0.16 | -0.26 | 0.4 | 0.36 | -0.01 | -1.12 | 0.53 | 0.08 | -0.045 | 0.775 | 0.4075 | 0.335058 | 0.868119 | 0.130505 | 0.996493 | 0.997988 | 0.994666 |

| | | | | | | | | | | | | | | | | | | | | | | | | | | |
|---|---|---|---|---|---|---|---|---|---|---|---|---|---|---|---|---|---|---|---|---|---|---|---|---|---|---|
| Hivep2_Q3UHF7_S865s_1 | Hivep2_Q3 | -0.19 | -1.16 | -2.09 | -2.1 | -0.52 | -1.8 | -4.6 | -3.12 | -1.24 | -0.41 | -1.24 | -2 | -0.35 | 0.15 | 0.02 | -2.31 | 0.075 | 0.735 | 0.52 | 0.264415 | 0.211091 | 0.756204 | 0.989223 | 0.997988 | 0.994666 |
| Hivep2_Q3UHF7_T2294t_1 | Hivep2_Q3 | -3.13 | 0.37 | -0.98 | -2.93 | -3.27 | -3.71 | -2.98 | -3.01 | -1.12 | -0.45 | -0.72 | -1.31 | 1.53 | -1.01 | -1.47 | -1.83 | -1.125 | 0.1625 | 0.7625 | 0.335707 | 0.058626 | 0.330774 | 0.989223 | 0.997988 | 0.994666 |
| Hmga1_Q3TE85_S101s_1 | Hmga1_Q3 | 0.64 | 0.25 | 0.75 | 0.41 | -0.54 | 0.37 | -0.89 | -0.24 | 1.16 | 0.75 | 0.56 | 0.03 | 0.54 | -0.39 | 0.72 | -0.04 | -1.575 | 0.7675 | 0.9725 | 0.35921 | 0.021504 | 0.708364 | 0.989223 | 0.997988 | 0.994666 |
| Hmga1_Q3TE85_S102s_1 | Hmga1_Q3 | 0.65 | 0.23 | -0.08 | 0.39 | -1.31 | 0.18 | -0.73 | -0.51 | 1.17 | -0.23 | 0.09 | -1.03 | -0.3 | -0.9 | 0.7 | 0.01 | -0.8375 | 0.1125 | -0.305 | 0.286067 | 0.027299 | 0.522682 | 0.989223 | 0.997988 | 0.994666 |
| Hmga1_Q3TE85_S36s_1 | Hmga1_Q3 | -0.11 | 0.15 | -0.43 | -0.86 | -0.03 | -0.62 | -0.2 | -0.76 | -0.73 | 0.09 | -1.01 | -1.49 | -0.49 | 0.34 | -0.42 | -0.26 | -0.89 | -0.2975 | -0.42 | 0.750501 | 0.765538 | 0.250868 | 0.995762 | 0.997988 | 0.9971 |
| Hmgcr_F2Z470_S384s_1 | Hmgcr_F22 | 0.3 | -0.06 | 0.32 | -0.76 | -1.08 | 1 | 0.53 | -0.52 | -0.54 | 0.55 | -1.16 | -1.19 | -0.22 | 0.35 | 0.75 | 1.9 | -0.09 | -0.4725 | 0.105 | 0.151854 | 0.947536 | 0.261607 | 0.999187 | 0.997988 | 0.994666 |
| Hmgn3_Q9DCB1_S6s_1 | Hmgn3_Q9 | -0.98 | -1.37 | -2.42 | -5 | -1.58 | -1.35 | -3.03 | -3.64 | 0.36 | -0.69 | -3.58 | -2.32 | 1.26 | -2.26 | -0.24 | -2.72 | 0.0325 | -0.535 | 0.745 | 0.215042 | 0.963177 | 0.43141 | 0.999187 | 0.997988 | 0.994666 |
| Hn1_P97825_S70s_1 | Hn1_P9782 | 0.91 | 0.11 | -0.03 | -0.2 | -0.56 | 1.95 | -0.05 | 0.4 | 3.16 | -0.18 | -1.41 | 0.15 | 2.32 | -0.92 | 0.3 | 0.59 | 0.0425 | 0.885 | 1.4525 | 0.563401 | 0.660948 | 0.793662 | 0.990434 | 0.997988 | 0.995259 |
| Hn1_P97825_S79s_1 | Hn1_P9782 | -3.97 | -2.9 | -2.58 | -4.58 | -3.34 | -3.64 | -5.36 | -4.54 | -1.02 | -3.54 | -2.01 | -2.92 | 0.52 | -2.99 | -2.12 | -3.6 | 0.2375 | 0.2325 | 0.375 | 0.12751 | 0.24194 | 0.108798 | 0.989223 | 0.997988 | 0.994666 |
| Hn1_P97825_S87s_1 | Hn1_P9782 | 0.05 | 0.58 | 0.18 | -0.26 | 0.6 | -0.05 | 0.26 | -0.23 | -0.33 | 0.19 | -0.05 | 0.2 | -0.37 | 0.61 | -0.11 | -0.1 | -0.7125 | 1.135 | 1.46 | 0.685917 | 0.978984 | 0.621404 | 0.999187 | 0.997988 | 0.996159 |
| Hnrnpa1_Q5EBP8_S257s_1 | Hnrnpa1_C | 1.22 | 0.77 | 1.89 | 0.99 | 0.6 | 0.71 | 1.65 | 1.87 | 0.66 | 0.86 | 1.76 | 1.97 | -0.29 | 0.49 | 1.6 | 1.77 | 0.0075 | -0.135 | -0.13 | 0.528995 | 0.979566 | 0.813245 | 0.999187 | 0.997988 | 0.994666 |
| Hnrnpa1_Q5EBP8_S369s_1 | Hnrnpa1_C | -1.44 | -0.34 | 0.12 | 0.28 | -0.73 | 1.96 | 0.58 | 0.3 | 0.86 | -1.2 | -0.92 | 0.35 | 1.88 | 0.66 | 0.59 | 0.65 | -0.01 | 0.095 | -0.325 | 0.021544 | 0.172057 | 0.839163 | 0.989223 | 0.997988 | 0.994666 |
| Hnrnpa1_Q5EBP8_S6s_1 | Hnrnpa1_C | 0.2 | -0.58 | -0.71 | -0.16 | -0.16 | -0.71 | -0.18 | -0.17 | -0.05 | -0.2 | 0.33 | -0.5 | 0.28 | 0.29 | 0.36 | -0.35 | 0.8725 | 0.1175 | 1.29 | 0.17033 | 0.978767 | 0.503064 | 0.999187 | 0.997988 | 0.994666 |
| Hnrnpa2b1_O88569_S259s_1 | Hnrnpa2b1 | 0.57 | 0.13 | 0.55 | 0.74 | -0.12 | 0.27 | 0.86 | 0.78 | 0 | -0.09 | 0.48 | 0.3 | 0.01 | -0.02 | 0.67 | 0.59 | 0.0075 | 0.2075 | 0.4575 | 0.531464 | 0.864301 | 0.223494 | 0.996493 | 0.997988 | 0.994666 |
| Hnrnpa2b1_O88569_S344s_1 | Hnrnpa2b1 | 0.43 | 0.25 | -0.31 | 0 | 0.04 | 0.54 | -0.27 | 0.46 | -0.23 | 0.47 | 0.38 | -0.38 | 0.5 | -0.4 | 0.07 | 0.37 | -0.05 | -0.325 | -0.185 | 0.890989 | 0.72409 | 0.914917 | 0.995437 | 0.997988 | 0.999507 |
| Hnrnpa3_Q8BG05_S359s_1 | Hnrnpa3_C | 0.31 | 0.23 | 0.56 | 0.15 | -0.26 | 0.49 | 0.14 | 0.07 | 0.94 | 0.02 | -0.39 | 0.21 | 2.09 | 0.36 | 1.05 | 0.77 | 0.1 | -0.0325 | 0.0425 | 0.078006 | 0.410832 | 0.714883 | 0.989223 | 0.997988 | 0.994666 |
| Hnrnpa3_Q8BG05_S376s_1 | Hnrnpa3_C | 0.29 | -0.12 | 0.3 | 0.42 | -0.03 | 0.16 | -0.75 | 0.14 | 0.98 | 0.57 | -1.29 | -0.07 | 1.4 | -1.83 | 0.67 | 0.5 | -0.2025 | -0.1175 | 0.755 | 0.953323 | 0.240466 | 0.717727 | 0.989223 | 0.997988 | 0.999507 |
| Hnrnpc_Q9Z204_S218s_1 | Hnrnpc_Q9 | 1.46 | 0.94 | 0.1 | -0.13 | 0.78 | 1.07 | 0.68 | 0.09 | 0.57 | 0.97 | -1.12 | -0.13 | -0.11 | 0.96 | 0.26 | -0.31 | -0.3425 | -0.175 | -0.0375 | 0.393581 | 0.877369 | 0.353194 | 0.996493 | 0.997988 | 0.994666 |
| Hnrnpc_Q9Z204_S228s_1 | Hnrnpc_Q9 | 1.52 | 0.53 | 0.63 | 0.67 | -0.51 | 1.42 | -0.61 | 0.45 | 2.07 | 0.88 | 1.09 | 0.61 | 1.84 | -0.59 | 0.67 | 0.02 | 0.0625 | -0.52 | -0.3925 | 0.511204 | 0.20228 | 0.416395 | 0.989223 | 0.997988 | 0.994666 |
| Hnrnpc_Q9Z204_S236s_1 | Hnrnpc_Q9 | -1.35 | -1.88 | -1.39 | -2.1 | -1.85 | -0.36 | -2.22 | -1.91 | 0.28 | -1.55 | -1.09 | -1.54 | 1.29 | -0.95 | -0.08 | -1.7 | -0.65 | 0.325 | -0.3525 | 0.0529 | 0.825179 | 0.143969 | 0.995762 | 0.997988 | 0.994666 |
| Hnrnpc_Q9Z204_S293s_1 | Hnrnpc_Q9 | -1.81 | -2 | -2.43 | -2.56 | -3.76 | -1.35 | -3.01 | -3.2 | -1.95 | -2.16 | -2.09 | -3.32 | -1.66 | -3.75 | -2.46 | -2.52 | 0.095 | 0.705 | 1.32 | 0.389953 | 0.228688 | 0.628871 | 0.989223 | 0.997988 | 0.994666 |
| Hnrnpd_Q60668_S80s_1 | Hnrnpd_Q6 | 1.46 | 0.19 | 0.45 | 1.23 | -1.65 | 2.05 | 0.18 | 0.48 | 1.06 | -0.14 | 0.33 | 0.43 | 1.69 | -2.74 | 2.68 | 0.7 | -0.63 | -0.18 | -0.3975 | 0.813146 | 0.43782 | 0.307744 | 0.989223 | 0.997988 | 0.999507 |
| Hnrnpd_Q60668_S80s_2 | Hnrnpd_Q6 | 2.85 | 0.38 | 0.63 | 0.43 | 0.19 | 1.91 | 0.79 | 0.71 | 2.04 | 1.7 | 0.53 | 0.52 | 0.8 | -2.45 | 3.34 | 1.47 | -0.5675 | -0.4125 | -0.25 | 0.808229 | 0.781287 | 0.847125 | 0.995762 | 0.997988 | 0.999507 |
| Hnrnpd_Q60668_S82s_1 | Hnrnpd_Q6 | 1.64 | -0.9 | -0.07 | 1.11 | -1.66 | 0.99 | -0.84 | 1.21 | 1.38 | 0.84 | 1.28 | 0.16 | 1.4 | -1.66 | 0.8 | 0.45 | -0.1725 | 0.125 | -0.2825 | 0.799675 | 0.51588 | 0.430436 | 0.989223 | 0.997988 | 0.999507 |
| Hnrnpd_Q60668_S83s_1 | Hnrnpd_Q6 | 1.23 | 0.21 | 0.66 | -1.26 | -0.6 | 1.27 | 0.54 | 0.75 | 0.24 | 0.42 | -1.08 | 0.98 | 1.06 | -1.4 | 2.37 | 0.79 | -0.52 | 0.47 | -0.1975 | 0.555987 | 0.638829 | 0.911099 | 0.989223 | 0.997988 | 0.995259 |
| Hnrnpd_Q60668_S83s_2 | Hnrnpd_Q6 | 2.85 | 0.38 | 0.63 | 0.43 | 0.19 | 1.91 | 0.79 | 0.71 | 2.04 | 1.7 | 0.53 | 0.52 | 0.8 | -2.45 | 3.34 | 1.47 | 0.28 | -0.07 | 0.495 | 0.808229 | 0.781287 | 0.847125 | 0.995762 | 0.997988 | 0.999507 |
| Hnrnpdl_D3YTQ3_S241s_1 | Hnrnpdl_D | -0.68 | -0.46 | -1.29 | -0.45 | -0.42 | -0.48 | 0.27 | -1.02 | -1.35 | -0.93 | -0.46 | -0.38 | -0.62 | -0.46 | -0.03 | -0.08 | -0.1725 | 0.125 | -0.2825 | 0.185685 | 0.37273 | 0.853904 | 0.989223 | 0.997988 | 0.994666 |
| Hnrnph1_Q8C2Q7_S104s_1 | Hnrnph1_C | 3.53 | 1.26 | 2.27 | 0.83 | 1.41 | 3.14 | 3.46 | 3.15 | 1.1 | 2.06 | 1.28 | 2.83 | 1.15 | 0.62 | 3.29 | 3.12 | 0.3075 | -0.06 | 0.4225 | 0.927796 | 0.242671 | 0.812537 | 0.989223 | 0.997988 | 0.999507 |
| Hnrnph2_P70333_S4s_1 | Hnrnph2_P | 0.21 | -0.92 | -3.92 | -0.38 | -0.91 | 0.59 | -1.33 | -3.67 | 2.44 | 0.37 | -1.41 | -4.98 | 3.04 | -0.65 | 0.24 | -0.62 | 0.8175 | -0.155 | 0.0725 | 0.134542 | 0.943672 | 0.821997 | 0.999187 | 0.997988 | 0.994666 |
| Hnrnpk_P61979_S284s_1 | Hnrnpk_P6 | 0.81 | -0.66 | -0.25 | -1.72 | -0.45 | -0.12 | -0.63 | -0.32 | -2.3 | 0.02 | -0.62 | -0.21 | -0.81 | -1.99 | 0.38 | 0.04 | -0.0775 | 0.3575 | 1.755 | 0.834146 | 0.878525 | 0.632626 | 0.996493 | 0.997988 | 0.999507 |
| Hnrnpl_G5E924_S174s_1 | Hnrnpl_G5 | 0.54 | 0.58 | -0.22 | 0.21 | -0.17 | -0.1 | -1 | -0.43 | 1.63 | 0.13 | -0.62 | -0.72 | 0.98 | 0.09 | 0.36 | -0.23 | 0.075 | -0.3225 | -0.14 | 0.948372 | 0.039731 | 0.747099 | 0.989223 | 0.997988 | 0.999507 |
| Hnrnpl_G5E924_S90s_1 | Hnrnpl_G5 | -0.38 | -2.21 | 1.69 | 0.4 | -0.84 | -1.39 | 1.33 | 0.6 | -2.11 | -1.6 | 0.2 | 1.45 | -0.04 | -0.22 | 1.98 | 1.88 | -0.7025 | -0.1725 | 0.0225 | 0.264381 | 0.955384 | 0.703379 | 0.999187 | 0.997988 | 0.994666 |
| Hnrnpll_Q921F4_S37s_1 | Hnrnpll_Q9 | -0.14 | 1.26 | 0.91 | -0.54 | -0.37 | 0.66 | 0.45 | 0.45 | 0 | -0.41 | 0.35 | -0.66 | -0.17 | 0.14 | -0.51 | -0.05 | 0.05 | -0.39 | 1.025 | 0.253586 | 0.867839 | 0.248638 | 0.996493 | 0.997988 | 0.994666 |
| Hnrnpm_B8JK33_S413s_1 | Hnrnpm_B | 0.48 | -1.54 | -1.09 | -0.9 | -1.38 | 0 | -0.91 | 0.3 | -1.04 | -1.25 | 2.46 | -0.29 | -0.29 | -0.68 | -1.02 | -1.53 | -0.075 | -0.5525 | -0.52 | 0.808603 | 0.62001 | 0.400593 | 0.989223 | 0.997988 | 0.999507 |
| Hnrnpm_B8JK33_S488s_1 | Hnrnpm_B | -0.98 | -2.88 | -2.2 | -2.35 | -2.04 | -1.01 | -0.93 | -1.12 | -2 | -2.38 | 1.44 | -1.61 | -2.8 | -0.98 | -2.21 | -3.77 | 0.265 | 0.7325 | -0.1175 | 0.60074 | 0.089457 | 0.276152 | 0.989223 | 0.997988 | 0.995259 |
| Hnrnpm_B8JK33_S535s_1 | Hnrnpm_B | -0.51 | -1.22 | -0.82 | -0.47 | -1.32 | -0.93 | -1.15 | -0.44 | -1.19 | -1.07 | 0.4 | -0.78 | -0.01 | -0.6 | -0.06 | -0.42 | 0.8275 | 0.965 | -0.3375 | 0.123643 | 0.483699 | 0.812581 | 0.989223 | 0.997988 | 0.994666 |
| Hnrnpm_B8JK33_S578s_1 | Hnrnpm_B | 0.35 | 0.05 | -1.26 | -1.54 | -0.91 | 0.66 | -1.93 | -0.18 | 0.76 | 0.02 | 0.78 | -1.07 | -0.26 | 0.98 | -2.47 | -1.89 | -0.205 | 0.095 | 0.4825 | 0.701813 | 0.987561 | 0.238692 | 0.999187 | 0.997988 | 0.996159 |
| Hnrnpm_B8JK33_S597s_1 | Hnrnpm_B | 0.24 | -0.31 | 0.62 | -0.02 | -0.5 | -0.58 | 0.84 | 0.83 | -0.59 | 0.42 | 0 | 0.28 | -1.32 | -0.9 | 0.87 | 1.24 | 0.01 | 0.7225 | -0.31 | 0.792068 | 0.971545 | 0.747174 | 0.999187 | 0.997988 | 0.999507 |
| Hnrnpu_Q8VEK3_S247s_1 | Hnrnpu_Q8 | 0.07 | -0.26 | -0.06 | -0.8 | 0.47 | 0.39 | -0.2 | -0.57 | -0.14 | 0.54 | -1.26 | -0.77 | -0.94 | 0.03 | -0.35 | -0.62 | 0.015 | -0.105 | -0.16 | 0.527916 | 0.391071 | 0.73337 | 0.989223 | 0.997988 | 0.994666 |
| Hnrnpul2_Q00PI9_S159s_1 | Hnrnpul2_( | 0.77 | 0.38 | 0.99 | 1.15 | -0.66 | 0.71 | 0 | 1.4 | 0.34 | -0.1 | 0.45 | 0.83 | 0.84 | -0.66 | 1.46 | 0.4 | 0.285 | -0.145 | -0.2075 | 0.502852 | 0.31533 | 0.159031 | 0.989223 | 0.997988 | 0.994666 |
| Hnrnpul2_Q00PI9_S159s_2 | Hnrnpul2_( | 1.12 | 0.54 | -0.78 | -0.8 | -0.81 | 1.29 | -1.62 | -1.08 | 0.5 | 0.03 | -1.48 | -1.55 | -2.48 | -1.84 | -0.41 | -1.06 | -0.46 | -0.4425 | -0.3125 | 0.033939 | 0.422862 | 0.33199 | 0.989223 | 0.997988 | 0.994666 |
| Hnrnpul2_Q00PI9_S166s_1 | Hnrnpul2_( | 1.36 | 0.21 | 0.41 | -0.01 | -0.71 | 1.3 | -0.4 | 0.15 | 0.71 | 0.68 | -1 | 0.2 | 1.14 | -0.89 | 0.34 | -0.12 | -0.575 | -0.645 | -1.4675 | 0.454998 | 0.416162 | 0.475188 | 0.989223 | 0.997988 | 0.994666 |
| Hnrnpul2_Q00PI9_S166s_2 | Hnrnpul2_( | 0.95 | 0.39 | -0.78 | -0.8 | -1.13 | 1.26 | -1.62 | -1.08 | 0.42 | 0.05 | -1.48 | -1.55 | -3.17 | -2.22 | -0.41 | -1.06 | -0.4075 | -0.345 | -0.375 | 0.033128 | 0.406507 | 0.357104 | 0.989223 | 0.997988 | 0.994666 |
| Hnrnpul2_Q00PI9_S183s_1 | Hnrnpul2_( | 0.29 | -0.94 | 1.18 | 2.18 | -2.88 | 0.08 | -0.62 | 1.53 | 0.88 | -1.2 | 0.87 | 0.59 | 0.64 | -1.78 | 0.73 | 0.55 | -0.5825 | -0.58 | -1.655 | 0.426635 | 0.258552 | 0.600958 | 0.989223 | 0.997988 | 0.994666 |
| Hnrnpul2_Q00PI9_S191s_1 | Hnrnpul2_( | 0.43 | -0.96 | 0.35 | -2.03 | -2.46 | 0.55 | -1.29 | -0.32 | 2.95 | -0.88 | -1.08 | -0.35 | 3.4 | -2.09 | 2.39 | -1.21 | -1.15 | -0.3925 | -0.6425 | 0.36305 | 0.669963 | 0.474111 | 0.992876 | 0.997988 | 0.994666 |
| Hnrnpul2_Q00PI9_S224s_1 | Hnrnpul2_( | -0.32 | -1 | -1.65 | -1.7 | -0.56 | -0.71 | -1.52 | -1.42 | -0.5 | -0.95 | -0.83 | -1.4 | -0.33 | 0.18 | -2.07 | -1.96 | -0.3275 | 0.7125 | 1.175 | 0.837945 | 0.769811 | 0.518374 | 0.995762 | 0.997988 | 0.999507 |
| Hnrnpul2_Q00PI9_S226s_1 | Hnrnpul2_( | -0.88 | -1.33 | -1.98 | -2.18 | -0.9 | -1.2 | -1.87 | -1.77 | -1.15 | -1.12 | -1.03 | -1.6 | -0.5 | -0.66 | -2.35 | -2.47 | 0.115 | 0.2475 | 0.1225 | 0.862119 | 0.673198 | 0.29993 | 0.992876 | 0.997988 | 0.999507 |
| Homer1_Q9Z2Y3_S286s_1 | Homer1_Q | 1.27 | 1.62 | 1.29 | 1.34 | 0.91 | 1.43 | 1.09 | 1.25 | 1.27 | 1.27 | 1.41 | 1.24 | 1.06 | 1.54 | 1.2 | 1 | 0.1575 | 0.3675 | 0.0975 | 0.484393 | 0.35631 | 0.700914 | 0.989223 | 0.997988 | 0.994666 |
| Homer3_Q99JP6_S159s_1 | Homer3_Q | 0.94 | 0.52 | 0.88 | -0.21 | -0.09 | 0.7 | -0.18 | -0.13 | 0.54 | 0.34 | -1.29 | -0.56 | 1.85 | 0.9 | -0.18 | -0.38 | -0.21 | -0.0825 | -0.18 | 0.97782 | 0.202593 | 0.128509 | 0.989223 | 0.997988 | 1 |
| Hook3_Q8BUK6_S238s_1 | Hook3_Q8I | 1.6 | 1 | 1.81 | 2.17 | 0.87 | 1.2 | 1.23 | 1.41 | 2.21 | 1.86 | 3.64 | 2.47 | 1.91 | 2.64 | 1.78 | 1.73 | -0.4575 | -0.775 | 0.015 | 0.308291 | 0.139152 | 0.066875 | 0.989223 | 0.997988 | 0.994666 |
| Hook3_Q8BUK6_S707s_1 | Hook3_Q8I | -1.57 | -2.58 | -2.29 | -2.73 | -2.05 | -2.26 | -1.78 | -2.7 | -0.15 | -1.07 | -2.23 | -3.45 | 1.09 | -1.15 | -2.18 | -1.66 | -0.4675 | 0.9 | 0.37 | 0.07976 | 0.775081 | 0.41762 | 0.995762 | 0.997988 | 0.994666 |
| Hopx_Q8R1H0_S61s_1 | Hopx_Q8R: | -1.52 | -0.94 | -0.87 | -1.75 | -0.47 | -1.4 | -0.57 | -0.99 | -1.44 | -1.43 | -0.8 | -1.8 | -0.58 | -1.66 | -1.79 | -0.63 | 0.095 | 0.5675 | 1.3175 | 0.791901 | 0.216221 | 0.765366 | 0.989223 | 0.997988 | 0.999507 |
| Hp1bp3_Z4YKA3_S475s_1 | Hp1bp3_Z4 | 1.54 | 1.18 | 1.29 | 0.95 | 0.27 | 0.39 | 0.2 | 0.61 | 0.7 | 0.21 | 1.22 | 1.08 | 1.22 | 0.28 | 0.85 | 0.78 | 0.4125 | -0.0975 | 0.105 | 0.144652 | 0.003555 | 0.165871 | 0.989223 | 0.997988 | 0.994666 |
| Hp1bp3_Z4YKA3_T13t_1 | Hp1bp3_Z4 | 2.15 | 0.95 | 1.94 | 1.93 | -0.92 | 0.76 | 0.05 | 1.03 | 0.7 | 0.66 | 0.51 | 1.11 | 1.62 | -1.01 | 1.07 | 0.89 | -0.8725 | -0.4375 | -0.4575 | 0.083345 | 0.009647 | 0.012104 | 0.989223 | 0.997988 | 0.994666 |
| Hpcal1_P62748_S175s_1 | Hpcal1_P6 | -0.13 | 0.34 | 0.68 | 0.3 | 0.76 | 0.11 | -0.13 | 0.01 | 0.11 | -0.22 | 0.74 | 0.29 | 0.55 | 0.6 | 0.1 | 0.11 | -1.5125 | -0.9975 | -1.1 | 0.88216 | 0.70401 | 0.821903 | 0.995437 | 0.997988 | 0.999507 |
| Hpcal1_P62748_T144t_1 | Hpcal1_P6 | 1.05 | 1.47 | 1.43 | 1.36 | 0.76 | 1.55 | 0.68 | 0.75 | 1.98 | 1.39 | 0.76 | 1.02 | 1.64 | 0.42 | 1.22 | 1.69 | -0.11 | -0.0675 | 0.0425 | 0.804251 | 0.168256 | 0.898419 | 0.989223 | 0.997988 | 0.999507 |
| Hpcal4_F2Z3Z1_T146t_1 | Hpcal4_F2 | -0.07 | 0.3 | 0.53 | 0.84 | 0.42 | 0.22 | 0.22 | 0.64 | 0.91 | 0.76 | 0.47 | 0.22 | 1.3 | 0.21 | 0.75 | 0.32 | -0.3925 | -0.04 | -0.085 | 0.484094 | 0.924276 | 0.518136 | 0.997335 | 0.997988 | 0.994666 |
| Hrh1_P70174_S230s_1 | Hrh1_P701 | -0.91 | -0.88 | 0.66 | 0.37 | -0.14 | -1.72 | 0.38 | 0.23 | -0.57 | -0.06 | -0.88 | -0.56 | -2.03 | 0.58 | 0.12 | 0.89 | -0.025 | 0.19 | 0.245 | 0.908373 | 0.829828 | 0.45616 | 0.995762 | 0.997988 | 0.999507 |
| Hrh1_P70174_S264s_1 | Hrh1_P701 | -0.16 | -0.63 | 0.71 | -0.1 | -0.13 | 0.11 | -0.35 | 0.53 | 0.05 | -0.55 | 0.89 | 0.34 | -0.29 | 0.58 | 0.03 | 0.33 | -0.1225 | -0.3275 | 0.08 | 0.568084 | 0.803583 | 0.577588 | 0.995762 | 0.997988 | 0.995259 |
| Hrh3_E9Q292_S302s_1 | Hrh3_E9Q2 | 0.63 | 0.9 | 1.55 | 1.42 | 0.85 | -0.17 | 1.47 | 1.61 | 0.67 | 1.06 | 3.93 | 0.49 | 0.92 | 0.15 | 1.17 | 1.61 | 0.085 | 0.2275 | 0.2075 | 0.675843 | 0.670534 | 0.584816 | 0.992876 | 0.997988 | 0.996159 |
| Hrh3_E9Q292_S332s_1 | Hrh3_E9Q2 | 0.57 | 1.05 | 2.25 | 2.29 | 0.64 | 0.73 | 1.69 | 1.57 | 1.01 | 0.7 | 2.88 | 2.64 | 1.58 | 0.74 | 2.67 | 1.73 | -0.185 | 0.4125 | -0.1625 | 0.797178 | 0.427892 | 0.677526 | 0.989223 | 0.997988 | 0.999507 |
| Hrh3_E9Q292_S343s_1 | Hrh3_E9Q2 | -0.27 | 0.23 | 0.78 | 0.05 | 0.63 | -0.49 | 0.1 | 0.03 | -0.33 | 0.54 | -0.15 | 0.32 | 1.02 | 0.6 | 1.13 | 1.21 | -0.3825 | 0.2675 | 0.14 | 0.028102 | 0.693381 | 0.753086 | 0.994893 | 0.997988 | 0.994666 |
| Hrh3_E9Q292_S439s_1 | Hrh3_E9Q2 | 0.1 | 0.25 | 0.21 | 0.72 | 1.01 | 0.08 | 0.81 | 0.44 | -0.12 | 0.43 | 0.36 | 0.55 | -0.03 | 0.9 | 1.34 | 0.53 | -0.13 | -0.1025 | 0.7925 | 0.311958 | 0.358725 | 0.954651 | 0.989223 | 0.997988 | 0.994666 |
| Hsd17b4_P51660_S308s_1 | Hsd17b4_F | -0.42 | 0.53 | 0.77 | 0.12 | 0.79 | 0.64 | 0.12 | 0.39 | 0.66 | 0.12 | 0.5 | 0.91 | 0.82 | 0.76 | 0.66 | 0.25 | 0.265 | -0.015 | 0.365 | 0.276864 | 0.463565 | 0.381071 | 0.989223 | 0.997988 | 0.994666 |
| Hsp90aa1_P07901_S231s_1 | Hsp90aa1_ | -1.68 | -1.13 | -0.46 | -2.86 | -1.97 | -0.84 | -2.03 | -1.85 | -0.3 | -1.41 | 0.44 | -0.56 | -1.15 | -2.31 | -0.91 | -1.43 | 0.235 | 0.2975 | 0.3725 | 0.880594 | 0.790775 | 0.092117 | 0.995762 | 0.997988 | 0.999507 |
| Hsp90aa1_P07901_S263s_1 | Hsp90aa1_ | -1.91 | -0.68 | -1.96 | -2.51 | -0.75 | -1.7 | -1.8 | -2.12 | -1.78 | -1.11 | -2.31 | -2.2 | -3 | -1.59 | -2.14 | -2.19 | -0.14 | 1.075 | 0.0825 | 0.331214 | 0.704503 | 0.851397 | 0.995437 | 0.997988 | 0.994666 |
| Hsp90aa1_P07901_S454s_1 | Hsp90aa1_ | 0.23 | 0.13 | -0.13 | 0.13 | 0.51 | -0.21 | -0.55 | 0.11 | 0.34 | 0.19 | 0.51 | -0.05 | 0.21 | 0.21 | -0.07 | 0.01 | 0.1725 | -0.085 | -0.465 | 1 | 0.653198 | 0.505866 | 0.990434 | 0.997988 | 1 |
| Hsp90ab1_P11499_S226s_1 | Hsp90ab1_ | -2.63 | -1.85 | -1.65 | -1.74 | -2.22 | -2.77 | -2.17 | -1.62 | -2.49 | -2.19 | -0.45 | -1.3 | -2.32 | -1.95 | -1.66 | -1.69 | -0.125 | 0.1575 | -4.9E-17 | 0.844909 | 0.500823 | 0.465397 | 0.989223 | 0.997988 | 0.999507 |
| Hsp90ab1_P11499_S255s_1 | Hsp90ab1_ | -1.46 | -0.71 | 1.72 | 1.45 | -0.95 | -1.5 | 1 | 1.66 | -1.92 | -0.87 | 1.28 | 1.7 | -1.83 | -1.8 | 1.82 | 1.9 | -0.2275 | 0.36 | 0.0625 | 0.842317 | 0.834895 | 0.844295 | 0.995762 | 0.997988 | 0.999507 |
| Hsp90ab1_P11499_S445s_1 | Hsp90ab1_ | -0.61 | 0.08 | 0.43 | 0.64 | 0.06 | -0.05 | -0.34 | 0.15 | -0.62 | -0.1 | 0.97 | 0.14 | 1.35 | 0.18 | 0.34 | 0.33 | -0.1975 | -0.2025 | -0.2275 | 0.305935 | 0.56768 | 0.928937 | 0.989223 | 0.997988 | 0.994666 |
| Hsp90ab1_P11499_T446t_1 | Hsp90ab1_ | -0.31 | 0.46 | 0.62 | 0.91 | 0.04 | -0.15 | 0.13 | 0.48 | 0.24 | 0.41 | 0.42 | 0.63 | 0.25 | 0.29 | 0.33 | 0.87 | -0.18 | -0.0375 | 0.415 | 0.964378 | 0.354722 | 0.987026 | 0.989223 | 0.997988 | 0.999507 |
| Hsp90b1_P08113_T44t_1 | Hsp90b1_F | -2.19 | 0.96 | -0.6 | -0.88 | -1.2 | -1.72 | -3.79 | -1.29 | -1.83 | -0.82 | -0.75 | -0.96 | -1.25 | -1.16 | -0.54 | -2.48 | -0.295 | 0.005 | 0.015 | 0.333624 | 0.114008 | 0.518576 | 0.989223 | 0.997988 | 0.994666 |
| Hspa12a_Q8K0U4_S23s_1 | Hspa12a_C | -0.85 | -0.93 | -0.55 | -0.57 | -1.38 | -1.3 | -0.52 | -0.48 | -0.86 | -1.25 | -0.34 | -0.62 | -0.45 | -1.03 | -1.49 | -0.52 | -1.3225 | -0.4125 | -0.68 | 0.63971 | 0.508109 | 0.876004 | 0.989223 | 0.997988 | 0.995259 |
| Hspa12a_Q8K0U4_S24s_1 | Hspa12a_C | -1.43 | -1.84 | -1.45 | -1.9 | -1.19 | -1.76 | -1.21 | -1.02 | -1.63 | -1.28 | -0.8 | -1.73 | -0.94 | -0.7 | -2.3 | -2.02 | -0.195 | -0.0425 | -0.1475 | 0.69139 | 0.181683 | 0.32276 | 0.989223 | 0.997988 | 0.996159 |
| Hspa12a_Q8K0U4_S24s_2 | Hspa12a_C | -0.1 | -0.91 | -0.44 | -0.6 | -0.99 | -0.7 | -1 | -0.32 | -1.01 | -0.59 | 0.08 | -0.48 | -1.05 | -0.3 | -0.98 | -0.72 | 0.36 | 0.295 | 0.165 | 0.413991 | 0.38907 | 0.968137 | 0.989223 | 0.997988 | 0.994666 |
| Hspa12a_Q8K0U4_S631s_1 | Hspa12a_C | -0.12 | 0.35 | -0.24 | 0.14 | 0.06 | -0.1 | 0.19 | -0.14 | -0.05 | 0.3 | -0.19 | 0.09 | -0.52 | 0.2 | -0.46 | 0 | -0.24 | 0.0125 | -0.25 | 0.439714 | 0.895739 | 0.983812 | 0.996493 | 0.997988 | 0.994666 |
| Hspa12b_Q9CZJ2_S276s_1 | Hspa12b_C | -0.49 | -0.57 | -0.11 | -0.43 | -0.68 | -1.05 | -0.45 | -0.75 | -0.68 | -0.36 | -0.18 | -0.06 | -0.34 | -1.11 | -0.77 | -0.5 | -0.03 | 0.005 | -0.2275 | 0.32601 | 0.175354 | 0.746167 | 0.989223 | 0.997988 | 0.994666 |
| Hspa12b_Q9CZJ2_S433s_1 | Hspa12b_C | -0.74 | -0.17 | -0.96 | -0.87 | -0.33 | -0.7 | -0.34 | -0.75 | -1.07 | -1.14 | -0.36 | -0.61 | -0.17 | -0.99 | -0.64 | -1.03 | -0.3325 | 0.08 | -0.28 | 0.943299 | 0.554432 | 0.713013 | 0.989223 | 0.997988 | 0.999507 |
| Hspa12b_Q9CZJ2_S44s_1 | Hspa12b_C | -0.63 | 0.34 | -0.19 | -0.04 | -0.39 | 0.08 | -0.14 | -0.31 | 0.6 | -0.39 | -1.66 | 0.33 | -0.12 | -0.96 | 0.66 | -0.09 | 0.155 | -0.11 | -0.0225 | 0.99495 | 0.823507 | 0.768988 | 0.995762 | 0.997988 | 1 |
| Hspa12b_Q9CZJ2_T42t_1 | Hspa12b_C | -0.11 | 0.14 | 0.37 | 0.04 | -0.27 | 0.22 | 0.78 | 0.19 | 0.39 | 0.68 | 0.33 | 0.37 | 0.22 | 0.77 | 0.24 | 0.4 | -0.06 | -0.15 | 0.0025 | 0.271275 | 0.664965 | 0.169061 | 0.991343 | 0.997988 | 0.994666 |
| Hspa1a_Q61696_T636t_1 | Hspa1a_Q6 | 0.6 | 0.29 | 0.2 | -0.27 | 0.73 | 0.39 | 0.87 | 0.04 | 0.38 | 1.26 | 0.48 | 0.15 | 0.4 | 0.36 | 0.29 | 0.54 | 0.12 | 0.3325 | 0.2975 | 0.486392 | 0.309793 | 0.283612 | 0.989223 | 0.997988 | 0.994666 |
| Hspa4_Q3U2G2_S503s_1 | Hspa4_Q3 | -2.19 | -1.1 | -0.86 | -2.16 | -1.38 | -1.87 | -2.11 | -1.61 | -1.82 | -1.79 | 0.18 | -1.37 | -2.47 | -1.2 | -1.76 | -1.94 | 0.3025 | 0.3625 | 0.1925 | 0.542211 | 0.661664 | 0.493637 | 0.990434 | 0.997988 | 0.994666 |
| Hspa4_Q3U2G2_S506s_1 | Hspa4_Q3 | -0.44 | 0.33 | -0.91 | -1.67 | -0.11 | -0.5 | -1.67 | -2.67 | -0.5 | 0.05 | -0.84 | -1.34 | -1.03 | 0.36 | -1.96 | -1.5 | -0.165 | 0.3775 | -0.265 | 0.550412 | 0.38534 | 0.975257 | 0.989223 | 0.997988 | 0.994666 |
| Hspa4_Q3U2G2_S548s_1 | Hspa4_Q3 | -1.01 | -0.77 | -1.49 | -1.71 | -1.95 | -2.94 | -2.75 | -1.55 | -0.54 | -1.82 | -0.6 | -1.6 | -1.3 | -2.79 | -2.46 | -1.86 | -0.565 | 0.015 | -0.36 | 0.055203 | 0.020106 | 0.790701 | 0.989223 | 0.997988 | 0.994666 |
| Hspa4_Q3U2G2_S694s_1 | Hspa4_Q3 | -0.55 | 0.22 | 1.24 | 0.6 | -0.31 | -0.17 | 0.15 | 0.25 | 0.98 | 0.92 | 1.92 | 1.88 | 1.26 | 0.14 | 0.67 | 0.11 | -1.0525 | 0.105 | -0.8575 | 0.710286 | 0.316537 | 0.04043 | 0.989223 | 0.997988 | 0.996966 |
| Hspa4_Q3U2G2_T468t_1 | Hspa4_Q3 | 0.44 | 0.25 | -0.15 | 0.14 | 0.25 | 0.48 | -0.11 | 0.33 | 1.48 | 0.25 | -0.35 | -1.17 | 1.86 | -0.63 | -0.34 | -0.18 | -0.3975 | 1.0475 | 0.1675 | 0.988923 | 0.779399 | 0.825302 | 0.995762 | 0.997988 | 1 |
| Hspa4_Q3U2G2_T540t_1 | Hspa4_Q3 | -1.96 | -1.42 | -1.48 | -1.46 | -1.53 | -1.93 | -1.73 | -2.27 | -1.56 | -2 | -0.99 | -1.44 | -1.07 | -0.52 | -1.84 | -1.79 | 0.0675 | -0.1175 | 0.0075 | 0.455888 | 0.280482 | 0.775813 | 0.989223 | 0.997988 | 0.994666 |
| Hspa4l_P48722_S53s_1 | Hspa4l_P4 | 1.65 | 1.14 | 1.58 | 0.34 | 1.51 | 1.56 | 1.74 | 1.27 | -1.06 | 1.72 | -1.08 | 1.43 | -0.5 | -1 | 1.28 | 1.69 | -0.285 | 0.0825 | 0.275 | 0.234111 | 0.310281 | 0.230799 | 0.989223 | 0.997988 | 0.994666 |
| Hspa4l_P48722_S805s_1 | Hspa4l_P4 | -0.65 | -0.66 | -1.09 | -0.75 | -0.36 | -0.47 | -1.86 | -0.31 | -1.02 | -1.7 | -0.12 | -2.18 | -1.35 | 0.41 | -1.4 | -0.46 | 0.3425 | -0.925 | -0.81 | 0.840232 | 0.920795 | 0.306597 | 0.997335 | 0.997988 | 0.999507 |
| Hspa4l_P48722_S811s_1 | Hspa4l_P4 | -0.22 | -0.52 | 0.04 | -0.17 | -0.9 | -0.31 | -1.01 | -0.53 | -0.16 | -0.46 | -0.52 | -0.72 | -0.56 | -0.59 | -1.18 | -0.52 | 0.0375 | -0.4675 | 0.0875 | 0.098634 | 0.088915 | 0.324724 | 0.989223 | 0.997988 | 0.994666 |
| Hspa4l_P48722_T524t_1 | Hspa4l_P4 | 5.62 | 1.01 | 1.74 | 0.27 | 1.78 | 5.58 | 3.94 | 2.74 | -0.55 | 3.99 | -0.94 | 2.2 | -0.39 | -1.64 | 3.81 | 3.78 | -0.47 | -0.2475 | -0.495 | 0.628943 | 0.292765 | 0.504884 | 0.989223 | 0.997988 | 0.995259 |
| Hspa4l_P48722_T572t_1 | Hspa4l_P4 | -0.83 | -0.9 | -0.18 | 0.03 | -1.25 | -1.3 | -0.78 | -2.09 | -0.88 | -0.73 | -1.31 | -1.09 | -1.54 | -1.41 | -2.18 | 0.38 | 1.35 | -0.985 | -0.77 | 0.214165 | 0.031496 | 0.104703 | 0.989223 | 0.997988 | 0.994666 |
| Hspa4l_P48722_T797t_1 | Hspa4l_P4 | 1 | 0.49 | 0.23 | -0.49 | 1.09 | 0.76 | 0.5 | 0.31 | 0.86 | 1.74 | 0.69 | 0.45 | 0.08 | 0.81 | 0.42 | 0.69 | -0.885 | -0.5325 | -0.7175 | 0.605232 | 0.325871 | 0.153088 | 0.989223 | 0.997988 | 0.995259 |
| Hsph1_E9Q0U7_S733s_1 | Hsph1_E9C | 0.22 | -0.28 | -0.25 | 0.64 | 1 | 0.86 | 0.46 | 0.31 | -0.43 | 0.1 | 0.64 | 0.1 | 0.57 | 1.28 | 1.26 | -0.4 | 0.3575 | 0.6275 | 0.1925 | 0.19958 | 0.078631 | 0.951938 | 0.989223 | 0.997988 | 0.994666 |
| Hsph1_E9Q0U7_S769s_1 | Hsph1_E9C | 1.27 | 0.93 | 1.23 | 1.46 | 0.91 | 1.12 | 0.96 | 1.51 | 1.73 | 1.81 | 2.1 | 1.26 | 1.25 | 1.47 | 1.39 | 1.23 | 0.575 | 0.02 | 0.595 | 0.648517 | 0.685773 | 0.085598 | 0.994626 | 0.997988 | 0.995259 |
| Hsph1_E9Q0U7_T445t_1 | Hsph1_E9C | -1.32 | -0.21 | -1.08 | -0.29 | -0.7 | -1.37 | 0.43 | -0.19 | -0.66 | -1.02 | -0.85 | -0.61 | -1.71 | -0.87 | -0.67 | -0.5 | -0.0975 | 0.5025 | 0.1125 | 0.59381 | 0.550155 | 0.852349 | 0.989223 | 0.997988 | 0.995259 |
| Hsph1_E9Q0U7_T462t_1 | Hsph1_E9C | -0.73 | 0.47 | -0.58 | -0.92 | -0.03 | -0.51 | -0.31 | -1.08 | -0.39 | -0.24 | -1.14 | -0.71 | -1.03 | 0.53 | -0.69 | -0.58 | 0.2675 | -0.06 | -0.2125 | 0.995541 | 0.909718 | 0.633578 | 0.997335 | 0.997988 | 1 |
| Htatsf1_Q8BGC0_S388s_1 | Htatsf1_Q8 | -2.56 | -3.44 | -2.45 | -7.03 | -3.4 | -3.03 | -2.58 | -3.31 | -1.42 | -2.9 | -0.3 | -3.06 | 0.18 | -0.32 | -2.82 | -2.51 | -0.0425 | -0.18 | -0.0025 | 0.050638 | 0.412719 | 0.104844 | 0.989223 | 0.997988 | 0.994666 |
| Htr2a_P35363_S280s_1 | Htr2a_P35 | -0.34 | -0.46 | -0.86 | -0.52 | -0.02 | 0.28 | -0.25 | -0.23 | -0.75 | -1.86 | -1.15 | -1.07 | -1.38 | -0.27 | -1.36 | -1.15 | 0.79 | 1.95 | 2.5025 | 0.154384 | 0.061352 | 0.049663 | 0.989223 | 0.997988 | 0.994666 |
| Htr2c_P34968_S408s_1 | Htr2c_P34 | -1.71 | -1.38 | -1.6 | -1.4 | -0.63 | -1.63 | -1.7 | -1.58 | -0.48 | -0.5 | -1.29 | -0.55 | -0.09 | 0.39 | 0.29 | 0.31 | 0.49 | -0.6625 | -0.495 | 5.26E-05 | 0.641105 | 0.012949 | 0.989223 | 0.997988 | 0.269064 |
| Htt_G3X9H5_S1180s_1 | Htt_G3X9H | -0.05 | -0.43 | -0.7 | -0.69 | -0.02 | -0.08 | -0.76 | -0.4 | -0.72 | -0.31 | 0.04 | -0.77 | -0.64 | -0.31 | -0.56 | -0.85 | 0.1375 | 0.8175 | 1.7475 | 0.655919 | 0.576025 | 0.924269 | 0.989223 | 0.997988 | 0.995259 |
| Htt_G3X9H5_S1842s_1 | Htt_G3X9H | 6.26 | 3.28 | 5.52 | 0.54 | 3.64 | 7.04 | 4.53 | 4.44 | -0.03 | 5.53 | -1.66 | 5.06 | 1.41 | -2.38 | 6.35 | 6.14 | 0.1525 | 0.0275 | -0.1225 | 0.628667 | 0.434426 | 0.395343 | 0.989223 | 0.997988 | 0.995259 |
| Htt_G3X9H5_S1854s_1 | Htt_G3X9H | -0.21 | 0.44 | -0.01 | -0.38 | -0.08 | 0.81 | -0.34 | 0.04 | -0.05 | -0.16 | 0.33 | -0.48 | 0.2 | 0.06 | -0.42 | -0.03 | 1.0125 | -1.675 | -1.02 | 0.979374 | 0.645729 | 0.862715 | 0.99033 | 0.997988 | 1 |
| Htt_G3X9H5_S638s_1 | Htt_G3X9H | 1.09 | 1.48 | 2.1 | 2.13 | 0.86 | 0.97 | 2.13 | 2.09 | 0.45 | 1.12 | 2.88 | 2.11 | -0.66 | 0.1 | 2.89 | 2.63 | 0.1475 | -0.05 | -0.0075 | 0.577001 | 0.648501 | 0.912927 | 0.99033 | 0.997988 | 0.995259 |
| Htt_G3X9H5_T2923t_1 | Htt_G3X9H | -0.53 | 0.03 | 0.25 | 0.62 | 0.83 | 0.26 | 0.54 | 0.53 | 0.01 | -0.12 | 0.2 | 0.5 | -0.5 | 0.43 | 0.44 | 0.52 | -0.1875 | -0.06 | -0.46 | 0.721588 | 0.152849 | 0.85911 | 0.989223 | 0.997988 | 0.9971 |
| Huwe1_Q7TMY8_S1298s_1 | Huwe1_Q7 | 1.01 | 0.13 | -0.97 | -0.15 | -1.55 | 1.39 | -2.02 | -0.96 | 2.45 | 0.44 | 0.56 | -0.68 | 1.58 | -1.85 | -0.78 | -0.66 | 0.4475 | 0.055 | 0.13 | 0.560815 | 0.309242 | 0.334043 | 0.989223 | 0.997988 | 0.995259 |
| Huwe1_Q7TMY8_S1368s_1 | Huwe1_Q7 | 1.69 | 1.66 | 2.38 | 1.38 | 1.54 | 1.57 | 1.87 | 2.41 | -0.3 | 1.45 | 1.3 | 2.03 | -0.02 | -0.25 | 1.87 | 2.04 | -0.79 | 0.6875 | -0.4325 | 0.163791 | 0.822424 | 0.217428 | 0.995762 | 0.997988 | 0.994666 |
| Huwe1_Q7TMY8_S1395s_1 | Huwe1_Q7 | 0.03 | 0.51 | 0.55 | 0.34 | 0.57 | 0.19 | 0.45 | 0.9 | 0.27 | 0.32 | 1.32 | 0.28 | 0.52 | 0.79 | 0.89 | 0.87 | 0.07 | -0.6575 | -0.8675 | 0.131355 | 0.497894 | 0.550448 | 0.989223 | 0.997988 | 0.994666 |
| Huwe1_Q7TMY8_S1907s_1 | Huwe1_Q7 | 0.01 | 0.15 | -0.21 | -0.12 | 0.29 | -0.4 | -0.26 | 0.02 | -0.32 | 0.24 | 0.28 | -0.6 | -0.66 | 0.1 | -0.58 | -0.22 | 0.17 | 0.19 | 0.41 | 0.29464 | 0.850499 | 0.837733 | 0.996349 | 0.997988 | 0.994666 |
| Huwe1_Q7TMY8_S2917s_1 | Huwe1_Q7 | -0.98 | -0.87 | -0.62 | -0.23 | -0.43 | -0.98 | -0.91 | 0.69 | -1.04 | -0.37 | -0.03 | -0.52 | 0.09 | -1.14 | 0.01 | 0.12 | -0.045 | -0.0575 | -0.2975 | 0.244565 | 0.511911 | 0.54791 | 0.989223 | 0.997988 | 0.994666 |
| Huwe1_Q7TMY8_S2918s_1 | Huwe1_Q7 | -0.42 | 0.02 | -1.47 | -1.27 | -1.36 | 0.84 | -2.56 | -1.6 | -0.37 | 0.5 | -2.53 | -1.59 | 0.56 | -0.33 | -1.69 | -2.18 | 0.2675 | 0.185 | 0.445 | 0.847278 | 0.587797 | 0.755641 | 0.989223 | 0.997988 | 0.999507 |
| Huwe1_Q7TMY8_S3374s_1 | Huwe1_Q7 | -1.54 | -1.68 | -0.9 | -1.14 | -1.5 | -1.29 | -0.34 | -1.56 | -1.5 | -0.81 | -1.52 | -2.06 | -1.33 | -1.08 | -1.2 | -0.75 | -0.385 | -0.2125 | -0.125 | 0.443384 | 0.677439 | 0.640001 | 0.99315 | 0.997988 | 0.994666 |
| Huwe1_Q7TMY8_S3663s_1 | Huwe1_Q7 | 0.38 | 0.58 | 0.33 | 0.88 | 0.09 | 0.3 | 0.53 | 0.68 | 0.09 | 0.56 | 1.07 | -0.07 | -0.22 | 0.53 | 0.16 | 0.48 | 0.1425 | -0.1575 | 0.225 | 0.301288 | 0.560817 | 0.682334 | 0.989223 | 0.997988 | 0.994666 |
| Huwe1_Q7TMY8_S3758s_1 | Huwe1_Q7 | 0.86 | 0.53 | 0.15 | 0.6 | 1.44 | 0.48 | 0.56 | 0.48 | 0.01 | 1.23 | 0.4 | 0.75 | 0.56 | -0.22 | 0.08 | 0.55 | -0.1425 | -0.13 | -0.305 | 0.343614 | 0.500531 | 0.847412 | 0.989223 | 0.997988 | 0.994666 |
| Huwe1_Q7TMY8_S3818s_1 | Huwe1_Q7 | -1.12 | -0.37 | 0.37 | 0.34 | -1.02 | -0.67 | 0.23 | 0.21 | -0.52 | -0.55 | -0.22 | -0.2 | -1.02 | -0.28 | -0.43 | 0.29 | 0.205 | 0.0625 | -0.2925 | 0.706052 | 0.791293 | 0.635267 | 0.995762 | 0.997988 | 0.996331 |
| Huwe1_Q7TMY8_S3922s_1 | Huwe1_Q7 | 0.76 | -0.04 | -0.11 | 0.73 | 0.32 | 0.62 | 0.52 | 0.7 | 0.02 | 0.35 | -0.57 | 0.47 | 0.01 | 0.52 | -0.29 | 0.29 | -0.1175 | -0.1775 | -0.165 | 0.548011 | 0.477115 | 0.450305 | 0.989223 | 0.997988 | 0.994666 |
| Huwe1_Q7TMY8_T3927t_1 | Huwe1_Q7 | -0.71 | 0.3 | -0.31 | -1.54 | 0.57 | -0.45 | 0.11 | -1.22 | -0.61 | -0.4 | -0.85 | -0.8 | 0.01 | 0.68 | -1.7 | -0.02 | 0.205 | -0.2675 | -0.2025 | 0.60117 | 0.529583 | 0.801508 | 0.989223 | 0.997988 | 0.995259 |
| Huwe1_Q7TMY8_T3930t_1 | Huwe1_Q7 | 1.3 | 0.25 | 0.84 | 1.03 | 0.9 | 0.83 | 1.02 | 1.94 | 0.86 | 1.35 | 1.44 | 0.67 | 1.05 | 1.24 | 1.2 | 1.49 | 0.3175 | -0.1 | 0.3075 | 0.215605 | 0.369013 | 0.487735 | 0.989223 | 0.997988 | 0.994666 |
| Ibtk_Q6ZPR6_S945s_1 | Ibtk_Q6ZPI | -2.81 | -1.55 | -1.22 | -2.22 | -1.52 | -1.17 | -0.76 | -1.2 | -4.48 | -1.77 | -1.35 | -1.9 | -0.67 | -1.3 | -2.41 | -1.44 | 0.3175 | 0.225 | 0.39 | 0.318204 | 0.061334 | 0.556246 | 0.989223 | 0.997988 | 0.994666 |
| Ica1l_Q3TY65_S294s_1 | Ica1l_Q3TY | -2.17 | -2.39 | 1.25 | 1.45 | -2.09 | -1.91 | 1.52 | 1.5 | -2.24 | -1.79 | 1.43 | 1.44 | -2.73 | -1.59 | 1.43 | 1.39 | 0.7875 | -0.425 | 0.495 | 0.944052 | 0.860675 | 0.890351 | 0.996493 | 0.997988 | 0.999507 |
| Ice1_E9Q286_T1607t_1 | Ice1_E9Q2 | -2.06 | -0.86 | -0.13 | -0.98 | -1.32 | -1.62 | -0.47 | -2.72 | 0.58 | -1.84 | 0.64 | -2.25 | -0.78 | -0.48 | 0.24 | -1.23 | 0.22 | 0.175 | 0.09 | 0.366081 | 0.353879 | 0.709608 | 0.989223 | 0.997988 | 0.994666 |
| Ifngr1_P15261_S296s_1 | Ifngr1_P15 | -1.2 | -2.06 | -2.42 | -2.63 | -0.31 | -1.4 | 0.01 | -0.73 | -3.45 | -2.5 | 0.23 | -0.96 | -2.62 | -0.89 | -1.64 | -2.53 | -0.525 | 0.29 | 0.445 | 0.748504 | 0.005664 | 0.604653 | 0.989223 | 0.997988 | 0.9971 |
| Ifngr1_P15261_T373t_1 | Ifngr1_P15 | -0.63 | -0.7 | 0.53 | 0.57 | -0.31 | -1.02 | 0.69 | 1.07 | 0 | 0.22 | 1.28 | 0.61 | 0.18 | -0.09 | 0.77 | 0.74 | 1.47 | 0.4075 | 0.1575 | 0.282782 | 0.758546 | 0.201828 | 0.995762 | 0.997988 | 0.994666 |
| Ifngr2_Q63953_T331t_1 | Ifngr2_Q63 | 0.15 | 0 | 0.27 | -0.36 | 0.89 | -0.43 | 0.36 | 0.25 | -0.76 | 0.62 | -0.33 | 0.84 | -0.13 | -1.69 | 1.27 | 0.85 | 0.165 | 0.585 | 0.4575 | 0.921731 | 0.436409 | 0.84706 | 0.989223 | 0.997988 | 0.999507 |
| Ift46_Q9DB07_T283t_1 | Ift46_Q9DE | -0.3 | 0.25 | -0.32 | -0.23 | 0.6 | -0.07 | 0.53 | -0.17 | -0.03 | 0.42 | 0.18 | -0.03 | 0.05 | 0.23 | 0.31 | -0.27 | 0.2525 | 0.0775 | 0.06 | 0.407373 | 0.201187 | 0.268852 | 0.989223 | 0.997988 | 0.994666 |
| Ift74_B1AWG1_S300s_1 | Ift74_B1AV | -0.28 | 0.16 | 0.27 | -0.41 | -0.27 | -0.4 | -0.71 | -0.57 | 0.81 | -0.14 | -0.15 | -0.43 | 0.98 | -0.15 | 0.15 | 0.1 | 0.3725 | 0.285 | 0.23 | 0.329831 | 0.114999 | 0.795988 | 0.989223 | 0.997988 | 0.994666 |
| Igbp1b_Q9QZ29_S282s_1 | Igbp1b_Q9 | 1.8 | 0.41 | 1.78 | 2.36 | 0.68 | 2.22 | 1.68 | 2.73 | 1.57 | 1.52 | 2.36 | 2.12 | -0.22 | 1.17 | 1.68 | 2.2 | -0.4225 | 0.0875 | 0.335 | 0.535116 | 0.661694 | 0.500457 | 0.990434 | 0.997988 | 0.994666 |
| Igsf5_D3YXH0_S284s_1 | Igsf5_D3YX | -1.49 | -1.37 | -1.09 | -1.84 | -0.44 | -1.67 | -2.33 | -1.7 | -1.69 | -1.23 | 0.95 | -2.12 | -0.93 | -0.32 | -3.1 | -1.66 | 0.24 | 0.305 | -0.38 | 0.923009 | 0.82971 | 0.509663 | 0.995762 | 0.997988 | 0.999507 |
| Igsf9b_E9PZ19_S1207s_1 | Igsf9b_E9P | 0.81 | 1.01 | 0.69 | 0.97 | 0.59 | 0.6 | 1.11 | 0.71 | 0.93 | 1.27 | 0.87 | 1.09 | 1.18 | 0.94 | 0.26 | 0.39 | -0.0875 | 0.425 | -0.055 | 0.552481 | 0.603302 | 0.453636 | 0.989223 | 0.997988 | 0.994666 |
| Igsf9b_E9PZ19_S1215s_1 | Igsf9b_E9P | 0.99 | 0.91 | 1.27 | 1.09 | 0.81 | 1.13 | 0.5 | 0.96 | 0.92 | 1.15 | 1.62 | 1.32 | 1.05 | 1.22 | 0.34 | 0.89 | -0.1175 | 0.17 | -0.1775 | 0.505833 | 0.361395 | 0.452709 | 0.989223 | 0.997988 | 0.994666 |
| Igsf9b_E9PZ19_S1258s_1 | Igsf9b_E9P | 0.86 | 1.59 | 0.96 | 0.88 | -0.31 | 0.96 | 0.36 | -0.27 | 3.98 | 1.03 | 0.83 | 0.72 | 4.24 | -0.64 | 0.66 | 1.05 | -0.215 | 0.1875 | -0.19 | 0.781591 | 0.028485 | 0.439348 | 0.989223 | 0.997988 | 0.998226 |
| Igsf9b_E9PZ19_S775s_1 | Igsf9b_E9P | 0.62 | 1.4 | 1.58 | -0.47 | 0.97 | 1.01 | 0.46 | 0.23 | 1.12 | 0.65 | 1.27 | 1.22 | -0.08 | 0.79 | 1.29 | 1.01 | -0.8875 | 0.5675 | 0.255 | 0.953785 | 0.805975 | 0.548122 | 0.995762 | 0.997988 | 0.999507 |
| Igsf9b_E9PZ19_S783s_1 | Igsf9b_E9P | 0.33 | 1.32 | 1.42 | 0.43 | 1 | 0.86 | 0.56 | 0.84 | 1.17 | 1.22 | 2.22 | 1.26 | 1.31 | 1.28 | 1.11 | 0.75 | -0.115 | 0.2825 | -0.03 | 0.498484 | 0.850142 | 0.148523 | 0.996349 | 0.997988 | 0.994666 |
| Ikbkap_Q7TT37_T1285t_1 | Ikbkap_Q7 | -1.12 | -0.07 | 0.45 | 0.11 | -0.06 | -1.46 | 0.56 | 0.11 | -0.94 | -0.51 | 0.82 | 0.24 | -0.91 | 0.14 | 0.04 | 0.18 | -0.06 | 0.5925 | 0.2375 | 0.962293 | 0.913193 | 0.902281 | 0.997335 | 0.997988 | 0.999507 |
| Ikbkb_Q5D0E0_S672s_1 | Ikbkb_Q5D | 0.93 | 0.83 | 0.79 | 0.63 | 0.26 | 0.29 | 0.37 | 0.21 | 0.71 | 0.73 | 0.8 | 0.45 | 0.79 | 1.18 | 0.39 | 0.33 | -0.055 | 0.06 | 0.02 | 0.66636 | 0.026409 | 0.575588 | 0.989223 | 0.997988 | 0.996159 |
| Ikbkg_Q8VC91_S379s_1 | Ikbkg_Q8V | 1.11 | 0.94 | 1.27 | 1.06 | 0.53 | 1.14 | 1 | 1.15 | 0.9 | 1.25 | 1.17 | 1.16 | 0.55 | 0.96 | 0.72 | 0.98 | -0.5125 | -0.1225 | -0.1225 | 0.251348 | 0.551947 | 0.908697 | 0.989223 | 0.997988 | 0.994666 |
| Ildr2_B5TVM2_S287s_1 | Ildr2_B5TV | -2.65 | -3.54 | -2.38 | -6.75 | -3.95 | -1.18 | -3.36 | -3.07 | -2.67 | -3.99 | 0.7 | -3.32 | -2 | -3.11 | -3.37 | -4.08 | -0.14 | 0.025 | -0.2925 | 0.475777 | 0.363697 | 0.25242 | 0.989223 | 0.997988 | 0.994666 |
| Ildr2_B5TVM2_S384s_1 | Ildr2_B5TV | -1.53 | -0.69 | -0.95 | -0.98 | -0.23 | -1.08 | -0.47 | -0.99 | -1.42 | -0.89 | -0.89 | -0.96 | -0.38 | -0.26 | -1.05 | -1.61 | 0.94 | 1.51 | 0.69 | 0.57605 | 0.262128 | 0.992707 | 0.989223 | 0.997988 | 0.995259 |
| Ildr2_B5TVM2_S435s_1 | Ildr2_B5TV | 0.61 | 0.61 | 0.94 | 0.78 | 0.3 | 0.97 | 0.27 | 0.32 | 0.92 | 0.87 | 0.81 | 0.85 | 0.79 | 0.3 | 0.56 | 0.81 | 0.345 | -0.0025 | 0.2125 | 0.637147 | 0.288528 | 0.55118 | 0.989223 | 0.997988 | 0.995259 |
| Ildr2_B5TVM2_S457s_1 | Ildr2_B5TV | 0.14 | 0.55 | 0.28 | 0.36 | 0.42 | 0.36 | 0.71 | 0.44 | -0.12 | 0.51 | -0.19 | 0.6 | -0.53 | 0.45 | 0.36 | 0.42 | -0.27 | 0.1275 | -0.12 | 0.610336 | 0.487172 | 0.634362 | 0.989223 | 0.997988 | 0.995259 |
| Ildr2_B5TVM2_T438t_1 | Ildr2_B5TV | 0.06 | 0.28 | 0.34 | -0.76 | 0.16 | 0.53 | -1.31 | -1.35 | 0.51 | 0.37 | -1.29 | -0.78 | 0.67 | -0.21 | -1.39 | -0.14 | 0.15 | -0.1325 | -0.1575 | 0.604029 | 0.360513 | 0.567636 | 0.989223 | 0.997988 | 0.995259 |
| Ilf3_Q9Z1X4_S382s_1 | Ilf3_Q9Z1X | 2.17 | 0.55 | 1.25 | 1.66 | -0.19 | 1.62 | 0.18 | 1.2 | 1.34 | 0.96 | 1.66 | 1.07 | 2.37 | 0.17 | 1.68 | 1.08 | -0.4725 | -0.2775 | -0.2475 | 0.87816 | 0.180699 | 0.694341 | 0.989223 | 0.997988 | 0.999507 |
| Ilf3_Q9Z1X4_T67t_1 | Ilf3_Q9Z1X | 0.7 | 0.03 | -0.17 | -0.23 | 0.53 | 0.08 | 1.52 | 0.54 | 0.39 | -0.05 | -0.31 | 0.05 | 0.12 | 0.89 | -0.15 | 0.55 | -0.705 | -0.15 | -0.0825 | 0.442404 | 0.13561 | 0.834055 | 0.989223 | 0.997988 | 0.994666 |
| Ina_P46660_S30s_1 | Ina_P4666 | 0.93 | 0.7 | 0.66 | 0.87 | 1 | 0.98 | 0.65 | 0.68 | 0.99 | 0.55 | 0.79 | 0.59 | 0.26 | 0.77 | 0.56 | 0.43 | 0.585 | -0.0625 | 0.27 | 0.264331 | 0.859984 | 0.789869 | 0.996493 | 0.997988 | 0.994666 |
| Ina_P46660_S335s_1 | Ina_P4666 | 1.27 | 0.7 | 1.13 | 0.81 | -0.33 | 1.27 | -0.2 | 0.4 | 2.08 | 0.73 | 0.08 | 0.62 | 1.78 | -0.53 | 0.99 | 1.06 | 0.0375 | -0.06 | -0.285 | 0.751666 | 0.093634 | 0.817473 | 0.989223 | 0.997988 | 0.9971 |
| Ina_P46660_S37s_1 | Ina_P4666 | 1.98 | 0.16 | 0.87 | 0.85 | 1.41 | 2.16 | 1.14 | 1.37 | 0.7 | 0.43 | 4.31 | 0.16 | -0.75 | 1 | -0.05 | 1.11 | -0.6925 | -0.1 | -0.1525 | 0.256599 | 0.203086 | 0.640637 | 0.989223 | 0.997988 | 0.994666 |
| Inf2_E9QLA5_S1170s_1 | Inf2_E9QL | 0.01 | 0.28 | -1.07 | -0.34 | -1.98 | -0.6 | -0.55 | -1.36 | 0.59 | 0.12 | -1 | -0.21 | -0.69 | -0.26 | -2.51 | -0.25 | 0.555 | 0.435 | -0.6375 | 0.268106 | 0.072188 | 0.720515 | 0.989223 | 0.997988 | 0.994666 |
| Inf2_E9QLA5_S1172s_1 | Inf2_E9QL | -0.45 | -0.4 | 0.63 | -0.03 | -1.41 | -0.15 | -0.07 | -0.37 | -0.27 | -0.1 | -0.32 | -0.1 | 0.01 | -0.89 | 0.09 | 0.18 | -0.8425 | 0.155 | -0.6475 | 0.808476 | 0.274085 | 0.650999 | 0.989223 | 0.997988 | 0.999507 |
| Inf2_E9QLA5_S1184s_1 | Inf2_E9QL | -0.9 | -1.15 | -1.17 | -1.77 | -0.78 | -1.79 | -2.28 | -1.27 | -0.39 | -0.99 | 0.23 | -1.41 | 0.38 | -0.05 | -0.41 | -1.28 | -0.4375 | -0.135 | -0.09 | 0.046058 | 0.450163 | 0.153838 | 0.989223 | 0.997988 | 0.994666 |
| Inpp4a_E9Q9A0_S186s_1 | Inpp4a_E9 | 4.62 | 3.22 | 0.11 | 0.44 | 3.75 | 4.91 | -0.27 | 0.69 | 3.45 | 3.33 | -0.18 | 0.19 | 2 | 2.79 | -0.25 | -0.2 | -0.2825 | 0.6075 | 0.9075 | 0.391688 | 0.902486 | 0.756108 | 0.996493 | 0.997988 | 0.994666 |
| Inpp4a_E9Q9A0_S516s_1 | Inpp4a_E9 | 0.43 | -0.32 | 0.14 | 1.09 | 0.35 | 0.08 | 0.85 | 0.68 | -0.57 | -0.31 | 0.3 | 0.12 | -1.25 | 0.56 | 0.1 | 0.37 | 0.1725 | -0.4 | -1.0125 | 0.426501 | 0.655213 | 0.237684 | 0.990434 | 0.997988 | 0.994666 |
| Inpp4a_E9Q9A0_S517s_1 | Inpp4a_E9 | 0.43 | -0.32 | 0.14 | 1.09 | 0.35 | 0.08 | 0.85 | 0.68 | -0.57 | -0.31 | 0.3 | 0.12 | -1.25 | 0.56 | 0.1 | 0.37 | 0.155 | -0.45 | -0.39 | 0.426501 | 0.655213 | 0.237684 | 0.990434 | 0.997988 | 0.994666 |
| Inpp4a_E9Q9A0_S916s_1 | Inpp4a_E9 | -1.44 | -2.1 | -2.72 | -2.39 | -0.37 | -2.06 | -1.56 | -1.8 | -2.67 | -1.01 | -1.83 | -1.9 | -1.75 | -0.82 | -2.29 | -1.82 | 0.155 | -0.45 | -0.39 | 0.250294 | 0.125427 | 0.472078 | 0.989223 | 0.997988 | 0.994666 |
| Inpp4b_E9Q2C4_S324s_1 | Inpp4b_E9 | -2.31 | 0.24 | -1.32 | -1.27 | -1.35 | 2.96 | -0.6 | -0.07 | -2.69 | -2.76 | -1.93 | -0.36 | -1.23 | -0.42 | -0.81 | 0.36 | 0.715 | 0.31 | 0.4925 | 0.281692 | 0.15933 | 0.282553 | 0.989223 | 0.997988 | 0.994666 |
| Inpp4b_E9Q2C4_T316t_1 | Inpp4b_E9 | 0.2 | -1 | -0.55 | 0.24 | -0.47 | -0.25 | -0.38 | 0.37 | 0.31 | -0.18 | 0.7 | -0.06 | -0.26 | -0.42 | -0.45 | 0.04 | 1.4 | -0.77 | 0.64 | 0.988628 | 0.789834 | 0.223777 | 0.995762 | 0.997988 | 1 |
| Inpp5b_F2Z437_S156s_1 | Inpp5b_F2 | 8.32 | 4.3 | 5.3 | 2.16 | 4.53 | 8.17 | 5.36 | 4.36 | -0.03 | 7.5 | 2.53 | 5.16 | 0.29 | -1.17 | 5.35 | 6.14 | 0.095 | 0.47 | 0.005 | 0.232194 | 0.662435 | 0.500759 | 0.990434 | 0.997988 | 0.994666 |
| Inpp5d_Q9ES52_S709s_1 | Inpp5d_Q9 | -0.13 | -0.52 | -0.5 | -0.03 | -0.09 | 0.31 | 0.24 | -0.24 | -0.03 | 0.36 | 0.47 | 0.07 | -0.01 | 0.47 | 0.45 | -1.05 | 0.585 | -1.23 | -2.3675 | 0.508776 | 0.174131 | 0.064019 | 0.989223 | 0.997988 | 0.994666 |
| Inpp5e_Q9JII1_S103s_1 | Inpp5e_Q9 | 0.22 | 0.33 | -0.23 | -0.08 | -0.06 | 0.64 | -0.28 | -0.3 | -0.41 | 0.41 | 0.49 | -0.06 | 0.11 | 0.15 | -0.4 | -0.54 | 0.35 | 0.5125 | 0.26 | 0.433778 | 0.834585 | 0.870681 | 0.995762 | 0.997988 | 0.994666 |
| Inpp5f_Q8CDA1_S1103s_1 | Inpp5f_Q8 | 0.04 | 0.1 | -0.82 | -0.87 | -0.15 | 0.87 | -0.69 | -1.29 | -0.76 | 0.5 | -0.52 | -1.22 | -0.64 | 0.82 | -1 | -0.11 | -0.06 | 0.0475 | -0.23 | 0.736826 | 0.881656 | 0.796627 | 0.996493 | 0.997988 | 0.9971 |
| Inpp5f_Q8CDA1_S879s_1 | Inpp5f_Q8 | -0.71 | 0.27 | 0.14 | -0.34 | -0.47 | -0.93 | -0.07 | 0.31 | -0.63 | 0.3 | -0.41 | 0.23 | -0.47 | -0.86 | -1.66 | 0.26 | 0.0725 | -0.1125 | 0.155 | 0.260622 | 0.711678 | 0.924242 | 0.995437 | 0.997988 | 0.994666 |
| Inpp5f_Q8CDA1_S908s_1 | Inpp5f_Q8 | -0.79 | 0.14 | 0.41 | 0.63 | 0.33 | 0.18 | 0.63 | 0.43 | -0.1 | 0.08 | 0.52 | 0.7 | -0.12 | 0.63 | 0.46 | 0.7 | -0.13 | 0.0325 | -0.5225 | 0.407721 | 0.38811 | 0.588316 | 0.989223 | 0.997988 | 0.994666 |
| Inpp5f_Q8CDA1_S936s_1 | Inpp5f_Q8 | -0.54 | 0.02 | -0.73 | -1.03 | 0.02 | -0.23 | -0.33 | -0.91 | -0.53 | -0.67 | -0.26 | -0.77 | -0.72 | 0.03 | -0.31 | -0.89 | 0.295 | 0.2025 | 0.32 | 0.773881 | 0.513197 | 0.965771 | 0.989223 | 0.997988 | 0.99798 |
| Inpp5f_Q8CDA1_S941s_1 | Inpp5f_Q8 | -0.62 | -0.61 | -1.4 | -0.86 | -0.45 | -1.35 | -1.59 | -1.27 | -1 | -1.15 | -0.96 | -1.07 | 0.04 | -0.52 | -0.91 | -0.79 | 0.2075 | 0.0125 | 0.0975 | 0.32709 | 0.375599 | 0.508642 | 0.989223 | 0.997988 | 0.994666 |
| Inpp5f_Q8CDA1_S941s_2 | Inpp5f_Q8 | 0.42 | 0.52 | -0.84 | 0.19 | -0.47 | -0.53 | -1.43 | -1.9 | 0.71 | 0.83 | -1.47 | -1.77 | -0.02 | 0.32 | 0.02 | -1.76 | -0.2925 | -0.1725 | 0.3275 | 0.421996 | 0.02473 | 0.47519 | 0.989223 | 0.997988 | 0.994666 |
| Inpp5f_Q8CDA1_S946s_2 | Inpp5f_Q8 | 0.42 | 0.52 | -0.84 | 0.19 | -0.47 | -0.53 | -1.43 | -1.9 | 0.71 | 0.83 | -1.47 | -1.77 | -0.02 | 0.32 | 0.02 | -1.76 | -1.155 | -0.4975 | -0.4325 | 0.421996 | 0.02473 | 0.47519 | 0.989223 | 0.997988 | 0.994666 |
| Inpp5j_P59644_S266s_1 | Inpp5j_P59 | 0 | -0.51 | 0.26 | 0.24 | 0.62 | 0.21 | 0.2 | 0.49 | -0.22 | -0.16 | 1.06 | 0.22 | 0.23 | 0.34 | 0.34 | 0.22 | -1.155 | -0.4975 | -0.4325 | 0.306262 | 0.161713 | 0.529232 | 0.989223 | 0.997988 | 0.994666 |
| Inpp5j_P59644_S691s_1 | Inpp5j_P59 | 1.92 | 0.89 | 0.77 | -0.25 | 1.04 | 2.41 | -0.8 | -0.26 | 2.38 | 1.67 | -1.43 | 0.44 | 2.14 | -0.61 | 0.4 | 0.74 | 0.3825 | 0.2275 | 0.285 | 0.799963 | 0.751372 | 0.935974 | 0.995762 | 0.997988 | 0.999507 |
| Inpp5j_P59644_S69s_1 | Inpp5j_P59 | 1.01 | -0.4 | -0.37 | -0.1 | 0.63 | -0.11 | 0.05 | 0.33 | 0.35 | 0.5 | 0.85 | -0.03 | 0.98 | 0.45 | 0.82 | 0.53 | -0.235 | -0.0675 | -0.165 | 0.102045 | 0.605917 | 0.330618 | 0.989223 | 0.997988 | 0.994666 |
| Ints1_K3W4P2_S131s_1 | Ints1_K3W | -0.9 | 0.8 | -0.9 | -1.08 | 0.15 | -1.15 | -0.9 | -0.81 | -0.61 | -1.56 | 0.01 | -1.09 | -0.3 | -0.97 | -0.43 | -1.14 | 0.19 | 0.3825 | 0.66 | 0.686077 | 0.745646 | 0.576437 | 0.995762 | 0.997988 | 0.996159 |
| Ints1_K3W4P2_S1354s_1 | Ints1_K3W | -0.59 | -0.87 | -0.11 | 0.41 | -0.39 | -0.58 | -1.37 | 0.25 | -0.35 | -0.37 | 0.73 | -0.51 | -0.16 | -0.49 | 0.28 | -0.15 | -0.1575 | -0.2925 | -0.19 | 0.650669 | 0.578753 | 0.680948 | 0.989223 | 0.997988 | 0.995259 |
| Ints10_E9Q361_S231s_1 | Ints10_E9C | -0.21 | -0.82 | -0.01 | -0.02 | -0.62 | -0.64 | -0.87 | 0.38 | -0.74 | -1.08 | -0.24 | -0.79 | 0.13 | -0.35 | 0.24 | -0.37 | -0.2325 | 0.165 | 0.16 | 0.56469 | 0.616751 | 0.158473 | 0.989223 | 0.997988 | 0.995259 |
| Ints12_Q9D168_S127s_1 | Ints12_Q9I | -0.03 | -0.33 | -0.06 | -0.48 | 0.99 | 0.22 | -0.32 | -0.18 | 0.17 | -0.42 | 0.61 | -0.06 | 0.4 | 1.35 | 0.27 | -0.66 | -0.1725 | -0.4475 | 0.1775 | 0.202512 | 0.235543 | 0.312261 | 0.989223 | 0.997988 | 0.994666 |
| Ip6k1_H3BIW9_S40s_1 | Ip6k1_H3B | 0.29 | 0.2 | 0.53 | 0.23 | 0.18 | 0.44 | 0.73 | 0.67 | -0.2 | 0.13 | -0.24 | -0.04 | -0.54 | 0.3 | 0.23 | 0.62 | 0.4025 | 0.3 | 0.565 | 0.609003 | 0.403363 | 0.097715 | 0.989223 | 0.997988 | 0.995259 |
| Ipcef1_Q5DU31_S192s_1 | Ipcef1_Q5I | 0.12 | -0.32 | 0.44 | 0 | 0.32 | 1.61 | 0.92 | 0.19 | 1.2 | 0.19 | 1.46 | 0.73 | 1.12 | 0.43 | 1.35 | -0.5 | 0.1925 | -0.4 | -0.16 | 0.235221 | 0.076461 | 0.033385 | 0.989223 | 0.997988 | 0.994666 |
| Ipo11_Q8K2V6_S343s_1 | Ipo11_Q8K | 2.25 | 2.68 | 2.03 | 2.01 | 0.72 | 1.27 | 1.57 | 1.83 | 1.12 | 2.09 | 2.37 | 2.1 | -0.16 | 1.24 | 2.11 | 1.44 | 0.7 | 0.835 | 0.54 | 0.045276 | 0.01496 | 0.350642 | 0.989223 | 0.997988 | 0.994666 |
| Iqgap2_Q3UQ44_S16s_1 | Iqgap2_Q3 | -1.1 | -1.15 | -1.83 | -2.63 | -0.68 | -1.05 | -1.72 | -2.19 | -0.68 | -1.45 | -0.23 | -2.32 | -0.95 | -0.27 | -3.16 | -2.21 | -0.895 | -0.3225 | -1.085 | 0.963924 | 0.562968 | 0.360102 | 0.989223 | 0.997988 | 0.999507 |
| Iqgap2_Q3UQ44_S396s_1 | Iqgap2_Q3 | 0.51 | -1.09 | -0.46 | -0.06 | -1.13 | -0.46 | -0.57 | 0.2 | 0.06 | 0.03 | -0.39 | -1.26 | -0.82 | -0.77 | -0.14 | -1 | 0.2675 | 0.5075 | 0.03 | 0.315393 | 0.605277 | 0.794513 | 0.989223 | 0.997988 | 0.994666 |
| Iqsec1_Q8R0S2_S104s_1 | Iqsec1_Q8I | -0.83 | -0.21 | -1.11 | -0.92 | 0.24 | -0.46 | -0.09 | -2.92 | -0.61 | -0.22 | -0.85 | -0.31 | -0.5 | 0.16 | -1.29 | -0.92 | -0.215 | -0.115 | -0.4075 | 0.733851 | 0.951594 | 0.361144 | 0.999187 | 0.997988 | 0.9971 |
| Iqsec1_Q8R0S2_S159s_1 | Iqsec1_Q8I | -1.74 | -0.67 | -1.42 | -6.22 | -0.61 | -1.09 | -1.47 | -1.43 | 0.76 | -1.24 | -2.92 | -1.22 | 1.6 | -1.46 | -1.31 | -1.47 | -0.04 | 0.27 | 0.13 | 0.165988 | 0.233236 | 0.305044 | 0.989223 | 0.997988 | 0.994666 |

| | | | | | | | | | | | | | | | | | | | | | | | | | | |
|---|---|---|---|---|---|---|---|---|---|---|---|---|---|---|---|---|---|---|---|---|---|---|---|---|---|---|
| Iqsec1_Q8R0S2_S179s_1 | Iqsec1_Q8I | 0.02 | 0.06 | 0.21 | -0.24 | 0.39 | -0.04 | 0.09 | 0.06 | -0.97 | 0.06 | -1.03 | 0.04 | -0.99 | -0.8 | -0.32 | -0.07 | 1.3625 | 1.3575 | 1.8525 | 0.084915 | 0.610809 | 0.173047 | 0.989223 | 0.997988 | 0.994666 |
| Iqsec1_Q8R0S2_S197s_1 | Iqsec1_Q8I | 0.25 | 0.42 | 0.32 | 0.39 | 0.59 | 0.25 | 0.65 | 0.21 | -0.14 | 0.02 | 0.37 | 0.25 | -0.1 | 0.77 | 0.24 | 0.23 | 0.1125 | -0.4875 | -0.5575 | 0.824925 | 0.710236 | 0.340927 | 0.995437 | 0.997988 | 0.999507 |
| Iqsec1_Q8R0S2_S362s_1 | Iqsec1_Q8I | 0.87 | 0.59 | 0.61 | 0.38 | 0.92 | 1.02 | -0.21 | 0.72 | 0.57 | 0.89 | 0.2 | 0.44 | 0.86 | 1.66 | 0.16 | -0.03 | 0.08 | -0.22 | -0.06 | 0.900992 | 1 | 0.727202 | 1 | 0.997988 | 0.999507 |
| Iqsec1_Q8R0S2_S415s_1 | Iqsec1_Q8I | 1 | 0.42 | 0.45 | -0.14 | 0.18 | 0.71 | -0.73 | -0.21 | 1.15 | 0.34 | -0.92 | 0.25 | 2.69 | 0.38 | 0.63 | 0.08 | -1.2E-16 | -0.0875 | 0.05 | 0.389837 | 0.254231 | 0.626095 | 0.989223 | 0.997988 | 0.994666 |
| Iqsec1_Q8R0S2_S513s_1 | Iqsec1_Q8I | 0.6 | 1.16 | 1.03 | 0.89 | 1.29 | 0.55 | 1.36 | 0.86 | 0.12 | 1.02 | 0.53 | 1.1 | -0.33 | 0.8 | 1.06 | 0.9 | -0.445 | -0.2275 | 0.5125 | 0.398181 | 0.724075 | 0.453426 | 0.995437 | 0.997988 | 0.994666 |
| Iqsec1_Q8R0S2_S795s_1 | Iqsec1_Q8I | -0.57 | 1.17 | 1.18 | -0.01 | 0.04 | -0.13 | 0.12 | 0.25 | -0.3 | 0.09 | -0.03 | 0.59 | -0.08 | 0.41 | 0.2 | 0.7 | 0.095 | -0.2275 | -0.3125 | 0.766686 | 0.387668 | 0.444637 | 0.989223 | 0.997988 | 0.9971 |
| Iqsec1_Q8R0S2_S923s_1 | Iqsec1_Q8I | -2.88 | -1.63 | -1.72 | -3.16 | -0.87 | -2.78 | -1.41 | -2.75 | -3.16 | -2.38 | -0.16 | -1.87 | -0.78 | -0.7 | -1.99 | -1.99 | -0.3725 | -0.355 | -0.135 | 0.076168 | 0.486767 | 0.506188 | 0.989223 | 0.997988 | 0.994666 |
| Iqsec1_Q8R0S2_S937s_1 | Iqsec1_Q8I | -0.17 | -0.08 | -0.38 | 0.05 | -0.09 | -0.37 | -0.1 | 0.27 | -0.33 | 0.29 | 0.26 | 0.35 | -0.29 | 0.37 | -0.07 | -0.1 | 0.395 | 0.455 | 0.9825 | 0.643002 | 0.755481 | 0.274634 | 0.995762 | 0.997988 | 0.995259 |
| Iqsec2_E9QAD8_S111s_1 | Iqsec2_E9C | -1.13 | -0.32 | -0.1 | -2.25 | -1.21 | -0.21 | -0.65 | -0.73 | 0.38 | -0.56 | -1.05 | -0.63 | 0.99 | 0.17 | -1.36 | -0.71 | 0.0725 | 0.2875 | 0.1225 | 0.27506 | 0.609457 | 0.374236 | 0.989223 | 0.997988 | 0.994666 |
| Iqsec2_E9QAD8_S1143s_1 | Iqsec2_E9C | -1.42 | -2 | -1.8 | -2.87 | -0.51 | -1.16 | -1.94 | -1.74 | -2.36 | -2.43 | 0.29 | -2.03 | -1.24 | -0.37 | -2.96 | -3.26 | 0.25 | 0.485 | 0.7225 | 0.923679 | 0.129384 | 0.552468 | 0.989223 | 0.997988 | 0.999507 |
| Iqsec2_E9QAD8_S1158s_1 | Iqsec2_E9C | 0.52 | 1.02 | 0.17 | -0.18 | 1.15 | 0.71 | 0.39 | -0.17 | 0.38 | 0.78 | -0.75 | -0.01 | 0.42 | 1.2 | -0.5 | 0.24 | 0.685 | 0.39 | 0.065 | 0.920889 | 0.712264 | 0.495712 | 0.995437 | 0.997988 | 0.999507 |
| Iqsec2_E9QAD8_S1161s_1 | Iqsec2_E9C | 0.46 | 0.37 | 1.45 | 1.32 | 1.11 | 0.77 | 1.15 | 1.28 | 0.6 | 0.32 | 1.33 | 1.09 | 0.92 | 1.95 | 0.9 | 0.79 | 0.1375 | -0.2825 | -0.0425 | 0.550774 | 0.579635 | 0.860849 | 0.989223 | 0.997988 | 0.994666 |
| Iqsec2_E9QAD8_S15s_1 | Iqsec2_E9C | -0.54 | -0.01 | 0.79 | 0.34 | -2.79 | -0.53 | 0.06 | 0.24 | 0.47 | -0.48 | -0.46 | 0.39 | 1.45 | -0.14 | 0.32 | 0.64 | 0.1775 | -0.065 | 0.24 | 0.340014 | 0.198151 | 0.670003 | 0.989223 | 0.997988 | 0.994666 |
| Iqsec2_E9QAD8_S214s_1 | Iqsec2_E9C | 0.06 | -0.41 | -0.74 | -0.95 | -0.38 | 0.04 | -0.83 | -0.98 | -0.16 | -0.2 | -1.15 | -1.08 | 0.78 | -0.53 | -0.41 | -0.61 | -0.9 | -0.165 | 0.4225 | 0.435555 | 0.933435 | 0.703617 | 0.998562 | 0.997988 | 0.994666 |
| Iqsec2_E9QAD8_S246s_1 | Iqsec2_E9C | 0.48 | 0.7 | 1.33 | 1.2 | 0.69 | 0.56 | 1.16 | 1.38 | 0.27 | 0.45 | 1 | 0.87 | -0.07 | 0.84 | 0.58 | 0.83 | -0.0275 | -0.1375 | 0.3175 | 0.26783 | 0.947264 | 0.366919 | 0.999187 | 0.997988 | 0.994666 |
| Iqsec2_E9QAD8_S262s_1 | Iqsec2_E9C | 0.46 | 1.2 | 1.47 | -1.09 | 0.5 | 0.59 | 0.21 | 0.08 | 0.37 | 0.84 | 1.69 | -0.52 | -0.14 | 0.8 | -0.68 | 0.14 | 0.02 | -0.28 | -0.3825 | 0.428488 | 0.757119 | 0.898574 | 0.995762 | 0.997988 | 0.994666 |
| Iqsec2_E9QAD8_S393s_1 | Iqsec2_E9C | 0.69 | 0.56 | 0.97 | 0.62 | 1.03 | 0.53 | 0.97 | 0.73 | -0.41 | 0.58 | -1.07 | 0.8 | -1.08 | -0.45 | 0.17 | 0.57 | -0.165 | 0.085 | -0.48 | 0.038641 | 0.644239 | 0.116103 | 0.99033 | 0.997988 | 0.994666 |
| Iqsec2_E9QAD8_S412s_1 | Iqsec2_E9C | 1.01 | 0.61 | 0.23 | 0.51 | 0.67 | 1.08 | 0.47 | 0.6 | 0.24 | 0.52 | 0.44 | 0.37 | 0.08 | 0.93 | 0.08 | 0.07 | 0.105 | -0.735 | -0.9075 | 0.355595 | 0.658018 | 0.43476 | 0.990434 | 0.997988 | 0.994666 |
| Iqsec2_E9QAD8_S528s_1 | Iqsec2_E9C | 0.21 | 0.83 | -0.09 | -0.8 | 0.83 | 0.15 | 0 | -0.36 | -0.56 | 0.32 | -0.09 | -0.29 | -0.69 | 0.19 | -0.46 | -0.15 | 0.115 | -0.1975 | -0.3 | 0.43496 | 0.771116 | 0.621696 | 0.995762 | 0.997988 | 0.994666 |
| Iqsec2_E9QAD8_S567s_1 | Iqsec2_E9C | 0.64 | 0.09 | 0.46 | -1.53 | -0.14 | 0.9 | -1.15 | -0.65 | 2.23 | -1.58 | -0.15 | -0.65 | 2.23 | -1.2 | 0.65 | -1.51 | 0.1175 | -0.1925 | -0.315 | 0.884605 | 0.767895 | 0.955186 | 0.995762 | 0.997988 | 0.999507 |
| Iqsec2_E9QAD8_S627s_1 | Iqsec2_E9C | -1.76 | -2.44 | -4.01 | -3.01 | -1.67 | -2.57 | -2.08 | -1.61 | -2.42 | -1.79 | -3.2 | -3.92 | -0.44 | -0.92 | -4.03 | -5.84 | -0.175 | 0.0475 | 0.1275 | 0.998315 | 0.114692 | 0.963861 | 0.989223 | 0.997988 | 1 |
| Iqsec2_E9QAD8_S741s_1 | Iqsec2_E9C | 0.51 | 0.68 | 0.81 | 1.36 | -0.05 | 0.2 | 1.33 | 0.98 | 0.46 | 0.55 | 1.21 | 1.27 | 0.57 | 0.93 | 0.62 | 0.79 | 0.8225 | -0.0275 | -0.0025 | 0.68882 | 0.544675 | 0.917456 | 0.989223 | 0.997988 | 0.996159 |
| Iqsec2_E9QAD8_S744s_1 | Iqsec2_E9C | 4.06 | 1.74 | 1.67 | 0.58 | 2.32 | 4.35 | 2.72 | 1.22 | 0.35 | 3.59 | 0.56 | 1.85 | 0.51 | 0.59 | 1.73 | 2.09 | -0.225 | 0.0325 | -0.1125 | 0.306176 | 0.459338 | 0.647821 | 0.989223 | 0.997988 | 0.994666 |
| Iqsec2_E9QAD8_S82s_1 | Iqsec2_E9C | 2.38 | 2.07 | 2.26 | 2.5 | 1.46 | 1.88 | 1.82 | 2.27 | 2.04 | 2.76 | 3.23 | 2.38 | 0.89 | 1.61 | 2.5 | 2.35 | 0.64 | -0.425 | -0.7825 | 0.253382 | 0.098142 | 0.341714 | 0.989223 | 0.997988 | 0.994666 |
| Iqsec2_E9QAD8_Y89y_1 | Iqsec2_E9C | 0.75 | 0.2 | 0.45 | 0.39 | 0.32 | 0.91 | -0.24 | -0.09 | 0.26 | 0.43 | -0.41 | 0.15 | 0.27 | -0.14 | 0.26 | -0.45 | -0.445 | 0.3 | -0.465 | 0.127294 | 0.470505 | 0.231015 | 0.989223 | 0.997988 | 0.994666 |
| Iqsec3_Q3TES0_S1068s_1 | Iqsec3_Q3T | 0.35 | 1.3 | 0.66 | 0.7 | 0.83 | 0.1 | -0.29 | 0.1 | 0.21 | 0.79 | 0.08 | 0.02 | 0.14 | 1.02 | 0.46 | 0.81 | -0.2225 | -0.34 | -0.4625 | 0.655118 | 0.102667 | 0.140573 | 0.989223 | 0.997988 | 0.995259 |
| Iqsec3_Q3TES0_S1152s_1 | Iqsec3_Q3T | 0.09 | 0.1 | -0.16 | -0.06 | -0.37 | 0.69 | -0.73 | -0.66 | 2.08 | 0.04 | -3.03 | 0 | 2.18 | -1.23 | 0.37 | 0.08 | -0.5675 | -0.4775 | -0.145 | 0.57827 | 0.452188 | 0.813845 | 0.989223 | 0.997988 | 0.995259 |
| Iqsec3_Q3TES0_S132s_1 | Iqsec3_Q3T | 0.8 | 1.43 | 0.22 | -0.13 | 1.55 | 0.87 | 0.58 | 0.84 | 0.82 | 1.01 | 0.51 | 0.66 | 0.64 | 1.79 | 0.47 | 0.31 | -0.26 | -0.22 | 0.3575 | 0.632671 | 0.340127 | 0.645357 | 0.989223 | 0.997988 | 0.995259 |
| Iqsec3_Q3TES0_S256s_1 | Iqsec3_Q3T | -0.21 | 0.57 | 0.72 | 0.46 | -0.44 | -0.07 | 0.82 | -0.78 | 0.36 | 0.54 | 1.69 | -0.32 | 0.33 | 0.18 | 1.66 | 1.19 | 0.38 | 0.17 | 0.2225 | 0.283355 | 0.216674 | 0.685045 | 0.989223 | 0.997988 | 0.994666 |
| Iqsec3_Q3TES0_S646s_1 | Iqsec3_Q3T | 1.23 | 0.31 | -0.33 | -0.05 | 1.04 | 1.1 | 0.58 | 0.36 | 0.16 | 0.76 | 0.3 | 0.14 | -0.33 | 0.51 | 0.22 | 0.26 | -0.5025 | 0.1825 | 0.455 | 0.750649 | 0.222125 | 0.893894 | 0.989223 | 0.997988 | 0.9971 |
| Iqsec3_Q3TES0_T357t_1 | Iqsec3_Q3T | 1 | 1.27 | 0.22 | 0.35 | 0.35 | 0.67 | -0.05 | 0.27 | 0.24 | 0.54 | 0.66 | 0.02 | 0.77 | 1.29 | -0.08 | 0.2 | 0.48 | 0.05 | -0.125 | 0.682517 | 0.21944 | 0.297188 | 0.989223 | 0.997988 | 0.996159 |
| Iqsec3_Q3TES0_T653t_1 | Iqsec3_Q3T | 1.06 | 1.61 | 0.81 | 0.65 | 1.33 | 0.29 | 0.49 | 0.01 | 0.64 | 0.84 | 0.32 | 0.49 | 0.1 | 1.53 | 0.72 | 0.98 | -0.4 | -0.345 | -0.165 | 0.600504 | 0.178546 | 0.132773 | 0.989223 | 0.997988 | 0.995259 |
| Ireb2_Q811J3_S178s_1 | Ireb2_Q81 | -0.89 | -1.33 | -1.21 | -2.1 | -1.99 | 0.55 | -1.77 | -0.93 | 0.43 | -1.26 | 0.33 | -1.43 | -0.54 | -1.83 | -1.85 | -2.22 | -0.5025 | -0.46 | -0.2 | 0.607271 | 0.544073 | 0.11254 | 0.989223 | 0.997988 | 0.995259 |
| Irf2bp1_Q8R3Y8_S365s_1 | Irf2bp1_Q8 | -1.52 | -1.68 | -1.68 | -1.77 | -1.68 | -2.4 | -1.65 | -3.03 | 0.18 | -1.2 | -2.02 | -1.95 | 0.29 | -1.47 | -1.5 | -2.8 | 0.3475 | 0.9 | -0.2275 | 0.618379 | 0.144927 | 0.402823 | 0.989223 | 0.997988 | 0.995259 |
| Irf2bp1_Q8R3Y8_S434s_1 | Irf2bp1_Q8 | -4.2 | -2.69 | -1.04 | -1.4 | -3.36 | -3.17 | -1.59 | -1.81 | -2.32 | -1.86 | -0.44 | -1.65 | -1.2 | -2.31 | -0.87 | -1.51 | -0.5275 | 0.415 | 0.2925 | 0.236783 | 0.840452 | 0.314082 | 0.995762 | 0.997988 | 0.994666 |
| Irf2bp2_E9Q1P8_S226s_1 | Irf2bp2_E9 | -3.22 | -2.51 | -2.37 | 0.12 | -2.87 | -5.68 | -2.47 | 0.02 | -2.94 | -4.69 | -0.63 | 0.45 | -1.4 | -1.83 | 0.29 | -2.42 | -0.15 | 0.765 | 0.86 | 0.433648 | 0.528129 | 0.971667 | 0.989223 | 0.997988 | 0.994666 |
| Irf2bp2_E9Q1P8_S343s_1 | Irf2bp2_E9 | -1.82 | -2.21 | -2.48 | -2.11 | -2.05 | -2.47 | -2.12 | -2.79 | -1.92 | -1.43 | -1.5 | -2.7 | -1.06 | -1.55 | -1.87 | -2.79 | -0.755 | 0.0425 | 0.655 | 0.404349 | 0.451184 | 0.440935 | 0.989223 | 0.997988 | 0.994666 |
| Irf2bp2_E9Q1P8_S440s_1 | Irf2bp2_E9 | -3.07 | -2.7 | -2.3 | -2.87 | -2.97 | -2.8 | -2.67 | -2.76 | -1.93 | -1.85 | -1.33 | -2.39 | -1.31 | -3.25 | -2.62 | -3.11 | -0.2025 | 0.2675 | 0.3375 | 0.722686 | 0.78706 | 0.018895 | 0.995762 | 0.997988 | 0.9971 |
| Irf2bp2_E9Q1P8_S443s_1 | Irf2bp2_E9 | -1.68 | -3.27 | -2.81 | -3.73 | -2.67 | -2.83 | -2.12 | -2.25 | -1.11 | -2.43 | -2.55 | -3.4 | -0.69 | -2.12 | -3.02 | -2.33 | -0.065 | 0.86 | 0.1625 | 0.187729 | 0.370186 | 0.408183 | 0.989223 | 0.997988 | 0.994666 |
| Irf2bpl_Q8K3X4_S195s_1 | Irf2bpl_Q8 | -7.44 | -9.57 | -0.4 | -0.91 | -6.48 | -6.76 | -1.37 | -1.25 | -4.97 | -5.33 | 0.84 | -2.29 | -4.35 | -5.94 | -1.94 | -1.06 | 0.405 | 0.5 | 0.8325 | 0.569019 | 0.794815 | 0.492177 | 0.995762 | 0.997988 | 0.995259 |
| Irf2bpl_Q8K3X4_S313s_1 | Irf2bpl_Q8 | -3.26 | -2.33 | -3.57 | -6.34 | -2.93 | -7.21 | -4.13 | -3.91 | -3.95 | -5.09 | -1.59 | -4.48 | -0.48 | -2.24 | -3.28 | -3.82 | 0.615 | 1.6425 | 1.2575 | 0.173481 | 0.543227 | 0.923594 | 0.989223 | 0.997988 | 0.994666 |
| Irf2bpl_Q8K3X4_S526s_1 | Irf2bpl_Q8 | -1.2 | -0.78 | -2.69 | -2.55 | -3.73 | -2.74 | -1.68 | -2.66 | -1.2 | -2.22 | -1.35 | -3.32 | -2.21 | -0.71 | -4.25 | -3 | -0.67 | 0.0975 | 1.42 | 0.356808 | 0.141357 | 0.727861 | 0.989223 | 0.997988 | 0.994666 |
| Irf2bpl_Q8K3X4_S636s_2 | Irf2bpl_Q8 | -0.69 | -0.3 | -1.02 | -0.86 | -0.48 | 0 | -1.25 | -2.28 | 0.05 | 0.26 | -1.01 | -1.54 | 0.5 | -0.58 | -2.41 | -1.19 | -0.8975 | -0.2175 | -0.7375 | 0.72576 | 0.556894 | 0.721692 | 0.989223 | 0.997988 | 0.9971 |
| Irf2bpl_Q8K3X4_S638s_1 | Irf2bpl_Q8 | -1.26 | -1.4 | -1.4 | -0.37 | -1.7 | -2.1 | -0.66 | -1.16 | -0.81 | -1.23 | -0.89 | -1.52 | -1.73 | -1.55 | -0.46 | -1.28 | -0.285 | 0.1575 | -0.2025 | 0.703805 | 0.45087 | 0.987681 | 0.989223 | 0.997988 | 0.996159 |
| Irf2bpl_Q8K3X4_S638s_2 | Irf2bpl_Q8 | -0.69 | -0.3 | -0.81 | -1.67 | -0.48 | 0 | -0.99 | -2.27 | 0.05 | 0.26 | -1.45 | -1.78 | 0.5 | -0.58 | -2.38 | -1.52 | -0.2975 | -0.005 | -0.1475 | 0.83723 | 0.896141 | 0.802432 | 0.996493 | 0.997988 | 0.999507 |
| Irf2bpl_Q8K3X4_S641s_1 | Irf2bpl_Q8 | -2.37 | -1.74 | -0.58 | -1.01 | -2.98 | -1.07 | -0.31 | -0.59 | -1.42 | -2.13 | -1.07 | -0.7 | -1.49 | -1.17 | -0.46 | -1.07 | -0.0675 | 0.1375 | -0.1275 | 0.401547 | 0.769784 | 0.841432 | 0.995762 | 0.997988 | 0.994666 |
| Irs1_Q543V3_S1072s_1 | Irs1_Q543\ | -0.81 | -0.34 | -0.72 | -1.38 | 0.1 | -0.35 | -0.01 | -0.26 | -0.06 | 0.55 | 1.63 | -0.64 | -0.74 | 0.64 | -0.55 | -1.69 | 0.1875 | 0.095 | 0.3775 | 0.64865 | 0.032148 | 0.038831 | 0.989223 | 0.997988 | 0.995259 |
| Irs1_Q543V3_S1094s_1 | Irs1_Q543\ | 7.01 | 2.87 | 2.4 | -0.27 | 4.77 | 7.52 | 3.37 | 3.62 | 0.79 | 5.76 | -0.97 | 3.11 | 1.18 | 0.39 | 3.39 | 3.17 | 0.6825 | 1.1825 | 0.2275 | 0.505776 | 0.250805 | 0.650781 | 0.989223 | 0.997988 | 0.994666 |
| Irs1_Q543V3_S325s_1 | Irs1_Q543\ | -1.19 | -1.28 | -1.27 | -1.81 | -1.29 | -0.75 | -1.75 | -2.1 | 0.48 | -1.04 | -0.86 | -0.85 | 1.26 | -1.15 | -1.34 | -1.6 | 1.8175 | -0.83 | -0.97 | 0.285464 | 0.799207 | 0.056236 | 0.995762 | 0.997988 | 0.994666 |
| Irs1_Q543V3_S336s_1 | Irs1_Q543\ | 1.12 | 0.84 | 0.46 | -0.01 | 1.05 | 0.75 | 0.18 | 0.39 | 1.05 | 1.17 | 0.48 | 0.68 | 0.08 | -0.52 | 0.98 | -0.72 | -0.085 | 0.82 | 0.68 | 0.167451 | 0.975381 | 0.457381 | 0.999187 | 0.997988 | 0.994666 |
| Irs1_Q543V3_S369s_1 | Irs1_Q543\ | -0.87 | -2.21 | -2.39 | -4.06 | -0.83 | -1.11 | -3.32 | -3.42 | -2.88 | -0.52 | -1.61 | -3.31 | -1.66 | -1.83 | -2.98 | -3.1 | -0.01 | 0.2425 | -0.6475 | 0.988209 | 0.798633 | 0.710304 | 0.995762 | 0.997988 | 1 |
| Irs1_Q543V3_T448t_1 | Irs1_Q543\ | -0.96 | -1.42 | -2.17 | -0.24 | -1.52 | -0.26 | -1.95 | -1.49 | -1.43 | -0.45 | -3.16 | -1.18 | 0.14 | -0.25 | -1.04 | -1.06 | 0.2125 | 0.3025 | -0.01 | 0.200767 | 0.829492 | 0.579049 | 0.995762 | 0.997988 | 0.994666 |
| Irs2_P81122_S1165s_1 | Irs2_P8112 | 2.15 | 0.44 | 1.18 | 0.72 | 0.63 | 2.58 | 1.1 | 1.22 | -0.53 | 1.52 | 0.93 | 1.12 | 1.05 | 0.05 | 0.76 | 1.31 | -0.1075 | -0.3575 | 0.645 | 0.470322 | 0.614623 | 0.508928 | 0.989223 | 0.997988 | 0.994666 |
| Irs2_P81122_S1190s_1 | Irs2_P8112 | 0.37 | 0.05 | 0.02 | 0.27 | 0.66 | 0.16 | 1 | 0.89 | 0.07 | -0.02 | 0.78 | 0.66 | -0.37 | 0.77 | 0.69 | 0.53 | 0.26 | -0.3625 | -0.33 | 0.486219 | 0.076179 | 0.48439 | 0.989223 | 0.997988 | 0.994666 |
| Irs2_P81122_S303s_1 | Irs2_P8112 | -2.43 | -2.82 | -1.46 | -0.74 | -2.01 | -2.76 | -0.75 | -1.99 | -1.69 | -2.36 | -1.94 | -0.49 | -2.09 | -2.25 | -1.29 | -0.95 | 0.5 | 0.195 | 0.2275 | 0.681587 | 0.97883 | 0.672116 | 0.999187 | 0.997988 | 0.996159 |
| Irs2_P81122_S362s_1 | Irs2_P8112 | -4.07 | -3.18 | -1.64 | -0.57 | -4.18 | -2.41 | -3.88 | -4.08 | -1 | -5.63 | 0.05 | -2.44 | -2.11 | -2.12 | -3.4 | -2.54 | -0.015 | 0.2425 | 0.2175 | 0.811684 | 0.1253 | 0.931457 | 0.989223 | 0.997988 | 0.999507 |
| Irs2_P81122_S388s_1 | Irs2_P8112 | -0.32 | -0.24 | -1.03 | -1.09 | -0.22 | -0.45 | -0.65 | -0.92 | -0.61 | -0.03 | -0.16 | -1.09 | -1.24 | -0.38 | -0.96 | -0.95 | -1.2725 | 0.11 | -0.1775 | 0.52456 | 0.711829 | 0.572496 | 0.995437 | 0.997988 | 0.994666 |
| Irs2_P81122_S573s_1 | Irs2_P8112 | -0.57 | -1.05 | -0.85 | -1.34 | -0.32 | -0.93 | -0.88 | -1.12 | -1.87 | -0.98 | -1.07 | -1.13 | -0.7 | -0.44 | -1.24 | -0.76 | 0.11 | 0.1975 | -0.2125 | 0.575583 | 0.613531 | 0.316785 | 0.989223 | 0.997988 | 0.995259 |
| Irs2_P81122_S604s_1 | Irs2_P8112 | 0.69 | 0.08 | 0.27 | 0.12 | -0.45 | 1.07 | -0.75 | -0.21 | 1.07 | 0.24 | 0.02 | -0.54 | 1.69 | 0.05 | 0.1 | -0.29 | 0.14 | -0.31 | 0.1675 | 0.829022 | 0.367022 | 0.80267 | 0.989223 | 0.997988 | 0.999507 |
| Irs2_P81122_S66s_1 | Irs2_P8112 | 0.07 | 0.73 | 1.33 | 0.26 | 0.52 | 0.82 | -1.1 | -2.1 | 0.49 | 0.61 | 1.57 | -1.13 | 1.31 | 0.65 | -0.74 | 1.62 | -0.375 | -0.0925 | 0.0975 | 0.838147 | 0.132041 | 0.713341 | 0.989223 | 0.997988 | 0.999507 |
| Irs2_P81122_S675s_1 | Irs2_P8112 | 0.61 | 0.03 | -0.07 | -0.49 | -0.18 | 0.7 | 0.03 | 0.4 | -0.01 | 0.8 | -0.07 | -0.22 | -0.18 | 0.45 | 0.2 | 0.5 | -1.0625 | -0.2125 | 0.1125 | 0.494228 | 0.496573 | 0.758952 | 0.989223 | 0.997988 | 0.994666 |
| Irs2_P81122_S723s_1 | Irs2_P8112 | -0.43 | -0.37 | -0.37 | -0.32 | -0.71 | -0.29 | -1.46 | -1.26 | 0.59 | -0.4 | -0.39 | -0.6 | 0.01 | -0.57 | -1.2 | -1.38 | 0.2175 | 0.105 | 0.2225 | 0.254192 | 0.083411 | 0.57533 | 0.989223 | 0.997988 | 0.994666 |
| Irs2_P81122_S723s_2 | Irs2_P8112 | 0.22 | 0.28 | -0.33 | 0.12 | 0.64 | 0.71 | -0.59 | -0.24 | -0.46 | 0.9 | -0.28 | -0.75 | 0.09 | 1.31 | -1.27 | -1.19 | -0.5575 | 0.1725 | -0.4125 | 0.560946 | 0.870368 | 0.574978 | 0.996493 | 0.997988 | 0.995259 |
| Irs2_P81122_S727s_1 | Irs2_P8112 | -0.72 | -0.49 | -0.09 | 0.42 | -0.22 | -1.64 | -0.88 | 0.1 | -0.54 | -0.28 | 1.29 | 0.33 | 0.45 | 0.13 | 0.06 | 0.94 | 0.0575 | -0.22 | -0.3375 | 0.104447 | 0.322899 | 0.370014 | 0.989223 | 0.997988 | 0.994666 |
| Irs2_P81122_S727s_2 | Irs2_P8112 | 0.22 | 0.28 | -0.33 | 0.12 | 0.64 | 0.71 | -0.59 | -0.24 | -0.46 | 0.9 | -0.28 | -0.75 | 0.09 | 1.31 | -1.27 | -1.19 | -0.44 | 0.42 | 0.615 | 0.560946 | 0.870368 | 0.574978 | 0.996493 | 0.997988 | 0.995259 |
| Irs2_P81122_S728s_1 | Irs2_P8112 | -0.04 | 0.47 | -0.42 | -1.15 | 0.01 | -0.23 | -0.58 | -0.74 | 0.3 | 0.14 | -0.21 | -0.22 | 0.88 | 0.32 | -0.25 | -0.32 | 0.0575 | -0.22 | -0.3375 | 0.322219 | 0.789361 | 0.447348 | 0.995762 | 0.997988 | 0.994666 |
| Irs2_P81122_S924s_1 | Irs2_P8112 | 1.02 | 0.62 | 0.51 | -0.04 | 1.04 | -0.12 | 0.57 | 0.35 | 0.05 | 0.49 | -0.15 | 0.02 | -0.41 | 0.48 | 0.16 | 0.81 | -0.1 | 0.2875 | 0.4425 | 0.467493 | 0.839868 | 0.177497 | 0.995762 | 0.997988 | 0.994666 |
| Irs2_P81122_T1189t_1 | Irs2_P8112 | 0.37 | 0.05 | 0.02 | 0.27 | 0.66 | 0.16 | 1 | 0.89 | 0.07 | -0.02 | 0.78 | 0.66 | -0.37 | 0.77 | 0.69 | 0.53 | -0.0675 | -0.425 | -0.2675 | 0.486219 | 0.076179 | 0.48439 | 0.989223 | 0.997988 | 0.994666 |
| Irs2_P81122_T347t_1 | Irs2_P8112 | -0.41 | -0.35 | 0.65 | 0.51 | -0.12 | -0.31 | 0.47 | 0.77 | -0.94 | -0.01 | 0.28 | 0.41 | -0.27 | 0.19 | 0.67 | 0.03 | 0.5 | 0.195 | 0.2275 | 0.879816 | 0.782092 | 0.687076 | 0.995762 | 0.997988 | 0.999507 |
| Irs2_P81122_T517t_1 | Irs2_P8112 | -1.01 | -0.09 | -1.03 | -0.52 | -0.66 | -0.34 | -0.99 | -0.13 | 0.71 | 0.17 | -1.16 | -1.27 | -0.07 | -0.68 | -1.09 | -0.39 | 0.1025 | -0.165 | 0.055 | 0.761094 | 0.671921 | 0.590327 | 0.992876 | 0.997988 | 0.9971 |
| Irs2_P81122_T524t_1 | Irs2_P8112 | 0.53 | -0.25 | 1.32 | -0.31 | 0.01 | 0.54 | -0.32 | 1.36 | 1.13 | 0.02 | -0.36 | 1.03 | 0.36 | 0.73 | -0.94 | 0.84 | 0.1325 | 0.275 | 0.105 | 0.886246 | 0.877835 | 0.791748 | 0.996493 | 0.997988 | 0.999507 |
| Iscu_D3Z7W0_S15s_1 | Iscu_D3Z7\ | -1.55 | -0.77 | -0.79 | -1.3 | -0.35 | -1.38 | -0.7 | -1.23 | -1.51 | -1 | -1.46 | -0.91 | -0.36 | -0.41 | -0.96 | -0.8 | 0.075 | 0.1325 | -0.075 | 0.144316 | 0.562606 | 0.688147 | 0.989223 | 0.997988 | 0.994666 |
| Iscu_D3Z7W0_S30s_1 | Iscu_D3Z7\ | -0.37 | -0.28 | 0.18 | -0.31 | -0.17 | -1.03 | -0.54 | -0.26 | -1.09 | -0.29 | 0.71 | -0.17 | 0.19 | 0.27 | 0.03 | -0.37 | 0.1875 | -0.1175 | 0.47 | 0.420514 | 0.278417 | 0.969308 | 0.989223 | 0.997988 | 0.994666 |
| Islr2_E9Q9N7_S694s_1 | Islr2_E9Q9 | 0.99 | 0.95 | 1.09 | 1.22 | 0.34 | 0.89 | 0.83 | 1.05 | 0.49 | 0.79 | 1.29 | 1.47 | 0.18 | 0.36 | 0.32 | 1.26 | -0.305 | -0.015 | 0.225 | 0.109831 | 0.243448 | 0.852772 | 0.989223 | 0.997988 | 0.994666 |
| Islr2_E9Q9N7_S698s_1 | Islr2_E9Q9 | -0.47 | 0.11 | 0.24 | -0.3 | 0.16 | -0.53 | -0.04 | -0.88 | -0.85 | -0.23 | -0.34 | -0.28 | -0.08 | -0.29 | 0.1 | -0.35 | -0.285 | -0.0525 | -0.5325 | 0.856933 | 0.487411 | 0.262897 | 0.989223 | 0.997988 | 0.999507 |
| Islr2_E9Q9N7_S739s_1 | Islr2_E9Q9 | -0.54 | 0.52 | 0.15 | 0.5 | 0.26 | -0.09 | 0.78 | 0.09 | -0.28 | 0.28 | 0.17 | 0.45 | -0.28 | 0.37 | 0.04 | 0.36 | -0.2175 | -0.32 | -0.05 | 0.915954 | 0.751927 | 0.993782 | 0.995762 | 0.997988 | 0.999507 |
| Isyna1_Q9JHU9_S524s_1 | Isyna1_Q9J | 0.14 | 0.36 | -0.28 | -0.21 | 0.6 | 0.05 | 0.64 | -0.11 | -0.1 | 0.3 | 0.11 | -0.19 | 0.15 | 0.84 | 0.37 | 0.3 | 0.1025 | -0.0025 | -0.035 | 0.171066 | 0.309254 | 0.914367 | 0.989223 | 0.997988 | 0.994666 |
| Isyna1_Q9JHU9_T523t_1 | Isyna1_Q9J | -1.41 | -1.17 | -0.85 | -1.11 | -0.86 | -1.4 | -0.59 | -0.67 | -1.98 | -1.06 | -0.31 | -0.94 | -1.26 | -0.06 | -0.71 | -1.21 | 0.2925 | 0.0275 | 0.4125 | 0.348738 | 0.344161 | 0.866065 | 0.989223 | 0.997988 | 0.994666 |
| Itfg2_Q91WI7_S219s_1 | Itfg2_Q91V | 0.38 | 0.26 | -0.05 | -0.21 | -0.53 | -0.17 | -1.11 | -0.66 | -0.07 | 0.22 | 0.6 | -0.06 | -0.04 | -1.16 | 0.08 | 0.05 | 0.255 | 0.0625 | 0.325 | 0.321252 | 0.026275 | 0.772896 | 0.989223 | 0.997988 | 0.994666 |
| Itfg2_Q91WI7_T440t_1 | Itfg2_Q91V | 0.1 | 0.3 | 0.28 | 0.3 | -0.83 | -0.77 | -0.91 | -0.25 | 2.03 | -0.03 | 0.62 | 0.14 | 2.3 | 0.55 | 0.16 | 0.36 | -0.7125 | 0.0775 | -0.3625 | 0.227368 | 0.0025 | 0.337916 | 0.989223 | 0.997988 | 0.994666 |
| Itga3_Q62470_S1013s_1 | Itga3_Q624 | 0.72 | 0 | 0.66 | 0.65 | -0.55 | 0.84 | -0.42 | -0.18 | 2.51 | 0.36 | -0.03 | -0.18 | 2.12 | -0.62 | 1.07 | 0.58 | -0.935 | 0.445 | 0.5975 | 0.612927 | 0.126828 | 0.791249 | 0.989223 | 0.997988 | 0.995259 |
| Itga6_F6VSK8_S658s_1 | Itga6_F6VS | -0.82 | -0.65 | 1.02 | 1.48 | -0.31 | -0.72 | 1.1 | 1.47 | -1.36 | -1.17 | 1.54 | 1.68 | 0.29 | -0.96 | 1.97 | 2.43 | -0.585 | 0.1575 | 0.28 | 0.437857 | 0.854465 | 0.924683 | 0.996349 | 0.997988 | 0.994666 |
| Itgav_F6RWM8_S57s_1 | Itgav_F6RV | -0.23 | 0.94 | 2.39 | 2.57 | 0.2 | -0.31 | 1.01 | 2.09 | 0.23 | -0.02 | 4.47 | 2.64 | 0.53 | 0.45 | 1.73 | 1.92 | 0.1275 | -0.085 | 0.675 | 0.705463 | 0.37551 | 0.708964 | 0.989223 | 0.997988 | 0.996159 |
| Itm2c_Q91VK4_T39t_1 | Itm2c_Q91 | 0.14 | 0.18 | 0.34 | 0.46 | 0.48 | 0.01 | 0.63 | 0.53 | 0.07 | 0.1 | 0.21 | 0.46 | 0.04 | 0.04 | 0.12 | 0.47 | -0.67 | 0.4125 | -0.26 | 0.649804 | 0.569336 | 0.753979 | 0.989223 | 0.997988 | 0.995259 |
| Itpka_Q8R071_S185s_1 | Itpka_Q8R( | -3.1 | -2.04 | -3.08 | -2.36 | -1.99 | -2.38 | -1.74 | -1.85 | -3.13 | -2.6 | -3.9 | -3.35 | -2.64 | -2.55 | -5.94 | -3.34 | 0.1325 | -0.07 | -0.1125 | 0.211919 | 0.06225 | 0.141314 | 0.989223 | 0.997988 | 0.994666 |
| Itpka_Q8R071_S20s_1 | Itpka_Q8R( | 0.56 | -2.97 | -3.13 | -2.35 | -3.15 | 1.01 | -1.59 | -0.12 | -1.79 | -2.54 | 3.68 | -2.17 | -3.18 | 0.26 | -5.17 | -5.79 | 0.655 | -0.6 | -0.9725 | 0.295228 | 0.359452 | 0.403692 | 0.989223 | 0.997988 | 0.994666 |
| Itpkb_B2RXC2_S269s_1 | Itpkb_B2R) | 0.36 | 0.32 | 0.38 | 0.13 | 0.32 | 0.98 | 0 | 0.27 | 1.09 | 0.37 | 0.24 | 0.09 | 1.44 | 0.2 | 0.54 | 0.1 | 1.01 | 1.2675 | -1.4975 | 0.435556 | 0.718579 | 0.595394 | 0.995437 | 0.997988 | 0.994666 |
| Itpr1_P11881_S1573s_1 | Itpr1_P118 | -0.71 | -0.66 | -0.15 | -0.35 | 0.1 | -0.59 | -0.26 | -0.01 | -1.04 | -0.97 | -0.64 | -0.66 | -0.07 | -0.06 | -0.18 | -0.21 | 0.095 | 0.15 | 0.2725 | 0.199717 | 0.292258 | 0.168015 | 0.989223 | 0.997988 | 0.994666 |
| Itpr1_P11881_S1700s_1 | Itpr1_P118 | 0.88 | 0.89 | -0.02 | -0.18 | 1.25 | 0.83 | 0.14 | -0.19 | 0.85 | 0.74 | -0.2 | -0.38 | 0.73 | 1.55 | -0.09 | -0.06 | 0.2775 | -0.36 | 0.3375 | 0.763838 | 0.78123 | 0.738441 | 0.995762 | 0.997988 | 0.9971 |
| Itpr3_P70227_S934s_1 | Itpr3_P702 | 0.04 | 0.18 | 0.43 | -0.16 | 0.34 | -0.06 | 0.75 | 0.59 | -0.09 | 0.32 | 0.92 | 0.89 | -0.26 | -0.06 | -0.35 | 0.31 | 0.115 | -0.14 | 0.14 | 0.447083 | 0.297498 | 0.228451 | 0.989223 | 0.997988 | 0.994666 |
| Itsn1_E9Q0N0_S1135s_1 | Itsn1_E9Q( | 0.37 | 0.29 | 0.67 | 0.34 | 0.53 | 0.25 | 0.39 | 0.3 | 0.1 | 0.28 | 0.87 | 0.31 | 0.64 | 0.84 | 0.45 | 0.39 | 0.2825 | 0.3875 | -0.2125 | 0.518739 | 0.811281 | 0.914566 | 0.995762 | 0.997988 | 0.994666 |
| Itsn1_E9Q0N0_S203s_1 | Itsn1_E9Q( | 0.11 | -0.08 | 0.11 | 0.2 | 0.47 | -0.2 | 0.26 | 0.51 | 0.03 | 0 | 0.61 | 0.12 | -0.56 | 0.17 | -0.09 | 0.28 | -0.05 | -0.0275 | 0.1625 | 0.628122 | 0.471 | 0.662671 | 0.989223 | 0.997988 | 0.995259 |
| Itsn1_E9Q0N0_S315s_1 | Itsn1_E9Q( | 0.6 | 0.13 | -0.15 | 0.5 | 0.31 | 0.58 | 0.36 | 0.35 | -0.05 | 0.57 | 0.18 | 0.03 | -0.51 | 0.47 | -0.32 | 0.07 | 0.175 | 0.105 | -0.135 | 0.303801 | 0.597641 | 0.751271 | 0.989223 | 0.997988 | 0.994666 |
| Itsn1_E9Q0N0_S324s_1 | Itsn1_E9Q( | -1.46 | -1.88 | -0.97 | -1.61 | -0.16 | -1.21 | -1.6 | -1.61 | -3.38 | -3.11 | -2.76 | -1.56 | -3.2 | -2.48 | -1.87 | -1.54 | 0.13 | -0.0875 | -0.3425 | 0.080305 | 0.389836 | 0.018155 | 0.989223 | 0.997988 | 0.994666 |
| Itsn1_E9Q0N0_S335s_1 | Itsn1_E9Q( | -0.32 | 0.49 | 0.56 | 0.57 | 0.1 | -0.12 | 0.06 | 0.54 | 0.29 | 0.07 | 0.81 | 0.31 | -0.38 | 0.72 | 0.33 | 0.26 | 0.335 | -1.2225 | -0.7925 | 0.789377 | 0.53659 | 0.882227 | 0.989223 | 0.997988 | 0.999507 |
| Itsn1_E9Q0N0_S558s_1 | Itsn1_E9Q( | -0.56 | -1.02 | 0.67 | -0.72 | -0.92 | -0.75 | -0.68 | -0.66 | -0.81 | -1.24 | 0.07 | -1.13 | 0.93 | -1.04 | 0.13 | 0.57 | -0.18 | 0.045 | -0.0925 | 0.308959 | 0.363245 | 0.426 | 0.989223 | 0.997988 | 0.994666 |
| Itsn1_E9Q0N0_S623s_1 | Itsn1_E9Q( | 0.95 | 0.09 | 0.84 | 1.86 | -0.27 | 0.11 | -0.18 | 0.92 | 2.45 | 1.97 | 1.76 | 0.71 | 2.64 | 1.03 | 0.72 | 0.86 | -0.345 | -0.37 | 0.555 | 0.487143 | 0.090302 | 0.132539 | 0.989223 | 0.997988 | 0.994666 |
| Itsn1_E9Q0N0_S902s_1 | Itsn1_E9Q( | 0.72 | 0.04 | -0.04 | 0.11 | 0.51 | 0.49 | 0.85 | 0.78 | -0.13 | 0.13 | 0 | 0.29 | -1.12 | 0.27 | -0.32 | 0.09 | -0.79 | 0.7875 | 0.3775 | 0.220335 | 0.099374 | 0.607411 | 0.989223 | 0.997988 | 0.994666 |
| Itsn1_E9Q0N0_S976s_1 | Itsn1_E9Q( | 0.22 | 0.42 | 0.31 | 0.47 | 0.2 | 0.44 | -0.36 | 0.54 | -0.65 | -0.24 | 0 | 0.89 | -0.71 | 0.63 | 0.22 | 0.15 | 0.45 | -0.135 | -0.4775 | 0.399813 | 0.565833 | 0.320698 | 0.989223 | 0.997988 | 0.994666 |
| Itsn1_E9Q0N0_S984s_1 | Itsn1_E9Q( | -1.22 | -1.73 | -0.24 | 0.89 | -0.76 | -0.83 | 0.64 | 1.23 | -1.41 | -1.19 | 1.49 | 0.21 | -0.74 | -1.16 | -0.17 | 0.67 | -0.15 | -0.355 | -0.2825 | 0.723493 | 0.356824 | 0.6608 | 0.989223 | 0.997988 | 0.9971 |
| Itsn1_E9Q0N0_T1142t_1 | Itsn1_E9Q( | 0.35 | -0.01 | 0.58 | 0.86 | 0.36 | 0.27 | 0.71 | 0.53 | 0 | 0.35 | 0.42 | 0.6 | -0.18 | 0.39 | 0.21 | 0.24 | 0.645 | 0.35 | 0.225 | 0.34646 | 0.930567 | 0.71193 | 0.997844 | 0.997988 | 0.994666 |
| Itsn2_E9QNG1_S110s_1 | Itsn2_E9QI | 0.68 | 0.04 | -0.11 | 0 | 0.31 | 0.2 | 0.77 | 0.44 | -1.38 | -0.63 | 0.48 | -0.11 | -1.19 | 0.24 | -0.38 | -0.9 | 0.0225 | -0.1025 | -0.28 | 0.085332 | 0.307155 | 0.206076 | 0.989223 | 0.997988 | 0.994666 |
| Iws1_Q8C1D8_S261s_1 | Iws1_Q8C1 | -2.03 | -2.21 | -2.81 | -4.9 | -2.19 | -0.69 | -5.4 | -3.96 | -1 | -4.14 | -1.18 | -2.84 | -1.58 | -2.11 | -4.68 | -2.45 | 0.2775 | -0.5625 | -0.71 | 0.735688 | 0.945074 | 0.43694 | 0.999187 | 0.997988 | 0.9971 |
| Iws1_Q8C1D8_S295s_1 | Iws1_Q8C1 | -3.67 | -3.65 | -3.82 | -3.76 | -4.16 | -3.73 | -4.94 | -4.83 | -1.61 | -3.85 | -2.89 | -3.69 | -1.48 | -4.25 | -2.99 | -4.63 | -0.0725 | 0.6975 | 0.2825 | 0.551079 | 0.047422 | 0.165463 | 0.989223 | 0.997988 | 0.994666 |
| Iws1_Q8C1D8_S667s_1 | Iws1_Q8C1 | -1.56 | -3.57 | -2.33 | -2.54 | -3.56 | -2.14 | -4.09 | -2.56 | -1.88 | -3.02 | -2.51 | -2.08 | 0.34 | -2.64 | -3.47 | -3.84 | -0.69 | 0.715 | 0.3875 | 0.9142 | 0.301203 | 0.783632 | 0.989223 | 0.997988 | 0.999507 |
| Jag2_Q9QYE5_S1193s_1 | Jag2_Q9QY | -1.24 | -1.21 | -0.46 | -1.47 | -0.86 | -1.9 | -0.84 | -1.65 | -1.02 | -0.71 | -0.22 | -0.3 | -1.68 | 0.07 | -0.47 | -2.47 | -0.5875 | 0.1275 | 0.0975 | 0.940236 | 0.539727 | 0.120059 | 0.989223 | 0.997988 | 0.999507 |
| Jag2_Q9QYE5_S1199s_1 | Jag2_Q9QY | -0.36 | -0.51 | -0.58 | -0.86 | -0.76 | 0.13 | -0.74 | -0.91 | 1.32 | -0.54 | 0.1 | -0.16 | 1.28 | -1.05 | -0.18 | -0.93 | -0.2175 | 0.5325 | -0.0425 | 0.491666 | 0.979286 | 0.091047 | 0.999187 | 0.997988 | 0.994666 |
| Jag2_Q9QYE5_T1164t_1 | Jag2_Q9QY | 0.38 | 0.45 | 0.68 | 0.45 | 0.98 | 0.38 | 0.89 | 0.63 | -0.5 | 0.36 | 0.23 | 0.89 | -0.03 | -0.63 | 0.7 | 0.85 | 0.0075 | 0.7575 | 0.3575 | 0.475167 | 0.326547 | 0.453967 | 0.989223 | 0.997988 | 0.994666 |
| Jak2_Q62120_S523s_1 | Jak2_Q621 | 0.04 | 0.33 | 0.25 | 0.41 | 0.18 | -0.39 | 0.45 | 0.09 | 0.01 | -0.04 | -0.04 | 0.42 | 0.82 | -0.28 | 0.1 | 0.3 | 0.23 | -0.245 | -0.2675 | 0.940607 | 0.488824 | 0.469874 | 0.989223 | 0.997988 | 0.999507 |
| Jak2_Q62120_Y570y_1 | Jak2_Q621 | 0.95 | 0 | 0.83 | 0.7 | 0.3 | 0.78 | -0.23 | 1.65 | 0.4 | 0.74 | 2.09 | 0.37 | 0.36 | 0.38 | 1.18 | 0.68 | -0.175 | -0.17 | -0.0225 | 0.927147 | 0.990659 | 0.531413 | 0.999187 | 0.997988 | 0.999507 |
| Jakmip1_G3UYZ9_T285t_1 | Jakmip1_G | -2.1 | -1.41 | -2.1 | -1.21 | -1.26 | -2.05 | -1.54 | -2.05 | -0.59 | -1.28 | 0.2 | -2.08 | -0.69 | 0.29 | -2.14 | -1.77 | 0.005 | 0.28 | 0.03 | 0.272733 | 0.94994 | 0.154255 | 0.999187 | 0.997988 | 0.994666 |
| Jakmip2_D3YXK0_S259s_1 | Jakmip2_D | 1.69 | 0.54 | 0.77 | -0.51 | 0.21 | 0.98 | -0.5 | 0.31 | 1.85 | 1.4 | 0.97 | 0 | 1.92 | 0.07 | 0.1 | 0.27 | -0.02 | 0.7675 | 0.6275 | 0.955429 | 0.461864 | 0.44396 | 0.989223 | 0.997988 | 0.999507 |
| Jakmip2_D3YXK0_S260s_1 | Jakmip2_D | 1.36 | 0.43 | 0.97 | 1.34 | -0.37 | 0.85 | 0.21 | 0.61 | 1.57 | 1.24 | 1.21 | 0.6 | 1.85 | -1.43 | 1.01 | 0.61 | -0.3725 | 0.4325 | -0.0325 | 0.441066 | 0.068411 | 0.689516 | 0.989223 | 0.997988 | 0.994666 |
| Jakmip2_D3YXK0_S278s_1 | Jakmip2_D | -0.47 | -0.34 | 0.3 | 0.03 | -0.64 | -0.3 | -0.03 | 0.45 | 0.2 | -0.2 | 1.42 | 0.3 | 0.85 | -0.22 | 0.71 | 0.19 | -0.7 | 0.13 | -0.515 | 0.1611 | 0.974181 | 0.18175 | 0.999187 | 0.997988 | 0.994666 |
| Jakmip3_E9Q7K7_S384s_1 | Jakmip3_E! | 0.01 | 0.44 | 0.35 | 0.91 | 0.41 | -0.21 | 0.43 | 0.99 | -0.24 | 0.2 | 0.64 | 0.43 | 0.19 | 0.39 | 1.08 | 1.41 | -0.01 | 0.55 | 0.5025 | 0.361879 | 0.944162 | 0.577535 | 0.999187 | 0.997988 | 0.994666 |
| Jazf1_Q80ZQ5_T98t_1 | Jazf1_Q80Z | -1.18 | -1.16 | -1.27 | -2.44 | -2.2 | -1.01 | -2.82 | -1.99 | -0.3 | -1.32 | -1.91 | -2.5 | 0.62 | -0.94 | -0.98 | -1.34 | -0.0225 | -0.17 | 0.34 | 0.116713 | 0.29366 | 0.992392 | 0.989223 | 0.997988 | 0.994666 |
| Jcad_Q5DTX6_S351s_1 | Jcad_Q5DT | 0.01 | 0.14 | -0.53 | -0.73 | 0.05 | -0.49 | -0.32 | -1.33 | -0.43 | 0.08 | -0.43 | -0.3 | -0.26 | 0.75 | -0.94 | -0.6 | -0.4925 | 0.005 | 0.8525 | 0.97144 | 0.499236 | 0.979206 | 0.989223 | 0.997988 | 0.999507 |
| Jcad_Q5DTX6_S405s_1 | Jcad_Q5DT | 0.34 | 0.47 | 0.49 | 0.48 | 0.81 | 0.59 | 0.57 | 0.34 | 0.35 | 0.69 | 0.24 | 0.06 | -0.29 | 0.29 | 0.16 | 0.34 | -0.245 | 0.0075 | 0.015 | 0.228626 | 0.528789 | 0.637138 | 0.989223 | 0.997988 | 0.994666 |
| Jcad_Q5DTX6_S498s_1 | Jcad_Q5DT | -0.93 | -0.64 | -0.21 | -0.35 | -0.28 | -0.63 | -0.31 | -0.28 | -1.22 | -0.55 | -0.03 | -0.1 | -0.46 | -0.04 | -0.69 | -0.56 | 0.1325 | -0.11 | -0.32 | 0.740224 | 0.52243 | 0.864827 | 0.989223 | 0.997988 | 0.9971 |
| Jcad_Q5DTX6_S543s_1 | Jcad_Q5DT | -0.36 | -0.07 | 0.33 | -0.24 | -1.4 | 0.92 | -1.04 | -1.08 | 1.9 | 0.08 | -0.17 | 0.38 | 1.73 | -1.11 | -0.51 | -0.26 | 0.1575 | 0.0575 | 0.095 | 0.934827 | 0.27846 | 0.19736 | 0.989223 | 0.997988 | 0.999507 |
| Jcad_Q5DTX6_S655s_1 | Jcad_Q5DT | 0.48 | -0.74 | 0.5 | 0.72 | -0.45 | 0.46 | -0.43 | 0.22 | 0.33 | 0.43 | 0.97 | 0.73 | -0.35 | 0.27 | 0.29 | -0.11 | -0.565 | 0.6325 | 0.0475 | 0.575305 | 0.464574 | 0.324514 | 0.989223 | 0.997988 | 0.995259 |
| Jcad_Q5DTX6_S888s_1 | Jcad_Q5DT | 0.89 | 0.24 | 0.41 | 0.84 | 1.03 | 0.45 | 1.09 | 0.77 | 0.34 | 0.82 | 0.44 | -0.01 | -0.05 | 0.59 | 0.39 | 0.85 | -0.29 | 0.375 | -0.215 | 0.625651 | 0.372896 | 0.491698 | 0.989223 | 0.997988 | 0.995259 |
| Jcad_Q5DTX6_S941s_1 | Jcad_Q5DT | -1.43 | -1.98 | -1.11 | -2.31 | -1.56 | -1.86 | -1.68 | -2.38 | -1.38 | -2.25 | -1.47 | -0.56 | -1.97 | -1.2 | -2.46 | -1.08 | 0.24 | -0.1975 | -0.15 | 0.943218 | 0.628005 | 0.499185 | 0.989223 | 0.997988 | 0.999507 |
| Jcad_Q5DTX6_T1001t_1 | Jcad_Q5DT | -0.21 | 0.49 | -0.42 | -0.44 | 0.1 | 0.29 | -0.17 | -0.74 | -0.97 | 0.07 | -0.59 | -0.27 | -0.52 | -0.33 | -0.35 | -0.02 | -0.1625 | 0.2925 | 0.03 | 0.598498 | 0.963166 | 0.387279 | 0.999187 | 0.997988 | 0.995259 |
| Jmjd1c_G3UZM1_S1970s_1 | Jmjd1c_G3 | -0.81 | -0.12 | -0.26 | -0.37 | -0.6 | -0.22 | 0.29 | -0.2 | -0.17 | -0.11 | 0.92 | -0.3 | -0.06 | -0.13 | -0.06 | 0.45 | 0.015 | -0.295 | -0.16 | 0.139879 | 0.457038 | 0.183748 | 0.989223 | 0.997988 | 0.994666 |
| Jmjd1c_G3UZM1_S617s_1 | Jmjd1c_G3 | -1.2 | -1.72 | -0.99 | -1.04 | -1.31 | -1.84 | -1.49 | -0.93 | -1.08 | -0.88 | -0.31 | -1.58 | -0.35 | -0.11 | -1.42 | -0.72 | 0.2075 | 0.475 | 0.44 | 0.124938 | 0.589732 | 0.418879 | 0.989223 | 0.997988 | 0.994666 |
| Jmy_Q9QXM1_S108s_1 | Jmy_Q9QX | -1.49 | -1.4 | -1.7 | -0.8 | -0.4 | -1.55 | -1.15 | -0.87 | -1.32 | -1.41 | -1.07 | -0.86 | -0.95 | -0.84 | -1.27 | -1.07 | -0.155 | 0.275 | 0.5875 | 0.287041 | 0.287333 | 0.520582 | 0.989223 | 0.997988 | 0.994666 |
| Jmy_Q9QXM1_S75s_1 | Jmy_Q9QX | -1.7 | -3.24 | -1.88 | -3.53 | -1.82 | -1.55 | -2.46 | -2.27 | 0.27 | -1.88 | -3.04 | -2.01 | 0.47 | -2.7 | -1.15 | -1.82 | 0.355 | 0.1825 | 0.315 | 0.10213 | 0.251405 | 0.238634 | 0.989223 | 0.997988 | 0.994666 |
| Jph1_Q9ET80_S157s_1 | Jph1_Q9ET | 1.91 | 1.18 | 0.97 | 1.88 | 1.68 | 2 | 1.45 | 1.95 | 1.04 | 1.92 | 1.49 | 1.72 | 0.39 | 2 | 0.78 | 0.8 | 0.5625 | 0.9225 | 1.2875 | 0.260261 | 0.350983 | 0.861777 | 0.989223 | 0.997988 | 0.994666 |
| Jph1_Q9ET80_S186s_1 | Jph1_Q9ET | -0.04 | 1.45 | -0.97 | -0.09 | -0.19 | -0.65 | 1.16 | 2.05 | -0.17 | 0.3 | 0.32 | 1.28 | -0.17 | -0.38 | 0.39 | 1.58 | 0.285 | 0.0575 | -0.4925 | 0.661282 | 0.478407 | 0.531073 | 0.989223 | 0.997988 | 0.995671 |
| Jph1_Q9ET80_S216s_1 | Jph1_Q9ET | -0.7 | -0.6 | -0.67 | -1.2 | 0.38 | -0.16 | 0.27 | -0.5 | -1.4 | -1.35 | -0.73 | -0.69 | -0.92 | 0.29 | -0.9 | -0.49 | 0.505 | 0.345 | 0.2675 | 0.415259 | 0.017817 | 0.390482 | 0.989223 | 0.997988 | 0.994666 |
| Jph1_Q9ET80_S448s_1 | Jph1_Q9ET | -0.29 | 1.14 | 1.08 | 0.02 | 0.26 | -0.25 | -0.24 | -0.65 | 0.53 | 0.05 | 0.63 | 0.18 | 0.71 | 0.74 | -0.39 | 0.47 | 0.79 | -0.25 | 0.2875 | 0.812104 | 0.099244 | 0.721043 | 0.989223 | 0.997988 | 0.999507 |
| Jph1_Q9ET80_S490s_1 | Jph1_Q9ET | 0.1 | -0.96 | 0.43 | -0.13 | -0.73 | 0.74 | -0.59 | -0.05 | 2.05 | -0.48 | -1.05 | 0.01 | 1.97 | -1.41 | 0.13 | -0.46 | -0.7075 | -0.14 | -0.105 | 0.775296 | 0.966944 | 0.684337 | 0.999187 | 0.997988 | 0.99798 |
| Jph3_Q9ET77_S166s_1 | Jph3_Q9ET | 0.41 | -0.14 | 0.67 | 0.27 | 0.74 | 0.38 | 0.64 | 0.73 | -0.59 | -0.4 | 0.3 | 0.25 | -0.83 | 0.28 | 0.31 | 0.04 | -0.0175 | 0.2725 | 0.1975 | 0.323695 | 0.216489 | 0.210512 | 0.989223 | 0.997988 | 0.994666 |
| Jph3_Q9ET77_S188s_1 | Jph3_Q9ET | 5.03 | 1.2 | 0.88 | 1.34 | 0.75 | 5.53 | 3.43 | 1.91 | 1.78 | 3.21 | 2.57 | 0.68 | 0.1 | 0.55 | 2.65 | 2.63 | 0.32 | -0.4125 | -0.3525 | 0.54553 | 0.521374 | 0.957587 | 0.989223 | 0.997988 | 0.994666 |
| Jph3_Q9ET77_S420s_1 | Jph3_Q9ET | 0.22 | 0.13 | 0.21 | 0.16 | 0.73 | -0.37 | 0.45 | 0.28 | 0.74 | -0.05 | -0.12 | 0.21 | 0.06 | 0.96 | 0.12 | 0.32 | 0.7925 | -0.0525 | -0.63 | 0.517745 | 0.736274 | 0.954187 | 0.995762 | 0.997988 | 0.994666 |
| Jph3_Q9ET77_S506s_1 | Jph3_Q9ET | 0.05 | -0.75 | 0.13 | -0.02 | -0.42 | -0.38 | -0.93 | -0.32 | -0.94 | -0.56 | -0.11 | -0.32 | 0.21 | -0.14 | -0.06 | 0.17 | 0.0925 | 0.015 | 0.185 | 0.511693 | 0.215669 | 0.290073 | 0.989223 | 0.997988 | 0.994666 |
| Jph3_Q9ET77_S578s_1 | Jph3_Q9ET | 0.2 | 1.23 | -0.29 | 0.12 | 0.45 | 1.33 | 0.03 | 0.19 | 0.75 | 0.73 | -0.92 | 0.61 | 0.76 | -0.19 | 0.57 | -0.05 | -0.365 | -0.335 | 0.1925 | 0.915952 | 0.656374 | 0.963337 | 0.990434 | 0.997988 | 0.999507 |
| Jph3_Q9ET77_T162t_1 | Jph3_Q9ET | 1.08 | 0.18 | 0.72 | 0.58 | 0.94 | 0.53 | 0.56 | 0.62 | 0.87 | 0.18 | 0.81 | 0.33 | -0.13 | 1.1 | 0.72 | 0.55 | 0.185 | -0.0225 | -0.0425 | 0.818394 | 0.930638 | 0.754699 | 0.997844 | 0.997988 | 0.999507 |
| Jph4_Q80WT0_T569t_1 | Jph4_Q80V | 1.26 | 0.99 | 0.08 | 0.88 | 0.12 | 1.34 | 1.36 | 1.79 | 0.51 | 0.3 | 1.1 | 0.44 | -1.49 | 0.59 | -0.15 | 1 | 0.0225 | -0.0925 | -0.08 | 0.166995 | 0.411538 | 0.522366 | 0.989223 | 0.997988 | 0.994666 |
| Jun_P05627_S73s_1 | Jun_P0562 | -2.66 | -3.49 | -2.46 | -3.08 | -2.56 | -2.83 | -3.11 | -2.56 | -2.79 | -2.09 | -0.8 | -1.94 | -1.58 | -1.55 | -1.58 | -2 | 0.35 | -0.215 | -0.815 | 0.002519 | 0.593248 | 0.047451 | 0.989223 | 0.997988 | 0.994666 |
| Kalrn_F6QYT9_S1586s_1 | Kalrn_F6Q\ | -0.05 | -0.38 | 1.59 | 1.16 | -0.73 | -1.05 | 2.01 | 2.09 | -0.35 | 0.55 | 2.22 | 0.76 | 2.06 | 0.39 | 1.71 | 1.67 | 0.1575 | 1.0175 | 1.245 | 0.137457 | 1 | 0.740075 | 1 | 0.997988 | 0.994666 |
| Kalrn_F6QYT9_S1785s_1 | Kalrn_F6Q\ | -0.26 | 0.26 | 0.61 | -0.26 | -1.09 | 0.03 | -1.13 | -0.31 | 0.29 | 0.37 | -4.18 | -0.18 | 0.35 | -1.04 | 0.79 | 0.33 | 3.14E-16 | 0.215 | 0.8775 | 0.963804 | 0.071154 | 0.317305 | 0.989223 | 0.997988 | 0.999507 |
| Kalrn_F6QYT9_S465s_1 | Kalrn_F6Q\ | 0.68 | 1 | 1 | 1.05 | 0.95 | 0.81 | 0.96 | 0.7 | 0.66 | 0.75 | -0.01 | 1.17 | 0.44 | 0.94 | 0.24 | 0.8 | -0.7125 | -1.0125 | 0.02 | 0.243907 | 0.711808 | 0.344516 | 0.995437 | 0.997988 | 0.994666 |
| Kank1_E9Q238_S887s_1 | Kank1_E9C | -0.35 | -0.29 | -0.61 | -0.32 | -1.21 | -0.44 | -0.72 | -1.1 | -0.07 | -0.33 | -0.45 | -1.05 | 0.25 | -1.08 | -0.65 | -0.02 | -0.0775 | -0.29 | -0.3275 | 0.959336 | 0.081671 | 0.764598 | 0.989223 | 0.997988 | 0.999507 |
| Kank3_Q9Z1P7_S166s_1 | Kank3_Q92 | -0.56 | -0.1 | -0.52 | 0.58 | 0.74 | -0.16 | 0.95 | 0.29 | 0.28 | 0.27 | 0.43 | 0.01 | -0.19 | 0.93 | -0.55 | 0.43 | -0.475 | -0.0825 | 0.0175 | 0.474058 | 0.118383 | 0.222883 | 0.989223 | 0.997988 | 0.994666 |
| Kank3_Q9Z1P7_S167s_1 | Kank3_Q92 | -0.56 | -0.1 | -0.52 | 0.58 | 0.74 | -0.16 | 0.95 | 0.29 | 0.28 | 0.27 | 0.43 | 0.01 | -0.19 | 0.93 | -0.55 | 0.43 | 0.605 | 0.3975 | 0.305 | 0.474058 | 0.118383 | 0.222883 | 0.989223 | 0.997988 | 0.994666 |
| Kank3_Q9Z1P7_S290s_1 | Kank3_Q92 | 1.14 | 0.79 | 0.98 | 1.25 | 0.58 | 0.93 | 0.7 | 1.49 | 1.11 | 1.1 | 0.92 | 1.19 | 0.49 | 0.8 | 0.86 | 1.08 | 0.605 | 0.3975 | 0.305 | 0.379824 | 0.669813 | 0.857536 | 0.992876 | 0.997988 | 0.994666 |
| Kank3_Q9Z1P7_T292t_1 | Kank3_Q92 | -3.06 | -2.2 | -2.6 | -4.53 | -4.84 | -1.7 | -4.03 | -3.41 | -0.03 | -2.84 | -2.05 | -1.34 | 1.13 | -2.16 | -1.14 | -3.49 | -0.115 | 0.04 | -0.2325 | 0.105138 | 0.592037 | 0.053779 | 0.989223 | 0.997988 | 0.994666 |
| Kank4_Q6P9J5_S74s_1 | Kank4_Q6F | 0.24 | 0.01 | -0.03 | -0.31 | 0.56 | -0.2 | 0.11 | 0.14 | 0.01 | -0.14 | -0.06 | 0.14 | 0.15 | 0.55 | 0.01 | 0.43 | -0.3975 | 1.5325 | 1.6825 | 0.260662 | 0.489067 | 0.96494 | 0.989223 | 0.997988 | 0.994666 |
| Kat7_Q5SVQ0_S124s_1 | Kat7_Q5SV | 0.95 | 1.28 | 1.12 | 0.64 | -0.49 | 1.59 | 0.28 | -0.54 | 1.82 | 1.08 | -0.68 | 0.48 | 1.54 | -0.66 | 1.1 | 1.33 | 0.175 | 0.01 | 0.3075 | 0.731901 | 0.123461 | 0.532991 | 0.989223 | 0.997988 | 0.9971 |
| Kat7_Q5SVQ0_T88t_1 | Kat7_Q5SV | -1.04 | -0.33 | -1.7 | -1.92 | -0.18 | -0.85 | -0.99 | -1.69 | -1.31 | -1.52 | -0.65 | -1.33 | -0.2 | -1.17 | -1 | -1.12 | -0.7875 | -0.3225 | -0.17 | 0.384434 | 0.477149 | 0.910927 | 0.989223 | 0.997988 | 0.994666 |
| Kat8_Q9D1P2_S42s_1 | Kat8_Q9D1 | 0.61 | 0.88 | 0.27 | 0.53 | 0.14 | 0.48 | 0.13 | 0.58 | 0.94 | 0.67 | 0.3 | 0 | 1.34 | 0.43 | 0.5 | 0.95 | 0.32 | 0.045 | 0.375 | 0.452441 | 0.326822 | 0.742077 | 0.989223 | 0.997988 | 0.994666 |
| Katnb1_Q8BG40_S360s_1 | Katnb1_Q8 | -0.37 | -0.4 | 0.65 | -0.51 | -1.8 | 0.15 | -1.05 | -0.71 | 2.13 | -0.78 | 1.17 | -0.04 | 1.5 | -1.2 | 0.76 | -0.65 | -0.24 | -0.095 | 0.2325 | 0.674716 | 0.150021 | 0.240973 | 0.989223 | 0.997988 | 0.996159 |
| Kazn_B7ZCA7_S210s_1 | Kazn_B7ZC | 0.25 | 0.66 | -1.35 | -1.57 | 0.73 | 0.32 | -0.52 | -0.76 | -0.74 | 0.39 | -0.46 | -0.83 | -1.03 | 0.73 | -1.31 | -1.32 | -0.695 | 0.7775 | 0.26 | 0.732293 | 0.459174 | 0.872474 | 0.989223 | 0.997988 | 0.9971 |
| Kazn_B7ZCA7_S231s_1 | Kazn_B7ZC | -0.68 | -1.71 | -1.74 | -2.96 | -1.6 | -1.51 | -1.58 | -1.57 | -1.9 | -0.94 | 0.5 | -1.93 | -0.39 | -0.49 | -1.64 | -1.64 | 0.445 | 0.0925 | -0.23 | 0.196866 | 0.637932 | 0.306132 | 0.989223 | 0.997988 | 0.994666 |
| Kazn_Q69ZS8_S8s_1 | Kazn_Q69Z | 0.07 | 0.1 | 0.3 | 0.73 | -0.82 | -0.77 | -0.43 | -0.03 | 0.11 | 0.29 | 1.25 | 0.15 | 0.18 | -0.46 | -0.06 | 0.1 | 0.2075 | 0.705 | 0.7325 | 0.223828 | 0.014437 | 0.653631 | 0.989223 | 0.997988 | 0.994666 |
| Kbtbd11_Q8BNW9_S107s_1 | Kbtbd11_C | -2.2 | -2.54 | 0.2 | -0.01 | -1.94 | -1.39 | 0.42 | -0.67 | -1.37 | -2.43 | 0.89 | 1.03 | -1.03 | -1.17 | 0.77 | -0.63 | -0.8125 | 0.15 | -0.36 | 0.414332 | 0.753208 | 0.503211 | 0.995762 | 0.997988 | 0.994666 |
| Kbtbd11_Q8BNW9_S316s_1 | Kbtbd11_C | 0.64 | 0 | -0.01 | 0.57 | -1.09 | 0.69 | -1.02 | -0.3 | 0.97 | -0.15 | -1.45 | -0.32 | 1.51 | -1.27 | -0.04 | 0.66 | 0.2425 | 0.6675 | 0.6225 | 0.880516 | 0.111933 | 0.294261 | 0.989223 | 0.997988 | 0.999507 |
| Kbtbd11_Q8BNW9_S320s_1 | Kbtbd11_C | 0.31 | 0.13 | 0.78 | 0.57 | -1.48 | 0.92 | -1.15 | -0.07 | 1.46 | 0.13 | -1.11 | 0.11 | 1.88 | -1.91 | -0.35 | 0.15 | -0.73 | -0.5375 | -0.085 | 0.482287 | 0.108352 | 0.560269 | 0.989223 | 0.997988 | 0.994666 |
| Kbtbd11_Q8BNW9_S62s_1 | Kbtbd11_C | 0.7 | -0.08 | -0.11 | -0.27 | 0.01 | 0.98 | 0.19 | 1.12 | 1.4 | 1.17 | -2.09 | 0.22 | 0.7 | 0.95 | 1.01 | -0.21 | -0.8925 | -0.3 | -0.505 | 0.163134 | 0.16867 | 0.876713 | 0.989223 | 0.997988 | 0.994666 |
| Kbtbd11_Q8BNW9_S70s_1 | Kbtbd11_C | 0.14 | 0.33 | 0.08 | 0.33 | -0.13 | -0.5 | 0.02 | -0.01 | 0.09 | 0.13 | 0.15 | 0.16 | 0.49 | 0.06 | 0.01 | -0.48 | 0.515 | 0.115 | 0.5525 | 0.486576 | 0.115594 | 0.674815 | 0.989223 | 0.997988 | 0.994666 |
| Kbtbd11_Q8BNW9_S70s_2 | Kbtbd11_C | 0.04 | 0.91 | 1.5 | 0.8 | 0.69 | 0.05 | 0.36 | 0.41 | 0.48 | 0.79 | 1.46 | 0.94 | 0.13 | 0.82 | 0.78 | 0.86 | -0.375 | -0.0875 | -0.2 | 0.654941 | 0.215034 | 0.777263 | 0.989223 | 0.997988 | 0.995259 |
| Kbtbd11_Q8BNW9_S73s_2 | Kbtbd11_C | 0.21 | 1.21 | 2.07 | 0.62 | 0.71 | 0.02 | 0.63 | 0.75 | 0.61 | 0.76 | 0.7 | 1.27 | 0.47 | 0.9 | 0.61 | 1.31 | -0.435 | 0.105 | -0.165 | 0.640451 | 0.248668 | 0.649178 | 0.989223 | 0.997988 | 0.995259 |
| Kcna1_P16388_S13s_1 | Kcna1_P16 | -0.51 | -0.07 | -0.69 | -1.01 | -0.06 | -0.38 | -0.59 | -0.56 | -0.24 | -0.5 | -1.53 | -1.57 | -0.06 | -0.69 | -1.53 | -0.63 | -0.5 | -0.1925 | -0.205 | 0.677532 | 0.530116 | 0.337567 | 0.989223 | 0.997988 | 0.996159 |
| Kcna1_P16388_S23s_1 | Kcna1_P16 | 0.01 | 0.5 | 0.51 | 0.02 | -0.1 | 0.58 | -0.09 | 0.33 | 0.35 | 0.5 | 0.31 | 0.13 | 0.81 | 0.05 | -0.24 | -0.36 | 0.1725 | -0.39 | -0.1575 | 0.5652 | 0.762781 | 0.797009 | 0.995762 | 0.997988 | 0.995259 |
| Kcna1_P16388_S5s_1 | Kcna1_P16 | -0.08 | 0.48 | -0.25 | 0.67 | -0.62 | -0.29 | 0.24 | 0.31 | -0.13 | 0 | 0.72 | 1.4 | 0.6 | 0.36 | 0.59 | -0.09 | -0.08 | 0.0625 | -0.195 | 0.619421 | 0.374549 | 0.482847 | 0.989223 | 0.997988 | 0.995259 |
| Kcna2_P63141_S434s_1 | Kcna2_P63 | 0.78 | 1.19 | 1.06 | 0.62 | 0.96 | 0.79 | 1.22 | 1.19 | 0.24 | 0.63 | 1.03 | 0.95 | 0.15 | 0.77 | 0.38 | 1.42 | -0.295 | 0.2925 | 0.16 | 0.501057 | 0.590267 | 0.475018 | 0.989223 | 0.997988 | 0.994666 |
| Kcna2_P63141_S440s_1 | Kcna2_P63 | -1.41 | 0.29 | -0.8 | -0.34 | 0.92 | -0.72 | -0.48 | -1.22 | -1.1 | -0.61 | -0.52 | -0.53 | 0.05 | -0.91 | -0.37 | -1.46 | 0.1275 | -0.2 | -0.2325 | 0.81846 | 0.720276 | 0.747213 | 0.995437 | 0.997988 | 0.999507 |
| Kcna2_P63141_S441s_1 | Kcna2_P63 | -0.71 | 0.07 | 0.2 | -0.43 | 0.28 | -0.69 | 0.21 | -0.26 | -0.47 | -0.7 | -0.14 | -0.32 | 0.07 | 0.27 | 0.02 | -0.22 | 0.19 | -0.125 | -0.1075 | 0.406471 | 0.751988 | 0.516544 | 0.995762 | 0.997988 | 0.994666 |
| Kcna2_P63141_S449s_1 | Kcna2_P63 | 0.53 | 0.61 | 1.58 | 1.28 | -0.46 | 0.92 | 0.79 | 0.97 | 3.29 | 1.31 | -1.56 | 1.03 | 3.03 | -0.79 | 2.66 | 0.75 | 0.1025 | -0.19 | 0.2525 | 0.615077 | 0.290603 | 0.98466 | 0.989223 | 0.997988 | 0.995259 |
| Kcna4_Q61423_S79s_1 | Kcna4_Q61 | -0.4 | 0.61 | -1.1 | 0.99 | -0.34 | 0.68 | -0.24 | 0.35 | 1.64 | 0.58 | -1.35 | -0.45 | 1.69 | -0.31 | 0.9 | 0.5 | -0.445 | 0.0175 | 0.4125 | 0.265526 | 0.858583 | 0.911682 | 0.996349 | 0.997988 | 0.994666 |
| Kcna4_Q61423_S82s_1 | Kcna4_Q61 | 0.41 | 0.3 | -0.41 | 0.83 | -1.11 | 1.26 | -0.38 | 0.29 | 1.7 | 0.62 | -0.35 | -0.27 | 1.76 | -0.46 | 0.45 | 0.63 | 0.0875 | 0.08 | 0.67 | 0.532992 | 0.606617 | 0.780117 | 0.989223 | 0.997988 | 0.994666 |
| Kcna5_Q61762_S29s_1 | Kcna5_Q61 | 1.05 | -0.93 | -1.31 | 0.08 | -1.74 | 0.94 | -0.38 | 0.19 | -0.09 | 0.96 | 1.27 | -0.78 | -0.34 | -0.11 | -0.08 | -0.66 | -0.2675 | 0.1425 | 0.3125 | 0.968954 | 0.965052 | 0.351769 | 0.999187 | 0.997988 | 0.999507 |
| Kcna5_Q61762_S535s_1 | Kcna5_Q61 | 1.15 | -0.38 | 0.24 | 1.19 | -1.33 | 0.68 | -0.54 | 0.28 | 1.77 | 1.12 | 0.9 | -0.3 | 1.71 | -1.87 | 0.98 | 0.36 | 0.03 | 0.6175 | -0.02 | 0.738923 | 0.167359 | 0.550925 | 0.989223 | 0.997988 | 0.9971 |
| Kcna6_Q61923_S511s_1 | Kcna6_Q61 | -1.53 | -2.21 | -2.29 | -4.08 | -1.38 | -1.5 | -1.31 | -2.1 | -1.65 | -1.9 | -0.33 | -1.71 | -0.22 | -0.64 | -1.06 | -3.02 | -0.7775 | 0.3225 | -0.255 | 0.103904 | 0.093031 | 0.085239 | 0.989223 | 0.997988 | 0.994666 |
| Kcna6_Q61923_S6s_1 | Kcna6_Q61 | 0.35 | 0.28 | -0.28 | 0.06 | 0.74 | 0.21 | 0.49 | 0 | -0.09 | -0.11 | -0.45 | 0.09 | -0.41 | 0.59 | -0.09 | 0.07 | 0.955 | 1.13 | 1.2925 | 0.839168 | 0.339016 | 0.350461 | 0.989223 | 0.997988 | 0.999507 |
| Kcnab2_E0CXI4_S111s_1 | Kcnab2_E0 | 0.94 | 1.12 | 1.15 | 0.16 | 1.35 | 0.71 | 0.56 | 0.12 | 1.42 | 1.19 | 1 | 0.99 | 1.12 | 1.42 | 0.32 | 0.72 | 0.2575 | -0.2425 | -0.0625 | 0.883627 | 0.652103 | 0.3125 | 0.990358 | 0.997988 | 0.999507 |
| Kcnab2_E0CXI4_S112s_1 | Kcnab2_E0 | 0.94 | 1.12 | 1.15 | 0.16 | 1.35 | 0.71 | 0.56 | 0.12 | 1.42 | 1.19 | 1 | 0.99 | 1.12 | 1.42 | 0.32 | 0.72 | -0.1575 | 0.3075 | 0.0525 | 0.883627 | 0.652103 | 0.3125 | 0.990358 | 0.997988 | 0.999507 |

| | | | | | | | | | | | | | | | | | | | | | | | | | | |
|---|---|---|---|---|---|---|---|---|---|---|---|---|---|---|---|---|---|---|---|---|---|---|---|---|---|---|
| Kcnab2_E0CXI4_S20s_1 | Kcnab2_E0 | -0.39 | -0.91 | -1.05 | -0.78 | -0.65 | -0.64 | -0.65 | -0.74 | -0.85 | -0.61 | -0.38 | -0.76 | -0.86 | -0.24 | -1.24 | -0.92 | -0.1575 | 0.3075 | 0.0525 | 0.915941 | 0.619308 | 0.59896 | 0.989223 | 0.997988 | 0.999507 |
| Kcnab2_E0CXI4_S9s_1 | Kcnab2_E0 | -0.83 | -0.85 | -0.92 | -1.2 | -0.36 | -1.44 | -1 | -1.14 | -1.15 | -0.74 | -0.36 | -1.09 | -0.28 | -0.28 | -1.04 | -1.22 | 0.1125 | 0.1325 | -0.0325 | 0.442602 | 0.900277 | 0.664898 | 0.996493 | 0.997988 | 0.994666 |
| Kcnab2_E0CXI4_Y25y_1 | Kcnab2_E0 | 2.24 | -1.35 | -0.05 | 1.08 | 0.43 | 1.73 | 0.11 | 1.19 | 0 | -0.81 | 0.64 | 0.12 | -2.4 | 0.42 | -0.7 | -0.13 | -0.035 | 0.115 | 0.245 | 0.191981 | 0.608768 | 0.511229 | 0.989223 | 0.997988 | 0.994666 |
| Kcnb1_Q03717_S15s_1 | Kcnb1_Q03 | 0.88 | 1.06 | 0.39 | 0.31 | 0.94 | 0.58 | 0.55 | 0.35 | 0.54 | 0.78 | -0.38 | 0.59 | 0.23 | 0.67 | 0.39 | 0.72 | 0.385 | -0.4925 | -1.1825 | 0.587546 | 0.836115 | 0.420795 | 0.995762 | 0.997988 | 0.995259 |
| Kcnb1_Q03717_S444s_1 | Kcnb1_Q03 | 0.61 | 1.49 | 2.38 | 1.99 | 1.1 | 0.67 | 1.92 | 1.46 | 1 | 1.25 | 3 | 2.04 | 0.72 | 1.63 | 1.49 | 2.12 | -0.055 | -0.2775 | -0.1575 | 0.783255 | 0.457576 | 0.709187 | 0.989223 | 0.997988 | 0.998491 |
| Kcnb1_Q03717_S484s_1 | Kcnb1_Q03 | 1.92 | 1.08 | 2.29 | 1.66 | 0.99 | 1.69 | 1.51 | 1.39 | 0.87 | 1.91 | 1.26 | 2.02 | 0.5 | 0.02 | 1.92 | 1.77 | -0.33 | 0.205 | -0.1275 | 0.195512 | 0.289102 | 0.55975 | 0.989223 | 0.997988 | 0.994666 |
| Kcnb1_Q03717_S511s_1 | Kcnb1_Q03 | -2.41 | -3.23 | -5.15 | -2.86 | -3.42 | -2.62 | -1.66 | -3.25 | 4.29 | -3.41 | -1.26 | -1.7 | 4.5 | -3.24 | 3.2 | -3.09 | -0.3425 | -0.2225 | -0.685 | 0.063376 | 0.30748 | 0.088101 | 0.989223 | 0.997988 | 0.994666 |
| Kcnb1_Q03717_S541s_1 | Kcnb1_Q03 | -0.67 | -1.21 | -2.09 | -2.25 | -1.07 | -1.07 | -1.1 | -1.14 | -1.75 | -1.64 | 0.12 | -1.21 | -2.01 | -0.6 | -2.21 | -2.14 | 0.675 | 2.8925 | 3.755 | 0.71448 | 0.231478 | 0.420871 | 0.989223 | 0.997988 | 0.996966 |
| Kcnb1_Q03717_S633s_1 | Kcnb1_Q03 | 2.46 | -1.71 | -0.92 | 0.95 | 0.14 | 2.59 | -0.1 | 1.28 | 1.26 | -0.43 | -0.09 | 0.54 | -2.01 | -0.65 | -1.51 | -0.14 | 0.46 | 0.435 | -0.185 | 0.180267 | 0.428902 | 0.888961 | 0.989223 | 0.997988 | 0.994666 |
| Kcnb1_Q03717_S655s_1 | Kcnb1_Q03 | 1.42 | -0.81 | 0.15 | 0.9 | 0.9 | 1.11 | 0.78 | 0.77 | 0.06 | 0.1 | -0.05 | 0.41 | -3.19 | 0.19 | -0.48 | -0.19 | 0.7825 | 0.125 | -1.2725 | 0.122362 | 0.313037 | 0.549068 | 0.989223 | 0.997988 | 0.994666 |
| Kcnb2_A6H8H5_S481s_2 | Kcnb2_A6H | 0.63 | 0.25 | 0.28 | 0.73 | 0.39 | 0.39 | 0.22 | 0.55 | 0.42 | 0.68 | 1.01 | 0.75 | -0.73 | 0.68 | 0.34 | 0.47 | 0.475 | -0.285 | -1.3325 | 0.441876 | 0.703807 | 0.341704 | 0.995437 | 0.997988 | 0.994666 |
| Kcnb2_A6H8H5_S488s_1 | Kcnb2_A6H | 0.83 | 0.53 | 1.03 | 0.96 | 0.81 | 0.33 | 0.61 | 0.47 | 1.29 | 0.61 | 1.99 | 0.97 | 1.17 | 2.25 | 0.99 | 1.23 | -0.085 | 0.2425 | -0.2825 | 0.117839 | 0.234754 | 0.277195 | 0.989223 | 0.997988 | 0.994666 |
| Kcnb2_A6H8H5_S488s_2 | Kcnb2_A6H | 0.63 | 0.25 | 0.28 | 0.73 | 0.39 | 0.39 | 0.22 | 0.55 | 0.42 | 0.68 | 1.01 | 0.75 | -0.73 | 0.68 | 0.34 | 0.47 | -0.2825 | 0.3775 | 0.5725 | 0.441876 | 0.703807 | 0.341704 | 0.995437 | 0.997988 | 0.994666 |
| Kcnb2_A6H8H5_S851s_1 | Kcnb2_A6H | -0.02 | -0.83 | 0.81 | 1.44 | -0.68 | -0.1 | 1.08 | 1.24 | -0.34 | -0.53 | 2.25 | 0.9 | 1.19 | -0.89 | 1.12 | 0.89 | -0.085 | 0.2425 | -0.2825 | 0.719778 | 0.953858 | 0.763262 | 0.999187 | 0.997988 | 0.9971 |
| Kcnb2_A6H8H5_T725t_1 | Kcnb2_A6H | -0.06 | -0.38 | 0.07 | 0.12 | 0.63 | -0.03 | -0.17 | 0 | -0.48 | -0.87 | 0.43 | 0 | -0.24 | 0.53 | -0.56 | -0.01 | 0.035 | 0.22 | 0.2275 | 0.980671 | 0.517742 | 0.613262 | 0.989223 | 0.997988 | 1 |
| Kcnc1_P15388_S468s_1 | Kcnc1_P15 | 5.27 | 4.54 | 3.48 | 2.3 | 4.89 | 5.54 | 3.41 | 3.59 | -0.51 | 5.1 | 1.41 | 3.37 | -1.25 | 0.56 | 3.15 | 3.64 | 0.17 | -0.1675 | -0.0075 | 0.061041 | 0.530941 | 0.218351 | 0.989223 | 0.997988 | 0.994666 |
| Kcnc1_P15388_T527t_1 | Kcnc1_P15 | 0.55 | 0.66 | 0.34 | 0.8 | 0.76 | 0.82 | 0.76 | 0.65 | -0.04 | 0.72 | 0.21 | 0.43 | -0.13 | 0.23 | 0.35 | 0.66 | 0.46 | -1.555 | -2.3725 | 0.27451 | 0.45005 | 0.330207 | 0.989223 | 0.997988 | 0.994666 |
| Kcnc2_Q14B80_S508s_1 | Kcnc2_Q14 | 4.36 | 3.31 | 5.43 | 3.62 | 3.98 | 4.66 | 5.44 | 5.75 | 0.65 | 4.26 | 0.14 | 5.9 | -0.06 | 0.34 | 5.42 | 6.47 | 0.16 | -0.2575 | -0.31 | 0.457342 | 0.184908 | 0.28075 | 0.989223 | 0.997988 | 0.994666 |
| Kcnc2_Q14B80_S509s_1 | Kcnc2_Q14 | 2.47 | 1.44 | 3.59 | 1.49 | 2.39 | 1.59 | 4.06 | 3.53 | 0.58 | 2.65 | 1.6 | 3.9 | -0.96 | 0.49 | 3.55 | 3.73 | 0.7775 | -1.4425 | -1.1375 | 0.620956 | 0.345286 | 0.933616 | 0.989223 | 0.997988 | 0.995259 |
| Kcnc2_Q14B80_S557s_1 | Kcnc2_Q14 | 0.21 | 0.45 | 0.77 | 0.73 | 0.24 | 0.16 | 0.85 | 0.18 | 0.25 | 0.26 | 0 | 0.85 | -0.2 | 0.53 | -0.18 | 0.82 | 0.645 | -0.065 | -0.545 | 0.3773 | 0.488373 | 0.476234 | 0.989223 | 0.997988 | 0.994666 |
| Kcnc2_Q14B80_S604s_1 | Kcnc2_Q14 | 0.02 | 0.08 | -0.42 | -1.36 | 0.29 | -0.62 | 0.58 | -0.25 | -1.95 | -0.86 | -0.68 | -1.01 | -0.74 | -0.12 | -1.85 | -0.27 | -0.1825 | -0.2 | -0.2975 | 0.510822 | 0.318179 | 0.124251 | 0.989223 | 0.997988 | 0.994666 |
| Kcnc2_Q14B80_S619s_1 | Kcnc2_Q14 | -0.42 | -0.04 | -0.12 | -0.33 | 1.09 | 0.62 | 0.9 | -0.68 | -0.66 | 0.05 | -0.43 | -0.7 | -0.11 | 0.3 | -1.15 | 0.66 | 0.42 | -0.705 | -0.325 | 0.707707 | 0.097757 | 0.433 | 0.989223 | 0.997988 | 0.996966 |
| Kcnc3_E9QKI2_S732s_1 | Kcnc3_E9Q | 0.02 | -1 | -0.84 | -0.39 | 0.41 | 0.42 | -0.51 | -0.46 | -1.05 | -0.32 | -1.22 | 0.07 | -1.14 | -0.15 | -1.4 | -0.46 | 0.71 | -0.2075 | 0.1525 | 0.544642 | 0.162506 | 0.840245 | 0.989223 | 0.997988 | 0.994666 |
| Kcnc3_E9QKI2_T751t_1 | Kcnc3_E9Q | 0.77 | 0.39 | 0.36 | 0.85 | 0.65 | 0.7 | 0.73 | 1.03 | 0.19 | 0.05 | 0.25 | 0.21 | -1.17 | -0.1 | 0.06 | 1.23 | 0.5175 | -0.0775 | -0.235 | 0.243193 | 0.427935 | 0.096292 | 0.989223 | 0.997988 | 0.994666 |
| Kcnd1_Q03719_S547s_1 | Kcnd1_Q03 | 0.48 | 0.1 | 1.62 | 0.3 | -0.27 | -0.02 | 0.65 | 0.01 | 0.62 | 0.45 | 1.31 | 1.15 | 1.4 | 0.27 | 1.21 | 0.65 | 0.185 | -0.4175 | -0.5875 | 0.548086 | 0.186625 | 0.521233 | 0.989223 | 0.997988 | 0.994666 |
| Kcnd1_Q03719_S550s_1 | Kcnd1_Q03 | 0.91 | 0.8 | 1.07 | 0.51 | 0.72 | 0.97 | 0.11 | 0.73 | 0.95 | -0.36 | -1.29 | 0.82 | 1.95 | -1.73 | 1.53 | 0.62 | -0.5325 | 0.2575 | 0.2575 | 0.756058 | 0.478305 | 0.144947 | 0.989223 | 0.997988 | 0.9971 |
| Kcnd1_Q03719_S555s_1 | Kcnd1_Q03 | 0.12 | 0.56 | 0.91 | 0.14 | 0.33 | -0.53 | 0.91 | 0.38 | 0.31 | 0.25 | 0.52 | 0.45 | 0.26 | 0.59 | 0.3 | 0.34 | -0.19 | -0.7925 | -0.23 | 0.830735 | 0.652401 | 0.84908 | 0.990358 | 0.997988 | 0.999507 |
| Kcnd1_Q03719_S568s_1 | Kcnd1_Q03 | 0.38 | -2.23 | -1.32 | 0 | -0.13 | 1.57 | -0.17 | 0 | 3.11 | -0.84 | -2.32 | 0.31 | 1.23 | -1.35 | 0.05 | -0.77 | -0.16 | -0.05 | -0.06 | 0.433857 | 0.113476 | 0.458929 | 0.989223 | 0.997988 | 0.994666 |
| Kcnd2_Q9Z0V2_S546s_1 | Kcnd2_Q9Z | -0.51 | 0.44 | -0.01 | -0.78 | 1.23 | -1.3 | 0.09 | -0.52 | -1.33 | 0.29 | -0.44 | -2.14 | 1.48 | 0.52 | -2.95 | -0.58 | 1.11 | 0.8575 | 0.5825 | 0.847338 | 0.868018 | 0.229963 | 0.996493 | 0.997988 | 0.999507 |
| Kcnd2_Q9Z0V2_S546s_2 | Kcnd2_Q9Z | 0.43 | 0.11 | -0.32 | 0.82 | 0.7 | 0.23 | -0.01 | 0.92 | 0.3 | 0.2 | -0.87 | 0.65 | -0.72 | 0.89 | -0.06 | 0.49 | 0.09 | -0.69 | -0.1675 | 0.794726 | 0.550297 | 0.639835 | 0.989223 | 0.997988 | 0.999507 |
| Kcnd2_Q9Z0V2_S548s_1 | Kcnd2_Q9Z | 0.21 | 0.11 | 0.54 | 0.17 | 0.58 | -0.05 | 0.15 | 0.03 | 0.05 | -0.23 | 0.77 | 0.07 | 0.57 | 1.06 | 0.1 | -0.38 | 0.2 | -0.19 | -0.11 | 0.820543 | 0.737133 | 0.744555 | 0.995762 | 0.997988 | 0.999507 |
| Kcnd2_Q9Z0V2_S548s_2 | Kcnd2_Q9Z | 0.41 | 0.3 | -0.56 | 0.36 | 0.69 | 0.19 | 0.16 | -0.06 | 0.19 | 0.31 | -1.2 | 0.62 | -0.31 | 0.68 | -0.09 | 0.14 | -0.08 | -0.0925 | 0.08 | 0.94805 | 0.69887 | 0.742384 | 0.994893 | 0.997988 | 0.999507 |
| Kcnd2_Q9Z0V2_S552s_1 | Kcnd2_Q9Z | -0.26 | 0.63 | 0.35 | 0.1 | 0.46 | -0.4 | 0.47 | -0.31 | -0.14 | 0.14 | -0.45 | 0.07 | 0.19 | 0.66 | -0.16 | 0.12 | 0.1175 | -0.1475 | -0.0225 | 0.993541 | 0.63984 | 0.301839 | 0.989223 | 0.997988 | 1 |
| Kcnd2_Q9Z0V2_S552s_2 | Kcnd2_Q9Z | 0.41 | 0.3 | -0.56 | 0.36 | 0.69 | 0.19 | 0.16 | -0.06 | 0.19 | 0.31 | -1.2 | 0.62 | -0.31 | 0.68 | -0.09 | 0.14 | -0.15 | -0.3 | -0.0025 | 0.94805 | 0.69887 | 0.742384 | 0.994893 | 0.997988 | 0.999507 |
| Kcnd2_Q9Z0V2_S572s_1 | Kcnd2_Q9Z | -0.1 | -1.08 | -0.78 | -0.73 | -1.15 | 0.44 | -0.92 | -0.68 | 0.23 | -1.02 | -1.56 | -0.72 | -0.7 | -1.28 | -1.41 | -1.27 | 0.1175 | -0.1475 | -0.0225 | 0.139238 | 0.809685 | 0.820893 | 0.995762 | 0.997988 | 0.994666 |
| Kcnd3_Q9Z0V1_S550s_1 | Kcnd3_Q9Z | -0.1 | 0.05 | -0.21 | -0.15 | 0.14 | -0.04 | 0.22 | -0.16 | -0.62 | 0.21 | -0.25 | -0.12 | -0.61 | -0.18 | -0.25 | 0.48 | 0.095 | -0.095 | -0.4925 | 0.899573 | 0.499014 | 0.715235 | 0.989223 | 0.997988 | 0.999507 |
| Kcnd3_Q9Z0V1_S566s_1 | Kcnd3_Q9Z | -0.52 | -0.38 | -0.51 | 0.31 | -0.68 | -0.51 | 0.18 | 0.01 | -0.46 | -0.05 | -0.92 | 0.15 | -0.91 | -0.54 | -0.02 | 0.19 | 0.1425 | -0.0925 | -0.0375 | 0.897288 | 0.934878 | 0.892107 | 0.99868 | 0.997988 | 0.999507 |
| Kcnh1_Q3UHC9_S831s_1 | Kcnh1_Q3U | 1 | -1.3 | -0.27 | 0.44 | 0.4 | 1.05 | -1.12 | 0.59 | 0.25 | -0.55 | 0.36 | 0.26 | -0.72 | -0.01 | -1.14 | -0.61 | 0.025 | -0.045 | -0.045 | 0.270666 | 0.668366 | 0.824281 | 0.992876 | 0.997988 | 0.994666 |
| Kcnh1_Q3UHC9_S872s_1 | Kcnh1_Q3U | 2.77 | 2.15 | 1.42 | 2.38 | 1.66 | 2.06 | 0.95 | 1.43 | 2.6 | 3.58 | 2.12 | 1.41 | 2.16 | 2.78 | 2.05 | 1.39 | 0.2625 | 0.1125 | -0.5875 | 0.834207 | 0.097041 | 0.625763 | 0.989223 | 0.997988 | 0.999507 |
| Kcnh1_Q3UHC9_S877s_1 | Kcnh1_Q3U | -0.49 | -1.17 | -0.98 | -0.88 | 0.2 | 0.33 | -0.38 | -1.8 | -0.61 | -0.46 | -0.96 | -1.87 | -0.98 | 0.01 | -2.13 | -1.06 | -0.655 | 0.2475 | -0.085 | 0.721917 | 0.333567 | 0.791564 | 0.989223 | 0.997988 | 0.9971 |
| Kcnh1_Q3UHC9_S951s_1 | Kcnh1_Q3U | -2.39 | -1.21 | 0.84 | 0.64 | -1.71 | -2.16 | 0.7 | 0.36 | -1.73 | -1.45 | 0.48 | 0.92 | -0.44 | -1.14 | 0.33 | 0.71 | 0.4675 | -0.095 | -0.16 | 0.612808 | 0.85079 | 0.925238 | 0.996349 | 0.997988 | 0.995259 |
| Kcnh1_Q3UHC9_S951s_2 | Kcnh1_Q3U | -2.68 | -0.31 | -2.22 | -1.33 | -1.03 | -1.78 | -1.92 | -2.99 | -0.67 | -0.29 | -0.72 | -2.07 | -0.59 | -1.18 | -2.71 | -2.66 | -0.1725 | 0.085 | 0.395 | 0.823309 | 0.620294 | 0.261377 | 0.989223 | 0.997988 | 0.999507 |
| Kcnh2_Q53Z09_S1140s_1 | Kcnh2_Q53 | 0.23 | -0.84 | -0.89 | -1.41 | 0.65 | -0.69 | -0.52 | -1.1 | -0.9 | 0.23 | -1.69 | -1.34 | 0.66 | 1.63 | -1.28 | 0.32 | -0.295 | 0.6975 | -0.15 | 0.118189 | 0.512388 | 0.698443 | 0.989223 | 0.997988 | 0.994666 |
| Kcnh2_Q53Z09_S241s_1 | Kcnh2_Q53 | 0.94 | 0.46 | 0.05 | 0.19 | 1.09 | 0.7 | 0.99 | 0.38 | 0.08 | 0.77 | 0.64 | 1.2 | 0.23 | 0.55 | 0.75 | 0.26 | 0.3125 | -0.1975 | 1.06 | 0.899229 | 0.204272 | 0.431991 | 0.989223 | 0.997988 | 0.999507 |
| Kcnh2_Q53Z09_S286s_1 | Kcnh2_Q53 | 0.26 | 0.19 | -0.33 | -0.05 | 0.8 | -0.45 | -0.15 | -0.95 | -0.03 | 0.26 | 0.58 | 0.09 | -0.01 | 1 | 0.39 | -0.17 | 0.38 | 0.2625 | 0.0375 | 0.400794 | 0.59415 | 0.427677 | 0.989223 | 0.997988 | 0.994666 |
| Kcnh7_Q9ER47_S1135s_1 | Kcnh7_Q9E | 0.2 | -0.34 | 0.36 | 0.67 | 0.63 | 0.19 | 0.62 | 0.22 | -0.52 | -0.97 | -0.16 | 0.37 | -0.99 | -0.37 | 0.12 | 0.32 | -0.205 | 0.2075 | 0.285 | 0.248129 | 0.496919 | 0.160364 | 0.989223 | 0.997988 | 0.994666 |
| Kcnh7_Q9ER47_S1147s_1 | Kcnh7_Q9E | 0.44 | 0.6 | 0.18 | 0.45 | 0.26 | -0.06 | -0.39 | 0.31 | -0.68 | 0.47 | -0.58 | -0.29 | -0.99 | -0.03 | -0.19 | -0.26 | 0.1925 | -0.5425 | -0.4525 | 0.023179 | 0.138448 | 0.048811 | 0.989223 | 0.997988 | 0.994666 |
| Kcnh7_Q9ER47_S174s_1 | Kcnh7_Q9E | 0.83 | 0.35 | 0.56 | 0.67 | 0.89 | 1.27 | 0.74 | 0.69 | -0.03 | 1.1 | 0.35 | 1.1 | 0 | 0.39 | 0.13 | 0.22 | -0.3875 | -0.6875 | -0.785 | 0.117209 | 0.229478 | 0.932703 | 0.989223 | 0.997988 | 0.994666 |
| Kcnh7_Q9ER47_S209s_1 | Kcnh7_Q9E | 0.76 | -1.64 | -0.35 | -0.66 | -0.34 | 0.91 | -0.91 | -0.61 | 2.29 | 0.37 | 0.13 | -0.82 | 1.62 | -0.28 | 0.04 | -1.19 | 0.295 | 0.0275 | -0.4175 | 0.455968 | 0.682189 | 0.210683 | 0.994114 | 0.997988 | 0.994666 |
| Kcnh7_Q9ER47_S350s_1 | Kcnh7_Q9E | -0.15 | -0.1 | 0.44 | -0.2 | -0.92 | 0 | -0.22 | -0.73 | 1.51 | 0.2 | -1.04 | 0.13 | 2 | -1.26 | 0.12 | -0.41 | 0.235 | 0.965 | 0.52 | 0.857121 | 0.134666 | 0.691647 | 0.989223 | 0.997988 | 0.999507 |
| Kcnh7_Q9ER47_S905s_1 | Kcnh7_Q9E | -1.6 | -1 | 1.92 | 0.99 | -0.87 | -1.03 | 1.29 | 1.8 | 0.28 | -0.84 | 3.39 | 1.67 | 0.65 | -0.51 | 1.12 | 1.11 | -0.465 | 0.2025 | 0.115 | 0.526621 | 0.817809 | 0.346619 | 0.995762 | 0.997988 | 0.994666 |
| Kcnh7_Q9ER47_S912s_1 | Kcnh7_Q9E | 0.65 | 1.44 | 1.41 | 1.58 | 0.44 | 1.12 | 0.87 | 1.85 | 1.13 | 1.02 | 2.07 | 1.73 | 1.11 | 0.16 | 1.1 | 1.13 | 0.22 | 1.0475 | 0.515 | 0.273502 | 0.582086 | 0.531604 | 0.989223 | 0.997988 | 0.994666 |
| Kcnh7_Q9ER47_T882t_1 | Kcnh7_Q9E | -0.46 | 0.07 | -2.17 | -3.06 | 0.16 | -0.6 | -1.63 | -2.22 | -0.69 | -0.84 | 0.55 | 0.1 | 0.4 | -1.59 | -0.63 | -2.42 | -0.2 | 0.2175 | -0.395 | 0.681219 | 0.674627 | 0.126083 | 0.992876 | 0.997988 | 0.996159 |
| Kcnip2_Q3YAB2_S19s_1 | Kcnip2_Q3Y | 1.68 | 1.16 | 1.68 | 1.57 | 1.47 | 1.08 | 1.79 | 1.54 | 0.34 | 1.39 | 1.1 | 1.77 | -0.37 | 0.49 | 1.53 | 1.8 | 0.3325 | 1.185 | 0.345 | 0.198217 | 0.833317 | 0.295787 | 0.995762 | 0.997988 | 0.994666 |
| Kcnip2_Q3YAB2_S9s_1 | Kcnip2_Q3Y | 1.05 | 0.36 | 0.71 | 0.8 | -0.74 | 0.57 | 0.01 | 0.28 | 1.84 | 0.35 | -0.35 | 0.37 | 1.99 | -0.81 | 0.21 | 0.52 | -0.0525 | -0.3725 | -0.66 | 0.648978 | 0.055413 | 0.701472 | 0.989223 | 0.997988 | 0.995259 |
| Kcnip4_Q6PHZ8_S8s_1 | Kcnip4_Q6 | 0.88 | 0.93 | 0.72 | 1.01 | 0.99 | 0.99 | 1.22 | 1.06 | 0.49 | 0.87 | 0.61 | 1.27 | 0.05 | 0.84 | 0.59 | 0.92 | -0.7 | -0.1775 | -0.2525 | 0.325655 | 0.382088 | 0.768365 | 0.989223 | 0.997988 | 0.994666 |
| Kcnj10_Q9JM63_S338s_1 | Kcnj10_Q9 | 0.82 | 0.85 | 0.26 | 0.55 | 1.42 | -0.69 | 0.05 | 0.02 | 1.18 | 0.61 | 1.49 | 0.03 | 0.75 | 0.9 | 0.96 | 1.44 | 0.18 | -0.075 | -0.285 | 0.184185 | 0.346622 | 0.569402 | 0.989223 | 0.997988 | 0.994666 |
| Kcnj12_B1ATI1_S502s_1 | Kcnj12_B1A | 1.65 | 1.83 | 2.29 | 2.09 | 1.42 | 1.83 | 1.07 | 2.01 | 0.67 | 1.83 | 1.95 | 2.31 | -1.04 | 0.42 | 1.44 | 1.38 | -0.42 | 0.2075 | 0.3925 | 0.027362 | 0.2022 | 0.48092 | 0.989223 | 0.997988 | 0.994666 |
| Kcnj12_B1ATI1_T456t_1 | Kcnj12_B1A | 0.73 | 1.33 | 1.12 | 0.65 | 1.31 | 1.02 | 0.11 | 0.68 | 0.36 | 0.52 | 0.61 | 0.53 | 0.87 | 1 | 0.38 | 0.06 | -0.3825 | -0.275 | -1.415 | 0.251715 | 0.581128 | 0.093234 | 0.989223 | 0.997988 | 0.994666 |
| Kcnj13_P86046_T327t_1 | Kcnj13_P86 | -1.4 | -0.89 | -0.66 | -0.71 | -0.26 | -1.63 | -0.93 | -1.54 | -0.43 | -0.42 | -1.33 | -3.8 | 0.36 | 0.27 | -1.34 | -0.02 | -0.1775 | -0.4525 | -0.38 | 0.110565 | 0.627418 | 0.436514 | 0.989223 | 0.997988 | 0.994666 |
| Kcnj16_Q9Z307_S374s_1 | Kcnj16_Q9 | 0.14 | 0.58 | 0.73 | 0.68 | 0.5 | 0.14 | 0.77 | 0.71 | 0.19 | 0 | -0.04 | 0.4 | 0.01 | 0.1 | 0.25 | 0.4 | -0.175 | -0.58 | 0.7325 | 0.208208 | 0.99206 | 0.134522 | 0.999187 | 0.997988 | 0.994666 |
| Kcnj3_P63250_S368s_1 | Kcnj3_P632 | 0.15 | 0.28 | 0.68 | 0.45 | 0.76 | 0.15 | 0.78 | 0.48 | -0.65 | -0.08 | -0.22 | 0.04 | -0.7 | 0 | 0.07 | 0.37 | -0.0025 | -0.395 | -0.3425 | 0.163866 | 0.540326 | 0.036594 | 0.989223 | 0.997988 | 0.994666 |
| Kcnj3_P63250_S369s_1 | Kcnj3_P632 | 0.45 | 0.19 | 0.46 | 0.49 | 0.83 | 0.28 | 0.6 | 0.22 | -0.42 | 0.47 | -0.35 | -0.1 | -1.04 | 0.47 | -0.19 | 0.41 | 0.1525 | -0.6175 | -0.455 | 0.216165 | 0.714984 | 0.092853 | 0.995437 | 0.997988 | 0.994666 |
| Kcnj3_P63250_S421s_1 | Kcnj3_P632 | -1.85 | -0.81 | -1.97 | -5 | -1.4 | -0.62 | -1.55 | -1.87 | 1.33 | -0.95 | -2.34 | -0.52 | 1.17 | -1.87 | -0.19 | -1.39 | 0.085 | -0.4975 | -0.485 | 0.087297 | 0.221912 | 0.11162 | 0.989223 | 0.997988 | 0.994666 |
| Kcnj3_P63250_S442s_1 | Kcnj3_P632 | 0.9 | -0.31 | -0.43 | 0.46 | -0.02 | 1.24 | 0.14 | 1.1 | -0.63 | -0.52 | 1.13 | 0.11 | -0.75 | 0 | -0.26 | -0.04 | 1.0475 | 1.7875 | 1.8375 | 0.284327 | 0.297619 | 0.785303 | 0.989223 | 0.997988 | 0.994666 |
| Kcnj3_P63250_S447s_1 | Kcnj3_P632 | 0.61 | 0.62 | 0.7 | 0.45 | 0.85 | 1.09 | -0.23 | 0.91 | 0.04 | 0.53 | 0.39 | 0.51 | -0.38 | 0.29 | 0.14 | 0.79 | 0.46 | -0.1325 | -0.4175 | 0.225691 | 0.850857 | 0.329244 | 0.996349 | 0.997988 | 0.994666 |
| Kcnj4_Q8R435_T344t_1 | Kcnj4_Q8R | 0.37 | 1.49 | 1.72 | 0.53 | 0.07 | 0.13 | 0.55 | 0.04 | 1.09 | 0.62 | 0.92 | -0.11 | 1.88 | 1.68 | 1.26 | 0.86 | 0.06 | -0.2275 | -0.385 | 0.350267 | 0.041133 | 0.35704 | 0.989223 | 0.997988 | 0.994666 |
| Kcnj6_P48542_S7s_1 | Kcnj6_P485 | 1.03 | 0.9 | 1.06 | 1.16 | 0.86 | 0.63 | 0.54 | 1.2 | 0.32 | 1.19 | 0.04 | 0.76 | 0.06 | 1.3 | 0.11 | 1.28 | -0.83 | -0.3975 | 0.3925 | 0.357493 | 0.33243 | 0.148065 | 0.989223 | 0.997988 | 0.994666 |
| Kcnj6_Q8C4T8_S386s_1 | Kcnj6_Q8C | -0.85 | -1.27 | -0.83 | -0.76 | -0.8 | -0.75 | -0.09 | -1.01 | -1.54 | -1.13 | 0.06 | -0.69 | -1.54 | -0.17 | -2 | -1.11 | -0.23 | -0.46 | -0.35 | 0.503124 | 0.341497 | 0.781686 | 0.989223 | 0.997988 | 0.994666 |
| Kcnk1_O08581_S326s_1 | Kcnk1_O08 | 0.15 | 0.68 | 0.29 | 0.43 | 0.51 | 0.59 | 0.7 | 0.55 | 0.29 | 0.65 | 0.28 | 0.51 | 0 | 1.01 | 0.5 | 0.3 | 0.265 | 0.1025 | -0.2775 | 0.829152 | 0.363215 | 0.8482 | 0.989223 | 0.997988 | 0.999507 |
| Kcnk12_Q76M80_S5s_1 | Kcnk12_Q7 | -0.21 | -0.2 | 0.59 | 0.75 | 0.3 | 0.44 | -0.15 | -0.94 | 0.37 | 0.09 | -3.68 | 0.61 | 1.27 | -1.19 | -0.14 | -0.1 | 0.2 | 0.045 | 0.065 | 0.6081 | 0.419834 | 0.351959 | 0.989223 | 0.997988 | 0.995259 |
| Kcnk2_Q6P6P9_S336s_1 | Kcnk2_Q6P | 0.52 | 0.15 | 0.68 | 0.82 | 1.07 | 1.56 | 1.17 | 0.65 | -0.57 | 0.95 | 0.9 | 0.77 | -0.88 | 1.15 | 0.44 | 0.58 | -0.32 | -0.885 | -0.2725 | 0.621563 | 0.062109 | 0.938789 | 0.989223 | 0.997988 | 0.995259 |
| Kcnk2_Q6P6P9_S354s_1 | Kcnk2_Q6P | -0.6 | -1.21 | -0.36 | 0.3 | -0.27 | -0.1 | 0.51 | 0.5 | -0.57 | -0.66 | 0.19 | 0.33 | -0.76 | 0.11 | 0.61 | 0.84 | 0.57 | -0.03 | -0.22 | 0.167866 | 0.113163 | 0.475441 | 0.989223 | 0.997988 | 0.994666 |
| Kcnk4_O88454_S324s_1 | Kcnk4_O88 | 0.43 | -0.32 | 0.5 | 0.26 | 1.01 | 0.11 | 1.05 | 0.54 | -0.11 | 0.08 | -0.09 | 0.4 | 0.05 | 1.01 | -0.16 | 0.47 | 0.6275 | 0.29 | 0.6675 | 0.720817 | 0.160652 | 0.594792 | 0.989223 | 0.997988 | 0.9971 |
| Kcnma1_J3QN27_S582s_1 | Kcnma1_J3 | 0.56 | 0.85 | 1.03 | 0.68 | 0.67 | 0.51 | 0.8 | 0.77 | 0.31 | 0.92 | 0.2 | 0.88 | 0.43 | 0.49 | 0.52 | 0.56 | 0.46 | -0.1475 | 0.125 | 0.259861 | 0.668925 | 0.462498 | 0.992876 | 0.997988 | 0.994666 |
| Kcnma1_J3QN27_S809s_1 | Kcnma1_J3 | -1.99 | -1.32 | -1.32 | -1.22 | 0.02 | -1.65 | -0.6 | -1.3 | -2.85 | -2.44 | -1.39 | -1.26 | -1.91 | -0.59 | -1.41 | -0.84 | -0.0925 | -0.2025 | -0.28 | 0.459747 | 0.167326 | 0.23403 | 0.989223 | 0.997988 | 0.994666 |
| Kcnma1_J3QN27_T580t_1 | Kcnma1_J3 | 0.37 | 0.71 | 0.89 | -0.12 | 0.33 | 0.67 | 0.35 | 0.89 | 0.39 | 0.97 | 0.83 | 0.84 | 0.34 | 0.93 | 0.8 | 0.45 | 0.58 | -0.5225 | 0.275 | 0.596365 | 0.73724 | 0.333782 | 0.995762 | 0.997988 | 0.995259 |
| Kcnma1_Q08460_S672s_1 | Kcnma1_Q | -0.24 | 0.22 | 0.62 | 1.23 | 0.59 | 0.76 | 1.03 | 0.22 | -0.09 | 0.46 | 0.17 | 0.28 | 0.03 | 0.89 | -0.04 | 0.8 | 0.0975 | 0.295 | 0.1675 | 0.925756 | 0.59047 | 0.475202 | 0.989223 | 0.997988 | 0.999507 |
| Kcnma1_Q08460_S782s_1 | Kcnma1_Q | -0.07 | -0.08 | 0.56 | 0.14 | -1.21 | -0.14 | -0.31 | -0.2 | 3.19 | 0.12 | -1.67 | 0.76 | 3.66 | -1.11 | 3.16 | -0.96 | 0.1925 | -0.2525 | -0.0375 | 0.361056 | 0.077773 | 0.610201 | 0.989223 | 0.997988 | 0.994666 |
| Kcnma1_Q08460_S966s_1 | Kcnma1_Q | -0.13 | -0.19 | -0.28 | -0.32 | 0.52 | 0.43 | 0.75 | 0.35 | -0.77 | 0.09 | -0.76 | 0.12 | -0.43 | 0.65 | -0.08 | 0.49 | -0.6025 | 0.4625 | 1.05 | 0.22943 | 0.004957 | 0.736464 | 0.989223 | 0.997988 | 0.994666 |
| Kcnma1_Q08460_S977s_1 | Kcnma1_Q | -0.19 | 0.23 | 0.38 | 0.08 | -0.07 | -0.2 | -0.15 | 0.15 | -0.14 | 0.04 | 0.08 | 0.3 | 0.13 | 0.23 | 0.07 | -0.27 | 0.7425 | -0.1 | 0.3875 | 0.745527 | 0.401776 | 0.817607 | 0.989223 | 0.997988 | 0.9971 |
| Kcnma1_Q08460_S978s_1 | Kcnma1_Q | -0.29 | 0.29 | 0.25 | 0.16 | -0.13 | -0.12 | 0.13 | 0.01 | -0.03 | 0.04 | -0.31 | -0.02 | 0.25 | 0.18 | -0.28 | -0.26 | -0.1925 | -0.055 | -0.085 | 0.639643 | 0.569278 | 0.454179 | 0.989223 | 0.997988 | 0.995259 |
| Kcnma1_Q08460_S978s_2 | Kcnma1_Q | 0.05 | 0.48 | 0.53 | -0.58 | 0.44 | 0.3 | -0.36 | 0.26 | 0.12 | 0.05 | 0.09 | -0.36 | -1.49 | 0.22 | -0.1 | -0.8 | -0.13 | -0.1825 | -0.13 | 0.162681 | 0.901967 | 0.645092 | 0.996493 | 0.997988 | 0.994666 |
| Kcnma1_Q08460_S982s_2 | Kcnma1_Q | 0.05 | 0.48 | 0.67 | -0.03 | 0.44 | 0.3 | 0.26 | 0.74 | 0.12 | 0.05 | 0.56 | 0.11 | -1.49 | 0.22 | 0.22 | -0.18 | 0.04 | -0.145 | -0.6625 | 0.187649 | 0.578011 | 0.757389 | 0.989223 | 0.997988 | 0.994666 |
| Kcnma1_Q08460_T1088t_1 | Kcnma1_Q | 0.73 | 1.01 | 0.86 | 0.71 | 1.22 | 0.61 | 0.44 | 0.54 | 0.56 | 0.36 | 0.68 | 0.9 | 0.96 | 0.89 | 0.66 | 0.4 | 0.1425 | -0.0825 | -0.6 | 0.694187 | 0.616212 | 0.388663 | 0.989223 | 0.997988 | 0.996159 |
| Kcnma1_Q08460_T734t_1 | Kcnma1_Q | 0.73 | 0.77 | 1.04 | 0.57 | -0.24 | 1.17 | 0.3 | 0.65 | 1.18 | -0.76 | 0.22 | 0.63 | 1.24 | 0.96 | 0.78 | 0.57 | -0.125 | -0.2025 | -0.1 | 0.679982 | 0.355558 | 0.283293 | 0.989223 | 0.997988 | 0.996159 |
| Kcnma1_Q08460_T970t_1 | Kcnma1_Q | 0.44 | 0.18 | 0.86 | 0.39 | 0.71 | 0.64 | 1.05 | 0.8 | 0.67 | 0.66 | 0.82 | 0.91 | 0.22 | 1.13 | 0.03 | 0.93 | -0.3075 | -0.46 | 0.11 | 0.745989 | 0.182554 | 0.234691 | 0.989223 | 0.997988 | 0.9971 |
| Kcnn2_P58390_S152s_1 | Kcnn2_P58 | 1.61 | -0.23 | 0.76 | 1.22 | 0.62 | 1.31 | 0.6 | 1.05 | 0.77 | 0.02 | 1.29 | 0.98 | -0.01 | 0.88 | 0.05 | -0.18 | 0.3325 | 0.2975 | 0.11 | 0.168524 | 0.893919 | 0.870371 | 0.996493 | 0.997988 | 0.994666 |
| Kcnn2_P58390_S827s_2 | Kcnn2_P58 | -0.64 | 0.29 | 0.56 | 0.51 | -0.3 | 1.28 | -0.16 | 0.5 | 2.18 | -0.04 | 0.7 | 0.44 | 1.6 | -0.62 | -0.37 | 0.4 | 0.055 | -0.075 | -0.655 | 0.891772 | 0.728543 | 0.236822 | 0.995762 | 0.997988 | 0.999507 |
| Kcnn2_P58390_S838s_1 | Kcnn2_P58 | -0.21 | 0.63 | 0.54 | 0.67 | -1.77 | 1.06 | -0.29 | -0.08 | 2.48 | 0.59 | -1.16 | 0.27 | 2.23 | -0.59 | 0.61 | 0.47 | 0.15 | 0.64 | 0.0725 | 0.633916 | 0.24244 | 0.844443 | 0.989223 | 0.997988 | 0.995259 |
| Kcnn2_P58390_S839s_1 | Kcnn2_P58 | -0.21 | 0.63 | 0.54 | 0.67 | -1.77 | 1.06 | -0.29 | -0.08 | 2.48 | 0.59 | -1.16 | 0.27 | 2.23 | -0.59 | 0.61 | 0.47 | -0.6775 | 0.1375 | 0.2725 | 0.633916 | 0.24244 | 0.844443 | 0.989223 | 0.997988 | 0.995259 |
| Kcnq2_B7ZBV3_S238s_1 | Kcnq2_B7Z | -0.7 | -0.11 | -1.01 | -0.62 | -0.41 | -0.91 | -0.5 | -2.13 | -0.71 | -0.61 | -1.92 | -1.43 | -0.97 | -0.6 | -1.63 | -1.39 | -0.6775 | 0.1375 | 0.2725 | 0.131924 | 0.375516 | 0.156584 | 0.989223 | 0.997988 | 0.994666 |
| Kcnq2_B7ZBV3_S302s_1 | Kcnq2_B7Z | 1.02 | 0.94 | 0.01 | -0.44 | 0.33 | 0.01 | -0.01 | -0.02 | 0.89 | -0.37 | 2 | 0.24 | 0.56 | 0.12 | 0.3 | 0.74 | -0.3775 | -0.5575 | -0.5375 | 0.903608 | 0.410991 | 0.593471 | 0.989223 | 0.997988 | 0.999507 |
| Kcnq2_B7ZBV3_S313s_1 | Kcnq2_B7Z | 0.08 | -0.31 | 0.93 | 0.91 | -1.11 | 0.23 | -0.16 | 0.17 | 1.31 | -0.1 | -0.47 | 0.47 | 1.56 | -1.65 | 0.39 | 0.52 | -0.305 | 0.3075 | 0.0475 | 0.766886 | 0.160182 | 0.832272 | 0.989223 | 0.997988 | 0.9971 |
| Kcnq2_B7ZBV3_S322s_1 | Kcnq2_B7Z | 0.39 | 0.74 | 0.2 | 0.38 | 0.66 | 0.22 | 0.39 | 0.55 | 0.25 | 0.72 | 1.17 | 0.35 | 0.56 | 0.67 | 0.55 | -0.08 | -0.62 | -0.1 | -0.1975 | 0.992906 | 0.903524 | 0.499967 | 0.996493 | 0.997988 | 1 |
| Kcnq2_B7ZBV3_S332s_1 | Kcnq2_B7Z | -0.33 | 1.51 | 0.8 | 0.21 | 0.57 | -0.17 | 0.13 | -0.22 | 0.49 | 0.54 | 0.22 | 0.13 | 1.45 | 0.82 | 1.09 | 0.56 | 0.0275 | 0.195 | -0.0025 | 0.33048 | 0.273268 | 0.618549 | 0.989223 | 0.997988 | 0.994666 |
| Kcnq2_B7ZBV3_S529s_1 | Kcnq2_B7Z | -0.37 | 0.45 | 0.29 | 0.2 | 0.19 | -0.4 | 0.14 | 0 | 0.07 | -0.15 | 0.44 | -0.04 | -0.34 | 0.63 | 0.1 | 0.23 | -0.47 | -0.2025 | 0.4325 | 0.968536 | 0.552971 | 0.820195 | 0.989223 | 0.997988 | 0.999507 |
| Kcnq2_B7ZBV3_S585s_1 | Kcnq2_B7Z | 7.6 | 2.35 | 0.8 | 1.39 | 3.85 | 7.98 | 1.49 | 1.52 | 0.94 | 5.04 | 1.19 | 1.95 | -2.76 | 0 | 1.31 | 1.85 | -0.16 | -0.0625 | 0.0125 | 0.091894 | 0.717499 | 0.636166 | 0.995437 | 0.997988 | 0.994666 |
| Kcnq2_B7ZBV8_S579s_1 | Kcnq2_B7Z | -0.34 | 0.86 | 0.11 | 0.75 | 0.1 | 0.22 | 0.08 | 0.21 | 0.29 | 0.16 | 0.73 | -0.31 | 0.53 | 0.28 | -0.1 | 0.08 | 0.675 | -0.755 | -2.935 | 0.670515 | 0.53337 | 0.726842 | 0.989223 | 0.997988 | 0.996159 |
| Kcnq3_Q8K3F6_S666s_1 | Kcnq3_Q8K | 0.66 | -0.01 | 0.5 | -0.57 | -1.79 | 0.7 | -1.01 | -0.11 | 2.89 | -0.03 | -1.55 | 0.35 | 2.06 | -1.31 | 0.1 | -0.33 | -0.1925 | -0.1275 | -0.1475 | 0.982437 | 0.223976 | 0.752869 | 0.989223 | 0.997988 | 1 |
| Kcnq3_Q8K3F6_T580t_1 | Kcnq3_Q8K | 0.36 | 0.54 | 0.82 | 0.95 | 0.9 | 0.07 | 1.03 | 0.51 | -0.19 | 0.29 | 0.38 | 0.6 | 0.21 | 0.8 | 0.8 | 0.85 | -0.6975 | 0.27 | -0.015 | 0.992878 | 0.88884 | 0.16668 | 0.996493 | 0.997988 | 1 |
| Kcnq3_Q8K3F6_T82t_1 | Kcnq3_Q8K | 0.52 | 1.1 | 1.45 | 0.9 | 0.94 | 1.08 | 1.32 | 1.2 | 0.78 | 0.82 | 1.31 | 0.78 | 0.32 | 1.1 | 0.73 | 1.24 | -0.04 | -0.3975 | -0.0025 | 0.65852 | 0.586334 | 0.804843 | 0.989223 | 0.997988 | 0.995259 |
| Kcnq5_E9Q9F1_S461s_1 | Kcnq5_E9C | -1.26 | -1.52 | -2.63 | -4.47 | -1.38 | 0.18 | -3.25 | -2.66 | 1.2 | -2.07 | -3.15 | -2.19 | 1.15 | -1.29 | -1.88 | -3.25 | 0.1425 | -0.07 | -0.145 | 0.275404 | 0.455971 | 0.39439 | 0.989223 | 0.997988 | 0.994666 |
| Kcnq5_E9Q9F1_S467s_2 | Kcnq5_E9C | -0.01 | -0.06 | 1.06 | 0.94 | 0.42 | -0.72 | 0.62 | 1.36 | -0.55 | 0.34 | 1.57 | 1.47 | -0.41 | -0.1 | 1.13 | 1.14 | 0.6925 | 0.9175 | 1.1525 | 0.929675 | 0.897084 | 0.68043 | 0.996493 | 0.997988 | 0.999507 |
| Kcnq5_E9Q9F1_S477s_1 | Kcnq5_E9C | 0.02 | 0.83 | 0.96 | 1.33 | 0.43 | 0.3 | 0.93 | 0.68 | 0.34 | 0.71 | 1.41 | 1.23 | 0.85 | 1.01 | 1.46 | 0.68 | -0.0625 | 0.225 | -0.0425 | 0.545135 | 0.539682 | 0.714579 | 0.989223 | 0.997988 | 0.994666 |
| Kcnq5_E9Q9F1_S477s_2 | Kcnq5_E9C | -0.01 | -0.06 | 1.06 | 0.94 | 0.42 | -0.72 | 0.62 | 1.36 | -0.55 | 0.34 | 1.57 | 1.47 | -0.41 | -0.1 | 1.13 | 1.14 | -0.2 | 0.1375 | 0.215 | 0.929675 | 0.897084 | 0.68043 | 0.996493 | 0.997988 | 0.999507 |
| Kcnq5_E9Q9F1_S79s_1 | Kcnq5_E9C | -0.52 | -1.2 | -0.18 | -0.79 | -0.51 | 0.25 | -0.7 | -0.13 | -0.33 | -0.36 | 0.17 | 0.37 | -0.12 | -0.46 | -0.97 | -0.34 | -0.0625 | 0.225 | -0.0425 | 0.542504 | 0.227233 | 0.068399 | 0.989223 | 0.997988 | 0.994666 |
| Kcns2_O35174_S441s_1 | Kcns2_O35 | 0.03 | 0.12 | 0.1 | 0.33 | 0.59 | -0.45 | 0.69 | 0.09 | -0.85 | -0.6 | 0.26 | 0.2 | -0.65 | 0.36 | -0.42 | 0.19 | 0.4 | 0.635 | 0.2 | 0.379684 | 0.774542 | 0.236717 | 0.995762 | 0.997988 | 0.994666 |
| Kcnt1_Q6ZPR4_S1028s_1 | Kcnt1_Q6Z | 0.75 | -0.04 | 1.09 | 0.03 | -1.16 | 0.26 | 1.54 | 2.72 | -0.26 | 0.71 | 3.56 | 0.87 | 0.74 | 0.15 | 1.94 | 3.08 | 0.085 | -0.3925 | -0.275 | 0.135672 | 0.621519 | 0.33751 | 0.989223 | 0.997988 | 0.994666 |
| Kcnt1_Q6ZPR4_S34s_1 | Kcnt1_Q6Z | 0.6 | 0 | 0.5 | 0.65 | 0.71 | 0.5 | 1.01 | 0.99 | 0.01 | 0.27 | 0.41 | 0.86 | -0.58 | 0.42 | 0.27 | 0.59 | 0.3825 | 0.7625 | 1.02 | 0.443367 | 0.166986 | 0.85941 | 0.989223 | 0.997988 | 0.994666 |
| Kcnt1_Q6ZPR4_S44s_1 | Kcnt1_Q6Z | 1.27 | 0.96 | 0.56 | 0.56 | 0.06 | 1.72 | -0.45 | 0.23 | 1.85 | 0.75 | 0.81 | 1.22 | 2.42 | 0.33 | 0.43 | 0.27 | 0.365 | -0.05 | -0.2625 | 0.961225 | 0.345113 | 0.34559 | 0.989223 | 0.997988 | 0.999507 |
| Kcnt1_Q6ZPR4_S47s_1 | Kcnt1_Q6Z | 0.67 | 0.74 | 1.6 | 1.82 | 0.28 | 0.48 | 0.3 | 0.87 | 1.19 | 0.32 | 1.81 | 1.41 | 1.63 | -0.34 | 1.34 | 1.26 | -0.4475 | 0.32 | 0.025 | 0.642328 | 0.052323 | 0.952659 | 0.989223 | 0.997988 | 0.995259 |
| Kcnt2_D3Z592_S1053s_1 | Kcnt2_D3Z | 0.41 | -0.05 | 0.86 | 2.46 | 0.18 | 1.31 | 1.65 | 2.92 | -0.12 | -0.79 | 1.88 | 1.36 | -1.49 | 0.73 | 0.79 | 0.45 | -0.725 | -0.025 | -0.235 | 0.263064 | 0.399771 | 0.651661 | 0.989223 | 0.997988 | 0.994666 |
| Kcnv1_Q8BZN2_S179s_1 | Kcnv1_Q8B | 1.93 | 1.93 | 1.6 | 1.91 | 1.6 | 2.21 | 1.12 | 1.72 | 0.96 | 2.34 | 0.39 | 1.48 | -0.32 | 0.8 | 1.39 | 2.28 | 0.595 | -0.3375 | -0.8 | 0.144121 | 0.519414 | 0.206678 | 0.989223 | 0.997988 | 0.994666 |
| Kctd1_E9Q6T9_S364s_1 | Kctd1_E9Q | -2.38 | -0.28 | -2.67 | -2.73 | -2.26 | -4.09 | -2.23 | -4.81 | -2.91 | -1.56 | -0.34 | -3.33 | 1.23 | 0.27 | -3.91 | -2.72 | -0.18 | -0.55 | -0.805 | 0.533756 | 0.107993 | 0.980025 | 0.989223 | 0.997988 | 0.994666 |
| Kctd1_E9Q6T9_S616s_1 | Kctd1_E9Q | -0.67 | 0.19 | -0.44 | -0.09 | -0.19 | -0.65 | -0.58 | 0.76 | -0.97 | -0.35 | -0.25 | 0.67 | -0.63 | -0.63 | -1.13 | -0.27 | -1.3325 | -0.02 | 0.7325 | 0.206913 | 0.813305 | 0.943579 | 0.995762 | 0.997988 | 0.994666 |
| Kctd12_Q6WVG3_S187s_1 | Kctd12_Q6 | 0.55 | 0.35 | 0.14 | 0.57 | 0.79 | 0.3 | 0.96 | 0.47 | 0.31 | 0.65 | 0.4 | 0.61 | -0.08 | 0.64 | 0.51 | 0.42 | 0.0875 | 0.0275 | -0.4125 | 0.912296 | 0.360756 | 0.694951 | 0.989223 | 0.997988 | 0.999507 |
| Kctd12_Q6WVG3_S189s_1 | Kctd12_Q6 | -0.21 | -0.33 | 2.11 | 0.78 | 0.48 | -0.13 | 1.37 | 1.15 | 0.8 | 0.34 | 1.46 | 0.1 | -0.26 | 0.24 | 0.99 | -0.1 | 0.2275 | 0.09 | -0.03 | 0.526774 | 0.825897 | 0.881534 | 0.995762 | 0.997988 | 0.994666 |
| Kctd12_Q6WVG3_S200s_1 | Kctd12_Q6 | 0.35 | -0.09 | -0.61 | -0.3 | 0.76 | 0.27 | -0.34 | 0 | 0.12 | 0.27 | 1.25 | -0.41 | 0.94 | 0.64 | -0.24 | 0.27 | 0.13 | 0.0875 | -0.37 | 0.133999 | 0.311726 | 0.256908 | 0.989223 | 0.997988 | 0.994666 |
| Kctd12_Q6WVG3_S202s_1 | Kctd12_Q6 | 1.15 | 1 | 0.47 | 0.62 | 0.51 | 0.32 | 0.4 | 0.97 | 0.85 | 0.94 | -0.16 | -0.08 | 0.92 | 1.2 | 0.19 | 0.47 | 0.335 | 0.47 | 0.565 | 0.721831 | 0.335091 | 0.246245 | 0.989223 | 0.997988 | 0.9971 |
| Kctd12_Q6WVG3_T198t_1 | Kctd12_Q6 | 0.61 | 0.1 | 0.17 | 0.75 | 0.91 | 0.74 | 0.86 | 0.34 | 0.21 | -0.18 | -0.13 | 0.54 | -0.49 | 0.45 | 0.06 | 0.19 | -0.26 | -0.4225 | -0.115 | 0.268273 | 0.253611 | 0.306776 | 0.989223 | 0.997988 | 0.994666 |
| Kctd16_Q5DTY9_S130s_1 | Kctd16_Q5 | -1.24 | -1 | -0.7 | -1.16 | -1.33 | -1.52 | -1 | -1.44 | -0.78 | -1.65 | -1.2 | -0.79 | -0.21 | 0.62 | -1.2 | -1.92 | 0.305 | -0.2975 | -0.355 | 0.517763 | 0.225649 | 0.780026 | 0.989223 | 0.997988 | 0.994666 |
| Kctd16_Q5DTY9_S137s_1 | Kctd16_Q5 | -0.34 | -0.15 | -0.34 | 0.48 | -0.47 | 0.01 | -0.19 | 0.42 | -0.27 | -0.07 | -0.7 | -0.16 | -0.69 | 0.29 | -0.97 | -0.22 | -0.2975 | -0.08 | 0.3475 | 0.40096 | 0.919316 | 0.465303 | 0.997335 | 0.997988 | 0.994666 |
| Kctd16_Q5DTY9_S143s_1 | Kctd16_Q5 | 0.52 | -0.01 | -0.13 | 0.41 | 0.46 | 0.7 | 0.42 | 0.31 | 0.59 | 0.61 | 0.18 | 0.38 | 0.44 | 0.93 | -0.17 | 0.52 | 0.03 | -0.2125 | -0.31 | 0.476999 | 0.272184 | 0.356701 | 0.989223 | 0.997988 | 0.994666 |
| Kctd16_Q5DTY9_S173s_1 | Kctd16_Q5 | 0.04 | -0.56 | 0.56 | -0.16 | -0.51 | 0.57 | -0.54 | -0.51 | 1.15 | -0.32 | -0.71 | 0.1 | 1.6 | -1.05 | 0.39 | 0.74 | 0.275 | 0.2425 | 0.2325 | 0.42521 | 0.547024 | 0.849119 | 0.989223 | 0.997988 | 0.994666 |
| Kctd16_Q5DTY9_S389s_1 | Kctd16_Q5 | 0.82 | 1.28 | 1.35 | 0.79 | 1.23 | 0.03 | 1.58 | 1.24 | 0.28 | 1.04 | 1.02 | 1.38 | -0.09 | 0.79 | 1.16 | 1.57 | -0.2175 | 0.085 | 0.45 | 0.608703 | 0.913034 | 0.675774 | 0.997335 | 0.997988 | 0.995259 |
| Kctd16_Q5DTY9_T340t_1 | Kctd16_Q5 | -1.33 | -1.28 | -1.48 | -2.22 | -0.45 | -1.56 | -1.22 | -1.96 | -2.06 | -1.49 | -1.91 | -1.49 | -0.98 | -1.24 | -2.42 | -1.5 | -0.04 | -0.13 | -0.2025 | 0.91359 | 0.466648 | 0.598302 | 0.989223 | 0.997988 | 0.999507 |
| Kctd17_F7B6A4_S166s_1 | Kctd17_F7B | 0.62 | -0.54 | -0.15 | 0.01 | -0.38 | 1.05 | -1.25 | -0.29 | -0.35 | -0.25 | 0.36 | 0.06 | -0.22 | -0.18 | -0.58 | -0.45 | 0.28 | -0.16 | 0.0425 | 0.287902 | 0.680368 | 0.925076 | 0.993464 | 0.997988 | 0.994666 |
| Kctd3_Q8BFX3_S712s_1 | Kctd3_Q8B | 0.34 | -0.1 | 0.38 | -0.4 | -0.34 | 1.12 | -0.9 | -0.53 | 2.1 | 0.62 | -0.85 | -0.25 | 1.47 | -1.01 | -0.05 | -0.38 | -0.2025 | -0.03 | -0.3425 | 0.92757 | 0.630351 | 0.569566 | 0.989223 | 0.997988 | 0.999507 |
| Kctd4_Q9D7X1_S20s_1 | Kctd4_Q9D | 8.64 | 3.74 | 4.42 | -1.04 | 3.85 | 10.21 | 4.72 | 1.77 | 0.52 | 6.83 | -0.28 | 4.05 | -0.94 | 0.2 | 3.89 | 5.11 | -0.2175 | 0.35 | -0.0475 | 0.381322 | 0.602356 | 0.606824 | 0.989223 | 0.997988 | 0.994666 |
| Kctd8_Q50H33_S413s_1 | Kctd8_Q50 | -0.98 | -1.22 | -0.33 | -0.27 | 0.14 | -1.02 | 0.31 | -0.2 | -1.73 | -1.01 | -0.71 | -0.54 | -1.18 | -0.31 | -0.55 | -0.7 | 1.1975 | -1.16 | -1.875 | 0.964421 | 0.194453 | 0.422091 | 0.989223 | 0.997988 | 0.999507 |
| Kdm1a_Q6ZQ88_S850s_1 | Kdm1a_Q6 | -0.2 | 0.23 | 1.43 | 0.57 | -0.08 | 0.31 | 0.7 | 0.75 | -0.11 | 0.38 | 1.84 | 1.15 | -0.84 | 0 | 1.25 | 0.95 | 0.5075 | -0.2975 | 0.015 | 0.75907 | 0.820353 | 0.554825 | 0.995762 | 0.997988 | 0.9971 |
| Kdm2a_F6YRW4_S422s_1 | Kdm2a_F6Y | -2.24 | -0.99 | 0.6 | -1.1 | -1.24 | -1.14 | -0.5 | -3.23 | 0.55 | -0.52 | -0.83 | -1.56 | 1.75 | -0.91 | -1.62 | -1.32 | -0.0875 | 0.3075 | -0.1675 | 0.633943 | 0.423378 | 0.60722 | 0.989223 | 0.997988 | 0.995259 |
| Kdm2a_F6YRW4_S739s_1 | Kdm2a_F6Y | -2.14 | -3.28 | -1.6 | -4.29 | -4.21 | -2.12 | -2.29 | -3.58 | 0.59 | -3.42 | -2.73 | -1.62 | 1.54 | -3.14 | -2.26 | -2.72 | -0.595 | 0.3425 | 0.4075 | 0.28491 | 0.748473 | 0.288702 | 0.995762 | 0.997988 | 0.994666 |
| Kdm2a_F6YRW4_S740s_1 | Kdm2a_F6Y | -2.14 | -3.28 | -1.6 | -4.29 | -4.21 | -2.12 | -2.29 | -3.58 | 0.59 | -3.42 | -2.73 | -1.62 | 1.54 | -3.14 | -2.26 | -2.72 | -0.2225 | 1.0325 | 1.1825 | 0.28491 | 0.748473 | 0.288702 | 0.995762 | 0.997988 | 0.994666 |
| Kdm5b_Q80Y84_S1405s_1 | Kdm5b_Q8 | 2.02 | 1.33 | 1.77 | 0.78 | 0.69 | 1.7 | 0.3 | 0.54 | 1.65 | 1.35 | 0.55 | 0.75 | 1.3 | -0.02 | 1.63 | 1.41 | -0.2225 | 1.0325 | 1.1825 | 0.389534 | 0.117572 | 0.305787 | 0.989223 | 0.997988 | 0.994666 |
| Kdm5c_P41230_S856s_1 | Kdm5c_P41 | -1.13 | -3.8 | -1.02 | 0.19 | -0.03 | -2.22 | -0.35 | -0.56 | -0.85 | -3.06 | 1.21 | -1.67 | -0.33 | 0.82 | -0.27 | -0.6 | -0.6675 | -0.4 | -0.395 | 0.115586 | 0.450456 | 0.748113 | 0.989223 | 0.997988 | 0.994666 |
| Keap1_Q9Z2X8_T43t_1 | Keap1_Q9Z | 0.62 | -0.13 | 1.48 | -0.31 | -0.39 | 0.33 | -0.3 | 0.39 | 2.34 | 0.27 | 2.11 | 0.17 | 2.19 | -1.11 | 2.21 | 2.13 | 0.65 | 0.3475 | 1.345 | 0.263007 | 0.35597 | 0.230964 | 0.989223 | 0.997988 | 0.994666 |
| Khdrbs1_Q60749_S38s_1 | Khdrbs1_Q | -0.17 | -3.32 | -2.66 | -4.17 | -1.3 | -0.44 | -1.66 | -4.42 | -4.34 | -0.29 | -2.97 | -3.62 | -0.98 | -2.6 | -2.24 | -2.27 | -0.4075 | 0.8075 | 0.94 | 0.502531 | 0.555622 | 0.835846 | 0.989223 | 0.997988 | 0.994666 |
| Khdrbs1_Q60749_S58s_1 | Khdrbs1_Q | -0.77 | -0.79 | -0.7 | -0.34 | -1.09 | -1.41 | -0.51 | -0.46 | -1.23 | -1.39 | -0.91 | -1.21 | -0.58 | -0.72 | -0.73 | -0.49 | 0.625 | -0.225 | 0.5575 | 0.934677 | 0.453766 | 0.0451 | 0.989223 | 0.997988 | 0.999507 |
| Khsrp_Q3U0V1_S182s_1 | Khsrp_Q3U | -2.44 | -2.01 | -1.7 | -1.92 | -2.05 | -3.45 | -1.53 | -2.26 | -2.34 | -2.06 | -0.6 | -1.76 | -1.55 | -1.11 | -2.25 | -2.04 | -0.2175 | -0.535 | 0.02 | 0.413921 | 0.467896 | 0.430126 | 0.989223 | 0.997988 | 0.994666 |
| Khsrp_Q3U0V1_S481s_1 | Khsrp_Q3U | -4.49 | -4.72 | -2.29 | -3.74 | -4.21 | -4.89 | -2.87 | -3.38 | -3.99 | -3.55 | -1.58 | -2.77 | -2.14 | -4.93 | -2.63 | -1.92 | -0.305 | 0.3275 | 0.28 | 0.263683 | 0.965097 | 0.242626 | 0.999187 | 0.997988 | 0.994666 |
| Kiaa0195_Q7TSH8_S445s_1 | Kiaa0195_( | 0.61 | 1.5 | 0.03 | 0.23 | 1.19 | 0.75 | 0.62 | 0.37 | 0.86 | 1.36 | 0.05 | 0.04 | 0.93 | 1.76 | 0.77 | 0.41 | -0.0275 | 0.8375 | 0.905 | 0.391318 | 0.701894 | 0.972991 | 0.995437 | 0.997988 | 0.994666 |
| Kiaa1045_Q80TL4_S18s_1 | Kiaa1045_( | -1.03 | -2.44 | -2.3 | -1.82 | -0.68 | -2.58 | -0.66 | -1.9 | -2.61 | -1.9 | -1.4 | -1.66 | -2.25 | -0.85 | -2.31 | -1.72 | 0.14 | -0.015 | 0.375 | 0.799038 | 0.404029 | 0.990175 | 0.989223 | 0.997988 | 0.999507 |
| Kiaa1045_Q80TL4_S309s_1 | Kiaa1045_( | -0.93 | -0.68 | -0.97 | -0.86 | -0.46 | -1.47 | -0.42 | -0.4 | -0.74 | -1.5 | 0.25 | -1.1 | -0.35 | -0.03 | 0.11 | -0.31 | 0.4425 | 0.005 | 0.115 | 0.015667 | 0.56321 | 0.819455 | 0.989223 | 0.997988 | 0.994666 |
| Kiaa1045_Q80TL4_S309s_2 | Kiaa1045_( | -0.86 | -1.36 | -0.93 | 0.09 | -0.12 | -0.08 | -0.19 | 0.05 | 0.38 | -1.52 | 0.25 | -0.25 | -2 | -0.88 | -0.6 | -0.68 | 0.1725 | 0.0875 | 0.715 | 0.534545 | 0.058945 | 0.349149 | 0.989223 | 0.997988 | 0.994666 |
| Kiaa1045_Q80TL4_S347s_1 | Kiaa1045_( | -0.82 | -1.1 | -0.96 | -0.73 | -1.25 | -0.9 | -0.58 | -0.84 | -1.24 | -0.87 | -1.14 | -2.1 | -1.21 | -1.49 | -0.92 | -1.15 | 0.68 | 0.48 | -0.275 | 0.268043 | 0.965683 | 0.185334 | 0.999187 | 0.997988 | 0.994666 |
| Kiaa1045_Q80TL4_S348s_1 | Kiaa1045_( | -0.82 | -1.1 | -2.72 | -2.12 | -1.25 | -0.9 | -1.76 | -1.42 | -1.24 | -0.87 | -2.91 | -2.14 | -1.21 | -1.49 | -2.62 | -1.93 | 0.01 | -0.435 | -0.29 | 0.809457 | 0.431658 | 0.864415 | 0.989223 | 0.997988 | 0.999507 |
| Kiaa1045_Q80TL4_S43s_1 | Kiaa1045_( | -0.64 | -0.91 | -0.51 | 0.18 | -0.71 | -0.5 | -0.24 | 0.28 | -0.72 | -1.27 | 0.06 | -0.31 | 0.09 | -0.2 | -0.25 | -0.01 | 0.3575 | -0.1 | -0.1225 | 0.233345 | 0.590188 | 0.810035 | 0.989223 | 0.997988 | 0.994666 |
| Kiaa1045_Q80TL4_S51s_1 | Kiaa1045_( | -0.04 | -1.08 | 0.56 | 0.71 | 0.14 | 0.24 | 0.26 | 0.75 | -1.18 | -0.08 | 0.99 | 0.65 | -0.83 | 1.23 | -0.36 | 0.08 | 0.1775 | -0.09 | 0.3775 | 0.9892 | 0.456078 | 0.920791 | 0.989223 | 0.997988 | 1 |
| Kiaa1045_Q80TL4_S73s_1 | Kiaa1045_( | -0.17 | -0.29 | -0.38 | -0.7 | -0.03 | -0.62 | 0.21 | 0.75 | 0.01 | -0.07 | 0.63 | -0.26 | -0.75 | 0.79 | -0.02 | 0.37 | 0.31 | 0.0575 | -0.0075 | 0.209823 | 0.171951 | 0.121099 | 0.989223 | 0.997988 | 0.994666 |
| Kiaa1045_Q80TL4_T100t_1 | Kiaa1045_( | -2.49 | -1.73 | -1.15 | -2.36 | -1.03 | -2.9 | -1.01 | -1.44 | -4.15 | -3.06 | -1.47 | -1.79 | -2.32 | -1.69 | -2.04 | -2.1 | 0.4625 | 0.4625 | 0.4825 | 0.771004 | 0.50299 | 0.291333 | 0.989223 | 0.997988 | 0.9971 |
| Kiaa1045_Q80TL4_T47t_1 | Kiaa1045_( | -0.46 | -0.33 | -0.36 | -0.96 | -0.53 | -0.61 | -0.66 | -1.04 | -0.79 | -0.61 | -0.59 | -0.28 | -0.64 | -0.28 | -1.72 | -0.65 | 0.3375 | -0.685 | -0.105 | 0.428318 | 0.464373 | 0.874397 | 0.989223 | 0.997988 | 0.994666 |
| Kiaa1107_Q80TK0_S1051s_1 | Kiaa1107_( | -0.49 | -0.96 | 0.01 | -0.13 | -0.97 | -0.1 | 0.26 | 0.32 | -1.51 | -0.83 | 0.94 | 0.45 | -1.67 | -0.2 | -0.11 | 0.05 | -0.1825 | -0.04 | -0.295 | 0.839339 | 0.465364 | 0.781959 | 0.989223 | 0.997988 | 0.999507 |
| Kiaa1107_Q80TK0_S353s_1 | Kiaa1107_( | -1.44 | -1.66 | -0.08 | 0.49 | -1.98 | -1 | -0.64 | -0.15 | 0.76 | -0.9 | -0.19 | 0.28 | 0.97 | -2.99 | -0.18 | -0.3 | 0.27 | 0.155 | -0.09 | 0.956272 | 0.644944 | 0.272502 | 0.99033 | 0.997988 | 0.999507 |
| Kiaa1107_Q80TK0_S481s_1 | Kiaa1107_( | -0.73 | -1.79 | -1.37 | -0.79 | -1.43 | 0.17 | -0.84 | -2.28 | -1.57 | -0.9 | -2.28 | -0.48 | -1.82 | -0.33 | -1.92 | -2.01 | -0.27 | 0.66 | 0.0475 | 0.450904 | 0.885664 | 0.7605 | 0.996493 | 0.997988 | 0.994666 |
| Kiaa1107_Q80TK0_S695s_1 | Kiaa1107_( | -1.75 | -0.59 | -1.68 | -1.05 | -1.41 | -2.01 | -1.03 | -0.42 | -0.98 | -1.53 | -1.54 | -0.72 | -1.74 | 0.5 | -1.05 | -1.46 | 0.075 | -0.1375 | -0.35 | 0.539245 | 0.903471 | 0.833269 | 0.996493 | 0.997988 | 0.994666 |
| Kiaa1107_Q80TK0_S714s_1 | Kiaa1107_( | 0.96 | 0.81 | 1.44 | 0.57 | -2.32 | 1.05 | -0.08 | 0.19 | 1.01 | 0.84 | 0.63 | 0.67 | 0.99 | -2.11 | 1.36 | 1.52 | 0.05 | 0.075 | 0.33 | 0.520035 | 0.085552 | 0.556376 | 0.989223 | 0.997988 | 0.994666 |
| Kiaa1107_Q80TK0_S714s_2 | Kiaa1107_( | 1.17 | 1.22 | 1.24 | 0.7 | 0.01 | 1.2 | -0.8 | 0.87 | 1.52 | 1.21 | 0.62 | 1.14 | 0.76 | -0.1 | 0.52 | 0.78 | -1.235 | -0.1575 | -0.505 | 0.074514 | 0.108205 | 0.885923 | 0.989223 | 0.997988 | 0.994666 |
| Kiaa1107_Q80TK0_S722s_2 | Kiaa1107_( | 1.17 | 1.22 | 1.24 | 0.7 | 0.01 | 1.2 | -0.8 | 0.87 | 1.52 | 1.21 | 0.62 | 1.14 | 0.76 | -0.1 | 0.52 | 0.78 | -0.7625 | 0.04 | -0.5925 | 0.074514 | 0.108205 | 0.885923 | 0.989223 | 0.997988 | 0.994666 |
| Kiaa1107_Q80TK0_S783s_2 | Kiaa1107_( | -0.1 | 0.26 | 0.3 | 0.01 | -0.31 | -0.34 | -0.76 | 0.14 | 0.21 | 0.12 | -0.7 | -0.46 | -0.56 | 0.07 | 0.15 | 0.25 | -0.7625 | 0.04 | -0.5925 | 0.622615 | 0.117313 | 0.276202 | 0.989223 | 0.997988 | 0.995259 |
| Kiaa1107_Q80TK0_S785s_1 | Kiaa1107_( | -0.29 | 0.15 | 0.6 | 1.13 | -0.16 | -0.08 | 0.09 | 0.44 | 0.15 | 0.23 | 0.12 | 0.97 | -0.03 | -0.25 | 0.44 | 0.53 | -0.435 | -0.325 | -0.14 | 0.55195 | 0.349573 | 0.935731 | 0.989223 | 0.997988 | 0.994666 |
| Kiaa1107_Q80TK0_S785s_2 | Kiaa1107_( | -0.1 | 0.26 | 0.3 | 0.01 | -0.31 | -0.34 | -0.76 | 0.14 | 0.21 | 0.12 | -0.7 | -0.46 | -0.56 | 0.07 | 0.15 | 0.25 | -0.325 | -0.03 | -0.225 | 0.622615 | 0.117313 | 0.276202 | 0.989223 | 0.997988 | 0.995259 |
| Kiaa1107_Q80TK0_S828s_1 | Kiaa1107_( | 2.46 | 1.9 | 0.35 | -0.66 | 1.17 | 0.57 | -1.21 | -3.76 | 1.09 | 1.39 | -2.14 | -0.72 | 2.07 | 1.61 | -0.97 | 0.83 | -0.435 | -0.325 | -0.14 | 0.882044 | 0.133469 | 0.268773 | 0.989223 | 0.997988 | 0.999507 |
| Kiaa1109_A2AAE1_S1294s_1 | Kiaa1109_/ | 1.05 | 1.15 | 1.05 | 0.99 | 0.34 | 0.96 | 0.66 | 0.31 | 1.12 | 1.13 | 0.74 | 0.75 | 0.45 | 0.35 | 0.28 | 1.36 | -1.82 | -1.1075 | -0.1275 | 0.168708 | 0.056091 | 0.577695 | 0.989223 | 0.997988 | 0.994666 |
| Kiaa1109_A2AAE1_S1418s_1 | Kiaa1109_/ | 1.16 | 0.82 | 1.26 | 1.11 | 0.75 | 0.92 | 1.04 | 0.89 | 0.84 | 1.07 | 2.18 | 0.85 | 0.76 | 0.21 | 0.32 | 0.68 | -0.4925 | -0.125 | -0.45 | 0.0444 | 0.385579 | 0.673978 | 0.989223 | 0.997988 | 0.994666 |
| Kiaa1109_A2AAE1_S1981s_1 | Kiaa1109_/ | 0.74 | 0.5 | -0.89 | 0 | 0.77 | 0.15 | -0.27 | 0.26 | 0.41 | 0.76 | 0.43 | -0.07 | 0.06 | -0.13 | -0.35 | -0.7 | -0.1875 | 0.1475 | -0.595 | 0.37095 | 0.728553 | 0.464746 | 0.995762 | 0.997988 | 0.994666 |
| Kiaa1109_A2AAE1_S2350s_1 | Kiaa1109_/ | 0.79 | 0.41 | 0.86 | 0.43 | -0.14 | 1.14 | 0.33 | 0.29 | 1.47 | 0.56 | -1 | 0.43 | 0.85 | -1.46 | -0.55 | 0.24 | 0.14 | 0.295 | -0.3675 | 0.106687 | 0.490007 | 0.604962 | 0.989223 | 0.997988 | 0.994666 |
| Kiaa1109_A2AAE1_S2368s_1 | Kiaa1109_/ | 0.22 | 0.66 | 0.49 | 0.27 | 0.56 | 0.25 | 0.22 | 0.39 | 0.3 | 0.49 | 0.01 | 0.51 | -0.44 | 0.58 | -0.24 | 0.51 | -0.2175 | -0.2575 | -0.8525 | 0.353754 | 0.801086 | 0.731378 | 0.995762 | 0.997988 | 0.994666 |
| Kiaa1109_A2AAE1_S2809s_1 | Kiaa1109_/ | 0.95 | 1.93 | 1.5 | 0.13 | 1.49 | 1.2 | -0.39 | 0.33 | 1.61 | 1.28 | 0.35 | 0.31 | 0.89 | 1.18 | -1.84 | 1.21 | -0.055 | -0.0825 | -0.3075 | 0.314284 | 0.381189 | 0.62108 | 0.989223 | 0.997988 | 0.994666 |
| Kiaa1109_A2AAE1_S903s_1 | Kiaa1109_/ | 1 | 0.83 | 1.49 | 0.47 | 1.48 | 0.96 | 1.68 | 1.23 | 0.48 | 0.83 | -0.82 | 1.14 | -0.42 | 0.52 | 1.6 | 0.68 | -0.47 | -0.24 | -0.7675 | 0.442782 | 0.202825 | 0.258511 | 0.989223 | 0.997988 | 0.994666 |
| Kiaa1109_A2AAE1_S925s_1 | Kiaa1109_/ | -2.51 | -2.97 | -2.52 | -4.03 | -2.77 | -0.81 | -3.15 | -2.88 | -1.47 | -2.92 | -2.21 | -2.98 | -0.79 | -5.21 | -3.48 | -3.38 | 0.39 | -0.54 | -0.3525 | 0.809364 | 0.310418 | 0.220904 | 0.989223 | 0.997988 | 0.999507 |
| Kiaa1109_A2AAE1_S936s_1 | Kiaa1109_/ | -1.43 | -1.58 | -1.37 | -0.9 | -0.53 | -0.45 | -0.95 | -0.38 | -2.48 | -1.08 | 1.07 | -1.24 | -1.92 | -0.98 | -0.22 | -0.31 | 0.605 | 0.6125 | -0.2075 | 0.28482 | 0.014034 | 0.57194 | 0.989223 | 0.997988 | 0.994666 |
| Kiaa1109_F6YAT5_S213s_1 | Kiaa1109_I | 0.95 | -0.22 | 0.64 | 0.78 | 0.77 | 1.62 | 0.01 | 0.6 | 0.44 | -0.13 | -0.3 | 0.01 | -0.99 | -1.67 | -0.38 | 0.28 | 0.7425 | 0.3875 | 0.4625 | 0.0257 | 0.602195 | 0.131516 | 0.989223 | 0.997988 | 0.994666 |
| Kiaa1109_F6YAT5_S475s_1 | Kiaa1109_I | 0.7 | -0.05 | 0.26 | -0.32 | -0.07 | 0.84 | -0.03 | 0.08 | 3.43 | 0.27 | -1.1 | 0.01 | 2.96 | -1.44 | 1.3 | 0.22 | 0.2125 | -0.5325 | -1.2275 | 0.470463 | 0.857682 | 0.571522 | 0.996349 | 0.997988 | 0.994666 |
| Kiaa1109_F6YAT5_S889s_1 | Kiaa1109_I | 1.34 | 0.55 | 1.16 | 1.09 | 0.72 | 0.5 | 0.91 | 1.38 | 0.15 | 0.55 | 0.12 | 1.53 | -0.76 | -1.67 | 0.29 | 0.84 | 0.0575 | 0.505 | 0.6125 | 0.029934 | 0.58423 | 0.252743 | 0.989223 | 0.997988 | 0.994666 |
| Kiaa1109_F6YAT5_T577t_1 | Kiaa1109_I | 0.25 | 0.54 | 0.65 | 0.83 | 0.26 | 0.07 | 0.91 | 0.42 | 0.08 | 0.14 | 0.21 | 0.38 | -0.22 | 0.13 | -0.1 | 0.21 | -0.1575 | -0.4475 | -1.36 | 0.052393 | 0.566343 | 0.141104 | 0.989223 | 0.997988 | 0.994666 |
| Kiaa1522_A2A7S8_S227s_1 | Kiaa1522_/ | 1.8 | 0.31 | 1.08 | 1.14 | 1.91 | 2.4 | 1.32 | 1.89 | 0.85 | 0.99 | 0.12 | 1.15 | 0.72 | -0.23 | 1.04 | 1.46 | -0.1525 | -0.365 | -0.5625 | 0.469663 | 0.05453 | 0.434614 | 0.989223 | 0.997988 | 0.994666 |
| Kiaa1522_A2A7S8_S264s_1 | Kiaa1522_/ | 0.43 | 0.14 | -1.24 | -1.68 | 0.2 | -0.46 | -1.86 | -1.52 | 0.81 | 0.73 | -1.09 | -0.42 | 0.37 | 1.37 | -0.87 | -0.36 | 0.7975 | -0.305 | -0.335 | 0.280453 | 0.607757 | 0.357052 | 0.989223 | 0.997988 | 0.994666 |

| | | | | | | | | | | | | | | | | | | | | | | | | | | |
|---|---|---|---|---|---|---|---|---|---|---|---|---|---|---|---|---|---|---|---|---|---|---|---|---|---|---|
| Kiaa1522_A2A7S8_S289s_1 | Kiaa1522_/ | 0.16 | 0.58 | 1.15 | 0.78 | 0.69 | 0.48 | 1.11 | 0.6 | 0.79 | 0.6 | 1.28 | 0.48 | 1.66 | 1.91 | 1.37 | 1.43 | -0.3225 | 0.595 | 0.715 | 0.011609 | 0.852422 | 0.696897 | 0.996349 | 0.997988 | 0.994666 |
| Kiaa1522_A2A7S8_S289s_2 | Kiaa1522_/ | -0.78 | -1.32 | -0.38 | -0.39 | 0.17 | 0.08 | -0.79 | 0 | -0.7 | -0.05 | -0.36 | 0.02 | -1.43 | 0.47 | 0.2 | 0.04 | 0.0525 | 0.12 | 0.925 | 0.263307 | 0.09886 | 0.174722 | 0.989223 | 0.997988 | 0.994666 |
| Kiaa1522_A2A7S8_S294s_2 | Kiaa1522_/ | -0.78 | -1.32 | -0.71 | -0.33 | 0.17 | 0.08 | -1.06 | 0.1 | -0.7 | -0.05 | -0.29 | -0.29 | -1.43 | 0.47 | -0.72 | -0.82 | 0.5825 | 0.445 | 0.5375 | 0.715846 | 0.114832 | 0.143687 | 0.989223 | 0.997988 | 0.996966 |
| Kiaa1522_A2A7S8_S306s_1 | Kiaa1522_/ | -1.33 | -1.33 | -0.72 | -1.4 | -0.75 | -0.23 | -1.3 | -0.89 | -0.2 | -1.84 | -2.22 | -0.89 | 0.22 | -2.39 | -0.62 | -0.75 | 0.6075 | 0.4525 | 0.16 | 0.562515 | 0.198126 | 0.841797 | 0.989223 | 0.997988 | 0.995259 |
| Kiaa1522_A2A7S8_S332s_1 | Kiaa1522_/ | -2.62 | -4.44 | -1.58 | -2.94 | -1 | -1.67 | -1.86 | -1.86 | -2.83 | -2.88 | -0.37 | -2.29 | -1.71 | -1.07 | -2.41 | -1.81 | 0.4025 | -0.0925 | 0.31 | 0.079722 | 0.042331 | 0.299511 | 0.989223 | 0.997988 | 0.994666 |
| Kiaa1522_A2A7S8_S430s_1 | Kiaa1522_/ | 0.66 | 0.77 | -0.11 | 0.23 | 1.01 | 1.42 | 0.58 | 0.29 | 0.77 | 0.22 | 0.21 | 0.91 | 0.3 | 1.53 | -0.09 | -0.03 | 1.2975 | 0.8025 | 1.145 | 0.924952 | 0.206081 | 0.651149 | 0.989223 | 0.997988 | 0.999507 |
| Kiaa1522_A2A7S8_S796s_1 | Kiaa1522_/ | 0.2 | 0.36 | 0.7 | 1.41 | 1.47 | 0.88 | 1.4 | 1.4 | 0.85 | 0.72 | 0.62 | 1.14 | 0.51 | 1.52 | 0.9 | 1.68 | 0.4375 | 0.14 | 0.04 | 0.235316 | 0.075704 | 0.608892 | 0.989223 | 0.997988 | 0.994666 |
| Kiaa1522_A2A7S8_S890s_1 | Kiaa1522_/ | 0.59 | 0.32 | 0.72 | 0.57 | 0.22 | 1.35 | 0.06 | 0.05 | 1.66 | 0.71 | 0.16 | 0.46 | 2.1 | 0.07 | 0.28 | 0.6 | 0.62 | 0.165 | 0.485 | 0.640866 | 0.696918 | 0.57618 | 0.994893 | 0.997988 | 0.995259 |
| Kiaa1598_Q8K2Q9_S3s_1 | Kiaa1598_( | 0.73 | 0.21 | 0.24 | 0.09 | -0.85 | 1.27 | -1.34 | -0.29 | 1.13 | 1.28 | 1.1 | -0.02 | 1.57 | 0.01 | -0.25 | -0.9 | -0.13 | 0.1975 | 0.2125 | 0.681974 | 0.259729 | 0.136354 | 0.989223 | 0.997988 | 0.996159 |
| Kiaa1671_Q8BRV5_S166s_1 | Kiaa1671_( | -5.03 | -5.01 | -4.75 | -12.27 | -3.08 | -3.26 | -6.34 | -6.06 | -2.48 | -4.28 | -1.31 | -5.14 | -1.5 | -2.53 | -5.05 | -5.21 | -0.62 | 0.555 | -0.21 | 0.095141 | 0.249765 | 0.075983 | 0.989223 | 0.997988 | 0.994666 |
| Kiaa1671_Q8BRV5_S175s_1 | Kiaa1671_( | -0.58 | -0.44 | -0.13 | -1.1 | -0.54 | 0.05 | -1.37 | -1.17 | -0.17 | -1.57 | -1.81 | -0.53 | -0.66 | -1.22 | -1.29 | -0.63 | 2.08 | 3.4625 | 3.1925 | 0.241113 | 0.604186 | 0.305725 | 0.989223 | 0.997988 | 0.994666 |
| Kiaa1671_Q8BRV5_S205s_1 | Kiaa1671_( | 1.12 | 1.14 | 1.17 | 1.02 | 0.8 | 1.05 | 0.64 | 0.99 | 1.26 | 1.22 | 0.79 | 0.95 | 0.32 | 1.08 | 0.84 | 0.85 | -0.195 | -0.4575 | -0.3875 | 0.214415 | 0.258753 | 0.797065 | 0.989223 | 0.997988 | 0.994666 |
| Kiaa1671_Q8BRV5_S277s_1 | Kiaa1671_( | 1.97 | 0.25 | -0.67 | -1.49 | 1.38 | 2.94 | -0.19 | -0.2 | 0.96 | 0.99 | 0.56 | -0.77 | -0.75 | 1.56 | -0.56 | -0.56 | -0.2425 | -0.0575 | -0.34 | 0.909421 | 0.305605 | 0.584179 | 0.989223 | 0.997988 | 0.999507 |
| Kiaa1671_Q8BRV5_S302s_1 | Kiaa1671_( | -2.47 | -1.7 | -2.71 | -4.28 | -1.2 | -1.88 | -1.75 | -2.58 | -1.98 | -1.96 | -0.19 | -1.64 | -0.59 | -0.31 | -2.33 | -2.56 | 0.9675 | 0.42 | -0.0925 | 0.083664 | 0.11492 | 0.055667 | 0.989223 | 0.997988 | 0.994666 |
| Kiaa1671_Q8BRV5_T215t_1 | Kiaa1671_( | -0.65 | -1.08 | -1.54 | -1.55 | -1.76 | 0.19 | -1.38 | -1.56 | 0.17 | -1.71 | 0.67 | -1.71 | 0.69 | -0.94 | -0.9 | -1.84 | 0.9375 | 1.3475 | 1.3425 | 0.397941 | 0.866312 | 0.365235 | 0.996493 | 0.997988 | 0.994666 |
| Kidins220_E9Q9B7_S1578s_1 | Kidins220_ | 0.2 | 0.68 | 1.46 | 1.91 | 1.23 | 0.18 | 1.33 | 1.79 | 0.58 | 0.33 | 1.36 | 0.91 | 0.42 | 1.12 | 1.71 | 1.04 | 0.0775 | 0.56 | 0.4575 | 0.982348 | 0.882588 | 0.539517 | 0.996493 | 0.997988 | 1 |
| Kidins220_E9Q9B7_S1704s_1 | Kidins220_ | 0.49 | -0.49 | 0.08 | -0.52 | -0.07 | 0.44 | 0.53 | 0.18 | 0.33 | 0.04 | 1.37 | -0.16 | 0.62 | 1.08 | -0.19 | 0.63 | 0.07 | -0.2675 | 0.01 | 0.111657 | 0.227614 | 0.239774 | 0.989223 | 0.997988 | 0.994666 |
| Kidins220_E9Q9B7_S885s_1 | Kidins220_ | 1.23 | 1.06 | 0.77 | -0.06 | 0.83 | 0.88 | 0.53 | -0.06 | 0.33 | 1.5 | 0.16 | 0.81 | 0.57 | 0.17 | 0.82 | 0.43 | 0.38 | 0.505 | 0.645 | 0.474158 | 0.569858 | 0.902714 | 0.989223 | 0.997988 | 0.994666 |
| Kif13b_E9Q4K7_S1380s_1 | Kif13b_E9C | -1.92 | -2.28 | -1.58 | -2.63 | -3.03 | -2.29 | -1.65 | -2.18 | -1.52 | -1.91 | -0.42 | -1.72 | 2.22 | -1.7 | -2.21 | -2.04 | -0.205 | -0.05 | -0.2525 | 0.234076 | 0.610941 | 0.102971 | 0.989223 | 0.997988 | 0.994666 |
| Kif13b_E9Q4K7_S1794s_1 | Kif13b_E9C | -0.67 | -0.5 | -0.23 | 0.12 | -0.4 | -0.45 | -0.05 | -0.51 | -0.08 | 0.32 | 0.48 | -0.2 | -0.11 | 0.01 | -0.38 | -0.04 | -0.185 | 0.71 | 1.17 | 0.496073 | 0.898275 | 0.139202 | 0.996493 | 0.997988 | 0.994666 |
| Kif13b_E9Q4K7_S1795s_1 | Kif13b_E9C | 0.11 | -0.14 | -0.01 | 0.47 | -0.24 | 0.63 | 0.11 | -0.27 | 0.34 | 0.63 | 0.22 | 0.23 | 0.44 | 0.08 | -0.24 | -0.02 | -0.0325 | 0.45 | 0.19 | 0.877433 | 0.859209 | 0.323542 | 0.996349 | 0.997988 | 0.999507 |
| Kif1a_Q6TA13_S1102s_1 | Kif1a_Q6T/ | -0.02 | 0.69 | 0.46 | 0.03 | 1.12 | -0.45 | 1.37 | -0.18 | -1.38 | -0.57 | -1.16 | -0.03 | -0.86 | 1.01 | -0.13 | 0.65 | -0.05 | 0.2475 | -0.0425 | 0.781658 | 0.701281 | 0.01409 | 0.99521 | 0.997988 | 0.998226 |
| Kif1a_Q6TA13_S1345s_1 | Kif1a_Q6T/ | 0.28 | 0.46 | -0.16 | 0.14 | 0.21 | -0.21 | 0.24 | -0.13 | -0.09 | 0.02 | -0.95 | -0.19 | -0.13 | 0.25 | -0.22 | -0.57 | 0.175 | -1.075 | -0.1225 | 0.242998 | 0.52952 | 0.130121 | 0.989223 | 0.997988 | 0.994666 |
| Kif1a_Q6TA13_S1378s_1 | Kif1a_Q6T/ | -0.05 | 0.7 | -0.11 | 0.81 | 0.68 | 0.03 | 0.83 | 0.72 | -0.45 | 0.55 | -0.14 | 0.55 | -1.47 | 0.58 | 0.51 | 0.52 | -0.1525 | -0.4825 | -0.3475 | 0.56563 | 0.480687 | 0.56424 | 0.989223 | 0.997988 | 0.995259 |
| Kif1a_Q6TA13_S1438s_1 | Kif1a_Q6T/ | 0.68 | 0.43 | 1.4 | 1.58 | 0.39 | 0.13 | 0.55 | 1.43 | -0.02 | 1.05 | 1 | 0.7 | -2.26 | 0.63 | 1.07 | 1.43 | 0.2275 | -0.21 | -0.3025 | 0.318504 | 0.315493 | 0.377479 | 0.989223 | 0.997988 | 0.994666 |
| Kif1a_Q6TA13_S1485s_1 | Kif1a_Q6T/ | -0.8 | -1.23 | -1.1 | -1.71 | -0.98 | -0.51 | -0.68 | -0.84 | -1.07 | -0.66 | -0.45 | -1.3 | -0.7 | -0.68 | -0.59 | -0.67 | -0.3975 | -0.34 | -0.805 | 0.069575 | 0.106569 | 0.283441 | 0.989223 | 0.997988 | 0.994666 |
| Kif1a_Q6TA13_S1503s_1 | Kif1a_Q6T/ | 0.05 | 0.64 | 0.96 | 0.75 | 0.99 | 0.13 | 1.27 | 1.34 | -0.2 | 0.59 | 0.85 | 1.11 | -0.56 | 0.6 | 0.31 | 0.54 | 0.4575 | 0.34 | 0.55 | 0.305653 | 0.346132 | 0.972015 | 0.989223 | 0.997988 | 0.994666 |
| Kif1a_Q6TA13_S1556s_1 | Kif1a_Q6T/ | 0.31 | 0.1 | 0.56 | 0.31 | 0.03 | 0.52 | -0.28 | -0.03 | 1.1 | 0.25 | -1.57 | 0.09 | 1.5 | -0.82 | 0.22 | 0.23 | 0.3325 | -0.0125 | -0.3775 | 0.93583 | 0.311336 | 0.509694 | 0.989223 | 0.997988 | 0.999507 |
| Kif1a_Q6TA13_S411s_1 | Kif1a_Q6T/ | 0.59 | 0.47 | 0.02 | 0.45 | 0.67 | 1.03 | 0.78 | -0.31 | 0.62 | 0.12 | -1.58 | 0.81 | 0.35 | 0.55 | -0.45 | 0.55 | -0.26 | -0.3525 | -0.0375 | 0.677854 | 0.629558 | 0.461657 | 0.989223 | 0.997988 | 0.996159 |
| Kif1a_Q6TA13_S425s_1 | Kif1a_Q6T/ | -0.1 | -1.22 | -1.94 | -2.13 | -0.26 | -0.02 | -0.78 | -0.41 | 0.38 | -1.26 | 1.21 | -0.29 | 0.62 | 0.92 | -1.07 | -1.4 | 0.16 | -0.39 | -0.1325 | 0.121189 | 0.053813 | 0.056288 | 0.989223 | 0.997988 | 0.994666 |
| Kif1a_Q6TA13_S428s_1 | Kif1a_Q6T/ | -0.08 | -1.19 | -1.1 | -0.5 | 0.02 | -0.52 | -0.19 | -2.01 | 0.09 | -0.94 | -0.38 | -2.33 | 0.5 | -0.5 | -0.76 | -1.36 | 0.98 | 1.3575 | 1.115 | 0.680877 | 0.930294 | 0.751623 | 0.997844 | 0.997988 | 0.996159 |
| Kif1a_Q6TA13_S941s_1 | Kif1a_Q6T/ | -0.13 | 0.05 | 1.37 | 0.55 | -0.1 | -0.35 | 1.18 | 1.36 | 0.28 | 0.43 | 1.87 | 0.33 | -0.01 | 0.08 | 0.69 | 2.1 | 0.0425 | -0.1725 | 0.1875 | 0.643732 | 0.90119 | 0.582177 | 0.996493 | 0.997988 | 0.995259 |
| Kif1a_Q6TA13_S946s_1 | Kif1a_Q6T/ | 0.8 | 0.21 | 0.71 | 0.51 | -0.56 | 1 | -0.22 | 0.4 | 0.97 | 0.3 | 0.68 | 0.18 | 0.42 | -0.4 | 0.59 | 0.35 | 0.0625 | 0.2675 | 0.255 | 0.31892 | 0.286071 | 0.927967 | 0.989223 | 0.997988 | 0.994666 |
| Kif1a_Q6TA13_T1511t_1 | Kif1a_Q6T/ | 0.31 | 0.7 | 0.37 | -0.64 | 0.37 | 0.53 | 0.91 | 0.33 | 0.05 | 0.49 | 1.1 | -0.35 | -0.46 | 0.43 | 0.22 | 0.28 | -0.4025 | -0.025 | -0.3175 | 0.854999 | 0.301061 | 0.741686 | 0.989223 | 0.997988 | 0.999507 |
| Kif1b_Q60575_S1165s_1 | Kif1b_Q60! | -0.33 | 0.12 | 0.83 | -0.25 | 0.26 | -0.67 | 1.04 | -0.23 | -0.66 | -0.16 | -1.38 | 0.62 | -1.27 | -0.1 | -0.68 | -0.22 | 0.35 | 0.1375 | -0.0675 | 0.113374 | 0.985981 | 0.317385 | 0.999187 | 0.997988 | 0.994666 |
| Kif1b_Q60575_S1408s_1 | Kif1b_Q60! | 0.54 | 0.12 | 0.58 | 0.43 | -0.33 | 0.88 | 0.06 | 0.38 | 0.72 | 0.22 | 0.22 | 0.46 | 0.91 | -1.77 | 0.27 | 0.55 | 0.0075 | -0.4875 | -0.66 | 0.454595 | 0.57491 | 0.9587 | 0.989223 | 0.997988 | 0.994666 |
| Kif1b_Q60575_S1567s_1 | Kif1b_Q60! | 1.4 | 0.84 | 1.1 | 0 | 1.23 | 1.09 | 1.48 | 1.15 | -0.84 | 1.03 | -0.75 | 1.18 | 0.1 | 0.41 | 0.72 | 0.55 | -0.17 | -0.0125 | -0.4275 | 0.288137 | 0.2351 | 0.25696 | 0.989223 | 0.997988 | 0.994666 |
| Kif1b_Q60575_S1613s_1 | Kif1b_Q60! | 0.05 | 0.36 | 0.41 | 0.63 | 0.46 | 0.38 | 0.69 | 0.49 | 0.36 | 0.56 | 0.53 | 0.45 | -0.4 | 0.3 | 0.32 | 0.38 | 0.4025 | -0.68 | -0.39 | 0.469256 | 0.524925 | 0.62362 | 0.989223 | 0.997988 | 0.994666 |
| Kif1b_Q60575_T1406t_1 | Kif1b_Q60! | 0.58 | -0.44 | -0.03 | -0.55 | -0.21 | 0.72 | -1 | -0.3 | 0.04 | -0.04 | -1.53 | 0.13 | 0.44 | -2.57 | -0.63 | -0.16 | 0.1425 | 0.1125 | -0.2125 | 0.340417 | 0.833109 | 0.598364 | 0.995762 | 0.997988 | 0.994666 |
| Kif1c_E9Q9B0_S1043s_1 | Kif1c_E9Q! | -0.51 | -0.18 | -0.4 | -0.99 | -0.63 | -0.11 | -1.02 | -0.66 | 1.77 | -0.23 | -2.7 | -0.14 | 1.88 | -1.22 | -0.25 | -0.62 | -0.0875 | -0.24 | -0.62 | 0.465049 | 0.766676 | 0.814419 | 0.995762 | 0.997988 | 0.994666 |
| Kif1c_E9Q9B0_T1092t_1 | Kif1c_E9Q! | 0.61 | -0.05 | 0.51 | 0.31 | 0.41 | 0.41 | 0.08 | 0.24 | -0.37 | 0.59 | 0.37 | 0.33 | 0.31 | 0.8 | 0.44 | 0.35 | -0.085 | 0.195 | 0.4675 | 0.633325 | 0.799121 | 0.698488 | 0.995762 | 0.997988 | 0.995259 |
| Kif21a_E9Q0J5_S1241s_1 | Kif21a_E9C | -1.39 | -0.74 | -0.65 | -1.15 | -0.85 | -1.2 | -1.41 | -1.41 | -1.9 | -0.96 | 2.66 | -1.41 | 0.14 | -0.04 | -0.81 | -2.2 | -0.06 | -0.115 | 0.13 | 0.628546 | 0.384466 | 0.538262 | 0.989223 | 0.997988 | 0.995259 |
| Kif21a_E9Q0J5_S258s_1 | Kif21a_E9C | -0.03 | 0.35 | 0.3 | 0.21 | 0.46 | 0.03 | 0.58 | 0.46 | 0.12 | 0.32 | 0.24 | 0.18 | 0.07 | 0.63 | 0.12 | 0.41 | -0.235 | 0.58 | 0.255 | 0.699559 | 0.447295 | 0.972213 | 0.989223 | 0.997988 | 0.996159 |
| Kif21a_E9Q0J5_S524s_1 | Kif21a_E9C | -2.82 | -1.52 | -4.2 | -3.77 | -1.21 | -2.03 | -2.6 | -5.09 | -1.9 | -1.99 | -2.06 | -3.79 | -2.36 | -0.89 | -4 | -3.56 | 0.175 | 0.0075 | 0.1 | 0.64422 | 0.699548 | 0.353944 | 0.994893 | 0.997988 | 0.995259 |
| Kif21a_E9Q0J5_S855s_1 | Kif21a_E9C | -2.1 | -1.54 | -0.91 | -1.58 | -0.62 | -2.02 | -0.74 | -1.22 | -3.66 | -2.16 | -0.41 | -1.18 | -1.25 | -1.14 | -2.25 | -1.36 | 0.345 | 0.6425 | 0.375 | 0.930171 | 0.337407 | 0.635718 | 0.989223 | 0.997988 | 0.999507 |
| Kif21b_Q9QXL1_S1150s_1 | Kif21b_Q9( | -0.67 | -0.79 | -0.82 | -0.16 | -0.96 | -0.93 | 0.7 | -0.5 | -0.57 | -0.5 | -0.52 | -0.03 | -0.56 | -1.29 | -0.4 | -0.55 | 0.3825 | -0.32 | 0.0325 | 0.770317 | 0.642223 | 0.442803 | 0.989327 | 0.997988 | 0.9971 |
| Kif21b_Q9QXL1_S1198s_1 | Kif21b_Q9( | -3.68 | -2.97 | -2.72 | -4.29 | -0.97 | -3.49 | -2.54 | -3.26 | -4.31 | -3.52 | -2.03 | -3.24 | -1.93 | -1.57 | -2.96 | -2.68 | 0.1875 | 0.205 | -0.09 | 0.033695 | 0.179186 | 0.798709 | 0.989223 | 0.997988 | 0.994666 |
| Kif21b_Q9QXL1_S1217s_1 | Kif21b_Q9( | 0.05 | -0.17 | 0.36 | 0.14 | 0.78 | 0.02 | 0.34 | 0.39 | -0.08 | 0.32 | 1.06 | 0.9 | 0.09 | 0.93 | 0.26 | -0.06 | 0.85 | 0.14 | 1.13 | 0.494239 | 0.264558 | 0.173824 | 0.989223 | 0.997988 | 0.994666 |
| Kif21b_Q9QXL1_S222s_1 | Kif21b_Q9( | 0.75 | 0.59 | 1.14 | -1.73 | 0.52 | -0.08 | -0.48 | -1.28 | 1.55 | 0.54 | 0.57 | -0.25 | 1.32 | 0.8 | 0.35 | 1.16 | 0.2875 | 0.455 | 0.21 | 0.263839 | 0.444019 | 0.5432 | 0.989223 | 0.997988 | 0.994666 |
| Kif21b_Q9QXL1_S223s_1 | Kif21b_Q9( | 0.53 | 1.03 | 1.27 | -0.33 | -1.69 | 0.09 | 0.44 | -0.25 | 1.6 | 0.17 | 1.15 | 0.5 | 1.75 | -0.89 | -0.19 | 0.95 | -0.5175 | 0.415 | 0.72 | 0.724373 | 0.091747 | 0.617768 | 0.989223 | 0.997988 | 0.9971 |
| Kif21b_Q9QXL1_S969s_1 | Kif21b_Q9( | 6.62 | 3.73 | 4.71 | 1.56 | 4.59 | 7 | 5.17 | 4.42 | -0.45 | 6.72 | -0.06 | 5.5 | -0.25 | -0.02 | 5.65 | 5.64 | -0.9775 | 0.23 | -0.22 | 0.416202 | 0.289722 | 0.513712 | 0.989223 | 0.997988 | 0.994666 |
| Kif21b_Q9QXL1_T1621t_1 | Kif21b_Q9( | -0.15 | -0.75 | -0.74 | -0.17 | -0.67 | -0.82 | -0.46 | -2.47 | -2.63 | -1.36 | -2.42 | -1.31 | -0.66 | -2 | -1.53 | -0.33 | 1.14 | -1.2275 | -1.4 | 0.13096 | 0.176421 | 0.003817 | 0.989223 | 0.997988 | 0.994666 |
| Kif2a_F8VQ42_S111s_1 | Kif2a_F8VC | -0.57 | -0.34 | 0.17 | 0.36 | -0.02 | -0.59 | -0.62 | -0.6 | -0.81 | -0.11 | 0.4 | -0.58 | -0.9 | 0.96 | 0.49 | 0.04 | -0.6525 | -1.4775 | -0.6775 | 0.586246 | 0.231832 | 0.616723 | 0.989223 | 0.997988 | 0.995259 |
| Kif2a_F8VQ42_S123s_1 | Kif2a_F8VC | 0.09 | 0.42 | 0.42 | 0.49 | 0.2 | -0.43 | 0.77 | 0.63 | 0.47 | 0.71 | 0.4 | 0.4 | 0.55 | 1.06 | 0.54 | 0.47 | -0.3625 | -0.18 | 0.2425 | 0.26811 | 0.838096 | 0.534893 | 0.995762 | 0.997988 | 0.994666 |
| Kif2a_F8VQ42_S190s_1 | Kif2a_F8VC | -0.04 | 0.28 | -0.44 | 0.2 | 0.37 | -0.09 | 0.5 | 0.51 | -1.37 | -0.19 | 1.04 | 0.06 | -0.07 | 1.25 | 0.57 | -0.28 | -0.0625 | 0.14 | 0.3 | 0.359252 | 0.237565 | 0.81532 | 0.989223 | 0.997988 | 0.994666 |
| Kif2a_F8VQ42_S410s_1 | Kif2a_F8VC | -2.98 | -1.94 | -2.05 | -0.67 | -2.39 | -2.54 | -0.9 | -1.33 | 1.26 | -0.98 | -2 | -1.48 | 2.46 | -1.91 | 0.27 | -1.42 | 0.3225 | -0.115 | 0.3675 | 0.091118 | 0.830343 | 0.173373 | 0.995762 | 0.997988 | 0.994666 |
| Kif2a_F8VQ42_S568s_1 | Kif2a_F8VC | -0.28 | -0.12 | -0.6 | -1.16 | -0.1 | -0.8 | -0.42 | -0.8 | -0.32 | -0.11 | -1.9 | -0.74 | -0.61 | 0.23 | -1.71 | -1.19 | 0.12 | 1.11 | 1.76 | 0.54621 | 0.973933 | 0.611403 | 0.999187 | 0.997988 | 0.994666 |
| Kif2a_F8VQ42_S569s_1 | Kif2a_F8VC | -0.39 | -0.23 | -1.02 | -0.29 | -0.09 | -0.92 | 0.18 | -0.53 | -0.99 | -0.31 | -0.6 | -0.46 | -0.88 | -0.24 | -0.54 | -0.24 | 0.01 | -0.2275 | -0.28 | 0.979995 | 0.656299 | 0.705028 | 0.990434 | 0.997988 | 1 |
| Kif2a_P28740_S100s_1 | Kif2a_P287 | -0.55 | -2.16 | -0.94 | -0.39 | 0.17 | -0.15 | -0.31 | -0.13 | -0.08 | -0.69 | 0.63 | -0.12 | -0.72 | 0.94 | -0.08 | -0.21 | 0.1425 | -0.1075 | 0.0075 | 0.072482 | 0.044268 | 0.065991 | 0.989223 | 0.997988 | 0.994666 |
| Kif3a_B1AQZ2_S713s_1 | Kif3a_B1A( | -2.89 | -2.53 | -0.71 | -1.58 | -1.93 | -1.7 | -0.73 | -1.58 | -2.55 | -1.36 | -0.45 | -1.25 | -1.67 | -1.03 | -1.33 | -1.21 | 0.905 | 0.945 | 0.9925 | 0.223031 | 0.393985 | 0.391627 | 0.989223 | 0.997988 | 0.994666 |
| Kif3b_Q61771_S681s_1 | Kif3b_Q61 | -1.21 | 0 | 0.78 | 0.24 | -0.71 | -1.07 | 0.54 | 0.46 | -0.88 | -0.07 | 0.27 | -0.02 | -0.2 | -0.53 | 0.14 | 0.2 | 0.4425 | 0.525 | 0.6175 | 0.910091 | 0.781811 | 0.784417 | 0.995762 | 0.997988 | 0.999507 |
| Kif3c_O35066_S768s_1 | Kif3c_O35C | -3.08 | -4.39 | -3.55 | -3.74 | -2.61 | -3.55 | -2.76 | -3.45 | -4.36 | -3.77 | -1.81 | -3.74 | -2.26 | -2.58 | -3.27 | -4.08 | -0.1475 | -0.1275 | -0.05 | 0.191701 | 0.121847 | 0.637657 | 0.989223 | 0.997988 | 0.994666 |
| Kif5a_P33175_S931s_1 | Kif5a_P331 | -1.08 | -1.38 | -0.44 | -0.44 | -1.39 | 0.23 | -0.4 | -0.67 | -0.65 | -0.68 | -0.45 | -0.87 | -0.7 | -0.31 | -0.91 | -2.21 | 0.5975 | 0.27 | 0.6425 | 0.668492 | 0.488936 | 0.561222 | 0.989223 | 0.997988 | 0.996159 |
| Kif5b_Q61768_S933s_1 | Kif5b_Q61 | -0.97 | -0.41 | -0.14 | -0.21 | -0.97 | -0.54 | 0.14 | -0.09 | -0.79 | -0.72 | -0.47 | -0.22 | 0.3 | -0.27 | 0.56 | -0.44 | 0.2775 | 0.1725 | -0.1975 | 0.186257 | 0.834582 | 0.676105 | 0.995762 | 0.997988 | 0.994666 |
| Kif5b_Q61768_S945s_1 | Kif5b_Q61 | -0.45 | -0.03 | -0.17 | -0.03 | 0.56 | 0.58 | 0.21 | 0.54 | -0.57 | 0.08 | 0.82 | 0.58 | 0.52 | 0.28 | -0.21 | 0.22 | 0.0675 | -0.1175 | 0.47 | 0.189447 | 0.014954 | 0.262801 | 0.989223 | 0.997988 | 0.994666 |
| Kif5c_P28738_S934s_1 | Kif5c_P287 | 0.97 | -0.12 | -1.87 | -1.56 | -0.66 | 1.4 | -0.35 | -1.55 | -0.44 | 0.42 | -2.67 | -1.35 | 0.03 | -0.41 | -1.21 | -2.18 | 0.6425 | 0.3975 | 0.3725 | 0.684611 | 0.654897 | 0.662285 | 0.990434 | 0.997988 | 0.996159 |
| Kif5c_P28738_T396t_1 | Kif5c_P287 | -0.11 | 0.44 | -0.64 | 0.07 | 0.4 | 0.43 | 0.21 | -1.29 | 0.29 | 0.55 | -0.69 | -0.54 | 0.06 | 0.8 | -0.35 | -0.19 | 0.355 | -0.365 | -0.2975 | 0.70051 | 0.995468 | 0.921801 | 0.999616 | 0.997988 | 0.996159 |
| Kif7_J3QNW9_S662s_1 | Kif7_J3QN\ | -0.18 | 0.33 | -0.21 | 0.38 | -0.02 | 0.49 | 0.08 | 0.35 | 0.39 | 0.09 | 0.75 | -0.56 | 0.73 | -0.29 | -0.63 | 0.23 | -0.0025 | -0.0375 | 0.14 | 0.846599 | 0.569504 | 0.797003 | 0.989223 | 0.997988 | 0.999507 |
| Kifap3_P70188_S102s_1 | Kifap3_P70 | 0.52 | -0.63 | 0.19 | 0.06 | -1.24 | 0.24 | -1.1 | -0.05 | 0.14 | -0.06 | 0.73 | -0.4 | 0.81 | -1.84 | -0.12 | 0.06 | 0.145 | 0.0875 | -0.07 | 0.587663 | 0.194987 | 0.848369 | 0.989223 | 0.997988 | 0.995259 |
| Kifap3_P70188_S60s_1 | Kifap3_P70 | -0.4 | -0.19 | 1.28 | 1.07 | 0.19 | -0.55 | 1.62 | 1.01 | -0.57 | -0.26 | 0.34 | 0.9 | -0.24 | -0.32 | 1.4 | 1.67 | -0.5725 | 0.0675 | -0.3075 | 0.761764 | 0.824358 | 0.509985 | 0.995762 | 0.997988 | 0.9971 |
| Kifc2_G3X8Q6_S643s_1 | Kifc2_G3X8 | -0.11 | -0.07 | 0.01 | -0.19 | 0.06 | -0.55 | 0.03 | -0.91 | -0.03 | -0.75 | 0.91 | -0.33 | 0.45 | 0.17 | -0.47 | -0.19 | 0.1275 | -0.3375 | 0.1875 | 0.777087 | 0.372789 | 0.912602 | 0.989223 | 0.997988 | 0.99798 |
| Kit_P05532_S185s_1 | Kit_P05532 | -0.32 | 0.24 | 0.47 | 0.35 | 0.43 | 0.28 | 1.41 | 1.46 | 0.84 | -0.68 | 0.32 | 0.89 | 0.19 | 0.26 | 0.91 | 1.02 | -0.2525 | 0.04 | 0.08 | 0.223869 | 0.072071 | 0.695745 | 0.989223 | 0.997988 | 0.994666 |
| Kiz_Q3UXL4_S454s_1 | Kiz_Q3UXL | -0.56 | -0.5 | -1.01 | -1.42 | -1.73 | -1.17 | -1.92 | -2.54 | 0.99 | -0.95 | -2.25 | -1.82 | 1.78 | -3.02 | -1.55 | -1.5 | 0.71 | 0.1575 | 0.41 | 0.825604 | 0.020925 | 0.842273 | 0.989223 | 0.997988 | 0.999507 |
| Klc1_Q5UE59_S162s_1 | Klc1_Q5UE | -1.7 | -1.69 | -0.9 | -1.14 | 1.98 | -2.66 | -0.71 | -0.67 | -2 | -1.89 | -1.15 | -0.76 | -1.99 | -1.28 | -0.72 | -0.51 | -0.9675 | -0.135 | -0.2 | 0.560199 | 0.333424 | 0.801553 | 0.989223 | 0.997988 | 0.995259 |
| Klc1_Q5UE59_S460s_1 | Klc1_Q5UE | 0.02 | 0.49 | 0.29 | -0.2 | 0.73 | -0.16 | 0.54 | 0.11 | -0.14 | 0.39 | 0.29 | 0.29 | 0.37 | 0.7 | 0.06 | 0.54 | 0.8425 | -0.0925 | 0.2325 | 0.35329 | 0.589698 | 0.824402 | 0.989223 | 0.997988 | 0.994666 |
| Klc1_Q5UE59_S521s_1 | Klc1_Q5UE | 0.65 | 0.29 | 0.38 | 0.27 | 0.37 | 0.51 | 0.19 | 0.08 | 0.11 | 0.85 | -0.48 | 0.28 | -0.05 | -0.07 | -0.58 | 0.12 | 0.155 | 0.0575 | 0.2675 | 0.06608 | 0.617091 | 0.51847 | 0.989223 | 0.997988 | 0.994666 |
| Klc1_Q5UE59_S524s_1 | Klc1_Q5UE | 0.14 | 0.25 | 1.03 | 1.02 | 0.32 | -0.04 | 0.75 | 1.2 | -0.28 | 0.52 | -0.01 | 1.54 | -0.16 | 0.5 | 0.52 | 1.01 | -0.11 | -0.2075 | -0.5425 | 0.695762 | 0.883678 | 0.710956 | 0.996493 | 0.997988 | 0.996159 |
| Klc1_Q5UE59_S534s_1 | Klc1_Q5UE | 1.38 | -0.49 | 1.41 | 1.23 | -0.81 | 1.95 | 0.44 | 1 | 1.79 | 0.05 | 2.71 | 0.76 | 1.58 | -0.47 | 0.84 | 0.07 | -0.0525 | -0.1675 | -0.1425 | 0.525062 | 0.717053 | 0.512827 | 0.995437 | 0.997988 | 0.994666 |
| Klc2_D3YXZ3_S151s_1 | Klc2_D3YX | -2 | -2.74 | -0.34 | -0.72 | -0.94 | -0.57 | -0.82 | -0.77 | -0.64 | -2.31 | -0.17 | -0.38 | -0.93 | -1.53 | -0.36 | -0.69 | -0.2375 | 0.445 | -0.3775 | 0.3226 | 0.209611 | 0.401124 | 0.989223 | 0.997988 | 0.994666 |
| Klc2_D3YXZ3_S443s_1 | Klc2_D3YX | -0.25 | -0.18 | -0.27 | -0.35 | 0.77 | -0.52 | 0 | -0.49 | -0.43 | -0.41 | -0.32 | -0.26 | -1.32 | 0.45 | -0.15 | 0.29 | 0.675 | 0.575 | 0.5725 | 0.843635 | 0.529541 | 0.653089 | 0.989223 | 0.997988 | 0.999507 |
| Klc2_D3YXZ3_S551s_1 | Klc2_D3YX | 0.46 | -1.54 | -0.97 | -0.55 | 0.45 | 0.65 | -0.36 | 0.15 | -0.08 | -0.65 | 0.18 | -0.27 | -2.03 | -0.15 | -0.89 | -0.53 | 0.2025 | -0.0925 | 0.08 | 0.647327 | 0.075135 | 0.327418 | 0.989223 | 0.997988 | 0.995259 |
| Klc2_D3YXZ3_S552s_1 | Klc2_D3YX | -3.06 | -2.68 | -0.72 | -0.36 | 1.11 | -2.17 | -0.03 | -2.59 | -2.1 | -0.75 | -0.64 | -1.77 | -0.5 | -0.66 | -0.12 | 0.08 | 0.8725 | 0.445 | -0.25 | 0.049343 | 0.423238 | 0.579015 | 0.989223 | 0.997988 | 0.994666 |
| Klc2_D3YXZ3_S577s_1 | Klc2_D3YX | -1.8 | -1.62 | 0.49 | 0.51 | 0.08 | -1.97 | 0.94 | -0.76 | -3.04 | -1.8 | 1.85 | -0.57 | -1.19 | -0.45 | 0.62 | 0.97 | 0.785 | 0.39 | 1.405 | 0.419726 | 0.819921 | 0.791495 | 0.995762 | 0.997988 | 0.994666 |
| Klc2_D3YXZ3_S584s_1 | Klc2_D3YX | 1.4 | 0.96 | -0.18 | 0.71 | 1.08 | 1.53 | 0.52 | 0.87 | 0.41 | 1.02 | 0.2 | 0.25 | -0.45 | 0.64 | -0.73 | -0.52 | 0.1775 | -0.285 | 0.5925 | 0.047903 | 0.475883 | 0.517125 | 0.989223 | 0.997988 | 0.994666 |
| Klc2_D3YXZ3_S605s_1 | Klc2_D3YX | -2.32 | -1.03 | -0.87 | -1.71 | -0.31 | -1.96 | -0.62 | -1.28 | -2.07 | -1.82 | -1 | -1.35 | 0.07 | -0.37 | -1.21 | -0.89 | 0.2775 | -0.2525 | -0.9875 | 0.064889 | 0.351639 | 0.848566 | 0.989223 | 0.997988 | 0.994666 |
| Klc2_D3YXZ3_S606s_1 | Klc2_D3YX | 0.53 | -0.23 | -0.09 | 0.41 | 0.65 | 0.32 | 0.12 | 0.2 | -0.21 | -0.28 | -0.59 | -0.01 | -0.48 | 0.18 | -0.01 | 0.34 | 0.44 | -0.0775 | 0.8825 | 0.636773 | 0.531622 | 0.146114 | 0.989223 | 0.997988 | 0.995259 |
| Klc4_Q9DBS5_S590s_1 | Klc4_Q9DB | -0.35 | 0 | 0.96 | 0.58 | 1.03 | -0.18 | 0.7 | 0.76 | -1.06 | -0.09 | -0.09 | 0.67 | -0.07 | 0.66 | 0.07 | 0.14 | 0.1675 | -0.4275 | -0.1475 | 0.78593 | 0.47104 | 0.334848 | 0.989223 | 0.997988 | 0.999507 |
| Klc4_Q9DBS5_S611s_1 | Klc4_Q9DB | -0.13 | 0.04 | -0.02 | -0.05 | 0.45 | 0.16 | 0.59 | 0.5 | -0.48 | 0.42 | -0.07 | 0.04 | -0.67 | -0.03 | 0.47 | 0.29 | 0.28 | -0.44 | -0.0975 | 0.858441 | 0.04647 | 0.945686 | 0.989223 | 0.997988 | 0.999507 |
| Klf16_P58334_S103s_1 | Klf16_P583 | 1.93 | 1.89 | -0.18 | -0.12 | 1.2 | 2.21 | -0.02 | 0.3 | 1.48 | 1.9 | -0.12 | -0.75 | 0.86 | 1.32 | 0.15 | -0.14 | 0.465 | 0.0175 | 0.055 | 0.593909 | 0.950597 | 0.74568 | 0.999187 | 0.997988 | 0.995259 |
| Klf3_Q60980_S249s_1 | Klf3_Q609 | -0.23 | -0.49 | 0.11 | 0.08 | -0.13 | -0.12 | -0.22 | -0.35 | 0.34 | 0.37 | 0.69 | 0.18 | 0.29 | 0.12 | -0.23 | 0.12 | 0.0425 | -0.2525 | -0.3325 | 0.447415 | 0.751547 | 0.06008 | 0.995762 | 0.997988 | 0.994666 |
| Klhdc7a_A2APT9_S361s_1 | Klhdc7a_A | -0.46 | -0.1 | -0.08 | 0.12 | 0.02 | -0.19 | 0.47 | 0.01 | -0.42 | -0.27 | 0.48 | 0.09 | -0.81 | 0.12 | -0.32 | 0.11 | -0.0725 | 0.5275 | 0.2075 | 0.757608 | 0.406636 | 0.72512 | 0.989223 | 0.997988 | 0.9971 |
| Klhl22_D3YZH5_S18s_1 | Klhl22_D3Y | -1.41 | -0.49 | -1.81 | -1.25 | -1.2 | -0.56 | -0.87 | -1.57 | -0.5 | -0.57 | -0.59 | -1.3 | -0.28 | -0.84 | -1.22 | -1.18 | 0.2075 | 0.1 | -0.095 | 0.344164 | 0.592672 | 0.176335 | 0.989223 | 0.997988 | 0.994666 |
| Klhl26_Q8BGY4_S13s_1 | Klhl26_Q8E | 0.78 | 0.86 | -1.2 | -0.56 | 0.07 | 1.06 | -2.99 | 0.45 | 1.23 | 0.39 | -2.14 | 0.57 | 0.26 | -0.01 | -0.43 | -0.06 | 0.19 | 0.5 | 0.36 | 0.952025 | 0.720664 | 0.957658 | 0.995437 | 0.997988 | 0.999507 |
| Klhl9_Q6ZPT1_S603s_1 | Klhl9_Q6ZF | -0.23 | -0.32 | -0.78 | -0.47 | 0.43 | -0.51 | -0.18 | -0.39 | 0.3 | 0.21 | 0.28 | -0.96 | -0.15 | 0.4 | -0.59 | -0.57 | -0.3225 | 0.0425 | -0.03 | 0.483877 | 0.315492 | 0.257682 | 0.989223 | 0.997988 | 0.994666 |
| Kmt2a_P55200_S2539s_1 | Kmt2a_P55 | -0.58 | 0.32 | 0.04 | 0.29 | -0.03 | -0.35 | -0.16 | 1.01 | -0.26 | -0.17 | 0.12 | 0.32 | -0.27 | 0.51 | 0.13 | -0.08 | 0.2875 | 0.4075 | 0.2225 | 0.862205 | 0.784653 | 0.95898 | 0.995762 | 0.997988 | 0.999507 |
| Kmt2a_P55200_S2931s_1 | Kmt2a_P55 | 0.08 | 0.13 | -0.2 | -0.44 | -0.1 | 0.14 | -0.71 | -0.98 | 1.11 | 0.23 | 0.36 | -0.08 | 1.35 | 0.8 | 0.14 | -0.71 | 0.1 | -0.015 | 0.055 | 0.280859 | 0.339552 | 0.130646 | 0.989223 | 0.997988 | 0.994666 |
| Kmt2a_P55200_S3029s_1 | Kmt2a_P55 | -0.04 | -0.43 | 1.51 | 0.68 | 0.32 | 0.14 | 0.81 | 0.67 | 0.06 | 0.43 | 0.02 | 1.23 | -0.29 | -0.08 | 0.24 | 0.17 | -0.305 | 0.5125 | 0.5025 | 0.348051 | 0.89792 | 0.99175 | 0.996493 | 0.997988 | 0.994666 |
| Kmt2b_F6W623_S158s_1 | Kmt2b_F6\ | 0.75 | -1.39 | 1.44 | 1.25 | -0.59 | -0.76 | 0.07 | 0.82 | -0.21 | -0.28 | 1.02 | 1.89 | 1.17 | -0.31 | 2.3 | 1.69 | 0.055 | 0.005 | -0.42 | 0.368224 | 0.352671 | 0.901558 | 0.989223 | 0.997988 | 0.994666 |
| Kmt2d_Q6PDK2_S2318s_1 | Kmt2d_Q6 | -0.77 | -0.88 | 0.27 | 0.37 | -0.51 | -0.7 | 0.74 | 0.19 | -0.95 | -0.95 | 1.59 | 1 | -0.65 | -0.1 | 0.28 | 0.26 | -0.6275 | 0.0925 | 0.7 | 0.621277 | 0.679583 | 0.529436 | 0.993464 | 0.997988 | 0.995259 |
| Kmt2d_Q6PDK2_S3618s_1 | Kmt2d_Q6 | 1.02 | 0.56 | -2.47 | -0.25 | -0.15 | -0.19 | -2.13 | -0.29 | 2.53 | 0.79 | -0.12 | 0.27 | 0.18 | 0.13 | -0.68 | -1.11 | 0.1825 | 0.425 | 0.2 | 0.908893 | 0.613444 | 0.203421 | 0.989223 | 0.997988 | 0.999507 |
| Kmt2d_Q6PDK2_S4410s_1 | Kmt2d_Q6 | -2.15 | -2.27 | -2.52 | -2.52 | -1.84 | -2.55 | -1.77 | -2.46 | -2.19 | -2.33 | -1.61 | -2.08 | -2.16 | -1.6 | -3.15 | -2.02 | -0.405 | 1.1525 | -0.085 | 0.716516 | 0.440043 | 0.237094 | 0.989223 | 0.997988 | 0.996966 |
| Kmt2d_Q6PDK2_S4789s_1 | Kmt2d_Q6 | -2.38 | -1.88 | -2.96 | -3.24 | -1.57 | -2.61 | -4.11 | -3.81 | -1.51 | -1.11 | -0.51 | -3.88 | -0.98 | -1.65 | -3.14 | -2.54 | 0.21 | 0.3125 | 0.1325 | 0.322437 | 0.492111 | 0.248847 | 0.989223 | 0.997988 | 0.994666 |
| Kmt2e_Q3UG20_S263s_1 | Kmt2e_Q3 | -3.08 | -2.72 | -1.91 | -1.7 | -2.93 | -2.48 | -2.58 | -2.48 | 0.19 | -2.29 | -0.99 | -2.24 | 0.62 | -3.3 | -1.27 | -1.46 | -0.41 | 0.8625 | 0.5375 | 0.212979 | 0.452792 | 0.124002 | 0.989223 | 0.997988 | 0.994666 |
| Kndc1_Q0KK55_S794s_1 | Kndc1_Q0 | -0.19 | 0.09 | -0.08 | -0.26 | 0.45 | -0.36 | 0.46 | 0.22 | 0.1 | 0.06 | 0 | 0.12 | -0.34 | 0.28 | 0.2 | 0.49 | -0.265 | 1.02 | 1 | 0.34365 | 0.258058 | 0.403966 | 0.989223 | 0.997988 | 0.994666 |
| Kndc1_Q0KK55_S843s_1 | Kndc1_Q0 | 0 | -2.02 | -0.75 | -0.37 | -1.25 | -0.95 | -1.58 | -1.22 | 1.02 | -0.23 | -2.08 | -1.05 | 1.74 | -2.62 | -1.25 | -1.88 | 0.3025 | 0.18 | 0.2675 | 0.813241 | 0.296664 | 0.778629 | 0.989223 | 0.997988 | 0.999507 |
| Kndc1_Q0KK55_S951s_1 | Kndc1_Q0 | -0.33 | 1 | -1.36 | -0.04 | 0.71 | 0.37 | -1.24 | -0.52 | 1.2 | 0.42 | -0.15 | -0.75 | 1.3 | -0.25 | 0.04 | -0.09 | -0.465 | 0.2 | -0.2175 | 0.444154 | 0.983013 | 0.541063 | 0.999187 | 0.997988 | 0.994666 |
| Kndc1_Q0KK55_S951s_2 | Kndc1_Q0 | -0.28 | -0.54 | 0.01 | 0.06 | -0.13 | -0.68 | 0.65 | 1.47 | -0.34 | -0.67 | 1.08 | -0.51 | -0.83 | 0.12 | -0.71 | 1.07 | 0.0125 | 0.3625 | 0.4325 | 0.823937 | 0.274882 | 0.852825 | 0.989223 | 0.997988 | 0.999507 |
| Kndc1_Q0KK55_S967s_1 | Kndc1_Q0 | -0.15 | -1.15 | -1.6 | -1.88 | -0.93 | -0.69 | -0.71 | -1.78 | -1.06 | -0.2 | -2.96 | -1.17 | 0.43 | 0.15 | -1.63 | -0.75 | 0.515 | 0.0775 | 0.1 | 0.201805 | 0.699127 | 0.809152 | 0.994893 | 0.997988 | 0.994666 |
| Kndc1_Q0KK55_S967s_2 | Kndc1_Q0 | -0.28 | -0.54 | 0.01 | 0.06 | -0.13 | -0.68 | 0.65 | 1.47 | -0.34 | -0.67 | 1.08 | -0.51 | -0.83 | 0.12 | -0.71 | 1.07 | 0.1675 | -0.1525 | 0.745 | 0.823937 | 0.274882 | 0.852825 | 0.989223 | 0.997988 | 0.999507 |
| Kpna6_O35345_S6s_1 | Kpna6_O35 | -1.88 | -1.75 | -1.83 | -2.81 | -1.6 | -2.25 | -2.53 | -2.31 | -1.11 | -1.71 | -0.99 | -2.31 | 0.21 | -1.21 | -1.79 | -1.98 | 0.515 | 0.0775 | 0.1 | 0.117629 | 0.750675 | 0.193813 | 0.995762 | 0.997988 | 0.994666 |
| Kri1_V9GXJ9_S24s_1 | Kri1_V9GX | 1.2 | 2.2 | 1.19 | 0.81 | -0.5 | 1.07 | 0.21 | -0.34 | 0.18 | 0.81 | 1.83 | 1.03 | 1.11 | -0.05 | -0.14 | 1.27 | -0.105 | 0.5375 | 0.875 | 0.107271 | 0.017044 | 0.386993 | 0.989223 | 0.997988 | 0.994666 |
| Krit1_Q6S5J6_S22s_1 | Krit1_Q6S5 | 0.51 | -1.01 | 0.03 | -0.49 | -1 | -0.25 | -1.03 | -0.77 | 0.39 | -1.06 | -0.67 | -0.48 | 0.71 | -1.68 | -0.64 | -0.9 | -1.24 | -0.3875 | -0.8025 | 0.48833 | 0.17963 | 0.623711 | 0.989223 | 0.997988 | 0.994666 |
| Krt8_P11679_S24s_1 | Krt8_P116 | -2.47 | -2.74 | -2.75 | -2.78 | -2.56 | -2.09 | -5.51 | -4.32 | -1.7 | -1.66 | -1.76 | -3.48 | -0.52 | -1.72 | -2.11 | -2.5 | -0.5225 | -0.215 | -0.3875 | 0.045015 | 0.203494 | 0.239493 | 0.989223 | 0.997988 | 0.994666 |
| Krtap9-1_Q64526_S4s_1 | Krtap9-1_C | -0.08 | 0.2 | -0.8 | 0.06 | -1.01 | -0.58 | -1.84 | -0.64 | 0.63 | 0.31 | -0.36 | -1.88 | 0.07 | -0.4 | -0.29 | -0.23 | -0.935 | 0.535 | 0.9725 | 0.849577 | 0.037394 | 0.76011 | 0.989223 | 0.997988 | 0.999507 |
| Ksr1_Q5SXE4_S320s_1 | Ksr1_Q5SX | -0.59 | -0.79 | -1.3 | -0.32 | -0.37 | -0.7 | -0.12 | -0.95 | -1.01 | -1.47 | 0.85 | -1.18 | -1.16 | -0.03 | -0.72 | -0.07 | -0.8625 | -0.17 | -0.0575 | 0.489946 | 0.480133 | 0.928179 | 0.989223 | 0.997988 | 0.994666 |
| Ksr1_Q5SXE4_S392s_1 | Ksr1_Q5SX | -0.04 | -0.38 | 0.22 | 0.15 | 0.18 | 0.16 | 0.23 | 0.32 | -0.21 | -0.41 | 0.36 | 0.1 | -0.17 | 0.52 | -0.19 | 0.19 | 0.215 | 0.0475 | 0.255 | 0.728397 | 0.306022 | 0.91966 | 0.989223 | 0.997988 | 0.9971 |
| Ksr2_M0QW59_T273t_1 | Ksr2_M0Q | 0.59 | 0.78 | -0.49 | -0.78 | 0.89 | 0.13 | 0.14 | -1.08 | 0.52 | 0.1 | 0.82 | -1.81 | 0.67 | 1.13 | -1.48 | 0.06 | 0.235 | -0.0275 | 0.1 | 0.910701 | 0.992218 | 0.854922 | 0.999187 | 0.997988 | 0.999507 |
| Ksr2_M0QW59_T291t_1 | Ksr2_M0Q | -1.09 | -2.55 | -1.16 | -1.05 | -0.61 | -1.13 | -1.1 | -1.12 | -1.01 | -1.08 | 0.97 | -0.61 | -0.98 | -0.55 | -1.8 | -0.23 | -0.005 | -0.1175 | 0.07 | 0.247782 | 0.22938 | 0.090418 | 0.989223 | 0.997988 | 0.994666 |
| Kti12_Q9D1R2_T156t_1 | Kti12_Q9D | -0.66 | -0.98 | -1.05 | -1.28 | 0.11 | -1.02 | -1.65 | -0.77 | -0.82 | -0.51 | -1.94 | -0.66 | -0.77 | -0.52 | -0.82 | -2.17 | 0.4725 | 1.03 | 0.5725 | 0.846478 | 0.673453 | 0.977873 | 0.992876 | 0.997988 | 0.999507 |
| Ktn1_Q61595_S124s_1 | Ktn1_Q615 | 0.05 | -0.27 | 0.14 | 0.25 | 0.25 | -0.05 | 0.6 | 0.67 | 0.12 | 0.05 | 0.17 | -0.23 | 0.03 | -0.04 | 0.98 | 0.29 | 0.16 | 0.01 | -0.0775 | 0.391391 | 0.221119 | 0.948956 | 0.989223 | 0.997988 | 0.994666 |
| Ktn1_Q61595_S75s_1 | Ktn1_Q615 | -1.6 | -1.09 | -1.89 | -4.83 | -2.13 | -1.18 | -2.11 | -2.73 | -0.09 | -1.14 | -1.61 | -2.05 | 0.81 | -0.55 | -1.32 | -0.88 | 0.325 | -0.015 | 0.2725 | 0.049213 | 0.690377 | 0.199869 | 0.994672 | 0.997988 | 0.994666 |
| L1cam_Q6PGJ3_S1165s_1 | L1cam_Q6 | -2.79 | -1.69 | -2.94 | -2.44 | -1.67 | -3.25 | -1.64 | -2.42 | -3.43 | -2.72 | -1.54 | -1.89 | -1.85 | -1.94 | -2.13 | -2.36 | 0.315 | 1.13 | 1.8675 | 0.259366 | 0.621133 | 0.884383 | 0.989223 | 0.997988 | 0.994666 |
| L1cam_Q6PGJ3_S1180s_1 | L1cam_Q6 | 0.5 | 1.41 | 1.59 | 1.11 | 0.02 | 1.29 | 0.27 | 0.54 | 2.09 | 0.93 | 1.84 | 0.94 | 0.73 | 0.66 | 1.79 | 1.13 | 0.22 | 0.07 | 0.395 | 0.840056 | 0.110778 | 0.449376 | 0.989223 | 0.997988 | 0.999507 |
| L1cam_Q6PGJ3_S1180s_2 | L1cam_Q6 | -0.16 | 0.87 | 0.58 | -0.18 | -0.39 | 1 | -0.33 | -0.25 | 1.43 | 0.08 | 0.76 | 0.33 | -0.74 | -0.05 | 0.86 | 0.46 | -0.6225 | 0.2975 | -0.075 | 0.736591 | 0.511792 | 0.358771 | 0.989223 | 0.997988 | 0.9971 |
| L1cam_Q6PGJ3_S1183s_1 | L1cam_Q6 | 0.47 | 0.81 | 1.04 | 0.99 | -0.49 | 0.96 | 0.02 | 0.7 | 1.82 | 0.86 | 0.67 | 0.88 | 0.95 | -0.47 | 0.51 | -0.13 | -0.27 | 0.3725 | -0.145 | 0.118893 | 0.15808 | 0.476831 | 0.989223 | 0.997988 | 0.994666 |
| L1cam_Q6PGJ3_S1183s_2 | L1cam_Q6 | -0.16 | 0.87 | 0.58 | -0.18 | -0.39 | 1 | -0.33 | -0.25 | 1.43 | 0.08 | 0.76 | 0.33 | -0.74 | -0.05 | 0.86 | 0.46 | -0.53 | 0.23 | -0.6125 | 0.736591 | 0.511792 | 0.358771 | 0.989223 | 0.997988 | 0.9971 |
| L1cam_Q6PGJ3_S1250s_1 | L1cam_Q6 | 0.15 | 0.85 | 0.24 | 0.39 | 0.82 | -0.09 | 1.16 | 0.73 | 0.11 | 0.3 | 0.28 | 0.05 | -0.42 | 0.42 | 0.83 | -0.11 | -0.27 | 0.3725 | -0.145 | 0.518908 | 0.448624 | 0.376954 | 0.989223 | 0.997988 | 0.994666 |
| L1cam_Q6PGJ3_T1174t_1 | L1cam_Q6 | -0.56 | 0.29 | 1.3 | -2.87 | 1.08 | -0.29 | 0.85 | 0.21 | -2.16 | -0.06 | 0.58 | 1.77 | -1.16 | -1.7 | -0.69 | 0.35 | 0.2475 | -0.2225 | -0.2275 | 0.696496 | 0.277864 | 0.646097 | 0.989223 | 0.997988 | 0.996159 |
| Lactb2_Q99KR3_T279t_1 | Lactb2_Q9 | -0.31 | -0.38 | -0.21 | 0.06 | -0.33 | -1.16 | 0.31 | 0.05 | -1.72 | -0.48 | 0.61 | 0.05 | -0.42 | -0.71 | -0.74 | -0.81 | 0.9225 | 0.4925 | -0.34 | 0.088393 | 0.83154 | 0.716649 | 0.995762 | 0.997988 | 0.994666 |
| Lad1_P57016_T19t_1 | Lad1_P570 | -0.88 | -1.16 | -1.57 | -2.08 | -1.7 | -1.22 | -2.27 | -1.82 | -1.65 | -0.86 | 0.24 | -1.7 | -1.15 | -0.77 | -2.23 | -2.14 | -0.0725 | -0.175 | -0.46 | 0.732828 | 0.34871 | 0.39407 | 0.989223 | 0.997988 | 0.9971 |
| Lamtor1_Q9CQ22_T28t_1 | Lamtor1_Q | 0.84 | 0.89 | 0.37 | 0.73 | 1.08 | 0.55 | 0.91 | 0.53 | 0.29 | 0.99 | 0.24 | 0.34 | 0.44 | 1.12 | 0.28 | 0.35 | -0.33 | 0.43 | -0.15 | 0.588107 | 0.80485 | 0.380129 | 0.995762 | 0.997988 | 0.995259 |
| Lancl2_F6RJV6_S302s_1 | Lancl2_F6R | 0.51 | 0.99 | 0.13 | 0.13 | 0.38 | 1.06 | -0.35 | 0.28 | 0.49 | 0.89 | -0.2 | -0.16 | 0.6 | 0.36 | -0.99 | 0.41 | 0.06 | -0.2425 | -0.16 | 0.417587 | 0.783247 | 0.599381 | 0.995762 | 0.997988 | 0.994666 |
| Larp1_Z4YJT3_S1023s_1 | Larp1_Z4YJ | 2.86 | 0.19 | 0.9 | -0.63 | 1.04 | 2.89 | 0.6 | 1.46 | 1.61 | 2.31 | 1.16 | 2.23 | 2.44 | 0.22 | 2.52 | 2.25 | -0.0975 | -0.185 | -0.345 | 0.227715 | 0.403494 | 0.18573 | 0.989223 | 0.997988 | 0.994666 |
| Larp1_Z4YJT3_S494s_1 | Larp1_Z4YJ | -0.17 | -0.25 | 0.16 | -0.26 | -0.83 | 0.58 | -1.11 | -0.62 | 0.9 | -0.35 | -1.22 | -0.01 | 1.35 | -1.77 | -0.28 | -0.1 | 0.6675 | 0.9975 | 1.0275 | 0.905606 | 0.345313 | 0.926809 | 0.989223 | 0.997988 | 0.999507 |
| Larp1_Z4YJT3_S494s_2 | Larp1_Z4YJ | -1.78 | -2.23 | -1.65 | -2 | -2.66 | -1.15 | -2 | -1.64 | 0.03 | -2.09 | -1.56 | -1.17 | -0.47 | -2.36 | -1.86 | -1.67 | -0.365 | -0.04 | -0.07 | 0.445636 | 0.87938 | 0.136935 | 0.996493 | 0.997988 | 0.994666 |
| Larp1_Z4YJT3_S498s_1 | Larp1_Z4YJ | 0.56 | 0.09 | 0.69 | 0.71 | -1.06 | 1.07 | -0.29 | 0.03 | 2.59 | 0.65 | -0.16 | 0.54 | 2.43 | -0.89 | 0.14 | 0.07 | 0.0525 | 0.7175 | 0.325 | 0.907959 | 0.208969 | 0.490138 | 0.989223 | 0.997988 | 0.999507 |
| Larp1_Z4YJT3_S498s_2 | Larp1_Z4YJ | -1.78 | -2.23 | -1.65 | -2 | -2.66 | -1.15 | -2 | -1.64 | 0.03 | -2.09 | -1.56 | -1.17 | -0.47 | -2.36 | -1.86 | -1.67 | -0.575 | 0.3925 | -0.075 | 0.445636 | 0.87938 | 0.136935 | 0.996493 | 0.997988 | 0.994666 |
| Larp1_Z4YJT3_S68s_1 | Larp1_Z4YJ | -2.57 | -2.69 | -2.86 | -2.57 | -2.04 | -2.59 | -2.01 | -2.87 | -1.15 | -1.99 | -1.25 | -2.43 | -0.63 | -2.01 | -2.12 | -2.28 | 0.0525 | 0.7175 | 0.325 | 0.04183 | 0.284637 | 0.016382 | 0.989223 | 0.997988 | 0.994666 |
| Larp1_Z4YJT3_S68s_2 | Larp1_Z4YJ | -2.46 | -1.96 | -2.17 | -1.79 | -1.94 | -2.3 | -1.63 | -2.33 | -1.4 | -1.38 | -0.57 | -1.29 | -1.4 | -0.93 | -1.73 | -1.21 | 0.295 | 0.9675 | 0.9125 | 0.022496 | 0.86525 | 0.009506 | 0.996493 | 0.997988 | 0.994666 |
| Larp1_Z4YJT3_S743s_1 | Larp1_Z4YJ | -0.76 | -0.71 | -1.69 | -0.93 | -0.94 | -1.28 | -1.03 | -0.99 | -1.48 | -1.05 | -0.78 | -1.38 | -1.43 | -0.59 | -1.23 | -0.92 | 0.045 | 0.935 | 0.7775 | 0.951955 | 0.892367 | 0.632051 | 0.996493 | 0.997988 | 0.999507 |
| Larp1_Z4YJT3_S751s_1 | Larp1_Z4YJ | 0.06 | -1.22 | -0.16 | 0.05 | -0.27 | -0.44 | 0.75 | 0.1 | -0.17 | -0.15 | -0.19 | -0.16 | -0.9 | 0.25 | 0.05 | -0.08 | -0.0375 | -0.15 | -0.02 | 0.714356 | 0.377312 | 0.650827 | 0.989223 | 0.997988 | 0.996966 |
| Larp1_Z4YJT3_S81s_1 | Larp1_Z4YJ | -0.69 | -0.08 | -0.15 | 0.02 | -0.41 | -0.36 | -0.12 | -0.06 | -0.06 | -0.09 | 0.58 | -0.1 | 0.37 | 0.34 | 0.06 | -0.05 | 0.3525 | 0.15 | 0.1475 | 0.162728 | 0.959002 | 0.289682 | 0.999187 | 0.997988 | 0.994666 |
| Larp1_Z4YJT3_S81s_2 | Larp1_Z4YJ | -2.46 | -1.96 | -2.17 | -1.79 | -1.94 | -2.3 | -1.63 | -2.33 | -1.4 | -1.38 | -0.57 | -1.29 | -1.4 | -0.93 | -1.73 | -1.21 | -0.0125 | 0.3075 | 0.405 | 0.022496 | 0.86525 | 0.009506 | 0.996493 | 0.997988 | 0.994666 |
| Larp1_Z4YJT3_S828s_1 | Larp1_Z4YJ | -0.33 | 0.05 | -0.18 | -0.02 | 0.25 | -0.15 | -0.15 | -0.44 | -0.16 | -0.04 | -0.22 | -0.32 | -0.23 | 0.25 | -0.18 | -0.11 | 0.045 | 0.935 | 0.7775 | 0.834289 | 0.991513 | 0.766414 | 0.999187 | 0.997988 | 0.999507 |
| Larp1_Z4YJT3_T503t_1 | Larp1_Z4YJ | -1.96 | -1.91 | -2.1 | -3.29 | -0.75 | -1.85 | -1.7 | -1.55 | -3.29 | -2.08 | 0.49 | -2.6 | -1.11 | -0.87 | -1.77 | -2.1 | -0.0025 | -0.065 | 0.0525 | 0.071754 | 0.053887 | 0.577822 | 0.989223 | 0.997988 | 0.994666 |
| Larp1_Z4YJT3_T626t_1 | Larp1_Z4YJ | 0.44 | 0.15 | 0.09 | -0.3 | 0.45 | -0.1 | 0.52 | -0.33 | -0.72 | 0.78 | -0.66 | -0.22 | -0.4 | -0.35 | -0.29 | 0 | 0.8525 | 0.445 | 0.8525 | 0.205587 | 0.88972 | 0.440445 | 0.996493 | 0.997988 | 0.994666 |
| Larp1b_Q3TQS1_S60s_1 | Larp1b_Q3 | 1.05 | 0.07 | 1.22 | -0.15 | 0.8 | 0.41 | 0.65 | 0.84 | 0.55 | 1.07 | 0.53 | 0.4 | 0.01 | 0.75 | 0.93 | 1.17 | 0.04 | -0.3 | -0.355 | 0.692986 | 0.721603 | 0.81242 | 0.995437 | 0.997988 | 0.996159 |
| Larp4_E9Q066_S578s_1 | Larp4_E9Q | -0.31 | -0.16 | 0.27 | 0.57 | 0.11 | -0.45 | 0.9 | -0.27 | -0.38 | -0.67 | 1.19 | 0.14 | -0.34 | -0.24 | 0.31 | 0.62 | 0.1275 | 0.09 | 0.1675 | 0.98824 | 0.955563 | 0.959264 | 0.999187 | 0.997988 | 1 |
| Larp4b_Q6A0A2_S517s_1 | Larp4b_Q6 | -0.32 | 0.08 | 0.75 | 0.33 | 0.15 | -0.06 | 0.67 | -0.27 | -0.61 | 0.08 | 0.85 | -0.58 | -0.12 | -0.76 | 0.73 | 0.44 | -0.02 | -0.0225 | -0.005 | 0.733697 | 0.783137 | 0.50396 | 0.995762 | 0.997988 | 0.9971 |
| Larp4b_Q6A0A2_S519s_1 | Larp4b_Q6 | -1.16 | -1.31 | -1.32 | -1.13 | -0.59 | -1.15 | -0.32 | -0.98 | -1.01 | -1.18 | -0.82 | -1.56 | -0.75 | -0.66 | -1.22 | -0.99 | -0.0875 | -0.275 | -0.1375 | 0.213773 | 0.085881 | 0.721624 | 0.989223 | 0.997988 | 0.994666 |
| Larp4b_Q6A0A2_S637s_1 | Larp4b_Q6 | -1.96 | -1.71 | -2.45 | -2.39 | -1.96 | -1.4 | -2.54 | -2.79 | -0.14 | -1.82 | -3.86 | -2.12 | 0.99 | -2.36 | -1.51 | -1.2 | 0.47 | 0.0875 | 0.325 | 0.119607 | 0.899607 | 0.839817 | 0.996493 | 0.997988 | 0.994666 |
| Larp4b_Q6A0A2_S655s_1 | Larp4b_Q6 | -1.1 | -2.28 | -0.8 | -0.92 | -1.11 | -0.26 | -0.73 | -0.83 | -0.76 | -1.08 | -2.52 | -0.77 | 0.19 | -3.22 | -0.75 | -0.42 | -0.045 | 0.1425 | 1.1075 | 0.759144 | 0.172279 | 0.988188 | 0.989223 | 0.997988 | 0.9971 |
| Larp4b_Q6A0A2_S655s_2 | Larp4b_Q6 | -1.49 | -1.59 | -1.81 | -1.81 | -1.03 | -1.41 | -1.57 | -1.37 | -0.1 | -2.52 | -1.58 | -1.31 | 0.21 | -1.37 | -2.08 | -2.04 | 0.5425 | -0.0075 | 0.225 | 0.491768 | 0.162211 | 0.539139 | 0.989223 | 0.997988 | 0.994666 |
| Larp4b_Q6A0A2_T651t_1 | Larp4b_Q6 | 0.09 | 0.77 | 0.71 | 0.68 | -0.55 | 0.32 | 0.33 | -0.09 | 1.17 | 0.99 | -0.26 | 0.4 | 2.44 | -0.44 | 1.12 | 1.22 | 0.33 | 0.2975 | 0.355 | 0.36527 | 0.079585 | 0.972912 | 0.989223 | 0.997988 | 0.994666 |
| Larp4b_Q6A0A2_T651t_2 | Larp4b_Q6 | -1.49 | -1.59 | -1.81 | -1.81 | -1.03 | -1.41 | -1.57 | -1.37 | -0.1 | -2.52 | -1.58 | -1.31 | 0.21 | -1.37 | -2.08 | -2.04 | -0.56 | 0.0125 | 0.5225 | 0.491768 | 0.162211 | 0.539139 | 0.989223 | 0.997988 | 0.994666 |
| Larp7_Q05CL8_S253s_2 | Larp7_Q05 | -0.04 | 0.18 | -0.19 | -0.15 | 1.67 | -0.63 | -2.12 | -0.48 | -0.67 | 0.41 | 0.13 | -1.72 | 0.78 | 1.05 | -0.17 | -0.34 | 0.33 | 0.2975 | 0.355 | 0.321275 | 0.625383 | 0.384027 | 0.989223 | 0.997988 | 0.994666 |
| Larp7_Q05CL8_S256s_2 | Larp7_Q05 | -0.04 | 0.18 | -0.19 | -0.15 | 1.67 | -0.63 | -2.12 | -0.48 | -0.67 | 0.41 | 0.13 | -1.72 | 0.78 | 1.05 | -0.17 | -0.34 | -0.34 | -0.4125 | 0.38 | 0.321275 | 0.625383 | 0.384027 | 0.989223 | 0.997988 | 0.994666 |
| Las1l_A2BE28_S20s_1 | Las1l_A2BE | -1.6 | -0.77 | -0.68 | -2.01 | -2.23 | -1.61 | -2.31 | -1.54 | 0.55 | -2.24 | -1.93 | -0.89 | 1.45 | -1.61 | -1.62 | -0.72 | -0.34 | -0.4125 | 0.38 | 0.376494 | 0.104467 | 0.830757 | 0.989223 | 0.997988 | 0.994666 |
| Las1l_A2BE28_S238s_1 | Las1l_A2BE | -2.2 | -1.35 | -2.86 | -2.85 | -2.41 | -1.74 | -4.13 | -3 | -1.51 | -1.97 | -1.34 | -3 | -0.82 | -1.43 | -2.37 | -2.7 | -0.6575 | 0.1375 | 0.64 | 0.365535 | 0.376892 | 0.467581 | 0.989223 | 0.997988 | 0.994666 |
| Lats1_Q8BYR2_S204s_1 | Lats1_Q8B | -3.77 | -2.14 | -1.75 | -3.06 | -1.87 | -2.14 | -3.23 | -2.36 | -1.69 | -1.47 | -0.91 | -2.16 | 0.46 | -1.76 | -4.47 | -2.08 | -0.505 | 0.36 | 0.485 | 0.46397 | 0.576532 | 0.045547 | 0.989223 | 0.997988 | 0.994666 |
| Lbr_Q3U9G9_S101s_1 | Lbr_Q3U9C | -2.03 | -1.77 | -0.89 | -0.52 | -2.53 | -1.24 | -1.53 | -1.82 | -0.49 | -1.57 | -0.63 | -1.38 | 0.21 | -2.11 | -0.74 | -0.43 | 0.28 | 1.1225 | 0.7175 | 0.350962 | 0.279824 | 0.516283 | 0.989223 | 0.997988 | 0.994666 |
| Lbr_Q3U9G9_S67s_1 | Lbr_Q3U9C | -0.71 | 0.13 | 0.84 | -0.31 | -0.11 | 0.19 | -0.44 | -0.48 | 1.33 | -0.17 | -1.4 | 0.29 | 0.99 | -0.81 | -0.26 | -0.15 | -0.4775 | 0.285 | 0.535 | 0.925313 | 0.590179 | 0.966776 | 0.989223 | 0.997988 | 0.999507 |
| Lbr_Q3U9G9_S67s_2 | Lbr_Q3U9C | -1.36 | -0.77 | -0.58 | -1.07 | -1.65 | -0.36 | -1.83 | -1.45 | 0.33 | -1.26 | -0.95 | -0.68 | 0.37 | -1.46 | -1.11 | -1.21 | -0.1975 | 0.025 | -0.045 | 0.833308 | 0.318049 | 0.438175 | 0.989223 | 0.997988 | 0.999507 |
| Lbr_Q3U9G9_S71s_2 | Lbr_Q3U9C | -1.36 | -0.77 | -0.58 | -1.07 | -1.65 | -0.36 | -1.83 | -1.45 | 0.33 | -1.26 | -0.95 | -0.68 | 0.37 | -1.46 | -1.11 | -1.21 | -0.3775 | 0.305 | 0.0925 | 0.833308 | 0.318049 | 0.438175 | 0.989223 | 0.997988 | 0.999507 |
| Lca5_G5E887_S7s_1 | Lca5_G5E8 | -0.55 | -0.14 | 0.38 | 0.09 | -1 | -0.4 | -0.9 | -0.67 | -0.23 | -0.09 | 0.45 | -0.37 | 0.38 | -0.69 | 0.48 | -0.09 | -0.3775 | 0.305 | 0.0925 | 0.833927 | 0.030668 | 0.986842 | 0.989223 | 0.997988 | 0.999507 |
| Lcp1_Q61233_S5s_1 | Lcp1_Q612 | -0.27 | -0.34 | -0.48 | 0.05 | -0.91 | 0.13 | -0.1 | -0.03 | 0.29 | -0.73 | -0.02 | -0.5 | 0.96 | -0.17 | -0.12 | -0.64 | -0.6875 | -0.005 | 0.075 | 0.480833 | 0.910375 | 0.946316 | 0.997335 | 0.997988 | 0.994666 |
| Ldb1_D3Z1C5_S266s_1 | Ldb1_D3Z1 | -4.2 | -3.29 | -3.03 | -3.76 | -4.13 | -3.65 | -3.13 | -3 | -3.03 | -2.72 | -2.24 | -2.72 | -1.34 | -3.53 | -3.31 | -3.31 | 0.0325 | 0.02 | 0.2675 | 0.213332 | 0.799065 | 0.022072 | 0.995762 | 0.997988 | 0.994666 |
| Ldb3_Q9JKS4_S179s_1 | Ldb3_Q9JK | 0.93 | -0.7 | -0.17 | -1.21 | 0.31 | 0.81 | -0.76 | -0.62 | 1.24 | 0.15 | -1.76 | 0.37 | -0.31 | 1.27 | -0.54 | 0.07 | 0.0925 | 0.8925 | 0.6975 | 0.472559 | 0.679935 | 0.684035 | 0.993464 | 0.997988 | 0.994666 |
| Lect1_G5E8Z6_S224s_3 | Lect1_G5E | 6.14 | 3.25 | 4.98 | 1.17 | 1.75 | 6.59 | 4.36 | 3.99 | 2.07 | 5.48 | 0.53 | 4.77 | 2.96 | -2.13 | 6.39 | 5.77 | 0.2225 | 0.2875 | 0.41 | 0.737845 | 0.819651 | 0.629441 | 0.995762 | 0.997988 | 0.9971 |
| Lect1_G5E8Z6_T228t_3 | Lect1_G5E | 6.14 | 3.25 | 4.98 | 1.17 | 1.75 | 6.59 | 4.36 | 3.99 | 2.07 | 5.48 | 0.53 | 4.77 | 2.96 | -2.13 | 6.39 | 5.77 | 0.2875 | -0.6725 | -0.6375 | 0.737845 | 0.819651 | 0.629441 | 0.995762 | 0.997988 | 0.9971 |
| Lect1_G5E8Z6_T229t_3 | Lect1_G5E | 6.14 | 3.25 | 4.98 | 1.17 | 1.75 | 6.59 | 4.36 | 3.99 | 2.07 | 5.48 | 0.53 | 4.77 | 2.96 | -2.13 | 6.39 | 5.77 | 0.2875 | -0.6725 | -0.6375 | 0.737845 | 0.819651 | 0.629441 | 0.995762 | 0.997988 | 0.9971 |
| Lemd3_E9QP59_S331s_1 | Lemd3_E9( | 1.11 | -1.49 | -1.61 | -0.71 | -1.07 | 1.34 | -1.24 | -0.7 | 0.42 | -1.25 | -0.77 | -0.96 | -0.45 | -1.12 | -1.38 | -1.21 | 0.2875 | -0.6725 | -0.6375 | 0.548174 | 0.735002 | 0.957611 | 0.995762 | 0.997988 | 0.994666 |
| Leng8_D3YWS8_S352s_1 | Leng8_D3Y | -0.46 | -0.75 | -0.83 | -0.37 | -0.97 | -0.28 | -2.13 | -1.56 | -0.07 | -0.11 | -1.46 | -1.12 | 0.88 | -1.45 | -0.91 | -1.47 | 0.2575 | 0.035 | -0.365 | 0.798464 | 0.136007 | 0.816125 | 0.989223 | 0.997988 | 0.999507 |
| Leo1_Q5XJE5_S162s_1 | Leo1_Q5XJ | -4.68 | -4.9 | -3.37 | -9.86 | -4 | -4.98 | -4.23 | -3.59 | -4.13 | -4.28 | -1.87 | -3.69 | -2.01 | -5.76 | -2.51 | -3.69 | -0.6325 | -0.0875 | -0.135 | 0.14417 | 0.247982 | 0.125031 | 0.989223 | 0.997988 | 0.994666 |
| Leo1_Q5XJE5_S198s_1 | Leo1_Q5XJ | -1.96 | -2.53 | -2.98 | -4.07 | -2.29 | -2.07 | -3.03 | -3.04 | -1.36 | -1.61 | -0.8 | -2.95 | -0.28 | -2.76 | -3.42 | -2.93 | 1.5025 | 2.21 | 2.21 | 0.474443 | 0.561676 | 0.064951 | 0.989223 | 0.997988 | 0.994666 |
| Leo1_Q5XJE5_S206s_1 | Leo1_Q5XJ | -3.88 | -4.49 | -3.2 | -4.89 | -3.35 | -4.55 | -4.61 | -4.35 | -4.34 | -3.93 | -3.9 | -4.07 | -2.44 | -4.47 | -6.09 | -4.66 | 0.2775 | 1.205 | 0.5375 | 0.687833 | 0.8212 | 0.886342 | 0.995762 | 0.997988 | 0.996159 |
| Leo1_Q5XJE5_T189t_1 | Leo1_Q5XJ | -0.79 | -2.51 | -1.1 | -1.72 | -3.73 | -1.37 | -1.67 | -1.43 | -0.63 | -2.96 | -0.02 | -1.53 | 0.84 | -2.74 | 0.65 | -1.69 | -0.1 | 0.055 | -0.3 | 0.35809 | 0.401044 | 0.7164 | 0.989223 | 0.997988 | 0.994666 |
| Letmd1_Q924L1_S35s_1 | Letmd1_Q | 3.42 | 1.27 | 0.7 | 0.64 | 0.97 | 3.21 | 0.67 | 0.61 | 0.74 | 2.46 | 0.27 | 0.46 | -0.51 | -0.43 | 1.13 | 0.81 | -0.52 | 0.245 | 0.795 | 0.096222 | 0.8564 | 0.483909 | 0.996349 | 0.997988 | 0.994666 |
| Lgalsl_Q8VED9_S25s_1 | Lgalsl_Q8V | -0.58 | -0.2 | -0.26 | -0.08 | -0.23 | -0.68 | 0.27 | -0.33 | -0.55 | -0.09 | -0.59 | -0.3 | -0.25 | 0.05 | -0.49 | -0.23 | -0.1425 | -0.525 | -1.2575 | 0.845943 | 0.88841 | 0.672676 | 0.996493 | 0.997988 | 0.999507 |

| | | | | | | | | | | | | | | | | | | | | | | | | | | |
|---|---|---|---|---|---|---|---|---|---|---|---|---|---|---|---|---|---|---|---|---|---|---|---|---|---|---|
| Lgalsl_Q8VED9_S4s_1 | Lgalsl_Q8V | -0.74 | -0.53 | -0.55 | -1.59 | -0.64 | -0.97 | -2.21 | -1.56 | -1.01 | -0.74 | 0.51 | -1.34 | -0.66 | -0.35 | -2.15 | -1.56 | 0.0375 | -0.1025 | 0.05 | 0.48732 | 0.245916 | 0.650808 | 0.989223 | 0.997988 | 0.994666 |
| Lima1_Q9ERG0_S200s_1 | Lima1_Q9E | 0.86 | 1.4 | 1.72 | 2.14 | 0.93 | 1.38 | 1.43 | 1.82 | 1.7 | 1.97 | 1.54 | 1.98 | 0.95 | 1.15 | 2.2 | 1.72 | -0.4925 | 0.2075 | -0.3275 | 0.949921 | 0.676504 | 0.41199 | 0.99315 | 0.997988 | 0.999507 |
| Lima1_Q9ERG0_S212s_1 | Lima1_Q9E | 1.2 | 1.05 | 0.49 | 0.45 | 1.26 | 0.68 | 0.33 | 0.4 | 1.02 | 1.05 | 0.92 | 0.85 | 1.08 | 1.02 | 0.88 | 0.34 | -0.14 | 0.2675 | -0.025 | 0.91636 | 0.6729 | 0.539188 | 0.992876 | 0.997988 | 0.999507 |
| Lima1_Q9ERG0_S328s_1 | Lima1_Q9E | 0.95 | 0.65 | 0.04 | 0.4 | 0.75 | 0.69 | 0.31 | 0.32 | 0.84 | 1.28 | -0.5 | 0.15 | 0.43 | 1.43 | -0.58 | -0.75 | -0.13 | 0.1625 | 0.0325 | 0.464903 | 0.977723 | 0.873901 | 0.999187 | 0.997988 | 0.994666 |
| Limch1_D3Z589_S10s_1 | Limch1_D3 | 0.53 | 0.55 | 0.47 | 0.11 | -0.77 | 0.56 | -0.29 | 0.5 | 0.95 | 0.63 | 2.04 | -0.99 | 0.96 | -1.3 | -0.01 | 1.23 | 0.0075 | -0.0675 | -0.3775 | 0.719085 | 0.244468 | 0.679111 | 0.989223 | 0.997988 | 0.9971 |
| Limch1_D3Z589_S144s_1 | Limch1_D3 | 0.71 | -0.48 | 0.1 | 0.6 | 0.24 | 0.46 | 0.54 | 0.66 | -0.87 | -0.4 | -1.24 | 0.1 | -1.25 | -0.2 | -0.12 | -0.94 | -0.415 | 0.2425 | -0.195 | 0.052906 | 0.437431 | 0.059767 | 0.989223 | 0.997988 | 0.994666 |
| Limch1_D3Z589_S218s_1 | Limch1_D3 | -0.36 | -0.87 | -0.57 | -1 | 0.36 | -2.12 | -0.01 | -1.07 | -0.82 | -0.64 | 0.06 | -0.75 | -1.23 | -0.07 | -0.83 | -0.47 | 0.2425 | -0.835 | -0.86 | 0.879142 | 0.984795 | 0.581872 | 0.999187 | 0.997988 | 0.999507 |
| Limch1_D3Z589_S364s_1 | Limch1_D3 | 0.39 | -0.44 | -0.84 | 0.55 | 0.71 | -0.04 | 0.9 | -0.42 | 0.77 | -0.91 | 0.12 | -1.2 | 0.22 | 0.09 | -0.2 | 0.49 | -0.01 | 0.1625 | 0.05 | 0.538247 | 0.395745 | 0.677601 | 0.989223 | 0.997988 | 0.994666 |
| Limch1_D3Z589_S560s_1 | Limch1_D3 | 0.43 | 0.09 | -0.52 | 0.16 | -0.13 | 0.09 | 0.73 | -0.09 | -0.22 | 0.28 | -0.26 | 0.01 | -1.23 | 0.44 | 0.04 | -0.11 | 0.3725 | -0.22 | 0.235 | 0.539831 | 0.718903 | 0.758933 | 0.995437 | 0.997988 | 0.994666 |
| Limch1_D3Z589_S58s_1 | Limch1_D3 | 0.16 | 0.4 | 0.23 | 0.39 | 0.43 | -0.02 | 0.66 | 0.11 | 0.13 | 0.61 | 0.35 | -0.01 | 0.02 | 0.6 | 0.43 | 0.4 | 0.11 | -0.0875 | -0.255 | 0.787524 | 1 | 0.915821 | 1 | 0.997988 | 0.999507 |
| Limch1_D3Z589_S598s_1 | Limch1_D3 | -1.5 | -0.82 | -1.64 | -1.98 | -0.54 | -1.09 | -1 | -1.36 | -1.07 | -0.87 | 0.18 | -2.05 | -1.43 | -0.59 | -0.56 | -1.32 | -3.9E-17 | -0.025 | 0.0675 | 0.179907 | 0.145625 | 0.292786 | 0.989223 | 0.997988 | 0.994666 |
| Limch1_D3Z589_S72s_1 | Limch1_D3 | -0.7 | -0.22 | -0.36 | -0.07 | -0.11 | -1.05 | -0.22 | -0.39 | 0.45 | -0.54 | -0.14 | -0.22 | 1.48 | -0.21 | 0.26 | -0.54 | 0.4875 | 0.5325 | 0.51 | 0.214139 | 0.712059 | 0.447142 | 0.995437 | 0.997988 | 0.994666 |
| Limch1_D3Z589_S74s_1 | Limch1_D3 | -0.24 | -0.1 | -1.05 | -0.81 | -0.65 | -0.62 | -1.02 | -0.48 | 0.18 | -0.26 | 0.08 | -0.87 | 0.82 | -0.12 | 0.11 | -0.09 | -0.105 | 0.225 | 0.585 | 0.058798 | 0.620966 | 0.348297 | 0.989223 | 0.997988 | 0.994666 |
| Limch1_D3Z589_S814s_1 | Limch1_D3 | -0.29 | -0.59 | -0.26 | -0.09 | -0.51 | -0.65 | -0.07 | -0.55 | 0.19 | -0.14 | 0.04 | -0.46 | 0.25 | -0.84 | -0.24 | -0.4 | -0.1425 | 0.3325 | 0.73 | 1 | 0.562385 | 0.398824 | 0.989223 | 0.997988 | 1 |
| Limd1_Q9QXD8_S659s_1 | Limd1_Q9C | 0.45 | 0.67 | 0.17 | -1.62 | 0.26 | -0.35 | -0.15 | -0.05 | 1.99 | -0.04 | 0.24 | -0.6 | 0.79 | 1.89 | 1.11 | -1.19 | -0.1375 | 0.215 | -7.9E-17 | 0.338274 | 0.983828 | 0.492972 | 0.999187 | 0.997988 | 0.994666 |
| Lin7a_Q8JZS0_S233s_1 | Lin7a_Q8JZ | 2.08 | 1.82 | 1.56 | 0.35 | 1.61 | 1.63 | 0.64 | 1.12 | 1.31 | 2.19 | -1.37 | 2.16 | 1.07 | 1.85 | 0.91 | -0.02 | 0.01 | 0.48 | 0.7325 | 0.339579 | 0.635623 | 0.645731 | 0.989223 | 0.997988 | 0.994666 |
| Lin7b_O88951_S115s_1 | Lin7b_O88 | 0.81 | 0.6 | 0.59 | 0.7 | 0.9 | 0.92 | 0.72 | 0.82 | 0.49 | 0.78 | 0.4 | 0.64 | -0.07 | 0.7 | 0.93 | 0.7 | -0.2025 | -0.38 | -0.5 | 0.707372 | 0.413456 | 0.654485 | 0.989223 | 0.997988 | 0.996966 |
| Lin7b_O88951_S199s_1 | Lin7b_O88 | 0.77 | -0.73 | 1.14 | 2.38 | 0.76 | 1.16 | 1.39 | 1.64 | 1.15 | 0.91 | 0.05 | 2.4 | 1.21 | -1.71 | 0.71 | 1.43 | 0.165 | -0.0975 | -0.11 | 0.574938 | 0.563892 | 0.743011 | 0.989223 | 0.997988 | 0.995259 |
| Lingo1_Q9D1T0_S596s_1 | Lingo1_Q9 | 0.34 | 0.79 | 0.57 | 0.61 | 0.95 | 0.5 | 0.44 | 0.64 | 0.2 | 0.65 | 1.02 | 0.6 | 0.72 | 0.87 | 0.89 | 0.66 | 0.3475 | 0.2375 | -0.48 | 0.39738 | 0.808108 | 0.877127 | 0.995762 | 0.997988 | 0.994666 |
| Lingo1_Q9D1T0_S602s_1 | Lingo1_Q9 | 1.05 | 1.37 | 0.96 | 0.61 | 1.19 | 0.72 | 0.59 | 0.62 | 1.09 | 1.64 | 0.49 | 0.82 | 1.54 | 0.25 | 1 | 0.61 | 0.055 | 0.04 | 0.2075 | 0.673231 | 0.410356 | 0.96869 | 0.989223 | 0.997988 | 0.996159 |
| Lipe_E9Q4M2_S963s_1 | Lipe_E9Q4 | -1.02 | -0.48 | -0.53 | -0.72 | -0.18 | -0.66 | 0 | -1.24 | -0.64 | -0.8 | -0.64 | -0.76 | -0.18 | -0.01 | -0.92 | -0.7 | -0.2175 | 0.0125 | -0.1475 | 0.447638 | 0.601685 | 0.921465 | 0.989223 | 0.997988 | 0.994666 |
| Lipe_E9Q4M2_S964s_1 | Lipe_E9Q4 | -1.02 | -0.48 | -0.53 | -0.72 | -0.18 | -0.66 | 0 | -1.24 | -0.64 | -0.8 | -0.64 | -0.76 | -0.18 | -0.01 | -0.92 | -0.7 | 0.1675 | -0.0225 | 0.235 | 0.447638 | 0.601685 | 0.921465 | 0.989223 | 0.997988 | 0.994666 |
| Llgl1_Q80Y17_S986s_1 | Llgl1_Q80Y | 0.72 | 0.55 | 0.75 | 0.48 | 0.65 | 0.2 | 0.99 | 0.62 | 1.1 | 1.06 | 0.42 | 0.45 | 0.19 | 0.96 | 0.6 | 0.9 | 0.1675 | -0.0225 | 0.235 | 0.890709 | 0.966731 | 0.61558 | 0.999187 | 0.997988 | 0.999507 |
| Lmf1_Q3U3R4_S19s_1 | Lmf1_Q3U | -0.75 | -0.55 | -0.67 | -0.48 | -0.27 | -0.88 | -0.09 | -0.75 | 0.16 | -0.54 | -0.47 | -0.59 | 0.6 | 0.38 | 0.83 | 0.21 | -0.01 | 0.1325 | 0.0375 | 0.001493 | 0.650842 | 0.335952 | 0.990358 | 0.997988 | 0.994666 |
| Lmf1_Q3U3R4_S7s_1 | Lmf1_Q3U | 0.04 | -0.09 | 0.74 | 1.52 | -0.45 | 0.37 | 0.49 | 0.2 | 1.7 | 0.31 | 1.22 | 0.85 | 1.8 | 1.46 | 1.52 | 1.79 | 0.115 | 0.2525 | 1.1175 | 0.018772 | 0.338715 | 0.31766 | 0.989223 | 0.997988 | 0.994666 |
| Lmna_P48678_S22s_1 | Lmna_P48 | -0.11 | -0.58 | -0.66 | 0 | -1 | -0.97 | -0.33 | -0.53 | -1.15 | -0.68 | -0.55 | -0.61 | 0.33 | 0.13 | -0.58 | -1 | -0.4 | 0.4675 | 1.09 | 0.876613 | 0.197937 | 0.156082 | 0.989223 | 0.997988 | 0.999507 |
| Lmna_P48678_S390s_1 | Lmna_P48 | -0.2 | -0.05 | 0.05 | -0.12 | -0.22 | -0.66 | -0.21 | 0.08 | -0.68 | -1.08 | -0.34 | -0.39 | 0.54 | -0.18 | 0.28 | -0.28 | -0.37 | -0.41 | 0.0575 | 0.54675 | 0.466491 | 0.054633 | 0.989223 | 0.997988 | 0.994666 |
| Lmna_P48678_S392s_1 | Lmna_P48 | 0.64 | 0.85 | 1 | 0.73 | 0.44 | 0.5 | 0.5 | 0.83 | 0.29 | 0.15 | 0.45 | 0.48 | 1.37 | 0.49 | 0.92 | 0.6 | -0.1725 | -0.5425 | 0.17 | 0.888521 | 0.28303 | 0.060357 | 0.989223 | 0.997988 | 0.999507 |
| Lmna_P48678_S395s_1 | Lmna_P48 | -2.14 | -2.14 | -1.38 | -4.81 | -1.35 | -2.77 | -2.57 | -3.21 | -3.36 | -1.97 | -1.19 | -3.05 | -1.54 | -1.76 | -0.81 | -3.08 | -0.2375 | -0.4625 | 0.04 | 0.311316 | 0.848711 | 0.780394 | 0.996349 | 0.997988 | 0.994666 |
| Lmna_P48678_S407s_1 | Lmna_P48 | -0.54 | -1.45 | -0.26 | -0.5 | -1.04 | -0.79 | -0.58 | -0.75 | 0.89 | -1.29 | -1.04 | -1.42 | 2.78 | -2.59 | 0.57 | -0.9 | 0.1425 | 0.225 | 0.82 | 0.525443 | 0.734823 | 0.960504 | 0.995762 | 0.997988 | 0.994666 |
| Lmna_P48678_S429s_1 | Lmna_P48 | -2.68 | 1.04 | 0.09 | -0.34 | 0.28 | -0.63 | -1.95 | -0.57 | 1.53 | -0.76 | -1.7 | -0.85 | 2.05 | 0.59 | -0.93 | -0.66 | -0.1025 | -0.0275 | 0.6525 | 0.429004 | 0.759569 | 0.976382 | 0.995762 | 0.997988 | 0.994666 |
| Lmna_P48678_T19t_1 | Lmna_P48 | 0.88 | -0.2 | 0.68 | 1.25 | 0.78 | -0.83 | 0.62 | 0.76 | -0.34 | -0.98 | -0.62 | 1 | 1.59 | 0.16 | 1.17 | 1.26 | -0.245 | 0.0275 | 0.735 | 0.372609 | 0.493419 | 0.102091 | 0.989223 | 0.997988 | 0.994666 |
| Lmna_P48678_T3t_1 | Lmna_P48 | 0.8 | -1.54 | 0.59 | 0.27 | 0.04 | 0.4 | -0.59 | -0.47 | 4.27 | 1.1 | 0.03 | -0.08 | 3.99 | 0.61 | 1.27 | 0.22 | -0.32 | -0.8875 | 0.3925 | 0.115683 | 0.727051 | 0.220061 | 0.995569 | 0.997988 | 0.994666 |
| Lmnb1_P14733_S17s_1 | Lmnb1_P1 | 0.61 | 0.52 | -0.24 | 0 | 0.71 | 0.23 | -0.02 | 0.21 | 0.14 | 0.27 | 0.36 | 0.43 | 0.69 | 0.75 | 0.34 | 0.43 | -0.185 | 1.3 | 1.4925 | 0.277469 | 0.834617 | 0.77555 | 0.995762 | 0.997988 | 0.994666 |
| Lmnb1_P14733_S24s_1 | Lmnb1_P1 | 0.68 | 0.3 | 0.75 | 0.75 | 0.57 | 0.65 | 0.49 | 0.87 | 0.09 | -0.06 | 0.96 | 0.5 | 0.79 | 0.9 | 1.2 | 0.62 | 0.06 | 0.0775 | 0.33 | 0.33718 | 0.909972 | 0.411062 | 0.997335 | 0.997988 | 0.994666 |
| Lmnb1_P14733_S394s_1 | Lmnb1_P1 | 0.71 | 0.38 | 0.88 | 0.76 | -0.04 | 0.27 | 0.26 | 0.55 | 0.3 | -0.24 | -0.1 | 0.06 | 1.46 | -0.06 | 1.18 | 1.08 | 0.025 | -0.2475 | 0.2575 | 0.534966 | 0.095411 | 0.018599 | 0.989223 | 0.997988 | 0.994666 |
| Lmnb1_P14733_T21t_1 | Lmnb1_P1 | 1.45 | 0.53 | 1.02 | 0.79 | 0.79 | 0.95 | 0.54 | 1.11 | 0.63 | 0.11 | 1 | 0.54 | 1.38 | 1.41 | 0.97 | 0.53 | -0.4225 | -0.6775 | 0.2325 | 0.703383 | 0.713233 | 0.234698 | 0.995437 | 0.997988 | 0.996159 |
| Lmnb2_P21619_S397s_1 | Lmnb2_P2 | -1.81 | 0.02 | -0.52 | -1.68 | 0.61 | -1.38 | -0.29 | 0.47 | -1.01 | -0.83 | 0.58 | -1.19 | 0.05 | -0.26 | 0.16 | -0.39 | -0.1 | -0.3775 | 0.125 | 0.074241 | 0.161902 | 0.495191 | 0.989223 | 0.997988 | 0.994666 |
| Lmnb2_P21619_S403s_1 | Lmnb2_P2 | -0.15 | -1.48 | 1.48 | 1.72 | 0.24 | -0.33 | 0.07 | 1.48 | -0.16 | 0.39 | 1.95 | 0.86 | -0.34 | 0.37 | 1.08 | 0.37 | 0.85 | 0.385 | 0.8875 | 0.974945 | 0.970408 | 0.6399 | 0.999187 | 0.997988 | 0.999668 |
| Lmnb2_P21619_S427s_1 | Lmnb2_P2 | 1.78 | 0.65 | 0.73 | 1.42 | -0.01 | 1.42 | 0.48 | 1.09 | 1.26 | 0.76 | 1.18 | 0.7 | 1.61 | 0.1 | 1.64 | 1.22 | -0.0275 | 0.3675 | -0.0225 | 0.995487 | 0.333501 | 0.610917 | 0.989223 | 0.997988 | 1 |
| Lmnb2_P21619_T494t_1 | Lmnb2_P2 | 0.3 | 0.52 | 0.4 | 0.33 | 0.45 | 0.46 | 0.6 | 0.72 | -0.28 | -0.04 | -0.74 | 0.12 | 0.35 | 0.77 | 0.55 | 0.96 | -0.4 | -0.17 | -0.0025 | 0.299204 | 0.407237 | 0.036794 | 0.989223 | 0.997988 | 0.994666 |
| Lmo7_E9PYI7_S1073s_1 | Lmo7_E9P | 0.37 | 0.79 | 1.15 | 0.78 | -0.82 | 0.37 | -0.76 | -0.24 | 1.79 | 0.71 | -0.38 | 0.33 | 2.47 | -0.53 | 0.61 | 0.15 | 0.17 | -0.6225 | 0.27 | 0.871652 | 0.006462 | 0.728506 | 0.989223 | 0.997988 | 0.999507 |
| Lmo7_E9PYI7_S122s_1 | Lmo7_E9P | 0.36 | -0.17 | 0.24 | 0.51 | 1.06 | -0.11 | 0.69 | 0.97 | 0.65 | 0.55 | 0.24 | 0.29 | 0.13 | 0.12 | 0.03 | 0.47 | -1.135 | -0.16 | -0.0975 | 0.858741 | 0.210994 | 0.438846 | 0.989223 | 0.997988 | 0.999507 |
| Lmo7_E9PYI7_S1269s_1 | Lmo7_E9P | 0.86 | -1.46 | -0.31 | 0.34 | 0.07 | 0.62 | 0.29 | 0.88 | -1.93 | -0.97 | -0.63 | 0.4 | -2.75 | -0.36 | -0.78 | 0.29 | 0.4175 | 0.1975 | -0.0475 | 0.315756 | 0.232994 | 0.325118 | 0.989223 | 0.997988 | 0.994666 |
| Lmo7_E9PYI7_S127s_1 | Lmo7_E9P | 0.63 | -1.12 | 0.5 | 0.81 | 1.35 | 0.82 | 0.96 | 0.86 | -0.19 | 0.42 | 0.42 | 0.12 | 0.25 | 0.04 | 0.43 | 0.77 | 0.6075 | -0.64 | -0.7575 | 0.71502 | 0.094194 | 0.977886 | 0.989223 | 0.997988 | 0.996966 |
| Lmo7_E9PYI7_S1300s_1 | Lmo7_E9P | 1.16 | -0.39 | -0.61 | 0.42 | 0.75 | 1.02 | 0.51 | 0.79 | -0.88 | 0.15 | -0.48 | 0.16 | -1.22 | 0.59 | -0.3 | -0.26 | 0.7925 | -0.0125 | 0.1675 | 0.399431 | 0.145296 | 0.38423 | 0.989223 | 0.997988 | 0.994666 |
| Lmo7_E9PYI7_S1369s_1 | Lmo7_E9P | 1.88 | -1.57 | -0.34 | 0.66 | -0.49 | 1.93 | -0.36 | 0.51 | 2.11 | -0.66 | 0.02 | 0.54 | 0.33 | -1.24 | -0.93 | -0.13 | 0.6225 | -0.4075 | -0.4425 | 0.382181 | 0.765744 | 0.681661 | 0.995762 | 0.997988 | 0.994666 |
| Lmo7_E9PYI7_S1369s_2 | Lmo7_E9P | 0.77 | -0.47 | 0.25 | 0.34 | -0.99 | 1.87 | -1.07 | 0.53 | 2.02 | -0.64 | 1.13 | 0.16 | 0.9 | -0.37 | 0.33 | 0.27 | 0.24 | 0.345 | -0.65 | 0.874655 | 0.835086 | 0.45187 | 0.995762 | 0.997988 | 0.999507 |
| Lmo7_E9PYI7_S1376s_1 | Lmo7_E9P | 6.57 | 1.55 | 2.31 | 1.05 | 2.68 | 6.85 | 4.08 | 3.56 | 0.57 | 4.34 | 1.41 | 3.12 | -0.16 | -1.36 | 3.6 | 3.82 | -0.1375 | 0.445 | 0.06 | 0.382099 | 0.29942 | 0.702054 | 0.989223 | 0.997988 | 0.994666 |
| Lmo7_E9PYI7_S1391s_1 | Lmo7_E9P | 1.93 | 0.47 | -1.05 | -0.46 | 0.94 | 0.98 | 0.36 | 1 | -0.42 | 0.29 | 0.91 | 0.15 | -0.89 | 0.48 | -0.47 | -1.31 | 1.4225 | -0.51 | -1.395 | 0.272173 | 0.33024 | 0.987534 | 0.989223 | 0.997988 | 0.994666 |
| Lmo7_E9PYI7_S206s_1 | Lmo7_E9P | 0.99 | 0.58 | 1.45 | 1.06 | 0.01 | 1.14 | 0.24 | 0.78 | 1.51 | 0.3 | 0.35 | 1.17 | 1.68 | 0.07 | 0.97 | 1.1 | 0.5975 | 0.01 | -0.77 | 0.867211 | 0.164955 | 0.606234 | 0.989223 | 0.997988 | 0.999507 |
| Lmo7_E9PYI7_S253s_1 | Lmo7_E9P | 0.07 | -0.35 | 0.67 | -0.57 | 0.14 | 0.28 | 0.15 | -0.03 | -0.16 | -0.71 | 0.51 | -0.07 | -1.01 | -0.11 | -0.84 | -1.11 | -0.4775 | -0.1875 | -0.065 | 0.077526 | 0.556262 | 0.867836 | 0.989223 | 0.997988 | 0.994666 |
| Lmo7_E9PYI7_S466s_1 | Lmo7_E9P | -0.34 | -0.46 | -0.66 | -1.8 | -0.52 | -0.25 | -0.78 | -0.88 | -1.29 | -0.31 | 0.86 | -0.52 | -0.72 | -0.85 | -1.27 | -0.56 | 0.18 | -0.0625 | -0.7225 | 0.926946 | 0.569027 | 0.349305 | 0.989223 | 0.997988 | 0.999507 |
| Lmo7_E9PYI7_S580s_1 | Lmo7_E9P | 0.83 | 1.03 | 1.48 | 1.16 | 0.92 | 0.46 | 0.91 | 0.9 | 0.43 | 1.34 | 1.82 | 0.85 | 0.17 | 1.07 | 0.95 | 1.5 | 0.2075 | 0.5 | -0.035 | 0.558296 | 0.196075 | 0.965506 | 0.989223 | 0.997988 | 0.995259 |
| Lmo7_E9PYI7_S589s_1 | Lmo7_E9P | 0.7 | -1.18 | -0.31 | 0.63 | 0.38 | 0.66 | 0.1 | 0.89 | -0.04 | -0.7 | 0 | 0.41 | -1.08 | 0.11 | -0.26 | 0.34 | -0.3275 | -0.015 | -0.2025 | 0.721493 | 0.238395 | 0.928715 | 0.989223 | 0.997988 | 0.9971 |
| Lmo7_E9PYI7_S787s_1 | Lmo7_E9P | 1.04 | 0.62 | 0.65 | 1.19 | 0.61 | 0.74 | 0.67 | 0.72 | 0.78 | 0.87 | 0.53 | 0.79 | 0.96 | 0.72 | 0.58 | 0.44 | 0.5475 | -0.0425 | -0.1825 | 0.46437 | 0.408525 | 0.587656 | 0.989223 | 0.997988 | 0.994666 |
| Lmo7_E9PYI7_T1365t_2 | Lmo7_E9P | 0.77 | -0.47 | 0.25 | 0.34 | -0.99 | 1.87 | -1.07 | 0.53 | 2.02 | -0.64 | 1.13 | 0.16 | 0.9 | -0.37 | 0.33 | 0.27 | -0.19 | -0.1325 | -0.2 | 0.874655 | 0.835086 | 0.45187 | 0.995762 | 0.997988 | 0.999507 |
| Lmtk2_Q3TYD6_S1104s_1 | Lmtk2_Q3T | -0.92 | -0.07 | -0.36 | 0.1 | 0.09 | -0.41 | 0.39 | 0.08 | 0.1 | -0.08 | -0.82 | 0.06 | 0.62 | 0.07 | 0.03 | -0.03 | -0.1375 | 0.445 | 0.06 | 0.151439 | 0.263563 | 0.702945 | 0.989223 | 0.997988 | 0.994666 |
| Lmtk2_Q3TYD6_S1359s_1 | Lmtk2_Q3T | 0.22 | -1.02 | -0.18 | -0.54 | 0.03 | -0.44 | 0.37 | 0.32 | -0.06 | -0.55 | -0.46 | 0.2 | 0.37 | -1.05 | 1.05 | 0.49 | 0.35 | 0.1275 | 0.485 | 0.245583 | 0.197005 | 0.632303 | 0.989223 | 0.997988 | 0.994666 |
| Lmtk2_Q3TYD6_S1418s_1 | Lmtk2_Q3T | 0.08 | -0.2 | -0.85 | -0.08 | 0.27 | -0.32 | -0.17 | -0.46 | -0.61 | 0.2 | -0.34 | -0.9 | -1.34 | 0.08 | -0.68 | -0.07 | 0.45 | 0.1625 | 0.595 | 0.54485 | 0.74932 | 0.654319 | 0.995762 | 0.997988 | 0.994666 |
| Lmtk2_Q3TYD6_S611s_1 | Lmtk2_Q3T | -1.22 | -1.73 | -2.06 | -1.64 | -0.26 | -1.25 | -1.59 | -2.34 | -2.27 | -2.03 | -1.9 | -2.56 | -2.28 | -0.79 | -3.64 | -1.31 | 0.0925 | -0.15 | -0.24 | 0.567603 | 0.494571 | 0.084118 | 0.989223 | 0.997988 | 0.995259 |
| Lmtk2_Q3TYD6_S742s_1 | Lmtk2_Q3T | 0.43 | -0.05 | 0.08 | 0.37 | 0.61 | 0.26 | 0.57 | 0.39 | -0.02 | 0.3 | 0.32 | 0.36 | -0.62 | 0.51 | -0.09 | 0.38 | 0.3025 | -0.5275 | -0.3425 | 0.619433 | 0.279757 | 0.890009 | 0.989223 | 0.997988 | 0.995259 |
| Lmtk3_Q5XJV6_S232s_1 | Lmtk3_Q5 | 0.12 | 0.43 | 0.41 | 0.29 | 0.29 | 0.07 | 0.4 | 0.32 | 0.37 | 0.43 | 0.4 | 0.3 | 0.81 | 0.75 | 0.36 | 0.5 | 0.25 | 0.0325 | -0.1625 | 0.253938 | 0.83769 | 0.766559 | 0.995762 | 0.997988 | 0.994666 |
| Lmtk3_Q5XJV6_S531s_1 | Lmtk3_Q5 | -0.05 | -0.77 | -0.02 | 0.13 | -0.1 | 0.16 | 0.41 | 0.59 | -0.23 | 0.05 | -0.01 | 0.02 | -0.6 | 0.22 | -0.31 | -0.32 | -0.0425 | 0.0625 | 0.2925 | 0.811949 | 0.144193 | 0.619175 | 0.989223 | 0.997988 | 0.999507 |
| Lmtk3_Q5XJV6_S947s_1 | Lmtk3_Q5 | 0.41 | 0.75 | -0.06 | -0.4 | 0.95 | 0.09 | 0.13 | -0.71 | 0.63 | 0.46 | -0.08 | 0.29 | 0.6 | 1.23 | -0.62 | 0.09 | 0.4425 | 0.135 | -0.075 | 0.741249 | 0.882334 | 0.644307 | 0.996493 | 0.997988 | 0.9971 |
| Lmtk3_Q5XJV6_S962s_1 | Lmtk3_Q5 | 0.08 | 0.38 | -0.31 | -0.06 | 0.42 | -0.04 | 0.46 | -0.24 | 0.2 | 0.24 | 0.13 | -0.07 | 0.35 | 0.24 | 0.62 | 0.02 | -0.06 | 0.15 | 0.15 | 0.312951 | 0.636327 | 0.673493 | 0.989223 | 0.997988 | 0.994666 |
| Lmtk3_Q5XJV6_S977s_1 | Lmtk3_Q5 | 0.25 | 0.29 | 0.61 | 0.26 | 0.81 | 0.33 | 0.96 | 0.67 | -0.01 | 0.26 | 0.09 | 0.41 | -0.22 | 0.54 | 0.38 | 0.35 | 0.1275 | 0.1025 | 0.285 | 0.741943 | 0.166969 | 0.474563 | 0.989223 | 0.997988 | 0.9971 |
| Lnp_Q7TQ95_S194s_1 | Lnp_Q7TQ | 0.29 | 1.19 | 1.44 | 0.48 | 0.46 | 1.39 | -0.16 | 0.91 | 0.6 | 1.05 | 1.44 | 1.37 | -0.12 | 0.05 | 0.51 | 1.07 | 0.34 | -0.165 | -0.09 | 0.247564 | 0.628384 | 0.457545 | 0.989223 | 0.997988 | 0.994666 |
| Lnp_Q7TQ95_S222s_1 | Lnp_Q7TQ | -0.03 | 0.19 | -0.19 | 0.26 | 0.51 | 0.33 | 0.2 | -0.01 | 0.32 | 0.22 | -0.01 | -0.81 | -0.09 | 0.65 | -0.02 | -0.73 | -0.2 | 0.265 | -0.4725 | 0.755988 | 0.390343 | 0.682241 | 0.989223 | 0.997988 | 0.9971 |
| Lnp_Q7TQ95_S411s_1 | Lnp_Q7TQ | 0.27 | -0.25 | -0.07 | -0.01 | 0.38 | 0.27 | 0.38 | 0.36 | -0.14 | 0.09 | -0.55 | -0.3 | -0.53 | 0.4 | 0.18 | 0.16 | 0.2 | -0.1275 | -0.105 | 0.81876 | 0.111529 | 0.407833 | 0.989223 | 0.997988 | 0.999507 |
| Loh12cr1_E9Q2W8_S76s_1 | Loh12cr1_ | 0.34 | 0.63 | 0.73 | 0.55 | 0.47 | 0.42 | 0.49 | 0.24 | 0.59 | 0.58 | 0.45 | 0.35 | 0.51 | 0.79 | 0.69 | 0.67 | 0.3625 | -0.21 | 0.0675 | 0.668108 | 0.454839 | 0.74817 | 0.989223 | 0.997988 | 0.996159 |
| Lpar1_P61793_S323s_1 | Lpar1_P61 | -1.66 | -0.67 | -0.87 | -0.94 | -1.97 | -0.93 | -2.55 | -2.17 | 0.77 | -1.89 | -0.81 | -1.49 | 0.95 | -3.6 | -1.38 | -1.36 | -0.1575 | -0.07 | 0.1025 | 0.710435 | 0.049776 | 0.754712 | 0.989223 | 0.997988 | 0.996966 |
| Lpgat1_E9QL80_S24s_1 | Lpgat1_E9 | 1.24 | 0.92 | 0.09 | 0.64 | 0.62 | 1.12 | 0.44 | 0.35 | 0.81 | 1.07 | 0.29 | 1 | 0.46 | 0.47 | 1.17 | 0.7 | -0.87 | 0.18 | -0.3125 | 0.946239 | 0.775621 | 0.830433 | 0.995762 | 0.997988 | 0.999507 |
| Lpgat1_E9QL80_S272s_1 | Lpgat1_E9 | -0.53 | -0.5 | 0.4 | -0.95 | -1.21 | 1 | 0.44 | 0.69 | 1.15 | -0.3 | -1 | 1.12 | 2.41 | -2.78 | 1.44 | 0.7 | -0.09 | 0.07 | -0.0225 | 0.417249 | 0.246729 | 0.272236 | 0.989223 | 0.997988 | 0.994666 |
| Lpin1_E9QKQ5_S328s_1 | Lpin1_E9Q | -0.97 | -1.18 | -1.56 | -1.49 | -1.18 | -1.08 | -1.75 | -1.7 | -0.66 | -1.26 | -2.23 | -1.61 | -0.11 | -1.28 | -0.92 | -1.46 | 0.625 | 0.6375 | 0.8375 | 0.328667 | 0.633764 | 0.702671 | 0.989223 | 0.997988 | 0.994666 |
| Lpin2_E9PWN0_S144s_1 | Lpin2_E9P | -1.23 | -0.51 | -1.18 | -2.17 | -0.6 | -1.59 | -1.52 | -1.57 | -1.09 | -1.57 | -1.5 | -1.57 | -0.85 | -0.44 | -2.17 | -1.79 | -0.1275 | -0.14 | 0.3575 | 0.936037 | 0.905676 | 0.665685 | 0.996513 | 0.997988 | 0.999507 |
| Lpin2_E9PWN0_T329t_1 | Lpin2_E9P | -0.85 | 0.17 | -0.65 | -1.22 | -0.37 | -0.17 | -0.89 | -0.65 | 0.36 | 0.25 | -0.9 | -1.24 | 0.45 | -1 | -1.47 | 0.31 | -0.0475 | -0.16 | -0.04 | 0.689725 | 0.730154 | 0.594054 | 0.995762 | 0.997988 | 0.996159 |
| Lppr2_Q8VCY8_S312s_1 | Lppr2_Q8V | 0.4 | 1.56 | -0.19 | 0.49 | 0.97 | 0.56 | 0.69 | 0.33 | 1.28 | 0.94 | 0.22 | 0.15 | -0.18 | 0.61 | 0.19 | 0.34 | 0.1175 | 0.255 | 0.21 | 0.429785 | 0.848387 | 0.851914 | 0.996349 | 0.997988 | 0.994666 |
| Lppr2_Q8VCY8_S330s_1 | Lppr2_Q8V | 0.13 | -3.32 | -0.27 | 0.01 | -0.29 | 0.66 | -0.22 | 0.37 | -0.43 | -1.36 | -0.18 | 0.18 | -1.52 | -0.46 | -0.64 | 0.04 | 0.0725 | 0.0825 | -0.325 | 0.781528 | 0.206796 | 0.603017 | 0.989223 | 0.997988 | 0.998226 |
| Lppr2_Q8VCY8_S330s_1_1 | Lppr2_Q8V | 6.69 | 1.59 | 4 | 0.62 | 2.55 | 8.37 | 6.04 | 3.61 | 2.52 | 4.28 | 0.57 | 4.88 | 3.44 | 1.93 | 6.95 | 5.44 | 0.9925 | 0.415 | 0.2175 | 0.426369 | 0.250866 | 0.911087 | 0.989223 | 0.997988 | 0.994666 |
| Lppr2_Q8VCY8_S364s_1 | Lppr2_Q8V | -0.28 | -0.93 | -0.12 | -0.27 | -0.48 | 0.15 | -0.39 | -0.32 | -0.38 | -0.43 | 0.26 | -0.17 | 0.19 | -0.75 | -0.66 | -0.43 | 1.9175 | -0.1625 | 1.215 | 0.969145 | 0.607301 | 0.450205 | 0.989223 | 0.997988 | 0.999507 |
| Lppr2_Q8VCY8_S398s_1 | Lppr2_Q8V | 0.6 | 0.29 | 0.63 | 1.27 | 0.71 | 0.35 | 0.54 | 0.84 | -0.41 | 0.81 | 0.56 | 1.03 | -1.34 | 0.44 | 0.1 | 1.46 | 0.14 | 0.22 | -0.0125 | 0.358617 | 0.748437 | 0.603083 | 0.995762 | 0.997988 | 0.994666 |
| Lppr3_Q7TPB0_S311s_1 | Lppr3_Q7T | 3.28 | 1.49 | 1.38 | 0.57 | 1.32 | 3.58 | 1.77 | 1.83 | 1.53 | 1.98 | 0.98 | 2.19 | 1.03 | 0.24 | 2.41 | 1.85 | -0.0875 | -0.2 | -0.5325 | 0.65763 | 0.511967 | 0.986243 | 0.989223 | 0.997988 | 0.995259 |
| Lppr3_Q7TPB0_S346s_1 | Lppr3_Q7T | -0.52 | -1.13 | -1.16 | -0.98 | -0.11 | -0.45 | -0.53 | -0.7 | -1.54 | -1.17 | -1.52 | -1.03 | -1.32 | -0.76 | -0.88 | -0.96 | 0.445 | -0.01 | -0.2975 | 0.90569 | 0.069838 | 0.181887 | 0.989223 | 0.997988 | 0.999507 |
| Lppr3_Q7TPB0_S351s_1 | Lppr3_Q7T | 0.68 | 0.56 | 0 | 1.04 | 0.18 | 0.57 | 0.85 | 0.16 | 1.03 | 0.79 | -0.06 | 0.64 | 0.04 | 0.61 | 0.17 | 0.52 | 0.5 | -0.3675 | -0.0325 | 0.454922 | 0.664001 | 0.929371 | 0.991206 | 0.997988 | 0.994666 |
| Lppr3_Q7TPB0_S382s_1 | Lppr3_Q7T | -0.56 | -0.39 | -1.55 | -1.09 | -0.49 | -1.01 | -0.35 | -0.15 | -0.78 | -0.71 | 1.14 | -0.98 | -0.06 | 0 | -1.92 | -0.26 | -0.13 | 0.03 | -0.235 | 0.504029 | 0.248364 | 0.296018 | 0.989223 | 0.997988 | 0.994666 |
| Lppr3_Q7TPB0_S426s_1 | Lppr3_Q7T | 0.76 | -0.52 | 0.05 | 0.69 | 0.58 | 0.66 | 1 | 1.11 | 0.47 | 0.44 | 0.47 | 0.34 | -0.51 | 0.8 | 0.33 | 0.52 | 0.3975 | 0.565 | 0.3375 | 0.922869 | 0.102816 | 0.575798 | 0.989223 | 0.997988 | 0.999507 |
| Lppr3_Q7TPB0_S505s_1 | Lppr3_Q7T | 0.21 | 0.58 | 0.46 | 0.31 | 0.79 | 0.37 | 0.67 | 0.47 | 0.24 | 0.72 | -0.5 | 0.54 | -0.26 | 0.72 | 0.02 | 0.09 | 0.5925 | 0.185 | 0.04 | 0.404163 | 0.402961 | 0.65666 | 0.989223 | 0.997988 | 0.994666 |
| Lppr3_Q7TPB0_S567s_1 | Lppr3_Q7T | 0.01 | 0.19 | 0.17 | 0.27 | -1.9 | 0.02 | -0.91 | 0.6 | 1.23 | 0.18 | 1.77 | 0.27 | 1.98 | -2.91 | 0.5 | 1.07 | 0.185 | -0.14 | -0.2475 | 1 | 0.182642 | 0.097755 | 0.989223 | 0.997988 | 1 |
| Lppr3_Q7TPB0_S575s_1 | Lppr3_Q7T | -1.42 | -1.48 | -2.32 | -1.46 | -2.09 | -1.34 | -1.42 | -1.67 | -0.21 | -1.41 | -1.47 | -1.29 | 0.3 | -1.19 | -0.91 | -0.7 | -0.7075 | 0.7025 | 0 | 0.024464 | 0.893757 | 0.149678 | 0.996493 | 0.997988 | 0.994666 |
| Lppr3_Q7TPB0_S641s_1 | Lppr3_Q7T | 0.19 | 0.31 | 0.06 | 0.2 | 0.13 | 0.39 | 0.43 | 0.43 | 0.35 | 0.3 | 0.16 | 0.5 | 0 | 0.38 | 0.48 | 0.45 | 0.04 | 0.575 | 1.045 | 0.577822 | 0.453493 | 0.523579 | 0.989223 | 0.997988 | 0.995259 |
| Lppr3_Q7TPB0_T342t_1 | Lppr3_Q7T | -0.31 | 0.12 | 0.32 | -0.08 | 0.47 | -0.44 | 0.18 | -0.43 | -0.4 | -0.04 | 0.3 | 0.05 | -0.14 | 0.29 | 0.43 | -0.1 | 0.155 | 0.1375 | 0.1375 | 0.698885 | 0.817798 | 0.893831 | 0.995762 | 0.997988 | 0.996159 |
| Lppr3_Q7TPB0_T374t_1 | Lppr3_Q7T | -0.14 | -1.12 | -1.1 | -0.41 | -0.36 | -0.69 | 0.19 | -0.98 | -0.79 | -0.54 | -1.35 | -0.1 | -1.3 | 0.17 | -0.69 | -0.16 | -0.0675 | -0.035 | 0.1075 | 0.630813 | 0.515231 | 0.994574 | 0.989223 | 0.997988 | 0.995259 |
| Lppr4_Q7TME0_S347s_1 | Lppr4_Q7T | 0.9 | 0.63 | 0.79 | 0.38 | 0.87 | 0.94 | 0.52 | 0.39 | 0.62 | 0.72 | 0.17 | 0.65 | 0.57 | 1.23 | 0.41 | 0.54 | 0.2325 | -0.0025 | 0.1975 | 0.965289 | 0.983369 | 0.587133 | 0.999187 | 0.997988 | 0.999507 |
| Lppr4_Q7TME0_S37s_1 | Lppr4_Q7T | 6.3 | -0.02 | 5.39 | 0.32 | 3.14 | 6.87 | 6.09 | 3.51 | 1.72 | 5.62 | 1.02 | 6.04 | 1.31 | -1.16 | 6.06 | 6.45 | 0.005 | -0.135 | 0.0125 | 0.937066 | 0.258178 | 0.742742 | 0.989223 | 0.997988 | 0.999507 |
| Lppr4_Q7TME0_S386s_1 | Lppr4_Q7T | -0.08 | -0.51 | -0.4 | 0.06 | 0.5 | 0.29 | 0.23 | -0.04 | -0.88 | -0.59 | -0.35 | 0.25 | -1.62 | -0.18 | -0.3 | -0.48 | 1.905 | 0.6025 | 0.1675 | 0.286365 | 0.070794 | 0.607352 | 0.989223 | 0.997988 | 0.994666 |
| Lppr4_Q7TME0_S435s_1 | Lppr4_Q7T | -0.22 | -2.35 | -0.16 | 0.29 | -1.73 | 0.64 | -0.38 | 0.2 | -0.97 | -1.96 | 0.76 | -0.39 | -1.02 | -1.43 | 0.19 | 0.04 | 0.4775 | -0.16 | -0.4125 | 0.931818 | 0.673735 | 0.967466 | 0.992876 | 0.997988 | 0.999507 |
| Lppr4_Q7TME0_S439s_1 | Lppr4_Q7T | -0.61 | -1.73 | -1.1 | -0.8 | -0.73 | -0.26 | -0.73 | -0.07 | -0.72 | -1.38 | -0.53 | -0.44 | -0.58 | -1.15 | -1.6 | -1.22 | 0.2925 | -0.03 | 0.055 | 0.825779 | 0.076295 | 0.403204 | 0.989223 | 0.997988 | 0.999507 |
| Lppr4_Q7TME0_S474s_1 | Lppr4_Q7T | 0.41 | 0.39 | 0.5 | 0.65 | 0.46 | 0.63 | 1.11 | 0.76 | 0.62 | 0.63 | 0.49 | 0.85 | 0.07 | 0.69 | 0.46 | 0.76 | 0.6125 | 0.2925 | -0.0775 | 0.97722 | 0.283673 | 0.465239 | 0.989223 | 0.997988 | 1 |
| Lppr4_Q7TME0_S549s_1 | Lppr4_Q7T | -0.08 | -0.34 | -0.02 | -0.24 | -0.25 | 0.08 | 0.21 | -0.01 | -0.22 | 0.35 | -0.12 | 0.38 | -0.3 | 0.05 | -0.31 | -0.5 | 0.2525 | 0.16 | 0.0075 | 0.705015 | 0.418473 | 0.297436 | 0.989223 | 0.997988 | 0.996159 |
| Lppr4_Q7TME0_S594s_1 | Lppr4_Q7T | 0.16 | 2.05 | 0.5 | -0.06 | 1.71 | 0.66 | 0.92 | 0.91 | 0.19 | 1.38 | 0.49 | 0.53 | 0.92 | 0.67 | 1.9 | 2.19 | 0.1775 | 0.2675 | -0.095 | 0.195723 | 0.433613 | 0.976408 | 0.989223 | 0.997988 | 0.994666 |
| Lppr4_Q7TME0_S608s_1 | Lppr4_Q7T | 1.06 | -0.29 | -2.75 | 0.27 | -1.24 | 1.62 | 0.12 | 1.17 | 0.19 | -0.22 | 0.22 | 0.2 | -0.16 | -1.97 | -2 | -1.27 | 0.3875 | -0.015 | 0.7575 | 0.276186 | 0.360592 | 0.486452 | 0.989223 | 0.997988 | 0.994666 |
| Lppr4_Q7TME0_S649s_1 | Lppr4_Q7T | -0.51 | -1.66 | -1.15 | -0.59 | 0.35 | -0.01 | -0.04 | -0.39 | -1.74 | -0.28 | -0.28 | -0.4 | -2.19 | -0.03 | -2.06 | -0.39 | 0.845 | 0.525 | -0.9225 | 0.739523 | 0.013968 | 0.490917 | 0.989223 | 0.997988 | 0.9971 |
| Lppr4_Q7TME0_S685s_1 | Lppr4_Q7T | 0.42 | 0.58 | 1.15 | 0.14 | 0.3 | 1.13 | -0.03 | 0 | 0.56 | 0.63 | 0.7 | 1.12 | -1.54 | -1.3 | 0.65 | 0.86 | 0.955 | 0.3025 | -0.19 | 0.159724 | 0.52634 | 0.541282 | 0.989223 | 0.997988 | 0.994666 |
| Lppr4_Q7TME0_S746s_1 | Lppr4_Q7T | 0.09 | -1.68 | 0.13 | -0.17 | -1.12 | 0.8 | -0.58 | -0.05 | 2.3 | -0.72 | -0.16 | 0.5 | 1.53 | -1.27 | 0.28 | 0 | -0.2225 | 0.18 | -0.905 | 0.411086 | 0.75223 | 0.229282 | 0.995762 | 0.997988 | 0.994666 |
| Lppr4_Q7TME0_T417t_1 | Lppr4_Q7T | 0.12 | -0.73 | -0.28 | -0.28 | 0.39 | -0.09 | 0.43 | 0.27 | -0.6 | -0.34 | -0.31 | -0.22 | -0.7 | -0.1 | -1.02 | -0.17 | 0.17 | 0.8875 | 0.5425 | 0.531982 | 0.060238 | 0.77244 | 0.989223 | 0.997988 | 0.994666 |
| Lppr4_Q7TME0_T548t_1 | Lppr4_Q7T | 0.13 | 0.15 | -0.1 | 0.21 | 0.22 | 0.48 | 0.59 | 0.2 | 0.35 | 0.62 | -0.28 | 0.49 | -0.28 | 0.28 | 0.12 | -0.23 | 0.5425 | -0.075 | -0.205 | 0.628113 | 0.21888 | 0.470401 | 0.989223 | 0.997988 | 0.995259 |
| Lppr5_Q8BJ52_S302s_1 | Lppr5_Q8B | 0.55 | 0.76 | 0.72 | 0.69 | 0.49 | 0.59 | 0.51 | 0.81 | 0.58 | 0.55 | 0.66 | 0.76 | 0.4 | 0.64 | 0.63 | 0.58 | 0.275 | 0.1975 | -0.125 | 0.612339 | 0.694379 | 0.83769 | 0.994893 | 0.997988 | 0.995259 |
| Lrba_E9Q3Y4_S994s_1 | Lrba_E9Q3 | -0.15 | -1.03 | -0.05 | -1.33 | -1.34 | -0.59 | -2.02 | -0.7 | -0.63 | 0.03 | -3.04 | -0.15 | 0.47 | -5.17 | -0.01 | 0.25 | -0.08 | -0.0425 | -0.1175 | 0.693841 | 0.245426 | 0.663109 | 0.989223 | 0.997988 | 0.996159 |
| Lrch1_P62046_S522s_1 | Lrch1_P62 | -0.02 | 0.57 | 1.05 | 1.06 | 0.88 | -0.48 | 1.54 | 1.01 | 0.65 | 0.38 | 2.18 | 1.85 | 0.38 | 1.3 | 1.24 | 1.56 | -0.5225 | -0.3075 | -0.475 | 0.246837 | 0.875922 | 0.234487 | 0.996493 | 0.997988 | 0.994666 |
| Lrch3_F6YZL6_S212s_1 | Lrch3_F6Y | -0.17 | 0.34 | -0.63 | -0.67 | -0.27 | -0.77 | 0.42 | 0.35 | -1.34 | -0.26 | 0.72 | -0.48 | -0.27 | -0.2 | -0.25 | 0.34 | 0.0725 | 0.6 | 0.455 | 0.565214 | 0.558265 | 0.901199 | 0.989223 | 0.997988 | 0.995259 |
| Lrfn1_Q2WF71_S669s_1 | Lrfn1_Q2W | 0.62 | 0.35 | -0.2 | 1.11 | 0.63 | 0.69 | 0.48 | 1.35 | 0.44 | 0.29 | -0.33 | 0.05 | -0.57 | -0.03 | 0.67 | 0.33 | 0.215 | -0.0575 | 0.1875 | 0.355697 | 0.362224 | 0.308457 | 0.989223 | 0.997988 | 0.994666 |
| Lrfn1_Q2WF71_S713s_1 | Lrfn1_Q2W | 1.04 | 0.43 | -1.05 | 0.1 | 0.01 | 0.24 | -0.22 | -0.53 | -0.67 | -0.05 | -1.16 | -0.99 | -1.16 | -1.14 | 0.14 | -0.9 | 0.3175 | -0.3575 | -0.37 | 0.102195 | 0.56569 | 0.102317 | 0.989223 | 0.997988 | 0.994666 |
| Lrfn1_Q2WF71_S726s_1 | Lrfn1_Q2W | 0.29 | 0.21 | -0.46 | -0.01 | 0.71 | 0.13 | 0.42 | -0.01 | 0.02 | -0.17 | -0.18 | -0.07 | -0.44 | 0.53 | -0.68 | 0.11 | -0.255 | -0.8475 | -0.895 | 0.716534 | 0.279472 | 0.668396 | 0.989223 | 0.997988 | 0.996966 |
| Lrfn2_Q80TG9_S742s_1 | Lrfn2_Q80 | 1.17 | 0.73 | 0.57 | 0.34 | 1.28 | 0.9 | 1.37 | 0.98 | 0.28 | 1.41 | 0.31 | 0.85 | -0.2 | 0.87 | 0.96 | 0.97 | 0.305 | -0.1075 | -0.1275 | 0.883715 | 0.121837 | 0.976466 | 0.989223 | 0.997988 | 0.999507 |
| Lrfn4_Q80XU8_S575s_1 | Lrfn4_Q80 | 1.03 | 1.35 | 1.66 | 1.35 | 0.84 | 1.1 | 1.15 | 1.33 | 1.01 | 1.17 | 0.16 | 1.46 | 1.49 | -0.32 | 1.7 | 1.63 | 0.43 | 0.01 | -0.0525 | 0.643735 | 0.315027 | 0.248911 | 0.989223 | 0.997988 | 0.995259 |
| Lrfn4_Q80XU8_S585s_1 | Lrfn4_Q80 | -0.48 | -0.59 | 0.02 | 0.66 | -0.32 | -0.37 | 0.37 | 0.34 | -0.52 | 0 | 0.5 | 0.6 | -0.8 | -0.63 | 0.37 | 0.53 | -0.2425 | -0.3975 | -0.2225 | 0.935975 | 0.770822 | 0.534731 | 0.995762 | 0.997988 | 0.999507 |
| Lrfn4_Q80XU8_S611s_1 | Lrfn4_Q80 | -1.71 | -1.63 | -1.46 | -3.73 | -0.88 | -1.46 | -1.02 | -1.78 | -3.46 | -1.63 | -1.63 | -2.16 | -1.68 | -2.96 | -1.68 | -1.42 | 0.1025 | 0.2425 | -0.035 | 0.736061 | 0.129486 | 0.888909 | 0.989223 | 0.997988 | 0.9971 |
| Lrfn4_Q80XU8_S627s_1 | Lrfn4_Q80 | 1.25 | 0.39 | 1.47 | 1.62 | 0.84 | 1.41 | 1.47 | 1.83 | 1.49 | 1.64 | 0.54 | 1.53 | 0.64 | 0.92 | 1.46 | 1.33 | 0.8475 | -0.0875 | 0.1975 | 0.791319 | 0.557508 | 0.757339 | 0.989223 | 0.997988 | 0.999507 |
| Lrp1_Q91ZX7_S4521s_1 | Lrp1_Q91Z | 7.12 | 2.41 | 5.17 | 2.07 | 2.82 | 8.52 | 6.04 | 3.98 | 0.57 | 5.25 | 2.4 | 5.69 | 0.28 | -1.15 | 5.84 | 6.91 | 0.205 | 0.1175 | -0.095 | 0.543259 | 0.446548 | 0.632907 | 0.989223 | 0.997988 | 0.994666 |
| Lrp4_Q8VI56_S1830s_1 | Lrp4_Q8VI | 1.25 | 1.8 | 0.26 | -1.25 | 1.82 | 0.26 | -0.06 | 0.3 | 1.2 | 1.94 | -0.72 | 0.32 | 1.01 | 2.07 | -0.79 | -0.25 | 1.1475 | -0.715 | -1.2225 | 0.995116 | 0.925658 | 0.829171 | 0.997335 | 0.997988 | 1 |
| Lrpap1_P55302_S53s_1 | Lrpap1_P5 | -0.14 | 0.16 | 0.12 | -0.14 | -0.52 | -0.35 | -0.28 | -0.42 | 0.94 | 0.66 | 0.78 | -0.43 | 1.54 | -0.15 | 1.22 | 0.85 | 0.065 | 0.17 | -0.005 | 0.047424 | 0.08179 | 0.176305 | 0.989223 | 0.997988 | 0.994666 |
| Lrrc16a_D3Z030_S1099s_1 | Lrrc16a_D | 1.35 | 0.3 | 0.76 | 1.12 | -0.25 | 1.29 | -0.09 | 1.19 | 2.56 | 1.63 | -0.06 | 0.51 | 2.11 | 0.19 | 0.94 | 0.54 | -0.3925 | 0.4875 | 0.865 | 0.891946 | 0.437462 | 0.632828 | 0.989223 | 0.997988 | 0.999507 |
| Lrrc16a_D3Z030_S1149s_1 | Lrrc16a_D | -2.91 | -2.1 | -3.76 | -4.41 | -1.83 | -2.83 | -0.79 | -1.92 | 3.36 | -3.45 | -1.56 | -0.89 | 4.3 | -2.46 | 3.23 | -3.61 | -0.3475 | 0.2775 | 0.0625 | 0.060981 | 0.031525 | 0.071543 | 0.989223 | 0.997988 | 0.994666 |
| Lrrc16b_Q3UFQ8_T131t_1 | Lrrc16b_Q | -0.03 | -0.25 | 0.86 | 1.03 | -1.34 | 0.86 | -0.35 | 0.52 | 1.97 | 0.11 | -0.19 | 0.56 | 1.7 | -0.52 | 0.31 | 0.42 | 1.4525 | 2.66 | 3.66 | 0.885802 | 0.377424 | 0.69549 | 0.989223 | 0.997988 | 0.999507 |
| Lrrc4b_P0C192_S698s_1 | Lrrc4b_P0 | 0.98 | -0.76 | 2.06 | 2.48 | -0.8 | 0.67 | 1.13 | 2.95 | 1.48 | 0.6 | 1.56 | 1.4 | 0.98 | 0.36 | -0.25 | 2.26 | -0.48 | 0.21 | 0.075 | 0.659132 | 0.825493 | 0.918346 | 0.995762 | 0.997988 | 0.995259 |
| Lrrc4b_P0C192_S700s_1 | Lrrc4b_P0 | 3.05 | -0.26 | 1.76 | 2.85 | -0.71 | 2.26 | 1.29 | 3.22 | 1.68 | 2.17 | 2.01 | 1.14 | 0.78 | 0.74 | 0.93 | 1.66 | -0.2025 | 0.07 | -0.3525 | 0.26052 | 0.732875 | 0.888382 | 0.995762 | 0.997988 | 0.994666 |
| Lrrc4c_Q8C031_S631s_1 | Lrrc4c_Q8 | 3.24 | 0.39 | 1.47 | 1.83 | 0.12 | 2.95 | 1.01 | 2.28 | 2.02 | 1.3 | 1.3 | 1.49 | 0.32 | 0.48 | 0.8 | 1.44 | -0.335 | -0.1 | -0.8225 | 0.121738 | 0.850911 | 0.717701 | 0.996349 | 0.997988 | 0.994666 |
| Lrrc7_E9Q6L9_S1123s_1 | Lrrc7_E9Q | -0.73 | -0.39 | -0.18 | -0.44 | 0.31 | -0.8 | -0.08 | -0.58 | -0.88 | -0.65 | -0.37 | -0.25 | -0.52 | 0.01 | -0.71 | -0.36 | -0.1425 | -0.205 | -0.9725 | 0.883978 | 0.624179 | 0.686904 | 0.989223 | 0.997988 | 0.999507 |
| Lrrc7_E9Q6L9_S1137s_1 | Lrrc7_E9Q | -0.37 | -0.88 | -0.28 | -0.01 | -0.34 | -0.14 | -0.41 | -0.31 | -0.45 | -0.46 | 0.08 | 0.04 | -0.5 | 0.22 | -0.91 | -0.78 | 0.1475 | -0.1025 | 0.04 | 0.755228 | 0.733068 | 0.514242 | 0.995762 | 0.997988 | 0.9971 |
| Lrrc7_E9Q6L9_S1162s_1 | Lrrc7_E9Q | -0.72 | 0.09 | 0.13 | 0.08 | 0.34 | -0.01 | 0.48 | 0.32 | -0.32 | -0.11 | 0.56 | 0.24 | -0.08 | 0.81 | -0.27 | -0.1 | 0.085 | 0.1875 | -0.1075 | 0.579494 | 0.176017 | 0.534211 | 0.989223 | 0.997988 | 0.995259 |
| Lrrc7_E9Q6L9_S1211s_1 | Lrrc7_E9Q | -0.21 | 0.35 | -0.85 | -0.89 | 0.41 | 0.09 | -1.54 | -2.32 | 0.42 | -0.02 | -0.55 | -1.85 | 0.66 | 0.82 | -0.97 | -1.17 | 0.3875 | 0.1975 | 0.195 | 0.674169 | 0.494795 | 0.851366 | 0.989223 | 0.997988 | 0.996159 |
| Lrrc7_E9Q6L9_S1395s_1 | Lrrc7_E9Q | 0.63 | -0.35 | 0.36 | 0.83 | 0.26 | 1.26 | 0.61 | 1.09 | -0.53 | 0.14 | 1.19 | 0.61 | -0.37 | 0.02 | 0.29 | 0.15 | -0.44 | -0.1 | 0.235 | 0.314882 | 0.227449 | 0.972364 | 0.989223 | 0.997988 | 0.994666 |
| Lrrc7_E9Q6L9_S1444s_1 | Lrrc7_E9Q | 0.23 | -0.4 | -0.09 | -0.22 | -0.12 | 0.5 | 0.2 | 0.33 | -0.6 | 0.22 | -0.41 | 0.03 | -0.7 | -0.56 | -0.42 | -0.19 | 0.4375 | -0.015 | -0.345 | 0.210864 | 0.180924 | 0.804169 | 0.989223 | 0.997988 | 0.994666 |
| Lrrc7_E9Q6L9_S713s_1 | Lrrc7_E9Q | 0.82 | 0.52 | 0.98 | 0.74 | -0.11 | 2.04 | -0.55 | 0.67 | 0.77 | 0.47 | 1.43 | 0.69 | 0.35 | 0.4 | -1.09 | -0.66 | 0.3475 | -0.07 | -0.3475 | 0.026407 | 0.631955 | 0.788756 | 0.989223 | 0.997988 | 0.994666 |
| Lrrc7_E9Q6L9_S855s_1 | Lrrc7_E9Q | 1.25 | -1.33 | -0.28 | 0.99 | 0.59 | 1.57 | 0.82 | 1.61 | 0.35 | -0.47 | 0.57 | 0.62 | -3.21 | 0.01 | -1.02 | -0.63 | -0.2525 | 0.075 | -1.015 | 0.116761 | 0.115873 | 0.853391 | 0.989223 | 0.997988 | 0.994666 |
| Lrrc7_E9Q6L9_S874s_1 | Lrrc7_E9Q | 0.58 | 0 | -0.79 | -0.13 | 0.93 | 0.19 | 0.09 | -0.7 | 0.03 | -0.13 | -0.49 | -0.1 | -0.95 | 0.44 | -1.35 | 0.22 | 0.99 | 0.11 | -1.37 | 0.514849 | 0.611181 | 0.789741 | 0.989223 | 0.997988 | 0.994666 |
| Lrrc7_E9Q6L9_S954s_1 | Lrrc7_E9Q | 0.23 | 0.68 | 0.86 | 0.65 | 0.66 | 0.22 | 0.75 | 0.51 | -0.1 | 0.1 | 0.33 | 0.76 | -0.07 | 0.71 | 0.31 | 0.49 | 0.2125 | -0.0875 | -0.325 | 0.400179 | 0.771957 | 0.252949 | 0.995762 | 0.997988 | 0.994666 |
| Lrrc7_E9Q6L9_T870t_1 | Lrrc7_E9Q | 2.59 | -0.23 | 0.38 | 0.78 | -0.03 | 1.94 | 1.1 | 1.21 | 2.31 | 1.97 | 0.14 | 0.6 | 0.05 | 0.78 | -0.39 | 0.32 | -0.07 | -0.3325 | -0.245 | 0.26544 | 0.787389 | 0.605657 | 0.995762 | 0.997988 | 0.994666 |
| Lrrc75a_Q7TSF4_S15s_1 | Lrrc75a_Q | 1.37 | 0.85 | 1.41 | 0.97 | 0.64 | 1.36 | 0.32 | 0.61 | 1.42 | 1.07 | 0.1 | 0.83 | 1.65 | 0.24 | 0.85 | 1.13 | 0.175 | 0.375 | -0.69 | 0.608334 | 0.174927 | 0.388777 | 0.989223 | 0.997988 | 0.995259 |
| Lrrc75a_Q7TSF4_S322s_1 | Lrrc75a_Q | -0.64 | -0.65 | -0.5 | -0.48 | 0.16 | 0.03 | -0.37 | 0.04 | -0.82 | -0.46 | 0.87 | -1.05 | -1.02 | -0.05 | -0.1 | -1.3 | -0.4175 | -0.295 | -0.1825 | 0.886698 | 0.032317 | 0.633475 | 0.989223 | 0.997988 | 0.999507 |
| Lrrc8b_Q5DU41_S215s_1 | Lrrc8b_Q5 | 0.91 | 0.5 | 0.4 | 0.54 | 1.14 | 0.88 | 1.36 | 0.5 | 0.59 | 0.69 | 0.89 | -0.26 | 0.22 | 1.1 | 0.29 | 0.27 | 0.5325 | 0.2025 | -0.05 | 0.696097 | 0.168818 | 0.724094 | 0.989223 | 0.997988 | 0.996159 |
| Lrrfip1_E9Q9T1_S236s_1 | Lrrfip1_E9 | 0.06 | 0.01 | -0.25 | 0.3 | -0.31 | 0 | -1.15 | 0.2 | 0.87 | 0.34 | 0.52 | 0.56 | 0.55 | 1.13 | -0.91 | -0.57 | 0.3825 | -0.11 | -0.1175 | 0.966045 | 0.308703 | 0.046922 | 0.989223 | 0.997988 | 0.999507 |
| Lrrfip1_E9Q9T1_S88s_1 | Lrrfip1_E9 | 1.28 | 1.73 | -0.98 | -2.9 | 1.71 | 1.14 | -1.38 | -1.87 | 1.5 | 1.98 | -0.11 | -1.09 | 0.54 | 1.08 | -1.09 | -2.54 | -0.345 | 0.5425 | 0.02 | 0.807363 | 0.921985 | 0.492205 | 0.997335 | 0.997988 | 0.999507 |
| Lrrfip1_E9Q9T1_S90s_1 | Lrrfip1_E9 | 0.13 | 0.3 | 1.29 | 0.78 | -0.43 | 0.24 | 0.63 | 0.68 | 1.42 | -0.18 | 1.71 | 1.18 | 0.11 | -1.08 | 1.57 | 0.7 | 0.1175 | 0.7875 | -0.285 | 0.598793 | 0.353651 | 0.39596 | 0.989223 | 0.997988 | 0.995259 |
| Lrrfip2_E9QN52_S103s_1 | Lrrfip2_E9 | -0.52 | -0.25 | 0.2 | -0.14 | 0.36 | -0.22 | -0.03 | -0.33 | -0.38 | 0.21 | 0.33 | 0.02 | 0.21 | -0.54 | -0.04 | 0.25 | -0.345 | 0.4075 | -0.3 | 0.622719 | 0.640634 | 0.422424 | 0.989223 | 0.997988 | 0.995259 |
| Lrrfip2_E9QN52_S103s_2 | Lrrfip2_E9 | -0.14 | 0.76 | 0.5 | 0.52 | -0.16 | 0.53 | 0.13 | 0.8 | -0.69 | 0.58 | 0.35 | -0.22 | -1.19 | -0.71 | 0.29 | 0.74 | 0.1225 | 0.2225 | 0.1475 | 0.199567 | 0.78244 | 0.274508 | 0.995762 | 0.997988 | 0.994666 |
| Lrrfip2_E9QN52_S111s_1 | Lrrfip2_E9 | 0.9 | 0.73 | 0.27 | -0.13 | 1.15 | 0.46 | 0.85 | -0.15 | 0.53 | 1.43 | -0.74 | 0.81 | 0.13 | 1.04 | 0.22 | 0.21 | -0.085 | -0.405 | -0.6275 | 0.902613 | 0.710053 | 0.893337 | 0.995437 | 0.997988 | 0.999507 |
| Lrrfip2_E9QN52_S95s_1 | Lrrfip2_E9 | 1.26 | 0.55 | 0.34 | 0.3 | 1.26 | 0.79 | 0.51 | 0.11 | 1.67 | 1.23 | 1.53 | 0.65 | 1.55 | 2.28 | 0.33 | 0.33 | 0.135 | 0.065 | -0.0425 | 0.322511 | 0.870079 | 0.077477 | 0.996493 | 0.997988 | 0.994666 |
| Lrrfip2_E9QN52_T99t_1 | Lrrfip2_E9 | 0.38 | -0.35 | -1.32 | 0.85 | 1.15 | 1.72 | -0.02 | -0.22 | 0.21 | 0.72 | 1.43 | 0.51 | 0.42 | 0.07 | -1.08 | -0.07 | 0.055 | 0.6575 | 0.51 | 0.917796 | 0.216705 | 0.128437 | 0.989223 | 0.997988 | 0.999507 |
| Lrrtm2_Q8BGA3_S458s_1 | Lrrtm2_Q8 | -0.46 | -1.7 | -1 | -1.48 | -1.52 | 0.35 | -1.48 | -1.02 | -0.26 | -2.22 | -0.33 | -1.56 | 0.53 | -1.38 | -1.1 | -1.28 | 0.7675 | 0.8275 | -0.055 | 0.485657 | 0.613812 | 0.896207 | 0.989223 | 0.997988 | 0.994666 |
| Lrrtm4_E9Q8Y9_S462s_1 | Lrrtm4_E9 | -0.91 | -1.51 | -1.47 | -1.3 | -0.47 | -0.74 | -1.04 | -1.02 | -1.13 | -1.4 | 0.08 | -0.57 | -1.01 | -0.61 | -1.51 | -1.76 | 0.2425 | 0.0675 | 0.3525 | 0.821073 | 0.079204 | 0.161507 | 0.989223 | 0.997988 | 0.999507 |
| Lsm11_Q8BUV6_S155s_1 | Lsm11_Q8 | -0.15 | 1.18 | 0.91 | 0.87 | 0.66 | -0.17 | 1.21 | 1.61 | 0.22 | 0.7 | 1.74 | 1.37 | -0.3 | 1.16 | 0.76 | 0.93 | 0.48 | 0.5425 | 0.075 | 0.879902 | 0.78221 | 0.48901 | 0.995762 | 0.997988 | 0.999507 |
| Lsm11_Q8BUV6_S15s_1 | Lsm11_Q8 | -0.16 | 0.32 | 0.04 | -1.78 | 0.11 | -0.38 | -0.45 | -0.17 | 0.09 | 0.41 | 0.15 | 0.5 | 0.9 | 0.08 | -0.33 | -0.39 | 0.125 | 0.305 | -0.065 | 0.388552 | 0.705712 | 0.163638 | 0.995437 | 0.997988 | 0.994666 |
| Lsm11_Q8BUV6_S15s_2 | Lsm11_Q8 | 0.15 | 0.69 | 1.07 | 0.58 | 0.49 | 0.68 | 0.35 | 0.34 | 0.75 | 0.51 | 0.33 | 0.98 | 0.54 | 0.34 | 0.06 | -0.06 | 0.1725 | 0.6825 | 0.46 | 0.196466 | 0.543647 | 0.943979 | 0.989223 | 0.997988 | 0.994666 |
| Lsm11_Q8BUV6_S18s_1 | Lsm11_Q8 | -0.7 | -0.08 | 0.45 | 0.01 | 0.14 | -0.46 | 0.01 | -0.29 | 0.71 | -0.11 | 0.69 | 0.28 | 0.94 | -1.19 | 0.79 | 0.97 | -0.1575 | 0.02 | -0.4025 | 0.402382 | 0.814934 | 0.176416 | 0.995762 | 0.997988 | 0.994666 |
| Lsm11_Q8BUV6_S21s_1 | Lsm11_Q8 | 0.08 | 0.82 | 0.7 | 0.89 | 0.84 | 0.11 | 0.7 | 0.45 | 0.63 | 0.96 | 1.01 | 0.84 | 1.29 | 1.2 | 0.93 | 0.21 | -0.07 | 0.4725 | 0.4575 | 0.412696 | 0.728714 | 0.381576 | 0.995762 | 0.997988 | 0.994666 |
| Lsm11_Q8BUV6_S21s_2 | Lsm11_Q8 | 0.2 | 0.64 | 1.07 | 0.58 | 0.42 | 0.58 | 0.35 | 0.34 | 0.77 | 0.39 | 0.33 | 0.98 | 0.59 | 0.27 | 0.06 | -0.06 | -0.0975 | 0.2375 | 0.285 | 0.187852 | 0.42524 | 0.985979 | 0.989223 | 0.997988 | 0.994666 |
| Lsm11_Q8BUV6_S285s_1 | Lsm11_Q8 | 0.98 | 2.03 | 1.85 | 1.53 | 1.84 | 1.58 | 1.5 | 1.46 | 1.41 | 1.37 | 1.81 | 2.07 | 1.82 | 1.94 | 1.74 | 1.88 | -0.2 | -0.005 | -0.4075 | 0.414293 | 0.992894 | 0.830888 | 0.999187 | 0.997988 | 0.994666 |
| Lsm11_Q8BUV6_T105t_1 | Lsm11_Q8 | -0.89 | -0.61 | -0.46 | -1.55 | -1.03 | 0.4 | -1.18 | -0.73 | -0.52 | -0.8 | -1.94 | -0.08 | 0.92 | -0.09 | -1.5 | -0.86 | -0.0025 | 0.0675 | 0.2475 | 0.366052 | 0.559255 | 0.924298 | 0.989223 | 0.997988 | 0.994666 |
| Lsm12_Q9D0R8_T75t_1 | Lsm12_Q9 | -2.2 | -1.77 | -2.15 | -2.65 | -1.27 | -2.96 | -2.53 | -3.32 | -3.51 | -2.13 | -1.16 | -2.96 | -1.97 | -1.77 | -3.03 | -2.14 | 0.2425 | 0.0425 | 0.495 | 0.921923 | 0.473276 | 0.629264 | 0.989223 | 0.997988 | 0.999507 |
| Lsm14a_Q8K2F8_S182s_1 | Lsm14a_Q | -2.57 | -2.39 | -1.87 | -2.18 | -1.59 | -3.15 | -1.31 | -2.12 | -3.23 | -2.47 | -1.34 | -1.79 | -2.02 | -1.19 | -1.86 | -1.69 | -0.3275 | -0.2475 | -0.035 | 0.083875 | 0.613026 | 0.916176 | 0.989223 | 0.997988 | 0.994666 |
| Lsm14a_Q8K2F8_S183s_1 | Lsm14a_Q | -2.05 | -1.79 | -1.61 | -1.45 | -1.22 | -2.34 | -1.08 | -1.39 | -2.81 | -1.91 | -1.36 | -1.42 | -1.72 | -0.82 | -1.56 | -1.53 | 0.21 | 0.045 | 0.5625 | 0.304031 | 0.508344 | 0.684503 | 0.989223 | 0.997988 | 0.994666 |
| Lsm14a_Q8K2F8_S192s_1 | Lsm14a_Q | -0.33 | -0.26 | -0.36 | 0.09 | -0.16 | -0.47 | 0.04 | -0.01 | -0.45 | -0.23 | -0.21 | -0.25 | -0.68 | -0.03 | -0.24 | 0.04 | 0.2175 | -0.15 | 0.3175 | 0.963758 | 0.777883 | 0.754941 | 0.995762 | 0.997988 | 0.999507 |
| Lsm14a_Q8K2F8_S216s_1 | Lsm14a_Q | -0.49 | -1.33 | -0.97 | -0.38 | -0.53 | -0.4 | -0.13 | -0.37 | -0.69 | -0.76 | -0.28 | -0.64 | -0.75 | -0.35 | -0.62 | -0.53 | 0.065 | -0.07 | -0.0125 | 0.448084 | 0.137893 | 0.496256 | 0.989223 | 0.997988 | 0.994666 |
| Lsr_D3YYU9_S258s_1 | Lsr_D3YYU | -0.78 | -0.77 | 0.05 | -0.62 | -3.42 | -0.11 | -0.87 | 0.13 | -0.64 | 0.77 | 0.15 | -0.28 | 0.48 | -0.18 | 0.35 | -1.3 | 0.435 | 0.2 | 0.23 | 0.41359 | 0.470734 | 0.175181 | 0.989223 | 0.997988 | 0.994666 |
| Ltbp4_D3Z598_S1491s_1 | Ltbp4_D3Z | -1.87 | -2.42 | -2.27 | -1.72 | -1.44 | -1.75 | -1.76 | -2.11 | -1.99 | -1.53 | -1.87 | -1.52 | -2.23 | -2.36 | -1.72 | -1.52 | -0.5375 | 0.53 | 0.3675 | 0.719373 | 0.261487 | 0.217648 | 0.989223 | 0.997988 | 0.9971 |
| Ltn1_Q14B86_S18s_1 | Ltn1_Q14B | -4.25 | -3.83 | -2.97 | -6.21 | -2.68 | -3.32 | -3.68 | -4.51 | -3.7 | -4.52 | -1.09 | -3.85 | -1.55 | -3.8 | -2.4 | -3 | 0.305 | 0.3425 | 0.1125 | 0.050789 | 0.283667 | 0.274164 | 0.989223 | 0.997988 | 0.994666 |
| Luc7l3_Q5SUF2_S425s_1 | Luc7l3_Q5 | -0.72 | 0.08 | 0.21 | -0.15 | -0.06 | -0.98 | 0.12 | -0.1 | -0.35 | -0.33 | 1.38 | 0.17 | -0.52 | 0.72 | 0.96 | 0.15 | 0.7675 | 1.025 | 1.6275 | 0.251352 | 0.740083 | 0.419073 | 0.995762 | 0.997988 | 0.994666 |
| Lurap1_Q9D6I9_S118s_1 | Lurap1_Q9 | -0.84 | -0.24 | -0.51 | -0.23 | -0.77 | -0.06 | -0.37 | -1.32 | -0.59 | 0.04 | -0.92 | -0.78 | -0.46 | -0.97 | -1.43 | -1.67 | -0.11 | 0.3625 | 0.4725 | 0.070231 | 0.589308 | 0.71835 | 0.989223 | 0.997988 | 0.994666 |
| Lurap1_Q9D6I9_S126s_1 | Lurap1_Q9 | -0.04 | 0.24 | -0.4 | 0.47 | 0.1 | -1.04 | 0.4 | -0.6 | 0.76 | -0.2 | -0.97 | -0.55 | 0.29 | 0.28 | -0.26 | -0.07 | -0.175 | -0.1075 | -0.6775 | 0.979762 | 0.354371 | 0.458732 | 0.989223 | 0.997988 | 1 |
| Lurap1l_Q8K2P1_S72s_1 | Lurap1l_Q | 0.24 | 1.17 | -0.72 | 0.06 | -0.98 | 0.68 | 0.12 | 0.24 | -1.32 | 0.23 | -1.21 | 0.19 | 0.55 | 0.3 | 0.76 | 1.24 | -0.3525 | -0.3075 | -0.0075 | 0.241688 | 0.721931 | 0.204542 | 0.995437 | 0.997988 | 0.994666 |
| Luzp1_Q8R4U7_S395s_1 | Luzp1_Q8 | 1.48 | 0.67 | -0.54 | -1.3 | -0.37 | 1.21 | -1.09 | -1.24 | 3.44 | 1.57 | -2.63 | -0.84 | 2.81 | -0.63 | -0.74 | -0.52 | -0.1725 | -0.715 | 0.525 | 0.869299 | 0.543657 | 0.811304 | 0.989223 | 0.997988 | 0.999507 |
| Luzp1_Q8R4U7_S612s_1 | Luzp1_Q8 | -0.83 | -0.57 | 0.84 | -0.42 | -0.78 | -1.25 | 0.76 | -0.68 | -2.27 | -0.06 | 1.43 | -0.31 | -1.52 | -0.56 | -0.62 | -1.95 | -0.45 | 0.3075 | 0.1525 | 0.082276 | 0.642871 | 0.939483 | 0.99 | 0.997988 | 0.994666 |
| Luzp1_Q8R4U7_S746s_1 | Luzp1_Q8 | -1.86 | -2.68 | -1.63 | -1.62 | -1.63 | -1.54 | -2.34 | -2.5 | -2.4 | -2.58 | 1 | -2.35 | -2.07 | -1.35 | -0.08 | -0.55 | -0.2425 | -0.0575 | -0.9175 | 0.077724 | 0.875515 | 0.650562 | 0.996493 | 0.997988 | 0.994666 |
| Luzp2_Q8BGY3_S279s_1 | Luzp2_Q8B | 1.14 | 0.98 | 0.78 | 0.65 | 0.61 | 0.41 | 0.49 | 0.54 | 0.5 | 0.9 | 0.76 | 0.62 | 0.25 | 0.69 | 0.42 | 0.25 | -0.055 | 0.365 | 0.935 | 0.083009 | 0.104001 | 0.416539 | 0.989223 | 0.997988 | 0.994666 |
| Lysmd1_Q9D0E3_S23s_1 | Lysmd1_Q | -0.55 | -0.58 | -1.05 | -1.02 | -1.31 | -1.24 | -0.57 | -0.71 | -1.81 | -0.22 | 0.18 | -1.39 | -2 | -0.3 | -0.63 | -1.39 | -0.375 | -0.1925 | -0.485 | 0.498627 | 0.565438 | 0.982929 | 0.989223 | 0.997988 | 0.994666 |
| Lysmd2_G3X915_S24s_1 | Lysmd2_G | 4.39 | 3.45 | 4.22 | 1.71 | 3.85 | 4.09 | 3.49 | 3.9 | -1.06 | 4.31 | -1.23 | 4.13 | -0.46 | -1.42 | 4.23 | 4.27 | -0.1575 | -0.01 | -0.28 | 0.222284 | 0.494276 | 0.212014 | 0.989223 | 0.997988 | 0.994666 |
| Lysmd2_G3X915_S31s_1 | Lysmd2_G | 0.16 | 0.04 | -0.08 | 0.03 | 0.18 | -0.19 | 0.22 | 0.36 | -0.29 | -0.25 | 0.28 | 0.41 | -0.6 | 0.36 | -0.35 | 0.12 | 0.39 | -1.905 | -1.7875 | 0.597753 | 0.631576 | 1 | 0.989223 | 1 | 0.995259 |

| | | | | | | | | | | | | | | | | | | | | | | | | | | |
|---|---|---|---|---|---|---|---|---|---|---|---|---|---|---|---|---|---|---|---|---|---|---|---|---|---|---|
| Lyst_G5E8Q0_S2098s_1 | Lyst_G5E8 | -0.72 | 0.13 | -0.23 | -0.39 | 0.29 | -0.84 | 0.33 | -0.57 | -1.07 | -0.6 | -0.99 | -0.56 | -0.92 | 0.08 | -0.51 | -0.44 | 0.105 | 1.18E-16 | -0.155 | 0.650974 | 0.763952 | 0.093823 | 0.995762 | 0.997988 | 0.995259 |
| Lyst_G5E8Q0_S2117s_1 | Lyst_G5E8 | -0.04 | 0.32 | 0.28 | -0.19 | 0.57 | -0.46 | 0.95 | -0.02 | -0.32 | -0.09 | -0.95 | 0.19 | -0.34 | 0.64 | -0.32 | 0.47 | 0.105 | -0.5025 | -0.145 | 0.951381 | 0.625761 | 0.23095 | 0.989223 | 0.997988 | 0.999507 |
| Lyst_G5E8Q0_S2202s_1 | Lyst_G5E8 | 1.08 | 1.48 | 1.48 | 1.92 | 1.14 | 1.18 | 1.2 | 1.87 | 1.66 | 1.47 | 1.65 | 1.43 | -0.34 | 0.82 | 2.44 | 1.91 | 0.1675 | -0.385 | 0.02 | 0.631768 | 0.613693 | 0.805389 | 0.989223 | 0.997988 | 0.995259 |
| Lyst_G5E8Q0_S2559s_1 | Lyst_G5E8 | -2.25 | -0.7 | -0.37 | -1.03 | -1.66 | -1.84 | -1.04 | -0.11 | 0.18 | -1.19 | -0.09 | -0.83 | 0.88 | -0.92 | -1.53 | -0.76 | -0.1425 | 0.0625 | -0.2825 | 0.409656 | 0.884454 | 0.237175 | 0.996493 | 0.997988 | 0.994666 |
| Lyst_G5E8Q0_S2559s_2 | Lyst_G5E8 | -1.28 | -1.34 | -2.99 | -0.87 | -1.43 | -1.44 | -2.39 | -1.11 | -0.98 | -1.31 | -1.48 | -1.79 | -0.68 | -1.11 | -1.04 | -1.13 | -0.075 | 0.605 | 0.505 | 0.195579 | 0.955983 | 0.629454 | 0.999187 | 0.997988 | 0.994666 |
| Lyst_G5E8Q0_S2563s_1 | Lyst_G5E8 | -1.04 | -1.99 | -2.01 | -1.4 | -1.36 | -1.16 | -1.32 | -1.23 | 0.09 | -1.44 | -1.55 | -2.27 | 1.1 | -0.36 | -0.93 | -1.1 | 0.0275 | 0.23 | 0.63 | 0.031357 | 0.237125 | 0.541758 | 0.989223 | 0.997988 | 0.994666 |
| Lyst_G5E8Q0_S2563s_2 | Lyst_G5E8 | -1.28 | -1.34 | -2.99 | -0.87 | -1.43 | -1.44 | -2.39 | -1.11 | -0.98 | -1.31 | -1.48 | -1.79 | -0.68 | -1.11 | -1.04 | -1.13 | 0.3425 | 0.3175 | 1.2875 | 0.195579 | 0.955983 | 0.629454 | 0.999187 | 0.997988 | 0.994666 |
| Lyst_G5E8Q0_S2613s_1 | Lyst_G5E8 | 0.3 | -0.11 | 0.21 | 0.47 | -0.04 | 0.42 | 0.11 | 0.3 | 0.51 | 0.38 | 0.07 | 0 | 0.38 | 0.32 | -0.41 | 0.03 | 0.0275 | 0.23 | 0.63 | 0.635392 | 0.931244 | 0.927846 | 0.998075 | 0.997988 | 0.995259 |
| Lyst_G5E8Q0_S669s_1 | Lyst_G5E8 | 0.4 | 0.03 | 1.33 | 0.5 | 0.46 | 0.21 | 1.3 | 1.85 | 0.49 | 0.65 | 1.39 | 1.33 | 0.05 | 0.67 | 0.35 | 0.6 | -0.02 | 0.0225 | -0.1375 | 0.667896 | 0.384787 | 0.292165 | 0.989223 | 0.997988 | 0.996159 |
| Lyst_G5E8Q0_S866s_1 | Lyst_G5E8 | -0.99 | 0.02 | -2.28 | -1.64 | -0.09 | -2.04 | -1.6 | -1.72 | -1.14 | -0.61 | -0.78 | -1.52 | 0.76 | -1.73 | -1.22 | -0.43 | 0.39 | 0.4 | -0.1475 | 0.398607 | 0.811715 | 0.675631 | 0.995762 | 0.997988 | 0.994666 |
| Lzts3_A2AHG0_T561t_1 | Lzts3_A2AH | -0.53 | -1.58 | -0.54 | 0.17 | -1.37 | -0.08 | -0.57 | 0.01 | 0.82 | -1.05 | -0.43 | 0.71 | 1.18 | -3.02 | -0.67 | -0.18 | -0.14 | 0.21 | 0.5675 | 0.949705 | 0.793464 | 0.25841 | 0.995762 | 0.997988 | 0.999507 |
| Lzts3_A2AHG0_T563t_1 | Lzts3_A2AH | -0.63 | -0.65 | 1.23 | 0.49 | -1.86 | -0.57 | 0 | -0.55 | 0.39 | -0.42 | 0.1 | 0.85 | 0.89 | -0.11 | -0.14 | 0.48 | 0.1175 | 0.6325 | -0.0525 | 0.73167 | 0.142598 | 0.810189 | 0.989223 | 0.997988 | 0.9971 |
| Macf1_Q9QXZ0_S3889s_1 | Macf1_Q9 | 1.1 | 0.58 | 0.21 | 0.6 | 0.6 | 0.9 | 0.42 | 0.7 | 0.73 | 1.17 | 0.28 | 0.31 | 0.45 | 0.84 | 0.1 | 0.3 | -0.855 | 0.12 | 0.17 | 0.511547 | 0.899914 | 1 | 0.996493 | 1 | 0.994666 |
| Macf1_Q9QXZ0_S4458s_1 | Macf1_Q9 | 0.84 | 0.98 | 0.99 | 1.02 | 0.79 | 0.57 | 0.78 | 1.08 | 0.55 | 1.02 | 1 | 1.06 | 1.08 | 0.48 | 0.92 | 0.67 | 0.0325 | -1.6E-16 | -0.2 | 0.503736 | 0.477506 | 0.825941 | 0.989223 | 0.997988 | 0.994666 |
| Macf1_Q9QXZ0_S4483s_1 | Macf1_Q9 | 0.13 | 1.05 | 0.07 | 0.17 | 0.56 | 0.61 | 0.68 | -0.22 | 0.45 | -0.86 | 0.21 | -0.22 | 0.27 | -1.02 | -0.48 | 0.11 | -0.1525 | -0.05 | -0.17 | 0.126145 | 0.87198 | 0.240077 | 0.996493 | 0.997988 | 0.994666 |
| Macf1_Q9QXZ0_S4611s_1 | Macf1_Q9 | 3.8 | 0.7 | 0.89 | 0.21 | 0.01 | 3.55 | 3.42 | 1.12 | 0.42 | 3.51 | -0.75 | 1.68 | -0.88 | 1.09 | 1.89 | 3.05 | 0.0525 | -0.46 | -0.635 | 0.911147 | 0.548047 | 0.863365 | 0.989223 | 0.997988 | 0.999507 |
| Macf1_Q9QXZ0_S5551s_1 | Macf1_Q9 | -1.57 | -0.81 | -0.63 | -0.4 | -0.72 | -0.94 | 0.17 | -0.11 | -0.88 | -0.74 | 0.13 | -0.32 | -0.77 | -0.22 | -0.59 | -0.22 | 0.625 | -0.185 | -0.1125 | 0.240942 | 0.228735 | 0.276187 | 0.989223 | 0.997988 | 0.994666 |
| Macf1_Q9QXZ0_S57s_1 | Macf1_Q9 | 0.97 | -0.26 | 0 | 0.53 | -0.08 | 0.79 | 0.33 | 0.73 | 0.36 | 0.56 | 0.18 | 0.91 | 0.05 | -0.89 | 0.53 | 0.52 | 0.4525 | 0.4 | 0.4025 | 0.550891 | 0.70184 | 0.571392 | 0.995437 | 0.997988 | 0.994666 |
| Macf1_Q9QXZ0_S5988s_1 | Macf1_Q9 | 2.34 | 2.35 | 2.56 | 1.98 | 2.64 | 2.02 | 2.59 | 2.28 | 0.56 | 2.62 | 1.32 | 2.42 | 0.39 | 1.25 | 2.66 | 2.73 | 0.1325 | 0.1925 | -0.2575 | 0.322822 | 0.762063 | 0.242499 | 0.995762 | 0.997988 | 0.994666 |
| Macf1_Q9QXZ0_S6923s_1 | Macf1_Q9 | -0.43 | -0.08 | -0.24 | -0.05 | -0.25 | 0.21 | 0.37 | -0.01 | -0.62 | 0.02 | 0.04 | -0.04 | -0.19 | 0.45 | -0.29 | -0.04 | 0.075 | -0.5775 | -0.55 | 0.507442 | 0.247576 | 0.84315 | 0.989223 | 0.997988 | 0.994666 |
| Macf1_Q9QXZ0_S7244s_1 | Macf1_Q9 | -4.62 | -3.86 | -4.32 | -6.42 | -3.27 | -4.88 | -4.9 | -5.93 | -3.32 | -3.89 | -1.39 | -4.17 | -2.14 | -2.62 | -5.29 | -5.69 | 0.28 | 0.05 | 0.1825 | 0.361818 | 0.930902 | 0.056475 | 0.998022 | 0.997988 | 0.994666 |
| Macf1_Q9QXZ0_S907s_1 | Macf1_Q9 | 0.25 | -0.19 | 0.33 | 0.44 | -0.67 | 0.89 | 0.7 | 0.39 | 0.76 | 0.34 | 1.99 | 0.19 | 0.08 | 0.13 | 0.74 | -0.32 | 0.06 | 1.6125 | 0.87 | 0.872508 | 0.745718 | 0.170453 | 0.995762 | 0.997988 | 0.999507 |
| Macrod2_Q3UYG8_S257s_1 | Macrod2_ | -2.32 | -2.48 | -2.07 | -4.42 | -1.9 | -2.36 | -6.93 | -3.56 | -1.65 | -2.04 | -2.3 | -2.84 | -1.37 | -2.15 | -2.59 | -4.35 | 0.12 | 0.6125 | -0.05 | 0.778939 | 0.433043 | 0.280809 | 0.989223 | 0.997988 | 0.998201 |
| Macrod2_Q3UYG8_S266s_1 | Macrod2_ | -2.25 | -1.56 | -1.41 | -1.83 | -1.36 | -2.58 | -1.49 | -2.39 | -1.9 | -1.76 | -0.66 | -1.8 | -0.81 | -0.69 | -1.52 | -1.57 | -0.865 | 0.615 | 0.2075 | 0.091056 | 0.594461 | 0.520273 | 0.989223 | 0.997988 | 0.994666 |
| Macrod2_Q3UYG8_S283s_1 | Macrod2_ | 0.16 | 0.97 | 1.01 | -0.56 | 0.09 | -0.46 | -1.62 | -1.72 | -1.88 | -0.82 | -1.77 | -1.4 | -0.06 | -2.66 | 0.12 | 1.16 | -0.1925 | 0.2325 | 0.615 | 0.351609 | 0.030004 | 0.001834 | 0.989223 | 0.997988 | 0.994666 |
| Macrod2_Q3UYG8_S284s_1 | Macrod2_ | 0.16 | 0.97 | 1.01 | -0.56 | 0.09 | -0.46 | -1.62 | -1.72 | -1.88 | -0.82 | -1.77 | -1.4 | -0.06 | -2.66 | 0.12 | 1.16 | -1.3225 | -1.8625 | -0.755 | 0.351609 | 0.030004 | 0.001834 | 0.989223 | 0.997988 | 0.994666 |
| Madd_E9QN47_S1038s_1 | Madd_E9Q | 0.51 | 0.42 | 0.4 | 0.67 | 0.81 | 0.23 | 0.61 | 0.8 | 0.49 | 0.61 | 1.13 | 0.77 | 0.63 | 1.01 | 0.49 | 0.33 | 0.1125 | 0.25 | 0.115 | 0.65864 | 0.622793 | 0.309605 | 0.989223 | 0.997988 | 0.995259 |
| Madd_E9QN47_S1089s_1 | Madd_E9Q | 0.42 | 1.26 | 0.88 | 1.07 | 1.99 | 0.86 | 1.77 | 0.88 | 0.31 | 0.53 | -0.24 | 1.32 | -0.38 | 1.9 | 1.37 | 1.22 | 0.4675 | -0.4275 | 0.12 | 0.808898 | 0.201003 | 0.272761 | 0.989223 | 0.997988 | 0.999507 |
| Madd_E9QN47_S1218s_1 | Madd_E9Q | -0.02 | 0.48 | 0.03 | 0.23 | 0.62 | -0.21 | 0.53 | 0.16 | 0.15 | 0.3 | -0.05 | 0.22 | 0.09 | 0.79 | 0.03 | 0.22 | 0.095 | -0.025 | 0.1025 | 0.71805 | 0.721833 | 0.913906 | 0.995437 | 0.997988 | 0.996966 |
| Madd_E9QN47_S1247s_1 | Madd_E9Q | 0.91 | 0.77 | 0.88 | 0.74 | 0.82 | 0.9 | 0.4 | 1.39 | 1.18 | 0.74 | 0.77 | 0.94 | 1.61 | 0.8 | 0.28 | 0.62 | 0.0525 | 0.0825 | 0.0025 | 0.993909 | 0.838713 | 0.709255 | 0.995762 | 0.997988 | 1 |
| Madd_E9QN47_S1480s_1 | Madd_E9Q | 1.39 | 1.17 | 1.5 | 1.1 | 0.24 | 1.82 | 0.42 | 0.68 | 2.35 | 1.22 | 0.65 | 1.24 | 2.4 | 0.13 | 1 | 1.1 | -0.5 | 0.075 | -0.1325 | 0.77414 | 0.191375 | 0.84113 | 0.989223 | 0.997988 | 0.99798 |
| Madd_E9QN47_S1627s_1 | Madd_E9Q | 0.04 | 0.1 | 0.92 | 0.51 | -0.19 | 0.25 | 0.65 | 0.49 | 0.61 | 0.49 | 0.35 | 0.95 | 1 | -0.18 | 0.92 | 0.87 | -0.0925 | 0.2075 | 0.26 | 0.484573 | 0.757706 | 0.477489 | 0.995762 | 0.997988 | 0.994666 |
| Madd_E9QN47_S817s_1 | Madd_E9Q | 0.41 | 0.7 | 1.12 | 0.66 | -0.36 | 1.2 | 0.05 | 0.3 | 2.05 | 0.73 | 0.56 | 0.38 | 2.47 | -0.57 | 1.19 | 0.79 | -0.425 | 0.2075 | 0.2475 | 0.675469 | 0.255396 | 0.609971 | 0.989223 | 0.997988 | 0.996159 |
| Madd_E9QN47_S819s_1 | Madd_E9Q | -0.85 | -0.7 | -0.74 | -0.99 | -2.08 | 0.1 | -1.71 | -1.21 | 1.02 | -0.53 | -0.41 | -0.71 | 1.41 | -2.01 | -1.05 | -1.09 | -0.405 | 0.6625 | 0.135 | 0.838491 | 0.377988 | 0.123943 | 0.989223 | 0.997988 | 0.999507 |
| Madd_E9QN47_S828s_1 | Madd_E9Q | 1.02 | 0.37 | 0.09 | 0.49 | 1.47 | 1.06 | 0.31 | 0.33 | 1.51 | 1.15 | -0.5 | 0.34 | -0.28 | 1.52 | -0.48 | 0.96 | 0.3 | 0.1325 | -0.0625 | 0.899277 | 0.398627 | 0.776529 | 0.989223 | 0.997988 | 0.999507 |
| Madd_E9QN47_S829s_1 | Madd_E9Q | 0.21 | 0.51 | 0.27 | 0.09 | 0.68 | 0.39 | 0.61 | 0.2 | 0.32 | 0.54 | 0.07 | 0.31 | -0.13 | 0.57 | -0.08 | 0.24 | 0.2 | 0.04 | -0.12 | 0.660004 | 0.381112 | 0.861307 | 0.989223 | 0.997988 | 0.995259 |
| Madd_E9QN47_S881s_1 | Madd_E9Q | -1.18 | -1 | -1.13 | -1.42 | -0.16 | -1.54 | -0.92 | -1.37 | -2.73 | -0.77 | 0.18 | -0.76 | -1.13 | -0.1 | -1.41 | -0.82 | 0.185 | 0.1625 | 0.3175 | 0.353798 | 0.578856 | 0.775917 | 0.989223 | 0.997988 | 0.994666 |
| Madd_E9QN47_S909s_1 | Madd_E9Q | -0.34 | -0.18 | -0.17 | -0.93 | -1.23 | 0.03 | -0.88 | -1.2 | 2.37 | -0.47 | -1.46 | -0.35 | 2.55 | -1.33 | 0.5 | -0.68 | -0.415 | 0.4275 | 0.665 | 0.398271 | 0.250668 | 0.573794 | 0.989223 | 0.997988 | 0.994666 |
| Madd_E9QN47_S968s_1 | Madd_E9Q | -1.56 | -1.42 | 0.07 | -1.45 | -1.56 | -0.01 | -0.95 | -1.01 | 0.59 | -1.63 | 0.29 | -0.12 | 1.26 | -0.95 | -0.54 | -1.05 | 0.2075 | 0.8725 | 0.77 | 0.222859 | 0.658492 | 0.155583 | 0.990434 | 0.997988 | 0.994666 |
| Madd_E9QN47_T1620t_1 | Madd_E9Q | -0.21 | 0.48 | 0.01 | 0.19 | 0.36 | 0.02 | 0.21 | 0.03 | 0.09 | 0.28 | 0 | 0.53 | 0.42 | 0.4 | 0.31 | 0.48 | 0.0375 | 0.1075 | 0.285 | 0.281312 | 0.873775 | 0.676334 | 0.996493 | 0.997988 | 0.994666 |
| Madd_E9QN47_T1620t_2 | Madd_E9Q | 1.01 | 0.79 | 0.23 | 1.2 | 0.84 | 0.44 | 0.79 | -0.12 | 0.96 | 1.32 | 0.03 | 1.66 | -0.65 | 0.22 | 1.32 | 1.18 | -0.32 | 0.185 | -0.29 | 0.551347 | 0.330224 | 0.649663 | 0.989223 | 0.997988 | 0.994666 |
| Mag_P20917_S547s_1 | Mag_P209 | -1.04 | -0.32 | -0.42 | -0.43 | -0.56 | -1.32 | -0.49 | -0.72 | -1.01 | -0.89 | -1.2 | -0.59 | -0.95 | -0.7 | -0.4 | -0.24 | -0.22 | -0.37 | -0.02 | 0.945719 | 0.445777 | 0.190262 | 0.989223 | 0.997988 | 0.999507 |
| Mag_P20917_S547s_1_1 | Mag_P209 | -1.04 | -0.32 | -0.42 | -0.43 | -0.56 | -1.32 | -0.49 | -0.72 | -1.01 | -0.89 | -1.2 | -0.59 | -0.95 | -0.7 | -0.4 | -0.24 | -0.22 | -0.37 | -0.02 | 0.945719 | 0.445777 | 0.190262 | 0.989223 | 0.997988 | 0.999507 |
| Mag_P20917_S565s_1 | Mag_P209 | -3.42 | -1.95 | 1.25 | 0.85 | -2.47 | -3.31 | 0.02 | -0.02 | -3.2 | -2.96 | 0.91 | 1.03 | -1.61 | -3.07 | 0.15 | 0.46 | -0.6275 | -0.2375 | -0.2 | 0.867766 | 0.606853 | 0.866857 | 0.989223 | 0.997988 | 0.999507 |
| Maged2_Q9ER67_S84s_1 | Maged2_Q | 0.21 | 1.19 | 0.67 | 0.28 | 0.65 | 0.63 | 1.12 | 0.74 | 0.35 | 0.86 | 1.02 | 0.28 | 0.83 | 0.66 | 1.11 | 0.91 | 0.1975 | 0.04 | 0.29 | 0.350202 | 0.494052 | 0.900385 | 0.989223 | 0.997988 | 0.994666 |
| Maged2_Q9ER67_T121t_1 | Maged2_Q | -0.18 | 0.17 | -0.99 | -0.73 | -0.71 | -0.21 | -1.38 | 0.31 | -0.51 | 0.04 | -1.35 | -0.57 | -0.05 | -0.39 | -0.07 | 0.38 | -0.065 | -0.165 | 0.4 | 0.258402 | 0.877704 | 0.672952 | 0.996493 | 0.997988 | 0.994666 |
| Magi1_Q4VBG2_S732s_1 | Magi1_Q4 | -2.19 | -3.36 | -2.84 | -3.16 | -3.91 | -1.55 | -3.26 | -2.88 | 0.15 | -2.62 | -4.34 | -2.83 | 1.4 | -3.56 | -2.84 | -3.45 | -0.0125 | 0.4775 | 0.775 | 0.466968 | 0.980465 | 0.582734 | 0.999187 | 0.997988 | 0.994666 |
| Magi1_Q6RHR9_T1235t_1 | Magi1_Q6 | 0.54 | 0.58 | 0.62 | 1.06 | 1 | 0.64 | 1.12 | 1.35 | 0.63 | 0.93 | 1.19 | 1.2 | 0.09 | 0.61 | -0.02 | 0.95 | 0.3275 | 0.2875 | -0.2925 | 0.358324 | 0.209128 | 0.2732 | 0.989223 | 0.997988 | 0.994666 |
| Magi2_Q9WVQ1_S522s_1 | Magi2_Q9 | 0.26 | -0.13 | -0.05 | 0.47 | 0.72 | 0.15 | 1.13 | 0.44 | -0.65 | 0.13 | -0.54 | 0.49 | -2.05 | 0.29 | -0.11 | -0.31 | 0.4725 | -0.28 | -0.6825 | 0.198121 | 0.121599 | 0.405786 | 0.989223 | 0.997988 | 0.994666 |
| Magi2_Q9WVQ1_S539s_1 | Magi2_Q9 | 0.55 | 0.42 | 1.04 | 0.3 | 0.85 | 0.42 | 0.96 | 0.72 | 0.42 | 0.66 | 1.52 | 1.13 | -0.37 | 0.58 | 0.55 | 1.18 | 0.16 | 0.355 | -0.0925 | 0.805701 | 0.532699 | 0.286694 | 0.989223 | 0.997988 | 0.999507 |
| Magi2_Q9WVQ1_S563s_1 | Magi2_Q9 | -0.09 | 0.21 | 0.54 | 0.6 | 0.76 | 0.2 | 1.13 | 0.6 | 0.37 | 0.24 | 0.33 | 0.81 | -0.29 | 0.6 | 0.51 | 0.4 | 0.3575 | 0.1225 | -0.01 | 0.974347 | 0.227559 | 0.646766 | 0.989223 | 0.997988 | 0.999507 |
| Magi2_Q9WVQ1_S56s_1 | Magi2_Q9 | -0.08 | -1.17 | -1.07 | -1.44 | -0.89 | -0.32 | -0.84 | -1.05 | -0.73 | -0.39 | 0.34 | -0.94 | -0.17 | -1 | -1.76 | -0.81 | 0.165 | 0.51 | 0.005 | 0.990819 | 0.631396 | 0.228037 | 0.989223 | 0.997988 | 1 |
| Magi2_Q9WVQ1_S711s_2 | Magi2_Q9 | -1.09 | -2.08 | 0.32 | 0.05 | -1.83 | -0.49 | -1.19 | -0.78 | 0.38 | -1.92 | -1.4 | -0.18 | 0.42 | -2.39 | -0.69 | 0.13 | -0.3725 | -0.08 | 0.0675 | 0.927785 | 0.512548 | 0.907883 | 0.989223 | 0.997988 | 0.999507 |
| Magi2_Q9WVQ1_S714s_1 | Magi2_Q9 | -0.69 | -1.1 | -0.51 | 0.11 | -1.14 | 0.01 | -0.79 | -0.09 | -0.7 | -1.55 | 0.18 | -0.1 | 0.28 | -2.04 | -0.8 | -0.42 | 0.045 | 0.005 | -0.1975 | 0.701887 | 0.902911 | 0.990975 | 0.996493 | 0.997988 | 0.996159 |
| Magi2_Q9WVQ1_S720s_1 | Magi2_Q9 | -1.76 | -0.41 | -0.91 | -0.29 | -1.7 | -0.55 | -0.84 | -1.3 | 0.52 | -1.21 | -1.16 | -0.74 | 3.13 | -1.71 | -1.31 | -1.21 | -0.255 | 0.195 | 0.5675 | 0.585507 | 0.530716 | 0.694107 | 0.989223 | 0.997988 | 0.995259 |
| Magi2_Q9WVQ1_S720s_2 | Magi2_Q9 | -0.85 | -0.61 | 0.03 | 0.65 | -1.35 | -0.18 | -0.74 | -0.2 | 1.18 | -0.87 | 0.44 | -0.23 | 0.8 | -1.72 | 0.1 | -0.11 | -0.4225 | 0.325 | -0.0375 | 0.948297 | 0.32263 | 0.535544 | 0.989223 | 0.997988 | 0.999507 |
| Magi2_Q9WVQ1_S720s_3 | Magi2_Q9 | 0.43 | 1.18 | 1.73 | 0.89 | -0.4 | 0.86 | 0.59 | 0.29 | 1.66 | 0.83 | 1.44 | 1.52 | 0.28 | 0 | 0.85 | 1.23 | -0.7225 | 0.305 | -0.4675 | 0.255975 | 0.080171 | 0.387845 | 0.989223 | 0.997988 | 0.994666 |
| Magi2_Q9WVQ1_S721s_1 | Magi2_Q9 | -1.47 | -1.21 | -1.33 | -0.44 | -1.66 | -0.8 | -0.86 | -1.34 | 0.06 | -1.21 | -1.43 | -1.06 | 1.28 | -1.55 | -0.76 | -1.53 | -0.0525 | 0.2025 | 0.4725 | 0.465239 | 0.870429 | 0.616236 | 0.996493 | 0.997988 | 0.994666 |
| Magi2_Q9WVQ1_S721s_2 | Magi2_Q9 | -0.52 | -0.13 | 0.32 | 0.05 | -1.11 | 0.15 | -1.19 | -0.78 | 1.27 | -0.14 | -1.4 | -0.18 | 0.96 | -1.41 | -0.69 | 0.13 | -0.6625 | -0.0425 | -0.1825 | 0.721114 | 0.087121 | 0.936499 | 0.989223 | 0.997988 | 0.9971 |
| Magi2_Q9WVQ1_S721s_3 | Magi2_Q9 | 0.43 | 1.18 | 1.73 | 0.89 | -0.4 | 0.86 | 0.59 | 0.29 | 1.66 | 0.83 | 1.44 | 1.52 | 0.28 | 0 | 0.85 | 1.23 | -0.7225 | 0.305 | -0.4675 | 0.255975 | 0.080171 | 0.387845 | 0.989223 | 0.997988 | 0.994666 |
| Magi2_Q9WVQ1_S850s_1 | Magi2_Q9 | 0.61 | 0.87 | 0.01 | 0.05 | 0.63 | 0.69 | 0.6 | 0.27 | 0.6 | 0.76 | 0.25 | 0.68 | 0.51 | 1.07 | 0.15 | -0.02 | 0.1625 | 0.1875 | 0.0425 | 0.90344 | 0.554702 | 0.518444 | 0.989223 | 0.997988 | 0.999507 |
| Magi2_Q9WVQ1_Y198y_1 | Magi2_Q9 | 0.25 | 1.41 | 0.03 | 0.8 | 0.91 | 0.83 | 1.38 | 0.93 | -0.1 | -0.05 | 0.29 | 0.93 | -0.65 | 0.88 | 0.4 | 0.82 | 0.39 | -0.355 | -0.26 | 0.571305 | 0.267178 | 0.373845 | 0.989223 | 0.997988 | 0.995259 |
| Magi3_G5E8T6_S236s_1 | Magi3_G5E | -2.21 | 0.02 | -3.16 | -2.36 | -1.77 | -2.66 | -3.74 | -1.75 | -1.38 | -1.32 | -1.65 | -0.36 | -1.28 | 0.11 | -2.05 | -5.28 | -0.5525 | 0.75 | -0.1975 | 0.863749 | 0.455082 | 0.278626 | 0.989223 | 0.997988 | 0.999507 |
| Magi3_G5E8T6_S237s_1 | Magi3_G5E | -0.93 | 1.76 | -2.11 | -3.44 | -0.33 | -2.09 | -2.16 | -2.21 | -0.89 | -0.32 | -0.45 | -1.18 | -1.01 | 0.26 | -2.49 | -3.37 | -0.5175 | 0.47 | -0.4725 | 0.690128 | 0.618753 | 0.637641 | 0.989223 | 0.997988 | 0.996159 |
| Magi3_G5E8T6_S568s_1 | Magi3_G5E | -0.1 | -0.05 | -0.4 | 0.58 | 0.68 | -0.57 | 0.76 | -0.08 | 0.68 | 0.01 | 0.56 | 0.09 | 0.18 | 0.48 | -0.38 | -0.73 | 0.19 | 0.3275 | -0.12 | 0.74223 | 0.612442 | 0.295537 | 0.989223 | 0.997988 | 0.9971 |
| Magi3_G5E8T6_S662s_1 | Magi3_G5E | 0.58 | -0.7 | 0.13 | 0.32 | -0.95 | 1.22 | -0.72 | -0.75 | 2.61 | -0.04 | -1.12 | -0.06 | 2.5 | -1.7 | 0.97 | 0.01 | -0.3825 | 0.265 | 0.3625 | 0.657411 | 0.474067 | 0.726221 | 0.989223 | 0.997988 | 0.995259 |
| Magi3_G5E8T6_S833s_1 | Magi3_G5E | 0.41 | 0.3 | -0.5 | 0.48 | 0.69 | 0.26 | 0.14 | 0.25 | 0.51 | 0.17 | 0.7 | -0.17 | 0.51 | 0.94 | 0.13 | -0.24 | 0.1625 | 0.13 | 0.1625 | 0.656214 | 0.576451 | 0.689717 | 0.989223 | 0.997988 | 0.995259 |
| Magi3_G5E8T6_S954s_1 | Magi3_G5E | 1.41 | 0.34 | 0.4 | 1.16 | 0.33 | 1 | 1.12 | 0.77 | 1.28 | 1.1 | 1.24 | 1.19 | 0.11 | 1.24 | 0.89 | 1.2 | -0.0225 | 0.375 | 0.0325 | 0.933102 | 0.945712 | 0.241897 | 0.999187 | 0.997988 | 0.999507 |
| Mal2_Q8BI08_S2s_1 | Mal2_Q8BI | 1.05 | 1.04 | 0.25 | 0.81 | 0.61 | 1.42 | 0.33 | 0.48 | 1.25 | 1.17 | 0.66 | 0.29 | 0 | 1.37 | -0.07 | 0.65 | -0.0775 | 0.055 | -0.3 | 0.453238 | 0.809608 | 0.864258 | 0.995762 | 0.997988 | 0.994666 |
| Maoa_Q64133_S388s_1 | Maoa_Q64 | 0.95 | 1.13 | 1.5 | 0.56 | 0.57 | 1.24 | 0.42 | 0.53 | 0.63 | 0.88 | 0.44 | 1.04 | 0.8 | 1.23 | 1.12 | 0.52 | -0.345 | -0.2875 | -0.1175 | 0.704079 | 0.261699 | 0.326489 | 0.989223 | 0.997988 | 0.996159 |
| Map1a_Q9QYR6_S1008s_2 | Map1a_Q9 | -0.2 | 1.29 | 0.13 | 0.25 | 0.23 | -0.69 | 0.44 | -0.21 | 0.07 | 0.54 | -0.03 | -0.34 | -0.54 | 0.6 | 0.88 | 0.31 | -0.425 | -0.3075 | -0.055 | 0.899793 | 0.296004 | 0.421857 | 0.989223 | 0.997988 | 0.999507 |
| Map1a_Q9QYR6_S1052s_1 | Map1a_Q9 | -0.67 | -0.95 | -0.58 | -0.98 | -0.58 | -1.11 | -0.59 | -0.88 | -0.91 | -0.78 | -0.85 | -0.9 | -1.8 | -0.19 | -0.91 | -0.87 | 0.005 | -0.065 | -0.1475 | 0.688439 | 0.982908 | 0.766849 | 0.999187 | 0.997988 | 0.996159 |
| Map1a_Q9QYR6_S1062s_1 | Map1a_Q9 | 0.53 | 0.76 | 0.82 | 0.48 | 0.58 | 0.16 | 0.81 | 0.52 | 0.03 | 0.56 | 0.33 | 0.08 | -0.41 | 0.63 | -0.17 | 0.3 | -0.13 | -0.3975 | -0.56 | 0.091976 | 0.57823 | 0.119343 | 0.989223 | 0.997988 | 0.994666 |
| Map1a_Q9QYR6_S1190s_1 | Map1a_Q9 | -0.34 | -0.16 | -0.51 | -0.28 | 0.16 | -1.29 | -0.07 | -0.56 | -0.41 | -0.22 | -0.6 | -0.39 | -1.26 | -0.72 | -0.42 | -0.34 | -0.1175 | -0.0825 | -0.3625 | 0.230725 | 0.727561 | 0.708099 | 0.995569 | 0.997988 | 0.994666 |
| Map1a_Q9QYR6_S1196s_1 | Map1a_Q9 | -1.12 | -0.97 | -1.45 | -1.28 | -0.77 | -1.51 | -1.08 | -2.05 | -1.24 | -0.98 | -0.31 | -1.58 | -0.99 | -1.1 | -1.13 | -1.42 | -0.1475 | 0.1775 | 0.045 | 0.857695 | 0.640214 | 0.579967 | 0.989223 | 0.997988 | 0.999507 |
| Map1a_Q9QYR6_S1205s_1 | Map1a_Q9 | 0.01 | 0.12 | -0.25 | -0.38 | -0.37 | -0.59 | -0.19 | -0.8 | -0.47 | 0.24 | -1.12 | -0.67 | -0.69 | 0.08 | -0.74 | -0.57 | -0.3625 | -0.38 | -0.355 | 0.240897 | 0.155505 | 0.267013 | 0.989223 | 0.997988 | 0.994666 |
| Map1a_Q9QYR6_S1305s_1 | Map1a_Q9 | -0.63 | -3.27 | -1.08 | -1.31 | -1.46 | -3.49 | -1.26 | -0.67 | -0.82 | -2.75 | -0.01 | -0.47 | -1.04 | -0.12 | -0.48 | -1.19 | -0.1475 | 0.56 | 0.865 | 0.160595 | 0.84264 | 0.463357 | 0.996349 | 0.997988 | 0.994666 |
| Map1a_Q9QYR6_S1306s_1 | Map1a_Q9 | -1.7 | -0.25 | -2.47 | -5.7 | -0.51 | -1.55 | -4.97 | -2.62 | -0.36 | -0.41 | -1.92 | -3.91 | -0.39 | 0 | -3.93 | -4.82 | 0.1175 | 0.88 | 0.245 | 0.865219 | 0.927091 | 0.486687 | 0.997335 | 0.997988 | 0.999507 |
| Map1a_Q9QYR6_S1310s_1 | Map1a_Q9 | -0.25 | 0.15 | -0.29 | -0.21 | 0.28 | -1.19 | -0.53 | -1.12 | 0.14 | 0.02 | -0.61 | -0.73 | -0.44 | -0.01 | -0.78 | -0.87 | -0.49 | -0.145 | -0.375 | 0.215565 | 0.189929 | 0.617017 | 0.989223 | 0.997988 | 0.994666 |
| Map1a_Q9QYR6_S1360s_1 | Map1a_Q9 | 1.06 | 0.67 | 0.61 | -0.19 | 1.03 | -0.11 | 0.6 | 0.82 | -0.07 | -0.25 | 1.25 | 0.5 | -0.88 | 0.73 | 1.12 | 0.76 | 0.0475 | -0.18 | -0.105 | 0.83092 | 0.894756 | 0.668996 | 0.996493 | 0.997988 | 0.999507 |
| Map1a_Q9QYR6_S1370s_1 | Map1a_Q9 | -0.17 | -0.19 | -1.35 | -1.76 | -0.41 | 0.07 | -2.1 | -1.39 | 1.08 | -0.85 | -2.76 | -1.16 | 0.38 | -2.1 | -2.08 | -1.71 | -0.09 | -0.055 | -0.51 | 0.439215 | 0.874541 | 0.944565 | 0.996493 | 0.997988 | 0.994666 |
| Map1a_Q9QYR6_S1580s_1 | Map1a_Q9 | 1.54 | 0.99 | 0.52 | 0.42 | 0.8 | 0.48 | 0.07 | 0.18 | 0.5 | 1.12 | 0.47 | 0.03 | 0.74 | 1.22 | 0.19 | 0.21 | -0.485 | -0.3375 | -0.2775 | 0.463854 | 0.152864 | 0.353094 | 0.989223 | 0.997988 | 0.994666 |
| Map1a_Q9QYR6_S1606s_1 | Map1a_Q9 | 0.06 | 0.26 | 0.57 | 0.5 | -0.03 | -0.07 | 0.52 | 0.19 | -0.03 | 0.45 | -0.02 | 0.15 | -0.36 | 0.37 | 0.31 | 0.13 | -0.195 | -0.21 | -0.235 | 0.410904 | 0.428649 | 0.397861 | 0.989223 | 0.997988 | 0.994666 |
| Map1a_Q9QYR6_S1648s_1 | Map1a_Q9 | -0.39 | 0 | -0.21 | 0.03 | 0.21 | -0.86 | 0.55 | -0.67 | -0.98 | -1.01 | -0.75 | -0.47 | -1.02 | -0.07 | -0.13 | -0.5 | -0.05 | -0.66 | -0.2875 | 0.350713 | 0.887721 | 0.021225 | 0.996493 | 0.997988 | 0.994666 |
| Map1a_Q9QYR6_S1747s_1 | Map1a_Q9 | 1 | 0.35 | 0.05 | 0.5 | 0.56 | 0.14 | 0.56 | 0.18 | -0.19 | 0.54 | -0.24 | 0.12 | -0.78 | 0.33 | -0.24 | -0.09 | -0.115 | -0.4175 | -0.67 | 0.072807 | 0.672842 | 0.192861 | 0.992876 | 0.997988 | 0.994666 |
| Map1a_Q9QYR6_S1768s_1 | Map1a_Q9 | 0.03 | -0.26 | -0.08 | -0.17 | -0.07 | -0.09 | -0.1 | -0.64 | -0.14 | 0.18 | -0.54 | -0.48 | -0.16 | 0.3 | -0.89 | 0 | -0.105 | -0.125 | -0.0675 | 0.829124 | 0.647801 | 0.621443 | 0.99033 | 0.997988 | 0.999507 |
| Map1a_Q9QYR6_S1772s_1 | Map1a_Q9 | -0.13 | 0.76 | 0.04 | 0.2 | 0.98 | -0.41 | 0.46 | -0.68 | -0.02 | 0.31 | 0.19 | -0.25 | 0.06 | 0.65 | 0.57 | -0.39 | -0.13 | -0.16 | 0.005 | 0.98837 | 0.751984 | 0.571515 | 0.995762 | 0.997988 | 1 |
| Map1a_Q9QYR6_S1789s_1 | Map1a_Q9 | -0.26 | -0.12 | -0.4 | -0.29 | -0.14 | -0.66 | -0.41 | -0.31 | -0.22 | -0.45 | -0.19 | -0.56 | -0.19 | 0.18 | -0.4 | -0.43 | -0.1125 | -0.0875 | 0.0575 | 0.822758 | 0.604976 | 0.691484 | 0.989223 | 0.997988 | 0.999507 |
| Map1a_Q9QYR6_S1993s_1 | Map1a_Q9 | -0.8 | -0.17 | -0.68 | -0.76 | -0.2 | -1.03 | -0.33 | -0.96 | -0.63 | -0.56 | -1.51 | -1.05 | -0.46 | -0.28 | -0.66 | -0.81 | -0.0275 | -0.335 | 0.05 | 0.854361 | 0.924131 | 0.283868 | 0.997335 | 0.997988 | 0.999507 |
| Map1a_Q9QYR6_S2029s_1 | Map1a_Q9 | 1.86 | 1.81 | -0.76 | -0.92 | 1.07 | 0.39 | -0.71 | -1.32 | 0.54 | 1.36 | -1.44 | -1.37 | 0.09 | -1.58 | -1.19 | -0.94 | -0.64 | -0.725 | -1.4025 | 0.088938 | 0.443309 | 0.440542 | 0.989223 | 0.997988 | 0.994666 |
| Map1a_Q9QYR6_S2048s_1 | Map1a_Q9 | 0.05 | 0.35 | 0.58 | 0.67 | 0.34 | -0.44 | 0.85 | 0.51 | -0.17 | -0.3 | 0.06 | 0.05 | -0.03 | 0.45 | 0.55 | 0.69 | -0.0975 | -0.5025 | 0.0025 | 0.992961 | 0.761443 | 0.064459 | 0.995762 | 0.997988 | 1 |
| Map1a_Q9QYR6_S2156s_1 | Map1a_Q9 | -0.35 | 0.04 | -0.1 | -0.69 | -0.25 | -0.51 | 0.33 | 0.01 | 0.31 | -0.41 | 0.52 | -0.14 | 0.13 | -0.61 | -0.46 | 0.38 | 0.17 | 0.345 | 0.135 | 0.681468 | 0.544691 | 0.271898 | 0.989223 | 0.997988 | 0.996159 |
| Map1a_Q9QYR6_S2162s_1 | Map1a_Q9 | -0.37 | -1.18 | -0.2 | -0.13 | -0.85 | -0.54 | -0.15 | 0.38 | -0.71 | -0.85 | 0.63 | -0.53 | -0.86 | -0.4 | -0.85 | -0.33 | 0.18 | 0.105 | -0.14 | 0.667965 | 0.617408 | 0.798112 | 0.989223 | 0.997988 | 0.996159 |
| Map1a_Q9QYR6_S2199s_1 | Map1a_Q9 | -0.28 | -0.42 | -0.17 | 0.08 | -0.12 | -0.65 | 0.14 | -0.08 | -0.7 | -0.37 | -0.15 | -0.32 | -0.94 | -0.24 | -0.43 | -0.02 | 0.02 | -0.1875 | -0.21 | 0.477076 | 0.936877 | 0.443575 | 0.998984 | 0.997988 | 0.994666 |
| Map1a_Q9QYR6_S2209s_1 | Map1a_Q9 | 0.19 | 0.26 | -0.59 | 0.04 | 0.13 | 0.27 | 0.32 | 0.14 | -0.23 | -0.4 | 0.46 | -0.32 | -0.56 | 0.12 | -0.7 | -0.31 | 0.24 | -0.0975 | -0.3375 | 0.299108 | 0.355576 | 0.75394 | 0.989223 | 0.997988 | 0.994666 |
| Map1a_Q9QYR6_S2226s_1 | Map1a_Q9 | 0.7 | 0.41 | 0.2 | 0.3 | 0.55 | 0.17 | 0.95 | 1.42 | -0.42 | 0.43 | -0.1 | 0.09 | 0.08 | 0.26 | -0.59 | 0.62 | 0.37 | -0.4025 | -0.31 | 0.347702 | 0.249478 | 0.15767 | 0.989223 | 0.997988 | 0.994666 |
| Map1a_Q9QYR6_S2230s_1 | Map1a_Q9 | -0.54 | 0.24 | 0.11 | -0.27 | 0.1 | 0.13 | 0.55 | -0.16 | -0.02 | 0.21 | -0.84 | -0.14 | -0.05 | 0.09 | 0.19 | 0.04 | 0.27 | -0.0825 | 0.1825 | 0.506926 | 0.333723 | 0.795511 | 0.989223 | 0.997988 | 0.994666 |
| Map1a_Q9QYR6_S2591s_1 | Map1a_Q9 | -1.96 | -3.42 | -2.32 | -4.48 | -2.56 | -2.58 | -2.86 | -3.28 | 0.52 | -2.96 | -5.43 | -2.44 | 1.38 | -3.07 | -1.14 | -2.25 | 0.225 | 0.4675 | 1.775 | 0.095373 | 0.677408 | 0.693757 | 0.99315 | 0.997988 | 0.994666 |
| Map1a_Q9QYR6_S2630s_1 | Map1a_Q9 | -1.06 | -0.39 | -0.34 | -1.92 | -1.33 | -1.67 | -4.22 | -0.92 | -0.63 | -1.16 | 0.93 | -0.75 | 0.48 | -2.02 | -0.46 | -0.99 | -1.1075 | 0.525 | 0.18 | 0.759021 | 0.153446 | 0.349043 | 0.989223 | 0.997988 | 0.9971 |
| Map1a_Q9QYR6_S2637s_1 | Map1a_Q9 | 0.43 | -0.76 | 0.62 | 1.39 | 0.61 | 0.39 | 0.29 | 1.79 | 0.65 | 0.4 | 0.22 | 0.25 | 0.4 | 0.36 | 0.08 | 0.93 | 0.35 | -0.04 | 0.0225 | 0.961145 | 0.501479 | 0.927597 | 0.989223 | 0.997988 | 0.999507 |
| Map1a_Q9QYR6_S384s_1 | Map1a_Q9 | -0.16 | 0.01 | 0.03 | -0.37 | -0.34 | -0.72 | -0.98 | 0.41 | -0.31 | -0.48 | 1.49 | -0.44 | 0.61 | -0.18 | -0.13 | -0.49 | -0.285 | 0.1875 | 0.075 | 0.807223 | 0.394323 | 0.687213 | 0.989223 | 0.997988 | 0.999507 |
| Map1a_Q9QYR6_S500s_1 | Map1a_Q9 | -2.99 | -2.7 | -3.64 | -4.2 | -2.32 | -3.74 | -4.25 | -5.29 | -3.51 | -2.49 | -2.2 | -4.85 | -2.07 | -3.78 | -2.01 | -2.36 | -0.5175 | 0.12 | 0.8275 | 0.125969 | 0.417042 | 0.848028 | 0.989223 | 0.997988 | 0.994666 |
| Map1a_Q9QYR6_S611s_1 | Map1a_Q9 | -0.11 | -0.48 | 0.39 | 0.74 | -1.38 | -0.09 | -0.96 | 0.2 | 0.23 | -0.62 | 1.25 | -0.63 | 0.35 | -2.03 | -0.11 | -0.45 | -0.6925 | -0.0775 | -0.695 | 0.219465 | 0.132839 | 0.874714 | 0.989223 | 0.997988 | 0.994666 |
| Map1a_Q9QYR6_S644s_1 | Map1a_Q9 | -2.29 | -1.8 | -4.51 | -7.09 | -2.51 | -2.4 | -7.53 | -5.84 | -1.91 | -2.31 | -2.77 | -4.22 | -0.66 | -2.54 | -4.1 | -4.44 | -0.6475 | 1.12 | 0.9875 | 0.448093 | 0.667509 | 0.343376 | 0.99282 | 0.997988 | 0.994666 |
| Map1a_Q9QYR6_S667s_1 | Map1a_Q9 | -3.16 | -2.17 | 0.26 | 0.23 | -2.1 | -3.45 | 0.43 | -0.04 | -2.78 | -2.45 | -0.01 | -0.02 | -2.45 | -1.53 | 0.07 | 0.07 | -0.08 | -0.105 | 0.25 | 0.788556 | 0.940874 | 0.917261 | 0.999187 | 0.997988 | 0.999507 |
| Map1a_Q9QYR6_S678s_1 | Map1a_Q9 | -4.12 | -3.88 | -4.13 | -7.09 | -2.71 | -5.01 | -4.63 | -4.66 | -5.36 | -3.77 | -0.42 | -4.5 | -3.24 | -2.38 | -5.07 | -4.82 | 0.5525 | 1.2925 | 0.9275 | 0.304373 | 0.499289 | 0.282797 | 0.989223 | 0.997988 | 0.994666 |
| Map1a_Q9QYR6_S852s_1 | Map1a_Q9 | 0.62 | 0.34 | 0.83 | 0.76 | 1.16 | 0.54 | 1.74 | 0.5 | 0.19 | 0.63 | 1.37 | 0.75 | 0.17 | 0.56 | 0.53 | 0.99 | 0.3475 | 0.0975 | -0.075 | 0.788302 | 0.300358 | 0.74884 | 0.989223 | 0.997988 | 0.999507 |
| Map1a_Q9QYR6_S895s_1 | Map1a_Q9 | 0.34 | 0.48 | -0.25 | 0.17 | 0.49 | 0.17 | 0.22 | 0.02 | -0.11 | 0.51 | -0.33 | -0.65 | -0.01 | 0.52 | 0.16 | -0.23 | 0.04 | -0.33 | -0.075 | 0.797313 | 0.870888 | 0.317421 | 0.996493 | 0.997988 | 0.999507 |
| Map1a_Q9QYR6_S908s_1 | Map1a_Q9 | -0.79 | -0.29 | -0.51 | -0.54 | -0.32 | -1.34 | -0.01 | -0.85 | -1.24 | -1.02 | -0.49 | -1.3 | -1.09 | -0.41 | -0.36 | -0.87 | -0.0975 | -0.48 | -0.15 | 0.597182 | 0.763698 | 0.101237 | 0.995762 | 0.997988 | 0.995259 |
| Map1a_Q9QYR6_S981s_1 | Map1a_Q9 | -0.38 | -0.05 | -0.31 | -0.41 | -0.42 | -0.68 | -0.29 | -0.29 | -0.79 | -0.02 | 0.1 | -0.49 | -0.2 | -0.07 | -0.33 | -0.44 | -0.1325 | -0.0125 | 0.0275 | 0.909596 | 0.543642 | 0.963918 | 0.989223 | 0.997988 | 0.999507 |
| Map1a_Q9QYR6_S991s_1 | Map1a_Q9 | -1.07 | -0.79 | -1.84 | -1.69 | -0.92 | -1.5 | -1.44 | -1.93 | -1.11 | -1.07 | -1.95 | -2.01 | -0.39 | -0.55 | -1.59 | -2.03 | -0.1 | -0.1875 | 0.2075 | 0.650645 | 0.764453 | 0.612196 | 0.995762 | 0.997988 | 0.995259 |
| Map1a_Q9QYR6_S991s_2 | Map1a_Q9 | -0.2 | 1.29 | 0.13 | 0.25 | 0.23 | -0.69 | 0.44 | -0.21 | 0.07 | 0.54 | -0.03 | -0.34 | -0.54 | 0.6 | 0.88 | 0.31 | -0.425 | -0.3075 | -0.055 | 0.899793 | 0.296004 | 0.421857 | 0.989223 | 0.997988 | 0.999507 |
| Map1a_Q9QYR6_T1712t_1 | Map1a_Q9 | -0.06 | -0.31 | -0.65 | 0.11 | 0.24 | -0.2 | -0.29 | -0.16 | -0.63 | -0.13 | -0.56 | 0.35 | -1.72 | 0.06 | -2.04 | -0.66 | 0.125 | -0.015 | -0.8625 | 0.101895 | 0.626439 | 0.961721 | 0.989223 | 0.997988 | 0.994666 |
| Map1a_Q9QYR6_T1806t_1 | Map1a_Q9 | -0.14 | -3.45 | -2.16 | -1.11 | -1.81 | -0.99 | -0.96 | -1.16 | -2.72 | -2.86 | 0.78 | -1.89 | -3.52 | -1.15 | -2.87 | -2.13 | 0.485 | 0.0425 | -0.7025 | 0.374841 | 0.464914 | 0.965211 | 0.989223 | 0.997988 | 0.994666 |
| Map1a_Q9QYR6_T1928t_1 | Map1a_Q9 | -2.11 | -3.1 | -3.32 | -10.82 | -2.31 | -3.27 | -2.78 | -2.79 | -2.24 | -1.79 | 2.43 | -2.31 | -2.39 | -0.67 | -3.39 | -3.49 | 2.05 | 3.86 | 2.3525 | 0.214086 | 0.253155 | 0.081078 | 0.989223 | 0.997988 | 0.994666 |
| Map1a_Q9QYR6_T2031t_1 | Map1a_Q9 | -0.36 | 0.05 | 0.02 | -0.21 | 0.13 | -0.49 | 0.38 | -0.13 | -0.96 | -0.4 | -0.49 | -0.48 | -0.71 | -0.08 | -0.04 | 0 | 0.0975 | -0.4575 | -0.0825 | 0.765659 | 0.708034 | 0.085361 | 0.995437 | 0.997988 | 0.9971 |
| Map1a_Q9QYR6_T2182t_1 | Map1a_Q9 | -0.46 | -2.32 | -1.68 | -2.04 | -1.49 | -1.13 | -0.99 | -1.17 | -2.44 | -2.27 | 1 | -1.35 | -2.35 | -0.48 | -2.22 | -1.96 | 0.43 | 0.36 | -0.1275 | 0.817218 | 0.303727 | 0.652946 | 0.989223 | 0.997988 | 0.999507 |
| Map1a_Q9QYR6_T633t_1 | Map1a_Q9 | 0.24 | 0.51 | -0.59 | -0.01 | 0.15 | -0.44 | 0.15 | -0.37 | -0.18 | 0.1 | -0.45 | -0.64 | -0.47 | 0.46 | -0.23 | -0.36 | -0.165 | -0.33 | -0.1875 | 0.590922 | 0.592443 | 0.310848 | 0.989223 | 0.997988 | 0.995259 |
| Map1b_P14873_S1013s_1 | Map1b_P1 | -2.17 | -0.82 | -1.41 | -2.16 | -1.38 | -1.81 | -1.65 | -1.91 | -1.5 | -1.68 | -0.55 | -1.7 | -1.32 | -0.99 | -1.58 | -1.77 | -0.0475 | 0.2825 | 0.225 | 0.559145 | 0.891588 | 0.503939 | 0.996493 | 0.997988 | 0.995259 |
| Map1b_P14873_S1073s_1 | Map1b_P1 | -2.1 | -1.42 | -0.7 | -1.28 | -1.14 | -0.98 | -1.2 | -1.01 | -1.79 | -1.39 | 0.09 | -1.18 | -0.43 | -1.55 | -0.73 | -0.9 | 0.2925 | 0.3075 | 0.4725 | 0.238823 | 0.358853 | 0.520897 | 0.989223 | 0.997988 | 0.994666 |
| Map1b_P14873_S1151s_1 | Map1b_P1 | 0.68 | 1.05 | 0.57 | 0.58 | 0.75 | 0.22 | 0.78 | 0.53 | 0.29 | 0.74 | 0.12 | 0.43 | -1.2 | 0.58 | 0.5 | 0.47 | -0.15 | -0.325 | -0.6325 | 0.170598 | 0.53363 | 0.212936 | 0.989223 | 0.997988 | 0.994666 |
| Map1b_P14873_S1204s_1 | Map1b_P1 | 1.41 | 1.82 | 1.46 | 1.15 | 1.36 | 1.09 | 1.7 | 1.23 | 0.71 | 1.94 | 0.17 | 1.3 | 0.65 | 0.83 | 1.54 | 1.71 | -0.115 | -0.43 | -0.2775 | 0.414393 | 0.644864 | 0.300563 | 0.99033 | 0.997988 | 0.994666 |
| Map1b_P14873_S1233s_1 | Map1b_P1 | -0.94 | -1.39 | -1.05 | -0.59 | -0.74 | -1.41 | -0.48 | -0.51 | -1.53 | -1.11 | -0.33 | -1.52 | -1.3 | -0.14 | -1.43 | -0.44 | 0.2075 | -0.13 | 0.165 | 0.661482 | 0.491508 | 0.706699 | 0.989223 | 0.997988 | 0.995671 |
| Map1b_P14873_S1247s_1 | Map1b_P1 | -0.27 | 0.12 | 0.1 | -0.41 | 0.31 | -1.2 | -0.08 | -0.18 | -0.5 | 0.12 | 0.88 | 0.16 | -0.57 | -0.52 | 0.25 | 0.24 | -0.1725 | 0.28 | -0.035 | 0.911464 | 0.624598 | 0.411526 | 0.989223 | 0.997988 | 0.999507 |
| Map1b_P14873_S1255s_1 | Map1b_P1 | -0.51 | 0.38 | -0.72 | 0.55 | 0.65 | -0.74 | 0.6 | -0.62 | -0.48 | 0.19 | 0.01 | -1.59 | -0.21 | 0.3 | -0.24 | 0.16 | 0.0475 | -0.3925 | 0.0775 | 0.832288 | 0.91746 | 0.425456 | 0.997335 | 0.997988 | 0.999507 |
| Map1b_P14873_S1260s_1 | Map1b_P1 | -0.98 | -0.29 | -0.35 | -0.28 | -0.48 | -1.33 | -0.14 | -0.31 | -0.85 | -0.48 | -0.93 | -0.6 | -0.73 | -0.09 | -0.54 | -0.57 | -0.09 | -0.24 | -0.0075 | 0.979238 | 0.782652 | 0.372617 | 0.995762 | 0.997988 | 1 |
| Map1b_P14873_S1307s_1 | Map1b_P1 | 0.85 | 1.82 | 0.32 | 0.63 | 1.37 | 0.99 | 0.66 | 0.41 | 1.19 | 1.09 | 0.23 | 0.34 | 0.72 | 1.74 | 0.47 | 0.31 | -0.0475 | -0.1925 | -0.095 | 0.831146 | 0.89965 | 0.636073 | 0.996493 | 0.997988 | 0.999507 |
| Map1b_P14873_S1317s_1 | Map1b_P1 | -0.59 | -0.34 | -0.8 | -0.55 | -0.1 | -0.86 | -0.22 | -0.41 | -0.67 | -0.27 | -1.56 | -0.85 | -0.23 | -0.24 | -0.54 | -0.67 | 0.1725 | -0.2675 | 0.15 | 0.557919 | 0.494267 | 0.407339 | 0.989223 | 0.997988 | 0.995259 |
| Map1b_P14873_S1334s_1 | Map1b_P1 | -0.35 | 0.1 | -0.87 | -0.84 | 0.02 | -0.58 | -0.04 | -0.38 | -0.45 | 0.24 | -1.45 | -1.01 | 0.27 | -0.04 | -0.08 | -0.65 | 0.245 | -0.1775 | 0.365 | 0.293589 | 0.417623 | 0.675491 | 0.989223 | 0.997988 | 0.994666 |
| Map1b_P14873_S1373s_1 | Map1b_P1 | 0.14 | 0.16 | -0.17 | -0.21 | 0.19 | -0.01 | 0.2 | -0.22 | 0.02 | 0.43 | -0.47 | -0.17 | -0.09 | 0.5 | -0.06 | -0.11 | 0.06 | -0.0275 | 0.08 | 0.765532 | 0.788304 | 0.919024 | 0.995762 | 0.997988 | 0.9971 |
| Map1b_P14873_S1382s_1 | Map1b_P1 | 0.27 | 0.53 | 0.25 | 0.3 | 0.49 | -0.15 | 1.1 | 0.4 | -0.08 | 0.66 | 0.99 | -0.14 | -0.4 | 0.49 | 1.03 | 1.49 | 0.1225 | 0.02 | 0.315 | 0.452275 | 0.676807 | 0.949596 | 0.99315 | 0.997988 | 0.994666 |
| Map1b_P14873_S1384s_1 | Map1b_P1 | 0.3 | 0.39 | -0.32 | -0.26 | 0.47 | -0.04 | -0.14 | -0.32 | 0 | 0.43 | -0.5 | -0.21 | -0.23 | 0.46 | -0.18 | -0.3 | -0.035 | -0.0975 | -0.09 | 0.771531 | 0.901886 | 0.750125 | 0.996493 | 0.997988 | 0.9971 |
| Map1b_P14873_S1391s_1 | Map1b_P1 | 0.38 | 0.59 | 0.4 | 0.66 | -0.12 | -0.11 | 0.21 | -0.04 | 0.53 | 1.19 | 0.41 | 0.27 | 0.37 | 0.65 | 0.36 | 0.7 | -0.5225 | 0.0925 | 0.0125 | 0.958779 | 0.030049 | 0.734612 | 0.989223 | 0.997988 | 0.999507 |
| Map1b_P14873_S1395s_1 | Map1b_P1 | -0.34 | 0.78 | 0.23 | 0.08 | 0.09 | -1.11 | 0.22 | -0.37 | -0.15 | 0.56 | -0.36 | -0.11 | -0.21 | 0.59 | 0.14 | -0.02 | -0.48 | -0.2025 | -0.0625 | 0.849448 | 0.218136 | 0.543419 | 0.989223 | 0.997988 | 0.999507 |
| Map1b_P14873_S1422s_1 | Map1b_P1 | 0.36 | -1.74 | -1.73 | -1.4 | -1.26 | 0.12 | -1.31 | -0.65 | -1.08 | -1.11 | 1.25 | -1.22 | -2.04 | -0.52 | -2.09 | -2.38 | 0.3525 | 0.5875 | -0.63 | 0.306299 | 0.522732 | 0.41286 | 0.989223 | 0.997988 | 0.994666 |
| Map1b_P14873_S1438s_1 | Map1b_P1 | 1.51 | 1.03 | -1.08 | -0.95 | 0.91 | 0.89 | -0.53 | -1.43 | 1.05 | 1.71 | -1.57 | -1.2 | 0 | 1.29 | -1.14 | -1.18 | -0.1675 | -0.13 | -0.385 | 0.625664 | 0.827914 | 0.88893 | 0.995762 | 0.997988 | 0.995259 |
| Map1b_P14873_S1497s_1 | Map1b_P1 | 0.08 | 0.2 | -0.34 | -0.14 | 0.46 | -0.04 | 0.32 | -0.28 | -0.18 | 0.19 | -0.52 | -0.38 | -0.54 | 0.49 | -0.29 | -0.05 | 0.165 | -0.1725 | -0.0475 | 0.8769 | 0.526357 | 0.513338 | 0.989223 | 0.997988 | 0.999507 |
| Map1b_P14873_S1523s_1 | Map1b_P1 | -1.65 | -0.33 | -0.07 | -0.7 | -0.75 | -0.84 | -0.44 | -0.1 | -0.5 | -0.71 | 0.31 | 0.76 | 0.53 | -0.48 | 0.02 | -0.53 | 0.155 | 0.6525 | 0.5725 | 0.197495 | 0.681792 | 0.184663 | 0.993835 | 0.997988 | 0.994666 |
| Map1b_P14873_S1616s_1 | Map1b_P1 | -1.88 | -0.84 | -0.01 | 0.04 | -1.77 | -1.38 | 0.3 | 0.1 | -1.76 | -1.66 | 0.6 | -0.32 | -0.54 | -1.42 | -0.57 | -0.64 | -0.015 | -0.1125 | -0.12 | 0.801076 | 0.980479 | 0.863897 | 0.999187 | 0.997988 | 0.999507 |
| Map1b_P14873_S1627s_1 | Map1b_P1 | -0.01 | -0.92 | -0.85 | -0.19 | -1.23 | 0.91 | -0.74 | -0.87 | 2.3 | -0.46 | -0.87 | -1.44 | 0.8 | -1.27 | -0.58 | -0.8 | 0.01 | 0.375 | 0.03 | 0.949988 | 0.983578 | 0.628942 | 0.999187 | 0.997988 | 0.999507 |
| Map1b_P14873_S1627s_2 | Map1b_P1 | 1.56 | 0.19 | 0.18 | 0.63 | -1.23 | 1.89 | -0.7 | 0.04 | 2.57 | 0.37 | 0.15 | 0.08 | 0.57 | -0.61 | -0.27 | 0.07 | -0.64 | 0.1525 | -0.7 | 0.114661 | 0.350117 | 0.805555 | 0.989223 | 0.997988 | 0.994666 |
| Map1b_P14873_S1649s_1 | Map1b_P1 | -0.19 | 0.38 | -0.41 | -0.8 | 0.12 | -0.13 | -0.25 | -0.61 | 0.07 | 0.26 | -1.3 | -0.53 | -0.45 | 0.18 | -0.92 | -0.6 | 0.0375 | -0.12 | -0.1925 | 0.597185 | 0.903438 | 0.775552 | 0.996493 | 0.997988 | 0.995259 |
| Map1b_P14873_S1775s_1 | Map1b_P1 | 0.88 | 1.2 | 1.07 | 0.79 | 0.49 | 0.27 | 0.75 | 0.24 | 0.35 | 1.29 | 0.6 | 0.64 | 0.45 | 0.66 | 0.95 | 0.7 | -0.5475 | -0.265 | -0.295 | 0.257325 | 0.035633 | 0.348579 | 0.989223 | 0.997988 | 0.994666 |
| Map1b_P14873_S1781s_1 | Map1b_P1 | 0.32 | 0.93 | 0.92 | 0.73 | 0.63 | -0.25 | 0.71 | 0.28 | 0.43 | 1.23 | 0.38 | 0.34 | -0.33 | 0.85 | 1.04 | 0.78 | -0.3825 | -0.13 | -0.14 | 0.701413 | 0.208857 | 0.663172 | 0.989223 | 0.997988 | 0.996159 |
| Map1b_P14873_S1781s_2 | Map1b_P1 | 0.41 | 0.35 | 0.4 | 1.11 | 0.04 | 0.56 | 0.66 | 0.99 | 0.76 | 0.71 | 0.76 | 0.49 | -0.17 | 0.71 | 0.18 | 0.28 | -0.005 | 0.1125 | -0.3175 | 0.319853 | 0.986463 | 0.665802 | 0.999187 | 0.997988 | 0.994666 |
| Map1b_P14873_S1789s_1 | Map1b_P1 | 0.39 | 0.64 | 0.35 | 0.44 | 0.4 | 0.01 | 0.72 | 0.08 | 0.19 | 0.68 | 0.1 | 0.1 | -0.09 | 0.75 | 0.48 | 0.34 | -0.1525 | -0.1875 | -0.085 | 0.755554 | 0.530204 | 0.440984 | 0.989223 | 0.997988 | 0.9971 |
| Map1b_P14873_S1789s_2 | Map1b_P1 | -0.21 | 0.58 | -0.17 | -0.15 | 0.21 | -0.19 | 0.3 | 0.02 | 0.27 | 0.61 | -0.56 | -0.23 | -0.66 | 0.28 | -0.4 | 0.1 | 0.0725 | 0.01 | -0.1825 | 0.583098 | 0.784251 | 0.976579 | 0.995762 | 0.997988 | 0.995259 |
| Map1b_P14873_S1814s_1 | Map1b_P1 | -0.38 | -0.02 | 0.12 | 0.18 | 0.17 | -0.52 | 0.16 | 0.09 | -0.22 | 0.17 | -0.35 | 0.17 | 0.01 | 0.3 | -0.08 | -0.03 | 7.85E-17 | -0.0325 | 0.075 | 0.770348 | 1 | 0.898331 | 1 | 0.997988 | 0.9971 |
| Map1b_P14873_S1877s_1 | Map1b_P1 | -1.42 | -2.16 | -0.49 | -4.3 | -0.85 | -1.9 | -3.01 | -0.75 | -0.57 | -1.12 | -1.38 | -4.47 | -1.26 | -0.38 | -2.45 | -0.66 | 0.465 | 0.2075 | 0.905 | 0.286519 | 0.584528 | 0.843816 | 0.989223 | 0.997988 | 0.994666 |
| Map1b_P14873_S1911s_1 | Map1b_P1 | -0.93 | -1.09 | 0.1 | 1.08 | -0.47 | -0.95 | 0.96 | 0.95 | -1.28 | -1.3 | 0.38 | 0.46 | -1.79 | -0.37 | 0.4 | 0.28 | 0.3325 | -0.225 | -0.16 | 0.803775 | 0.598163 | 0.726636 | 0.989223 | 0.997988 | 0.999507 |
| Map1b_P14873_S2030s_1 | Map1b_P1 | 1.12 | 0.73 | 0.11 | 0.34 | 0.04 | 0.29 | 0.3 | 0.1 | 0.31 | 1.23 | 0.04 | 0.02 | -0.3 | 0.82 | 0.29 | 0.11 | -0.3925 | -0.175 | -0.345 | 0.338119 | 0.172611 | 0.637387 | 0.989223 | 0.997988 | 0.994666 |
| Map1b_P14873_S2054s_1 | Map1b_P1 | 1.82 | -1.31 | 0.85 | 1.43 | -0.02 | 1.75 | 0.31 | 1.97 | -0.45 | 0.25 | 2.09 | 1.06 | -2.17 | 0.56 | 0.3 | 1.02 | 0.305 | 0.04 | -0.77 | 0.389713 | 0.686929 | 0.959681 | 0.994672 | 0.997988 | 0.994666 |
| Map1b_P14873_S2122s_1 | Map1b_P1 | -3.02 | -1.03 | -1.12 | -1.4 | -2.08 | -1.81 | -2 | -1.81 | -2.22 | -0.91 | -1.94 | -1.68 | -0.41 | -1.09 | -2.18 | -1.54 | -0.2825 | -0.045 | 0.3375 | 0.545192 | 0.526993 | 0.92971 | 0.989223 | 0.997988 | 0.994666 |
| Map1b_P14873_S2267s_1 | Map1b_P1 | -1.79 | -1.45 | -1.22 | -1.23 | -1.12 | -2.01 | -0.69 | -1.38 | -1.76 | -1.19 | -0.44 | -0.98 | -0.83 | -1.12 | -1.1 | -1.45 | 0.1225 | 0.33 | 0.2975 | 0.288383 | 0.70368 | 0.32907 | 0.995437 | 0.997988 | 0.994666 |
| Map1b_P14873_S2410s_1 | Map1b_P1 | 0.16 | 0.45 | 0.25 | 0.15 | 1.15 | 0.09 | 1.15 | -0.23 | 0.05 | 0.38 | -0.34 | 0.46 | -0.52 | -0.4 | 0.09 | 0.02 | 0.2875 | -0.115 | -0.455 | 0.108957 | 0.435306 | 0.660683 | 0.989223 | 0.997988 | 0.994666 |
| Map1b_P14873_S25s_1 | Map1b_P1 | 0.36 | 0.68 | 0.1 | 0.13 | 0.64 | 0.23 | 0.53 | 0.21 | 0.38 | 0.56 | -0.33 | 0.11 | -0.39 | 0.78 | -0.16 | 0.07 | 0.085 | -0.1375 | -0.2425 | 0.466924 | 0.722508 | 0.630672 | 0.995437 | 0.997988 | 0.994666 |
| Map1b_P14873_S336s_1 | Map1b_P1 | 0.79 | 0.17 | -0.15 | 0.23 | 0.31 | 0.11 | 0.28 | 0.35 | 0.66 | 1.23 | 0.68 | 0.19 | 0.9 | 0.51 | 0.36 | 0.38 | 0.0025 | 0.43 | 0.2775 | 0.360496 | 0.992175 | 0.199108 | 0.999187 | 0.997988 | 0.994666 |
| Map1b_P14873_S339s_1 | Map1b_P1 | 0.01 | 0.3 | 0.27 | -0.12 | 0.22 | -0.51 | -0.19 | 0.13 | 0.12 | 0.92 | -0.01 | -0.24 | 0.19 | 0.66 | -0.05 | -0.23 | -0.2025 | 0.0825 | 0.0275 | 0.924032 | 0.427779 | 0.78863 | 0.989223 | 0.997988 | 0.999507 |
| Map1b_P14873_S341s_1 | Map1b_P1 | -0.08 | 0.16 | 0.1 | 0.42 | 0.26 | -0.37 | 0.48 | 0.12 | -0.13 | 0.06 | 0.6 | 0.14 | -0.39 | 0.64 | -0.42 | -1.07 | -0.0275 | 0.0175 | -0.46 | 0.247163 | 0.915121 | 0.945432 | 0.997335 | 0.997988 | 0.994666 |
| Map1b_P14873_S343s_1 | Map1b_P1 | -0.38 | 0.03 | -0.37 | 0.11 | 0.21 | -0.11 | 0.09 | -0.02 | -0.18 | 0.14 | 0.04 | -0.32 | 0.18 | -0.4 | 0.09 | -0.09 | 0.195 | 0.0725 | 0.0975 | 0.719002 | 0.398524 | 0.768134 | 0.989223 | 0.997988 | 0.9971 |
| Map1b_P14873_S601s_1 | Map1b_P1 | 0.72 | -1.07 | -0.05 | -0.24 | -1.01 | 0.77 | -1.35 | -0.5 | -0.12 | -0.74 | -0.99 | -0.44 | -0.01 | -1.45 | -0.45 | -1.11 | -0.3625 | -0.4125 | -0.595 | 0.225788 | 0.504991 | 0.325931 | 0.989223 | 0.997988 | 0.994666 |
| Map1b_P14873_S614s_1 | Map1b_P1 | -0.45 | -0.38 | 0.83 | -0.03 | -0.37 | -0.52 | 0.09 | 0.32 | 0.35 | -0.19 | 0.21 | 0.57 | 0.97 | -0.45 | 0.9 | 0.42 | -0.1125 | 0.2425 | 0.4675 | 0.295817 | 0.75096 | 0.494286 | 0.995762 | 0.997988 | 0.994666 |
| Map1b_P14873_S934s_1 | Map1b_P1 | -0.45 | -0.3 | -1.5 | -1.57 | -0.01 | -0.45 | -1.27 | -1.7 | -0.7 | -0.38 | -1.76 | -1.47 | -0.99 | -0.1 | -1.82 | -1.62 | 0.0975 | -0.1225 | -0.1775 | 0.716787 | 0.836447 | 0.785195 | 0.995762 | 0.997988 | 0.996966 |
| Map1b_P14873_S960s_1 | Map1b_P1 | -1.55 | -0.62 | -0.87 | -1.07 | -1.34 | -1.28 | -1.24 | -1.1 | -0.87 | -0.91 | -0.26 | -1.17 | -0.35 | -0.73 | -0.97 | -0.97 | -0.2125 | 0.225 | 0.2725 | 0.380303 | 0.414834 | 0.474144 | 0.989223 | 0.997988 | 0.994666 |
| Map1b_P14873_S989s_1 | Map1b_P1 | -2.11 | -1.68 | -0.69 | -1.45 | -2.43 | -1.91 | -1.49 | -1.53 | -1.2 | -2.11 | -0.46 | -1.03 | -0.23 | -2.46 | -1.04 | -1.05 | -0.3575 | 0.2825 | 0.2875 | 0.581012 | 0.340306 | 0.524605 | 0.989223 | 0.997988 | 0.995259 |
| Map1b_P14873_S989s_2 | Map1b_P1 | -3.13 | -2.96 | -2.59 | -3.26 | -3.23 | -2.69 | -3.44 | -2.95 | -1.84 | -2.39 | -1.39 | -2.35 | -1.34 | -2.95 | -2.46 | -2.77 | -0.0925 | 0.9925 | 0.605 | 0.15156 | 0.727372 | 0.010002 | 0.995569 | 0.997988 | 0.994666 |
| Map1b_P14873_S992s_1 | Map1b_P1 | -2.76 | -2.53 | -1.45 | -2.88 | -3.1 | -2.78 | -2.18 | -2.21 | -2.35 | -2.45 | -0.75 | -1.92 | -1.08 | -3.7 | -2.03 | -2.1 | -0.1625 | 0.5375 | 0.1775 | 0.760532 | 0.67461 | 0.28167 | 0.992876 | 0.997988 | 0.9971 |
| Map1b_P14873_S992s_2 | Map1b_P1 | -3.13 | -2.96 | -2.59 | -3.26 | -3.23 | -2.69 | -3.44 | -2.95 | -1.84 | -2.39 | -1.39 | -2.35 | -1.34 | -2.95 | -2.46 | -2.77 | -0.0925 | 0.9925 | 0.605 | 0.15156 | 0.727372 | 0.010002 | 0.995569 | 0.997988 | 0.994666 |
| Map1b_P14873_T1202t_1 | Map1b_P1 | 0.1 | 0.29 | -0.14 | -0.64 | 0.63 | -0.01 | 0.36 | -0.68 | 0 | 0.52 | -1.13 | -0.44 | -0.43 | 0.14 | 0.35 | 0.22 | 0.1725 | -0.165 | 0.1675 | 0.597861 | 0.62418 | 0.681636 | 0.989223 | 0.997988 | 0.995259 |
| Map1b_P14873_T1499t_1 | Map1b_P1 | 0.9 | 0.5 | -0.27 | -0.3 | 0.9 | 0.39 | 0.27 | -0.5 | 0.24 | 0.76 | -0.53 | -0.62 | -0.21 | 0.82 | -0.47 | -0.33 | 0.0575 | -0.245 | -0.255 | 0.544681 | 0.885189 | 0.57245 | 0.996493 | 0.997988 | 0.994666 |
| Map1b_P14873_T1629t_1 | Map1b_P1 | 0.5 | -1.6 | -0.66 | -0.37 | -1.43 | 1.01 | -0.72 | -0.64 | 1.67 | -0.59 | -1.17 | -0.79 | -0.07 | -2.48 | -0.88 | -1.01 | 0.0875 | 0.3125 | -0.5775 | 0.351206 | 0.884444 | 0.655967 | 0.996493 | 0.997988 | 0.994666 |
| Map1b_P14873_T1629t_2 | Map1b_P1 | 1.56 | 0.19 | 0.18 | 0.63 | -1.23 | 1.89 | -0.7 | 0.04 | 2.57 | 0.37 | 0.15 | 0.08 | 0.57 | -0.61 | -0.27 | 0.07 | -0.64 | 0.1525 | -0.7 | 0.114661 | 0.350117 | 0.805555 | 0.989223 | 0.997988 | 0.994666 |
| Map1b_P14873_T1784t_1 | Map1b_P1 | 0.78 | -0.52 | 0.45 | 0.37 | -0.94 | 0.45 | 1.67 | 0.55 | 0.07 | 0.03 | 1.46 | 0.22 | -0.7 | 0.59 | 0.53 | -0.33 | 0.1625 | 0.175 | -0.2475 | 0.560569 | 0.765944 | 0.684135 | 0.995762 | 0.997988 | 0.995259 |
| Map1b_P14873_T1784t_2 | Map1b_P1 | 0.41 | 0.35 | 0.4 | 1.11 | 0.04 | 0.56 | 0.66 | 0.99 | 0.76 | 0.71 | 0.76 | 0.49 | -0.17 | 0.71 | 0.18 | 0.28 | -0.005 | 0.1125 | -0.3175 | 0.319853 | 0.986463 | 0.665802 | 0.999187 | 0.997988 | 0.994666 |
| Map1b_P14873_T1925t_1 | Map1b_P1 | 0.91 | 0.93 | 0.36 | 0.17 | 0.44 | 0.3 | 0.8 | 0.31 | 0.58 | 0.9 | 0.08 | 0.21 | 0.09 | 0.43 | 0.43 | 0.42 | -0.13 | -0.15 | -0.25 | 0.38974 | 0.630606 | 0.624756 | 0.989223 | 0.997988 | 0.994666 |
| Map1b_P14873_T1928t_1 | Map1b_P1 | 0.82 | 0.97 | 0.06 | 0.48 | 0.86 | 0.43 | 0.89 | 0.22 | 0.36 | 0.56 | -0.02 | 0.05 | -0.33 | 1.12 | 0.27 | 0.19 | 0.0175 | -0.345 | -0.27 | 0.481042 | 0.951915 | 0.251421 | 0.999187 | 0.997988 | 0.994666 |
| Map1b_P14873_T1945t_1 | Map1b_P1 | -0.21 | 0.83 | 0 | -0.47 | 0.06 | 0.11 | 0.02 | -0.42 | 0.05 | 0.48 | -0.31 | -0.69 | -0.78 | 0.54 | -0.5 | -0.79 | -0.095 | -0.155 | -0.42 | 0.330864 | 0.767545 | 0.684812 | 0.995762 | 0.997988 | 0.994666 |
| Map1b_P14873_T1962t_1 | Map1b_P1 | -0.12 | -0.6 | 0.57 | 0.46 | -0.43 | -0.05 | 0.45 | 0.29 | -0.15 | -0.53 | 0.46 | 0.47 | -0.48 | 0.44 | -0.34 | 0.22 | -0.0125 | -0.015 | -0.1175 | 0.751125 | 0.970692 | 0.967872 | 0.999187 | 0.997988 | 0.9971 |
| Map1b_P14873_T2047t_1 | Map1b_P1 | -0.62 | -4.12 | -2.12 | -1.93 | -2.8 | -1.47 | -2.18 | -1.63 | -1.95 | -3.46 | 0.08 | -2.3 | -2.52 | -0.2 | -2.91 | -2.18 | 0.1775 | 0.29 | 0.245 | 0.767851 | 0.797372 | 0.751628 | 0.995762 | 0.997988 | 0.9971 |

| | | | | | | | | | | | | | | | | | | | | | | | | | | |
|---|---|---|---|---|---|---|---|---|---|---|---|---|---|---|---|---|---|---|---|---|---|---|---|---|---|---|
| Map1b_P14873_T2301t_1 | Map1b_P1 | -0.9 | -0.46 | -0.35 | -1 | -0.8 | -0.2 | -1.98 | -0.83 | -0.5 | -0.19 | -1.18 | -1.07 | 0.32 | -1.4 | -1.2 | -1.77 | -0.275 | -0.0575 | -0.335 | 0.480402 | 0.488548 | 0.855447 | 0.989223 | 0.997988 | 0.994666 |
| Map1b_P14873_T527t_1 | Map1b_P1 | -0.73 | -0.58 | -0.42 | -0.82 | -0.3 | -1.22 | -0.11 | -0.44 | -0.81 | -0.55 | -0.21 | -0.77 | -0.6 | -0.18 | -1.25 | -0.86 | 0.12 | 0.0525 | -0.085 | 0.778335 | 0.679685 | 0.829666 | 0.993464 | 0.997988 | 0.99798 |
| Map1b_P14873_T945t_1 | Map1b_P1 | -2.27 | -1.22 | -2.34 | -1.29 | -2.46 | -2.22 | -1.93 | -1.58 | -1.12 | -1.86 | -1.19 | -2.42 | -1.31 | -1.12 | -1.42 | -1.35 | -0.2675 | 0.1325 | 0.48 | 0.184796 | 0.461028 | 0.754173 | 0.989223 | 0.997988 | 0.994666 |
| Map1s_Q8C052_S458s_1 | Map1s_Q8 | -1.12 | -1.41 | -2.83 | -4.12 | -1.13 | -1.89 | -2.77 | -3.41 | -0.93 | -1.17 | -2.12 | -3.08 | -0.48 | -0.75 | -3.36 | -3.21 | 0.07 | 0.545 | 0.42 | 0.645644 | 0.925487 | 0.480022 | 0.997335 | 0.997988 | 0.995259 |
| Map1s_Q8C052_S462s_1 | Map1s_Q8 | 0.66 | 0.15 | 0.46 | 0.41 | -1.03 | 0.68 | -0.44 | 0.1 | 1.33 | 0.37 | 0.22 | 0.31 | 1.4 | -0.98 | 0.12 | 0.05 | -0.5925 | 0.1375 | -0.2725 | 0.57118 | 0.137833 | 0.66154 | 0.989223 | 0.997988 | 0.995259 |
| Map1s_Q8C052_S462s_2 | Map1s_Q8 | -0.93 | -2.07 | -2.76 | -1.83 | -2.19 | -1.55 | -2.65 | -1.46 | -1.17 | -1.34 | -0.71 | -1.19 | -0.4 | -2.96 | -2.31 | -0.71 | -0.065 | 0.795 | 0.3025 | 0.645142 | 0.882983 | 0.072218 | 0.996493 | 0.997988 | 0.995259 |
| Map1s_Q8C052_S465s_1 | Map1s_Q8 | -0.29 | -0.28 | 0.33 | -0.26 | -2.59 | -0.17 | -0.6 | 0.1 | 0.38 | -0.74 | -1.65 | 0.46 | 0.53 | -3.09 | -0.71 | -0.44 | -0.69 | -0.2625 | -0.8025 | 0.268981 | 0.240885 | 0.598491 | 0.989223 | 0.997988 | 0.994666 |
| Map1s_Q8C052_S465s_2 | Map1s_Q8 | -0.93 | -2.07 | -2.76 | -1.83 | -2.19 | -1.55 | -2.65 | -1.46 | -1.17 | -1.34 | -0.71 | -1.19 | -0.4 | -2.96 | -2.31 | -0.71 | -0.065 | 0.795 | 0.3025 | 0.645142 | 0.882983 | 0.072218 | 0.996493 | 0.997988 | 0.995259 |
| Map1s_Q8C052_S684s_1 | Map1s_Q8 | 0.34 | -0.53 | -1.29 | -0.56 | -0.04 | 0.42 | -1.18 | -0.76 | -0.16 | -0.4 | 0 | -0.36 | -0.31 | -0.17 | -1.09 | -0.6 | 0.12 | 0.28 | -0.0325 | 0.934838 | 0.792627 | 0.442404 | 0.995762 | 0.997988 | 0.999507 |
| Map2_P20357_S1013s_1 | Map2_P20 | 2.53 | 2.17 | 3.45 | 3.12 | 1.15 | 2.27 | 1.33 | 2.55 | 3.51 | 2.6 | 0.38 | 3.24 | 3.29 | 0.57 | 4.45 | 2.19 | -0.9925 | -0.385 | -0.1925 | 0.803871 | 0.040075 | 0.581006 | 0.989223 | 0.997988 | 0.999507 |
| Map2_P20357_S1023s_1 | Map2_P20 | -0.67 | 0.13 | 0.13 | -0.39 | -0.02 | -0.37 | -0.23 | -0.41 | -0.91 | 0.06 | -0.36 | 0.17 | -0.84 | 0.42 | -0.28 | -0.31 | -0.0575 | -0.06 | -0.0525 | 0.882046 | 0.827594 | 0.858584 | 0.995762 | 0.997988 | 0.999507 |
| Map2_P20357_S1161s_1 | Map2_P20 | -0.64 | -0.5 | -0.68 | -0.51 | -0.58 | -1.12 | -0.3 | -0.62 | -0.71 | -0.5 | -0.81 | -0.88 | -0.68 | 0.13 | -0.82 | -0.81 | -0.0725 | -0.1425 | 0.0375 | 0.899185 | 0.763972 | 0.513526 | 0.995762 | 0.997988 | 0.999507 |
| Map2_P20357_S1352s_1 | Map2_P20 | -2.12 | -1.42 | -0.18 | -1.75 | -1.84 | -2.21 | -0.69 | -0.51 | -2.25 | -1.57 | -1.36 | -0.56 | -0.48 | -2.14 | 0.09 | 0.22 | 0.055 | -0.0675 | 0.79 | 0.224776 | 0.918609 | 0.895052 | 0.997335 | 0.997988 | 0.994666 |
| Map2_P20357_S1485s_1 | Map2_P20 | -2.4 | -3.02 | -3.23 | -2.95 | -2.67 | -2.15 | -3.39 | -3.34 | -0.31 | -2.13 | -3.42 | -3.23 | 0.87 | -3.35 | -2.41 | -2.61 | 0.0125 | 0.6275 | 1.025 | 0.240704 | 0.971283 | 0.358031 | 0.999187 | 0.997988 | 0.994666 |
| Map2_P20357_S1539s_1 | Map2_P20 | 1.34 | 1.61 | 0.43 | 0.3 | 1.15 | 1.28 | 0.17 | 0.62 | 1.15 | 1.32 | 0.3 | 0.2 | 1.3 | 1.51 | 0.27 | 0.14 | -0.115 | -0.1775 | -0.115 | 0.803608 | 0.773332 | 0.677153 | 0.995762 | 0.997988 | 0.999507 |
| Map2_P20357_S1560s_1 | Map2_P20 | -1.64 | -1.75 | -2.13 | -1.4 | -1.54 | -1.6 | -1.11 | -1.37 | -1.46 | -1.61 | -0.22 | -1.55 | -1.58 | -0.93 | -1.98 | -1.96 | 0.325 | 0.52 | 0.1175 | 0.722462 | 0.208776 | 0.185078 | 0.989223 | 0.997988 | 0.9971 |
| Map2_P20357_S1625s_1 | Map2_P20 | -1.23 | -1.83 | -1.44 | -1.56 | -1.46 | -2.02 | -1.5 | -1.56 | -1.97 | -1.17 | -1.38 | -2.28 | -1.03 | -0.92 | -1.22 | -1.38 | -0.12 | -0.185 | 0.3775 | 0.170565 | 0.623721 | 0.562633 | 0.989223 | 0.997988 | 0.994666 |
| Map2_P20357_S1635s_1 | Map2_P20 | 0.26 | -0.36 | -3.07 | -5.84 | 0.52 | 0.22 | -2.59 | -4.56 | -2.21 | 0.41 | -0.48 | -3.3 | -0.93 | -2.1 | -3.83 | -4.21 | 0.65 | 0.8575 | -0.515 | 0.707995 | 0.682113 | 0.550716 | 0.994114 | 0.997988 | 0.996966 |
| Map2_P20357_S1654s_2 | Map2_P20 | -0.79 | -0.8 | -3.14 | -4.1 | -0.86 | -0.67 | -3.05 | -2.49 | -0.84 | -0.47 | -0.37 | -3.2 | -0.17 | -1.33 | -2.23 | -3.82 | 0.44 | 0.9875 | 0.32 | 0.747372 | 0.62376 | 0.312589 | 0.989223 | 0.997988 | 0.9971 |
| Map2_P20357_S1743s_1 | Map2_P20 | -0.99 | -1.31 | -3.41 | -1.77 | -2.11 | -0.8 | -2.4 | -1.21 | -2.44 | -1.71 | 0.13 | -3.32 | 0.02 | -0.8 | -4.23 | -4.14 | 0.24 | 0.035 | -0.4175 | 0.697111 | 0.684109 | 0.965541 | 0.994379 | 0.997988 | 0.996159 |
| Map2_P20357_S1783s_1 | Map2_P20 | 3.55 | 0.13 | 0.71 | -0.09 | 0.24 | 3.93 | 5.85 | 1.65 | 0.25 | 2.06 | 1.34 | 2 | 0.63 | 0.27 | 1.99 | 2.66 | 1.8425 | 0.3375 | 0.3125 | 0.72555 | 0.174777 | 0.688268 | 0.989223 | 0.997988 | 0.9971 |
| Map2_P20357_S1791s_1 | Map2_P20 | -1.62 | -2.08 | -1.94 | -2.55 | -2.27 | -0.97 | -3.24 | -2.34 | 0.24 | -1.13 | -3.26 | -2.18 | 0.23 | -2.23 | -2.26 | -2.21 | -0.1575 | 0.465 | 0.43 | 0.472959 | 0.737349 | 0.50944 | 0.995762 | 0.997988 | 0.994666 |
| Map2_P20357_S238s_1 | Map2_P20 | -0.07 | -1.19 | -1.16 | -0.38 | -0.31 | -0.36 | 0.21 | -0.5 | -0.99 | -1.1 | 0.57 | -0.63 | -1.23 | 1.86 | -1.23 | -1.13 | 0.46 | 0.1625 | 0.2675 | 0.713326 | 0.187169 | 0.722173 | 0.989223 | 0.997988 | 0.996966 |
| Map2_P20357_S283s_1 | Map2_P20 | -0.21 | -1.75 | -1.88 | -1.14 | -0.76 | -0.07 | -0.48 | -0.44 | -1.76 | -1.77 | 1.24 | -1.71 | -1.94 | -0.24 | -2.4 | -2.23 | 0.8075 | 0.245 | -0.4575 | 0.434328 | 0.063974 | 0.745009 | 0.989223 | 0.997988 | 0.994666 |
| Map2_P20357_S28s_1 | Map2_P20 | -3.58 | -4.69 | -2 | -1.34 | -1.55 | -5.55 | -1.12 | -1.79 | -2.58 | -3.44 | -2.83 | -1.57 | 0.16 | -1.85 | -1.85 | -2.39 | 0.4 | 0.2975 | 1.42 | 0.113188 | 0.716776 | 0.697973 | 0.995437 | 0.997988 | 0.994666 |
| Map2_P20357_S42s_1 | Map2_P20 | 0.21 | -0.09 | -1.03 | -1.85 | -0.38 | 0.54 | -1.68 | -0.61 | 0.6 | -0.52 | 0.58 | -0.99 | -0.87 | -1.47 | -1.38 | -1.17 | 0.1575 | 0.6075 | -0.5325 | 0.272302 | 0.788341 | 0.300513 | 0.995762 | 0.997988 | 0.994666 |
| Map2_P20357_S447s_1 | Map2_P20 | 0.1 | -0.2 | -0.88 | -0.53 | -0.71 | -0.36 | -0.71 | -0.69 | -0.95 | -0.24 | -1.38 | -1.31 | -0.49 | -0.43 | -2.08 | -1 | -0.24 | -0.5925 | -0.6225 | 0.171653 | 0.384554 | 0.117442 | 0.989223 | 0.997988 | 0.994666 |
| Map2_P20357_S472s_1 | Map2_P20 | 0 | 0.43 | -0.4 | -0.36 | 0.34 | 0.23 | 0.13 | -0.15 | -0.52 | -0.18 | -0.12 | -0.47 | -0.52 | 0.43 | -0.5 | -0.05 | 0.22 | -0.24 | -0.0775 | 0.816958 | 0.415546 | 0.390805 | 0.989223 | 0.997988 | 0.999507 |
| Map2_P20357_S476s_1 | Map2_P20 | -1.49 | -2.15 | -1.1 | -1.27 | -1.78 | -1.43 | -0.61 | -0.51 | -2.09 | -1.72 | -1.14 | -0.91 | -2.45 | -0.77 | -1.4 | -1.08 | 0.42 | 0.0375 | 0.0775 | 0.856063 | 0.282863 | 0.917918 | 0.989223 | 0.997988 | 0.999507 |
| Map2_P20357_S599s_1 | Map2_P20 | -0.25 | -0.24 | -1.03 | -1.55 | -0.19 | -0.78 | -1.4 | -2.03 | -0.03 | -1.07 | -1.93 | -1.23 | 0.1 | 0.03 | -1.8 | -2.55 | -0.3325 | -0.2975 | -0.2875 | 0.666362 | 0.486541 | 0.539848 | 0.989223 | 0.997988 | 0.996159 |
| Map2_P20357_S608s_1 | Map2_P20 | -0.67 | -0.35 | -0.49 | -0.83 | -0.45 | -1.14 | -0.62 | -0.8 | -0.76 | -0.36 | -1.07 | -0.86 | -0.53 | -0.18 | -0.68 | -0.79 | -0.1675 | -0.1775 | 0.04 | 0.8796 | 0.496783 | 0.489958 | 0.989223 | 0.997988 | 0.999507 |
| Map2_P20357_S63s_1 | Map2_P20 | -2 | -3.28 | -4.02 | -3.95 | -3.99 | -3.6 | -2.32 | -2.43 | -1.03 | -3.59 | -1.76 | -2.25 | -0.44 | -2.11 | -2.4 | -3.27 | 0.2275 | 1.155 | 1.2575 | 0.08899 | 0.688426 | 0.101497 | 0.994672 | 0.997988 | 0.994666 |
| Map2_P20357_S654s_1 | Map2_P20 | 0.91 | 0.79 | -0.64 | 0.32 | 1.07 | 0.31 | -0.06 | 0.13 | 0.96 | 0.81 | -0.63 | -0.17 | -0.76 | 0.75 | -0.25 | -0.02 | 0.0175 | -0.1025 | -0.415 | 0.374719 | 0.966 | 0.834979 | 0.999187 | 0.997988 | 0.994666 |
| Map2_P20357_S655s_1 | Map2_P20 | -0.49 | -0.48 | -2.09 | -1.43 | -0.55 | -0.94 | -1.66 | -1.99 | -0.96 | -0.5 | -2.83 | -2.62 | -0.47 | -0.04 | -2.33 | -2.35 | -0.1625 | -0.605 | -0.175 | 0.788818 | 0.731953 | 0.357006 | 0.995762 | 0.997988 | 0.999507 |
| Map2_P20357_S823s_1 | Map2_P20 | -3.19 | -4.46 | -4.91 | -7.51 | -3.76 | -2.95 | -3.88 | -3.75 | -4.48 | -4.08 | 3.36 | -4.28 | -3.49 | 0.02 | -6.8 | -7.37 | 1.4325 | 2.6475 | 0.6075 | 0.714482 | 0.103861 | 0.174423 | 0.989223 | 0.997988 | 0.996966 |
| Map2_P20357_S825s_1 | Map2_P20 | -0.41 | -2.63 | -3.86 | -5.66 | -0.99 | -0.66 | -3.06 | -3.06 | -1.32 | -1.51 | 3.06 | -3.61 | -2.39 | 0.27 | -5.58 | -5.53 | 1.1975 | 2.295 | -0.1675 | 0.912954 | 0.292856 | 0.164487 | 0.989223 | 0.997988 | 0.999507 |
| Map2_P20357_S938s_1 | Map2_P20 | -1.21 | -1.03 | -1.6 | -1.64 | -2.05 | -1.13 | -2.07 | -2.08 | -0.18 | -1.03 | -2.26 | -2.19 | 0.77 | -2.22 | -1.94 | -1.91 | -0.4625 | -0.045 | 0.045 | 0.944566 | 0.149201 | 0.927039 | 0.989223 | 0.997988 | 0.999507 |
| Map2_P20357_T1160t_1 | Map2_P20 | -0.36 | -0.1 | -1.41 | -1.66 | 0.13 | -0.49 | -0.26 | -0.4 | -0.53 | -0.4 | -1 | -1.44 | -0.79 | 0.16 | -0.6 | -1.49 | 0.6275 | 0.04 | 0.2025 | 0.679048 | 0.135813 | 0.927059 | 0.989223 | 0.997988 | 0.996159 |
| Map2_P20357_T1260t_1 | Map2_P20 | -1.88 | -1.53 | -1.63 | -1.56 | -0.93 | -1.4 | -1.17 | -1.18 | -1.91 | -1.63 | -0.16 | -1.27 | -1.39 | -0.63 | -1.22 | -1.49 | 0.48 | 0.4075 | 0.4675 | 0.124021 | 0.048812 | 0.313461 | 0.989223 | 0.997988 | 0.994666 |
| Map2_P20357_T1358t_1 | Map2_P20 | 1.83 | 2.06 | 0.57 | -0.06 | 1.47 | 1.26 | 0.8 | 0.73 | 0.18 | 1.51 | 0.05 | 0.61 | -0.71 | 0.18 | 0.86 | 0.72 | -0.035 | -0.5125 | -0.8375 | 0.165182 | 0.943406 | 0.370715 | 0.999187 | 0.997988 | 0.994666 |
| Map2_P20357_T1599t_2 | Map2_P20 | -0.29 | -0.76 | -1.1 | -1.68 | -0.17 | -1.53 | -1.28 | -2.78 | 1.2 | -0.05 | -0.24 | -0.63 | -0.05 | 0.52 | -0.17 | -0.33 | -0.4825 | 1.0275 | 0.95 | 0.026146 | 0.391334 | 0.05194 | 0.989223 | 0.997988 | 0.994666 |
| Map2_P20357_T1609t_1 | Map2_P20 | -2.73 | -2.57 | -2.81 | -2.44 | -2.4 | -2.89 | -1.76 | -2.6 | -2.59 | -2.25 | -2.16 | -2.55 | -1.67 | -1.78 | -2.56 | -2.39 | 0.225 | 0.25 | 0.5375 | 0.096826 | 0.438975 | 0.295058 | 0.989223 | 0.997988 | 0.994666 |
| Map2_P20357_T1609t_2 | Map2_P20 | 0.31 | 0.49 | 0.25 | 0.16 | 0.65 | 0.31 | 0.22 | 0.3 | 0.47 | 0.61 | 0.29 | 0.56 | -0.41 | -0.15 | 0.61 | 0.36 | 0.0675 | 0.18 | -0.2 | 0.513427 | 0.752969 | 0.415896 | 0.995762 | 0.997988 | 0.994666 |
| Map2_P20357_T1617t_1 | Map2_P20 | -2.48 | -2.14 | -1.11 | -1.36 | -1.97 | -2.59 | -1.6 | -1.5 | -2.51 | -2.27 | 0.35 | -1.48 | -1.6 | -1.5 | -0.88 | -1.57 | -0.1425 | 0.295 | 0.385 | 0.324068 | 0.7172 | 0.654671 | 0.995437 | 0.997988 | 0.994666 |
| Map2_P20357_T1617t_2 | Map2_P20 | -2.14 | -2.02 | -1.99 | -2.35 | -1.84 | -1.14 | -1.86 | -2.41 | -2.86 | -1.59 | -0.79 | -1.88 | -0.02 | -1.36 | -2.4 | -1.9 | 0.3125 | 0.345 | 0.705 | 0.175396 | 0.309119 | 0.425513 | 0.989223 | 0.997988 | 0.994666 |
| Map2_P20357_T1620t_1 | Map2_P20 | -3.09 | -3.6 | -2.92 | -4.04 | -2.56 | -3.73 | -2.68 | -3.15 | -3.76 | -2.68 | -1.31 | -3.5 | -1.81 | -2.17 | -2.4 | -3.49 | 0.3825 | 0.6 | 0.945 | 0.052637 | 0.308377 | 0.300378 | 0.989223 | 0.997988 | 0.994666 |
| Map2_P20357_T1623t_1 | Map2_P20 | -1.33 | -1.51 | -1.81 | -1.13 | -1 | -2.21 | -1.42 | -1.73 | -2.27 | -1.91 | -0.29 | -1.61 | -0.82 | -0.96 | -1.72 | -1.67 | -0.145 | -0.075 | 0.1525 | 0.637249 | 0.643619 | 0.864718 | 0.990258 | 0.997988 | 0.995259 |
| Map2_P20357_T1623t_2 | Map2_P20 | -2.14 | -2.02 | -2.42 | -2.48 | -1.84 | -1.14 | -2.1 | -2.94 | -2.86 | -1.59 | -1.25 | -2.22 | -0.02 | -1.36 | -2.42 | -2.33 | 0.26 | 0.285 | 0.7325 | 0.189824 | 0.498198 | 0.4574 | 0.989223 | 0.997988 | 0.994666 |
| Map2_P20357_T1650t_1 | Map2_P20 | -1.92 | -2.48 | -3.08 | -2.66 | -1.84 | -3.05 | -2.45 | -2.81 | -2.41 | -1.49 | -1.87 | -3.21 | -0.87 | -1.44 | -2.54 | -2.87 | -0.0025 | 0.29 | 0.605 | 0.242648 | 0.994394 | 0.506915 | 0.999278 | 0.997988 | 0.994666 |
| Map2_P20357_T1650t_2 | Map2_P20 | -0.79 | -0.8 | -3.14 | -4.1 | -0.86 | -0.67 | -3.05 | -2.49 | -0.84 | -0.47 | -0.37 | -3.2 | -0.17 | -1.33 | -2.23 | -3.82 | 0.44 | 0.9875 | 0.32 | 0.747372 | 0.62376 | 0.312589 | 0.989223 | 0.997988 | 0.9971 |
| Map2k1_P31938_S231s_1 | Map2k1_P3 | -0.43 | -0.03 | -0.09 | 0.26 | 0.38 | -0.7 | -0.39 | 0.01 | -0.28 | -0.01 | 0.45 | 0.33 | 0.02 | -0.34 | 0.09 | -0.25 | -0.1025 | 0.195 | -0.0475 | 0.85884 | 0.732757 | 0.482549 | 0.995762 | 0.997988 | 0.999507 |
| Map2k1_P31938_T286t_1 | Map2k1_P3 | -0.06 | 0.18 | -2.57 | -1.53 | -0.01 | 0.15 | -0.96 | -3.13 | -0.31 | 0.53 | -1.22 | -0.89 | -0.85 | 0.12 | -1.28 | -2.99 | 0.0075 | 0.5225 | -0.255 | 0.753062 | 0.993075 | 0.449751 | 0.999187 | 0.997988 | 0.9971 |
| Map2k2_Q91YS7_S295s_1 | Map2k2_Q | -0.09 | -0.38 | -1.22 | -1.33 | 0.25 | 0.11 | -0.62 | -1.24 | -0.99 | 0.4 | -0.75 | -1.48 | -1.3 | -0.27 | -1.27 | -1.44 | 0.38 | 0.05 | -0.315 | 0.450014 | 0.392353 | 0.916565 | 0.989223 | 0.997988 | 0.994666 |
| Map2k2_Q91YS7_S306s_1 | Map2k2_Q | -1.85 | -1.92 | -0.84 | -1.18 | -1.57 | -1.6 | -0.53 | -0.9 | -1.06 | -2.53 | -0.74 | -1.02 | -0.33 | -1.52 | -0.49 | -1.34 | 0.2975 | 0.11 | 0.5275 | 0.210924 | 0.42554 | 0.811484 | 0.989223 | 0.997988 | 0.994666 |
| Map2k2_Q91YS7_T394t_1 | Map2k2_Q | 0.61 | 0.52 | -0.91 | -1.96 | 0.67 | 0.46 | -1.66 | -1.9 | 0.71 | 0.59 | -2.17 | -1.44 | 1.37 | 0.55 | -1.15 | -0.99 | -0.1725 | -0.1425 | 0.38 | 0.623142 | 0.829683 | 0.866259 | 0.995762 | 0.997988 | 0.995259 |
| Map2k4_P47809_S392s_1 | Map2k4_P4 | 0.62 | 0.68 | 0.56 | 0.45 | 0.76 | 0.38 | 0.75 | 0.46 | 0.02 | 0.64 | -0.01 | 0.36 | -0.44 | 0.27 | 0.25 | 0.71 | 0.01 | -0.325 | -0.38 | 0.228133 | 0.962127 | 0.203272 | 0.999187 | 0.997988 | 0.994666 |
| Map2k4_P47809_S88s_1 | Map2k4_P4 | -0.02 | 0.49 | -0.07 | 0.03 | 0.61 | -0.28 | 0.59 | 0.16 | -0.31 | 0.15 | -0.22 | 0.17 | -1.24 | 0.77 | 0 | 0.26 | 0.1625 | -0.16 | -0.16 | 0.7156 | 0.567658 | 0.530204 | 0.989223 | 0.997988 | 0.996966 |
| Map3k1_P53349_S915s_1 | Map3k1_P5 | 0.5 | -1.01 | -0.04 | 0.25 | -0.07 | 0.61 | -0.21 | 0.22 | -0.11 | -0.07 | 1.89 | 0.45 | 0.54 | 0.85 | 0.6 | 0.39 | 0.2125 | 0.615 | 0.67 | 0.092473 | 0.570215 | 0.266656 | 0.989223 | 0.997988 | 0.994666 |
| Map3k10_D3YXM8_S502s_1 | Map3k10_ | 0.52 | 0.66 | 0.25 | 0.59 | 0.62 | 0.15 | 0.52 | 0.4 | 0.29 | 0.54 | 0.16 | 0.46 | 0.29 | 0.69 | 0.27 | 0.56 | -0.0825 | -0.1425 | -0.0525 | 0.833972 | 0.710011 | 0.533176 | 0.995437 | 0.997988 | 0.999507 |
| Map3k10_D3YXM8_S714s_1 | Map3k10_ | -0.85 | -2.75 | 0.36 | 1.93 | -0.27 | -0.6 | 2.04 | 2.75 | -2.7 | -2.79 | 1.92 | 1.71 | -2.48 | -1.26 | 0.99 | 0.97 | 1.3075 | -0.1375 | -0.1175 | 0.917377 | 0.257857 | 0.923782 | 0.989223 | 0.997988 | 0.999507 |
| Map3k10_D3YXM8_S818s_1 | Map3k10_ | -0.37 | -1.07 | -0.22 | 0.21 | -0.16 | -0.38 | -0.12 | 0.19 | -0.72 | -0.85 | 0.5 | -0.01 | 0.02 | -0.13 | 0.3 | 0.21 | 0.245 | 0.0925 | 0.4625 | 0.178882 | 0.437304 | 0.820785 | 0.989223 | 0.997988 | 0.994666 |
| Map3k10_D3YXM8_T552t_1 | Map3k10_ | -2.13 | -1.34 | -2.74 | -1.31 | -0.89 | -2.29 | -1.37 | -1.86 | -2.5 | -2.01 | -1.08 | -1.5 | -1.26 | -1.61 | -1.87 | -1.84 | 0.2775 | 0.1075 | 0.235 | 0.545165 | 0.525217 | 0.809417 | 0.989223 | 0.997988 | 0.994666 |
| Map3k12_Q60700_S569s_1 | Map3k12_ | -0.33 | -0.85 | 0.15 | -0.1 | -0.13 | 0.26 | -0.24 | -0.24 | 1.81 | -0.64 | -1.7 | 0.18 | 1.8 | -1.06 | 0.51 | 0.23 | 0.195 | 0.195 | 0.6525 | 0.27225 | 0.491622 | 0.779369 | 0.989223 | 0.997988 | 0.994666 |
| Map3k12_Q60700_T43t_1 | Map3k12_ | 2.3 | 2.58 | 2.67 | 1.63 | 3.34 | 2.55 | 2.65 | 2.59 | 0.26 | 2.49 | 0.78 | 2.41 | -0.01 | 0.27 | 2.82 | 3.03 | 0.4875 | -0.81 | -0.7675 | 0.319528 | 0.148086 | 0.176711 | 0.989223 | 0.997988 | 0.994666 |
| Map3k13_Q1HKZ5_S38s_1 | Map3k13_ | 0.2 | -0.1 | -0.7 | 0.73 | 0.24 | 0.28 | -0.4 | -0.61 | -0.01 | -0.31 | 0.43 | 0.03 | -0.78 | 0.35 | -0.69 | -0.08 | -0.155 | 0.0025 | -0.3325 | 0.419882 | 0.675526 | 0.9943 | 0.992876 | 0.997988 | 0.994666 |
| Map3k13_Q1HKZ5_S680s_1 | Map3k13_ | -0.17 | 0.24 | -0.15 | -1.47 | 0 | 0.73 | -0.79 | -0.61 | 1.1 | -0.6 | 0.87 | -0.6 | 1.48 | 0 | -1.46 | -0.24 | 0.22 | 0.58 | 0.3325 | 0.60654 | 0.641388 | 0.305962 | 0.989223 | 0.997988 | 0.995259 |
| Map3k13_Q1HKZ5_S914s_1 | Map3k13_ | -1.18 | -0.75 | -0.09 | -0.78 | -0.54 | -1.09 | -0.46 | -0.96 | -1.41 | -1.29 | 0.45 | -0.47 | -1.2 | -0.35 | -1.06 | -0.4 | -0.0625 | 0.02 | -0.0525 | 0.879649 | 0.834398 | 0.965626 | 0.995762 | 0.997988 | 0.999507 |
| Map3k2_G5E8L8_S163s_1 | Map3k2_G | 1.63 | 1.84 | 1.25 | 0.95 | 1.95 | 1.71 | 1.57 | 1.15 | 0.93 | 1.46 | 0.35 | 1.45 | -1.84 | 1.63 | 1.17 | 1.41 | 0.1775 | -0.37 | -0.825 | 0.28498 | 0.544133 | 0.301045 | 0.989223 | 0.997988 | 0.994666 |
| Map3k4_O08648_S116s_1 | Map3k4_O | -2.61 | -2.39 | -1.54 | -1.84 | -2.09 | -2.8 | -1.51 | -2.47 | -1.2 | -1.73 | -1.79 | -1.94 | -0.02 | -2.5 | -1.32 | -1.53 | -0.1225 | 0.43 | 0.7525 | 0.177599 | 0.738548 | 0.203574 | 0.995762 | 0.997988 | 0.994666 |
| Map3k4_O08648_S424s_1 | Map3k4_O | 1.44 | 0.4 | 0.43 | 1.05 | 0.84 | 0.95 | 0.92 | 1.56 | 0.16 | 0.72 | -0.16 | 0.7 | -0.85 | 1.07 | 0.6 | 0.53 | 0.2375 | -0.475 | -0.4925 | 0.305913 | 0.461741 | 0.194818 | 0.989223 | 0.997988 | 0.994666 |
| Map3k4_O08648_S454s_1 | Map3k4_O | 0.54 | 0.8 | -0.59 | -0.26 | -0.05 | -0.75 | 0.04 | -0.19 | 0.69 | 0.16 | 0.24 | -0.68 | 0.18 | 0.59 | 0.3 | -0.21 | -0.36 | -0.02 | 0.0925 | 0.808568 | 0.340279 | 0.962386 | 0.989223 | 0.997988 | 0.999507 |
| Map3k7_Q923A8_S389s_1 | Map3k7_Q | 0.7 | -0.7 | -0.21 | -0.31 | -0.64 | 0.15 | -0.21 | 0.17 | 0.74 | -0.37 | 0.03 | -0.02 | 0.25 | 0.46 | -0.79 | -0.45 | -0.0025 | 0.225 | -0.0025 | 0.995206 | 0.994343 | 0.558236 | 0.999278 | 0.997988 | 1 |
| Map3k9_Q3U1V8_S540s_1 | Map3k9_Q | 0.42 | 0.63 | 0.87 | 0.44 | 0.65 | 0.18 | 0.7 | 0.34 | 0.52 | 0.8 | 0.54 | 0.71 | 0.76 | 1.08 | 0.27 | 0.47 | -0.1225 | 0.0525 | 0.055 | 0.845272 | 0.603147 | 0.81685 | 0.989223 | 0.997988 | 0.999507 |
| Map4_E9PZ43_S298s_1 | Map4_E9P | -0.89 | -0.24 | -0.55 | -0.44 | -0.83 | -0.68 | 0.56 | -0.17 | -1.05 | -0.74 | 1.14 | -0.16 | -1.16 | 0.01 | -1.02 | -0.66 | 0.25 | 0.3275 | -0.1775 | 0.597394 | 0.475804 | 0.498496 | 0.989223 | 0.997988 | 0.995259 |
| Map4_E9PZ43_S366s_1 | Map4_E9P | -0.79 | -0.97 | -1.66 | -0.66 | 0.29 | -1.26 | -0.41 | -1.64 | -0.97 | -0.97 | -0.32 | -1.36 | -1.85 | -0.93 | -0.95 | -1.7 | 0.265 | 0.115 | -0.3375 | 0.355135 | 0.562948 | 0.730414 | 0.989223 | 0.997988 | 0.994666 |
| Map4_E9PZ43_S514s_1 | Map4_E9P | -1.49 | -1.01 | -0.69 | -0.89 | -0.94 | -1.65 | -0.62 | -0.92 | -1 | -0.99 | -0.94 | -0.65 | -0.09 | -1.33 | -0.9 | -0.77 | -0.0125 | 0.125 | 0.2475 | 0.475835 | 0.966835 | 0.629352 | 0.999187 | 0.997988 | 0.994666 |
| Map4_E9PZ43_S632s_1 | Map4_E9P | -0.39 | -1.05 | -0.94 | -0.75 | -0.02 | -0.43 | -0.25 | -0.51 | -0.85 | -0.34 | 0.12 | -0.74 | -0.51 | -0.12 | -0.73 | -0.85 | 0.48 | 0.33 | 0.23 | 0.431799 | 0.073301 | 0.290365 | 0.989223 | 0.997988 | 0.994666 |
| Map4_E9PZ43_S645s_1 | Map4_E9P | -0.06 | -1.16 | -2.31 | -3.54 | -1.09 | -0.84 | -2.02 | -1.76 | -0.76 | -0.49 | 0.29 | -2.02 | -0.45 | -0.35 | -2.42 | -2.94 | 0.34 | 1.0225 | 0.2275 | 0.796009 | 0.630753 | 0.218872 | 0.989223 | 0.997988 | 0.999507 |
| Map4_E9PZ43_S675s_1 | Map4_E9P | -0.7 | -1.67 | -0.8 | -1.82 | -1.62 | -0.19 | -2.12 | -1.04 | -0.01 | -0.78 | 0.08 | -2.24 | 0.67 | -0.45 | -1.85 | -1.9 | 0.005 | 0.51 | 0.365 | 0.559131 | 0.991478 | 0.376356 | 0.999187 | 0.997988 | 0.995259 |
| Map4_E9PZ43_S723s_1 | Map4_E9P | -0.54 | -1.25 | -2.22 | -2.51 | -1.11 | -1.08 | -2.21 | -2.6 | -2.03 | -0.95 | 0.5 | -2.48 | -1.75 | -0.41 | -1.31 | -2.25 | -0.12 | 0.39 | 0.2 | 0.71839 | 0.823897 | 0.591901 | 0.995762 | 0.997988 | 0.9971 |
| Map4_E9PZ43_S855s_1 | Map4_E9P | -0.3 | -0.44 | -0.17 | -0.4 | -0.59 | -0.35 | 0.23 | 0.09 | -0.54 | -1.06 | 0.47 | -0.46 | -1.06 | -0.12 | -0.73 | -0.22 | 0.1725 | -0.07 | -0.205 | 0.492045 | 0.501813 | 0.83815 | 0.989223 | 0.997988 | 0.994666 |
| Map4_E9PZ43_S93s_1 | Map4_E9P | 0.92 | 0.28 | -0.45 | 0.26 | 0.74 | 1.18 | 0.41 | 1.11 | 0.56 | 0.19 | 0.98 | 0.16 | -0.07 | 0.65 | -0.03 | 0.64 | 0.6075 | 0.22 | 0.045 | 0.902129 | 0.098394 | 0.538102 | 0.989223 | 0.997988 | 0.999507 |
| Map4_E9PZ43_T505t_1 | Map4_E9P | -2.59 | -1.36 | -1.83 | -1.36 | -1.32 | -2.61 | -1.33 | -1.36 | -2.08 | -1.86 | -1.28 | -1.72 | -1.65 | -0.83 | -1.71 | -1.22 | 0.13 | 0.05 | 0.4325 | 0.264319 | 0.752385 | 0.886674 | 0.995762 | 0.997988 | 0.994666 |
| Map4_E9PZ43_T694t_1 | Map4_E9P | -0.65 | -0.22 | -0.8 | -0.91 | -0.85 | -1 | -0.44 | -0.8 | -0.26 | -0.27 | 0.12 | -0.46 | 0.38 | 0 | -0.27 | -0.34 | -0.1275 | 0.4275 | 0.5875 | 0.067837 | 0.611757 | 0.126193 | 0.989223 | 0.997988 | 0.994666 |
| Map4_P27546_S506s_1 | Map4_P27 | 0.11 | 0.57 | -0.03 | 0.48 | 0.24 | 0.32 | 0.55 | -0.06 | -0.22 | 0.33 | 0.23 | 0.26 | 0.35 | 0.32 | 0.07 | 0.3 | -0.02 | -0.1325 | -0.0225 | 0.930737 | 0.935935 | 0.610542 | 0.99868 | 0.997988 | 0.999507 |
| Map4_P27546_S517s_1 | Map4_P27 | 1.42 | 1.83 | 1.29 | 1.15 | 1.46 | 1.12 | 1.1 | 1.17 | -0.15 | 1.8 | 0.67 | 1.31 | -0.22 | 0.51 | 1.48 | 1.55 | -0.21 | -0.515 | -0.5925 | 0.198596 | 0.385886 | 0.25357 | 0.989223 | 0.997988 | 0.994666 |
| Map4k2_D3YUH9_S284s_1 | Map4k2_D | 0.21 | -0.14 | -0.09 | -0.66 | -0.13 | 0.26 | 0.64 | 0.07 | -0.31 | -0.51 | -0.73 | -1.18 | 0.03 | 0.4 | -0.4 | -0.25 | 0.38 | -0.5125 | 0.115 | 0.709567 | 0.196165 | 0.113243 | 0.989223 | 0.997988 | 0.996966 |
| Map4k4_E9PVG7_S661s_1 | Map4k4_E | 0.26 | 0.2 | 0.57 | 0.13 | -0.27 | 0.52 | -0.46 | 0.24 | 0.13 | 0.35 | 0.16 | 0.45 | 0.06 | -0.78 | 0.47 | 0.33 | -0.2825 | -0.0175 | -0.27 | 0.42797 | 0.32817 | 0.938272 | 0.989223 | 0.997988 | 0.994666 |
| Map4k4_E9PVG7_S676s_1 | Map4k4_E | 1.72 | 0.17 | 1.04 | 1.78 | -0.19 | 1.82 | 0.89 | 1.85 | 1.72 | 1.65 | 1.51 | 1.42 | 1.55 | 1.08 | 1.38 | 0.94 | -0.085 | 0.3975 | 0.06 | 0.882115 | 0.877541 | 0.315549 | 0.996493 | 0.997988 | 0.999507 |
| Map4k4_E9PVG7_S684s_1 | Map4k4_E | 1.36 | 0.56 | -0.29 | 0.55 | 0.99 | 0.9 | 0.53 | -0.5 | 0.84 | 1 | 1.22 | 1.2 | 0.89 | 0.81 | 1.12 | 0.99 | -0.065 | 0.52 | 0.4075 | 0.280922 | 0.884725 | 0.17263 | 0.996493 | 0.997988 | 0.994666 |
| Map4k4_E9PVG7_S761s_1 | Map4k4_E | -0.96 | -1.15 | -1.17 | -2.28 | -1.93 | -0.49 | -1.35 | -1.54 | 1.67 | -1.21 | -1.7 | -0.49 | 1.77 | -2.08 | 0.17 | -1.23 | 0.0625 | 0.9575 | 1.0475 | 0.206587 | 0.878412 | 0.206257 | 0.996493 | 0.997988 | 0.994666 |
| Map4k4_E9PVG7_S763s_1 | Map4k4_E | -1.39 | -1.72 | -0.49 | -1.78 | -2.2 | -0.97 | -2.16 | -1.72 | 1.52 | -1.39 | -1.22 | -0.31 | 1.54 | -2.95 | 0.19 | -1.32 | -0.4175 | 0.995 | 0.71 | 0.431739 | 0.308048 | 0.157577 | 0.989223 | 0.997988 | 0.994666 |
| Map4k4_E9PVG7_S849s_1 | Map4k4_E | -0.09 | 0.03 | 1.14 | 0.49 | 0.08 | -0.23 | 0.35 | 1.1 | 0.12 | 0.33 | 1.79 | 0.56 | 0.03 | -0.48 | 1.11 | 0.84 | -0.0675 | 0.3075 | -0.0175 | 0.968825 | 0.8614 | 0.499317 | 0.996493 | 0.997988 | 0.999507 |
| Map4k4_E9PVG7_S850s_1 | Map4k4_E | 0.66 | 0.81 | 0.05 | 0.55 | 1.3 | 0.88 | 0.69 | 0.63 | 0.96 | 0.86 | 0.5 | 0.1 | 1.54 | 1.25 | 0.49 | 0.32 | 0.3575 | 0.0875 | 0.3825 | 0.304131 | 0.202593 | 0.768449 | 0.989223 | 0.997988 | 0.994666 |
| Map4k4_E9PVG7_S891s_1 | Map4k4_E | 0.57 | 0.49 | -0.27 | -0.64 | -0.82 | 0.79 | -0.12 | -0.66 | 0.88 | -0.32 | -0.17 | 0.23 | 0.78 | -0.7 | 0.12 | -0.31 | -0.24 | 0.1175 | -0.065 | 0.879304 | 0.587697 | 0.76758 | 0.989223 | 0.997988 | 0.999507 |
| Map4k4_E9PVG7_S949s_1 | Map4k4_E | 0.04 | -1.21 | -0.56 | -0.4 | -0.24 | -0.26 | 0.39 | -0.73 | -0.41 | -0.65 | -0.36 | -0.72 | -0.14 | -0.58 | -0.82 | -0.7 | 0.3225 | -0.0025 | -0.0275 | 0.934736 | 0.366446 | 0.993529 | 0.989223 | 0.997988 | 0.999507 |
| Map4k5_E9Q1T3_S268s_1 | Map4k5_E | -1.3 | -1.81 | -1.57 | -1.49 | -0.78 | -2.1 | -0.87 | -1.56 | -1.58 | -0.92 | -1.42 | -1.25 | -1.81 | -0.83 | -1.69 | -1.27 | 0.215 | 0.25 | 0.1425 | 0.641491 | 0.52687 | 0.332118 | 0.989223 | 0.997988 | 0.995259 |
| Map4k5_E9Q1T3_S366s_1 | Map4k5_E | 0 | -0.93 | -0.75 | -0.18 | -0.02 | 0.29 | 0.18 | -0.2 | -0.25 | 0.73 | 0.1 | -0.41 | -0.57 | -0.67 | 0.3 | -0.64 | 0.5275 | 0.5075 | 0.07 | 0.842064 | 0.086881 | 0.172741 | 0.989223 | 0.997988 | 0.999507 |
| Map6_Q7TSJ2_S207s_1 | Map6_Q7T | -1.11 | -1.9 | -0.03 | -0.58 | -0.39 | -1.61 | 0.61 | 0.17 | -1.08 | -1.16 | 1.35 | 0.4 | -1.3 | -0.36 | 0.32 | -0.23 | 0.6 | 0.7825 | 0.5125 | 0.314094 | 0.301031 | 0.25373 | 0.989223 | 0.997988 | 0.994666 |
| Map6_Q7TSJ2_S294s_1 | Map6_Q7T | 2.27 | 2.12 | 1.02 | 0.9 | 2.12 | 2.91 | 1.03 | 1.54 | 1.05 | 1.06 | -0.35 | 0.72 | 0.29 | 1.54 | 1.18 | 0.9 | 0.3225 | -0.9575 | -0.6 | 0.191987 | 0.519863 | 0.065413 | 0.989223 | 0.997988 | 0.994666 |
| Map6_Q7TSJ2_S386s_1 | Map6_Q7T | -0.93 | -0.96 | -1.17 | -0.83 | -0.32 | -0.93 | -0.95 | -0.55 | -0.52 | -1.31 | -0.04 | -1.11 | -0.1 | -0.96 | -1.15 | -1.03 | 0.285 | 0.2275 | 0.1625 | 0.598895 | 0.247741 | 0.489019 | 0.989223 | 0.997988 | 0.995259 |
| Map6_Q7TSJ2_S424s_1 | Map6_Q7T | 0.37 | 0.82 | 0.36 | 0.2 | 0.33 | 0.39 | 0.4 | 0.3 | -0.95 | 0.22 | 1.09 | 0.72 | 0.15 | -0.01 | 0.56 | 0.73 | -0.0825 | -0.1675 | -0.08 | 0.781954 | 0.710078 | 0.708692 | 0.995437 | 0.997988 | 0.998226 |
| Map6_Q7TSJ2_S632s_1 | Map6_Q7T | 0.51 | 0.36 | 1.59 | 1.41 | 0.57 | 0.22 | 0.31 | 1.28 | -0.18 | 0.48 | 1.21 | 1.03 | -0.2 | 0.16 | 1.23 | 1.44 | -0.3725 | -0.3325 | -0.31 | 0.525954 | 0.34312 | 0.446653 | 0.989223 | 0.997988 | 0.994666 |
| Map6_Q7TSJ2_S687s_1 | Map6_Q7T | 0.05 | -0.48 | -0.42 | -0.04 | 0.5 | 0.03 | 0.63 | 0.23 | -0.78 | -0.48 | -0.11 | -0.47 | -1.35 | 0.04 | 0.1 | -0.12 | 0.57 | -0.2375 | -0.11 | 0.773518 | 0.042438 | 0.369644 | 0.989223 | 0.997988 | 0.99798 |
| Map6_Q7TSJ2_S832s_1 | Map6_Q7T | -0.99 | -0.74 | 0.41 | -0.3 | 0.19 | -0.17 | 0.61 | 1.29 | -0.24 | -0.73 | 0.14 | 0.17 | -1.09 | 0.05 | -0.38 | 0.16 | 0.885 | 0.24 | 0.09 | 0.829273 | 0.057337 | 0.529178 | 0.989223 | 0.997988 | 0.999507 |
| Map6_Q7TSJ2_S86s_1 | Map6_Q7T | -0.96 | -1.41 | -0.22 | -0.38 | -0.36 | -0.16 | -0.35 | 0.51 | -0.93 | -0.75 | 1.97 | -0.2 | -0.54 | 0.64 | -0.98 | -0.55 | 0.6525 | 0.765 | 0.385 | 0.386636 | 0.084477 | 0.260684 | 0.989223 | 0.997988 | 0.994666 |
| Map6_Q7TSJ2_S888s_1 | Map6_Q7T | -0.26 | 0.19 | 1.04 | 0.59 | 0.33 | 0.37 | 0 | 0.88 | -0.39 | 0.2 | 0.37 | 0.43 | -0.32 | 0.22 | 0.35 | 0.34 | 0.005 | -0.2375 | -0.2425 | 0.493689 | 0.988204 | 0.50327 | 0.999187 | 0.997988 | 0.994666 |
| Map6_Q7TSJ2_S905s_1 | Map6_Q7T | 0.21 | 0.59 | 0.18 | 0.28 | 0.43 | -0.03 | 0.52 | 0.19 | 0.12 | 0.44 | 0.08 | 0.14 | -0.44 | 0.58 | 0.11 | 0.21 | -0.0375 | -0.12 | -0.2 | 0.503926 | 0.870731 | 0.599412 | 0.996493 | 0.997988 | 0.994666 |
| Map6_Q7TSJ2_T634t_1 | Map6_Q7T | 1.72 | 0.95 | 0.14 | 0.24 | 0.61 | 1.97 | 0.21 | 0.17 | 0.97 | 1.62 | -0.54 | 0.31 | 1.58 | 0.58 | 0.32 | 0.31 | -0.0225 | -0.1725 | -0.065 | 0.887295 | 0.964803 | 0.752952 | 0.999187 | 0.997988 | 0.999507 |
| Map6d1_Q14BB9_S154s_1 | Map6d1_Q | -0.28 | -0.45 | 0.45 | -0.52 | -0.66 | -0.98 | 0.45 | -0.12 | -0.06 | -0.08 | 0.73 | -0.44 | -1.39 | -0.89 | -0.98 | 0.04 | -0.1275 | 0.2375 | -0.605 | 0.142466 | 0.735744 | 0.501038 | 0.995762 | 0.997988 | 0.994666 |
| Map6d1_Q14BB9_S53s_1 | Map6d1_Q | -3.52 | -3.7 | -4.14 | -6.14 | -2.61 | -4.41 | -5.25 | -6.51 | -3.47 | -3.37 | -3.55 | -5.44 | -1.67 | -5.08 | -3.73 | -4.19 | -0.32 | 0.4175 | 0.7075 | 0.402534 | 0.717851 | 0.556537 | 0.995437 | 0.997988 | 0.994666 |
| Map7d1_A8Y5P4_S118s_1 | Map7d1_A | -0.17 | 1.77 | 0.94 | 0.53 | 0.34 | 1.1 | 0.41 | 0.45 | 1.78 | 1.43 | 0.38 | 0.45 | 0.81 | 0.96 | 0.05 | 0.71 | -0.1925 | 0.2425 | -0.135 | 0.760664 | 0.648112 | 0.632471 | 0.99033 | 0.997988 | 0.9971 |
| Map7d1_A8Y5P4_S256s_1 | Map7d1_A | -0.18 | 0.23 | -1.38 | -0.46 | -0.51 | -0.76 | -0.53 | -0.33 | -0.62 | 0 | -0.18 | -0.75 | -0.21 | 0.27 | 0.27 | -0.36 | -0.085 | 0.06 | 0.44 | 0.275102 | 0.810113 | 0.876724 | 0.995762 | 0.997988 | 0.994666 |
| Map7d1_A8Y5P4_S275s_1 | Map7d1_A | -1.45 | -0.7 | -1.62 | -2.48 | -1.09 | -1.78 | -1.75 | -2.24 | -2.19 | -0.85 | -0.73 | -2.49 | -1.65 | -0.86 | -2.3 | -2.66 | -0.1525 | -0.0025 | -0.305 | 0.551581 | 0.713781 | 0.996304 | 0.995437 | 0.998451 | 0.994666 |
| Map7d1_A8Y5P4_S78s_1 | Map7d1_A | -0.35 | -0.28 | -0.18 | 0.6 | -0.09 | 0.09 | 0.4 | 0.21 | -0.53 | 0.09 | 1.18 | 0.3 | -0.34 | 0.97 | -0.26 | -0.02 | 0.205 | 0.3125 | 0.14 | 0.717695 | 0.469628 | 0.454204 | 0.989223 | 0.997988 | 0.996966 |
| Map7d1_A8Y5P4_S95s_2 | Map7d1_A | -1.11 | -2.37 | -2.48 | -0.88 | -1.79 | 0.3 | -1.32 | -2.03 | 0.82 | -1.27 | -1.03 | -1.97 | -0.5 | 0.39 | -1.46 | -2.18 | 0.5 | 0.8475 | 0.7725 | 0.240626 | 0.412595 | 0.219132 | 0.989223 | 0.997988 | 0.994666 |
| Map7d1_A8Y5P4_T99t_1 | Map7d1_A | -0.64 | 0.44 | 0.33 | 0.09 | 0.27 | 0.19 | 0.05 | 0.22 | -0.6 | -0.05 | 0.76 | -0.02 | 0.03 | -0.21 | 0.22 | -0.03 | 0.1275 | -0.0325 | -0.0525 | 0.866179 | 0.65292 | 0.931076 | 0.990434 | 0.997988 | 0.999507 |
| Map7d1_A8Y5P4_T99t_2 | Map7d1_A | -1.11 | -2.37 | -2.48 | -0.88 | -1.79 | 0.3 | -1.32 | -2.03 | 0.82 | -1.27 | -1.03 | -1.97 | -0.5 | 0.39 | -1.46 | -2.18 | 0.5 | 0.8475 | 0.7725 | 0.240626 | 0.412595 | 0.219132 | 0.989223 | 0.997988 | 0.994666 |
| Map7d1_F6WF36_S146s_1 | Map7d1_F | -1.95 | -1.38 | -1.57 | -2.7 | -0.93 | -2.69 | -1.81 | -2.6 | -2.07 | -1.59 | -0.46 | -2.04 | -0.83 | -0.75 | -2.52 | -2.53 | -0.1075 | 0.36 | 0.2425 | 0.654569 | 0.817942 | 0.438613 | 0.995762 | 0.997988 | 0.995259 |
| Map7d1_F6WF36_S189s_1 | Map7d1_F | -4.23 | -4.26 | -3.26 | -3.78 | -3.34 | -2.81 | -3.98 | -4.07 | -3.49 | -3.76 | -1.26 | -3.88 | -2.03 | -3.09 | -3.37 | -3.41 | 0.3325 | 0.785 | 0.9075 | 0.046459 | 0.38101 | 0.215632 | 0.989223 | 0.997988 | 0.994666 |
| Map7d1_F6WF36_S205s_1 | Map7d1_F | -0.81 | -0.57 | -1.35 | -1.89 | -0.2 | -1.49 | -0.97 | -1.6 | -0.9 | -0.83 | -2.17 | -1.44 | -0.46 | -0.29 | -1.29 | -0.91 | 0.09 | -0.18 | 0.4175 | 0.292628 | 0.827899 | 0.6689 | 0.995762 | 0.997988 | 0.994666 |
| Map7d1_F6WF36_S214s_1 | Map7d1_F | -5.03 | -3.79 | -4.87 | -4.6 | -3.08 | -4.8 | -3.82 | -4.44 | -3.03 | -3.73 | -2.76 | -3.98 | -1.98 | -3.16 | -3.8 | -4.62 | 0.5375 | 1.1975 | 1.1825 | 0.062611 | 0.239559 | 0.013316 | 0.989223 | 0.997988 | 0.994666 |
| Map7d1_F6WF36_S264s_1 | Map7d1_F | -1.61 | -0.28 | -1.13 | -1.36 | -1.3 | -1.03 | -1.31 | -2.3 | 0.02 | -0.34 | -2.3 | -2.85 | 0.78 | -0.95 | -1.98 | -1.93 | -0.39 | -0.2725 | 0.075 | 0.906546 | 0.329825 | 0.694747 | 0.989223 | 0.997988 | 0.999507 |
| Map7d1_F6WF36_S289s_1 | Map7d1_F | 0.47 | 0.73 | 1.18 | 1 | 0.62 | 0.33 | 0.91 | 1.14 | 0.07 | 0.99 | 1.5 | 0.84 | 0.6 | 1.03 | 1.16 | 1.08 | -0.095 | 0.005 | 0.1225 | 0.662083 | 0.729677 | 0.988587 | 0.995762 | 0.997988 | 0.995804 |
| Map7d1_F6WF36_S293s_1 | Map7d1_F | 0.41 | 0.59 | 0.08 | 0.64 | 0.79 | -0.11 | 0.88 | 0.9 | -0.15 | 0.41 | 0.72 | 0.39 | -0.2 | 0.95 | 1.09 | 0.85 | 0.185 | -0.0875 | 0.2425 | 0.494688 | 0.540233 | 0.751193 | 0.989223 | 0.997988 | 0.994666 |
| Map7d1_F6WF36_S60s_1 | Map7d1_F | -3.13 | -3.61 | -3.28 | -5.27 | -2.41 | -1.52 | -2.48 | -3.6 | -1.88 | -3.1 | -1.01 | -3.82 | -2.34 | -1.86 | -2.95 | -2.81 | 1.32 | 1.37 | 1.3325 | 0.026804 | 0.045559 | 0.082142 | 0.989223 | 0.997988 | 0.994666 |
| Map7d2_A2AG50_S131s_1 | Map7d2_A | -2.44 | -2.75 | -2.96 | -3.7 | -2.27 | -2.55 | -3.21 | -3.45 | -2.23 | -2.3 | -1.38 | -2.96 | -1.5 | -1.81 | -3.46 | -3.21 | 0.0925 | 0.745 | 0.4675 | 0.382667 | 0.806222 | 0.098264 | 0.995762 | 0.997988 | 0.994666 |
| Map7d2_A2AG50_S162s_1 | Map7d2_A | -3.94 | -4.28 | -4.53 | -12.42 | -3.87 | -5.4 | -3.45 | -4.46 | -4.09 | -3.65 | -2.01 | -4.29 | -1.37 | -3.06 | -4.7 | -3.75 | 1.9975 | 2.7825 | 3.0725 | 0.122059 | 0.279291 | 0.154777 | 0.989223 | 0.997988 | 0.994666 |
| Map7d2_A2AG50_S173s_1 | Map7d2_A | -1.05 | -1.27 | -1.26 | -1.46 | -0.63 | -1.17 | -1 | -1.09 | -1.18 | -1.06 | -0.7 | -1.22 | -1.4 | -0.84 | -1.34 | -1.27 | 0.2875 | 0.22 | 0.0475 | 0.85318 | 0.22292 | 0.36166 | 0.989223 | 0.997988 | 0.999507 |
| Map7d2_A2AG50_S515s_1 | Map7d2_A | -0.89 | 0.09 | -0.22 | -0.04 | -0.34 | -1.29 | 0.1 | 0.25 | -0.94 | -0.7 | -0.22 | -0.68 | -0.61 | -0.74 | 0.08 | -0.07 | -0.055 | -0.37 | -0.07 | 0.834299 | 0.889247 | 0.240898 | 0.996493 | 0.997988 | 0.999507 |
| Map7d2_A2AG50_S567s_1 | Map7d2_A | 2.4 | -0.73 | -0.28 | 1.2 | 0.26 | 1.47 | 0.52 | 1.27 | 0.74 | 0.32 | 1.57 | -0.07 | -1.56 | -0.2 | -0.66 | -0.14 | 0.2325 | -0.0075 | -1.2875 | 0.092675 | 0.733728 | 0.991645 | 0.995762 | 0.997988 | 0.994666 |
| Map7d2_A2AG50_S572s_1 | Map7d2_A | 0.19 | 0.2 | -0.7 | -0.35 | -0.01 | 0.58 | 0.08 | -1.73 | 0.06 | 0.27 | 0.27 | -0.6 | 0.42 | 0.86 | -1.27 | -0.38 | -0.105 | 0.165 | 0.0725 | 0.882747 | 0.834695 | 0.616193 | 0.995762 | 0.997988 | 0.999507 |
| Map7d2_Q8C0A6_S34s_1 | Map7d2_Q | -3.91 | -3.69 | -2.69 | -4.13 | -3.25 | -3.96 | -2.3 | -2.3 | -5.92 | -5.27 | 0.47 | -2.84 | -4.67 | -1.94 | -2.63 | -2.55 | 0.6525 | 0.215 | 0.6575 | 0.298794 | 0.191527 | 0.86835 | 0.989223 | 0.997988 | 0.994666 |
| Map9_Q3TRR0_S257s_1 | Map9_Q3T | -0.62 | -1.2 | 0.16 | 0.09 | -2.45 | 0.63 | -0.27 | 1.38 | 1.1 | -0.71 | -0.27 | 0.16 | 1.72 | -2.21 | -0.13 | -0.37 | 0.215 | 0.4625 | 0.145 | 0.850117 | 0.782633 | 0.345726 | 0.995762 | 0.997988 | 0.999507 |
| Map9_Q3TRR0_S260s_1 | Map9_Q3T | -2.02 | 0.24 | -0.91 | -0.44 | -1.41 | -0.12 | -1.15 | -0.88 | 0.74 | -0.36 | -0.94 | -0.14 | 1.39 | -1.09 | -0.52 | -0.11 | -0.1075 | 0.6075 | 0.7 | 0.292205 | 0.831174 | 0.283189 | 0.995762 | 0.997988 | 0.994666 |
| Mapk10_Q3TQZ7_T221t_1 | Mapk10_Q | 0 | -0.09 | 0.96 | 1.36 | 0.23 | 0.04 | 0.51 | 1.21 | 0.9 | -0.15 | 0.93 | 1.03 | 1.6 | 0.38 | 1.34 | 0.81 | -0.06 | 0.12 | 0.475 | 0.29629 | 0.885884 | 0.78454 | 0.996493 | 0.997988 | 0.994666 |
| Mapk10_Q3TQZ7_Y223y_1 | Mapk10_Q | 0.77 | 0.95 | 0.61 | 0.88 | 0.87 | 0.85 | 1.15 | 1.03 | 0.93 | 0.97 | 0.43 | 0.8 | 0.51 | 0.68 | 0.73 | 0.8 | 0.1725 | -0.02 | -0.1225 | 0.607156 | 0.416078 | 0.932037 | 0.989223 | 0.997988 | 0.995259 |
| Mapk14_B2KF34_Y182y_1 | Mapk14_B | 5.45 | 0.42 | 3.5 | 1.95 | 0.97 | 6.49 | 5.01 | 3.46 | 0.88 | 4.1 | 0.32 | 4.14 | 1.85 | -1.95 | 3.83 | 4.77 | 1.1525 | -0.47 | -0.705 | 0.65599 | 0.408925 | 0.718413 | 0.989223 | 0.997988 | 0.995259 |
| Mapk3_D3Z3G6_T203t_1 | Mapk3_D3 | -0.21 | -0.49 | -0.04 | 0.18 | 0.12 | -0.3 | 0.54 | 0.15 | -0.8 | -0.26 | -0.56 | 0 | -0.24 | 0.01 | 0.1 | 0.24 | 0.2675 | -0.265 | 0.1675 | 0.533593 | 0.329168 | 0.352223 | 0.989223 | 0.997988 | 0.994666 |
| Mapk4_Q6P5G0_S382s_1 | Mapk4_Q6 | 0.74 | 0.43 | 0.57 | -0.47 | 1.08 | 0.63 | -0.24 | -0.2 | 0.05 | 0.75 | 1.18 | -0.39 | -0.67 | 0.7 | -0.32 | 0.24 | -2E-16 | 0.08 | -0.33 | 0.427392 | 1 | 0.852667 | 1 | 0.997988 | 0.994666 |
| Mapk4_Q6P5G0_S430s_1 | Mapk4_Q6 | 0.34 | -0.18 | -0.26 | 0.21 | 0.23 | -0.14 | -0.23 | -0.43 | 0.44 | -0.27 | 0.49 | -0.16 | 0.02 | 0.51 | -1.3 | -0.44 | -0.17 | 0.0975 | -0.33 | 0.432255 | 0.509163 | 0.738049 | 0.989223 | 0.997988 | 0.994666 |
| Mapk4_Q6P5G0_S504s_1 | Mapk4_Q6 | -0.56 | -0.1 | -0.31 | -1.03 | -0.04 | -0.75 | -0.68 | -0.91 | -0.38 | -0.32 | 0.13 | -0.76 | 0.48 | 0.22 | -1.44 | -1.04 | -0.095 | 0.1675 | 0.055 | 0.909808 | 0.752667 | 0.587872 | 0.995762 | 0.997988 | 0.999507 |
| Mapk8_D3Z1Z4_S301s_1 | Mapk8_D3 | 0.63 | 0.25 | 0.06 | 0.61 | -0.15 | 1.04 | 0.67 | 0.8 | 0.51 | -0.32 | 0.46 | -0.33 | -1.31 | 0.32 | 0.95 | 0.79 | 0.2025 | -0.3075 | -0.2 | 0.69314 | 0.521196 | 0.3314 | 0.989223 | 0.997988 | 0.996159 |
| Mapk8ip1_Q9WVI9_S14s_1 | Mapk8ip1_ | -0.08 | 0.23 | -0.13 | 0.04 | -0.12 | -0.64 | 0.1 | -0.11 | -0.43 | 0.02 | -0.22 | -0.06 | 0.12 | 0.15 | -0.18 | 0.31 | -0.2075 | -0.1875 | 0.085 | 0.732194 | 0.399022 | 0.419205 | 0.989223 | 0.997988 | 0.9971 |
| Mapk8ip1_Q9WVI9_S189s_1 | Mapk8ip1_ | 7.98 | 3.86 | 3.42 | 1.82 | 5.7 | 8.08 | 4.4 | 4.15 | 0.51 | 6.11 | 1.58 | 4.03 | -6.36 | -0.37 | 3.5 | 4.07 | 1.3125 | -1.2125 | -4.06 | 0.107557 | 0.348107 | 0.451537 | 0.989223 | 0.997988 | 0.994666 |
| Mapk8ip1_Q9WVI9_S231s_1 | Mapk8ip1_ | 0.47 | 0.12 | 0.43 | 0.51 | -1.36 | 0.19 | 0.04 | 0.04 | 2.61 | 0.02 | -0.59 | 0.25 | 2.37 | -1.35 | 1.73 | 0.49 | -0.655 | 0.19 | 0.4275 | 0.563374 | 0.10198 | 0.768034 | 0.989223 | 0.997988 | 0.995259 |
| Mapk8ip1_Q9WVI9_S351s_1 | Mapk8ip1_ | 1.88 | 0.6 | 0.06 | 0.78 | 1.19 | 1.28 | -0.46 | 0.54 | 1.78 | 0.05 | 1.41 | 0.45 | 1 | 0.98 | 0 | 0.4 | -0.1925 | 0.0925 | -0.235 | 0.598014 | 0.704831 | 0.85874 | 0.995437 | 0.997988 | 0.995259 |
| Mapk8ip2_Q9ERE9_S128s_1 | Mapk8ip2_ | 0.73 | 1.02 | 0.76 | 0.65 | 1.17 | 0.61 | 1.27 | 0.86 | 0.13 | 1.16 | 0.84 | 0.6 | 0.18 | 0.73 | 0.8 | 1.88 | 0.1875 | -0.1075 | 0.1075 | 0.777423 | 0.438281 | 0.703756 | 0.989223 | 0.997988 | 0.99798 |
| Mapk8ip2_Q9ERE9_S175s_1 | Mapk8ip2_ | 0.46 | -0.4 | 0.49 | 0.43 | 0.79 | 1.5 | 0.73 | 0.72 | 0.3 | 0.28 | 1.93 | 0.18 | 0.12 | 0.57 | 0.09 | -0.15 | 0.69 | 0.4275 | -0.0875 | 0.780748 | 0.046534 | 0.357876 | 0.989223 | 0.997988 | 0.998226 |
| Mapk8ip2_Q9ERE9_S248s_1 | Mapk8ip2_ | -0.2 | -0.77 | -0.39 | -0.23 | -0.6 | -0.6 | -1.49 | -0.72 | -0.02 | -0.1 | -0.49 | -0.06 | 0.78 | 0.3 | 0.24 | -0.24 | -0.455 | 0.23 | 0.6675 | 0.050689 | 0.13445 | 0.364417 | 0.989223 | 0.997988 | 0.994666 |
| Mapk8ip2_Q9ERE9_S249s_1 | Mapk8ip2_ | -0.2 | -0.77 | -0.39 | -0.23 | -0.6 | -0.6 | -1.49 | -0.72 | -0.02 | -0.1 | -0.49 | -0.06 | 0.78 | 0.3 | 0.24 | -0.24 | -0.455 | 0.23 | 0.6675 | 0.050689 | 0.13445 | 0.364417 | 0.989223 | 0.997988 | 0.994666 |
| Mapk8ip2_Q9ERE9_S249s_2 | Mapk8ip2_ | -0.52 | -0.3 | -0.3 | -0.55 | -1.91 | -1.56 | 0.02 | -0.46 | -0.35 | 0.84 | -0.35 | -0.11 | -0.24 | -0.45 | -0.3 | -0.15 | -0.56 | 0.425 | 0.1325 | 0.576978 | 0.215902 | 0.20607 | 0.989223 | 0.997988 | 0.995259 |
| Mapk8ip2_Q9ERE9_S370s_1 | Mapk8ip2_ | 0.42 | -0.2 | -0.43 | 0.16 | 0.79 | 0.24 | 0.58 | 0 | 0.08 | 0.04 | 0.25 | -0.09 | -0.35 | 0.36 | 0.32 | -0.06 | 0.415 | 0.0825 | 0.08 | 0.795338 | 0.173208 | 0.755193 | 0.989223 | 0.997988 | 0.999507 |
| Mapk8ip2_Q9ERE9_S380s_1 | Mapk8ip2_ | 0.9 | -1 | -1.31 | -0.86 | -0.08 | 0.72 | -0.61 | -0.16 | -0.2 | -0.6 | 1.07 | -0.37 | -1.06 | 0.46 | -0.88 | -0.6 | 0.535 | 0.5425 | 0.0475 | 0.932036 | 0.316674 | 0.356711 | 0.989223 | 0.997988 | 0.999507 |
| Mapk8ip3_K3W4S4_S611s_1 | Mapk8ip3_ | 0.66 | 0.54 | 1.6 | -0.23 | 1.06 | 0.83 | 1.44 | 0.76 | 0.98 | 0.42 | 2.54 | 0.69 | 1.07 | 1.34 | 1.77 | 0.35 | 0.38 | 0.515 | 0.49 | 0.304206 | 0.344736 | 0.368801 | 0.989223 | 0.997988 | 0.994666 |
| Mapk8ip3_K3W4S4_S685s_1 | Mapk8ip3_ | 0.85 | 0.52 | 0.21 | 0.96 | 0.06 | 1.07 | -0.19 | 0.4 | 0.84 | 0.77 | 0.12 | 0.35 | 0.95 | 0.36 | 0.21 | 0.16 | -0.3 | -0.115 | -0.215 | 0.487739 | 0.375992 | 0.690763 | 0.989223 | 0.997988 | 0.994666 |
| Mapk8ip3_K3W4S4_S763s_1 | Mapk8ip3_ | 6.86 | 1.05 | 2.16 | -0.23 | -0.02 | 7.55 | 6.11 | 0.92 | 0.58 | 5.75 | -0.69 | 2.75 | 1.98 | 0.26 | 3.23 | 4.66 | 1.18 | -0.3625 | 0.0725 | 0.962645 | 0.572342 | 0.842233 | 0.989223 | 0.997988 | 0.999507 |
| Mapk8ip3_K3W4S4_T1199t_1 | Mapk8ip3_ | -1.87 | -0.57 | -0.29 | -1.95 | -1.62 | -1.23 | -1.24 | -1.91 | 0.17 | -1.88 | -1.58 | -0.54 | 1.44 | -2.48 | -0.78 | -1.25 | -0.33 | 0.2125 | 0.4025 | 0.623878 | 0.454685 | 0.718538 | 0.989223 | 0.997988 | 0.995259 |
| Mapkap1_Q8BKH7_S318s_1 | Mapkap1_ | -1.76 | -1.19 | -0.28 | -1.56 | -0.79 | -2.13 | -0.22 | -1.43 | -2.15 | -1 | 0.39 | -0.56 | -0.54 | -0.86 | -0.28 | -1.12 | 0.055 | 0.3675 | 0.4975 | 0.219259 | 0.909597 | 0.525318 | 0.997335 | 0.997988 | 0.994666 |
| Mapkbp1_Q6NS57_S756s_1 | Mapkbp1_ | -0.35 | -1.72 | 0.08 | -0.49 | -2 | 0.66 | -1.03 | -0.65 | 1.91 | 0.07 | -1.82 | 0.09 | 2.1 | -4.71 | -0.86 | -1.31 | -0.135 | 0.6825 | -0.575 | 0.646771 | 0.822429 | 0.382846 | 0.995762 | 0.997988 | 0.995259 |
| Mapre2_Q8R001_S199s_1 | Mapre2_Q | -3.15 | -1.92 | -2.77 | -1.97 | -2.81 | -1.97 | -3.85 | -2.93 | -0.06 | -2.39 | -4.97 | -2.3 | 0.77 | -3.16 | -1.73 | -2.63 | -0.4375 | 0.0225 | 0.765 | 0.358509 | 0.349885 | 0.980645 | 0.989223 | 0.997988 | 0.994666 |
| Mapre2_Q8R001_S222s_1 | Mapre2_Q | -5.12 | -4.85 | 0.46 | -0.04 | -3.86 | -3.37 | 0.78 | 1.11 | -3.2 | -3.9 | 0.9 | 0.71 | -1.64 | -4.68 | 0.25 | 0.21 | 1.0525 | 1.015 | 0.9225 | 0.573978 | 0.542531 | 0.557201 | 0.989223 | 0.997988 | 0.995259 |
| Mapre2_Q8R001_S9s_1 | Mapre2_Q | 1.75 | 0.87 | 0.59 | 0.5 | 0.79 | 2.08 | 1.53 | 0.55 | 2.22 | 2.52 | -0.07 | 0.76 | 1.23 | 1.41 | 0.91 | 0.82 | 0.31 | 0.43 | 0.165 | 0.637091 | 0.473953 | 0.491309 | 0.989223 | 0.997988 | 0.995259 |
| Mapre3_D3Z6G3_S147s_1 | Mapre3_D | -0.16 | -0.85 | -1.14 | -1.65 | -0.76 | 0.48 | -1.41 | -1.5 | 2.73 | -0.18 | -3.76 | -0.42 | 2.89 | -1.13 | 1.05 | -1.42 | 0.1525 | 0.5425 | 1.2975 | 0.183035 | 0.763191 | 0.651788 | 0.995762 | 0.997988 | 0.994666 |
| Mapre3_Q6PER3_S162s_1 | Mapre3_Q | -0.16 | -0.85 | -1.14 | -1.65 | -0.76 | 0.48 | -1.41 | -1.5 | 2.73 | -0.18 | -3.76 | -0.42 | 2.89 | -1.13 | 1.05 | -1.42 | 0.1525 | 0.5425 | 1.2975 | 0.183035 | 0.763191 | 0.651788 | 0.995762 | 0.997988 | 0.994666 |
| Mapt_B1AQW2_T42t_1 | Mapt_B1A | -1.38 | -1.29 | -1.01 | -0.86 | -1.3 | -1.57 | -0.86 | -0.89 | -1.3 | -1.15 | -0.47 | -1.17 | -0.47 | -1.5 | -1.32 | -0.97 | -0.02 | 0.1125 | 0.07 | 0.821813 | 0.938427 | 0.685443 | 0.999187 | 0.997988 | 0.999507 |
| Mapt_B1AQW2_T54t_1 | Mapt_B1A | -0.63 | -0.74 | 0.08 | -0.04 | -0.45 | -1.03 | -0.17 | -0.62 | -1 | -0.46 | 0.17 | -0.25 | -1.01 | -0.3 | -1.11 | 0.12 | -0.235 | -0.0525 | -0.2425 | 0.523347 | 0.439373 | 0.87693 | 0.989223 | 0.997988 | 0.994666 |
| Mapt_P10637_S167s_2 | Mapt_P10 | 0.95 | 0.87 | 0.24 | 0.4 | 0.54 | 0.14 | 0.24 | 0.19 | 0.81 | 1.03 | 0.79 | 0.36 | 0.34 | 1.17 | 0.91 | 0.1 | -0.3375 | 0.1325 | 0.015 | 0.964647 | 0.200931 | 0.634537 | 0.989223 | 0.997988 | 0.999507 |
| Mapt_P10637_S180s_1 | Mapt_P10 | -0.12 | -0.07 | 0.1 | 0.3 | -1.01 | -0.22 | -0.24 | -0.54 | 0.61 | 0.31 | -0.05 | -0.15 | 0.72 | -0.34 | 0.57 | 0.06 | -0.555 | 0.1275 | 0.2 | 0.527143 | 0.054362 | 0.629277 | 0.989223 | 0.997988 | 0.994666 |
| Mapt_P10637_S180s_2 | Mapt_P10 | 0.63 | 0.38 | 0.37 | 0.52 | -0.11 | 0.32 | -0.25 | 0.17 | 0.56 | 0.59 | 0.72 | 0.38 | -0.1 | -0.6 | 0.42 | 0.55 | -0.4425 | 0.0875 | -0.4075 | 0.220843 | 0.073847 | 0.685506 | 0.989223 | 0.997988 | 0.994666 |
| Mapt_P10637_S184s_2 | Mapt_P10 | -1.58 | -0.41 | -0.21 | -0.38 | -0.36 | -0.64 | -1.68 | -0.45 | -1.86 | -1.19 | 0.66 | -0.29 | -0.61 | -0.32 | -0.42 | -0.88 | -0.1375 | -0.025 | 0.0875 | 0.809048 | 0.741991 | 0.965645 | 0.995762 | 0.997988 | 0.999507 |
| Mapt_P10637_S187s_1 | Mapt_P10 | 0.29 | 0.83 | -0.52 | -0.53 | 0.52 | 0.32 | -0.86 | -0.63 | 0.18 | 0.93 | -0.76 | 0.01 | 0.8 | 0.88 | -0.14 | 0.32 | -0.18 | 0.0725 | 0.4475 | 0.290683 | 0.687771 | 0.874652 | 0.994672 | 0.997988 | 0.994666 |
| Mapt_P10637_S187s_2 | Mapt_P10 | 0.44 | 1.81 | -0.17 | 0.13 | 1.3 | -0.14 | -0.2 | -0.27 | 0.26 | 1.42 | 0.76 | -0.56 | -0.44 | 0.92 | 1.35 | 0.15 | -0.38 | -0.0825 | -0.0575 | 0.916435 | 0.475372 | 0.882621 | 0.989223 | 0.997988 | 0.999507 |
| Mapt_P10637_S187s_3 | Mapt_P10 | 0.32 | 1.06 | 0.9 | 1.11 | 0.09 | -0.05 | 0.45 | 1.16 | 0.24 | 0.86 | 1.81 | 1.57 | -0.39 | 0.42 | 1.6 | 1.39 | -0.435 | 0.2725 | -0.0925 | 0.845337 | 0.213665 | 0.498991 | 0.989223 | 0.997988 | 0.999507 |
| Mapt_P10637_S188s_1 | Mapt_P10 | 0.22 | 0.8 | 0.81 | 0.31 | 0.6 | 0.06 | 0 | 0.58 | 0.45 | 0.43 | 0.93 | 0.45 | 1.2 | 0.56 | 0.4 | 0.74 | -0.225 | 0.03 | 0.19 | 0.527018 | 0.411408 | 0.909029 | 0.989223 | 0.997988 | 0.994666 |
| Mapt_P10637_S188s_2 | Mapt_P10 | 0.74 | 1.23 | 0.93 | 0.67 | 0.53 | 0.2 | 0.47 | 0.44 | 0.59 | 1.38 | 1.58 | 0.68 | -0.98 | 0.84 | 1.23 | 0.51 | -0.4825 | 0.165 | -0.4925 | 0.155429 | 0.936271 | 0.598776 | 0.989223 | 0.997988 | 0.994666 |
| Mapt_P10637_S188s_3 | Mapt_P10 | 0.37 | 1.15 | 0.9 | 1.14 | 0.08 | -0.09 | 0.46 | 1.02 | 0.37 | 0.85 | 1.97 | 1.38 | -0.42 | 0.49 | 1.65 | 1.45 | -0.5225 | 0.2525 | -0.0975 | 0.841118 | 0.128466 | 0.522734 | 0.989223 | 0.997988 | 0.994666 |
| Mapt_P10637_S191s_1 | Mapt_P10 | 0.52 | 0.63 | 0.09 | 0.29 | 0.34 | 0.09 | -0.31 | 0.05 | 0.88 | 0.7 | 0.71 | 0.22 | 1.35 | 1.37 | 0.22 | 0.42 | -0.34 | 0.245 | 0.4575 | 0.216366 | 0.183899 | 0.350743 | 0.989223 | 0.997988 | 0.994666 |
| Mapt_P10637_S191s_2 | Mapt_P10 | 0.6 | 0.92 | 1.53 | 1.18 | 0.25 | -0.02 | 0.82 | 0.96 | 0.43 | 1.15 | 2.15 | 1.16 | -1.27 | 0.75 | 1.59 | 0.97 | -0.555 | 0.165 | -0.5475 | 0.367978 | 0.107949 | 0.682408 | 0.989223 | 0.997988 | 0.994666 |

| | | | | | | | | | | | | | | | | | | | | | | | | | | |
|---|---|---|---|---|---|---|---|---|---|---|---|---|---|---|---|---|---|---|---|---|---|---|---|---|---|---|
| Mapt_P10637_S191s_3 | Mapt_P106 | 0.54 | 1.16 | 0.99 | 0.92 | 0.05 | 0.02 | 0.6 | 0.92 | 0.27 | 0.97 | 2.05 | 1.15 | -0.46 | 0.42 | 1.65 | 1.15 | -0.505 | 0.2075 | -0.2125 | 0.646702 | 0.104035 | 0.597357 | 0.989223 | 0.997988 | 0.995259 |
| Mapt_P10637_S197s_1 | Mapt_P106 | -1.48 | -2.98 | 0.71 | 1.27 | -0.24 | 0.24 | 0.81 | 0.8 | 0.12 | -0.1 | 0.77 | 0.55 | 0.11 | 0.88 | 1.31 | 0.38 | 1.0225 | 0.955 | 1.29 | 0.172215 | 0.264668 | 0.299829 | 0.989223 | 0.997988 | 0.994666 |
| Mapt_P10637_S224s_2 | Mapt_P106 | -0.72 | -0.9 | 1.29 | 0.79 | -1.15 | -0.31 | 0.96 | 1.21 | 0.45 | -0.83 | -0.02 | 1.24 | 0.22 | -1.1 | 1.51 | 1.29 | 0.0625 | 0.095 | 0.365 | 0.614959 | 0.927344 | 0.880814 | 0.997344 | 0.997988 | 0.995259 |
| Mapt_P10637_S313s_1 | Mapt_P106 | -4.45 | -1.98 | -4.04 | -8.78 | -2.5 | -4.04 | -10.33 | -4.87 | -1.5 | -2.9 | 0.91 | -5.38 | -0.07 | 1 | -5.32 | -6.66 | -0.6225 | 2.595 | 2.05 | 0.325534 | 0.743443 | 0.150272 | 0.995762 | 0.997988 | 0.994666 |
| Mapt_P10637_S345s_1 | Mapt_P106 | -0.46 | -0.75 | -0.39 | -0.48 | -0.05 | -0.34 | -0.08 | -0.22 | -0.42 | -0.43 | 0.39 | -0.15 | -1.17 | 0.11 | -0.96 | -0.43 | 0.3475 | 0.3675 | -0.0925 | 0.783005 | 0.120869 | 0.192521 | 0.989223 | 0.997988 | 0.998491 |
| Mapt_P10637_S35s_1 | Mapt_P106 | -1.05 | 0.54 | 0.25 | -0.34 | -0.66 | -0.4 | 0.21 | -0.05 | 0.23 | 0.24 | -0.52 | -0.59 | 0.31 | 0.78 | 0.19 | 0.52 | -0.075 | -0.01 | 0.6 | 0.145073 | 0.846825 | 0.98065 | 0.996349 | 0.997988 | 0.994666 |
| Mapt_P10637_S385s_1 | Mapt_P106 | 1.73 | -0.11 | -0.74 | -1.22 | -0.29 | 1.3 | -0.91 | -1.06 | -0.85 | 0.31 | -0.17 | -1.14 | -0.34 | -0.26 | -0.94 | -1.07 | -0.155 | -0.3775 | -0.5675 | 0.368451 | 0.833984 | 0.569256 | 0.995762 | 0.997988 | 0.994666 |
| Mapt_P10637_S385s_2 | Mapt_P106 | -2.46 | -1.43 | -1.21 | -0.88 | -1.15 | -1.95 | -1.2 | -0.46 | -2.4 | -2.18 | -0.22 | -1.09 | -2.53 | -1.29 | -1.26 | -1.54 | 0.305 | 0.0225 | -0.16 | 0.718746 | 0.485232 | 0.968128 | 0.989223 | 0.997988 | 0.9971 |
| Mapt_P10637_S385s_3 | Mapt_P106 | -1.36 | -0.89 | -0.89 | -1.84 | -1.05 | -1.45 | -1.95 | -1 | -0.46 | -1.01 | 0.25 | -0.92 | -0.33 | -0.67 | -1.35 | -1.08 | -0.1175 | 0.71 | 0.3875 | 0.283607 | 0.720667 | 0.083413 | 0.995437 | 0.997988 | 0.994666 |
| Mapt_P10637_S389s_1 | Mapt_P106 | 0.82 | 1.51 | 1.01 | 0.88 | 0.58 | -0.75 | -0.17 | -0.08 | 0.94 | 1.05 | 0.81 | 0.83 | 1.11 | 1.16 | 1.23 | 0.76 | -1.16 | -0.1475 | 0.01 | 0.970788 | 0.005384 | 0.551913 | 0.989223 | 0.997988 | 0.999507 |
| Mapt_P10637_S389s_2 | Mapt_P106 | 0.32 | 1.12 | 0.54 | 0.19 | 0.55 | -0.09 | -0.15 | 0.05 | 0.21 | 0.94 | -0.1 | 0.03 | -0.92 | 0.5 | 0.8 | 0.7 | -0.4525 | -0.2725 | -0.2725 | 0.541277 | 0.143038 | 0.421984 | 0.989223 | 0.997988 | 0.994666 |
| Mapt_P10637_S389s_3 | Mapt_P106 | -1.36 | -0.89 | -0.89 | -1.84 | -1.05 | -1.45 | -1.95 | -1 | -0.46 | -1.01 | 0.25 | -0.92 | -0.33 | -0.67 | -1.35 | -1.08 | -0.1175 | 0.71 | 0.3875 | 0.283607 | 0.720667 | 0.083413 | 0.995437 | 0.997988 | 0.994666 |
| Mapt_P10637_S393s_1 | Mapt_P106 | -0.08 | 0.45 | 0.37 | 0.25 | 0.34 | -0.03 | 0.14 | 0.24 | 0 | 0.37 | 0.52 | 0.15 | 0.83 | 0.74 | 0.42 | 0.2 | -0.075 | 0.0125 | 0.3 | 0.286419 | 0.737912 | 0.959169 | 0.995762 | 0.997988 | 0.994666 |
| Mapt_P10637_S393s_2 | Mapt_P106 | -0.87 | -0.02 | 0.09 | -0.28 | -0.23 | -0.88 | -0.45 | -0.28 | -1.01 | -0.33 | -0.09 | -0.38 | -1.83 | -0.23 | 0.32 | 0.23 | -0.19 | -0.1825 | -0.1075 | 0.83341 | 0.517644 | 0.571728 | 0.989223 | 0.997988 | 0.999507 |
| Mapt_P10637_S393s_3 | Mapt_P106 | -1.36 | -0.89 | -0.89 | -1.84 | -1.05 | -1.45 | -1.95 | -1 | -0.46 | -1.01 | 0.25 | -0.92 | -0.33 | -0.67 | -1.35 | -1.08 | -0.1175 | 0.71 | 0.3875 | 0.283607 | 0.720667 | 0.083413 | 0.995437 | 0.997988 | 0.994666 |
| Mapt_P10637_S402s_1 | Mapt_P106 | 4.46 | 2.25 | 2.65 | 1.15 | 2.75 | 4.83 | 3.07 | 2.53 | -0.84 | 4.3 | 0.07 | 2.96 | -1.2 | -1.16 | 2.99 | 3.49 | 0.6675 | -1.005 | -1.5975 | 0.222515 | 0.388622 | 0.417962 | 0.989223 | 0.997988 | 0.994666 |
| Mapt_P10637_S405s_1 | Mapt_P106 | 3.28 | 1.74 | 1.53 | 0.61 | 2.23 | 3.98 | 1.79 | 1.76 | -0.3 | 3.09 | 0.36 | 1.68 | -0.73 | 0.02 | 1.37 | 1.7 | 0.65 | -0.5825 | -1.2 | 0.116564 | 0.347479 | 0.489233 | 0.989223 | 0.997988 | 0.994666 |
| Mapt_P10637_S57s_1 | Mapt_P106 | 0.17 | 0.78 | 0.27 | 0.03 | 0.69 | 0.04 | 0.52 | 0.32 | -0.09 | 0.05 | 0.18 | 0.11 | -0.33 | 0.27 | 0.12 | 0 | 0.08 | -0.25 | -0.2975 | 0.307469 | 0.760885 | 0.329099 | 0.995762 | 0.997988 | 0.994666 |
| Mapt_P10637_T170t_1 | Mapt_P106 | 0.54 | 0.73 | 0.57 | 0.5 | -0.1 | 0.35 | -0.27 | -0.01 | 0.96 | 0.81 | 0.12 | 0.18 | 1.35 | 0.15 | 0.37 | 0.15 | -0.5925 | -0.0675 | -0.08 | 0.809297 | 0.023413 | 0.806483 | 0.989223 | 0.997988 | 0.999507 |
| Mapt_P10637_T170t_2 | Mapt_P106 | 0.86 | 0.74 | 0.31 | 0.46 | 0.37 | 0.19 | 0.02 | 0.18 | 0.74 | 0.91 | 0.76 | 0.37 | 0.23 | 0.77 | 0.68 | 0.33 | -0.4025 | 0.1025 | -0.09 | 0.739658 | 0.09967 | 0.680214 | 0.989223 | 0.997988 | 0.9971 |
| Mapt_P10637_T194t_1 | Mapt_P106 | -1.16 | -1.31 | -2.39 | -1.95 | -0.95 | -1.87 | -0.86 | -1.09 | -2.87 | -1.64 | -0.2 | -0.76 | -1.83 | -0.87 | -1.36 | -1.19 | 0.51 | 0.335 | 0.39 | 0.305553 | 0.183853 | 0.576498 | 0.989223 | 0.997988 | 0.994666 |
| Mapt_P10637_T194t_2 | Mapt_P106 | -0.28 | 0.11 | -0.11 | -0.38 | -0.04 | -0.87 | -0.11 | -0.64 | -0.61 | 0.09 | 0.71 | 0.04 | -1.95 | 0.29 | 0.54 | -0.46 | -0.25 | 0.2225 | -0.23 | 0.66688 | 0.366951 | 0.49205 | 0.989223 | 0.997988 | 0.996159 |
| Mapt_P10637_T194t_3 | Mapt_P106 | 0.15 | 0.68 | 0.72 | 1.55 | 0.11 | 0.12 | 0.2 | 0.9 | -0.33 | 0.89 | 2.14 | 1.38 | -0.29 | 0.13 | 1.77 | 2.04 | -0.4425 | 0.245 | 0.1375 | 0.817066 | 0.224363 | 0.657607 | 0.989223 | 0.997988 | 0.999507 |
| Mapt_P10637_T201t_1 | Mapt_P106 | -1.95 | -2.24 | -2.27 | -3.55 | -1.47 | -2.02 | -1.7 | -3.39 | -2.84 | -1.82 | 0.29 | -2.21 | -1.65 | -0.91 | -2.57 | -2.61 | 0.3575 | 0.8575 | 0.5675 | 0.281631 | 0.489249 | 0.235068 | 0.989223 | 0.997988 | 0.994666 |
| Mapt_P10637_T206t_1 | Mapt_P106 | -0.9 | -0.17 | -0.68 | -1.09 | -0.51 | -1.83 | -0.42 | -1.42 | -1.16 | -0.02 | -0.55 | -0.72 | -0.51 | 0 | -0.08 | -0.7 | -0.335 | 0.0975 | 0.3875 | 0.23427 | 0.396443 | 0.769146 | 0.989223 | 0.997988 | 0.994666 |
| Mapt_P10637_T220t_1 | Mapt_P106 | 2.23 | 1.68 | 1.73 | 2.13 | -0.24 | 1.99 | 0.52 | 1.16 | 4.01 | 2.45 | 0.29 | 1.7 | 3.66 | 0.73 | 1.89 | 1.72 | -1.085 | 0.17 | 0.0575 | 0.920243 | 0.038352 | 0.81068 | 0.989223 | 0.997988 | 0.999507 |
| Mapt_P10637_T220t_2 | Mapt_P106 | -1.03 | -1.1 | 1.29 | 0.79 | -1.29 | -0.74 | 0.96 | 1.21 | 0.23 | -1.1 | -0.02 | 1.24 | 0 | -1.47 | 1.51 | 1.29 | 0.0475 | 0.1 | 0.345 | 0.672991 | 0.950447 | 0.88705 | 0.999187 | 0.997988 | 0.996159 |
| Mapt_P10637_T252t_1 | Mapt_P106 | -0.98 | -2.08 | -1.15 | -3.82 | -0.76 | -0.87 | -0.4 | -1.08 | -1.33 | -1.58 | 2.12 | -1.71 | -1.63 | -0.19 | -1.06 | -2.59 | 1.23 | 1.3825 | 0.64 | 0.392208 | 0.063137 | 0.18723 | 0.989223 | 0.997988 | 0.994666 |
| Mapt_P10637_T392t_1 | Mapt_P106 | 0.15 | 1.15 | 0.75 | 0.51 | 0.76 | -0.15 | -0.15 | 0.32 | 0.31 | 0.71 | 0.85 | 0.27 | 1 | 1.18 | 0.71 | 0.69 | -0.445 | -0.105 | 0.255 | 0.4067 | 0.184693 | 0.723934 | 0.989223 | 0.997988 | 0.994666 |
| Mapt_P10637_T392t_2 | Mapt_P106 | -2.3 | -1.13 | 0.92 | 2.65 | -1.09 | -2.21 | 1.36 | 2.93 | -2.02 | -1.82 | 1.74 | 1.82 | -2.1 | -1.78 | 1.18 | 0.6 | 0.2125 | -0.105 | -0.56 | 0.639038 | 0.876803 | 0.93748 | 0.996493 | 0.997988 | 0.995259 |
| Mapt_P10637_T403t_1 | Mapt_P106 | 3.26 | 1.98 | 2.25 | -0.07 | 2.42 | 3.3 | 2.96 | 3.2 | 0.11 | 3.27 | 0.12 | 2.14 | -1.17 | 0.31 | 2.26 | 3.22 | 1.115 | -0.445 | -0.7 | 0.509047 | 0.109004 | 0.633822 | 0.989223 | 0.997988 | 0.994666 |
| March11_Q8CBH7_S115s_1 | March11_Q | 0.85 | 1.49 | 0.71 | 0.85 | -0.09 | 0.33 | -0.67 | -0.64 | 0.81 | 0.85 | 0.28 | 0.19 | 1.24 | -0.56 | 0.59 | 1.03 | -1.2425 | -0.4425 | -0.4 | 0.36588 | 0.002887 | 0.152764 | 0.989223 | 0.997988 | 0.994666 |
| Marcks_P26645_S112s_1 | Marcks_P2 | -1.49 | -0.03 | 0.28 | -0.21 | -0.19 | -1.21 | -0.35 | 0.1 | -0.82 | -0.86 | -0.33 | 0.01 | -0.41 | -0.13 | 0.05 | -0.1 | -0.05 | -0.1375 | 0.215 | 0.598329 | 0.911295 | 0.749521 | 0.997335 | 0.997988 | 0.995259 |
| Marcks_P26645_S113s_1 | Marcks_P2 | -0.73 | 0.17 | 0.22 | -0.15 | -0.22 | -0.79 | -0.33 | -0.58 | -0.02 | -0.21 | -1.78 | 0 | -0.72 | 0.37 | -0.62 | -0.69 | -0.3575 | -0.38 | -0.2925 | 0.429923 | 0.230151 | 0.418146 | 0.989223 | 0.997988 | 0.994666 |
| Marcks_P26645_S122s_1 | Marcks_P2 | -1.03 | -0.41 | 0.66 | 0.52 | 0.31 | -1.46 | 0.43 | 0.16 | 0.23 | -0.63 | 0.54 | 0.76 | 0.49 | -0.47 | 0.97 | 0.21 | -0.075 | 0.29 | 0.365 | 0.452707 | 0.889627 | 0.547794 | 0.996493 | 0.997988 | 0.994666 |
| Marcks_P26645_S138s_1 | Marcks_P2 | 0.23 | 0.46 | 0.44 | 0.32 | 0.47 | 0.5 | 0.35 | 0.51 | 0.23 | 0.67 | 0.41 | 0.4 | 0.72 | 0.29 | 0.46 | 0.43 | 0.095 | 0.065 | 0.1125 | 0.639835 | 0.63194 | 0.76729 | 0.989223 | 0.997988 | 0.995259 |
| Marcks_P26645_S138s_2 | Marcks_P2 | -0.32 | -0.41 | -0.57 | -0.42 | -0.26 | -0.17 | -0.49 | -0.66 | -0.17 | -0.11 | -0.19 | -0.49 | -0.01 | -0.49 | -1.04 | -0.4 | 0.035 | 0.19 | -0.055 | 0.849015 | 0.871135 | 0.391899 | 0.996493 | 0.997988 | 0.999507 |
| Marcks_P26645_S140s_1 | Marcks_P2 | 0.17 | 0.64 | -0.24 | 0.37 | 0.12 | -0.45 | 0.01 | 0.28 | 0.51 | 0.46 | 0.95 | 0.05 | 0.78 | 0.5 | 0.4 | -0.03 | -0.245 | 0.2575 | 0.1775 | 0.56607 | 0.390096 | 0.401122 | 0.989223 | 0.997988 | 0.995259 |
| Marcks_P26645_S141s_2 | Marcks_P2 | 0.31 | 0.05 | -0.55 | -0.67 | 0.9 | 0.22 | -0.61 | -0.78 | -0.22 | 0.3 | -0.9 | -0.26 | 0.64 | 0.58 | -1.76 | -0.11 | 0.1475 | -0.055 | 0.0525 | 0.925263 | 0.732015 | 0.876554 | 0.995762 | 0.997988 | 0.999507 |
| Marcks_P26645_S160s_1 | Marcks_P2 | -1.51 | -1.75 | -1.18 | -2.47 | -0.35 | -1.07 | -1.25 | -1.63 | -2.32 | -1.69 | -0.46 | -0.76 | -1.76 | -0.67 | -2.27 | -1.76 | 0.6525 | 0.42 | 0.1125 | 0.793471 | 0.108049 | 0.392928 | 0.989223 | 0.997988 | 0.999507 |
| Marcks_P26645_S163s_1 | Marcks_P2 | -4.73 | -4.52 | -3.82 | -6.33 | -2.03 | -4.72 | -3.19 | -4.34 | -6.89 | -4.52 | -1.47 | -3.37 | -3.48 | -2.19 | -3.87 | -3.36 | 1.28 | 0.7875 | 1.625 | 0.01957 | 0.097122 | 0.480494 | 0.989223 | 0.997988 | 0.994666 |
| Marcks_P26645_S171s_1 | Marcks_P2 | -1.09 | -0.1 | -0.23 | -0.66 | -0.74 | -0.73 | -0.85 | -0.89 | -1 | -1.01 | -0.58 | -0.56 | 0.08 | -0.46 | -0.53 | -0.61 | -0.2825 | -0.2675 | 0.14 | 0.663818 | 0.310059 | 0.380099 | 0.989223 | 0.997988 | 0.996159 |
| Marcks_P26645_S26s_1 | Marcks_P2 | -0.04 | 1.17 | 1.51 | 0.49 | 0.26 | 0.79 | 0.58 | 0.83 | 0.05 | 0.65 | 2.16 | 1.24 | 0.36 | -0.17 | 0.81 | 1.78 | -0.1675 | 0.2425 | -0.0875 | 0.863277 | 0.649074 | 0.648554 | 0.99033 | 0.997988 | 0.999507 |
| Marcks_P26645_S27s_1 | Marcks_P2 | 0.13 | 0.71 | 0.39 | -0.01 | 0.68 | 0.33 | 0.26 | 0.02 | 0.36 | 0.55 | 0.69 | 0.32 | 0.9 | 0.96 | 0.08 | -0.07 | 0.0175 | 0.175 | 0.1625 | 0.63936 | 0.946075 | 0.494415 | 0.999187 | 0.997988 | 0.995259 |
| Marcks_P26645_S29s_1 | Marcks_P2 | -1.71 | 0.04 | 0.7 | 0.11 | -0.44 | -1.08 | 0.18 | 0.16 | -0.8 | -0.56 | 1.2 | 0.44 | -0.49 | -0.55 | 0.58 | 0.03 | -0.08 | 0.285 | 0.1075 | 0.842694 | 0.882747 | 0.654467 | 0.996493 | 0.997988 | 0.999507 |
| Marcks_P26645_S46s_1 | Marcks_P2 | 0.1 | 0.43 | 0.46 | 0.12 | 0.5 | 0.28 | 0.25 | 0.28 | 0.23 | 0.59 | 0.78 | 0.67 | 0.74 | 0.62 | 0.56 | 0.43 | 0.05 | 0.29 | 0.31 | 0.221889 | 0.813693 | 0.244663 | 0.995762 | 0.997988 | 0.994666 |
| Marcks_P26645_T143t_1 | Marcks_P2 | -0.55 | -0.47 | -1.05 | -0.78 | -0.41 | -0.89 | -0.6 | -1.04 | -0.63 | -0.08 | 0.07 | -0.17 | 0.29 | -0.84 | -0.44 | -0.31 | -0.0225 | 0.51 | 0.3875 | 0.240382 | 0.928123 | 0.078261 | 0.997447 | 0.997988 | 0.994666 |
| Marcks_P26645_T143t_2 | Marcks_P2 | -0.57 | -0.69 | -0.64 | -0.42 | -0.89 | -0.32 | -0.69 | -0.9 | -0.28 | -0.36 | -0.39 | -0.76 | -0.16 | -0.86 | -1.04 | -0.5 | -0.12 | 0.1325 | -0.06 | 0.831114 | 0.59935 | 0.560358 | 0.989223 | 0.997988 | 0.999507 |
| Marcks_P26645_T79t_1 | Marcks_P2 | -1.75 | -1.81 | -2.09 | -2.08 | -1.39 | -1.71 | -2.52 | -2.81 | -2.93 | -1.87 | -2.1 | -2.47 | -1.71 | -1.89 | -2.53 | -2.92 | -0.175 | -0.41 | -0.33 | 0.335599 | 0.617493 | 0.183456 | 0.989223 | 0.997988 | 0.994666 |
| Marcksl1_P28667_S101s_1 | Marcksl1_P | -0.75 | -0.67 | -0.4 | -1.26 | 0.41 | 0.12 | 0.45 | 0.11 | 0.32 | -0.11 | -0.12 | 0.66 | -0.88 | 0.78 | -0.03 | -0.48 | 1.0425 | 0.9575 | 0.6175 | 0.15027 | 0.002317 | 0.009912 | 0.989223 | 0.997988 | 0.994666 |
| Marcksl1_P28667_S104s_1 | Marcksl1_P | -3.76 | -3.58 | -3.35 | -5.73 | -1.66 | -3.83 | -2.86 | -3.99 | -4.83 | -3.34 | -1.83 | -3.22 | -2.4 | -1.49 | -3.43 | -3.32 | 1.02 | 0.8 | 1.445 | 0.046408 | 0.156406 | 0.294569 | 0.989223 | 0.997988 | 0.994666 |
| Marcksl1_P28667_S120s_1 | Marcksl1_P | 0.63 | 1.26 | -0.03 | -0.28 | 0.9 | 0.35 | 0.17 | -0.28 | 0.96 | 1.52 | -0.17 | -0.25 | 0.31 | 1.53 | -0.04 | 0.12 | -0.11 | 0.12 | 0.085 | 0.858375 | 0.786598 | 0.817394 | 0.995762 | 0.997988 | 0.999507 |
| Marcksl1_P28667_S132s_1 | Marcksl1_P | 0.23 | 0.24 | -0.26 | 0.31 | 0.13 | 0.01 | 0.44 | 0.18 | -0.06 | 1.02 | 0.88 | 0.79 | -0.2 | 0.37 | 0.24 | 0.52 | 0.06 | 0.5275 | 0.1025 | 0.715837 | 0.796464 | 0.11558 | 0.995762 | 0.997988 | 0.996966 |
| Marcksl1_P28667_S135s_1 | Marcksl1_P | -0.3 | 0.39 | 0.26 | 0.56 | 0.24 | 0.6 | 0.59 | 0.75 | 0.86 | 1.45 | 0.17 | 0.7 | 0.43 | 0.98 | 1.08 | 0.42 | 0.3175 | 0.5675 | 0.5 | 0.131736 | 0.24505 | 0.122737 | 0.989223 | 0.997988 | 0.994666 |
| Marcksl1_P28667_S151s_1 | Marcksl1_P | 0.67 | 0.91 | 0.06 | -0.01 | -0.1 | 1.16 | -0.69 | -0.02 | 1.43 | 1.25 | -0.27 | 0.11 | 1.83 | 0.19 | -0.06 | -0.11 | -0.32 | 0.2225 | 0.055 | 0.910251 | 0.458316 | 0.628246 | 0.989223 | 0.997988 | 0.999507 |
| Marcksl1_P28667_S151s_2 | Marcksl1_P | 0.44 | -0.05 | -0.63 | 0.37 | -0.57 | 0.96 | -0.55 | 0.23 | 1.89 | 0.94 | -1.06 | 0.58 | 0.96 | -0.61 | -0.43 | 0.29 | -0.015 | 0.555 | 0.02 | 0.962891 | 0.971371 | 0.371033 | 0.999187 | 0.997988 | 0.999507 |
| Marcksl1_P28667_S22s_1 | Marcksl1_P | 0.15 | 0.38 | 0.29 | 0.23 | 0.65 | 0.28 | 0.13 | 0.53 | 0.54 | 0.81 | 1.09 | 0.76 | 0.81 | 1.26 | 0.49 | 0.37 | 0.135 | 0.5375 | 0.47 | 0.120181 | 0.539521 | 0.03778 | 0.989223 | 0.997988 | 0.994666 |
| Marcksl1_P28667_S36s_1 | Marcksl1_P | 0.35 | 0.13 | 0.35 | 0.19 | -0.18 | 1.04 | -0.01 | 0.41 | 1.21 | 1.13 | -0.39 | 0.68 | 1.66 | 0.15 | 0.49 | 0.23 | 0.06 | 0.4025 | 0.3775 | 0.324917 | 0.842177 | 0.30182 | 0.996212 | 0.997988 | 0.994666 |
| Marcksl1_P28667_T122t_1 | Marcksl1_P | 0.41 | 0.99 | 0.8 | 0.68 | 0.86 | 0.13 | 0.71 | 0.29 | 0.71 | 1.2 | 0.63 | 0.67 | -0.17 | 0.85 | 0.81 | 1.09 | -0.2225 | 0.0825 | -0.075 | 0.825268 | 0.401143 | 0.74474 | 0.989223 | 0.997988 | 0.999507 |
| Marcksl1_P28667_T148t_1 | Marcksl1_P | 1.13 | 0.61 | 0.35 | 0.5 | 0 | 1.33 | -0.21 | 0.14 | 1.66 | 1.41 | 0.35 | 0.69 | 1.63 | 0.12 | 0.41 | 0.52 | -0.3325 | 0.38 | 0.0225 | 0.953327 | 0.388027 | 0.307081 | 0.989223 | 0.997988 | 0.999507 |
| Marcksl1_P28667_T148t_2 | Marcksl1_P | 0.44 | -0.05 | -0.63 | 0.37 | -0.57 | 0.96 | -0.55 | 0.23 | 1.89 | 0.94 | -1.06 | 0.58 | 0.96 | -0.61 | -0.43 | 0.29 | -0.015 | 0.555 | 0.02 | 0.962891 | 0.971371 | 0.371033 | 0.999187 | 0.997988 | 0.999507 |
| Marcksl1_P28667_T41t_1 | Marcksl1_P | 0.11 | -0.04 | 0.57 | 0.32 | 0.15 | 0.85 | 0.22 | 0.44 | 1.59 | 0.96 | -0.27 | 1.03 | 1.75 | -0.26 | 0.7 | 0.67 | 0.175 | 0.5875 | 0.475 | 0.282364 | 0.501469 | 0.174382 | 0.989223 | 0.997988 | 0.994666 |
| Marcksl1_P28667_T85t_1 | Marcksl1_P | 0.6 | 0.41 | 0 | -0.64 | 0.02 | 1.36 | -0.36 | -1.09 | 1.9 | 1.3 | -1.73 | -0.8 | 1.67 | 0.21 | 0.32 | 0.68 | -0.11 | 0.075 | 0.6275 | 0.164914 | 0.835318 | 0.925572 | 0.995762 | 0.997988 | 0.994666 |
| Mark1_Q8VHJ5_S382s_1 | Mark1_Q8V | 0.19 | 0.11 | -0.7 | -0.18 | 0.52 | -0.29 | 0.23 | -0.22 | -0.45 | 0.1 | -0.43 | -0.63 | -0.43 | -0.04 | -0.14 | 0.01 | 0.205 | -0.2075 | -0.005 | 0.986321 | 0.502201 | 0.490436 | 0.989223 | 0.997988 | 1 |
| Mark1_Q8VHJ5_S403s_1 | Mark1_Q8V | -2.32 | -3.29 | -2.61 | -2.37 | -2.11 | -2.38 | -1.54 | -1.97 | -2.6 | -3.32 | -0.57 | -2.1 | -1.75 | -1.25 | -2.58 | -2.59 | 0.6475 | 0.5 | 0.605 | 0.15723 | 0.057491 | 0.394336 | 0.989223 | 0.997988 | 0.994666 |
| Mark1_Q8VHJ5_S414s_1 | Mark1_Q8V | -2.4 | -1.65 | -1.2 | -3.66 | -1.72 | -2.1 | -2.52 | -2.29 | -1.31 | -1.95 | -4.7 | -2.03 | 0.12 | -2.15 | -1.68 | -2.03 | 0.07 | -0.27 | 0.7925 | 0.259139 | 0.891684 | 0.743618 | 0.996493 | 0.997988 | 0.994666 |
| Mark1_Q8VHJ5_S423s_1 | Mark1_Q8V | 0.22 | 0.35 | 1.24 | 0.7 | 0.91 | 0.15 | 0.88 | 0.98 | 0.15 | 0.47 | 0.41 | 0.89 | -0.2 | 0.51 | -0.78 | 0.43 | 0.1025 | -0.1475 | -0.6375 | 0.126504 | 0.746297 | 0.635336 | 0.995762 | 0.997988 | 0.994666 |
| Mark1_Q8VHJ5_S444s_1 | Mark1_Q8V | -1.91 | -2.29 | -3.36 | -4.59 | -2.86 | -1.9 | -3.93 | -3.43 | -1.38 | -1.49 | -1.86 | -3.51 | -0.39 | -2.15 | -4.09 | -3.09 | 0.0075 | 0.9775 | 0.6075 | 0.490443 | 0.990898 | 0.186836 | 0.999187 | 0.997988 | 0.994666 |
| Mark1_Q8VHJ5_S468s_1 | Mark1_Q8V | -0.98 | -0.21 | 0.31 | 0.37 | -0.53 | -1.12 | 0.63 | 0.47 | -0.46 | -0.35 | 0.72 | 0.34 | -0.73 | -0.35 | 0.31 | 0.55 | -0.01 | 0.19 | 0.0725 | 0.864793 | 0.983367 | 0.648006 | 0.999187 | 0.997988 | 0.999507 |
| Mark1_Q8VHJ5_S46s_1 | Mark1_Q8V | -0.62 | -0.55 | -0.2 | -0.01 | 0.15 | -0.03 | 0.58 | 0.38 | -0.33 | -0.45 | 1.27 | -0.18 | -0.95 | 0.12 | 0.59 | 0.73 | 0.615 | 0.4225 | 0.4675 | 0.270572 | 0.033381 | 0.326505 | 0.989223 | 0.997988 | 0.994666 |
| Mark1_Q8VHJ5_S488s_1 | Mark1_Q8V | 0.41 | 0.33 | 0.02 | 0.32 | -0.53 | 0.36 | 0.07 | -0.06 | 0.28 | 0.27 | 0.51 | 0.17 | 0.72 | -0.42 | 0.22 | 0.32 | -0.31 | 0.0375 | -0.06 | 0.845204 | 0.244753 | 0.865601 | 0.989223 | 0.997988 | 0.999507 |
| Mark1_Q8VHJ5_S588s_1 | Mark1_Q8V | -0.76 | -0.24 | -1.78 | -0.95 | -0.31 | -1.01 | -1.89 | -0.67 | -0.31 | -1.37 | 0.45 | -0.95 | -0.29 | 0.53 | -0.87 | -0.88 | -0.0375 | 0.3875 | 0.555 | 0.235786 | 0.931667 | 0.430822 | 0.998317 | 0.997988 | 0.994666 |
| Mark1_Q8VHJ5_S590s_1 | Mark1_Q8V | -1.21 | -1.49 | -0.58 | -1.16 | -1.62 | -0.59 | -0.39 | -0.76 | -0.83 | -1.51 | 0.62 | -1.09 | 1.11 | -0.39 | -0.71 | -0.25 | 0.27 | 0.4075 | 1.05 | 0.035762 | 0.432712 | 0.401462 | 0.989223 | 0.997988 | 0.994666 |
| Mark1_Q8VHJ5_S593s_1 | Mark1_Q8V | -1.22 | -1.55 | -1.2 | -0.83 | -0.94 | -1.65 | -1.4 | -0.74 | -1.89 | -0.67 | -0.44 | -0.85 | -0.43 | -1.34 | -1.2 | -1.66 | 0.0175 | 0.2375 | 0.0425 | 0.89952 | 0.951314 | 0.517881 | 0.999187 | 0.997988 | 0.999507 |
| Mark1_Q8VHJ5_S633s_1 | Mark1_Q8V | -2.74 | -1.99 | -1.79 | -1.65 | -2.39 | -2.8 | -1.39 | -2.12 | -1.98 | -1.72 | -2.23 | -1.44 | -1.2 | -1.59 | -2.06 | -1.72 | -0.1325 | 0.2 | 0.4 | 0.253866 | 0.725057 | 0.54076 | 0.995437 | 0.997988 | 0.994666 |
| Mark1_Q8VHJ5_S649s_1 | Mark1_Q8V | -2.54 | -2.03 | -1.6 | -2.27 | -1.24 | -2.93 | -0.8 | -0.97 | -2.86 | -1.92 | 0.85 | -1.61 | -1.49 | -0.55 | -1.66 | -1.43 | 0.625 | 0.725 | 0.8275 | 0.037694 | 0.219541 | 0.33592 | 0.989223 | 0.997988 | 0.994666 |
| Mark1_Q8VHJ5_S666s_1 | Mark1_Q8V | 0.65 | -0.12 | -0.39 | -0.76 | 0 | 1.43 | -1.04 | -0.33 | 1.85 | 0.47 | 0.07 | 0.14 | 1.21 | -3.18 | -0.3 | -0.57 | 0.17 | 0.7875 | -0.555 | 0.515596 | 0.754317 | 0.129973 | 0.995762 | 0.997988 | 0.994666 |
| Mark2_Q05512_S390s_1 | Mark2_Q05 | -1.46 | -1.51 | -1.11 | -1.27 | -1.59 | -0.34 | -0.55 | -0.52 | -1.44 | -1.38 | 1.11 | -0.32 | -2.3 | -1.07 | -1.61 | -1.85 | 0.5875 | 0.83 | -0.37 | 0.264274 | 0.087771 | 0.162033 | 0.989223 | 0.997988 | 0.994666 |
| Mark2_Q05512_S408s_1 | Mark2_Q05 | 0.99 | 0.4 | 0.89 | 0.28 | 0.41 | 0.46 | 0.78 | 1.29 | -0.02 | 0.52 | 0.25 | 1.31 | -0.28 | 0.6 | 0.31 | 0.44 | 0.095 | -0.125 | -0.3725 | 0.251534 | 0.748212 | 0.722984 | 0.995762 | 0.997988 | 0.994666 |
| Mark2_Q05512_S409s_1 | Mark2_Q05 | 0.99 | 0.4 | 0.89 | 0.28 | 0.41 | 0.46 | 0.78 | 1.29 | -0.02 | 0.52 | 0.25 | 1.31 | -0.28 | 0.6 | 0.31 | 0.44 | 0.095 | -0.125 | -0.3725 | 0.251534 | 0.748212 | 0.722984 | 0.995762 | 0.997988 | 0.994666 |
| Mark2_Q05512_S40s_1 | Mark2_Q05 | -0.08 | -1.92 | -0.86 | -0.21 | -1.09 | -0.19 | -0.5 | -0.17 | -1.5 | -1.74 | 1.23 | -0.08 | -1.09 | -0.58 | -1.23 | -0.7 | 0.28 | 0.245 | -0.1325 | 0.76331 | 0.531323 | 0.736386 | 0.989223 | 0.997988 | 0.9971 |
| Mark2_Q05512_S453s_1 | Mark2_Q05 | -0.88 | -2.7 | -2.26 | -2.74 | -1.61 | -1.16 | -1.34 | -1.16 | -1.85 | -2.13 | 2.93 | -2.31 | -2.01 | -0.58 | -2.54 | -2.18 | 0.8275 | 1.305 | 0.3175 | 0.576933 | 0.076249 | 0.281705 | 0.989223 | 0.997988 | 0.995259 |
| Mark2_Q05512_S483s_1 | Mark2_Q05 | -1.07 | -1.77 | -1.49 | -1.42 | -0.96 | -0.67 | -0.78 | -0.91 | -1.09 | -1.32 | -0.13 | -1.08 | -1.92 | -0.93 | -1.49 | -1.34 | 0.6075 | 0.5325 | 0.0175 | 0.954437 | 0.024686 | 0.128886 | 0.989223 | 0.997988 | 0.999507 |
| Mark2_Q05512_S566s_1 | Mark2_Q05 | -3.04 | -3.29 | -1.44 | -1.58 | -3.26 | -2.29 | -0.5 | -0.95 | -3.04 | -3.47 | -0.6 | -2.47 | -2.34 | -2.06 | -1.38 | -0.97 | 0.5875 | -0.0575 | 0.65 | 0.242878 | 0.410012 | 0.935881 | 0.989223 | 0.997988 | 0.994666 |
| Mark2_Q05512_S616s_1 | Mark2_Q05 | 0.42 | -1.5 | -0.43 | 0.31 | -0.48 | 0.08 | 0.35 | 0.08 | -0.59 | -0.46 | 0.54 | -0.24 | -0.28 | -0.39 | 0.06 | 0.05 | 0.3075 | 0.1125 | 0.16 | 0.720517 | 0.495376 | 0.815892 | 0.989223 | 0.997988 | 0.9971 |
| Mark3_Q03141_S215s_1 | Mark3_Q03 | -0.76 | -0.02 | 0.36 | 0.23 | 0.01 | -0.28 | 0.98 | 0.36 | -0.04 | -0.15 | 1.34 | 1.58 | -0.66 | -0.9 | 0.25 | 0.47 | 0.315 | 0.73 | -0.1625 | 0.698433 | 0.398053 | 0.159819 | 0.989223 | 0.997988 | 0.996159 |
| Mark3_Q03141_S419s_1 | Mark3_Q03 | -0.7 | -0.14 | 0.05 | -0.14 | -0.09 | -0.83 | 0.59 | 0 | -0.45 | -0.16 | 0.03 | -0.05 | -0.24 | -0.01 | -0.06 | 0.01 | 0.15 | 0.075 | 0.1575 | 0.556537 | 0.660231 | 0.773045 | 0.990434 | 0.997988 | 0.995259 |
| Mark3_Q03141_S42s_1 | Mark3_Q03 | -1.76 | -1.59 | 0.69 | 1.22 | -1.33 | -1.6 | 1.46 | 1.13 | -1.8 | -1.2 | 1.14 | 0.91 | -1.45 | -0.55 | 1.05 | 1.71 | 0.275 | 0.1225 | 0.55 | 0.554764 | 0.775079 | 0.896677 | 0.995762 | 0.997988 | 0.99525 |
| Mark3_Q03141_S469s_1 | Mark3_Q03 | -0.94 | -1 | -0.81 | 0.03 | -0.6 | -1.14 | -0.3 | -0.69 | -1.45 | -0.58 | 0.56 | 0.08 | -1.01 | 0.03 | -1.4 | -0.69 | -0.0025 | 0.3325 | -0.0875 | 0.824757 | 0.993639 | 0.488271 | 0.999278 | 0.997988 | 0.999507 |
| Mark3_Q03141_S489s_1 | Mark3_Q03 | -0.95 | -2.3 | -0.67 | -0.25 | -0.8 | -1.26 | 0.13 | 0.05 | -1.26 | -0.8 | 2.01 | 0.86 | -0.83 | -0.08 | -0.05 | 0.36 | 0.5725 | 1.245 | 0.8925 | 0.090748 | 0.276942 | 0.137492 | 0.989223 | 0.997988 | 0.994666 |
| Mark3_Q03141_S499s_1 | Mark3_Q03 | -0.69 | -0.51 | -0.99 | -0.82 | -1.91 | -0.56 | -1.38 | -0.75 | 0 | -0.35 | 0.32 | -1.04 | -0.31 | -1.34 | -0.75 | -0.86 | -0.3975 | 0.485 | -0.0625 | 0.83379 | 0.251095 | 0.168123 | 0.989223 | 0.997988 | 0.999507 |
| Mark3_Q03141_S543s_1 | Mark3_Q03 | -2.87 | -2.23 | -1.75 | -0.38 | -1.71 | -1.8 | -0.94 | -0.51 | -1.25 | -1.86 | -0.46 | -1.56 | 0.03 | -0.78 | -0.65 | -1.27 | 0.5675 | 0.525 | 1.14 | 0.061803 | 0.318888 | 0.361951 | 0.989223 | 0.997988 | 0.994666 |
| Mark3_Q03141_S563s_1 | Mark3_Q03 | -2.3 | -2.78 | -3.25 | -7.53 | -2.22 | -2.31 | -4.57 | -3.86 | -0.88 | -2.74 | -1.52 | -3.22 | -0.1 | -1.88 | -3.65 | -2.86 | 0.725 | 1.875 | 1.8425 | 0.158365 | 0.533121 | 0.131514 | 0.989223 | 0.997988 | 0.994666 |
| Mark3_Q03141_S583s_1 | Mark3_Q03 | 0.08 | -0.14 | 0.11 | 0.12 | -0.24 | -0.89 | 0.45 | 0.21 | -0.58 | -0.03 | -0.39 | -0.06 | -0.6 | 0.23 | -0.42 | -0.58 | -0.16 | -0.3075 | -0.385 | 0.192998 | 0.616048 | 0.213711 | 0.989223 | 0.997988 | 0.994666 |
| Mark3_Q03141_S619s_1 | Mark3_Q03 | 0.86 | 0.52 | 1.98 | 2.08 | -0.11 | 1.98 | 1.21 | 2.62 | 2.7 | 0.22 | 2.83 | 2.36 | 1.93 | -0.55 | 2.34 | 1.91 | 0.065 | 0.6675 | 0.0475 | 0.944954 | 0.917639 | 0.324976 | 0.997335 | 0.997988 | 0.999507 |
| Mark3_Q03141_T211t_1 | Mark3_Q03 | -0.3 | 0.01 | -0.29 | -0.14 | -0.13 | -0.13 | 0.38 | -0.17 | -0.33 | 0.12 | -0.32 | -0.43 | -0.03 | 0.3 | -0.11 | -0.04 | 0.1675 | -0.06 | 0.21 | 0.397585 | 0.4689 | 0.798092 | 0.989223 | 0.997988 | 0.994666 |
| Mark4_Q8CIP4_S438s_1 | Mark4_Q8C | 1.81 | 1.35 | -1.77 | -1.94 | 1.26 | 1.59 | -1.37 | -1.38 | 1.43 | 1.88 | -0.88 | -1.34 | 1.18 | 1.87 | -0.76 | -1.59 | 0.1625 | 0.41 | 0.3125 | 0.77906 | 0.883288 | 0.717086 | 0.996493 | 0.997988 | 0.998217 |
| Mark4_Q8CIP4_S458s_1 | Mark4_Q8C | -0.72 | -1.26 | -3.36 | -4.59 | -2.15 | -0.53 | -0.7 | -1.32 | 0.33 | -1.45 | -0.17 | 0.08 | 1.62 | -1.28 | -0.41 | -0.13 | 1.3075 | 2.18 | 2.4325 | 0.02764 | 0.148223 | 0.031853 | 0.989223 | 0.997988 | 0.994666 |
| Mark4_Q8CIP4_S45s_1 | Mark4_Q8C | -0.95 | -2.18 | -0.34 | -2.11 | -1.4 | -1.61 | -1.22 | -0.48 | -1.5 | -1.1 | -0.25 | -0.89 | -0.89 | -0.48 | -0.85 | -3.02 | 0.2175 | 0.46 | 0.085 | 0.897438 | 0.648518 | 0.361623 | 0.99033 | 0.997988 | 0.999507 |
| Mark4_Q8CIP4_S464s_1 | Mark4_Q8C | -2.31 | -1.32 | -1.02 | -2.91 | -1.69 | -1.02 | -2.46 | -2.07 | 0.9 | -2.28 | -0.78 | -0.22 | 0.47 | -3.09 | -1.23 | -1.5 | 0.08 | 1.295 | 0.5525 | 0.470597 | 0.870574 | 0.096224 | 0.996493 | 0.997988 | 0.994666 |
| Mark4_Q8CIP4_S567s_1 | Mark4_Q8C | -0.98 | -1 | -0.47 | -0.93 | -1.08 | -1.06 | -1.12 | -1.67 | -0.83 | -1.17 | -1.75 | -1.67 | 0.04 | -0.95 | -0.18 | -0.38 | -0.3875 | -0.51 | 0.4775 | 0.141487 | 0.145306 | 0.108847 | 0.989223 | 0.997988 | 0.994666 |
| Mark4_Q8CIP4_S594s_1 | Mark4_Q8C | 0.48 | -0.25 | 0.13 | 0.24 | 0.51 | 0.05 | 0.19 | 0.31 | 0.17 | 0.46 | -0.24 | 0.32 | -0.5 | -0.83 | 0.15 | 0.15 | 0.115 | 0.0275 | -0.4075 | 0.235288 | 0.638144 | 0.919334 | 0.989223 | 0.997988 | 0.994666 |
| Mark4_Q8CIP4_S654s_1 | Mark4_Q8C | -2.51 | -1.95 | -2.02 | -3.47 | -0.85 | -2.17 | -1.59 | -2.31 | -2.9 | -1.79 | -0.3 | -1.15 | -0.29 | -1.1 | -1.11 | -2.73 | 0.7575 | 0.9525 | 1.18 | 0.062696 | 0.118403 | 0.135918 | 0.989223 | 0.997988 | 0.994666 |
| Mast1_Q9R1L5_S1138s_1 | Mast1_Q9R | -1.93 | -3.2 | -2.67 | -3.82 | -2.77 | -0.66 | -1.87 | -1.2 | 0.67 | -2.19 | -1.11 | -1.36 | 0.11 | -1.71 | -0.94 | -1.34 | 1.28 | 1.9075 | 1.935 | 0.004404 | 0.040128 | 0.015985 | 0.989223 | 0.997988 | 0.994666 |
| Mast1_Q9R1L5_S1244s_1 | Mast1_Q9R | 2 | 2.02 | 1.32 | 0.36 | 1.9 | 1.83 | 1.22 | 1.03 | -0.62 | 2.79 | 0.4 | 1.16 | 0.12 | -0.62 | 1.43 | 1.42 | 0.07 | -0.4925 | -0.8375 | 0.175848 | 0.868683 | 0.507892 | 0.996493 | 0.997988 | 0.994666 |
| Mast1_Q9R1L5_S1414s_1 | Mast1_Q9R | -0.52 | -1.07 | -0.9 | -0.73 | 0.03 | 0.15 | 0.27 | 0.37 | 0.29 | -0.38 | -0.26 | -0.36 | 0.57 | -0.09 | -0.27 | -0.71 | 1.01 | 0.6275 | 0.68 | 0.063981 | 0.001207 | 0.036683 | 0.989223 | 0.997988 | 0.994666 |
| Mast1_Q9R1L5_S346s_1 | Mast1_Q9R | 0.01 | 1.16 | -0.62 | 0.41 | -0.84 | -1.14 | -0.47 | -0.65 | -0.19 | -0.3 | 0.24 | -0.35 | 0.61 | 0.42 | -1.09 | -1.07 | -1.015 | -0.39 | -0.5225 | 0.354068 | 0.024965 | 0.337148 | 0.989223 | 0.997988 | 0.994666 |
| Mast1_Q9R1L5_S718s_1 | Mast1_Q9R | 0.55 | 1.26 | 0.11 | 0.53 | 0.4 | 1.07 | -0.06 | -0.16 | 0.43 | 1.03 | 1.14 | -0.34 | 0.6 | 0.42 | -0.37 | -0.02 | -0.3 | -0.0475 | -0.455 | 0.216752 | 0.419219 | 0.907603 | 0.989223 | 0.997988 | 0.994666 |
| Mast1_Q9R1L5_S791s_1 | Mast1_Q9R | 0.22 | -0.16 | 0.05 | 0.29 | 0.34 | 0.19 | 1.03 | 0.16 | 0.05 | 0.35 | 0.19 | 0.59 | -0.4 | -1.39 | 0.52 | 0.68 | 0.33 | 0.195 | -0.2475 | 0.600478 | 0.239739 | 0.423526 | 0.989223 | 0.997988 | 0.995259 |
| Mast1_Q9R1L5_S895s_1 | Mast1_Q9R | 5.36 | 2.93 | 4.69 | 2.4 | 2.92 | 6.64 | 5.38 | 4.7 | 1.52 | 4.48 | 2.89 | 4.5 | 1.6 | 1.22 | 3.7 | 5.62 | 1.065 | -0.4975 | -0.81 | 0.45792 | 0.258874 | 0.579708 | 0.989223 | 0.997988 | 0.994666 |
| Mast1_Q9R1L5_S90s_1 | Mast1_Q9R | 1.42 | -0.56 | 0.2 | -0.12 | -0.45 | 0.05 | 0.41 | -0.38 | -0.22 | 0.22 | -0.31 | -0.11 | -0.27 | 1.55 | -0.34 | 0.08 | -0.3275 | -0.34 | 0.02 | 0.971717 | 0.464195 | 0.435919 | 0.989223 | 0.997988 | 0.999507 |
| Mast1_Q9R1L5_S968s_1 | Mast1_Q9R | -0.5 | -0.01 | 0.57 | -0.33 | 0.04 | -0.28 | 0.5 | 0.28 | 0.07 | -0.41 | -0.02 | 0.24 | 0.02 | 0.48 | 0.45 | 0.07 | 0.2025 | 0.0375 | 0.3225 | 0.320419 | 0.516917 | 0.903034 | 0.989223 | 0.997988 | 0.994666 |
| Mast1_Q9R1L5_T1371t_1 | Mast1_Q9R | -0.04 | -0.13 | -1.17 | -0.43 | -0.72 | -0.16 | -1.27 | -1.13 | 0.16 | 0.24 | 0.23 | -0.51 | 0.47 | -0.64 | 0.15 | -0.56 | -0.3775 | 0.4725 | 0.2975 | 0.447923 | 0.3049 | 0.182251 | 0.989223 | 0.997988 | 0.994666 |
| Mast1_Q9R1L5_T351t_1 | Mast1_Q9R | 0.13 | 0.44 | -1.14 | -1.54 | -0.31 | -0.29 | -0.96 | -0.8 | 0.43 | -0.19 | -0.62 | -0.82 | -0.99 | 0.17 | -1.2 | -1.48 | -0.0625 | 0.2275 | -0.3475 | 0.536458 | 0.894565 | 0.662339 | 0.996493 | 0.997988 | 0.994666 |
| Mast2_B1AST8_S1177s_1 | Mast2_B1A | -2.82 | -1.13 | -3.48 | -2.99 | -2.22 | -1.18 | -2.32 | -1.91 | -0.16 | -0.87 | -1.03 | -2.62 | 1.22 | -1.84 | 0.06 | -1.25 | 0.6975 | 1.435 | 2.1525 | 0.017268 | 0.202092 | 0.053486 | 0.989223 | 0.997988 | 0.994666 |
| Mast3_Q3U214_S107s_1 | Mast3_Q3U | 0.03 | -0.08 | -0.17 | 0 | 0.13 | -0.06 | -0.01 | -0.18 | 0 | -0.44 | -0.13 | 0.03 | -1.03 | 0.31 | -0.56 | 0.41 | 0.025 | -0.08 | -0.1625 | 0.662133 | 0.900735 | 0.720921 | 0.996493 | 0.997988 | 0.995804 |
| Mast3_Q3U214_S156s_1 | Mast3_Q3U | -0.44 | -1.13 | -1.18 | -1.39 | -1.32 | -0.52 | -1.54 | -1.01 | -0.8 | -0.64 | 0.68 | -1.16 | -0.63 | -0.63 | -1.47 | -1.54 | -0.0625 | 0.555 | -0.0325 | 0.92682 | 0.844199 | 0.225178 | 0.996349 | 0.997988 | 0.999507 |
| Mast3_Q3U214_S159s_1 | Mast3_Q3U | -2.71 | 1.06 | 0.4 | -1.09 | -0.83 | -1.94 | -2.01 | -2.02 | -0.86 | 0.2 | 0.99 | -0.36 | 0.38 | 0.55 | -0.38 | -0.76 | -1.115 | 0.5775 | 0.5325 | 0.505946 | 0.174322 | 0.490866 | 0.989223 | 0.997988 | 0.994666 |
| Mast3_Q3U214_S168s_1 | Mast3_Q3U | 0.17 | -0.22 | -0.45 | 0.45 | 0.21 | 0.7 | 0.51 | 1.02 | 0.29 | -0.16 | 0.58 | -0.33 | 0.51 | -0.05 | -0.09 | 0.02 | 0.6225 | 0.1075 | 0.11 | 0.717456 | 0.055999 | 0.736383 | 0.989223 | 0.997988 | 0.996966 |
| Mast3_Q3U214_S61s_1 | Mast3_Q3U | -0.99 | -0.33 | -0.13 | -0.65 | -0.09 | -0.45 | 0.28 | -0.54 | -0.45 | -0.65 | -0.39 | -0.43 | -0.68 | -0.53 | -0.53 | -0.51 | 0.325 | 0.045 | -0.0375 | 0.891615 | 0.284292 | 0.86354 | 0.989223 | 0.997988 | 0.999507 |
| Mast3_Q3U214_S803s_1 | Mast3_Q3U | -2.48 | 0.54 | -0.29 | -0.28 | -0.48 | -1.75 | -1.52 | -0.71 | -1.47 | -0.35 | -0.46 | -1.25 | -0.45 | -0.45 | -1.12 | -0.98 | -0.4875 | -0.255 | -0.1225 | 0.841019 | 0.451287 | 0.692063 | 0.989223 | 0.997988 | 0.999507 |
| Mast3_Q3U214_S942s_1 | Mast3_Q3U | -1.97 | 1.03 | -1.02 | -1.27 | -0.99 | -1.58 | -1.07 | -1.55 | 1.13 | -0.08 | -0.28 | -0.45 | 3.17 | -0.03 | -1.36 | -1.11 | -0.49 | 0.8875 | 0.975 | 0.370117 | 0.417802 | 0.205747 | 0.989223 | 0.997988 | 0.994666 |
| Mast3_Q3U214_S947s_1 | Mast3_Q3U | 0.16 | 0.03 | -0.75 | -1.89 | -1.47 | 1.01 | -1.26 | -1.2 | 1.66 | -0.79 | 0.6 | -0.91 | 1 | -0.88 | -1.56 | -1.16 | -0.1175 | 0.7525 | -0.0375 | 0.954896 | 0.859417 | 0.295777 | 0.996349 | 0.997988 | 0.999507 |
| Mast4_E9PWX8_S1176s_1 | Mast4_E9P | -0.83 | -1.5 | -1.42 | -1.3 | -0.39 | -0.88 | -0.04 | -0.96 | -0.95 | -0.5 | -1.09 | -1.42 | -1.45 | 0.15 | -1.04 | -1.13 | 0.695 | 0.2725 | 0.395 | 0.326121 | 0.03708 | 0.35726 | 0.989223 | 0.997988 | 0.994666 |
| Mast4_E9PWX8_S2161s_1 | Mast4_E9P | -0.87 | -2.18 | -2.23 | -2.88 | -2.89 | -2.29 | -2.89 | -3.41 | 0.69 | -0.85 | -1.35 | -2.76 | 0.84 | -2.34 | -3.44 | -2.05 | -0.83 | 0.9725 | 0.2925 | 0.740633 | 0.089666 | 0.211704 | 0.989223 | 0.997988 | 0.9971 |
| Mat2b_Q99LB6_S271s_1 | Mat2b_Q99 | -2.13 | -1.06 | -0.71 | -1.76 | -1.54 | -1.54 | -0.45 | -1.68 | -2.34 | -0.82 | -0.84 | -0.81 | -0.94 | -0.41 | -1.3 | -1.91 | 0.1125 | 0.2125 | 0.275 | 0.537814 | 0.784945 | 0.65555 | 0.995762 | 0.997988 | 0.994666 |
| Matr3_Q8K310_S188s_1 | Matr3_Q8K | -2.37 | -2.34 | -2.42 | -4.67 | -3.49 | -2.11 | -2.56 | -3.42 | -1.86 | -2.99 | -2.6 | -3.05 | -0.54 | -2.73 | -2.67 | -3.29 | 0.055 | 0.325 | 0.6425 | 0.395982 | 0.926306 | 0.581141 | 0.997335 | 0.997988 | 0.994666 |
| Matr3_Q8K310_S211s_1 | Matr3_Q8K | 1.06 | 0.22 | 1.48 | 0.01 | -1.48 | 1.14 | -0.48 | 0.81 | 0.6 | 0.32 | 0.3 | 0.26 | 1.24 | -0.07 | 1.27 | 1.27 | -0.695 | -0.3225 | 0.235 | 0.614557 | 0.279089 | 0.38761 | 0.989223 | 0.997988 | 0.995259 |
| Matr3_Q8K310_S4s_1 | Matr3_Q8K | -0.95 | -1.91 | -2.07 | -2.49 | -0.67 | -1.52 | -2.01 | -1.68 | -0.81 | -1.26 | -0.87 | -1.72 | -0.31 | -0.77 | -2.15 | -1.68 | 0.385 | 0.69 | 0.6275 | 0.230265 | 0.362467 | 0.102378 | 0.989223 | 0.997988 | 0.994666 |
| Matr3_Q8K310_S598s_1 | Matr3_Q8K | 0.14 | -0.48 | -0.31 | -0.81 | -1.33 | -0.18 | -0.94 | -0.74 | 0.42 | -1.03 | -1.78 | -0.77 | 1.29 | -1.39 | 0.3 | -0.68 | -0.4325 | -0.425 | 0.245 | 0.667519 | 0.202735 | 0.380301 | 0.989223 | 0.997988 | 0.996159 |
| Matr3_Q8K310_S619s_1 | Matr3_Q8K | 2.84 | 1.02 | -0.45 | -0.87 | -1.1 | 2.92 | -1.44 | -0.71 | 3.8 | 1.3 | -1.74 | -0.15 | 2.88 | -0.78 | 0.59 | -1.45 | -0.7175 | 0.1675 | -0.325 | 0.768486 | 0.53067 | 0.894676 | 0.989223 | 0.997988 | 0.9971 |
| Matr3_Q8K310_S746s_1 | Matr3_Q8K | -4.83 | -4.43 | -2.36 | -2.3 | -4.95 | -5.06 | -1.64 | -1.31 | -4.22 | -3.92 | -0.85 | -1.75 | -3.55 | -3.06 | -2.72 | -1.31 | 0.24 | 0.795 | 0.82 | 0.279927 | 0.819956 | 0.406532 | 0.995762 | 0.997988 | 0.994666 |
| Matr3_Q8K310_S9s_1 | Matr3_Q8K | 0.25 | -0.67 | -0.26 | -1.43 | -0.01 | -0.39 | -0.51 | -0.05 | -1.32 | -0.39 | -0.44 | -0.4 | 0.56 | -1.61 | -0.35 | -1.19 | 0.2875 | -0.11 | -0.12 | 0.828178 | 0.444959 | 0.790983 | 0.989223 | 0.997988 | 0.999507 |
| Matr3_Q8K310_T150t_1 | Matr3_Q8K | -1.68 | -0.88 | -1.05 | -1.12 | -1.44 | -2.1 | -1.26 | -2.53 | -1.12 | -1.47 | -1.52 | -1.02 | 0.17 | -0.54 | -0.68 | -1.63 | -0.65 | -0.1 | 0.5125 | 0.231511 | 0.085332 | 0.7132 | 0.989223 | 0.997988 | 0.994666 |
| Mb21d2_D3Z742_S433s_1 | Mb21d2_D | -1.79 | -1.49 | -2.1 | -1.54 | -1.76 | -1.82 | -3.3 | -1.97 | -1.15 | -1.16 | -1.28 | -0.83 | -0.46 | -0.52 | -1.95 | -2.36 | -0.4825 | 0.625 | 0.4075 | 0.408877 | 0.225654 | 0.02952 | 0.989223 | 0.997988 | 0.994666 |
| Mb21d2_D3Z742_S433s_2 | Mb21d2_D | -0.63 | -0.8 | -1.16 | -1.48 | -0.58 | -0.2 | -0.89 | -0.81 | 0.17 | -0.37 | -0.23 | -1.31 | 0.13 | 0.29 | -2.44 | -1.33 | 0.3975 | 0.5825 | 0.18 | 0.7694 | 0.17885 | 0.143737 | 0.989223 | 0.997988 | 0.9971 |
| Mb21d2_D3Z742_S439s_1 | Mb21d2_D | 0.37 | 0.51 | 0.96 | 1.04 | 0.17 | -0.04 | 0.64 | 0.95 | 0.67 | 0.75 | 0.94 | 0.76 | 0.58 | 0.36 | 0.64 | 0.81 | -0.29 | 0.06 | -0.1225 | 0.656121 | 0.348906 | 0.81027 | 0.989223 | 0.997988 | 0.995259 |
| Mb21d2_D3Z742_S439s_2 | Mb21d2_D | -0.51 | -0.62 | 0.1 | -0.1 | -0.46 | -0.35 | -0.32 | 0.39 | 0.32 | -0.52 | 0.27 | -0.12 | 0.26 | 0.16 | -0.29 | 0.01 | 0.0975 | 0.27 | 0.3175 | 0.278367 | 0.736084 | 0.377483 | 0.995762 | 0.997988 | 0.994666 |
| Mb21d2_D3Z742_S439s_3 | Mb21d2_D | -0.86 | 0.01 | -0.05 | -2.29 | 0.21 | -0.54 | -0.69 | -0.54 | -0.54 | -0.51 | -0.3 | 0.64 | -1.22 | -0.22 | -0.45 | 0.66 | 0.4075 | 0.62 | 0.49 | 0.424549 | 0.442096 | 0.283192 | 0.989223 | 0.997988 | 0.994666 |
| Mb21d2_D3Z742_S442s_2 | Mb21d2_D | -0.2 | -0.22 | 0.63 | 0.19 | -0.18 | -0.79 | 0.31 | 0.93 | 0.66 | -0.95 | 0.41 | 0.2 | 0.58 | -0.18 | 0.57 | 0.46 | -0.0325 | -0.02 | 0.2575 | 0.427004 | 0.935242 | 0.960534 | 0.99868 | 0.997988 | 0.994666 |
| Mbp_P04370_S139s_1 | Mbp_P043 | 0.44 | 0.55 | 0.78 | 0.63 | 1.04 | -0.25 | 1.06 | 0.51 | -0.22 | 0.55 | -0.15 | 0.85 | -0.59 | 0.5 | 1.27 | 0.87 | -0.01 | -0.3425 | -0.0875 | 0.830608 | 0.975602 | 0.282508 | 0.999187 | 0.997988 | 0.999507 |
| Mbp_P04370_S145s_1 | Mbp_P043 | 1.66 | 1.23 | 1.03 | 1.22 | 0.98 | 0.84 | 0.72 | 1.09 | -0.22 | 1.15 | -1.7 | 1.57 | -0.82 | -1.54 | 1.09 | 0.88 | -0.3775 | -1.085 | -1.3825 | 0.042192 | 0.126598 | 0.136122 | 0.989223 | 0.997988 | 0.994666 |
| Mbp_P04370_S151s_1 | Mbp_P043 | 1.24 | 1.04 | 0.59 | 0.61 | 1.41 | 0.5 | 0.45 | 0.51 | 0.88 | 1.03 | -0.59 | 0.91 | 1.09 | 1.45 | 1.72 | 1.42 | -0.1525 | -0.3125 | 0.55 | 0.076348 | 0.618421 | 0.453946 | 0.989223 | 0.997988 | 0.994666 |
| Mbp_P04370_S172s_1 | Mbp_P043 | 0.38 | 0.47 | -0.66 | -0.66 | 0.87 | 0.3 | 0.49 | -0.28 | 0.12 | 0.18 | -0.48 | -0.2 | -0.71 | 0.55 | -0.28 | 0.04 | 0.4625 | 0.0225 | 0.0175 | 0.966126 | 0.247122 | 0.950188 | 0.989223 | 0.997988 | 0.999507 |
| Mbp_P04370_S176s_1 | Mbp_P043 | 1.02 | 0.97 | 0.55 | 0.54 | 1.26 | 0.82 | 0.53 | 0.88 | 0.9 | 1.14 | 1.16 | 0.83 | 0.71 | 0.82 | 1.2 | 1.03 | 0.1025 | 0.2375 | 0.17 | 0.525036 | 0.686635 | 0.33552 | 0.994672 | 0.997988 | 0.994666 |
| Mbp_P04370_S201s_1 | Mbp_P043 | 7.61 | 0.33 | 2.08 | 0.81 | 0.16 | 7.09 | 5.94 | 1.25 | -0.03 | 5.74 | 2.11 | 2.33 | -0.23 | 0.22 | 3.77 | 4.74 | 0.9025 | -0.17 | -0.5825 | 0.745098 | 0.65982 | 0.924302 | 0.990434 | 0.997988 | 0.9971 |
| Mbp_P04370_S243s_1 | Mbp_P043 | 0.2 | -0.34 | 0.96 | 0.21 | -0.41 | 0.08 | -0.48 | -0.33 | 1.57 | 0.49 | -1.07 | 0.77 | 1.99 | -0.72 | 0.92 | 0.87 | -0.5425 | 0.1825 | 0.5075 | 0.383414 | 0.108634 | 0.747921 | 0.989223 | 0.997988 | 0.994666 |
| Mbp_P04370_S245s_1 | Mbp_P043 | 0.53 | 0.33 | 0.6 | 0.26 | -0.47 | 0.13 | -0.43 | -0.86 | 1.79 | 0.49 | -1.65 | 0.23 | 1.92 | -1.17 | 1.04 | 0.54 | -0.8375 | -0.215 | 0.1525 | 0.798939 | 0.010054 | 0.740716 | 0.989223 | 0.997988 | 0.999507 |
| Mbp_P04370_S66s_1 | Mbp_P043 | 1.02 | 0.77 | 0.83 | 0.96 | 0.58 | 0.34 | 0.17 | 0.63 | 1.52 | 1.68 | 1.47 | 0.9 | 1.73 | 0.89 | 1.73 | 1.2 | -0.465 | 0.4975 | 0.4925 | 0.111697 | 0.053753 | 0.074179 | 0.989223 | 0.997988 | 0.994666 |
| Mbp_P04370_S70s_1 | Mbp_P043 | 0.55 | 1.19 | 1.23 | 1.18 | 0.33 | 0.46 | 0.9 | 1.1 | 2.18 | 0.59 | 1.61 | 0.69 | 2.14 | 1.25 | 1.29 | 1.64 | -0.34 | 0.23 | 0.5425 | 0.109451 | 0.242806 | 0.577218 | 0.989223 | 0.997988 | 0.994666 |
| Mbp_P04370_T122t_1 | Mbp_P043 | -0.04 | 0.09 | 0.18 | 0.2 | 0.18 | -1.28 | -0.28 | -0.33 | 2.63 | -0.31 | -0.39 | 0.14 | 2.62 | -0.56 | 1.23 | 0.13 | -0.535 | 0.41 | 0.7475 | 0.254073 | 0.124081 | 0.532705 | 0.989223 | 0.997988 | 0.994666 |
| Mbp_P04370_T149t_1 | Mbp_P043 | 0.13 | 0.09 | 1.07 | 1.01 | 0.23 | -0.06 | 0.36 | 0.38 | 0.62 | 0.47 | 0.59 | 1.01 | 1.1 | 0.58 | 1.02 | 0.66 | -0.3475 | 0.0975 | 0.265 | 0.437945 | 0.275571 | 0.762147 | 0.989223 | 0.997988 | 0.994666 |
| Mbp_P04370_T152t_1 | Mbp_P043 | 0.54 | 0.48 | 1.02 | 0.87 | 0.61 | 0.27 | 0.45 | 0.82 | 1.25 | 0.66 | 0.45 | 1.38 | 1.57 | 0.86 | 1.7 | 1.42 | -0.19 | 0.2075 | 0.66 | 0.045807 | 0.436588 | 0.49431 | 0.989223 | 0.997988 | 0.994666 |
| Mbp_P04370_T167t_1 | Mbp_P043 | 6.65 | 3.3 | 4.47 | 1.92 | 4.45 | 5.86 | 4.76 | 3.8 | -0.16 | 5.71 | 0.96 | 4.92 | -1.81 | 0.14 | 4.69 | 5.24 | 0.6325 | -1.2275 | -2.02 | 0.254727 | 0.50814 | 0.432194 | 0.989223 | 0.997988 | 0.994666 |
| Mbp_P04370_T174t_1 | Mbp_P043 | 0.64 | 0.3 | 0.51 | -0.02 | 0.39 | 0.34 | 0.02 | 0.04 | 0.52 | 0.76 | 0.41 | 0.75 | 0.53 | 0.39 | 0.27 | 0.43 | -0.16 | 0.2525 | 0.0475 | 0.853565 | 0.509377 | 0.319778 | 0.989223 | 0.997988 | 0.999507 |
| Mbp_P04370_T211t_1 | Mbp_P043 | -1.36 | -0.83 | 0.14 | -2.26 | -0.04 | -0.79 | -0.64 | -1.04 | -0.52 | -0.35 | 1.29 | 0.06 | 0.59 | 1.55 | -1.16 | -0.44 | 0.45 | 1.1975 | 1.2125 | 0.106739 | 0.378161 | 0.069058 | 0.989223 | 0.997988 | 0.994666 |
| Mbp_P04370_T229t_1 | Mbp_P043 | 0.03 | -0.27 | 0.54 | 0.41 | -0.63 | -0.6 | -0.05 | 0.1 | 3 | -0.09 | 0.08 | 0.82 | 3.3 | -1.34 | 1.85 | 0.35 | -0.4725 | 0.775 | 0.8625 | 0.343744 | 0.130325 | 0.26197 | 0.989223 | 0.997988 | 0.994666 |
| Mbp_P04370_Y146y_1 | Mbp_P043 | -0.34 | -0.66 | 0.76 | 1.28 | -1.81 | -0.85 | -0.42 | 1.35 | -0.28 | -0.56 | 0.48 | 0.94 | 0.09 | -0.45 | 0.38 | 0.42 | -0.6925 | -0.115 | -0.15 | 0.753586 | 0.340466 | 0.829003 | 0.989223 | 0.997988 | 0.9971 |
| Mbp_P04370_Y169y_1 | Mbp_P043 | 1.95 | 1.09 | 0.89 | 1.59 | 1.23 | 1.96 | 0.31 | 1.42 | 1.25 | 1.15 | 1.26 | 1.05 | 0.8 | 1.22 | 0.53 | 0.5 | -0.15 | -0.2025 | -0.6175 | 0.088415 | 0.710191 | 0.491458 | 0.995437 | 0.997988 | 0.994666 |
| Mcf2l_E9PXE2_S1146s_1 | Mcf2l_E9PX | -1.93 | -2.66 | -1.73 | -1.91 | -1.43 | -0.5 | -1.34 | -1.66 | -0.62 | -1.95 | -1.86 | -1.13 | -0.63 | -1.66 | -1.23 | -0.9 | 0.825 | 0.6675 | 0.9525 | 0.017949 | 0.032266 | 0.106032 | 0.989223 | 0.997988 | 0.994666 |
| Mcf2l_E9PXE2_S558s_1 | Mcf2l_E9PX | 1.15 | -0.73 | -0.14 | -0.2 | -0.77 | 1.14 | -0.84 | 0.53 | 0.22 | -0.46 | -0.92 | 0.14 | 0.12 | -1.28 | -0.35 | -0.45 | -0.005 | -0.275 | -0.51 | 0.297013 | 0.992976 | 0.554383 | 0.999187 | 0.997988 | 0.994666 |
| Mcf2l_E9PXE2_S936s_1 | Mcf2l_E9PX | 0.33 | 0.28 | 0.67 | 0.93 | -0.3 | -0.05 | 1.62 | 1.03 | -0.88 | 0.88 | 1.4 | 1.32 | 0.14 | 0.9 | 2 | 0.75 | 0.0225 | 0.1275 | 0.395 | 0.354575 | 0.959798 | 0.805791 | 0.999187 | 0.997988 | 0.994666 |
| Mcf2l_E9PXE2_S946s_1 | Mcf2l_E9PX | 1.72 | 1.44 | 1.18 | 1.46 | 1 | 0.37 | 1.39 | 1.7 | 1.79 | 2.06 | 1.57 | 1.14 | 0.45 | 1.17 | 1.9 | 1.29 | -0.335 | 0.19 | -0.2475 | 0.483057 | 0.311455 | 0.498532 | 0.989223 | 0.997988 | 0.994666 |
| Mchr1_Q8JZL2_S325s_1 | Mchr1_Q8J | 1.51 | 0.07 | 1.83 | 1.44 | 2.47 | 1.01 | 1.77 | 1.08 | 0.68 | 0.79 | 0.98 | 1.32 | 0.58 | 2.1 | 1.54 | 1.79 | 0.37 | -0.27 | 0.29 | 0.554122 | 0.447156 | 0.514517 | 0.989223 | 0.997988 | 0.994677 |
| Mcm2_P97310_S40s_1 | Mcm2_P97 | -1 | -1.14 | -0.91 | -1.58 | -1.42 | -1.04 | -1.54 | -1.01 | -0.55 | -0.44 | -0.98 | -1.47 | 0.22 | -1.1 | -0.43 | -1.11 | -0.095 | 0.2975 | 0.5525 | 0.159354 | 0.708987 | 0.351215 | 0.995437 | 0.997988 | 0.994666 |
| Mcmbp_H9KV04_S154s_1 | Mcmbp_H9 | -0.39 | -1.76 | -0.87 | -0.86 | -1.11 | -1.69 | -0.96 | -0.79 | -0.16 | -2.03 | -0.54 | -0.98 | 0.05 | -1.23 | -0.78 | -0.87 | -0.1675 | 0.0425 | 0.2625 | 0.516689 | 0.633228 | 0.928105 | 0.989223 | 0.997988 | 0.994666 |
| Mcoln1_Q99J21_S547s_1 | Mcoln1_Q9 | 0.21 | 0.51 | 0.35 | 0.07 | 0.35 | 0.38 | 1.04 | 0.62 | 0.07 | 0.67 | 0.69 | 0.33 | -0.14 | 0.46 | 0.31 | 0.33 | 0.3125 | 0.155 | -0.045 | 0.862926 | 0.223126 | 0.540172 | 0.989223 | 0.997988 | 0.999507 |
| Mcoln1_Q99J21_T34t_1 | Mcoln1_Q9 | 0.25 | 0.59 | 0.41 | 0.52 | 0.73 | 0.22 | 0.65 | 0.87 | 0.34 | 0.84 | 0.54 | 0.84 | -0.03 | 0.67 | 0.51 | 0.71 | 0.175 | 0.1975 | 0.0225 | 0.934049 | 0.457403 | 0.409012 | 0.989223 | 0.997988 | 0.999507 |
| Mctp1_E9PV86_S56s_1 | Mctp1_E9P | 0.72 | 0.53 | 0.7 | 0.53 | 0.31 | 0.41 | 0.66 | 0.66 | 0.35 | 0.58 | -0.37 | 0.22 | 0.43 | 1.12 | 0.18 | 0.01 | -0.11 | -0.425 | -0.185 | 0.549699 | 0.600234 | 0.138992 | 0.989223 | 0.997988 | 0.994666 |
| Mctp1_E9PV86_S941s_1 | Mctp1_E9P | -0.07 | -0.57 | -0.21 | 0 | 0.08 | -0.92 | 0.67 | 0.06 | -0.3 | -0.11 | 0.32 | -0.01 | 0 | 0.38 | 0.01 | 0.11 | 0.185 | 0.1875 | 0.3375 | 0.211019 | 0.603468 | 0.466114 | 0.989223 | 0.997988 | 0.994666 |
| Mctp1_E9PV86_T145t_1 | Mctp1_E9P | 0.45 | -0.1 | -0.19 | -0.18 | 0.53 | 0 | 0.08 | -0.01 | 0.03 | 0 | -0.12 | -0.1 | 0.24 | 0.18 | -0.02 | -0.02 | 0.155 | -0.0425 | 0.1 | 0.705 | 0.545039 | 0.860096 | 0.989223 | 0.997988 | 0.996159 |
| Mctp1_E9PYR7_S174s_1 | Mctp1_E9P | 0.74 | 0.86 | -0.7 | -0.1 | 0.84 | 0.44 | 0.21 | -0.23 | 0.8 | 0.74 | -1.47 | -0.55 | 0.17 | 1.09 | -0.12 | -0.49 | 0.115 | -0.32 | -0.0375 | 0.937385 | 0.779972 | 0.598856 | 0.995762 | 0.997988 | 0.999507 |
| Mdc1_E9QK89_T624t_1 | Mdc1_E9Q | -1.58 | -3.06 | -1.66 | -4.53 | -2.92 | -0.9 | -3.46 | -3.59 | 0.35 | -0.7 | -1.39 | -2.48 | 0.7 | -2.84 | -1.86 | -3.44 | -0.01 | 1.6525 | 0.8475 | 0.405764 | 0.990203 | 0.068287 | 0.999187 | 0.997988 | 0.994666 |
| Mdh1_P14152_S241s_1 | Mdh1_P14 | 1.08 | 0.96 | 0.84 | 0.94 | 0.56 | 0.96 | 0.33 | 1.11 | 1.29 | 1.56 | 0.76 | 0.59 | 1.37 | 0.77 | 0.99 | 1.04 | -0.215 | 0.095 | 0.0875 | 0.726083 | 0.392057 | 0.736659 | 0.989223 | 0.997988 | 0.9971 |
| Mdh1_P14152_S242s_1 | Mdh1_P14 | 0.73 | 0.6 | 0.38 | 0.65 | 0.69 | 0.24 | 0.13 | 0.15 | 0.43 | 0.52 | 1.3 | 0.44 | 0.67 | 1.19 | 0.34 | 0.01 | -0.2875 | 0.0825 | -0.0375 | 0.904866 | 0.227688 | 0.766073 | 0.989223 | 0.997988 | 0.999507 |
| Mdn1_J3QMC5_S4822s_1 | Mdn1_J3Q | -3.06 | -4.02 | -3.47 | -2.16 | -1.69 | -3.19 | -2.09 | -2.16 | -2.32 | -1.69 | -2.85 | -1.32 | -1.8 | -3.09 | -2.36 | -2.15 | 0.895 | 1.1325 | 0.8275 | 0.098405 | 0.081542 | 0.042426 | 0.989223 | 0.997988 | 0.994666 |
| Mecp2_Q9Z2D6_S13s_1 | Mecp2_Q9 | -3.62 | -1.96 | -1.45 | -3.03 | -2.23 | -3.61 | -1.31 | -3.37 | -3.96 | -2.09 | -1.46 | -0.91 | -1.72 | -2.02 | -1.4 | -2.05 | -0.115 | 0.41 | 0.7175 | 0.167277 | 0.858515 | 0.583925 | 0.996349 | 0.997988 | 0.994666 |
| Mecp2_Q9Z2D6_S229s_1 | Mecp2_Q9 | 0.87 | 0.45 | 0.5 | 0.79 | 0.57 | 0.39 | 0.88 | 0.56 | 0.33 | 0.08 | -0.8 | 0.32 | 0.66 | 0.68 | 0.44 | 0.37 | -0.0525 | -0.67 | -0.115 | 0.644478 | 0.816364 | 0.058824 | 0.995762 | 0.997988 | 0.995259 |
| Mecp2_Q9Z2D6_S421s_1 | Mecp2_Q9 | 0.67 | 0.51 | 0.61 | 0.66 | -0.29 | 1.66 | 0.19 | 0.68 | 1.15 | 0.77 | 1.01 | 0.2 | 0.58 | 0.21 | 0.61 | 0.77 | -0.0525 | 0.17 | -0.07 | 0.775947 | 0.89598 | 0.534349 | 0.996493 | 0.997988 | 0.99798 |

| | | | | | | | | | | | | | | | | | | | | | | | | | | |
|---|---|---|---|---|---|---|---|---|---|---|---|---|---|---|---|---|---|---|---|---|---|---|---|---|---|---|
| Mecp2_Q9Z2D6_S80s_1 | Mecp2_Q9 | 0.66 | 1.07 | 0.83 | 0.94 | 0.43 | 0.47 | 0.43 | 0.87 | 0.43 | 0.8 | 1.44 | 0.95 | 0.76 | 0.66 | 0.47 | 1.34 | -0.325 | 0.03 | -0.0675 | 0.811292 | 0.166955 | 0.914265 | 0.989223 | 0.997988 | 0.999507 |
| Med1_Q925J9_T1036t_1 | Med1_Q92 | -1.17 | 0.26 | -3.16 | -1.7 | -0.54 | -1.09 | -1.66 | -0.71 | 0.06 | -0.69 | -1.54 | -2.38 | 0.23 | -0.21 | -1.85 | -0.83 | 0.4425 | 0.305 | 0.7775 | 0.310948 | 0.509609 | 0.699721 | 0.989223 | 0.997988 | 0.994666 |
| Med15_E9Q7C1_T564t_1 | Med15_E9 | -0.85 | -1.24 | 0.76 | 0.37 | -1.71 | -1.05 | 0.74 | 0.25 | -1.19 | -0.49 | 0.54 | 0.52 | -0.49 | -1.4 | 0.91 | 0.2 | -0.2025 | 0.085 | 0.045 | 0.942396 | 0.758372 | 0.884462 | 0.995762 | 0.997988 | 0.999507 |
| Med26_Q7TN02_S435s_1 | Med26_Q7 | 0.62 | 0.32 | 0.37 | -0.23 | 0.77 | -0.78 | -0.14 | -0.13 | -0.73 | 0.96 | -0.15 | -0.55 | -0.38 | -0.45 | 0.57 | 0.72 | -0.34 | -0.3875 | -0.155 | 0.679816 | 0.360131 | 0.359864 | 0.989223 | 0.997988 | 0.996159 |
| Med7_Q9CZB6_S195s_1 | Med7_Q9C | -2.41 | -1.38 | -3.76 | -3.69 | -1.78 | -1.1 | -3.47 | -2.35 | -1.5 | -1.33 | -1.36 | -3.69 | -1.65 | -1.04 | -2.56 | -2.01 | 0.635 | 0.84 | 0.995 | 0.120289 | 0.355668 | 0.265634 | 0.989223 | 0.997988 | 0.994666 |
| Mef2a_Q60929_S255s_1 | Mef2a_Q6 | -0.54 | -0.36 | -1.4 | -0.83 | -1.45 | -0.79 | -1.65 | -1.3 | -0.06 | -0.05 | -0.43 | -1.2 | 0.18 | 0.02 | -0.77 | -1.44 | -0.515 | 0.3475 | 0.28 | 0.522086 | 0.123217 | 0.351261 | 0.989223 | 0.997988 | 0.994666 |
| Mef2a_Q60929_S400s_1 | Mef2a_Q6 | -4.23 | -3.46 | -3.33 | -1.7 | -2.77 | -4.3 | -2.51 | -3.21 | -4.26 | -4.65 | -0.86 | -4.25 | -1.71 | -1.31 | -2.22 | -3.25 | -0.0175 | -0.325 | 1.0575 | 0.111447 | 0.976434 | 0.723496 | 0.999187 | 0.997988 | 0.994666 |
| Mef2c_Q8CFN5_S222s_1 | Mef2c_Q8 | -0.94 | -0.72 | -2.51 | -2.58 | -0.55 | -0.66 | -2.08 | -2.93 | -0.83 | 0.11 | -1.29 | -2.21 | -0.84 | -0.61 | -2.58 | -2.64 | 0.1325 | 0.6325 | 0.02 | 0.975975 | 0.843898 | 0.330134 | 0.996349 | 0.997988 | 1 |
| Mef2c_Q8CFN5_S240s_1 | Mef2c_Q8 | -0.89 | -0.66 | -1.73 | -3.64 | -0.18 | -0.43 | -2.06 | -2.88 | 0.05 | -0.33 | -1.74 | -2.29 | 0.62 | -0.03 | -2.83 | -2.51 | 0.3425 | 0.6525 | 0.5425 | 0.575424 | 0.676798 | 0.414642 | 0.99315 | 0.997988 | 0.995259 |
| Mef2c_Q8CFN5_S396s_1 | Mef2c_Q8 | -5.23 | -4.6 | -5.04 | -5.74 | -4.34 | -6 | -6.09 | -4.3 | -3.43 | -4.5 | -2.05 | -4.12 | -0.86 | -3.61 | -3.78 | -5.47 | -0.03 | 1.6275 | 1.7225 | 0.070634 | 0.952543 | 0.014208 | 0.999187 | 0.997988 | 0.994666 |
| Mef2c_Q8CFN5_S446s_1 | Mef2c_Q8 | -0.59 | -0.47 | -2.18 | -0.62 | -0.58 | -0.8 | -0.62 | -1.92 | -1.43 | -0.51 | -1.27 | -1.94 | -1.27 | 0.18 | -1.94 | -2.57 | -0.015 | -0.3225 | -0.435 | 0.506344 | 0.974845 | 0.504801 | 0.999187 | 0.997988 | 0.994666 |
| Mef2d_E9QKT0_S250s_1 | Mef2d_E9 | -0.86 | -0.34 | -1.85 | -2.09 | -0.28 | -0.89 | -1.71 | -1.59 | -0.21 | -0.47 | -1.52 | -1.12 | -0.21 | -0.35 | -1.93 | -1.82 | 0.1675 | 0.455 | 0.2075 | 0.716596 | 0.731798 | 0.358151 | 0.995762 | 0.997988 | 0.996966 |
| Mef2d_E9QKT0_S429s_1 | Mef2d_E9 | -5.26 | -3.53 | -3.26 | -4.87 | -3.28 | -5.77 | -3.12 | -3.18 | -3.28 | -4.1 | -1.74 | -3.01 | -1.59 | -3.14 | -3.24 | -3.35 | 0.3925 | 1.1975 | 1.4 | 0.037477 | 0.585424 | 0.084253 | 0.989223 | 0.997988 | 0.994666 |
| Mef2d_E9QKT0_S435s_1 | Mef2d_E9 | -0.49 | 0.74 | -1.18 | -0.31 | -0.31 | -0.48 | -0.62 | -0.41 | -0.34 | -0.76 | -0.42 | 0.03 | -0.35 | -1.23 | -1.69 | -0.44 | -0.145 | -0.0625 | -0.6175 | 0.224382 | 0.710773 | 0.88169 | 0.995437 | 0.997988 | 0.994666 |
| Men1_F7CJP6_S41s_1 | Men1_F7C | 0.86 | 1.67 | -0.41 | -0.12 | 0.98 | 1.04 | 0.78 | 0.77 | -0.28 | 1.37 | -0.16 | -0.11 | 0.6 | 0.65 | -0.51 | 0.29 | 0.3925 | -0.295 | -0.2425 | 0.638225 | 0.391926 | 0.605642 | 0.989223 | 0.997988 | 0.995259 |
| Mepce_Q8K3A9_S192s_1 | Mepce_Q8 | -1.63 | -1.14 | -1.97 | -2.56 | -0.69 | -1.32 | -2.15 | -2.5 | -1.45 | -0.9 | -1.54 | -2.07 | -0.3 | -0.86 | -1.35 | -1.07 | 0.16 | 0.335 | 0.93 | 0.034919 | 0.733498 | 0.394555 | 0.995762 | 0.997988 | 0.994666 |
| Mepce_Q8K3A9_T188t_1 | Mepce_Q8 | -0.52 | -1.44 | -1.69 | -1.75 | -2.4 | -0.18 | -2.31 | -2.28 | 0.26 | -1.24 | -0.84 | -2.85 | -1.39 | -1.66 | -1.58 | -2.21 | -0.4425 | 0.1825 | -0.36 | 0.329955 | 0.429265 | 0.776021 | 0.989223 | 0.997988 | 0.994666 |
| Met_F8VQL0_S988s_1 | Met_F8VQ | 0.18 | -0.21 | 0.82 | 0.37 | 0.13 | -0.28 | 0.16 | 0.94 | -0.18 | 1.07 | 0.26 | 0.88 | 0.86 | 1.36 | 0.42 | 0.69 | -0.0525 | 0.2175 | 0.5425 | 0.126811 | 0.876948 | 0.557209 | 0.996493 | 0.997988 | 0.994666 |
| Mettl22_D3Z0D7_S83s_1 | Mettl22_D | -0.65 | -1.09 | -1.01 | -1 | -1.47 | -1.38 | -1.95 | -0.81 | -0.75 | -0.61 | 0.52 | -0.7 | 0.58 | -0.94 | -0.81 | 0.05 | -0.465 | 0.5525 | 0.6575 | 0.114194 | 0.128771 | 0.128741 | 0.989223 | 0.997988 | 0.994666 |
| Mex3c_Q05A36_S530s_1 | Mex3c_Q0 | -1.77 | -1.07 | -3.42 | -5.11 | -1.3 | -1.79 | -3.77 | -3.05 | -0.21 | -1.02 | -3.37 | -4.71 | 1.53 | -1.91 | -2.71 | -1.68 | 0.365 | 0.515 | 1.65 | 0.165694 | 0.693938 | 0.671011 | 0.994893 | 0.997988 | 0.994666 |
| Mff_Q6PCP5_S129s_1 | Mff_Q6PCI | -3.71 | -3.36 | -2.85 | -3.84 | -1.88 | -3.14 | -1.19 | -3.02 | -4.02 | -3.39 | -0.44 | -2.71 | -1.38 | -2.63 | -2.36 | -2.59 | 1.1325 | 0.8 | 1.2 | 0.010345 | 0.037981 | 0.288929 | 0.989223 | 0.997988 | 0.994666 |
| Mff_Q6PCP5_S129s_2 | Mff_Q6PCI | 0.89 | -1.64 | -1.35 | -0.03 | -0.26 | 1.74 | -0.32 | 0.35 | -0.48 | -1.08 | -0.02 | -0.57 | -2.27 | -0.07 | -1.24 | -0.16 | 0.91 | -0.005 | -0.4025 | 0.569696 | 0.197142 | 0.993094 | 0.989223 | 0.997988 | 0.995259 |
| Mff_Q6PCP5_S131s_1 | Mff_Q6PCI | -1.1 | -0.25 | 0.07 | -0.65 | -0.35 | -0.92 | -0.2 | -1.25 | -0.61 | -0.63 | -0.67 | -0.12 | 1.11 | -0.73 | -0.76 | -0.62 | -0.1975 | -0.025 | 0.2325 | 0.639327 | 0.57919 | 0.936821 | 0.989223 | 0.997988 | 0.995259 |
| Mff_Q6PCP5_S131s_2 | Mff_Q6PCI | 0.89 | -1.64 | -1.35 | -0.03 | -0.26 | 1.74 | -0.32 | 0.35 | -0.48 | -1.08 | -0.02 | -0.57 | -2.27 | -0.07 | -1.24 | -0.16 | 0.91 | -0.005 | -0.4025 | 0.569696 | 0.197142 | 0.993094 | 0.989223 | 0.997988 | 0.995259 |
| Mff_Q6PCP5_S146s_1 | Mff_Q6PCI | -0.48 | 0.81 | 0.4 | 0.03 | 1.11 | 1.83 | 0.69 | 1.13 | -1.42 | 0.11 | 0.28 | 0.7 | -0.96 | -0.63 | -0.21 | 0.48 | 1 | -0.2725 | -0.52 | 0.229514 | 0.019189 | 0.592674 | 0.989223 | 0.997988 | 0.994666 |
| Mff_Q6PCP5_S146s_1_1 | Mff_Q6PCI | 0.63 | 0.52 | 0.09 | 0.21 | 0.72 | 0.31 | 1.05 | 0.74 | -0.95 | 0.42 | 0.07 | 0.72 | -0.18 | -0.48 | 0.33 | 0.92 | 0.3425 | -0.2975 | -0.215 | 0.551925 | 0.196726 | 0.449531 | 0.989223 | 0.997988 | 0.994666 |
| Mff_Q6PCP5_S151s_1 | Mff_Q6PCI | 0.41 | 1.94 | 0.26 | -0.01 | 0.5 | -0.47 | -0.37 | -0.73 | 1.14 | 1.63 | 1.34 | 0.35 | 2.38 | 1.03 | 0.21 | -0.11 | -0.9175 | 0.465 | 0.2275 | 0.722949 | 0.078428 | 0.354017 | 0.989223 | 0.997988 | 0.9971 |
| Mff_Q6PCP5_S151s_2 | Mff_Q6PCI | 1.13 | 0.98 | 1.18 | 1.38 | 1.48 | 1.08 | 1.31 | 1.28 | 1.15 | 1.13 | 0.24 | 1.17 | -1.24 | 1.05 | 1.81 | 1.59 | 0.12 | -0.245 | -0.365 | 0.570037 | 0.576926 | 0.405166 | 0.989223 | 0.997988 | 0.995259 |
| Mff_Q6PCP5_T149t_2 | Mff_Q6PCI | 1 | 1.07 | 1.18 | 1.38 | 1.39 | 0.88 | 1.31 | 1.28 | 0.91 | 1.15 | 0.24 | 1.17 | -1.18 | 0.83 | 1.81 | 1.59 | 0.0575 | -0.29 | -0.395 | 0.53011 | 0.797338 | 0.319008 | 0.995762 | 0.997988 | 0.994666 |
| Mfsd5_Q921Y4_S440s_1 | Mfsd5_Q9 | 0.64 | -0.31 | 0.13 | 1.25 | 0.16 | -0.15 | 0.48 | -0.11 | 1.57 | 0.36 | 0.64 | 0.01 | -0.24 | 0.24 | -1.03 | 0.21 | -0.3325 | 0.2175 | -0.6325 | 0.172815 | 0.370882 | 0.634522 | 0.989223 | 0.997988 | 0.994666 |
| Mfsd6_D3Z183_S646s_1 | Mfsd6_D3 | -0.25 | 0.72 | 0.37 | 0.38 | 0.5 | -0.72 | 0.32 | -0.18 | 0.16 | -0.09 | -0.08 | 0.05 | -0.6 | 0.05 | -0.15 | 0.54 | -0.325 | -0.295 | -0.345 | 0.329186 | 0.357526 | 0.289786 | 0.989223 | 0.997988 | 0.994666 |
| Mfsd6_D3Z183_S647s_1 | Mfsd6_D3 | 0.16 | 0.22 | 0.52 | 0.56 | 0.61 | -0.1 | 0.59 | 0.72 | -0.11 | 0.28 | 0.64 | 0.37 | -0.24 | 0.62 | 0.14 | 0.33 | 0.09 | -0.07 | -0.1525 | 0.592522 | 0.73125 | 0.78448 | 0.995762 | 0.997988 | 0.995259 |
| Mfsd6_D3Z183_S736s_1 | Mfsd6_D3 | 0.12 | 0.12 | 0.51 | 0.18 | -0.46 | 0.87 | -0.46 | -0.03 | 2.86 | 0.31 | 0.14 | -0.12 | 2.81 | -0.72 | 1.35 | 0.88 | -0.2525 | 0.565 | 0.8475 | 0.219519 | 0.458287 | 0.385349 | 0.989223 | 0.997988 | 0.994666 |
| Mfsd8_Q8BH31_S18s_1 | Mfsd8_Q8 | -1.45 | -0.86 | -1 | -1.06 | -1.11 | -1.22 | -0.79 | -0.69 | -1.76 | -0.7 | 0.22 | -0.63 | -0.68 | -1.1 | -0.41 | -1.1 | 0.14 | 0.375 | 0.27 | 0.355228 | 0.566593 | 0.37942 | 0.989223 | 0.997988 | 0.994666 |
| Mga_H7BX50_S287s_1 | Mga_H7BX | -1.44 | -1.67 | -0.12 | -0.04 | -1.99 | -0.18 | -0.68 | -1.08 | 0.76 | -1.27 | -2.6 | -0.63 | 1.36 | -3.27 | -0.12 | -0.52 | -0.165 | -0.1175 | 0.18 | 0.845574 | 0.752862 | 0.873099 | 0.995762 | 0.997988 | 0.999507 |
| Mgea5_Q9EQQ9_T709t_1 | Mgea5_Q9 | -0.55 | -0.1 | -0.64 | -0.17 | 0.05 | -0.58 | 0.11 | -0.34 | 0.06 | 0.06 | 0 | 0.03 | -1.22 | -0.27 | -0.66 | -0.03 | 0.175 | 0.4025 | -0.18 | 0.590971 | 0.507295 | 0.107841 | 0.989223 | 0.997988 | 0.995259 |
| Mgll_E9Q3B9_S205s_1 | Mgll_E9Q3 | 0.03 | 0.33 | -0.26 | -0.07 | 0.58 | 0.15 | 0.05 | -0.4 | -0.6 | -0.2 | -0.51 | -0.1 | -0.34 | 0.21 | 0.03 | 0.09 | 0.0875 | -0.36 | -0.01 | 0.96987 | 0.750742 | 0.171083 | 0.995762 | 0.997988 | 0.999507 |
| Mgll_E9Q3B9_S22s_1 | Mgll_E9Q3 | 0.15 | -0.07 | 0.43 | -0.06 | -0.22 | -0.14 | -0.3 | -0.24 | 0.4 | -0.3 | -0.49 | -0.08 | 0.6 | -0.83 | -0.14 | 0.02 | -0.3375 | -0.23 | -0.2 | 0.568581 | 0.141816 | 0.416772 | 0.989223 | 0.997988 | 0.995259 |
| Mgrn1_Q9D074_S106s_1 | Mgrn1_Q9 | -4.08 | -2.83 | -2.92 | -4.51 | -3.7 | -3.08 | -3.76 | -4.71 | -2.4 | -3.13 | -2.69 | -4.2 | -0.47 | -6.76 | -4.48 | -3.86 | -0.2275 | 0.48 | -0.3075 | 0.794715 | 0.646702 | 0.381308 | 0.99033 | 0.997988 | 0.999507 |
| Mgrn1_Q9D074_S428s_1 | Mgrn1_Q9 | 0.41 | 0.37 | 0.79 | -0.01 | -0.08 | 0.85 | -0.47 | -0.19 | 1.05 | 0.43 | -0.21 | 0.13 | 1.36 | -0.35 | 0.32 | 0.2 | -0.3625 | -0.04 | -0.0075 | 0.985002 | 0.297249 | 0.905001 | 0.989223 | 0.997988 | 1 |
| Mgrn1_Q9D074_S437s_1 | Mgrn1_Q9 | -1.65 | -1.6 | -0.97 | -3.27 | -1.44 | -2.21 | -0.71 | -3.35 | -1.94 | -1.51 | -0.28 | -1.57 | -3.26 | -0.73 | -1.71 | -2.77 | -0.055 | 0.5475 | -0.245 | 0.716271 | 0.933775 | 0.343652 | 0.998562 | 0.997988 | 0.996966 |
| Mia3_Q8BI84_S1733s_1 | Mia3_Q8BI | -0.33 | -2.06 | 0.02 | -0.54 | -0.16 | -0.33 | -0.14 | -0.05 | -0.87 | -0.15 | -0.61 | -0.56 | -2.59 | -0.87 | 0.38 | -0.01 | 0.5575 | 0.18 | -0.045 | 0.949909 | 0.220552 | 0.697613 | 0.989223 | 0.997988 | 0.999507 |
| Mia3_Q8BI84_S1915s_1 | Mia3_Q8BI | -6.17 | -3.87 | -1.97 | -1.59 | -4.49 | -4.7 | -1.56 | -1.47 | -5.16 | -3.88 | -1.55 | -1.72 | -2.28 | -3.86 | -0.98 | -1.3 | 0.345 | 0.3225 | 1.295 | 0.24461 | 0.77105 | 0.787995 | 0.995762 | 0.997988 | 0.994666 |
| Mia3_Q8BI84_S820s_1 | Mia3_Q8BI | 0.05 | 0.57 | 0.31 | 0.69 | 0.46 | 0.19 | 0.96 | 1 | 0.06 | 0.69 | 1.09 | 0.81 | -0.02 | 0.46 | 1.96 | 1.6 | 0.2475 | 0.2575 | 0.595 | 0.224571 | 0.385947 | 0.400072 | 0.989223 | 0.997988 | 0.994666 |
| Mib2_Q8R516_S251s_1 | Mib2_Q8R | -0.37 | -0.04 | -0.13 | -0.2 | 0.8 | -0.06 | -0.06 | -0.28 | 0.01 | 0.12 | 0.28 | -0.05 | 0.39 | 0.49 | 0.02 | -0.23 | 0.285 | 0.275 | 0.3525 | 0.211514 | 0.327361 | 0.226488 | 0.989223 | 0.997988 | 0.994666 |
| Mical1_E9PUI4_S777s_1 | Mical1_E9 | 0.47 | 0.88 | 0.17 | -0.06 | 0.51 | 0.31 | -0.06 | -0.08 | -0.12 | 0.04 | 0.47 | 0.07 | 0.45 | 0.48 | 0.38 | 0.12 | -0.195 | -0.25 | -0.0075 | 0.979284 | 0.496632 | 0.39268 | 0.989223 | 0.997988 | 1 |
| Mical2_Q8BML1_S965s_1 | Mical2_Q8 | 0.49 | 0.98 | 0.57 | 0.93 | 1.02 | 0.28 | 0.63 | 0.67 | -0.12 | 0.17 | 1.45 | 0.92 | -0.05 | 0.57 | 1.29 | 1.42 | -0.0925 | -0.1375 | 0.065 | 0.863808 | 0.713744 | 0.719365 | 0.995437 | 0.997988 | 0.999507 |
| Mical3_Q8CJ19_S1131s_1 | Mical3_Q8 | 0.23 | -0.59 | -2.46 | -1.52 | -1.02 | -0.01 | -1.31 | -1.26 | -0.79 | -1.04 | -1.97 | -2.31 | -0.8 | -0.08 | -2.16 | -1.13 | 0.185 | -0.4425 | 0.0425 | 0.948077 | 0.753336 | 0.485268 | 0.995762 | 0.997988 | 0.999507 |
| Mical3_Q8CJ19_S1167s_1 | Mical3_Q8 | -0.54 | -0.54 | -0.62 | -0.89 | -0.01 | -0.69 | -0.24 | -0.45 | -1.48 | -0.92 | -0.61 | -0.88 | -0.83 | -0.16 | -0.71 | -0.34 | 0.3 | -0.325 | 0.1375 | 0.610126 | 0.223996 | 0.237491 | 0.989223 | 0.997988 | 0.995259 |
| Mical3_Q8CJ19_S1187s_1 | Mical3_Q8 | 0.32 | 0.67 | 0.55 | 0.65 | 0.79 | 0.34 | 0.97 | 0.53 | 0.17 | 0.66 | 0.2 | 0.38 | -0.13 | 0.8 | 0.41 | 0.35 | 0.11 | -0.195 | -0.19 | 0.506629 | 0.638643 | 0.410489 | 0.989223 | 0.997988 | 0.994666 |
| Mical3_Q8CJ19_S1645s_1 | Mical3_Q8 | 1.6 | -0.38 | -0.19 | 1.85 | 0.27 | 1.6 | 0.91 | 1.87 | 1.66 | 1.21 | 0.83 | 0.98 | 0.59 | 0.29 | 0.95 | 0.45 | 0.4425 | 0.45 | -0.15 | 0.787315 | 0.475787 | 0.433521 | 0.989223 | 0.997988 | 0.999507 |
| Mical3_Q8CJ19_S1691s_1 | Mical3_Q8 | -0.62 | -1.52 | -2.29 | -3.36 | -0.01 | -0.85 | -1.64 | -3.81 | -1.64 | -1.21 | -0.69 | -2.98 | -2.1 | -0.98 | -2.32 | -1.9 | 0.37 | 0.3175 | 0.1225 | 0.837075 | 0.672271 | 0.645735 | 0.992876 | 0.997988 | 0.999507 |
| Mical3_Q8CJ19_S1692s_1 | Mical3_Q8 | 0.12 | -1.4 | -2.29 | -3.36 | 0.44 | -0.1 | -1.64 | -3.81 | -0.95 | -0.59 | -0.69 | -2.98 | -2.08 | -0.56 | -2.32 | -1.9 | 0.455 | 0.43 | 0.0175 | 0.981177 | 0.662996 | 0.605438 | 0.990434 | 0.997988 | 1 |
| Mical3_Q8CJ19_S1695s_1 | Mical3_Q8 | -0.58 | -1.95 | -0.66 | -0.36 | -0.24 | -0.66 | 0.22 | -0.38 | -1.43 | -0.92 | -0.55 | -0.05 | -1.92 | -0.05 | -0.4 | -0.66 | 0.6225 | 0.15 | 0.13 | 0.799421 | 0.136282 | 0.737535 | 0.989223 | 0.997988 | 0.999507 |
| Mical3_Q8CJ19_S1716s_1 | Mical3_Q8 | -2.56 | -3.05 | -2.95 | -2.78 | -1.98 | -3.96 | -3.22 | -3.57 | -3.14 | -3.7 | -2.45 | -3.42 | -3.2 | -1.91 | -4.76 | -3.15 | -0.3475 | -0.3425 | -0.42 | 0.452155 | 0.416066 | 0.297868 | 0.989223 | 0.997988 | 0.994666 |
| Mical3_Q8CJ19_S1736s_1 | Mical3_Q8 | -2.56 | -2.24 | -2.5 | -6.05 | 0.22 | -2.09 | -2.29 | -2.81 | -2.94 | -2.03 | -0.65 | -3.17 | -1.9 | -1.13 | -3.65 | -2.86 | 1.595 | 1.14 | 0.9525 | 0.318248 | 0.127579 | 0.248322 | 0.989223 | 0.997988 | 0.994666 |
| Mical3_Q8CJ19_S1769s_1 | Mical3_Q8 | 0.75 | 0.41 | 0.32 | 0.02 | 0.25 | 0.81 | 0.24 | 0.1 | 1.74 | 0.71 | 0.42 | -0.2 | 1.11 | 0.26 | 0.3 | -0.77 | -0.025 | 0.2925 | -0.15 | 0.717994 | 0.924451 | 0.494337 | 0.997335 | 0.997988 | 0.996966 |
| Mical3_Q8CJ19_S685s_1 | Mical3_Q8 | 0.79 | 0.55 | 0.63 | 0.29 | -0.53 | 1.01 | -0.91 | -0.41 | 1.32 | 0.41 | 0.18 | -0.02 | 1.31 | -0.45 | -0.19 | 0.23 | -0.775 | -0.0925 | -0.34 | 0.410996 | 0.085984 | 0.783058 | 0.989223 | 0.997988 | 0.994666 |
| Mical3_Q8CJ19_S687s_1 | Mical3_Q8 | 1.54 | 0.27 | 1.11 | 1.54 | -0.42 | 1.61 | -0.22 | 1.05 | 1.62 | 0.71 | 0.39 | 0.82 | 0.96 | 0.09 | 0.47 | 0.93 | -0.61 | -0.23 | -0.5025 | 0.20627 | 0.261464 | 0.565141 | 0.989223 | 0.997988 | 0.994666 |
| Mical3_Q8CJ19_S867s_1 | Mical3_Q8 | 0.75 | 0.18 | 0.96 | 0.91 | 0.4 | 0.43 | 0.88 | 1.61 | 0.93 | -0.53 | 2.48 | 1.03 | 0.18 | 1.42 | 0.69 | 0.55 | 0.13 | 0.2775 | 0.01 | 0.976994 | 0.703329 | 0.638907 | 0.995437 | 0.997988 | 1 |
| Mical3_Q8CJ19_T684t_1 | Mical3_Q8 | 0.58 | -0.11 | 0.57 | 0.57 | -1.24 | 0.79 | -0.85 | -0.43 | 1.13 | 0.61 | 0.08 | 0.04 | 0.88 | -0.85 | -0.24 | 0.02 | -0.835 | 0.0625 | -0.45 | 0.280134 | 0.084511 | 0.85067 | 0.989223 | 0.997988 | 0.994666 |
| Micall1_Q8BGT6_S628s_1 | Micall1_Q8 | -0.24 | -0.06 | -0.93 | -1.5 | -0.66 | -0.38 | -0.06 | -0.8 | -0.84 | -0.58 | -1.35 | -0.87 | -1.38 | -0.55 | -1.55 | -0.86 | 0.2075 | -0.2275 | -0.4025 | 0.335565 | 0.573209 | 0.54709 | 0.989223 | 0.997988 | 0.994666 |
| Micall1_Q8BGT6_S647s_1 | Micall1_Q8 | 0.42 | 0.46 | 0.52 | 0.63 | 0.87 | 0.09 | 0.15 | 0.49 | 0.29 | 0.58 | 1.13 | 0.74 | 0.83 | 0.62 | 0.89 | 0.44 | -0.1075 | 0.1775 | 0.1875 | 0.445232 | 0.66329 | 0.489068 | 0.990434 | 0.997988 | 0.994666 |
| Micall2_E9PZD2_S67s_1 | Micall2_E9 | -0.24 | 0.3 | 0.83 | 0.29 | 0.11 | -0.06 | -0.1 | 0.01 | 0.21 | 0.83 | 0.15 | 0.29 | 0.38 | -0.19 | 0.87 | -0.54 | -0.305 | 0.075 | -0.165 | 0.673993 | 0.27057 | 0.805984 | 0.989223 | 0.997988 | 0.996159 |
| Mid1_B1AV01_S96s_1 | Mid1_B1A | -0.53 | -0.56 | 0.42 | 0.83 | -2 | -1.54 | -0.37 | 0.77 | 0.47 | -0.47 | 2.64 | 1.55 | 0.74 | -1.81 | 0.49 | 0.42 | -0.825 | 1.0075 | -0.08 | 0.898029 | 0.213391 | 0.1652 | 0.989223 | 0.997988 | 0.999507 |
| Mid1ip1_Q9CQ20_S71s_1 | Mid1ip1_Q | 0.02 | 0.54 | -0.24 | 0.4 | 0.48 | -0.6 | 1.33 | 0.68 | 0.41 | 0.2 | 0.31 | 0.32 | 0.42 | 0.48 | 0.57 | 1 | 0.2925 | 0.13 | 0.4375 | 0.155512 | 0.488911 | 0.611522 | 0.989223 | 0.997988 | 0.994666 |
| Mid2_B1AVF6_S96s_1 | Mid2_B1A | 1.94 | 0.62 | 0.47 | 0.99 | -0.07 | 1.23 | -0.18 | 0.24 | 2.15 | 2.04 | 0.89 | 0.67 | 2.04 | 0.31 | 0.58 | 0.35 | -0.7 | 0.4325 | -0.185 | 0.711588 | 0.131609 | 0.378397 | 0.989223 | 0.997988 | 0.996966 |
| Mief1_Q8BGV8_S59s_1 | Mief1_Q8 | -0.04 | 0.36 | -0.18 | -0.17 | 0.29 | 0.11 | -0.44 | 0.06 | -0.22 | -0.15 | 0.03 | 0.18 | 0.73 | 0.55 | -0.43 | -0.42 | 0.0125 | -0.0325 | 0.115 | 0.749537 | 0.960784 | 0.892341 | 0.999187 | 0.997988 | 0.9971 |
| Mief1_Q8BGV8_S94s_1 | Mief1_Q8 | -0.1 | 0.14 | -0.16 | -0.03 | 0.11 | -0.54 | -0.48 | -0.13 | -0.44 | -0.23 | 0.21 | -0.45 | -0.95 | -0.25 | -0.43 | 0.13 | -0.2225 | -0.19 | -0.3375 | 0.272688 | 0.356804 | 0.447709 | 0.989223 | 0.997988 | 0.994666 |
| Mier1_Q5UAK0_S195s_1 | Mier1_Q5 | -2.74 | -1.69 | -2.06 | -1.11 | -4.9 | -4.86 | -1.33 | -1.11 | -1.57 | -2.03 | -0.58 | -1.07 | -2.48 | -2.16 | -0.48 | -1.48 | -1.15 | 0.5875 | 0.25 | 0.634562 | 0.250286 | 0.209999 | 0.989223 | 0.997988 | 0.995259 |
| Mink1_Q9JM52_S537s_1 | Mink1_Q9 | -0.6 | -2.03 | -1.34 | -0.48 | -1.63 | -0.66 | -1.46 | -0.89 | -0.84 | -1.91 | -1.07 | -1.64 | -0.47 | -2.62 | -1.43 | -1.94 | -0.0475 | -0.2525 | -0.5025 | 0.362041 | 0.907364 | 0.557454 | 0.997038 | 0.997988 | 0.994666 |
| Mink1_Q9JM52_S559s_1 | Mink1_Q9 | 0.19 | 0.11 | -0.64 | 0 | 0.03 | -0.48 | 0.83 | 0.32 | -0.62 | -0.15 | -0.07 | -0.08 | -0.68 | 0.78 | -0.27 | -0.2 | 0.26 | -0.145 | -0.0075 | 0.984175 | 0.450667 | 0.608422 | 0.989223 | 0.997988 | 1 |
| Mink1_Q9JM52_S603s_1 | Mink1_Q9 | -0.83 | -1.6 | -0.86 | -1.01 | -0.68 | -0.69 | -0.98 | -0.51 | -1.33 | -1.56 | -0.06 | -0.79 | -0.82 | -1.27 | -0.98 | -0.63 | 0.36 | 0.14 | 0.15 | 0.610181 | 0.182256 | 0.714889 | 0.989223 | 0.997988 | 0.995259 |
| Mink1_Q9JM52_S644s_1 | Mink1_Q9 | 0.57 | 0.46 | -0.11 | 0.34 | 0.67 | 0.21 | 0.55 | 0.24 | -0.16 | 0.47 | 0.11 | 0.31 | -0.13 | 0.78 | 0.29 | 0.38 | 0.1025 | -0.1325 | 0.015 | 0.960117 | 0.679444 | 0.618691 | 0.993464 | 0.997988 | 0.999507 |
| Mink1_Q9JM52_S699s_1 | Mink1_Q9 | 0.5 | -1.39 | -1.12 | -0.35 | -0.91 | 0.85 | -1.11 | -0.06 | 0.27 | -0.82 | -0.42 | -0.39 | -0.09 | -1.38 | -2.3 | -1.66 | 0.2825 | 0.25 | -0.7675 | 0.205297 | 0.614308 | 0.590194 | 0.989223 | 0.997988 | 0.994666 |
| Mink1_Q9JM52_S729s_1 | Mink1_Q9 | -1.1 | -1.06 | -1.64 | -0.35 | -0.6 | -0.87 | -0.14 | -0.44 | -0.82 | -1.1 | -1.16 | -0.94 | -1 | -0.38 | -1.01 | -0.54 | 0.525 | 0.0325 | 0.305 | 0.384336 | 0.126168 | 0.9164 | 0.989223 | 0.997988 | 0.994666 |
| Mink1_Q9JM52_S745s_1 | Mink1_Q9 | 0.47 | 0.72 | -0.03 | 0.63 | 0.84 | 1.13 | 0.39 | 1.07 | -0.22 | 0.96 | 1.31 | 0.67 | -0.57 | 0.23 | -0.37 | 0.66 | 0.41 | 0.2325 | -0.46 | 0.21504 | 0.160473 | 0.53848 | 0.989223 | 0.997988 | 0.994666 |
| Mink1_Q9JM52_S769s_1 | Mink1_Q9 | -1.2 | -2.69 | -1.42 | -2.6 | -2.71 | -1.22 | -2.16 | -1.08 | -2.59 | -3.18 | 0.52 | -2.08 | -2.14 | -2.85 | -2.09 | -2.35 | 0.185 | 0.145 | -0.38 | 0.379306 | 0.713815 | 0.857266 | 0.995437 | 0.997988 | 0.994666 |
| Mink1_Q9JM52_S894s_1 | Mink1_Q9 | -0.25 | -0.1 | 0.37 | 0.39 | 0.11 | -0.32 | 0.82 | 0.43 | 0.17 | -0.31 | 0.71 | 0.19 | -0.46 | 0.22 | -0.09 | 0.24 | 0.1575 | 0.0875 | -0.125 | 0.674111 | 0.614616 | 0.772836 | 0.989223 | 0.997988 | 0.996159 |
| Mink1_Q9JM52_S903s_1 | Mink1_Q9 | 0.06 | -0.62 | 0.08 | -0.01 | -1.16 | 0.32 | -0.43 | -0.46 | 2.08 | 0.02 | -1.46 | 0.08 | 2.44 | -1.1 | 0.51 | -0.31 | -0.31 | 0.3025 | 0.5075 | 0.472418 | 0.383551 | 0.655456 | 0.989223 | 0.997988 | 0.994666 |
| Mink1_Q9JM52_T675t_1 | Mink1_Q9 | -0.19 | 0.19 | -0.32 | -0.39 | 0.32 | -1.49 | 0.26 | -1.04 | -0.58 | -0.03 | 0.62 | -0.77 | -0.58 | 0.7 | -0.87 | -0.73 | -0.31 | -0.0125 | -0.1925 | 0.62657 | 0.492538 | 0.971687 | 0.989223 | 0.997988 | 0.995259 |
| Mios_Q8VE19_S766s_1 | Mios_Q8V | 0.07 | -0.24 | 0.11 | 0.19 | -0.14 | -0.02 | 0 | 0.14 | -0.55 | 0.3 | -0.08 | -0.04 | -0.16 | -0.53 | 0.01 | 0.28 | -0.0375 | -0.125 | -0.1325 | 0.633214 | 0.858943 | 0.636052 | 0.996349 | 0.997988 | 0.995259 |
| Misp_Q9D279_S416s_3 | Misp_Q9D | -1.7 | -0.14 | -0.41 | -0.19 | -1.37 | -2.17 | -0.12 | 0.26 | 0.84 | -0.66 | -0.14 | -0.52 | 1.66 | -2.41 | 1.42 | 0.23 | -0.24 | 0.49 | 0.835 | 0.354069 | 0.690238 | 0.317117 | 0.994672 | 0.997988 | 0.994666 |
| Misp_Q9D279_S419s_3 | Misp_Q9D | -1.7 | -0.14 | -0.41 | -0.19 | -1.37 | -2.17 | -0.12 | 0.26 | 0.84 | -0.66 | -0.14 | -0.52 | 1.66 | -2.41 | 1.42 | 0.23 | -0.24 | 0.49 | 0.835 | 0.354069 | 0.690238 | 0.317117 | 0.994672 | 0.997988 | 0.994666 |
| Misp_Q9D279_T420t_3 | Misp_Q9D | -1.7 | -0.14 | -0.41 | -0.19 | -1.37 | -2.17 | -0.12 | 0.26 | 0.84 | -0.66 | -0.14 | -0.52 | 1.66 | -2.41 | 1.42 | 0.23 | -0.24 | 0.49 | 0.835 | 0.354069 | 0.690238 | 0.317117 | 0.994672 | 0.997988 | 0.994666 |
| Misp_Q9D279_T423t_3 | Misp_Q9D | -1.7 | -0.14 | -0.41 | -0.19 | -1.37 | -2.17 | -0.12 | 0.26 | 0.84 | -0.66 | -0.14 | -0.52 | 1.66 | -2.41 | 1.42 | 0.23 | -0.24 | 0.49 | 0.835 | 0.354069 | 0.690238 | 0.317117 | 0.994672 | 0.997988 | 0.994666 |
| Mkl1_D3YUI2_S621s_1 | Mkl1_D3Y | 1.31 | 0.08 | -0.01 | 0.29 | 0.79 | 1.5 | -0.43 | 0.63 | 0.64 | 0.26 | 0.7 | 0.3 | 0.63 | 0.64 | -0.28 | -0.69 | 0.205 | 0.0575 | -0.3425 | 0.445844 | 0.660973 | 0.866871 | 0.990434 | 0.997988 | 0.994666 |
| Mkl1_D3YUI2_S855s_1 | Mkl1_D3Y | -1.19 | 0.16 | 1.07 | 0.15 | -0.34 | -1.1 | 0.48 | 0.59 | 0.04 | -0.37 | 0.83 | 0.01 | -0.74 | -1.3 | 1.01 | 0.53 | -0.14 | 0.08 | -0.1725 | 0.788728 | 0.799697 | 0.87249 | 0.995762 | 0.997988 | 0.999507 |
| Mkl1_D3YUI2_T503t_1 | Mkl1_D3Y | 0.36 | 1.28 | 0.53 | 0.19 | 0.09 | 0.34 | 0.29 | -0.19 | 0.14 | 1.01 | 0.99 | 0.65 | 0.86 | 0.67 | -0.19 | 0.26 | -0.4575 | 0.1075 | -0.19 | 0.599916 | 0.146002 | 0.74969 | 0.989223 | 0.997988 | 0.995259 |
| Mkl2_Q5DTZ3_S207s_1 | Mkl2_Q5D | -0.24 | -0.06 | 0.01 | -1.1 | -0.07 | -0.2 | -0.17 | -0.08 | 0.42 | 0.44 | -0.87 | -0.28 | -0.1 | 0.28 | -0.14 | -0.84 | 0.2175 | 0.275 | 0.1475 | 0.689335 | 0.458574 | 0.499825 | 0.989223 | 0.997988 | 0.996159 |
| Mkl2_Q5DTZ3_S209s_1 | Mkl2_Q5D | -0.43 | 0.6 | 0.17 | 0.56 | -0.19 | -0.24 | 0.27 | 0.35 | -0.43 | 0.34 | -0.17 | 0.25 | 0.48 | 0.16 | -0.19 | -0.55 | -0.1775 | -0.2275 | -0.25 | 0.485913 | 0.564521 | 0.490911 | 0.989223 | 0.997988 | 0.994666 |
| Mlc1_E9QP87_S189s_1 | Mlc1_E9Q | -0.11 | -0.15 | 0.8 | 0.78 | 0.31 | 0.46 | 0.94 | 0.2 | -0.66 | 0.37 | 0.51 | -0.53 | -0.24 | -0.65 | 0.42 | -0.3 | 0.1475 | -0.4075 | -0.5225 | 0.178118 | 0.651107 | 0.321948 | 0.990358 | 0.997988 | 0.994666 |
| Mlc1_E9QP87_T41t_1 | Mlc1_E9Q | 2.14 | 1.42 | 3.9 | 3.42 | 0.91 | 2.3 | 2.13 | 3.58 | 1 | 2.52 | 5.64 | 3.99 | 1.28 | 0.99 | 4.11 | 4.02 | -0.49 | 0.5675 | -0.12 | 0.893341 | 0.487665 | 0.577829 | 0.989223 | 0.997988 | 0.999507 |
| Mlf2_Q99KX1_S237s_1 | Mlf2_Q99K | -0.28 | -0.17 | -0.47 | -0.65 | -0.3 | -0.84 | -0.55 | -0.7 | -0.83 | -0.41 | -0.8 | -0.84 | -0.18 | -0.29 | -0.48 | -0.51 | -0.205 | -0.3275 | 0.0275 | 0.911838 | 0.385081 | 0.189333 | 0.989223 | 0.997988 | 0.999507 |
| Mlf2_Q99KX1_S32s_1 | Mlf2_Q99K | -0.79 | -0.58 | -1.15 | -1.24 | -0.17 | -0.81 | -0.32 | -0.67 | -0.52 | -0.78 | -1.17 | -0.9 | -0.46 | -0.99 | -0.64 | -0.65 | 0.4475 | 0.0975 | 0.255 | 0.363468 | 0.113112 | 0.715131 | 0.989223 | 0.997988 | 0.994666 |
| Mlh3_Q68FG1_S1283s_2 | Mlh3_Q68 | -1.86 | 1.21 | 1.1 | 0.83 | 0.98 | -1.76 | -0.12 | 0.33 | 1.18 | -1.24 | 1.65 | 0.51 | 1.35 | 1.78 | 0.95 | 0.25 | -0.4625 | 0.205 | 0.7625 | 0.300635 | 0.574441 | 0.811661 | 0.989223 | 0.997988 | 0.994666 |
| Mlip_V9GWW6_S61s_1 | Mlip_V9GV | 0.3 | 0.33 | 0.57 | 1.03 | 0.18 | -0.19 | 0.77 | 0.89 | -0.24 | -0.4 | 0.37 | 0.46 | -0.53 | 0.2 | 0.68 | 0.44 | -0.145 | -0.51 | -0.36 | 0.309982 | 0.651744 | 0.124204 | 0.990358 | 0.997988 | 0.994666 |
| Mllt4_E9Q9C3_S1090s_1 | Mllt4_E9Q | -1.04 | -0.24 | -0.18 | -0.41 | -0.06 | -0.71 | 0.37 | -0.59 | -0.62 | -0.74 | -1.29 | -0.21 | -0.33 | -0.21 | -0.62 | -0.51 | 0.22 | -0.2475 | 0.05 | 0.862215 | 0.508526 | 0.452737 | 0.989223 | 0.997988 | 0.999507 |
| Mllt4_E9Q9C3_S1114s_1 | Mllt4_E9Q | -0.21 | 0.03 | -0.47 | -0.2 | 0.36 | -0.15 | 0.28 | -0.25 | -0.47 | 0.07 | -0.3 | -0.1 | -0.7 | 0.23 | -0.48 | -0.39 | 0.2725 | 0.0125 | -0.1225 | 0.675917 | 0.28154 | 0.958487 | 0.989223 | 0.997988 | 0.996159 |
| Mllt4_E9Q9C3_S1179s_1 | Mllt4_E9Q | -0.47 | -0.11 | 0 | 0.36 | -0.22 | -0.83 | 0.14 | 0.25 | -0.08 | -0.44 | 0.8 | 0.44 | -0.29 | 0.11 | 0.65 | 0.17 | -0.11 | 0.235 | 0.215 | 0.494833 | 0.727296 | 0.497874 | 0.995569 | 0.997988 | 0.994666 |
| Mllt4_E9Q9C3_S1180s_2 | Mllt4_E9Q | 0.45 | -0.09 | -0.64 | 0.3 | 0.1 | 0.91 | -0.6 | 0.01 | 0.33 | 0.43 | 0.09 | -0.77 | 0.51 | 0.15 | -0.49 | -0.29 | 0.1 | 0.015 | -0.035 | 0.921857 | 0.7946 | 0.967844 | 0.995762 | 0.997988 | 0.999507 |
| Mllt4_E9Q9C3_S1189s_1 | Mllt4_E9Q | 0.59 | 0.45 | 0.37 | 0.39 | 0.81 | 0.59 | 0.5 | 0.62 | 0.01 | 0.57 | 0.83 | 0.67 | 1.01 | 1.14 | 0.38 | 0.28 | 0.18 | 0.07 | 0.2525 | 0.395913 | 0.382221 | 0.784164 | 0.989223 | 0.997988 | 0.994666 |
| Mllt4_E9Q9C3_S1189s_2 | Mllt4_E9Q | 0.45 | -0.09 | -0.64 | 0.3 | 0.1 | 0.91 | -0.6 | 0.01 | 0.33 | 0.43 | 0.09 | -0.77 | 0.51 | 0.15 | -0.49 | -0.29 | 0.1 | 0.015 | -0.035 | 0.921857 | 0.7946 | 0.967844 | 0.995762 | 0.997988 | 0.999507 |
| Mllt4_E9Q9C3_S1218s_1 | Mllt4_E9Q | 0.36 | 0.7 | 0.2 | 0.48 | 0.53 | 0.26 | 0.74 | 0.7 | 0.45 | 0.79 | 0.66 | 0.53 | 0.16 | 0.67 | 0.75 | 0.57 | 0.1225 | 0.1725 | 0.1025 | 0.698504 | 0.594805 | 0.457269 | 0.989223 | 0.997988 | 0.996159 |
| Mllt4_E9Q9C3_S1282s_1 | Mllt4_E9Q | 2.54 | 1.26 | 1.18 | 1.32 | -0.09 | 3.02 | 0.25 | 1.17 | 2.32 | 1.59 | 0.68 | 1.12 | 0.83 | -0.15 | 0.54 | 1.15 | -0.4875 | -0.1475 | -0.9825 | 0.04083 | 0.478825 | 0.747474 | 0.989223 | 0.997988 | 0.994666 |
| Mllt4_E9Q9C3_S1461s_1 | Mllt4_E9Q | -1.68 | -0.85 | -1.07 | -1.16 | -1.26 | -1.73 | -0.64 | -1.57 | -1.62 | -1.38 | -1.11 | -1.2 | -1.47 | -1.18 | -0.6 | -2.22 | -0.11 | -0.1375 | -0.1775 | 0.651053 | 0.727687 | 0.61094 | 0.995569 | 0.997988 | 0.995259 |
| Mllt4_E9Q9C3_S216s_1 | Mllt4_E9Q | -2.38 | -1.7 | -0.7 | -0.6 | -0.74 | -1.65 | -0.53 | -0.86 | -1.94 | -1.82 | 0.01 | -0.92 | -0.53 | -0.34 | -0.43 | -1 | 0.4 | 0.1775 | 0.77 | 0.106061 | 0.391231 | 0.757138 | 0.989223 | 0.997988 | 0.994666 |
| Mllt4_E9Q9C3_S246s_1 | Mllt4_E9Q | 0.68 | 0.68 | 0.64 | 0 | 0.24 | 0.48 | -0.77 | 0.11 | 0.16 | 0.82 | -1.62 | 0.94 | 0.95 | -1.85 | 0.89 | 0.5 | -0.485 | -0.425 | -0.3775 | 0.548065 | 0.165131 | 0.460091 | 0.989223 | 0.997988 | 0.994666 |
| Mllt4_E9Q9C3_S2s_1 | Mllt4_E9Q | -2 | -0.71 | 0.06 | -2.01 | -2.26 | -1.1 | -2.19 | -1.37 | -0.69 | -1.65 | -0.28 | -0.43 | -0.34 | -2.3 | -0.55 | -0.9 | -0.565 | 0.4025 | 0.1425 | 0.816378 | 0.303811 | 0.472241 | 0.989223 | 0.997988 | 0.999507 |
| Mllt4_E9Q9C3_S391s_1 | Mllt4_E9Q | -0.02 | -0.09 | 1.21 | 0.83 | -0.27 | -0.93 | 1.1 | 0.74 | 0.46 | -0.26 | 0.27 | 0.96 | -0.19 | 0.55 | 0.43 | 0.73 | -0.3225 | -0.125 | -0.1025 | 0.792206 | 0.535281 | 0.757372 | 0.989223 | 0.997988 | 0.999507 |
| Mllt4_E9Q9C3_S562s_1 | Mllt4_E9Q | -0.24 | -1.2 | -0.43 | -1.03 | -1.13 | 0.22 | -0.97 | -1.31 | 1.25 | -1.01 | -0.95 | -0.16 | 1.39 | -1.14 | -0.61 | -1.41 | -0.0725 | 0.5075 | 0.2825 | 0.646665 | 0.85624 | 0.354956 | 0.996349 | 0.997988 | 0.995259 |
| Mllt4_E9Q9C3_S655s_1 | Mllt4_E9Q | -1.53 | -0.78 | -1.1 | -0.94 | -0.46 | -1.48 | -0.58 | -1.15 | -1.25 | -0.86 | -1.17 | -0.36 | 0.59 | -0.15 | -0.8 | -0.56 | 0.17 | 0.1775 | 0.8575 | 0.039498 | 0.585573 | 0.555981 | 0.989223 | 0.997988 | 0.994666 |
| Mllt4_E9Q9C3_T214t_1 | Mllt4_E9Q | -4.6 | -1.97 | -2.03 | -0.94 | -0.57 | -1.27 | -0.88 | -2.72 | -2.73 | -1.78 | 1.46 | -4.13 | -1.46 | 0.06 | -1.24 | -2.78 | 1.025 | 0.59 | 1.03 | 0.246549 | 0.219141 | 0.63751 | 0.989223 | 0.997988 | 0.994666 |
| Mllt6_B1AR09_S273s_1 | Mllt6_B1A | -0.01 | -0.1 | -0.68 | -0.28 | 0.4 | -0.04 | 0.45 | -0.26 | -0.19 | 0.41 | -0.8 | 0.39 | -0.23 | -0.23 | -0.13 | 0.14 | 0.405 | 0.22 | 0.155 | 0.563015 | 0.157061 | 0.524729 | 0.989223 | 0.997988 | 0.995259 |
| Mmrn2_A6H6E2_S403s_1 | Mmrn2_A6 | -3.58 | -1.03 | -2.26 | -1.33 | -0.44 | -3.46 | -4.62 | -1.94 | -1.64 | -2.82 | -0.69 | -2.86 | -0.1 | 0.29 | -2.44 | -3.44 | -0.565 | 0.0475 | 0.6275 | 0.507092 | 0.548256 | 0.945748 | 0.989223 | 0.997988 | 0.994666 |
| Mobp_Q9D2P8_S107s_1 | Mobp_Q9D | -1.05 | -0.98 | -0.16 | -0.53 | -1.21 | -1.59 | -1.26 | -0.69 | -0.21 | -1 | -1.02 | -0.37 | 0.63 | -2.17 | -0.01 | -0.24 | -0.5075 | 0.03 | 0.2325 | 0.691914 | 0.118618 | 0.926037 | 0.989223 | 0.997988 | 0.996159 |
| Mobp_Q9D2P8_S74s_1 | Mobp_Q9D | 0.97 | 0.06 | 1.14 | 0.22 | -0.2 | 1.42 | -0.62 | -0.53 | 3.8 | 1.09 | -2.1 | 0.46 | 3.34 | -1.39 | 0.73 | 0.41 | -0.58 | 0.215 | 0.175 | 0.843482 | 0.264022 | 0.843881 | 0.989223 | 0.997988 | 0.999507 |
| Mobp_Q9D2P8_S85s_1 | Mobp_Q9D | 5.43 | 3.67 | 3.62 | 0.86 | 4.18 | 4.88 | 3.17 | 3.02 | -0.01 | 5.05 | -1.77 | 3.87 | -0.15 | -0.42 | 4.14 | 4.51 | 0.4175 | -1.61 | -1.375 | 0.340411 | 0.645931 | 0.334539 | 0.99033 | 0.997988 | 0.994666 |
| Mobp_Q9D2P8_S98s_1 | Mobp_Q9D | -0.49 | -0.21 | -0.1 | -0.59 | 0.12 | -1.05 | -0.77 | -0.55 | -0.17 | -0.38 | 0.33 | -0.16 | 0.27 | 0.01 | -0.04 | -0.39 | -0.215 | 0.2525 | 0.31 | 0.265016 | 0.479072 | 0.340124 | 0.989223 | 0.997988 | 0.994666 |
| Mon1a_Q6PDG8_S56s_1 | Mon1a_Q6 | 0.06 | 0 | 0.47 | 0.32 | 0.57 | 0.03 | 0.71 | 0.24 | -0.14 | 0.02 | 0.63 | 0.25 | -0.49 | 0.45 | 0.35 | 0.42 | 0.175 | -0.0225 | -0.03 | 0.922099 | 0.486669 | 0.931927 | 0.989223 | 0.997988 | 0.999507 |
| Mon2_B9EKJ3_S1175s_1 | Mon2_B9E | -0.83 | -0.36 | -0.26 | -0.71 | -0.26 | -0.64 | 0 | -0.61 | -1.5 | -0.59 | -1.04 | -0.66 | -1.32 | -1.28 | -0.2 | -0.15 | 0.1625 | -0.4075 | -0.1975 | 0.597728 | 0.53076 | 0.187262 | 0.989223 | 0.997988 | 0.995259 |
| Mon2_B9EKJ3_S1213s_1 | Mon2_B9E | 0.4 | -0.35 | -0.13 | -0.52 | -0.77 | 0.9 | -0.76 | -0.51 | 0.78 | 0.09 | -2.22 | -0.34 | 0.68 | -1.93 | -0.51 | -0.37 | -0.135 | -0.2725 | -0.3825 | 0.479458 | 0.750267 | 0.658927 | 0.995762 | 0.997988 | 0.994666 |
| Mon2_B9EKJ3_S534s_1 | Mon2_B9E | -1.03 | -0.84 | -0.23 | -0.63 | -1.04 | -0.71 | -1.06 | -1.59 | -0.1 | -1.04 | 0.34 | -0.53 | -0.21 | -0.95 | -0.51 | -0.83 | -0.4175 | 0.35 | 0.0575 | 0.848165 | 0.164544 | 0.337382 | 0.989223 | 0.997988 | 0.999507 |
| Mon2_Q80TL7_S534s_1 | Mon2_Q8C | -1.27 | -1.18 | -1.67 | -1.45 | -1.03 | -1.21 | -1.37 | -2.16 | -0.56 | -1.01 | -0.27 | -0.92 | -0.09 | -0.32 | -1.32 | -1.19 | -0.05 | 0.7025 | 0.6625 | 0.086554 | 0.866498 | 0.023889 | 0.996493 | 0.997988 | 0.994666 |
| Morc2a_Q69ZX6_S614s_1 | Morc2a_Q | -0.54 | -0.59 | -0.83 | -0.51 | -0.29 | -1.52 | 0.13 | -0.7 | -1.17 | -0.76 | -0.42 | -0.68 | -0.84 | -0.61 | -1.06 | -0.87 | 0.0225 | -0.14 | -0.2275 | 0.361219 | 0.949865 | 0.576043 | 0.999187 | 0.997988 | 0.994666 |
| Morc2a_Q69ZX6_S775s_1 | Morc2a_Q | -0.98 | -4.13 | -1.3 | -1.26 | -1.24 | -2.65 | -1.16 | -2.15 | -1.51 | -0.41 | -2.28 | -1.51 | -0.72 | -1.02 | -0.93 | -1.32 | 0.1175 | 0.49 | 0.92 | 0.19457 | 0.871199 | 0.517843 | 0.996493 | 0.997988 | 0.994666 |
| Morc2a_Q69ZX6_S854s_1 | Morc2a_Q | -1.89 | -0.92 | -2.15 | -1.54 | -1.54 | -0.61 | -0.57 | -1.83 | -0.44 | -0.14 | -1.73 | -0.45 | -1.08 | -1.04 | -1.38 | -0.96 | 0.4875 | 0.935 | 0.51 | 0.141563 | 0.242986 | 0.053749 | 0.989223 | 0.997988 | 0.994666 |
| Mpdz_D3YUX2_S230s_1 | Mpdz_D3Y | 0.13 | 0.38 | 0.03 | 0.53 | 0.33 | 0.12 | -0.29 | -0.35 | 0.55 | 0.43 | 0.7 | 0.24 | 0.19 | 0.4 | 0.35 | -0.03 | -0.315 | 0.2125 | -0.04 | 0.875658 | 0.233242 | 0.382001 | 0.989223 | 0.997988 | 0.999507 |
| Mpdz_D3YUX2_S231s_1 | Mpdz_D3Y | 0.33 | 0.05 | -0.08 | 0.18 | 0.25 | -0.13 | -0.1 | -0.17 | 0.42 | 0.54 | 0.76 | 0.02 | 0.02 | 0.62 | 0.11 | -0.16 | -0.1575 | 0.315 | 0.0275 | 0.919866 | 0.4786 | 0.230752 | 0.989223 | 0.997988 | 0.999507 |
| Mphosph8_Q3TYA6_S25s_1 | Mphosph8 | -2.53 | -0.62 | -0.51 | -0.42 | -2.16 | -0.86 | -1.36 | -1.23 | -1.24 | -1.77 | -0.14 | -1.09 | -0.42 | -1.48 | -1.06 | -1.91 | -0.3825 | -0.04 | -0.1975 | 0.721343 | 0.470766 | 0.943228 | 0.989223 | 0.997988 | 0.9971 |
| Mpp1_B7ZCM1_S57s_1 | Mpp1_B7Z | -0.13 | -0.77 | 0.51 | 0.54 | -0.1 | -0.29 | -0.03 | 0.03 | -0.23 | -0.29 | 0.06 | 0.5 | -1.2 | -0.23 | -0.03 | 0.12 | -0.135 | -0.0275 | -0.3725 | 0.391681 | 0.682609 | 0.940323 | 0.994175 | 0.997988 | 0.994666 |
| Mpp1_B7ZCM1_S93s_1 | Mpp1_B7Z | 0.34 | 0.71 | -0.98 | -0.54 | -0.2 | 0.36 | 0.12 | -0.21 | 0.18 | 0.74 | -2.57 | -1.06 | 0.3 | 0.44 | -1.31 | -1.35 | 0.135 | -0.56 | -0.3625 | 0.533486 | 0.73476 | 0.459797 | 0.995762 | 0.997988 | 0.994666 |
| Mpp2_Q9WV34_S121s_1 | Mpp2_Q9V | 0.51 | 0.83 | 1.12 | 0.6 | 0.77 | 0.58 | 1.44 | 1.53 | 0.7 | 1.09 | 0.2 | 1.78 | 0.39 | 1.03 | 1.2 | 0.31 | 0.315 | 0.1775 | -0.0325 | 0.917451 | 0.306753 | 0.632055 | 0.989223 | 0.997988 | 0.999507 |
| Mpp2_Q9WV34_S42s_1 | Mpp2_Q9V | 1.21 | -0.82 | 0.32 | 0.98 | 0.79 | 0.98 | 0.94 | 1.06 | 0.08 | -0.23 | 0.11 | 0.65 | -2.81 | 0.54 | -0.45 | 0.11 | 0.52 | -0.27 | -1.075 | 0.18749 | 0.247442 | 0.567361 | 0.989223 | 0.997988 | 0.994666 |
| Mpp2_Q9WV34_T117t_1 | Mpp2_Q9V | 0.99 | 1.81 | 0.47 | 0.33 | 1.03 | 0.92 | 0.76 | 0.42 | 1.08 | 1.43 | 0.48 | 0.49 | 1.23 | 1.13 | 0.62 | 0.73 | -0.1175 | -0.03 | 0.0275 | 0.942432 | 0.743941 | 0.940817 | 0.995762 | 0.997988 | 0.999507 |
| Mpp3_Q6XE40_S307s_1 | Mpp3_Q6X | -0.15 | 0.14 | 0.7 | 0.42 | 0.51 | -0.23 | 1.08 | 0.74 | -0.5 | -0.01 | 0.25 | 0.42 | 0.19 | 0.49 | 0.83 | 0.53 | 0.2475 | -0.2375 | 0.2325 | 0.434183 | 0.471593 | 0.446742 | 0.989223 | 0.997988 | 0.994666 |
| Mpp3_Q6XE40_S354s_1 | Mpp3_Q6X | 0.46 | 0.28 | 0.59 | 0.77 | 0.12 | 0.59 | -1.02 | -0.36 | 1.57 | 0.44 | -1.77 | 0.27 | 1.89 | -0.33 | 1.37 | -0.47 | -0.6925 | -0.3975 | 0.09 | 0.872053 | 0.078329 | 0.538413 | 0.989223 | 0.997988 | 0.999507 |
| Mpp6_Q9JLB0_S199s_1 | Mpp6_Q9J | -0.2 | 0.68 | 0.64 | 0.28 | 0.24 | -0.2 | 0.3 | 0.29 | -0.24 | -0.19 | -0.09 | 0.08 | -0.56 | -0.74 | -0.18 | 0.43 | -0.1925 | -0.46 | -0.6125 | 0.110846 | 0.490409 | 0.117646 | 0.989223 | 0.997988 | 0.994666 |
| Mprip_F6RND9_S350s_1 | Mprip_F6R | 0.53 | 0.4 | -0.05 | 0.12 | 0.59 | 1.23 | 0.31 | -0.01 | 0.44 | 0.51 | -0.1 | 0.69 | -0.14 | 0.41 | 0.14 | -0.27 | 0.28 | 0.135 | -0.215 | 0.448951 | 0.381065 | 0.621969 | 0.989223 | 0.997988 | 0.994666 |
| Mprip_F6RND9_S89s_2 | Mprip_F6R | 0.47 | 0.75 | 0.22 | -0.05 | 0.74 | 0.49 | 0.25 | 0.32 | 0.91 | -0.02 | 2.42 | 0.8 | 0.73 | 1.05 | 0.3 | 0.17 | 0.1025 | 0.68 | 0.215 | 0.500453 | 0.689305 | 0.199741 | 0.994672 | 0.997988 | 0.994666 |
| Mprip_F6RND9_S93s_2 | Mprip_F6R | 0.47 | 0.75 | 0.22 | -0.05 | 0.74 | 0.49 | 0.25 | 0.32 | 0.91 | -0.02 | 2.42 | 0.8 | 0.73 | 1.05 | 0.3 | 0.17 | 0.1025 | 0.68 | 0.215 | 0.500453 | 0.689305 | 0.199741 | 0.994672 | 0.997988 | 0.994666 |
| Mprip_F6RND9_S962s_1 | Mprip_F6R | -1.7 | -0.65 | 1.42 | 0.78 | -1.01 | -0.83 | 0.51 | 0.34 | -1.15 | -1.07 | 0.02 | 0.37 | -0.93 | -1.78 | 0.41 | 0.94 | -0.21 | -0.42 | -0.3025 | 0.715052 | 0.767363 | 0.563597 | 0.995762 | 0.997988 | 0.996966 |
| Mprip_F6RND9_S976s_1 | Mprip_F6R | -0.15 | -1.06 | -0.81 | -0.78 | 0.56 | -0.31 | -0.74 | -0.11 | -3.06 | -1.79 | -0.78 | -0.59 | -1.96 | 0.12 | -1.15 | -0.53 | 0.55 | -0.855 | -0.18 | 0.700427 | 0.130127 | 0.148219 | 0.989223 | 0.997988 | 0.996159 |
| Mprip_F6RND9_S978s_1 | Mprip_F6R | -1.32 | -2.64 | -1.43 | -1.65 | -0.66 | -0.7 | -1.19 | -0.94 | -2 | -2.08 | -0.84 | -1.51 | -2.12 | -1.02 | -1.81 | -1.43 | 0.8875 | 0.1525 | 0.165 | 0.676045 | 0.023355 | 0.710404 | 0.989223 | 0.997988 | 0.996159 |
| Mprip_Q5SWZ5_S998s_1 | Mprip_Q5S | -1.21 | -2.39 | -0.61 | -1.56 | -0.61 | -1.13 | -0.28 | -0.4 | -2.32 | -3 | 0.82 | -0.92 | -2.18 | -0.76 | -1.26 | -0.3 | 0.8375 | 0.0875 | 0.3175 | 0.54144 | 0.06028 | 0.915204 | 0.989223 | 0.997988 | 0.994666 |
| Msantd1_Q8BIL2_S161s_1 | Msantd1_( | 1.11 | 1.11 | 0.51 | 0.59 | 0.79 | 0.63 | 1.14 | 0.8 | 0.67 | 0.98 | 1.11 | 0.6 | 0.01 | 0.46 | 0.54 | 1.32 | 0.01 | 0.01 | -0.2475 | 0.482827 | 0.968172 | 0.969903 | 0.999187 | 0.997988 | 0.994666 |
| Msh6_P54276_S63s_1 | Msh6_P54 | -2.37 | -1.64 | -2.37 | -3.3 | -2.57 | -2.29 | -3.57 | -3.16 | -0.95 | -1.79 | -2.76 | -2.89 | -0.43 | -1.51 | -3.04 | -3.26 | -0.4775 | 0.3225 | 0.36 | 0.595783 | 0.275105 | 0.546325 | 0.989223 | 0.997988 | 0.995259 |
| Msi2_Q920Q6_S279s_1 | Msi2_Q92C | 0.05 | -0.37 | 0.63 | 0.36 | -0.07 | -0.55 | 0.96 | 0.54 | 0.08 | -0.2 | 0.79 | 0.77 | -0.06 | 0.31 | 0.48 | 1.25 | 0.0525 | 0.1925 | 0.3275 | 0.385664 | 0.891488 | 0.581274 | 0.996493 | 0.997988 | 0.994666 |
| Msi2_Q920Q6_S6s_1 | Msi2_Q92C | 0.25 | -0.03 | 0 | 0.21 | 0.22 | -0.07 | -0.25 | 0.06 | 0.41 | -0.01 | 1.14 | 0.38 | 0.93 | 0.37 | 0.39 | 0.35 | -0.1175 | 0.3725 | 0.4025 | 0.144365 | 0.588962 | 0.224791 | 0.989223 | 0.997988 | 0.994666 |
| Msl3_Q9WVG9_S345s_1 | Msl3_Q9W | 0.66 | 0.84 | 0.65 | 0.52 | 0.34 | 0.5 | 0.6 | 0.79 | 0.46 | 1.15 | 0.79 | 0.56 | 0.27 | 0.95 | 0.68 | 1 | -0.11 | 0.0725 | 0.0575 | 0.830832 | 0.607586 | 0.768317 | 0.989223 | 0.997988 | 0.999507 |
| Msrb3_D3YUC9_S105s_1 | Msrb3_D3 | -2.14 | -1.56 | -1.56 | -2.21 | -3.19 | -1.21 | -2.6 | -3.53 | -0.32 | -2.19 | -2.35 | -1.97 | 0.26 | -2.18 | -1.36 | -2.37 | -0.765 | 0.16 | 0.455 | 0.436191 | 0.148013 | 0.737742 | 0.989223 | 0.997988 | 0.994666 |
| Mt3_P28184_S33s_1 | Mt3_P281 | 0.06 | -0.25 | -0.36 | -0.31 | -1.29 | 1.02 | -0.27 | -0.35 | 0.48 | 0.24 | -0.71 | -0.61 | 1.52 | -1.21 | -0.44 | -0.01 | -0.0075 | 0.065 | 0.18 | 0.740036 | 0.986739 | 0.846412 | 0.999187 | 0.997988 | 0.9971 |
| Mta1_F8WHY8_S432s_1 | Mta1_F8W | -0.01 | -0.22 | -0.46 | -0.44 | -0.2 | -0.13 | -0.02 | -0.9 | 0.07 | 0.06 | -0.13 | -0.49 | 0.8 | -0.89 | 0.31 | -0.12 | -0.03 | 0.16 | 0.3075 | 0.433598 | 0.911063 | 0.521199 | 0.997335 | 0.997988 | 0.994666 |
| Mta1_F8WHY8_S505s_1 | Mta1_F8W | -0.35 | -0.18 | 0.42 | -0.16 | -0.09 | -0.41 | 0.5 | -0.52 | -0.86 | 0.53 | 0.09 | 0.29 | -0.08 | 0.27 | 0.25 | 0.19 | -0.0625 | 0.08 | 0.225 | 0.417386 | 0.838009 | 0.823979 | 0.995762 | 0.997988 | 0.994666 |
| Mta1_F8WHY8_S622s_1 | Mta1_F8W | -3.04 | -2.19 | -2.32 | -2.99 | -1.97 | -3.78 | -2.19 | -3.24 | -4.47 | -2.77 | -2.28 | -2.66 | -2.35 | -1.92 | -2.69 | -2.17 | -0.16 | -0.41 | 0.3525 | 0.287756 | 0.72363 | 0.423391 | 0.995437 | 0.997988 | 0.994666 |
| Mta3_H3BIY7_S176s_1 | Mta3_H3B | 3.35 | 2 | 3.65 | 1.12 | 2.86 | 4.02 | 4.51 | 2.64 | -1.13 | 3.34 | -0.47 | 3.22 | -2.45 | 0.35 | 3.95 | 5.69 | 0.9775 | -1.29 | -0.645 | 0.695532 | 0.161584 | 0.284155 | 0.989223 | 0.997988 | 0.996159 |
| Mta3_H3BIY7_S207s_1 | Mta3_H3B | -0.07 | 0.45 | -1.1 | -2.62 | -0.14 | -0.5 | -0.2 | -0.13 | -0.06 | 1.21 | -2.79 | -2.79 | -0.33 | -0.01 | -0.1 | 0.23 | 0.5925 | -0.2725 | 0.7825 | 0.228707 | 0.343099 | 0.79861 | 0.989223 | 0.997988 | 0.994666 |
| Mta3_H3BIY7_S85s_1 | Mta3_H3B | -3.31 | -3.29 | -3.24 | -4.9 | -3.84 | -3.52 | -4.34 | -4.36 | -3.44 | -2.75 | -1.53 | -3.98 | -1.7 | -3.17 | -3.55 | -3.46 | -0.33 | 0.76 | 0.715 | 0.212962 | 0.448712 | 0.232306 | 0.989223 | 0.997988 | 0.994666 |
| Mtch1_Q791T5_S364s_1 | Mtch1_Q7 | 0.09 | -0.01 | 0.19 | -0.01 | 0.43 | -0.31 | 0.35 | 0.01 | -0.24 | -0.44 | -0.05 | -0.05 | -0.11 | 0.87 | 0.15 | 0.73 | 0.055 | -0.26 | 0.345 | 0.266567 | 0.819567 | 0.253574 | 0.995762 | 0.997988 | 0.994666 |
| Mtcl1_H3BL83_S1095s_1 | Mtcl1_H3B | -2.63 | -4.51 | -2.1 | -1.88 | -1.9 | -3.2 | -2 | -2.19 | -3.3 | -3.48 | -0.62 | -2.82 | -3.26 | -1.92 | -2.34 | -2.41 | 0.4575 | 0.225 | 0.2975 | 0.623555 | 0.451251 | 0.777531 | 0.989223 | 0.997988 | 0.995259 |
| Mtcl1_H3BL83_S1369s_2 | Mtcl1_H3B | -1.89 | -2.46 | -2.38 | -4.25 | -2.53 | -2.57 | -3.83 | -3 | -0.55 | -2.23 | -1 | -2.34 | -0.37 | -3.06 | -3.24 | -3.04 | -0.2375 | 1.215 | 0.3175 | 0.678491 | 0.662793 | 0.077075 | 0.990434 | 0.997988 | 0.996159 |
| Mtcl1_H3BL83_T1366t_2 | Mtcl1_H3B | -1.89 | -2.46 | -2.38 | -4.25 | -2.53 | -2.57 | -3.83 | -3 | -0.55 | -2.23 | -1 | -2.34 | -0.37 | -3.06 | -3.24 | -3.04 | -0.2375 | 1.215 | 0.3175 | 0.678491 | 0.662793 | 0.077075 | 0.990434 | 0.997988 | 0.996159 |
| Mtcl1_Q3UHU5_S255s_1 | Mtcl1_Q3L | 0.5 | -0.2 | 0.46 | 0.39 | 0.29 | 0.13 | 0.64 | 0.7 | -0.05 | 0.79 | 0.77 | -0.04 | -0.21 | -0.08 | -0.11 | 1.15 | 0.1525 | 0.08 | -0.1 | 0.791378 | 0.563693 | 0.802079 | 0.989223 | 0.997988 | 0.999507 |
| Mtcl1_Q3UHU5_S312s_1 | Mtcl1_Q3L | -0.48 | -1.22 | -0.23 | -1.05 | -0.8 | -1.83 | -0.52 | -0.16 | -0.73 | -0.63 | 0.94 | -0.75 | -0.85 | 0.4 | 0.23 | -0.05 | -0.0825 | 0.4525 | 0.6775 | 0.098604 | 0.840243 | 0.332926 | 0.995762 | 0.997988 | 0.994666 |
| Mtdh_Q80WJ7_S297s_1 | Mtdh_Q80 | -2.42 | -0.61 | -2.6 | -2.57 | -0.04 | -2.27 | -0.88 | -1.57 | -3.37 | -3.48 | -0.78 | -3.1 | -2.88 | -1.49 | -1.05 | -1.23 | 0.86 | -0.6325 | 0.3875 | 0.511519 | 0.178256 | 0.390076 | 0.989223 | 0.997988 | 0.994666 |
| Mtdh_Q80WJ7_S423s_1 | Mtdh_Q80 | -1.01 | -2.05 | 0.36 | 0.06 | -0.76 | -2.8 | -0.45 | -0.43 | -2.27 | -2.36 | -0.16 | -0.52 | -1.79 | -1.34 | 0.67 | -0.39 | -0.45 | -0.6675 | -0.0525 | 0.939546 | 0.523001 | 0.362739 | 0.989223 | 0.997988 | 0.999507 |
| Mtdh_Q80WJ7_S565s_1 | Mtdh_Q80 | -0.84 | -0.77 | -0.48 | -0.38 | 0 | -1.3 | 0.28 | -0.31 | -1.52 | -0.56 | -0.2 | -0.32 | -0.92 | -0.46 | 0.27 | -0.03 | 0.285 | -0.0325 | 0.3325 | 0.321163 | 0.436384 | 0.923675 | 0.989223 | 0.997988 | 0.994666 |
| Mtfr1l_Q9CWE0_S100s_1 | Mtfr1l_Q9 | -1.97 | -1.87 | -2.12 | -2.19 | -1.36 | -2.6 | -1.45 | -2.95 | -2.68 | -1.94 | -2.01 | -2.09 | -1.15 | -1.49 | -2.66 | -2.67 | -0.0525 | -0.1425 | 0.045 | 0.911532 | 0.894103 | 0.579198 | 0.996493 | 0.997988 | 0.999507 |
| Mtfr1l_Q9CWE0_S107s_1 | Mtfr1l_Q9 | 1.08 | 0.58 | 0.43 | 0.93 | 0.83 | 0.48 | 0.41 | 0.92 | 1.02 | 0.95 | 0.85 | 1.28 | 1.21 | 0.46 | 1.88 | 1.33 | -0.095 | 0.27 | 0.465 | 0.211713 | 0.706973 | 0.29719 | 0.995437 | 0.997988 | 0.994666 |
| Mtfr1l_Q9CWE0_S235s_1 | Mtfr1l_Q9 | -1.05 | -0.65 | -0.75 | -0.7 | 0.03 | -1.24 | 0.01 | -0.75 | -1.52 | -0.62 | -0.88 | -0.91 | -1.14 | 0 | -1.04 | -0.46 | 0.3 | -0.195 | 0.1275 | 0.695963 | 0.376732 | 0.475897 | 0.989223 | 0.997988 | 0.996159 |
| Mtif2_Q5M6W9_S83s_1 | Mtif2_Q5N | 0.31 | 0.22 | 0.72 | 0.52 | -0.93 | 0.06 | -0.77 | -0.14 | 3.79 | 0.13 | -0.33 | 0.28 | 4.41 | 0.32 | 0.15 | 0.35 | -0.8875 | 0.525 | 0.865 | 0.354245 | 0.012503 | 0.541367 | 0.989223 | 0.997988 | 0.994666 |
| Mtmr1_I7HJQ9_S659s_1 | Mtmr1_I7I | -1.34 | -0.77 | -0.53 | -0.55 | -0.82 | -0.96 | -0.41 | -0.72 | -0.02 | -0.97 | -0.93 | -0.9 | 0.23 | -0.25 | -0.52 | -1.05 | 0.07 | 0.0925 | 0.4 | 0.275743 | 0.793054 | 0.775491 | 0.995762 | 0.997988 | 0.994666 |
| Mtmr1_I7HJQ9_T665t_2 | Mtmr1_I7I | -0.1 | -2.11 | -2.55 | -1.89 | -1.46 | -0.34 | -1.5 | -2.19 | 0.06 | -1.22 | -1.06 | -2.75 | -0.53 | -1.21 | -3.13 | -3.56 | 0.29 | 0.42 | -0.445 | 0.582545 | 0.626156 | 0.55857 | 0.989223 | 0.997988 | 0.995259 |
| Mtmr2_Q9Z2D1_S21s_1 | Mtmr2_Q9 | 0.52 | 0.59 | -0.74 | 0.07 | 0.18 | -0.75 | 0.08 | 0.22 | -0.15 | 0.22 | -1.24 | -0.92 | 0.65 | 1.35 | -0.48 | -0.4 | -0.1775 | -0.6325 | 0.17 | 0.737155 | 0.637856 | 0.174679 | 0.989223 | 0.997988 | 0.9971 |
| Mtmr3_M0QW74_S633s_1 | Mtmr3_M0 | 0.12 | -0.05 | 0.76 | 0.47 | -1.2 | 0.74 | 0.02 | 0.32 | 0.38 | -0.05 | 0.54 | 0.64 | 0.87 | -0.03 | 0.41 | 0.67 | -0.355 | 0.0525 | 0.155 | 0.625578 | 0.416937 | 0.853806 | 0.989223 | 0.997988 | 0.995259 |
| Mtmr4_Q91XS1_S572s_1 | Mtmr4_Q9 | -0.32 | -0.48 | 0.29 | 0.32 | 0.19 | -0.3 | 0.12 | -0.01 | -0.46 | -0.41 | 0.22 | -0.1 | 0.45 | 0.36 | -0.32 | -0.66 | 0.0475 | -0.14 | 0.005 | 0.988969 | 0.861914 | 0.641852 | 0.996493 | 0.997988 | 1 |
| Mtmr6_Q8VE11_S585s_1 | Mtmr6_Q8 | 0.27 | 0.15 | -0.93 | -0.07 | -0.71 | 0.57 | -0.77 | -0.9 | 0.13 | -0.82 | 0.12 | -0.87 | 0.29 | -0.4 | -0.75 | -0.71 | -0.3075 | -0.215 | -0.2475 | 0.517702 | 0.466453 | 0.584954 | 0.989223 | 0.997988 | 0.994666 |
| Mtmr7_Q9Z2C9_S593s_1 | Mtmr7_Q9 | 0 | 0.26 | 0.71 | 0.19 | -0.48 | 0.38 | 1.06 | 0.03 | 0.14 | -0.33 | 0.98 | -0.78 | -0.09 | 0.87 | 0.13 | 0.4 | -0.0425 | -0.2875 | 0.0375 | 0.90362 | 0.904977 | 0.480776 | 0.996493 | 0.997988 | 0.999507 |
| Mtmr7_Q9Z2C9_S595s_1 | Mtmr7_Q9 | -0.41 | -0.08 | 0.35 | -0.1 | -0.17 | -0.39 | 0.33 | -0.05 | -0.3 | -0.18 | -0.02 | -0.82 | -0.45 | 0.29 | -0.67 | -0.09 | -0.01 | -0.27 | -0.17 | 0.590691 | 0.969713 | 0.352154 | 0.999187 | 0.997988 | 0.995259 |

| | | | | | | | | | | | | | | | | | | | | | | | | | | |
|---|---|---|---|---|---|---|---|---|---|---|---|---|---|---|---|---|---|---|---|---|---|---|---|---|---|---|
| Mtmr7_Q9Z2C9_S634s_1 | Mtmr7_Q9 | 0.33 | 0.09 | 1.56 | 1.49 | -0.75 | 1.41 | 0.22 | 1.01 | 1.98 | 0.29 | 1.3 | 1.49 | 1.8 | -0.77 | 1.15 | 1.75 | -0.395 | 0.3975 | 0.115 | 0.858468 | 0.480471 | 0.428607 | 0.989223 | 0.997988 | 0.999507 |
| Mtmr7_Q9Z2C9_T578t_1 | Mtmr7_Q9 | -2.8 | -0.45 | -0.35 | 0.78 | -0.65 | -3.42 | -1.63 | 0.12 | 0.4 | -2.02 | 1.61 | -0.64 | 0.8 | 1.21 | -0.72 | 0.03 | -0.69 | 0.5425 | 1.035 | 0.198467 | 0.46464 | 0.572427 | 0.989223 | 0.997988 | 0.994666 |
| Mtor_Q9JLN9_S567s_1 | Mtor_Q9JL | -0.47 | -0.29 | 0.23 | -0.58 | 0.08 | -0.85 | 0.07 | -0.26 | -0.11 | -0.37 | -0.49 | 0.42 | -0.05 | -0.05 | -0.74 | 0.04 | 0.0375 | 0.14 | 0.0775 | 0.802438 | 0.901972 | 0.649339 | 0.996493 | 0.997988 | 0.999507 |
| Mtpn_P62774_T31t_1 | Mtpn_P62 | -0.19 | 0.12 | 0.55 | 0.3 | 0.19 | -0.47 | -0.23 | 0.88 | -0.82 | 0.11 | 0.95 | 0.53 | 0.6 | 0.07 | 0.23 | 0.14 | -0.1025 | -0.0025 | 0.065 | 0.814435 | 0.763633 | 0.995072 | 0.995762 | 0.998021 | 0.999507 |
| Mtss1l_D3YWC9_S199s_1 | Mtss1l_D3 | 1.48 | 1.77 | 1.59 | 1.67 | 0.99 | 1.34 | 2.13 | 1.29 | 1.33 | 1.42 | -0.34 | 1.75 | 0.68 | 0.78 | 1.48 | 1.58 | -0.19 | -0.5875 | -0.4975 | 0.123634 | 0.509106 | 0.216822 | 0.989223 | 0.997988 | 0.994666 |
| Mtss1l_D3YWC9_S329s_1 | Mtss1l_D3 | 0.19 | 0.21 | -0.13 | -0.13 | 0.25 | -0.01 | -0.85 | -0.27 | 0.32 | 0.36 | -0.13 | 0.2 | 0.58 | 0.08 | 0.08 | -0.26 | -0.255 | 0.1525 | 0.085 | 0.760099 | 0.383158 | 0.522954 | 0.989223 | 0.997988 | 0.9971 |
| Mtss1l_D3YWC9_S357s_1 | Mtss1l_D3 | 0.28 | -0.23 | 0.54 | 1.17 | 0.25 | 0.56 | 1.65 | 3.14 | 0.15 | -0.14 | 2.91 | 1.34 | 0.01 | 0.97 | 1.45 | 1.35 | 0.96 | 0.625 | 0.505 | 0.260594 | 0.154991 | 0.369449 | 0.989223 | 0.997988 | 0.994666 |
| Mtss1l_D3YWC9_S502s_1 | Mtss1l_D3 | 0.87 | 0.05 | -0.45 | -0.21 | 0.97 | 0.94 | -0.16 | -0.36 | 0.69 | -0.38 | 0.16 | -0.81 | 0.29 | 1.1 | -0.63 | 0.14 | 0.2825 | -0.15 | 0.16 | 0.720179 | 0.515741 | 0.724358 | 0.989223 | 0.997988 | 0.9971 |
| Mtss1l_D3YWC9_S540s_1 | Mtss1l_D3 | 0.23 | 0.32 | 0.51 | 0.44 | 0.59 | -0.25 | 0.87 | 0.33 | 0.08 | 0.08 | 0.11 | 0.25 | -0.89 | 0.15 | 0.76 | 1 | 0.01 | -0.245 | -0.12 | 0.777095 | 0.971646 | 0.260458 | 0.999187 | 0.997988 | 0.99798 |
| Mtss1l_D3YWC9_T195t_1 | Mtss1l_D3 | 0.69 | 0.84 | 0.56 | 0.63 | 0.47 | 0.55 | 0.91 | 0.76 | 0.01 | 0.97 | 0.12 | 0.53 | -0.27 | -0.08 | 0.76 | 0.28 | -0.0075 | -0.2725 | -0.5075 | 0.113394 | 0.971932 | 0.341242 | 0.999187 | 0.997988 | 0.994666 |
| Mtss1l_F7D291_S157s_1 | Mtss1l_F7D | 0 | -0.23 | -0.65 | -1.04 | -0.93 | 0.27 | -0.26 | -0.86 | 0.51 | -0.77 | -0.11 | -1.02 | -0.24 | -1.22 | 0.49 | 0.09 | 0.035 | 0.1325 | 0.26 | 0.546861 | 0.922808 | 0.746415 | 0.997335 | 0.997988 | 0.994666 |
| Mtus2_Q3UHD3_S109s_1 | Mtus2_Q3 | -1 | -0.45 | 0.07 | 1.2 | -0.63 | 0 | 0.5 | 1.55 | -1.49 | -1.52 | 1.78 | 0.78 | -1.16 | -0.24 | 0.02 | 0.18 | 0.4 | -0.0675 | -0.255 | 0.626738 | 0.502332 | 0.936484 | 0.989223 | 0.997988 | 0.995259 |
| Mtus2_Q3UHD3_S1324s_1 | Mtus2_Q3 | 0.03 | -0.52 | 0.32 | 0.32 | 0.49 | 0.34 | 0.54 | 0.88 | -1.04 | -0.26 | -0.48 | 0.65 | -0.84 | -0.21 | 0.15 | 0.57 | 0.525 | -0.32 | -0.12 | 0.749768 | 0.07711 | 0.43291 | 0.989223 | 0.997988 | 0.9971 |
| Mtus2_Q3UHD3_S1340s_2 | Mtus2_Q3 | 0.42 | -2.81 | -2.56 | -2.39 | -2.84 | 0.61 | -1.1 | -0.25 | 0.47 | -1.7 | -0.16 | -1.28 | -0.57 | -1.98 | -2.23 | -1.59 | 0.94 | 1.1675 | 0.2425 | 0.745839 | 0.318762 | 0.172836 | 0.989223 | 0.997988 | 0.9971 |
| Mtus2_Q3UHD3_S1344s_1 | Mtus2_Q3 | 0.34 | -3.76 | -1.5 | -1.61 | -2.01 | 0.96 | -1.12 | -0.07 | 0.49 | -2.54 | -2.75 | -0.93 | -0.41 | -3.16 | -1.89 | -2.61 | 1.0725 | 0.2 | -0.385 | 0.670352 | 0.259745 | 0.841228 | 0.989223 | 0.997988 | 0.996159 |
| Mtus2_Q3UHD3_S1344s_2 | Mtus2_Q3 | 0.42 | -2.81 | -2.56 | -2.39 | -2.84 | 0.61 | -1.1 | -0.25 | 0.47 | -1.7 | -0.16 | -1.28 | -0.57 | -1.98 | -2.23 | -1.59 | 0.94 | 1.1675 | 0.2425 | 0.745839 | 0.318762 | 0.172836 | 0.989223 | 0.997988 | 0.9971 |
| Mtus2_Q3UHD3_S141s_1 | Mtus2_Q3 | 1.02 | -0.29 | 0.17 | 1.15 | 0.26 | 1.53 | -0.4 | 0.84 | -0.07 | -0.32 | 0.03 | -1.04 | -0.3 | -0.63 | 0.39 | -0.27 | 0.045 | -0.8625 | -0.715 | 0.105534 | 0.9272 | 0.061962 | 0.997335 | 0.997988 | 0.994666 |
| Mtus2_Q3UHD3_S332s_1 | Mtus2_Q3 | 0.75 | 0.08 | 0.68 | 0.88 | 0.54 | 0.43 | 0.36 | 0.35 | 0.2 | 0.35 | 0.92 | 0.43 | 0.22 | 0.15 | 0.03 | -0.06 | -0.1775 | -0.1225 | -0.5125 | 0.085777 | 0.473825 | 0.668417 | 0.989223 | 0.997988 | 0.994666 |
| Mtus2_Q3UHD3_S435s_2 | Mtus2_Q3 | 1.36 | -0.17 | 0.3 | -0.1 | -1.1 | 0.68 | -0.64 | 0.97 | 1.09 | -0.03 | 1.53 | 1.16 | 0.6 | 0.5 | 0.26 | -0.31 | -0.37 | 0.59 | -0.085 | 0.835795 | 0.509137 | 0.225803 | 0.989223 | 0.997988 | 0.999507 |
| Mtus2_Q3UHD3_S951s_1 | Mtus2_Q3 | 0.94 | -1.5 | -1.21 | -0.85 | 0.83 | 0.61 | 0.14 | 0.67 | -1.37 | -0.71 | -0.83 | -0.66 | -1.17 | 0.01 | -0.28 | -0.53 | 1.2175 | -0.2375 | 0.1625 | 0.770912 | 0.038938 | 0.657221 | 0.989223 | 0.997988 | 0.9971 |
| Mtus2_Q3UHD3_T1331t_1 | Mtus2_Q3 | 0.99 | -0.63 | -0.09 | 0.67 | 0.44 | 1.17 | 1.1 | 1.16 | -0.54 | -0.1 | -0.16 | 0.18 | -2.15 | -0.99 | -0.37 | 0.33 | 0.7325 | -0.39 | -1.03 | 0.105031 | 0.087973 | 0.336474 | 0.989223 | 0.997988 | 0.994666 |
| Mtus2_Q3UHD3_T433t_2 | Mtus2_Q3 | 1.36 | -0.17 | 0.3 | -0.1 | -1.1 | 0.68 | -0.64 | 0.97 | 1.09 | -0.03 | 1.53 | 1.16 | 0.6 | 0.5 | 0.26 | -0.31 | -0.37 | 0.59 | -0.085 | 0.835795 | 0.509137 | 0.225803 | 0.989223 | 0.997988 | 0.999507 |
| Mtx3_D3YTP3_S249s_1 | Mtx3_D3Y | -0.6 | -0.26 | -0.99 | -0.46 | -0.4 | -1.09 | 0.13 | -0.48 | -0.72 | -0.18 | -0.02 | -1.64 | -0.44 | -0.2 | -1.87 | -0.84 | 0.1175 | -0.0625 | -0.26 | 0.524695 | 0.707417 | 0.874425 | 0.995437 | 0.997988 | 0.994666 |
| Mtx3_D3YTP3_S284s_1 | Mtx3_D3Y | 0.97 | 0.2 | 0.51 | 0.91 | 0.23 | 0.51 | 0.54 | 0.91 | 0.73 | 0.59 | 0.36 | 0.83 | 0.03 | 0.46 | 0.32 | 0.62 | -0.1 | -0.02 | -0.29 | 0.3298 | 0.712305 | 0.940395 | 0.995437 | 0.997988 | 0.994666 |
| Mtx3_D3YTP3_S311s_1 | Mtx3_D3Y | 0.42 | 0.5 | -0.18 | 0.05 | 0.3 | 0.15 | -0.21 | -0.36 | 0.48 | 0.46 | 0.15 | -0.34 | 0.06 | 0.41 | -0.63 | -0.51 | -0.2275 | -0.01 | -0.365 | 0.287121 | 0.402034 | 0.972726 | 0.989223 | 0.997988 | 0.994666 |
| Mvb12a_Q78HU3_S168s_1 | Mvb12a_Q | -2.23 | -1.58 | -2.47 | -1.29 | -1.14 | -1.74 | -1.01 | -1.57 | -2.03 | -1.57 | -0.88 | -1.74 | -1.37 | -0.77 | -1.88 | -1.79 | 0.5275 | 0.3375 | 0.44 | 0.271345 | 0.138692 | 0.378061 | 0.989223 | 0.997988 | 0.994666 |
| Mvb12b_Q6KAU4_S307s_1 | Mvb12b_Q | 0.03 | 0.14 | 0.18 | 0.05 | 0.57 | -0.17 | 0.28 | 0.08 | -0.21 | 0.17 | 0.46 | -0.03 | 0.63 | 0.59 | 0.18 | -0.11 | 0.09 | -0.0025 | 0.2225 | 0.417217 | 0.700218 | 0.991565 | 0.994893 | 0.997988 | 0.994666 |
| Mxra7_Q9D1T2_S118s_1 | Mxra7_Q9 | 0.23 | 0.49 | 0.42 | 0.1 | 0.43 | 0.21 | 0.41 | -0.27 | 0.76 | 0.25 | 0.53 | 0.11 | 0.48 | 0.92 | 0.83 | 0.79 | -0.115 | 0.1025 | 0.445 | 0.098214 | 0.641697 | 0.680019 | 0.989223 | 0.997988 | 0.994666 |
| Mxra7_Q9D1T2_S79s_1 | Mxra7_Q9 | 1.2 | 2.38 | 1.18 | 1.36 | 2.18 | 1.7 | 1.14 | 0.82 | 1.62 | 1.6 | 2.95 | 0.94 | 1.56 | 2.2 | 1.33 | 0.42 | -0.07 | 0.2475 | -0.1525 | 0.737015 | 0.860921 | 0.610737 | 0.996493 | 0.997988 | 0.9971 |
| Mybbp1a_Q7TPV4_S1164s_1 | Mybbp1a_ | 1.6 | 0.69 | 0.87 | 1.24 | 0.73 | 0.63 | 0.37 | 0.49 | 0.01 | 1.34 | 1.24 | 1.18 | 0.85 | 0.6 | 0.89 | 1.03 | -0.545 | -0.1575 | -0.2575 | 0.385225 | 0.062538 | 0.677766 | 0.989223 | 0.997988 | 0.994666 |
| Mybbp1a_Q7TPV4_S1253s_1 | Mybbp1a_ | 0.92 | 0.37 | 0.8 | 0.92 | 0.44 | 0.87 | -0.38 | -0.14 | 0.23 | 1.31 | -1.91 | 0.86 | 0.78 | 0.62 | 0.12 | 0.8 | -0.555 | -0.63 | -0.1725 | 0.544585 | 0.112376 | 0.350362 | 0.989223 | 0.997988 | 0.994666 |
| Mybbp1a_Q7TPV4_S722s_1 | Mybbp1a_ | -4.34 | -3.87 | -2.43 | -7.38 | -3.4 | -2.65 | -3.22 | -2.08 | -1.49 | -3.3 | -0.33 | -3.05 | -2.21 | -2 | -3.43 | -2.9 | 1.6675 | 2.4625 | 1.87 | 0.074499 | 0.101296 | 0.048299 | 0.989223 | 0.997988 | 0.994666 |
| Mycbp2_F6SMY7_S2298s_1 | Mycbp2_F | -0.08 | 0.16 | 0.08 | 0.36 | 0.63 | 0.13 | 1.2 | 0.74 | 0.01 | -0.15 | 0.21 | 0.34 | -0.16 | 0.24 | 0.4 | 0.71 | 0.545 | -0.0275 | 0.1675 | 0.554233 | 0.073128 | 0.906321 | 0.989223 | 0.997988 | 0.994702 |
| Mycbp2_F6SMY7_S2801s_1 | Mycbp2_F | -1.43 | -1.99 | -1.25 | -1.45 | -1.81 | -0.98 | -0.86 | -1.16 | 1.44 | -1.26 | -0.44 | -0.74 | 0.95 | -3.57 | 0.65 | -0.35 | 0.3275 | 1.28 | 0.95 | 0.313783 | 0.280474 | 0.045311 | 0.989223 | 0.997988 | 0.994666 |
| Mycbp2_F6SMY7_S3893s_1 | Mycbp2_F | 1.17 | 1.23 | 0.77 | 0.33 | 0.64 | 1.08 | 0.57 | 0.56 | 0.96 | 1.01 | -0.46 | 0.51 | 0.25 | 0.95 | 0.29 | 0.82 | -0.1625 | -0.37 | -0.2975 | 0.366041 | 0.56366 | 0.364003 | 0.989223 | 0.997988 | 0.994666 |
| Mycbp2_F6SMY7_T1589t_1 | Mycbp2_F | 0.19 | 0.15 | 1.23 | 1.31 | 0.4 | 0.13 | -0.4 | 1.44 | 1.49 | -0.13 | 1.37 | -1.37 | 0.87 | 0.99 | 0.98 | 1.38 | -0.3275 | -0.38 | 0.335 | 0.364926 | 0.486894 | 0.57832 | 0.989223 | 0.997988 | 0.994666 |
| Mycbp2_F6SMY7_T3882t_1 | Mycbp2_F | 0.8 | 0.38 | 0.63 | 1.41 | 1.44 | -0.02 | 0.79 | 0.92 | 0.83 | 0.82 | 1 | 0.63 | 0.33 | 0.51 | 0.91 | 1.32 | -0.0225 | 0.015 | -0.0375 | 0.913161 | 0.951257 | 0.957568 | 0.999187 | 0.997988 | 0.999507 |
| Myef2_Q8C854_S16s_1 | Myef2_Q8 | -1.45 | -0.04 | -0.34 | -1.58 | -1.96 | -0.88 | -1.17 | -2.83 | 0.3 | -1.04 | -1.14 | -1.49 | 0.15 | -0.41 | 0.19 | 0.85 | -0.8575 | 0.01 | 1.0475 | 0.04155 | 0.131426 | 0.98455 | 0.989223 | 0.997988 | 0.994666 |
| Myef2_Q8C854_S403s_1 | Myef2_Q8 | -0.29 | -0.95 | -0.93 | -0.08 | -0.96 | 0 | -1.28 | 0.44 | -0.56 | -1.11 | 1.1 | -0.63 | 0.22 | -0.15 | -0.02 | -1.08 | 0.1125 | 0.2625 | 0.305 | 0.427216 | 0.795274 | 0.602238 | 0.995762 | 0.997988 | 0.994666 |
| Myh10_Q5SV64_S1970s_1 | Myh10_Q5 | -2.53 | -2.82 | -2.31 | -3.97 | -1.05 | -2.07 | -1.8 | -1.71 | -3.8 | -3.1 | -0.14 | -2.02 | -2.62 | -0.61 | -2.22 | -1.97 | 1.25 | 0.6425 | 1.0525 | 0.073045 | 0.012098 | 0.422169 | 0.989223 | 0.997988 | 0.994666 |
| Myh10_Q5SV64_S1996s_1 | Myh10_Q5 | -0.93 | -2.17 | -2.75 | -1.41 | -2.7 | -2.14 | -1.61 | -1.41 | -1.71 | -1.55 | -1.33 | -0.8 | -0.52 | -1.88 | -1.12 | -1.29 | -0.15 | 0.4675 | 0.6125 | 0.214041 | 0.745494 | 0.29889 | 0.995762 | 0.997988 | 0.994666 |
| Myh10_Q5SV64_S2006s_1 | Myh10_Q5 | 1.62 | 1.06 | 1.64 | -0.03 | -0.06 | 1.62 | 0.27 | 0.8 | -0.75 | 1.58 | -1.07 | 1.15 | -0.09 | -2.31 | 1.33 | 1.53 | -0.415 | -0.845 | -0.9575 | 0.276841 | 0.408513 | 0.244469 | 0.989223 | 0.997988 | 0.994666 |
| Myh10_Q5SV64_T1995t_1 | Myh10_Q5 | -0.93 | -2.17 | -2.75 | -1.41 | -2.7 | -2.14 | -1.61 | -1.41 | -1.71 | -1.55 | -1.33 | -0.8 | -0.52 | -1.88 | -1.12 | -1.29 | -0.15 | 0.4675 | 0.6125 | 0.214041 | 0.745494 | 0.29889 | 0.995762 | 0.997988 | 0.994666 |
| Myh11_E9QPE7_S1954s_1 | Myh11_E9 | -2.28 | -2.13 | -4.25 | -4.37 | -2 | -2.24 | -3.11 | -3.17 | -2.42 | -1.69 | -3.07 | -4.04 | -1.5 | -2.74 | -3.72 | -3.68 | 0.6275 | 0.4525 | 0.3475 | 0.629242 | 0.314489 | 0.528045 | 0.989223 | 0.997988 | 0.995259 |
| Myh9_Q8VDD5_S1943s_1 | Myh9_Q8V | -1.22 | -1.7 | 1.25 | 1.72 | -2.92 | -1.15 | 0.47 | 1.59 | -0.6 | -1.52 | 2.42 | 1.35 | -0.57 | -2.22 | 1.59 | 1.44 | -0.515 | 0.4 | 0.0475 | 0.965003 | 0.648935 | 0.715539 | 0.99033 | 0.997988 | 0.999507 |
| Myl12a_Q6ZWQ9_S20s_1 | Myl12a_Q | -1.85 | -2.2 | -2.11 | -2.1 | -1.35 | -1.96 | -1.13 | -1.7 | -2.1 | -1.57 | -0.54 | -1.45 | -1.74 | -1.2 | -1.78 | -1.97 | 0.53 | 0.65 | 0.3925 | 0.169074 | 0.059799 | 0.088495 | 0.989223 | 0.997988 | 0.994666 |
| Myo18a_E9PUR2_S279s_1 | Myo18a_E | -1.17 | -0.62 | -0.53 | -0.96 | 0.29 | -1.2 | -0.58 | -1.5 | 1.02 | -0.45 | -1.72 | -1.19 | 1.63 | -0.2 | -0.55 | -1.31 | 0.0725 | 0.235 | 0.7125 | 0.243718 | 0.857876 | 0.67934 | 0.996349 | 0.997988 | 0.994666 |
| Myo18a_E9PUR2_S52s_1 | Myo18a_E | 0 | 0.04 | 0.66 | 1 | 0.17 | 0.12 | 0.31 | 0.91 | 0.33 | -0.9 | 1.4 | 1.39 | 0.13 | 0.05 | 1.04 | 0.95 | -0.0475 | 0.13 | 0.1175 | 0.754814 | 0.881754 | 0.814524 | 0.996493 | 0.997988 | 0.9971 |
| Myo18a_E9QA74_S1595s_1 | Myo18a_E | -3.64 | -3.17 | -3.1 | -2.92 | -2.24 | -4.76 | -3.16 | -2.22 | -4.42 | -3.48 | -1.49 | -2.02 | -1.84 | -2.8 | -2.33 | -3.13 | 0.1125 | 0.355 | 0.6825 | 0.076102 | 0.839659 | 0.574722 | 0.995762 | 0.997988 | 0.994666 |
| Myo18a_E9QA74_S472s_1 | Myo18a_E | 0.39 | -2.63 | -1.05 | -1.9 | -1.47 | 0.14 | -1.97 | -1 | -1.2 | -1.45 | 0.39 | -1.09 | -1.5 | -1.39 | -2.07 | -1.97 | 0.2225 | 0.46 | -0.435 | 0.481878 | 0.749889 | 0.51334 | 0.995762 | 0.997988 | 0.994666 |
| Myo18a_Q9JMH9_S164s_1 | Myo18a_Q | -0.27 | -1.55 | -0.54 | -1.32 | -0.84 | -0.12 | 0.7 | -0.35 | -1.65 | -0.29 | 0.49 | 0.44 | -0.88 | -1 | -1.48 | 0.39 | 0.7675 | 0.6675 | 0.1775 | 0.712608 | 0.093889 | 0.238894 | 0.989223 | 0.997988 | 0.996966 |
| Myo18a_Q9JMH9_S340s_1 | Myo18a_Q | 2.62 | 0.95 | 1.27 | 1.73 | 0.25 | 1.43 | 1.05 | 1.69 | 1.71 | 2.91 | 1.16 | 1.08 | 0.99 | 0.98 | 0.97 | 1.19 | -0.5375 | 0.0725 | -0.61 | 0.13443 | 0.249202 | 0.889175 | 0.989223 | 0.997988 | 0.994666 |
| Myo5a_D3YZ62_S1425s_1 | Myo5a_D3 | 0.52 | 0.43 | 0.46 | 0.58 | 0.74 | 0.37 | 0.89 | 0.46 | 0.44 | 0.82 | 0.47 | 0.57 | -0.26 | 0.84 | 0.35 | 0.68 | 0.1175 | 0.0775 | -0.095 | 0.755072 | 0.590892 | 0.719121 | 0.989223 | 0.997988 | 0.9971 |
| Myo5a_D3YZ62_S1625s_1 | Myo5a_D3 | 0.3 | -0.24 | 0.04 | 0.39 | 0.32 | 0.54 | 0.57 | 0.53 | -0.11 | -0.04 | 0.51 | 0.33 | -1.19 | 0.55 | -0.26 | -0.01 | 0.3675 | 0.05 | -0.35 | 0.388207 | 0.133647 | 0.851109 | 0.989223 | 0.997988 | 0.994666 |
| Myo5a_D3YZ62_S600s_1 | Myo5a_D3 | 0.23 | 0.31 | 0.89 | 0.72 | 0.49 | 0.22 | 0.36 | 0.42 | 0.4 | 0.65 | 1.11 | 0.48 | 1.13 | 0.57 | 0.57 | 0.49 | -0.165 | 0.1225 | 0.1525 | 0.599291 | 0.492354 | 0.660844 | 0.989223 | 0.997988 | 0.995259 |
| Myo5a_D3YZ62_T1623t_1 | Myo5a_D3 | 0.56 | 0.4 | 0.06 | 0.27 | 0.9 | 0.11 | 0.31 | 0.45 | 0.75 | 1.03 | 0.12 | 0.38 | -0.02 | 0.93 | -0.19 | 0.2 | 0.12 | 0.2475 | -0.0925 | 0.771064 | 0.636912 | 0.385503 | 0.989223 | 0.997988 | 0.9971 |
| Myo6_E9Q3L1_S1123s_1 | Myo6_E9Q | 0.5 | 0.74 | 0.11 | 0.58 | 1.17 | 0.39 | 0.63 | 0.96 | 0.64 | 0.75 | 0.32 | -0.21 | 0.73 | 0.83 | 0.77 | 0.78 | 0.305 | -0.1075 | 0.295 | 0.255494 | 0.266039 | 0.716846 | 0.989223 | 0.997988 | 0.994666 |
| Myo9a_D3Z3A8_S1259s_1 | Myo9a_D3 | 0.6 | 0.71 | 0.77 | 0.8 | 0.73 | -0.48 | 0.78 | 0.27 | 0.31 | -0.22 | 1.23 | 0.5 | 0.1 | 0.84 | 0.84 | 0.89 | -0.395 | -0.265 | -0.0525 | 0.849363 | 0.226826 | 0.427962 | 0.989223 | 0.997988 | 0.999507 |
| Myo9a_D3Z3A8_S1300s_1 | Myo9a_D3 | -0.17 | -0.26 | 0.57 | 0.55 | -0.08 | 0.07 | 0.67 | 0.86 | -0.02 | -0.15 | 0.53 | 0.61 | -0.43 | -0.55 | 0.71 | 0.21 | 0.2075 | 0.07 | -0.1875 | 0.626772 | 0.533874 | 0.828428 | 0.989223 | 0.997988 | 0.995259 |
| Myo9b_E9QKV6_S1113s_1 | Myo9b_E9 | 1.29 | 2.05 | 0.5 | 0.86 | 2.05 | 1.48 | 0.41 | 0.35 | 1.77 | 1.85 | 0.55 | 0.89 | 0.91 | 2.05 | 0.82 | 1.04 | -0.1025 | 0.09 | 0.03 | 0.944659 | 0.834499 | 0.840506 | 0.995762 | 0.997988 | 0.999507 |
| Myo9b_E9QKV6_S1264s_1 | Myo9b_E9 | 0.81 | 0.65 | 0.38 | -0.01 | 0.35 | 0.38 | 0.25 | 0.55 | -0.09 | 0.99 | -0.51 | 0.73 | 1.12 | 0.12 | 1.25 | 0.82 | -0.075 | -0.1775 | 0.37 | 0.296177 | 0.762886 | 0.653098 | 0.995762 | 0.997988 | 0.994666 |
| Myo9b_E9QKV6_S1335s_1 | Myo9b_E9 | 0.28 | 0.84 | 0.94 | 0.88 | 0.66 | 0.19 | 0.76 | 1.1 | 0.98 | 1.29 | -0.3 | 0.78 | 0.84 | -0.27 | 0.87 | 1.22 | -0.0575 | -0.0475 | -0.07 | 0.851982 | 0.836693 | 0.900877 | 0.995762 | 0.997988 | 0.999507 |
| Myof_E9Q390_S174s_1 | Myof_E9Q | 0.17 | -0.32 | 0.43 | 0.33 | 0.22 | -1.42 | -0.05 | 0.1 | -0.55 | -0.21 | -0.43 | 0.52 | -0.51 | 0.18 | 0.01 | 0.74 | -0.44 | -0.32 | -0.0475 | 0.88926 | 0.28739 | 0.331703 | 0.989223 | 0.997988 | 0.999507 |
| Mypop_Q8R4U1_S182s_1 | Mypop_Q8 | 0.93 | 0.7 | 1.08 | 0.37 | 0.89 | 0.5 | 0.58 | 0.02 | 0.63 | 0.45 | 0.04 | 0 | 0.1 | 0.82 | 0.93 | 0.73 | -0.2725 | -0.49 | -0.125 | 0.680087 | 0.335914 | 0.100189 | 0.989223 | 0.997988 | 0.996159 |
| Myrip_Q8K3I4_S540s_1 | Myrip_Q8K | -0.11 | -0.45 | 0.16 | -0.04 | -0.14 | -0.2 | -0.14 | -0.46 | 0.5 | -0.28 | 0.4 | -0.45 | 0.62 | -0.02 | -0.5 | 0.01 | -0.125 | 0.1525 | 0.1375 | 0.662693 | 0.584758 | 0.620914 | 0.989223 | 0.997988 | 0.995804 |
| Myrip_Q8K3I4_S618s_1 | Myrip_Q8K | 0.59 | 1.31 | 1.32 | 0.86 | 1.04 | 0.67 | 1.21 | 0.43 | 0.8 | 0.67 | 0.6 | 0.94 | 0.24 | 1.75 | 1.6 | 0.91 | -0.1825 | -0.2675 | 0.105 | 0.79207 | 0.525882 | 0.318198 | 0.989223 | 0.997988 | 0.999507 |
| Myt1l_P97500_S391s_1 | Myt1l_P97 | 0.05 | -0.58 | -0.74 | 1.1 | -0.14 | 0.15 | 0.86 | 0.37 | 0.18 | -0.32 | 0.41 | 0.34 | -0.53 | 0.02 | 0.97 | 1.09 | 0.3525 | 0.195 | 0.43 | 0.426921 | 0.430321 | 0.655774 | 0.989223 | 0.997988 | 0.994666 |
| Myt1l_P97500_S470s_1 | Myt1l_P97 | -0.75 | -1.92 | 0.13 | 0.58 | -2.33 | -0.68 | 0.02 | -0.63 | 0.52 | -0.85 | 1.34 | 0.52 | 1.05 | -2.24 | 2.05 | 1.7 | -0.415 | 0.8725 | 1.13 | 0.266581 | 0.533615 | 0.200628 | 0.989223 | 0.997988 | 0.994666 |
| Myt1l_P97500_S685s_1 | Myt1l_P97 | -0.2 | -0.72 | -1.42 | -1.29 | -0.07 | 0.24 | 0.08 | -1.66 | 0.79 | 0.6 | -2.18 | -1.43 | -0.22 | 0.73 | 1.13 | 0.32 | 0.555 | 0.3525 | 1.3975 | 0.006461 | 0.266278 | 0.622597 | 0.989223 | 0.997988 | 0.994666 |
| Myt1l_P97500_T372t_1 | Myt1l_P97 | -0.39 | -0.08 | -0.53 | -0.73 | -0.33 | -0.2 | -0.73 | -1.16 | -0.56 | -0.29 | -1.15 | -0.5 | -0.02 | -0.7 | -1.04 | 0.02 | -0.1725 | -0.1925 | -0.0025 | 0.994012 | 0.552414 | 0.49836 | 0.989223 | 0.997988 | 1 |
| N4bp1_Q6A037_S560s_1 | N4bp1_Q6 | -0.86 | -0.07 | 0.05 | -0.37 | -0.4 | -0.25 | 0.15 | -0.16 | -0.59 | 0.31 | -0.64 | -0.2 | -0.29 | -0.7 | -0.35 | 0.15 | 0.1475 | 0.0325 | 0.015 | 0.962212 | 0.592525 | 0.920415 | 0.989223 | 0.997988 | 0.999507 |
| Nab2_Q61127_S147s_1 | Nab2_Q61 | -2.62 | -3.66 | -2.12 | -1.86 | -3.97 | -4.54 | -1.86 | -2.16 | -1.71 | -2.56 | -0.91 | -2.71 | 1.02 | -4.32 | -3.21 | -2.66 | -0.5675 | 0.5925 | 0.2725 | 0.797631 | 0.413972 | 0.285601 | 0.989223 | 0.997988 | 0.999507 |
| Nab2_Q61127_S6s_1 | Nab2_Q61 | -2.22 | -2.23 | -1.89 | -2.14 | -1.95 | -3.66 | -1.45 | -2.04 | -1.72 | -2.45 | 0.52 | -2.52 | 0.18 | -1.49 | -1.15 | -1.47 | -0.155 | 0.5775 | 1.1375 | 0.018483 | 0.733105 | 0.384636 | 0.995762 | 0.997988 | 0.994666 |
| Nab2_Q61127_T434t_1 | Nab2_Q61 | -0.94 | -0.63 | -1.62 | -2.72 | -0.34 | -0.36 | -1.2 | -1.85 | 0.42 | -0.83 | 0.25 | -2.04 | -0.41 | -0.91 | -2.84 | -1.03 | 0.54 | 0.9275 | 0.18 | 0.777989 | 0.32567 | 0.185819 | 0.989223 | 0.997988 | 0.99798 |
| Nacad_Q5SWP3_S1077s_1 | Nacad_Q5 | -0.61 | -1.79 | -1.55 | -1.69 | -0.33 | -0.74 | -1.6 | -1.7 | -1.77 | -1.27 | -1.25 | -2.29 | -1.11 | -0.62 | -1.08 | -1.13 | 0.3175 | -0.235 | 0.425 | 0.224786 | 0.446424 | 0.533262 | 0.989223 | 0.997988 | 0.994666 |
| Nacad_Q5SWP3_S1214s_1 | Nacad_Q5 | -3.07 | -1.85 | -2.62 | -4.04 | -2.86 | -3.33 | -1.64 | -2.52 | -3.19 | -2.16 | -2.32 | -2.83 | -0.95 | -2.02 | -2.93 | -2.28 | 0.3075 | 0.27 | 0.85 | 0.157816 | 0.563159 | 0.582438 | 0.989223 | 0.997988 | 0.994666 |
| Nacad_Q5SWP3_S1268s_1 | Nacad_Q5 | 0.46 | -0.03 | -0.16 | -1.1 | 0.47 | -1.32 | -0.6 | -1.72 | -1.82 | 0.23 | -0.04 | -1.16 | -0.18 | -1.22 | 0.13 | 0.39 | -0.585 | -0.49 | -0.0125 | 0.978471 | 0.283597 | 0.374903 | 0.989223 | 0.997988 | 1 |
| Nadk_P58058_S46s_1 | Nadk_P58 | 0.06 | -0.41 | -0.68 | -1.29 | 1.02 | -1.28 | -0.36 | -0.64 | -1.87 | 0.08 | -0.93 | -1.09 | 0.23 | 0.87 | -1.67 | -0.78 | 0.265 | -0.3725 | 0.2425 | 0.67471 | 0.607142 | 0.43375 | 0.989223 | 0.997988 | 0.996159 |
| Nadk_P58058_S48s_1 | Nadk_P58 | -0.33 | -1.18 | -0.54 | -0.55 | -0.3 | 0.17 | 0.28 | 0.28 | 0.18 | -0.47 | 0.14 | -0.14 | -0.39 | -0.15 | 0.41 | 0.21 | 0.7575 | 0.5775 | 0.67 | 0.053732 | 0.018574 | 0.068193 | 0.989223 | 0.997988 | 0.994666 |
| Nans_Q99J77_S275s_1 | Nans_Q99J | 0.72 | 0.17 | 0.65 | 0.39 | -0.04 | 0.83 | -0.66 | -0.23 | 1.87 | 0.52 | -1.51 | 0.12 | 2.02 | -0.09 | -0.12 | -0.17 | -0.5075 | -0.2325 | -0.0725 | 0.888519 | 0.162183 | 0.71842 | 0.989223 | 0.997988 | 0.999507 |
| Napa_Q9DB05_S26s_1 | Napa_Q9D | 0.38 | 0.34 | -0.28 | -0.16 | 1 | -0.27 | 0.45 | -0.76 | -3.01 | -0.61 | -7.15 | -0.78 | -1.34 | -3.34 | -0.27 | -0.24 | 0.035 | -2.9575 | -1.3675 | 0.065937 | 0.931203 | 0.05057 | 0.998075 | 0.997988 | 0.994666 |
| Napepld_Q8BH82_S27s_1 | Napepld_Q | -1.38 | -1.37 | -0.31 | -0.9 | -0.84 | 0.23 | -1.9 | -0.38 | 0.29 | -1.21 | -1.4 | -0.01 | 1.52 | -1.65 | -0.79 | -0.3 | 0.2675 | 0.4075 | 0.685 | 0.303859 | 0.577672 | 0.395967 | 0.989223 | 0.997988 | 0.994666 |
| Napg_Q9CWZ7_S284s_1 | Napg_Q9C | -1.03 | -0.52 | 0.48 | -0.14 | -0.79 | -1.02 | 0.61 | 0.75 | -0.63 | -0.49 | 0.52 | 1.28 | -0.01 | -0.92 | -0.05 | 0.59 | 0.19 | 0.4725 | 0.205 | 0.640803 | 0.711037 | 0.371211 | 0.995437 | 0.997988 | 0.995259 |
| Napg_Q9CWZ7_T287t_1 | Napg_Q9C | -2.33 | -1.44 | -1.46 | -1.45 | -2.06 | -2.1 | -1.2 | -1.34 | -1.14 | -1.65 | -2.53 | -1.54 | 0.3 | -2.19 | -1.4 | -1.31 | -0.005 | -0.045 | 0.52 | 0.338462 | 0.98797 | 0.90394 | 0.999187 | 0.997988 | 0.994666 |
| Nasp_B1AU75_S712s_1 | Nasp_B1AU | -3.92 | -3.6 | -3.28 | -8.12 | -3.72 | -3.3 | -2.79 | -4.52 | -1.95 | -3.04 | -2.86 | -3.51 | -0.73 | -3.42 | -2.84 | -3.52 | 1.1475 | 1.89 | 2.1025 | 0.089226 | 0.282347 | 0.096472 | 0.989223 | 0.997988 | 0.994666 |
| Nav1_Q8CH77_S1010s_1 | Nav1_Q8C | -1.37 | -0.22 | -0.26 | -0.4 | -0.42 | -1.39 | 0.21 | -0.46 | -0.91 | -0.72 | -1.02 | -0.04 | -0.65 | 0.23 | -0.61 | -0.27 | 0.0475 | -0.11 | 0.2375 | 0.517697 | 0.907373 | 0.760923 | 0.997038 | 0.997988 | 0.994666 |
| Nav1_Q8CH77_S1251s_1 | Nav1_Q8C | -0.21 | 0 | 0.45 | 0.23 | 0.47 | -0.07 | 0.73 | 0.48 | -0.06 | -0.04 | -0.47 | 0.58 | -0.1 | 0.09 | 0.4 | 0.56 | 0.285 | -0.115 | 0.12 | 0.672153 | 0.298753 | 0.70172 | 0.989223 | 0.997988 | 0.996159 |
| Nav1_Q8CH77_S291s_1 | Nav1_Q8C | -0.2 | 0.16 | -1.18 | -0.17 | 0.47 | -0.35 | -0.08 | 0.08 | 0.81 | 0.35 | -0.07 | -0.81 | -0.85 | 0.92 | -0.38 | -0.95 | 0.3775 | 0.4175 | 0.0325 | 0.947285 | 0.285176 | 0.350715 | 0.989223 | 0.997988 | 0.999507 |
| Nav1_Q8CH77_S454s_1 | Nav1_Q8C | -1.28 | -1.24 | -0.55 | -1.81 | -0.23 | -0.94 | -1.4 | -1.84 | -1.02 | -1.66 | 1.03 | -3.08 | -1.42 | -0.47 | -3.13 | -1.53 | 0.1175 | 0.0375 | -0.4175 | 0.464239 | 0.775128 | 0.962423 | 0.995762 | 0.997988 | 0.994666 |
| Nav1_Q8CH77_S455s_1 | Nav1_Q8C | -3.13 | -4.51 | -3.37 | -5.14 | -2.63 | -1.96 | -2.59 | -4.11 | -2.88 | -2.77 | -0.36 | -3.14 | -1.23 | -0.85 | -2.6 | -2.69 | 1.215 | 1.75 | 2.195 | 0.004987 | 0.063503 | 0.035357 | 0.989223 | 0.997988 | 0.994666 |
| Nav1_Q8CH77_S510s_1 | Nav1_Q8C | -0.17 | -0.18 | -0.44 | -1.3 | -1.49 | -0.44 | -1.94 | -1.11 | 2.74 | -0.2 | 0.79 | -1.12 | 2.92 | -0.33 | -0.41 | -2.16 | -0.7225 | 1.075 | 0.5275 | 0.583053 | 0.095735 | 0.189534 | 0.989223 | 0.997988 | 0.995259 |
| Nav1_Q8CH77_S544s_1 | Nav1_Q8C | -0.02 | -0.26 | -0.37 | -0.48 | -0.26 | 0.27 | 0.14 | -0.07 | -0.05 | -0.15 | -0.84 | 0.25 | -0.21 | 0.16 | -0.35 | -0.44 | 0.3025 | 0.085 | 0.0725 | 0.782383 | 0.207469 | 0.772574 | 0.989223 | 0.997988 | 0.998226 |
| Nav1_Q8CH77_S762s_1 | Nav1_Q8C | -0.41 | 0.35 | -0.25 | -0.44 | 0.05 | 0.44 | -0.59 | -0.3 | -0.26 | 0.4 | 0.22 | -0.2 | 0.07 | 0.36 | -0.04 | -0.34 | 0.0875 | 0.2275 | 0.2 | 0.506417 | 0.777204 | 0.440181 | 0.995762 | 0.997988 | 0.994666 |
| Nav1_Q8CH77_S810s_1 | Nav1_Q8C | -1.37 | -0.96 | -0.56 | -1.47 | -1.06 | -1.07 | -0.6 | -0.49 | -0.7 | -0.55 | -1.13 | -0.76 | -0.12 | -1.38 | -0.66 | -1.05 | 0.285 | 0.305 | 0.2875 | 0.43776 | 0.336199 | 0.305236 | 0.989223 | 0.997988 | 0.994666 |
| Nav1_Q8CH77_S93s_1 | Nav1_Q8C | -1.47 | -0.89 | -1.07 | -1.25 | -0.94 | -1.64 | -0.41 | -0.93 | -1.61 | -1.1 | -0.38 | -0.77 | -1.1 | -0.85 | -1.06 | -1.1 | 0.19 | 0.205 | 0.1425 | 0.573174 | 0.536013 | 0.524437 | 0.989223 | 0.997988 | 0.995259 |
| Nav1_Q8CH77_S952s_1 | Nav1_Q8C | -0.5 | 0.24 | -0.67 | -0.8 | -0.45 | 0.22 | -0.54 | -0.72 | -0.4 | -0.09 | 0.96 | 0.5 | 0.14 | 0.11 | -0.46 | -0.36 | 0.06 | 0.675 | 0.29 | 0.38177 | 0.852798 | 0.10587 | 0.996349 | 0.997988 | 0.994666 |
| Nav1_Q8CH77_S998s_1 | Nav1_Q8C | -0.01 | 0 | -0.82 | 0.44 | 0.08 | -0.14 | 0.66 | 0.67 | -0.24 | -0.29 | 1.21 | 0.81 | -0.58 | 0.03 | 0.28 | 0.34 | 0.415 | 0.47 | 0.115 | 0.750385 | 0.240792 | 0.304881 | 0.989223 | 0.997988 | 0.9971 |
| Nav1_Q8CH77_T360t_1 | Nav1_Q8C | -1.81 | -1.95 | -0.86 | -1.13 | -1.57 | -1.49 | -1.29 | -1.38 | -0.58 | -2.11 | -1.36 | -0.95 | 0.44 | -1.5 | -0.16 | -0.33 | 0.005 | 0.1875 | 1.05 | 0.045982 | 0.986531 | 0.651991 | 0.999187 | 0.997988 | 0.994666 |
| Nav2_V9GXT3_S1240s_1 | Nav2_V9G | 0.52 | 0.03 | 0.18 | 0.06 | 0.6 | 0.26 | 0.36 | 0.31 | -1.01 | 0.68 | -0.23 | 0.43 | -0.4 | -0.55 | 0.87 | 1.01 | 0.185 | -0.23 | 0.035 | 0.934003 | 0.411701 | 0.562992 | 0.989223 | 0.997988 | 0.999507 |
| Nav3_E9QMF5_S1539s_1 | Nav3_E9Q | -0.96 | -0.7 | 1.03 | -0.12 | 0.07 | -0.22 | 0.04 | -0.04 | 1.04 | 0.07 | 1.73 | 0.91 | 1.63 | 0.07 | 1.08 | 0.33 | 0.15 | 1.125 | 0.965 | 0.094104 | 0.723858 | 0.054908 | 0.995437 | 0.997988 | 0.994666 |
| Nav3_E9QMF5_S1848s_1 | Nav3_E9Q | -0.77 | -1.18 | 0.75 | 0.26 | -1.36 | 0.61 | 0.71 | -1.19 | 0.69 | 0.05 | -0.17 | 0.87 | 1.35 | -1.01 | -0.75 | -0.26 | -0.0725 | 0.595 | 0.0675 | 0.914326 | 0.909282 | 0.239072 | 0.997335 | 0.997988 | 0.999507 |
| Nav3_E9QMF5_S275s_1 | Nav3_E9Q | -0.04 | 0.13 | -0.96 | -1.07 | 0.12 | 0.32 | -0.25 | -0.29 | -0.28 | 0.6 | -0.08 | -0.48 | 0.34 | 0.62 | -0.55 | -0.48 | 0.46 | 0.425 | 0.4675 | 0.285085 | 0.20472 | 0.290774 | 0.989223 | 0.997988 | 0.994666 |
| Nav3_E9QMF5_S354s_1 | Nav3_E9Q | -0.47 | -0.8 | -1.66 | -0.13 | 0.18 | 0.51 | -0.12 | -2.54 | 0.16 | -1.18 | 1.02 | 0 | -0.63 | -0.57 | -0.78 | -0.42 | 0.2725 | 0.765 | 0.165 | 0.64916 | 0.689213 | 0.167291 | 0.994672 | 0.997988 | 0.995259 |
| Nbas_E9Q411_S472s_1 | Nbas_E9Q | 0.39 | 0.33 | 0.83 | 0.06 | -0.23 | -0.22 | 0.18 | 0.54 | -0.05 | 0.64 | 0.22 | 0.42 | -0.56 | -0.33 | 1.37 | 1.4 | -0.335 | -0.095 | 0.0675 | 0.896514 | 0.249244 | 0.728659 | 0.989223 | 0.997988 | 0.999507 |
| Nbas_E9Q411_S474s_1 | Nbas_E9Q | 0.27 | -0.47 | 0.74 | 0.24 | -0.32 | -1.23 | 0.29 | -0.24 | 0.15 | 0.49 | 1.45 | 0.09 | 0.09 | -0.76 | 0.43 | 1.08 | -0.57 | 0.35 | 0.015 | 0.97315 | 0.166999 | 0.390855 | 0.989223 | 0.997988 | 0.999507 |
| Nbea_Q9EPN1_S1311s_1 | Nbea_Q9E | -1.16 | -0.66 | -0.53 | -0.4 | -0.39 | -1.13 | -0.5 | -0.22 | -1.14 | -0.89 | 0.2 | -0.45 | -0.26 | -0.12 | -0.77 | -0.61 | 0.1275 | 0.1175 | 0.2475 | 0.4059 | 0.660986 | 0.739203 | 0.990434 | 0.997988 | 0.994666 |
| Nbea_Q9EPN1_S1519s_1 | Nbea_Q9E | -0.11 | 0.41 | 0.32 | 0.08 | 0.48 | -0.14 | 0.67 | -0.08 | 0.23 | 0.43 | -0.26 | -0.1 | 0.06 | 0.86 | 0.39 | 0.11 | 0.0575 | -0.1 | 0.18 | 0.536929 | 0.833608 | 0.702634 | 0.995762 | 0.997988 | 0.994666 |
| Nbea_Q9EPN1_S1596s_1 | Nbea_Q9E | -1.54 | -2.15 | -0.96 | -0.04 | -1.11 | -2.01 | -0.13 | -0.59 | -1.67 | -1.32 | -1.37 | -0.81 | -1.56 | -0.57 | -0.93 | -0.65 | 0.2125 | -0.12 | 0.245 | 0.611868 | 0.697983 | 0.795295 | 0.994893 | 0.997988 | 0.995259 |
| Nbea_Q9EPN1_S1724s_1 | Nbea_Q9E | -1.41 | -1.97 | -1.95 | -1.81 | -0.26 | -1.17 | -1.98 | -1.75 | -1.91 | -1.68 | -1.13 | -2.13 | -1.32 | 0.78 | -2.31 | -3.32 | 0.495 | 0.0725 | 0.2425 | 0.75717 | 0.226006 | 0.805375 | 0.989223 | 0.997988 | 0.9971 |
| Nbea_Q9EPN1_S2565s_1 | Nbea_Q9E | -1.24 | -1.24 | -1.25 | -1.16 | -0.85 | -0.84 | -0.76 | -1.01 | -1.25 | -1.3 | -1.54 | -1.62 | -1.39 | -1.18 | -1.33 | -1.25 | 0.3575 | -0.205 | -0.065 | 0.774176 | 0.092409 | 0.352405 | 0.989223 | 0.997988 | 0.99798 |
| Ncam1_A0A0A6YY91_S1019s_1 | Ncam1_A0 | 1.03 | 0.45 | 0.39 | 0.74 | 0.45 | 0.4 | 0.68 | 0.55 | 0.19 | 0.47 | -0.19 | 0.23 | -0.18 | 0.81 | -0.11 | 0.4 | -0.1325 | -0.4775 | -0.4225 | 0.209605 | 0.571978 | 0.096573 | 0.989223 | 0.997988 | 0.994666 |
| Ncam1_P13595_S1005s_1 | Ncam1_P1 | 0.51 | 0.98 | 0.75 | 0.9 | 0.72 | 0.22 | 0.57 | 0.64 | 0.56 | 0.91 | 1.4 | 0.57 | 1.01 | 0.86 | 0.77 | 0.52 | -0.2475 | 0.075 | 0.005 | 0.984279 | 0.293713 | 0.786621 | 0.989223 | 0.997988 | 1 |
| Ncam1_P13595_S1016s_1 | Ncam1_P1 | -0.27 | 0.05 | 0.92 | 0.63 | 0.2 | 0.28 | 0.7 | 0.02 | 0.39 | 0.95 | 0.88 | 0.16 | 0.02 | 1.99 | 1.54 | 1.37 | -0.0325 | 0.2625 | 0.8975 | 0.087134 | 0.919217 | 0.457336 | 0.997335 | 0.997988 | 0.994666 |
| Ncam1_P13595_S774s_1 | Ncam1_P1 | 2.63 | 2.03 | 3.53 | 2.84 | 1.83 | 2.3 | 3.24 | 3.42 | 1.7 | 3.01 | 1.73 | 3.61 | 0.88 | 1.03 | 3.4 | 3.35 | -0.06 | -0.245 | -0.5925 | 0.397145 | 0.895297 | 0.645454 | 0.996493 | 0.997988 | 0.994666 |
| Ncam1_P13595_S817s_1 | Ncam1_P1 | 0.66 | 1.27 | 1.13 | 1 | 1.39 | 0.99 | 1.03 | 1.25 | 0.31 | 0.71 | 1.41 | 1.5 | 1.03 | 1.46 | 1.6 | 1.19 | 0.15 | -0.0325 | 0.305 | 0.276509 | 0.524581 | 0.92279 | 0.989223 | 0.997988 | 0.994666 |
| Ncam1_P13595_S887s_1 | Ncam1_P1 | 1.05 | 1.39 | 0.62 | 0.57 | 1 | 0.79 | 0.13 | 0.43 | 0.76 | 1.08 | 0.7 | 0.7 | 1.06 | 1.27 | 0.94 | 0.28 | -0.32 | -0.0975 | -0.02 | 0.951586 | 0.298727 | 0.720224 | 0.989223 | 0.997988 | 0.999507 |
| Ncam1_P13595_S946s_1 | Ncam1_P1 | 1 | 0.27 | -0.01 | 0.44 | 1.24 | 1.03 | 0.96 | 1.05 | 0.58 | 1.12 | 0.06 | 0.34 | -0.21 | 1.09 | 0.36 | 0.49 | 0.645 | 0.1 | 0.0075 | 0.983546 | 0.034386 | 0.764214 | 0.989223 | 0.997988 | 1 |
| Ncam1_P13595_S946s_2 | Ncam1_P1 | 1.16 | -0.29 | -0.72 | 0.74 | 1.84 | 0.97 | 1.17 | 0.21 | -0.1 | 0.87 | 0.53 | 0.53 | 0.65 | 1.21 | -0.11 | -0.35 | 0.825 | 0.235 | 0.1275 | 0.809213 | 0.12715 | 0.612894 | 0.989223 | 0.997988 | 0.999507 |
| Ncam1_P13595_S958s_1 | Ncam1_P1 | 0.54 | 0.43 | 0.49 | 0.53 | 0.77 | 0.3 | 0.8 | 0.62 | 0.34 | 0.73 | 0.03 | 0.38 | 0.01 | 0.77 | 0.38 | 0.33 | 0.125 | -0.1275 | -0.125 | 0.631469 | 0.562008 | 0.591174 | 0.989223 | 0.997988 | 0.995259 |
| Ncam1_P13595_S958s_2 | Ncam1_P1 | 1.16 | -0.29 | -0.72 | 0.74 | 1.84 | 0.97 | 1.17 | 0.21 | -0.1 | 0.87 | 0.53 | 0.53 | 0.65 | 1.21 | -0.11 | -0.35 | 0.825 | 0.235 | 0.1275 | 0.809213 | 0.12715 | 0.612894 | 0.989223 | 0.997988 | 0.999507 |
| Ncam1_P13595_T1030t_1 | Ncam1_P1 | -0.96 | -0.81 | -0.09 | 0.13 | -0.42 | -1.08 | 0.48 | 0.06 | -1.54 | -0.83 | -0.08 | -0.11 | -0.74 | -0.34 | 0.15 | -0.09 | 0.1925 | -0.2075 | 0.1775 | 0.619235 | 0.63993 | 0.628782 | 0.989223 | 0.997988 | 0.995259 |
| Ncam1_P13595_T783t_1 | Ncam1_P1 | -3.63 | -2.45 | 1.15 | 1.07 | -2.32 | -5.34 | 0.36 | 0.99 | -4.67 | -2.25 | 0.87 | 0.96 | -3.19 | -2.58 | 1.14 | 0.81 | -0.6125 | -0.3075 | 0.01 | 0.994401 | 0.705952 | 0.846858 | 0.995437 | 0.997988 | 1 |
| Ncam1_P13595_T783t_1_1 | Ncam1_P1 | -3.63 | -2.45 | 0.99 | 0.88 | -2.32 | -5.34 | 0.18 | 0.81 | -4.67 | -2.25 | 0.61 | 0.84 | -3.19 | -2.58 | 1.24 | 0.67 | -0.615 | -0.315 | 0.0875 | 0.949951 | 0.694687 | 0.837097 | 0.994893 | 0.997988 | 0.999507 |
| Ncam1_P13595_T788t_1 | Ncam1_P1 | -0.23 | -0.16 | 0.2 | 0.09 | -1.17 | 0.52 | -0.49 | -0.65 | 3.01 | 0.33 | 0.05 | 0.6 | 2.7 | -1.21 | 1.06 | -0.32 | -0.4225 | 1.0225 | 0.5825 | 0.451869 | 0.263764 | 0.128895 | 0.989223 | 0.997988 | 0.994666 |
| Ncam1_P13595_T788t_1_1 | Ncam1_P1 | -0.23 | -0.16 | 0.2 | 0.09 | -1.17 | 0.52 | -0.49 | -0.65 | 3.01 | 0.33 | 0.05 | 0.6 | 2.7 | -1.21 | 1.06 | -0.32 | -0.4225 | 1.0225 | 0.5825 | 0.451869 | 0.263764 | 0.128895 | 0.989223 | 0.997988 | 0.994666 |
| Ncam1_P13595_T803t_1 | Ncam1_P1 | 5.5 | 2.87 | 5.58 | 1.54 | 2.34 | 5.77 | 6.61 | 4.54 | 1.11 | 5.06 | -0.1 | 5.91 | -0.14 | 1.27 | 6.21 | 6.55 | 0.9425 | -0.8775 | -0.4 | 0.813293 | 0.430228 | 0.574764 | 0.989223 | 0.997988 | 0.999507 |
| Ncam2_O35136_S765s_1 | Ncam2_O3 | 5.98 | 5.37 | 6.26 | 4.69 | 5.49 | 6.16 | 5.58 | 5.99 | 1.62 | 6.21 | 2.87 | 6.38 | 0.23 | 1.69 | 6.74 | 6.91 | 0.23 | -1.305 | -1.6825 | 0.280151 | 0.544908 | 0.25286 | 0.989223 | 0.997988 | 0.994666 |
| Ncam2_O35136_S786s_1 | Ncam2_O3 | 0.03 | 0.65 | -1.87 | -1.23 | 0.93 | 0.15 | -0.42 | -1.59 | 0.42 | 0.43 | -2.11 | -0.75 | 0.17 | 0.89 | -0.08 | -1.21 | 0.3725 | 0.1025 | 0.5475 | 0.409396 | 0.592318 | 0.890894 | 0.989223 | 0.997988 | 0.994666 |
| Ncbp1_Q3UYV9_S7s_1 | Ncbp1_Q3 | 3.83 | -0.29 | 1.55 | -7.65 | -0.41 | 4.07 | 0.67 | 1.09 | -4.96 | 1.22 | -2.36 | 1.35 | 1.05 | -4.44 | 0.64 | 0.85 | 1.995 | -0.5475 | 0.165 | 0.945146 | 0.389331 | 0.828683 | 0.989223 | 0.997988 | 0.999507 |
| Ncdn_Q9Z0E0_S566s_1 | Ncdn_Q9Z | 0.62 | 1 | -0.44 | -0.2 | 0.4 | -0.06 | 0.07 | -0.36 | -0.33 | 0.96 | -0.36 | -0.41 | 0.25 | 1.54 | -0.36 | -0.33 | -0.2325 | -0.28 | 0.03 | 0.954222 | 0.532594 | 0.542793 | 0.989223 | 0.997988 | 0.999507 |
| Ncf2_O70145_S332s_1 | Ncf2_O701 | 0.28 | -0.1 | 0.16 | -0.31 | -0.52 | -0.22 | -1.2 | -2.1 | -0.24 | -0.6 | 0.79 | 0 | 0.08 | 0.07 | 0.02 | -1.51 | -1.0175 | -0.02 | -0.3425 | 0.417086 | 0.033446 | 0.953307 | 0.989223 | 0.997988 | 0.994666 |
| Nckap1_P28660_S337s_1 | Nckap1_P2 | -0.37 | 0.17 | -0.49 | -0.38 | 0.28 | -0.4 | -0.58 | -0.68 | 1.27 | -0.38 | 0.42 | -0.44 | 1.28 | -0.51 | -0.4 | -0.72 | -0.0775 | 0.485 | 0.18 | 0.700346 | 0.79056 | 0.26575 | 0.995762 | 0.997988 | 0.996159 |
| Nckap5l_Q6GQX2_S1284s_1 | Nckap5l_Q | -1.43 | 0.22 | -0.46 | -1.54 | -1.86 | -1.29 | -1.34 | -1.83 | -0.09 | -0.69 | 1.11 | -0.74 | 0.82 | -0.55 | -0.98 | -1.9 | -0.7775 | 0.7 | 0.15 | 0.813884 | 0.091229 | 0.227963 | 0.989223 | 0.997988 | 0.999507 |
| Nckipsd_Q9ESJ4_S122s_1 | Nckipsd_Q | -0.05 | 0.02 | 0.09 | 0.54 | -0.08 | -0.1 | -0.13 | 0.31 | 0.41 | 0.21 | 0.37 | 0.15 | 0.76 | -0.13 | 0.39 | 0.12 | -0.15 | 0.135 | 0.135 | 0.650605 | 0.531267 | 0.571119 | 0.989223 | 0.997988 | 0.995259 |
| Nckipsd_Q9ESJ4_T174t_1 | Nckipsd_Q | -0.4 | -0.44 | -0.2 | 0.24 | 0.2 | -0.6 | -0.12 | 0.46 | -1.03 | -1.09 | -0.24 | -0.05 | -0.24 | 0.05 | -0.06 | 0.23 | 0.185 | -0.4025 | 0.195 | 0.478428 | 0.542323 | 0.245118 | 0.989223 | 0.997988 | 0.994666 |
| Ncl_P09405_S303s_1 | Ncl_P0940 | -1.39 | -0.74 | -1.79 | -2.55 | -0.61 | -0.7 | -0.78 | -2.01 | -0.79 | -0.51 | -1.65 | -1.97 | -1.41 | -0.75 | -0.77 | -1.9 | 0.5925 | 0.3875 | 0.41 | 0.378318 | 0.223638 | 0.43314 | 0.989223 | 0.997988 | 0.994666 |
| Ncl_P09405_S403s_1 | Ncl_P0940 | -2.55 | -1.97 | -2.3 | -1.62 | -0.77 | -2.61 | -1.37 | -1.82 | -2.87 | -2.33 | -2.35 | -1.63 | -2.12 | -0.98 | -2.04 | -1.97 | 0.4675 | -0.185 | 0.3325 | 0.366779 | 0.277478 | 0.592801 | 0.989223 | 0.997988 | 0.994666 |
| Ncoa1_P70365_S570s_1 | Ncoa1_P7 | -4.01 | -2.85 | -4.3 | -5 | -5.06 | -4.01 | -4.02 | -4.45 | -2.22 | -3.51 | -3.43 | -3.7 | -0.49 | -6.79 | -3.88 | -3.61 | -0.345 | 0.825 | 0.3475 | 0.768386 | 0.472363 | 0.140405 | 0.989223 | 0.997988 | 0.9971 |
| Ncoa2_Q61026_S493s_1 | Ncoa2_Q6 | -2.99 | -2 | -2.62 | -3.72 | -1.53 | -2.65 | -2.94 | -3.82 | -2.42 | -1.59 | -1.32 | -2.92 | -0.71 | -1.4 | -2.86 | -2.64 | 0.0975 | 0.77 | 0.93 | 0.130508 | 0.856175 | 0.139899 | 0.996349 | 0.997988 | 0.994666 |
| Ncoa2_Q61026_S493s_2 | Ncoa2_Q6 | -2.73 | -2.43 | -0.97 | -0.2 | -2.52 | -2.96 | -0.84 | -0.11 | -1.57 | -2.01 | 0.46 | -0.4 | -0.92 | -2.05 | 0.59 | -0.63 | -0.025 | 0.7025 | 0.83 | 0.266901 | 0.974751 | 0.353374 | 0.999187 | 0.997988 | 0.994666 |
| Ncoa2_Q61026_S499s_2 | Ncoa2_Q6 | -2.73 | -2.43 | -0.97 | -0.2 | -2.52 | -2.96 | -0.84 | -0.11 | -1.57 | -2.01 | 0.46 | -0.4 | -0.92 | -2.05 | 0.59 | -0.63 | -0.025 | 0.7025 | 0.83 | 0.266901 | 0.974751 | 0.353374 | 0.999187 | 0.997988 | 0.994666 |
| Ncoa2_Q61026_S565s_1 | Ncoa2_Q6 | 0.09 | 0.09 | -0.91 | -0.18 | 0.23 | -0.21 | -0.43 | -0.7 | 0.23 | 0.63 | -0.4 | -0.58 | -0.34 | 0.15 | -0.93 | -0.64 | -0.05 | 0.1975 | -0.2125 | 0.554924 | 0.876354 | 0.59941 | 0.996493 | 0.997988 | 0.99525 |
| Ncoa5_B7ZC24_S126s_1 | Ncoa5_B7Z | 0.23 | -1.02 | -0.29 | -0.88 | -2.3 | 0.52 | -1.18 | 0.04 | 1.75 | -0.83 | -3.01 | -0.42 | 2.13 | -2.55 | 1.86 | -0.45 | -0.24 | -0.1375 | 0.7375 | 0.462264 | 0.699765 | 0.878743 | 0.994893 | 0.997988 | 0.994666 |
| Ncoa5_B7ZC24_S162s_1 | Ncoa5_B7Z | -0.69 | -0.33 | -0.61 | -1.34 | -0.92 | -0.37 | -1.49 | -1.54 | -0.83 | -1.69 | -2.39 | -1.12 | 0.45 | -2.19 | -0.44 | -1.26 | -0.3375 | -0.765 | -0.1175 | 0.83315 | 0.347677 | 0.083233 | 0.989223 | 0.997988 | 0.999507 |
| Ncoa5_B7ZC24_S34s_1 | Ncoa5_B7Z | 2.27 | 0.58 | 2.12 | 2.16 | -0.04 | 1.28 | 0.52 | 1.65 | 2.21 | 1.57 | 0.83 | 1 | 1.68 | 0.56 | 2.46 | 1.89 | -0.93 | -0.38 | -0.135 | 0.798561 | 0.091365 | 0.439006 | 0.989223 | 0.997988 | 0.999507 |
| Ncoa5_B7ZC24_S378s_1 | Ncoa5_B7Z | 0.76 | 0.33 | 0.97 | 0.63 | -0.14 | 0.52 | 0.05 | 0.88 | 0.37 | 0.69 | 1.46 | 0.83 | 0.67 | 1.04 | 0.49 | -0.33 | -0.345 | 0.165 | -0.205 | 0.560743 | 0.25864 | 0.588946 | 0.989223 | 0.997988 | 0.995259 |
| Ncoa5_B7ZC24_S6s_1 | Ncoa5_B7Z | -0.02 | 0.52 | 0.19 | -0.74 | -0.57 | 0.93 | -1.47 | -1.45 | 2.05 | 0.65 | 1.13 | -0.3 | 1.76 | 0.74 | -0.02 | -0.36 | -0.6275 | 0.895 | 0.5425 | 0.301433 | 0.280454 | 0.112342 | 0.989223 | 0.997988 | 0.994666 |
| Ncoa5_B7ZC24_S9s_1 | Ncoa5_B7Z | -0.62 | 0.12 | 0.06 | 0.42 | -0.45 | 0.04 | -0.36 | -0.81 | 1.19 | 0.09 | -1.08 | -0.3 | 1.26 | -0.97 | 0.39 | 0.58 | -0.39 | -0.02 | 0.32 | 0.51894 | 0.218739 | 0.967537 | 0.989223 | 0.997988 | 0.994666 |
| Ncoa5_Q91W39_S377s_1 | Ncoa5_Q9 | 0.86 | 0.64 | -0.2 | 0.06 | 0.25 | -0.44 | 0.35 | -0.09 | -0.12 | -0.22 | 0 | 0.14 | 0.22 | 1.04 | 0.23 | -0.06 | -0.3225 | -0.39 | 0.0175 | 0.96169 | 0.326654 | 0.212995 | 0.989223 | 0.997988 | 0.999507 |
| Ncoa7_B9EIW5_S342s_1 | Ncoa7_B9E | 1.55 | 0.31 | 0.97 | 1.08 | -0.4 | 1.59 | -0.45 | 0.66 | 1.99 | 1.6 | 0.9 | 0.65 | 1.24 | -0.24 | 0.93 | 0.64 | -0.6275 | 0.3075 | -0.335 | 0.422995 | 0.231457 | 0.448596 | 0.989223 | 0.997988 | 0.994666 |
| Ncor1_Q5RIM6_S1123s_1 | Ncor1_Q5 | -2.85 | -4.42 | -3.72 | -3.08 | -3.14 | -3.63 | -4.41 | -3.31 | -3.31 | -2.27 | -2.01 | -2.67 | -0.31 | -2.05 | -2.67 | -2.76 | -0.105 | 0.9525 | 1.57 | 0.026707 | 0.80576 | 0.05334 | 0.995762 | 0.997988 | 0.994666 |
| Ncor1_Q5RIM6_S1334s_1 | Ncor1_Q5 | -1.2 | -0.98 | -1.96 | -2.37 | -1.72 | -1.45 | -1.76 | -3.28 | -2.92 | -1.18 | -1.4 | -2.3 | 0.22 | -1.87 | -1.56 | -2.55 | -0.425 | -0.3225 | 0.1875 | 0.760155 | 0.390962 | 0.515294 | 0.989223 | 0.997988 | 0.9971 |
| Ncor1_Q5RIM6_S2117s_1 | Ncor1_Q5 | -0.06 | -1.13 | -0.47 | -0.93 | 0.38 | 0.24 | -1.19 | -0.44 | 0.96 | -0.33 | -1.86 | -0.02 | -0.14 | 0.41 | 0.39 | -1.35 | 0.395 | 0.335 | 0.475 | 0.316956 | 0.350435 | 0.567754 | 0.989223 | 0.997988 | 0.994666 |
| Ncor1_Q5RIM6_S2135s_1 | Ncor1_Q5 | -0.77 | -0.58 | -0.2 | -0.61 | -0.38 | -1.3 | 0.19 | -0.12 | -1.34 | -0.47 | -1.15 | 0.34 | -0.5 | -0.35 | -0.73 | -0.56 | 0.1375 | -0.115 | 0.005 | 0.984238 | 0.692551 | 0.772764 | 0.994893 | 0.997988 | 1 |
| Ncor2_E9PY55_S1818s_1 | Ncor2_E9P | -2.28 | -2.45 | -1.91 | -5.27 | -2.47 | -4.4 | -2.9 | -2.94 | -0.93 | -2.83 | -1.99 | -1.6 | 0.34 | -2.79 | -2.18 | -2.95 | -0.2 | 1.14 | 1.0825 | 0.270383 | 0.795365 | 0.166525 | 0.995762 | 0.997988 | 0.994666 |
| Ncor2_E9PY55_S2211s_1 | Ncor2_E9P | -1.05 | -1.27 | -0.23 | -0.83 | -0.88 | -0.93 | -0.54 | -0.83 | -1.26 | -1.44 | -0.71 | -0.23 | -0.32 | -0.54 | -1.64 | -0.95 | 0.05 | -0.065 | -0.0175 | 0.96334 | 0.85708 | 0.858388 | 0.996349 | 0.997988 | 0.999507 |
| Ncor2_E9PY55_T1349t_1 | Ncor2_E9P | -1.29 | -1.66 | -0.86 | -2.34 | -1.6 | -2.05 | -1.16 | -1.67 | -0.23 | -1.38 | 0.27 | -2.03 | 0.63 | -0.68 | -0.69 | -1.69 | -0.0825 | 0.695 | 0.93 | 0.105718 | 0.819166 | 0.238233 | 0.995762 | 0.997988 | 0.994666 |
| Ncor2_E9PY55_T1405t_1 | Ncor2_E9P | -1.98 | -2.13 | -1.34 | -1.92 | -2.01 | -3.23 | -0.8 | -0.81 | -3.35 | -3.02 | -0.81 | -1.73 | -0.97 | -1.06 | -0.48 | -1.87 | 0.13 | -0.385 | 0.7475 | 0.061924 | 0.812383 | 0.501595 | 0.995762 | 0.997988 | 0.994666 |
| Ndel1_Q9ERR1_S194s_1 | Ndel1_Q9E | 0.19 | 0.86 | 1.3 | 0.68 | 1.1 | 0.62 | 1.21 | 0.9 | 0.25 | 0.73 | 1.21 | 0.8 | 0.95 | 1.09 | 0.63 | 1.22 | 0.2 | -0.01 | 0.215 | 0.49867 | 0.499357 | 0.975637 | 0.989223 | 0.997988 | 0.994666 |
| Ndel1_Q9ERR1_S194s_2 | Ndel1_Q9E | 0.03 | 0.47 | -0.1 | 0.38 | 0.68 | -0.22 | 0.56 | 0.15 | -0.46 | -0.23 | -0.36 | -0.39 | -2.84 | 0.53 | 0.33 | 0.13 | 0.0975 | -0.555 | -0.6575 | 0.372554 | 0.729708 | 0.039084 | 0.995762 | 0.997988 | 0.994666 |
| Ndel1_Q9ERR1_S198s_1 | Ndel1_Q9E | 1.12 | 1.61 | 1.14 | 1.02 | 0.96 | 0.42 | 0.64 | 0.54 | 1.23 | 1.25 | 1.34 | 0.66 | 0.82 | 0.94 | 1.07 | 0.84 | -0.5825 | -0.1025 | -0.305 | 0.246595 | 0.03449 | 0.700444 | 0.989223 | 0.997988 | 0.994666 |
| Ndel1_Q9ERR1_S198s_2 | Ndel1_Q9E | -0.04 | 0.36 | -0.1 | 0.38 | 0.65 | -0.33 | 0.56 | 0.15 | -0.48 | -0.29 | -0.36 | -0.39 | -3.23 | 0.48 | 0.33 | 0.13 | 0.1075 | -0.53 | -0.7225 | 0.373399 | 0.710468 | 0.042996 | 0.995437 | 0.997988 | 0.994666 |
| Ndel1_Q9ERR1_S215s_1 | Ndel1_Q9E | -0.46 | 0.02 | 0.29 | 0.13 | 0.34 | -0.35 | 0.14 | 0.09 | -0.58 | -0.69 | -0.98 | 0.16 | -0.3 | 0 | -0.29 | -0.01 | 0.06 | -0.5175 | -0.145 | 0.594178 | 0.82015 | 0.131861 | 0.995762 | 0.997988 | 0.995259 |
| Ndel1_Q9ERR1_S231s_1 | Ndel1_Q9E | 0.5 | 0.3 | 0.24 | 0.5 | 0.78 | 0.4 | 0.58 | 0.42 | 0.19 | 0.25 | 0.26 | 0.17 | 0.45 | 0.79 | 0.74 | 0.51 | 0.16 | -0.1675 | 0.2375 | 0.335321 | 0.455856 | 0.43067 | 0.989223 | 0.997988 | 0.994666 |
| Ndel1_Q9ERR1_S336s_1 | Ndel1_Q9E | 0.07 | 0.89 | -0.9 | 0 | 0.16 | 0.24 | 0.59 | -0.07 | 1.2 | 0.19 | -0.12 | 0.11 | -0.06 | -0.06 | 0.62 | -0.65 | 0.215 | 0.33 | -0.0525 | 0.90491 | 0.575822 | 0.47065 | 0.989223 | 0.997988 | 0.999507 |
| Ndel1_Q9ERR1_T219t_1 | Ndel1_Q9E | 0.11 | 1.15 | 0.24 | 0.75 | 0.62 | 0.99 | 0.66 | 0.14 | 0.66 | 1.01 | 0.86 | 0.69 | 0.86 | 0.56 | 0.31 | 1.45 | 0.04 | 0.2425 | 0.2325 | 0.528839 | 0.898648 | 0.420253 | 0.996493 | 0.997988 | 0.994666 |
| Ndfip2_M0QWK1_S12s_1 | Ndfip2_M0 | 0.08 | 0.79 | -0.4 | -0.03 | -0.18 | 0.74 | 0.11 | -0.12 | 0.36 | 0.76 | -1.15 | -0.04 | 0.16 | 0.66 | -0.15 | -0.17 | 0.0275 | -0.1275 | 0.015 | 0.965477 | 0.934385 | 0.782023 | 0.998658 | 0.997988 | 0.999507 |
| Ndfip2_M0QWK1_T216t_1 | Ndfip2_M0 | -0.47 | -0.37 | -0.95 | -0.98 | -2.19 | -0.89 | -1.46 | -1.9 | 0.49 | -1.02 | -1.11 | -1.26 | 0.65 | -3.05 | -1.71 | -1.69 | -0.9175 | -0.0325 | -0.7575 | 0.295902 | 0.019971 | 0.939269 | 0.989223 | 0.997988 | 0.994666 |
| Ndrg1_Q62433_S330s_1 | Ndrg1_Q62 | 0.53 | 0.24 | 0.5 | 0.06 | 0.18 | -0.54 | 0.12 | -0.14 | 0.07 | 0.14 | 0.3 | 0.13 | 0.34 | 0 | 1.03 | 0.84 | -0.4275 | -0.1725 | 0.22 | 0.487108 | 0.116122 | 0.451768 | 0.989223 | 0.997988 | 0.994666 |
| Ndrg1_Q62433_S330s_2 | Ndrg1_Q62 | 0.96 | 0.34 | 0.19 | -0.81 | 0.89 | -0.05 | -0.07 | -0.37 | 0.69 | 1.16 | 1.18 | -0.38 | -0.33 | 0.52 | 0.18 | 0.77 | -0.07 | 0.4925 | 0.115 | 0.790018 | 0.87106 | 0.328107 | 0.996493 | 0.997988 | 0.999507 |
| Ndrg1_Q62433_S333s_2 | Ndrg1_Q62 | 0.96 | 0.34 | 0.11 | -0.25 | 0.89 | -0.05 | -0.18 | 0.58 | 0.69 | 1.16 | 0.72 | -0.05 | -0.33 | 0.52 | 0.86 | 0.5 | 0.02 | 0.34 | 0.0975 | 0.795461 | 0.955487 | 0.365017 | 0.999187 | 0.997988 | 0.999507 |
| Ndrg1_Q62433_T328t_2 | Ndrg1_Q62 | 1.08 | 1.05 | 0.14 | -0.47 | 1.3 | 0.2 | -0.13 | 0.21 | 0.75 | 1.5 | 0.92 | -0.18 | -0.69 | 0.62 | 0.59 | 0.62 | -0.055 | 0.2975 | -0.165 | 0.729604 | 0.903837 | 0.544166 | 0.996493 | 0.997988 | 0.9971 |
| Ndrg1_Q62433_T375t_1 | Ndrg1_Q62 | 0.35 | 1.29 | 0.29 | 0.66 | 0.86 | -0.27 | 0.65 | -0.01 | 0.71 | 0.37 | 0.88 | 0.66 | 0.52 | 1.32 | 0.66 | 0.22 | -0.34 | 0.0075 | 0.0325 | 0.92679 | 0.347556 | 0.979684 | 0.989223 | 0.997988 | 0.999507 |
| Ndrg2_Q9QYG0_S314s_2 | Ndrg2_Q9 | -0.3 | 0.13 | 0.38 | 0.23 | -0.1 | -0.29 | 0.14 | 0.32 | -0.04 | -0.13 | -0.16 | -0.28 | -1.23 | 0.07 | -0.66 | -0.22 | -0.0925 | -0.2625 | -0.62 | 0.100999 | 0.71514 | 0.289428 | 0.995437 | 0.997988 | 0.994666 |
| Ndrg2_Q9QYG0_S318s_1 | Ndrg2_Q9 | 0.92 | 0.78 | 0.32 | 0.44 | 0.55 | 0.63 | 0.71 | 0.51 | 0.55 | 0.82 | 0.19 | 0.37 | 0.49 | 0.49 | 0.22 | 0.16 | -0.015 | -0.1325 | -0.275 | 0.309392 | 0.947207 | 0.613379 | 0.999187 | 0.997988 | 0.994666 |
| Ndrg2_Q9QYG0_S318s_2 | Ndrg2_Q9 | 0.1 | 0.1 | 0.55 | 0.56 | -0.11 | -0.17 | 0.51 | 0.48 | -0.21 | 0.23 | 0.5 | 0.56 | -0.88 | 0.59 | 0.13 | 0.18 | -0.15 | -0.0575 | -0.3225 | 0.383835 | 0.580022 | 0.83411 | 0.989223 | 0.997988 | 0.994666 |
| Ndrg2_Q9QYG0_S324s_1 | Ndrg2_Q9 | 0.58 | 0.68 | 0.64 | 0.69 | 0 | 0.22 | 0.58 | 0.51 | 0.43 | 0.65 | -0.07 | 0.69 | 0.41 | 0.17 | -0.08 | 0.06 | -0.32 | -0.2225 | -0.5075 | 0.057522 | 0.171788 | 0.385304 | 0.989223 | 0.997988 | 0.994666 |
| Ndrg2_Q9QYG0_S324s_2 | Ndrg2_Q9 | 0.64 | 0.74 | 0.4 | 0.57 | 0.3 | 0.47 | 0.46 | 0.63 | 0.54 | 0.25 | 0 | 0.45 | -1.14 | 0.9 | -0.26 | 0.23 | -0.1225 | -0.2775 | -0.655 | 0.151804 | 0.557134 | 0.252018 | 0.989223 | 0.997988 | 0.994666 |
| Ndrg2_Q9QYG0_S336s_1 | Ndrg2_Q9 | -0.6 | -0.19 | -1.48 | -1.09 | -0.7 | -0.47 | -0.67 | -1.18 | -0.85 | 0.08 | -1.37 | -1.63 | -0.8 | -0.74 | -1.75 | -1.36 | 0.085 | -0.1025 | -0.3225 | 0.409756 | 0.796808 | 0.820928 | 0.995762 | 0.997988 | 0.994666 |

| | | | | | | | | | | | | | | | | | | | | | | | | | | |
|---|---|---|---|---|---|---|---|---|---|---|---|---|---|---|---|---|---|---|---|---|---|---|---|---|---|---|
| Ndrg2_Q9QYG0_S336s_2 | Ndrg2_Q9( | -1.04 | -1.13 | 0.06 | 0.62 | -1.93 | -1 | 0.79 | 0.62 | -0.37 | -0.63 | 0.2 | 0.7 | -1.04 | -1.36 | -0.52 | -0.28 | -0.0075 | 0.3475 | -0.4275 | 0.376401 | 0.991325 | 0.485823 | 0.999187 | 0.997988 | 0.994666 |
| Ndrg2_Q9QYG0_S338s_1 | Ndrg2_Q9( | -0.41 | -0.08 | -0.44 | 0 | -0.61 | -0.5 | 0.08 | -0.15 | -0.73 | 0.03 | -0.84 | -0.3 | -0.64 | -0.54 | -0.39 | -0.11 | -0.0625 | -0.2275 | -0.1875 | 0.477929 | 0.803387 | 0.426639 | 0.995762 | 0.997988 | 0.994666 |
| Ndrg2_Q9QYG0_S339s_1 | Ndrg2_Q9( | 0.41 | 0.47 | 0.29 | 0.63 | 0.34 | 0.12 | 0.61 | 0.52 | 0.49 | 0.35 | -0.22 | 0.55 | 0.38 | 0.36 | 0.18 | 0.28 | -0.0525 | -0.1575 | -0.15 | 0.524714 | 0.810311 | 0.545187 | 0.995762 | 0.997988 | 0.994666 |
| Ndrg2_Q9QYG0_T316t_1 | Ndrg2_Q9( | -0.4 | 0.03 | -0.41 | -0.38 | -0.38 | -0.99 | 0.14 | -0.23 | -0.38 | -0.12 | -0.56 | -0.65 | -0.3 | -0.41 | -0.55 | -0.33 | -0.075 | -0.1375 | -0.1075 | 0.661542 | 0.795452 | 0.572655 | 0.995762 | 0.997988 | 0.995671 |
| Ndrg2_Q9QYG0_T316t_2 | Ndrg2_Q9( | 0.29 | 0.4 | 0.47 | 0.54 | 0.1 | 0.07 | 0.44 | 0.52 | 0.13 | 0.09 | 0.05 | 0.41 | -0.94 | 0.61 | -0.3 | 0.17 | -0.1425 | -0.255 | -0.54 | 0.157893 | 0.517448 | 0.257116 | 0.989223 | 0.997988 | 0.994666 |
| Ndrg2_Q9QYG0_T334t_2 | Ndrg2_Q9( | -0.49 | -0.81 | -1.59 | -1.02 | -1.44 | -0.9 | -1.02 | -1.27 | -0.42 | -0.25 | -1.79 | -1.84 | -1.48 | -1.16 | -1.2 | -0.76 | -0.18 | -0.0975 | -0.1725 | 0.594147 | 0.540304 | 0.83408 | 0.989223 | 0.997988 | 0.995259 |
| Ndrg3_Q9QYF9_S335s_1 | Ndrg3_Q9( | 1.98 | 2.1 | 3.98 | 2.38 | 2.49 | 2.36 | 2.92 | 3.53 | 1.44 | 2.78 | 1.98 | 4.09 | 1.14 | 0.43 | 5.26 | 4.31 | 0.215 | -0.0375 | 0.175 | 0.873427 | 0.662787 | 0.955304 | 0.990434 | 0.997988 | 0.999507 |
| Ndrg3_Q9QYF9_S352s_1 | Ndrg3_Q9( | 1.96 | 1.45 | 0.9 | 0.58 | 0.94 | 1.56 | 0.44 | 0.47 | 1.84 | 1.74 | 0.63 | 0.9 | 0.71 | 1.38 | 0.39 | 0.37 | -0.37 | 0.055 | -0.51 | 0.215884 | 0.353931 | 0.895735 | 0.989223 | 0.997988 | 0.994666 |
| Ndrg3_Q9QYF9_S361s_1 | Ndrg3_Q9( | 0.24 | 0.66 | 0.45 | 0.44 | 0.43 | 0.44 | -0.08 | 0.28 | 0.57 | 0.87 | 0.37 | 0.4 | -0.18 | 0.22 | 0.2 | -0.12 | -0.18 | 0.105 | -0.4175 | 0.119902 | 0.436094 | 0.655972 | 0.989223 | 0.997988 | 0.994666 |
| Ndrg3_Q9QYF9_S374s_1 | Ndrg3_Q9( | 2.94 | 0.69 | 2.61 | -0.54 | 0.72 | 3.29 | 1.6 | 1.64 | 0.85 | 3.07 | 2.09 | 1.99 | 1.24 | -0.91 | 2.86 | 3.41 | 0.3875 | 0.575 | 0.225 | 0.838162 | 0.651721 | 0.497243 | 0.990358 | 0.997988 | 0.999507 |
| Ndrg3_Q9QYF9_T355t_1 | Ndrg3_Q9( | -0.02 | 0.91 | -0.34 | 0.24 | 0.17 | 0.08 | -0.13 | 0.05 | 0.73 | 0.95 | 0.17 | 0.38 | 0.48 | 0.89 | -0.14 | -0.19 | -0.155 | 0.36 | 0.0625 | 0.870801 | 0.605836 | 0.30253 | 0.989223 | 0.997988 | 0.999507 |
| Ndrg4_Q8BTG7_S298s_1 | Ndrg4_Q8E | -0.56 | -0.26 | -0.13 | -0.29 | 0.04 | -0.61 | 0.19 | -0.23 | -0.36 | -0.15 | -0.02 | 0.01 | 0.49 | 0.27 | 0 | 0.02 | 0.1575 | 0.18 | 0.505 | 0.071786 | 0.53671 | 0.434081 | 0.989223 | 0.997988 | 0.994666 |
| Ndrg4_Q8BTG7_S298s_2 | Ndrg4_Q8E | -0.74 | -0.03 | -0.71 | -1.04 | -0.14 | -0.81 | -0.77 | -0.75 | -0.47 | -2.25 | -1.59 | 0.21 | -0.19 | -0.38 | -0.41 | 0.13 | 0.0125 | -0.395 | 0.4175 | 0.19241 | 0.966083 | 0.477843 | 0.999187 | 0.997988 | 0.994666 |
| Ndrg4_Q8BTG7_S306s_1 | Ndrg4_Q8E | -1.54 | -0.99 | -0.88 | -2.32 | -1.19 | -1.95 | -1.57 | -1.76 | -2.23 | -1.05 | -2.09 | -1.3 | -0.86 | -2.25 | -1.35 | -1.13 | -0.185 | -0.235 | 0.035 | 0.936263 | 0.613409 | 0.58582 | 0.989223 | 0.997988 | 0.999507 |
| Ndrg4_Q8BTG7_S306s_2 | Ndrg4_Q8E | -0.74 | -0.03 | -0.71 | -1.04 | -0.14 | -0.81 | -0.77 | -0.75 | -0.47 | -2.25 | -1.59 | 0.21 | -0.19 | -0.38 | -0.41 | 0.13 | 0.0125 | -0.395 | 0.4175 | 0.19241 | 0.966083 | 0.477843 | 0.999187 | 0.997988 | 0.994666 |
| Ndrg4_Q8BTG7_S317s_1 | Ndrg4_Q8E | -1.83 | -1.03 | 0.29 | -0.01 | -1.1 | -1.37 | -0.14 | 0.07 | -0.7 | -0.68 | 0.05 | 0.56 | -0.06 | -0.56 | 0.36 | 1.09 | 0.01 | 0.4525 | 0.8525 | 0.147857 | 0.985305 | 0.406435 | 0.999187 | 0.997988 | 0.994666 |
| Ndrg4_Q8BTG7_S317s_2 | Ndrg4_Q8E | -2.01 | 0.48 | -1.63 | -1.38 | -0.57 | -1.34 | -1.16 | -1.35 | -1.29 | -0.85 | -1.35 | -0.38 | -0.47 | -0.73 | -2.93 | -0.29 | 0.03 | 0.1675 | 0.03 | 0.967468 | 0.954799 | 0.76258 | 0.999187 | 0.997988 | 0.999507 |
| Ndrg4_Q8BTG7_S337s_1 | Ndrg4_Q8E | -1.24 | -1.01 | -1.62 | -1.66 | -1.32 | -1.3 | -2.15 | -2.59 | -1.48 | -1.03 | -2.02 | -1.75 | -0.59 | -0.76 | -2.19 | -2.19 | -0.4575 | -0.1875 | -0.05 | 0.911945 | 0.218009 | 0.538063 | 0.989223 | 0.997988 | 0.999507 |
| Ndrg4_Q8BTG7_T315t_2 | Ndrg4_Q8E | -2.01 | 0.48 | -1.63 | -1.38 | -0.57 | -1.34 | -1.16 | -1.35 | -1.29 | -0.85 | -1.35 | -0.38 | -0.47 | -0.73 | -2.93 | -0.29 | 0.03 | 0.1675 | 0.03 | 0.967468 | 0.954799 | 0.76258 | 0.999187 | 0.997988 | 0.999507 |
| Ndufb10_Q9DCS9_S21s_1 | Ndufb10_C | -0.26 | 0.62 | 0.67 | 0.36 | 0.49 | -0.01 | 0.03 | 0.21 | 0.18 | 0.03 | -0.03 | -0.01 | 0.03 | 0.31 | 0.44 | 0.61 | -0.1675 | -0.305 | 6.87E-17 | 1 | 0.55168 | 0.282703 | 0.989223 | 0.997988 | 1 |
| Ndufs5_Q99LY9_T93t_1 | Ndufs5_Q9 | -1.11 | -0.69 | -0.04 | -0.3 | -1.5 | -1.34 | -1.61 | -1.33 | -0.38 | -1.57 | -1.05 | -1.43 | -0.04 | -1.23 | -0.37 | -0.49 | -0.91 | -0.5725 | 0.0025 | 0.994533 | 0.008623 | 0.141385 | 0.989223 | 0.997988 | 1 |
| Ndufs8_Q8K3J1_S108s_1 | Ndufs8_Q8 | -0.16 | -0.2 | -0.41 | -0.6 | -1.86 | 0.4 | -1.9 | -1.21 | 1.8 | -0.34 | 0.47 | -0.23 | 1.82 | -0.24 | -0.35 | -0.33 | -0.8 | 0.7675 | 0.5675 | 0.282006 | 0.135492 | 0.133787 | 0.989223 | 0.997988 | 0.994666 |
| Ndufv3_Q3U422_S465s_1 | Ndufv3_Q3 | 0.59 | 1.34 | 0.91 | 0.72 | 0.56 | 0.8 | 0.04 | 0.36 | 0.75 | 0.81 | 0.71 | 0.97 | 1.15 | 0.96 | 0.47 | 0.65 | -0.45 | -0.08 | -0.0825 | 0.77764 | 0.122008 | 0.74867 | 0.989223 | 0.997988 | 0.99798 |
| Nedd4_P46935_S309s_1 | Nedd4_P4( | -0.3 | -0.68 | -0.46 | -0.88 | -0.39 | -0.29 | -0.49 | -0.04 | -0.57 | -0.63 | -0.01 | -0.71 | -0.57 | -0.22 | -0.98 | -0.68 | 0.2775 | 0.1 | -0.0325 | 0.907545 | 0.250367 | 0.707316 | 0.989223 | 0.997988 | 0.999507 |
| Nedd4_P46935_T287t_1 | Nedd4_P4( | -0.1 | 0.35 | -0.12 | -0.12 | -0.03 | -0.09 | 0.26 | 0.3 | 0.28 | 0.27 | 0.67 | -0.6 | -0.25 | 0.06 | 0.14 | 0.72 | 0.1075 | 0.1525 | 0.165 | 0.581542 | 0.640696 | 0.636621 | 0.989223 | 0.997988 | 0.995259 |
| Nedd4l_Q8CFI0_S504s_1 | Nedd4l_Q8 | -0.37 | -0.55 | 0.43 | 0.55 | 0 | -0.14 | 0.88 | 0.53 | -0.53 | -0.38 | 0.19 | 0.63 | -0.71 | -0.14 | 0.9 | 0.68 | 0.3025 | -0.0375 | 0.1675 | 0.711509 | 0.412603 | 0.922781 | 0.989223 | 0.997988 | 0.996966 |
| Nedd4l_Q8CFI0_S508s_1 | Nedd4l_Q8 | 1.81 | 0.2 | 0.32 | 1.19 | 1.14 | 1.75 | 0.84 | 1.31 | 1.16 | 0.65 | 0.97 | 0.85 | -0.73 | 0.9 | 0.55 | 0.59 | 0.38 | 0.0275 | -0.5525 | 0.282222 | 0.361959 | 0.944379 | 0.989223 | 0.997988 | 0.994666 |
| Nefh_P19246_S122s_1 | Nefh_P192 | -0.19 | 0.52 | 0.78 | 0.57 | -0.15 | 0.63 | 1.1 | -0.21 | -2.04 | -0.05 | -0.22 | -0.22 | 0.11 | -0.55 | 2.16 | -0.57 | -0.0775 | -1.0525 | -0.1325 | 0.829736 | 0.835773 | 0.054919 | 0.995762 | 0.997988 | 0.999507 |
| Nefh_P19246_S345s_1 | Nefh_P192 | 0.24 | 0.02 | 0.36 | -0.43 | -0.48 | 0.36 | -0.64 | 0.47 | 2.04 | -0.08 | -0.54 | -0.4 | 1.41 | -0.67 | 0.78 | 1 | -0.12 | 0.2075 | 0.5825 | 0.231543 | 0.724511 | 0.720135 | 0.995437 | 0.997988 | 0.994666 |
| Nefh_P19246_S416s_1 | Nefh_P192 | 1.34 | 0.48 | 0.32 | 0.39 | 1.3 | 0.39 | 0.96 | 0.51 | -0.06 | 0.25 | -0.72 | 0.26 | -0.26 | -0.61 | 0.82 | 0.88 | 0.1575 | -0.7 | -0.425 | 0.34468 | 0.633358 | 0.069194 | 0.989223 | 0.997988 | 0.994666 |
| Nefh_P19246_S500s_1 | Nefh_P192 | -1.79 | -0.51 | -3.53 | -2.47 | -0.49 | -3.15 | -3.43 | -3.43 | 0.31 | -1.8 | -1.96 | -3.16 | -0.62 | -0.1 | -3.3 | -3.59 | -0.55 | 0.4225 | 0.1725 | 0.857226 | 0.513577 | 0.622698 | 0.989223 | 0.997988 | 0.999507 |
| Nefh_P19246_S535s_1 | Nefh_P192 | 2.02 | 0.63 | 2.42 | 2.79 | 0.02 | 2.14 | 0.51 | 1.95 | 3.95 | 2.16 | 0.08 | 1.76 | 3.13 | -0.8 | 2.36 | 2.27 | -0.81 | 0.0225 | -0.225 | 0.795564 | 0.217588 | 0.978178 | 0.989223 | 0.997988 | 0.999507 |
| Nefh_P19246_S54s_1 | Nefh_P192 | 1.26 | 0.62 | 0.36 | 0.72 | 1.27 | 1.33 | 0.24 | 0.92 | 1.11 | -0.4 | 0.85 | 0.3 | 0.8 | 0.79 | 0.3 | 0.99 | 0.2 | -0.275 | -0.02 | 0.946924 | 0.543379 | 0.482578 | 0.989223 | 0.997988 | 0.999507 |
| Nefh_P19246_S577s_2 | Nefh_P192 | 0.56 | 0.62 | 1.1 | 0.45 | 1.39 | 0.71 | 0.6 | 0.33 | 0.66 | -0.02 | 0.39 | 0.58 | -0.38 | 0.83 | 1.04 | 1.19 | 0.075 | -0.28 | -0.0125 | 0.974684 | 0.799302 | 0.311784 | 0.995762 | 0.997988 | 0.999507 |
| Nefh_P19246_S583s_2 | Nefh_P192 | 0.56 | 0.62 | 1.1 | 0.45 | 1.39 | 0.71 | 0.6 | 0.33 | 0.66 | -0.02 | 0.39 | 0.58 | -0.38 | 0.83 | 1.04 | 1.19 | 0.075 | -0.28 | -0.0125 | 0.974684 | 0.799302 | 0.311784 | 0.995762 | 0.997988 | 0.999507 |
| Nefh_P19246_S607s_1 | Nefh_P192 | 2.38 | 0.89 | 1.24 | 2.49 | 0.2 | 1.94 | 0.53 | 1.05 | 3.1 | 2.17 | -1.69 | 1.26 | 2.39 | -0.57 | 1.13 | 1.2 | -0.82 | -0.54 | -0.7125 | 0.294041 | 0.130105 | 0.585672 | 0.989223 | 0.997988 | 0.994666 |
| Nefh_P19246_S61s_1 | Nefh_P192 | 1.17 | 0.85 | 0.73 | 0.81 | 1.2 | 0.72 | 0.73 | 0.67 | 0.66 | 0.79 | 0.76 | 0.78 | 1.35 | 0.98 | 1.36 | 1.11 | -0.06 | -0.1425 | 0.31 | 0.233177 | 0.795517 | 0.517743 | 0.995762 | 0.997988 | 0.994666 |
| Nefh_P19246_S637s_1 | Nefh_P192 | 3.68 | 1.01 | 2.28 | 0.54 | 0.71 | 2.4 | 1.52 | -0.22 | 2.28 | 3.27 | 2.66 | 2.22 | 2.07 | 0.29 | 2.38 | 2.43 | -0.775 | 0.73 | -0.085 | 0.911937 | 0.337462 | 0.294521 | 0.989223 | 0.997988 | 0.999507 |
| Nefh_P19246_S727s_1 | Nefh_P192 | 1.74 | 1.15 | 1.72 | 2.5 | 0.99 | 1.12 | 0.94 | 1.92 | 1.13 | 1.75 | 1.77 | 1.21 | 1.15 | 0.84 | 2.1 | 2.05 | -0.535 | -0.3125 | -0.2425 | 0.567317 | 0.15917 | 0.374442 | 0.989223 | 0.997988 | 0.995259 |
| Nefh_P19246_S769s_1 | Nefh_P192 | 1.44 | 1.38 | 1.49 | 1.42 | 1.36 | 1.07 | 1.36 | 1.38 | 1.24 | 1.64 | 1.42 | 1.37 | 0.93 | 1.06 | 1.36 | 1.2 | -0.14 | -0.015 | -0.295 | 0.231116 | 0.49047 | 0.943807 | 0.989223 | 0.997988 | 0.994666 |
| Nefh_P19246_S795s_1 | Nefh_P192 | 1.33 | 1.23 | 2.54 | 1.32 | 0.68 | 1.82 | 1.18 | 1.57 | 1.44 | 2.27 | 1.74 | 1.54 | 1.94 | 0.25 | 1.52 | 2.25 | -0.2925 | 0.1425 | -0.115 | 0.821018 | 0.457381 | 0.701675 | 0.989223 | 0.997988 | 0.999507 |
| Nefh_P19246_S834s_1 | Nefh_P192 | 0.65 | 0.86 | 1.04 | 1.08 | 0.72 | 0.24 | 1.08 | 0.84 | 0.46 | 0.72 | 0.9 | 0.85 | 0.68 | 0.59 | 1.61 | 1.63 | -0.1875 | -0.175 | 0.22 | 0.519786 | 0.468659 | 0.459118 | 0.989223 | 0.997988 | 0.994666 |
| Nefh_P19246_S834s_2 | Nefh_P192 | -0.36 | 0.36 | 0.37 | -0.22 | 0.36 | -0.59 | 0.16 | -0.31 | -0.85 | -0.72 | 0.86 | 0.76 | -1.91 | 0.09 | 0.76 | 0.15 | -0.1325 | -0.025 | -0.265 | 0.640282 | 0.669422 | 0.957995 | 0.992876 | 0.997988 | 0.995259 |
| Nefh_P19246_S867s_1 | Nefh_P192 | 1.3 | 1.39 | 1.76 | 0.97 | -0.62 | 1.86 | 0.02 | 0.24 | 3.2 | 0.68 | -2.46 | 1.24 | 3.57 | -0.38 | 3.3 | 0.79 | -0.98 | -0.69 | 0.465 | 0.59102 | 0.077375 | 0.513727 | 0.989223 | 0.997988 | 0.995259 |
| Nefh_P19246_S888s_1 | Nefh_P192 | 0.14 | 0.66 | 0.1 | -0.72 | 0.74 | 0.21 | -0.54 | -1.1 | 1.24 | 0.07 | -1.93 | -0.45 | 1.57 | -0.56 | 0.4 | -0.3 | -0.2175 | -0.3125 | 0.2325 | 0.656411 | 0.638914 | 0.632967 | 0.989223 | 0.997988 | 0.995259 |
| Nefh_P19246_S947s_1 | Nefh_P192 | 0.2 | 0.7 | 0.82 | 0.55 | 0.77 | 0.28 | 0.62 | 0.48 | -0.3 | 0.54 | 0.22 | 0.52 | 0.34 | 1.09 | 0.22 | 0.19 | -0.03 | -0.3225 | -0.1075 | 0.727217 | 0.899519 | 0.275742 | 0.996493 | 0.997988 | 0.9971 |
| Nefh_P19246_S965s_1 | Nefh_P192 | 0.64 | 1.24 | 1.45 | 1.16 | 1.02 | 0.9 | 0.24 | 0.88 | 1.52 | 0.69 | 0.27 | 0.29 | 1.23 | -0.07 | 0.87 | 1.54 | -0.3625 | -0.43 | -0.23 | 0.565532 | 0.217696 | 0.242918 | 0.989223 | 0.997988 | 0.995259 |
| Nefh_P19246_T839t_1 | Nefh_P192 | 1.18 | 1.08 | 0.76 | 0.44 | 1.29 | 0.66 | 0.56 | 0.4 | 0.74 | 0.92 | 0.76 | -0.52 | 0.74 | 0.92 | 1.05 | 0.68 | -0.1375 | -0.39 | -0.0175 | 0.948842 | 0.635237 | 0.317397 | 0.989223 | 0.997988 | 0.999507 |
| Nefh_P19246_T839t_2 | Nefh_P192 | -0.36 | 0.36 | 0.37 | -0.22 | 0.36 | -0.59 | 0.16 | -0.31 | -0.85 | -0.72 | 0.86 | 0.76 | -1.91 | 0.09 | 0.76 | 0.15 | -0.1325 | -0.025 | -0.265 | 0.640282 | 0.669422 | 0.957995 | 0.992876 | 0.997988 | 0.995259 |
| Nefl_P08551_S473s_1 | Nefl_P085! | -2.19 | -0.84 | -0.85 | -1.43 | -0.94 | -2.21 | -1.24 | -1.21 | -1.79 | -1.29 | 0.12 | -0.43 | -1.35 | -0.69 | 0.01 | -0.92 | -0.0725 | 0.48 | 0.59 | 0.186362 | 0.858077 | 0.35133 | 0.996349 | 0.997988 | 0.994666 |
| Nefl_P08551_S532s_1 | Nefl_P085! | -2.19 | -2.08 | -2.34 | -2.9 | -2.6 | -2.13 | -2.53 | -3.13 | -1.63 | -1.94 | -1.32 | -2.51 | -0.77 | -3.09 | -2.46 | -2.66 | -0.22 | 0.5275 | 0.1325 | 0.794821 | 0.469039 | 0.140246 | 0.989223 | 0.997988 | 0.999507 |
| Nefl_P08551_S56s_1 | Nefl_P085! | 0.14 | 0.68 | 0.16 | 0.17 | 0.63 | -0.19 | 0.79 | 0.51 | -0.01 | -0.14 | -0.47 | 0.26 | -0.18 | -0.1 | 0.4 | 0.2 | 0.1475 | -0.3775 | -0.2075 | 0.45418 | 0.607729 | 0.175695 | 0.989223 | 0.997988 | 0.994666 |
| Nefm_P08553_S28s_1 | Nefm_P08! | 0.56 | 0.76 | 0.5 | 0.05 | 0.75 | 0.77 | 0.2 | 0.66 | 0.36 | 0.47 | 1.48 | 0.19 | 0.68 | 0.82 | 0.31 | 0.87 | 0.1275 | 0.1575 | 0.2025 | 0.471401 | 0.618358 | 0.649595 | 0.989223 | 0.997988 | 0.994666 |
| Nefm_P08553_S30s_1 | Nefm_P08! | 0.56 | 0.18 | 0.2 | 0.43 | -0.07 | 0.42 | 0.54 | 0.73 | -0.07 | 0.36 | 0.25 | 0.89 | 0.3 | 0.43 | 1.17 | 1.06 | 0.0625 | 0.015 | 0.3975 | 0.205427 | 0.803078 | 0.956437 | 0.995762 | 0.997988 | 0.994666 |
| Nefm_P08553_S428s_1 | Nefm_P08! | 0.41 | -0.22 | -0.7 | -0.04 | 0.34 | 0.45 | -0.15 | 0.72 | 0.38 | 0.25 | 1.49 | -0.44 | -1.02 | 0.38 | -0.5 | 0.32 | 0.4775 | 0.5575 | -0.0675 | 0.869382 | 0.149579 | 0.230186 | 0.989223 | 0.997988 | 0.999507 |
| Nefm_P08553_S44s_1 | Nefm_P08! | 1.33 | 0.73 | 1.78 | 1.05 | 0.79 | 0.52 | 0.45 | 0.72 | 1.05 | 1.16 | 1.55 | 1.49 | 1.65 | 0.57 | 1.96 | 2.12 | -0.6025 | 0.09 | 0.3525 | 0.404365 | 0.051183 | 0.761435 | 0.989223 | 0.997988 | 0.994666 |
| Nefm_P08553_S610s_1 | Nefm_P08! | -2.85 | -2.18 | -2.26 | -3.67 | -1.14 | -2.72 | -3.25 | -2.91 | -2.59 | -1.97 | -1.77 | -2.68 | -1.29 | -2.01 | -2.38 | -2.08 | 0.235 | 0.4875 | 0.8 | 0.078921 | 0.657711 | 0.24966 | 0.990434 | 0.997988 | 0.994666 |
| Nefm_P08553_S645s_1 | Nefm_P08! | 0.38 | 0.48 | -0.46 | -1.05 | 0.04 | 0.86 | -1.48 | -1.6 | 2.56 | 0.07 | -2.28 | -0.71 | 2.49 | -0.7 | -0.14 | -0.95 | -0.3825 | 0.0725 | 0.3375 | 0.662798 | 0.543746 | 0.939053 | 0.989223 | 0.997988 | 0.995804 |
| Nefm_P08553_S669s_1 | Nefm_P08! | 2.63 | -0.55 | -1.11 | -1.25 | -0.74 | 2.68 | 0.03 | -0.22 | 3.06 | 1.94 | -4.08 | -0.24 | 3.2 | -1.14 | 1.24 | -1.01 | 0.5075 | 0.24 | 0.6425 | 0.592176 | 0.623798 | 0.879713 | 0.989223 | 0.997988 | 0.995259 |
| Nefm_P08553_S715s_1 | Nefm_P08! | 0.62 | -0.45 | 0.68 | 1.11 | -2.05 | 0.67 | -0.69 | 0.42 | 2.5 | 0.81 | 1.37 | 0.82 | 1.51 | -1.86 | 0.89 | 0.31 | -0.9025 | 0.885 | -0.2775 | 0.699999 | 0.172803 | 0.096954 | 0.989223 | 0.997988 | 0.996159 |
| Nefm_P08553_S715s_2 | Nefm_P08! | 0.45 | 0.07 | 1.08 | 0.91 | -1.68 | 1.06 | -0.8 | 0.06 | 2.91 | 0.3 | 1.29 | 1.09 | 0.77 | -1.66 | 1.09 | 1.21 | -0.9675 | 0.77 | -0.275 | 0.671461 | 0.113205 | 0.184689 | 0.989223 | 0.997988 | 0.996159 |
| Nefm_P08553_S715s_3 | Nefm_P08! | -1.51 | -0.44 | -0.39 | -1.36 | -2.54 | -0.14 | -1.38 | -1.12 | 1.76 | -1.33 | 0.6 | -0.12 | -0.16 | -1.84 | -0.01 | -0.03 | -0.37 | 1.1525 | 0.415 | 0.419056 | 0.485658 | 0.101397 | 0.989223 | 0.997988 | 0.994666 |
| Nefm_P08553_S720s_1 | Nefm_P08! | 0.61 | -0.42 | 0.9 | 0.71 | -1.37 | 0.72 | -0.61 | -0.4 | 2.26 | 0.9 | -0.28 | 0.83 | 2.05 | -1.64 | 1.06 | 1.03 | -0.865 | 0.4775 | 0.175 | 0.815709 | 0.09877 | 0.398654 | 0.989223 | 0.997988 | 0.999507 |
| Nefm_P08553_S720s_3 | Nefm_P08! | -1.51 | -0.44 | -0.39 | -1.36 | -2.54 | -0.14 | -1.38 | -1.12 | 1.76 | -1.33 | 0.6 | -0.12 | -0.16 | -1.84 | -0.01 | -0.03 | -0.37 | 1.1525 | 0.415 | 0.419056 | 0.485658 | 0.101397 | 0.989223 | 0.997988 | 0.994666 |
| Nefm_P08553_S723s_1 | Nefm_P08! | -0.81 | -1.12 | 0.94 | 0.19 | -2.1 | 0.08 | -0.41 | -0.33 | 1.79 | -0.69 | 0.41 | 0.45 | 1.37 | -2.07 | 0.95 | 1.39 | -0.49 | 0.69 | 0.61 | 0.475068 | 0.424652 | 0.290464 | 0.989223 | 0.997988 | 0.994666 |
| Nefm_P08553_S723s_2 | Nefm_P08! | 0.45 | 0.07 | 0.9 | 0.68 | -1.68 | 1.06 | -0.85 | -0.15 | 2.91 | 0.3 | 1.42 | 0.93 | 0.77 | -1.66 | 0.99 | 1.04 | -0.93 | 0.865 | -0.24 | 0.696752 | 0.115225 | 0.136345 | 0.989223 | 0.997988 | 0.996159 |
| Nefm_P08553_S723s_3 | Nefm_P08! | -1.51 | -0.44 | -0.39 | -1.36 | -2.54 | -0.14 | -1.38 | -1.12 | 1.76 | -1.33 | 0.6 | -0.12 | -0.16 | -1.84 | -0.01 | -0.03 | -0.37 | 1.1525 | 0.415 | 0.419056 | 0.485658 | 0.101397 | 0.989223 | 0.997988 | 0.994666 |
| Nefm_P08553_S769s_1 | Nefm_P08! | 0.28 | 0.54 | 0.35 | 0.4 | 0.7 | 0.22 | 0.8 | 0.45 | 0.56 | 0.84 | 0.36 | 0.73 | 0.55 | 1.01 | 1 | 0.98 | 0.15 | 0.23 | 0.4925 | 0.069565 | 0.507956 | 0.318883 | 0.989223 | 0.997988 | 0.994666 |
| Nefm_P08553_T565t_1 | Nefm_P08! | -1.39 | -0.1 | -4.48 | -3.09 | -1.07 | -0.79 | -3.17 | -2 | -1.13 | -0.84 | -0.47 | -2.99 | -2.69 | 0.06 | -2.46 | -1.9 | 0.5075 | 0.9075 | 0.5175 | 0.60607 | 0.596793 | 0.366753 | 0.989223 | 0.997988 | 0.995259 |
| Nek1_B2RXQ4_S972s_1 | Nek1_B2R) | -2.93 | -2.76 | -1.84 | -2 | -1.37 | -2.23 | -1.32 | -1.73 | -0.86 | -2.33 | -0.52 | -2.28 | -1.85 | -2.42 | -1.58 | -1.62 | 0.72 | 0.885 | 0.515 | 0.173808 | 0.061682 | 0.109298 | 0.989223 | 0.997988 | 0.994666 |
| Nek4_F8WGA9_S423s_1 | Nek4_F8W | -3.66 | -2.96 | -0.72 | -0.79 | -0.92 | -3.77 | -1.3 | -0.65 | -5.09 | -4.16 | 1.59 | -0.51 | -3.96 | -3.53 | -1.45 | -0.78 | 0.3725 | -0.01 | -0.3975 | 0.674811 | 0.680566 | 0.994723 | 0.993464 | 0.998021 | 0.996159 |
| Nelfa_Q8BG30_S363s_1 | Nelfa_Q8B( | -0.29 | -0.2 | -0.72 | -0.21 | -0.88 | -0.83 | -0.2 | -0.69 | -0.8 | 0.26 | 0.74 | -0.04 | -0.3 | -0.61 | -0.87 | 0.26 | -0.295 | 0.395 | -0.025 | 0.937676 | 0.260723 | 0.286196 | 0.989223 | 0.997988 | 0.999507 |
| Nelfb_Q8C4Y3_S557s_1 | Nelfb_Q8C | -0.23 | -0.15 | -0.98 | -0.69 | -0.2 | -0.65 | -0.16 | -1.15 | -0.56 | 0.01 | -0.52 | -1.02 | -0.3 | 0.27 | -0.72 | -0.86 | -0.0275 | -0.01 | 0.11 | 0.754026 | 0.931209 | 0.97488 | 0.998075 | 0.997988 | 0.9971 |
| Nelfb_Q8C4Y3_T559t_1 | Nelfb_Q8C | -0.75 | -0.67 | -0.97 | -0.31 | -0.36 | -1.04 | -0.16 | -0.6 | -0.83 | -0.96 | -0.3 | -0.56 | -0.51 | -0.18 | -0.25 | -0.2 | 0.135 | 0.0125 | 0.39 | 0.155569 | 0.623967 | 0.962196 | 0.989223 | 0.997988 | 0.994666 |
| Nelfe_P19426_S115s_1 | Nelfe_P19 | -1.79 | -1.42 | -1.89 | -3.06 | -1.85 | -1.36 | -2.63 | -2.07 | -1.47 | -1.58 | -0.43 | -1.71 | -1.8 | -1 | -1.94 | -2.87 | 0.0625 | 0.7425 | 0.1375 | 0.781455 | 0.881511 | 0.120951 | 0.996493 | 0.997988 | 0.998226 |
| Nelfe_P19426_S181s_1 | Nelfe_P19 | -1.43 | -2.57 | -1.26 | -2.16 | -1.8 | -1.87 | -1.74 | -1.97 | 0.43 | -1.93 | -3.39 | -1.79 | 1.06 | -2.28 | -1.46 | -1.86 | 0.01 | 0.185 | 0.72 | 0.332562 | 0.975404 | 0.807361 | 0.999187 | 0.997988 | 0.994666 |
| Nelfe_P19426_S49s_1 | Nelfe_P19 | -0.14 | -0.02 | 0.42 | 0.58 | 0.24 | -0.23 | 0.63 | 0.12 | -2.6 | 0.44 | 0.04 | 0.01 | -0.26 | -0.66 | 0.49 | 0.65 | -0.02 | -0.7375 | -0.155 | 0.679037 | 0.943414 | 0.275486 | 0.999187 | 0.997988 | 0.996159 |
| Neo1_Q7TQG5_S1182s_1 | Neo1_Q7T | 0.72 | 0.66 | -0.1 | 0.53 | 1.15 | 0.75 | 0.66 | 0.32 | 0.6 | 0.92 | 0.18 | -0.1 | -0.22 | 1.28 | 0.13 | 0 | 0.2675 | -0.0525 | -0.155 | 0.694864 | 0.361613 | 0.870389 | 0.989223 | 0.997988 | 0.996159 |
| Neo1_Q7TQG5_S1198s_2 | Neo1_Q7T | 0.86 | 0.41 | 1.66 | 1.3 | 0.7 | 1.2 | 1.47 | 1.94 | -1.54 | 1.46 | 2.11 | 1.5 | 1.06 | -1.77 | 0.58 | 1.56 | 0.27 | -0.175 | -0.7 | 0.331812 | 0.471351 | 0.821136 | 0.989223 | 0.997988 | 0.994666 |
| Neo1_Q7TQG5_S1224s_1 | Neo1_Q7T | 3.36 | 2.14 | 2.42 | 1.1 | 0.91 | 5.05 | 1.56 | 2.25 | 3.05 | 3.5 | 0.02 | 1.82 | 2.06 | 0.83 | 2.09 | 2.29 | 0.1875 | -0.1575 | -0.4375 | 0.420529 | 0.832985 | 0.845628 | 0.995762 | 0.997988 | 0.994666 |
| Neo1_Q7TQG5_S1405s_1 | Neo1_Q7T | 1.44 | -0.83 | -0.87 | 1.9 | -0.93 | -0.48 | -0.5 | 0.92 | 0.46 | 2.54 | 1.3 | 0.51 | 0.37 | -0.14 | 0.53 | -0.36 | -0.6575 | 0.7925 | -0.31 | 0.652089 | 0.381521 | 0.32743 | 0.989223 | 0.997988 | 0.995259 |
| Neo1_Q7TQG5_T1202t_1 | Neo1_Q7T | -0.55 | -0.71 | -0.64 | -0.54 | -0.58 | 0.01 | 0 | 0.27 | -0.23 | 0.1 | -0.65 | -0.44 | -0.92 | -0.75 | -0.44 | -0.27 | 0.535 | 0.305 | 0.015 | 0.953377 | 0.051499 | 0.232716 | 0.989223 | 0.997988 | 0.999507 |
| Neo1_Q7TQG5_T1202t_2 | Neo1_Q7T | 0.86 | 0.41 | 1.66 | 1.3 | 0.7 | 1.2 | 1.47 | 1.94 | -1.54 | 1.46 | 2.11 | 1.5 | 1.06 | -1.77 | 0.58 | 1.56 | 0.27 | -0.175 | -0.7 | 0.331812 | 0.471351 | 0.821136 | 0.989223 | 0.997988 | 0.994666 |
| Nes_Q6P5H2_S1793s_1 | Nes_Q6P5I | -0.61 | -0.4 | -0.93 | -1.01 | -0.17 | -1.64 | -0.45 | -1.51 | -0.05 | 0.38 | -2.07 | -0.79 | 0.49 | 0.27 | -0.75 | -0.06 | -0.205 | 0.105 | 0.725 | 0.056422 | 0.597735 | 0.839736 | 0.989223 | 0.997988 | 0.994666 |
| Neto1_Q8R4I7_S520s_1 | Neto1_Q8I | -0.91 | -1.55 | -1.7 | -1.95 | -0.97 | -0.48 | -0.54 | -0.7 | -2.18 | -1.8 | -0.31 | -1.15 | -1.18 | -1.82 | -2.47 | -2.26 | 0.855 | 0.1675 | -0.405 | 0.298316 | 0.012468 | 0.709703 | 0.989223 | 0.997988 | 0.994666 |
| Neurl4_H9H9T3_S247s_1 | Neurl4_H9 | -0.08 | 0.15 | 0.23 | -0.14 | 0.2 | -0.22 | 0.07 | -0.42 | 0.02 | -0.02 | 0.1 | 0.06 | -0.35 | 0.72 | 0.22 | 0.1 | -0.1325 | 0 | 0.1325 | 0.659405 | 0.577 | 1 | 0.989223 | 1 | 0.995259 |
| Neurl4_H9H9T3_S647s_1 | Neurl4_H9 | -1.15 | -2.83 | -1.62 | 0.68 | -0.92 | -2.35 | 0.2 | 0.44 | -1.1 | -1.74 | 0.75 | 0.53 | 0.11 | -1.59 | -0.12 | 0.33 | 0.5725 | 0.84 | 0.9125 | 0.244078 | 0.50243 | 0.334093 | 0.989223 | 0.997988 | 0.994666 |
| Neurl4_H9H9T3_S652s_1 | Neurl4_H9 | -0.96 | -0.4 | -0.44 | 0.91 | -0.64 | -0.38 | 0.54 | 0.11 | -0.3 | -0.14 | 0.32 | 0.15 | -0.06 | 0.34 | 0.52 | 0.1 | 0.13 | 0.23 | 0.4475 | 0.299242 | 0.771185 | 0.582603 | 0.995762 | 0.997988 | 0.994666 |
| Nexn_Q7TPW1_S16s_1 | Nexn_Q7TI | 0.46 | -0.05 | 0.73 | 0.38 | 0.27 | -0.49 | 0.39 | 0.81 | 0.33 | 0.11 | 0.23 | -0.44 | 0.07 | 0.17 | 0.58 | 0.33 | -0.135 | -0.3225 | -0.0925 | 0.739213 | 0.681041 | 0.274235 | 0.993613 | 0.997988 | 0.9971 |
| Nexn_Q7TPW1_S296s_1 | Nexn_Q7TI | 1.15 | 0.31 | 0.1 | 0.42 | 0.38 | 0.26 | 0.59 | 0.01 | 0.55 | 0.55 | 0.04 | 0.07 | 0.19 | 1.02 | 0.22 | 0.48 | -0.185 | -0.1925 | -0.0175 | 0.958426 | 0.52656 | 0.533343 | 0.989223 | 0.997988 | 0.999507 |
| Nf1_Q04690_S2169s_1 | Nf1_Q046! | 0 | 0.43 | 0.28 | 0.26 | 0.26 | -0.12 | 0.1 | 0.2 | 0.04 | 0.43 | 0.1 | 0.3 | -0.18 | 0.33 | 0.27 | 0.29 | -0.1325 | -0.025 | -0.065 | 0.799391 | 0.542987 | 0.912421 | 0.989223 | 0.997988 | 0.999507 |
| Nf1_Q04690_S2477s_1 | Nf1_Q046! | -0.49 | -0.84 | -0.64 | -0.39 | -0.15 | -1.17 | -0.57 | -0.47 | -0.71 | -0.21 | -0.89 | -0.65 | -0.81 | 0.1 | -0.65 | -0.5 | -7.9E-17 | -0.025 | 0.125 | 0.668343 | 1 | 0.920176 | 1 | 0.997988 | 0.996159 |
| Nf1_Q04690_S2496s_1 | Nf1_Q046! | -0.27 | -0.76 | 0.15 | 0.19 | -0.21 | -0.24 | 0.4 | 0.38 | -1 | -0.58 | -1.04 | 0.33 | -1.24 | -0.8 | -0.06 | -0.23 | 0.255 | -0.4 | -0.41 | 0.283281 | 0.413151 | 0.318361 | 0.989223 | 0.997988 | 0.994666 |
| Nf1_Q04690_S2502s_1 | Nf1_Q046! | 0.59 | 0.46 | 0.45 | 0.22 | 0.27 | 0.67 | 0.19 | 0.15 | 0.95 | 0.65 | 0.22 | 0.02 | 1.22 | 0.73 | 0.37 | 0.56 | -0.11 | 0.03 | 0.29 | 0.311077 | 0.625666 | 0.913685 | 0.989223 | 0.997988 | 0.994666 |
| Nf1_Q04690_S2504s_1 | Nf1_Q046! | 0.5 | 0.4 | 0.44 | 0.29 | 0.47 | 0.36 | 0.24 | 0.13 | 0.57 | 0.74 | 0.03 | -0.05 | 0.63 | 0.7 | 0.01 | 0.19 | -0.1075 | -0.085 | -0.025 | 0.925119 | 0.598592 | 0.74786 | 0.989223 | 0.997988 | 0.999507 |
| Nf1_Q04690_S2524s_1 | Nf1_Q046! | 1.71 | 0.58 | 1.59 | 0.04 | 1.07 | 1.87 | 2.09 | 1.31 | -1.06 | 1.83 | -0.46 | 1.62 | -0.77 | -0.75 | 1.39 | 1.79 | 0.605 | -0.4975 | -0.565 | 0.433238 | 0.190537 | 0.510762 | 0.989223 | 0.997988 | 0.994666 |
| Nf1_Q04690_S2578s_1 | Nf1_Q046! | 0.34 | 0.62 | 0.05 | 0.15 | 0.94 | -0.02 | 0.67 | 0.65 | -0.1 | 0.57 | 0.13 | 0.96 | -0.21 | 0.72 | 0.04 | 0.1 | 0.27 | 0.1 | -0.1275 | 0.669494 | 0.342652 | 0.742766 | 0.989223 | 0.997988 | 0.996159 |
| Nf1_Q04690_S2783s_1 | Nf1_Q046! | -1.98 | -1.95 | -1.43 | -0.92 | -0.88 | -1.91 | -0.75 | -1.48 | -1.25 | -1.43 | -0.53 | -1.47 | -0.6 | -0.95 | -0.87 | -2.8 | 0.315 | 0.4 | 0.265 | 0.616783 | 0.39766 | 0.268935 | 0.989223 | 0.997988 | 0.995259 |
| Nf1_Q04690_S2798s_1 | Nf1_Q046! | 7.1 | 2.19 | 3.25 | -2.5 | 3.04 | 8.18 | 4.82 | 3.52 | 0.76 | 5.76 | 1.21 | 3.98 | 0.96 | -0.21 | 5.77 | 5.36 | 2.38 | 0.4175 | 0.46 | 0.828871 | 0.242025 | 0.834843 | 0.989223 | 0.997988 | 0.999507 |
| Nf1_Q04690_S659s_1 | Nf1_Q046! | -0.42 | -2.25 | -2.42 | -1.21 | -1.02 | -0.17 | -1.11 | -1.07 | -0.77 | -1.64 | 0.69 | -0.99 | -0.98 | 0.06 | -1.49 | -1.47 | 0.7325 | 0.8975 | 0.605 | 0.286707 | 0.151052 | 0.171766 | 0.989223 | 0.997988 | 0.994666 |
| Nf1_Q04690_S866s_1 | Nf1_Q046! | 0.7 | 0.69 | 0.83 | 0.67 | 0.84 | 0.56 | 0.94 | 0.78 | -1.23 | 1.14 | -1.97 | 0.84 | -1.01 | -1.83 | 0.85 | 0.79 | 0.0575 | -1.0275 | -1.0225 | 0.118676 | 0.777788 | 0.161901 | 0.995762 | 0.997988 | 0.994666 |
| Nf1_Q04690_S878s_1 | Nf1_Q046! | 0.81 | 0.03 | -0.72 | -0.72 | 1.5 | 0.52 | 0.46 | -0.33 | 0.3 | 0.43 | 0.52 | -0.31 | 0.03 | 1.68 | -0.34 | -0.23 | 0.6875 | 0.385 | 0.435 | 0.438662 | 0.177117 | 0.355602 | 0.989223 | 0.997988 | 0.994666 |
| Nf1_Q04690_T2545t_1 | Nf1_Q046! | 0.37 | -0.03 | -0.37 | 0.24 | 0.26 | 0.11 | -0.25 | -0.16 | -0.16 | 0.49 | -0.02 | -0.12 | 0.1 | 0.3 | -0.03 | -0.24 | -0.0625 | -0.005 | -0.02 | 0.942734 | 0.806395 | 0.985549 | 0.995762 | 0.997988 | 0.999507 |
| Nf1_Q04690_T2546t_1 | Nf1_Q046! | 0.18 | -0.11 | -0.26 | 0.22 | 0.25 | 0 | -0.23 | -0.21 | -0.27 | 0.4 | -0.15 | -0.19 | 0.14 | 0.14 | -0.07 | -0.27 | -0.055 | -0.06 | -0.0225 | 0.930051 | 0.813597 | 0.816798 | 0.995762 | 0.997988 | 0.999507 |
| Nfat5_Q80TR0_S117s_1 | Nfat5_Q80 | 0.34 | 0.41 | -0.26 | 0.29 | 0.22 | 0.08 | 0.67 | 0.59 | 0.08 | -0.02 | 0.55 | -0.05 | -0.4 | 0.82 | -0.26 | 0.14 | 0.195 | -0.055 | -0.12 | 0.729237 | 0.458775 | 0.837271 | 0.989223 | 0.997988 | 0.9971 |
| Nfic_P70255_S194s_1 | Nfic_P702! | -2.07 | -3.41 | -2.18 | -5.04 | -3.71 | -2.15 | -3.06 | -4.1 | 0.24 | -2.75 | -2.61 | -1.74 | 1.21 | -4.32 | 0.22 | -2.41 | -0.08 | 1.46 | 1.85 | 0.158917 | 0.910882 | 0.118723 | 0.997335 | 0.997988 | 0.994666 |
| Nfic_P70255_S305s_1 | Nfic_P702! | -0.57 | 1.29 | -2.49 | -0.82 | -0.3 | 0.59 | -1.65 | -2.18 | -0.7 | 0.36 | -2.56 | -2.01 | 0.22 | -0.13 | 1.23 | -0.94 | -0.2375 | -0.58 | 0.7425 | 0.359538 | 0.784765 | 0.524262 | 0.995762 | 0.997988 | 0.994666 |
| Nfic_P70255_S339s_1 | Nfic_P702! | -0.55 | -0.17 | -0.01 | 0.26 | -0.38 | -0.52 | -0.39 | 0 | 0.17 | -0.01 | 0.1 | 1.19 | -0.1 | -0.06 | 0.42 | 0.22 | -0.205 | 0.48 | 0.2375 | 0.411645 | 0.430454 | 0.18572 | 0.989223 | 0.997988 | 0.994666 |
| Nfix_D3YZ00_S301s_1 | Nfix_D3YZ( | 0.18 | 0.05 | -0.08 | -0.05 | 0.04 | 0 | 0.26 | 0.05 | 0.45 | 0.33 | -0.32 | 0.14 | -0.57 | 0.62 | -0.14 | -0.31 | 0.0625 | 0.125 | -0.125 | 0.691511 | 0.757264 | 0.62221 | 0.995762 | 0.997988 | 0.996159 |
| Nfx1_B1AY10_S197s_1 | Nfx1_B1AY | -2.02 | -0.13 | -2.93 | -4.69 | -4.11 | -1.11 | -1.94 | -2.93 | -2.08 | -1.26 | -1.27 | -2.39 | -1.18 | -1.21 | -2.73 | -2.34 | -0.08 | 0.6925 | 0.5775 | 0.525818 | 0.93577 | 0.439626 | 0.99868 | 0.997988 | 0.994666 |
| Ngef_E9QK62_S606s_1 | Ngef_E9QK | -1.44 | -2.31 | -1.95 | -1.59 | -0.94 | -1.49 | -0.91 | -1.17 | -2.6 | -2 | -1.26 | -1.74 | -2.21 | -1.21 | -2.14 | -1.82 | 0.695 | -0.0775 | -0.0225 | 0.946709 | 0.028997 | 0.826861 | 0.989223 | 0.997988 | 0.999507 |
| Ngef_E9QK62_S696s_1 | Ngef_E9QK | 1.5 | -0.13 | 0.42 | 1.55 | -0.2 | 2.09 | -0.03 | 0.97 | 2.74 | 1.04 | -0.45 | 0.93 | 2.09 | -0.32 | 0.61 | 0.2 | -0.1275 | 0.23 | -0.19 | 0.753726 | 0.831785 | 0.742182 | 0.995762 | 0.997988 | 0.9971 |
| Nhp2l1_Q9D0T1_S122s_1 | Nhp2l1_Q9 | 2.02 | 2.61 | 0.56 | 0.22 | 1.6 | 2.08 | 0.61 | 0.97 | 1.14 | 2.61 | -0.97 | 0.92 | 0.63 | 1.6 | 1.39 | 1.17 | -0.0375 | -0.4275 | -0.155 | 0.783131 | 0.949338 | 0.608964 | 0.999187 | 0.997988 | 0.998491 |
| Nhs_B9EJX2_S715s_1 | Nhs_B9EJX | -0.26 | -0.45 | -0.94 | -0.68 | 0 | -0.47 | 0.16 | -0.63 | 0.23 | 0.4 | 0.35 | -0.84 | 0.6 | 0.24 | 0 | -0.15 | 0.3475 | 0.6175 | 0.755 | 0.025269 | 0.228177 | 0.100457 | 0.989223 | 0.997988 | 0.994666 |
| Nhsl1_Q8CAF4_S1075s_1 | Nhsl1_Q8C | -0.15 | -0.75 | 0.08 | -0.07 | 0.01 | -0.76 | 0.5 | 0.39 | -0.62 | -0.52 | -0.16 | 0.05 | -0.37 | -0.46 | -0.25 | 0.22 | 0.2575 | -0.09 | 0.0075 | 0.979967 | 0.459857 | 0.754317 | 0.989223 | 0.997988 | 1 |
| Nhsl1_Q8CAF4_S1360s_1 | Nhsl1_Q8C | -1.73 | -2.84 | -1.99 | -7.42 | -3.01 | -1.58 | -1.36 | -1.37 | 2.84 | -2.22 | -0.7 | -0.72 | 2.13 | -2.11 | 1.8 | -2.98 | 1.665 | 3.295 | 3.205 | 0.070296 | 0.185278 | 0.05046 | 0.989223 | 0.997988 | 0.994666 |
| Nhsl1_Q8CAF4_S535s_1 | Nhsl1_Q8C | 0.2 | 0.02 | 0.9 | 0.25 | -1.42 | 0.38 | -0.4 | -0.48 | 2.11 | 0.98 | 0.78 | 0.37 | 1.67 | -1.1 | -0.31 | -0.83 | -0.8225 | 0.7175 | -0.485 | 0.425681 | 0.063796 | 0.108541 | 0.989223 | 0.997988 | 0.994666 |
| Nhsl2_B1AXH1_S1145s_1 | Nhsl2_B1A | -1.55 | -0.11 | 0.68 | 2 | -1 | -1.55 | 1.9 | 1.54 | 0.19 | -0.92 | 2.96 | 2.17 | 0.39 | 1.69 | 1.87 | -0.32 | -0.0325 | 0.845 | 0.6525 | 0.424899 | 0.973853 | 0.417564 | 0.999187 | 0.997988 | 0.994666 |
| Nhsl2_B1AXH1_S1208s_1 | Nhsl2_B1A | -2.29 | -0.61 | -1.13 | -2.61 | -1.04 | -1.92 | -1.38 | -1.97 | -1.99 | -1.2 | -0.16 | -1.5 | -0.7 | -0.53 | -2.25 | -1.48 | 0.0825 | 0.4475 | 0.42 | 0.465963 | 0.864121 | 0.435773 | 0.996493 | 0.997988 | 0.994666 |
| Nhsl2_B1AXH1_S690s_1 | Nhsl2_B1A | -0.16 | 0.04 | 0.37 | 0.43 | -0.12 | -0.81 | 0.74 | 0.6 | -0.57 | -0.28 | 0.78 | 0.33 | -0.52 | 0.09 | 0.03 | 0.35 | -0.0675 | -0.105 | -0.1825 | 0.540449 | 0.857608 | 0.764286 | 0.996349 | 0.997988 | 0.994666 |
| Nhsl2_B1AXH1_T284t_1 | Nhsl2_B1A | -3.56 | -2.28 | -1.79 | -1.77 | -1.51 | -3.46 | -1.06 | -2.01 | -4.3 | -2.89 | -0.54 | -1.84 | -2.85 | -3.15 | -2.44 | -1.21 | 0.34 | -0.0425 | -0.0625 | 0.910108 | 0.573231 | 0.957811 | 0.989223 | 0.997988 | 0.999507 |
| Nicn1_Q9CQM0_S142s_1 | Nicn1_Q9C | 0.13 | 0.32 | -0.06 | -0.12 | 0.7 | 0.01 | 0.56 | -0.13 | -0.26 | 0.1 | 0.19 | 0.21 | 0.16 | 0.43 | -0.1 | 0.14 | 0.2175 | -0.0075 | 0.09 | 0.724427 | 0.427224 | 0.974695 | 0.989223 | 0.997988 | 0.9971 |
| Nifk_Q91VE6_S256s_1 | Nifk_Q91V | 0.6 | 0.61 | 0.08 | -0.76 | -0.11 | 0.9 | -0.94 | -0.95 | 1.84 | 0.02 | -1.59 | -0.41 | 1.95 | -0.49 | -0.66 | -1.43 | -0.4075 | -0.1675 | -0.29 | 0.685605 | 0.422314 | 0.81209 | 0.989223 | 0.997988 | 0.996159 |
| Nipbl_Q6KCD5_S2652s_1 | Nipbl_Q6K | 0.91 | -0.13 | 0.61 | 0.61 | 0.32 | -0.2 | 0.86 | 0.45 | 0.49 | 0.49 | -0.18 | -0.12 | 0.39 | 0.73 | 0.32 | 0.34 | -0.1425 | -0.33 | -0.055 | 0.855333 | 0.661685 | 0.314755 | 0.990434 | 0.997988 | 0.999507 |
| Nipbl_Q6KCD5_T599t_1 | Nipbl_Q6K | -0.74 | -0.53 | -0.25 | -0.04 | -1.33 | -1.05 | -0.22 | -0.55 | -0.28 | -0.64 | -0.57 | 0.09 | 0.39 | 0 | 0.24 | 0.11 | -0.3975 | 0.04 | 0.575 | 0.054173 | 0.22158 | 0.885881 | 0.989223 | 0.997988 | 0.994666 |
| Nisch_Q80TM9_S1128s_1 | Nisch_Q80 | -2.64 | -1.5 | -0.08 | -0.22 | -0.76 | -2.38 | -0.56 | -0.09 | -2.73 | -1.58 | -0.14 | -0.08 | -1.19 | -1.64 | 0.04 | -0.51 | 0.1625 | -0.0225 | 0.285 | 0.657163 | 0.813742 | 0.977037 | 0.995762 | 0.997988 | 0.995259 |
| Nisch_Q80TM9_S492s_1 | Nisch_Q80 | 0.12 | 0.3 | 0.59 | 0.66 | 0.36 | 0.5 | 1.09 | 0.71 | 0.24 | 0.48 | 1.15 | 0.32 | -0.24 | 0.1 | 0.65 | 1.43 | 0.2475 | 0.13 | 0.0675 | 0.86411 | 0.344848 | 0.653972 | 0.989223 | 0.997988 | 0.999507 |
| Nkd2_Q8VE28_S259s_1 | Nkd2_Q8V | 0.68 | 1.3 | 0.76 | 1.28 | 0.82 | 1.39 | 0.81 | 1.07 | 1.9 | 1.73 | 0.75 | 1.23 | 0.77 | 0.23 | 0.94 | 1.18 | 0.0175 | 0.3975 | -0.225 | 0.478268 | 0.946718 | 0.249389 | 0.999187 | 0.997988 | 0.994666 |
| Nktr_P30415_S1139s_1 | Nktr_P304 | -1.33 | -1.32 | 0.1 | 0.05 | -1.5 | -0.89 | -0.83 | 0.15 | -1.04 | -1.72 | -0.5 | -1.01 | 0.07 | 0.01 | 1.1 | -1.46 | -0.1425 | -0.4425 | 0.555 | 0.370074 | 0.770117 | 0.344881 | 0.995762 | 0.997988 | 0.994666 |
| Nktr_P30415_S468s_1 | Nktr_P304 | 0.99 | 0.65 | 0.69 | 0.55 | -0.53 | 0.67 | 0.73 | -0.12 | 0.19 | 0.46 | 1.16 | 0.4 | 0.33 | 0.44 | 0.82 | -0.12 | -0.5325 | -0.1675 | -0.3525 | 0.23846 | 0.133253 | 0.555667 | 0.989223 | 0.997988 | 0.994666 |
| Nlgn2_Q69ZK9_S714s_1 | Nlgn2_Q69 | 0.5 | 0.88 | 0.76 | 0.74 | 0.92 | 0.51 | 1.01 | 0.58 | 0.56 | 0.77 | 0.43 | 0.89 | 0.61 | 0.96 | 0.62 | 0.81 | 0.035 | -0.0575 | 0.03 | 0.90156 | 0.877102 | 0.801855 | 0.996493 | 0.997988 | 0.999507 |
| Nlgn3_Q8BYM5_S722s_1 | Nlgn3_Q8E | 1.67 | 1.25 | 1.58 | 1.93 | 1.13 | 1.49 | 1.3 | 1.9 | 1.86 | 2 | 2.22 | 1.68 | 1.76 | 1.55 | 1.79 | 1.58 | -0.1525 | 0.3325 | 0.0625 | 0.808076 | 0.566343 | 0.210149 | 0.989223 | 0.997988 | 0.999507 |
| Nlgn4l_B0F2B4_S880s_1 | Nlgn4l_B0F | 0.94 | -0.09 | 0.48 | 0.75 | -0.25 | 1.22 | 0.07 | 1.04 | 0.79 | 0.11 | 0.2 | 0.48 | 0.29 | -0.79 | 0.05 | -0.31 | 2.36E-16 | -0.125 | -0.71 | 0.068655 | 1 | 0.685294 | 1 | 0.997988 | 0.994666 |
| Nlk_O54949_T298t_1 | Nlk_O5494 | 2.06 | 1.25 | 0.35 | 0.51 | 0.22 | 2.1 | 1.27 | 0.83 | 0.09 | 1.13 | -0.6 | 0.02 | 0.93 | -0.45 | -0.25 | 0.7 | 0.0625 | -0.8825 | -0.81 | 0.125841 | 0.902138 | 0.104071 | 0.996493 | 0.997988 | 0.994666 |
| Nmt1_O70310_S47s_1 | Nmt1_O70 | -2.32 | -2.68 | -2.47 | -3.7 | -2 | -2.42 | -2.48 | -2.53 | -3.15 | -2.37 | -1.35 | -3.03 | -1.61 | -1.6 | -2.58 | -2.34 | 0.435 | 0.3175 | 0.76 | 0.08576 | 0.221125 | 0.520663 | 0.989223 | 0.997988 | 0.994666 |
| Nmt2_O70311_S38s_1 | Nmt2_O70 | -2.11 | -1.64 | -2.23 | -2.29 | -2.03 | -2.63 | -2.72 | -2.12 | -1.42 | -1.67 | -1.54 | -2.01 | -0.61 | -1.44 | -2.18 | -2.49 | -0.3075 | 0.4075 | 0.3875 | 0.38505 | 0.272563 | 0.142982 | 0.989223 | 0.997988 | 0.994666 |
| Nmt2_O70311_S66s_1 | Nmt2_O70 | -1.76 | -1.28 | -2.83 | -1.87 | -2.3 | -1.72 | -1.61 | -1.39 | -0.34 | -1.67 | -0.91 | -2.47 | -0.49 | -2.85 | -2.33 | -3.02 | 0.18 | 0.5875 | -0.2375 | 0.695937 | 0.630536 | 0.282041 | 0.989223 | 0.997988 | 0.996159 |
| No66_Q9JJF3_S105s_1 | No66_Q9JJ | -1.2 | -1.9 | -2.51 | -0.6 | -1.06 | -0.97 | -2.89 | 0.32 | -0.96 | -1.77 | -0.85 | 0.27 | -0.46 | -0.29 | -1.77 | -2.83 | 0.4025 | 0.725 | 0.215 | 0.743542 | 0.562274 | 0.207118 | 0.989223 | 0.997988 | 0.9971 |
| Noc2l_J3QK52_S60s_1 | Noc2l_J3Q | 1.86 | 0.75 | 1.54 | 1.95 | -0.37 | 1.79 | 0.37 | 1.02 | 3.1 | 1.19 | 0.57 | 1.6 | 2.7 | 0.39 | 0.94 | 0.89 | -0.8225 | 0.09 | -0.295 | 0.584407 | 0.119815 | 0.871874 | 0.989223 | 0.997988 | 0.995259 |
| Noc2l_J3QK52_T59t_1 | Noc2l_J3Q | -1.36 | -1.26 | 1.46 | 2.72 | -2.22 | -1.05 | 0.73 | 1.68 | 0.74 | -2.72 | 1.33 | 1.95 | 1.36 | -2.42 | 1.83 | 2.1 | -0.605 | -0.065 | 0.3275 | 0.795669 | 0.602588 | 0.959321 | 0.989223 | 0.997988 | 0.999507 |
| Nol3_Q9D1X0_S142s_1 | Nol3_Q9D1 | -0.7 | -0.64 | 0.99 | 1.42 | -0.3 | -0.36 | 0.31 | 0.2 | -0.95 | -0.76 | 2.07 | 0.47 | 0.24 | 1.06 | 0.15 | 2.92 | -0.305 | -0.06 | 0.825 | 0.287403 | 0.56306 | 0.939538 | 0.989223 | 0.997988 | 0.994666 |
| Nol3_Q9D1X0_T114t_1 | Nol3_Q9D1 | -0.15 | 0.12 | 0.98 | 1.47 | -0.4 | -0.93 | 1.52 | 1.37 | -0.49 | 0.56 | 0.79 | 1.14 | -0.14 | 0.1 | 0.45 | 1.17 | -0.215 | -0.105 | -0.21 | 0.647617 | 0.738772 | 0.829179 | 0.995762 | 0.997988 | 0.995259 |
| Nol4_F6XSA1_S135s_1 | Nol4_F6XS | -3.15 | -2.71 | -2.15 | -1.01 | -2.64 | -2.97 | -2.87 | -2.67 | -3.04 | -2.85 | -2.44 | -1.79 | -1.15 | -2.45 | -2.07 | -2.32 | -0.5325 | -0.275 | 0.2575 | 0.619021 | 0.245407 | 0.589996 | 0.989223 | 0.997988 | 0.995259 |
| Nol4l_Q3UH85_S17s_1 | Nol4l_Q3U | -1.86 | -1.75 | -2.16 | -1.54 | -1.89 | -1.55 | -1.81 | -2.21 | -1.59 | -0.94 | 0.39 | -1.4 | 0.09 | -1.91 | -0.52 | -0.51 | -0.0375 | 0.9425 | 1.115 | 0.027536 | 0.879256 | 0.059801 | 0.996493 | 0.997988 | 0.994666 |
| Nol9_Q3TZX8_S88s_1 | Nol9_Q3TZ | 0.85 | 0.54 | 0.22 | 0.57 | 0.88 | 0.52 | 0.86 | 0.34 | 0.68 | 0.73 | 0.84 | 0.27 | -0.51 | 1.28 | 0.26 | 0.62 | 0.105 | 0.085 | -0.1325 | 0.741506 | 0.670064 | 0.736683 | 0.992876 | 0.997988 | 0.9971 |
| Nolc1_E9Q5C9_S563s_1 | Nolc1_E9Q | 0.45 | 0.44 | 0.9 | 0.87 | -1.18 | -0.02 | -0.15 | 0.08 | -0.22 | 0.52 | 0.52 | 0.76 | -1.02 | -0.27 | 0.77 | 1 | -0.9825 | -0.27 | -0.545 | 0.26188 | 0.013486 | 0.36656 | 0.989223 | 0.997988 | 0.994666 |
| Nolc1_E9Q5C9_S626s_1 | Nolc1_E9Q | 1.99 | 2.32 | 2.63 | 2.2 | 1.39 | 1.52 | 1.36 | 1.54 | 1.72 | 1.9 | 1.63 | 2.07 | 2.16 | 1.59 | 2.29 | 1.86 | -0.8325 | -0.455 | -0.31 | 0.287727 | 0.004114 | 0.089598 | 0.989223 | 0.997988 | 0.994666 |
| Nono_Q99K48_T452t_1 | Nono_Q99 | -0.4 | -0.69 | -1.08 | -0.87 | -0.54 | -1.61 | -0.76 | -1.41 | -0.9 | -0.87 | -0.87 | -1.21 | -1.58 | -0.65 | -1.04 | -0.64 | -0.32 | -0.2025 | -0.2175 | 0.495255 | 0.318814 | 0.418282 | 0.989223 | 0.997988 | 0.994666 |
| Nop56_Q9D6Z1_S536s_1 | Nop56_Q9 | 0.7 | -0.28 | -0.13 | -0.1 | 0.52 | 0.63 | 0.17 | -0.32 | -0.09 | 0.38 | -0.64 | -0.32 | 0.34 | 0.85 | 0 | 0.29 | 0.2025 | -0.215 | 0.3225 | 0.335889 | 0.533201 | 0.521572 | 0.989223 | 0.997988 | 0.994666 |
| Nop58_Q6DFW4_S509s_1 | Nop58_Q6 | 0.15 | 0.45 | 0.3 | 0.3 | 0.51 | 0.11 | 0.91 | -0.03 | 0.22 | 0.3 | -0.1 | 0.18 | -0.74 | 0.66 | 0.45 | 0.55 | 0.075 | -0.15 | -0.07 | 0.844888 | 0.778018 | 0.500065 | 0.995762 | 0.997988 | 0.999507 |
| Nop58_Q6DFW4_S521s_1 | Nop58_Q6 | -0.57 | 0.28 | 0.88 | 0.71 | -0.2 | -0.49 | 0.37 | 0.8 | -0.59 | 0.3 | 0.86 | 0.67 | -0.65 | 0.11 | 0.55 | 0.83 | -0.205 | -0.015 | -0.115 | 0.796801 | 0.622203 | 0.972864 | 0.989223 | 0.997988 | 0.999507 |
| Nos1_S4R255_S1412s_1 | Nos1_S4R2 | -0.26 | -0.27 | -0.91 | -0.19 | 0.15 | -0.18 | 0.85 | 0.04 | -0.9 | -0.59 | -0.38 | 0.03 | -0.93 | 0.41 | -0.32 | 0.18 | 0.6225 | -0.0525 | 0.2425 | 0.510458 | 0.063531 | 0.860272 | 0.989223 | 0.997988 | 0.994666 |
| Nos1_S4R255_S287s_1 | Nos1_S4R2 | 1.43 | 0.94 | 1.12 | 1.41 | 0.64 | 0.97 | 1.53 | 1.49 | 1.08 | 1.47 | 1.81 | 1.41 | 1.01 | 1.12 | 1.32 | 0.89 | -0.0675 | 0.2175 | -0.14 | 0.58679 | 0.810051 | 0.409198 | 0.995762 | 0.997988 | 0.995259 |
| Nos1_S4R255_S847s_1 | Nos1_S4R2 | -0.49 | 0.19 | -0.93 | -1.24 | -0.28 | -0.31 | -0.94 | -0.8 | -0.64 | -0.2 | -0.04 | -1.35 | -0.57 | 0.56 | -0.16 | -1.19 | 0.035 | 0.06 | 0.2775 | 0.553223 | 0.92109 | 0.885916 | 0.997335 | 0.997988 | 0.994666 |
| Nos1_S4R255_S857s_1 | Nos1_S4R2 | 0.37 | 0.49 | -0.18 | -0.32 | -0.2 | 0.58 | -0.46 | -0.14 | 1.02 | 0.24 | -0.32 | -0.41 | 1.63 | -0.79 | -0.09 | -0.32 | -0.145 | 0.0425 | 0.0175 | 0.973507 | 0.647968 | 0.91245 | 0.99033 | 0.997988 | 0.999507 |
| Nos1_S4R255_T116t_1 | Nos1_S4R2 | 1.02 | 1.27 | 0.69 | 0.48 | 1.17 | 0.89 | 0.9 | 0.46 | 0.13 | 0.61 | 0 | 0.8 | 0.32 | 0.59 | 0.75 | 0.32 | -0.01 | -0.48 | -0.37 | 0.209356 | 0.970463 | 0.133787 | 0.999187 | 0.997988 | 0.994666 |
| Nos1_S4R255_T128t_1 | Nos1_S4R2 | 0.2 | 0 | 0.89 | 0.85 | -0.05 | -0.46 | 0.8 | 0.77 | -0.4 | -0.18 | 1.41 | 0.74 | -0.08 | 0.38 | 0.72 | 0.31 | -0.22 | -0.0925 | -0.1525 | 0.639877 | 0.563083 | 0.839625 | 0.989223 | 0.997988 | 0.995259 |
| Nos1ap_Q9D3A8_S465s_1 | Nos1ap_Q9 | -0.25 | 0.09 | 0.38 | -0.85 | 0.23 | -1.14 | -0.64 | 0.47 | -1.3 | -0.37 | -0.01 | 0.83 | -0.1 | -0.17 | 0.95 | 0.52 | -0.1125 | -0.055 | 0.4575 | 0.255163 | 0.795076 | 0.90995 | 0.995762 | 0.997988 | 0.994666 |
| Nostrin_Q6WKZ7_S429s_1 | Nostrin_Q6 | -0.3 | -0.18 | -1.1 | -0.41 | 0.69 | -0.9 | -1.4 | -1.24 | -0.14 | 0.27 | -1.12 | -0.4 | 0.47 | 0.32 | -0.65 | -0.94 | -0.215 | 0.15 | 0.2975 | 0.473291 | 0.656455 | 0.683683 | 0.990434 | 0.997988 | 0.994666 |
| Npdc1_Q64322_S236s_2 | Npdc1_Q6 | -0.3 | -0.16 | -0.42 | -0.8 | -0.83 | 0.22 | -1 | -0.98 | 1.15 | -0.81 | -1.26 | -0.73 | 0.46 | -1.23 | -0.63 | -0.42 | -0.2275 | 0.0075 | -0.035 | 0.928007 | 0.498471 | 0.98839 | 0.989223 | 0.997988 | 0.999507 |
| Npdc1_Q64322_S243s_1 | Npdc1_Q6 | 0.54 | 0.48 | 0.18 | 0.29 | -0.41 | 0.99 | -0.4 | -0.16 | 1.74 | 0.46 | -1.26 | 0.32 | 1.81 | -0.72 | 0.56 | 0.46 | -0.3675 | -0.0575 | 0.155 | 0.755147 | 0.304295 | 0.919821 | 0.989223 | 0.997988 | 0.9971 |
| Npdc1_Q64322_T239t_1 | Npdc1_Q6 | -0.56 | -0.95 | -0.5 | -0.16 | -0.96 | -1.02 | -0.05 | -0.39 | -0.59 | -0.48 | -1.77 | -0.41 | 0.94 | -2.16 | 0.7 | -0.28 | -0.0625 | -0.27 | 0.3425 | 0.602011 | 0.837974 | 0.470157 | 0.995762 | 0.997988 | 0.995259 |
| Npdc1_Q64322_T239t_2 | Npdc1_Q6 | -0.3 | -0.16 | -0.42 | -0.8 | -0.83 | 0.22 | -1 | -0.98 | 1.15 | -0.81 | -1.26 | -0.73 | 0.46 | -1.23 | -0.63 | -0.42 | -0.2275 | 0.0075 | -0.035 | 0.928007 | 0.498471 | 0.98839 | 0.989223 | 0.997988 | 0.999507 |
| Npepps_E9Q039_S746s_1 | Npepps_E9 | -0.26 | 0.53 | 0.05 | 0.22 | 0.34 | -0.15 | -0.08 | -0.21 | -0.06 | -0.05 | -0.34 | 0 | -0.48 | 0.49 | -0.35 | 0.03 | -0.16 | -0.2475 | -0.2125 | 0.512184 | 0.538666 | 0.342073 | 0.989223 | 0.997988 | 0.994666 |
| Npy1r_Q04573_S367s_1 | Npy1r_Q04 | 0.66 | 0.05 | -0.1 | -0.03 | 1.42 | 1.07 | -0.39 | -0.22 | 0.21 | 0.39 | -1.13 | -0.25 | 0.69 | 1.7 | 0.06 | 0.08 | 0.325 | -0.34 | 0.4875 | 0.263708 | 0.480155 | 0.387016 | 0.989223 | 0.997988 | 0.994666 |
| Nr2f1_Q32NY6_T48t_1 | Nr2f1_Q32 | -3.17 | -0.89 | 0.05 | -1.19 | -0.93 | -0.98 | -0.69 | -0.7 | -1.76 | -1.92 | -0.54 | -0.32 | 0.02 | -0.03 | 1.12 | 0.53 | 0.475 | 0.165 | 1.71 | 0.025515 | 0.442412 | 0.81709 | 0.989223 | 0.997988 | 0.994666 |
| Nr3c1_E9Q310_T168t_1 | Nr3c1_E9Q | -0.59 | -0.84 | -2.36 | -3.05 | -2.01 | -0.43 | -2.26 | -1.46 | -1.41 | 0.3 | -0.29 | -0.85 | 0.59 | -0.94 | -3.94 | -2.28 | 0.17 | 1.1475 | 0.0675 | 0.9454 | 0.790426 | 0.097445 | 0.995762 | 0.997988 | 0.999507 |
| Nr3c2_D6RIL1_S299s_1 | Nr3c2_D6R | -1.77 | -1.47 | -2.5 | -3.1 | -1.11 | -2.2 | -2.4 | -2.32 | -1.31 | -1.41 | -1.22 | -2.56 | 0.97 | -1.27 | -3.03 | -2.26 | 0.2025 | 0.585 | 0.8125 | 0.340034 | 0.650862 | 0.226077 | 0.990358 | 0.997988 | 0.994666 |
| Nrbp1_D3YUV1_S2s_1 | Nrbp1_D3Y | -0.5 | -0.25 | 0.96 | -0.69 | -0.16 | -0.72 | 0.46 | -0.47 | -0.3 | -0.09 | 0.54 | 0.23 | 0.19 | -0.19 | 0.26 | -0.4 | -0.1025 | 0.215 | 0.085 | 0.834454 | 0.809909 | 0.601755 | 0.995762 | 0.997988 | 0.999507 |
| Nrbp1_D3YUV1_S430s_1 | Nrbp1_D3Y | 0.49 | 0.77 | 0.31 | 0.41 | 0.62 | 0.69 | 0.73 | 0.44 | 0.32 | 0.88 | 0.07 | 0.31 | 0.35 | 0.67 | 0.3 | 0.23 | 0.125 | -0.1 | -0.1075 | 0.670011 | 0.562445 | 0.704069 | 0.989223 | 0.997988 | 0.996159 |
| Nrbp1_D3YUV1_T439t_1 | Nrbp1_D3Y | -1.21 | -0.9 | -1.24 | -1.43 | -0.61 | -1.13 | -2.05 | -1.31 | -1.5 | -1.43 | -0.5 | -1.17 | -0.7 | -0.63 | -1.75 | -1.54 | -0.08 | 0.045 | 0.04 | 0.906672 | 0.807716 | 0.878821 | 0.995762 | 0.997988 | 0.999507 |
| Nrbp1_D3YUV1_T441t_1 | Nrbp1_D3Y | -2.48 | -2.39 | -0.2 | -1.16 | -1.01 | -3.09 | -2.91 | -1.16 | -3.18 | -2.95 | -0.26 | -1.65 | -1.18 | -2.14 | -1.17 | -1.09 | -0.485 | -0.4525 | 0.1625 | 0.769345 | 0.485373 | 0.562662 | 0.989223 | 0.997988 | 0.9971 |
| Nrbp2_Q91V36_T409t_1 | Nrbp2_Q9 | -0.08 | 0.4 | 1.04 | 0.94 | 0.73 | 0.96 | 1.21 | 0.32 | 0.97 | 0.36 | 0.49 | 1.06 | -0.78 | 0.51 | 0.9 | 1.78 | 0.23 | 0.145 | 0.0275 | 0.959986 | 0.492849 | 0.666111 | 0.989223 | 0.997988 | 0.999507 |
| Nrcam_Q810U4_S1178s_1 | Nrcam_Q8 | -0.57 | -0.56 | -4.9 | -4.68 | -0.41 | -1.09 | -4.57 | -5.85 | -1.11 | -0.39 | -2.23 | -4.7 | -1.6 | -0.32 | -5.47 | -4.49 | -0.3025 | 0.57 | -0.2925 | 0.842676 | 0.844194 | 0.674072 | 0.996349 | 0.997988 | 0.999507 |
| Nrcam_Q810U4_S1179s_1 | Nrcam_Q8 | -0.28 | 0.15 | 1.11 | 1.16 | 0.42 | 0.05 | 0.41 | 1.62 | -0.8 | -0.16 | 1.13 | 1.81 | -0.93 | 0.22 | 2.02 | 1.77 | 0.09 | -0.04 | 0.235 | 0.736728 | 0.84507 | 0.949458 | 0.996349 | 0.997988 | 0.9971 |
| Nrcam_Q810U4_S1182s_1 | Nrcam_Q8 | 0.96 | 1.8 | 0.96 | 0.69 | 1.26 | 0.64 | 0.8 | 0.65 | 0.85 | 0.94 | 0.56 | 0.66 | -0.41 | 1.25 | 0.41 | 0.5 | -0.265 | -0.35 | -0.665 | 0.134968 | 0.393244 | 0.257218 | 0.989223 | 0.997988 | 0.994666 |
| Nrcam_Q810U4_T1173t_1 | Nrcam_Q8 | -3.45 | -0.56 | -1.96 | -2.62 | -1.47 | -2.54 | -2.32 | -2.01 | -1.44 | -1.94 | -1.26 | -2.99 | -1.2 | -1.91 | -2.75 | -2.78 | 0.0625 | 0.24 | -0.0125 | 0.984604 | 0.914959 | 0.715516 | 0.997335 | 0.997988 | 1 |
| Nrd1_Q8BHG1_S85s_1 | Nrd1_Q8BI | 0.51 | 0.7 | 0.89 | 0.42 | -0.36 | 0.79 | -0.95 | 0.18 | 0.65 | 0.34 | 1.28 | 0.54 | 0.06 | -0.3 | 0.31 | 0.25 | -0.715 | 0.0725 | -0.55 | 0.062298 | 0.08387 | 0.795828 | 0.989223 | 0.997988 | 0.994666 |
| Nrg2_D3YZR2_S464s_1 | Nrg2_D3YZ | 1.26 | 0.8 | -0.6 | -0.64 | 1.77 | 1.3 | -0.25 | -0.22 | 0.92 | 0.91 | -1.35 | -0.63 | 1.59 | 1.09 | 0.12 | -0.7 | 0.445 | -0.2425 | 0.32 | 0.617658 | 0.487639 | 0.720854 | 0.989223 | 0.997988 | 0.995259 |

| | | | | | | | | | | | | | | | | | | | | | | | | | | |
|---|---|---|---|---|---|---|---|---|---|---|---|---|---|---|---|---|---|---|---|---|---|---|---|---|---|---|
| Nrg2_D3YZR2_S470s_1 | Nrg2_D3YZ | -1.47 | -0.86 | -0.08 | -0.2 | -2.68 | 0.22 | -0.66 | -0.52 | 1.04 | 0.17 | 0.03 | -0.72 | 1.87 | -0.33 | 1.01 | -0.25 | -0.2575 | 0.7825 | 1.2275 | 0.054502 | 0.680512 | 0.117 | 0.993464 | 0.997988 | 0.994666 |
| Nrgn_P60761_S76s_1 | Nrgn_P607 | 0.19 | -2.28 | -1.83 | -2.58 | -3.99 | -0.25 | -0.59 | -0.34 | 2.29 | -0.49 | -2.1 | -0.74 | 2.98 | -1.03 | 2.11 | -1.8 | 0.3325 | 1.365 | 2.19 | 0.081149 | 0.727133 | 0.187795 | 0.995569 | 0.997988 | 0.994666 |
| Nrxn1_E0CY11_S1450s_1 | Nrxn1_E0C | 2.32 | 2.66 | 3.01 | 1.81 | 1.61 | 2.39 | 1.41 | 2.24 | 0.77 | 2.64 | 0.99 | 3.01 | 0.3 | -0.61 | 2.67 | 2.22 | -0.5375 | -0.5975 | -1.305 | 0.099373 | 0.149904 | 0.312779 | 0.989223 | 0.997988 | 0.994666 |
| Nrxn1_E0CY11_S1466s_1 | Nrxn1_E0C | 0.76 | 1.11 | 0.77 | 1.01 | 0.68 | 0.37 | 1.12 | 1.29 | 0.64 | 0.99 | 2.02 | 0.96 | 0.1 | 0.95 | 1.25 | 1.72 | -0.0475 | 0.24 | 0.0925 | 0.803325 | 0.86003 | 0.479474 | 0.996493 | 0.997988 | 0.999507 |
| Nsd1_E9QAE4_S2232s_1 | Nsd1_E9Q | 0.82 | 0.9 | 0.74 | 0.98 | 0.69 | 0.7 | 1.22 | 0.98 | 0.6 | 1.13 | 1.05 | 1.22 | 0.36 | 1 | 1.16 | 0.83 | 0.0375 | 0.14 | -0.0225 | 0.933392 | 0.865764 | 0.557339 | 0.996493 | 0.997988 | 0.999507 |
| Nsd1_E9QAE4_S2370s_1 | Nsd1_E9Q | 1.1 | 0.53 | -0.2 | 0.38 | 0.74 | 1.16 | -0.89 | -0.32 | 1.5 | 1.38 | -0.69 | 0.21 | 1.78 | 0.12 | 0.21 | -0.16 | -0.28 | 0.1475 | 0.035 | 0.942778 | 0.576394 | 0.785866 | 0.989223 | 0.997988 | 0.999507 |
| Nsf_P46460_S207s_1 | Nsf_P4646 | 0.11 | 0.48 | 0.17 | 0.03 | 0.75 | 0.03 | 0.49 | 0.38 | -0.81 | -0.51 | -0.74 | 0.45 | -0.46 | 0.42 | -0.07 | 0.66 | 0.215 | -0.6 | -0.06 | 0.850308 | 0.384944 | 0.096295 | 0.989223 | 0.997988 | 0.999507 |
| Nsf_P46460_S298s_1 | Nsf_P4646 | -0.31 | 0.26 | 0.98 | 0.58 | -0.37 | 0.44 | -0.04 | 0.22 | -0.87 | -0.03 | 1.37 | -0.1 | 0.07 | 0.72 | 0.22 | 0.97 | -0.315 | -0.285 | 0.1175 | 0.748723 | 0.355447 | 0.577416 | 0.989223 | 0.997988 | 0.9971 |
| Nsf_P46460_S31s_1 | Nsf_P4646 | 0.61 | 0.45 | 0.78 | -1.12 | -0.6 | -0.04 | 0.74 | -0.23 | 1.33 | 0.47 | 1.24 | -0.02 | 1.15 | 0.25 | 0.36 | 0.35 | -0.2125 | 0.575 | 0.3475 | 0.464022 | 0.660242 | 0.277148 | 0.990434 | 0.997988 | 0.994666 |
| Nsfl1c_Q9CZ44_S114s_1 | Nsfl1c_Q9C | -1.14 | -0.5 | -0.44 | -0.61 | 0.04 | -1.44 | -0.26 | -0.99 | -0.94 | -0.77 | -0.9 | -0.45 | -1.12 | -0.63 | -0.57 | -0.48 | 0.01 | -0.0925 | -0.0275 | 0.923554 | 0.978328 | 0.72322 | 0.999187 | 0.997988 | 0.999507 |
| Nsfl1c_Q9CZ44_S140s_1 | Nsfl1c_Q9C | -3.29 | -3.1 | -0.59 | -1.07 | -2.15 | -2.42 | -0.7 | -0.25 | -0.76 | -2.32 | -0.93 | -0.94 | 0.04 | -1.13 | 0.25 | -0.18 | 0.6325 | 0.775 | 1.7575 | 0.026046 | 0.416839 | 0.287073 | 0.989223 | 0.997988 | 0.994666 |
| Nsfl1c_Q9CZ44_S176s_1 | Nsfl1c_Q9C | 4.33 | 1.82 | 3.47 | 0.35 | 2.54 | 4.68 | 4.07 | 3.32 | -0.29 | 3.47 | 0.53 | 3.48 | 0.02 | -0.41 | 3.85 | 4.22 | 1.16 | -0.695 | -0.5725 | 0.661737 | 0.202775 | 0.553223 | 0.989223 | 0.997988 | 0.995804 |
| Nsmf_Q99NF2_S148s_1 | Nsmf_Q99 | -0.5 | -0.8 | -0.68 | -1.43 | 0.05 | -0.33 | -1.39 | -0.95 | -0.45 | -1.24 | -0.96 | -0.77 | 0.36 | -0.09 | -2.08 | -1.22 | 0.1975 | -0.0025 | 0.095 | 0.86174 | 0.598407 | 0.993359 | 0.989223 | 0.997988 | 0.999507 |
| Nsmf_Q99NF2_S153s_1 | Nsmf_Q99 | -1.23 | -1.04 | -1.86 | -0.73 | -1.04 | -0.97 | -1.37 | -2.02 | -1.43 | -0.78 | -1.81 | -1.69 | -0.17 | -0.81 | -1.25 | -0.92 | -0.135 | -0.2125 | 0.4275 | 0.247451 | 0.694925 | 0.545253 | 0.994893 | 0.997988 | 0.994666 |
| Nsmf_Q99NF2_S158s_1 | Nsmf_Q99 | 0.41 | 0.23 | 0.39 | 0.17 | 0.29 | 0.66 | 0.21 | 0.61 | -0.02 | 0.18 | -0.14 | 0.05 | -0.16 | -0.08 | 0.49 | 1.13 | 0.1425 | -0.2825 | 0.045 | 0.894693 | 0.517679 | 0.208023 | 0.989223 | 0.997988 | 0.999507 |
| Nsmf_Q99NF2_S292s_1 | Nsmf_Q99 | -0.11 | -0.13 | 0.53 | 0.6 | 0.43 | -0.31 | 0.59 | 0.37 | -0.14 | 0.15 | 0.24 | 0.85 | -0.29 | 0.73 | 1.02 | 0.35 | 0.0475 | 0.0525 | 0.23 | 0.534952 | 0.875622 | 0.868714 | 0.996493 | 0.997988 | 0.994666 |
| Nsmf_Q99NF2_S36s_1 | Nsmf_Q99 | 0.24 | 0.14 | -0.8 | -0.44 | 0.28 | 0.28 | -1.24 | -1.9 | 1.36 | 0.56 | -1.35 | -0.82 | 0.66 | 1.94 | -2.01 | -0.85 | -0.43 | 0.1525 | 0.15 | 0.850048 | 0.438371 | 0.803183 | 0.989223 | 0.997988 | 0.999507 |
| Nt5c_Q9JM14_T102t_1 | Nt5c_Q9JN | -1.09 | -2.17 | -2.12 | -1.93 | -0.95 | -1.69 | -2.55 | -1.06 | -1.91 | -1.3 | -1.02 | -3 | -2.34 | -0.72 | -3.67 | -1.17 | 0.265 | 0.02 | -0.1475 | 0.8175 | 0.534273 | 0.966715 | 0.989223 | 0.997988 | 0.999507 |
| Ntrk3_Q6VNS1_S561s_1 | Ntrk3_Q6V | -0.03 | -0.31 | -2.24 | -2.05 | -0.85 | 0.99 | -0.74 | -1.69 | -0.09 | 0.17 | -0.29 | -3 | -0.31 | -0.86 | -2.2 | -1.15 | 0.585 | 0.355 | 0.0275 | 0.965276 | 0.417345 | 0.671443 | 0.989223 | 0.997988 | 0.999507 |
| Ntsr1_O88319_S407s_1 | Ntsr1_O88 | -1.26 | -1.42 | -1.71 | -0.52 | -1.14 | -1.13 | -0.59 | -0.58 | -0.64 | -0.98 | -1.05 | -0.65 | -0.4 | -0.72 | -0.85 | -0.86 | 0.3675 | 0.3975 | 0.52 | 0.130809 | 0.262025 | 0.220081 | 0.989223 | 0.997988 | 0.994666 |
| Nuak1_Q641K5_T212t_1 | Nuak1_Q6 | 0.99 | 0.05 | 0.68 | -0.02 | 1.24 | -0.25 | 0.44 | 0.17 | 0.99 | 0.16 | -1.82 | 0.13 | -1.38 | 1.52 | -0.29 | 0.62 | -0.025 | -0.56 | -0.3075 | 0.615447 | 0.94844 | 0.355989 | 0.999187 | 0.997988 | 0.995259 |
| Nucb1_Q02819_S368s_1 | Nucb1_Q0 | -0.48 | -0.31 | -0.2 | -0.86 | 0.49 | -0.84 | -0.19 | -0.11 | -0.25 | 0.09 | -0.08 | -0.45 | 0.27 | 0.47 | 0.58 | 0.42 | 0.3 | 0.29 | 0.8975 | 0.006187 | 0.362798 | 0.273096 | 0.989223 | 0.997988 | 0.994666 |
| Nucb1_Q02819_S456s_1 | Nucb1_Q0 | 0.36 | 1.1 | 0.09 | 0.43 | 1.03 | -0.28 | 1 | 0.72 | -0.29 | 0.66 | 0.36 | 0.52 | -0.1 | 0.65 | 1.14 | 0.37 | 0.1225 | -0.1825 | 0.02 | 0.955775 | 0.740752 | 0.579124 | 0.995762 | 0.997988 | 0.999507 |
| Nucb1_Q02819_S85s_1 | Nucb1_Q0 | 0.62 | 0.45 | 0.79 | 0.37 | 1.03 | 0.06 | 0.86 | 0.41 | 0.27 | 1 | 0.28 | 0.52 | -0.27 | 0.63 | 1.05 | 0.93 | 0.0325 | -0.04 | 0.0275 | 0.936447 | 0.906462 | 0.877947 | 0.996975 | 0.997988 | 0.999507 |
| Nucks1_Q80XU3_S181s_1 | Nucks1_Q8 | -2.47 | -1.8 | -3.33 | -1.96 | -2.41 | -2.24 | -2.05 | -3.24 | -0.71 | -0.99 | -1.4 | -2.28 | -1.49 | -0.63 | -3.04 | -2.84 | -0.095 | 1.045 | 0.39 | 0.525804 | 0.818037 | 0.046941 | 0.995762 | 0.997988 | 0.994666 |
| Nucks1_Q80XU3_S214s_1 | Nucks1_Q8 | -3.58 | -3.05 | -2.68 | -3.22 | -3.79 | -4.04 | -2.92 | -2.99 | -3.59 | -2.86 | -1.73 | -2.6 | -2.56 | -3.56 | -2.74 | -2.5 | -0.3025 | 0.4375 | 0.2925 | 0.402382 | 0.388283 | 0.310973 | 0.989223 | 0.997988 | 0.994666 |
| Nucks1_Q80XU3_S58s_2 | Nucks1_Q8 | -3.13 | -2.78 | -1.91 | -1.81 | -2.61 | -3.54 | -3.48 | -3.44 | -1.17 | -2.54 | -0.67 | -1.58 | -1.24 | -3.34 | -2.37 | -2.28 | -0.86 | 0.9175 | 0.1 | 0.84372 | 0.046337 | 0.084875 | 0.989223 | 0.997988 | 0.999507 |
| Nucks1_Q80XU3_S61s_2 | Nucks1_Q8 | -3.13 | -2.78 | -1.91 | -1.81 | -2.61 | -3.54 | -3.48 | -3.44 | -1.17 | -2.54 | -0.67 | -1.58 | -1.24 | -3.34 | -2.37 | -2.28 | -0.86 | 0.9175 | 0.1 | 0.84372 | 0.046337 | 0.084875 | 0.989223 | 0.997988 | 0.999507 |
| Nucks1_Q80XU3_S79s_1 | Nucks1_Q8 | 4.39 | 2.65 | -1.79 | -4.82 | 1.46 | 4.4 | -2.76 | -2.82 | 3.54 | 4.53 | -1 | -2.36 | 0.86 | -2.34 | -1.82 | -2.75 | -0.0375 | 1.07 | -1.62 | 0.408211 | 0.987146 | 0.648563 | 0.999187 | 0.997988 | 0.994666 |
| Nudc_O35685_S136s_1 | Nudc_O35 | 2.28 | 3.46 | 5 | 3.63 | 3.33 | 2.21 | 3.96 | 4.33 | 1.03 | 2.64 | 4.39 | 4.2 | -1.92 | 1.04 | 5.3 | 5.35 | -0.135 | -0.5275 | -1.15 | 0.476116 | 0.833944 | 0.541426 | 0.995762 | 0.997988 | 0.994666 |
| Nudc_O35685_S139s_1 | Nudc_O35 | -2.23 | -1.49 | 1.63 | 0.7 | -0.88 | -2.47 | 1.21 | 1.04 | -2.44 | -1.54 | 0.47 | 1.15 | -2.05 | -1.55 | 1.8 | 1.86 | 0.0725 | -0.2425 | 0.3625 | 0.762968 | 0.946646 | 0.823639 | 0.999187 | 0.997988 | 0.9971 |
| Nudc_O35685_T108t_1 | Nudc_O35 | -0.1 | -0.3 | 0.11 | -0.03 | -0.95 | -0.13 | -0.19 | -0.07 | -0.08 | -0.5 | 1.62 | 0.4 | 0.27 | -0.16 | 1 | 0.87 | -0.255 | 0.44 | 0.575 | 0.10364 | 0.352155 | 0.340274 | 0.989223 | 0.997988 | 0.994666 |
| Nudt11_P0C028_T150t_1 | Nudt11_P0 | 1.26 | 1.16 | -0.01 | 0.81 | 1.4 | 1.05 | 0.4 | 0.98 | 0.43 | 1.56 | 0.64 | -0.11 | -0.26 | 0.77 | 1.19 | -0.02 | 0.1525 | -0.175 | -0.385 | 0.387964 | 0.669017 | 0.690457 | 0.992876 | 0.997988 | 0.994666 |
| Nufip2_Q5F2E7_S213s_1 | Nufip2_Q5 | -1.49 | -1.7 | -0.48 | -0.26 | -3.46 | -1.33 | -0.77 | -0.4 | -1.28 | -1.55 | -0.19 | -0.78 | -0.08 | -0.84 | -0.3 | 0.1 | -0.5075 | 0.0325 | 0.7025 | 0.114906 | 0.463743 | 0.942358 | 0.989223 | 0.997988 | 0.994666 |
| Nufip2_Q5F2E7_S376s_1 | Nufip2_Q5 | 1.44 | 1.08 | -0.55 | -1.49 | 0.24 | -0.04 | -0.81 | -0.37 | -0.03 | -0.09 | 0.4 | -2.59 | 0.95 | 0.76 | 0.37 | -0.08 | -0.365 | -0.6975 | 0.38 | 0.565083 | 0.574173 | 0.426297 | 0.989223 | 0.997988 | 0.995259 |
| Nufip2_Q5F2E7_S626s_1 | Nufip2_Q5 | 0.1 | 0.49 | -0.05 | -0.02 | 0.82 | -0.33 | 0.2 | 0.06 | -1 | 0.33 | 1.1 | 0.49 | -0.74 | 0.23 | 0.32 | 0.7 | 0.0575 | 0.1 | -0.0025 | 0.994408 | 0.845972 | 0.821745 | 0.996349 | 0.997988 | 1 |
| Numa1_E9Q7G0_S1739s_1 | Numa1_E9 | 1.15 | 0.86 | 1.7 | 1.98 | 0.08 | 0.03 | 0.9 | 0.63 | 0.5 | 0.18 | 1.08 | 1.51 | 0.76 | 0.52 | 2.25 | 2.19 | -1.0125 | -0.605 | 0.0075 | 0.987948 | 0.013444 | 0.147344 | 0.989223 | 0.997988 | 1 |
| Numa1_E9Q7G0_S398s_1 | Numa1_E9 | -0.69 | -0.19 | 0.51 | 0.41 | -0.64 | -0.99 | -1 | 0.44 | -0.37 | 0.42 | 0.08 | 0.46 | 0.4 | -0.18 | -0.57 | 0.42 | -0.5575 | 0.1375 | 0.0075 | 0.98435 | 0.203738 | 0.698495 | 0.989223 | 0.997988 | 1 |
| Numa1_E9Q7G0_T1982t_1 | Numa1_E9 | -0.46 | -0.12 | 1.04 | 0.43 | -1.48 | 0.42 | -0.61 | -0.19 | 0.24 | -0.03 | -0.51 | 0.87 | 1.21 | -1.43 | 0 | -0.42 | -0.6875 | -0.08 | -0.3825 | 0.51756 | 0.172844 | 0.851083 | 0.989223 | 0.997988 | 0.994666 |
| Numb_Q9QZS3_S230s_1 | Numb_Q9 | 0.61 | 0.51 | 0.78 | 1.29 | 0.36 | 0.59 | 1.11 | 1.08 | 0.48 | 1.21 | 0.78 | 1.71 | 0.19 | 1.07 | 1.22 | 1.28 | -0.0125 | 0.2475 | 0.1425 | 0.677983 | 0.965107 | 0.471975 | 0.999187 | 0.997988 | 0.996159 |
| Numb_Q9QZS3_S351s_1 | Numb_Q9 | -0.83 | -0.99 | -3.58 | -3.33 | -0.8 | -0.75 | -1.53 | -1.55 | -2.14 | -0.75 | -2.97 | -2.53 | -1.71 | -0.21 | -2.66 | -1.88 | 1.025 | 0.085 | 0.5675 | 0.480423 | 0.155738 | 0.913858 | 0.989223 | 0.997988 | 0.994666 |
| Numb_Q9QZS3_T378t_1 | Numb_Q9 | -1.59 | -1.76 | -1.33 | -1.7 | -2.21 | -0.45 | -1.41 | -1.86 | 0.63 | -1.54 | -0.8 | -0.87 | 0.97 | -1.25 | -0.97 | -1.21 | 0.1125 | 0.95 | 0.98 | 0.079305 | 0.769154 | 0.058765 | 0.995762 | 0.997988 | 0.994666 |
| Numbl_O08919_S263s_1 | Numbl_O0 | -1.02 | -0.16 | 0.26 | -0.92 | -0.14 | -1.03 | 0.43 | 0.08 | -1.09 | -1.01 | -0.37 | 0.16 | -0.65 | 0.09 | -0.14 | -0.08 | 0.295 | -0.1175 | 0.265 | 0.476788 | 0.484414 | 0.778595 | 0.989223 | 0.997988 | 0.994666 |
| Numbl_O08919_S270s_1 | Numbl_O0 | -1.23 | 0.03 | -0.42 | -0.58 | -0.6 | -0.92 | 0.45 | 0.25 | -0.17 | -0.34 | -0.92 | -0.2 | -0.05 | -0.01 | -0.81 | 0 | 0.345 | 0.1425 | 0.3325 | 0.360113 | 0.401702 | 0.672607 | 0.989223 | 0.997988 | 0.994666 |
| Numbl_O08919_S411s_1 | Numbl_O0 | -0.79 | -1.29 | 0.26 | -0.46 | -2.22 | 0.09 | -0.76 | -0.29 | 1.35 | -0.97 | -0.07 | -0.45 | 1.37 | -1.87 | 0.32 | -0.54 | -0.225 | 0.535 | 0.39 | 0.570003 | 0.680387 | 0.343986 | 0.993464 | 0.997988 | 0.995259 |
| Numbl_O08919_T279t_1 | Numbl_O0 | -0.46 | -0.67 | -1.07 | -1.23 | -0.31 | -0.85 | -0.96 | -0.81 | -1.22 | -0.57 | -0.35 | -0.44 | -1.5 | -0.36 | -0.82 | -0.58 | 0.125 | 0.2125 | 0.0425 | 0.900345 | 0.645755 | 0.488858 | 0.99033 | 0.997988 | 0.999507 |
| Numbl_O08919_T409t_1 | Numbl_O0 | -0.32 | -1.38 | -0.61 | 0.05 | -1.45 | 0.28 | -0.44 | -0.56 | 1.1 | -0.89 | -1.22 | -0.59 | 1.26 | -2.03 | 0.09 | -0.47 | 0.0225 | 0.165 | 0.2775 | 0.680878 | 0.959052 | 0.766343 | 0.999187 | 0.997988 | 0.996159 |
| Nup107_E9Q4V9_S11s_1 | Nup107_E9 | 0.88 | 1.33 | 0.27 | 0.76 | 1.12 | 0.64 | 0.99 | 0.78 | -0.21 | 0.31 | 0.12 | 0.64 | -0.07 | 0.31 | 1.03 | 0.57 | 0.0725 | -0.595 | -0.35 | 0.328224 | 0.795266 | 0.083632 | 0.995762 | 0.997988 | 0.994666 |
| Nup160_Q9Z0W3_S1123s_1 | Nup160_Q | 0.55 | 0.42 | 0.11 | 0.43 | 0.63 | -0.2 | 1.07 | 1.18 | -0.79 | -0.06 | -0.48 | -0.1 | -0.29 | 0.62 | 0.53 | 0.14 | 0.2925 | -0.735 | -0.1275 | 0.666112 | 0.392813 | 0.018487 | 0.989223 | 0.997988 | 0.996159 |
| Nup188_Q6ZQH8_S1719s_1 | Nup188_Q | 0.79 | 0.95 | 0.23 | 0.35 | 0.57 | 0.59 | 0.66 | 0.63 | 0.09 | 0.49 | 0.51 | -0.02 | 0.97 | 0.72 | 0.69 | 0.23 | 0.0325 | -0.3125 | 0.0725 | 0.806564 | 0.892 | 0.27262 | 0.996493 | 0.997988 | 0.999507 |
| Nup205_B9EJ54_S1165s_1 | Nup205_B9 | -0.8 | -0.74 | -0.8 | -1.9 | -0.77 | -1.52 | -1.01 | -0.64 | -1.53 | -1.1 | -1.61 | -0.59 | -0.75 | -0.42 | -0.33 | -0.91 | 0.075 | -0.1475 | 0.4575 | 0.205289 | 0.827994 | 0.693111 | 0.995762 | 0.997988 | 0.994666 |
| Nup210_Q9QY81_S1848s_1 | Nup210_Q | -0.63 | 0.03 | -0.17 | 0.13 | -0.32 | -0.39 | 0.21 | 0.02 | 0.16 | 0.01 | -0.64 | -0.29 | 0.38 | 0.05 | 0.1 | 0.62 | 0.04 | -0.03 | 0.4475 | 0.142898 | 0.880236 | 0.917482 | 0.996493 | 0.997988 | 0.994666 |
| Nup35_Q8R4R6_S258s_1 | Nup35_Q8 | 0.5 | 0.51 | 0.94 | 0.41 | 0.04 | 0.17 | 1.05 | 0.9 | -0.06 | 0.15 | 0.98 | 0.43 | 0.07 | 0.75 | 0.97 | 0.59 | -0.05 | -0.215 | 0.005 | 0.986354 | 0.869132 | 0.474532 | 0.996493 | 0.997988 | 1 |
| Nup35_Q8R4R6_T265t_1 | Nup35_Q8 | 0.33 | 0.35 | 0.37 | 0.09 | 0.57 | 0.13 | 0.77 | 0.53 | 0.22 | -0.31 | 0.24 | 0.06 | -0.1 | 0.18 | 0.63 | 0.37 | 0.215 | -0.2325 | -0.015 | 0.954538 | 0.356242 | 0.334867 | 0.989223 | 0.997988 | 0.999507 |
| Nup35_Q8R4R6_T274t_1 | Nup35_Q8 | 0.08 | 0.28 | 0.55 | 0.18 | 0.06 | 0.36 | 0.03 | 0.58 | 0.28 | -0.26 | -0.41 | -0.19 | 1.13 | 0.87 | 0.17 | 0.75 | -0.015 | -0.4175 | 0.4575 | 0.143138 | 0.949125 | 0.124211 | 0.999187 | 0.997988 | 0.994666 |
| Nup50_Q9JIH2_S322s_1 | Nup50_Q9 | 0.09 | -0.62 | -0.38 | -0.66 | -1.86 | 0.65 | -1.51 | -0.3 | 1.99 | -0.21 | 0.48 | -0.27 | 2.57 | -1.07 | -0.15 | -0.23 | -0.3625 | 0.89 | 0.6725 | 0.362968 | 0.509842 | 0.112043 | 0.989223 | 0.997988 | 0.994666 |
| Nup88_Q8CEC0_S699s_1 | Nup88_Q8 | 1.08 | 0.95 | 0.43 | 0.06 | 0.86 | 0.8 | 1.17 | 0.77 | 0.74 | 0.66 | 0.79 | 0.22 | 0.87 | 1.09 | 1.59 | 0.69 | 0.27 | -0.0275 | 0.43 | 0.226819 | 0.356679 | 0.92834 | 0.989223 | 0.997988 | 0.994666 |
| Nup93_Q8BJ71_S644s_1 | Nup93_Q8 | 1.18 | 1.2 | 1.17 | 1.04 | -0.08 | 1.17 | -0.02 | 0.52 | 1.3 | 0.49 | -0.03 | 0.44 | 2.32 | -0.36 | 1.26 | 1.16 | -0.75 | -0.5975 | -0.0525 | 0.919259 | 0.035517 | 0.081082 | 0.989223 | 0.997988 | 0.999507 |
| Nwd2_Q6P5U7_S1662s_1 | Nwd2_Q6F | -1.84 | -0.84 | -1.45 | -2.64 | -0.64 | -1.18 | -1.11 | -1.56 | 0.15 | -2.15 | -2.76 | -1.02 | 1.26 | -1.73 | 0.41 | -1.51 | 0.57 | 0.2475 | 1.3 | 0.10144 | 0.181107 | 0.714255 | 0.989223 | 0.997988 | 0.994666 |
| Nwd2_Q6P5U7_S792s_1 | Nwd2_Q6F | -0.44 | 0.26 | 1.59 | 0.52 | 0.83 | -0.31 | 0.51 | 0.29 | -1.07 | -0.73 | 1.78 | 1.98 | -1.02 | 0.38 | 0.47 | 0.99 | -0.1525 | 0.0075 | -0.2775 | 0.620148 | 0.736742 | 0.992619 | 0.995762 | 0.997988 | 0.995259 |
| Nyap1_E9PXJ0_T705t_1 | Nyap1_E9F | 0.04 | -0.26 | 0.05 | -0.1 | 0.38 | -3.45 | 0.48 | 0.01 | 1.05 | 0.29 | 0.44 | -0.02 | -1.06 | -0.44 | -1.69 | -0.36 | -0.5775 | 0.5075 | -0.82 | 0.038887 | 0.488551 | 0.100951 | 0.989223 | 0.997988 | 0.994666 |
| Nyap2_Q8BM65_S184s_1 | Nyap2_Q8 | -0.17 | 0.01 | 0.34 | -0.35 | -0.48 | -0.18 | 0.31 | 0.39 | 0.51 | 0.24 | 0.67 | 0.38 | 0.2 | -0.29 | 0.78 | 0.32 | 0.0525 | 0.4925 | 0.295 | 0.360663 | 0.85428 | 0.073514 | 0.996349 | 0.997988 | 0.994666 |
| Nyap2_Q8BM65_S238s_1 | Nyap2_Q8 | -0.93 | -0.89 | -1.43 | -1.24 | -0.16 | -0.98 | -0.62 | -1.61 | -1.61 | -1.09 | -0.76 | -1.33 | -1.1 | -0.03 | -1 | -1.17 | 0.28 | -0.075 | 0.2975 | 0.384697 | 0.417095 | 0.786107 | 0.989223 | 0.997988 | 0.994666 |
| Nyap2_Q8BM65_S732s_1 | Nyap2_Q8 | 1.02 | 1.01 | 1.18 | 0.31 | 1.15 | 0.8 | 0.75 | 0.33 | 0.89 | 1.3 | 0.44 | 0.81 | 1.31 | 1.18 | 0.68 | 0.24 | -0.1225 | -0.02 | -0.0275 | 0.936445 | 0.671793 | 0.946991 | 0.992876 | 0.997988 | 0.999507 |
| Nyap2_Q8BM65_Y300y_1 | Nyap2_Q8 | -0.85 | -1.91 | -0.7 | -1.16 | -1.11 | -1.16 | -1.54 | -1.62 | -1.35 | -1.13 | -0.1 | -1.09 | -1.07 | -0.91 | -2.01 | -1.35 | -0.2025 | 0.2375 | -0.18 | 0.636165 | 0.525808 | 0.545248 | 0.989223 | 0.997988 | 0.995259 |
| O35375-5_O35375_S869s_1 | O35375-5_ | 1.54 | 1 | 0.29 | 1.12 | 0.62 | 1.51 | 0.75 | 1.38 | 0.45 | 0.78 | 1.06 | 1.05 | 0.67 | -0.59 | 0.83 | 0.72 | 0.0775 | -0.1525 | -0.58 | 0.190481 | 0.822751 | 0.639782 | 0.995762 | 0.997988 | 0.994666 |
| Ociad1_Q9CRD0_T146t_1 | Ociad1_Q9 | 1.33 | 0.96 | 0.63 | 0.57 | 1.09 | 1.43 | 0.96 | 0.99 | 0.44 | 1.02 | 0.38 | 0.81 | 0.25 | 0.96 | 0.4 | 0.69 | 0.245 | -0.21 | -0.2975 | 0.331483 | 0.352193 | 0.463717 | 0.989223 | 0.997988 | 0.994666 |
| Ocln_Q61146_S368s_1 | Ocln_Q611 | 1 | 0.73 | -0.3 | -1.18 | 0.49 | 0.51 | -0.95 | 0.68 | 0.1 | 0.91 | 2.34 | 0.75 | 0.42 | 0.08 | -0.39 | 1.51 | 0.12 | 0.9625 | 0.3425 | 0.564054 | 0.831955 | 0.149786 | 0.995762 | 0.997988 | 0.995259 |
| Ogfr_Q99PG2_S403s_1 | Ogfr_Q99P | 0.45 | 0.45 | 0.49 | 0.67 | 0.85 | 0.36 | 0.95 | 0.67 | 0.37 | 0.79 | 0.34 | 0.88 | -0.18 | 0.94 | 0.5 | 0.75 | 0.1925 | 0.08 | -0.0125 | 0.96746 | 0.397863 | 0.736739 | 0.989223 | 0.997988 | 0.999507 |
| Ogfrl1_Q8VE52_S406s_1 | Ogfrl1_Q8 | 0.26 | 0.38 | 1.08 | 1.02 | 0.18 | -0.01 | 0.24 | 0.66 | 2.24 | 0.04 | 1.09 | 1.22 | 1.31 | 0.03 | 1.21 | 1.07 | -0.4175 | 0.4625 | 0.22 | 0.565573 | 0.168834 | 0.341673 | 0.989223 | 0.997988 | 0.995259 |
| Ogfrl1_Q8VE52_S65s_1 | Ogfrl1_Q8 | -0.99 | -0.63 | 0.67 | 0.66 | -0.34 | -1.8 | 0.83 | 0.8 | -0.78 | -0.85 | 0.78 | 0.79 | -1.16 | 0.05 | 1.05 | 0.81 | -0.055 | 0.0575 | 0.26 | 0.666724 | 0.934423 | 0.921211 | 0.998658 | 0.997988 | 0.996159 |
| Opalin_Q7M750_S100s_1 | Opalin_Q7 | 1.08 | 1.11 | 1.76 | 1.16 | 0.85 | 0.76 | 0.84 | 1 | 0.39 | 0.54 | 0.84 | 0.53 | 0.76 | 0.73 | 1.25 | 0.89 | -0.415 | -0.7025 | -0.37 | 0.206589 | 0.105692 | 0.020837 | 0.989223 | 0.997988 | 0.994666 |
| Opalin_Q7M750_S90s_1 | Opalin_Q7 | 1.34 | 1.24 | 1.58 | 1.46 | 0.77 | 0.37 | 1 | 0.42 | 0.78 | 0.73 | 1.19 | 0.76 | 1.42 | -0.01 | 1.44 | 1.23 | -0.765 | -0.54 | -0.385 | 0.315706 | 0.008823 | 0.039375 | 0.989223 | 0.997988 | 0.994666 |
| Ophn1_B1AXS4_T52t_1 | Ophn1_B1 | -0.14 | 0.21 | 0.35 | 0.18 | 0.38 | -0.22 | 0.22 | 0.34 | 0.21 | 0.59 | 1.15 | 0.54 | 0.85 | 0.84 | 0.67 | 0.22 | 0.03 | 0.4725 | 0.495 | 0.091419 | 0.899971 | 0.112412 | 0.996493 | 0.997988 | 0.994666 |
| Oprl1_B0R0C0_S329s_1 | Oprl1_B0R | 1.85 | 0.59 | 0.73 | 0.98 | 0.43 | 1.46 | 0.03 | 1.07 | 0.89 | 0.23 | 0.77 | 0.96 | 1.2 | -0.37 | 0.82 | 0.28 | -0.29 | -0.325 | -0.555 | 0.22262 | 0.483337 | 0.358352 | 0.989223 | 0.997988 | 0.994666 |
| Oprm1_E9Q2X5_S311s_1 | Oprm1_E9 | 1.1 | 1.03 | 0.14 | 0.09 | 0.44 | 1.47 | -0.23 | -0.36 | 0.63 | 0.84 | -1.27 | -1.2 | 1 | -2.38 | 0.93 | 1.03 | -0.26 | -0.84 | -0.445 | 0.573313 | 0.578951 | 0.173044 | 0.989223 | 0.997988 | 0.995259 |
| Oprm1_E9Q2X5_T306t_1 | Oprm1_E9 | 2.11 | -0.26 | -0.15 | 1.16 | -0.46 | 2.15 | -0.27 | 1.52 | 0.96 | 0.8 | 1.12 | 0.85 | 1.27 | 1 | 1.28 | 0.51 | 0.02 | 0.2175 | 0.3 | 0.588487 | 0.978847 | 0.68351 | 0.999187 | 0.997988 | 0.995259 |
| Osbp_Q3B7Z2_S236s_1 | Osbp_Q3B | -0.98 | -1 | -0.53 | -0.27 | 0.4 | -0.71 | 0.04 | -1.1 | -1.22 | -1.01 | -0.3 | -0.36 | -1.02 | -0.09 | -0.88 | -0.65 | 0.3525 | -0.0275 | 0.035 | 0.912685 | 0.36221 | 0.931596 | 0.989223 | 0.997988 | 0.999507 |
| Osbp_Q3B7Z2_S238s_1 | Osbp_Q3B | 0.44 | 0.38 | 0.58 | 0.54 | 0.53 | 0.32 | 0.76 | 0.51 | 0.47 | 0.56 | 0.51 | 0.3 | -0.27 | 0.94 | 0.16 | 0.49 | 0.045 | -0.025 | -0.155 | 0.62203 | 0.828525 | 0.904805 | 0.995762 | 0.997988 | 0.995259 |
| Osbp_Q3B7Z2_S380s_1 | Osbp_Q3B | 0.53 | 1.57 | 0.85 | 0.47 | 1.29 | 0.63 | 0.75 | 0.83 | 0.22 | 0.95 | -1.58 | 1.08 | 0.31 | 0.6 | 0.35 | 0.94 | 0.02 | -0.6875 | -0.305 | 0.368525 | 0.948551 | 0.274022 | 0.999187 | 0.997988 | 0.994666 |
| Osbp_Q3B7Z2_S383s_1 | Osbp_Q3B | 0.32 | -0.3 | 0.21 | 0.27 | 0.49 | -0.03 | 0.45 | 1.21 | 0.08 | 0.31 | -0.6 | 0.04 | -0.08 | 0.19 | 0.05 | 0.5 | 0.405 | -0.1675 | 0.04 | 0.883923 | 0.214357 | 0.564976 | 0.989223 | 0.997988 | 0.999507 |
| Osbp_Q3B7Z2_S384s_1 | Osbp_Q3B | 1.35 | 0.99 | 0.69 | 0.18 | 0.93 | 0.2 | 0.41 | 0.31 | 0.67 | 0.95 | -2.01 | 0.04 | -0.56 | 0.39 | -0.78 | 0.61 | -0.34 | -0.89 | -0.8875 | 0.058748 | 0.293432 | 0.192058 | 0.989223 | 0.997988 | 0.994666 |
| Osbp2_Q5QNQ6_S759s_1 | Osbp2_Q5 | 3.73 | 1.07 | 6.91 | -1.41 | 0.57 | 3.99 | 5.63 | 4.16 | 1.97 | 3.16 | 0.85 | 6.74 | 2.44 | -0.05 | 5.72 | 6.5 | 1.0125 | 0.605 | 1.0775 | 0.593001 | 0.572428 | 0.751919 | 0.989223 | 0.997988 | 0.995259 |
| Osbpl10_S4R1M9_S28s_1 | Osbpl10_S | -0.71 | 0.55 | -1.19 | -0.8 | 0.65 | -0.13 | -1.1 | -1.3 | -0.52 | -0.46 | 0.93 | -0.36 | 0.51 | -1.04 | 0.03 | 0.05 | 0.0675 | 0.435 | 0.425 | 0.38452 | 0.899404 | 0.380241 | 0.996493 | 0.997988 | 0.994666 |
| Osbpl10_S4R1M9_S58s_1 | Osbpl10_S | -1.2 | -0.39 | -0.7 | -0.77 | -0.45 | -0.69 | -1.27 | -0.96 | -1.57 | -1.33 | -0.13 | -0.62 | 0.12 | -0.53 | -0.78 | -1.05 | -0.0775 | -0.1475 | 0.205 | 0.547559 | 0.781658 | 0.695451 | 0.995762 | 0.997988 | 0.994666 |
| Osbpl10_S4R1M9_S58s_2 | Osbpl10_S | -0.32 | -0.14 | -1.52 | -1.22 | -0.32 | -0.1 | -1.63 | -1.61 | 0.08 | 0.11 | -1.82 | -1.44 | -0.03 | -0.54 | -1.28 | -1.34 | -0.115 | 0.0325 | 0.0025 | 0.995548 | 0.813745 | 0.953698 | 0.995762 | 0.997988 | 1 |
| Osbpl10_S4R1M9_S66s_1 | Osbpl10_S | 1.29 | 0.59 | -0.21 | 0.84 | 0.37 | 1.14 | 0.09 | 0.5 | 1.19 | 0.91 | 0.83 | 0.63 | 0.95 | 0.35 | 0.27 | 0.79 | -0.1025 | 0.2625 | -0.0375 | 0.919989 | 0.785953 | 0.461026 | 0.995762 | 0.997988 | 0.999507 |
| Osbpl10_S4R1M9_S66s_2 | Osbpl10_S | -0.32 | -0.14 | -1.08 | -0.83 | -0.32 | -0.1 | -1.18 | -1.18 | 0.08 | 0.11 | -0.97 | -0.93 | -0.03 | -0.54 | -0.96 | -0.95 | -0.1025 | 0.165 | -0.0275 | 0.936113 | 0.774509 | 0.663731 | 0.995762 | 0.997988 | 0.999507 |
| Osbpl3_D3Z194_S250s_1 | Osbpl3_D3 | -0.45 | -0.87 | -0.63 | -0.59 | 0.37 | -0.39 | -0.11 | -0.65 | -1.53 | -1.97 | -0.18 | -0.38 | -1.28 | -0.76 | -2.02 | -1.53 | 0.44 | -0.38 | -0.7625 | 0.037927 | 0.133613 | 0.38945 | 0.989223 | 0.997988 | 0.994666 |
| Osbpl3_D3Z194_S400s_1 | Osbpl3_D3 | 0.82 | 0.4 | 0.78 | 0.7 | 0.43 | 0.43 | 0.21 | 0.53 | 0.35 | 0.56 | 0.43 | 0.52 | 0.31 | 0.52 | 0.48 | 0.42 | -0.275 | -0.21 | -0.2425 | 0.324461 | 0.217712 | 0.350956 | 0.989223 | 0.997988 | 0.994666 |
| Osbpl3_D3Z194_S478s_1 | Osbpl3_D3 | -0.6 | -0.04 | 0.55 | 0.4 | 0.26 | -1.06 | 0.75 | -0.16 | -0.24 | -0.63 | -0.62 | 0.33 | -0.72 | 0.23 | -0.11 | 0.15 | -0.13 | -0.3675 | -0.19 | 0.600783 | 0.765634 | 0.316523 | 0.995762 | 0.997988 | 0.995259 |
| Osbpl3_D3Z194_T477t_1 | Osbpl3_D3 | -0.6 | -0.04 | 0.55 | 0.4 | 0.26 | -1.06 | 0.75 | -0.16 | -0.24 | -0.63 | -0.62 | 0.33 | -0.72 | 0.23 | -0.11 | 0.15 | -0.13 | -0.3675 | -0.19 | 0.600783 | 0.765634 | 0.316523 | 0.995762 | 0.997988 | 0.995259 |
| Osbpl5_Q9ER64_S311s_1 | Osbpl5_Q9 | -0.84 | -0.83 | -1.8 | -1.06 | -2.53 | -0.35 | -1.81 | -1.95 | 1.7 | -0.66 | -1 | -1.72 | 1.44 | -2.64 | -1.01 | -1.53 | -0.5275 | 0.7125 | 0.1975 | 0.802315 | 0.285247 | 0.322223 | 0.989223 | 0.997988 | 0.999507 |
| Osbpl6_F8WIZ7_S290s_1 | Osbpl6_F8 | 0.48 | -0.06 | 0.29 | 0.51 | 0.37 | -0.06 | 0.59 | 0.5 | -0.94 | 0.54 | 0.43 | 0.05 | -0.71 | 0.37 | 0.47 | 0.46 | 0.045 | -0.285 | -0.1575 | 0.651403 | 0.857609 | 0.448049 | 0.996349 | 0.997988 | 0.995259 |
| Osbpl6_F8WIZ7_S9s_1 | Osbpl6_F8 | -0.46 | -1.57 | -2.39 | -1.68 | -0.83 | -2.19 | -2.72 | -1.95 | -0.57 | -1.67 | -4.15 | -3.12 | -2.47 | -2.68 | -3.72 | -1.77 | -0.3975 | -0.8525 | -1.135 | 0.055272 | 0.446767 | 0.296157 | 0.989223 | 0.997988 | 0.994666 |
| Osbpl6_F8WIZ7_T288t_1 | Osbpl6_F8 | 0.09 | -0.28 | -0.11 | -0.22 | -0.19 | 0.37 | 1.35 | 0.62 | -0.54 | 0.66 | -1.44 | -0.97 | -0.02 | 0.21 | 0.84 | -0.42 | 0.6675 | -0.4425 | 0.2825 | 0.390179 | 0.072409 | 0.330527 | 0.989223 | 0.997988 | 0.994666 |
| Osbpl8_B9EJ86_S342s_1 | Osbpl8_B9 | -2.14 | -1.56 | -2.25 | -3.23 | -1.58 | -2.44 | -2.35 | -3.33 | -3.35 | -1.84 | 0.03 | -1.86 | -2.15 | -1.99 | -5.2 | -3.75 | -0.13 | 0.54 | -0.9775 | 0.207041 | 0.779122 | 0.446918 | 0.995762 | 0.997988 | 0.994666 |
| Osbpl8_B9EJ86_S65s_1 | Osbpl8_B9 | 0.37 | -0.64 | -1.11 | -0.26 | -0.13 | -0.19 | 0.26 | -0.2 | -0.86 | -0.55 | 0.14 | -0.52 | -0.1 | 0.09 | -1.19 | -0.27 | 0.345 | -0.0375 | 0.0425 | 0.919457 | 0.321454 | 0.921512 | 0.989223 | 0.997988 | 0.999507 |
| Osbpl8_B9EJ86_T39t_1 | Osbpl8_B9 | 0.6 | -0.18 | -0.03 | 0.83 | 0.99 | 0.38 | 1.19 | 0.98 | -0.08 | 0.25 | 0.47 | 0.58 | -1.18 | 0.68 | 1.22 | 0.76 | 0.58 | 5.89E-17 | 0.065 | 0.904484 | 0.092139 | 1 | 0.989223 | 1 | 0.999507 |
| Otud5_Q3U2S4_S165s_1 | Otud5_Q3 | -1.5 | -1.97 | -2.18 | -1.45 | -1.75 | -1.98 | -1.93 | -1.71 | -1.95 | -0.56 | -0.72 | -0.7 | -0.73 | -2.4 | -0.94 | -0.9 | -0.0675 | 0.7925 | 0.5325 | 0.229139 | 0.786226 | 0.059497 | 0.995762 | 0.997988 | 0.994666 |
| Otud5_Q3U2S4_S64s_1 | Otud5_Q3 | -0.64 | 1.04 | -1 | -0.08 | 0.25 | -0.65 | 0.26 | 0.29 | -0.6 | 0.53 | 0.45 | -0.28 | -0.22 | -0.05 | 0.28 | -0.13 | 0.2075 | 0.195 | 0.14 | 0.754709 | 0.657046 | 0.694545 | 0.990434 | 0.997988 | 0.9971 |
| Otud7a_Q8R554_S593s_1 | Otud7a_Q8 | -0.53 | -0.53 | 0.32 | -0.32 | -1.9 | 0.01 | -0.64 | -0.4 | 1.48 | -0.05 | -1.95 | -0.03 | 1.9 | -2.38 | -0.35 | -0.61 | -0.4675 | 0.1275 | -0.095 | 0.904927 | 0.294562 | 0.847337 | 0.989223 | 0.997988 | 0.999507 |
| Otud7a_Q8R554_S711s_1 | Otud7a_Q8 | -0.7 | -0.4 | -0.77 | 0.37 | 0.16 | -0.79 | -0.34 | 0.34 | -0.06 | -0.62 | -0.04 | -0.93 | -0.21 | -0.14 | -0.96 | -0.34 | 0.2175 | -0.0375 | -0.0375 | 0.914786 | 0.552145 | 0.915653 | 0.989223 | 0.997988 | 0.999507 |
| Otud7a_Q8R554_S749s_1 | Otud7a_Q8 | -0.53 | -0.75 | 0.21 | -0.88 | -0.99 | 0.36 | -1.92 | -0.79 | 0.28 | -0.74 | -1.72 | -0.31 | 1.23 | -2.93 | -1.77 | -1.17 | -0.3475 | -0.135 | -0.6725 | 0.411344 | 0.480607 | 0.771838 | 0.989223 | 0.997988 | 0.994666 |
| Otud7a_Q8R554_S771s_1 | Otud7a_Q8 | -0.74 | -0.64 | -0.21 | -0.31 | -0.88 | -0.15 | -0.3 | 0.21 | 0.02 | -0.6 | -0.26 | -0.21 | 0.89 | -0.42 | -0.32 | -0.17 | 0.195 | 0.2125 | 0.47 | 0.206943 | 0.506711 | 0.410035 | 0.989223 | 0.997988 | 0.994666 |
| Otud7a_Q8R554_T804t_1 | Otud7a_Q8 | -0.19 | -0.66 | -0.37 | -0.2 | -0.05 | -0.03 | 0.1 | -0.45 | -0.79 | -0.06 | -0.67 | -0.66 | -0.3 | -0.22 | -0.74 | -0.66 | 0.2475 | -0.19 | -0.125 | 0.638108 | 0.303946 | 0.475014 | 0.989223 | 0.997988 | 0.995259 |
| Otud7a_Q8R554_T829t_1 | Otud7a_Q8 | 0.97 | 1.27 | 1.42 | 0.83 | 1.22 | 0.57 | 1.49 | 0.79 | 0.69 | 0.2 | 0.19 | 1.63 | 0.43 | 1.05 | 0.97 | 1.43 | -0.105 | -0.445 | -0.1525 | 0.61869 | 0.710655 | 0.249881 | 0.995437 | 0.997988 | 0.995259 |
| Oxr1_Q4KMM3_S104s_1 | Oxr1_Q4K | -0.25 | 0.63 | -0.03 | -0.94 | 0 | -0.03 | -0.97 | -0.39 | -0.44 | -0.45 | -1.05 | -1.51 | -0.04 | 0.22 | -1.02 | -0.55 | -0.2 | -0.715 | -0.2 | 0.637778 | 0.605002 | 0.106966 | 0.989223 | 0.997988 | 0.995259 |
| Oxr1_Q4KMM3_S13s_1 | Oxr1_Q4K | -1.79 | -1 | -1.69 | -1.83 | -0.75 | -1.06 | -1.52 | -1.9 | -0.98 | -0.85 | -0.35 | -2.01 | -0.36 | -0.45 | -1.81 | -1.63 | 0.27 | 0.53 | 0.515 | 0.244307 | 0.421371 | 0.204381 | 0.989223 | 0.997988 | 0.994666 |
| Oxr1_Q4KMM3_S15s_1 | Oxr1_Q4K | -2.47 | -2.14 | -0.93 | -1.09 | -1.43 | -1.97 | -1.06 | -0.95 | -2.53 | -1.58 | -0.4 | -1.08 | -1.97 | -0.66 | -1.35 | -0.84 | 0.305 | 0.26 | 0.4525 | 0.34224 | 0.476231 | 0.635913 | 0.989223 | 0.997988 | 0.994666 |
| Oxr1_Q4KMM3_S202s_1 | Oxr1_Q4K | 0.12 | 0.08 | 0.75 | 0.5 | -0.16 | 0.06 | -0.07 | 0.11 | -0.44 | 0.27 | 0.37 | 0.72 | -0.6 | 0.24 | -0.29 | 0.09 | -0.3775 | -0.1325 | -0.5025 | 0.125001 | 0.137958 | 0.680292 | 0.989223 | 0.997988 | 0.994666 |
| Oxr1_Q4KMM3_S323s_1 | Oxr1_Q4K | 1.37 | 1.13 | 1.87 | 1.38 | 0 | 2.12 | 0.11 | 1.22 | 3.15 | 1.53 | 1.53 | 1.81 | 2.49 | -0.62 | 1.74 | 0.92 | -0.575 | 0.5675 | -0.305 | 0.625563 | 0.252711 | 0.190514 | 0.989223 | 0.997988 | 0.995259 |
| Oxr1_Q4KMM3_S488s_1 | Oxr1_Q4K | 1.85 | 1.25 | 2.2 | 1.48 | 2.58 | 0.68 | 0.55 | 1.69 | 1.31 | 1.41 | 1.57 | 1.04 | 0.22 | 1.83 | 1.15 | 0.76 | -0.32 | -0.3625 | -0.705 | 0.105788 | 0.509544 | 0.223829 | 0.989223 | 0.997988 | 0.994666 |
| Oxr1_Q4KMM3_S518s_1 | Oxr1_Q4K | -0.8 | -1.44 | 0.13 | 0.06 | 1.65 | 0.1 | 0.18 | 0.08 | -0.74 | -0.66 | -0.39 | 1.02 | -0.13 | 1.32 | -0.04 | 0.91 | 1.015 | 0.32 | 1.0275 | 0.060287 | 0.063294 | 0.542465 | 0.989223 | 0.997988 | 0.994666 |
| Oxr1_Q4KMM3_S568s_1 | Oxr1_Q4K | -1.51 | -0.87 | 0.29 | 0.52 | 0.15 | 0.94 | 0.69 | 0.54 | -1.89 | -0.67 | 0.46 | 0.87 | -1.74 | -0.2 | -0.18 | 0.55 | 0.9725 | 0.085 | 3.93E-16 | 1 | 0.062977 | 0.904232 | 0.989223 | 0.997988 | 1 |
| Oxtr_P97926_S360s_1 | Oxtr_P979 | 0.24 | 0.27 | 0.45 | 0.5 | 0.15 | 1.06 | -0.22 | 0.11 | 0.61 | 0.59 | 0.48 | 0.41 | 1.24 | 0.08 | 0.36 | 0.21 | -0.09 | 0.1575 | 0.1075 | 0.736267 | 0.767787 | 0.462645 | 0.995762 | 0.997988 | 0.9971 |
| P0C913_S28s_1 | P0C913_S2 | 2.89 | 2.64 | 3.15 | 3.03 | 1.83 | 2.62 | 2.62 | 3.25 | 1.77 | 2.48 | 3.04 | 2.98 | -1.55 | 1.39 | 2.91 | 2.99 | -0.3475 | -0.36 | -1.4925 | 0.134199 | 0.298279 | 0.29884 | 0.989223 | 0.997988 | 0.994666 |
| P2ry12_Q9CPV9_S342s_1 | P2ry12_Q9 | 0.19 | -0.05 | -0.51 | -0.13 | -0.28 | 0.39 | -0.19 | 0.02 | 0.05 | 0.08 | 0.21 | -0.11 | 0.06 | 0.15 | -0.39 | -0.34 | 0.11 | 0.1825 | -0.005 | 0.985691 | 0.671617 | 0.458172 | 0.992876 | 0.997988 | 1 |
| P32507-2_P32507_S414s_1 | P32507-2_ | -0.61 | -0.25 | -0.26 | -1.37 | -0.17 | -0.61 | -0.99 | -0.48 | -0.25 | -0.42 | -0.47 | -0.75 | 0.09 | 0.2 | -0.75 | -0.6 | 0.06 | 0.15 | 0.3575 | 0.350816 | 0.853656 | 0.635315 | 0.996349 | 0.997988 | 0.994666 |
| Pabpc4_V9GXG3_S96s_1 | Pabpc4_V9 | -1.55 | -1.27 | -0.23 | -0.58 | 0.1 | -1.68 | -0.13 | -0.65 | -1.17 | -0.72 | -0.62 | -0.19 | -0.9 | -0.22 | -0.4 | -0.04 | 0.3175 | 0.2325 | 0.5175 | 0.188466 | 0.498858 | 0.536187 | 0.989223 | 0.997988 | 0.994666 |
| Pacrgl_Q9D3X5_S47s_1 | Pacrgl_Q9 | -0.94 | -1.83 | -1.25 | -2.54 | -1.75 | -1.35 | -1.74 | -3.23 | -1.43 | -0.68 | -2.37 | -1.04 | -0.54 | -0.97 | -0.78 | -1.13 | -0.3775 | 0.26 | 0.785 | 0.066532 | 0.456129 | 0.591236 | 0.989223 | 0.997988 | 0.994666 |
| Pacs1_Q8K212_S23s_1 | Pacs1_Q8K | 0.37 | 1.4 | 0.09 | 0.1 | 1.13 | 0.92 | 0.97 | 0.43 | 0.94 | 1.29 | 1.09 | 1.05 | -0.65 | 1.24 | 0.74 | 1.25 | 0.3725 | 0.6025 | 0.155 | 0.762577 | 0.299391 | 0.101752 | 0.989223 | 0.997988 | 0.9971 |
| Pacs1_Q8K212_S28s_1 | Pacs1_Q8K | 0.81 | 0.72 | 0.35 | 0.26 | 0.86 | 0.68 | 0.25 | 0.36 | 0.66 | 0.97 | 0.27 | 0.27 | 0.51 | 1 | 0.48 | 1 | 0.0025 | 0.0075 | 0.2125 | 0.452748 | 0.992053 | 0.978076 | 0.999187 | 0.997988 | 0.994666 |
| Pacs1_Q8K212_S493s_1 | Pacs1_Q8K | 2.94 | 0.41 | -0.3 | 0.04 | 1.52 | 3.04 | 0.02 | 0.68 | 1.48 | 0.4 | 0.4 | 0.35 | 0.35 | 0.58 | -0.39 | -0.4 | 0.5425 | -0.115 | -0.7375 | 0.304834 | 0.53081 | 0.871003 | 0.989223 | 0.997988 | 0.994666 |
| Pacs1_Q8K212_S526s_1 | Pacs1_Q8K | -0.6 | -2.94 | -1.62 | -1.96 | -2.75 | 0.07 | -1.05 | 0.65 | 3.12 | -1.09 | -0.38 | -0.02 | 3.04 | -1.74 | 3.11 | -2.55 | 1.01 | 2.1875 | 2.245 | 0.126157 | 0.215422 | 0.039567 | 0.989223 | 0.997988 | 0.994666 |
| Pacs1_Q8K212_S532s_1 | Pacs1_Q8K | -1.28 | -0.92 | -0.61 | -0.35 | -1.4 | -2.04 | 0.68 | -0.31 | -2.17 | -0.75 | -1.9 | -0.98 | -0.74 | -1.32 | 0.02 | -0.33 | 0.0225 | -0.66 | 0.1975 | 0.596685 | 0.968445 | 0.123247 | 0.999187 | 0.997988 | 0.995259 |
| Pacs1_Q8K212_T44t_1 | Pacs1_Q8K | 0.63 | 0.87 | 0.43 | 0.35 | -0.02 | 0.42 | 0.33 | 0.45 | 0.48 | 0.7 | 0.21 | 0.47 | 0.48 | 0.45 | 0.06 | 0.32 | -0.275 | -0.105 | -0.2425 | 0.355135 | 0.253622 | 0.662234 | 0.989223 | 0.997988 | 0.994666 |
| Pacs2_E9Q7E9_S349s_1 | Pacs2_E9Q | 0.28 | -0.91 | 0.43 | -0.18 | 0.33 | -0.53 | 0.23 | -0.14 | -0.6 | -0.92 | -0.27 | 0.43 | -0.42 | 0.28 | 0.98 | 0.54 | 0.0675 | -0.245 | 0.44 | 0.308169 | 0.850461 | 0.555212 | 0.996349 | 0.997988 | 0.994666 |
| Pacs2_E9Q7E9_S417s_1 | Pacs2_E9Q | -1.49 | -0.85 | -0.34 | -0.83 | 0.09 | -1.61 | -0.26 | -0.93 | -2.12 | -1.44 | -0.59 | -1.05 | -1.19 | -0.56 | -0.67 | -0.11 | 0.2 | -0.4225 | 0.245 | 0.492489 | 0.636677 | 0.30451 | 0.989223 | 0.997988 | 0.994666 |
| Pacs2_E9Q7E9_S454s_1 | Pacs2_E9Q | 0.51 | 0.3 | 0.76 | 0.46 | -0.6 | 0.82 | -0.57 | 0.33 | 1.87 | 0.64 | 0.54 | 0.75 | 1.79 | -0.59 | 0.95 | 0.19 | -0.5125 | 0.4425 | 0.0775 | 0.874924 | 0.177796 | 0.217479 | 0.989223 | 0.997988 | 0.999507 |
| Pacsin1_Q61644_S326s_1 | Pacsin1_Q6 | 0.66 | 0.57 | 0.74 | 1.22 | 0.49 | 0.2 | 1.05 | 1.31 | -0.39 | 0.61 | -0.56 | 1.34 | -0.21 | -0.75 | 1.26 | -0.21 | -0.035 | -0.5475 | -0.775 | 0.107016 | 0.910406 | 0.243909 | 0.997335 | 0.997988 | 0.994666 |
| Pacsin1_Q61644_S343s_1 | Pacsin1_Q6 | 1.6 | 0.54 | -0.1 | 0.48 | 0.68 | 1.19 | 0.5 | 0.63 | 0.5 | 1.22 | -0.66 | 0.65 | -0.28 | -0.91 | -0.02 | 0.24 | 0.12 | -0.2025 | -0.8725 | 0.065324 | 0.750771 | 0.686092 | 0.995762 | 0.997988 | 0.994666 |
| Pacsin1_Q61644_S343s_2 | Pacsin1_Q6 | -4 | -4.03 | -3.04 | -4.86 | -4.66 | -3.88 | -3.54 | -3.72 | -3.87 | -4.38 | -1.13 | -3.06 | -3.23 | -3.65 | -3.88 | -3.58 | 0.0325 | 0.8725 | 0.3975 | 0.335022 | 0.938702 | 0.247156 | 0.999187 | 0.997988 | 0.994666 |
| Pacsin1_Q61644_S345s_2 | Pacsin1_Q6 | 0.57 | 0.2 | -1.62 | -1.06 | -0.14 | 0.21 | -1.18 | -2.4 | -0.03 | 0 | -1.13 | -1.24 | -0.7 | -0.35 | -2.18 | -1.91 | -0.4 | -0.1225 | -0.8075 | 0.213925 | 0.565595 | 0.82975 | 0.989223 | 0.997988 | 0.994666 |
| Pacsin1_Q61644_S346s_1 | Pacsin1_Q6 | 0.36 | 0.15 | 0.55 | 0.48 | -0.71 | 0.63 | -0.69 | 0.28 | 1.1 | 0.1 | 0.34 | 0.26 | 1.06 | -0.75 | -0.4 | -0.37 | -0.5075 | 0.065 | -0.5 | 0.243529 | 0.173569 | 0.82059 | 0.989223 | 0.997988 | 0.994666 |
| Pacsin1_Q61644_S427s_1 | Pacsin1_Q6 | 0.07 | 0.2 | -0.19 | -0.31 | 0.44 | -0.07 | 0.41 | 0.19 | 0.19 | -0.01 | -0.24 | -0.55 | -0.47 | 0.82 | -0.42 | 0.14 | 0.3 | -0.095 | 0.075 | 0.831285 | 0.222951 | 0.717665 | 0.989223 | 0.997988 | 0.999507 |
| Pacsin1_Q61644_S76s_1 | Pacsin1_Q6 | -0.59 | -0.25 | -0.29 | -1.06 | -0.82 | -1.11 | -0.63 | -0.7 | -0.39 | -0.35 | -0.13 | -0.3 | 0.03 | 0.6 | -0.98 | -0.92 | -0.2675 | 0.255 | 0.23 | 0.590045 | 0.321483 | 0.341393 | 0.989223 | 0.997988 | 0.995259 |
| Pacsin1_Q61644_T181t_1 | Pacsin1_Q6 | 0.88 | 0.5 | 0.9 | 0.78 | -0.17 | 1.72 | -0.42 | 0.23 | 2.11 | 0.87 | -0.81 | 0.42 | 2.4 | -1.14 | 0.02 | 0.23 | -0.425 | -0.1175 | -0.3875 | 0.565903 | 0.361585 | 0.834854 | 0.989223 | 0.997988 | 0.995259 |
| Pacsin3_F6W8Y8_S44s_1 | Pacsin3_F6 | 0.68 | 0.5 | 0.63 | 0.54 | 0.58 | 0.79 | 1.29 | 0.65 | 0.2 | 0.93 | 0.65 | 0.86 | 0.11 | 0.38 | 0.47 | 0.65 | 0.24 | 0.0725 | -0.185 | 0.455569 | 0.321478 | 0.769631 | 0.989223 | 0.997988 | 0.994666 |
| Pacsin3_F6W8Y8_S79s_1 | Pacsin3_F6 | 0.77 | 1.35 | 0.77 | 0.51 | 0.8 | 0.38 | 0.8 | -0.31 | 1.6 | 1.38 | 1.71 | -0.1 | -0.82 | 1.77 | 0.04 | 0.94 | -0.4325 | 0.2975 | -0.3675 | 0.505326 | 0.207879 | 0.506291 | 0.989223 | 0.997988 | 0.994666 |
| Pafah1b2_Q61206_S64s_1 | Pafah1b2_ | -0.68 | -0.07 | -0.34 | 0.1 | 0.41 | -0.64 | 0.18 | -0.06 | -0.62 | -0.13 | -0.31 | -0.35 | -0.85 | 0.11 | -0.44 | -0.23 | 0.22 | -0.105 | -0.105 | 0.73898 | 0.476561 | 0.690147 | 0.989223 | 0.997988 | 0.9971 |
| Pag1_Q3U1F9_S379s_1 | Pag1_Q3U | 0.26 | 0.47 | 0.25 | 0.11 | 1.03 | 0.84 | 0.81 | 0.38 | 0.42 | 0.05 | 0.18 | 0.41 | -0.64 | 1.4 | 0.07 | 0.35 | 0.4925 | -0.0075 | 0.0225 | 0.957828 | 0.055901 | 0.973213 | 0.989223 | 0.997988 | 0.999507 |
| Pag1_Q3U1F9_Y224y_1 | Pag1_Q3U | 0.18 | 0.09 | -0.67 | -0.13 | 0.87 | 0.38 | 0.1 | 0.33 | -0.34 | 0.54 | -0.35 | -0.51 | -0.53 | 0.34 | 0.23 | 0.41 | 0.5525 | -0.0325 | 0.245 | 0.464716 | 0.076388 | 0.921398 | 0.989223 | 0.997988 | 0.994666 |
| Pagr1a_Q99L02_S147s_2 | Pagr1a_Q9 | -2.08 | -1.05 | -1.8 | -1.79 | -1.72 | -1.35 | -2.18 | -1.37 | -2.01 | -2.07 | -2.7 | -2.23 | -0.16 | -1.78 | -1.77 | -3.12 | 0.025 | -0.5725 | -0.0275 | 0.962771 | 0.93616 | 0.087663 | 0.99868 | 0.997988 | 0.999507 |
| Pagr1a_Q99L02_T137t_2 | Pagr1a_Q9 | -2.08 | -1.05 | -1.8 | -1.79 | -1.72 | -1.35 | -2.18 | -1.37 | -2.01 | -2.07 | -2.7 | -2.23 | -0.16 | -1.78 | -1.77 | -3.12 | 0.025 | -0.5725 | -0.0275 | 0.962771 | 0.93616 | 0.087663 | 0.99868 | 0.997988 | 0.999507 |
| Paics_Q9DCL9_S107s_1 | Paics_Q9D | 1.11 | 0.84 | 1.17 | 0.23 | 1.04 | 0.39 | 0.15 | 0.5 | 0.34 | 0.89 | -1.35 | 0.66 | 0.13 | -0.06 | -0.48 | -1.27 | -0.3175 | -0.7025 | -1.2575 | 0.008983 | 0.31465 | 0.195954 | 0.989223 | 0.997988 | 0.994666 |
| Paip1_G3UZS6_S52s_1 | Paip1_G3U | 5.57 | 1.17 | 0.3 | 0.23 | 0.65 | 5.65 | 5.49 | 1.49 | 1.82 | 3.13 | 0.04 | 1.48 | 1.4 | -0.9 | 1.29 | 1.41 | 1.5025 | -0.2 | -1.0175 | 0.40595 | 0.347953 | 0.872195 | 0.989223 | 0.997988 | 0.994666 |
| Pak1_G5E884_S144s_1 | Pak1_G5E8 | -0.22 | 0.4 | 0.47 | -0.41 | 1.14 | -0.28 | -0.31 | -0.35 | -1.19 | -0.6 | -0.91 | -0.11 | -0.46 | 0.7 | -0.32 | 0.01 | -0.01 | -0.7625 | -0.0775 | 0.831018 | 0.980376 | 0.047197 | 0.999187 | 0.997988 | 0.999507 |
| Pak1_G5E884_S222s_1 | Pak1_G5E8 | 0.14 | 0.69 | 0.53 | 0.86 | 0.62 | 0.2 | 0.73 | 0.84 | 0.13 | 0.29 | 0.4 | 0.67 | 0.14 | 0.74 | 0.85 | 0.54 | 0.0425 | -0.1825 | 0.0125 | 0.965504 | 0.869301 | 0.486432 | 0.996493 | 0.997988 | 0.999507 |
| Pak1_G5E884_S28s_1 | Pak1_G5E8 | -0.78 | -1.28 | 0.35 | 0.13 | -0.89 | -0.71 | -0.45 | -0.13 | -0.38 | -1.61 | 0.95 | -0.86 | -1.14 | -0.87 | -1.62 | -0.94 | -0.15 | -0.08 | -0.7475 | 0.099302 | 0.709218 | 0.894691 | 0.995437 | 0.997988 | 0.994666 |
| Pak1_G5E884_T146t_1 | Pak1_G5E8 | -0.13 | 0.5 | 0.35 | 0.1 | 1.26 | -0.26 | 0.32 | -0.01 | -1.2 | -0.48 | -1.28 | 0.15 | -0.58 | 0.82 | 0.27 | 0.74 | 0.1225 | -0.9075 | 0.1075 | 0.771656 | 0.733634 | 0.034071 | 0.995762 | 0.997988 | 0.9971 |
| Pak1_G5E884_T211t_1 | Pak1_G5E8 | -0.25 | -1.3 | -1.58 | -1.17 | 0.12 | 0.1 | -0.1 | -0.31 | -1.19 | -1.18 | 0.47 | -0.77 | -1.56 | 0 | -2 | -1.26 | 1.0275 | 0.4075 | -0.13 | 0.791453 | 0.009341 | 0.390617 | 0.989223 | 0.997988 | 0.999507 |
| Pak1_G5E884_T228t_1 | Pak1_G5E8 | 0.34 | 0.51 | 0.61 | 0.7 | 0.77 | 0.49 | 0.82 | 0.77 | -0.37 | 1.18 | -0.21 | 0.85 | 0.01 | 0.72 | 0.97 | 0.79 | 0.1725 | -0.1775 | 0.0825 | 0.777925 | 0.420012 | 0.652322 | 0.989223 | 0.997988 | 0.99798 |
| Pak1_G5E884_T229t_1 | Pak1_G5E8 | 0.02 | 0.91 | 0.76 | 0.7 | 0.4 | 0.96 | 0.58 | 0.55 | 0.6 | 0.65 | 1.07 | 0.46 | -0.12 | 0.49 | 0.44 | 0.56 | 0.025 | 0.0975 | -0.255 | 0.415767 | 0.926651 | 0.733229 | 0.997335 | 0.997988 | 0.994666 |
| Pak1ip1_Q9DCE5_S361s_1 | Pak1ip1_Q | -2.37 | -1.92 | -2.19 | -1.01 | -2.04 | -1.42 | -3.1 | -0.54 | 0.08 | -1.57 | -0.79 | -1.73 | 0.08 | -1.39 | 0.58 | -2.73 | 0.0975 | 0.87 | 1.0075 | 0.184297 | 0.860972 | 0.099922 | 0.996493 | 0.997988 | 0.994666 |
| Pak2_Q8CIN4_S141s_1 | Pak2_Q8CI | -1.03 | -0.77 | -0.97 | -1.01 | -0.04 | -1.36 | -0.45 | -1.06 | -1.45 | -1 | -1.22 | -1.02 | -0.98 | -0.53 | -1.27 | -0.75 | 0.2175 | -0.2275 | 0.0625 | 0.813157 | 0.500169 | 0.325826 | 0.989223 | 0.997988 | 0.999507 |
| Pak2_Q8CIN4_T143t_1 | Pak2_Q8CI | 0.69 | 0.88 | 0.76 | 0.35 | 1.06 | 0.31 | 0.58 | 0.38 | 0.76 | 0.47 | 0.79 | 0.47 | 0.74 | 0.88 | 0.19 | -0.12 | -0.0875 | -0.0475 | -0.2475 | 0.436372 | 0.733069 | 0.839751 | 0.995762 | 0.997988 | 0.994666 |
| Pak3_Q61036_S4s_1 | Pak3_Q610 | 0.79 | 0.42 | -0.4 | 0.68 | 0.54 | 0.51 | 0.18 | 0.92 | 0.27 | 0.41 | -0.25 | 0.79 | -0.47 | 0.17 | -0.04 | 0.18 | 0.165 | -0.0675 | -0.4125 | 0.247054 | 0.611075 | 0.850092 | 0.989223 | 0.997988 | 0.994666 |
| Pak4_Q8BTW9_S104s_1 | Pak4_Q8B | -0.08 | -0.15 | 0.39 | 0.17 | -0.77 | 0.3 | -0.8 | -0.34 | 0.23 | 0.47 | -0.18 | 0.16 | 0.88 | -0.96 | 0.01 | 0.12 | -0.485 | 0.0875 | -0.07 | 0.861931 | 0.137851 | 0.730949 | 0.989223 | 0.997988 | 0.999507 |
| Pak4_Q8BTW9_S181s_1 | Pak4_Q8B | -0.59 | -0.49 | -1.55 | -0.92 | -0.22 | -0.63 | -1.11 | -1.34 | -0.25 | -0.31 | -0.8 | -1.5 | -0.86 | 0.35 | -2.05 | -1.22 | 0.0625 | 0.1725 | -0.0575 | 0.911629 | 0.857728 | 0.651298 | 0.996349 | 0.997988 | 0.999507 |
| Pak4_Q8BTW9_S266s_1 | Pak4_Q8B | -0.27 | 0.15 | -0.21 | 0.3 | 1.31 | 0.04 | 0.27 | -0.01 | 0.22 | 0.14 | 0.2 | -0.26 | -0.67 | 1.04 | 0.16 | -0.01 | 0.41 | 0.0825 | 0.1375 | 0.724672 | 0.250196 | 0.743732 | 0.989223 | 0.997988 | 0.9971 |
| Pak4_Q8BTW9_S476s_1 | Pak4_Q8B | 0.6 | 0.59 | 0.17 | 0.56 | 0.47 | 0.2 | 0.67 | 0.09 | -0.29 | 0.63 | 0.38 | -0.02 | -0.42 | 0.37 | 0.15 | -0.02 | -0.1225 | -0.305 | -0.46 | 0.12232 | 0.606592 | 0.293106 | 0.989223 | 0.997988 | 0.994666 |
| Pak6_Q3ULB5_S348s_2 | Pak6_Q3U | -1.89 | -2.51 | -1.96 | -3.41 | -2.17 | 0.04 | -2.06 | -0.88 | 1.95 | -1.46 | -3.38 | 0.3 | 0.89 | -1.22 | -0.28 | -1.37 | 1.175 | 1.795 | 1.9475 | 0.007197 | 0.062602 | 0.115581 | 0.989223 | 0.997988 | 0.994666 |
| Pak6_Q3ULB5_S352s_1 | Pak6_Q3U | -0.61 | -1.52 | -0.13 | -0.5 | -0.91 | -0.07 | -1.2 | -0.8 | 1.94 | -0.65 | -0.27 | 0.36 | 2.51 | -0.84 | 0.57 | 0 | -0.055 | 1.035 | 1.25 | 0.095692 | 0.88295 | 0.105923 | 0.996493 | 0.997988 | 0.994666 |
| Pak6_Q3ULB5_S352s_2 | Pak6_Q3U | -1.89 | -2.51 | -1.96 | -3.41 | -2.17 | 0.04 | -2.06 | -0.88 | 1.95 | -1.46 | -3.38 | 0.3 | 0.89 | -1.22 | -0.28 | -1.37 | 1.175 | 1.795 | 1.9475 | 0.007197 | 0.062602 | 0.115581 | 0.989223 | 0.997988 | 0.994666 |
| Pak6_Q3ULB5_S561s_1 | Pak6_Q3U | -0.05 | 0.38 | 0.36 | 0.31 | 0.53 | -0.22 | 0.75 | 0.29 | -0.31 | 0.23 | -0.13 | 0.47 | -0.43 | 0.29 | 0.09 | 0.41 | 0.0875 | -0.185 | -0.16 | 0.577596 | 0.748398 | 0.490844 | 0.995762 | 0.997988 | 0.995259 |
| Pak7_Q8C015_S104s_1 | Pak7_Q8C | 0.03 | 1.11 | 0.77 | 0.65 | 0.64 | -0.04 | 0.36 | 0.64 | 0.32 | 0.5 | 0.61 | 0.6 | 1.33 | 0.83 | 0.72 | 0.51 | -0.24 | -0.1325 | 0.2075 | 0.530725 | 0.433219 | 0.642885 | 0.989223 | 0.997988 | 0.994666 |
| Pak7_Q8C015_S104s_2 | Pak7_Q8C | 0.1 | -0.29 | 1.25 | 0.91 | 0.43 | 0.46 | 0.24 | 0.6 | -0.04 | 0.45 | 1.41 | 0.62 | -0.84 | 0.14 | 0.45 | 0.39 | -0.06 | 0.1175 | -0.4575 | 0.324558 | 0.867901 | 0.793553 | 0.996493 | 0.997988 | 0.994666 |
| Pak7_Q8C015_S277s_1 | Pak7_Q8C | -2.2 | -2.16 | -1.02 | -0.25 | -1.44 | -1.78 | -0.9 | -0.63 | -1.65 | -1.71 | -1.42 | -0.62 | -0.15 | -1.06 | -0.78 | -0.57 | 0.22 | 0.0575 | 0.7675 | 0.139034 | 0.658328 | 0.909021 | 0.990434 | 0.997988 | 0.994666 |
| Pak7_Q8C015_S288s_1 | Pak7_Q8C | -0.57 | -0.15 | -0.97 | -1.5 | -0.06 | -0.19 | -0.41 | -0.45 | -0.63 | 0.35 | -0.95 | -1 | -0.17 | 0.12 | -1.62 | -0.76 | 0.52 | 0.24 | 0.19 | 0.682782 | 0.127095 | 0.568917 | 0.989223 | 0.997988 | 0.996159 |
| Pak7_Q8C015_S290s_1 | Pak7_Q8C | 0.59 | -0.5 | -1.41 | -1.69 | -0.01 | 0.21 | -0.49 | -1.31 | -0.24 | -0.09 | -0.83 | -1.77 | -0.89 | -0.01 | -0.9 | -1.41 | 0.3525 | 0.02 | -0.05 | 0.92724 | 0.529731 | 0.972836 | 0.989223 | 0.997988 | 0.999507 |
| Pak7_Q8C015_S341s_1 | Pak7_Q8C | 0.66 | 0.6 | 0.66 | -0.13 | 0.26 | 0.65 | -0.23 | 0.77 | 0.71 | 1.3 | -1.19 | 0.99 | 0.26 | 0.04 | -1.06 | 0.21 | -0.085 | 0.005 | -0.585 | 0.148673 | 0.787041 | 0.992733 | 0.995762 | 0.997988 | 0.994666 |
| Pak7_Q8C015_T107t_1 | Pak7_Q8C | 0.17 | 0.33 | 0.6 | 0.15 | 0.07 | -0.02 | -0.05 | -0.2 | -0.11 | 0.25 | 0.73 | 0.31 | 0.13 | -0.17 | -0.5 | -0.24 | -0.3625 | -0.0175 | -0.5075 | 0.078359 | 0.115585 | 0.947116 | 0.989223 | 0.997988 | 0.994666 |
| Pak7_Q8C015_T107t_2 | Pak7_Q8C | 0.1 | -0.29 | 1.25 | 0.91 | 0.43 | 0.46 | 0.24 | 0.6 | -0.04 | 0.45 | 1.41 | 0.62 | -0.84 | 0.14 | 0.45 | 0.39 | -0.06 | 0.1175 | -0.4575 | 0.324558 | 0.867901 | 0.793553 | 0.996493 | 0.997988 | 0.994666 |
| Pakap_F7AA26_S1043s_1 | Pakap_F7A | 1.38 | -1.14 | 0.39 | 1.38 | 0.41 | 1.47 | 1.58 | 1.95 | -0.21 | -0.13 | 0.27 | 1.07 | -1.24 | 0.14 | 0.66 | 0.47 | 0.85 | -0.2525 | -0.495 | 0.460713 | 0.184416 | 0.678611 | 0.989223 | 0.997988 | 0.994666 |
| Pakap_F7AA26_S239s_1 | Pakap_F7A | 0.59 | 0.36 | 0.45 | 0.31 | 0.66 | -0.21 | 0.55 | -0.17 | 0.57 | 0.31 | 0.69 | -0.04 | 0.55 | 0.93 | 0.59 | 0.16 | -0.22 | -0.045 | 0.13 | 0.624714 | 0.434467 | 0.856518 | 0.989223 | 0.997988 | 0.995259 |
| Pald1_P70261_S347s_1 | Pald1_P70 | -1.24 | -0.32 | -1.12 | 0.26 | 0.26 | -1.89 | 0.43 | -0.99 | -1.76 | -0.95 | 0.08 | -0.24 | -0.86 | 0.2 | 0.47 | -1.37 | 0.0575 | -0.1125 | 0.215 | 0.68286 | 0.921615 | 0.824572 | 0.997335 | 0.997988 | 0.996159 |
| Palm_Q9Z0P4_S116s_1 | Palm_Q9Z | -0.88 | -1.47 | 0.01 | 0.13 | -0.84 | 1.07 | 0.1 | -0.52 | 1.55 | 0.16 | -0.59 | 0.01 | 2.18 | -3.13 | 0.03 | -0.15 | 0.505 | 0.835 | 0.285 | 0.777106 | 0.340875 | 0.152512 | 0.989223 | 0.997988 | 0.99798 |
| Palm_Q9Z0P4_S116s_2 | Palm_Q9Z | -0.39 | -0.3 | -2.75 | -3.24 | 0.2 | -0.74 | -2.68 | -3.72 | -0.86 | -0.53 | -1.61 | -2.99 | -0.64 | -0.35 | -3.25 | -3.58 | -0.065 | 0.1725 | -0.285 | 0.775495 | 0.949131 | 0.837638 | 0.999187 | 0.997988 | 0.99798 |

| | | | | | | | | | | | | | | | | | | | | | | | | | | |
|---|---|---|---|---|---|---|---|---|---|---|---|---|---|---|---|---|---|---|---|---|---|---|---|---|---|---|
| Palm_Q9Z0P4_S116s_3 | Palm_Q9Z0 | -4.99 | -5.04 | -2.56 | -3.78 | -2.95 | -4.98 | -2.27 | -3.37 | -6.64 | -4.96 | -1.41 | -3.2 | -1.71 | -3.24 | -2.96 | -3.29 | 0.7 | 0.04 | 1.2925 | 0.064353 | 0.34574 | 0.971408 | 0.989223 | 0.997988 | 0.994666 |
| Palm_Q9Z0P4_S122s_1 | Palm_Q9Z0 | 0.28 | 0.81 | 0.61 | 0.62 | 0.89 | 0.16 | 0.54 | 0.87 | -0.14 | -0.14 | 0.19 | 0.6 | 0.6 | 0.03 | 0.96 | 0.68 | 0.035 | -0.4525 | -0.0125 | 0.965805 | 0.891201 | 0.116901 | 0.996493 | 0.997988 | 0.999507 |
| Palm_Q9Z0P4_S122s_2 | Palm_Q9Z0 | -0.44 | 0.52 | 1.36 | 0.99 | 0.78 | -0.63 | 0.99 | 1.01 | -0.77 | -0.15 | 0.75 | 1.21 | -0.47 | -0.11 | 1.21 | 1.11 | -0.07 | -0.3475 | -0.1725 | 0.748434 | 0.889837 | 0.530488 | 0.996493 | 0.997988 | 0.9971 |
| Palm_Q9Z0P4_S122s_3 | Palm_Q9Z0 | -4.99 | -5.04 | -2.56 | -3.78 | -2.95 | -4.98 | -2.27 | -3.37 | -6.64 | -4.96 | -1.41 | -3.2 | -1.71 | -3.24 | -2.96 | -3.29 | 0.7 | 0.04 | 1.2925 | 0.064353 | 0.34574 | 0.971408 | 0.989223 | 0.997988 | 0.994666 |
| Palm_Q9Z0P4_S124s_1 | Palm_Q9Z0 | 0.48 | 1.32 | 1.67 | 1.28 | 1.06 | 0.38 | 0.95 | 1.07 | 0.07 | 0.95 | 0.72 | 1.46 | 1.18 | 0.43 | 1.58 | 1.25 | -0.3225 | -0.3875 | -0.0775 | 0.833986 | 0.322068 | 0.328189 | 0.989223 | 0.997988 | 0.999507 |
| Palm_Q9Z0P4_S124s_2 | Palm_Q9Z0 | 0.33 | 1.04 | 1.18 | 0.84 | 1.3 | 0.05 | 0.86 | 0.84 | -0.22 | 0.44 | 0.59 | 1.04 | -0.21 | 0.44 | 1.03 | 0.93 | -0.085 | -0.385 | -0.3 | 0.416642 | 0.796986 | 0.274752 | 0.995762 | 0.997988 | 0.994666 |
| Palm_Q9Z0P4_S124s_3 | Palm_Q9Z0 | -4.99 | -5.04 | -2.56 | -3.78 | -2.95 | -4.98 | -2.27 | -3.37 | -6.64 | -4.96 | -1.41 | -3.2 | -1.71 | -3.24 | -2.96 | -3.29 | 0.7 | 0.04 | 1.2925 | 0.064353 | 0.34574 | 0.971408 | 0.989223 | 0.997988 | 0.994666 |
| Palm_Q9Z0P4_S157s_1 | Palm_Q9Z0 | 0.62 | 0.05 | 0.21 | 0.35 | 0.47 | 0.32 | 0.2 | 0.2 | 0.27 | 0.28 | -0.3 | -0.22 | 0.61 | 0.4 | 0.15 | 0.06 | -0.01 | -0.3 | -0.0025 | 0.992499 | 0.96401 | 0.26904 | 0.999187 | 0.997988 | 1 |
| Palm_Q9Z0P4_S157s_2 | Palm_Q9Z0 | 1.04 | 0.33 | 0.67 | 1.14 | 1.07 | 0.82 | 0.81 | 0.89 | 0.25 | 0.56 | 0.68 | 0.71 | -0.63 | 1.04 | 0.54 | 0.76 | 0.1025 | -0.245 | -0.3675 | 0.383225 | 0.683909 | 0.376559 | 0.994379 | 0.997988 | 0.994666 |
| Palm_Q9Z0P4_S161s_1 | Palm_Q9Z0 | 0.1 | -0.98 | -0.99 | -0.76 | 0.12 | -0.45 | -0.52 | -0.69 | -0.58 | -0.95 | 0.08 | -0.58 | -0.37 | 0.48 | -0.83 | -0.97 | 0.2725 | 0.15 | 0.235 | 0.575677 | 0.410441 | 0.670292 | 0.989223 | 0.997988 | 0.995259 |
| Palm_Q9Z0P4_S161s_2 | Palm_Q9Z0 | 1.29 | 0.26 | 0.83 | 1.57 | 1.28 | 0.7 | 1.35 | 0.64 | -0.02 | 0.12 | 1.18 | 0.61 | -1.07 | 1.53 | 0.82 | 0.54 | 0.005 | -0.515 | -0.5325 | 0.361884 | 0.988456 | 0.21287 | 0.999187 | 0.997988 | 0.994666 |
| Palm_Q9Z0P4_S200s_1 | Palm_Q9Z0 | -0.88 | -0.48 | 0.13 | -0.98 | -1.52 | -1.19 | -1.19 | -1.7 | -0.07 | -0.31 | -1.37 | -0.96 | -0.62 | 0.01 | -1.39 | -0.9 | -0.8475 | -0.125 | -0.1725 | 0.663356 | 0.018817 | 0.749435 | 0.989223 | 0.997988 | 0.996159 |
| Palm_Q9Z0P4_S204s_1 | Palm_Q9Z0 | 0.1 | -0.15 | 1.37 | 1.71 | 0.6 | -0.12 | 0.47 | 0.83 | -0.26 | -0.31 | 1.26 | 0.13 | -0.31 | -0.17 | -0.13 | 0.45 | -0.3125 | -0.5525 | -0.7975 | 0.115449 | 0.507798 | 0.324763 | 0.989223 | 0.997988 | 0.994666 |
| Palm_Q9Z0P4_S301s_1 | Palm_Q9Z0 | 0.88 | 0.32 | 0.41 | 0.74 | 0.96 | 1.02 | 1.31 | 1.16 | 0.55 | 0.58 | 0.45 | 1.23 | -0.05 | 1 | 0.68 | 1.07 | 0.525 | 0.115 | 0.0875 | 0.791017 | 0.043468 | 0.678493 | 0.989223 | 0.997988 | 0.999507 |
| Palm_Q9Z0P4_S321s_1 | Palm_Q9Z0 | 0.78 | 1.12 | 1.07 | 0.8 | 0.93 | 0.29 | 0.44 | 0.94 | 0.55 | 0.86 | -0.06 | 0.88 | 0.07 | 0.64 | 0.67 | 0.91 | -0.2925 | -0.385 | -0.37 | 0.205074 | 0.255833 | 0.198732 | 0.989223 | 0.997988 | 0.994666 |
| Palm_Q9Z0P4_T141t_1 | Palm_Q9Z0 | 0.94 | 0.56 | 0.59 | 0.71 | 1.02 | 1.09 | 0.74 | 0.55 | 0.51 | 0.86 | -0.11 | 0.7 | 0.4 | 0.63 | -0.13 | 0.01 | 0.15 | -0.21 | -0.4725 | 0.112866 | 0.517045 | 0.461131 | 0.989223 | 0.997988 | 0.994666 |
| Palm_Q9Z0P4_T141t_2 | Palm_Q9Z0 | 0.76 | 0.35 | 0.65 | 1.3 | 0.87 | 0.47 | 1.06 | 0.86 | 0.18 | 0.49 | 0.82 | 1.13 | -0.91 | 0.55 | 0.79 | 0.68 | 0.05 | -0.11 | -0.4875 | 0.281254 | 0.85502 | 0.728704 | 0.996349 | 0.997988 | 0.994666 |
| Palm_Q9Z0P4_T145t_1 | Palm_Q9Z0 | 0.35 | 0.71 | 1.02 | 0.6 | 0.82 | -0.21 | 1.38 | 0.43 | 0.25 | 0.65 | -0.26 | 1.04 | 0.36 | 0.84 | 0.78 | 1.05 | -0.065 | -0.25 | 0.0875 | 0.754575 | 0.856791 | 0.460641 | 0.996349 | 0.997988 | 0.9971 |
| Palm_Q9Z0P4_T145t_2 | Palm_Q9Z0 | 0.76 | 0.35 | 0.65 | 1.3 | 0.87 | 0.47 | 1.06 | 0.86 | 0.18 | 0.49 | 0.82 | 1.13 | -0.91 | 0.55 | 0.79 | 0.68 | 0.05 | -0.11 | -0.4875 | 0.281254 | 0.85502 | 0.728704 | 0.996349 | 0.997988 | 0.994666 |
| Palm_Q9Z0P4_T153t_2 | Palm_Q9Z0 | 0.97 | 0.26 | 0.61 | 1.27 | 1.09 | 0.86 | 0.89 | 0.79 | 0.21 | 0.5 | 0.87 | 0.69 | -0.67 | 0.97 | 0.58 | 0.71 | 0.13 | -0.21 | -0.38 | 0.379443 | 0.632509 | 0.489844 | 0.989223 | 0.997988 | 0.994666 |
| Palm_Q9Z0P4_T291t_1 | Palm_Q9Z0 | 1.26 | 1.06 | -0.06 | 0.59 | 1.02 | 1.5 | 1.28 | 0.79 | -0.12 | 1.32 | 1.26 | 0.29 | -0.91 | -0.28 | 0.66 | -0.23 | 0.435 | -0.025 | -0.9025 | 0.059794 | 0.21872 | 0.955281 | 0.989223 | 0.997988 | 0.994666 |
| Palm_Q9Z0P4_T319t_1 | Palm_Q9Z0 | 0.59 | 1.19 | 0.78 | 0.77 | 1.11 | 0.86 | 0.66 | 0.58 | 0.88 | 1.17 | 0.6 | 0.8 | 0.37 | 0.71 | 0.96 | 0.81 | -0.03 | 0.03 | -0.12 | 0.656381 | 0.900237 | 0.904108 | 0.996493 | 0.997988 | 0.995259 |
| Palm2_Q8BR92_S225s_1 | Palm2_Q8B | -0.39 | -0.71 | -0.54 | 0.5 | 1.42 | -0.14 | 0.14 | 0.16 | 0.08 | -0.51 | 1.09 | -0.53 | 0.03 | 0.75 | 0.43 | 0.21 | 0.68 | 0.3175 | 0.64 | 0.088182 | 0.129411 | 0.485256 | 0.989223 | 0.997988 | 0.994666 |
| Palm2_Q8BR92_S355s_1 | Palm2_Q8B | -0.33 | -0.65 | -1.65 | -1.94 | 0.1 | -0.86 | -1.91 | -1.3 | -1.18 | 0.17 | -0.27 | -2.39 | 0.44 | 0.47 | -1.59 | -2.19 | 0.15 | 0.225 | 0.425 | 0.550669 | 0.77406 | 0.719562 | 0.995762 | 0.997988 | 0.994666 |
| Palm3_A2TJV2_S270s_1 | Palm3_A2T | 0.12 | -0.65 | -0.46 | -0.33 | 0.49 | -0.32 | -0.1 | -0.18 | -0.43 | 0.26 | -0.59 | -0.76 | 0.16 | 0.05 | -0.76 | 0.8 | 0.3025 | -0.05 | 0.3925 | 0.311308 | 0.293273 | 0.872204 | 0.989223 | 0.997988 | 0.994666 |
| Palm3_A2TJV2_S524s_1 | Palm3_A2T | 3.58 | 3.42 | 3.23 | 2.08 | 2.25 | 4.8 | 2.05 | 2.41 | 2.72 | 3.54 | 0.85 | 3.39 | 0.79 | 0.35 | 3.49 | 3.17 | -0.2 | -0.4525 | -1.1275 | 0.168288 | 0.7577 | 0.486558 | 0.995762 | 0.997988 | 0.994666 |
| Palm3_A2TJV2_S565s_1 | Palm3_A2T | 1.2 | 0.9 | 1.09 | 1.58 | 0.65 | 1.15 | 1.09 | 0.92 | 0.81 | 1.07 | 1.3 | 1.28 | -0.15 | 1.08 | 1.36 | 0.93 | -0.24 | -0.0775 | -0.3875 | 0.317703 | 0.337709 | 0.760947 | 0.989223 | 0.997988 | 0.994666 |
| Palm3_A2TJV2_S601s_1 | Palm3_A2T | 1.59 | 1.41 | 1.56 | 1.72 | 0.29 | 1.96 | 0.71 | 1.12 | 2.43 | 1.9 | 1.48 | 1.67 | 1.46 | -0.2 | 1.88 | 1.57 | -0.55 | 0.3 | -0.3925 | 0.400163 | 0.150856 | 0.286546 | 0.989223 | 0.997988 | 0.994666 |
| Palm3_A2TJV2_S620s_1 | Palm3_A2T | -0.17 | 0.32 | 0.35 | 0.48 | 0.28 | -0.14 | 0.63 | -0.74 | -0.3 | 0.24 | 0.47 | -0.29 | -0.32 | 0.18 | -0.72 | 0.18 | -0.2375 | -0.215 | -0.415 | 0.205388 | 0.48483 | 0.461248 | 0.989223 | 0.997988 | 0.994666 |
| Palm3_A2TJV2_S87s_1 | Palm3_A2T | -0.57 | 0.09 | -0.09 | -1.06 | -0.91 | 0.15 | -1.41 | -0.81 | -0.03 | -0.04 | -0.67 | -0.74 | 0.28 | -2.18 | -0.46 | -0.06 | -0.3375 | 0.0375 | -0.1975 | 0.724245 | 0.407854 | 0.91262 | 0.989223 | 0.997988 | 0.9971 |
| Palmd_Q3UVT7_S364s_1 | Palmd_Q3U | 0.11 | -0.92 | -1.08 | -0.83 | -1.52 | 1.12 | -1.73 | -0.62 | 1.51 | -0.94 | -1.4 | -0.09 | 0.13 | -2.42 | -0.32 | -0.5 | -0.0075 | 0.45 | -0.0975 | 0.865025 | 0.990418 | 0.482847 | 0.999187 | 0.997988 | 0.999507 |
| Palmd_Q3UVT7_S370s_1 | Palmd_Q3U | -1.04 | -1.79 | -1.55 | -2.23 | -0.68 | -1.3 | -1.39 | -1.64 | -1.55 | -1.49 | -0.92 | -1.44 | -1.33 | -0.76 | -2.04 | -1.82 | 0.4 | 0.3025 | 0.165 | 0.672433 | 0.245347 | 0.353114 | 0.989223 | 0.997988 | 0.996159 |
| Palmd_Q3UVT7_S385s_1 | Palmd_Q3U | -0.16 | -1.13 | -0.17 | 0.03 | -0.05 | 0.39 | -0.37 | 0.65 | 0.72 | 0.05 | 0.71 | -0.37 | 0.13 | 0.29 | -0.16 | -0.23 | 0.5125 | 0.635 | 0.365 | 0.2845 | 0.165289 | 0.119575 | 0.989223 | 0.997988 | 0.994666 |
| Pank2_V9GXP9_S43s_1 | Pank2_V9G | -2.43 | -2.14 | -2.81 | -2.7 | -1.18 | -1.84 | -1.97 | -3.03 | -2.18 | -2.24 | -0.74 | -2.53 | -0.63 | -0.73 | -2.13 | -2.39 | 0.515 | 0.5975 | 1.05 | 0.045916 | 0.214424 | 0.178297 | 0.989223 | 0.997988 | 0.994666 |
| Pank2_V9GXP9_S44s_1 | Pank2_V9G | -2.79 | -2.65 | -1.35 | -1.47 | -1.88 | -2.52 | -1.33 | -0.88 | -4.15 | -3.39 | -0.27 | -1.36 | -1.66 | -1.82 | -1.36 | -1.31 | 0.4125 | -0.2275 | 0.5275 | 0.212145 | 0.399124 | 0.792587 | 0.989223 | 0.997988 | 0.994666 |
| Pank4_Q80YV4_S63s_1 | Pank4_Q80 | 2 | 1.17 | 1.58 | 0.62 | 1.07 | 2 | 0.89 | 0.68 | 0 | 1.29 | 0.63 | 0.96 | 0.63 | 0.31 | 1.02 | 1.02 | -0.1825 | -0.6225 | -0.5975 | 0.125436 | 0.648734 | 0.145452 | 0.99033 | 0.997988 | 0.994666 |
| Papd5_E9QJT6_S97s_1 | Papd5_E9Q | 0.05 | 0.24 | 0.95 | 0.97 | -0.95 | -0.34 | -0.26 | -0.13 | 2.03 | 1.82 | 0.35 | -0.66 | 1.07 | -0.72 | 0.37 | -0.2 | -0.9725 | 0.3325 | -0.4225 | 0.351661 | 0.011703 | 0.594663 | 0.989223 | 0.997988 | 0.994666 |
| Papola_F6VTN8_S234s_1 | Papola_F6V | -0.65 | -0.23 | -1.56 | -1.52 | -1.17 | -2.04 | -1.58 | -1.77 | 0.92 | -2.48 | 0.04 | -1.01 | 1.62 | -0.42 | -1.05 | -1.14 | -0.65 | 0.3575 | 0.7425 | 0.271217 | 0.105124 | 0.622421 | 0.989223 | 0.997988 | 0.994666 |
| Paqr9_Q6TCG2_S33s_1 | Paqr9_Q6T | -0.4 | -0.96 | -0.33 | -0.94 | -0.92 | -0.54 | -0.78 | -1.04 | -0.4 | -0.28 | -1.44 | -0.97 | 0.43 | -1.55 | -0.34 | -1.08 | -0.1625 | -0.115 | 0.0225 | 0.960295 | 0.52712 | 0.734493 | 0.989223 | 0.997988 | 0.999507 |
| Pard3_B7ZNY3_S1203s_1 | Pard3_B7Z | -1.82 | -1.44 | -1.46 | -2.08 | -1.31 | -1.72 | -0.4 | -1.79 | -2.2 | -1.97 | -1.32 | -1.93 | -1.71 | -1.75 | -1.63 | -1.51 | 0.395 | -0.155 | 0.05 | 0.848236 | 0.281958 | 0.594417 | 0.989223 | 0.997988 | 0.999507 |
| Pard3_B7ZNY3_S383s_1 | Pard3_B7Z | 0.43 | 0.15 | -0.13 | 1.02 | 0.13 | -0.17 | 0.53 | 0.77 | 0.4 | 0.76 | 0.99 | 0.5 | 0.22 | -0.51 | 0.61 | 0.67 | -0.0525 | 0.295 | -0.12 | 0.753182 | 0.874275 | 0.356908 | 0.996493 | 0.997988 | 0.9971 |
| Pard3_B7ZNY3_S849s_1 | Pard3_B7Z | -0.78 | -0.73 | 0.35 | 0.08 | 0.11 | -0.39 | 0.78 | -1.31 | -0.81 | -0.67 | -0.37 | -0.63 | -0.09 | -1.8 | 0.95 | 0.57 | 0.0675 | -0.35 | 0.1775 | 0.771974 | 0.888917 | 0.298692 | 0.996493 | 0.997988 | 0.997259 |
| Pard3_B7ZNY3_S854s_1 | Pard3_B7Z | -0.72 | -0.68 | -1.22 | -0.76 | -0.6 | -0.63 | -0.62 | -1.66 | -0.95 | -0.88 | -2.18 | -1.62 | -0.24 | -1.12 | -0.56 | -1.18 | -0.0325 | -0.5625 | 0.07 | 0.822541 | 0.916315 | 0.131786 | 0.997335 | 0.997988 | 0.999507 |
| Pard3b_Q5SV54_S338s_1 | Pard3b_Q5 | -0.15 | 1.02 | -1.52 | 1.19 | 0.43 | 0.26 | -1.22 | -0.74 | 0.82 | 0.56 | 0.14 | -1.29 | 1.69 | 0.09 | -0.42 | 0.43 | -0.4525 | -0.0775 | 0.3125 | 0.652713 | 0.496691 | 0.912566 | 0.989223 | 0.997988 | 0.995259 |
| Pard3b_Q5SV54_S746s_1 | Pard3b_Q5 | 1.2 | 1.27 | 0.78 | 0.59 | 0.96 | 0.69 | 1.1 | 1.26 | 0.67 | 0.61 | -0.04 | 1.04 | 0.39 | 1.01 | 0.86 | 1.46 | 0.0425 | -0.39 | -0.03 | 0.92558 | 0.868091 | 0.230439 | 0.996493 | 0.997988 | 0.999507 |
| Parg_G3X8U8_S297s_1 | Parg_G3X8 | -0.15 | -0.8 | -1.04 | -0.72 | -0.31 | -0.64 | -0.47 | 0.25 | -0.19 | -0.03 | -0.11 | -1.03 | 0.52 | 0.44 | -0.87 | -1.97 | 0.385 | 0.3375 | 0.2075 | 0.717682 | 0.214267 | 0.314455 | 0.989223 | 0.997988 | 0.996966 |
| Parn_Q8VDG3_S556s_1 | Parn_Q8VD | -1.13 | -1.88 | -0.12 | -0.97 | -2.29 | -2.55 | -1 | -0.44 | -1.47 | -1.51 | -0.72 | 0.11 | -0.29 | -0.72 | -0.73 | -0.51 | -0.545 | 0.1275 | 0.4625 | 0.250724 | 0.343541 | 0.797133 | 0.989223 | 0.997988 | 0.994666 |
| Pbrm1_D3Z1N4_S1468s_1 | Pbrm1_D3Z | 0.55 | -0.36 | 1.72 | 1.43 | -1.12 | -0.12 | -0.36 | 0.96 | 2.44 | 0.02 | -0.37 | 1.81 | 2.37 | -0.88 | 0.87 | 1.52 | -0.995 | 0.14 | 0.135 | 0.855497 | 0.107723 | 0.850429 | 0.989223 | 0.997988 | 0.999507 |
| Pbx2_Q3UR63_S248s_1 | Pbx2_Q3U | -2.5 | -1 | -0.19 | -2.07 | -1.86 | -1.59 | -1.81 | -0.78 | -0.73 | -1.18 | -0.26 | -0.66 | 0.54 | -1.16 | -1.48 | -1.26 | -0.07 | 0.7325 | 0.6 | 0.355254 | 0.894204 | 0.181898 | 0.996493 | 0.997988 | 0.994666 |
| Pcbp1_P60335_S189s_1 | Pcbp1_P60 | 0.93 | 0.98 | 0.31 | 0.38 | 0.72 | 0.72 | 0.79 | 0.47 | 0.61 | 1.17 | 0.03 | 0.47 | 0.39 | 1.08 | 0.4 | 0.55 | 0.025 | -0.08 | -0.045 | 0.881096 | 0.919804 | 0.804097 | 0.997335 | 0.997988 | 0.999507 |
| Pcbp1_P60335_S190s_1 | Pcbp1_P60 | 0.79 | 0.98 | 0.47 | 0.3 | 0.78 | 0.48 | 0.86 | 0.51 | 0.44 | 0.71 | 0.36 | 0.48 | 0.09 | 0.96 | 0.39 | 0.75 | 0.0225 | -0.1375 | -0.0875 | 0.774075 | 0.926245 | 0.582125 | 0.997335 | 0.997988 | 0.99798 |
| Pcbp2_B2M1R7_S185s_1 | Pcbp2_B2M | -0.76 | -1.29 | -2.44 | -2.6 | -1.18 | -0.95 | -1.3 | -1.89 | -1.58 | -0.77 | -1.22 | -1.95 | -1.1 | -0.42 | -2.25 | -2.92 | 0.4425 | 0.3925 | 0.1 | 0.877245 | 0.344022 | 0.427085 | 0.989223 | 0.997988 | 0.999507 |
| Pcdh1_E0CXD4_S343s_1 | Pcdh1_E0C | -0.23 | -1.33 | -0.7 | 0.35 | -1.17 | 0.29 | 0.41 | 0.59 | 0.41 | -1.09 | 0.39 | 0.43 | -0.73 | 0.03 | -1.22 | -1.11 | 0.5075 | 0.5125 | -0.28 | 0.532887 | 0.318981 | 0.309615 | 0.989223 | 0.997988 | 0.994666 |
| Pcdh1_F7BJK1_S1010s_1 | Pcdh1_F7B | 0.3 | -0.04 | -0.17 | 0.47 | 0.18 | 0.4 | -0.52 | 0.41 | 0.12 | 0.17 | 0.19 | -0.07 | -0.31 | 0.4 | -0.18 | -0.51 | -0.0225 | -0.0375 | -0.29 | 0.35169 | 0.93871 | 0.876911 | 0.999187 | 0.997988 | 0.994666 |
| Pcdh1_F7BJK1_S1010s_2 | Pcdh1_F7B | 0.27 | 0.65 | 0.82 | 0.54 | 0.34 | 0.2 | 0.2 | 0.49 | 0.48 | 0.67 | 1.14 | 0.75 | -0.87 | 0.83 | 0.66 | 0.44 | -0.2625 | 0.19 | -0.305 | 0.460523 | 0.252512 | 0.459425 | 0.989223 | 0.997988 | 0.994666 |
| Pcdh1_F7BJK1_S1012s_1 | Pcdh1_F7B | 0.21 | 0.76 | 0.8 | 0.49 | 0.5 | -0.05 | 0.45 | 0.42 | 0.97 | 0.55 | 1.39 | 0.71 | 1.06 | 1.43 | 0.98 | 0.6 | -0.235 | 0.34 | 0.4525 | 0.141603 | 0.352777 | 0.244171 | 0.989223 | 0.997988 | 0.994666 |
| Pcdh1_F7BJK1_S1012s_2 | Pcdh1_F7B | 0.27 | 0.65 | 0.82 | 0.54 | 0.34 | 0.2 | 0.2 | 0.49 | 0.48 | 0.67 | 1.14 | 0.75 | -0.87 | 0.83 | 0.66 | 0.44 | -0.2625 | 0.19 | -0.305 | 0.460523 | 0.252512 | 0.459425 | 0.989223 | 0.997988 | 0.994666 |
| Pcdh1_F7BJK1_S741s_1 | Pcdh1_F7B | 0.19 | -1.04 | -0.44 | 0.32 | 0.13 | -0.49 | 0.59 | 0.87 | -0.27 | -0.95 | 0.72 | 0.37 | -1.29 | 0.54 | -0.27 | -0.17 | 0.5175 | 0.21 | -0.055 | 0.906929 | 0.228675 | 0.65141 | 0.989223 | 0.997988 | 0.999507 |
| Pcdh1_F7BJK1_S788s_1 | Pcdh1_F7B | 0.84 | 0.32 | 0.44 | 0.88 | 1.02 | 0.74 | 0.69 | 0.7 | 0.46 | 0.64 | 0.54 | 0.75 | -0.16 | 0.67 | 0.4 | 0.44 | 0.1675 | -0.0225 | -0.2825 | 0.346734 | 0.478798 | 0.925151 | 0.989223 | 0.997988 | 0.994666 |
| Pcdh1_F7BJK1_S801s_1 | Pcdh1_F7B | 2.89 | 1.47 | 1.79 | 0.37 | 2.49 | 2.96 | 2.74 | 2.41 | -0.62 | 2.51 | -0.67 | 2.42 | -1.54 | -0.82 | 2.17 | 2.18 | 1.02 | -0.72 | -1.1325 | 0.258938 | 0.061412 | 0.441152 | 0.989223 | 0.997988 | 0.994666 |
| Pcdh1_F7BJK1_S812s_1 | Pcdh1_F7B | 0.04 | -0.2 | -0.21 | 0.64 | 0.06 | 0.77 | 0.64 | 0.73 | 0.18 | 0.43 | -1.08 | 0.69 | -0.32 | 0.19 | -0.2 | 0.06 | 0.4825 | -0.0125 | -0.135 | 0.649757 | 0.120766 | 0.976717 | 0.989223 | 0.997988 | 0.995259 |
| Pcdh1_F7BJK1_S823s_1 | Pcdh1_F7B | 0.57 | 0.17 | -0.22 | 0.23 | 0.82 | 0.72 | 0.73 | 0.36 | 0.24 | 0.58 | -0.34 | 0.29 | -0.41 | 0.81 | -0.16 | 0.13 | 0.47 | 0.005 | -0.095 | 0.782019 | 0.08414 | 0.986435 | 0.989223 | 0.997988 | 0.998226 |
| Pcdh1_F7BJK1_S823s_2 | Pcdh1_F7B | -0.03 | 0.16 | 0.96 | 0.83 | 0.29 | 0.46 | 1 | 0.76 | -0.09 | 0.18 | 0.64 | 0.9 | -0.64 | 0.33 | -0.06 | 0.26 | 0.1475 | -0.0725 | -0.5075 | 0.176766 | 0.636217 | 0.834679 | 0.989223 | 0.997988 | 0.994666 |
| Pcdh1_F7BJK1_S978s_1 | Pcdh1_F7B | 0.73 | -0.11 | -0.62 | -0.73 | -0.37 | 0.57 | -0.19 | -0.15 | 0.08 | 0.71 | -0.1 | -0.91 | 0.18 | -0.08 | -0.77 | -1 | 0.1475 | 0.1275 | -0.235 | 0.587015 | 0.700509 | 0.778595 | 0.99503 | 0.997988 | 0.995259 |
| Pcdh1_F7BJK1_S979s_1 | Pcdh1_F7B | -0.59 | -0.76 | -0.96 | -0.87 | -0.34 | -0.53 | -0.08 | -1 | -0.91 | -0.28 | -1.29 | -1.24 | -1.22 | -0.26 | -0.1 | -1.14 | 0.3075 | -0.135 | 0.115 | 0.734384 | 0.253714 | 0.64414 | 0.989223 | 0.997988 | 0.9971 |
| Pcdh10_Q925I8_S841s_1 | Pcdh10_Q9 | 0.86 | -0.54 | 0.33 | 0.82 | 0.08 | 1.44 | 0.97 | 0.74 | 0.3 | 0.23 | 1.42 | 0.28 | -0.87 | 0.22 | 0.23 | 0.3 | 0.44 | 0.19 | -0.3975 | 0.362211 | 0.300088 | 0.656087 | 0.989223 | 0.997988 | 0.994666 |
| Pcdh10_Q925I8_S898s_1 | Pcdh10_Q9 | -0.01 | 0.84 | -1.36 | -1.12 | -0.24 | -0.2 | 0.13 | -0.91 | 0.28 | -0.33 | -0.98 | 0.61 | 0.3 | 0.68 | -0.35 | -2.6 | 0.1075 | 0.3075 | -0.08 | 0.919328 | 0.832303 | 0.593127 | 0.995762 | 0.997988 | 0.999507 |
| Pcdh11x_Q2TJH7_S333s_1 | Pcdh11x_Q | -0.44 | -1.57 | -1.18 | -0.07 | -0.37 | 0.33 | -0.56 | -0.48 | 0.01 | -0.07 | 0.09 | -0.44 | 0.56 | -0.51 | -0.72 | -1.29 | 0.545 | 0.7125 | 0.325 | 0.512381 | 0.180913 | 0.079543 | 0.989223 | 0.997988 | 0.994666 |
| Pcdh17_E9PXF0_S1024s_1 | Pcdh17_E9 | 0.19 | 0.53 | -1.06 | 0.05 | 0.36 | 1.09 | 0.69 | 0.37 | -0.06 | 0.92 | -1.13 | -0.83 | 0.03 | 0.32 | -1.18 | -0.24 | 0.7 | -0.2025 | -0.195 | 0.671789 | 0.088772 | 0.705069 | 0.989223 | 0.997988 | 0.996159 |
| Pcdh17_E9PXF0_S1036s_1 | Pcdh17_E9 | -0.5 | -0.33 | 0.66 | 0.27 | 0.53 | -0.47 | 0.97 | 0.56 | 0.02 | 0.44 | 0.47 | 0.47 | 0.11 | 0.58 | 0.11 | 0.67 | 0.3725 | 0.325 | 0.3425 | 0.329491 | 0.35552 | 0.323107 | 0.989223 | 0.997988 | 0.994666 |
| Pcdh17_E9PXF0_S1109s_1 | Pcdh17_E9 | -0.12 | -0.21 | 0.2 | 0.52 | -0.51 | 0.33 | 0.93 | 0.75 | 0.4 | -0.12 | 0.11 | 0.47 | -0.33 | 0.25 | 0.18 | -0.01 | 0.2775 | 0.1175 | -0.075 | 0.792425 | 0.447569 | 0.66709 | 0.989223 | 0.997988 | 0.999507 |
| Pcdh17_E9PXF0_S1110s_1 | Pcdh17_E9 | 0.03 | 0.01 | -0.06 | 0.28 | -0.59 | 0.66 | -0.51 | -0.19 | 0.7 | 0.15 | -0.55 | -0.11 | -0.05 | 0.78 | -1.27 | -2.06 | -0.2225 | -0.0175 | -0.715 | 0.238691 | 0.4835 | 0.954514 | 0.989223 | 0.997988 | 0.994666 |
| Pcdh18_Q8VHR0_S781s_2 | Pcdh18_Q8 | 0.09 | 0.33 | 0.23 | 0.2 | -0.6 | -0.6 | -0.37 | 0.4 | 0.86 | -0.18 | 0.08 | 0.69 | -0.02 | -0.71 | 0.69 | 0.54 | -0.505 | 0.15 | -0.0875 | 0.803192 | 0.095513 | 0.612555 | 0.989223 | 0.997988 | 0.999507 |
| Pcdh7_E9Q2S0_S1000s_2 | Pcdh7_E9Q | 0.23 | -0.41 | -0.04 | 0.6 | 0.78 | 0.48 | 0.75 | 0.31 | 0.02 | 0.15 | 0.53 | 0.09 | -1.28 | 0.4 | -0.21 | -0.28 | 0.485 | 0.1025 | -0.4375 | 0.301212 | 0.106694 | 0.722925 | 0.989223 | 0.997988 | 0.994666 |
| Pcdh7_E9Q2S0_S1011s_1 | Pcdh7_E9Q | 0.44 | 0.59 | 0.6 | 0.79 | 0.58 | 0.53 | 0.87 | 0.8 | 0.54 | 0.89 | 0.32 | 0.75 | 0.55 | 1.01 | 0.58 | 0.77 | 0.09 | 0.02 | 0.1225 | 0.622206 | 0.670638 | 0.932054 | 0.992876 | 0.997988 | 0.995259 |
| Pcdh7_E9Q2S0_S1011s_2 | Pcdh7_E9Q | 0.23 | -0.41 | -0.19 | 0.58 | 0.78 | 0.48 | 0.67 | 0.14 | 0.02 | 0.15 | 0.37 | 0.01 | -1.28 | 0.4 | -0.31 | -0.38 | 0.465 | 0.085 | -0.445 | 0.294066 | 0.134545 | 0.765465 | 0.989223 | 0.997988 | 0.994666 |
| Pcdh7_E9Q2S0_S1235s_1 | Pcdh7_E9Q | 0.43 | -0.33 | -0.29 | 0.72 | 0.24 | 0.55 | 0.69 | 0.7 | -0.2 | -0.14 | 0.96 | 0.48 | -0.92 | -0.14 | 0 | 0.01 | 0.4125 | 0.1425 | -0.395 | 0.295105 | 0.198108 | 0.71099 | 0.989223 | 0.997988 | 0.994666 |
| Pcdh7_E9Q2S0_S974s_1 | Pcdh7_E9Q | 0.51 | 0.37 | 0.46 | 0.47 | 0.57 | 0.28 | 0.16 | 0.2 | -0.17 | 0.32 | 0.13 | 0.65 | 0.14 | -0.94 | 0.07 | 0.17 | -0.15 | -0.22 | -0.5925 | 0.087585 | 0.473606 | 0.388551 | 0.989223 | 0.997988 | 0.994666 |
| Pcdh7_E9Q2S0_S989s_1 | Pcdh7_E9Q | 0.82 | 0.97 | 0.89 | 0.65 | 1.19 | 1.12 | 0.52 | 0.54 | 0.4 | 1.07 | -1.51 | 0.81 | 0.42 | -0.56 | 0.69 | 0.8 | 0.01 | -0.64 | -0.495 | 0.177514 | 0.96808 | 0.25907 | 0.999187 | 0.997988 | 0.994666 |
| Pcdh8_Q7TSK3_S798s_1 | Pcdh8_Q7T | 0.4 | 0.88 | 0.03 | 0.47 | 1.32 | 0.06 | 1.29 | 0.54 | 0.68 | 0.35 | 1.08 | 0.5 | 1.83 | 1.96 | 0.71 | 1.42 | 0.3575 | 0.2075 | 1.035 | 0.015527 | 0.324991 | 0.471569 | 0.989223 | 0.997988 | 0.994666 |
| Pcdh8_Q7TSK3_S802s_1 | Pcdh8_Q7T | 0.64 | 1.38 | 0.38 | 0.76 | 0.8 | 0.05 | 0.59 | 0.39 | 0.75 | 1.08 | 1.28 | 0.4 | 1.77 | 1.14 | 0.71 | 1.12 | -0.3325 | 0.0875 | 0.395 | 0.262605 | 0.273831 | 0.782951 | 0.989223 | 0.997988 | 0.994666 |
| Pcdh8_Q7TSK3_S904s_1 | Pcdh8_Q7T | 1.55 | 0.17 | 1.69 | 0.21 | 1.52 | 2.39 | 0.69 | 1 | 2.04 | 1.07 | 1.6 | 0.84 | 1.74 | 0.65 | 1.84 | 1.92 | 0.495 | 0.4825 | 0.6325 | 0.213514 | 0.342745 | 0.319565 | 0.989223 | 0.997988 | 0.994666 |
| Pcdh9_F8VPK8_S1109s_1 | Pcdh9_F8V | 2.24 | 2.24 | 1.24 | 1.06 | 2.11 | 2.27 | 1.07 | 1.09 | 0.64 | 1.33 | 0.92 | 1.34 | 2.17 | 1.64 | 1.11 | 1.16 | -0.06 | -0.6375 | -0.175 | 0.667807 | 0.888234 | 0.109681 | 0.996493 | 0.997988 | 0.996159 |
| Pcdh9_F8VPK8_S1156s_1 | Pcdh9_F8V | 1.06 | 0.4 | 0.12 | 0.6 | 1.16 | 1.2 | 0.55 | 0.88 | -0.49 | -0.14 | 2.34 | 0.79 | -1.2 | 0.28 | -0.58 | -0.15 | 0.4025 | 0.08 | -0.9575 | 0.031108 | 0.177295 | 0.894946 | 0.989223 | 0.997988 | 0.994666 |
| Pcdh9_F8VPK8_S1231s_1 | Pcdh9_F8V | 0.36 | 0.65 | 0.63 | 0.72 | 0.38 | 0.64 | -0.33 | 0.53 | 0.36 | 0.9 | -1.58 | 0.19 | 0.42 | -0.29 | 0.31 | 0.59 | -0.285 | -0.6225 | -0.3325 | 0.257257 | 0.31021 | 0.241508 | 0.989223 | 0.997988 | 0.994666 |
| Pcdh9_F8VPK8_S932s_1 | Pcdh9_F8V | 0.3 | 0.65 | 0.52 | 0.8 | 1.05 | 0.23 | 0.94 | 0.56 | 0.2 | 0.45 | 0.15 | 0.81 | -0.21 | 0.82 | 0.28 | 0.11 | 0.1275 | -0.165 | -0.3175 | 0.305817 | 0.629159 | 0.522192 | 0.989223 | 0.997988 | 0.994666 |
| Pcdha9_Q91Y11_S789s_1 | Pcdha9_Q9 | -0.08 | 0.7 | 1.08 | 1 | 0.06 | 1.24 | -0.03 | 0.29 | 1.02 | 0.43 | 0.42 | -0.13 | 0.97 | -0.49 | 0.41 | 0.39 | -0.285 | -0.24 | -0.355 | 0.391171 | 0.463821 | 0.514632 | 0.989223 | 0.997988 | 0.994666 |
| Pcdha9_Q91Y11_S922s_1 | Pcdha9_Q9 | -0.26 | 0.14 | -1.22 | -0.8 | 0.54 | -0.41 | -0.42 | -0.87 | 0.05 | -0.76 | -0.54 | -1.16 | -0.29 | 0.77 | -0.83 | -0.46 | 0.245 | -0.0675 | 0.3325 | 0.461247 | 0.547946 | 0.863084 | 0.989223 | 0.997988 | 0.994666 |
| Pcdhga12_Q91XY7_S136s_1 | Pcdhga12_ | 0.22 | 0.58 | 0.77 | 0.76 | 0.21 | -0.31 | -0.01 | 1.31 | -0.14 | 0.26 | 1.67 | 1.08 | -0.32 | 0.12 | 0.79 | 0.79 | -0.2825 | 0.135 | -0.2375 | 0.48762 | 0.452575 | 0.747348 | 0.989223 | 0.997988 | 0.994666 |
| Pcdhgc3_Q91XX1_S780s_1 | Pcdhgc3_Q | 0.27 | -0.1 | 1.18 | 0.82 | 0.73 | 0.23 | 0.83 | 0.3 | 0.53 | 0.84 | 3.19 | 0.23 | 0.7 | 0.91 | 0.5 | 1.08 | -0.02 | 0.655 | 0.255 | 0.465297 | 0.951851 | 0.337436 | 0.999187 | 0.997988 | 0.994666 |
| Pcdhgc5_Q91XW9_S746s_1 | Pcdhgc5_Q | 0.05 | 0.94 | 0.25 | 0.61 | 0.27 | 0.44 | 0.6 | 0.58 | -0.41 | 0.99 | 0.42 | 0.71 | 0.84 | 0.19 | 0.55 | 0.45 | 0.01 | -0.035 | 0.045 | 0.880662 | 0.969311 | 0.924388 | 0.999187 | 0.997988 | 0.999507 |
| Pcif1_P59114_S116s_1 | Pcif1_P591 | -0.4 | -1.18 | -0.22 | -0.22 | -1.32 | -1.23 | -0.49 | -0.36 | 0.27 | -0.63 | 0.06 | 0.27 | -0.4 | -0.95 | 0.43 | 0.21 | -0.345 | 0.4975 | 0.3275 | 0.415978 | 0.326982 | 0.161798 | 0.989223 | 0.997988 | 0.994666 |
| Pclo_Q9QYX7_S1116s_1 | Pclo_Q9QY | -0.34 | 0.19 | 1.3 | 1.22 | 0.38 | 0.29 | 1.62 | 0.63 | 0.07 | 0.18 | 1.05 | 1.51 | 0.57 | 0.42 | 1.74 | 1.51 | 0.1375 | 0.11 | 0.4675 | 0.35491 | 0.768818 | 0.825658 | 0.995762 | 0.997988 | 0.994666 |
| Pclo_Q9QYX7_S1292s_1 | Pclo_Q9QY | 0.07 | 0.49 | 0.27 | 0.16 | 0.48 | 0.16 | 0.52 | 0.02 | -0.13 | 0.38 | -0.21 | -0.18 | -0.54 | 0.66 | 0.1 | 0.42 | 0.0475 | -0.2825 | -0.0875 | 0.785858 | 0.835569 | 0.267354 | 0.995762 | 0.997988 | 0.999507 |
| Pclo_Q9QYX7_S131s_1 | Pclo_Q9QY | -3.89 | -4.19 | -3.72 | -2.81 | -2.26 | -3.22 | -2.43 | -4.67 | -5.32 | -4.03 | -0.83 | -3.56 | -2.92 | -2.29 | -3.42 | -2.93 | 0.5075 | 0.2175 | 0.7625 | 0.074294 | 0.377893 | 0.804791 | 0.989223 | 0.997988 | 0.994666 |
| Pclo_Q9QYX7_S1332s_1 | Pclo_Q9QY | 0.72 | 0.86 | 0.88 | 1.12 | 0.46 | 0.76 | 0.44 | 0.88 | 0.47 | 0.67 | 0.92 | 1.02 | 0.07 | 0.83 | 0.98 | 0.8 | -0.26 | -0.125 | -0.225 | 0.444824 | 0.259336 | 0.601117 | 0.989223 | 0.997988 | 0.994666 |
| Pclo_Q9QYX7_S1341s_1 | Pclo_Q9QY | 1 | 1.21 | 1.1 | 0.99 | 1.13 | 0.81 | 1.05 | 0.98 | 0.66 | 1.02 | 0.56 | 1.08 | 0.2 | 0.95 | 0.9 | 1.26 | -0.0825 | -0.245 | -0.2475 | 0.410364 | 0.684985 | 0.307369 | 0.994624 | 0.997988 | 0.994666 |
| Pclo_Q9QYX7_S1378s_1 | Pclo_Q9QY | 0.54 | 1.04 | 0.65 | 0.57 | 0.96 | 0.18 | 0.14 | 0.34 | 0.3 | 0.63 | 0.69 | 0.7 | 0.79 | 0.84 | 0.49 | 0.76 | -0.295 | -0.12 | 0.02 | 0.936591 | 0.28474 | 0.615579 | 0.989223 | 0.997988 | 0.999507 |
| Pclo_Q9QYX7_S1439s_1 | Pclo_Q9QY | 0.35 | 0.45 | 0.26 | 0.17 | 0.91 | 0.19 | 1.06 | 0.05 | -0.56 | 0.21 | -0.3 | 0.19 | -1.11 | 0.39 | 0.34 | 0.36 | 0.245 | -0.4225 | -0.3125 | 0.426248 | 0.407619 | 0.133451 | 0.989223 | 0.997988 | 0.994666 |
| Pclo_Q9QYX7_S1444s_1 | Pclo_Q9QY | 0.51 | 0.17 | -0.06 | 0.2 | 0.05 | 0.65 | 0.02 | 0.07 | 0.73 | 0.4 | 0.57 | 0.41 | 0.56 | 1.3 | 0.35 | 0.04 | -0.0075 | 0.3225 | 0.3575 | 0.297103 | 0.97594 | 0.18967 | 0.999187 | 0.997988 | 0.994666 |
| Pclo_Q9QYX7_S1444s_2 | Pclo_Q9QY | 0.66 | 0.5 | -0.08 | 0.91 | 0.24 | -0.56 | 0.13 | 0.33 | 0.78 | 0.81 | 1.37 | 0.57 | 0.12 | 1.08 | 1.02 | 0.72 | -0.4625 | 0.385 | 0.2375 | 0.490009 | 0.160516 | 0.229555 | 0.989223 | 0.997988 | 0.994666 |
| Pclo_Q9QYX7_S1457s_1 | Pclo_Q9QY | 1.84 | 1.87 | 0.92 | 1.34 | 1.62 | 1.1 | 1.1 | 1.47 | 1.42 | 1.99 | 0.6 | 0.1 | 1.25 | 1.02 | 1.48 | 1.33 | -0.17 | -0.465 | -0.2225 | 0.47078 | 0.56301 | 0.323912 | 0.989223 | 0.997988 | 0.994666 |
| Pclo_Q9QYX7_S1457s_2 | Pclo_Q9QY | 0.66 | 0.5 | -0.08 | 0.91 | 0.24 | -0.56 | 0.13 | 0.33 | 0.78 | 0.81 | 1.37 | 0.57 | 0.12 | 1.08 | 1.02 | 0.72 | -0.4625 | 0.385 | 0.2375 | 0.490009 | 0.160516 | 0.229555 | 0.989223 | 0.997988 | 0.994666 |
| Pclo_Q9QYX7_S1484s_1 | Pclo_Q9QY | -1.34 | -0.17 | 1.56 | 1.37 | -0.71 | -0.82 | 1.97 | 1.39 | -0.74 | -0.21 | 0.71 | 1.83 | -0.9 | -0.25 | 2.23 | 2.05 | 0.1025 | 0.0425 | 0.4275 | 0.643327 | 0.905319 | 0.957191 | 0.996493 | 0.997988 | 0.995259 |
| Pclo_Q9QYX7_S1550s_1 | Pclo_Q9QY | 0.07 | 0.52 | 2.09 | 2.35 | 0.16 | 0.5 | 0.84 | 2.21 | -0.17 | -0.1 | 2.7 | 1.82 | -0.24 | -0.08 | 1.86 | 1.85 | -0.33 | -0.195 | -0.41 | 0.574299 | 0.609492 | 0.810539 | 0.989223 | 0.997988 | 0.995259 |
| Pclo_Q9QYX7_S1551s_1 | Pclo_Q9QY | 2 | 1.4 | 0.41 | 0.61 | 0.88 | 1.95 | 0.12 | 0.34 | 1.69 | 1.47 | 0.58 | 0.65 | 1.26 | 1.51 | 0.75 | 0.51 | -0.2825 | -0.0075 | -0.0975 | 0.819925 | 0.57721 | 0.986585 | 0.989223 | 0.997988 | 0.999507 |
| Pclo_Q9QYX7_S1553s_1 | Pclo_Q9QY | -1.75 | 0.07 | -0.17 | -1.24 | -0.49 | -2.33 | -1.44 | -1.6 | -1.09 | -1.26 | -1.83 | -0.67 | -0.16 | -1.4 | -1.07 | -1.19 | -0.6925 | -0.44 | -0.1825 | 0.70886 | 0.2065 | 0.362481 | 0.989223 | 0.997988 | 0.996966 |
| Pclo_Q9QYX7_S1563s_1 | Pclo_Q9QY | 3.23 | 2.61 | 1.26 | 1.11 | 2.47 | 2.87 | 1.89 | 1.41 | -0.15 | 2.71 | -0.64 | 1.54 | -1.6 | 0.01 | 1.67 | 2.08 | 0.1075 | -1.1875 | -1.5125 | 0.106868 | 0.84509 | 0.175382 | 0.996349 | 0.997988 | 0.994666 |
| Pclo_Q9QYX7_S1585s_1 | Pclo_Q9QY | 1.4 | 0.9 | 0.21 | 0.43 | 1.46 | 1.6 | 0.66 | 0.26 | 0.19 | 0.9 | -0.42 | 0.35 | -0.07 | 1.25 | -0.2 | 0.04 | 0.26 | -0.48 | -0.48 | 0.274112 | 0.520147 | 0.229613 | 0.989223 | 0.997988 | 0.994666 |
| Pclo_Q9QYX7_S1766s_1 | Pclo_Q9QY | 0.4 | 1.54 | 1.08 | 0.91 | 0.31 | 0.45 | 1.08 | 0.42 | 1.23 | 1.53 | 1.06 | 1.23 | 0.69 | 1.38 | 0.81 | 1.02 | -0.4175 | 0.28 | -0.0075 | 0.981537 | 0.201559 | 0.3569 | 0.989223 | 0.997988 | 1 |
| Pclo_Q9QYX7_S1772s_1 | Pclo_Q9QY | 0.18 | 0.96 | 1.19 | 0.93 | 0.25 | 0.45 | 0.81 | 0.35 | 0.56 | 1.02 | 1.4 | 1.39 | 0.21 | 1.18 | 0.97 | 0.87 | -0.35 | 0.2775 | -0.0075 | 0.982341 | 0.237377 | 0.400443 | 0.989223 | 0.997988 | 1 |
| Pclo_Q9QYX7_S1829s_1 | Pclo_Q9QY | 0.57 | 0.68 | 0.08 | 0.79 | 0.91 | 0.35 | 0.5 | 0.42 | 0.05 | 0.42 | -0.07 | 0.33 | -0.37 | 0.67 | 0.36 | 0.19 | 0.015 | -0.3475 | -0.3175 | 0.330456 | 0.952898 | 0.203741 | 0.999187 | 0.997988 | 0.994666 |
| Pclo_Q9QYX7_S212s_1 | Pclo_Q9QY | 0.33 | 0.41 | -0.01 | 0.5 | 1.14 | 0.67 | 1.09 | 0.65 | 0.04 | 0.09 | -0.24 | 0.43 | -0.09 | 0.62 | 0.34 | 0.26 | 0.58 | -0.2275 | -0.025 | 0.92646 | 0.034227 | 0.375666 | 0.989223 | 0.997988 | 0.999507 |
| Pclo_Q9QYX7_S2288s_1 | Pclo_Q9QY | -1.31 | -1.5 | -1.19 | -0.88 | -1.01 | -0.82 | -0.77 | -0.95 | -1.19 | -0.96 | -0.16 | -1.92 | -0.96 | -0.7 | -1.58 | -1.32 | 0.3325 | 0.1625 | 0.08 | 0.788023 | 0.161098 | 0.67585 | 0.989223 | 0.997988 | 0.999507 |
| Pclo_Q9QYX7_S2495s_1 | Pclo_Q9QY | 0.22 | 0.02 | -2.38 | -0.3 | 0.25 | 0.57 | 0.02 | -0.42 | 0.28 | 0.25 | -1.04 | -1.41 | 0.17 | 0.29 | -1.01 | -0.3 | 0.715 | 0.13 | 0.3975 | 0.518399 | 0.229889 | 0.846497 | 0.989223 | 0.997988 | 0.994666 |
| Pclo_Q9QYX7_S258s_1 | Pclo_Q9QY | -0.47 | 0.53 | 0.39 | 0.22 | -0.15 | 0.21 | -0.21 | 0.1 | -0.39 | 0 | 0.4 | 1.02 | 0.29 | -0.5 | 0.14 | 0.16 | -0.18 | 0.09 | -0.145 | 0.658942 | 0.524146 | 0.812236 | 0.989223 | 0.997988 | 0.995259 |
| Pclo_Q9QYX7_S3358s_1 | Pclo_Q9QY | -1.21 | -0.3 | -1.07 | -0.02 | 0.23 | -0.65 | 0.22 | -0.45 | -1.39 | -0.44 | 1.85 | -0.58 | -0.38 | -0.13 | -0.36 | 1.52 | 0.4875 | 0.51 | 0.8125 | 0.135278 | 0.202869 | 0.461162 | 0.989223 | 0.997988 | 0.994666 |
| Pclo_Q9QYX7_S3514s_1 | Pclo_Q9QY | 1.24 | 0.79 | 1 | 1.45 | 0.94 | 1 | 1.51 | 1.16 | 1.29 | 1.25 | 1.55 | 1.14 | 1.09 | 0.98 | 1.46 | 1.21 | 0.0325 | 0.1875 | 0.065 | 0.808212 | 0.896274 | 0.453621 | 0.996493 | 0.997988 | 0.999507 |
| Pclo_Q9QYX7_S3545s_1 | Pclo_Q9QY | 0.49 | -0.1 | 0.34 | 0.15 | 1.04 | 0.42 | 0.88 | 0.58 | -0.17 | -0.07 | 0.24 | 0.33 | -0.61 | 0.52 | -0.08 | 0.09 | 0.51 | -0.1375 | -0.24 | 0.454738 | 0.063762 | 0.585338 | 0.989223 | 0.997988 | 0.994666 |
| Pclo_Q9QYX7_S3561s_1 | Pclo_Q9QY | -0.81 | -0.76 | -2.02 | -2.46 | -0.25 | -1.48 | -1.49 | -2.54 | -0.58 | -1.01 | -0.08 | -2.52 | -0.75 | -0.62 | -2.19 | -3.05 | 0.0725 | 0.465 | -0.14 | 0.830837 | 0.898972 | 0.460395 | 0.996493 | 0.997988 | 0.999507 |
| Pclo_Q9QYX7_S3598s_1 | Pclo_Q9QY | 0.39 | -0.16 | 0.45 | 0.77 | 0.37 | 0.92 | 0.27 | 0.06 | 0.83 | 0.23 | -1.58 | 0.1 | 1.06 | 0.17 | 0.22 | -0.11 | 0.0425 | -0.4675 | -0.0275 | 0.937049 | 0.885061 | 0.37638 | 0.996493 | 0.997988 | 0.999507 |
| Pclo_Q9QYX7_S3608s_1 | Pclo_Q9QY | 0.8 | 1.65 | 1.19 | 0.94 | 0.67 | 0.46 | 0.5 | 0.74 | 0.86 | 1.35 | 1.05 | 0.92 | 0.92 | 1.26 | 1.17 | 0.94 | -0.5525 | -0.1 | -0.0725 | 0.797037 | 0.051379 | 0.714741 | 0.989223 | 0.997988 | 0.999507 |
| Pclo_Q9QYX7_S3610s_1 | Pclo_Q9QY | 0.38 | 1.46 | 1.43 | 1.33 | 0.32 | 0.22 | 0.82 | 0.9 | 0.93 | 1.11 | 1.45 | 1.3 | 1.17 | 1 | 1.52 | 1.41 | -0.585 | 0.0475 | 0.125 | 0.702958 | 0.096131 | 0.8797 | 0.989223 | 0.997988 | 0.996159 |
| Pclo_Q9QYX7_S3610s_2 | Pclo_Q9QY | 1.08 | 1.01 | 0.35 | 0.73 | 0.98 | 1.15 | 0.14 | 0.89 | 0.74 | 0.89 | 0.34 | 0.4 | -0.64 | 1.17 | 0.09 | 0.92 | -0.0025 | -0.2 | -0.4075 | 0.362147 | 0.993383 | 0.466432 | 0.999244 | 0.997988 | 0.994666 |
| Pclo_Q9QYX7_S3616s_1 | Pclo_Q9QY | 0.26 | -0.05 | 0.09 | 0.4 | -0.23 | -0.08 | 0.09 | -0.35 | -0.55 | 0.4 | -0.77 | 0.29 | -0.04 | -0.87 | -0.21 | -0.09 | -0.3175 | -0.3325 | -0.4775 | 0.121911 | 0.175058 | 0.331858 | 0.989223 | 0.997988 | 0.994666 |
| Pclo_Q9QYX7_S3616s_2 | Pclo_Q9QY | 0.87 | 0.9 | 0.35 | 0.73 | 0.94 | 0.92 | 0.14 | 0.89 | 1.06 | 1.19 | 0.34 | 0.4 | -0.55 | 1.17 | 0.09 | 0.92 | 0.01 | 0.035 | -0.305 | 0.46819 | 0.9707 | 0.9057 | 0.999187 | 0.997988 | 0.994666 |
| Pclo_Q9QYX7_S3763s_1 | Pclo_Q9QY | -1.4 | -1.04 | -1.87 | -2.59 | -1.06 | -1.6 | -2.19 | -2.28 | -2 | -1.5 | -0.62 | -1.78 | -1.47 | -1.42 | -2.34 | -2.17 | -0.0575 | 0.25 | -0.125 | 0.761758 | 0.890383 | 0.571013 | 0.996493 | 0.997988 | 0.9971 |
| Pclo_Q9QYX7_S3777s_1 | Pclo_Q9QY | -4.01 | -1.14 | -0.29 | -1.62 | -2.53 | -3.65 | -0.41 | -1.46 | -0.77 | -1.65 | 0 | -0.23 | 0.94 | -0.46 | 0.03 | -0.16 | -0.2475 | 1.1025 | 1.8525 | 0.033156 | 0.787813 | 0.183216 | 0.995762 | 0.997988 | 0.994666 |
| Pclo_Q9QYX7_S4016s_1 | Pclo_Q9QY | 0.39 | 0.95 | 0.51 | 0.5 | 0.6 | 0.32 | 0.73 | 0.53 | 0.14 | 0.51 | 1.25 | 0.59 | -0.06 | 0.72 | 0.37 | 0.19 | -0.0425 | 0.035 | -0.2825 | 0.329596 | 0.852216 | 0.907386 | 0.996349 | 0.997988 | 0.994666 |
| Pclo_Q9QYX7_S4066s_1 | Pclo_Q9QY | -1.83 | -1.06 | 0.15 | -0.39 | -0.8 | -1.39 | -0.22 | -0.13 | -0.95 | -0.48 | -0.22 | -0.02 | 0.01 | 0.61 | -0.54 | 0.15 | 0.1475 | 0.365 | 0.84 | 0.099527 | 0.758272 | 0.429892 | 0.995762 | 0.997988 | 0.994666 |
| Pclo_Q9QYX7_S4132s_1 | Pclo_Q9QY | -0.92 | -0.26 | -0.02 | -0.57 | -0.15 | -0.91 | -0.63 | -1.23 | -1.04 | -0.65 | -0.39 | 0.02 | -0.08 | -0.03 | -1.05 | -0.51 | -0.2875 | -0.0725 | 0.025 | 0.941564 | 0.375005 | 0.822739 | 0.989223 | 0.997988 | 0.999507 |
| Pclo_Q9QYX7_S4325s_1 | Pclo_Q9QY | 1.07 | 1.53 | 2.49 | 2.21 | 1.3 | 1.41 | 2.46 | 2.74 | 0.13 | 1.22 | 1.74 | 2.89 | 0.34 | 0.63 | 2.62 | 2.79 | 0.1525 | -0.33 | -0.23 | 0.72428 | 0.737503 | 0.586795 | 0.995762 | 0.997988 | 0.9971 |
| Pclo_Q9QYX7_S4544s_1 | Pclo_Q9QY | -0.51 | -0.99 | -0.92 | -0.56 | -0.83 | 0.15 | -0.73 | -0.26 | -0.08 | -1.09 | -1.86 | -0.71 | -0.34 | -1.28 | -0.69 | -0.74 | 0.3275 | -0.19 | -0.0175 | 0.952638 | 0.272712 | 0.629906 | 0.989223 | 0.997988 | 0.999507 |
| Pclo_Q9QYX7_S4598s_1 | Pclo_Q9QY | -1.05 | 0.36 | -0.85 | -0.34 | -1.07 | -1.27 | -2.98 | -0.5 | 0.19 | -0.61 | -0.21 | -0.54 | 1.02 | 0.49 | -0.47 | -0.91 | -0.985 | 0.1775 | 0.5025 | 0.336721 | 0.1035 | 0.634538 | 0.989223 | 0.997988 | 0.994666 |
| Pclo_Q9QYX7_S4706s_1 | Pclo_Q9QY | -0.27 | 0.45 | 1.04 | 0.83 | 1 | -0.02 | 0.52 | 0.56 | -0.15 | 0.42 | 0.04 | 0.4 | -0.1 | 1.15 | 0.55 | 0.76 | 0.0025 | -0.335 | 0.0775 | 0.845152 | 0.994386 | 0.337601 | 0.999278 | 0.997988 | 0.999507 |
| Pclo_Q9QYX7_S4814s_1 | Pclo_Q9QY | -0.81 | -1.49 | -0.64 | -2.32 | -1.55 | -0.1 | -1.12 | -0.89 | 0.8 | -1.41 | -5.16 | -0.75 | 0.85 | -1.54 | -0.07 | -0.75 | 0.4 | -0.315 | 0.9375 | 0.132553 | 0.389565 | 0.786212 | 0.989223 | 0.997988 | 0.994666 |
| Pclo_Q9QYX7_S5063s_1 | Pclo_Q9QY | 0.27 | 0.36 | 0 | 0.14 | 0.72 | -1.25 | 0.69 | 0.59 | 0.59 | -0.16 | -1.01 | 0.82 | 0.43 | 0.54 | 0.2 | -0.53 | -0.005 | -0.1325 | -0.0325 | 0.916011 | 0.991197 | 0.749439 | 0.999187 | 0.997988 | 0.999507 |
| Pclo_Q9QYX7_S599s_1 | Pclo_Q9QY | 0.61 | -0.37 | -0.07 | 0.03 | 0.86 | 0.26 | 0.35 | 0.43 | -1.17 | 0.81 | 1.75 | 0.16 | 0.07 | 0.49 | -1.51 | 0.28 | 0.425 | 0.3375 | -0.2175 | 0.650719 | 0.153256 | 0.57275 | 0.989223 | 0.997988 | 0.995259 |
| Pclo_Q9QYX7_S625s_1 | Pclo_Q9QY | 1.73 | 1.3 | 1.6 | 1.3 | 1.85 | 1.94 | 0.54 | 1.33 | 1.22 | 1.45 | 0.93 | 1.37 | 1.31 | 1.78 | 1.01 | 1.2 | -0.0675 | -0.24 | -0.1575 | 0.573397 | 0.844306 | 0.333523 | 0.996349 | 0.997988 | 0.995259 |
| Pclo_Q9QYX7_S709s_1 | Pclo_Q9QY | 0 | -0.08 | -0.26 | 0.15 | -0.12 | -0.29 | -0.45 | -0.05 | 0.13 | -0.05 | -0.1 | -0.35 | 0.09 | 0.17 | 0.21 | 0.7 | -0.18 | -0.045 | 0.34 | 0.213285 | 0.414113 | 0.844303 | 0.989223 | 0.997988 | 0.994666 |
| Pclo_Q9QYX7_S709s_3 | Pclo_Q9QY | -0.9 | 0.11 | 0.18 | 0.03 | -0.25 | -0.46 | 0.59 | 0.37 | -1.5 | -0.28 | 0.73 | -0.24 | -0.94 | -0.1 | 0.18 | 0.64 | 0.2075 | -0.1775 | 0.09 | 0.829313 | 0.562729 | 0.719613 | 0.989223 | 0.997988 | 0.999507 |
| Pclo_Q9QYX7_S75s_1 | Pclo_Q9QY | -0.43 | -0.56 | -0.36 | -0.11 | -0.2 | -0.26 | -0.25 | -0.08 | 0.21 | -0.33 | 0.04 | 0.11 | 0.64 | 0.06 | 0.04 | -0.35 | 0.1675 | 0.3725 | 0.4625 | 0.138149 | 0.429708 | 0.142962 | 0.989223 | 0.997988 | 0.994666 |
| Pclo_Q9QYX7_S769s_1 | Pclo_Q9QY | 2.82 | 1.49 | 1.72 | 2.09 | 1.12 | 2.07 | 0.94 | 1.75 | 2.32 | 2.24 | 1.05 | 1.52 | 2.05 | 1.18 | 1.58 | 1.7 | -0.56 | -0.2475 | -0.4025 | 0.285103 | 0.167285 | 0.55338 | 0.989223 | 0.997988 | 0.994666 |
| Pclo_Q9QYX7_S813s_1 | Pclo_Q9QY | -0.93 | -1.23 | -0.23 | -0.06 | -0.44 | -0.91 | 0.02 | 0.14 | -0.63 | -0.87 | 0.48 | 0.19 | -0.84 | -0.06 | 0.08 | -0.12 | 0.315 | 0.405 | 0.3775 | 0.31837 | 0.396785 | 0.345674 | 0.989223 | 0.997988 | 0.994666 |
| Pclo_Q9QYX7_S844s_1 | Pclo_Q9QY | -3.5 | -2.79 | -2.7 | -2.62 | -1.79 | -3.73 | -1.6 | -2.19 | -4.28 | -3.44 | -0.28 | -2.2 | -2.12 | -1.46 | -2.5 | -1.82 | 0.575 | 0.3525 | 0.9275 | 0.019747 | 0.251909 | 0.659747 | 0.989223 | 0.997988 | 0.994666 |
| Pclo_Q9QYX7_T1120t_1 | Pclo_Q9QY | 0.28 | 1.33 | 0.68 | 0.31 | 1.18 | 0.44 | 0.6 | 0.14 | 0.62 | 0.52 | 0.26 | 0.5 | 0.52 | 1.3 | 0.55 | 0.72 | -0.06 | -0.175 | 0.1225 | 0.719147 | 0.85812 | 0.559475 | 0.996349 | 0.997988 | 0.9971 |
| Pclo_Q9QYX7_T2998t_1 | Pclo_Q9QY | -0.95 | -0.69 | 0.79 | 0.81 | -0.16 | -0.84 | 0.8 | 0.55 | -1.29 | -0.57 | 0.64 | 0.75 | -1.05 | -0.53 | 0.14 | 0.36 | 0.0975 | -0.1075 | -0.26 | 0.626921 | 0.857229 | 0.862609 | 0.996349 | 0.997988 | 0.995259 |
| Pclo_Q9QYX7_T3376t_1 | Pclo_Q9QY | -0.06 | 0.51 | 0.64 | 0.29 | 0 | 0.06 | -0.14 | 0.65 | -0.68 | -0.06 | -0.05 | 0.4 | -0.39 | -0.66 | -0.33 | -0.23 | -0.2025 | -0.4425 | -0.7475 | 0.018817 | 0.464162 | 0.171669 | 0.989223 | 0.997988 | 0.994666 |
| Pclo_Q9QYX7_T815t_1 | Pclo_Q9QY | -0.05 | -0.42 | -0.53 | -0.23 | 0.02 | -0.15 | -0.19 | -0.02 | -1.17 | -0.28 | -0.28 | 0.33 | -0.35 | 0.64 | -0.04 | -0.41 | 0.2225 | -0.0425 | 0.2675 | 0.403832 | 0.31196 | 0.901734 | 0.989223 | 0.997988 | 0.994666 |
| Pclo_Q9QYX7_T860t_1 | Pclo_Q9QY | -0.04 | -0.1 | 0.11 | 0.09 | 0.32 | 0.13 | -0.33 | 0.11 | -0.73 | 0.41 | 1.23 | 0.22 | 0.13 | 0.78 | -0.19 | 0 | 0.0425 | 0.2675 | 0.165 | 0.569746 | 0.851017 | 0.511392 | 0.996349 | 0.997988 | 0.995259 |
| Pcm1_Q9R0L6_S1471s_1 | Pcm1_Q9R | 3.71 | 0.58 | 1.85 | -1.1 | 0.44 | 4.37 | 2.55 | 0.74 | -0.26 | 2.6 | 1.71 | 2.51 | 0.93 | -2.67 | 3.73 | 3.7 | 0.765 | 0.38 | 0.1625 | 0.917167 | 0.519204 | 0.722877 | 0.989223 | 0.997988 | 0.999507 |
| Pcm1_Q9R0L6_S65s_1 | Pcm1_Q9R | -0.13 | -0.43 | 0.18 | 0.74 | -0.4 | -0.31 | -0.11 | -0.12 | -0.54 | -0.09 | 0.13 | -0.05 | 0.33 | -0.4 | 0.34 | 0.22 | -0.325 | -0.2275 | 0.0325 | 0.923964 | 0.278749 | 0.480343 | 0.989223 | 0.997988 | 0.999507 |
| Pcm1_Q9R0L6_S996s_1 | Pcm1_Q9R | -0.21 | -0.6 | -1.99 | -2.31 | -0.56 | -0.37 | -2.1 | -2.24 | -0.3 | -0.06 | -0.99 | -2.25 | -0.05 | 0.11 | -1.62 | -1.99 | -0.04 | 0.3775 | 0.39 | 0.56283 | 0.94967 | 0.562682 | 0.999187 | 0.997988 | 0.995259 |
| Pcnp_D3YWA4_S80s_1 | Pcnp_D3YW | 0.32 | 0.49 | -0.54 | -0.7 | -0.11 | -0.65 | -0.44 | 0.74 | 0.37 | -0.48 | 1.93 | -0.29 | -0.12 | 0.72 | -0.32 | 0.62 | -0.0075 | 0.49 | 0.3325 | 0.417655 | 0.985364 | 0.403465 | 0.999187 | 0.997988 | 0.994666 |
| Pcnx_Q9QYC1_S574s_1 | Pcnx_Q9Q | 0.56 | 0.6 | 0.84 | 0.57 | -0.25 | 1.41 | 0.04 | 0.15 | 1.61 | 0.73 | -0.5 | 0.56 | 1.76 | -0.26 | 0.41 | -0.25 | -0.305 | -0.0425 | -0.2275 | 0.625175 | 0.416426 | 0.920665 | 0.989223 | 0.997988 | 0.995259 |
| Pcnxl2_Q5DU28_S660s_1 | Pcnxl2_Q5 | -1.21 | 0.15 | -1.01 | -0.77 | 0.11 | -1.83 | -1.17 | -0.42 | -0.19 | 0.27 | -0.47 | -1.08 | 0.34 | -0.34 | -1.47 | -1.39 | -0.1175 | 0.3425 | -0.005 | 0.99202 | 0.806834 | 0.409956 | 0.995762 | 0.997988 | 1 |
| Pcnxl3_Q8VI59_S1955s_1 | Pcnxl3_Q8 | 0.8 | 0.96 | 0.56 | 0.76 | 0.93 | 0.81 | 0.76 | 0.62 | 0.42 | 0.59 | 0.85 | 0.97 | 0.63 | 0.68 | 0.76 | 0.81 | 0.01 | -0.0625 | -0.05 | 0.831743 | 0.961774 | 0.792436 | 0.999187 | 0.997988 | 0.999507 |
| Pcnxl3_Q8VI59_S287s_1 | Pcnxl3_Q8 | 1.48 | 0.57 | 0.52 | 1.44 | 0.63 | 1.05 | 0.34 | 0.73 | 1.4 | 0.92 | 1.16 | 0.9 | 1 | 0.47 | 0.2 | 0.1 | -0.315 | 0.0925 | -0.56 | 0.142335 | 0.334814 | 0.772176 | 0.989223 | 0.997988 | 0.994666 |
| Pcnxl3_Q8VI59_S297s_1 | Pcnxl3_Q8 | 0.66 | 0.7 | 0.24 | -0.32 | 0.1 | 0.59 | 0.29 | 0.5 | 0.87 | 1.26 | 0.67 | 0.39 | 0.75 | -0.38 | 0.61 | 0.45 | 0.05 | 0.4775 | 0.0375 | 0.918771 | 0.863573 | 0.166862 | 0.996493 | 0.997988 | 0.999507 |
| Pcnxl3_Q8VI59_S96s_1 | Pcnxl3_Q8 | 0.11 | 0.47 | 0.69 | 0.79 | 0.05 | 0.44 | 0.71 | 0.31 | 0.2 | 0.43 | 0.08 | 0.37 | 0.05 | 0.96 | 0.3 | 0.21 | -0.1375 | -0.245 | -0.135 | 0.661436 | 0.593892 | 0.335808 | 0.989223 | 0.997988 | 0.995671 |
| Pcnxl3_Q8VI59_T199t_1 | Pcnxl3_Q8 | 0.52 | 0.95 | 1.24 | 0.83 | 0.74 | 0.67 | 0.85 | 0.13 | -0.49 | 0.29 | 0.65 | 1 | -0.05 | 0.74 | 0.27 | 0.76 | -0.2875 | -0.5225 | -0.455 | 0.158559 | 0.292691 | 0.173432 | 0.989223 | 0.997988 | 0.994666 |
| Pcnxl3_Q8VI59_T370t_1 | Pcnxl3_Q8 | 1.83 | 1.47 | 2.68 | 1.2 | 1.63 | 1.63 | 2.86 | 2.47 | -0.15 | 1.75 | -1.63 | 2.38 | -0.7 | -0.9 | 2.38 | 2.65 | 0.3525 | -1.2075 | -0.9375 | 0.305278 | 0.413549 | 0.184248 | 0.989223 | 0.997988 | 0.994666 |
| Pcnxl3_Q8VI59_T97t_1 | Pcnxl3_Q8 | -0.34 | -0.23 | 0.86 | 0.57 | 0.86 | -0.02 | 0.71 | 0.23 | 0.39 | -0.24 | 0.08 | 0.29 | 0.66 | 0.94 | 0.2 | 0.23 | 0.23 | -0.085 | 0.2925 | 0.432932 | 0.525749 | 0.805299 | 0.989223 | 0.997988 | 0.994666 |
| Pcp4l1_Q6W8Q3_S16s_1 | Pcp4l1_Q6 | 0.28 | 0.93 | 0.31 | 0.53 | -0.36 | -0.64 | -0.01 | -0.11 | 0.69 | 1.04 | -0.09 | -0.73 | 0.39 | -0.05 | 0.21 | 0.66 | -0.7925 | -0.285 | -0.21 | 0.468094 | 0.011446 | 0.499608 | 0.989223 | 0.997988 | 0.994666 |
| Pcsk2_P21661_S526s_2 | Pcsk2_P21 | -0.45 | -0.45 | -1.23 | -1.82 | 0.12 | -0.58 | -0.7 | -1.65 | 0.21 | -0.16 | -0.56 | -1.43 | 0.8 | 0.31 | -1.32 | -2.79 | 0.285 | 0.5025 | 0.2375 | 0.761404 | 0.538795 | 0.29399 | 0.989223 | 0.997988 | 0.9971 |
| Pcsk2_P21661_T525t_2 | Pcsk2_P21 | -0.45 | -0.45 | -1.23 | -1.82 | 0.12 | -0.58 | -0.7 | -1.65 | 0.21 | -0.16 | -0.56 | -1.43 | 0.8 | 0.31 | -1.32 | -2.79 | 0.285 | 0.5025 | 0.2375 | 0.761404 | 0.538795 | 0.29399 | 0.989223 | 0.997988 | 0.9971 |
| Pcyt1a_P49586_S319s_1 | Pcyt1a_P4 | -2.47 | -2.67 | -2.06 | -2.02 | -1.86 | -2.69 | -2.88 | -1.93 | 0.47 | -1.58 | -0.66 | -2.62 | 1.4 | -3.15 | -0.67 | -2.19 | -0.035 | 1.2075 | 1.1525 | 0.211586 | 0.912753 | 0.076668 | 0.997335 | 0.997988 | 0.994666 |
| Pcyt1a_P49586_S331s_1 | Pcyt1a_P4 | -3.81 | -1.97 | -1.31 | -2.1 | -1.96 | -6.43 | -1.51 | -4.41 | -7.39 | -3.59 | 1.15 | -2.31 | -1.41 | -1.19 | -3.16 | -1.5 | -1.28 | -0.7375 | 0.4825 | 0.453356 | 0.257429 | 0.647904 | 0.989223 | 0.997988 | 0.994666 |
| Pcyt1a_P49586_S343s_1 | Pcyt1a_P4 | -2.1 | -1.96 | -2.06 | -2.35 | -1.82 | -2.68 | -2.19 | -2.24 | -2.23 | -1.35 | -0.64 | -2.39 | -0.63 | -1 | -1.94 | -2.01 | -0.115 | 0.465 | 0.7225 | 0.077108 | 0.648294 | 0.27436 | 0.99033 | 0.997988 | 0.994666 |
| Pcyt1a_P49586_S343s_2 | Pcyt1a_P4 | -2.61 | -1.18 | -1.77 | -3.57 | -1.6 | -2.44 | -1.86 | -4.46 | -0.83 | -1.66 | 0.24 | -1.77 | -1.2 | -1.35 | -2.07 | -2.25 | -0.3075 | 1.2775 | 0.565 | 0.310171 | 0.674804 | 0.069659 | 0.992876 | 0.997988 | 0.994666 |
| Pcyt1b_Q811Q9_S289s_1 | Pcyt1b_Q8 | -1.61 | -1.04 | -0.35 | -1.13 | -1.51 | -1.43 | -1.17 | -1.82 | -1.03 | -0.81 | -1.69 | -0.8 | 1.34 | -1.95 | -1.16 | -1.57 | -0.45 | -0.05 | 0.1975 | 0.779156 | 0.171099 | 0.88615 | 0.989223 | 0.997988 | 0.998217 |
| Pcyt1b_Q811Q9_S289s_2 | Pcyt1b_Q8 | -0.58 | -0.45 | -0.14 | -0.22 | -1.65 | 0.16 | -1.22 | -0.32 | 1.26 | -1.02 | -0.69 | 0.06 | 0.4 | -1.58 | -0.68 | -0.55 | -0.41 | 0.25 | -0.255 | 0.545033 | 0.326954 | 0.610848 | 0.989223 | 0.997988 | 0.994666 |
| Pcyt1b_Q811Q9_S293s_1 | Pcyt1b_Q8 | 0.42 | -0.02 | 0.69 | 0.06 | -0.66 | 0 | 0.15 | -0.19 | 1.15 | 0.55 | -0.1 | 0.39 | 1.25 | -1.24 | 0.42 | 0.23 | -0.4625 | 0.21 | -0.1225 | 0.810804 | 0.121491 | 0.529319 | 0.989223 | 0.997988 | 0.999507 |
| Pcyt1b_Q811Q9_S293s_2 | Pcyt1b_Q8 | -0.58 | -0.45 | -0.14 | -0.22 | -1.65 | 0.16 | -1.22 | -0.32 | 1.26 | -1.02 | -0.69 | 0.06 | 0.4 | -1.58 | -0.68 | -0.55 | -0.41 | 0.25 | -0.255 | 0.545033 | 0.326954 | 0.610848 | 0.989223 | 0.997988 | 0.994666 |
| Pcyt1b_Q811Q9_S316s_1 | Pcyt1b_Q8 | -1.95 | -0.94 | -1.07 | -1.68 | -0.82 | -1.01 | -2.07 | -2.15 | 0.09 | -0.19 | -1.63 | -1.61 | 1.21 | -0.92 | -1.07 | -1.42 | -0.1025 | 0.575 | 0.86 | 0.167448 | 0.800622 | 0.257712 | 0.995762 | 0.997988 | 0.994666 |
| Pdap1_Q3UHX2_S176s_1 | Pdap1_Q3U | -1.71 | -2.21 | -0.28 | -0.72 | -1.51 | -2.32 | -0.13 | -0.16 | -2.15 | -1.39 | 0.54 | -0.41 | -1.78 | -0.63 | -0.54 | -0.54 | 0.2 | 0.3775 | 0.3575 | 0.48583 | 0.747617 | 0.573488 | 0.995762 | 0.997988 | 0.994666 |
| Pdap1_Q3UHX2_S19s_1 | Pdap1_Q3U | -0.31 | 0.46 | 0.59 | 0.19 | 0.6 | -0.09 | 0.86 | 0.57 | 0.21 | 0.5 | -0.67 | -0.08 | -0.33 | 0.24 | -0.49 | 1.15 | 0.2525 | -0.2425 | -0.09 | 0.830135 | 0.417345 | 0.481284 | 0.989223 | 0.997988 | 0.999507 |
| Pdap1_Q3UHX2_S48s_1 | Pdap1_Q3U | -0.93 | -0.93 | -1.16 | -2.55 | -1.68 | -0.3 | -6.14 | -3.69 | 1.21 | -1.68 | -1.39 | -1.99 | 0.94 | -0.35 | -2.48 | -1.31 | -1.56 | 0.43 | 0.5925 | 0.427099 | 0.194428 | 0.567801 | 0.989223 | 0.997988 | 0.994666 |
| Pdap1_Q3UHX2_S63s_1 | Pdap1_Q3U | 1.79 | 2.17 | 1.14 | 1.09 | 1.66 | 2 | 1.19 | 1.5 | 2.35 | 2.71 | 0.96 | 1.15 | 0.13 | 1.42 | 1.43 | 1.62 | 0.04 | 0.245 | -0.3975 | 0.363519 | 0.901738 | 0.61264 | 0.996493 | 0.997988 | 0.994666 |

| | | | | | | | | | | | | | | | | | | | | | | | | | | |
|---|---|---|---|---|---|---|---|---|---|---|---|---|---|---|---|---|---|---|---|---|---|---|---|---|---|---|
| Pdap1_Q3UHX2_T18t_1 | Pdap1_Q3U | -0.82 | -1.32 | -2.01 | -0.92 | -0.55 | -0.72 | -0.81 | -2.77 | -1.26 | -1.16 | -0.5 | -2.46 | -1.22 | -0.24 | -3.13 | -2.94 | 0.055 | -0.0775 | -0.615 | 0.370351 | 0.917787 | 0.868165 | 0.997335 | 0.997988 | 0.994666 |
| Pdcd4_Q61823_S71s_1 | Pdcd4_Q61 | -0.27 | 1.2 | -0.76 | -0.99 | -0.52 | -0.16 | -1.39 | -2.98 | -1.04 | -0.35 | -0.03 | 0.08 | 0.08 | -0.5 | -1.5 | -0.12 | -1.0575 | -0.13 | -0.305 | 0.587786 | 0.156457 | 0.8017 | 0.989223 | 0.997988 | 0.995259 |
| Pdcd4_Q61823_S76s_1 | Pdcd4_Q61 | -0.48 | -1.7 | -0.94 | -0.54 | -0.42 | -2.06 | -0.13 | -0.5 | -0.64 | -1.49 | 0.83 | -0.19 | -1.13 | -0.37 | 0.21 | 1.21 | 0.1375 | 0.5425 | 0.895 | 0.11665 | 0.773923 | 0.313506 | 0.995762 | 0.997988 | 0.994666 |
| Pdcd5_D3Z7Q5_S198s_1 | Pdcd5_D3Z | -0.52 | 0.62 | -0.41 | -0.35 | -0.02 | -1.06 | -0.59 | -0.35 | -0.57 | 0.01 | 0.37 | -0.07 | 0.17 | 0.56 | -0.42 | -0.18 | -0.34 | 0.1 | 0.1975 | 0.589084 | 0.339099 | 0.772304 | 0.989223 | 0.997988 | 0.995259 |
| Pdcl_Q9DBX2_S296s_1 | Pdcl_Q9DB | -0.09 | -0.03 | 0.19 | 0.23 | -0.14 | -0.69 | 0.01 | 0.18 | -0.52 | 0.13 | 0.56 | 0.34 | -0.69 | -0.21 | 0.12 | 0.01 | -0.235 | 0.0525 | -0.2675 | 0.346738 | 0.370125 | 0.856777 | 0.989223 | 0.997988 | 0.994666 |
| Pde10a_F8WHK3_T706t_1 | Pde10a_F8 | 2.74 | 0.6 | 0.85 | 2.33 | 0 | 1.2 | 2.18 | 1.86 | 1.61 | 2.13 | 1.27 | 1.24 | 0.36 | 1.8 | 1.76 | 1.54 | -0.32 | -0.0675 | -0.265 | 0.648969 | 0.618033 | 0.898878 | 0.989223 | 0.997988 | 0.995259 |
| Pde11a_P0C1Q2_S163s_1 | Pde11a_P0 | -1.81 | -0.47 | -1.68 | -1.43 | -1.04 | -2.33 | -1.63 | -1.78 | -1.4 | -1.12 | -1.91 | -1.23 | -0.47 | -1.15 | -2 | -1.92 | -0.3475 | -0.0675 | -0.0375 | 0.934436 | 0.383277 | 0.851665 | 0.989223 | 0.997988 | 0.999507 |
| Pde11a_P0C1Q2_T136t_1 | Pde11a_P0 | -1.28 | -0.71 | -1.09 | -1.51 | -0.91 | -1.32 | -1.63 | -1.26 | -2.32 | -1.66 | -0.49 | -1.43 | -0.49 | -1.43 | -1.29 | -0.74 | -0.1325 | -0.3275 | 0.16 | 0.624021 | 0.623252 | 0.432025 | 0.989223 | 0.997988 | 0.995259 |
| Pde1b_Q01065_S7s_1 | Pde1b_Q01 | 0.38 | 0.57 | -0.06 | 0.53 | 0.65 | 0.08 | 0.8 | 0.29 | 0.04 | 0.47 | 0.13 | -0.37 | -0.14 | 0.38 | 0.4 | 0.08 | 0.1 | -0.2875 | -0.175 | 0.530378 | 0.70645 | 0.315527 | 0.995437 | 0.997988 | 0.994666 |
| Pde1c_Q8CDV2_S440s_1 | Pde1c_Q8C | 0.65 | 0.28 | 0.14 | 0.6 | 0.43 | 0.46 | 0.67 | 0.91 | 0.8 | 1.24 | 1.65 | 0.2 | 1.07 | 1.24 | 1.31 | 0.83 | 0.2 | 0.555 | 0.695 | 0.023231 | 0.405427 | 0.138122 | 0.989223 | 0.997988 | 0.994666 |
| Pde1c_Q8CDV2_S441s_1 | Pde1c_Q8C | 0.65 | 0.28 | 0.14 | 0.6 | 0.43 | 0.46 | 0.67 | 0.91 | 0.8 | 1.24 | 1.65 | 0.2 | 1.07 | 1.24 | 1.31 | 0.83 | 0.2 | 0.555 | 0.695 | 0.023231 | 0.405427 | 0.138122 | 0.989223 | 0.997988 | 0.994666 |
| Pde2a_Q5PR72_S116s_1 | Pde2a_Q5P | -0.65 | 0.19 | -1.15 | -0.81 | 0.02 | -0.1 | 0.07 | -0.4 | -1.71 | -0.91 | -1 | -1.22 | 0.07 | 0.34 | -0.6 | 0.45 | 0.5025 | -0.605 | 0.67 | 0.105856 | 0.13975 | 0.111314 | 0.989223 | 0.997988 | 0.994666 |
| Pde2a_Q5PR72_S13s_1 | Pde2a_Q5P | 1.34 | -0.73 | -0.34 | 0.3 | 0.44 | 2.42 | 0.28 | 1.55 | 0.3 | -0.67 | -0.12 | 0.22 | 1.19 | -0.65 | 1.11 | 1.23 | 1.03 | -0.21 | 0.5775 | 0.339347 | 0.113683 | 0.661785 | 0.989223 | 0.997988 | 0.994666 |
| Pde2a_Q5PR72_S204s_1 | Pde2a_Q5P | -2.3 | -0.49 | 1.1 | 1.46 | 0.23 | -0.52 | 1.27 | 1.41 | -0.86 | -0.73 | 0.92 | 1.27 | -0.42 | -0.12 | 1.12 | 0.93 | 0.655 | 0.2075 | 0.435 | 0.601806 | 0.446852 | 0.818729 | 0.989223 | 0.997988 | 0.995259 |
| Pde4a_O89084_S13s_1 | Pde4a_O89 | -3.89 | -1.58 | -2.11 | -5.04 | -0.93 | -3.08 | -2.03 | -2.82 | -4.47 | -2.93 | -0.2 | -3.42 | -2.14 | -0.98 | -1.66 | -3.93 | 0.94 | 0.4 | 0.9775 | 0.288938 | 0.265985 | 0.708544 | 0.989223 | 0.997988 | 0.994666 |
| Pde4a_O89084_S147s_1 | Pde4a_O89 | -0.77 | 0.55 | 0.57 | 0.17 | -1.29 | 0.05 | -0.38 | -0.23 | 0.79 | 0.62 | 0.21 | -0.24 | 1.3 | 0 | 0.14 | 0.14 | -0.5925 | 0.215 | 0.265 | 0.542629 | 0.170119 | 0.584381 | 0.989223 | 0.997988 | 0.994666 |
| Pde4a_O89084_S204s_1 | Pde4a_O89 | 0.15 | -0.41 | -1.19 | 0.4 | -0.31 | -0.06 | 0.61 | 0.64 | -0.89 | 0.1 | 1.66 | -0.53 | -1.03 | -0.02 | 0.38 | 1.03 | 0.4825 | 0.3475 | 0.3525 | 0.504547 | 0.254488 | 0.570989 | 0.989223 | 0.997988 | 0.994666 |
| Pde4a_O89084_S333s_1 | Pde4a_O89 | 0.93 | 0.33 | 1.25 | 1.3 | 0.99 | 0.97 | 1.06 | 2 | -0.18 | 1.26 | 0.49 | 1.57 | -0.06 | 1.08 | 0.97 | 1.4 | 0.3025 | -0.1675 | -0.105 | 0.790714 | 0.384426 | 0.703304 | 0.989223 | 0.997988 | 0.999507 |
| Pde4a_O89084_S49s_1 | Pde4a_O89 | -0.37 | -0.8 | 0.03 | -0.75 | -1.45 | 1.13 | -1.23 | -0.17 | 1.34 | -0.11 | 1.1 | -0.38 | 1.04 | -1.46 | -0.03 | -0.74 | 0.0425 | 0.96 | 0.175 | 0.741426 | 0.939437 | 0.058163 | 0.999187 | 0.997988 | 0.9971 |
| Pde4a_O89084_S786s_1 | Pde4a_O89 | -0.2 | 0.04 | 0.11 | -0.31 | 0.16 | -0.15 | 0.78 | 0.35 | 0.07 | -0.04 | 0.44 | 0.46 | -0.34 | 0.58 | -0.64 | 0.88 | 0.375 | 0.3225 | 0.21 | 0.590809 | 0.178532 | 0.206304 | 0.989223 | 0.997988 | 0.995259 |
| Pde4b_Q8VBU5_S139s_1 | Pde4b_Q8V | 0.32 | 0.5 | 0.15 | -0.08 | 0.92 | 0.66 | 0.37 | -0.07 | -0.12 | 0.12 | 0.03 | 0.4 | -1.87 | 0.45 | -0.45 | 0.21 | 0.2475 | -0.115 | -0.6375 | 0.22665 | 0.388308 | 0.640537 | 0.989223 | 0.997988 | 0.994666 |
| Pde4b_Q8VBU5_S275s_1 | Pde4b_Q8V | 0.51 | 0.97 | 0.37 | 0.34 | 0.39 | 0.05 | 0.33 | 0.58 | -0.05 | 1.22 | 0.17 | 0.53 | 0.07 | -0.36 | 0.47 | 0.39 | -0.21 | -0.08 | -0.405 | 0.198038 | 0.399089 | 0.811673 | 0.989223 | 0.997988 | 0.994666 |
| Pde4b_Q8VBU5_S294s_1 | Pde4b_Q8V | -2.63 | -0.47 | -1.56 | -2.35 | -1.1 | -2.7 | -1.23 | -1.87 | -0.87 | -0.02 | 0.56 | -2.54 | -0.68 | -0.44 | -3.48 | -0.71 | 0.0275 | 1.035 | 0.425 | 0.583363 | 0.959962 | 0.187775 | 0.999187 | 0.997988 | 0.995259 |
| Pde4b_Q8VBU5_S304s_1 | Pde4b_Q8V | -0.09 | 0.8 | 2.37 | 0.69 | 0.59 | 0.33 | 2.86 | 1.18 | 0.12 | -0.04 | -0.11 | 1.45 | 0.61 | 0.93 | 2.23 | 2.9 | 0.2975 | -0.5875 | 0.725 | 0.2952 | 0.662385 | 0.327037 | 0.990434 | 0.997988 | 0.994666 |
| Pde4d_E9PVU7_S105s_1 | Pde4d_E9P | -0.35 | -0.31 | 0.09 | -0.78 | -1.68 | 0.35 | -1.42 | -0.65 | 1.55 | -0.14 | 0.28 | 0.22 | 1.81 | -0.63 | 0.15 | -0.46 | -0.5125 | 0.815 | 0.555 | 0.320352 | 0.277485 | 0.070028 | 0.989223 | 0.997988 | 0.994666 |
| Pde4d_E9PVU7_S137s_1 | Pde4d_E9P | -0.16 | 0.04 | -1.23 | -1.17 | 1.08 | -1.04 | 0.16 | -0.59 | -0.88 | -0.09 | -1.09 | -0.71 | -0.78 | 0.29 | -1.13 | -0.7 | 0.5325 | -0.0625 | 0.05 | 0.909702 | 0.319286 | 0.874181 | 0.989223 | 0.997988 | 0.999507 |
| Pde4d_E9PVU7_S14s_1 | Pde4d_E9P | 0.9 | 1.14 | 0.97 | 0.98 | 1.29 | 0.9 | 0.59 | 0.85 | 0.28 | 0.78 | 0.7 | 0.74 | 0.67 | 1.7 | 0.44 | 0.78 | -0.09 | -0.3725 | -0.1 | 0.758991 | 0.695738 | 0.12723 | 0.994893 | 0.997988 | 0.9971 |
| Pde4d_E9PVU7_S185s_1 | Pde4d_E9P | -0.55 | 0.51 | -1.15 | -0.89 | 0.46 | -0.37 | -1.18 | -1.77 | -0.41 | -0.09 | -0.36 | -0.97 | 0.02 | 0.96 | -1.03 | -1.31 | -0.195 | 0.0625 | 0.18 | 0.757545 | 0.723057 | 0.87726 | 0.995437 | 0.997988 | 0.9971 |
| Pde4d_E9PVU7_S257s_1 | Pde4d_E9P | -0.78 | 0 | -1.25 | -0.95 | 0 | -0.25 | 0.91 | -0.11 | -0.08 | 0.13 | -0.24 | 0.57 | 0.74 | -0.15 | -0.29 | -0.73 | 0.8825 | 0.84 | 0.6375 | 0.1444 | 0.036528 | 0.032244 | 0.989223 | 0.997988 | 0.994666 |
| Pde4d_E9PVU7_S343s_1 | Pde4d_E9P | 5.69 | 2.76 | 1.79 | 0.27 | 2.85 | 5.74 | 2.5 | 1.61 | 0.69 | 5.08 | -0.49 | 1.85 | -0.73 | 0.38 | 2.27 | 2.47 | 0.5475 | -0.845 | -1.53 | 0.218833 | 0.661871 | 0.562658 | 0.990434 | 0.997988 | 0.994666 |
| Pde7b_E9Q0W7_S478s_1 | Pde7b_E9Q | -0.38 | -0.47 | 1.18 | 0.74 | -0.3 | -0.12 | 0.85 | 0.9 | -0.32 | -0.17 | 0.67 | -0.29 | -0.92 | 0.13 | 0.58 | -0.15 | 0.065 | -0.295 | -0.3575 | 0.473046 | 0.891757 | 0.520628 | 0.996493 | 0.997988 | 0.994666 |
| Pde8a_O88502_S452s_1 | Pde8a_O88 | 0.24 | 0.52 | 1.38 | 0.78 | 0.29 | 0.12 | 0.31 | 0.74 | -0.05 | 0.3 | -0.28 | 0.28 | 0.15 | 0.22 | 0.53 | 0.5 | -0.365 | -0.6675 | -0.38 | 0.243731 | 0.243534 | 0.05681 | 0.989223 | 0.997988 | 0.994666 |
| Pde8b_E9PUL4_S304s_1 | Pde8b_E9P | 2.09 | 1.15 | 1.09 | 1.31 | 1.01 | 1.79 | 0.96 | 1.37 | 0.77 | 1.15 | -0.25 | 1.63 | 0.87 | -0.07 | 0.79 | 0.74 | -0.1275 | -0.585 | -0.8275 | 0.037613 | 0.688614 | 0.210389 | 0.994672 | 0.997988 | 0.994666 |
| Pde8b_E9PUL4_S381s_1 | Pde8b_E9P | 0.01 | 0.16 | 0.8 | 0.24 | 0.35 | -0.08 | 0.27 | 0.84 | 0.26 | 0.3 | 0.98 | 1.1 | 0.41 | 0.01 | -0.12 | -0.3 | 0.0425 | 0.3575 | -0.3025 | 0.318429 | 0.882533 | 0.270597 | 0.996493 | 0.997988 | 0.994666 |
| Pde8b_E9PUL4_S385s_1 | Pde8b_E9P | 1.15 | -0.01 | 0.91 | 0.94 | 0.15 | 1.01 | 0.38 | 1.12 | 0.5 | 0.32 | 1.26 | 0.78 | 0.08 | 0.57 | -0.02 | -0.08 | -0.0825 | -0.0325 | -0.61 | 0.094341 | 0.814625 | 0.92522 | 0.995762 | 0.997988 | 0.994666 |
| Pde8b_E9PUL4_S470s_1 | Pde8b_E9P | 0.91 | -0.12 | 0.17 | 0.59 | 0.31 | 0.48 | 0.68 | 0.7 | -0.11 | -0.06 | 0.15 | 0.64 | -0.51 | 0.46 | 0.2 | -0.11 | 0.155 | -0.2325 | -0.3775 | 0.286155 | 0.58366 | 0.468094 | 0.989223 | 0.997988 | 0.994666 |
| Pde9a_F6ZAU1_S177s_1 | Pde9a_F6Z | 1.56 | 1.34 | 3.28 | 2.14 | -0.35 | 2.37 | 0.17 | 2.12 | 2.3 | 1.34 | 1.22 | 2.63 | 1.46 | -0.99 | 3.01 | 2.57 | -1.0025 | -0.2075 | -0.5675 | 0.520516 | 0.182453 | 0.691499 | 0.989223 | 0.997988 | 0.994666 |
| Pdha1_P35486_S232s_1 | Pdha1_P35 | 0.81 | -0.62 | 0.42 | 0.13 | 0.2 | 1.08 | 0.65 | 0.75 | -1.71 | -0.13 | 0.6 | 0.89 | -2.69 | -2.09 | -0.71 | -0.35 | 0.485 | -0.2725 | -1.645 | 0.01743 | 0.191943 | 0.651739 | 0.989223 | 0.997988 | 0.994666 |
| Pdha1_P35486_S293s_1 | Pdha1_P35 | 2.47 | 0.66 | 1.63 | -0.64 | 1.43 | 2.59 | 2.63 | 1.31 | -1.29 | 2.56 | -2 | 2.42 | -0.11 | -2.59 | 2.66 | 2.55 | 0.96 | -0.6075 | -0.4025 | 0.742581 | 0.175459 | 0.617062 | 0.989223 | 0.997988 | 0.9971 |
| Pdha1_P35486_S293s_2 | Pdha1_P35 | 2.48 | 0.09 | 1.23 | -1.02 | 1.59 | 2.73 | 2.65 | 1.58 | -1.51 | 2.13 | -1.86 | 1.65 | -1.49 | -1.6 | 1.54 | 1.56 | 1.4425 | -0.5925 | -0.6925 | 0.500734 | 0.069304 | 0.601797 | 0.989223 | 0.997988 | 0.994666 |
| Pdha1_P35486_S295s_1 | Pdha1_P35 | 0.5 | 0.35 | 1.3 | -0.28 | -0.12 | -0.34 | 1.98 | 1.25 | -1.2 | 0.7 | -0.8 | 1.7 | 3.25 | -2.31 | 1.68 | 1.76 | 0.225 | -0.3675 | 0.6275 | 0.561515 | 0.697784 | 0.58927 | 0.994893 | 0.997988 | 0.995259 |
| Pdha1_P35486_S300s_1 | Pdha1_P35 | 0.75 | 0.96 | 1.55 | 0.49 | 0.45 | 1.11 | 2.01 | 2.17 | 0.32 | 0.98 | -1.2 | 1.92 | 0.99 | -0.73 | 1.85 | 1.4 | 0.4975 | -0.4325 | -0.06 | 0.914723 | 0.269383 | 0.499406 | 0.989223 | 0.997988 | 0.999507 |
| Pdha1_P35486_S300s_2 | Pdha1_P35 | 2.48 | 0.09 | 1.23 | -1.02 | 1.59 | 2.73 | 2.65 | 1.58 | -1.51 | 2.13 | -1.86 | 1.65 | -1.49 | -1.6 | 1.54 | 1.56 | 1.4425 | -0.5925 | -0.6925 | 0.500734 | 0.069304 | 0.601797 | 0.989223 | 0.997988 | 0.994666 |
| Pdha1_P35486_Y301y_1 | Pdha1_P35 | 1.51 | 1.65 | 3.08 | 0.63 | 0.62 | 1.79 | 2.05 | 2.85 | -0.35 | 1.45 | -0.19 | 2.73 | 1 | -2.02 | 3.13 | 2.24 | 0.11 | -0.8075 | -0.63 | 0.559602 | 0.8573 | 0.323481 | 0.996349 | 0.997988 | 0.995259 |
| Pdlim1_S4R1V0_S185s_1 | Pdlim1_S4I | -3.83 | -3.94 | -2.17 | -2.28 | -1.83 | -2.99 | -1.35 | -2.06 | -3.47 | -3.4 | -2.02 | -2.29 | -2.88 | -1.08 | -1.55 | -1.79 | 0.9975 | 0.26 | 1.23 | 0.052363 | 0.088105 | 0.645591 | 0.989223 | 0.997988 | 0.994666 |
| Pdlim4_Q5SWV3_S111s_1 | Pdlim4_Q5 | 2.18 | 1.36 | 1.34 | 0.33 | 0.61 | 1.25 | 0.56 | 0.29 | 1.01 | 0.65 | -1.46 | 1.02 | 1.07 | 0.43 | -0.6 | 1.74 | -0.625 | -0.9975 | -0.6425 | 0.280058 | 0.151979 | 0.145068 | 0.989223 | 0.997988 | 0.994666 |
| Pdlim4_Q5SWV3_S118s_1 | Pdlim4_Q5 | -1.15 | -0.48 | -0.47 | -0.32 | -0.56 | -1.05 | -0.52 | -0.67 | -1.4 | -0.66 | -0.45 | -0.87 | -0.71 | -0.33 | -0.66 | -0.25 | -0.095 | -0.24 | 0.1175 | 0.685288 | 0.721833 | 0.446115 | 0.995437 | 0.997988 | 0.996159 |
| Pdlim5_D9J2Z9_S119s_1 | Pdlim5_D9 | -0.98 | -1.76 | -1.05 | -1.98 | -2.39 | -0.57 | -2.06 | -1.83 | 1.23 | -1.05 | -5.23 | -1.44 | 1.45 | -2.69 | -0.11 | -1.58 | -0.27 | -0.18 | 0.71 | 0.39774 | 0.544785 | 0.880246 | 0.989223 | 0.997988 | 0.994666 |
| Pdlim5_D9J2Z9_S223s_1 | Pdlim5_D9 | -0.01 | 0.18 | 0.49 | 0.71 | 0.24 | 0.21 | 0.27 | 0.7 | 0.18 | 1.18 | 0.73 | -0.01 | 0.07 | 0.01 | 0.73 | 0.15 | 0.0125 | 0.1775 | -0.1025 | 0.729291 | 0.960464 | 0.599758 | 0.999187 | 0.997988 | 0.9971 |
| Pdlim5_D9J2Z9_S250s_1 | Pdlim5_D9 | -0.45 | -0.88 | -1.97 | -0.27 | -1.6 | -0.68 | -1.37 | -0.56 | -0.34 | -0.35 | 0.11 | -0.8 | -0.09 | -0.08 | -1.77 | -1.61 | -0.16 | 0.5475 | 0.005 | 0.992813 | 0.712254 | 0.210236 | 0.995437 | 0.997988 | 1 |
| Pdpk1_F2Z3Z9_S117s_1 | Pdpk1_F2Z | -0.16 | 0.04 | -0.19 | -0.14 | 0.43 | -0.37 | 0.46 | -0.01 | -0.17 | -0.01 | -0.57 | -0.34 | 0.17 | 0.64 | 0.15 | -0.1 | 0.24 | -0.16 | 0.3275 | 0.229525 | 0.360388 | 0.490703 | 0.989223 | 0.997988 | 0.994666 |
| Pds5b_F8WHU5_S1162s_2 | Pds5b_F8W | -1.58 | -0.07 | -1.27 | -0.86 | -2.66 | -1.27 | -1.49 | 0.16 | 0.46 | -1.07 | -0.24 | -0.15 | 0.05 | 0.19 | -0.06 | 0 | -0.37 | 0.695 | 0.99 | 0.019203 | 0.538055 | 0.139107 | 0.989223 | 0.997988 | 0.994666 |
| Pds5b_F8WHU5_S1166s_1 | Pds5b_F8W | 0.17 | 0.62 | -0.51 | -0.37 | 0.22 | 0.34 | -0.71 | -0.56 | 1.02 | 0.24 | -0.37 | -0.13 | 1.2 | -0.1 | 0.41 | -0.14 | -0.155 | 0.2125 | 0.365 | 0.382225 | 0.674666 | 0.59604 | 0.992876 | 0.997988 | 0.994666 |
| Pds5b_F8WHU5_S1166s_2 | Pds5b_F8W | -1.58 | -0.07 | -1.27 | -0.86 | -2.66 | -1.27 | -1.49 | 0.16 | 0.46 | -1.07 | -0.24 | -0.15 | 0.05 | 0.19 | -0.06 | 0 | -0.37 | 0.695 | 0.99 | 0.019203 | 0.538055 | 0.139107 | 0.989223 | 0.997988 | 0.994666 |
| Pds5b_F8WHU5_S1359s_1 | Pds5b_F8W | -2.99 | -3.27 | -1.18 | -0.78 | -3.3 | -2.79 | -0.32 | -0.84 | -2.63 | -3.32 | -0.13 | -1.12 | -2.41 | -2.5 | 0.41 | -0.44 | 0.2425 | 0.255 | 0.82 | 0.343934 | 0.772484 | 0.764917 | 0.995762 | 0.997988 | 0.994666 |
| Pds5b_F8WHU5_S1384s_1 | Pds5b_F8W | 0.07 | -0.55 | -1.06 | -1.59 | -1.01 | -0.39 | -1.24 | -2.31 | 0.24 | -0.6 | -1.18 | -0.69 | 0.51 | -1.74 | -0.12 | -0.69 | -0.455 | 0.225 | 0.2725 | 0.62308 | 0.365664 | 0.615531 | 0.989223 | 0.997988 | 0.995259 |
| Pds5b_F8WHU5_S1386s_1 | Pds5b_F8W | 0.75 | 0.31 | -0.45 | -0.23 | 0.02 | 0.62 | -0.57 | -0.78 | 0.56 | 0.6 | -1.4 | -0.22 | 1.4 | -0.14 | 0.03 | 0.3 | -0.2725 | -0.21 | 0.3025 | 0.490498 | 0.500574 | 0.680317 | 0.989223 | 0.997988 | 0.994666 |
| Pds5b_F8WHU5_T1255t_1 | Pds5b_F8W | -0.53 | -0.06 | -0.89 | -1.21 | -0.83 | -0.76 | -0.51 | -0.95 | -1.04 | -0.01 | -1.29 | -0.63 | -1.2 | -0.4 | -0.77 | -1.05 | -0.09 | -0.07 | -0.1825 | 0.593172 | 0.758736 | 0.853062 | 0.995762 | 0.997988 | 0.995259 |
| Pdxdc1_Q99K01_T687t_1 | Pdxdc1_Q9 | -2.75 | -2.55 | -2.1 | -2.03 | -0.98 | -2.66 | -1.45 | -2 | -2.89 | -1.7 | -1.05 | -1.81 | -2.18 | -1.51 | -2.38 | -1.79 | 0.585 | 0.495 | 0.3925 | 0.229712 | 0.156882 | 0.249442 | 0.989223 | 0.997988 | 0.994666 |
| Pdzd2_E9Q1M1_S1061s_1 | Pdzd2_E9C | 1.17 | 1.19 | 0.3 | 0.93 | 0.7 | 1.27 | 0.83 | 0.21 | 0.9 | 1.03 | 0.51 | 0.12 | 0.64 | 1.44 | -0.73 | 0.25 | -0.145 | -0.2575 | -0.4975 | 0.309843 | 0.649306 | 0.430611 | 0.990343 | 0.997988 | 0.994666 |
| Pdzd2_E9Q1M1_S323s_1 | Pdzd2_E9C | -0.14 | 0.27 | 0 | -0.27 | 0.08 | -0.41 | -0.05 | 0.15 | -0.15 | -0.05 | -0.48 | -0.64 | -0.03 | 0.36 | -0.04 | 0.04 | -0.0225 | -0.295 | 0.1175 | 0.647208 | 0.924522 | 0.261039 | 0.997335 | 0.997988 | 0.995259 |
| Pdzd2_E9Q1M1_S677s_1 | Pdzd2_E9C | -0.25 | 1.42 | 0.72 | 1.51 | 0.89 | 1.12 | 1.08 | 0.4 | 1.11 | 0.7 | 1.42 | 1.36 | 0.32 | 0.38 | 1.08 | 1.31 | 0.0225 | 0.2975 | -0.0775 | 0.866443 | 0.956988 | 0.492422 | 0.999187 | 0.997988 | 0.999507 |
| Pdzd2_E9Q1M1_S947s_1 | Pdzd2_E9C | 0.19 | 0.27 | 0.17 | 0 | 0.41 | 0.21 | 0.66 | 0.16 | 0.13 | 0.43 | 0.11 | 0.21 | -0.25 | 0.39 | 0.5 | 0.49 | 0.2025 | 0.0625 | 0.125 | 0.648858 | 0.363451 | 0.771565 | 0.989223 | 0.997988 | 0.995259 |
| Pdzd4_Q9QY39_S278s_1 | Pdzd4_Q9Q | -0.14 | 0.56 | 1.33 | 1.02 | 0.29 | 1.02 | 1.19 | 0.92 | 0.74 | 0.95 | 2.47 | 1.63 | 0.72 | 0.52 | 1.41 | 0.86 | 0.1625 | 0.755 | 0.185 | 0.632756 | 0.66167 | 0.140362 | 0.990434 | 0.997988 | 0.995259 |
| Pdzd8_B9EJ80_S515s_1 | Pdzd8_B9E | -1.2 | -1.03 | -1.14 | -1.64 | -0.62 | -1.02 | -0.93 | -1.31 | -0.89 | -1.19 | -0.44 | -0.87 | -0.32 | -0.33 | -0.91 | -0.93 | 0.2825 | 0.405 | 0.63 | 0.051466 | 0.276971 | 0.152442 | 0.989223 | 0.997988 | 0.994666 |
| Pdzd8_B9EJ80_S532s_1 | Pdzd8_B9E | -0.79 | -1.96 | -0.81 | -0.42 | -0.62 | -0.71 | -0.03 | -0.42 | -1.64 | -1.35 | -0.32 | -0.99 | -1.76 | -0.74 | -1 | -1.39 | 0.55 | -0.08 | -0.2275 | 0.578212 | 0.154253 | 0.851621 | 0.989223 | 0.997988 | 0.995259 |
| Pdzrn4_E9Q5P6_S520s_1 | Pdzrn4_E9 | -1.02 | 0.07 | -0.15 | -0.28 | -0.62 | -1.04 | -0.24 | -0.56 | -0.09 | -0.01 | -0.13 | -0.31 | 0.98 | 0.05 | 0.21 | -0.21 | -0.27 | 0.21 | 0.6025 | 0.12722 | 0.390976 | 0.475416 | 0.989223 | 0.997988 | 0.994666 |
| Pea15_Q62048_S94s_1 | Pea15_Q62 | -0.21 | 0.19 | 0.25 | 0.37 | -0.33 | -0.21 | 0.05 | -0.21 | -0.24 | 0.02 | -0.56 | -0.11 | -0.38 | 0.26 | 0.16 | -0.03 | -0.325 | -0.3725 | -0.1475 | 0.592497 | 0.175691 | 0.161934 | 0.989223 | 0.997988 | 0.995259 |
| Peak1_Q69Z38_S1363s_1 | Peak1_Q69 | -2.53 | -1.42 | -0.95 | -2.94 | -1.74 | -2.96 | -2.13 | -2.11 | -1.79 | -1.23 | -2.23 | -1.58 | -1.23 | -1.28 | -2.72 | -3.15 | -0.275 | 0.2525 | -0.135 | 0.826643 | 0.577092 | 0.60394 | 0.989223 | 0.997988 | 0.999507 |
| Peak1_Q69Z38_S282s_1 | Peak1_Q69 | 0.19 | 0.18 | 0.35 | 0.45 | 0.62 | 0.27 | 0.29 | 0.59 | 0.29 | 0.55 | 0.33 | 0.08 | 0.99 | 0.59 | 0.22 | -0.17 | 0.15 | 0.02 | 0.115 | 0.712076 | 0.486392 | 0.928577 | 0.989223 | 0.997988 | 0.996966 |
| Peak1_Q69Z38_T1141t_1 | Peak1_Q69 | 0.16 | 0.55 | 0.07 | 0.7 | 0.23 | 0.16 | 0.97 | 0.45 | -0.16 | 0.45 | 0.11 | 0.79 | -0.02 | 0.51 | 0.44 | 0.11 | 0.0825 | -0.0725 | -0.11 | 0.693582 | 0.765447 | 0.807329 | 0.995762 | 0.997988 | 0.996159 |
| Peak1_Q69Z38_T851t_1 | Peak1_Q69 | -0.64 | -0.05 | 0.55 | 0.44 | 0.12 | -1.09 | 0.33 | 0.95 | 0.83 | -0.59 | 0.76 | 0.21 | 1.08 | 1.46 | 0.85 | 0.89 | 0.0025 | 0.2275 | 0.995 | 0.014922 | 0.995744 | 0.589194 | 0.999616 | 0.997988 | 0.994666 |
| Pebp1_P70296_S113s_1 | Pebp1_P70 | 0.3 | 0.5 | 0.31 | -0.13 | 0.23 | 0.1 | 0.34 | 0.09 | 0.31 | 0.03 | -0.06 | -0.02 | -0.17 | -0.5 | 0.06 | 0.41 | -0.055 | -0.18 | -0.295 | 0.333583 | 0.807775 | 0.462334 | 0.995762 | 0.997988 | 0.994666 |
| Pebp1_P70296_S185s_1 | Pebp1_P70 | 0.38 | 0.21 | 0.2 | 0.11 | 0.56 | 0.02 | 0.13 | 0.5 | -0.39 | 0.4 | -0.7 | 0.63 | 0.4 | -1.19 | 0.22 | 0.69 | 0.0775 | -0.24 | -0.195 | 0.644361 | 0.73185 | 0.486985 | 0.995762 | 0.997988 | 0.995259 |
| Pebp1_P70296_S52s_1 | Pebp1_P70 | 0.04 | 0.62 | -0.51 | -0.91 | 0.57 | -0.47 | 0.41 | -1.1 | -0.27 | 0.07 | -0.65 | 0.15 | -0.28 | 0.68 | -0.36 | -0.35 | 0.0425 | 0.015 | 0.1125 | 0.787859 | 0.928637 | 0.968724 | 0.99758 | 0.997988 | 0.999507 |
| Pebp1_P70296_T44t_1 | Pebp1_P70 | -0.08 | 0.4 | 0.54 | -1.97 | 0.51 | -0.67 | -0.49 | 0.4 | -0.45 | 0.32 | -0.34 | -0.45 | 0.02 | 0.63 | -1.62 | 0.76 | 0.215 | 0.0475 | 0.225 | 0.75228 | 0.714437 | 0.9326 | 0.995437 | 0.997988 | 0.9971 |
| Peg10_Q7TN75_S30s_1 | Peg10_Q7T | -0.39 | 0.06 | -0.29 | -0.68 | -0.72 | -1.73 | -1.41 | -0.83 | -0.81 | -0.98 | -0.02 | -0.66 | -0.6 | -0.06 | -0.53 | -0.12 | -0.8475 | -0.2925 | -0.0025 | 0.992926 | 0.019252 | 0.341288 | 0.989223 | 0.997988 | 1 |
| Pelo_Q80X73_S55s_1 | Pelo_Q80X | -0.2 | -1.68 | -0.33 | -1.05 | -0.86 | -0.98 | -0.82 | -1.31 | 0.86 | -1.04 | 0.35 | 0.32 | 0.34 | -1.2 | -0.07 | -1.23 | -0.1775 | 0.9375 | 0.275 | 0.583887 | 0.623671 | 0.087727 | 0.989223 | 0.997988 | 0.995259 |
| Penk_P22005_S252s_1 | Penk_P220 | 0.24 | 0.41 | 0.21 | 0.17 | 0.15 | -0.08 | 0.09 | 0.17 | 0.17 | 0.49 | -0.01 | -0.44 | -0.35 | 0.1 | 0.79 | 0.09 | -0.175 | -0.205 | -0.1 | 0.740743 | 0.392192 | 0.445234 | 0.989223 | 0.997988 | 0.9971 |
| Pex1_Q5BL07_S1142s_1 | Pex1_Q5BL | 0.01 | -1.23 | -0.23 | -0.32 | -1.62 | 0.17 | -0.81 | -1.06 | 0.47 | -0.58 | 0.06 | -1.03 | 0.76 | -0.21 | -0.15 | -0.31 | -0.3875 | 0.1725 | 0.465 | 0.24212 | 0.383381 | 0.683012 | 0.989223 | 0.997988 | 0.994666 |
| Pex1_Q5BL07_S1172s_1 | Pex1_Q5BL | 0.99 | 0.4 | -0.02 | -0.47 | 0.63 | 0.84 | 0.45 | 0.9 | 0.58 | 0.47 | -0.43 | 0.59 | 0.36 | 0.51 | 0.82 | 1.08 | 0.48 | 0.0775 | 0.4675 | 0.225787 | 0.175281 | 0.844559 | 0.989223 | 0.997988 | 0.994666 |
| Pex14_Q9R0A0_S370s_1 | Pex14_Q9R | 0.35 | 0.25 | 0.83 | 0.75 | -0.55 | 1.71 | -0.39 | 0.63 | 2.08 | -0.12 | -0.11 | 0.09 | 1.09 | -0.93 | 0.23 | 0.39 | -0.195 | -0.06 | -0.35 | 0.429814 | 0.696168 | 0.907616 | 0.994893 | 0.997988 | 0.994666 |
| Pex14_Q9R0A0_T48t_1 | Pex14_Q9R | 1.14 | 1.03 | 1.82 | 1.38 | 0.89 | 0.3 | 1.24 | 0.71 | 2.29 | 1.85 | -0.69 | 0.85 | 1.83 | 0.74 | 1.13 | 1.45 | -0.5575 | -0.2675 | -0.055 | 0.868053 | 0.081008 | 0.66938 | 0.989223 | 0.997988 | 0.999507 |
| Pex16_Q91XC9_S181s_1 | Pex16_Q91 | -0.69 | -0.82 | -0.67 | -0.73 | 0.02 | -0.63 | 0.3 | -0.01 | 0.97 | 0.38 | -0.25 | 0.05 | 0.62 | 0.58 | 1.29 | 0.65 | 0.6475 | 1.015 | 1.5125 | 0.00025 | 0.027507 | 0.007595 | 0.989223 | 0.997988 | 0.851508 |
| Pex19_Q8VCI5_S35s_1 | Pex19_Q8V | 0.23 | 0.35 | 0.04 | 0.06 | -0.12 | -0.08 | 0.4 | 0.08 | 0.71 | 1.41 | 0 | -0.26 | 0.29 | 1.24 | 0.2 | 0.28 | -0.1 | 0.295 | 0.3325 | 0.299778 | 0.655283 | 0.451462 | 0.990434 | 0.997988 | 0.994666 |
| Pex5l_F8SLQ1_S240s_1 | Pex5l_F8SL | -0.77 | -0.68 | -0.08 | 0.02 | 0.47 | -0.49 | 0.16 | 0.06 | -0.64 | -0.7 | 0.02 | -0.33 | -0.26 | 0.14 | -0.02 | -0.03 | 0.4275 | -0.035 | 0.335 | 0.263432 | 0.184786 | 0.90726 | 0.989223 | 0.997988 | 0.994666 |
| Pex5l_F8SLQ1_S277s_1 | Pex5l_F8SL | 0.11 | -1.64 | -0.17 | 0.4 | 0.38 | -0.31 | -0.23 | -0.7 | -1.84 | -1.22 | 0.04 | -0.19 | -2.16 | 0.08 | -1.27 | -1.54 | 0.11 | -0.4775 | -0.8975 | 0.157323 | 0.814496 | 0.420703 | 0.995762 | 0.997988 | 0.994666 |
| Pex5l_F8SLQ1_S285s_1 | Pex5l_F8SL | 6.5 | 1.3 | 2.54 | -0.29 | 1.19 | 6.65 | 4.98 | 2.77 | -0.01 | 4.26 | -0.03 | 2.73 | 0.35 | 0.64 | 3.12 | 3.97 | 1.385 | -0.775 | -0.4925 | 0.737374 | 0.399417 | 0.623236 | 0.989223 | 0.997988 | 0.9971 |
| Pex5l_F8SLQ1_S461s_1 | Pex5l_F8SL | 0.99 | 1.02 | 1.17 | 0.75 | 0.41 | 1.3 | -0.04 | 0.19 | 1.76 | 0.91 | -0.64 | 0.51 | 2.13 | 0.21 | 0.7 | 0.75 | -0.5175 | -0.3475 | -0.035 | 0.933543 | 0.131179 | 0.475721 | 0.989223 | 0.997988 | 0.999507 |
| Pex5l_F8SLQ1_S471s_1 | Pex5l_F8SL | -0.3 | 0.26 | 1.23 | 1.2 | 0.32 | -0.32 | 1.61 | 1.38 | -0.13 | 0.22 | 1.09 | 1.38 | 0.02 | 0.51 | 1.47 | 1.58 | 0.15 | 0.0425 | 0.2975 | 0.557447 | 0.778853 | 0.930603 | 0.995762 | 0.997988 | 0.995259 |
| Pfkfb2_B2Z892_S447s_1 | Pfkfb2_B2Z | -0.3 | -1.15 | 0.15 | -0.03 | -0.11 | -0.29 | 0.67 | -0.15 | -0.94 | -1.2 | 0.79 | -0.83 | -0.93 | 0.27 | 0.29 | 0.04 | 0.3625 | -0.2125 | 0.25 | 0.545694 | 0.326001 | 0.674368 | 0.989223 | 0.997988 | 0.994666 |
| Pfkfb2_B2Z892_S464s_1 | Pfkfb2_B2Z | -3.18 | -3.88 | -3.42 | -7.09 | -2.57 | -3.22 | -3.99 | -4.67 | -4.96 | -4.17 | -0.72 | -3.75 | -2.64 | -2.68 | -2.96 | -3.77 | 0.78 | 0.9925 | 1.38 | 0.123414 | 0.388449 | 0.395042 | 0.989223 | 0.997988 | 0.994666 |
| Pfkfb2_B2Z892_T449t_1 | Pfkfb2_B2Z | -0.45 | -0.49 | 0.31 | -0.28 | -0.12 | -0.08 | 0.36 | 0.07 | -0.95 | -0.57 | -0.13 | -0.73 | -1.33 | -0.19 | -0.13 | -0.43 | 0.285 | -0.3675 | -0.2925 | 0.423401 | 0.292327 | 0.233486 | 0.989223 | 0.997988 | 0.994666 |
| Pfkfb4_Q6DTY7_S176s_1 | Pfkfb4_Q6I | 0.34 | 0.07 | -0.2 | -0.04 | 0.94 | 0.27 | 0.36 | -0.11 | 0.2 | 0.82 | 0.66 | -0.35 | -0.59 | 1.2 | -0.77 | -0.26 | 0.3225 | 0.29 | -0.1475 | 0.743382 | 0.267329 | 0.371329 | 0.989223 | 0.997988 | 0.9971 |
| Pfkl_P12382_S775s_1 | Pfkl_P1238 | 0.51 | 1.01 | 2.19 | 2.5 | 1.26 | 0.64 | 1.81 | 2.61 | 0.64 | 0.86 | 1.44 | 1.98 | 0.95 | 1.58 | 1.58 | 1.83 | 0.0275 | -0.3225 | -0.0675 | 0.889256 | 0.96136 | 0.542374 | 0.999187 | 0.997988 | 0.999507 |
| Pfkl_P12382_T304t_1 | Pfkl_P1238 | -0.38 | -0.07 | 0.4 | 0.42 | -0.11 | -0.64 | 0.12 | 0.72 | -0.17 | -0.38 | 0.98 | 0.39 | 0.18 | 0.22 | 0.01 | 0.12 | -0.07 | 0.1125 | 0.04 | 0.885859 | 0.839615 | 0.761255 | 0.995762 | 0.997988 | 0.999507 |
| Pfkm_P47857_S377s_1 | Pfkm_P478 | -1.03 | 0.53 | -0.57 | -0.2 | 0.4 | -0.96 | 0.05 | 0.2 | -1.45 | -0.66 | -1.47 | -0.01 | -1.86 | -1.2 | -0.06 | -0.56 | 0.24 | -0.58 | -0.6025 | 0.234844 | 0.574465 | 0.228448 | 0.989223 | 0.997988 | 0.994666 |
| Pfkm_P47857_S667s_1 | Pfkm_P478 | 0.67 | 1.03 | 0.28 | 0.44 | 1.32 | 1.02 | 1.03 | -0.13 | 0.56 | 1.14 | 0.08 | 0.17 | 0.4 | 1.04 | 0.85 | 1.12 | 0.205 | -0.1175 | 0.2475 | 0.409809 | 0.570851 | 0.714314 | 0.989223 | 0.997988 | 0.994666 |
| Pfkp_Q8C605_S2s_1 | Pfkp_Q8C6 | 0.32 | 0.47 | 0.47 | 0.43 | 0.81 | 0.36 | -0.09 | 0.45 | 0.13 | 0.52 | 0.07 | 0.22 | 0.5 | 0.67 | 0.4 | 0.4 | -0.04 | -0.1875 | 0.07 | 0.761808 | 0.871533 | 0.402494 | 0.996493 | 0.997988 | 0.9971 |
| Pgam1_Q9DBJ1_S14s_1 | Pgam1_Q9 | 2.73 | -0.27 | 1.52 | 0.39 | -0.9 | 2.14 | 3.04 | 1.92 | -1.26 | 1.27 | 0.84 | 1.28 | -2.39 | -1.44 | 1.77 | 2.65 | 0.4575 | -0.56 | -0.945 | 0.437872 | 0.62628 | 0.489684 | 0.989223 | 0.997988 | 0.994666 |
| Pgam1_Q9DBJ1_Y92y_1 | Pgam1_Q9 | 5.12 | 2.5 | 4.53 | 1.72 | 2.68 | 4.52 | 4.2 | 3 | -0.79 | 4.34 | 0.62 | 3.62 | -1.07 | 0.91 | 3.76 | 5.01 | 0.1325 | -1.52 | -1.315 | 0.349968 | 0.869635 | 0.253655 | 0.996493 | 0.997988 | 0.994666 |
| Pgbd5_D3YZI9_S519s_1 | Pgbd5_D3Y | 0.71 | 0.76 | -0.33 | -0.42 | 0.17 | 0.45 | 0.23 | -0.35 | 0.46 | 0.37 | -0.91 | 0.08 | -0.04 | 0.63 | -0.64 | -0.32 | -0.055 | -0.18 | -0.2725 | 0.519857 | 0.878842 | 0.680737 | 0.996493 | 0.997988 | 0.994666 |
| Pgbd5_D3YZI9_S520s_1 | Pgbd5_D3Y | 0.75 | 0.76 | 0.04 | 0.11 | 0.82 | 0.55 | 0.62 | 0.07 | 0.4 | 0.73 | -0.37 | 0.14 | -0.07 | 0.91 | -0.07 | 0.22 | 0.1 | -0.19 | -0.1675 | 0.623725 | 0.726931 | 0.567232 | 0.995569 | 0.997988 | 0.995259 |
| Pgk1_P09411_S203s_1 | Pgk1_P094 | -0.3 | 0.05 | 0.11 | 0.11 | 0.12 | -0.53 | 0.2 | -0.02 | 0.01 | 0.07 | -0.31 | 0.04 | -0.43 | 0.25 | -0.44 | -0.18 | -0.05 | -0.04 | -0.1925 | 0.48769 | 0.840848 | 0.861783 | 0.995818 | 0.997988 | 0.994666 |
| Pgls_F6X8L5_S43s_1 | Pgls_F6X8L | -0.98 | -0.27 | -0.46 | -0.07 | 0.2 | -0.66 | 0.51 | -0.13 | -0.72 | -0.53 | -0.36 | -0.5 | -0.4 | -0.27 | -0.42 | 0.07 | 0.425 | -0.0825 | 0.19 | 0.521118 | 0.216004 | 0.759082 | 0.989223 | 0.997988 | 0.994666 |
| Pgm2_Q7TSV4_S173s_1 | Pgm2_Q7T | -0.38 | -1.27 | -0.43 | -1.02 | -1.01 | 0.11 | 0.03 | -1.4 | -2.74 | -0.38 | -2.3 | -0.53 | -0.84 | -3.36 | -1.01 | -0.59 | 0.2075 | -0.7125 | -0.675 | 0.289761 | 0.619614 | 0.246174 | 0.989223 | 0.997988 | 0.994666 |
| Pgm2_Q9D0F9_S117s_1 | Pgm2_Q9D | 1.42 | 1.18 | 0.56 | 0.59 | 1.28 | 1.36 | 1.03 | 0.63 | 0.57 | 0.45 | -0.38 | 1.12 | -0.56 | 0.48 | 0.59 | 1 | 0.1375 | -0.4975 | -0.56 | 0.185201 | 0.644932 | 0.213014 | 0.99033 | 0.997988 | 0.994666 |
| Pgm2_Q9D0F9_T507t_1 | Pgm2_Q9D | 0.75 | 0.72 | 0.89 | 1.05 | 0.45 | 0.21 | -0.1 | 0.2 | 0.69 | 0.24 | 0.86 | -0.02 | 0.88 | 0.71 | 1.05 | 0.6 | -0.6625 | -0.41 | -0.0425 | 0.862656 | 0.013277 | 0.156999 | 0.989223 | 0.997988 | 0.999507 |
| Pgm2l1_Q8CAA7_S175s_1 | Pgm2l1_Q8 | 0.31 | -0.17 | -0.24 | -0.08 | -0.04 | 0.09 | 0.07 | 0.36 | 0.38 | 0.51 | -0.62 | 0.35 | 0.06 | -0.26 | -0.48 | -0.36 | 0.165 | 0.2 | -0.215 | 0.423726 | 0.474582 | 0.533987 | 0.989223 | 0.997988 | 0.994666 |
| Pgm3_Q8BWW3_S64s_1 | Pgm3_Q8B | 0.45 | 0.64 | 0.46 | 0.81 | 0.55 | 0.39 | 0.71 | 0.64 | 0.1 | 0.09 | -0.02 | 0.49 | -0.19 | 0.68 | 0.14 | 0.54 | -0.0175 | -0.425 | -0.2975 | 0.313997 | 0.933797 | 0.09475 | 0.998562 | 0.997988 | 0.994666 |
| Pgrmc1_O55022_S181s_1 | Pgrmc1_O5 | -0.01 | 0.18 | -0.12 | 0.11 | 0.15 | -0.24 | 0.06 | -0.04 | 0.19 | 0.31 | 0.36 | -0.12 | -0.76 | 0.43 | 0.26 | 0.14 | -0.0575 | 0.145 | -0.0225 | 0.944051 | 0.78419 | 0.52815 | 0.995762 | 0.997988 | 0.999507 |
| Pgrmc2_Q80UU9_S84s_1 | Pgrmc2_Q8 | 0.03 | 0.03 | -0.16 | 0.29 | 0.02 | 0.16 | 0.62 | -0.1 | -0.06 | 0.1 | 0.51 | 0.06 | -0.23 | -0.17 | -0.36 | 0.43 | 0.1275 | 0.105 | -0.13 | 0.642118 | 0.604234 | 0.663015 | 0.989223 | 0.997988 | 0.995259 |
| Phactr1_Q2M3X8_S230s_1 | Phactr1_Q2 | -2.2 | -1.53 | -1.39 | -1.9 | -1.16 | -2.5 | -0.98 | -1.63 | -2.62 | -1.99 | -0.97 | -1.43 | -1.3 | -1.05 | -1.84 | -1.49 | 0.1875 | 0.0025 | 0.335 | 0.287929 | 0.62175 | 0.994993 | 0.989223 | 0.998021 | 0.994666 |
| Phactr1_Q2M3X8_S498s_1 | Phactr1_Q2 | -2.51 | -0.68 | -1.17 | -1.83 | -1.47 | -3.49 | -1.14 | -1.39 | -2.7 | -2 | 0.77 | -1.4 | 0.25 | -0.51 | -1.64 | -1.7 | -0.325 | 0.215 | 0.6475 | 0.271149 | 0.592249 | 0.777645 | 0.989223 | 0.997988 | 0.994666 |
| Phactr1_Q2M3X8_S60s_1 | Phactr1_Q2 | -3.6 | -3.98 | -2.64 | -3.72 | -3.06 | -3.72 | -1.55 | -2.24 | -3.77 | -2.71 | 1.24 | -2.59 | -1.86 | -1.14 | -2.58 | -2.34 | 0.8425 | 1.5275 | 1.505 | 0.00568 | 0.122946 | 0.152849 | 0.989223 | 0.997988 | 0.994666 |
| Phactr1_Q2M3X8_T97t_1 | Phactr1_Q2 | 0.19 | -0.73 | -0.02 | -0.51 | 0.64 | 0.32 | 0.52 | 0.06 | -0.41 | -0.33 | 0.57 | 0.17 | -0.18 | -0.12 | -0.42 | 0.08 | 0.6525 | 0.2675 | 0.1075 | 0.719714 | 0.041551 | 0.432654 | 0.989223 | 0.997988 | 0.9971 |
| Phactr2_B1AVN9_S204s_1 | Phactr2_B1 | -0.61 | -1.93 | -2.02 | -0.6 | -0.03 | 0.66 | -0.51 | 0.61 | -0.83 | -0.28 | 1.2 | -0.58 | -2.59 | 0.43 | -0.48 | 0.06 | 1.4725 | 1.1675 | 0.645 | 0.367809 | 0.008727 | 0.060997 | 0.989223 | 0.997988 | 0.994666 |
| Phactr2_B1AVN9_S444s_1 | Phactr2_B1 | -0.8 | -1.3 | -1.37 | -0.68 | 0.02 | -1.49 | -1.05 | -1.26 | 0.74 | -0.08 | 0.8 | -0.5 | 0.06 | 0.26 | -1.12 | -1.6 | 0.0925 | 1.2775 | 0.4375 | 0.359478 | 0.803175 | 0.006688 | 0.995762 | 0.997988 | 0.994666 |
| Phactr2_B1AVN9_T25t_1 | Phactr2_B1 | 0.31 | -0.51 | -0.01 | -0.65 | -0.42 | -0.32 | -0.01 | 0.38 | -0.18 | -0.23 | 1.06 | 0.98 | -0.77 | 0.66 | -0.24 | -0.29 | 0.1225 | 0.6225 | 0.055 | 0.88636 | 0.691072 | 0.155684 | 0.994893 | 0.997988 | 0.999507 |
| Phactr3_Q8BYK5_S189s_1 | Phactr3_Q8 | -0.11 | 0.16 | 0.69 | -1.13 | 0.49 | -0.18 | 0.28 | 0.46 | 0.2 | 0.48 | 0.46 | -0.45 | -0.12 | 0.36 | -0.12 | 0.95 | 0.36 | 0.27 | 0.365 | 0.423067 | 0.375539 | 0.532906 | 0.989223 | 0.997988 | 0.994666 |
| Phactr3_Q8BYK5_T18t_1 | Phactr3_Q8 | -1.07 | -1.03 | -1.29 | -1.75 | -0.8 | -1.3 | -0.74 | -1.33 | -2 | -0.71 | -1.1 | -1.75 | -0.47 | 0.08 | -1.95 | -1.01 | 0.2425 | -0.105 | 0.4475 | 0.333478 | 0.380361 | 0.766476 | 0.989223 | 0.997988 | 0.994666 |
| Phactr3_Q8BYK5_T195t_1 | Phactr3_Q8 | -1.03 | 0.3 | -0.64 | -0.72 | 0.79 | 0.25 | -0.82 | 0.05 | 1.03 | -0.76 | 0.22 | 1.01 | -0.23 | -0.18 | -0.42 | -0.04 | 0.59 | 0.8975 | 0.305 | 0.373857 | 0.18133 | 0.090054 | 0.989223 | 0.997988 | 0.994666 |
| Phactr4_Q501J7_S147s_1 | Phactr4_Q5 | -0.3 | -0.51 | -1.5 | -0.68 | -0.51 | -0.2 | -0.74 | -0.78 | -0.33 | -0.59 | -0.72 | -1.23 | 0.53 | -0.32 | 0.16 | -0.36 | 0.19 | 0.03 | 0.75 | 0.061707 | 0.547295 | 0.930122 | 0.989223 | 0.997988 | 0.994666 |
| Phactr4_Q501J7_S430s_1 | Phactr4_Q5 | -1.01 | -0.94 | -1.55 | -1.83 | -0.05 | -1.21 | -0.63 | -1.5 | -1.5 | -0.86 | -0.6 | -2.09 | -1.34 | -0.03 | -1.17 | -0.54 | 0.485 | 0.07 | 0.5625 | 0.16543 | 0.219827 | 0.859735 | 0.989223 | 0.997988 | 0.994666 |
| Phb2_O35129_S267s_1 | Phb2_O351 | 0.07 | 0.55 | -0.22 | 0.23 | -0.01 | 0.84 | -0.18 | -0.22 | 1.83 | 0.5 | -1.14 | 0.13 | 2.14 | -0.7 | 0.44 | 0.22 | -0.05 | 0.1725 | 0.3675 | 0.520574 | 0.87327 | 0.766936 | 0.996493 | 0.997988 | 0.994666 |
| Phc3_F8WIN6_S281s_1 | Phc3_F8WI | -0.68 | -1.03 | -0.28 | -0.08 | -1.95 | -0.64 | -1.23 | -0.72 | -0.1 | -0.71 | -1.14 | -0.45 | 0.96 | -1.69 | -0.31 | -0.43 | -0.6175 | -0.0825 | 0.15 | 0.781626 | 0.115903 | 0.802316 | 0.989223 | 0.997988 | 0.998226 |
| Phf10_K4DI61_S326s_1 | Phf10_K4D | -0.12 | -0.83 | -0.19 | -0.43 | -0.17 | 1 | -0.15 | -0.31 | 0.82 | 0.05 | 0.52 | -0.49 | 0.05 | -0.51 | 0.86 | -0.96 | 0.485 | 0.6175 | 0.2525 | 0.552902 | 0.184964 | 0.100157 | 0.989223 | 0.997988 | 0.994666 |
| Phf2_Q9WTU0_S1057s_1 | Phf2_Q9W | -0.02 | -0.28 | 0.92 | 0.27 | 0.02 | 0.93 | -0.24 | -0.45 | 1.72 | -0.57 | -2.05 | 0.32 | 2.33 | -0.32 | 0.03 | 0.06 | -0.1575 | -0.3675 | 0.3025 | 0.617791 | 0.685323 | 0.624618 | 0.994624 | 0.997988 | 0.995259 |
| Phf2_Q9WTU0_S1060s_1 | Phf2_Q9W | 1.05 | -0.13 | 0.4 | 0.53 | -1.02 | 1.24 | -0.6 | -0.9 | 2.66 | 0.41 | -0.81 | -0.06 | 2.75 | -1.02 | 0.42 | 0.59 | -0.7825 | 0.0875 | 0.2225 | 0.7595 | 0.161638 | 0.901502 | 0.989223 | 0.997988 | 0.9971 |
| Phf20_Q8BLG0_S894s_1 | Phf20_Q8B | -0.33 | -0.29 | -0.11 | -0.67 | -0.69 | -0.97 | -0.22 | 0.3 | -1.24 | -0.89 | -1.62 | -0.54 | -0.39 | -0.89 | 0.18 | -0.1 | -0.045 | -0.7225 | 0.05 | 0.872028 | 0.887372 | 0.035719 | 0.996493 | 0.997988 | 0.999507 |
| Phf3_B2RQG2_S377s_1 | Phf3_B2RC | -0.73 | -0.52 | -1.1 | -1.34 | -0.19 | -1.08 | 0.01 | -0.54 | -1.78 | -0.27 | -1.14 | -0.18 | -0.81 | -0.93 | -0.57 | 0.13 | 0.4725 | 0.08 | 0.3775 | 0.278884 | 0.160128 | 0.847152 | 0.989223 | 0.997988 | 0.994666 |
| Phf3_B2RQG2_S660s_1 | Phf3_B2RC | -1.33 | -0.94 | -0.49 | -1.16 | -0.3 | -1.49 | -0.12 | -1.67 | -1.19 | -1.18 | -1.63 | -1.93 | 0.03 | -0.03 | -0.07 | 0.01 | 0.085 | -0.5025 | 0.965 | 0.005254 | 0.838474 | 0.117876 | 0.995762 | 0.997988 | 0.994666 |
| Phf6_E9QAG2_S203s_1 | Phf6_E9QA | -5.92 | -7.48 | -1.36 | -2.41 | -4.97 | -6.16 | -1.45 | -1.15 | -3.78 | -4.81 | -3.17 | -0.86 | -1.04 | -5.89 | 0.45 | -0.68 | 0.86 | 1.1375 | 2.5025 | 0.169513 | 0.601519 | 0.442537 | 0.989223 | 0.997988 | 0.994666 |
| Phip_F8VQ93_S1783s_1 | Phip_F8VQ | 0.53 | 0.46 | 0.05 | 0.46 | 1.24 | 0.49 | -0.04 | 1.1 | 0.87 | 0.54 | 1.12 | 0.98 | -0.69 | 1.28 | 1.68 | 0.78 | 0.3225 | 0.5025 | 0.3875 | 0.446257 | 0.336057 | 0.065034 | 0.989223 | 0.997988 | 0.994666 |
| Phkb_Q7TSH2_S19s_1 | Phkb_Q7TS | -0.3 | -0.42 | 0.05 | -0.4 | 0.09 | -0.48 | -0.26 | -0.05 | -0.76 | -0.26 | -0.39 | -0.6 | -0.49 | -0.71 | -0.79 | -0.06 | 0.0925 | -0.235 | -0.245 | 0.387215 | 0.69538 | 0.340912 | 0.994893 | 0.997988 | 0.994666 |
| Phldb1_D3Z4N0_S382s_1 | Phldb1_D3 | 0.9 | -0.44 | 1.32 | -0.4 | 0.11 | 0.28 | 0.39 | -0.07 | 0.19 | 0.48 | 1.4 | 0.35 | 2.05 | 0.77 | 0.64 | 0.47 | -0.1675 | 0.26 | 0.6375 | 0.253231 | 0.700651 | 0.602503 | 0.995094 | 0.997988 | 0.994666 |
| Phldb1_D3Z4N0_S445s_1 | Phldb1_D3 | -0.3 | 0.17 | -0.08 | -0.04 | -0.41 | -0.68 | -0.07 | -0.55 | 0.03 | -0.35 | -0.28 | -0.11 | 0.51 | 0.05 | 0 | -0.38 | -0.365 | -0.115 | 0.1075 | 0.704295 | 0.143248 | 0.616777 | 0.989223 | 0.997988 | 0.996159 |
| Phldb1_D3Z4N0_S463s_1 | Phldb1_D3 | 0.21 | 0.72 | 0.08 | -0.06 | 0.63 | 0.34 | 0.84 | -0.04 | 0.05 | 0.44 | 0.58 | 0.16 | -0.32 | 0.62 | 0.09 | 0.26 | 0.205 | 0.07 | -0.075 | 0.810001 | 0.481355 | 0.794805 | 0.989223 | 0.997988 | 0.999507 |
| Phldb1_D3Z4N0_S472s_1 | Phldb1_D3 | 0.03 | -0.64 | 0.25 | 0.09 | -0.25 | -0.3 | -0.35 | -0.43 | -0.77 | 0.01 | 0.23 | -0.26 | -0.21 | -0.72 | -1.33 | -1.51 | -0.265 | -0.13 | -0.875 | 0.039625 | 0.310831 | 0.686161 | 0.989223 | 0.997988 | 0.994666 |
| Phldb1_D3Z4N0_S491s_1 | Phldb1_D3 | 0.38 | -0.4 | -0.12 | 0.04 | -0.65 | -0.49 | -1.67 | -1.62 | 0.46 | -0.88 | 0 | -0.75 | 1.17 | -0.89 | -0.9 | -0.68 | -1.0825 | -0.2675 | -0.3 | 0.551064 | 0.011885 | 0.471258 | 0.989223 | 0.997988 | 0.994666 |
| Phldb1_D3Z4N0_S51s_1 | Phldb1_D3 | -2.19 | -1.51 | -1.85 | -0.2 | 0.11 | -1.4 | -0.86 | -0.99 | -1.55 | -1.98 | -1.12 | -0.9 | -1.56 | -1.48 | -0.97 | -0.69 | 0.6525 | 0.05 | 0.2625 | 0.575522 | 0.208838 | 0.915701 | 0.989223 | 0.997988 | 0.995259 |
| Phldb1_D3Z4N0_S541s_1 | Phldb1_D3 | 1.3 | 0.97 | 0.29 | 0.39 | 0.86 | 0.19 | 0.54 | 0.51 | 0.31 | 0.83 | 0.27 | 0.39 | -0.34 | 1 | 0.74 | 0.12 | -0.2125 | -0.2875 | -0.3575 | 0.376037 | 0.48551 | 0.362076 | 0.989223 | 0.997988 | 0.994666 |
| Phldb1_D3Z4N0_S950s_1 | Phldb1_D3 | 1.06 | 0.92 | 1.05 | 0.79 | 0.71 | 0.76 | 0.92 | 0.8 | 0.92 | 1.13 | 1.21 | 0.76 | 0.68 | 0.96 | 1.01 | 1.02 | -0.1575 | 0.05 | -0.0375 | 0.874984 | 0.43956 | 0.824063 | 0.989223 | 0.997988 | 0.999507 |
| Phldb2_D3Z069_S456s_1 | Phldb2_D3 | -2.13 | -1.45 | -2.06 | -3.12 | -1.41 | -2.3 | -2.41 | -2.45 | -2.46 | -1.61 | -1.18 | -1.86 | -1.05 | -1.79 | -2.34 | -1.67 | 0.0475 | 0.4125 | 0.4775 | 0.282875 | 0.906869 | 0.346001 | 0.99702 | 0.997988 | 0.994666 |
| Phlpp1_Q8CHE4_S304s_1 | Phlpp1_Q8 | 0.69 | 0.51 | -0.47 | 1.34 | 0.72 | 0.83 | 0.98 | 0.96 | 0.28 | 0.7 | 0.5 | -0.45 | -0.82 | 1.15 | 0.81 | 1.26 | 0.355 | -0.26 | 0.0825 | 0.883568 | 0.352843 | 0.555568 | 0.989223 | 0.997988 | 0.999507 |
| Phlpp2_J3QM82_S1214s_1 | Phlpp2_J3C | -0.49 | -0.5 | -0.58 | -0.13 | -1.04 | -0.5 | -0.45 | -0.6 | -0.28 | -0.34 | -0.33 | -0.45 | 0.14 | 0.12 | -0.33 | -0.62 | -0.2225 | 0.075 | 0.2525 | 0.384687 | 0.35849 | 0.733407 | 0.989223 | 0.997988 | 0.994666 |
| Phlpp2_J3QM82_S1242s_1 | Phlpp2_J3C | -1.12 | -0.25 | -0.38 | -0.38 | -1.32 | -0.32 | 0.25 | 0.24 | -0.4 | -0.27 | 0.34 | -0.39 | 0.32 | -0.86 | -0.39 | -0.19 | 0.245 | 0.3525 | 0.2525 | 0.472462 | 0.546382 | 0.26237 | 0.989223 | 0.997988 | 0.994666 |
| Phospho2_Q9D9M5_T203t_1 | Phospho2_ | -1.15 | -2.33 | 0.01 | -0.02 | -0.66 | -1.63 | -0.13 | -0.39 | 1.3 | -1.8 | -0.64 | -0.95 | 0.74 | -1.54 | 0.4 | 0.44 | 0.17 | 0.35 | 0.8825 | 0.217275 | 0.769454 | 0.650762 | 0.995762 | 0.997988 | 0.994666 |
| Phrf1_A6H619_S622s_1 | Phrf1_A6H | 1.15 | -0.04 | 0.47 | -0.12 | -0.3 | 0.23 | -0.49 | 0.44 | 0.1 | -0.76 | 1.52 | -0.95 | 1.38 | -0.6 | 1.4 | 0.7 | -0.395 | -0.3875 | 0.355 | 0.498922 | 0.291333 | 0.511445 | 0.989223 | 0.997988 | 0.994666 |
| Phrf1_F7ANA1_S247s_1 | Phrf1_F7AI | 0.06 | -0.44 | -0.23 | -1.09 | -0.38 | -0.51 | -0.02 | 0.04 | 0.08 | -1.2 | -0.39 | -0.55 | 0.93 | 0.81 | 0.52 | -0.03 | 0.2075 | -0.09 | 0.9825 | 0.018812 | 0.497828 | 0.807185 | 0.989223 | 0.997988 | 0.994666 |
| Phrf1_F7ANA1_S277s_1 | Phrf1_F7AI | 1.34 | 0.18 | 0.44 | 0.6 | 0.13 | 0.33 | -0.59 | 0.85 | 0.24 | -0.2 | 0.02 | -0.45 | 1.4 | 0.59 | 0.96 | -0.01 | -0.46 | -0.7375 | 0.095 | 0.810774 | 0.244039 | 0.042263 | 0.989223 | 0.997988 | 0.999507 |
| Phyhipl_Q8BGT8_T116t_1 | Phyhipl_Q8 | -0.38 | 0.24 | 0.23 | -0.53 | 0.05 | -4.05 | -1.19 | -0.82 | 0.22 | 0.45 | -2.35 | -0.49 | 0.79 | 0.53 | -0.34 | -0.07 | -1.3925 | -0.4325 | 0.3375 | 0.355414 | 0.105774 | 0.483841 | 0.989223 | 0.997988 | 0.994666 |
| Pi4k2a_Q2TBE6_S47s_1 | Pi4k2a_Q2 | -0.06 | 0.47 | -0.06 | 0.09 | 0.61 | -1.11 | 0.17 | -0.08 | 0.31 | 0 | 0.46 | 0.1 | 1.47 | 1.09 | 0.25 | 0.23 | -0.2125 | 0.1075 | 0.65 | 0.095899 | 0.576648 | 0.660517 | 0.989223 | 0.997988 | 0.994666 |
| Pi4k2a_Q2TBE6_S47s_2 | Pi4k2a_Q2 | 0.63 | 0.36 | -1.87 | -1.95 | -0.17 | 0.82 | -0.38 | -2.59 | 1.14 | 1.05 | -1.83 | -1.81 | 1.82 | 0.11 | -0.53 | -1.61 | 0.1275 | 0.345 | 0.655 | 0.462186 | 0.883559 | 0.721684 | 0.996493 | 0.997988 | 0.994666 |
| Pi4k2a_Q2TBE6_S51s_2 | Pi4k2a_Q2 | 0.63 | 0.36 | -1.87 | -1.95 | -0.17 | 0.82 | -0.38 | -2.59 | 1.14 | 1.05 | -1.83 | -1.81 | 1.82 | 0.11 | -0.53 | -1.61 | 0.1275 | 0.345 | 0.655 | 0.462186 | 0.883559 | 0.721684 | 0.996493 | 0.997988 | 0.994666 |
| Pi4k2a_Q2TBE6_S9s_1 | Pi4k2a_Q2 | -0.01 | 0.37 | 0.46 | 1.21 | 0.04 | 1.09 | 0.6 | 0.42 | 0.44 | 0.56 | 1.24 | 0.57 | 0.02 | 0.68 | 0.9 | 1.1 | 0.03 | 0.195 | 0.1675 | 0.650361 | 0.929911 | 0.564376 | 0.997796 | 0.997988 | 0.995259 |
| Pi4ka_E9Q3L2_S1427s_1 | Pi4ka_E9Q | 0.24 | 0.3 | 0.88 | 0.79 | 0.65 | 0.77 | 0.94 | 0.9 | 0.33 | 0.84 | 0.37 | 0.39 | 0.61 | 1.35 | 0.57 | 0.95 | 0.2625 | -0.07 | 0.3175 | 0.309939 | 0.292339 | 0.792257 | 0.989223 | 0.997988 | 0.994666 |
| Pi4ka_E9Q3L2_S259s_1 | Pi4ka_E9Q | 0.13 | 0.63 | 0.02 | 0.22 | 1.45 | 0.32 | 0.51 | 0.23 | -0.3 | 0.07 | 0.36 | 1.04 | 0.52 | 1.69 | -0.15 | 0.08 | 0.3775 | 0.0425 | 0.285 | 0.509022 | 0.260712 | 0.899025 | 0.989223 | 0.997988 | 0.994666 |
| Pi4ka_E9Q3L2_S263s_2 | Pi4ka_E9Q | -0.01 | 0.28 | 1 | 0.89 | 1.11 | 0.51 | 0.93 | 1.52 | 0.02 | -0.09 | 2.21 | 1.28 | -0.64 | 0.92 | 1.09 | 1.24 | 0.4775 | 0.315 | 0.1125 | 0.813711 | 0.171487 | 0.573534 | 0.989223 | 0.997988 | 0.999507 |
| Pi4ka_E9Q3L2_S268s_1 | Pi4ka_E9Q | 0.57 | 0.3 | 0.76 | 0.91 | 0.85 | 0.36 | 0.7 | 0.77 | 0.28 | 0.43 | 0.39 | 0.73 | 0.17 | 0.47 | 0.81 | 0.84 | 0.035 | -0.1775 | -0.0625 | 0.82475 | 0.882901 | 0.473278 | 0.996493 | 0.997988 | 0.999507 |
| Pi4ka_E9Q3L2_S268s_2 | Pi4ka_E9Q | -0.01 | 0.28 | 1 | 0.89 | 1.11 | 0.51 | 0.93 | 1.52 | 0.02 | -0.09 | 2.21 | 1.28 | -0.64 | 0.92 | 1.09 | 1.24 | 0.4775 | 0.315 | 0.1125 | 0.813711 | 0.171487 | 0.573534 | 0.989223 | 0.997988 | 0.999507 |
| Pi4kb_E9Q8A3_S523s_1 | Pi4kb_E9Q | -0.57 | -0.33 | -1.2 | -0.9 | -0.73 | -1.08 | -0.21 | -0.93 | -0.96 | -0.04 | -0.36 | -0.79 | -1.43 | 0.01 | -1.23 | -1.26 | 0.0125 | 0.2125 | -0.2275 | 0.565451 | 0.966244 | 0.504126 | 0.999187 | 0.997988 | 0.995259 |
| Pianp_Q6P1B3_S32s_1 | Pianp_Q6P | -1.45 | -1.22 | -0.52 | -1.21 | -1.58 | -0.52 | -1.3 | -0.49 | -0.12 | -1.78 | -2.53 | -0.98 | 0.84 | -1.53 | -1.01 | -1.14 | 0.1275 | -0.2525 | 0.39 | 0.466789 | 0.712851 | 0.629389 | 0.995437 | 0.997988 | 0.994666 |
| Pias1_Q3U556_S485s_1 | Pias1_Q3U | -1.37 | -0.53 | -1.47 | -1.95 | -0.77 | -1.67 | -1.38 | -2.62 | -0.66 | -1.41 | 0.48 | -2.1 | 0.1 | -0.44 | -1.54 | -0.41 | -0.28 | 0.4075 | 0.7575 | 0.113479 | 0.540588 | 0.485814 | 0.989223 | 0.997988 | 0.994666 |
| Pik3c3_E9Q824_S282s_1 | Pik3c3_E9Q | 1.19 | 1.36 | 0.89 | 0.76 | 1.22 | 1.02 | -0.25 | 0.38 | 1.44 | 1.18 | 0.63 | 0.14 | 1.24 | 1.34 | 0.63 | 0.91 | -0.4575 | -0.2025 | -0.02 | 0.944095 | 0.222374 | 0.55575 | 0.989223 | 0.997988 | 0.999507 |
| Pik3cd_B0QZL5_S516s_1 | Pik3cd_B0C | -0.16 | 0.34 | 0.5 | -0.38 | -0.83 | 0.1 | -0.53 | 0.02 | -0.34 | 0.26 | 1.29 | 0.54 | 0.37 | 0.05 | 0.75 | 0.36 | -0.385 | 0.3625 | 0.3075 | 0.330367 | 0.246111 | 0.37075 | 0.989223 | 0.997988 | 0.994666 |
| Pikfyve_D3Z5N5_S318s_1 | Pikfyve_D3 | 0.13 | 0.39 | -0.44 | -0.53 | 0.6 | 0.38 | -0.15 | -0.11 | -0.15 | 0.29 | -0.17 | 0.05 | 0.13 | 1.02 | -0.47 | -0.03 | 0.2925 | 0.1175 | 0.275 | 0.489857 | 0.355599 | 0.687839 | 0.989223 | 0.997988 | 0.994666 |
| Pip4k2b_Q80XI4_S131s_1 | Pip4k2b_Q | 0.55 | 0.3 | 1.16 | 0.76 | -0.9 | 1.31 | -0.83 | 0.4 | 1.15 | -0.23 | 1 | 1.08 | 1.71 | -1.38 | 1.14 | 0.45 | -0.6975 | 0.0575 | -0.2125 | 0.736607 | 0.194728 | 0.879561 | 0.989223 | 0.997988 | 0.9971 |
| Pip4k2c_Q91XU3_S129s_1 | Pip4k2c_Q | 0.16 | -0.31 | 0.6 | 0.1 | -1.24 | -0.24 | -0.67 | -0.4 | 2.31 | 0.03 | 0.47 | 0.32 | 2.17 | -0.86 | 0 | -0.03 | -0.775 | 0.645 | 0.1825 | 0.766419 | 0.029634 | 0.230543 | 0.989223 | 0.997988 | 0.9971 |
| Pip5k1a_F8WI18_S457s_1 | Pip5k1a_F8 | -0.52 | -0.7 | -0.06 | 0.09 | -0.61 | -0.58 | 0.46 | 0.22 | -1.01 | -0.35 | 0.81 | 0.38 | 0.49 | -1.14 | 0.22 | 0.35 | 0.17 | 0.255 | 0.2775 | 0.513352 | 0.617838 | 0.557741 | 0.989223 | 0.997988 | 0.994666 |
| Pip5k1c_F8WHW6_S475s_1 | Pip5k1c_F8 | 1.66 | 0.42 | 0.16 | 0 | 0.33 | 1.07 | 0.41 | 0.67 | -0.15 | 0.07 | -0.22 | 0.34 | -0.48 | 0.67 | -0.03 | 0.04 | 0.06 | -0.55 | -0.51 | 0.260711 | 0.879826 | 0.187973 | 0.996493 | 0.997988 | 0.994666 |
| Pip5k1c_F8WHW6_S538s_1 | Pip5k1c_F8 | -1.66 | -1.97 | -1.34 | -3.14 | -2.63 | -0.81 | -1.65 | -1.57 | 0.06 | -2.28 | -1.2 | -1.08 | 0.21 | -2.65 | -1.46 | -1.79 | 0.3625 | 0.9025 | 0.605 | 0.361103 | 0.471984 | 0.139317 | 0.989223 | 0.997988 | 0.994666 |
| Pip5k1c_F8WHW6_S540s_1 | Pip5k1c_F8 | -1.41 | -1.48 | -0.11 | -0.81 | -2.05 | -0.48 | -1.44 | -0.69 | 0.74 | -1.42 | -0.84 | -0.24 | 0.77 | -2.54 | -0.33 | -0.63 | -0.2125 | 0.5125 | 0.27 | 0.692657 | 0.638041 | 0.340707 | 0.989223 | 0.997988 | 0.996159 |
| Pip5k1c_F8WHW6_S554s_1 | Pip5k1c_F8 | 0.42 | -0.02 | -0.32 | 0.19 | 0.18 | 0.09 | 0.04 | 0.1 | -0.93 | -0.06 | -0.14 | -0.24 | -0.45 | -0.16 | -0.25 | -0.15 | 0.035 | -0.41 | -0.32 | 0.246376 | 0.880378 | 0.188429 | 0.996493 | 0.997988 | 0.994666 |
| Pip5k1c_F8WHW6_S671s_1 | Pip5k1c_F8 | 0.26 | -1.2 | -1.38 | -2.08 | -0.48 | 0.28 | -0.65 | -0.67 | -1.54 | -0.94 | -1.95 | -0.48 | -1.24 | -0.69 | -1.16 | -1.04 | 0.72 | -0.1275 | 0.0675 | 0.888783 | 0.169567 | 0.815534 | 0.989223 | 0.997988 | 0.999507 |
| Pip5k1c_F8WHW6_S681s_1 | Pip5k1c_F8 | 0.13 | -0.33 | 0.98 | 1.11 | -1.82 | 0.75 | -0.12 | 0.07 | 2.61 | 0.41 | 1.02 | 0.47 | 2.44 | -0.82 | 2.03 | 0.23 | -0.7525 | 0.655 | 0.4975 | 0.510007 | 0.214806 | 0.268055 | 0.989223 | 0.997988 | 0.994666 |
| Pip5k1c_F8WHW6_T661t_1 | Pip5k1c_F8 | 4.46 | 0.44 | -1.49 | -0.85 | 1.75 | 4.38 | -0.31 | -0.22 | -0.61 | 4.05 | -0.96 | -0.55 | -1.33 | -0.83 | -1.94 | -1.56 | 0.76 | -0.1575 | -2.055 | 0.104996 | 0.610421 | 0.919605 | 0.989223 | 0.997988 | 0.994666 |
| Pitpnc1_Q8K4R4_S274s_1 | Pitpnc1_Q8 | -1.01 | -1.03 | -0.93 | -0.72 | -0.22 | -1.22 | -0.25 | -0.78 | -1.38 | -0.86 | -0.4 | -0.92 | -1.15 | -0.27 | -0.93 | -0.68 | 0.305 | 0.0325 | 0.165 | 0.55918 | 0.296224 | 0.904362 | 0.989223 | 0.997988 | 0.995259 |
| Pitpnc1_Q8K4R4_S299s_1 | Pitpnc1_Q8 | -0.09 | 0.64 | 0.59 | 0.17 | 1.29 | 0.5 | 1.31 | 0.69 | 0.18 | 0.41 | -0.16 | 0.8 | 0.21 | 1.44 | 0.67 | 0.86 | 0.62 | -0.02 | 0.4675 | 0.194051 | 0.060572 | 0.9475 | 0.989223 | 0.997988 | 0.994666 |

| | | | | | | | | | | | | | | | | | | | | | | | | | | |
|---|---|---|---|---|---|---|---|---|---|---|---|---|---|---|---|---|---|---|---|---|---|---|---|---|---|---|
| Pitpnm1_O35954_S326s_1 | Pitpnm1_C | -0.83 | -0.5 | -0.95 | 0.11 | -0.49 | -0.62 | 0.06 | -0.45 | -0.25 | -0.33 | -1.08 | -0.62 | -0.35 | 0.1 | 0.02 | -0.02 | 0.1675 | -0.0275 | 0.48 | 0.146673 | 0.583831 | 0.933112 | 0.989223 | 0.997988 | 0.994666 |
| Pitpnm1_O35954_S593s_1 | Pitpnm1_C | -0.18 | 0.07 | 0.38 | 0.45 | 0.28 | 0.05 | 0.68 | 0.47 | 0 | 0.15 | 0.34 | 0.41 | 0 | 0.46 | 0.47 | 0.5 | 0.19 | 0.045 | 0.1775 | 0.520715 | 0.459107 | 0.856487 | 0.989223 | 0.997988 | 0.994666 |
| Pitpnm1_O35954_S621s_1 | Pitpnm1_C | 1.24 | -1.63 | -1.4 | -0.21 | -0.3 | 0.98 | -0.19 | 0.86 | -0.76 | -1.18 | 1.53 | -0.72 | -1.44 | -0.6 | -2.01 | -1.79 | 0.8375 | 0.2175 | -0.96 | 0.166589 | 0.22165 | 0.786718 | 0.989223 | 0.997988 | 0.994666 |
| Pitpnm1_O35954_S664s_1 | Pitpnm1_C | -0.16 | 0.53 | -0.23 | 0.03 | 0.28 | 0.73 | 0.21 | 0.02 | 0.42 | -0.11 | -0.35 | 0.08 | 1.03 | 0.74 | 0.26 | 0.39 | 0.2675 | -0.0325 | 0.5625 | 0.088967 | 0.335065 | 0.909095 | 0.989223 | 0.997988 | 0.994666 |
| Pitpnm1_O35954_S895s_1 | Pitpnm1_C | 0.21 | -0.22 | -0.61 | -0.14 | 0.01 | 0.44 | 0.72 | 0.42 | 0.34 | 0.49 | -0.32 | -0.08 | 0.17 | 0.3 | -0.1 | -0.03 | 0.5875 | 0.2975 | 0.275 | 0.330273 | 0.050138 | 0.325729 | 0.989223 | 0.997988 | 0.994666 |
| Pitpnm1_O35954_T270t_1 | Pitpnm1_C | -0.88 | -0.81 | -0.62 | -0.47 | -1.06 | -0.73 | -1.25 | -0.7 | -2.01 | -0.94 | -0.87 | -0.64 | 0.03 | -1.7 | -0.39 | -0.33 | -0.24 | -0.42 | 0.0975 | 0.806465 | 0.318376 | 0.235301 | 0.989223 | 0.997988 | 0.999507 |
| Pitpnm2_Q6ZPQ6_S1209s_1 | Pitpnm2_C | -1.31 | -1.28 | -1.08 | -1.13 | -0.61 | -0.79 | -0.04 | -0.5 | -1.21 | -1.36 | 0.82 | -0.58 | -1.99 | -0.16 | -1.21 | -0.73 | 0.715 | 0.6175 | 0.1775 | 0.657557 | 0.012885 | 0.215375 | 0.989223 | 0.997988 | 0.995259 |
| Pitpnm2_Q6ZPQ6_S1230s_1 | Pitpnm2_C | -2.23 | -2.1 | -2.79 | -6.56 | -0.6 | -2.68 | -2.18 | -2.53 | 1.11 | -2.43 | -1.41 | -1.94 | 4.09 | -0.97 | -3.35 | -3.99 | 1.4225 | 2.2525 | 2.365 | 0.212219 | 0.178862 | 0.077642 | 0.989223 | 0.997988 | 0.994666 |
| Pitpnm2_Q6ZPQ6_S1234s_1 | Pitpnm2_C | 0.35 | 0.38 | -0.34 | -0.43 | 0.75 | -0.01 | -0.19 | -0.28 | 1.35 | 0.51 | 0.81 | 0.63 | 1.16 | 0.62 | 0.62 | 0.41 | 0.0775 | 0.835 | 0.7125 | 0.047065 | 0.814413 | 0.024612 | 0.995762 | 0.997988 | 0.994666 |
| Pitpnm2_Q6ZPQ6_S302s_1 | Pitpnm2_C | 1.75 | 0.93 | 1.55 | 1.68 | 0.53 | 1.91 | 0.89 | 1.51 | 1.95 | 0.97 | 0.48 | 1.19 | 1.55 | 0.88 | 1.19 | 0.43 | -0.2675 | -0.33 | -0.465 | 0.191433 | 0.463604 | 0.379959 | 0.989223 | 0.997988 | 0.994666 |
| Pitpnm2_Q6ZPQ6_S365s_1 | Pitpnm2_C | 0.05 | -0.4 | -0.36 | -0.17 | 0.44 | 0.02 | -0.17 | -0.93 | -0.44 | -0.38 | -0.01 | 0.04 | -1.3 | 0.21 | -0.4 | -0.79 | 0.06 | 0.0225 | -0.35 | 0.343022 | 0.851047 | 0.926147 | 0.996349 | 0.997988 | 0.994666 |
| Pitpnm2_Q6ZPQ6_S397s_1 | Pitpnm2_C | -0.06 | 0.07 | -0.75 | 0.32 | -0.08 | -0.17 | 0.03 | -0.36 | -0.25 | -0.31 | 0.02 | -0.35 | -1.03 | 0.33 | -0.7 | -0.47 | -0.04 | -0.1175 | -0.3625 | 0.355621 | 0.886194 | 0.685909 | 0.996493 | 0.997988 | 0.994666 |
| Pitpnm3_Q3UHE1_S279s_1 | Pitpnm3_C | 0.91 | 0.92 | 0.63 | 0.75 | 0.94 | 0.3 | 0.7 | 0.64 | 0.54 | 0.94 | 0.47 | 0.69 | 1.01 | 0.79 | 0.69 | 0.3 | -0.1575 | -0.1425 | -0.105 | 0.689429 | 0.494203 | 0.534112 | 0.989223 | 0.997988 | 0.996159 |
| Pitpnm3_Q3UHE1_S305s_1 | Pitpnm3_C | -1.27 | -1.51 | -0.46 | -0.45 | -1.02 | -1.53 | 0.26 | 0.1 | -2.21 | -1.22 | 0.88 | 0.25 | -1.8 | -0.26 | 0.25 | -0.1 | 0.375 | 0.3475 | 0.445 | 0.384672 | 0.437337 | 0.611822 | 0.989223 | 0.997988 | 0.994666 |
| Pitpnm3_Q3UHE1_S479s_1 | Pitpnm3_C | 0.58 | 0.59 | 0.34 | 0.35 | 0.56 | 0.17 | 0.65 | 0.49 | -0.21 | 0.38 | 0.4 | 0.4 | 0.42 | 0.49 | 0.7 | 0.39 | 0.0025 | -0.2225 | 0.035 | 0.882668 | 0.990797 | 0.375655 | 0.999187 | 0.997988 | 0.999507 |
| Pitpnm3_Q3UHE1_S488s_1 | Pitpnm3_C | 0.77 | 1.31 | 0.91 | 0.54 | 0.97 | 0.9 | 0.69 | 0.14 | 0.71 | 0.95 | 0.19 | -0.1 | 0.88 | 1.37 | 0.2 | 0 | -0.2075 | -0.445 | -0.27 | 0.47493 | 0.468961 | 0.185332 | 0.989223 | 0.997988 | 0.994666 |
| Pitpnm3_Q3UHE1_S891s_1 | Pitpnm3_C | -1.85 | -2.04 | -1.64 | -1.11 | -0.59 | -1.87 | -0.68 | -1.1 | -2.33 | -2.17 | -1.1 | -1.09 | -2.35 | -0.75 | -1.65 | -0.57 | 0.6 | -0.0125 | 0.33 | 0.468201 | 0.116105 | 0.974472 | 0.989223 | 0.997988 | 0.994666 |
| Pitpnm3_Q3UHE1_S912s_1 | Pitpnm3_C | -0.37 | -0.17 | 0.24 | 0.13 | 0.85 | 0 | 0.68 | 0.23 | -1.33 | -1.18 | 0.29 | 0.2 | -0.09 | 1.11 | 0.14 | 0.84 | 0.4825 | -0.4625 | 0.5425 | 0.142701 | 0.108047 | 0.309537 | 0.989223 | 0.997988 | 0.994666 |
| Pitpnm3_Q3UHE1_S93s_1 | Pitpnm3_C | 0.37 | 0.63 | -0.41 | 0.06 | 0.18 | 0.25 | 0.05 | -0.69 | 0.99 | 0.67 | -0.94 | -1.45 | 0.01 | 0.6 | -0.84 | -0.34 | -0.215 | -0.345 | -0.305 | 0.439452 | 0.512098 | 0.55976 | 0.989223 | 0.997988 | 0.994666 |
| Pja1_O55176_S137s_1 | Pja1_O551 | -2.92 | -0.8 | -1.87 | -4.14 | -1.18 | -3.03 | -3.31 | -4.41 | -2.57 | -1.51 | -2.1 | -3.43 | -0.02 | -2.54 | -1.79 | -1.83 | -0.55 | 0.03 | 0.8875 | 0.277916 | 0.523892 | 0.967542 | 0.989223 | 0.997988 | 0.994666 |
| Pja1_O55176_S32s_1 | Pja1_O551 | -2.65 | -2.92 | -3 | -4.51 | -2.17 | -2.84 | -1.84 | -2.07 | -2.25 | -2.05 | 0.87 | -2.32 | -1.41 | -0.71 | -2.84 | -2.49 | 1.04 | 1.8325 | 1.4075 | 0.037017 | 0.039096 | 0.04099 | 0.989223 | 0.997988 | 0.994666 |
| Pja2_Q80U04_S320s_1 | Pja2_Q80U | -0.64 | -0.34 | -0.31 | 0.16 | -0.45 | -1.08 | 0.27 | -0.36 | -0.74 | -0.62 | -0.06 | -0.13 | -1.12 | -0.42 | -0.16 | 0.02 | -0.1225 | -0.105 | -0.1375 | 0.684385 | 0.712563 | 0.714203 | 0.995437 | 0.997988 | 0.996159 |
| Pkd2_O35245_S754s_1 | Pkd2_O352 | -0.84 | -0.14 | 0.57 | 0.06 | -2.36 | -0.41 | -0.48 | -0.52 | 0.56 | -0.32 | 1.46 | 0.19 | -0.55 | -2.02 | 0.65 | -0.7 | -0.855 | 0.56 | -0.5675 | 0.332576 | 0.117819 | 0.239249 | 0.989223 | 0.997988 | 0.994666 |
| Pkib_F7D7A9_S24s_1 | Pkib_F7D7 | -0.86 | -0.8 | -0.21 | -0.51 | -1.62 | -0.26 | -0.56 | 0.05 | 0.03 | -0.52 | 0.44 | -1.68 | 0.55 | -1.13 | -0.97 | 0.13 | -0.0025 | 0.1625 | 0.24 | 0.581462 | 0.994779 | 0.726041 | 0.99937 | 0.997988 | 0.995259 |
| Pkn1_P70268_S540s_1 | Pkn1_P702 | -1.18 | -0.82 | -0.95 | 0.18 | 0.77 | -0.29 | 0.21 | -1.6 | -1.4 | -0.43 | 0.97 | 0.37 | 0.04 | -0.43 | -0.63 | -0.02 | 0.465 | 0.57 | 0.4325 | 0.252386 | 0.393567 | 0.315563 | 0.989223 | 0.997988 | 0.994666 |
| Pkn1_P70268_S920s_1 | Pkn1_P702 | 0.26 | 0.67 | 0.27 | 0.09 | 0.48 | 0.1 | 0.33 | 0.07 | 0.17 | 0.35 | 0.24 | 0.43 | -0.52 | 0.63 | -0.17 | 0.21 | -0.0775 | -0.025 | -0.285 | 0.386603 | 0.73801 | 0.913859 | 0.995762 | 0.997988 | 0.994666 |
| Pkn2_G3UXH4_S534s_1 | Pkn2_G3U) | 0.41 | 0.14 | 1.07 | 0.45 | 1.47 | 0.35 | -0.68 | 0.29 | 0.27 | 0.19 | 0.56 | 0.48 | 0.85 | 0.61 | 0.35 | 0.67 | -0.16 | -0.1425 | 0.1025 | 0.725223 | 0.722864 | 0.603002 | 0.995437 | 0.997988 | 0.9971 |
| Pkp2_Q9CQ73_S82s_1 | Pkp2_Q9C( | -0.59 | -0.73 | -0.89 | -0.99 | -0.07 | -0.62 | -0.43 | 0.23 | -0.97 | -1.28 | 0.29 | -0.4 | -0.97 | -0.18 | -0.79 | -0.46 | 0.5775 | 0.21 | 0.2 | 0.476817 | 0.047472 | 0.56936 | 0.989223 | 0.997988 | 0.994666 |
| Pkp4_Q68FH0_S1036s_1 | Pkp4_Q68I | 1.41 | 0.87 | 1.14 | 1.5 | 1.47 | 1.53 | 1.13 | 1.25 | 0.9 | 1.13 | 0.48 | 1.05 | -0.72 | 0.01 | 0.42 | 1.12 | 0.115 | -0.34 | -1.0225 | 0.031256 | 0.630709 | 0.220386 | 0.989223 | 0.997988 | 0.994666 |
| Pkp4_Q68FH0_S1090s_1 | Pkp4_Q68I | 0.37 | 0.95 | 1.17 | 1.11 | 0.8 | 0.29 | 1.58 | 0.76 | 0.68 | 0.66 | 0.69 | 0.91 | 0.88 | 1.1 | 0.92 | 1.45 | -0.0425 | -0.165 | 0.1875 | 0.52526 | 0.898329 | 0.528594 | 0.996493 | 0.997988 | 0.994666 |
| Pkp4_Q68FH0_S1090s_2 | Pkp4_Q68I | 0.61 | 0.45 | 0.21 | 0.56 | 0.65 | 0.19 | 1.08 | 0.29 | 0.36 | 0.38 | 0.03 | 0.58 | -0.44 | 0.69 | 0.41 | 0.36 | 0.095 | -0.12 | -0.2025 | 0.520453 | 0.721348 | 0.612258 | 0.995437 | 0.997988 | 0.994666 |
| Pkp4_Q68FH0_S1099s_1 | Pkp4_Q68I | -0.15 | 0.34 | 0.14 | -0.04 | 0.08 | 0.03 | 0.2 | -0.49 | -0.21 | -0.08 | -0.72 | 0.15 | -0.29 | 0.44 | -0.25 | -0.97 | -0.1175 | -0.2875 | -0.34 | 0.332724 | 0.635333 | 0.303817 | 0.989223 | 0.997988 | 0.994666 |
| Pkp4_Q68FH0_S1133s_1 | Pkp4_Q68I | 0.47 | 0.47 | 0.71 | 0.68 | 0.53 | 0.34 | 0.52 | 0.27 | 0.04 | 0.4 | 0.18 | 0.7 | 0.24 | 0.51 | 0.31 | -0.03 | -0.1675 | -0.2525 | -0.325 | 0.209888 | 0.421681 | 0.310445 | 0.989223 | 0.997988 | 0.994666 |
| Pkp4_Q68FH0_S1136s_1 | Pkp4_Q68I | -0.54 | 0.18 | -0.27 | -0.26 | -0.47 | -0.18 | -0.74 | -0.93 | -0.22 | -0.15 | 0.33 | -0.24 | -1.45 | -0.56 | -0.43 | -0.96 | -0.3575 | 0.1525 | -0.6275 | 0.075139 | 0.200243 | 0.566939 | 0.989223 | 0.997988 | 0.994666 |
| Pkp4_Q68FH0_S213s_1 | Pkp4_Q68I | -1.31 | -2.56 | -2.23 | -2.5 | -2.49 | -0.73 | -1.84 | -1.72 | -1.66 | -2.08 | -1.7 | -1.7 | -0.46 | -1.94 | -2.82 | -1.35 | 0.455 | 0.365 | 0.5075 | 0.354857 | 0.311819 | 0.284905 | 0.989223 | 0.997988 | 0.994666 |
| Pkp4_Q68FH0_S220s_1 | Pkp4_Q68I | 0.97 | 1.14 | 0.23 | 0.57 | 0.25 | 0.78 | -0.1 | -0.31 | 0.18 | 0.48 | 0.1 | 1.09 | 0.49 | 1.12 | 0.13 | -0.01 | -0.5725 | -0.265 | -0.295 | 0.412147 | 0.104575 | 0.428448 | 0.989223 | 0.997988 | 0.994666 |
| Pkp4_Q68FH0_S246s_1 | Pkp4_Q68I | 0.82 | 1.78 | 0.25 | 1.07 | 1.53 | 0.9 | 0.67 | 1.06 | 0.89 | 1.2 | 0.12 | 0.32 | 0.52 | 1.13 | 0.93 | 0.96 | 0.06 | -0.3475 | -0.095 | 0.794751 | 0.868597 | 0.396151 | 0.996493 | 0.997988 | 0.999507 |
| Pkp4_Q68FH0_S272s_1 | Pkp4_Q68I | -3.04 | -2.96 | 1.66 | 1.29 | -2.7 | -2.43 | 1.38 | 1.25 | -1.14 | -3 | 1.7 | 1.86 | 0.13 | -1.79 | 1.63 | 1.93 | 0.1375 | 0.6175 | 1.2375 | 0.364766 | 0.925247 | 0.686782 | 0.997335 | 0.997988 | 0.994666 |
| Pkp4_Q68FH0_S280s_1 | Pkp4_Q68I | 1.82 | 1.29 | 0.94 | 1 | 0.89 | 1.53 | 0.16 | 0.28 | 1.82 | 1.98 | 0.57 | 0.86 | 1.73 | 1.58 | 0.63 | 0.29 | -0.5475 | 0.045 | -0.205 | 0.618525 | 0.161315 | 0.910369 | 0.989223 | 0.997988 | 0.995259 |
| Pkp4_Q68FH0_S313s_1 | Pkp4_Q68I | 0.57 | 0.71 | 0.67 | 0.79 | 0.54 | 0.35 | 0.82 | 0.68 | 0.47 | 0.68 | 0.65 | 0.95 | 0.7 | 0.98 | 0.8 | 0.62 | -0.0875 | 0.0025 | 0.09 | 0.702489 | 0.679618 | 0.990947 | 0.993464 | 0.997988 | 0.996159 |
| Pkp4_Q68FH0_S326s_2 | Pkp4_Q68I | 0.58 | 0.16 | 1.35 | 1.43 | 0.15 | 0.28 | 0.97 | 1.72 | 0.14 | 0.66 | 1.77 | 1.61 | -0.07 | 0.64 | 1.27 | 0.87 | -0.1 | 0.165 | -0.2025 | 0.628671 | 0.822002 | 0.727173 | 0.995762 | 0.997988 | 0.995259 |
| Pkp4_Q68FH0_S336s_1 | Pkp4_Q68I | 0.15 | 0.81 | 2.08 | 1.88 | 0.36 | 0.16 | 1.27 | 1.46 | 0.53 | 0.32 | 1.84 | 1.93 | 0.97 | 0.98 | 1.65 | 1.26 | -0.4175 | -0.075 | -0.015 | 0.974246 | 0.421787 | 0.895858 | 0.989223 | 0.997988 | 0.999507 |
| Pkp4_Q68FH0_S336s_2 | Pkp4_Q68I | -1.57 | 0.17 | -0.29 | -1.48 | -0.72 | -0.22 | 0.42 | -0.09 | 0.21 | 0.26 | 1.37 | 0.26 | -0.34 | -1.91 | -0.18 | 0.21 | 0.64 | 1.3175 | 0.2375 | 0.685229 | 0.185269 | 0.022625 | 0.989223 | 0.997988 | 0.996159 |
| Pkp4_Q68FH0_S405s_1 | Pkp4_Q68I | -2.43 | -1.38 | -0.49 | -0.07 | -1.87 | -2.71 | 0.29 | -0.27 | -1.25 | -1.77 | -0.68 | 0.19 | -0.52 | -0.97 | -0.66 | -0.61 | -0.0475 | 0.215 | 0.4025 | 0.431106 | 0.950265 | 0.726359 | 0.999187 | 0.997988 | 0.994666 |
| Pkp4_Q68FH0_S421s_1 | Pkp4_Q68I | 0.03 | 0.6 | 0.65 | 0.13 | 0.26 | -0.08 | 0 | 0.01 | -0.12 | 0.06 | 0.07 | 0.44 | -0.16 | 0.43 | 0.1 | 0.18 | -0.305 | -0.24 | -0.215 | 0.448525 | 0.224188 | 0.371264 | 0.989223 | 0.997988 | 0.994666 |
| Pkp4_Q68FH0_S426s_1 | Pkp4_Q68I | -0.69 | -0.66 | -0.63 | -0.44 | -0.45 | -0.94 | -0.56 | -0.66 | -0.21 | -0.32 | -0.77 | -0.58 | 0.43 | 0.17 | -0.91 | -0.76 | -0.4775 | 0.135 | 0.3375 | 0.362325 | 0.824771 | 0.564949 | 0.995762 | 0.997988 | 0.994666 |
| Pkp4_Q68FH0_S437s_1 | Pkp4_Q68I | -0.1 | -0.67 | -0.29 | -0.31 | -0.44 | -0.93 | 0 | -0.19 | 0.02 | -0.36 | 0.23 | -0.51 | 0.57 | 0.09 | 0.3 | 0.04 | -0.0475 | 0.1875 | 0.5925 | 0.046692 | 0.862197 | 0.489628 | 0.996493 | 0.997988 | 0.994666 |
| Pkp4_Q68FH0_S446s_1 | Pkp4_Q68I | 0.05 | -0.53 | -0.37 | -0.6 | -0.3 | -0.01 | -0.63 | -0.16 | -0.14 | -0.41 | 0.31 | -0.3 | -0.05 | 0.7 | -1.31 | -1.12 | 0.0875 | 0.0025 | -0.0825 | 0.861665 | 0.7291 | 0.994034 | 0.995762 | 0.997988 | 0.999507 |
| Pkp4_Q68FH0_S460s_1 | Pkp4_Q68I | -1.53 | -1 | -0.76 | -1.17 | -1.17 | -1.89 | -1.22 | -1.04 | -1.95 | -2.21 | 0.31 | -0.5 | -0.46 | -0.48 | -0.81 | -1.32 | -0.215 | 0.0275 | 0.3475 | 0.280352 | 0.455547 | 0.961597 | 0.989223 | 0.997988 | 0.994666 |
| Pkp4_Q68FH0_S75s_1 | Pkp4_Q68I | 0.55 | 0.03 | -0.16 | 0.61 | 0.31 | 0.14 | 0.39 | 0.59 | 0.31 | -0.47 | 1.08 | -1.19 | 0.28 | 0.22 | -1.23 | -0.41 | 0.1 | -0.325 | -0.5425 | 0.20112 | 0.702348 | 0.517101 | 0.995437 | 0.997988 | 0.994666 |
| Pkp4_Q68FH0_T1012t_1 | Pkp4_Q68I | 1.19 | 1.2 | 1.02 | 0.93 | 1.1 | 0.61 | 1.28 | 1.03 | 0.32 | 0.94 | 0.55 | 1.44 | -0.02 | 0.64 | 1.17 | 0.67 | -0.08 | -0.2725 | -0.47 | 0.151292 | 0.729685 | 0.368402 | 0.995762 | 0.997988 | 0.994666 |
| Pkp4_Q68FH0_T1016t_1 | Pkp4_Q68I | 0.65 | 1.11 | 1.38 | 0.97 | 1.41 | 0.84 | 1.43 | 0.71 | -0.41 | 0.39 | 0.41 | 1.49 | -1.38 | 1.09 | 1.27 | 1.17 | 0.07 | -0.5575 | -0.49 | 0.422941 | 0.801772 | 0.198561 | 0.995762 | 0.997988 | 0.994666 |
| Pkp4_Q68FH0_T282t_1 | Pkp4_Q68I | -0.81 | 0.01 | -1.42 | -1.89 | -1.03 | -0.37 | -1.29 | -2.02 | -0.6 | -0.33 | -2.79 | -0.96 | 0.6 | -0.18 | -2.01 | -2.23 | -0.15 | -0.1425 | 0.0725 | 0.919534 | 0.760183 | 0.820889 | 0.995762 | 0.997988 | 0.999507 |
| Plcb1_Q9Z1B3_S1197s_1 | Plcb1_Q9Z | -0.82 | -0.87 | -0.71 | -1.54 | -0.71 | -0.89 | -1.41 | -1.24 | -1.27 | -1.35 | 0.08 | -0.99 | -0.92 | -0.78 | -1.54 | -1.77 | -0.0775 | 0.1025 | -0.2675 | 0.438506 | 0.783447 | 0.789229 | 0.995762 | 0.997988 | 0.994666 |
| Plcb1_Q9Z1B3_S1200s_1 | Plcb1_Q9Z | -0.01 | -0.32 | 0.15 | -0.2 | -0.53 | 0 | -0.54 | -0.08 | -0.85 | -0.86 | 0.19 | -0.24 | -0.81 | -0.38 | -0.36 | -0.64 | -0.1525 | -0.345 | -0.69 | 0.03784 | 0.54858 | 0.281885 | 0.989223 | 0.997988 | 0.994666 |
| Plcb1_Q9Z1B3_S848s_1 | Plcb1_Q9Z | 0.27 | 0.72 | 0.74 | 0.52 | -0.05 | 0.24 | -0.69 | 0.46 | 0.06 | -0.03 | 0.34 | -0.21 | 0.08 | -0.27 | -0.36 | -0.07 | -0.5725 | -0.5225 | -0.7175 | 0.017809 | 0.07984 | 0.054178 | 0.989223 | 0.997988 | 0.994666 |
| Plcb1_Q9Z1B3_S981s_1 | Plcb1_Q9Z | -0.87 | -0.38 | 0.23 | 1.08 | -0.44 | -0.74 | 0.36 | 0.63 | -1.06 | -0.8 | 0.27 | 0.04 | -0.9 | -0.26 | -0.18 | -0.45 | -0.0625 | -0.4025 | -0.4625 | 0.308631 | 0.898063 | 0.427737 | 0.996493 | 0.997988 | 0.994666 |
| Plcb1_Q9Z1B3_S982s_1 | Plcb1_Q9Z | -0.66 | -0.56 | -1.24 | -0.81 | -0.5 | -0.53 | -0.36 | -0.96 | -0.86 | -0.47 | -1.3 | -1.14 | -0.79 | 0.1 | -1.2 | -1.21 | 0.23 | -0.125 | 0.0425 | 0.90716 | 0.373618 | 0.661849 | 0.989223 | 0.997988 | 0.999507 |
| Plcb1_Q9Z1B3_T1164t_1 | Plcb1_Q9Z | -0.62 | -0.87 | -1.61 | -0.75 | -0.73 | -1.02 | 0.05 | -0.64 | -1.08 | -0.82 | -1.58 | -0.8 | -1.23 | -0.75 | -0.96 | -1.21 | 0.3775 | -0.1075 | -0.075 | 0.806615 | 0.267171 | 0.735252 | 0.989223 | 0.997988 | 0.999507 |
| Plcb3_P51432_S1105s_1 | Plcb3_P51 | -0.97 | -1.13 | -1.3 | -0.6 | -0.31 | 0.4 | -1.91 | -2.12 | -0.74 | -1.37 | 0.23 | 0.35 | -0.08 | -3.63 | 0.54 | -0.82 | 0.015 | 0.6175 | 0.0025 | 0.99757 | 0.978918 | 0.17092 | 0.999187 | 0.997988 | 1 |
| Plcg1_Q62077_S1248s_1 | Plcg1_Q62 | -0.92 | -2.19 | -0.58 | -1.53 | -1.74 | -0.1 | -1.63 | -0.6 | -0.8 | -1.26 | 0.45 | -0.3 | -1.09 | -1.56 | -0.76 | -1 | 0.2875 | 0.8275 | 0.2025 | 0.614093 | 0.561519 | 0.11304 | 0.989223 | 0.997988 | 0.995259 |
| Plcg1_Q62077_S1263s_1 | Plcg1_Q62 | 0.62 | 0.4 | -1.53 | -0.48 | -0.1 | -0.08 | -1.54 | -1.92 | -0.21 | 0.22 | -0.8 | -1.8 | -0.6 | 0.59 | -1.38 | -0.81 | -0.6625 | -0.4 | -0.3025 | 0.608761 | 0.293961 | 0.510883 | 0.989223 | 0.997988 | 0.995259 |
| Plch2_H3BIW6_S1024s_1 | Plch2_H3B | 0.03 | -0.01 | -0.61 | 0.24 | -0.66 | -0.52 | -0.1 | -0.14 | 0.08 | 0.35 | 0.21 | 0.4 | -0.28 | 0.8 | -0.25 | -0.11 | -0.25 | -0.2675 | 0.2575 | 0.135 | 0.696369 | 0.336387 | 0.380229 | 0.989223 | 0.997988 |
| Plch2_H3BIW6_S1044s_1 | Plch2_H3B | -0.98 | -1.04 | -0.23 | 0.01 | -1.65 | -0.43 | -0.28 | 0.39 | -1.67 | -0.46 | 0.9 | -0.66 | -1.04 | -0.95 | 0.19 | 0.06 | 0.0675 | 0.0875 | 0.125 | 0.765403 | 0.884484 | 0.873137 | 0.996493 | 0.997988 | 0.9971 |
| Plch2_H3BIW6_S558s_1 | Plch2_H3B | -1.21 | -1.41 | -1.06 | -1.68 | -0.52 | -0.77 | -0.22 | -0.64 | -2.38 | -1.28 | -2.41 | -0.68 | -0.63 | -2.09 | -0.99 | -0.62 | 0.5525 | -0.5975 | 0.0075 | 0.986058 | 0.098294 | 0.229601 | 0.989223 | 0.997988 | 1 |
| Plch2_H3BIW6_S559s_1 | Plch2_H3B | -0.68 | -0.63 | -0.41 | -2.63 | 0.73 | -1.14 | -0.44 | -1.4 | -4.47 | -1.25 | -3 | -0.34 | -1.33 | -1.09 | -0.09 | -0.13 | 0.525 | -1.1775 | 0.4275 | 0.454064 | 0.410689 | 0.22774 | 0.989223 | 0.997988 | 0.994666 |
| Plch2_H3BIW6_S575s_1 | Plch2_H3B | 0.61 | 0.61 | 0.42 | 0.44 | 0.13 | 0.22 | 0.04 | 0.23 | 0.36 | 0.48 | 0.13 | 0.15 | -0.01 | 0.62 | 0.26 | 0.38 | -0.365 | -0.24 | -0.2075 | 0.418979 | 0.090536 | 0.285378 | 0.989223 | 0.997988 | 0.994666 |
| Plch2_H3BIW6_S585s_1 | Plch2_H3B | -0.26 | -0.61 | -0.64 | -0.12 | 1.37 | 0.48 | 0.72 | 0.64 | -0.07 | 0.56 | -0.52 | 1.03 | -0.28 | 1.18 | 0.98 | 0.16 | 1.21 | 0.6575 | 0.9175 | 0.03551 | 0.001433 | 0.103688 | 0.989223 | 0.997988 | 0.994666 |
| Plch2_H3BIW6_S886s_1 | Plch2_H3B | 1.54 | 1.54 | 0.52 | 0.7 | 1.45 | 1.91 | 0.18 | 0.53 | 1.86 | 1.73 | 0.68 | 0.53 | 1.56 | 1.01 | 0.53 | 0.5 | -0.0575 | 0.125 | -0.175 | 0.648726 | 0.898575 | 0.770682 | 0.996493 | 0.997988 | 0.995259 |
| Plch2_H3BIW6_S911s_1 | Plch2_H3B | -0.21 | -1.23 | -0.9 | -2.81 | -1.23 | -0.39 | -2.16 | -2.1 | 3.58 | -1.13 | -3.82 | -2.07 | 2.9 | -0.96 | 0.74 | -2.02 | -0.1825 | 0.4275 | 1.4525 | 0.188035 | 0.767425 | 0.770373 | 0.995762 | 0.997988 | 0.994666 |
| Plch2_H3BIW6_S947s_1 | Plch2_H3B | -0.33 | -0.05 | -0.42 | -0.45 | 0.43 | -0.07 | 0.38 | -1.05 | -0.05 | 0.03 | -0.91 | -0.28 | -0.78 | -0.05 | -1.03 | -0.72 | 0.235 | 0.01 | -0.3325 | 0.274954 | 0.513073 | 0.971678 | 0.989223 | 0.997988 | 0.994666 |
| Plcl1_Q3USB7_S48s_1 | Plcl1_Q3US | 1.91 | 0.34 | -0.24 | 0.05 | 0.87 | 0.1 | 0.3 | 0.17 | -0.19 | 1.23 | 0.63 | -0.47 | 0.63 | 1.42 | 0.18 | 0.1 | -0.155 | -0.215 | 0.0675 | 0.898517 | 0.74347 | 0.706535 | 0.995762 | 0.997988 | 0.999507 |
| Plcl1_Q3USB7_S570s_1 | Plcl1_Q3US | 0.96 | -0.31 | 0.19 | 0.27 | -1.23 | 0.89 | -0.48 | 0.26 | 1.41 | 0.54 | 0.16 | 0.34 | 1.43 | -1.72 | 0.35 | 0.06 | -0.4175 | 0.335 | -0.2475 | 0.698535 | 0.399806 | 0.392825 | 0.989223 | 0.997988 | 0.996159 |
| Plcl1_Q3USB7_S77s_1 | Plcl1_Q3US | -0.05 | -0.03 | -0.53 | 0.22 | 1.11 | 0.31 | 0.14 | -0.4 | -0.57 | -0.26 | -0.59 | -0.31 | 0.05 | 1.35 | -0.65 | 0 | 0.3875 | -0.335 | 0.435 | 0.337715 | 0.285765 | 0.205191 | 0.989223 | 0.997988 | 0.994666 |
| Plcl1_Q3USB7_S78s_1 | Plcl1_Q3US | 0.31 | 0.21 | -0.53 | 0.22 | 1.29 | 0.74 | 0.14 | -0.4 | 0.05 | 0.22 | -0.59 | -0.31 | 0.52 | 1.47 | -0.65 | 0.6 | 0.39 | -0.21 | 0.4325 | 0.359754 | 0.341284 | 0.495521 | 0.989223 | 0.997988 | 0.994666 |
| Plcl1_Q3USB7_S96s_1 | Plcl1_Q3US | -1.33 | -1.98 | -0.65 | -0.54 | -0.46 | -0.41 | -1.73 | -0.76 | -2.42 | -0.89 | -0.88 | -2.78 | -0.96 | 0.14 | -2.32 | -1.61 | 0.285 | -0.6175 | -0.0625 | 0.912745 | 0.510978 | 0.284078 | 0.989223 | 0.997988 | 0.999507 |
| Plcl2_Q8K394_S1114s_1 | Plcl2_Q8K3 | 0.06 | -0.11 | 0.39 | 0.15 | 0.15 | -0.25 | 0 | -0.14 | 0.13 | 0.24 | 0.11 | -0.35 | 0.28 | -0.02 | -0.11 | -0.1475 | -0.0375 | -0.1725 | 0.516814 | 0.508122 | 0.870135 | 0.989223 | 0.997988 | 0.994666 | |
| Plcl2_Q8K394_S124s_1 | Plcl2_Q8K3 | -0.83 | -0.23 | -0.62 | -0.63 | 0.12 | -0.31 | -0.55 | -0.81 | -0.73 | -0.47 | 0.57 | -0.27 | -1.15 | 0.3 | -0.75 | -0.56 | 0.19 | 0.3525 | 0.0375 | 0.91626 | 0.493368 | 0.30299 | 0.989223 | 0.997988 | 0.999507 |
| Plcl2_Q8K394_S18s_1 | Plcl2_Q8K3 | -1.94 | -2.39 | -1.77 | -0.4 | -1.53 | -1.84 | -0.72 | -1.47 | -2.28 | -2.06 | 0.38 | -1.62 | -1.54 | -1.02 | -0.87 | -1.12 | 0.235 | 0.23 | 0.4875 | 0.286343 | 0.609236 | 0.733112 | 0.989223 | 0.997988 | 0.994666 |
| Plcl2_Q8K394_T585t_1 | Plcl2_Q8K3 | -0.22 | 0.15 | 0.74 | 1.15 | 0.63 | -0.7 | 0.95 | 1.06 | 0.27 | 0.22 | 0.64 | 0.94 | 1.46 | 0.36 | 1.39 | 1.36 | 0.03 | 0.0625 | 0.6875 | 0.115458 | 0.949029 | 0.862141 | 0.999187 | 0.997988 | 0.994666 |
| Plcl2_Q8K394_T8st_1 | Plcl2_Q8K3 | 0.08 | -0.61 | -2.9 | -4.39 | -1.41 | 0.39 | -1.61 | -1.69 | -0.54 | -0.54 | 1.55 | -1.9 | -0.26 | 0.82 | -3.81 | -3.08 | 0.875 | 1.5975 | 0.3725 | 0.775448 | 0.386617 | 0.171377 | 0.989223 | 0.997988 | 0.99798 |
| Plec_Q9QXS1_S1324s_1 | Plec_Q9QX | 0.38 | 0.79 | 0.67 | 0.29 | 0.75 | -0.11 | 0.27 | 0.26 | 0.39 | 0.27 | 0.75 | 0.45 | 0.52 | 0.97 | 0.25 | 0.5 | -0.24 | -0.0675 | 0.0275 | 0.920137 | 0.368413 | 0.779295 | 0.989223 | 0.997988 | 0.999507 |
| Plec_Q9QXS1_S4274s_1 | Plec_Q9QX | 0.63 | -0.61 | -0.28 | 0.16 | 0.51 | -0.46 | -0.25 | 0.52 | -0.39 | -0.5 | 0.1 | 0.2 | -1.08 | -0.19 | -0.12 | -0.29 | 0.08 | -0.1225 | -0.395 | 0.300035 | 0.827799 | 0.719795 | 0.995762 | 0.997988 | 0.994666 |
| Plec_Q9QXS1_S4530s_1 | Plec_Q9QX | 0.92 | 0.08 | 0.33 | 0.64 | 0.79 | 0.56 | 0.54 | 0.8 | 0.57 | 0.75 | 0.26 | 0.61 | 0.08 | 1.01 | 0.54 | 0.35 | 0.18 | 0.055 | 0.0025 | 0.993705 | 0.480409 | 0.838121 | 0.989223 | 0.997988 | 1 |
| Plec_Q9QXS1_S609s_1 | Plec_Q9QX | 0.46 | 1.08 | 1.29 | 0.48 | 0.66 | 0.69 | 0.6 | 0.16 | 0.06 | 0.56 | 0.57 | 0.9 | 0.72 | 0.08 | 0.77 | 0.52 | -0.3 | -0.305 | -0.305 | 0.343998 | 0.299466 | 0.335378 | 0.989223 | 0.997988 | 0.994666 |
| Plekha1_Q8BUL6_S308s_1 | Plekha1_Q | -1.9 | -3.97 | -3.9 | -3.18 | -3.73 | -1.61 | -3.75 | -3.46 | -1.35 | -2.4 | -0.61 | -3.76 | -1.81 | -1.68 | -3.09 | -2.76 | 0.1 | 1.2075 | 0.9025 | 0.125831 | 0.873024 | 0.132979 | 0.996493 | 0.997988 | 0.994666 |
| Plekha1_Q8BUL6_S372s_1 | Plekha1_Q | 0.29 | -0.87 | -0.04 | -0.58 | -0.58 | -0.85 | -0.28 | -1.86 | 0.06 | 0.2 | 0.18 | 0.23 | -1.42 | -0.15 | -0.75 | 0.08 | -0.5925 | 0.4675 | -0.26 | 0.542676 | 0.172854 | 0.146866 | 0.989223 | 0.997988 | 0.994666 |
| Plekha2_F8WIK5_S312s_1 | Plekha2_F | -0.97 | -1.17 | -0.28 | -0.4 | -0.39 | -0.49 | 0.87 | -0.35 | -1.14 | -0.6 | 0.08 | 0.1 | -0.63 | -0.1 | -0.8 | -0.29 | 0.615 | 0.315 | 0.25 | 0.438495 | 0.129412 | 0.409664 | 0.989223 | 0.997988 | 0.994666 |
| Plekha5_E9Q6H8_S1020s_1 | Plekha5_E | 0.14 | 0.09 | -0.06 | 0.53 | 0.39 | 0.49 | 0.81 | 0.24 | -0.02 | -0.03 | 0.05 | 0.63 | -0.67 | 0.54 | 0.12 | 0.62 | 0.3125 | -0.0075 | -0.0225 | 0.948779 | 0.211032 | 0.977249 | 0.989223 | 0.997988 | 0.999507 |
| Plekha5_E9Q6H8_S141s_1 | Plekha5_E | 0.52 | 1.19 | -0.01 | 0.51 | 0.24 | 0.54 | 0.87 | 0.49 | 0.96 | 0.89 | 0.64 | 0.38 | -0.61 | 0.57 | 0.26 | 0.84 | -0.0175 | 0.165 | -0.2875 | 0.482884 | 0.953698 | 0.599687 | 0.999187 | 0.997988 | 0.994666 |
| Plekha5_E9Q6H8_S417s_1 | Plekha5_E | 0.61 | 0.26 | -0.43 | -0.35 | 0.89 | 1.02 | 0.21 | 0.18 | -0.17 | 0.35 | -0.66 | 0.09 | -0.26 | 0.99 | -0.91 | -1.44 | 0.5525 | -0.12 | -0.4275 | 0.435995 | 0.128744 | 0.729756 | 0.989223 | 0.997988 | 0.994666 |
| Plekha5_E9Q6H8_S484s_1 | Plekha5_E | 0.65 | 0.84 | -1.4 | -0.76 | 0.57 | -0.02 | -0.74 | -1.1 | 1.27 | 0.83 | -1.93 | -0.93 | 1.39 | 0.83 | -0.82 | -1.18 | -0.155 | -0.0225 | 0.2225 | 0.76302 | 0.793237 | 0.978235 | 0.995762 | 0.997988 | 0.9971 |
| Plekha5_E9Q6H8_S532s_1 | Plekha5_E | 8.15 | 4.25 | 7.39 | 2.51 | 3.96 | 9.3 | 7.16 | 6.36 | 2.58 | 7.52 | 1.04 | 7.23 | 1.02 | 0.73 | 7.55 | 8.59 | 1.12 | 0.9825 | -1.1025 | 0.604157 | 0.455118 | 0.594971 | 0.989223 | 0.997988 | 0.995259 |
| Plekha5_E9Q6H8_S565s_1 | Plekha5_E | 0.05 | -0.21 | 0.68 | 0.34 | 0.39 | 0.05 | 1.12 | 0.75 | -0.67 | 0.1 | 0.85 | 1.55 | -0.68 | 0.16 | 0.56 | 1.03 | 0.3625 | 0.2425 | 0.0525 | 0.898453 | 0.268529 | 0.621104 | 0.989223 | 0.997988 | 0.999507 |
| Plekha5_E9Q6H8_S574s_1 | Plekha5_E | 0.92 | 0.76 | 0.63 | 0.74 | 0.86 | 0.73 | 0.43 | 0.73 | 0.94 | 1.32 | 1.17 | 1.03 | 0.98 | 1.01 | 0.69 | 0.9 | -0.075 | 0.3525 | 0.1325 | 0.577379 | 0.722392 | 0.131784 | 0.995437 | 0.997988 | 0.995259 |
| Plekha6_Q7TQG1_S336s_1 | Plekha6_Q | 0.32 | 0.28 | 0.38 | 0.66 | 0.73 | 0.29 | 0.75 | 0.69 | 0.4 | 0.31 | 0.11 | 0.63 | 0.47 | 0.54 | 0.34 | 0.57 | 0.205 | -0.0475 | 0.07 | 0.768703 | 0.368348 | 0.837906 | 0.989223 | 0.997988 | 0.9971 |
| Plekha6_Q7TQG1_S426s_1 | Plekha6_Q | 0.02 | 0.48 | 0 | 0.51 | 1.12 | 0.06 | 0.98 | 0.38 | -0.27 | 0.27 | -0.28 | 0.36 | -0.55 | 0.73 | 0.36 | 0.36 | 0.3825 | -0.2325 | -0.0275 | 0.935722 | 0.232649 | 0.408589 | 0.989223 | 0.997988 | 0.999507 |
| Plekha6_Q7TQG1_S573s_1 | Plekha6_Q | -0.29 | 0 | 0.13 | 0.4 | -0.95 | 0.95 | 0.06 | -0.46 | 2.05 | -0.12 | -0.16 | 0.13 | 2.21 | -1.83 | 0.46 | 0.21 | -0.16 | 0.415 | 0.2025 | 0.786321 | 0.698155 | 0.427496 | 0.994893 | 0.997988 | 0.999507 |
| Plekha6_Q7TQG1_S574s_1 | Plekha6_Q | 0.16 | -0.05 | 0.6 | -0.24 | -0.24 | 0.21 | -0.44 | -0.33 | 1.47 | 0.22 | -1.82 | 0.27 | 2.14 | -0.75 | 0.32 | 0.63 | 0.3175 | -0.0825 | 0.4675 | 0.423184 | 0.259425 | 0.897666 | 0.989223 | 0.997988 | 0.994666 |
| Plekha6_Q7TQG1_S816s_1 | Plekha6_Q | 0.81 | 1.34 | 0.62 | 0.65 | 0.76 | 1.13 | 0.48 | 0.88 | 0.38 | 0.88 | 0.05 | 0.5 | 0.68 | 1.04 | 0.24 | 0.83 | -0.0425 | -0.4025 | -0.1575 | 0.601642 | 0.87123 | 0.183256 | 0.996493 | 0.997988 | 0.995259 |
| Plekha6_Q7TQG1_S864s_1 | Plekha6_Q | 1.18 | 1.17 | 1.47 | 0.76 | 1.32 | 1 | 1.01 | 1.09 | 1.18 | 1.11 | 1.32 | 1.29 | 1.2 | 1.68 | 0.72 | 0.97 | -0.04 | 0.08 | -0.0025 | 0.993499 | 0.864578 | 0.73894 | 0.996493 | 0.997988 | 1 |
| Plekha6_Q7TQG1_S901s_1 | Plekha6_Q | 0.31 | 0.38 | 0.06 | 0.26 | 0.62 | 0.13 | 0.59 | 0.1 | 0.15 | 0.62 | -0.21 | 0.37 | 0.17 | 0.94 | -0.07 | 0.32 | 0.1075 | -0.02 | 0.0875 | 0.765642 | 0.644053 | 0.938004 | 0.99033 | 0.997988 | 0.9971 |
| Plekha6_Q7TQG1_S967s_1 | Plekha6_Q | 1.56 | 1.12 | 0.93 | 1.18 | 0.97 | 1.35 | 1.26 | 1.4 | 0.45 | 1.8 | 0.92 | 0.84 | 0.44 | 0.68 | 0.88 | 1.19 | 0.0475 | -0.195 | -0.4 | 0.17928 | 0.839386 | 0.564961 | 0.995762 | 0.997988 | 0.994666 |
| Plekha6_Q7TQG1_T1045t_1 | Plekha6_Q | -0.13 | -0.27 | 1.01 | 0.68 | 0.21 | 0.15 | 0.57 | 0.68 | 0.61 | 0.15 | 1.39 | 1.14 | 0.42 | 0.53 | 0.72 | 0.75 | 0.08 | 0.5 | 0.2825 | 0.42784 | 0.815239 | 0.242407 | 0.995762 | 0.997988 | 0.994666 |
| Plekha7_Q3UIL6_S373s_1 | Plekha7_Q | 0.88 | 0.97 | 0.57 | 0.9 | 0.58 | 0.64 | 0.82 | 0.61 | 0.37 | 1.14 | -0.56 | 0.76 | 0.1 | 0.43 | -0.13 | 0.75 | -0.1675 | -0.4025 | -0.5425 | 0.082289 | 0.430225 | 0.304508 | 0.989223 | 0.997988 | 0.994666 |
| Plekha7_Q3UIL6_S668s_1 | Plekha7_Q | 0.64 | 0.78 | 0.92 | 0.84 | 1.22 | 0.3 | 1.34 | 0.66 | 0.09 | 0.84 | 1.5 | 1.32 | 0.49 | 0.99 | 0.41 | 1.12 | 0.085 | 0.1425 | -0.0425 | 0.87608 | 0.765185 | 0.67662 | 0.995762 | 0.997988 | 0.999507 |
| Plekhf2_Q91WB4_S16s_1 | Plekhf2_Q | -0.5 | -0.38 | -0.95 | -0.58 | -1.04 | -0.3 | -1.37 | -1.1 | -1.36 | -1.92 | -3.92 | -2.3 | -0.55 | -1.1 | -1.22 | -3.16 | -0.35 | -1.7725 | -0.905 | 0.120139 | 0.245841 | 0.007622 | 0.989223 | 0.997988 | 0.994666 |
| Plekhf2_Q91WB4_S226s_1 | Plekhf2_Q | 1.13 | 0.74 | -1.03 | -0.51 | 0.36 | 0.65 | -0.76 | -0.48 | 0.22 | 0.77 | -0.03 | -0.71 | 0.26 | 0.44 | -0.67 | -0.74 | -0.14 | -0.02 | -0.26 | 0.63897 | 0.799604 | 0.97103 | 0.995762 | 0.997988 | 0.995259 |
| Plekhg1_Q3UX37_S107s_1 | Plekhg1_Q | -0.02 | 1.03 | 0.3 | 0.74 | 0.55 | 0.24 | 0.67 | 0.39 | 0.74 | 0.7 | 0.52 | 0.98 | 2.17 | 0.54 | 1.02 | 0.88 | -0.1 | 0.2225 | 0.64 | 0.152772 | 0.72029 | 0.456278 | 0.995437 | 0.997988 | 0.994666 |
| Plekhg1_Q3UX37_S613s_1 | Plekhg1_Q | -0.36 | -0.62 | -0.75 | -0.71 | -0.34 | -0.46 | -0.52 | -0.73 | -0.47 | 0.39 | 0.63 | -0.05 | 0.27 | 0.14 | -0.41 | -0.32 | 0.0975 | 0.735 | 0.53 | 0.077709 | 0.651336 | 0.033022 | 0.990358 | 0.997988 | 0.994666 |
| Plekhg5_B1AS67_S925s_1 | Plekhg5_B | -3.55 | -2.41 | -3.58 | -6.38 | -1.99 | -2.96 | -3.07 | -3.6 | -3.61 | -2.75 | -2.49 | -4.23 | -2.52 | -1.73 | -4.96 | -3.75 | 1.075 | 0.71 | 0.74 | 0.44892 | 0.197733 | 0.403476 | 0.989223 | 0.997988 | 0.994666 |
| Pllp_Q9DCU2_S13s_1 | Pllp_Q9DC | 0.52 | 0.97 | 2.13 | 2.34 | 0.95 | 0.79 | 2.62 | 0.96 | 0.48 | 0.12 | 3.64 | 2.54 | 0.41 | 1.18 | 1.73 | 2.54 | -0.16 | 0.205 | -0.025 | 0.965495 | 0.774328 | 0.808261 | 0.995762 | 0.997988 | 0.999507 |
| Pllp_Q9DCU2_S14s_1 | Pllp_Q9DC | -0.01 | 0.41 | 0.56 | 0.08 | 0.3 | 0.2 | 0.72 | 0.11 | 0.13 | 0 | 0.31 | 0 | 0.33 | 0.42 | 0.3 | 0.05 | 0.0725 | -0.15 | 0.015 | 0.953677 | 0.770902 | 0.535091 | 0.995762 | 0.997988 | 0.999507 |
| Pllp_Q9DCU2_S182s_1 | Pllp_Q9DC | 0.91 | 0.77 | 0.67 | 0.68 | -1.83 | -0.05 | 0.15 | 0.46 | 0.74 | 0.62 | 0.84 | 1.36 | 0.73 | 0.45 | 0.97 | 0.89 | -0.075 | 0.1325 | 0.0025 | 0.991913 | 0.046441 | 0.596713 | 0.989223 | 0.997988 | 1 |
| Pln_P61014_S16s_1 | Pln_P6101 | -0.29 | -1.18 | -0.7 | -0.08 | -1.65 | -1.01 | -1.19 | -0.82 | -0.47 | -0.72 | -2.21 | -1.38 | 0.37 | -0.38 | -0.35 | -1.21 | -0.605 | -0.6325 | 0.17 | 0.677754 | 0.081534 | 0.176999 | 0.989223 | 0.997988 | 0.996159 |
| Plp1_P60202_S114s_1 | Plp1_P602 | 1.43 | 0.49 | 0.7 | 0.93 | 0.81 | 0.79 | 0.16 | 0.2 | 1.24 | 1.08 | 1.04 | 0.72 | 0.66 | 0.81 | 0.58 | 0.58 | -0.3975 | 0.1325 | -0.23 | 0.425993 | 0.201207 | 0.638738 | 0.989223 | 0.997988 | 0.994666 |
| Plp1_P60202_T118t_1 | Plp1_P602 | 0.33 | -0.48 | 0.95 | 1.1 | 0.89 | 0.22 | -0.19 | 0.5 | 0.03 | 1.18 | 1.92 | 0.88 | 0.65 | 0 | 0.8 | 0.89 | -0.12 | 0.5275 | 0.11 | 0.790198 | 0.768526 | 0.306531 | 0.995762 | 0.997988 | 0.999507 |
| Pls3_B1AX58_S549s_1 | Pls3_B1AX | -0.3 | -0.18 | -0.05 | -0.39 | 0.28 | -0.48 | -0.2 | -0.15 | 0.1 | -0.51 | -0.25 | 0.08 | -0.01 | 0.18 | -0.14 | -0.1 | 0.1025 | 0.095 | 0.2225 | 0.363791 | 0.671041 | 0.699141 | 0.992876 | 0.997988 | 0.994666 |
| Plxdc2_B1AY86_S456s_1 | Plxdc2_B1A | -0.06 | -0.02 | 0.17 | 0.2 | 0.18 | 0.66 | 0.39 | 0.41 | 0.52 | 0.42 | 0.1 | 0.56 | 0.21 | 0.41 | 0.57 | 0.54 | 0.3375 | 0.3275 | 0.36 | 0.156121 | 0.139397 | 0.171063 | 0.989223 | 0.997988 | 0.994666 |
| Plxna1_P70206_S1630s_1 | Plxna1_P7C | 0.2 | 0.06 | 0.91 | 0.29 | 0.15 | -0.06 | 0.42 | -0.05 | 0.27 | 0.22 | 0.71 | 0.6 | 0.35 | -0.13 | 0.74 | 0.6 | -0.25 | 0.085 | 0.025 | 0.937127 | 0.356618 | 0.759334 | 0.989223 | 0.997988 | 0.999507 |
| Plxna2_P70207_S1630s_1 | Plxna2_P7C | 0.82 | 0.96 | 1.13 | 0.81 | 1.05 | 0.66 | 0.63 | 0.62 | 0.63 | 1.01 | 1.11 | 0.88 | 0.73 | 1.31 | 0.43 | 0.45 | -0.19 | -0.0225 | -0.2475 | 0.494143 | 0.393258 | 0.921317 | 0.989223 | 0.997988 | 0.994666 |
| Plxna2_P70207_S1633s_1 | Plxna2_P7C | 0.9 | -0.31 | 0.33 | 0.57 | 0.8 | 0.92 | 0.52 | 1.15 | -0.1 | -0.06 | 1.26 | 1.05 | -0.22 | 0.81 | 0.83 | 0.85 | 0.475 | 0.165 | 0.195 | 0.611314 | 0.147211 | 0.701734 | 0.989223 | 0.997988 | 0.995259 |
| Plxna3_P70208_S1531s_1 | Plxna3_P7C | -0.03 | -0.52 | 0.42 | 0.45 | -0.78 | 0.17 | -0.27 | 0.95 | 0.27 | 0.2 | 0.28 | -1.2 | -0.41 | -0.35 | -0.1 | -0.13 | -0.4025 | -0.0575 | 0.5925 | 0.506545 | 0.244349 | 0.888086 | 0.989223 | 0.997988 | 0.994666 |
| Plxnc1_Q9QZC2_S984s_1 | Plxnc1_Q9 | 0.26 | 0.58 | -0.07 | -0.47 | 0.55 | 0.26 | 1.07 | 0.38 | -0.95 | -0.02 | -0.95 | -0.1 | -0.09 | 0.1 | 0.17 | -0.19 | 0.49 | -0.58 | -0.0775 | 0.796813 | 0.136133 | 0.128449 | 0.989223 | 0.997988 | 0.999507 |
| Plxnd1_Q3UH93_S1384s_1 | Plxnd1_Q3 | 0.7 | 0.15 | 0.63 | -0.75 | -0.07 | 0.49 | 0.16 | 0.22 | -0.08 | 0.44 | -1.46 | -1.84 | -0.74 | 0.07 | -0.44 | 0.49 | 0.0175 | -0.9175 | -0.3375 | 0.436946 | 0.960519 | 0.14334 | 0.999187 | 0.997988 | 0.994666 |
| Pml_F7BTZ2_S9s_1 | Pml_F7BTZ | 0.48 | 0.12 | -0.27 | -0.16 | -0.57 | -0.74 | -1.1 | -0.78 | -0.14 | -0.5 | -0.02 | -1.24 | 0 | 0.2 | -0.7 | -0.86 | -0.84 | -0.1575 | 0.2 | 0.280345 | 0.007972 | 0.153837 | 0.989223 | 0.997988 | 0.994666 |
| Pml_F7BTZ2_T473t_1 | Pml_F7BTZ | -1.7 | -0.48 | 0.2 | -0.13 | -2.14 | -1.67 | -0.32 | -0.22 | -1.61 | -2.08 | -0.37 | 0.08 | -1.66 | -0.92 | -0.2 | -0.2 | -0.56 | -0.4675 | -0.2175 | 0.67122 | 0.340049 | 0.444402 | 0.989223 | 0.997988 | 0.996159 |
| Pmm1_Q91W01_S182s_1 | Pmm1_Q9 | 0.27 | -0.17 | 0.52 | 0.37 | 0.12 | -0.19 | 0.21 | 0.12 | 1.14 | 0.86 | 1.36 | -0.16 | 1.17 | 0.09 | 0.81 | -0.5 | -0.1325 | 0.6025 | 0.195 | 0.637784 | 0.598291 | 0.139463 | 0.989223 | 0.997988 | 0.995259 |
| Pmpcb_Q9CXT8_T45t_1 | Pmpcb_Q9 | 0.56 | 0.55 | 0.29 | 0.22 | 0.44 | 0.66 | 1.03 | -0.04 | -0.42 | 0.38 | 0.05 | 0.75 | 0.2 | 0.08 | 0.88 | 0.88 | 0.1175 | -0.215 | 0.165 | 0.538585 | 0.673383 | 0.482695 | 0.992876 | 0.997988 | 0.994666 |
| Pnck_Q9QYK9_S316s_1 | Pnck_Q9QY | -0.12 | 0.73 | 0.52 | 0.34 | -0.93 | 0.79 | -0.94 | -0.01 | 0.27 | 0.31 | 1.27 | -0.04 | 1.56 | -0.06 | 0.71 | 0.88 | -0.64 | 0.085 | 0.405 | 0.312806 | 0.159088 | 0.808991 | 0.989223 | 0.997988 | 0.994666 |
| Pnisr_A2AJT4_S211s_1 | Pnisr_A2AJ | 0.43 | 0.48 | -0.25 | -0.28 | -0.32 | -0.35 | -0.57 | -0.03 | -0.59 | -0.65 | -0.69 | -0.5 | 0.52 | 0.68 | -0.32 | -1.34 | -0.4125 | -0.7025 | -0.21 | 0.665998 | 0.157225 | 0.026304 | 0.989223 | 0.997988 | 0.996159 |
| Pnisr_A2AJT4_S601s_1 | Pnisr_A2AJ | 4.26 | 2.41 | 1.81 | 2.13 | 1.63 | 4.15 | 0.24 | 2.2 | 5.26 | 3.45 | -0.19 | 1.48 | 3.14 | 2.06 | 2.03 | 0.83 | -0.5975 | -0.1525 | -0.6375 | 0.341739 | 0.489255 | 0.894079 | 0.989223 | 0.997988 | 0.994666 |
| Pnn_Q3TUQ5_S100s_1 | Pnn_Q3TU | -4.35 | -4.25 | -3.36 | -6.17 | -4.33 | -4.04 | -5.56 | -3.74 | -4.85 | -4.21 | -2.19 | -3.44 | -3.11 | -5.35 | -3.67 | -4.34 | 0.14 | 0.885 | 0.44 | 0.51787 | 0.822083 | 0.243424 | 0.995762 | 0.997988 | 0.994666 |
| Pnn_Q3TUQ5_S347s_1 | Pnn_Q3TU | -2.21 | -3.29 | -4.92 | -6.85 | -2.68 | -4.43 | -5.7 | -6.66 | -2.51 | -4.09 | -2.36 | -5.06 | -1.08 | -2.55 | -4.02 | -6.37 | -0.55 | 0.8125 | 0.8125 | 0.537741 | 0.63242 | 0.450301 | 0.989223 | 0.997988 | 0.994666 |
| Pnn_Q3TUQ5_S443s_1 | Pnn_Q3TU | 2.08 | 1.99 | 2.43 | 2.19 | 0.62 | 1.76 | 1.15 | 2.41 | 1.18 | 0.85 | 2.23 | 2.36 | 1.28 | 1.91 | 2.58 | 1.85 | -0.6875 | -0.5175 | -0.2675 | 0.420515 | 0.100931 | 0.205684 | 0.989223 | 0.997988 | 0.994666 |
| Pnn_Q3TUQ5_S66s_1 | Pnn_Q3TU | -0.47 | 0.29 | -0.15 | -0.33 | -0.1 | -0.5 | -0.13 | -0.39 | -0.2 | -0.07 | -0.49 | -0.19 | 0.6 | 0.32 | 0.05 | -0.22 | -0.115 | -0.0725 | 0.3525 | 0.260039 | 0.646463 | 0.777927 | 0.99033 | 0.997988 | 0.994666 |
| Pnn_Q3TUQ5_S699s_1 | Pnn_Q3TU | 0.5 | 0.17 | 0.1 | -0.18 | -1.74 | 0.82 | -1.26 | -0.29 | 0.99 | -1.14 | -2.56 | -0.8 | 2.07 | -1.38 | -0.45 | -1.3 | -0.765 | -1.025 | -0.4125 | 0.574613 | 0.171756 | 0.152552 | 0.989223 | 0.997988 | 0.995259 |
| Pnpla6_Q3TRM4_S306s_1 | Pnpla6_Q3 | 1.05 | 0.53 | 0.61 | 0.17 | 1.11 | 0.07 | 1.62 | 0.02 | 0.93 | 0.5 | 0.48 | -0.27 | 0.24 | 1.05 | -1.23 | 0.42 | 0.115 | -0.18 | -0.47 | 0.349242 | 0.781444 | 0.590102 | 0.995762 | 0.997988 | 0.994666 |
| Pnpo_Q91XF0_S164s_1 | Pnpo_Q91 | -0.8 | -0.82 | -0.7 | -1.17 | -0.68 | -0.76 | -0.73 | -1.69 | -0.24 | -1.01 | -0.56 | -0.87 | -0.23 | -0.83 | -1.19 | -0.84 | -0.0925 | 0.1025 | 0.1 | 0.732557 | 0.751896 | 0.702084 | 0.995762 | 0.997988 | 0.9971 |
| Pnpo_Q91XF0_S165s_1 | Pnpo_Q91 | -0.72 | -0.2 | -0.11 | -0.65 | 0.42 | -0.51 | -0.39 | -0.46 | -0.93 | -0.29 | -0.21 | -0.64 | 0.16 | -0.28 | 0.11 | -0.24 | 0.185 | -0.0975 | 0.3575 | 0.214621 | 0.535744 | 0.727653 | 0.989223 | 0.997988 | 0.994666 |
| Pnpo_Q91XF0_S241s_1 | Pnpo_Q91 | 0.2 | 0.34 | 0.06 | 0.1 | 0.5 | 0.1 | 1.14 | 1.05 | 0.53 | 0.45 | 0.52 | 0.94 | 0.48 | -0.34 | 0.48 | -0.19 | 0.35 | 0.5225 | 0.4225 | -0.1 | 0.72571 | 0.092146 | 0.09004 | 0.989223 | 0.997988 |
| Pogz_D3YUW8_S1265s_1 | Pogz_D3YL | -0.46 | -0.07 | -0.5 | -0.06 | -0.04 | -0.7 | -0.16 | -0.22 | -0.14 | -0.77 | 0.39 | -0.28 | 0.33 | 0.42 | 0.64 | 0.13 | -0.0075 | 0.0725 | 0.6525 | 0.029805 | 0.97578 | 0.811793 | 0.999187 | 0.997988 | 0.994666 |
| Poldip3_F6VR84_S127s_1 | Poldip3_F6 | -0.27 | -2.26 | -1.37 | -1.61 | -3.37 | -1.81 | -2.94 | -3.5 | -2.19 | -1.33 | -3.19 | -1.63 | -1.7 | -2.01 | -1.19 | -2.94 | -1.5275 | -0.7075 | -0.5825 | 0.278412 | 0.014046 | 0.212379 | 0.989223 | 0.997988 | 0.994666 |
| Pole4_E0CYA9_T11t_1 | Pole4_E0C | -1.22 | -0.37 | -2.69 | -1.59 | -0.3 | -2.19 | -1.19 | -1.88 | 1.73 | 0.23 | -1.11 | -0.85 | 2.07 | -0.46 | -0.18 | -1.05 | 0.0775 | 1.4675 | 1.5625 | 0.058999 | 0.892428 | 0.067958 | 0.996493 | 0.997988 | 0.994666 |
| Poll_Q9QXE2_S175s_1 | Poll_Q9QX | -0.28 | -0.29 | 0.33 | 0.36 | -0.21 | -0.36 | -0.44 | 0.34 | 0.26 | -0.3 | 0.52 | 0.25 | -0.54 | -0.27 | 0.24 | 0.24 | -0.1975 | 0.1525 | -0.1125 | 0.722636 | 0.494773 | 0.606299 | 0.989223 | 0.997988 | 0.9971 |
| Polr2a_P08775_S1913s_1 | Polr2a_P0 | 1.12 | 0.76 | 0.19 | -0.55 | 1.15 | 0.51 | 0.52 | -0.29 | 0.54 | 0.77 | 0.44 | 0.42 | 0.2 | 0.94 | 0.95 | -0.79 | 0.24 | 0.0925 | -0.17 | 0.71373 | 0.833583 | 0.780595 | 0.995762 | 0.997988 | 0.996966 |
| Polr2a_P08775_S1920s_1 | Polr2a_P0 | 0.05 | -0.25 | -0.26 | -0.23 | -0.27 | -0.44 | 0.03 | -0.34 | -0.25 | -0.28 | -0.35 | -0.03 | 0.67 | -0.16 | -0.43 | -0.03 | -0.1725 | -0.145 | 0.095 | 0.759402 | 0.449662 | 0.525158 | 0.989223 | 0.997988 | 0.9971 |
| Polr3c_F6QK97_S78s_1 | Polr3c_F6C | -1.42 | -2.12 | -2.66 | -13.05 | -4.17 | -0.93 | -2.1 | -2.34 | -1.47 | -0.64 | -2.79 | -2.47 | -0.94 | -1.78 | -1.13 | -1.85 | 2.4275 | 2.97 | 3.3875 | 0.174132 | 0.328643 | 0.24143 | 0.989223 | 0.997988 | 0.994666 |
| Polr3e_Q9CZT4_S136s_1 | Polr3e_Q9 | -0.67 | -0.3 | 0.36 | -0.57 | -1.29 | -0.6 | -0.19 | 0.04 | -0.21 | -0.32 | 1.47 | -0.07 | -1.13 | 0.35 | 0.22 | -0.59 | -0.215 | 0.5125 | 0.2575 | 0.462657 | 0.562505 | 0.281918 | 0.989223 | 0.997988 | 0.995259 |
| Pom121_Q8K3Z9_S159s_1 | Pom121_Q | 0.62 | -0.99 | 0.55 | 0.81 | 0.23 | 0.49 | 0.35 | 1.07 | 0.2 | -0.16 | 0.64 | 0.39 | 0.69 | 0.88 | 0.32 | 0.37 | 0.2875 | 0.02 | 0.3175 | 0.468098 | 0.509048 | 0.963339 | 0.989223 | 0.997988 | 0.994666 |
| Pom121_Q8K3Z9_T421t_1 | Pom121_Q | -0.65 | 0.16 | 0.21 | -0.08 | -0.68 | -0.1 | -0.8 | -0.35 | 1.14 | -1.49 | -1.9 | 0.09 | 1.84 | -1.15 | 0.38 | 0.11 | -0.3925 | -0.45 | 0.385 | 0.518645 | 0.191425 | 0.503243 | 0.989223 | 0.997988 | 0.994666 |
| Ppap2a_Q61469_T279t_1 | Ppap2a_Q6 | 2.83 | 0.68 | 0.9 | 1.6 | 0.52 | 2.98 | 0.63 | 1.83 | 1.11 | 1.34 | 1.46 | 0.96 | 1.04 | -0.05 | 1.46 | 1.51 | -0.0125 | -0.285 | -0.5125 | 0.3699 | 0.985045 | 0.550722 | 0.999187 | 0.997988 | 0.994666 |
| Ppap2b_Q99JY8_S19s_1 | Ppap2b_Q9 | 0.54 | 1.26 | 0.9 | 0.76 | 0.93 | 0.56 | 0.41 | 0.44 | 0.99 | 1.15 | 1.02 | 1.34 | 1.09 | 0.39 | 0.73 | 0.48 | -0.28 | 0.26 | -0.1925 | 0.509494 | 0.278387 | 0.308959 | 0.989223 | 0.997988 | 0.994666 |
| Ppap2c_G3XA61_S209s_1 | Ppap2c_G3 | 0.13 | 0.42 | -0.51 | -0.35 | 0.53 | -0.42 | 0.52 | -0.31 | -0.4 | 0.22 | -0.17 | -0.13 | -0.85 | 1.05 | 0.53 | -0.17 | 0.1575 | -0.1925 | 0.2175 | 0.632978 | 0.645996 | 0.558572 | 0.99033 | 0.997988 | 0.995259 |
| Ppapdc2_Q9D4F2_S24s_1 | Ppapdc2_C | -3.17 | -1.96 | -2.95 | -1.26 | -1.81 | -2.12 | -0.86 | -1.95 | -2.18 | -2.87 | -2.37 | -2.02 | -1.03 | -1.02 | -1.38 | -1.47 | 0.65 | -0.025 | 1.11 | 0.031571 | 0.201519 | 0.95668 | 0.989223 | 0.997988 | 0.994666 |
| Ppapdc2_Q9D4F2_S60s_1 | Ppapdc2_C | -0.75 | -0.72 | 0.23 | 0.42 | -0.92 | -0.17 | -0.38 | 0.18 | 1.46 | -0.57 | -1.35 | 0.08 | 1.51 | -2.24 | 1.24 | 0.91 | -0.1175 | 0.11 | 0.56 | 0.498467 | 0.755687 | 0.857472 | 0.995762 | 0.997988 | 0.994666 |
| Ppapdc2_Q9D4F2_S67s_1 | Ppapdc2_C | -0.36 | -0.13 | 0.08 | 0.35 | 0.33 | 0.47 | 0.49 | 0.54 | 0.1 | 0.08 | 0.09 | 0.63 | -0.41 | 0.2 | 0.08 | 0.45 | 0.2925 | 0.06 | -0.145 | 0.646809 | 0.199476 | 0.812057 | 0.989223 | 0.997988 | 0.995259 |
| Ppapdc3_Q91WB2_S43s_1 | Ppapdc3_C | 0.39 | 0.88 | -0.53 | -0.6 | 1.78 | -0.02 | -0.5 | -0.24 | 0.69 | 0.9 | -0.74 | -0.16 | -0.13 | 1.38 | 1.18 | -0.83 | -0.54 | 0.22 | -0.0675 | 0.2625 | 0.670769 | 0.699415 | 0.886381 | 0.994893 | 0.997988 |
| Ppapdc3_Q91WB2_S62s_1 | Ppapdc3_C | 0.76 | 0.52 | -0.4 | 0.12 | 0.32 | 1.05 | 0.43 | 0.1 | 0.15 | 0.53 | -0.29 | -0.66 | -0.54 | 0.59 | -0.51 | 0.18 | 0.225 | -0.3175 | -0.32 | 0.416756 | 0.505632 | 0.401001 | 0.989223 | 0.997988 | 0.994666 |
| Ppfia1_B2RXQ2_S150s_1 | Ppfia1_B2F | 0.4 | 0.99 | -0.14 | 0.38 | 0.75 | 0 | 0.86 | -0.84 | 0.96 | 0.33 | -1.21 | 0.03 | -0.47 | 1.28 | 0.15 | -0.64 | -0.215 | -0.38 | -0.3275 | 0.493605 | 0.619789 | 0.440326 | 0.989223 | 0.997988 | 0.994666 |
| Ppfia1_B2RXQ2_S239s_1 | Ppfia1_B2F | 0.47 | 0.32 | -0.01 | 0.55 | 0.41 | -0.03 | 0.21 | -0.34 | 0.28 | 0.62 | 0.47 | 0.3 | -0.03 | 0.16 | -0.02 | 0.6 | -0.27 | -0.055 | -0.155 | 0.576592 | 0.305701 | 0.845691 | 0.989223 | 0.997988 | 0.995259 |
| Ppfia1_B2RXQ2_S732s_1 | Ppfia1_B2F | 0.04 | -0.24 | -0.57 | -0.45 | -0.7 | 1.85 | -1.11 | -0.9 | 0.09 | -0.72 | -0.75 | 1.9 | -0.67 | -1.04 | 0.03 | 0.09 | -0.1 | 0.36 | 0.557335 | 0.886072 | 0.833946 | 0.996493 | 0.997988 | 0.995259 | |
| Ppfia1_B2RXQ2_S775s_1 | Ppfia1_B2F | 0.37 | 0.06 | -0.2 | -0.33 | -0.89 | -0.01 | -0.83 | -0.88 | 0.43 | 0.63 | 0.55 | -0.6 | 0.74 | -0.89 | 0.23 | -0.13 | -0.6275 | 0.2775 | 0.0125 | 0.974251 | 0.055205 | 0.428478 | 0.989223 | 0.997988 | 0.999507 |
| Ppfia1_B2RXQ2_S797s_1 | Ppfia1_B2F | -0.62 | -0.34 | 0.69 | 0.15 | -0.24 | -0.63 | -0.11 | 0.22 | -0.66 | -0.92 | 1.54 | 0.07 | -0.44 | -0.18 | -0.47 | -0.17 | -0.16 | 0.0375 | -0.285 | 0.406188 | 0.64215 | 0.947856 | 0.989327 | 0.997988 | 0.994666 |
| Ppfia2_F7CYX4_S236s_1 | Ppfia2_F7C | -1.24 | -1.08 | -0.65 | -0.11 | -0.99 | -0.97 | 0.07 | -0.08 | 0.17 | -1.64 | -0.53 | -0.38 | 0.18 | -0.73 | -0.58 | -0.13 | 0.2775 | 0.175 | 0.455 | 0.219937 | 0.463624 | 0.692556 | 0.989223 | 0.997988 | 0.994666 |
| Ppfia2_F7CYX4_S689s_1 | Ppfia2_F7C | -0.68 | 0.05 | -0.54 | -0.61 | 0.2 | -0.52 | 0.05 | -0.52 | -0.86 | -0.69 | -0.81 | -0.62 | -0.73 | 0.31 | -0.9 | -0.78 | 0.2475 | -0.3 | -0.08 | 0.821489 | 0.395103 | 0.249947 | 0.989223 | 0.997988 | 0.999507 |
| Ppfia2_F7CYX4_S774s_1 | Ppfia2_F7C | 0.21 | 0.26 | 0.01 | 0.23 | -0.21 | -0.04 | -0.3 | -0.23 | 0.55 | 0.65 | -0.17 | 0.6 | 0.08 | 0.37 | 0.02 | -0.315 | 0.0225 | 0.09 | 0.723968 | 0.147207 | 0.937346 | 0.989223 | 0.997988 | 0.9971 | |
| Ppfia3_P60469_S1164s_1 | Ppfia3_P6C | -0.38 | -0.58 | -0.31 | 0.1 | -0.08 | -0.41 | 0.63 | 0.32 | -0.42 | -0.74 | -0.17 | 0.67 | -0.51 | -0.27 | -0.73 | 0.2 | 0.4075 | 0.1275 | -0.035 | 0.90822 | 0.189881 | 0.716159 | 0.989223 | 0.997988 | 0.999507 |
| Ppfia3_P60469_S142s_1 | Ppfia3_P6C | -0.12 | 0.32 | 0.17 | 0.04 | 0.55 | 0.1 | 0.54 | 0.22 | 0.06 | 0.38 | -0.69 | 0.2 | 0.22 | 0.36 | 0.44 | 0.5 | 0.25 | -0.115 | 0.2775 | 0.267151 | 0.285786 | 0.697972 | 0.989223 | 0.997988 | 0.994666 |
| Ppfia3_P60469_S17s_1 | Ppfia3_P6C | -0.46 | -0.79 | -0.07 | 0.02 | 0.25 | 0.01 | -0.29 | 0.38 | -0.3 | -0.52 | 0.36 | 0.55 | -0.28 | 0.44 | 0.12 | -0.5 | 0.4125 | 0.3475 | 0.27 | 0.414461 | 0.158882 | 0.318332 | 0.989223 | 0.997988 | 0.994666 |
| Ppfia3_P60469_S508s_1 | Ppfia3_P6C | -4.18 | -3.77 | -3.46 | -3.81 | -1.67 | -4.85 | -2.88 | -3.94 | -7.36 | -4.52 | -2.13 | -3.16 | -3.61 | -2.41 | -3.66 | -3.33 | 0.47 | -0.4875 | 0.5525 | 0.142419 | 0.458581 | 0.63094 | 0.989223 | 0.997988 | 0.994666 |
| Ppfia3_P60469_S537s_1 | Ppfia3_P6C | 0.43 | 0.55 | -1.49 | -0.27 | 1.48 | 1.12 | 0.06 | -0.13 | -0.23 | -0.77 | -0.66 | -0.69 | 0.44 | 0.6 | -0.34 | -1.02 | 0.8275 | -0.3925 | 0.115 | 0.835755 | 0.158103 | 0.405572 | 0.989223 | 0.997988 | 0.999507 |
| Ppfia3_P60469_S645s_1 | Ppfia3_P6C | -0.14 | -0.6 | -0.75 | -0.42 | 0.19 | -0.55 | -0.26 | -0.54 | -1.71 | -0.66 | -1.65 | 0.46 | -0.83 | -2.12 | -0.18 | 0.24 | 0.1875 | -0.4125 | -0.245 | 0.627717 | 0.483354 | 0.415362 | 0.989223 | 0.997988 | 0.995259 |
| Ppfia3_P60469_S683s_1 | Ppfia3_P6C | 0.11 | -1.25 | -1.19 | -0.98 | 0.53 | 0.29 | -0.14 | -0.01 | -1.04 | -0.71 | 0.26 | -0.45 | -1.28 | 0.37 | -1.4 | -0.8 | 0.995 | 0.3425 | 0.05 | 0.918455 | 0.017829 | 0.417227 | 0.989223 | 0.997988 | 0.999507 |
| Ppfia3_P60469_S69s_1 | Ppfia3_P6C | 0.89 | 1.18 | 1.88 | 1.53 | 0.66 | 0.84 | 0.87 | 1.38 | 1.16 | 1.06 | 2.23 | 1.32 | 1.09 | 1.36 | 1.81 | 1.48 | -0.4325 | 0.0725 | 0.065 | 0.83565 | 0.163193 | 0.838779 | 0.989223 | 0.997988 | 0.999507 |
| Ppfia3_P60469_S75s_1 | Ppfia3_P6C | -1.85 | -2.38 | -2.18 | -1.47 | -0.91 | -1.21 | -0.17 | -1.15 | -2.56 | -2.75 | -1.5 | -2.34 | -1.68 | 0.11 | -3.56 | -1.44 | 1.11 | -0.3175 | 0.3275 | 0.640018 | 0.006625 | 0.380592 | 0.989223 | 0.997988 | 0.995259 |
| Ppfia3_P60469_T714t_1 | Ppfia3_P6C | 0.32 | 0.53 | 0.41 | 0.58 | 0.7 | 0.01 | 0.69 | 0.54 | 0.16 | 0.45 | 0.72 | 0.49 | 0.44 | 1.07 | 0.67 | 0.52 | 0.025 | -0.005 | 0.215 | 0.410933 | 0.916672 | 0.982516 | 0.997335 | 0.997988 | 0.994666 |
| Ppfia3_P60469_T714t_2 | Ppfia3_P6C | -0.81 | -0.67 | -0.56 | 0.16 | -0.37 | -1.22 | -0.11 | -0.85 | -1.4 | -0.08 | 3.93 | -0.37 | -1.02 | -1.01 | -1.08 | -0.45 | -0.1675 | 0.99 | -0.42 | 0.201395 | 0.620508 | 0.357651 | 0.989223 | 0.997988 | 0.994666 |
| Ppfia3_P60469_T718t_2 | Ppfia3_P6C | -0.81 | -0.67 | -0.56 | 0.16 | -0.37 | -1.22 | -0.11 | -0.85 | -1.4 | -0.08 | 3.93 | -0.37 | -1.02 | -1.01 | -1.08 | -0.45 | -0.1675 | 0.99 | -0.42 | 0.201395 | 0.620508 | 0.357651 | 0.989223 | 0.997988 | 0.994666 |

| | | | | | | | | | | | | | | | | | | | | | | | | | | |
|---|---|---|---|---|---|---|---|---|---|---|---|---|---|---|---|---|---|---|---|---|---|---|---|---|---|---|
| Ppfia4_B8QI36_S140s_1 | Ppfia4_B8C | 0.06 | 0.01 | 0.36 | 0.91 | 0.55 | -0.24 | 0.42 | -0.01 | -0.21 | 0.33 | 0.23 | 0.62 | 0.4 | 0.42 | 0.27 | 0.76 | -0.155 | -0.0925 | 0.1275 | 0.66811 | 0.608918 | 0.762404 | 0.989223 | 0.997988 | 0.996159 |
| Ppfia4_B8QI36_S641s_1 | Ppfia4_B8C | 0.35 | 0.88 | -0.33 | -0.33 | 0.34 | 0.13 | 0.16 | -1.06 | 0.54 | 0.25 | -0.47 | -0.59 | -0.48 | -0.03 | -1 | -0.09 | -0.25 | -0.21 | -0.5425 | 0.178949 | 0.549998 | 0.601895 | 0.989223 | 0.997988 | 0.994666 |
| Ppfia4_B8QI36_S701s_1 | Ppfia4_B8C | 0.08 | -0.11 | -0.2 | -0.33 | -1.13 | -0.27 | -0.83 | -0.88 | 0.16 | 0.41 | 0.55 | -0.6 | 0.74 | -1.22 | 0.23 | -0.13 | -0.6375 | 0.27 | 0.045 | 0.914997 | 0.029914 | 0.388606 | 0.989223 | 0.997988 | 0.999507 |
| Ppfibp1_F6S1C4_S128s_1 | Ppfibp1_F6 | -0.6 | -1.05 | -1.44 | -2.46 | -0.35 | -1.01 | -2.64 | -2.04 | -0.96 | -0.29 | -0.93 | -1.31 | -0.33 | 0.21 | -1.78 | -1.56 | -0.1225 | 0.515 | 0.5225 | 0.372333 | 0.833208 | 0.256295 | 0.995762 | 0.997988 | 0.994666 |
| Ppfibp1_F6S1C4_S48s_1 | Ppfibp1_F6 | -0.19 | 0.37 | -0.41 | -0.01 | -0.7 | -0.51 | -0.89 | -0.19 | 0.19 | 0.4 | -0.21 | -0.13 | 0.71 | -1.12 | -0.36 | -0.37 | -0.5125 | 0.1225 | -0.225 | 0.588221 | 0.079788 | 0.655952 | 0.989223 | 0.997988 | 0.995259 |
| Ppfibp1_F6YZ95_S99s_1 | Ppfibp1_F6 | -0.22 | 0.16 | 0.32 | -0.48 | -0.5 | -0.37 | 0.02 | -0.67 | 0.1 | 0.07 | 0.2 | -0.29 | -0.16 | -0.11 | -0.73 | 0.15 | -0.325 | 0.075 | -0.1575 | 0.616024 | 0.252424 | 0.78106 | 0.989223 | 0.997988 | 0.995259 |
| Ppig_A2AR02_S413s_1 | Ppig_A2AR | 7.91 | 3.6 | 7.02 | 2.72 | 4.13 | 8.36 | 6.5 | 6.08 | 3.08 | 6.09 | -0.06 | 6.75 | 3.07 | -2.17 | 8.12 | 7.33 | 0.955 | -1.3475 | -1.225 | 0.595005 | 0.475299 | 0.449435 | 0.989223 | 0.997988 | 0.995259 |
| Ppig_A2AR02_S714s_1 | Ppig_A2AR | 0.9 | -0.4 | 0.36 | -0.06 | -1.46 | 0.64 | -1.27 | -0.37 | 1.5 | -0.67 | -4.47 | -0.03 | 2.22 | -1.4 | 0.72 | -1.05 | -0.815 | -1.1175 | -0.0775 | 0.921075 | 0.134147 | 0.340695 | 0.989223 | 0.997988 | 0.999507 |
| Ppig_A2AR02_T356t_1 | Ppig_A2AR | 0.52 | 0.33 | -0.02 | 0.02 | -0.27 | -0.62 | -0.1 | 0 | -0.7 | -1.03 | -0.94 | -0.67 | -0.72 | 0.08 | -0.15 | -0.83 | -0.46 | -1.0475 | -0.6175 | 0.070431 | 0.087683 | 0.001698 | 0.989223 | 0.997988 | 0.994666 |
| Ppip5k1_F6Q5C6_S140s_1 | Ppip5k1_F6 | 0.08 | 0.7 | -0.8 | -0.93 | -0.07 | 0.23 | -0.54 | -0.46 | 0.37 | 0.02 | -0.85 | -0.45 | -0.22 | 0.68 | -0.44 | -0.47 | 0.0275 | 0.01 | 0.125 | 0.784108 | 0.945984 | 0.982271 | 0.999187 | 0.997988 | 0.998711 |
| Ppip5k1_F6Q5C6_S160s_1 | Ppip5k1_F6 | 0.37 | 0.3 | 0.47 | 0.32 | -0.24 | -0.03 | 0.54 | 0.5 | -0.39 | -0.16 | 0.87 | -0.07 | -0.61 | 0.45 | -0.09 | 0.24 | -0.1725 | -0.3025 | -0.3675 | 0.235629 | 0.499914 | 0.346099 | 0.989223 | 0.997988 | 0.994666 |
| Ppip5k2_E9Q9J4_S1005s_1 | Ppip5k2_E9 | 0.07 | 1.46 | 1.63 | 0.55 | -0.67 | -0.03 | 0.7 | 0.07 | -0.49 | 0.82 | 1.41 | 0.05 | -1.55 | -0.48 | 0.48 | 0.33 | -0.91 | -0.48 | -1.2325 | 0.04743 | 0.061433 | 0.369912 | 0.989223 | 0.997988 | 0.994666 |
| Ppm1a_Q9EQE2_S135s_1 | Ppm1a_Q9 | -0.65 | -0.09 | -1.34 | -0.52 | 0.28 | -0.97 | -0.25 | -1.64 | -0.78 | -0.51 | -1.66 | -0.87 | -0.52 | -1.15 | -0.47 | -0.46 | 0.005 | -0.305 | 7.85E-17 | 1 | 0.991293 | 0.415162 | 0.999187 | 0.997988 | 1 |
| Ppm1a_Q9EQE2_S216s_1 | Ppm1a_Q9 | 0.78 | -0.65 | 0.72 | 0.73 | 0.07 | 0.42 | 1.47 | 1.19 | -1.16 | 0.89 | 2.86 | 0.1 | -0.91 | -0.56 | 0.65 | 0.87 | 0.3925 | 0.2775 | -0.3825 | 0.472509 | 0.38973 | 0.733571 | 0.989223 | 0.997988 | 0.994666 |
| Ppm1e_Q80TL0_S395s_1 | Ppm1e_Q8 | -1.23 | -1.81 | -0.02 | -0.44 | 0.11 | -0.57 | -0.18 | 0 | -2.12 | -2 | 0.01 | 0 | -1.92 | -0.63 | -0.62 | -0.8 | 0.715 | -0.1525 | -0.1175 | 0.808202 | 0.105507 | 0.815191 | 0.989223 | 0.997988 | 0.999507 |
| Ppm1e_Q80TL0_S532s_1 | Ppm1e_Q8 | -2 | -1.06 | -2.38 | -1.67 | -0.74 | -1.57 | -1.08 | -2.17 | -2.26 | -1.81 | -2.34 | -1.63 | -1.95 | -0.7 | -2.14 | -2.46 | 0.3875 | -0.2325 | -0.035 | 0.939351 | 0.34695 | 0.506679 | 0.989223 | 0.997988 | 0.999507 |
| Ppm1e_Q80TL0_S545s_1 | Ppm1e_Q8 | -2.51 | -2.71 | -2.19 | -2.87 | -1.9 | -2.39 | -2.41 | -2.12 | -3.73 | -3.15 | -0.78 | -2.19 | -2.36 | -1.39 | -2.92 | -2.63 | 0.365 | 0.1075 | 0.245 | 0.522222 | 0.164975 | 0.858927 | 0.989223 | 0.997988 | 0.994666 |
| Ppm1h_Q3UYC0_S118s_1 | Ppm1h_Q3 | 0.26 | 0.7 | 1.07 | 1.31 | 0.49 | 0.8 | 0.19 | 1.7 | 0.44 | 0.35 | 1.17 | 1.15 | 1.35 | 0.76 | 1.32 | 1.09 | -0.04 | -0.0575 | 0.295 | 0.36243 | 0.917647 | 0.865292 | 0.997335 | 0.997988 | 0.994666 |
| Ppm1h_Q3UYC0_S123s_1 | Ppm1h_Q3 | 2 | 0.55 | 0.47 | 0.9 | 1.14 | 1.8 | 1.01 | 1.44 | 1.73 | 2.29 | 0.89 | 0.84 | 1.21 | 2.3 | 0.85 | 0.89 | 0.3675 | 0.4575 | 0.3325 | 0.485161 | 0.350121 | 0.346021 | 0.989223 | 0.997988 | 0.994666 |
| Ppm1h_Q3UYC0_S197s_1 | Ppm1h_Q3 | -0.01 | 0.53 | -0.38 | -0.53 | 0.48 | 0.33 | 0.05 | 0.21 | 0.19 | 0.4 | -0.36 | -0.52 | -0.38 | 0.36 | -0.43 | -0.09 | 0.365 | 0.025 | -0.0375 | 0.910947 | 0.221159 | 0.941539 | 0.989223 | 0.997988 | 0.999507 |
| Ppm1h_Q3UYC0_S220s_1 | Ppm1h_Q3 | -0.8 | -2.02 | -1.34 | -1.08 | -1 | -1.01 | -1.52 | -0.72 | -1.04 | -1.4 | -0.42 | -0.94 | -1.03 | -0.6 | -1.5 | -1.32 | 0.2475 | 0.36 | 0.1975 | 0.5803 | 0.450887 | 0.314291 | 0.989223 | 0.997988 | 0.995259 |
| Ppm1h_Q3UYC0_S220s_2 | Ppm1h_Q3 | 0.18 | -2 | -2.05 | -2.38 | -2.44 | -0.05 | -2.13 | -1.92 | 0.61 | 0.42 | -2.49 | -3.37 | 0.48 | 0.67 | -1.74 | -0.99 | -0.0725 | 0.355 | 1.1675 | 0.137222 | 0.917723 | 0.731229 | 0.997335 | 0.997988 | 0.994666 |
| Ppm1h_Q3UYC0_T109t_1 | Ppm1h_Q3 | -0.74 | -0.81 | -1.24 | -1.13 | -0.04 | -0.33 | -1.15 | -1.22 | -0.08 | -0.94 | -1.29 | -0.98 | 1.63 | 1.21 | -0.44 | -0.47 | 0.295 | 0.1575 | 1.4625 | 0.018884 | 0.381107 | 0.621592 | 0.989223 | 0.997988 | 0.994666 |
| Ppm1h_Q3UYC0_T113t_1 | Ppm1h_Q3 | -0.29 | -0.6 | -1.68 | -1.14 | 0.38 | 0.37 | -0.9 | -1.14 | -0.19 | 0.25 | -1.12 | -1.02 | 0.34 | 0.3 | -0.86 | -1.15 | 0.605 | 0.4075 | 0.585 | 0.235683 | 0.218906 | 0.362756 | 0.989223 | 0.997988 | 0.994666 |
| Ppm1h_Q3UYC0_T223t_1 | Ppm1h_Q3 | -0.96 | -2.14 | -1.73 | -1.96 | -1.11 | -1.77 | -1.49 | -1.82 | -1.89 | -1.52 | -1.01 | -1.57 | -0.61 | -0.44 | -1.32 | -1.66 | 0.15 | 0.2 | 0.69 | 0.107335 | 0.642035 | 0.557681 | 0.989327 | 0.997988 | 0.994666 |
| Ppm1h_Q3UYC0_T223t_2 | Ppm1h_Q3 | 0.18 | -2 | -2.05 | -2.38 | -2.44 | -0.05 | -2.13 | -1.92 | 0.61 | 0.42 | -2.49 | -3.37 | 0.48 | 0.67 | -1.74 | -0.99 | -0.0725 | 0.355 | 1.1675 | 0.137222 | 0.917723 | 0.731229 | 0.997335 | 0.997988 | 0.994666 |
| Ppm1h_Q3UYC0_T361t_1 | Ppm1h_Q3 | 0.19 | 1.05 | 1.24 | -0.19 | 0.07 | 0.7 | 0.88 | 0.99 | -0.13 | 0.75 | -0.1 | 1.43 | -0.15 | -0.3 | 0.51 | 0.74 | 0.0875 | -0.085 | -0.3725 | 0.387575 | 0.821287 | 0.859689 | 0.995762 | 0.997988 | 0.994666 |
| Ppm1j_Q149T7_S92s_1 | Ppm1j_Q14 | 0.61 | 0.19 | 0.33 | 0.92 | -0.82 | 0.49 | -0.24 | 0.15 | 0.53 | 0.49 | 0.06 | -0.11 | 0.7 | -0.36 | 0.29 | 0.31 | -0.6175 | -0.27 | -0.2775 | 0.395515 | 0.089308 | 0.345397 | 0.989223 | 0.997988 | 0.994666 |
| Ppm1k_Q8BXN7_S248s_1 | Ppm1k_Q8 | 0.08 | -0.63 | -1.16 | -0.27 | 0.06 | -0.3 | 0.13 | 0.24 | -0.87 | -1.04 | -1.81 | 0.13 | -0.92 | -0.13 | -0.77 | 0.65 | 0.5275 | -0.4025 | 0.2025 | 0.645025 | 0.113519 | 0.390623 | 0.989223 | 0.997988 | 0.995259 |
| Ppm1m_Q8BU27_S42s_1 | Ppm1m_Q8 | -0.85 | -0.54 | -0.5 | -1.37 | -2.87 | 0.52 | -1.41 | -0.52 | 1.74 | 0.34 | -0.23 | -2.14 | 2.01 | -1.31 | -0.86 | -0.62 | -0.255 | 0.7425 | 0.62 | 0.382119 | 0.700373 | 0.331672 | 0.994975 | 0.997988 | 0.994666 |
| Ppme1_Q8BVQ5_S243s_1 | Ppme1_Q8 | 0.18 | 0.42 | 0.54 | 0.22 | -0.15 | -0.07 | 0.1 | 0.28 | -1.74 | 0.34 | -3.02 | 0.43 | -0.71 | -4 | 0.68 | 0.7 | -0.3 | -1.3375 | -1.1725 | 0.244112 | 0.190498 | 0.103678 | 0.989223 | 0.997988 | 0.994666 |
| Ppp1ca_P62137_T320t_1 | Ppp1ca_P6 | -0.79 | -0.31 | -1.1 | -0.89 | 0.38 | -0.71 | -0.04 | -0.62 | -1.17 | -0.6 | -1.32 | -0.72 | -0.51 | -0.67 | -0.29 | -0.79 | 0.525 | -0.18 | 0.2075 | 0.46318 | 0.126671 | 0.535773 | 0.989223 | 0.997988 | 0.994666 |
| Ppp1cb_P62141_T316t_1 | Ppp1cb_P6 | -0.25 | 0.22 | -0.3 | -0.14 | -0.18 | -0.48 | -0.03 | -0.09 | -1.22 | -0.16 | -2.41 | -0.24 | -0.26 | -2.15 | -0.56 | -0.55 | -0.0775 | -0.89 | -0.7625 | 0.106509 | 0.736517 | 0.105227 | 0.995762 | 0.997988 | 0.994666 |
| Ppp1r10_Q80W00_S313s_1 | Ppp1r10_C | -3.43 | -4.81 | -4.05 | -12.8 | -2.86 | -3.65 | -3.23 | -4.31 | -3.81 | -3.48 | -1.3 | -3.85 | -1.89 | -2.07 | -2.9 | -3.33 | 2.76 | 3.1625 | 3.725 | 0.074654 | 0.16786 | 0.13605 | 0.989223 | 0.997988 | 0.994666 |
| Ppp1r11_G3UZ30_T103t_1 | Ppp1r11_G | -0.1 | 0.68 | 0.57 | 0.26 | 0.09 | -0.03 | 0.83 | 0.44 | 0.08 | 0.4 | -0.11 | 0.82 | 0.1 | 0.35 | 0.85 | 0.7 | -0.02 | -0.055 | 0.1475 | 0.628358 | 0.945097 | 0.856955 | 0.999187 | 0.997988 | 0.995259 |
| Ppp1r12a_Q9DBR7_S299s_1 | Ppp1r12a_ | 0.13 | 0.1 | -0.05 | 0.02 | 0.01 | -0.25 | 0.23 | -0.19 | 0.05 | 0.23 | -0.21 | 0 | -0.41 | 0.25 | -0.14 | -0.25 | -0.1 | -0.0325 | -0.1875 | 0.46736 | 0.64083 | 0.880949 | 0.989223 | 0.997988 | 0.994666 |
| Ppp1r12a_Q9DBR7_S336s_1 | Ppp1r12a_ | -0.48 | -0.38 | -1.19 | -1.19 | -0.28 | -0.85 | -1.65 | -0.95 | -1.85 | -0.01 | 0.21 | -0.85 | -0.92 | 0.12 | -1.27 | -0.9 | -0.1225 | 0.185 | 0.0675 | 0.860796 | 0.731855 | 0.706067 | 0.995762 | 0.997988 | 0.999507 |
| Ppp1r12a_Q9DBR7_S422s_1 | Ppp1r12a_ | 1.68 | 1.31 | 1.79 | 2.07 | 0.05 | 1.86 | 0.72 | 1.53 | 2.22 | 1.91 | 2.08 | 1.7 | 1.89 | 0.39 | 1.77 | 1.9 | -0.6725 | 0.265 | -0.225 | 0.580697 | 0.130928 | 0.32158 | 0.989223 | 0.997988 | 0.995259 |
| Ppp1r12a_Q9DBR7_S445s_1 | Ppp1r12a_ | -0.73 | -0.28 | 0.29 | 0.12 | -0.28 | -0.53 | 0.34 | 0.22 | -0.89 | -0.03 | 0.81 | 0.43 | -0.66 | -0.59 | 0.21 | 0.87 | 0.0875 | 0.23 | 0.1075 | 0.800316 | 0.785342 | 0.588422 | 0.995762 | 0.997988 | 0.999507 |
| Ppp1r12a_Q9DBR7_T596t_1 | Ppp1r12a_ | -0.5 | -0.84 | -1.09 | -1.09 | -1.12 | -0.12 | -1.75 | -1.21 | 1.32 | -0.56 | -1.47 | -0.08 | 1.13 | -0.88 | -0.3 | -0.77 | -0.17 | 0.6825 | 0.675 | 0.170248 | 0.642337 | 0.237907 | 0.989327 | 0.997988 | 0.994666 |
| Ppp1r12b_Q8BG95_S29s_1 | Ppp1r12b_ | -0.26 | -0.58 | -0.82 | 0.12 | -0.79 | -0.68 | -0.96 | -0.28 | -0.68 | 0.68 | -0.7 | -1.15 | -0.49 | -0.44 | -0.72 | -1.2 | -0.2925 | -0.0775 | -0.3275 | 0.315512 | 0.316215 | 0.857848 | 0.989223 | 0.997988 | 0.994666 |
| Ppp1r12b_Q8BG95_S446s_1 | Ppp1r12b_ | 0.33 | -0.43 | 0.17 | 0.37 | -0.7 | 0.07 | 0.39 | 0.09 | 0.11 | -0.35 | 0.61 | 0.13 | 0.19 | 0.41 | 0.44 | -0.48 | -0.1475 | 0.015 | 0.03 | 0.926711 | 0.640982 | 0.960854 | 0.989223 | 0.997988 | 0.999507 |
| Ppp1r12b_Q8BG95_S729s_1 | Ppp1r12b_ | -1.33 | -1.04 | -1.4 | -1.86 | -0.83 | -1.33 | -0.82 | -1.1 | -1.14 | -1.18 | 1.11 | -1.22 | -0.2 | -0.17 | -0.52 | -1.22 | 0.3875 | 0.8 | 0.88 | 0.024556 | 0.158233 | 0.172092 | 0.989223 | 0.997988 | 0.994666 |
| Ppp1r12b_Q8BG95_T444t_1 | Ppp1r12b_ | 0.5 | -0.01 | 0.23 | 0.5 | -0.12 | 0.39 | 0.47 | 0.48 | 0.24 | 0.3 | 0.79 | 0.5 | 0.35 | 0.39 | 0.36 | -0.02 | 5.89E-17 | 0.1525 | -0.035 | 0.892267 | 1 | 0.545475 | 1 | 0.997988 | 0.999507 |
| Ppp1r12c_F6XWD4_S324s_1 | Ppp1r12c_I | 0.64 | -0.42 | 0.14 | 0.68 | 0.61 | 0.5 | 0.4 | 0.03 | -0.15 | 0.61 | -0.69 | 0.29 | -0.69 | -0.59 | 0.33 | 0.04 | 0.125 | -0.245 | -0.4875 | 0.213196 | 0.6856 | 0.528838 | 0.994624 | 0.997988 | 0.994666 |
| Ppp1r12c_F6XWD4_S569s_1 | Ppp1r12c_I | -0.16 | 0.13 | 0.65 | 0.63 | -0.16 | -0.29 | 0.26 | 0.45 | 0.09 | 0.07 | 1.1 | 0.09 | 0.36 | 0.29 | 0.65 | -0.28 | -0.2475 | 0.025 | -0.0575 | 0.858769 | 0.406451 | 0.941591 | 0.989223 | 0.997988 | 0.999507 |
| Ppp1r13b_Q62415_S335s_1 | Ppp1r13b_ | -0.22 | 0.17 | -0.56 | 0.03 | -0.3 | -0.34 | -0.6 | -2.22 | -0.44 | -0.43 | 0.47 | 0.58 | -1.03 | -0.52 | -0.29 | -0.31 | -0.72 | 0.19 | -0.3925 | 0.208587 | 0.135821 | 0.579981 | 0.989223 | 0.997988 | 0.994666 |
| Ppp1r14a_Q91VC7_S136s_1 | Ppp1r14a_ | 0.29 | 0.35 | 0.13 | 0.23 | 0.38 | 0.09 | 0.12 | 0.16 | -0.22 | 0.14 | -0.41 | 0.09 | -0.44 | 0.35 | 0.18 | 0.27 | -0.0625 | -0.35 | -0.16 | 0.560154 | 0.756561 | 0.156748 | 0.995762 | 0.997988 | 0.995259 |
| Ppp1r14a_Q91VC7_S26s_1 | Ppp1r14a_ | 0.5 | 0.19 | 0.65 | 0.73 | 0.6 | 0.05 | -1.24 | -0.22 | 0.77 | 0.59 | 1.54 | 0.39 | 1.22 | 0.49 | 0.39 | 0.77 | -0.72 | 0.305 | 0.2 | 0.495984 | 0.090933 | 0.34057 | 0.989223 | 0.997988 | 0.994666 |
| Ppp1r16a_Q923M0_S353s_1 | Ppp1r16a_ | -4.54 | -4.76 | -3.71 | -4.96 | -2.37 | -4.88 | -4.05 | -5.3 | -5.12 | -4.08 | 0.6 | -4.2 | -2.59 | -2.22 | -3.68 | -4.09 | 0.3425 | 1.2925 | 1.3475 | 0.020708 | 0.587332 | 0.280563 | 0.989223 | 0.997988 | 0.994666 |
| Ppp1r18_Q8BQ30_S146s_1 | Ppp1r18_C | 0.79 | 1.38 | 0.81 | 0.78 | -0.25 | 1.32 | 0.26 | 0.88 | 1.65 | 1.77 | 1.45 | 1.02 | 1.55 | 0 | 1.37 | 0.85 | -0.3875 | 0.5325 | 0.0025 | 0.994861 | 0.308686 | 0.078847 | 0.989223 | 0.997988 | 1 |
| Ppp1r18_Q8BQ30_S194s_1 | Ppp1r18_C | -0.32 | 0.46 | 0.99 | -0.12 | 1 | 0.68 | -0.55 | 0.23 | -0.71 | 0.2 | -0.59 | 0.84 | -0.21 | 0.16 | 0.33 | 0.25 | 0.0875 | -0.3175 | -0.12 | 0.732018 | 0.836638 | 0.487228 | 0.995762 | 0.997988 | 0.9971 |
| Ppp1r18_Q8BQ30_S234s_1 | Ppp1r18_C | -1.68 | -0.31 | -2.25 | -0.57 | -0.83 | -0.72 | -1.22 | -3.06 | 0.67 | -1.29 | -4.45 | -2.05 | 0.72 | -2.74 | -0.49 | -2.33 | -0.255 | -0.5775 | -0.0075 | 0.992687 | 0.688256 | 0.573651 | 0.994672 | 0.997988 | 1 |
| Ppp1r1a_Q9ERT9_S47s_1 | Ppp1r1a_Q | 0.23 | 1.04 | 0.11 | 0.58 | 0.5 | 0.77 | 0.85 | 0.3 | -0.07 | 0.15 | 0.07 | 0.28 | -0.12 | 0.72 | 0.55 | 1.02 | 0.115 | -0.3825 | 0.0525 | 0.880483 | 0.68236 | 0.188059 | 0.994114 | 0.997988 | 0.999507 |
| Ppp1r1a_Q9ERT9_S67s_1 | Ppp1r1a_Q | -2.2 | -4.01 | -3.17 | -4.01 | -1.67 | -1.86 | -2 | -2 | -3.15 | -2.96 | -0.19 | -2.59 | -2.69 | -1.61 | -3.08 | -2.93 | 1.465 | 1.125 | 0.77 | 0.156316 | 0.005737 | 0.148073 | 0.989223 | 0.997988 | 0.994666 |
| Ppp1r1a_Q9ERT9_S6s_1 | Ppp1r1a_Q | -1.23 | -1.7 | -1.08 | -1.41 | -1.18 | -1.49 | -1.67 | -1.16 | -0.55 | -1.03 | 0.13 | -1.24 | 0.37 | -0.87 | -0.89 | -1.5 | -0.02 | 0.6825 | 0.6325 | 0.151496 | 0.934614 | 0.075655 | 0.99868 | 0.997988 | 0.994666 |
| Ppp1r1b_Q60829_S10s_1 | Ppp1r1b_C | -0.51 | -0.7 | -0.02 | -0.47 | -1.42 | -1.17 | -0.95 | -0.39 | 0.17 | -0.47 | 0.51 | -0.89 | 1.02 | -0.73 | 0.7 | -0.09 | -0.5575 | 0.255 | 0.65 | 0.145186 | 0.081062 | 0.482669 | 0.989223 | 0.997988 | 0.994666 |
| Ppp1r1b_Q60829_S10s_2 | Ppp1r1b_C | -0.58 | -1.25 | -2.57 | -0.8 | -1.98 | -0.97 | -1.84 | -1.25 | -0.21 | -1.14 | -0.47 | -2.11 | -1.18 | -1.93 | -1.53 | -1.88 | -0.21 | 0.3175 | -0.33 | 0.481252 | 0.655856 | 0.579037 | 0.990434 | 0.997988 | 0.994666 |
| Ppp1r1b_Q60829_S135s_1 | Ppp1r1b_C | 4.69 | 2.48 | 3.08 | 1.91 | 0.98 | 5.33 | 2.08 | 3.07 | 1.76 | 2.39 | 1.74 | 3.06 | -2.42 | -1.84 | 2.11 | 2.49 | -0.175 | -0.8025 | -2.955 | 0.035281 | 0.854674 | 0.215037 | 0.996349 | 0.997988 | 0.994666 |
| Ppp1r1b_Q60829_S156s_1 | Ppp1r1b_C | -0.52 | -0.59 | -1.03 | -0.58 | -0.85 | -1.43 | 0.22 | -0.43 | -0.77 | -0.5 | -0.67 | -1.21 | -0.12 | -0.6 | -0.03 | -0.35 | 0.0575 | -0.1075 | 0.405 | 0.152104 | 0.874394 | 0.679279 | 0.996493 | 0.997988 | 0.994666 |
| Ppp1r1b_Q60829_S61s_1 | Ppp1r1b_C | -0.02 | 0.1 | 0.09 | 0.22 | 0.31 | 0.13 | 0.4 | 0.17 | -0.27 | 0.51 | -0.01 | 0.2 | -0.46 | 0.37 | -0.27 | 0 | 0.155 | 0.01 | -0.1875 | 0.496756 | 0.447642 | 0.967872 | 0.989223 | 0.997988 | 0.994666 |
| Ppp1r1b_Q60829_S9s_1 | Ppp1r1b_C | -1.53 | -2.49 | -1.81 | -1.33 | -2.97 | -2.81 | -2.51 | -1.94 | -0.64 | -2.23 | -0.87 | -2.09 | 0 | -1.81 | 0.36 | -2.07 | -0.7675 | 0.3325 | 0.91 | 0.158802 | 0.047934 | 0.476901 | 0.989223 | 0.997988 | 0.994666 |
| Ppp1r1b_Q60829_S9s_2 | Ppp1r1b_C | -0.38 | -1.22 | -2.57 | -0.8 | -1.96 | -0.97 | -1.84 | -1.25 | -0.1 | -0.99 | -0.47 | -2.11 | -1.14 | -1.96 | -1.53 | -1.88 | -0.2625 | 0.325 | -0.385 | 0.436048 | 0.593251 | 0.586168 | 0.989223 | 0.997988 | 0.994666 |
| Ppp1r1b_Q60829_T39t_1 | Ppp1r1b_C | -0.85 | 0.51 | -0.9 | -0.78 | -0.5 | -0.46 | -0.78 | -1.36 | -1.06 | 0.44 | -0.62 | -0.83 | -0.61 | 0.81 | -0.89 | -1.08 | -0.27 | -0.0125 | 0.0625 | 0.903088 | 0.490398 | 0.978072 | 0.989223 | 0.997988 | 0.999507 |
| Ppp1r2_D3Z3A0_S122s_1 | Ppp1r2_D3 | 1.28 | 1.42 | -0.24 | -0.42 | 0.86 | 0.63 | -0.35 | -0.42 | 0.96 | 1.46 | -1.74 | -0.91 | 0.6 | 1.38 | 0.09 | 0.36 | -0.33 | -0.5675 | 0.0975 | 0.852555 | 0.540529 | 0.486519 | 0.989223 | 0.997988 | 0.999507 |
| Ppp1r2_D3Z3A0_S122s_2 | Ppp1r2_D3 | -2.03 | 0.2 | -0.6 | -0.49 | -0.19 | -0.6 | -0.84 | -0.56 | -0.6 | 0.63 | -0.02 | -1.03 | -1.02 | -1.21 | -0.73 | -0.71 | 0.1825 | 0.475 | -0.1875 | 0.688342 | 0.688896 | 0.395963 | 0.994672 | 0.997988 | 0.996159 |
| Ppp1r2_D3Z3A0_S123s_1 | Ppp1r2_D3 | -0.41 | 0.38 | 0.29 | 0.72 | -0.59 | 0.18 | 0.51 | 0.76 | -0.6 | -0.5 | 1.41 | -0.07 | -0.95 | 0.17 | 1.34 | 1.12 | -0.03 | -0.185 | 0.175 | 0.743934 | 0.935552 | 0.708013 | 0.99868 | 0.997988 | 0.9971 |
| Ppp1r2_D3Z3A0_S123s_2 | Ppp1r2_D3 | -2.03 | 0.2 | -0.6 | -0.49 | -0.19 | -0.6 | -0.84 | -0.56 | -0.6 | 0.63 | -0.02 | -1.03 | -1.02 | -1.21 | -0.73 | -0.71 | 0.1825 | 0.475 | -0.1875 | 0.688342 | 0.688896 | 0.395963 | 0.994672 | 0.997988 | 0.996159 |
| Ppp1r2_D3Z3A0_S23s_1 | Ppp1r2_D3 | -0.92 | -0.54 | -0.73 | -0.55 | -0.67 | -0.87 | -0.22 | -0.96 | -0.94 | -0.15 | -1.21 | -0.79 | -0.62 | -0.15 | -0.4 | -0.49 | 0.005 | -0.0875 | 0.27 | 0.293979 | 0.983833 | 0.762024 | 0.999187 | 0.997988 | 0.994666 |
| Ppp1r21_Q3TDD9_S101s_1 | Ppp1r21_C | 1.13 | -0.84 | 0.77 | -0.18 | -1.16 | 2.17 | 0.13 | 0.03 | 1.23 | 0.45 | -0.23 | 0.96 | 0.39 | -1.36 | -0.42 | -0.57 | 0.0725 | 0.3825 | -0.71 | 0.205521 | 0.920221 | 0.465763 | 0.997335 | 0.997988 | 0.994666 |
| Ppp1r21_Q3TDD9_T652t_1 | Ppp1r21_C | 1.42 | 0.06 | -0.15 | 0.02 | 0.4 | 0.91 | 0.07 | 0.34 | 1.54 | 1.49 | 0.27 | 0.26 | 0.26 | 1.08 | -0.21 | -0.46 | 0.0925 | 0.5525 | -0.17 | 0.721707 | 0.813047 | 0.272234 | 0.995762 | 0.997988 | 0.9971 |
| Ppp1r3d_A2AJW4_S54s_2 | Ppp1r3d_A | 0.13 | -0.41 | 0.01 | 0.09 | 0.3 | 0.09 | -0.08 | 0.23 | -0.93 | -0.11 | 0.1 | 0.04 | -1.76 | 0.37 | 0.09 | -0.05 | 0.18 | -0.18 | -0.2925 | 0.543431 | 0.437176 | 0.559786 | 0.989223 | 0.997988 | 0.994666 |
| Ppp1r3d_A2AJW4_S58s_1 | Ppp1r3d_A | 0.87 | 1.49 | 1.17 | 0.76 | 1.17 | 0.85 | 0.47 | 0.98 | 1.47 | 1.54 | 1.7 | 1.1 | 1.7 | 1.38 | 1.47 | 0.92 | -0.205 | 0.38 | 0.295 | 0.332104 | 0.447916 | 0.178943 | 0.989223 | 0.997988 | 0.994666 |
| Ppp1r3d_A2AJW4_S58s_2 | Ppp1r3d_A | -1.67 | -1.63 | -1.09 | -1.28 | -0.83 | -1.7 | -1.17 | -1.68 | -2.56 | -1.7 | -0.94 | -1.3 | -2.28 | -0.99 | -1.05 | -1.61 | 0.0725 | -0.2075 | -0.065 | 0.855665 | 0.799868 | 0.587677 | 0.995762 | 0.997988 | 0.999507 |
| Ppp1r3d_A2AJW4_T57t_2 | Ppp1r3d_A | 1.98 | 1.04 | 0.64 | 0.8 | 1.64 | 1.96 | 0.51 | 1.13 | 0.77 | 1.55 | 0.76 | 0.77 | -1.03 | 1.92 | 0.73 | 0.76 | 0.195 | -0.1525 | -0.52 | 0.408112 | 0.640436 | 0.679074 | 0.989223 | 0.997988 | 0.994666 |
| Ppp1r3e_H3BJB2_S16s_1 | Ppp1r3e_H | -1.67 | -2.03 | -1.52 | -8.62 | -0.48 | -2.04 | -1.24 | -3.52 | -1.97 | -2.29 | -0.94 | -3.03 | -1.69 | -1.31 | -1.83 | -1.43 | 1.64 | 1.4025 | 1.895 | 0.221438 | 0.312221 | 0.376756 | 0.989223 | 0.997988 | 0.994666 |
| Ppp1r3g_Q9CW07_S79s_1 | Ppp1r3g_Q | -1.4 | -2.03 | -0.92 | -1.83 | -0.94 | -1.9 | -1.21 | -1.82 | -1.67 | -2.11 | -1.86 | -1.91 | -1.32 | -0.41 | -2.14 | -0.97 | 0.0775 | -0.3425 | 0.335 | 0.445547 | 0.821871 | 0.272714 | 0.995762 | 0.997988 | 0.994666 |
| Ppp1r3g_Q9CW07_S81s_1 | Ppp1r3g_Q | -0.54 | -0.46 | -1.59 | -1.67 | -0.38 | -1.08 | -2.01 | -1.67 | -1.26 | -0.44 | -1.53 | -1.86 | -0.6 | -1 | -2.22 | -0.46 | -0.22 | -0.2075 | -0.005 | 0.991864 | 0.62874 | 0.634057 | 0.989223 | 0.997988 | 1 |
| Ppp1r7_Q3UM45_S12s_1 | Ppp1r7_Q3 | 1.24 | -0.29 | 0.48 | 0.87 | 0.17 | 1.09 | 0.43 | 1 | 0.65 | 0.06 | 0.51 | 0.57 | 0.26 | 0.72 | -0.07 | 0.05 | 0.0975 | -0.1275 | -0.335 | 0.39287 | 0.800207 | 0.726438 | 0.995762 | 0.997988 | 0.994666 |
| Ppp1r7_Q3UM45_S24s_1 | Ppp1r7_Q3 | 0.28 | 0.34 | -0.5 | -0.54 | -0.83 | 1.16 | -0.75 | -1.25 | 2.87 | 1.02 | 0.25 | -0.84 | 2.29 | -0.55 | -0.11 | -0.68 | -0.3125 | 0.93 | 0.3425 | 0.607328 | 0.561752 | 0.225759 | 0.989223 | 0.997988 | 0.995259 |
| Ppp1r7_Q3UM45_S24s_2 | Ppp1r7_Q3 | -1.3 | -0.68 | -1.55 | -1.96 | -2.31 | -0.73 | -2.09 | -2.2 | 0.87 | -1.05 | -0.35 | -1.35 | -0.2 | -2.16 | -1.29 | -1.8 | -0.46 | 0.9025 | 0.01 | 0.983398 | 0.300956 | 0.111591 | 0.989223 | 0.997988 | 1 |
| Ppp1r7_Q3UM45_S27s_1 | Ppp1r7_Q3 | -1.08 | -1.41 | -1.26 | -3 | -1.78 | -0.79 | -2.45 | -2.01 | 1.36 | -1.14 | 0.93 | -1.14 | 0.51 | -1.47 | -0.48 | -2.27 | -0.07 | 1.69 | 0.76 | 0.274843 | 0.891887 | 0.039916 | 0.996493 | 0.997988 | 0.994666 |
| Ppp1r7_Q3UM45_S27s_2 | Ppp1r7_Q3 | -1.3 | -0.68 | -1.55 | -1.96 | -2.31 | -0.73 | -2.09 | -2.2 | 0.87 | -1.05 | -0.35 | -1.35 | -0.2 | -2.16 | -1.29 | -1.8 | -0.46 | 0.9025 | 0.01 | 0.983398 | 0.300956 | 0.111591 | 0.989223 | 0.997988 | 1 |
| Ppp1r7_Q3UM45_S45s_1 | Ppp1r7_Q3 | 5.29 | 2.81 | 3.28 | 1.75 | 3.22 | 4.8 | 3.47 | 2.85 | 0.31 | 5.05 | 0.73 | 3.08 | -0.29 | 0.11 | 3.04 | 3.51 | 0.3025 | -0.99 | -1.69 | 0.137751 | 0.687706 | 0.405802 | 0.994672 | 0.997988 | 0.994666 |
| Ppp1r9a_H3BKQ7_S160s_1 | Ppp1r9a_H | 2.15 | 2.73 | 4.14 | 3.16 | 1.69 | 2.02 | 3.29 | 2.91 | 2.9 | 2.1 | 3.45 | 3.55 | 2.34 | 2.07 | 3.76 | 3.93 | -0.5675 | -0.045 | -0.02 | 0.972611 | 0.28446 | 0.928692 | 0.989223 | 0.997988 | 0.999507 |
| Ppp1r9a_H3BKQ7_S187s_1 | Ppp1r9a_H | 1.09 | 0.63 | 1.25 | 1.35 | 0.62 | 0.65 | 0.65 | 1.26 | 1.05 | 1.2 | 1.92 | 1.31 | 0.08 | 1.03 | 1.03 | 1.42 | -0.285 | 0.29 | -0.19 | 0.594511 | 0.300268 | 0.334731 | 0.989223 | 0.997988 | 0.995259 |
| Ppp1r9a_H3BKQ7_S192s_1 | Ppp1r9a_H | 0.99 | 0.34 | 0.68 | 0.88 | 1.25 | 0.88 | 0.65 | 1.11 | 0.31 | 0.3 | -0.13 | 0.4 | 0.34 | 0.43 | 0.09 | -0.02 | 0.25 | -0.5025 | -0.5125 | 0.080801 | 0.331553 | 0.074839 | 0.989223 | 0.997988 | 0.994666 |
| Ppp1r9a_H3BKQ7_S199s_1 | Ppp1r9a_H | 0.58 | 0.36 | 0.06 | 0.41 | 0.76 | 0.57 | 0.52 | 0.62 | 0.43 | 0.62 | -0.48 | 0.27 | 0.03 | 0.76 | -0.17 | 0.28 | 0.265 | -0.1425 | -0.1275 | 0.666016 | 0.236197 | 0.639285 | 0.989223 | 0.997988 | 0.996159 |
| Ppp1r9a_H3BKQ7_S372s_1 | Ppp1r9a_H | 1.09 | 1.29 | 1.37 | 1.98 | 0.9 | 1.04 | 1.32 | 1.7 | 0.64 | 1.22 | 1.62 | 1.26 | 0.26 | 0.81 | 1.58 | 1.21 | -0.1925 | -0.2475 | -0.4675 | 0.219358 | 0.512412 | 0.435834 | 0.989223 | 0.997988 | 0.994666 |
| Ppp1r9a_H3BKQ7_S841s_1 | Ppp1r9a_H | 0.74 | 0.62 | 0.69 | 0.44 | 0.51 | 0.82 | 0.28 | 0.42 | 0.78 | 0.71 | 0.74 | 0.43 | 0.89 | 0.49 | 0.48 | 0.49 | -0.115 | 0.0425 | -0.035 | 0.886048 | 0.603198 | 0.845764 | 0.989223 | 0.997988 | 0.999507 |
| Ppp1r9a_H3BKQ7_T371t_1 | Ppp1r9a_H | 1.07 | 1.24 | 1.51 | 1.52 | 1.03 | 0.92 | 0.77 | 1.81 | 0.7 | 0.99 | 1.96 | 1.27 | -0.25 | 0.68 | 1.18 | 1.48 | -0.2025 | -0.105 | -0.5625 | 0.182541 | 0.487608 | 0.743484 | 0.989223 | 0.997988 | 0.994666 |
| Ppp1r9b_Q6R891_S100s_1 | Ppp1r9b_C | -1.44 | -1.6 | -1.62 | -1.2 | -0.49 | -1.34 | -0.68 | -0.89 | -1.78 | -1.62 | -0.52 | -1.35 | -1.93 | -0.65 | -1.29 | -1.15 | 0.615 | 0.1475 | 0.21 | 0.522926 | 0.036623 | 0.650466 | 0.989223 | 0.997988 | 0.994666 |
| Ppp1r9b_Q6R891_S192s_1 | Ppp1r9b_C | -2.91 | -2.25 | -1.14 | -1.41 | -1.29 | -3.24 | -0.82 | -1 | -3.66 | -2.82 | 0.21 | -0.84 | -2.95 | -1.18 | -1.02 | -1.02 | 0.34 | 0.15 | 0.385 | 0.505174 | 0.583485 | 0.862583 | 0.989223 | 0.997988 | 0.994666 |
| Ppp1r9b_Q6R891_S205s_1 | Ppp1r9b_C | -1.71 | -1.45 | -2.22 | -2.62 | -1.34 | -1.73 | -1.52 | -2.79 | -2.23 | -2.27 | -1.24 | -2.17 | -1.94 | -0.72 | -2.45 | -2.86 | 0.155 | 0.0225 | 0.0075 | 0.988089 | 0.699893 | 0.951196 | 0.994893 | 0.997988 | 1 |
| Ppp1r9b_Q6R891_S382s_1 | Ppp1r9b_C | -2.73 | -2.49 | -1.53 | -0.74 | -2.54 | -2.57 | -0.7 | -1.97 | -1.88 | -3.07 | -0.32 | -0.35 | -1.32 | -3.23 | -0.64 | -1.16 | -0.0725 | 0.4675 | 0.285 | 0.665602 | 0.89866 | 0.523899 | 0.996493 | 0.997988 | 0.996159 |
| Ppp1r9b_Q6R891_S99s_1 | Ppp1r9b_C | -0.3 | -0.44 | -1.86 | -1.27 | 0.37 | -0.06 | -0.6 | -0.88 | -0.09 | -0.21 | -0.73 | -1.1 | -0.72 | 0.35 | -1.27 | -1.45 | 0.675 | 0.435 | 0.195 | 0.704995 | 0.144207 | 0.320348 | 0.989223 | 0.997988 | 0.996159 |
| Ppp1r9b_Q6R891_T193t_1 | Ppp1r9b_C | -4.43 | -3.2 | -2.62 | -2.56 | -1.92 | -4.22 | -1.81 | -2.5 | -6.27 | -4.5 | -0.87 | -2.19 | -4.04 | -1.33 | -2.99 | -2.8 | 0.59 | -0.255 | 0.4125 | 0.5233 | 0.359232 | 0.820331 | 0.989223 | 0.997988 | 0.994666 |
| Ppp2r5a_Q6PD03_S41s_1 | Ppp2r5a_Q | -3.19 | -2.99 | -1.56 | -1.28 | -3.61 | -4.61 | -2.1 | -0.93 | -1.07 | -3.28 | -1 | -0.59 | -0.76 | -2.94 | -2.23 | -1.82 | -0.5575 | 0.77 | 0.3175 | 0.603835 | 0.504975 | 0.289112 | 0.989223 | 0.997988 | 0.995259 |
| Ppp2r5d_Q91V89_S44s_1 | Ppp2r5d_C | -0.32 | 0.53 | 0.06 | -0.35 | -0.43 | -0.08 | -0.55 | -0.42 | -0.47 | 0.16 | 0.41 | 0.18 | 0.33 | -1.06 | 0.31 | -0.14 | -0.35 | 0.09 | -0.12 | 0.761037 | 0.21654 | 0.773443 | 0.989223 | 0.997988 | 0.9971 |
| Ppp2r5d_Q91V89_S565s_1 | Ppp2r5d_C | -2.22 | -1.96 | -3.87 | -4.69 | -2.12 | -3.09 | -2.2 | -3.32 | -2.58 | -1.55 | -0.48 | -3.23 | -2.44 | -1.47 | -3.66 | -3.29 | 0.5025 | 1.225 | 0.47 | 0.523177 | 0.441967 | 0.148853 | 0.989223 | 0.997988 | 0.994666 |
| Ppp2r5d_Q91V89_S80s_1 | Ppp2r5d_C | -0.18 | -0.73 | -0.41 | -0.38 | -0.2 | -0.71 | 0.42 | -0.28 | -0.89 | -0.39 | -0.74 | -0.07 | -0.99 | -1.51 | -0.17 | -0.22 | 0.2325 | -0.0975 | -0.2975 | 0.422887 | 0.430022 | 0.721406 | 0.989223 | 0.997988 | 0.994666 |
| Ppp2r5d_Q91V89_S81s_1 | Ppp2r5d_C | -0.43 | -0.97 | -1.66 | -2 | -0.41 | -1.08 | -0.9 | -1.8 | -1.14 | -0.53 | -1.25 | -1.54 | -0.95 | -1.24 | -1.36 | -1.38 | 0.2175 | 0.15 | 0.0325 | 0.931801 | 0.61446 | 0.712732 | 0.989223 | 0.997988 | 0.999507 |
| Ppp2r5e_Q3UZJ4_S32s_1 | Ppp2r5e_Q | -1.3 | -1.38 | -0.32 | -0.7 | -1.17 | -0.46 | -0.94 | -0.62 | -0.81 | -1.43 | -2.16 | -0.26 | 0.05 | -1.33 | -0.53 | -0.3 | 0.1275 | -0.24 | 0.3975 | 0.327708 | 0.687115 | 0.604457 | 0.994672 | 0.997988 | 0.994666 |
| Ppp2r5e_Q3UZJ4_T7t_1 | Ppp2r5e_Q | -2.04 | -0.95 | 1.32 | 1.6 | -0.27 | -2.53 | 1.33 | 1.04 | -1.81 | -1.58 | 1.21 | 1.33 | 0.62 | -0.33 | 0.82 | 1.75 | -0.09 | -0.195 | 0.7325 | 0.404509 | 0.933231 | 0.857162 | 0.998562 | 0.997988 | 0.994666 |
| Ppp3ca_P63328_S462s_1 | Ppp3ca_P6 | -1.14 | -0.64 | -0.63 | -0.89 | 0.06 | 0.45 | -1.16 | -1.03 | -0.83 | -0.43 | -1.38 | -1.45 | -0.38 | -1.24 | -0.22 | -0.69 | 0.405 | -0.1975 | 0.1925 | 0.538297 | 0.325407 | 0.523552 | 0.989223 | 0.997988 | 0.994666 |
| Ppp3ca_P63328_S469s_1 | Ppp3ca_P6 | 0.15 | -0.33 | -0.09 | 0.17 | 1.06 | 0.75 | 0.36 | 0.6 | -1.03 | -0.54 | -0.46 | -0.43 | -1.21 | 0.5 | -0.33 | -0.42 | 0.7175 | -0.59 | -0.34 | 0.384559 | 0.015401 | 0.041598 | 0.989223 | 0.997988 | 0.994666 |
| Ppp3ca_P63328_T427t_1 | Ppp3ca_P6 | -0.08 | -0.06 | -0.87 | -1.14 | 0.5 | -0.03 | 0.22 | -0.11 | -0.88 | -1.33 | -0.52 | -0.89 | -0.75 | -0.16 | -0.49 | -0.76 | 0.6825 | -0.3675 | -0.0025 | 0.994182 | 0.05739 | 0.296755 | 0.989223 | 0.997988 | 1 |
| Ppp3cb_E0CZ78_S116s_1 | Ppp3cb_E0 | 0.87 | 1.06 | 0.6 | 0.05 | 1.52 | 0.43 | 0.69 | 0.56 | 0.84 | 0.86 | 0.95 | 0.24 | 0.56 | 1 | 0.48 | 0.62 | 0.155 | 0.0775 | 0.02 | 0.947745 | 0.647263 | 0.801692 | 0.99033 | 0.997988 | 0.999507 |
| Ppp3cb_E0CZ78_S471s_1 | Ppp3cb_E0 | 0.13 | 0.42 | 0.48 | 0.2 | 0.19 | -0.28 | 0.25 | 0.2 | 0.32 | 0.38 | 0.83 | 0.33 | 0.01 | 0.61 | 0.04 | -0.09 | -0.2175 | 0.1575 | -0.165 | 0.543538 | 0.351732 | 0.511531 | 0.989223 | 0.997988 | 0.994666 |
| Ppp3cb_E0CZ78_S478s_1 | Ppp3cb_E0 | -1.26 | -2.02 | -2.3 | -2.04 | -0.49 | -0.6 | -1.22 | -1.32 | -2.03 | -1.25 | -0.73 | -2.38 | -1.89 | -1.01 | -1.86 | -1.95 | 0.9975 | 0.3075 | 0.2275 | 0.517616 | 0.011269 | 0.476052 | 0.989223 | 0.997988 | 0.994666 |
| Ppp4r4_Q8C0Y0_S777s_1 | Ppp4r4_Q8 | -1.89 | -1.83 | -0.24 | -0.16 | -0.92 | -0.78 | -0.47 | -0.13 | -1.36 | -0.89 | -0.14 | -0.85 | -0.84 | -0.58 | -1.15 | -0.55 | 0.455 | 0.22 | 0.25 | 0.603375 | 0.34794 | 0.66664 | 0.989223 | 0.997988 | 0.995259 |
| Ppp6r1_Q7TSI3_S739s_1 | Ppp6r1_Q7 | -0.32 | -0.07 | 0.34 | 0.03 | -0.48 | -0.84 | 0.31 | 0.09 | -0.9 | -0.22 | -0.33 | 0.62 | -0.57 | -0.15 | 0.23 | 0.27 | -0.225 | -0.2025 | -0.05 | 0.86768 | 0.479458 | 0.572156 | 0.989223 | 0.997988 | 0.999507 |
| Ppp6r1_Q7TSI3_S817s_1 | Ppp6r1_Q7 | 2.67 | 1.1 | 2.14 | 0.86 | 1.41 | 2.63 | 1.3 | 1.64 | 1.2 | 2.38 | 0.65 | 1.85 | 1.29 | 0.23 | 1.96 | 2.05 | 0.0525 | -0.1725 | -0.31 | 0.579366 | 0.913484 | 0.746102 | 0.997335 | 0.997988 | 0.995259 |
| Ppp6r1_Q7TSI3_S826s_1 | Ppp6r1_Q7 | 0.64 | 0.62 | 0.81 | 0.72 | 0.73 | 0.54 | 1.02 | 0.82 | 0.18 | 1.04 | 0.02 | 0.81 | 0.05 | 0.42 | 0.67 | 0.51 | 0.08 | -0.185 | -0.285 | 0.272878 | 0.704719 | 0.53161 | 0.995437 | 0.997988 | 0.994666 |
| Ppp6r2_G3X9K4_S435s_1 | Ppp6r2_G3 | -0.27 | -0.56 | -0.6 | -0.02 | 0.2 | 0.2 | 0.37 | 0.17 | -0.61 | -0.15 | 0.23 | -0.09 | -1.07 | -0.19 | 0.12 | 0.78 | 0.5975 | 0.2075 | 0.2725 | 0.511192 | 0.023181 | 0.45766 | 0.989223 | 0.997988 | 0.994666 |
| Ppp6r2_G3X9K4_S670s_1 | Ppp6r2_G3 | 0.67 | 0.74 | 0.61 | 0.6 | 0.41 | 0.3 | 0.44 | 0.6 | 0.28 | 1.08 | 0.61 | 0.66 | 0.46 | 0.84 | 0.2 | 0.14 | -0.2175 | 0.0025 | -0.245 | 0.35969 | 0.288376 | 0.991884 | 0.989223 | 0.997988 | 0.994666 |
| Ppp6r2_G3X9K4_S745s_1 | Ppp6r2_G3 | -0.67 | -1.16 | -1.7 | -2.58 | -1.53 | -1.96 | -3.43 | -1.41 | -0.11 | -0.35 | -1.98 | -2.52 | -1.56 | 0.29 | -2.63 | -2.9 | -0.555 | 0.2875 | -0.1725 | 0.815626 | 0.333085 | 0.662448 | 0.989223 | 0.997988 | 0.999507 |
| Ppp6r2_G3X9K4_S746s_1 | Ppp6r2_G3 | -4.23 | -3.25 | -3.94 | -3.04 | -2.62 | -1.19 | -3.2 | -2.35 | -2.44 | -2.14 | -0.52 | -3.16 | -1.25 | -2.7 | -3.15 | -2.67 | 1.275 | 1.55 | 1.1725 | 0.032783 | 0.021322 | 0.022615 | 0.989223 | 0.997988 | 0.994666 |
| Ppp6r2_G3X9K4_S760s_1 | Ppp6r2_G3 | -0.41 | -1.63 | 0.04 | -0.17 | -1.04 | -1.08 | -0.64 | -0.19 | 0.28 | -0.18 | -0.21 | -0.93 | -0.4 | -0.72 | -1.49 | -0.22 | -0.195 | 0.2825 | -0.165 | 0.716951 | 0.635377 | 0.522043 | 0.989223 | 0.997988 | 0.996966 |
| Ppp6r2_G3X9K4_S760s_2 | Ppp6r2_G3 | -1.98 | -2.23 | -2.32 | -1.26 | -1.65 | -1.68 | -1.66 | -2.57 | -1.85 | -1.64 | -0.14 | -1.78 | -0.44 | -0.74 | -2.14 | -2.47 | 0.0575 | 0.595 | 0.5 | 0.350431 | 0.864686 | 0.211093 | 0.996493 | 0.997988 | 0.994666 |
| Ppp6r2_G3X9K4_S820s_1 | Ppp6r2_G3 | 0.71 | 0.91 | 0.28 | 0.35 | 0.78 | 0.54 | 0.92 | 0.05 | 0.57 | 0.81 | 0.5 | 0.37 | 0.58 | 0.84 | 0.81 | 0.32 | 0.01 | -5.9E-17 | 0.075 | 0.78538 | 0.97138 | 1 | 0.999187 | 1 | 0.999507 |
| Prcc_Q9EQC8_S157s_1 | Prcc_Q9EQ | -3.45 | -2.55 | 0.72 | -1.3 | -2.67 | -2.77 | -0.66 | -1.49 | 0.39 | -2.05 | 0.2 | -2.37 | 0.1 | -3.46 | -1 | -0.69 | -0.2525 | 0.6875 | 0.3825 | 0.711362 | 0.778976 | 0.506727 | 0.995762 | 0.997988 | 0.996966 |
| Prex1_I7HPV9_S1021s_1 | Prex1_I7HF | 0.12 | 0.52 | 0.05 | 0.86 | 0.72 | 0.23 | 0.8 | 0.16 | 0.93 | 0.88 | 0.11 | 0.72 | 0.96 | 1.19 | 1.02 | 0.43 | 0.09 | 0.2725 | 0.5125 | 0.119636 | 0.751648 | 0.380829 | 0.995762 | 0.997988 | 0.994666 |
| Prex1_I7HPV9_S144s_1 | Prex1_I7HF | -2.87 | -2.21 | -4.15 | -1.94 | -1.09 | -2.6 | -1.64 | -2.94 | -4.59 | -1.97 | -4.15 | -2.45 | -3.09 | -2.41 | -2.57 | -2.28 | 0.725 | -0.4975 | 0.205 | 0.680782 | 0.2341 | 0.498014 | 0.989223 | 0.997988 | 0.996159 |
| Prex1_I7HPV9_S973s_1 | Prex1_I7HF | -2.85 | -2.49 | -2.7 | -2.14 | -3 | -2.12 | -2.61 | -2.36 | -2.35 | -1.94 | -2.46 | -1.97 | -0.41 | -2.44 | -2.93 | -3.12 | 0.0225 | 0.365 | 0.32 | 0.588323 | 0.935739 | 0.190947 | 0.99868 | 0.997988 | 0.995259 |
| Prickle1_Q3U5C7_S592s_1 | Prickle1_Q | 0.13 | 0.3 | 0.29 | -0.02 | 0.19 | 0.22 | 0.74 | 0.16 | 0 | 0.19 | 0.68 | 0.62 | -0.63 | 0.36 | 0.17 | -0.12 | 0.1525 | 0.1975 | -0.23 | 0.442102 | 0.514781 | 0.443756 | 0.989223 | 0.997988 | 0.994666 |
| Prickle1_Q3U5C7_S595s_1 | Prickle1_Q | -0.24 | 0.24 | 0.82 | 1.06 | 0.52 | -0.39 | 1.27 | 0.14 | 1.12 | -0.29 | 0.1 | 0.6 | 0.97 | -0.06 | -0.23 | 1.53 | -0.085 | -0.0875 | 0.0825 | 0.865514 | 0.843299 | 0.833414 | 0.996349 | 0.997988 | 0.999507 |
| Prickle2_Q80Y24_S319s_1 | Prickle2_Q | 1.71 | 0.4 | 0.34 | 0.32 | 0.32 | 1.55 | 1.22 | 0.04 | 0.09 | 1.44 | 0.25 | -0.1 | 0.4 | -0.2 | 1.44 | 0.34 | -0.2225 | 0.14 | 0.42 | 0.964349 | 0.593866 | 0.789174 | 0.989223 | 0.997988 | 0.994666 |
| Prickle2_Q80Y24_S321s_1 | Prickle2_Q | 0.32 | 0.4 | 0.78 | 0.67 | 0.05 | 0.32 | 1.01 | 0.5 | -0.81 | 0.56 | 0.64 | 0.07 | -0.57 | -1.13 | 0.76 | 0.76 | -0.0725 | -0.4275 | -0.5875 | 0.231982 | 0.789484 | 0.255395 | 0.995762 | 0.997988 | 0.994666 |
| Prickle2_Q80Y24_S322s_1 | Prickle2_Q | 0 | 0.56 | -0.01 | 0.02 | 0.91 | 0.41 | -0.56 | -1.19 | 1.07 | 0.65 | -1.26 | -1.05 | 0.89 | 0.12 | -1 | -0.11 | -0.25 | -0.29 | -0.1675 | 0.686924 | 0.588493 | 0.607085 | 0.989223 | 0.997988 | 0.996159 |
| Prickle2_Q80Y24_S322s_2 | Prickle2_Q | 0.32 | 0.4 | 0.78 | 0.67 | 0.05 | 0.32 | 1.01 | 0.5 | -0.81 | 0.56 | 0.64 | 0.07 | -0.57 | -1.13 | 0.76 | 0.76 | -0.0725 | -0.4275 | -0.5875 | 0.231982 | 0.789484 | 0.255395 | 0.995762 | 0.997988 | 0.994666 |
| Prickle2_Q80Y24_S396s_1 | Prickle2_Q | -1.83 | 0.43 | -0.16 | -0.73 | -0.03 | -0.16 | -1.23 | -0.59 | -0.73 | -0.29 | 1.41 | -0.56 | -0.75 | 0.68 | -0.22 | -0.67 | 0.07 | 0.53 | 0.3325 | 0.543178 | 0.889726 | 0.407962 | 0.996493 | 0.997988 | 0.994666 |
| Prickle2_Q80Y24_S414s_1 | Prickle2_Q | 0.23 | 0.28 | 0.13 | 0.74 | 0.25 | 0.6 | 0.62 | 0.05 | 0.09 | 0.6 | -0.33 | 0.22 | -0.1 | 0.35 | -0.16 | 0.62 | 0.035 | -0.2 | -0.1675 | 0.573773 | 0.888808 | 0.48704 | 0.996493 | 0.997988 | 0.995259 |
| Prickle2_Q80Y24_S463s_1 | Prickle2_Q | 0.08 | 0.91 | 1.59 | 0.06 | 0.18 | 2.38 | 0.45 | 1.79 | -0.21 | 1.46 | 1.85 | 1.43 | 1.01 | -0.44 | 2.15 | 2.36 | 0.54 | 0.4725 | 0.61 | 0.370753 | 0.361448 | 0.395947 | 0.989223 | 0.997988 | 0.994666 |
| Prickle2_Q80Y24_S544s_1 | Prickle2_Q | 0.14 | 0.59 | 0.48 | 0.47 | 0.69 | 0.31 | 0.71 | 0.4 | 0.2 | 0.61 | 0.21 | 0.94 | -0.03 | 0.87 | -0.08 | 0.5 | 0.1075 | 0.07 | -0.105 | 0.731018 | 0.632413 | 0.791854 | 0.989223 | 0.997988 | 0.9971 |
| Prickle2_Q80Y24_S572s_1 | Prickle2_Q | -1.67 | -2.02 | -1.54 | -2.8 | -0.57 | -1.76 | -1.09 | -1.88 | -2.92 | -2.03 | -0.81 | -1.95 | -2.03 | -0.55 | -2.94 | -2.18 | 0.6825 | 0.08 | 0.0825 | 0.877405 | 0.11388 | 0.869927 | 0.989223 | 0.997988 | 0.999507 |
| Prickle2_Q80Y24_S667s_1 | Prickle2_Q | 0.65 | 0.54 | 0.74 | 0.34 | 0.3 | 0.62 | 0.36 | 0.33 | 0.28 | 0.91 | 1.43 | 0.55 | 0.6 | 0.52 | 0.48 | 0.61 | -0.165 | 0.225 | -0.015 | 0.949126 | 0.444161 | 0.462732 | 0.989223 | 0.997988 | 0.999507 |
| Prickle2_Q80Y24_S696s_1 | Prickle2_Q | -0.62 | -0.74 | -0.55 | -0.08 | -0.41 | -0.54 | -0.59 | -0.36 | -0.7 | -1.19 | 0.26 | -0.43 | -0.65 | -0.66 | -0.28 | -0.14 | 0.0225 | -0.0175 | 0.065 | 0.814495 | 0.921981 | 0.96015 | 0.997335 | 0.997988 | 0.999507 |
| Prickle2_Q80Y24_S732s_1 | Prickle2_Q | -1.04 | -0.47 | -0.75 | -1.28 | -0.64 | -0.61 | -1.09 | -1.01 | -1.69 | -1.01 | -0.35 | -1.02 | -0.82 | -0.75 | -1.24 | -1.44 | 0.0475 | -0.1325 | -0.1775 | 0.559942 | 0.856363 | 0.700641 | 0.996349 | 0.997988 | 0.995259 |
| Prickle2_Q80Y24_S807s_1 | Prickle2_Q | -0.16 | 0.52 | 0.61 | -0.39 | -0.06 | 0.13 | 0.17 | -0.02 | -0.28 | 0.04 | -0.34 | -0.58 | -0.66 | 1.03 | -0.65 | -0.35 | -0.09 | -0.435 | -0.3025 | 0.513473 | 0.753623 | 0.18588 | 0.995762 | 0.997988 | 0.994666 |
| Prickle2_Q80Y24_T535t_1 | Prickle2_Q | -0.26 | 0.46 | -0.25 | 0.29 | 0.53 | 0 | 0.56 | 0.54 | 0.7 | 0.57 | 1.35 | 0.55 | 1.06 | 0.87 | -0.21 | 0.1 | 0.3475 | 0.7325 | 0.395 | 0.305132 | 0.220054 | 0.035233 | 0.989223 | 0.997988 | 0.994666 |
| Prickle2_Q80Y24_T535t_2 | Prickle2_Q | -1.69 | -0.25 | 0.11 | 0.95 | -0.79 | -0.94 | -0.52 | -1.46 | -0.01 | -0.73 | 1.84 | 1.24 | -0.3 | -0.1 | 0.85 | -0.1 | -0.7075 | 0.805 | 0.3075 | 0.586849 | 0.204073 | 0.281539 | 0.989223 | 0.997988 | 0.995259 |
| Prickle2_Q80Y24_T540t_2 | Prickle2_Q | -1.69 | -0.25 | 0.11 | 0.95 | -0.79 | -0.94 | -0.52 | -1.46 | -0.01 | -0.73 | 1.84 | 1.24 | -0.3 | -0.1 | 0.85 | -0.1 | -0.7075 | 0.805 | 0.3075 | 0.586849 | 0.204073 | 0.281539 | 0.989223 | 0.997988 | 0.995259 |
| Prickle2_Q80Y24_T806t_1 | Prickle2_Q | -0.16 | 0.52 | 0.83 | -0.42 | -0.14 | 0.18 | 0.23 | 0.23 | -0.16 | -0.05 | 0.42 | -0.6 | -0.77 | 1.05 | -0.13 | -0.23 | -0.0675 | -0.29 | -0.2125 | 0.648633 | 0.832519 | 0.437608 | 0.995762 | 0.997988 | 0.995259 |
| Prkaa1_Q3TUQ7_S487s_1 | Prkaa1_Q3 | -0.61 | -0.55 | 0.06 | -0.09 | -0.65 | -0.58 | -0.07 | 0.18 | -0.74 | -0.71 | 0.5 | -0.3 | 0.08 | -0.61 | -0.02 | -0.08 | 0.0175 | -0.015 | 0.14 | 0.635275 | 0.951911 | 0.965707 | 0.999187 | 0.997988 | 0.995259 |
| Prkaa1_Q3TUQ7_T174t_1 | Prkaa1_Q3 | -0.66 | 0.62 | -0.02 | -0.77 | -0.3 | -0.73 | 0.28 | -0.17 | -1.64 | -0.55 | -2.13 | -0.28 | -1.21 | -1.17 | -0.04 | 0.46 | -0.0225 | -0.9425 | -0.2825 | 0.573992 | 0.95217 | 0.091408 | 0.999187 | 0.997988 | 0.995259 |
| Prkaa1_Q3TUQ7_T481t_1 | Prkaa1_Q3 | -0.23 | -0.29 | 0.21 | -0.01 | -0.48 | -0.11 | -0.34 | -0.28 | 2.66 | -0.23 | -2.73 | -0.15 | 2.79 | -0.96 | 1.62 | -0.29 | -0.2225 | -0.0325 | 0.87 | 0.275387 | 0.329471 | 0.973466 | 0.989223 | 0.997988 | 0.994666 |
| Prkaa2_Q8BRK8_S481s_1 | Prkaa2_Q8 | -0.69 | -2.27 | -0.8 | -2.13 | -1.45 | -0.89 | -0.62 | -0.8 | 2.31 | -0.59 | -2.7 | -0.86 | 3.12 | -2.02 | 2.31 | -1.04 | 0.5325 | 1.0125 | 2.065 | 0.096559 | 0.23668 | 0.319542 | 0.989223 | 0.997988 | 0.994666 |
| Prkab1_Q9R078_S182s_1 | Prkab1_Q9 | 1.69 | 1.29 | 1.72 | 1.21 | 1.28 | 1.37 | 1.22 | 1.7 | -0.67 | 1.93 | 0.68 | 1.52 | -0.19 | -0.44 | 1.79 | 2.03 | -0.085 | -0.6125 | -0.68 | 0.275089 | 0.721537 | 0.279327 | 0.995437 | 0.997988 | 0.994666 |
| Prkab2_Q6PAM0_S183s_1 | Prkab2_Q6 | 0.59 | 0.87 | 0.96 | 0.68 | 0.89 | 0.48 | 1.14 | 0.87 | 0.8 | 0.73 | 0.8 | 0.85 | 0.02 | 0.58 | 0.76 | 0.69 | 0.07 | 0.02 | -0.2625 | 0.348979 | 0.764168 | 0.925347 | 0.995762 | 0.997988 | 0.994666 |
| Prkaca_P05132_T188t_1 | Prkaca_P05 | -0.48 | -0.27 | -0.76 | -0.98 | -0.01 | -0.27 | 0.05 | -0.77 | -0.38 | 0.25 | -0.83 | -0.46 | -0.78 | -0.03 | -0.14 | -0.14 | 0.3725 | 0.2675 | 0.35 | 0.254733 | 0.203582 | 0.395328 | 0.989223 | 0.997988 | 0.994666 |
| Prkaca_P05132_T190t_1 | Prkaca_P05 | -0.32 | -0.42 | -0.66 | -0.25 | 0.07 | -0.1 | 0.49 | -0.64 | -0.63 | 0.12 | -1.08 | -0.67 | -0.91 | 0.07 | -0.13 | -0.3 | 0.3675 | -0.1525 | 0.095 | 0.747909 | 0.21593 | 0.617355 | 0.989223 | 0.997988 | 0.9971 |
| Prkag2_Q91WG5_S122s_1 | Prkag2_Q9 | -2.45 | -0.59 | 0.44 | -1.78 | 0.09 | -1.92 | -0.16 | -0.4 | -4.93 | -1.7 | -1.59 | -2.59 | -2.95 | -0.3 | -0.55 | -0.31 | 0.4975 | -1.6075 | 0.0675 | 0.932824 | 0.478023 | 0.098103 | 0.989223 | 0.997988 | 0.999507 |
| Prkag2_Q91WG5_S196s_1 | Prkag2_Q9 | 0.17 | 0.97 | 0.49 | 0.46 | 0.68 | 0.11 | 0.36 | 0.33 | 0.25 | 0.92 | 0.16 | 0.35 | 0.6 | 0.21 | 0.49 | 0.28 | -0.1525 | -0.1025 | -0.1275 | 0.642902 | 0.554369 | 0.720801 | 0.989223 | 0.997988 | 0.995259 |
| Prkag2_Q91WG5_S87s_1 | Prkag2_Q9 | 0.35 | 0.14 | 0.12 | 0.36 | -0.19 | 0.62 | 0.47 | 0.62 | 0.43 | 0.37 | 0.91 | 0.8 | -0.35 | 0.99 | 0.16 | -0.22 | 0.1375 | 0.385 | -0.0975 | 0.776263 | 0.593974 | 0.130115 | 0.989223 | 0.997988 | 0.99798 |
| Prkar1a_Q9DBC7_S5s_1 | Prkar1a_Q | 1.15 | 1.51 | -0.82 | 0.34 | -1.71 | 0.87 | -1.39 | -0.24 | 3.17 | 0.33 | -0.78 | -0.68 | 2.58 | 0.76 | 0.57 | -0.45 | -1.1625 | -0.035 | 0.32 | 0.660951 | 0.117734 | 0.970112 | 0.989223 | 0.997988 | 0.995671 |
| Prkar1a_Q9DBC7_S77s_1 | Prkar1a_Q | 0.48 | 0.16 | 1 | 0.58 | 0.36 | 0.48 | 0.58 | 0.06 | 0.19 | 0.37 | 0.36 | 0.63 | -0.07 | 0.23 | 0.75 | 0.51 | -0.185 | -0.1675 | -0.2 | 0.51713 | 0.47878 | 0.52574 | 0.989223 | 0.997988 | 0.994666 |
| Prkar1a_Q9DBC7_S83s_1 | Prkar1a_Q | 0.44 | 0.86 | 0.3 | 0.42 | 0.54 | 0.37 | 0.52 | 0.45 | 0.29 | 0.74 | 0.27 | 0.4 | 0.36 | 0.7 | 0.51 | 0.57 | -0.035 | -0.08 | 0.03 | 0.905246 | 0.872499 | 0.743816 | 0.996493 | 0.997988 | 0.999507 |
| Prkar1a_Q9DBC7_T75t_1 | Prkar1a_Q | 0.78 | 1.24 | 0.77 | 1.33 | 1.73 | 1.15 | 0.73 | 0.22 | 0.7 | 1.47 | 0.06 | 0.99 | -0.16 | 0.45 | 1.13 | 1.31 | -0.0725 | -0.225 | -0.3475 | 0.373655 | 0.837545 | 0.521394 | 0.995762 | 0.997988 | 0.994666 |
| Prkar1b_P12849_S3s_1 | Prkar1b_P1 | -1.09 | -0.19 | -1.69 | -0.41 | -0.02 | -1.15 | -1.07 | -1.33 | -0.63 | -0.64 | -0.98 | -0.67 | -0.78 | 0.4 | -0.47 | -0.65 | -0.0475 | 0.115 | 0.47 | 0.287647 | 0.911294 | 0.751029 | 0.997335 | 0.997988 | 0.994666 |
| Prkar1b_P12849_S83s_1 | Prkar1b_P1 | -0.89 | -0.4 | 0.2 | 0.46 | -0.55 | -0.14 | 0.56 | 0.69 | -0.93 | -0.72 | 0.02 | 0.15 | -0.79 | -0.34 | 0.45 | 0.68 | 0.2975 | -0.2125 | 0.1575 | 0.724811 | 0.468693 | 0.599377 | 0.989223 | 0.997988 | 0.9971 |
| Prkar1b_P12849_S9s_1 | Prkar1b_P1 | -0.95 | -0.01 | -0.13 | -0.65 | -0.4 | -0.25 | 0.45 | -0.43 | -0.9 | -0.24 | -0.88 | -0.43 | -0.64 | -0.14 | -0.11 | -0.01 | 0.2775 | -0.1775 | 0.21 | 0.508233 | 0.39258 | 0.5701 | 0.989223 | 0.997988 | 0.994666 |
| Prkar2a_Q8K1M3_S381s_1 | Prkar2a_Q8 | 0.9 | 1.1 | -0.33 | -1.15 | 1.06 | -0.57 | -0.37 | -1.2 | -0.06 | 0.51 | 0.97 | 0.58 | -0.25 | 0.08 | -0.52 | 1.03 | -0.4 | 0.37 | -0.045 | 0.937895 | 0.532935 | 0.493099 | 0.989223 | 0.997988 | 0.999507 |
| Prkar2a_Q8K1M3_S97s_1 | Prkar2a_Q8 | -1.01 | -0.44 | -1.07 | -1.55 | -0.45 | -1.13 | -1.19 | -1.54 | -1.03 | -0.91 | -0.7 | -2.05 | -1.43 | -0.49 | -1.45 | -1.3 | -0.06 | -0.155 | -0.15 | 0.670759 | 0.856366 | 0.685079 | 0.996349 | 0.997988 | 0.996159 |
| Prkar2b_P31324_S112s_1 | Prkar2b_P3 | -0.96 | -0.74 | -1.13 | -1.24 | -0.97 | -0.82 | -0.75 | -1.22 | -1.42 | -1.05 | -0.68 | -1.58 | -1.24 | -1.34 | -0.95 | -0.84 | 0.0775 | -0.165 | -0.075 | 0.773595 | 0.734694 | 0.559522 | 0.995762 | 0.997988 | 0.99798 |
| Prkca_Q4VA93_S13s_1 | Prkca_Q4V | 0.9 | 0.84 | -0.13 | -0.69 | 0.23 | 0.23 | 0.15 | -0.66 | -0.43 | 0.53 | 0.63 | -0.01 | -0.84 | 0.22 | 0.01 | -0.48 | -0.2425 | -0.05 | -0.5025 | 0.274881 | 0.567797 | 0.910166 | 0.989223 | 0.997988 | 0.994666 |
| Prkca_Q4VA93_S226s_1 | Prkca_Q4V | 0.34 | 0.55 | 0.5 | 0.71 | 0.77 | 0.53 | 1.02 | 0.73 | -0.04 | 0.16 | -0.42 | 0.51 | -0.37 | 0.48 | 0.33 | 0.56 | 0.2375 | -0.4725 | -0.275 | 0.358834 | 0.289961 | 0.104519 | 0.989223 | 0.997988 | 0.994666 |
| Prkca_Q4VA93_S319s_1 | Prkca_Q4V | 1.12 | 0.9 | 1.19 | 1.4 | 0.55 | 1.04 | 0.24 | 0.89 | 1.17 | 0.91 | 1.52 | 0.83 | 1.23 | 0.22 | 0.81 | 0.95 | -0.4725 | -0.045 | -0.35 | 0.25898 | 0.091847 | 0.860488 | 0.989223 | 0.997988 | 0.994666 |
| Prkcb_P68404_S11s_1 | Prkcb_P68 | 0.3 | 0.29 | -0.23 | -0.08 | -0.15 | 0.58 | -0.41 | -0.61 | 0.46 | 0.33 | -1.38 | 0.18 | 0.54 | 0.59 | -0.84 | 0.43 | -0.2175 | -0.1725 | 0.11 | 0.773304 | 0.490412 | 0.693636 | 0.989223 | 0.997988 | 0.99798 |
| Prkcb_P68404_S206s_1 | Prkcb_P68 | 2.02 | 1.16 | 1.47 | 1.72 | 1.39 | 1.45 | 1.5 | 1.34 | 1.53 | 1.88 | 2.07 | 1.84 | 1.03 | 2.12 | 1.99 | 0.92 | -0.1725 | 0.2375 | -0.0775 | 0.837917 | 0.489154 | 0.392157 | 0.989223 | 0.997988 | 0.999507 |
| Prkcb_P68404_S660s_1 | Prkcb_P68 | -1.78 | -1.14 | -1.26 | -1.18 | -0.39 | -1.73 | -0.33 | -1.13 | -3.17 | -1.72 | -2.92 | -1.41 | -2.86 | -1.82 | -1.49 | -0.98 | 0.445 | -0.965 | -0.4475 | 0.303489 | 0.237751 | 0.053356 | 0.989223 | 0.997988 | 0.994666 |
| Prkcb_P68404_T641t_1 | Prkcb_P68 | 8.26 | 4.82 | 6.15 | 2.86 | 5.62 | 8.14 | 6.48 | 5.54 | -0.07 | 7.19 | 1.21 | 6.21 | -1.27 | 0.56 | 6 | 6.7 | 0.9225 | -1.8875 | -2.525 | 0.215347 | 0.414012 | 0.322379 | 0.989223 | 0.997988 | 0.994666 |
| Prkcb_P68404_T642t_1 | Prkcb_P68 | -0.56 | -0.44 | -1.4 | -1.45 | -0.27 | -0.85 | -0.96 | -1.37 | -0.96 | -0.67 | -2.4 | -1.49 | -0.5 | -0.16 | -1.21 | -1.29 | 0.1 | -0.4175 | 0.1725 | 0.662531 | 0.776979 | 0.363345 | 0.995762 | 0.997988 | 0.995804 |
| Prkcb_P68404_T644t_1 | Prkcb_P68 | 0.28 | 0.37 | -0.02 | 0.02 | 0.75 | -0.13 | 0.56 | -0.01 | 0.51 | 0.28 | -0.27 | 0.24 | 0.46 | 0.37 | 0.21 | 0.36 | 0.13 | 0.0275 | 0.1875 | 0.443137 | 0.637365 | 0.91511 | 0.989223 | 0.997988 | 0.994666 |
| Prkcd_Q1MX43_S389s_1 | Prkcd_Q1N | -0.73 | -0.07 | -0.1 | 0.19 | -0.05 | -0.06 | 0.23 | -0.12 | -0.55 | -0.13 | -0.34 | -0.38 | -0.09 | 0.04 | -0.29 | 0.14 | 0.1775 | -0.1275 | 0.1275 | 0.659115 | 0.49975 | 0.529327 | 0.989223 | 0.997988 | 0.995259 |
| Prkcd_Q1MX43_S528s_1 | Prkcd_Q1N | -2.66 | -2.12 | -3.88 | -4.48 | -1.84 | -3.22 | -3.01 | -3.9 | -3.08 | -2.09 | -3.06 | -3.99 | -2.3 | -1.93 | -4.19 | -3.65 | 0.2925 | 0.23 | 0.2675 | 0.697131 | 0.636972 | 0.706926 | 0.989223 | 0.997988 | 0.994666 |
| Prkcd_Q1MX43_S547s_1 | Prkcd_Q1N | -1.91 | -1.25 | -1.31 | -1.53 | -0.77 | -2.51 | -0.72 | -1.7 | -2.59 | -1.44 | -1.58 | -1.59 | -2.12 | -1.06 | -1.48 | -1.27 | 0.075 | -0.3 | 0.0175 | 0.956425 | 0.86053 | 0.373444 | 0.996493 | 0.997988 | 0.994666 |
| Prkcd_Q1MX43_T390t_1 | Prkcd_Q1N | -0.33 | 0.03 | 0.07 | 0.09 | 0.16 | -0.04 | 0.57 | -0.06 | 0 | -0.08 | -0.7 | 0.2 | 0.18 | -0.04 | 0.04 | -0.02 | 0.1925 | -0.11 | 0.075 | 0.756189 | 0.432577 | 0.689009 | 0.989223 | 0.997988 | 0.9971 |
| Prkcd_Q1MX43_Y196y_1 | Prkcd_Q1N | 0.83 | 1.33 | 1.35 | 1.25 | 1.08 | 0.14 | 1.42 | 0.92 | -0.11 | 1.18 | -0.52 | 1.39 | -0.18 | -0.44 | 1.07 | 1.4 | -0.3 | -0.705 | -0.7275 | 0.13545 | 0.351971 | 0.154467 | 0.989223 | 0.997988 | 0.994666 |

| | | | | | | | | | | | | | | | | | | | | | | | | | | |
|---|---|---|---|---|---|---|---|---|---|---|---|---|---|---|---|---|---|---|---|---|---|---|---|---|---|---|
| Prkce_P16054_S234s_1 | Prkce_P16 | -0.11 | -0.38 | -0.21 | -0.17 | -0.43 | 0.4 | -0.19 | 0.31 | 0.11 | -0.07 | 0.28 | 0.1 | 0.73 | -0.53 | 0.2 | 0.62 | 0.24 | 0.3225 | 0.4725 | 0.176787 | 0.363396 | 0.157605 | 0.989223 | 0.997988 | 0.994666 |
| Prkce_P16054_S329s_1 | Prkce_P16 | 0.94 | 0.66 | 1.01 | 1.27 | 0.52 | 0.53 | 1.02 | 1.23 | 0.64 | 0.97 | 0.9 | 0.97 | 0.71 | 0.38 | 1.06 | 0.74 | -0.145 | -0.1 | -0.2475 | 0.374772 | 0.58608 | 0.673982 | 0.989223 | 0.997988 | 0.994666 |
| Prkce_P16054_S329s_2 | Prkce_P16 | -0.26 | -0.28 | -0.27 | 0 | 0.94 | 0.34 | 0.08 | -0.21 | -0.3 | -0.29 | -1 | 0 | -1.18 | -0.81 | -0.45 | -0.31 | 0.49 | -0.195 | -0.485 | 0.111269 | 0.112214 | 0.487131 | 0.989223 | 0.997988 | 0.994666 |
| Prkce_P16054_S337s_1 | Prkce_P16 | 1.61 | 1.17 | 1.64 | 1.87 | 0.81 | 1.21 | 1.35 | 1.97 | 1.14 | 2 | 1.37 | 1.63 | 1.26 | -0.19 | 1.59 | 1.47 | -0.2375 | -0.0375 | -0.54 | 0.230913 | 0.442268 | 0.894996 | 0.989223 | 0.997988 | 0.994666 |
| Prkce_P16054_S337s_2 | Prkce_P16 | -0.26 | -0.28 | -0.56 | 0.14 | 0.94 | 0.34 | -0.01 | -0.21 | -0.3 | -0.29 | -0.61 | -0.17 | -1.18 | -0.81 | -0.34 | -0.15 | 0.505 | -0.1025 | -0.38 | 0.253691 | 0.128193 | 0.681096 | 0.989223 | 0.997988 | 0.994666 |
| Prkce_P16054_S346s_1 | Prkce_P16 | -0.06 | 0.58 | -0.4 | 0.03 | 0.59 | -0.39 | 0.22 | -0.43 | -1.08 | -0.51 | 0.06 | 0.11 | -0.45 | 0.77 | 0.22 | 0.32 | -0.04 | -0.3925 | 0.1775 | 0.616632 | 0.90353 | 0.289107 | 0.996493 | 0.997988 | 0.995259 |
| Prkce_P16054_S346s_2 | Prkce_P16 | -0.2 | 0.42 | -0.19 | -0.06 | 0.49 | -0.3 | -0.33 | 0.03 | -0.51 | -0.04 | -0.35 | -0.68 | -1.42 | 0.81 | -0.03 | -0.09 | -0.02 | -0.3875 | -0.175 | 0.70798 | 0.942526 | 0.165146 | 0.999187 | 0.997988 | 0.996966 |
| Prkce_P16054_S350s_1 | Prkce_P16 | 1.04 | 1.3 | 1.74 | 1.5 | 0.85 | 0.9 | 1.07 | 1.2 | 1.31 | 1.23 | 1.97 | 1.23 | 0.9 | 1.35 | 1.41 | 0.9 | -0.39 | 0.04 | -0.255 | 0.374793 | 0.125047 | 0.887453 | 0.989223 | 0.997988 | 0.994666 |
| Prkce_P16054_S350s_2 | Prkce_P16 | -0.2 | 0.42 | -0.19 | -0.06 | 0.49 | -0.3 | -0.33 | 0.03 | -0.51 | -0.04 | -0.35 | -0.68 | -1.42 | 0.81 | -0.03 | -0.09 | -0.02 | -0.3875 | -0.175 | 0.70798 | 0.942526 | 0.165146 | 0.999187 | 0.997988 | 0.996966 |
| Prkce_P16054_S368s_1 | Prkce_P16 | -0.33 | -1.07 | -0.69 | -0.58 | -0.1 | -0.31 | -0.33 | -0.34 | -1.03 | -0.76 | 0.24 | -0.91 | -1.25 | 0.1 | -0.87 | -0.83 | 0.3975 | 0.0525 | -0.045 | 0.898673 | 0.11556 | 0.87931 | 0.989223 | 0.997988 | 0.999507 |
| Prkce_P16054_S388s_1 | Prkce_P16 | 0.54 | 0.55 | 0.74 | 0.62 | 0.46 | 0.32 | 0.45 | 0.62 | 0.4 | 0.81 | 0.28 | 0.61 | 0.68 | -0.07 | 0.63 | 0.53 | -0.15 | -0.0875 | -0.17 | 0.531861 | 0.460347 | 0.700879 | 0.989223 | 0.997988 | 0.994666 |
| Prkce_P16054_S62s_1 | Prkce_P16 | -0.24 | -0.22 | 1.1 | 0.02 | -0.45 | 0.08 | 0.4 | 0.47 | -1.03 | 0.51 | 0.58 | 0.08 | -0.39 | -0.19 | 0.42 | 1.07 | -0.04 | -0.13 | 0.0625 | 0.888572 | 0.914646 | 0.780782 | 0.997335 | 0.997988 | 0.999507 |
| Prkce_P16054_S729s_1 | Prkce_P16 | -0.93 | -0.33 | -0.82 | -1.37 | 0.05 | -2.31 | -0.32 | -1.05 | -0.89 | -0.93 | -1.05 | -3.21 | -0.47 | -0.12 | -0.81 | -0.2 | -0.045 | -0.6575 | 0.4625 | 0.16538 | 0.930077 | 0.25716 | 0.997844 | 0.997988 | 0.994666 |
| Prkce_P16054_T228t_1 | Prkce_P16 | 0.79 | 1.04 | 0.51 | 0.72 | 0.08 | 0.51 | -0.06 | 0.19 | 1.22 | 1 | 1.11 | 0.36 | 1.27 | 0.45 | 0.6 | 0.68 | -0.585 | 0.1575 | -0.015 | 0.957917 | 0.03042 | 0.571543 | 0.989223 | 0.997988 | 0.999507 |
| Prkce_P16054_T710t_1 | Prkce_P16 | -4.28 | -3.1 | -3.26 | -2.15 | -2.59 | -4.72 | -2.98 | -2.95 | -4.14 | -3.82 | -4.37 | -3.18 | -2.79 | -1.9 | -3.41 | -2.36 | -0.1125 | -0.68 | 0.5825 | 0.270216 | 0.846441 | 0.180761 | 0.996349 | 0.997988 | 0.994666 |
| Prkcg_P63318_S132s_1 | Prkcg_P63 | -3.18 | -2.23 | -1.97 | -3.47 | -3.01 | -0.83 | -2.85 | -2.96 | 0.19 | -1.45 | -2.57 | -2.8 | 0.53 | -0.93 | -2.16 | -2.84 | 0.3 | 1.055 | 1.3625 | 0.088939 | 0.604834 | 0.155205 | 0.989223 | 0.997988 | 0.994666 |
| Prkcg_P63318_S148s_1 | Prkcg_P63 | -1.37 | -0.56 | -1.09 | -0.94 | 0.15 | -0.22 | 0.67 | -0.86 | -1.81 | 0.6 | 0.35 | 0.44 | 0.01 | 0.21 | -0.79 | -0.55 | 0.925 | 0.885 | 0.71 | 0.05422 | 0.027477 | 0.134299 | 0.989223 | 0.997988 | 0.994666 |
| Prkcg_P63318_S326s_1 | Prkcg_P63 | 0.59 | 0.48 | 0.26 | 0.22 | 0.54 | 0.24 | 0.24 | 0.25 | 0.37 | 0.67 | -0.27 | 0.35 | 0.16 | 0.88 | -0.29 | -0.39 | -0.07 | -0.1075 | -0.2975 | 0.389571 | 0.742948 | 0.694625 | 0.995762 | 0.997988 | 0.994666 |
| Prkcg_P63318_S328s_1 | Prkcg_P63 | 0.35 | 0.37 | 1.66 | 1.61 | 0.22 | -0.01 | 1.91 | 1.52 | 1.19 | 0.92 | 1.7 | 1.86 | 1.37 | 0.63 | 1.87 | 2.05 | -0.0875 | 0.42 | 0.4825 | 0.316549 | 0.871976 | 0.330704 | 0.996493 | 0.997988 | 0.994666 |
| Prkcg_P63318_S330s_1 | Prkcg_P63 | 0.41 | 1.12 | 0.81 | 0.93 | 0.88 | 0.19 | 1.39 | 0.8 | 0.74 | 1.33 | 1.04 | 1 | 0.81 | 0.97 | 1.29 | 1.03 | -0.0025 | 0.21 | 0.2075 | 0.448353 | 0.993525 | 0.426661 | 0.999278 | 0.997988 | 0.994666 |
| Prkcg_P63318_S373s_1 | Prkcg_P63 | -0.2 | 0.94 | 0.8 | 0.05 | -0.05 | 0.07 | 0.24 | 0.1 | 0.66 | 0.45 | 0.58 | 0.48 | 1.55 | 1.45 | 0.65 | 0.45 | -0.3075 | 0.145 | 0.6275 | 0.141444 | 0.329011 | 0.646826 | 0.989223 | 0.997988 | 0.994666 |
| Prkcg_P63318_S510s_1 | Prkcg_P63 | 0.23 | -0.38 | -0.18 | -0.27 | 0.27 | 0.07 | 0.26 | 0.02 | -0.13 | 0.08 | 0.12 | -0.28 | 0.15 | 0.36 | -0.16 | 0.1 | 0.305 | 0.0975 | 0.2625 | 0.334552 | 0.200027 | 0.690284 | 0.989223 | 0.997988 | 0.994666 |
| Prkcg_P63318_S639s_1 | Prkcg_P63 | 0.92 | 0.17 | 0.24 | 0.92 | -0.66 | 1.31 | -0.02 | 0.41 | 1.49 | 0.61 | 0.13 | 1 | 1.28 | -0.87 | 0.28 | 0.8 | -0.3025 | 0.245 | -0.19 | 0.694906 | 0.492526 | 0.507364 | 0.989223 | 0.997988 | 0.996159 |
| Prkcg_P63318_S687s_1 | Prkcg_P63 | 0.22 | 0.44 | 0.41 | 0.22 | 0.6 | -0.13 | 0.77 | 0.51 | 0.37 | 0.46 | 0.64 | 0.29 | 0.07 | 0.48 | 0.13 | 0.12 | 0.115 | 0.1175 | -0.1225 | 0.614003 | 0.656006 | 0.589339 | 0.990434 | 0.997988 | 0.995259 |
| Prkcg_P63318_S690s_1 | Prkcg_P63 | 0.83 | -0.3 | 0.43 | 0.93 | 0.29 | 1.03 | 1.11 | 1.21 | 0.34 | 0.12 | 0.05 | 0.76 | -0.09 | 0.73 | -0.06 | 0.1 | 0.4375 | -0.155 | -0.3025 | 0.41238 | 0.232041 | 0.651253 | 0.989223 | 0.997988 | 0.994666 |
| Prkcg_P63318_T514t_1 | Prkcg_P63 | 0.16 | -0.27 | -0.12 | -0.18 | -0.03 | 0.11 | 0.43 | -0.04 | -0.78 | -0.03 | -1.27 | -0.28 | -0.98 | -1.29 | -0.12 | -0.14 | 0.22 | -0.4875 | -0.53 | 0.147731 | 0.340505 | 0.151947 | 0.989223 | 0.997988 | 0.994666 |
| Prkcg_P63318_T655t_1 | Prkcg_P63 | 0.72 | 0.47 | 0.76 | 0.54 | 0.53 | 0.45 | 0.68 | 0.59 | 0.55 | 0.71 | 0.7 | 0.5 | 0.47 | 0.72 | 0.51 | 0.33 | -0.06 | -0.0075 | -0.115 | 0.633415 | 0.767353 | 0.971933 | 0.995762 | 0.997988 | 0.995259 |
| Prkch_P23298_S675s_1 | Prkch_P23 | 0.21 | -0.33 | -0.48 | -1.69 | -0.14 | -0.4 | 0.24 | -2.2 | -0.6 | 0.3 | -0.16 | 0.6 | -0.21 | 0.13 | -0.64 | 0.45 | -0.0525 | 0.6075 | 0.505 | 0.279331 | 0.930448 | 0.207683 | 0.997844 | 0.997988 | 0.994666 |
| Prkci_Q62074_T411t_1 | Prkci_Q62 | 1.6 | 1.65 | 0.98 | 0.37 | 1.45 | 1.53 | 0.82 | 0.71 | 0.7 | 1.54 | -0.06 | -0.35 | 0.59 | 1.08 | -0.14 | 0.72 | -0.0225 | -0.6925 | -0.5875 | 0.16656 | 0.950733 | 0.181731 | 0.999187 | 0.997988 | 0.994666 |
| Prkcq_Q02111_T536t_1 | Prkcq_Q02 | 0.94 | 0.95 | 1.27 | 1.14 | 1.17 | 0.88 | 1.27 | 0.97 | 0.64 | 0.83 | 0.59 | 0.52 | 0.79 | 0.69 | 0.8 | 1.33 | -0.0025 | -0.43 | -0.1725 | 0.515667 | 0.9907 | 0.074712 | 0.999187 | 0.997988 | 0.994666 |
| Prkcz_Q02956_T227t_1 | Prkcz_Q02 | 0.17 | 0.59 | -0.05 | 0.18 | 0.43 | -0.19 | -0.04 | 0.06 | 0.09 | 0.59 | -0.04 | -0.71 | -0.27 | 0.39 | -0.12 | 0.39 | -0.1575 | -0.24 | -0.125 | 0.665996 | 0.528235 | 0.467835 | 0.989223 | 0.997988 | 0.996159 |
| Prkd1_Q62101_S255s_1 | Prkd1_Q62 | -1.3 | -0.01 | -1.22 | -2.55 | -0.84 | -1.51 | -1.54 | -2.08 | -0.4 | -0.91 | -0.13 | -1.14 | -0.38 | -0.39 | -1.12 | -1.12 | -0.2225 | 0.625 | 0.5175 | 0.336529 | 0.673124 | 0.25626 | 0.992876 | 0.997988 | 0.994666 |
| Prkd2_Q8BZ03_S197s_1 | Prkd2_Q8B | -0.53 | -0.05 | -0.66 | -1.04 | -0.6 | -1.65 | -1.07 | -0.65 | -0.54 | 0.82 | -0.68 | -0.41 | 0.39 | -0.35 | -1.17 | -1.45 | -0.4225 | 0.3675 | -0.075 | 0.867939 | 0.218718 | 0.368633 | 0.989223 | 0.997988 | 0.999507 |
| Prkd3_Q5FWX6_S735s_1 | Prkd3_Q5F | -1.32 | 0.06 | -0.4 | -0.93 | 0.32 | -0.17 | -0.07 | -0.05 | -0.56 | -0.29 | -0.02 | -0.12 | -0.68 | -0.16 | -0.39 | -0.41 | 0.655 | 0.4 | 0.2375 | 0.502688 | 0.072303 | 0.261271 | 0.989223 | 0.997988 | 0.994666 |
| Prkg1_P0C605_T532t_1 | Prkg1_P0C | 0.11 | 0.54 | -0.94 | -0.12 | -0.16 | -0.34 | 0.63 | 0.44 | -0.16 | -0.84 | -1.18 | -0.08 | -0.83 | 0.62 | -0.63 | 0.09 | 0.245 | -0.4625 | -0.085 | 0.848667 | 0.523157 | 0.27143 | 0.989223 | 0.997988 | 0.999507 |
| Prkg2_E9QPH0_S431s_1 | Prkg2_E9Q | 0.4 | 0.53 | 0.5 | 0.41 | 0.87 | 0.24 | 0.79 | 0.25 | 0.02 | 0.22 | 0.2 | 0.46 | -0.22 | 0.77 | 0.07 | 0.39 | 0.0775 | -0.235 | -0.2075 | 0.476041 | 0.746327 | 0.292528 | 0.995762 | 0.997988 | 0.994666 |
| Prkg2_E9QPH0_S97s_1 | Prkg2_E9Q | -1.77 | -0.55 | -0.35 | -0.47 | -0.43 | -1.78 | -0.57 | -0.85 | -2.59 | -0.57 | -1.02 | -1.49 | -1.33 | -1.34 | -0.35 | -0.06 | -0.1225 | -0.6325 | 0.015 | 0.973646 | 0.773518 | 0.236039 | 0.995762 | 0.997988 | 0.999507 |
| Prkg2_E9QPH0_T109t_1 | Prkg2_E9Q | -0.36 | -0.53 | 0.88 | 0.61 | 0.52 | 0.13 | 0.55 | 0.93 | -0.95 | -0.66 | 0.45 | 0.89 | -0.74 | -0.4 | 0.32 | 1.09 | 0.3825 | -0.2175 | -0.0825 | 0.870325 | 0.324733 | 0.679448 | 0.989223 | 0.997988 | 0.999507 |
| Prkg2_E9QPH0_T580t_1 | Prkg2_E9Q | 0.56 | 0.49 | 0.62 | 0.23 | 0.6 | 0.61 | 1.28 | -0.33 | 0.04 | 0.58 | -0.9 | 0.38 | -0.34 | 0.99 | -0.07 | 0.27 | 0.065 | -0.45 | -0.2625 | 0.443737 | 0.850816 | 0.223191 | 0.996349 | 0.997988 | 0.994666 |
| Prkra_Q9WTX2_S18s_1 | Prkra_Q9W | -0.3 | -0.24 | -0.66 | -0.62 | 0.18 | -0.33 | -0.1 | -0.71 | -0.44 | -0.08 | -0.66 | -0.81 | -0.55 | 0.15 | -0.44 | -0.44 | 0.215 | -0.0425 | 0.135 | 0.625654 | 0.422968 | 0.869576 | 0.989223 | 0.997988 | 0.995259 |
| Prmt7_Q922X9_T351t_1 | Prmt7_Q92 | -0.72 | -0.57 | -0.8 | -1.01 | -0.17 | -1.03 | -0.81 | -1.69 | -2.08 | -0.66 | -0.24 | -1.78 | 0.19 | -0.39 | -1.16 | -0.56 | -0.15 | -0.415 | 0.295 | 0.38488 | 0.65579 | 0.353505 | 0.990434 | 0.997988 | 0.994666 |
| Prnp_P04925_S43s_1 | Prnp_P049 | -0.05 | -0.29 | 1 | 0.5 | -0.67 | 0.48 | -0.2 | 0.19 | 0.07 | -0.93 | 0.65 | -0.14 | 0.64 | -0.94 | -0.78 | 0.7 | -0.34 | -0.3775 | -0.385 | 0.448673 | 0.374525 | 0.385692 | 0.989223 | 0.997988 | 0.994666 |
| Proser1_Q5PRE5_S596s_1 | Proser1_Q | -0.82 | -0.84 | -0.24 | -0.77 | -0.99 | -1.95 | -0.67 | -0.42 | -0.39 | -0.89 | -0.12 | -1.23 | -0.29 | -0.37 | -1.52 | -0.66 | -0.34 | 0.01 | -0.0425 | 0.902527 | 0.359162 | 0.97484 | 0.989223 | 0.997988 | 0.999507 |
| Proser1_Q5PRE5_T610t_1 | Proser1_Q | -0.17 | 0.3 | -0.06 | -1.75 | -0.79 | 0.54 | -1.43 | -1.33 | 2.08 | -0.04 | 0.2 | -1.17 | 2.36 | 1.2 | -0.84 | -0.04 | -0.3325 | 0.6875 | 1.09 | 0.166203 | 0.567502 | 0.358724 | 0.989223 | 0.997988 | 0.994666 |
| Prox1_P48437_S179s_1 | Prox1_P48 | -0.2 | 1.23 | -0.41 | -0.35 | 0.7 | -0.03 | -0.12 | -0.39 | 1.45 | 0.85 | -0.06 | -0.38 | 2.48 | 1.54 | 1.91 | 0.73 | -0.0275 | 0.3975 | 1.5975 | 0.010146 | 0.948843 | 0.462915 | 0.999187 | 0.997988 | 0.994666 |
| Prox1_P48437_S199s_1 | Prox1_P48 | -0.56 | 0.15 | 0.29 | -0.72 | -0.17 | -0.08 | 0.16 | 0.1 | 0.01 | -0.19 | 0 | -0.57 | 1.59 | 0.87 | 1.26 | 0.81 | 0.2125 | 0.0225 | 1.3425 | 0.0031 | 0.473519 | 0.943333 | 0.989223 | 0.997988 | 0.994666 |
| Prox1_P48437_S220s_1 | Prox1_P48 | -0.5 | -0.04 | 0.23 | 0.23 | 0 | -0.19 | 0.84 | 0.39 | -0.87 | 0.23 | 0.72 | 0.51 | 0.18 | -0.65 | 0.85 | -0.2 | 0.28 | 0.1675 | 0.065 | 0.863256 | 0.372238 | 0.671476 | 0.989223 | 0.997988 | 0.999507 |
| Prpf3_Q922U1_S619s_1 | Prpf3_Q92 | -1.62 | -1.42 | -0.09 | -0.6 | -2.56 | -1.45 | -1.23 | -0.08 | -1.58 | -1.88 | 0.81 | -0.71 | -1.74 | -2.34 | 0.06 | -0.35 | -0.3975 | 0.0925 | -0.16 | 0.793275 | 0.483739 | 0.884655 | 0.989223 | 0.997988 | 0.999507 |
| Prpf38a_Q4FK66_S209s_1 | Prpf38a_Q | 1.56 | -0.7 | -0.08 | 0.25 | -1.63 | 0.97 | -0.9 | -0.32 | 2.22 | 0.09 | -2.27 | -0.39 | 2.19 | -1.08 | 0.01 | -0.44 | -0.7275 | -0.345 | -0.0875 | 0.908094 | 0.276518 | 0.708527 | 0.989223 | 0.997988 | 0.999507 |
| Prpf40a_Q9R1C7_S884s_1 | Prpf40a_Q | -2.11 | -2.65 | -3.02 | -10.52 | -4.36 | -1.41 | -3.7 | -3.85 | -1.24 | -2.5 | -1.26 | -3.4 | -0.09 | -6.61 | -2.53 | -3.38 | 1.245 | 2.475 | 1.4225 | 0.493884 | 0.492305 | 0.190134 | 0.989223 | 0.997988 | 0.994666 |
| Prpf40b_F6XW22_S218s_1 | Prpf40b_F6 | -3.16 | -3.7 | -2.17 | -2.16 | -2.72 | -4.3 | -1.87 | -2.74 | -4.48 | -3.39 | -1.13 | -2.19 | -3.17 | -2.6 | -2.75 | -2.18 | -0.11 | 5.5E-16 | 0.1225 | 0.775106 | 0.847057 | 1 | 0.996349 | 1 | 0.99798 |
| Prpf4b_Q61136_S21s_1 | Prpf4b_Q6 | -0.6 | -1.28 | -3.63 | -5.6 | -2.36 | -2.26 | -4.99 | -4.13 | -2.06 | -1.4 | -1.52 | -4.59 | -1.48 | -3.44 | -3.99 | -4.04 | -0.6575 | 0.385 | -0.46 | 0.682075 | 0.568194 | 0.748371 | 0.989223 | 0.997988 | 0.996159 |
| Prpf4b_Q61136_S24s_2 | Prpf4b_Q6 | -2.83 | -2.02 | -1.14 | -3.64 | -2.19 | -1.76 | -4.29 | -2.8 | -2.71 | -1.01 | -0.21 | -3.63 | -0.14 | -0.92 | -4.01 | -3.2 | -0.3525 | 0.5175 | 0.34 | 0.71502 | 0.607006 | 0.543212 | 0.989223 | 0.997988 | 0.996966 |
| Prpf4b_Q61136_S258s_1 | Prpf4b_Q6 | 0.66 | 0.11 | -0.59 | -1.13 | -0.61 | 1.34 | -1.09 | -0.41 | 2.57 | -0.62 | -1.37 | -0.45 | 3.4 | -0.75 | 2.25 | -0.79 | 0.045 | 0.27 | 1.265 | 0.220446 | 0.939242 | 0.750777 | 0.999187 | 0.997988 | 0.994666 |
| Prpf4b_Q61136_S278s_1 | Prpf4b_Q6 | 0.72 | 0.05 | -0.37 | -0.07 | -0.93 | -0.2 | -0.98 | 0.42 | 0.09 | -1.07 | -0.4 | -0.06 | 1.11 | -0.16 | 0.73 | -0.3 | -0.505 | -0.4425 | 0.2625 | 0.529824 | 0.217326 | 0.236167 | 0.989223 | 0.997988 | 0.994666 |
| Prpf4b_Q61136_S33s_1 | Prpf4b_Q6 | 1.2 | 0.31 | -1.03 | 0.89 | -0.95 | 1.13 | -0.69 | 0.12 | 1.18 | -0.07 | 0.66 | -0.85 | 0.9 | 0.58 | 0.94 | 0.06 | -0.44 | -0.1125 | 0.2775 | 0.584611 | 0.475098 | 0.852723 | 0.989223 | 0.997988 | 0.995259 |
| Prpf4b_Q61136_S33s_2 | Prpf4b_Q6 | -2.83 | -2.02 | -1.14 | -3.64 | -2.19 | -1.76 | -4.29 | -2.8 | -2.71 | -1.01 | -0.21 | -3.63 | -0.14 | -0.92 | -4.01 | -3.2 | -0.3525 | 0.5175 | 0.34 | 0.71502 | 0.607006 | 0.543212 | 0.989223 | 0.997988 | 0.996966 |
| Prpf4b_Q61136_S341s_1 | Prpf4b_Q6 | -0.49 | -3.31 | -3.08 | -2.45 | -3.1 | -1.31 | -2.2 | -1.33 | 3.59 | -2.69 | -1.34 | -1.48 | 4.83 | -2.18 | 4.32 | -3.34 | 0.3475 | 1.8525 | 3.24 | 0.114289 | 0.611475 | 0.189465 | 0.989223 | 0.997988 | 0.994666 |
| Prpf4b_Q61136_S431s_1 | Prpf4b_Q6 | 0.62 | 0.51 | 1.05 | 2.17 | 0 | 1.04 | -0.93 | 0.94 | -0.72 | -0.85 | 3.03 | 2.9 | 0.74 | -1.78 | 1.66 | 1.18 | -0.825 | 0.0025 | -0.6375 | 0.40992 | 0.151334 | 0.998028 | 0.989223 | 0.999885 | 0.994666 |
| Prpf4b_Q61136_S94s_1 | Prpf4b_Q6 | -1.27 | -1.5 | -1.38 | -2.39 | -1.6 | -1.96 | -2.15 | -2.16 | -2.44 | -2.31 | -1.47 | -2.09 | -0.13 | -1.59 | -1.72 | -2.35 | -0.3325 | -0.4425 | 0.1875 | 0.711033 | 0.295599 | 0.22636 | 0.989223 | 0.997988 | 0.996966 |
| Prpf4b_Q61136_Y849y_1 | Prpf4b_Q6 | 0.23 | 0.52 | 0.51 | -0.03 | 0.63 | -0.18 | 0.57 | -0.23 | -0.05 | 0.29 | -0.11 | 0.17 | -0.01 | 0.67 | -0.09 | 0.59 | -0.11 | -0.2325 | -0.0175 | 0.953314 | 0.709827 | 0.350922 | 0.995437 | 0.997988 | 0.999507 |
| Prpsap2_Q8R574_S198s_1 | Prpsap2_Q | 1.35 | 1.06 | 0.56 | 0.39 | 0.64 | 0.81 | -0.62 | -0.27 | 0.84 | 0.92 | -0.21 | -1.02 | 1.59 | -1.37 | 0.15 | -0.05 | -0.7 | -0.7075 | -0.76 | 0.218217 | 0.103342 | 0.169172 | 0.989223 | 0.997988 | 0.994666 |
| Prpsap2_Q8R574_S219s_1 | Prpsap2_Q | 0.83 | 0.23 | 0.65 | 0.62 | 0.96 | -0.77 | 1.01 | -0.66 | 0.92 | 0.71 | -0.22 | -0.2 | 0 | -0.02 | -1.09 | 0.65 | -0.4475 | -0.28 | -0.6975 | 0.100493 | 0.35295 | 0.423108 | 0.989223 | 0.997988 | 0.994666 |
| Prpsap2_Q8R574_S227s_1 | Prpsap2_Q | 1.31 | 0.4 | 0.31 | 0.38 | 0.38 | 1.2 | 0.5 | 0.07 | 0.29 | 0.67 | 0.16 | 0.36 | -0.28 | -0.23 | 0.07 | 0.5 | -0.0625 | -0.23 | -0.585 | 0.106541 | 0.854986 | 0.451161 | 0.996349 | 0.997988 | 0.994666 |
| Prr12_E9PYL2_S648s_1 | Prr12_E9P | -1.57 | -3.33 | -1.61 | -5.24 | -2.42 | -2.29 | -1.45 | -1.42 | -3.98 | -1.01 | -2.73 | -1.26 | -2.8 | -4.32 | -1.62 | -2.17 | 1.0425 | 0.6925 | 0.21 | 0.818809 | 0.210823 | 0.486735 | 0.989223 | 0.997988 | 0.999507 |
| Prr18_Q6PAN7_T91t_1 | Prr18_Q6P | 0.39 | 0.54 | -0.07 | 0.62 | 0.4 | -0.48 | -0.42 | 0.44 | -0.71 | -0.07 | 0.1 | -0.03 | -0.23 | -0.47 | 0.06 | 0.04 | -0.385 | -0.5475 | -0.52 | 0.087137 | 0.237766 | 0.081252 | 0.989223 | 0.997988 | 0.994666 |
| Prr36_E9PV26_S197s_1 | Prr36_E9P | 0.61 | 0.51 | 0.4 | -0.58 | 0.53 | 0.64 | 0.27 | 0 | -0.06 | 0.57 | -0.12 | 0.14 | 0.02 | 0.52 | -0.09 | -0.16 | 0.125 | -0.1025 | -0.1625 | 0.641116 | 0.69993 | 0.761598 | 0.994893 | 0.997988 | 0.995259 |
| Prr36_E9PV26_S519s_1 | Prr36_E9P | 0 | 0.24 | 0.04 | -0.01 | 0.28 | -0.81 | 0.2 | -0.03 | -0.44 | 0.38 | 0.25 | 0.01 | 0.05 | 0.49 | -0.03 | -0.13 | -0.1575 | -0.0175 | 0.0275 | 0.914144 | 0.586411 | 0.945822 | 0.989223 | 0.997988 | 0.999507 |
| Prr5_Q812A5_S232s_1 | Prr5_Q812 | 0.13 | -1.28 | 0.08 | 0.09 | 0.09 | 0.38 | 0.75 | 0.54 | -1.18 | -0.91 | 0.56 | 0.52 | -0.6 | -0.71 | 0.62 | 0.35 | 0.685 | -0.0075 | 0.16 | 0.730671 | 0.087837 | 0.98882 | 0.989223 | 0.997988 | 0.9971 |
| Prr5_Q812A5_S302s_1 | Prr5_Q812 | -0.75 | -1.54 | -0.62 | 0.12 | -0.77 | -0.5 | -0.23 | -0.54 | 0.14 | -0.3 | 0.19 | -0.76 | 1.17 | -0.33 | 0.29 | 0.35 | 0.1875 | 0.515 | 1.0675 | 0.036744 | 0.601933 | 0.221681 | 0.989223 | 0.997988 | 0.994666 |
| Prr7_Q3V0I2_S185s_1 | Prr7_Q3V0 | 1.66 | 2.27 | 1.79 | 0.94 | 1.96 | 3.28 | 0.96 | 0.77 | 1.27 | 1.71 | 0.66 | 1.46 | 2.45 | 2.51 | 1.38 | 1.59 | 0.0775 | -0.39 | 0.3175 | 0.44033 | 0.89236 | 0.300174 | 0.996493 | 0.997988 | 0.994666 |
| Prr7_Q3V0I2_S68s_1 | Prr7_Q3V0 | 0.67 | 0.68 | -0.32 | 0.11 | 0.72 | -0.15 | 0.28 | 0.09 | 0.77 | 0.8 | 0.09 | 0.13 | -0.46 | 0.77 | -1.3 | 0.85 | -0.05 | 0.1625 | -0.32 | 0.551663 | 0.875415 | 0.628074 | 0.996493 | 0.997988 | 0.994666 |
| Prrc2a_G3UX48_S1049s_1 | Prrc2a_G3 | -1.06 | -2.9 | -1.22 | -1.3 | -2.2 | -2.01 | -1.23 | -1.09 | -1.99 | -0.54 | -0.65 | -1.52 | 0.15 | -0.44 | -0.97 | -1.14 | -0.0125 | 0.445 | 1.02 | 0.062353 | 0.978892 | 0.400056 | 0.999187 | 0.997988 | 0.994666 |
| Prrc2a_G3UX48_S111s_1 | Prrc2a_G3 | -0.36 | -1.27 | -1.98 | -0.87 | -0.49 | -0.69 | -0.34 | -0.58 | -0.38 | -1.27 | -1.22 | -0.29 | -1.94 | -0.32 | -1.71 | -1.08 | 0.595 | 0.33 | -0.1425 | 0.76557 | 0.115814 | 0.442336 | 0.989223 | 0.997988 | 0.9971 |
| Prrc2a_G3UX48_S1162s_1 | Prrc2a_G3 | -0.71 | -1.18 | -1.03 | -0.82 | -0.11 | -0.74 | 0.11 | -0.9 | -0.8 | -0.92 | -0.54 | -0.9 | -0.69 | -0.91 | -0.19 | -0.26 | 0.525 | 0.145 | 0.4225 | 0.155393 | 0.098799 | 0.535785 | 0.989223 | 0.997988 | 0.994666 |
| Prrc2a_G3UX48_S295s_1 | Prrc2a_G3 | -1.04 | 0.42 | 0.01 | -0.77 | -0.36 | -1.23 | -0.41 | -1.34 | -0.99 | -0.5 | -0.65 | -0.38 | -0.44 | -1.58 | 0.41 | 0.11 | -0.49 | -0.285 | -0.03 | 0.95389 | 0.248773 | 0.449765 | 0.989223 | 0.997988 | 0.999507 |
| Prrc2a_G3UX48_S323s_1 | Prrc2a_G3 | 0.09 | 0.55 | -0.08 | -0.59 | 0.46 | -0.05 | 0.04 | 0.18 | 0.15 | 0.49 | -0.62 | 0.27 | 0.36 | 0.31 | -0.57 | 0.13 | 0.165 | 0.08 | 0.065 | 0.852306 | 0.573036 | 0.820265 | 0.989223 | 0.997988 | 0.999507 |
| Prrc2a_G3UX48_S399s_1 | Prrc2a_G3 | -2.5 | -5 | -2.41 | -1.13 | -2.91 | -3.04 | -1.59 | 1.5 | -3.32 | -2.85 | 0.31 | -0.72 | -2.58 | -2.61 | -0.97 | 0.31 | 1.25 | 1.115 | 1.2975 | 0.188396 | 0.290681 | 0.3005 | 0.989223 | 0.997988 | 0.994666 |
| Prrc2a_G3UX48_S706s_1 | Prrc2a_G3 | -1.92 | -2.26 | -1 | -1.02 | -1.27 | -1.21 | -1.05 | -1.13 | -1.71 | -1.56 | 0.36 | -0.9 | -0.76 | -1.12 | -0.5 | -0.07 | 0.385 | 0.5975 | 0.9375 | 0.038041 | 0.263948 | 0.277344 | 0.989223 | 0.997988 | 0.994666 |
| Prrc2a_G3UX48_S834s_1 | Prrc2a_G3 | -0.04 | -1.18 | 0.02 | 0.62 | -0.21 | 0.09 | 0.34 | 0.57 | -0.49 | -0.74 | 0.26 | -0.48 | -1.65 | 0.13 | -0.6 | -0.11 | 0.3425 | -0.2175 | -0.4125 | 0.428928 | 0.397469 | 0.610147 | 0.989223 | 0.997988 | 0.994666 |
| Prrc2a_G3UX48_T480t_1 | Prrc2a_G3 | 0.44 | 0.94 | 0.22 | -0.04 | 0.76 | 0.73 | 0.33 | 0.02 | -1.03 | 0.5 | 0.67 | -1.48 | -0.14 | 0.64 | -0.22 | -0.24 | 0.07 | -0.725 | -0.38 | 0.272856 | 0.814536 | 0.200176 | 0.995762 | 0.997988 | 0.994666 |
| Prrc2a_G3UX48_T554t_1 | Prrc2a_G3 | -0.81 | -0.3 | -0.2 | 0.51 | 0.01 | -0.73 | 0.68 | 0.74 | -0.57 | -0.08 | 1.2 | -0.05 | 0.03 | 0.61 | 0.02 | 0.02 | 0.375 | 0.325 | 0.37 | 0.295472 | 0.379019 | 0.474872 | 0.989223 | 0.997988 | 0.994666 |
| Prrc2a_G3UX48_T727t_1 | Prrc2a_G3 | -1.35 | -1.67 | -1.11 | -0.86 | -0.97 | -1.74 | -0.58 | -1.33 | -1.31 | -0.85 | -0.18 | -0.99 | -0.56 | -0.62 | -0.04 | -0.13 | 0.0925 | 0.415 | 0.91 | 0.011134 | 0.771409 | 0.218323 | 0.995762 | 0.997988 | 0.994666 |
| Prrc2b_F8WHT3_S1171s_1 | Prrc2b_F8 | -3.64 | -3.62 | -3.74 | -3.71 | -3.75 | -2.91 | -2.82 | -3.3 | -1.19 | -2.91 | -1.87 | -4.07 | -0.53 | -1.89 | -3 | -2.84 | 0.4825 | 1.1675 | 1.6125 | 0.012372 | 0.090807 | 0.069269 | 0.989223 | 0.997988 | 0.994666 |
| Prrc2b_F8WHT3_S1681s_1 | Prrc2b_F8 | -1.6 | -0.64 | -2.69 | -2.78 | -0.46 | -1.1 | -2.07 | -2.31 | -1.3 | -0.64 | -3.18 | -2.57 | -1.59 | -1.84 | -0.29 | -1.72 | 0.4425 | 0.005 | 0.5675 | 0.333778 | 0.463443 | 0.994249 | 0.989223 | 0.997988 | 0.994666 |
| Prrc2b_F8WHT3_S415s_1 | Prrc2b_F8 | -1.61 | -2.82 | -2.27 | -4.43 | -1.34 | -2.42 | -1.88 | -2.97 | -1.67 | -1.88 | -2.53 | -2.03 | -0.72 | -0.58 | -3.37 | -2.03 | 0.63 | 0.755 | 1.1075 | 0.181142 | 0.323867 | 0.213769 | 0.989223 | 0.997988 | 0.994666 |
| Prrc2b_F8WHT3_S621s_1 | Prrc2b_F8 | -1.28 | -0.63 | -1.3 | -0.86 | -0.36 | -1.86 | -0.6 | -1.52 | -1 | -0.77 | -1.39 | -0.94 | 0.6 | -0.22 | -0.51 | -0.82 | -0.0675 | -0.0075 | 0.78 | 0.056954 | 0.86064 | 0.977753 | 0.996493 | 0.997988 | 0.994666 |
| Prrc2c_S4R294_S1965s_1 | Prrc2c_S4R | -2.9 | -0.7 | -1.63 | -0.96 | -1.49 | -1.94 | -0.48 | -1.11 | -3.5 | -1.76 | -0.29 | -1.71 | -2.58 | -1.39 | -1.41 | -2.31 | 0.2925 | -0.2675 | -0.375 | 0.492203 | 0.582336 | 0.717372 | 0.989223 | 0.997988 | 0.994666 |
| Prrc2c_S4R294_S2646s_1 | Prrc2c_S4R | -1.09 | -2.01 | -2.42 | -3.96 | -2.19 | -1.51 | -1.26 | -2.07 | 0.66 | -1.73 | -1.17 | -0.24 | 1.63 | -2.11 | 0.11 | -0.88 | 0.6125 | 1.75 | 2.0575 | 0.038232 | 0.300789 | 0.034363 | 0.989223 | 0.997988 | 0.994666 |
| Prrc2c_S4R294_S761s_1 | Prrc2c_S4R | 0.24 | 0.67 | -1.07 | -0.98 | 0.23 | 0.31 | 1.04 | 0.67 | 0.95 | 0.42 | 2.1 | -0.42 | 0.79 | 0.37 | -0.52 | -1.19 | 0.8475 | 1.0475 | 0.1475 | 0.797045 | 0.081833 | 0.119731 | 0.989223 | 0.997988 | 0.999507 |
| Prrc2c_S4R294_S899s_1 | Prrc2c_S4R | -5.4 | -6.2 | -3.56 | -1.96 | -5.28 | -4.52 | -1.86 | -3.44 | -1.16 | -5.06 | -1.22 | -2.33 | 0.06 | -3.75 | -1.8 | -2.46 | 0.505 | 1.8375 | 2.2925 | 0.056131 | 0.62961 | 0.138081 | 0.989223 | 0.997988 | 0.994666 |
| Prrc2c_S4R294_S901s_2 | Prrc2c_S4R | -1.82 | -2.35 | -2.04 | -2.28 | -3.5 | -1.7 | -4.57 | -3.82 | -0.74 | -1.88 | -1.83 | -2.96 | -1.97 | -2.46 | -0.72 | -1.02 | -1.275 | 0.27 | 0.58 | 0.190441 | 0.044062 | 0.553408 | 0.989223 | 0.997988 | 0.994666 |
| Prrc2c_S4R294_S904s_1 | Prrc2c_S4R | -4.98 | -3.03 | -3.58 | -4.01 | -3.96 | -3.49 | -3.01 | -5.89 | -0.21 | -2.96 | -3.18 | -4 | 0.97 | -3.03 | -2.54 | -2.83 | -0.1875 | 1.3125 | 2.0425 | 0.045685 | 0.778989 | 0.135619 | 0.995762 | 0.997988 | 0.994666 |
| Prrc2c_S4R294_S904s_2 | Prrc2c_S4R | -1.7 | -2.43 | -2.04 | -2.28 | -3.64 | -2.24 | -4.57 | -3.82 | -1.31 | -2.09 | -1.83 | -2.96 | -2.18 | -3.55 | -0.72 | -1.02 | -1.455 | 0.065 | 0.245 | 0.686197 | 0.011733 | 0.865193 | 0.989223 | 0.997988 | 0.996159 |
| Prrc2c_S4R294_T2625t_1 | Prrc2c_S4R | -1.35 | -1.48 | -3.09 | -2.08 | -0.8 | -1.67 | -1.27 | -2.49 | -1.93 | -1.32 | -2.03 | -2.57 | -1.63 | -1.5 | -2.63 | -2.08 | 0.4425 | 0.0375 | 0.04 | 0.930209 | 0.378142 | 0.934044 | 0.989223 | 0.997988 | 0.999507 |
| Prrc2c_S4R294_T358t_1 | Prrc2c_S4R | 0.06 | -0.78 | -1.6 | -1.83 | -1.17 | 1.15 | -1.67 | -1.76 | 1.05 | 0.3 | -2.01 | -1.71 | 1.33 | -2.62 | -0.32 | -1.76 | 0.175 | 0.445 | 0.195 | 0.81857 | 0.805651 | 0.568596 | 0.995762 | 0.997988 | 0.999507 |
| Prrt2_E9PUL5_S100s_1 | Prrt2_E9PU | 1.04 | 0.68 | 0.25 | 0.18 | -0.29 | 1.54 | -1.13 | -0.25 | 1.13 | 0.68 | -1.17 | -0.4 | 0.71 | -0.82 | -0.32 | 0.25 | -0.57 | -0.4775 | -0.5825 | 0.1655 | 0.30627 | 0.371255 | 0.989223 | 0.997988 | 0.994666 |
| Prrt2_E9PUL5_S102s_1 | Prrt2_E9PU | 0.39 | 0.89 | 0.93 | 0.93 | 0.34 | 0.2 | 0.14 | 0.42 | 0.81 | 0.46 | 0.93 | 0.83 | 1.24 | 0.26 | 0.9 | 1.15 | -0.51 | -0.0275 | 0.1025 | 0.742263 | 0.045792 | 0.910927 | 0.989223 | 0.997988 | 0.9971 |
| Prrt2_E9PUL5_S102s_2 | Prrt2_E9PU | 0.42 | -0.42 | 0.32 | -0.16 | -0.58 | 0.93 | -0.21 | -0.38 | 0.51 | -0.9 | -0.37 | 0 | -1.07 | -1.18 | -0.27 | 0.15 | -0.1 | -0.23 | -0.6325 | 0.128225 | 0.79401 | 0.535145 | 0.995762 | 0.997988 | 0.994666 |
| Prrt2_E9PUL5_S104s_1 | Prrt2_E9PU | 0.31 | -0.25 | 0.66 | -0.09 | -0.04 | 0.15 | -0.4 | -0.33 | 0.43 | -0.44 | 0.37 | 0.19 | 0.22 | -0.34 | -0.45 | 0.26 | -0.3125 | -0.02 | -0.235 | 0.471856 | 0.277971 | 0.949562 | 0.989223 | 0.997988 | 0.994666 |
| Prrt2_E9PUL5_S104s_2 | Prrt2_E9PU | 0.07 | -1.23 | -0.42 | 0 | -1.23 | 0.92 | -0.73 | -0.61 | -0.03 | -1.5 | 0.42 | -1.09 | -1.15 | -2.21 | -0.38 | -0.9 | -0.0175 | -0.155 | -0.765 | 0.127721 | 0.972318 | 0.759601 | 0.999187 | 0.997988 | 0.994666 |
| Prrt2_E9PUL5_S208s_1 | Prrt2_E9PU | 0.66 | 1.06 | 1.4 | 0.8 | 0.56 | 0.73 | 1.23 | 1 | -1.83 | 0.91 | -0.6 | 1.37 | -0.53 | -2.51 | 0.87 | 1.21 | -0.1 | -1.0175 | -1.22 | 0.134838 | 0.707015 | 0.157699 | 0.995437 | 0.997988 | 0.994666 |
| Prrt2_E9PUL5_S244s_1 | Prrt2_E9PU | 3.4 | 1.81 | 2.68 | 0.66 | 1.55 | 4.23 | 1.75 | 2.5 | 1.36 | 2.59 | -2.62 | 2.8 | 1.57 | -3.21 | 3.58 | 2.06 | 0.37 | -1.105 | -1.1375 | 0.412088 | 0.621398 | 0.375694 | 0.989223 | 0.997988 | 0.994666 |
| Prrt2_E9PUL5_S254s_1 | Prrt2_E9PU | 2.8 | 1.16 | 1.83 | 0.7 | 2.37 | 4.28 | 2.18 | 1.2 | -0.08 | 2.94 | 1 | 1.89 | -0.48 | -0.26 | 1.57 | 2.29 | 0.885 | -0.185 | -0.8425 | 0.265771 | 0.223455 | 0.794562 | 0.989223 | 0.997988 | 0.994666 |
| Prrt2_E9PUL5_S6s_1 | Prrt2_E9PU | 0.8 | 0.33 | 0.46 | -0.25 | 0.61 | 0.59 | -0.68 | -0.37 | 0.82 | 0.48 | 0.11 | -0.69 | 1.2 | 0.49 | -0.28 | 0 | -0.2975 | -0.155 | 0.0175 | 0.96492 | 0.447755 | 0.693211 | 0.989223 | 0.997988 | 0.999507 |
| Prrt2_E9PUL5_S98s_1 | Prrt2_E9PU | 0.34 | 1.38 | 0.82 | 0.42 | 0.32 | 0.05 | -0.58 | -0.05 | 0.64 | 0.13 | -0.51 | 0.11 | 1.2 | -0.06 | 0.08 | 0.54 | -0.805 | -0.6475 | -0.3 | 0.44193 | 0.029138 | 0.090568 | 0.989223 | 0.997988 | 0.994666 |
| Prrt2_E9PUL5_S98s_2 | Prrt2_E9PU | 0.09 | -0.25 | 0.12 | -0.11 | -0.66 | 0.91 | -0.35 | -0.44 | 0.72 | -0.78 | -0.13 | -0.28 | -0.76 | -1.26 | -0.3 | -0.11 | -0.0975 | -0.08 | -0.57 | 0.099492 | 0.788081 | 0.815452 | 0.995762 | 0.997988 | 0.994666 |
| Prrt2_E9PUL5_T214t_1 | Prrt2_E9PU | 0.61 | 1.31 | 1.24 | 0.64 | 0.69 | 0.75 | 0.24 | 0.33 | 0.75 | 0.67 | 0.52 | 0.73 | 1.4 | 0.39 | 0.61 | 0.58 | -0.4475 | -0.2825 | -0.205 | 0.541292 | 0.122268 | 0.294618 | 0.989223 | 0.997988 | 0.994666 |
| Prrt3_Q6PE13_S783s_1 | Prrt3_Q6P | 1.05 | 1.15 | 0.96 | 0.98 | 0.4 | 0.69 | 0.65 | 1.38 | 0.66 | 1.06 | 1.28 | 1.64 | 0.99 | 1.13 | 0.53 | 1.49 | -0.255 | 0.125 | -8.8E-17 | 1 | 0.342933 | 0.643847 | 0.989223 | 0.997988 | 1 |
| Prrt3_Q6PE13_S783s_2 | Prrt3_Q6P | 0.03 | -0.01 | 0.22 | 0.71 | 0.14 | -0.01 | 0.89 | 0.69 | -0.28 | -0.02 | 0.35 | 0.52 | -1.23 | 0.32 | 0.4 | 0.83 | 0.19 | -0.095 | -0.1575 | 0.734503 | 0.527491 | 0.743767 | 0.989223 | 0.997988 | 0.9971 |
| Prrt3_Q6PE13_S791s_2 | Prrt3_Q6P | 0.05 | -1.08 | -0.49 | -0.13 | 0.67 | 0 | 0.3 | 0.28 | -0.21 | -0.31 | -0.51 | -0.16 | -1.56 | 0.02 | -0.87 | -0.12 | 0.725 | 0.115 | -0.22 | 0.615599 | 0.037811 | 0.702722 | 0.989223 | 0.997988 | 0.995259 |
| Prrt3_Q6PE13_S799s_1 | Prrt3_Q6P | 0.52 | 1.23 | 1.14 | 0.94 | 1.21 | 0.67 | 1.26 | 1.11 | 0.56 | 1.04 | 0.43 | 0.74 | 0.57 | 1.07 | 0.95 | 0.77 | 0.105 | -0.265 | -0.1175 | 0.670594 | 0.685213 | 0.333959 | 0.994624 | 0.997988 | 0.996159 |
| Prrt3_Q6PE13_S799s_2 | Prrt3_Q6P | 0.03 | -0.26 | -0.15 | 0.27 | 0.41 | 0.13 | 0.58 | 0.47 | -0.36 | -0.11 | -0.11 | 0.16 | -1.17 | 0.24 | -0.24 | 0.33 | 0.425 | -0.0775 | -0.1825 | 0.631595 | 0.087568 | 0.748311 | 0.989223 | 0.997988 | 0.995259 |
| Prrt3_Q6PE13_S806s_1 | Prrt3_Q6P | -2.63 | -3.37 | -3.75 | -4.49 | -1.64 | -3.85 | -2.41 | -3.4 | -5.11 | -4.07 | -0.51 | -3.45 | -4.35 | -2.53 | -5.1 | -3.71 | 0.735 | 0.275 | -0.3625 | 0.555025 | 0.215232 | 0.769732 | 0.989223 | 0.997988 | 0.99525 |
| Prrt3_Q6PE13_S833s_1 | Prrt3_Q6P | 0.42 | 0.29 | 0.34 | 0.52 | 0.57 | 0.35 | 0.65 | 0.36 | -0.15 | 0.05 | -0.2 | 0.37 | -0.24 | 0.67 | 0.02 | 0.08 | 0.09 | -0.375 | -0.26 | 0.361762 | 0.660919 | 0.132433 | 0.990434 | 0.997988 | 0.994666 |
| Prrt3_Q6PE13_S845s_1 | Prrt3_Q6P | 0.29 | 0.57 | 0.37 | 0.47 | 0.76 | 0.2 | 0.66 | 0.4 | 0.08 | 0.45 | 0.15 | 0.43 | -0.29 | 0.67 | 0.12 | 0.37 | 0.08 | -0.1475 | -0.2075 | 0.473473 | 0.72118 | 0.511419 | 0.995437 | 0.997988 | 0.994666 |
| Prrt3_Q6PE13_S865s_1 | Prrt3_Q6P | -2.11 | -2.54 | -2.11 | -2.68 | -1.4 | -1.97 | -1.79 | -2.34 | -3.52 | -2.47 | -1.14 | -2.97 | -2.61 | -1.76 | -2.6 | -2.6 | 0.485 | -0.165 | -0.0325 | 0.916642 | 0.108966 | 0.741471 | 0.989223 | 0.997988 | 0.999507 |
| Prune2_Q52KR3_S2850s_1 | Prune2_Q5 | -1.18 | -0.96 | 0.25 | -0.46 | -0.42 | -1.24 | 0 | 0.56 | -1.25 | -0.97 | 0.29 | -1.18 | -1.91 | -0.52 | -0.88 | -0.65 | 0.3125 | -0.19 | -0.4025 | 0.370123 | 0.50211 | 0.680809 | 0.989223 | 0.997988 | 0.994666 |
| Prune2_Q52KR3_S3070s_1 | Prune2_Q5 | 0.89 | 0.94 | 0.56 | 0.66 | 1.23 | 0.43 | 0.52 | 1.11 | 0.69 | 0.71 | 0.2 | 1.52 | 0.12 | -0.44 | 0.98 | 0.89 | 0.06 | 0.0175 | -0.375 | 0.323046 | 0.822148 | 0.956091 | 0.995762 | 0.997988 | 0.994666 |
| Psd_Q5DTT2_S1011s_1 | Psd_Q5DT | 0.49 | -3.75 | -1.43 | -1.81 | -2.81 | 1.52 | -2.11 | -0.31 | 0.47 | -2.76 | -0.31 | -1.88 | -0.09 | -2.22 | -2.23 | -2.38 | 0.6975 | 0.505 | -0.105 | 0.907024 | 0.538516 | 0.617112 | 0.989223 | 0.997988 | 0.999507 |
| Psd_Q5DTT2_S126s_1 | Psd_Q5DT | 2.97 | 1 | 1.3 | -0.49 | 1.16 | 3.1 | 2.29 | 1.69 | -2.5 | 1.62 | -2.92 | 1.81 | -1.97 | -1.87 | 0.97 | 1.38 | 0.865 | -1.6925 | -1.5675 | 0.140246 | 0.250031 | 0.208559 | 0.989223 | 0.997988 | 0.994666 |
| Psd_Q5DTT2_S17s_1 | Psd_Q5DT | 0.03 | -0.82 | -0.78 | -0.66 | -1.14 | -0.43 | -0.16 | -0.44 | -1.8 | -0.32 | -1.65 | -0.99 | -1 | -2.76 | -0.66 | -0.74 | 0.015 | -0.6325 | -0.7325 | 0.169235 | 0.961226 | 0.133346 | 0.999187 | 0.997988 | 0.994666 |
| Psd_Q5DTT2_S283s_1 | Psd_Q5DT | -1.18 | -2.48 | -1.8 | -3.51 | -1.62 | -1.58 | -1.32 | -1.16 | -1.13 | -1.16 | 1.62 | -2.62 | -0.81 | -0.19 | -2.46 | -2.51 | 0.8225 | 1.42 | 0.75 | 0.292652 | 0.107418 | 0.140691 | 0.989223 | 0.997988 | 0.994666 |
| Psd_Q5DTT2_S34s_1 | Psd_Q5DT | 0.08 | 0.1 | 0.05 | 0.19 | 0.42 | 0.13 | 0.33 | 0.22 | -0.05 | 0.04 | 0.02 | -0.01 | -0.73 | 0.21 | -0.39 | 0.06 | 0.17 | -0.105 | -0.3175 | 0.28579 | 0.400858 | 0.605777 | 0.989223 | 0.997988 | 0.994666 |
| Psd_Q5DTT2_S720s_1 | Psd_Q5DT | 0.65 | -0.46 | -0.36 | 0.46 | -0.04 | 0.66 | 0.39 | 0.7 | -0.1 | -0.23 | 0.13 | 0.29 | -0.66 | -0.13 | -0.99 | -0.82 | 0.355 | -0.05 | -0.7225 | 0.070213 | 0.306608 | 0.879361 | 0.989223 | 0.997988 | 0.994666 |
| Psd_Q5DTT2_S736s_1 | Psd_Q5DT | -0.03 | -0.7 | 0.21 | 1.12 | 0.37 | -0.08 | 1.17 | 1.35 | -0.52 | -0.26 | -0.41 | 1.11 | -0.41 | 0.97 | 0.29 | 0.76 | 0.5525 | -0.17 | 0.2525 | 0.591443 | 0.255758 | 0.736087 | 0.989223 | 0.997988 | 0.995259 |
| Psd2_Q6P1I6_S473s_1 | Psd2_Q6P1 | 1.1 | 0.96 | -0.89 | -1.35 | 0.75 | 1.6 | -0.12 | -0.98 | 1.26 | 0.98 | -0.25 | -1.66 | 0.1 | 0.9 | -0.9 | -0.65 | 0.3575 | 0.1275 | -0.0925 | 0.890502 | 0.629538 | 0.875959 | 0.989223 | 0.997988 | 0.999507 |
| Psd3_E9PUC5_S337s_1 | Psd3_E9PU | 5.4 | 1.41 | 1.28 | -0.45 | 1.69 | 6.11 | 3.04 | 1.63 | -0.06 | 3.97 | -0.35 | 2.17 | -2.02 | -1.46 | 2.44 | 2.4 | 1.2075 | -0.4775 | -1.57 | 0.304895 | 0.39515 | 0.733685 | 0.989223 | 0.997988 | 0.994666 |
| Psd3_E9PUC5_S339s_1 | Psd3_E9PU | 3.52 | 1.63 | 2.29 | 1.09 | 1.7 | 3.69 | 2.49 | 1.97 | 0.02 | 2.98 | 0.77 | 2.46 | -0.06 | 0.7 | 2.04 | 2.48 | 0.33 | -0.575 | -0.8425 | 0.249533 | 0.5918 | 0.467458 | 0.989223 | 0.997988 | 0.994666 |
| Psd3_E9PUC5_S340s_2 | Psd3_E9PU | 3.09 | 1.26 | 1.87 | 1.07 | 1.92 | 3.51 | 2.06 | 1.73 | -0.76 | 2.41 | 0.39 | 1.38 | 0.34 | -0.15 | 2.06 | 2.1 | 0.4825 | -0.9675 | -0.735 | 0.284967 | 0.39193 | 0.209065 | 0.989223 | 0.997988 | 0.994666 |
| Psd3_E9PUC5_S342s_1 | Psd3_E9PU | 4.2 | 2.26 | 2.04 | 0.88 | 1.67 | 4.08 | 1.39 | 1.33 | 1.01 | 3.68 | -0.29 | 1.29 | 0.56 | 0.67 | 1.56 | 2.03 | -0.2275 | -0.9225 | -1.14 | 0.125382 | 0.784388 | 0.345429 | 0.995762 | 0.997988 | 0.994666 |
| Psd3_E9PUC5_S348s_1 | Psd3_E9PU | 2.2 | 2.25 | 2.32 | 1.68 | 1.65 | 2.15 | 1.66 | 2.36 | 0.63 | 2.42 | 1 | 2.39 | 1.43 | 0.77 | 2.48 | 2.3 | -0.1575 | -0.5025 | -0.3675 | 0.393197 | 0.565669 | 0.296388 | 0.989223 | 0.997988 | 0.994666 |
| Psd3_E9PUC5_S348s_2 | Psd3_E9PU | 2.4 | 1.01 | 1.52 | 1.02 | 1.64 | 2.89 | 2.04 | 1.59 | 0 | 2.09 | 0.44 | 1.45 | 0.11 | 0.26 | 1.78 | 1.91 | 0.5525 | -0.4925 | -0.4725 | 0.391892 | 0.210362 | 0.369736 | 0.989223 | 0.997988 | 0.994666 |
| Psd3_E9PUC5_S44s_1 | Psd3_E9PU | 0.64 | 0.59 | -1.2 | -0.93 | 0.7 | 1 | -0.79 | 0.04 | 0.61 | 0.93 | -0.86 | -0.88 | -0.1 | 0.21 | -1.58 | -1.66 | 0.4625 | 0.175 | -0.5575 | 0.383514 | 0.422587 | 0.779125 | 0.989223 | 0.997988 | 0.994666 |
| Psd3_E9PUC5_S76s_1 | Psd3_E9PU | -1.3 | -1.04 | -0.1 | -1.14 | -2.3 | -0.66 | -1.04 | -0.88 | 1.07 | -1.46 | -1.47 | -0.14 | 2.09 | -2.19 | 1.57 | -0.65 | -0.325 | 0.395 | 1.1 | 0.241826 | 0.457793 | 0.521871 | 0.989223 | 0.997988 | 0.994666 |
| Psd3_E9PUC5_S78s_1 | Psd3_E9PU | -1.54 | -2.54 | -0.83 | -2.47 | -1.9 | -1.52 | -1.03 | -1.41 | -0.33 | -1.62 | -1.66 | -1.13 | -0.45 | -2.05 | 0.78 | -1.27 | 0.38 | 0.66 | 1.0975 | 0.120409 | 0.378794 | 0.195942 | 0.989223 | 0.997988 | 0.994666 |
| Psd3_E9PUC5_S98s_1 | Psd3_E9PU | -2.24 | -2.53 | -2.02 | -1.58 | -1.38 | -1.81 | -0.98 | -1.38 | -3.05 | -2.89 | -0.49 | -1.43 | -2.65 | -0.94 | -2.35 | -1.74 | 0.705 | 0.1275 | 0.1725 | 0.685439 | 0.034739 | 0.829252 | 0.989223 | 0.997988 | 0.996159 |
| Psen2_Q3U4P5_S25s_1 | Psen2_Q3U | 0.42 | 0.38 | 0.6 | 0.08 | 0.11 | 0.26 | 0.13 | 0.05 | 0.6 | 0.83 | 0.92 | 0.16 | 0.93 | 0.61 | 0.32 | 0.17 | -0.2325 | 0.2575 | 0.1375 | 0.624099 | 0.291426 | 0.342433 | 0.989223 | 0.997988 | 0.995259 |
| Psen2_Q3U4P5_S52s_1 | Psen2_Q3U | -1.48 | -2.34 | -0.89 | -1.55 | -2.88 | -0.44 | -2.88 | -0.7 | 0.31 | -1.61 | 0.29 | -0.39 | -0.88 | -3.5 | -1.69 | -1.27 | -0.16 | 1.215 | -0.27 | 0.651678 | 0.805581 | 0.036978 | 0.995762 | 0.997988 | 0.995259 |
| Psip1_Q99JF8_S106s_1 | Psip1_Q99 | -2.28 | -3 | -1.57 | -1.32 | -2.58 | -3.22 | -1.82 | -1.57 | -1.94 | -1.82 | -0.15 | -1.41 | -1.76 | -2.28 | -0.68 | -1.59 | -0.255 | 0.7125 | 0.465 | 0.346387 | 0.605183 | 0.194124 | 0.989223 | 0.997988 | 0.994666 |
| Psip1_Q99JF8_S176s_1 | Psip1_Q99 | 2.07 | 0.35 | 1.65 | 1.84 | 0.78 | 0.89 | 1.32 | 2.37 | 0.51 | 1.91 | 1.42 | 1.45 | 0.54 | -0.45 | 1.16 | 1.6 | -0.1375 | -0.155 | -0.765 | 0.183499 | 0.777993 | 0.738408 | 0.995762 | 0.997988 | 0.994666 |
| Psip1_Q99JF8_T122t_1 | Psip1_Q99 | -7.41 | -2.12 | -0.06 | -0.41 | -5.56 | -7.77 | -0.46 | -0.68 | -3.87 | -6.52 | -0.89 | 0.18 | -2.25 | -4.64 | -0.28 | 0.39 | -1.1175 | -0.275 | 0.805 | 0.64639 | 0.600729 | 0.889385 | 0.989223 | 0.997988 | 0.995259 |
| Psma2_P49722_S7s_1 | Psma2_P49 | 0.09 | 0.42 | 1.01 | -0.1 | -0.34 | -0.24 | 0.27 | -0.54 | -0.14 | 0.17 | -2.09 | 0.37 | -0.87 | 0.33 | 0.9 | 0.45 | -0.5675 | -0.7775 | -0.1525 | 0.729808 | 0.09736 | 0.193756 | 0.989223 | 0.997988 | 0.9971 |
| Psma3_E0CX62_S175s_1 | Psma3_E0C | -0.16 | -0.1 | -0.02 | 0.07 | -0.6 | -0.71 | -0.23 | 0.19 | -0.31 | -0.15 | 0.66 | 0.09 | -1.24 | 0.05 | -0.04 | 0.09 | -0.285 | 0.125 | -0.2325 | 0.513986 | 0.287419 | 0.649859 | 0.989223 | 0.997988 | 0.994666 |
| Psma5_Q9Z2U1_S16s_1 | Psma5_Q9 | 2.6 | 2.86 | -0.19 | 0.18 | 2.9 | 3.24 | -0.13 | -0.02 | 2.98 | 2.42 | -0.11 | -0.06 | 1.45 | 3.1 | -0.04 | -0.11 | 0.135 | -0.055 | -0.2625 | 0.784325 | 0.897032 | 0.956223 | 0.996493 | 0.997988 | 0.998711 |
| Psma5_Q9Z2U1_S56s_1 | Psma5_Q9 | 0.11 | 0.68 | 0.64 | 0.05 | 0.44 | -0.17 | 0.4 | -0.09 | 0.36 | 0.45 | 0.28 | 0.55 | -0.15 | 0.57 | 0.18 | -0.15 | -0.225 | 0.04 | -0.2575 | 0.401257 | 0.417647 | 0.873579 | 0.989223 | 0.997988 | 0.994666 |
| Psmd1_Q3TXS7_T273t_1 | Psmd1_Q3 | -0.05 | 0.51 | 0.94 | 1.5 | 0.61 | -0.02 | 0.95 | 1.09 | -0.01 | 0.23 | 0.86 | 1.2 | 0.37 | 0.94 | 0.77 | 1.04 | -0.0675 | -0.155 | 0.055 | 0.883861 | 0.864739 | 0.713948 | 0.996493 | 0.997988 | 0.999507 |
| Psmd1_Q3TXS7_T311t_1 | Psmd1_Q3 | 0.6 | 0.37 | 2.28 | 0.79 | 0.18 | 0.83 | 1.48 | 2.08 | 0.54 | 1.08 | 2.46 | 1.07 | 1.39 | 0.64 | 1.11 | 1.06 | 0.1325 | 0.2775 | 0.04 | 0.9287 | 0.806125 | 0.617544 | 0.995762 | 0.997988 | 0.999507 |
| Psmd11_G3UX67_S7s_1 | Psmd11_G | -0.51 | -1.1 | -0.06 | -0.03 | 0.14 | -0.35 | 0.12 | 0.32 | -1.03 | -0.62 | 0.28 | 0.21 | -0.85 | 0.03 | -0.29 | 0.03 | 0.4825 | 0.135 | 0.155 | 0.662361 | 0.142384 | 0.738322 | 0.989223 | 0.997988 | 0.995804 |
| Psmd2_Q8VDM4_S361s_1 | Psmd2_Q8 | -0.21 | 0.07 | 0.03 | -0.29 | 0.03 | -0.31 | -0.3 | -0.21 | -0.24 | -0.2 | -0.27 | -0.31 | 0.1 | 0.41 | -0.29 | -0.23 | -0.0975 | -0.155 | 0.0975 | 0.720223 | 0.650711 | 0.476471 | 0.990358 | 0.997988 | 0.9971 |
| Psmd2_Q8VDM4_T20t_1 | Psmd2_Q8 | 1.01 | 0.71 | 0.53 | 0.66 | 1.19 | 1.01 | 0.53 | 0.22 | 0.97 | 1.17 | 0.71 | 0.35 | 1.08 | 1.56 | 0.72 | 0.62 | 0.01 | 0.0725 | 0.2675 | 0.380311 | 0.971469 | 0.785296 | 0.999187 | 0.997988 | 0.994666 |
| Psmd2_Q8VDM4_T9t_1 | Psmd2_Q8 | 1.23 | 1.46 | 0.76 | 0.91 | 1.03 | 1.23 | -0.31 | 0.88 | -0.04 | 1.54 | -1.63 | -0.04 | 0.05 | 0.29 | 0.63 | 0.72 | -0.3825 | -1.1325 | -0.6675 | 0.042321 | 0.320105 | 0.089344 | 0.989223 | 0.997988 | 0.994666 |
| Psmd4_F7AI87_T65t_1 | Psmd4_F7 | -1.05 | -3.08 | 0.08 | 0.83 | -1.44 | -1.13 | 0.23 | 0.38 | 0.15 | -0.77 | -2.11 | 0.43 | -0.44 | 0 | 0.78 | 1.27 | 0.315 | 0.23 | 1.2075 | 0.16564 | 0.71001 | 0.800444 | 0.995437 | 0.997988 | 0.994666 |
| Psmd9_Q9CR00_S128s_1 | Psmd9_Q9 | 0.99 | 0.79 | 0.94 | 0.44 | 0.98 | 1.07 | 0.94 | 0.31 | 0.72 | 1.22 | 0.12 | 0.54 | 0.54 | 0.59 | 0.68 | 0.39 | 0.035 | -0.14 | -0.24 | 0.350918 | 0.893519 | 0.641858 | 0.996493 | 0.997988 | 0.994666 |
| Psmd9_Q9CR00_S2s_1 | Psmd9_Q9 | -0.44 | -0.04 | 0.39 | -0.36 | -0.69 | 0.13 | -0.72 | -0.47 | 2.64 | -0.16 | 0.26 | -0.88 | 2.94 | -0.21 | 0.5 | 0.44 | -0.325 | 0.5775 | 1.03 | 0.136963 | 0.291401 | 0.423075 | 0.989223 | 0.997988 | 0.994666 |
| Psmf1_Q8BHL8_S153s_1 | Psmf1_Q8 | 0.42 | 0.3 | 1.51 | 1.92 | 0.1 | 0.8 | 1.27 | 2.04 | 1.29 | 1.27 | 1.92 | 1.44 | 1.06 | 0.7 | 1.36 | 1.09 | 0.015 | 0.4425 | 0.015 | 0.971556 | 0.976984 | 0.306997 | 0.999187 | 0.997988 | 0.999507 |
| Psmg1_Q9JK23_T18t_1 | Psmg1_Q9 | 1.65 | 1.03 | 0.21 | -0.38 | 0.77 | 1.51 | -1.19 | -0.22 | 1.71 | 1.23 | -0.15 | -0.51 | 0.31 | 1.49 | -0.44 | -0.7 | -0.41 | -0.0575 | -0.4625 | 0.451853 | 0.535422 | 0.927659 | 0.989223 | 0.997988 | 0.994666 |
| Pspc1_Q8R326_S477s_1 | Pspc1_Q8R | -0.33 | -0.47 | -0.48 | -0.4 | -0.23 | -0.69 | -0.17 | -0.45 | -0.47 | -0.32 | -0.51 | -0.45 | 0.02 | -0.34 | -0.36 | -0.22 | 0.035 | -0.0175 | 0.195 | 0.417178 | 0.871282 | 0.931895 | 0.996493 | 0.997988 | 0.994666 |
| Ptbp1_F7DCW4_S91s_1 | Ptbp1_F7D | -1.81 | -3.22 | -1.92 | -4.18 | -2.51 | -2.52 | -2.78 | -2.68 | 0.34 | -3.28 | -3.78 | -1.7 | 1.63 | -2.88 | -0.78 | -3.17 | 0.16 | 0.6775 | 1.4825 | 0.192574 | 0.758018 | 0.486673 | 0.995762 | 0.997988 | 0.994666 |
| Ptbp2_Q91Z31_S308s_1 | Ptbp2_Q91 | -1.13 | -1.48 | -3.09 | -1.73 | -1.2 | -1.9 | -1.65 | -2.14 | -1.95 | -0.67 | -0.02 | -1.53 | -1.37 | -0.46 | -1.02 | -1.18 | 0.135 | 0.815 | 0.85 | 0.088296 | 0.76138 | 0.17152 | 0.995762 | 0.997988 | 0.994666 |
| Ptdss1_Q99LH2_S442s_1 | Ptdss1_Q9 | -0.89 | -0.83 | 0.31 | 0.22 | -1.03 | -0.37 | -0.04 | 0.49 | 1.05 | -0.89 | 0.33 | 0.55 | 2 | -1.69 | 0.64 | 0.51 | 0.06 | 0.5575 | 0.6625 | 0.380001 | 0.888885 | 0.277258 | 0.996493 | 0.997988 | 0.994666 |
| Ptdss2_G3UYQ3_S16s_1 | Ptdss2_G3 | 0.26 | -0.44 | 0.46 | 0.22 | 0.19 | -0.08 | 0.65 | 0.14 | -1.09 | 0.39 | -0.57 | 0.39 | 0.2 | -0.66 | 0.82 | 0.81 | 0.1 | -0.345 | 0.1675 | 0.68011 | 0.724291 | 0.410293 | 0.995437 | 0.997988 | 0.996159 |
| Ptgdr2_Q9Z2J6_S375s_1 | Ptgdr2_Q9 | -1.8 | -0.67 | -0.93 | -1.5 | -0.88 | -0.08 | -1.54 | -1.38 | 0.38 | 0.03 | 0.55 | -1.02 | 0.98 | -0.43 | -1.14 | -0.51 | 0.255 | 1.21 | 0.95 | 0.078051 | 0.52965 | 0.017847 | 0.989223 | 0.997988 | 0.994666 |
| Ptgfrn_Q9WV91_S875s_1 | Ptgfrn_Q9 | 0.09 | 0.19 | 0.27 | -0.22 | 1.05 | 0.17 | -0.27 | -0.37 | 1.27 | 0.07 | 0.33 | -0.97 | -0.36 | 0.63 | -0.47 | -0.38 | 0.0625 | 0.0925 | -0.2275 | 0.489654 | 0.85626 | 0.838621 | 0.996349 | 0.997988 | 0.994666 |
| Ptk2_P34152_S910s_1 | Ptk2_P341 | 0.09 | 0.47 | 0.3 | 0.51 | 0.43 | -0.01 | 0.65 | 0.49 | 0.02 | 0.25 | 0.28 | 0.37 | 0.09 | 0.64 | 0.17 | 0.22 | 0.0475 | -0.1125 | -0.0625 | 0.808898 | 0.841807 | 0.619683 | 0.996064 | 0.997988 | 0.999507 |
| Ptk2_P34152_Y576y_1 | Ptk2_P341 | 0.64 | -0.27 | 0.53 | 0.92 | -0.03 | 0.36 | 0.36 | 0.33 | -0.46 | 0.1 | 0.83 | 0.47 | -0.36 | -0.1 | 0.43 | 0.04 | -0.2 | -0.22 | -0.4525 | 0.203605 | 0.507335 | 0.566009 | 0.989223 | 0.997988 | 0.994666 |
| Ptk2b_E9Q2A6_S375s_1 | Ptk2b_E9Q | -0.11 | -0.09 | -0.49 | -0.99 | 0.69 | -0.3 | 0.09 | -0.49 | -0.14 | 0.14 | -0.41 | -0.49 | -0.05 | 0 | -0.85 | -0.38 | 0.4175 | 0.195 | 0.1 | 0.761865 | 0.239731 | 0.515658 | 0.989223 | 0.997988 | 0.9971 |
| Ptma_P26350_S2s_1 | Ptma_P263 | 0.78 | 1.34 | -0.71 | -1.57 | 1.13 | 0.08 | -0.88 | -0.86 | 0.71 | 1.15 | -0.89 | -1.33 | -0.85 | 0.67 | -1.05 | -0.79 | -0.0925 | -0.05 | -0.465 | 0.50884 | 0.898044 | 0.950328 | 0.996493 | 0.997988 | 0.994666 |
| Ptms_Q9D0J8_S5s_1 | Ptms_Q9D | -2.26 | -2.27 | -2.25 | -2.39 | -1.96 | -2.9 | -2.25 | -2.66 | -2.52 | -1.49 | -1.79 | -2.46 | -1.86 | -1.81 | -2.69 | -2.69 | -0.15 | 0.2275 | 0.03 | 0.921867 | 0.568341 | 0.450646 | 0.989223 | 0.997988 | 0.999507 |
| Ptov1_Q91VU8_S53s_1 | Ptov1_Q91 | -2.22 | -2.09 | -1.37 | -2.33 | -1.95 | -2.56 | -2.36 | -2.08 | -1.56 | -1.89 | -1.41 | -2.33 | -0.2 | -2 | -1.66 | -1.53 | -0.235 | 0.205 | 0.655 | 0.161275 | 0.422511 | 0.531838 | 0.989223 | 0.997988 | 0.994666 |
| Ptpdc1_Q6NZK8_S475s_1 | Ptpdc1_Q6 | -0.23 | -0.11 | 1.39 | 0.69 | -0.36 | -1.08 | -0.38 | 0.74 | -0.28 | 0.56 | 2.69 | -0.04 | 0.41 | 0.17 | 0.97 | 0.69 | -0.705 | 0.2975 | 0.125 | 0.764169 | 0.173459 | 0.671609 | 0.989223 | 0.997988 | 0.9971 |
| Ptpn1_P35821_S364s_1 | Ptpn1_P35 | -1.02 | -2.73 | -0.02 | -0.34 | -2.43 | -1.89 | -0.63 | -0.63 | -0.87 | -0.54 | -0.41 | 0.69 | 0.98 | -2.07 | -0.08 | 0.38 | -0.3675 | 0.745 | 0.83 | 0.308965 | 0.586039 | 0.256255 | 0.989223 | 0.997988 | 0.994666 |
| Ptpn11_P35235_Y542y_1 | Ptpn11_P3 | 3.76 | 1.47 | 1.86 | 0.52 | 0.99 | 3.93 | 2 | 2.22 | -1.3 | 2.61 | -0.25 | 2.03 | -0.79 | -0.29 | 2.3 | 2.63 | 0.3825 | -1.13 | -0.94 | 0.343523 | 0.633828 | 0.281825 | 0.989223 | 0.997988 | 0.994666 |
| Ptpn11_P35235_Y62y_1 | Ptpn11_P3 | 0.8 | 0.94 | 0.68 | 0.78 | 0.4 | -0.07 | 0.75 | 0.51 | 0.27 | 1.21 | -0.59 | 0.41 | 0.11 | -0.46 | 0.36 | 0.32 | -0.4025 | -0.475 | -0.7175 | 0.02612 | 0.122743 | 0.228708 | 0.989223 | 0.997988 | 0.994666 |
| Ptpn12_P35831_S19s_1 | Ptpn12_P3 | -1.28 | -0.08 | 0.26 | -0.68 | -0.43 | -0.56 | -0.84 | -0.63 | -0.33 | -0.55 | 1.05 | -0.07 | 0.63 | 0.43 | 0.07 | -0.32 | -0.17 | 0.47 | 0.6475 | 0.133542 | 0.630713 | 0.32973 | 0.989223 | 0.997988 | 0.994666 |
| Ptpn12_P35831_S331s_1 | Ptpn12_P3 | -2.01 | -1.41 | -1.24 | -1.36 | -1.44 | -1.93 | -2.21 | -2.2 | -1.5 | -1.13 | -0.56 | -1.69 | -0.5 | -1.3 | -1.52 | -1.62 | -0.44 | 0.285 | 0.27 | 0.436755 | 0.14466 | 0.394734 | 0.989223 | 0.997988 | 0.994666 |
| Ptpn12_P35831_S368s_1 | Ptpn12_P3 | 0.23 | 0.61 | -0.91 | -0.15 | 0.34 | -0.38 | -0.16 | -1.58 | -0.78 | -0.38 | -1.13 | -1.37 | -1.42 | 0 | -1.66 | -0.04 | -0.39 | -0.86 | -0.725 | 0.181539 | 0.425212 | 0.050807 | 0.989223 | 0.997988 | 0.994666 |
| Ptpn12_P35831_S396s_1 | Ptpn12_P3 | 0.92 | 0.73 | 0.68 | 1.14 | 0.08 | 0.67 | 1.08 | 0.53 | -0.17 | 0.85 | 1.13 | 0.83 | -1.15 | -0.34 | 1.53 | 1.53 | -0.2775 | -0.2075 | -0.475 | 0.452299 | 0.321387 | 0.532182 | 0.989223 | 0.997988 | 0.994666 |
| Ptpn12_P35831_S434s_1 | Ptpn12_P3 | -1.05 | -0.2 | -0.91 | -0.93 | -0.48 | -1.36 | -0.43 | -1.05 | -1.08 | -0.77 | -0.24 | -0.61 | -1.69 | -0.37 | -1.06 | -0.9 | -0.0575 | 0.0975 | -0.2325 | 0.521697 | 0.854951 | 0.745777 | 0.996349 | 0.997988 | 0.994666 |

| | | | | | | | | | | | | | | | | | | | | | | | | | | |
|---|---|---|---|---|---|---|---|---|---|---|---|---|---|---|---|---|---|---|---|---|---|---|---|---|---|---|
| Ptpn12_P35831_S632s_1 | Ptpn12_P3 | -0.47 | -0.4 | -0.33 | -0.56 | -0.78 | 0.29 | 0.21 | 0.43 | 0.12 | 0.32 | 0.89 | 0.53 | -0.41 | -0.26 | 0.27 | -0.09 | 0.4775 | 0.905 | 0.3175 | 0.236653 | 0.140111 | 0.004876 | 0.989223 | 0.997988 | 0.994666 |
| Ptpn12_P35831_T519t_1 | Ptpn12_P3 | 0.11 | -0.52 | -0.12 | 0.37 | -0.98 | -0.08 | -0.57 | -0.36 | -0.06 | 0.41 | -0.11 | -0.53 | -0.42 | -0.57 | -0.18 | -0.27 | -0.4575 | -0.0325 | -0.32 | 0.274489 | 0.145079 | 0.915298 | 0.989223 | 0.997988 | 0.994666 |
| Ptpn13_G5E8B1_S2059s_1 | Ptpn13_G5 | 0.3 | 0.5 | 0.24 | 0.25 | 0.26 | 0.09 | -0.32 | 0.4 | 0.28 | 0.64 | 0.47 | -0.05 | 0.3 | 0.46 | 0.23 | -0.07 | -0.215 | 0.0125 | -0.0925 | 0.708369 | 0.373667 | 0.958826 | 0.989223 | 0.997988 | 0.996966 |
| Ptpn2_Q06180_S268s_1 | Ptpn2_Q06 | -1.03 | -0.44 | -1.41 | -1.08 | -1.14 | -1.07 | -0.28 | -0.56 | -1.3 | -1.97 | -0.9 | -1.17 | -0.55 | 0.02 | -1.6 | -1.03 | 0.2275 | -0.345 | 0.2 | 0.622947 | 0.467729 | 0.309088 | 0.989223 | 0.997988 | 0.995259 |
| Ptpn2_Q06180_S298s_1 | Ptpn2_Q06 | -1.49 | -1.23 | -1.43 | -1.01 | -2.04 | -1.23 | -0.04 | -1.87 | -1.68 | -1.45 | -0.02 | -1.37 | -1.41 | -1.23 | -0.75 | -0.84 | -0.005 | 0.16 | 0.2325 | 0.406243 | 0.990883 | 0.683976 | 0.999187 | 0.997988 | 0.994666 |
| Ptpn23_Q6PB44_T744t_1 | Ptpn23_Q6 | -0.46 | -0.16 | 0.46 | 0.83 | -0.37 | -0.77 | -0.1 | -0.34 | -0.4 | -0.88 | -0.31 | -0.11 | -0.89 | -0.61 | 0.23 | 0.02 | -0.5625 | -0.5925 | -0.48 | 0.247879 | 0.116381 | 0.116905 | 0.989223 | 0.997988 | 0.994666 |
| Ptpn3_A2ALK8_S372s_1 | Ptpn3_A2A | 0.59 | 0.77 | 0.79 | 0.71 | 1.35 | 0.5 | 1.22 | 1.03 | 0.41 | 0.32 | 0.15 | 1.12 | -0.14 | 1.37 | 0.5 | 0.51 | 0.31 | -0.215 | -0.155 | 0.655248 | 0.233211 | 0.440482 | 0.989223 | 0.997988 | 0.995259 |
| Ptpn3_A2ALK8_S434s_1 | Ptpn3_A2A | 1.75 | 1.57 | 2.83 | 0.11 | 0.32 | 2.98 | -0.05 | 1.59 | -0.35 | 1.89 | -0.66 | 1.23 | 0.58 | -2.05 | 2.49 | 2.26 | -0.355 | -1.0375 | -0.745 | 0.476441 | 0.648632 | 0.187027 | 0.99033 | 0.997988 | 0.994666 |
| Ptpn3_A2ALK8_S459s_1 | Ptpn3_A2A | 0.23 | -0.11 | -0.83 | -1 | -0.2 | -0.11 | -0.04 | 0.9 | -0.55 | 0.96 | 0.59 | -0.59 | 0.7 | -0.42 | 0.38 | 0.22 | 0.565 | 0.53 | 0.6475 | 0.119596 | 0.160341 | 0.27549 | 0.989223 | 0.997988 | 0.994666 |
| Ptpn4_E9Q0S8_S392s_1 | Ptpn4_E9Q | -0.53 | -0.11 | 0.27 | -0.01 | -0.88 | -0.08 | -0.68 | -0.25 | 1.51 | -0.51 | -1.02 | 0.07 | 2.15 | -1.3 | 0.69 | 0.49 | -0.3775 | 0.1075 | 0.6025 | 0.368551 | 0.203192 | 0.839902 | 0.989223 | 0.997988 | 0.994666 |
| Ptpn4_E9Q0S8_S403s_1 | Ptpn4_E9Q | 1.57 | -0.24 | 0.39 | 1.11 | 0.99 | 1.52 | 0.62 | 1.52 | 0.15 | 0.26 | 0.65 | 0.86 | -0.05 | 0.63 | -0.06 | 0.36 | 0.455 | -0.2275 | -0.4875 | 0.271372 | 0.304283 | 0.592849 | 0.989223 | 0.997988 | 0.994666 |
| Ptpn5_P54830_S319s_1 | Ptpn5_P54 | -0.46 | 0.3 | 1.47 | 0.52 | 0.29 | -0.57 | 0.78 | 0.87 | 0.16 | 0.28 | 1.36 | -0.13 | -0.23 | 0.65 | 0.33 | 0.99 | -0.115 | -0.04 | -0.0225 | 0.960881 | 0.809641 | 0.934392 | 0.995762 | 0.997988 | 0.999507 |
| Ptpn5_P54830_S39s_1 | Ptpn5_P54 | 0.33 | 0.67 | 0.93 | 0.56 | 0.81 | 0.84 | 0.8 | 1.03 | -0.46 | -0.11 | 0.45 | 1.12 | -0.07 | 1 | 0.63 | 1.17 | 0.2475 | -0.3725 | 0.06 | 0.859205 | 0.279768 | 0.332325 | 0.989223 | 0.997988 | 0.999507 |
| Ptpra_Q91V35_S176s_2 | Ptpra_Q91 | 1.28 | 0.41 | 0.72 | 0.89 | -0.28 | 1.69 | -0.38 | 0.74 | 1.88 | 0.47 | 0.06 | 0.49 | 1.07 | -1.01 | 0.6 | 0.5 | -0.3825 | -0.1 | -0.535 | 0.269527 | 0.432942 | 0.814645 | 0.989223 | 0.997988 | 0.994666 |
| Ptpra_Q91V35_S180s_1 | Ptpra_Q91 | 0.65 | 0.61 | 0.23 | 0.37 | -0.4 | 0.95 | -0.31 | -0.04 | 1.75 | 0.51 | -0.11 | 0.17 | 2.1 | -1.18 | 1.02 | 0.07 | -0.415 | 0.115 | 0.0375 | 0.953046 | 0.233243 | 0.781919 | 0.989223 | 0.997988 | 0.999507 |
| Ptpra_Q91V35_S180s_2 | Ptpra_Q91 | 1.28 | 0.41 | 0.72 | 0.89 | -0.28 | 1.69 | -0.38 | 0.74 | 1.88 | 0.47 | 0.06 | 0.49 | 1.07 | -1.01 | 0.6 | 0.5 | -0.3825 | -0.1 | -0.535 | 0.269527 | 0.432942 | 0.814645 | 0.989223 | 0.997988 | 0.994666 |
| Ptprc_S4R1M0_S801s_1 | Ptprc_S4R1 | -0.11 | -0.15 | 0.38 | -0.13 | 0.02 | 0.13 | 0.09 | -1.04 | 0.7 | -0.14 | -0.15 | -0.06 | 0.76 | -0.63 | -0.36 | -0.27 | -0.1975 | 0.09 | -0.1225 | 0.731991 | 0.544048 | 0.755231 | 0.989223 | 0.997988 | 0.9971 |
| Ptprd_Q64487_S1296s_1 | Ptprd_Q64 | -2.53 | -3.55 | -3.66 | -3.92 | -3.57 | -1.87 | -2.16 | -2.35 | 3.68 | -3.03 | -0.59 | -1.86 | 4 | -3.24 | 3.83 | -3.72 | 0.9275 | 2.965 | 3.6325 | 0.074039 | 0.063808 | 0.045944 | 0.989223 | 0.997988 | 0.994666 |
| Ptprk_B2RRF0_S856s_1 | Ptprk_B2RI | 0.81 | 0.8 | 0.79 | 0.82 | 0.69 | 0.75 | 0.99 | 1.26 | 0.93 | 1.28 | 1.85 | 1.19 | 0.1 | 0.42 | 0.88 | 1.27 | 0.1175 | 0.5075 | -0.1375 | 0.6595 | 0.594131 | 0.0767 | 0.989223 | 0.997988 | 0.995259 |
| Ptprn_Q60673_S307s_1 | Ptprn_Q60 | 0.31 | 0.51 | -0.13 | 0.15 | 0.53 | -0.12 | -0.16 | -0.14 | 0.59 | 0.7 | 0.54 | -0.14 | -0.18 | 0.71 | 0.83 | 0.52 | -0.1825 | 0.2125 | 0.26 | 0.416832 | 0.492751 | 0.459978 | 0.989223 | 0.997988 | 0.994666 |
| Ptprn2_Q3UU93_S184s_1 | Ptprn2_Q3 | 0.26 | 0.99 | -0.09 | 0.03 | 0.54 | -0.35 | 0.67 | 0.05 | 0.16 | 0.81 | -0.23 | 0.11 | -0.48 | 0.31 | 0.31 | 0.26 | -0.07 | -0.085 | -0.1975 | 0.568566 | 0.83789 | 0.804706 | 0.995762 | 0.997988 | 0.995259 |
| Ptprn2_Q3UU93_S185s_1 | Ptprn2_Q3 | -0.25 | 0.34 | -0.17 | -0.04 | 0.56 | -0.1 | 0.49 | 0.21 | 0.52 | 0.52 | -0.07 | 0.12 | -0.15 | 0.59 | 0.35 | 0.31 | 0.32 | 0.3025 | 0.305 | 0.292069 | 0.226225 | 0.266015 | 0.989223 | 0.997988 | 0.994666 |
| Ptprn2_Q3UU93_S339s_1 | Ptprn2_Q3 | 0.27 | 0.42 | 0.75 | 1.11 | 0.47 | 0.05 | 0.09 | 0.86 | 0.67 | 0.73 | 0.93 | 0.99 | 0.19 | 1.03 | 1.18 | 1.15 | -0.27 | 0.1925 | 0.25 | 0.464833 | 0.369271 | 0.473384 | 0.989223 | 0.997988 | 0.994666 |
| Ptprn2_Q3UU93_S423s_1 | Ptprn2_Q3 | -0.14 | 0.38 | -0.45 | -0.44 | 0.11 | -0.87 | -0.45 | -0.81 | -0.41 | -0.13 | -0.63 | -0.74 | -0.17 | -0.14 | -0.38 | -0.42 | -0.3425 | -0.315 | -0.115 | 0.685531 | 0.291962 | 0.285443 | 0.989223 | 0.997988 | 0.996159 |
| Ptprn2_Q3UU93_S678s_1 | Ptprn2_Q3 | -0.04 | 0.58 | -0.03 | 0.63 | 0.42 | 0.18 | 0.74 | 0.33 | 0.39 | 0.24 | 1.41 | -0.31 | 0.25 | -0.06 | 0.13 | 0.25 | 0.1325 | 0.1475 | -0.1425 | 0.611394 | 0.619493 | 0.71386 | 0.989223 | 0.997988 | 0.995259 |
| Ptprn2_Q3UU93_S690s_1 | Ptprn2_Q3 | -0.7 | 0.09 | -0.36 | 0.38 | 0.05 | -0.16 | -0.09 | 0.08 | -0.61 | -0.25 | 0.39 | -0.69 | -1.43 | 0.39 | -0.45 | -0.76 | 0.1175 | -0.1425 | -0.415 | 0.355722 | 0.677073 | 0.690148 | 0.99315 | 0.997988 | 0.994666 |
| Ptprr_Q62132_S27s_1 | Ptprr_Q62 | 0.82 | 1.14 | 1.69 | 0.45 | 0.14 | -0.29 | 1.61 | 0.66 | 1.52 | 0.49 | 0.8 | 0.9 | -0.47 | 0.79 | 1.33 | 0.35 | -0.495 | -0.0975 | -0.525 | 0.260766 | 0.290679 | 0.783262 | 0.989223 | 0.997988 | 0.994666 |
| Ptpru_Q3V360_S781s_1 | Ptpru_Q3V | 1.05 | 1.75 | 0.36 | 0.33 | 1.58 | 1.25 | 0.79 | 0.39 | 1.27 | 1.57 | -0.1 | -0.14 | 0.69 | 1 | 0.37 | 0.34 | 0.13 | -0.2225 | -0.2725 | 0.483714 | 0.75004 | 0.672383 | 0.995762 | 0.997988 | 0.994666 |
| Ptpru_Q3V360_S791s_1 | Ptpru_Q3V | -0.48 | 0.28 | 0.11 | 0.31 | -0.05 | 0.02 | -0.04 | -0.11 | -0.04 | -0.37 | -0.82 | 0.17 | 0.14 | -0.44 | 0.13 | -0.04 | -0.1 | -0.32 | -0.1075 | 0.71564 | 0.685361 | 0.323634 | 0.994624 | 0.997988 | 0.996966 |
| Ptprz1_B9EKR1_S1457s_1 | Ptprz1_B9E | -1.95 | -2.01 | -2.08 | -2.24 | -1.41 | -2.63 | -2.17 | -2.18 | -5.4 | -1.52 | -4.03 | -1.83 | -2.43 | -4.59 | -1.09 | -1.27 | -0.0275 | -1.125 | -0.275 | 0.703473 | 0.924522 | 0.195056 | 0.997335 | 0.997988 | 0.996159 |
| Ptprz1_B9EKR1_S647s_1 | Ptprz1_B9E | -0.69 | -0.29 | -2.51 | -2.14 | -0.71 | -0.71 | -1.94 | -2.51 | -0.58 | -0.53 | -1.6 | -2.18 | 0.18 | -0.3 | -1.86 | -1.25 | -0.06 | 0.185 | 0.6 | 0.361423 | 0.923815 | 0.764466 | 0.997335 | 0.997988 | 0.994666 |
| Ptprz1_B9EKR1_T1681t_1 | Ptprz1_B9E | 0.05 | 0.85 | 0.38 | 0.55 | 0.57 | -0.03 | 0.46 | 0.28 | 0.51 | 0.82 | 0.39 | -0.16 | 0.76 | 0.46 | 0.26 | 0.18 | -0.1375 | -0.0675 | -0.0425 | 0.881388 | 0.600693 | 0.823381 | 0.989223 | 0.997988 | 0.999507 |
| Pum1_Q80U78_S709s_1 | Pum1_Q80 | -1.25 | -0.55 | -2.05 | -1.51 | -0.79 | -1.11 | -1.52 | -2.76 | -1.27 | -0.75 | -0.76 | -2.28 | -0.72 | -0.39 | -2.07 | -1.89 | -0.205 | 0.075 | 0.0725 | 0.88343 | 0.676657 | 0.869417 | 0.99315 | 0.997988 | 0.999507 |
| Pum1_Q80U78_S98s_1 | Pum1_Q80 | 2.52 | 2.14 | 2.43 | 1.19 | 2.17 | 2.8 | 1.82 | 2.26 | 1.46 | 2.38 | 0.75 | 2.77 | 1.45 | 0 | 2.55 | 2.3 | 0.1925 | -0.23 | -0.495 | 0.414193 | 0.598388 | 0.655704 | 0.989223 | 0.997988 | 0.994666 |
| Pum1_Q80U78_T112t_1 | Pum1_Q80 | 0.2 | -0.19 | -1.14 | -1.86 | 0.44 | -0.06 | -1.32 | -1.33 | 0.7 | 0.15 | -0.99 | -0.71 | 1.56 | 0.77 | -1.02 | -1.11 | 0.18 | 0.535 | 0.7975 | 0.285799 | 0.756932 | 0.352422 | 0.995762 | 0.997988 | 0.994666 |
| Pum2_Q3UR91_S136s_1 | Pum2_Q3L | -0.53 | -0.54 | -0.89 | -0.86 | -0.18 | -0.82 | -0.79 | -0.9 | -0.51 | -0.12 | -0.03 | -0.71 | -0.17 | -0.13 | -0.63 | -0.63 | 0.0325 | 0.3625 | 0.315 | 0.251821 | 0.896426 | 0.181712 | 0.996493 | 0.997988 | 0.994666 |
| Pum2_Q3UR91_S181s_1 | Pum2_Q3L | -1.24 | -1.13 | -1.49 | -1.78 | -0.84 | -1.52 | -2 | -2.48 | -1.28 | -0.87 | -0.96 | -1.21 | -0.74 | -0.38 | -0.89 | -0.9 | -0.3 | 0.33 | 0.6825 | 0.030487 | 0.428571 | 0.207932 | 0.989223 | 0.997988 | 0.994666 |
| Pura_P42669_S181s_1 | Pura_P426 | -0.13 | 0.04 | -0.26 | -0.28 | 0.02 | 0.15 | 0.15 | -0.05 | -0.39 | -0.17 | -0.42 | -0.12 | -0.31 | -0.05 | -0.27 | 0 | 0.225 | -0.1175 | 3.93E-17 | 1 | 0.285174 | 0.595085 | 0.989223 | 0.997988 | 1 |
| Pura_P42669_T182t_1 | Pura_P426 | -0.09 | 0.1 | -0.46 | -0.41 | -0.17 | 0.17 | 0.15 | -0.17 | -0.32 | -0.25 | -0.55 | -0.37 | -0.3 | -0.16 | -0.39 | -0.1 | 0.21 | -0.1575 | -0.0225 | 0.929689 | 0.380977 | 0.510853 | 0.989223 | 0.997988 | 0.999507 |
| Purb_O35295_S316s_1 | Purb_O352 | -0.83 | -0.81 | -0.99 | -0.62 | -2.19 | -0.29 | -1.96 | -1.14 | 0.89 | -0.13 | 0.63 | -0.77 | -0.04 | -1.78 | -0.75 | -0.84 | -0.5825 | 0.9675 | -0.04 | 0.916152 | 0.184751 | 0.030897 | 0.989223 | 0.997988 | 0.999507 |
| Purb_O35295_S6s_2 | Purb_O352 | -0.55 | -0.45 | -0.42 | 1.12 | -0.65 | -0.22 | 1.7 | 0.68 | -0.82 | 0.01 | 2.33 | 2.21 | 0.54 | -0.26 | 1.08 | 1.78 | 0.4525 | 1.0075 | 0.86 | 0.13947 | 0.448899 | 0.223804 | 0.989223 | 0.997988 | 0.994666 |
| Purb_O35295_S8s_2 | Purb_O352 | -0.55 | -0.45 | -0.42 | 1.12 | -0.65 | -0.22 | 1.7 | 0.68 | -0.82 | 0.01 | 2.33 | 2.21 | 0.54 | -0.26 | 1.08 | 1.78 | 0.4525 | 1.0075 | 0.86 | 0.13947 | 0.448899 | 0.223804 | 0.989223 | 0.997988 | 0.994666 |
| Purg_Q8R4E6_S342s_1 | Purg_Q8R4 | -0.65 | -1.24 | -0.71 | -0.63 | -1.49 | -0.57 | -0.91 | -0.92 | -0.5 | -0.45 | -0.16 | -1.13 | -0.26 | -1.28 | -0.47 | -0.78 | -0.165 | 0.2475 | 0.11 | 0.727914 | 0.555469 | 0.408731 | 0.989223 | 0.997988 | 0.9971 |
| Pvr_Q8K094_S384s_1 | Pvr_Q8K09 | -0.25 | -0.2 | 0.89 | 0.46 | -0.47 | -0.01 | 0.32 | 0.78 | -0.25 | -0.64 | 0.69 | 0.62 | 0.85 | -1.91 | 0.65 | 0.23 | -0.07 | -0.12 | -0.27 | 0.668233 | 0.851465 | 0.774888 | 0.996349 | 0.997988 | 0.996159 |
| Pvr_Q8K094_S393s_1 | Pvr_Q8K09 | 0.74 | 0.42 | 0.17 | 0.19 | 0.57 | 0.11 | 0.08 | 0.66 | -0.11 | -0.65 | 1.87 | -0.84 | 0.56 | 0.75 | 0.03 | 0.43 | -0.025 | -0.3125 | 0.0625 | 0.823576 | 0.921706 | 0.594938 | 0.997335 | 0.997988 | 0.999507 |
| Pvrl1_Q9JKF6_S421s_1 | Pvrl1_Q9JK | 1.2 | 0.87 | 0.36 | 0.78 | 0.98 | 0.38 | 0.7 | 0.45 | 0.51 | 1.02 | 0.38 | -0.07 | 0.17 | 1.23 | 0.7 | 0.39 | -0.175 | -0.3425 | -0.18 | 0.587484 | 0.514534 | 0.292551 | 0.989223 | 0.997988 | 0.995259 |
| Pvrl1_Q9JKF6_S433s_1 | Pvrl1_Q9JK | -0.48 | 0.55 | 1.4 | 1.39 | -0.04 | -0.87 | 1.37 | -0.22 | 0.66 | 0.28 | 1.23 | 0.57 | -0.36 | 1.09 | 1.37 | -0.23 | -0.655 | -0.03 | -0.2475 | 0.669837 | 0.276908 | 0.948593 | 0.989223 | 0.997988 | 0.996159 |
| Pvrl1_Q9JKF6_S434s_1 | Pvrl1_Q9JK | 0.32 | 1.03 | 0.21 | 0.86 | 0.73 | 0.17 | 0.56 | 0.13 | 0.36 | 0.64 | -0.35 | -0.14 | -0.09 | 0.91 | -0.15 | 0.68 | -0.2075 | -0.4775 | -0.2675 | 0.464056 | 0.46986 | 0.169237 | 0.989223 | 0.997988 | 0.994666 |
| Pwp1_Q99LL5_S493s_1 | Pwp1_Q99 | 0.34 | -0.18 | 0.16 | 0.07 | -1.1 | 0.05 | -0.3 | -0.68 | 0.82 | 0.24 | 0.55 | -0.01 | 0.75 | -1.13 | -0.83 | 0.21 | -0.605 | 0.3025 | -0.3475 | 0.440465 | 0.065448 | 0.278649 | 0.989223 | 0.997988 | 0.994666 |
| Pxn_F8VQ28_S258s_1 | Pxn_F8VQ2 | -1.77 | -2.13 | -1.98 | -2.71 | -2.65 | -1.64 | -2.67 | -2.7 | -2.04 | -1.66 | -1.59 | -2 | -0.22 | -3.26 | -2.55 | -1.75 | -0.2675 | 0.325 | 0.2025 | 0.744567 | 0.433502 | 0.26712 | 0.989223 | 0.997988 | 0.9971 |
| Pygo2_Q3V113_T264t_1 | Pygo2_Q3\ | -0.73 | -0.35 | 0.28 | 1.18 | -0.32 | -1.01 | 0.73 | 0.85 | -0.39 | -0.35 | 0.31 | 0.41 | -0.52 | 0.02 | 0.79 | 0.2 | -0.0325 | -0.1 | 0.0275 | 0.953886 | 0.952862 | 0.824347 | 0.999187 | 0.997988 | 0.999507 |
| Q3TQI7_S261s_1 | Q3TQI7_S2 | -2.61 | -2.05 | -2.04 | -1.9 | -1.79 | -2.9 | -1.74 | -2.21 | -1.91 | -1.94 | -1.27 | -1.86 | -0.74 | -2.54 | -1.77 | -2.32 | -0.01 | 0.405 | 0.3075 | 0.477661 | 0.975343 | 0.167891 | 0.999187 | 0.997988 | 0.994666 |
| Q3TYS2_S168s_1 | Q3TYS2_S1 | -1.2 | -0.92 | -0.41 | -1.62 | -1.25 | -1.08 | -1.04 | -1.95 | -0.04 | -1.27 | -0.41 | -2.01 | 0.8 | -2.12 | -1.32 | -1.65 | -0.2925 | 0.105 | -0.035 | 0.955559 | 0.395977 | 0.827978 | 0.989223 | 0.997988 | 0.999507 |
| Q3UPC7_S376s_1 | Q3UPC7_S | 0.1 | 0.39 | 0.02 | 0.32 | 0.74 | 0.56 | 0.77 | 0.1 | 0.84 | 1.43 | -0.66 | 0.36 | -0.05 | 0.25 | 0.34 | 0.35 | 0.335 | 0.285 | 0.015 | 0.951578 | 0.187815 | 0.519158 | 0.989223 | 0.997988 | 0.999507 |
| Q3UTZ3_S517s_1 | Q3UTZ3_S | -1.16 | -1.25 | -0.77 | -1.86 | -0.1 | -0.83 | -1.16 | -0.86 | -0.34 | -1.26 | -2.02 | -1.58 | 0.27 | -0.02 | -0.87 | -1.2 | 0.5225 | -0.04 | 0.805 | 0.076965 | 0.137365 | 0.923034 | 0.989223 | 0.997988 | 0.994666 |
| Q3UTZ3_S533s_1 | Q3UTZ3_S | -1.87 | -1.76 | -1.13 | -2.09 | -1.03 | -1.33 | -1.8 | -1.6 | -2.58 | -1.94 | -1.35 | -2.27 | -0.65 | -1.34 | -0.98 | -1.35 | 0.2725 | -0.3225 | 0.6325 | 0.069434 | 0.364305 | 0.367979 | 0.989223 | 0.997988 | 0.994666 |
| Q3UTZ3_S546s_1 | Q3UTZ3_S | -1.26 | -1.3 | -1.41 | -1.74 | -1.09 | -1.53 | -0.99 | -0.62 | 2.24 | -2.32 | 7.08 | -1.76 | 1.83 | 0.7 | -0.87 | -1.33 | 0.37 | 2.7375 | 1.51 | 0.04383 | 0.182302 | 0.172468 | 0.989223 | 0.997988 | 0.994666 |
| Q3UTZ3_T541t_1 | Q3UTZ3_T | -0.49 | 0.11 | -0.23 | -0.51 | 0.8 | -0.5 | 0.38 | -0.26 | -0.68 | -0.11 | -0.12 | -0.29 | 0.49 | 0.86 | 0.16 | 0.36 | 0.385 | -0.02 | 0.7475 | 0.023693 | 0.270846 | 0.939021 | 0.989223 | 0.997988 | 0.994666 |
| Q3UY34_S139s_1 | Q3UY34_S | -3.36 | -2.68 | -2.42 | -3.64 | -3.35 | -2.02 | -3.33 | -3.46 | -0.74 | -2.87 | -2.99 | -3.22 | 0.6 | -3.66 | -2.63 | -1.94 | -0.015 | 0.57 | 1.1175 | 0.203137 | 0.971595 | 0.3469 | 0.999187 | 0.997988 | 0.994666 |
| Q6DFW0-2_Q6DFW0_S9s_1 | Q6DFW0-2 | 1.24 | 1.43 | 0.81 | 1.23 | 0.78 | 0.74 | 1.16 | 0.95 | 0.32 | 0.35 | 0.49 | 0.6 | -0.3 | 1.32 | 0.91 | 1.18 | -0.27 | -0.7375 | -0.4 | 0.328909 | 0.265057 | 0.011056 | 0.989223 | 0.997988 | 0.994666 |
| Q6NV72-2_Q6NV72_S535s_1 | Q6NV72-2_ | 1.16 | 0.98 | 1.05 | 1.44 | 0.65 | 1.46 | 1.24 | 1.6 | 1.12 | 0.96 | 2.05 | 1.08 | 0.37 | 1.06 | 1.5 | 1.15 | 0.08 | 0.145 | -0.1375 | 0.659723 | 0.769755 | 0.638459 | 0.995762 | 0.997988 | 0.995259 |
| Q7TNG5-3_Q7TNG5_S151s_1 | Q7TNG5-3_ | 0.16 | -0.18 | -0.04 | 0.31 | 0.04 | -0.93 | 0.4 | -0.39 | -0.66 | -0.03 | 0.48 | -1.26 | 0.34 | 0.47 | 0.21 | -0.04 | -0.2825 | -0.43 | 0.1825 | 0.483867 | 0.388385 | 0.290068 | 0.989223 | 0.997988 | 0.994666 |
| Q7TNG5-3_Q7TNG5_S153s_1 | Q7TNG5-3_ | -2.38 | -1.17 | -0.82 | -0.8 | -1.62 | -2.43 | 0.06 | -0.89 | -1.64 | -0.91 | -0.27 | -0.38 | -0.77 | -1.12 | -0.41 | -0.88 | 0.0725 | 0.4925 | 0.4975 | 0.23815 | 0.90068 | 0.304497 | 0.996493 | 0.997988 | 0.994666 |
| Q7TNG5-3_Q7TNG5_S154s_1 | Q7TNG5-3_ | 0.98 | 1.09 | -0.18 | -0.03 | 0.83 | 0.77 | -0.62 | -0.78 | 0.82 | 1.46 | 0.15 | -0.38 | 1.57 | 1.61 | -0.23 | -0.76 | -0.415 | 0.0475 | 0.0825 | 0.895801 | 0.415765 | 0.922822 | 0.989223 | 0.997988 | 0.999507 |
| Q7TNG5-3_Q7TNG5_S170s_1 | Q7TNG5-3_ | 0.69 | 0.72 | 0.37 | 0.96 | 0.49 | 0.84 | 0.88 | 1.11 | 1.68 | 1.64 | 2.48 | 0.94 | 1.78 | 1.56 | 1.26 | 1.05 | 0.145 | 1 | 0.7275 | 0.025627 | 0.551015 | 0.017673 | 0.989223 | 0.997988 | 0.994666 |
| Q8BP99_S139s_1 | Q8BP99_S | -0.98 | -1.21 | 0.06 | -0.2 | 0.37 | -0.12 | 0.79 | 0.41 | -1.05 | -0.58 | 0.33 | -0.03 | -1.38 | 0.76 | 1.15 | 0.54 | 0.945 | 0.25 | 0.85 | 0.170516 | 0.023658 | 0.556312 | 0.989223 | 0.997988 | 0.994666 |
| Q8BY89-2_Q8BY89_T12t_1 | Q8BY89-2_ | 1.17 | 1.39 | 1.57 | 1.11 | 0.81 | 1.14 | 1.28 | 1.4 | 1.13 | 1.43 | 1.83 | 1.53 | 0.87 | 1.37 | 1.27 | 1.57 | -0.1525 | 0.17 | -0.04 | 0.882084 | 0.521676 | 0.505195 | 0.989223 | 0.997988 | 0.999507 |
| Q8BZR9_S563s_1 | Q8BZR9_S | -0.68 | -1.28 | 1.99 | 0.97 | -2.33 | -1.03 | 0.96 | -0.07 | 1.23 | -1.28 | 3.18 | 1.81 | 0.16 | -0.97 | 2.65 | 1.41 | -0.8675 | 0.985 | 0.5625 | 0.555782 | 0.343144 | 0.362067 | 0.989223 | 0.997988 | 0.995259 |
| Q8C3W1_S130s_1 | Q8C3W1_S | 0.08 | 1.84 | 0.37 | 0.34 | 0.37 | 0.13 | 0.58 | 0.33 | -0.28 | 0.95 | -0.17 | 0.26 | -0.59 | -0.71 | 0.54 | 0.77 | -0.305 | -0.4675 | -0.655 | 0.22547 | 0.448016 | 0.328852 | 0.989223 | 0.997988 | 0.994666 |
| Q8C3W1_S176s_1 | Q8C3W1_S | 0.6 | 0.24 | 0.32 | 0.44 | 0.2 | 0.51 | 0.25 | 0.35 | 0.02 | 0.34 | -0.08 | 0.5 | -0.56 | 0.47 | 0.05 | 0.5 | -0.0725 | -0.205 | -0.285 | 0.371994 | 0.728972 | 0.404026 | 0.995762 | 0.997988 | 0.994666 |
| Q8C3W1_S278s_1 | Q8C3W1_S | 0.1 | 0.61 | 0.89 | 0.11 | 0.89 | 0.35 | -0.02 | 0.16 | 0.22 | 0.72 | -0.32 | 0.04 | 0.74 | 1.29 | 0.02 | 0.63 | -0.0825 | -0.2625 | 0.2425 | 0.497909 | 0.784241 | 0.421019 | 0.995762 | 0.997988 | 0.994666 |
| Q8C3W1_T274t_1 | Q8C3W1_T | -0.15 | 0.41 | -0.01 | 0.46 | 0.55 | 0.1 | 0.1 | 0.26 | 0.17 | 0.61 | 0.22 | -0.36 | -0.08 | -0.49 | 0.23 | 0.23 | 0.075 | -0.0175 | -0.205 | 0.491611 | 0.760506 | 0.952432 | 0.995762 | 0.997988 | 0.994666 |
| Q8C3W1_T276t_1 | Q8C3W1_T | 0.63 | 0.81 | 1.25 | 0.71 | 0.22 | 0.42 | 0.89 | 0.5 | 0.61 | 1.12 | 0.64 | 0.88 | 0.86 | 0.74 | 0.91 | 1.08 | -0.3425 | -0.0375 | 0.0475 | 0.853933 | 0.195948 | 0.882531 | 0.989223 | 0.997988 | 0.999507 |
| Q8C5K5-2_Q8C5K5_T196t_1 | Q8C5K5-2_ | -1.2 | -2.12 | -0.46 | 0.6 | -0.85 | -0.27 | -1.42 | -0.58 | 1.46 | -1.25 | -2 | -0.28 | 1.45 | -2.62 | 0.12 | -0.42 | 0.015 | 0.2775 | 0.4275 | 0.635685 | 0.978733 | 0.741629 | 0.999187 | 0.997988 | 0.995259 |
| Q8K207_S115s_1 | Q8K207_S1 | 2.15 | 1.62 | 0.94 | 0.98 | 1.33 | 1.94 | 1.19 | 0.96 | 2.05 | 2.24 | 0.7 | 0.41 | -0.1 | 2.05 | 0.73 | 0.99 | -0.0675 | -0.0725 | -0.505 | 0.325995 | 0.849625 | 0.887415 | 0.996349 | 0.997988 | 0.994666 |
| Q8K207_S95s_1 | Q8K207_S | -0.3 | -0.05 | 0.18 | 0.35 | 0.25 | -0.1 | 0.5 | 0.26 | -0.24 | -0.22 | -0.32 | 0.08 | -0.08 | 0.12 | -0.01 | 0.46 | 0.1825 | -0.22 | 0.0775 | 0.775815 | 0.466473 | 0.380942 | 0.989223 | 0.997988 | 0.99798 |
| Q91Z58_S529s_1 | Q91Z58_S | -1.59 | -1.88 | -2.06 | -2.35 | -1.81 | -2.3 | -2.24 | -2.25 | -2.5 | -1.49 | -0.48 | -1.86 | -1.2 | -1.72 | -2.41 | -1.99 | -0.18 | 0.3875 | 0.14 | 0.678792 | 0.480379 | 0.386477 | 0.989223 | 0.997988 | 0.996159 |
| Q99K99_S268s_1 | Q99K99_S2 | -2.47 | -2.03 | -2.83 | -3.4 | -2.82 | -1.75 | -2.31 | -3.32 | -1.82 | -1.97 | -1.8 | -3.04 | -0.67 | -1.49 | -2.48 | -2.49 | 0.1325 | 0.525 | 0.9 | 0.096187 | 0.753735 | 0.220477 | 0.995762 | 0.997988 | 0.994666 |
| Q99M80-2_Q99M80_S1005s_1 | Q99M80-2_ | 1.3 | 0.49 | 0.77 | 1.04 | 1.61 | 0.95 | 0.53 | 0.64 | 1.47 | 1.04 | 2.2 | 1 | 0.71 | 1.31 | 0.97 | 0.86 | 0.0325 | 0.5275 | 0.0625 | 0.827968 | 0.917926 | 0.152113 | 0.997335 | 0.997988 | 0.999507 |
| Q9CPS8_S40s_1 | Q9CPS8_S | 0.75 | 1.08 | -0.6 | 0.1 | 0.43 | 1.25 | -0.61 | -0.84 | 1.39 | 0.89 | 0.57 | -0.97 | 1.18 | 0.35 | -0.39 | -1.12 | -0.275 | 0.1375 | -0.3275 | 0.568104 | 0.620036 | 0.812614 | 0.989223 | 0.997988 | 0.995259 |
| Q9CPS8_T38t_1 | Q9CPS8_T | 0.64 | 1.36 | -0.6 | 0.1 | 0.84 | 0.53 | -0.61 | -0.84 | 0.89 | 0.92 | 0.57 | -0.97 | 0.93 | 0.89 | -0.39 | -1.12 | -0.395 | -0.0225 | -0.2975 | 0.619873 | 0.464787 | 0.968138 | 0.989223 | 0.997988 | 0.995259 |
| Q9CQF4_S106s_1 | Q9CQF4_S | 0.34 | 0.75 | 0.54 | 0.26 | -0.91 | 0.37 | 0.14 | 0.63 | 0.27 | 0.5 | 1.06 | 0.62 | -1.01 | -0.22 | 0.52 | 0.56 | -0.415 | 0.14 | -0.51 | 0.216836 | 0.260135 | 0.597022 | 0.989223 | 0.997988 | 0.994666 |
| Q9CZL2_S38s_1 | Q9CZL2_S3 | 0 | -0.56 | 0.47 | 0.13 | -0.67 | 0.01 | 0.4 | 0.03 | 0.33 | -0.11 | 0.47 | -0.22 | 0.57 | -0.01 | 0.42 | 0.99 | -0.0675 | 0.1075 | 0.4825 | 0.173216 | 0.834391 | 0.727163 | 0.995762 | 0.997988 | 0.994666 |
| Q9D2Q3_S423s_1 | Q9D2Q3_S | 0.43 | -0.31 | 0.14 | 0.28 | 0.4 | -0.12 | 0.61 | 0 | -0.02 | 0.14 | 0.1 | 0.18 | -0.05 | 0.61 | 0.13 | -0.02 | 0.0875 | -0.035 | 0.0325 | 0.91077 | 0.749526 | 0.886412 | 0.995762 | 0.997988 | 0.999507 |
| Q9D5Q8-3_Q9D5Q8_S157s_1 | Q9D5Q8-3_ | 1.07 | 1.12 | 1.13 | 1.46 | 0.6 | 1.09 | 0.8 | 1.15 | 1.21 | 1.58 | 1.81 | 0.97 | 0.33 | 1.25 | 0.82 | 0.96 | -0.285 | 0.1975 | -0.355 | 0.232593 | 0.236151 | 0.467497 | 0.989223 | 0.997988 | 0.994666 |
| Q9D6W8_S173s_1 | Q9D6W8_S | 0.16 | 0.48 | 0.84 | 1.05 | -0.53 | 0.53 | 0.87 | 1.49 | -0.43 | 0.08 | 1.24 | 0.84 | 0.27 | 0.08 | 1.39 | 0.86 | -0.0425 | -0.2 | 0.0175 | 0.962671 | 0.922694 | 0.632598 | 0.997335 | 0.997988 | 0.999507 |
| Q9D6W8_S43s_1 | Q9D6W8_S | 1.62 | 0.91 | -0.01 | 0.96 | 0.22 | 0.48 | 0.94 | 0.92 | 0.46 | 1.04 | -0.04 | -0.15 | 1.04 | 0.76 | 0.18 | 0.33 | -0.23 | -0.5425 | -0.2925 | 0.467337 | 0.540533 | 0.219727 | 0.989223 | 0.997988 | 0.994666 |
| Q9D6W8_T201t_1 | Q9D6W8_T | 1.02 | 1.37 | 0.24 | -0.15 | 1.19 | 1.31 | 0.51 | -0.24 | 0.09 | 0.22 | 0.57 | 0.24 | 0.5 | 1.33 | -0.24 | -0.01 | 0.0725 | -0.34 | -0.225 | 0.635961 | 0.875718 | 0.370755 | 0.996493 | 0.997988 | 0.995259 |
| Q9D9H8_S44s_1 | Q9D9H8_S | -0.55 | -1.99 | -1.49 | -8.28 | -3.2 | 1.21 | -0.29 | -0.53 | 1.35 | -0.46 | -3.45 | 0.33 | 1.61 | -2.37 | 1.16 | -1.29 | 2.375 | 2.52 | 2.855 | 0.121399 | 0.183794 | 0.179617 | 0.989223 | 0.997988 | 0.994666 |
| Q9D9H8_S77s_1 | Q9D9H8_S | -0.71 | -0.15 | -0.42 | -0.71 | -2.08 | 0.79 | -1.77 | -1.29 | 2.47 | -0.27 | -0.33 | 0.16 | 2.43 | -1.33 | 1.4 | -1.28 | -0.59 | 1.005 | 0.8025 | 0.354896 | 0.331661 | 0.129805 | 0.989223 | 0.997988 | 0.994666 |
| Qars_D3Z158_S70s_1 | Qars_D3Z1 | -0.67 | -0.43 | -0.61 | -1.35 | -0.07 | -0.23 | -0.81 | -0.63 | -2.14 | -0.64 | -0.75 | -1.06 | -1.56 | -0.08 | -1.17 | -0.82 | 0.33 | -0.3825 | -0.1425 | 0.712828 | 0.276647 | 0.347106 | 0.989223 | 0.997988 | 0.996966 |
| R3hcc1_E9PUE1_S167s_1 | R3hcc1_E9 | -0.96 | -1.16 | -1.71 | -1.57 | -1.21 | -1.01 | -1.95 | -2.58 | -0.61 | -1.62 | -1.06 | -2.59 | -0.61 | -1.52 | -2.71 | -1.69 | -0.3375 | -0.12 | -0.2825 | 0.535793 | 0.394161 | 0.787613 | 0.989223 | 0.997988 | 0.994666 |
| R3hdm1_E9Q9Q2_S1007s_1 | R3hdm1_E | 0.37 | 0.23 | -0.32 | -0.31 | 0 | 0.27 | 0.03 | -0.3 | 0.01 | 0.83 | -0.42 | -0.2 | 0.28 | 0.35 | -0.69 | -0.02 | 0.0075 | 0.0625 | -0.0125 | 0.970263 | 0.97715 | 0.856078 | 0.999187 | 0.997988 | 0.999507 |
| R3hdm1_E9Q9Q2_S280s_1 | R3hdm1_E | 0.86 | 1.89 | 0.38 | -0.17 | 1.68 | 0.9 | -0.19 | -0.16 | 0.85 | 1.65 | 0.38 | 0.37 | 0.46 | 2.26 | 0.42 | -0.05 | -0.1825 | 0.0725 | 0.0325 | 0.957533 | 0.747656 | 0.884593 | 0.995762 | 0.997988 | 0.999507 |
| R3hdm1_E9Q9Q2_S282s_1 | R3hdm1_E | 0.03 | 0.99 | 1.81 | 1.57 | 1.59 | 0.58 | 0.98 | 1.43 | 0.85 | 0.58 | 2.22 | 2.02 | 0.47 | 2.01 | 1.76 | 1.65 | 0.045 | 0.3175 | 0.3725 | 0.459614 | 0.916825 | 0.554426 | 0.997335 | 0.997988 | 0.994666 |
| R3hdm1_E9Q9Q2_S347s_1 | R3hdm1_E | 0.44 | 0.4 | 0.27 | -0.52 | -0.93 | 0.09 | 0 | -0.17 | 0.89 | -0.06 | 1.1 | 0.09 | 1.25 | -0.15 | 1.39 | 0.5 | -0.4 | 0.3575 | 0.6 | 0.17657 | 0.247692 | 0.349734 | 0.989223 | 0.997988 | 0.994666 |
| R3hdm1_E9Q9Q2_S381s_1 | R3hdm1_E | -1.09 | -0.8 | -0.68 | -0.73 | -0.31 | -0.8 | -0.33 | -0.66 | -1.26 | -0.63 | -0.61 | -0.65 | -0.85 | -0.61 | -0.73 | -0.81 | 0.3 | 0.0375 | 0.075 | 0.754613 | 0.210806 | 0.882526 | 0.989223 | 0.997988 | 0.9971 |
| R3hdm2_E9Q9D6_S263s_1 | R3hdm2_E | -5.27 | -4.43 | -3.21 | -5.04 | -4.34 | -6.48 | -3.62 | -3.76 | -4.35 | -5.03 | -3.18 | -3.01 | -2.83 | -4.58 | -3.44 | -2.94 | -0.0625 | 0.595 | 1.04 | 0.090349 | 0.929841 | 0.342418 | 0.997796 | 0.997988 | 0.994666 |
| R3hdm2_E9Q9D6_S349s_1 | R3hdm2_E | -1.42 | -0.12 | -0.79 | -1.81 | 0.28 | -1.77 | -0.91 | -1.55 | -2.11 | -0.37 | -2.46 | -1.65 | -0.69 | 0.29 | -1.79 | -1.97 | 0.0475 | -0.6125 | -0.005 | 0.993218 | 0.92928 | 0.278756 | 0.997639 | 0.997988 | 1 |
| R3hdm2_E9Q9D6_S361s_1 | R3hdm2_E | -2.9 | -2.35 | -1.54 | -3.73 | -2.97 | -1.93 | -2.41 | -2.5 | -0.47 | -2.29 | -4.45 | -1.34 | 1.23 | -2.34 | -0.97 | -2.08 | 0.1775 | 0.4925 | 1.59 | 0.078629 | 0.70676 | 0.57199 | 0.995437 | 0.997988 | 0.994666 |
| R3hdm2_E9Q9D6_S857s_1 | R3hdm2_E | -0.64 | -0.31 | 0.1 | -0.27 | -0.06 | -0.97 | 0.12 | -0.05 | -0.9 | -0.48 | -0.18 | -0.13 | -0.19 | 0.46 | -0.4 | -0.02 | 0.04 | -0.1425 | 0.2425 | 0.426248 | 0.897068 | 0.616567 | 0.996493 | 0.997988 | 0.994666 |
| R3hdm2_E9Q9D6_T35t_1 | R3hdm2_E | 0.14 | 0.9 | 1.44 | 0.78 | -0.48 | 1.4 | -0.13 | 0.39 | 2.43 | 0.2 | 0.3 | 1.27 | 2.08 | -0.49 | 0.71 | 0.69 | -0.52 | 0.235 | -0.0675 | 0.901659 | 0.270515 | 0.666266 | 0.989223 | 0.997988 | 0.999507 |
| R3hdm2_Q80TM6_S263s_1 | R3hdm2_Q | -1.37 | -0.7 | -1.52 | -1.23 | -1.67 | -1.55 | -2.31 | -2.07 | -0.55 | -0.39 | -0.95 | -1.33 | 0.24 | -0.97 | -1.97 | -2.08 | -0.695 | 0.4 | 0.01 | 0.984942 | 0.033169 | 0.218133 | 0.989223 | 0.997988 | 1 |
| Rab11a_P62492_S190s_1 | Rab11a_P6 | 0.95 | 0.93 | 0.87 | 0.48 | 1.12 | 0.8 | 1.14 | 0.85 | -0.31 | 1.32 | -0.61 | 0.93 | 0.19 | -0.61 | 0.92 | 0.99 | 0.17 | -0.475 | -0.435 | 0.289499 | 0.454954 | 0.317049 | 0.989223 | 0.997988 | 0.994666 |
| Rab11b_P46638_T77t_1 | Rab11b_P4 | 0.4 | 0.15 | 0.19 | 0.66 | 0.71 | 0.24 | -0.52 | 0.32 | -0.07 | 0.1 | 0.14 | 0 | 0.76 | 1.75 | 0.42 | 0.88 | -0.1625 | -0.3075 | 0.6025 | 0.102722 | 0.596509 | 0.198125 | 0.989223 | 0.997988 | 0.994666 |
| Rab11fip2_G3XA57_S277s_1 | Rab11fip2_ | -1.68 | -1.86 | -1.27 | -1.33 | -0.75 | -1.09 | -0.66 | -0.97 | -1.73 | -1.72 | -0.47 | -0.99 | -1.44 | -0.45 | -0.99 | -0.78 | 0.6675 | 0.3075 | 0.62 | 0.067699 | 0.018454 | 0.392463 | 0.989223 | 0.997988 | 0.994666 |
| Rab11fip5_Q8R361_S296s_1 | Rab11fip5_ | 1.11 | -1.5 | -0.48 | 0.01 | -0.01 | 1.41 | -0.4 | 0.29 | 0.67 | -0.95 | -0.67 | -0.15 | -1.29 | -0.93 | -0.52 | -0.19 | 0.5375 | -0.06 | -0.5175 | 0.357871 | 0.378548 | 0.919588 | 0.989223 | 0.997988 | 0.994666 |
| Rab11fip5_Q8R361_S307s_1 | Rab11fip5_ | 1.8 | 0.96 | 0.53 | 0.65 | 1.47 | 1.92 | 0.52 | 0.88 | 2.44 | 1.56 | 0.82 | 0.85 | 0.83 | 1.51 | 0.19 | 0.45 | 0.2125 | 0.4325 | -0.24 | 0.560288 | 0.602162 | 0.356401 | 0.989223 | 0.997988 | 0.995259 |
| Rab11fip5_Q8R361_S480s_2 | Rab11fip5_ | -0.71 | -0.44 | -0.55 | -0.79 | -0.04 | -0.43 | -0.67 | -0.97 | -0.52 | -1.08 | -0.14 | -0.69 | -1.1 | -0.69 | -0.99 | 0.16 | 0.095 | 0.015 | -0.0325 | 0.92252 | 0.716104 | 0.955541 | 0.995437 | 0.997988 | 0.999507 |
| Rab11fip5_Q8R361_S486s_1 | Rab11fip5_ | -1.15 | -1.7 | -1.91 | -1.63 | -0.45 | -1.36 | -1.05 | -1.23 | -0.4 | -2.48 | -1.93 | -1.78 | -1.39 | -0.14 | -1.55 | -1.82 | 0.575 | -0.05 | 0.3725 | 0.372696 | 0.070675 | 0.911901 | 0.989223 | 0.997988 | 0.994666 |
| Rab11fip5_Q8R361_S530s_1 | Rab11fip5_ | -2.74 | -1.76 | -0.1 | -0.5 | -1.88 | -2.86 | 0.05 | -0.55 | -2.01 | -2.04 | -0.17 | -0.26 | -2.04 | -1.93 | -0.76 | 0.19 | -0.035 | 0.155 | 0.14 | 0.844844 | 0.964142 | 0.82918 | 0.999187 | 0.997988 | 0.999507 |
| Rab11fip5_Q8R361_S539s_1 | Rab11fip5_ | -0.31 | -0.35 | 0.57 | -0.34 | 1.07 | 0.2 | 0.48 | 0.12 | -1.11 | 0 | 0.04 | 0.24 | -1.76 | 0.67 | 0.04 | 0.27 | 0.575 | -0.1 | -0.0875 | 0.871695 | 0.10212 | 0.794282 | 0.989223 | 0.997988 | 0.999507 |
| Rab11fip5_Q8R361_S640s_1 | Rab11fip5_ | 0.38 | 0.06 | 0.17 | 0.28 | 1.13 | 0.7 | 0.48 | 0.65 | -0.19 | 0.18 | -1.62 | -0.27 | -0.75 | -0.24 | -0.58 | 0.06 | 0.5175 | -0.6975 | -0.6 | 0.052066 | 0.046085 | 0.10635 | 0.989223 | 0.997988 | 0.994666 |
| Rab12_P35283_S15s_1 | Rab12_P35 | -0.07 | 0.11 | 0.09 | -0.03 | 0 | 0.1 | 0.63 | -0.15 | -0.29 | -0.32 | 0.77 | 0.86 | -0.41 | 0.43 | 0.35 | 0.33 | 0.12 | 0.23 | 0.15 | 0.594487 | 0.619871 | 0.50983 | 0.989223 | 0.997988 | 0.995259 |
| Rab12_P35283_S20s_1 | Rab12_P35 | 0.16 | -0.34 | 0.47 | 0.47 | -0.51 | -0.63 | 0.2 | 0.45 | -0.81 | -0.7 | 2.07 | 0.06 | -0.16 | -0.03 | -0.66 | 0.37 | -0.3125 | -0.035 | -0.31 | 0.356424 | 0.361904 | 0.955785 | 0.989223 | 0.997988 | 0.994666 |
| Rab14_Q91V41_S204s_1 | Rab14_Q9 | 0.92 | 1.05 | 2.06 | 1.86 | 0.84 | 0.95 | 1.44 | 1.11 | 0.79 | 0.76 | 1.72 | 1.99 | 1.32 | 0.97 | 1.66 | 1.74 | -0.3875 | -0.1575 | -0.05 | 0.88956 | 0.252756 | 0.707415 | 0.989223 | 0.997988 | 0.999507 |
| Rab1A_P62821_S194s_1 | Rab1A_P62 | 1.37 | 0.92 | 1.35 | 1 | 0.48 | 1.39 | 0.14 | 0.61 | 1.99 | 1.12 | 0.3 | 0.87 | 2.41 | 0.4 | 0.87 | 0.59 | -0.505 | -0.09 | -0.0925 | 0.839816 | 0.12717 | 0.810935 | 0.989223 | 0.997988 | 0.999507 |
| Rab1A_P62821_T195t_1 | Rab1A_P62 | 1.37 | 0.95 | 1.39 | 1.05 | 0.51 | 1.36 | 0.17 | 0.66 | 1.95 | 1.07 | 0.31 | 0.94 | 2.39 | 0.44 | 0.91 | 0.68 | -0.515 | -0.1225 | -0.085 | 0.84783 | 0.11051 | 0.737979 | 0.989223 | 0.997988 | 0.999507 |
| Rab1b_Q9D1G1_S179s_1 | Rab1b_Q9 | 0.51 | 0.61 | 0.84 | 1.07 | 0.74 | 0.31 | 0.71 | 0.26 | 0.12 | 0.38 | 1.07 | 0.51 | 0.69 | 1.27 | 0.88 | 0.86 | -0.2525 | -0.2375 | 0.1675 | 0.53458 | 0.311279 | 0.413399 | 0.989223 | 0.997988 | 0.994666 |
| Rab22a_P35285_S187s_1 | Rab22a_P3 | -0.88 | -0.58 | 0.95 | 0.97 | -1.59 | 0.36 | -0.68 | 0.12 | 1.59 | -1.2 | -0.53 | 0.97 | 2.43 | -1.8 | 0.32 | 0.5 | -0.5625 | 0.0925 | 0.2475 | 0.777032 | 0.353539 | 0.899115 | 0.989223 | 0.997988 | 0.99798 |
| Rab23_Q9D4I9_S197s_1 | Rab23_Q9 | -2.3 | -2.75 | -1.58 | -2.12 | -0.97 | -2.15 | -0.96 | -1.48 | -3.02 | -2.71 | -0.37 | -1.52 | -2.04 | -0.88 | -1.98 | -1.37 | 0.7975 | 0.2825 | 0.62 | 0.12844 | 0.052387 | 0.63825 | 0.989223 | 0.997988 | 0.994666 |
| Rab31_Q3TXV4_S36s_1 | Rab31_Q3 | 0.08 | 0.46 | 0.66 | 0.79 | 0.11 | -0.12 | 0.67 | 0.45 | 0.14 | 0.37 | 0.48 | 0.81 | -0.35 | 0.25 | 0.41 | 0.74 | -0.22 | -0.0475 | -0.235 | 0.471807 | 0.429827 | 0.859403 | 0.989223 | 0.997988 | 0.994666 |
| Rab37_Q544E8_T4t_1 | Rab37_Q5 | 0.3 | 0.78 | 0.9 | -0.59 | 0.58 | 0.08 | 1.13 | 1.09 | -0.48 | 0.02 | 0.72 | 0.8 | 0.26 | 0.02 | 0.63 | 1.11 | 0.3725 | -0.0825 | 0.1575 | 0.704621 | 0.365807 | 0.85128 | 0.989223 | 0.997988 | 0.996159 |
| Rab39b_Q8BHC1_S201s_1 | Rab39b_Q | 0.07 | 0.49 | 0.36 | 0.53 | 0.91 | -0.19 | 1.13 | 0.52 | 0.25 | 0.36 | 0.26 | 0.47 | 0.41 | 0.97 | 0.73 | 0.86 | 0.23 | -0.0275 | 0.38 | 0.166967 | 0.480816 | 0.901872 | 0.989223 | 0.997988 | 0.994666 |
| Rab3a_P63011_S188s_1 | Rab3a_P63 | 0.52 | 0.71 | 0.38 | 0.33 | 0.73 | 0.33 | 0.55 | 0.63 | -0.18 | 0.03 | -0.04 | 0.32 | -0.17 | 0.37 | 0.3 | 0.66 | 0.075 | -0.4525 | -0.195 | 0.484234 | 0.727759 | 0.075783 | 0.995569 | 0.997988 | 0.994666 |
| Rab3a_P63011_S190s_1 | Rab3a_P63 | 1.27 | 0.52 | 0.97 | 1.16 | 1.11 | 1.42 | 1.44 | 1.42 | 0.54 | 0.4 | 0.85 | 1.05 | -0.06 | 1.1 | 0.98 | 1.26 | 0.3675 | -0.27 | -0.16 | 0.66192 | 0.156852 | 0.340645 | 0.989223 | 0.997988 | 0.995804 |
| Rab3gap2_E9QKE4_S449s_1 | Rab3gap2_ | 0.2 | -0.36 | 0.29 | 0.7 | 0.46 | 0.25 | 0.82 | 0.55 | -0.4 | -0.07 | 0.13 | 0.76 | -0.73 | 0.26 | 0.29 | 0.59 | 0.3125 | -0.1025 | -0.105 | 0.781011 | 0.284592 | 0.766825 | 0.989223 | 0.997988 | 0.998226 |
| Rab3ip_Q68EF0_S218s_1 | Rab3ip_Q6 | 0.2 | 0.32 | -0.13 | 0.27 | 0.53 | 0.08 | 0.26 | 0.16 | 0.16 | 0.24 | 0.29 | 0.34 | -0.11 | 0.7 | 0.03 | 0 | 0.0925 | 0.0925 | -0.01 | 0.971927 | 0.680455 | 0.675895 | 0.993464 | 0.997988 | 0.999507 |
| Rab3ip_Q68EF0_S240s_1 | Rab3ip_Q6 | -1.6 | -1.8 | -1.2 | -0.99 | -1.46 | -1.82 | -0.62 | -1.16 | -1.74 | -1.88 | -0.59 | -0.92 | -2.06 | -0.8 | -1.44 | -1.25 | 0.1325 | 0.115 | 0.01 | 0.977171 | 0.685437 | 0.757007 | 0.994624 | 0.997988 | 1 |
| Rab40c_E9Q016_S249s_1 | Rab40c_E9 | -0.55 | -0.36 | -0.79 | -0.86 | -0.26 | -0.52 | -0.35 | -1.25 | -0.08 | 0.02 | -0.27 | -1.17 | 0.49 | -0.41 | -0.39 | -0.09 | 0.045 | 0.265 | 0.54 | 0.097486 | 0.874599 | 0.42029 | 0.996493 | 0.997988 | 0.994666 |
| Rab5a_Q9CQD1_S193s_1 | Rab5a_Q9 | 0.52 | -0.62 | 0.42 | 0.52 | -0.31 | 1.6 | -0.75 | 0.57 | 1.75 | -0.29 | -0.33 | 0.75 | 2.21 | -0.79 | 0.6 | 1.1 | 0.0675 | 0.26 | 0.57 | 0.368206 | 0.899381 | 0.625121 | 0.996493 | 0.997988 | 0.994666 |
| Rab5b_P61021_S205s_1 | Rab5b_P61 | -0.88 | 0.61 | -0.29 | 0.02 | -0.04 | -0.19 | -0.51 | -0.48 | 0.3 | -0.42 | -0.2 | -0.37 | 0.95 | 0.32 | -0.11 | -0.05 | -0.17 | -0.0375 | 0.4125 | 0.317356 | 0.617168 | 0.917485 | 0.989223 | 0.997988 | 0.994666 |
| Rab8a_P55258_S181s_1 | Rab8a_P55 | 0.88 | 0.93 | 1.37 | 0.9 | -0.12 | 0.93 | 0.2 | 0.5 | 1.02 | 1.08 | 1.31 | 1.01 | 1.19 | -0.7 | 0.7 | 0.53 | -0.6425 | 0.085 | -0.59 | 0.180483 | 0.045836 | 0.714389 | 0.989223 | 0.997988 | 0.994666 |
| Rab8a_P55258_S185s_1 | Rab8a_P55 | -0.72 | -0.32 | -0.18 | -0.91 | -0.22 | -0.51 | -0.67 | -0.86 | -0.61 | -0.8 | -0.65 | -1.08 | 0.67 | -1.02 | -0.32 | -0.27 | -0.0325 | -0.2525 | 0.2975 | 0.458146 | 0.901908 | 0.350932 | 0.996493 | 0.997988 | 0.994666 |
| Rab8b_P61028_S182s_1 | Rab8b_P61 | 1.53 | 0.64 | 0.52 | 0.44 | 0.65 | 1.1 | -0.23 | 0.45 | 1.18 | 1.6 | -0.11 | 0.61 | 1.41 | 0.12 | 0.14 | 0.59 | -0.29 | 0.0375 | -0.2175 | 0.589463 | 0.439949 | 0.931256 | 0.989223 | 0.997988 | 0.995259 |
| Rab8b_P61028_S183s_1 | Rab8b_P61 | -0.34 | 0.19 | 0.58 | 0 | -0.36 | 1.06 | -0.3 | 0.39 | 0.69 | 0.97 | -0.79 | 0.68 | 1.61 | -1.19 | 0.21 | 0.71 | 0.09 | 0.28 | 0.2275 | 0.689469 | 0.811524 | 0.520052 | 0.995762 | 0.997988 | 0.996159 |
| Rab9a_Q9R0M6_S179s_1 | Rab9a_Q9 | 0.18 | 0.1 | 0.32 | 0.16 | 0.19 | 0.27 | -0.61 | 1.44 | 0.02 | 0.44 | -0.39 | 0.45 | -0.68 | 0.36 | 1.26 | 0.61 | 0.1325 | -0.06 | 0.1975 | 0.630976 | 0.745033 | 0.822027 | 0.995762 | 0.997988 | 0.995259 |
| Rabep1_J3QJV7_S323s_1 | Rabep1_J3 | 0.33 | -0.31 | -0.07 | 0.14 | 0.28 | 0.31 | 0.44 | 0.38 | -0.34 | -0.19 | 0.88 | 0.02 | -0.49 | 0.67 | -0.07 | -0.17 | 0.33 | 0.07 | -0.0375 | 0.908177 | 0.164531 | 0.832004 | 0.989223 | 0.997988 | 0.999507 |
| Rabep1_J3QJV7_S326s_1 | Rabep1_J3 | 0.04 | 0.26 | -0.86 | -0.73 | -0.29 | 0.03 | -0.39 | -0.35 | -0.49 | 0.08 | -0.1 | -0.21 | -0.75 | -0.05 | -0.93 | -0.9 | 0.0725 | 0.1425 | -0.335 | 0.372369 | 0.816451 | 0.665158 | 0.995762 | 0.997988 | 0.994666 |
| Rabgap1_A2AWA9_S42s_1 | Rabgap1_A | 1.52 | 0.87 | -0.11 | -0.63 | -0.38 | 2.12 | -1.13 | -1.05 | 1.72 | 1.45 | -0.82 | -0.32 | 1.79 | -0.26 | -0.14 | -0.79 | -0.5225 | 0.095 | -0.2625 | 0.695371 | 0.512689 | 0.894436 | 0.989223 | 0.997988 | 0.996159 |
| Rabgap1_A2AWA9_S486s_1 | Rabgap1_A | -1.44 | -1.17 | -0.3 | -1.04 | -0.88 | -0.66 | -1.52 | -1.03 | 0.02 | -1.01 | -1.08 | -0.68 | 0.65 | -1.69 | -0.87 | -0.54 | -0.035 | 0.3 | 0.375 | 0.469531 | 0.912713 | 0.415471 | 0.997335 | 0.997988 | 0.994666 |
| Rabgef1_Q9JM13_S124s_1 | Rabgef1_Q | -2.95 | -3.24 | -3.57 | -2.75 | -1.18 | -2.73 | -2.94 | -3.3 | -3.59 | -2.59 | -0.12 | -2.38 | -2.49 | -1.86 | -2.28 | -1.7 | 0.59 | 0.9575 | 1.045 | 0.007131 | 0.224053 | 0.183351 | 0.989223 | 0.997988 | 0.994666 |
| Rabgef1_Q9JM13_S132s_1 | Rabgef1_Q | -0.32 | -1.02 | -0.18 | -0.54 | -1.71 | -0.02 | -2.1 | -0.71 | 0.08 | -0.73 | -0.8 | -0.9 | 0.38 | -1.71 | -0.99 | -0.69 | -0.62 | -0.0725 | -0.2375 | 0.605545 | 0.208188 | 0.820805 | 0.989223 | 0.997988 | 0.995259 |
| Rabgef1_Q9JM13_S390s_1 | Rabgef1_Q | 0.15 | -0.2 | 0.13 | -0.18 | -1.09 | 0.12 | -0.79 | -0.57 | -3.94 | 0.11 | -5.37 | -0.3 | -0.68 | -6.45 | -0.61 | -0.4 | -0.5575 | -2.35 | -2.01 | 0.138926 | 0.087856 | 0.073734 | 0.989223 | 0.997988 | 0.994666 |
| Rabl6_Q5U3K5_S377s_1 | Rabl6_Q5U | 0.16 | 0.04 | 0.36 | 0.51 | 0.61 | 0.44 | 0.32 | 0.28 | 0.13 | -0.41 | 0.53 | 0.19 | -0.77 | -0.19 | 0.91 | 0.75 | 0.145 | -0.1575 | -0.0925 | 0.822636 | 0.511026 | 0.570763 | 0.989223 | 0.997988 | 0.999507 |
| Rabl6_Q5U3K5_S482s_1 | Rabl6_Q5U | 0.94 | 1.2 | 0.43 | 0.66 | 1.02 | 1.2 | 0.76 | 1.29 | 0.91 | 1.64 | 1.8 | 1.1 | -1.24 | 0.47 | 0.37 | 0.71 | 0.26 | 0.555 | -0.73 | 0.137116 | 0.32354 | 0.096348 | 0.989223 | 0.997988 | 0.994666 |
| Rabl6_Q5U3K5_S502s_1 | Rabl6_Q5U | 0.91 | 0.58 | 1.36 | 1.13 | 0.31 | 0.62 | 0.7 | 0.9 | 0.25 | 0.68 | 1.19 | 0.62 | 0.14 | 0.12 | 1.05 | 0.8 | -0.3625 | -0.31 | -0.4675 | 0.178389 | 0.181481 | 0.310404 | 0.989223 | 0.997988 | 0.994666 |
| Rad23a_Q3TN85_S123s_1 | Rad23a_Q3 | 0.59 | 0.77 | 1.88 | 1.73 | 0.1 | 0.62 | 2 | 0.52 | 1.08 | 0.55 | 2.95 | 1.69 | 0.01 | 0.87 | 2.01 | 1.65 | -0.4325 | 0.325 | -0.1075 | 0.835635 | 0.383232 | 0.568548 | 0.989223 | 0.997988 | 0.999507 |
| Rad23a_Q3TN85_S125s_1 | Rad23a_Q3 | 0.02 | 0.16 | 0.14 | -0.21 | 0.04 | 0.44 | -0.22 | 0.4 | 0.02 | 0.64 | 0.01 | 0.89 | 0.79 | 0.58 | 0.39 | -0.2 | 0.1375 | 0.3625 | 0.3625 | 0.237635 | 0.573073 | 0.225224 | 0.989223 | 0.997988 | 0.994666 |
| Rad23a_Q3TN85_S92s_1 | Rad23a_Q3 | 0.15 | 0.14 | 0.44 | 0.15 | 0.46 | 0.03 | 0.84 | 0.31 | 0.32 | -0.14 | 0.78 | 0.06 | 0.05 | -0.66 | 0.67 | 0.2 | 0.19 | 0.035 | -0.155 | 0.63801 | 0.445485 | 0.896889 | 0.989223 | 0.997988 | 0.995259 |
| Rad23b_P54728_S160s_1 | Rad23b_P5 | -1.32 | -0.88 | -1.17 | -1.1 | -0.8 | -1.33 | -0.72 | -0.95 | -1.16 | -0.85 | -0.79 | -0.87 | -1.1 | -0.45 | -1.36 | -1.22 | 0.1675 | 0.2 | 0.085 | 0.770002 | 0.480749 | 0.386655 | 0.989223 | 0.997988 | 0.9971 |
| Rad23b_P54728_S160s_2 | Rad23b_P5 | 0.23 | 0.35 | 0.01 | 0.19 | -0.36 | 0.21 | 0.88 | 0.5 | 1.23 | 0.17 | 0.16 | -0.38 | 0.65 | 1.77 | 0.03 | 0.21 | 0.1125 | 0.1 | 0.47 | 0.260308 | 0.705687 | 0.779746 | 0.995437 | 0.997988 | 0.994666 |
| Rad23b_P54728_T155t_1 | Rad23b_P5 | 0.31 | 0.06 | 0.62 | 0.41 | -0.14 | -0.1 | 0.32 | 0.01 | -0.19 | -0.01 | -0.07 | 0.51 | -0.31 | -0.48 | 0.4 | 0.88 | -0.3275 | -0.29 | -0.2275 | 0.533068 | 0.17836 | 0.281001 | 0.989223 | 0.997988 | 0.994666 |
| Rad23b_P54728_T155t_2 | Rad23b_P5 | 0.23 | 0.35 | 0.08 | 1 | -0.36 | 0.21 | 0.84 | 0.83 | 1.23 | 0.17 | 0.3 | 0.21 | 0.65 | 1.77 | 0.11 | 0.52 | -0.035 | 0.0625 | 0.3475 | 0.406782 | 0.920884 | 0.855162 | 0.997335 | 0.997988 | 0.994666 |
| Radil_D3YWJ9_S206s_1 | Radil_D3Y | -2.12 | -2.31 | -1.71 | -1.62 | -2.25 | -2.81 | -1.28 | -1.38 | -2.29 | -1.96 | -2.18 | -2.15 | -0.95 | -1.97 | -1.89 | -1.49 | 0.01 | -0.205 | 0.365 | 0.281696 | 0.979448 | 0.423895 | 0.999187 | 0.997988 | 0.994666 |
| Raf1_Q99N57_S233s_1 | Raf1_Q99N | -0.63 | -1.09 | -1.86 | -2.08 | -0.33 | -0.54 | -0.41 | -2.06 | -1.39 | -0.21 | -1.26 | -0.44 | -0.72 | -1.87 | -0.73 | -0.62 | 0.58 | 0.59 | 0.43 | 0.340695 | 0.25319 | 0.196466 | 0.989223 | 0.997988 | 0.994666 |
| Rai1_Q61818_S899s_1 | Rai1_Q618 | 0.41 | 0.6 | 0.29 | 1.46 | 0.37 | 0.26 | 1.02 | 0.7 | 0.15 | 0.1 | 1.04 | 0.38 | -0.28 | -0.45 | 0.96 | 0.74 | -0.1025 | -0.2725 | -0.4475 | 0.318193 | 0.754391 | 0.45039 | 0.995762 | 0.997988 | 0.994666 |
| Rala_P63321_S11s_1 | Rala_P633 | 0.36 | 1.44 | 1.07 | 0.73 | 1.28 | 0.63 | 0.92 | 0.72 | 0.31 | 0.56 | 0.54 | -0.02 | 0.99 | 1.3 | 1.12 | 0.86 | -0.0125 | -0.5525 | 0.1675 | 0.587305 | 0.966515 | 0.095921 | 0.999187 | 0.997988 | 0.995259 |
| Ralbp1_Q62172_S11s_1 | Ralbp1_Q6 | 0.24 | 0.84 | 1.01 | 0.41 | 0.91 | 0.25 | 1.14 | 1.09 | 0.02 | 0.71 | -0.41 | 1.42 | -0.7 | -0.39 | 1.17 | 0.98 | 0.2225 | -0.19 | -0.36 | 0.46297 | 0.462637 | 0.659034 | 0.989223 | 0.997988 | 0.994666 |
| Ralbp1_Q62172_S34s_1 | Ralbp1_Q6 | 0.47 | 1.34 | 1.56 | 1.42 | 0.68 | -0.27 | 1.47 | 1.27 | 0.6 | 0.81 | 1.7 | 1.56 | 1.09 | 0.88 | 1.22 | 0.77 | -0.41 | -0.03 | -0.2075 | 0.516839 | 0.357369 | 0.935961 | 0.989223 | 0.997988 | 0.994666 |
| Ralbp1_Q62172_S34s_2 | Ralbp1_Q6 | -5.02 | -2.36 | -1.21 | -2.87 | -2.94 | -4.16 | -3.08 | -0.82 | -6.22 | -3.27 | -0.42 | -2.07 | -3.81 | -2.11 | -1.87 | -2.27 | 0.115 | -0.13 | 0.35 | 0.664796 | 0.900472 | 0.918994 | 0.996493 | 0.997988 | 0.996159 |
| Ralbp1_Q62172_S463s_1 | Ralbp1_Q6 | -0.5 | -0.46 | -0.91 | -0.92 | 0.15 | -0.68 | -0.38 | -0.73 | -0.65 | -0.56 | -0.66 | -0.94 | -0.81 | -0.23 | -0.64 | -1.04 | 0.2875 | -0.005 | 0.0175 | 0.951084 | 0.312206 | 0.98321 | 0.989223 | 0.997988 | 0.999507 |
| Ralbp1_Q62172_S601s_1 | Ralbp1_Q6 | 0.68 | 0.03 | 0.32 | 0.27 | 0.27 | 1.14 | -0.22 | 0.86 | -0.76 | -0.65 | -0.05 | 0.01 | -0.99 | -0.84 | 0.38 | 0.33 | 0.1875 | -0.6875 | -0.605 | 0.153168 | 0.583155 | 0.036924 | 0.989223 | 0.997988 | 0.994666 |
| Ralgapa1_Q6GYP7_S1003s_1 | Ralgapa1_( | 0.24 | -0.18 | 0.35 | 0.54 | 0.54 | -0.04 | 0.57 | 0.32 | 0.43 | 0.62 | 1.1 | 0.21 | 0.45 | 0.78 | 0.39 | 0.49 | 0.11 | 0.3525 | 0.29 | 0.292755 | 0.671173 | 0.241873 | 0.992876 | 0.997988 | 0.994666 |
| Ralgapa1_Q6GYP7_S1003s_2 | Ralgapa1_( | 0.28 | 0.07 | 0.43 | 0.25 | 0.52 | 0.18 | 0.6 | 0.63 | -0.06 | 0.3 | 0.06 | 0.5 | -2.36 | 0.6 | 0.83 | -0.31 | 0.225 | -0.0575 | -0.5675 | 0.397901 | 0.315321 | 0.807175 | 0.989223 | 0.997988 | 0.994666 |
| Ralgapa1_Q6GYP7_S1477s_1 | Ralgapa1_( | -2.51 | -1.95 | -2.25 | -3.13 | -2.06 | -1.78 | -2 | -1.62 | -1.98 | -1.81 | -0.77 | -1.9 | 0.54 | -1.16 | -1.6 | -2.56 | 0.595 | 0.845 | 1.265 | 0.069142 | 0.069529 | 0.050229 | 0.989223 | 0.997988 | 0.994666 |
| Ralgapa1_Q6GYP7_S368s_1 | Ralgapa1_( | -0.95 | -0.76 | -0.35 | -0.2 | -0.96 | -0.62 | -0.46 | -0.41 | 0.2 | -0.49 | 0 | -0.4 | 0.28 | 0.11 | -0.41 | -1.56 | -0.0475 | 0.3925 | 0.17 | 0.700862 | 0.856117 | 0.193278 | 0.996349 | 0.997988 | 0.996159 |
| Ralgapa1_Q6GYP7_S371s_1 | Ralgapa1_( | -1.2 | -1.49 | -0.94 | -1.41 | -1.09 | -0.5 | -0.59 | -2.28 | -0.04 | -0.1 | -0.85 | -2.23 | -0.1 | -0.25 | -1.39 | -3.42 | 0.145 | 0.455 | -0.03 | 0.96552 | 0.723455 | 0.368954 | 0.995437 | 0.997988 | 0.999507 |
| Ralgapa1_Q6GYP7_S739s_1 | Ralgapa1_( | 0.44 | -0.76 | 0.73 | -0.19 | -1.27 | 0.78 | -0.08 | -0.13 | 2.25 | 0.1 | -1.05 | 0.58 | 2.08 | -0.94 | -0.66 | -0.31 | -0.23 | 0.415 | -0.0125 | 0.985481 | 0.642315 | 0.550154 | 0.989327 | 0.997988 | 1 |
| Ralgapa1_Q6GYP7_S859s_1 | Ralgapa1_( | -0.28 | -1.44 | -0.49 | -0.21 | -0.38 | -0.25 | 0.45 | -0.13 | -0.69 | -0.23 | -0.32 | -0.17 | -0.41 | 0.19 | -0.51 | -0.5 | 0.5275 | 0.2525 | 0.2975 | 0.412987 | 0.14866 | 0.453325 | 0.989223 | 0.997988 | 0.994666 |
| Ralgapa1_Q6GYP7_S863s_1 | Ralgapa1_( | 0.96 | 0.78 | 0.68 | 0.6 | 0.95 | 0.29 | 0.86 | 0.91 | 0.25 | 0.72 | 1.05 | 0.73 | -0.31 | 1.1 | 0.67 | 1.2 | -0.0025 | -0.0675 | -0.09 | 0.809041 | 0.991664 | 0.790419 | 0.999187 | 0.997988 | 0.999507 |
| Ralgapa1_Q6GYP7_T1001t_2 | Ralgapa1_( | 0.28 | 0.07 | 0.43 | 0.25 | 0.52 | 0.18 | 0.6 | 0.63 | -0.06 | 0.3 | 0.06 | 0.5 | -2.36 | 0.6 | 0.83 | -0.31 | 0.225 | -0.0575 | -0.5675 | 0.397901 | 0.315321 | 0.807175 | 0.989223 | 0.997988 | 0.994666 |
| Ralgapb_F8WHN4_S1450s_1 | Ralgapb_F | 0.51 | -1.24 | -0.09 | -0.89 | -1.43 | -0.7 | -0.51 | -0.54 | -1.07 | 0.27 | -0.71 | -0.47 | 0.77 | -0.36 | -1.41 | 0.19 | -0.3675 | -0.0675 | 0.225 | 0.689948 | 0.397323 | 0.884298 | 0.989223 | 0.997988 | 0.996159 |
| Ralgapb_F8WHN4_S359s_1 | Ralgapb_F | -0.08 | 0.57 | 0.47 | 0.26 | 0.69 | -0.24 | -0.15 | 0.32 | -0.13 | 0.19 | 0.44 | 0.27 | 0.93 | 0.66 | 0.38 | 0.28 | -0.15 | -0.1125 | 0.2575 | 0.371725 | 0.607386 | 0.662145 | 0.989223 | 0.997988 | 0.994666 |
| Ralgapb_F8WHN4_T363t_1 | Ralgapb_F | 0.6 | 0.07 | 0.65 | 0.51 | 0.42 | 0.51 | 0.2 | 0.67 | 0.49 | 0.5 | 0.64 | 0.55 | 0.31 | 0.5 | 0.07 | 0.29 | -0.0075 | 0.0875 | -0.165 | 0.529454 | 0.974516 | 0.70658 | 0.999187 | 0.997988 | 0.994666 |
| Ralgapb_F8WHN4_T363t_2 | Ralgapb_F | -0.37 | -0.47 | -0.12 | -0.18 | -0.21 | -0.62 | -0.65 | -0.53 | -1.09 | -0.62 | -0.41 | -0.13 | -1.99 | -0.71 | -0.75 | -0.23 | -0.2175 | -0.2775 | -0.635 | 0.131571 | 0.332999 | 0.324995 | 0.989223 | 0.997988 | 0.994666 |
| Ralgapb_F8WHN4_T379t_1 | Ralgapb_F | -0.35 | -0.27 | -0.28 | -0.24 | -0.34 | -0.57 | 0.06 | -0.19 | -0.5 | -0.19 | 0.01 | -0.24 | 0.24 | 0.04 | 0.09 | -0.15 | 0.025 | 0.055 | 0.34 | 0.167927 | 0.909764 | 0.802744 | 0.997335 | 0.997988 | 0.994666 |
| Ralgapb_F8WHN4_T734t_1 | Ralgapb_F | 1.61 | 1.47 | 0.99 | 0.96 | 1.38 | 1.49 | 1.07 | 1.2 | -0.38 | 1.8 | -1.85 | 1.21 | 0.15 | -0.34 | 1.21 | 1.26 | 0.0275 | -1.0625 | -0.6875 | 0.131684 | 0.911774 | 0.181298 | 0.997335 | 0.997988 | 0.994666 |
| Ralgps1_A2AR50_S292s_1 | Ralgps1_A | 0.91 | -0.29 | 0.49 | -0.08 | -0.89 | 0.68 | -0.44 | -0.42 | 1.22 | 0.09 | -0.05 | 0.34 | 1.5 | -1.13 | 0.59 | 0.3 | -0.525 | 0.1425 | 0.0575 | 0.918533 | 0.222081 | 0.718415 | 0.989223 | 0.997988 | 0.999507 |

| | | | | | | | | | | | | | | | | | | | | | | | | | | |
|---|---|---|---|---|---|---|---|---|---|---|---|---|---|---|---|---|---|---|---|---|---|---|---|---|---|---|
| Ralgps1_A2AR50_S298s_1 | Ralgps1_A | 2.01 | 0.31 | 0.57 | 1.2 | 0.85 | 1.83 | 0.89 | 1.49 | 0.65 | 0.81 | -0.07 | 1.36 | -0.77 | 0.84 | -0.05 | 1.15 | 0.2425 | -0.335 | -0.73 | 0.196014 | 0.570448 | 0.472538 | 0.989223 | 0.997988 | 0.994666 |
| Ralgps1_A2AR50_S358s_1 | Ralgps1_A | -0.48 | -0.73 | -0.73 | -0.73 | -0.25 | -0.21 | -0.49 | -0.34 | -1.2 | -0.93 | 0.3 | -0.61 | -1.51 | -0.42 | -1.94 | -0.46 | 0.345 | 0.0575 | -0.415 | 0.307847 | 0.115169 | 0.868938 | 0.989223 | 0.997988 | 0.994666 |
| Ralgps1_A2AR50_T319t_1 | Ralgps1_A | 0.88 | -0.07 | 0.69 | 0.03 | -0.03 | 0.32 | 0.08 | -0.24 | -0.39 | 0.29 | -1.15 | 1.04 | -1.31 | -0.17 | -0.69 | -0.16 | -0.35 | -0.435 | -0.965 | 0.027426 | 0.249426 | 0.389622 | 0.989223 | 0.997988 | 0.994666 |
| Ralgps1_A2AR50_T331t_1 | Ralgps1_A | 0.4 | 0.43 | -0.14 | 0.62 | 0.5 | -0.1 | 1.16 | 0.92 | -0.24 | 0.23 | 1.52 | 0.46 | -0.65 | 0.66 | 0.47 | 0.86 | 0.2925 | 0.165 | 0.0075 | 0.984529 | 0.386627 | 0.683468 | 0.989223 | 0.997988 | 1 |
| Ralgps1_A2AR50_T360t_1 | Ralgps1_A | -0.15 | -0.18 | 0.11 | 0.58 | 0.08 | -0.25 | 0.38 | -0.4 | -0.73 | -0.28 | -0.27 | -0.04 | -0.16 | -0.41 | -0.06 | -0.82 | -0.1375 | -0.42 | -0.4525 | 0.159119 | 0.628084 | 0.157392 | 0.989223 | 0.997988 | 0.994666 |
| Raly_Q64012_S119s_1 | Raly_Q640 | -0.2 | -0.91 | 1.24 | 1.13 | -0.28 | -0.82 | 0.98 | 1.29 | -0.51 | -0.32 | 0.25 | 0.88 | 0.77 | 0.03 | 0.5 | 0.21 | -0.0225 | -0.24 | 0.0625 | 0.903298 | 0.972111 | 0.671422 | 0.999187 | 0.997988 | 0.999507 |
| Raly_Q64012_S254s_1 | Raly_Q640 | 0.38 | -0.28 | 1 | 0.61 | 0.15 | 0.2 | 0.56 | 0.37 | 0.03 | 0.33 | 1.5 | 0.26 | 0.16 | -0.58 | 0.03 | -0.05 | -0.1075 | 0.1025 | -0.5375 | 0.144437 | 0.72571 | 0.806058 | 0.995437 | 0.997988 | 0.994666 |
| Raly_Q64012_T258t_1 | Raly_Q640 | -0.45 | -0.28 | -1.41 | -0.14 | 0.11 | 0.05 | 0.26 | -0.67 | -0.25 | -0.05 | -0.46 | -0.53 | -0.17 | 0.19 | -0.56 | -0.4 | 0.5075 | 0.2475 | 0.335 | 0.358905 | 0.175139 | 0.46143 | 0.989223 | 0.997988 | 0.994666 |
| Ranbp2_Q9ERU9_S2113s_1 | Ranbp2_Q | 0.8 | -0.32 | 0.75 | 1.12 | -0.32 | 0.27 | 0.64 | 1.3 | 0.44 | -0.49 | 0.31 | 0.91 | 0.71 | -0.48 | 2.14 | 1.5 | -0.115 | -0.295 | 0.38 | 0.524514 | 0.791647 | 0.487604 | 0.995762 | 0.997988 | 0.994666 |
| Ranbp2_Q9ERU9_S781s_1 | Ranbp2_Q | 7.2 | 2.92 | 6.22 | 1.11 | 3.52 | 8.17 | 5.69 | 5.29 | 2.07 | 6.19 | -2.07 | 5.98 | 2.31 | 0.97 | 7.59 | 6.99 | 1.305 | -1.32 | 0.1025 | 0.956158 | 0.383774 | 0.533349 | 0.989223 | 0.997988 | 0.999507 |
| Ranbp2_Q9ERU9_S788s_1 | Ranbp2_Q | 1.24 | 0.25 | 0.92 | 0.77 | 0.64 | 0.66 | 0.75 | 1 | 0.17 | 0.58 | 1.39 | 0.84 | 1.07 | 0.36 | 1.65 | 1.31 | -0.0325 | -0.05 | 0.3025 | 0.415585 | 0.902983 | 0.884974 | 0.996493 | 0.997988 | 0.994666 |
| Ranbp2_Q9ERU9_T898t_1 | Ranbp2_Q | -0.47 | -0.22 | -0.86 | -0.82 | -0.6 | -0.43 | -1.26 | -1.39 | -0.23 | -0.6 | -0.69 | -0.67 | 0.03 | -1.22 | -0.21 | -0.9 | -0.3275 | 0.045 | 0.0175 | 0.960733 | 0.297289 | 0.860384 | 0.989223 | 0.997988 | 0.999507 |
| Ranbp3_Q9CT10_S277s_1 | Ranbp3_Q | -4.79 | -2.87 | -3.44 | -3.22 | -4.66 | -1.43 | -3.34 | -2.74 | -2.02 | -2.82 | -3.88 | -2.89 | 0.26 | -3.89 | -4.44 | -4.93 | 0.5375 | 0.6775 | 0.33 | 0.76314 | 0.448449 | 0.220938 | 0.989223 | 0.997988 | 0.9971 |
| Rangap1_P46061_S405s_1 | Rangap1_P | -0.59 | 0.02 | 0.32 | -0.46 | -0.85 | -0.51 | -0.55 | -0.96 | 0.3 | -0.62 | -0.61 | -0.27 | 1.31 | -1.16 | -0.18 | -0.05 | -0.54 | -0.1225 | 0.1575 | 0.760747 | 0.075796 | 0.709376 | 0.989223 | 0.997988 | 0.9971 |
| Rangap1_P46061_S444s_1 | Rangap1_P | -0.33 | -0.62 | -0.86 | -0.06 | -0.97 | -1.05 | 0.24 | -0.21 | -0.48 | -0.11 | -0.08 | -0.44 | -0.18 | 0.32 | -0.31 | -0.45 | -0.03 | 0.19 | 0.3125 | 0.31423 | 0.932698 | 0.480335 | 0.998562 | 0.997988 | 0.994666 |
| Rangap1_P46061_T411t_1 | Rangap1_P | -0.33 | -1.29 | -1.29 | -1.35 | -1.53 | -0.53 | -2.17 | -1.55 | 0.63 | -0.92 | -1.65 | -1.3 | 0.74 | -1.76 | -1.11 | -1.35 | -0.38 | 0.255 | 0.195 | 0.727884 | 0.356365 | 0.627672 | 0.989223 | 0.997988 | 0.9971 |
| Rap1gap_A2ALS5_S439s_1 | Rap1gap_A | 0.01 | -0.03 | -0.51 | -0.24 | -0.02 | 0.01 | -0.64 | -0.77 | -0.18 | -0.08 | 0.32 | -0.78 | 0.48 | -0.15 | -0.7 | -0.69 | -0.1625 | 0.0125 | -0.0725 | 0.83099 | 0.558357 | 0.966458 | 0.989223 | 0.997988 | 0.999507 |
| Rap1gap_A2ALS5_S441s_1 | Rap1gap_A | -0.45 | -0.47 | -0.61 | -0.37 | -0.24 | -0.26 | -0.6 | -0.63 | -0.86 | -0.5 | 0.04 | -0.55 | -0.1 | -0.22 | -0.72 | -0.57 | 0.0425 | 0.0075 | 0.0725 | 0.778448 | 0.842252 | 0.976952 | 0.996212 | 0.997988 | 0.99798 |
| Rap1gap_A2ALS5_S460s_1 | Rap1gap_A | -0.86 | -0.7 | -0.4 | 0.64 | -0.34 | 0.41 | 0.55 | 0.72 | -0.85 | -0.19 | 0.35 | 0.12 | -0.87 | 0.44 | 0.18 | 0.07 | 0.665 | 0.1875 | 0.285 | 0.517033 | 0.118263 | 0.655328 | 0.989223 | 0.997988 | 0.994666 |
| Rap1gap_A2ALS5_S484s_1 | Rap1gap_A | 0.19 | -1.39 | -0.95 | 0.68 | -0.05 | 0.49 | 0.79 | 0.2 | -0.78 | -0.73 | 1.03 | -0.43 | -1.67 | -0.35 | -0.61 | -0.36 | 0.725 | 0.14 | -0.38 | 0.484087 | 0.152133 | 0.812816 | 0.989223 | 0.997988 | 0.994666 |
| Rap1gap_A2ALS5_S498s_1 | Rap1gap_A | 1.51 | 0.67 | -0.26 | -0.11 | 0.65 | 1.33 | 0.34 | -0.02 | 0.3 | 1.73 | -0.85 | -0.02 | 0.06 | 0.52 | -0.15 | 0.28 | 0.1225 | -0.1625 | -0.275 | 0.524665 | 0.791371 | 0.792146 | 0.995762 | 0.997988 | 0.994666 |
| Rap1gap_A2ALS5_S499s_1 | Rap1gap_A | 1.51 | 0.67 | -0.26 | -0.11 | 0.65 | 1.33 | 0.34 | -0.02 | 0.3 | 1.73 | -0.85 | -0.02 | 0.06 | 0.52 | -0.15 | 0.28 | 0.1225 | -0.1625 | -0.275 | 0.524665 | 0.791371 | 0.792146 | 0.995762 | 0.997988 | 0.994666 |
| Rap1gap_A2ALS5_S515s_1 | Rap1gap_A | -1.1 | -1.08 | -0.61 | -1.25 | -1.82 | -0.8 | -1.89 | -1.69 | 1.57 | -1.01 | -2.2 | -1.35 | 1.81 | -2.14 | -0.93 | -1.59 | -0.54 | 0.2625 | 0.2975 | 0.70558 | 0.105921 | 0.723372 | 0.989223 | 0.997988 | 0.996159 |
| Rap1gap_A2ALS5_S541s_1 | Rap1gap_A | -0.43 | 0.09 | 2.04 | 2.41 | -0.29 | -0.64 | 0.61 | 1.6 | 0.12 | 0.1 | 2.35 | 2.22 | 1.61 | -0.36 | 2.69 | 2.71 | -0.7075 | 0.17 | 0.635 | 0.478465 | 0.361682 | 0.839603 | 0.989223 | 0.997988 | 0.994666 |
| Rap1gap_A2ALS5_S541s_2 | Rap1gap_A | -0.56 | -0.1 | 0.22 | 0.32 | -0.66 | -0.72 | 0.36 | 0.3 | -0.3 | 0.15 | 0.12 | 0.35 | -0.25 | -0.21 | 0.5 | 0.62 | -0.15 | 0.11 | 0.195 | 0.568556 | 0.674206 | 0.702931 | 0.992876 | 0.997988 | 0.995259 |
| Rap1gap_A2ALS5_S542s_1 | Rap1gap_A | 0.22 | 1.37 | 0.64 | 0.34 | 0.29 | -0.21 | 0.39 | 0.42 | 0.95 | 1.07 | 0.27 | 0.3 | 1.76 | 0.15 | 1.02 | 0.62 | -0.42 | 0.005 | 0.245 | 0.567101 | 0.202697 | 0.988563 | 0.989223 | 0.997988 | 0.995259 |
| Rap1gap_A2ALS5_S542s_2 | Rap1gap_A | -0.56 | -0.1 | 0.22 | 0.32 | -0.66 | -0.72 | 0.36 | 0.3 | -0.3 | 0.15 | 0.12 | 0.35 | -0.25 | -0.21 | 0.5 | 0.62 | -0.15 | 0.11 | 0.195 | 0.568556 | 0.674206 | 0.702931 | 0.992876 | 0.997988 | 0.995259 |
| Rap1gap_A2ALS5_S566s_1 | Rap1gap_A | 0.32 | -0.03 | -0.49 | -0.72 | -0.03 | 0.39 | -0.22 | -0.05 | -0.25 | 0.42 | 0.62 | -0.21 | -0.46 | 0.29 | -1.07 | -0.59 | 0.2525 | 0.375 | -0.2275 | 0.553454 | 0.400197 | 0.286788 | 0.989223 | 0.997988 | 0.994666 |
| Rap1gap_A2ALS5_T522t_1 | Rap1gap_A | 0.44 | -0.93 | -2.14 | -1.84 | 0.22 | -0.29 | -2.52 | -1.74 | 0.48 | 0.28 | -1.5 | -1.9 | -0.35 | -0.4 | -2.56 | -2 | 0.035 | 0.4575 | -0.21 | 0.770787 | 0.9629 | 0.547323 | 0.999187 | 0.997988 | 0.9971 |
| Rap1gap2_Q5SVL6_S26s_1 | Rap1gap2_ | 0.15 | 0.34 | 0.31 | 0.24 | 0.4 | 0.16 | 0.62 | 0.44 | 0.03 | 0.38 | 0 | 0.3 | 0.04 | 0.64 | -0.01 | 0.09 | 0.145 | -0.0825 | -0.07 | 0.787054 | 0.49259 | 0.707124 | 0.989223 | 0.997988 | 0.999507 |
| Rap1gap2_Q5SVL6_S399s_1 | Rap1gap2_ | 0.09 | 0.26 | 0.09 | -0.08 | 0.64 | -0.37 | 0.69 | 0.82 | 0.04 | 0.35 | 0.54 | 0.23 | -0.01 | 0.71 | 0.27 | 0.29 | 0.355 | 0.2 | 0.225 | 0.399496 | 0.261765 | 0.388737 | 0.989223 | 0.997988 | 0.994666 |
| Rap1gap2_Q5SVL6_S488s_1 | Rap1gap2_ | 3.06 | 0.09 | 1.55 | -2.5 | 0.6 | 3.9 | 2.96 | 2.43 | 1.06 | 1.68 | 0.84 | 2.56 | 1.35 | -0.95 | 1.81 | 2.05 | 1.9225 | 0.985 | 0.515 | 0.664713 | 0.128547 | 0.379347 | 0.989223 | 0.997988 | 0.996159 |
| Rap1gap2_Q5SVL6_S525s_1 | Rap1gap2_ | -0.53 | -1.31 | -2.43 | -1.15 | 0 | -0.64 | -1.86 | -1.4 | -0.85 | -0.96 | -2.17 | -1.14 | -2.08 | -0.08 | -1.47 | -0.76 | 0.38 | 0.075 | 0.2575 | 0.638314 | 0.47107 | 0.87424 | 0.989223 | 0.997988 | 0.995259 |
| Rap1gap2_Q5SVL6_S581s_1 | Rap1gap2_ | -0.15 | -1.37 | -0.5 | -0.23 | -1.34 | 0.2 | -1.14 | -0.4 | 0.69 | -0.24 | -0.55 | -0.27 | 0.45 | -0.61 | -0.74 | -0.59 | -0.1075 | 0.47 | 0.19 | 0.636271 | 0.801027 | 0.246693 | 0.995762 | 0.997988 | 0.995259 |
| Rap1gap2_Q5SVL6_S594s_1 | Rap1gap2_ | 0.62 | 0.56 | 0.49 | 0.65 | 0.35 | 0.82 | 0.46 | 0.36 | 0.93 | 1.42 | 0.41 | 0.55 | 0.8 | 0.65 | 0.86 | 0.81 | -0.0825 | 0.2475 | 0.2 | 0.389113 | 0.699675 | 0.387505 | 0.994893 | 0.997988 | 0.994666 |
| Rap1gap2_Q5SVL6_S660s_1 | Rap1gap2_ | 0.65 | 0.48 | 0.33 | 0.44 | 0.7 | 1.19 | 0.9 | 0.7 | 0.42 | 0.99 | 0.32 | 0.6 | -0.26 | 0.93 | -0.07 | 1.06 | 0.3975 | 0.1075 | -0.06 | 0.869841 | 0.096703 | 0.660239 | 0.989223 | 0.997988 | 0.999507 |
| Rap1gds1_E9Q912_S548s_1 | Rap1gds1_ | -0.32 | 0.61 | 0.82 | 0.37 | -0.34 | 0.18 | 0.05 | -0.13 | 1.23 | 0.31 | 0.37 | 0.29 | 1.76 | -0.01 | -0.03 | -0.14 | -0.43 | 0.18 | 0.025 | 0.959474 | 0.171894 | 0.610799 | 0.989223 | 0.997988 | 0.999507 |
| Rapgef1_Q3UHC1_S308s_1 | Rapgef1_Q | -1.5 | -0.33 | 0.1 | -0.57 | -1.89 | -0.83 | -1.34 | -1.34 | 0.44 | -1.01 | -1.8 | -0.42 | 1.54 | -1.89 | -1.09 | -0.8 | -0.775 | -0.1225 | 0.015 | 0.983425 | 0.071144 | 0.820699 | 0.989223 | 0.997988 | 1 |
| Rapgef1_Q3UHC1_S326s_1 | Rapgef1_Q | 0.85 | -0.71 | 0.49 | 0.1 | 0.91 | 0.61 | 0.84 | 0.15 | -0.24 | 0.2 | 1.17 | 0.9 | 0.2 | 0.66 | -0.67 | 0.6 | 0.445 | 0.325 | 0.015 | 0.973001 | 0.24794 | 0.474356 | 0.989223 | 0.997988 | 0.999507 |
| Rapgef1_Q3UHC1_S375s_1 | Rapgef1_Q | 0.15 | -0.21 | 0.3 | 0.03 | 0.05 | 0.44 | -0.34 | 0.45 | -0.26 | -0.28 | 0.19 | 0.01 | -1.13 | -0.41 | -0.34 | -0.19 | 0.0825 | -0.1525 | -0.585 | 0.074485 | 0.754581 | 0.530655 | 0.995762 | 0.997988 | 0.994666 |
| Rapgef1_Q3UHC1_S381s_1 | Rapgef1_Q | -1.64 | -2.07 | -1.05 | -0.86 | -1.42 | -1.39 | -1.57 | -1.33 | -1.56 | -1.44 | -0.37 | -0.35 | -1.64 | -0.73 | -0.54 | -1.39 | -0.0225 | 0.475 | 0.33 | 0.408189 | 0.940968 | 0.277017 | 0.999187 | 0.997988 | 0.994666 |
| Rapgef2_E9QNQ4_S1086s_1 | Rapgef2_E | -2.2 | -2.55 | -1.6 | -2.11 | -1.74 | -3.19 | -0.69 | -0.9 | -2.72 | -1.97 | 0.37 | -1.31 | -2.47 | -0.56 | -1.65 | -1.1 | 0.485 | 0.7075 | 0.67 | 0.15408 | 0.382366 | 0.276006 | 0.989223 | 0.997988 | 0.994666 |
| Rapgef2_E9QNQ4_S1113s_1 | Rapgef2_E | 0.56 | 0.74 | 0.97 | 1.04 | 0.7 | 0.51 | 1.06 | 1.09 | 0.41 | 0.95 | 1.35 | 0.96 | 0.08 | -0.48 | 0.55 | 0.38 | 0.0125 | 0.09 | -0.695 | 0.045189 | 0.958817 | 0.745133 | 0.999187 | 0.997988 | 0.994666 |
| Rapgef2_E9QNQ4_S1220s_1 | Rapgef2_E | 0.54 | 0.56 | 0.57 | 1.11 | 1.24 | -0.24 | 0.86 | 0.72 | 0.2 | 0.52 | 0.8 | 0.42 | -0.81 | 0.38 | 0.04 | 0.24 | -0.05 | -0.21 | -0.7325 | 0.05259 | 0.885424 | 0.420589 | 0.996493 | 0.997988 | 0.994666 |
| Rapgef2_E9QNQ4_S1235s_1 | Rapgef2_E | -0.45 | -1.46 | -1.49 | -3.21 | -1.39 | 0.39 | -2.22 | -1.88 | -1.01 | -0.87 | 0.74 | -1.89 | -0.73 | -1 | -2.95 | -2.68 | 0.3775 | 0.895 | -0.1875 | 0.794605 | 0.601406 | 0.229978 | 0.989223 | 0.997988 | 0.999507 |
| Rapgef2_E9QNQ4_S1313s_1 | Rapgef2_E | 0.88 | -0.34 | 1.11 | 1.99 | -0.74 | 1.23 | 0.45 | 2.15 | -0.01 | 0.19 | 0.68 | 1.99 | -0.58 | -1.26 | 0.65 | 1.07 | -0.1375 | -0.1975 | -0.94 | 0.172151 | 0.841414 | 0.743656 | 0.995932 | 0.997988 | 0.994666 |
| Rapgef2_E9QNQ4_S928s_1 | Rapgef2_E | -0.7 | -0.44 | -0.65 | -0.69 | -0.13 | -1.01 | -0.01 | -0.74 | -0.69 | -0.53 | -0.91 | -0.69 | -0.78 | -0.06 | -0.67 | -0.48 | 0.1475 | -0.085 | 0.1225 | 0.644949 | 0.603808 | 0.696356 | 0.989223 | 0.997988 | 0.995259 |
| Rapgef2_E9QNQ4_S931s_1 | Rapgef2_E | -0.47 | -1.26 | -0.37 | 0.06 | 0.05 | -0.36 | 0.19 | -0.34 | -0.84 | -0.5 | -0.59 | 0.02 | -0.72 | 0.45 | -0.55 | 0.17 | 0.395 | 0.0325 | 0.3475 | 0.394445 | 0.238842 | 0.925179 | 0.989223 | 0.997988 | 0.994666 |
| Rapgef2_E9QNQ4_S933s_1 | Rapgef2_E | -0.37 | -0.35 | 0.66 | -0.4 | 0.27 | -0.04 | 0.59 | 0.09 | -0.14 | 0.01 | -0.73 | 0.19 | -0.39 | 0.47 | -0.59 | 0.18 | 0.3425 | -0.0525 | 0.0325 | 0.930798 | 0.286966 | 0.878788 | 0.989223 | 0.997988 | 0.999507 |
| Rapgef2_E9QNQ4_S957s_1 | Rapgef2_E | 0.44 | 0.41 | -0.19 | -0.75 | 0.91 | 0.37 | -0.17 | -0.71 | -0.49 | 0.45 | -0.7 | -0.42 | -0.32 | 0.5 | -0.46 | -0.11 | 0.1225 | -0.2675 | -0.075 | 0.840245 | 0.773368 | 0.490846 | 0.995762 | 0.997988 | 0.999507 |
| Rapgef2_E9QNQ4_S958s_1 | Rapgef2_E | -0.17 | -0.03 | 0 | -0.58 | 0.46 | 0.01 | -0.24 | -0.4 | -0.81 | -0.19 | -0.02 | -0.6 | -0.36 | 0.19 | -0.26 | -0.19 | 0.1525 | -0.21 | 0.04 | 0.881787 | 0.577451 | 0.45767 | 0.989223 | 0.997988 | 0.999507 |
| Rapgef3_H3BK39_S489s_1 | Rapgef3_H | 0.75 | 0.45 | 1.46 | 0.46 | 0.56 | 0.46 | 0.42 | 0.87 | 0.17 | 0.7 | -1.14 | 1.39 | 0.43 | -0.84 | 1.37 | 1.26 | -0.2025 | -0.5 | -0.225 | 0.670138 | 0.488955 | 0.369058 | 0.989223 | 0.997988 | 0.996159 |
| Rapgef6_Q5NCJ4_S809s_1 | Rapgef6_Q | 1.37 | 0.33 | 0.19 | -0.21 | 0.75 | 0.88 | -0.33 | 0.01 | 1.25 | 0.4 | 0.61 | 0.52 | 0.7 | 1.48 | -0.01 | -0.17 | -0.0925 | 0.275 | 0.08 | 0.868302 | 0.82657 | 0.484694 | 0.995762 | 0.997988 | 0.999507 |
| Raph1_F2Z3U3_S1027s_1 | Raph1_F2Z | -0.74 | -0.06 | -0.59 | -0.94 | 0.2 | -0.75 | -0.25 | -0.37 | -0.26 | -0.07 | -0.08 | -0.27 | -0.45 | -0.06 | -0.45 | -0.47 | 0.29 | 0.4125 | 0.225 | 0.438908 | 0.342196 | 0.139791 | 0.989223 | 0.997988 | 0.994666 |
| Raph1_F2Z3U3_S1169s_1 | Raph1_F2Z | 0.22 | 0.38 | -0.05 | 0.5 | 0.58 | 0.48 | 0.8 | 0.19 | 0.18 | 0.57 | -0.21 | 0.53 | -0.11 | 0.46 | 0.14 | 0.56 | 0.25 | 0.005 | 7.85E-17 | 1 | 0.310943 | 0.985389 | 0.989223 | 0.997988 | 1 |
| Raph1_F2Z3U3_S192s_1 | Raph1_F2Z | 0.17 | -1.08 | -0.41 | -0.21 | -0.02 | 0.12 | -0.58 | 0.41 | -0.55 | -0.63 | 0.93 | -0.12 | -0.02 | 0.31 | -0.45 | -0.61 | 0.365 | 0.29 | 0.19 | 0.599945 | 0.298514 | 0.507175 | 0.989223 | 0.997988 | 0.995259 |
| Raph1_F2Z3U3_S850s_1 | Raph1_F2Z | -0.91 | -0.28 | -1.09 | -0.94 | -0.12 | -0.38 | -0.63 | -0.5 | -0.76 | -0.7 | -0.49 | -1.23 | -0.91 | -0.62 | -1.07 | -0.84 | 0.3975 | 0.01 | -0.055 | 0.844444 | 0.14941 | 0.972054 | 0.989223 | 0.997988 | 0.999507 |
| Raph1_F2Z3U3_S899s_1 | Raph1_F2Z | 0.3 | 0.25 | 0.59 | 0.41 | 0.43 | -0.24 | 0.33 | 0.45 | -0.13 | 0.23 | 0.21 | 0.51 | 0.3 | 0.33 | 0.38 | 0.18 | -0.145 | -0.1825 | -0.09 | 0.701323 | 0.553801 | 0.450954 | 0.989223 | 0.997988 | 0.996159 |
| Rasa2_P58069_S554s_1 | Rasa2_P58 | -0.01 | -0.09 | -0.04 | 0.22 | 0.02 | 0.25 | 0.65 | 0.42 | -0.4 | -0.33 | 1.24 | 0.66 | -1.02 | 0.2 | 0.2 | -0.25 | 0.315 | 0.2725 | -0.2375 | 0.48352 | 0.188652 | 0.50232 | 0.989223 | 0.997988 | 0.994666 |
| Rasa3_Q60790_S809s_1 | Rasa3_Q60 | 1.01 | 1.04 | 1.74 | 1.6 | 1.06 | 1.23 | 0.96 | 1.52 | 0.39 | 0.85 | 1.75 | 1.52 | 0.32 | 0.54 | 1.1 | 1.32 | -0.155 | -0.22 | -0.5275 | 0.14226 | 0.566751 | 0.557237 | 0.989223 | 0.997988 | 0.994666 |
| Rasa3_Q60790_S833s_1 | Rasa3_Q60 | -0.42 | 0.08 | -0.64 | 0.96 | -0.51 | -0.26 | -0.1 | 1.47 | -0.72 | -0.45 | 0.32 | 1.32 | -0.28 | -0.64 | 0.7 | 0.49 | 0.155 | 0.1225 | 0.0725 | 0.874694 | 0.766251 | 0.820113 | 0.995762 | 0.997988 | 0.999507 |
| Rasal1_Q9Z268_S194s_1 | Rasal1_Q9 | -0.23 | 0.48 | 0.38 | -0.6 | 0.34 | 0.13 | -0.14 | -0.15 | -0.03 | -0.11 | -0.16 | 0.24 | 0.62 | 0.55 | 0.22 | -0.07 | 0.0375 | -0.0225 | 0.3225 | 0.352081 | 0.901869 | 0.941516 | 0.996493 | 0.997988 | 0.994666 |
| Rasal1_Q9Z268_T400t_1 | Rasal1_Q9 | -0.13 | 0.6 | 0.71 | -0.82 | -0.14 | -0.47 | -0.72 | -1.17 | 0.63 | -0.71 | -0.43 | -0.16 | 1 | -0.4 | -0.92 | -0.89 | -0.715 | -0.2575 | -0.3925 | 0.469459 | 0.100024 | 0.564434 | 0.989223 | 0.997988 | 0.994666 |
| Rasal2_D3YUS5_S1068s_1 | Rasal2_D3 | 0.57 | 0.85 | -0.97 | -0.96 | 0.81 | 0.2 | -0.65 | -1.12 | 0.71 | 1.26 | -3.54 | -1.63 | 0.96 | 0.4 | -1.57 | -1.14 | -0.0625 | -0.6725 | -0.21 | 0.762943 | 0.914603 | 0.532391 | 0.997335 | 0.997988 | 0.9971 |
| Rasal2_D3YUS5_S1068s_2 | Rasal2_D3 | -0.16 | 0.09 | 0.06 | -0.05 | -0.09 | -0.46 | 0.59 | 0.12 | -0.46 | 0.24 | 0.28 | 0.36 | -0.12 | -0.09 | 0.24 | 0.85 | 0.055 | 0.12 | 0.235 | 0.435806 | 0.838115 | 0.649334 | 0.995762 | 0.997988 | 0.994666 |
| Rasal2_D3YUS5_S1076s_1 | Rasal2_D3 | 0.14 | 1.07 | 0.51 | 1.07 | 0.85 | 0.77 | 0.88 | 0.83 | 0.88 | 1.07 | 1.15 | 0.87 | 0.4 | 0.85 | 0.51 | 1.17 | 0.135 | 0.295 | 0.035 | 0.915199 | 0.620334 | 0.316876 | 0.989223 | 0.997988 | 0.999507 |
| Rasal2_D3YUS5_S1076s_2 | Rasal2_D3 | -0.16 | 0.09 | 0.06 | -0.05 | -0.09 | -0.46 | 0.59 | 0.12 | -0.46 | 0.24 | 0.28 | 0.36 | -0.12 | -0.09 | 0.24 | 0.85 | 0.055 | 0.12 | 0.235 | 0.435806 | 0.838115 | 0.649334 | 0.995762 | 0.997988 | 0.994666 |
| Rasal2_D3YUS5_S115s_1 | Rasal2_D3 | -0.11 | 0.61 | -0.44 | -0.34 | 0.9 | -0.03 | 0.12 | -0.2 | -0.13 | 0.32 | -0.72 | -0.47 | -0.15 | 0.81 | -0.02 | -0.03 | 0.2675 | -0.18 | 0.2225 | 0.532392 | 0.444297 | 0.604145 | 0.989223 | 0.997988 | 0.994666 |
| Rasal2_D3YUS5_S140s_1 | Rasal2_D3 | 1.92 | -0.85 | -0.33 | 1.67 | 0.93 | 1.91 | 1.27 | 1.86 | 1.59 | 1.02 | 0.51 | 1.2 | 0.55 | 0.94 | 0.85 | 0.46 | 0.89 | 0.4775 | 0.0975 | 0.878906 | 0.195306 | 0.478871 | 0.989223 | 0.997988 | 0.999507 |
| Rasal2_D3YUS5_S164s_1 | Rasal2_D3 | -0.48 | 0.84 | -0.11 | -1.08 | 0.64 | 0.52 | 0.11 | -0.89 | -0.41 | 0.26 | -0.16 | -0.15 | 0.83 | 0 | -0.21 | 0.26 | 0.3025 | 0.0925 | 0.4275 | 0.352879 | 0.54019 | 0.824765 | 0.989223 | 0.997988 | 0.994666 |
| Rasal2_D3YUS5_S785s_1 | Rasal2_D3 | -0.76 | -0.59 | -1.3 | -2.16 | -1.86 | -1.94 | -1.78 | -1.13 | -0.48 | -2.27 | 0.13 | -1.82 | -1.33 | -2.09 | -0.95 | -2.71 | -0.475 | 0.0925 | -0.5675 | 0.272805 | 0.238962 | 0.878821 | 0.989223 | 0.997988 | 0.994666 |
| Rasal2_D3YUS5_S822s_1 | Rasal2_D3 | -0.93 | 0.09 | -0.69 | -1.79 | -0.53 | -0.99 | -1.47 | -1.28 | -0.77 | -1.28 | 0.24 | -0.88 | 0.32 | -0.59 | -1.07 | -1.28 | -0.2375 | 0.1575 | 0.175 | 0.725772 | 0.571749 | 0.74273 | 0.989223 | 0.997988 | 0.9971 |
| Rasal2_D3YUS5_S925s_1 | Rasal2_D3 | 0.74 | 0.06 | 0.61 | 0.11 | 1.06 | 0.43 | 1.02 | 0.92 | 0.68 | 0.52 | -0.54 | 0.76 | 0.69 | 0.68 | 0.81 | 0.93 | 0.4775 | -0.025 | 0.3975 | 0.164835 | 0.101105 | 0.944574 | 0.989223 | 0.997988 | 0.994666 |
| Rasal2_D3YUS5_S986s_1 | Rasal2_D3 | -1.02 | -0.48 | 0.06 | -0.19 | -1.78 | 0.21 | -1.87 | -0.56 | 0.99 | -0.26 | -0.92 | -0.09 | 1.87 | -1.58 | -0.52 | -1.02 | -0.5925 | 0.3375 | 0.095 | 0.893351 | 0.25901 | 0.453902 | 0.989223 | 0.997988 | 0.999507 |
| Rasal2_D3YUS5_T742t_1 | Rasal2_D3 | 0.01 | -0.07 | -0.34 | 0.13 | 0.74 | -0.03 | 0.74 | 0.61 | -0.16 | 0.11 | 0.75 | 0.87 | -0.14 | 0.55 | -0.3 | -0.51 | 0.5825 | 0.46 | -0.0325 | 0.915553 | 0.04629 | 0.155677 | 0.989223 | 0.997988 | 0.999507 |
| Rasgef1a_A6H650_T157t_1 | Rasgef1a_ | -0.15 | -0.08 | 0.27 | 0.34 | -0.76 | 0.42 | -0.26 | -0.02 | 0.93 | -0.21 | 0.3 | 0.45 | 1.58 | -1.69 | 0.95 | 0.54 | -0.25 | 0.2725 | 0.25 | 0.701987 | 0.413047 | 0.37914 | 0.989223 | 0.997988 | 0.996159 |
| Rasgrf1_P27671_S485s_1 | Rasgrf1_P2 | 0.65 | 0.02 | -0.26 | -1.08 | 0.54 | 0.98 | -0.19 | -0.63 | -0.28 | 0.35 | 0.45 | 0.02 | 0.16 | -0.04 | -0.34 | -0.62 | 0.3425 | 0.3025 | -0.0425 | 0.915904 | 0.473591 | 0.450861 | 0.989223 | 0.997988 | 0.999507 |
| Rasgrf1_P27671_S726s_1 | Rasgrf1_P2 | 0.23 | 0.57 | 0 | 0.98 | 0.62 | -0.13 | 0.72 | 0.79 | 0.05 | 0.43 | 0.97 | 0.64 | -0.18 | 0.71 | 1.06 | 0.84 | 0.055 | 0.0775 | 0.1625 | 0.65952 | 0.862392 | 0.807436 | 0.996493 | 0.997988 | 0.995259 |
| Rasgrf1_P27671_S727s_1 | Rasgrf1_P2 | 1.03 | 1.47 | 0.66 | 0.42 | 1.01 | 0.33 | 1.12 | 0.52 | 0.68 | 1.63 | 0.44 | 0.38 | -0.38 | 1.45 | 0.48 | 0.81 | -0.15 | -0.1125 | -0.305 | 0.490656 | 0.635683 | 0.764775 | 0.989223 | 0.997988 | 0.994666 |
| Rasgrf1_P27671_S737s_1 | Rasgrf1_P2 | -0.13 | -0.25 | -0.94 | -0.49 | 0.45 | -0.05 | 0.11 | -0.52 | -0.69 | -0.06 | 0.07 | -0.7 | -0.95 | -0.18 | -0.7 | -0.44 | 0.45 | 0.1075 | -0.115 | 0.705265 | 0.152611 | 0.726866 | 0.989223 | 0.997988 | 0.996159 |
| Rasgrf1_P27671_S837s_1 | Rasgrf1_P2 | -3.35 | -0.4 | -3.68 | -2.8 | -1.81 | -3.43 | -2.27 | -3.81 | -2.8 | -2.18 | -2.2 | -2.36 | -2.01 | -1.03 | -2.26 | -3.3 | -0.2725 | 0.1725 | 0.4075 | 0.601951 | 0.724102 | 0.800738 | 0.995437 | 0.997988 | 0.995259 |
| Rasgrf2_P70392_S484s_1 | Rasgrf2_P7 | -3.01 | -3.68 | -0.76 | -4.43 | -0.87 | -2.4 | 0.3 | -1.99 | -3.68 | -3.15 | 0.81 | -0.25 | -2.91 | -0.69 | -0.65 | -1.75 | 1.73 | 1.4025 | 1.47 | 0.105875 | 0.070881 | 0.255132 | 0.989223 | 0.997988 | 0.994666 |
| Rasgrf2_P70392_S726s_1 | Rasgrf2_P7 | -0.18 | 0.06 | -0.09 | -0.33 | 0.09 | -0.49 | 0.11 | -0.3 | -1.14 | 0.24 | -1.01 | -0.41 | 0.05 | -0.56 | -0.43 | -0.22 | -0.0125 | -0.445 | -0.155 | 0.552437 | 0.957941 | 0.217339 | 0.999187 | 0.997988 | 0.994666 |
| Rasgrf2_P70392_S801s_1 | Rasgrf2_P7 | 0.02 | -0.68 | 0.03 | -0.06 | -1.31 | -0.5 | -1.18 | -0.59 | 0.36 | -0.4 | -0.03 | -0.48 | 0.58 | -0.91 | -0.8 | -0.3 | -0.7225 | 0.035 | -0.185 | 0.637461 | 0.032562 | 0.906406 | 0.989223 | 0.997988 | 0.995259 |
| Rasgrf2_P70392_S801s_2 | Rasgrf2_P7 | -0.22 | -1.36 | -1.93 | -0.21 | -0.27 | 0.11 | 0.06 | -0.54 | -0.69 | -1 | -0.47 | -1.06 | -1.09 | -0.61 | -1.62 | -0.7 | 0.77 | 0.125 | -0.075 | 0.87269 | 0.098715 | 0.775045 | 0.989223 | 0.997988 | 0.999507 |
| Rasgrf2_P70392_S805s_1 | Rasgrf2_P7 | 1.71 | 0.86 | -0.26 | 1.35 | 0.7 | 1.93 | 0.86 | 1.57 | 0.77 | 1.32 | 0.68 | 0.6 | 1.47 | 1.07 | 0.8 | 0.61 | 0.35 | -0.0725 | 0.0725 | 0.873026 | 0.470942 | 0.869961 | 0.989223 | 0.997988 | 0.999507 |
| Rasgrf2_P70392_S805s_2 | Rasgrf2_P7 | -0.15 | -1.26 | -1.83 | -0.56 | -0.14 | -0.02 | -0.54 | -0.83 | -0.61 | -0.86 | 0.28 | -1.08 | -1.05 | -0.6 | -1.65 | -0.98 | 0.5675 | 0.3825 | -0.12 | 0.778886 | 0.178832 | 0.412243 | 0.989223 | 0.997988 | 0.998201 |
| Rasgrp1_G3UYC3_S562s_1 | Rasgrp1_G | 0.51 | -0.64 | -0.63 | -0.2 | 0.65 | -0.34 | 1.09 | 0.42 | -0.08 | 0.69 | 0.81 | 0.56 | 0.39 | 0.54 | 0.8 | 0.6 | 0.695 | 0.735 | 0.8225 | 0.029311 | 0.100701 | 0.060411 | 0.989223 | 0.997988 | 0.994666 |
| Rasgrp1_G3UYC3_S759s_1 | Rasgrp1_G | 1.78 | 0.87 | 1.14 | -0.59 | 0.83 | 1.69 | 0 | 1.44 | 1.51 | 1.56 | 1.55 | 1.34 | 1.48 | 0.71 | 0.04 | 0.85 | 0.19 | 0.69 | -0.03 | 0.955712 | 0.736915 | 0.17303 | 0.995762 | 0.997988 | 0.999507 |
| Rasgrp2_Q9QUG9_S585s_1 | Rasgrp2_Q | -2.14 | -1.99 | -1.9 | -2.43 | -0.54 | -2.33 | -1.22 | -1.96 | -1.77 | -1.73 | 0.38 | -1.34 | -0.33 | -0.38 | -0.98 | -1 | 0.6025 | 1 | 1.4425 | 0.000621 | 0.154967 | 0.067278 | 0.989223 | 0.997988 | 0.994666 |
| Rasgrp2_Q9QUG9_S586s_1 | Rasgrp2_Q | -2.14 | -1.99 | -0.6 | -0.07 | -0.54 | -2.33 | -0.34 | -0.57 | -1.77 | -1.73 | 0.67 | -0.11 | -0.33 | -0.38 | -0.24 | 0.05 | 0.255 | 0.465 | 0.975 | 0.073275 | 0.680274 | 0.520358 | 0.993464 | 0.997988 | 0.994666 |
| Rasip1_Q3U0S6_S286s_1 | Rasip1_Q3 | -2.77 | -2.84 | -2.16 | -1.65 | -2.05 | -3.44 | -1.97 | -2.26 | -0.76 | -2.32 | -0.05 | -2.44 | -0.8 | -0.57 | -2.81 | -3.18 | -0.075 | 0.9625 | 0.515 | 0.440631 | 0.858224 | 0.133177 | 0.996349 | 0.997988 | 0.994666 |
| Rasip1_Q3U0S6_T960t_1 | Rasip1_Q3 | 3.12 | 1.13 | 1.91 | 0.76 | 1.12 | 3.23 | 2.11 | 1.97 | 0.57 | 2.89 | 1.71 | 2.01 | 1.08 | 0.03 | 2.4 | 2.66 | 0.3775 | 0.065 | -0.1875 | 0.79391 | 0.537389 | 0.919506 | 0.989223 | 0.997988 | 0.999507 |
| Rassf5_D3Z6G2_S177s_1 | Rassf5_D3 | -0.7 | -0.27 | 0.18 | 0.1 | 0.03 | -0.37 | 0.47 | 0.3 | -0.69 | -0.69 | 0.83 | 0.1 | -0.32 | -0.69 | 0.22 | -0.73 | 0.28 | 0.06 | -0.2075 | 0.540984 | 0.358785 | 0.883903 | 0.989223 | 0.997988 | 0.994666 |
| Raver1_Q9CW46_S14s_1 | Raver1_Q9 | 0.25 | 0.71 | 0.44 | 0.02 | 0.21 | 0.2 | 0.56 | 0.24 | 0.04 | 0.55 | 0.38 | 0.74 | -0.68 | 0.95 | 0.29 | 0.85 | -0.0525 | 0.0725 | -0.0025 | 0.995075 | 0.825157 | 0.787224 | 0.995762 | 0.997988 | 1 |
| Raver1_Q9CW46_S576s_1 | Raver1_Q9 | -0.09 | -2.28 | -1.79 | -1.49 | -0.75 | -3.33 | -1.74 | -2.06 | -1.56 | -0.3 | -1 | -1.25 | -0.77 | -0.13 | -2.55 | -1.12 | -0.5575 | 0.385 | 0.27 | 0.669497 | 0.387485 | 0.455771 | 0.989223 | 0.997988 | 0.996159 |
| Raver1_Q9CW46_S626s_1 | Raver1_Q9 | 0.17 | 0.05 | -0.12 | -0.39 | 0.13 | -0.48 | 0.09 | 0.05 | -0.53 | 0.01 | 0.5 | -0.41 | -0.11 | 0.02 | -0.08 | -0.96 | 0.02 | -0.035 | -0.21 | 0.50497 | 0.935444 | 0.907554 | 0.99868 | 0.997988 | 0.994666 |
| Rb1_P13405_S243s_1 | Rb1_P134 | -2.87 | -2.54 | -2.05 | -1.94 | -1.78 | -3.2 | -1.34 | -2.03 | -3.06 | -2.57 | -2.68 | -2.55 | -0.93 | -1.51 | -2.18 | -1.6 | 0.2625 | -0.365 | 0.795 | 0.049434 | 0.543491 | 0.23058 | 0.989223 | 0.997988 | 0.994666 |
| Rb1cc1_E9PX56_S222s_1 | Rb1cc1_E9 | -1.12 | -0.8 | -1.1 | -0.94 | -1.98 | -0.66 | -0.96 | -1.08 | -2.22 | -0.09 | 0.16 | -1 | -0.03 | -1.29 | -0.9 | -1.48 | -0.18 | 0.2025 | 0.065 | 0.855451 | 0.567657 | 0.692496 | 0.989223 | 0.997988 | 0.999507 |
| Rb1cc1_E9PX56_S257s_1 | Rb1cc1_E9 | -1.02 | -0.39 | -0.18 | -1.4 | 0.06 | -0.65 | -0.22 | -0.12 | -2.82 | -1.81 | -0.26 | -1.03 | -0.73 | -0.85 | -0.63 | 0.3 | 0.515 | -0.7325 | 0.27 | 0.498683 | 0.142791 | 0.218098 | 0.989223 | 0.997988 | 0.994666 |
| Rb1cc1_E9PX56_S261s_1 | Rb1cc1_E9 | -0.09 | -0.35 | 0.46 | 0.94 | 0.49 | 0.15 | -0.56 | -0.57 | -0.17 | -0.09 | 1.38 | -1.05 | 0.12 | 0.26 | -0.28 | 0.46 | -0.3625 | -0.2225 | -0.1 | 0.778483 | 0.353984 | 0.681121 | 0.989223 | 0.997988 | 0.99798 |
| Rb1cc1_E9PX56_S629s_1 | Rb1cc1_E9 | 0.53 | 0.99 | 0.28 | 0.27 | 0.21 | -0.31 | 0.31 | 0.18 | 0.83 | 1.3 | -0.1 | 0.33 | 0.97 | -0.5 | 0.43 | 0.06 | -0.42 | 0.0725 | -0.2775 | 0.461807 | 0.136733 | 0.840042 | 0.989223 | 0.997988 | 0.994666 |
| Rb1cc1_E9PX56_S629s_2 | Rb1cc1_E9 | 0.97 | 0.88 | -0.49 | 0.39 | -0.27 | 0.57 | -0.32 | -0.16 | 0.98 | 0.95 | -0.05 | -0.33 | 1.03 | 0.62 | -1.01 | -0.28 | -0.4825 | -0.05 | -0.3475 | 0.516176 | 0.227854 | 0.912685 | 0.989223 | 0.997988 | 0.994666 |
| Rb1cc1_E9PX56_S635s_2 | Rb1cc1_E9 | 0.97 | 0.88 | -0.49 | 0.39 | -0.27 | 0.57 | -0.32 | -0.16 | 0.98 | 0.95 | -0.05 | -0.33 | 1.03 | 0.62 | -1.01 | -0.28 | -0.4825 | -0.05 | -0.3475 | 0.516176 | 0.227854 | 0.912685 | 0.989223 | 0.997988 | 0.994666 |
| Rb1cc1_E9PX56_S649s_1 | Rb1cc1_E9 | 0.44 | 0.53 | 1.53 | 0.36 | 0.69 | -0.18 | 0.75 | 0.76 | 0.22 | 0.36 | 1.32 | 1.12 | 0.92 | 0.51 | 1.95 | 1.46 | -0.21 | 0.04 | 0.495 | 0.252701 | 0.559391 | 0.917869 | 0.989223 | 0.997988 | 0.994666 |
| Rbbp6_P97868_S1244s_1 | Rbbp6_P97 | 1.04 | 0.78 | 1.22 | 1.16 | 0.67 | 0.89 | 0.83 | 0.92 | 0.55 | 0.97 | 0.98 | 0.99 | 0.69 | 0.37 | 1.34 | 0.75 | -0.2225 | -0.1775 | -0.2625 | 0.379282 | 0.307853 | 0.458016 | 0.989223 | 0.997988 | 0.994666 |
| Rbl2_Q64700_S37s_1 | Rbl2_Q647 | -1.08 | -1.69 | -0.44 | -0.81 | -1.2 | -2.08 | -1.15 | -1.59 | -0.64 | -0.98 | -0.19 | -1.33 | 0.09 | -1.25 | -1.12 | -0.82 | -0.5 | 0.22 | 0.23 | 0.57341 | 0.169927 | 0.553116 | 0.989223 | 0.997988 | 0.995259 |
| Rbl2_Q64700_S946s_1 | Rbl2_Q647 | 0.97 | 0.98 | 2.22 | 0.48 | 0.44 | 2.17 | 2.79 | 1.99 | 0.96 | 1.89 | 1.4 | 1.61 | 2 | 0.17 | 2.79 | 2.32 | 0.685 | 0.3025 | 0.6575 | 0.304038 | 0.240812 | 0.47084 | 0.989223 | 0.997988 | 0.994666 |
| Rbl2_Q64700_S960s_1 | Rbl2_Q647 | -0.27 | -1.22 | 0.11 | -0.29 | -1.43 | -1.15 | -0.89 | 0.12 | -0.17 | -0.39 | -0.56 | -0.42 | 0.65 | -2.08 | -0.29 | 0.43 | -0.42 | 0.0325 | 0.095 | 0.877898 | 0.328929 | 0.919594 | 0.989223 | 0.997988 | 0.999507 |
| Rbm10_Q99KG3_S646s_1 | Rbm10_Q9 | -0.6 | -2.23 | -2.02 | -2.08 | -1.7 | -2.62 | -1.5 | -1.62 | -2.32 | -1.83 | -1.5 | -1.67 | -0.35 | -1.42 | -0.27 | -1.36 | -0.1275 | -0.0975 | 0.8825 | 0.087073 | 0.768406 | 0.813349 | 0.995762 | 0.997988 | 0.994666 |
| Rbm14_E9QL13_S582s_1 | Rbm14_E9 | -0.23 | -2.4 | -1.73 | -1.63 | -2.07 | -2.08 | -1.54 | -1.31 | -0.88 | -1.82 | -1.05 | -1.17 | -0.08 | -1.26 | -0.74 | -1.71 | -0.2525 | 0.2675 | 0.55 | 0.31835 | 0.586362 | 0.576847 | 0.989223 | 0.997988 | 0.994666 |
| Rbm14_E9QL13_T206t_1 | Rbm14_E9 | 0.2 | -1.68 | -1.01 | -0.92 | -1.13 | -0.88 | -0.68 | -0.48 | -1.47 | -1.23 | -1.29 | -0.54 | -0.69 | -0.73 | -0.4 | -0.62 | 0.06 | -0.28 | 0.2425 | 0.550295 | 0.880218 | 0.518425 | 0.996493 | 0.997988 | 0.994666 |
| Rbm15_Q0VBL3_S293s_1 | Rbm15_Q0 | -0.11 | -0.46 | 1.01 | -1.01 | -0.04 | -0.88 | 0.51 | -0.7 | -0.67 | -0.43 | -0.04 | 0.32 | -0.07 | 0.24 | 0.85 | 0.46 | -0.135 | -0.0625 | 0.5125 | 0.275948 | 0.78297 | 0.891594 | 0.995762 | 0.997988 | 0.994666 |
| Rbm15_Q0VBL3_S655s_1 | Rbm15_Q0 | 5.14 | 1.07 | 1.97 | -0.48 | 2.02 | 4.59 | 3.13 | 1.84 | 1.23 | 3.62 | -1.1 | 1.76 | 1.53 | -1.58 | 3.33 | 3.2 | 0.97 | -0.5475 | -0.305 | 0.829405 | 0.410587 | 0.684244 | 0.989223 | 0.997988 | 0.999507 |
| Rbm15_Q0VBL3_S674s_1 | Rbm15_Q0 | -1.39 | -2.06 | -4.02 | -7.87 | -1.64 | -1.53 | -4.62 | -3.88 | -0.04 | -1.69 | -2.15 | -3.9 | 1.01 | -1.36 | -2.76 | -4.63 | 0.9175 | 1.89 | 1.9 | 0.25607 | 0.522041 | 0.213361 | 0.989223 | 0.997988 | 0.994666 |
| Rbm15_Q0VBL3_S782s_1 | Rbm15_Q0 | 0.2 | 0.96 | -0.26 | -0.27 | 0.29 | -0.69 | -0.76 | -0.49 | -0.45 | 0 | -2.79 | -1.09 | 1 | 0.43 | -0.6 | -0.05 | -0.57 | -1.24 | 0.0375 | 0.931813 | 0.148445 | 0.07015 | 0.989223 | 0.997988 | 0.999507 |
| Rbm15_Q0VBL3_T567t_1 | Rbm15_Q0 | 2.62 | 3.37 | 0.91 | 0.34 | 2.87 | 1.8 | 2.49 | 1.32 | -2.39 | 2.39 | 2.06 | 1.1 | -0.94 | 2.15 | 1.81 | 2.31 | 0.31 | -1.02 | -0.4775 | 0.603624 | 0.65815 | 0.384337 | 0.990434 | 0.997988 | 0.995259 |
| Rbm15b_Q6PHZ5_S107s_1 | Rbm15b_Q | 0.77 | -1.19 | -0.05 | -1 | -1.1 | 0.34 | -0.77 | -0.59 | 1.35 | -0.94 | -1.8 | -0.41 | 2.06 | -1.65 | 0.58 | -0.22 | -0.1625 | -0.0825 | 0.56 | 0.486231 | 0.746884 | 0.909201 | 0.995762 | 0.997988 | 0.994666 |
| Rbm15b_Q6PHZ5_S111s_1 | Rbm15b_Q | 0.19 | 1.36 | -0.35 | 0.08 | 1.02 | 0.49 | 1.04 | -0.4 | 0.42 | 0.66 | -0.35 | 0.85 | 0.08 | 1.64 | 0.26 | 0.36 | 0.2175 | 0.075 | 0.265 | 0.588637 | 0.639925 | 0.863682 | 0.989223 | 0.997988 | 0.995259 |
| Rbm15b_Q6PHZ5_S261s_1 | Rbm15b_Q | -0.76 | -0.97 | 2.23 | 0.72 | 0.05 | -1.69 | 2.26 | 2.66 | -1.13 | -1.45 | 2.06 | 2.7 | -0.2 | -0.5 | 1.66 | 0.89 | 0.515 | 0.24 | 0.1575 | 0.842284 | 0.636852 | 0.833927 | 0.989223 | 0.997988 | 0.999507 |
| Rbm15b_Q6PHZ5_S263s_1 | Rbm15b_Q | 2.47 | -0.58 | 1.16 | 0.2 | -1.24 | 0.3 | 1.12 | 1.21 | 0.27 | -0.04 | 1.02 | 0.96 | 0.41 | 1.14 | 1.09 | 0.5 | -0.465 | -0.26 | -0.0275 | 0.964673 | 0.545092 | 0.686978 | 0.989223 | 0.997988 | 0.999507 |
| Rbm15b_Q6PHZ5_S606s_1 | Rbm15b_Q | 7.45 | -1.01 | 0.21 | -1.92 | -0.6 | 7.7 | 6.09 | -0.44 | 0.9 | 3.78 | -0.14 | 0.7 | 2.59 | -1.4 | 1.34 | 2.12 | 2.005 | 0.1275 | -0.02 | 0.991918 | 0.446174 | 0.949093 | 0.989223 | 0.997988 | 1 |
| Rbm15b_Q6PHZ5_T234t_1 | Rbm15b_Q | 0.58 | 0.7 | 0.75 | 0.18 | 0.32 | 0.34 | 0.5 | 0.27 | -1.05 | 0.55 | -2.06 | 0.05 | -0.06 | -1.13 | 1.2 | 0.07 | -0.195 | -1.18 | -0.5325 | 0.277141 | 0.390812 | 0.05698 | 0.989223 | 0.997988 | 0.994666 |
| Rbm15b_Q6PHZ5_T529t_1 | Rbm15b_Q | 1.27 | 1.22 | 1.45 | 1.03 | 0.9 | 0.82 | 1.15 | 0.93 | -0.5 | 0.69 | 1.28 | 1.46 | -0.79 | 0.29 | 1.45 | 1.51 | -0.2925 | -0.51 | -0.6275 | 0.2447 | 0.188603 | 0.261359 | 0.989223 | 0.997988 | 0.994666 |
| Rbm17_Q8JZX4_S155s_1 | Rbm17_Q8 | -0.45 | -0.36 | -1.17 | -1.17 | -1.05 | -1.06 | -1.17 | -0.93 | -0.81 | -0.85 | -0.3 | -1.21 | -0.23 | -0.43 | -0.15 | -0.63 | -0.265 | -0.005 | 0.4275 | 0.18253 | 0.338181 | 0.987499 | 0.989223 | 0.997988 | 0.994666 |
| Rbm17_Q8JZX4_S222s_1 | Rbm17_Q8 | -1.1 | -0.83 | -1.07 | -0.86 | -0.93 | -1.56 | -0.38 | -0.65 | -1.19 | -1.04 | -1.25 | -1.03 | -0.46 | -0.14 | -1.02 | -0.99 | 0.085 | -0.1625 | 0.3125 | 0.300044 | 0.771014 | 0.45434 | 0.995762 | 0.997988 | 0.994666 |
| Rbm17_Q8JZX4_T71t_1 | Rbm17_Q8 | -2.63 | -1.82 | -2.91 | -2.16 | -2.18 | -3.07 | -2.08 | -2.1 | -2.93 | -3.04 | -1.58 | -1.63 | -1.19 | -0.88 | -1.58 | -2.49 | 0.0225 | 0.085 | 0.845 | 0.070063 | 0.947865 | 0.850027 | 0.999187 | 0.997988 | 0.994666 |
| Rbm20_Q3UQS8_S1184s_1 | Rbm20_Q3 | -2.66 | -2.66 | -3.63 | -4.58 | -3.26 | -2.85 | -2.92 | -3.61 | -1.96 | -2.23 | -2.31 | -2.97 | -0.2 | -6.16 | -2.78 | -3.24 | 0.2225 | 1.015 | 0.2875 | 0.799286 | 0.6292 | 0.059418 | 0.989223 | 0.997988 | 0.999507 |
| Rbm25_B2RY56_S578s_1 | Rbm25_B2 | -0.01 | -0.28 | -3.23 | -4.18 | -1.89 | -0.15 | -3.39 | -5.31 | -0.4 | -0.89 | -2.6 | -3.42 | -0.52 | -0.11 | -3.9 | -3.82 | -0.76 | 0.0975 | -0.1625 | 0.897707 | 0.562857 | 0.930104 | 0.989223 | 0.997988 | 0.999507 |
| Rbm25_B2RY56_S672s_1 | Rbm25_B2 | 1.28 | 0.03 | 1.32 | 1.17 | 0.52 | 0.37 | 1.01 | 1.08 | 0.02 | 0.7 | 1.35 | 0.51 | 0.41 | 1.01 | 1.1 | 0.86 | -0.205 | -0.305 | -0.105 | 0.774524 | 0.567311 | 0.462031 | 0.989223 | 0.997988 | 0.99798 |
| Rbm26_E9PUF4_S518s_1 | Rbm26_E9 | -2.74 | -3.53 | -2.76 | -3.01 | -2.99 | -3.69 | -3.1 | -3.25 | -3.1 | -3.08 | -1.67 | -2 | -1.58 | -2.55 | -3.3 | -3.58 | -0.2475 | 0.5475 | 0.2575 | 0.583432 | 0.382364 | 0.200919 | 0.989223 | 0.997988 | 0.995259 |
| Rbm26_E9PUF4_S616s_1 | Rbm26_E9 | -0.62 | -1.65 | -1.2 | -0.31 | -1.05 | -1.02 | -0.88 | -1.41 | -1.27 | -1.2 | -0.63 | -1.04 | -0.32 | -0.55 | -0.95 | -1.65 | -0.145 | -0.09 | 0.0775 | 0.85257 | 0.661643 | 0.796027 | 0.990434 | 0.997988 | 0.999507 |
| Rbm27_Q5SFM8_S120s_1 | Rbm27_Q5 | -2.9 | -3.65 | -2.78 | -3.67 | -2.52 | -4.42 | -3.92 | -4.65 | -2.12 | -2.1 | -1.94 | -3.61 | -1.75 | -4.38 | -1.5 | -2.53 | -0.6275 | 0.8075 | 0.71 | 0.275302 | 0.220831 | 0.094511 | 0.989223 | 0.997988 | 0.994666 |
| Rbm28_Q8CGC6_S252s_1 | Rbm28_Q8 | -1.64 | -1.68 | -0.44 | 0.1 | -0.44 | -0.4 | -0.43 | -0.07 | -0.23 | -1.64 | -0.52 | -0.7 | -0.6 | 0.36 | 0.12 | -0.82 | 0.58 | 0.1425 | 0.68 | 0.193952 | 0.196961 | 0.778549 | 0.989223 | 0.997988 | 0.994666 |
| Rbm33_D3Z5I9_S203s_1 | Rbm33_D3 | -1.54 | -1.01 | -0.37 | -0.19 | -2.16 | -1.23 | -1.38 | -1.45 | -0.15 | -1.44 | 3.54 | 0.25 | -1.01 | 0.89 | -0.54 | -1.11 | -0.7775 | 1.3275 | 0.335 | 0.524253 | 0.057845 | 0.196068 | 0.989223 | 0.997988 | 0.994666 |
| Rbm33_D3Z5I9_S992s_1 | Rbm33_D3 | -4.91 | -5.29 | -2.79 | -3.29 | -5.17 | -4.17 | -2.97 | -2.65 | -4.24 | -3.96 | -0.97 | -3.15 | -1.85 | -3.36 | -2.85 | -2.78 | 0.33 | 0.99 | 1.36 | 0.051357 | 0.655918 | 0.263407 | 0.990434 | 0.997988 | 0.994666 |
| Rbm39_B7ZD61_S135s_1 | Rbm39_B7 | 0.6 | 0.75 | 0.95 | 1.02 | 0.52 | 0.44 | 0.91 | 1.18 | -0.36 | 0.31 | 0.53 | 0.99 | -0.83 | 0 | 1.15 | 1.01 | -0.0675 | -0.4625 | -0.4975 | 0.293756 | 0.789424 | 0.176729 | 0.995762 | 0.997988 | 0.994666 |
| Rbm39_B7ZD61_S97s_1 | Rbm39_B7 | 0.47 | -0.13 | -0.67 | -1.38 | -0.14 | -0.06 | -1.61 | -1.2 | -0.1 | -0.66 | -1.08 | -0.53 | 1.02 | 0.53 | -0.94 | -1.28 | -0.325 | -0.165 | 0.26 | 0.67621 | 0.524133 | 0.702484 | 0.989223 | 0.997988 | 0.996159 |
| Rbm45_Q8BHN5_S199s_1 | Rbm45_Q8 | 0.8 | -0.99 | -0.29 | 0.08 | -0.94 | 0.48 | -0.72 | -1.11 | 0.03 | -0.63 | 0.38 | -0.69 | 0.86 | -0.51 | 0.73 | 0.14 | -0.4725 | -0.1275 | 0.405 | 0.397218 | 0.336345 | 0.772343 | 0.989223 | 0.997988 | 0.994666 |
| Rbm5_S4R1N4_S59s_1 | Rbm5_S4R | 1.21 | 0.67 | 0.85 | -2.64 | -0.5 | 1.16 | -0.87 | -0.11 | -2.71 | 0.44 | -1.3 | 0.46 | -0.28 | -4.36 | 1.52 | 0.7 | -0.1025 | -0.8 | -0.6275 | 0.646497 | 0.905918 | 0.448247 | 0.996513 | 0.997988 | 0.995259 |
| Rbm8a_Q9CWZ3_S42s_1 | Rbm8a_Q9 | 0.46 | -0.44 | 0.09 | -0.28 | -1.43 | -0.26 | -0.47 | 0.04 | 0.51 | -1.27 | 0.36 | -0.09 | 0.82 | -0.52 | 0.16 | -0.39 | -0.4875 | -0.08 | 0.06 | 0.874666 | 0.208913 | 0.854645 | 0.989223 | 0.997988 | 0.999507 |
| Rbm8a_Q9CWZ3_S42s_1_1 | Rbm8a_Q9 | 0.86 | -0.31 | 0.18 | -0.18 | -1.55 | 0.11 | -1.04 | 0.09 | -0.24 | -0.59 | 0.22 | -0.11 | 0.34 | -0.84 | 0.23 | -0.54 | -0.735 | -0.3175 | -0.34 | 0.401853 | 0.133358 | 0.353654 | 0.989223 | 0.997988 | 0.994666 |
| Rbm8a_Q9CWZ3_S56s_1 | Rbm8a_Q9 | 0.08 | 0.11 | -0.06 | 0.12 | -0.17 | -0.41 | -0.1 | -0.77 | -0.35 | -0.18 | -0.06 | -0.3 | -0.26 | -0.49 | -0.9 | 0.7 | -0.425 | -0.285 | -0.3 | 0.418962 | 0.091251 | 0.197588 | 0.989223 | 0.997988 | 0.994666 |
| Rbmxl1_Q91VM5_S205s_1 | Rbmxl1_Q | 0.66 | 0.11 | 0.21 | -0.06 | 0.1 | 0.08 | 0.06 | 0.3 | -0.21 | -0.28 | -0.39 | -0.04 | 0.01 | 0.44 | 0.52 | -0.42 | -0.095 | -0.46 | -0.0925 | 0.770064 | 0.686401 | 0.08929 | 0.994672 | 0.997988 | 0.9971 |
| Rbmxl1_Q91VM5_S246s_1 | Rbmxl1_Q | 3.32 | 2.37 | 3.34 | 2.61 | 1.27 | 2.47 | 1.95 | 3.32 | 0.63 | 2.25 | 0.89 | 3.13 | 0.92 | -0.23 | 4.42 | 3.15 | -0.6575 | -1.185 | -0.845 | 0.381321 | 0.178212 | 0.068502 | 0.989223 | 0.997988 | 0.994666 |
| Rbmxl1_Q91VM5_S323s_1 | Rbmxl1_Q | 1.49 | 0.5 | 1.53 | 1.72 | -0.78 | 1.78 | 0.27 | 1.63 | 1.01 | 0.53 | 1.83 | 1.28 | 0.83 | -0.8 | 1.92 | 1.3 | -0.585 | -0.1475 | -0.4975 | 0.407525 | 0.338491 | 0.7045 | 0.989223 | 0.997988 | 0.994666 |
| Rbmxl1_Q91VM5_S349s_1 | Rbmxl1_Q | 0.56 | 0.15 | -0.08 | -0.14 | -0.21 | -0.37 | -0.26 | -0.07 | -0.04 | -0.14 | 0.19 | -0.23 | 0.34 | 0.06 | 0.59 | -0.31 | -0.35 | -0.1775 | 0.0475 | 0.876823 | 0.164548 | 0.490482 | 0.989223 | 0.997988 | 0.999507 |
| Rbmxl1_Q91VM5_S88s_1 | Rbmxl1_Q | 0.81 | -0.39 | 0.72 | -0.38 | 0.61 | 0.15 | -0.32 | 0.08 | -0.37 | 0.05 | 1.22 | -0.06 | 0.93 | 0.65 | 0.91 | -0.63 | -0.06 | 0.02 | 0.275 | 0.567301 | 0.873073 | 0.965345 | 0.996493 | 0.997988 | 0.995259 |
| Rbmxl1_Q91VM5_T163t_1 | Rbmxl1_Q | 0.98 | 0.21 | 1.1 | 0.79 | -0.62 | 0.3 | 0.18 | 0.7 | 0.2 | -0.01 | -1.7 | 0.51 | 1.65 | 0.15 | 0.56 | 0.35 | -0.63 | -1.02 | -0.0925 | 0.815647 | 0.092764 | 0.067168 | 0.989223 | 0.997988 | 0.999507 |
| Rbsn_Q80Y56_S214s_1 | Rbsn_Q80 | -1.23 | -0.55 | 0.11 | -0.26 | -1.41 | -0.55 | -1.42 | 0.07 | -0.49 | -0.65 | -0.5 | -0.7 | 0.38 | -0.52 | -0.55 | 0.52 | -0.345 | -0.1025 | 0.44 | 0.294244 | 0.433787 | 0.747923 | 0.989223 | 0.997988 | 0.994666 |
| Rbsn_Q80Y56_S214s_2 | Rbsn_Q80 | -0.1 | 0.3 | 0.31 | -0.06 | -0.2 | -0.42 | -0.4 | 0.76 | -0.61 | -0.21 | 0.42 | 0.69 | -2.6 | -0.03 | -0.06 | -0.11 | -0.1775 | -0.04 | -0.8125 | 0.18974 | 0.578627 | 0.905451 | 0.989223 | 0.997988 | 0.994666 |
| Rbsn_Q80Y56_S216s_2 | Rbsn_Q80 | -0.1 | 0.34 | -1.33 | -1.21 | -0.18 | -0.68 | -1.57 | -0.72 | -0.53 | -0.3 | -1.17 | -1.18 | -3.16 | -0.27 | -1.58 | -2.04 | -0.2125 | -0.22 | -1.1875 | 0.095271 | 0.650339 | 0.627914 | 0.990358 | 0.997988 | 0.994666 |
| Rbsn_Q80Y56_S218s_1 | Rbsn_Q80 | 0.19 | 1.39 | -0.32 | -0.4 | -0.19 | -0.01 | 0.32 | 0.67 | 0.89 | 0.29 | 0.22 | 0.01 | 1.2 | 0.83 | 0.41 | 0.19 | -0.0175 | 0.1375 | 0.4425 | 0.343941 | 0.967434 | 0.754814 | 0.999187 | 0.997988 | 0.994666 |
| Rcan1_Q9JHG6_S32s_1 | Rcan1_Q9J | 0.58 | 0.89 | 0.58 | 1.02 | 0.82 | 0.12 | 1.12 | 0.87 | 0.14 | 0.53 | 0.31 | 0.67 | -0.02 | 0.94 | 0.6 | 0.44 | -0.035 | -0.355 | -0.2775 | 0.357344 | 0.899945 | 0.168411 | 0.996493 | 0.997988 | 0.994666 |
| Rcc1_Q6PFB2_S11s_1 | Rcc1_Q6PF | 0.74 | -0.24 | 0.21 | 0.26 | -0.51 | -0.11 | -0.32 | -0.05 | 0.05 | -0.12 | 0.05 | -0.12 | -0.05 | -0.43 | -0.01 | 0.01 | -0.49 | -0.2775 | -0.3625 | 0.234777 | 0.093928 | 0.313291 | 0.989223 | 0.997988 | 0.994666 |
| Rcor1_Q8CFE3_S454s_1 | Rcor1_Q8C | -1.15 | -1.43 | -1.49 | -1.22 | -1.6 | -1.4 | -1.48 | -0.96 | -0.6 | -0.53 | -1.27 | -1.42 | 0.02 | -1.08 | -1.74 | -1.44 | -0.0375 | 0.3675 | 0.2625 | 0.516108 | 0.872356 | 0.222821 | 0.996493 | 0.997988 | 0.994666 |
| Rcsd1_Q3UZA1_S149s_1 | Rcsd1_Q3U | -1.67 | -1.59 | -1.98 | -1.82 | -2.52 | -1.44 | -2.46 | -2.17 | -1.22 | -1.63 | -1.16 | -2.28 | 0.16 | -2.54 | -2.2 | -2.51 | -0.3825 | 0.1925 | -0.0075 | 0.989977 | 0.210259 | 0.535541 | 0.989223 | 0.997988 | 1 |
| Rcsd1_Q3UZA1_S303s_1 | Rcsd1_Q3U | 0.17 | 0.37 | -0.11 | -0.2 | 0.43 | 0.11 | -0.3 | -0.56 | 0.54 | 0.64 | -0.44 | -0.08 | 0.12 | 0.56 | -0.67 | -0.63 | -0.1375 | 0.1075 | -0.2125 | 0.552593 | 0.633588 | 0.736107 | 0.989223 | 0.997988 | 0.994666 |
| Rcsd1_Q3UZA1_S90s_1 | Rcsd1_Q3U | -1.37 | -0.8 | -0.68 | -1.3 | -0.58 | -1.1 | -1.48 | -0.24 | -0.76 | -0.19 | 0.5 | -0.69 | -0.52 | 0.09 | -0.91 | -1.21 | 0.1875 | 0.7525 | 0.4 | 0.278338 | 0.577419 | 0.057152 | 0.989223 | 0.997988 | 0.994666 |
| Reep1_Q8BGH4_S152s_1 | Reep1_Q8 | 0.04 | -0.19 | -0.36 | -0.43 | 0.31 | -0.03 | -0.09 | -0.06 | -0.91 | -0.36 | -0.53 | -0.6 | -0.33 | 0.17 | -0.65 | -0.57 | 0.2675 | -0.365 | -0.11 | 0.700972 | 0.249131 | 0.153353 | 0.989223 | 0.997988 | 0.996159 |
| Reep1_Q8BGH4_S192s_1 | Reep1_Q8 | 0.06 | -0.04 | -0.59 | -1.57 | -0.69 | -0.31 | -0.77 | -0.17 | 1.24 | 0.59 | 0.44 | -1.08 | 2.34 | -0.76 | 0.7 | -0.46 | 0.05 | 0.8325 | 0.99 | 0.185909 | 0.897559 | 0.167603 | 0.996493 | 0.997988 | 0.994666 |
| Reep2_Q8VCD6_S201s_1 | Reep2_Q8 | 1.82 | 0.29 | -0.74 | -1.52 | -1.03 | 3.16 | -1.56 | -0.78 | 1.44 | 0.7 | -1.98 | -1.1 | 0.81 | -0.63 | -1.05 | -0.47 | -0.015 | -0.1975 | -0.2975 | 0.686633 | 0.989315 | 0.834685 | 0.999187 | 0.997988 | 0.996159 |
| Rell2_E9Q812_S104s_1 | Rell2_E9Q | 0.62 | 1.13 | -0.16 | 0.99 | 0.27 | 0.4 | 0.39 | 0.87 | 0.93 | 0.44 | 0.45 | 0 | 0.76 | 0.2 | 0.71 | 0.59 | -0.1625 | -0.19 | -0.08 | 0.817833 | 0.623326 | 0.598546 | 0.989223 | 0.997988 | 0.999507 |
| Rell2_E9Q812_S96s_1 | Rell2_E9Q | 0.47 | 0.99 | 0.42 | 0.73 | 0.65 | 0.69 | 0.76 | 0.1 | 1.32 | 0.89 | -0.63 | 0.15 | 0.72 | 0.82 | 1.24 | 0.97 | -0.1025 | -0.22 | 0.285 | 0.299196 | 0.687911 | 0.615513 | 0.994672 | 0.997988 | 0.994666 |
| Rem2_E9Q4D5_S27s_1 | Rem2_E9Q | -0.8 | -1.08 | 0 | 0.2 | -1.03 | -0.49 | 0.57 | -0.07 | -1.78 | -0.46 | -0.76 | -0.07 | -1.53 | -0.71 | -0.15 | 0.21 | 0.165 | -0.3475 | -0.125 | 0.790317 | 0.703083 | 0.455773 | 0.995437 | 0.997988 | 0.999507 |
| Rem2_E9Q4D5_S296s_1 | Rem2_E9Q | 1.52 | 1.69 | 3.54 | 1.92 | 1.45 | 1.22 | 3.12 | 3.39 | -0.85 | 2.01 | 0.47 | 3.35 | 0.41 | -2.39 | 4.64 | 4.17 | 0.1275 | -0.9225 | -0.46 | 0.756624 | 0.843389 | 0.323921 | 0.996349 | 0.997988 | 0.9971 |
| Rem2_E9Q4D5_S69s_1 | Rem2_E9Q | -1.59 | -2.07 | -1.7 | -2.01 | -1.71 | -1.94 | -1.29 | -1.57 | -2.17 | -1.75 | -0.47 | -1.74 | -1.08 | -1.45 | -1.69 | -1.64 | 0.215 | 0.31 | 0.3775 | 0.183923 | 0.385405 | 0.432346 | 0.989223 | 0.997988 | 0.994666 |
| Reps1_O54916_S509s_1 | Reps1_O54 | 1.5 | 0.85 | 0.65 | 0.82 | 1.1 | 1.7 | 0.15 | 0.35 | 1.34 | 0.92 | 1.08 | 0.27 | 0.84 | 0.98 | 0.18 | 0.09 | -0.13 | -0.0525 | -0.4325 | 0.214298 | 0.739299 | 0.87069 | 0.995762 | 0.997988 | 0.994666 |
| Reps1_O54916_S93s_1 | Reps1_O54 | 1.02 | 0.71 | 1.14 | -0.03 | 0.14 | 0.97 | -0.01 | 0.05 | 0.61 | 0.4 | 1.26 | -0.13 | 0.73 | 0.21 | 0.24 | 0.4 | -0.4225 | -0.175 | -0.315 | 0.351502 | 0.246797 | 0.655461 | 0.989223 | 0.997988 | 0.994666 |
| Reps2_B9EI38_S481s_1 | Reps2_B9E | 1.81 | 2.09 | -0.03 | 0.15 | 1.97 | 1.49 | -0.05 | -0.11 | 1.52 | 1.33 | 0.5 | -0.64 | 1.21 | 2.25 | -0.17 | -0.39 | -0.18 | -0.3275 | -0.28 | 0.704887 | 0.790302 | 0.626134 | 0.995762 | 0.997988 | 0.996159 |
| Reps2_B9EI38_S538s_1 | Reps2_B9E | -0.34 | -0.71 | 0.04 | 0.22 | -0.51 | 0.09 | -0.19 | -0.51 | 1.1 | -0.56 | -2 | -0.65 | 1.41 | -0.72 | -0.17 | -0.39 | -0.0825 | -0.33 | 0.23 | 0.640899 | 0.77272 | 0.591588 | 0.995762 | 0.997988 | 0.995259 |
| Reps2_B9EI38_S619s_1 | Reps2_B9E | 0.09 | -0.35 | 0.81 | 0.2 | -0.22 | -0.41 | 0.62 | 0.66 | -0.16 | -0.05 | 0.88 | 1.09 | -0.39 | 0.78 | 0.63 | 0.76 | -0.025 | 0.2525 | 0.2575 | 0.505932 | 0.945172 | 0.529134 | 0.999187 | 0.997988 | 0.994666 |
| Reps2_B9EI38_T467t_1 | Reps2_B9E | 0.57 | 0.61 | -0.4 | -0.07 | 0.12 | 0.26 | -0.39 | -0.34 | 0.28 | 0.3 | -0.52 | -0.48 | 0.21 | 0.02 | -0.54 | -1.31 | -0.265 | -0.2825 | -0.5825 | 0.187571 | 0.407877 | 0.430162 | 0.989223 | 0.997988 | 0.994666 |
| Rer1_Q9CQU3_S2s_1 | Rer1_Q9C | -2.16 | -0.14 | -1.29 | -1.21 | -0.69 | -0.64 | -2.76 | -1.98 | -0.59 | -0.66 | -1.05 | -1.43 | 0.47 | -0.5 | -1.18 | -1.19 | -0.3175 | 0.2675 | 0.6 | 0.275388 | 0.595132 | 0.548854 | 0.989223 | 0.997988 | 0.994666 |
| Rftn2_Q8CHX7_S228s_1 | Rftn2_Q8C | 0.83 | 0.92 | 1.31 | 1.14 | 0.34 | 1.25 | -0.35 | 0.35 | 0.66 | 1.24 | 0.6 | 1.05 | 1.31 | -0.14 | 0.57 | 0.57 | -0.6525 | -0.1625 | -0.4725 | 0.194678 | 0.086413 | 0.531822 | 0.989223 | 0.997988 | 0.994666 |
| Rftn2_Q8CHX7_S429s_1 | Rftn2_Q8C | 0.72 | 0.36 | -0.27 | -0.19 | 0.83 | 0.21 | 0.06 | -0.37 | 0.18 | 0.22 | 0.38 | -0.57 | -0.04 | 0.7 | 0.18 | 0.19 | 0.0275 | -0.1025 | 0.1025 | 0.753593 | 0.936519 | 0.761598 | 0.998821 | 0.997988 | 0.9971 |
| Rftn2_Q8CHX7_T408t_1 | Rftn2_Q8C | 1.09 | 1.09 | 1.24 | 1.42 | 0.36 | 0.82 | 0.82 | 1.01 | 0.87 | 1.57 | 0.95 | 1.66 | 0.25 | 1.22 | 1.28 | 0.5 | -0.4575 | 0.0525 | -0.3975 | 0.230897 | 0.073439 | 0.848441 | 0.989223 | 0.997988 | 0.994666 |
| Rfx3_G3UWP0_S155s_1 | Rfx3_G3UV | -1.86 | -1.33 | -1.72 | -1.18 | -1.39 | -2.98 | -1.46 | -2.88 | -3.01 | -1.54 | -2.61 | -2.34 | -1.58 | -1.29 | -2.45 | -1.9 | -0.655 | -0.8525 | -0.2825 | 0.407093 | 0.157119 | 0.038888 | 0.989223 | 0.997988 | 0.994666 |
| Rgcc_Q9DBX1_S97s_1 | Rgcc_Q9DE | 0.26 | -0.2 | -0.06 | 0.55 | -0.73 | -0.43 | 0.11 | -0.1 | -1.59 | -0.57 | -1.34 | 0.01 | -1.82 | 1.24 | -0.23 | 0.05 | -0.425 | -1.01 | -0.3275 | 0.585383 | 0.157454 | 0.029854 | 0.989223 | 0.997988 | 0.995259 |

| | | | | | | | | | | | | | | | | | | | | | | | | | | |
|---|---|---|---|---|---|---|---|---|---|---|---|---|---|---|---|---|---|---|---|---|---|---|---|---|---|---|
| Rgl1_E9Q8N0_S518s_1 | Rgl1_E9Q8 | -1.19 | -1.68 | -1.38 | -0.99 | -1.63 | -1.58 | -0.83 | -1.11 | -1.45 | -2.32 | -0.59 | -1.52 | -1.65 | -0.6 | -2.03 | -0.7 | 0.0225 | -0.16 | 0.065 | 0.868259 | 0.935595 | 0.679193 | 0.99868 | 0.997988 | 0.999507 |
| Rgl3_Q3UYI5_S707s_1 | Rgl3_Q3UY | -0.44 | 0.24 | -1.24 | -0.35 | -0.05 | -1 | -0.25 | -0.39 | 1.27 | -0.28 | -0.85 | 0.07 | 0.75 | 0.05 | 0.48 | -0.66 | 0.025 | 0.5 | 0.6025 | 0.18172 | 0.945127 | 0.33846 | 0.999187 | 0.997988 | 0.994666 |
| Rgma_Q8C4C4_S26s_1 | Rgma_Q8C | -0.39 | 0.19 | 0.07 | 0.01 | 0.1 | -0.07 | -0.24 | -0.08 | 0.09 | -0.09 | 0.08 | 0.46 | 0.27 | 0.11 | 0.04 | -0.31 | -0.0425 | 0.165 | 0.0575 | 0.829597 | 0.850112 | 0.510868 | 0.996349 | 0.997988 | 0.999507 |
| Rgs12_E9Q652_S1125s_1 | Rgs12_E9Q | -3.53 | -3.18 | -2.46 | -4.47 | -2.61 | -2.09 | -2.44 | -2.99 | -0.54 | -2.73 | -1.87 | -1.52 | -0.64 | -3.16 | -2.64 | -2.53 | 0.8775 | 1.745 | 1.1675 | 0.088203 | 0.066407 | 0.01229 | 0.989223 | 0.997988 | 0.994666 |
| Rgs12_E9Q652_S1130s_1 | Rgs12_E9Q | -1.12 | -1.02 | -0.13 | -1.98 | -0.39 | -0.55 | -1.04 | -1.39 | 0.74 | 0.06 | -2.37 | -0.37 | 1.71 | -2.18 | 0.41 | 0.02 | 0.22 | 0.5775 | 1.0525 | 0.203555 | 0.602747 | 0.414031 | 0.989223 | 0.997988 | 0.994666 |
| Rgs12_E9Q652_S171s_1 | Rgs12_E9Q | -0.12 | -0.59 | 0.73 | 0.4 | 0.57 | 0.14 | 0.69 | 0.44 | -0.44 | 0.05 | 0.48 | -0.05 | -0.37 | 0.42 | 0.97 | 0.46 | 0.355 | -0.095 | 0.265 | 0.51797 | 0.291242 | 0.791269 | 0.989223 | 0.997988 | 0.994666 |
| Rgs12_E9Q652_S850s_1 | Rgs12_E9Q | 0.41 | 0.72 | 0.92 | 0.64 | 0.67 | 0.32 | 0.68 | 0.83 | 0.78 | 0.75 | 0.18 | 0.67 | 0.59 | 1.02 | 0.25 | 0.37 | -0.0475 | -0.0775 | -0.115 | 0.681329 | 0.835237 | 0.757218 | 0.995762 | 0.997988 | 0.996159 |
| Rgs12_E9Q652_S879s_1 | Rgs12_E9Q | -0.87 | -0.23 | -0.77 | -0.37 | -0.11 | -0.38 | -0.51 | -0.3 | -0.65 | -0.29 | -0.2 | -0.32 | -1.89 | 0.24 | 0.06 | -0.41 | 0.235 | 0.195 | 0.06 | 0.901206 | 0.340969 | 0.450294 | 0.989223 | 0.997988 | 0.999507 |
| Rgs14_P97492_S203s_1 | Rgs14_P97 | -1.2 | -0.37 | -0.54 | -0.71 | -0.39 | -1.28 | -0.81 | -0.88 | -0.68 | -0.85 | -0.16 | -0.83 | -0.05 | -0.31 | -0.31 | -0.75 | -0.135 | 0.075 | 0.35 | 0.253961 | 0.640102 | 0.794499 | 0.989223 | 0.997988 | 0.994666 |
| Rgs14_P97492_S218s_1 | Rgs14_P97 | -1.6 | -0.4 | -1.17 | -0.79 | -0.87 | -1.29 | -0.33 | -1.53 | -1.2 | -0.98 | -0.73 | -1.54 | -0.22 | -0.41 | -1.3 | -0.89 | -0.015 | -0.1225 | 0.285 | 0.451404 | 0.967131 | 0.713307 | 0.999187 | 0.997988 | 0.994666 |
| Rgs14_P97492_S289s_1 | Rgs14_P97 | -0.91 | -1.02 | 0.38 | -0.64 | -1.35 | -0.43 | -1.44 | -0.92 | -0.14 | -1.12 | 0.67 | -0.68 | -0.19 | -1.23 | -0.28 | -0.92 | -0.4875 | 0.23 | -0.1075 | 0.792919 | 0.224019 | 0.630834 | 0.989223 | 0.997988 | 0.999507 |
| Rgs19_Q8BFU4_S58s_1 | Rgs19_Q8E | 0.07 | -0.3 | 0.03 | -0.17 | -0.09 | -0.02 | 0.35 | 0.11 | -0.31 | 0.15 | 0.29 | 0.02 | -0.18 | 0.37 | 0.19 | -0.01 | 0.18 | 0.13 | 0.185 | 0.474101 | 0.419468 | 0.590902 | 0.989223 | 0.997988 | 0.994666 |
| Rgs20_E9Q720_S219s_1 | Rgs20_E9Q | 2.72 | 1.72 | 2.21 | 3.09 | 1 | 2.01 | 1.83 | 3.25 | 2 | 3.22 | 3.1 | 2.9 | 1.29 | 1.06 | 2.43 | 2.15 | -0.4125 | 0.37 | -0.7025 | 0.133195 | 0.422728 | 0.370606 | 0.989223 | 0.997988 | 0.994666 |
| Rgs20_Q9QZB1_S13s_1 | Rgs20_Q9C | -0.18 | 0.69 | 0.75 | 0.54 | -0.03 | -1.19 | 0.74 | 0.96 | 2.68 | 0.7 | 3.36 | 0.15 | 1.74 | 2.65 | 0.68 | 0.67 | -0.33 | 1.2725 | 0.985 | 0.071484 | 0.504406 | 0.103401 | 0.989223 | 0.997988 | 0.994666 |
| Rgs3_Q9DC04_S287s_1 | Rgs3_Q9DC | 1.14 | 0.52 | 0.18 | 0.91 | 0.66 | 0.59 | 1.4 | 1.32 | 0.88 | 0.94 | 0.97 | 0.88 | 0.1 | 0.81 | 1.21 | 1.16 | 0.305 | 0.23 | 0.1325 | 0.712217 | 0.348113 | 0.405646 | 0.989223 | 0.997988 | 0.996966 |
| Rgs3_Q9DC04_S288s_1 | Rgs3_Q9DC | 1.14 | 0.52 | -0.04 | 0.04 | 0.66 | 0.59 | 0.98 | 0.65 | 0.88 | 0.94 | 0.47 | 0.29 | 0.1 | 0.81 | 0.38 | 0.8 | 0.305 | 0.23 | 0.1075 | 0.759894 | 0.332836 | 0.498875 | 0.989223 | 0.997988 | 0.9971 |
| Rgs6_Q9Z2H2_S244s_1 | Rgs6_Q9Z2 | 1.1 | 0.33 | 0.67 | 1.01 | 0.86 | 0.63 | 1.3 | 1.02 | 0.7 | 0.89 | 0.53 | 1.13 | 0.36 | 1.01 | 0.45 | 0.56 | 0.175 | 0.035 | -0.1825 | 0.538393 | 0.519081 | 0.898188 | 0.989223 | 0.997988 | 0.994666 |
| Rgs7_Q80XD3_S229s_1 | Rgs7_Q80X | -1.43 | -1.82 | -1.48 | -1.66 | -0.85 | -1.88 | -0.7 | -1.46 | -2.21 | -1.61 | -1.07 | -1.47 | -1.72 | -0.76 | -1.71 | -1.61 | 0.375 | 0.0075 | 0.1475 | 0.63103 | 0.242576 | 0.979689 | 0.989223 | 0.997988 | 0.995259 |
| Rgs7_Q80XD3_S239s_1 | Rgs7_Q80X | 6.32 | 2.09 | 3.16 | 2.15 | 2.29 | 5.72 | 4.53 | 3.45 | 2.3 | 3.46 | 4.09 | 3.23 | 1.51 | 2.29 | 3.48 | 3.45 | 0.5675 | -0.16 | -0.7475 | 0.444863 | 0.597214 | 0.864688 | 0.989223 | 0.997988 | 0.994666 |
| Rgs7bp_Q8BQP9_S18s_1 | Rgs7bp_Q8 | 0.33 | -1.95 | -1.11 | -0.31 | -0.53 | 0.34 | 0.08 | 0.28 | -1.8 | -2.36 | 0.52 | -0.4 | -1.67 | -0.25 | -1.56 | -0.67 | 0.8025 | -0.25 | -0.2775 | 0.620526 | 0.12644 | 0.735165 | 0.989223 | 0.997988 | 0.995259 |
| Rgs8_D3Z5I4_S46s_1 | Rgs8_D3Z5 | 1.83 | 0.75 | 1.43 | 1.3 | -0.12 | 0.36 | 0.98 | 1.13 | 0.98 | 1.37 | 1.87 | 0.73 | 1.34 | 0.21 | 1.46 | 1.24 | -0.74 | -0.09 | -0.265 | 0.490623 | 0.065382 | 0.797102 | 0.989223 | 0.997988 | 0.994666 |
| Rgs8_D3Z5I4_T47t_1 | Rgs8_D3Z5 | 0.31 | -0.1 | 0.31 | 0.5 | -0.88 | -0.17 | -0.47 | 0 | 0.5 | -0.25 | 0.08 | -1.04 | 1.31 | -0.95 | -0.79 | -0.51 | -0.635 | -0.4325 | -0.49 | 0.346235 | 0.03985 | 0.249546 | 0.989223 | 0.997988 | 0.994666 |
| Rhbdd3_Q8BP97_S365s_1 | Rhbdd3_Q8 | -1.55 | -1.53 | -2.84 | -3.96 | -1.53 | -1.17 | -1.96 | -2.59 | 1.13 | -1.43 | -4.31 | -2.33 | 1.4 | -1.36 | -1.23 | -1.14 | 0.6575 | 0.735 | 1.8875 | 0.035028 | 0.281341 | 0.515601 | 0.989223 | 0.997988 | 0.994666 |
| Rhbdf1_F2Z3W3_S11s_1 | Rhbdf1_F2 | -1.79 | -2.5 | -0.15 | -0.91 | -1.29 | -0.98 | -0.9 | -0.4 | -0.16 | -1.41 | 0.12 | -0.02 | 0.2 | -1.15 | -0.68 | -0.29 | 0.445 | 0.97 | 0.8575 | 0.140277 | 0.382907 | 0.116767 | 0.989223 | 0.997988 | 0.994666 |
| Rhbdf1_F2Z3W3_S61s_1 | Rhbdf1_F2 | -0.6 | -0.05 | 0.24 | 0 | -0.01 | -0.42 | -0.06 | -0.28 | 0.8 | -0.38 | 0.08 | 0.03 | 1.63 | -0.46 | 0.68 | 0.76 | -0.09 | 0.235 | 0.755 | 0.119604 | 0.724612 | 0.479949 | 0.995437 | 0.997988 | 0.994666 |
| Rhbdf1_F2Z3W3_S8s_1 | Rhbdf1_F2 | 0.15 | -0.56 | -0.49 | -0.89 | 0.16 | -0.01 | -0.82 | -1.05 | 0.61 | -0.73 | -0.27 | -0.07 | 1.42 | -0.52 | 0 | -0.15 | 0.0175 | 0.3325 | 0.635 | 0.190686 | 0.961701 | 0.371636 | 0.999187 | 0.997988 | 0.994666 |
| Rhbdl3_P58873_S6s_1 | Rhbdl3_P5 | 1 | -0.45 | 0.22 | 0.2 | 0.35 | 0.56 | -0.54 | 0.52 | 0.07 | 0.04 | -1.31 | 0.21 | 4.21 | -1.52 | 0.34 | 0.29 | -0.02 | -0.49 | 0.5875 | 0.588613 | 0.958322 | 0.289059 | 0.999187 | 0.997988 | 0.995259 |
| Rhob_P62746_S123s_1 | Rhob_P627 | 5.19 | 4.56 | 4.87 | 3.84 | 3.76 | 4.93 | 3.18 | 4.24 | 1.79 | 4.47 | 4.05 | 3.99 | -0.62 | 1.25 | 4.52 | 4.25 | -0.5875 | -1.04 | -2.265 | 0.063691 | 0.203925 | 0.115742 | 0.989223 | 0.997988 | 0.994666 |
| Rhog_P84096_T125t_1 | Rhog_P84C | -0.2 | 0.21 | 1.33 | 1.03 | -0.31 | -0.35 | -1.65 | 0.28 | 1.38 | -0.37 | 0.11 | -0.02 | 1.56 | -1.11 | 0.12 | 0.54 | -1.1 | -0.3175 | -0.315 | 0.601781 | 0.048654 | 0.523675 | 0.989223 | 0.997988 | 0.995259 |
| Ric8a_Q3TIR3_S435s_1 | Ric8a_Q3TI | -0.37 | 0.37 | 0.49 | 0.39 | 0.16 | -0.58 | 0.68 | 0.36 | 0.21 | -0.15 | 0.71 | 0.52 | -0.02 | 0.15 | 0.57 | -0.14 | -0.065 | 0.1025 | -0.08 | 0.795041 | 0.848108 | 0.740219 | 0.996349 | 0.997988 | 0.999507 |
| Ric8a_Q3TIR3_S435s_2 | Ric8a_Q3TI | -0.64 | 0.3 | 0.25 | -0.62 | 0.41 | 0.27 | -0.25 | -0.58 | -0.26 | 0.83 | -0.67 | -0.09 | 0.32 | 1.52 | -0.46 | -0.2 | 0.14 | 0.13 | 0.4725 | 0.3441 | 0.690406 | 0.749527 | 0.994672 | 0.997988 | 0.994666 |
| Ric8a_Q3TIR3_T440t_2 | Ric8a_Q3TI | -0.64 | 0.3 | 0.25 | -0.62 | 0.41 | 0.27 | -0.25 | -0.58 | -0.26 | 0.83 | -0.67 | -0.09 | 0.32 | 1.52 | -0.46 | -0.2 | 0.14 | 0.13 | 0.4725 | 0.3441 | 0.690406 | 0.749527 | 0.994672 | 0.997988 | 0.994666 |
| Rictor_Q6QI06_S1234s_1 | Rictor_Q6C | 0.26 | 0.29 | 0.64 | -0.06 | 0.25 | -0.05 | 0.04 | -0.1 | 0.56 | 0.21 | -0.79 | 0.12 | 0.45 | -0.41 | 0.49 | 0.17 | -0.2475 | -0.2575 | -0.1075 | 0.727568 | 0.304739 | 0.457136 | 0.989223 | 0.997988 | 0.9971 |
| Rictor_Q6QI06_S1591s_1 | Rictor_Q6C | -2.76 | -2.84 | -3.24 | -3.55 | -1.24 | -3.01 | -1.62 | -2.31 | -2.93 | -2.94 | -2.42 | -1.87 | -2.87 | -2.44 | -2.38 | -1.12 | 1.0525 | 0.5575 | 0.895 | 0.055608 | 0.027723 | 0.122511 | 0.989223 | 0.997988 | 0.994666 |
| Rilpl1_Q9JJC6_S349s_1 | Rilpl1_Q9JJ | -1.43 | -1.13 | -0.52 | -0.27 | -0.21 | -0.44 | -0.52 | -0.06 | -0.29 | -0.17 | 0.58 | -0.24 | -0.18 | -0.43 | -0.17 | -0.14 | 0.53 | 0.8075 | 0.6075 | 0.084391 | 0.110821 | 0.043509 | 0.989223 | 0.997988 | 0.994666 |
| Rilpl1_Q9JJC6_S7s_1 | Rilpl1_Q9JJ | -0.78 | -1.63 | -0.06 | -0.38 | -0.02 | 0.1 | 0.15 | 0.11 | -0.05 | -0.14 | 0.54 | 0.32 | -0.04 | 0.7 | 0.27 | -0.16 | 0.7975 | 0.88 | 0.905 | 0.044093 | 0.041909 | 0.041919 | 0.989223 | 0.997988 | 0.994666 |
| Rimbp2_D3YXR8_S279s_1 | Rimbp2_D3 | -0.33 | -0.09 | 0.09 | 0.31 | 0.87 | 0.44 | 0.64 | 0.18 | -1.2 | -0.44 | 0.24 | 0.04 | -1.22 | 0.21 | 0.07 | 0.38 | 0.5375 | -0.335 | -0.135 | 0.734713 | 0.057435 | 0.363249 | 0.989223 | 0.997988 | 0.9971 |
| Rimbp2_D3YXR8_S978s_1 | Rimbp2_D3 | 1.11 | -0.5 | 0.18 | -7.77 | 0.1 | 1.72 | -0.45 | 0.44 | -1.66 | 0.61 | -3.94 | -0.07 | 1.24 | -1.31 | 0.55 | 0.96 | 2.1975 | 0.48 | 2.105 | 0.262037 | 0.236663 | 0.808237 | 0.989223 | 0.997988 | 0.994666 |
| Rims1_E9PWP6_S1077s_1 | Rims1_E9P | 0.14 | -0.7 | -0.85 | -0.18 | -0.89 | 0.66 | -0.44 | -0.27 | 0.04 | 0.44 | -0.48 | -0.96 | 0.21 | -1.36 | -0.15 | -0.09 | 0.1625 | 0.1575 | 0.05 | 0.904232 | 0.675819 | 0.683316 | 0.992909 | 0.997988 | 0.999507 |
| Rims1_E9PWP6_S1105s_1 | Rims1_E9P | 0.05 | 0.53 | -0.32 | -0.01 | 0.47 | 0.1 | 0.04 | -0.02 | 0.9 | 0.6 | 0.07 | 0.19 | 1.01 | 0.96 | 0.18 | 0.47 | 0.085 | 0.3775 | 0.5925 | 0.086013 | 0.742365 | 0.22732 | 0.995762 | 0.997988 | 0.994666 |
| Rims1_E9PWP6_S1166s_1 | Rims1_E9P | -1.98 | -1.71 | -1.19 | -1.34 | -2.52 | -1.24 | -1.96 | -1.19 | 0.21 | -2.71 | -1.67 | -1.59 | -0.15 | -4.05 | -0.67 | -1.64 | -0.1725 | 0.115 | -0.0725 | 0.926005 | 0.635479 | 0.843213 | 0.989223 | 0.997988 | 0.999507 |
| Rims1_E9PWP6_S1201s_2 | Rims1_E9P | 0.37 | 0.19 | 0.83 | 0.78 | 0.4 | 0.11 | 0.37 | 0.37 | 0.42 | 0.42 | 0.89 | 0.83 | 0.46 | 0.84 | 0.61 | 0.23 | -0.23 | 0.0975 | -0.0075 | 0.978628 | 0.345825 | 0.713211 | 0.989223 | 0.997988 | 1 |
| Rims1_E9PWP6_S1205s_1 | Rims1_E9P | 0.67 | 0.89 | 1.19 | 1.21 | 0.45 | 0.11 | 0.32 | 0.8 | 0.45 | 1.39 | 0.89 | 0.72 | 0.95 | 0.95 | 0.69 | 1.08 | -0.57 | -0.1275 | -0.0725 | 0.778207 | 0.044163 | 0.656045 | 0.989223 | 0.997988 | 0.99798 |
| Rims1_E9PWP6_S1205s_2 | Rims1_E9P | 0.37 | 0.19 | 0.76 | 0.74 | 0.4 | 0.11 | 0.29 | 0.27 | 0.42 | 0.42 | 0.82 | 0.78 | 0.46 | 0.84 | 0.5 | 0.18 | -0.2475 | 0.095 | -0.02 | 0.94232 | 0.295362 | 0.706895 | 0.989223 | 0.997988 | 0.999507 |
| Rims1_E9PWP6_S1229s_1 | Rims1_E9P | 0.78 | 1 | 0.99 | 0.98 | 0.82 | 1.09 | 1.11 | 0.97 | 0.67 | 0.95 | -0.01 | 0.76 | -0.09 | 1.07 | -0.07 | 0.62 | 0.06 | -0.345 | -0.555 | 0.116877 | 0.767353 | 0.22571 | 0.995762 | 0.997988 | 0.994666 |
| Rims1_E9PWP6_S1259s_2 | Rims1_E9P | 0.67 | -0.19 | 0.3 | 0.15 | -0.87 | 0.12 | -0.24 | 0.31 | -0.07 | -0.33 | 0.41 | -0.75 | -0.81 | 0.09 | -0.03 | -0.36 | -0.4025 | -0.4175 | -0.51 | 0.133272 | 0.237032 | 0.222182 | 0.989223 | 0.997988 | 0.994666 |
| Rims1_E9PWP6_S1260s_2 | Rims1_E9P | 0.67 | -0.19 | 0.3 | 0.15 | -0.87 | 0.12 | -0.24 | 0.31 | -0.07 | -0.33 | 0.41 | -0.75 | -0.81 | 0.09 | -0.03 | -0.36 | -0.4025 | -0.4175 | -0.51 | 0.133272 | 0.237032 | 0.222182 | 0.989223 | 0.997988 | 0.994666 |
| Rims1_E9PWP6_S1288s_1 | Rims1_E9P | 1.52 | 2.29 | 0.64 | 0.28 | 2.39 | 0.82 | 0.25 | 0.75 | 1.96 | 2.65 | 0.36 | 1.25 | 2.31 | 2.61 | 0.96 | 0.9 | -0.13 | 0.3725 | 0.5125 | 0.388833 | 0.822993 | 0.545278 | 0.995762 | 0.997988 | 0.994666 |
| Rims1_E9PWP6_S1397s_1 | Rims1_E9P | -1.99 | -1.99 | -5.53 | -3.45 | -1.28 | -0.97 | -3.23 | -3.52 | -1.69 | -1.56 | -1.68 | -2.74 | 0.01 | -1.61 | -1.42 | -1.83 | 0.99 | 1.3225 | 2.0275 | 0.032485 | 0.298258 | 0.119542 | 0.989223 | 0.997988 | 0.994666 |
| Rims1_E9PWP6_S1402s_1 | Rims1_E9P | -2.79 | -2.97 | -2.06 | -1.91 | -2.37 | -2.15 | -1.92 | -1.82 | -1.62 | -2.21 | -0.25 | -1.19 | -0.66 | -1.73 | -1.64 | -1.22 | 0.3675 | 1.115 | 1.12 | 0.013581 | 0.253335 | 0.037493 | 0.989223 | 0.997988 | 0.994666 |
| Rims1_E9PWP6_S1418s_1 | Rims1_E9P | -0.99 | -0.48 | -0.6 | -0.88 | -0.31 | -0.05 | -1.35 | -0.19 | -0.61 | -0.32 | 0.68 | -0.64 | -0.03 | 0.22 | -1.03 | -1.03 | 0.2625 | 0.515 | 0.27 | 0.471405 | 0.43399 | 0.163009 | 0.989223 | 0.997988 | 0.994666 |
| Rims1_E9PWP6_S1626s_1 | Rims1_E9P | 0.35 | 0.42 | 0.2 | 0.5 | 0.2 | 1.05 | 0.63 | 1.06 | 0.22 | 0.82 | 0.82 | 0.9 | 0.66 | 0.62 | 1.04 | 0.91 | 0.3675 | 0.3225 | 0.44 | 0.096587 | 0.182972 | 0.21365 | 0.989223 | 0.997988 | 0.994666 |
| Rims1_E9PWP6_S1626s_2 | Rims1_E9P | 0.27 | 0.9 | 1.22 | 1.83 | 0.32 | 0.59 | 0.86 | 1.82 | 0.51 | 0.2 | 0.58 | 1.59 | -0.09 | 0.74 | 0.58 | 0.48 | -0.1575 | -0.335 | -0.6275 | 0.127972 | 0.717477 | 0.44503 | 0.995437 | 0.997988 | 0.994666 |
| Rims1_E9PWP6_S1629s_1 | Rims1_E9P | 0.36 | -0.67 | -0.15 | -0.8 | -0.29 | -0.79 | -0.25 | 0.2 | -0.54 | -0.51 | 0.21 | -2.91 | -0.03 | -0.96 | 0.45 | -0.65 | 0.0325 | -0.6225 | 0.0175 | 0.966185 | 0.923748 | 0.358814 | 0.997335 | 0.997988 | 0.999507 |
| Rims1_E9PWP6_S1629s_2 | Rims1_E9P | 0.27 | 0.9 | 1.22 | 1.83 | 0.32 | 0.59 | 0.86 | 1.82 | 0.51 | 0.2 | 0.58 | 1.59 | -0.09 | 0.74 | 0.58 | 0.48 | -0.1575 | -0.335 | -0.6275 | 0.127972 | 0.717477 | 0.44503 | 0.995437 | 0.997988 | 0.994666 |
| Rims1_E9PWP6_S1641s_1 | Rims1_E9P | 0.93 | 1.02 | 0.38 | 0.64 | 0.8 | 0.63 | 0.46 | 0.11 | 0.91 | 1.16 | -0.97 | 0.34 | 0.4 | 0.79 | 0.55 | 0.38 | -0.2425 | -0.3825 | -0.2125 | 0.43266 | 0.358518 | 0.427379 | 0.989223 | 0.997988 | 0.994666 |
| Rims1_E9PWP6_S211s_1 | Rims1_E9P | 0.92 | 0.76 | 0.23 | 0.92 | 0.65 | 0.65 | 1.02 | 0.65 | 0.19 | 0.75 | 0.26 | 0.66 | -0.19 | 0.66 | 0.43 | 0.6 | 0.035 | -0.2425 | -0.3325 | 0.296982 | 0.887326 | 0.383971 | 0.996493 | 0.997988 | 0.994666 |
| Rims1_E9PWP6_S241s_1 | Rims1_E9P | -1.89 | -1.51 | -2.1 | -1.36 | -0.89 | -1.22 | -0.9 | -1.24 | -1.58 | -1.09 | -0.67 | -1.94 | -1.27 | -1.03 | -2.37 | -2 | 0.6525 | 0.395 | 0.0475 | 0.89875 | 0.025775 | 0.268354 | 0.989223 | 0.997988 | 0.999507 |
| Rims1_E9PWP6_S252s_1 | Rims1_E9P | 0.06 | 0.41 | 0.87 | 0.78 | 0.02 | 0.22 | 0.17 | 0.58 | 0.15 | 0.1 | 0.9 | 0.44 | 0.43 | 0.2 | 0.71 | 0.65 | -0.2825 | -0.1325 | -0.0325 | 0.910426 | 0.302204 | 0.660628 | 0.989223 | 0.997988 | 0.999507 |
| Rims1_E9PWP6_S346s_1 | Rims1_E9P | 0.51 | -0.13 | -0.37 | -1.09 | -0.32 | 0.93 | -1.2 | -1.23 | -0.15 | 0.43 | -1.9 | -0.97 | 0.54 | -1.14 | -0.58 | -0.55 | -0.185 | -0.3775 | -0.1625 | 0.727402 | 0.736069 | 0.505124 | 0.995762 | 0.997988 | 0.9971 |
| Rims1_E9PWP6_S443s_1 | Rims1_E9P | 1.47 | 0.74 | 1.24 | 1.24 | 0.94 | 0.68 | 0.79 | 1.12 | 1.43 | 2.13 | 1.77 | 1.18 | 1.01 | 0.66 | 0.96 | 0.73 | -0.29 | 0.455 | -0.3325 | 0.232904 | 0.251641 | 0.151827 | 0.989223 | 0.997988 | 0.994666 |
| Rims1_E9PWP6_S443s_2 | Rims1_E9P | 0.79 | 0.68 | 1.08 | 0.8 | 0.76 | 0.67 | 0.39 | 0.67 | 0.31 | 0.38 | 0.81 | 0.8 | -0.87 | 0.94 | 0.49 | 0.37 | -0.215 | -0.2625 | -0.605 | 0.156399 | 0.327555 | 0.293291 | 0.989223 | 0.997988 | 0.994666 |
| Rims1_E9PWP6_S447s_2 | Rims1_E9P | 0.79 | 0.68 | 1.08 | 0.8 | 0.76 | 0.67 | 0.39 | 0.67 | 0.31 | 0.38 | 0.81 | 0.8 | -0.87 | 0.94 | 0.49 | 0.37 | -0.215 | -0.2625 | -0.605 | 0.156399 | 0.327555 | 0.293291 | 0.989223 | 0.997988 | 0.994666 |
| Rims1_E9PWP6_S514s_1 | Rims1_E9P | 0.19 | 1.74 | 0.93 | 0.09 | 1.31 | 0.79 | 0.09 | 0.51 | -2.79 | -0.53 | -0.12 | 0.08 | -1.39 | -0.89 | 0.23 | 0.8 | -0.0625 | -1.5775 | -1.05 | 0.095696 | 0.885427 | 0.044541 | 0.996493 | 0.997988 | 0.994666 |
| Rims1_E9PWP6_S726s_1 | Rims1_E9P | -0.53 | 0.33 | 0.55 | 0.1 | -0.38 | -0.16 | 0.9 | 0.3 | -0.29 | -0.08 | 0.7 | 0.49 | 0 | 0.55 | 0.85 | 0.78 | 0.0525 | 0.0925 | 0.4325 | 0.221326 | 0.885117 | 0.789824 | 0.996493 | 0.997988 | 0.994666 |
| Rims1_E9PWP6_S741s_1 | Rims1_E9P | 0.49 | 0.76 | 1.01 | 1.07 | 0.67 | 0.48 | 1.28 | 0.88 | 0.64 | 1.03 | 1.29 | 1.18 | 0.45 | 0.96 | 0.99 | 0.89 | -0.005 | 0.2025 | -0.01 | 0.970488 | 0.984842 | 0.444212 | 0.999187 | 0.997988 | 0.999507 |
| Rims1_E9PWP6_S744s_1 | Rims1_E9P | 1.26 | 1.16 | 0.49 | 0.5 | 1.6 | 1.18 | 0.78 | 0.55 | 0.71 | 1.17 | 0.15 | 0.32 | 0.45 | 1.56 | -0.01 | 0.32 | 0.175 | -0.265 | -0.2725 | 0.504959 | 0.591438 | 0.43183 | 0.989223 | 0.997988 | 0.994666 |
| Rims1_E9PWP6_S894s_1 | Rims1_E9P | -0.33 | -0.12 | -0.3 | 0.09 | 0.59 | -0.58 | 0.51 | 0.21 | -0.51 | -0.33 | -0.6 | -0.03 | -0.23 | 0.65 | -0.05 | 0.01 | 0.3475 | -0.2025 | 0.26 | 0.375544 | 0.272308 | 0.411515 | 0.989223 | 0.997988 | 0.994666 |
| Rims1_E9PWP6_S966s_2 | Rims1_E9P | 1.21 | 1.39 | 0.33 | 0.58 | 0.38 | 2.1 | 0.37 | 1.2 | 1.29 | 1.02 | 0.55 | 0.48 | 0.72 | 0.99 | 1.46 | 1.75 | 0.135 | -0.0425 | 0.3525 | 0.345631 | 0.764849 | 0.899933 | 0.995762 | 0.997988 | 0.994666 |
| Rims1_E9PWP6_S968s_2 | Rims1_E9P | 1.21 | 1.39 | 0.33 | 0.58 | 0.38 | 2.1 | 0.37 | 1.2 | 1.29 | 1.02 | 0.55 | 0.48 | 0.72 | 0.99 | 1.46 | 1.75 | 0.135 | -0.0425 | 0.3525 | 0.345631 | 0.764849 | 0.899933 | 0.995762 | 0.997988 | 0.994666 |
| Rims1_E9PWP6_S990s_1 | Rims1_E9P | -0.16 | -0.44 | -1.39 | -0.57 | -0.38 | -0.53 | -1.09 | -0.43 | 0 | -0.24 | -1.39 | -1.26 | 0.18 | -0.18 | -1.07 | -1.28 | 0.0325 | -0.0825 | 0.0525 | 0.903265 | 0.920016 | 0.847483 | 0.997335 | 0.997988 | 0.999507 |
| Rims1_E9PWP6_T1203t_1 | Rims1_E9P | 0.4 | 0.15 | 0.49 | -0.01 | 0.36 | -0.03 | -0.17 | 0.26 | 0.57 | 0.42 | 0.75 | 0.49 | 0.49 | 0.59 | 0.48 | -0.31 | -0.1525 | 0.3 | 0.055 | 0.854459 | 0.524255 | 0.214766 | 0.989223 | 0.997988 | 0.999507 |
| Rims1_E9PWP6_T326t_1 | Rims1_E9P | 0.22 | 1.63 | 0.81 | 0.24 | 1.49 | -1.51 | -0.37 | 0.18 | -0.19 | 1.23 | 1.28 | 0.4 | 0.54 | 1.82 | 0.97 | 0.38 | -0.7775 | -0.045 | 0.2025 | 0.654265 | 0.234893 | 0.922515 | 0.989223 | 0.997988 | 0.995259 |
| Rims1_E9PWP6_T743t_1 | Rims1_E9P | 0.47 | 0.38 | -0.28 | -0.18 | 0.91 | 0.72 | 0.16 | -0.77 | 0.67 | 0.31 | -1.06 | 0.08 | 0.31 | 1.23 | -0.65 | -0.71 | 0.1575 | -0.0975 | -0.0525 | 0.912124 | 0.698331 | 0.813516 | 0.994893 | 0.997988 | 0.999507 |
| Rims2_Q0VF51_S1075s_1 | Rims2_Q0\ | -0.87 | -0.81 | -0.75 | -0.83 | -0.44 | -0.94 | -0.12 | -1.26 | -1.28 | -1.09 | -0.8 | -0.94 | -0.72 | -0.78 | -1 | -0.56 | 0.125 | -0.2125 | 0.05 | 0.832309 | 0.664694 | 0.34548 | 0.991343 | 0.997988 | 0.999507 |
| Rims2_Q0VF51_S1099s_1 | Rims2_Q0\ | 0.62 | 0.26 | 0.17 | 0.78 | 1.01 | 0.77 | 0.58 | 0.91 | 0.54 | 0.38 | 0.71 | 0.29 | 0.7 | 1.01 | 1.08 | 0.79 | 0.36 | 0.0225 | 0.4375 | 0.123489 | 0.155568 | 0.927767 | 0.989223 | 0.997988 | 0.994666 |
| Rims2_Q0VF51_S1115s_2 | Rims2_Q0\ | -0.11 | 0.24 | 0.34 | 0.03 | 0.24 | 0.03 | 0.14 | 0.84 | -0.36 | 0.21 | 0.9 | 0.74 | -0.51 | 0.17 | 1.36 | 0.71 | 0.1875 | 0.2475 | 0.3075 | 0.461961 | 0.473815 | 0.457683 | 0.989223 | 0.997988 | 0.994666 |
| Rims2_Q0VF51_S1163s_1 | Rims2_Q0\ | -0.66 | -1.32 | -1.05 | -3.16 | -1.74 | -1.79 | -1.84 | -2.32 | 0.44 | -1.42 | -1.02 | -1.17 | 0.66 | -1.4 | -1.35 | -1.8 | -0.375 | 0.755 | 0.575 | 0.420324 | 0.476549 | 0.251584 | 0.989223 | 0.997988 | 0.994666 |
| Rims2_Q0VF51_S1535s_1 | Rims2_Q0\ | -0.07 | 0.36 | 0.83 | 0.72 | 0.69 | 0.73 | 0.82 | 1 | 0.05 | 1.27 | 1.38 | 1.5 | 1.34 | 1.06 | 1.37 | 1.13 | 0.35 | 0.59 | 0.765 | 0.023322 | 0.202939 | 0.158026 | 0.989223 | 0.997988 | 0.994666 |
| Rims2_Q0VF51_S1535s_2 | Rims2_Q0\ | 1.08 | 0.22 | 0.2 | 0.47 | 1.07 | 0.53 | 0.06 | 0.46 | 0.66 | 0.53 | 0.8 | 0.81 | 1.57 | 1.08 | 0.21 | 0.49 | 0.0375 | 0.2075 | 0.345 | 0.37632 | 0.903963 | 0.453594 | 0.996493 | 0.997988 | 0.994666 |
| Rims2_Q0VF51_S1538s_1 | Rims2_Q0\ | 0.59 | 0.63 | 0.81 | 0.17 | 0.33 | 0.21 | 0.27 | 0.55 | 0.84 | 0.86 | -0.27 | 0.96 | 0.88 | 0.12 | 0.17 | -0.03 | -0.21 | 0.0475 | -0.265 | 0.391659 | 0.371917 | 0.888889 | 0.989223 | 0.997988 | 0.994666 |
| Rims2_Q0VF51_S1538s_2 | Rims2_Q0\ | 1.08 | 0.22 | 0.2 | 0.47 | 1.07 | 0.53 | 0.06 | 0.46 | 0.66 | 0.53 | 0.8 | 0.81 | 1.57 | 1.08 | 0.21 | 0.49 | 0.0375 | 0.2075 | 0.345 | 0.37632 | 0.903963 | 0.453594 | 0.996493 | 0.997988 | 0.994666 |
| Rims2_Q0VF51_S236s_1 | Rims2_Q0\ | 0.14 | -0.5 | -1.43 | -0.41 | -1.07 | 0.16 | -0.94 | -1.29 | 2.49 | 0.2 | -3.46 | -1.36 | 2.38 | -0.88 | -0.3 | -1.73 | -0.235 | 0.0175 | 0.4175 | 0.618178 | 0.589704 | 0.987738 | 0.989223 | 0.997988 | 0.995259 |
| Rims2_Q0VF51_S250s_1 | Rims2_Q0\ | -1.75 | -1.75 | -1.72 | -5.68 | -2.67 | -1.04 | -2.79 | -2.77 | -0.21 | -2.52 | -0.82 | -3.23 | -0.16 | -2.47 | -2.13 | -2.61 | 0.4075 | 1.03 | 0.8825 | 0.383518 | 0.662999 | 0.347493 | 0.990434 | 0.997988 | 0.994666 |
| Rims2_Q0VF51_S269s_1 | Rims2_Q0\ | -0.29 | -0.25 | -0.31 | -0.07 | -0.91 | 0.13 | -0.62 | -0.31 | 0.6 | -0.24 | -0.18 | 0.17 | 0.84 | -1.04 | -0.1 | 0.04 | -0.1975 | 0.3175 | 0.165 | 0.678881 | 0.472286 | 0.24788 | 0.989223 | 0.997988 | 0.996159 |
| Rims2_Q0VF51_S271s_1 | Rims2_Q0\ | -0.7 | -1.18 | -0.79 | -0.18 | -1.66 | -0.12 | -0.53 | -1.45 | -1.02 | -1.53 | -0.12 | -0.95 | -1.21 | -1.41 | -2.39 | -1.18 | -0.2275 | -0.1925 | -0.835 | 0.046081 | 0.576338 | 0.601906 | 0.989223 | 0.997988 | 0.994666 |
| Rims2_Q0VF51_S335s_1 | Rims2_Q0\ | -1.49 | -1 | -1.68 | -2.04 | -0.83 | -1.44 | -1.49 | -1.97 | -0.72 | -1.63 | 0.05 | -1.64 | 0.21 | -0.72 | -1.37 | -1.22 | 0.12 | 0.5675 | 0.7775 | 0.087525 | 0.716315 | 0.222771 | 0.995437 | 0.997988 | 0.994666 |
| Rims2_Q0VF51_S432s_1 | Rims2_Q0\ | 0.92 | 1.08 | 1.21 | 0.73 | 0.23 | 0.91 | 0.61 | 0.86 | 0.45 | 1.08 | -0.78 | 1.05 | 1.52 | -1.01 | 1.45 | 1.51 | -0.3325 | -0.535 | -0.1175 | 0.840033 | 0.198711 | 0.237544 | 0.989223 | 0.997988 | 0.999507 |
| Rims2_Q0VF51_S439s_1 | Rims2_Q0\ | 0.02 | 0.04 | 0.15 | 0.06 | 0.31 | -0.25 | -0.18 | -0.05 | -0.23 | 0 | 0.06 | 0.14 | -0.24 | 0.41 | 0.05 | -0.06 | -0.11 | -0.075 | -0.0275 | 0.912945 | 0.616093 | 0.724791 | 0.989223 | 0.997988 | 0.999507 |
| Rims2_Q0VF51_S482s_1 | Rims2_Q0\ | 1.37 | 2.16 | 2.84 | 2.02 | 1.49 | 1.57 | 1.77 | 2.17 | -0.9 | 1.27 | 0.93 | 2.31 | -0.39 | -0.62 | 2.43 | 2.74 | -0.3475 | -1.195 | -1.0575 | 0.223606 | 0.324054 | 0.098684 | 0.989223 | 0.997988 | 0.994666 |
| Rims2_Q0VF51_S925s_1 | Rims2_Q0\ | 4.93 | 0.34 | 1.81 | 1.02 | 0.12 | 5.46 | 2.47 | 0.84 | 1.88 | 4.11 | 1.49 | 1.73 | 2.49 | -0.27 | 2.05 | 2.64 | 0.1975 | 0.2775 | -0.2975 | 0.778948 | 0.882632 | 0.79017 | 0.996493 | 0.997988 | 0.998201 |
| Rims2_Q0VF51_T1114t_1 | Rims2_Q0\ | 0.66 | 1.51 | 0.62 | 0.56 | -0.24 | 0.8 | 0.42 | 0.24 | 0.98 | 1.3 | 1.07 | 0.19 | 1.76 | 0.67 | 0.73 | 0.93 | -0.5325 | 0.0475 | 0.185 | 0.611453 | 0.126059 | 0.891026 | 0.989223 | 0.997988 | 0.995259 |
| Rims3_Q80U57_S22s_1 | Rims3_Q80 | 0.42 | -0.61 | 0.74 | 0.96 | -0.7 | -1.07 | 0.26 | 1.16 | -2.09 | -0.76 | 1 | 0.66 | -0.36 | -0.02 | 0.71 | 0.62 | -0.465 | -0.675 | -0.14 | 0.743938 | 0.407857 | 0.354616 | 0.989223 | 0.997988 | 0.9971 |
| Rims3_Q80U57_S297s_1 | Rims3_Q80 | 0.53 | 0.13 | -0.04 | 0.84 | 1.07 | 0.57 | 0.81 | 0.7 | 0.44 | 0.49 | 0.42 | 0.41 | -0.47 | 0.61 | 0.8 | 0.77 | 0.4225 | 0.075 | 0.0625 | 0.868509 | 0.139866 | 0.775817 | 0.989223 | 0.997988 | 0.999507 |
| Rims4_P60191_S257s_1 | Rims4_P60 | 0 | 0.13 | 0.37 | 0.4 | 0.11 | 0.21 | 0.46 | 0.86 | 0.45 | 0.88 | -1.04 | 1.01 | -0.07 | -0.23 | 0.52 | 0.53 | 0.185 | 0.1 | -0.0375 | 0.896908 | 0.465582 | 0.827909 | 0.989223 | 0.997988 | 0.999507 |
| Rin1_Q921Q7_S198s_1 | Rin1_Q921 | -0.11 | -0.61 | -1.57 | -0.7 | -0.52 | 0.7 | -0.33 | -0.57 | -0.53 | -0.56 | -0.96 | -0.63 | -1.89 | -0.18 | -1.23 | -0.1 | 0.5675 | 0.0775 | -0.1025 | 0.837349 | 0.185437 | 0.819379 | 0.989223 | 0.997988 | 0.999507 |
| Rin1_Q921Q7_S246s_1 | Rin1_Q921 | 0.48 | 0.85 | 0.7 | -0.65 | 0.59 | -0.07 | 0.9 | -0.36 | 0.59 | 0.71 | -2.74 | -0.86 | 0.74 | 1 | 0.43 | 0.56 | -0.08 | -0.92 | 0.3375 | 0.382265 | 0.850243 | 0.257535 | 0.996349 | 0.997988 | 0.994666 |
| Rin1_Q921Q7_S280s_1 | Rin1_Q921 | -2.7 | -2.54 | -2.19 | -4.63 | -2.12 | -2.13 | -2.93 | -2.7 | -1.88 | -2.96 | -0.5 | -2.65 | -0.75 | -2.55 | -3.09 | -2.5 | 0.545 | 1.0175 | 0.7925 | 0.256351 | 0.319186 | 0.169951 | 0.989223 | 0.997988 | 0.994666 |
| Rin1_Q921Q7_S340s_1 | Rin1_Q921 | -1.01 | -0.44 | -0.49 | -0.52 | -0.43 | -0.98 | -0.25 | -0.61 | -1.09 | -0.34 | -0.63 | -0.56 | -1.15 | -0.56 | -0.87 | -0.81 | 0.0475 | -0.04 | -0.2325 | 0.397131 | 0.853079 | 0.880819 | 0.996349 | 0.997988 | 0.994666 |
| Rin1_Q921Q7_S446s_1 | Rin1_Q921 | -0.41 | -0.34 | -1.16 | 0.75 | 0.46 | -0.65 | -0.24 | -0.33 | 0.94 | -0.32 | 1.21 | 0.05 | 0.25 | 0.87 | 0.26 | -0.31 | 0.1 | 0.76 | 0.5575 | 0.233729 | 0.816822 | 0.155287 | 0.995762 | 0.997988 | 0.994666 |
| Rin1_Q921Q7_S590s_1 | Rin1_Q921 | -0.02 | -0.41 | -0.28 | 0.3 | 0.8 | -0.12 | 0.56 | 0.37 | -0.39 | -0.23 | -0.38 | -0.13 | -0.6 | 0.42 | -0.03 | 0.3 | 0.505 | -0.18 | 0.125 | 0.699413 | 0.100678 | 0.472364 | 0.989223 | 0.997988 | 0.996159 |
| Rin1_Q921Q7_Y35y_1 | Rin1_Q921 | 0.78 | 0.29 | 0.96 | 1.19 | 0.44 | 0.72 | 0.53 | 1.11 | 0.53 | 0.56 | 1.01 | 0.56 | 0.2 | 0.81 | 0.99 | 0.45 | -0.105 | -0.14 | -0.1925 | 0.542457 | 0.707292 | 0.615434 | 0.995437 | 0.997988 | 0.994666 |
| Ring1_O35730_S219s_1 | Ring1_O35 | 0.32 | 0.64 | -0.56 | 0.64 | 0.64 | 0.17 | 0.82 | 0.23 | -0.37 | 0.33 | 0.71 | 0.52 | -0.85 | 0.07 | 0.34 | 0.7 | 0.205 | 0.0375 | -0.195 | 0.652222 | 0.542667 | 0.92024 | 0.989223 | 0.997988 | 0.995259 |
| Ripk2_P58801_S364s_1 | Ripk2_P58 | 0.35 | 0.62 | 0.16 | -0.01 | 0.17 | 0.52 | -1.09 | -0.03 | 0.18 | -0.06 | -0.06 | 0.31 | 0.03 | -0.01 | 0.43 | -0.05 | -0.3875 | -0.1875 | -0.18 | 0.505167 | 0.306328 | 0.449903 | 0.989223 | 0.997988 | 0.994666 |
| Rltpr_S0DHL8_S1377s_1 | Rltpr_S0DF | 0.19 | 0.17 | -0.09 | -0.19 | 0.13 | 0.08 | -0.22 | -0.47 | -0.46 | 0.25 | -2.53 | -1.34 | -0.72 | 0.13 | -2.21 | -1.1 | -0.14 | -1.04 | -0.995 | 0.059537 | 0.558541 | 0.09003 | 0.989223 | 0.997988 | 0.994666 |
| Rltpr_S0DHL8_S1382s_1 | Rltpr_S0DF | -2.42 | -0.46 | -0.72 | -1.28 | -1.47 | -2.18 | -1.79 | -1.46 | -0.8 | -1.41 | -1.08 | -1.35 | 0.72 | -2.08 | -0.71 | -1.03 | -0.505 | 0.06 | 0.445 | 0.500561 | 0.266924 | 0.891902 | 0.989223 | 0.997988 | 0.994666 |
| Rltpr_S0DHL8_S1382s_2 | Rltpr_S0DF | -2.41 | -2.77 | -1.9 | -2.68 | -3.12 | -1.57 | -2.53 | -3 | -1.8 | -1.59 | -0.97 | -2.02 | -0.84 | -1.43 | -3.12 | -2.17 | -0.115 | 0.845 | 0.55 | 0.287689 | 0.768538 | 0.026338 | 0.995762 | 0.997988 | 0.994666 |
| Rltpr_S0DHL8_S1385s_2 | Rltpr_S0DF | -2.41 | -2.77 | -1.9 | -2.68 | -3.12 | -1.57 | -2.53 | -3 | -1.8 | -1.59 | -0.97 | -2.02 | -0.84 | -1.43 | -3.12 | -2.17 | -0.115 | 0.845 | 0.55 | 0.287689 | 0.768538 | 0.026338 | 0.995762 | 0.997988 | 0.994666 |
| Rltpr_S0DHL8_S955s_1 | Rltpr_S0DF | -0.1 | 0.94 | 0.32 | -0.23 | 0.29 | 0.8 | -0.08 | 0.05 | 0.46 | 0.33 | -0.22 | -0.09 | 1.17 | 0.37 | 0.05 | 0.36 | 0.0325 | -0.1125 | 0.255 | 0.500537 | 0.922721 | 0.735999 | 0.997335 | 0.997988 | 0.994666 |
| Rltpr_S0DHL8_T1287t_1 | Rltpr_S0DF | 0.32 | 0.15 | -0.02 | 0.35 | 0.06 | 0.22 | 1.18 | 0.56 | -0.55 | 0.05 | -0.25 | 0.44 | -2.61 | 0.47 | 0.21 | 0.12 | 0.305 | -0.2775 | -0.6525 | 0.333099 | 0.309827 | 0.334771 | 0.989223 | 0.997988 | 0.994666 |
| Rmdn2_Q8BSE0_S51s_1 | Rmdn2_Q8 | -0.47 | 0.55 | 0.61 | 0.33 | 0.01 | -0.25 | 0.18 | 0.89 | -1.01 | -0.04 | 0.41 | -0.48 | -0.83 | -0.22 | 0.16 | 1.2 | -0.0475 | -0.535 | -0.1775 | 0.708541 | 0.892172 | 0.195564 | 0.996493 | 0.997988 | 0.996966 |
| Rmdn3_Q3UJU9_S46s_1 | Rmdn3_Q3 | 3.15 | 0.01 | -0.02 | 0.4 | 0.38 | 3.08 | 2.22 | 0.79 | -0.98 | 2.18 | -1.03 | 0.84 | -1.27 | 0.18 | 1.38 | 1.69 | 0.7325 | -0.6325 | -0.39 | 0.662683 | 0.402778 | 0.51545 | 0.989223 | 0.997988 | 0.995804 |
| Rnf113a1_Q8R3P8_S6s_1 | Rnf113a1_ | 0.85 | 1.02 | -0.05 | 0.02 | 1.75 | 0.37 | -0.31 | -1.67 | 0.28 | 0.05 | 1.32 | 0.92 | 1.76 | -0.29 | 0.59 | -0.89 | -0.425 | 0.1825 | -0.1675 | 0.775111 | 0.53339 | 0.649593 | 0.989223 | 0.997988 | 0.99798 |
| Rnf123_Q05CH9_S675s_1 | Rnf123_Q0 | -0.23 | 0.14 | -0.38 | -0.31 | -0.31 | -0.01 | -0.27 | 0.33 | -0.52 | -0.74 | -0.19 | -0.21 | -1.77 | -1.22 | -0.11 | 0.74 | 0.13 | -0.22 | -0.395 | 0.46654 | 0.601113 | 0.389724 | 0.989223 | 0.997988 | 0.994666 |
| Rnf139_Q7TMV1_S627s_1 | Rnf139_Q7 | -1.62 | -0.92 | -0.97 | -4.14 | -1.45 | -0.46 | -0.09 | -1.01 | 1.95 | -1.05 | -1.28 | -0.83 | 2.17 | 0 | 1.75 | -0.69 | 1.16 | 1.61 | 2.72 | 0.012504 | 0.131373 | 0.115916 | 0.989223 | 0.997988 | 0.994666 |
| Rnf14_Q9JI90_S349s_1 | Rnf14_Q9J | -2.71 | -2.91 | -2.47 | -4.79 | -2.01 | -3.14 | -3.54 | -3.65 | -4.25 | -2.18 | -2.14 | -3.47 | -3.42 | -1.25 | -3.97 | -2.94 | 0.135 | 0.21 | 0.325 | 0.647632 | 0.816975 | 0.754519 | 0.995762 | 0.997988 | 0.995259 |
| Rnf157_F7D0T8_S239s_1 | Rnf157_F7I | 0.3 | -0.35 | 0.06 | 0.53 | 0.67 | 0.12 | 0.5 | 0.3 | -0.21 | 0.22 | 0.15 | 0.46 | -0.46 | 0.46 | 0.05 | -0.15 | 0.2625 | 0.02 | -0.16 | 0.617075 | 0.338207 | 0.943638 | 0.989223 | 0.997988 | 0.995259 |
| Rnf157_F7D0T8_S281s_1 | Rnf157_F7I | 0.15 | 0.84 | 0.46 | 0.1 | -0.27 | 0.07 | -0.52 | -0.55 | 0.12 | 0.49 | -0.03 | -0.83 | 1.53 | 0.04 | -0.04 | -0.06 | -0.705 | -0.45 | -0.02 | 0.962139 | 0.023962 | 0.212432 | 0.989223 | 0.997988 | 0.999507 |
| Rnf169_E9Q7F2_S279s_1 | Rnf169_E9 | -6.72 | -4.57 | -1.76 | -0.58 | -6.66 | -6.65 | -0.7 | -1.23 | -6.2 | -4.19 | -0.25 | -0.52 | -2.68 | -7.29 | 0.28 | -0.94 | -0.4025 | 0.6175 | 0.75 | 0.686091 | 0.82663 | 0.724316 | 0.995762 | 0.997988 | 0.996159 |
| Rnf169_E9Q7F2_S390s_1 | Rnf169_E9 | -2.93 | -4.68 | -2.41 | -3.13 | -1.41 | -4.34 | -2.91 | -4.61 | -2.48 | -2.19 | -3.01 | -3.36 | -2.51 | -1.45 | -5.1 | -3.5 | -0.03 | 0.5275 | 0.1475 | 0.855314 | 0.969054 | 0.323184 | 0.999187 | 0.997988 | 0.999507 |
| Rnf214_Q8BFU3_S48s_1 | Rnf214_Q8 | -0.53 | -0.2 | 0.22 | 0.14 | -0.16 | -0.39 | 0.34 | 0.32 | -0.09 | 0.03 | -0.3 | 0.03 | 0.11 | -0.15 | -0.26 | 0.12 | 0.12 | 0.01 | 0.0475 | 0.86399 | 0.673329 | 0.96896 | 0.992876 | 0.997988 | 0.999507 |
| Rnf214_Q8BFU3_S500s_1 | Rnf214_Q8 | -0.26 | -0.12 | -0.83 | -0.96 | -0.54 | -0.39 | -0.36 | -0.13 | -2.07 | -0.46 | -0.22 | -0.81 | -0.92 | -0.82 | -0.83 | -0.33 | 0.1875 | -0.3475 | -0.1825 | 0.552751 | 0.489336 | 0.442426 | 0.989223 | 0.997988 | 0.994666 |
| Rnf217_D3YYI7_S27s_1 | Rnf217_D3 | -0.13 | -0.49 | 0.34 | 0.52 | -0.59 | 0.79 | -0.71 | 0.28 | -1.32 | 0.58 | -2.44 | 0.5 | 0.63 | -4.73 | 0.35 | 0.36 | -0.1175 | -0.73 | -0.9075 | 0.433469 | 0.773123 | 0.308524 | 0.995762 | 0.997988 | 0.994666 |
| Rnf6_Q9DBU5_S183s_1 | Rnf6_Q9DE | 1.36 | 0.96 | 0.87 | 0.73 | 0.6 | 0.56 | 0.15 | 0.4 | 0.81 | 0.84 | 0.05 | 0.09 | 1.74 | 0.72 | 1.23 | 1.23 | -0.5525 | -0.5325 | 0.25 | 0.421666 | 0.040451 | 0.099755 | 0.989223 | 0.997988 | 0.994666 |
| Rnf6_Q9DBU5_S301s_1 | Rnf6_Q9DE | 1.02 | 0.99 | 0.14 | 0.15 | -0.26 | 1.3 | -0.18 | -0.53 | 2.38 | 0.61 | -0.98 | -0.8 | 2.62 | 0.69 | 0.64 | 0.36 | -0.4925 | -0.2725 | 0.5025 | 0.360071 | 0.289499 | 0.711302 | 0.989223 | 0.997988 | 0.994666 |
| Rnmt_D3Z4K1_S100s_1 | Rnmt_D3Z4 | -5.18 | -6.32 | -4.39 | -6.74 | -3.71 | -6.87 | -4.78 | -4.1 | -7.51 | -3.92 | -1.55 | -3.47 | -3.93 | -2.37 | -3.5 | -4.96 | 0.7925 | 1.545 | 1.9675 | 0.01582 | 0.319999 | 0.214814 | 0.989223 | 0.997988 | 0.994666 |
| Rnmt_D3Z4K1_S11s_1 | Rnmt_D3Z4 | -2.08 | -0.79 | -1.66 | -1.2 | -1.03 | -0.97 | -1.81 | -2.45 | -0.97 | 0.2 | -1.07 | -2.37 | -0.29 | -0.11 | -2.4 | -2.1 | -0.1325 | 0.38 | 0.2075 | 0.730457 | 0.755734 | 0.496385 | 0.995762 | 0.997988 | 0.9971 |
| Rnmt_D3Z4K1_S11s_2 | Rnmt_D3Z4 | -1.92 | -1.43 | -1.89 | -1.83 | -1.5 | -2.12 | -2.29 | -2.35 | -0.96 | -1.49 | -0.62 | -1.92 | -0.41 | -0.96 | -1.56 | -2.06 | -0.2975 | 0.52 | 0.52 | 0.201202 | 0.284021 | 0.142585 | 0.989223 | 0.997988 | 0.994666 |
| Rnmt_D3Z4K1_S15s_1 | Rnmt_D3Z4 | -1.95 | -1.28 | -3.46 | -3.67 | -2.31 | -1.94 | -3.2 | -2.8 | -1.14 | -1.44 | -0.73 | -3.63 | -0.46 | -1.45 | -2.88 | -3.7 | 0.0275 | 0.855 | 0.4675 | 0.570885 | 0.961985 | 0.288469 | 0.999187 | 0.997988 | 0.995259 |
| Rnmt_D3Z4K1_S15s_2 | Rnmt_D3Z4 | -1.92 | -1.43 | -1.89 | -1.83 | -1.5 | -2.12 | -2.29 | -2.35 | -0.96 | -1.49 | -0.62 | -1.92 | -0.41 | -0.96 | -1.56 | -2.06 | -0.2975 | 0.52 | 0.52 | 0.201202 | 0.284021 | 0.142585 | 0.989223 | 0.997988 | 0.994666 |
| Robo1_G5E843_S1403s_1 | Robo1_G5I | -1.3 | -0.37 | -0.57 | -0.72 | -0.32 | -0.85 | -0.17 | -0.56 | -0.57 | -0.67 | -0.66 | 0.19 | -0.93 | -0.24 | 0.09 | -1.3 | 0.265 | 0.3125 | 0.145 | 0.708595 | 0.360773 | 0.338748 | 0.989223 | 0.997988 | 0.996966 |
| Robo1_G5E843_S901s_1 | Robo1_G5I | 1.48 | 0.87 | 0.15 | 0.63 | 1.2 | 1.33 | 1.13 | 0.7 | 0.97 | 0.58 | 0.87 | 0.45 | -0.15 | 1.24 | 0.44 | 0.57 | 0.3075 | -0.065 | -0.2575 | 0.526953 | 0.351783 | 0.842725 | 0.989223 | 0.997988 | 0.994666 |
| Robo2_Q19AB2_S1197s_1 | Robo2_Q1 | -0.1 | 0.38 | 0.28 | 0.62 | 0.32 | 0.14 | 1.06 | 0.05 | -0.35 | 0.42 | 0.57 | -0.13 | -0.7 | -0.01 | 0.08 | 0.62 | 0.0975 | -0.1675 | -0.2975 | 0.395642 | 0.744959 | 0.583615 | 0.995762 | 0.997988 | 0.994666 |
| Rock2_F8VPK5_S1052s_1 | Rock2_F8V | -0.44 | -0.59 | -1.16 | -1.08 | -1.37 | -0.75 | -1.88 | -2.15 | 1.57 | -1.64 | -2.74 | -1.17 | 2.14 | -1.24 | -0.3 | -1.1 | -0.72 | -0.1775 | 0.6925 | 0.346033 | 0.067509 | 0.830967 | 0.989223 | 0.997988 | 0.994666 |
| Rock2_F8VPK5_S439s_1 | Rock2_F8V | 0.75 | 0.46 | 0.23 | 0.68 | -0.14 | 1.15 | -0.47 | 0.29 | 1.71 | 0.59 | -0.44 | -0.02 | 1.72 | -0.53 | 0.3 | -0.44 | -0.3225 | -0.07 | -0.2675 | 0.597873 | 0.388093 | 0.879285 | 0.989223 | 0.997988 | 0.995259 |
| Rogdi_Q3TDK6_S227s_1 | Rogdi_Q3T | -0.75 | -0.73 | 0 | -0.62 | -0.7 | -0.28 | -0.41 | -0.02 | -0.84 | -0.74 | -1.5 | -0.7 | -0.98 | -1.44 | -0.98 | -0.74 | 0.1725 | -0.42 | -0.51 | 0.10931 | 0.526496 | 0.182321 | 0.989223 | 0.997988 | 0.994666 |
| Ror1_Q9Z139_S436s_1 | Ror1_Q9Z1 | 0.43 | 0.87 | 0.86 | 1.07 | -1.7 | -1.02 | -1.19 | -1.15 | 1.99 | 0.76 | 0.31 | 0.42 | 1.96 | -1.51 | 0.96 | 1.5 | -2.0725 | 0.0625 | -0.08 | 0.909201 | 1.63E-05 | 0.877158 | 0.166877 | 0.997988 | 0.999507 |
| Rpap3_Q9D706_S429s_1 | Rpap3_Q9I | 0.68 | -0.02 | 0.76 | 0.94 | -0.39 | 0.83 | -0.11 | 0.95 | 0.14 | 0.44 | -1.7 | 0.38 | 1.07 | -1.82 | 0.6 | 0.96 | -0.27 | -0.775 | -0.3875 | 0.551454 | 0.488657 | 0.157742 | 0.989223 | 0.997988 | 0.994666 |
| Rph3a_P47708_S245s_1 | Rph3a_P47 | 0.74 | -0.66 | 2.39 | 0.67 | -0.09 | 0.71 | 2.59 | 4 | 0.96 | 0.26 | 2.28 | 3.78 | -0.11 | 0.76 | 2.49 | 3.96 | 1.0175 | 1.035 | 0.99 | 0.316693 | 0.306327 | 0.260009 | 0.989223 | 0.997988 | 0.994666 |
| Rph3a_P47708_S271s_1 | Rph3a_P47 | -1.86 | -1.72 | -1.46 | -3.08 | -0.87 | -0.95 | -1.43 | -1.63 | -0.58 | -1.9 | -2.33 | -1.28 | 1.03 | -1.37 | -1.49 | -1.24 | 0.81 | 0.5075 | 1.2625 | 0.071304 | 0.062399 | 0.31898 | 0.989223 | 0.997988 | 0.994666 |
| Rph3a_P47708_S679s_1 | Rph3a_P47 | 2.42 | 1.05 | 2.63 | 1.46 | 1.28 | 2.03 | 3.71 | 2.91 | 0.96 | 2.1 | 1.13 | 3.01 | 0.82 | 0.49 | 3.35 | 3.65 | 0.5925 | -0.09 | 0.1875 | 0.815364 | 0.322761 | 0.872577 | 0.989223 | 0.997988 | 0.999507 |
| Rph3a_P47708_S680s_1 | Rph3a_P47 | 4.07 | 1.78 | 4 | 1.26 | 1.97 | 4.01 | 5.19 | 4.06 | 0.47 | 3.41 | 1.01 | 4.07 | 0.48 | 0.24 | 4.81 | 4.86 | 1.03 | -0.5375 | -0.18 | 0.888228 | 0.251748 | 0.598491 | 0.989223 | 0.997988 | 0.999507 |
| Rph3a_P47708_T188t_1 | Rph3a_P47 | -0.85 | -1.34 | -1.63 | -0.25 | -0.68 | -0.14 | -0.53 | -0.47 | -2.22 | -2.15 | 0.45 | -0.24 | -1.07 | -0.49 | -1.7 | -2.07 | 0.5625 | -0.0225 | -0.315 | 0.488938 | 0.115958 | 0.973199 | 0.989223 | 0.997988 | 0.994666 |
| Rpl13_P47963_S139s_1 | Rpl13_P47 | -1.23 | -2.07 | -1.27 | -1.46 | -1.77 | -1.28 | -1.56 | -1.89 | -0.97 | -1.3 | -2.35 | -1.59 | 0.24 | -1.31 | -1.37 | -1.11 | -0.1175 | -0.045 | 0.62 | 0.166711 | 0.670328 | 0.90131 | 0.992876 | 0.997988 | 0.994666 |
| Rpl15_Q9CZM2_T197t_1 | Rpl15_Q9C | -1.66 | -2.91 | -2.24 | -4.48 | -1.77 | -4.39 | -3.46 | -3.13 | -1.88 | -4.15 | -0.82 | -4.33 | -1.12 | -1.49 | -3.1 | -5.91 | -0.365 | 0.0275 | -0.0825 | 0.939205 | 0.613291 | 0.976502 | 0.989223 | 0.997988 | 0.999507 |
| Rpl17_Q6ZWZ7_S141s_1 | Rpl17_Q6Z | -2.29 | -1.88 | -1.11 | -1.24 | -2.06 | -3.1 | -0.72 | -1.33 | -2.6 | -2.65 | -0.9 | -1.82 | -1.06 | -2 | -1.28 | -1.81 | -0.1725 | -0.3625 | 0.0925 | 0.804053 | 0.744764 | 0.449806 | 0.995762 | 0.997988 | 0.999507 |
| Rpl24_Q8BP67_T83t_1 | Rpl24_Q8B | -1.22 | -1.54 | -1.14 | -0.29 | -1.69 | -2.25 | -0.59 | -0.35 | -1.5 | -1.22 | 0.42 | -0.67 | -2.1 | -1.63 | -0.6 | -0.01 | -0.1725 | 0.305 | -0.0375 | 0.941668 | 0.721539 | 0.526577 | 0.995437 | 0.997988 | 0.999507 |
| Rpl6_P47911_S21s_1 | Rpl6_P479 | -1.11 | -1.7 | -1.16 | -2.28 | -2.32 | -0.69 | -2.29 | -2.53 | 1.22 | -1.1 | -2.76 | -1.95 | 2.07 | -2.53 | -1.08 | -1.89 | -0.395 | 0.415 | 0.705 | 0.452316 | 0.40895 | 0.60852 | 0.989223 | 0.997988 | 0.994666 |
| Rprd2_Q6NXI6_T332t_1 | Rprd2_Q6N | -0.82 | -0.92 | -1.99 | -1.83 | -1.5 | -2.77 | -1.14 | -1.03 | -0.2 | 0.1 | -0.03 | -1 | 0.89 | -0.45 | -0.47 | -1.28 | -0.22 | 1.1075 | 1.0625 | 0.061125 | 0.638128 | 0.01804 | 0.989223 | 0.997988 | 0.994666 |
| Rprd2_Q6NXI6_T698t_1 | Rprd2_Q6N | 0.54 | 0.38 | -0.51 | -1.17 | -0.38 | 0.21 | -1.08 | -0.79 | -0.15 | 0.01 | -0.4 | -0.62 | 0.55 | 0.35 | -1.37 | -1.56 | -0.32 | -0.1 | -0.3175 | 0.611897 | 0.487733 | 0.810131 | 0.989223 | 0.997988 | 0.995259 |
| Rps17_P63276_S113s_1 | Rps17_P63 | -0.98 | -0.18 | -0.33 | -0.87 | -0.6 | -0.87 | 0.5 | 0.02 | -1.94 | -0.18 | -2.81 | -0.11 | -0.72 | -1.92 | 0 | 0.46 | 0.3525 | -0.67 | 0.045 | 0.930983 | 0.344098 | 0.305863 | 0.989223 | 0.997988 | 0.999507 |
| Rps2_P25444_S264s_1 | Rps2_P254 | -2.54 | -1.95 | -2.46 | -2.24 | -1.68 | -2.73 | -1.45 | -2.33 | -1.99 | -1.74 | -2.51 | -2.79 | -1.63 | -0.52 | -1.25 | -2.23 | 0.25 | 0.04 | 0.89 | 0.044427 | 0.458196 | 0.896707 | 0.989223 | 0.997988 | 0.994666 |
| Rps26_P62855_T96t_1 | Rps26_P62 | -1.38 | -0.73 | 0.51 | -2.24 | -0.82 | -1.12 | -1.04 | -1.06 | -0.04 | -1.12 | -1.84 | -0.79 | 1.26 | -2.32 | 0.93 | 1.28 | -0.05 | 0.0125 | 1.2475 | 0.195156 | 0.924723 | 0.984109 | 0.997335 | 0.997988 | 0.994666 |
| Rps27_Q6ZWU9_S11s_1 | Rps27_Q6Z | 1.64 | 2.08 | 1.29 | 0.25 | 2.25 | 1.52 | 1.37 | 1.2 | -1.14 | 2.04 | 0.17 | 0.8 | -1.24 | 0.22 | 1.98 | 2.1 | 0.27 | -0.8475 | -0.55 | 0.488276 | 0.5328 | 0.241082 | 0.989223 | 0.997988 | 0.994666 |
| Rps28_G3UYV7_S10s_1 | Rps28_G3U | -1.74 | -2.7 | -2.26 | -2.76 | -2.13 | -0.38 | -2.14 | -2.1 | -0.38 | -0.77 | -1.66 | -1.99 | 0.2 | -3.36 | -1.15 | -1.89 | 0.6775 | 1.165 | 0.815 | 0.259984 | 0.165176 | 0.022434 | 0.989223 | 0.997988 | 0.994666 |
| Rps3_P62908_T221t_1 | Rps3_P629 | -0.87 | -0.3 | -0.45 | -0.69 | 0.13 | -0.93 | 0.15 | -0.58 | -1.51 | -0.41 | -0.43 | -0.41 | -0.55 | -0.27 | 0.17 | 0.28 | 0.27 | -0.1125 | 0.485 | 0.12593 | 0.399305 | 0.731418 | 0.989223 | 0.997988 | 0.994666 |
| Rps6ka2_Q9WUT3_S218s_1 | Rps6ka2_Q | 0.97 | -0.54 | -0.81 | -0.88 | 0.04 | 0.64 | 0.53 | 0.6 | -0.13 | -1.07 | -2.3 | -2.57 | -0.55 | 0.34 | -1.31 | 0.65 | 0.7675 | -1.2025 | 0.0975 | 0.864556 | 0.100313 | 0.089986 | 0.989223 | 0.997988 | 0.999507 |
| Rps6ka2_Q9WUT3_S360s_1 | Rps6ka2_Q | 1.34 | 0.59 | 0.69 | 0.77 | 0.8 | 1.4 | 0.65 | 0.85 | 0.67 | 1.28 | 0.66 | 1.05 | 0.59 | 0.32 | 1.23 | 0.98 | 0.0775 | 0.0675 | -0.0675 | 0.829965 | 0.777858 | 0.808923 | 0.995762 | 0.997988 | 0.999507 |
| Rps6ka2_Q9WUT3_S377s_1 | Rps6ka2_Q | -1.59 | -1.48 | -0.97 | -1.04 | -0.7 | -1.86 | -0.4 | -0.91 | -1.7 | -1.59 | -0.5 | -0.83 | -1.76 | -0.76 | -1.12 | -0.74 | 0.3025 | 0.115 | 0.175 | 0.595515 | 0.400443 | 0.741047 | 0.989223 | 0.997988 | 0.995259 |
| Rps6ka3_B1AXN9_S199s_1 | Rps6ka3_B | 0.3 | 0.04 | -0.58 | -0.51 | 0 | 0.51 | -0.42 | -0.13 | 0.12 | 0.3 | -0.5 | -0.53 | -0.36 | 0.44 | -0.17 | -0.81 | 0.1775 | 0.035 | -0.0375 | 0.917064 | 0.568793 | 0.91472 | 0.989223 | 0.997988 | 0.999507 |
| Rps6ka3_B1AXN9_S341s_1 | Rps6ka3_B | 1.45 | 1.07 | 1.77 | 2.18 | 0.76 | 0.99 | 1.42 | 1.2 | 0.82 | 1.22 | 0.93 | 2.26 | 0.38 | 1.73 | 1.75 | 2.04 | -0.525 | -0.31 | -0.1425 | 0.743012 | 0.10528 | 0.44769 | 0.989223 | 0.997988 | 0.9971 |
| Rps6ka3_B1AXN9_S387s_1 | Rps6ka3_B | 0.29 | 0.68 | 0.77 | 0.62 | 0.63 | 0.24 | 1.03 | 0.69 | 0.34 | 0.26 | 0.23 | 0.58 | 0 | 0.74 | 0.18 | 0.28 | 0.0575 | -0.2375 | -0.29 | 0.304065 | 0.818029 | 0.314891 | 0.995762 | 0.997988 | 0.994666 |
| Rps6ka3_B1AXN9_S687s_1 | Rps6ka3_B | -0.42 | -0.07 | 0.54 | 0.8 | -0.88 | -0.6 | 0.55 | 1 | -0.2 | -0.2 | -0.34 | 0.53 | -0.5 | 0 | -0.52 | 0.37 | -0.195 | -0.265 | -0.375 | 0.325511 | 0.690236 | 0.46223 | 0.994672 | 0.997988 | 0.994666 |
| Rps6kb1_Q8BSK8_S424s_1 | Rps6kb1_C | 1.84 | -1.2 | 0.18 | -0.26 | 0.96 | 1.62 | -0.7 | 0.4 | 1.04 | 1.19 | 0.35 | 0.34 | 0.06 | 0.96 | -0.39 | -0.5 | 0.43 | 0.59 | -0.1075 | 0.868094 | 0.547308 | 0.350405 | 0.989223 | 0.997988 | 0.999507 |
| Rps6kc1_E9QMX4_S448s_1 | Rps6kc1_E | 0.8 | 0.35 | 1.07 | 0.21 | 0.54 | 0.24 | 0.2 | 0.11 | 0.86 | 0.95 | -0.21 | 0.2 | -0.28 | 0.67 | 0.72 | 0.62 | -0.335 | -0.1575 | -0.175 | 0.613067 | 0.226169 | 0.65828 | 0.989223 | 0.997988 | 0.995259 |
| Rps6kc1_E9QMX4_S558s_1 | Rps6kc1_E | -0.54 | -0.17 | -0.42 | 0.51 | 0.09 | -0.21 | 0.48 | 0.15 | 0.07 | -0.38 | -0.35 | 0 | 0.15 | -0.66 | 0.15 | 0.21 | 0.2825 | -0.01 | 0.1175 | 0.7346 | 0.356584 | 0.973469 | 0.989223 | 0.997988 | 0.9971 |
| Rps6kc1_E9QMX4_S779s_1 | Rps6kc1_E | -0.53 | -0.46 | -0.11 | 0.03 | -0.13 | -1.4 | 0.13 | -0.06 | -1.37 | -0.56 | -0.33 | -0.06 | -1.39 | -0.54 | -0.31 | -0.38 | -0.0975 | -0.3125 | -0.3875 | 0.253778 | 0.791997 | 0.362815 | 0.995762 | 0.997988 | 0.994666 |
| Rps8_P62242_T130t_1 | Rps8_P622 | -0.04 | 0.47 | -0.25 | -0.48 | 0.22 | -0.66 | 0.29 | -1 | 0.43 | 0.29 | -0.7 | -0.73 | 0.43 | 1.47 | 0.22 | 0.2 | -0.2125 | -0.1025 | 0.655 | 0.109115 | 0.572248 | 0.786012 | 0.989223 | 0.997988 | 0.994666 |
| Rptor_Q8K4Q0_S863s_1 | Rptor_Q8K | 0.73 | 0.86 | 0.29 | 0.57 | 0.63 | 0.6 | 0.64 | 0.63 | 0.05 | 0.71 | -0.54 | 0.28 | 0.08 | -0.2 | 0.24 | 0.52 | 0.0125 | -0.4875 | -0.4525 | 0.126356 | 0.953813 | 0.15014 | 0.999187 | 0.997988 | 0.994666 |
| Rptor_Q8K4Q0_T706t_1 | Rptor_Q8K | -1.74 | -1.43 | -2.44 | -1.42 | 0.43 | 0.14 | -1.16 | -1.35 | 0.2 | -0.12 | 2.9 | 0.3 | 0.66 | -0.8 | -1.71 | -1.96 | 1.2725 | 2.5775 | 0.805 | 0.191149 | 0.021918 | 0.003629 | 0.989223 | 0.997988 | 0.994666 |
| Rrad_Q6PGA2_S38s_1 | Rrad_Q6PC | -2.61 | -2.59 | -0.82 | -0.87 | -2.89 | -2.61 | -2.57 | -1.98 | -1.94 | -3.05 | 0.79 | -0.64 | -0.38 | -2.87 | -0.49 | -0.6 | -0.79 | 0.5125 | 0.6375 | 0.37343 | 0.136641 | 0.556607 | 0.989223 | 0.997988 | 0.994666 |
| Rragb_G3UYP1_S4s_1 | Rragb_G3U | -0.83 | -1.43 | -0.81 | -0.44 | -2.5 | -1.6 | -2.21 | -1.38 | 0.72 | -1.07 | 1.34 | -1.74 | 0.72 | -1.9 | -1.37 | -2.34 | -1.045 | 0.69 | -0.345 | 0.592489 | 0.011449 | 0.326214 | 0.989223 | 0.997988 | 0.995259 |
| Rragc_Q99K70_S15s_1 | Rragc_Q99 | -0.55 | -0.47 | -0.38 | -0.06 | -0.08 | -0.6 | -0.13 | -0.1 | -0.89 | -0.71 | 0.07 | 0.1 | -1.01 | -0.27 | -0.79 | -0.53 | 0.1375 | 0.0075 | -0.285 | 0.314989 | 0.561807 | 0.980832 | 0.989223 | 0.997988 | 0.994666 |
| Rragd_B1AWT4_T270t_1 | Rragd_B1A | -1.43 | -2.13 | -2.17 | -3.95 | -2.1 | -1.31 | -2.75 | -2.44 | -0.65 | -1.48 | -2.43 | -2.96 | 0.94 | -2.26 | -2.81 | -2.46 | 0.27 | 0.54 | 0.7725 | 0.398086 | 0.631311 | 0.429901 | 0.989223 | 0.997988 | 0.994666 |
| Rras2_P62071_S186s_1 | Rras2_P62( | 0.85 | 0.37 | 0.84 | 1.55 | 0.11 | 0.74 | 0.76 | 1.49 | 0.6 | 1.22 | 1.15 | 0.99 | 0.64 | -0.24 | 1.23 | 0.94 | -0.1275 | 0.0875 | -0.26 | 0.525153 | 0.72947 | 0.77987 | 0.995762 | 0.997988 | 0.994666 |
| Rrbp1_Q99PL5_S1162s_1 | Rrbp1_Q99 | 0.45 | 1.05 | -2.03 | 0.82 | 1.68 | -0.01 | -0.07 | 0.5 | 0.17 | 0.68 | -1.88 | 0.31 | -0.36 | 0.36 | 0.45 | 0.82 | 0.4525 | -0.2525 | 0.245 | 0.71795 | 0.534319 | 0.757657 | 0.989223 | 0.997988 | 0.996966 |
| Rrbp1_Q99PL5_S786s_1 | Rrbp1_Q99 | 0.04 | 0.42 | -0.81 | -2.79 | -1.31 | 0.82 | -0.27 | -0.53 | 2.34 | 0.36 | -0.72 | 0.42 | 3.22 | -1.35 | 5.62 | -0.61 | 0.4625 | 1.385 | 2.505 | 0.128078 | 0.535233 | 0.130074 | 0.989223 | 0.997988 | 0.994666 |
| Rrp1_P56183_S383s_1 | Rrp1_P561 | -1.64 | -1.16 | -1.74 | -1.99 | -3.89 | -1.39 | -3.18 | -1.8 | -2 | -1.24 | -1.7 | -2.07 | -0.83 | -2.86 | -1.73 | -1.49 | -0.9325 | -0.12 | -0.095 | 0.831407 | 0.115954 | 0.687851 | 0.989223 | 0.997988 | 0.999507 |
| Rrp36_Q3UFY0_T64t_1 | Rrp36_Q3U | 0.28 | 0.17 | -1.28 | -0.81 | 0.51 | 0.48 | -0.92 | -0.86 | -0.08 | 0.7 | -0.79 | -1.39 | 1.05 | 0.01 | -0.91 | -0.6 | 0.2125 | 0.02 | 0.2975 | 0.581655 | 0.674935 | 0.970803 | 0.992876 | 0.997988 | 0.995259 |

| | | | | | | | | | | | | | | | | | | | | | | | | | | |
|---|---|---|---|---|---|---|---|---|---|---|---|---|---|---|---|---|---|---|---|---|---|---|---|---|---|---|
| Rsf1_E9PWW9_S1312s_1 | Rsf1_E9PW | 1.35 | 1.54 | 1.56 | 1.55 | 0.62 | 0.69 | 1.01 | 1.01 | 0.45 | 1.57 | 3.3 | 1.48 | 0.6 | 0.86 | 1.79 | 1.24 | -0.6675 | 0.2 | -0.3775 | 0.247995 | 0.01049 | 0.716707 | 0.989223 | 0.997988 | 0.994666 |
| Rsf1_E9PWW9_T1308t_1 | Rsf1_E9PW | 1.13 | 1.08 | 0.31 | 0.26 | 0.32 | 0.9 | 0.31 | -0.12 | 1.01 | 1.28 | 0.51 | -0.46 | 0.51 | 1.45 | 0.51 | 0.02 | -0.3425 | -0.11 | -0.0725 | 0.85314 | 0.310294 | 0.801336 | 0.989223 | 0.997988 | 0.999507 |
| Rsph3b_Q9DA80_T100t_1 | Rsph3b_Q9 | -0.75 | -0.49 | -0.13 | -0.38 | -1.3 | -1.45 | -0.36 | -1.51 | 0.32 | -0.81 | -0.87 | -1.29 | 0.32 | -1.57 | -0.59 | -0.33 | -0.7175 | -0.225 | -0.105 | 0.799845 | 0.044063 | 0.551431 | 0.989223 | 0.997988 | 0.999507 |
| Rspry1_Q8BVR6_S50s_1 | Rspry1_Q8 | 2.03 | 1.7 | 1.8 | 0.25 | 1.36 | 1.92 | 1.76 | 1.51 | -0.3 | 1.67 | 0.01 | 1.67 | -0.68 | -0.83 | 1.99 | 2.34 | 0.1925 | -0.6825 | -0.74 | 0.380995 | 0.636839 | 0.278683 | 0.989223 | 0.997988 | 0.994666 |
| Rsrc1_Q3TGY1_S6s_1 | Rsrc1_Q3T | -0.33 | -1.59 | -0.12 | -3.83 | -2.73 | -0.33 | -1.57 | -0.48 | 1.9 | -1.17 | -1.09 | -1.11 | 2.36 | -2.07 | 1.91 | -1.21 | 0.19 | 1.1 | 1.715 | 0.178003 | 0.83001 | 0.290011 | 0.995762 | 0.997988 | 0.994666 |
| Rsrc2_S4R2L4_S103s_1 | Rsrc2_S4R | -1.53 | -2.24 | -1.26 | -2.51 | -2.54 | -1.76 | -1.93 | -1.8 | -0.23 | -2.25 | -1.28 | -1.94 | 1 | -2.42 | -0.25 | -1.33 | -0.1225 | 0.46 | 1.135 | 0.132502 | 0.72539 | 0.371519 | 0.995437 | 0.997988 | 0.994666 |
| Rsrc2_S4R2L4_S104s_1 | Rsrc2_S4R | -1.71 | -2.33 | -1.44 | -3 | -2.75 | -1.95 | -2.07 | -1.97 | -0.77 | -2.71 | -1.62 | -2.21 | 0.73 | -2.67 | -0.37 | -1.59 | -0.065 | 0.2925 | 1.145 | 0.13962 | 0.865972 | 0.569216 | 0.996493 | 0.997988 | 0.994666 |
| Rsrc2_S4R2L4_S32s_1 | Rsrc2_S4R | -1.86 | -1.66 | -2.82 | -3.25 | -1.05 | -2.23 | -2.09 | -2.36 | -1.82 | -1.61 | -2.17 | -1.67 | -1.07 | -2.27 | -0.62 | -1.68 | 0.465 | 0.58 | 0.9875 | 0.071267 | 0.317557 | 0.169872 | 0.989223 | 0.997988 | 0.994666 |
| Rtkn_D3Z4N1_S125s_1 | Rtkn_D3Z4 | -0.95 | -0.96 | -1.6 | -2.88 | -0.94 | -0.59 | -2.15 | -1.17 | -1.75 | -0.85 | -0.4 | -2.2 | -0.22 | -0.55 | -2.12 | -2.04 | 0.385 | 0.2975 | 0.365 | 0.553791 | 0.460849 | 0.601234 | 0.989223 | 0.997988 | 0.994666 |
| Rtkn_D3Z4N1_S134s_1 | Rtkn_D3Z4 | -0.53 | 0.27 | -1.57 | -1.32 | -0.87 | -1.63 | -0.17 | -1.43 | -1.12 | -1.59 | 0.17 | -1.1 | 0.87 | -1.61 | 0.17 | -1.4 | -0.2375 | -0.1225 | 0.295 | 0.656549 | 0.62807 | 0.815484 | 0.989223 | 0.997988 | 0.995259 |
| Rtkn_D3Z4N1_S56s_1 | Rtkn_D3Z4 | 0.2 | 0.88 | 0.75 | -0.57 | -0.5 | 1.13 | -0.83 | -0.68 | 0.62 | 0.08 | 0.44 | 0.3 | 1.26 | -0.17 | -0.46 | 0.25 | -0.535 | 0.045 | -0.095 | 0.842589 | 0.311204 | 0.90057 | 0.989223 | 0.997988 | 0.999507 |
| Rtkn_Q8C6B2_S232s_1 | Rtkn_Q8C6 | 0.62 | 0.65 | 0.04 | 0.92 | 0.86 | 1.1 | 0.43 | 0.77 | 0.5 | 1 | 0.31 | 0.49 | 1.03 | 0.76 | -0.34 | 0.89 | 0.2325 | 0.0175 | 0.0275 | 0.942158 | 0.402384 | 0.95106 | 0.989223 | 0.997988 | 0.999507 |
| Rtkn_Q8C6B2_S520s_1 | Rtkn_Q8C6 | -0.77 | -0.39 | -0.41 | -0.24 | -0.24 | 1.4 | 1.17 | -0.72 | -1.17 | -1.26 | -0.63 | 0.27 | -0.1 | -0.71 | -0.14 | 0.23 | 0.855 | -0.245 | 0.2725 | 0.36288 | 0.106596 | 0.518341 | 0.989223 | 0.997988 | 0.994666 |
| Rtn1_Q7M6W1_S7s_1 | Rtn1_Q7M | -0.47 | -0.48 | -0.11 | -0.97 | -1.02 | -0.71 | -1.02 | -1.12 | -0.16 | 0.06 | -0.41 | -0.68 | 1.04 | -0.05 | -0.79 | -0.83 | -0.46 | 0.21 | 0.35 | 0.451115 | 0.093888 | 0.469253 | 0.989223 | 0.997988 | 0.994666 |
| Rtn1_Q8K0T0_S16s_1 | Rtn1_Q8K0 | 0.05 | 0.15 | 0.37 | 0.69 | 0.96 | -0.07 | 0.61 | 0.39 | 0.51 | 0.38 | -1 | 0.33 | 0.54 | 0.46 | 0.36 | 0.51 | 0.1575 | -0.26 | 0.1525 | 0.552903 | 0.588348 | 0.503765 | 0.989223 | 0.997988 | 0.994666 |
| Rtn1_Q8K0T0_S352s_1 | Rtn1_Q8K0 | 1.65 | 1.61 | 0.13 | 0.96 | 2.2 | 2.1 | 1.12 | 0.7 | 1.02 | 0.86 | 0.64 | 0.3 | -0.01 | 2.07 | -0.26 | 0.2 | 0.4425 | -0.3825 | -0.5875 | 0.329588 | 0.36207 | 0.339528 | 0.989223 | 0.997988 | 0.994666 |
| Rtn1_Q8K0T0_S487s_1 | Rtn1_Q8K0 | -0.25 | 0.37 | 0.32 | 0.11 | 0.1 | -0.27 | 0.54 | 0.23 | -0.29 | -0.04 | -0.05 | 0.08 | -0.14 | -0.72 | 0.24 | 0.08 | 0.0125 | -0.2125 | -0.2725 | 0.386747 | 0.962313 | 0.391709 | 0.999187 | 0.997988 | 0.994666 |
| Rtn1_Q8K0T0_S70s_1 | Rtn1_Q8K0 | -0.02 | 0.14 | -0.08 | 0.19 | 0.19 | -0.17 | 0.38 | -0.11 | -0.28 | -0.12 | -0.7 | -0.18 | -0.07 | 0.34 | -0.11 | -0.18 | 0.015 | -0.3775 | -0.0625 | 0.802132 | 0.946845 | 0.134744 | 0.999187 | 0.997988 | 0.999507 |
| Rtn1_Q8K0T0_S92s_1 | Rtn1_Q8K0 | 0.14 | -0.27 | 0.1 | 0.34 | 0.16 | 0 | 0.61 | 0.44 | -0.34 | -0.25 | -0.13 | 0.06 | -0.9 | 0.14 | -0.17 | 0.06 | 0.225 | -0.2425 | -0.295 | 0.364358 | 0.372358 | 0.325049 | 0.989223 | 0.997988 | 0.994666 |
| Rtn3_Q9ES97_S113s_1 | Rtn3_Q9ES | 0.4 | -0.21 | 0.28 | 0.2 | -0.64 | 0.45 | -0.48 | -0.01 | 0.98 | 0.5 | -1.47 | 0.22 | 0.96 | -0.77 | 0.02 | 0.38 | -0.3375 | -0.11 | -0.02 | 0.959506 | 0.280965 | 0.830679 | 0.989223 | 0.997988 | 0.999507 |
| Rtn3_Q9ES97_S31s_1 | Rtn3_Q9ES | 0.26 | 0.62 | 0.47 | 0.24 | 0.64 | 0.14 | 0.75 | 0 | 0.48 | 0.75 | 0.42 | 0.39 | 0.42 | 0.95 | 0.44 | 0.51 | -0.015 | 0.1125 | 0.1825 | 0.484497 | 0.953364 | 0.620309 | 0.999187 | 0.997988 | 0.994666 |
| Rtn3_Q9ES97_S510s_1 | Rtn3_Q9ES | 0.92 | 0.29 | -0.71 | 0.65 | 0.5 | 0.52 | 0.65 | 0.17 | 0.22 | 0.86 | -0.58 | -0.52 | -0.41 | 0.86 | -0.07 | -0.34 | 0.1725 | -0.2925 | -0.2775 | 0.540866 | 0.639922 | 0.538196 | 0.989223 | 0.997988 | 0.994666 |
| Rtn3_Q9ES97_S654s_1 | Rtn3_Q9ES | 0.34 | 0.82 | 0.47 | 0.39 | 0.67 | 0.09 | 0.66 | 0.38 | 0.17 | 0.56 | -0.13 | 0.53 | 0.05 | 0.35 | 0.26 | 0.25 | -0.055 | -0.2225 | -0.2775 | 0.276032 | 0.819057 | 0.40414 | 0.995762 | 0.997988 | 0.994666 |
| Rtn3_Q9ES97_S735s_1 | Rtn3_Q9ES | 0.27 | -0.56 | 0.09 | 0.16 | 0.46 | 0.41 | 0.6 | 0.52 | -0.64 | -0.35 | -0.35 | 0.19 | -0.66 | 0.38 | -0.14 | 0.16 | 0.5075 | -0.2775 | -0.055 | 0.868573 | 0.065704 | 0.361049 | 0.989223 | 0.997988 | 0.999507 |
| Rtn3_Q9ES97_T505t_1 | Rtn3_Q9ES | 0.41 | 0.88 | 0.34 | 0.42 | 0.4 | 0.06 | 0.42 | 0.45 | -0.15 | 0.17 | -0.02 | 0.18 | 0.26 | 0.18 | 0.31 | 0.47 | -0.18 | -0.4675 | -0.2075 | 0.416935 | 0.440835 | 0.073361 | 0.989223 | 0.997988 | 0.994666 |
| Rtn4_Q99P72_S145s_1 | Rtn4_Q99P | -0.85 | -0.74 | -0.98 | -0.96 | 0.09 | -0.64 | 0.05 | -0.84 | -1.34 | -0.55 | -0.59 | -0.96 | -0.65 | -0.04 | -0.89 | -0.52 | 0.5475 | 0.0225 | 0.3575 | 0.210566 | 0.075228 | 0.930849 | 0.989223 | 0.997988 | 0.994666 |
| Rtn4_Q99P72_S165s_1 | Rtn4_Q99P | 0.1 | 0.45 | -0.28 | -0.32 | 0.52 | 0.05 | 0.36 | -0.17 | 0.14 | 0.32 | -0.79 | -0.1 | -0.24 | 0.59 | -0.11 | -0.02 | 0.2025 | -0.095 | 0.0675 | 0.828378 | 0.469951 | 0.772609 | 0.989223 | 0.997988 | 0.999507 |
| Rtn4_Q99P72_S167s_1 | Rtn4_Q99P | 0.85 | 0.9 | 0.2 | 0.37 | 0.75 | 0.35 | 0.63 | 0.31 | 0.9 | 1.08 | 0.23 | -0.14 | 0.45 | 0.8 | 0.13 | 0.12 | -0.07 | -0.0625 | -0.205 | 0.498637 | 0.785017 | 0.858248 | 0.995762 | 0.997988 | 0.994666 |
| Rtn4_Q99P72_S16s_1 | Rtn4_Q99P | 0.29 | 0.68 | 0.37 | 0.33 | 0.67 | -0.02 | 0.82 | 0.33 | 0.19 | 0.63 | 0.2 | 0.42 | 0.27 | 0.8 | 0.46 | 0.5 | 0.0325 | -0.0575 | 0.09 | 0.721912 | 0.899687 | 0.804329 | 0.996493 | 0.997988 | 0.9971 |
| Rtn4_Q99P72_S223s_1 | Rtn4_Q99P | 0.9 | -0.81 | -0.69 | 0.44 | 0.66 | 0.7 | 0 | 1.08 | -1.56 | -0.46 | -1.02 | 0.33 | -2.29 | 0.53 | -1.3 | 0.09 | 0.65 | -0.6375 | -0.7025 | 0.32316 | 0.168875 | 0.257974 | 0.989223 | 0.997988 | 0.994666 |
| Rtn4_Q99P72_S329s_1 | Rtn4_Q99P | -0.36 | 0 | -0.13 | -0.51 | 0.11 | -1.23 | 0.16 | -0.6 | -0.81 | -0.51 | 0.25 | -0.32 | -1.26 | -0.21 | -0.51 | -0.1 | -0.14 | -0.0975 | -0.27 | 0.418721 | 0.690694 | 0.740593 | 0.994893 | 0.997988 | 0.994666 |
| Rtn4_Q99P72_S344s_1 | Rtn4_Q99P | 0.49 | 0.77 | 1.41 | 1.39 | 0.63 | -0.04 | 1.11 | 1.15 | 0.17 | 0.52 | 1.59 | 1.12 | 0.4 | 0.77 | 1.22 | 1.11 | -0.3025 | -0.165 | -0.14 | 0.676176 | 0.40815 | 0.674008 | 0.989223 | 0.997988 | 0.996159 |
| Rtn4_Q99P72_S690s_1 | Rtn4_Q99P | 0.55 | 0 | 0.29 | 0.51 | 0.79 | 0.49 | 0.87 | 0.43 | 0.23 | 0.33 | 0.31 | 0.29 | -0.48 | 0.73 | 0.46 | 0.47 | 0.3075 | -0.0475 | -0.0425 | 0.898513 | 0.212911 | 0.834986 | 0.989223 | 0.997988 | 0.999507 |
| Rtn4_Q99P72_S768s_1 | Rtn4_Q99P | -1.39 | -1.36 | -1.49 | -0.81 | -0.71 | -1.6 | -0.12 | -0.47 | -1.34 | -1.19 | -1.3 | -1.09 | -0.76 | -0.74 | -0.75 | -0.92 | 0.5375 | 0.0325 | 0.47 | 0.095714 | 0.151286 | 0.894034 | 0.989223 | 0.997988 | 0.994666 |
| Rtn4_Q99P72_S857s_1 | Rtn4_Q99P | 4.49 | 3.85 | 3.49 | 2.22 | 3.68 | 4.47 | 3.2 | 3.35 | 0.47 | 4.62 | 1.32 | 3.18 | 0.37 | 0.47 | 3.57 | 4 | 0.1625 | -1.115 | -1.41 | 0.160769 | 0.749434 | 0.248411 | 0.995762 | 0.997988 | 0.994666 |
| Rtn4_Q99P72_S953s_1 | Rtn4_Q99P | -3.75 | -3.99 | -3.52 | -5.33 | -2.71 | -5.02 | -3.78 | -4.21 | -4.19 | -2.77 | -2.72 | -3.54 | -2.15 | -4.03 | -3.23 | -3.73 | 0.2175 | 0.8425 | 0.8625 | 0.134116 | 0.701943 | 0.12057 | 0.995437 | 0.997988 | 0.994666 |
| Rtn4_Q99P72_S961s_1 | Rtn4_Q99P | -0.48 | -0.29 | -0.24 | -0.79 | 0.14 | -1.86 | -0.91 | -1.79 | -1.37 | -0.16 | -2.3 | -0.96 | 0.08 | -0.08 | -1.32 | -0.72 | -0.655 | -0.7475 | -0.06 | 0.869681 | 0.170528 | 0.119919 | 0.989223 | 0.997988 | 0.999507 |
| Rtn4_Q99P72_T430t_1 | Rtn4_Q99P | 2.79 | 0.86 | 0.95 | 1.61 | 1.06 | 1.73 | 1.41 | 1.58 | 0.64 | 2.03 | 0.8 | 1.17 | -0.24 | 0.43 | 1.17 | 1.1 | -0.1075 | -0.3925 | -0.9375 | 0.096732 | 0.806859 | 0.448457 | 0.995762 | 0.997988 | 0.994666 |
| Rtn4_Q99P72_T488t_1 | Rtn4_Q99P | 2.91 | 1.14 | 1.93 | 2.15 | 0.48 | 1.96 | 1.51 | 2.51 | 2.7 | 3.09 | 3.03 | 1.76 | 1.64 | 1.42 | 1.93 | 0.98 | -0.4175 | 0.6125 | -0.54 | 0.214371 | 0.425025 | 0.202127 | 0.989223 | 0.997988 | 0.994666 |
| Rufy3_Q9D394_S27s_1 | Rufy3_Q9D | -0.49 | -0.59 | -0.31 | -0.98 | -0.41 | -0.28 | -1.26 | -0.09 | -0.05 | -0.22 | 0.3 | 0.48 | -0.38 | 0.42 | -1.33 | -1.71 | 0.0825 | 0.72 | -0.1575 | 0.742565 | 0.792209 | 0.023905 | 0.995762 | 0.997988 | 0.9971 |
| Rufy3_Q9D394_T51t_1 | Rufy3_Q9D | -0.14 | 0.23 | 0 | 0.19 | 0.48 | -0.16 | 0.39 | 0.26 | -0.46 | 0.34 | -0.37 | 0.17 | -0.57 | 0.34 | 0.35 | 0.55 | 0.1725 | -0.15 | 0.0975 | 0.757633 | 0.470275 | 0.584703 | 0.989223 | 0.997988 | 0.9971 |
| Rufy3_Q9D394_T5t_1 | Rufy3_Q9D | -0.05 | 0.53 | 0.57 | 0.24 | 0.51 | -0.09 | 0.7 | 0.19 | 0.09 | 0.24 | 0.21 | 0.4 | 0.42 | 0.65 | 0.5 | 0.52 | 0.005 | -0.0875 | 0.2 | 0.443236 | 0.985166 | 0.717751 | 0.999187 | 0.997988 | 0.994666 |
| Rundc3a_O08576_S361s_1 | Rundc3a_O | 0.97 | 0.57 | 0.48 | 0.29 | 1.3 | 1.14 | 0.85 | 0.6 | -0.14 | 0.73 | -0.01 | 0.65 | -0.21 | 0.28 | 1.69 | 1.02 | 0.395 | -0.27 | 0.1175 | 0.786906 | 0.152946 | 0.383418 | 0.989223 | 0.997988 | 0.999507 |
| Rusc1_Q8BG26_S367s_1 | Rusc1_Q8B | -0.58 | -0.98 | 0.49 | 0.56 | -0.96 | -0.79 | 1.04 | 0.47 | -1.23 | -0.42 | 0.69 | 0.3 | 0.38 | 0.12 | -0.25 | 0.64 | 0.0675 | -0.0375 | 0.35 | 0.419998 | 0.90389 | 0.943862 | 0.996493 | 0.997988 | 0.994666 |
| Rusc1_Q8BG26_T355t_1 | Rusc1_Q8B | 1.4 | 1.09 | -0.08 | 0.68 | 0.57 | 0.41 | 0.46 | 0.78 | 0.62 | 0.82 | 0.71 | 0.46 | 0.78 | 1.47 | 0.53 | 0.23 | -0.2175 | -0.12 | -0.02 | 0.961563 | 0.524029 | 0.72957 | 0.989223 | 0.997988 | 0.999507 |
| Rusc2_Q3V1Z0_S242s_1 | Rusc2_Q3V | -0.48 | -0.47 | -0.09 | 0.58 | -0.97 | -0.55 | 0.06 | 0.06 | 0.16 | -0.3 | 0.28 | 0.28 | 0.68 | -1.76 | 0.65 | 1.09 | -0.235 | 0.22 | 0.28 | 0.658368 | 0.512204 | 0.491721 | 0.989223 | 0.997988 | 0.995259 |
| Rusc2_Q3V1Z0_S542s_1 | Rusc2_Q3V | 0.92 | 1 | 1.23 | 0.6 | 0.5 | 0.46 | 0.8 | 0.92 | 0.17 | 0.75 | -0.12 | 0.78 | -0.17 | -0.16 | 1.18 | 1.41 | -0.2675 | -0.5425 | -0.3725 | 0.40242 | 0.278922 | 0.09477 | 0.989223 | 0.997988 | 0.994666 |
| Rusc2_Q3V1Z0_S565s_1 | Rusc2_Q3V | 0.06 | 0.4 | 0.76 | 0.36 | 0.2 | -0.31 | 0.21 | 0.03 | 0.04 | 0.19 | 0.86 | 0.36 | 1.01 | 0.04 | 0.67 | 0.49 | -0.3625 | -0.0325 | 0.1575 | 0.608277 | 0.165643 | 0.907614 | 0.989223 | 0.997988 | 0.995259 |
| Rusc2_Q3V1Z0_S580s_1 | Rusc2_Q3V | 0.66 | 0.64 | 0.52 | 0.17 | 0.76 | -0.07 | 0.48 | 0.44 | 0.15 | 0.48 | 0.95 | 0.89 | 0.14 | 0.36 | 0.41 | -0.12 | -0.095 | 0.12 | -0.3 | 0.270186 | 0.71315 | 0.663544 | 0.995437 | 0.997988 | 0.994666 |
| Rusc2_Q3V1Z0_S779s_1 | Rusc2_Q3V | 0.3 | 0.63 | 0.94 | 0.95 | 0.64 | 0.52 | 0.65 | 0.95 | 1.09 | 0.77 | 0.3 | 0.11 | 0.5 | 0.55 | 1.28 | 0.83 | -0.015 | -0.1375 | 0.085 | 0.776149 | 0.950628 | 0.655499 | 0.999187 | 0.997988 | 0.99798 |
| Rusc2_Q3V1Z0_S785s_1 | Rusc2_Q3V | 0.77 | 0.94 | 1.27 | 1.14 | 0.92 | 0.58 | 0.95 | 1 | 0.65 | 1.16 | 1.42 | 1.22 | 0.84 | 1.38 | 1.44 | 1.39 | -0.1675 | 0.0825 | 0.2325 | 0.396777 | 0.465004 | 0.753341 | 0.989223 | 0.997988 | 0.994666 |
| Ryr2_E9Q401_S1495s_1 | Ryr2_E9Q4 | 0.16 | 0.12 | 0.07 | 0.15 | -0.82 | 0.16 | -0.37 | 0.55 | 0.31 | -0.29 | 0.57 | 0.02 | 0.62 | -0.57 | 0.13 | 0.35 | -0.245 | 0.0275 | 0.0075 | 0.980665 | 0.446794 | 0.914411 | 0.989223 | 0.997988 | 1 |
| Ryr2_E9Q401_S2362s_1 | Ryr2_E9Q4 | 2.42 | 2.77 | 2.19 | 1.91 | 0.89 | 3.48 | 0.99 | 1.76 | 2.86 | 2.76 | 0.9 | 2.25 | 2.82 | -0.66 | 2.19 | 1.51 | -0.5425 | -0.13 | -0.8575 | 0.23754 | 0.348229 | 0.77994 | 0.989223 | 0.997988 | 0.994666 |
| Ryr2_E9Q401_S2367s_1 | Ryr2_E9Q4 | 0.01 | -0.17 | 1.06 | 0.72 | 0.14 | 0.73 | -0.02 | -0.04 | 1.45 | -0.63 | -0.3 | 0.11 | 1.45 | -0.97 | 1.22 | 0.4 | -0.2025 | -0.2475 | 0.12 | 0.833344 | 0.561826 | 0.628355 | 0.989223 | 0.997988 | 0.999507 |
| Ryr2_E9Q401_S2693s_1 | Ryr2_E9Q4 | 0.02 | 0.46 | 0.25 | -0.15 | 0.21 | 0.03 | 0.52 | -0.1 | -0.08 | 0.12 | -0.6 | -0.13 | 0.18 | 0.62 | -0.16 | 0.09 | 0.02 | -0.3175 | 0.0375 | 0.895058 | 0.935621 | 0.249285 | 0.99868 | 0.997988 | 0.999507 |
| Ryr2_E9Q401_S2807s_1 | Ryr2_E9Q4 | 0.18 | -0.09 | -1.41 | -0.24 | 0.17 | -0.36 | -0.5 | 0.45 | 0.4 | -0.06 | -1.88 | -0.69 | 0.3 | 0.22 | -1.14 | -0.61 | 0.33 | -0.1675 | 0.0825 | 0.861373 | 0.417882 | 0.765341 | 0.989223 | 0.997988 | 0.999507 |
| Ryr2_E9Q401_S2813s_1 | Ryr2_E9Q4 | 0.11 | 0.09 | -0.98 | -0.5 | -0.16 | -0.18 | -0.54 | -0.59 | -1.05 | -0.03 | -2.13 | -0.46 | -0.08 | 0.46 | -0.89 | -0.45 | -0.0475 | -0.5975 | 0.08 | 0.839868 | 0.876881 | 0.246572 | 0.996493 | 0.997988 | 0.999507 |
| Ryr2_E9Q401_S4549s_1 | Ryr2_E9Q4 | 0.55 | 0.18 | -0.05 | -0.26 | -1.2 | 1.51 | -0.63 | -0.53 | 0.88 | 0.36 | -0.21 | -0.44 | 1.17 | 0.38 | -0.62 | 0.13 | -0.3175 | 0.0425 | 0.16 | 0.697391 | 0.572025 | 0.905033 | 0.989223 | 0.997988 | 0.996159 |
| S1pr2_P52592_S332s_1 | S1pr2_P52 | -0.3 | -0.23 | 0.58 | 0.32 | -1.12 | 0.65 | -0.55 | -0.47 | 1.64 | 0.39 | -1.88 | 0.37 | 1.46 | -1.68 | 1.12 | 0.07 | -0.465 | 0.0375 | 0.15 | 0.821282 | 0.271986 | 0.956572 | 0.989223 | 0.997988 | 0.999507 |
| S1pr5_Q91X56_S340s_1 | S1pr5_Q91 | 1.13 | 1.16 | 1.34 | 0.91 | 0.7 | 0.18 | 0.29 | 0.62 | 0.67 | 1.07 | 1.59 | 0.74 | 1.22 | 0.71 | 1.04 | 0.83 | -0.6875 | -0.1175 | -0.185 | 0.471038 | 0.013249 | 0.675546 | 0.989223 | 0.997988 | 0.994666 |
| S1pr5_Q91X56_S342s_1 | S1pr5_Q91 | 0.09 | 0.42 | 0 | -0.26 | 0.2 | -0.34 | -0.29 | -0.58 | -0.43 | -0.44 | -0.48 | -0.77 | 0.33 | -0.29 | -0.93 | -0.31 | -0.315 | -0.5925 | -0.3625 | 0.291288 | 0.248728 | 0.034463 | 0.989223 | 0.997988 | 0.994666 |
| Sacs_E9QNY8_S1781s_1 | Sacs_E9QN | 0.04 | 0.12 | 0.22 | -0.1 | 1.05 | 0.26 | 0.39 | 0.3 | -0.44 | 0.22 | -1.12 | 0.08 | -1.2 | -0.23 | -0.06 | 0.75 | 0.43 | -0.385 | -0.255 | 0.536934 | 0.113585 | 0.266184 | 0.989223 | 0.997988 | 0.994666 |
| Sacs_E9QNY8_S2513s_1 | Sacs_E9QN | -2.05 | -1.58 | -3.02 | -2.65 | -1.03 | -1.05 | -2.21 | -3.87 | -2.11 | -0.94 | -2 | -3.36 | -1.76 | -1.22 | -2.94 | -3.24 | 0.285 | 0.2225 | 0.035 | 0.947859 | 0.665904 | 0.685033 | 0.991876 | 0.997988 | 0.999507 |
| Sacs_E9QNY8_S4266s_1 | Sacs_E9QN | -0.53 | -0.62 | 0.76 | 0.95 | 0.39 | 0.25 | 0.54 | 0.76 | -0.13 | 0.81 | 0.23 | 0.82 | 0.45 | 0.37 | 0.32 | 0.47 | 0.345 | 0.2925 | 0.2625 | 0.532741 | 0.409079 | 0.526316 | 0.989223 | 0.997988 | 0.994666 |
| Safb_S4R1M2_S298s_1 | Safb_S4R1 | -1 | -2 | -2.28 | -3.34 | -2.62 | -0.75 | -1.68 | -1.59 | 2.18 | -2.06 | -3.07 | -1.71 | 3.38 | -2.2 | 2.2 | -2.84 | 0.495 | 0.99 | 2.29 | 0.127367 | 0.382565 | 0.378218 | 0.989223 | 0.997988 | 0.994666 |
| Safb_S4R1M2_S366s_1 | Safb_S4R1 | -0.03 | -0.12 | 0.73 | 0.77 | -0.06 | -0.2 | 0.39 | 0.8 | -0.52 | -0.27 | 0.37 | 0.51 | -0.2 | -0.03 | 0.96 | 0.47 | -0.105 | -0.315 | -0.0375 | 0.91986 | 0.755861 | 0.388202 | 0.995762 | 0.997988 | 0.999507 |
| Safb_S4R1M2_S626s_1 | Safb_S4R1 | -1.01 | -0.76 | -1.41 | -2.04 | -0.91 | -2.32 | -1.54 | -2.16 | -2.49 | -0.89 | -2.07 | -2.13 | -1.72 | -1.7 | -0.01 | -1.49 | -0.4275 | -0.59 | 0.075 | 0.874555 | 0.308644 | 0.195674 | 0.989223 | 0.997988 | 0.999507 |
| Safb_S4R1M2_S783s_1 | Safb_S4R1 | -0.8 | -0.18 | 0.93 | 0.12 | -0.93 | 0.56 | -0.63 | 1.39 | 1.03 | -0.96 | 1.16 | -0.26 | 1.06 | -0.88 | 1.23 | 1.27 | 0.08 | 0.225 | 0.6525 | 0.276496 | 0.890333 | 0.697528 | 0.996493 | 0.997988 | 0.994666 |
| Samd1_D3YXK1_S150s_1 | Samd1_D3 | -1.22 | -0.46 | -1 | 0.54 | -0.45 | -0.99 | -0.7 | -1.19 | -0.41 | -0.75 | 0.67 | -0.84 | 0.11 | -0.68 | 0.03 | -0.5 | -0.2975 | 0.2025 | 0.275 | 0.528822 | 0.470685 | 0.683089 | 0.989223 | 0.997988 | 0.994666 |
| Samd1_D3YXK1_S242s_1 | Samd1_D3 | -2.32 | -1.36 | -0.31 | -1.81 | -1.77 | -2.27 | -2.43 | -1.65 | -1.97 | -1.3 | -1.01 | -2.07 | 0.94 | -2.22 | -1.11 | -1.21 | -0.58 | -0.1375 | 0.55 | 0.44277 | 0.20763 | 0.772398 | 0.989223 | 0.997988 | 0.994666 |
| Samd1_D3YXK1_T107t_1 | Samd1_D3 | -0.28 | -0.78 | -0.65 | -0.3 | -0.91 | -0.35 | -0.5 | -0.76 | -0.15 | -0.41 | 0.2 | -0.03 | 0.91 | -0.73 | -0.32 | -0.55 | -0.1275 | 0.405 | 0.33 | 0.416116 | 0.600544 | 0.133374 | 0.989223 | 0.997988 | 0.994666 |
| Samd14_Q8K070_S108s_1 | Samd14_Q | -0.4 | 0.42 | -0.35 | -0.21 | 1.22 | -0.43 | 0.09 | -0.04 | 0.71 | 0.37 | -0.34 | -0.6 | 0.64 | 0.35 | -0.67 | -0.38 | 0.345 | 0.17 | 0.12 | 0.750488 | 0.385815 | 0.645282 | 0.989223 | 0.997988 | 0.9971 |
| Samd14_Q8K070_S140s_1 | Samd14_Q | 0.65 | 1.13 | -1.23 | -0.51 | 0.79 | 0.56 | -0.17 | -0.94 | 1 | 1.08 | -1.18 | -1.09 | 0.76 | 1 | -0.33 | -0.57 | 0.05 | -0.0575 | 0.205 | 0.735589 | 0.932996 | 0.938207 | 0.998562 | 0.997988 | 0.9971 |
| Samd14_Q8K070_S151s_1 | Samd14_Q | 0.03 | -0.68 | -0.22 | -0.19 | -0.12 | -0.14 | 0.27 | -0.1 | -0.13 | 0.14 | 0.04 | -0.01 | -0.88 | -1.71 | -0.59 | -0.26 | 0.2425 | 0.275 | -0.595 | 0.128952 | 0.329507 | 0.273182 | 0.989223 | 0.997988 | 0.994666 |
| Samd14_Q8K070_S165s_1 | Samd14_Q | 0.27 | -0.79 | 0.37 | -0.43 | -0.07 | 0.68 | 0.28 | -0.03 | 1.69 | 0.23 | -0.27 | 0.37 | 1.53 | -0.15 | 0.72 | -0.51 | 0.36 | 0.65 | 0.5425 | 0.298206 | 0.299505 | 0.195991 | 0.989223 | 0.997988 | 0.994666 |
| Samd14_Q8K070_S173s_1 | Samd14_Q | 1.87 | 2.33 | 1.96 | 1.11 | 2.13 | 1.79 | 2.45 | 1.51 | 0.85 | 1.85 | 1.64 | 1.97 | 0 | -0.16 | 2.18 | 1.74 | 0.1525 | -0.24 | -0.8775 | 0.162911 | 0.651002 | 0.518984 | 0.990358 | 0.997988 | 0.994666 |
| Samd14_Q8K070_S256s_1 | Samd14_Q | 1.55 | -0.13 | -0.19 | 0.3 | 0.14 | 1.33 | -0.73 | -0.18 | 2.76 | 0.43 | -0.73 | 0.19 | 1.56 | -0.2 | -0.27 | -0.16 | -0.2425 | 0.28 | -0.15 | 0.787097 | 0.653981 | 0.712547 | 0.990434 | 0.997988 | 0.999507 |
| Samd14_Q8K070_S260s_1 | Samd14_Q | 0.91 | 0.89 | 0.42 | 0.9 | 0.5 | 0.29 | 0.07 | 0.09 | 2.49 | 1.03 | -0.97 | 0.59 | 2.72 | 0.48 | 1.25 | 0.69 | -0.5425 | 0.005 | 0.505 | 0.31933 | 0.039065 | 0.993905 | 0.989223 | 0.997988 | 0.994666 |
| Samd14_Q8K070_S77s_1 | Samd14_Q | -0.88 | -2.9 | -0.24 | -0.04 | -1.29 | -2.52 | -0.5 | -0.04 | -2.86 | -1.02 | -0.71 | -0.53 | -1.06 | -3.23 | -0.31 | 0.03 | -0.0725 | -0.265 | -0.1275 | 0.882338 | 0.922462 | 0.727172 | 0.997335 | 0.997988 | 0.999507 |
| Samd4b_G5E8A7_S271s_1 | Samd4b_G | 0.1 | -0.21 | 0.26 | 0.46 | 0.23 | -0.08 | 0.6 | 0.3 | -0.05 | 0.34 | -0.15 | 0.1 | -0.18 | 0.39 | 0.15 | 0.78 | 0.11 | -0.0925 | 0.1325 | 0.665078 | 0.665264 | 0.713408 | 0.991354 | 0.997988 | 0.996159 |
| Samd8_D6REI9_S15s_1 | Samd8_D6 | -0.19 | -0.7 | -2.01 | -1.19 | -0.13 | -0.48 | -0.47 | -1.03 | -0.31 | -1.36 | 0.27 | -1.15 | -0.18 | -0.1 | -1.39 | -1.26 | 0.495 | 0.385 | 0.29 | 0.558851 | 0.246227 | 0.455908 | 0.989223 | 0.997988 | 0.995259 |
| Sap130_J3QNK5_S274s_1 | Sap130_J3 | 0.54 | 0.5 | -1.71 | -0.99 | 0.09 | -0.3 | -1.09 | -1.29 | -0.87 | -0.21 | -1.75 | -1.43 | 0.08 | 0.13 | -0.53 | -1.06 | -0.2325 | -0.65 | 0.07 | 0.903151 | 0.690119 | 0.293872 | 0.994672 | 0.997988 | 0.999507 |
| Sarm1_Q6PDS3_S548s_1 | Sarm1_Q6 | 0.11 | 0.56 | -0.21 | -0.06 | 0.44 | 0.83 | 0.78 | 0.09 | 0.75 | 0.75 | 1.07 | -0.52 | 0.76 | 0.4 | 0.84 | 1.24 | 0.435 | 0.4125 | 0.71 | 0.038188 | 0.140889 | 0.305965 | 0.989223 | 0.997988 | 0.994666 |
| Sarm1_Q6PDS3_S558s_1 | Sarm1_Q6 | -0.06 | 0.73 | -0.05 | -0.15 | 0.17 | -0.13 | -0.25 | -0.08 | 0.81 | 0.56 | 0.56 | -0.17 | 1.01 | 0.57 | -0.31 | 0.39 | -0.19 | 0.3225 | 0.2975 | 0.423466 | 0.483473 | 0.331183 | 0.989223 | 0.997988 | 0.994666 |
| Sarm1_Q6PDS3_S572s_1 | Sarm1_Q6 | -0.05 | 0.67 | -0.06 | -0.62 | 1.13 | -1.08 | 0.28 | -0.32 | -1.1 | 0.26 | -0.91 | 0.66 | 0.2 | -0.3 | -0.18 | -0.17 | 0.0175 | -0.2575 | -0.0975 | 0.76666 | 0.971672 | 0.595057 | 0.999187 | 0.997988 | 0.9971 |
| Sash1_F8VQK5_S1020s_1 | Sash1_F8V | -0.09 | -0.29 | 0.39 | -0.38 | -0.11 | 0.12 | 0.35 | 0.06 | -0.44 | 0.08 | 0.75 | 1.03 | 0.09 | -0.43 | -0.08 | 0.01 | 0.1975 | 0.4475 | -0.01 | 0.971737 | 0.440454 | 0.255378 | 0.989223 | 0.997988 | 0.999507 |
| Sash1_F8VQK5_S241s_1 | Sash1_F8V | -0.67 | 0.74 | 0.38 | 0.13 | -1.91 | 0.17 | -1.32 | -0.78 | 2.56 | 1.58 | -0.02 | -0.29 | 0.46 | 1.75 | -0.37 | -0.08 | -1.105 | 0.8125 | 0.295 | 0.574657 | 0.046016 | 0.243592 | 0.989223 | 0.997988 | 0.995259 |
| Sash1_F8VQK5_S500s_1 | Sash1_F8V | 0.63 | 0.57 | 1.09 | 0.77 | 0.96 | 0.85 | 0.74 | 1.2 | 0.55 | 0.71 | 1.5 | 0.84 | 0.52 | 1.16 | 1.21 | 1.1 | 0.1725 | 0.135 | 0.2325 | 0.412289 | 0.45796 | 0.6392 | 0.989223 | 0.997988 | 0.994666 |
| Sash1_F8VQK5_S723s_1 | Sash1_F8V | 0.66 | -0.01 | 0.31 | 0.27 | 0.12 | 0.37 | -0.39 | 0.71 | -0.41 | -0.07 | 1.17 | 0.2 | -0.38 | -1.08 | 0.29 | 0.67 | -0.105 | -0.085 | -0.4325 | 0.307212 | 0.723497 | 0.819864 | 0.995437 | 0.997988 | 0.994666 |
| Sash1_F8VQK5_S805s_1 | Sash1_F8V | -0.41 | -1.69 | -2.52 | -1.96 | 0.21 | -1.38 | 0.42 | -1.31 | -1.36 | -0.76 | -0.58 | -3.28 | -1.35 | -1.01 | -0.01 | -2.18 | 1.13 | 0.15 | 0.5075 | 0.392258 | 0.079977 | 0.82759 | 0.989223 | 0.997988 | 0.994666 |
| Sash1_F8VQK5_T279t_1 | Sash1_F8V | 0.12 | 0.79 | 0.21 | 0.71 | 0.2 | 0.56 | 0.19 | -0.39 | 1.12 | 0.19 | 0.77 | -1.01 | -0.22 | 0.95 | 0.09 | 0.42 | -0.3175 | -0.19 | -0.1475 | 0.664786 | 0.28955 | 0.689159 | 0.989223 | 0.997988 | 0.996159 |
| Satb1_E9PVB7_S669s_1 | Satb1_E9P | -3.31 | -1.98 | -3.1 | -2.07 | -2.56 | -2.63 | -2.89 | -2.2 | -2.6 | -1.86 | -2.64 | -3.21 | 0.72 | -2.48 | -2.41 | -3.09 | 0.045 | 0.0375 | 0.8 | 0.339387 | 0.90249 | 0.930398 | 0.996493 | 0.997988 | 0.994666 |
| Satb2_H3BKH3_S13s_1 | Satb2_H3B | -4.87 | -3.67 | -3.46 | -5.34 | -4.6 | -5.88 | -3.98 | -3.86 | -3.66 | -3.61 | -2.08 | -2.81 | -0.47 | -2.74 | -2.54 | -3.27 | -0.245 | 1.295 | 2.08 | 0.012612 | 0.675699 | 0.039017 | 0.992876 | 0.997988 | 0.994666 |
| Satb2_H3BKH3_S13s_2 | Satb2_H3B | -4.16 | -1.98 | -3.6 | -5.3 | -2.3 | -3.3 | -6.48 | -5.56 | -0.47 | -2.3 | -1.21 | -4.2 | 0.02 | -3.31 | -2.57 | -2.48 | -0.65 | 1.715 | 1.675 | 0.081935 | 0.531829 | 0.095309 | 0.989223 | 0.997988 | 0.994666 |
| Satb2_H3BKH3_S7s_1 | Satb2_H3B | -3.04 | -2.95 | -4.49 | -9.73 | -2.1 | -3.32 | -4.16 | -7.43 | -2.09 | -2.49 | -2.66 | -4.75 | -1.4 | -2.31 | -5.02 | -4.03 | 0.8 | 2.055 | 1.8625 | 0.24522 | 0.635519 | 0.190569 | 0.989223 | 0.997988 | 0.994666 |
| Satb2_H3BKH3_S7s_2 | Satb2_H3B | -4.16 | -1.98 | -3.6 | -5.3 | -2.3 | -3.3 | -6.48 | -5.56 | -0.47 | -2.3 | -1.21 | -4.2 | 0.02 | -3.31 | -2.57 | -2.48 | -0.65 | 1.715 | 1.675 | 0.081935 | 0.531829 | 0.095309 | 0.989223 | 0.997988 | 0.994666 |
| Saysd1_Q8K190_S16s_1 | Saysd1_Q8 | -0.74 | -0.51 | 0.07 | 0.3 | -0.68 | -0.25 | -0.14 | -0.2 | -0.32 | -1.28 | 0.1 | -0.53 | -0.49 | -0.31 | -0.79 | -0.34 | -0.0975 | -0.2875 | -0.2625 | 0.415528 | 0.744085 | 0.458099 | 0.995762 | 0.997988 | 0.994666 |
| Sbf1_Q6ZPE2_S1141s_1 | Sbf1_Q6ZP | 0.04 | -0.28 | -0.42 | -0.8 | -0.06 | -0.44 | -0.17 | -0.93 | -1.42 | -0.64 | -1.22 | -0.51 | -0.9 | 0.2 | -0.8 | -0.87 | -0.035 | -0.5825 | -0.2275 | 0.518058 | 0.904015 | 0.089103 | 0.996493 | 0.997988 | 0.994666 |
| Sbf1_Q6ZPE2_S1228s_1 | Sbf1_Q6ZP | 0.08 | 0.06 | -0.4 | 0.04 | 0.07 | -0.27 | 0.16 | 0.51 | -0.29 | -0.01 | 0.28 | -0.28 | -0.69 | -0.36 | -0.64 | 0.57 | 0.1725 | -0.02 | -0.225 | 0.520543 | 0.499641 | 0.936504 | 0.989223 | 0.997988 | 0.994666 |
| Sbf1_Q6ZPE2_S1746s_1 | Sbf1_Q6ZP | -0.4 | 0.42 | 1.29 | 0.96 | 0.62 | 0.53 | 2.06 | 1.25 | 0.3 | -0.27 | 0.85 | -1.93 | -0.12 | 0.68 | -0.56 | 2.42 | 0.5475 | -0.83 | 0.0375 | 0.955768 | 0.264124 | 0.216717 | 0.989223 | 0.997988 | 0.999507 |
| Sbf1_Q6ZPE2_S559s_1 | Sbf1_Q6ZP | -1.04 | -0.83 | -0.18 | -1.58 | -2.12 | -0.91 | -0.63 | -1.58 | -1.1 | -0.47 | -0.64 | -0.86 | 0.33 | -1.96 | 0.08 | -0.21 | -0.4025 | 0.14 | 0.4675 | 0.40475 | 0.350589 | 0.681653 | 0.989223 | 0.997988 | 0.994666 |
| Sbf1_Q6ZPE2_T1137t_1 | Sbf1_Q6ZP | -1.49 | -0.35 | -1.32 | -1.01 | -0.59 | -2.79 | -0.28 | -2.02 | -2.85 | -2.02 | -0.91 | -0.76 | -1.14 | -1.81 | -1.45 | -1.13 | -0.3775 | -0.5925 | -0.34 | 0.324972 | 0.51887 | 0.269994 | 0.989223 | 0.997988 | 0.994666 |
| Sbf1_Q6ZPE2_T1819t_1 | Sbf1_Q6ZP | 0.29 | -0.16 | 0.42 | 0.4 | -0.42 | -0.19 | 0.44 | -0.47 | 0.71 | 0.06 | -0.32 | 0.44 | -0.01 | 1 | 0.59 | 0.15 | -0.3975 | -0.015 | 0.195 | 0.540544 | 0.182741 | 0.960246 | 0.989223 | 0.997988 | 0.994666 |
| Sbf2_E9Q0D4_T1077t_1 | Sbf2_E9Q0 | 1.25 | 1.58 | 0.82 | 1.74 | 1.52 | 0.87 | 1.47 | 1.23 | 0.7 | 1.14 | 1.01 | 1.58 | 1 | 1.11 | 1.44 | 0.75 | -0.075 | -0.24 | -0.2725 | 0.383043 | 0.792467 | 0.443954 | 0.995762 | 0.997988 | 0.994666 |
| Scaf1_Q5U4C3_S240s_1 | Scaf1_Q5U | 0.45 | 0.24 | -0.47 | 0.08 | 0.5 | -0.16 | 0.62 | 0.27 | -0.6 | 0.11 | -1.2 | -0.35 | -0.44 | -0.28 | 0.28 | -0.06 | 0.2325 | -0.585 | -0.2 | 0.519694 | 0.432157 | 0.122072 | 0.989223 | 0.997988 | 0.994666 |
| Scaf1_Q5U4C3_S491s_1 | Scaf1_Q5U | -1.88 | -2.33 | -1.63 | -1.99 | -1.56 | -3.21 | -2.66 | -1.66 | -1.63 | -1.93 | -1.06 | -1.56 | -0.42 | -1.75 | -1.66 | -1.23 | -0.315 | 0.4125 | 0.6925 | 0.079987 | 0.445961 | 0.167458 | 0.989223 | 0.997988 | 0.994666 |
| Scaf1_Q5U4C3_S491s_2 | Scaf1_Q5U | 0.68 | -0.16 | -0.15 | -0.16 | -0.79 | 0.33 | -0.56 | 0.25 | -0.18 | -0.34 | -0.99 | -0.77 | -0.62 | -0.09 | -0.09 | -0.05 | -0.245 | -0.6225 | -0.265 | 0.396576 | 0.493115 | 0.072275 | 0.989223 | 0.997988 | 0.994666 |
| Scaf1_Q5U4C3_S493s_1 | Scaf1_Q5U | 0.5 | 0.13 | 1.49 | 1.62 | -0.15 | 0.24 | 0.69 | 1.53 | -0.2 | 0.41 | -0.01 | 1.34 | 1.25 | -0.01 | 1.6 | 1.71 | -0.3575 | -0.55 | 0.2025 | 0.691125 | 0.459154 | 0.266717 | 0.989223 | 0.997988 | 0.996159 |
| Scaf1_Q5U4C3_S493s_2 | Scaf1_Q5U | 0.68 | -0.16 | -0.15 | -0.16 | -0.79 | 0.33 | -0.56 | 0.25 | -0.18 | -0.34 | -0.99 | -0.77 | -0.62 | -0.09 | -0.09 | -0.05 | -0.245 | -0.6225 | -0.265 | 0.396576 | 0.493115 | 0.072275 | 0.989223 | 0.997988 | 0.994666 |
| Scaf1_Q5U4C3_S577s_2 | Scaf1_Q5U | 0.75 | 0.07 | -0.51 | -0.49 | -1.48 | 0.85 | -0.6 | -0.43 | 0.99 | -0.56 | -0.35 | -0.95 | -0.04 | -1.57 | 0.39 | -0.33 | -0.37 | -0.1725 | -0.3425 | 0.490096 | 0.478716 | 0.724245 | 0.989223 | 0.997988 | 0.994666 |
| Scaf1_Q5U4C3_S579s_2 | Scaf1_Q5U | 0.75 | 0.07 | -0.51 | -0.49 | -1.48 | 0.85 | -0.6 | -0.43 | 0.99 | -0.56 | -0.35 | -0.95 | -0.04 | -1.57 | 0.39 | -0.33 | -0.37 | -0.1725 | -0.3425 | 0.490096 | 0.478716 | 0.724245 | 0.989223 | 0.997988 | 0.994666 |
| Scaf1_Q5U4C3_S682s_1 | Scaf1_Q5U | 0.07 | -0.31 | -0.55 | -0.01 | 0.22 | -0.57 | -0.28 | 0.14 | 0.14 | 0.32 | 0.69 | -0.57 | 0.83 | 0.52 | -0.46 | -0.22 | 0.0775 | 0.345 | 0.3675 | 0.321184 | 0.777233 | 0.305571 | 0.995762 | 0.997988 | 0.994666 |
| Scaf1_Q5U4C3_S912s_1 | Scaf1_Q5U | 0.99 | -0.02 | 1 | 2.48 | 0.15 | 0.49 | 2.02 | 2.74 | -0.2 | -0.22 | 4.24 | 2.29 | -0.81 | 0.77 | 2.83 | 1.53 | 0.2375 | 0.415 | -0.0325 | 0.967962 | 0.737677 | 0.694657 | 0.995762 | 0.997988 | 0.999507 |
| Scaf1_Q5U4C3_T328t_1 | Scaf1_Q5U | 1.86 | 1.8 | 1.65 | 1.56 | 1.6 | 1.06 | 1.51 | 1.44 | 1.16 | 1.53 | 1.61 | 1.65 | -0.75 | 1.53 | 2.45 | 2.1 | -0.315 | -0.23 | -0.385 | 0.558498 | 0.178462 | 0.329548 | 0.989223 | 0.997988 | 0.995259 |
| Scaf1_Q5U4C3_T923t_1 | Scaf1_Q5U | -0.36 | -0.93 | -0.96 | -0.72 | -1.2 | -0.03 | -1.13 | -1.08 | 0.74 | -1.06 | -1.93 | -0.95 | 1.26 | -2.18 | -1.44 | 0.08 | -0.1175 | -0.0575 | 0.1725 | 0.805499 | 0.71718 | 0.914371 | 0.995437 | 0.997988 | 0.999507 |
| Scaf11_E9PZM7_S857s_1 | Scaf11_E9 | 0.25 | 0.41 | 0.41 | 0.51 | -0.01 | 0.19 | 0.49 | 0.29 | 0.44 | -0.15 | 0.77 | 0.27 | 0.76 | 0.32 | 1.17 | 0.43 | -0.155 | -0.0625 | 0.275 | 0.336002 | 0.474411 | 0.812151 | 0.989223 | 0.997988 | 0.994666 |
| Scaf11_E9PZM7_S859s_1 | Scaf11_E9 | -0.78 | -0.43 | -0.76 | -0.38 | -0.76 | -1.53 | -0.28 | -0.73 | -1.27 | -0.75 | -0.6 | -0.7 | -0.04 | 0.07 | -0.25 | -0.25 | -0.2375 | -0.2425 | 0.47 | 0.084034 | 0.440961 | 0.352908 | 0.989223 | 0.997988 | 0.994666 |
| Scaf4_Q7TSH6_S154s_1 | Scaf4_Q7T | -0.65 | -0.9 | -0.79 | -0.23 | -0.56 | -1.32 | -0.14 | -0.25 | -1.18 | -0.51 | 0.06 | -0.47 | -0.95 | -0.49 | -0.62 | -0.41 | 0.075 | 0.1175 | 0.025 | 0.927128 | 0.814278 | 0.715705 | 0.995762 | 0.997988 | 0.999507 |
| Scaf8_Q6DID3_S617s_1 | Scaf8_Q6D | 1.71 | 1.36 | 1 | 0.66 | 1.04 | 1.37 | 0.57 | 0.2 | 1.51 | 1.87 | 0.81 | 0.64 | 1.59 | 1.16 | 0.83 | 0.92 | -0.3875 | 0.025 | -0.0575 | 0.860209 | 0.279687 | 0.946715 | 0.989223 | 0.997988 | 0.999507 |
| Scai_B0R0E3_S2s_1 | Scai_B0R0 | 2.16 | 0.46 | -0.3 | -0.17 | 0.75 | 0.92 | -0.24 | -1.18 | 1.5 | 1 | 0.14 | 0.14 | 2.37 | 0.71 | 0.09 | 0.76 | -0.475 | 0.1575 | 0.445 | 0.512142 | 0.478563 | 0.794012 | 0.989223 | 0.997988 | 0.994666 |
| Scai_B0R0E3_S3s_1 | Scai_B0R0 | 2.16 | 0.46 | -0.3 | -0.17 | 0.75 | 0.92 | -0.24 | -1.18 | 1.5 | 1 | 0.14 | 0.14 | 2.37 | 0.71 | 0.09 | 0.76 | -0.475 | 0.1575 | 0.445 | 0.512142 | 0.478563 | 0.794012 | 0.989223 | 0.997988 | 0.994666 |
| Scamp1_Q8K021_S112s_1 | Scamp1_Q | 0.09 | 0.51 | 0.25 | 0.05 | 0.95 | 0.22 | -0.18 | 0.21 | 0.02 | 0.08 | -0.88 | -0.04 | 0.94 | -0.31 | 0.72 | 0.46 | 0.075 | -0.43 | 0.2275 | 0.498391 | 0.795239 | 0.166044 | 0.995762 | 0.997988 | 0.994666 |
| Scamp1_Q8K021_S295s_1 | Scamp1_Q | 1.19 | 0.83 | 0.23 | 0.28 | 0.8 | 0.75 | 0.4 | 0.82 | 0.54 | 0.66 | 0.83 | 0.5 | 0.61 | 0.64 | 0.68 | 0.87 | 0.06 | 5.89E-17 | 0.0675 | 0.821714 | 0.832732 | 1 | 0.995762 | 1 | 0.999507 |
| Scamp1_Q8K021_S50s_1 | Scamp1_Q | -1.56 | -0.55 | -0.82 | -1.5 | -1.09 | -0.45 | -1 | -1.22 | -1.4 | -1.57 | -2.03 | -2.11 | 0.69 | -0.71 | -1.14 | -0.47 | 0.1675 | -0.67 | 0.7 | 0.145514 | 0.600987 | 0.066742 | 0.989223 | 0.997988 | 0.994666 |
| Scamp1_Q8K021_T45t_1 | Scamp1_Q | -0.68 | -0.54 | -0.54 | -0.89 | -0.44 | -0.34 | -0.69 | -0.96 | -0.24 | -0.99 | -1.81 | -1.04 | 0.65 | -0.84 | -0.73 | -0.5 | 0.055 | -0.3575 | 0.3075 | 0.416367 | 0.813796 | 0.318196 | 0.995762 | 0.997988 | 0.994666 |
| Scamp3_E9Q855_S78s_1 | Scamp3_E9 | 0.31 | 0.38 | 0.4 | 0.58 | 0.8 | 0.3 | 0.47 | 0.81 | 0.18 | 0.27 | 1.01 | 0.44 | 0.52 | 0.7 | 0.88 | 1.02 | 0.1775 | 0.0575 | 0.3625 | 0.162777 | 0.433223 | 0.825491 | 0.989223 | 0.997988 | 0.994666 |
| Scap_Q6GQT6_S1051s_1 | Scap_Q6G | -0.16 | 0.23 | -0.71 | -0.28 | -0.59 | 0.21 | -0.16 | -0.26 | -0.62 | 0.9 | -0.65 | -0.36 | 0.69 | 0.13 | 0.04 | -1.17 | 0.03 | 0.0475 | 0.1525 | 0.724222 | 0.916492 | 0.907536 | 0.997335 | 0.997988 | 0.9971 |
| Scap_Q6GQT6_S821s_1 | Scap_Q6G | -0.15 | -0.67 | -2.59 | -1.05 | -0.14 | -0.4 | -1.76 | -2.16 | -0.11 | -0.36 | 0.54 | -2.56 | -0.2 | 0.56 | -3.23 | -3.08 | -1.6E-16 | 0.4925 | -0.3725 | 0.700796 | 1 | 0.524249 | 1 | 0.997988 | 0.996159 |
| Scap_Q6GQT6_S850s_1 | Scap_Q6G | -0.09 | -0.08 | -0.08 | -0.21 | 0.64 | -0.41 | 0.02 | -0.25 | 0 | 0.21 | 0.15 | -0.41 | 0.44 | 0.75 | 0.3 | 0.18 | 0.115 | 0.1025 | 0.5325 | 0.053693 | 0.675211 | 0.663604 | 0.992876 | 0.997988 | 0.994666 |
| Scap_Q6GQT6_S934s_1 | Scap_Q6G | 0.4 | 0.1 | -0.42 | 0.34 | 0.77 | -0.06 | 1.22 | -0.28 | 0.58 | 0.51 | 0.03 | 0.13 | 0.32 | 0.59 | 1.13 | -0.56 | 0.3075 | 0.2075 | 0.265 | 0.516612 | 0.434292 | 0.468124 | 0.989223 | 0.997988 | 0.994666 |
| Scaper_Q8BIZ8_S184s_1 | Scaper_Q8 | 1.49 | 0.98 | 1.3 | 0.29 | 0.8 | 2.6 | 0.15 | 1.14 | 1.47 | 1.71 | 1.64 | 1.3 | 1.62 | 0.5 | 1.37 | 0.7 | 0.1575 | 0.515 | 0.0325 | 0.933 | 0.765951 | 0.124795 | 0.995762 | 0.997988 | 0.999507 |
| Sccpdh_Q8R127_S215s_1 | Sccpdh_Q8 | -0.28 | -0.09 | -0.08 | -0.82 | -0.59 | 0.67 | -0.59 | -0.73 | -0.71 | 0.16 | -0.94 | -1.22 | -0.03 | 0.01 | -0.98 | 0.02 | 0.0075 | -0.36 | 0.0725 | 0.829918 | 0.983681 | 0.327734 | 0.999187 | 0.997988 | 0.999507 |
| Sccpdh_Q8R127_S217s_1 | Sccpdh_Q8 | -0.66 | -0.37 | -0.82 | -0.46 | 0.13 | 0.89 | 0.25 | -0.48 | -0.45 | -0.23 | 1.65 | -0.52 | -0.16 | -0.75 | -1.44 | -0.28 | 0.775 | 0.69 | -0.08 | 0.815018 | 0.032603 | 0.187198 | 0.989223 | 0.997988 | 0.999507 |
| Scfd1_Q8BRF7_S300s_1 | Scfd1_Q8B | 0.07 | -0.2 | -0.05 | 0.37 | 0.11 | 0.03 | 0.7 | 0.12 | 0.19 | 0.28 | 0.44 | 0.55 | 0.49 | 0.5 | 0.52 | 0.49 | 0.1925 | 0.3175 | 0.4525 | 0.089583 | 0.451581 | 0.199503 | 0.989223 | 0.997988 | 0.994666 |
| Scg2_Q4W8U9_S270s_1 | Scg2_Q4W | -1.22 | -0.29 | -0.86 | -0.39 | 0.11 | -1.62 | -0.65 | -0.89 | -1.2 | -0.66 | 0.11 | -0.29 | -0.23 | -0.32 | 0.47 | -0.09 | -0.0725 | 0.18 | 0.6475 | 0.070242 | 0.856403 | 0.622203 | 0.996349 | 0.997988 | 0.994666 |
| Scg2_Q4W8U9_S492s_1 | Scg2_Q4W | -0.33 | 0.4 | -0.25 | 0.01 | -0.64 | -0.34 | 0.67 | 0.11 | -0.58 | 0.45 | 0 | -0.1 | -1.22 | 0.09 | 0.61 | 0.62 | -0.0075 | -0.015 | 0.0675 | 0.880882 | 0.98221 | 0.960694 | 0.999187 | 0.997988 | 0.999507 |
| Scg2_Q4W8U9_S515s_1 | Scg2_Q4W | 7.26 | 3.37 | 3.6 | 0.85 | 3.99 | 7.11 | 5.07 | 3.74 | 0.58 | 6.52 | 1.25 | 3.71 | -0.58 | -0.29 | 5.85 | 5.96 | 1.2075 | -0.755 | -1.035 | 0.594352 | 0.367773 | 0.647903 | 0.989223 | 0.997988 | 0.995259 |
| Scg2_Q4W8U9_S516s_1 | Scg2_Q4W | 5.4 | 1.3 | 4.5 | 0.57 | 2.07 | 4.94 | 5.86 | 4.63 | 0.66 | 4.95 | 0.07 | 3.59 | 1.63 | 0.9 | 6.95 | 6.75 | 1.4325 | -0.625 | 1.115 | 0.520826 | 0.262625 | 0.668208 | 0.989223 | 0.997988 | 0.994666 |
| Scg5_P12961_S205s_1 | Scg5_P129 | 0.13 | 0.12 | 0.58 | 0.85 | 0.23 | -0.31 | 0.94 | 0.82 | 0.02 | 0.56 | 0.27 | 0.61 | -0.83 | 0.53 | 0.89 | 1.22 | 0 | -0.055 | 0.0325 | 0.944408 | 1 | 0.843374 | 1 | 0.997988 | 0.999507 |
| Scn1a_A0A1D5RLV0_S757s_1 | Scn1a_A0A | -0.34 | -1.39 | -0.88 | -0.91 | -0.4 | 0.04 | -1.26 | -0.84 | -1.54 | -1.69 | 0 | -0.65 | -1.88 | -0.13 | -1.99 | -0.85 | 0.265 | -0.09 | -0.3325 | 0.487226 | 0.46044 | 0.836868 | 0.989223 | 0.997988 | 0.994666 |
| Scn1a_A2APX8_S1801s_1 | Scn1a_A2A | -2.76 | 0.04 | -2.39 | -4.14 | -1.37 | -1.67 | -3.53 | -3.35 | -2.34 | -1.26 | -1.5 | -3.16 | -0.98 | -2.56 | -3.71 | -3.27 | -0.1675 | 0.2475 | -0.3175 | 0.731585 | 0.852024 | 0.774675 | 0.996349 | 0.997988 | 0.9971 |
| Scn1a_A2APX8_S1911s_1 | Scn1a_A2A | 0.32 | -1.26 | 0.96 | 1.32 | 0.34 | 0.82 | 0.96 | 0.82 | -1.06 | -0.78 | 2.16 | 0.42 | -0.3 | 0.6 | 0.96 | -0.15 | 0.4 | -0.15 | -0.0575 | 0.922328 | 0.459189 | 0.856088 | 0.989223 | 0.997988 | 0.999507 |
| Scn1a_A2APX8_S467s_1 | Scn1a_A2A | -0.01 | 0.78 | 1.22 | 1.05 | 0.77 | 0.28 | 0.87 | 0.94 | 0.56 | 0.65 | 0.72 | 0.85 | 1.25 | 0.64 | 1.37 | 1.66 | -0.045 | -0.065 | 0.47 | 0.220994 | 0.889328 | 0.835038 | 0.996493 | 0.997988 | 0.994666 |
| Scn1a_A2APX8_S467s_2 | Scn1a_A2A | -0.4 | 0.55 | -0.79 | -0.3 | 0.12 | -0.21 | -1.08 | -1.46 | -0.07 | -0.08 | -0.74 | -0.72 | 0.53 | -0.64 | -0.24 | -0.56 | -0.4225 | -0.1675 | 0.0075 | 0.984889 | 0.345367 | 0.637819 | 0.989223 | 0.997988 | 1 |
| Scn1a_A2APX8_S470s_2 | Scn1a_A2A | -0.47 | -0.3 | -1.67 | -0.71 | -0.29 | -0.76 | -1.3 | -2.86 | -0.26 | -0.85 | -1.29 | -1.29 | -0.33 | -1.16 | -1.13 | -1.86 | -0.515 | -0.135 | -0.3325 | 0.448411 | 0.378991 | 0.730958 | 0.989223 | 0.997988 | 0.994666 |
| Scn1a_A2APX8_S623s_1 | Scn1a_A2A | 0.46 | -1.28 | 0.01 | -0.75 | -0.43 | 2.27 | -1.25 | -0.23 | 0.67 | -0.7 | -1.44 | 0.2 | 0.63 | -2.74 | -1.46 | -0.63 | 0.48 | 0.0725 | -0.66 | 0.370464 | 0.525681 | 0.897432 | 0.989223 | 0.997988 | 0.994666 |
| Scn1a_A2APX8_T464t_2 | Scn1a_A2A | -0.14 | 0.94 | -0.12 | -0.11 | 0.49 | 0.5 | -1.61 | -0.37 | 0.94 | 0.6 | -0.51 | -0.41 | 1.17 | -0.05 | -0.38 | -0.11 | -0.39 | 0.0125 | 0.015 | 0.972162 | 0.455709 | 0.9771 | 0.989223 | 0.997988 | 0.999507 |
| Scn1a_A2APX8_T465t_2 | Scn1a_A2A | -0.18 | 0.65 | -0.02 | 0.11 | 0.3 | 0.53 | -0.85 | -0.41 | -0.05 | 0.29 | -0.19 | -0.18 | 0.86 | -0.87 | 0.54 | 0.47 | -0.2475 | -0.1725 | 0.11 | 0.794133 | 0.500496 | 0.528077 | 0.989223 | 0.997988 | 0.999507 |
| Scn1b_P97952_S207s_1 | Scn1b_P97 | -0.49 | 0.54 | -0.36 | -0.21 | 0.4 | -0.39 | 1.08 | 0.42 | -0.03 | -0.71 | 0.29 | -0.2 | -0.51 | 0.24 | 0.04 | 0.31 | 0.5075 | -0.0325 | 0.15 | 0.655455 | 0.194821 | 0.92236 | 0.989223 | 0.997988 | 0.995259 |
| Scn2a1_B1AWN6_S468s_1 | Scn2a1_B1 | 0.19 | 0.89 | 1.4 | 1.13 | 0.22 | 0.86 | 0.86 | 0.88 | 0.42 | 1.25 | 0.07 | 1.33 | 0.75 | -0.03 | 1.39 | 1.08 | -0.1975 | -0.135 | -0.105 | 0.7956 | 0.541539 | 0.737341 | 0.989223 | 0.997988 | 0.999507 |
| Scn2a1_B1AWN6_S468s_2 | Scn2a1_B1 | 0.37 | -0.34 | -0.62 | 0.33 | -0.06 | -0.3 | 0.3 | -0.58 | 0.07 | 0.03 | 0.86 | -0.08 | -1.19 | -0.29 | -0.27 | 0.26 | -0.095 | 0.285 | -0.3075 | 0.445667 | 0.768774 | 0.417851 | 0.995762 | 0.997988 | 0.994666 |
| Scn2a1_B1AWN6_S471s_2 | Scn2a1_B1 | 0.37 | -0.34 | -0.62 | 0.33 | -0.06 | -0.3 | 0.3 | -0.58 | 0.07 | 0.03 | 0.86 | -0.08 | -1.19 | -0.29 | -0.27 | 0.26 | -0.095 | 0.285 | -0.3075 | 0.445667 | 0.768774 | 0.417851 | 0.995762 | 0.997988 | 0.994666 |
| Scn2a1_B1AWN6_S526s_1 | Scn2a1_B1 | 0.98 | 1.09 | 0.73 | 0.12 | -0.04 | 1.48 | 0.1 | -0.38 | 2.12 | 1.16 | -0.21 | -0.41 | 2.07 | 0.13 | 0.01 | -0.19 | -0.44 | -0.065 | -0.225 | 0.672924 | 0.326162 | 0.911343 | 0.989223 | 0.997988 | 0.996159 |
| Scn2a1_B1AWN6_S528s_1 | Scn2a1_B1 | 0.2 | 0.52 | 0.85 | 0.7 | -0.37 | 0.48 | -0.11 | 0.28 | 1.29 | 0.26 | -0.31 | 0.63 | 1.51 | -0.5 | 0.59 | 0.67 | -0.4975 | -0.1 | -4.7E-16 | 1 | 0.096341 | 0.78773 | 0.989223 | 0.997988 | 1 |
| Scn2a1_B1AWN6_S531s_1 | Scn2a1_B1 | 0.21 | 0.23 | 0.32 | 0.3 | -0.73 | 1.22 | -0.34 | 0.48 | 1.26 | -0.42 | -0.53 | 0.32 | 0.95 | -0.76 | 0.47 | 0.16 | -0.1075 | -0.1075 | -0.06 | 0.873716 | 0.795692 | 0.792775 | 0.995762 | 0.997988 | 0.999507 |
| Scn2a1_B1AWN6_S540s_1 | Scn2a1_B1 | -0.86 | -0.93 | -0.39 | -1.19 | -0.33 | -0.7 | -0.89 | -0.92 | -1.47 | -1.26 | -0.09 | -0.59 | -1.97 | -0.13 | -1.31 | -0.94 | 0.1325 | -0.01 | -0.245 | 0.561066 | 0.615861 | 0.978203 | 0.989223 | 0.997988 | 0.995259 |
| Scn2a1_B1AWN6_S568s_1 | Scn2a1_B1 | 0.64 | -1.88 | -0.54 | 0.26 | -0.06 | 0.42 | 0.48 | 0.4 | -0.49 | -1.54 | 0.37 | 0.3 | -2.93 | 0.12 | -1.32 | -1.2 | 0.69 | 0.04 | -0.9525 | 0.220545 | 0.205575 | 0.950711 | 0.989223 | 0.997988 | 0.994666 |
| Scn2a1_B1AWN6_S579s_1 | Scn2a1_B1 | -1.87 | -2.46 | -1.87 | -2.29 | -0.42 | -1.67 | -1.75 | -1.11 | -2.83 | -2.2 | 0.32 | -2.4 | -1.98 | -0.56 | -2.33 | -1.87 | 0.885 | 0.345 | 0.4375 | 0.307165 | 0.027567 | 0.603376 | 0.989223 | 0.997988 | 0.994666 |
| Scn2a1_B1AWN6_S623s_1 | Scn2a1_B1 | -1.55 | -1.68 | 0.37 | 0.28 | -2.28 | 0.04 | -0.37 | -0.4 | 1.32 | -1.45 | -0.26 | 0.41 | 1.73 | -0.68 | -0.8 | 0.15 | -0.1075 | 0.65 | 0.745 | 0.315773 | 0.873544 | 0.382309 | 0.996493 | 0.997988 | 0.994666 |
| Scn2a1_B1AWN6_T1944t_1 | Scn2a1_B1 | -0.46 | 0.72 | -0.96 | -2 | 0.99 | -0.06 | -1.22 | -1.26 | -0.04 | 0.09 | -0.59 | -1.3 | -0.48 | 0.72 | -1.18 | -1.54 | 0.2875 | 0.215 | 0.055 | 0.93512 | 0.677279 | 0.718126 | 0.99315 | 0.997988 | 0.999507 |
| Scn2a1_B1AWN6_T1960t_1 | Scn2a1_B1 | 0.75 | 1.61 | 2.16 | 1.53 | 1.61 | 0.79 | 1.94 | 1.57 | 0.78 | 0.83 | 0.85 | 2.11 | 1.07 | 1.05 | 1.68 | 1.79 | -0.035 | -0.37 | -0.115 | 0.756418 | 0.925088 | 0.393193 | 0.997335 | 0.997988 | 0.9971 |
| Scn2a1_B1AWN6_T1967t_1 | Scn2a1_B1 | -1.02 | -0.92 | -0.38 | -0.24 | -0.37 | -0.79 | 0.17 | -0.41 | -1.12 | -1.1 | -0.13 | -0.07 | -1.16 | 0.09 | -0.68 | -0.44 | 0.29 | 0.035 | 0.0925 | 0.793312 | 0.347224 | 0.922784 | 0.989223 | 0.997988 | 0.999507 |
| Scn2b_Q56A07_S192s_1 | Scn2b_Q56 | -1.8 | -0.95 | -1.09 | -1.91 | -0.48 | -1.61 | -1.12 | -1.86 | -1.62 | -1.56 | -0.76 | -1.62 | -0.93 | -0.61 | -1.34 | -1.69 | 0.17 | 0.0475 | 0.295 | 0.424222 | 0.656612 | 0.889269 | 0.990434 | 0.997988 | 0.994666 |
| Scn2b_Q56A07_T214t_1 | Scn2b_Q56 | -1.27 | -1.51 | 0.21 | -1.07 | -2.65 | -1.49 | -1.02 | -1.91 | 1.11 | -1.15 | -0.31 | -0.29 | 1.3 | -0.74 | -0.74 | -0.75 | -0.8575 | 0.75 | 0.6775 | 0.264557 | 0.098025 | 0.202478 | 0.989223 | 0.997988 | 0.994666 |
| Scn3a_A2ASI5_S466s_1 | Scn3a_A2A | 0.58 | 1.4 | -0.87 | -0.17 | -0.13 | 0.22 | -0.68 | -0.29 | -0.02 | -0.13 | -2.09 | -0.47 | -0.66 | -0.25 | 0.44 | -0.13 | -0.455 | -0.9125 | -0.385 | 0.455104 | 0.356923 | 0.168273 | 0.989223 | 0.997988 | 0.994666 |
| Scn3a_A2ASI5_S466s_2 | Scn3a_A2A | 0.25 | 0.86 | -0.02 | -1.04 | 0.92 | 0.98 | -0.35 | -1.55 | 0.94 | 0.3 | -3.18 | -0.21 | 0.8 | 1.84 | -2.39 | -0.67 | -0.0125 | -0.55 | -0.1175 | 0.893492 | 0.984386 | 0.537486 | 0.999187 | 0.997988 | 0.999507 |
| Scn3a_A2ASI5_S469s_1 | Scn3a_A2A | 0.58 | 1.4 | -0.56 | 0.57 | -0.13 | 0.22 | -0.07 | 0.06 | -0.02 | -0.13 | -1.56 | 0.35 | -0.66 | -0.25 | 0.82 | 0.83 | -0.4775 | -0.8375 | -0.3125 | 0.55049 | 0.245835 | 0.146423 | 0.989223 | 0.997988 | 0.994666 |
| Scn3a_A2ASI5_S469s_2 | Scn3a_A2A | 0.25 | 0.86 | -0.02 | -1.04 | 0.92 | 0.98 | -0.35 | -1.55 | 0.94 | 0.3 | -3.18 | -0.21 | 0.8 | 1.84 | -2.39 | -0.67 | -0.0125 | -0.55 | -0.1175 | 0.893492 | 0.984386 | 0.537486 | 0.999187 | 0.997988 | 0.999507 |
| Scn3a_A2ASI5_S623s_1 | Scn3a_A2A | -0.01 | -0.71 | 0.6 | 0.82 | -1.38 | 1.06 | -0.08 | 0.49 | 1.13 | 0.08 | 0.6 | 1.21 | 0.7 | -2.28 | -0.17 | -0.19 | -0.1525 | 0.58 | -0.66 | 0.323568 | 0.78671 | 0.192688 | 0.995762 | 0.997988 | 0.994666 |
| Scn8a_F7D6K4_S678s_1 | Scn8a_F7D | 1.19 | 0.85 | 0.34 | 0.53 | 1.81 | 1.23 | 0.41 | 0.23 | 1.17 | 0.49 | -2.27 | 0.26 | -0.31 | 0.99 | -0.7 | -0.4 | 0.1925 | -0.815 | -0.8325 | 0.070878 | 0.63016 | 0.262005 | 0.989223 | 0.997988 | 0.994666 |
| Scrib_Q80U72_S1263s_1 | Scrib_Q80U | -0.56 | -0.75 | -1.52 | -2.01 | -0.9 | -1.31 | -1.55 | -1.71 | -0.52 | -0.36 | -1.02 | -0.64 | 0.56 | -0.39 | -0.97 | -1 | -0.1575 | 0.575 | 0.76 | 0.135317 | 0.67503 | 0.148153 | 0.992876 | 0.997988 | 0.994666 |
| Scrib_Q80U72_S1464s_1 | Scrib_Q80U | 0.33 | -0.26 | 0.26 | 0.67 | 0.4 | 0.58 | 0.84 | 0.27 | -0.02 | 0.88 | 0.76 | 1.16 | -0.58 | 1.35 | 1.47 | 0.26 | 0.2725 | 0.445 | 0.375 | 0.45552 | 0.326851 | 0.209481 | 0.989223 | 0.997988 | 0.994666 |
| Scrib_Q80U72_S506s_1 | Scrib_Q80U | 0.27 | 0.44 | 0.17 | 0.83 | -0.12 | 0.22 | 0.99 | 0.03 | -0.11 | 0.6 | 0.92 | 0.12 | -0.33 | -0.12 | 0.66 | 0.66 | -0.1475 | -0.045 | -0.21 | 0.535181 | 0.633053 | 0.884084 | 0.989223 | 0.997988 | 0.994666 |
| Scrn1_Q9CZC8_S322s_1 | Scrn1_Q9C | 0.45 | 0.36 | -0.81 | 0.44 | 0.13 | 0.33 | -0.53 | -0.31 | -0.11 | 0.05 | -0.61 | -2.66 | -0.11 | 1.23 | -0.77 | 0.36 | -0.205 | -0.9425 | 0.0675 | 0.891254 | 0.574874 | 0.161791 | 0.989223 | 0.997988 | 0.999507 |
| Scrn1_Q9CZC8_S360s_1 | Scrn1_Q9C | 0.27 | -0.16 | -0.41 | -0.18 | -0.73 | -0.59 | -0.78 | -0.64 | 0.08 | 0.05 | 0.82 | -0.68 | 0.15 | 0.19 | -0.51 | -1.65 | -0.565 | 0.1875 | -0.335 | 0.455976 | 0.030833 | 0.597222 | 0.989223 | 0.997988 | 0.994666 |
| Scyl2_G5E8J9_S676s_1 | Scyl2_G5E8 | 0.57 | 0.03 | 0.51 | 0.14 | 0.59 | 0.3 | 0.39 | 0.69 | 0.73 | 0.41 | 1.14 | 0.61 | 0.09 | 0.92 | 0.57 | 0.66 | 0.18 | 0.41 | 0.2475 | 0.401353 | 0.447394 | 0.147831 | 0.989223 | 0.997988 | 0.994666 |
| Sdc3_Q64519_S418s_1 | Sdc3_Q645 | -0.38 | 0.76 | -0.12 | -0.35 | 0.95 | 0.54 | -0.05 | -0.43 | 0.41 | 0.18 | 0.74 | 0.18 | 0.08 | 0.41 | 0.21 | -0.58 | 0.275 | 0.4 | 0.0525 | 0.885549 | 0.489785 | 0.237829 | 0.989223 | 0.997988 | 0.999507 |
| Sdccag8_Q05DD9_S92s_1 | Sdccag8_Q | -0.36 | -0.16 | -0.63 | -1.69 | -1.1 | 0.09 | -1.76 | -2.15 | 1.33 | -0.36 | -1.16 | -0.84 | 1.92 | -2.39 | -0.64 | -0.72 | -0.52 | 0.4525 | 0.2525 | 0.763546 | 0.348417 | 0.455255 | 0.989223 | 0.997988 | 0.9971 |
| Sde2_Q8K1J5_S269s_1 | Sde2_Q8K1 | -0.92 | -1.42 | -1.19 | -0.35 | -2.22 | -1.61 | -0.79 | -1.8 | -1.14 | -1.02 | 0.1 | -1.2 | -1.31 | -0.48 | -1.4 | -1.62 | -0.635 | 0.155 | -0.2325 | 0.526296 | 0.11345 | 0.689353 | 0.989223 | 0.997988 | 0.994666 |

| | | | | | | | | | | | | | | | | | | | | | | | | | | |
|---|---|---|---|---|---|---|---|---|---|---|---|---|---|---|---|---|---|---|---|---|---|---|---|---|---|---|
| Sdk1_Q3UH53_S297s_2 | Sdk1_Q3UI | -0.48 | 0.13 | 0.88 | 0.11 | -0.18 | -0.97 | -0.1 | 0.1 | -0.85 | -0.16 | 1.22 | 0.8 | -0.7 | -0.14 | 0.17 | 0.52 | -0.4475 | 0.0925 | -0.1975 | 0.615432 | 0.235507 | 0.856168 | 0.989223 | 0.997988 | 0.995259 |
| Sdk1_Q3UH53_T296t_2 | Sdk1_Q3UI | -0.48 | 0.13 | 0.88 | 0.11 | -0.18 | -0.97 | -0.1 | 0.1 | -0.85 | -0.16 | 1.22 | 0.8 | -0.7 | -0.14 | 0.17 | 0.52 | -0.4475 | 0.0925 | -0.1975 | 0.615432 | 0.235507 | 0.856168 | 0.989223 | 0.997988 | 0.995259 |
| Sdpr_Q63918_S27s_1 | Sdpr_Q639 | 0.25 | 0.56 | -0.76 | 0.87 | 0.47 | 0.05 | 0.5 | 0.49 | -0.15 | 0.47 | -1.38 | 0.3 | -0.02 | 1.24 | 0.4 | 0.75 | 0.1475 | -0.42 | 0.3625 | 0.414211 | 0.688021 | 0.421575 | 0.994672 | 0.997988 | 0.994666 |
| Sdpr_Q63918_S293s_1 | Sdpr_Q639 | 0.02 | -0.25 | 0.43 | 0.52 | 0.19 | 0.27 | 0.55 | 0.24 | 0.57 | 0.87 | 0.29 | 0.79 | 0.35 | 0.31 | 0.33 | 0.7 | 0.1325 | 0.45 | 0.2425 | 0.396746 | 0.602004 | 0.128905 | 0.989223 | 0.997988 | 0.994666 |
| Sec14l2_Q99J08_T401t_1 | Sec14l2_Q | 0.09 | -1.6 | -0.56 | -0.38 | -1.46 | -1.4 | -0.69 | -0.87 | 0.1 | -1.04 | -0.59 | -0.53 | 0.9 | -4.96 | 0.08 | -0.93 | -0.4925 | 0.0975 | -0.615 | 0.600161 | 0.22819 | 0.815876 | 0.989223 | 0.997988 | 0.995259 |
| Sec16a_E9QAT4_S1028s_1 | Sec16a_E9 | 0.56 | 0.22 | 0.07 | 0.56 | 0.13 | 0.68 | 0.02 | 0.46 | 0.98 | 0.23 | 0.73 | 0.35 | 0.37 | 1.24 | -0.23 | 0.16 | -0.03 | 0.22 | 0.0325 | 0.927926 | 0.904921 | 0.424662 | 0.996493 | 0.997988 | 0.999507 |
| Sec16a_E9QAT4_S2101s_1 | Sec16a_E9 | -2.27 | -1.93 | -1.72 | -1.67 | -1.17 | -0.53 | -1.39 | -1.86 | -2.47 | -2.27 | -0.43 | -1.93 | -1.69 | -1.03 | -1.86 | -1.46 | 0.66 | 0.1225 | 0.3875 | 0.206454 | 0.064521 | 0.790664 | 0.989223 | 0.997988 | 0.994666 |
| Sec16a_E9QAT4_S554s_1 | Sec16a_E9 | -0.53 | -0.29 | -1.37 | -0.86 | -0.28 | -1.06 | -0.38 | -1.26 | -0.65 | -0.69 | -1.35 | -1.21 | -1.16 | 0.34 | -1.25 | -1.04 | 0.0175 | -0.2125 | -0.015 | 0.972455 | 0.95925 | 0.514603 | 0.999187 | 0.997988 | 0.999507 |
| Sec22b_O08547_S137s_1 | Sec22b_OC | -0.29 | -0.35 | 0.26 | 0.37 | 0.34 | 0.12 | 0.95 | 0.37 | -0.03 | 0.11 | 0.57 | 0.38 | -0.22 | 0.51 | 0.53 | 0.66 | 0.4475 | 0.26 | 0.3725 | 0.261302 | 0.144181 | 0.365126 | 0.989223 | 0.997988 | 0.994666 |
| Sec22b_O08547_S175s_1 | Sec22b_OC | 0.27 | 0.12 | 1.12 | 1.35 | 0.54 | 0.46 | 1.87 | 0.66 | -0.16 | 0.39 | 2.43 | 1.03 | 0.8 | 0.74 | 2.05 | 0.46 | 0.1675 | 0.2075 | 0.2975 | 0.516788 | 0.695497 | 0.723455 | 0.994893 | 0.997988 | 0.994666 |
| Sec22b_O08547_S177s_1 | Sec22b_OC | -1.18 | -1.14 | -0.33 | -0.96 | -0.55 | -0.89 | -0.56 | -0.25 | -1.45 | -0.77 | 0.76 | -0.53 | -0.1 | -0.7 | -0.15 | -0.05 | 0.34 | 0.405 | 0.6525 | 0.055784 | 0.235563 | 0.405664 | 0.989223 | 0.997988 | 0.994666 |
| Sec23ip_G3X928_S600s_1 | Sec23ip_G: | 1.58 | 1.64 | -0.05 | -0.07 | 0.84 | 0.43 | 0.66 | 0.05 | 0.46 | 0.41 | 0.19 | 0 | 1.21 | 2.44 | -0.65 | 0.28 | -0.28 | -0.51 | 0.045 | 0.95068 | 0.557669 | 0.294336 | 0.989223 | 0.997988 | 0.999507 |
| Sec23ip_G3X928_S748s_1 | Sec23ip_G: | -1.76 | -0.92 | -0.98 | 1.63 | -0.24 | -2.98 | -0.4 | 0.01 | -3.51 | -1.55 | 0.12 | 1.8 | -0.88 | -0.92 | 0.29 | -0.52 | -0.395 | -0.2775 | 7.85E-17 | 1 | 0.656027 | 0.815612 | 0.990434 | 0.997988 | 1 |
| Sec24b_Q80ZX0_S1203s_1 | Sec24b_Q8 | -0.4 | -0.01 | 0.92 | 0.46 | 0.2 | -1.18 | 0.53 | -0.06 | -0.75 | 0.34 | 1.02 | 0.73 | -0.97 | 0.11 | 0.59 | 1.08 | -0.37 | 0.0925 | -0.04 | 0.935653 | 0.409432 | 0.841017 | 0.989223 | 0.997988 | 0.999507 |
| Sec24c_G3X972_S864s_1 | Sec24c_G3 | -0.14 | -0.39 | 0.75 | 0.41 | 0.01 | 0.01 | -0.23 | -0.04 | -0.34 | 0.17 | -0.06 | -0.17 | 0.97 | 1.13 | -0.17 | -0.75 | -0.22 | -0.2575 | 0.1375 | 0.781245 | 0.459895 | 0.419014 | 0.989223 | 0.997988 | 0.998226 |
| Sec31a_S4R2A9_S502s_1 | Sec31a_S4 | 0.37 | 0.55 | -0.76 | -0.31 | 0.19 | 0.89 | -0.49 | -0.62 | 0.3 | 0.49 | 0.26 | -1.24 | -0.05 | 0.19 | -0.69 | -0.38 | 0.03 | -0.01 | -0.195 | 0.606757 | 0.944946 | 0.983276 | 0.999187 | 0.997988 | 0.995259 |
| Sec31a_S4R2A9_S877s_1 | Sec31a_S4 | -0.75 | 0.31 | 0.85 | 0.83 | 0.67 | 0.23 | 0.52 | 0.9 | -0.03 | 0.29 | 0.62 | 1.07 | 0.14 | 0.51 | 0.55 | 0.46 | 0.27 | 0.1775 | 0.105 | 0.790533 | 0.492441 | 0.681379 | 0.989223 | 0.997988 | 0.999507 |
| Sec61b_Q9CQS8_S13s_1 | Sec61b_Q9 | -1.47 | 0.07 | -1.79 | -1.69 | -1.47 | -0.14 | -0.67 | -1.28 | -1.09 | -0.46 | 0.07 | -1.24 | 0.03 | 0.31 | 0 | -0.88 | 0.33 | 0.54 | 1.085 | 0.046411 | 0.503913 | 0.294885 | 0.989223 | 0.997988 | 0.994666 |
| Sec61b_Q9CQS8_S17s_1 | Sec61b_Q9 | 2.47 | 0.56 | 1.19 | 2.03 | -0.06 | 1.84 | 1.08 | 1.78 | 1.68 | 2.49 | 1.75 | 1.75 | 1.63 | 1.83 | 2.46 | 1.3 | -0.4025 | 0.355 | 0.2425 | 0.609785 | 0.474147 | 0.438392 | 0.989223 | 0.997988 | 0.995259 |
| Sec62_Q8BU14_S335s_2 | Sec62_Q8E | -1.28 | 0.02 | 0.23 | -0.82 | -2.42 | -1.12 | -1.71 | -4.07 | -0.62 | -0.76 | 0.02 | -2.48 | 0.09 | -1.23 | -0.03 | -0.81 | -1.8675 | -0.4975 | -0.0325 | 0.94355 | 0.016069 | 0.407667 | 0.989223 | 0.997988 | 0.999507 |
| Sec62_Q8BU14_S341s_1 | Sec62_Q8E | -1.33 | -1.25 | 0.25 | -0.37 | -1.9 | -1.36 | -1.84 | -1.42 | -0.51 | -1.58 | -0.05 | -0.7 | 0.28 | -2.6 | 0.04 | 0.53 | -0.955 | -0.035 | 0.2375 | 0.745439 | 0.033107 | 0.940769 | 0.989223 | 0.997988 | 0.9971 |
| Sec62_Q8BU14_S341s_2 | Sec62_Q8E | -2.52 | -0.86 | -0.6 | -2.04 | -2.49 | -1.42 | -3.34 | -3.62 | -0.99 | -2.46 | -0.5 | -2.96 | -0.03 | -1.61 | -0.76 | -1.18 | -1.2125 | -0.2225 | 0.61 | 0.268839 | 0.069409 | 0.741867 | 0.989223 | 0.997988 | 0.994666 |
| Sec62_Q8BU14_S341s_3 | Sec62_Q8E | -1.4 | 0.25 | 1.31 | -0.04 | -2.45 | -0.05 | -0.98 | -0.3 | 0.25 | -0.85 | 0.4 | -0.75 | -1.47 | -0.07 | -0.71 | 0.3 | -0.975 | -0.2675 | -0.5175 | 0.411184 | 0.177437 | 0.65372 | 0.989223 | 0.997988 | 0.994666 |
| Sec62_Q8BU14_S345s_3 | Sec62_Q8E | -1.4 | 0.25 | 1.31 | -0.04 | -2.45 | -0.05 | -0.98 | -0.3 | 0.25 | -0.85 | 0.4 | -0.75 | -1.47 | -0.07 | -0.71 | 0.3 | -0.975 | -0.2675 | -0.5175 | 0.411184 | 0.177437 | 0.65372 | 0.989223 | 0.997988 | 0.994666 |
| Sec62_Q8BU14_T158t_1 | Sec62_Q8E | 0.64 | -0.15 | 0.42 | 0.18 | -0.49 | 1.33 | -0.83 | 0.01 | 2.22 | 0.34 | -0.79 | 0.05 | 2.52 | -0.76 | 0.71 | 0.53 | -0.2675 | 0.1825 | 0.4775 | 0.455765 | 0.570049 | 0.762178 | 0.989223 | 0.997988 | 0.994666 |
| Sec62_Q8BU14_T343t_2 | Sec62_Q8E | -2.52 | -0.86 | -1.43 | -3.63 | -2.49 | -1.42 | -6.52 | -3.32 | -0.99 | -2.46 | -0.95 | -3.39 | -0.03 | -1.61 | -1.47 | -1.49 | -1.3275 | 0.1625 | 0.96 | 0.162255 | 0.239895 | 0.831851 | 0.989223 | 0.997988 | 0.994666 |
| Sec62_Q8BU14_T343t_3 | Sec62_Q8E | -1.4 | 0.25 | 1.31 | -0.04 | -2.45 | -0.05 | -0.98 | -0.3 | 0.25 | -0.85 | 0.4 | -0.75 | -1.47 | -0.07 | -0.71 | 0.3 | -0.975 | -0.2675 | -0.5175 | 0.411184 | 0.177437 | 0.65372 | 0.989223 | 0.997988 | 0.994666 |
| Secisbp2l_Q6A098_S897s_1 | Secisbp2l_( | -3.25 | -1.71 | -0.09 | 1.99 | -3.2 | -1.99 | 1.35 | 1.36 | -0.96 | -1.57 | 2.14 | 1.52 | 0 | -1.31 | 2.27 | 1.77 | 0.145 | 1.0475 | 1.4475 | 0.247016 | 0.916721 | 0.417136 | 0.997335 | 0.997988 | 0.994666 |
| Selplg_Q3TA56_S365s_1 | Selplg_Q3T | -0.3 | -0.69 | -0.2 | -0.18 | -0.58 | -0.27 | 0.47 | -0.06 | -1.09 | -0.51 | -0.5 | -0.29 | -0.9 | -0.28 | -0.25 | -0.22 | 0.2325 | -0.255 | -0.07 | 0.802734 | 0.421747 | 0.35445 | 0.989223 | 0.997988 | 0.999507 |
| Sema4b_Q62179_S780s_1 | Sema4b_Q | -1.72 | -0.61 | -0.28 | -0.43 | -1.29 | -1.73 | -0.7 | 0.67 | -0.57 | 0.02 | -0.75 | 0.4 | 0.44 | -0.62 | 0.46 | -0.99 | -0.0025 | 0.535 | 0.5825 | 0.237736 | 0.996407 | 0.217616 | 0.99973 | 0.997988 | 0.994666 |
| Sema4c_Q64151_S761s_1 | Sema4c_Q | 1.07 | -0.12 | 0.95 | 0.99 | 0.44 | 0.99 | 1.5 | 1.25 | 0.54 | 1.71 | 1.27 | 1.3 | 0.18 | 0.28 | 0.48 | 1.14 | 0.3225 | 0.4825 | -0.2025 | 0.590337 | 0.380967 | 0.221991 | 0.989223 | 0.997988 | 0.995259 |
| Sema4d_O09126_S832s_1 | Sema4d_O | 1.24 | 1.38 | 0.51 | 0.51 | 1.61 | 1.59 | 0.9 | 0.72 | 1.4 | 1.5 | 1.5 | 1.36 | 0.47 | 1.85 | 1.19 | 1.81 | 0.295 | 0.53 | 0.42 | 0.312526 | 0.389657 | 0.088328 | 0.989223 | 0.997988 | 0.994666 |
| Sema6a_K3W4S5_S672s_1 | Sema6a_K: | 0.63 | 0.46 | 0.7 | 1.28 | -0.2 | 0.44 | 0.46 | 0.52 | 1.22 | 0.42 | 0.44 | 0.47 | 1.75 | -0.59 | 0.68 | 0.65 | -0.4625 | -0.13 | -0.145 | 0.765827 | 0.124805 | 0.668344 | 0.989223 | 0.997988 | 0.9971 |
| Sema6b_O54951_S647s_1 | Sema6b_O | -0.41 | -1.94 | -0.14 | 1.18 | -0.61 | -0.48 | 0.53 | 1.35 | -1.06 | -0.68 | 0.41 | 0.38 | -1.61 | -0.75 | -0.7 | 1.03 | 0.525 | 0.09 | -0.18 | 0.810789 | 0.457399 | 0.893201 | 0.989223 | 0.997988 | 0.999507 |
| Senp3_Q9EP97_S206s_1 | Senp3_Q9E | -0.77 | -4.01 | 0.06 | -0.32 | -1.28 | -2.19 | 0.24 | -1.17 | -0.93 | -0.8 | -0.78 | 0.21 | -1.07 | -0.99 | -0.25 | -0.46 | 0.16 | 0.685 | 0.5675 | 0.504028 | 0.861696 | 0.434611 | 0.996493 | 0.997988 | 0.994666 |
| Senp3_Q9EP97_S226s_1 | Senp3_Q9E | 0.14 | 0.13 | 0.06 | 0.14 | 0.58 | 0.04 | 0.27 | 0.54 | 0.32 | 0.36 | -0.08 | 0.08 | -0.51 | 0.02 | -0.12 | 0.69 | 0.24 | 0.0525 | -0.0975 | 0.751215 | 0.286531 | 0.810914 | 0.989223 | 0.997988 | 0.9971 |
| Senp7_Q9CX65_S12s_1 | Senp7_Q9( | 1.11 | -0.48 | 0.99 | 0.81 | -0.29 | 0.24 | 0.14 | 0.93 | 0.21 | -0.32 | 0.65 | 1.12 | 0.61 | 0.47 | -0.75 | -0.66 | -0.3525 | -0.1925 | -0.69 | 0.181457 | 0.413338 | 0.676335 | 0.989223 | 0.997988 | 0.994666 |
| Senp7_Q9CX65_S214s_1 | Senp7_Q9( | 0.47 | 0.65 | 0.81 | 0.93 | 0.5 | 0.32 | 0.55 | 1.02 | 0.27 | 0.02 | 0.89 | 0.94 | 0.93 | 1.14 | 0.89 | 1.1 | -0.1175 | -0.185 | 0.3 | 0.236456 | 0.630185 | 0.531977 | 0.989223 | 0.997988 | 0.994666 |
| Sepn1_D3Z2R5_S13s_1 | Sepn1_D3Z | 0.11 | 0.84 | 0.65 | 0.49 | 0.5 | -0.05 | 0.53 | 0.38 | 0.6 | 0.06 | 0.98 | 0.65 | 1.13 | 0.53 | 1.39 | 0.02 | -0.1825 | 0.05 | 0.245 | 0.50834 | 0.482846 | 0.863358 | 0.989223 | 0.997988 | 0.994666 |
| Sept11_Q8C1B7_S9s_1 | Sept11_Q8 | 1.58 | 0.94 | -2.42 | -1.78 | 1.47 | 0.57 | -1.62 | -1.65 | 0.38 | 1.27 | -0.96 | -0.99 | 0.1 | 1.14 | -3.05 | -1.03 | 0.1125 | 0.345 | -0.29 | 0.80173 | 0.917684 | 0.730222 | 0.997335 | 0.997988 | 0.999507 |
| Sept4_P28661_S100s_1 | Sept4_P28 | 0.81 | 0.57 | 0.26 | 1.04 | 0.37 | 0.11 | 0.83 | 0.65 | 0.17 | 0.39 | 0.6 | 0.96 | -0.66 | 0.72 | 0.25 | 0.58 | -0.18 | -0.14 | -0.4475 | 0.246571 | 0.512038 | 0.625664 | 0.989223 | 0.997988 | 0.994666 |
| Sept5_Q9Z2Q6_S327s_1 | Sept5_Q9Z | -0.15 | -0.05 | -0.11 | 0.13 | 0.28 | 0.23 | 0.47 | 0.23 | -0.04 | -0.09 | -0.32 | 0.1 | -0.83 | 0.25 | -0.52 | -0.07 | 0.3475 | -0.0425 | -0.2475 | 0.424838 | 0.11107 | 0.846696 | 0.989223 | 0.997988 | 0.994666 |
| Sept5_Q9Z2Q6_T13t_1 | Sept5_Q9Z | -2.36 | -1.91 | -0.56 | -1.06 | -1.77 | -2.23 | -0.69 | -0.91 | -2.57 | -2.38 | -0.81 | -0.83 | -0.97 | -2.47 | -0.88 | -0.96 | 0.0725 | -0.175 | 0.1525 | 0.77102 | 0.884366 | 0.762612 | 0.996493 | 0.997988 | 0.9971 |
| Sept5_Q9Z2Q6_T336t_1 | Sept5_Q9Z | 0 | -1.06 | -0.55 | 0.08 | -0.25 | 0.11 | 0.04 | 0.19 | -0.78 | -0.94 | 0.34 | -0.33 | -1.44 | -0.47 | -1.24 | -0.65 | 0.405 | -0.045 | -0.5675 | 0.150761 | 0.20476 | 0.908365 | 0.989223 | 0.997988 | 0.994666 |
| Sept6_Q9R1T4_S411s_1 | Sept6_Q9R | -0.42 | 0.36 | -0.43 | -0.44 | 0.65 | 0.32 | -0.27 | 0.28 | -0.16 | -0.07 | 0.58 | 0.51 | -0.42 | 0.74 | -0.07 | 0.48 | 0.4775 | 0.4475 | 0.415 | 0.26041 | 0.1355 | 0.172173 | 0.989223 | 0.997988 | 0.994666 |
| Sept7_E9Q9F5_S334s_1 | Sept7_E9Q | -0.11 | -0.16 | 0.42 | 0.46 | 0.19 | 0.02 | 0.86 | 0.58 | -0.48 | -0.22 | 0.29 | 0.19 | -0.22 | 0.51 | 0.22 | 0.26 | 0.26 | -0.2075 | 0.04 | 0.891197 | 0.37282 | 0.480712 | 0.989223 | 0.997988 | 0.999507 |
| Sept7_E9Q9F5_S423s_1 | Sept7_E9Q | -0.68 | -0.57 | -0.61 | -0.72 | -0.13 | 0.23 | -0.32 | -0.06 | -0.49 | -0.8 | -0.33 | -0.08 | 0.32 | -0.34 | -0.13 | 0.06 | 0.575 | 0.22 | 0.6225 | 0.032577 | 0.022293 | 0.370225 | 0.989223 | 0.997988 | 0.994666 |
| Sept7_E9Q9F5_S77s_1 | Sept7_E9Q | -0.36 | 0.01 | -0.11 | 0.06 | 0.03 | -0.03 | 0.29 | -0.06 | -0.35 | -0.19 | -0.93 | -0.03 | -0.89 | -0.21 | -0.53 | -0.09 | 0.1575 | -0.275 | -0.33 | 0.256699 | 0.472243 | 0.328325 | 0.989223 | 0.997988 | 0.994666 |
| Sept8_B1AQZ0_S10s_1 | Sept8_B1A | -0.25 | 0.11 | 0.52 | -0.1 | -0.35 | 0.07 | -0.5 | 0 | -0.26 | 0.03 | -0.76 | 0.48 | 0.66 | -1.23 | 0.21 | 0.1 | -0.265 | -0.1975 | -0.135 | 0.755822 | 0.327142 | 0.556163 | 0.989223 | 0.997988 | 0.9971 |
| Sept8_B1AQZ0_S442s_1 | Sept8_B1A | -0.11 | 0.11 | -0.68 | -0.36 | 0.19 | 0.01 | -0.13 | -0.64 | 0.11 | 0.24 | -0.06 | -0.58 | -0.13 | 0.31 | -1.06 | -0.35 | 0.1175 | 0.1875 | -0.0475 | 0.894444 | 0.677436 | 0.525062 | 0.99315 | 0.997988 | 0.999507 |
| Sept8_B1AQZ0_S444s_1 | Sept8_B1A | -0.1 | 0.52 | -0.21 | 0.35 | -0.22 | -0.04 | 0.21 | 0.48 | -0.49 | 0.57 | -0.25 | 0.27 | -0.26 | -0.01 | 0.18 | 0.39 | -0.0325 | -0.115 | -0.065 | 0.823608 | 0.905072 | 0.724076 | 0.996493 | 0.997988 | 0.999507 |
| Sept8_B1AQZ0_S446s_1 | Sept8_B1A | 0.12 | 0.12 | -0.38 | -0.21 | 0.23 | 0.15 | -0.09 | 0.04 | -0.03 | 0.27 | 0.02 | -0.12 | -0.18 | 0.41 | 0.03 | -0.1 | 0.17 | 0.1225 | 0.1275 | 0.638072 | 0.455884 | 0.608812 | 0.989223 | 0.997988 | 0.995259 |
| Sept9_Q80UG5_S30s_1 | Sept9_Q80 | 0.15 | -0.39 | -0.66 | -0.35 | 2.16 | 0.93 | -0.05 | -0.12 | 1.24 | 0.38 | -0.26 | -0.41 | 0.46 | 0.54 | -0.7 | -0.66 | 1.0425 | 0.55 | 0.2225 | 0.572488 | 0.065639 | 0.198962 | 0.989223 | 0.997988 | 0.995259 |
| Sept9_Q80UG5_S330s_1 | Sept9_Q80 | 0.2 | 0.69 | 1.51 | 0.72 | -0.64 | 0.85 | -0.61 | 0.19 | 1.5 | -0.11 | -0.67 | 0.51 | 1.58 | -0.94 | 0.31 | 0.01 | -0.8325 | -0.4725 | -0.54 | 0.334247 | 0.074744 | 0.361852 | 0.989223 | 0.997988 | 0.994666 |
| Serbp1_Q9CY58_S234s_1 | Serbp1_Q9 | -1.49 | -2.2 | -1.27 | -1.67 | -1.23 | -1.4 | -3.94 | -1.59 | -0.99 | -1.72 | 0.07 | -2.28 | -1.07 | -1.36 | -0.8 | -1.03 | -0.3825 | 0.4275 | 0.5925 | 0.0688 | 0.526502 | 0.412081 | 0.989223 | 0.997988 | 0.994666 |
| Serbp1_Q9CY58_S327s_1 | Serbp1_Q9 | -1.09 | -2.35 | -1.7 | -1.6 | -1.93 | -1.04 | -1.38 | -1.34 | -3.16 | -1.15 | -1.16 | -1.61 | -2.69 | -1.83 | -1.92 | -1.73 | 0.2625 | -0.085 | -0.3575 | 0.336943 | 0.433034 | 0.867145 | 0.989223 | 0.997988 | 0.994666 |
| Serbp1_Q9CY58_S329s_1 | Serbp1_Q9 | -1.74 | -1.94 | -2.03 | -1.8 | -2.11 | -1.42 | -1.51 | -1.68 | -1.99 | -1.52 | -1.38 | -1.73 | -1.55 | -1.63 | -1.92 | -1.79 | 0.1975 | 0.2225 | 0.155 | 0.522031 | 0.411556 | 0.35942 | 0.989223 | 0.997988 | 0.994666 |
| Serinc1_Q9QZI8_S364s_1 | Serinc1_Q9 | 0.84 | 0.34 | 0.36 | 0.08 | 0.69 | 0.68 | -0.26 | 0.03 | 1.13 | 0.47 | -0.06 | 0.25 | -0.4 | 0.59 | 0.95 | 0.04 | -0.12 | 0.0425 | -0.11 | 0.761212 | 0.697028 | 0.895786 | 0.994893 | 0.997988 | 0.9971 |
| Sertm1_Q8CD78_S106s_1 | Sertm1_Q8 | -0.06 | 1.32 | 0.92 | 0.71 | 0.83 | 0.08 | 0.66 | 0.69 | 0.62 | 1 | 0.69 | 0.96 | 1.13 | 0.83 | 1.25 | 0.58 | -0.1575 | 0.095 | 0.225 | 0.529959 | 0.645024 | 0.773532 | 0.99033 | 0.997988 | 0.994666 |
| Sertm1_Q8CD78_S107s_1 | Sertm1_Q8 | -0.06 | 1.32 | 0.92 | 0.71 | 0.83 | 0.08 | 0.66 | 0.69 | 0.62 | 1 | 0.69 | 0.96 | 1.13 | 0.83 | 1.25 | 0.58 | -0.1575 | 0.095 | 0.225 | 0.529959 | 0.645024 | 0.773532 | 0.99033 | 0.997988 | 0.994666 |
| Set_Q9EQU5_S30s_1 | Set_Q9EQU | -3.05 | -2.72 | -3.97 | -8.61 | -2.64 | -3.61 | -5.68 | -5.19 | -3.26 | -2.03 | -2 | -4.85 | -1.56 | -1.43 | -4.83 | -5.83 | 0.3075 | 1.5525 | 1.175 | 0.446795 | 0.815713 | 0.262156 | 0.995762 | 0.997988 | 0.994666 |
| Setd1a_E9PYH6_S1159s_1 | Setd1a_E9I | -1.92 | -2.08 | -1.99 | -1.3 | -2.05 | -2.83 | -1.42 | -3.65 | -2.08 | -1.58 | -0.33 | -2.75 | -1.45 | -1.55 | -1.3 | -1.23 | -0.665 | 0.1375 | 0.44 | 0.134033 | 0.184399 | 0.786888 | 0.989223 | 0.997988 | 0.994666 |
| Setd1a_E9PYH6_S1160s_1 | Setd1a_E9I | -1.41 | -1.78 | -1.99 | -1.3 | -1.91 | -1.72 | -1.42 | -3.65 | -2.03 | -0.74 | -0.33 | -2.75 | -0.82 | -1.37 | -1.3 | -1.23 | -0.555 | 0.1575 | 0.44 | 0.140996 | 0.2697 | 0.771865 | 0.989223 | 0.997988 | 0.994666 |
| Setd1a_E9PYH6_T931t_1 | Setd1a_E9I | -4.6 | -5.25 | -5.11 | -4.99 | -4.42 | -5.34 | -9.77 | -4.19 | -4.3 | -3.81 | -2.63 | -4.31 | -3.12 | -2.86 | -3.86 | -3.83 | -0.9425 | 1.225 | 1.57 | 0.000874 | 0.412915 | 0.014472 | 0.989223 | 0.997988 | 0.994666 |
| Setd1b_Q8CFT2_S985s_2 | Setd1b_Q8 | -2.77 | -1.86 | -1.85 | -3.13 | -1.28 | -1.81 | -1.68 | -0.57 | -1.32 | -2.48 | -2 | -1.02 | -0.15 | -1.55 | -2.77 | -2.4 | 1.0675 | 0.6975 | 0.685 | 0.275536 | 0.025314 | 0.143453 | 0.989223 | 0.997988 | 0.994666 |
| Setd2_E9Q5F9_T1827t_1 | Setd2_E9Q | 0.09 | -0.04 | -0.75 | -1.73 | 0.09 | -0.53 | 0.41 | -0.48 | -0.11 | 0.11 | -0.25 | -0.22 | -0.91 | 0.09 | -0.18 | 0.26 | 0.48 | 0.49 | 0.4225 | 0.380328 | 0.296212 | 0.258586 | 0.989223 | 0.997988 | 0.994666 |
| Setd3_Q91WC0_S12s_1 | Setd3_Q91 | -3.15 | -3.86 | -2.67 | -1.78 | -2.75 | -4.33 | -3.79 | -1.5 | -3.12 | -3.68 | -0.93 | -1.22 | -1.77 | -3.51 | -0.55 | -3.58 | -0.2275 | 0.6275 | 0.5125 | 0.503434 | 0.736074 | 0.399181 | 0.995762 | 0.997988 | 0.994666 |
| Setd3_Q91WC0_S21s_1 | Setd3_Q91 | 0.19 | 0.33 | -0.7 | -0.65 | -0.13 | -0.35 | -1.15 | -1.32 | -0.76 | -0.19 | -0.57 | -1.6 | 0.21 | -0.72 | -1.65 | -0.63 | -0.53 | -0.5725 | -0.49 | 0.29545 | 0.193378 | 0.17661 | 0.989223 | 0.997988 | 0.994666 |
| Setd3_Q91WC0_S571s_1 | Setd3_Q91 | -0.92 | 0.2 | -1.09 | -1.41 | 0.44 | 0.12 | 0.17 | -2.91 | 0.48 | -0.12 | -0.55 | -1.25 | -0.2 | 1.16 | -1.24 | -2.25 | 0.26 | 0.445 | 0.1725 | 0.811139 | 0.732419 | 0.36502 | 0.995762 | 0.997988 | 0.999507 |
| Sf1_D3YVH4_S54s_1 | Sf1_D3YVH | -0.51 | 0.16 | 0.44 | -0.05 | -0.18 | -0.97 | 0.44 | 0.46 | 0.01 | -0.32 | 0.65 | 0.68 | -3.23 | -0.44 | 0.66 | 0.81 | -0.0725 | 0.245 | -0.56 | 0.510941 | 0.849545 | 0.475076 | 0.996349 | 0.997988 | 0.994666 |
| Sf1_D3YVH4_S56s_1 | Sf1_D3YVH | -0.51 | 0.16 | 0.44 | -0.05 | -0.18 | -0.97 | 0.44 | 0.46 | 0.01 | -0.32 | 0.65 | 0.68 | -3.23 | -0.44 | 0.66 | 0.81 | -0.0725 | 0.245 | -0.56 | 0.510941 | 0.849545 | 0.475076 | 0.996349 | 0.997988 | 0.994666 |
| Sf3b1_Q99NB9_S129s_1 | Sf3b1_Q99 | 0 | -0.57 | -0.74 | -1.15 | -0.69 | -0.63 | -1.05 | -0.8 | -1.02 | -1.16 | -1.37 | -0.81 | -0.66 | -0.38 | -0.65 | -1.48 | -0.1775 | -0.475 | -0.1775 | 0.625177 | 0.540914 | 0.142778 | 0.989223 | 0.997988 | 0.995259 |
| Sf3b1_Q99NB9_T211t_1 | Sf3b1_Q99 | 2.67 | 1.13 | 0.89 | 1.33 | 1.79 | 1.08 | 0.84 | 1.3 | -2.64 | 1.14 | -0.58 | 1.04 | -1.46 | -0.11 | 1.05 | 1.16 | -0.2525 | -1.765 | -1.345 | 0.06515 | 0.554716 | 0.066084 | 0.989223 | 0.997988 | 0.994666 |
| Sf3b1_Q99NB9_T223t_1 | Sf3b1_Q99 | 0.47 | 0.67 | 0.25 | 0.15 | 0.79 | 0.85 | 0.62 | 0.39 | 0.49 | 0.84 | -0.04 | -0.54 | -0.02 | 0.45 | -0.23 | -0.84 | 0.2775 | -0.1975 | -0.545 | 0.125469 | 0.24664 | 0.567894 | 0.989223 | 0.997988 | 0.994666 |
| Sf3b1_Q99NB9_T235t_1 | Sf3b1_Q99 | 0.68 | -0.19 | 0.04 | -0.73 | 0 | -0.26 | 0.14 | -0.03 | -1.07 | -1.22 | -1 | 0.11 | -0.24 | 0.33 | 0.33 | -0.6 | 0.0125 | -0.745 | 0.005 | 0.989604 | 0.968695 | 0.099229 | 0.999187 | 0.997988 | 1 |
| Sf3b1_Q99NB9_T248t_1 | Sf3b1_Q99 | 0.32 | 1.07 | 0.19 | -0.44 | -1.59 | 0.78 | -1.74 | 0.16 | 1.36 | -0.12 | -1.4 | -0.67 | 1.66 | -1.22 | -1.73 | -0.56 | -0.8825 | -0.4925 | -0.7475 | 0.314063 | 0.180612 | 0.424642 | 0.989223 | 0.997988 | 0.994666 |
| Sf3b1_Q99NB9_T267t_1 | Sf3b1_Q99 | -0.43 | 0.36 | -1.3 | -0.16 | -0.15 | 0.46 | -0.1 | -0.24 | -0.54 | -0.16 | -0.14 | -0.9 | 0.06 | -0.38 | -1.08 | -0.73 | 0.375 | -0.0525 | -0.15 | 0.722863 | 0.329791 | 0.893237 | 0.989223 | 0.997988 | 0.9971 |
| Sf3b1_Q99NB9_T273t_1 | Sf3b1_Q99 | -2.41 | -2.91 | -3.27 | -2.55 | -2.85 | -4.82 | -3.12 | -2.81 | -1.76 | -4.73 | -3.99 | -3.22 | -0.41 | -3.34 | -2.56 | -4.44 | -0.615 | -0.64 | 0.0975 | 0.89961 | 0.217202 | 0.306906 | 0.989223 | 0.997988 | 0.999507 |
| Sf3b1_Q99NB9_T278t_1 | Sf3b1_Q99 | -2.7 | -2.19 | -1.63 | -1.94 | -2.08 | -2.06 | -2.34 | -1.91 | 0.68 | -2.62 | -2.22 | -2.06 | 3.08 | -2.05 | -1.36 | -2.55 | 0.0175 | 0.56 | 1.395 | 0.237492 | 0.950017 | 0.437885 | 0.999187 | 0.997988 | 0.994666 |
| Sf3b1_Q99NB9_T303t_1 | Sf3b1_Q99 | 1.8 | 1.11 | 0.45 | 0.84 | 1.12 | 0.71 | 0.48 | 0.44 | -0.59 | 0.33 | 1.52 | 0.29 | 0.64 | -0.3 | 1.32 | -0.14 | -0.3625 | -0.6625 | -0.67 | 0.16521 | 0.292197 | 0.198419 | 0.989223 | 0.997988 | 0.994666 |
| Sf3b1_Q99NB9_T313t_1 | Sf3b1_Q99 | 0.96 | -0.2 | 0.08 | 0 | -0.44 | 0.31 | -0.05 | 0.21 | 0.44 | -0.91 | -0.11 | 0.85 | -0.3 | 0.4 | 0.18 | 0.8 | -0.2025 | -0.1425 | 0.06 | 0.869766 | 0.532364 | 0.74844 | 0.989223 | 0.997988 | 0.999507 |
| Sf3b1_Q99NB9_T426t_1 | Sf3b1_Q99 | -1.65 | -1.99 | -0.58 | -1.1 | 0.1 | -1.8 | -0.46 | -1.3 | -3.87 | -1.59 | 0.2 | -0.3 | -1.44 | -1.05 | -1.58 | -0.73 | 0.465 | -0.06 | 0.13 | 0.732964 | 0.348897 | 0.943924 | 0.989223 | 0.997988 | 0.9971 |
| Sfn_O70456_S45s_1 | Sfn_O7045 | -1.02 | -1.42 | -0.27 | -0.68 | 0.36 | -1.23 | -0.42 | -0.38 | -1.34 | -1.54 | -0.99 | -0.69 | -0.87 | 0.34 | -0.8 | -0.29 | 0.43 | -0.2925 | 0.4425 | 0.267218 | 0.289695 | 0.388819 | 0.989223 | 0.997988 | 0.994666 |
| Sfr1_Q8BP27_S115s_1 | Sfr1_Q8BP | -1.14 | -1.67 | -2.7 | -2.92 | -1.82 | -2.31 | -2.45 | -2.57 | -0.18 | -1.42 | -1.3 | -2.44 | 0.15 | -1.77 | -2.12 | -1.94 | -0.18 | 0.7725 | 0.6875 | 0.279821 | 0.675458 | 0.202259 | 0.992876 | 0.997988 | 0.994666 |
| Sfr1_Q8BP27_S67s_1 | Sfr1_Q8BP | -1.34 | -0.94 | -1 | -0.69 | -1.43 | -1.84 | -1.7 | -1.18 | -1.24 | -1.11 | 0.27 | -0.93 | -1.29 | -1.12 | -0.98 | -1.33 | -0.545 | 0.24 | -0.1875 | 0.47445 | 0.054029 | 0.52898 | 0.989223 | 0.997988 | 0.994666 |
| Sfr1_Q8BP27_S67s_2 | Sfr1_Q8BP | -2.48 | -1.91 | -3.51 | -3.63 | -2.08 | -2.65 | -5.1 | -3.84 | -2.39 | -1.64 | -2.44 | -3.63 | -1.28 | -1.53 | -3.88 | -4.95 | -0.535 | 0.3575 | -0.0275 | 0.97469 | 0.448691 | 0.514764 | 0.989223 | 0.997988 | 0.999507 |
| Sfr1_Q8BP27_S71s_2 | Sfr1_Q8BP | -1.18 | -0.48 | -3.51 | -3.63 | -1.16 | -1.02 | -5.1 | -3.84 | -0.45 | -0.34 | -2.44 | -3.63 | -0.29 | -0.67 | -3.88 | -4.95 | -0.58 | 0.485 | -0.2475 | 0.839016 | 0.604504 | 0.630198 | 0.989223 | 0.997988 | 0.999507 |
| Sfswap_Q3USH5_S831s_1 | Sfswap_Q3 | 0.26 | -1.47 | -1.25 | -1.51 | -1.55 | -0.44 | -1.77 | -1.71 | -0.49 | -1.47 | -2.34 | -1.64 | 1.18 | -1.67 | -0.2 | -0.5 | -0.375 | -0.4925 | 0.695 | 0.300723 | 0.445468 | 0.364316 | 0.989223 | 0.997988 | 0.994666 |
| Sgip1_F6VMS8_S27s_1 | Sgip1_F6VI | 1.42 | 1.4 | 0.32 | -2.08 | -0.05 | 1.7 | -0.74 | -0.34 | 2.47 | 0.79 | -1.26 | -0.16 | 1.59 | -1.53 | 1.19 | -1.49 | -0.1225 | 0.195 | -0.325 | 0.751206 | 0.886133 | 0.846411 | 0.996493 | 0.997988 | 0.9971 |
| Sgip1_Q8VD37_S160s_1 | Sgip1_Q8V | -1.46 | -2.29 | -2.23 | -2.97 | -1.86 | -0.53 | -2.29 | -2.49 | -0.61 | -1.83 | -3.82 | -2.91 | -0.43 | -2.04 | -2.5 | -2.5 | 0.445 | -0.055 | 0.37 | 0.498355 | 0.378412 | 0.935971 | 0.989223 | 0.997988 | 0.994666 |
| Sgip1_Q8VD37_S169s_1 | Sgip1_Q8V | 0.3 | 0.02 | 1.31 | 1.22 | 0.51 | 0.43 | 0.75 | 1.22 | 0.24 | 0.19 | 0.51 | 0.79 | 0.19 | 0.38 | 0.45 | 0.82 | 0.015 | -0.28 | -0.2525 | 0.50046 | 0.967278 | 0.448356 | 0.999187 | 0.997988 | 0.994666 |
| Sgip1_Q8VD37_S170s_1 | Sgip1_Q8V | -1.27 | -0.99 | -0.51 | -1.15 | -0.28 | -0.79 | -1.06 | -1.04 | -2 | -1.32 | -0.06 | -1.05 | -1.1 | -0.49 | -0.99 | -1.25 | 0.1875 | -0.1275 | 0.0225 | 0.939692 | 0.510448 | 0.764675 | 0.989223 | 0.997988 | 0.999507 |
| Sgip1_Q8VD37_S24s_2 | Sgip1_Q8V | 4.21 | 4.18 | 2.15 | 1.75 | 3.87 | 3.71 | 1.44 | 1.99 | 2.53 | 3.9 | 1.18 | 2.7 | -2.76 | 1.61 | 1.68 | 2.04 | -0.32 | -0.495 | -2.43 | 0.055629 | 0.683597 | 0.524216 | 0.994272 | 0.997988 | 0.994666 |
| Sgip1_Q8VD37_S264s_1 | Sgip1_Q8V | 0 | 0.81 | 0.65 | 0.43 | 0.57 | -0.25 | 0.85 | 0.96 | 0.04 | 0.29 | 0.45 | 0.57 | 0.38 | 0.9 | 0.08 | 0.31 | 0.06 | -0.135 | -0.055 | 0.856715 | 0.857319 | 0.61809 | 0.996349 | 0.997988 | 0.999507 |
| Sgip1_Q8VD37_S265s_1 | Sgip1_Q8V | 0.34 | 0.73 | 0.69 | 0.4 | 0.34 | -0.01 | 0.71 | 0.59 | -0.17 | 0.52 | -0.1 | 0.26 | 0.11 | 0.59 | 0.36 | 0.41 | -0.1325 | -0.4125 | -0.1725 | 0.498656 | 0.594226 | 0.134776 | 0.989223 | 0.997988 | 0.994666 |
| Sgip1_Q8VD37_S27s_1 | Sgip1_Q8V | -0.13 | -0.91 | 0.32 | -2.08 | -1.02 | -0.16 | -0.74 | -0.34 | 1.24 | -0.28 | -1.26 | -0.16 | 0.3 | -2.58 | 1.19 | -1.49 | 0.135 | 0.585 | 0.055 | 0.950021 | 0.792024 | 0.390236 | 0.995762 | 0.997988 | 0.999507 |
| Sgip1_Q8VD37_S27s_1_1 | Sgip1_Q8V | 0.83 | 0.8 | 1.48 | 0.27 | 0.78 | 0.83 | 0.9 | 1.34 | 1.33 | 0.83 | 0.19 | 1.08 | 0 | -0.76 | 2.01 | 1.23 | 0.1175 | 0.0125 | -0.225 | 0.711774 | 0.698484 | 0.972264 | 0.994893 | 0.997988 | 0.996966 |
| Sgip1_Q8VD37_S300s_1 | Sgip1_Q8V | 0.28 | 1.31 | 0.78 | 0.84 | 0.79 | 0.42 | 0.31 | 0.88 | 0.82 | 0.91 | 0.51 | 0.24 | 0.09 | 1.22 | 0.36 | 0.55 | -0.2025 | -0.1825 | -0.2475 | 0.485286 | 0.483501 | 0.546849 | 0.989223 | 0.997988 | 0.994666 |
| Sgip1_Q8VD37_S319s_1 | Sgip1_Q8V | 2.6 | 0.17 | 3.19 | -0.95 | 0.43 | 2.2 | 4.34 | 1.15 | -0.88 | 2.41 | -2.93 | 3.76 | 1.37 | 0.14 | 3.52 | 4.37 | 0.7775 | -0.6625 | 1.0975 | 0.368734 | 0.493286 | 0.67678 | 0.989223 | 0.997988 | 0.994666 |
| Sgip1_Q8VD37_S371s_1 | Sgip1_Q8V | 5.98 | 3.27 | 4.58 | 1.78 | 4.8 | 6.33 | 5.31 | 4.77 | -0.82 | 4.85 | 0.18 | 4.62 | -2.38 | 0.06 | 4.72 | 5.69 | 1.4 | -1.695 | -1.88 | 0.312857 | 0.122579 | 0.277698 | 0.989223 | 0.997988 | 0.994666 |
| Sgip1_Q8VD37_S377s_1 | Sgip1_Q8V | -1.01 | -1.74 | -2.08 | -2.87 | 0.43 | -0.54 | -1.19 | -1.02 | -2.48 | -2.19 | -0.7 | -2.72 | -1.68 | -1.16 | -2.78 | -2.37 | 1.345 | -0.0975 | -0.0725 | 0.884484 | 0.019845 | 0.859779 | 0.989223 | 0.997988 | 0.999507 |
| Sgip1_Q8VD37_S398s_1 | Sgip1_Q8V | -0.55 | 0.18 | 0.41 | 0.26 | 0.37 | -0.61 | 0.71 | 0.12 | -1.09 | -0.29 | -0.2 | 0.36 | -1.09 | -0.28 | -0.04 | 0.16 | 0.0725 | -0.38 | -0.3875 | 0.307288 | 0.837349 | 0.323321 | 0.995762 | 0.997988 | 0.994666 |
| Sgip1_Q8VD37_S423s_1 | Sgip1_Q8V | -2.2 | -1.09 | -0.55 | -0.49 | -0.76 | -2.13 | -1.34 | -2.17 | -1.55 | -0.57 | -1.59 | 0.57 | -0.52 | 0.31 | -0.97 | -1.25 | -0.5175 | 0.2975 | 0.475 | 0.349691 | 0.29715 | 0.617753 | 0.989223 | 0.997988 | 0.994666 |
| Sgip1_Q8VD37_S424s_1 | Sgip1_Q8V | -0.23 | -0.42 | -0.08 | -0.43 | -0.71 | -1.21 | -0.64 | -0.26 | -1.75 | -1.7 | -0.22 | -0.69 | -0.65 | 0.49 | 0.03 | 0.31 | -0.415 | -0.8 | 0.335 | 0.301984 | 0.13636 | 0.064889 | 0.989223 | 0.997988 | 0.994666 |
| Sgip1_Q8VD37_S424s_2 | Sgip1_Q8V | 0.31 | 0.17 | 1.48 | 1.03 | 0.02 | -0.16 | -0.79 | -0.24 | 0.9 | -0.12 | 0.44 | 0.29 | 0.41 | -0.62 | 0.58 | 0.59 | -1.04 | -0.37 | -0.5075 | 0.246523 | 0.014815 | 0.3411 | 0.989223 | 0.997988 | 0.994666 |
| Sgip1_Q8VD37_S429s_1 | Sgip1_Q8V | -0.51 | 0.38 | 1.71 | 0.52 | -0.95 | 0.49 | -0.62 | 0.52 | 0.85 | 0.9 | 1.66 | 0.03 | 1.07 | 1.44 | 0.19 | -0.71 | -0.665 | 0.335 | -0.0275 | 0.963648 | 0.234968 | 0.52949 | 0.989223 | 0.997988 | 0.999507 |
| Sgip1_Q8VD37_S437s_2 | Sgip1_Q8V | 0.12 | -0.53 | 1.7 | 1.2 | 0.25 | 0.07 | -0.96 | -0.02 | 0.27 | -0.1 | 0.8 | 0.42 | -0.42 | -0.69 | 0.8 | 0.61 | -0.7875 | -0.275 | -0.5475 | 0.353813 | 0.155742 | 0.590185 | 0.989223 | 0.997988 | 0.994666 |
| Sgip1_Q8VD37_S441s_1 | Sgip1_Q8V | -0.56 | -0.13 | -0.02 | -0.28 | 0.12 | -0.23 | -0.46 | -0.39 | -0.1 | -0.33 | 0.02 | -0.07 | 0.11 | 0.3 | -0.51 | 0.1 | 0.0075 | 0.1275 | 0.2475 | 0.394984 | 0.975038 | 0.58648 | 0.999187 | 0.997988 | 0.994666 |
| Sgip1_Q8VD37_S465s_1 | Sgip1_Q8V | -0.8 | 1.33 | 0.24 | 0.21 | -0.06 | 0.4 | 0.05 | -0.05 | 0.2 | 1.48 | -0.44 | -1.08 | -0.86 | 0.52 | 0.23 | -0.41 | -0.16 | -0.205 | -0.375 | 0.464197 | 0.707025 | 0.747606 | 0.995437 | 0.997988 | 0.994666 |
| Sgip1_Q8VD37_S484s_2 | Sgip1_Q8V | 0.11 | 0.38 | -0.14 | 0.01 | 0.26 | 0.15 | 0.24 | 0.13 | -0.2 | 0.42 | -0.49 | 0.11 | -0.51 | 0.48 | -0.26 | -0.09 | 0.105 | -0.13 | -0.185 | 0.540295 | 0.623307 | 0.641676 | 0.989223 | 0.997988 | 0.994666 |
| Sgip1_Q8VD37_S490s_1 | Sgip1_Q8V | -0.99 | 0.38 | 0.1 | 0.12 | -0.52 | -0.54 | 0.34 | -0.43 | -1.1 | -0.49 | -0.71 | 0.07 | -0.36 | -0.14 | -0.27 | -0.91 | -0.19 | -0.46 | -0.3225 | 0.391591 | 0.606136 | 0.257475 | 0.989223 | 0.997988 | 0.994666 |
| Sgip1_Q8VD37_S490s_2 | Sgip1_Q8V | -1.3 | -0.4 | -3.48 | -3.11 | -0.36 | -1.14 | -1.33 | -2.75 | -1.05 | -0.49 | -2.74 | -2.52 | -1.04 | -0.45 | -2.87 | -2.68 | 0.6775 | 0.3725 | 0.3125 | 0.708674 | 0.39242 | 0.650422 | 0.989223 | 0.997988 | 0.996966 |
| Sgip1_Q8VD37_S491s_1 | Sgip1_Q8V | -0.49 | 0.13 | 0.19 | 1.41 | -0.5 | -0.8 | 1.14 | 1.04 | -0.36 | 0.13 | 0.78 | 0.86 | 0.03 | 0.34 | 0.23 | 0.18 | -0.09 | 0.0425 | -0.115 | 0.777558 | 0.876248 | 0.92764 | 0.996493 | 0.997988 | 0.99798 |
| Sgip1_Q8VD37_S491s_2 | Sgip1_Q8V | -0.37 | -0.07 | -0.5 | -0.28 | 0.07 | 0.06 | -0.3 | 0.04 | -0.48 | 0.04 | -0.44 | -0.1 | -0.52 | 0.2 | -0.41 | -0.36 | 0.2725 | 0.06 | 0.0325 | 0.904655 | 0.230003 | 0.803312 | 0.989223 | 0.997988 | 0.999507 |
| Sgip1_Q8VD37_S503s_1 | Sgip1_Q8V | 0.07 | 0.37 | 0.4 | 0.71 | 0.41 | 0.38 | 0.12 | -0.23 | 0.41 | 0.47 | 0.53 | 1.15 | 0.32 | 0.41 | 0.58 | 0.84 | -0.2175 | 0.2525 | 0.15 | 0.576354 | 0.398662 | 0.365814 | 0.989223 | 0.997988 | 0.995259 |
| Sgip1_Q8VD37_S78s_1 | Sgip1_Q8V | 0.38 | 0.43 | 1.27 | 1.14 | 0.3 | -0.07 | 0.95 | 1.02 | 0.21 | 0.59 | 0.3 | 0.65 | -0.3 | 0.69 | 0.94 | 0.38 | -0.255 | -0.3675 | -0.3775 | 0.325308 | 0.475264 | 0.236138 | 0.989223 | 0.997988 | 0.994666 |
| Sgip1_Q8VD37_T180t_1 | Sgip1_Q8V | 0.89 | -0.2 | -1.12 | -1.17 | 1.09 | 1.37 | -0.9 | -1.13 | -0.04 | 0.62 | 0.08 | -0.89 | -0.04 | 1.03 | -1.42 | -1.86 | 0.5075 | 0.3425 | -0.1725 | 0.8131 | 0.483849 | 0.527673 | 0.989223 | 0.997988 | 0.999507 |
| Sgip1_Q8VD37_T180t_2 | Sgip1_Q8V | 0.44 | 0.59 | -0.14 | 0.32 | 0.45 | 0.21 | 0.12 | 0.45 | -0.03 | -0.19 | 0.28 | 0.51 | -0.89 | 0.5 | 0.43 | -0.08 | 0.005 | -0.16 | -0.3125 | 0.413132 | 0.98351 | 0.565969 | 0.999187 | 0.997988 | 0.994666 |
| Sgip1_Q8VD37_T182t_1 | Sgip1_Q8V | -0.5 | -0.87 | 0.45 | 0.22 | 0.24 | -0.19 | 0.09 | 0.58 | -1.05 | -0.89 | 0.76 | 0.33 | -1 | -0.06 | 0.08 | -0.09 | 0.355 | -0.0375 | -0.0925 | 0.817513 | 0.323091 | 0.941253 | 0.989223 | 0.997988 | 0.999507 |
| Sgip1_Q8VD37_T182t_2 | Sgip1_Q8V | 0.44 | 0.59 | -0.14 | 0.32 | 0.45 | 0.21 | 0.12 | 0.45 | -0.03 | -0.19 | 0.28 | 0.51 | -0.89 | 0.5 | 0.43 | -0.08 | 0.005 | -0.16 | -0.3125 | 0.413132 | 0.98351 | 0.565969 | 0.999187 | 0.997988 | 0.994666 |
| Sgip1_Q8VD37_T247t_1 | Sgip1_Q8V | -1.11 | -0.44 | -0.78 | -0.78 | -0.68 | -1.06 | -0.35 | -1.08 | -1.39 | -0.88 | -1.17 | -0.72 | -1.05 | -0.04 | -1 | -1.45 | -0.015 | -0.2625 | -0.1075 | 0.762929 | 0.955003 | 0.33493 | 0.999187 | 0.997988 | 0.9971 |
| Sgip1_Q8VD37_T258t_1 | Sgip1_Q8V | 0.21 | 0.8 | 0.46 | 0.2 | 0.81 | -0.16 | 0.1 | 0.04 | 0 | 0.25 | 0.6 | 0.1 | 0.74 | 0.82 | 0.26 | 0.02 | -0.22 | -0.18 | 0.0425 | 0.887184 | 0.449635 | 0.493482 | 0.989223 | 0.997988 | 0.999507 |
| Sgip1_Q8VD37_T259t_1 | Sgip1_Q8V | 0.21 | 0.8 | 0.46 | 0.2 | 0.81 | -0.16 | 0.1 | 0.04 | 0 | 0.25 | 0.6 | 0.1 | 0.74 | 0.82 | 0.26 | 0.02 | -0.22 | -0.18 | 0.0425 | 0.887184 | 0.449635 | 0.493482 | 0.989223 | 0.997988 | 0.999507 |
| Sgip1_Q8VD37_T259t_2 | Sgip1_Q8V | 2.21 | 1.28 | 1.42 | 2.24 | 1.16 | 1.66 | 1.99 | 2.2 | 1.32 | 2.43 | 1.35 | 2.08 | 0.71 | 1.66 | 1.65 | 1.45 | -0.035 | 0.0075 | -0.42 | 0.263901 | 0.919085 | 0.984202 | 0.997335 | 0.997988 | 0.994666 |
| Sgip1_Q8VD37_T25t_2 | Sgip1_Q8V | 4.21 | 4.18 | 2.72 | 2.41 | 3.87 | 3.71 | 2.05 | 2.51 | 2.53 | 3.9 | 1.53 | 3.17 | -2.76 | 1.61 | 2.22 | 2.48 | -0.345 | -0.5975 | -2.4925 | 0.05153 | 0.558846 | 0.355128 | 0.989223 | 0.997988 | 0.994666 |
| Sgip1_Q8VD37_T263t_1 | Sgip1_Q8V | 1.14 | 0.81 | 0.95 | -0.27 | 1.21 | 1.01 | -0.48 | -0.58 | 0.93 | 0.78 | 0.12 | -1.1 | 0.11 | 1.4 | -0.42 | -0.37 | -0.3675 | -0.475 | -0.4775 | 0.351757 | 0.485803 | 0.375169 | 0.989223 | 0.997988 | 0.994666 |
| Sgip1_Q8VD37_T263t_2 | Sgip1_Q8V | 2.21 | 1.28 | 1.42 | 2.24 | 1.16 | 1.66 | 1.99 | 2.2 | 1.32 | 2.43 | 1.35 | 2.08 | 0.71 | 1.66 | 1.65 | 1.45 | -0.035 | 0.0075 | -0.42 | 0.263901 | 0.919085 | 0.984202 | 0.997335 | 0.997988 | 0.994666 |
| Sgip1_Q8VD37_T335t_1 | Sgip1_Q8V | 0.19 | 0.76 | 0.39 | 0.46 | 0.43 | 0.57 | 0.56 | 0.35 | 0.03 | 0.64 | -0.05 | 0.11 | 0.08 | 0.66 | 0.37 | 0.22 | 0.0275 | -0.2675 | -0.1175 | 0.659228 | 0.899875 | 0.319686 | 0.996493 | 0.997988 | 0.995259 |
| Sgip1_Q8VD37_T409t_1 | Sgip1_Q8V | 0.01 | 0.25 | 0.12 | 0.16 | 0.36 | 0 | -0.33 | 0 | 0.21 | -0.03 | -0.06 | -0.2 | 0.14 | 0.27 | -0.49 | -0.55 | -0.1275 | -0.155 | -0.2925 | 0.323497 | 0.57841 | 0.481044 | 0.989223 | 0.997988 | 0.994666 |
| Sgip1_Q8VD37_T455t_1 | Sgip1_Q8V | 0.19 | 0.89 | -0.52 | 0.07 | 0.59 | 0.63 | 0.77 | -0.22 | 1.7 | 0.39 | 0.03 | -0.23 | 0.71 | 1.12 | -0.24 | -0.97 | 0.285 | 0.315 | -0.0025 | 0.996148 | 0.440145 | 0.524049 | 0.989223 | 0.997988 | 1 |
| Sgk2_Q9QZS5_S10s_1 | Sgk2_Q9Q | 0.88 | 1.45 | 1.84 | 0.39 | 0.97 | 0.45 | 1.2 | -0.16 | 1.42 | 1.48 | 0.64 | 0.72 | 0.6 | 1.15 | 1.36 | 1.38 | -0.525 | -0.075 | -0.0175 | 0.963348 | 0.228078 | 0.847409 | 0.989223 | 0.997988 | 0.999507 |
| Sgk223_Q571I4_S219s_1 | Sgk223_Q5 | 0.74 | 0.41 | 0.04 | 0.27 | 0.31 | 0.87 | -0.36 | 0.81 | 0.15 | 0.06 | -0.1 | 0.15 | -0.73 | 0.64 | -0.05 | 0.49 | 0.0425 | -0.3 | -0.2775 | 0.454334 | 0.897623 | 0.233705 | 0.996493 | 0.997988 | 0.994666 |
| Sgk223_Q571I4_S40s_1 | Sgk223_Q5 | -0.6 | -0.84 | -0.86 | -1.33 | -0.37 | -0.44 | -1.56 | -1.19 | -1.37 | -0.82 | 0.03 | -0.83 | -0.13 | -0.01 | -1.43 | -1.28 | 0.0175 | 0.16 | 0.195 | 0.633592 | 0.95841 | 0.644478 | 0.999187 | 0.997988 | 0.995259 |
| Sgk223_Q571I4_S720s_1 | Sgk223_Q5 | 0.16 | -0.48 | -0.45 | 0.69 | 0.74 | -0.01 | -0.19 | 0.46 | -0.61 | 0.7 | -0.06 | 0.08 | -0.68 | -0.12 | -0.08 | 0.21 | 0.27 | 0.0475 | -0.1475 | 0.683132 | 0.450594 | 0.902698 | 0.989223 | 0.997988 | 0.996159 |
| Sgpp1_Q9JI99_S101s_1 | Sgpp1_Q9J | 0.39 | 0.5 | 0.39 | 0.29 | 0.51 | -0.13 | 0.4 | 0.22 | 0.6 | 0.85 | -0.09 | 0.14 | 0.28 | 0.93 | 0.57 | 0.56 | -0.1425 | -0.0175 | 0.1925 | 0.451491 | 0.533025 | 0.948984 | 0.989223 | 0.997988 | 0.994666 |
| Sgpp1_Q9JI99_S65s_1 | Sgpp1_Q9J | 0.81 | 0.51 | 0.17 | 0.24 | 0.17 | 0.82 | -0.4 | -0.02 | 2.58 | 1.14 | 0.1 | -0.19 | 2.54 | 1.08 | 0.88 | 0.57 | -0.29 | 0.475 | 0.835 | 0.088168 | 0.364154 | 0.429457 | 0.989223 | 0.997988 | 0.994666 |
| Sgsm1_B2RQR5_S229s_1 | Sgsm1_B2F | 6.31 | 2.92 | 3.95 | 0.85 | 4.21 | 6.94 | 4.48 | 3.85 | 0.62 | 5.57 | 0.59 | 3.7 | 0.33 | -0.2 | 5.18 | 4.82 | 1.3625 | -0.8875 | -0.975 | 0.537763 | 0.253621 | 0.546831 | 0.989223 | 0.997988 | 0.994666 |
| Sgsm1_B2RQR5_S397s_1 | Sgsm1_B2F | 0.1 | 1.33 | 1 | 0.63 | 0.39 | 0.45 | 1.14 | 0.79 | 0.81 | 0.43 | 1.34 | 1.1 | -0.15 | 1.13 | 1.37 | 1.19 | -0.0725 | 0.155 | 0.12 | 0.781187 | 0.824682 | 0.655713 | 0.995762 | 0.997988 | 0.998226 |
| Sgsm1_B2RQR5_S633s_1 | Sgsm1_B2F | -1.37 | -2.05 | -0.51 | -0.35 | -3.2 | -1.05 | -0.93 | -0.54 | -1.34 | -1.1 | -0.38 | -0.47 | -0.39 | -2.27 | -0.07 | -0.55 | -0.36 | 0.2475 | 0.25 | 0.668171 | 0.576188 | 0.580954 | 0.989223 | 0.997988 | 0.996159 |
| Sgsm1_B2RQR5_T647t_1 | Sgsm1_B2F | 0.06 | 0.01 | 0.41 | 0.51 | -0.29 | 1.01 | 0.22 | -0.19 | 2.09 | 0.02 | 0.32 | 1.08 | 2.03 | -0.63 | 0.48 | 1.35 | -0.06 | 0.63 | 0.56 | 0.318665 | 0.856055 | 0.191973 | 0.996349 | 0.997988 | 0.994666 |
| Sgsm2_Q80U12_S682s_1 | Sgsm2_Q8( | 0.42 | 0.56 | -0.28 | 0.31 | 0.53 | 0.3 | 0.16 | 0.03 | -1.44 | 0.45 | 3.58 | -0.15 | -0.27 | -1.49 | 0.4 | 0.25 | 0.0025 | 0.3575 | -0.53 | 0.260004 | 0.992365 | 0.709447 | 0.999187 | 0.997988 | 0.994666 |
| Sgta_Q8BJU0_S306s_1 | Sgta_Q8BJ | 0.16 | -1.77 | -0.9 | -0.81 | -2.5 | 1.28 | -1.22 | -0.5 | 1.04 | -0.27 | -1.03 | -0.44 | 0.54 | -2.37 | -1.12 | -1.17 | 0.095 | 0.655 | -0.2 | 0.757585 | 0.902034 | 0.249224 | 0.996493 | 0.997988 | 0.9971 |
| Sgta_Q8BJU0_S306s_2 | Sgta_Q8BJ | 0.06 | 0.62 | 0.88 | 0.78 | -1.86 | 0.25 | -1.19 | 0.5 | 2.9 | 0.49 | 0.2 | 0.83 | 0.18 | -0.75 | -0.57 | -1.44 | -1.16 | 0.52 | -1.23 | 0.010284 | 0.054896 | 0.386129 | 0.989223 | 0.997988 | 0.994666 |
| Sgta_Q8BJU0_S308s_1 | Sgta_Q8BJ | -0.21 | -0.92 | -0.04 | -0.21 | -2.43 | 0.66 | -1.8 | -0.38 | 0.72 | -0.4 | -0.15 | 0.06 | 0.73 | -1.8 | -1.3 | -0.75 | -0.6425 | 0.4025 | -0.435 | 0.428895 | 0.330621 | 0.245494 | 0.989223 | 0.997988 | 0.994666 |
| Sgta_Q8BJU0_T304t_1 | Sgta_Q8BJ | -1.76 | -2.86 | -2.22 | -1.74 | -3.03 | -1.08 | -3.16 | -1.71 | -0.36 | -2.23 | -0.58 | -1.72 | -0.36 | -3.39 | -3.8 | -2.83 | -0.1 | 0.9225 | -0.45 | 0.53944 | 0.847731 | 0.086808 | 0.996349 | 0.997988 | 0.994666 |
| Sgta_Q8BJU0_T304t_2 | Sgta_Q8BJ | 0.06 | 0.62 | 0.88 | 0.78 | -1.86 | 0.25 | -1.19 | 0.5 | 2.9 | 0.49 | 0.2 | 0.83 | 0.18 | -0.75 | -0.57 | -1.44 | -1.16 | 0.52 | -1.23 | 0.010284 | 0.054896 | 0.386129 | 0.989223 | 0.997988 | 0.994666 |
| Sgta_Q8BJU0_T81t_1 | Sgta_Q8BJ | -0.66 | 0.1 | -0.3 | -0.76 | -0.72 | -0.52 | -0.81 | -0.68 | 0.77 | -0.34 | 0.01 | -0.34 | 1.09 | -0.19 | -0.38 | -0.5 | -0.2775 | 0.43 | 0.41 | 0.337073 | 0.294174 | 0.231761 | 0.989223 | 0.997988 | 0.994666 |
| Sgta_Q8BJU0_T81t_2 | Sgta_Q8BJ | -0.73 | 0.14 | 2.04 | 0.43 | -0.64 | 0.22 | -0.94 | -0.9 | 0.31 | 0.05 | -0.92 | 0.17 | -0.11 | -0.22 | 0.14 | 0.58 | -1.035 | -0.5675 | -0.3725 | 0.510184 | 0.096836 | 0.348535 | 0.989223 | 0.997988 | 0.994666 |
| Sgtb_Q8VD33_S295s_1 | Sgtb_Q8VD | -0.55 | -1.65 | -1.15 | -1.38 | -1.13 | -0.58 | -1.38 | -0.77 | -1.12 | -1.86 | 0.32 | -0.93 | 0.06 | -0.98 | -1.68 | -1.38 | 0.2175 | 0.285 | 0.1875 | 0.669962 | 0.492801 | 0.558924 | 0.989223 | 0.997988 | 0.996159 |
| Sgtb_Q8VD33_S295s_2 | Sgtb_Q8VD | -1.2 | -0.99 | 0.39 | -0.8 | -1.37 | -0.41 | -0.82 | 0.15 | 0.06 | -0.8 | 0.02 | -0.21 | -2.1 | -1.14 | -0.27 | 0.33 | 0.0375 | 0.4175 | -0.145 | 0.804318 | 0.933316 | 0.315982 | 0.998562 | 0.997988 | 0.999507 |
| Sgtb_Q8VD33_S299s_1 | Sgtb_Q8VD | 0.41 | 0.15 | 0.37 | 0.06 | -0.89 | 0.44 | -0.96 | -0.48 | 0.98 | 0.09 | -0.04 | 0.32 | 0.88 | -0.43 | -0.72 | -0.02 | -0.72 | 0.09 | -0.32 | 0.403644 | 0.057348 | 0.755405 | 0.989223 | 0.997988 | 0.994666 |
| Sgtb_Q8VD33_S304s_2 | Sgtb_Q8VD | -1.2 | -0.99 | 1.05 | -1.25 | -1.37 | -0.41 | -1.13 | 0.1 | 0.06 | -0.8 | -1.58 | -0.65 | -2.1 | -1.14 | -0.32 | -1.35 | -0.105 | -0.145 | -0.63 | 0.311233 | 0.856385 | 0.806982 | 0.996349 | 0.997988 | 0.994666 |
| Sh2b1_Q91ZM2_S126s_1 | Sh2b1_Q91 | 1.08 | 0.31 | 0.09 | -0.13 | 1.07 | 1.05 | 0.59 | 0.29 | 0.58 | 0.97 | 0.46 | 0.08 | -0.1 | 0.92 | 0.1 | 0.46 | 0.4125 | 0.185 | 0.0075 | 0.983677 | 0.234886 | 0.589788 | 0.989223 | 0.997988 | 1 |
| Sh2b1_Q91ZM2_S127s_1 | Sh2b1_Q91 | 0.59 | 0.19 | 0.23 | 0.24 | 0.49 | 0.61 | 0.69 | 0.53 | 0.1 | 0.3 | 0.77 | 0.3 | -0.27 | 0.65 | -0.02 | 0.48 | 0.2675 | 0.055 | -0.1025 | 0.730723 | 0.219376 | 0.824155 | 0.989223 | 0.997988 | 0.9971 |
| Sh2b1_Q91ZM2_S219s_1 | Sh2b1_Q91 | 0.04 | -0.39 | -0.56 | -0.55 | -0.79 | -0.23 | -0.66 | 0.24 | 0.3 | 0.14 | 0.68 | -0.35 | -0.06 | -0.72 | -0.84 | -0.23 | 0.005 | 0.5575 | -0.0975 | 0.743955 | 0.986594 | 0.08634 | 0.999187 | 0.997988 | 0.9971 |
| Sh2b3_D3Z0T9_S102s_1 | Sh2b3_D3Z | -1.94 | -0.91 | -1.01 | -2.16 | -1.09 | -1.51 | -1.83 | -2.22 | -2.01 | -2.2 | -1.57 | -1.41 | -0.98 | -1.75 | -2.12 | -1.67 | -0.1575 | -0.2925 | -0.125 | 0.756735 | 0.68524 | 0.442509 | 0.994624 | 0.997988 | 0.9971 |
| Sh2d3c_Q9QZS8_S189s_1 | Sh2d3c_Q9 | 0.56 | 0.13 | 0.19 | 0.31 | 0.05 | 1.17 | -0.42 | -0.39 | 2.11 | 0.5 | -0.59 | 0.71 | 2.31 | -0.25 | 0.51 | 0.44 | -0.195 | 0.385 | 0.455 | 0.391582 | 0.606512 | 0.470275 | 0.989223 | 0.997988 | 0.994666 |
| Sh2d3c_Q9QZS8_S283s_1 | Sh2d3c_Q9 | 0.25 | 0.94 | 0.57 | 0.86 | 0.62 | 0.97 | 1.45 | 1.13 | 0.36 | 1.04 | 0.82 | 1.36 | 1.2 | -0.83 | 1.11 | 1.03 | 0.3875 | 0.24 | -0.0275 | 0.954927 | 0.178679 | 0.433381 | 0.989223 | 0.997988 | 0.999507 |
| Sh2d3c_Q9QZS8_S289s_1 | Sh2d3c_Q9 | 0.01 | 0.28 | 0.35 | 0.14 | -0.22 | -0.43 | 0.48 | -0.2 | -0.21 | 0.43 | 0.96 | -0.01 | 0.46 | 0.34 | 0.27 | 0.53 | -0.2875 | 0.0975 | 0.205 | 0.395453 | 0.285591 | 0.750795 | 0.989223 | 0.997988 | 0.994666 |
| Sh2d3c_Q9QZS8_S290s_1 | Sh2d3c_Q9 | 0.01 | 0.28 | 0.35 | 0.14 | -0.22 | -0.43 | 0.48 | -0.2 | -0.21 | 0.43 | 0.96 | -0.01 | 0.46 | 0.34 | 0.27 | 0.53 | -0.2875 | 0.0975 | 0.205 | 0.395453 | 0.285591 | 0.750795 | 0.989223 | 0.997988 | 0.994666 |
| Sh2d5_Q8JZW5_S12s_1 | Sh2d5_Q8J | -1.81 | -2.5 | -3.25 | -3.31 | -1.78 | -1.35 | -1.93 | -1.2 | -2.19 | -1.73 | 0.62 | -2.66 | -1.07 | -1.12 | -2.3 | -3.4 | 1.1525 | 1.2275 | 0.745 | 0.233858 | 0.013186 | 0.118161 | 0.989223 | 0.997988 | 0.994666 |
| Sh2d5_Q8JZW5_S237s_1 | Sh2d5_Q8J | -1.35 | -2.03 | -0.53 | -0.41 | -1.35 | -1.39 | 0.23 | 0.08 | -2.04 | -1.27 | -0.56 | -0.64 | -1.76 | -1.88 | -0.43 | -0.3 | 0.4725 | -0.0475 | -0.0125 | 0.981145 | 0.381895 | 0.921829 | 0.989223 | 0.997988 | 1 |
| Sh3bgrl2_Q8BG73_S103s_1 | Sh3bgrl2_( | -1.07 | -2.3 | 0 | 0.63 | -2.15 | -1.53 | -0.34 | -0.78 | -1.56 | -1.59 | -0.8 | -0.35 | -0.75 | -1.58 | -1.43 | -2.91 | -0.515 | -0.39 | -0.9825 | 0.18426 | 0.4494 | 0.549747 | 0.989223 | 0.997988 | 0.994666 |
| Sh3bp1_P55194_S535s_1 | Sh3bp1_P5 | -1.03 | -0.91 | -2.05 | -1.91 | -0.39 | -1.12 | -1.77 | -2.09 | -1.71 | -0.92 | -1.77 | -1.95 | -1.17 | -1.03 | -1.9 | -2.08 | 0.1325 | -0.1125 | -0.07 | 0.860982 | 0.767158 | 0.766001 | 0.995762 | 0.997988 | 0.999507 |
| Sh3bp4_Q921I6_S245s_1 | Sh3bp4_Q9 | -2.59 | -2.89 | -0.64 | -0.47 | -1.85 | -2.68 | -0.13 | -1.28 | -4.04 | -2.97 | -1.1 | -1.47 | -2.09 | -1.41 | -0.55 | -0.65 | 0.1625 | -0.7475 | 0.4725 | 0.478617 | 0.823856 | 0.378474 | 0.995762 | 0.997988 | 0.994666 |
| Sh3bp4_Q921I6_S42s_1 | Sh3bp4_Q9 | -0.56 | 0.15 | -1.38 | -0.16 | -1.08 | -1.17 | -0.81 | -0.83 | -0.52 | -0.05 | 0.17 | 0.06 | -0.14 | 0.55 | -0.03 | -0.74 | -0.485 | 0.4025 | 0.3975 | 0.357345 | 0.184114 | 0.294556 | 0.989223 | 0.997988 | 0.994666 |
| Sh3bp5l_Q99LH9_S30s_1 | Sh3bp5l_Q | -0.86 | -0.33 | 0.07 | 0.6 | -0.48 | -1.07 | 1.13 | 0.55 | -0.15 | 0.06 | 0.7 | 0.74 | -0.38 | 0.55 | 0.72 | 0.08 | 0.1625 | 0.4675 | 0.3725 | 0.365099 | 0.759771 | 0.243804 | 0.995762 | 0.997988 | 0.994666 |
| Sh3bp5l_Q99LH9_S361s_1 | Sh3bp5l_Q | -0.29 | 0.79 | 0.27 | 0.53 | 1.1 | 0.68 | -0.22 | 0.62 | 0.37 | 0.68 | 1.22 | -0.25 | 0.66 | 0.04 | 0.21 | 0.73 | 0.22 | 0.18 | 0.085 | 0.795858 | 0.543831 | 0.642975 | 0.989223 | 0.997988 | 0.999507 |
| Sh3bp5l_Q99LH9_S377s_1 | Sh3bp5l_Q | -0.11 | -0.18 | -0.1 | 0.06 | -0.32 | -0.09 | -0.14 | -0.03 | 0.17 | 0.41 | 0.43 | 0.59 | 0.46 | -0.17 | 0.33 | -0.42 | -0.0625 | 0.4825 | 0.1325 | 0.645071 | 0.756474 | 0.049309 | 0.995762 | 0.997988 | 0.995259 |
| Sh3bp5l_Q99LH9_T13t_1 | Sh3bp5l_Q | -3.16 | -2.16 | -0.71 | 0.24 | -2.49 | -3.34 | -1.2 | -0.6 | -3.03 | -2.17 | -1.03 | -0.9 | -1.09 | -3.09 | -0.84 | -0.9 | -0.46 | -0.335 | -0.0325 | 0.968264 | 0.591192 | 0.680313 | 0.989223 | 0.997988 | 0.999507 |
| Sh3d19_Q91X43_S195s_1 | Sh3d19_Q9 | -1.44 | -1.63 | -2.22 | -1.92 | -2.45 | -0.72 | -3.41 | -2.72 | -1.02 | -0.88 | 0.33 | -3.16 | -0.21 | -1.29 | -2.46 | -3.38 | -0.5225 | 0.62 | -0.0325 | 0.959618 | 0.346022 | 0.369358 | 0.989223 | 0.997988 | 0.999507 |
| Sh3gl1_Q62419_S288s_1 | Sh3gl1_Q6 | -2.63 | -2.25 | -1.67 | -3.77 | -0.44 | -2.43 | -0.17 | -2.59 | -4.05 | -1.4 | 0.34 | -0.07 | -0.13 | 0.07 | -2.08 | -1.74 | 1.1725 | 1.285 | 1.61 | 0.029251 | 0.113411 | 0.202572 | 0.989223 | 0.997988 | 0.994666 |
| Sh3gl2_Q62420_S262s_1 | Sh3gl2_Q6 | -0.52 | -0.25 | -0.8 | -0.47 | -0.09 | -0.33 | -0.22 | -0.85 | -0.47 | -0.46 | -0.82 | -0.81 | -1.01 | -0.08 | -0.79 | -0.53 | 0.1375 | -0.13 | -0.0925 | 0.754283 | 0.591848 | 0.588551 | 0.989223 | 0.997988 | 0.9971 |
| Sh3gl3_J3QQ44_T243t_1 | Sh3gl3_J3C | -1.66 | -2.35 | -4.31 | -1.39 | -1.85 | -2.12 | -2.11 | -1.53 | -1.16 | -2.54 | 1.28 | -1.67 | -0.5 | -2.29 | -2.47 | -2.39 | 0.525 | 1.405 | 0.515 | 0.482228 | 0.393161 | 0.155993 | 0.989223 | 0.997988 | 0.994666 |
| Sh3gl3_J3QQ44_T248t_1 | Sh3gl3_J3C | -0.72 | -0.87 | -0.94 | -1.2 | -1.37 | -0.63 | -1.16 | -3.93 | 0.1 | -1.12 | -0.33 | -1.73 | 1.05 | -0.88 | -1.03 | -1.26 | -0.84 | 0.1625 | 0.4025 | 0.437748 | 0.22183 | 0.695198 | 0.989223 | 0.997988 | 0.994666 |
| Sh3glb2_Q8R3V5_S379s_1 | Sh3glb2_Q | 0.27 | 0.75 | 0.4 | 0.09 | 0.44 | 0.27 | 0.45 | 0.11 | 0.66 | 0.64 | 0.13 | 0.1 | -0.01 | 0.58 | 0.19 | 0.47 | -0.06 | 0.005 | -0.07 | 0.799807 | 0.797132 | 0.985108 | 0.995762 | 0.997988 | 0.999507 |
| Sh3kbp1_Q8R550_S373s_1 | Sh3kbp1_Q | -0.95 | -1.25 | -0.78 | -0.88 | -1.44 | -0.68 | -0.77 | -0.8 | -1.33 | -0.63 | -0.46 | -0.67 | -0.78 | -0.75 | -0.63 | -0.88 | 0.0425 | 0.1925 | 0.205 | 0.406075 | 0.867463 | 0.486593 | 0.996493 | 0.997988 | 0.994666 |
| Sh3pxd2a_O89032_S377s_1 | Sh3pxd2a_ | -1.98 | -0.83 | -1.38 | -1.33 | -0.58 | -1.7 | -0.59 | -1.14 | -2.21 | -1.48 | -0.96 | -1.64 | -1.38 | -0.79 | -1.44 | -1.89 | 0.3775 | -0.1925 | 0.005 | 0.988719 | 0.303372 | 0.595912 | 0.989223 | 0.997988 | 1 |
| Sh3pxd2a_O89032_S503s_1 | Sh3pxd2a_ | 1.18 | 1.51 | 1.1 | 1.09 | 0.63 | 1.26 | 1.15 | 1.26 | 1.49 | 1.31 | 1.2 | 0.27 | 1 | 1.29 | 0.85 | 1.05 | -0.145 | -0.1525 | -0.1725 | 0.494927 | 0.554262 | 0.634869 | 0.989223 | 0.997988 | 0.994666 |

| | | | | | | | | | | | | | | | | | | | | | | | | |
|---|---|---|---|---|---|---|---|---|---|---|---|---|---|---|---|---|---|---|---|---|---|---|---|---|
| Sh3pxd2a_O89032_S769s_1 | Sh3pxd2a_ | 0.18 | 0.19 | 0.32 | 0.57 | 0.1 | 0.43 | 0.79 | 0.62 | -0.13 | 0.46 | 0.67 | 0.24 | -0.82 | 0.51 | 0.18 | 0.54 | 0.17 | -0.005 | -0.2125 | 0.555422 | 0.483728 | 0.984634 | 0.989223 | 0.997988 | 0.995259 |
| Sh3pxd2a_O89032_S950s_1 | Sh3pxd2a_ | 0.14 | 0.05 | -0.31 | 0.07 | 0.39 | 0.51 | -0.8 | 0.21 | -0.42 | -0.39 | 0.23 | 0.11 | -0.89 | -0.18 | -0.45 | -0.18 | 0.09 | -0.105 | -0.4125 | 0.159892 | 0.783469 | 0.688838 | 0.995762 | 0.997988 | 0.994666 |
| Sh3pxd2a_O89032_S965s_1 | Sh3pxd2a_ | -0.38 | -0.86 | -0.04 | -0.28 | -0.24 | 0.52 | -0.72 | -0.29 | -1.39 | -1.17 | -0.68 | -0.46 | -0.23 | -1.58 | -0.93 | -0.62 | 0.2075 | -0.535 | -0.45 | 0.228923 | 0.525476 | 0.110033 | 0.989223 | 0.997988 | 0.994666 |
| Sh3pxd2a_O89032_S986s_1 | Sh3pxd2a_ | 0.44 | 0.83 | 0.24 | 0.17 | 0.83 | -0.02 | 0.26 | 0.64 | -0.1 | -0.19 | -0.18 | -0.06 | 0.51 | 0.74 | 0.16 | 0.12 | 0.0075 | -0.5525 | -0.0375 | 0.894892 | 0.978479 | 0.041807 | 0.999187 | 0.997988 | 0.999507 |
| Sh3rf1_E9QQ33_S740s_1 | Sh3rf1_E9Q | 0.13 | 0.47 | -0.06 | -0.26 | 0.54 | -0.42 | -0.11 | 0.37 | -0.14 | -0.03 | -1.91 | -0.47 | -0.75 | -0.01 | 0.15 | 0.48 | 0.025 | -0.7075 | -0.1025 | 0.762683 | 0.932644 | 0.137636 | 0.998562 | 0.997988 | 0.9971 |
| Sh3rf1_E9QQ33_S804s_1 | Sh3rf1_E9Q | 1.08 | 0.54 | 0.62 | 1.03 | 0.57 | 0.83 | 0.66 | 1.02 | 0.61 | 0.46 | 0.33 | 0.56 | 0.19 | 0.96 | -0.01 | 0.21 | -0.0475 | -0.3275 | -0.48 | 0.144729 | 0.841741 | 0.192041 | 0.996064 | 0.997988 | 0.994666 |
| Shank1_D3YZU4_S1283s_1 | Shank1_D3 | -1.3 | -0.67 | -0.5 | -0.87 | -0.4 | -1.5 | -0.25 | -0.61 | -1.19 | -0.76 | 0.08 | -0.58 | -0.73 | -0.78 | -0.91 | -0.61 | 0.145 | 0.2225 | 0.0775 | 0.775059 | 0.667817 | 0.513673 | 0.992842 | 0.997988 | 0.99798 |
| Shank1_D3YZU4_S1434s_1 | Shank1_D3 | 0.59 | -0.12 | -0.6 | 0.45 | 0.97 | 0.7 | 0.17 | 0.88 | 0.09 | -0.43 | 0.85 | 0.47 | -0.34 | -0.12 | -0.8 | 0.04 | 0.6 | 0.165 | -0.385 | 0.294536 | 0.099068 | 0.672232 | 0.989223 | 0.997988 | 0.994666 |
| Shank1_D3YZU4_S1531s_1 | Shank1_D3 | 1.13 | -1.84 | -0.66 | -0.01 | 0.75 | 1.71 | 0.48 | 0.84 | -0.92 | -1.58 | 0.33 | 0.19 | -2.24 | -0.02 | -1.52 | -1.52 | 1.29 | -0.15 | -0.98 | 0.181231 | 0.055819 | 0.8293 | 0.989223 | 0.997988 | 0.994666 |
| Shank1_D3YZU4_S1899s_1 | Shank1_D3 | -0.19 | -0.02 | -1.61 | -0.64 | -0.04 | 0.4 | -1.27 | -1.32 | 0.99 | 0 | -0.59 | -1.74 | 0.79 | -0.68 | -1.67 | -1.73 | 0.0575 | 0.28 | -0.2075 | 0.740296 | 0.910762 | 0.650423 | 0.997335 | 0.997988 | 0.9971 |
| Shank1_D3YZU4_S1998s_1 | Shank1_D3 | 0.18 | -1.72 | -2.67 | -3.83 | 0.96 | -0.38 | -0.86 | -3.52 | -1.27 | -1.43 | 0.47 | -0.43 | -1.05 | -0.28 | -0.93 | -1.51 | 1.06 | 1.345 | 1.0675 | 0.194605 | 0.343714 | 0.138722 | 0.989223 | 0.997988 | 0.994666 |
| Shank1_D3YZU4_S2017s_1 | Shank1_D3 | -0.25 | 0.06 | 0.54 | -0.64 | -0.66 | -0.59 | -0.35 | 0.32 | 0.04 | -1.05 | -1.05 | 0.34 | -0.77 | -0.65 | 0.47 | 0.84 | -0.2475 | -0.3575 | 0.045 | 0.921673 | 0.473863 | 0.413907 | 0.989223 | 0.997988 | 0.999507 |
| Shank1_D3YZU4_S413s_1 | Shank1_D3 | 1.66 | 1.14 | 1.35 | 0.81 | 1.32 | 1.66 | 1.95 | 1.51 | -0.48 | 1.5 | -0.8 | 1.95 | 0 | -1.11 | 1.72 | 1.41 | 0.37 | -0.6975 | -0.735 | 0.252076 | 0.187857 | 0.298256 | 0.989223 | 0.997988 | 0.994666 |
| Shank1_D3YZU4_S519s_1 | Shank1_D3 | -0.23 | -1.88 | -0.85 | -3.49 | -2.16 | 0.54 | -1.7 | -1.31 | 0.76 | -2.65 | -1.89 | -1.02 | 0.3 | -2.66 | -4.12 | -2.1 | 0.455 | 0.4125 | -0.5325 | 0.601294 | 0.576822 | 0.650735 | 0.989223 | 0.997988 | 0.995259 |
| Shank1_D3YZU4_S890s_1 | Shank1_D3 | -2.01 | -0.65 | -1.19 | -1.84 | -0.76 | -2.32 | -0.9 | -1.75 | -2.43 | -1.7 | 0.19 | -1.06 | -1.66 | -0.2 | -1.87 | -1.57 | -0.01 | 0.1725 | 0.0975 | 0.836456 | 0.982282 | 0.768696 | 0.999187 | 0.997988 | 0.999507 |
| Shank1_D3YZU5_S638s_1 | Shank1_D3 | 3.29 | 2.5 | 2.36 | 0.79 | 2.44 | 3.09 | 2.68 | 2.04 | 1.63 | 2.66 | 1.77 | 1.16 | 0.18 | 1.54 | 2.66 | 2.82 | 0.3275 | -0.43 | -0.435 | 0.548149 | 0.531098 | 0.452782 | 0.989223 | 0.997988 | 0.994666 |
| Shank2_D3Z5K8_S1013s_1 | Shank2_D3 | 0.04 | 1.33 | 1 | 0.6 | -0.49 | 0.52 | -0.75 | -0.36 | 1.98 | 1.13 | 1.26 | 1.03 | 1.74 | -1.17 | 1.93 | 1.3 | -1.0125 | 0.6075 | 0.2075 | 0.764124 | 0.023544 | 0.119564 | 0.989223 | 0.997988 | 0.9971 |
| Shank2_D3Z5K8_S1043s_1 | Shank2_D3 | 0.4 | 0.64 | 0.55 | 0.13 | 0.47 | 0.5 | 0.65 | 0.93 | -0.62 | 0.58 | -0.98 | 0.16 | -0.36 | 0.07 | -0.13 | 0.56 | 0.2075 | -0.645 | -0.395 | 0.198382 | 0.37637 | 0.114385 | 0.989223 | 0.997988 | 0.994666 |
| Shank2_D3Z5K8_S1370s_1 | Shank2_D3 | 1.62 | 1.11 | -0.13 | 0.34 | 1.92 | 1.37 | 0.63 | 0.75 | 0.3 | 0.46 | -0.12 | -0.2 | -0.08 | 1.66 | -0.71 | 0.02 | 0.4325 | -0.625 | -0.5125 | 0.391217 | 0.35639 | 0.156674 | 0.989223 | 0.997988 | 0.994666 |
| Shank2_D3Z5K8_S1603s_1 | Shank2_D3 | 0.25 | -0.38 | -0.31 | -1.06 | 0.58 | 0.38 | 0.27 | -0.16 | -1.8 | -0.21 | 0.14 | -0.56 | -1.13 | -0.64 | 0.13 | -0.28 | 0.6425 | -0.2325 | -0.105 | 0.787901 | 0.071841 | 0.627161 | 0.989223 | 0.997988 | 0.999507 |
| Shank2_D3Z5K8_S1699s_1 | Shank2_D3 | 0.44 | -0.84 | -1.53 | 0.44 | 0.52 | 1.06 | 0.53 | 1.16 | -0.6 | -0.6 | 1.14 | 0.19 | -1.61 | 0.11 | -1.47 | -1.26 | 1.19 | 0.405 | -0.685 | 0.254083 | 0.03116 | 0.49709 | 0.989223 | 0.997988 | 0.994666 |
| Shank2_D3Z5K8_S1703s_1 | Shank2_D3 | -0.19 | 0.77 | 0.93 | 0.72 | 0.96 | 0.08 | 0.96 | 0.38 | -0.23 | 0.26 | -0.4 | 0.84 | -0.2 | 0.81 | 0.21 | 0.33 | 0.0375 | -0.44 | -0.27 | 0.45385 | 0.912328 | 0.265347 | 0.997335 | 0.997988 | 0.994666 |
| Shank2_D3Z5K8_S446s_1 | Shank2_D3 | 1.36 | 0.15 | 0.02 | 0.6 | 0.69 | 1.69 | 0.6 | 0.47 | -0.15 | 0.77 | 1.82 | 0.63 | -0.13 | 0.99 | -0.81 | -0.56 | 0.33 | 0.235 | -0.66 | 0.189614 | 0.414601 | 0.626388 | 0.989223 | 0.997988 | 0.994666 |
| Shank2_D3Z5K8_S821s_1 | Shank2_D3 | 0.78 | -0.06 | -1.34 | -0.67 | 0.47 | 0.79 | 0.31 | 0.68 | -0.8 | -0.03 | -0.4 | -0.15 | -0.27 | 0.15 | -1.04 | -0.4 | 0.885 | -0.0225 | -0.0675 | 0.890096 | 0.066836 | 0.961053 | 0.989223 | 0.997988 | 0.999507 |
| Shank2_D3Z5K8_S951s_1 | Shank2_D3 | 0.01 | 0.45 | 1.15 | 0.95 | 0.85 | 0.47 | 1.22 | 0.92 | -0.58 | 0.31 | 0.74 | 1.28 | -1.09 | -0.77 | 0.7 | 0.84 | 0.225 | -0.2025 | -0.72 | 0.190355 | 0.482313 | 0.655254 | 0.989223 | 0.997988 | 0.994666 |
| Shank2_Q80Z38_S372s_1 | Shank2_Q8 | -2.94 | -1.87 | -1.94 | -1.67 | -2.57 | -3.75 | -1.51 | -2.02 | -3.03 | -2.95 | -0.72 | -1.63 | -4.18 | -1.8 | -1.84 | -2.35 | -0.3575 | 0.0225 | -0.4375 | 0.455248 | 0.489073 | 0.968764 | 0.989223 | 0.997988 | 0.994666 |
| Shank3_Q4ACU6_S1032s_1 | Shank3_Q4 | 0.22 | -0.16 | -0.4 | -0.46 | -0.2 | 0.25 | 0.33 | -0.47 | -0.84 | 0.77 | -0.27 | -0.69 | -0.32 | 0.34 | -0.25 | 0.26 | 0.1775 | -0.0575 | 0.2075 | 0.487639 | 0.530666 | 0.883929 | 0.989223 | 0.997988 | 0.994666 |
| Shank3_Q4ACU6_S1134s_1 | Shank3_Q4 | 0.46 | -0.66 | -0.28 | 0.91 | 0.19 | 0.65 | 0.86 | 1.13 | -0.09 | -0.1 | -0.59 | 0.27 | -0.61 | 0.28 | -0.34 | -0.19 | 0.6 | -0.235 | -0.3225 | 0.433706 | 0.150801 | 0.555812 | 0.989223 | 0.997988 | 0.994666 |
| Shank3_Q4ACU6_S1510s_1 | Shank3_Q4 | -0.77 | -1.39 | -0.53 | -0.53 | -0.23 | 0.03 | -0.43 | -0.32 | -1.02 | -1.48 | 0.6 | -0.91 | -1.98 | 0.05 | -1.36 | -1.17 | 0.5675 | 0.1025 | -0.31 | 0.502446 | 0.058003 | 0.827924 | 0.989223 | 0.997988 | 0.994666 |
| Shank3_Q4ACU6_S1529s_1 | Shank3_Q4 | 0.86 | -0.78 | -1.68 | -0.81 | 0.38 | 1.71 | 0.05 | 0.47 | 0.74 | -1.12 | 1.29 | -0.52 | 0.04 | -0.33 | -1.59 | -0.17 | 1.255 | 0.7 | 0.09 | 0.878756 | 0.052624 | 0.326377 | 0.989223 | 0.997988 | 0.999507 |
| Shank3_Q4ACU6_S1577s_1 | Shank3_Q4 | 2.7 | -0.58 | -1.12 | -0.3 | 0.96 | 2.79 | -0.3 | 0.11 | 1.29 | -0.02 | -0.3 | -0.53 | 0.44 | -0.04 | -1.93 | -1.22 | 0.715 | -0.065 | -0.8625 | 0.344693 | 0.45946 | 0.93867 | 0.989223 | 0.997988 | 0.994666 |
| Shank3_Q4ACU6_S351s_1 | Shank3_Q4 | 0.59 | 0.46 | 0.51 | 0.38 | 0.71 | 0.69 | 0.08 | -0.34 | 0.88 | 0.08 | -0.16 | -0.31 | 1.19 | -0.09 | -0.59 | 0.33 | -0.2 | -0.3625 | -0.275 | 0.486708 | 0.494418 | 0.25346 | 0.989223 | 0.997988 | 0.994666 |
| Shank3_Q4ACU6_S375s_1 | Shank3_Q4 | -1.6 | -0.91 | -1.24 | -1.68 | -0.18 | -1.43 | -1.1 | -1.32 | -1.35 | -1.52 | -0.37 | -1.71 | -0.29 | 0.69 | -2.02 | -2.1 | 0.35 | 0.12 | 0.4275 | 0.507912 | 0.318319 | 0.738783 | 0.989223 | 0.997988 | 0.994666 |
| Shank3_Q4ACU6_S434s_1 | Shank3_Q4 | 0.67 | -0.07 | 0.08 | 0.26 | 0.53 | 0.59 | 0.74 | 0.75 | 0.16 | 0.59 | -0.64 | -0.03 | -0.39 | 0.72 | -0.11 | -0.48 | 0.4175 | -0.215 | -0.3 | 0.397837 | 0.103776 | 0.515908 | 0.989223 | 0.997988 | 0.994666 |
| Shank3_Q4ACU6_S482s_1 | Shank3_Q4 | 4.59 | 1.95 | 3.68 | 1.73 | 2.5 | 4.53 | 4.39 | 3.78 | 0.53 | 4.02 | 1.38 | 3.39 | 1 | -0.92 | 4.38 | 4.64 | 0.8125 | -0.6575 | -0.7125 | 0.588119 | 0.28181 | 0.494469 | 0.989223 | 0.997988 | 0.995259 |
| Shank3_Q4ACU6_S685s_1 | Shank3_Q4 | -3.09 | -2.28 | -1.91 | -2.5 | -1.34 | -2.38 | -2.13 | -1.93 | -3.14 | -2.39 | -0.31 | -2.05 | -2.2 | -0.43 | -2.97 | -2.14 | 0.5 | 0.4725 | 0.51 | 0.362903 | 0.163399 | 0.434849 | 0.989223 | 0.997988 | 0.994666 |
| Shank3_Q4ACU6_S694s_1 | Shank3_Q4 | -2.9 | -2.84 | -1.95 | -1.33 | -1.57 | -2.9 | -1.2 | -1.53 | -2.64 | -2.41 | 0.01 | -1.39 | -1.81 | -1.93 | -1.9 | -1.32 | 0.455 | 0.6475 | 0.515 | 0.225231 | 0.364097 | 0.330445 | 0.989223 | 0.997988 | 0.994666 |
| Shank3_Q4ACU6_S801s_1 | Shank3_Q4 | -0.77 | -1.59 | -1.16 | -2.01 | -0.36 | 0.27 | -1.38 | -0.5 | -1.71 | -1.58 | 0.6 | -1.7 | -0.96 | -0.42 | -2.71 | -1.79 | 0.89 | 0.285 | -0.0875 | 0.86877 | 0.054098 | 0.623569 | 0.989223 | 0.997988 | 0.999507 |
| Shank3_Q4ACU6_S838s_1 | Shank3_Q4 | 0.55 | 0.07 | 0.3 | 0.81 | 0.14 | -0.88 | 0.45 | 1.18 | -0.28 | -0.25 | 0.14 | 0.35 | -1.06 | 1.2 | 0.95 | 1.22 | -0.21 | -0.4425 | 0.145 | 0.785907 | 0.627662 | 0.134146 | 0.989223 | 0.997988 | 0.999507 |
| Shank3_Q4ACU6_S995s_1 | Shank3_Q4 | 0.97 | 0.74 | 0.35 | 0.63 | 0.89 | 0.8 | 0.95 | 0.86 | 0.25 | 0.74 | 0.35 | 0.44 | -0.33 | 1.02 | 0.08 | 0.62 | 0.2025 | -0.2275 | -0.325 | 0.367266 | 0.368255 | 0.36599 | 0.989223 | 0.997988 | 0.994666 |
| Shank3_Q4ACU6_T1234t_1 | Shank3_Q4 | 1.09 | 0.07 | 0.16 | 0.93 | 0.58 | 0.62 | 0.62 | 1.16 | 0.26 | 0.29 | 0.12 | 0.71 | -0.42 | 0.41 | 0.19 | 0.62 | 0.1825 | -0.2175 | -0.3625 | 0.334903 | 0.563863 | 0.501713 | 0.989223 | 0.997988 | 0.994666 |
| Shc2_Q8BMC3_S286s_1 | Shc2_Q8BM | 0.43 | 1.31 | 0.58 | 0.29 | 0.48 | 0.23 | 0.21 | 0.82 | 0.69 | 0.38 | 1.25 | 0.62 | 1.07 | -0.14 | 1 | 0.96 | -0.2175 | 0.0825 | 0.07 | 0.854411 | 0.467683 | 0.797306 | 0.989223 | 0.997988 | 0.999507 |
| Shc3_Q61120_S282s_1 | Shc3_Q611 | 0.01 | -0.03 | -0.16 | -0.31 | 0.15 | -0.54 | 0.37 | -0.01 | -0.22 | 0.11 | 0.1 | -0.23 | -0.3 | 0.27 | -0.61 | -0.12 | 0.115 | 0.0625 | -0.0675 | 0.80817 | 0.656977 | 0.781043 | 0.990434 | 0.997988 | 0.999507 |
| Shc3_Q61120_S354s_1 | Shc3_Q611 | -1.18 | -1.26 | -1.22 | -0.81 | -1.08 | -0.57 | -0.79 | -1.04 | -0.13 | -0.69 | 0.18 | -1.62 | -0.63 | -0.41 | -1.36 | -1.35 | 0.2475 | 0.5525 | 0.18 | 0.572392 | 0.300069 | 0.194673 | 0.989223 | 0.997988 | 0.995259 |
| Shc3_Q61120_S359s_1 | Shc3_Q611 | 0.95 | 0.05 | -0.52 | 0.06 | 0.35 | 0.9 | 0.77 | 0.89 | 0.16 | 0.61 | 0.52 | -0.62 | 0.07 | 0.13 | -0.16 | -0.33 | 0.5925 | 0.0325 | -0.2075 | 0.558022 | 0.104527 | 0.936453 | 0.989223 | 0.997988 | 0.995259 |
| Shisa6_F6VQZ6_S191s_1 | Shisa6_F6V | -0.35 | 0.08 | 0.49 | 0.1 | 0.09 | -0.84 | 0.23 | 0.22 | -0.77 | -0.29 | 0.6 | 0.12 | -0.34 | -0.09 | 0.36 | 0.25 | -0.155 | -0.165 | -0.035 | 0.906388 | 0.632852 | 0.642041 | 0.989223 | 0.997988 | 0.999507 |
| Shisa6_F6VQZ6_S410s_1 | Shisa6_F6V | 4.66 | 3.63 | 3.27 | 1.92 | 3.89 | 4.96 | 3 | 3.33 | -0.62 | 3.91 | 0.11 | 3.22 | -1.05 | -0.51 | 2.85 | 3.4 | 0.425 | -1.715 | -2.1975 | 0.07075 | 0.506444 | 0.145979 | 0.989223 | 0.997988 | 0.994666 |
| Shisa6_F6VQZ6_T190t_1 | Shisa6_F6V | -0.49 | 0.72 | 0.06 | -0.8 | 0.89 | -1.58 | 0.1 | -0.15 | -0.89 | -0.23 | 0.19 | -0.63 | 0.38 | 1.41 | 0.28 | -0.05 | -0.0575 | -0.2625 | 0.6325 | 0.180279 | 0.917284 | 0.521686 | 0.997335 | 0.997988 | 0.994666 |
| Shisa6_F6VQZ6_T264t_1 | Shisa6_F6V | -3.62 | -3.99 | -4.33 | -4.11 | -3.23 | -5.67 | -3.43 | -4.41 | -4.15 | -3.55 | -4.11 | -3.66 | -4.54 | -3.81 | -5.68 | -3.67 | -0.1725 | 0.145 | -0.4125 | 0.385817 | 0.743097 | 0.595705 | 0.995762 | 0.997988 | 0.994666 |
| Shisa7_F6T9T6_S104s_1 | Shisa7_F6T | 1.29 | 0.76 | 0.82 | 0.86 | 1.33 | 1.06 | 1.31 | 0.59 | 0.81 | 1.15 | -0.04 | 0.86 | 0.87 | 1.61 | 1.13 | 1.07 | 0.14 | -0.2375 | 0.2375 | 0.402477 | 0.592645 | 0.457889 | 0.989223 | 0.997988 | 0.994666 |
| Shisa7_F6T9T6_S121s_1 | Shisa7_F6T | -0.12 | 0.08 | -0.63 | -0.27 | 0.36 | -0.02 | -0.57 | -0.35 | 0.22 | 0.13 | -1.07 | -0.17 | 0.31 | 0.19 | -0.96 | -0.87 | 0.09 | 0.0125 | -0.0975 | 0.799324 | 0.752687 | 0.971255 | 0.995762 | 0.997988 | 0.999507 |
| Shisa7_F6T9T6_S222s_1 | Shisa7_F6T | 0.04 | -0.74 | -0.13 | 0.58 | -0.54 | 0.23 | 0.09 | 0.03 | -0.67 | -0.48 | 0.64 | 0.21 | -1.6 | 0.2 | -0.48 | -0.36 | 0.015 | -0.0125 | -0.4975 | 0.286668 | 0.96376 | 0.975305 | 0.999187 | 0.997988 | 0.994666 |
| Shisa7_F6T9T6_S255s_1 | Shisa7_F6T | -0.28 | 0.31 | -0.87 | -0.17 | 0.67 | 0.3 | -0.43 | -0.01 | -0.08 | 0.35 | -0.91 | 0.59 | 0.7 | 1.06 | -1.56 | -0.94 | 0.385 | 0.24 | 0.0675 | 0.912614 | 0.276623 | 0.55741 | 0.989223 | 0.997988 | 0.999507 |
| Shisa7_F6T9T6_S268s_1 | Shisa7_F6T | 6.97 | 3.78 | 6.03 | 2.37 | 4.65 | 6.96 | 5.68 | 5.35 | -1.18 | 5.88 | -0.65 | 5.89 | -2.76 | -0.2 | 5.39 | 5.81 | 0.8725 | -2.3025 | -2.7275 | 0.197116 | 0.391756 | 0.253432 | 0.989223 | 0.997988 | 0.994666 |
| Shisa7_F6T9T6_S272s_1 | Shisa7_F6T | 0.12 | -0.54 | 1.68 | 0.14 | 0.09 | -0.13 | 1.06 | 1.56 | -0.92 | 0.07 | 2.35 | 0.51 | -0.23 | 0.36 | 1.32 | 0.8 | 0.295 | 0.1525 | 0.2125 | 0.692266 | 0.599023 | 0.837925 | 0.989223 | 0.997988 | 0.996159 |
| Shisa7_F6T9T6_S67s_1 | Shisa7_F6T | 0.12 | -0.24 | 0.25 | 0.23 | 0.5 | 0.12 | 0.26 | 0.29 | -0.46 | -0.12 | -0.04 | 0.19 | -0.43 | 0.57 | -0.35 | 0.12 | 0.2025 | -0.1975 | -0.1125 | 0.718605 | 0.37334 | 0.438565 | 0.989223 | 0.997988 | 0.9971 |
| Shisa7_F6T9T6_T349t_1 | Shisa7_F6T | -0.92 | 0.2 | 0.46 | 0.47 | 0.59 | -1.1 | 0.49 | 0.31 | -0.56 | 0.06 | -0.3 | 0.96 | -2.38 | -0.68 | -0.57 | 1.02 | 0.02 | -0.0125 | -0.705 | 0.320088 | 0.966441 | 0.977838 | 0.999187 | 0.997988 | 0.994666 |
| Shisa9_E9QN38_S321s_1 | Shisa9_E9Q | 0.57 | 0.86 | 0.37 | 0.81 | 0.66 | 0.61 | 0.48 | 0.7 | 0.75 | 0.51 | 1.31 | 0.65 | 0.6 | 1.27 | 0.38 | 0.14 | -0.04 | 0.1525 | -0.055 | 0.862398 | 0.85311 | 0.573396 | 0.996349 | 0.997988 | 0.999507 |
| Shroom1_D3Z3D5_S293s_1 | Shroom1_D | -0.3 | -0.72 | -0.66 | -0.4 | -0.73 | -0.71 | -0.53 | -0.41 | -0.44 | -0.2 | -0.35 | -0.3 | -0.11 | -0.37 | -0.21 | -0.06 | -0.075 | 0.1975 | 0.3325 | 0.196052 | 0.730819 | 0.384038 | 0.995762 | 0.997988 | 0.994666 |
| Shroom2_A7TU71_S1047s_1 | Shroom2_A | 0.15 | 0.39 | 0.67 | 0.6 | 0.43 | -0.25 | -0.27 | 0.67 | 0.4 | 0.1 | 1.42 | 0.45 | 0.69 | 0.59 | 0.09 | 0.16 | -0.3075 | 0.14 | -0.07 | 0.798916 | 0.30986 | 0.675128 | 0.989223 | 0.997988 | 0.999507 |
| Shroom2_A7TU71_S1095s_1 | Shroom2_A | 6.49 | 2.1 | 4.85 | -4.46 | 3.4 | 7.51 | 4.28 | 3.1 | -2.3 | 5.58 | -2.6 | 5.25 | 1.47 | -1.75 | 4.85 | 5.83 | 2.3275 | -0.7625 | 0.355 | 0.888229 | 0.310723 | 0.791001 | 0.989223 | 0.997988 | 0.999507 |
| Shroom2_A7TU71_S1159s_1 | Shroom2_A | 0.48 | 0.68 | -1.18 | 0.57 | 0.46 | 0.49 | -0.05 | -0.27 | 0.64 | 0.36 | 1.03 | -0.4 | 0.49 | 0.68 | 0.28 | 0.03 | 0.02 | 0.27 | 0.2325 | 0.607993 | 0.964405 | 0.594133 | 0.999187 | 0.997988 | 0.995259 |
| Shroom2_A7TU71_S1171s_1 | Shroom2_A | 0.36 | 0.73 | 0.3 | 0.16 | 0.47 | 0.41 | 0.27 | 0.04 | 0.06 | 0.73 | 0.71 | -0.1 | -0.18 | -0.3 | 0.54 | 0.09 | -0.09 | -0.0375 | -0.35 | 0.247334 | 0.696642 | 0.897786 | 0.994893 | 0.997988 | 0.994666 |
| Shroom2_A7TU71_S318s_1 | Shroom2_A | 0.53 | 0.93 | -0.18 | 0.02 | 1.1 | -0.24 | 0.63 | 0.14 | 0.9 | 0.91 | 0.38 | -0.11 | 0.62 | 1.01 | -0.02 | 0.14 | 0.0825 | 0.195 | 0.1125 | 0.758606 | 0.826738 | 0.592048 | 0.995762 | 0.997988 | 0.9971 |
| Sik2_H3BJC9_S342s_1 | Sik2_H3BJC | -1.31 | -0.74 | -0.29 | -1.48 | -0.43 | -0.9 | -0.15 | -0.73 | -0.82 | -0.32 | 0.33 | -1.41 | -0.17 | 0 | 0.23 | -0.06 | 0.4025 | 0.4 | 0.955 | 0.01513 | 0.240507 | 0.378039 | 0.989223 | 0.997988 | 0.994666 |
| Sik2_H3BJC9_T359t_1 | Sik2_H3BJC | -1.64 | -0.34 | -0.77 | -0.76 | -0.69 | -0.42 | -1.69 | -0.39 | -0.45 | -0.71 | -1.37 | -2.02 | -0.35 | -0.18 | -1 | -1.47 | 0.08 | -0.26 | 0.1275 | 0.754888 | 0.839597 | 0.552163 | 0.995762 | 0.997988 | 0.9971 |
| Sik3_F6U6U5_S456s_1 | Sik3_F6U6U | -0.76 | -1.3 | -0.5 | -0.58 | -1.01 | -0.91 | -0.21 | -0.31 | -1.35 | -1.08 | 0.1 | -0.61 | -1.44 | -0.46 | -0.86 | -0.96 | 0.175 | 0.05 | -0.145 | 0.650611 | 0.560467 | 0.893063 | 0.989223 | 0.997988 | 0.995259 |
| Sik3_F6U6U5_S684s_1 | Sik3_F6U6U | -0.81 | -0.25 | 0.09 | -0.44 | -0.22 | -0.91 | 0.01 | -0.3 | -0.71 | -0.39 | -1.18 | -0.22 | -0.06 | -0.75 | -0.48 | -0.17 | -0.0025 | -0.2725 | -0.0125 | 0.967086 | 0.99329 | 0.396038 | 0.999244 | 0.997988 | 0.999507 |
| Sin3a_Q60520_S941s_1 | Sin3a_Q605 | -0.58 | -0.1 | -0.18 | -0.42 | 0.8 | 0.19 | -1.16 | -0.23 | 0.52 | -0.11 | -0.22 | 0.45 | 0.12 | -0.02 | 0.17 | -0.32 | 0.22 | 0.48 | 0.3075 | 0.251656 | 0.592282 | 0.106583 | 0.989223 | 0.997988 | 0.994666 |
| Sin3b_Q62141_S698s_1 | Sin3b_Q62 | 1.15 | 0.93 | 1.1 | 0.52 | -0.16 | 0.32 | 1.15 | 1.07 | 1.41 | 1.19 | 0.15 | 0.6 | 0.86 | -0.19 | 0.44 | 0.37 | -0.33 | -0.0875 | -0.555 | 0.10041 | 0.354317 | 0.796902 | 0.989223 | 0.997988 | 0.994666 |
| Sipa1_P46062_S72s_1 | Sipa1_P460 | 1.96 | 0.38 | -0.01 | 0.49 | -0.49 | 1.12 | 0.14 | 1.24 | 0.6 | 1.14 | 0.28 | 0.91 | 1.58 | -2.42 | 1.27 | 0.64 | -0.2025 | 0.0275 | -0.4375 | 0.62447 | 0.708935 | 0.951477 | 0.995437 | 0.997988 | 0.995259 |
| Sipa1l1_Q8C0T5_S1127s_1 | Sipa1l1_Q8 | 0.04 | 0.58 | -0.17 | 0.61 | 1.31 | 0.98 | 0.37 | 0.57 | 0.08 | 0.53 | -0.7 | -0.68 | -0.4 | 0.87 | -1.12 | 0.01 | 0.5425 | -0.4575 | -0.425 | 0.354048 | 0.103321 | 0.233659 | 0.989223 | 0.997988 | 0.994666 |
| Sipa1l1_Q8C0T5_S114s_1 | Sipa1l1_Q8 | 0.44 | 0.43 | 0.26 | 0.04 | 0.78 | 0.14 | 0.6 | 0.04 | 0.32 | 0.17 | -0.27 | 0.16 | 0.02 | 1.15 | -0.12 | 0.21 | 0.0975 | -0.1975 | 0.0225 | 0.94687 | 0.702676 | 0.422 | 0.995437 | 0.997988 | 0.999507 |
| Sipa1l1_Q8C0T5_S1170s_1 | Sipa1l1_Q8 | -0.72 | -0.6 | -2.26 | -3.05 | -0.33 | -1.07 | -1.44 | -2.52 | -0.72 | -0.05 | 0.41 | -2.22 | -0.41 | 0.41 | -2.27 | -2.14 | 0.3175 | 1.0125 | 0.555 | 0.486607 | 0.635502 | 0.196144 | 0.989223 | 0.997988 | 0.994666 |
| Sipa1l1_Q8C0T5_S1184s_1 | Sipa1l1_Q8 | 3.29 | -0.08 | -0.43 | -0.04 | 0.79 | 3.47 | -0.71 | 0.59 | 2.37 | 2.23 | 0.74 | -0.66 | 0.44 | 1.6 | -1.67 | -1.21 | 0.35 | 0.485 | -0.895 | 0.383036 | 0.743548 | 0.628017 | 0.995762 | 0.997988 | 0.994666 |
| Sipa1l1_Q8C0T5_S1249s_1 | Sipa1l1_Q8 | 0.33 | 0.46 | -0.4 | 0.15 | 0.6 | 0.88 | 0.58 | 0.64 | -0.24 | 0.05 | -0.52 | 0.32 | -0.43 | 0.38 | 0.15 | 0.66 | 0.54 | -0.2325 | 0.055 | 0.87012 | 0.057299 | 0.447586 | 0.989223 | 0.997988 | 0.999507 |
| Sipa1l1_Q8C0T5_S1372s_1 | Sipa1l1_Q8 | -0.01 | -0.21 | -2.01 | -2.71 | 0.19 | 0.52 | -1.86 | -1.63 | -0.71 | -0.32 | -0.54 | -3.27 | -1.46 | 0.57 | -2.89 | -4.28 | 0.54 | 0.025 | -0.78 | 0.471995 | 0.5001 | 0.976618 | 0.989223 | 0.997988 | 0.994666 |
| Sipa1l1_Q8C0T5_S1384s_1 | Sipa1l1_Q8 | 1.44 | -1.7 | -0.44 | 0.6 | -0.13 | 1.93 | 0.81 | 1.86 | -1.53 | -0.41 | -4.51 | 0.26 | -1.25 | -2.33 | -0.99 | -1.12 | 1.1425 | -1.5225 | -1.3975 | 0.060575 | 0.144118 | 0.19014 | 0.989223 | 0.997988 | 0.994666 |
| Sipa1l1_Q8C0T5_S148s_1 | Sipa1l1_Q8 | -3.88 | -1.48 | -1.53 | -5.44 | -2.3 | -3.39 | -2.86 | -3.26 | -2.8 | -2.18 | 0.53 | -1.84 | -1.39 | -2.15 | -3.01 | -2.74 | 0.13 | 1.51 | 0.76 | 0.408284 | 0.880639 | 0.179618 | 0.996493 | 0.997988 | 0.994666 |
| Sipa1l1_Q8C0T5_S1528s_1 | Sipa1l1_Q8 | 0.67 | 0.2 | 0.45 | 0.52 | 0.61 | 0.43 | 0.8 | 0.66 | -0.27 | 0.47 | -0.47 | 0.53 | -0.28 | -0.39 | 0.11 | 0.25 | 0.165 | -0.395 | -0.5375 | 0.071089 | 0.453131 | 0.220459 | 0.989223 | 0.997988 | 0.994666 |
| Sipa1l1_Q8C0T5_S1564s_1 | Sipa1l1_Q8 | 0.78 | 0.93 | 1.41 | 0.53 | 0.99 | 0.6 | 0.86 | 1.77 | 0.2 | 0.72 | 1.55 | 1.39 | 0.23 | 0.4 | 1.17 | 1.74 | 0.1425 | 0.0525 | -0.0275 | 0.945442 | 0.66255 | 0.887268 | 0.990434 | 0.997988 | 0.999507 |
| Sipa1l1_Q8C0T5_S1582s_1 | Sipa1l1_Q8 | -1.7 | -0.29 | -0.86 | 0.44 | 0.41 | -0.92 | 0.54 | -0.44 | -1.48 | -1.1 | -0.24 | -0.47 | -0.98 | 0.34 | -1.25 | -0.44 | 0.5 | -0.22 | 0.02 | 0.970063 | 0.348493 | 0.663152 | 0.989223 | 0.997988 | 0.999507 |
| Sipa1l1_Q8C0T5_S1600s_1 | Sipa1l1_Q8 | -1.42 | -0.63 | -2.07 | -1.63 | -0.48 | -1.21 | -2.35 | -2.22 | -0.9 | -1.07 | -1.38 | -2.28 | -1.4 | -0.43 | -2.71 | -1.79 | -0.1275 | 0.03 | -0.145 | 0.782292 | 0.795628 | 0.943177 | 0.995762 | 0.997988 | 0.998226 |
| Sipa1l1_Q8C0T5_S1712s_1 | Sipa1l1_Q8 | 1.95 | 2.2 | 3.41 | 1.95 | 2.27 | 2.23 | 2.52 | 2.44 | 0.69 | 2.33 | 1.09 | 3.01 | 0.88 | 0.5 | 3.14 | 3.23 | -0.0125 | -0.5975 | -0.44 | 0.543783 | 0.971943 | 0.324659 | 0.999187 | 0.997988 | 0.994666 |
| Sipa1l1_Q8C0T5_S208s_1 | Sipa1l1_Q8 | 0.06 | 0.49 | 0.29 | 0.02 | 0.69 | 0.71 | 0.17 | 0.33 | -0.11 | 0.38 | 0.53 | 0.1 | 0.07 | 0.54 | 0.25 | 0.07 | 0.26 | 0.01 | 0.0175 | 0.945918 | 0.292383 | 0.968379 | 0.989223 | 0.997988 | 0.999507 |
| Sipa1l1_Q8C0T5_S211s_1 | Sipa1l1_Q8 | -0.42 | -0.33 | 0.14 | -0.24 | -0.21 | 0 | -0.07 | -0.03 | -0.44 | -0.74 | 0.32 | -0.06 | -1.21 | 0 | -0.1 | -0.55 | 0.135 | -0.0175 | -0.2525 | 0.462107 | 0.542674 | 0.953492 | 0.989223 | 0.997988 | 0.994666 |
| Sipa1l1_Q8C0T5_S288s_1 | Sipa1l1_Q8 | -2.34 | -1.4 | -2.62 | -5.21 | -1.24 | -3.59 | -1.93 | -2.89 | -5.95 | -2.92 | 0.33 | -3.21 | -2.52 | -1.3 | -2.77 | -2.21 | 0.48 | -0.045 | 0.6925 | 0.382531 | 0.571763 | 0.973019 | 0.989223 | 0.997988 | 0.994666 |
| Sipa1l1_Q8C0T5_S310s_1 | Sipa1l1_Q8 | -0.7 | -1.21 | 0.16 | 0.44 | -1.51 | 0.95 | 0.19 | 0.71 | -1.98 | -0.83 | 0.32 | 0.64 | -0.86 | -3.87 | -0.03 | 0.39 | 0.4125 | -0.135 | -0.765 | 0.40734 | 0.498466 | 0.833822 | 0.989223 | 0.997988 | 0.994666 |
| Sipa1l1_Q8C0T5_T1190t_1 | Sipa1l1_Q8 | -1.33 | -0.96 | -0.28 | -1.2 | -1.05 | -1.04 | -1.86 | -2.19 | 0.58 | -0.75 | 0.1 | -0.91 | 0.98 | -0.61 | -0.84 | -1.48 | -0.5925 | 0.6975 | 0.455 | 0.403217 | 0.132628 | 0.119928 | 0.989223 | 0.997988 | 0.994666 |
| Sipa1l2_Q80TE4_S1406s_1 | Sipa1l2_Q8 | 0.23 | 0.94 | 0.5 | 0.31 | 0.42 | 0.21 | 0.59 | 0.23 | 0.53 | 0.61 | 0.47 | 0.8 | 0.56 | 0.22 | 0.57 | 0.31 | -0.1325 | 0.1075 | -0.08 | 0.767443 | 0.589998 | 0.668107 | 0.989223 | 0.997988 | 0.9971 |
| Sipa1l2_Q80TE4_S1488s_1 | Sipa1l2_Q8 | 1.34 | 1.31 | 1.17 | 1.87 | 0.85 | 1.32 | 0.99 | 1.16 | 1.05 | 1.45 | 2.26 | 1.16 | 0.92 | 0.46 | 1.36 | 1.14 | -0.3425 | 0.0575 | -0.4525 | 0.160561 | 0.184782 | 0.863878 | 0.989223 | 0.997988 | 0.994666 |
| Sipa1l2_Q80TE4_S195s_1 | Sipa1l2_Q8 | 0.12 | 0.33 | 0.15 | -0.19 | 0.43 | 0.12 | -0.52 | -0.11 | 0.04 | 0.11 | 0.58 | -0.42 | 0.91 | 0.24 | -0.6 | 0.02 | -0.1225 | -0.025 | 0.04 | 0.910568 | 0.651362 | 0.929253 | 0.990358 | 0.997988 | 0.999507 |
| Sipa1l3_G3X9J0_S1358s_1 | Sipa1l3_G3 | -0.06 | -0.5 | 0.36 | 0.11 | 0.37 | -0.61 | 0.04 | 0.61 | -0.13 | -0.28 | 1.02 | 0.06 | -0.01 | -0.32 | -0.08 | -0.27 | 0.125 | 0.19 | -0.1475 | 0.596965 | 0.706326 | 0.597198 | 0.995437 | 0.997988 | 0.995259 |
| Sipa1l3_G3X9J0_S94s_1 | Sipa1l3_G3 | 0.04 | 0.61 | -1.04 | -0.14 | 0.08 | -0.82 | 0.76 | 0.31 | 0.55 | 0.33 | 0.67 | -0.02 | 0.66 | -0.68 | 1.52 | 0.55 | 0.215 | 0.515 | 0.645 | 0.242132 | 0.632063 | 0.196449 | 0.989223 | 0.997988 | 0.994666 |
| Sirpa_P97797_S462s_1 | Sirpa_P977 | 0.57 | -1.19 | 1.03 | 0.27 | 1.38 | 0.17 | 0.96 | 2 | 0.37 | -0.16 | 2.33 | 1.63 | -0.43 | 1.3 | -0.01 | 1.11 | 0.9575 | 0.8725 | 0.3225 | 0.584607 | 0.109763 | 0.216706 | 0.989223 | 0.997988 | 0.995259 |
| Sirpa_P97797_Y460y_1 | Sirpa_P977 | 1.26 | 0.03 | 1.35 | 1.42 | 0.26 | 0.36 | 1.11 | 1.86 | 0.14 | -0.29 | 1.13 | 0.69 | 0.53 | -0.09 | 1.71 | 1.32 | -0.1175 | -0.5975 | -0.1475 | 0.765517 | 0.799636 | 0.195928 | 0.995762 | 0.997988 | 0.9971 |
| Sirt2_Q8VDQ8_S331s_1 | Sirt2_Q8VD | 1.94 | 1.79 | 2.5 | 1.17 | 1.87 | 1.94 | 2.61 | 1.69 | 0.94 | 2.05 | 0.08 | 2.06 | 0.58 | 1.61 | 2.33 | 2.6 | 0.1775 | -0.5675 | -0.07 | 0.888417 | 0.608221 | 0.288241 | 0.989223 | 0.997988 | 0.999507 |
| Sirt2_Q8VDQ8_S335s_1 | Sirt2_Q8VD | 5.34 | 3.88 | 4.43 | 2.77 | 3.95 | 4.04 | 4.29 | 4.07 | 0.75 | 4.86 | 1.17 | 4.54 | 1.19 | -0.71 | 5.06 | 5.14 | -0.0175 | -1.275 | -1.435 | 0.297867 | 0.971892 | 0.250589 | 0.999187 | 0.997988 | 0.994666 |
| Skiv2l_Q6NZR5_S240s_2 | Skiv2l_Q6N | -0.59 | 0.66 | -0.19 | 0.14 | -0.22 | -0.81 | 0.41 | 0.18 | 0.62 | 0.2 | 0.94 | 0.67 | -0.37 | 1.29 | 0.65 | 0.14 | -0.115 | 0.6025 | 0.4225 | 0.344278 | 0.756577 | 0.093971 | 0.995762 | 0.997988 | 0.994666 |
| Skiv2l_Q6NZR5_S242s_2 | Skiv2l_Q6N | -0.59 | 0.66 | -0.19 | 0.14 | -0.22 | -0.81 | 0.41 | 0.18 | 0.62 | 0.2 | 0.94 | 0.67 | -0.37 | 1.29 | 0.65 | 0.14 | -0.115 | 0.6025 | 0.4225 | 0.344278 | 0.756577 | 0.093971 | 0.995762 | 0.997988 | 0.994666 |
| Skiv2l_Q6NZR5_S253s_1 | Skiv2l_Q6N | -0.08 | 0.11 | -0.15 | -0.21 | 0.29 | 0.23 | -0.07 | -0.36 | -0.03 | 0.09 | -0.37 | -0.4 | 0.02 | 0.22 | -0.2 | 0.04 | 0.105 | -0.095 | 0.1025 | 0.671947 | 0.656868 | 0.685228 | 0.990434 | 0.997988 | 0.996159 |
| Skp1_Q9WTX5_T131t_1 | Skp1_Q9W | -1.43 | -0.67 | -1.03 | -1.28 | -1.19 | -1.31 | -1.35 | -1.56 | -1.22 | -1.24 | -0.48 | -1.01 | -0.22 | -1.18 | -1.38 | -1.44 | -0.25 | 0.115 | 0.0475 | 0.893734 | 0.320312 | 0.691564 | 0.989223 | 0.997988 | 0.999507 |
| Slain1_F7B673_S270s_1 | Slain1_F7B | -1.3 | -0.5 | -1.07 | -0.62 | -0.65 | -0.37 | -1.24 | -0.95 | -0.78 | -0.06 | -3.86 | -0.44 | -0.03 | -0.53 | -0.51 | -0.87 | 0.07 | -0.4125 | 0.3875 | 0.230136 | 0.811787 | 0.60674 | 0.995762 | 0.997988 | 0.994666 |
| Slain2_Q8CI08_S354s_1 | Slain2_Q8C | -2 | -2.74 | -2.38 | -3.74 | -2.46 | -1.65 | -3.14 | -3.05 | -0.44 | -2.76 | -4.66 | -2.35 | -0.51 | -3.47 | -3.39 | -2.73 | 0.14 | 0.1625 | 0.19 | 0.78717 | 0.766133 | 0.846466 | 0.995762 | 0.997988 | 0.999507 |
| Slain2_Q8CI08_S87s_1 | Slain2_Q8C | -0.36 | -0.21 | 0.45 | -0.04 | -1.07 | 0.61 | -0.67 | -0.54 | 1.35 | 0.4 | -0.24 | 0.11 | 1.68 | -0.66 | 0.5 | -0.22 | -0.3775 | 0.445 | 0.365 | 0.479403 | 0.344249 | 0.266724 | 0.989223 | 0.997988 | 0.994666 |
| Slbp_Q3V2L1_S7s_1 | Slbp_Q3V2 | -1.75 | -2.22 | -1.99 | -2.28 | -2.73 | -1.89 | -2.22 | -3.01 | -1.72 | -1.58 | -1.3 | -2.16 | -0.09 | -2.47 | -2.26 | -2.24 | -0.4025 | 0.37 | 0.295 | 0.583957 | 0.204019 | 0.197122 | 0.989223 | 0.997988 | 0.995259 |
| Slc12a2_E9QM38_S75s_1 | Slc12a2_E9 | -1.65 | 0.79 | -0.14 | -1.1 | 0.11 | -1.73 | 1.27 | -1.39 | -1.41 | -0.64 | -1.38 | -1.27 | 1.42 | 0.11 | -0.41 | -1.17 | 0.09 | -0.65 | 0.5125 | 0.462142 | 0.906843 | 0.238984 | 0.99702 | 0.997988 | 0.994666 |
| Slc12a2_E9QM38_T211t_1 | Slc12a2_E9 | -0.06 | -0.4 | -1.63 | -2.09 | -0.66 | -0.07 | -2.27 | -1.67 | -1.32 | -0.8 | -1.17 | -2.45 | -1.57 | -0.36 | -2.65 | -2.12 | -0.1225 | -0.39 | -0.63 | 0.327375 | 0.84268 | 0.489225 | 0.996349 | 0.997988 | 0.994666 |
| Slc12a5_Q91V14_S1044s_1 | Slc12a5_Q9 | 0.45 | -0.09 | 1.14 | 1.36 | 0.79 | 0.79 | 1.12 | 1.54 | -0.07 | -0.22 | 1.59 | 1.59 | -0.36 | 0.74 | 0.91 | 1.01 | 0.345 | 0.0075 | -0.14 | 0.754468 | 0.362819 | 0.98923 | 0.989223 | 0.997988 | 0.9971 |
| Slc12a5_Q91V14_S1047s_1 | Slc12a5_Q9 | -0.4 | 0.3 | -0.5 | -0.48 | 0.9 | -0.44 | -0.77 | -0.59 | 0.26 | -0.01 | -0.65 | -0.96 | 0.29 | 0.5 | -0.9 | -0.31 | 0.045 | -0.07 | 0.165 | 0.667928 | 0.912071 | 0.843298 | 0.997335 | 0.997988 | 0.996159 |
| Slc12a5_Q91V14_S1048s_1 | Slc12a5_Q9 | -1.31 | 0.78 | 0.08 | -0.75 | 0.95 | -1.59 | -0.26 | -1.35 | -0.83 | -0.58 | -1.26 | -0.28 | 0.18 | 0.34 | 0.17 | -0.28 | -0.2625 | -0.4375 | 0.4025 | 0.393488 | 0.690437 | 0.371972 | 0.994672 | 0.997988 | 0.994666 |
| Slc12a5_Q91V14_S48s_1 | Slc12a5_Q9 | 0.68 | 0.48 | 0.08 | 1.15 | 0.55 | 0.4 | 0.98 | 1.09 | 0.87 | 0.88 | 0.16 | 1 | 0.29 | 0.88 | 0.63 | 0.85 | 0.1575 | 0.13 | 0.065 | 0.835475 | 0.603836 | 0.687319 | 0.989223 | 0.997988 | 0.999507 |
| Slc12a5_Q91V14_S49s_1 | Slc12a5_Q9 | 0.53 | -0.51 | 0.08 | 0.73 | 0.96 | 0.82 | 0.9 | 0.89 | 0.19 | 0.07 | 0.45 | 0.84 | -1.74 | 0.6 | 0.03 | 0.41 | 0.685 | 0.18 | -0.3825 | 0.4971 | 0.044044 | 0.601736 | 0.989223 | 0.997988 | 0.994666 |
| Slc12a5_Q91V14_S54s_1 | Slc12a5_Q9 | 0.06 | 1.03 | 1.79 | 1.48 | 0.46 | 0.34 | 1.02 | 1.19 | 0.75 | 0.44 | 2.24 | 1.92 | 0.03 | -0.22 | 1.44 | 1.62 | -0.3375 | 0.2475 | -0.3725 | 0.510216 | 0.420171 | 0.647054 | 0.989223 | 0.997988 | 0.994666 |
| Slc12a5_Q91V14_S955s_1 | Slc12a5_Q9 | 0.28 | -3.15 | -1.04 | -0.68 | 0.14 | 0.42 | -0.3 | -0.1 | -1.63 | -1.88 | -0.75 | -0.8 | -4.31 | -0.22 | -2.18 | -1.06 | 1.1875 | -0.1175 | -0.795 | 0.432494 | 0.095748 | 0.867003 | 0.989223 | 0.997988 | 0.994666 |
| Slc12a5_Q91V14_S963s_1 | Slc12a5_Q9 | 2.3 | 0.01 | 1.18 | 2.22 | 1.46 | 2.35 | 1.56 | 2.57 | 0.56 | 0.44 | 0.16 | 2.1 | -1.04 | 1.06 | 0.09 | 1.12 | 0.5575 | -0.6125 | -1.12 | 0.117456 | 0.321822 | 0.344121 | 0.989223 | 0.997988 | 0.994666 |
| Slc12a5_Q91V14_T1029t_1 | Slc12a5_Q9 | -2.51 | -2.06 | -2.69 | -2.41 | -0.46 | -2.75 | -1.74 | -3.17 | -2.79 | -2.61 | -2.5 | -2.11 | -2.06 | -0.98 | -2.65 | -2.95 | 0.3875 | -0.085 | 0.2575 | 0.565384 | 0.492361 | 0.745207 | 0.989223 | 0.997988 | 0.995259 |
| Slc12a6_Q6P6P5_S50s_1 | Slc12a6_Q6 | -1.88 | -1.13 | -2.16 | -2 | -1.43 | -0.12 | -2.38 | -5.32 | -0.62 | -1.09 | -1.23 | -3.59 | -0.44 | -1.74 | -1.16 | -1.43 | -0.52 | 0.16 | 0.6 | 0.135348 | 0.596854 | 0.802685 | 0.989223 | 0.997988 | 0.994666 |
| Slc14a1_Q8VHL0_S4s_1 | Slc14a1_Q8 | 0.02 | 0.5 | 0.27 | 0.23 | 0.7 | 0.41 | 0.13 | 0.36 | 0.97 | 0.83 | 0.5 | 0.5 | 1.06 | 0.57 | 0.66 | 0.65 | 0.145 | 0.445 | 0.48 | 0.084768 | 0.531399 | 0.089488 | 0.989223 | 0.997988 | 0.994666 |
| Slc16a1_P53986_S213s_1 | Slc16a1_P5 | 0.25 | 0.46 | 2.28 | 1.76 | -0.33 | 0.67 | 0.48 | 1.15 | 0.92 | 0.3 | 1.44 | 1.51 | 1.41 | -0.7 | 1.67 | 0.45 | -0.695 | -0.145 | -0.48 | 0.472001 | 0.209752 | 0.784479 | 0.989223 | 0.997988 | 0.994666 |
| Slc16a1_P53986_S491s_1 | Slc16a1_P5 | 0.15 | 0.3 | 0.33 | 0.94 | 0.64 | 0.12 | -0.1 | 0.39 | 0.31 | 0.58 | 1.74 | 0.21 | 1.09 | 1.35 | 0.34 | 0.5 | -0.1675 | 0.28 | 0.39 | 0.2611 | 0.547564 | 0.482259 | 0.989223 | 0.997988 | 0.994666 |
| Slc16a1_P53986_S491s_2 | Slc16a1_P5 | -0.02 | 0.85 | -0.09 | 0.38 | 0.76 | 0.06 | 0.3 | 0.08 | 0.14 | 0.54 | 0.42 | 0.31 | 0.29 | 0.85 | 0.46 | 0.07 | 0.02 | 0.0725 | 0.1375 | 0.668442 | 0.946254 | 0.797632 | 0.999187 | 0.997988 | 0.996159 |
| Slc16a8_O35308_S207s_1 | Slc16a8_O3 | -3.42 | -1.22 | -1.53 | -0.69 | -1.05 | -0.82 | -1.84 | -2.56 | 0.65 | -1.64 | -1.74 | -3.15 | 1.5 | -1.86 | -1.77 | -1.6 | 0.1475 | 0.245 | 0.7825 | 0.385563 | 0.816425 | 0.779836 | 0.995762 | 0.997988 | 0.994666 |
| Slc17a7_Q3TXX4_S36s_1 | Slc17a7_Q3 | 0.7 | 0.93 | 0.66 | 1.12 | 1.13 | 0.56 | 1.58 | 1.6 | 1.02 | -0.32 | -0.71 | 1.04 | 0.58 | 1.5 | 1.29 | 1.16 | 0.365 | -0.595 | 0.28 | 0.349669 | 0.234382 | 0.206384 | 0.989223 | 0.997988 | 0.994666 |
| Slc17a8_D3YTT3_S376s_1 | Slc17a8_D3 | 0.31 | -0.02 | -0.99 | -0.16 | 0.4 | 0.13 | 0.07 | -0.45 | -0.31 | 0.48 | 0.22 | -0.1 | -0.38 | 1.09 | -0.4 | 0.27 | 0.2525 | 0.2875 | 0.36 | 0.420355 | 0.459619 | 0.413249 | 0.989223 | 0.997988 | 0.994666 |
| Slc1a1_P51906_S516s_1 | Slc1a1_P51 | 0.04 | -0.92 | 1.2 | 1.12 | 0.69 | 0.61 | 1.08 | 0.96 | -0.63 | -0.4 | -0.44 | -0.03 | -1.13 | 0.17 | 0.38 | 1.41 | 0.475 | -0.735 | -0.1525 | 0.815437 | 0.330739 | 0.157709 | 0.989223 | 0.997988 | 0.999507 |
| Slc1a2_P43006_S24s_1 | Slc1a2_P43 | 2.65 | 2.06 | 2.3 | 1.44 | 2.03 | 2.46 | 2.61 | 1.92 | 1.12 | 2.23 | 1.94 | 2.12 | 0.94 | 1.53 | 2.37 | 2.33 | 0.1425 | -0.26 | -0.32 | 0.458012 | 0.656888 | 0.484701 | 0.990434 | 0.997988 | 0.994666 |
| Slc1a2_P43006_S25s_1 | Slc1a2_P43 | -1.12 | 2.26 | 1.44 | 1.8 | 1.92 | -1.2 | -0.08 | 0.91 | 1.07 | 0.67 | 2.57 | 1.18 | 0.83 | 2.39 | 1.01 | 0.47 | -0.7075 | 0.2775 | 0.08 | 0.916928 | 0.428568 | 0.720121 | 0.989223 | 0.997988 | 0.999507 |
| Slc1a2_P43006_S3s_1 | Slc1a2_P43 | 0.84 | 0.94 | 1.05 | 0.78 | 1.04 | 0.77 | 0.38 | 0.57 | 1.14 | 0.87 | 0.88 | 0.61 | 1.44 | 1.21 | 0.59 | 0.49 | -0.2125 | -0.0275 | 0.03 | 0.920461 | 0.365344 | 0.903115 | 0.989223 | 0.997988 | 0.999507 |
| Slc1a2_P43006_S505s_1 | Slc1a2_P43 | 0.19 | 0.69 | 0.47 | 0.76 | 0.83 | 0.14 | 0.32 | 0.26 | -0.07 | -0.01 | -0.16 | 0.46 | -0.2 | 1.15 | 0.41 | 0.94 | -0.14 | -0.4725 | 0.0475 | 0.893525 | 0.583416 | 0.092442 | 0.989223 | 0.997988 | 0.999507 |
| Slc1a2_P43006_S520s_1 | Slc1a2_P43 | -1.24 | -0.9 | -0.47 | -1.01 | -0.45 | -0.82 | -0.8 | -0.88 | -1.61 | -1.49 | -0.55 | -0.84 | -0.96 | -0.81 | -0.95 | -0.45 | 0.1675 | -0.2175 | 0.1125 | 0.688364 | 0.503632 | 0.511552 | 0.989223 | 0.997988 | 0.996159 |
| Slc1a2_P43006_S530s_1 | Slc1a2_P43 | 4.73 | 2.35 | 2.1 | 1.32 | 2.49 | 5.28 | 3.03 | 2.08 | 0.39 | 3.35 | 0.71 | 1.95 | 0.36 | 1.51 | 1.94 | 2.33 | 0.595 | -1.025 | -1.09 | 0.171557 | 0.508976 | 0.264667 | 0.989223 | 0.997988 | 0.994666 |
| Slc1a2_P43006_S532s_1 | Slc1a2_P43 | 0.36 | 0.99 | 1.03 | 0.7 | 0.71 | 0.26 | 1.14 | 0.83 | 0.03 | 0.65 | -0.01 | 0.74 | 0.13 | 1.04 | 0.75 | 1.04 | -0.035 | -0.4175 | -0.03 | 0.924138 | 0.899459 | 0.179811 | 0.996493 | 0.997988 | 0.999507 |
| Slc1a2_P43006_S542s_1 | Slc1a2_P43 | -0.16 | 0.72 | 0.08 | 0.11 | 1.47 | -0.14 | 0.96 | 0.4 | -0.86 | 0.19 | -1.41 | 0.72 | -0.64 | -0.3 | 0.43 | 0.58 | 0.485 | -0.5275 | -0.17 | 0.645469 | 0.226986 | 0.298368 | 0.989223 | 0.997988 | 0.995259 |
| Slc1a2_P43006_S558s_1 | Slc1a2_P43 | 2.22 | 1.06 | 0.54 | 0.55 | 1.33 | 1.69 | 0.68 | 1.21 | 0.16 | 0.96 | -0.85 | 0.97 | -0.6 | -0.02 | -0.34 | 0.35 | 0.135 | -0.7825 | -1.245 | 0.017003 | 0.750302 | 0.173238 | 0.995762 | 0.997988 | 0.994666 |
| Slc1a2_P43006_S562s_1 | Slc1a2_P43 | 1.89 | 1.64 | 1.64 | 2.02 | 1.53 | 1.9 | 1.46 | 2.15 | 1.09 | 1.85 | 1.31 | 1.76 | -0.15 | 0.57 | 1.51 | 1.69 | -0.0375 | -0.295 | -0.8925 | 0.063281 | 0.879262 | 0.283917 | 0.996493 | 0.997988 | 0.994666 |
| Slc1a2_P43006_T4t_1 | Slc1a2_P43 | 0.87 | 0.96 | 1.33 | 0.77 | 1.07 | 0.78 | 0.35 | 0.56 | 1.16 | 0.88 | 0.67 | 0.82 | 1.49 | 1.28 | 0.66 | 0.78 | -0.2925 | -0.1 | 0.07 | 0.813636 | 0.262876 | 0.6809 | 0.989223 | 0.997988 | 0.999507 |
| Slc1a2_P43006_Y522y_1 | Slc1a2_P43 | -1.06 | -0.68 | 0.73 | 1.81 | 0.04 | -0.04 | 0.49 | 1.84 | -0.22 | -0.83 | 2.2 | 0.64 | -0.15 | -0.24 | 0.19 | 1.09 | 0.3825 | 0.2475 | 0.0225 | 0.972583 | 0.586322 | 0.765964 | 0.989223 | 0.997988 | 0.999507 |
| Slc1a2_P43006_Y537y_1 | Slc1a2_P43 | 0.29 | 0.02 | -0.06 | -0.39 | 0.44 | 0.96 | -0.5 | 0.59 | 0 | 0.78 | -0.02 | 0.06 | -0.07 | -0.03 | -0.57 | 0.89 | 0.4075 | 0.24 | 0.09 | 0.802625 | 0.255299 | 0.410384 | 0.989223 | 0.997988 | 0.999507 |
| Slc1a3_P56564_S4s_1 | Slc1a3_P56 | -3.34 | -1.81 | 2.16 | 1.35 | -3.73 | -2.23 | 0.87 | 1.21 | -0.36 | -2.6 | 1.24 | 2.29 | 0.81 | -3.12 | 3.75 | 0.41 | -0.56 | 0.5525 | 0.8725 | 0.597601 | 0.712458 | 0.707267 | 0.995437 | 0.997988 | 0.995259 |
| Slc1a3_P56564_S512s_1 | Slc1a3_P56 | 0.55 | 1.34 | 1.17 | 0.82 | 1.21 | 0.54 | 1.1 | 0.91 | 0.46 | 0.67 | 0.53 | 1 | -0.57 | 0.69 | 1.05 | 1.29 | -0.03 | -0.305 | -0.355 | 0.4288 | 0.911927 | 0.277833 | 0.997335 | 0.997988 | 0.994666 |
| Slc1a4_O35874_S507s_1 | Slc1a4_O35 | 0.6 | 1 | 0.77 | 0.61 | 0.88 | 0.65 | 0.77 | 0.47 | 0.47 | 0.97 | 0.15 | 0.51 | -0.19 | 1.1 | 0.17 | 0.39 | -0.0525 | -0.22 | -0.3775 | 0.268351 | 0.809952 | 0.406127 | 0.995762 | 0.997988 | 0.994666 |
| Slc1a4_O35874_T24t_1 | Slc1a4_O35 | 0.28 | 0.68 | 1.17 | 1.11 | -0.12 | 0.78 | 0.2 | 0.6 | 0.93 | 0.33 | 0.25 | 1.23 | 1.25 | -0.25 | 0.61 | 0.55 | -0.445 | -0.125 | -0.27 | 0.487923 | 0.173407 | 0.711008 | 0.989223 | 0.997988 | 0.994666 |
| Slc20a1_Q61609_S269s_1 | Slc20a1_Q6 | 0.41 | 0.38 | 0.79 | 1.16 | 0.66 | 0.44 | 0.73 | 0.54 | 0.6 | 0.77 | 0.54 | 0.76 | 0.52 | 0.5 | 1.81 | 1.16 | -0.0925 | -0.0175 | 0.3125 | 0.416604 | 0.713115 | 0.946158 | 0.995437 | 0.997988 | 0.994666 |
| Slc20a2_Q80UP8_S256s_1 | Slc20a2_Q8 | 0.76 | 0.98 | 0.92 | 0.73 | -0.42 | 1.19 | -0.17 | 0.39 | 2.28 | 1.12 | 0.79 | 0.56 | 2.53 | 0.28 | 1.27 | 0.25 | -0.6 | 0.34 | 0.235 | 0.646252 | 0.121101 | 0.391342 | 0.989223 | 0.997988 | 0.995259 |
| Slc20a2_Q80UP8_S268s_1 | Slc20a2_Q8 | 1.46 | 0.72 | 0.92 | 0.96 | 0.52 | 1.39 | 0.91 | 1.18 | 1.16 | 0.97 | 1.08 | 0.8 | 0.25 | 1.3 | 0.77 | 0.91 | -0.015 | -0.0125 | -0.2075 | 0.517868 | 0.957323 | 0.960128 | 0.999187 | 0.997988 | 0.994666 |
| Slc20a2_Q80UP8_S316s_1 | Slc20a2_Q8 | -3.27 | -3.39 | -2.69 | -3.52 | -1.97 | -2.74 | -2.58 | -3.28 | -3 | -3.44 | -1.75 | -3.08 | -1.58 | -2.11 | -3.42 | -2.91 | 0.575 | 0.4 | 0.7125 | 0.130212 | 0.111004 | 0.338767 | 0.989223 | 0.997988 | 0.994666 |
| Slc23a2_Q9EPR4_S18s_1 | Slc23a2_Q9 | 0.82 | -0.16 | 0.35 | 0.8 | 0.52 | 0.68 | -0.13 | 0.68 | 0.06 | 0.09 | 0.11 | 0.76 | -1.35 | 0.55 | 0.33 | 0.04 | -0.015 | -0.1975 | -0.56 | 0.249194 | 0.962244 | 0.537365 | 0.999187 | 0.997988 | 0.994666 |
| Slc23a2_Q9EPR4_S81s_1 | Slc23a2_Q9 | 0.68 | 0.86 | 1.34 | 1.05 | 0.29 | 0.3 | 1.06 | 0.6 | 0.42 | 0.7 | 0.42 | 0.76 | 0.03 | 0.39 | 0.45 | 1.52 | -0.42 | -0.4075 | -0.385 | 0.312778 | 0.145367 | 0.123719 | 0.989223 | 0.997988 | 0.994666 |
| Slc24a2_F6RT95_S43s_1 | Slc24a2_F6 | -1.8 | -1.08 | -0.86 | -2.1 | -1.85 | -1.14 | -2.16 | -1.88 | -2.2 | -1.07 | -1.42 | -1.22 | -0.45 | -4.37 | -1.35 | -1.62 | -0.2975 | -0.0175 | -0.4875 | 0.541712 | 0.419961 | 0.96395 | 0.989223 | 0.997988 | 0.994666 |
| Slc24a2_F6RT95_S65s_1 | Slc24a2_F6 | -0.17 | -0.27 | 0.43 | -0.02 | 0.35 | 0 | 0.02 | 0.18 | -0.03 | -0.18 | -0.08 | 0.11 | 0.57 | 0.58 | -0.04 | -0.11 | 0.145 | -0.0375 | 0.2575 | 0.404383 | 0.54978 | 0.878499 | 0.989223 | 0.997988 | 0.994666 |
| Slc24a2_F6RT95_S68s_1 | Slc24a2_F6 | -1.97 | -0.17 | -0.16 | -0.28 | -0.39 | -1.48 | -0.51 | -0.7 | -1.04 | -0.48 | -0.9 | -0.57 | 0.29 | -0.13 | -0.62 | -0.87 | -0.125 | -0.1025 | 0.3125 | 0.526068 | 0.78997 | 0.818191 | 0.995762 | 0.997988 | 0.994666 |
| Slc24a2_Q8BUN9_S312s_1 | Slc24a2_Q8 | -0.25 | -0.17 | -1.54 | -2.4 | -0.17 | -0.29 | -1.61 | -1.65 | 1.52 | -0.02 | -2.15 | -1.23 | 1.89 | 0.11 | 0.84 | -1.83 | 0.16 | 0.62 | 1.3425 | 0.133674 | 0.790545 | 0.473801 | 0.995762 | 0.997988 | 0.994666 |
| Slc24a4_E9PUL6_S296s_1 | Slc24a4_E9 | -0.71 | 0.04 | 0.5 | -0.36 | 0.15 | -0.54 | 0.25 | -0.84 | -0.83 | -0.04 | -0.25 | 0.05 | -0.51 | -0.46 | 0.03 | 0.37 | -0.1125 | -0.135 | -0.01 | 0.97779 | 0.759703 | 0.696513 | 0.995762 | 0.997988 | 1 |
| Slc25a35_Q5SWT3_T99t_1 | Slc25a35_Q | -0.96 | -0.51 | -0.26 | 0.1 | -0.08 | 0.01 | -0.21 | -0.21 | 0.29 | -0.55 | 0.95 | 0.46 | 0.26 | 0.55 | 0.07 | 0.33 | 0.285 | 0.695 | 0.71 | 0.039347 | 0.307097 | 0.098103 | 0.989223 | 0.997988 | 0.994666 |
| Slc25a4_P48962_S149s_1 | Slc25a4_P4 | 1.53 | 1.39 | 1.47 | 1.75 | 1.06 | 1.48 | 0.67 | 1.53 | 1.13 | 1.51 | 1.44 | 1.4 | 1.42 | 0.4 | 1.17 | 1 | -0.35 | -0.165 | -0.5375 | 0.094142 | 0.204006 | 0.464583 | 0.989223 | 0.997988 | 0.994666 |
| Slc25a4_P48962_T84t_1 | Slc25a4_P4 | 1.4 | 1.95 | 1.81 | 1.73 | 1.44 | 1.51 | 1.47 | 1.3 | 1.09 | 1.28 | 0.41 | 1.4 | 0.78 | 1.37 | 1.26 | 1.51 | -0.2925 | -0.6775 | -0.4925 | 0.101337 | 0.198882 | 0.042396 | 0.989223 | 0.997988 | 0.994666 |
| Slc25a4_P48962_Y191y_1 | Slc25a4_P4 | 0.19 | 0.61 | 0.23 | 0.85 | 0.65 | 0.26 | 0.47 | 0.27 | 0.02 | -0.1 | -0.8 | -0.03 | -0.03 | 0.01 | 0.47 | 0.33 | -0.0575 | -0.6975 | -0.275 | 0.33702 | 0.814294 | 0.037554 | 0.995762 | 0.997988 | 0.994666 |
| Slc25a46_Q9CQS4_S61s_1 | Slc25a46_Q | 0.42 | 0.26 | 0.49 | 0.33 | 0.75 | 0.82 | 0.62 | 0.69 | -0.05 | 0.38 | 0.63 | 0.43 | -0.49 | 0.49 | 0.54 | 0.81 | 0.345 | -0.0275 | -0.0375 | 0.909518 | 0.105845 | 0.90813 | 0.989223 | 0.997988 | 0.999507 |
| Slc25a46_Q9CQS4_S94s_1 | Slc25a46_Q | -1.43 | -0.87 | -0.2 | -0.66 | -0.46 | -0.59 | -1.35 | -1.05 | -1 | -0.8 | -0.41 | -0.42 | -0.03 | -0.15 | -1.08 | -1.57 | -0.0725 | 0.1325 | 0.0825 | 0.851612 | 0.8291 | 0.681931 | 0.995762 | 0.997988 | 0.999507 |
| Slc25a46_Q9CQS4_T44t_1 | Slc25a46_Q | -1.62 | -0.87 | -0.32 | -0.35 | -1.17 | -1.93 | 0.4 | -0.54 | -0.81 | -1.47 | -2.22 | -0.43 | -0.97 | -0.57 | -0.78 | -0.63 | -0.02 | -0.4425 | 0.0525 | 0.880024 | 0.969684 | 0.361436 | 0.999187 | 0.997988 | 0.999507 |
| Slc25a46_Q9CQS4_T45t_1 | Slc25a46_Q | -1.42 | -0.71 | -1.06 | -0.56 | -0.86 | -1.57 | -0.38 | -0.62 | -0.6 | -1.21 | -1.24 | -0.97 | -0.91 | -0.35 | -1.13 | -0.98 | 0.08 | -0.0675 | 0.095 | 0.760032 | 0.808986 | 0.815139 | 0.995762 | 0.997988 | 0.9971 |
| Slc26a6_Q8CIW6_S728s_1 | Slc26a6_Q8 | 0.27 | 0.52 | 0.62 | 0.41 | 0.55 | -0.18 | 0.34 | 0.56 | 0.18 | 0.45 | 0.49 | 0.46 | 0.26 | 0.28 | 0.47 | 0.3 | -0.1375 | -0.06 | -0.1275 | 0.589655 | 0.581834 | 0.784172 | 0.989223 | 0.997988 | 0.995259 |
| Slc29a1_E9PXM6_S273s_1 | Slc29a1_E9 | -0.08 | 0.3 | 0.53 | 0.15 | 0.41 | 0.65 | 0.34 | 0.71 | 0.61 | 0.13 | -0.95 | 1.16 | 1.17 | -0.26 | 0.49 | 1.07 | 0.3025 | 0.0125 | 0.3925 | 0.305647 | 0.211 | 0.977764 | 0.989223 | 0.997988 | 0.994666 |
| Slc2a13_Q3UHK1_S629s_1 | Slc2a13_Q3 | -0.06 | 0.43 | -0.47 | 0.13 | 0.51 | 0.06 | -0.25 | 0.16 | 0.77 | 0.68 | -0.1 | -0.77 | 0.75 | 0.89 | -0.76 | -0.05 | 0.1125 | 0.1375 | 0.2 | 0.639005 | 0.689543 | 0.734507 | 0.994672 | 0.997988 | 0.995259 |
| Slc2a13_Q3UHK1_S634s_1 | Slc2a13_Q3 | -1.04 | 0.08 | -0.65 | -0.25 | -0.9 | -0.88 | -0.18 | -0.11 | -0.09 | -0.28 | -0.13 | 0.2 | -0.53 | -0.29 | -0.11 | 0.71 | -0.0525 | 0.39 | 0.41 | 0.293895 | 0.874903 | 0.216741 | 0.996493 | 0.997988 | 0.994666 |
| Slc2a3_P32037_S471s_1 | Slc2a3_P32 | 2.32 | -0.17 | 0.92 | 0.87 | 0.5 | 2.89 | 0.52 | 1.82 | 1.23 | 0.33 | -0.07 | 1.09 | 0.59 | -0.65 | 0.73 | 0.27 | 0.4475 | -0.34 | -0.75 | 0.196887 | 0.515458 | 0.543071 | 0.989223 | 0.997988 | 0.994666 |
| Slc30a1_Q60738_S501s_1 | Slc30a1_Q6 | 1.99 | 1.97 | 1.31 | 1.3 | 1.97 | 2.3 | 1.04 | 1.46 | 1.93 | 2.7 | 1.27 | 1.17 | 2.16 | 2.88 | 1.46 | 1.16 | 0.05 | 0.125 | 0.2725 | 0.527402 | 0.884347 | 0.755711 | 0.996493 | 0.997988 | 0.994666 |
| Slc30a10_Q3UVU3_S157s_1 | Slc30a10_Q | 1.22 | 0.15 | 0.43 | 1.17 | 0.1 | 1.9 | 0.9 | 2.41 | 2 | 0.66 | 1.31 | 1.48 | 1.81 | -0.11 | 1.16 | 2.18 | 0.585 | 0.62 | 0.5175 | 0.342726 | 0.283752 | 0.134624 | 0.989223 | 0.997988 | 0.994666 |
| Slc30a3_P97441_S10s_1 | Slc30a3_P9 | 0.66 | 0.18 | 0.81 | 1.11 | 0.25 | 1.66 | -0.23 | 1.5 | 1.26 | 1.47 | -0.84 | 1.56 | 1.91 | 0.41 | 0.97 | 0.82 | 0.105 | 0.1725 | 0.3375 | 0.390303 | 0.822236 | 0.757548 | 0.995762 | 0.997988 | 0.994666 |
| Slc30a3_P97441_S22s_1 | Slc30a3_P9 | -1.43 | -1.01 | -0.02 | -3.47 | -0.13 | 0.57 | -0.12 | -0.95 | -0.15 | -2.15 | -1.5 | -0.25 | 1.31 | -0.39 | -0.89 | -0.79 | 1.325 | 0.47 | 1.2925 | 0.124972 | 0.082274 | 0.551645 | 0.989223 | 0.997988 | 0.994666 |
| Slc30a3_P97441_S23s_1 | Slc30a3_P9 | -0.18 | -0.51 | 1.03 | -0.05 | 0.29 | 1.29 | 1.01 | 0.47 | 0.67 | -0.84 | 1.84 | 0.78 | 1.85 | 0.23 | 2.43 | 2.19 | 0.6925 | 0.54 | 1.6025 | 0.01572 | 0.103631 | 0.372485 | 0.989223 | 0.997988 | 0.994666 |
| Slc32a1_O35633_S95s_1 | Slc32a1_O3 | 0.74 | 1.2 | 1.55 | 1.38 | 0.68 | 0.6 | 1.31 | 1.63 | 0.91 | 0.52 | 1.42 | 1.46 | 1.03 | 1.21 | 1.49 | 1.28 | -0.1625 | -0.14 | 0.035 | 0.899925 | 0.612937 | 0.658668 | 0.989223 | 0.997988 | 0.999507 |
| Slc32a1_O35633_S97s_1 | Slc32a1_O3 | -0.9 | -0.01 | -0.04 | -0.19 | 0.17 | -1.26 | 0.08 | -0.17 | -0.93 | -0.79 | 0.36 | -0.11 | 0.14 | 0.35 | 0.27 | -0.28 | -0.01 | -0.0825 | 0.405 | 0.20839 | 0.979027 | 0.825175 | 0.999187 | 0.997988 | 0.994666 |
| Slc33a1_Q99J27_S42s_1 | Slc33a1_Q9 | -2.9 | -3.45 | -2.81 | -4.54 | -3.04 | -2.8 | -2.87 | -2.42 | -1.47 | -3.25 | -2.17 | -2.51 | -1.36 | -3.03 | -2.05 | -1.15 | 0.6425 | 1.075 | 1.5275 | 0.01762 | 0.134926 | 0.059363 | 0.989223 | 0.997988 | 0.994666 |
| Slc35a5_M0QWE5_S129s_1 | Slc35a5_M | -0.34 | -0.39 | 0.41 | -0.16 | -0.69 | -0.06 | -1.14 | -0.16 | -0.09 | -0.65 | 0.54 | -0.33 | -0.01 | -1.02 | -0.01 | -0.26 | -0.3925 | -0.0125 | -0.205 | 0.547927 | 0.24392 | 0.970119 | 0.989223 | 0.997988 | 0.994666 |
| Slc35d3_Q8BGF8_S344s_1 | Slc35d3_Q8 | -1.21 | -1.45 | -4.04 | -0.69 | -0.88 | -2.33 | 0.51 | -0.84 | -0.63 | -0.58 | -0.76 | -0.84 | -0.19 | -1.23 | 0.3 | -0.33 | 0.9625 | 1.145 | 1.485 | 0.064647 | 0.261071 | 0.117799 | 0.989223 | 0.997988 | 0.994666 |
| Slc35e1_Q8CD26_S363s_1 | Slc35e1_Q8 | 0.65 | -1.14 | 0.5 | 0.81 | 0.11 | 1 | 1.17 | -0.53 | -1.77 | 0.08 | -0.02 | 0.47 | -0.68 | -1.31 | 0.25 | 0.39 | 0.2325 | -0.515 | -0.5425 | 0.344363 | 0.671031 | 0.411545 | 0.992876 | 0.997988 | 0.994666 |
| Slc35g2_D3YVE8_S4s_1 | Slc35g2_D3 | 0.35 | 0.46 | 0.97 | 1.11 | 0.02 | 0.6 | 0.36 | 0.27 | 1.13 | 0.37 | 0.9 | 0.92 | 1.78 | 0.06 | 1 | 1.27 | -0.41 | 0.1075 | 0.305 | 0.462632 | 0.148149 | 0.713243 | 0.989223 | 0.997988 | 0.994666 |

| | | | | | | | | | | | | | | | | | | | | | | | | | | |
|---|---|---|---|---|---|---|---|---|---|---|---|---|---|---|---|---|---|---|---|---|---|---|---|---|---|---|
| Slc38a1_Q8K2P7_S52s_1 | Slc38a1_Q8 | 0.4 | 0.31 | 1.26 | 0.61 | 0.99 | 0.48 | 0.68 | 0.92 | 0.14 | 0.14 | 2.28 | 1.56 | 0.39 | 0.95 | 1.33 | 0.81 | 0.1225 | 0.385 | 0.225 | 0.501002 | 0.663196 | 0.4788 | 0.990434 | 0.997988 | 0.994666 |
| Slc38a7_Q8BWH0_S28s_1 | Slc38a7_Q8 | 0.77 | 0.28 | 0.48 | 0.79 | 1.48 | 1.19 | 1.15 | 1.12 | 0.39 | 0.98 | -0.53 | -0.32 | -0.02 | 1.03 | -0.05 | 0.84 | 0.655 | -0.45 | -0.13 | 0.704994 | 0.015846 | 0.246608 | 0.989223 | 0.997988 | 0.996159 |
| Slc39a10_Q6P5F6_S537s_1 | Slc39a10_( | 1.01 | 0.31 | 0.02 | 0.08 | 0.3 | 1.71 | -0.18 | 0.01 | 0.65 | 0.34 | 0.33 | -0.07 | 1.08 | 0.29 | 0.37 | 0 | 0.105 | -0.0425 | 0.08 | 0.820148 | 0.81656 | 0.889738 | 0.995762 | 0.997988 | 0.999507 |
| Slc39a10_Q6P5F6_S593s_1 | Slc39a10_( | -0.08 | 0.53 | -1.18 | -0.89 | 0.55 | -0.16 | -0.55 | -0.97 | -0.08 | 0.14 | -2.44 | -1.32 | -0.46 | 0.5 | -2.1 | -1.13 | 0.1225 | -0.52 | -0.3925 | 0.525603 | 0.793984 | 0.43094 | 0.995762 | 0.997988 | 0.994666 |
| Slc39a10_Q6P5F6_T538t_1 | Slc39a10_( | 2.58 | 0.22 | 0.25 | 1.11 | -0.08 | 2.73 | -0.33 | 1.26 | 1.35 | 0.77 | 1.1 | 0.36 | 1.35 | -0.2 | 0.35 | 0.36 | -0.145 | -0.145 | -0.575 | 0.339734 | 0.85316 | 0.792226 | 0.996349 | 0.997988 | 0.994666 |
| Slc39a10_Q6P5F6_T542t_1 | Slc39a10_( | -0.45 | -1.29 | -1.01 | -1.49 | -0.72 | 0.47 | -1.32 | -0.84 | 0.12 | -1.5 | -0.97 | -0.74 | 0.68 | -1.46 | -0.5 | -0.47 | 0.4575 | 0.2875 | 0.6225 | 0.208691 | 0.291491 | 0.481575 | 0.989223 | 0.997988 | 0.994666 |
| Slc39a6_D3Z7N4_S204s_1 | Slc39a6_D3 | -0.1 | 0.66 | 0.63 | 0.25 | 0.39 | 0.55 | -0.06 | -0.1 | 0.77 | 0.84 | 1.05 | 0.5 | 0.74 | 0.46 | 0.83 | 0.32 | -0.165 | 0.43 | 0.2275 | 0.436027 | 0.557458 | 0.137365 | 0.989223 | 0.997988 | 0.994666 |
| Slc39a7_Q31125_S312s_1 | Slc39a7_Q3 | 0.81 | 1.3 | 0.56 | 0.84 | 1.48 | 0.48 | 0.61 | 0.4 | 0.92 | 1.37 | 1.21 | 0.96 | 0.01 | 1.18 | 1.14 | 0.91 | -0.135 | 0.2375 | -0.0675 | 0.845283 | 0.666592 | 0.366064 | 0.992177 | 0.997988 | 0.999507 |
| Slc3a2_P10852_S2s_1 | Slc3a2_P10 | -0.33 | 0.08 | -0.11 | -0.18 | 0.17 | 0.04 | -0.4 | 0.18 | -0.48 | -0.86 | 0.15 | -0.27 | 0.31 | 0.36 | -0.1 | -0.26 | 0.1325 | -0.23 | 0.2125 | 0.433534 | 0.572786 | 0.419053 | 0.989223 | 0.997988 | 0.994666 |
| Slc43a2_Q8CGA3_S274s_1 | Slc43a2_Q8 | -0.12 | 0.35 | -0.09 | 0.18 | 0.29 | 0.55 | 0.82 | -0.24 | 0.62 | 0.54 | 0.07 | -0.01 | 0.98 | 0.54 | 0.4 | 0.19 | 0.275 | 0.225 | 0.4475 | 0.134513 | 0.347124 | 0.398839 | 0.989223 | 0.997988 | 0.994666 |
| Slc45a4_F7BEQ1_S344s_1 | Slc45a4_F7 | 0.41 | 0.9 | 0.32 | 0.22 | 0.82 | 0.15 | 0.21 | 0.18 | 0.3 | 0.64 | -0.2 | 0.34 | 0.84 | 1.4 | 0.54 | 0.43 | -0.1225 | -0.1925 | 0.34 | 0.295757 | 0.646278 | 0.499191 | 0.99033 | 0.997988 | 0.994666 |
| Slc45a4_F7BEQ1_S365s_1 | Slc45a4_F7 | -0.36 | 0.3 | -0.34 | -0.85 | 0.33 | 0.41 | -0.03 | -0.91 | -1.03 | -0.71 | -0.33 | -1.01 | 0.09 | 0.68 | -0.68 | -0.33 | 0.2625 | -0.4575 | 0.2525 | 0.519424 | 0.490499 | 0.173199 | 0.989223 | 0.997988 | 0.994666 |
| Slc47a1_D6RGQ3_S18s_1 | Slc47a1_D6 | 0.61 | 0.12 | 0.01 | 1.42 | -0.1 | 0.49 | 0.32 | 1.55 | 0.5 | 0.18 | 1.58 | 0.92 | 0.78 | 0.73 | 1.15 | 0.53 | 0.025 | 0.255 | 0.2575 | 0.48892 | 0.955196 | 0.556423 | 0.999187 | 0.997988 | 0.994666 |
| Slc4a10_Q5DTL9_S1056s_1 | Slc4a10_Q5 | -0.15 | 0.47 | -0.15 | 0.1 | 0.41 | -0.15 | 0.32 | 0.14 | -0.13 | 0.07 | 0.27 | 0.16 | -0.36 | 0.65 | 0.09 | 0.31 | 0.1125 | 0.025 | 0.105 | 0.736153 | 0.653214 | 0.919332 | 0.990434 | 0.997988 | 0.9971 |
| Slc4a10_Q5DTL9_S246s_1 | Slc4a10_Q5 | -0.67 | -1.07 | -0.37 | 0.47 | -1.73 | -0.55 | -0.9 | -0.6 | 0.62 | -1.26 | -1.98 | -0.16 | 1.09 | -1.45 | 0.52 | 0.03 | -0.535 | -0.285 | 0.4575 | 0.439549 | 0.209698 | 0.640087 | 0.989223 | 0.997988 | 0.994666 |
| Slc4a10_Q5DTL9_S260s_1 | Slc4a10_Q5 | 0.41 | 1.12 | 0.53 | 0.95 | 0.7 | 1.02 | 0.44 | 0.69 | 0.37 | 0.67 | 1.24 | 0.79 | 0.26 | 0.67 | 0.72 | 0.57 | -0.04 | 0.015 | -0.1975 | 0.483232 | 0.876443 | 0.958915 | 0.996493 | 0.997988 | 0.994666 |
| Slc4a10_Q5DTL9_S275s_1 | Slc4a10_Q5 | 2.93 | 0.85 | 1.44 | 2.36 | 2.16 | 3 | 1.86 | 2.62 | 1.88 | 1.65 | 1.29 | 2.14 | -0.75 | 1.28 | 0.62 | 1.35 | 0.515 | -0.155 | -1.27 | 0.061997 | 0.304088 | 0.744122 | 0.989223 | 0.997988 | 0.994666 |
| Slc4a10_Q5DTL9_S279s_1 | Slc4a10_Q5 | -1.93 | -2.72 | -1.81 | -2.04 | -1.03 | -1.61 | -1.18 | -1.31 | -2.37 | -2.39 | -0.15 | -1.32 | -2.17 | -0.9 | -2.44 | -1.71 | 0.8425 | 0.5675 | 0.32 | 0.432104 | 0.012124 | 0.300359 | 0.989223 | 0.997988 | 0.994666 |
| Slc4a10_Q5DTL9_S296s_1 | Slc4a10_Q5 | 4.02 | 2.78 | 3.32 | 0.72 | 3.14 | 4.9 | 3.23 | 2.31 | -0.63 | 3.76 | -1.63 | 2.74 | 1.35 | -3.68 | 3.81 | 3.96 | 0.685 | -1.65 | -1.35 | 0.420311 | 0.391016 | 0.223383 | 0.989223 | 0.997988 | 0.994666 |
| Slc4a10_Q5DTL9_S80s_2 | Slc4a10_Q5 | -0.62 | -0.04 | 2.02 | 1.7 | 0.09 | -0.45 | 1.32 | 0.38 | -0.54 | -0.73 | -0.12 | -0.05 | -0.48 | -0.69 | 1.02 | 0.37 | -0.43 | -1.125 | -0.71 | 0.31073 | 0.519549 | 0.091519 | 0.989223 | 0.997988 | 0.994666 |
| Slc4a10_Q5DTL9_S88s_1 | Slc4a10_Q5 | 1.44 | 1.74 | 1.38 | 0.87 | 1.42 | 1.27 | 1.01 | 0.93 | 1.07 | 1.67 | 1.24 | 1.09 | 0.13 | 1.09 | 0.91 | 1.31 | -0.2 | -0.09 | -0.4975 | 0.173248 | 0.451493 | 0.74827 | 0.989223 | 0.997988 | 0.994666 |
| Slc4a10_Q5DTL9_S90s_1 | Slc4a10_Q5 | 1.37 | 1.8 | 0.97 | 0.86 | 2.17 | 1.63 | 0.81 | 0.73 | 1.03 | 1.35 | 0.33 | 0.48 | 0.7 | 1.78 | 0.56 | 0.78 | 0.085 | -0.4525 | -0.295 | 0.433984 | 0.828477 | 0.204488 | 0.995762 | 0.997988 | 0.994666 |
| Slc4a2_P13808_S156s_2 | Slc4a2_P13 | 0.62 | 0.93 | 0.12 | 0.65 | 0.89 | 0.42 | -0.18 | -0.15 | 0.68 | 0.48 | 0.54 | -0.51 | -0.6 | -0.64 | 0.29 | 0.44 | -0.335 | -0.2825 | -0.7075 | 0.072643 | 0.310367 | 0.415202 | 0.989223 | 0.997988 | 0.994666 |
| Slc4a2_P13808_S158s_1 | Slc4a2_P13 | 0.09 | 0.26 | -0.06 | -0.79 | 0.34 | -0.59 | -1.42 | -0.55 | 0.46 | -0.4 | -0.29 | -0.52 | 1.11 | 0 | -0.73 | 0.05 | -0.43 | -0.0625 | 0.2325 | 0.596669 | 0.307192 | 0.854006 | 0.989223 | 0.997988 | 0.995259 |
| Slc4a2_P13808_S158s_2 | Slc4a2_P13 | 0.62 | 0.93 | 0.12 | 0.65 | 0.89 | 0.42 | -0.18 | -0.15 | 0.68 | 0.48 | 0.54 | -0.51 | -0.6 | -0.64 | 0.29 | 0.44 | -0.335 | -0.2825 | -0.7075 | 0.072643 | 0.310367 | 0.415202 | 0.989223 | 0.997988 | 0.994666 |
| Slc4a3_F8WHM3_S198s_1 | Slc4a3_F8W | 0.6 | 0.39 | 0.13 | 0.13 | 1.14 | 1 | 0.84 | 0.14 | 0.75 | 0.89 | 0.33 | 0.82 | 0.31 | 0.03 | 0.34 | 0.19 | 0.4675 | 0.385 | -0.095 | 0.704037 | 0.124011 | 0.144487 | 0.989223 | 0.997988 | 0.996159 |
| Slc4a3_F8WHM3_S296s_1 | Slc4a3_F8W | 0.48 | 0.42 | 0.42 | 0.3 | 0.96 | 0.3 | 0.96 | 0.35 | 0.43 | 0.74 | 0.52 | 0.39 | 0.49 | 1.01 | 1.17 | 0.71 | 0.2375 | 0.115 | 0.44 | 0.114088 | 0.348127 | 0.592358 | 0.989223 | 0.997988 | 0.994666 |
| Slc4a4_D3Z7G8_S192s_1 | Slc4a4_D3Z | -1.98 | -0.52 | -1.6 | -0.89 | -1.08 | -2.34 | -0.64 | -1.19 | -0.19 | -0.77 | -1.06 | -0.92 | 0.96 | -0.58 | -0.44 | -0.87 | -0.065 | 0.5125 | 1.015 | 0.065546 | 0.886946 | 0.205083 | 0.996493 | 0.997988 | 0.994666 |
| Slc4a4_D3Z7G8_S204s_1 | Slc4a4_D3Z | -1.37 | -0.24 | -0.1 | -0.02 | -0.12 | -1.2 | 1.01 | -0.06 | -1.14 | -0.37 | -0.14 | 0.14 | -0.31 | 0.53 | 0.1 | 0.25 | 0.34 | 0.055 | 0.575 | 0.151652 | 0.505159 | 0.89381 | 0.989223 | 0.997988 | 0.994666 |
| Slc4a4_E9Q8N8_S1029s_1 | Slc4a4_E9C | -0.33 | 1.09 | -0.21 | -0.84 | 0.16 | -0.16 | -0.77 | -0.64 | -0.06 | 0.07 | 0.23 | -1.44 | 0.37 | 0.46 | -1.01 | -0.18 | -0.28 | -0.2275 | -0.0175 | 0.972183 | 0.525657 | 0.666414 | 0.989223 | 0.997988 | 0.999507 |
| Slc4a4_E9Q8N8_S1060s_1 | Slc4a4_E9C | -0.43 | 0.63 | 0.7 | 0.26 | 0.47 | -0.36 | 0.52 | -0.88 | -0.2 | -0.16 | -0.2 | -0.68 | -1.02 | 0.14 | 0.44 | 0.5 | -0.3525 | -0.6 | -0.275 | 0.52908 | 0.397164 | 0.084508 | 0.989223 | 0.997988 | 0.994666 |
| Slc4a4_E9Q8N8_S239s_1 | Slc4a4_E9C | 1.06 | 1.35 | 1.2 | 0.59 | 1.47 | 1.37 | 1.07 | 0.82 | 0.23 | 1.92 | 1.39 | 0.99 | 0.64 | 0.3 | 1.45 | 1.45 | 0.1325 | 0.0825 | -0.09 | 0.802465 | 0.620527 | 0.833459 | 0.995762 | 0.997988 | 0.999507 |
| Slc4a4_E9Q8N8_S245s_1 | Slc4a4_E9C | -0.27 | 0.21 | -0.58 | -0.49 | 0.04 | -0.44 | -0.14 | -0.66 | -0.37 | 0.5 | -1.46 | -0.07 | 0.67 | -1.59 | 0.15 | -0.16 | -0.0175 | -0.0675 | 0.05 | 0.918595 | 0.949144 | 0.876573 | 0.999187 | 0.997988 | 0.999507 |
| Slc4a4_E9Q8N8_S86s_1 | Slc4a4_E9C | -2.92 | -3.13 | -2.84 | -2.69 | -2.78 | -2.75 | -2.87 | -2.56 | -2.54 | -2.58 | -1.34 | -3.2 | -1.46 | -1.35 | -3.04 | -2.79 | 0.155 | 0.48 | 0.735 | 0.120126 | 0.470794 | 0.246722 | 0.989223 | 0.997988 | 0.994666 |
| Slc4a4_E9Q8N8_T1064t_1 | Slc4a4_E9C | -3.9 | -2.57 | -2.58 | -2.97 | -0.33 | -2.99 | -0.41 | -3.23 | -2.41 | -2.23 | -2.15 | -1.45 | -2.22 | -1.69 | -0.72 | -1.61 | 1.265 | 0.945 | 1.445 | 0.007735 | 0.116087 | 0.032063 | 0.989223 | 0.997988 | 0.994666 |
| Slc4a7_H6WZF5_S1072s_1 | Slc4a7_H6\ | -0.54 | -1.41 | -0.1 | -0.57 | 0.8 | 0.22 | 0.41 | 0.05 | -1.4 | -1.56 | 0.54 | 0.02 | -1.83 | 0.61 | -0.59 | -0.24 | 1.025 | 0.055 | 0.1425 | 0.790543 | 0.010867 | 0.919601 | 0.989223 | 0.997988 | 0.999507 |
| Slc4a7_H6WZF5_S233s_1 | Slc4a7_H6\ | -3.17 | -4.26 | -3.55 | -5.16 | -2.92 | -3.26 | -2.64 | -3.53 | -6.32 | -3.64 | -0.42 | -2.65 | -3.42 | -2.4 | -3.98 | -3.47 | 0.9475 | 0.7775 | 0.7175 | 0.186131 | 0.057978 | 0.501213 | 0.989223 | 0.997988 | 0.994666 |
| Slc4a7_H6WZF5_S255s_1 | Slc4a7_H6\ | 1.37 | 0.72 | -0.09 | 0.3 | 1.38 | 1.27 | 0.65 | 1.3 | 0.49 | 1.16 | 0.28 | -0.05 | 0.59 | 1.37 | 0.58 | 0.55 | 0.575 | -0.105 | 0.1975 | 0.609363 | 0.130338 | 0.793559 | 0.989223 | 0.997988 | 0.995259 |
| Slc4a7_H6WZF5_S283s_1 | Slc4a7_H6\ | -1.93 | -2.72 | -1.81 | -2.04 | -1.03 | -1.61 | -1.18 | -1.31 | -2.37 | -2.39 | -0.15 | -1.32 | -2.17 | -0.9 | -2.44 | -1.71 | 0.8425 | 0.5675 | 0.32 | 0.432104 | 0.012124 | 0.300359 | 0.989223 | 0.997988 | 0.994666 |
| Slc4a7_H6WZF5_S84s_1 | Slc4a7_H6\ | -2.85 | -2.19 | 0.07 | -0.61 | -1.71 | -2.91 | 0 | -0.36 | -3.55 | -2.71 | -0.91 | 0.4 | -2.39 | -1.89 | 0.15 | -0.28 | 0.15 | -0.2975 | 0.2925 | 0.717799 | 0.856319 | 0.763483 | 0.996349 | 0.997988 | 0.996966 |
| Slc4a8_Q8JZR6_S194s_1 | Slc4a8_Q8J | 0.66 | 0.41 | 0.42 | 0.75 | -0.25 | 1.4 | 0.22 | 0.12 | 2.11 | 0.58 | 0.11 | 0.61 | 2.32 | -0.68 | 0.87 | -0.67 | -0.1875 | 0.2925 | -0.1 | 0.878259 | 0.608882 | 0.502722 | 0.989223 | 0.997988 | 0.999507 |
| Slc4a8_Q8JZR6_S207s_1 | Slc4a8_Q8J | 0.43 | 1.31 | 1.58 | 1.4 | 0.97 | 0.39 | 1.63 | 1.3 | 0.56 | 1.16 | 1.22 | 1.79 | 0.42 | 0.4 | 1.75 | 1.34 | -0.1075 | 0.0025 | -0.2025 | 0.633272 | 0.768494 | 0.994569 | 0.995762 | 0.997988 | 0.995259 |
| Slc4a8_Q8JZR6_S994s_1 | Slc4a8_Q8J | 0.15 | 0.04 | 0.61 | 0.12 | 0.78 | -0.01 | 0.37 | 0.09 | 0.13 | 0.47 | 0.52 | 0.48 | 0.65 | 0.85 | 0.36 | 0.13 | 0.0775 | 0.17 | 0.2675 | 0.353462 | 0.769971 | 0.486948 | 0.995762 | 0.997988 | 0.994666 |
| Slc5a3_Q9JKZ2_S37s_2 | Slc5a3_Q9J | -0.85 | 0.05 | -1.72 | -2.03 | -0.54 | -1.42 | -2.37 | -4.61 | -2.47 | -2.11 | -0.52 | -1.19 | 0.1 | -1.3 | -1.41 | -1.53 | -1.0975 | -0.435 | 0.1025 | 0.854203 | 0.22378 | 0.468539 | 0.989223 | 0.997988 | 0.999507 |
| Slc5a3_Q9JKZ2_Y39y_2 | Slc5a3_Q9J | -0.85 | 0.05 | -1.72 | -2.03 | -0.54 | -1.42 | -2.37 | -4.61 | -2.47 | -2.11 | -0.52 | -1.19 | 0.1 | -1.3 | -1.41 | -1.53 | -1.0975 | -0.435 | 0.1025 | 0.854203 | 0.22378 | 0.468539 | 0.989223 | 0.997988 | 0.999507 |
| Slc5a7_Q8BGY9_S550s_1 | Slc5a7_Q8I | 0.59 | -0.02 | 9.48 | 8.97 | -0.35 | 0.29 | 7.87 | 11.22 | -0.69 | -0.27 | 14.47 | 9.52 | -0.28 | 1.01 | 9.41 | 10.02 | 0.0025 | 1.0025 | 0.285 | 0.928777 | 0.99939 | 0.799366 | 1 | 0.997988 | 0.999507 |
| Slc6a1_P31648_S14s_1 | Slc6a1_P31 | 1.13 | 1.35 | 0.85 | 0.76 | 0.96 | 1.21 | 1.1 | 1.01 | 0.48 | 0.88 | 0.42 | 0.85 | 0.39 | -0.18 | 0.87 | 0.85 | 0.0475 | -0.365 | -0.54 | 0.122099 | 0.833365 | 0.172708 | 0.995762 | 0.997988 | 0.994666 |
| Slc6a1_P31648_S591s_1 | Slc6a1_P31 | -0.44 | 0.06 | 0.44 | 0.21 | 0.72 | -0.29 | 0.86 | 0.58 | 0.11 | 0.36 | 0.15 | 0.37 | -0.54 | 0.56 | -0.07 | 0.49 | 0.4 | 0.18 | 0.0425 | 0.90304 | 0.243404 | 0.498671 | 0.989223 | 0.997988 | 0.999507 |
| Slc6a1_P31648_S592s_1 | Slc6a1_P31 | 0.75 | 0.35 | 1.13 | 1.21 | 1.17 | 0.49 | 1.85 | 0.86 | 0.66 | 1.09 | 0.54 | 1.61 | -0.04 | 0.76 | 1.51 | 1.24 | 0.2325 | 0.115 | 0.0075 | 0.985014 | 0.512724 | 0.731845 | 0.989223 | 0.997988 | 1 |
| Slc6a1_P31648_T15t_1 | Slc6a1_P31 | 1.09 | 1.68 | 0.86 | 0.87 | 1.12 | 0.58 | 0.99 | 1.34 | 0.54 | 0.46 | 0.4 | 0.77 | 0.14 | -1.19 | 1.02 | 1.01 | -0.1175 | -0.5825 | -0.88 | 0.116258 | 0.680067 | 0.053484 | 0.993464 | 0.997988 | 0.994666 |
| Slc6a11_P31650_S21s_1 | Slc6a11_P3 | 2.26 | 1.87 | 1.19 | 0.76 | 1.5 | 1.53 | 0.49 | 0.76 | 2.06 | 2.19 | 1.56 | 0.44 | 1.88 | 1.53 | 1.27 | 0.98 | -0.45 | 0.0425 | -0.105 | 0.790759 | 0.286123 | 0.931104 | 0.989223 | 0.997988 | 0.999507 |
| Slc6a13_P31649_T587t_1 | Slc6a13_P3 | -0.08 | 0.31 | 0.04 | 0.43 | 0.39 | -1.73 | 0.85 | 0.54 | 0.41 | 0.35 | 0.5 | 0.05 | -0.14 | 0.57 | -0.06 | 0.51 | -0.1625 | 0.1525 | 0.045 | 0.876526 | 0.765234 | 0.528072 | 0.995762 | 0.997988 | 0.999507 |
| Slc6a15_Q8BG16_S55s_1 | Slc6a15_Q8 | 0.24 | 1.55 | 0.79 | 0.91 | 1.8 | 0.7 | 0.19 | 1.15 | -0.97 | 1.4 | 0.97 | -0.09 | -2.3 | -0.02 | 1.15 | 1.22 | 0.0875 | -0.545 | -0.86 | 0.278163 | 0.832765 | 0.338074 | 0.995762 | 0.997988 | 0.994666 |
| Slc6a15_Q8BG16_S675s_1 | Slc6a15_Q8 | -0.82 | -0.83 | -0.59 | -1.08 | 0.38 | -0.53 | -0.31 | -0.74 | -1.18 | -1.32 | -0.39 | -0.54 | -1.44 | -0.05 | -1.09 | -0.18 | 0.53 | -0.0275 | 0.14 | 0.70848 | 0.0948 | 0.925449 | 0.989223 | 0.997988 | 0.996966 |
| Slc6a15_Q8BG16_S686s_1 | Slc6a15_Q8 | -0.24 | -0.43 | -0.02 | -0.34 | -0.45 | -1.23 | 0.06 | -0.2 | -0.47 | 0.24 | -0.03 | -0.02 | 0.02 | 0.32 | 0.01 | 0.06 | -0.1975 | 0.1875 | 0.36 | 0.160959 | 0.529949 | 0.457222 | 0.989223 | 0.997988 | 0.994666 |
| Slc6a15_Q8BG16_S701s_1 | Slc6a15_Q8 | -0.33 | 0.33 | 0.19 | -0.19 | 0.31 | -0.04 | -1.18 | -0.45 | 0.44 | 0.17 | 0.11 | -0.26 | 1.15 | 0.35 | 0.23 | 0.19 | -0.34 | 0.115 | 0.48 | 0.15846 | 0.351287 | 0.67189 | 0.989223 | 0.997988 | 0.994666 |
| Slc6a17_Q8BJI1_S665s_1 | Slc6a17_Q8 | 2.79 | 1.79 | 1.93 | 2.17 | 2.93 | 3.41 | 2.01 | 2.69 | -0.18 | 1.81 | 0.65 | 2.6 | -2.81 | 0.21 | 1.97 | 2.48 | 0.59 | -0.95 | -1.7075 | 0.130458 | 0.129094 | 0.138289 | 0.989223 | 0.997988 | 0.994666 |
| Slc6a17_Q8BJI1_S682s_1 | Slc6a17_Q8 | -1.19 | -1.54 | -1.61 | -0.69 | -0.46 | -0.6 | -0.61 | -0.77 | -1.27 | -1.71 | -0.32 | -0.53 | -0.94 | -0.39 | -1.55 | -0.69 | 0.6475 | 0.3 | 0.365 | 0.314467 | 0.033369 | 0.44628 | 0.989223 | 0.997988 | 0.994666 |
| Slc6a17_Q8BJI1_S682s_2 | Slc6a17_Q8 | -0.15 | -0.91 | 0.12 | -0.67 | -0.81 | -0.91 | -0.59 | -0.59 | -0.34 | -2.6 | -0.04 | -0.87 | -1.02 | 0.48 | 0.85 | -0.85 | -0.3225 | -0.56 | 0.2675 | 0.593346 | 0.271275 | 0.339491 | 0.989223 | 0.997988 | 0.995259 |
| Slc6a17_Q8BJI1_S686s_1 | Slc6a17_Q8 | 0.4 | 0.29 | -0.38 | -0.12 | -0.04 | 1.24 | -0.23 | 0.43 | 0.71 | 0.41 | 0.76 | -0.09 | 0.55 | 0.36 | -0.37 | -0.22 | 0.3025 | 0.4 | 0.0325 | 0.921165 | 0.420994 | 0.209233 | 0.989223 | 0.997988 | 0.999507 |
| Slc6a17_Q8BJI1_S686s_2 | Slc6a17_Q8 | -0.15 | -0.91 | 0.12 | -0.67 | -0.81 | -0.91 | -0.59 | -0.59 | -0.34 | -2.6 | -0.04 | -0.87 | -1.02 | 0.48 | 0.85 | -0.85 | -0.3225 | -0.56 | 0.2675 | 0.593346 | 0.271275 | 0.339491 | 0.989223 | 0.997988 | 0.995259 |
| Slc6a17_Q8BJI1_S701s_1 | Slc6a17_Q8 | 0.76 | 0.81 | 0.08 | 0.71 | 0.94 | 0.58 | 0.78 | 0.6 | 0.24 | 0.62 | 0.02 | 0.49 | 0.3 | 0.44 | 0.01 | 0.5 | 0.135 | -0.2475 | -0.2775 | 0.335689 | 0.590213 | 0.376025 | 0.989223 | 0.997988 | 0.994666 |
| Slc6a17_Q8BJI1_S715s_1 | Slc6a17_Q8 | 0.05 | 0.53 | 0 | 0.78 | 0.11 | -0.5 | 0.9 | 0.89 | -0.19 | -0.71 | 0.24 | 0.24 | 0.12 | 0.33 | 0.26 | 0.9 | 0.01 | -0.445 | 0.0625 | 0.840084 | 0.978926 | 0.190122 | 0.999187 | 0.997988 | 0.999507 |
| Slc6a17_Q8BJI1_T722t_1 | Slc6a17_Q8 | 0.79 | -0.01 | 0.34 | 0.67 | 0.44 | 0.75 | 0.55 | 0.62 | -0.15 | 0.19 | -0.34 | 0.51 | -0.97 | -0.8 | -0.32 | 0.17 | 0.1425 | -0.395 | -0.9275 | 0.022231 | 0.570207 | 0.208865 | 0.989223 | 0.997988 | 0.994666 |
| Slc6a6_O35316_T596t_1 | Slc6a6_O3! | 0.19 | -0.43 | -0.3 | -0.68 | 0.09 | -0.27 | 0.03 | -0.08 | 0.54 | -0.44 | 0.91 | -0.63 | 0.83 | 0.35 | -0.24 | -0.44 | 0.2475 | 0.4 | 0.43 | 0.255583 | 0.341183 | 0.342512 | 0.989223 | 0.997988 | 0.994666 |
| Slc6a7_Q6PGE7_S598s_1 | Slc6a7_Q6I | 0.4 | 0.57 | 0.4 | 0.65 | 0.11 | 0.94 | 1.27 | 0.68 | 1.11 | 1.01 | -0.14 | 0.79 | 0.38 | -0.07 | 0.87 | 0.86 | 0.245 | 0.1875 | 0.005 | 0.986553 | 0.400447 | 0.562631 | 0.989223 | 0.997988 | 1 |
| Slc6a7_Q6PGE7_S600s_1 | Slc6a7_Q6I | -0.04 | -0.4 | -0.34 | 0.35 | 0.49 | 0.12 | 0.99 | 0.65 | -0.34 | 0.16 | -1.33 | 0.3 | -0.75 | 0.16 | 0.17 | 0.18 | 0.67 | -0.195 | 0.0475 | 0.885118 | 0.03792 | 0.630939 | 0.989223 | 0.997988 | 0.999507 |
| Slc6a7_Q6PGE7_S616s_1 | Slc6a7_Q6I | -0.22 | -1.09 | 1.06 | 1.34 | 0.34 | 0.12 | 1.62 | 1.86 | -1.43 | -0.94 | -0.49 | 1.73 | -1.85 | -0.12 | 0.64 | 1.86 | 0.7125 | -0.555 | -0.14 | 0.868469 | 0.280563 | 0.495021 | 0.989223 | 0.997988 | 0.999507 |
| Slc6a8_F6UKB4_T561t_1 | Slc6a8_F6U | 1.11 | 1.87 | 1.09 | 1.38 | 0.52 | 2.19 | 1.04 | 1.62 | 0.14 | 0.85 | 0.35 | 1.22 | -0.21 | 0.83 | 0.94 | 0.65 | -0.02 | -0.7225 | -0.81 | 0.040736 | 0.959177 | 0.051192 | 0.999187 | 0.997988 | 0.994666 |
| Slc7a10_E9Q1V6_S5s_1 | Slc7a10_E9 | -0.23 | 0.49 | -0.14 | 0.43 | 0.08 | -0.93 | 0.61 | -0.02 | -0.58 | -0.93 | 0.45 | 0.11 | 0.48 | 0.7 | 0.58 | 0.54 | -0.2025 | -0.375 | 0.4375 | 0.137105 | 0.583549 | 0.328789 | 0.989223 | 0.997988 | 0.994666 |
| Slc7a10_E9Q1V6_S9s_1 | Slc7a10_E9 | -0.03 | 0.23 | -0.13 | 0.35 | 0.21 | -0.24 | -0.35 | 0.02 | 0.36 | 0.59 | 0.69 | 0.28 | 0.4 | 0.11 | 0.36 | -0.26 | -0.195 | 0.375 | 0.0475 | 0.861981 | 0.41914 | 0.137943 | 0.989223 | 0.997988 | 0.999507 |
| Slc7a14_Q8BXR1_S465s_1 | Slc7a14_Q8 | 0.41 | -0.31 | 0.06 | -0.37 | 0.54 | 0.2 | 1.64 | 0.41 | 0.19 | -0.25 | 2.61 | -0.48 | -0.07 | -0.43 | 0.07 | 1.66 | 0.75 | 0.57 | 0.36 | 0.45652 | 0.064374 | 0.401133 | 0.989223 | 0.997988 | 0.994666 |
| Slc7a14_Q8BXR1_S691s_1 | Slc7a14_Q8 | -0.25 | -0.66 | -1.02 | -1.3 | 0.14 | -0.12 | -0.27 | -0.74 | 0.21 | -0.1 | -1.22 | -1.01 | 0.45 | -0.41 | 0.13 | -0.94 | 0.56 | 0.2775 | 0.615 | 0.140553 | 0.097855 | 0.502929 | 0.989223 | 0.997988 | 0.994666 |
| Slc7a2_E9QJY0_S662s_1 | Slc7a2_E9C | 0.07 | -0.12 | 0.42 | 0.22 | 1.2 | 0.2 | -0.52 | 0.43 | -0.57 | -0.37 | 0.01 | 0.66 | -3.91 | 0.09 | 0.45 | 0.38 | 0.18 | -0.215 | -0.895 | 0.3477 | 0.626514 | 0.509524 | 0.989223 | 0.997988 | 0.994666 |
| Slc7a4_Q8BLQ7_S427s_1 | Slc7a4_Q8I | 0.79 | 0.19 | 0.54 | 0.49 | 0.63 | 0.44 | 0.72 | 0.23 | 0.73 | 0.11 | 0.11 | 1.23 | -0.06 | 0.71 | 1.04 | 0.65 | 0.0025 | 0.0425 | 0.0825 | 0.792785 | 0.991494 | 0.89578 | 0.999187 | 0.997988 | 0.999507 |
| Slc7a4_Q8BLQ7_S626s_1 | Slc7a4_Q8I | -0.39 | 0.06 | 0.32 | 0.25 | -0.01 | -0.58 | 0.74 | 0.46 | -0.3 | 0.1 | 0.16 | 0.77 | -0.28 | 0.11 | 0.25 | 0.35 | 0.0925 | 0.1225 | 0.0475 | 0.867601 | 0.784241 | 0.691767 | 0.995762 | 0.997988 | 0.999507 |
| Slc7a6_Q8BGK6_S30s_1 | Slc7a6_Q8I | 0.39 | 0.58 | 0.23 | 0.24 | 0.66 | -0.2 | 0.96 | 0.2 | 0.23 | 0.82 | 1.61 | 0.37 | 0.35 | 1.14 | 0.07 | 0.7 | 0.045 | 0.3975 | 0.205 | 0.504855 | 0.878791 | 0.261511 | 0.996493 | 0.997988 | 0.994666 |
| Slc7a6os_Q7TPE5_S256s_1 | Slc7a6os_C | -1.75 | -2.1 | -2.57 | -3.38 | -3.66 | -2.53 | -3.17 | -3.14 | -2.91 | -1.52 | -0.94 | -5.22 | -0.09 | -1.93 | -2.92 | -2.04 | -0.675 | -0.1975 | 0.705 | 0.277153 | 0.120407 | 0.826419 | 0.989223 | 0.997988 | 0.994666 |
| Slc7a6os_Q7TPE5_S257s_1 | Slc7a6os_C | -1.75 | -2.1 | -2.57 | -3.38 | -3.66 | -2.53 | -3.17 | -3.14 | -2.91 | -1.52 | -0.94 | -5.22 | -0.09 | -1.93 | -2.92 | -2.04 | -0.675 | -0.1975 | 0.705 | 0.277153 | 0.120407 | 0.826419 | 0.989223 | 0.997988 | 0.994666 |
| Slc7a8_Q9QXW9_S21s_1 | Slc7a8_Q9( | 3.82 | 2.33 | 2.62 | 0.62 | 2.49 | 4.47 | 2.96 | 2.41 | 0.48 | 3.94 | 0.53 | 3.13 | 0.17 | -0.1 | 3.28 | 3.54 | 0.735 | -0.3275 | -0.625 | 0.546062 | 0.319308 | 0.738582 | 0.989223 | 0.997988 | 0.994666 |
| Slc7a8_Q9QXW9_S21s_2 | Slc7a8_Q9( | 3.21 | 2.21 | 1.51 | 0.87 | 2.92 | 3.71 | 2.4 | 2.25 | 0.49 | 3.32 | 1.81 | 2.25 | 0.13 | 0.75 | 1.86 | 2.54 | 0.87 | 0.0175 | -0.63 | 0.354711 | 0.134022 | 0.979907 | 0.989223 | 0.997988 | 0.994666 |
| Slc7a8_Q9QXW9_S527s_1 | Slc7a8_Q9( | 0.97 | 0.47 | 0.46 | -0.22 | 0.45 | 2.07 | -0.16 | 0.36 | 0.18 | 1.1 | 0.61 | 0.44 | 0.17 | -0.97 | 0.19 | 0.31 | 0.26 | 0.1625 | -0.495 | 0.229883 | 0.602789 | 0.628941 | 0.989223 | 0.997988 | 0.994666 |
| Slc8a1_Q68FL0_S282s_1 | Slc8a1_Q68 | -1.24 | -0.38 | -0.85 | -1.11 | -0.2 | -1.05 | -0.74 | -1.33 | -0.65 | -0.62 | -0.02 | -0.71 | -0.54 | 0.06 | -0.71 | -0.97 | 0.065 | 0.395 | 0.355 | 0.297795 | 0.839891 | 0.199197 | 0.995762 | 0.997988 | 0.994666 |
| Slc8a2_Q8K596_S266s_1 | Slc8a2_Q8I | -0.93 | -0.77 | -0.08 | -0.01 | -0.48 | -1.31 | 0.2 | 0.02 | -1.04 | -0.74 | -1.32 | -0.45 | -0.6 | 0.03 | -0.14 | 0.39 | 0.055 | -0.44 | 0.3675 | 0.301503 | 0.88971 | 0.200124 | 0.996493 | 0.997988 | 0.994666 |
| Slc8a2_Q8K596_S394s_1 | Slc8a2_Q8I | -2.05 | -0.33 | -0.46 | 0.31 | -0.93 | -1.06 | -1.22 | 0.31 | -0.49 | -1.04 | -0.39 | -0.43 | -0.82 | -2.23 | 0.16 | 1.19 | -0.0925 | 0.045 | 0.2075 | 0.791071 | 0.86697 | 0.927428 | 0.996493 | 0.997988 | 0.999507 |
| Slc8a2_Q8K596_S622s_1 | Slc8a2_Q8I | -0.33 | -0.09 | 0.28 | 0.39 | 0.73 | -0.04 | 0.88 | 0.45 | -0.35 | -0.34 | -0.1 | 0.49 | -0.41 | 0.5 | -0.06 | 0.35 | 0.4425 | -0.1375 | 0.0325 | 0.917811 | 0.152671 | 0.646596 | 0.989223 | 0.997988 | 0.999507 |
| Slc8a3_E9PX50_S629s_1 | Slc8a3_E9F | 1.87 | 1.64 | -0.3 | 0.53 | 1.95 | 0.43 | 1.36 | 0.27 | 1.11 | 1.49 | -0.05 | -0.72 | 0.16 | 1.75 | -0.03 | 0.08 | 0.0675 | -0.4775 | -0.445 | 0.465313 | 0.906792 | 0.470821 | 0.99702 | 0.997988 | 0.994666 |
| Slc9a1_Q61165_S697s_1 | Slc9a1_Q61 | 0.74 | 0.97 | 0.85 | 0.68 | 0.98 | 0.33 | 0.62 | 0.6 | 0.52 | 0.46 | 0.88 | 0.37 | 0.53 | 0.82 | 0.67 | 0.44 | -0.1775 | -0.2525 | -0.195 | 0.423262 | 0.441542 | 0.285076 | 0.989223 | 0.997988 | 0.994666 |
| Slc9a1_Q61165_S790s_1 | Slc9a1_Q61 | 1.5 | 1.31 | 1.16 | 0.83 | 1.86 | 1.29 | 0.75 | 0.65 | 1.01 | 1.95 | 0.58 | 1.02 | -1.21 | 0.75 | 1.06 | 0.81 | -0.0625 | -0.06 | -0.8475 | 0.122887 | 0.847591 | 0.860333 | 0.996349 | 0.997988 | 0.994666 |
| Slc9a3r1_P70441_S117s_1 | Slc9a3r1_P | -0.76 | 1.14 | 0.55 | -0.06 | 0.47 | -0.21 | -1.47 | -0.09 | -1.07 | -0.27 | -0.23 | -0.17 | -0.19 | 0.51 | 0.47 | 0.03 | -0.5425 | -0.6525 | -0.0125 | 0.977047 | 0.315846 | 0.165928 | 0.989223 | 0.997988 | 1 |
| Slc9a3r1_P70441_S273s_1 | Slc9a3r1_P | 1.09 | 1.34 | 0.8 | 0.96 | 0.75 | 0.38 | 0.57 | 0.61 | 0.94 | 1.11 | 1.03 | 1.02 | 0.21 | 1.23 | 0.35 | 0.5 | -0.47 | -0.0225 | -0.475 | 0.148296 | 0.057595 | 0.920086 | 0.989223 | 0.997988 | 0.994666 |
| Slc9a3r1_P70441_S275s_1 | Slc9a3r1_P | 0.24 | 0.95 | 0.57 | 0.45 | 0.29 | 0.07 | 0.22 | 0.18 | 0.51 | 0.88 | 0.63 | 0.74 | 0.55 | 0.96 | 0.33 | 0.33 | -0.3625 | 0.1375 | -0.01 | 0.971937 | 0.140568 | 0.580424 | 0.989223 | 0.997988 | 0.999507 |
| Slc9a3r1_P70441_S285s_2 | Slc9a3r1_P | 0.24 | 0.64 | 0.58 | 0.95 | 1.03 | 0.26 | 0.13 | 0.36 | 0.09 | 0.9 | 0.53 | 0.69 | -2.65 | 0.37 | -0.35 | 0.22 | -0.1575 | -0.05 | -1.205 | 0.087449 | 0.580014 | 0.85728 | 0.989223 | 0.997988 | 0.994666 |
| Slc9a3r1_P70441_S286s_1 | Slc9a3r1_P | 1.3 | 1.27 | 0.9 | 1.43 | 1.08 | 1.43 | 0.53 | 1.83 | 1.16 | 0.86 | 1.1 | 1.6 | 0.71 | 1.37 | 0.68 | 0.65 | -0.0075 | -0.045 | -0.3725 | 0.208418 | 0.980998 | 0.861975 | 0.999187 | 0.997988 | 0.994666 |
| Slc9a3r1_P70441_S289s_2 | Slc9a3r1_P | -0.98 | 0.73 | 0.88 | 0.5 | -0.39 | -0.55 | 0.72 | 0.53 | -0.3 | 0.47 | 1.31 | 1.07 | -1.53 | 0.36 | 1.08 | 0.22 | -0.205 | 0.355 | -0.25 | 0.69428 | 0.677676 | 0.501952 | 0.993224 | 0.997988 | 0.996159 |
| Slc9a3r1_P70441_S297s_1 | Slc9a3r1_P | 0.93 | 1.46 | -1.24 | -0.43 | 0.83 | 0.46 | -0.58 | 0.68 | 0.43 | 1.6 | -1.56 | -0.51 | 0.89 | 0.61 | -0.19 | -0.65 | 0.1675 | -0.19 | -0.015 | 0.981407 | 0.787213 | 0.815813 | 0.995762 | 0.997988 | 1 |
| Slc9a3r1_P70441_S297s_2 | Slc9a3r1_P | 0.65 | 0.75 | 0.58 | 0.95 | 1.12 | 0.58 | 0.13 | 0.36 | 0.47 | 1.09 | 0.53 | 0.69 | -2.76 | 0.87 | -0.35 | 0.22 | -0.185 | -0.0375 | -1.2375 | 0.107265 | 0.497511 | 0.877312 | 0.989223 | 0.997988 | 0.994666 |
| Slc9a3r1_P70441_S2s_1 | Slc9a3r1_P | -0.75 | -0.73 | 0.5 | 0.49 | -1.69 | -0.73 | 0.41 | 0.78 | 0.36 | -0.14 | 0.42 | 0.53 | 0.72 | -0.09 | 0.37 | 0.44 | -0.185 | 0.415 | 0.4825 | 0.246565 | 0.756465 | 0.300119 | 0.995762 | 0.997988 | 0.994666 |
| Slc9a3r2_Q9JHL1_S192s_1 | Slc9a3r2_C | -0.92 | -0.11 | 0.41 | 0.01 | 0.93 | -0.33 | 1.43 | 0.08 | -1.07 | 0 | -0.39 | 0.6 | -0.47 | -0.02 | 1.32 | 1 | 0.68 | -0.0625 | 0.61 | 0.225455 | 0.158188 | 0.885603 | 0.989223 | 0.997988 | 0.994666 |
| Slc9a6_B0QZV3_S668s_1 | Slc9a6_B0( | 0.47 | 0.47 | 0.25 | 0.47 | 0.95 | 0.06 | 0.61 | 0.44 | 0.22 | 0.35 | 0.11 | 0.35 | -0.27 | 0.75 | 0.15 | 0.42 | 0.1 | -0.1575 | -0.1525 | 0.602934 | 0.690158 | 0.462585 | 0.994672 | 0.997988 | 0.995259 |
| Slc9a6_B0QZV3_T665t_1 | Slc9a6_B0( | 0.37 | 0.88 | 0.15 | 0.59 | 0.76 | 0.41 | 0.46 | 0.46 | 0.29 | 0.57 | -0.18 | 0.25 | 0.2 | 0.42 | -0.02 | -0.05 | 0.025 | -0.265 | -0.36 | 0.210077 | 0.91712 | 0.347547 | 0.997335 | 0.997988 | 0.994666 |
| Slc9a7_Q8BLV3_S696s_1 | Slc9a7_Q8I | 1.14 | -0.23 | 0.47 | 0.23 | 0.81 | 1.29 | 0.09 | 0.9 | 0.19 | -0.13 | 0.91 | 0.56 | -0.58 | 0.83 | 0.16 | -0.04 | 0.37 | -0.02 | -0.31 | 0.456777 | 0.334277 | 0.956987 | 0.989223 | 0.997988 | 0.994666 |
| Slc9b2_Q5BKR2_S62s_1 | Slc9b2_Q5I | 0.34 | 1.31 | 0.92 | 0 | 0.86 | 0.43 | 0.63 | 0.5 | 0.39 | 1 | 0.4 | 0.71 | 0.5 | 1.39 | 1.63 | 1 | -0.0375 | -0.0175 | 0.4875 | 0.233345 | 0.90709 | 0.959452 | 0.997038 | 0.997988 | 0.994666 |
| Slco2b1_D3YY31_S35s_1 | Slco2b1_D3 | 0.03 | -0.62 | -0.26 | -0.49 | 0.04 | 0.11 | -0.68 | 0.04 | -0.67 | 0.37 | -0.2 | -0.04 | 0.85 | 0.57 | -0.03 | -0.24 | 0.2125 | 0.2 | 0.6225 | 0.085718 | 0.445907 | 0.507952 | 0.989223 | 0.997988 | 0.994666 |
| Slitrk5_Q810B7_S829s_1 | Slitrk5_Q81 | 0.47 | 0 | 0.2 | 0.19 | 0.73 | 0.72 | 0.65 | 0.41 | 0.13 | 0 | 0.45 | 0.24 | -0.36 | 1.11 | -0.48 | -0.41 | 0.4125 | -0.01 | -0.25 | 0.536859 | 0.080783 | 0.96547 | 0.989223 | 0.997988 | 0.994666 |
| Slk_O54988_S189s_1 | Slk_O5498 | 0.4 | 0.24 | 0.75 | 0.5 | 0.88 | 0.58 | 1.09 | 0.47 | 0.33 | 0.12 | 0.68 | 0.31 | -0.5 | 1.01 | 0.83 | 0.78 | 0.2825 | -0.1125 | 0.0575 | 0.879275 | 0.259359 | 0.643266 | 0.989223 | 0.997988 | 0.999507 |
| Slmap_F8WIH0_S27s_1 | Slmap_F8V | -0.14 | -0.32 | 0.08 | 1.15 | 0.39 | -0.09 | 0.93 | 0.53 | -0.63 | -0.24 | 1.06 | 0.53 | -0.14 | 0.42 | 0.38 | 0.25 | 0.2475 | -0.0125 | 0.035 | 0.925018 | 0.521068 | 0.979092 | 0.989223 | 0.997988 | 0.999507 |
| Slmap_F8WIH0_S314s_1 | Slmap_F8V | 0.02 | 0.31 | 0.81 | 0.2 | 0.45 | 0.5 | 0.66 | 0.78 | -0.45 | 0.4 | -0.4 | 0.23 | -0.13 | -0.28 | 0.7 | 0.62 | 0.2625 | -0.39 | -0.1075 | 0.751895 | 0.3003 | 0.227628 | 0.989223 | 0.997988 | 0.9971 |
| Sltm_Q8CH25_S199s_1 | Sltm_Q8CH | -1.29 | -1.68 | -1.56 | -2.96 | -1.64 | -1.69 | -2 | -2.21 | -1.21 | -1.17 | -1.24 | -3.02 | -1.05 | -1.16 | -1.79 | -3.04 | -0.0125 | 0.2125 | 0.1125 | 0.836863 | 0.974037 | 0.697237 | 0.999187 | 0.997988 | 0.999507 |
| Sltm_Q8CH25_S534s_1 | Sltm_Q8CH | 0.76 | 0.7 | 0.31 | 0.24 | 0.19 | 0.3 | -0.15 | 0.26 | 0.07 | 0.06 | 0.43 | 0.1 | 0.43 | 0.44 | 0.35 | 0.31 | -0.3525 | -0.3375 | -0.12 | 0.633534 | 0.15968 | 0.186398 | 0.989223 | 0.997988 | 0.995259 |
| Sltm_Q8CH25_S794s_1 | Sltm_Q8CH | -0.54 | -1.56 | -0.62 | -2.07 | -1.62 | -1.35 | -1.72 | -0.33 | -3.35 | -1.1 | -0.38 | -2.19 | -0.98 | -0.71 | -1.74 | -2.44 | -0.0575 | -0.5575 | -0.27 | 0.597775 | 0.899583 | 0.418812 | 0.996493 | 0.997988 | 0.995259 |
| Sltm_Q8CH25_S902s_1 | Sltm_Q8CH | -0.01 | -0.61 | 0 | -0.59 | -2.19 | -0.23 | -1.83 | -0.69 | 0.52 | -1.21 | -1.54 | -0.08 | 1.67 | -2.02 | 0.21 | -0.99 | -0.9325 | -0.275 | 0.02 | 0.977952 | 0.066846 | 0.573267 | 0.989223 | 0.997988 | 1 |
| Sltm_Q8CH25_S981s_1 | Sltm_Q8CH | 1.11 | 0.31 | 0.09 | 0.04 | -0.71 | 1.34 | -0.08 | 1.15 | 0.04 | 0.68 | -1.22 | 1.08 | 1.62 | -0.04 | 0.53 | -0.1 | 0.0375 | -0.2425 | 0.115 | 0.80076 | 0.940659 | 0.645071 | 0.999187 | 0.997988 | 0.999507 |
| Slx4_Q6P1D7_S909s_1 | Slx4_Q6P1I | -1.42 | -2.51 | -0.65 | -0.04 | -2.04 | -1.79 | -1.39 | -0.94 | -0.21 | -2.48 | -1.36 | -1.27 | 1 | -2.01 | -0.36 | -0.47 | -0.385 | -0.175 | 0.695 | 0.350035 | 0.474466 | 0.785462 | 0.989223 | 0.997988 | 0.994666 |
| Smad1_Q8C3Y6_T322t_1 | Smad1_Q8 | -1.15 | -0.97 | 0.36 | -0.37 | -2.38 | -0.62 | -1.28 | -0.67 | -0.22 | -0.96 | -0.11 | -0.53 | 0.18 | -2.99 | -0.02 | -0.5 | -0.705 | 0.0775 | -0.3 | 0.678586 | 0.173143 | 0.842922 | 0.989223 | 0.997988 | 0.996159 |
| Smad2_E9PXA8_T8t_1 | Smad2_E9I | -0.73 | -0.41 | 0.13 | -0.43 | -0.5 | -0.56 | -0.33 | -0.53 | -0.18 | 0.04 | -0.07 | -0.68 | 0.13 | 0.28 | -0.43 | 0.18 | -0.12 | 0.1375 | 0.4 | 0.204465 | 0.628024 | 0.63354 | 0.989223 | 0.997988 | 0.994666 |
| Smad3_Q8BUN5_T8t_1 | Smad3_Q8 | -0.34 | -0.23 | -0.73 | -0.94 | -0.09 | -0.93 | -0.56 | -0.94 | 0.01 | -0.15 | -0.29 | -0.71 | 0.21 | -0.38 | -0.37 | 0.12 | -0.07 | 0.275 | 0.455 | 0.146315 | 0.809568 | 0.337398 | 0.995762 | 0.997988 | 0.994666 |
| Smap_Q9R0P4_S145s_1 | Smap_Q9R | -8.37 | -9.06 | -4.95 | -4.13 | -6.04 | -10.02 | -6.13 | -3.57 | -7.51 | -6.47 | -2.59 | -4.67 | -5.42 | -3.31 | -3.49 | -5.34 | 0.1875 | 1.3175 | 2.2375 | 0.080887 | 0.903579 | 0.367394 | 0.996493 | 0.997988 | 0.994666 |
| Smap2_Q7TN29_S219s_1 | Smap2_Q7 | -0.07 | -0.56 | -0.67 | -0.53 | 0.22 | -0.14 | -0.3 | -0.4 | 0.03 | -0.45 | -0.03 | -0.26 | -1.48 | -0.15 | -0.75 | -0.4 | 0.3025 | 0.28 | -0.2375 | 0.501135 | 0.241597 | 0.277072 | 0.989223 | 0.997988 | 0.994666 |
| Smap2_Q7TN29_S239s_1 | Smap2_Q7 | -0.2 | 0.24 | 0.05 | 0.2 | 0.15 | -0.12 | 0.44 | 0.09 | -0.27 | 0.06 | -0.02 | -0.24 | -0.4 | 0.47 | -0.02 | 0.06 | 0.0675 | -0.19 | -0.045 | 0.872898 | 0.768581 | 0.414117 | 0.995762 | 0.997988 | 0.999507 |
| Smarca2_Q6DIC0_S1382s_1 | Smarca2_C | -2.23 | -1.81 | -2.69 | -3.8 | -1.68 | -2.27 | -3.11 | -3.49 | -2.16 | -1.46 | -1.5 | -3.26 | -1.3 | -1.34 | -3.35 | -3.42 | -0.005 | 0.5375 | 0.28 | 0.673302 | 0.992559 | 0.346047 | 0.999187 | 0.997988 | 0.996159 |
| Smarca2_Q6DIC0_S166s_1 | Smarca2_C | -1.39 | 0.87 | -0.09 | -0.58 | 1.22 | -1.19 | -0.29 | -0.05 | -1.52 | -1 | -0.7 | -0.48 | -0.19 | -0.9 | 0.68 | -0.16 | 0.22 | -0.6275 | 0.155 | 0.771762 | 0.720013 | 0.224531 | 0.995437 | 0.997988 | 0.997111 |
| Smarca2_Q6DIC0_S321s_1 | Smarca2_C | -0.73 | -0.15 | -0.65 | -1.34 | -0.87 | -0.22 | -1.5 | -1.5 | -0.22 | -0.7 | -1.05 | -1.31 | -1.09 | -0.39 | -2.08 | -0.74 | -0.305 | -0.1025 | -0.3575 | 0.416909 | 0.432218 | 0.772179 | 0.989223 | 0.997988 | 0.994666 |
| Smarca4_Q3TKT4_T1390t_1 | Smarca4_C | 0.12 | -1.41 | 0.34 | -0.53 | -1.89 | -0.15 | -0.34 | -0.57 | 0.94 | -0.59 | 0.59 | 0.2 | 1.12 | -3.9 | 0.3 | 0.66 | -0.3675 | 0.655 | -0.085 | 0.936234 | 0.475846 | 0.199052 | 0.989223 | 0.997988 | 0.999507 |
| Smarcc1_Q3UZD0_S309s_1 | Smarcc1_C | -1.1 | -1.43 | -0.93 | -0.51 | -1.93 | -1.58 | -1.55 | -1.19 | -0.89 | -1.52 | 0.01 | -0.97 | -0.65 | -1.69 | -0.87 | -0.43 | -0.57 | 0.15 | 0.0825 | 0.81879 | 0.065846 | 0.690944 | 0.989223 | 0.997988 | 0.999507 |
| Smarcc1_Q3UZD0_S327s_1 | Smarcc1_C | -0.18 | 0.17 | -0.63 | -0.59 | -0.36 | -0.38 | -0.24 | 0.19 | 0.05 | -0.36 | 1.39 | 0.04 | 1.52 | -0.11 | -0.68 | 0.24 | 0.11 | 0.5875 | 0.55 | 0.269255 | 0.686926 | 0.183288 | 0.994672 | 0.997988 | 0.994666 |
| Smarcc1_Q3UZD0_S327s_2 | Smarcc1_C | -0.21 | -0.11 | -1.08 | -1.07 | -0.08 | -0.35 | -1.17 | -0.4 | 0.14 | -0.15 | 1.48 | -1 | 2.16 | 0.5 | -0.76 | -0.23 | 0.1175 | 0.735 | 1.035 | 0.122902 | 0.740553 | 0.194601 | 0.995762 | 0.997988 | 0.994666 |
| Smarcc1_Q3UZD0_S329s_2 | Smarcc1_C | -0.21 | -0.11 | -1.08 | -1.07 | -0.08 | -0.35 | -1.17 | -0.4 | 0.14 | -0.15 | 1.48 | -1 | 2.16 | 0.5 | -0.76 | -0.23 | 0.1175 | 0.735 | 1.035 | 0.122902 | 0.740553 | 0.194601 | 0.995762 | 0.997988 | 0.994666 |
| Smarcc2_Q3UID0_S283s_1 | Smarcc2_C | -0.71 | -1.62 | -1.54 | -1.45 | -1.36 | -1.36 | -1.53 | -1.34 | -0.95 | -1.24 | -1.3 | -1.36 | -1.22 | -0.98 | -0.9 | -1.45 | -0.0675 | 0.1175 | 0.1925 | 0.529511 | 0.796853 | 0.676081 | 0.995762 | 0.997988 | 0.994666 |
| Smarcc2_Q3UID0_S302s_1 | Smarcc2_C | 0.62 | -0.24 | 0.37 | 0.6 | -0.29 | 0.36 | -0.12 | 0.32 | 0.59 | 0.92 | -0.24 | -0.09 | 0.61 | 0.36 | 0.04 | -0.01 | -0.27 | -0.0425 | -0.0875 | 0.774698 | 0.3607 | 0.904565 | 0.989223 | 0.997988 | 0.99798 |
| Smarcc2_Q3UID0_S302s_2 | Smarcc2_C | -0.04 | -0.68 | -1.3 | -0.91 | -1.41 | -0.85 | -1.83 | -1.49 | -0.06 | -1.02 | -1.47 | -1.6 | -0.45 | -0.71 | -1.37 | -1.59 | -0.6625 | -0.305 | -0.2975 | 0.450747 | 0.075998 | 0.480926 | 0.989223 | 0.997988 | 0.994666 |
| Smarcc2_Q3UID0_S304s_2 | Smarcc2_C | 0.15 | -0.82 | -1.22 | -0.84 | -1.39 | -0.72 | -1.73 | -1.44 | 0.04 | -0.91 | -1.39 | -1.5 | -0.3 | -0.55 | -1.3 | -1.49 | -0.6375 | -0.2575 | -0.2275 | 0.583437 | 0.102384 | 0.563385 | 0.989223 | 0.997988 | 0.995259 |
| Smarcc2_Q3UID0_S306s_2 | Smarcc2_C | -0.82 | -0.23 | -2.13 | -1.66 | -1.46 | -1.33 | -2.93 | -2 | -0.46 | -1.42 | -2.19 | -2.7 | -1.09 | -1.38 | -2.04 | -2.7 | -0.72 | -0.4825 | -0.5925 | 0.272114 | 0.180908 | 0.424666 | 0.989223 | 0.997988 | 0.994666 |
| Smc1a_Q9CU62_S956s_1 | Smc1a_Q9( | -0.86 | 0.79 | 0.12 | 0.68 | -0.25 | -1.56 | 0.94 | 0.28 | -1.5 | 0.35 | -1.24 | 1.68 | 1.11 | -0.15 | 0.97 | 0.53 | -0.33 | -0.36 | 0.4325 | 0.355802 | 0.574889 | 0.631784 | 0.989223 | 0.997988 | 0.994666 |
| Smc3_Q9CW03_S1067s_1 | Smc3_Q9C | 0.2 | -1.25 | -0.23 | 0.38 | -2.76 | 0.54 | -1.71 | -0.6 | 1.41 | 0.3 | 0.63 | -0.03 | 1.62 | -2.55 | 0.08 | -0.5 | -0.9075 | 0.8025 | -0.1125 | 0.891495 | 0.217727 | 0.106208 | 0.989223 | 0.997988 | 0.999507 |
| Smchd1_Q6P5D8_S1975s_1 | Smchd1_Q | -0.87 | -1.76 | -1.52 | -1.37 | -2.5 | -0.5 | -1.28 | -1.9 | -0.95 | -0.63 | -1.52 | -2.02 | -0.81 | -1.48 | -0.91 | -0.76 | -0.165 | 0.1 | 0.39 | 0.223895 | 0.70807 | 0.786584 | 0.995437 | 0.997988 | 0.994666 |
| Smcr8_Q3UMB5_S393s_1 | Smcr8_Q3( | 0.22 | -0.89 | 0.21 | 0.16 | -0.68 | -0.05 | 0.48 | 0.95 | -1.77 | -0.68 | -1.05 | 0.31 | -0.71 | -1.5 | 0.37 | 0.04 | 0.25 | -0.7225 | -0.375 | 0.438869 | 0.556299 | 0.160384 | 0.989223 | 0.997988 | 0.994666 |
| Smcr8_Q3UMB5_S788s_1 | Smcr8_Q3( | -0.65 | -0.32 | 0.67 | 0.31 | 0.06 | -0.43 | 0.8 | 0.64 | -0.49 | -0.16 | 0.13 | 0.4 | -0.07 | 0.07 | 0.72 | 0.63 | 0.265 | -0.0325 | 0.335 | 0.38251 | 0.508356 | 0.928893 | 0.989223 | 0.997988 | 0.994666 |
| Smek1_E9Q481_S780s_1 | Smek1_E9( | -0.88 | -0.48 | -0.91 | -0.45 | -0.75 | -1.65 | -0.15 | -0.88 | -0.45 | -0.03 | -1.47 | -0.72 | -0.67 | -1.32 | -1.79 | -1.25 | -0.1775 | 0.0125 | -0.5775 | 0.090436 | 0.603116 | 0.971073 | 0.989223 | 0.997988 | 0.994666 |
| Smg1_Q8BKX6_S3553s_1 | Smg1_Q8B | -1.55 | -1.84 | -0.55 | -0.6 | -1.65 | -3.42 | -0.81 | -1.14 | -1.11 | -1.09 | -0.71 | -1.61 | -0.47 | -0.55 | -2.03 | -0.73 | -0.62 | 0.005 | 0.19 | 0.688429 | 0.3136 | 0.989519 | 0.989223 | 0.997988 | 0.996159 |
| Smg1_Q8BKX6_S3585s_1 | Smg1_Q8B | -1.05 | -2.16 | -1.54 | -2.8 | -1.31 | -0.79 | -1.49 | -1.79 | 0.11 | -1.24 | -1.63 | -1.73 | -0.11 | -2.49 | -1.74 | -1.63 | 0.5425 | 0.765 | 0.395 | 0.498293 | 0.210212 | 0.173203 | 0.989223 | 0.997988 | 0.994666 |
| Smg6_P61406_S331s_1 | Smg6_P61- | 1.22 | 0.7 | 0.33 | 0.38 | -0.5 | 0.79 | -0.22 | 0.69 | 0.25 | 0.78 | 0.58 | -0.57 | -0.02 | -0.27 | 0.86 | -0.64 | -0.4675 | -0.3975 | -0.675 | 0.108766 | 0.232173 | 0.297359 | 0.989223 | 0.997988 | 0.994666 |
| Smg9_Q9DB90_S32s_1 | Smg9_Q9D | -0.98 | -1.01 | -1.38 | -2.73 | -0.78 | -0.94 | -1.75 | -1.5 | -0.06 | -0.42 | 0.38 | -1.4 | -0.78 | -0.06 | -0.59 | -0.61 | 0.2825 | 1.15 | 1.015 | 0.039729 | 0.527241 | 0.051139 | 0.989223 | 0.997988 | 0.994666 |
| Smim13_E9Q942_S70s_1 | Smim13_E! | -0.51 | -0.23 | -0.24 | -0.32 | 0.13 | -0.61 | 0.33 | -0.59 | 0.32 | -0.1 | 0.5 | -0.05 | -0.3 | 0.44 | 0.36 | -0.06 | 0.14 | 0.4925 | 0.435 | 0.135139 | 0.625021 | 0.066703 | 0.989223 | 0.997988 | 0.994666 |
| Smpd3_Q9JJY3_S173s_1 | Smpd3_Q9 | 0.04 | 0.41 | -0.51 | 0.04 | 0.3 | 0.07 | 0.74 | 0.37 | 0.13 | 0.26 | -0.47 | 0.14 | -0.03 | 0.28 | -0.64 | 0.15 | 0.375 | 0.02 | -0.055 | 0.864935 | 0.193479 | 0.945697 | 0.989223 | 0.997988 | 0.999507 |
| Smpd3_Q9JJY3_S202s_1 | Smpd3_Q9 | -1.39 | -1.79 | -1 | -0.91 | -0.64 | -0.8 | 0.16 | -0.06 | -1.96 | -2.62 | 1.19 | -1.38 | -1.07 | -0.5 | -1.03 | -1.19 | 0.9375 | 0.08 | 0.325 | 0.306351 | 0.014827 | 0.916955 | 0.989223 | 0.997988 | 0.994666 |
| Smpd3_Q9JJY3_S208s_1 | Smpd3_Q9 | 0 | -0.47 | 0.06 | 0.76 | -0.22 | 1.48 | 0.54 | 1.39 | 0.31 | -0.48 | 0.74 | -0.1 | 0.03 | -0.29 | -0.24 | 0.21 | 0.71 | 0.03 | -0.16 | 0.62377 | 0.134506 | 0.935708 | 0.989223 | 0.997988 | 0.995259 |
| Smpd3_Q9JJY3_S289s_2 | Smpd3_Q9 | 4.23 | 1.62 | 3.05 | 2.93 | 1.35 | 4.26 | 2.14 | 3.31 | 0.52 | 2.4 | 1.72 | 3.12 | -2.89 | -0.93 | 2.29 | 2.86 | -0.1925 | -1.0175 | -2.625 | 0.061084 | 0.792933 | 0.166634 | 0.995762 | 0.997988 | 0.994666 |
| Smpd3_Q9JJY3_S292s_1 | Smpd3_Q9 | 2.46 | 1.23 | 2.45 | 2.59 | 1.7 | 2.73 | 1.94 | 2.79 | 0.93 | 1.61 | 1.75 | 1.77 | -0.83 | -0.33 | 1.91 | 2.53 | 0.1075 | -0.6675 | -1.3625 | 0.107139 | 0.790625 | 0.105394 | 0.995762 | 0.997988 | 0.994666 |
| Smpd3_Q9JJY3_S292s_2 | Smpd3_Q9 | 4.23 | 1.62 | 3.34 | 3.25 | 1.35 | 4.26 | 2.41 | 3.69 | 0.52 | 2.4 | 1.91 | 3.44 | -2.89 | -0.93 | 2.61 | 3.19 | -0.1825 | -1.0425 | -2.615 | 0.074752 | 0.806753 | 0.177932 | 0.995762 | 0.997988 | 0.994666 |
| Smtn_D3Z3Q3_S357s_1 | Smtn_D3Z3 | -0.26 | -0.91 | -1.26 | -0.49 | -1.17 | -0.29 | -0.42 | -0.55 | -0.54 | -0.47 | -1.33 | 0.04 | 0.52 | -1.56 | -0.99 | -0.52 | 0.1225 | 0.155 | 0.0925 | 0.845033 | 0.696832 | 0.675096 | 0.994893 | 0.997988 | 0.999507 |
| Snap23_B0R030_S110s_1 | Snap23_B0 | 0.64 | 0.94 | 0.19 | 0.54 | 1.19 | 0.15 | 0.36 | 0.45 | 0.44 | 1.01 | 0.14 | 0.24 | 0.4 | 0.97 | 0.31 | 0.02 | -0.04 | -0.12 | -0.1525 | 0.62231 | 0.893594 | 0.682672 | 0.996493 | 0.997988 | 0.995259 |
| Snap25_P60879_S115s_1 | Snap25_P6 | 1.01 | 1.33 | 0.79 | 0.82 | 0.71 | 0.75 | 0.6 | 0.74 | 1.35 | 1.44 | -0.7 | 0.52 | 1.38 | 0.52 | 1.1 | 0.73 | -0.2875 | -0.335 | -0.055 | 0.851662 | 0.208727 | 0.494942 | 0.989223 | 0.997988 | 0.999507 |
| Snap25_P60879_S130s_1 | Snap25_P6 | 1.27 | 1.19 | -0.15 | 0 | 1.91 | 1.4 | 0.64 | -0.03 | 0.24 | 0.84 | -0.26 | 0.2 | 0.2 | 1.18 | -0.29 | 0.35 | 0.4025 | -0.3225 | -0.2175 | 0.643828 | 0.445438 | 0.45936 | 0.989223 | 0.997988 | 0.995259 |
| Snap25_P60879_S25s_1 | Snap25_P6 | -0.11 | -0.01 | 0.9 | 0.97 | 0.01 | -0.13 | 0.32 | 0.57 | -0.36 | -0.14 | 1.1 | 0.92 | -0.34 | 0.31 | 0.77 | 0.75 | -0.245 | -0.0575 | -0.065 | 0.869696 | 0.472371 | 0.898291 | 0.989223 | 0.997988 | 0.999507 |
| Snap25_P60879_S28s_1 | Snap25_P6 | -0.37 | -0.34 | -0.98 | -0.57 | -0.22 | -0.11 | -0.37 | -0.26 | -0.81 | -0.85 | 1.71 | -0.07 | -0.44 | 0.44 | -0.21 | 0.17 | 0.325 | 0.56 | 0.555 | 0.092965 | 0.182076 | 0.338352 | 0.989223 | 0.997988 | 0.994666 |
| Snap29_Q9ERB0_S153s_1 | Snap29_Q9 | 2.85 | 0.85 | 0.19 | -0.86 | 0.48 | 3.59 | -0.53 | 0.34 | 2.21 | 1.37 | -0.83 | 0.05 | 1.27 | 0.13 | -0.03 | -0.77 | 0.2125 | -0.0575 | -0.6075 | 0.446605 | 0.836648 | 0.949888 | 0.995762 | 0.997988 | 0.994666 |
| Snap29_Q9ERB0_S163s_1 | Snap29_Q9 | -0.71 | 0.82 | 0.54 | -1.04 | 0.28 | -0.83 | -0.99 | -0.38 | 0.01 | -0.13 | -0.06 | -0.48 | 0.79 | -0.05 | -0.69 | -0.7 | -0.3825 | -0.0675 | -0.065 | 0.903652 | 0.446605 | 0.881259 | 0.989223 | 0.997988 | 0.999507 |
| Snap91_E9Q9A3_S16s_1 | Snap91_E9 | -0.53 | -0.95 | -0.31 | -0.27 | -0.08 | -1.13 | -0.35 | 0.07 | -0.32 | -0.01 | 0.98 | -0.48 | -0.98 | -0.4 | -0.91 | 0.04 | 0.1425 | 0.5575 | -0.0475 | 0.884725 | 0.660308 | 0.156926 | 0.990434 | 0.997988 | 0.999507 |
| Snap91_E9Q9A3_S298s_1 | Snap91_E9 | -0.4 | 0.27 | -0.73 | -0.37 | 0.73 | -0.08 | -0.49 | -0.38 | 0.31 | 0.52 | 0.15 | -0.39 | 1.05 | 0.95 | 0.05 | -0.23 | 0.2525 | 0.455 | 0.7625 | 0.076881 | 0.47515 | 0.17402 | 0.989223 | 0.997988 | 0.994666 |
| Snap91_E9Q9A3_S303s_1 | Snap91_E9 | -0.13 | 0.4 | -0.24 | -0.78 | 0.79 | -0.04 | -0.12 | 0.11 | -0.61 | 0.96 | 0.39 | -0.54 | 1.29 | 0.69 | -0.06 | 0.16 | 0.3725 | 0.2375 | 0.7075 | 0.098634 | 0.274461 | 0.589182 | 0.989223 | 0.997988 | 0.994666 |
| Snap91_E9Q9A3_S311s_1 | Snap91_E9 | 0.36 | 0.78 | 0.49 | 0.45 | 0.83 | 0.32 | 0.36 | 0.52 | 0.54 | 0.8 | 0.92 | 0.5 | 0.48 | 1.22 | 0.59 | 0.52 | -0.0125 | 0.17 | 0.1825 | 0.515205 | 0.955958 | 0.469293 | 0.999187 | 0.997988 | 0.994666 |
| Snap91_E9Q9A3_S323s_1 | Snap91_E9 | 0.14 | 0.87 | 0.43 | 0.19 | 0.78 | 0.05 | 0.56 | -0.01 | -0.53 | 0.26 | -0.71 | 0.34 | -0.09 | -0.02 | 0.22 | 0.24 | -0.0625 | -0.5675 | -0.32 | 0.257583 | 0.827693 | 0.11841 | 0.995762 | 0.997988 | 0.994666 |
| Snap91_E9Q9A3_S333s_1 | Snap91_E9 | 0.58 | 0.92 | 1.23 | 0.61 | 1.09 | 0.51 | 0.83 | 0.87 | -0.74 | 0.93 | 0.62 | 0.64 | -0.32 | -0.34 | 1.12 | 0.67 | -0.01 | -0.4725 | -0.5525 | 0.190883 | 0.968102 | 0.257406 | 0.999187 | 0.997988 | 0.994666 |
| Snap91_E9Q9A3_S587s_1 | Snap91_E9 | -1.23 | -0.19 | -0.38 | -0.78 | -0.49 | -0.75 | -0.38 | -1.15 | -0.82 | -0.79 | -0.95 | -0.49 | -0.7 | -0.02 | -0.98 | -1.34 | -0.0475 | -0.1175 | -0.115 | 0.761425 | 0.877284 | 0.689985 | 0.996493 | 0.997988 | 0.9971 |
| Snap91_E9Q9A3_S593s_1 | Snap91_E9 | 0.65 | 0.09 | -0.45 | 0.02 | 0.82 | 0.46 | 0.57 | 0.68 | -0.22 | 0.04 | -0.13 | -0.16 | -0.91 | 0.3 | -0.34 | -0.14 | 0.555 | -0.195 | -0.35 | 0.345092 | 0.068976 | 0.495396 | 0.989223 | 0.997988 | 0.994666 |
| Snap91_E9Q9A3_T310t_1 | Snap91_E9 | 0.28 | 0.64 | 0.43 | 0.88 | 0.64 | 0.48 | 0.59 | 0.4 | 0.49 | 0.4 | 0.89 | 1.06 | 1.06 | 0.92 | 0.53 | 0.53 | -0.03 | 0.1525 | 0.2025 | 0.465268 | 0.893305 | 0.57013 | 0.996493 | 0.997988 | 0.994666 |
| Snap91_E9Q9A3_T310t_2 | Snap91_E9 | 0.57 | 0.83 | 0.47 | 0.47 | 0.74 | 0.63 | 0.94 | 0.09 | 0.46 | 0.92 | 0.62 | 0.98 | -0.3 | 0.78 | 0.06 | 1.04 | 0.015 | 0.16 | -0.19 | 0.591523 | 0.952893 | 0.505985 | 0.999187 | 0.997988 | 0.995259 |
| Snap91_E9Q9A3_T322t_1 | Snap91_E9 | 0.16 | 0.91 | 0.68 | 0.27 | 0.73 | 0.06 | 0.68 | 0.1 | -0.67 | 0.22 | -0.72 | 0.41 | 0.02 | -0.09 | -0.08 | 0.49 | -0.1125 | -0.695 | -0.42 | 0.17202 | 0.693999 | 0.076502 | 0.994893 | 0.997988 | 0.994666 |
| Snap91_Q61548_S296s_1 | Snap91_Q6 | 0.94 | 0.98 | 1.13 | 1.11 | 0.81 | 0.82 | 0.88 | 1.32 | 0.85 | 0.88 | 1.34 | 1.56 | 0.86 | 1.64 | 1.46 | 0.87 | -0.0825 | 0.1175 | 0.1675 | 0.556907 | 0.707454 | 0.644355 | 0.995437 | 0.997988 | 0.995259 |
| Snap91_Q61548_S296s_2 | Snap91_Q6 | -0.4 | 0.7 | 1.11 | 0.95 | 1.18 | -0.99 | 0.52 | 0.57 | -0.98 | -0.23 | 1.31 | 1.11 | -0.66 | 1.01 | 0.41 | 0.5 | -0.27 | -0.2875 | -0.275 | 0.561924 | 0.607558 | 0.629107 | 0.989223 | 0.997988 | 0.995259 |
| Snap91_Q61548_S300s_1 | Snap91_Q6 | 0.6 | 0.68 | 0.44 | 0.34 | 0.73 | 0.54 | 0.29 | 0.5 | 0.32 | 0.78 | 0.52 | 0.26 | 0.43 | 0.74 | 0.36 | 0.11 | -1.5E-16 | -0.045 | -0.105 | 0.682946 | 1 | 0.846998 | 1 | 0.997988 | 0.996159 |
| Snap91_Q61548_S300s_2 | Snap91_Q6 | -0.4 | 0.7 | 1.11 | 0.95 | 1.18 | -0.99 | 0.52 | 0.57 | -0.98 | -0.23 | 1.31 | 1.11 | -0.66 | 1.01 | 0.41 | 0.5 | -0.27 | -0.2875 | -0.275 | 0.561924 | 0.607558 | 0.629107 | 0.989223 | 0.997988 | 0.995259 |
| Snapin_Q9Z266_S10s_1 | Snapin_Q9 | -0.17 | 0.07 | 0.06 | 1.61 | 0.01 | 0.04 | 0.77 | 1.12 | -0.25 | -0.34 | 0.59 | 0.6 | -0.43 | 0.67 | 1.06 | -0.79 | 0.0925 | -0.2425 | -0.265 | 0.635242 | 0.84036 | 0.602849 | 0.995762 | 0.997988 | 0.995259 |
| Snapin_Q9Z266_S133s_1 | Snapin_Q9 | -0.04 | 0.24 | -0.55 | -0.56 | 0.57 | -0.21 | 0.17 | -0.29 | 0.31 | 0.24 | -0.4 | -0.37 | -0.11 | 0.45 | -0.38 | -0.28 | 0.2875 | 0.1725 | 0.1475 | 0.645465 | 0.353802 | 0.580259 | 0.989223 | 0.997988 | 0.995259 |
| Snapin_Q9Z266_T14t_1 | Snapin_Q9 | 0.22 | 0.46 | 1.2 | 1.04 | -0.21 | 0.13 | -0.01 | 0.99 | 1.5 | 1.04 | 0.42 | 1.56 | 0.76 | 0.73 | 1.04 | 1.14 | -0.505 | 0.4 | 0.1875 | 0.547764 | 0.175585 | 0.286074 | 0.989223 | 0.997988 | 0.994666 |
| Snapin_Q9Z266_T14t_2 | Snapin_Q9 | 1.47 | 2.19 | -0.48 | -1.14 | 1.24 | 0.57 | -0.83 | 0.32 | 1.45 | 2.01 | -0.58 | -0.3 | 1.58 | 0.49 | -0.42 | -0.35 | -0.185 | 0.135 | -0.185 | 0.818986 | 0.81391 | 0.880688 | 0.995762 | 0.997988 | 0.999507 |
| Snca_O55042_S42s_1 | Snca_O550 | -0.58 | -0.57 | -0.57 | -0.65 | 0.82 | -0.42 | 0.81 | 0.32 | -0.54 | -0.23 | -0.07 | 1.24 | -0.5 | 0.94 | 1.03 | 1.2 | 0.975 | 0.6925 | 1.26 | 0.010289 | 0.010631 | 0.104331 | 0.989223 | 0.997988 | 0.994666 |
| Snca_O55042_T81t_1 | Snca_O550 | -0.65 | -1.13 | -0.08 | -1.32 | 0.22 | -0.87 | -0.42 | -1.85 | -1.62 | -0.37 | 0.69 | -0.01 | -2.42 | 0.39 | -1.74 | -0.21 | 0.065 | 0.4675 | -0.2 | 0.755599 | 0.891458 | 0.380441 | 0.996493 | 0.997988 | 0.9971 |

| | | | | | | | | | | | | | | | | | | | | | | | | | | |
|---|---|---|---|---|---|---|---|---|---|---|---|---|---|---|---|---|---|---|---|---|---|---|---|---|---|---|
| Sncb_Q91ZZ3_S45s_1 | Sncb_Q912 | -0.13 | 0.51 | -0.75 | -1.11 | 1.09 | -0.66 | 0.84 | -0.49 | -0.58 | 0.59 | 1.05 | 1 | 0.58 | 1.68 | -0.35 | 0.86 | 0.565 | 0.885 | 1.0625 | 0.063634 | 0.29417 | 0.098291 | 0.989223 | 0.997988 | 0.994666 |
| Sncb_Q91ZZ3_S63s_1 | Sncb_Q912 | -0.27 | -0.33 | -0.27 | -0.56 | 0.12 | -0.43 | 0 | -0.26 | -0.57 | 0.02 | -0.25 | 0.39 | -1.41 | 0.71 | -0.62 | -0.13 | 0.215 | 0.255 | -0.005 | 0.990933 | 0.349736 | 0.360497 | 0.989223 | 0.997988 | 1 |
| Sncb_Q91ZZ3_S71s_1 | Sncb_Q912 | -0.6 | -0.04 | 0.04 | -0.09 | 0.12 | -0.62 | 0.18 | -0.14 | -0.69 | -0.2 | -0.99 | -0.11 | -0.39 | 0.73 | -0.67 | -0.01 | 0.0575 | -0.325 | 0.0875 | 0.808469 | 0.833256 | 0.2876 | 0.995762 | 0.997988 | 0.999507 |
| Sncb_Q91ZZ3_T44t_1 | Sncb_Q912 | -0.13 | 0.51 | -0.75 | -1.11 | 1.09 | -0.66 | 0.84 | -0.49 | -0.58 | 0.59 | 1.05 | 1 | 0.58 | 1.68 | -0.35 | 0.86 | 0.565 | 0.885 | 1.0625 | 0.063634 | 0.29417 | 0.098291 | 0.989223 | 0.997988 | 0.994666 |
| Snd1_Q3TJ56_S426s_1 | Snd1_Q3TJ | -0.19 | 0.4 | 0.16 | 0.83 | 0.7 | -0.01 | 0.76 | 0.58 | 0.26 | 0.3 | 1.06 | -0.07 | 0.2 | 0.67 | 1.11 | 0.95 | 0.2075 | 0.0875 | 0.4325 | 0.213634 | 0.496669 | 0.797414 | 0.989223 | 0.997988 | 0.994666 |
| Snip1_Q8BIZ6_S144s_1 | Snip1_Q8B | -1.01 | -1.47 | -0.09 | -0.5 | -1.24 | -0.19 | -1.19 | -0.79 | 0.49 | -1.45 | -1.64 | -1.24 | 1.27 | -2.37 | -0.07 | -0.69 | -0.085 | -0.1925 | 0.3025 | 0.677515 | 0.822098 | 0.719787 | 0.995762 | 0.997988 | 0.996159 |
| Snip1_Q8BIZ6_S145s_1 | Snip1_Q8B | -0.75 | -1.16 | -0.28 | -0.54 | -1.47 | -0.71 | -1.53 | -0.86 | 1.17 | -1.55 | -1.56 | -1.47 | 1.28 | -1.94 | -0.32 | -0.76 | -0.46 | -0.17 | 0.2475 | 0.694136 | 0.152788 | 0.789679 | 0.989223 | 0.997988 | 0.996159 |
| Snip1_Q8BIZ6_S18s_1 | Snip1_Q8B | -0.46 | 0.26 | -0.22 | -0.6 | 0.49 | -0.68 | -0.26 | -0.35 | -1.43 | -0.46 | -0.72 | -1.12 | 0.26 | -0.22 | -1.11 | -0.77 | 0.055 | -0.6775 | -0.205 | 0.586544 | 0.865147 | 0.056058 | 0.996493 | 0.997988 | 0.995259 |
| Snip1_Q8BIZ6_S33s_1 | Snip1_Q8B | -0.44 | -0.72 | -1.08 | -0.19 | 0.18 | -0.48 | -0.1 | 0.15 | -0.75 | -1.05 | 0.6 | -0.72 | 0.34 | 0.68 | 0.95 | -0.39 | 0.545 | 0.1275 | 1.0025 | 0.02077 | 0.076738 | 0.755714 | 0.989223 | 0.997988 | 0.994666 |
| Snip1_Q8BIZ6_S50s_1 | Snip1_Q8B | 0.99 | -0.3 | 0.4 | 0.47 | -0.28 | 0.36 | 0.61 | 0.96 | -0.23 | 0.24 | -0.38 | 0.6 | 1.09 | 1.03 | 1.45 | 0.8 | 0.0225 | -0.3325 | 0.7025 | 0.059673 | 0.951161 | 0.365868 | 0.999187 | 0.997988 | 0.994666 |
| Snn_P61807_S49s_1 | Snn_P6180 | -0.45 | 0.77 | 2.02 | 2.17 | -0.64 | -0.55 | 1.1 | 1.32 | 0.88 | 0.17 | 1.19 | 0.27 | -0.21 | 1.82 | 1.14 | 2 | -0.82 | -0.5 | 0.06 | 0.93232 | 0.264969 | 0.416527 | 0.989223 | 0.997988 | 0.999507 |
| Snn_P61807_S83s_1 | Snn_P6180 | 0.17 | 0.39 | 0.34 | 0.34 | 0.5 | 0.2 | 0.19 | 0.3 | 0.55 | 0.55 | 0.86 | 0.62 | 0.54 | 0.46 | 0.46 | 0.34 | -0.0125 | 0.335 | 0.14 | 0.544119 | 0.950845 | 0.141779 | 0.999187 | 0.997988 | 0.994666 |
| Snph_Q80U23_S200s_1 | Snph_Q80U | -0.86 | 0.06 | 0.93 | -0.39 | -2.25 | -0.16 | -1.23 | -0.45 | 2.28 | -0.41 | -0.23 | 0.03 | 2.68 | -1.92 | 0.06 | -0.09 | -0.9575 | 0.4825 | 0.2475 | 0.782114 | 0.104826 | 0.474683 | 0.989223 | 0.997988 | 0.998226 |
| Snph_Q80U23_S200s_2 | Snph_Q80U | -1.1 | -0.18 | -0.11 | -0.75 | -1.34 | -0.49 | -0.8 | -0.97 | 1.82 | -0.92 | 0.81 | -0.2 | 0.49 | -1.15 | 0 | -0.39 | -0.365 | 0.9125 | 0.2725 | 0.519683 | 0.261647 | 0.147254 | 0.989223 | 0.997988 | 0.994666 |
| Snph_Q80U23_S204s_1 | Snph_Q80U | 0.35 | 0.38 | 0.73 | -0.25 | -0.51 | 1.16 | -0.82 | -0.34 | 2.9 | 0.3 | 0.03 | 0.43 | 2.94 | -1.04 | 0.06 | 0.03 | -0.43 | 0.6125 | 0.195 | 0.80194 | 0.354156 | 0.347235 | 0.989223 | 0.997988 | 0.999507 |
| Snph_Q80U23_S204s_2 | Snph_Q80U | -1.1 | -0.18 | -0.11 | -0.75 | -1.34 | -0.49 | -0.8 | -0.97 | 1.82 | -0.92 | 0.81 | -0.2 | 0.49 | -1.15 | 0 | -0.39 | -0.365 | 0.9125 | 0.2725 | 0.519683 | 0.261647 | 0.147254 | 0.989223 | 0.997988 | 0.994666 |
| Snph_Q80U23_S20s_1 | Snph_Q80U | 0.24 | 0.39 | 0.41 | 0.2 | 0.66 | -0.04 | 0.23 | 0.21 | 0.65 | 0.44 | 0.79 | 0.3 | 1.34 | 1.02 | 0.53 | 0.25 | -0.045 | 0.235 | 0.475 | 0.145275 | 0.844915 | 0.311188 | 0.996349 | 0.997988 | 0.994666 |
| Snph_Q80U23_S219s_1 | Snph_Q80U | -0.32 | 0.79 | 1.53 | 1.62 | 0.46 | -0.51 | 1 | 1.37 | -0.27 | -0.36 | 1.3 | 1.71 | 0.19 | 0.44 | 1.51 | 1.4 | -0.325 | -0.31 | -0.02 | 0.969486 | 0.557124 | 0.626919 | 0.989223 | 0.997988 | 0.999507 |
| Snph_Q80U23_T19t_1 | Snph_Q80U | -0.05 | 0.42 | 0.93 | 0.3 | 0.74 | -0.23 | 0.42 | 0.34 | 0.23 | 0.2 | 0.91 | 0.46 | 1.03 | 0.96 | 0.5 | 0.44 | -0.0825 | 0.05 | 0.3325 | 0.29677 | 0.788461 | 0.867705 | 0.995762 | 0.997988 | 0.994666 |
| Snph_Q80U23_T292t_1 | Snph_Q80U | 0.88 | 0.44 | 0.45 | 0.72 | 0.81 | 1.44 | 0.63 | 0.81 | 0.28 | 0.38 | 0.39 | 0.62 | -1.72 | 0.85 | -0.44 | 0.5 | 0.3 | -0.205 | -0.825 | 0.15273 | 0.262501 | 0.380474 | 0.989223 | 0.997988 | 0.994666 |
| Snrk_Q8VDU5_S518s_1 | Snrk_Q8VD | -1.67 | -1.35 | -1.22 | -1.57 | -0.83 | -1.5 | -1.18 | -1.04 | -1.19 | -1.32 | 0.04 | -1.41 | -0.88 | -0.54 | -1.16 | -1.47 | 0.315 | 0.4825 | 0.44 | 0.154849 | 0.209442 | 0.20739 | 0.989223 | 0.997988 | 0.994666 |
| Snrk_Q8VDU5_S569s_1 | Snrk_Q8VD | -1.54 | -1.25 | -0.96 | -1.38 | -1.91 | -1.31 | -1.58 | -1.51 | -0.65 | -1.24 | -1.34 | -1.39 | 0.09 | -1.69 | -1.1 | -1.55 | -0.295 | 0.1275 | 0.22 | 0.603541 | 0.238322 | 0.638119 | 0.989223 | 0.997988 | 0.995259 |
| Snrk_Q8VDU5_S596s_1 | Snrk_Q8VD | -2.19 | 0.75 | -1 | -1.29 | -0.69 | 0.24 | -1.11 | -1.48 | -0.28 | 0.2 | -1.52 | 0.05 | 0.38 | -3.69 | -0.58 | -1.2 | 0.1725 | 0.545 | -0.34 | 0.715604 | 0.787173 | 0.418458 | 0.995762 | 0.997988 | 0.996966 |
| Snrnp200_Q6P4T2_S225s_1 | Snrnp200_ | 0.31 | 0.01 | -0.2 | -0.19 | -0.36 | 0.17 | -0.08 | 0.12 | 0.07 | -0.31 | 0.11 | 0.2 | 0.28 | 0.06 | -0.05 | -0.03 | -0.02 | 0.035 | 0.0825 | 0.744118 | 0.932812 | 0.88609 | 0.998562 | 0.997988 | 0.9971 |
| Snrnp70_Q62376_S226s_1 | Snrnp70_Q | -0.13 | -0.76 | -0.87 | -0.77 | -0.4 | -0.77 | -0.66 | -0.54 | -0.34 | -0.98 | -0.86 | -0.89 | 0.38 | -0.36 | 0.22 | -0.32 | 0.04 | -0.135 | 0.6125 | 0.071438 | 0.871148 | 0.627672 | 0.996493 | 0.997988 | 0.994666 |
| Snrnp70_Q62376_S408s_1 | Snrnp70_Q | -0.06 | -0.8 | -1.74 | -1.25 | 0.03 | -0.34 | -0.83 | -0.3 | -0.96 | -0.96 | -0.61 | 0.13 | 0.79 | 0.28 | 0.43 | -0.87 | 0.6025 | 0.3625 | 1.12 | 0.041152 | 0.143978 | 0.407015 | 0.989223 | 0.997988 | 0.994666 |
| Snta1_Q61234_S183s_1 | Snta1_Q61 | 0.65 | 0.64 | 0.92 | 0.72 | 0.65 | 0.51 | 1.05 | 0.97 | -0.1 | 0.88 | 0.23 | 0.78 | -0.15 | -0.32 | 0.95 | 0.98 | 0.0625 | -0.285 | -0.3675 | 0.336889 | 0.781482 | 0.334906 | 0.995762 | 0.997988 | 0.994666 |
| Snta1_Q61234_S194s_1 | Snta1_Q61 | -1.04 | -1.53 | -1.45 | -0.57 | -0.6 | -0.48 | -0.6 | -0.73 | -0.95 | -0.93 | 0.27 | -0.91 | -0.35 | -0.96 | -0.35 | -0.74 | 0.545 | 0.5175 | 0.5475 | 0.10926 | 0.067062 | 0.193036 | 0.989223 | 0.997988 | 0.994666 |
| Snta1_Q61234_S195s_1 | Snta1_Q61 | 0.03 | -0.32 | -0.58 | -0.04 | -0.57 | 0.61 | -0.6 | -0.81 | 1.17 | 0.11 | -0.61 | -0.69 | 1.03 | -0.75 | -0.03 | -0.24 | -0.115 | 0.2225 | 0.23 | 0.572664 | 0.744556 | 0.614448 | 0.995762 | 0.997988 | 0.995259 |
| Sntb1_Q99L88_S86s_1 | Sntb1_Q99 | -0.4 | -0.53 | -0.16 | -0.25 | -0.63 | -0.89 | 0.25 | 0.54 | 0.06 | -0.03 | 0.73 | -1.24 | -0.29 | -0.59 | -0.18 | -0.03 | 0.1525 | 0.215 | 0.0625 | 0.805238 | 0.667316 | 0.604291 | 0.99272 | 0.997988 | 0.999507 |
| Sntb2_B7ZNU9_S373s_1 | Sntb2_B7Z | 0.65 | 0.26 | 0.54 | 0.72 | 1.1 | 0.36 | 0.89 | 0.65 | 0.12 | 0.85 | 0.69 | 0.92 | -0.4 | 1.11 | -0.28 | 0.85 | 0.2075 | 0.1025 | -0.2225 | 0.584356 | 0.410788 | 0.702967 | 0.989223 | 0.997988 | 0.995259 |
| Sntb2_B7ZNU9_S75s_1 | Sntb2_B7Z | 0.33 | 0.36 | 0.61 | 0.43 | 0.68 | -0.05 | 0.65 | 0.45 | -0.09 | 0.37 | 0.48 | 0.27 | 0.14 | 0.7 | 0.18 | 0.41 | -1.6E-16 | -0.175 | -0.075 | 0.767239 | 1 | 0.458673 | 1 | 0.997988 | 0.9971 |
| Sntb2_B7ZNU9_S90s_1 | Sntb2_B7Z | -0.09 | 0.09 | 0.14 | 0.28 | 0.2 | -0.36 | -0.25 | -0.17 | -0.34 | -0.04 | 0.61 | 0.13 | 0.32 | 0.49 | -0.03 | 0.03 | -0.25 | -0.015 | 0.0975 | 0.701453 | 0.282167 | 0.955791 | 0.989223 | 0.997988 | 0.996159 |
| Snurf_Q9WU12_S32s_1 | Snurf_Q9W | -0.53 | -1.18 | -2.84 | -1.47 | -1.38 | -0.54 | -2.18 | -1.58 | -1.04 | -1.69 | -1.04 | -2.35 | -1.01 | -0.73 | -2.15 | -1.75 | 0.085 | -0.025 | 0.095 | 0.861432 | 0.874288 | 0.962903 | 0.996493 | 0.997988 | 0.999507 |
| Snw1_Q9CSN1_S224s_1 | Snw1_Q9C | -0.76 | -0.73 | -0.66 | -0.56 | -0.81 | -0.2 | -1.32 | -0.73 | 0.15 | -0.64 | -0.74 | -0.63 | 0.4 | 0.05 | -0.26 | -1.46 | -0.0875 | 0.2125 | 0.36 | 0.388934 | 0.749487 | 0.438967 | 0.995762 | 0.997988 | 0.994666 |
| Snw1_Q9CSN1_S224s_2 | Snw1_Q9C | -0.5 | -0.86 | -0.53 | -0.4 | -1.35 | -1.19 | -0.67 | -0.54 | -0.93 | -1.45 | -0.64 | -0.48 | -0.47 | -0.55 | -0.44 | -0.55 | -0.365 | -0.3025 | 0.07 | 0.769773 | 0.191182 | 0.302435 | 0.989223 | 0.997988 | 0.9971 |
| Snw1_Q9CSN1_S232s_2 | Snw1_Q9C | -0.5 | -0.86 | -0.53 | -0.4 | -1.35 | -1.19 | -0.67 | -0.54 | -0.93 | -1.45 | -0.64 | -0.48 | -0.47 | -0.55 | -0.44 | -0.55 | -0.365 | -0.3025 | 0.07 | 0.769773 | 0.191182 | 0.302435 | 0.989223 | 0.997988 | 0.9971 |
| Snx14_G3UXB6_S695s_1 | Snx14_G3U | -1.11 | -0.72 | -0.6 | -0.25 | -0.66 | -1.76 | 0.35 | -0.05 | -1.26 | -1.32 | -1.8 | 1 | -0.48 | -0.04 | -0.55 | -0.62 | 0.14 | -0.175 | 0.2475 | 0.402443 | 0.760515 | 0.769794 | 0.995762 | 0.997988 | 0.994666 |
| Snx15_D3YVT2_S215s_1 | Snx15_D3Y | -1.02 | -1.01 | -0.59 | -0.52 | -0.55 | -2.22 | 0.46 | -0.32 | -1.65 | -1.57 | -0.26 | -0.12 | -2.13 | -0.9 | -0.29 | -0.36 | 0.1275 | -0.115 | -0.135 | 0.758333 | 0.808861 | 0.785874 | 0.995762 | 0.997988 | 0.9971 |
| Snx16_Q8BLP0_S29s_1 | Snx16_Q8E | -0.39 | -0.35 | -0.44 | -0.16 | -0.16 | -0.03 | 0.08 | 0.76 | -0.79 | -0.65 | 0.66 | -0.29 | 0 | 0.03 | -0.1 | -0.6 | 0.4975 | 0.0675 | 0.1675 | 0.523456 | 0.082559 | 0.846435 | 0.989223 | 0.997988 | 0.994666 |
| Snx16_Q8BLP0_S70s_1 | Snx16_Q8E | 0.31 | 0.36 | 0.78 | 0.96 | 0.23 | 0.42 | 0.88 | 0.37 | 0.82 | 0.05 | 1.29 | 0.47 | -0.29 | -0.25 | 0.8 | 0.84 | -0.1275 | 0.055 | -0.3275 | 0.388341 | 0.627185 | 0.86809 | 0.989223 | 0.997988 | 0.994666 |
| Snx17_Q8BVL3_S409s_1 | Snx17_Q8E | -2.34 | -2.94 | -1.95 | -2.88 | -1.89 | -2.99 | -2.43 | -1.98 | -1.05 | -2.8 | -5.74 | -1.61 | -0.46 | -2.53 | -0.49 | -3 | 0.205 | -0.2725 | 0.9075 | 0.179113 | 0.558952 | 0.774135 | 0.989223 | 0.997988 | 0.994666 |
| Snx17_Q8BVL3_S415s_1 | Snx17_Q8E | -0.73 | 0.35 | -0.15 | -1.27 | 0.24 | -0.58 | 0.03 | -0.49 | -0.41 | -0.13 | -0.22 | -0.22 | 0.39 | 0 | -0.21 | -0.53 | 0.25 | 0.205 | 0.3625 | 0.381286 | 0.526923 | 0.578267 | 0.989223 | 0.997988 | 0.994666 |
| Snx2_Q9CWK8_S93s_1 | Snx2_Q9C\ | -0.18 | 0.17 | 0.82 | 0.12 | -0.14 | -0.3 | 0.49 | 0.45 | -0.33 | -0.28 | 0.1 | 0.09 | -1.43 | 0.27 | -0.36 | 0.28 | -0.1075 | -0.3375 | -0.5425 | 0.240298 | 0.730129 | 0.258251 | 0.995762 | 0.997988 | 0.994666 |
| Snx2_Q9CWK8_S97s_1 | Snx2_Q9C\ | 0.18 | 0.36 | 0.01 | 0.13 | 0.31 | -0.5 | 0.12 | -0.02 | 0.37 | 0.21 | -0.37 | 0.09 | -0.08 | 0.55 | 0.25 | -0.05 | -0.1925 | -0.095 | -0.0025 | 0.992382 | 0.443169 | 0.704772 | 0.989223 | 0.997988 | 1 |
| Snx2_Q9CWK8_T101t_1 | Snx2_Q9C\ | -0.76 | -0.25 | 0.1 | 0.74 | 0.38 | 0.17 | 0.84 | 1.2 | -0.21 | 0.24 | 0.33 | 1.28 | -0.29 | -0.44 | 0.23 | 1.33 | 0.69 | 0.4525 | 0.25 | 0.609055 | 0.095911 | 0.309699 | 0.989223 | 0.997988 | 0.995259 |
| Snx2_Q9CWK8_T104t_1 | Snx2_Q9C\ | -0.16 | 0.58 | 0.23 | -0.36 | 0.92 | 0.31 | 0.83 | -0.18 | -0.72 | 0.57 | -0.07 | -0.3 | 0.31 | 0.96 | -0.15 | 0.24 | 0.3975 | -0.2025 | 0.2675 | 0.44418 | 0.255981 | 0.571312 | 0.989223 | 0.997988 | 0.994666 |
| Snx25_E9Q1K0_S126s_1 | Snx25_E9Q | 0.43 | 0.44 | -0.1 | 0.25 | -0.01 | 0.3 | 0.61 | 0.38 | 0.48 | 0.68 | 0.38 | 0.26 | 0.08 | 0.41 | 0.48 | 0.89 | 0.065 | 0.195 | 0.21 | 0.465721 | 0.78912 | 0.424484 | 0.995762 | 0.997988 | 0.994666 |
| Snx27_Q3UHD6_S49s_1 | Snx27_Q3U | -0.58 | -0.02 | -0.2 | -0.49 | 0.42 | -0.77 | -0.1 | -0.36 | -1.5 | -0.71 | -1.11 | -0.57 | -0.86 | -0.43 | -0.25 | 0.08 | 0.12 | -0.65 | -0.0425 | 0.886484 | 0.693607 | 0.048163 | 0.994893 | 0.997988 | 0.999507 |
| Snx29_D3YZ90_S371s_1 | Snx29_D3Y | 2.38 | -0.02 | 0.99 | -2.99 | 0 | 2.8 | 1.37 | 0.8 | 1.98 | 2.18 | -4.31 | 1.33 | 2.13 | -2.02 | 2.54 | 1.16 | 1.1525 | 0.205 | 0.8625 | 0.519284 | 0.311401 | 0.90241 | 0.989223 | 0.997988 | 0.994666 |
| Snx30_Q8CE50_S67s_1 | Snx30_Q8C | -0.13 | 0.48 | -1.37 | 0.61 | 0.77 | 0.17 | 1.12 | 0.49 | 0.72 | 0.27 | -0.36 | 0.58 | -0.18 | 0.76 | -0.57 | 1.12 | 0.74 | 0.405 | 0.385 | 0.49364 | 0.133175 | 0.412765 | 0.989223 | 0.997988 | 0.994666 |
| Snx7_F8WI30_S8s_1 | Snx7_F8WI | 0.95 | 0.12 | 0.83 | 0.56 | 0.91 | 0.69 | 0.82 | 0.96 | 0.63 | 1.07 | 0.7 | 0.93 | 0.18 | 1.42 | 0.33 | 0.83 | 0.23 | 0.2175 | 0.075 | 0.83501 | 0.36877 | 0.427835 | 0.989223 | 0.997988 | 0.999507 |
| Sod1_P08228_S99s_1 | Sod1_P082 | 1.4 | 1.25 | 1.71 | 0.69 | 0.77 | 1.24 | 1.38 | 1.28 | -0.29 | 1.3 | 0.27 | 1.18 | -2.04 | 0.47 | 1.48 | 1.73 | -0.095 | -0.6475 | -0.8525 | 0.292745 | 0.739986 | 0.152672 | 0.995762 | 0.997988 | 0.994666 |
| Soga1_A2ACV6_S173s_1 | Soga1_A2A | 0.78 | 0.75 | 0.54 | 0.38 | 0.13 | 0.11 | 0.81 | 0.63 | 0.8 | 0.8 | 0.68 | 0.69 | 0.02 | 0.41 | 1.08 | 0.57 | -0.1925 | 0.13 | -0.0925 | 0.75835 | 0.45573 | 0.553508 | 0.989223 | 0.997988 | 0.9971 |
| Soga1_E1U8D0_S1015s_1 | Soga1_E1U | 0.01 | 0.4 | 0.58 | -0.07 | 0.09 | -0.21 | -0.49 | -0.53 | 0.42 | 0.02 | -0.07 | -0.12 | 0.64 | 0.13 | 0.3 | 0.07 | -0.515 | -0.1675 | 0.055 | 0.844188 | 0.072707 | 0.527754 | 0.989223 | 0.997988 | 0.999507 |
| Soga3_Q6NZL0_S637s_1 | Soga3_Q6N | -0.44 | -1.37 | -0.78 | -0.68 | 0.42 | 0.01 | -0.31 | -1.31 | -1.81 | -1.33 | -1.19 | -1.97 | -1.91 | -0.14 | -0.97 | -0.59 | 0.52 | -0.7575 | -0.085 | 0.840565 | 0.21623 | 0.03282 | 0.989223 | 0.997988 | 0.999507 |
| Soga3_Q6NZL0_S639s_1 | Soga3_Q6N | 0.83 | 0.61 | 0.42 | 0.35 | 0.81 | 0.7 | 0.29 | -0.16 | 0.58 | 0.43 | -0.06 | 0.1 | 0.66 | 0.76 | -0.24 | 0.26 | -0.1425 | -0.29 | -0.1925 | 0.535423 | 0.614115 | 0.270382 | 0.989223 | 0.997988 | 0.994666 |
| Soga3_Q6NZL0_S760s_1 | Soga3_Q6N | -2.37 | -1.71 | -0.75 | -2.5 | -2.88 | -2.12 | -2.69 | -2.56 | -0.01 | -2.56 | -1.55 | -1.04 | 0.18 | -3.14 | -2.1 | -2.14 | -0.73 | 0.5425 | 0.0325 | 0.964005 | 0.101825 | 0.383011 | 0.989223 | 0.997988 | 0.999507 |
| Soga3_Q6NZL0_S779s_1 | Soga3_Q6N | 0.05 | -0.13 | 0.42 | 0.18 | 0.36 | -0.02 | -0.19 | -0.15 | 0.1 | 0.5 | 0.72 | -0.2 | -0.45 | 0.42 | -0.28 | -0.43 | -0.13 | 0.15 | -0.315 | 0.305526 | 0.587742 | 0.600464 | 0.989223 | 0.997988 | 0.994666 |
| Soga3_Q6NZL0_T514t_1 | Soga3_Q6N | -0.93 | -0.18 | -0.56 | -0.47 | -0.75 | -1.01 | -0.6 | -0.47 | -0.74 | -0.07 | -0.31 | -0.28 | 0.15 | -0.05 | -0.13 | -0.33 | -0.1725 | 0.185 | 0.445 | 0.125632 | 0.495771 | 0.496552 | 0.989223 | 0.997988 | 0.994666 |
| Son_H9KV01_S1723s_1 | Son_H9KV( | 0.85 | 0.46 | -0.71 | 0.17 | 0.23 | 0.39 | -0.51 | 0.97 | -0.29 | -0.4 | -1.04 | -0.61 | 0.98 | -0.58 | 0.41 | 0.53 | 0.0775 | -0.7775 | 0.1425 | 0.753722 | 0.855628 | 0.063591 | 0.996349 | 0.997988 | 0.9971 |
| Sorbs1_Q62417_S1209s_1 | Sorbs1_Q6 | -0.74 | 0.24 | 0.22 | 0.49 | -0.5 | -0.71 | 0.28 | 0.55 | -0.06 | -0.24 | 1.39 | 0.78 | 1.07 | 0.03 | 1.46 | 0.63 | -0.1475 | 0.415 | 0.745 | 0.095627 | 0.708169 | 0.367008 | 0.995437 | 0.997988 | 0.994666 |
| Sorbs1_Q62417_S250s_1 | Sorbs1_Q6 | 1.86 | 1.05 | 0.48 | 0.51 | 1.03 | 2.01 | 0.07 | 0.52 | -0.05 | 0.61 | 0.4 | 0.7 | 0.2 | -0.61 | 0.98 | 1.15 | -0.0675 | -0.56 | -0.545 | 0.283053 | 0.889324 | 0.155969 | 0.996493 | 0.997988 | 0.994666 |
| Sorbs1_Q62417_S254s_1 | Sorbs1_Q6 | -0.1 | -0.11 | -0.01 | -0.25 | 0.49 | -0.07 | 0.3 | -0.36 | -0.6 | -0.47 | 0.2 | -0.21 | -0.54 | 0.44 | 0.01 | 0 | 0.2075 | -0.1525 | 0.095 | 0.737333 | 0.418211 | 0.552584 | 0.989223 | 0.997988 | 0.9971 |
| Sorbs1_Q62417_S270s_1 | Sorbs1_Q6 | -0.46 | 0.41 | 0.32 | 0.41 | 0.88 | 0.04 | 0.1 | -0.07 | -0.54 | -0.52 | -0.56 | 0.44 | -0.24 | 0.38 | 1.02 | 0.87 | 0.0675 | -0.465 | 0.3375 | 0.375658 | 0.832106 | 0.19658 | 0.995762 | 0.997988 | 0.994666 |
| Sorbs1_Q62417_S2s_1 | Sorbs1_Q6 | 0.37 | -0.23 | 0.19 | 0.89 | -1.16 | 0.32 | -0.19 | 0.45 | 1.23 | -0.28 | 0.4 | 0.47 | 1.62 | -0.28 | 0.61 | -0.05 | -0.45 | 0.15 | 0.17 | 0.716512 | 0.29118 | 0.699912 | 0.989223 | 0.997988 | 0.996966 |
| Sorbs1_Q62417_S3s_1 | Sorbs1_Q6 | 0.37 | -0.23 | 0.19 | 0.89 | -1.16 | 0.32 | -0.19 | 0.45 | 1.23 | -0.28 | 0.4 | 0.47 | 1.62 | -0.28 | 0.61 | -0.05 | -0.45 | 0.15 | 0.17 | 0.716512 | 0.29118 | 0.699912 | 0.989223 | 0.997988 | 0.996966 |
| Sorbs1_Q62417_S533s_1 | Sorbs1_Q6 | -0.15 | -0.32 | -0.67 | 0.12 | 0.46 | 0.09 | 0.19 | -0.23 | -0.72 | -0.31 | -0.67 | 0.08 | -0.66 | 0.37 | 0.98 | -0.03 | 0.3825 | -0.15 | 0.42 | 0.299042 | 0.170902 | 0.610503 | 0.989223 | 0.997988 | 0.994666 |
| Sorbs1_Q62417_S554s_1 | Sorbs1_Q6 | 0.02 | -0.16 | -0.15 | 0.1 | 0.31 | 0.04 | 0.18 | -0.32 | 0.05 | -0.09 | 0.4 | -0.83 | -0.05 | 0.25 | -0.49 | -0.58 | 0.1 | -0.07 | -0.17 | 0.549647 | 0.66238 | 0.818008 | 0.990434 | 0.997988 | 0.994666 |
| Sorbs1_Q62417_S576s_1 | Sorbs1_Q6 | 0.46 | -0.94 | -0.35 | 0.47 | 0.64 | 0.59 | 0.6 | -0.37 | -0.46 | -0.79 | 0.58 | -0.01 | -2.01 | 0.6 | -0.36 | 0.12 | 0.455 | -0.08 | -0.3225 | 0.596195 | 0.276428 | 0.85529 | 0.989223 | 0.997988 | 0.995259 |
| Sorbs1_Q62417_S58s_1 | Sorbs1_Q6 | -1.6 | -1.65 | -0.51 | -0.48 | -1.24 | -1.51 | -0.11 | -0.28 | -1.42 | -1.44 | -0.35 | -0.52 | -0.35 | -0.68 | -0.65 | -0.63 | 0.275 | 0.1275 | 0.4825 | 0.201324 | 0.541483 | 0.764949 | 0.989223 | 0.997988 | 0.994666 |
| Sorbs1_Q62417_S58s_2 | Sorbs1_Q6 | -1.78 | -1.33 | 1.97 | 2.31 | -0.77 | -2.01 | 1.81 | 2.12 | -1.61 | -1.45 | 2.05 | 1.99 | -0.8 | -0.67 | 1.17 | 2.24 | -0.005 | -0.0475 | 0.1925 | 0.864392 | 0.996834 | 0.970807 | 0.999864 | 0.997988 | 0.999507 |
| Sorbs1_Q62417_S62s_1 | Sorbs1_Q6 | 0.26 | 0.94 | 0.09 | -0.58 | 0.28 | -0.6 | -0.87 | -1.11 | 0.95 | 1.53 | -0.05 | -0.73 | 2.42 | 1.3 | 0.35 | -0.13 | -0.7525 | 0.2475 | 0.8075 | 0.19253 | 0.095142 | 0.654355 | 0.989223 | 0.997988 | 0.994666 |
| Sorbs1_Q62417_S635s_1 | Sorbs1_Q6 | -0.84 | -2.38 | -1.13 | -0.52 | -0.65 | -0.52 | -0.21 | -0.16 | -1.04 | -1.52 | -0.09 | -0.38 | -1.68 | -0.66 | -0.89 | -1.24 | 0.8325 | 0.46 | 0.1 | 0.824596 | 0.064575 | 0.360511 | 0.989223 | 0.997988 | 0.999507 |
| Sorbs1_Q62417_S638s_1 | Sorbs1_Q6 | 1.11 | -0.17 | 0.22 | 0.94 | 0.72 | 0.9 | 0.46 | -0.02 | 0.25 | 0.27 | 0.05 | 0.52 | -0.84 | 0.43 | 0.44 | 0.01 | -0.01 | -0.2525 | -0.515 | 0.242063 | 0.977814 | 0.461276 | 0.999187 | 0.997988 | 0.994666 |
| Sorbs1_Q62417_S666s_1 | Sorbs1_Q6 | 1.87 | 0.4 | -0.55 | -0.13 | 0.69 | 1.56 | -0.73 | 0.26 | 0.24 | 0.47 | -1.07 | 0.8 | 0.3 | 0.8 | -2 | -0.32 | 0.0475 | -0.2875 | -0.7025 | 0.341322 | 0.940051 | 0.639824 | 0.999187 | 0.997988 | 0.994666 |
| Sorbs1_Q62417_S694s_1 | Sorbs1_Q6 | -0.61 | -0.62 | -1.73 | -1.53 | -0.21 | -0.68 | -1.02 | -1.77 | -0.78 | -0.4 | -1.07 | -2.53 | -0.78 | 0.03 | -2.76 | -1.64 | 0.2025 | -0.0725 | -0.165 | 0.786161 | 0.631423 | 0.888299 | 0.989223 | 0.997988 | 0.999507 |
| Sorbs1_Q62417_S707s_1 | Sorbs1_Q6 | -0.19 | -0.8 | -0.33 | -0.28 | 0.67 | 0.24 | 0.22 | -0.11 | 0.02 | -0.38 | 0.42 | -0.27 | -0.25 | 0.87 | -0.44 | 0.06 | 0.655 | 0.3475 | 0.46 | 0.209016 | 0.029136 | 0.231015 | 0.989223 | 0.997988 | 0.994666 |
| Sorbs1_Q62417_S722s_1 | Sorbs1_Q6 | 0.23 | 0.03 | 0.02 | -0.06 | 0.47 | 0.05 | 0.19 | -0.01 | 0.17 | 0.16 | -0.55 | 0.02 | -0.35 | 0.44 | -0.38 | 0.09 | 0.12 | -0.105 | -0.105 | 0.710203 | 0.581796 | 0.679519 | 0.989223 | 0.997988 | 0.996966 |
| Sorbs1_Q62417_S791s_1 | Sorbs1_Q6 | -0.86 | 0.17 | 0.35 | -0.01 | -0.21 | -0.13 | 0 | -0.05 | -0.27 | 0.15 | -0.25 | 0.27 | 0.9 | -0.09 | -0.19 | 0.24 | -0.01 | 0.0625 | 0.3025 | 0.432972 | 0.973168 | 0.848421 | 0.999187 | 0.997988 | 0.994666 |
| Sorbs1_Q62417_S969s_1 | Sorbs1_Q6 | 0.18 | 1.13 | 0.78 | 0.22 | 0.3 | 0.91 | 0.17 | 0.2 | 0.77 | 1.24 | -0.59 | -0.21 | 0.53 | 0.82 | 0.17 | 0.67 | -0.1825 | -0.275 | -0.03 | 0.924652 | 0.557501 | 0.55492 | 0.989223 | 0.997988 | 0.999507 |
| Sorbs1_Q62417_S96s_1 | Sorbs1_Q6 | -0.13 | -0.68 | -2.29 | -0.84 | 0.19 | -0.28 | -0.04 | -0.81 | -0.62 | -0.47 | -1.7 | -1.25 | -1.17 | -0.57 | -1.29 | -1.67 | 0.75 | -0.025 | -0.19 | 0.698543 | 0.135908 | 0.960764 | 0.989223 | 0.997988 | 0.996159 |
| Sorbs1_Q62417_T286t_1 | Sorbs1_Q6 | -0.72 | -0.11 | -0.52 | -1 | 0.8 | -0.93 | 0.11 | -0.79 | -1.3 | -0.88 | -1.49 | -0.48 | -0.17 | 0.09 | -0.44 | 0.07 | 0.385 | -0.45 | 0.475 | 0.128879 | 0.374058 | 0.184363 | 0.989223 | 0.997988 | 0.994666 |
| Sorbs2_B7ZWM6_S109s_1 | Sorbs2_B7 | -0.5 | -0.82 | 0.39 | -0.32 | -0.32 | -0.44 | 0.89 | 1.06 | -1.3 | -0.17 | 0.38 | 0.46 | -1.25 | -0.31 | 0.57 | 1.23 | 0.61 | 0.155 | 0.3725 | 0.505128 | 0.188376 | 0.736471 | 0.989223 | 0.997988 | 0.994666 |
| Sorbs2_B7ZWM6_S1111s_1 | Sorbs2_B7 | -0.42 | -0.22 | -1.03 | -1.32 | -0.02 | 0.08 | -0.51 | -1.45 | -1.23 | 0.01 | -2 | -0.06 | -0.2 | 0.36 | -1.4 | -0.44 | 0.2725 | -0.0725 | 0.3275 | 0.462883 | 0.514863 | 0.887877 | 0.989223 | 0.997988 | 0.994666 |
| Sorbs2_B7ZWM6_S1113s_1 | Sorbs2_B7 | 0.26 | 0.31 | -0.11 | 0.19 | 0.21 | 0.15 | 0.14 | 0.27 | -0.01 | 0.05 | 0.04 | 0.41 | -0.38 | 0.36 | -0.19 | -0.14 | 0.03 | -0.04 | -0.25 | 0.367441 | 0.884677 | 0.862448 | 0.996493 | 0.997988 | 0.994666 |
| Sorbs2_B7ZWM6_S130s_1 | Sorbs2_B7 | -0.09 | 0.2 | 0 | 0.19 | 0.38 | -0.22 | 0.53 | 0.25 | -0.45 | -0.08 | -0.4 | -0.02 | -0.5 | -0.14 | 0.07 | 0.16 | 0.16 | -0.3125 | -0.1775 | 0.502544 | 0.51244 | 0.194921 | 0.989223 | 0.997988 | 0.994666 |
| Sorbs2_B7ZWM6_S143s_1 | Sorbs2_B7 | 0.43 | 0.49 | 0.27 | 0.56 | 0.13 | 0.4 | 0.9 | 0.41 | -0.3 | 0.31 | -0.45 | 0.56 | -0.59 | 0.32 | 0.6 | 0.73 | 0.0225 | -0.4075 | -0.1725 | 0.613752 | 0.924825 | 0.186749 | 0.997335 | 0.997988 | 0.995259 |
| Sorbs2_B7ZWM6_S27s_1 | Sorbs2_B7 | -2.41 | -4.48 | 0.13 | -0.35 | -1.23 | -2.45 | 0.45 | 0.17 | -5.95 | -3.37 | 0.21 | -0.31 | -2.9 | -1.45 | -0.14 | 0.03 | 1.0125 | -0.5775 | 0.6625 | 0.546956 | 0.359688 | 0.71112 | 0.989223 | 0.997988 | 0.994666 |
| Sorbs2_B7ZWM6_S354s_1 | Sorbs2_B7 | -3.49 | -3.97 | -2.4 | -2.98 | -2.1 | -3.83 | -2.31 | -2.88 | -6.06 | -3.72 | -1.26 | -2.59 | -3.51 | -2.84 | -2.86 | -2.73 | 0.43 | -0.1975 | 0.225 | 0.568545 | 0.375611 | 0.835004 | 0.989223 | 0.997988 | 0.995259 |
| Sorbs2_B7ZWM6_S397s_1 | Sorbs2_B7 | 0.81 | 0.79 | 1.65 | 1.79 | 0.98 | 1.08 | 1.6 | 1.1 | 0.43 | 0.72 | 2.24 | 1.31 | -0.04 | 0.97 | 1.78 | 0.15 | -0.07 | -0.085 | -0.545 | 0.268016 | 0.825441 | 0.85342 | 0.995762 | 0.997988 | 0.994666 |
| Sorbs2_B7ZWM6_S40s_1 | Sorbs2_B7 | 0.29 | 0.37 | 0.25 | 0.46 | 0.48 | -0.62 | 0.26 | 0.42 | -0.13 | 0.03 | 0.32 | 0.46 | 0.4 | 0.44 | 0.12 | -0.5 | -0.2075 | -0.1725 | -0.2275 | 0.44146 | 0.480518 | 0.467775 | 0.989223 | 0.997988 | 0.994666 |
| Sorbs2_B7ZWM6_S592s_1 | Sorbs2_B7 | -0.68 | 0.42 | 0.55 | 0.66 | 0.47 | -0.72 | 0.64 | 0.46 | 0.23 | -0.3 | -0.14 | 0.39 | -1.13 | 0.81 | 0.19 | 0.53 | -0.025 | -0.1925 | -0.1375 | 0.783733 | 0.952345 | 0.595748 | 0.999187 | 0.997988 | 0.998578 |
| Sorbs2_B7ZWM6_S645s_1 | Sorbs2_B7 | 1.71 | -0.68 | -0.04 | 1 | 0.57 | 1.65 | 0.89 | 1.09 | 0.36 | -0.75 | 0.36 | 0.93 | -2.78 | -0.1 | 0.23 | 0.75 | 0.5525 | -0.2725 | -0.9725 | 0.262587 | 0.308024 | 0.643773 | 0.989223 | 0.997988 | 0.994666 |
| Sorbs2_B7ZWM6_S820s_1 | Sorbs2_B7 | -0.92 | -0.82 | -0.61 | -1.94 | 0.34 | -0.23 | -0.81 | -2.2 | -0.94 | -0.77 | -1.52 | -1.28 | -0.93 | 0.71 | -1.4 | -2.61 | 0.3475 | -0.055 | 0.015 | 0.982187 | 0.538456 | 0.876682 | 0.989223 | 0.997988 | 1 |
| Sorbs2_B7ZWM6_S82s_1 | Sorbs2_B7 | -0.59 | 0.18 | -1.66 | -0.18 | -0.53 | -2.02 | 0.02 | -0.29 | -2.03 | -0.79 | -1.37 | -1.29 | -0.1 | -0.01 | -0.27 | 0.37 | -0.1425 | -0.8075 | 0.56 | 0.202895 | 0.79403 | 0.101806 | 0.995762 | 0.997988 | 0.994666 |
| Sorbs2_B7ZWM6_S844s_1 | Sorbs2_B7 | 0.28 | 0.1 | 0.05 | 0.13 | 0.15 | 0.33 | 0.32 | 0.14 | -0.03 | 0.09 | -0.51 | 0.25 | -0.2 | 0.44 | -0.31 | -0.01 | 0.095 | -0.19 | -0.16 | 0.550948 | 0.634676 | 0.450903 | 0.989223 | 0.997988 | 0.994666 |
| Sorbs2_B7ZWM6_S99s_1 | Sorbs2_B7 | -0.27 | -0.29 | 0.96 | 0.72 | 0.88 | 0.35 | 1.28 | 1.28 | -0.73 | 0.25 | 1.46 | 1.59 | -0.74 | -0.19 | 1.07 | 1.46 | 0.6675 | 0.3625 | 0.12 | 0.8318 | 0.107658 | 0.540654 | 0.989223 | 0.997988 | 0.999507 |
| Sorbs2_D3Z080_S330s_1 | Sorbs2_D3 | 0.52 | 1.55 | 1.02 | 0.55 | 0.57 | 0.35 | -0.35 | -0.29 | 1.09 | 1.01 | -0.65 | 0.57 | 1.94 | 0.01 | 0.36 | 0.62 | -0.84 | -0.405 | -0.1775 | 0.70522 | 0.031987 | 0.38013 | 0.989223 | 0.997988 | 0.996159 |
| Sorbs2_D3Z080_S350s_1 | Sorbs2_D3 | 0.31 | -0.03 | 0.38 | 0.42 | 0.11 | 0.55 | 0.27 | 0.27 | 0.13 | 0.07 | 0.13 | 0.78 | 0.09 | 0.93 | -0.22 | -0.71 | 0.03 | 0.0075 | -0.2475 | 0.515483 | 0.892506 | 0.977117 | 0.996493 | 0.997988 | 0.994666 |
| Sorbs2_D3Z080_S40s_1 | Sorbs2_D3 | -0.72 | -2.05 | -1.73 | -3.14 | -3 | 0.08 | -0.74 | -0.59 | 2.23 | -0.86 | -1.37 | -1.05 | 2.75 | -2.63 | 1.84 | -0.73 | 0.8475 | 1.6475 | 2.2175 | 0.077813 | 0.265233 | 0.083346 | 0.989223 | 0.997988 | 0.994666 |
| Sorbs2_Q3UTJ2_S373s_1 | Sorbs2_Q3 | 0.35 | -0.01 | 0.42 | 0.45 | 0.17 | 0.57 | 0.3 | 0.3 | 0.2 | 0.15 | 0.19 | 0.82 | 0.13 | 0.97 | -0.17 | -0.66 | 0.0325 | 0.0375 | -0.235 | 0.535575 | 0.883184 | 0.884937 | 0.996493 | 0.997988 | 0.994666 |
| Sorcs1_E9PYT6_S1133s_1 | Sorcs1_E9F | 0.36 | 1 | -0.64 | -0.2 | 0.73 | 0.16 | 0.05 | -0.35 | 0.2 | 0.64 | -0.27 | -0.44 | 0.32 | 0.75 | -0.25 | -0.19 | 0.0175 | -0.0975 | 0.0275 | 0.94813 | 0.965276 | 0.817093 | 0.999187 | 0.997988 | 0.999507 |
| Sorcs1_E9PYT6_T1152t_1 | Sorcs1_E9F | -0.06 | 1.6 | 0.95 | 0.04 | -0.26 | 0.44 | -0.24 | -1.09 | 0.87 | 0.65 | 0.78 | 0.93 | 1.37 | 0.17 | 0.92 | 0.12 | -0.92 | 0.175 | 0.0125 | 0.979042 | 0.073773 | 0.661326 | 0.989223 | 0.997988 | 1 |
| Sorcs3_Q8VI51_T175t_1 | Sorcs3_Q8' | 0.25 | 0.9 | -0.16 | 0.16 | -0.02 | 1 | -0.16 | -0.37 | 0.79 | 0.66 | 0.43 | -0.3 | 0.7 | 0.16 | -0.16 | 0.38 | -0.175 | 0.1075 | -0.0175 | 0.957509 | 0.639069 | 0.756884 | 0.989223 | 0.997988 | 0.999507 |
| Sort1_Q6PHU5_S791s_1 | Sort1_Q6P | -1.01 | 0.68 | 0.72 | -0.43 | 0.71 | -1.24 | 0.7 | -0.16 | -1.28 | 0.05 | 1.07 | 0.45 | 0.24 | 1.29 | 0.5 | 0.79 | 0.0125 | 0.0825 | 0.715 | 0.148489 | 0.98236 | 0.890629 | 0.999187 | 0.997988 | 0.994666 |
| Sos1_Q62245_S1082s_1 | Sos1_Q622 | 0.48 | 0.78 | -0.36 | 0.7 | 0.62 | 0.2 | -0.06 | 0.53 | 1.1 | 0.03 | 0.64 | 0.24 | 0.84 | 0.51 | 1.09 | -1 | -0.0775 | 0.1025 | -0.04 | 0.936927 | 0.808399 | 0.777707 | 0.995762 | 0.997988 | 0.999507 |
| Sos1_Q62245_S1147s_1 | Sos1_Q622 | -0.33 | 0.01 | -0.03 | -0.79 | -0.76 | 0.44 | -1.31 | -1.07 | 0.13 | -0.55 | -1.41 | -1.05 | 1.22 | -1.33 | -0.78 | -0.38 | -0.39 | -0.435 | -0.0325 | 0.951808 | 0.35406 | 0.274785 | 0.989223 | 0.997988 | 0.999507 |
| Sos1_Q62245_S1196s_1 | Sos1_Q622 | 0.76 | 0.96 | 1.06 | 1.12 | 0.22 | 0.35 | 0.93 | 1.14 | 0.34 | 0.61 | 1 | 1.1 | 0.4 | 0.29 | 0.81 | 0.71 | -0.315 | -0.2125 | -0.4225 | 0.12182 | 0.268791 | 0.421676 | 0.989223 | 0.997988 | 0.994666 |
| Sos2_S4R2C4_S1003s_1 | Sos2_S4R2 | -0.66 | 0.13 | -0.11 | -0.37 | -0.58 | -0.63 | -0.03 | -0.39 | 0.75 | -1.2 | 0.36 | -0.27 | 0.73 | 0.26 | 0.28 | -0.33 | -0.155 | 0.1625 | 0.4875 | 0.153908 | 0.560643 | 0.714386 | 0.989223 | 0.997988 | 0.994666 |
| Sowaha_Q8BLS7_S242s_1 | Sowaha_Q | -2.11 | -1.46 | -0.89 | -2.3 | -1.91 | -0.15 | -2.31 | -1.8 | 0.6 | -1.4 | -0.82 | -1.07 | 1.32 | -2.23 | -2.13 | -1.93 | 0.1475 | 1.0175 | 0.4475 | 0.582408 | 0.778351 | 0.072273 | 0.995762 | 0.997988 | 0.995259 |
| Sowaha_Q8BLS7_S258s_1 | Sowaha_Q | -0.16 | 0.33 | -1.08 | -0.23 | 0.61 | -0.56 | -0.62 | -0.91 | 0.03 | 0.46 | -0.84 | -1.48 | -0.59 | -0.13 | -1.71 | -1.18 | -0.085 | -0.1725 | -0.6175 | 0.184993 | 0.840459 | 0.727998 | 0.995762 | 0.997988 | 0.994666 |
| Sox1_P53783_S325s_1 | Sox1_P537 | 0.35 | -0.53 | -1.95 | -0.53 | -0.84 | -0.49 | -2.65 | -1.01 | 0.54 | -0.36 | 0.2 | -2.41 | 1.2 | -0.48 | -2.85 | -1.53 | -0.5825 | 0.1575 | -0.25 | 0.771599 | 0.347037 | 0.82962 | 0.989223 | 0.997988 | 0.9971 |
| Sox10_Q04888_S24s_1 | Sox10_Q04 | 0.23 | 0.36 | 0.7 | 1.16 | 0.26 | 0.96 | 0.44 | 1.12 | -0.57 | 0.73 | 1.28 | 0.53 | 0.42 | 0.88 | 1.05 | 1.5 | 0.0825 | -0.12 | 0.35 | 0.317363 | 0.790984 | 0.779839 | 0.995762 | 0.997988 | 0.994666 |
| Sox10_Q04888_S45s_1 | Sox10_Q04 | -1.75 | -1.49 | -2.62 | -3.1 | -1.14 | -2.33 | -1.64 | -1.79 | -1.93 | -1.48 | -1.64 | -1.68 | -1.15 | -1.43 | -1.5 | -1.18 | 0.515 | 0.5575 | 0.925 | 0.039181 | 0.243258 | 0.174406 | 0.989223 | 0.997988 | 0.994666 |
| Sox5_B2KFM4_S327s_1 | Sox5_B2KF | -1.43 | -0.12 | -2.36 | -1.27 | -0.21 | 0.01 | -0.76 | -0.94 | -0.6 | -0.28 | -0.39 | -0.81 | 0.57 | -1.11 | -1.08 | -1.52 | 0.82 | 0.775 | 0.51 | 0.401464 | 0.108576 | 0.115024 | 0.989223 | 0.997988 | 0.994666 |
| Spag9_F8WH31_S1036s_1 | Spag9_F8W | -1.71 | -1.34 | -1.08 | -1.2 | -1.31 | -1.05 | -1.53 | -1.7 | -1.4 | -1.35 | -1.37 | -1.24 | -0.67 | -1.5 | -1.62 | -1.28 | -0.065 | -0.0075 | 0.065 | 0.832621 | 0.796339 | 0.974355 | 0.995762 | 0.997988 | 0.999507 |
| Spag9_F8WH31_S1086s_1 | Spag9_F8W | -0.09 | -0.3 | 0.2 | -0.01 | -0.01 | -0.39 | 0.49 | 0.38 | -0.08 | 0.24 | 0.45 | 0.56 | -0.37 | 0.37 | 0.52 | 0.31 | 0.1675 | 0.3425 | 0.2575 | 0.386729 | 0.537054 | 0.19278 | 0.989223 | 0.997988 | 0.994666 |
| Spag9_F8WH31_S175s_1 | Spag9_F8W | -1.06 | 0.22 | -0.76 | -1.44 | -1.3 | -0.4 | -2.66 | -1.45 | 1.19 | -0.97 | -1.4 | -0.65 | 0.41 | -1.02 | -1.21 | -1.42 | -0.6925 | 0.3025 | -0.05 | 0.922328 | 0.212678 | 0.624303 | 0.989223 | 0.997988 | 0.999507 |
| Spag9_F8WH31_S578s_1 | Spag9_F8W | -2.37 | -1.12 | -1 | -2.64 | -1.88 | -3.41 | -4.42 | -3.07 | -3.01 | -2.7 | -0.96 | -1.48 | -0.97 | -2.09 | -3.21 | -1.74 | -1.4125 | -0.255 | -0.22 | 0.703843 | 0.039125 | 0.667908 | 0.989223 | 0.997988 | 0.996159 |
| Sparcl1_P70663_S155s_1 | Sparcl1_P7 | -1.57 | -1.24 | -1.32 | -1.63 | -1.58 | -2.33 | -2.09 | -1.48 | -2.14 | -1.07 | -1.28 | -1.39 | -0.38 | -1.49 | -1.26 | -1.31 | -0.43 | -0.03 | 0.33 | 0.310394 | 0.133251 | 0.918737 | 0.989223 | 0.997988 | 0.994666 |
| Sparcl1_P70663_S171s_1 | Sparcl1_P7 | -3.01 | -2.88 | -1.17 | -2.44 | -1.95 | -3.78 | -1.68 | -2.24 | -3.69 | -2.94 | -0.92 | -1.49 | -1.98 | -2.06 | -2.37 | -1.79 | -0.0375 | 0.115 | 0.325 | 0.457877 | 0.947231 | 0.867623 | 0.999187 | 0.997988 | 0.994666 |
| Sparcl1_P70663_S406s_1 | Sparcl1_P7 | -0.9 | -1.06 | -2.69 | -1.56 | -2.7 | -0.41 | -2.39 | -1.62 | 0.9 | -0.6 | -0.94 | -2.52 | 1.66 | -1.95 | -0.43 | -1.69 | -0.2275 | 0.7625 | 0.95 | 0.258431 | 0.697734 | 0.309873 | 0.994893 | 0.997988 | 0.994666 |
| Sparcl1_P70663_S411s_1 | Sparcl1_P7 | -1.63 | -1.46 | -1.56 | -1.34 | -2.6 | -1.12 | -1.71 | -1.78 | -0.26 | -1.53 | -1.71 | -1.58 | 0.63 | -2.57 | -1.64 | -1.31 | -0.305 | 0.2275 | 0.275 | 0.656026 | 0.357842 | 0.528872 | 0.989223 | 0.997988 | 0.995259 |
| Sparcl1_P70663_S70s_1 | Sparcl1_P7 | -4.3 | -2.72 | -6.58 | -4.98 | -3.33 | -3.44 | -4.49 | -5.39 | -2.74 | -2.84 | -3.28 | -4.61 | -3.1 | -2.02 | -6.08 | -5.16 | 0.4825 | 1.2775 | 0.555 | 0.604592 | 0.558277 | 0.140567 | 0.989223 | 0.997988 | 0.995259 |
| Sparcl1_P70663_S70s_2 | Sparcl1_P7 | -3.05 | -3.04 | -2.9 | -2.22 | -2.68 | -3.8 | -2.8 | -3.15 | -2.6 | -3.3 | -2.67 | -3.04 | -1.63 | -1.34 | -1.88 | -3.43 | -0.305 | -0.1 | 0.7325 | 0.153562 | 0.366413 | 0.737285 | 0.989223 | 0.997988 | 0.994666 |
| Sparcl1_P70663_S78s_1 | Sparcl1_P7 | -0.68 | -0.72 | -5.27 | -6.8 | -0.17 | -1.02 | -6.33 | -5.76 | -1.2 | -0.12 | -3.79 | -6.79 | -0.03 | -0.06 | -4.95 | -5.38 | 0.0475 | 0.3925 | 0.7625 | 0.680848 | 0.980099 | 0.834867 | 0.999187 | 0.997988 | 0.996159 |
| Sparcl1_P70663_S78s_2 | Sparcl1_P7 | -2.68 | -2.16 | -2.9 | -2.22 | -2.02 | -3.43 | -2.8 | -3.15 | -2.55 | -2.85 | -2.67 | -3.04 | -1.53 | -1.03 | -1.88 | -3.43 | -0.36 | -0.2875 | 0.5225 | 0.323639 | 0.323346 | 0.299728 | 0.989223 | 0.997988 | 0.994666 |
| Spast_Q9QYY8_S15s_1 | Spast_Q9Q | -0.16 | -0.64 | 0.35 | 0.94 | -0.37 | -0.21 | -0.43 | 0.07 | 1.11 | 0.06 | 1.34 | 1.42 | -0.03 | 0.11 | 0.42 | 1.57 | -0.3575 | 0.86 | 0.395 | 0.41538 | 0.329507 | 0.079949 | 0.989223 | 0.997988 | 0.994666 |
| Spast_Q9QYY8_S171s_1 | Spast_Q9Q | -1.26 | -0.95 | -0.76 | -0.91 | -1.13 | -1.42 | -0.85 | -1.54 | 1.22 | -0.41 | -0.62 | -0.82 | 1.59 | -0.57 | -0.23 | -0.34 | -0.265 | 0.8125 | 1.0825 | 0.047544 | 0.297454 | 0.102852 | 0.989223 | 0.997988 | 0.994666 |
| Spast_Q9QYY8_T217t_1 | Spast_Q9Q | -1.98 | -3.07 | -0.61 | -0.65 | -1.74 | -2.99 | -2.4 | -1.41 | -3.19 | -2.12 | 0.53 | -0.04 | -1.53 | -1.89 | -1.08 | -1.55 | -0.5575 | 0.3725 | 0.065 | 0.908438 | 0.37405 | 0.69086 | 0.989223 | 0.997988 | 0.999507 |
| Spata18_J3QPR5_S132s_1 | Spata18_J3 | 0.57 | 0.99 | 0.85 | 0.14 | -0.26 | 0.61 | 0.15 | -0.05 | 2.71 | 0.43 | -0.53 | -0.25 | 2.38 | -0.5 | 0.65 | 0.99 | -0.525 | -0.0475 | 0.2425 | 0.672955 | 0.098125 | 0.944679 | 0.989223 | 0.997988 | 0.996159 |
| Spata2_Q8K004_S247s_1 | Spata2_Q8 | -0.01 | 0.57 | -0.22 | 0.15 | 0.24 | -0.09 | 0.38 | -0.08 | -0.27 | 0.54 | 0.08 | 0.32 | -0.59 | 0.74 | -0.06 | 0.03 | -0.01 | 0.045 | -0.0925 | 0.791922 | 0.968853 | 0.875676 | 0.999187 | 0.997988 | 0.999507 |
| Spata2l_Q8BNN1_S318s_1 | Spata2l_Q8 | -1.22 | -1.34 | 0.66 | -0.54 | -1.61 | -1.91 | -0.19 | -0.91 | -0.42 | -3.05 | -0.89 | -0.96 | 0.32 | -1.34 | -0.45 | 0.4 | -0.545 | -0.72 | 0.3425 | 0.547909 | 0.327525 | 0.300392 | 0.989223 | 0.997988 | 0.994666 |
| Spata2l_Q8BNN1_S326s_1 | Spata2l_Q8 | 0.66 | -0.04 | -1.15 | 0.77 | 0.15 | 0.42 | -0.77 | -0.39 | 0.42 | 0.47 | 2.43 | -0.86 | 0.59 | 0.23 | -0.72 | -1.58 | -0.2075 | 0.555 | -0.43 | 0.480425 | 0.664799 | 0.453145 | 0.991343 | 0.997988 | 0.994666 |
| Spats2l_E9Q106_S179s_1 | Spats2l_E9 | -1.26 | -1.8 | -1.59 | -1.01 | -0.98 | -1.62 | -1.27 | -1.2 | -0.92 | -1.15 | -0.69 | -1.4 | -0.97 | -0.86 | -1.38 | -1.26 | 0.1475 | 0.375 | 0.2975 | 0.312228 | 0.580965 | 0.205452 | 0.989223 | 0.997988 | 0.994666 |
| Specc1_Q5SXY1_S133s_1 | Specc1_Q5 | -0.1 | -1.41 | 0.14 | 0.06 | -0.53 | -1.32 | -0.28 | -0.95 | -1.26 | 0 | 0.43 | -0.62 | -0.3 | 0.33 | -0.71 | -0.5 | -0.4425 | -0.035 | 0.0325 | 0.93894 | 0.2969 | 0.942983 | 0.989223 | 0.997988 | 0.999507 |
| Specc1_Q5SXY1_S355s_1 | Specc1_Q5 | -0.98 | 0.58 | 0.32 | -0.28 | 0.29 | -0.74 | 0.11 | -0.44 | -0.35 | -0.34 | -0.92 | 0.11 | -0.2 | 0.24 | -0.23 | 0.31 | -0.105 | -0.285 | 0.12 | 0.756967 | 0.795161 | 0.485368 | 0.995762 | 0.997988 | 0.9971 |
| Specc1_Q5SXY1_S55s_1 | Specc1_Q5 | -1.06 | -0.72 | -1.47 | -0.55 | -0.52 | -1.42 | -1.28 | -1.19 | -1.1 | -0.77 | 0.09 | -0.02 | -0.76 | -0.33 | -1.15 | -1.59 | -0.1525 | 0.5 | -0.0075 | 0.983425 | 0.620781 | 0.191592 | 0.989223 | 0.997988 | 1 |
| Specc1_Q5SXY1_S847s_1 | Specc1_Q5 | -0.91 | -0.46 | -0.36 | 0.12 | 0.08 | -0.16 | 0.7 | 0.12 | -0.01 | -0.53 | 2.21 | -0.25 | -0.55 | 0.74 | 0.56 | -0.29 | 0.5875 | 0.7575 | 0.5175 | 0.205854 | 0.077025 | 0.231305 | 0.989223 | 0.997988 | 0.994666 |
| Specc1_Q5SXY1_S9s_1 | Specc1_Q5 | -2.02 | -0.79 | 0.91 | 0.71 | -1.16 | -2.55 | 1.62 | 0.81 | -2.36 | -1.46 | 1.02 | 0.92 | -1.35 | -1.74 | 1.54 | 1.69 | -0.0225 | -0.1725 | 0.3325 | 0.739454 | 0.982218 | 0.85852 | 0.999187 | 0.997988 | 0.9971 |
| Specc1l_Q2KN98_S171s_1 | Specc1l_Q | 0.52 | 0.14 | 0.47 | -0.57 | -0.72 | 0.48 | 0.09 | -0.96 | 1.72 | 0.49 | -1.88 | -0.09 | 2.03 | -0.34 | -0.28 | -1.02 | -0.4175 | -0.08 | -0.0425 | 0.947251 | 0.31536 | 0.910377 | 0.989223 | 0.997988 | 0.999507 |
| Speg_D3Z7T0_S34s_1 | Speg_D3Z7 | 0.83 | 1.21 | 0.22 | -0.29 | 1.24 | 1.22 | 0.53 | 0.75 | -0.16 | 0.93 | -0.1 | 1.1 | -0.12 | 1.36 | -0.39 | -1.07 | 0.4425 | -0.05 | -0.5475 | 0.343074 | 0.249543 | 0.911718 | 0.989223 | 0.997988 | 0.994666 |
| Speg_E9QQ25_S1177s_1 | Speg_E9Q( | 0.48 | 0.59 | 0.24 | -0.39 | 0.78 | 0.16 | 0.34 | 0.37 | 0.78 | 0.15 | -0.03 | 0.01 | 0.27 | 1.18 | 0.26 | 0.46 | 0.1825 | -0.0025 | 0.3125 | 0.372565 | 0.529409 | 0.993734 | 0.989223 | 0.997988 | 0.994666 |
| Speg_E9QQ25_S2020s_1 | Speg_E9Q( | -0.72 | 0.05 | 0.41 | 0.11 | 0.04 | -1.14 | 0.07 | 0.04 | -0.13 | -0.23 | 0.87 | 0.13 | 0.79 | 0.51 | 0.19 | -0.06 | -0.21 | 0.1975 | 0.395 | 0.261971 | 0.578919 | 0.584521 | 0.989223 | 0.997988 | 0.994666 |
| Speg_E9QQ25_S2042s_1 | Speg_E9Q( | 0.04 | 0.14 | -0.26 | -0.62 | 0.46 | -0.72 | -0.35 | -0.82 | -0.46 | -0.06 | -1.01 | -0.87 | 0.61 | 0.77 | -0.7 | -0.43 | -0.1825 | -0.425 | 0.2375 | 0.564531 | 0.59667 | 0.191483 | 0.989223 | 0.997988 | 0.995259 |
| Speg_E9QQ25_S2114s_1 | Speg_E9Q( | -0.55 | -0.12 | -0.92 | 0.14 | 0.03 | -0.46 | -0.49 | -0.04 | 0.15 | -0.29 | 0.14 | -0.36 | 0.95 | 0 | -0.24 | 0.08 | 0.1225 | 0.2725 | 0.56 | 0.1537 | 0.681313 | 0.38488 | 0.993613 | 0.997988 | 0.994666 |
| Speg_E9QQ25_S2115s_1 | Speg_E9Q( | -1.18 | 1.14 | -0.27 | -0.44 | -1.41 | -0.45 | -0.71 | -0.07 | 0.02 | -0.29 | -0.74 | -0.43 | 0.49 | -0.18 | 0.02 | -0.11 | -0.4725 | -0.1725 | 0.2425 | 0.618488 | 0.367246 | 0.721713 | 0.989223 | 0.997988 | 0.995259 |
| Speg_E9QQ25_S2135s_1 | Speg_E9Q( | -0.11 | -0.16 | 0.33 | 0.26 | 0.24 | -0.12 | -0.06 | 0.23 | -0.61 | -0.14 | 0.6 | 0.36 | 0.43 | 0.48 | 0.14 | 0.6 | -0.0075 | -0.0275 | 0.3325 | 0.220181 | 0.9741 | 0.932334 | 0.999187 | 0.997988 | 0.994666 |
| Speg_E9QQ25_S2171s_1 | Speg_E9Q( | 1.15 | 0.04 | 0.62 | 0.85 | 0.17 | 0.81 | 0.91 | 1.12 | 0.56 | 1.06 | 0.79 | 0.79 | 0.66 | 0.97 | 0.48 | 0.32 | 0.0875 | 0.135 | -0.0575 | 0.857538 | 0.787558 | 0.651397 | 0.995762 | 0.997988 | 0.999507 |
| Speg_E9QQ25_S2327s_1 | Speg_E9Q( | -0.32 | 0.68 | 0.31 | 0.48 | 0.7 | 0.65 | 1.27 | 1.01 | -0.33 | -0.57 | 0.67 | 0.51 | -0.7 | 0 | 0.07 | -0.54 | 0.62 | -0.2175 | -0.58 | 0.10453 | 0.055685 | 0.569662 | 0.989223 | 0.997988 | 0.994666 |
| Speg_E9QQ25_S2347s_1 | Speg_E9Q( | 0.08 | -0.83 | 0.45 | 0.76 | 0.04 | 0.61 | 0.29 | 1 | 0.76 | 0.19 | 0.76 | 0.51 | 0.95 | -0.31 | 0.57 | 0.7 | 0.37 | 0.44 | 0.3625 | 0.411981 | 0.354135 | 0.259715 | 0.989223 | 0.997988 | 0.994666 |
| Speg_E9QQ25_S2396s_1 | Speg_E9Q( | 0.56 | 0.49 | 1.06 | 1.23 | 0.52 | -0.59 | 0.17 | 1.14 | 0.91 | 0.83 | 3.12 | 0.41 | 1.2 | 1.63 | 1.03 | 0.69 | -0.525 | 0.4825 | 0.3025 | 0.351874 | 0.201256 | 0.419068 | 0.989223 | 0.997988 | 0.994666 |
| Speg_E9QQ25_S2451s_1 | Speg_E9Q( | 0.99 | -0.18 | -0.25 | 0.2 | 1.3 | 1.05 | 0.18 | 0.14 | 0.25 | 0.45 | 0 | -0.07 | -1.06 | 0.37 | -0.21 | 0.2 | 0.4775 | -0.0325 | -0.365 | 0.399192 | 0.247157 | 0.921929 | 0.989223 | 0.997988 | 0.994666 |
| Speg_E9QQ25_S2461s_1 | Speg_E9Q( | -0.79 | -0.46 | -0.01 | -0.42 | 0.3 | -0.33 | 0.3 | 0.47 | -0.75 | -1.73 | 1.56 | 0.59 | -0.17 | -0.16 | -0.28 | 0.06 | 0.605 | 0.3375 | 0.2825 | 0.304441 | 0.050923 | 0.617866 | 0.989223 | 0.997988 | 0.994666 |
| Speg_E9QQ25_S2499s_1 | Speg_E9Q( | -1.59 | -0.66 | -1.16 | -3.27 | -0.57 | -1.58 | -1.96 | -2.35 | -1.87 | -1.7 | -1.02 | -1.7 | 0.08 | -0.68 | -1.4 | -0.69 | 0.055 | 0.0975 | 0.9975 | 0.11511 | 0.928068 | 0.859942 | 0.997447 | 0.997988 | 0.994666 |
| Speg_E9QQ25_S2777s_1 | Speg_E9Q( | 0.44 | -0.76 | -0.4 | 0.42 | 0.83 | 0.71 | 0.16 | 1.04 | 0.52 | -0.62 | 1.32 | 0.53 | -0.26 | 0.43 | 0.02 | 0.56 | 0.76 | 0.5125 | 0.2625 | 0.487551 | 0.055398 | 0.296653 | 0.989223 | 0.997988 | 0.994666 |
| Speg_E9QQ25_S2777s_2 | Speg_E9Q( | -1.58 | 0.22 | 0.49 | 0.42 | -0.07 | -2 | 0.05 | 0.44 | -0.93 | -0.53 | 0.52 | 0.41 | -0.3 | -0.21 | 1.47 | 0.37 | -0.2825 | -0.02 | 0.445 | 0.454357 | 0.66625 | 0.971539 | 0.991957 | 0.997988 | 0.994666 |
| Speg_E9QQ25_S490s_1 | Speg_E9Q( | 0.21 | -0.49 | -0.44 | -0.45 | -0.05 | 0.14 | 0.21 | -0.07 | -0.57 | -0.41 | -0.41 | -0.37 | -0.57 | 0.33 | -0.65 | -0.59 | 0.35 | -0.1475 | -0.0775 | 0.814143 | 0.17385 | 0.557743 | 0.989223 | 0.997988 | 0.999507 |
| Speg_E9QQ25_S511s_1 | Speg_E9Q( | 0.01 | 0.19 | 0.3 | 0.4 | 0.26 | 0.36 | -0.16 | 0.73 | 0.56 | 0.35 | 0.32 | 1.03 | 0.51 | 0.35 | 0.36 | 0.84 | 0.0725 | 0.34 | 0.29 | 0.267389 | 0.77599 | 0.203649 | 0.995762 | 0.997988 | 0.994666 |
| Speg_E9QQ25_S554s_1 | Speg_E9Q( | 0.53 | -0.41 | 0.44 | 1.04 | 0.17 | 0.36 | 0.61 | 1.36 | -0.65 | 0.31 | 0.86 | 0.6 | -1.11 | -0.22 | -0.12 | 0.48 | 0.225 | -0.12 | -0.6425 | 0.164927 | 0.564114 | 0.782138 | 0.989223 | 0.997988 | 0.994666 |
| Speg_E9QQ25_S858s_1 | Speg_E9Q( | -1.1 | -1.22 | -0.77 | -1.15 | -2.39 | -0.64 | -1.27 | -0.44 | 1.05 | -1.04 | -0.49 | -0.82 | 1.21 | -1.78 | -1.37 | -1.03 | -0.125 | 0.735 | 0.3175 | 0.607985 | 0.769431 | 0.137066 | 0.995762 | 0.997988 | 0.995259 |
| Speg_E9QQ25_T2303t_1 | Speg_E9Q( | 0.71 | -0.67 | -0.86 | 0.52 | 0.1 | 0.98 | 0.55 | 1.28 | 0.42 | -0.61 | 0.05 | 1.06 | -1.33 | 0.78 | 0.26 | 0.81 | 0.8025 | 0.305 | 0.205 | 0.728601 | 0.098063 | 0.54687 | 0.989223 | 0.997988 | 0.9971 |
| Speg_E9QQ25_T2774t_1 | Speg_E9Q( | -0.99 | 0.59 | 0.35 | 0.48 | -0.68 | -0.36 | 0.89 | 0.34 | -0.15 | -0.19 | 0.92 | 0.6 | 0.74 | 0.74 | 0.87 | 0.08 | -0.06 | 0.1875 | 0.5 | 0.243141 | 0.898856 | 0.674816 | 0.996493 | 0.997988 | 0.994666 |
| Speg_E9QQ25_T2774t_2 | Speg_E9Q( | -1.58 | 0.22 | 0.49 | 0.42 | -0.07 | -2 | 0.05 | 0.44 | -0.93 | -0.53 | 0.52 | 0.41 | -0.3 | -0.21 | 1.47 | 0.37 | -0.2825 | -0.02 | 0.445 | 0.454357 | 0.66625 | 0.971539 | 0.991957 | 0.997988 | 0.994666 |
| Speg_E9QQ25_T2847t_1 | Speg_E9Q( | -0.07 | -0.2 | 0 | 0 | 0.09 | 0.05 | 0 | 0.46 | 0.2 | 0.04 | 0.82 | 0.18 | 0.8 | 0.35 | -0.07 | 0.35 | 0.2175 | 0.3775 | 0.425 | 0.141324 | 0.320511 | 0.159395 | 0.989223 | 0.997988 | 0.994666 |
| Speg_E9QQ25_T453t_1 | Speg_E9Q( | 0.49 | 0.89 | 0.4 | 0.54 | 0.59 | 0.23 | 0.69 | 0.58 | -0.09 | 0.98 | 0.1 | 0.02 | -0.67 | 0.34 | 1.12 | 0.57 | -0.0575 | -0.3275 | -0.24 | 0.549409 | 0.799772 | 0.298267 | 0.995762 | 0.997988 | 0.994666 |
| Spen_Q62504_S1267s_1 | Spen_Q62! | -1.51 | -1.9 | -0.92 | -1.62 | -2.56 | -1.25 | -2.14 | -1.54 | -0.18 | -1.75 | -1.24 | -1.82 | 0.44 | -2.1 | -1.13 | -1.45 | -0.385 | 0.24 | 0.4275 | 0.434182 | 0.298024 | 0.572744 | 0.989223 | 0.997988 | 0.994666 |
| Spen_Q62504_S1276s_1 | Spen_Q62! | 0.72 | -0.5 | 0.06 | 0.24 | -1.6 | 0.47 | -0.92 | -0.19 | 1.37 | -0.32 | -0.56 | -1.14 | 1.9 | -1.61 | 0.08 | -0.38 | -0.69 | -0.2925 | -0.1325 | 0.847698 | 0.169742 | 0.598455 | 0.989223 | 0.997988 | 0.999507 |
| Spen_Q62504_S188s_1 | Spen_Q62! | 1.29 | 0.89 | 1.13 | 1.19 | 0.36 | 0.61 | 0.36 | 0.99 | 2.14 | 0.79 | 0.87 | 0.25 | 1.2 | 0.57 | 1.28 | 1.52 | -0.545 | -0.1125 | 0.0175 | 0.951756 | 0.043354 | 0.781529 | 0.989223 | 0.997988 | 0.999507 |
| Spen_Q62504_S2128s_1 | Spen_Q62! | 0.37 | 0.8 | 1.02 | 0.93 | 0.47 | -0.36 | 0.42 | 0.62 | 1.68 | -0.31 | 1.43 | 0.69 | 1.48 | 2.09 | 0.57 | 0.05 | -0.4925 | 0.0925 | 0.2675 | 0.565953 | 0.116807 | 0.837644 | 0.989223 | 0.997988 | 0.995259 |
| Spen_Q62504_S647s_1 | Spen_Q62! | 0.68 | 0.05 | -0.09 | 0.36 | -1.53 | 1.06 | -0.74 | 0.03 | 0.96 | -0.51 | -0.03 | -0.49 | 0.84 | -0.14 | 0.31 | -0.4 | -0.545 | -0.2675 | -0.0975 | 0.781434 | 0.314341 | 0.494671 | 0.989223 | 0.997988 | 0.998226 |
| Spen_Q62504_S758s_2 | Spen_Q62! | 0.83 | -0.07 | 0.58 | 0.6 | -0.11 | 0.47 | 0.03 | 0.75 | 0.77 | 0.32 | 0.79 | 0.15 | -0.13 | 0.64 | 0.99 | 0.88 | -0.2 | 0.0225 | 0.11 | 0.752874 | 0.511462 | 0.939045 | 0.989223 | 0.997988 | 0.9971 |
| Spg20_D3Z3F8_S126s_1 | Spg20_D3Z | 0.98 | 0.91 | 0.52 | 0.73 | 0.94 | 0.6 | 0.2 | 0.67 | 0.61 | 0.57 | 1.29 | 0.52 | 0.64 | 1.04 | 0.96 | 0.79 | -0.1825 | -0.0375 | 0.0725 | 0.772319 | 0.463281 | 0.888872 | 0.989223 | 0.997988 | 0.997453 |
| Sphkap_E9PUC4_S1067s_1 | Sphkap_E9 | 0.41 | -1.4 | -1.48 | -0.14 | 0.25 | 1.59 | -0.75 | 0.08 | 0.65 | 0.65 | 1.08 | -0.42 | -0.37 | 0.95 | -0.58 | -1.34 | 0.945 | 1.1425 | 0.3175 | 0.604373 | 0.141398 | 0.05464 | 0.989223 | 0.997988 | 0.995259 |
| Sphkap_E9PUC4_S1070s_1 | Sphkap_E9 | -1.64 | -0.5 | -0.08 | 0.16 | -0.41 | -1.69 | -0.81 | -0.84 | -0.99 | -0.23 | -0.26 | -0.12 | -0.06 | -1.15 | 0.06 | -0.63 | -0.4225 | 0.115 | 0.07 | 0.881154 | 0.359236 | 0.790857 | 0.989223 | 0.997988 | 0.999507 |
| Sphkap_E9PUC4_S1223s_1 | Sphkap_E9 | -2.31 | -2.39 | -1.62 | -1.92 | -2.09 | -1.02 | -1.71 | -2.07 | 0.41 | -1.39 | -1.41 | -1.4 | 0.34 | -1.28 | -1.45 | -2.04 | 0.3375 | 1.1125 | 0.9525 | 0.086441 | 0.30833 | 0.037178 | 0.989223 | 0.997988 | 0.994666 |
| Sphkap_E9PUC4_S1244s_1 | Sphkap_E9 | 0.16 | -0.21 | 0.4 | 0.54 | 0.28 | 0.34 | 0.92 | 0.65 | 0.26 | 0.3 | 0.99 | 0.62 | 0.07 | 0.59 | 0.2 | 0.4 | 0.325 | 0.32 | 0.0925 | 0.74085 | 0.240594 | 0.277728 | 0.989223 | 0.997988 | 0.9971 |
| Sphkap_E9PUC4_S1273s_1 | Sphkap_E9 | -0.64 | -1.95 | -1.77 | -1.83 | -0.33 | -0.75 | -0.91 | -2.24 | -0.52 | -1.07 | 0.29 | -1.31 | -0.87 | 0.13 | -2.24 | -1.15 | 0.49 | 0.895 | 0.515 | 0.348291 | 0.31577 | 0.072489 | 0.989223 | 0.997988 | 0.994666 |

| | | | | | | | | | | | | | | | | | | | | | | | | | | |
|---|---|---|---|---|---|---|---|---|---|---|---|---|---|---|---|---|---|---|---|---|---|---|---|---|---|---|
| Sphkap_E9PUC4_S1369s_1 | Sphkap_E9 | 0.32 | -0.15 | 0.5 | 0.59 | 0.61 | 0.75 | 1.06 | 1.49 | -0.53 | 0.16 | 1.01 | 1.2 | -0.43 | -0.28 | 0.4 | 0.17 | 0.6625 | 0.145 | -0.35 | 0.273615 | 0.041824 | 0.732411 | 0.989223 | 0.997988 | 0.994666 |
| Sphkap_E9PUC4_S1403s_1 | Sphkap_E9 | -0.44 | -1.68 | -3.3 | -2.04 | -0.45 | 0.19 | -1.46 | -1.34 | 0.33 | -0.32 | -1.52 | -2.21 | -0.06 | -1.01 | -2.52 | -2.53 | 1.1 | 0.935 | 0.335 | 0.656842 | 0.105551 | 0.225598 | 0.989223 | 0.997988 | 0.995259 |
| Sphkap_E9PUC4_S841s_1 | Sphkap_E9 | 0.05 | 0.74 | 0.06 | -0.27 | 0.03 | 1.3 | -0.85 | 0.43 | 1.96 | 0.8 | -1.5 | 0.11 | 1.94 | 1.13 | 0 | -0.86 | 0.0825 | 0.1975 | 0.4075 | 0.499393 | 0.857636 | 0.772239 | 0.996349 | 0.997988 | 0.994666 |
| Spire1_D3YTL8_S229s_1 | Spire1_D3Y | 0.01 | 0.35 | 0.12 | -0.02 | 0.56 | -0.22 | 0.45 | 0.17 | -0.99 | -0.71 | -0.38 | -0.69 | -0.87 | 0.17 | -0.43 | 0.42 | 0.125 | -0.8075 | -0.2925 | 0.398181 | 0.619234 | 0.007196 | 0.989223 | 0.997988 | 0.994666 |
| Spire1_D3YTL8_S275s_1 | Spire1_D3Y | -0.5 | -0.49 | -0.18 | 0.41 | -0.16 | -1.06 | -0.71 | -0.15 | -0.16 | -0.98 | -0.56 | -1.41 | 0.55 | 0.57 | 0.1 | -0.91 | -0.33 | -0.5875 | 0.2675 | 0.518419 | 0.319411 | 0.125283 | 0.989223 | 0.997988 | 0.994666 |
| Spire1_D3YTL8_S565s_1 | Spire1_D3Y | 1.24 | 1.79 | 2 | 1.82 | 2.14 | 1.81 | 1.89 | 2.2 | 0.24 | 0.52 | 1.31 | 1.3 | 0.3 | 1.72 | 1.94 | 2.46 | 0.2975 | -0.87 | -0.1075 | 0.819475 | 0.248426 | 0.027705 | 0.989223 | 0.997988 | 0.999507 |
| Spock2_F6SLR4_S72s_1 | Spock2_F6 | -0.4 | 0.03 | 1.4 | 0.99 | 0.2 | -0.88 | 1.77 | 1.28 | -0.46 | -1.2 | 2.21 | 1.34 | -1.82 | -0.62 | 0.73 | 2.43 | 0.0875 | -0.0325 | -0.325 | 0.712743 | 0.891469 | 0.967378 | 0.996493 | 0.997988 | 0.996966 |
| Sppl2b_Q3TD49_S534s_1 | Sppl2b_Q3 | -4.04 | -2.11 | -1.73 | -2.71 | 0.04 | -1.95 | -4.19 | -1.92 | -0.37 | -2.18 | -4.07 | -1.74 | -1.02 | -0.8 | -5.48 | -1.62 | 0.6425 | 0.5575 | 0.4175 | 0.691762 | 0.467397 | 0.500738 | 0.989223 | 0.997988 | 0.996159 |
| Spred1_Q924S8_S318s_1 | Spred1_Q9 | -1.43 | -2.03 | -1.06 | -1.07 | -0.99 | -1.41 | -0.67 | -0.82 | -2.14 | -1.89 | 0.46 | -0.87 | -1.53 | -0.91 | -1.77 | -1.32 | 0.425 | 0.2875 | 0.015 | 0.963912 | 0.18155 | 0.623762 | 0.989223 | 0.997988 | 0.999507 |
| Spred2_Q5SRF8_S195s_1 | Spred2_Q5 | -0.38 | -0.21 | 0.23 | 0.78 | -0.81 | -0.1 | 0.14 | 0.43 | 1.09 | 0.14 | 1.56 | 0.14 | 1.64 | 0 | 0.83 | 0.48 | -0.19 | 0.6275 | 0.6325 | 0.162623 | 0.606505 | 0.167277 | 0.989223 | 0.997988 | 0.994666 |
| Spred3_Q6P6N5_S286s_1 | Spred3_Q6 | 0.14 | 0.25 | 0.71 | 0.36 | -0.68 | 0.92 | -1.08 | -0.34 | 1.43 | 0.24 | 0.4 | 0.49 | 1.57 | -1.15 | 0.18 | -0.01 | -0.66 | 0.275 | -0.2175 | 0.684181 | 0.145308 | 0.404142 | 0.989223 | 0.997988 | 0.996159 |
| Spry1_Q9QXV9_S133s_1 | Spry1_Q9C | 0.4 | 1.29 | 0.6 | 0.58 | 0.41 | 2.24 | -0.29 | 0.5 | 0.77 | 0.29 | 0.53 | 0.9 | 1.61 | 0.75 | 1.06 | 0.89 | -0.0025 | -0.095 | 0.36 | 0.276401 | 0.99617 | 0.740166 | 0.99973 | 0.997988 | 0.994666 |
| Spry2_Q9QXV8_S117s_1 | Spry2_Q9C | 0.09 | 0.03 | -0.12 | -0.68 | -0.12 | -0.5 | -0.52 | 0.63 | -0.55 | -0.6 | -0.05 | 0.31 | 0.15 | 0.14 | 0.13 | 0.11 | 0.0425 | -0.0525 | 0.3025 | 0.274227 | 0.897765 | 0.866341 | 0.996493 | 0.997988 | 0.994666 |
| Sptan1_P16546_S1029s_1 | Sptan1_P1 | 0.79 | 0.31 | 0.34 | 0.57 | 0.81 | 0.55 | 0.97 | 0.55 | 0.07 | 0.05 | 0.07 | 0.71 | -0.02 | 0.49 | 0.3 | 0.73 | 0.2175 | -0.2775 | -0.1275 | 0.645941 | 0.354082 | 0.304076 | 0.989223 | 0.997988 | 0.995259 |
| Sptan1_P16546_S1031s_1 | Sptan1_P1 | 1.12 | 0.12 | 0.18 | 0.69 | 1.18 | 0.91 | 1.11 | 0.64 | 0.13 | -0.09 | -0.09 | 0.7 | -0.89 | 0.47 | 0.28 | 0.64 | 0.4325 | -0.365 | -0.4025 | 0.347667 | 0.163322 | 0.280032 | 0.989223 | 0.997988 | 0.994666 |
| Sptan1_P16546_S1190s_1 | Sptan1_P1 | 0.46 | -0.49 | -1.05 | 0.41 | 1.29 | -2.83 | 0.85 | 0.41 | 0.21 | -1.02 | -3.57 | -1.12 | -3.33 | -0.39 | -2.07 | -1.36 | 0.0975 | -1.2075 | -1.62 | 0.030637 | 0.911228 | 0.14698 | 0.997335 | 0.997988 | 0.994666 |
| Sptan1_P16546_S1217s_1 | Sptan1_P1 | -0.85 | 0.04 | 1.14 | 0.47 | 0.39 | -1.08 | 0.74 | 0.4 | -1.41 | -1.24 | 0.66 | 0.39 | -0.96 | -0.07 | 0.48 | 0.88 | -0.0875 | -0.6 | -0.1175 | 0.82678 | 0.868458 | 0.346702 | 0.996493 | 0.997988 | 0.999507 |
| Sptan1_P16546_S328s_1 | Sptan1_P1 | 0.58 | 1.15 | 0.45 | 0.17 | 1.06 | 0.71 | 1.05 | 0.35 | -0.35 | 0.72 | -0.27 | 0.54 | 0.14 | -0.02 | 0.87 | 0.44 | 0.205 | -0.4275 | -0.23 | 0.487751 | 0.491301 | 0.248908 | 0.989223 | 0.997988 | 0.994666 |
| Sptan1_P16546_S330s_1 | Sptan1_P1 | 0.18 | 0.27 | 0.69 | 0.86 | 0.88 | 0.17 | 1.02 | 0.24 | -0.5 | 0.5 | -0.33 | 1.46 | 0 | -0.55 | 0.34 | 1 | 0.0775 | -0.2175 | -0.3025 | 0.432576 | 0.794896 | 0.636922 | 0.995762 | 0.997988 | 0.994666 |
| Sptan1_P16546_S809s_1 | Sptan1_P1 | 0.67 | 0.49 | 0.49 | 0.73 | 0.74 | 0.59 | 0.31 | 0.41 | -0.19 | -0.41 | 0.37 | 0.7 | 0.98 | 0.42 | 0.55 | 1.19 | -0.0825 | -0.4775 | 0.19 | 0.494291 | 0.698573 | 0.137783 | 0.994893 | 0.997988 | 0.994666 |
| Sptb_E9Q397_S1226s_1 | Sptb_E9Q3 | 1.04 | 0.33 | 0.54 | -0.39 | 0.98 | 0.82 | 0.59 | -0.43 | 0.52 | 0.67 | -0.36 | 0.04 | 0.9 | 1.18 | -0.34 | 0.69 | 0.11 | -0.1625 | 0.2275 | 0.605148 | 0.790549 | 0.671899 | 0.995762 | 0.997988 | 0.995259 |
| Sptb_E9Q397_S2114s_1 | Sptb_E9Q3 | 1.7 | -0.11 | 0.26 | 1.51 | 0.39 | 2.46 | 0.22 | 0.91 | 0.73 | 0.36 | -0.16 | 0.25 | 0.94 | 1.43 | -0.78 | -0.03 | 0.155 | -0.545 | -0.45 | 0.466215 | 0.798684 | 0.258571 | 0.995762 | 0.997988 | 0.994666 |
| Sptb_E9Q397_T2113t_1 | Sptb_E9Q3 | 1.89 | 0.42 | 1.03 | 2.27 | 1.41 | 2.18 | 1.12 | 1.98 | 2.31 | 0.49 | 0.34 | 0.82 | 2.18 | 1.44 | 0.25 | 0.46 | 0.27 | -0.4125 | -0.32 | 0.574352 | 0.554428 | 0.47381 | 0.989223 | 0.997988 | 0.995259 |
| Sptb_Q3UGX2_S2300s_1 | Sptb_Q3U | -0.45 | 0.23 | -0.05 | -1.1 | 0.48 | -0.26 | 0.56 | -0.25 | -0.54 | -0.69 | -0.43 | -0.91 | -0.2 | -0.22 | -0.41 | -0.38 | 0.475 | -0.3 | 0.04 | 0.904315 | 0.21103 | 0.375286 | 0.989223 | 0.997988 | 0.999507 |
| Sptb_Q3UGX2_T2113t_1 | Sptb_Q3U | 0.81 | -0.06 | 0.78 | 0.21 | -1.44 | 1.52 | -0.85 | 0.06 | 2.54 | 0.23 | -0.94 | 0.43 | 2.22 | -0.91 | 1.16 | -0.13 | -0.6125 | 0.13 | 0.15 | 0.818592 | 0.326242 | 0.849025 | 0.989223 | 0.997988 | 0.999507 |
| Sptbn1_Q62261_S1662s_1 | Sptbn1_Q6 | -1.55 | 0.39 | -0.09 | -0.68 | -0.88 | -1.48 | -2.37 | -1.26 | 1.1 | -0.35 | -0.15 | -1 | 0.58 | -0.08 | -1.44 | -1.41 | -1.015 | 0.3825 | -0.105 | 0.860277 | 0.059229 | 0.500368 | 0.989223 | 0.997988 | 0.999507 |
| Sptbn1_Q62261_S2102s_1 | Sptbn1_Q6 | 0.98 | 0.7 | 0.89 | 0.95 | 0.59 | 0.66 | 0.19 | 0.61 | 1.13 | 1.29 | 0.28 | 0.58 | 1.68 | 0.3 | 0.83 | 0.68 | -0.3675 | -0.06 | -0.0075 | 0.982165 | 0.11555 | 0.835848 | 0.989223 | 0.997988 | 1 |
| Sptbn1_Q62261_S2127s_1 | Sptbn1_Q6 | 1.32 | 0.99 | 0.87 | 0.96 | 0.86 | 0.67 | 0.71 | 0.77 | 0.91 | 1.61 | 0.78 | 0.8 | 0.32 | 0.88 | 0.97 | 1 | -0.2825 | -0.01 | -0.2425 | 0.384307 | 0.19922 | 0.97098 | 0.989223 | 0.997988 | 0.994666 |
| Sptbn1_Q62261_S2127s_2 | Sptbn1_Q6 | -0.47 | 0.02 | 1.11 | 1.53 | 0.4 | -0.13 | 0.4 | 1.17 | -0.09 | -0.34 | 0.33 | 1.07 | 0.41 | 0.48 | 1 | 1.17 | -0.0875 | -0.305 | 0.2175 | 0.651803 | 0.85906 | 0.56232 | 0.996349 | 0.997988 | 0.995259 |
| Sptbn1_Q62261_S2137s_1 | Sptbn1_Q6 | 1.19 | 0.69 | 0.94 | 1.35 | 0.83 | 0.69 | 1.1 | 1.26 | 1.07 | 1.34 | 1.21 | 1.31 | 0.83 | 1.14 | 1.13 | 1.2 | -0.0725 | 0.19 | 0.0325 | 0.90164 | 0.772382 | 0.43834 | 0.995762 | 0.997988 | 0.999507 |
| Sptbn1_Q62261_S2137s_2 | Sptbn1_Q6 | -0.47 | 0.02 | 1.05 | 1.6 | 0.4 | -0.13 | 0.74 | 1.38 | -0.09 | -0.34 | 0 | 1.14 | 0.41 | 0.48 | 1.01 | 1.22 | 0.0475 | -0.3725 | 0.23 | 0.638834 | 0.926851 | 0.492397 | 0.997335 | 0.997988 | 0.995259 |
| Sptbn1_Q62261_S2164s_1 | Sptbn1_Q6 | -1.34 | 0 | -0.92 | -1.58 | -1.01 | -2.07 | -1.19 | -1.53 | -0.38 | -0.34 | -0.69 | -0.98 | 0.24 | -0.67 | -0.9 | -1.14 | -0.49 | 0.3625 | 0.3425 | 0.452224 | 0.241736 | 0.355122 | 0.989223 | 0.997988 | 0.994666 |
| Sptbn1_Q62261_S2164s_2 | Sptbn1_Q6 | -3.8 | -3.07 | -2.89 | -3.59 | -2.42 | -4.62 | -2.82 | -3.37 | -4.71 | -3.33 | -1.18 | -2.77 | -3.47 | -2.51 | -2.68 | -3.2 | 0.03 | 0.34 | 0.3725 | 0.292832 | 0.950466 | 0.623029 | 0.999187 | 0.997988 | 0.994666 |
| Sptbn1_Q62261_S2168s_1 | Sptbn1_Q6 | -0.16 | 0.94 | 0.04 | -1.04 | 0.28 | -0.78 | -0.39 | -1.68 | 0.2 | 0.27 | -0.32 | -0.16 | 0.98 | 0.33 | 0.22 | -0.75 | -0.5875 | 0.0525 | 0.25 | 0.625507 | 0.278996 | 0.900734 | 0.989223 | 0.997988 | 0.995259 |
| Sptbn1_Q62261_S2168s_2 | Sptbn1_Q6 | -3.8 | -3.07 | -2.89 | -3.59 | -2.42 | -4.62 | -2.82 | -3.37 | -4.71 | -3.33 | -1.18 | -2.77 | -3.47 | -2.51 | -2.68 | -3.2 | 0.03 | 0.34 | 0.3725 | 0.292832 | 0.950466 | 0.623029 | 0.999187 | 0.997988 | 0.994666 |
| Sptbn1_Q62261_S2315s_1 | Sptbn1_Q6 | 1.92 | -0.12 | 1.25 | 0.45 | -0.11 | 2.46 | 0.42 | 0.47 | 1.12 | 0.65 | -2.09 | 1.09 | 1.64 | -2.91 | 1.03 | 1.02 | -0.065 | -0.6825 | -0.68 | 0.497678 | 0.919113 | 0.400069 | 0.997335 | 0.997988 | 0.994666 |
| Sptbn1_Q62261_S2315s_2 | Sptbn1_Q6 | 3.42 | 0.88 | 3.18 | -1.74 | 1.16 | 4.27 | 1.66 | 1.43 | 1.67 | 1.8 | 0.38 | 3.15 | 1.68 | -1.75 | 1.08 | 2.48 | 0.695 | 0.315 | -0.5625 | 0.667785 | 0.567762 | 0.787547 | 0.989223 | 0.997988 | 0.996159 |
| Sptbn1_Q62261_S2318s_1 | Sptbn1_Q6 | 2.22 | 2.76 | 3.05 | 0.73 | 2.38 | 2.44 | 1.64 | 1.76 | 1.25 | 2.66 | -0.02 | 3.09 | 2.05 | -0.94 | 3.09 | 3.93 | -0.135 | -0.445 | -0.1575 | 0.878084 | 0.790709 | 0.573181 | 0.995762 | 0.997988 | 0.999507 |
| Sptbn1_Q62261_S2318s_2 | Sptbn1_Q6 | 3.44 | 0.91 | 2.4 | -1.03 | 1.3 | 4.31 | 1.54 | 0.85 | 2.23 | 1.68 | 0.26 | 2.31 | 1.75 | -1.37 | 1.57 | 2.42 | 0.57 | 0.19 | -0.3375 | 0.761975 | 0.598824 | 0.842382 | 0.989223 | 0.997988 | 0.9971 |
| Sptbn1_Q62261_S2339s_1 | Sptbn1_Q6 | 1.03 | 0.01 | 0.58 | 0.67 | 0.97 | 0.88 | 1.24 | 0.82 | 0.48 | 0.36 | -0.02 | 0.77 | -0.06 | 0.74 | 0.43 | 0.65 | 0.405 | -0.175 | -0.1325 | 0.682439 | 0.159744 | 0.568691 | 0.989223 | 0.997988 | 0.996159 |
| Sptbn1_Q62261_S2340s_1 | Sptbn1_Q6 | 0.69 | 0.15 | 0.29 | 0.7 | 0.81 | 0.74 | 1.01 | 0.87 | 0.29 | 0.46 | 0.25 | 0.77 | -0.2 | 0.5 | 0.22 | 0.45 | 0.4 | -0.015 | -0.215 | 0.458404 | 0.105406 | 0.95293 | 0.989223 | 0.997988 | 0.994666 |
| Sptbn1_Q62261_S2357s_1 | Sptbn1_Q6 | -2.3 | -3.6 | -3.06 | -3.42 | -1.32 | -1.14 | -1.56 | -3.03 | -3.76 | -3.15 | 0.85 | -2.2 | -2.19 | -1.24 | -3.09 | -3.12 | 1.3325 | 1.03 | 0.685 | 0.19369 | 0.018942 | 0.289839 | 0.989223 | 0.997988 | 0.994666 |
| Sptbn1_Q62261_T2186t_1 | Sptbn1_Q6 | -0.21 | -0.65 | 0.24 | -0.07 | 0.15 | -1.34 | 0.27 | 0.34 | -1 | -0.38 | 0.25 | -0.37 | -1.11 | 0.05 | 0.21 | -0.09 | 0.0275 | -0.2025 | -0.0625 | 0.865967 | 0.9476 | 0.551207 | 0.999187 | 0.997988 | 0.999507 |
| Sptbn1_Q62261_T2194t_1 | Sptbn1_Q6 | 0.15 | 0.26 | -0.01 | -0.03 | 0.83 | 0.07 | 0.52 | 0.35 | -0.14 | 0 | -0.53 | 0.32 | -0.25 | 0.39 | -0.17 | 0.28 | 0.35 | -0.18 | -0.03 | 0.910278 | 0.166885 | 0.490339 | 0.989223 | 0.997988 | 0.999507 |
| Sptbn1_Q62261_T2316t_2 | Sptbn1_Q6 | 5.39 | 2.67 | 3.72 | -2.93 | 3.24 | 6.8 | 3.93 | 2.72 | 1.61 | 4.93 | 1.87 | 4.82 | 1.98 | 0.19 | 4.95 | 4.87 | 1.96 | 1.095 | 0.785 | 0.670215 | 0.272856 | 0.537688 | 0.989223 | 0.997988 | 0.996159 |
| Sptbn1_Q62261_T2319t_1 | Sptbn1_Q6 | 2.72 | 3.06 | 3.64 | 1.11 | 2.77 | 2.97 | 1.5 | 2.22 | 3.23 | 3.8 | 0.45 | 3.56 | 2.74 | -3.73 | 3.72 | 3.66 | -0.2675 | 0.1275 | -1.035 | 0.523163 | 0.640781 | 0.879924 | 0.989223 | 0.997988 | 0.994666 |
| Sptbn1_Q62261_T2319t_2 | Sptbn1_Q6 | 3.18 | 1.95 | 3.72 | -2.93 | 1.47 | 4.07 | 3.93 | 2.72 | 2.85 | 2.79 | 1.87 | 4.82 | 3.03 | -0.41 | 4.95 | 4.87 | 1.5675 | 1.6025 | 1.63 | 0.345746 | 0.279056 | 0.279809 | 0.989223 | 0.997988 | 0.994666 |
| Sptbn2_Q68FG2_S1060s_1 | Sptbn2_Q6 | 1.12 | -0.53 | 0.1 | -0.6 | 0.36 | 0.66 | 0.12 | 0.11 | 0.67 | -0.18 | -0.89 | -1.99 | 1.32 | 0.27 | -0.39 | -0.42 | 0.29 | -0.62 | 0.1725 | 0.746012 | 0.476998 | 0.337229 | 0.989223 | 0.997988 | 0.9971 |
| Sptbn2_Q68FG2_S2199s_1 | Sptbn2_Q6 | 0.1 | -0.6 | 0.23 | 0.75 | 0.27 | 0.5 | 0.65 | 0.84 | 0.17 | -0.5 | -0.38 | 0.33 | -0.11 | 0.43 | -0.02 | 0.55 | 0.445 | -0.215 | 0.0925 | 0.792635 | 0.18467 | 0.551138 | 0.989223 | 0.997988 | 0.999507 |
| Sptbn2_Q68FG2_S2200s_1 | Sptbn2_Q6 | -0.43 | -1.51 | -0.75 | -0.51 | -0.02 | -0.03 | -0.42 | 0.26 | -0.2 | -1.15 | 0.54 | -0.91 | -0.31 | -0.02 | 0.14 | -2.08 | 0.7475 | 0.37 | 0.2325 | 0.662789 | 0.032981 | 0.408985 | 0.989223 | 0.997988 | 0.995804 |
| Sptbn2_Q68FG2_S2254s_1 | Sptbn2_Q6 | -0.73 | -0.38 | 0.07 | -1.02 | 0.23 | -0.63 | -0.26 | -0.42 | -1.65 | -1.9 | -0.85 | -0.62 | -1.1 | -0.22 | -1.6 | -1.02 | 0.245 | -0.74 | -0.47 | 0.23892 | 0.444049 | 0.083086 | 0.989223 | 0.997988 | 0.994666 |
| Sptbn2_Q68FG2_T2354t_1 | Sptbn2_Q6 | -0.32 | -0.09 | -0.49 | -0.19 | 0.15 | -0.23 | 0.38 | -0.77 | -0.07 | 0.37 | -0.1 | 0.34 | -0.3 | -0.25 | -0.15 | -0.06 | 0.155 | 0.4075 | 0.0825 | 0.729712 | 0.600282 | 0.114929 | 0.989223 | 0.997988 | 0.9971 |
| Sptbn4_Q91ZE6_S1445s_1 | Sptbn4_Q9 | -0.27 | -0.17 | 0.41 | 0.62 | 0.04 | -0.15 | 1.01 | 0.77 | -0.8 | 0.16 | 0.25 | 0 | -1.12 | -0.25 | 0.18 | -0.02 | 0.27 | -0.245 | -0.45 | 0.249882 | 0.453619 | 0.480607 | 0.989223 | 0.997988 | 0.994666 |
| Sptbn4_Q91ZE6_S1577s_1 | Sptbn4_Q9 | 0.12 | 0.73 | 0.81 | -0.09 | 0.08 | 0.57 | -0.05 | -0.23 | 0.53 | -0.18 | -0.03 | 0.19 | 1.34 | -0.18 | 0.57 | 0.44 | -0.3 | -0.265 | 0.15 | 0.70356 | 0.336078 | 0.397507 | 0.989223 | 0.997988 | 0.996159 |
| Sptbn4_Q91ZE6_S1863s_1 | Sptbn4_Q9 | 0.11 | 0.4 | 0.47 | -0.09 | -0.87 | 0.6 | -0.37 | 0.06 | 0.9 | -0.29 | -0.5 | 0.12 | 0.89 | -0.89 | 0.27 | -0.87 | -0.3675 | -0.165 | -0.3725 | 0.413942 | 0.299621 | 0.639774 | 0.989223 | 0.997988 | 0.994666 |
| Sptbn4_Q91ZE6_S2156s_1 | Sptbn4_Q9 | 1.07 | -0.64 | 0.18 | 0.18 | -0.2 | 0.9 | -0.65 | 0.22 | -0.14 | -0.11 | 0.08 | -0.4 | -0.2 | 0.28 | -0.78 | -0.47 | -0.13 | -0.34 | -0.49 | 0.255988 | 0.772417 | 0.370234 | 0.995762 | 0.997988 | 0.994666 |
| Sptbn4_Q91ZE6_S2245s_1 | Sptbn4_Q9 | 1.45 | 0.62 | 1.19 | 1.91 | 0.1 | 1.24 | 0.59 | 1.87 | 1.13 | 1.25 | 2.11 | 1.62 | 0.29 | 1.07 | 0.88 | 0.9 | -0.3425 | 0.235 | -0.5075 | 0.168943 | 0.444968 | 0.51835 | 0.989223 | 0.997988 | 0.994666 |
| Sptbn4_Q91ZE6_S2263s_1 | Sptbn4_Q9 | -0.11 | -0.6 | -1.24 | -0.08 | -0.24 | -0.4 | -0.29 | 0.05 | 0.06 | -0.17 | -0.13 | -0.48 | -1.23 | 0.87 | -0.61 | -0.85 | 0.2875 | 0.3275 | 0.0525 | 0.916983 | 0.362654 | 0.323077 | 0.989223 | 0.997988 | 0.999507 |
| Sptbn4_Q91ZE6_S2296s_1 | Sptbn4_Q9 | 2.29 | 1.4 | 1.46 | 1.76 | 0.64 | 1.8 | 1.06 | 1.59 | 2.34 | 2.25 | 1.49 | 1.87 | 1.52 | 0.86 | 1.07 | 1.53 | -0.455 | 0.26 | -0.4825 | 0.148965 | 0.200306 | 0.41619 | 0.989223 | 0.997988 | 0.994666 |
| Sptbn4_Q91ZE6_S2297s_1 | Sptbn4_Q9 | 2.79 | 1.73 | 0.33 | 0.03 | 0.57 | 2.2 | -0.86 | -0.64 | 2.54 | 2.76 | -0.85 | 0.29 | 1.66 | 1.11 | -0.27 | -0.34 | -0.9025 | -0.035 | -0.68 | 0.359868 | 0.291799 | 0.970932 | 0.989223 | 0.997988 | 0.994666 |
| Sptbn4_Q91ZE6_S2391s_1 | Sptbn4_Q9 | 0.28 | 1.29 | 0.31 | 0.22 | 0.26 | 0.41 | 0.43 | 0.48 | 0.72 | 0.09 | 0.43 | 0.76 | 0.11 | 0.7 | 0.44 | 0.56 | -0.13 | -0.025 | -0.0725 | 0.824951 | 0.65562 | 0.938732 | 0.990434 | 0.997988 | 0.999507 |
| Sqstm1_D3YZJ1_T272t_1 | Sqstm1_D3 | 0.91 | 1.2 | 0.12 | 0.05 | 1.21 | 0.98 | 0.24 | 0.22 | 0.63 | 0.83 | -0.52 | -0.05 | -0.25 | 1.18 | 0.5 | 0.36 | 0.0925 | -0.3475 | -0.1225 | 0.766477 | 0.805569 | 0.412285 | 0.995762 | 0.997988 | 0.9971 |
| Src_P05480_S17s_1 | Src_P0548 | -1.25 | -0.51 | -0.57 | -0.54 | -0.51 | -1.15 | -0.08 | -0.52 | -0.87 | -0.95 | -0.62 | -0.33 | -0.67 | -0.21 | -0.57 | -0.42 | 0.1525 | 0.025 | 0.25 | 0.384269 | 0.619428 | 0.928475 | 0.989223 | 0.997988 | 0.994666 |
| Src_P05480_S74s_1 | Src_P0548 | 0.73 | 0.26 | 0.83 | 0.58 | 0.77 | 0.89 | 1.14 | 0.93 | -1.01 | 0.74 | -1.27 | 0.77 | -1.17 | -1.86 | 0.21 | 0.72 | 0.3325 | -0.7925 | -1.125 | 0.070468 | 0.164811 | 0.156669 | 0.989223 | 0.997988 | 0.994666 |
| Srcap_E9Q9V7_S559s_1 | Srcap_E9Q | 1.96 | 3.04 | 0.63 | 3.15 | 2.13 | 0.94 | 2.21 | 1.3 | 3.84 | 4.34 | 4.09 | 1.09 | 0.91 | 3.56 | 0.18 | 1.75 | -0.55 | 1.145 | -0.595 | 0.476325 | 0.366401 | 0.201069 | 0.989223 | 0.997988 | 0.994666 |
| Srcin1_B1AQX9_S1021s_1 | Srcin1_B1A | 0.31 | 0.26 | 0.61 | 0.35 | 0.06 | 0.52 | 0.46 | 0.65 | 0.51 | 0.87 | 0.42 | 0.72 | 0.49 | 0.68 | 0.2 | 0.2 | 0.04 | 0.2475 | 0.01 | 0.96834 | 0.860405 | 0.293612 | 0.996493 | 0.997988 | 0.999507 |
| Srcin1_B1AQX9_S1037s_1 | Srcin1_B1A | 0.39 | -0.3 | -0.14 | 0.1 | 0.72 | 0.6 | 0.34 | 0.72 | 0.44 | -0.01 | 0.99 | 0.13 | 0.66 | 0.47 | 0.34 | 0.09 | 0.5825 | 0.375 | 0.3775 | 0.191813 | 0.034353 | 0.23803 | 0.989223 | 0.997988 | 0.994666 |
| Srcin1_B1AQX9_S1077s_1 | Srcin1_B1A | 0.85 | 0.49 | 0.7 | 0.46 | 0.69 | 0.75 | 0.54 | 0.4 | 0.37 | 0.66 | 0.51 | 0.57 | 0 | 1.12 | 0.19 | 0.29 | -0.03 | -0.0975 | -0.225 | 0.480819 | 0.889077 | 0.660674 | 0.996493 | 0.997988 | 0.994666 |
| Srcin1_B1AQX9_S1094s_1 | Srcin1_B1A | -1.93 | -1.87 | -2.03 | -2.32 | -1.26 | -1.77 | -1.46 | -2.27 | -2.2 | -1.49 | -0.87 | -2.3 | -1.69 | -0.71 | -2.98 | -1.75 | 0.3475 | 0.3225 | 0.255 | 0.58223 | 0.231277 | 0.381679 | 0.989223 | 0.997988 | 0.995259 |
| Srcin1_B1AQX9_S1101s_1 | Srcin1_B1A | -2.4 | -2.79 | -2.71 | 0.84 | -0.33 | -1.75 | -1.91 | 0.71 | -1.39 | -1.31 | 1.51 | 0.72 | -1.98 | 0.02 | -1.93 | -1.22 | 0.945 | 1.6475 | 0.4875 | 0.577569 | 0.322654 | 0.126843 | 0.989223 | 0.997988 | 0.995259 |
| Srcin1_B1AQX9_S1152s_1 | Srcin1_B1A | -0.11 | -0.04 | 0 | -0.32 | -0.48 | 0.27 | -0.83 | -0.6 | 1.14 | 0.1 | -2.75 | -0.03 | 1.86 | -2.39 | -0.23 | 0.1 | -0.2925 | -0.2675 | -0.0475 | 0.951073 | 0.31491 | 0.721337 | 0.989223 | 0.997988 | 0.999507 |
| Srcin1_B1AQX9_S1189s_1 | Srcin1_B1A | -0.04 | 3.39 | -3.91 | -6.21 | 3.69 | 0.35 | -4.31 | -4.05 | 2.25 | 0.89 | -0.62 | -3.7 | 1.19 | 2.94 | -3.7 | -4.42 | 0.6125 | 1.3975 | 0.695 | 0.770277 | 0.801452 | 0.520641 | 0.995762 | 0.997988 | 0.9971 |
| Srcin1_B1AQX9_S12s_1 | Srcin1_B1A | -0.63 | 0.16 | -0.49 | -0.62 | 0.07 | -0.63 | -0.36 | -0.95 | -0.52 | -0.28 | -0.4 | -0.27 | -0.49 | 0.2 | -0.59 | -0.8 | -0.0725 | 0.0275 | -0.025 | 0.939308 | 0.81348 | 0.916232 | 0.995762 | 0.997988 | 0.999507 |
| Srcin1_B1AQX9_S13s_1 | Srcin1_B1A | -0.44 | 0.72 | -0.06 | -0.68 | 0.12 | -0.43 | -0.1 | -0.77 | 0.04 | 0.3 | -0.27 | -0.41 | 0.22 | 0.35 | -0.75 | -0.57 | -0.18 | 0.03 | -0.0725 | 0.860827 | 0.619983 | 0.933085 | 0.989223 | 0.997988 | 0.999507 |
| Srcin1_B1AQX9_S199s_1 | Srcin1_B1A | -1.25 | -0.73 | -0.7 | -0.88 | -0.51 | -1.46 | -0.27 | -0.79 | -1.59 | -1.13 | -0.63 | -0.59 | -0.52 | -0.86 | -0.83 | -0.69 | 0.1325 | -0.095 | 0.165 | 0.522328 | 0.667743 | 0.75567 | 0.992842 | 0.997988 | 0.994666 |
| Srcin1_B1AQX9_S213s_1 | Srcin1_B1A | -0.12 | 0.28 | 0.42 | -0.12 | 0.64 | -0.5 | 0.39 | -0.2 | 0.38 | 0.13 | -0.46 | 0.14 | -0.52 | 0.37 | -0.32 | 0.54 | -0.0325 | -0.0675 | -0.0975 | 0.76981 | 0.917524 | 0.808775 | 0.997335 | 0.997988 | 0.9971 |
| Srcin1_B1AQX9_S259s_1 | Srcin1_B1A | 0.44 | 0.9 | 0.98 | 0.53 | 1.13 | 0.33 | 0.86 | 0.51 | 0.48 | 0.58 | 0.67 | 0.89 | 0.5 | 1.26 | 0.46 | 0.68 | -0.005 | -0.0575 | 0.0125 | 0.966056 | 0.985058 | 0.812523 | 0.999187 | 0.997988 | 0.999507 |
| Srcin1_B1AQX9_S332s_1 | Srcin1_B1A | -0.51 | -0.65 | 0.78 | 1.29 | -0.51 | -0.87 | 0.61 | 0.61 | 0.28 | -0.25 | 3.26 | 0.9 | 0.02 | -0.45 | -1.17 | 1.43 | -0.2675 | 0.82 | -0.27 | 0.68186 | 0.630618 | 0.32639 | 0.989223 | 0.997988 | 0.996159 |
| Srcin1_B1AQX9_S340s_1 | Srcin1_B1A | 0.48 | 0.52 | -0.33 | -0.03 | 0.37 | -0.49 | 0.7 | 0.52 | -0.9 | 0.32 | -1.02 | 0.36 | -0.56 | -0.31 | 0.39 | 0.97 | 0.115 | -0.47 | -0.0375 | 0.926506 | 0.736141 | 0.280073 | 0.995762 | 0.997988 | 0.999507 |
| Srcin1_B1AQX9_S373s_1 | Srcin1_B1A | -1.3 | 0.42 | -0.57 | -0.66 | -0.87 | -0.54 | -0.96 | -1.09 | -0.21 | -0.04 | -1.06 | -1.04 | 1.63 | 0.11 | -1.1 | -0.95 | -0.3375 | -0.06 | 0.45 | 0.495678 | 0.371297 | 0.889752 | 0.989223 | 0.997988 | 0.994666 |
| Srcin1_B1AQX9_S377s_1 | Srcin1_B1A | -0.04 | 0.06 | 2.03 | 1.98 | -0.35 | -0.75 | 1.61 | 1.53 | 0.19 | 0.51 | 2.78 | 2.05 | 0.62 | -0.49 | 2.49 | 2.78 | -0.4975 | 0.375 | 0.3425 | 0.688468 | 0.507465 | 0.622693 | 0.989223 | 0.997988 | 0.996159 |
| Srcin1_B1AQX9_S396s_1 | Srcin1_B1A | -0.56 | -0.42 | 0.97 | 0.34 | 0.12 | -0.73 | 0.42 | 0.52 | -1.7 | -0.26 | 0.09 | 0.73 | -0.47 | -1.07 | -1.01 | -0.45 | 0 | -0.3675 | -0.8325 | 0.061159 | 1 | 0.528019 | 1 | 0.997988 | 0.994666 |
| Srcin1_B1AQX9_S398s_1 | Srcin1_B1A | 0 | 0.8 | 1.49 | -1.25 | 0.67 | 0.71 | 1.78 | 0.75 | 0.04 | 0.77 | 1.98 | 0.79 | 0.1 | 0.4 | 0.35 | 0.6 | 0.7175 | 0.635 | 0.1025 | 0.852884 | 0.235377 | 0.339569 | 0.989223 | 0.997988 | 0.999507 |
| Srcin1_B1AQX9_S39s_1 | Srcin1_B1A | 2.73 | 0.95 | 0.94 | 1 | 0.77 | 2.73 | 0.3 | 1.04 | 3.39 | 1.27 | -1.12 | 0.99 | 3.12 | -0.37 | 1.41 | 0.55 | -0.195 | -0.2725 | -0.2275 | 0.766813 | 0.751605 | 0.764154 | 0.995762 | 0.997988 | 0.9971 |
| Srcin1_B1AQX9_S45s_1 | Srcin1_B1A | -0.17 | 0.25 | 0.18 | -0.12 | 0.29 | 0.22 | 0.2 | -0.34 | -0.37 | 0.37 | 0.03 | 0.55 | 0.23 | 1.06 | -0.39 | -0.55 | 0.0575 | 0.11 | 0.0525 | 0.89302 | 0.812821 | 0.695536 | 0.995762 | 0.997988 | 0.999507 |
| Srcin1_B1AQX9_S468s_1 | Srcin1_B1A | -1.27 | -0.1 | -0.79 | -1.03 | -0.27 | -1.86 | -0.37 | -1.35 | -2.9 | -0.66 | -2.09 | -0.27 | -0.29 | 0.04 | -0.67 | -0.74 | -0.165 | -0.6825 | 0.3825 | 0.281923 | 0.704817 | 0.278127 | 0.995437 | 0.997988 | 0.994666 |
| Srcin1_B1AQX9_S506s_1 | Srcin1_B1A | 0.64 | 0.26 | 0.56 | 0.63 | 0.16 | 0.69 | 0.08 | 0.42 | 0.35 | 0.48 | 0.19 | 0.47 | 0.76 | 0.02 | -0.01 | 0.24 | -0.185 | -0.15 | -0.27 | 0.345012 | 0.438579 | 0.503948 | 0.989223 | 0.997988 | 0.994666 |
| Srcin1_B1AQX9_S507s_1 | Srcin1_B1A | 0.64 | 0.34 | 0.6 | 0.71 | 0.27 | 0.88 | 0.15 | 0.55 | 0.49 | 0.41 | 0.28 | 0.4 | 0.96 | -0.32 | 0.02 | 0.22 | -0.11 | -0.1775 | -0.3525 | 0.29489 | 0.65323 | 0.417134 | 0.990434 | 0.997988 | 0.994666 |
| Srcin1_B1AQX9_S526s_1 | Srcin1_B1A | 0.25 | 0.24 | 0.58 | 0.22 | 0.15 | 0.38 | 0.59 | 0.6 | -0.31 | 0.65 | -0.29 | 0.53 | 0.02 | -0.57 | 0.46 | 0.11 | 0.1075 | -0.1775 | -0.3175 | 0.297622 | 0.629905 | 0.567282 | 0.989223 | 0.997988 | 0.994666 |
| Srcin1_B1AQX9_S529s_1 | Srcin1_B1A | 0.17 | 0.14 | 0.12 | 0.1 | 0.44 | -0.19 | 0.82 | 0.35 | -0.14 | 0.41 | -0.1 | -0.04 | -1.09 | -0.4 | -0.02 | 1.11 | 0.2225 | -0.1 | -0.2325 | 0.606658 | 0.399442 | 0.662611 | 0.989223 | 0.997988 | 0.995259 |
| Srcin1_B1AQX9_S53s_1 | Srcin1_B1A | -0.7 | 0.64 | -0.3 | -0.42 | 0.43 | -0.07 | 0.15 | -0.69 | -0.85 | 0.55 | -0.02 | 0.1 | 0.95 | -0.88 | 0.05 | 0.12 | 0.15 | 0.14 | 0.255 | 0.58117 | 0.686457 | 0.731702 | 0.994672 | 0.997988 | 0.995259 |
| Srcin1_B1AQX9_S541s_1 | Srcin1_B1A | 1.05 | 1.27 | 0.87 | 0.8 | 0.99 | 1.43 | 0.58 | 1 | 1.24 | 1.53 | 0.93 | 1.43 | 2.68 | 2.48 | 1.12 | 0.27 | 0.0025 | 0.285 | 0.64 | 0.254791 | 0.992182 | 0.265694 | 0.999187 | 0.997988 | 0.994666 |
| Srcin1_B1AQX9_S546s_1 | Srcin1_B1A | -0.28 | 0.45 | 0.63 | 0.37 | 0.21 | -0.5 | 0.57 | 0.19 | 0.07 | 0.22 | 0.61 | 0.75 | 0.78 | 0.96 | 0.36 | 0.41 | -0.175 | 0.12 | 0.335 | 0.286432 | 0.582363 | 0.686786 | 0.989223 | 0.997988 | 0.994666 |
| Srcin1_B1AQX9_S555s_1 | Srcin1_B1A | -0.08 | 0.55 | -0.02 | -0.06 | 0.21 | -0.07 | 0 | -0.15 | 0.34 | 0.21 | 0.07 | 0.36 | 0.36 | 0.81 | 0.16 | -0.06 | -0.1 | 0.1475 | 0.22 | 0.470272 | 0.675146 | 0.551355 | 0.992876 | 0.997988 | 0.994666 |
| Srcin1_B1AQX9_S625s_1 | Srcin1_B1A | -0.82 | -0.45 | 1.07 | 0.75 | -0.54 | 0.33 | 0.65 | 0.14 | 0.67 | -0.93 | 0.47 | 1.04 | 1.17 | -0.91 | 1.87 | 1.03 | 0.0075 | 0.175 | 0.6525 | 0.345674 | 0.98755 | 0.762467 | 0.999187 | 0.997988 | 0.994666 |
| Srcin1_B1AQX9_S64s_1 | Srcin1_B1A | 2.78 | 1.72 | 2.57 | 0.7 | 2.21 | 3.06 | 3.09 | 1.82 | -0.29 | 2.55 | 0.14 | 2.96 | -0.61 | 0.19 | 1.92 | 3.07 | 0.6025 | -0.6025 | -0.8 | 0.353808 | 0.262726 | 0.483079 | 0.989223 | 0.997988 | 0.994666 |
| Srcin1_B1AQX9_S654s_1 | Srcin1_B1A | 0.31 | 0.86 | 0.95 | 1.08 | 0.9 | 0.44 | 1.16 | 1.27 | 0.42 | 0.29 | 1.26 | 0.79 | 0.05 | 0.77 | 1.23 | 1.48 | 0.1425 | -0.11 | 0.0825 | 0.825723 | 0.618162 | 0.723375 | 0.989223 | 0.997988 | 0.999507 |
| Srcin1_B1AQX9_S667s_1 | Srcin1_B1A | 0.82 | 0.85 | 1.18 | 1.12 | 0.15 | 0.82 | 0.5 | 1.05 | 0.49 | 0.3 | 0.75 | 0.8 | -0.25 | 0.67 | 0.32 | 1.32 | -0.3625 | -0.4075 | -0.4775 | 0.210123 | 0.190234 | 0.111725 | 0.989223 | 0.997988 | 0.994666 |
| Srcin1_B1AQX9_S670s_1 | Srcin1_B1A | 0.19 | 0.2 | 0.29 | 0.04 | -0.03 | -0.39 | 0.64 | 0.39 | 0.32 | 0.04 | 0.23 | 0.57 | 0.24 | -0.13 | 0.23 | 0.49 | -0.0275 | 0.11 | 0.0275 | 0.912626 | 0.919836 | 0.627603 | 0.997335 | 0.997988 | 0.999507 |
| Srcin1_B1AQX9_S878s_1 | Srcin1_B1A | 0.12 | 0.6 | 0.32 | 0.41 | 0.51 | 0.12 | 0.78 | 0.25 | -0.22 | 0.43 | -0.79 | 0.43 | -0.24 | 0.13 | 0.1 | -0.12 | 0.0525 | -0.4 | -0.395 | 0.13749 | 0.82781 | 0.248766 | 0.995762 | 0.997988 | 0.994666 |
| Srcin1_B1AQX9_S900s_1 | Srcin1_B1A | -0.23 | 0.19 | 0.38 | 0.36 | 0.28 | -0.03 | 0.57 | 0.32 | -0.5 | 0.34 | -0.69 | 0.44 | 0.07 | -0.57 | 0.19 | 0.09 | 0.11 | -0.2775 | -0.23 | 0.438402 | 0.657526 | 0.423507 | 0.990434 | 0.997988 | 0.994666 |
| Srcin1_B1AQX9_S941s_1 | Srcin1_B1A | -0.27 | -1.7 | -1.09 | -1.5 | 0.41 | 0.11 | -0.86 | -0.16 | -1.45 | -1.74 | 2.31 | -0.22 | -1.32 | -0.32 | -1.74 | -0.68 | 1.015 | 0.865 | 0.125 | 0.776573 | 0.028803 | 0.331888 | 0.989223 | 0.997988 | 0.99798 |
| Srcin1_B1AQX9_T629t_1 | Srcin1_B1A | -0.39 | -0.11 | 0.13 | -0.95 | -1.9 | 0.06 | -1.5 | -1.69 | 2.27 | -0.15 | -0.58 | -0.46 | 1.12 | -2.2 | -1.77 | -1.54 | -0.9275 | 0.6 | -0.7675 | 0.290827 | 0.071799 | 0.36521 | 0.989223 | 0.997988 | 0.994666 |
| Srcin1_B1AQX9_T642t_1 | Srcin1_B1A | 0 | 0.36 | 1.11 | 1.55 | 0.44 | -0.15 | 0.91 | 0.55 | -0.36 | 0.22 | 1.8 | 0.84 | 0.65 | 0.23 | 0.89 | 1.03 | -0.3175 | -0.13 | -0.055 | 0.890466 | 0.434316 | 0.80976 | 0.989223 | 0.997988 | 0.999507 |
| Srcin1_B1AQX9_T658t_1 | Srcin1_B1A | 0.86 | 0.77 | 0.86 | 0.57 | 0.37 | 0.66 | 0.8 | 1.07 | 0.84 | 0.8 | 0.71 | 1.35 | 0.13 | 0.93 | 1.01 | 0.73 | -0.04 | 0.16 | -0.065 | 0.819378 | 0.863759 | 0.514011 | 0.996493 | 0.997988 | 0.999507 |
| Srcin1_B1AQX9_T658t_2 | Srcin1_B1A | 2.55 | 2.78 | 0.56 | 0.64 | 2.35 | 3.01 | 0.68 | 1.03 | 0.2 | 1.99 | 0.35 | 0.79 | 1.01 | 0 | 0.41 | 1.14 | 0.135 | -0.8 | -0.9925 | 0.122212 | 0.85011 | 0.2419 | 0.996349 | 0.997988 | 0.994666 |
| Srcin1_B1AQX9_T671t_2 | Srcin1_B1A | 1.33 | 0.77 | 2.24 | 2.36 | 0.74 | 1 | 1.74 | 1.27 | 1.27 | 0.9 | 3.39 | 2.15 | 0.85 | 1.05 | 2.32 | 2.83 | -0.4875 | 0.2525 | 0.0875 | 0.877003 | 0.257399 | 0.681712 | 0.989223 | 0.997988 | 0.999507 |
| Srcin1_B1AQX9_Y430y_1 | Srcin1_B1A | 0.64 | 1.44 | 1.12 | 1.08 | -0.02 | -1.23 | 1.15 | 1.8 | -1.68 | 0.26 | 1.4 | 1.02 | -1.52 | 0.64 | 1.25 | 1.59 | -0.645 | -0.82 | -0.58 | 0.380937 | 0.307749 | 0.221695 | 0.989223 | 0.997988 | 0.994666 |
| Srcin1_Q9QWI6_S46s_1 | Srcin1_Q9 | -0.36 | 0.33 | 0.67 | 0.76 | 0.3 | -0.43 | 0.85 | 0.66 | -0.37 | -0.03 | 0.64 | 0.95 | -0.45 | 0.33 | 0.57 | 0.64 | -0.005 | -0.0525 | -0.0775 | 0.83596 | 0.9893 | 0.894011 | 0.999187 | 0.997988 | 0.999507 |
| Srcin1_Q9QWI6_S47s_1 | Srcin1_Q9 | 0.73 | 1.12 | 0.41 | 0.19 | 0.95 | 0.58 | 0.49 | 0.11 | 0.88 | 1.11 | 0.26 | 0.38 | 0.5 | 1.15 | 0.02 | 0.17 | -0.08 | 0.045 | -0.1525 | 0.665742 | 0.785622 | 0.886917 | 0.995762 | 0.997988 | 0.996159 |
| Srgap1_D3YZW1_S835s_1 | Srgap1_D3 | 1.05 | 0.55 | 1.29 | 1.15 | -0.11 | 1.11 | 0.46 | 0.3 | 2.35 | 0.97 | 0.19 | 0.73 | 1.25 | -1.05 | 1.42 | 0.34 | -0.57 | 0.05 | -0.52 | 0.352494 | 0.097399 | 0.914375 | 0.989223 | 0.997988 | 0.994666 |
| Srgap1_D3YZW1_S906s_1 | Srgap1_D3 | 0.15 | -1.16 | -0.55 | 0.05 | -1.47 | 0.05 | -0.96 | -0.51 | 0.54 | -0.46 | 0.57 | 0.16 | 0.86 | -2.42 | 0.51 | -0.45 | -0.345 | 0.58 | 0.0025 | 0.99719 | 0.420822 | 0.159606 | 0.989223 | 0.997988 | 1 |
| Srgap1_D3YZW1_S917s_1 | Srgap1_D3 | -1.08 | -1.73 | -1.66 | -1.12 | -0.49 | -0.4 | -0.42 | -0.24 | -0.81 | -1.11 | -0.54 | -0.89 | -0.6 | 0.11 | -0.72 | -1.14 | 1.01 | 0.56 | 0.81 | 0.038909 | 0.002113 | 0.061447 | 0.989223 | 0.997988 | 0.994666 |
| Srgap2_Q91Z67_S206s_1 | Srgap2_Q9 | -0.19 | -0.23 | 0.61 | 0.79 | -1.2 | 0.3 | -0.83 | 0.55 | 1.59 | -0.03 | 0.5 | 0.77 | 1.37 | -1.1 | -0.01 | -0.06 | -0.54 | 0.4625 | -0.195 | 0.715369 | 0.263336 | 0.287976 | 0.989223 | 0.997988 | 0.996966 |
| Srgap2_Q91Z67_S427s_1 | Srgap2_Q9 | 1.41 | 0.83 | 1.19 | 0.74 | 1.96 | 0.84 | 1.76 | 1.34 | 0.4 | 1.12 | 0.1 | 1.03 | 0.03 | 1.27 | 0.78 | 0.9 | 0.4325 | -0.38 | -0.2975 | 0.390499 | 0.187528 | 0.254485 | 0.989223 | 0.997988 | 0.994666 |
| Srgap2_Q91Z67_S500s_1 | Srgap2_Q9 | 0.56 | -0.42 | -1.1 | -0.49 | -0.13 | 2.45 | -0.26 | -0.26 | -0.36 | 0.58 | -0.82 | -0.93 | 0.12 | -0.55 | -1.04 | 0.22 | 0.8125 | -0.02 | 0.05 | 0.910192 | 0.240262 | 0.965602 | 0.989223 | 0.997988 | 0.999507 |
| Srgap2_Q91Z67_S860s_1 | Srgap2_Q9 | -0.58 | 0.25 | -0.55 | -0.64 | -0.25 | -0.51 | -0.22 | -0.34 | -0.66 | -0.19 | -0.96 | -0.26 | -0.73 | -0.07 | -0.18 | -0.58 | 0.05 | -0.1375 | -0.01 | 0.9746 | 0.85078 | 0.659927 | 0.996349 | 0.997988 | 0.999507 |
| Srgap2_Q91Z67_S983s_1 | Srgap2_Q9 | -1.33 | -1.05 | -0.91 | -0.85 | -0.75 | -1.23 | -1.04 | -0.76 | -1.31 | -1.11 | -1.27 | -1.54 | -1.08 | -0.66 | -1.29 | -1.43 | 0.09 | -0.2725 | -0.08 | 0.774326 | 0.698655 | 0.259492 | 0.994893 | 0.997988 | 0.99798 |
| Srgap2_Q91Z67_S994s_1 | Srgap2_Q9 | -0.81 | -1.31 | -0.91 | -0.37 | -0.25 | -0.54 | -0.04 | -0.25 | -0.74 | -0.74 | -0.53 | -0.09 | -0.89 | -0.14 | -0.55 | -0.66 | 0.58 | 0.325 | 0.29 | 0.354768 | 0.050765 | 0.280898 | 0.989223 | 0.997988 | 0.994666 |
| Srgap2_Q91Z67_T982t_1 | Srgap2_Q9 | -1.33 | -1.05 | -0.91 | -0.85 | -0.75 | -1.23 | -1.04 | -0.76 | -1.31 | -1.11 | -1.27 | -1.54 | -1.08 | -0.66 | -1.29 | -1.43 | 0.09 | -0.2725 | -0.08 | 0.774326 | 0.698655 | 0.259492 | 0.994893 | 0.997988 | 0.99798 |
| Srgap3_E9QN14_S1054s_1 | Srgap3_E9 | 0.61 | 0.32 | 0.79 | 0.71 | -0.7 | 0.18 | 0.16 | 1.26 | 0.23 | 0.47 | 1.03 | 1.01 | -0.74 | -1.56 | 0.54 | 0.67 | -0.3825 | 0.0775 | -0.88 | 0.11099 | 0.34957 | 0.780412 | 0.989223 | 0.997988 | 0.994666 |
| Srgap3_E9QN14_S413s_1 | Srgap3_E9 | -1.03 | -0.76 | -0.36 | -0.48 | -0.84 | -0.62 | 0.23 | -0.5 | -1.28 | -1.3 | 0.89 | -1.09 | -0.43 | -0.99 | -0.92 | -0.59 | 0.225 | -0.0375 | -0.075 | 0.7886 | 0.460628 | 0.942014 | 0.989223 | 0.997988 | 0.999507 |
| Srgap3_E9QN14_S813s_1 | Srgap3_E9 | 0.54 | 0.83 | 0.64 | 0.81 | 1.09 | 0.78 | 1.04 | 0.97 | 0.62 | 1.12 | 0.67 | 0.86 | -0.19 | 0.97 | 0.54 | 0.91 | 0.265 | 0.1125 | -0.1475 | 0.648432 | 0.219199 | 0.626946 | 0.989223 | 0.997988 | 0.995259 |
| Srgap3_E9QN14_S834s_1 | Srgap3_E9 | -1.75 | -1.72 | -0.11 | -0.65 | -0.99 | -1.19 | -0.18 | 0.17 | -1.1 | -1.63 | 0.78 | -0.28 | -0.23 | -0.86 | 0.06 | -0.16 | 0.51 | 0.5 | 0.76 | 0.111109 | 0.302489 | 0.419412 | 0.989223 | 0.997988 | 0.994666 |
| Srgap3_E9QN14_S844s_1 | Srgap3_E9 | 1.85 | 0.71 | 2.59 | 2 | 0.53 | 3.02 | 1.3 | 2.4 | 2.57 | 0.7 | -0.15 | 2.08 | 2.17 | -1.73 | 2.47 | 1.93 | 0.025 | -0.4875 | -0.5775 | 0.537896 | 0.967171 | 0.472415 | 0.999187 | 0.997988 | 0.994666 |
| Srgap3_E9QN14_S844s_2 | Srgap3_E9 | 3.07 | 1.61 | 2.12 | 1.13 | 1.19 | 3.75 | 1.29 | 2.28 | 1.65 | 1.99 | -0.61 | 2.16 | 0.56 | -1.46 | 1.75 | 1.75 | 0.145 | -0.685 | -1.3325 | 0.107322 | 0.821805 | 0.335683 | 0.995762 | 0.997988 | 0.994666 |
| Srgap3_E9QN14_S850s_1 | Srgap3_E9 | 0.08 | 0.32 | 1.26 | 0.94 | -0.78 | 0.45 | 0.04 | 0.1 | 2.33 | 0.39 | -0.83 | 0.91 | 2.53 | -1.47 | 1.15 | 0.4 | -0.6975 | 0.05 | 0.0025 | 0.997429 | 0.085464 | 0.938075 | 0.989223 | 0.997988 | 1 |
| Srgap3_E9QN14_S850s_2 | Srgap3_E9 | 3.07 | 1.61 | 2.12 | 1.13 | 1.19 | 3.75 | 1.29 | 2.28 | 1.65 | 1.99 | -0.61 | 2.16 | 0.56 | -1.46 | 1.75 | 1.75 | 0.145 | -0.685 | -1.3325 | 0.107322 | 0.821805 | 0.335683 | 0.995762 | 0.997988 | 0.994666 |
| Srgap3_E9QN14_S884s_1 | Srgap3_E9 | 0.32 | -0.28 | 0.08 | 0.58 | -0.32 | 0.46 | -0.83 | 0.2 | 1.09 | -0.07 | -0.47 | -0.49 | 0.97 | -0.9 | -0.32 | -0.15 | -0.2975 | -0.16 | -0.275 | 0.524556 | 0.396901 | 0.696405 | 0.989223 | 0.997988 | 0.994666 |
| Srgap3_E9QN14_S895s_1 | Srgap3_E9 | 0.82 | 0.48 | 0.66 | 1.04 | 0.63 | 0.74 | 1.08 | 1.29 | -0.01 | 0.27 | 0.08 | 0.85 | -0.16 | 0.51 | 0.54 | 0.86 | 0.185 | -0.4525 | -0.3125 | 0.317445 | 0.465953 | 0.130645 | 0.989223 | 0.997988 | 0.994666 |
| Srgap3_E9QN14_S924s_1 | Srgap3_E9 | -0.62 | 0.9 | 3.34 | 0.45 | 1.25 | -0.06 | 4.93 | 3.24 | 2.53 | -0.04 | -2.05 | 3.76 | 2 | 0.73 | 5.47 | 4.04 | 1.3225 | 0.0325 | 2.0425 | 0.104822 | 0.281655 | 0.980826 | 0.989223 | 0.997988 | 0.994666 |
| Srgap3_E9QN14_S987s_1 | Srgap3_E9 | -0.67 | -0.89 | -0.23 | -0.22 | -0.83 | -0.98 | -0.5 | -0.28 | -1.37 | -1.41 | 2.5 | 0.01 | -0.48 | -0.27 | -0.16 | -0.45 | -0.145 | 0.435 | 0.1625 | 0.551945 | 0.59552 | 0.603043 | 0.989223 | 0.997988 | 0.994666 |
| Srp68_Q8BMA6_S12s_1 | Srp68_Q8B | 0.3 | -0.04 | 0.3 | 0.4 | -0.68 | -0.5 | -0.12 | -1.63 | -0.12 | -0.37 | 0.11 | 0.89 | 0.12 | -0.77 | -0.21 | -0.33 | -0.9725 | -0.1125 | -0.5375 | 0.082703 | 0.016924 | 0.724783 | 0.989223 | 0.997988 | 0.994666 |
| Srp68_Q8BMA6_S26s_1 | Srp68_Q8B | -0.8 | -1.36 | -0.2 | -0.49 | -0.47 | 0.34 | -0.99 | -0.63 | 0.58 | 0.79 | 0.41 | 0.66 | 1.21 | -1.19 | -0.91 | 0.55 | 0.275 | 1.3225 | 0.6275 | 0.291746 | 0.463565 | 0.001567 | 0.989223 | 0.997988 | 0.994666 |
| Srp72_E9Q740_S560s_1 | Srp72_E9Q | -2.49 | -1.81 | -1.94 | -2.11 | -1.82 | -2.62 | -2.8 | -2.78 | -1.91 | -2.07 | -2.07 | -2.06 | -0.37 | -2.02 | -1.94 | -2.03 | -0.4175 | 0.06 | 0.4975 | 0.262858 | 0.185677 | 0.801883 | 0.989223 | 0.997988 | 0.994666 |
| Srpk2_O54781_S487s_1 | Srpk2_O54 | -1.42 | 0.76 | 0.72 | 0.27 | -0.08 | 0.68 | 0.9 | 0.53 | -0.63 | -0.72 | 0.56 | 0.76 | -0.21 | 0.33 | 0.04 | 0.23 | 0.425 | -0.09 | 0.015 | 0.975964 | 0.410715 | 0.878795 | 0.989223 | 0.997988 | 1 |
| Srpk2_O54781_S490s_1 | Srpk2_O54 | -0.23 | -0.46 | 0.07 | 0.19 | -0.08 | -0.26 | -0.2 | -0.35 | -0.33 | -0.15 | 0.25 | -0.35 | 0.21 | -0.11 | -0.23 | 0.3 | -0.115 | -0.0375 | 0.15 | 0.589682 | 0.621439 | 0.887343 | 0.989223 | 0.997988 | 0.995259 |
| Srpk2_O54781_T331t_1 | Srpk2_O54 | -1.6 | -1.24 | -0.81 | -1.87 | -1 | -2.21 | -1.49 | -1.3 | -1.44 | -0.92 | -0.77 | -1.42 | -1.03 | -0.9 | -1.75 | -1.07 | -0.12 | 0.2425 | 0.1925 | 0.57026 | 0.730738 | 0.451964 | 0.995762 | 0.997988 | 0.995259 |
| Srpk2_O54781_T332t_1 | Srpk2_O54 | -1.16 | -0.75 | -1.69 | -3.74 | -1.12 | -1.59 | -0.51 | -2.4 | -1 | -1.23 | -1.48 | -1.98 | -0.59 | -0.28 | -2.57 | -2.29 | 0.43 | 0.4125 | 0.4025 | 0.608797 | 0.533674 | 0.519958 | 0.989223 | 0.997988 | 0.995259 |
| Srrm1_E9PUK6_S220s_1 | Srrm1_E9P | 0.84 | 0.28 | 0.17 | 0.2 | -0.04 | 0.24 | 0.45 | 0.45 | -0.04 | -0.05 | 0.14 | 0.05 | 0.37 | 0.66 | 0.82 | 0.16 | -0.0975 | -0.3475 | 0.13 | 0.652601 | 0.699118 | 0.17815 | 0.994893 | 0.997988 | 0.995259 |
| Srrm1_E9PUK6_S387s_2 | Srrm1_E9P | 0.79 | 0.34 | 0.64 | 0.35 | 0.53 | 0.18 | 0.53 | 0.72 | 0.13 | -0.2 | 0.87 | 0.74 | 0.43 | 1.16 | 1.46 | 0.54 | -0.04 | -0.145 | 0.3675 | 0.266189 | 0.86297 | 0.642187 | 0.996493 | 0.997988 | 0.994666 |
| Srrm1_E9PUK6_S391s_2 | Srrm1_E9P | 0.79 | 0.34 | 0.57 | 0.53 | 0.53 | 0.18 | 0.41 | 1 | 0.13 | -0.2 | 0.84 | 1.34 | 0.43 | 1.16 | 1.58 | 0.68 | -0.0275 | -0.03 | 0.405 | 0.224593 | 0.91286 | 0.93493 | 0.997335 | 0.997988 | 0.994666 |
| Srrm1_E9PUK6_S412s_1 | Srrm1_E9P | 8.1 | 3.12 | 3.28 | -1.53 | 3.59 | 8.8 | 3.55 | 2.25 | 4.35 | 5.18 | 0.22 | 3.05 | 4.03 | 2.34 | 4.24 | 3.95 | 1.305 | -0.0425 | 0.3975 | 0.817789 | 0.535445 | 0.982625 | 0.989223 | 0.997988 | 0.999507 |
| Srrm1_E9PUK6_S429s_1 | Srrm1_E9P | 1.97 | 1.56 | 1.7 | 1.05 | 0.96 | 1.7 | 0.46 | 1.15 | 1.18 | 0.71 | 0.88 | 0.98 | 2.42 | 1.34 | 1.85 | 1.11 | -0.5025 | -0.6325 | 0.11 | 0.766251 | 0.152823 | 0.041897 | 0.989223 | 0.997988 | 0.9971 |
| Srrm1_E9PUK6_S448s_1 | Srrm1_E9P | 1.25 | 0.27 | 0.36 | 0.55 | -0.45 | 0.86 | -0.59 | 0.65 | 1.19 | 0 | -0.32 | 0.47 | 1.85 | -0.43 | 0.81 | -0.08 | -0.49 | -0.2725 | -0.07 | 0.892995 | 0.253194 | 0.496132 | 0.989223 | 0.997988 | 0.999507 |
| Srrm1_E9PUK6_S461s_1 | Srrm1_E9P | 4.29 | 2.61 | -0.09 | -0.43 | 1.86 | 3.12 | -0.68 | -0.14 | 2.23 | 2.61 | -0.41 | -0.03 | 2.33 | 3.02 | -0.16 | -1.06 | -0.555 | -0.495 | -0.5625 | 0.662977 | 0.653445 | 0.680532 | 0.990434 | 0.997988 | 0.995926 |
| Srrm1_E9PUK6_S550s_1 | Srrm1_E9P | 0.05 | -0.18 | -0.09 | -0.36 | 0 | -0.65 | -0.08 | 0.09 | -0.96 | -0.77 | -0.12 | -0.45 | 0.36 | 0.52 | 0.62 | -0.36 | -0.015 | -0.43 | 0.43 | 0.173814 | 0.951541 | 0.130967 | 0.999187 | 0.997988 | 0.994666 |
| Srrm1_E9PUK6_S561s_2 | Srrm1_E9P | 0.83 | -0.23 | 0.15 | -0.17 | -1.11 | 0.53 | -1.01 | -0.02 | 0.86 | -0.67 | -1.41 | -0.39 | 0.14 | -1.15 | 0.43 | 0.04 | -0.5475 | -0.5475 | -0.28 | 0.512284 | 0.230009 | 0.29006 | 0.989223 | 0.997988 | 0.994666 |
| Srrm1_E9PUK6_S563s_2 | Srrm1_E9P | 0.83 | -0.23 | 0.15 | -0.17 | -1.11 | 0.53 | -1.01 | -0.02 | 0.86 | -0.67 | -1.41 | -0.39 | 0.14 | -1.15 | 0.43 | 0.04 | -0.5475 | -0.5475 | -0.28 | 0.512284 | 0.230009 | 0.29006 | 0.989223 | 0.997988 | 0.994666 |
| Srrm1_E9PUK6_S572s_2 | Srrm1_E9P | 0.45 | 0.16 | 0.71 | 0.51 | -0.16 | -0.15 | 0.67 | 0.78 | 0.04 | 0.09 | 0.13 | 0.88 | 0.36 | 0.19 | 1.02 | 1.07 | -0.1725 | -0.1725 | 0.2025 | 0.515553 | 0.57178 | 0.542848 | 0.989223 | 0.997988 | 0.994666 |
| Srrm1_E9PUK6_S574s_2 | Srrm1_E9P | 0.45 | 0.16 | 0.71 | 0.51 | -0.16 | -0.15 | 0.67 | 0.78 | 0.04 | 0.09 | 0.13 | 0.88 | 0.36 | 0.19 | 1.02 | 1.07 | -0.1725 | -0.1725 | 0.2025 | 0.515553 | 0.57178 | 0.542848 | 0.989223 | 0.997988 | 0.994666 |
| Srrm1_E9PUK6_S591s_1 | Srrm1_E9P | 1.45 | 1.38 | 1.57 | 1.29 | 0.32 | 1.42 | 0.21 | 1.3 | 1.98 | 0.3 | -0.64 | 0.78 | 2.48 | 0.33 | 1.46 | 1.54 | -0.61 | -0.8175 | 0.03 | 0.945141 | 0.092115 | 0.137195 | 0.989223 | 0.997988 | 0.999507 |
| Srrm1_E9PUK6_S591s_2 | Srrm1_E9P | 0.15 | -0.9 | -0.27 | -0.45 | -1.65 | -0.06 | -1.5 | -0.29 | -0.07 | -1.66 | -1.47 | -0.8 | 0.22 | -1.66 | 0.29 | -0.54 | -0.5075 | -0.6325 | -0.055 | 0.908773 | 0.261076 | 0.151886 | 0.989223 | 0.997988 | 0.999507 |
| Srrm1_E9PUK6_S602s_2 | Srrm1_E9P | 0.4 | 0.02 | -0.48 | -0.2 | -0.8 | -0.1 | -0.61 | -0.07 | -0.2 | -0.7 | -0.26 | -1 | 0.31 | -0.27 | 0.6 | -0.8 | -0.33 | -0.475 | 0.025 | 0.947271 | 0.272808 | 0.142354 | 0.989223 | 0.997988 | 0.999507 |
| Srrm1_E9PUK6_S616s_1 | Srrm1_E9P | 0.59 | -0.06 | -0.29 | -0.51 | -0.17 | -0.27 | -0.33 | 0.09 | -0.11 | -0.5 | 0.19 | -0.51 | 0.57 | 0.29 | 0.53 | -0.07 | -0.1025 | -0.165 | 0.3975 | 0.238906 | 0.721657 | 0.609062 | 0.995437 | 0.997988 | 0.994666 |
| Srrm1_E9PUK6_S624s_1 | Srrm1_E9P | 1.02 | 0.78 | 0.22 | -0.36 | 0.28 | 0.05 | -0.04 | 0.32 | 0.06 | 0.65 | 0.19 | 0.3 | 1.26 | 1.11 | 1.16 | -0.7 | -0.2625 | -0.115 | 0.2925 | 0.580877 | 0.434698 | 0.742296 | 0.989223 | 0.997988 | 0.995259 |
| Srrm1_E9PUK6_S635s_1 | Srrm1_E9P | 0.23 | 0.13 | -0.09 | -0.16 | -0.08 | -0.36 | -0.46 | -0.02 | -0.62 | -0.43 | 0.53 | -0.82 | 0.6 | 0.33 | 0.51 | -0.55 | -0.2575 | -0.3625 | 0.195 | 0.550957 | 0.266251 | 0.294843 | 0.989223 | 0.997988 | 0.994666 |
| Srrm1_E9PUK6_S635s_2 | Srrm1_E9P | 0.52 | 0.74 | -0.58 | -0.34 | -0.17 | 0.15 | 0.09 | 0.55 | 0.85 | 0.21 | 0.31 | -0.06 | 0.82 | 1 | 0.63 | -0.15 | 0.07 | 0.2425 | 0.49 | 0.251453 | 0.843553 | 0.526643 | 0.996349 | 0.997988 | 0.994666 |
| Srrm1_E9PUK6_S655s_2 | Srrm1_E9P | 0.24 | -0.57 | -1.18 | -0.75 | -1.82 | 0.1 | -1.23 | -0.63 | 0.93 | -0.92 | -1.55 | -0.87 | 0.83 | -1.12 | 0.08 | -1.05 | -0.33 | -0.0375 | 0.25 | 0.633441 | 0.488353 | 0.946954 | 0.989223 | 0.997988 | 0.995259 |
| Srrm1_E9PUK6_S657s_2 | Srrm1_E9P | 0.24 | -0.57 | -1.18 | -0.75 | -1.82 | 0.1 | -1.23 | -0.63 | 0.93 | -0.92 | -1.55 | -0.87 | 0.83 | -1.12 | 0.08 | -1.05 | -0.33 | -0.0375 | 0.25 | 0.633441 | 0.488353 | 0.946954 | 0.989223 | 0.997988 | 0.995259 |
| Srrm1_E9PUK6_S664s_1 | Srrm1_E9P | 1.22 | -0.06 | -0.07 | 0.63 | -0.98 | 0.6 | -0.33 | -0.2 | 2.06 | -0.43 | -0.61 | -0.31 | 2.67 | -0.84 | 0.83 | -0.1 | -0.6575 | -0.2525 | 0.21 | 0.773733 | 0.14612 | 0.693877 | 0.989223 | 0.997988 | 0.99798 |
| Srrm1_E9PUK6_S672s_1 | Srrm1_E9P | 0.28 | -0.26 | 0.5 | -0.3 | -0.56 | 0.39 | -0.38 | -0.02 | 0.62 | -0.57 | -1.71 | 0.07 | 1.41 | -0.98 | 0.67 | -0.55 | -0.1975 | -0.4525 | 0.0825 | 0.879159 | 0.527327 | 0.381899 | 0.989223 | 0.997988 | 0.999507 |
| Srrm1_E9PUK6_S714s_1 | Srrm1_E9P | 0.63 | 0.09 | 0.05 | 0.02 | -0.09 | 0.03 | -0.16 | 0.43 | -0.03 | -0.2 | 0.36 | -0.16 | 1.07 | 0.39 | 0.76 | -0.12 | -0.145 | -0.205 | 0.3275 | 0.338545 | 0.567594 | 0.437834 | 0.989223 | 0.997988 | 0.994666 |
| Srrm1_E9PUK6_S723s_1 | Srrm1_E9P | -1.96 | -1.12 | -2.38 | -4.71 | -2.11 | -4.15 | -1.79 | -1.63 | 0.49 | -1.81 | -3.31 | -1.21 | 0.9 | -1.65 | -0.31 | -1.62 | 0.1225 | 1.0825 | 1.8725 | 0.05255 | 0.884225 | 0.281882 | 0.996493 | 0.997988 | 0.994666 |
| Srrm1_E9PUK6_S725s_1 | Srrm1_E9P | 1.02 | -0.14 | 0.15 | 0.41 | -1.3 | 1.03 | 0.01 | 0.59 | 3.17 | -0.14 | -1.78 | 0.54 | 4.09 | -0.97 | 2.81 | -0.57 | -0.2775 | 0.0875 | 0.98 | 0.384519 | 0.591839 | 0.925572 | 0.989223 | 0.997988 | 0.994666 |
| Srrm1_E9PUK6_S731s_1 | Srrm1_E9P | 0.81 | 0.16 | -1.1 | -2.21 | -0.16 | 0.09 | -1.63 | -1.52 | 0.04 | -0.11 | -1.48 | -1.48 | 0.85 | 0.64 | -0.5 | -1.02 | -0.22 | -0.1725 | 0.5775 | 0.430784 | 0.757307 | 0.808749 | 0.995762 | 0.997988 | 0.994666 |
| Srrm1_E9PUK6_S756s_1 | Srrm1_E9P | 2.3 | 0.94 | 0.1 | -0.25 | 0.28 | 1.36 | -0.22 | 0.02 | 1.27 | 1.05 | -0.31 | -0.52 | 1.76 | 0.95 | 0.55 | 0.11 | -0.4125 | -0.4 | 0.07 | 0.908193 | 0.494225 | 0.54926 | 0.989223 | 0.997988 | 0.999507 |
| Srrm1_E9PUK6_S756s_2 | Srrm1_E9P | 1.35 | 0.44 | 0.81 | 0.47 | -0.51 | 0.8 | -0.29 | 0.4 | 0.67 | -0.08 | 0.31 | 0.16 | 0.88 | -0.02 | 1.04 | 0.49 | -0.6675 | -0.5025 | -0.17 | 0.627191 | 0.093646 | 0.121971 | 0.989223 | 0.997988 | 0.995259 |
| Srrm1_E9PUK6_S758s_2 | Srrm1_E9P | 1.35 | 0.44 | 0.81 | 0.47 | -0.51 | 0.8 | -0.29 | 0.4 | 0.67 | -0.08 | 0.31 | 0.16 | 0.88 | -0.02 | 1.04 | 0.49 | -0.6675 | -0.5025 | -0.17 | 0.627191 | 0.093646 | 0.121971 | 0.989223 | 0.997988 | 0.995259 |
| Srrm1_E9PUK6_S774s_1 | Srrm1_E9P | 1.22 | 0.35 | 1.03 | 0.91 | -0.49 | 1.3 | 0.41 | 1.06 | 1.08 | -0.01 | -0.24 | 0.72 | 2.39 | -2.91 | 1.11 | 0.58 | -0.3075 | -0.49 | -0.585 | 0.561933 | 0.470025 | 0.205964 | 0.989223 | 0.997988 | 0.995259 |
| Srrm1_E9PUK6_S787s_1 | Srrm1_E9P | 0.73 | 0.08 | -0.04 | -0.34 | -0.8 | 0.49 | 0.24 | 0.47 | 0.07 | 0.68 | -2.04 | 0.06 | 0.49 | 1.34 | -0.02 | -0.09 | -0.0075 | -0.415 | 0.3225 | 0.432904 | 0.983941 | 0.484313 | 0.999187 | 0.997988 | 0.994666 |
| Srrm1_E9PUK6_T593t_2 | Srrm1_E9P | 0.15 | -0.9 | -0.27 | -0.45 | -1.65 | -0.06 | -1.5 | -0.29 | -0.07 | -1.66 | -1.47 | -0.8 | 0.22 | -1.66 | 0.29 | -0.54 | -0.5075 | -0.6325 | -0.055 | 0.908773 | 0.261076 | 0.151886 | 0.989223 | 0.997988 | 0.999507 |
| Srrm1_E9PUK6_T633t_2 | Srrm1_E9P | 0.52 | 0.74 | -0.58 | -0.34 | -0.17 | 0.15 | 0.09 | 0.55 | 0.85 | 0.21 | 0.31 | -0.06 | 0.82 | 1 | 0.63 | -0.15 | 0.07 | 0.2425 | 0.49 | 0.251453 | 0.843553 | 0.526643 | 0.996349 | 0.997988 | 0.994666 |
| Srrm2_Q8BTI8_S1068s_1 | Srrm2_Q8E | 0.92 | 0.38 | 0.12 | 0.19 | -0.16 | 0.31 | 0.03 | 0.39 | -0.49 | 0.02 | -0.21 | -0.04 | 0.27 | 0.33 | 0.11 | -0.21 | -0.26 | -0.5825 | -0.2775 | 0.34812 | 0.340934 | 0.055665 | 0.989223 | 0.997988 | 0.994666 |
| Srrm2_Q8BTI8_S1097s_1 | Srrm2_Q8E | 0.93 | 0.35 | 0.25 | 0.29 | -0.75 | 0.9 | -1.18 | 0.12 | 1.45 | 0.09 | -1.44 | 0.04 | 2.15 | -0.42 | 0.64 | -0.69 | -0.6825 | -0.42 | -0.035 | 0.95382 | 0.158571 | 0.463902 | 0.989223 | 0.997988 | 0.999507 |
| Srrm2_Q8BTI8_S1151s_1 | Srrm2_Q8E | 1.66 | 1.3 | 0.19 | 0.68 | 0.4 | 0.23 | 0.57 | 0.88 | -0.37 | 0.17 | 0.49 | 0.46 | 0.57 | 0.98 | 0.92 | 0.59 | -0.4375 | -0.77 | -0.1925 | 0.600987 | 0.236511 | 0.070937 | 0.989223 | 0.997988 | 0.995259 |
| Srrm2_Q8BTI8_S1179s_1 | Srrm2_Q8E | 0.18 | -0.42 | 0.36 | 0.38 | -1.42 | 0.15 | -0.55 | -0.07 | 0.93 | -0.61 | -2.94 | 0.13 | 2 | -2.22 | 0.88 | -0.4 | -0.5975 | -0.7475 | -0.06 | 0.941307 | 0.144676 | 0.34185 | 0.989223 | 0.997988 | 0.999507 |
| Srrm2_Q8BTI8_S1216s_1 | Srrm2_Q8E | 0.49 | 0.65 | -0.22 | 0.02 | -0.18 | -0.29 | 0.08 | -0.07 | -0.12 | 0.11 | -0.35 | -0.59 | 0 | 0.34 | 0.09 | -0.82 | -0.35 | -0.4725 | -0.3325 | 0.356021 | 0.205246 | 0.134263 | 0.989223 | 0.997988 | 0.994666 |
| Srrm2_Q8BTI8_S1269s_1 | Srrm2_Q8E | 1.31 | 0.99 | -0.56 | 0.11 | 0.83 | 0.26 | 0.11 | 0.07 | -0.08 | 0.2 | -0.42 | -0.51 | 0.3 | 0.95 | 0.18 | -0.84 | -0.145 | -0.665 | -0.315 | 0.553487 | 0.738417 | 0.155809 | 0.995762 | 0.997988 | 0.994666 |

| | | | | | | | | | | | | | | | | | | | | | | | | | | |
|---|---|---|---|---|---|---|---|---|---|---|---|---|---|---|---|---|---|---|---|---|---|---|---|---|---|---|
| Srrm2_Q8BTI8_S1278s_1 | Srrm2_Q8E | -0.71 | -0.55 | -0.78 | -1.25 | -0.98 | -1.37 | -1.43 | -0.96 | -1.34 | -1.51 | -0.83 | -1.44 | 0.01 | -0.56 | -0.85 | -1.2 | -0.3625 | -0.4575 | 0.1725 | 0.609101 | 0.172016 | 0.118836 | 0.989223 | 0.997988 | 0.995259 |
| Srrm2_Q8BTI8_S1305s_1 | Srrm2_Q8E | -0.02 | 0.1 | -0.42 | -0.19 | 0.15 | -0.73 | 0.22 | -0.08 | -0.61 | -0.36 | -0.33 | 0.01 | -0.19 | 0.07 | 0.28 | -0.14 | 0.0225 | -0.19 | 0.1375 | 0.597123 | 0.935893 | 0.450101 | 0.99868 | 0.997988 | 0.995259 |
| Srrm2_Q8BTI8_S1359s_1 | Srrm2_Q8E | 0.54 | 0.32 | 0.85 | -1.04 | 0.56 | -0.29 | 1.06 | 0.01 | -0.03 | 0.21 | 1.44 | 0.18 | 0.05 | 0.49 | -1.16 | -0.11 | 0.1675 | 0.2825 | -0.35 | 0.499439 | 0.724727 | 0.57732 | 0.995437 | 0.997988 | 0.994666 |
| Srrm2_Q8BTI8_S1360s_1 | Srrm2_Q8E | 0.86 | 0.41 | 0.3 | 0.21 | -0.07 | -0.09 | 0.11 | 0.49 | -0.38 | -0.24 | -0.07 | 0.01 | 0.41 | 0.46 | 0.29 | -0.19 | -0.335 | -0.615 | -0.2025 | 0.479967 | 0.205056 | 0.031279 | 0.989223 | 0.997988 | 0.994666 |
| Srrm2_Q8BTI8_S1378s_1 | Srrm2_Q8E | 0.97 | 0.45 | -0.23 | 0.44 | -0.03 | 0.4 | -0.25 | 0.03 | 0.23 | 0.04 | -0.51 | -0.14 | 0.95 | -0.05 | 0.17 | 0.11 | -0.37 | -0.5025 | -0.1125 | 0.753726 | 0.241285 | 0.142449 | 0.989223 | 0.997988 | 0.9971 |
| Srrm2_Q8BTI8_S1380s_1 | Srrm2_Q8E | 1.07 | 0.77 | 0.23 | -0.05 | -0.06 | 0.12 | -0.52 | 0.32 | 0.03 | 0.11 | -1.2 | -0.36 | 1.55 | -0.59 | 0.02 | -0.05 | -0.54 | -0.86 | -0.2725 | 0.586649 | 0.120685 | 0.051975 | 0.989223 | 0.997988 | 0.995259 |
| Srrm2_Q8BTI8_S1397s_1 | Srrm2_Q8E | 0.33 | -0.14 | -0.72 | -0.53 | -1.43 | -0.06 | -1.25 | -0.33 | 0.13 | -1.31 | -1.75 | -0.81 | 1.08 | -1.46 | -0.19 | -0.59 | -0.5025 | -0.67 | -0.025 | 0.962829 | 0.223319 | 0.159831 | 0.989223 | 0.997988 | 0.999507 |
| Srrm2_Q8BTI8_S1400s_1 | Srrm2_Q8E | 0.55 | 0.14 | -1.09 | -0.24 | -0.85 | 0.35 | -1.81 | -0.96 | 0.74 | -0.55 | -4.75 | -0.56 | 1.73 | -0.65 | -0.69 | -0.5 | -0.6575 | -1.12 | 0.1325 | 0.830983 | 0.222162 | 0.321259 | 0.989223 | 0.997988 | 0.999507 |
| Srrm2_Q8BTI8_S1419s_1 | Srrm2_Q8E | 4.96 | 1.86 | 1.82 | 1.6 | 2.45 | 4.83 | 2.79 | 2.1 | 1.65 | 3.5 | 2.16 | 3.35 | 1.54 | 0.74 | 3.78 | 2.66 | 0.4825 | 0.105 | -0.38 | 0.677678 | 0.585269 | 0.898134 | 0.989223 | 0.997988 | 0.996159 |
| Srrm2_Q8BTI8_S1453s_3 | Srrm2_Q8E | 1.58 | 2.53 | 0.96 | -0.4 | -3.02 | 2.41 | -1.06 | 0.38 | 3.06 | 0.35 | 1.53 | 0.11 | 0.77 | 1.65 | 1.2 | 0.94 | -1.49 | 0.095 | -0.0275 | 0.962722 | 0.204813 | 0.907004 | 0.989223 | 0.997988 | 0.999507 |
| Srrm2_Q8BTI8_S1455s_3 | Srrm2_Q8E | 1.58 | 2.53 | 0.96 | -0.4 | -3.02 | 2.41 | -1.06 | 0.38 | 3.06 | 0.35 | 1.53 | 0.11 | 0.77 | 1.65 | 1.2 | 0.94 | -1.49 | 0.095 | -0.0275 | 0.962722 | 0.204813 | 0.907004 | 0.989223 | 0.997988 | 0.999507 |
| Srrm2_Q8BTI8_S1557s_1 | Srrm2_Q8E | 0.19 | -0.74 | -0.57 | -0.94 | -1.85 | -0.42 | -1.34 | -0.79 | 1.59 | -1.04 | -3.04 | -0.71 | 2.11 | -1.79 | 0.7 | -1.12 | -0.585 | -0.285 | 0.49 | 0.549363 | 0.154957 | 0.743968 | 0.989223 | 0.997988 | 0.994666 |
| Srrm2_Q8BTI8_S1647s_2 | Srrm2_Q8E | 1.06 | 0.87 | 0.9 | 1.39 | -0.47 | 0.64 | 0.06 | 1.86 | 0.49 | 0.19 | 0.58 | 1.1 | 0.3 | -0.04 | 1.36 | 0.6 | -0.5325 | -0.465 | -0.5 | 0.176772 | 0.278903 | 0.11946 | 0.989223 | 0.997988 | 0.994666 |
| Srrm2_Q8BTI8_S1649s_1 | Srrm2_Q8E | 0.91 | 0.18 | 0.08 | -0.2 | -0.51 | 0.14 | -0.74 | -0.05 | 0.17 | -0.4 | -0.32 | -0.11 | 0.83 | -0.2 | 0.4 | -0.53 | -0.5325 | -0.4075 | -0.1175 | 0.765903 | 0.125823 | 0.203683 | 0.989223 | 0.997988 | 0.9971 |
| Srrm2_Q8BTI8_S1650s_1 | Srrm2_Q8E | 0.91 | 0.18 | -0.11 | 0.5 | -0.51 | 0.14 | -0.94 | 0.26 | 0.17 | -0.4 | 0.12 | 0.26 | 0.83 | -0.2 | 0.43 | -0.01 | -0.6325 | -0.3325 | -0.1075 | 0.75823 | 0.101755 | 0.288137 | 0.989223 | 0.997988 | 0.9971 |
| Srrm2_Q8BTI8_S1825s_1 | Srrm2_Q8E | -0.13 | -1.6 | -1.12 | -3.59 | -2.3 | 0.38 | -0.82 | -0.37 | 2.75 | -1.21 | -2.19 | -0.18 | 4.23 | -2.46 | 2.86 | -1.73 | 0.8325 | 1.4025 | 2.335 | 0.156769 | 0.314808 | 0.236542 | 0.989223 | 0.997988 | 0.994666 |
| Srrm2_Q8BTI8_S2075s_1 | Srrm2_Q8E | -0.32 | -0.3 | -0.53 | -0.69 | -1.14 | -0.34 | -0.73 | -0.53 | -1.34 | -1.15 | -1.17 | -0.54 | -0.16 | -0.36 | -0.41 | -1.53 | -0.225 | -0.59 | -0.155 | 0.661266 | 0.380203 | 0.046983 | 0.989223 | 0.997988 | 0.995671 |
| Srrm2_Q8BTI8_S2084s_1 | Srrm2_Q8E | 0.17 | 0.81 | 0.05 | -0.28 | 0.32 | -0.07 | -0.16 | -0.47 | 0.44 | 0.59 | 0.41 | 0.14 | 0.59 | 1.05 | -0.63 | -0.7 | -0.2825 | 0.2075 | -0.11 | 0.816716 | 0.362883 | 0.482417 | 0.989223 | 0.997988 | 0.999507 |
| Srrm2_Q8BTI8_S2224s_1 | Srrm2_Q8E | 0.65 | 1.08 | 0.73 | 1.04 | -0.24 | 0.58 | 0.95 | 1.15 | 0.15 | 0.48 | 1.41 | 0.92 | 0.12 | -0.3 | 0.93 | 0.98 | -0.265 | -0.135 | -0.4425 | 0.235208 | 0.43554 | 0.676472 | 0.989223 | 0.997988 | 0.994666 |
| Srrm2_Q8BTI8_S2351s_1 | Srrm2_Q8E | 0.59 | 0.07 | 0.43 | 0.02 | -0.6 | 0.15 | -0.35 | 0.48 | 0.23 | -0.5 | 0.13 | 0.12 | 0.54 | 0.22 | 0.03 | -0.59 | -0.3575 | -0.2825 | -0.2275 | 0.485234 | 0.255723 | 0.315502 | 0.989223 | 0.997988 | 0.994666 |
| Srrm2_Q8BTI8_S2360s_2 | Srrm2_Q8E | 1.01 | 0.35 | 0.17 | -0.07 | -0.49 | 0.38 | -0.58 | 0.59 | -0.31 | -0.53 | 0.68 | 0.03 | -0.43 | -0.13 | 0.07 | -0.58 | -0.39 | -0.3975 | -0.6325 | 0.073314 | 0.308304 | 0.288423 | 0.989223 | 0.997988 | 0.994666 |
| Srrm2_Q8BTI8_S2381s_1 | Srrm2_Q8E | 0.27 | -0.14 | -1.29 | -0.82 | -1.37 | -1.93 | -1.39 | -0.37 | 2.08 | -0.77 | -2.62 | -0.72 | 2.94 | -0.35 | -0.76 | -0.99 | -0.77 | -0.0125 | 0.705 | 0.422553 | 0.109608 | 0.989058 | 0.989223 | 0.997988 | 0.994666 |
| Srrm2_Q8BTI8_S2404s_1 | Srrm2_Q8E | 0.84 | 0.23 | 0.44 | 0.46 | -0.5 | 0.84 | -0.07 | 0.65 | -0.32 | 0.67 | 0.32 | 0.26 | 0.02 | -0.23 | 0.23 | 0.1 | -0.2625 | -0.26 | -0.4625 | 0.100582 | 0.450542 | 0.377373 | 0.989223 | 0.997988 | 0.994666 |
| Srrm2_Q8BTI8_S2535s_1 | Srrm2_Q8E | 1.35 | 0.66 | -0.06 | 0.05 | 0.24 | 0.79 | -0.32 | 0.13 | 0.45 | 0.39 | 0.17 | -0.32 | 0.93 | 0.69 | 0.01 | -0.28 | -0.29 | -0.3275 | -0.1625 | 0.704319 | 0.458983 | 0.392447 | 0.989223 | 0.997988 | 0.996159 |
| Srrm2_Q8BTI8_S2642s_1 | Srrm2_Q8E | 0.35 | 0.53 | -0.04 | -0.27 | -0.29 | -0.27 | -0.55 | 0.78 | 0.12 | -0.48 | 0.72 | 0.09 | 0.59 | 0.68 | 0.47 | -1.14 | -0.225 | -0.03 | 0.0075 | 0.986831 | 0.523392 | 0.927431 | 0.989223 | 0.997988 | 1 |
| Srrm2_Q8BTI8_S2646s_1 | Srrm2_Q8E | 0.07 | 0.12 | -1.17 | -1.57 | -0.09 | 0.03 | -1.05 | -1.05 | -1 | -0.7 | 0.02 | -1.38 | -0.46 | 0.57 | -0.83 | -1.5 | 0.0975 | -0.1275 | 0.0825 | 0.883466 | 0.839636 | 0.796319 | 0.995762 | 0.997988 | 0.999507 |
| Srrm2_Q8BTI8_S2648s_1 | Srrm2_Q8E | 1.31 | 0.66 | 0.42 | 0.47 | 0.04 | 0.76 | 0.01 | 0.7 | 0.1 | 0.38 | 0.02 | 0.22 | 0.62 | 0.53 | 0.42 | -0.21 | -0.3375 | -0.535 | -0.375 | 0.263316 | 0.290465 | 0.077063 | 0.989223 | 0.997988 | 0.994666 |
| Srrm2_Q8BTI8_S2656s_1 | Srrm2_Q8E | 0.53 | 0.28 | 0.4 | 0.02 | -0.84 | 0.27 | -1.16 | 0.02 | 0.81 | -0.54 | -0.74 | -0.28 | 1.56 | -1.01 | -0.11 | -0.54 | -0.735 | -0.495 | -0.3325 | 0.535864 | 0.063655 | 0.202998 | 0.989223 | 0.997988 | 0.994666 |
| Srrm2_Q8BTI8_S2656s_2 | Srrm2_Q8E | 0.75 | 0.27 | 0.6 | 0.25 | -0.81 | 0.34 | -1.06 | -0.01 | 1.51 | -0.83 | -0.82 | 0.1 | 1.43 | -0.86 | 0.11 | 0.12 | -0.8525 | -0.4775 | -0.2675 | 0.57066 | 0.035056 | 0.376297 | 0.989223 | 0.997988 | 0.995259 |
| Srrm2_Q8BTI8_S2660s_2 | Srrm2_Q8E | 0.75 | 0.27 | 0.6 | 0.25 | -0.81 | 0.34 | -1.06 | -0.01 | 1.51 | -0.83 | -0.82 | 0.1 | 1.43 | -0.86 | 0.11 | 0.12 | -0.8525 | -0.4775 | -0.2675 | 0.57066 | 0.035056 | 0.376297 | 0.989223 | 0.997988 | 0.995259 |
| Srrm2_Q8BTI8_S2681s_1 | Srrm2_Q8E | -1.48 | -3.15 | -2.91 | -2.33 | -2.59 | -2.02 | -2 | -1.55 | 3.85 | -3.55 | -1.56 | -1.35 | 5.18 | -2.74 | 3.23 | -3.33 | 0.4275 | 1.815 | 3.0525 | 0.124169 | 0.310573 | 0.221631 | 0.989223 | 0.997988 | 0.994666 |
| Srrm2_Q8BTI8_S333s_1 | Srrm2_Q8E | 0.04 | -0.49 | -0.53 | -0.67 | -1.74 | -0.27 | -1.31 | -0.82 | 0.66 | -1.24 | -0.88 | -0.47 | 1.68 | -1.46 | 0.14 | -1.18 | -0.6225 | -0.07 | 0.2075 | 0.754401 | 0.103976 | 0.870351 | 0.989223 | 0.997988 | 0.9971 |
| Srrm2_Q8BTI8_S349s_1 | Srrm2_Q8E | 1.03 | 0.42 | 0.6 | -0.08 | 0.1 | 0.46 | -0.08 | 0.67 | 0.18 | -0.06 | 0.46 | 0.18 | 1.11 | 0.57 | 0.68 | 0.04 | -0.205 | -0.3025 | 0.1075 | 0.758298 | 0.5091 | 0.319904 | 0.989223 | 0.997988 | 0.9971 |
| Srrm2_Q8BTI8_S349s_2 | Srrm2_Q8E | 0.81 | 0.32 | 0.38 | 0.19 | -0.35 | -0.24 | -0.44 | 0.59 | -0.38 | -0.54 | 0.15 | -0.18 | -1.67 | 0.1 | 0.06 | -0.39 | -0.535 | -0.6625 | -0.9 | 0.059891 | 0.098036 | 0.030276 | 0.989223 | 0.997988 | 0.994666 |
| Srrm2_Q8BTI8_S351s_1 | Srrm2_Q8E | 0.64 | 0.26 | 0.44 | 0.29 | 0.4 | 0.28 | -0.1 | 0.5 | 0.85 | -0.1 | 0.32 | 0.27 | 1.35 | 1.01 | 0.69 | -0.06 | -0.1375 | -0.0725 | 0.34 | 0.337891 | 0.556046 | 0.789658 | 0.989223 | 0.997988 | 0.994666 |
| Srrm2_Q8BTI8_S351s_2 | Srrm2_Q8E | 0.81 | 0.32 | 0.38 | 0.19 | -0.35 | -0.24 | -0.44 | 0.59 | -0.38 | -0.54 | 0.15 | -0.18 | -1.67 | 0.1 | 0.06 | -0.39 | -0.535 | -0.6625 | -0.9 | 0.059891 | 0.098036 | 0.030276 | 0.989223 | 0.997988 | 0.994666 |
| Srrm2_Q8BTI8_S396s_1 | Srrm2_Q8E | -0.8 | -0.57 | -0.57 | 0.21 | -0.6 | -1.02 | -1.25 | -1.55 | -1.34 | -0.75 | -2.04 | -0.89 | -0.66 | -0.4 | -0.33 | -0.64 | -0.6725 | -0.8225 | -0.075 | 0.801878 | 0.056114 | 0.05012 | 0.989223 | 0.997988 | 0.999507 |
| Srrm2_Q8BTI8_S454s_1 | Srrm2_Q8E | 1.53 | 0.6 | 0.54 | 0.39 | -0.67 | 1.38 | -0.78 | 0.08 | 2.03 | 0.33 | -0.53 | 0.88 | 2.8 | -0.6 | 0.67 | -0.07 | -0.7625 | -0.0875 | -0.065 | 0.92662 | 0.160029 | 0.873901 | 0.989223 | 0.997988 | 0.999507 |
| Srrm2_Q8BTI8_S778s_1 | Srrm2_Q8E | 1.69 | 0.56 | -0.46 | 0.25 | -0.23 | 1.74 | -0.69 | -0.09 | 1.28 | 0.67 | -1.07 | -0.13 | 0.74 | -0.18 | 0.8 | -0.41 | -0.3275 | -0.3225 | -0.2725 | 0.59761 | 0.600676 | 0.604768 | 0.989223 | 0.997988 | 0.995259 |
| Srrm2_Q8BTI8_S829s_2 | Srrm2_Q8E | 0.38 | -0.29 | -0.52 | -0.31 | -1.03 | 0.34 | 0 | 0.66 | 0.43 | -0.57 | 0.06 | -0.54 | -1.26 | 0.21 | 0.08 | -0.25 | 0.1775 | 0.03 | -0.12 | 0.761435 | 0.658376 | 0.928424 | 0.990434 | 0.997988 | 0.9971 |
| Srrm2_Q8BTI8_S829s_3 | Srrm2_Q8E | 0.49 | 0.79 | -1.01 | -0.46 | -0.46 | 0.01 | -0.94 | -0.39 | -0.64 | -0.01 | -0.37 | -0.58 | -2.68 | 0.05 | -0.45 | -0.92 | -0.3975 | -0.3525 | -0.9525 | 0.168792 | 0.370296 | 0.420285 | 0.989223 | 0.997988 | 0.994666 |
| Srrm2_Q8BTI8_S870s_1 | Srrm2_Q8E | 0.65 | -1.26 | -1.34 | -4.58 | -1.35 | 0.06 | -2.94 | -1.61 | 2.51 | -0.96 | -2.18 | -1.68 | 4.46 | -2.05 | 2.97 | -3 | 0.1725 | 1.055 | 2.2275 | 0.241023 | 0.872541 | 0.434728 | 0.996493 | 0.997988 | 0.994666 |
| Srrm2_Q8BTI8_S928s_1 | Srrm2_Q8E | 1.37 | 0 | -1.64 | -1.64 | -0.56 | 0.68 | -1.36 | -0.87 | 0.45 | 0.51 | -1.14 | -1.25 | 0.94 | 0.08 | -0.91 | -1.37 | -0.05 | 0.12 | 0.1625 | 0.83676 | 0.946344 | 0.877873 | 0.999187 | 0.997988 | 0.999507 |
| Srrm2_Q8BTI8_T1390t_1 | Srrm2_Q8E | 3.06 | 0.4 | 0.03 | 2.29 | -0.58 | 1.71 | -0.21 | 2.3 | 1.3 | 2.43 | 1.41 | 0.34 | 1.33 | 0.55 | 0.49 | -0.21 | -0.64 | -0.075 | -0.905 | 0.222986 | 0.475429 | 0.921214 | 0.989223 | 0.997988 | 0.994666 |
| Srrm2_Q8BTI8_T1428t_1 | Srrm2_Q8E | -0.53 | -0.46 | 0.49 | -0.54 | -1.77 | -0.21 | -1.51 | -0.67 | 0.32 | -1.49 | -1.09 | -0.59 | 1.43 | -2.85 | -0.16 | -0.78 | -0.78 | -0.4525 | -0.33 | 0.685519 | 0.088064 | 0.325503 | 0.989223 | 0.997988 | 0.996159 |
| Srrm2_Q8BTI8_T1448t_1 | Srrm2_Q8E | 0.59 | 0.22 | 0.16 | -0.2 | 0.14 | -0.05 | -0.58 | 0.17 | -0.98 | -0.81 | 0.24 | 0.02 | 0.73 | 0.31 | 0.43 | -0.32 | -0.2725 | -0.575 | 0.095 | 0.76744 | 0.335949 | 0.131393 | 0.989223 | 0.997988 | 0.9971 |
| Srrm2_Q8BTI8_T1812t_1 | Srrm2_Q8E | -0.2 | -0.49 | -0.34 | -1.18 | -0.19 | -0.84 | -0.34 | -0.37 | -1.25 | -1.38 | -0.35 | -0.34 | -0.37 | 0.11 | 0.02 | -1.16 | 0.1175 | -0.2775 | 0.2025 | 0.594868 | 0.685835 | 0.454595 | 0.994626 | 0.997988 | 0.995259 |
| Srrm2_Q8BTI8_T1880t_1 | Srrm2_Q8E | 2.4 | 2.1 | 1.91 | 1.54 | 1.48 | 1.17 | 0.95 | 1.67 | 1.33 | 0.73 | 0.23 | 1.24 | 2.39 | 0.89 | 2.15 | 1.89 | -0.67 | -1.105 | -0.1575 | 0.685457 | 0.035235 | 0.008299 | 0.989223 | 0.997988 | 0.996159 |
| Srrm2_Q8BTI8_T2056t_1 | Srrm2_Q8E | 0.36 | -0.1 | 0.12 | -0.23 | -0.18 | -0.27 | -0.27 | 0.25 | -0.49 | -0.97 | 0.01 | -0.24 | 0.46 | 0.09 | 0.15 | -0.72 | -0.155 | -0.46 | -0.0425 | 0.896369 | 0.527392 | 0.139031 | 0.989223 | 0.997988 | 0.999507 |
| Srrm2_Q8BTI8_T2350t_1 | Srrm2_Q8E | 0.48 | -0.11 | -0.13 | -0.1 | -0.68 | -0.22 | -0.46 | -0.03 | -0.4 | -0.68 | -0.44 | -0.32 | -0.13 | 0.38 | -0.24 | -1.16 | -0.3825 | -0.495 | -0.3225 | 0.395809 | 0.159782 | 0.069463 | 0.989223 | 0.997988 | 0.994666 |
| Srrm2_Q8BTI8_T2362t_1 | Srrm2_Q8E | -0.45 | -0.6 | -0.03 | -0.44 | -1.43 | -0.53 | -1.32 | 0.09 | -0.89 | -1.31 | -1.47 | -0.66 | 0.57 | -1.75 | -0.68 | -1.16 | -0.4175 | -0.7025 | -0.375 | 0.44601 | 0.27766 | 0.029026 | 0.989223 | 0.997988 | 0.994666 |
| Srrm2_Q8BTI8_T2362t_2 | Srrm2_Q8E | 0.04 | -0.41 | 0.08 | 0.02 | -0.59 | -0.56 | -0.73 | 0.42 | 0.73 | -1.01 | 0.49 | 0.05 | 0.18 | -0.27 | 0.03 | -0.3 | -0.2975 | 0.1325 | -0.0225 | 0.931448 | 0.346993 | 0.740719 | 0.989223 | 0.997988 | 0.999507 |
| Srrm2_Q8BTI8_T823t_1 | Srrm2_Q8E | 0.67 | 0.12 | 0.08 | 0.22 | -0.8 | 0.37 | 0.52 | 0.69 | 0.07 | -0.59 | 0.25 | -0.2 | 0.78 | -0.32 | 0.96 | -0.06 | -0.0775 | -0.39 | 0.0675 | 0.852794 | 0.830448 | 0.185786 | 0.995762 | 0.997988 | 0.999507 |
| Srrm2_Q8BTI8_T823t_2 | Srrm2_Q8E | 0.66 | -0.07 | -0.07 | -0.62 | -0.6 | 0.43 | -0.51 | 0.23 | -0.05 | -0.32 | -0.56 | -0.31 | -1.29 | 0.2 | -0.08 | -0.59 | -0.0875 | -0.285 | -0.415 | 0.334348 | 0.810822 | 0.37564 | 0.995762 | 0.997988 | 0.994666 |
| Srrm2_Q8BTI8_T823t_3 | Srrm2_Q8E | 0.49 | 0.79 | -1.01 | -0.46 | -0.46 | 0.01 | -0.94 | -0.39 | -0.64 | -0.01 | -0.37 | -0.58 | -2.68 | 0.05 | -0.45 | -0.92 | -0.3975 | -0.3525 | -0.9525 | 0.168792 | 0.370296 | 0.420285 | 0.989223 | 0.997988 | 0.994666 |
| Srrm2_Q8BTI8_T841t_2 | Srrm2_Q8E | 1.04 | 0.24 | 0.13 | -0.79 | -0.06 | 0.58 | -0.8 | 0 | -1 | 0.03 | -0.89 | -0.21 | -1.33 | 0.19 | -0.16 | -0.77 | -0.225 | -0.6725 | -0.6725 | 0.18404 | 0.611975 | 0.149673 | 0.989223 | 0.997988 | 0.994666 |
| Srrm2_Q8BTI8_T841t_3 | Srrm2_Q8E | 0.49 | 0.79 | -1.01 | -0.46 | -0.46 | 0.01 | -0.94 | -0.39 | -0.64 | -0.01 | -0.37 | -0.58 | -2.68 | 0.05 | -0.45 | -0.92 | -0.3975 | -0.3525 | -0.9525 | 0.168792 | 0.370296 | 0.420285 | 0.989223 | 0.997988 | 0.994666 |
| Srrm2_Q8BTI8_T877t_2 | Srrm2_Q8E | 0.49 | 0.43 | 0 | 0.09 | -1.27 | -0.02 | -1.24 | -0.36 | 2.56 | -1.27 | -1.89 | -0.81 | 1.15 | -1.56 | -0.56 | -0.25 | -0.975 | -0.605 | -0.5575 | 0.310776 | 0.01717 | 0.502235 | 0.989223 | 0.997988 | 0.994666 |
| Srrm2_Q8BTI8_T955t_1 | Srrm2_Q8E | -0.42 | -0.7 | -0.51 | -0.67 | -0.92 | -1.5 | -0.31 | -0.5 | -1.6 | -0.81 | -0.47 | -0.31 | -0.7 | -0.55 | -0.9 | -0.82 | -0.2325 | -0.2225 | -0.1675 | 0.487384 | 0.442387 | 0.495964 | 0.989223 | 0.997988 | 0.994666 |
| Srrm2_Q8BTI8_T973t_1 | Srrm2_Q8E | 0.81 | 1.09 | -0.05 | 0.04 | 0.19 | 0.34 | -0.02 | 0.23 | -0.04 | 0.14 | -0.23 | -0.26 | 0.26 | 0.45 | 0.1 | 0.2 | -0.2875 | -0.57 | -0.22 | 0.512182 | 0.366481 | 0.104881 | 0.989223 | 0.997988 | 0.994666 |
| Srrm3_Q80WV7_S562s_1 | Srrm3_Q80 | 0.02 | -0.44 | -0.36 | -0.18 | -1.4 | -0.86 | -0.64 | -0.2 | -0.4 | -1.05 | -0.42 | -0.6 | 0.37 | 0.2 | 0.23 | -0.87 | -0.535 | -0.3775 | 0.2225 | 0.517365 | 0.096337 | 0.161287 | 0.989223 | 0.997988 | 0.994666 |
| Srrm3_Q80WV7_T352t_1 | Srrm3_Q80 | -0.13 | -0.4 | -0.65 | -0.93 | -1.13 | -1.08 | -1.65 | -0.48 | -1.21 | -0.77 | -0.18 | -1.28 | 0.67 | 0.04 | -0.4 | -1.24 | -0.5575 | -0.3325 | 0.295 | 0.498786 | 0.099932 | 0.327351 | 0.989223 | 0.997988 | 0.994666 |
| Srrm4_Q8BKA3_S271s_1 | Srrm4_Q8E | 0.48 | -0.02 | 0.07 | 0.29 | -0.82 | -0.17 | -1.19 | 0.49 | -0.64 | -0.03 | 0.98 | 0.18 | 0.59 | 1.02 | 0.39 | -0.68 | -0.6275 | -0.0825 | 0.125 | 0.748721 | 0.122475 | 0.820452 | 0.989223 | 0.997988 | 0.9971 |
| Srrm4_Q8BKA3_S285s_1 | Srrm4_Q8E | 0.19 | -0.66 | -1.12 | -2.01 | -0.91 | -1.57 | -2.69 | -1.17 | 0.96 | 1.16 | -2.82 | -3.21 | 1.15 | 0.39 | -1.9 | -1.69 | -0.685 | -0.0775 | 0.3875 | 0.623128 | 0.229273 | 0.944434 | 0.989223 | 0.997988 | 0.995259 |
| Srrm4_Q8BKA3_S496s_1 | Srrm4_Q8E | 0.03 | -0.21 | 0.51 | -0.07 | -1.36 | 0.22 | -1.09 | 0.73 | 0.01 | -0.16 | 0.53 | -0.07 | 0.66 | -0.25 | 0.31 | -0.17 | -0.44 | 0.0125 | 0.0725 | 0.818194 | 0.376338 | 0.963706 | 0.989223 | 0.997988 | 0.999507 |
| Srrt_Q99MR6_S4s_1 | Srrt_Q99M | -0.17 | 0.39 | 0.67 | 0.75 | -2.46 | 1.09 | -1.5 | -0.45 | 2.23 | -0.06 | 0.26 | 0.28 | 2.31 | -1.38 | -0.52 | 0.42 | -1.24 | 0.2675 | -0.2025 | 0.7814 | 0.10014 | 0.613246 | 0.989223 | 0.997988 | 0.998226 |
| Srrt_Q99MR6_S67s_1 | Srrt_Q99M | -2.26 | -2.7 | -2.53 | -3.78 | -2.48 | -3.67 | -2.79 | -3.35 | -3.69 | -3.13 | -1.7 | -2.81 | -1.54 | -1.98 | -2.13 | -2.64 | -0.255 | -0.015 | 0.745 | 0.092528 | 0.537268 | 0.976206 | 0.989223 | 0.997988 | 0.994666 |
| Srrt_Q99MR6_T543t_1 | Srrt_Q99M | -0.57 | -0.46 | -0.61 | -0.16 | -0.77 | -0.69 | -0.21 | 0.02 | -0.7 | -0.5 | -1.26 | -0.3 | -0.54 | 0.34 | -0.38 | 0.04 | 0.0375 | -0.24 | 0.315 | 0.296007 | 0.886441 | 0.403109 | 0.996493 | 0.997988 | 0.994666 |
| Srsf1_H7BX95_S249s_1 | Srsf1_H7B | 8.93 | 3.04 | 3.73 | 0.73 | 3.42 | 8.08 | 5.04 | 3.69 | -1.06 | 5.8 | -0.78 | 3.56 | -4.48 | -0.23 | 5.68 | 5.27 | 0.95 | -2.2275 | -2.5475 | 0.328742 | 0.587431 | 0.303569 | 0.989223 | 0.997988 | 0.994666 |
| Srsf10_Q3TFP0_S133s_1 | Srsf10_Q3T | 3.67 | 1.43 | 0.64 | 1.28 | 1.85 | 3.36 | 0.82 | 1.81 | -2.96 | 2.25 | 2.66 | 1.21 | -1.78 | -0.87 | 0.98 | 0.81 | 0.205 | -0.965 | -1.97 | 0.037219 | 0.782159 | 0.453732 | 0.995762 | 0.997988 | 0.994666 |
| Srsf11_E9Q6E5_S462s_1 | Srsf11_E9C | 0.4 | -0.01 | 0.86 | -1.09 | 0.4 | 0.07 | 0.09 | 0.23 | -1.44 | -0.25 | -0.93 | 0.75 | -1.11 | -1.52 | 1.55 | 1.08 | 0.1575 | -0.5075 | -0.04 | 0.958802 | 0.698763 | 0.391917 | 0.994893 | 0.997988 | 0.999507 |
| Srsf11_E9Q6E5_S477s_1 | Srsf11_E9C | -0.35 | -2.27 | -1.22 | -1.84 | -2.17 | -1.22 | -1.54 | -1.26 | -1.53 | -2.12 | -0.21 | -1.13 | 0.25 | -1.16 | -0.49 | -0.98 | -0.1275 | 0.1725 | 0.825 | 0.121092 | 0.773198 | 0.748942 | 0.995762 | 0.997988 | 0.994666 |
| Srsf11_E9Q6E5_S510s_1 | Srsf11_E9C | 0.17 | -1.02 | -0.33 | -0.95 | -1.19 | -0.88 | -0.46 | -0.6 | -1.32 | -0.54 | -0.35 | -0.51 | -1.39 | -2.02 | 0.73 | -0.29 | -0.25 | -0.1475 | -0.21 | 0.730948 | 0.45967 | 0.687205 | 0.989223 | 0.997988 | 0.9971 |
| Srsf2_Q62093_S191s_1 | Srsf2_Q62 | -0.1 | -0.12 | 0.9 | 0.38 | -1.23 | -0.18 | -0.05 | 0.85 | 0.38 | -1.2 | -0.22 | 0.5 | 1.53 | -0.96 | 1.38 | 1.1 | -0.4175 | -0.4 | 0.4975 | 0.398741 | 0.37098 | 0.376446 | 0.989223 | 0.997988 | 0.994666 |
| Srsf2_Q62093_S212s_1 | Srsf2_Q62 | -0.16 | 0.01 | -1.49 | -2.34 | -1.76 | 0.09 | -1.59 | -1.39 | 0.58 | -0.1 | -1 | -0.86 | 1.42 | 0.26 | -0.16 | 0.07 | -0.1675 | 0.65 | 1.3925 | 0.041864 | 0.789063 | 0.303229 | 0.995762 | 0.997988 | 0.994666 |
| Srsf2_Q62093_S220s_1 | Srsf2_Q62 | -3.23 | -2.67 | -0.7 | -1.67 | -2.43 | -2.8 | -1.06 | -1.31 | -1.91 | -2.51 | 0.62 | -0.82 | -0.59 | -3.08 | -0.69 | 0 | 0.1675 | 0.9125 | 0.9775 | 0.229259 | 0.788518 | 0.266621 | 0.995762 | 0.997988 | 0.994666 |
| Srsf2_Q62093_S221s_1 | Srsf2_Q62 | -3.23 | -2.67 | -0.7 | -1.67 | -2.43 | -2.8 | -1.06 | -1.31 | -1.91 | -2.51 | 0.62 | -0.82 | -0.59 | -3.08 | -0.69 | 0 | 0.1675 | 0.9125 | 0.9775 | 0.229259 | 0.788518 | 0.266621 | 0.995762 | 0.997988 | 0.994666 |
| Srsf2_Q62093_S26s_1 | Srsf2_Q62 | 0.85 | -0.36 | 0.18 | -0.2 | -1.24 | -0.04 | -1.48 | -0.45 | 0.24 | -0.07 | 0.04 | -0.73 | 0.68 | -0.52 | 0.08 | -0.85 | -0.92 | -0.2475 | -0.27 | 0.532017 | 0.047196 | 0.491515 | 0.989223 | 0.997988 | 0.994666 |
| Srsf2_Q62093_T25t_1 | Srsf2_Q62 | 0.72 | -0.4 | 1.04 | 1.64 | -1 | -0.15 | -0.15 | 1.12 | 0.17 | -0.21 | 0.95 | 0.31 | 0.64 | -0.47 | 1.59 | 0.47 | -0.795 | -0.445 | -0.1925 | 0.729746 | 0.173165 | 0.354702 | 0.989223 | 0.997988 | 0.9971 |
| Srsf3_P84104_S108s_1 | Srsf3_P841 | 0.52 | -0.94 | -0.39 | -0.85 | -1.27 | -0.55 | -1.15 | -1.28 | 2.01 | -0.86 | -4.18 | -2.47 | 2.5 | -2.28 | 0.04 | -1.2 | -0.6475 | -0.96 | 0.18 | 0.848698 | 0.106278 | 0.428927 | 0.989223 | 0.997988 | 0.999507 |
| Srsf4_Q542V3_S455s_1 | Srsf4_Q542 | 0.35 | 1.02 | 0.57 | -0.22 | -1.13 | -0.12 | -0.56 | -0.18 | -0.3 | -0.71 | 0.32 | -0.31 | 0.18 | 0.38 | -0.09 | -0.2 | -0.9275 | -0.68 | -0.3625 | 0.28796 | 0.023471 | 0.077613 | 0.989223 | 0.997988 | 0.994666 |
| Srsf7_Q8BL97_S181s_1 | Srsf7_Q8BI | 1.7 | 1.76 | 1.23 | 0.86 | 1.5 | 1.62 | 0.89 | 1.32 | 1.02 | 0.71 | 0.82 | 0.89 | 1.46 | 1.7 | 1.36 | 0.96 | -0.055 | -0.5275 | -0.0175 | 0.955475 | 0.851412 | 0.081865 | 0.996349 | 0.997988 | 0.999507 |
| Srsf7_Q8BL97_S233s_1 | Srsf7_Q8BI | 2.84 | 0.89 | 2.01 | 2.81 | -0.24 | 2.1 | 1.34 | 2.49 | 2.51 | 1.58 | 1.15 | 1.94 | 2.35 | 0.53 | 2.56 | 1.66 | -0.715 | -0.3425 | -0.3625 | 0.544368 | 0.301056 | 0.505499 | 0.989223 | 0.997988 | 0.994666 |
| Srsf9_F6RTE1_S141s_1 | Srsf9_F6RT | 0.26 | -0.58 | -0.41 | -0.78 | -2.55 | 0.25 | -2.07 | -1.04 | 1.29 | -0.72 | -1.42 | -1.21 | 1.71 | -2.02 | -0.16 | -1.31 | -0.975 | -0.1375 | -0.0675 | 0.928262 | 0.124259 | 0.819826 | 0.989223 | 0.997988 | 0.999507 |
| Ssbp2_Q9CYZ8_S291s_1 | Ssbp2_Q9C | -1.56 | -0.92 | -1.84 | -1.98 | -1.88 | -1.62 | -1.56 | -2.2 | -1.11 | -1.2 | -1.22 | -1.74 | -0.61 | -1.14 | -2.01 | -1.81 | -0.24 | 0.2575 | 0.1825 | 0.652069 | 0.433014 | 0.414497 | 0.989223 | 0.997988 | 0.995259 |
| Ssbp2_Q9CYZ8_T303t_1 | Ssbp2_Q9C | -0.8 | -0.51 | -0.27 | -0.05 | -0.89 | -1.28 | -0.08 | -0.41 | -0.02 | -0.82 | 0.56 | 0.2 | 0.32 | -1.15 | -0.07 | 0.26 | -0.2575 | 0.3875 | 0.2475 | 0.52848 | 0.432931 | 0.284025 | 0.989223 | 0.997988 | 0.994666 |
| Ssbp3_B1AS37_S327s_1 | Ssbp3_B1A | -0.7 | -0.1 | -0.06 | -0.66 | -0.98 | -1.64 | 0.31 | 0.04 | -1.32 | -1.04 | -0.67 | -0.39 | -0.73 | -1.06 | 0.16 | -0.01 | -0.1875 | -0.475 | -0.03 | 0.933875 | 0.67984 | 0.144397 | 0.993464 | 0.997988 | 0.999507 |
| Ssbp4_Q3U4B1_S320s_1 | Ssbp4_Q3U | -0.88 | -0.53 | -1.04 | -1.38 | -0.99 | -0.57 | 0.26 | 0.52 | -0.33 | -1.17 | 0.16 | -0.23 | -0.59 | 0.21 | -0.95 | -0.31 | 0.7625 | 0.565 | 0.5475 | 0.130273 | 0.071725 | 0.129777 | 0.989223 | 0.997988 | 0.994666 |
| Ssbp4_Q3U4B1_T333t_1 | Ssbp4_Q3U | -0.23 | -0.29 | 0.8 | 0.19 | 0.27 | 0.81 | 0.1 | 0.65 | 0.48 | 0.22 | -0.72 | 1.21 | 0.34 | 0.28 | 0.16 | -0.19 | 0.34 | 0.18 | 0.03 | 0.925998 | 0.298028 | 0.692111 | 0.989223 | 0.997988 | 0.999507 |
| Ssfa2_Q922B9_S667s_1 | Ssfa2_Q92 | 0.31 | 0.42 | 0.11 | -0.45 | 1.31 | 0.41 | -0.33 | -0.04 | 0.18 | 0.4 | -1.45 | -0.36 | -0.01 | 0.24 | -0.85 | -0.36 | 0.24 | -0.405 | -0.3425 | 0.32769 | 0.545705 | 0.369881 | 0.989223 | 0.997988 | 0.994666 |
| Ssh2_Q5SW75_S1216s_1 | Ssh2_Q5SV | -1.01 | -0.24 | -1.36 | -0.78 | -0.28 | 0.2 | -0.44 | -1.08 | -0.28 | -1.39 | -0.99 | -1.85 | 0.07 | 0.34 | -1.79 | -0.91 | 0.4475 | -0.28 | 0.275 | 0.591733 | 0.226158 | 0.493769 | 0.989223 | 0.997988 | 0.995259 |
| Ssh2_Q5SW75_S17s_1 | Ssh2_Q5SV | 0.37 | 0.26 | -1.85 | 0.49 | -0.41 | 0.15 | -1.61 | -0.37 | 0.83 | 0.08 | 1.6 | 0.04 | 0.93 | 0.36 | 0.25 | -0.69 | -0.3775 | 0.82 | 0.395 | 0.511774 | 0.533416 | 0.201857 | 0.989223 | 0.997988 | 0.994666 |
| Ssh2_Q5SW75_S25s_1 | Ssh2_Q5SV | 0.17 | -0.04 | -0.48 | 0.07 | -0.7 | 1.07 | -1.15 | 0.24 | 0.91 | 0.66 | 1.76 | -0.31 | -0.2 | 1.07 | -0.1 | 0.43 | -0.065 | 0.825 | 0.37 | 0.309053 | 0.891396 | 0.084659 | 0.996493 | 0.997988 | 0.994666 |
| Ssh2_Q5SW75_S25s_2 | Ssh2_Q5SV | 0.86 | 0.63 | 1.23 | 1.65 | 1.04 | -0.7 | 0.44 | 2.12 | 0.14 | 0.46 | 3.1 | 0.01 | 0.54 | 0.03 | 1.85 | 2.55 | -0.3675 | -0.165 | 0.15 | 0.793349 | 0.521272 | 0.811009 | 0.989223 | 0.997988 | 0.999507 |
| Ssh2_Q5SW75_S461s_1 | Ssh2_Q5SV | -1.97 | -2.78 | -1.71 | -0.87 | -1.92 | -1.6 | -2.2 | -1.55 | -2.37 | -2.57 | -0.26 | -2.3 | -0.26 | -1.11 | -1.8 | -1.41 | 0.015 | -0.0425 | 0.6875 | 0.180677 | 0.970375 | 0.944558 | 0.999187 | 0.997988 | 0.994666 |
| Ssh2_Q5SW75_S9s_1 | Ssh2_Q5SV | -1.09 | -2.39 | -1 | -1.24 | -1.24 | -2.32 | -1.41 | -2.44 | -1.94 | -2.05 | -1.42 | -2.39 | -0.66 | -4.01 | -0.57 | -1.54 | -0.4225 | -0.52 | -0.265 | 0.730271 | 0.331208 | 0.198015 | 0.989223 | 0.997988 | 0.9971 |
| Ssh2_Q5SW75_T1422t_1 | Ssh2_Q5SV | -1.43 | 0.41 | -0.01 | -1.09 | 0.15 | -1.36 | -0.23 | -0.23 | -0.85 | 0.35 | -0.3 | -0.75 | -0.46 | -0.42 | -0.01 | -0.35 | 0.1125 | 0.1425 | 0.22 | 0.618505 | 0.821742 | 0.770024 | 0.995762 | 0.997988 | 0.995259 |
| Ssh3_Q8K330_S643s_1 | Ssh3_Q8K3 | 0 | -0.32 | 0.63 | 1.06 | -1.92 | 0.4 | -0.81 | 0 | 1.44 | -0.15 | 0.81 | 0.38 | 1.41 | -1.21 | 0.05 | 0.37 | -0.925 | 0.2775 | -0.1875 | 0.743813 | 0.113323 | 0.534762 | 0.989223 | 0.997988 | 0.9971 |
| Ssh3_Q8K330_S9s_1 | Ssh3_Q8K3 | -0.92 | 0.23 | -0.64 | -0.09 | 0.41 | -0.68 | -1.27 | 0.18 | -0.22 | -0.71 | 0.14 | -0.97 | 1.08 | -0.06 | 0.19 | -1.2 | 0.015 | -0.085 | 0.3575 | 0.485934 | 0.97274 | 0.817491 | 0.999187 | 0.997988 | 0.994666 |
| Ssr3_Q9DCF9_S105s_1 | Ssr3_Q9DC | -1.41 | -0.92 | -1.19 | -1.94 | -1.08 | 0.26 | -1.06 | -1.6 | 0.12 | -1.77 | -2.08 | -1.63 | 1.44 | -1.24 | -0.92 | -1.62 | 0.495 | 0.025 | 0.78 | 0.24996 | 0.263958 | 0.960688 | 0.989223 | 0.997988 | 0.994666 |
| Sst_P60041_S92s_1 | Sst_P6004 | 0.58 | 0.31 | 0.67 | -0.11 | 0.77 | 0.17 | -0.57 | 0.14 | 0.48 | 0.46 | 1.71 | 0.26 | 1.1 | 0.64 | 1.46 | 1.34 | -0.235 | 0.365 | 0.7725 | 0.02972 | 0.487353 | 0.348432 | 0.989223 | 0.997988 | 0.994666 |
| Sstr3_P30935_S245s_1 | Sstr3_P309 | -1.15 | -0.09 | -0.35 | -1.1 | -0.87 | -0.24 | -0.29 | -0.34 | -0.54 | -0.8 | -0.8 | -1.15 | -1.33 | -0.46 | -0.85 | -0.6 | 0.2375 | -0.15 | -0.1375 | 0.699751 | 0.463801 | 0.643722 | 0.989223 | 0.997988 | 0.996159 |
| Stac2_Q8R1B0_S221s_1 | Stac2_Q8R | -0.42 | 0.06 | -0.55 | -0.59 | 0.93 | 0.17 | -0.61 | -0.46 | 0.29 | 0.64 | 0.53 | 0.45 | 1 | 1.22 | -0.5 | 0.01 | 0.3825 | 0.8525 | 0.8075 | 0.085094 | 0.320644 | 0.006408 | 0.989223 | 0.997988 | 0.994666 |
| Stac2_Q8R1B0_S228s_1 | Stac2_Q8R | -0.15 | 0.45 | -1.26 | -0.86 | -0.11 | -0.52 | -1.06 | -1.05 | -1.35 | -0.67 | -2.04 | -0.84 | -0.21 | -1.26 | -1.15 | -1.01 | -0.23 | -0.77 | -0.4525 | 0.31669 | 0.587283 | 0.124699 | 0.989223 | 0.997988 | 0.994666 |
| Stac2_Q8R1B0_S274s_1 | Stac2_Q8R | -0.77 | -0.79 | 0.23 | -0.24 | -0.59 | -0.28 | -0.02 | -0.38 | 0.5 | -0.86 | -0.97 | 0.18 | 1.11 | -0.91 | 0 | 0.06 | 0.075 | 0.105 | 0.4575 | 0.335763 | 0.800763 | 0.807282 | 0.995762 | 0.997988 | 0.994666 |
| Stam_Q3UGN9_S156s_1 | Stam_Q3U | -0.3 | -0.51 | -0.63 | -0.71 | -0.07 | -0.52 | -0.82 | -0.83 | -0.84 | -0.77 | 0.61 | -0.87 | -0.74 | 0.26 | -1.64 | -1.6 | -0.0225 | 0.07 | -0.3925 | 0.388526 | 0.92923 | 0.85241 | 0.997639 | 0.997988 | 0.994666 |
| Stambpl1_Q76N33_S76s_1 | Stambpl1_ | -0.12 | 0.07 | -0.08 | -0.02 | 0.09 | -0.3 | 0.18 | -0.11 | -0.23 | 0.07 | -0.2 | -0.16 | -0.25 | 0.06 | -0.18 | -0.07 | 0.0025 | -0.0925 | -0.0725 | 0.75516 | 0.990614 | 0.662719 | 0.999187 | 0.997988 | 0.9971 |
| Stard10_G3UY87_S276s_1 | Stard10_G | -0.06 | 0.09 | -0.24 | -0.24 | -1.11 | -0.58 | -0.45 | -0.36 | 0.01 | 0.01 | 0.74 | -0.66 | -0.01 | -0.31 | 0.1 | 0.09 | -0.5125 | 0.1375 | 0.08 | 0.745402 | 0.060884 | 0.672629 | 0.989223 | 0.997988 | 0.9971 |
| Stard13_F8WIY7_S572s_1 | Stard13_F8 | -0.72 | -0.4 | -0.94 | -1.06 | -0.53 | -0.39 | -1.35 | -1.17 | 0.17 | -0.06 | -0.35 | -0.89 | 0.48 | -0.58 | -1.87 | -1.05 | -0.08 | 0.4975 | 0.025 | 0.959003 | 0.790478 | 0.130414 | 0.995762 | 0.997988 | 0.999507 |
| Stard7_F7BBA2_S392s_1 | Stard7_F7E | -0.31 | -0.73 | -0.7 | -1.14 | -0.98 | -0.1 | -1.11 | -1.42 | 1.42 | -0.27 | -1.89 | -1.1 | 1.81 | -1.48 | -0.71 | -0.77 | -0.1825 | 0.26 | 0.4325 | 0.520826 | 0.591132 | 0.694919 | 0.989223 | 0.997988 | 0.994666 |
| Stau2_E2QRQ3_S447s_1 | Stau2_E2Q | -0.67 | -1.47 | -0.64 | 0.1 | -0.58 | -0.24 | -0.51 | 0.49 | -0.75 | -0.89 | 1.6 | -0.11 | 0.55 | 0.53 | 0.6 | 0 | 0.46 | 0.6325 | 1.09 | 0.014358 | 0.256628 | 0.307269 | 0.989223 | 0.997988 | 0.994666 |
| Steap2_Q8BWB6_S482s_1 | Steap2_Q8 | 0.71 | 1.06 | 0.62 | 0.71 | 1.09 | 0.52 | 0.86 | 0.66 | 0.22 | 0.75 | 0.47 | 0.1 | 0.1 | 0.68 | 0.87 | 0.52 | 0.0075 | -0.39 | -0.2325 | 0.406012 | 0.97413 | 0.143009 | 0.999187 | 0.997988 | 0.994666 |
| Stim1_P70302_S575s_1 | Stim1_P70 | 1.06 | -0.1 | -0.92 | -0.09 | 0.52 | 1.01 | -0.29 | -0.02 | 0.22 | 0.28 | 0.33 | -0.41 | -0.42 | 0.22 | -0.97 | -0.87 | 0.3175 | 0.1175 | -0.4975 | 0.304534 | 0.498888 | 0.785097 | 0.989223 | 0.997988 | 0.994666 |
| Stim1_P70302_S660s_1 | Stim1_P70 | -1.07 | -2.49 | -0.24 | -0.25 | -1.18 | -1.68 | -0.47 | -0.43 | -0.93 | -2.42 | 0.07 | -0.59 | -2.8 | -0.96 | -0.08 | -0.51 | 0.0725 | 0.045 | -0.075 | 0.9162 | 0.895114 | 0.94685 | 0.996493 | 0.997988 | 0.999507 |
| Stim1_P70302_S660s_2 | Stim1_P70 | -0.02 | -0.49 | 0.05 | 1 | -1.02 | -0.79 | 0.25 | 0.64 | -0.51 | -1.12 | 0.49 | -0.12 | 0.43 | 0.5 | 0.64 | 1.06 | -0.365 | -0.45 | 0.5225 | 0.174953 | 0.445229 | 0.325188 | 0.989223 | 0.997988 | 0.994666 |
| Stim1_P70302_S667s_2 | Stim1_P70 | -0.02 | -0.49 | 0.11 | 1.19 | -1.02 | -0.79 | 0.69 | 1.14 | -0.51 | -1.12 | 0.6 | 0.24 | 0.43 | 0.5 | 0.99 | 1.48 | -0.1925 | -0.395 | 0.6525 | 0.150483 | 0.739228 | 0.432236 | 0.995762 | 0.997988 | 0.994666 |
| Stim1_P70302_S668s_1 | Stim1_P70 | -0.38 | 0.07 | -0.28 | 0.02 | -0.07 | -1.46 | 0.02 | -0.12 | -1.04 | -0.48 | -0.91 | -0.25 | -0.15 | 0.66 | 0.63 | -0.08 | -0.265 | -0.5275 | 0.4075 | 0.201739 | 0.474443 | 0.078124 | 0.989223 | 0.997988 | 0.994666 |
| Stim2_P83093_S523s_1 | Stim2_P83 | 0.94 | -2.2 | -0.98 | 0.3 | 0.5 | 0.61 | -0.04 | 0.56 | -0.71 | -1.37 | -0.2 | -0.38 | -1.82 | -0.26 | -1.46 | -0.59 | 0.8925 | -0.18 | -0.5475 | 0.443874 | 0.181917 | 0.789246 | 0.989223 | 0.997988 | 0.994666 |
| Stim2_P83093_S609s_1 | Stim2_P83 | 0.32 | 0.76 | 0.08 | 0.02 | 0.57 | -0.27 | 0.54 | -0.12 | -0.23 | 0.22 | 0.08 | -0.07 | 0.05 | 0.65 | 0.41 | 0.31 | -0.115 | -0.295 | 0.06 | 0.832592 | 0.702665 | 0.273819 | 0.995437 | 0.997988 | 0.999507 |
| Stim2_P83093_S631s_1 | Stim2_P83 | 0.73 | 0.36 | 1.36 | 0.76 | 1.17 | 0.86 | 1.06 | 1.46 | -1.21 | 0.4 | 1.46 | 0.9 | -3.75 | 0.46 | 1.49 | 1.24 | 0.335 | -0.415 | -0.9425 | 0.391058 | 0.247437 | 0.46873 | 0.989223 | 0.997988 | 0.994666 |
| Stim2_P83093_S640s_1 | Stim2_P83 | -1.91 | -1.05 | -0.94 | -1.97 | -0.43 | -2.14 | -1.63 | -1.59 | -2.65 | -1.95 | -0.04 | -1.05 | -1.06 | -0.45 | -0.97 | -1 | 0.02 | 0.045 | 0.5975 | 0.106286 | 0.962738 | 0.937909 | 0.999187 | 0.997988 | 0.994666 |
| Stim2_P83093_S650s_1 | Stim2_P83 | -0.08 | 0.07 | -0.01 | -0.17 | 0.43 | -0.23 | 0.01 | -0.46 | -0.34 | -0.04 | -0.04 | -0.11 | 0.1 | 0.3 | -0.06 | -0.08 | -0.015 | -0.085 | 0.1125 | 0.638626 | 0.952504 | 0.691426 | 0.999187 | 0.997988 | 0.995259 |
| Stip1_Q60864_S481s_1 | Stip1_Q60 | 6.79 | 1.67 | 4.22 | 1.3 | 1.01 | 7.97 | 4.5 | 3.58 | 0.58 | 5 | 0.47 | 4.6 | 0.89 | -0.9 | 4.35 | 5.15 | 0.77 | -0.8325 | -1.1225 | 0.49959 | 0.64134 | 0.59413 | 0.989223 | 0.997988 | 0.994666 |
| Stk10_O55098_T950t_1 | Stk10_O55 | -0.28 | -0.44 | -0.11 | 0 | -0.68 | -0.84 | -0.24 | -0.54 | 0.07 | -0.28 | -0.25 | -0.4 | -0.24 | 0.46 | -0.66 | 0.03 | -0.3675 | -0.0075 | 0.105 | 0.734633 | 0.138633 | 0.974272 | 0.989223 | 0.997988 | 0.9971 |
| Stk11ip_Q3TAA7_S388s_1 | Stk11ip_Q3 | 0.16 | -0.22 | -1.83 | -0.03 | -0.65 | -0.56 | 0.13 | -0.4 | -0.16 | 0.04 | -0.89 | -1.15 | -0.4 | -0.02 | -0.27 | -0.99 | 0.11 | -0.06 | 0.06 | 0.900226 | 0.809343 | 0.905502 | 0.995762 | 0.997988 | 0.999507 |
| Stk24_Q99KH8_S4s_1 | Stk24_Q99 | 0.22 | 1.04 | 0.08 | 0.31 | 1.2 | 0.02 | 0.69 | 0.23 | -0.07 | 0.38 | 0.46 | 0.09 | -0.13 | 0.66 | 0.08 | 0.5 | 0.1225 | -0.1975 | -0.135 | 0.679673 | 0.722028 | 0.503991 | 0.995437 | 0.997988 | 0.996159 |
| Stk26_Q99JT2_S4s_1 | Stk26_Q99 | 0.35 | -0.04 | 0.91 | 0.03 | 0.81 | 0.34 | 0.84 | 0.34 | 0.57 | 0.72 | 0.48 | 0.15 | 0.09 | 0.38 | 0.77 | -0.54 | 0.27 | 0.1675 | -0.1375 | 0.711338 | 0.361125 | 0.569602 | 0.989223 | 0.997988 | 0.996966 |
| Stk32c_Q8QZV4_S10s_2 | Stk32c_Q8 | -1.11 | -0.23 | -1.24 | -1.56 | -0.25 | -0.93 | -1.09 | -0.75 | -0.52 | -0.62 | 0.82 | -0.28 | -0.18 | 0.4 | -1.02 | -0.97 | 0.28 | 0.885 | 0.5925 | 0.19593 | 0.422475 | 0.062575 | 0.989223 | 0.997988 | 0.994666 |
| Stk32c_Q8QZV4_S18s_1 | Stk32c_Q8 | -0.17 | 0.14 | 0.21 | 0.21 | 0.16 | -0.34 | -0.05 | 0.05 | -0.34 | -0.1 | 0.01 | -0.14 | 0.71 | -0.28 | 0.12 | -0.16 | -0.1425 | -0.24 | -3.9E-16 | 1 | 0.528129 | 0.298087 | 0.989223 | 0.997988 | 1 |
| Stk32c_Q8QZV4_S18s_2 | Stk32c_Q8 | -1.11 | -0.23 | -1.24 | -1.56 | -0.25 | -0.93 | -1.09 | -0.75 | -0.52 | -0.62 | 0.82 | -0.28 | -0.18 | 0.4 | -1.02 | -0.97 | 0.28 | 0.885 | 0.5925 | 0.19593 | 0.422475 | 0.062575 | 0.989223 | 0.997988 | 0.994666 |
| Stk32c_Q8QZV4_S32s_1 | Stk32c_Q8 | -0.18 | 0.5 | -0.21 | -0.4 | -0.78 | 0.8 | -0.07 | 0 | -0.14 | 0.64 | -0.55 | -0.83 | 0.46 | 0.04 | -0.43 | -0.28 | 0.06 | -0.1475 | 0.02 | 0.950681 | 0.871872 | 0.698817 | 0.996493 | 0.997988 | 0.999507 |
| Stk32c_Q8QZV4_S69s_1 | Stk32c_Q8 | -2.43 | -1.73 | -1.5 | -1.91 | -1.77 | -1.81 | -1.28 | -1.77 | -1.35 | -1.97 | -0.1 | -0.75 | -0.21 | -1.49 | -1.91 | -1.7 | 0.235 | 0.85 | 0.565 | 0.206195 | 0.400809 | 0.076091 | 0.989223 | 0.997988 | 0.994666 |
| Stk38_Q91VJ4_S281s_1 | Stk38_Q91 | -1.03 | -0.68 | -1.05 | -0.52 | -0.11 | -1.11 | 0.52 | -0.49 | -1.11 | -0.25 | -0.32 | -0.83 | -1.32 | -0.2 | -0.34 | -0.38 | 0.5225 | 0.1925 | 0.26 | 0.437705 | 0.17316 | 0.511804 | 0.989223 | 0.997988 | 0.994666 |
| Stk39_Q9Z1W9_S382s_1 | Stk39_Q9Z | -1.85 | -1.72 | -1.19 | -1.01 | -1.58 | -2.4 | -0.86 | -0.67 | -1.02 | -2.25 | -0.02 | -0.89 | -0.51 | -0.56 | -0.78 | -0.52 | 0.065 | 0.3975 | 0.85 | 0.013671 | 0.876963 | 0.413618 | 0.996493 | 0.997988 | 0.994666 |
| Stk39_Q9Z1W9_S383s_1 | Stk39_Q9Z | -1.83 | -1.64 | -1.19 | -1.01 | -1.56 | -2.14 | -0.86 | -0.67 | -1.31 | -2.49 | -0.02 | -0.89 | -0.5 | -0.74 | -0.78 | -0.52 | 0.11 | 0.24 | 0.7825 | 0.018883 | 0.771812 | 0.641932 | 0.995762 | 0.997988 | 0.994666 |
| Stk39_Q9Z1W9_T366t_1 | Stk39_Q9Z | -0.22 | -0.08 | -0.26 | 0.08 | -0.22 | -0.21 | -0.37 | -0.07 | -0.3 | -0.22 | -0.79 | -0.28 | 0.11 | 0.21 | -0.24 | 0.05 | -0.0975 | -0.2775 | 0.1525 | 0.538593 | 0.639129 | 0.263331 | 0.989223 | 0.997988 | 0.994666 |
| Stmn1_P54227_S16s_1 | Stmn1_P54 | -1.39 | -1.69 | -0.46 | -0.08 | -0.35 | -1.18 | 0.25 | 0.34 | -0.9 | -0.33 | -0.26 | 0.3 | -1.14 | -0.78 | -0.18 | -0.11 | 0.67 | 0.6075 | 0.3525 | 0.434057 | 0.182781 | 0.188398 | 0.989223 | 0.997988 | 0.994666 |
| Stmn1_P54227_S25s_1 | Stmn1_P54 | -0.71 | -1.04 | -0.41 | -0.74 | -0.19 | -0.49 | -0.02 | -0.26 | -1.54 | -0.75 | -1.7 | -0.79 | -1.02 | -0.49 | -1.28 | -1 | 0.485 | -0.47 | -0.2225 | 0.441064 | 0.061403 | 0.156678 | 0.989223 | 0.997988 | 0.994666 |
| Stmn1_P54227_S38s_1 | Stmn1_P54 | -0.11 | 0.28 | 0.13 | 0.01 | 0.44 | -0.21 | 0.38 | 0.09 | -0.04 | 0.45 | -0.14 | 0.12 | 0.19 | 0.7 | 0.1 | 0.08 | 0.0975 | 0.02 | 0.19 | 0.47746 | 0.683496 | 0.933368 | 0.994272 | 0.997988 | 0.994666 |
| Stmn1_P54227_S3s_1 | Stmn1_P54 | 1.45 | 0.34 | 0.58 | -0.19 | 0.28 | 0.7 | -0.07 | 0.19 | 0.28 | 1.33 | 0.33 | -0.06 | 0.07 | 0.35 | -0.26 | -0.1 | -0.27 | -0.075 | -0.53 | 0.186019 | 0.473595 | 0.864572 | 0.989223 | 0.997988 | 0.994666 |
| Stmn1_P54227_S46s_1 | Stmn1_P54 | 3.41 | 3.85 | 3.38 | 2.41 | 3.68 | 3.48 | 2.8 | 3.12 | 0.03 | 3.75 | 0.67 | 3.1 | -0.9 | 0.5 | 3.2 | 3.34 | 0.0075 | -1.375 | -1.7275 | 0.093857 | 0.983333 | 0.132015 | 0.999187 | 0.997988 | 0.994666 |
| Stmn1_P54227_S4s_1 | Stmn1_P54 | 1.45 | 0.34 | 0.58 | -0.19 | 0.28 | 0.7 | -0.07 | 0.19 | 0.28 | 1.33 | 0.33 | -0.06 | 0.07 | 0.35 | -0.26 | -0.1 | -0.27 | -0.075 | -0.53 | 0.186019 | 0.473595 | 0.864572 | 0.989223 | 0.997988 | 0.994666 |
| Stmn1_P54227_S63s_1 | Stmn1_P54 | 1.1 | 0.21 | 0.59 | 0.87 | 0.33 | 1.38 | 0.32 | 1.05 | 0.65 | 0.61 | 1.01 | 0.48 | 0.58 | 0.86 | 0.8 | 0.41 | 0.0775 | -0.005 | -0.03 | 0.917123 | 0.817427 | 0.985564 | 0.995762 | 0.997988 | 0.999507 |
| Stmn2_P55821_S62s_1 | Stmn2_P55 | 0.37 | 0.77 | 0.63 | 0.52 | 0.73 | 0.21 | 0.94 | 0.67 | 0.34 | 0.86 | 0.09 | 0.72 | 0.23 | 0.16 | 0.56 | 0.68 | 0.065 | -0.07 | -0.165 | 0.524623 | 0.787236 | 0.788669 | 0.995762 | 0.997988 | 0.994666 |
| Stmn2_P55821_S97s_1 | Stmn2_P55 | -0.63 | -0.34 | 0.85 | 0.2 | 0.53 | -0.13 | 1.36 | 0.8 | -0.9 | -0.76 | 1.33 | 0.29 | -0.3 | -0.35 | 1.21 | 0.09 | 0.62 | -0.03 | 0.1425 | 0.761569 | 0.168157 | 0.957692 | 0.989223 | 0.997988 | 0.9971 |
| Stmn3_O70166_S50s_1 | Stmn3_O7 | -0.29 | 0.97 | 0.65 | 0.19 | 0.3 | -0.17 | 0.59 | 0.28 | -0.14 | 0.37 | 0.06 | 0.52 | 0.14 | 0.11 | 0.35 | 0.41 | -0.13 | -0.1775 | -0.1275 | 0.698036 | 0.692659 | 0.598654 | 0.994893 | 0.997988 | 0.996159 |
| Stmn3_O70166_S53s_1 | Stmn3_O7 | 0.62 | 1.45 | 0.45 | 0.18 | 0.96 | 0.56 | 0.73 | -0.07 | 0.92 | 1.38 | -0.38 | 0.42 | 1.16 | 0.85 | 0 | 0.1 | -0.13 | -0.09 | -0.1475 | 0.713434 | 0.713268 | 0.840975 | 0.995437 | 0.997988 | 0.996966 |
| Stmn3_O70166_S81s_1 | Stmn3_O7 | 1.82 | 1.64 | 1.91 | 1.69 | 1.84 | 1.45 | 1.71 | 1.82 | 0.67 | 1.76 | 0.95 | 1.64 | 0.01 | 0.83 | 1.77 | 1.95 | -0.06 | -0.51 | -0.625 | 0.181381 | 0.775849 | 0.122642 | 0.995762 | 0.997988 | 0.994666 |
| Stmn4_Q05DI3_S90s_1 | Stmn4_Q0 | 0.61 | 0.92 | 1.27 | 1.3 | 0.57 | 0.65 | 0.81 | 0.96 | 0.22 | 0.65 | 1.22 | 0.65 | 0.49 | 0.82 | 0.96 | 1.08 | -0.2775 | -0.34 | -0.1875 | 0.512131 | 0.274515 | 0.275625 | 0.989223 | 0.997988 | 0.994666 |
| Stom_P54116_S7s_1 | Stom_P541 | -1.56 | -1.99 | -2.18 | -2.94 | -3.57 | -1.33 | -1.84 | -2.22 | 1.37 | -1.57 | -4.81 | -3.07 | 2.53 | -3.19 | -1.04 | -1.89 | -0.0725 | 0.1475 | 1.27 | 0.26186 | 0.887081 | 0.900216 | 0.996493 | 0.997988 | 0.994666 |
| Stox2_Q499E5_S110s_1 | Stox2_Q49 | 0.5 | 0.4 | -0.73 | 0.26 | -0.27 | 1.39 | -0.67 | 0.43 | 0.86 | 0.61 | 0 | -0.24 | 0.55 | 0.06 | 0.55 | 0.22 | 0.1125 | 0.2 | 0.2375 | 0.493927 | 0.818519 | 0.606014 | 0.995762 | 0.997988 | 0.994666 |
| Strada_Q3UUJ4_S276s_1 | Strada_Q3 | 0.11 | 0.25 | 0.35 | 0.49 | 0.48 | 0.25 | 0.77 | 0.34 | -0.39 | 0.37 | 0.49 | 0.49 | -0.24 | 0.45 | 0.42 | 0.96 | 0.16 | -0.06 | 0.0975 | 0.754863 | 0.478881 | 0.829735 | 0.989223 | 0.997988 | 0.9971 |
| Strip1_Q8C079_S335s_1 | Strip1_Q8C | 0.58 | 0.45 | 0.65 | 0.33 | 0.6 | 0.43 | 0.89 | 0.45 | 0.43 | 0.72 | 0.57 | 0.73 | 0.28 | 0.77 | 0.48 | 0.45 | 0.09 | 0.11 | -0.0075 | 0.975617 | 0.680791 | 0.615332 | 0.993508 | 0.997988 | 1 |
| Strip2_E9QN41_S328s_1 | Strip2_E9Q | 0.46 | 0.39 | 0.99 | 0.37 | 0.59 | -0.08 | 0.92 | 0.73 | 0.51 | 0.56 | 0.92 | 0.37 | 0.53 | 0.73 | 0.44 | 0.6 | -0.0125 | 0.0375 | 0.0225 | 0.930923 | 0.965786 | 0.883887 | 0.999187 | 0.997988 | 0.999507 |
| Strip2_E9QN41_S364s_1 | Strip2_E9Q | -0.01 | -0.51 | 0.35 | -0.45 | -0.46 | -1.1 | 0.12 | -0.12 | -0.73 | -0.95 | -0.54 | -0.25 | -1.03 | 0.02 | -0.4 | -0.49 | -0.235 | -0.4625 | -0.32 | 0.350332 | 0.494301 | 0.140586 | 0.989223 | 0.997988 | 0.994666 |
| Strn_F6Z700_S227s_1 | Strn_F6Z7 | 0.74 | 0.4 | 0.25 | 0.24 | 0.4 | 0.74 | 0.75 | 0.86 | -0.27 | 0.15 | 0.27 | 0.41 | -0.78 | 0.53 | 0.19 | 0.65 | 0.28 | -0.2675 | -0.26 | 0.483614 | 0.241057 | 0.311769 | 0.989223 | 0.997988 | 0.994666 |
| Strn_F6Z700_S81s_1 | Strn_F6Z7 | -2 | -2.12 | -1.21 | -1.95 | -0.75 | -1.86 | -0.15 | -1.69 | -2.73 | -1.49 | 0.74 | -1.31 | -1.48 | -0.8 | -0.83 | -1.38 | 0.7075 | 0.6225 | 0.6975 | 0.052008 | 0.123583 | 0.368981 | 0.989223 | 0.997988 | 0.994666 |
| Strn3_B2RQS1_S229s_1 | Strn3_B2R | -0.12 | 0.09 | 0.96 | -0.17 | -0.27 | 0.13 | 0.68 | 0.07 | 0 | 0.88 | 0.91 | 0.76 | 0 | 0.94 | -0.08 | 0.38 | -0.0375 | 0.4475 | 0.12 | 0.746576 | 0.911073 | 0.226045 | 0.997335 | 0.997988 | 0.9971 |
| Strn4_P58404_S206s_1 | Strn4_P58 | 1.04 | 0.21 | 0.31 | 0.04 | 0.22 | 1.04 | 0.18 | -0.15 | 0.11 | 0.74 | 0.88 | 0.08 | -0.64 | 0.62 | 0.3 | -0.07 | -0.0775 | 0.0525 | -0.3475 | 0.358124 | 0.820449 | 0.872986 | 0.995762 | 0.997988 | 0.994666 |
| Strn4_P58404_S223s_1 | Strn4_P58 | -0.18 | -0.35 | 0.37 | -0.01 | 0.35 | -0.02 | 0.71 | 0.46 | -0.29 | 0.11 | 0.1 | -0.05 | 0.39 | 0.38 | -0.39 | 0.88 | 0.4175 | 0.01 | 0.3575 | 0.307182 | 0.137173 | 0.968393 | 0.989223 | 0.997988 | 0.994666 |
| Stt3b_Q3TDQ1_S495s_1 | Stt3b_Q3T | -1.07 | -0.64 | -0.11 | 0.28 | -1.01 | -0.74 | -0.62 | 0.03 | -0.72 | -0.33 | 0.77 | 0.36 | -1.2 | -1.03 | 1.17 | 1.08 | -0.2 | 0.405 | 0.39 | 0.548425 | 0.587087 | 0.362819 | 0.989223 | 0.997988 | 0.994666 |
| Stub1_Q9WUD1_S20s_1 | Stub1_Q9V | 1.46 | 1.74 | 1.62 | 1.7 | 1.15 | 1.59 | 1.54 | 1.83 | 1.35 | 1.55 | 1.53 | 1.72 | 1.37 | 1.68 | 2.03 | 1.77 | -0.1025 | -0.0925 | 0.0825 | 0.747371 | 0.656948 | 0.670727 | 0.990434 | 0.997988 | 0.9971 |
| Stub1_Q9WUD1_S274s_1 | Stub1_Q9V | -0.24 | 0.05 | 0 | -0.04 | 0.25 | -0.31 | 0.94 | 0.39 | -0.47 | 0.07 | 0.05 | 0.66 | -0.05 | 0.7 | 0.17 | -0.01 | 0.375 | 0.135 | 0.26 | 0.349608 | 0.220493 | 0.639727 | 0.989223 | 0.997988 | 0.994666 |
| Stx12_Q9ER00_S142s_1 | Stx12_Q9E | 0.77 | 0.44 | 1 | 0.88 | -0.57 | 0.85 | 0.02 | 0.99 | 1.48 | 0.58 | 1.02 | 0.29 | 1.34 | -0.34 | 0.66 | 0.86 | -0.45 | 0.07 | -0.1425 | 0.712724 | 0.251425 | 0.825384 | 0.989223 | 0.997988 | 0.996966 |
| Stx17_Q9D0I4_S288s_1 | Stx17_Q9D | 1.01 | 0.62 | 0.98 | 1.12 | 1.08 | 0.77 | 1.15 | 0.74 | 0.93 | 0.75 | 1.08 | 0.66 | 0.17 | 1.1 | 1.14 | 1.24 | 0.0025 | -0.0775 | -0.02 | 0.950048 | 0.991255 | 0.741679 | 0.999187 | 0.997988 | 0.999507 |
| Stx1a_O35526_S14s_1 | Stx1a_O35 | 1.14 | 1.33 | 1.46 | 1.35 | 1.07 | 1.15 | 0.97 | 1.18 | 1.05 | 1.32 | 1.19 | 1.38 | -0.88 | 1.2 | 1.04 | 1.16 | -0.2275 | -0.085 | -0.69 | 0.177526 | 0.275162 | 0.696319 | 0.989223 | 0.997988 | 0.994666 |
| Stx1a_O35526_S59s_1 | Stx1a_O35 | 5.53 | 2.66 | 3.65 | 1.35 | 3.22 | 5.59 | 3.7 | 3.3 | 0.84 | 4.62 | 0.32 | 3.07 | -0.15 | -0.37 | 3.55 | 3.6 | 0.655 | -1.085 | -1.64 | 0.200888 | 0.475102 | 0.36535 | 0.989223 | 0.997988 | 0.994666 |
| Stx1a_O35526_S64s_1 | Stx1a_O35 | 4.76 | 3.01 | 4.3 | 1.72 | 3.3 | 4.86 | 3.88 | 3.24 | 0.08 | 4.26 | 0.79 | 3.88 | -0.37 | 0.77 | 4.01 | 4.51 | 0.3725 | -1.195 | -1.2175 | 0.321244 | 0.591681 | 0.29792 | 0.989223 | 0.997988 | 0.994666 |
| Stx1a_O35526_T10t_1 | Stx1a_O35 | 0.76 | 0.49 | 1.07 | 0.15 | 0.86 | 0.52 | 0.61 | 0.14 | 0.86 | 0.75 | 0.72 | -0.18 | -0.52 | 1.45 | 0.19 | 0.87 | -0.085 | -0.08 | -0.12 | 0.792062 | 0.762933 | 0.810463 | 0.995762 | 0.997988 | 0.999507 |
| Stx1b_P61264_S14s_1 | Stx1b_P61 | 1.09 | 1.6 | 1.42 | 1.18 | 1.43 | 1.01 | 0.88 | 0.72 | 0.67 | 0.97 | 0.82 | 1.05 | -1.6 | 1.14 | 1.11 | 1.17 | -0.3125 | -0.445 | -0.8675 | 0.190343 | 0.228668 | 0.083634 | 0.989223 | 0.997988 | 0.994666 |
| Stx4_P70452_S117s_1 | Stx4_P704 | 1.58 | 1.41 | 1.57 | 0.63 | 0.93 | 1.34 | 0.65 | 1.03 | 1.42 | 1.39 | 1.11 | 0.79 | 0.97 | 2.26 | 1.47 | 1.24 | -0.31 | -0.12 | 0.1875 | 0.619588 | 0.307108 | 0.695771 | 0.989223 | 0.997988 | 0.995259 |
| Stx4_P70452_S15s_1 | Stx4_P704 | 0.48 | 0.64 | 0.46 | -0.03 | 0.29 | -0.04 | -0.2 | -0.31 | 0.27 | 0.54 | 0.01 | 0.08 | -0.54 | 0.04 | 0.13 | 0.22 | -0.4525 | -0.1625 | -0.425 | 0.168789 | 0.096892 | 0.530755 | 0.989223 | 0.997988 | 0.994666 |
| Stx6_Q9JKK1_S2s_1 | Stx6_Q9JK | 0.07 | -0.37 | 0.12 | 0.14 | 0.14 | 0.07 | 0.25 | 0.24 | 0.03 | -0.36 | 0.2 | -0.18 | -0.57 | 0.25 | -0.39 | 0.19 | 0.185 | -0.0675 | -0.12 | 0.690263 | 0.40563 | 0.786023 | 0.989223 | 0.997988 | 0.996159 |
| Stx7_Q8BH40_S129s_1 | Stx7_Q8BH | -0.71 | -2.16 | 0.06 | 0.38 | 0.58 | 0.61 | 0.42 | 0.21 | -0.1 | 0.27 | 0.74 | 0.15 | 0.71 | 0 | 0.05 | -0.42 | 1.0625 | 0.8725 | 0.6925 | 0.238775 | 0.066039 | 0.137891 | 0.989223 | 0.997988 | 0.994666 |
| Stx7_Q8BH40_S75s_1 | Stx7_Q8BH | -0.11 | 0.05 | -0.47 | -0.52 | 0.58 | -0.03 | -0.13 | -0.06 | 0.43 | 0.09 | -1.08 | 0.35 | -0.17 | 0.4 | -0.64 | 0.12 | 0.3525 | 0.21 | 0.19 | 0.548312 | 0.200599 | 0.584173 | 0.989223 | 0.997988 | 0.994666 |
| Stx7_Q8BH40_T41t_1 | Stx7_Q8BH | 0.61 | 1.26 | -0.25 | 0.25 | 1.67 | 1.24 | 0.65 | 0.19 | 0.81 | 1.32 | -0.07 | 0.29 | 0.27 | 1.35 | 0.55 | 0.36 | 0.47 | 0.12 | 0.165 | 0.685504 | 0.289277 | 0.779585 | 0.989223 | 0.997988 | 0.996159 |
| Stx7_Q8BH40_T79t_1 | Stx7_Q8BH | -0.88 | -0.18 | -0.16 | -0.45 | 0.39 | -0.91 | 0.13 | -0.46 | -1.37 | -0.49 | -0.98 | -0.45 | -0.04 | -0.16 | -0.48 | -0.27 | 0.205 | -0.405 | 0.18 | 0.517701 | 0.553208 | 0.212338 | 0.989223 | 0.997988 | 0.994666 |
| Stxbp1_O08599_S507s_1 | Stxbp1_O0 | 0.12 | -0.19 | -0.09 | 0.25 | 0 | 0.02 | -0.47 | -0.06 | -0.47 | -0.09 | -0.15 | 0.57 | -0.48 | -0.16 | -0.91 | -0.21 | -0.15 | -0.0575 | -0.4625 | 0.121719 | 0.517266 | 0.841252 | 0.989223 | 0.997988 | 0.994666 |
| Stxbp1_O08599_S509s_1 | Stxbp1_O0 | 1.2 | 0.9 | -0.29 | -0.27 | 0.74 | 0.78 | -0.21 | 0.39 | 0.28 | 1.04 | -0.46 | -0.01 | 0.26 | 0.19 | -0.52 | 0.12 | 0.04 | -0.1725 | -0.3725 | 0.39069 | 0.92526 | 0.717687 | 0.997335 | 0.997988 | 0.994666 |

| | | | | | | | | | | | | | | | | | | | | | | | | | | |
|---|---|---|---|---|---|---|---|---|---|---|---|---|---|---|---|---|---|---|---|---|---|---|---|---|---|---|
| Stxbp1_O08599_S516s_1 | Stxbp1_O0 | 0.01 | -1.51 | -2.52 | -1.57 | -1.16 | -0.1 | -0.88 | 0.14 | -1.76 | -1.42 | -0.41 | -1.45 | -1.27 | -1.99 | -2.73 | -1.99 | 0.8975 | 0.1375 | -0.5975 | 0.298171 | 0.12742 | 0.80454 | 0.989223 | 0.997988 | 0.994666 |
| Stxbp1_O08599_S593s_1 | Stxbp1_O0 | 1.66 | 0.92 | 1.67 | 2.07 | -0.03 | 1.63 | 0.9 | 1.84 | 1.69 | 1.61 | 1.79 | 1.49 | 0.89 | 0.27 | 1.15 | 1.23 | -0.495 | 0.065 | -0.695 | 0.073228 | 0.2906 | 0.824 | 0.989223 | 0.997988 | 0.994666 |
| Stxbp1_O08599_S594s_1 | Stxbp1_O0 | 0.39 | 0.68 | 0.34 | 0.2 | 0.58 | 0.31 | 0.55 | 0.03 | 0.32 | 0.45 | 0.24 | 0.23 | 0.16 | 0.42 | -0.26 | 0.36 | -0.035 | -0.0925 | -0.2325 | 0.400507 | 0.881063 | 0.678192 | 0.996493 | 0.997988 | 0.994666 |
| Stxbp1_O08599_S594s_1_1 | Stxbp1_O0 | 2.25 | 1.16 | 1.87 | 2.44 | 0.21 | 1.91 | 1.18 | 2.26 | 2.02 | 2.22 | 1.99 | 1.7 | 1.34 | 0.79 | 1.37 | 1.4 | -0.54 | 0.0525 | -0.705 | 0.067384 | 0.287135 | 0.872691 | 0.989223 | 0.997988 | 0.994666 |
| Stxbp1_O08599_Y473y_1 | Stxbp1_O0 | 0.79 | -0.6 | 0.67 | 1.27 | -0.22 | 0.69 | 0.62 | 1.61 | 0.11 | -0.04 | 0.59 | 0.92 | -0.35 | -0.09 | -0.44 | -0.08 | 0.1425 | -0.1375 | -0.7725 | 0.085657 | 0.776379 | 0.755664 | 0.995762 | 0.997988 | 0.994666 |
| Stxbp4_B1ATH0_S11s_1 | Stxbp4_B1 | -5.01 | -3 | -3.04 | -4.92 | -2.36 | -3.18 | -4.66 | -3.71 | -3.29 | -4.11 | -2.97 | -3.75 | -1.16 | -3.77 | -4.42 | -3.83 | 0.515 | 0.4625 | 0.6975 | 0.398947 | 0.440236 | 0.42399 | 0.989223 | 0.997988 | 0.994666 |
| Stxbp5_D3Z079_S1023s_1 | Stxbp5_D3 | -0.15 | -0.08 | 0.07 | -0.15 | 0.21 | -0.23 | 0.04 | -0.09 | -0.5 | -0.22 | -0.63 | -0.34 | -0.44 | 0.33 | -0.57 | -0.9 | 0.06 | -0.345 | -0.3175 | 0.327809 | 0.775255 | 0.139905 | 0.995762 | 0.997988 | 0.994666 |
| Stxbp5_D3Z079_S693s_1 | Stxbp5_D3 | 0.61 | 1.04 | 0.28 | 0.14 | 1 | 0.77 | 0.76 | 0.41 | 0.72 | 0.29 | -0.08 | 0.18 | -0.3 | 0.74 | 0.32 | 0.69 | 0.2175 | -0.24 | -0.155 | 0.654806 | 0.435011 | 0.431892 | 0.989223 | 0.997988 | 0.995259 |
| Stxbp5_D3Z079_S750s_1 | Stxbp5_D3 | 0.4 | 0.67 | 1.61 | 1.8 | 0.67 | 0.25 | 1.28 | 1.85 | 0.46 | 0.61 | 1.66 | 1.89 | 0.9 | 0.68 | 1.25 | 1.77 | -0.1075 | 0.035 | 0.03 | 0.942706 | 0.814303 | 0.941242 | 0.995762 | 0.997988 | 0.999507 |
| Stxbp5_D3Z079_S811s_1 | Stxbp5_D3 | 0.74 | 0.65 | 0.29 | 0.4 | 0.94 | 0.13 | 0.49 | 0.34 | 0.21 | 0.98 | 0.41 | 0.64 | 0.24 | 1.02 | 0.68 | 0.25 | -0.045 | 0.04 | 0.0275 | 0.923653 | 0.859666 | 0.878234 | 0.996417 | 0.997988 | 0.999507 |
| Stxbp5_D3Z079_S867s_1 | Stxbp5_D3 | 0.91 | 0.69 | 0.67 | 0.85 | 0.99 | 0.3 | 1.19 | 0.59 | 0.94 | 1.49 | 0.74 | 0.76 | 0.39 | 0.83 | 0.46 | 1.14 | -0.0125 | 0.2025 | -0.075 | 0.782347 | 0.961392 | 0.435434 | 0.999187 | 0.997988 | 0.998226 |
| Stxbp5_D3Z079_T749t_1 | Stxbp5_D3 | -0.57 | -0.13 | -0.58 | -1.44 | 0.19 | -0.57 | -1.17 | -1.25 | -0.62 | -0.16 | -0.83 | -0.64 | 0.82 | -0.02 | -0.65 | -1.02 | -0.02 | 0.1175 | 0.4625 | 0.336526 | 0.96122 | 0.724305 | 0.999187 | 0.997988 | 0.994666 |
| Stxbp5l_Q5DQR4_S1120s_1 | Stxbp5l_Q5 | 0.14 | 0.16 | 0.37 | 0.03 | 0.48 | 0.09 | 0.58 | 0.19 | -0.17 | -0.05 | 0.08 | -0.38 | -0.5 | -0.06 | 0.22 | -0.1 | 0.16 | -0.305 | -0.285 | 0.292072 | 0.476406 | 0.197436 | 0.989223 | 0.997988 | 0.994666 |
| Stxbp5l_Q5DQR4_S588s_1 | Stxbp5l_Q5 | -0.26 | -0.38 | 0.28 | -1.22 | 0.72 | 0.02 | 0.07 | -0.26 | -1.05 | 0.6 | 1.15 | 0.28 | -0.27 | -0.38 | 1.02 | 1.25 | 0.5325 | 0.64 | 0.8 | 0.133029 | 0.171323 | 0.241396 | 0.989223 | 0.997988 | 0.994666 |
| Stxbp5l_Q5DQR4_S597s_1 | Stxbp5l_Q5 | -0.03 | 1.13 | 0.39 | 0.72 | 1.08 | 0.68 | 0.68 | 0.86 | 0.21 | 0.69 | 0.35 | 0.45 | 0.21 | -0.46 | 0.62 | 0.47 | 0.2725 | -0.1275 | -0.3425 | 0.359768 | 0.362931 | 0.675877 | 0.989223 | 0.997988 | 0.994666 |
| Stxbp5l_Q5DQR4_S598s_1 | Stxbp5l_Q5 | 0.49 | 1.05 | 0.66 | 0.36 | 1.08 | 0.67 | 0.64 | 0.45 | 0.31 | 0.79 | -0.82 | 0.73 | 0.38 | 0.52 | 0.69 | 0.72 | 0.07 | -0.3875 | -0.0625 | 0.813196 | 0.783002 | 0.34519 | 0.995762 | 0.997988 | 0.999507 |
| Stxbp5l_Q5DQR4_S792s_1 | Stxbp5l_Q5 | 0.51 | 0.58 | 0.13 | 0.18 | 1 | 0.85 | 0.48 | 0.51 | 0.3 | 0.58 | -0.3 | -0.06 | -0.28 | 0.9 | -0.14 | 0.3 | 0.36 | -0.22 | -0.155 | 0.640467 | 0.154809 | 0.436846 | 0.989223 | 0.997988 | 0.995259 |
| Stxbp5l_Q5DQR4_S819s_2 | Stxbp5l_Q5 | -1.9 | -0.88 | -1.43 | -1.94 | -0.64 | -1.43 | -1.9 | -3.13 | -1.72 | -1.01 | 0.09 | -1.6 | -1.51 | -0.39 | -1.18 | -1.71 | -0.2375 | 0.4775 | 0.34 | 0.39521 | 0.652494 | 0.315309 | 0.990358 | 0.997988 | 0.994666 |
| Stxbp5l_Q5DQR4_S822s_1 | Stxbp5l_Q5 | -0.46 | 0.27 | -2.16 | -3.52 | 0.1 | -0.39 | -3.28 | -3.67 | -0.06 | 0.33 | -3.1 | -2.75 | 0 | 0.18 | -2.09 | -2.98 | -0.3425 | 0.0725 | 0.245 | 0.807673 | 0.758644 | 0.946653 | 0.995762 | 0.997988 | 0.999507 |
| Stxbp5l_Q5DQR4_S822s_2 | Stxbp5l_Q5 | -1.9 | -0.88 | -1.43 | -1.94 | -0.64 | -1.43 | -1.9 | -3.13 | -1.72 | -1.01 | 0.09 | -1.6 | -1.51 | -0.39 | -1.18 | -1.71 | -0.2375 | 0.4775 | 0.34 | 0.39521 | 0.652494 | 0.315309 | 0.990358 | 0.997988 | 0.994666 |
| Sucla2_Q9Z2I9_S279s_1 | Sucla2_Q9Z | 0.8 | 1.12 | 0.67 | 0.72 | 0.19 | 0.53 | 0.41 | 0.48 | 0.43 | 0.39 | -0.33 | 0.54 | 0.12 | 0.29 | 0 | 0.49 | -0.425 | -0.57 | -0.6025 | 0.037792 | 0.075115 | 0.064001 | 0.989223 | 0.997988 | 0.994666 |
| Suds3_Q8BR65_S234s_2 | Suds3_Q8B | 0.28 | -0.4 | 0.72 | -1.03 | 0.63 | -0.06 | 0.47 | -0.04 | 0.5 | 0.72 | -0.44 | -1.28 | 0.37 | -1.04 | 0.44 | 0.5 | 0.3575 | -0.0175 | 0.175 | 0.728889 | 0.387628 | 0.974805 | 0.989223 | 0.997988 | 0.9971 |
| Suds3_Q8BR65_S236s_2 | Suds3_Q8B | 0.28 | -0.4 | 0.72 | -1.03 | 0.63 | -0.06 | 0.47 | -0.04 | 0.5 | 0.72 | -0.44 | -1.28 | 0.37 | -1.04 | 0.44 | 0.5 | 0.3575 | -0.0175 | 0.175 | 0.728889 | 0.387628 | 0.974805 | 0.989223 | 0.997988 | 0.9971 |
| Sugp2_Q8CH09_S206s_1 | Sugp2_Q8C | -0.62 | -0.79 | -1.29 | -1.17 | -0.49 | -0.84 | -0.51 | -0.98 | -1.38 | -0.96 | -0.12 | -1.01 | -1.18 | -0.34 | -1.03 | -1.34 | 0.2625 | 0.1 | -0.005 | 0.987474 | 0.314061 | 0.763958 | 0.989223 | 0.997988 | 1 |
| Sugp2_Q8CH09_S733s_1 | Sugp2_Q8C | 1.21 | 1.33 | 0.81 | 1.04 | 0.5 | 0.81 | 1.15 | 1.14 | 1.13 | 0.75 | 1.96 | 1.09 | -0.49 | 1.28 | 0.9 | 0.98 | -0.1975 | 0.135 | -0.43 | 0.310783 | 0.435817 | 0.667865 | 0.989223 | 0.997988 | 0.994666 |
| Sugp2_Q8CH09_S838s_1 | Sugp2_Q8C | 0.4 | 0.18 | 0.08 | 0.11 | 0.13 | 0.61 | 0.39 | 0.23 | -0.13 | -0.22 | -0.38 | -0.5 | -0.11 | 0.46 | -0.41 | -0.17 | 0.1475 | -0.5 | -0.25 | 0.379109 | 0.50336 | 0.045657 | 0.989223 | 0.997988 | 0.994666 |
| Sulf2_Q8CFG0_S843s_1 | Sulf2_Q8CF | 1.36 | 1.31 | 2.61 | 1.67 | 1.14 | 0.97 | 1.55 | 1.39 | 0.06 | 1.36 | 2.06 | 1.79 | -0.66 | -0.67 | 2.67 | 1.99 | -0.475 | -0.42 | -0.905 | 0.283796 | 0.179858 | 0.413817 | 0.989223 | 0.997988 | 0.994666 |
| Sumo1_P63166_S2s_1 | Sumo1_P63 | -0.78 | -0.76 | -1.11 | -1.25 | -0.43 | -0.81 | -1.08 | -1.01 | -2.05 | -1.07 | -0.5 | -0.54 | -0.59 | -0.4 | -0.66 | -1.05 | 0.1425 | -0.065 | 0.3 | 0.283627 | 0.569314 | 0.865499 | 0.989223 | 0.997988 | 0.994666 |
| Susd5_G3UW60_S237s_1 | Susd5_G3U | 0.2 | -0.06 | 0.39 | 0.62 | -1.09 | 1.12 | 0.09 | -0.37 | 2.71 | 0.85 | 0.4 | 1.77 | 2.15 | -0.75 | -0.61 | 0.36 | -0.35 | 1.145 | -3.1E-16 | 1 | 0.446688 | 0.044412 | 0.989223 | 0.997988 | 1 |
| Sv2a_Q9JIS5_S127s_1 | Sv2a_Q9JIS | 1.17 | 0.99 | 1.3 | 1.4 | 0.76 | 0.65 | 0.46 | 1.23 | 1.05 | 1.34 | 1.91 | 1.14 | 0.72 | 1.03 | 0.86 | 0.78 | -0.44 | 0.145 | -0.3675 | 0.151595 | 0.099327 | 0.594816 | 0.989223 | 0.997988 | 0.994666 |
| Svil_Q8K4L2_S666s_1 | Svil_Q8K4L | -1.21 | -1.48 | -2.46 | -1.8 | -0.68 | -1.48 | -1 | -2.12 | -2.94 | -0.86 | -3.24 | -0.74 | -1.03 | -2.65 | -0.58 | -1.78 | 0.4175 | -0.2075 | 0.2275 | 0.650119 | 0.308559 | 0.75036 | 0.989223 | 0.997988 | 0.995259 |
| Svil_Q8K4L2_S857s_1 | Svil_Q8K4L | -0.78 | -0.46 | -0.29 | -0.01 | -0.52 | -1.18 | 0.02 | 0.01 | -0.3 | -0.37 | 0.19 | -0.03 | -0.5 | -0.08 | 0.08 | 0.06 | -0.0325 | 0.2575 | 0.275 | 0.345708 | 0.922568 | 0.347364 | 0.997335 | 0.997988 | 0.994666 |
| Svop_Q8BFT9_S25s_1 | Svop_Q8BF | 0.83 | 0.43 | 0.69 | 0.68 | 0.69 | 0.53 | 0.66 | 0.77 | 0.52 | 0.92 | 0.59 | 0.75 | -0.25 | 0.89 | 0.63 | 0.83 | 0.005 | 0.0375 | -0.1325 | 0.682566 | 0.980646 | 0.867883 | 0.999187 | 0.997988 | 0.996159 |
| Svop_Q8BFT9_S31s_1 | Svop_Q8BF | 0.82 | 1.27 | 0.99 | 0.96 | 0.67 | 0.89 | 1.16 | 1.27 | 0.23 | 1.05 | 0.24 | 1.31 | -0.41 | 0.37 | 1.18 | 1.29 | -0.0125 | -0.3025 | -0.4025 | 0.338764 | 0.957551 | 0.359008 | 0.999187 | 0.997988 | 0.994666 |
| Svop_Q8BFT9_S31s_2 | Svop_Q8BF | 1.16 | 0.64 | 1.18 | 1.18 | 0.47 | 0.8 | 1.39 | 0.94 | 1.02 | 1.41 | 1.37 | 1.8 | -2.07 | 0.73 | 1.46 | 1.27 | -0.14 | 0.36 | -0.6925 | 0.360504 | 0.610513 | 0.201073 | 0.989223 | 0.997988 | 0.994666 |
| Svop_Q8BFT9_T534t_1 | Svop_Q8BF | -0.24 | -0.12 | -0.72 | 0.07 | -0.49 | 0 | -0.38 | 0.26 | 0.51 | -0.48 | -0.79 | -0.23 | 0.75 | 0.32 | 0.12 | -0.08 | 0.1 | 0.005 | 0.53 | 0.106057 | 0.720147 | 0.988356 | 0.995437 | 0.997988 | 0.994666 |
| Syap1_Q9D5V6_T262t_1 | Syap1_Q9D | -0.4 | -0.01 | -0.04 | -0.39 | 0.12 | -0.34 | -0.24 | -0.53 | -0.37 | 0.08 | -0.42 | -0.37 | -0.13 | 0.33 | -0.47 | -0.39 | -0.0375 | -0.06 | 0.045 | 0.874125 | 0.875507 | 0.804203 | 0.996493 | 0.997988 | 0.999507 |
| Sybu_Q8BHS8_S197s_1 | Sybu_Q8BH | -0.24 | 0.43 | -0.23 | -0.53 | -0.67 | 0.07 | -1.07 | -1.32 | 1.58 | 0.28 | -1.95 | -0.52 | 2.13 | -1.32 | -0.24 | -0.12 | -0.605 | -0.01 | 0.255 | 0.706824 | 0.120531 | 0.988456 | 0.989223 | 0.997988 | 0.996966 |
| Sybu_Q8BHS8_S201s_2 | Sybu_Q8BH | -0.2 | -0.12 | 0.24 | 0.05 | -1.4 | 0.36 | -0.71 | -0.19 | 1.84 | -0.08 | -1.01 | 0.2 | -0.19 | -1.23 | 0.05 | -0.1 | -0.4775 | 0.245 | -0.36 | 0.306897 | 0.22732 | 0.661273 | 0.989223 | 0.997988 | 0.994666 |
| Sybu_Q8BHS8_S244s_1 | Sybu_Q8BH | 0.03 | -0.98 | -2.25 | -2.13 | -1.03 | 0.47 | -2.61 | -1.68 | -0.17 | -0.23 | -0.73 | -1.95 | 0.55 | -1.66 | -1.74 | -1.99 | 0.12 | 0.5625 | 0.1225 | 0.863508 | 0.870935 | 0.37415 | 0.996493 | 0.997988 | 0.999507 |
| Sybu_Q8BHS8_S548s_1 | Sybu_Q8BH | -0.61 | -0.43 | -0.52 | 0.29 | 0.11 | -0.11 | 0.14 | 0.49 | 0.56 | 0.24 | -0.18 | 0.08 | 0.26 | 0.37 | 0.07 | -0.35 | 0.475 | 0.4925 | 0.405 | 0.214967 | 0.112447 | 0.124372 | 0.989223 | 0.997988 | 0.994666 |
| Sympk_Q80X82_S1260s_1 | Sympk_Q8 | -1 | -1.27 | -1.11 | -0.8 | 0.21 | -0.74 | 0.39 | -1.02 | -0.87 | -0.2 | -0.6 | -0.84 | -1.12 | -0.68 | -0.08 | -1.08 | 0.755 | 0.4175 | 0.305 | 0.342134 | 0.059427 | 0.126669 | 0.989223 | 0.997988 | 0.994666 |
| Sympk_Q80X82_S1276s_1 | Sympk_Q8 | -0.95 | -2.77 | -1.49 | -2.52 | -1.48 | 0.47 | -1.55 | -1.57 | 1.79 | -1.78 | -4.36 | -0.93 | 2.51 | -2.82 | 1.06 | -0.7 | 0.9 | 0.6125 | 1.945 | 0.093377 | 0.151886 | 0.604647 | 0.989223 | 0.997988 | 0.994666 |
| Sympk_Q80X82_S494s_1 | Sympk_Q8 | -1.14 | -0.98 | 0.27 | 0.19 | -0.61 | -1.15 | -0.62 | -0.01 | -0.9 | -1.03 | -2.05 | -0.91 | -1.46 | -0.15 | -0.34 | 0.39 | -0.1825 | -0.8075 | 0.025 | 0.960732 | 0.664382 | 0.098368 | 0.991343 | 0.997988 | 0.999507 |
| Syn1_O88935_S11s_1 | Syn1_O889 | -1.7 | -1.16 | -0.05 | -0.21 | -0.61 | -1.22 | 0.41 | -0.14 | -1.51 | -1.32 | -0.23 | 0.15 | -1.68 | -0.52 | -0.34 | -0.21 | 0.39 | 0.0525 | 0.0925 | 0.850648 | 0.427234 | 0.920585 | 0.989223 | 0.997988 | 0.999507 |
| Syn1_O88935_S330s_1 | Syn1_O889 | 0.61 | 1.03 | 0.38 | 0.49 | 0.83 | 0.48 | 0.35 | 0.35 | 0.29 | -0.03 | 0.52 | 0.37 | 0.09 | 0.99 | 0.08 | -0.08 | -0.125 | -0.34 | -0.3575 | 0.28781 | 0.610598 | 0.203249 | 0.989223 | 0.997988 | 0.994666 |
| Syn1_O88935_S332s_1 | Syn1_O889 | 0.09 | 0.69 | 1.23 | 0.81 | 0.82 | 0 | 0.78 | 1.19 | -0.35 | 0.49 | 0.98 | 1.18 | 0.26 | 0.64 | 0.46 | 0.49 | -0.0075 | -0.13 | -0.2425 | 0.435051 | 0.982731 | 0.751016 | 0.999187 | 0.997988 | 0.994666 |
| Syn1_O88935_S341s_1 | Syn1_O889 | 0.07 | 0.42 | 0.56 | 0.53 | 0.56 | 0.38 | 0.5 | 0.58 | 0 | 0.3 | -0.1 | 0.58 | 0.09 | 0.27 | 0.14 | 0.74 | 0.11 | -0.2 | -0.085 | 0.755252 | 0.611628 | 0.446333 | 0.989223 | 0.997988 | 0.9971 |
| Syn1_O88935_S39s_1 | Syn1_O889 | -1.23 | -1.46 | -1.51 | -1.28 | -1.3 | -1.04 | -1.04 | -1.59 | -1.6 | -1.55 | -1.08 | -1.24 | -1.4 | -0.44 | -1.59 | -1.99 | 0.1275 | 0.0025 | 0.015 | 0.966766 | 0.577009 | 0.991465 | 0.989223 | 0.997988 | 0.999507 |
| Syn1_O88935_S427s_1 | Syn1_O889 | 1.2 | 0.5 | 1.31 | 1.61 | 0.71 | 0.79 | 0.47 | 0.96 | 2.74 | 1.34 | -0.18 | 0.8 | 2.27 | 0.83 | 1.39 | 0.9 | -0.4225 | 0.02 | 0.1925 | 0.639882 | 0.165022 | 0.973306 | 0.989223 | 0.997988 | 0.995259 |
| Syn1_O88935_S437s_1 | Syn1_O889 | 3.34 | 0.78 | 1.57 | 1.25 | 0.91 | 3.38 | 5.33 | 0.39 | 1.38 | 2.3 | 1.34 | 2.27 | 0.92 | 1.55 | 3.22 | 4.08 | 0.7675 | 0.0875 | 0.7075 | 0.393069 | 0.490742 | 0.878211 | 0.989223 | 0.997988 | 0.994666 |
| Syn1_O88935_S449s_1 | Syn1_O889 | -2.93 | -2.36 | -2.17 | -0.86 | -1.5 | -2.73 | -1.5 | -0.98 | -2.45 | -2.33 | -0.73 | -0.31 | -1.35 | -0.52 | -0.57 | -1.06 | 0.4025 | 0.625 | 1.205 | 0.025861 | 0.448369 | 0.339998 | 0.989223 | 0.997988 | 0.994666 |
| Syn1_O88935_S510s_1 | Syn1_O889 | 0.72 | 0.29 | 0.17 | 0.66 | 0.24 | 0.66 | 0.66 | 0.86 | 0.06 | 0.32 | 1.48 | 0.62 | -0.53 | 0.67 | 0.46 | 0.47 | 0.145 | 0.16 | -0.1925 | 0.572824 | 0.561112 | 0.651067 | 0.989223 | 0.997988 | 0.995259 |
| Syn1_O88935_S520s_1 | Syn1_O889 | 0.03 | 0.93 | 0.91 | 0.65 | 0.16 | 0.25 | 0.97 | 0.82 | 0.28 | 0.9 | 1.36 | 0.72 | 0.34 | 0.29 | 1.2 | 0.89 | -0.08 | 0.185 | 0.05 | 0.88297 | 0.797008 | 0.579015 | 0.995762 | 0.997988 | 0.999507 |
| Syn1_O88935_S551s_1 | Syn1_O889 | 1.15 | 0.4 | 0.97 | 1.23 | 0.18 | 1.46 | 1.6 | 1.54 | 0.19 | 0.91 | 0.8 | 1.25 | 0.73 | 1.01 | 1.32 | 1.04 | 0.2575 | -0.15 | 0.0875 | 0.764147 | 0.502192 | 0.640242 | 0.989223 | 0.997988 | 0.9971 |
| Syn1_O88935_S551s_2 | Syn1_O889 | -0.41 | -0.52 | -0.87 | -0.21 | -0.59 | -0.35 | 0.13 | -0.43 | -0.94 | -0.58 | 0.17 | -0.46 | -1.01 | -0.75 | -0.04 | -1.37 | 0.1925 | 0.05 | -0.29 | 0.410384 | 0.462597 | 0.870078 | 0.989223 | 0.997988 | 0.994666 |
| Syn1_O88935_S553s_1 | Syn1_O889 | 0.12 | 0.45 | 0.21 | 0.33 | 0.05 | 0.44 | 0.75 | 0.59 | -0.5 | 0.07 | -0.57 | 0.44 | 0.42 | 0.7 | 0.35 | 0.1 | 0.18 | -0.4175 | 0.115 | 0.650823 | 0.452237 | 0.178951 | 0.989223 | 0.997988 | 0.995259 |
| Syn1_O88935_S553s_2 | Syn1_O889 | -0.41 | -0.52 | -0.87 | -0.21 | -0.59 | -0.35 | 0.13 | -0.43 | -0.94 | -0.58 | 0.17 | -0.46 | -1.01 | -0.75 | -0.04 | -1.37 | 0.1925 | 0.05 | -0.29 | 0.410384 | 0.462597 | 0.870078 | 0.989223 | 0.997988 | 0.994666 |
| Syn1_O88935_S55s_1 | Syn1_O889 | 1.48 | 2.01 | 0.45 | 0.93 | 1.26 | 1.1 | 1.21 | 1.05 | 0.4 | 1.28 | 1.82 | 2 | 0.78 | 0.88 | 0.51 | 0.42 | -0.0625 | 0.1575 | -0.57 | 0.150658 | 0.856311 | 0.73808 | 0.996349 | 0.997988 | 0.994666 |
| Syn1_O88935_S605s_1 | Syn1_O889 | -2.02 | -1.72 | -0.7 | -1.34 | -2.19 | -1.57 | -2.08 | -1.53 | -0.4 | -1.86 | -1.36 | -0.75 | 0.58 | -1.91 | -1.25 | -1 | -0.3975 | 0.3525 | 0.55 | 0.334426 | 0.258061 | 0.412561 | 0.989223 | 0.997988 | 0.994666 |
| Syn1_O88935_S624s_1 | Syn1_O889 | -0.04 | -0.16 | 0.08 | 1.24 | -0.71 | 0.16 | 0.04 | -0.02 | 0.46 | -0.11 | 1.11 | -0.13 | 1.86 | -0.2 | 1.08 | 0.99 | -0.4125 | 0.0525 | 0.6525 | 0.216063 | 0.283796 | 0.901875 | 0.989223 | 0.997988 | 0.994666 |
| Syn1_O88935_S62s_1 | Syn1_O889 | -0.1 | -0.38 | 0.04 | 0.25 | -0.74 | 0.26 | 0.47 | 0.66 | -0.95 | -0.91 | 1.19 | 0.53 | -1.34 | -0.62 | -0.72 | 0.02 | 0.21 | 0.0125 | -0.6175 | 0.096572 | 0.544762 | 0.980622 | 0.989223 | 0.997988 | 0.994666 |
| Syn1_O88935_S642s_1 | Syn1_O889 | -3.78 | -3.13 | -3.54 | -3.32 | -2.7 | -4.62 | -2.99 | -4.39 | -3.36 | -3.09 | -0.94 | -3.66 | -2.95 | -1.75 | -3.72 | -4.34 | -0.2325 | 0.68 | 0.2525 | 0.639255 | 0.621603 | 0.261678 | 0.989223 | 0.997988 | 0.995259 |
| Syn1_O88935_S642s_1_1 | Syn1_O889 | -3.54 | -3.33 | -4.5 | -4.44 | -2.81 | -3.87 | -3.03 | -4.01 | -3.69 | -2.99 | -0.93 | -4.23 | -3.18 | -1.57 | -3.78 | -4.57 | 0.5225 | 0.9925 | 0.6775 | 0.302269 | 0.219373 | 0.183025 | 0.989223 | 0.997988 | 0.994666 |
| Syn1_O88935_S664s_1 | Syn1_O889 | 0.17 | 0.19 | 0.94 | 1.06 | -1.35 | 0 | -0.03 | 0.68 | 1.76 | -0.12 | -0.29 | 0.84 | 2.06 | -0.59 | 0.8 | 0.71 | -0.765 | -0.0425 | 0.155 | 0.777656 | 0.117407 | 0.932182 | 0.989223 | 0.997988 | 0.99798 |
| Syn1_O88935_S666s_1 | Syn1_O889 | -0.32 | -0.2 | -0.28 | -1.05 | -0.62 | -0.67 | -0.95 | -0.68 | -0.16 | -1.09 | 0.14 | -0.08 | 0.82 | -0.25 | -0.35 | 0.13 | -0.2675 | 0.165 | 0.55 | 0.147453 | 0.317597 | 0.640188 | 0.989223 | 0.997988 | 0.994666 |
| Syn1_O88935_S666s_1_1 | Syn1_O889 | 0.61 | 1.24 | -0.98 | -1.09 | 2.07 | 1.46 | 0.16 | -0.34 | 0.89 | 0.23 | 0.32 | -0.13 | 0.5 | 2.57 | -0.73 | -0.36 | 0.8925 | 0.3825 | 0.55 | 0.511145 | 0.228314 | 0.507365 | 0.989223 | 0.997988 | 0.994666 |
| Syn1_O88935_S67s_1 | Syn1_O889 | 0.04 | -0.03 | 0.03 | 0.38 | 0.18 | 0.23 | 0.49 | 0.33 | -0.33 | 0.01 | -0.5 | 0.04 | -0.84 | -0.01 | -0.38 | 0.08 | 0.2025 | -0.3 | -0.3925 | 0.203823 | 0.353489 | 0.236686 | 0.989223 | 0.997988 | 0.994666 |
| Syn1_O88935_S682s_1 | Syn1_O889 | 0.77 | 0.72 | 0.36 | 0.46 | 1.18 | 1.29 | 0.84 | 0.73 | 0.2 | 0.44 | -0.43 | 0.49 | -0.36 | 0.63 | -0.19 | 0.36 | 0.4325 | -0.4025 | -0.4675 | 0.152648 | 0.092137 | 0.178467 | 0.989223 | 0.997988 | 0.994666 |
| Syn1_O88935_S684s_1 | Syn1_O889 | 0.89 | 0.34 | 1.64 | 1.97 | 1.17 | 1.43 | 2.84 | 2.49 | 0.06 | 0.34 | 1.85 | 2.98 | -0.6 | 0.5 | 2.23 | 2.75 | 0.7725 | 0.0975 | 0.01 | 0.989494 | 0.147052 | 0.888996 | 0.989223 | 0.997988 | 1 |
| Syn1_O88935_S9s_1 | Syn1_O889 | -0.76 | -0.38 | -0.38 | -0.27 | -0.19 | -0.8 | 0.21 | -0.28 | -0.83 | -0.37 | -0.32 | -0.24 | -0.77 | 0.05 | -0.38 | -0.23 | 0.1825 | 0.0075 | 0.115 | 0.682531 | 0.509885 | 0.975827 | 0.989223 | 0.997988 | 0.996159 |
| Syn1_O88935_T339t_1 | Syn1_O889 | -0.08 | 0.63 | 0.02 | 0.46 | 0.66 | -0.17 | 0.58 | 0.3 | -0.47 | -0.09 | -0.55 | 0.19 | -0.33 | 0.2 | 0.3 | 0.53 | 0.085 | -0.4875 | -0.0825 | 0.787901 | 0.766664 | 0.11732 | 0.995762 | 0.997988 | 0.999507 |
| Syn1_O88935_T448t_1 | Syn1_O889 | 0 | -0.01 | -2.73 | -2.78 | -0.8 | -0.9 | -0.68 | -2.75 | 0.3 | -0.07 | 1.49 | -1.89 | 0.34 | -0.73 | -1.64 | -1.24 | 0.0975 | 1.3375 | 0.5625 | 0.486201 | 0.904697 | 0.177089 | 0.996493 | 0.997988 | 0.994666 |
| Syn2_Q64332_S108s_1 | Syn2_Q643 | -2.73 | -1.48 | -3.36 | -2.38 | -2.36 | -0.62 | -0.78 | 1.32 | -0.71 | -2.07 | -3.4 | 0.18 | -2.36 | -2.31 | -3.65 | -1.1 | 1.8775 | 0.9875 | 0.1325 | 0.824065 | 0.02954 | 0.22821 | 0.989223 | 0.997988 | 0.999507 |
| Syn2_Q64332_S111s_1 | Syn2_Q643 | -2.13 | -1.24 | -2.26 | -0.61 | -1.73 | -0.74 | -0.17 | -0.29 | -0.85 | -1.68 | -1.02 | -0.36 | -2.04 | -1.87 | -0.17 | -0.44 | 0.8275 | 0.5825 | 0.43 | 0.457246 | 0.113464 | 0.222346 | 0.989223 | 0.997988 | 0.994666 |
| Syn2_Q64332_S426s_1 | Syn2_Q643 | 1.57 | 0.83 | 0.07 | 0.17 | 0.84 | 1.31 | 0.52 | 0.01 | 1.05 | 1.3 | 0.04 | 0.1 | 0.86 | 1.37 | 0.21 | 0.34 | 0.01 | -0.0375 | 0.035 | 0.93524 | 0.980962 | 0.934286 | 0.999187 | 0.997988 | 0.999507 |
| Syn2_Q64332_S426s_2 | Syn2_Q643 | -1.07 | -0.09 | -0.99 | -0.8 | -0.02 | -0.97 | 0.3 | -0.5 | -1.61 | -1.32 | -1.06 | -0.73 | -1.34 | -0.01 | -0.64 | -0.26 | 0.44 | -0.4425 | 0.175 | 0.647334 | 0.235826 | 0.189773 | 0.989223 | 0.997988 | 0.995259 |
| Syn2_Q64332_S480s_1 | Syn2_Q643 | -0.92 | -0.21 | 0.9 | 0.5 | 0.16 | -0.4 | 0.92 | 0.66 | -0.49 | -0.48 | 1.19 | 0.75 | -0.68 | 0.17 | 1.07 | 0.58 | 0.2675 | 0.175 | 0.2175 | 0.673105 | 0.565234 | 0.748834 | 0.989223 | 0.997988 | 0.996159 |
| Syn2_Q64332_S483s_1 | Syn2_Q643 | 0.19 | 1.32 | 0.55 | 0.38 | 0.35 | 0.37 | 0.21 | 0.88 | 0.63 | 0.36 | 0.97 | 1.02 | 0.26 | 0.42 | 0.43 | 1.04 | -0.1575 | 0.135 | -0.0725 | 0.83027 | 0.611897 | 0.675283 | 0.989223 | 0.997988 | 0.999507 |
| Syn2_Q64332_S53s_1 | Syn2_Q643 | 0.56 | 1.82 | 0.81 | 0.98 | 0.81 | 1.3 | 1.1 | 0.93 | 0.22 | 1.2 | 1.13 | 0.39 | -1.08 | 1.32 | 0.87 | 1.25 | -0.0075 | -0.3075 | -0.4525 | 0.440411 | 0.98079 | 0.422834 | 0.999187 | 0.997988 | 0.994666 |
| Syn2_Q64332_S546s_1 | Syn2_Q643 | -1.64 | -2.4 | -1.14 | -0.65 | -1.49 | -1.34 | -0.13 | -0.27 | -0.09 | -1.22 | -0.95 | -0.57 | -0.41 | 1.32 | -1.13 | -0.64 | 0.65 | 0.75 | 1.2425 | 0.060602 | 0.193265 | 0.110385 | 0.989223 | 0.997988 | 0.994666 |
| Syn2_Q64332_S552s_1 | Syn2_Q643 | -1.53 | -0.48 | -0.5 | -0.82 | 0.18 | -1.61 | -0.2 | -0.28 | -0.54 | -0.99 | -0.95 | -0.65 | -1.5 | -0.77 | -0.97 | -0.34 | 0.355 | 0.05 | -0.0625 | 0.864233 | 0.422177 | 0.869994 | 0.989223 | 0.997988 | 0.999507 |
| Syn2_Q64332_T422t_1 | Syn2_Q643 | -0.97 | -0.14 | -0.6 | -0.93 | -0.06 | -1.25 | -0.31 | -1 | -1.37 | -1.04 | -0.95 | -0.75 | -0.39 | -0.13 | -0.49 | -0.45 | 0.005 | -0.3675 | 0.295 | 0.312316 | 0.988409 | 0.213503 | 0.999187 | 0.997988 | 0.994666 |
| Syn2_Q64332_T422t_2 | Syn2_Q643 | -0.93 | 0.06 | -0.99 | -0.8 | 0.03 | -0.82 | 0.3 | -0.5 | -1.45 | -1.16 | -1.06 | -0.73 | -1.17 | 0.09 | -0.64 | -0.26 | 0.4175 | -0.435 | 0.17 | 0.655926 | 0.254883 | 0.192421 | 0.989223 | 0.997988 | 0.995259 |
| Syn2_Q64332_T449t_1 | Syn2_Q643 | -2.49 | -0.07 | -1.76 | -2.35 | -0.52 | -1.26 | -0.99 | -0.84 | -1.13 | -1.12 | -0.42 | -1.54 | 0.77 | 0.21 | -2.17 | -2.9 | 0.765 | 0.615 | 0.645 | 0.48895 | 0.167939 | 0.286089 | 0.989223 | 0.997988 | 0.994666 |
| Syn2_Q64332_T449t_1_1 | Syn2_Q643 | -0.97 | -0.97 | -1.5 | -1.1 | -0.72 | -0.25 | -0.81 | -1.63 | -0.78 | -0.95 | -1.94 | -1.14 | -1.01 | -0.65 | -1.77 | -1.25 | 0.2825 | -0.0675 | -0.035 | 0.911816 | 0.394991 | 0.831153 | 0.989223 | 0.997988 | 0.999507 |
| Syn3_Q8JZP2_S404s_1 | Syn3_Q8JZ | -0.53 | -1.31 | -1.51 | -0.4 | -0.35 | -0.13 | -0.03 | -0.21 | -1.62 | -0.8 | -0.29 | -0.78 | -1.95 | -0.88 | -1.04 | -0.92 | 0.7575 | 0.065 | -0.26 | 0.507063 | 0.031821 | 0.868029 | 0.989223 | 0.997988 | 0.994666 |
| Syn3_Q8JZP2_S418s_1 | Syn3_Q8JZ | 0.23 | -0.29 | 0.6 | 1.22 | 0.19 | 0.23 | 2 | 1.48 | -0.9 | 0.16 | 2.49 | 1.31 | -0.67 | -0.32 | 0.8 | 1.36 | 0.535 | 0.325 | -0.1475 | 0.781964 | 0.305856 | 0.652144 | 0.989223 | 0.997988 | 0.998226 |
| Syn3_Q8JZP2_S446s_1 | Syn3_Q8JZ | -1.2 | -1.15 | -1.21 | -2.56 | -2.24 | -1.02 | -2.28 | -0.93 | -1.6 | -1.77 | 1.32 | -0.82 | -0.1 | -1.03 | -1.37 | -1.8 | -0.0875 | 0.8125 | 0.455 | 0.352204 | 0.851979 | 0.270279 | 0.996349 | 0.997988 | 0.994666 |
| Syn3_Q8JZP2_S461s_1 | Syn3_Q8JZ | 0.54 | 0.3 | 0.3 | 0.43 | 0.16 | 0.11 | 0.19 | 0.22 | -0.03 | 0.43 | 0.37 | 0 | 0.37 | 0.65 | 0.38 | 0.29 | -0.2225 | -0.2 | 0.03 | 0.899202 | 0.274137 | 0.395337 | 0.989223 | 0.997988 | 0.999507 |
| Syn3_Q8JZP2_S461s_2 | Syn3_Q8JZ | 1.63 | 0.53 | 0.1 | 0.15 | -0.23 | 1.36 | 0.22 | 0.72 | 1.55 | 1.01 | -0.02 | 0.9 | 0.65 | -0.07 | 0.07 | 0.65 | -0.085 | 0.2575 | -0.2775 | 0.500521 | 0.853019 | 0.580108 | 0.996349 | 0.997988 | 0.994666 |
| Syn3_Q8JZP2_S469s_1 | Syn3_Q8JZ | -0.05 | 0.11 | -0.47 | -0.13 | -0.44 | 0.15 | -0.23 | -0.1 | -0.15 | -0.51 | -0.37 | 0.67 | 0.95 | -0.25 | -0.47 | 0.23 | -0.02 | 0.045 | 0.25 | 0.494586 | 0.933428 | 0.888116 | 0.998562 | 0.997988 | 0.994666 |
| Syn3_Q8JZP2_S469s_2 | Syn3_Q8JZ | 1.34 | 0.31 | -1.03 | -1.52 | 0.01 | 1 | 0.01 | -0.02 | 1.21 | 0.81 | -1.37 | 0.24 | 0.72 | 0.33 | -0.72 | -0.24 | 0.475 | 0.4475 | 0.2475 | 0.704999 | 0.450904 | 0.565462 | 0.989223 | 0.997988 | 0.996159 |
| Syn3_Q8JZP2_S480s_1 | Syn3_Q8JZ | 0.25 | -0.71 | -0.59 | -0.06 | 0.37 | 0.33 | 0.24 | 0.51 | -1.16 | -0.78 | 0.66 | 0.23 | -1.11 | 0.21 | -0.45 | -0.42 | 0.64 | 0.015 | -0.165 | 0.657909 | 0.03922 | 0.973988 | 0.989223 | 0.997988 | 0.995259 |
| Syn3_Q8JZP2_S480s_2 | Syn3_Q8JZ | 0.82 | -0.19 | 0.06 | 0.29 | -0.03 | 0.82 | 0.29 | 0.67 | -0.04 | -0.2 | 0.68 | 0.09 | -1.55 | 0.09 | 0.31 | 0.14 | 0.1925 | -0.1125 | -0.4975 | 0.301904 | 0.5368 | 0.724489 | 0.989223 | 0.997988 | 0.994666 |
| Syn3_Q8JZP2_S483s_1 | Syn3_Q8JZ | 0.4 | -0.11 | 0.64 | -0.37 | 0.29 | -0.02 | 0.26 | 0.04 | 0.66 | -0.18 | 0.38 | 0.92 | 0.38 | 0.62 | 0.42 | -1.69 | 0.0025 | 0.305 | -0.2075 | 0.7056 | 0.992863 | 0.389593 | 0.999187 | 0.997988 | 0.996159 |
| Syn3_Q8JZP2_S483s_2 | Syn3_Q8JZ | 0.88 | -0.31 | 0.06 | 0.29 | -0.08 | 0.88 | 0.29 | 0.67 | 0.01 | -0.22 | 0.68 | 0.09 | -1.72 | 0.1 | 0.31 | 0.14 | 0.21 | -0.09 | -0.5225 | 0.317636 | 0.535484 | 0.789101 | 0.989223 | 0.997988 | 0.994666 |
| Syn3_Q8JZP2_S494s_1 | Syn3_Q8JZ | 1.3 | -0.38 | -0.75 | 0.1 | 0.75 | 2.09 | -0.29 | 0.31 | 0.91 | 0.96 | -1.68 | -0.05 | 0.28 | -0.13 | 0.35 | 0.04 | 0.6475 | -0.0325 | 0.0675 | 0.880185 | 0.297753 | 0.962272 | 0.989223 | 0.997988 | 0.999507 |
| Syn3_Q8JZP2_S519s_1 | Syn3_Q8JZ | 0.37 | -0.58 | -0.09 | 0.37 | -0.55 | 0.8 | -0.3 | 0.12 | 1.62 | -0.26 | 1.08 | 0.21 | 1.44 | -0.77 | -0.04 | 0.31 | 3.14E-16 | 0.645 | 0.2175 | 0.657478 | 1 | 0.183786 | 1 | 0.997988 | 0.995259 |
| Syn3_Q8JZP2_S540s_1 | Syn3_Q8JZ | -0.69 | -0.42 | 0.47 | 0.66 | -0.15 | 0.76 | -0.15 | 0.09 | -0.18 | -0.61 | 1.21 | 0.86 | -0.38 | 0.28 | -0.14 | -0.32 | 0.1325 | 0.315 | -0.145 | 0.702403 | 0.73051 | 0.53918 | 0.995762 | 0.997988 | 0.996159 |
| Syn3_Q8JZP2_S544s_2 | Syn3_Q8JZ | -0.75 | 0.39 | -1.26 | 0.61 | -1.77 | 0.36 | -1.11 | 0.51 | 0.37 | -2.61 | 0.14 | -0.97 | 0.72 | 0.35 | -0.7 | 0.41 | -0.25 | -0.515 | 0.4475 | 0.392487 | 0.695716 | 0.487938 | 0.994893 | 0.997988 | 0.994666 |
| Syn3_Q8JZP2_S9s_1 | Syn3_Q8JZ | 0.25 | 0.44 | 0.45 | 0.36 | 0.7 | 0.39 | 0.47 | 0.59 | 0.25 | 0.41 | 0.53 | 0.85 | -0.26 | 0.7 | 0.29 | 0.34 | 0.1625 | 0.135 | -0.1075 | 0.702888 | 0.42851 | 0.562538 | 0.989223 | 0.997988 | 0.996159 |
| Syncrip_G3UZ48_S151s_1 | Syncrip_G3 | -0.99 | -0.53 | -0.57 | -1.03 | 0.23 | -1.04 | -0.47 | -1 | -0.93 | -0.26 | -2.11 | -0.35 | -0.37 | -0.67 | -0.75 | -0.87 | 0.21 | -0.1325 | 0.115 | 0.665568 | 0.53366 | 0.760261 | 0.989223 | 0.997988 | 0.996159 |
| Syndig1l_Q3USQ7_S15s_1 | Syndig1l_Q | -1.3 | -3.38 | -2.21 | -2.46 | -2.64 | -1.59 | -1.63 | -2.51 | -0.88 | -2.75 | -1.44 | -2.99 | -0.34 | -2.7 | -1.34 | -1.46 | 0.245 | 0.3225 | 0.8775 | 0.161094 | 0.606876 | 0.598611 | 0.989223 | 0.997988 | 0.994666 |
| Syne1_Q6ZWR6_S1360s_1 | Syne1_Q6Z | 0.74 | -2.25 | -1.37 | -2.56 | -1.02 | -0.39 | -1.58 | 0.33 | 0.06 | -1.6 | 0.37 | -0.67 | -1.03 | -0.48 | -1.65 | -0.33 | 0.695 | 0.9 | 0.4875 | 0.501137 | 0.362905 | 0.262193 | 0.989223 | 0.997988 | 0.994666 |
| Syne1_Q6ZWR6_S1369s_1 | Syne1_Q6Z | 0.13 | 1.34 | 0.23 | 0.48 | 0.87 | 0.15 | 0.18 | -0.14 | 0.3 | 0.09 | 0.09 | 0.59 | -0.03 | 0.34 | 0.63 | 1.83 | -0.28 | -0.2775 | 0.1475 | 0.753493 | 0.432389 | 0.403846 | 0.989223 | 0.997988 | 0.9971 |
| Syne1_Q6ZWR6_S6239s_1 | Syne1_Q6Z | 0.02 | -0.94 | -0.1 | 0.01 | -1.98 | -0.01 | -1.6 | -0.51 | 1.84 | -0.54 | -3.05 | -0.26 | 1.9 | -1.18 | -0.3 | -1.05 | -0.7725 | -0.25 | 0.095 | 0.887712 | 0.129414 | 0.784011 | 0.989223 | 0.997988 | 0.999507 |
| Syne1_Q6ZWR6_S8252s_1 | Syne1_Q6Z | -0.11 | 1.34 | 0.58 | 0.74 | 0.79 | -0.11 | 1.02 | 0.54 | -0.12 | 0.71 | -0.22 | 0.4 | 0.04 | 0.08 | 0.46 | -0.19 | -0.0775 | -0.445 | -0.54 | 0.151738 | 0.837192 | 0.255026 | 0.995762 | 0.997988 | 0.994666 |
| Syne1_Q6ZWR6_S8306s_1 | Syne1_Q6Z | -1.07 | -0.63 | -0.69 | -0.84 | -0.72 | -1.28 | -0.8 | -1.86 | -1 | -0.67 | -0.2 | -1.04 | -1.68 | 0 | -0.08 | -0.77 | -0.3575 | 0.08 | 0.175 | 0.667686 | 0.25611 | 0.770367 | 0.989223 | 0.997988 | 0.996159 |
| Syne2_E9QP46_S6349s_1 | Syne2_E9Q | -3.1 | -1.99 | -2.57 | -2.81 | -1.6 | -3.93 | -2.56 | -1.9 | -3.12 | -2.67 | -0.48 | -1.76 | -1.59 | -0.99 | -1.52 | -1.81 | 0.12 | 0.61 | 1.14 | 0.006523 | 0.817024 | 0.306664 | 0.995762 | 0.997988 | 0.994666 |
| Syngap1_F6SEU4_S1027s_1 | Syngap1_F | 0.01 | 0.07 | -0.45 | 0.06 | 0.34 | -1.01 | 0.82 | 0.46 | -0.4 | -0.17 | -0.17 | 0.13 | -0.72 | 0.13 | 0.16 | 0.35 | 0.23 | -0.075 | 0.0575 | 0.856453 | 0.571289 | 0.760077 | 0.989223 | 0.997988 | 0.999507 |
| Syngap1_F6SEU4_S1033s_1 | Syngap1_F | -0.66 | -0.19 | -0.98 | -1.01 | -0.09 | -0.47 | -0.53 | -0.83 | 0.07 | 0.04 | -1.75 | -0.99 | -0.2 | -0.29 | -1.1 | -0.47 | 0.23 | 0.0525 | 0.195 | 0.550509 | 0.419605 | 0.908885 | 0.989223 | 0.997988 | 0.994666 |
| Syngap1_F6SEU4_S1053s_1 | Syngap1_F | 0.02 | -0.25 | -0.07 | -0.01 | -0.31 | -0.36 | -0.03 | -0.4 | -0.6 | 0.36 | -0.42 | -0.12 | 0.21 | -0.2 | -0.26 | -0.57 | -0.1975 | -0.1175 | -0.1275 | 0.632615 | 0.355173 | 0.669487 | 0.989223 | 0.997988 | 0.995259 |
| Syngap1_F6SEU4_S1070s_1 | Syngap1_F | -4.89 | -5.04 | -3.33 | -4.24 | -3.78 | -6.33 | -2.76 | -4.04 | -5.44 | -4.76 | 0.98 | -4.53 | -3.56 | -3.67 | -3.97 | -3.65 | 0.1475 | 0.9375 | 0.6625 | 0.126163 | 0.842904 | 0.491088 | 0.996349 | 0.997988 | 0.994666 |
| Syngap1_F6SEU4_S1091s_1 | Syngap1_F | -1.4 | -1.74 | 0.17 | 0.18 | -0.86 | -0.49 | 0.62 | -0.08 | -1.18 | -1.13 | 1.25 | 0.32 | -0.94 | 0.67 | -0.33 | -0.45 | 0.495 | 0.5125 | 0.435 | 0.447191 | 0.372391 | 0.472666 | 0.989223 | 0.997988 | 0.994666 |
| Syngap1_F6SEU4_S1111s_1 | Syngap1_F | -0.18 | 0.93 | 0.85 | 1.01 | 0.79 | 1.41 | 0.91 | 1.96 | 0.82 | 1.06 | 1.89 | 1.26 | 0.21 | 1.93 | 0.17 | 0.13 | 0.615 | 0.605 | -0.0425 | 0.93145 | 0.128937 | 0.127494 | 0.989223 | 0.997988 | 0.999507 |
| Syngap1_F6SEU4_S1115s_1 | Syngap1_F | 0.02 | 0.11 | 0.05 | 0.13 | -0.01 | -0.18 | -0.13 | -0.04 | 0.13 | 0.03 | -0.31 | 0.15 | -0.01 | 0.56 | -0.31 | 0.03 | -0.1675 | -0.0775 | -0.01 | 0.970483 | 0.396145 | 0.726262 | 0.989223 | 0.997988 | 0.999507 |
| Syngap1_F6SEU4_S1118s_1 | Syngap1_F | -0.76 | -0.04 | -0.58 | -0.92 | -0.39 | -0.87 | -0.89 | -0.49 | -0.69 | -0.1 | -0.32 | -0.94 | -0.53 | -0.78 | -1.3 | -0.33 | -0.085 | 0.0625 | -0.16 | 0.626608 | 0.754879 | 0.83755 | 0.995762 | 0.997988 | 0.995259 |
| Syngap1_F6SEU4_S1135s_1 | Syngap1_F | 1.34 | 0.12 | 1.65 | 0.97 | 0.62 | 1.5 | 0.89 | 0.4 | 1.25 | 1.84 | 0.87 | 1.67 | 0.38 | 0.74 | 1.37 | 1.1 | -0.1675 | 0.3875 | -0.1225 | 0.760263 | 0.671733 | 0.339856 | 0.992876 | 0.997988 | 0.9971 |
| Syngap1_F6SEU4_S1162s_1 | Syngap1_F | -1.42 | -1.06 | -1.21 | -0.91 | -0.79 | -1.04 | -0.44 | -0.86 | -1.71 | -0.88 | -1.84 | -0.87 | -1.2 | -0.85 | -1.79 | -0.97 | 0.3675 | -0.175 | -0.0525 | 0.860267 | 0.143452 | 0.581251 | 0.989223 | 0.997988 | 0.999507 |
| Syngap1_F6SEU4_S117s_1 | Syngap1_F | -0.01 | -0.15 | -0.1 | 0.41 | 0.42 | 0.73 | 0.32 | 0.77 | -0.22 | -0.13 | 0.18 | 0.3 | -0.88 | 0.5 | 0 | -0.04 | 0.5225 | -0.005 | -0.1425 | 0.68122 | 0.050086 | 0.984112 | 0.989223 | 0.997988 | 0.996159 |
| Syngap1_F6SEU4_S1201s_1 | Syngap1_F | -2.46 | -1.74 | -0.55 | -3.55 | -1.11 | -3.3 | -1.27 | -2.15 | -4.58 | -2.94 | -0.71 | -0.51 | -1.91 | -0.94 | -1.82 | -2.42 | 0.1175 | -0.11 | 0.3025 | 0.635466 | 0.868461 | 0.914056 | 0.996493 | 0.997988 | 0.995259 |
| Syngap1_F6SEU4_S165s_1 | Syngap1_F | 1.67 | 0.13 | 1.15 | 1.1 | -0.11 | 2.07 | 0.17 | 1.15 | 3.25 | 0.81 | 1.99 | 1.22 | 1.96 | 0.28 | 0.75 | 1.09 | -0.1925 | 0.805 | 0.0075 | 0.987035 | 0.720307 | 0.185279 | 0.995437 | 0.997988 | 1 |
| Syngap1_F6SEU4_S534s_1 | Syngap1_F | 0.7 | 0.85 | -0.23 | -0.78 | 1.1 | -0.76 | 0.02 | -1.05 | -0.29 | -0.36 | -0.94 | 0.14 | 0.41 | 0.84 | -0.06 | -0.56 | -0.3075 | -0.4975 | 0.0225 | 0.961895 | 0.583764 | 0.268913 | 0.989223 | 0.997988 | 0.999507 |
| Syngap1_F6SEU4_S535s_1 | Syngap1_F | -0.77 | -0.36 | -0.31 | -0.08 | -0.23 | -0.36 | 0.21 | -0.04 | -0.68 | -0.77 | -1.25 | -0.28 | 0.06 | -0.22 | -0.32 | 0.49 | 0.275 | -0.365 | 0.3825 | 0.216314 | 0.283528 | 0.229088 | 0.989223 | 0.997988 | 0.994666 |
| Syngap1_F6SEU4_S752s_1 | Syngap1_F | 1.25 | -0.02 | 0.09 | 0.81 | 0.72 | 1.18 | 0.73 | 1.07 | 0.32 | 0.7 | 0.74 | 0.46 | -0.91 | 1.01 | -0.1 | 0.05 | 0.3925 | 0.0225 | -0.52 | 0.289565 | 0.25627 | 0.946939 | 0.989223 | 0.997988 | 0.994666 |
| Syngap1_F6SEU4_S766s_1 | Syngap1_F | 0.55 | 0.03 | 0.06 | 0.57 | 0.7 | 1.09 | 0.83 | 1.51 | -0.21 | -0.04 | 0.99 | 0.74 | -1.73 | 0.49 | -0.52 | 0.06 | 0.73 | 0.0675 | -0.7275 | 0.154731 | 0.022561 | 0.845139 | 0.989223 | 0.997988 | 0.994666 |
| Syngap1_F6SEU4_S779s_1 | Syngap1_F | 0.42 | -1.2 | -0.35 | 0.37 | 0.25 | 1.2 | 0.12 | 1.09 | 0.24 | -0.59 | 1.1 | 0.13 | -1.18 | 0.52 | -0.82 | -0.64 | 0.855 | 0.41 | -0.34 | 0.501717 | 0.078314 | 0.408733 | 0.989223 | 0.997988 | 0.994666 |
| Syngap1_F6SEU4_S780s_1 | Syngap1_F | 0.56 | -1.18 | -0.34 | 0.21 | 0.37 | 1.08 | 0.05 | 1.08 | 0.03 | -0.62 | 1.11 | 0.12 | -1.56 | 0.8 | -1.05 | -0.94 | 0.8325 | 0.3475 | -0.5 | 0.402297 | 0.079716 | 0.485916 | 0.989223 | 0.997988 | 0.994666 |
| Syngap1_F6SEU4_S824s_1 | Syngap1_F | 1.48 | -1.03 | 0.43 | 0.92 | -0.24 | 2.33 | 0.36 | 1.24 | 0.36 | -0.11 | 0.99 | 0.58 | -1.19 | 0.74 | 0.39 | 0.15 | 0.4725 | 0.005 | -0.4275 | 0.496581 | 0.495917 | 0.992644 | 0.989223 | 0.997988 | 0.994666 |
| Syngap1_F6SEU4_S87s_1 | Syngap1_F | -1.15 | -1.08 | -5.12 | -10.31 | -0.65 | -0.89 | -6.77 | -6.24 | -1.4 | -1.56 | -0.79 | -5.95 | -0.8 | -0.48 | -5.58 | -5.32 | 0.7775 | 1.99 | 1.37 | 0.538887 | 0.739683 | 0.367794 | 0.995762 | 0.997988 | 0.994666 |
| Syngap1_F6SEU4_S892s_1 | Syngap1_F | 0.22 | 0.12 | 0.84 | -0.13 | -0.29 | 0.01 | 0.43 | 0.36 | 0.31 | 0.23 | -0.4 | 0.5 | 0.74 | 0.3 | -0.24 | -0.34 | -0.135 | -0.1025 | -0.1475 | 0.677501 | 0.647061 | 0.745588 | 0.99033 | 0.997988 | 0.996159 |
| Syngap1_F6SEU4_S895s_1 | Syngap1_F | 1.07 | 1.3 | 1.26 | 0.69 | 1.03 | 1.1 | 1.66 | 1.38 | -0.99 | 1.62 | -1.23 | 1.83 | 0 | -1.62 | 1.32 | 0.42 | 0.2125 | -0.7725 | -1.05 | 0.095319 | 0.411705 | 0.315721 | 0.989223 | 0.997988 | 0.994666 |
| Syngap1_F6SEU4_S985s_1 | Syngap1_F | -0.72 | -0.31 | -0.9 | -0.57 | -0.06 | -1.29 | 0.3 | -0.32 | -1.26 | -0.7 | -1.15 | -0.53 | -1.77 | 0.3 | -0.5 | -0.62 | 0.2825 | -0.285 | -0.0225 | 0.958851 | 0.441037 | 0.309819 | 0.989223 | 0.997988 | 0.999507 |
| Syngap1_F6SEU4_Y100y_1 | Syngap1_F | 3.52 | 0.48 | 2.54 | -0.14 | 0.76 | 3.75 | 3.04 | 1.32 | 1.07 | 1.28 | 1.09 | 1.83 | 1.21 | 0.23 | 2.96 | 2.46 | 0.6175 | -0.2825 | 0.115 | 0.900941 | 0.525468 | 0.719393 | 0.989223 | 0.997988 | 0.999507 |
| Syngap1_F6SEU4_Y34y_1 | Syngap1_F | 0.29 | 0.22 | -0.36 | 0.25 | 0.11 | 0.34 | -0.2 | 0.28 | 1.07 | 0.48 | -0.24 | -0.34 | 0.66 | -0.25 | 0.23 | 0.12 | 0.0325 | 0.1425 | 0.09 | 0.766221 | 0.897131 | 0.702657 | 0.996493 | 0.997988 | 0.9971 |
| Syngap1_F6SEU4_Y39y_1 | Syngap1_F | 0.15 | 0.44 | 0.59 | 0.34 | 0.19 | -0.04 | 1.53 | -0.12 | 0.36 | -0.86 | 0.05 | 0.32 | 0.18 | 0.17 | -0.5 | 0.12 | 0.01 | -0.4125 | -0.3875 | 0.17904 | 0.979284 | 0.225239 | 0.999187 | 0.997988 | 0.994666 |
| Syngap1_J3QQ18_S1298s_1 | Syngap1_J3 | 0.08 | 0.58 | -1.04 | -0.24 | 0.92 | 0.76 | -0.19 | 0.14 | 0.09 | -0.36 | -0.39 | -0.91 | -0.51 | 0.94 | -0.84 | -0.24 | 0.5625 | -0.2375 | -0.0075 | 0.987771 | 0.191329 | 0.552173 | 0.989223 | 0.997988 | 1 |
| Syngr3_Q8R191_S5s_1 | Syngr3_Q8 | 0.44 | -1.05 | -0.53 | -0.45 | -1.64 | -0.48 | -0.72 | 0.32 | 0.93 | 0.55 | 0.2 | -0.21 | 0.93 | 0.17 | 1.03 | -0.22 | -0.2325 | 0.765 | 0.875 | 0.064868 | 0.623498 | 0.077695 | 0.989223 | 0.997988 | 0.994666 |
| Syngr3_Q8R191_T133t_1 | Syngr3_Q8 | 0.89 | 1.65 | 1.52 | 1.17 | 1.38 | 1.02 | 0.17 | 0.53 | 0.75 | 1.25 | 0.55 | 0.93 | 1.11 | 1.13 | 0.76 | 0.77 | -0.5325 | -0.4375 | -0.365 | 0.211834 | 0.129866 | 0.142587 | 0.989223 | 0.997988 | 0.994666 |
| Synj1_F6XIX0_S44s_1 | Synj1_F6XI | -1.2 | -0.79 | -1.19 | 0.03 | 0.26 | -1.13 | 0.24 | -0.24 | -1.32 | -1.08 | -0.27 | 0.61 | -1.14 | 0.23 | -0.82 | -0.35 | 0.57 | 0.2725 | 0.2675 | 0.524067 | 0.190962 | 0.584473 | 0.989223 | 0.997988 | 0.994666 |
| Synj1_F7BQW7_S1064s_1 | Synj1_F7B | 0.06 | -0.99 | -0.51 | -0.14 | 0.08 | -0.04 | 0.54 | -0.11 | -0.39 | -0.58 | 0.27 | -0.49 | -1.26 | -0.05 | -0.97 | -0.21 | 0.5125 | 0.0975 | -0.2275 | 0.558402 | 0.111443 | 0.765804 | 0.989223 | 0.997988 | 0.995259 |
| Synj1_F7BQW7_S1068s_1 | Synj1_F7B | 0.57 | -1.25 | -0.34 | 0.45 | 0.1 | 0.54 | 0.49 | 0.44 | -0.09 | -0.85 | -0.56 | 0.04 | -2.56 | -0.33 | -0.31 | -0.11 | 0.535 | -0.2225 | -0.685 | 0.304318 | 0.214993 | 0.62391 | 0.989223 | 0.997988 | 0.994666 |
| Synj1_F7BQW7_S1099s_1 | Synj1_F7B | 0.37 | 0.49 | 0.6 | 0.67 | 0.4 | 0.16 | 0.1 | 0.49 | 0.24 | 0.42 | 0.64 | 0.48 | 0.5 | 0.9 | 0.49 | 0.12 | -0.245 | -0.0875 | -0.03 | 0.910015 | 0.266056 | 0.690965 | 0.989223 | 0.997988 | 0.999507 |
| Synj1_F7BQW7_S1162s_1 | Synj1_F7B | -1.87 | -2.58 | -2.59 | -4.06 | -2.04 | -1.79 | -1.68 | -1.62 | -5.03 | -2.91 | -0.64 | -2.4 | -3.15 | -3.05 | -3.5 | -3.27 | 0.9925 | 0.03 | -0.4675 | 0.319175 | 0.046118 | 0.973363 | 0.989223 | 0.997988 | 0.994666 |
| Synj1_F7BQW7_S1175s_1 | Synj1_F7B | 3.19 | -0.27 | 1.06 | 2.93 | -0.18 | 2.7 | 1.09 | 3.59 | 1.46 | 2.28 | 1.11 | 1.53 | 0.1 | 0.13 | 0.85 | 1.08 | 0.0725 | -0.1325 | -1.1875 | 0.142065 | 0.94285 | 0.862405 | 0.999187 | 0.997988 | 0.994666 |
| Synj1_F7BQW7_S1190s_1 | Synj1_F7B | -0.95 | -0.71 | -1.53 | -0.48 | -1.21 | -1.48 | -0.23 | -0.39 | -1.59 | -0.38 | -1.21 | -0.17 | -1.41 | -0.31 | -0.94 | -0.96 | 0.09 | 0.08 | 0.0125 | 0.971445 | 0.809546 | 0.842329 | 0.995762 | 0.997988 | 0.999507 |
| Synj1_F7BQW7_S11s_1 | Synj1_F7B | -0.68 | -0.95 | -0.05 | -0.98 | -1.34 | 0.11 | -0.77 | 0.03 | 3.12 | -1.03 | -1.22 | 0.71 | 2.76 | -1.98 | 2.24 | -1.47 | 0.1725 | 1.06 | 1.0525 | 0.343123 | 0.662566 | 0.26342 | 0.990434 | 0.997988 | 0.994666 |
| Synj1_F7BQW7_S1308s_1 | Synj1_F7B | 1.66 | 1.4 | 0.69 | 0.3 | 1.23 | 1.16 | 0 | 0.86 | 0.21 | 1.09 | 0.94 | -0.01 | -0.06 | 0.99 | -0.01 | -0.26 | -0.2 | -0.455 | -0.8475 | 0.067682 | 0.62327 | 0.282188 | 0.989223 | 0.997988 | 0.994666 |
| Synj1_F7BQW7_T1063t_1 | Synj1_F7B | -0.05 | -0.74 | -1.15 | -1.79 | -0.02 | -0.3 | -0.07 | -1.65 | -0.61 | -0.83 | -0.04 | -1.93 | -0.94 | -0.6 | -1.61 | -1.05 | 0.4225 | 0.08 | -0.1175 | 0.779736 | 0.39629 | 0.874295 | 0.989223 | 0.997988 | 0.998226 |
| Synm_Q70IV5_S1075s_1 | Synm_Q70 | -2.66 | -1.2 | 0.14 | -0.11 | -1.79 | -2.11 | 0.14 | -0.08 | -1.61 | -1.55 | -0.57 | 1.01 | -0.95 | -1.73 | -0.11 | 0.58 | -0.0025 | 0.2775 | 0.405 | 0.57866 | 0.997352 | 0.725979 | 1 | 0.997988 | 0.995259 |
| Synm_Q70IV5_S172s_1 | Synm_Q70 | -0.33 | 0.08 | 0.03 | -0.09 | -0.1 | -0.62 | 0.27 | 0.02 | 0.41 | 0.19 | -0.17 | 0.53 | 1.03 | -0.95 | 0.14 | 0.53 | -0.03 | 0.3175 | 0.265 | 0.538533 | 0.90761 | 0.227827 | 0.997192 | 0.997988 | 0.994666 |
| Synpo_Q8CC35_S19s_1 | Synpo_Q8C | 0.82 | 0.5 | 0.8 | 0.72 | 0.87 | 0.77 | 0.74 | 0.63 | 0.29 | 0.33 | 0.25 | 0.73 | -0.75 | 0.76 | 0.35 | 0.38 | 0.0425 | -0.31 | -0.525 | 0.166626 | 0.834623 | 0.19994 | 0.995762 | 0.997988 | 0.994666 |
| Synpo_Q8CC35_S328s_1 | Synpo_Q8C | 0.2 | 0.61 | 0.45 | 0.53 | 0.63 | 0.3 | 0.7 | 0.57 | 0.15 | 0.09 | 0.03 | 0.46 | 0.23 | 0.73 | 0.39 | 0.41 | 0.1025 | -0.265 | -0.0075 | 0.976103 | 0.637967 | 0.264877 | 0.989223 | 0.997988 | 1 |
| Synpo_Q8CC35_S331s_1 | Synpo_Q8C | 1.3 | 1.02 | 1.9 | 1.63 | 1.07 | 1.25 | 1.62 | 1.83 | 0.87 | 0.69 | 0.78 | 1.37 | 0.48 | 1.6 | 0.77 | 0.67 | -0.02 | -0.535 | -0.5825 | 0.116963 | 0.944747 | 0.091242 | 0.999187 | 0.997988 | 0.994666 |
| Synpo_Q8CC35_S433s_1 | Synpo_Q8C | 0.6 | 0.47 | -0.25 | 0.2 | 0.3 | 0.45 | 0.65 | 0.2 | -0.02 | 0.29 | -0.27 | 0.23 | -1.03 | 0.01 | -0.09 | 0.51 | 0.145 | -0.1975 | -0.405 | 0.307412 | 0.580899 | 0.485687 | 0.989223 | 0.997988 | 0.994666 |
| Synpo_Q8CC35_S501s_1 | Synpo_Q8C | 0.42 | -0.08 | 0.28 | 0.52 | 0.29 | 0.55 | 0.9 | 0.65 | -0.55 | 0.38 | -0.3 | 0.03 | -0.69 | 0.19 | 0.2 | 0.42 | 0.3125 | -0.395 | -0.255 | 0.438534 | 0.22022 | 0.192126 | 0.989223 | 0.997988 | 0.994666 |
| Synpo_Q8CC35_S566s_1 | Synpo_Q8C | 0.61 | 0.06 | -0.02 | 0.41 | 0.99 | 1.13 | 0.77 | 0.71 | -0.15 | 0.62 | 0.01 | 0.1 | -0.8 | 0.62 | 0.19 | 0.04 | 0.635 | -0.12 | -0.2525 | 0.486448 | 0.024341 | 0.665997 | 0.989223 | 0.997988 | 0.994666 |
| Synpo_Q8CC35_S594s_1 | Synpo_Q8C | 1.09 | -0.11 | 1.46 | 1.32 | 1.16 | 1.18 | 1.63 | 1.89 | 0.56 | 0.69 | 0.69 | 1.2 | 0.17 | 1.09 | 0.9 | 1.54 | 0.525 | -0.155 | -0.015 | 0.973184 | 0.197864 | 0.689948 | 0.989223 | 0.997988 | 0.999507 |
| Synpo_Q8CC35_S615s_1 | Synpo_Q8C | 0.11 | -1.28 | -0.73 | -0.14 | 0.58 | 0.5 | 0.56 | 0.51 | -0.85 | -0.87 | 0.46 | -0.07 | -1.37 | -0.08 | -0.63 | -0.21 | 1.0475 | 0.1775 | -0.0625 | 0.88252 | 0.009112 | 0.684688 | 0.989223 | 0.997988 | 0.999507 |
| Synpo_Q8CC35_S643s_1 | Synpo_Q8C | 0.78 | 0.77 | 0.9 | 1.6 | 0.49 | 0.79 | 1.2 | 1.3 | 0.54 | 0.62 | -0.04 | 1.19 | 0.47 | 0.92 | -0.08 | -0.02 | -0.0675 | -0.435 | -0.69 | 0.06748 | 0.821059 | 0.221404 | 0.995762 | 0.997988 | 0.994666 |
| Synpo_Q8CC35_S654s_1 | Synpo_Q8C | 0.82 | -0.25 | -0.54 | 1.16 | 0.96 | 0.75 | 1.05 | 1.04 | 0.43 | 0.02 | -0.09 | 0.75 | -0.9 | 0.26 | 0.27 | 0.44 | 0.6525 | -0.02 | -0.28 | 0.569616 | 0.127991 | 0.963618 | 0.989223 | 0.997988 | 0.995259 |
| Synpo_Q8CC35_S819s_1 | Synpo_Q8C | -3.1 | -1.63 | -0.93 | -1.11 | -2.97 | -2.07 | -1.49 | -1.1 | -2.97 | -3.08 | -0.79 | -1.09 | -0.87 | -5.36 | -0.88 | -0.39 | -0.215 | -0.29 | -0.1825 | 0.86795 | 0.708643 | 0.681316 | 0.995437 | 0.997988 | 0.999507 |
| Synpo_Q8CC35_S892s_1 | Synpo_Q8C | -0.56 | -0.58 | -1.75 | -1.09 | -0.53 | 0.4 | -1.26 | -1.05 | 0.44 | -0.08 | -0.11 | -1.52 | 0.7 | -0.66 | -1.39 | -1.05 | 0.385 | 0.6775 | 0.395 | 0.442877 | 0.387807 | 0.180932 | 0.989223 | 0.997988 | 0.994666 |
| Synpo_Q8CC35_T308t_1 | Synpo_Q8C | 0.08 | 0.2 | 0.09 | 0.47 | 0.35 | -0.46 | 0.25 | 0.34 | 0.83 | -0.4 | 0.57 | -0.13 | 0.82 | 1.83 | -0.39 | 0.09 | -0.09 | 0.0075 | 0.3775 | 0.431757 | 0.73214 | 0.981743 | 0.995762 | 0.997988 | 0.994666 |
| Synpo_Q8CC35_T503t_1 | Synpo_Q8C | -0.43 | 0.1 | 1.23 | 1.28 | -0.95 | 0.19 | 1.99 | 2.19 | -0.22 | -0.21 | 1.23 | 1.72 | -0.79 | 0.1 | -0.17 | 0.72 | 0.31 | 0.085 | -0.58 | 0.262318 | 0.683048 | 0.887162 | 0.994272 | 0.997988 | 0.994666 |
| Synpo_Q8CC35_T530t_1 | Synpo_Q8C | -0.56 | -0.33 | -0.48 | 0.13 | -0.42 | -1.23 | 0.06 | -0.68 | -0.96 | -0.53 | -1.01 | -0.86 | -0.79 | 0.09 | -0.2 | 0.41 | -0.2575 | -0.53 | 0.1875 | 0.579643 | 0.433581 | 0.063844 | 0.989223 | 0.997988 | 0.995259 |
| Synrg_V9GXD9_S743s_1 | Synrg_V9G | -1.28 | -0.67 | -0.98 | -1.23 | -0.16 | -0.81 | -1.53 | -0.99 | -1.84 | -1.35 | -2.84 | -1.17 | -1.69 | -0.84 | -1.27 | -1.01 | 0.1675 | -0.76 | -0.1625 | 0.586151 | 0.610709 | 0.082299 | 0.989223 | 0.997988 | 0.995259 |
| Synrg_V9GXD9_S803s_1 | Synrg_V9G | -0.66 | -1.06 | -1.42 | -0.35 | -0.12 | -0.06 | -0.69 | -0.84 | -0.94 | -0.6 | -1.68 | -0.65 | -0.54 | -0.81 | -0.73 | -0.96 | 0.445 | -0.095 | 0.1125 | 0.713998 | 0.186988 | 0.789156 | 0.989223 | 0.997988 | 0.996966 |
| Synrg_V9GXD9_S920s_1 | Synrg_V9G | -0.79 | -1.22 | -1.36 | -0.41 | -1.42 | -0.92 | -1.58 | -1.35 | -1.15 | -0.86 | -1.59 | -1.38 | -0.94 | -0.81 | -1.2 | -1.5 | -0.3725 | -0.3 | -0.1675 | 0.59744 | 0.216938 | 0.336777 | 0.989223 | 0.997988 | 0.995259 |
| Synrg_V9GXD9_S989s_1 | Synrg_V9G | -1.86 | -1.82 | -0.82 | -1.91 | -0.31 | -2.06 | -1.4 | -0.9 | -2.74 | -1.67 | 0.12 | -0.46 | -1.06 | -0.33 | -1.78 | -1.5 | 0.435 | 0.415 | 0.435 | 0.305504 | 0.324316 | 0.51396 | 0.989223 | 0.997988 | 0.994666 |
| Syp_Q62277_Y81y_1 | Syp_Q6227 | 5.91 | 3.2 | 3.04 | 1.24 | 3.62 | 5.88 | 3.31 | 3.08 | 0.3 | 5.28 | 0.92 | 3.31 | -0.61 | 0.67 | 3.24 | 3.34 | 0.625 | -0.895 | -1.6875 | 0.177331 | 0.537077 | 0.499757 | 0.989223 | 0.997988 | 0.994666 |
| Syt1_P46096_S264s_1 | Syt1_P460 | 2.76 | 2.98 | -1.46 | -3.36 | 2.51 | 2.81 | -2.55 | -2.69 | 1.51 | 2.78 | -1.89 | -2.16 | -0.71 | 1.64 | -3.07 | -2.82 | -0.21 | -0.17 | -1.47 | 0.381406 | 0.910568 | 0.922147 | 0.997335 | 0.997988 | 0.994666 |
| Syt1_P46096_T128t_1 | Syt1_P460 | -3.1 | -2.56 | -2.62 | -2.79 | -1.93 | -3.11 | -2.51 | -3.89 | -2.74 | -2.5 | -1.08 | -2.71 | -2.52 | -2.01 | -3.3 | -2.87 | -0.0925 | 0.51 | 0.0925 | 0.783499 | 0.823908 | 0.232446 | 0.995762 | 0.997988 | 0.998491 |
| Syt1_P46096_T201t_1 | Syt1_P460 | 0.19 | 0.52 | 0.82 | 0.91 | 0.33 | 0.25 | 0.4 | 0.55 | 0.32 | 0.4 | 0.19 | 0.4 | -0.33 | 0.93 | -0.36 | 0.19 | -0.2275 | -0.2825 | -0.5025 | 0.190622 | 0.355344 | 0.27112 | 0.989223 | 0.997988 | 0.994666 |
| Syt1_P46096_T285t_1 | Syt1_P460 | 1.7 | 1.61 | 1.4 | 1.1 | 1.19 | 1.64 | 0.51 | 0.53 | 1.49 | 1.5 | 0.69 | 0.97 | 1.06 | 0.67 | 0.52 | 1.1 | -0.485 | -0.29 | -0.615 | 0.048433 | 0.153378 | 0.326825 | 0.989223 | 0.997988 | 0.994666 |
| Syt1_P46096_T334t_1 | Syt1_P460 | 0.64 | 0.29 | -0.57 | 0.63 | 0.49 | 0.18 | 0.69 | 0.4 | -0.22 | 0.94 | -0.02 | 0.08 | -0.3 | 0.34 | 0.5 | -0.11 | 0.1925 | -0.0525 | -0.14 | 0.701118 | 0.549738 | 0.891525 | 0.989223 | 0.997988 | 0.996159 |
| Syt10_Q9R0N4_S169s_1 | Syt10_Q9R | -0.56 | -1.12 | -1.72 | -1.34 | -1.35 | 0.15 | -0.5 | -0.8 | 0.93 | -1.17 | -1 | -1.16 | 2.01 | -2.31 | 0.14 | -0.75 | 0.56 | 0.585 | 0.9575 | 0.261876 | 0.168869 | 0.284415 | 0.989223 | 0.997988 | 0.994666 |
| Syt10_Q9R0N4_S218s_1 | Syt10_Q9R | -1.93 | -1.82 | -1.53 | -4.08 | -1.81 | -1.37 | -1.45 | -1.61 | 0.94 | -1.8 | -2.8 | -1.99 | 2.23 | -2.74 | 0.78 | -1.37 | 0.78 | 0.9275 | 2.065 | 0.082291 | 0.170561 | 0.31209 | 0.989223 | 0.997988 | 0.994666 |

| | | | | | | | | | | | | | | | | | | | | | | | | | | |
|---|---|---|---|---|---|---|---|---|---|---|---|---|---|---|---|---|---|---|---|---|---|---|---|---|---|---|
| Syt11_Q9R0N3_S113s_1 | Syt11_Q9R | 0.33 | -0.05 | 0.57 | 0.44 | 0.11 | 0.2 | 1.14 | 0.61 | 0.12 | 0.33 | 0.24 | 0.41 | 0.15 | 0.37 | 0.77 | 0.38 | 0.1925 | -0.0475 | 0.095 | 0.727505 | 0.521105 | 0.840789 | 0.989223 | 0.997988 | 0.9971 |
| Syt11_Q9R0N3_S133s_1 | Syt11_Q9R | 0.43 | 1.06 | 0.41 | 0.58 | 0.9 | 0.27 | 1 | 0.47 | 0.36 | 0.89 | 0.58 | 0.48 | -0.37 | 0.33 | 0.76 | 0.87 | 0.04 | -0.0425 | -0.2225 | 0.528805 | 0.882863 | 0.868901 | 0.996493 | 0.997988 | 0.994666 |
| Syt11_Q9R0N3_S416s_1 | Syt11_Q9R | -1.45 | -2.71 | -0.74 | -1.25 | -3.3 | -0.71 | -1.35 | -2.02 | -0.64 | -1.98 | 0.53 | -0.91 | 0.15 | -3.45 | -1.12 | -1.72 | -0.3075 | 0.7875 | 0.0025 | 0.997374 | 0.621346 | 0.215976 | 0.989223 | 0.997988 | 1 |
| Syt11_Q9R0N3_T139t_1 | Syt11_Q9R | -0.52 | -0.94 | -0.62 | -0.44 | -0.8 | -0.57 | -0.36 | -0.39 | 0.09 | -0.32 | -0.04 | 0.18 | -0.68 | -0.19 | -0.25 | -1.42 | 0.1 | 0.6075 | -0.005 | 0.988241 | 0.661742 | 0.029268 | 0.990434 | 0.997988 | 1 |
| Syt14_B7ZP31_S195s_2 | Syt14_B7ZI | 1.1 | 0.94 | 1.67 | 0.81 | -0.55 | 1.51 | 0.52 | 0.75 | 2.48 | 0.86 | 1.46 | 0.36 | 1.64 | -0.1 | 1.09 | 1.28 | -0.5725 | 0.16 | -0.1525 | 0.717438 | 0.211706 | 0.734423 | 0.989223 | 0.997988 | 0.996966 |
| Syt14_B7ZP31_S197s_2 | Syt14_B7ZI | 1.1 | 0.94 | 1.67 | 0.81 | -0.55 | 1.51 | 0.52 | 0.75 | 2.48 | 0.86 | 1.46 | 0.36 | 1.64 | -0.1 | 1.09 | 1.28 | -0.5725 | 0.16 | -0.1525 | 0.717438 | 0.211706 | 0.734423 | 0.989223 | 0.997988 | 0.996966 |
| Syt17_Q920M7_S100s_1 | Syt17_Q92 | -0.18 | -1.22 | -1.04 | -1.33 | 0.19 | 0.25 | -0.29 | -0.06 | -0.29 | -0.82 | -0.29 | -0.53 | -0.42 | 0.59 | -0.47 | -0.29 | 0.965 | 0.46 | 0.795 | 0.058433 | 0.010777 | 0.173308 | 0.989223 | 0.997988 | 0.994666 |
| Syt17_Q920M7_S114s_1 | Syt17_Q92 | 0.38 | -1.96 | 0.69 | 1.17 | -0.18 | 0.72 | 0.73 | 1.19 | -0.21 | -0.27 | -0.89 | 1.41 | -1.96 | 0.68 | -0.61 | -0.14 | 0.545 | -0.06 | -0.5775 | 0.469023 | 0.421828 | 0.937268 | 0.989223 | 0.997988 | 0.994666 |
| Syt17_Q920M7_S115s_1 | Syt17_Q92 | 0.38 | -1.96 | 0.69 | 1.17 | -0.18 | 0.72 | 0.73 | 1.19 | -0.21 | -0.27 | -0.89 | 1.41 | -1.96 | 0.68 | -0.61 | -0.14 | 0.545 | -0.06 | -0.5775 | 0.469023 | 0.421828 | 0.937268 | 0.989223 | 0.997988 | 0.994666 |
| Syt17_Q920M7_S84s_1 | Syt17_Q92 | -1.5 | -1.35 | -1.14 | -0.63 | -1.67 | -1.01 | 0.14 | -0.26 | 3.36 | -1.19 | -0.34 | 0.24 | 4.12 | -1.94 | 3.79 | -1.05 | 0.455 | 1.6725 | 2.385 | 0.108026 | 0.295617 | 0.088299 | 0.989223 | 0.997988 | 0.994666 |
| Syt3_G3X9Y1_S290s_1 | Syt3_G3X9 | 0.4 | -0.11 | -1.22 | -0.27 | -1.42 | 1.68 | -1.02 | -1.48 | 1.75 | -0.15 | -0.48 | -0.91 | 2.15 | -2.84 | -0.08 | -0.22 | -0.26 | 0.3525 | 0.0525 | 0.955374 | 0.721076 | 0.572022 | 0.995437 | 0.997988 | 0.999507 |
| Syt3_G3X9Y1_S586s_1 | Syt3_G3X9 | 1.84 | 0.62 | 1.54 | 1.61 | -0.79 | 1.77 | 0.46 | 1.18 | 2.85 | 1.36 | 0.96 | 0.88 | 1.47 | 0.55 | 1.11 | 0.36 | -0.7475 | 0.11 | -0.53 | 0.189226 | 0.198388 | 0.825901 | 0.989223 | 0.997988 | 0.994666 |
| Syt4_P40749_S135s_1 | Syt4_P407 | 0.68 | 1.16 | 0.02 | 0.71 | 1.27 | 1.31 | 0.6 | -0.17 | 0.88 | 0.89 | 0.56 | 0.21 | 1.66 | 0.07 | 0.89 | 0.52 | 0.11 | -0.0075 | 0.1425 | 0.729999 | 0.785051 | 0.981036 | 0.995762 | 0.997988 | 0.9971 |
| Syt5_Q9R0N5_S95s_1 | Syt5_Q9R0 | -0.85 | -0.85 | -1.4 | -0.86 | -0.59 | -0.47 | -0.74 | -0.45 | -0.85 | -0.79 | -1.3 | -0.85 | -0.54 | -0.18 | -0.52 | -0.74 | 0.4275 | 0.0425 | 0.495 | 0.09099 | 0.086994 | 0.866576 | 0.989223 | 0.997988 | 0.994666 |
| Syt7_Q9R0N7_S107s_1 | Syt7_Q9R0 | -0.42 | -0.1 | -0.25 | -0.08 | -0.09 | -0.21 | -0.24 | 0.23 | -0.35 | -0.25 | -0.66 | -0.42 | 0.09 | 0.47 | -1.13 | -0.23 | 0.135 | -0.2075 | 0.0125 | 0.973035 | 0.544034 | 0.365385 | 0.989223 | 0.997988 | 0.999507 |
| Syt7_Q9R0N7_S129s_1 | Syt7_Q9R0 | 1.07 | 0.29 | -0.12 | -0.47 | 0.87 | 0.84 | 0.6 | 0 | -0.11 | 1.2 | -0.77 | -0.69 | 0 | 0.53 | -0.23 | -0.58 | 0.385 | -0.285 | -0.2625 | 0.524559 | 0.323395 | 0.590469 | 0.989223 | 0.997988 | 0.994666 |
| Syt7_Q9R0N7_S172s_1 | Syt7_Q9R0 | -0.84 | -0.85 | 0.3 | 0.31 | -1.26 | -0.79 | 0.78 | 0.98 | -0.47 | -0.6 | -0.01 | 0.39 | -0.33 | 0.74 | 0.26 | -0.36 | 0.1975 | 0.0975 | 0.3475 | 0.418232 | 0.735685 | 0.807321 | 0.995762 | 0.997988 | 0.994666 |
| Syt7_Q9R0N7_S52s_1 | Syt7_Q9R0 | 1.22 | 1.11 | 0.72 | -0.17 | 0.9 | 1.45 | -0.28 | 0.16 | 0.94 | 1.01 | 1.22 | 0.42 | 0.98 | 1.37 | 1.22 | -1.25 | -0.1625 | 0.1775 | -0.14 | 0.823386 | 0.72581 | 0.630887 | 0.995437 | 0.997988 | 0.999507 |
| Syt7_Q9R0N7_S61s_1 | Syt7_Q9R0 | 1.71 | 1.33 | 0.84 | 1.13 | 1.68 | 1.36 | 0.44 | 0.93 | 1.64 | 1.47 | 0.94 | 1.17 | 1.24 | 1.76 | 1.11 | 0.89 | -0.15 | 0.0525 | -0.0025 | 0.993607 | 0.654808 | 0.854909 | 0.990434 | 0.997988 | 1 |
| Syt7_Q9R0N7_S61s_2 | Syt7_Q9R0 | 1.63 | 1.63 | 1.37 | 1.49 | 1.58 | 1.31 | 1.31 | 1.14 | 1.7 | 1.72 | 2.17 | 1.62 | -2.13 | 1.86 | 1.53 | 1.2 | -0.195 | 0.2725 | -0.915 | 0.279247 | 0.366732 | 0.259818 | 0.989223 | 0.997988 | 0.994666 |
| Syt7_Q9R0N7_T253t_1 | Syt7_Q9R0 | -0.05 | 0.22 | -0.84 | 0.3 | 0.34 | -0.49 | 0.12 | 0.23 | -0.51 | 0.13 | 0.07 | -0.06 | 0.38 | -0.08 | -0.1 | -0.5 | 0.1425 | 1.18E-16 | 0.0175 | 0.959803 | 0.667179 | 1 | 0.99272 | 1 | 0.999507 |
| Syt7_Q9R0N7_T262t_1 | Syt7_Q9R0 | 0.51 | -0.06 | 0.35 | 0.82 | 1.13 | 0.31 | 0.43 | 0.5 | 1.08 | 0.32 | 0.13 | 0.16 | 0.98 | 0.97 | 0.21 | -0.02 | 0.1875 | 0.0175 | 0.13 | 0.70941 | 0.521912 | 0.956138 | 0.989223 | 0.997988 | 0.996966 |
| Syt7_Q9R0N7_T58t_1 | Syt7_Q9R0 | 0.24 | 0.68 | 0.73 | 0.82 | 0.81 | 0.29 | 0.7 | 0.7 | 0.62 | 0.36 | 1.16 | 1.18 | 0.48 | 1.03 | 0.66 | 0.69 | 0.0075 | 0.2125 | 0.0975 | 0.714593 | 0.974946 | 0.466851 | 0.999187 | 0.997988 | 0.996966 |
| Syt7_Q9R0N7_T58t_2 | Syt7_Q9R0 | 1.63 | 1.63 | 1.37 | 1.49 | 1.58 | 1.31 | 1.31 | 1.14 | 1.7 | 1.72 | 2.17 | 1.62 | -2.13 | 1.86 | 1.53 | 1.2 | -0.195 | 0.2725 | -0.915 | 0.279247 | 0.366732 | 0.259818 | 0.989223 | 0.997988 | 0.994666 |
| Syt9_Q9R0N9_S207s_1 | Syt9_Q9R0 | 0.63 | 0.53 | 0.16 | 0.19 | 0.07 | 1.42 | -0.79 | -0.39 | 2.14 | 0.5 | -0.94 | 0.06 | 1.77 | -1.77 | 0.25 | -0.36 | -0.3 | 0.0625 | -0.405 | 0.54831 | 0.519532 | 0.916718 | 0.989223 | 0.997988 | 0.994666 |
| Syt9_Q9R0N9_T159t_1 | Syt9_Q9R0 | -2.6 | -2.04 | -1.64 | -2.22 | -2.09 | -2.56 | -1.56 | -2.71 | -1.53 | -2.25 | -1.64 | -1.92 | -0.22 | -1.02 | -1.92 | -2.04 | -0.105 | 0.29 | 0.825 | 0.095975 | 0.754603 | 0.342146 | 0.995762 | 0.997988 | 0.994666 |
| Sytl5_Q80T23_S347s_1 | Sytl5_Q80T | -0.23 | -0.25 | -0.79 | -1.41 | 0.78 | -0.49 | -0.22 | -0.76 | -0.26 | -0.33 | -0.04 | -0.95 | 0.02 | 0.22 | 0.05 | -0.1 | 0.4975 | 0.275 | 0.7175 | 0.05143 | 0.249505 | 0.445072 | 0.989223 | 0.997988 | 0.994666 |
| Syvn1_E9QQ99_S557s_1 | Syvn1_E9Q | 0.52 | -0.2 | -0.62 | -0.31 | 1.45 | 0.28 | -0.6 | -0.39 | 0.11 | -0.78 | -0.39 | -0.22 | 0.67 | 0.11 | 0.7 | 0.31 | 0.3375 | -0.1675 | 0.6 | 0.090017 | 0.488159 | 0.612963 | 0.989223 | 0.997988 | 0.994666 |
| Szrd1_Q6NXN1_S18s_1 | Szrd1_Q6N | -0.02 | 0.62 | 0.93 | 0.86 | 0.49 | 0.13 | 0.38 | 1.04 | 0.22 | 0.91 | 0.09 | -0.32 | -0.08 | 0.21 | 0.84 | 0.75 | -0.0875 | -0.3725 | -0.1675 | 0.626703 | 0.777447 | 0.302958 | 0.995762 | 0.997988 | 0.995259 |
| Szrd1_Q6NXN1_S20s_1 | Szrd1_Q6N | 0.66 | 0.55 | 0.72 | 0.73 | 0.59 | 0.46 | 0.69 | 1.14 | 0.97 | 1 | 0.06 | -0.03 | 0.31 | 0.56 | 0.61 | 0.61 | 0.055 | -0.165 | -0.1425 | 0.544669 | 0.810687 | 0.603242 | 0.995762 | 0.997988 | 0.994666 |
| Szt2_A2A9C3_T2432t_1 | Szt2_A2A9 | 0.46 | 0.33 | 0.2 | 0.17 | 1.07 | 0.74 | 0.34 | 0.39 | 1.05 | 1.03 | 0.72 | -0.65 | 0.36 | 1.5 | 0.16 | 0.59 | 0.345 | 0.2475 | 0.3625 | 0.299987 | 0.180307 | 0.544823 | 0.989223 | 0.997988 | 0.994666 |
| Tab3_Q571K4_S103s_1 | Tab3_Q571 | -0.14 | 0.7 | -0.49 | 0.09 | 0.06 | -0.41 | -0.6 | -0.38 | 0.96 | 0.12 | -0.38 | -0.7 | 0.45 | 0.98 | -0.37 | -0.67 | -0.3725 | -0.04 | 0.0575 | 0.896665 | 0.24405 | 0.925573 | 0.989223 | 0.997988 | 0.999507 |
| Tacc1_F8VQ95_S119s_1 | Tacc1_F8V | 0.64 | 0.93 | -0.15 | 0.07 | 0.65 | 0.53 | 0.02 | -0.33 | 0.85 | 0.63 | 0.13 | 0.2 | 0.15 | 0.76 | 0.36 | -0.32 | -0.155 | 0.08 | -0.135 | 0.708815 | 0.652849 | 0.80769 | 0.990434 | 0.997988 | 0.996966 |
| Tacc1_F8VQ95_S403s_1 | Tacc1_F8V | 0.29 | 0.97 | 1.4 | 1.56 | 0.93 | 0.35 | 1.5 | 0.89 | -0.3 | 0.59 | -0.09 | 1.19 | -0.46 | -0.1 | 1.08 | 1.57 | -0.1375 | -0.7075 | -0.5325 | 0.321339 | 0.70714 | 0.126241 | 0.995437 | 0.997988 | 0.994666 |
| Tacc1_F8VQ95_S45s_1 | Tacc1_F8V | 0.63 | 1.72 | 0.53 | 1.32 | 0.57 | 1.2 | 0.63 | 1.29 | 1.13 | 0.59 | 2.55 | 1.03 | -0.09 | 1.1 | 1.67 | 0.82 | -0.1275 | 0.275 | -0.175 | 0.699177 | 0.712246 | 0.573355 | 0.995437 | 0.997988 | 0.996159 |
| Tacc2_E9QL08_S213s_1 | Tacc2_E9Q | -0.95 | -0.45 | -0.4 | -0.63 | -0.18 | -0.99 | -0.08 | -0.48 | -0.96 | -0.74 | -0.4 | -0.81 | 0.16 | 0.12 | -0.67 | -0.87 | 0.175 | -0.12 | 0.2925 | 0.38971 | 0.531874 | 0.630944 | 0.989223 | 0.997988 | 0.994666 |
| Tacc2_E9QL08_S240s_1 | Tacc2_E9Q | -0.08 | 0.49 | 0.7 | 0.12 | 0.26 | -0.37 | -0.01 | -0.21 | 0.41 | 0.15 | -0.79 | 0.4 | 0.2 | 0.79 | 1.44 | 0.28 | -0.39 | -0.265 | 0.37 | 0.318304 | 0.16709 | 0.455994 | 0.989223 | 0.997988 | 0.994666 |
| Tacc2_E9QL08_S305s_1 | Tacc2_E9Q | 0.18 | 0.21 | 0.32 | 0.02 | -0.1 | -0.22 | -0.11 | -0.21 | -0.4 | 0.39 | -0.54 | 0.05 | 0.77 | -0.58 | -0.05 | -0.31 | -0.3425 | -0.3075 | -0.225 | 0.508022 | 0.10943 | 0.281819 | 0.989223 | 0.997988 | 0.994666 |
| Tacc2_E9QL08_S396s_1 | Tacc2_E9Q | -0.31 | 1.41 | 0.98 | -0.06 | 0.59 | -1.27 | 0 | -0.8 | 0.39 | 0.59 | 0.53 | 0.36 | 1.62 | 1.01 | 0.7 | 0.42 | -0.875 | -0.0375 | 0.4325 | 0.365622 | 0.123813 | 0.926955 | 0.989223 | 0.997988 | 0.994666 |
| Tacc2_E9QL08_S600s_1 | Tacc2_E9Q | 0.08 | 1.54 | 1.4 | 0.37 | 1.18 | -0.61 | 0.09 | 0.13 | 0.97 | 0.64 | 1.02 | 0.21 | 0.26 | 0.58 | 0.22 | 0.55 | -0.65 | -0.1375 | -0.445 | 0.269095 | 0.196663 | 0.735483 | 0.989223 | 0.997988 | 0.994666 |
| Tada3_Q8R0L9_S279s_1 | Tada3_Q8R | -1.83 | -1.62 | -0.75 | -1.7 | -2.79 | -2.2 | -2.28 | -1.38 | 1.32 | -2.46 | -1.02 | -1.09 | 2 | -2.85 | 0.19 | -0.65 | -0.6875 | 0.6625 | 1.1475 | 0.225449 | 0.091759 | 0.38022 | 0.989223 | 0.997988 | 0.994666 |
| Taf10_Q8K0H5_S44s_1 | Taf10_Q8K | -0.01 | -0.17 | 1.02 | 0.08 | 0.6 | 0.19 | 0.5 | -0.48 | -0.24 | 0.35 | 0.63 | 0.12 | 0.27 | 0.88 | 0.73 | 0.7 | -0.0275 | -0.015 | 0.415 | 0.236016 | 0.939143 | 0.965151 | 0.999187 | 0.997988 | 0.994666 |
| Taf15_Q8BQ46_S373s_1 | Taf15_Q8B | -3.98 | -1.64 | -2.67 | -3.39 | -1.42 | -3.17 | -2.57 | -4.66 | -1.25 | -2.38 | -3.2 | -2.71 | 0.42 | -3.98 | -2.5 | -2.44 | -0.035 | 0.535 | 0.795 | 0.395908 | 0.962196 | 0.381566 | 0.999187 | 0.997988 | 0.994666 |
| Taf3_Q5HZG4_T56t_1 | Taf3_Q5HZ | 0.47 | 1.52 | -0.3 | -1.01 | 1.08 | 1.05 | 0.78 | -0.16 | -0.68 | 0.93 | -0.45 | -0.36 | 0.89 | 0.92 | -0.23 | 0.48 | 0.5175 | -0.31 | 0.345 | 0.540904 | 0.362519 | 0.606671 | 0.989223 | 0.997988 | 0.994666 |
| Taf6_Q62311_S653s_1 | Taf6_Q623 | 0.81 | 0.89 | 0.12 | 1.22 | 0.46 | 0.66 | -0.01 | 0.95 | 1.09 | 1.05 | 2.13 | 0.6 | 0.48 | 0.51 | 1 | 0.59 | -0.245 | 0.4575 | -0.115 | 0.713954 | 0.452576 | 0.266354 | 0.989223 | 0.997988 | 0.996966 |
| Tagln3_Q9R1Q8_S163s_1 | Tagln3_Q9 | -0.05 | -0.42 | -0.15 | 0.2 | 0.43 | 0.22 | 0 | 0.27 | -0.03 | -0.17 | 0 | -0.17 | -0.51 | 0.64 | -0.2 | 0.1 | 0.335 | 0.0125 | 0.1125 | 0.727748 | 0.169519 | 0.956865 | 0.989223 | 0.997988 | 0.9971 |
| Taldo1_Q93092_S227s_1 | Taldo1_Q9 | -1.89 | -1.11 | -1.01 | -2.09 | -0.33 | -2.21 | -1.02 | -1.53 | -2.8 | -1.73 | -0.95 | -1.36 | -1.85 | -1 | -1.52 | -1.08 | 0.2525 | -0.185 | 0.1625 | 0.654394 | 0.578062 | 0.689077 | 0.989223 | 0.997988 | 0.995259 |
| Tanc1_E9QAF9_S211s_1 | Tanc1_E9Q | 0.95 | 0.38 | 0.12 | 0.13 | 0.54 | 0.93 | 0.08 | 0.26 | 0.61 | 0.32 | -0.51 | 0.15 | -1.25 | 0.84 | 0.48 | -0.13 | 0.0575 | -0.2525 | -0.41 | 0.398845 | 0.845764 | 0.452664 | 0.996349 | 0.997988 | 0.994666 |
| Tanc1_E9QAF9_S438s_1 | Tanc1_E9Q | 0.43 | 0.18 | 1.33 | 1.09 | 0.85 | 0.56 | 0.47 | 1.12 | 0.17 | 0.36 | 0.56 | 0.97 | 0.58 | 0.98 | 1.48 | 1.14 | -0.0075 | -0.2425 | 0.2875 | 0.427195 | 0.981404 | 0.482022 | 0.999187 | 0.997988 | 0.994666 |
| Tanc1_E9QAF9_S455s_1 | Tanc1_E9Q | 0.65 | 0.47 | 1.04 | 0.73 | 0.66 | 0.81 | 0.93 | 0.35 | 0.08 | 0.87 | 0.32 | 0.18 | -0.38 | 0.55 | 0.02 | 0.69 | -0.035 | -0.36 | -0.5025 | 0.141611 | 0.883583 | 0.205139 | 0.996493 | 0.997988 | 0.994666 |
| Tanc2_A2A690_S220s_1 | Tanc2_A2A | 0.54 | -0.56 | 0.23 | 0.43 | -0.19 | 0.94 | -0.27 | 0.52 | 0.25 | -0.13 | 1.43 | 0.01 | -0.31 | 0.5 | 0.13 | 0.04 | 0.09 | 0.23 | -0.07 | 0.835218 | 0.810441 | 0.590236 | 0.995762 | 0.997988 | 0.999507 |
| Tanc2_A2A690_S238s_1 | Tanc2_A2A | -0.45 | 0.46 | -1.39 | -1.78 | 0.26 | -0.79 | -0.79 | -0.88 | -0.43 | -0.43 | -0.94 | -1.55 | 0.31 | -0.36 | -1.23 | -0.84 | 0.24 | -0.0475 | 0.26 | 0.641844 | 0.64562 | 0.928435 | 0.99033 | 0.997988 | 0.995259 |
| Tanc2_A2A690_S254s_1 | Tanc2_A2A | -0.48 | -0.74 | 0.38 | 0.64 | 0.28 | -1.01 | 0.79 | -0.13 | -0.07 | -0.24 | 0.71 | 0.23 | -0.75 | -0.45 | 0.08 | 0.94 | 0.0325 | 0.2075 | 0.005 | 0.991615 | 0.944603 | 0.599281 | 0.999187 | 0.997988 | 1 |
| Tanc2_A2A690_S268s_1 | Tanc2_A2A | -1.39 | -0.95 | -0.42 | -0.85 | -0.4 | -1.23 | -0.69 | -0.61 | -0.71 | -0.75 | -2.96 | -0.76 | -0.86 | -0.34 | -1.95 | -0.68 | 0.17 | -0.3925 | -0.055 | 0.891978 | 0.566536 | 0.479374 | 0.989223 | 0.997988 | 0.999507 |
| Tanc2_A2A690_S400s_1 | Tanc2_A2A | -0.7 | 0.16 | -0.11 | -0.18 | -0.83 | -0.46 | -0.72 | -0.54 | -0.02 | 0.1 | -0.08 | -0.14 | 1.12 | -0.18 | -0.11 | -0.09 | -0.43 | 0.1725 | 0.3925 | 0.311892 | 0.114426 | 0.506214 | 0.989223 | 0.997988 | 0.994666 |
| Tanc2_A2A690_T434t_1 | Tanc2_A2A | -0.17 | -0.25 | -0.26 | 0.09 | 0.12 | 0 | 0.14 | 0.15 | -0.05 | 0.38 | -0.4 | -0.04 | -0.26 | 0.09 | 0.08 | 0.26 | 0.25 | 0.12 | 0.19 | 0.45464 | 0.238387 | 0.636148 | 0.989223 | 0.997988 | 0.994666 |
| Tanc2_E9Q2K2_S21s_1 | Tanc2_E9Q | 0.23 | -0.61 | -0.01 | -0.41 | -1.72 | 1.42 | -1.22 | -0.29 | 0.51 | -0.37 | -1.24 | 0.28 | 0.48 | -3.26 | -0.78 | -1.02 | -0.2525 | -0.005 | -0.945 | 0.207091 | 0.6928 | 0.990611 | 0.994893 | 0.997988 | 0.994666 |
| Tanc2_E9Q2K2_S22s_1 | Tanc2_E9Q | 0.85 | -0.03 | 0.45 | 0.69 | -2.37 | 2.19 | -1.57 | -0.27 | 2.84 | 0.81 | -0.65 | 0.05 | 2.48 | -0.34 | -1.06 | -0.93 | -0.995 | 0.2725 | -0.4525 | 0.552902 | 0.276123 | 0.698667 | 0.989223 | 0.997988 | 0.994666 |
| Tanc2_F6VP19_S1124s_1 | Tanc2_F6V | -0.63 | -0.99 | 0.86 | 0.64 | -0.52 | 0.04 | 0.05 | 0.5 | -0.62 | -0.19 | -0.04 | 0.77 | 0.86 | -0.94 | 0.4 | 0.61 | 0.0475 | 0.01 | 0.2625 | 0.642577 | 0.918962 | 0.984319 | 0.997335 | 0.997988 | 0.995259 |
| Tanc2_F6VP19_S589s_1 | Tanc2_F6V | 0.01 | 0.48 | 0.54 | -0.23 | -0.02 | 0.18 | 0.29 | 0.2 | -1.48 | 0.75 | -2.63 | 0.62 | -0.71 | -2.11 | 0.26 | 0.03 | -0.0375 | -0.885 | -0.8325 | 0.139789 | 0.881753 | 0.261315 | 0.996493 | 0.997988 | 0.994666 |
| Tanc2_F6VP19_S623s_1 | Tanc2_F6V | 0.14 | -0.77 | 0.22 | -0.24 | -0.77 | 1.04 | -0.4 | -0.12 | 0.75 | -0.25 | -3.04 | 0.39 | 1.76 | -2.65 | -0.26 | -0.3 | 0.1 | -0.375 | -0.2 | 0.807272 | 0.814897 | 0.638563 | 0.995762 | 0.997988 | 0.999507 |
| Tanc2_F6VP19_S664s_1 | Tanc2_F6V | -0.55 | 0.43 | 0.07 | -0.46 | 0.21 | -0.8 | 0.05 | -0.37 | -0.55 | -0.3 | -0.58 | -0.1 | 0.53 | 0.58 | -0.02 | 0.04 | -0.1 | -0.255 | 0.41 | 0.225846 | 0.76407 | 0.401482 | 0.995762 | 0.997988 | 0.994666 |
| Tanc2_F6VP19_S677s_1 | Tanc2_F6V | 0.56 | 0.57 | 0.86 | 0.61 | 0.7 | 0.08 | 0.8 | 0.51 | 0.57 | 0.77 | 0.39 | 0.75 | 0.8 | 1.11 | 0.6 | 0.7 | -0.1275 | -0.03 | 0.1525 | 0.543038 | 0.59774 | 0.892578 | 0.989223 | 0.997988 | 0.994666 |
| Tanc2_F6VP19_S677s_2 | Tanc2_F6V | -0.11 | 0.43 | 0 | -0.06 | 0.68 | 0.55 | 0.39 | -0.08 | -0.62 | -0.78 | 0.59 | -0.17 | -0.34 | -0.19 | -0.45 | 0.12 | 0.32 | -0.31 | -0.28 | 0.308473 | 0.233609 | 0.382969 | 0.989223 | 0.997988 | 0.994666 |
| Tanc2_F6VP19_S692s_1 | Tanc2_F6V | -0.34 | -0.01 | 0.46 | 0.13 | 0.2 | -0.35 | 0.53 | 0.36 | 0.08 | 0.17 | 0.59 | 0.76 | 0.46 | 0.48 | 0.53 | 0.75 | 0.125 | 0.34 | 0.495 | 0.090651 | 0.662835 | 0.247963 | 0.990434 | 0.997988 | 0.994666 |
| Tanc2_F6VP19_S971s_1 | Tanc2_F6V | -0.62 | -0.29 | -0.05 | 0.13 | 0.34 | 0.05 | 0.43 | 0.47 | -1.14 | -0.53 | 0.01 | 0.33 | -2.4 | -0.45 | 0.35 | -0.01 | 0.53 | -0.125 | -0.42 | 0.476509 | 0.054942 | 0.735479 | 0.989223 | 0.997988 | 0.994666 |
| Tanc2_F6VP19_S974s_1 | Tanc2_F6V | -0.85 | -0.29 | 1.01 | 0.17 | 0.13 | 0.19 | 0.68 | 1.11 | -1.67 | -0.61 | 0.15 | 1.43 | -1.96 | -1.86 | 0.95 | 0.79 | 0.5175 | -0.185 | -0.53 | 0.507827 | 0.24714 | 0.78821 | 0.989223 | 0.997988 | 0.994666 |
| Tanc2_F6VP19_T1123t_1 | Tanc2_F6V | 0.86 | 0.92 | 1.08 | 0.73 | 0.56 | 1.44 | -0.06 | 0.83 | 0.85 | 0.91 | 0.08 | 1.16 | 1.81 | 0.19 | 0.26 | 0.68 | -0.205 | -0.1475 | -0.1625 | 0.679233 | 0.538449 | 0.613092 | 0.989223 | 0.997988 | 0.996159 |
| Taok1_Q5F2E8_S421s_1 | Taok1_Q5F | 0.58 | 0.09 | 0.65 | 0.94 | 0.52 | 0.26 | 0.58 | 0.99 | 0.17 | 0.36 | 1.27 | 0.94 | 0.44 | 0.55 | 0.65 | 0.45 | 0.0225 | 0.12 | -0.0425 | 0.875246 | 0.934204 | 0.719228 | 0.998658 | 0.997988 | 0.999507 |
| Taok1_Q5F2E8_S9s_1 | Taok1_Q5F | -1.19 | -0.53 | -0.85 | -1.46 | -0.04 | -1.64 | -0.41 | -1.55 | -1.61 | -1.48 | -1.92 | -1.26 | -1.73 | -0.18 | -1.32 | -0.84 | 0.0975 | -0.56 | -0.01 | 0.979901 | 0.819258 | 0.079292 | 0.995762 | 0.997988 | 1 |
| Taok2_Q6ZQ29_S1018s_1 | Taok2_Q6Z | 0.09 | -1.14 | 0.45 | 0.32 | 0.28 | 0.56 | 0.12 | 0.69 | 0.32 | 0.02 | 0.49 | -0.14 | 0.14 | 1.06 | -1.3 | -0.15 | 0.4825 | 0.2425 | 0.0075 | 0.989323 | 0.222149 | 0.539942 | 0.989223 | 0.997988 | 1 |
| Taok2_Q6ZQ29_S493s_1 | Taok2_Q6Z | -3.32 | -2.85 | -2.23 | -2.57 | -2.23 | -3.01 | -2.76 | -2.72 | -3.15 | -3.02 | -0.52 | -1.85 | -2.18 | -1.69 | -2.82 | -2.29 | 0.0625 | 0.6075 | 0.4975 | 0.182752 | 0.837458 | 0.325052 | 0.995762 | 0.997988 | 0.994666 |
| Taok3_Q8BYC6_S324s_1 | Taok3_Q8E | 0.75 | -0.31 | 0.48 | 0.26 | -1.02 | -0.05 | 0.83 | 0.4 | -0.29 | 0.34 | 1 | 1.18 | -1.02 | -0.27 | 0.61 | 1.82 | -0.255 | 0.2625 | -0.01 | 0.986571 | 0.55704 | 0.517793 | 0.989223 | 0.997988 | 1 |
| Tardbp_H3BJV1_S35s_1 | Tardbp_H3 | -1.71 | -1.81 | -1.27 | -0.57 | -1.54 | -1.64 | -0.99 | -1.31 | -1.15 | -1.49 | -0.88 | -1.28 | -0.53 | -1.09 | -1.61 | -0.82 | -0.03 | 0.14 | 0.3275 | 0.397458 | 0.927087 | 0.675284 | 0.997335 | 0.997988 | 0.994666 |
| Tbc1d1_Q60949_S660s_1 | Tbc1d1_Q6 | -0.86 | -0.54 | -0.11 | -0.18 | -0.61 | -1.22 | 0.58 | 0.06 | -1.18 | -0.53 | -0.1 | -0.72 | -0.42 | -0.07 | -0.38 | 0.09 | 0.125 | -0.21 | 0.2275 | 0.433923 | 0.76074 | 0.51006 | 0.995762 | 0.997988 | 0.994666 |
| Tbc1d10a_Q5SPX8_S441s_1 | Tbc1d10a_ | -0.13 | -0.84 | -0.95 | -1.37 | -0.35 | 0.46 | -0.75 | -1.47 | -0.05 | 0.37 | 0.89 | -0.93 | 0.28 | 0.3 | -0.97 | -0.67 | 0.295 | 0.8925 | 0.5575 | 0.20124 | 0.514315 | 0.071156 | 0.989223 | 0.997988 | 0.994666 |
| Tbc1d10a_Q5SPX8_S52s_1 | Tbc1d10a_ | 0.08 | -0.4 | 0.13 | -0.08 | -0.37 | -0.25 | -0.41 | -0.05 | -0.19 | -0.23 | 0.43 | -0.08 | -0.27 | 0.14 | -0.39 | -0.29 | -0.2025 | 0.05 | -0.135 | 0.6106 | 0.379031 | 0.847517 | 0.989223 | 0.997988 | 0.995259 |
| Tbc1d10b_Q8BHL3_S22s_1 | Tbc1d10b_ | 6.06 | 4.08 | 4.37 | 2.57 | 4.18 | 5.98 | 4.38 | 4.37 | 0.59 | 5.22 | 1.04 | 4.74 | 1.25 | -0.05 | 4.78 | 4.79 | 0.4575 | -1.3725 | -1.5775 | 0.22088 | 0.534616 | 0.282079 | 0.989223 | 0.997988 | 0.994666 |
| Tbc1d10b_Q8BHL3_S644s_1 | Tbc1d10b_ | 0.99 | 0.23 | -1.12 | -0.67 | 0.54 | 1.64 | -0.01 | -0.5 | -0.46 | -0.91 | 0.2 | -1.24 | -0.06 | 0.48 | -1.96 | -1.27 | 0.56 | -0.46 | -0.56 | 0.403092 | 0.353777 | 0.392513 | 0.989223 | 0.997988 | 0.994666 |
| Tbc1d10b_Q8BHL3_S647s_1 | Tbc1d10b_ | -1.84 | -2.16 | -2.82 | -1.51 | -1.5 | -0.92 | -0.08 | -0.67 | -2.02 | -2.64 | 0.5 | -0.98 | -2.64 | -0.73 | -2.29 | -1.27 | 1.29 | 0.7975 | 0.35 | 0.486341 | 0.008296 | 0.25327 | 0.989223 | 0.997988 | 0.994666 |
| Tbc1d10b_Q8BHL3_S664s_1 | Tbc1d10b_ | 0.84 | -1.06 | -0.34 | 0.01 | -0.58 | 0.69 | 0.51 | 0.62 | -0.73 | -0.66 | 0.31 | 0.13 | -0.98 | -0.36 | -0.45 | -0.19 | 0.4475 | -0.1 | -0.3575 | 0.410444 | 0.343612 | 0.826846 | 0.989223 | 0.997988 | 0.994666 |
| Tbc1d10b_Q8BHL3_S673s_1 | Tbc1d10b_ | -0.1 | -0.52 | -0.86 | -0.5 | -0.2 | -0.44 | 0.32 | 0.28 | -1.27 | 0 | 0.16 | -0.53 | 0.28 | 0.16 | -0.68 | 0.29 | 0.485 | 0.085 | 0.5075 | 0.14216 | 0.107344 | 0.81689 | 0.989223 | 0.997988 | 0.994666 |
| Tbc1d10b_Q8BHL3_S693s_1 | Tbc1d10b_ | 0.46 | -0.44 | -0.74 | -0.09 | 0.28 | 0.51 | 0.39 | 0.36 | -1.18 | -0.15 | -3.02 | 0.26 | -1.07 | -0.84 | -0.69 | -0.31 | 0.5875 | -0.82 | -0.525 | 0.145741 | 0.068098 | 0.259328 | 0.989223 | 0.997988 | 0.994666 |
| Tbc1d10b_Q8BHL3_T136t_1 | Tbc1d10b_ | -0.27 | -1.82 | -0.24 | 0.33 | -0.55 | -0.24 | 0.15 | 0.55 | -0.99 | -1.27 | 0.5 | 0.22 | -1.4 | -0.38 | -0.19 | -0.37 | 0.4775 | 0.115 | -0.085 | 0.866647 | 0.332176 | 0.843997 | 0.989223 | 0.997988 | 0.999507 |
| Tbc1d13_Q8R3D1_S184s_1 | Tbc1d13_C | -3.08 | -3.23 | -2.54 | -2.82 | -2.54 | -3.4 | -2.64 | -2.28 | -2.34 | -2.63 | -1.9 | -2.61 | -0.97 | -2.13 | -2.43 | -2.56 | 0.2025 | 0.5475 | 0.895 | 0.046862 | 0.512985 | 0.075862 | 0.989223 | 0.997988 | 0.994666 |
| Tbc1d14_E9Q2V7_S296s_1 | Tbc1d14_E | -2.07 | -2.32 | -1.14 | -1.8 | -1.92 | -1.36 | -2.03 | -2.74 | -1.73 | -2.18 | -2.77 | -0.94 | 0.12 | -1.79 | -1.57 | -2.82 | -0.18 | -0.0725 | 0.3175 | 0.600626 | 0.632074 | 0.870711 | 0.989223 | 0.997988 | 0.995259 |
| Tbc1d14_E9Q2V7_S613s_1 | Tbc1d14_E | 0.73 | 0.38 | -0.32 | 0.54 | -0.19 | 1.24 | -0.67 | -0.44 | 2.88 | 1.3 | -2.19 | -0.26 | 2.38 | -0.72 | 0.32 | -0.22 | -0.3475 | 0.1 | 0.1075 | 0.868262 | 0.451308 | 0.918559 | 0.989223 | 0.997988 | 0.999507 |
| Tbc1d15_Q9CXF4_S205s_1 | Tbc1d15_C | -0.27 | -0.82 | 0.04 | 1.07 | 0.05 | -0.75 | 0.64 | 0.58 | -0.67 | -0.1 | 1.22 | -0.03 | -1.01 | 0.36 | -0.59 | 0.4 | 0.125 | 0.1 | -0.215 | 0.669248 | 0.791746 | 0.848944 | 0.995762 | 0.997988 | 0.996159 |
| Tbc1d15_Q9CXF4_S655s_1 | Tbc1d15_C | 3.19 | 2.49 | 3.79 | 2.26 | 1.54 | 2.6 | 2.64 | 3.42 | 2.49 | 2.23 | 1.4 | 3.42 | 1.19 | -0.54 | 3.15 | 3.75 | -0.3825 | -0.5475 | -1.045 | 0.266508 | 0.432913 | 0.297609 | 0.989223 | 0.997988 | 0.994666 |
| Tbc1d15_Q9CXF4_S662s_1 | Tbc1d15_C | 0.45 | -0.05 | 0.83 | 0.97 | -0.73 | 1.22 | 0.13 | 0.19 | 0.84 | -0.85 | 1.14 | 0.32 | -0.26 | -0.76 | -0.12 | 0.46 | -0.3475 | -0.1875 | -0.72 | 0.071617 | 0.430462 | 0.691484 | 0.989223 | 0.997988 | 0.994666 |
| Tbc1d22a_Q8R5A6_S166s_1 | Tbc1d22a_ | -2.73 | -2.22 | -2.01 | -2.19 | -1.6 | -2.48 | -1.34 | -1.96 | -1.81 | -2.7 | -2.19 | -1.95 | -0.79 | -1.49 | -1.84 | -2.12 | 0.4425 | 0.125 | 0.7275 | 0.063803 | 0.176932 | 0.670943 | 0.989223 | 0.997988 | 0.994666 |
| Tbc1d22b_Q80VE5_S141s_1 | Tbc1d22b_ | -0.88 | -3.22 | -2.33 | -3.76 | -1.72 | -1.3 | -2.65 | -2.46 | -1.02 | -2.21 | -1.93 | -2.06 | 0.18 | -2.25 | -1.9 | -2.33 | 0.515 | 0.7425 | 0.9725 | 0.224169 | 0.41981 | 0.252559 | 0.989223 | 0.997988 | 0.994666 |
| Tbc1d22b_Q80VE5_S58s_1 | Tbc1d22b_ | -2.02 | -2.79 | -1.12 | -1.96 | -1.29 | -2.18 | -1.5 | -1.67 | -3 | -2.08 | -0.9 | -1.4 | -1.66 | -0.85 | -2.02 | -0.89 | 0.3125 | 0.1275 | 0.6175 | 0.18204 | 0.420743 | 0.809973 | 0.989223 | 0.997988 | 0.994666 |
| Tbc1d22b_Q80VE5_T117t_1 | Tbc1d22b_ | -1.81 | -1.48 | -1.38 | -0.53 | -0.79 | -0.71 | -0.44 | -0.63 | -1.15 | -1.37 | -0.63 | -0.74 | -0.82 | -0.39 | -0.94 | -0.47 | 0.6575 | 0.3275 | 0.645 | 0.082333 | 0.05392 | 0.350731 | 0.989223 | 0.997988 | 0.994666 |
| Tbc1d23_Q8K0F1_S468s_1 | Tbc1d23_C | 0.29 | -0.39 | -0.47 | -0.1 | 0.16 | -1 | 0.1 | -0.21 | 0.97 | -0.71 | 0.4 | -0.26 | 1.2 | -0.19 | -0.16 | 0.26 | -0.07 | 0.2675 | 0.445 | 0.260777 | 0.831266 | 0.512475 | 0.995762 | 0.997988 | 0.994666 |
| Tbc1d23_Q8K0F1_S469s_2 | Tbc1d23_C | -0.84 | -0.71 | -0.27 | -0.33 | -0.87 | -1.65 | 0.12 | -0.05 | -0.51 | -1.27 | 1.53 | -0.06 | -0.7 | -0.8 | -0.06 | -1.07 | -0.075 | 0.46 | -0.12 | 0.699606 | 0.855373 | 0.421407 | 0.996349 | 0.997988 | 0.996159 |
| Tbc1d25_S4R2S6_S378s_1 | Tbc1d25_S | -0.22 | -0.06 | -0.38 | 0.21 | 0.14 | -0.24 | 0.39 | 0.16 | -0.57 | 0.2 | 0.59 | 0.02 | -0.06 | 0.38 | -0.31 | 0.03 | 0.225 | 0.1725 | 0.1225 | 0.65665 | 0.366653 | 0.578102 | 0.989223 | 0.997988 | 0.995259 |
| Tbc1d30_Q69ZT9_S624s_1 | Tbc1d30_C | 0.07 | -1.33 | -0.78 | -1.57 | -2.08 | -0.47 | -4.25 | -0.82 | 1.72 | 0.84 | -1.23 | -0.89 | 2.01 | 1.79 | -0.26 | -1.25 | -1.0025 | 1.0125 | 1.475 | 0.081811 | 0.23556 | 0.178786 | 0.989223 | 0.997988 | 0.994666 |
| Tbc1d5_Q80XQ2_S546s_1 | Tbc1d5_Q8 | -1.13 | -2.25 | -0.8 | -0.63 | -0.38 | -0.82 | -0.34 | -0.42 | -2.08 | -1.75 | -0.11 | -0.19 | -0.56 | -0.79 | -0.45 | -0.72 | 0.7125 | 0.17 | 0.5725 | 0.16072 | 0.082022 | 0.769703 | 0.989223 | 0.997988 | 0.994666 |
| Tbc1d5_Q80XQ2_S565s_1 | Tbc1d5_Q8 | -0.5 | -0.21 | 0.41 | 0.98 | -0.52 | -0.41 | 0.61 | 0.83 | -0.59 | -0.36 | -0.12 | 1.01 | -0.35 | -0.34 | 0.26 | 0.22 | -0.0425 | -0.185 | -0.2225 | 0.565814 | 0.924228 | 0.690772 | 0.997335 | 0.997988 | 0.995259 |
| Tbc1d5_Q80XQ2_S578s_1 | Tbc1d5_Q8 | -0.03 | 0.36 | 0.99 | 1.05 | 0.31 | -0.41 | 0.78 | 1.11 | -1.01 | 0.08 | 0.4 | 1 | 0.15 | -1.26 | 1.06 | 0.54 | -0.145 | -0.475 | -0.47 | 0.380755 | 0.719559 | 0.327553 | 0.995437 | 0.997988 | 0.994666 |
| Tbc1d7_Q9D0K0_S124s_1 | Tbc1d7_Q9 | -0.21 | 0.81 | 0.22 | 0.22 | 0.33 | -0.15 | -0.1 | -0.06 | -0.23 | 0.27 | -0.74 | -0.65 | -0.34 | 0.3 | 0.12 | 0.81 | -0.255 | -0.5975 | -0.0375 | 0.914103 | 0.365583 | 0.100065 | 0.989223 | 0.997988 | 0.999507 |
| Tbc1d9_Q3UYK3_S238s_1 | Tbc1d9_Q3 | 0.69 | 0.42 | 0.4 | 0.49 | 1.16 | 0.16 | 0.72 | 0.65 | 0.7 | 1.32 | 1.01 | 0.74 | 1.11 | 1.72 | 0.95 | 1 | 0.1725 | 0.4425 | 0.695 | 0.028512 | 0.51597 | 0.093005 | 0.989223 | 0.997988 | 0.994666 |
| Tbc1d9b_Q5SVR0_S464s_1 | Tbc1d9b_C | 0.57 | 0.61 | 1.11 | 0.74 | 1.01 | 0.49 | 0.86 | 0.54 | 0.74 | 0.86 | 0.4 | 0.79 | 0.24 | 0.97 | 1.16 | 1.19 | -0.0325 | -0.06 | 0.1325 | 0.669235 | 0.892503 | 0.80464 | 0.996493 | 0.997988 | 0.996159 |
| Tbc1d9b_Q5SVR0_T1082t_1 | Tbc1d9b_C | 2.19 | 0.87 | 0.92 | -0.03 | 1.26 | 2.26 | 1.01 | 1.32 | 0.55 | 2.53 | 1.96 | 0.64 | 0.27 | 0.18 | 1.06 | 1.26 | 0.475 | 0.4325 | -0.295 | 0.560778 | 0.344362 | 0.486145 | 0.989223 | 0.997988 | 0.995259 |
| Tbcb_Q9D1E6_S110s_1 | Tbcb_Q9D1 | -0.16 | -0.37 | -0.06 | 0.03 | -0.02 | -0.44 | 0.53 | 0.13 | -0.63 | 0.15 | 0.33 | 0.19 | -0.97 | 0.7 | 0.18 | 0.34 | 0.19 | 0.15 | 0.2025 | 0.600774 | 0.478168 | 0.598705 | 0.989223 | 0.997988 | 0.995259 |
| Tbcb_Q9D1E6_S173s_1 | Tbcb_Q9D1 | -2.38 | -1.79 | -1.64 | -1.98 | -2.28 | -1.83 | -2.38 | -2.74 | -0.91 | -2.09 | -2.1 | -2.06 | 0.3 | -2.28 | -1.76 | -1.61 | -0.36 | 0.1575 | 0.61 | 0.279014 | 0.220821 | 0.654271 | 0.989223 | 0.997988 | 0.994666 |
| Tbcel_D3YVP4_S18s_1 | Tbcel_D3Y | -0.7 | -0.22 | -0.18 | -0.73 | -0.11 | -0.85 | -0.39 | -0.65 | -0.61 | -0.25 | 0.14 | -0.6 | -0.5 | 0.38 | -0.58 | -0.28 | -0.0425 | 0.1275 | 0.2125 | 0.504775 | 0.872454 | 0.652963 | 0.996493 | 0.997988 | 0.994666 |
| Tbcel_D3YVP4_S41s_1 | Tbcel_D3Y | 0.06 | 0.26 | -0.51 | -0.28 | 0 | 0.28 | -0.62 | -0.66 | -0.27 | 0.27 | -0.45 | -0.26 | -0.35 | 0.19 | -1.14 | -0.26 | -0.1325 | -0.06 | -0.2725 | 0.448187 | 0.66949 | 0.83158 | 0.992876 | 0.997988 | 0.994666 |
| Tbck_E9Q1W7_S890s_1 | Tbck_E9Q1 | -0.63 | 0.08 | -0.41 | -0.05 | 0.26 | -0.2 | 0.45 | -0.14 | -0.43 | 0.07 | -0.71 | 0.18 | -0.58 | 0.44 | -0.45 | 0.14 | 0.345 | 0.03 | 0.14 | 0.674316 | 0.219843 | 0.921166 | 0.989223 | 0.997988 | 0.996159 |
| Tbr1_Q64336_S549s_1 | Tbr1_Q643 | -3.1 | -2.4 | -1.66 | -1.92 | -2.83 | -4.61 | -1.93 | -1.76 | -2.8 | -3.18 | -1.37 | -2.29 | -1.83 | -1.25 | -2.06 | -1.49 | -0.5125 | -0.14 | 0.6125 | 0.131039 | 0.434119 | 0.769991 | 0.989223 | 0.997988 | 0.994666 |
| Tbr1_Q64336_S640s_1 | Tbr1_Q643 | -0.76 | -0.4 | -0.01 | 0.42 | -0.55 | -1.44 | 0.05 | 0.09 | -0.48 | -0.16 | 0.59 | -0.37 | -0.74 | -0.09 | 0.08 | -0.46 | -0.275 | 0.0825 | -0.115 | 0.740077 | 0.514305 | 0.819422 | 0.989223 | 0.997988 | 0.9971 |
| Tbr1_Q64336_S84s_1 | Tbr1_Q643 | -0.28 | -0.91 | -0.21 | -0.02 | -0.14 | -0.44 | 0.93 | 0.19 | -0.06 | 0 | 0.25 | 0.21 | -0.24 | -0.61 | 0.12 | -0.24 | 0.49 | 0.455 | 0.1125 | 0.711246 | 0.187401 | 0.115357 | 0.989223 | 0.997988 | 0.996966 |
| Tcaf3_Q6QR59_Y458y_2 | Tcaf3_Q6Q | 0.5 | 1.01 | 0.64 | 1.04 | 0.51 | 0.81 | 0.54 | 0.88 | 0.8 | 0.64 | 0.92 | 0.48 | 0.96 | 0.58 | 1.04 | 1.04 | -0.1125 | -0.0875 | 0.1075 | 0.687567 | 0.633519 | 0.721705 | 0.989223 | 0.997988 | 0.996159 |
| Tcea1_E9PYD5_S111s_1 | Tcea1_E9P | -1.65 | -1.62 | -1.97 | -2.57 | -1.59 | -1.97 | -2.04 | -2.41 | -1.37 | -1.2 | -0.53 | -2.66 | -0.14 | -1.09 | -2.18 | -1.84 | -0.05 | 0.5125 | 0.64 | 0.204176 | 0.868133 | 0.294463 | 0.996493 | 0.997988 | 0.994666 |
| Tceal3_Q8R0A5_S120s_1 | Tceal3_Q8I | -4.8 | -4.77 | -1.71 | -1.56 | -4.79 | -5.38 | -1.83 | -1.45 | -2.28 | -5.02 | -2.74 | -1.91 | -0.38 | -2.56 | -1.68 | -2.13 | -0.1525 | 0.2225 | 1.5225 | 0.115283 | 0.895927 | 0.826041 | 0.996493 | 0.997988 | 0.994666 |
| Tceal3_Q8R0A5_S124s_1 | Tceal3_Q8I | -4.55 | -4.21 | -3.51 | -5.74 | -5.63 | -5.63 | -5.54 | -5.51 | -5.18 | -5.27 | -2.8 | -4.18 | -2.86 | -3.15 | -5.21 | -4.96 | -1.075 | 0.145 | 0.4575 | 0.509292 | 0.033071 | 0.828707 | 0.989223 | 0.997988 | 0.994666 |
| Tceal5_Q8CCT4_S23s_1 | Tceal5_Q8( | -4.6 | -4.77 | -3.31 | -5.19 | -4.51 | -5.53 | -4.57 | -4.19 | -4.78 | -4.01 | -1.61 | -4.23 | -2.51 | -3.16 | -4.13 | -3.89 | -0.2325 | 0.81 | 1.045 | 0.066424 | 0.61742 | 0.28271 | 0.989223 | 0.997988 | 0.994666 |
| Tceal5_Q8CCT4_T24t_1 | Tceal5_Q8( | -4.43 | -5 | -3.31 | -5.19 | -4.26 | -5.53 | -4.57 | -4.19 | -5.06 | -3.95 | -1.61 | -4.23 | -2.7 | -3.46 | -4.13 | -3.89 | -0.155 | 0.77 | 0.9375 | 0.085368 | 0.74859 | 0.32681 | 0.995762 | 0.997988 | 0.994666 |
| Tceb3_Q8CB77_S125s_1 | Tceb3_Q8( | -0.87 | -1.29 | -0.58 | -0.63 | -1.81 | -0.92 | -1.43 | -0.62 | 0.21 | -0.91 | -0.53 | -1.13 | 0.62 | -1.71 | 0.17 | -1.33 | -0.3525 | 0.2525 | 0.28 | 0.610471 | 0.291305 | 0.478193 | 0.989223 | 0.997988 | 0.995259 |
| Tceb3_Q8CB77_S195s_1 | Tceb3_Q8( | -2.68 | -3.4 | -1.44 | -0.82 | -4.57 | -3.24 | -1.21 | -1.24 | -2.71 | -2.61 | 0.41 | -0.14 | -1.35 | -0.95 | -0.46 | -1.05 | -0.48 | 0.8225 | 1.1325 | 0.069698 | 0.586451 | 0.366958 | 0.989223 | 0.997988 | 0.994666 |
| Tcf12_Q61286_T313t_1 | Tcf12_Q61 | -0.11 | 0.66 | 0.84 | 0.8 | 0.51 | 0.52 | 0.67 | 0.05 | 0.33 | 0.8 | 0.38 | 0.46 | -0.66 | 0.19 | 1.15 | 0.3 | -0.11 | -0.055 | -0.3025 | 0.485124 | 0.70536 | 0.850156 | 0.995437 | 0.997988 | 0.994666 |
| Tcf20_Q9EPQ8_S261s_1 | Tcf20_Q9E | -0.91 | -0.77 | -0.05 | 0.09 | -0.39 | -0.97 | 0.01 | -0.17 | -0.63 | -0.81 | 0.31 | 0.34 | -0.51 | -0.35 | -0.44 | 0.15 | 0.03 | 0.2125 | 0.1225 | 0.713271 | 0.929002 | 0.592556 | 0.997639 | 0.997988 | 0.996966 |
| Tcf20_Q9EPQ8_S587s_1 | Tcf20_Q9E | 1.63 | 0.75 | -0.49 | -0.31 | 0.4 | 1.06 | 0.01 | -0.74 | 1.14 | 1.23 | -0.04 | -0.47 | 1.56 | 1.25 | 0.75 | 0.77 | -0.2125 | 0.07 | 0.6875 | 0.19309 | 0.705199 | 0.906692 | 0.995437 | 0.997988 | 0.994666 |
| Tcf20_Q9EPQ8_S588s_1 | Tcf20_Q9E | -0.2 | -0.27 | 0.53 | 0.72 | -0.99 | 0.44 | -0.53 | 0.59 | -0.03 | 0.08 | -0.18 | 0.2 | 0.84 | -0.21 | 0.57 | 0.29 | -0.3175 | -0.1775 | 0.1775 | 0.625254 | 0.467749 | 0.560987 | 0.989223 | 0.997988 | 0.995259 |
| Tcf20_Q9EPQ8_S603s_1 | Tcf20_Q9E | 0.82 | 0.34 | 0.25 | 0.08 | -0.05 | 0.61 | -0.35 | 0 | 0.66 | 0.72 | 0.08 | -0.07 | 1.35 | 0.04 | 0.45 | 0.33 | -0.32 | -0.025 | 0.17 | 0.631682 | 0.282006 | 0.932839 | 0.989223 | 0.997988 | 0.995259 |
| Tcf20_Q9EPQ8_S669s_1 | Tcf20_Q9E | 0.7 | 0.11 | 0.89 | 1.13 | 0.45 | 0.13 | 0.53 | 0.74 | 0.53 | 0.58 | 0.74 | 0.7 | 0.99 | 0.6 | 1.15 | 0.6 | -0.245 | -0.07 | 0.1275 | 0.683591 | 0.399264 | 0.800624 | 0.989223 | 0.997988 | 0.996159 |
| Tcf20_Q9EPQ8_T1699t_1 | Tcf20_Q9E | 0.58 | 0.47 | 0.67 | 0.93 | 0.47 | 0.27 | 0.73 | 1.13 | 0.31 | 0.47 | 0.97 | 0.67 | 0.73 | 0.87 | 1.39 | 1.13 | -0.0125 | -0.0575 | 0.3675 | 0.190865 | 0.961545 | 0.817351 | 0.999187 | 0.997988 | 0.994666 |
| Tcf4_Q60722_S212s_1 | Tcf4_Q607 | 0.18 | 0.98 | -0.14 | 0.44 | 0.56 | -0.02 | 0.25 | 0.02 | -0.6 | 0.9 | 0.82 | -0.62 | 0.43 | 0.9 | -0.03 | -0.44 | -0.1625 | -0.24 | -0.15 | 0.698731 | 0.587914 | 0.607965 | 0.989223 | 0.997988 | 0.996159 |
| Tcf4_Q60722_S355s_1 | Tcf4_Q607 | -3.81 | -4.8 | -2.46 | -4.27 | -3.04 | -4.68 | -3.41 | -2.97 | -4.43 | -1.77 | -1.28 | -2.85 | -1.29 | -2.32 | -3.1 | -3.25 | 0.31 | 1.2525 | 1.345 | 0.050524 | 0.592163 | 0.12936 | 0.989223 | 0.997988 | 0.994666 |
| Tcof1_H3BL37_S1252s_1 | Tcof1_H3B | 0.28 | -0.8 | 0.06 | 0.38 | -1.03 | 0.14 | -0.44 | 0.16 | 0.58 | -0.27 | -0.22 | 0.43 | 0.67 | -0.81 | 0.16 | 0.39 | -0.2725 | 0.15 | 0.1225 | 0.769861 | 0.479922 | 0.676989 | 0.989223 | 0.997988 | 0.9971 |
| Tcof1_H3BL37_S593s_1 | Tcof1_H3B | 0.91 | 0.87 | 1.85 | 0.97 | -0.17 | -0.02 | 1.1 | 0.59 | 0.39 | 0.45 | 1.5 | 1.54 | -0.41 | 0.68 | 0.41 | 1.11 | -0.775 | -0.18 | -0.7025 | 0.106178 | 0.059573 | 0.649735 | 0.989223 | 0.997988 | 0.994666 |
| Tcof1_H3BL37_S855s_1 | Tcof1_H3B | -5.03 | -3.83 | -2.52 | -4.47 | -2.9 | -4.27 | -2.15 | -3.41 | -0.84 | -3.9 | -3.51 | -1.99 | -0.24 | -3.37 | -1.76 | -2.42 | 0.78 | 1.4025 | 2.015 | 0.023009 | 0.230239 | 0.103802 | 0.989223 | 0.997988 | 0.994666 |
| Tcof1_H3BL37_S879s_1 | Tcof1_H3B | -1 | -2.88 | -1.6 | -1.59 | -2.62 | -1.77 | -2.31 | -2.36 | -2.12 | -1.97 | -1.04 | -1 | -0.82 | -1.8 | -1.63 | -1.11 | -0.4975 | 0.235 | 0.4275 | 0.349861 | 0.248045 | 0.620836 | 0.989223 | 0.997988 | 0.994666 |
| Tcp11l1_Q8BTG3_S44s_1 | Tcp11l1_Q | -1.64 | -1.41 | -1.38 | -4.16 | -2.1 | -1.57 | -1.45 | -1.56 | 1.76 | -1.11 | -2.23 | -1.12 | 2.42 | -1.33 | 1.56 | -1.01 | 0.4775 | 1.4725 | 2.5575 | 0.027675 | 0.444618 | 0.150571 | 0.989223 | 0.997988 | 0.994666 |
| Tcp11l1_Q8BTG3_S55s_1 | Tcp11l1_Q | -0.44 | -0.87 | -0.98 | -1.24 | -0.47 | 0 | -1.08 | -1.04 | 0.78 | -0.57 | -1.98 | -0.76 | 1.53 | -2.31 | -0.92 | -0.36 | 0.235 | 0.25 | 0.3675 | 0.614198 | 0.470111 | 0.649071 | 0.989223 | 0.997988 | 0.995259 |
| Tcp11l2_Q8K1H7_S16s_1 | Tcp11l2_Q | -0.84 | 0.44 | 0.06 | -0.52 | -0.42 | -0.42 | -1.16 | -1.44 | 0.78 | 0.45 | -0.89 | -0.3 | 1.98 | -0.82 | -0.61 | -0.08 | -0.645 | 0.225 | 0.3325 | 0.603068 | 0.114137 | 0.622367 | 0.989223 | 0.997988 | 0.995259 |
| Tdrd3_Q91W18_S349s_1 | Tdrd3_Q91 | -2.07 | -2.97 | -1.97 | -1 | -1.68 | -2.2 | -0.83 | -1.13 | -3.64 | -2.9 | -1.52 | -0.59 | -1.16 | -1.23 | -1.15 | -0.7 | 0.5425 | -0.16 | 0.9425 | 0.04637 | 0.26313 | 0.822889 | 0.989223 | 0.997988 | 0.994666 |
| Tecpr1_Q80VP0_S272s_1 | Tecpr1_Q8 | 1.42 | 1.24 | 1.06 | 0.7 | 0.75 | 0.86 | -0.18 | 0.59 | 0.66 | 1.23 | -1.22 | 0.49 | 1.43 | 1.34 | 0.07 | -0.02 | -0.6 | -0.815 | -0.4 | 0.353513 | 0.07277 | 0.138983 | 0.989223 | 0.997988 | 0.994666 |
| Tecpr1_Q80VP0_S303s_1 | Tecpr1_Q8 | 2.57 | 1.39 | 1 | 1.3 | 2.91 | 2.61 | 1.87 | 2.2 | 1.5 | 1.82 | 1.24 | 1.1 | 1.01 | 3.43 | 1.63 | 1.68 | 0.8325 | -0.15 | 0.3725 | 0.521106 | 0.060477 | 0.696403 | 0.989223 | 0.997988 | 0.994666 |
| Tenm1_Q9WTS4_S145s_1 | Tenm1_Q9 | -0.76 | -0.58 | -0.69 | 0.21 | -1.26 | -0.66 | 0.02 | 0.78 | -0.51 | 1.2 | 0.02 | 0.32 | 0.49 | 0.83 | 1 | 0.52 | 0.175 | 0.7125 | 1.165 | 0.003896 | 0.704133 | 0.112727 | 0.995437 | 0.997988 | 0.994666 |
| Tenm1_Q9WTS4_S59s_1 | Tenm1_Q9 | -0.12 | -2.69 | -0.92 | -1.79 | -2.25 | 0.34 | -1.33 | -0.09 | 0 | -1.54 | -0.72 | -0.61 | -0.17 | -3.19 | -1.15 | -1.68 | 0.5475 | 0.6625 | -0.1675 | 0.822403 | 0.449704 | 0.275652 | 0.989223 | 0.997988 | 0.999507 |
| Tenm2_Q9WTS5_S41s_1 | Tenm2_Q9 | 0.39 | -0.24 | -0.69 | -0.34 | 0.39 | 0.54 | -0.51 | 0.2 | -0.47 | 0.38 | 0.31 | -0.34 | -0.38 | 0.24 | -0.28 | -0.35 | 0.375 | 0.19 | 0.0275 | 0.930877 | 0.27613 | 0.574526 | 0.989223 | 0.997988 | 0.999507 |
| Tenm2_Q9WTS5_S42s_1 | Tenm2_Q9 | 0.66 | -0.29 | 0.6 | 0.6 | -0.65 | 0.08 | -0.04 | 0.81 | 0 | 0.61 | 0.43 | 0.84 | -0.23 | -0.46 | -0.25 | 0.17 | -0.3425 | 0.0775 | -0.585 | 0.087975 | 0.36668 | 0.808036 | 0.989223 | 0.997988 | 0.994666 |
| Tenm4_F6UP10_S21s_1 | Tenm4_F6I | 0.03 | 0.26 | 0.59 | -0.37 | -1.42 | 0.77 | -0.42 | 0.03 | 2.03 | 0.25 | -1.11 | 0.61 | 1.86 | -2.14 | 1.19 | -0.64 | -0.3875 | 0.3175 | -0.06 | 0.941488 | 0.4131 | 0.609488 | 0.989223 | 0.997988 | 0.999507 |
| Tenm4_Q3UHK6_T2737t_2 | Tenm4_Q3 | -5.85 | 0.2 | -0.57 | -0.06 | -0.32 | -6.98 | -1.49 | -0.87 | 0.7 | -2.47 | 1.41 | 0.09 | 0.62 | 0.98 | -0.56 | -1.18 | -0.845 | 1.5025 | 1.535 | 0.259093 | 0.640237 | 0.316354 | 0.989223 | 0.997988 | 0.994666 |
| Tenm4_Q3UHK6_T2738t_2 | Tenm4_Q3 | -5.85 | 0.2 | -0.57 | -0.06 | -0.32 | -6.98 | -1.49 | -0.87 | 0.7 | -2.47 | 1.41 | 0.09 | 0.62 | 0.98 | -0.56 | -1.18 | -0.845 | 1.5025 | 1.535 | 0.259093 | 0.640237 | 0.316354 | 0.989223 | 0.997988 | 0.994666 |
| Tenm4_Q3UHK6_Y2735y_2 | Tenm4_Q3 | -5.85 | 0.2 | -0.57 | -0.06 | -0.32 | -6.98 | -1.49 | -0.87 | 0.7 | -2.47 | 1.41 | 0.09 | 0.62 | 0.98 | -0.56 | -1.18 | -0.845 | 1.5025 | 1.535 | 0.259093 | 0.640237 | 0.316354 | 0.989223 | 0.997988 | 0.994666 |
| Terf2_D3YZ08_S398s_1 | Terf2_D3Y | -0.45 | -1.64 | -0.46 | -0.36 | -2.32 | -0.85 | -0.58 | -0.31 | -0.74 | -2.35 | -0.28 | -1.06 | 0.36 | -1.95 | 0.78 | -0.27 | -0.2875 | -0.38 | 0.4575 | 0.461895 | 0.566777 | 0.459818 | 0.989223 | 0.997988 | 0.994666 |
| Terf2ip_Q91VL8_S43s_1 | Terf2ip_Q9 | -1.69 | -0.58 | 0.25 | -0.36 | -0.17 | -1.64 | 0.09 | -0.88 | -0.83 | -2.03 | -1.77 | -0.11 | -0.95 | -0.9 | -0.94 | -0.07 | -0.055 | -0.59 | -0.12 | 0.788694 | 0.914435 | 0.302786 | 0.997335 | 0.997988 | 0.999507 |
| Tex2_B1ATR2_S165s_1 | Tex2_B1AT | -0.09 | 0.33 | 0.6 | 0.18 | 1.09 | 0.07 | 0.5 | -0.84 | -1.3 | 0.55 | -0.91 | 0.73 | -0.35 | -2.23 | 0.4 | 0.49 | -0.05 | -0.4875 | -0.6775 | 0.269557 | 0.903203 | 0.342658 | 0.996493 | 0.997988 | 0.994666 |
| Tex2_B1ATR2_S221s_1 | Tex2_B1AT | -0.07 | -1.27 | -1.07 | -1.36 | -1.37 | 0.26 | -1.91 | -1.24 | -0.14 | -0.96 | -0.32 | -1 | -0.4 | -1.03 | -2.3 | -1.78 | -0.1225 | 0.3375 | -0.435 | 0.382545 | 0.808366 | 0.37955 | 0.995762 | 0.997988 | 0.994666 |
| Tex2_B1ATR2_S265s_1 | Tex2_B1AT | 1.22 | -0.77 | 0.1 | 0.84 | 0.52 | 1.51 | 0.54 | 0.61 | 0.51 | 0.17 | 1.44 | 0.85 | -0.32 | 2.27 | -0.26 | 0.12 | 0.4475 | 0.395 | 0.105 | 0.876785 | 0.347352 | 0.426774 | 0.989223 | 0.997988 | 0.999507 |
| Tex2_B1ATR2_S265s_2 | Tex2_B1AT | 0.26 | -0.12 | 0.26 | 0.11 | 0.21 | 0.91 | 0.6 | 0.6 | 0.47 | 0.16 | 0.65 | 0.19 | -0.54 | 0.6 | -0.19 | -0.49 | 0.4525 | 0.24 | -0.2825 | 0.392188 | 0.081531 | 0.324264 | 0.989223 | 0.997988 | 0.994666 |
| Tex2_B1ATR2_S269s_1 | Tex2_B1AT | 0.77 | 0.41 | 0.43 | 0.73 | 0.5 | -0.01 | 0.59 | 0.71 | 0.91 | 0.49 | 0.08 | 0.64 | -0.07 | 0.91 | 0.83 | 0.36 | -0.1375 | -0.055 | -0.0775 | 0.799419 | 0.578667 | 0.833999 | 0.989223 | 0.997988 | 0.999507 |
| Tex2_B1ATR2_S269s_2 | Tex2_B1AT | 0.26 | -0.12 | 0.26 | 0.11 | 0.21 | 0.91 | 0.6 | 0.6 | 0.47 | 0.16 | 0.65 | 0.19 | -0.54 | 0.6 | -0.19 | -0.49 | 0.4525 | 0.24 | -0.2825 | 0.392188 | 0.081531 | 0.324264 | 0.989223 | 0.997988 | 0.994666 |
| Tex2_B1ATR2_S294s_1 | Tex2_B1AT | -0.85 | -2 | -2.56 | -1.79 | -1.3 | 0.07 | -0.24 | -0.56 | -1.87 | -1.99 | 0.02 | -1.45 | -1.6 | -0.77 | -3.55 | -1.51 | 1.2925 | 0.4775 | -0.0575 | 0.92697 | 0.013831 | 0.388665 | 0.989223 | 0.997988 | 0.999507 |
| Tex2_B1ATR2_S791s_1 | Tex2_B1AT | -0.58 | -0.58 | -1.02 | -0.7 | -2.32 | 0.37 | -1.4 | -0.77 | 0.6 | -0.16 | -0.74 | -1.88 | 1.39 | -3 | 0.06 | -0.37 | -0.31 | 0.175 | 0.24 | 0.769053 | 0.556731 | 0.727819 | 0.989223 | 0.997988 | 0.9971 |
| Tex2_B1ATR2_S795s_1 | Tex2_B1AT | -0.77 | 0.3 | 1.43 | 0.26 | 0.06 | 0.02 | 0.51 | 0.09 | 0.56 | 0.65 | 1.92 | 1.29 | 0.96 | 0.57 | 0.63 | 0.4 | -0.135 | 0.8 | 0.335 | 0.464784 | 0.757374 | 0.145563 | 0.995762 | 0.997988 | 0.994666 |
| Tex2_B1ATR2_S803s_1 | Tex2_B1AT | 1.26 | 1.65 | 1.41 | 1.29 | 0.59 | 0.5 | 0.3 | 1.58 | 1.5 | 1.41 | 1.67 | 1.59 | 1.84 | 0.68 | 1.69 | 1.11 | -0.66 | 0.14 | -0.0725 | 0.823997 | 0.060366 | 0.527404 | 0.989223 | 0.997988 | 0.999507 |
| Tex264_E9Q137_S200s_1 | Tex264_E9 | 0.28 | 0.51 | 0.94 | 0.24 | 0.48 | 0.35 | 0.6 | -0.1 | 0.93 | -0.04 | 1.24 | 0.79 | 0.89 | 0.03 | 0.82 | 0.78 | -0.16 | 0.2375 | 0.1375 | 0.659859 | 0.551896 | 0.488 | 0.989223 | 0.997988 | 0.995259 |
| Tex264_E9Q137_S205s_1 | Tex264_E9 | 0.26 | 0.01 | 0.13 | -0.08 | 0.05 | 0.59 | 0.78 | 0.8 | 0.28 | 0.65 | 0.64 | 0.62 | 0.67 | 0.48 | -0.36 | 0.43 | 0.475 | 0.4675 | 0.225 | 0.460631 | 0.080631 | 0.060477 | 0.989223 | 0.997988 | 0.994666 |
| Tfeb_Q3UKG7_S180s_1 | Tfeb_Q3UI | -2.08 | -1.79 | -2.05 | -2.28 | -1.67 | -1.71 | -1.06 | -2.9 | -1.61 | -1.44 | -1 | -1.76 | -1.12 | -0.83 | -1.07 | -2.26 | 0.215 | 0.5975 | 0.73 | 0.066714 | 0.581029 | 0.042856 | 0.989223 | 0.997988 | 0.994666 |
| Tgfb1i1_E9Q1D5_S155s_1 | Tgfb1i1_E9 | -0.35 | -0.47 | -0.41 | -0.18 | -1.2 | -0.89 | -0.73 | -0.85 | -0.56 | -0.46 | -0.48 | -0.4 | 0.12 | -1.68 | -0.53 | -0.49 | -0.565 | -0.1225 | -0.2925 | 0.461294 | 0.023916 | 0.560834 | 0.989223 | 0.997988 | 0.994666 |
| Tgfbrap1_Q3UR70_S853s_1 | Tgfbrap1_( | 7.85 | 5.42 | 2.25 | 0.5 | 4.75 | 8.11 | 2 | 1.64 | 3.67 | 6.5 | -0.84 | 2.66 | 3.03 | 1.99 | 2.65 | 3.39 | 0.12 | -1.0075 | -1.24 | 0.395573 | 0.949445 | 0.606184 | 0.999187 | 0.997988 | 0.994666 |
| Tgfbrap1_Q3UR70_S853s_2 | Tgfbrap1_( | 1.43 | 1.06 | 2.23 | 1.26 | -0.61 | 2.37 | 1.2 | 1.37 | 1.85 | 1.3 | 0.79 | 1.41 | 1.89 | 0.77 | 2.16 | 2.75 | -0.4125 | -0.1575 | 0.3975 | 0.405974 | 0.497401 | 0.65564 | 0.989223 | 0.997988 | 0.994666 |
| Tgfbrap1_Q3UR70_S855s_2 | Tgfbrap1_( | 1.43 | 1.06 | 2.23 | 1.26 | -0.61 | 2.37 | 1.2 | 1.37 | 1.85 | 1.3 | 0.79 | 1.41 | 1.89 | 0.77 | 2.16 | 2.75 | -0.4125 | -0.1575 | 0.3975 | 0.405974 | 0.497401 | 0.65564 | 0.989223 | 0.997988 | 0.994666 |
| Th_P24529_S19s_1 | Th_P24529 | -0.22 | -1.93 | 0.18 | 0.19 | -2.42 | -0.75 | -0.23 | -0.66 | 1.39 | -1.3 | -1.96 | -1.51 | 2.01 | -1.16 | 1.78 | 0.18 | -0.57 | -0.4 | 1.1475 | 0.171085 | 0.36996 | 0.624374 | 0.989223 | 0.997988 | 0.994666 |
| Th_P24529_S31s_1 | Th_P24529 | 0.81 | -0.37 | 0.73 | 0.88 | -1.95 | 0.72 | 0.41 | 0.56 | 1.61 | 0.51 | -0.37 | -0.24 | 2.22 | -0.94 | 0.84 | 0.43 | -0.5775 | -0.135 | 0.125 | 0.846437 | 0.36396 | 0.790954 | 0.989223 | 0.997988 | 0.999507 |
| Th_P24529_S40s_1 | Th_P24529 | 0.8 | -0.08 | -0.75 | -0.51 | 0.32 | 0.41 | 0.3 | -0.09 | 0.19 | 0.26 | -0.17 | -1.24 | 0.39 | 0.73 | 0.11 | -0.67 | 0.37 | -0.105 | 0.275 | 0.53897 | 0.315167 | 0.821237 | 0.989223 | 0.997988 | 0.994666 |
| Thada_A8C756_S78s_1 | Thada_A8C | 0.9 | 0.91 | 0.26 | 0.3 | 0.44 | 0 | -0.13 | 0.01 | 1.55 | 1.02 | 1.11 | 0.59 | 1.27 | 0.8 | 1.01 | 0.99 | -0.5125 | 0.475 | 0.425 | 0.15505 | 0.077927 | 0.143676 | 0.989223 | 0.997988 | 0.994666 |
| Thoc1_Q8R3N6_S560s_1 | Thoc1_Q8F | -0.38 | -1.02 | 0.8 | 0.73 | -1.72 | -0.56 | -0.07 | 1.1 | 0.68 | -1.23 | 0.26 | 0.69 | 1.22 | -1.63 | 1.91 | 1.16 | -0.345 | 0.0675 | 0.6325 | 0.434422 | 0.598404 | 0.907825 | 0.989223 | 0.997988 | 0.994666 |
| Thoc2_B1AZI6_S1417s_1 | Thoc2_B1A | -2.01 | -2.32 | -3.32 | -3.04 | -1.47 | -3.15 | -2.28 | -2.82 | -3.33 | -2.92 | -1.84 | -2.94 | -0.32 | -1.48 | -0.84 | -1.91 | 0.2425 | -0.085 | 1.535 | 0.00691 | 0.589947 | 0.843445 | 0.989223 | 0.997988 | 0.994666 |
| Thra_P63058_S12s_1 | Thra_P630 | -1.56 | -1.68 | 0.25 | -0.99 | -3.42 | -0.37 | -1.62 | -1.74 | 0.22 | -1.05 | -1.01 | -0.57 | 0.71 | -3.52 | -1.32 | -0.98 | -0.7925 | 0.3925 | -0.2825 | 0.742111 | 0.259497 | 0.440274 | 0.989223 | 0.997988 | 0.9971 |
| Thrap3_Q569Z6_S190s_1 | Thrap3_Q5 | 0.16 | 0.09 | 1.13 | 0.68 | -2 | 0.38 | -1.79 | 0.14 | 0.68 | -0.65 | 0.53 | 0.93 | 0.85 | -1.81 | 0.48 | 0.41 | -1.3325 | -0.1425 | -0.5325 | 0.382093 | 0.048639 | 0.734214 | 0.989223 | 0.997988 | 0.994666 |
| Thrap3_Q569Z6_S238s_2 | Thrap3_Q5 | 0.11 | -0.25 | -0.61 | -0.58 | -0.51 | -0.19 | -0.99 | 0.1 | -0.37 | -0.96 | 0.65 | -0.87 | -0.68 | 0.69 | -0.56 | -1.57 | -0.065 | -0.055 | -0.1975 | 0.678202 | 0.833072 | 0.891316 | 0.995762 | 0.997988 | 0.996159 |
| Thrap3_Q569Z6_S243s_1 | Thrap3_Q5 | 0.81 | 0.63 | -0.07 | -0.23 | 0 | 0.11 | 0.24 | 0.24 | -0.37 | -0.12 | 0.33 | 0.58 | 0.68 | 0.29 | 0.56 | 0.38 | -0.1375 | -0.18 | 0.1925 | 0.549745 | 0.639097 | 0.609224 | 0.989223 | 0.997988 | 0.994666 |
| Thrap3_Q569Z6_S243s_2 | Thrap3_Q5 | 0.11 | -0.25 | -0.61 | -0.58 | -0.51 | -0.19 | -0.99 | 0.1 | -0.37 | -0.96 | 0.65 | -0.87 | -0.68 | 0.69 | -0.56 | -1.57 | -0.065 | -0.055 | -0.1975 | 0.678202 | 0.833072 | 0.891316 | 0.995762 | 0.997988 | 0.996159 |
| Thrap3_Q569Z6_S248s_1 | Thrap3_Q5 | 0.76 | 0.2 | 0.5 | 0.23 | -1.15 | 0.4 | -1 | 0.1 | 0.9 | -0.42 | -1.91 | -0.01 | 1.48 | -0.88 | 0.1 | -0.47 | -0.835 | -0.7825 | -0.365 | 0.47355 | 0.058396 | 0.182271 | 0.989223 | 0.997988 | 0.994666 |

| | | | | | | | | | | | | | | | | | | | | | | | | | | |
|---|---|---|---|---|---|---|---|---|---|---|---|---|---|---|---|---|---|---|---|---|---|---|---|---|---|---|
| Thrap3_Q569Z6_S264s_1 | Thrap3_Q5 | 1.5 | 1.17 | 0.96 | 0.48 | 0.59 | 0.95 | 0.49 | 0.7 | 1.35 | 1 | 0.77 | 0.65 | 1.86 | 1.03 | 0.57 | 0.15 | -0.345 | -0.085 | -0.125 | 0.767326 | 0.228578 | 0.778567 | 0.989223 | 0.997988 | 0.9971 |
| Thrap3_Q569Z6_S320s_1 | Thrap3_Q5 | 0.35 | 0.56 | 0.01 | -0.38 | -0.16 | -0.04 | -0.95 | -0.8 | -0.09 | -0.02 | -0.07 | -0.48 | 0.61 | -0.24 | -0.13 | -0.73 | -0.6225 | -0.3 | -0.2575 | 0.487976 | 0.077367 | 0.302402 | 0.989223 | 0.997988 | 0.994666 |
| Thrap3_Q569Z6_S532s_1 | Thrap3_Q5 | -0.46 | -0.78 | -0.92 | -1.76 | -1.64 | -0.66 | -1.23 | -1.04 | 2.08 | -0.75 | -3.51 | -1.01 | 3.14 | -1 | 0.3 | -1.49 | -0.1625 | 0.1825 | 1.2175 | 0.215203 | 0.641646 | 0.860274 | 0.989223 | 0.997988 | 0.994666 |
| Thrap3_Q569Z6_S55s_1 | Thrap3_Q5 | 0.56 | 0.85 | 0.5 | -0.87 | 0.18 | 0.46 | -0.12 | -0.18 | 1.62 | -0.96 | -3 | 0.25 | 2.58 | -0.12 | 1.11 | 0.61 | -0.175 | -0.7825 | 0.785 | 0.228193 | 0.660728 | 0.410358 | 0.990434 | 0.997988 | 0.994666 |
| Thrap3_Q569Z6_S572s_1 | Thrap3_Q5 | -0.41 | -0.23 | -0.62 | -1.07 | -0.64 | -1.22 | -1.04 | -0.68 | -2.52 | -1.23 | -0.77 | -0.98 | -2.14 | -0.92 | -1.03 | -1.44 | -0.3125 | -0.7925 | -0.8 | 0.046586 | 0.265803 | 0.087386 | 0.989223 | 0.997988 | 0.994666 |
| Thrap3_Q569Z6_S669s_1 | Thrap3_Q5 | 2.86 | 0.67 | 0.47 | 0.61 | -0.45 | 2.08 | -0.99 | 0.33 | 4.4 | 1.96 | -2.21 | -0.01 | 2.19 | -0.24 | 0.35 | -0.47 | -0.91 | -0.1175 | -0.695 | 0.358239 | 0.255471 | 0.929526 | 0.989223 | 0.997988 | 0.994666 |
| Thrap3_Q569Z6_S679s_1 | Thrap3_Q5 | 0.74 | 0.45 | 0.35 | 0 | -0.07 | -0.23 | 0.04 | 0 | -0.57 | -0.36 | -0.59 | -0.04 | -0.07 | 0.24 | 0.04 | -0.45 | -0.45 | -0.775 | -0.445 | 0.142306 | 0.080069 | 0.014795 | 0.989223 | 0.997988 | 0.994666 |
| Thrap3_Q569Z6_S695s_1 | Thrap3_Q5 | 9.94 | 5.14 | 5.4 | 2.04 | 5.62 | 9.9 | 5.99 | 5.12 | 0.77 | 7.17 | 1.89 | 5.27 | -1 | 1.54 | 5.77 | 5.99 | 1.0275 | -1.855 | -2.555 | 0.22374 | 0.543909 | 0.345393 | 0.989223 | 0.997988 | 0.994666 |
| Thrap3_Q569Z6_S860s_1 | Thrap3_Q5 | 0.57 | 0.32 | -0.16 | 0.13 | -0.05 | 0.13 | -0.25 | 0.43 | -0.7 | -0.41 | -1.48 | 0.02 | -1.31 | 0.09 | -0.05 | -1.52 | -0.15 | -0.8575 | -0.9125 | 0.060601 | 0.567582 | 0.038673 | 0.989223 | 0.997988 | 0.994666 |
| Thrap3_Q569Z6_S935s_1 | Thrap3_Q5 | 1.5 | 1.83 | 1.12 | 0.81 | 0.77 | 1.07 | 0.42 | 0.87 | 1.22 | 0.89 | 0.73 | 1.11 | -0.04 | 1.52 | 1.11 | 0.49 | -0.5325 | -0.3275 | -0.545 | 0.205626 | 0.092035 | 0.278839 | 0.989223 | 0.997988 | 0.994666 |
| Tiam1_G3UWG2_S1407s_1 | Tiam1_G3U | -0.2 | 0.27 | -1.23 | -0.9 | 0.05 | -0.57 | -0.45 | -0.74 | -0.51 | -0.41 | -0.53 | -0.63 | -0.48 | -0.33 | -0.28 | 0.01 | 0.0875 | -0.005 | 0.245 | 0.515411 | 0.814338 | 0.988752 | 0.995762 | 0.997988 | 0.994666 |
| Tiam1_G3UWG2_S1519s_1 | Tiam1_G3U | 0.57 | 0.86 | 0.9 | 0.76 | 0.72 | 1.26 | 0.95 | 0.95 | 0.66 | 0.86 | 0.65 | 1.27 | 0.3 | 1.02 | -0.31 | 0.31 | 0.1975 | 0.0875 | -0.4425 | 0.195617 | 0.380193 | 0.720337 | 0.989223 | 0.997988 | 0.994666 |
| Tiam1_G3UWG2_S358s_1 | Tiam1_G3U | -0.11 | 0.02 | -0.46 | -0.19 | -0.62 | -0.3 | -0.41 | -0.46 | -0.24 | -0.09 | 0.08 | -0.41 | 0.39 | -0.17 | -0.15 | -0.63 | -0.2625 | 0.02 | 0.045 | 0.879218 | 0.241316 | 0.93231 | 0.989223 | 0.997988 | 0.999507 |
| Tiam1_G3UWG2_T1405t_1 | Tiam1_G3U | -1.31 | -0.75 | 0 | 0.84 | 0.11 | -0.94 | 1.2 | 0.69 | -0.74 | -0.72 | 1.16 | 0.71 | -1.87 | -0.04 | 0.99 | 1 | 0.57 | 0.4075 | 0.325 | 0.658169 | 0.343839 | 0.514293 | 0.989223 | 0.997988 | 0.995259 |
| Tiam2_Q6ZPF3_S20s_1 | Tiam2_Q6Z | 8.55 | 3.93 | 5.09 | 0.57 | 3.64 | 9.89 | 5.29 | 4.32 | 2.18 | 6.59 | 0.43 | 4.93 | 1.29 | 1.64 | 5.15 | 5.62 | 1.25 | -1.0025 | -1.11 | 0.521754 | 0.504521 | 0.594271 | 0.989223 | 0.997988 | 0.994666 |
| Tiam2_Q6ZPF3_S440s_1 | Tiam2_Q6Z | 0.65 | 0.83 | 0.6 | 1.56 | 0.72 | 0.65 | 0.84 | 1.19 | 0.58 | 0.18 | 0.86 | 0.87 | -0.28 | 0.66 | 0.3 | 0.88 | -0.06 | -0.2875 | -0.52 | 0.172759 | 0.833482 | 0.364496 | 0.995762 | 0.997988 | 0.994666 |
| Tiam2_Q6ZPF3_S600s_1 | Tiam2_Q6Z | 0.24 | -0.07 | -0.2 | 0.47 | 0.46 | -0.17 | 0.5 | 0.17 | -0.57 | -0.16 | -0.89 | 0.16 | -1.1 | 0.06 | -0.39 | -0.23 | 0.13 | -0.475 | -0.525 | 0.136618 | 0.623846 | 0.149961 | 0.989223 | 0.997988 | 0.994666 |
| Tjp1_B9EHJ3_S1534s_1 | Tjp1_B9EH | -0.03 | 0.19 | 0.28 | 0.6 | 0.53 | -0.25 | 0.43 | 0.25 | 0.59 | 0.26 | -0.02 | 0.34 | -0.53 | 0.43 | -0.59 | 0.09 | -0.02 | 0.0325 | -0.41 | 0.225611 | 0.939478 | 0.897842 | 0.999187 | 0.997988 | 0.994666 |
| Tjp1_B9EHJ3_S175s_1 | Tjp1_B9EH | -0.22 | -0.27 | -0.52 | -0.25 | -0.46 | -0.49 | -1.03 | -0.7 | -0.02 | -0.37 | -0.87 | -0.66 | 0.36 | -0.72 | -0.89 | -0.73 | -0.355 | -0.165 | -0.18 | 0.593154 | 0.142409 | 0.532932 | 0.989223 | 0.997988 | 0.995259 |
| Tjp1_B9EHJ3_S212s_1 | Tjp1_B9EH | 1.3 | 0.94 | -0.45 | -0.33 | 1.01 | 1.4 | -0.77 | -0.03 | -0.24 | 0.92 | 1.03 | -0.46 | 0.23 | 0.78 | -0.88 | -1.48 | 0.0375 | -0.0525 | -0.7025 | 0.271436 | 0.949589 | 0.923193 | 0.999187 | 0.997988 | 0.994666 |
| Tjp1_B9EHJ3_S353s_1 | Tjp1_B9EH | 1.62 | 0.95 | -2.21 | -1.95 | 1.21 | 1.03 | -2.29 | -2.41 | 1.37 | 0.89 | -1.02 | -2.32 | 1.03 | 1.23 | -1.83 | -2.18 | -0.2175 | 0.1275 | -0.04 | 0.972403 | 0.856975 | 0.911264 | 0.996349 | 0.997988 | 0.999507 |
| Tjp1_B9EHJ3_S617s_1 | Tjp1_B9EH | 0.3 | 0.62 | 0.47 | 0.07 | 0.29 | 0.16 | 0.2 | 0.24 | -0.03 | 0.7 | 0.5 | 0.14 | 0.43 | -0.03 | -0.16 | -0.01 | -0.1425 | -0.0375 | -0.3075 | 0.265885 | 0.51297 | 0.887695 | 0.989223 | 0.997988 | 0.994666 |
| Tjp1_B9EHJ3_S622s_1 | Tjp1_B9EH | 0.59 | 0.02 | 0.84 | 0.73 | 0.06 | 0.87 | 0.8 | 0.77 | 0.45 | 0 | 0.38 | 0.47 | 0.7 | -0.15 | 0.4 | -0.25 | 0.08 | -0.22 | -0.37 | 0.279732 | 0.784335 | 0.425233 | 0.995762 | 0.997988 | 0.994666 |
| Tjp1_B9EHJ3_S899s_1 | Tjp1_B9EH | -2.22 | -2.61 | 0.42 | 0.04 | -2.13 | -0.41 | 1.06 | 0.41 | -1.27 | -0.02 | 1.09 | 0.33 | 0.09 | -0.22 | -0.6 | 0.13 | 0.825 | 1.125 | 0.9425 | 0.20307 | 0.369752 | 0.190869 | 0.989223 | 0.997988 | 0.994666 |
| Tjp1_B9EHJ3_S912s_1 | Tjp1_B9EH | -0.9 | -0.55 | -1.54 | -1.1 | -0.56 | -1.27 | -1.37 | -1.34 | -0.99 | -0.4 | -0.79 | -0.72 | -0.47 | -0.49 | -1 | -1.67 | -0.1125 | 0.2975 | 0.115 | 0.756184 | 0.712805 | 0.314778 | 0.995437 | 0.997988 | 0.9971 |
| Tjp1_B9EHJ3_T354t_1 | Tjp1_B9EH | 2 | 1.12 | 1.94 | 2.25 | 1.48 | 1.31 | 1.66 | 1.87 | 1.51 | 1.27 | 2.28 | 2.15 | 0.61 | 1.75 | 1.5 | 1.55 | -0.2475 | -0.025 | -0.475 | 0.221086 | 0.415723 | 0.944356 | 0.989223 | 0.997988 | 0.994666 |
| Tjp1_P39447_S912s_1 | Tjp1_P394 | -3.13 | -2.84 | -4.14 | -3.27 | -2.5 | -4.17 | -4.18 | -2.98 | -2.99 | -2.66 | -1.89 | -2.41 | -1.64 | -2.34 | -2.7 | -4.34 | -0.1125 | 0.8575 | 0.59 | 0.325822 | 0.81155 | 0.042414 | 0.995762 | 0.997988 | 0.994666 |
| Tjp2_Q9Z0U1_S107s_1 | Tjp2_Q9Z0 | 0.93 | 0.6 | 0.22 | 0.22 | 0.68 | 0.35 | 0.86 | 0.03 | 0.71 | 0.82 | 0.23 | 0.24 | 0.11 | 0.92 | 0.33 | 0.18 | -0.0125 | 0.0075 | -0.1075 | 0.727209 | 0.964893 | 0.978714 | 0.999187 | 0.997988 | 0.9971 |
| Tjp2_Q9Z0U1_S1136s_1 | Tjp2_Q9Z0 | -0.11 | 0.42 | 0.64 | 0.58 | 0.02 | -0.62 | 0.62 | 0.49 | -0.8 | 0.29 | 0.52 | 0.26 | -1.01 | 0.28 | 0.32 | 0.88 | -0.255 | -0.315 | -0.265 | 0.54152 | 0.455048 | 0.384797 | 0.989223 | 0.997988 | 0.994666 |
| Tjp2_Q9Z0U1_S213s_1 | Tjp2_Q9Z0 | 1.77 | 1.83 | 2.49 | 2.01 | 1.38 | 1.84 | 1.83 | 1.86 | 1.46 | 1.77 | 1.04 | 2.2 | 0.56 | 0.75 | 2.27 | 2.4 | -0.2975 | -0.4075 | -0.53 | 0.29355 | 0.25885 | 0.226468 | 0.989223 | 0.997988 | 0.994666 |
| Tjp2_Q9Z0U1_S393s_1 | Tjp2_Q9Z0 | -1.79 | 0.26 | -0.52 | -0.94 | -0.24 | -1.08 | -0.84 | -0.92 | -0.6 | -0.65 | 0.34 | -0.3 | 0.2 | -0.24 | -0.01 | -0.45 | -0.0225 | 0.445 | 0.6225 | 0.180354 | 0.958921 | 0.349029 | 0.999187 | 0.997988 | 0.994666 |
| Tjp2_Q9Z0U1_S395s_1 | Tjp2_Q9Z0 | 0.79 | 1.38 | 1.18 | 0.75 | 1.12 | 1.19 | 0.94 | 0.57 | 1.35 | 1.32 | 1.08 | 0.98 | 0.92 | 1.46 | 1.03 | 0.78 | -0.07 | 0.1575 | 0.0225 | 0.937104 | 0.785995 | 0.535422 | 0.995762 | 0.997988 | 0.999507 |
| Tjp2_Q9Z0U1_S404s_1 | Tjp2_Q9Z0 | -0.16 | -0.15 | 0.43 | 0.48 | -2.09 | -0.04 | -1.06 | -0.01 | 0.69 | -0.37 | 1.28 | 0.29 | 0.95 | -1.76 | 0.13 | -0.01 | -0.95 | 0.3225 | -0.3225 | 0.562796 | 0.074472 | 0.417152 | 0.989223 | 0.997988 | 0.995259 |
| Tjp2_Q9Z0U1_S421s_1 | Tjp2_Q9Z0 | 1.49 | 1.07 | 0.66 | 1.17 | 0.4 | 2.06 | 0.05 | 0.59 | 4.3 | 2.04 | 0.91 | 0.72 | 3.84 | 0.41 | 1.66 | 0.14 | -0.3225 | 0.895 | 0.415 | 0.589474 | 0.474549 | 0.253413 | 0.989223 | 0.997988 | 0.995259 |
| Tjp2_Q9Z0U1_S684s_1 | Tjp2_Q9Z0 | 0.23 | -0.12 | 0.11 | 0.4 | -0.15 | -0.15 | 0.49 | 0.15 | -0.35 | 0.06 | -0.57 | 0.02 | -0.53 | -0.33 | -0.41 | 0.25 | -0.07 | -0.365 | -0.41 | 0.168548 | 0.777086 | 0.17755 | 0.995762 | 0.997988 | 0.994666 |
| Tjp2_Q9Z0U1_S988s_1 | Tjp2_Q9Z0 | 0.56 | 0.39 | -0.92 | -1.21 | 0.41 | 0.55 | -1.07 | -1.01 | 0.21 | 0.38 | 0.01 | -1.28 | 0.04 | 0.35 | -0.96 | -0.95 | 0.015 | 0.125 | -0.085 | 0.871588 | 0.978848 | 0.818705 | 0.999187 | 0.997988 | 0.999507 |
| Tkt_P40142_T287t_1 | Tkt_P4014 | 1.35 | 1.86 | 0.46 | 0.39 | 0.96 | 0.92 | 0.38 | 0.03 | 1.14 | 2.71 | 0.57 | 0.49 | 1.2 | 1.32 | -1.14 | 0.87 | -0.4425 | 0.2125 | -0.4525 | 0.468719 | 0.288898 | 0.71332 | 0.989223 | 0.997988 | 0.994666 |
| Tle1_Q5SQA7_S31s_1 | Tle1_Q5SQ | -1.86 | -2 | -1.14 | -1.09 | -2.03 | -1.98 | -1.17 | -1.28 | -2.36 | -1.57 | -1.57 | -1.27 | -1.55 | -1.09 | -2.28 | -1.34 | -0.0925 | -0.17 | -0.0425 | 0.908327 | 0.783241 | 0.628182 | 0.995762 | 0.997988 | 0.999507 |
| Tle3_E0CY76_S202s_1 | Tle3_E0CY | -2.33 | -3.31 | -4.4 | -6.26 | -3.01 | -3.46 | -3.88 | -3.84 | -2.68 | -2.17 | -2.55 | -4.73 | -0.91 | -2.4 | -3.76 | -4.16 | 0.5275 | 1.0425 | 1.2675 | 0.214104 | 0.49281 | 0.266944 | 0.989223 | 0.997988 | 0.994666 |
| Tle3_E0CY76_S262s_1 | Tle3_E0CY | -0.48 | -1.02 | -0.28 | -0.15 | -0.23 | -0.78 | -0.16 | -0.48 | 0.2 | -1 | 0.57 | -0.03 | 0.02 | 0.16 | -0.54 | -0.23 | 0.07 | 0.4175 | 0.335 | 0.286258 | 0.799936 | 0.297366 | 0.995762 | 0.997988 | 0.994666 |
| Tle3_E0CY76_S285s_1 | Tle3_E0CY | -0.85 | -1.44 | -0.33 | -0.36 | -0.62 | -1.17 | -0.67 | -0.02 | -1.46 | -0.25 | -0.28 | -0.37 | -0.38 | -0.3 | -0.61 | -1.06 | 0.125 | 0.155 | 0.1575 | 0.649106 | 0.723457 | 0.692976 | 0.995437 | 0.997988 | 0.995259 |
| Tle4_Q62441_S292s_1 | Tle4_Q624 | 0.01 | -0.16 | 0.04 | 0.03 | -0.43 | -0.05 | 0.71 | 0.02 | -0.39 | -0.05 | 0.53 | 0.15 | -0.49 | 0.36 | 0.05 | 0.1 | 0.0825 | 0.08 | 0.025 | 0.92656 | 0.767593 | 0.760803 | 0.995762 | 0.997988 | 0.999507 |
| Tln1_P26039_S2040s_1 | Tln1_P260 | -1.85 | -2.2 | -1.97 | -2.03 | -1.7 | -2.1 | -1.16 | -1.42 | -1.52 | -1.3 | -0.19 | -1.25 | -2.12 | -1.46 | -1.03 | -1.31 | 0.4175 | 0.9475 | 0.5325 | 0.103404 | 0.135503 | 0.016864 | 0.989223 | 0.997988 | 0.994666 |
| Tln2_E9PUM4_S473s_1 | Tln2_E9PU | 1.42 | 1.5 | -0.1 | 0.02 | 0.24 | 1.78 | 0.73 | 0.07 | 1.29 | 1.35 | -0.87 | -0.65 | 0.02 | 0.39 | 0.39 | 0.37 | -0.005 | -0.43 | -0.4175 | 0.350328 | 0.992423 | 0.526872 | 0.999187 | 0.997988 | 0.994666 |
| Tln2_E9PUM4_T1843t_1 | Tln2_E9PU | 0.76 | 1.15 | 0.95 | 0.96 | 1.01 | 0.84 | 0.55 | 0.74 | 0.83 | 1.42 | 0.95 | 0.75 | 0.64 | 0.88 | 0.92 | 0.61 | -0.17 | 0.0325 | -0.1925 | 0.433831 | 0.440434 | 0.895383 | 0.989223 | 0.997988 | 0.994666 |
| Tmcc1_F8WJ98_S131s_1 | Tmcc1_F8V | -0.84 | -2.54 | -1.99 | -2.19 | -1.15 | -0.88 | -1.5 | -1.31 | -0.78 | -2.35 | -0.99 | -1.12 | -1.03 | -1.33 | -1.9 | -1.29 | 0.68 | 0.58 | 0.5025 | 0.241823 | 0.100052 | 0.249323 | 0.989223 | 0.997988 | 0.994666 |
| Tmcc1_F8WJ98_S419s_1 | Tmcc1_F8V | 0.08 | 0.77 | 0.1 | -0.35 | 0.83 | -0.36 | 0.56 | 0.04 | -0.08 | 0.1 | -0.21 | 0.12 | 0.3 | 0.16 | -0.32 | 0.06 | 0.1175 | -0.1675 | -0.1 | 0.752446 | 0.739907 | 0.566618 | 0.995762 | 0.997988 | 0.9971 |
| Tmcc1_F8WJ98_S450s_1 | Tmcc1_F8V | 1.63 | 1.57 | -0.26 | 0.79 | 1.52 | 0.1 | -0.03 | -0.96 | 0.43 | 1.53 | 0.1 | -0.49 | 0.89 | 1.96 | -0.07 | 0.13 | -0.775 | -0.54 | -0.205 | 0.72633 | 0.219495 | 0.35171 | 0.989223 | 0.997988 | 0.9971 |
| Tmcc1_F8WJ98_S451s_1 | Tmcc1_F8V | 0.19 | -0.37 | -0.31 | 0.7 | 0.25 | 0.02 | 0.48 | -0.83 | 0.76 | 0.09 | -0.18 | -0.31 | -0.52 | 0.08 | 0.17 | 0.45 | -0.0725 | 0.0375 | -0.0075 | 0.982964 | 0.845886 | 0.916444 | 0.996349 | 0.997988 | 1 |
| Tmcc2_Q80W04_S119s_1 | Tmcc2_Q8 | -1.13 | -0.38 | 0.12 | -0.79 | -1.35 | -0.8 | 0.13 | 0.15 | -0.53 | 0.51 | 1.03 | -1.15 | 0.45 | -0.19 | 0.51 | 0.62 | 0.0775 | 0.51 | 0.8925 | 0.029048 | 0.857164 | 0.344017 | 0.996349 | 0.997988 | 0.994666 |
| Tmcc2_Q80W04_S126s_1 | Tmcc2_Q8 | -0.63 | -1.14 | -0.98 | -1.34 | -1.01 | -0.67 | -0.6 | -2.11 | 0.5 | -0.85 | -1.19 | -1.15 | 0.6 | -2.43 | -0.51 | -1.26 | -0.075 | 0.35 | 0.1225 | 0.837686 | 0.84082 | 0.411205 | 0.995818 | 0.997988 | 0.999507 |
| Tmcc2_Q80W04_S189s_1 | Tmcc2_Q8 | -0.73 | -2.01 | -1.25 | -1.4 | 0.11 | -0.8 | -0.05 | -0.75 | -1.41 | -1.64 | -0.18 | -0.83 | -1.77 | -0.79 | -0.28 | -0.46 | 0.975 | 0.3325 | 0.5225 | 0.234547 | 0.019808 | 0.429103 | 0.989223 | 0.997988 | 0.994666 |
| Tmcc2_Q80W04_S435s_1 | Tmcc2_Q8 | -1.29 | -0.85 | -0.03 | -0.27 | 0.07 | -0.7 | 0.43 | 0.38 | -1.5 | -0.65 | 0.42 | 0.16 | -1.93 | -0.14 | -0.24 | -0.12 | 0.655 | 0.2175 | 0.0025 | 0.995986 | 0.108491 | 0.658873 | 0.989223 | 0.997988 | 1 |
| Tmcc2_Q80W04_S75s_1 | Tmcc2_Q8 | -0.25 | -0.7 | -0.94 | -0.85 | 0.87 | 0.07 | -0.05 | -0.01 | -1.47 | -0.9 | -0.61 | -1.06 | -1.25 | -0.39 | -2.17 | -0.74 | 0.905 | -0.325 | -0.4525 | 0.292244 | 0.011606 | 0.270761 | 0.989223 | 0.997988 | 0.994666 |
| Tmcc3_Q8R310_S216s_1 | Tmcc3_Q8 | -1.29 | -0.85 | -0.59 | -1.03 | 0.07 | -0.7 | -0.17 | -0.63 | -1.5 | -0.65 | 0.13 | -0.57 | -1.93 | -0.14 | -1.03 | -0.94 | 0.5825 | 0.2925 | -0.07 | 0.861355 | 0.057649 | 0.439755 | 0.989223 | 0.997988 | 0.999507 |
| Tmed8_Q3UHI4_S21s_1 | Tmed8_Q3 | -1.82 | -1.7 | -1.34 | -2.16 | -2.18 | -1.42 | -1.88 | -1.9 | 0.48 | -1.72 | -2.7 | -1.08 | 1.19 | -2.43 | -0.3 | -1.61 | -0.09 | 0.5 | 0.9675 | 0.201478 | 0.741279 | 0.433825 | 0.995762 | 0.997988 | 0.994666 |
| Tmeff2_Q9QYM9_T358t_1 | Tmeff2_Q9 | -0.47 | -0.94 | -0.46 | 0.03 | -2.01 | 0.5 | -0.3 | -0.51 | 1.75 | -0.42 | -1.02 | 0.15 | 2.07 | -2.34 | 0.56 | 0.03 | -0.12 | 0.575 | 0.54 | 0.517527 | 0.814588 | 0.332834 | 0.995762 | 0.997988 | 0.994666 |
| Tmem100_Q9CQG9_S121s_1 | Tmem100_ | -0.25 | -0.03 | 0.29 | 0.14 | 0.28 | -0.05 | 0.42 | 0.1 | 0.35 | 0.31 | 0.3 | 0.18 | 0.26 | -0.01 | 0.45 | 0.45 | 0.15 | 0.2475 | 0.25 | 0.347761 | 0.519386 | 0.288286 | 0.989223 | 0.997988 | 0.994666 |
| Tmem100_Q9CQG9_S15s_1 | Tmem100_ | -0.19 | -0.06 | 0.09 | 0.07 | -0.4 | -0.19 | 0.2 | -0.08 | 0.56 | 0.42 | 0.79 | -0.03 | 0.67 | 0.31 | 0.38 | 0.29 | -0.095 | 0.4575 | 0.435 | 0.096003 | 0.672285 | 0.099017 | 0.992876 | 0.997988 | 0.994666 |
| Tmem11_Q8BK08_S16s_1 | Tmem11_( | 1.43 | 0.76 | 1.6 | 1.51 | 0.1 | 1.74 | 0.28 | 0.95 | 1.46 | 0.23 | -0.25 | 0.68 | 1.11 | -0.4 | 0.55 | 0.96 | -0.5575 | -0.795 | -0.77 | 0.077656 | 0.187663 | 0.07635 | 0.989223 | 0.997988 | 0.994666 |
| Tmem121_Q80XA0_S310s_1 | Tmem121_ | 1.08 | 0.9 | 0.73 | 0.63 | 1.07 | 1.18 | 0.23 | 0.46 | 0.8 | 0.62 | -1.01 | -0.15 | 1.06 | 0.86 | 0.14 | -1.35 | -0.1 | -0.77 | -0.6575 | 0.225968 | 0.725976 | 0.091347 | 0.995437 | 0.997988 | 0.994666 |
| Tmem127_G3UXI2_S17s_1 | Tmem127_ | -0.25 | 0.96 | 0.41 | 0.49 | 0.11 | 0.29 | 0.62 | 0.63 | -0.02 | 0.57 | 1.05 | 0.9 | 0.33 | 0.97 | 0.77 | 0.24 | 0.01 | 0.2225 | 0.175 | 0.608899 | 0.973651 | 0.537226 | 0.999187 | 0.997988 | 0.995259 |
| Tmem132a_Q922P8_S883s_1 | Tmem132a | -0.13 | 0.47 | 0.26 | 0.26 | 0.53 | 0.06 | 1.42 | 0.36 | -0.01 | 0.41 | 0.48 | 0.53 | -0.42 | 0.86 | 0.56 | 0.04 | 0.3775 | 0.1375 | 0.045 | 0.895595 | 0.268112 | 0.585864 | 0.989223 | 0.997988 | 0.999507 |
| Tmem132a_Q922P8_S938s_1 | Tmem132a | -0.5 | -0.95 | 1.51 | 1.14 | -0.04 | -0.11 | 1.22 | 1.05 | -0.02 | 0.09 | 1.16 | 2.05 | -0.51 | -0.01 | 1.63 | 2.74 | 0.23 | 0.52 | 0.6625 | 0.440931 | 0.712433 | 0.464747 | 0.995437 | 0.997988 | 0.994666 |
| Tmem132c_E9QP73_S563s_1 | Tmem132c | -1.17 | -0.86 | -0.48 | -0.35 | -3.15 | -0.34 | -0.94 | -1.22 | 0.25 | -0.45 | -0.04 | -0.77 | 0.15 | -2.13 | -0.83 | -0.5 | -0.6975 | 0.4625 | -0.1125 | 0.818184 | 0.241015 | 0.172685 | 0.989223 | 0.997988 | 0.999507 |
| Tmem132d_Q76HP3_S1035s_1 | Tmem132c | 0.54 | 0.55 | 1.04 | 1.37 | -0.28 | 0.86 | 0.02 | 0.8 | 1.41 | 0.37 | 0.11 | 0.86 | 1.79 | -0.06 | 0.51 | 0.66 | -0.525 | -0.1875 | -0.15 | 0.72862 | 0.158068 | 0.606732 | 0.989223 | 0.997988 | 0.9971 |
| Tmem132d_Q76HP3_S554s_1 | Tmem132c | -2.28 | -1.32 | -0.24 | -0.93 | -1.81 | -0.06 | -1.49 | -1.89 | 1.52 | -0.78 | -0.12 | -0.09 | 0.37 | -1.6 | -0.1 | -2.07 | -0.12 | 1.325 | 0.3425 | 0.602132 | 0.825808 | 0.048835 | 0.995762 | 0.997988 | 0.995259 |
| Tmem135_Q8C8G3_S198s_1 | Tmem135_ | 0.07 | 0.13 | -0.11 | -1.31 | 0.79 | 0.66 | 0.25 | 0.45 | 0.41 | 0.36 | 0.36 | -0.28 | 0.44 | 1.39 | 0.74 | 0.23 | 0.8425 | 0.5175 | 1.005 | 0.036631 | 0.038933 | 0.196493 | 0.989223 | 0.997988 | 0.994666 |
| Tmem145_E9PUE6_S511s_1 | Tmem145_ | 1.3 | 1.54 | 1.12 | 0.85 | 1.17 | 1.27 | 0.82 | 0.83 | 1.09 | 1.37 | 1.06 | 0.59 | 0.85 | 1.48 | 0.86 | 1.27 | -0.18 | -0.175 | -0.0875 | 0.760199 | 0.470979 | 0.527022 | 0.989223 | 0.997988 | 0.9971 |
| Tmem151a_Q6GQT5_S323s_1 | Tmem151a | 0.85 | 0.52 | 0.98 | 0.98 | 0.52 | 0.88 | 0.83 | 0.45 | 0.51 | 0.9 | 0.41 | 0.56 | 0.52 | 1.19 | 0.9 | 0.54 | -0.1625 | -0.2375 | -0.045 | 0.870365 | 0.484642 | 0.332939 | 0.989223 | 0.997988 | 0.999507 |
| Tmem151b_Q68FE7_S486s_1 | Tmem151b | -0.49 | 0.2 | -0.2 | -0.45 | 0.2 | 0.87 | -0.64 | -0.98 | -0.21 | 0.11 | 0.1 | -0.4 | 0.64 | 0.23 | -0.48 | -0.37 | 0.0975 | 0.135 | 0.24 | 0.488037 | 0.817857 | 0.612053 | 0.995762 | 0.997988 | 0.994666 |
| Tmem151b_Q68FE7_S488s_1 | Tmem151b | 0.57 | 0.99 | 0.34 | 0.29 | 0.06 | 0.94 | -0.35 | 0.44 | 0.06 | 0.98 | 0.36 | 0.26 | 1 | 1.4 | 0.05 | 0.42 | -0.275 | -0.1325 | 0.17 | 0.641364 | 0.411731 | 0.656307 | 0.989223 | 0.997988 | 0.995259 |
| Tmem163_Q8C996_S11s_1 | Tmem163_ | 0.46 | 0.68 | 0.66 | 0.76 | 0.49 | 0.46 | 0.8 | 0.61 | 0.74 | 1.01 | 0.8 | 0.75 | 0.83 | 0.92 | 0.62 | 0.44 | -0.05 | 0.185 | 0.0625 | 0.799674 | 0.809583 | 0.397223 | 0.995762 | 0.997988 | 0.999507 |
| Tmem163_Q8C996_S45s_1 | Tmem163_ | 2.12 | 2.01 | 2.56 | 1.51 | 1.48 | 1.96 | 1.36 | 1.57 | 0.88 | 1.77 | 1.25 | 1.81 | 0.93 | -0.15 | 1.5 | 1.76 | -0.4575 | -0.6225 | -1.04 | 0.045356 | 0.13297 | 0.087603 | 0.989223 | 0.997988 | 0.994666 |
| Tmem163_Q8C996_S56s_1 | Tmem163_ | 0.4 | 0.68 | 0.74 | 0.62 | 0.39 | 0.13 | 0.85 | 0.64 | 0.34 | 0.52 | 0.46 | 0.69 | -0.28 | 0.81 | 0.55 | 0.59 | -0.1075 | -0.1075 | -0.1925 | 0.534794 | 0.654388 | 0.625241 | 0.990434 | 0.997988 | 0.994666 |
| Tmem163_Q8C996_S7s_1 | Tmem163_ | -0.96 | -0.85 | 0 | -0.79 | -1.07 | -0.12 | -1.09 | -2.11 | 1.64 | -0.26 | -0.49 | -1.94 | 1.6 | -0.75 | -0.42 | -0.33 | -0.4475 | 0.3875 | 0.675 | 0.227939 | 0.31739 | 0.57899 | 0.989223 | 0.997988 | 0.994666 |
| Tmem169_D3YUR6_S15s_1 | Tmem169_ | 2.19 | 1.28 | -0.29 | 0.45 | 1.54 | 2.09 | 0.61 | 0.01 | 0.29 | 2.19 | -1.78 | 0.26 | 0.52 | 1.3 | 0.34 | 0.51 | 0.155 | -0.6675 | -0.24 | 0.656067 | 0.805888 | 0.446924 | 0.995762 | 0.997988 | 0.995259 |
| Tmem169_D3YUR6_S20s_1 | Tmem169_ | -4.1 | -2.21 | -1.05 | -3.38 | -2.54 | -4.34 | -1.77 | -1.27 | -3.08 | -2.32 | -1.23 | -0.51 | -0.91 | -3.94 | -2.25 | -1.03 | 0.205 | 0.9 | 0.6525 | 0.452066 | 0.804572 | 0.269843 | 0.995762 | 0.997988 | 0.994666 |
| Tmem169_D3YUR6_S20s_2 | Tmem169_ | 0.14 | 1.98 | -1.28 | 0.42 | 0.31 | 1.1 | 0.32 | 1.46 | 1.5 | 0.56 | 0.54 | 1.94 | 1.13 | 0.96 | 0.77 | 1.22 | 0.4825 | 0.82 | 0.705 | 0.268208 | 0.461827 | 0.24802 | 0.989223 | 0.997988 | 0.994666 |
| Tmem169_D3YUR6_S69s_1 | Tmem169_ | 0.49 | 0.94 | -0.46 | -0.04 | -1.04 | 0.78 | -2.29 | -1.34 | 0.47 | 0.2 | 0.48 | -0.08 | -0.61 | -1.88 | -0.51 | 0.02 | -1.205 | 0.035 | -0.9775 | 0.067175 | 0.082015 | 0.920089 | 0.989223 | 0.997988 | 0.994666 |
| Tmem169_D3YUR6_T62t_1 | Tmem169_ | -3.21 | -2.08 | -1.1 | -1.32 | -2.35 | -1.98 | -2.18 | -1.92 | -1.92 | -2.08 | -0.32 | -1.22 | -3.41 | -2.15 | -1.67 | -1.57 | -0.18 | 0.5425 | -0.2725 | 0.642449 | 0.692551 | 0.35653 | 0.994893 | 0.997988 | 0.995259 |
| Tmem184b_Q8BG09_S382s_1 | Tmem184b | -4.46 | -3.74 | -2.67 | -5.44 | -2.53 | -2.64 | -2.85 | -3.44 | -1.19 | -6.1 | -4.09 | -3.89 | 0.7 | -3.29 | -1.86 | -3.01 | 1.2125 | 0.26 | 2.2125 | 0.039502 | 0.052577 | 0.80038 | 0.989223 | 0.997988 | 0.994666 |
| Tmem19_Q91W52_S294s_1 | Tmem19_( | 0.8 | -0.3 | 0.84 | 0.8 | 0.57 | 0.46 | 0.9 | 0.95 | 0.77 | 0.34 | 0.89 | 0.44 | 0.3 | 1.34 | 1.04 | 0.8 | 0.185 | 0.075 | 0.335 | 0.379694 | 0.565244 | 0.821061 | 0.989223 | 0.997988 | 0.994666 |
| Tmem200b_D3Z0P8_S177s_1 | Tmem200b | 2.35 | 0.7 | 0.23 | 1.34 | 1.19 | 2.74 | 0.48 | 0.61 | 2.09 | 0.94 | 1.02 | -0.05 | 1.14 | 0.62 | -0.21 | 0.09 | 0.1 | -0.155 | -0.745 | 0.170318 | 0.87125 | 0.790456 | 0.996493 | 0.997988 | 0.994666 |
| Tmem200b_D3Z0P8_S5s_1 | Tmem200b | -0.22 | 0.58 | 0.41 | -0.27 | -0.36 | 0.37 | -1.04 | -1.05 | 2.46 | 0.01 | -3.45 | 0 | 2.66 | -0.73 | 1 | -1.06 | -0.645 | -0.37 | 0.3425 | 0.663615 | 0.122636 | 0.733967 | 0.989223 | 0.997988 | 0.996159 |
| Tmem200c_J3QK46_S487s_1 | Tmem200c | -0.59 | -0.21 | -0.87 | -1.09 | -1.05 | -0.49 | -0.67 | -0.78 | -0.75 | -0.62 | 0.15 | -0.62 | -0.23 | -1.26 | -1.88 | -0.88 | -0.0575 | 0.23 | -0.3725 | 0.363758 | 0.82971 | 0.468669 | 0.995762 | 0.997988 | 0.994666 |
| Tmem201_A2A8U2_S454s_1 | Tmem201_ | -0.53 | -0.22 | 0.51 | -1.06 | 0.34 | -0.01 | -0.55 | -0.92 | -0.8 | -0.33 | 0.49 | -0.39 | 0.03 | 0.28 | -0.94 | -1.49 | 0.04 | 0.0675 | -0.205 | 0.682807 | 0.922487 | 0.871394 | 0.997335 | 0.997988 | 0.996159 |
| Tmem229a_B9EJI9_S4s_1 | Tmem229a | -0.29 | 0.26 | 0.84 | 1.22 | 0.2 | 0.3 | 0.71 | 1.38 | 0.24 | 0.46 | 1.89 | 1.04 | 0.94 | 0.18 | 1.51 | 1.47 | 0.14 | 0.4 | 0.5175 | 0.260765 | 0.73181 | 0.406001 | 0.995762 | 0.997988 | 0.994666 |
| Tmem245_D3YWD3_S12s_1 | Tmem245_ | -0.31 | -0.31 | 0.2 | -0.2 | -0.12 | -0.39 | -0.12 | 0.28 | -0.02 | -0.12 | 0.14 | 0.27 | 0.8 | -0.5 | 0.51 | 0.64 | 0.0675 | 0.2225 | 0.5175 | 0.160902 | 0.783034 | 0.359771 | 0.995762 | 0.997988 | 0.994666 |
| Tmem245_D3YWD3_S329s_1 | Tmem245_ | -1.66 | -0.37 | 0.64 | -0.25 | 0.43 | 0.21 | 0.58 | 0.75 | 0.36 | -0.16 | 0.94 | 0.01 | 1.32 | 0.4 | 0.68 | 0.62 | 0.9025 | 0.6975 | 1.165 | 0.03683 | 0.071811 | 0.186912 | 0.989223 | 0.997988 | 0.994666 |
| Tmem25_Q9DCF1_S297s_1 | Tmem25_( | -0.08 | -0.17 | 0.12 | 0.4 | 0.14 | 0.2 | 0.84 | 0.98 | -1.57 | 1.01 | 0.48 | -0.27 | 1.03 | -0.1 | 1.82 | 1.43 | 0.4725 | -0.155 | 0.9775 | 0.043826 | 0.121048 | 0.77197 | 0.989223 | 0.997988 | 0.994666 |
| Tmem261_Q9CQ00_S36s_1 | Tmem261_ | -1.64 | -1.48 | -1 | -0.73 | -1.11 | -0.93 | -1.37 | -0.46 | -1.92 | -1.01 | 0.89 | -0.94 | 0.2 | -1.12 | -0.21 | -1.13 | 0.245 | 0.4675 | 0.6475 | 0.131219 | 0.431677 | 0.425598 | 0.989223 | 0.997988 | 0.994666 |
| Tmem38a_Q3TMP8_S289s_1 | Tmem38a_ | -1.47 | -0.31 | -0.03 | -0.84 | -0.86 | -1.11 | -1.11 | -1.05 | -0.78 | -1.31 | -0.76 | -0.54 | 1.17 | -1.83 | 0.05 | -0.32 | -0.37 | -0.185 | 0.43 | 0.500732 | 0.281467 | 0.615764 | 0.989223 | 0.997988 | 0.994666 |
| Tmem40_E9Q6I9_S104s_1 | Tmem40_E | -0.43 | -0.77 | 0.05 | 0.28 | -0.99 | -0.28 | -0.17 | 0.57 | 0.58 | -0.4 | 0.06 | 0.23 | 0.22 | 0.8 | -0.29 | 1.03 | 1.18E-16 | 0.335 | 0.6575 | 0.117243 | 1 | 0.329846 | 1 | 0.997988 | 0.994666 |
| Tmem47_Q9JJG6_S6s_1 | Tmem47_( | 0.76 | 0.86 | 1.44 | 1.2 | 1.22 | 0.68 | 1.12 | 0.99 | 0.82 | 0.77 | 0.35 | 0.93 | 1.31 | -0.69 | 0.59 | 1.19 | -0.0625 | -0.3475 | -0.465 | 0.331645 | 0.803785 | 0.209965 | 0.995762 | 0.997988 | 0.994666 |
| Tmem51_Q99LG1_S114s_1 | Tmem51_( | 0.79 | 1.39 | 0.94 | 1.33 | 0.94 | 1.07 | -0.19 | 0.92 | 0.54 | 1.27 | 0.41 | 0.76 | 0.22 | 0.41 | 1.33 | 0.79 | -0.4275 | -0.3675 | -0.425 | 0.215141 | 0.223139 | 0.220649 | 0.989223 | 0.997988 | 0.994666 |
| Tmem51_Q99LG1_S151s_1 | Tmem51_( | -1.04 | 1.08 | -0.23 | -1.28 | 1 | 0.18 | 0.84 | -0.49 | -1.8 | 0.32 | -0.89 | -2.63 | 0.3 | -0.37 | -0.46 | 0.13 | 0.75 | -0.8825 | 0.2675 | 0.613988 | 0.208062 | 0.252998 | 0.989223 | 0.997988 | 0.995259 |
| Tmem51_Q99LG1_S153s_1 | Tmem51_( | 0.16 | 0.38 | 0.08 | 0.26 | 0.19 | -0.1 | 0.24 | 0.01 | 0.35 | 0.54 | -0.28 | -0.21 | 0.03 | 1.15 | 0.34 | 0.05 | -0.135 | -0.12 | 0.1725 | 0.590912 | 0.521147 | 0.659685 | 0.989223 | 0.997988 | 0.995259 |
| Tmem51_Q99LG1_T230t_1 | Tmem51_( | 0.07 | -0.67 | -0.3 | 0.07 | 0.92 | -0.3 | 0.53 | 0.12 | -0.52 | -0.12 | -0.96 | 0.49 | -1.06 | 0.55 | -0.1 | -0.53 | 0.525 | -0.07 | -0.0775 | 0.843777 | 0.13441 | 0.84793 | 0.989223 | 0.997988 | 0.999507 |
| Tmem55a_Q9CZX7_S47s_1 | Tmem55a_ | -0.61 | -0.66 | 0.1 | 0.29 | -0.6 | -0.28 | 0.43 | 0.28 | -0.97 | -0.03 | -0.15 | 0.32 | -0.74 | -0.78 | -0.08 | 0.13 | 0.1775 | 0.0125 | -0.1475 | 0.683077 | 0.610039 | 0.973197 | 0.989223 | 0.997988 | 0.996159 |
| Tmem55a_Q9CZX7_T22t_1 | Tmem55a_ | -0.95 | 0.3 | 0.55 | 0.18 | -0.51 | -0.52 | 0.61 | 0.48 | 0 | -0.16 | -0.75 | -0.12 | -0.04 | 0.43 | 0.05 | 0.69 | -0.005 | -0.2775 | 0.2625 | 0.501539 | 0.990663 | 0.468958 | 0.999187 | 0.997988 | 0.994666 |
| Tmem55b_E0CYE6_S169s_1 | Tmem55b_ | 0.71 | 0.83 | 0.35 | 0.59 | 0.64 | 0.45 | 0.58 | 0.39 | 0.33 | 0.59 | -0.32 | 0.4 | -0.02 | 0.41 | -0.02 | 0.26 | -0.105 | -0.37 | -0.4625 | 0.095072 | 0.625413 | 0.202846 | 0.989223 | 0.997988 | 0.994666 |
| Tmem63c_D6RI74_S94s_2 | Tmem63c_ | -0.92 | 0.06 | 0.52 | -0.87 | 0.02 | -0.37 | 0.38 | -0.32 | -0.81 | -0.63 | -0.39 | -0.89 | -2.33 | -0.04 | -0.36 | -0.3 | 0.23 | -0.3775 | -0.455 | 0.443294 | 0.5535 | 0.330479 | 0.989223 | 0.997988 | 0.994666 |
| Tmem74b_A2AU34_S198s_1 | Tmem74b_ | -0.91 | -0.04 | -0.21 | -0.11 | 0.1 | -0.85 | 0.15 | -0.32 | -1.64 | -1.13 | -0.71 | -0.44 | -0.87 | 0.33 | 0.22 | 0.23 | 0.0875 | -0.6625 | 0.295 | 0.430892 | 0.785403 | 0.081989 | 0.995762 | 0.997988 | 0.994666 |
| Tmem74b_A2AU34_S3s_1 | Tmem74b_ | 0.43 | 0.9 | 0.65 | 0.36 | 0.94 | 0.22 | 1.16 | -0.01 | 0.97 | 0.75 | 0.15 | 0.23 | 0.97 | 1.18 | 0.52 | 0.42 | -0.0075 | -0.06 | 0.1875 | 0.520691 | 0.981303 | 0.832221 | 0.999187 | 0.997988 | 0.994666 |
| Tmem74b_A2AU34_S62s_1 | Tmem74b_ | 0.41 | 0.64 | 0.27 | 0.52 | 0.8 | 0.51 | 1.26 | 0.45 | 0.79 | 0.84 | 0.36 | 0.78 | 1.13 | 0.9 | 0.63 | 1.05 | 0.295 | 0.2325 | 0.4675 | 0.086568 | 0.263351 | 0.329144 | 0.989223 | 0.997988 | 0.994666 |
| Tmem88_Q9D0N8_S145s_1 | Tmem88_( | -0.32 | 0 | -0.78 | -0.24 | 0.41 | 0.24 | 0.7 | 0.72 | 0.58 | 0.22 | -0.04 | 0.71 | 0.48 | 0.77 | 1.06 | 1.02 | 0.8525 | 0.7025 | 1.1675 | 0.002296 | 0.007352 | 0.032666 | 0.989223 | 0.997988 | 0.994666 |
| Tmem88_Q9D0N8_S148s_1 | Tmem88_( | -0.76 | -0.99 | -0.29 | -0.25 | 1.42 | 0.29 | 0.42 | 0.27 | -0.09 | -0.37 | -0.22 | 0.08 | 0.01 | 1.21 | 1.01 | 1.24 | 1.1725 | 0.4225 | 1.44 | 0.00288 | 0.006088 | 0.13781 | 0.989223 | 0.997988 | 0.994666 |
| Tmpo_Q61033_S155s_1 | Tmpo_Q61 | 0.69 | 0.23 | 0.33 | -0.31 | -0.14 | 0.99 | -0.29 | -0.33 | 2.4 | -0.47 | -0.97 | -0.36 | 2.77 | -0.92 | 1.36 | 0.6 | -0.1775 | -0.085 | 0.7175 | 0.326508 | 0.633415 | 0.904751 | 0.989223 | 0.997988 | 0.994666 |
| Tmpo_Q61033_T159t_1 | Tmpo_Q61 | 0.14 | 0.64 | 0.56 | 0.05 | 0.38 | -0.12 | 0.12 | -0.03 | 0.18 | 0.08 | -0.17 | 0.48 | -0.16 | 0.18 | 0.4 | 0.45 | -0.26 | -0.205 | -0.13 | 0.644865 | 0.303204 | 0.444066 | 0.989223 | 0.997988 | 0.995259 |
| Tmtc1_D6RG80_S168s_1 | Tmtc1_D6F | -1.5 | -2.39 | -3.01 | -2.29 | -0.96 | -0.39 | -1.68 | -2.15 | -1.14 | -0.37 | -1.48 | -4.12 | -0.16 | -1.57 | -0.11 | -1.36 | 1.0025 | 0.52 | 1.4975 | 0.010286 | 0.052828 | 0.509962 | 0.989223 | 0.997988 | 0.994666 |
| Tmub1_D3Z5X7_S71s_1 | Tmub1_D3 | -0.05 | -0.13 | -0.25 | -0.91 | 0.51 | -0.67 | -0.12 | -0.46 | -0.67 | -0.41 | 0.5 | 0.15 | -0.13 | 0.62 | 0.66 | -0.18 | 0.15 | 0.2275 | 0.5775 | 0.113178 | 0.653966 | 0.516647 | 0.990434 | 0.997988 | 0.994666 |
| Tmx1_Q8VBT0_S245s_1 | Tmx1_Q8V | 0.35 | 0.46 | -0.51 | -0.74 | -0.01 | 0.27 | -0.39 | -0.86 | 0.55 | 0.6 | -0.01 | -0.79 | -1.85 | 0.85 | -0.28 | -0.06 | -0.1375 | 0.1975 | -0.225 | 0.701016 | 0.718031 | 0.648196 | 0.995437 | 0.997988 | 0.996159 |
| Tmx2_D3Z2J6_S205s_1 | Tmx2_D3Z | 0.61 | 1.1 | 0.96 | 1.18 | 0.93 | -0.69 | 0.6 | 0.91 | 0.16 | 0.65 | 0.15 | 0.83 | -0.61 | 0.49 | 0.48 | 0.59 | -0.525 | -0.515 | -0.725 | 0.058259 | 0.200455 | 0.083688 | 0.989223 | 0.997988 | 0.994666 |
| Tmx4_Q8C0L0_S304s_1 | Tmx4_Q8C | -0.58 | -0.49 | -0.36 | -1.89 | -1.51 | 0.09 | -1.26 | -1.41 | 0.96 | -1.05 | -4.42 | -0.63 | 1.79 | -1.33 | 0.37 | -1.46 | -0.1925 | -0.455 | 0.6725 | 0.383015 | 0.687838 | 0.664736 | 0.994672 | 0.997988 | 0.994666 |
| Tnc_Q80YX0_S72s_1 | Tnc_Q80Y) | -0.72 | -0.46 | -0.82 | -0.74 | -0.04 | -0.83 | 0.04 | -0.6 | -0.63 | -0.01 | -0.67 | -0.48 | -0.11 | 0 | -0.13 | -0.58 | 0.3275 | 0.2375 | 0.48 | 0.085803 | 0.242111 | 0.349901 | 0.989223 | 0.997988 | 0.994666 |
| Tnfrsf19_Q9JLL3_S273s_1 | Tnfrsf19_Q | -0.94 | 0.12 | -0.24 | -0.87 | 0.13 | -0.21 | -0.78 | -0.2 | -0.62 | -1 | 0.72 | -0.45 | -0.02 | -0.42 | 0.11 | 0.16 | 0.2175 | 0.145 | 0.44 | 0.202066 | 0.512849 | 0.740277 | 0.989223 | 0.997988 | 0.994666 |
| Tnfrsf21_Q9EPU5_S565s_1 | Tnfrsf21_Q | 0.35 | -0.46 | -0.11 | 0.26 | -0.26 | 0.51 | 0 | 0.39 | 1.02 | 0.57 | -0.38 | 0.76 | 0.79 | -0.18 | 1.12 | 0.84 | 0.15 | 0.4825 | 0.6325 | 0.106344 | 0.604933 | 0.209345 | 0.989223 | 0.997988 | 0.994666 |
| Tnik_E9PUL9_S519s_1 | Tnik_E9PU | -0.09 | -1.27 | -3.03 | -2.72 | -1.47 | -0.06 | -1.96 | -1.84 | -1.42 | -1.22 | -0.01 | -2.2 | -1.83 | 0.7 | -3.37 | -3.37 | 0.445 | 0.565 | -0.19 | 0.852733 | 0.536304 | 0.449407 | 0.989223 | 0.997988 | 0.999507 |
| Tnik_E9PUL9_S581s_1 | Tnik_E9PU | -1.93 | -2.28 | -1.68 | -1.01 | -2.19 | -1.89 | -1.01 | -1.05 | -1.94 | -2.96 | 0.52 | -1.47 | -1.51 | -0.9 | -1.57 | -2.25 | 0.19 | 0.2625 | 0.1675 | 0.671993 | 0.627126 | 0.709147 | 0.989223 | 0.997988 | 0.996159 |
| Tnik_E9PUL9_S611s_1 | Tnik_E9PU | 0.8 | 1.12 | 0.04 | 1.03 | 0.78 | 0.46 | 0.08 | 0.56 | 0.59 | 0.85 | 0.28 | 0.43 | -0.15 | 0.55 | 0.67 | 0.17 | -0.2775 | -0.21 | -0.4375 | 0.221164 | 0.376196 | 0.501796 | 0.989223 | 0.997988 | 0.994666 |
| Tnik_E9PUL9_S649s_1 | Tnik_E9PU | -0.69 | -0.86 | -1.63 | -0.67 | -0.93 | -0.73 | -1.07 | -2.3 | -1.51 | -1.02 | -0.18 | -1.82 | -0.57 | -0.4 | -1.92 | -2.79 | -0.295 | -0.17 | -0.4575 | 0.426321 | 0.471176 | 0.683938 | 0.989223 | 0.997988 | 0.994666 |
| Tnik_E9PUL9_S651s_1 | Tnik_E9PU | 0.53 | -1.82 | -1.3 | -0.32 | -0.79 | 0.42 | -0.43 | 0.3 | -1.31 | -1.21 | 0.05 | -0.66 | -1.94 | -0.37 | -1.57 | -1.51 | 0.6025 | -0.055 | -0.62 | 0.294309 | 0.281528 | 0.921961 | 0.989223 | 0.997988 | 0.994666 |
| Tnik_E9PUL9_S659s_1 | Tnik_E9PU | 0.4 | 0.1 | 0.2 | 0.57 | 0.39 | 0.33 | 0 | 0.55 | 0.48 | 0.46 | 1.13 | 0.68 | 0.63 | 0.79 | 0.38 | 0.03 | -5.9E-17 | 0.37 | 0.14 | 0.615637 | 1 | 0.173021 | 1 | 0.997988 | 0.995259 |
| Tnik_E9PUL9_S678s_1 | Tnik_E9PU | 0.5 | -1.39 | -1 | 0.01 | -0.91 | 0.85 | -0.68 | 0.5 | 0.27 | -0.82 | -0.19 | -0.02 | -0.09 | -1.38 | -1.83 | -1.54 | 0.41 | 0.28 | -0.74 | 0.193635 | 0.467031 | 0.556128 | 0.989223 | 0.997988 | 0.994666 |
| Tnik_E9PUL9_S740s_1 | Tnik_E9PU | -0.43 | -0.08 | 0.43 | 0.5 | 0.18 | -0.8 | 0.44 | 0.5 | -0.11 | 0.47 | 0.88 | 0.7 | -0.25 | 0.22 | 0.52 | 0.48 | -0.025 | 0.38 | 0.1375 | 0.674632 | 0.94585 | 0.269259 | 0.999187 | 0.997988 | 0.996159 |
| Tnik_E9PUL9_S925s_1 | Tnik_E9PU | 0.78 | 1.03 | 0.46 | 0.89 | 0.39 | 0.63 | 0.68 | 0.99 | 0.59 | 1.02 | 1.07 | 0.98 | -0.16 | 0.49 | 0.53 | 0.7 | -0.1175 | 0.125 | -0.4 | 0.192616 | 0.625365 | 0.611243 | 0.989223 | 0.997988 | 0.994666 |
| Tnik_E9PUL9_T552t_1 | Tnik_E9PU | -0.19 | -0.69 | -1.08 | -0.48 | 0.15 | -0.29 | -0.22 | -0.53 | -0.84 | -0.66 | -0.24 | -0.71 | -1.01 | -0.23 | -1.31 | -0.8 | 0.3875 | -0.0025 | -0.2275 | 0.500936 | 0.179692 | 0.99286 | 0.989223 | 0.997988 | 0.994666 |
| Tnk2_D3Z3U5_S757s_1 | Tnk2_D3Z3 | 0.96 | 0 | -0.01 | 0.53 | 0.68 | 1.1 | 0.31 | 0.28 | -0.12 | 0.41 | 1.62 | 0.14 | -1.1 | 0.68 | -1.25 | -1.2 | 0.2225 | 0.1425 | -1.0875 | 0.050518 | 0.489974 | 0.744566 | 0.989223 | 0.997988 | 0.994666 |
| Tnk2_D3Z3U5_S785s_1 | Tnk2_D3Z3 | -1.53 | -2.28 | -1.43 | -2.23 | -0.97 | -0.9 | -1.58 | -1.11 | -1.69 | -1.93 | 0.06 | -1.26 | -1 | -0.48 | -1.09 | -2.21 | 0.7275 | 0.6625 | 0.6725 | 0.137965 | 0.033344 | 0.184935 | 0.989223 | 0.997988 | 0.994666 |
| Tnks1bp1_P58871_S1063s_1 | Tnks1bp1_ | 0.16 | 0.38 | -1.4 | -1.16 | 0 | 0.03 | -0.93 | -1.04 | -0.29 | 0.32 | -0.73 | -0.95 | -0.29 | 0.25 | -1.09 | -1.37 | 0.02 | 0.0925 | -0.12 | 0.825225 | 0.967625 | 0.853437 | 0.999187 | 0.997988 | 0.999507 |
| Tnks1bp1_P58871_S1131s_1 | Tnks1bp1_ | -0.43 | -2.12 | -1.85 | -1.56 | 0.06 | -0.4 | -1.15 | -0.91 | -1.44 | -1.31 | -0.78 | -1.34 | -1.73 | -0.54 | -1.98 | -1.88 | 0.89 | 0.2725 | -0.0425 | 0.929126 | 0.063854 | 0.49931 | 0.989223 | 0.997988 | 0.999507 |
| Tnks1bp1_P58871_S1290s_1 | Tnks1bp1_ | 0.15 | -1.11 | -0.26 | 0.08 | -0.52 | 0.5 | 0.04 | -0.07 | 0.6 | 0.05 | 1.02 | -0.15 | 0.97 | 0.24 | -0.66 | -0.22 | 0.2725 | 0.665 | 0.3675 | 0.415291 | 0.451961 | 0.116727 | 0.989223 | 0.997988 | 0.994666 |
| Tnks1bp1_P58871_S1375s_1 | Tnks1bp1_ | -0.07 | -0.1 | 0.09 | 0.24 | 0.43 | 0.29 | 0.54 | 0.25 | -0.15 | 0.16 | -0.23 | 0.22 | -0.91 | 0.18 | -0.07 | 0 | 0.3375 | -0.04 | -0.24 | 0.445356 | 0.130152 | 0.862842 | 0.989223 | 0.997988 | 0.994666 |
| Tnks1bp1_P58871_S1437s_1 | Tnks1bp1_ | 0.07 | -0.05 | -0.29 | 0 | 0.02 | -0.4 | 0.38 | -0.16 | 0.11 | -0.32 | -0.64 | -0.24 | 0.12 | 0.43 | -0.16 | -0.55 | 0.0275 | -0.205 | 0.0275 | 0.92467 | 0.910176 | 0.41881 | 0.997335 | 0.997988 | 0.999507 |
| Tnks1bp1_P58871_S1611s_1 | Tnks1bp1_ | -0.96 | -0.81 | -0.63 | -0.08 | -0.46 | -0.97 | -0.14 | 0 | -1.17 | -0.41 | -0.01 | -0.29 | -1.01 | -0.05 | -0.06 | -0.35 | 0.2275 | 0.15 | 0.2525 | 0.457343 | 0.467702 | 0.656444 | 0.989223 | 0.997988 | 0.994666 |
| Tnks1bp1_P58871_S1657s_1 | Tnks1bp1_ | 1.48 | -1.32 | 0.65 | 1.74 | -1.5 | 2.08 | -0.1 | 1.29 | 1.16 | 0.43 | 0.67 | 1.04 | -0.59 | -0.88 | -0.08 | 0.73 | -0.195 | 0.1875 | -0.8425 | 0.244507 | 0.830585 | 0.772451 | 0.995762 | 0.997988 | 0.994666 |
| Tnks1bp1_P58871_S220s_1 | Tnks1bp1_ | 0 | -0.93 | -0.38 | 0.04 | -0.31 | -0.25 | 1.1 | 0.58 | -0.66 | -1.19 | -0.18 | -0.44 | -1.59 | 0.35 | 1.27 | 1.4 | 0.5975 | -0.3 | 0.675 | 0.316318 | 0.153643 | 0.379348 | 0.989223 | 0.997988 | 0.994666 |
| Tnks1bp1_P58871_S429s_1 | Tnks1bp1_ | -0.17 | -0.76 | -0.73 | -0.74 | -0.02 | -0.19 | 0.17 | -0.17 | -1.01 | -0.05 | 0.06 | -0.12 | -0.39 | 0.11 | -0.27 | -0.34 | 0.5475 | 0.32 | 0.3775 | 0.185232 | 0.040694 | 0.325075 | 0.989223 | 0.997988 | 0.994666 |
| Tnks1bp1_P58871_S539s_1 | Tnks1bp1_ | -0.08 | -0.26 | -1.05 | -1.05 | -0.15 | -0.41 | 0.09 | -0.16 | -0.43 | -0.03 | -0.4 | -0.79 | -0.41 | 0.21 | -0.85 | -0.94 | 0.4525 | 0.1975 | 0.1125 | 0.768528 | 0.155995 | 0.548776 | 0.989223 | 0.997988 | 0.9971 |
| Tnks1bp1_P58871_S539s_2 | Tnks1bp1_ | -0.41 | -0.26 | -0.3 | 0.14 | 0 | -0.47 | -0.02 | 0.04 | -0.44 | -0.33 | -0.36 | -0.42 | -1.05 | -0.29 | -0.05 | 0.6 | 0.095 | -0.18 | 0.01 | 0.978847 | 0.690226 | 0.433586 | 0.994672 | 0.997988 | 1 |
| Tnks1bp1_P58871_S568s_1 | Tnks1bp1_ | 0.31 | -0.71 | -0.96 | -0.29 | -0.29 | 0.73 | -0.14 | 0.5 | -0.61 | -0.93 | 0.83 | -0.36 | -1.01 | -0.15 | -1.42 | -0.68 | 0.6125 | 0.145 | -0.4025 | 0.321661 | 0.120256 | 0.750393 | 0.989223 | 0.997988 | 0.994666 |
| Tnks1bp1_P58871_S602s_1 | Tnks1bp1_ | 0.7 | 1.12 | 0.47 | 0.91 | 1.06 | 0.7 | 1.04 | 0.79 | 0.64 | 1.11 | 0.71 | 0.93 | -0.78 | 1.43 | 0.8 | 0.83 | 0.0975 | 0.0475 | -0.23 | 0.628746 | 0.680259 | 0.849564 | 0.993464 | 0.997988 | 0.995259 |
| Tnks1bp1_P58871_S763s_1 | Tnks1bp1_ | 1.87 | 0.76 | 0.27 | 0.37 | 0.62 | 1.51 | 0.25 | 1 | 0.28 | 1.62 | 1.74 | 0.5 | 0.46 | -0.06 | 0.39 | 0.27 | 0.0275 | 0.2175 | -0.5525 | 0.18253 | 0.94889 | 0.661958 | 0.999187 | 0.997988 | 0.994666 |
| Tnks1bp1_P58871_S866s_1 | Tnks1bp1_ | 0.05 | -0.54 | 0.52 | 0.21 | -0.22 | -0.11 | -0.22 | 0.28 | -0.62 | -0.19 | 0.35 | 0.86 | -0.84 | -0.42 | 0.03 | 0.04 | -0.1275 | 0.04 | -0.3575 | 0.308807 | 0.656287 | 0.918569 | 0.990434 | 0.997988 | 0.994666 |
| Tnks1bp1_P58871_S887s_1 | Tnks1bp1_ | 0.07 | 0.45 | -1.79 | -1.17 | -0.51 | 0.37 | -2.26 | -0.96 | 0.35 | -0.91 | -0.38 | -1 | -0.36 | 1.59 | -2.33 | -1.34 | -0.23 | 0.125 | 0 | 1 | 0.73225 | 0.824166 | 0.995762 | 0.997988 | 1 |
| Tnks1bp1_P58871_S911s_1 | Tnks1bp1_ | -3.25 | -1.18 | 0.13 | 1.14 | -4.77 | -2.64 | 0.13 | 1.97 | -2.94 | -3.31 | 1.6 | 0.89 | -1.95 | -5.37 | -0.4 | 0.11 | -0.5375 | -0.15 | -1.1125 | 0.415418 | 0.722597 | 0.9138 | 0.995437 | 0.997988 | 0.994666 |
| Tnks1bp1_P58871_S912s_1 | Tnks1bp1_ | -3.46 | -3.67 | -1.75 | -2.36 | -3.4 | -2.47 | -3.14 | -2.42 | -2.57 | -3.69 | -0.52 | -1.53 | -1.87 | -2.79 | -1.83 | -1.85 | -0.0475 | 0.7325 | 0.725 | 0.160804 | 0.920663 | 0.333116 | 0.997335 | 0.997988 | 0.994666 |
| Tnks1bp1_P58871_S927s_1 | Tnks1bp1_ | -2.68 | -1.21 | -1.09 | -2.21 | -1.91 | -3.74 | -0.94 | -1.85 | -1.09 | -1.7 | 0.11 | -0.48 | -1.58 | -1.53 | -2.03 | -1.03 | -0.3125 | 1.0075 | 0.255 | 0.558262 | 0.619698 | 0.076033 | 0.989223 | 0.997988 | 0.995259 |
| Tnks1bp1_P58871_T131t_1 | Tnks1bp1_ | -3.07 | -0.78 | -3.65 | -1.95 | -1.17 | -0.96 | -2.11 | -2.59 | -1.72 | -0.6 | -1.48 | -3.47 | -0.76 | 0 | -3.75 | -3.21 | 0.655 | 0.545 | 0.4325 | 0.657468 | 0.332754 | 0.491549 | 0.989223 | 0.997988 | 0.995259 |
| Tnks1bp1_P58871_T533t_1 | Tnks1bp1_ | 1.07 | 1.42 | -0.4 | 0.29 | 0.34 | -0.28 | 0.36 | 0.68 | 0.33 | 1.01 | 0.46 | 0.96 | 0.54 | 1.3 | 0.38 | 0.09 | -0.32 | 0.095 | -0.0175 | 0.969935 | 0.462247 | 0.82537 | 0.989223 | 0.997988 | 0.999507 |
| Tnks1bp1_P58871_T533t_2 | Tnks1bp1_ | -0.41 | -0.26 | -0.3 | 0.14 | 0 | -0.47 | -0.02 | 0.04 | -0.44 | -0.33 | -0.36 | -0.42 | -1.05 | -0.29 | -0.05 | 0.6 | 0.095 | -0.18 | 0.01 | 0.978847 | 0.690226 | 0.433586 | 0.994672 | 0.997988 | 1 |
| Tnpo1_J3QMX2_S24s_1 | Tnpo1_J3Q | -0.18 | 0.1 | -0.06 | -0.47 | 0.07 | -0.51 | -0.28 | -0.38 | 0.41 | -0.07 | 0.92 | -0.31 | 2.45 | -0.05 | 0.16 | -0.25 | -0.1225 | 0.39 | 0.73 | 0.233121 | 0.610959 | 0.248367 | 0.989223 | 0.997988 | 0.994666 |
| Tnrc18_F6Z7L1_T1852t_1 | Tnrc18_F6 | -1.35 | -1.97 | -1.29 | -1.76 | -0.62 | -1.12 | -2.17 | -2.09 | -2.5 | -2.12 | -0.91 | -1.36 | -0.41 | -1.42 | -0.79 | -2.1 | 0.0925 | -0.13 | 0.4125 | 0.32573 | 0.815874 | 0.742369 | 0.995762 | 0.997988 | 0.994666 |
| Tnrc6b_Q8BKI2_S1313s_1 | Tnrc6b_Q8 | 0.96 | -1.47 | -1.44 | -0.58 | 1.04 | 2.23 | 1.24 | -1.81 | -1.88 | 0.45 | -2.19 | -0.09 | -0.81 | -1.98 | 0.76 | 0.92 | 1.3075 | -0.295 | 0.355 | 0.654806 | 0.17024 | 0.704256 | 0.989223 | 0.997988 | 0.995259 |
| Tnrc6b_Q8BKI2_S1793s_1 | Tnrc6b_Q8 | 0.18 | -0.34 | 0.09 | 0.43 | 0.16 | 0.03 | -0.08 | -0.11 | -0.49 | -0.04 | 0.21 | 0.22 | -0.06 | -0.32 | 0.16 | -0.46 | -0.09 | -0.115 | -0.26 | 0.372524 | 0.706819 | 0.684174 | 0.995437 | 0.997988 | 0.994666 |
| Tnrc6c_B1ATC3_T986t_1 | Tnrc6c_B1 | -1.4 | -0.93 | -0.37 | 0.08 | -0.41 | -1.61 | -0.14 | -0.11 | -0.92 | -0.83 | 0.38 | -0.45 | 0.06 | -0.55 | 0.09 | -0.74 | 0.0875 | 0.2 | 0.37 | 0.360123 | 0.845218 | 0.641696 | 0.996349 | 0.997988 | 0.994666 |
| Tns1_E9Q0S6_S1054s_1 | Tns1_E9Q0 | -0.36 | -0.28 | 1.24 | 0.85 | -0.16 | 0.47 | 0.46 | 0.49 | 0.47 | -0.13 | 0.93 | 1.71 | -0.52 | -0.19 | 0.52 | 1.51 | -0.0475 | 0.3825 | -0.0325 | 0.953456 | 0.908243 | 0.470603 | 0.997335 | 0.997988 | 0.999507 |
| Tns1_E9Q0S6_S1062s_1 | Tns1_E9Q0 | -0.26 | -0.36 | 0.85 | 0.51 | 0.15 | 0.05 | 0.51 | 0.02 | 0.88 | 0.46 | 2.5 | 0.38 | 0.86 | -2.15 | 0.99 | 0.08 | -0.0025 | 0.87 | -0.24 | 0.732803 | 0.993897 | 0.130125 | 0.999278 | 0.997988 | 0.9971 |
| Tns1_E9Q0S6_S1067s_1 | Tns1_E9Q0 | 0.05 | -1.36 | 0.77 | 0.62 | -0.87 | -0.37 | 0.09 | 0.84 | 0.93 | -0.11 | 2.47 | 0.66 | 0.8 | -0.91 | 0.86 | 0.86 | -0.0975 | 0.9675 | 0.3825 | 0.52629 | 0.858787 | 0.166451 | 0.996349 | 0.997988 | 0.994666 |
| Tns1_E9Q0S6_S1118s_1 | Tns1_E9Q0 | -0.06 | -0.69 | -0.09 | 0.55 | -1.77 | 0.29 | -0.01 | 0.43 | 1.19 | 0.09 | 0.71 | 0.7 | 0.11 | 0.14 | 0.03 | 0.71 | -0.1925 | 0.745 | 0.32 | 0.351906 | 0.711387 | 0.05919 | 0.995437 | 0.997988 | 0.994666 |
| Tns1_E9Q0S6_S1126s_1 | Tns1_E9Q0 | 0.71 | 0.92 | 0.34 | -0.09 | 0.82 | -0.2 | 0.41 | 0.09 | -0.03 | 0.59 | 0.56 | -0.36 | 0.9 | 1 | 0.18 | -0.05 | -0.19 | -0.28 | 0.0375 | 0.918032 | 0.561635 | 0.420052 | 0.989223 | 0.997988 | 0.999507 |
| Tns1_E9Q0S6_S1346s_1 | Tns1_E9Q0 | -0.2 | -0.18 | -0.16 | -0.3 | -0.51 | -0.46 | -0.39 | -0.63 | -0.57 | -0.02 | -0.23 | 0.08 | -0.04 | -0.25 | -0.35 | -0.32 | -0.2875 | 0.025 | -0.03 | 0.896786 | 0.165392 | 0.915673 | 0.989223 | 0.997988 | 0.999507 |
| Tns1_E9Q0S6_S1363s_1 | Tns1_E9Q0 | 0.02 | -0.09 | 0.61 | 0.25 | -0.5 | -0.14 | 0.07 | 0.42 | 0.25 | 0.03 | 0.63 | 0.3 | 0.3 | 0.25 | 0.1 | -0.08 | -0.235 | 0.105 | -0.055 | 0.837246 | 0.413236 | 0.690225 | 0.989223 | 0.997988 | 0.999507 |
| Tns1_E9Q0S6_S1397s_1 | Tns1_E9Q0 | 0.1 | -1.36 | -1.07 | -0.74 | -0.18 | -0.61 | -0.23 | -0.58 | 0.04 | -0.8 | -0.96 | -0.4 | -0.06 | -1.06 | -0.93 | -0.78 | 0.3675 | 0.2375 | 0.06 | 0.879275 | 0.296514 | 0.544851 | 0.989223 | 0.997988 | 0.999507 |
| Tns1_E9Q0S6_S1468s_1 | Tns1_E9Q0 | 1.9 | 2.05 | 2.19 | 1.76 | 2.37 | 1.96 | 2.09 | 1.72 | 1.52 | 2.4 | 1.3 | 2.12 | 1.68 | 1.95 | 1.96 | 1.59 | 0.06 | -0.14 | -0.18 | 0.475476 | 0.798467 | 0.650757 | 0.995762 | 0.997988 | 0.994666 |
| Tns1_E9Q0S6_S154s_1 | Tns1_E9Q0 | 0.24 | -0.21 | -0.97 | -2.7 | 0.37 | -0.35 | -0.81 | -0.44 | -1.66 | 1.43 | -1.99 | -1.63 | 0.78 | 0.62 | 0.79 | -0.18 | 0.6025 | -0.0525 | 1.4125 | 0.045018 | 0.341547 | 0.954093 | 0.989223 | 0.997988 | 0.994666 |
| Tns1_E9Q0S6_S792s_1 | Tns1_E9Q0 | 0.12 | 0.2 | -0.61 | -0.02 | 0.28 | 0.59 | 1.1 | 0.06 | -0.09 | 0.8 | -0.33 | 1.37 | 0.01 | 0.71 | 0.65 | 0.94 | 0.585 | 0.515 | 0.655 | 0.063892 | 0.084422 | 0.244695 | 0.989223 | 0.997988 | 0.994666 |
| Tns1_E9Q0S6_S847s_1 | Tns1_E9Q0 | 0.15 | 0.3 | 1.54 | 0.36 | -0.49 | 0.84 | 0.62 | 0.51 | 0.68 | 0.55 | 2.12 | 0.78 | 0.47 | 0.7 | 1.02 | 1.09 | -0.2175 | 0.445 | 0.2325 | 0.535009 | 0.60252 | 0.350607 | 0.989223 | 0.997988 | 0.994666 |
| Tns2_Q8CGB6_S1064s_1 | Tns2_Q8C | 2.46 | 2.35 | 1.51 | 0.19 | 1.87 | 2.23 | 1.25 | 1.41 | 1.57 | 2.67 | 0.75 | 2.25 | 1.79 | 0.97 | 1.95 | 2.1 | 0.0625 | 0.1825 | 0.075 | 0.889617 | 0.904044 | 0.766469 | 0.996493 | 0.997988 | 0.999507 |
| Tns2_Q8CGB6_S802s_1 | Tns2_Q8C | -0.7 | -0.79 | -0.09 | -0.02 | -0.92 | -0.99 | -0.65 | -0.79 | 0.42 | -0.53 | 0.54 | -0.12 | 0.82 | -0.2 | 0.04 | -0.38 | -0.4375 | 0.4775 | 0.47 | 0.209545 | 0.11998 | 0.182397 | 0.989223 | 0.997988 | 0.994666 |
| Tns2_Q8CGB6_S908s_1 | Tns2_Q8C | -0.58 | 0 | -0.61 | -0.87 | 0.55 | -1.33 | -0.99 | -1.66 | 0.5 | -0.53 | -0.57 | -0.12 | 0.39 | 0.17 | -0.5 | -1.21 | -0.3425 | 0.335 | 0.2275 | 0.580338 | 0.482828 | 0.327055 | 0.989223 | 0.997988 | 0.995259 |
| Tns2_Q8CGB6_Y460y_1 | Tns2_Q8C | 0.46 | -0.02 | 0.21 | 0.03 | 0.49 | 0.09 | 0.37 | 0.33 | 0.46 | 0.72 | -0.29 | 0.13 | 0.35 | -0.09 | 0.6 | 0.34 | 0.15 | 0.085 | 0.13 | 0.631022 | 0.504444 | 0.769294 | 0.989223 | 0.997988 | 0.995259 |
| Tns3_Q5SSZ5_S934s_1 | Tns3_Q5SS | -0.14 | -0.03 | -0.04 | 0.04 | -1.49 | 0.04 | -0.86 | -1.02 | 1.21 | -0.11 | -0.83 | -0.43 | 1.31 | -1.51 | -0.82 | -0.07 | -0.79 | 0.0025 | -0.23 | 0.681882 | 0.037121 | 0.99537 | 0.989223 | 0.998102 | 0.996159 |
| Tollip_Q9QZ06_T25t_1 | Tollip_Q9Q | 0.26 | 0.95 | 1.23 | 0.98 | 0.75 | -0.02 | 1.09 | 1.27 | -0.63 | 0.69 | 0.76 | 0.94 | -0.42 | 0.35 | 0.93 | 1.05 | -0.0825 | -0.415 | -0.3775 | 0.358264 | 0.815692 | 0.326016 | 0.995762 | 0.997988 | 0.994666 |
| Tom1_Q3UDC3_S160s_1 | Tom1_Q3U | -0.45 | -0.41 | -0.3 | -0.24 | -0.15 | -0.66 | 0.04 | -0.23 | -0.46 | -0.19 | -0.12 | 0.02 | -0.9 | 0.02 | -0.32 | 0.03 | 0.1 | 0.1625 | 0.0575 | 0.843706 | 0.665886 | 0.469974 | 0.991876 | 0.997988 | 0.999507 |
| Tom1_Q3UDC3_S176s_1 | Tom1_Q3U | -0.42 | -0.24 | -0.03 | -0.37 | 0.21 | -0.2 | -0.25 | -0.06 | -0.14 | 0.01 | 0.67 | 0.16 | 0.05 | 0.39 | -0.32 | -0.07 | 0.19 | 0.44 | 0.2775 | 0.309643 | 0.399469 | 0.118378 | 0.989223 | 0.997988 | 0.994666 |
| Tom1_Q3UDC3_S180s_1 | Tom1_Q3U | -1.31 | -2.57 | -1.69 | -2.46 | -2.51 | -0.24 | -2.54 | -1.48 | 1.26 | -1.81 | -1.1 | -1.56 | 1.87 | -1.77 | -0.88 | -1.2 | 0.315 | 1.205 | 1.5125 | 0.073329 | 0.577583 | 0.10788 | 0.989223 | 0.997988 | 0.994666 |
| Tom1_Q3UDC3_S183s_1 | Tom1_Q3U | 0.95 | -1.14 | 0.91 | 0.2 | -0.98 | 0.86 | -0.95 | 0.63 | 2.52 | 0.14 | 0.91 | -0.35 | 2.69 | -0.29 | 0.96 | 1.25 | -0.34 | 0.575 | 0.9225 | 0.208468 | 0.586358 | 0.430174 | 0.989223 | 0.997988 | 0.994666 |

| | | | | | | | | | | | | | | | | | | | | | | | | | | |
|---|---|---|---|---|---|---|---|---|---|---|---|---|---|---|---|---|---|---|---|---|---|---|---|---|---|---|
| Tom1_Q3UDC3_S400s_1 | Tom1_Q3U | -1.57 | -2.08 | -0.53 | -1.96 | -0.61 | -0.12 | -0.22 | -0.77 | -0.44 | -1.22 | 0.66 | 0.18 | 0.64 | -0.79 | 0.27 | 0.07 | 1.105 | 1.33 | 1.5825 | 0.005857 | 0.014825 | 0.025076 | 0.989223 | 0.997988 | 0.994666 |
| Tom1_Q3UDC3_S486s_1 | Tom1_Q3U | 0.21 | -0.09 | -0.45 | -0.1 | 0.21 | 0.34 | 0.44 | -0.15 | 0.1 | 0.14 | -0.44 | -0.24 | -0.35 | 0.38 | -0.47 | 0.02 | 0.3175 | -0.0025 | 0.0025 | 0.993299 | 0.217721 | 0.992321 | 0.989223 | 0.997988 | 1 |
| Tom1_Q3UDC3_S496s_1 | Tom1_Q3U | -0.04 | 0.8 | 0.04 | -0.05 | 0.33 | 0.27 | 0.28 | 0.05 | -0.01 | 0.71 | 0.6 | 0.48 | 0.38 | 0.46 | 0.19 | 0.34 | 0.045 | 0.2575 | 0.155 | 0.590426 | 0.863646 | 0.399421 | 0.996493 | 0.997988 | 0.995259 |
| Tom1l1_Q8BZR6_S93s_1 | Tom1l1_Q8 | -0.08 | 0.06 | -0.07 | -0.11 | 0.42 | 0.01 | 0.15 | 0.2 | -0.82 | 0.29 | -1.9 | 0.07 | -0.79 | -0.8 | -0.3 | -0.37 | 0.245 | -0.54 | -0.515 | 0.064346 | 0.250101 | 0.273078 | 0.989223 | 0.997988 | 0.994666 |
| Tom1l2_Q5SXA5_S110s_1 | Tom1l2_Q5 | -0.4 | 0 | -0.29 | -0.15 | 0.31 | -0.28 | 0.43 | -0.04 | -0.3 | -0.01 | -0.25 | -0.28 | -0.72 | 0.33 | -0.36 | -0.21 | 0.315 | -2E-17 | -0.03 | 0.919695 | 0.218815 | 1 | 0.989223 | 1 | 0.999507 |
| Tom1l2_Q5SXA5_S344s_1 | Tom1l2_Q5 | 0.72 | -0.66 | 0.03 | 0.45 | -0.11 | 1.5 | -0.01 | 0.95 | 0.97 | -0.29 | 0.52 | 1.09 | 0.7 | 0.23 | 0.06 | 0.27 | 0.4475 | 0.4375 | 0.18 | 0.615921 | 0.340351 | 0.316493 | 0.989223 | 0.997988 | 0.995259 |
| Tom1l2_Q5SXA5_S429s_1 | Tom1l2_Q5 | 0.11 | 0.49 | 0.19 | 0.17 | 0.43 | -0.02 | 0.54 | 0.06 | 0.08 | 0.46 | -0.01 | 0.36 | 0.12 | 0.59 | 0 | 0.09 | 0.0125 | -0.0175 | -0.04 | 0.8772 | 0.957233 | 0.940148 | 0.999187 | 0.997988 | 0.999507 |
| Tomm20_Q9DCC8_S138s_1 | Tomm20_Q | -0.5 | 0.63 | 1.61 | 0.73 | 0.61 | 0.36 | 1.84 | 1.94 | 1.09 | 0.84 | -0.07 | 1.86 | -0.28 | -0.14 | 1.55 | 1.57 | 0.57 | 0.3125 | 0.0575 | 0.924858 | 0.305464 | 0.570091 | 0.989223 | 0.997988 | 0.999507 |
| Tomm22_Q9CPQ3_S15s_1 | Tomm22_Q | 0.23 | 0.79 | 0.58 | 0.8 | 0.67 | 0.01 | 0.86 | 0.68 | 0.46 | 0.6 | 0.46 | 0.69 | 0.42 | 0.82 | 0.43 | 0.54 | -0.045 | -0.0475 | -0.0475 | 0.855729 | 0.868132 | 0.840032 | 0.996493 | 0.997988 | 0.999507 |
| Tomm34_Q9CYG7_S8s_1 | Tomm34_Q | -0.68 | -0.25 | -0.09 | -0.29 | -1 | -1.92 | -0.28 | -0.47 | -1.22 | -0.96 | 0.26 | -0.38 | -1.03 | -0.14 | -0.64 | -0.18 | -0.59 | -0.2475 | -0.17 | 0.578705 | 0.144255 | 0.500509 | 0.989223 | 0.997988 | 0.995259 |
| Tomm70a_Q9CZW5_S105s_2 | Tomm70a_ | -0.44 | 0.65 | -1.54 | -0.94 | -0.11 | 0.26 | -1.26 | -1.56 | 0.53 | -0.14 | -1.75 | -1.21 | -1.17 | 0.61 | -1.82 | -1.48 | -0.1 | -0.075 | -0.3975 | 0.541321 | 0.861755 | 0.905325 | 0.996493 | 0.997988 | 0.994666 |
| Tomm70a_Q9CZW5_S108s_1 | Tomm70a_ | 0.41 | -1.51 | -1.27 | -0.29 | -0.32 | 0.39 | -0.19 | -0.06 | 0.22 | -0.83 | -1.89 | 0.03 | -0.81 | 0.43 | -0.73 | -1.14 | 0.62 | 0.0475 | 0.1025 | 0.845268 | 0.183049 | 0.936919 | 0.989223 | 0.997988 | 0.999507 |
| Tomm70a_Q9CZW5_S94s_1 | Tomm70a_ | -0.31 | 0.14 | -1.3 | -0.63 | -0.09 | -0.09 | -0.45 | -0.87 | 0.07 | -0.06 | -1.26 | -0.97 | -0.09 | 0.06 | -1.11 | -1.16 | 0.15 | -0.03 | -0.05 | 0.908743 | 0.67409 | 0.944909 | 0.992876 | 0.997988 | 0.999507 |
| Tomm70a_Q9CZW5_S94s_2 | Tomm70a_ | -0.44 | 0.65 | -0.8 | -0.24 | -0.11 | 0.26 | -0.58 | -0.93 | 0.53 | -0.14 | -0.88 | -0.53 | -1.17 | 0.61 | -0.97 | -0.54 | -0.1325 | -0.0475 | -0.31 | 0.523821 | 0.735418 | 0.910388 | 0.995762 | 0.997988 | 0.994666 |
| Top1_Q04750_S114s_1 | Top1_Q047 | -0.97 | -0.57 | -0.5 | -0.64 | 0.01 | -0.76 | -0.19 | -0.51 | -1.06 | -1.11 | -0.01 | -0.23 | -0.18 | -0.1 | -0.18 | -0.51 | 0.3075 | 0.0675 | 0.4275 | 0.113826 | 0.243744 | 0.836382 | 0.989223 | 0.997988 | 0.994666 |
| Top2b_Q64511_S1537s_1 | Top2b_Q64 | 0.69 | -0.02 | 1.17 | 1.3 | -1.24 | 0.98 | -0.47 | 0.31 | 0.48 | 0.57 | 0.24 | 0.95 | 1.51 | -0.68 | 0.9 | 0.93 | -0.89 | -0.225 | -0.12 | 0.818078 | 0.110041 | 0.52396 | 0.989223 | 0.997988 | 0.999507 |
| Top2b_Q64511_S1539s_1 | Top2b_Q64 | 1.64 | 0.69 | 0.44 | 0.32 | -0.48 | 1.18 | -0.56 | -0.5 | 0.66 | 1.48 | 0.61 | 0.38 | 1.29 | -0.45 | 0.56 | 1.05 | -0.8625 | 0.01 | -0.16 | 0.733672 | 0.097272 | 0.979312 | 0.989223 | 0.997988 | 0.9971 |
| Topors_Q80Z37_S1025s_1 | Topors_Q8 | -0.81 | -0.88 | -1.85 | -2.7 | -1.07 | -1.15 | -2.38 | -2.27 | -0.79 | -1.12 | -1.09 | -2.06 | 0.12 | -0.89 | -1.14 | -2.28 | -0.1575 | 0.295 | 0.5125 | 0.407556 | 0.761887 | 0.555668 | 0.995762 | 0.997988 | 0.994666 |
| Tor1aip1_Q921T2_S140s_1 | Tor1aip1_Q | 0.51 | -0.15 | 0.42 | 0.66 | 0.56 | 0.17 | 0.71 | 0.58 | -0.03 | 0.66 | 0.63 | 0.43 | 0.5 | -0.16 | 1.1 | 0.61 | 0.145 | 0.0625 | 0.1525 | 0.660928 | 0.581029 | 0.827051 | 0.989223 | 0.997988 | 0.995671 |
| Tor1aip1_Q921T2_S151s_1 | Tor1aip1_Q | -1.57 | -0.54 | -0.31 | -1.25 | -1 | -1.42 | -1.73 | -1.84 | -0.61 | -1.25 | -0.44 | -1.1 | 0.04 | -0.98 | -0.11 | -0.48 | -0.58 | 0.0675 | 0.535 | 0.187207 | 0.12442 | 0.852644 | 0.989223 | 0.997988 | 0.994666 |
| Tor1aip1_Q921T2_S253s_1 | Tor1aip1_Q | 0.12 | -0.4 | 1.02 | 0.58 | -0.86 | 0.71 | -0.24 | 0.64 | 0.85 | -0.51 | -0.09 | 0.89 | 1.39 | -0.78 | 0.72 | 0.36 | -0.2675 | -0.045 | 0.0925 | 0.857156 | 0.557477 | 0.919659 | 0.989223 | 0.997988 | 0.999507 |
| Tox2_E9Q0B9_S199s_1 | Tox2_E9Q0 | -2.09 | -1.57 | -0.98 | -1.67 | -1.18 | -0.95 | -1.68 | -0.29 | -1.21 | -0.59 | -0.84 | -2.08 | 0.51 | -0.52 | -0.55 | -0.99 | 0.5525 | 0.3975 | 1.19 | 0.013512 | 0.153869 | 0.330206 | 0.989223 | 0.997988 | 0.994666 |
| Tox2_E9Q0B9_S506s_1 | Tox2_E9Q0 | -0.1 | -1.15 | -0.29 | -1.66 | -1.26 | 0.39 | -0.96 | -0.02 | -0.14 | 0.18 | -1.03 | -0.43 | 1.04 | -0.18 | -0.88 | -0.66 | 0.3375 | 0.445 | 0.63 | 0.250323 | 0.496975 | 0.320241 | 0.989223 | 0.997988 | 0.994666 |
| Tp53bp1_P70399_S1622s_1 | Tp53bp1_P | -0.13 | 0.68 | -0.46 | -0.39 | -0.18 | -0.23 | -0.5 | -0.72 | -1.65 | 0.1 | -0.29 | -0.68 | -0.05 | -0.88 | -0.62 | -1.34 | -0.3325 | -0.555 | -0.6475 | 0.119654 | 0.298154 | 0.229694 | 0.989223 | 0.997988 | 0.994666 |
| Tp53bp1_P70399_S1669s_1 | Tp53bp1_P | -1.11 | -1.63 | -1.86 | -1.63 | -1.99 | -1.51 | -2.68 | -1.41 | -2.16 | -1.6 | -1.17 | -1.96 | -0.72 | -1.42 | -2.53 | -2.68 | -0.34 | -0.165 | -0.28 | 0.557365 | 0.327325 | 0.592246 | 0.989223 | 0.997988 | 0.995259 |
| Tp53bp1_P70399_S332s_1 | Tp53bp1_P | -0.85 | -0.65 | 0.03 | -0.54 | -1.27 | -1 | 0.17 | -0.81 | -1.73 | -0.7 | -0.9 | -1.03 | -1.85 | -0.88 | -0.85 | -0.19 | -0.225 | -0.5875 | -0.44 | 0.284653 | 0.539665 | 0.093622 | 0.989223 | 0.997988 | 0.994666 |
| Tp53bp1_P70399_S381s_1 | Tp53bp1_P | 0.19 | 0.31 | 0.31 | -0.05 | -0.34 | 0.06 | 0.09 | -0.07 | 0.15 | -0.18 | -1.48 | -0.47 | -0.72 | 0.12 | -0.66 | -0.45 | -0.255 | -0.685 | -0.6175 | 0.0523 | 0.261071 | 0.09036 | 0.989223 | 0.997988 | 0.994666 |
| Tp53bp1_P70399_S429s_1 | Tp53bp1_P | 1.14 | 0.55 | -1.65 | -1.14 | -0.28 | 0.37 | -0.99 | -1.34 | 0.15 | 0.56 | -1.42 | -1.59 | -0.32 | -0.01 | -0.49 | -1.55 | -0.285 | -0.3 | -0.3175 | 0.637401 | 0.675441 | 0.697857 | 0.992876 | 0.997988 | 0.995259 |
| Tp53bp1_P70399_S533s_1 | Tp53bp1_P | -0.14 | -1.11 | -1.31 | -1.43 | -2.07 | -1.66 | -2.88 | -1 | -2.3 | -1.71 | -1.15 | -1.66 | -1.38 | -1.74 | -2.07 | -2.39 | -0.905 | -0.7075 | -0.8975 | 0.038206 | 0.072515 | 0.089098 | 0.989223 | 0.997988 | 0.994666 |
| Tp53bp1_P70399_S552s_1 | Tp53bp1_P | 0.49 | -0.03 | 0.23 | 0.18 | -0.45 | -0.29 | -0.49 | 0.19 | -0.24 | -0.1 | 0.06 | -0.36 | -0.82 | 0.18 | -0.81 | -0.38 | -0.4775 | -0.3775 | -0.675 | 0.053158 | 0.078957 | 0.131058 | 0.989223 | 0.997988 | 0.994666 |
| Tp53bp1_P70399_T323t_1 | Tp53bp1_P | -2.4 | -3.28 | -4.93 | -7.51 | -4.88 | -3.18 | -5.6 | -4.19 | -3.58 | -3.62 | -3.23 | -5 | -1.59 | -6.74 | -5.62 | -4.59 | 0.0675 | 0.6725 | -0.105 | 0.938243 | 0.949477 | 0.525729 | 0.999187 | 0.997988 | 0.999507 |
| Tpbgl_Q8C013_S375s_1 | Tpbgl_Q8C | 0.62 | -0.45 | -0.2 | 0.32 | 0.43 | 0.74 | 0.16 | 0.69 | 0.57 | 0.58 | 0.51 | 0.26 | 0.74 | 0.84 | 0.43 | 0.12 | 0.4325 | 0.4075 | 0.46 | 0.188242 | 0.173905 | 0.191172 | 0.989223 | 0.997988 | 0.994666 |
| Tpd52_F8WHQ1_S155s_1 | Tpd52_F8W | -0.36 | -0.9 | 0.52 | 0.8 | 0.86 | 0.68 | 0.54 | 1.11 | -1.78 | 0.01 | -1.44 | 0.42 | -0.29 | -1.35 | -0.22 | 0.98 | 0.7825 | -0.7125 | -0.235 | 0.680487 | 0.073351 | 0.260233 | 0.989223 | 0.997988 | 0.996159 |
| Tpd52_F8WHQ1_S186s_1 | Tpd52_F8W | -0.76 | -1.23 | -0.55 | -0.46 | -0.27 | -0.65 | -0.31 | -0.27 | -1.36 | -0.78 | -0.62 | -0.28 | -0.38 | 0.36 | -0.32 | -0.54 | 0.375 | -0.01 | 0.53 | 0.116774 | 0.158852 | 0.974657 | 0.989223 | 0.997988 | 0.994666 |
| Tpd52_F8WHQ1_S198s_1 | Tpd52_F8W | 2.67 | 1.34 | 1.81 | 1.93 | 1.45 | 2.73 | 1.89 | 2.17 | 2.16 | 2.59 | 1.2 | 1.75 | 0.9 | 2.06 | 1.28 | 1.72 | 0.1225 | -0.0125 | -0.4475 | 0.264042 | 0.745378 | 0.975142 | 0.995762 | 0.997988 | 0.994666 |
| Tpd52l1_B2RUC1_S142s_1 | Tpd52l1_B2 | 2.28 | 1.68 | 3.48 | 0.44 | -0.1 | 2.73 | 3.17 | 2.57 | 0.15 | 1.75 | 1.78 | 2.85 | 1.24 | 0.41 | 3.01 | 3.19 | 0.1225 | -0.3375 | -0.0075 | 0.992679 | 0.885218 | 0.656548 | 0.996493 | 0.997988 | 1 |
| Tpd52l2_F6VQ81_S140s_1 | Tpd52l2_F6 | -1.68 | -1.86 | -1.03 | -0.52 | -0.77 | -1.13 | -0.59 | -0.74 | -1.41 | -1.23 | -1.21 | -0.73 | -0.99 | -0.55 | -0.92 | -0.61 | 0.465 | 0.1275 | 0.505 | 0.176768 | 0.188598 | 0.719676 | 0.989223 | 0.997988 | 0.994666 |
| Tpd52l2_Q8BKP1_S12s_1 | Tpd52l2_Q | -0.22 | 0.13 | 0.3 | 0.22 | 0.29 | -0.2 | 0.22 | 0.13 | -0.49 | -0.22 | 1.21 | 0.46 | -0.35 | 0.1 | -0.31 | 0.16 | 0.0025 | 0.1325 | -0.2075 | 0.445068 | 0.991377 | 0.738446 | 0.999187 | 0.997988 | 0.994666 |
| Tpd52l2_Q8BKP1_S148s_1 | Tpd52l2_Q | 4.66 | 2.19 | 3.08 | -0.02 | 3.11 | 4.84 | 4.14 | 2.72 | 2.54 | 3.64 | 0.37 | 2.51 | 3.33 | -0.2 | 3.46 | 4.82 | 1.225 | -0.2125 | 0.375 | 0.764689 | 0.215035 | 0.839731 | 0.989223 | 0.997988 | 0.9971 |
| Tpd52l2_Q8BKP1_S186s_1 | Tpd52l2_Q | -0.71 | -0.22 | -0.09 | -0.03 | -0.12 | -0.53 | 0.34 | -0.07 | -0.17 | -0.11 | -0.43 | 0.09 | 0.24 | 0.52 | -0.23 | -0.1 | 0.1675 | 0.1075 | 0.37 | 0.228773 | 0.545975 | 0.676814 | 0.989223 | 0.997988 | 0.994666 |
| Tpgs1_Q99MS8_S279s_1 | Tpgs1_Q99 | -1.01 | -0.73 | -0.33 | -1.14 | -0.33 | -1.33 | -0.61 | -0.88 | -0.59 | -0.92 | -0.59 | -1.1 | 0.37 | -0.22 | -0.23 | -0.57 | 0.015 | 0.0025 | 0.64 | 0.066656 | 0.960334 | 0.992731 | 0.999187 | 0.997988 | 0.994666 |
| Tph2_Q8CGV2_S19s_1 | Tph2_Q8C | -0.45 | -0.32 | -0.58 | -0.06 | -0.27 | -0.18 | -0.09 | -0.28 | -0.31 | -0.4 | -0.66 | -0.46 | -0.42 | -0.04 | -0.99 | -0.54 | 0.1475 | -0.105 | -0.145 | 0.622127 | 0.497326 | 0.650129 | 0.989223 | 0.997988 | 0.995259 |
| Tpi1_P17751_S245s_1 | Tpi1_P177 | -1.78 | -1.86 | -0.6 | -1.9 | -1.89 | -1.78 | -1.8 | -1.91 | -1.34 | -1.6 | 0.52 | -2.12 | 0.85 | 0.06 | -1.07 | -1.04 | -0.31 | 0.4 | 1.235 | 0.038712 | 0.353922 | 0.509971 | 0.989223 | 0.997988 | 0.994666 |
| Tpi1_P17751_S262s_1 | Tpi1_P177 | 0.11 | 0.6 | -1.2 | -0.24 | 0.57 | -0.2 | -0.3 | -0.45 | -0.79 | 0.06 | 0.45 | -1.36 | -0.02 | 0.07 | 0.06 | -0.9 | 0.0875 | -0.2275 | -0.015 | 0.972697 | 0.83525 | 0.664406 | 0.995762 | 0.997988 | 0.999507 |
| Tpi1_P17751_T228t_1 | Tpi1_P177 | -0.34 | -0.06 | 0.38 | 0.32 | -0.79 | -0.78 | -0.68 | -0.05 | -0.58 | -0.14 | 0.58 | -0.3 | 0.43 | -0.42 | 0.38 | -0.17 | -0.65 | -0.185 | -0.02 | 0.949746 | 0.041082 | 0.573591 | 0.989223 | 0.997988 | 0.999507 |
| Tpm1_G5E8R0_S6s_1 | Tpm1_G5E | 0.66 | -0.2 | 0.97 | 1.68 | -0.01 | 1.56 | 0.43 | 0.72 | 0.38 | -0.05 | 0.61 | 1.16 | 0.57 | -1.98 | 1.5 | 0.66 | -0.1025 | -0.2525 | -0.59 | 0.440495 | 0.828478 | 0.575585 | 0.995762 | 0.997988 | 0.994666 |
| Tpm3_D3Z6I8_S51s_1 | Tpm3_D3Z | -0.83 | -0.13 | -0.71 | -0.52 | -0.44 | -1.41 | -1.18 | -1.24 | -1.79 | -0.42 | -0.75 | -0.99 | -0.88 | -0.34 | -0.76 | -0.07 | -0.52 | -0.44 | 0.035 | 0.907674 | 0.100817 | 0.224714 | 0.989223 | 0.997988 | 0.999507 |
| Tpm3_P21107_S207s_1 | Tpm3_P21 | -0.38 | -0.26 | 0.88 | 0.76 | -0.9 | -0.13 | 0.23 | 0.4 | -0.58 | -0.37 | 0.14 | 0.24 | 0.29 | -0.99 | 0.74 | 0.96 | -0.35 | -0.3925 | 0 | 1 | 0.410815 | 0.324854 | 0.989223 | 0.997988 | 1 |
| Tppp_Q7TQD2_S151s_1 | Tppp_Q7T | -1.43 | -1 | -0.64 | -1.28 | -0.36 | -0.8 | -0.33 | -1.25 | -3.06 | -0.98 | -0.18 | -1.03 | -0.16 | -2.43 | -1.55 | -0.28 | 0.4025 | -0.225 | -0.0175 | 0.97369 | 0.203146 | 0.702174 | 0.989223 | 0.997988 | 0.999507 |
| Tppp_Q7TQD2_S159s_1 | Tppp_Q7T | -4.4 | -6.93 | -4.21 | -6.35 | -2.48 | -5.78 | -4.71 | -5.61 | -7.5 | -6.7 | -1.51 | -4.33 | -3.64 | -3.15 | -3.51 | -4.94 | 0.8275 | 0.4625 | 1.6625 | 0.038631 | 0.36312 | 0.727265 | 0.989223 | 0.997988 | 0.994666 |
| Tppp_Q7TQD2_S182s_1 | Tppp_Q7T | -2.28 | -3.23 | -2.75 | -1.15 | -2.17 | -2.91 | -1.55 | -2.14 | -1.44 | -3.16 | -0.85 | -1.87 | -0.5 | -2.13 | -1.92 | -1.82 | 0.16 | 0.5225 | 0.76 | 0.180775 | 0.741467 | 0.398474 | 0.995762 | 0.997988 | 0.994666 |
| Tppp_Q7TQD2_S19s_1 | Tppp_Q7T | 2.42 | 1.57 | 1.79 | 1.77 | 1.79 | 2.41 | 0.95 | 1.86 | 1.8 | 2.02 | 1.07 | 1.32 | 1.38 | 1.07 | 1.89 | 1.84 | -0.135 | -0.335 | -0.3425 | 0.296103 | 0.703704 | 0.304267 | 0.995437 | 0.997988 | 0.994666 |
| Tppp_Q7TQD2_S19s_2 | Tppp_Q7T | -4.44 | -3.14 | -2.92 | -4.56 | -3.18 | -3.93 | -2.94 | -3.31 | -3.3 | -3.3 | -0.31 | -4.41 | -3.22 | -2.16 | -3.94 | -3.38 | 0.425 | 0.935 | 0.59 | 0.280961 | 0.352833 | 0.297743 | 0.989223 | 0.997988 | 0.994666 |
| Tppp_Q7TQD2_S31s_1 | Tppp_Q7T | 0.34 | 0.52 | 0.47 | 0.35 | 0.64 | 0.34 | 0.45 | 0.23 | 0.26 | 0.6 | -0.24 | 0.18 | -0.13 | 0.63 | 0 | 0 | -0.005 | -0.22 | -0.295 | 0.286263 | 0.98068 | 0.392205 | 0.999187 | 0.997988 | 0.994666 |
| Tppp_Q7TQD2_T15t_1 | Tppp_Q7T | 1.3 | 1.79 | 2 | 1.41 | 1.2 | 1.32 | 1.28 | 1.36 | 1.93 | 1.59 | 1.75 | 1.55 | 2.48 | 1.39 | 1.71 | 1.65 | -0.335 | 0.08 | 0.1825 | 0.581153 | 0.17836 | 0.754293 | 0.989223 | 0.997988 | 0.995259 |
| Tppp_Q7TQD2_T15t_2 | Tppp_Q7T | -4.44 | -3.14 | -2.92 | -4.56 | -3.18 | -3.93 | -2.94 | -3.31 | -3.3 | -3.3 | -0.31 | -4.41 | -3.22 | -2.16 | -3.94 | -3.38 | 0.425 | 0.935 | 0.59 | 0.280961 | 0.352833 | 0.297743 | 0.989223 | 0.997988 | 0.994666 |
| Tpr_Q7M739_S2031s_1 | Tpr_Q7M7 | 0.28 | 1.92 | 0.89 | 0.99 | 0.24 | -0.66 | 0.27 | 0.71 | 0.25 | -0.34 | -0.34 | 0.78 | 0.33 | 1.55 | 0.1 | 0.45 | -0.88 | -0.9325 | -0.4125 | 0.374188 | 0.060552 | 0.050789 | 0.989223 | 0.997988 | 0.994666 |
| Tpr_Q7M739_S2067s_1 | Tpr_Q7M7 | -1.54 | -1.33 | -0.52 | -1.07 | -1.42 | -2.22 | -0.75 | -0.68 | -1.12 | -1.74 | -0.54 | -0.71 | 0.62 | -0.62 | -0.5 | -0.65 | -0.1525 | 0.0875 | 0.8275 | 0.055855 | 0.706536 | 0.807244 | 0.995437 | 0.997988 | 0.994666 |
| Tpr_Q7M739_S379s_1 | Tpr_Q7M7 | 0.21 | 0.53 | -0.34 | -0.02 | 0.02 | 0.13 | 0.33 | -0.18 | 0.1 | -0.12 | -0.64 | 0.12 | 0.28 | 0.52 | 0.08 | 0.03 | -0.02 | -0.23 | 0.1325 | 0.645934 | 0.938837 | 0.445583 | 0.999187 | 0.997988 | 0.995259 |
| Tprgl_B1AX95_S10s_1 | Tprgl_B1A | 1.22 | 1.46 | 1.73 | 1.41 | 0.89 | 1 | 1.28 | 1.22 | 0.75 | 1.2 | 1.51 | 0.96 | 0.1 | 0.71 | 1.27 | 2.01 | -0.3575 | -0.35 | -0.4325 | 0.315973 | 0.133745 | 0.20068 | 0.989223 | 0.997988 | 0.994666 |
| Tprgl_B1AX95_S14s_1 | Tprgl_B1A | 0.64 | 0.85 | -0.11 | 0.21 | 1.04 | 1.47 | 0.74 | 0.47 | 0.07 | 1.05 | 0.37 | 0.23 | -0.47 | 0.79 | 0.23 | 0.02 | 0.5325 | 0.0325 | -0.255 | 0.486997 | 0.12039 | 0.921329 | 0.989223 | 0.997988 | 0.994666 |
| Tprgl_B1AX95_T34t_1 | Tprgl_B1A | -1.13 | -0.55 | -1.01 | -1.76 | -0.46 | -0.94 | -0.73 | -1.47 | -1.67 | -1.25 | -2.18 | -1.11 | -0.56 | -1.51 | -0.98 | -1.35 | 0.2125 | -0.44 | 0.0125 | 0.971847 | 0.53176 | 0.239414 | 0.989223 | 0.997988 | 0.999507 |
| Tprn_A2AI08_S457s_1 | Tprn_A2AI | -0.24 | 0.53 | 1.02 | 0.17 | 0.64 | -0.08 | 1.24 | 0.88 | 0.18 | 0.23 | 1.63 | 0.33 | -0.35 | 0.16 | 0.81 | 0.4 | 0.3 | 0.2225 | -0.115 | 0.760973 | 0.435396 | 0.605302 | 0.989223 | 0.997988 | 0.9971 |
| Tra2a_E9QP00_S14s_1 | Tra2a_E9Q | -0.03 | 0.04 | 0.12 | -0.27 | -0.14 | -0.05 | -0.42 | 0.28 | -0.73 | 0.27 | -0.49 | -0.55 | 0.55 | -0.83 | 0.42 | 0.77 | -0.0475 | -0.34 | 0.2625 | 0.49874 | 0.840606 | 0.251307 | 0.995762 | 0.997988 | 0.994666 |
| Tra2a_E9QP00_S14s_2 | Tra2a_E9Q | 0.68 | 0.25 | 0.52 | 0.41 | -0.34 | 1.13 | 2.13 | 1.11 | 0.98 | 1.05 | -0.27 | 1.33 | 0.56 | -0.95 | 1.63 | 1.98 | 0.5425 | 0.3075 | 0.34 | 0.577686 | 0.272211 | 0.419315 | 0.989223 | 0.997988 | 0.995259 |
| Tra2a_E9QP00_S20s_1 | Tra2a_E9Q | -2.66 | -2.7 | -2.9 | -4.48 | -3.33 | -2.01 | -3.16 | -2.14 | -0.32 | -2.32 | -1.88 | -2.73 | 0.6 | -4.3 | -1.8 | -3.3 | 0.525 | 1.3725 | 0.985 | 0.338145 | 0.313004 | 0.050922 | 0.989223 | 0.997988 | 0.994666 |
| Tra2a_E9QP00_S236s_1 | Tra2a_E9Q | -2.49 | -2.16 | -1.58 | -4.86 | -2.77 | -1.91 | -3.98 | -4.54 | -1.66 | -2.39 | -0.89 | -2.33 | 0.16 | -3.34 | -3.28 | -3.79 | -0.5275 | 0.955 | 0.21 | 0.835807 | 0.521839 | 0.207143 | 0.989223 | 0.997988 | 0.999507 |
| Tra2a_E9QP00_S260s_1 | Tra2a_E9Q | 1.68 | 1.54 | 0.6 | 0.6 | 0.47 | 0.95 | 0.16 | 0.62 | 1.1 | 0.59 | 0.13 | 0.77 | 1.35 | 1.28 | 1.28 | 0.86 | -0.555 | -0.4575 | 0.0875 | 0.800424 | 0.129119 | 0.230802 | 0.989223 | 0.997988 | 0.999507 |
| Tra2a_E9QP00_S260s_2 | Tra2a_E9Q | 0.77 | -0.01 | -0.26 | -0.12 | -1.76 | -0.15 | -0.83 | 0.25 | -0.31 | -0.41 | 0.91 | -1.04 | -1.47 | -0.72 | 0.31 | -1.22 | -0.7175 | -0.3075 | -0.87 | 0.075875 | 0.144672 | 0.500519 | 0.989223 | 0.997988 | 0.994666 |
| Tra2a_E9QP00_S262s_2 | Tra2a_E9Q | 0.75 | 0 | -0.26 | -0.12 | -1.83 | -0.1 | -0.83 | 0.25 | -0.22 | -0.39 | 0.91 | -1.04 | -1.52 | -0.72 | 0.31 | -1.22 | -0.72 | -0.2775 | -0.88 | 0.074655 | 0.152576 | 0.539211 | 0.989223 | 0.997988 | 0.994666 |
| Tra2a_E9QP00_S2s_1 | Tra2a_E9Q | 0.03 | 0.27 | -0.57 | -0.3 | -0.03 | 0.4 | -0.47 | 0.27 | 0.31 | 0.27 | 0.29 | 0.46 | 0.71 | 0.75 | -0.06 | 0.15 | 0.185 | 0.475 | 0.53 | 0.123494 | 0.534153 | 0.091269 | 0.989223 | 0.997988 | 0.994666 |
| Tra2a_E9QP00_S2s_2 | Tra2a_E9Q | 0.68 | 0.25 | 0.52 | 0.41 | -0.34 | 1.13 | 2.13 | 1.11 | 0.98 | 1.05 | -0.27 | 1.33 | 0.56 | -0.95 | 1.63 | 1.98 | 0.5425 | 0.3075 | 0.34 | 0.577686 | 0.272211 | 0.419315 | 0.989223 | 0.997988 | 0.995259 |
| Tra2a_E9QP00_T200t_1 | Tra2a_E9Q | 4.8 | 1.11 | 1.53 | -2.35 | 1.4 | 4.51 | 2.93 | 1.61 | -3.19 | 3.06 | -1.85 | 0.7 | -1.29 | -2.35 | 2.42 | 1.74 | 1.34 | -1.5925 | -1.1425 | 0.480534 | 0.349281 | 0.375212 | 0.989223 | 0.997988 | 0.994666 |
| Tra2b_P62996_S29s_2 | Tra2b_P62 | 2.58 | 0.79 | 0.7 | 1.16 | -0.46 | 1.27 | -0.52 | 0.71 | 3.85 | 1.44 | -2.09 | 0.22 | 1.89 | -0.69 | 2.67 | 0.64 | -1.0575 | -0.4525 | -0.18 | 0.812628 | 0.084235 | 0.695678 | 0.989223 | 0.997988 | 0.999507 |
| Tra2b_P62996_S2s_1 | Tra2b_P62 | -0.83 | 0.33 | 0.2 | 0.18 | -0.71 | -0.2 | -0.21 | -0.11 | 1.78 | 0.51 | 1.19 | 0.55 | 2.48 | 0.27 | 1.11 | 0.6 | -0.2775 | 1.0375 | 1.145 | 0.050434 | 0.391623 | 0.026916 | 0.989223 | 0.997988 | 0.994666 |
| Tra2b_P62996_S99s_2 | Tra2b_P62 | 2.76 | -0.07 | 0.28 | 0.55 | -0.8 | 1.49 | 0.27 | 0.42 | 0.26 | 1.11 | -0.99 | -0.65 | -0.02 | -2.11 | 2.06 | 0.6 | -0.535 | -0.9475 | -0.7475 | 0.434286 | 0.451173 | 0.207396 | 0.989223 | 0.997988 | 0.994666 |
| Tra2b_P62996_T201t_1 | Tra2b_P62 | -1.52 | -1.49 | -1.94 | -1.9 | -1.47 | -2.36 | -1.79 | -1.84 | -3.1 | -2.59 | -1.4 | -2.08 | -1.51 | -1.69 | -2 | -2.82 | -0.1525 | -0.58 | -0.2925 | 0.406925 | 0.568833 | 0.156761 | 0.989223 | 0.997988 | 0.994666 |
| Tra2b_P62996_T33t_2 | Tra2b_P62 | 2.58 | 0.79 | 0.7 | 1.16 | -0.46 | 1.27 | -0.52 | 0.71 | 3.85 | 1.44 | -2.09 | 0.22 | 1.89 | -0.69 | 2.67 | 0.64 | -1.0575 | -0.4525 | -0.18 | 0.812628 | 0.084235 | 0.695678 | 0.989223 | 0.997988 | 0.999507 |
| Traf3_Q3UHJ1_S243s_1 | Traf3_Q3U | 0.35 | 0.49 | 2.73 | 2.52 | 0.75 | -0.06 | 2.56 | 2.72 | -0.3 | 0.38 | 3.55 | 3.12 | -0.13 | 0.33 | 2.88 | 2.82 | -0.03 | 0.165 | -0.0475 | 0.957694 | 0.970657 | 0.87155 | 0.999187 | 0.997988 | 0.999507 |
| Traf3_Q3UHJ1_S330s_1 | Traf3_Q3U | -0.21 | 0.67 | 0.99 | 0.36 | 0.38 | 0.17 | 0.2 | 0.65 | 0.1 | -0.16 | 1.12 | 0.29 | 1.28 | 1.08 | 0.73 | 1.21 | -0.1025 | -0.115 | 0.6225 | 0.08168 | 0.73484 | 0.762972 | 0.995762 | 0.997988 | 0.994666 |
| Trafd1_Q3UDK1_S450s_1 | Trafd1_Q3U | -1.41 | -0.77 | -1.06 | -1.57 | -0.1 | -1.52 | -0.14 | -1.04 | -0.64 | -0.78 | 0.1 | -0.76 | 0.19 | -0.53 | -1.68 | -1.06 | 0.5025 | 0.6825 | 0.4325 | 0.328105 | 0.209653 | 0.050493 | 0.989223 | 0.997988 | 0.994666 |
| Tram1_Q91V04_S365s_1 | Tram1_Q91 | 0.78 | 1.36 | 0.71 | 0.84 | 0.72 | 0.64 | 0.53 | 1.22 | 1.08 | 0.79 | 1.44 | 0.67 | 1.35 | 1.28 | 1.75 | 1.35 | -0.145 | 0.0725 | 0.51 | 0.084644 | 0.581655 | 0.79516 | 0.989223 | 0.997988 | 0.994666 |
| Trappc10_Q3TLI0_S707s_1 | Trappc10_Q | 0.32 | 0.13 | -0.06 | 0.43 | 0.55 | 0.29 | 0.69 | 0.68 | -0.23 | 0.52 | 0 | 0.31 | -0.22 | 0.39 | 0.27 | 0.47 | 0.3475 | -0.055 | 0.0225 | 0.934364 | 0.14573 | 0.833896 | 0.989223 | 0.997988 | 0.999507 |
| Trappc12_Q8C0C3_S10s_1 | Trappc12_Q | 0.55 | 0.46 | 1.32 | 1.65 | 0.31 | 0.74 | 1.02 | 1.26 | 0.25 | 1.18 | 1.34 | 1.53 | -0.21 | 0.09 | 0.89 | 1.45 | -0.1625 | 0.08 | -0.44 | 0.352032 | 0.649382 | 0.84285 | 0.990343 | 0.997988 | 0.994666 |
| Trappc12_Q8C0C3_S140s_1 | Trappc12_Q | 1.35 | 1.89 | 0.72 | 0.36 | 1.91 | 1.56 | 0.79 | 0.44 | 0.19 | 1.46 | -0.41 | 0.92 | -0.09 | -0.55 | 0.84 | 0.95 | 0.095 | -0.54 | -0.7925 | 0.121268 | 0.832127 | 0.296455 | 0.995762 | 0.997988 | 0.994666 |
| Trappc12_Q8C0C3_S309s_1 | Trappc12_Q | 1.07 | 0.96 | -0.16 | -0.04 | 1 | 0.85 | 0.12 | -0.21 | 0.52 | 1.12 | -0.02 | -0.1 | 0.16 | 0.89 | -0.02 | -0.06 | -0.0175 | -0.0775 | -0.215 | 0.592606 | 0.966234 | 0.853811 | 0.999187 | 0.997988 | 0.995259 |
| Trappc12_Q8C0C3_T143t_1 | Trappc12_Q | 1.24 | 1.65 | 1.75 | 0.92 | 1.23 | 0.96 | 0.71 | 1.09 | 0.5 | 1.47 | 2.01 | 1.27 | 0.35 | 0.43 | 1.79 | 1.67 | -0.3925 | -0.0775 | -0.33 | 0.448143 | 0.163581 | 0.835604 | 0.989223 | 0.997988 | 0.994666 |
| Trappc4_Q9ES56_S219s_1 | Trappc4_Q | 0.63 | 1.24 | 0.31 | 0.11 | 0.1 | 1.38 | -0.66 | -1.49 | 0.13 | 0.85 | 0.84 | -0.23 | 0.58 | 0.13 | 0.3 | -0.59 | -0.74 | -0.175 | -0.4675 | 0.226695 | 0.228743 | 0.639943 | 0.989223 | 0.997988 | 0.994666 |
| Trappc8_E9PY51_S657s_1 | Trappc8_E9 | 0.31 | 0.71 | 0.45 | 0.74 | 1.04 | 0.74 | 0.9 | 0.75 | 0.7 | 0.94 | 1.13 | 0.36 | -0.13 | 0.81 | 0.8 | 0.94 | 0.305 | 0.23 | 0.0525 | 0.868524 | 0.182401 | 0.388778 | 0.989223 | 0.997988 | 0.999507 |
| Trappc9_Q3U0M1_S646s_1 | Trappc9_Q | -1.7 | -2.19 | -1.2 | -2.14 | -1.86 | -1.46 | -1.37 | -1.34 | -1.65 | -1.36 | -2.02 | -1.59 | -0.55 | -1.8 | -1.11 | -1.19 | 0.3 | 0.1525 | 0.645 | 0.103434 | 0.314998 | 0.619199 | 0.989223 | 0.997988 | 0.994666 |
| Trex1_Q91XB0_S78s_1 | Trex1_Q91 | 0.1 | 1.39 | -0.71 | -0.33 | -0.7 | 1.54 | 0.67 | -0.62 | -0.1 | 1.09 | 0.71 | -0.23 | 1.25 | 0.37 | 1.57 | -0.08 | 0.11 | 0.255 | 0.665 | 0.246391 | 0.861209 | 0.626318 | 0.996493 | 0.997988 | 0.994666 |
| Trhde_Q8K093_S11s_1 | Trhde_Q8K | 2.4 | 0.83 | 1.18 | 1.69 | 0.94 | 2.4 | 0.92 | 2.27 | 1.42 | 1.11 | 1.35 | 1.79 | 1.11 | 1.33 | 1.67 | 1.85 | 0.1075 | -0.1075 | -0.035 | 0.928443 | 0.825746 | 0.774648 | 0.995762 | 0.997988 | 0.999507 |
| Trim2_Q9ESN6_S424s_1 | Trim2_Q9E | -0.41 | 0.15 | 0.48 | -1.54 | 0.32 | -0.64 | 0.11 | -1.49 | -1.73 | -0.1 | -1.66 | -1.59 | -0.64 | -1.06 | 0.46 | -1.26 | -0.095 | -0.94 | -0.295 | 0.590729 | 0.861919 | 0.112756 | 0.996493 | 0.997988 | 0.995259 |
| Trim2_Q9ESN6_S424s_2 | Trim2_Q9E | -0.5 | 0.11 | 1.04 | 1.12 | 0.36 | -0.2 | 0.81 | 0.66 | -0.86 | 0.06 | -0.01 | 0.66 | -0.9 | -0.57 | -0.04 | 1.28 | -0.035 | -0.48 | -0.5 | 0.389319 | 0.93423 | 0.32583 | 0.998658 | 0.997988 | 0.994666 |
| Trim2_Q9ESN6_S428s_1 | Trim2_Q9E | 0.61 | 0.59 | 0.64 | 0.3 | 0.34 | 0.5 | 0.08 | 0.41 | 0.06 | 1.05 | 0.6 | 0.34 | 0.75 | -0.43 | 0.61 | 0.45 | -0.2025 | -0.0225 | -0.19 | 0.559573 | 0.357362 | 0.935394 | 0.989223 | 0.997988 | 0.995259 |
| Trim2_Q9ESN6_S440s_1 | Trim2_Q9E | -3.8 | -4.59 | -4.07 | -1 | -3.82 | -3.54 | -4.41 | -5.19 | -0.04 | -3.44 | -5.41 | -3.24 | 1.34 | -4.66 | -1.45 | -3.1 | -0.875 | 0.3325 | 1.3975 | 0.299151 | 0.274837 | 0.782644 | 0.989223 | 0.997988 | 0.994666 |
| Trim2_Q9ESN6_S456s_1 | Trim2_Q9E | -3.29 | -3.58 | -2.94 | -4.69 | -2.78 | -2.93 | -2.91 | -2.88 | -3.38 | -3.22 | 0.02 | -2.59 | -1.68 | -1.74 | -2.95 | -3.03 | 0.75 | 1.3325 | 1.275 | 0.028081 | 0.069029 | 0.115095 | 0.989223 | 0.997988 | 0.994666 |
| Trim24_E9Q1U8_S675s_1 | Trim24_E9 | -0.4 | -0.15 | -0.61 | -0.35 | -1.69 | -0.55 | -0.91 | -0.38 | 0.43 | -0.28 | 0.27 | -1.62 | 1.06 | 0.95 | 0.43 | -0.97 | -0.505 | 0.0775 | 0.745 | 0.130532 | 0.139716 | 0.864952 | 0.989223 | 0.997988 | 0.994666 |
| Trim24_E9Q1U8_S742s_1 | Trim24_E9 | -1.98 | -0.92 | -0.79 | -1.57 | -0.94 | -3.09 | -0.85 | -1.75 | -1.76 | -1.7 | -0.2 | -0.95 | -0.45 | -1.11 | -0.3 | -0.52 | -0.3425 | 0.1625 | 0.72 | 0.06828 | 0.526169 | 0.715964 | 0.989223 | 0.997988 | 0.994666 |
| Trim28_Q62318_S23s_1 | Trim28_Q6 | -0.14 | -0.16 | -0.12 | -0.07 | 0.03 | -0.39 | 0.28 | -0.06 | -0.37 | 0.05 | -0.29 | -0.19 | -0.38 | 0.25 | -0.19 | 0.02 | 0.0875 | -0.0775 | 0.0475 | 0.849635 | 0.696014 | 0.719388 | 0.994893 | 0.997988 | 0.999507 |
| Trim28_Q62318_S473s_1 | Trim28_Q6 | -0.6 | -0.42 | -1.42 | -1.59 | -0.44 | -1.51 | -0.61 | -1.81 | -0.67 | -0.54 | -0.98 | -1.73 | -0.42 | -0.01 | -1.34 | -1.61 | -0.085 | 0.0275 | 0.1625 | 0.72508 | 0.840163 | 0.944363 | 0.995762 | 0.997988 | 0.9971 |
| Trim28_Q62318_S51s_1 | Trim28_Q6 | -0.39 | -0.22 | -0.26 | 0.15 | -0.23 | -0.43 | 0.17 | -0.29 | -0.2 | -0.18 | -0.03 | -0.4 | -0.39 | 0 | -0.13 | -0.39 | -0.015 | -0.0225 | -0.0475 | 0.85301 | 0.949971 | 0.922833 | 0.999187 | 0.997988 | 0.999507 |
| Trim28_Q62318_S594s_1 | Trim28_Q6 | -0.05 | 0.13 | -0.14 | -0.35 | -0.44 | -0.11 | 0.19 | -0.12 | -0.42 | 0.01 | -0.83 | -0.96 | -0.55 | 0.06 | -0.18 | 0.07 | -0.0175 | -0.4475 | -0.0475 | 0.859037 | 0.940346 | 0.144886 | 0.999187 | 0.997988 | 0.999507 |
| Trim28_Q62318_S697s_1 | Trim28_Q6 | -0.54 | -1.87 | -1.08 | -0.91 | -2.59 | -0.64 | -1.35 | -0.79 | 0.59 | -1.04 | -0.58 | -1.57 | 1.14 | -0.73 | -0.1 | -0.39 | -0.2425 | 0.45 | 1.08 | 0.044167 | 0.61798 | 0.384626 | 0.989223 | 0.997988 | 0.994666 |
| Trim3_Q9R1R2_S437s_1 | Trim3_Q9R | 1.04 | 0.39 | -2.4 | -3.04 | 1.01 | 0.89 | -2.23 | -3.18 | 0.77 | 1.01 | -2.39 | -2.46 | 1.13 | 1.16 | -1.58 | -2.37 | 0.125 | 0.235 | 0.5875 | 0.619958 | 0.921224 | 0.847287 | 0.997335 | 0.997988 | 0.995259 |
| Trim3_Q9R1R2_S529s_1 | Trim3_Q9R | -0.29 | -0.2 | -0.05 | -0.31 | -0.21 | -1.12 | -0.84 | -0.77 | 1.24 | -0.67 | 0.3 | -0.36 | 2.13 | -0.58 | 0.05 | -0.49 | -0.5225 | 0.34 | 0.49 | 0.409648 | 0.063268 | 0.425014 | 0.989223 | 0.997988 | 0.994666 |
| Trim3_Q9R1R2_S7s_1 | Trim3_Q9R | -0.55 | -0.32 | -0.36 | -0.2 | 0.22 | -0.89 | -0.26 | -0.15 | -0.39 | 0.1 | -0.25 | -0.24 | -0.02 | 0.36 | -0.14 | -0.56 | 0.0875 | 0.1625 | 0.2675 | 0.352358 | 0.753972 | 0.481291 | 0.995762 | 0.997988 | 0.994666 |
| Trim32_Q8CH72_S330s_1 | Trim32_Q8 | 0.95 | 0.57 | 1.29 | 0.78 | 0.67 | 0.84 | 1.23 | 0.88 | 0.18 | 1.01 | -0.31 | 1.12 | 0.61 | -0.53 | 1.05 | 1.11 | 0.0075 | -0.3975 | -0.3375 | 0.420362 | 0.975996 | 0.309154 | 0.999187 | 0.997988 | 0.994666 |
| Trim46_D3YXA6_S604s_1 | Trim46_D3 | 1 | 0.73 | 0.67 | 1.13 | 0.79 | 1.06 | 0.9 | 0.67 | 0.65 | 1.13 | 0.51 | 0.14 | 0.7 | 0.66 | 0.44 | 0.97 | -0.0275 | -0.275 | -0.19 | 0.467226 | 0.901371 | 0.342735 | 0.996493 | 0.997988 | 0.994666 |
| Trim56_Q80VI1_S454s_1 | Trim56_Q8 | 0.31 | 0.82 | -0.36 | -0.05 | 0.34 | 0.24 | 0.66 | 0.09 | 0.04 | 0.64 | 0.84 | -0.55 | 0.21 | 0.98 | 0.28 | -0.44 | 0.1525 | 0.0625 | 0.0775 | 0.844182 | 0.61758 | 0.876076 | 0.989223 | 0.997988 | 0.999507 |
| Trim66_E9PZP2_S767s_1 | Trim66_E9 | 0.66 | 1.35 | 1.24 | 0.71 | 0.75 | 0.57 | 0.41 | 0.74 | 0.99 | 0.97 | 0.84 | -0.33 | 1.05 | 1.11 | 0.91 | 0.68 | -0.3725 | -0.3725 | -0.0525 | 0.851314 | 0.161484 | 0.329529 | 0.989223 | 0.997988 | 0.999507 |
| Trim67_Q505D9_S275s_1 | Trim67_Q5 | -0.13 | -0.53 | -0.3 | 0.31 | -0.89 | 0.34 | -1.07 | -0.42 | 1.73 | 0.28 | -0.09 | 0.14 | 1.46 | -0.72 | -0.14 | -0.27 | -0.3475 | 0.6775 | 0.245 | 0.615051 | 0.346318 | 0.14507 | 0.989223 | 0.997988 | 0.995259 |
| Trim67_Q505D9_T38t_1 | Trim67_Q5 | 0.38 | 1.49 | -0.6 | -0.25 | 0.57 | 0.86 | 1.27 | 0.13 | 0.22 | 0.91 | -1.01 | 0.19 | 0.9 | 0.94 | 0.3 | -1.47 | 0.4525 | -0.1775 | -0.0875 | 0.893857 | 0.355833 | 0.752496 | 0.989223 | 0.997988 | 0.999507 |
| Trim9_E9Q524_S46s_1 | Trim9_E9Q | 0.86 | 0.64 | 0.01 | 0.41 | 0.63 | 0.44 | 0.61 | 0.12 | 0.61 | 1.01 | 0.61 | 0.12 | 0.33 | 0.68 | 0.1 | -0.01 | -0.03 | 0.1075 | -0.205 | 0.498582 | 0.909245 | 0.718992 | 0.997335 | 0.997988 | 0.994666 |
| Trim9_E9Q524_T41t_1 | Trim9_E9Q | 1.26 | 0.42 | 1.13 | 1.49 | 0.86 | 1.03 | 1.56 | 1.53 | 0.63 | 0.88 | 0.85 | 1.41 | -0.08 | 1.03 | 0.72 | 0.72 | 0.17 | -0.1325 | -0.4775 | 0.202343 | 0.586424 | 0.675641 | 0.989223 | 0.997988 | 0.994666 |
| Trim9_E9Q524_T443t_1 | Trim9_E9Q | -1.32 | 0.17 | -1.87 | -0.7 | -1.55 | -1.03 | -0.02 | -0.59 | -0.09 | -0.42 | -2.09 | -0.89 | -2.26 | -0.52 | -1.06 | -2.16 | 0.1325 | 0.0575 | -0.57 | 0.324718 | 0.791053 | 0.919687 | 0.995762 | 0.997988 | 0.994666 |
| Trio_Q0KL02_S1724s_1 | Trio_Q0KL | 0.48 | -0.76 | 0.35 | 0.38 | 0.03 | 0.68 | 0.42 | 1.23 | 0.78 | 0.3 | 0.52 | 0.97 | -1.46 | 0.06 | 0.23 | 0.2 | 0.4775 | 0.53 | -0.355 | 0.465529 | 0.225644 | 0.149161 | 0.989223 | 0.997988 | 0.994666 |
| Trio_Q0KL02_S1818s_1 | Trio_Q0KL | 0.5 | 0.3 | -0.53 | 0.22 | -0.22 | 0.49 | 0.25 | 0.59 | 0.17 | 1.12 | 0.65 | 0.5 | -0.45 | -0.1 | 0.09 | 0.28 | 0.155 | 0.4875 | -0.1675 | 0.604086 | 0.617961 | 0.159013 | 0.989223 | 0.997988 | 0.995259 |
| Trio_Q0KL02_S1845s_1 | Trio_Q0KL | -0.75 | -0.7 | -1.03 | -0.64 | 0.37 | -1.1 | -0.6 | -0.44 | -0.81 | -0.03 | -1.16 | -0.46 | -0.32 | 0.07 | -0.67 | -0.73 | 0.3375 | 0.165 | 0.3675 | 0.211503 | 0.317756 | 0.582349 | 0.989223 | 0.997988 | 0.994666 |
| Trio_Q0KL02_S2282s_1 | Trio_Q0KL | 0.03 | 0.01 | 0.4 | 0.47 | 0.59 | 0.07 | 0.61 | 0.55 | -0.31 | 0.13 | -0.18 | 0.13 | -0.84 | -0.11 | 0.15 | 0.64 | 0.2275 | -0.285 | -0.2675 | 0.461039 | 0.357014 | 0.26176 | 0.989223 | 0.997988 | 0.994666 |
| Trio_Q0KL02_S2458s_1 | Trio_Q0KL | -0.67 | -0.4 | -0.62 | -0.62 | 0.08 | 0.04 | -0.03 | -0.45 | -0.53 | -0.07 | 0.43 | -0.68 | -0.9 | 0 | -2.11 | -0.94 | 0.4875 | 0.365 | -0.41 | 0.353436 | 0.050091 | 0.240192 | 0.989223 | 0.997988 | 0.994666 |
| Trio_Q0KL02_T2560t_1 | Trio_Q0KL | 0.3 | 0.17 | 0.15 | 0.29 | 0.51 | 0.09 | 0.24 | 0.3 | -0.34 | 0.27 | 0.31 | 0.01 | 0.13 | 0.62 | -0.03 | -0.08 | 0.0575 | -0.165 | -0.0675 | 0.796923 | 0.78021 | 0.497257 | 0.995762 | 0.997988 | 0.999507 |
| Triobp_Q99KW3_S1101s_1 | Triobp_Q99 | 0.37 | 0.78 | -0.35 | 0.23 | 1.2 | 1.09 | -0.18 | -0.5 | 1.18 | 0.44 | -1.21 | 0.21 | 1.64 | 0.59 | 0.75 | -0.45 | 0.145 | -0.1025 | 0.375 | 0.432181 | 0.752487 | 0.842709 | 0.995762 | 0.997988 | 0.994666 |
| Trip12_G5E870_S1049s_1 | Trip12_G5E | 1.77 | 1.71 | 2 | 2 | 1 | 1.36 | 1.65 | 1.87 | 1.81 | 1.81 | 2.41 | 2.04 | 1.67 | 1.76 | 1.99 | 1.99 | -0.4 | 0.1475 | -0.0175 | 0.942138 | 0.141924 | 0.547536 | 0.989223 | 0.997988 | 0.999507 |
| Trip12_G5E870_S239s_1 | Trip12_G5E | -0.21 | -2.68 | 0.14 | -0.69 | -2.92 | 0.31 | -1.09 | -1.11 | 2.41 | -0.33 | -3.28 | 0.08 | 2.96 | -3.38 | 1.58 | -0.33 | -0.3425 | 0.58 | 1.0675 | 0.420605 | 0.669359 | 0.620897 | 0.992876 | 0.997988 | 0.994666 |
| Trip12_G5E870_S312s_1 | Trip12_G5E | 0.25 | 0.23 | 0.33 | 0.28 | 0.09 | -0.12 | 0.32 | -0.03 | 0.25 | -0.04 | -0.01 | 0.03 | 0.02 | 0.5 | 0.21 | 0.25 | -0.2075 | -0.215 | -0.0275 | 0.907963 | 0.328316 | 0.316451 | 0.989223 | 0.997988 | 0.999507 |
| Trip12_G5E870_S77s_1 | Trip12_G5E | 0.58 | 0.14 | 0.34 | 0.02 | -1.34 | 0.28 | -1.02 | -0.51 | 1.89 | 0 | -0.93 | -0.13 | 1.85 | -1.53 | 0.01 | -0.57 | -0.9175 | -0.0625 | -0.33 | 0.614694 | 0.031091 | 0.911579 | 0.989223 | 0.997988 | 0.995259 |
| Trmt10a_Q8C1Z8_S22s_1 | Trmt10a_Q | -4.18 | -4.06 | -1.74 | -1.52 | -4.44 | -6.09 | -1.75 | -1.63 | -8.97 | -3.64 | 0.02 | -1.49 | -1.98 | -4.14 | -1.27 | -1.25 | -0.6025 | -0.645 | 0.715 | 0.419978 | 0.59423 | 0.724138 | 0.989223 | 0.997988 | 0.994666 |
| Trmt10a_Q8C1Z8_S324s_1 | Trmt10a_Q | -2.3 | -0.05 | -0.78 | -1.37 | -1.1 | -1.87 | -1.39 | -2.59 | -1.89 | -1.87 | -1.55 | -1.55 | -0.54 | -0.49 | -1.99 | -2.42 | -0.6125 | -0.59 | -0.235 | 0.707146 | 0.260243 | 0.223907 | 0.989223 | 0.997988 | 0.996966 |
| Trpc5_Q9QX29_S770s_1 | Trpc5_Q9Q | 0.43 | 0.37 | 0.89 | 0.62 | 1.22 | 0.62 | 0.47 | 0.49 | 0.6 | -0.08 | 0.39 | 0.62 | 0.72 | 0.63 | 0.55 | 0.7 | 0.1225 | -0.195 | 0.0725 | 0.767958 | 0.640042 | 0.466403 | 0.989223 | 0.997988 | 0.9971 |
| Trpc6_Q3UZG1_S736s_1 | Trpc6_Q3U | -0.02 | -0.55 | 0.2 | -0.03 | -0.5 | -0.55 | -0.26 | 0.14 | -0.2 | 0.15 | 0.5 | 0.69 | 0.03 | 0.11 | -0.73 | -0.42 | -0.1925 | 0.385 | -0.1525 | 0.623309 | 0.478137 | 0.213237 | 0.989223 | 0.997988 | 0.995259 |
| Trpm3_Q5F4S6_S1292s_1 | Trpm3_Q5F | 0.32 | 0.57 | 1.17 | 0.92 | 0.23 | -0.01 | 0.23 | 0.21 | 0.31 | 0.44 | 1 | 0.78 | 0.27 | 0.72 | 0.47 | 1.33 | -0.58 | -0.1125 | -0.0475 | 0.887224 | 0.04217 | 0.699278 | 0.989223 | 0.997988 | 0.999507 |
| Trpm3_Q5F4S6_S1629s_1 | Trpm3_Q5F | -0.23 | 1.38 | 0.4 | 0.55 | -0.09 | -1.68 | 0.03 | 0.93 | 0.05 | 0.05 | 0.09 | 0.8 | 0.72 | 0.52 | -0.88 | 0.07 | -0.7275 | -0.2775 | -0.4175 | 0.382708 | 0.223077 | 0.474761 | 0.989223 | 0.997988 | 0.994666 |
| Trpm3_Q5F4S6_T102t_1 | Trpm3_Q5F | 0.24 | -0.26 | -0.04 | -0.13 | 1.69 | 1.26 | 0.29 | 0.28 | 0.5 | 1.09 | -0.5 | -0.71 | -0.55 | 1.41 | -0.72 | 0.13 | 0.9275 | 0.1425 | 0.115 | 0.808661 | 0.02876 | 0.738751 | 0.989223 | 0.997988 | 0.999507 |
| Trpm4_Q7TN37_S527s_1 | Trpm4_Q7T | 0.18 | 1.2 | 0.37 | 0.69 | -0.32 | -0.31 | 0.28 | 0.05 | 0.35 | -0.06 | 1.13 | 0.57 | 0.67 | -0.14 | 0.74 | 0.05 | -0.685 | -0.1125 | -0.28 | 0.426285 | 0.040513 | 0.747964 | 0.989223 | 0.997988 | 0.994666 |
| Trpm7_Q923J1_T555t_1 | Trpm7_Q92 | 0.48 | -0.52 | 0.59 | 0.39 | 1.08 | 0.49 | 0.55 | -0.09 | 0.69 | 0.67 | 0.03 | 0.79 | 1.03 | -0.29 | 0.91 | 0.95 | 0.2725 | 0.31 | 0.415 | 0.322626 | 0.445331 | 0.362101 | 0.989223 | 0.997988 | 0.994666 |
| Trps1_Q80V18_S584s_1 | Trps1_Q80 | -2.35 | -2.55 | -1.21 | 0.33 | -3.11 | -1.85 | -0.21 | -0.27 | -1.18 | -1.39 | 1.03 | -0.36 | -0.21 | -1.59 | 0.5 | -0.64 | 0.085 | 0.97 | 0.96 | 0.196758 | 0.919019 | 0.227678 | 0.997335 | 0.997988 | 0.994666 |
| Trpv2_Q9WTR1_S15s_1 | Trpv2_Q9W | 0.87 | 1.15 | 1.23 | 0.81 | 0.62 | 0.63 | 0.21 | 0.86 | 0.52 | 0.95 | 1.22 | 0.87 | -0.17 | 1.06 | 0.87 | 0.86 | -0.435 | -0.125 | -0.36 | 0.298037 | 0.092574 | 0.620974 | 0.989223 | 0.997988 | 0.994666 |
| Trpv2_Q9WTR1_S47s_1 | Trpv2_Q9W | 1.06 | 1.25 | 0.53 | 0.87 | 0.67 | 0.95 | 0.33 | 0.68 | 0.36 | 0.85 | 0.26 | 0.76 | -0.38 | 0.85 | 0.33 | 0.73 | -0.27 | -0.37 | -0.545 | 0.141764 | 0.30144 | 0.19306 | 0.989223 | 0.997988 | 0.994666 |
| Trpv2_Q9WTR1_T14t_1 | Trpv2_Q9W | 0.85 | 1.14 | 1.84 | 1.57 | 1.63 | 0.59 | 0.51 | 1.88 | 0.82 | 0.41 | 1.79 | 1.11 | -0.23 | 0.4 | 1.74 | 1.51 | -0.1975 | -0.3175 | -0.495 | 0.325257 | 0.623153 | 0.402817 | 0.989223 | 0.997988 | 0.994666 |
| Trrap_F7CGG2_S1350s_1 | Trrap_F7C | -2.11 | -2.66 | -2.76 | -3.69 | -2.21 | -1.89 | -2.41 | -2.74 | -2.34 | -1.88 | -1.75 | -1.85 | -0.36 | -1.7 | -1.57 | -1.84 | 0.4925 | 0.85 | 1.4375 | 0.010511 | 0.202398 | 0.040535 | 0.989223 | 0.997988 | 0.994666 |
| Tsc1_Q9EP53_S1068s_1 | Tsc1_Q9EP | -0.2 | 0.08 | -0.78 | -0.94 | -0.72 | -0.68 | -0.24 | -0.47 | -0.94 | -0.18 | -1.31 | -1.09 | -0.29 | 0.13 | -1.03 | -0.49 | -0.0675 | -0.42 | 0.04 | 0.912262 | 0.817792 | 0.256877 | 0.995762 | 0.997988 | 0.999507 |
| Tsc1_Q9EP53_S1133s_1 | Tsc1_Q9EP | -0.55 | -0.29 | -0.33 | -0.01 | -0.16 | -0.32 | -0.01 | -0.04 | -0.11 | -0.07 | -0.39 | -0.37 | -0.4 | -0.03 | -0.14 | -0.24 | 0.1625 | 0.06 | 0.0925 | 0.7123 | 0.46562 | 0.796777 | 0.989223 | 0.997988 | 0.996966 |
| Tsc1_Q9EP53_S321s_1 | Tsc1_Q9EP | -0.01 | -0.17 | 0.22 | 0.25 | 0.16 | 0.05 | 0.14 | 0.03 | -0.18 | -0.2 | -0.08 | -0.01 | -0.15 | 0.03 | -0.41 | 0.25 | 0.0225 | -0.19 | -0.1425 | 0.59389 | 0.914193 | 0.398749 | 0.997335 | 0.997988 | 0.995259 |
| Tsc1_Q9EP53_S511s_1 | Tsc1_Q9EP | -1.12 | -0.26 | -1.23 | -3.11 | -1.67 | 0.08 | -1.81 | -1.85 | 1.39 | -0.26 | -1.41 | -1.59 | 1.6 | -0.98 | -0.82 | -0.47 | 0.1175 | 0.9625 | 1.2625 | 0.118965 | 0.861214 | 0.255012 | 0.996493 | 0.997988 | 0.994666 |
| Tsc1_Q9EP53_S580s_1 | Tsc1_Q9EP | -0.63 | -0.29 | 1.26 | 1.41 | -0.39 | 0.01 | 0.98 | 1.78 | -0.89 | 0.18 | 1.23 | 1.34 | -0.83 | 0.93 | -0.16 | 0.6 | 0.1575 | 0.0275 | -0.3025 | 0.615599 | 0.804738 | 0.967197 | 0.995762 | 0.997988 | 0.995259 |
| Tsc2_Q3UHB2_S1132s_1 | Tsc2_Q3U | -2.12 | -1.97 | -1.5 | -1.82 | -1.41 | -2.19 | -1.87 | -1.75 | -1.82 | -2.2 | -0.49 | -1.6 | -0.07 | -1.8 | -1.65 | -1.52 | 0.0475 | 0.325 | 0.5925 | 0.181236 | 0.854322 | 0.415 | 0.996349 | 0.997988 | 0.994666 |
| Tsc2_Q3UHB2_S1254s_1 | Tsc2_Q3U | -1.14 | 0.25 | -0.78 | -0.17 | 0.36 | -1.19 | -0.68 | -0.7 | -1.11 | -1.32 | -0.68 | -0.32 | 0.64 | 0.1 | -0.52 | 0.65 | -0.0925 | -0.3975 | 0.6775 | 0.128424 | 0.828067 | 0.316019 | 0.995762 | 0.997988 | 0.994666 |
| Tsc2_Q3UHB2_S1366s_1 | Tsc2_Q3U | 1.93 | 0.22 | -0.22 | -0.73 | 1.29 | 1.74 | 0.18 | 0.12 | 0.41 | 0.55 | 0.71 | 0.75 | -0.69 | 1.3 | -0.14 | -0.4 | 0.5325 | 0.305 | -0.2825 | 0.667741 | 0.405223 | 0.576142 | 0.989223 | 0.997988 | 0.996159 |
| Tsc2_Q3UHB2_S1413s_1 | Tsc2_Q3U | -0.19 | 0.27 | 0.45 | 0.24 | 0.43 | -0.53 | 0.15 | -0.37 | -0.94 | -0.46 | -0.51 | 0.35 | -0.6 | 0.55 | 0.2 | 0.08 | -0.2725 | -0.5825 | -0.135 | 0.676587 | 0.360526 | 0.101174 | 0.989223 | 0.997988 | 0.996159 |
| Tsc2_Q3UHB2_S664s_1 | Tsc2_Q3U | 0.39 | 0.3 | 0.56 | 0.78 | 0.66 | 0.38 | 0.82 | 0.75 | 0.39 | 0.61 | 0.26 | 0.73 | 0.56 | 0.64 | 0.47 | 0.73 | 0.145 | -0.01 | 0.0925 | 0.70549 | 0.523249 | 0.966378 | 0.989223 | 0.997988 | 0.996159 |
| Tsc2_Q3UHB2_S981s_1 | Tsc2_Q3U | -1.91 | -1.57 | -0.97 | -0.73 | -0.82 | -1 | -1.15 | -0.72 | -1.83 | -1.44 | -0.2 | -0.27 | -1.93 | -1.36 | -0.8 | -1.03 | 0.3725 | 0.36 | 0.015 | 0.968502 | 0.244422 | 0.451104 | 0.989223 | 0.997988 | 0.999507 |
| Tsc2_Q3UHB2_T1271t_1 | Tsc2_Q3U | 0.97 | 0.81 | 0.16 | 0.66 | 0.68 | 1.13 | 0.75 | -0.35 | 0.48 | 0.98 | -1.21 | 0.95 | -0.17 | 0.59 | 0.3 | 0.59 | -0.0975 | -0.35 | -0.3225 | 0.307881 | 0.786801 | 0.499133 | 0.995762 | 0.997988 | 0.994666 |
| Tsc22d2_E9Q7M2_S587s_1 | Tsc22d2_E9 | -1.4 | -0.28 | -1.05 | -1.36 | -0.01 | -1.37 | -0.62 | -1.8 | -1.89 | -2.08 | -2.05 | -0.6 | -0.45 | 0.34 | -0.99 | -1.53 | 0.0725 | -0.6325 | 0.365 | 0.435554 | 0.870374 | 0.163838 | 0.996493 | 0.997988 | 0.994666 |
| Tsc22d4_D3YZZ4_S38s_1 | Tsc22d4_D | -0.56 | -0.52 | -1.2 | -1.32 | -0.56 | -0.54 | 0.19 | -1.3 | -0.53 | 0.41 | 0.62 | -1.38 | 0.63 | -1.29 | -0.24 | -0.82 | 0.3475 | 0.68 | 0.47 | 0.311476 | 0.353622 | 0.180066 | 0.989223 | 0.997988 | 0.994666 |
| Tsc22d4_Q9EQN3_S62s_1 | Tsc22d4_Q | 0.92 | 0.89 | 1.16 | 0.69 | -0.35 | 0.71 | 0.26 | 0.72 | 1.28 | 1.12 | 0.5 | 0.73 | 1.56 | 0.2 | 1.24 | 0.97 | -0.58 | -0.0075 | 0.0775 | 0.820225 | 0.075262 | 0.977395 | 0.989223 | 0.997988 | 0.999507 |
| Tsnax_Q9QZE7_T165t_1 | Tsnax_Q9Q | 1 | 0.36 | 0.65 | 0.91 | -0.91 | 1.05 | -0.53 | 0.02 | 2.53 | 0.85 | -0.29 | 0.11 | 2.64 | -1.49 | 0.17 | -0.04 | -0.8225 | 0.07 | -0.41 | 0.597025 | 0.078371 | 0.905083 | 0.989223 | 0.997988 | 0.995259 |
| Tspan13_Q9D8C2_S143s_1 | Tspan13_Q | -0.29 | -0.61 | -0.9 | -0.18 | -0.81 | 0.04 | 0.62 | 1.05 | 0.11 | 0.34 | 0.98 | -0.39 | 0.2 | -0.14 | 0.58 | 0.12 | 0.72 | 0.755 | 0.685 | 0.038179 | 0.107325 | 0.05203 | 0.989223 | 0.997988 | 0.994666 |
| Tspyl1_O88852_S51s_1 | Tspyl1_O88 | 0.16 | -0.06 | -0.44 | 0 | 0.28 | 0.24 | -1.07 | -0.46 | 0.33 | 0.06 | 0.56 | -0.63 | 0.82 | 0.38 | -0.39 | -0.87 | -0.1675 | 0.165 | 0.07 | 0.862588 | 0.63244 | 0.606019 | 0.989223 | 0.997988 | 0.999507 |
| Tspyl2_Q7TQI8_S504s_1 | Tspyl2_Q7T | 0.54 | -0.32 | -1.52 | -2.9 | 0.37 | 0.68 | -0.77 | -0.82 | -1.35 | 0.63 | -1.07 | -2 | -0.52 | -0.89 | -1.24 | -2.94 | 0.915 | 0.1025 | -0.3475 | 0.669592 | 0.234788 | 0.901921 | 0.989223 | 0.997988 | 0.996159 |
| Tspyl4_Q8VD63_S116s_1 | Tspyl4_Q8V | -2.16 | -2.54 | -1.03 | -2.41 | -1.76 | -1.51 | -2.5 | -0.88 | -1.91 | -1.66 | -1.07 | -1.37 | -0.46 | -1.47 | -0.83 | -0.81 | 0.3725 | 0.5325 | 1.1425 | 0.018147 | 0.4155 | 0.194628 | 0.989223 | 0.997988 | 0.994666 |
| Tspyl4_Q8VD63_S397s_1 | Tspyl4_Q8V | -1.97 | -3.35 | -3.43 | -4.1 | -2.52 | -2.13 | -3.24 | -2.97 | -0.76 | -2.64 | -2.98 | -2.54 | -0.06 | -2.97 | -1.95 | -2.86 | 0.4975 | 0.9825 | 1.2525 | 0.107788 | 0.306393 | 0.134386 | 0.989223 | 0.997988 | 0.994666 |
| Tspyl4_Q8VD63_S87s_1 | Tspyl4_Q8V | -1.07 | -0.72 | -1.88 | -1.59 | -2.11 | -1.74 | -2.75 | -2.03 | -1.26 | -2.9 | -1.43 | -1.69 | -1.24 | -0.84 | -1.56 | -2.52 | -0.8425 | -0.505 | -0.225 | 0.607681 | 0.032107 | 0.26706 | 0.989223 | 0.997988 | 0.995259 |
| Tssc4_D3Z6S3_T124t_1 | Tssc4_D3Z | -1.88 | -1.43 | -1.79 | -2.33 | -1.39 | -2.21 | -1.74 | -2.08 | -1.78 | -1.53 | -0.44 | -1.86 | -0.5 | -1.21 | -1.55 | -1.9 | 0.0025 | 0.455 | 0.5675 | 0.149929 | 0.993131 | 0.251112 | 0.999187 | 0.997988 | 0.994666 |
| Ttbk1_Q6PCN3_S1222s_1 | Ttbk1_Q6P | -2.41 | -3.41 | -2.55 | -4.19 | -1.66 | -1.99 | -2.88 | -2.44 | -3.02 | -2.89 | -0.55 | -3.52 | -1.89 | -1.49 | -3.81 | -3.26 | 0.8975 | 0.645 | 0.5275 | 0.407678 | 0.075041 | 0.370997 | 0.989223 | 0.997988 | 0.994666 |
| Ttbk1_Q6PCN3_S433s_1 | Ttbk1_Q6P | -0.24 | -1.22 | -1 | -0.89 | -0.05 | -0.28 | -0.34 | -0.58 | -0.67 | -0.93 | -0.22 | -0.61 | -1.64 | -0.35 | -1.06 | -1.12 | 0.525 | 0.23 | -0.205 | 0.57457 | 0.081968 | 0.447138 | 0.989223 | 0.997988 | 0.995259 |
| Ttbk1_Q6PCN3_S441s_1 | Ttbk1_Q6P | 0.4 | 1.16 | 0.35 | 0.07 | 0.66 | 0.15 | 0.02 | 0.05 | 2 | 1.29 | 0.16 | -0.09 | 1.77 | 1.39 | 0.08 | -0.09 | -0.275 | 0.345 | 0.2925 | 0.557241 | 0.371801 | 0.50326 | 0.989223 | 0.997988 | 0.995259 |
| Ttbk1_Q6PCN3_S441s_2 | Ttbk1_Q6P | -0.69 | -1.82 | -1.27 | -1.15 | -0.51 | -0.91 | -0.7 | -0.46 | -1.14 | -1.76 | 0.54 | -0.76 | -1.57 | -0.26 | -1.98 | -0.86 | 0.5875 | 0.4525 | 0.065 | 0.881961 | 0.064044 | 0.381916 | 0.989223 | 0.997988 | 0.999507 |
| Ttbk1_Q6PCN3_S456s_1 | Ttbk1_Q6P | 0.05 | 0.06 | 0.01 | -0.09 | -1.31 | 0.51 | -0.89 | -0.46 | 2.49 | 0.24 | -1.31 | -0.42 | 2.01 | -1.93 | -0.2 | -0.53 | -0.545 | 0.2425 | -0.17 | 0.815254 | 0.176233 | 0.740267 | 0.989223 | 0.997988 | 0.999507 |
| Ttbk1_Q6PCN3_S484s_1 | Ttbk1_Q6P | -0.47 | -0.45 | -0.02 | -0.19 | -0.26 | -0.31 | -0.21 | -0.36 | -0.43 | -0.23 | 0 | -0.1 | -0.33 | 0 | -0.16 | -0.23 | -0.0025 | 0.0925 | 0.1025 | 0.679767 | 0.990583 | 0.694035 | 0.999187 | 0.997988 | 0.996159 |
| Ttbk1_Q6PCN3_S638s_1 | Ttbk1_Q6P | 0.5 | 0.46 | -0.28 | -0.66 | -0.84 | 0.1 | -1.04 | -0.41 | 0.66 | 0.76 | -0.55 | -0.57 | 1.25 | 0.17 | -0.27 | -0.97 | -0.5525 | 0.07 | 0.04 | 0.937787 | 0.163233 | 0.875766 | 0.989223 | 0.997988 | 0.999507 |
| Ttbk1_Q6PCN3_S655s_1 | Ttbk1_Q6P | -1.21 | -1.98 | -1.4 | -1.7 | -2.13 | -1.05 | -2.11 | -1.36 | 0.75 | -1.62 | -1.19 | -1.14 | 0.77 | -2.08 | -1.11 | -1.03 | -0.09 | 0.7725 | 0.71 | 0.231427 | 0.785458 | 0.160829 | 0.995762 | 0.997988 | 0.994666 |
| Ttbk1_Q6PCN3_S827s_1 | Ttbk1_Q6P | -0.13 | 0.02 | 0.06 | -0.21 | 0.76 | -0.37 | 0.24 | 1.48 | 0.27 | 0.19 | -0.6 | -0.51 | 0.43 | 0.5 | -1.08 | 0.21 | 0.5925 | -0.0975 | 0.08 | 0.836436 | 0.149421 | 0.732901 | 0.989223 | 0.997988 | 0.999507 |
| Ttbk1_Q6PCN3_T325t_1 | Ttbk1_Q6P | 0.63 | 0.79 | 0.16 | 0.6 | 0.71 | 0.59 | 0.18 | 0.6 | 0.55 | 0.79 | -0.2 | 0.41 | 0.38 | 0.32 | 0.16 | 0.35 | -0.025 | -0.1575 | -0.2425 | 0.350069 | 0.917657 | 0.59447 | 0.997335 | 0.997988 | 0.994666 |
| Ttbk1_Q6PCN3_T444t_2 | Ttbk1_Q6P | -0.69 | -1.82 | -1.27 | -1.15 | -0.51 | -0.91 | -0.7 | -0.46 | -1.14 | -1.76 | 0.54 | -0.76 | -1.57 | -0.26 | -1.98 | -0.86 | 0.5875 | 0.4525 | 0.065 | 0.881961 | 0.064044 | 0.381916 | 0.989223 | 0.997988 | 0.999507 |
| Ttbk2_Q3UVR3_S1181s_1 | Ttbk2_Q3U | -0.68 | -0.55 | -1.4 | -1.28 | -0.51 | -1.55 | -1.75 | -1.59 | 0.35 | -1.43 | -0.9 | -0.76 | 0.96 | -0.5 | -0.68 | -1.19 | -0.3725 | 0.2925 | 0.625 | 0.217037 | 0.307852 | 0.493055 | 0.989223 | 0.997988 | 0.994666 |
| Ttc14_E9PXW3_S570s_1 | Ttc14_E9P | -0.76 | 0.32 | -0.22 | -0.45 | -1.99 | -0.21 | -1.3 | -0.34 | 1.36 | -0.21 | -0.87 | -1.25 | 1.6 | -0.28 | 0.08 | -0.59 | -0.6825 | 0.035 | 0.48 | 0.354479 | 0.151447 | 0.951064 | 0.989223 | 0.997988 | 0.994666 |
| Ttc26_E0CXQ7_S14s_1 | Ttc26_E0C | 0.5 | 0.14 | -0.45 | -1.23 | 0.06 | 0.32 | -1.14 | -1.08 | 0.22 | 1.01 | -0.38 | -1.06 | 0.61 | -0.02 | -0.64 | -0.3 | -0.2 | 0.2075 | 0.1725 | 0.702011 | 0.685473 | 0.701418 | 0.994624 | 0.997988 | 0.996159 |
| Ttc3_G5E8T2_S1903s_1 | Ttc3_G5E8 | -1.1 | -0.61 | -1.3 | -0.27 | -1.24 | -1.14 | -0.59 | -2.1 | -0.86 | -0.28 | -0.85 | -1.11 | -0.07 | -1.42 | 0.03 | -1.84 | -0.4475 | 0.045 | -0.005 | 0.991999 | 0.257443 | 0.888449 | 0.989223 | 0.997988 | 1 |

| | | | | | | | | | | | | | | | | | | | | | | | | | | |
|---|---|---|---|---|---|---|---|---|---|---|---|---|---|---|---|---|---|---|---|---|---|---|---|---|---|---|
| Ttc7b_E9Q6P5_S158s_1 | Ttc7b_E9Q | -0.52 | 0.6 | -0.46 | -2.6 | 0.41 | -1.89 | 0.46 | 0.12 | -0.78 | -0.41 | -0.71 | -0.88 | -0.53 | -0.16 | -1.63 | -1.07 | 0.52 | 0.05 | -0.1025 | 0.877978 | 0.501898 | 0.935491 | 0.989223 | 0.997988 | 0.999507 |
| Ttc7b_E9Q6P5_S160s_1 | Ttc7b_E9Q | -1.34 | -0.57 | -0.41 | -0.24 | -0.25 | -0.9 | -0.12 | 0.12 | -1.1 | -0.88 | -0.46 | 0.31 | 0.16 | 0.21 | 0.38 | 0.38 | 0.3525 | 0.1075 | 0.9225 | 0.012867 | 0.306898 | 0.78515 | 0.989223 | 0.997988 | 0.994666 |
| Ttc7b_E9Q6P5_S202s_1 | Ttc7b_E9Q | 1.06 | 0.76 | 1.3 | 1.37 | 1.61 | 1.05 | 1.56 | 1.57 | 0.6 | 1.55 | 0.69 | 1.3 | 0.01 | 1.59 | 1.09 | 1.32 | 0.325 | -0.0875 | -0.12 | 0.755959 | 0.212457 | 0.775129 | 0.989223 | 0.997988 | 0.9971 |
| Ttc7b_E9Q6P5_S629s_1 | Ttc7b_E9Q | 0.03 | 0.49 | 0.32 | -0.07 | 0.77 | 0.03 | 0.43 | 0.58 | 0.65 | 0.04 | 0.79 | 0.49 | -0.19 | 0.72 | 0.29 | 0.08 | 0.26 | 0.3 | 0.0325 | 0.912499 | 0.322792 | 0.279516 | 0.989223 | 0.997988 | 0.999507 |
| Ttc7b_E9Q6P5_S630s_1 | Ttc7b_E9Q | -0.04 | 0.11 | -0.16 | 0.17 | 0.87 | -0.08 | 0.11 | 0.32 | 0.29 | -0.22 | 0.33 | 0.29 | -0.53 | 0.64 | 0.38 | 0.45 | 0.285 | 0.1525 | 0.215 | 0.505759 | 0.296247 | 0.526435 | 0.989223 | 0.997988 | 0.994666 |
| Ttc7b_E9Q6P5_S678s_1 | Ttc7b_E9Q | 0.66 | 0.98 | 1.51 | 1.44 | 1 | 0.9 | 1.55 | 1.53 | 0.96 | 1.42 | 1.67 | 1.45 | 0.88 | 0.96 | 1.31 | 1.1 | 0.0975 | 0.2275 | -0.085 | 0.77019 | 0.739526 | 0.445589 | 0.995762 | 0.997988 | 0.9971 |
| Ttc9_Q3V038_S15s_1 | Ttc9_Q3V0 | 0.45 | -0.05 | 0.39 | -0.06 | -0.35 | -0.53 | 0.08 | -0.14 | -0.6 | 0.01 | -0.03 | 0.02 | -0.27 | -0.64 | -0.23 | 0.04 | -0.4175 | -0.3325 | -0.4575 | 0.123897 | 0.118259 | 0.230695 | 0.989223 | 0.997988 | 0.994666 |
| Ttc9b_Q9D6E4_S226s_1 | Ttc9b_Q9D | -0.09 | -0.3 | 0.24 | -0.44 | -0.49 | 0.3 | -0.8 | -0.64 | 1.61 | -0.15 | -0.44 | -0.28 | 1.38 | -1.52 | 0.27 | -0.81 | -0.26 | 0.3325 | -0.0225 | 0.969816 | 0.40567 | 0.490905 | 0.989223 | 0.997988 | 0.999507 |
| Ttc9b_Q9D6E4_S27s_1 | Ttc9b_Q9D | 0.34 | 0 | -0.29 | -1.17 | 0.06 | 0.26 | 0.34 | -0.12 | 0.21 | 0.15 | 0.64 | -0.28 | -0.46 | 0.91 | -0.16 | 0.01 | 0.415 | 0.46 | 0.355 | 0.419472 | 0.246158 | 0.243931 | 0.989223 | 0.997988 | 0.994666 |
| Tti1_Q91V83_S783s_1 | Tti1_Q91V | 0.45 | -0.13 | 0.83 | 0.36 | -1.96 | 0.64 | -0.42 | 0.24 | 1.28 | 0.14 | -0.37 | 0.72 | 0.57 | -0.35 | 0.6 | 0.59 | -0.7525 | 0.065 | -0.025 | 0.941529 | 0.190871 | 0.872228 | 0.989223 | 0.997988 | 0.999507 |
| Ttll12_F2Z423_S11s_1 | Ttll12_F2Z4 | -2.1 | -0.55 | -0.82 | -1.17 | -1.13 | -1.89 | -1.09 | -1.22 | 3.64 | -2.26 | -0.68 | -0.88 | 2.57 | 1.93 | -0.74 | -0.54 | -0.1725 | 1.115 | 1.965 | 0.033397 | 0.650021 | 0.349259 | 0.990358 | 0.997988 | 0.994666 |
| Ttll7_E9Q6C6_S410s_1 | Ttll7_E9Q6 | 0.5 | -0.11 | 0.6 | -0.5 | -0.25 | -0.23 | 0.19 | 0.13 | -0.41 | 0.17 | -0.3 | 0.51 | 0.65 | 0.49 | -0.41 | 0.21 | -0.1625 | -0.13 | 0.1125 | 0.761205 | 0.59835 | 0.712353 | 0.989223 | 0.997988 | 0.9971 |
| Ttll9_D6RG93_S81s_2 | Ttll9_D6RG | 1.14 | -0.42 | -1.21 | 2.53 | 0.16 | 0.78 | -1.8 | -0.29 | -1.1 | 0.32 | 5.15 | -0.8 | -0.71 | -1.04 | -1.2 | -2.32 | -0.7975 | 0.3825 | -1.8275 | 0.043044 | 0.368921 | 0.7937 | 0.989223 | 0.997988 | 0.994666 |
| Ttll9_D6RG93_Y87y_2 | Ttll9_D6RG | 1.14 | -0.42 | -1.21 | 2.53 | 0.16 | 0.78 | -1.8 | -0.29 | -1.1 | 0.32 | 5.15 | -0.8 | -0.71 | -1.04 | -1.2 | -2.32 | -0.7975 | 0.3825 | -1.8275 | 0.043044 | 0.368921 | 0.7937 | 0.989223 | 0.997988 | 0.994666 |
| Ttyh1_Q9D3A9_S449s_1 | Ttyh1_Q9D | 0.27 | 0.4 | 0.4 | 0.47 | 1.22 | 0.02 | 0.92 | 0.34 | -0.04 | 0.43 | 0.28 | 0.29 | -0.94 | 0.82 | 0.61 | 0.62 | 0.24 | -0.145 | -0.1075 | 0.794494 | 0.431578 | 0.515052 | 0.989223 | 0.997988 | 0.999507 |
| Ttyh2_Q3TH73_S442s_1 | Ttyh2_Q3T | -1.23 | -0.37 | -0.46 | -0.63 | -0.74 | -0.42 | -0.43 | -0.48 | -1.31 | -0.91 | -0.3 | -0.34 | -0.77 | -0.32 | 0.29 | -0.01 | 0.155 | -0.0425 | 0.47 | 0.183738 | 0.55199 | 0.898407 | 0.989223 | 0.997988 | 0.994666 |
| Ttyh3_Q6P5F7_S496s_1 | Ttyh3_Q6P | 0.23 | 0.74 | 0.53 | 0.13 | 0.7 | 0.23 | 0.79 | 0.25 | -0.52 | 0.97 | -0.91 | 0.58 | 0.23 | -0.64 | 0.67 | 0.77 | 0.085 | -0.3775 | -0.15 | 0.686414 | 0.73981 | 0.409647 | 0.995762 | 0.997988 | 0.996159 |
| Ttyh3_Q6P5F7_S519s_1 | Ttyh3_Q6P | -0.11 | -0.18 | 0.42 | 0.28 | 0.33 | -0.38 | -0.19 | -0.07 | -0.13 | 0.17 | -0.09 | -0.03 | 1.01 | -0.05 | -0.39 | -0.28 | -0.18 | -0.1225 | -0.03 | 0.935573 | 0.493198 | 0.616298 | 0.989223 | 0.997988 | 0.999507 |
| Ttyh3_Q6P5F7_S522s_1 | Ttyh3_Q6P | 0.54 | 0.05 | -0.95 | -0.67 | 0.85 | -0.12 | -0.37 | -0.79 | 0.58 | 0.51 | 0.39 | -0.57 | 0.87 | 0.71 | -0.6 | -0.42 | 0.15 | 0.485 | 0.3975 | 0.420427 | 0.741233 | 0.26923 | 0.995762 | 0.997988 | 0.994666 |
| Ttyh3_Q6P5F7_T427t_1 | Ttyh3_Q6P | -0.35 | 0.72 | -0.31 | 1.06 | 1.21 | 0.51 | -1.53 | 0.59 | 0.19 | -0.56 | 0.35 | -0.32 | 0.54 | 0.62 | 0.16 | -0.1 | -0.085 | -0.365 | 0.025 | 0.950379 | 0.890893 | 0.385713 | 0.996493 | 0.997988 | 0.999507 |
| Tuba4a_P68368_S158s_1 | Tuba4a_P6 | 0.27 | 0.93 | -0.03 | -0.5 | 0.79 | 0.58 | -0.56 | -0.29 | 1.32 | 1.04 | 0.41 | -0.03 | 0.77 | 1.96 | -0.25 | -0.48 | -0.0375 | 0.5175 | 0.3325 | 0.570149 | 0.929015 | 0.236641 | 0.997639 | 0.997988 | 0.995259 |
| Tuba4a_P68368_S277s_1 | Tuba4a_P6 | 0.92 | 2.64 | 2.24 | 0.6 | 2.15 | 1.75 | 2.23 | 1.52 | -0.02 | 3.1 | 0.42 | 2.15 | -0.04 | 0.31 | 2.17 | 2.56 | 0.3125 | -0.1875 | -0.35 | 0.633571 | 0.522076 | 0.812538 | 0.989223 | 0.997988 | 0.995259 |
| Tuba4a_P68368_S340s_1 | Tuba4a_P6 | -1.65 | -1.85 | -1.47 | -0.89 | -1.22 | -0.65 | -0.78 | -1.01 | -2.11 | -1.04 | -1.3 | -1.65 | -1.46 | -0.93 | -1.46 | -1.23 | 0.55 | -0.06 | 0.195 | 0.52296 | 0.073191 | 0.857132 | 0.989223 | 0.997988 | 0.994666 |
| Tuba4a_P68368_T223t_1 | Tuba4a_P6 | -0.56 | 0.07 | 1.05 | 0.91 | -0.59 | -0.63 | 0.78 | 0.56 | -1.38 | -0.19 | 0.54 | 0.9 | 0.13 | -0.69 | 0.36 | 0.75 | -0.3375 | -0.4 | -0.23 | 0.624225 | 0.494552 | 0.495258 | 0.989223 | 0.997988 | 0.995259 |
| Tuba4a_P68368_Y103y_1 | Tuba4a_P6 | -1.97 | -0.27 | -0.37 | -2.42 | -2.13 | -0.12 | -1.94 | -1.29 | 1.5 | -1.13 | 1.22 | -0.99 | 2.46 | 1.63 | -1.77 | -1.52 | -0.1125 | 1.4075 | 1.4575 | 0.188082 | 0.859062 | 0.103487 | 0.996349 | 0.997988 | 0.994666 |
| Tuba4a_P68368_Y357y_1 | Tuba4a_P6 | 0.21 | 1 | 0.62 | -0.73 | 0.54 | 0.86 | 0.83 | 0.01 | 0.79 | 1.4 | -0.03 | 0.36 | 0.45 | 0.38 | 0.5 | 1.21 | 0.285 | 0.355 | 0.36 | 0.398864 | 0.486006 | 0.44895 | 0.989223 | 0.997988 | 0.994666 |
| Tubb3_Q9ERD7_S48s_1 | Tubb3_Q9I | -0.94 | -0.34 | -0.09 | 0.22 | -1.01 | -0.54 | -0.42 | -0.34 | -0.8 | -0.78 | -0.76 | -0.65 | -0.13 | -0.15 | -0.6 | -0.19 | -0.29 | -0.46 | 0.02 | 0.949869 | 0.358446 | 0.14062 | 0.989223 | 0.997988 | 0.999507 |
| Tubb4b_P68372_S338s_1 | Tubb4b_P6 | -1.92 | -0.85 | 0.5 | -0.2 | -0.88 | -1.96 | -0.35 | -0.81 | -0.29 | -0.49 | -0.67 | -0.92 | -0.97 | -0.1 | -1.82 | -0.38 | -0.3825 | 0.025 | -0.2 | 0.733471 | 0.49743 | 0.960102 | 0.989223 | 0.997988 | 0.9971 |
| Tubb4b_P68372_T219t_1 | Tubb4b_P6 | -1.61 | -0.85 | -1.02 | -0.75 | -0.16 | -1.29 | -0.83 | -0.55 | -0.9 | -0.7 | -1.04 | -0.6 | -1.02 | -0.53 | -0.5 | -0.8 | 0.35 | 0.2475 | 0.345 | 0.2584 | 0.290388 | 0.375348 | 0.989223 | 0.997988 | 0.994666 |
| Tubgcp3_P58854_Y542y_1 | Tubgcp3_P | 1.52 | 1.39 | 0.17 | 0.56 | 1.46 | 1.16 | 0.61 | 0.57 | 1.07 | 1.19 | 1.05 | 0.78 | 1.45 | 1.32 | 0.77 | 0.59 | 0.04 | 0.1125 | 0.1225 | 0.756991 | 0.916454 | 0.749638 | 0.997335 | 0.997988 | 0.9971 |
| Tubgcp5_Q8BKN5_S182s_1 | Tubgcp5_C | -0.18 | 1.3 | -0.74 | -0.88 | 0.08 | -2.39 | -1.29 | -0.87 | -2.2 | -1.23 | 0.21 | 0.06 | 0.13 | -0.58 | -0.43 | -0.74 | -0.9925 | -0.665 | -0.28 | 0.581104 | 0.142756 | 0.344186 | 0.989223 | 0.997988 | 0.995259 |
| Tusc1_Q673H1_S146s_1 | Tusc1_Q67 | -0.49 | -0.57 | -0.18 | -1.02 | -1.79 | 0.2 | -1.19 | -1.15 | 1.36 | -0.51 | -1.1 | -0.71 | 1.8 | -1.17 | -0.91 | -2.01 | -0.4175 | 0.325 | -0.0075 | 0.992001 | 0.343432 | 0.547115 | 0.989223 | 0.997988 | 1 |
| Tusc5_Q8C838_S74s_1 | Tusc5_Q8C | 0.92 | 0.77 | 0.28 | -0.4 | 0.32 | 0.08 | -0.34 | -0.64 | 1.32 | 2.51 | 0.88 | 0.26 | 1.97 | -0.45 | 0.6 | 1.1 | -0.5375 | 0.85 | 0.4125 | 0.45549 | 0.162742 | 0.13044 | 0.989223 | 0.997988 | 0.994666 |
| Twf1_Q91YR1_T349t_1 | Twf1_Q91Y | -0.44 | -0.53 | 0.41 | 0.42 | -1.14 | 0.19 | -1.2 | 0.21 | 0.77 | 0.19 | 1.07 | 0.53 | 0.68 | -1.27 | 0.46 | 0.82 | -0.45 | 0.675 | 0.2075 | 0.689641 | 0.324326 | 0.072551 | 0.989223 | 0.997988 | 0.996159 |
| Twf2_Q9Z0P5_S165s_1 | Twf2_Q9ZC | 1 | -0.58 | 0.21 | 0.51 | 0.5 | 1.02 | 0.37 | 0.63 | -0.37 | -0.17 | 0.58 | 0.79 | -1.21 | -0.17 | 0.02 | 0.27 | 0.345 | -0.0775 | -0.5575 | 0.235815 | 0.348068 | 0.855208 | 0.989223 | 0.997988 | 0.994666 |
| Txlna_Q6PAM1_S515s_1 | Txlna_Q6P. | 0.2 | 0.66 | 0.68 | 0.57 | 0.29 | 0.34 | -0.27 | 0.38 | 0.86 | 0.7 | 1.34 | 0.72 | 0.88 | 0.73 | 0.23 | -0.58 | -0.3425 | 0.3775 | -0.2125 | 0.567247 | 0.188978 | 0.164499 | 0.989223 | 0.997988 | 0.995259 |
| Txlna_Q6PAM1_S515s_2 | Txlna_Q6P. | -0.47 | -0.04 | -0.21 | 0.22 | -0.01 | -1.48 | 0.03 | -0.38 | -0.52 | 0.14 | 0.87 | 0.46 | 0.15 | 1.21 | 0.47 | 0.67 | -0.335 | 0.3625 | 0.75 | 0.037321 | 0.380913 | 0.3083 | 0.989223 | 0.997988 | 0.994666 |
| Txnrd3_Q99MD6_S16s_1 | Txnrd3_Q9 | 0.03 | -0.07 | 0.01 | -0.36 | -0.77 | -0.75 | -0.52 | -0.95 | -0.84 | -0.21 | -0.27 | -0.93 | 0.56 | -0.27 | -0.37 | -0.33 | -0.65 | -0.465 | -0.005 | 0.98671 | 0.013311 | 0.108937 | 0.989223 | 0.997988 | 1 |
| Txnrd3_Q99MD6_S50s_1 | Txnrd3_Q9 | -0.08 | -0.41 | 1.03 | 1.55 | -1.31 | 0.43 | 0.56 | 1.64 | -0.85 | -0.5 | 2.51 | 1.48 | 0.49 | -1.55 | 1.28 | 0.93 | -0.1925 | 0.1375 | -0.235 | 0.737847 | 0.776094 | 0.867491 | 0.995762 | 0.997988 | 0.9971 |
| Txnrd3_Q99MD6_T15t_1 | Txnrd3_Q9 | 0.03 | -0.07 | 0.01 | -0.36 | -0.77 | -0.75 | -0.52 | -0.95 | -0.84 | -0.21 | -0.27 | -0.93 | 0.56 | -0.27 | -0.37 | -0.33 | -0.65 | -0.465 | -0.005 | 0.98671 | 0.013311 | 0.108937 | 0.989223 | 0.997988 | 1 |
| Tyro3_P55144_S804s_1 | Tyro3_P55 | 0.25 | 0.48 | 1.07 | 0.9 | 0.32 | 0.29 | 0.2 | 0.55 | 0.4 | 0.26 | 1.38 | 1.04 | 0.71 | 0.4 | 0.34 | 0.31 | -0.335 | 0.095 | -0.235 | 0.416732 | 0.209925 | 0.782297 | 0.989223 | 0.997988 | 0.994666 |
| Tyro3_P55144_S855s_1 | Tyro3_P55 | 0.41 | 0.81 | 1.09 | 0.78 | 0.91 | 0.24 | 1.43 | 1.03 | 0.51 | 1.03 | 0.75 | 0.97 | -0.33 | 0.81 | 0.95 | 1.22 | 0.13 | 0.0425 | -0.11 | 0.77418 | 0.671872 | 0.867137 | 0.992876 | 0.997988 | 0.99798 |
| U2af1_Q9D883_S61s_1 | U2af1_Q9I | -0.02 | -1.5 | -0.94 | -1 | -1.95 | -1.21 | -3.29 | -1.97 | -1.87 | -2.42 | -0.69 | -1.77 | 0.75 | -2.43 | -1.08 | -1.12 | -1.24 | -0.8225 | -0.105 | 0.871994 | 0.028615 | 0.097868 | 0.989223 | 0.997988 | 0.999507 |
| U2surp_Q6NV83_S202s_1 | U2surp_Q6 | -3.32 | -4.38 | -3.19 | -2.64 | -2.85 | -5.15 | -3.57 | -3.86 | -6.89 | -2.5 | -3.6 | -3.5 | -2.59 | -4.21 | -4.17 | -3.8 | -0.475 | -0.74 | -0.31 | 0.537256 | 0.393287 | 0.422952 | 0.989223 | 0.997988 | 0.994666 |
| U2surp_Q6NV83_S67s_1 | U2surp_Q6 | -0.77 | -1.17 | -0.79 | -1.86 | -0.39 | -1.65 | -0.82 | -2.03 | -0.78 | -1.73 | -0.6 | -0.71 | 0.38 | -1.54 | -0.94 | -1.13 | -0.075 | 0.1925 | 0.34 | 0.474302 | 0.861023 | 0.607223 | 0.996493 | 0.997988 | 0.994666 |
| U2surp_Q6NV83_S810s_1 | U2surp_Q6 | -1.02 | -1.47 | -1.79 | -1.94 | -1.64 | -2.03 | -1.32 | -1.66 | 0.14 | -0.88 | -0.95 | -1.69 | 0.24 | -1.31 | -1.19 | -1.61 | -0.1075 | 0.71 | 0.5875 | 0.210409 | 0.705638 | 0.116604 | 0.995437 | 0.997988 | 0.994666 |
| Uba1_Q02053_S13s_1 | Uba1_Q02 | -4.5 | -4.14 | -3.45 | -4.38 | -3.33 | -5.21 | -3.56 | -5.08 | -4.93 | -4.01 | -2.64 | -4.42 | -2.86 | -3.14 | -4.41 | -3.84 | -0.1775 | 0.1175 | 0.555 | 0.207001 | 0.723707 | 0.818002 | 0.995437 | 0.997988 | 0.994666 |
| Uba1_Q02053_S810s_1 | Uba1_Q02 | 0.67 | 1.22 | 0.6 | -0.15 | 0.73 | 0.67 | 1.06 | 0.95 | 0.38 | 0.75 | 0.09 | 0.64 | -0.17 | 0.36 | 0.78 | 0.57 | 0.2675 | -0.12 | -0.2 | 0.589875 | 0.403209 | 0.723485 | 0.989223 | 0.997988 | 0.995259 |
| Uba1_Q02053_S816s_1 | Uba1_Q02 | 1.01 | 0.57 | 0.05 | 1.41 | 1.03 | -0.37 | 0.93 | 0.16 | -1.06 | -0.4 | 1.89 | 0.95 | -0.22 | 1.07 | 0.41 | 0.66 | -0.3225 | -0.415 | -0.28 | 0.492947 | 0.449763 | 0.532469 | 0.989223 | 0.997988 | 0.994666 |
| Uba1_Q02053_S835s_1 | Uba1_Q02 | -1.79 | -1.26 | 0.9 | 0.9 | -0.7 | -2.43 | 0.68 | 0.61 | -2.25 | -1.37 | 0.78 | 0.61 | -1.05 | -0.8 | 0.49 | 0.55 | -0.1475 | -0.245 | 0.11 | 0.880704 | 0.867405 | 0.788374 | 0.996493 | 0.997988 | 0.999507 |
| Uba1_Q02053_S876s_1 | Uba1_Q02 | 0.48 | 1.04 | 0.24 | 0.58 | 0.3 | -0.05 | 0.15 | 0.09 | -0.54 | 0.48 | -0.56 | 0.09 | 0.05 | -0.17 | 0.26 | 0.04 | -0.4625 | -0.7175 | -0.54 | 0.073148 | 0.083435 | 0.052502 | 0.989223 | 0.997988 | 0.994666 |
| Uba3_Q3TL72_S363s_1 | Uba3_Q3TI | -0.17 | 0.23 | 0.09 | -0.02 | 0.36 | -0.79 | 0.29 | -0.23 | -0.04 | -0.18 | -0.11 | -0.44 | -0.95 | -0.25 | -0.22 | -0.24 | -0.125 | -0.225 | -0.4475 | 0.132074 | 0.681335 | 0.331586 | 0.993613 | 0.997988 | 0.994666 |
| Ubac1_F6Z1E7_S18s_1 | Ubac1_F6Z | 0.64 | 0.2 | 0.01 | 0.12 | 0.18 | 0.46 | -0.27 | 0.04 | 0.45 | 0.25 | 0.65 | 0.34 | 1.26 | 1.04 | -0.01 | -0.39 | -0.14 | 0.18 | 0.2325 | 0.583838 | 0.588439 | 0.467078 | 0.989223 | 0.997988 | 0.995259 |
| Ubap1_Q8BH48_S146s_1 | Ubap1_Q8I | -0.46 | -0.07 | 0.03 | 0.24 | 0.32 | -0.43 | 0.3 | 0.35 | 0.2 | 0.29 | 0.2 | 0.01 | 0.51 | 1 | 0.4 | 0.44 | 0.2 | 0.24 | 0.6525 | 0.04042 | 0.476176 | 0.333679 | 0.989223 | 0.997988 | 0.994666 |
| Ubap2l_Q80X50_S467s_1 | Ubap2l_Q8 | -1.21 | -1.02 | 0.42 | -0.48 | -1.26 | -1.05 | 0 | -0.1 | -1.57 | -0.66 | -0.2 | 0.39 | -0.11 | -0.45 | 0.11 | 0.5 | -0.03 | 0.0625 | 0.585 | 0.182095 | 0.947305 | 0.903611 | 0.999187 | 0.997988 | 0.994666 |
| Ubap2l_Q80X50_S609s_1 | Ubap2l_Q8 | -1.46 | -1.59 | -0.32 | -0.71 | -1.1 | -1.74 | -0.55 | -0.43 | -1.49 | -1.11 | -0.43 | -0.58 | -0.31 | -0.22 | -0.2 | -0.45 | 0.065 | 0.1175 | 0.725 | 0.057636 | 0.873334 | 0.763717 | 0.996493 | 0.997988 | 0.994666 |
| Ubap2l_Q80X50_S992s_1 | Ubap2l_Q8 | -0.43 | -1.35 | -0.54 | -1.52 | -1.33 | 0.14 | -1.19 | -0.91 | -1.65 | -0.71 | -1.54 | -0.82 | -0.57 | -2.86 | -1.02 | -1.42 | 0.1375 | -0.22 | -0.5075 | 0.350428 | 0.739645 | 0.560283 | 0.995762 | 0.997988 | 0.994666 |
| Ube2e2_H3BJB5_S11s_1 | Ube2e2_H: | -0.99 | -1.68 | -1.48 | -0.78 | -0.45 | -0.63 | -3.21 | -1.39 | -0.96 | -0.75 | 0.66 | -1.37 | -0.39 | -0.78 | -1.51 | -1.79 | -0.1875 | 0.6275 | 0.115 | 0.770529 | 0.752942 | 0.200281 | 0.995762 | 0.997988 | 0.9971 |
| Ube2e3_B2FDH7_S8s_1 | Ube2e3_B: | -5.21 | -5.79 | -4.57 | -2.99 | -4.65 | -3.64 | -3.44 | -2.3 | -3.15 | -2.79 | -2.18 | -5.58 | -1.12 | -3.02 | -7.04 | -5.35 | 1.1325 | 1.215 | 0.5075 | 0.682564 | 0.122298 | 0.17813 | 0.989223 | 0.997988 | 0.996159 |
| Ube2j1_F6SPK0_S210s_1 | Ube2j1_F6: | -0.38 | -0.69 | -0.31 | -0.05 | 0.4 | -0.21 | 0.5 | 0.12 | -0.65 | -0.36 | -0.24 | -0.18 | -0.01 | 0.26 | 0.29 | 0.47 | 0.56 | 9.81E-17 | 0.61 | 0.040478 | 0.05232 | 1 | 0.989223 | 1 | 0.994666 |
| Ube2o_Q6ZPJ3_S110s_1 | Ube2o_Q6. | -0.9 | -1.26 | -1.57 | -1.09 | -0.09 | -0.87 | -0.13 | -0.86 | 0.09 | -0.94 | -1.53 | -0.73 | 1.14 | -0.48 | 0.31 | -0.27 | 0.7175 | 0.4275 | 1.38 | 0.006082 | 0.031745 | 0.267346 | 0.989223 | 0.997988 | 0.994666 |
| Ube2o_Q6ZPJ3_S1196s_1 | Ube2o_Q6. | -0.35 | -0.53 | -0.21 | -1.03 | 0 | -0.59 | -1.01 | -0.49 | -0.53 | -0.43 | 0.73 | -0.88 | -0.78 | 0.53 | -0.55 | -0.54 | 0.0075 | 0.2525 | 0.195 | 0.596628 | 0.979976 | 0.524875 | 0.999187 | 0.997988 | 0.995259 |
| Ube2o_Q6ZPJ3_S317s_1 | Ube2o_Q6. | 0.47 | 0.4 | 0.45 | -0.08 | 0.5 | -0.38 | 0.98 | 0.39 | -0.05 | 0.52 | -0.62 | 0.42 | -0.97 | 0.63 | 0.09 | 0.58 | 0.0625 | -0.2425 | -0.2275 | 0.572905 | 0.846888 | 0.456483 | 0.996349 | 0.997988 | 0.995259 |
| Ube2o_Q6ZPJ3_S376s_1 | Ube2o_Q6. | -0.29 | -1.81 | -1.26 | 0.4 | -2.13 | -0.72 | -0.57 | -0.71 | 0.3 | -2.18 | -1.2 | -0.91 | 0.44 | -0.74 | -1.06 | 0.41 | -0.2925 | -0.2575 | 0.5025 | 0.393213 | 0.60073 | 0.690899 | 0.989223 | 0.997988 | 0.994666 |
| Ube2o_Q6ZPJ3_S510s_1 | Ube2o_Q6. | -3.44 | -3.85 | -4.31 | -6.25 | -3.07 | -4.24 | -4.12 | -4.06 | -5.32 | -3.07 | -1.47 | -4.15 | -2.78 | -1.91 | -4.74 | -4.55 | 0.59 | 0.96 | 0.9675 | 0.254706 | 0.342437 | 0.307188 | 0.989223 | 0.997988 | 0.994666 |
| Ube2o_Q6ZPJ3_S833s_1 | Ube2o_Q6. | 0.71 | 1 | 0.22 | 0.14 | 0.09 | 0.92 | 0.7 | 0.33 | 0.39 | 0.92 | 0.46 | 0.26 | 0.09 | 0.74 | 0.59 | 0.79 | -0.0075 | -0.01 | 0.035 | 0.910657 | 0.98006 | 0.972761 | 0.999187 | 0.997988 | 0.999507 |
| Ube2o_Q6ZPJ3_S836s_1 | Ube2o_Q6. | 0.04 | 0.39 | 0.21 | 0.2 | 0.33 | 0.26 | 0.42 | 0.35 | 0.11 | 0.49 | -0.16 | 0.04 | -0.11 | 0.41 | 0.06 | 0.02 | 0.13 | -0.09 | -0.115 | 0.645447 | 0.52174 | 0.707824 | 0.989223 | 0.997988 | 0.995259 |
| Ube2o_Q6ZPJ3_S893s_1 | Ube2o_Q6. | -0.03 | 0.48 | -0.61 | -1.06 | 0.37 | -0.11 | -0.87 | -0.87 | -0.22 | -0.01 | -0.55 | -0.64 | 0.33 | 0.83 | -1.08 | -1.31 | -0.065 | -0.05 | -0.0025 | 0.996515 | 0.879408 | 0.893355 | 0.996493 | 0.997988 | 1 |
| Ube2ql1_E9QMD2_S66s_1 | Ube2ql1_E | -1.44 | -3.13 | -1.11 | -1.2 | -1.09 | -1.93 | -1.64 | -1.72 | -1.97 | -1.94 | -1.74 | -0.84 | -0.98 | -3.8 | -2.87 | -0.88 | 0.125 | 0.0975 | -0.4125 | 0.592443 | 0.790615 | 0.84857 | 0.995762 | 0.997988 | 0.995259 |
| Ube2v1_Q9CZY3_S104s_1 | Ube2v1_Q! | 0.34 | 0.31 | 0.6 | 0.46 | 0.42 | 0.5 | 0.27 | 0.42 | 0.23 | 0.5 | 0.6 | -0.06 | 0.32 | 0.57 | 0.12 | 0.41 | -0.025 | -0.11 | -0.0725 | 0.76524 | 0.901202 | 0.653102 | 0.996493 | 0.997988 | 0.9971 |
| Ube3a_O08759_S8s_1 | Ube3a_O0 | -0.76 | -0.04 | -0.97 | -0.48 | -0.06 | -0.77 | -0.38 | -0.48 | -0.12 | -0.29 | 0.14 | -0.96 | 0.25 | 0.45 | -0.12 | -0.41 | 0.14 | 0.255 | 0.605 | 0.086346 | 0.622815 | 0.450359 | 0.989223 | 0.997988 | 0.994666 |
| Ube3b_Q9ES34_S421s_1 | Ube3b_Q9 | 0.77 | -0.65 | 0 | -0.12 | 0.55 | 0.49 | 1.06 | 0.46 | 0.56 | 0.19 | 0.11 | 0.28 | -0.1 | 1 | -0.27 | 0.35 | 0.64 | 0.285 | 0.245 | 0.553945 | 0.080468 | 0.400702 | 0.989223 | 0.997988 | 0.994666 |
| Ube4b_G3UZD6_S101s_1 | Ube4b_G3 | 0.71 | 0.48 | 1.18 | 1.07 | 0.95 | 0.63 | 1.13 | 0.96 | 0.02 | 0.46 | 0.97 | 1.08 | -1.45 | 0.02 | 0.42 | 1.04 | 0.0575 | -0.2275 | -0.8525 | 0.12557 | 0.817876 | 0.485411 | 0.995762 | 0.997988 | 0.994666 |
| Ube4b_G3UZD6_S105s_1 | Ube4b_G3 | -0.4 | -0.28 | -0.44 | -0.35 | 0.04 | -0.15 | -0.6 | -0.22 | -0.36 | -0.35 | -0.78 | -0.62 | -0.58 | 0.01 | -1.18 | -0.59 | 0.135 | -0.16 | -0.2175 | 0.480173 | 0.548136 | 0.475568 | 0.989223 | 0.997988 | 0.994666 |
| Ube4b_G3UZD6_S31s_1 | Ube4b_G3 | -0.05 | 1.17 | -0.81 | -0.87 | -0.43 | -0.19 | -0.88 | -0.73 | 0.14 | 1.06 | -2.49 | -0.74 | 0.47 | 0.5 | -1.53 | -1.19 | -0.4175 | -0.3675 | -0.2975 | 0.647523 | 0.378491 | 0.646659 | 0.989223 | 0.997988 | 0.995259 |
| Ubn1_Q4G0F8_S493s_1 | Ubn1_Q4G | 0.28 | -0.11 | 0.07 | 0.69 | -1.58 | -0.73 | -1.11 | -0.51 | 1.09 | 0.84 | 0.4 | -0.32 | 0.47 | -0.44 | 0.73 | -0.11 | -1.215 | 0.27 | -0.07 | 0.840733 | 0.003061 | 0.464941 | 0.989223 | 0.997988 | 0.999507 |
| Ubn2_Q80WC1_S777s_2 | Ubn2_Q80 | -0.3 | 0.78 | 0.08 | -0.5 | 0.45 | -0.3 | 0.11 | -0.69 | -0.32 | 0.04 | -0.78 | -1.12 | 0.75 | 0.67 | 0.03 | -0.48 | -0.1225 | -0.56 | 0.2275 | 0.57983 | 0.740938 | 0.168032 | 0.995762 | 0.997988 | 0.995259 |
| Ubn2_Q80WC1_S780s_2 | Ubn2_Q80 | -0.3 | 0.78 | 0.08 | -0.5 | 0.45 | -0.3 | 0.11 | -0.69 | -0.32 | 0.04 | -0.78 | -1.12 | 0.75 | 0.67 | 0.03 | -0.48 | -0.1225 | -0.56 | 0.2275 | 0.57983 | 0.740938 | 0.168032 | 0.995762 | 0.997988 | 0.995259 |
| Ubqln2_Q9QZM0_S25s_1 | Ubqln2_Q9 | -0.01 | 0.2 | 0.18 | 0.32 | 0.18 | -0.03 | 0.06 | 0.22 | 0.32 | 0.63 | 0.63 | 0.34 | 0.54 | 0.62 | -0.06 | 0.27 | -0.065 | 0.3075 | 0.17 | 0.523715 | 0.750128 | 0.186941 | 0.995762 | 0.997988 | 0.994666 |
| Ubqln4_Q99NB8_S5s_1 | Ubqln4_Q9 | 2.23 | 0.23 | 1.76 | 1.87 | 0.14 | 2.84 | 0.71 | 2.01 | 2.62 | 1.08 | 1.38 | 1.26 | 1.38 | -0.51 | 0.72 | 1.39 | -0.0975 | 0.0625 | -0.7775 | 0.199602 | 0.884043 | 0.905459 | 0.996493 | 0.997988 | 0.994666 |
| Ubr4_A2AN08_S178s_1 | Ubr4_A2AI | 0.04 | 0.96 | 1.02 | 0.48 | 1.35 | 1.1 | 1.16 | 0.42 | 0.41 | 0.6 | 0.47 | 1.17 | 0.67 | 1.34 | 0.28 | 0.93 | 0.3825 | 0.0375 | 0.18 | 0.609258 | 0.251127 | 0.90604 | 0.989223 | 0.997988 | 0.995259 |
| Ubr4_A2AN08_S181s_1 | Ubr4_A2AI | -0.26 | -0.15 | 0.47 | 0.42 | 0.21 | -0.32 | 0.42 | 0.21 | -0.19 | -0.2 | -0.1 | 0.08 | 0.04 | 0.55 | 0.16 | 0.3 | 0.01 | -0.2225 | 0.1425 | 0.624025 | 0.971637 | 0.407825 | 0.999187 | 0.997988 | 0.995259 |
| Ubr4_A2AN08_S457s_1 | Ubr4_A2AI | -1.42 | -1.79 | -1.01 | -1.94 | -1.75 | -1.51 | -1.66 | -1.15 | -0.26 | -1.66 | -3.43 | -1.14 | 0.85 | -2.48 | -0.45 | -1.03 | 0.0225 | -0.0825 | 0.7625 | 0.258754 | 0.936202 | 0.89685 | 0.99868 | 0.997988 | 0.994666 |
| Ubr4_F6SSP6_S1439s_1 | Ubr4_F6SS | 1.09 | 4.62 | 3.97 | 4.31 | 3.69 | 0.41 | -1.35 | 4.3 | 2.06 | 1.36 | 3.59 | 2.9 | 1.49 | 0.92 | 3.78 | 3.06 | -1.735 | -1.02 | -1.185 | 0.217813 | 0.218121 | 0.245749 | 0.989223 | 0.997988 | 0.994666 |
| Ubr4_F6SSP6_S1442s_1 | Ubr4_F6SS | -0.23 | -0.48 | 0.85 | 0.7 | -1.61 | 0.09 | -0.7 | -0.35 | 0.18 | -0.5 | 1.24 | 1.07 | -0.52 | -1.27 | 0.61 | 0.8 | -0.8525 | 0.2875 | -0.305 | 0.580371 | 0.08728 | 0.564407 | 0.989223 | 0.997988 | 0.995259 |
| Ubr4_F6SSP6_S1464s_1 | Ubr4_F6SS | 0.02 | 0.77 | -0.48 | 0.61 | -0.7 | 0.8 | -0.77 | 0.19 | 0.41 | 0.63 | -1.05 | 0.89 | 0.78 | -0.54 | 0.58 | 1.28 | -0.35 | -0.01 | 0.295 | 0.528196 | 0.437727 | 0.983751 | 0.989223 | 0.997988 | 0.994666 |
| Ubr4_F6SSP6_S1570s_1 | Ubr4_F6SS | -1.26 | -0.79 | -2.06 | -0.21 | -1.18 | -0.94 | -1.44 | -0.71 | -0.14 | -0.4 | -1.14 | -0.96 | 0.21 | 0.06 | -2.18 | -1.08 | 0.0125 | 0.42 | 0.3325 | 0.593738 | 0.97529 | 0.353056 | 0.999187 | 0.997988 | 0.995259 |
| Ubr4_F6SSP6_T1444t_1 | Ubr4_F6SS | 0.26 | -0.04 | 1.04 | 1.13 | -1.45 | 0.59 | -0.53 | 0.19 | 0.91 | 0.23 | 1.06 | 1.19 | -2.27 | -1.61 | 0.48 | 0.64 | -0.8975 | 0.25 | -1.2875 | 0.093611 | 0.092519 | 0.502139 | 0.989223 | 0.997988 | 0.994666 |
| Ubr4_F6SSP6_T1569t_1 | Ubr4_F6SS | -1.26 | -0.79 | -2.06 | -0.21 | -1.18 | -0.94 | -1.44 | -0.71 | -0.14 | -0.4 | -1.14 | -0.96 | 0.21 | 0.06 | -2.18 | -1.08 | 0.0125 | 0.42 | 0.3325 | 0.593738 | 0.97529 | 0.353056 | 0.999187 | 0.997988 | 0.995259 |
| Ubr5_E9Q2H1_S1543s_1 | Ubr5_E9Q2 | -2.25 | -2.07 | -3.66 | -1.67 | -1.48 | -2.15 | -2.01 | -1.37 | -1.98 | -1.53 | -1.17 | -0.91 | -0.27 | -1.75 | -0.93 | -3.21 | 0.66 | 1.015 | 0.8725 | 0.224526 | 0.161002 | 0.054088 | 0.989223 | 0.997988 | 0.994666 |
| Ubxn1_Q922Y1_T65t_1 | Ubxn1_Q9. | 0.2 | 2.33 | 1.59 | 0.98 | 1.61 | 1.88 | 0.33 | 0.65 | 2.74 | 1.79 | 1.04 | 0.29 | 0.75 | 1.24 | 1.57 | 0.47 | -0.1575 | 0.19 | -0.2675 | 0.587567 | 0.767418 | 0.763513 | 0.995762 | 0.997988 | 0.995259 |
| Ubxn2b_Q0KL01_S216s_1 | Ubxn2b_Q( | 0.22 | 0.44 | 0.37 | -0.22 | -0.13 | 0 | -0.75 | 0.09 | -0.32 | -0.4 | -0.14 | 0.12 | 0.3 | -0.6 | -0.3 | 0.24 | -0.4 | -0.3875 | -0.2925 | 0.363006 | 0.172859 | 0.155725 | 0.989223 | 0.997988 | 0.994666 |
| Ubxn4_Q8VCH8_T487t_1 | Ubxn4_Q8' | -0.24 | -0.61 | -1.12 | -3.27 | -0.18 | -1.13 | 0.05 | -0.64 | 0.17 | -0.75 | -0.53 | 0.05 | -1.27 | -0.02 | 0.41 | -1.55 | 0.835 | 1.045 | 0.7025 | 0.352113 | 0.215791 | 0.132699 | 0.989223 | 0.997988 | 0.994666 |
| Ubxn6_Q99PL6_S36s_1 | Ubxn6_Q9! | 0.05 | -1.4 | -1.22 | -0.97 | -0.06 | -0.33 | -0.94 | -1.12 | -0.96 | -0.69 | 0.41 | -1.14 | -1.35 | -0.26 | -0.56 | -0.75 | 0.2725 | 0.29 | 0.155 | 0.701198 | 0.495699 | 0.529238 | 0.989223 | 0.997988 | 0.996159 |
| Ubxn6_Q99PL6_S75s_1 | Ubxn6_Q9! | 0.36 | -0.61 | 0.13 | 0.06 | -0.01 | 0.59 | -0.3 | 0.08 | -0.74 | -0.78 | -0.26 | -0.1 | -1.01 | 0.06 | -0.54 | 0.03 | 0.105 | -0.455 | -0.35 | 0.338623 | 0.729003 | 0.161079 | 0.995762 | 0.997988 | 0.994666 |
| Ubxn6_Q99PL6_S78s_1 | Ubxn6_Q9! | -0.61 | -2.08 | -0.24 | 0.21 | -0.41 | 0.1 | 0.14 | 0.66 | -0.75 | -1.26 | -0.01 | 1.59 | -0.37 | -0.03 | -0.02 | 0.47 | 0.8025 | 0.5725 | 0.6925 | 0.186014 | 0.131425 | 0.431987 | 0.989223 | 0.997988 | 0.994666 |
| Uck2_Q99PM9_S254s_1 | Uck2_Q99I | -2.88 | -3.87 | -2.17 | -3.88 | -3.83 | -3.09 | -2.9 | -3.72 | -0.64 | -2.72 | -2.26 | -2.65 | 0.33 | -2.17 | -2.18 | -2.53 | -0.185 | 1.1325 | 1.5625 | 0.047343 | 0.678763 | 0.080084 | 0.993464 | 0.997988 | 0.994666 |
| Ufd1l_P70362_S209s_1 | Ufd1l_P70: | -1.52 | -1.22 | -1.32 | -2.58 | -0.67 | -1.35 | -1.52 | -0.07 | -0.18 | -1.9 | 0.33 | -0.84 | -1.02 | 1.14 | -1.2 | -0.84 | 0.7575 | 1.0125 | 1.18 | 0.065186 | 0.104396 | 0.08439 | 0.989223 | 0.997988 | 0.994666 |
| Ufd1l_P70362_S247s_1 | Ufd1l_P70: | -1.58 | -2.08 | -2.21 | -1.87 | -0.38 | -0.78 | -1.28 | -1.78 | -1.93 | -0.68 | -0.89 | -1.73 | -0.78 | -1.43 | -2.52 | -2.78 | 0.88 | 0.6275 | 0.0575 | 0.90242 | 0.02617 | 0.100672 | 0.989223 | 0.997988 | 0.999507 |
| Ufd1l_P70362_S299s_1 | Ufd1l_P70: | -0.84 | -0.66 | -1.89 | -2.42 | -0.15 | -1.44 | -0.46 | -2.53 | -1.55 | -0.53 | -0.79 | -1.53 | -0.61 | -0.43 | -2.23 | -0.38 | 0.3075 | 0.3525 | 0.54 | 0.349829 | 0.6162 | 0.46161 | 0.989223 | 0.997988 | 0.994666 |
| Uhrf1bp1_B2KF50_S929s_1 | Uhrf1bp1_ | 2.98 | 2.88 | 2.99 | 1.87 | 2.36 | 2.67 | 1.84 | 2.37 | 0.84 | 3.05 | 1.27 | 2.33 | 0.74 | 1.26 | 2.6 | 2.44 | -0.37 | -0.8075 | -0.92 | 0.09083 | 0.2801 | 0.153731 | 0.989223 | 0.997988 | 0.994666 |
| Uhrf1bp1l_A2RSJ4_S1080s_1 | Uhrf1bp1l_ | -2.35 | -4.34 | -2.3 | -2.57 | -1.74 | -0.89 | -2.82 | -2.59 | -0.53 | -1.99 | -2.62 | -1.22 | -1.21 | -2.32 | -2.18 | -2.64 | 0.88 | 1.3 | 0.8025 | 0.158463 | 0.158301 | 0.05703 | 0.989223 | 0.997988 | 0.994666 |
| Uhrf1bp1l_A2RSJ4_S1336s_1 | Uhrf1bp1l_ | 0.21 | 0.91 | 0.38 | 0.44 | 0.27 | 0.28 | 0.6 | 0.52 | 0.66 | 0.13 | 0.38 | 1.21 | 0.17 | 0.48 | 0.64 | 0.19 | -0.0675 | 0.11 | -0.115 | 0.675297 | 0.777494 | 0.723527 | 0.995762 | 0.997988 | 0.996159 |
| Uhrf1bp1l_A2RSJ4_S774s_1 | Uhrf1bp1l_ | 0.22 | 0.05 | 0.1 | 0.26 | 0.12 | 0.51 | 0.7 | 0.63 | -0.28 | 0.42 | -0.15 | 0.55 | -0.47 | 0.24 | 0.18 | 0.54 | 0.3325 | -0.0225 | -0.035 | 0.903467 | 0.158871 | 0.933521 | 0.989223 | 0.997988 | 0.999507 |
| Uhrf1bp1l_A2RSJ4_S879s_1 | Uhrf1bp1l_ | -0.97 | -0.25 | -0.51 | -1.83 | -0.18 | -0.51 | 0.07 | -1.65 | 0.04 | -0.65 | -1.94 | -0.65 | -0.63 | 0.93 | -2.83 | -1.8 | 0.3225 | 0.09 | -0.1925 | 0.804718 | 0.502905 | 0.859025 | 0.989223 | 0.997988 | 0.999507 |
| Uhrf1bp1l_A2RSJ4_S890s_1 | Uhrf1bp1l_ | 0.34 | 0.83 | 0.33 | 0.67 | -0.01 | -0.46 | 0.05 | 0.11 | 0.27 | 1.03 | 0.58 | 0.02 | -0.25 | 0.79 | 0.02 | -0.54 | -0.62 | -0.0675 | -0.5375 | 0.142906 | 0.027615 | 0.818308 | 0.989223 | 0.997988 | 0.994666 |
| Uhrf1bp1l_A2RSJ4_S914s_1 | Uhrf1bp1l_ | -0.17 | -1.44 | -0.23 | -0.41 | -1 | 0.1 | 0.36 | -0.81 | 1.67 | -0.97 | -2.71 | 0.47 | 1.85 | -1.48 | 0.75 | -0.17 | 0.225 | 0.1775 | 0.8 | 0.261329 | 0.597866 | 0.839997 | 0.989223 | 0.997988 | 0.994666 |
| Uhrf1bp1l_A2RSJ4_S934s_1 | Uhrf1bp1l_ | -3.31 | -3.35 | -2.94 | -3.2 | -2.25 | -3.1 | -2.1 | -3.59 | -3.53 | -3.06 | -1.86 | -3.52 | -2.37 | -2.17 | -2.97 | -3.3 | 0.44 | 0.2075 | 0.4975 | 0.147736 | 0.243658 | 0.607404 | 0.989223 | 0.997988 | 0.994666 |
| Uhrf1bp1l_A2RSJ4_S952s_1 | Uhrf1bp1l_ | -0.13 | -0.31 | 0.02 | 0.06 | -0.72 | 0.01 | -0.43 | -0.32 | 0.41 | -0.14 | -0.72 | -0.36 | 0.93 | -0.72 | -0.35 | -0.46 | -0.275 | -0.1125 | -0.06 | 0.877348 | 0.266888 | 0.702931 | 0.989223 | 0.997988 | 0.999507 |
| Uhrf1bp1l_A2RSJ4_S988s_1 | Uhrf1bp1l_ | 0.35 | -0.31 | 0.28 | 0.35 | -1.51 | 0.74 | -1.17 | -0.34 | 2.45 | 0.29 | 0.28 | 0.28 | 2.14 | -1.25 | -0.39 | -1.15 | -0.7375 | 0.6575 | -0.33 | 0.648333 | 0.152145 | 0.232221 | 0.989223 | 0.997988 | 0.995259 |
| Uhrf1bp1l_Q8C318_S326s_1 | Uhrf1bp1l_ | 0.14 | 1.17 | 0.79 | 0.09 | 0.5 | -0.34 | -0.23 | -0.6 | 0.6 | 0.69 | 1 | 0.44 | 0.98 | 0.81 | 0.12 | 0 | -0.715 | 0.135 | -0.07 | 0.852122 | 0.067289 | 0.67192 | 0.989223 | 0.997988 | 0.999507 |
| Uhrf1bp1l_Q8C318_S326s_2 | Uhrf1bp1l_ | -0.61 | -0.76 | -0.57 | -0.39 | -0.38 | -0.48 | -0.76 | -0.2 | -1.08 | -0.47 | -0.58 | -0.02 | -2.55 | 0.41 | -0.83 | -0.41 | 0.1275 | 0.045 | -0.2625 | 0.650974 | 0.570879 | 0.872882 | 0.989223 | 0.997988 | 0.995259 |
| Uhrf1bp1l_Q8C318_S330s_1 | Uhrf1bp1l_ | -1.98 | -1.51 | -1.88 | -2.3 | -3.8 | -0.82 | -2.54 | -2.03 | -0.34 | -1.51 | -0.68 | -3.58 | 0.3 | -1.54 | -3.06 | -1.98 | -0.38 | 0.39 | 0.3475 | 0.59505 | 0.512655 | 0.566396 | 0.989223 | 0.997988 | 0.995259 |
| Uhrf1bp1l_Q8C318_S330s_2 | Uhrf1bp1l_ | -1.26 | -1.21 | -1.05 | -1.02 | -0.97 | -1.1 | -1.37 | -0.88 | -1.21 | -1.1 | -1.23 | -0.68 | -2.23 | -0.14 | -1.65 | -1.05 | 0.055 | 0.08 | -0.1325 | 0.764203 | 0.798764 | 0.732657 | 0.995762 | 0.997988 | 0.9971 |
| Uhrf1bp1l_Q8C318_S335s_1 | Uhrf1bp1l_ | -2.43 | -1.96 | 0.15 | -0.96 | -2.62 | -2.6 | -1.43 | -1.62 | 0.75 | -1.51 | -2.73 | -1.08 | 1.45 | -3.96 | -0.67 | -1.05 | -0.7675 | 0.1575 | 0.2425 | 0.823245 | 0.211002 | 0.847818 | 0.989223 | 0.997988 | 0.999507 |
| Uhrf1bp1l_Q8C318_T205t_1 | Uhrf1bp1l_ | -1.38 | -2.3 | -1.74 | -1.89 | -0.5 | -2.17 | -0.61 | -1.64 | -0.34 | -1.25 | -1.36 | -0.94 | -1.4 | 0.12 | -3.05 | -2.61 | 0.5975 | 0.855 | 0.0925 | 0.888788 | 0.181286 | 0.02475 | 0.989223 | 0.997988 | 0.999507 |
| Uhrf2_Q7TMI3_S668s_1 | Uhrf2_Q7T | -2.49 | -2.93 | -3.44 | -4.66 | -1.85 | 0.01 | -4.37 | -3.52 | -0.07 | -2.17 | -2.81 | -4.06 | 0.07 | -1.54 | -2.99 | -4.11 | 0.9475 | 1.1025 | 1.2375 | 0.188254 | 0.323213 | 0.216293 | 0.989223 | 0.997988 | 0.994666 |
| Ulk1_Q6PB82_S450s_1 | Ulk1_Q6PB | 0.37 | 1.09 | 0.76 | 0.18 | 0.7 | 0.88 | 0.59 | 1.29 | -0.08 | 0.57 | 0.82 | 0.47 | -0.08 | 0.75 | 0.2 | 0.44 | 0.265 | -0.155 | -0.2725 | 0.40066 | 0.366473 | 0.620919 | 0.989223 | 0.997988 | 0.994666 |
| Ulk1_Q6PB82_S628s_1 | Ulk1_Q6PB | 0.46 | -0.07 | 1.01 | 0.48 | 0.64 | -0.12 | 0.45 | 0.75 | 0.1 | 0.11 | 0.77 | 1.13 | -0.08 | 0.81 | -0.18 | 0.3 | -0.04 | 0.0575 | -0.2575 | 0.46396 | 0.897911 | 0.870135 | 0.996493 | 0.997988 | 0.994666 |
| Ulk1_Q6PB82_S643s_1 | Ulk1_Q6PB | -0.83 | 0.22 | -0.06 | -1.1 | 0.8 | -0.08 | 0.14 | -0.31 | -0.21 | -0.5 | 0 | 0.35 | 0.26 | 0.41 | -0.08 | 0.13 | 0.58 | 0.3525 | 0.6225 | 0.105476 | 0.153962 | 0.350794 | 0.989223 | 0.997988 | 0.994666 |
| Ulk1_Q6PB82_S763s_1 | Ulk1_Q6PB | 0.85 | 0.78 | 1.07 | 1.23 | 0.91 | 0.5 | 1.4 | 1.58 | 0.42 | 0.99 | 1.24 | 1.04 | -0.56 | 1.74 | 1.41 | 1.24 | 0.115 | -0.06 | -0.025 | 0.959951 | 0.696095 | 0.821431 | 0.994893 | 0.997988 | 0.999507 |
| Ulk2_Q9QY01_S322s_1 | Ulk2_Q9Q\ | -1.21 | 0.14 | -0.2 | 0.29 | 0.81 | 0.02 | 0.46 | 0.39 | 0.74 | 0.05 | 0.12 | 0.41 | 0.45 | 0.55 | 0.14 | 0.27 | 0.665 | 0.575 | 0.5975 | 0.130907 | 0.097295 | 0.15274 | 0.989223 | 0.997988 | 0.994666 |
| Ulk2_Q9QY01_S430s_1 | Ulk2_Q9Q\ | -0.27 | 0.78 | 0.27 | 0.58 | 0.37 | 0.4 | -0.34 | 0.57 | 0.28 | 0.3 | 1.88 | 1.02 | -0.26 | 0.83 | 0.45 | 0.22 | -0.09 | 0.53 | -0.03 | 0.931997 | 0.77865 | 0.237637 | 0.995762 | 0.997988 | 0.999507 |
| Ulk2_Q9QY01_S600s_1 | Ulk2_Q9Q\ | -0.01 | -0.02 | 0.5 | -0.01 | 0 | 0.04 | 0.47 | 0.02 | -0.01 | 0.02 | 0.48 | 0.68 | -0.08 | -0.07 | 0.05 | -0.27 | 0.0175 | 0.1775 | -0.2075 | 0.421416 | 0.941365 | 0.518007 | 0.999187 | 0.997988 | 0.994666 |
| Ulk2_Q9QY01_S772s_1 | Ulk2_Q9Q\ | 0.51 | 0.91 | 0.83 | 0.03 | 0.72 | 1.38 | 1.42 | 1.16 | 1.12 | 1.31 | 1.11 | -0.2 | 0.62 | 0.48 | 0.47 | 0.7 | 0.6 | 0.265 | -0.0025 | 0.992943 | 0.060624 | 0.511895 | 0.989223 | 0.997988 | 1 |
| Ulk2_Q9QY01_S781s_1 | Ulk2_Q9Q\ | 0.76 | 0.41 | 0.62 | 1.22 | 1.55 | -0.17 | 1.61 | 0.94 | -0.74 | 0.79 | 1.2 | 0.47 | -0.39 | 1.29 | 1.11 | 0.38 | 0.23 | -0.3225 | -0.155 | 0.712261 | 0.589976 | 0.467867 | 0.989223 | 0.997988 | 0.996966 |
| Ulk3_Q3U3Q1_S464s_1 | Ulk3_Q3U3 | -2.13 | -1.96 | -0.87 | -2.09 | -0.53 | -2.79 | -1.94 | -1.6 | -2.64 | -1.87 | 0.71 | -1.49 | -1.06 | -0.82 | -2.39 | -1.46 | 0.0475 | 0.44 | 0.33 | 0.466204 | 0.925393 | 0.535154 | 0.997335 | 0.997988 | 0.994666 |
| Unc119_B1AQD9_S37s_1 | Unc119_B1 | 0.64 | 0.4 | 3.01 | 1.95 | -0.15 | 0.36 | 0.77 | 1.42 | 0.78 | 0.52 | 1.49 | 1.96 | -0.09 | 0.58 | 1.6 | 1.87 | -0.9 | -0.3125 | -0.51 | 0.460768 | 0.168188 | 0.62193 | 0.989223 | 0.997988 | 0.994666 |
| Unc13a_Q4KUS2_S241s_1 | Unc13a_Q4 | 0.94 | -0.53 | 1.12 | 0.6 | 1.48 | 0.61 | 1.37 | 1.34 | 0.75 | 0.47 | -0.01 | 1.61 | 0.49 | 0.62 | 1.14 | 1.82 | 0.6675 | 0.1725 | 0.485 | 0.307937 | 0.123159 | 0.71868 | 0.989223 | 0.997988 | 0.994666 |
| Unc13a_Q4KUS2_S244s_1 | Unc13a_Q4 | -0.81 | -0.42 | -0.89 | -0.52 | -0.26 | -0.96 | -0.42 | -0.58 | -0.35 | -0.63 | -0.65 | -0.66 | -0.23 | -0.06 | -0.48 | -0.52 | 0.105 | 0.0875 | 0.3375 | 0.21223 | 0.672051 | 0.705733 | 0.992876 | 0.997988 | 0.994666 |
| Unc13a_Q4KUS2_S249s_1 | Unc13a_Q4 | 0.28 | 0.16 | 0.45 | -0.28 | 1.29 | 1.11 | 0.02 | -1.02 | 1.17 | -0.12 | -2.27 | 0.48 | 1.35 | 0.95 | -0.06 | 0.81 | 0.1975 | -0.3375 | 0.61 | 0.115574 | 0.699454 | 0.624959 | 0.994893 | 0.997988 | 0.994666 |
| Unc13a_Q4KUS2_S255s_1 | Unc13a_Q4 | 0.99 | 1.11 | 1.61 | 1.39 | 0.31 | 1.2 | 0.48 | 0.92 | 1.7 | 1.06 | 0.09 | 0.99 | 2 | -0.15 | 1.06 | 1.11 | -0.5475 | -0.315 | -0.27 | 0.55265 | 0.076709 | 0.401672 | 0.989223 | 0.997988 | 0.994666 |
| Unc13a_Q4KUS2_S255s_2 | Unc13a_Q4 | 1.55 | 0.57 | 1.61 | 1.81 | 0.92 | 2.14 | 0.36 | 1.56 | 1.45 | 0.17 | -1.05 | 1.63 | 0.71 | 0.68 | -0.11 | 1.17 | -0.14 | -0.835 | -0.7725 | 0.074304 | 0.75362 | 0.202428 | 0.995762 | 0.997988 | 0.994666 |
| Unc13a_Q4KUS2_S259s_1 | Unc13a_Q4 | 2.46 | 1 | 1.61 | 1.91 | 1.66 | 2.42 | 1.45 | 1.82 | 1.39 | 0.99 | 0.5 | 1.39 | 1.15 | 1.39 | 1.91 | 2.12 | 0.0925 | -0.6775 | -0.1025 | 0.792545 | 0.800203 | 0.098129 | 0.995762 | 0.997988 | 0.999507 |
| Unc13a_Q4KUS2_S385s_1 | Unc13a_Q4 | -1.73 | -1.95 | -2.1 | -0.15 | -0.53 | -1.02 | -1.15 | -1.25 | -1.51 | -0.97 | -1.22 | -0.86 | -1.6 | 0.7 | -2.19 | -0.43 | 0.495 | 0.3425 | 0.6025 | 0.39953 | 0.284426 | 0.458181 | 0.989223 | 0.997988 | 0.994666 |
| Unc13a_Q4KUS2_S440s_1 | Unc13a_Q4 | 0.29 | 0.86 | 0.94 | 0.29 | -0.08 | 1.08 | 0.05 | 0.15 | 1.04 | -0.13 | 0.12 | 0.56 | 0.33 | -0.13 | 0.45 | 0.74 | -0.295 | -0.1975 | -0.2475 | 0.43017 | 0.380097 | 0.558608 | 0.989223 | 0.997988 | 0.994666 |
| Unc13a_Q4KUS2_T416t_1 | Unc13a_Q4 | 0.08 | -0.46 | 0.83 | -0.51 | -2.59 | 1.15 | -1.28 | -0.26 | 2.59 | -0.33 | -3.7 | 0.88 | 3.01 | -2.41 | 2.65 | -0.63 | -0.73 | -0.125 | 0.67 | 0.568183 | 0.340707 | 0.916825 | 0.989223 | 0.997988 | 0.995259 |
| Unc13b_E9Q263_S1250s_1 | Unc13b_E9 | 0.23 | -0.4 | -0.83 | -1.49 | -2.02 | -0.01 | -1.23 | -1.81 | 2.02 | -0.78 | -2.97 | -0.23 | 2.31 | -1.6 | 0.09 | -1.11 | -0.645 | 0.1325 | 0.545 | 0.516701 | 0.238953 | 0.890272 | 0.989223 | 0.997988 | 0.994666 |
| Unc13c_Q8K0T7_T511t_1 | Unc13c_Q8 | 0.45 | 0.49 | 0.52 | 1.1 | -0.16 | 0.29 | -0.05 | 0.56 | 0.48 | -0.32 | 0.23 | 0.72 | 0.54 | -0.14 | 1.13 | 0.86 | -0.48 | -0.3625 | -0.0425 | 0.902225 | 0.099121 | 0.255613 | 0.989223 | 0.997988 | 0.999507 |
| Unc5a_F7CVI0_S143s_1 | Unc5a_F7C | -0.78 | -1.45 | -1.05 | -1.03 | -1.97 | 0.42 | -0.79 | -1.02 | 1.22 | -1.23 | -3.29 | -0.46 | 1.43 | -1.79 | -0.07 | -1.69 | 0.2375 | 0.1375 | 0.5475 | 0.438124 | 0.617519 | 0.870212 | 0.989223 | 0.997988 | 0.994666 |
| Unc5a_F7CVI0_S155s_1 | Unc5a_F7C | 0.01 | -0.45 | -0.4 | 0.1 | 1.27 | -0.13 | 0.48 | -0.16 | -1.3 | -0.98 | -1.15 | -0.12 | -1.28 | -0.33 | -0.97 | -0.26 | 0.55 | -0.7025 | -0.525 | 0.133823 | 0.152315 | 0.054449 | 0.989223 | 0.997988 | 0.994666 |
| Unc5c_E9PVI4_S447s_1 | Unc5c_E9P | -0.34 | 0.06 | -0.16 | -0.04 | 0.36 | -0.26 | 0.18 | -0.31 | -0.3 | -0.15 | -0.28 | -0.02 | -0.17 | 0.55 | 0 | 0.06 | 0.1125 | -0.0675 | 0.23 | 0.399785 | 0.649087 | 0.759509 | 0.99033 | 0.997988 | 0.994666 |
| Unc5c_E9PVI4_S638s_1 | Unc5c_E9P | -0.06 | 0.23 | -0.72 | -0.73 | 0.15 | 0.21 | 0.18 | -0.75 | -0.06 | 0.47 | -0.74 | -0.83 | -0.65 | 0.19 | -0.49 | -0.2 | 0.2675 | 0.03 | 0.0325 | 0.923651 | 0.440413 | 0.938698 | 0.989223 | 0.997988 | 0.999507 |
| Unc79_E5CYJ9_S1759s_1 | Unc79_E5C | 1.78 | 1.41 | 0.26 | 1.03 | 1.68 | 1.98 | 1.04 | 1.16 | 0.42 | 1.26 | 0.42 | 1.1 | -0.26 | 0.15 | 0.12 | 0.78 | 0.345 | -0.32 | -0.8725 | 0.050523 | 0.378089 | 0.424452 | 0.989223 | 0.997988 | 0.994666 |
| Unc79_E5CYJ9_S689s_1 | Unc79_E5C | 1.85 | 0.4 | 0.91 | 1.46 | 1.26 | 2.36 | 0.91 | 1.32 | 1.73 | 0.87 | 1.08 | 1.78 | 1.3 | 1.22 | 0.41 | 1.1 | 0.3075 | 0.21 | -0.1475 | 0.704851 | 0.472661 | 0.594631 | 0.989223 | 0.997988 | 0.996159 |
| Unc79_G8JL63_S1078s_1 | Unc79_G8J | 0.86 | 0.4 | 0.02 | -0.26 | -0.55 | 0.68 | -0.27 | -0.52 | 0.81 | -0.32 | -0.37 | -0.37 | 0.67 | 0.27 | 0.08 | 0.7 | -0.42 | -0.3175 | 0.175 | 0.597074 | 0.27451 | 0.415568 | 0.989223 | 0.997988 | 0.995259 |
| Unc80_Q8BLN6_S1259s_1 | Unc80_Q8I | -1.09 | -1.14 | 0.23 | -0.11 | -0.72 | 0.14 | -0.42 | -0.63 | 0.43 | -1.15 | -1.6 | 0.53 | 0.82 | -1.16 | -0.15 | 0.5 | 0.12 | 0.08 | 0.53 | 0.323629 | 0.756016 | 0.892375 | 0.995762 | 0.997988 | 0.994666 |
| Unc80_Q8BLN6_S166s_1 | Unc80_Q8I | -2.05 | -1.82 | -0.66 | -0.07 | -2.07 | -1.77 | -2.13 | -2.61 | 2.24 | -2.83 | -1.11 | -0.64 | 2.33 | -2.9 | 1.53 | -0.81 | -0.995 | 0.565 | 1.1875 | 0.297475 | 0.056079 | 0.5818 | 0.989223 | 0.997988 | 0.994666 |
| Unc80_Q8BLN6_S246s_1 | Unc80_Q8I | 0.66 | 1.39 | 0.74 | 0.54 | 0.4 | 0.95 | 0.55 | 0.47 | 0.12 | 0.85 | 1.37 | 0.33 | 1.14 | 1.62 | 1.02 | 0.73 | -0.24 | -0.165 | 0.295 | 0.361647 | 0.38314 | 0.64198 | 0.989223 | 0.997988 | 0.994666 |
| Unc80_Q8BLN6_S246s_2 | Unc80_Q8I | 1.06 | 0.88 | 0.86 | 1.17 | 0.34 | 1.17 | 0.67 | 0.98 | 0.56 | 0.94 | 1.56 | 1.27 | 0.63 | 1.02 | 0.74 | 0.86 | -0.2025 | 0.09 | -0.18 | 0.462657 | 0.429519 | 0.748099 | 0.989223 | 0.997988 | 0.994666 |
| Unc80_Q8BLN6_S257s_1 | Unc80_Q8I | 0 | -0.03 | 0.09 | 0.68 | 0.06 | -0.18 | -0.03 | 0.19 | 0.23 | 0.35 | -0.17 | 0.37 | 0.15 | 0.35 | 0.1 | 0.29 | -0.175 | 0.01 | 0.0375 | 0.888579 | 0.481255 | 0.970263 | 0.989223 | 0.997988 | 0.999507 |
| Unc80_Q8BLN6_S257s_2 | Unc80_Q8I | 1.06 | 0.88 | 0.86 | 1.17 | 0.34 | 1.17 | 0.67 | 0.98 | 0.56 | 0.94 | 1.56 | 1.27 | 0.63 | 1.02 | 0.74 | 0.86 | -0.2025 | 0.09 | -0.18 | 0.462657 | 0.429519 | 0.748099 | 0.989223 | 0.997988 | 0.994666 |
| Unc80_Q8BLN6_S295s_1 | Unc80_Q8I | -0.32 | -0.66 | 0.25 | 0.61 | -0.27 | -0.59 | 0.18 | 0.43 | -0.3 | -0.04 | 1.1 | 0.44 | -0.65 | -0.15 | -0.19 | 0.22 | -0.0325 | 0.33 | -0.1625 | 0.653422 | 0.928359 | 0.431934 | 0.99758 | 0.997988 | 0.995259 |
| Unc80_Q8BLN6_S3110s_1 | Unc80_Q8I | 0.86 | 0.94 | 0.66 | 0.29 | 0.25 | -0.27 | 1.1 | 0.19 | -0.22 | 0.31 | 1.21 | -0.1 | 0.24 | 0.86 | 0.31 | 1.45 | -0.37 | -0.3875 | 0.0275 | 0.93692 | 0.279084 | 0.30329 | 0.989223 | 0.997988 | 0.999507 |
| Unc80_Q8BLN6_S316s_1 | Unc80_Q8I | -0.15 | 0.17 | 1.25 | -0.4 | 0.28 | 0.2 | 0.48 | 1.15 | -0.04 | -0.28 | 0.09 | -0.71 | 0.24 | 0.42 | 0.24 | 0.66 | 0.31 | -0.4525 | 0.1725 | 0.658343 | 0.451217 | 0.276269 | 0.989223 | 0.997988 | 0.995259 |
| Unc80_Q8BLN6_S3203s_1 | Unc80_Q8I | -0.04 | 0.55 | 0.03 | 0.52 | 0.32 | -0.04 | 0.41 | -0.66 | 0.27 | 1.08 | -0.2 | 0.07 | -0.11 | 0.64 | 0.01 | -0.14 | -0.2575 | 0.04 | -0.165 | 0.586792 | 0.413454 | 0.9057 | 0.989223 | 0.997988 | 0.995259 |
| Unc80_Q8BLN6_S415s_1 | Unc80_Q8I | 0.78 | 0.24 | 0.75 | 1.08 | 0.48 | 0.99 | 1.62 | 1.28 | 0.84 | 0.79 | 1.45 | 1.34 | -0.54 | 0.39 | 1.47 | 1.22 | 0.38 | 0.3925 | -0.0775 | 0.868535 | 0.245884 | 0.196462 | 0.989223 | 0.997988 | 0.999507 |
| Unc80_Q8BLN6_S526s_1 | Unc80_Q8I | 0.24 | 0.22 | 0.22 | 0.28 | -0.12 | 0.43 | -0.32 | 0.3 | 0.78 | 0.07 | 0.76 | 0.42 | 0.33 | 0.55 | 0.07 | -0.07 | -0.1675 | 0.2675 | -0.02 | 0.93646 | 0.492541 | 0.293664 | 0.989223 | 0.997988 | 0.999507 |
| Unc80_Q8BLN6_T1014t_1 | Unc80_Q8I | -0.16 | -0.15 | -0.13 | -0.63 | -0.48 | -0.12 | -0.48 | -0.67 | -0.43 | -0.34 | -1 | -0.6 | 0.45 | 0.16 | -0.04 | -1.76 | -0.17 | -0.325 | -0.03 | 0.950882 | 0.477613 | 0.227589 | 0.989223 | 0.997988 | 0.999507 |
| Unk_Q8BL48_S240s_1 | Unk_Q8BL | -0.75 | -0.05 | -1.69 | -1.3 | -0.28 | -0.9 | -1.28 | -1.95 | -0.35 | 0.26 | -0.94 | -1.35 | -0.42 | -0.17 | -0.54 | -1.15 | -0.155 | 0.3525 | 0.3775 | 0.371997 | 0.738604 | 0.465291 | 0.995762 | 0.997988 | 0.994666 |
| Unk_Q8BL48_S371s_1 | Unk_Q8BL | -1.64 | -1.75 | -1.15 | -1.41 | -2.01 | -1.71 | -2.11 | -3.42 | -1.87 | -1.34 | -1.54 | -1.76 | -0.63 | -0.99 | -2.43 | -1.93 | -0.825 | -0.14 | -0.0075 | 0.986082 | 0.0575 | 0.580681 | 0.989223 | 0.997988 | 1 |
| Unkl_Q5FWH2_S15s_1 | Unkl_Q5FV | 1.09 | 1.83 | 0.02 | -0.17 | 1.47 | 1.06 | 0.15 | 0.36 | 0.84 | 1.58 | 0.86 | -0.21 | 1.29 | 1.64 | 0.06 | 0.52 | 0.0675 | 0.075 | 0.185 | 0.736356 | 0.895046 | 0.892045 | 0.996493 | 0.997988 | 0.9971 |
| Uqcrc2_Q9DB77_S169s_1 | Uqcrc2_Q9 | 0.91 | -0.36 | 0.65 | 0.02 | 1.08 | 0.35 | -0.43 | 0.12 | 0.4 | 0.27 | 0.94 | -0.03 | 0.87 | 0 | 0.6 | -0.01 | -0.025 | 0.09 | 0.06 | 0.87438 | 0.951089 | 0.804809 | 0.999187 | 0.997988 | 0.999507 |
| Urb1_E9PU96_S17s_1 | Urb1_E9PL | -0.38 | 1.5 | 0.82 | 0.82 | 0.31 | 0.51 | 0.37 | 0.38 | 0.56 | -0.04 | 1.15 | 0.58 | 1.27 | -0.4 | 0.39 | 1.09 | -0.2975 | -0.1275 | -0.1025 | 0.841536 | 0.444903 | 0.774491 | 0.989223 | 0.997988 | 0.999507 |
| Uri1_Q3TLD5_S341s_1 | Uri1_Q3TLI | 0.01 | -0.19 | 0.51 | -2.68 | -1.33 | -1.38 | -1.17 | -0.91 | -0.63 | 0.49 | 0.29 | 0.17 | 0.71 | 0.6 | -0.52 | 0.25 | -0.61 | 0.6675 | 0.8475 | 0.235764 | 0.352639 | 0.340527 | 0.989223 | 0.997988 | 0.994666 |
| Usp10_P52479_S209s_1 | Usp10_P52 | 0.91 | 0.81 | 0.75 | 0.81 | 0.7 | 0.23 | 1.23 | 0.55 | 0.55 | 0.59 | 0.97 | 0.75 | -0.91 | 0.54 | 0.66 | 0.43 | -0.1425 | -0.105 | -0.64 | 0.119112 | 0.586868 | 0.631594 | 0.989223 | 0.997988 | 0.994666 |
| Usp11_Q99K46_S906s_1 | Usp11_Q9! | -0.16 | 0.2 | 1.21 | 0.36 | -0.31 | -0.33 | 0.99 | 0.41 | -0.45 | -0.33 | 0.91 | 0.05 | -0.8 | -0.14 | 0.44 | -0.16 | -0.2125 | -0.3575 | -0.5675 | 0.172887 | 0.607298 | 0.399444 | 0.989223 | 0.997988 | 0.994666 |
| Usp14_E9PYI8_S108s_1 | Usp14_E9P | -0.18 | 0.87 | -0.31 | -0.11 | 0.71 | 0.41 | -0.16 | -0.01 | 0.7 | 0.77 | -0.07 | 0.11 | 0.16 | 1.19 | -0.2 | -0.11 | 0.17 | 0.31 | 0.1925 | 0.646767 | 0.620534 | 0.393838 | 0.989223 | 0.997988 | 0.995259 |
| Usp14_E9PYI8_S358s_1 | Usp14_E9P | 1.11 | 1.3 | 0.73 | 0.84 | 0.61 | 0.86 | 0.19 | 0.58 | 0.76 | 1.06 | 0.59 | 0.68 | 1.3 | 0.45 | 0.25 | 0.69 | -0.435 | -0.2225 | -0.3225 | 0.317507 | 0.10471 | 0.3743 | 0.989223 | 0.997988 | 0.994666 |
| Usp14_E9PYI8_T200t_1 | Usp14_E9P | -2.62 | -1.45 | -0.19 | 0.18 | -1.34 | -2.29 | -0.25 | -0.32 | -2.47 | -2.6 | 0.34 | 0.48 | -1.07 | -1.79 | -0.05 | -0.48 | -0.03 | -0.0425 | 0.1725 | 0.795903 | 0.965995 | 0.963936 | 0.999187 | 0.997988 | 0.999507 |
| Usp15_Q8R5H1_S229s_1 | Usp15_Q8I | 0.15 | -0.21 | -0.83 | -0.04 | -0.04 | 0.06 | 0.25 | -0.06 | -0.06 | -0.68 | 0.2 | -0.28 | -0.7 | -0.12 | -0.86 | -0.87 | 0.285 | 0.0275 | -0.405 | 0.228316 | 0.301755 | 0.930229 | 0.989223 | 0.997988 | 0.994666 |
| Usp15_Q8R5H1_S621s_1 | Usp15_Q8I | 0.25 | 0.07 | -0.12 | -0.38 | -0.24 | -0.63 | -0.75 | 0.04 | -0.03 | 0.13 | 0.09 | -0.06 | 0.35 | 0.16 | -0.09 | -0.7 | -0.35 | 0.0775 | -0.025 | 0.936813 | 0.213487 | 0.741262 | 0.989223 | 0.997988 | 0.999507 |
| Usp15_Q8R5H1_T226t_1 | Usp15_Q8I | 0.11 | -0.8 | -0.69 | -0.52 | -0.42 | 0.35 | -0.67 | -0.41 | -1.19 | -1.16 | -1.47 | -1.43 | -0.63 | -1.1 | -0.71 | -0.73 | 0.1875 | -0.8375 | -0.3175 | 0.296072 | 0.557276 | 0.012416 | 0.989223 | 0.997988 | 0.994666 |
| Usp15_Q8R5H1_T622t_1 | Usp15_Q8I | 0.46 | -0.21 | -0.2 | -0.16 | -0.12 | -0.95 | -1.83 | 0.01 | -0.18 | 0.2 | 0.61 | 0.44 | -0.2 | -0.07 | 0.25 | -1.1 | -0.695 | 0.295 | -0.2525 | 0.486182 | 0.131396 | 0.314845 | 0.989223 | 0.997988 | 0.994666 |
| Usp20_Q8C6M1_S112s_1 | Usp20_Q8( | 0.3 | 0.7 | 0.35 | 0.42 | 0.68 | -0.05 | 0.23 | 0.41 | 0.4 | 0.61 | 1 | 0.54 | 0.64 | 0.84 | 0.62 | 0.73 | -0.125 | 0.195 | 0.265 | 0.282392 | 0.607319 | 0.426398 | 0.989223 | 0.997988 | 0.994666 |
| Usp20_Q8C6M1_S369s_1 | Usp20_Q8( | 0.41 | 0.01 | 0.38 | 0.44 | -0.04 | 0.7 | -0.24 | 0.2 | 1.03 | 0.16 | -0.92 | 0.14 | 1.38 | -0.43 | 0.28 | 0.64 | -0.155 | -0.2075 | 0.1575 | 0.692966 | 0.56806 | 0.612532 | 0.989223 | 0.997988 | 0.996159 |
| Usp20_Q8C6M1_S374s_2 | Usp20_Q8( | 0.63 | -0.9 | -0.23 | 1.32 | -0.48 | -1.56 | 0.12 | 1 | 0.76 | 0.6 | 1.27 | 0.79 | 0.11 | 0.56 | 0.16 | 0.78 | -0.435 | 0.65 | 0.1975 | 0.68638 | 0.504223 | 0.19881 | 0.989223 | 0.997988 | 0.996159 |
| Usp20_Q8C6M1_S410s_1 | Usp20_Q8( | 0.14 | 0.51 | 0.67 | -0.11 | 0.03 | -0.07 | -0.02 | -0.08 | 0.35 | 0.33 | 0.27 | 0.01 | 1.14 | -0.05 | 0.56 | 0.18 | -0.3375 | -0.0625 | 0.155 | 0.655964 | 0.185426 | 0.81001 | 0.989223 | 0.997988 | 0.995259 |
| Usp20_Q8C6M1_T378t_1 | Usp20_Q8( | -1.55 | -0.41 | -0.35 | -0.26 | 0.37 | -0.76 | -1.08 | 0.53 | 0.28 | -0.11 | 0.01 | -0.9 | 1.24 | 0.75 | 1.16 | -0.01 | 0.4075 | 0.4625 | 1.4275 | 0.006631 | 0.393244 | 0.25989 | 0.989223 | 0.997988 | 0.994666 |

| | | | | | | | | | | | | | | | | | | | | | | | | | | |
|---|---|---|---|---|---|---|---|---|---|---|---|---|---|---|---|---|---|---|---|---|---|---|---|---|---|---|
| Usp20_Q8C6M1_T378t_2 | Usp20_Q8C | 0.63 | -0.9 | -0.23 | 1.32 | -0.48 | -1.56 | 0.12 | 1 | 0.76 | 0.6 | 1.27 | 0.79 | 0.11 | 0.56 | 0.16 | 0.78 | -0.435 | 0.65 | 0.1975 | 0.68638 | 0.504223 | 0.19881 | 0.989223 | 0.997988 | 0.996159 |
| Usp22_Q5DU02_T147t_1 | Usp22_Q5I | -1.6 | -1.89 | -0.78 | -1.81 | -0.93 | -1.26 | -0.5 | -2.19 | -0.9 | -1.07 | -1.37 | -1.3 | -1.06 | -0.43 | -1.69 | -1.73 | 0.3 | 0.36 | 0.2925 | 0.474779 | 0.478989 | 0.263488 | 0.989223 | 0.997988 | 0.994666 |
| Usp24_E9PV45_S1139s_1 | Usp24_E9P | 0.3 | -0.23 | 0.5 | -0.3 | -0.56 | 0.36 | -0.38 | -0.02 | 0.6 | -0.66 | -1.71 | 0.07 | 1.37 | -0.93 | 0.67 | -0.55 | -0.2175 | -0.4925 | 0.0725 | 0.891262 | 0.482 | 0.342522 | 0.989223 | 0.997988 | 0.999507 |
| Usp24_E9PV45_S1297s_1 | Usp24_E9P | -1.35 | -2.13 | -1.6 | -1.9 | -1.3 | -1.14 | -1.69 | -0.69 | -2.05 | -2.19 | 0.79 | -1.35 | -1.42 | -0.05 | -1.77 | -1.7 | 0.54 | 0.545 | 0.51 | 0.254742 | 0.092833 | 0.407265 | 0.989223 | 0.997988 | 0.994666 |
| Usp24_E9PV45_S1923s_1 | Usp24_E9P | -1.11 | -1.74 | 0.9 | 0.83 | -2.36 | -0.67 | -2.64 | -0.9 | 1.7 | -2.17 | 1.81 | 0 | 1.67 | -2.07 | -0.07 | -0.72 | -1.3625 | 0.615 | -0.0175 | 0.984478 | 0.090043 | 0.548415 | 0.989223 | 0.997988 | 1 |
| Usp24_E9PV45_S1941s_1 | Usp24_E9P | 1.37 | 0.76 | 1.47 | -0.8 | 0.19 | 1.67 | 0.26 | 0.62 | 0.93 | 1.22 | -2.18 | 1.01 | 1.81 | -3.43 | 1.46 | 0.58 | -0.015 | -0.455 | -0.595 | 0.603712 | 0.978739 | 0.598736 | 0.999187 | 0.997988 | 0.995259 |
| Usp24_E9PV45_S2045s_1 | Usp24_E9P | 0.34 | 0.59 | 0.51 | 0.1 | 0.79 | 0.27 | 0.52 | 0.19 | -0.36 | 0.59 | -0.12 | 0.41 | -0.12 | -0.61 | 0.31 | 0.2 | 0.0575 | -0.255 | -0.44 | 0.161719 | 0.810367 | 0.392013 | 0.995762 | 0.997988 | 0.994666 |
| Usp24_E9PV45_S2559s_1 | Usp24_E9P | -1.21 | -0.74 | -1.11 | -1.5 | -0.77 | -1.16 | 0 | -1.49 | -0.37 | -0.84 | -0.45 | -1.7 | -0.94 | -0.57 | -0.83 | -1.42 | 0.285 | 0.3 | 0.2 | 0.508858 | 0.432186 | 0.408505 | 0.989223 | 0.997988 | 0.994666 |
| Usp25_P57080_T61t_1 | Usp25_P57 | -0.23 | 1.3 | 1.41 | 0.83 | 0.24 | -0.3 | 0.23 | -0.06 | 0.71 | 0.27 | 0.58 | 0.37 | -0.7 | 0.74 | 1.11 | 0.2 | -0.8 | -0.345 | -0.49 | 0.349471 | 0.061725 | 0.385266 | 0.989223 | 0.997988 | 0.994666 |
| Usp31_E9Q6Y8_S1103s_1 | Usp31_E9C | -0.62 | -0.64 | -0.51 | -0.29 | -1.23 | -1.22 | -0.58 | -0.71 | -0.09 | -0.39 | -0.22 | -0.04 | 0.36 | -0.77 | -1.14 | -0.7 | -0.42 | 0.33 | -0.0475 | 0.894373 | 0.114251 | 0.160839 | 0.989223 | 0.997988 | 0.999507 |
| Usp31_E9Q6Y8_S1127s_1 | Usp31_E9C | -1.47 | -1.65 | -0.34 | -0.8 | -1.43 | -0.18 | -0.87 | -1.08 | 0.63 | -1.47 | -1.56 | -0.48 | -0.02 | -1.47 | -0.87 | -0.52 | 0.175 | 0.345 | 0.345 | 0.426331 | 0.654551 | 0.535904 | 0.990434 | 0.997988 | 0.994666 |
| Usp31_E9Q6Y8_S49s_1 | Usp31_E9C | -0.73 | 0.02 | -0.68 | -0.46 | 0.13 | -0.9 | -0.64 | -0.62 | -0.99 | -0.68 | -0.14 | -0.42 | 0.37 | -0.49 | -0.6 | -0.58 | -0.045 | -0.095 | 0.1375 | 0.678303 | 0.881962 | 0.746301 | 0.996493 | 0.997988 | 0.996159 |
| Usp31_E9Q6Y8_S793s_1 | Usp31_E9C | -0.54 | -1.26 | -0.87 | -0.27 | -0.57 | 0.42 | -1.13 | -0.9 | -0.11 | -1.02 | -0.29 | -0.59 | -0.99 | -0.44 | -1.26 | -1.05 | 0.19 | 0.2325 | -0.2 | 0.538866 | 0.628403 | 0.474917 | 0.989223 | 0.997988 | 0.994666 |
| Usp32_F8VPZ3_S1372s_1 | Usp32_F8V | 0.73 | 0.24 | 0.26 | 0.31 | 0.42 | 0.58 | 0.06 | 0.01 | 0.85 | 0.74 | 0.62 | 0.33 | -0.08 | 0.13 | 0.41 | 0.01 | -0.1175 | 0.25 | -0.2675 | 0.315595 | 0.631002 | 0.317921 | 0.989223 | 0.997988 | 0.994666 |
| Usp32_F8VPZ3_S1394s_1 | Usp32_F8V | 0.16 | 0.85 | -0.38 | -1.12 | 0.36 | 1.13 | -0.79 | -0.72 | 3.14 | 0.02 | -2.24 | 0.61 | 3.49 | -0.23 | 2.34 | -0.56 | 0.1175 | 0.505 | 1.3825 | 0.163041 | 0.833919 | 0.629768 | 0.995762 | 0.997988 | 0.994666 |
| Usp32_F8VPZ3_T1326t_1 | Usp32_F8V | 0.98 | 0.64 | 2.02 | 1.04 | 1.71 | 1.19 | 2.29 | 1.34 | 0.1 | 0.95 | 3.03 | 1.26 | -0.84 | 0.41 | 2.41 | 2.39 | 0.4625 | 0.165 | -0.0775 | 0.91816 | 0.238589 | 0.791512 | 0.989223 | 0.997988 | 0.999507 |
| Usp35_M0QWN7_S929s_1 | Usp35_M0 | 1.24 | 0.27 | 0.34 | -0.3 | -0.36 | 0.28 | -1.04 | -1.3 | 0.98 | 0.74 | -0.2 | -0.93 | 0.93 | 1.26 | -0.96 | -0.24 | -0.9925 | -0.24 | -0.14 | 0.802537 | 0.048419 | 0.639621 | 0.989223 | 0.997988 | 0.999507 |
| Usp35_M0QWN7_T738t_1 | Usp35_M0 | -0.4 | -0.26 | -0.22 | -0.66 | -0.24 | -0.27 | -0.92 | -0.84 | 0.34 | -0.19 | -0.59 | -0.84 | 1.16 | -0.22 | -0.4 | -0.07 | -0.1825 | 0.065 | 0.5025 | 0.209016 | 0.484626 | 0.834033 | 0.989223 | 0.997988 | 0.994666 |
| Usp36_B0QZF8_S264s_1 | Usp36_B0C | 0.18 | 0.4 | 0.32 | 0.14 | -0.22 | 0.17 | -0.58 | -0.01 | 0.81 | -0.13 | -0.73 | 0.1 | 1.17 | -0.26 | -0.42 | -0.15 | -0.42 | -0.2475 | -0.175 | 0.650357 | 0.103943 | 0.476491 | 0.989223 | 0.997988 | 0.995259 |
| Usp39_Q3TIX9_S81s_1 | Usp39_Q3T | -0.41 | -1.14 | -1.25 | -1.41 | -2.07 | -1.06 | -1.69 | -1.05 | 0.31 | -0.98 | -0.31 | -1.26 | 0.62 | -2.71 | 0.5 | -0.74 | -0.415 | 0.4925 | 0.47 | 0.517104 | 0.241333 | 0.248701 | 0.989223 | 0.997988 | 0.994666 |
| Usp4_P35123_S655s_1 | Usp4_P351 | -2.73 | -1.37 | -2.36 | -2.39 | -2.03 | -2.06 | -3.26 | -3.16 | -2.46 | -1.95 | -1.2 | -2.35 | -3.03 | -1.42 | -2.02 | -2.19 | -0.415 | 0.2225 | 0.0475 | 0.913131 | 0.339456 | 0.585763 | 0.989223 | 0.997988 | 0.999507 |
| Usp4_P35123_S680s_1 | Usp4_P351 | -1.39 | -0.54 | -1.38 | -1.19 | -0.74 | -1.7 | -0.9 | -1.97 | -1.26 | -1.23 | -0.81 | -2.14 | -0.89 | -0.36 | -1.47 | -0.92 | -0.2025 | -0.235 | 0.215 | 0.529927 | 0.576106 | 0.515157 | 0.989223 | 0.997988 | 0.994666 |
| Usp42_B2RQC2_S936s_1 | Usp42_B2F | -0.29 | -0.67 | 0.5 | 0.43 | -0.91 | -0.6 | -0.63 | -0.29 | 0.89 | -0.54 | -1.92 | -0.31 | 1.9 | -1.18 | 0.1 | 0.35 | -0.6 | -0.4625 | 0.3 | 0.634844 | 0.089456 | 0.440776 | 0.989223 | 0.997988 | 0.995259 |
| Usp43_Q8BUM9_S967s_1 | Usp43_Q8I | 0.66 | 1.02 | 1.28 | 1.22 | 0.95 | 1 | 1.03 | 0.94 | 0.42 | 1.11 | 0.92 | 1.57 | 0.55 | 0.72 | 1.22 | 0.74 | -0.065 | -0.04 | -0.2375 | 0.404671 | 0.771915 | 0.897315 | 0.995762 | 0.997988 | 0.994666 |
| Usp43_Q8BUM9_S970s_1 | Usp43_Q8I | 1.57 | 1.55 | 0.8 | 0.57 | 1.44 | 1.55 | 0.52 | 0.59 | 1.88 | 1.53 | 1.31 | 0.37 | 0.43 | 1.06 | 0.53 | 0.16 | -0.0975 | 0.15 | -0.5775 | 0.123157 | 0.792024 | 0.713826 | 0.995762 | 0.997988 | 0.994666 |
| Usp47_Q8BY87_S812s_1 | Usp47_Q8I | 0.42 | -0.17 | -2.04 | -1.67 | 0.04 | -0.53 | 0.3 | -1.18 | -0.29 | -0.66 | -0.58 | -0.46 | -0.29 | -0.82 | -1.64 | -0.19 | 0.5225 | 0.3675 | 0.13 | 0.832574 | 0.395753 | 0.509894 | 0.989223 | 0.997988 | 0.999507 |
| Usp47_Q8BY87_S829s_1 | Usp47_Q8I | 1.66 | -4.31 | -0.47 | 0.74 | -0.97 | 2.41 | -0.43 | 0.12 | 2.33 | -0.44 | 0.66 | -0.77 | 1.05 | -1.39 | -0.73 | -0.9 | 0.8775 | 1.04 | 0.1025 | 0.933155 | 0.503305 | 0.43274 | 0.989223 | 0.997988 | 0.999507 |
| Usp5_Q3U4W8_S728s_1 | Usp5_Q3U | 0.87 | 0.28 | 0.95 | 0.49 | -0.81 | 1.27 | -0.7 | -0.04 | 1.84 | 0.86 | -1.45 | 0.34 | 2.03 | -0.8 | 0.47 | -0.01 | -0.7175 | -0.25 | -0.225 | 0.693198 | 0.148733 | 0.698616 | 0.989223 | 0.997988 | 0.996159 |
| Usp5_Q3U4W8_S756s_1 | Usp5_Q3U | 0.23 | 0.17 | 0 | -0.1 | -0.09 | 0.34 | 0.33 | 0.15 | 0.33 | 0.68 | 0.37 | -0.22 | 0.12 | 0.63 | -0.48 | 0.09 | 0.1075 | 0.215 | 0.015 | 0.960132 | 0.622949 | 0.425004 | 0.989223 | 0.997988 | 0.999507 |
| Usp5_Q3U4W8_S760s_1 | Usp5_Q3U | 0 | 0.38 | 0.36 | 0.42 | 0.21 | 0.18 | 0.48 | 0.55 | 0.38 | 0.65 | 0.16 | 0.3 | 0.63 | 0.69 | 0.19 | 0.15 | 0.065 | 0.0825 | 0.125 | 0.639818 | 0.769139 | 0.725061 | 0.995762 | 0.997988 | 0.995259 |
| Usp5_Q3U4W8_S762s_1 | Usp5_Q3U | 0.81 | 1.34 | 0.67 | 0.6 | 1.12 | 0.61 | 0.75 | 0.65 | -0.27 | 1.29 | 0.48 | 0.51 | 0.13 | -0.02 | 0.87 | 0.69 | -0.0725 | -0.3525 | -0.4375 | 0.192991 | 0.777315 | 0.351017 | 0.995762 | 0.997988 | 0.994666 |
| Usp5_Q3U4W8_T623t_1 | Usp5_Q3U | -0.04 | 0.6 | 0.09 | 0.17 | 0.35 | -0.04 | 0.33 | 0.14 | -0.1 | -0.08 | -0.43 | -0.03 | -0.03 | 0.42 | 0.16 | 0.28 | -0.01 | -0.365 | 0.0025 | 0.99243 | 0.966124 | 0.16113 | 0.999187 | 0.997988 | 1 |
| Usp53_P15975_S490s_1 | Usp53_P15 | -0.12 | -0.02 | 0.14 | 0.57 | -0.37 | -0.97 | -0.1 | 0.29 | -0.37 | -0.54 | 1.11 | 0.56 | 0.45 | 0.01 | -0.45 | 0.03 | -0.43 | 0.0475 | -0.1325 | 0.660282 | 0.200554 | 0.908301 | 0.989223 | 0.997988 | 0.995533 |
| Usp54_F6ZXV1_S74s_1 | Usp54_F6Z | 0.91 | 0.73 | 0.59 | 1.2 | 0.05 | 0.99 | 0.44 | 0.42 | 1.42 | 1.3 | 1.99 | 0.79 | 0.39 | 0.36 | -0.61 | -0.1 | -0.3825 | 0.5175 | -0.8475 | 0.022964 | 0.184737 | 0.123344 | 0.989223 | 0.997988 | 0.994666 |
| Usp6nl_Q80XC3_S405s_1 | Usp6nl_Q8 | -1.05 | -3.03 | -1.53 | -2.43 | -3.12 | -0.14 | -1.27 | -1.94 | 2.6 | -2.23 | -1.51 | -1 | 3.21 | -3.64 | 2.39 | -1.28 | 0.3925 | 1.475 | 2.18 | 0.15119 | 0.565381 | 0.174275 | 0.989223 | 0.997988 | 0.994666 |
| Usp6nl_Q80XC3_S544s_1 | Usp6nl_Q8 | -0.59 | -1.46 | -1.42 | -0.58 | -0.94 | -0.58 | -0.71 | -0.68 | -0.6 | -1.07 | -1.1 | -1.09 | -0.02 | -0.25 | -0.6 | -0.43 | 0.285 | 0.0475 | 0.6875 | 0.056061 | 0.336894 | 0.878054 | 0.989223 | 0.997988 | 0.994666 |
| Usp8_Q80U87_S603s_1 | Usp8_Q80U | 0.35 | -0.02 | -1.75 | -2.07 | -2.13 | 0.06 | -1.8 | -1.46 | 1.63 | -0.33 | -0.91 | -1.8 | 1.93 | -0.71 | -1.28 | -1.72 | -0.46 | 0.52 | 0.4275 | 0.633977 | 0.507427 | 0.541502 | 0.989223 | 0.997988 | 0.995259 |
| Usp8_Q80U87_S680s_1 | Usp8_Q80U | -0.45 | 0.06 | -0.62 | -0.19 | 0.35 | -0.22 | 0.91 | 1.41 | -0.94 | -0.42 | -1.49 | -0.67 | -0.49 | -0.1 | -0.8 | 0.65 | 0.9125 | -0.58 | 0.115 | 0.754874 | 0.034015 | 0.086088 | 0.989223 | 0.997988 | 0.9971 |
| Usp9x_P70398_S2443s_1 | Usp9x_P70 | 0.41 | 0.75 | 0.23 | 0.26 | 0.73 | -0.1 | 0.65 | 0.26 | -0.08 | 0.59 | -0.19 | 0.3 | -0.33 | 0.69 | 0.16 | 0.39 | -0.0275 | -0.2575 | -0.185 | 0.54703 | 0.918423 | 0.356045 | 0.997335 | 0.997988 | 0.994666 |
| Utrn_E9Q6R7_S933s_1 | Utrn_E9Q6 | -0.07 | 0.57 | 0.32 | -0.06 | 0.32 | -0.51 | 0.11 | 0.32 | 0.17 | 0.05 | -0.12 | 0.16 | 0.1 | 0.83 | -0.09 | 0.88 | -0.13 | -0.125 | 0.24 | 0.47711 | 0.648966 | 0.615589 | 0.99033 | 0.997988 | 0.994666 |
| Uvrag_Q8K245_T517t_1 | Uvrag_Q8K | 0.72 | 0.51 | 0.49 | 1.04 | 0.16 | 1.23 | 0.75 | 0.3 | 0.29 | 0.94 | 0.47 | 0.52 | -0.04 | 0.69 | 0.54 | 0.76 | -0.08 | -0.135 | -0.2025 | 0.491878 | 0.789155 | 0.601583 | 0.995762 | 0.997988 | 0.994666 |
| Vamp2_B0QZN5_S75s_1 | Vamp2_B0 | 0.9 | 0.18 | 0.62 | 0.46 | 1.62 | 1.36 | 1.37 | 1.13 | 0.16 | 0.49 | 0.28 | 1.03 | 0.34 | 1.34 | 0.71 | 1.46 | 0.83 | -0.05 | 0.4225 | 0.233115 | 0.006713 | 0.863004 | 0.989223 | 0.997988 | 0.994666 |
| Vamp2_B0QZN5_S80s_1 | Vamp2_B0 | 0.53 | 0.81 | 0.58 | -0.18 | 1.19 | 1.18 | 0.47 | 0.04 | 0.68 | 1.73 | -0.43 | 0.72 | 0.72 | 1.46 | -0.27 | 1.39 | 0.285 | 0.24 | 0.39 | 0.390045 | 0.429694 | 0.610373 | 0.989223 | 0.997988 | 0.994666 |
| Vapa_Q9WV55_S166s_1 | Vapa_Q9W | 0.2 | 0.54 | 0.13 | 0.67 | 0.72 | 0.96 | 1.07 | 0.98 | 0.35 | 0.05 | 0.28 | 0.67 | -0.86 | 0.89 | 0.44 | 1.23 | 0.5475 | -0.0475 | 0.04 | 0.930694 | 0.035765 | 0.851709 | 0.989223 | 0.997988 | 0.999507 |
| Vapa_Q9WV55_S3s_1 | Vapa_Q9W | 0.75 | 0.12 | 0.87 | 1.36 | 0.12 | 0.73 | 1.11 | 1.68 | 1.32 | -0.04 | 0.2 | -0.43 | 1.42 | 0.77 | 1.14 | -0.5 | 0.135 | -0.5125 | -0.0675 | 0.886775 | 0.73605 | 0.261698 | 0.995762 | 0.997988 | 0.999507 |
| Vapb_Q8BH80_S144s_1 | Vapb_Q8BI | 0.99 | 0.21 | 0.6 | 0.68 | 0.9 | 0.58 | 0.41 | 0.61 | 0.3 | 0.12 | 0.36 | 0.46 | 0.49 | 1.21 | 0.22 | 0.39 | 0.005 | -0.31 | -0.0425 | 0.893926 | 0.983915 | 0.235206 | 0.999187 | 0.997988 | 0.999507 |
| Vapb_Q8BH80_S206s_1 | Vapb_Q8BI | 0.09 | 0.5 | 0.5 | 0.06 | 0.88 | 0.41 | 0.34 | 0.05 | 0.52 | 0.04 | 0.06 | 0.48 | 0.39 | 0.98 | -0.37 | -0.53 | 0.1325 | -0.0125 | -0.17 | 0.659961 | 0.613063 | 0.960411 | 0.989223 | 0.997988 | 0.995259 |
| Vash1_Q8C1W1_S323s_1 | Vash1_Q8C | -0.07 | 0.28 | 1.75 | 3.09 | -0.19 | 1.58 | 0.7 | 2.93 | 1.36 | 0.57 | 2.21 | 1.24 | 1.93 | -0.61 | 0.35 | 1.79 | -0.0075 | 0.0825 | -0.3975 | 0.635232 | 0.993008 | 0.908674 | 0.999187 | 0.997988 | 0.995259 |
| Vash1_Q8C1W1_S334s_1 | Vash1_Q8C | -1.65 | -1.54 | -1.58 | -2.71 | -2.29 | -1.1 | -2.05 | -1.38 | 1 | -1.24 | -2.84 | -1.29 | 0.69 | -2.09 | -0.69 | -1.65 | 0.165 | 0.7775 | 0.935 | 0.152677 | 0.669569 | 0.315793 | 0.992876 | 0.997988 | 0.994666 |
| Vasp_P70460_S317s_1 | Vasp_P704 | -1.95 | -3.47 | -4.2 | -3.8 | -3.04 | -3.24 | -2.6 | -3.29 | -2.31 | -3.25 | -2.94 | -2.44 | -1.89 | -2.47 | -2.38 | -2.44 | 0.3125 | 0.62 | 1.06 | 0.051839 | 0.516972 | 0.239652 | 0.989223 | 0.997988 | 0.994666 |
| Vat1l_Q80TB8_S31s_1 | Vat1l_Q80T | -2.17 | -0.71 | -2.29 | -4.03 | -1.76 | -1.9 | -1.98 | -2.95 | -0.66 | -0.82 | -4.31 | -3.41 | 2.17 | -1.93 | -0.85 | -3.66 | 0.1525 | -1.6E-16 | 1.2325 | 0.321816 | 0.814337 | 1 | 0.995762 | 1 | 0.994666 |
| Vat1l_Q80TB8_S394s_1 | Vat1l_Q80T | -1.22 | 0.19 | 1 | 1.07 | -0.42 | -0.98 | 1.45 | 0.64 | -0.92 | -1.07 | 2.17 | 1.61 | -2.01 | -0.42 | 1.27 | 1.93 | -0.0875 | 0.1875 | -0.0675 | 0.940448 | 0.896361 | 0.832007 | 0.996493 | 0.997988 | 0.999507 |
| Vcam1_Q3UPN1_S166s_1 | Vcam1_Q3 | 0.16 | 0.99 | 1.38 | 0.91 | 0.47 | 0.6 | -0.16 | 0.8 | 0.33 | 0.84 | 0.99 | 1.06 | 0.75 | 0.49 | 0.96 | 1.03 | -0.4325 | -0.055 | -0.0525 | 0.872027 | 0.218569 | 0.867041 | 0.989223 | 0.997988 | 0.999507 |
| Vcl_Q64727_S290s_1 | Vcl_Q6472 | -1.56 | -1.18 | 0.12 | 0.32 | -1.53 | -1.86 | 0.27 | 0.59 | -1.69 | -1.3 | 1.45 | 0.65 | -0.51 | 0.34 | 0.17 | -0.01 | -0.0575 | 0.3525 | 0.5725 | 0.25092 | 0.933138 | 0.658853 | 0.998562 | 0.997988 | 0.994666 |
| Vcl_Q64727_S346s_1 | Vcl_Q6472 | -1.39 | -0.95 | -0.82 | -0.64 | -0.71 | -1.39 | -1 | -0.53 | -0.81 | -0.91 | -0.81 | -1.03 | -0.07 | -0.71 | -1.24 | -1.89 | 0.0425 | 0.06 | -0.0275 | 0.94749 | 0.879977 | 0.807824 | 0.996493 | 0.997988 | 0.999507 |
| Vcp_Q01853_S702s_1 | Vcp_Q0185 | -0.05 | 0.24 | 0.34 | 0.16 | 0.42 | -0.15 | 0.42 | 0.25 | 0.08 | 0.11 | -0.18 | -0.11 | 0.06 | 0.91 | -0.05 | 0.43 | 0.0625 | -0.1975 | 0.165 | 0.580613 | 0.787528 | 0.381264 | 0.995762 | 0.997988 | 0.995259 |
| Vcp_Q01853_S748s_1 | Vcp_Q0185 | 0.88 | 0.73 | 1.29 | 0.58 | -0.23 | 1.02 | -0.43 | 0.19 | 0.93 | 0.77 | 0.93 | 0.31 | 1.07 | 0.5 | 0.88 | 0.22 | -0.7325 | -0.135 | -0.2025 | 0.509263 | 0.063569 | 0.619576 | 0.989223 | 0.997988 | 0.994666 |
| Vcpip1_Q8CDG3_S1007s_1 | Vcpip1_Q8 | 0.14 | -0.04 | -1.06 | -1.18 | 1.15 | 0.06 | -0.28 | -0.23 | -1.27 | -0.1 | -0.15 | -0.33 | 0.3 | 0.14 | -0.73 | -0.84 | 0.71 | 0.0725 | 0.2525 | 0.569548 | 0.136448 | 0.864928 | 0.989223 | 0.997988 | 0.995259 |
| Vcpip1_Q8CDG3_S1196s_1 | Vcpip1_Q8 | -0.34 | -0.57 | -0.81 | -0.23 | -0.45 | -0.52 | -0.76 | -0.11 | -0.05 | -0.09 | 0.47 | -0.71 | -0.61 | 0.28 | -0.58 | -0.66 | 0.0275 | 0.3925 | 0.095 | 0.761053 | 0.910962 | 0.22373 | 0.997335 | 0.997988 | 0.9971 |
| Vcpip1_Q8CDG3_S746s_1 | Vcpip1_Q8 | -1.55 | -1.62 | -0.98 | -1.13 | -0.59 | -1.75 | -0.94 | -0.98 | -1.94 | -1.34 | 0.04 | -0.88 | -0.93 | -0.58 | -0.89 | -1.09 | 0.255 | 0.29 | 0.4475 | 0.1272 | 0.41927 | 0.508515 | 0.989223 | 0.997988 | 0.994666 |
| Vcpip1_Q8CDG3_T760t_1 | Vcpip1_Q8 | -2.3 | -1.53 | -1.83 | -1.36 | -1.37 | -1.62 | -3.73 | -2.11 | -0.97 | -1.81 | 0.22 | -0.36 | -1.38 | -2.44 | -0.78 | -1.59 | -0.4525 | 1.025 | 0.2075 | 0.610726 | 0.392519 | 0.048894 | 0.989223 | 0.997988 | 0.995259 |
| Vdac1_Q60932_S104s_1 | Vdac1_Q60 | 0.65 | 0.86 | 0.41 | 0.28 | 0.49 | 0.44 | 0.72 | 0.07 | 0.82 | 0.94 | -0.28 | 0.37 | 0.67 | 0.73 | 0.12 | 0.22 | -0.12 | -0.0875 | -0.115 | 0.682254 | 0.62799 | 0.790723 | 0.989223 | 0.997988 | 0.996159 |
| Vdac1_Q60932_S215s_1 | Vdac1_Q60 | 1.02 | 1.47 | 1.16 | 1.19 | 0.06 | 0.69 | 1.7 | 0.85 | 0.33 | 0.32 | 1.32 | 1.54 | -0.1 | 0.54 | 0.7 | 0.31 | -0.385 | -0.3325 | -0.8475 | 0.012097 | 0.289807 | 0.355476 | 0.989223 | 0.997988 | 0.994666 |
| Vdac1_Q60932_S240s_1 | Vdac1_Q60 | -2.06 | -1.72 | 0.19 | 0.32 | -1.7 | -1.43 | 0.55 | 0.01 | -4.46 | -2.52 | -0.62 | 0.52 | -1.46 | -1.25 | -0.36 | 0.03 | 0.175 | -0.9525 | 0.0575 | 0.92923 | 0.810395 | 0.399654 | 0.995762 | 0.997988 | 0.999507 |
| Vdac1_Q60932_S57s_1 | Vdac1_Q60 | -0.13 | -0.01 | -0.13 | 0.06 | -0.5 | 1.09 | -0.73 | 0.34 | 1.4 | 0.62 | -0.55 | 0.16 | 1.49 | -0.22 | 0.21 | -0.06 | 0.1025 | 0.46 | 0.4075 | 0.320265 | 0.798923 | 0.2757 | 0.995762 | 0.997988 | 0.994666 |
| Vdac1_Q60932_T107t_1 | Vdac1_Q60 | 0.82 | 1.08 | -0.5 | -0.38 | 1.25 | 0.06 | 0.7 | -1.37 | 0.86 | 0.3 | -0.85 | -0.56 | 0.3 | 1.43 | -0.7 | 0.09 | -0.095 | -0.3175 | 0.025 | 0.963937 | 0.878127 | 0.550558 | 0.996493 | 0.997988 | 0.999507 |
| Vdac1_Q60932_T250t_1 | Vdac1_Q60 | 0.9 | 1.3 | 1.08 | 1.17 | 0.99 | 1.41 | 0.93 | 1.07 | 0.6 | 1.33 | 0.55 | 0.78 | 0.85 | 0.56 | 0.64 | 0.97 | -0.0125 | -0.2975 | -0.3575 | 0.169897 | 0.954972 | 0.273335 | 0.999187 | 0.997988 | 0.994666 |
| Vdac2_G3UX26_S104s_1 | Vdac2_G3U | 0.19 | 0.97 | 0.94 | 0.81 | 1.05 | -0.07 | 0.92 | 1.09 | 0.64 | 0.6 | 0.77 | 0.81 | 0.54 | 0.75 | 0.74 | 1.08 | 0.02 | -0.0225 | 0.05 | 0.861428 | 0.952664 | 0.930336 | 0.999187 | 0.997988 | 0.999507 |
| Vdac2_G3UX26_S240s_1 | Vdac2_G3U | 1.23 | 1.62 | 0.92 | 1.36 | 0.56 | 0.89 | 1.09 | 1.65 | 0.73 | 0.96 | 0.27 | 0.44 | 0.12 | 0.72 | 1.22 | 1.13 | -0.235 | -0.6825 | -0.485 | 0.165687 | 0.436795 | 0.029443 | 0.989223 | 0.997988 | 0.994666 |
| Vdac2_G3UX26_T250t_1 | Vdac2_G3U | 1.22 | 1.7 | 1.43 | 0.97 | 0.96 | 1.7 | 1.22 | 1.41 | 1.2 | 1.72 | 0.54 | 1.25 | 0.78 | 0.77 | 1.02 | 1.32 | -0.0075 | -0.1525 | -0.3575 | 0.221805 | 0.977429 | 0.633571 | 0.999187 | 0.997988 | 0.994666 |
| Vdac2_G3UX26_T57t_1 | Vdac2_G3U | -0.01 | 0.34 | 1.5 | 1.21 | 0.48 | 0.84 | 0.15 | 0.26 | 0.73 | 0.44 | 1.41 | 0.81 | 1.18 | 0.96 | 0.84 | 1.04 | -0.3275 | 0.0875 | 0.245 | 0.522255 | 0.39653 | 0.829317 | 0.989223 | 0.997988 | 0.994666 |
| Vgf_Q0VGU4_S423s_1 | Vgf_Q0VGU | 0.62 | 0 | 0.19 | 0.4 | -0.97 | 0.72 | -0.39 | 0 | 2.95 | 0.86 | -0.42 | 0.2 | 2.34 | -1.18 | 0.8 | 0.18 | -0.4625 | 0.595 | 0.2325 | 0.728275 | 0.233727 | 0.387623 | 0.989223 | 0.997988 | 0.9971 |
| Vill_G5E8H0_S232s_1 | Vill_G5E8H | -0.02 | 1.24 | 0.13 | 0.21 | 0.59 | 1.27 | 0.76 | -0.15 | 0.29 | 0.71 | 0.1 | 0.09 | 0.58 | 0.26 | 0.15 | -0.05 | 0.2275 | -0.0925 | -0.155 | 0.656998 | 0.569207 | 0.786534 | 0.989223 | 0.997988 | 0.995259 |
| Vim_P20152_S10s_1 | Vim_P2015 | -1.48 | -2.7 | -3.07 | -4.74 | -3.21 | -0.34 | -3.36 | -2.32 | -1.29 | -2.24 | -1.33 | -1.94 | 0.01 | -3.67 | -1.5 | -3.67 | 0.69 | 1.2975 | 0.79 | 0.427667 | 0.421383 | 0.07044 | 0.989223 | 0.997988 | 0.994666 |
| Vim_P20152_S39s_1 | Vim_P2015 | -3.08 | -3.96 | -2.37 | -3.74 | -2.48 | -3.64 | -2.02 | -2.21 | -4.28 | -3.68 | 0.33 | -2.58 | -1.38 | -2.88 | -2.7 | -2.19 | 0.7 | 0.735 | 1 | 0.057449 | 0.162001 | 0.450805 | 0.989223 | 0.997988 | 0.994666 |
| Vim_P20152_S73s_1 | Vim_P2015 | -2.08 | -1.86 | -1.4 | -1.95 | -0.95 | 0.39 | -1.44 | -2.12 | 0.74 | -2.1 | -0.51 | -1.37 | 0.68 | -2.01 | -0.27 | -1.88 | 0.7925 | 1.0125 | 0.9525 | 0.141776 | 0.140594 | 0.106345 | 0.989223 | 0.997988 | 0.994666 |
| Vipas39_Q8BGQ1_S119s_1 | Vipas39_Q | -0.02 | -0.37 | 0 | 0.12 | 0.31 | 0.1 | 0.36 | 0.2 | -0.09 | -0.05 | 0.48 | 0.39 | -0.76 | 0.26 | -0.41 | -0.19 | 0.31 | 0.25 | -0.2075 | 0.494291 | 0.172129 | 0.336553 | 0.989223 | 0.997988 | 0.994666 |
| Vipas39_Q8BGQ1_S77s_1 | Vipas39_Q | -0.64 | -0.73 | -1.07 | -0.79 | -1.14 | -0.57 | -1.75 | -1.43 | 0.9 | -0.9 | -0.65 | 0.98 | 1.07 | -2.92 | -0.35 | -0.64 | -0.415 | 0.89 | 0.0975 | 0.89502 | 0.181494 | 0.090422 | 0.989223 | 0.997988 | 0.999507 |
| Vps13c_Q8BX70_S1891s_1 | Vps13c_Q8 | 0.5 | 0.92 | 0.19 | 0.63 | 0.51 | 0.07 | 0.75 | 0.06 | 0.1 | 0.84 | -0.5 | 0.32 | -0.59 | 0.57 | -0.16 | 0.66 | -0.2125 | -0.37 | -0.44 | 0.24058 | 0.439052 | 0.288609 | 0.989223 | 0.997988 | 0.994666 |
| Vps13c_Q8BX70_S2442s_1 | Vps13c_Q8 | -0.15 | -0.42 | 1.28 | 1.07 | -0.45 | -0.24 | 0.8 | 0.65 | 1 | -0.02 | 0.74 | 0.64 | 1.09 | -1.12 | 0.93 | 0.88 | -0.255 | 0.145 | 3.14E-16 | 1 | 0.60347 | 0.752561 | 0.989223 | 0.997988 | 1 |
| Vps13c_Q8BX70_S736s_1 | Vps13c_Q8 | 0.68 | 0.68 | 1.01 | 1.07 | 0.83 | 0.48 | 1.53 | 0.86 | 0.32 | 0.67 | 0.88 | 0.6 | -0.3 | 0.85 | 1.28 | 1.48 | 0.065 | -0.2425 | -0.0325 | 0.937211 | 0.816113 | 0.327026 | 0.995762 | 0.997988 | 0.999507 |
| Vps13c_Q8BX70_S841s_1 | Vps13c_Q8 | 0.39 | 0.07 | 0.78 | 0.56 | -0.71 | 0.87 | -0.32 | 0.13 | 2.59 | 0.13 | -0.42 | 0.56 | 2.35 | -1.06 | 0.36 | 0.58 | -0.4575 | 0.265 | 0.1075 | 0.86779 | 0.23136 | 0.667969 | 0.989223 | 0.997988 | 0.999507 |
| Vps13c_Q8BX70_S841s_2 | Vps13c_Q8 | 0.42 | 0.19 | 1.13 | 0.67 | -0.32 | 1.09 | 0.2 | 0.18 | 2.3 | 0.23 | -0.62 | 0.89 | -0.66 | -0.64 | 1.12 | 0.86 | -0.315 | 0.0975 | -0.4325 | 0.388673 | 0.385966 | 0.869734 | 0.989223 | 0.997988 | 0.994666 |
| Vps13d_B1ART2_S1043s_1 | Vps13d_B1 | 0.9 | -0.06 | -0.67 | 0.42 | 1.04 | 0.46 | 0.28 | -0.13 | 0.23 | 1.38 | -2.5 | -0.39 | 0.27 | 1.01 | -0.42 | 0.69 | 0.265 | -0.4675 | 0.24 | 0.592754 | 0.51134 | 0.556267 | 0.989223 | 0.997988 | 0.995259 |
| Vps13d_B1ART2_S1725s_1 | Vps13d_B1 | 0.7 | 0.6 | 0.22 | 0.34 | 0.86 | 0.15 | 0.64 | 0.4 | 0.03 | 0.21 | 0.31 | 0.41 | -0.15 | 1.04 | 0.01 | 0.79 | 0.0475 | -0.225 | -0.0425 | 0.901717 | 0.848057 | 0.345131 | 0.996349 | 0.997988 | 0.999507 |
| Vps13d_B1ART2_S2080s_1 | Vps13d_B1 | 1.13 | 0.4 | 0.25 | 0.05 | 1.13 | 1.32 | 0.27 | 0.05 | 0.3 | 0.91 | 0.82 | -0.03 | 0.57 | 1.67 | 0.31 | 0.92 | 0.235 | 0.0425 | 0.41 | 0.306219 | 0.542413 | 0.901145 | 0.989223 | 0.997988 | 0.994666 |
| Vps13d_B1ART2_S2435s_1 | Vps13d_B1 | 0.56 | 0.75 | 1.11 | 0.27 | 0.94 | 0.45 | 1.16 | 0.84 | 0.62 | 0.11 | 0.69 | 0.61 | 0.03 | 1.45 | 1.16 | 0.77 | 0.175 | -0.165 | 0.18 | 0.631227 | 0.523832 | 0.553294 | 0.989223 | 0.997988 | 0.994666 |
| Vps13d_B1ART2_S2862s_1 | Vps13d_B1 | -2.14 | -1.21 | -2.42 | -2.15 | -1.13 | -1.83 | -1.22 | -2.05 | -2.32 | -1.62 | -0.62 | -0.39 | -1 | -0.78 | -2.58 | -1.82 | 0.4225 | 0.7425 | 0.435 | 0.36591 | 0.246091 | 0.156159 | 0.989223 | 0.997988 | 0.994666 |
| Vps13d_B1ART2_S2865s_1 | Vps13d_B1 | -0.11 | 0.54 | -0.02 | -0.59 | 0.13 | 0.26 | 0.86 | -0.52 | -0.68 | 0.76 | -0.9 | 0.05 | 0.17 | -0.6 | -1.01 | -0.13 | 0.2275 | -0.1475 | -0.3475 | 0.357373 | 0.534999 | 0.732512 | 0.989223 | 0.997988 | 0.994666 |
| Vps13d_B1ART2_S2920s_1 | Vps13d_B1 | 0.97 | 0.58 | 0.33 | 0.51 | 1.57 | 1.29 | 0.94 | 0.48 | 0.1 | 0.88 | -1.74 | 0.62 | -0.46 | 1.35 | 0.05 | 0.42 | 0.4725 | -0.6325 | -0.2575 | 0.532464 | 0.136507 | 0.27614 | 0.989223 | 0.997988 | 0.994666 |
| Vps13d_B1ART2_S3011s_1 | Vps13d_B1 | -0.3 | 0.11 | 0.17 | 0.23 | -0.55 | 0.1 | -0.42 | -0.12 | -1.41 | 0.12 | -0.67 | 0 | 0.79 | -4.62 | 0.31 | 0.06 | -0.3 | -0.5425 | -0.9175 | 0.412168 | 0.244826 | 0.174466 | 0.989223 | 0.997988 | 0.994666 |
| Vps26a_P40336_S315s_1 | Vps26a_P4 | 0.71 | 1.99 | 0.52 | 0.37 | 0.76 | 1.29 | 0.6 | 0.34 | 0.77 | 1.28 | 0.37 | 0.2 | 2.02 | 0.86 | 0.37 | 0.44 | -0.15 | -0.2425 | 0.025 | 0.96028 | 0.711363 | 0.576049 | 0.995437 | 0.997988 | 0.999507 |
| Vps26b_Q8C0E2_S302s_1 | Vps26b_Q8 | -1.08 | -0.26 | 1.13 | 1.25 | -0.41 | -0.31 | 0.28 | 1.55 | -0.17 | -1.06 | 0 | 1.55 | 0.46 | -0.09 | 0.64 | 0.76 | 0.0175 | -0.18 | 0.1825 | 0.740533 | 0.978182 | 0.798525 | 0.999187 | 0.997988 | 0.9971 |
| Vps26b_Q8C0E2_S304s_1 | Vps26b_Q8 | -0.73 | -0.77 | -0.41 | -0.47 | -0.14 | -0.55 | -0.57 | -0.67 | -0.24 | -0.76 | -0.63 | 0.01 | 0.4 | -0.15 | -0.11 | -0.47 | 0.1125 | 0.19 | 0.5125 | 0.0924 | 0.622298 | 0.4762 | 0.989223 | 0.997988 | 0.994666 |
| Vps26b_Q8C0E2_S330s_1 | Vps26b_Q8 | 1.08 | 0.41 | 1.2 | 1.31 | -0.04 | 1.05 | 0.08 | 0.71 | 1.17 | 1.08 | 1.61 | 0.88 | 1.1 | 0.21 | 0.55 | 0.53 | -0.55 | 0.185 | -0.4025 | 0.228986 | 0.12703 | 0.53632 | 0.989223 | 0.997988 | 0.994666 |
| Vps26b_Q8C0E2_S334s_1 | Vps26b_Q8 | -0.27 | -1.59 | -0.33 | 0.31 | -1.9 | 0.25 | -0.78 | 0.21 | -0.06 | 0.09 | 0.54 | 0.33 | -0.1 | -0.24 | 0.26 | 0.14 | -0.085 | 0.695 | 0.485 | 0.261099 | 0.883487 | 0.119891 | 0.996493 | 0.997988 | 0.994666 |
| Vps26b_Q8C0E2_T325t_1 | Vps26b_Q8 | -1.42 | -3.37 | -1.9 | -3.15 | -2.12 | -2.46 | -1.43 | -1.73 | -2.92 | -2.32 | 1.28 | -1.48 | -2.26 | -1.56 | -1.83 | -3.55 | 0.525 | 1.1 | 0.16 | 0.787276 | 0.293198 | 0.252263 | 0.989223 | 0.997988 | 0.999507 |
| Vps35_Q9EQH3_S783s_1 | Vps35_Q9E | -0.02 | 0.07 | -0.06 | -0.17 | 0.23 | 0.12 | 0.77 | 0.07 | -0.17 | 0.14 | 0.39 | 0.34 | -0.41 | 0.37 | 0.21 | 0.37 | 0.3425 | 0.22 | 0.18 | 0.517741 | 0.171236 | 0.354225 | 0.989223 | 0.997988 | 0.994666 |
| Vps35_Q9EQH3_S7s_1 | Vps35_Q9E | 0.05 | 0.55 | 0.87 | 0.5 | -0.9 | 1.08 | -0.8 | -0.05 | 2.44 | 0.28 | -0.42 | 0.95 | 2.11 | -0.55 | 0.33 | -0.94 | -0.66 | 0.32 | -0.255 | 0.688178 | 0.169569 | 0.585666 | 0.989223 | 0.997988 | 0.996159 |
| Vps45_P97390_S441s_1 | Vps45_P97 | 0.07 | 0.22 | 0.34 | 0.47 | -0.21 | -0.57 | 0.6 | 0.17 | 0.25 | 0.11 | -0.02 | 0.53 | 0.51 | 0.78 | 0.38 | 0.56 | -0.2775 | -0.0575 | 0.2825 | 0.263995 | 0.356127 | 0.807311 | 0.989223 | 0.997988 | 0.994666 |
| Vps4a_Q8VEJ9_S97s_1 | Vps4a_Q8\ | -1.46 | -1.17 | -1.49 | -2.5 | -2.37 | -1.09 | -2.13 | -1.88 | 0.52 | -0.71 | -0.05 | -2.31 | 1.49 | -1.76 | -0.75 | -1.42 | -0.2125 | 1.0175 | 1.045 | 0.160942 | 0.588399 | 0.125891 | 0.989223 | 0.997988 | 0.994666 |
| Vps4b_P46467_S102s_1 | Vps4b_P46 | -1.12 | -0.41 | 0 | -0.93 | -2.19 | -0.34 | -1.53 | -0.95 | 1.48 | -0.37 | 0.51 | -0.35 | 0.59 | -1.24 | -0.24 | -0.57 | -0.6375 | 0.9325 | 0.25 | 0.578442 | 0.171732 | 0.078379 | 0.989223 | 0.997988 | 0.995259 |
| Vps51_Q3UVL4_S640s_1 | Vps51_Q3U | -2.32 | -1.76 | 0.13 | -0.48 | -2 | -1.51 | 2.04 | 1.75 | 3.41 | -2.17 | -1.11 | 1.44 | 3.56 | -1.68 | 3.17 | -1.36 | 1.1775 | 1.5 | 2.03 | 0.146031 | 0.272946 | 0.2351 | 0.989223 | 0.997988 | 0.994666 |
| Vps51_Q3UVL4_S652s_1 | Vps51_Q3U | -2.47 | -3.54 | -3.58 | -4.14 | -2.23 | -2.58 | -2.92 | -3.08 | -3.05 | -2.26 | 0.29 | -2.14 | -2.01 | -1.75 | -3.19 | -4.02 | 0.73 | 1.6425 | 0.69 | 0.253086 | 0.083707 | 0.04505 | 0.989223 | 0.997988 | 0.994666 |
| Vsnl1_P62761_T146t_1 | Vsnl1_P62 | 0.21 | 1.08 | 0.69 | 0.08 | 0.78 | 0.56 | 0.5 | -0.01 | 0.75 | 0.78 | 0.88 | 0.38 | 1.19 | 0.65 | 0.51 | 0.5 | -0.0575 | 0.1825 | 0.1975 | 0.547766 | 0.850749 | 0.541755 | 0.996349 | 0.997988 | 0.994666 |
| Vti1b_F6UHS3_S80s_1 | Vti1b_F6UI | 0.59 | -0.26 | -0.43 | 0.25 | 0.53 | 0.51 | 0.42 | 0.25 | -0.56 | -0.3 | 0.03 | -0.02 | -0.76 | 0.29 | -0.77 | -0.18 | 0.39 | -0.25 | -0.3925 | 0.300379 | 0.184508 | 0.423853 | 0.989223 | 0.997988 | 0.994666 |
| Vwa5b1_A9Z1V5_S664s_1 | Vwa5b1_A | 1.67 | -0.33 | 0.81 | 1.63 | -1.24 | 2.06 | 0.75 | 0.9 | 1.49 | -0.1 | 0.81 | -0.03 | 1.57 | -0.16 | 1.53 | 1.28 | -0.3275 | -0.4025 | 0.11 | 0.847743 | 0.655056 | 0.476304 | 0.990434 | 0.997988 | 0.999507 |
| Vwa5b1_A9Z1V5_S804s_1 | Vwa5b1_A | 0.48 | 0.32 | 0.41 | 0.47 | 1.46 | -0.04 | 0.46 | 0.2 | 0.08 | -0.25 | 1.19 | -0.48 | 0.69 | 0.96 | 1.06 | 0.42 | 0.1 | -0.285 | 0.3625 | 0.177972 | 0.767944 | 0.457001 | 0.995762 | 0.997988 | 0.994666 |
| Vwa5b2_E0CZ01_S1017s_1 | Vwa5b2_E( | 0.13 | 0.82 | -0.83 | -0.02 | -0.01 | -0.03 | 0.61 | -0.11 | 0.17 | 0.7 | 0.78 | -0.76 | 0.62 | 0.53 | 0.04 | -0.53 | 0.09 | 0.1975 | 0.14 | 0.742904 | 0.808625 | 0.674115 | 0.995762 | 0.997988 | 0.9971 |
| Vwa5b2_E0CZ01_S759s_1 | Vwa5b2_E( | 1.35 | 0.48 | 0.23 | -0.04 | 0.99 | 1.78 | -0.37 | 0.03 | 2.24 | 0.63 | -0.77 | 0.03 | 2.39 | 0.78 | -0.05 | 0.05 | 0.1025 | 0.0275 | 0.2875 | 0.625528 | 0.843491 | 0.965741 | 0.996349 | 0.997988 | 0.995259 |
| Wasf1_Q8R5H6_S310s_1 | Wasf1_Q8I | -1.53 | -1.15 | -0.86 | -1.56 | -1.07 | -0.73 | -1.48 | -1.12 | -1.19 | -1.65 | -0.99 | -0.97 | -1.01 | -0.51 | -1.66 | -1.61 | 0.175 | 0.075 | 0.0775 | 0.824689 | 0.520583 | 0.789672 | 0.989223 | 0.997988 | 0.999507 |
| Wasf1_Q8R5H6_S321s_1 | Wasf1_Q8I | -0.02 | 0.75 | 1.02 | 1.12 | 0.34 | 0.31 | 0.93 | 0.81 | 0.09 | -0.27 | 0.93 | 0.45 | -0.11 | 0.29 | -0.13 | 1.19 | -0.12 | -0.4175 | -0.4075 | 0.329113 | 0.707322 | 0.277585 | 0.995437 | 0.997988 | 0.994666 |
| Wasf1_Q8R5H6_T323t_1 | Wasf1_Q8I | 1.02 | -0.13 | 0.15 | -0.03 | 0.75 | -0.16 | 0.28 | 0.34 | 0.01 | 0.54 | -0.79 | -0.61 | -0.42 | -0.31 | -0.44 | 0.12 | 0.05 | -0.465 | -0.515 | 0.145828 | 0.880031 | 0.262161 | 0.996493 | 0.997988 | 0.994666 |
| Wasf3_Q8VHI6_S229s_1 | Wasf3_Q8\ | -0.78 | -0.69 | -0.04 | 0.12 | -0.16 | -1.44 | -0.59 | -0.28 | -0.17 | -0.45 | 1.01 | 0.83 | 0.41 | -0.21 | -0.14 | -0.43 | -0.27 | 0.6525 | 0.255 | 0.446911 | 0.464834 | 0.144404 | 0.989223 | 0.997988 | 0.994666 |
| Wbp11_Q923D5_S237s_1 | Wbp11_Q9 | -1.37 | -0.68 | -0.79 | -0.29 | -1.58 | -1.01 | 0.22 | 0.65 | -1.67 | -0.81 | 0.23 | -0.62 | -0.64 | -0.87 | 0.35 | -0.16 | 0.3525 | 0.065 | 0.4525 | 0.24058 | 0.500016 | 0.881368 | 0.989223 | 0.997988 | 0.994666 |
| Wbp4_Q61048_S97s_1 | Wbp4_Q61 | -0.86 | -0.57 | -1.82 | -1.37 | -0.95 | -1.48 | -1.2 | -1.63 | -2.69 | -0.76 | -2.31 | -1.04 | -0.87 | -1 | -1.48 | -1.79 | -0.16 | -0.545 | -0.13 | 0.725596 | 0.626425 | 0.30309 | 0.989223 | 0.997988 | 0.9971 |
| Wdfy3_G3UYW1_S1942s_1 | Wdfy3_G3U | 0.33 | 0.73 | 0.58 | 0.53 | -0.01 | 0.3 | 0.59 | 0.57 | 0.07 | 0.6 | 0.66 | 0.87 | 0.61 | 1.07 | 1.06 | 0.67 | -0.18 | 0.0075 | 0.31 | 0.242837 | 0.449404 | 0.976738 | 0.989223 | 0.997988 | 0.994666 |
| Wdfy3_G3UYW1_S2277s_1 | Wdfy3_G3U | -1.86 | -1.79 | -2.14 | -1.91 | -0.78 | -1.5 | -0.63 | -2.02 | -3.33 | -1.34 | -2.41 | -1.04 | -1.57 | -2.67 | -1.97 | -2.06 | 0.6925 | -0.105 | -0.1425 | 0.637309 | 0.065482 | 0.83319 | 0.989223 | 0.997988 | 0.995259 |
| Wdfy3_G3UYW1_S2434s_1 | Wdfy3_G3U | -2 | -1.49 | -0.79 | -1.48 | -0.83 | -1.58 | -1.33 | -1.67 | -1.1 | -0.96 | -0.19 | -1.05 | -0.61 | -0.65 | -1.38 | -2.16 | 0.0875 | 0.615 | 0.24 | 0.584071 | 0.787658 | 0.101103 | 0.995762 | 0.997988 | 0.995259 |
| Wdr11_Q8K1X1_S405s_1 | Wdr11_Q8 | 0.44 | 0.1 | 0.59 | 0.4 | 0.43 | -0.63 | 0.93 | 1.19 | -0.37 | 0.11 | -0.05 | 0.1 | -0.85 | 0.08 | -0.32 | 0.79 | 0.0975 | -0.435 | -0.4575 | 0.243431 | 0.807258 | 0.09441 | 0.995762 | 0.997988 | 0.994666 |
| Wdr13_Q91V09_S70s_1 | Wdr13_Q9 | -0.85 | -0.07 | -0.18 | -0.59 | -0.32 | -0.93 | -0.41 | -0.47 | -1.18 | -0.66 | -0.51 | -0.52 | -0.77 | -0.48 | -0.67 | 0.14 | -0.11 | -0.295 | -0.0225 | 0.94393 | 0.684073 | 0.318788 | 0.994379 | 0.997988 | 0.999507 |
| Wdr13_Q91V09_S79s_1 | Wdr13_Q9 | -2.03 | -2.26 | -1.57 | -2.43 | -1.55 | -2.58 | -2.06 | -2.05 | -2.47 | -1.6 | 0.13 | -1.36 | -1.54 | -1.37 | -1.81 | -2.23 | 0.0125 | 0.7475 | 0.335 | 0.302454 | 0.967139 | 0.183654 | 0.999187 | 0.997988 | 0.994666 |
| Wdr17_H3BLI8_S309s_1 | Wdr17_H3 | 0.71 | 0.67 | 0.22 | 0.06 | 0.09 | 0.42 | 0.6 | 0.12 | 0.29 | 0.17 | 0.64 | 0.24 | -0.67 | 0.23 | 0.46 | -0.23 | -0.1075 | -0.08 | -0.4675 | 0.186549 | 0.67554 | 0.758415 | 0.992876 | 0.997988 | 0.994666 |
| Wdr20_Q3UWE6_S357s_1 | Wdr20_Q3 | 0.24 | -0.12 | 0.48 | 0.05 | 0.75 | 0.79 | 0.32 | 0.85 | 0.85 | -0.15 | 0.33 | 0.75 | 0.5 | 0.37 | 0.51 | -0.11 | 0.515 | 0.2825 | 0.155 | 0.57814 | 0.055801 | 0.359782 | 0.989223 | 0.997988 | 0.995259 |
| Wdr20_Q3UWE6_S434s_1 | Wdr20_Q3 | -1.07 | -0.68 | -0.59 | 0.15 | -0.14 | -1.1 | -0.31 | -0.45 | 0.02 | -0.56 | -0.88 | -0.49 | 0.14 | 0.01 | 0 | 0.28 | 0.0475 | 0.07 | 0.655 | 0.060803 | 0.887952 | 0.835418 | 0.996493 | 0.997988 | 0.994666 |
| Wdr20_Q3UWE6_S465s_1 | Wdr20_Q3 | -1.61 | -1.93 | -1.67 | -2 | -0.8 | -2.63 | -1.05 | -1.26 | -2.42 | -2.17 | -0.66 | -1.5 | -2.78 | -1.36 | -1.55 | -1.48 | 0.3675 | 0.115 | 0.01 | 0.978213 | 0.373017 | 0.77519 | 0.989223 | 0.997988 | 1 |
| Wdr25_E9Q349_S60s_1 | Wdr25_E9( | 0.35 | -0.18 | -0.67 | 0.38 | 0.63 | 0.21 | -1.13 | 0.21 | 0.93 | 0.3 | 0.95 | 0.15 | 1.29 | 0.14 | 0.09 | -0.08 | 0.01 | 0.6125 | 0.39 | 0.347637 | 0.981471 | 0.100721 | 0.999187 | 0.997988 | 0.994666 |
| Wdr26_E0CYH4_S101s_1 | Wdr26_E0( | 1.04 | 1.06 | 0.54 | 0.44 | 1.1 | 0.69 | 0.08 | -0.08 | 1.01 | 0.5 | 0.45 | 0.71 | 1.06 | 1.56 | 1.1 | 0.01 | -0.3225 | -0.1025 | 0.1625 | 0.670923 | 0.339815 | 0.702375 | 0.989223 | 0.997988 | 0.996159 |
| Wdr33_Q8K4P0_S1204s_1 | Wdr33_Q8 | 1.27 | -0.21 | 0.05 | -0.47 | 0.75 | 0.48 | 0.32 | 1 | 0.17 | 0.22 | 0.3 | 0.14 | 0.91 | -0.39 | 2.13 | 1.64 | 0.4775 | 0.0475 | 0.9125 | 0.158417 | 0.248389 | 0.90251 | 0.989223 | 0.997988 | 0.994666 |
| Wdr37_Q8CBE3_S30s_1 | Wdr37_Q8 | -1.57 | -1.93 | -1.62 | -2.23 | -3.2 | -0.92 | -0.49 | -0.47 | 3.18 | -3.07 | -2.7 | -0.22 | 2.92 | -3.13 | 2.38 | -2.22 | 0.5675 | 1.135 | 1.825 | 0.195759 | 0.355024 | 0.381921 | 0.989223 | 0.997988 | 0.994666 |
| Wdr37_Q8CBE3_S31s_1 | Wdr37_Q8 | -1.86 | -2.22 | -1.56 | -6.54 | -2.93 | -0.36 | -1.26 | -1.21 | 3.35 | -2.36 | -0.99 | -0.61 | 3.35 | -3.42 | 3.09 | -2.91 | 1.605 | 2.8925 | 3.0725 | 0.125702 | 0.171972 | 0.07684 | 0.989223 | 0.997988 | 0.994666 |
| Wdr4_E9Q156_S440s_1 | Wdr4_E9Q | -0.65 | -0.96 | -1.26 | -2.11 | 0.34 | -1.5 | -1.07 | -1.66 | -0.38 | -0.33 | -0.39 | -1.55 | -0.72 | -0.92 | -1.16 | -1.58 | 0.2725 | 0.5825 | 0.15 | 0.693887 | 0.592673 | 0.190585 | 0.989223 | 0.997988 | 0.996159 |
| Wdr44_Q6NVE8_S405s_1 | Wdr44_Q6 | -0.97 | -0.3 | -0.63 | -1.25 | -1.22 | -1.2 | -1.37 | -1.35 | -1.08 | -0.73 | -0.29 | -0.81 | -0.94 | -1.37 | -0.46 | -1.19 | -0.4975 | 0.06 | -0.2025 | 0.540916 | 0.080628 | 0.842457 | 0.989223 | 0.997988 | 0.994666 |
| Wdr44_Q6NVE8_S50s_1 | Wdr44_Q6 | 0.07 | 0.17 | 0.5 | -0.95 | -0.46 | 0.09 | -0.74 | -0.7 | -0.51 | 0.11 | -0.3 | -0.21 | -0.36 | -0.23 | -0.06 | 0.25 | -0.4 | -0.175 | -0.0475 | 0.895807 | 0.287101 | 0.622168 | 0.989223 | 0.997988 | 0.999507 |
| Wdr44_Q6NVE8_S563s_1 | Wdr44_Q6 | 0.27 | 0.87 | 0.58 | 0.61 | 1.1 | -0.12 | 0.45 | 0.27 | 0.06 | 0.72 | 0.58 | 0.3 | 0.36 | 1.24 | 0.65 | 0.57 | -0.1575 | -0.1675 | 0.1225 | 0.676528 | 0.607711 | 0.522308 | 0.989223 | 0.997988 | 0.996159 |
| Wdr44_Q6NVE8_S576s_1 | Wdr44_Q6 | -2.37 | -1.09 | -0.21 | -2.7 | -1.59 | -2.91 | -1.31 | -1.47 | -1.62 | -1.46 | 0.27 | -0.88 | -0.9 | -2.34 | -0.71 | -0.82 | -0.2275 | 0.67 | 0.4 | 0.528955 | 0.710344 | 0.3187 | 0.995437 | 0.997988 | 0.994666 |
| Wdr47_Q8CGF6_S292s_1 | Wdr47_Q8 | 1.57 | 0.34 | 1.24 | -0.99 | 0.71 | 0.64 | -0.03 | 0.8 | -0.08 | 0.15 | 1.18 | 1.05 | 0.62 | -0.04 | -0.07 | 0.44 | -0.01 | 0.035 | -0.3025 | 0.587516 | 0.985356 | 0.953379 | 0.999187 | 0.997988 | 0.995259 |
| Wdr47_Q8CGF6_S297s_1 | Wdr47_Q8 | -0.11 | -0.51 | -0.68 | -0.69 | -0.27 | -0.37 | -0.44 | -0.3 | -0.4 | -0.4 | 0.2 | -0.81 | -0.74 | -0.12 | -1.09 | -0.76 | 0.1525 | 0.145 | -0.18 | 0.555973 | 0.500603 | 0.622021 | 0.989223 | 0.997988 | 0.995259 |
| Wdr47_Q8CGF6_S304s_1 | Wdr47_Q8 | -0.21 | -0.44 | -0.43 | -0.37 | -0.05 | -0.22 | -0.13 | 0.07 | 0.04 | 0.09 | 0.17 | -0.74 | -0.36 | -0.03 | -0.94 | -0.48 | 0.28 | 0.2525 | -0.09 | 0.745696 | 0.186536 | 0.368147 | 0.989223 | 0.997988 | 0.9971 |
| Wdr47_Q8CGF6_S359s_1 | Wdr47_Q8 | -0.79 | -0.66 | 0.71 | 0.91 | -0.27 | -1.37 | 0.88 | 0.59 | -0.53 | -0.18 | -0.51 | 0.18 | -0.3 | 0.33 | 0.36 | 0.78 | -0.085 | -0.3025 | 0.25 | 0.602686 | 0.887725 | 0.512814 | 0.996493 | 0.997988 | 0.995259 |
| Wdr47_Q8CGF6_S374s_1 | Wdr47_Q8 | -0.93 | 0.27 | -1.18 | -0.95 | 0.14 | -0.6 | -0.85 | -0.55 | -0.73 | 0.61 | -1.11 | -0.67 | -0.09 | 0.65 | -0.66 | -0.68 | 0.2325 | 0.2225 | 0.5025 | 0.273377 | 0.545425 | 0.640251 | 0.989223 | 0.997988 | 0.994666 |
| Wdr47_Q8CGF6_T285t_1 | Wdr47_Q8 | -1.5 | -1.95 | -0.76 | -0.16 | -0.08 | -1.21 | -0.32 | -0.68 | -1.84 | -2.52 | -0.4 | -0.42 | -1.94 | -1.35 | -0.84 | -0.82 | 0.52 | -0.2025 | -0.145 | 0.753195 | 0.253148 | 0.738473 | 0.989223 | 0.997988 | 0.9971 |
| Wdr59_Q8C0M0_S603s_1 | Wdr59_Q8 | -0.1 | -0.37 | 0.07 | -1 | -0.47 | -0.62 | -0.91 | -0.93 | -3.04 | -0.1 | -3.6 | -0.63 | -0.93 | -4.22 | -0.2 | -0.11 | -0.3825 | -1.4925 | -1.015 | 0.263353 | 0.208433 | 0.089212 | 0.989223 | 0.997988 | 0.994666 |
| Wdr7_Q920I9_S1151s_1 | Wdr7_Q92 | 0.03 | 0.38 | 0.43 | 0.31 | 0.59 | 0.34 | 0.45 | 0.45 | -0.5 | 0.23 | 0.18 | 0.88 | -0.18 | 0.58 | -0.04 | 0.46 | 0.17 | -0.09 | -0.0825 | 0.770411 | 0.422913 | 0.780961 | 0.989223 | 0.997988 | 0.9971 |
| Wdr7_Q920I9_S1455s_1 | Wdr7_Q92 | 0.24 | 0.76 | 0.61 | 0.65 | 0.71 | 0.28 | 0.85 | 0.59 | 0.42 | 0.56 | 0.3 | 0.64 | 0.49 | 0.94 | 0.66 | 0.73 | 0.0425 | -0.085 | 0.14 | 0.584834 | 0.856797 | 0.713984 | 0.996349 | 0.997988 | 0.995259 |
| Wdr7_Q920I9_S716s_1 | Wdr7_Q92 | -1.92 | -1.06 | -1.76 | -1.5 | -1.81 | -1.1 | -1.11 | -1.79 | -2.25 | -2.05 | -4.29 | -1.11 | -0.96 | -1.07 | -1.36 | -3.26 | 0.1075 | -0.865 | -0.1025 | 0.847336 | 0.720515 | 0.194466 | 0.995437 | 0.997988 | 0.999507 |
| Wdr7_Q920I9_S772s_1 | Wdr7_Q92 | 2.83 | 0.55 | 2.08 | 2.57 | -0.07 | 3.4 | -0.02 | 2.41 | 2.21 | 1.41 | 1.82 | 2.34 | 1.53 | 0.15 | 0 | 1.11 | -0.5775 | -0.0625 | -1.31 | 0.045276 | 0.516813 | 0.903589 | 0.989223 | 0.997988 | 0.994666 |
| Wdr7_Q920I9_S772s_2 | Wdr7_Q92 | 2.62 | 1.05 | 1.55 | 2.06 | -0.56 | 2.99 | -0.4 | 1.5 | 2.66 | 0.94 | 0.74 | 1.81 | -0.31 | -0.62 | 0.41 | 0.79 | -0.9375 | -0.2825 | -1.7525 | 0.003331 | 0.255781 | 0.588991 | 0.989223 | 0.997988 | 0.994666 |
| Wdr7_Q920I9_S776s_1 | Wdr7_Q92 | 0.6 | 0.27 | 0.8 | 0.81 | -0.88 | 0.93 | -0.47 | 0.26 | 1.62 | 0.6 | 0.5 | 0.63 | 0.74 | -1.5 | 0.3 | 0.2 | -0.66 | 0.2175 | -0.685 | 0.180125 | 0.126458 | 0.502351 | 0.989223 | 0.997988 | 0.994666 |
| Wdr7_Q920I9_S776s_2 | Wdr7_Q92 | 2.62 | 1.05 | 1.55 | 2.06 | -0.56 | 2.99 | -0.4 | 1.5 | 2.66 | 0.94 | 0.74 | 1.81 | -0.31 | -0.62 | 0.41 | 0.79 | -0.9375 | -0.2825 | -1.7525 | 0.003331 | 0.255781 | 0.588991 | 0.989223 | 0.997988 | 0.994666 |
| Wdr7_Q920I9_S935s_1 | Wdr7_Q92 | -0.06 | 1.06 | -1.51 | 0.25 | 0.33 | -0.41 | 0.21 | -0.27 | 0.13 | 0.6 | -0.97 | 0.02 | 0.29 | 0.75 | -0.12 | 0.17 | 0.03 | 0.01 | 0.3375 | 0.527711 | 0.953584 | 0.986216 | 0.999187 | 0.997988 | 0.994666 |
| Wdr7_Q920I9_T1195t_1 | Wdr7_Q92 | 1.39 | 0.49 | 0.7 | 0.93 | 0.86 | 0.38 | 0.65 | 0.67 | 1.04 | 1.09 | 0.56 | 0.67 | 1.57 | 0.56 | 0.83 | 0.83 | -0.2375 | -0.0375 | 0.07 | 0.832582 | 0.377922 | 0.894673 | 0.989223 | 0.997988 | 0.999507 |
| Wdr7_Q920I9_T926t_1 | Wdr7_Q92 | -1.55 | -1.12 | -0.84 | -1.14 | -1.14 | -1.05 | -0.78 | -0.93 | -1.38 | -1.34 | -0.05 | -0.21 | -1.6 | -0.85 | -1.39 | -1.1 | 0.1875 | 0.4175 | -0.0725 | 0.80249 | 0.433785 | 0.296766 | 0.989223 | 0.997988 | 0.999507 |
| Wdr77_D6RIL4_T5t_1 | Wdr77_D6 | -0.13 | 0.33 | 0.05 | 0 | 0.43 | -0.07 | 0.43 | -0.14 | -0.45 | 0.25 | 0.07 | -0.15 | 0.11 | 0.25 | 0.1 | -0.1 | 0.1 | -0.1325 | 0.0275 | 0.910379 | 0.683596 | 0.601935 | 0.994272 | 0.997988 | 0.999507 |
| Wdr81_K4DI77_S1570s_1 | Wdr81_K4I | 0.64 | 0.54 | 0.71 | 0.44 | 0.94 | 0.5 | 0.74 | 0.69 | 0.31 | 0.87 | 0.51 | 0.46 | 0.02 | 0.88 | 0.71 | 0.77 | 0.135 | -0.045 | 0.0125 | 0.964482 | 0.525138 | 0.844805 | 0.989223 | 0.997988 | 0.999507 |
| Wdr91_S4R1X1_S257s_1 | Wdr91_S4I | -0.55 | 0 | -0.88 | -0.48 | -0.71 | -0.45 | -1.63 | -0.33 | -0.42 | -0.01 | -0.72 | -0.65 | 0.5 | -0.65 | -0.37 | -0.82 | -0.3025 | 0.0275 | 0.1425 | 0.698396 | 0.395042 | 0.923956 | 0.989223 | 0.997988 | 0.996159 |
| Wdtc1_Q80ZK9_S227s_1 | Wdtc1_Q8( | -1 | 0.81 | 0.35 | 0.38 | -1.68 | 0.68 | -0.73 | 0.6 | 0.31 | -0.85 | 1.31 | -0.98 | 0.34 | 0.21 | 0.4 | -0.45 | -0.4175 | -0.1875 | -0.01 | 0.981536 | 0.502598 | 0.75884 | 0.989223 | 0.997988 | 1 |
| Wfs1_P56695_S158s_1 | Wfs1_P566 | 0.56 | 0.3 | 0.84 | 0.9 | 0.15 | 0.16 | 0.7 | 1.16 | 0.24 | 0.87 | 1.34 | 0.72 | 0.81 | 1.23 | 1.26 | 0.14 | -0.1075 | 0.1425 | 0.21 | 0.533956 | 0.723046 | 0.640239 | 0.995437 | 0.997988 | 0.994666 |
| Wfs1_P56695_S236s_1 | Wfs1_P566 | -0.05 | -1.04 | -1.09 | -2.39 | -0.94 | 0.06 | -1.13 | -1.91 | 0.47 | -0.64 | -2.28 | -1.72 | 0.89 | -1.13 | -0.67 | -0.92 | 0.1625 | 0.1 | 0.685 | 0.27435 | 0.774134 | 0.886154 | 0.995762 | 0.997988 | 0.994666 |
| Wfs1_P56695_S32s_1 | Wfs1_P566 | 0.86 | 0.54 | -1.18 | -1.08 | 0.83 | 0.63 | -0.66 | -0.93 | 0.41 | 0.35 | -0.87 | -1.42 | 0.34 | 0.87 | -0.79 | -0.98 | 0.1825 | -0.1675 | 0.075 | 0.905064 | 0.768445 | 0.792843 | 0.995762 | 0.997988 | 0.999507 |
| Wfs1_P56695_S3s_1 | Wfs1_P566 | 0.33 | 0.7 | -0.43 | -0.04 | 0.57 | 0.32 | -0.41 | -0.61 | 0.61 | 0.84 | 0.47 | -0.11 | -0.19 | 0.92 | 0.11 | 0.43 | -0.1725 | 0.3125 | 0.1775 | 0.626761 | 0.641272 | 0.365551 | 0.989223 | 0.997988 | 0.995259 |
| Wfs1_P56695_S3s_2 | Wfs1_P566 | -0.31 | -0.03 | -0.32 | -0.15 | -0.54 | -0.44 | -0.19 | -0.12 | -0.17 | -0.1 | 0.92 | 0.55 | -0.08 | 0.26 | 0.31 | -0.13 | -0.12 | 0.5025 | 0.2925 | 0.257673 | 0.58055 | 0.127708 | 0.989223 | 0.997988 | 0.994666 |
| Wfs1_P56695_S9s_2 | Wfs1_P566 | -0.31 | -0.03 | -0.32 | -0.15 | -0.54 | -0.44 | -0.19 | -0.12 | -0.17 | -0.1 | 0.92 | 0.55 | -0.08 | 0.26 | 0.31 | -0.13 | -0.12 | 0.5025 | 0.2925 | 0.257673 | 0.58055 | 0.127708 | 0.989223 | 0.997988 | 0.994666 |
| Wfs1_P56695_T30t_1 | Wfs1_P566 | 1.51 | 1.11 | 1.44 | 1.67 | 1.63 | 1.26 | 1.13 | 1.53 | 0.96 | 0.98 | 1.17 | 0.43 | 1.04 | 1.69 | 1.54 | 1.66 | -0.045 | -0.5475 | 0.05 | 0.855646 | 0.84836 | 0.061211 | 0.996349 | 0.997988 | 0.999507 |
| Wfs1_P56695_T5t_1 | Wfs1_P566 | 0.33 | 0.7 | -0.43 | -0.04 | 0.57 | 0.32 | -0.41 | -0.61 | 0.61 | 0.84 | 0.47 | -0.11 | -0.19 | 0.92 | 0.11 | 0.43 | -0.1725 | 0.3125 | 0.1775 | 0.626761 | 0.641272 | 0.365551 | 0.989223 | 0.997988 | 0.995259 |
| Wfs1_P56695_T5t_2 | Wfs1_P566 | -0.31 | -0.03 | -0.32 | -0.15 | -0.54 | -0.44 | -0.19 | -0.12 | -0.17 | -0.1 | 0.92 | 0.55 | -0.08 | 0.26 | 0.31 | -0.13 | -0.12 | 0.5025 | 0.2925 | 0.257673 | 0.58055 | 0.127708 | 0.989223 | 0.997988 | 0.994666 |
| Wipf2_Q6PEV3_S155s_1 | Wipf2_Q6F | 0.22 | 0.24 | -0.05 | 0.13 | 0.01 | -0.44 | 0.13 | 0.33 | 0.01 | 0.01 | -0.21 | 0.25 | 0.33 | 0.95 | -0.21 | -0.29 | -0.1275 | -0.12 | 0.06 | 0.857221 | 0.599057 | 0.592894 | 0.989223 | 0.997988 | 0.999507 |
| Wipf2_Q6PEV3_S175s_1 | Wipf2_Q6F | 3.6 | 2.1 | 2.49 | 0.56 | 2.1 | 3.92 | 3.03 | 2.15 | 0.47 | 3.38 | -0.45 | 2.35 | 1.36 | -0.81 | 2.54 | 3.16 | 0.6125 | -0.75 | -0.625 | 0.51141 | 0.374027 | 0.438026 | 0.989223 | 0.997988 | 0.994666 |
| Wipf3_D3Z5R4_S161s_1 | Wipf3_D3Z | -0.9 | -0.67 | -0.6 | -1.04 | -1.11 | -2.08 | 0.03 | -0.44 | -3 | -1.19 | 1.41 | -0.89 | -1.54 | -0.25 | -0.48 | -0.22 | -0.0975 | -0.115 | 0.18 | 0.614312 | 0.824766 | 0.887374 | 0.995762 | 0.997988 | 0.995259 |
| Wipf3_D3Z5R4_S211s_1 | Wipf3_D3Z | 0.57 | 0.98 | 0.38 | 0.46 | 0.78 | 0.59 | 0.66 | 0.54 | -0.03 | 0.25 | -0.26 | 0.04 | 0.4 | 1.06 | 0.38 | 0.27 | 0.045 | -0.5975 | -0.07 | 0.810516 | 0.84127 | 0.035528 | 0.995932 | 0.997988 | 0.999507 |
| Wipf3_D3Z5R4_T106t_1 | Wipf3_D3Z | -0.72 | 0.04 | 0.15 | 0.1 | -0.07 | -0.46 | 0.24 | -0.29 | -0.7 | -0.06 | -0.36 | -0.51 | -0.65 | 0.23 | -0.37 | -0.48 | -0.0375 | -0.3 | -0.21 | 0.522784 | 0.89595 | 0.316539 | 0.996493 | 0.997988 | 0.994666 |
| Wipf3_D3Z5R4_T240t_1 | Wipf3_D3Z | -0.11 | -0.34 | -0.31 | -0.43 | 0.02 | 0.47 | 0.03 | 0.29 | -0.82 | -0.53 | -0.67 | -0.92 | -1.27 | 0.25 | -0.74 | -0.2 | 0.5 | -0.4375 | -0.1925 | 0.596238 | 0.042894 | 0.072623 | 0.989223 | 0.997988 | 0.995259 |
| Wipf3_D3Z5R4_T338t_1 | Wipf3_D3Z | 0.92 | 0.99 | -0.1 | 0.14 | 0.13 | 0.52 | 0.51 | -0.34 | 0.21 | 0.87 | -1.45 | -0.01 | -0.38 | 0.27 | -0.1 | 0.22 | -0.2825 | -0.5825 | -0.485 | 0.183104 | 0.422039 | 0.282713 | 0.989223 | 0.997988 | 0.994666 |
| Wipf3_D3Z5R4_T357t_1 | Wipf3_D3Z | -0.16 | -0.96 | -0.26 | -0.26 | 0.37 | -0.88 | -0.14 | 0.13 | -0.61 | -0.15 | -0.23 | -0.92 | -1.03 | -0.16 | -1.1 | -0.1 | 0.28 | -0.0675 | -0.1875 | 0.59996 | 0.413213 | 0.820719 | 0.989223 | 0.997988 | 0.995259 |
| Wipi2_Q80W47_S395s_1 | Wipi2_Q80 | -0.37 | -0.51 | 0.05 | 0.68 | 0.3 | -0.26 | -0.14 | 0.4 | -0.04 | -0.68 | 0.8 | -0.04 | 0.79 | 0.2 | 0.41 | 0.81 | 0.1125 | 0.0475 | 0.59 | 0.108825 | 0.729817 | 0.905743 | 0.995762 | 0.997988 | 0.994666 |
| Wnk1_P83741_S1037s_1 | Wnk1_P83 | 0.17 | 1 | 0.45 | 0.69 | 1.58 | 1.12 | 1.67 | 0.49 | 0.57 | 0.96 | 2.23 | 1.18 | -0.08 | 0.36 | 0.46 | 1.99 | 0.6375 | 0.6575 | 0.105 | 0.822047 | 0.080014 | 0.122464 | 0.989223 | 0.997988 | 0.999507 |
| Wnk1_P83741_S165s_1 | Wnk1_P83 | -0.57 | -0.93 | -0.69 | -0.35 | 0.68 | -0.79 | 0.84 | -0.95 | -1.62 | -0.76 | -0.48 | 0.36 | -1.01 | -0.37 | -0.4 | -2.42 | 0.58 | 0.01 | -0.415 | 0.391566 | 0.222393 | 0.98088 | 0.989223 | 0.997988 | 0.994666 |
| Wnk1_P83741_S168s_1 | Wnk1_P83 | -2.85 | -3.38 | -4.41 | -0.72 | -1.21 | -3.83 | -2.22 | -4.08 | -3.33 | -3.14 | 0.66 | -3.87 | -3.51 | -1.71 | -4.07 | -2.84 | 0.005 | 0.42 | -0.1925 | 0.81415 | 0.99555 | 0.713294 | 0.999616 | 0.997988 | 0.999507 |
| Wnk1_P83741_S2222s_1 | Wnk1_P83 | -0.15 | 0.47 | 0.39 | -0.26 | 0.23 | -0.6 | 0.08 | 0.02 | -0.17 | -0.18 | 0.22 | 0.18 | -0.12 | 0.58 | -0.01 | -0.26 | -0.18 | -0.1 | -0.065 | 0.83577 | 0.539411 | 0.713948 | 0.989223 | 0.997988 | 0.999507 |

| | | | | | | | | | | | | | | | | | | | | | | | | | | |
|---|---|---|---|---|---|---|---|---|---|---|---|---|---|---|---|---|---|---|---|---|---|---|---|---|---|---|
| Wnk1_P83741_S2273s_1 | Wnk1_P83 | -0.65 | 0.32 | -0.35 | -0.61 | -0.17 | -0.72 | -0.46 | -1.11 | -0.58 | 0.05 | -0.41 | -0.69 | 0.62 | -0.31 | -0.51 | -0.29 | -0.2925 | -0.085 | 0.2 | 0.583167 | 0.366685 | 0.784806 | 0.989223 | 0.997988 | 0.995259 |
| Wnk1_P83741_S2276s_1 | Wnk1_P83 | -2.54 | -2.08 | -2.01 | -1.23 | -1.93 | -1.68 | -1.21 | -1.85 | -1 | -1.6 | -2.04 | -2.57 | -0.61 | -1.67 | -2.08 | -1.8 | 0.2975 | 0.1625 | 0.425 | 0.324672 | 0.3743 | 0.700372 | 0.989223 | 0.997988 | 0.994666 |
| Wnk1_P83741_S2616s_1 | Wnk1_P83 | -0.42 | -0.2 | -0.06 | 0.12 | 0.45 | -0.73 | 0.08 | 0.12 | -1.18 | -0.73 | 0.5 | -0.66 | -0.78 | -0.32 | -0.43 | -0.03 | 0.12 | -0.3775 | -0.25 | 0.374512 | 0.690283 | 0.334146 | 0.994672 | 0.997988 | 0.994666 |
| Wnk1_P83741_T71t_1 | Wnk1_P83 | -0.4 | -0.45 | -1.49 | -1.26 | -0.84 | 0.14 | -2.19 | -1.94 | 1.13 | -0.47 | -1.2 | -0.8 | 1.72 | -1.52 | -0.26 | -0.88 | -0.3075 | 0.565 | 0.665 | 0.338645 | 0.576804 | 0.310752 | 0.989223 | 0.997988 | 0.994666 |
| Wnk2_E9Q6Q2_S125s_1 | Wnk2_E9Q | 0.81 | 0.57 | 0.23 | 0.37 | 1.18 | 1.39 | 0.75 | 0.94 | 0.99 | 1.42 | -0.54 | 1 | 0.67 | 1.7 | 0.65 | 0.59 | 0.57 | 0.2225 | 0.4075 | 0.240686 | 0.04195 | 0.611949 | 0.989223 | 0.997988 | 0.994666 |
| Wnk2_E9Q6Q2_S1456s_1 | Wnk2_E9Q | 0.24 | -0.03 | 0.31 | 0.4 | -0.1 | 0.22 | 0.62 | 0.54 | -0.65 | -0.45 | 0.67 | 0.69 | -1.2 | 0.16 | 0.71 | -0.35 | 0.09 | -0.165 | -0.4 | 0.348695 | 0.717027 | 0.661649 | 0.995437 | 0.997988 | 0.994666 |
| Wnk2_E9Q6Q2_S1660s_1 | Wnk2_E9Q | -0.13 | -0.5 | 0.59 | 0.18 | 0.56 | 0.18 | 0.92 | 0.79 | -0.17 | 0.16 | -0.04 | 0.34 | -0.57 | 0.76 | -0.81 | 0.06 | 0.5775 | 0.0375 | -0.175 | 0.677861 | 0.083289 | 0.899748 | 0.989223 | 0.997988 | 0.996159 |
| Wnk2_E9Q6Q2_S1687s_1 | Wnk2_E9Q | -0.18 | -0.5 | 0.13 | 0.18 | 0.32 | 0.82 | 0.18 | 0.59 | -0.33 | -0.09 | 1.39 | -0.11 | 0.06 | 0.93 | -0.22 | -0.33 | 0.57 | 0.3075 | 0.2025 | 0.570406 | 0.051352 | 0.46833 | 0.989223 | 0.997988 | 0.995259 |
| Wnk2_E9Q6Q2_S988s_1 | Wnk2_E9Q | 2.32 | 2.16 | 1.75 | 1.6 | 1.27 | 3.03 | 1.4 | 2.2 | 0.31 | 1.77 | 2.58 | 1.21 | -1.21 | 0.18 | 1.29 | 1.1 | 0.0175 | -0.49 | -1.6175 | 0.014767 | 0.966649 | 0.322069 | 0.999187 | 0.997988 | 0.994666 |
| Wnk2_E9Q6Q2_S990s_1 | Wnk2_E9Q | 1.86 | 1.99 | -0.37 | 1 | 1.33 | 2.1 | 0.64 | 1.5 | 1.25 | 1.85 | -0.02 | -0.07 | -0.5 | 1.21 | 0.43 | 0.31 | 0.2725 | -0.3675 | -0.7575 | 0.220141 | 0.628246 | 0.578275 | 0.989223 | 0.997988 | 0.994666 |
| Wnk4_F6Z709_S18s_1 | Wnk4_F6Z | -0.86 | -2.38 | -2.44 | -0.33 | -2.09 | -1.19 | -1.98 | -2.43 | 0.17 | -2.07 | -1.08 | -1.31 | 0.58 | -2.42 | -0.91 | -1.4 | -0.42 | 0.43 | 0.465 | 0.52969 | 0.444056 | 0.509802 | 0.989223 | 0.997988 | 0.994666 |
| Wrnip1_Q91XU0_S153s_1 | Wrnip1_Q9 | -3.49 | -3.24 | -2.37 | -1.6 | -2.66 | -3.92 | -1.11 | -1.77 | -3.21 | -1.84 | -1.12 | -2.32 | -0.95 | -3.17 | -1.19 | -1.34 | 0.31 | 0.5525 | 1.0125 | 0.121006 | 0.640519 | 0.343195 | 0.989223 | 0.997988 | 0.994666 |
| Wrnip1_Q91XU0_S465s_1 | Wrnip1_Q9 | -3.04 | -3.54 | -2.19 | -5.05 | -4.22 | -1.91 | -1.81 | -3.24 | -1.68 | -1.65 | -1.81 | -1.87 | -0.76 | -3.98 | -1.12 | -1.67 | 0.66 | 1.7025 | 1.5725 | 0.082915 | 0.3773 | 0.012596 | 0.989223 | 0.997988 | 0.994666 |
| Wrnip1_Q91XU0_S65s_1 | Wrnip1_Q9 | -3.56 | -3.07 | -3.05 | -4.54 | -2.17 | -4.49 | -4.42 | -4.3 | -3.08 | -2.95 | -3.34 | -3.86 | -1.21 | -3.73 | -3.45 | -3.18 | -0.29 | 0.2475 | 0.6625 | 0.292896 | 0.625583 | 0.540584 | 0.989223 | 0.997988 | 0.994666 |
| Wrnip1_Q91XU0_S75s_1 | Wrnip1_Q9 | -3.5 | -3.36 | -2.54 | -4.08 | -1.58 | -4.27 | -2.98 | -3.09 | -4.54 | -2.92 | -0.83 | -3.26 | -1.01 | -1.49 | -2.39 | -4.03 | 0.39 | 0.4825 | 1.14 | 0.111101 | 0.500289 | 0.522993 | 0.989223 | 0.997988 | 0.994666 |
| Wtap_E0CYH0_S297s_1 | Wtap_E0C | 0.19 | 1.17 | 1.05 | 0.41 | 1 | 0.97 | 1.73 | 1.42 | -0.18 | -0.21 | 1.34 | 1.64 | 0.2 | -0.59 | 1.82 | 1.33 | 0.575 | -0.0575 | -0.015 | 0.978271 | 0.095049 | 0.910441 | 0.989223 | 0.997988 | 1 |
| Wtip_Q7TQJ8_S29s_1 | Wtip_Q7T | -1.99 | -3.12 | -1.74 | -1.68 | -3.41 | -1.96 | -3.39 | -2.4 | -0.52 | -2.21 | -1.51 | -1.88 | -0.27 | -5.02 | -1.46 | -1.51 | -0.6575 | 0.6025 | 0.0675 | 0.942862 | 0.176091 | 0.223184 | 0.989223 | 0.997988 | 0.999507 |
| Wwc1_Q5SXA9_S891s_1 | Wwc1_Q5 | 0.57 | 0.71 | 0.03 | 0.55 | 0.76 | 0.72 | 0.75 | 0.63 | 0.02 | 0.74 | 0.51 | 0.62 | 0.28 | 0.32 | 0.52 | 0.9 | 0.25 | 0.0075 | 0.04 | 0.887223 | 0.290255 | 0.978096 | 0.989223 | 0.997988 | 0.999507 |
| Wwc2_Q6NXJ0_T999t_1 | Wwc2_Q6 | -2.06 | -1.44 | -0.8 | -1.9 | -0.37 | -1.47 | -0.95 | -1.23 | -1.24 | -2.17 | -1.73 | -1.73 | -0.53 | -0.46 | -1.72 | -1.07 | 0.545 | -0.1675 | 0.605 | 0.162648 | 0.159168 | 0.638252 | 0.989223 | 0.997988 | 0.994666 |
| Wwp2_Q9DBH0_S211s_1 | Wwp2_Q9 | -0.38 | -0.81 | 0.37 | -0.16 | -1.08 | -0.13 | -0.93 | -0.1 | 1.32 | -0.3 | -0.21 | -0.08 | 1.51 | -2.84 | 0.04 | -0.06 | -0.315 | 0.4275 | -0.0925 | 0.911042 | 0.386845 | 0.343526 | 0.989223 | 0.997988 | 0.999507 |
| Xkr4_Q5GH67_S207s_1 | Xkr4_Q5G | 0.45 | 0.26 | 1.05 | 0.76 | 0.16 | 0.84 | -1.66 | 0.02 | 0.5 | 0.8 | 0.8 | 0.76 | 0.71 | 0.95 | 0.17 | -2.04 | -0.79 | 0.085 | -0.6825 | 0.299116 | 0.146084 | 0.741406 | 0.989223 | 0.997988 | 0.994666 |
| Xkr4_Q5GH67_S520s_1 | Xkr4_Q5G | 0.3 | 0.19 | 0.14 | 0.06 | 0.64 | 0.51 | 0.2 | -0.03 | -0.05 | 0.59 | 0.26 | -0.12 | -0.29 | 0.64 | -0.16 | 0.07 | 0.1575 | -0.0025 | -0.1075 | 0.707112 | 0.503218 | 0.991931 | 0.989223 | 0.997988 | 0.996966 |
| Xkr4_Q5GH67_S522s_1 | Xkr4_Q5G | 0.14 | 0.13 | 0.17 | 0.49 | 0.36 | 0.25 | 0.51 | 0.32 | -0.23 | 0.49 | 0.28 | 0.22 | -0.02 | 0.49 | 0.22 | 0.76 | 0.1275 | -0.0425 | 0.13 | 0.636773 | 0.544086 | 0.864789 | 0.989223 | 0.997988 | 0.995259 |
| Xkr4_Q5GH67_S87s_1 | Xkr4_Q5G | -0.65 | -0.94 | -1.54 | -1.68 | -0.82 | -0.17 | -1.25 | -1.33 | -1.01 | -0.4 | -0.5 | -1.47 | -0.47 | -0.77 | -1.53 | -1.62 | 0.31 | 0.3575 | 0.105 | 0.785962 | 0.397915 | 0.333492 | 0.989223 | 0.997988 | 0.999507 |
| Xkr4_Q5GH67_T208t_1 | Xkr4_Q5G | 0.2 | 0.71 | 0.89 | 1.28 | 0.12 | 0.5 | 1.05 | 0.66 | 0.22 | 0.77 | 0.8 | 0.99 | 0.89 | 0.65 | 1.12 | 1.04 | -0.1875 | -0.075 | 0.155 | 0.613307 | 0.55341 | 0.810098 | 0.989223 | 0.997988 | 0.995259 |
| Xkr4_Q5GH67_T579t_1 | Xkr4_Q5G | -0.57 | -0.8 | -2.27 | -2.25 | -0.39 | -0.35 | -1.34 | -2.38 | -0.27 | -0.74 | -1.53 | -1.54 | 0.1 | 0.07 | -1.92 | -2.51 | 0.3575 | 0.4525 | 0.4075 | 0.577832 | 0.550583 | 0.392617 | 0.989223 | 0.997988 | 0.995259 |
| Xkr6_E9Q6C8_T106t_1 | Xkr6_E9Q6 | 4.9 | 2.6 | 3.25 | 0.75 | 2.74 | 5.37 | 3.03 | 1.82 | 2.57 | 3.72 | 1.29 | 2.9 | 3.6 | -0.21 | 3.82 | 3.6 | 0.365 | -0.255 | -0.1725 | 0.877853 | 0.712854 | 0.773245 | 0.995437 | 0.997988 | 0.999507 |
| Xkr6_E9Q6C8_T539t_1 | Xkr6_E9Q6 | 0.91 | 0.78 | 1.52 | 1.37 | 0.21 | 1.41 | 0.61 | 0.91 | 2.63 | 1.07 | -0.16 | 1.26 | 2.69 | 0.13 | 1.32 | 0.92 | -0.36 | 0.055 | 0.12 | 0.820046 | 0.280834 | 0.920838 | 0.989223 | 0.997988 | 0.999507 |
| Xkr6_E9Q6C8_T569t_1 | Xkr6_E9Q6 | -2.19 | -1.46 | -1.04 | -1.54 | 0.03 | 0.16 | -1.2 | -1.64 | -0.77 | -1.34 | -0.92 | -0.84 | 0.26 | -0.32 | -1.21 | -1.61 | 0.895 | 0.59 | 0.8375 | 0.098781 | 0.081787 | 0.079663 | 0.989223 | 0.997988 | 0.994666 |
| Xkr7_Q5GH64_S154s_1 | Xkr7_Q5G | 0.22 | 0.66 | 0.88 | -0.29 | 0.69 | 0.2 | 0.24 | 0.05 | 0.12 | 0.24 | 0.57 | 0.46 | 0.22 | 0.8 | 0.17 | -0.09 | -0.0725 | -0.02 | -0.0925 | 0.791563 | 0.816181 | 0.94875 | 0.995762 | 0.997988 | 0.999507 |
| Xkr7_Q5GH64_T496t_1 | Xkr7_Q5G | 0.44 | 0.64 | 0.43 | 0.1 | 0.92 | 0.34 | 0.61 | 0.47 | 0.18 | 0.66 | 0.06 | 0.19 | 0.46 | 0.86 | -0.17 | 0.25 | 0.1825 | -0.13 | -0.0525 | 0.861944 | 0.447322 | 0.604405 | 0.989223 | 0.997988 | 0.999507 |
| Xpo4_Q9ESJ0_S521s_1 | Xpo4_Q9E | 0.87 | 0.49 | 2.26 | 1.61 | 1.03 | 1.24 | 3.03 | 2.7 | -0.46 | 1.52 | 0.51 | 2.76 | -0.45 | 0.15 | 2.64 | 2.56 | 0.6925 | -0.225 | -0.0825 | 0.916859 | 0.24741 | 0.753472 | 0.989223 | 0.997988 | 0.999507 |
| Xpr1_D3Z2B4_S600s_1 | Xpr1_D3Z2 | -2.41 | -3.06 | -3.5 | -4.03 | -1.56 | -2.7 | -3.63 | -3.02 | -3.35 | -1.89 | -1.81 | -2.45 | -0.95 | -1.2 | -2.02 | -2.08 | 0.5225 | 0.875 | 1.6875 | 0.003406 | 0.316016 | 0.089207 | 0.989223 | 0.997988 | 0.994666 |
| Xrcc1_Q60596_S484s_1 | Xrcc1_Q60 | -1.69 | -1.76 | -1.14 | -0.49 | -1.54 | 0.05 | -0.99 | -1.52 | -0.4 | -2.55 | -0.7 | -2.84 | 1.27 | -1.77 | -0.3 | -0.88 | 0.27 | -0.3525 | 0.85 | 0.203277 | 0.547782 | 0.578751 | 0.989223 | 0.997988 | 0.994666 |
| Xrn1_F8VQ87_T1041t_1 | Xrn1_F8VC | -1.45 | -0.95 | -0.72 | -1.25 | 0.04 | -1.78 | -0.72 | -0.45 | -0.85 | -1.58 | -1.53 | -1.37 | 0.86 | -0.17 | -1.44 | -1.12 | 0.365 | -0.24 | 0.625 | 0.239854 | 0.37321 | 0.404639 | 0.989223 | 0.997988 | 0.994666 |
| Xrn2_Q9DBR1_S475s_1 | Xrn2_Q9D | -0.21 | -0.5 | -0.27 | 0.19 | -0.09 | -0.48 | 0.61 | 0.17 | -0.31 | -0.52 | -0.55 | 0.01 | -0.78 | -1.03 | -0.07 | -0.26 | 0.25 | -0.145 | -0.3375 | 0.299733 | 0.40828 | 0.5796 | 0.989223 | 0.997988 | 0.994666 |
| Xrn2_Q9DBR1_S499s_1 | Xrn2_Q9D | -0.92 | -1.53 | -1.66 | -1.17 | -1.8 | -0.89 | -1.12 | -1.68 | -0.83 | -0.88 | -0.61 | -2.25 | -0.82 | -1.65 | -1.65 | -0.67 | -0.0525 | 0.1775 | 0.1225 | 0.723344 | 0.861334 | 0.663389 | 0.996493 | 0.997988 | 0.9971 |
| Yaf2_Q99LW6_S173s_1 | Yaf2_Q99L | -2.04 | -1.57 | -2.52 | -2.97 | -2.64 | -1.52 | -4.34 | -3.16 | -0.92 | -1.74 | -2.1 | -3.35 | -0.12 | -3.29 | -2.86 | -3.07 | -0.64 | 0.2475 | -0.06 | 0.933172 | 0.293575 | 0.652121 | 0.989223 | 0.997988 | 0.999507 |
| Yap1_G3UYA6_S94s_1 | Yap1_G3U | -0.68 | 0.2 | 0.05 | -0.06 | 0.15 | -0.5 | -0.08 | -0.04 | -0.48 | -0.04 | 0.36 | 0.19 | 0.41 | -0.01 | 0.36 | 0.01 | 0.005 | 0.13 | 0.315 | 0.295001 | 0.985511 | 0.669545 | 0.999187 | 0.997988 | 0.994666 |
| Yars_Q91WQ3_S6s_1 | Yars_Q91V | -0.77 | 0.29 | -0.37 | -0.45 | -0.85 | 0 | -0.85 | -1.14 | 0.95 | -0.55 | -0.12 | -1.1 | 1.26 | -0.84 | 0.34 | -0.89 | -0.385 | 0.12 | 0.2925 | 0.580861 | 0.272342 | 0.796874 | 0.989223 | 0.997988 | 0.995259 |
| Ybx1_P62960_S163s_1 | Ybx1_P629 | -0.06 | -0.24 | -0.36 | -1.84 | 0.19 | -1.2 | -1.09 | -1.31 | -0.95 | 0.55 | -0.43 | -0.61 | 0.24 | -0.93 | -0.45 | -0.69 | -0.2275 | 0.265 | 0.1675 | 0.718885 | 0.647374 | 0.593005 | 0.99033 | 0.997988 | 0.9971 |
| Ybx1_P62960_S172s_1 | Ybx1_P629 | -2.92 | -3.36 | -2.33 | -3.16 | -4.94 | -2.74 | -1.58 | -3.68 | -1.98 | -2.44 | -2.38 | -3.12 | -0.69 | -3.56 | -1.97 | -2.46 | -0.2925 | 0.4625 | 0.7725 | 0.205946 | 0.659707 | 0.200137 | 0.990434 | 0.997988 | 0.994666 |
| Ybx1_P62960_S174s_1 | Ybx1_P629 | -2.61 | -1.64 | -2.95 | -0.91 | -2.55 | -1.82 | -2.82 | -3.12 | -1.38 | -2.42 | -1.55 | -0.56 | 0.24 | -3.8 | -0.16 | -0.82 | -0.55 | 0.55 | 0.8925 | 0.333615 | 0.284098 | 0.336033 | 0.989223 | 0.997988 | 0.994666 |
| Ybx1_P62960_S312s_1 | Ybx1_P629 | -1.49 | -0.49 | -1.1 | -0.84 | -1.6 | -0.58 | -0.35 | -0.57 | -0.3 | 0.25 | 0.06 | -1.44 | 0.82 | -1.33 | -0.86 | -1.52 | 0.205 | 0.6225 | 0.2575 | 0.63155 | 0.563756 | 0.1659 | 0.989223 | 0.997988 | 0.995259 |
| Ybx3_Q9JKB3_S259s_1 | Ybx3_Q9JK | -2.85 | -1.26 | -2.77 | -4.27 | -1.02 | -3.2 | -2.15 | -3.38 | -3.24 | -1.5 | -1.31 | -3.29 | -1.54 | -0.82 | -2.71 | -2.78 | 0.35 | 0.4525 | 0.825 | 0.253567 | 0.630066 | 0.541855 | 0.989223 | 0.997988 | 0.994666 |
| Yeats2_Q3TUF7_S467s_1 | Yeats2_Q3 | 0.66 | 0.37 | 0.52 | 0.33 | 0.91 | -0.08 | 0.61 | 0.52 | -0.62 | 0.74 | 0.3 | 0.5 | -0.11 | 0.26 | -0.09 | 0.55 | 0.02 | -0.24 | -0.3175 | 0.251714 | 0.939974 | 0.474468 | 0.999187 | 0.997988 | 0.994666 |
| Yes1_Q04736_S26s_1 | Yes1_Q047 | -1.2 | -0.94 | -1.84 | -1.38 | -1.51 | -1.47 | -1 | -1.09 | -1.54 | -1.31 | -3.19 | -1.44 | -0.17 | -1.2 | -1.62 | -1.35 | 0.0725 | -0.53 | 0.255 | 0.510732 | 0.7895 | 0.267761 | 0.995762 | 0.997988 | 0.994666 |
| Ythdc1_D6RG95_S321s_1 | Ythdc1_D6 | 0.01 | -0.44 | -0.49 | -0.91 | -0.49 | -0.98 | -0.37 | -0.54 | -1.02 | -1.3 | -0.44 | -1.07 | -0.94 | -0.29 | -0.42 | -1.18 | -0.1375 | -0.5 | -0.25 | 0.450627 | 0.615045 | 0.122862 | 0.989223 | 0.997988 | 0.994666 |
| Ythdc2_B2RR83_S1104s_1 | Ythdc2_B2 | -0.62 | -0.31 | -1.16 | -1.4 | -0.36 | -0.9 | -1.8 | -1.96 | -0.8 | 0.01 | -1.42 | -2.09 | -0.46 | 0.45 | -1.2 | -1.64 | -0.3825 | -0.2025 | 0.16 | 0.746488 | 0.381923 | 0.677386 | 0.989223 | 0.997988 | 0.9971 |
| Ythdc2_B2RR83_S1105s_1 | Ythdc2_B2 | -0.66 | -0.34 | -1.09 | -0.29 | -0.79 | -0.66 | -0.66 | -1.12 | -0.12 | -0.5 | 0.34 | -0.99 | 1.45 | 0.46 | -0.87 | -1.27 | -0.2125 | 0.2775 | 0.5375 | 0.376679 | 0.42516 | 0.438423 | 0.989223 | 0.997988 | 0.994666 |
| Ythdc2_B2RR83_S1284s_1 | Ythdc2_B2 | -0.15 | 0.77 | 1.4 | 0.09 | 0.48 | 0.66 | 0.61 | 1.11 | 0.22 | 0.88 | 1.65 | -0.41 | 0.17 | 0.68 | -0.24 | 0.08 | 0.1875 | 0.0575 | -0.355 | 0.389086 | 0.614924 | 0.912915 | 0.989223 | 0.997988 | 0.994666 |
| Ywhae_P62259_S65s_1 | Ywhae_P62 | -0.61 | -0.02 | 1.97 | 2.05 | -0.63 | -1.26 | 1.45 | 2.07 | -1.4 | -1.11 | 1.63 | 2.48 | -0.45 | -0.36 | 1.78 | 2.18 | -0.44 | -0.4475 | -0.06 | 0.943991 | 0.631949 | 0.670463 | 0.989223 | 0.997988 | 0.999507 |
| Ywhaz_P63101_S28s_1 | Ywhaz_P63 | -1.79 | -0.54 | -0.26 | -0.88 | -0.96 | -0.95 | -0.37 | 0.14 | -0.84 | -1.42 | 0.98 | -0.2 | 0.18 | 0.96 | -0.85 | -0.28 | 0.3325 | 0.4975 | 0.87 | 0.097472 | 0.421849 | 0.389366 | 0.989223 | 0.997988 | 0.994666 |
| Ywhaz_P63101_S37s_1 | Ywhaz_P63 | -0.15 | 0.24 | 0.44 | -0.4 | -0.33 | -0.35 | 0.19 | 0.03 | 0.01 | -0.45 | 0.11 | -0.12 | 2.56 | 1.04 | 0.57 | 0.86 | -0.1475 | -0.145 | 1.225 | 0.024474 | 0.59107 | 0.604576 | 0.989223 | 0.997988 | 0.994666 |
| Ywhaz_P63101_S64s_1 | Ywhaz_P63 | 0.22 | 0.59 | 0.38 | 0.38 | 0.3 | 0.53 | -0.13 | 0.24 | -0.06 | 0.38 | 0.57 | 0.27 | 0.64 | 0.49 | 0.28 | 0 | -0.1575 | -0.1025 | -0.04 | 0.877349 | 0.500515 | 0.669109 | 0.989223 | 0.997988 | 0.999507 |
| Zbed3_Q9D0L1_S168s_1 | Zbed3_Q9D | -1.16 | -2.41 | -1.88 | -0.92 | -2.09 | 0 | -3.96 | -1.06 | -0.91 | -1.64 | -1.27 | -1.74 | -1.24 | -1.99 | -2.11 | -2.04 | -0.185 | 0.2025 | -0.2525 | 0.534235 | 0.816068 | 0.606365 | 0.995762 | 0.997988 | 0.994666 |
| Zbtb11_G5E8B9_S473s_1 | Zbtb11_G5 | -2.76 | -2.77 | 3.7 | 0.9 | -2.3 | -1.93 | 3.21 | 2.14 | -1.1 | -1.94 | -0.37 | 4.07 | 0.63 | -2.22 | 3.79 | 4.82 | 0.5125 | 0.3975 | 1.9875 | 0.313347 | 0.775845 | 0.825039 | 0.995762 | 0.997988 | 0.994666 |
| Zbtb11_G5E8B9_S493s_1 | Zbtb11_G5 | 1.2 | 1.26 | -1.01 | -0.07 | -0.51 | 1.81 | -0.66 | -1.15 | 0.32 | 0.83 | -0.25 | -0.51 | 0 | -0.43 | -0.2 | -0.48 | -0.4725 | -0.2475 | -0.6225 | 0.250938 | 0.533749 | 0.667596 | 0.989223 | 0.997988 | 0.994666 |
| Zbtb16_Q3UQ17_S307s_1 | Zbtb16_Q3 | 0.21 | 0.76 | 0.25 | 0.93 | -0.13 | -0.12 | 0.34 | 0.58 | 1.24 | 1.07 | 0.28 | -0.36 | 0.22 | -0.84 | 0.02 | 0.35 | -0.37 | 0.02 | -0.6 | 0.113678 | 0.2149 | 0.960822 | 0.989223 | 0.997988 | 0.994666 |
| Zbtb18_H7BX69_S223s_1 | Zbtb18_H7 | 1 | -1.52 | 1.68 | 1.28 | 0.34 | 1 | -0.79 | -0.04 | 1.96 | 0.59 | 2.03 | 1.31 | 0.23 | 0.5 | 0.57 | 1.19 | -0.4825 | 0.8625 | 0.0125 | 0.985209 | 0.505982 | 0.248575 | 0.989223 | 0.997988 | 1 |
| Zbtb20_Q8K0L9_S359s_1 | Zbtb20_Q8 | 0 | 0.68 | -0.42 | -0.45 | -0.18 | -0.24 | 0.48 | -0.74 | 0.3 | 0.32 | 0.89 | 0.79 | -0.18 | 0.06 | 0.93 | 1.23 | -0.1225 | 0.6225 | 0.5575 | 0.210556 | 0.735087 | 0.085024 | 0.995762 | 0.997988 | 0.994666 |
| Zbtb20_Q8K0L9_T232t_1 | Zbtb20_Q8 | -0.3 | -0.7 | 0.23 | -0.31 | -0.7 | -0.85 | -0.4 | -0.75 | -0.91 | -0.23 | 1 | -0.92 | 0.05 | -0.96 | 0.41 | -0.2 | -0.405 | 0.005 | 0.095 | 0.796497 | 0.146211 | 0.991461 | 0.989223 | 0.997988 | 0.999507 |
| Zbtb22_Q9Z0G7_S602s_1 | Zbtb22_Q9 | -0.29 | -1.08 | -3.45 | -2.77 | -2.62 | -1.57 | -2.06 | -2.19 | -0.53 | -0.84 | -0.84 | -2.39 | -1.4 | -1.71 | -2.48 | -2.34 | -0.2125 | 0.7475 | -0.085 | 0.902478 | 0.753205 | 0.336193 | 0.995762 | 0.997988 | 0.999507 |
| Zbtb44_Q8R0A2_S161s_1 | Zbtb44_Q8 | -0.28 | 0.43 | -0.63 | -0.91 | -0.81 | -0.27 | -1.62 | -0.48 | -0.87 | 0.7 | 0.66 | -0.21 | -0.51 | 0.47 | 0.66 | -0.58 | -0.4475 | 0.4175 | 0.3575 | 0.413423 | 0.278281 | 0.371362 | 0.989223 | 0.997988 | 0.994666 |
| Zbtb45_Q52KG4_S435s_1 | Zbtb45_Q5 | -1.39 | -0.22 | 0.31 | -0.84 | 0.32 | -0.79 | 0.24 | -0.01 | 0.01 | -0.31 | 0.89 | -0.05 | -1.1 | 0.91 | 0.57 | 0.57 | 0.475 | 0.67 | 0.7725 | 0.17756 | 0.279099 | 0.151322 | 0.989223 | 0.997988 | 0.994666 |
| Zbtb7a_D3Z4C3_S331s_1 | Zbtb7a_D3 | -3.21 | -4.04 | -3.28 | -8.64 | -3.92 | -3.37 | -3.88 | -4.04 | -0.85 | -3.27 | -3.25 | -3.36 | -0.63 | -3.23 | -1.96 | -4.43 | 0.99 | 2.11 | 2.23 | 0.116572 | 0.388449 | 0.119212 | 0.989223 | 0.997988 | 0.994666 |
| Zc2hc1a_Q8BJH1_S222s_1 | Zc2hc1a_Q | -1.66 | -1.84 | -3.33 | -2.67 | -1.46 | -2.02 | -1.95 | -1.86 | -1.86 | -2.1 | -0.61 | -2.16 | -1.49 | -1.24 | -1.98 | -2.77 | 0.5525 | 0.6925 | 0.505 | 0.315022 | 0.182349 | 0.188786 | 0.989223 | 0.997988 | 0.994666 |
| Zc2hc1a_Q8BJH1_S251s_1 | Zc2hc1a_Q | -0.87 | -0.25 | -1.28 | -1.25 | -0.64 | -0.58 | -0.6 | -0.75 | -1.28 | -0.3 | -0.5 | -2.55 | -0.95 | -0.92 | -0.8 | -0.39 | 0.27 | -0.245 | 0.1475 | 0.646386 | 0.345708 | 0.643177 | 0.989223 | 0.997988 | 0.995259 |
| Zc2hc1a_Q8BJH1_T242t_1 | Zc2hc1a_Q | -1.7 | -1.7 | -1.7 | -2.04 | -0.84 | -2.58 | -1.2 | -1.91 | -1.93 | -1.3 | -1.04 | -1.77 | -0.77 | -1.14 | -1.7 | -1.92 | 0.1525 | 0.275 | 0.4025 | 0.230258 | 0.692855 | 0.333605 | 0.994893 | 0.997988 | 0.994666 |
| Zc2hc1a_Q8BJH1_T243t_1 | Zc2hc1a_Q | -1.61 | -1.56 | -1.7 | -2.04 | -0.9 | -2.44 | -1.2 | -1.91 | -1.92 | -1.33 | -1.04 | -1.77 | -0.99 | -0.73 | -1.7 | -1.92 | 0.115 | 0.2125 | 0.3925 | 0.263942 | 0.751021 | 0.455709 | 0.995762 | 0.997988 | 0.994666 |
| Zc3h12a_Q5D1E7_S394s_1 | Zc3h12a_Q | -0.99 | -0.08 | -1 | -0.38 | -2.36 | -0.95 | -1.53 | -0.16 | 0.4 | -0.37 | 0.95 | -0.41 | 0.31 | -2.91 | -1.14 | -0.48 | -0.6375 | 0.755 | -0.4425 | 0.502728 | 0.203718 | 0.083987 | 0.989223 | 0.997988 | 0.994666 |
| Zc3h12b_G3X9I7_S702s_1 | Zc3h12b_G | -1.78 | 0.84 | -0.65 | -1.53 | -0.96 | -0.77 | -1.94 | -2.61 | 0.24 | -0.17 | -1.7 | -1.84 | 1.62 | -1.54 | -1.01 | -1.29 | -0.79 | -0.0875 | 0.225 | 0.786491 | 0.242641 | 0.902451 | 0.989223 | 0.997988 | 0.999507 |
| Zc3h12c_E9Q1I3_S511s_1 | Zc3h12c_E | -0.13 | 0.18 | 1.11 | 0.2 | -0.31 | -0.21 | 0.13 | -0.05 | 0.62 | 0.07 | 0.87 | 0.5 | 1.51 | -0.75 | 0.82 | 0.48 | -0.45 | 0.175 | 0.175 | 0.732731 | 0.164663 | 0.605845 | 0.989223 | 0.997988 | 0.9971 |
| Zc3h13_E9Q784_S110s_1 | Zc3h13_E9 | -1.41 | -1.13 | -1.19 | -2.1 | -1.82 | -1.95 | -1.77 | -1.83 | -2.31 | -1.88 | -1.38 | -2.19 | 0.13 | -2.41 | -0.56 | -1.17 | -0.385 | -0.4825 | 0.455 | 0.410135 | 0.174786 | 0.166506 | 0.989223 | 0.997988 | 0.994666 |
| Zc3h13_E9Q784_S1273s_1 | Zc3h13_E9 | -0.25 | -0.9 | -1.98 | -1.4 | -3.35 | 0.45 | -2.33 | -1.48 | 1.3 | -0.95 | -2.1 | -1.19 | 2.1 | -1.78 | -0.26 | -0.57 | -0.545 | 0.3975 | 1.005 | 0.222191 | 0.487836 | 0.588238 | 0.989223 | 0.997988 | 0.994666 |
| Zc3h13_E9Q784_S1499s_1 | Zc3h13_E9 | -0.9 | -0.97 | -0.54 | -0.67 | -1 | -0.99 | 0.46 | -0.74 | -0.06 | -0.63 | 2.26 | 0.01 | -0.86 | -0.94 | -0.43 | 0.16 | 0.2025 | 1.165 | 0.2525 | 0.436597 | 0.57695 | 0.075086 | 0.989223 | 0.997988 | 0.994666 |
| Zc3h13_E9Q784_S198s_1 | Zc3h13_E9 | -0.07 | -0.12 | -0.73 | -1.05 | 0.19 | -1.06 | 0.5 | 0.55 | -0.84 | -0.72 | -0.16 | -1.42 | 0.08 | 0.44 | 0.12 | -0.15 | 0.5375 | -0.2925 | 0.615 | 0.077373 | 0.223589 | 0.428197 | 0.989223 | 0.997988 | 0.994666 |
| Zc3h13_E9Q784_S242s_1 | Zc3h13_E9 | -0.16 | -0.68 | -0.14 | -0.08 | -0.34 | -0.9 | 0.8 | 0.92 | -0.32 | -0.33 | 0.16 | -0.3 | -0.15 | 0.39 | 0.3 | 0.15 | 0.385 | 0.0675 | 0.4375 | 0.130753 | 0.388122 | 0.791002 | 0.989223 | 0.997988 | 0.994666 |
| Zc3h13_E9Q784_S316s_3 | Zc3h13_E9 | 2.18 | 2.63 | 1.06 | -0.32 | 2.95 | 2.69 | 0.64 | 0.62 | 1.81 | 1.5 | 0.86 | 0.34 | 0.82 | 2.23 | 1.15 | 1.95 | 0.3375 | -0.26 | 0.15 | 0.821088 | 0.673891 | 0.697372 | 0.992876 | 0.997988 | 0.999507 |
| Zc3h13_E9Q784_S317s_3 | Zc3h13_E9 | 2.18 | 2.63 | 1.06 | -0.32 | 2.95 | 2.69 | 0.64 | 0.62 | 1.81 | 1.5 | 0.86 | 0.34 | 0.82 | 2.23 | 1.15 | 1.95 | 0.3375 | -0.26 | 0.15 | 0.821088 | 0.673891 | 0.697372 | 0.992876 | 0.997988 | 0.999507 |
| Zc3h13_E9Q784_S324s_3 | Zc3h13_E9 | 2.18 | 2.63 | 1.06 | -0.32 | 2.95 | 2.69 | 0.64 | 0.62 | 1.81 | 1.5 | 0.86 | 0.34 | 0.82 | 2.23 | 1.15 | 1.95 | 0.3375 | -0.26 | 0.15 | 0.821088 | 0.673891 | 0.697372 | 0.992876 | 0.997988 | 0.999507 |
| Zc3h13_E9Q784_S703s_1 | Zc3h13_E9 | 1.17 | -0.06 | 0.85 | 0.26 | -0.06 | 0.22 | 0.7 | 1.35 | -0.39 | 0.11 | -0.2 | 0.04 | 0.98 | -0.49 | 2.15 | 1.48 | -0.0025 | -0.665 | 0.475 | 0.417587 | 0.995006 | 0.066865 | 0.999402 | 0.997988 | 0.994666 |
| Zc3h13_E9Q784_S704s_1 | Zc3h13_E9 | 0.74 | -0.45 | 0.77 | 0.24 | -0.11 | 0.23 | 0.2 | 1.01 | -0.06 | -0.17 | 1.38 | -0.19 | 0.99 | 0 | 1.9 | 1.49 | 0.0075 | -0.085 | 0.77 | 0.132003 | 0.9837 | 0.852338 | 0.999187 | 0.997988 | 0.994666 |
| Zc3h13_E9Q784_S77s_1 | Zc3h13_E9 | -0.33 | -1.27 | -1.12 | -0.81 | -0.1 | -0.7 | -0.68 | -0.28 | -0.85 | -0.99 | 0.23 | -1.44 | 0.23 | 0.25 | 0.27 | -0.47 | 0.4425 | 0.12 | 0.9525 | 0.01394 | 0.14756 | 0.767929 | 0.989223 | 0.997988 | 0.994666 |
| Zc3h13_E9Q784_S907s_1 | Zc3h13_E9 | -1.04 | -0.35 | -1.32 | -2.58 | -1.61 | -0.25 | -1.58 | -1.46 | 1.01 | -1.3 | -2.72 | -1.15 | 2.32 | -2.41 | 0.89 | -0.69 | 0.0975 | 0.2825 | 1.35 | 0.188913 | 0.850847 | 0.725703 | 0.996349 | 0.997988 | 0.994666 |
| Zc3h13_E9Q784_T1109t_1 | Zc3h13_E9 | -2.28 | -2.01 | -1.67 | -1.5 | -0.99 | -2.45 | -0.36 | -1.29 | -2.3 | -2.59 | -1.49 | -1.73 | -0.73 | -0.8 | -0.11 | -1.4 | 0.5925 | -0.1625 | 1.105 | 0.009751 | 0.201394 | 0.625686 | 0.989223 | 0.997988 | 0.994666 |
| Zc3h13_E9Q784_T363t_1 | Zc3h13_E9 | -0.67 | -0.88 | -0.51 | -1.06 | -0.19 | -1.34 | -0.05 | 0.08 | -1.91 | -1.89 | -0.93 | -1 | 0.01 | -0.2 | 0.79 | 0.08 | 0.405 | -0.6525 | 0.95 | 0.010765 | 0.264402 | 0.068676 | 0.989223 | 0.997988 | 0.994666 |
| Zc3h14_Q8BJ05_S309s_1 | Zc3h14_Q8 | -1.53 | -0.91 | -1.35 | -0.39 | -0.88 | -1.2 | -2.47 | -0.51 | -0.42 | -1.89 | 0.04 | -1.17 | -1.21 | 0.46 | -0.7 | -1.68 | -0.22 | 0.185 | 0.2625 | 0.600401 | 0.635125 | 0.695629 | 0.989223 | 0.997988 | 0.995259 |
| Zc3h14_Q8BJ05_S515s_1 | Zc3h14_Q8 | 0.8 | 0.54 | -0.43 | -0.38 | -0.08 | 0.16 | -0.44 | -0.03 | 0.02 | 0.15 | -0.44 | -0.03 | 0.53 | 0.27 | -0.07 | -0.26 | -0.23 | -0.2075 | -0.015 | 0.968247 | 0.508419 | 0.561234 | 0.989223 | 0.997988 | 0.999507 |
| Zc3h18_H3BIW0_S507s_1 | Zc3h18_H3 | -0.08 | -0.78 | -0.6 | -0.69 | -2.16 | -0.55 | -1.94 | -1.01 | 0.1 | -1.12 | -2.1 | -1.34 | 1.19 | -2.2 | -0.9 | -1.29 | -0.8775 | -0.5775 | -0.2625 | 0.692322 | 0.049504 | 0.231032 | 0.989223 | 0.997988 | 0.996159 |
| Zc3h18_H3BIW0_S64s_1 | Zc3h18_H3 | -1.88 | -1.77 | -3.04 | -1.76 | -2.15 | -1.79 | -2.9 | -3.24 | 0.63 | -2.86 | -1.84 | -2.57 | 0.66 | -1.87 | -2.98 | -2.85 | -0.4075 | 0.4525 | 0.3525 | 0.658814 | 0.354982 | 0.556792 | 0.989223 | 0.997988 | 0.995259 |
| Zc3h18_H3BIW0_S64s_2 | Zc3h18_H3 | -0.78 | -1.05 | 0.28 | 0.21 | -2.23 | -0.32 | -1.24 | 0.07 | -0.33 | -0.85 | 0.06 | -0.3 | 1.22 | -3.09 | -0.25 | -0.46 | -0.595 | -0.02 | -0.31 | 0.713985 | 0.298989 | 0.958959 | 0.989223 | 0.997988 | 0.996966 |
| Zc3h18_H3BIW0_S71s_2 | Zc3h18_H3 | -0.21 | -0.37 | 0.28 | 0.21 | -2.57 | -0.13 | -1.24 | 0.07 | -0.47 | -0.78 | 0.06 | -0.3 | 0.5 | -2.65 | -0.25 | -0.46 | -0.945 | -0.35 | -0.6925 | 0.28761 | 0.119691 | 0.238455 | 0.989223 | 0.997988 | 0.994666 |
| Zc3h18_H3BIW0_S75s_1 | Zc3h18_H3 | -2.85 | -2.93 | -2.49 | -3.08 | -3.31 | -3.45 | -1.92 | -1.52 | -4.71 | -4.17 | -2.17 | -2.63 | -2.1 | -4.69 | -2.23 | -2.73 | 0.2875 | -0.5825 | -0.1 | 0.859468 | 0.541717 | 0.321824 | 0.989223 | 0.997988 | 0.999507 |
| Zc3h18_H3BIW0_S80s_1 | Zc3h18_H3 | 0.74 | 0.67 | -0.13 | 0.24 | -0.06 | 0.03 | -0.46 | 0.8 | -0.55 | 0.49 | -0.23 | -0.04 | 0.15 | -0.42 | 0.38 | 0.12 | -0.3025 | -0.4625 | -0.3225 | 0.319612 | 0.382494 | 0.177526 | 0.989223 | 0.997988 | 0.994666 |
| Zc3h18_H3BIW0_S840s_1 | Zc3h18_H3 | -2.86 | -3.08 | -2.26 | -2.97 | -3.5 | -3.07 | -3.59 | -3.1 | -0.95 | -3.76 | -5.87 | -2.07 | 0.55 | -3.89 | -1.26 | -2.6 | -0.5225 | -0.37 | 0.9925 | 0.260585 | 0.077466 | 0.701225 | 0.989223 | 0.997988 | 0.994666 |
| Zc3h18_H3BIW0_S849s_1 | Zc3h18_H3 | 0.66 | -0.26 | -0.94 | -1.09 | -1.71 | -0.08 | -1.3 | -1.1 | 0.08 | -0.88 | -0.39 | -0.89 | 0.87 | -1.6 | -0.35 | -0.73 | -0.64 | -0.1125 | -0.045 | 0.939547 | 0.209218 | 0.800646 | 0.989223 | 0.997988 | 0.999507 |
| Zc3h18_H3BIW0_S890s_1 | Zc3h18_H3 | 1.06 | 0.12 | -0.45 | -0.22 | -0.87 | 0.33 | -1.61 | -0.77 | -0.23 | -0.35 | -1.96 | -0.18 | 1.18 | -0.41 | -0.25 | -0.74 | -0.8575 | -0.8075 | -0.1825 | 0.719945 | 0.099485 | 0.13804 | 0.989223 | 0.997988 | 0.9971 |
| Zc3h3_Q8CHP0_S212s_1 | Zc3h3_Q8 | -0.3 | -1.05 | -1.43 | -3.59 | -1.89 | -1.81 | -2.53 | -2.38 | 0.63 | -1.07 | -3.05 | -2.11 | 0.91 | -1.31 | -0.22 | -1.36 | -0.56 | 0.1925 | 1.0975 | 0.184341 | 0.395899 | 0.837207 | 0.989223 | 0.997988 | 0.994666 |
| Zc3h4_E9Q8K8_S1221s_1 | Zc3h4_E9Q | -0.55 | -1.44 | -1.3 | -0.31 | -1.9 | -2.2 | -0.83 | -2.63 | -1.61 | -0.78 | -0.65 | -2.28 | -0.6 | -3.21 | -0.62 | -0.41 | -0.99 | -0.43 | -0.31 | 0.636664 | 0.047747 | 0.354854 | 0.989223 | 0.997988 | 0.995259 |
| Zc3h4_E9Q8K8_S1227s_1 | Zc3h4_E9Q | -0.34 | -0.88 | -0.65 | -0.41 | -1.48 | -1.23 | -0.35 | -0.63 | -1.77 | -1.37 | 0.97 | 0 | -0.6 | -0.59 | 0.14 | -0.52 | -0.3525 | 0.0275 | 0.1775 | 0.541496 | 0.270407 | 0.962778 | 0.989223 | 0.997988 | 0.994666 |
| Zc3h4_E9Q8K8_S868s_1 | Zc3h4_E9Q | -1.94 | -2.89 | -3.24 | -2.19 | -2.78 | -2.37 | -1.77 | -3.05 | -1.22 | -0.94 | -5.49 | -2.89 | -1.3 | -1.66 | -2.86 | -3.54 | 0.0725 | -0.07 | 0.225 | 0.687065 | 0.85472 | 0.941893 | 0.996349 | 0.997988 | 0.996159 |
| Zc3h6_Q8BYK8_S1150s_1 | Zc3h6_Q8 | -1.35 | -1.47 | -0.18 | -1.75 | -2.02 | -1.48 | -1.04 | -0.7 | -1.21 | -1.23 | 0.03 | -0.82 | -1 | -0.2 | -0.84 | -0.79 | -0.1225 | 0.38 | 0.48 | 0.243962 | 0.773346 | 0.397664 | 0.995762 | 0.997988 | 0.994666 |
| Zc3h7b_F8VPP8_S372s_1 | Zc3h7b_F8 | -2.11 | -1.47 | -2.44 | -1.78 | -1.42 | -1.68 | -1.2 | -2.04 | -1.06 | -1.8 | -1.58 | -2.61 | -1.7 | -1.68 | -2.22 | -1.81 | 0.365 | 0.1875 | 0.0975 | 0.748217 | 0.243246 | 0.62952 | 0.989223 | 0.997988 | 0.9971 |
| Zc3hc1_H3BKM2_S301s_1 | Zc3hc1_H3 | -2.5 | -2.96 | -3.61 | -5.53 | -2.42 | -3.48 | -4.76 | -4.21 | -3.17 | -3.44 | -0.47 | -2.81 | -0.83 | -2.07 | -3.53 | -3.58 | -0.0675 | 1.1775 | 1.1475 | 0.186756 | 0.92678 | 0.187629 | 0.997335 | 0.997988 | 0.994666 |
| Zc3hc1_H3BKM2_S351s_1 | Zc3hc1_H3 | 0.7 | 0.47 | -0.95 | -0.18 | -0.89 | 0.59 | -0.45 | -0.31 | 0.79 | 0.91 | -0.12 | -0.03 | 0.05 | 0.72 | -0.45 | -0.56 | -0.275 | 0.3775 | -0.07 | 0.878188 | 0.546274 | 0.402996 | 0.989223 | 0.997988 | 0.999507 |
| Zc3hc1_H3BKM2_S352s_1 | Zc3hc1_H3 | 0.6 | 0.36 | -0.95 | -0.18 | -0.48 | 0.28 | -0.45 | -0.31 | 0.24 | 0.7 | -0.12 | -0.03 | 0.07 | 0.72 | -0.45 | -0.56 | -0.1975 | 0.24 | -0.0125 | 0.977405 | 0.604069 | 0.543046 | 0.989223 | 0.997988 | 1 |
| Zc3hc1_H3BKM2_S367s_1 | Zc3hc1_H3 | -1.85 | -2.71 | -2.79 | -2.28 | -3.33 | -2 | -3.03 | -1.89 | -2.57 | -2.09 | -3.27 | -2.3 | -0.56 | -3.72 | -2.76 | -1.83 | -0.155 | -0.15 | 0.19 | 0.766966 | 0.702757 | 0.670846 | 0.995437 | 0.997988 | 0.9971 |
| Zcchc6_E9PUA2_S705s_1 | Zcchc6_E9 | -0.14 | -0.8 | 0.11 | -0.05 | -0.23 | 0.1 | -1.57 | -0.66 | 0.72 | 0.09 | 0.32 | -0.32 | 0.92 | -1.26 | -1.28 | 0.77 | -0.37 | 0.4225 | 0.0075 | 0.989828 | 0.364077 | 0.212181 | 0.989223 | 0.997988 | 1 |
| Zdbf2_Q5SS00_S144s_1 | Zdbf2_Q5S | -1.47 | -1.59 | -1.03 | -1.06 | -0.9 | -0.67 | -1.78 | -1.67 | -0.29 | -0.24 | -0.24 | -2.39 | 0.27 | 0.24 | -1.58 | -1.98 | 0.0325 | 0.4975 | 0.525 | 0.362291 | 0.919995 | 0.348345 | 0.997335 | 0.997988 | 0.994666 |
| Zdhhc14_Q8BQQ1_S351s_1 | Zdhhc14_C | 0.16 | -1.15 | 0.28 | 0.72 | -0.1 | 0.43 | -0.02 | 0.92 | -0.41 | -0.76 | 0.68 | -0.09 | 0.06 | 0.06 | -0.23 | -0.36 | 0.305 | -0.1475 | -0.12 | 0.773424 | 0.492039 | 0.759235 | 0.989223 | 0.997988 | 0.99798 |
| Zdhhc14_Q8BQQ1_T19t_1 | Zdhhc14_C | 0.41 | 0.55 | 0.47 | 0.92 | 0.43 | -0.33 | 0.87 | 1.4 | 0.49 | 0.88 | 1.55 | 0.74 | 0.81 | 0.16 | 1.07 | 0.41 | 0.005 | 0.3275 | 0.025 | 0.932826 | 0.989373 | 0.284536 | 0.999187 | 0.997988 | 0.999507 |
| Zdhhc18_Q5Y5T2_S34s_1 | Zdhhc18_C | 0.88 | 0.36 | 0.57 | 1.19 | 0.83 | 1.08 | 1.56 | 1.2 | 0.29 | 0.29 | 1.1 | 1.01 | 0.04 | 0.51 | 0.49 | 0.25 | 0.4175 | -0.0775 | -0.4275 | 0.158333 | 0.153387 | 0.806993 | 0.989223 | 0.997988 | 0.994666 |
| Zdhhc18_Q5Y5T2_S358s_1 | Zdhhc18_C | 0.11 | 0.59 | -0.13 | 0.27 | 0.77 | 0.16 | 0.15 | 0.15 | 0.56 | 0.37 | -0.79 | -0.28 | 0.18 | 1.08 | -0.24 | -0.06 | 0.0975 | -0.245 | 0.03 | 0.932829 | 0.711571 | 0.498138 | 0.995437 | 0.997988 | 0.999507 |
| Zdhhc23_B7ZNY9_S242s_1 | Zdhhc23_B | -2.29 | -1.87 | -1.97 | -1.1 | -1.21 | -2.2 | -1.71 | -2.07 | -2.22 | -2.07 | -0.55 | -1.72 | -0.91 | -1.66 | -1.15 | -1.31 | 0.01 | 0.1675 | 0.55 | 0.125971 | 0.976604 | 0.704305 | 0.999187 | 0.997988 | 0.994666 |
| Zdhhc5_Q8VDZ4_S372s_1 | Zdhhc5_Q8 | -1.86 | -2.29 | -0.65 | -0.2 | -1.11 | -2.17 | -1.44 | -0.54 | -2.17 | -2.29 | -2.08 | -1.99 | -0.74 | -1.54 | -0.86 | -1.36 | -0.065 | -0.8825 | 0.125 | 0.802913 | 0.904523 | 0.088503 | 0.996493 | 0.997988 | 0.999507 |
| Zdhhc5_Q8VDZ4_S380s_1 | Zdhhc5_Q8 | 0.19 | 0.61 | 0.5 | 0.83 | 0.79 | 0.59 | 0.83 | 0.92 | 0.19 | 0.36 | 0.52 | 0.8 | 0.15 | 0.85 | 0.7 | 0.95 | 0.25 | -0.065 | 0.13 | 0.656732 | 0.28849 | 0.799541 | 0.989223 | 0.997988 | 0.995259 |
| Zdhhc5_Q8VDZ4_S398s_1 | Zdhhc5_Q8 | -1.68 | -0.77 | -1.05 | -1.46 | 0.18 | -1.82 | -0.34 | -1.12 | -2.15 | -1.62 | -1.62 | -1.34 | -1.62 | -0.11 | -0.83 | -0.4 | 0.465 | -0.4425 | 0.5 | 0.22545 | 0.317446 | 0.167979 | 0.989223 | 0.997988 | 0.994666 |
| Zdhhc5_Q8VDZ4_S409s_1 | Zdhhc5_Q8 | 1.36 | 1.26 | 1.16 | 1.39 | 1.24 | 1.88 | 1.14 | 1.76 | 1.02 | 1.62 | 1.23 | 0.88 | 0.02 | 1.33 | 0.77 | 0.53 | 0.2125 | -0.105 | -0.63 | 0.075647 | 0.403254 | 0.67223 | 0.989223 | 0.997988 | 0.994666 |
| Zdhhc5_Q8VDZ4_S554s_1 | Zdhhc5_Q8 | -2.3 | -1.04 | -1.51 | -3.44 | -0.67 | -2.82 | -2.26 | -2.4 | -2.97 | -2.18 | -0.95 | -1.87 | -1.1 | -1.08 | -1.84 | -1.58 | 0.035 | 0.08 | 0.6725 | 0.217638 | 0.955487 | 0.896079 | 0.999187 | 0.997988 | 0.994666 |
| Zdhhc5_Q8VDZ4_S621s_1 | Zdhhc5_Q8 | 0.93 | 0.78 | 0.91 | 0.89 | 0.57 | 1.05 | 0.83 | 1.33 | 0.57 | 1.07 | 0.87 | 1.08 | -0.31 | 1.19 | 0.96 | 0.6 | 0.0675 | 0.02 | -0.2675 | 0.4611 | 0.774361 | 0.929525 | 0.995762 | 0.997988 | 0.994666 |
| Zdhhc5_Q8VDZ4_S694s_1 | Zdhhc5_Q8 | -0.06 | 0.28 | 0.48 | 0.85 | 0.18 | 0.02 | 0.98 | 0.69 | -0.24 | 0.18 | 0.57 | 0.82 | -0.01 | 0.65 | 0.82 | 0.66 | 0.08 | -0.055 | 0.1425 | 0.652904 | 0.797566 | 0.866034 | 0.995762 | 0.997988 | 0.995259 |
| Zdhhc8_Q5Y5T5_S335s_1 | Zdhhc8_Q5 | 0.82 | 0.11 | 0.84 | 0.86 | 0.6 | 0.74 | 0.77 | 0.57 | 0.72 | 0.62 | 0.77 | 1.11 | 0.33 | 0.98 | 0.32 | 0.57 | 0.0125 | 0.1475 | -0.1075 | 0.721177 | 0.959749 | 0.587768 | 0.999187 | 0.997988 | 0.9971 |
| Zdhhc8_Q5Y5T5_S672s_1 | Zdhhc8_Q5 | 0.52 | 0.54 | 0.87 | 0.76 | 0.54 | 0.32 | 0.77 | 0.53 | 0.46 | 0.7 | 0.55 | 0.81 | 0.56 | 0.91 | 0.52 | 0.71 | -0.1325 | -0.0425 | 0.0025 | 0.991858 | 0.545393 | 0.848836 | 0.989223 | 0.997988 | 1 |
| Zdhhc8_Q5Y5T5_S679s_1 | Zdhhc8_Q5 | 0.18 | -0.62 | -0.53 | -0.12 | -0.18 | 0.51 | -0.68 | 0.08 | 0.47 | -1.23 | -0.9 | -0.19 | 0.04 | -1.27 | -1.39 | -1.42 | 0.205 | -0.19 | -0.7375 | 0.092629 | 0.530969 | 0.648853 | 0.989223 | 0.997988 | 0.994666 |
| Zfand2b_E0CY18_S163s_1 | Zfand2b_E | -0.44 | -0.68 | -1.85 | -0.97 | -0.66 | -0.84 | -1.01 | -0.88 | -1.03 | -2.56 | 1.47 | -1.17 | -1.23 | 0.33 | -1.39 | -2.5 | 0.1375 | 0.1625 | -0.2125 | 0.724374 | 0.676057 | 0.838622 | 0.992974 | 0.997988 | 0.9971 |
| Zfand2b_E0CY18_S173s_1 | Zfand2b_E | 0.17 | -0.56 | 0.58 | -0.05 | 0.13 | -0.26 | -0.2 | 0.32 | 0.15 | -0.59 | 0.8 | 0.09 | 0.23 | 0.93 | -0.02 | -0.13 | -0.0375 | 0.0775 | 0.2175 | 0.550257 | 0.900249 | 0.8367 | 0.996493 | 0.997988 | 0.994666 |
| Zfand2b_E0CY18_T185t_1 | Zfand2b_E | 0.06 | 0.42 | 0.67 | 0.3 | 0.71 | 0.22 | 0.71 | -0.09 | 0.13 | 0.33 | -1.38 | -0.96 | 0.33 | -0.64 | 0.32 | -0.46 | 0.025 | -0.8325 | -0.475 | 0.170174 | 0.927199 | 0.07551 | 0.997335 | 0.997988 | 0.994666 |
| Zfand5_O88878_S128s_1 | Zfand5_O8 | 0.64 | 0.7 | 0.89 | 0.39 | 0.69 | 0.63 | 0.77 | 0.85 | -0.04 | 0.71 | 1.25 | 0.6 | -0.44 | 0.49 | -3.77 | 0.75 | 0.08 | -0.025 | -1.3975 | 0.150935 | 0.707211 | 0.936773 | 0.995437 | 0.997988 | 0.994666 |
| Zfand5_O88878_S48s_1 | Zfand5_O8 | -0.4 | 0.01 | 0 | -0.18 | -0.7 | -0.51 | -0.3 | 0.16 | 0.43 | -0.21 | -0.58 | 0.32 | 0.65 | -0.3 | -0.04 | 0.23 | -0.195 | 0.1325 | 0.2775 | 0.354035 | 0.457741 | 0.656385 | 0.989223 | 0.997988 | 0.994666 |
| Zfand5_O88878_S48s_2 | Zfand5_O8 | -1.32 | 0.08 | -1.18 | -1.22 | -0.03 | -0.51 | -0.58 | -0.12 | 1.56 | -1.46 | 0.58 | -0.91 | -0.09 | 0.18 | -0.81 | -0.01 | 0.6 | 0.8525 | 0.7275 | 0.095573 | 0.118986 | 0.237195 | 0.989223 | 0.997988 | 0.994666 |
| Zfand5_O88878_S58s_2 | Zfand5_O8 | -0.08 | -0.36 | -1.18 | -1.22 | -0.85 | -0.71 | -0.58 | -0.12 | 1.07 | -1 | 0.58 | -0.91 | 0.11 | -0.25 | -0.81 | -0.01 | 0.145 | 0.645 | 0.47 | 0.226433 | 0.668131 | 0.262216 | 0.992876 | 0.997988 | 0.994666 |
| Zfml_E9QML5_S1571s_1 | Zfml_E9Q | 0.82 | -0.08 | -0.58 | -1.31 | 0.18 | 0.04 | -0.62 | -1.33 | -0.56 | -0.21 | -0.97 | -0.41 | 0.14 | 0.87 | -2.18 | -1.19 | -0.145 | -0.25 | -0.3025 | 0.677937 | 0.778886 | 0.586589 | 0.995762 | 0.997988 | 0.996159 |
| Zfml_E9QML5_S606s_1 | Zfml_E9Q | -3.61 | -3.4 | -4.02 | -4.96 | -3.2 | -3.88 | -3.74 | -4.59 | -5.81 | -3.73 | -3.38 | -3.96 | -2.79 | -2.09 | -3.43 | -3.48 | 0.145 | -0.2225 | 1.05 | 0.043681 | 0.733495 | 0.707462 | 0.995762 | 0.997988 | 0.994666 |
| Zfml_E9QML5_S881s_1 | Zfml_E9Q | -1.17 | -0.38 | 0.78 | -0.54 | -1.11 | -1.83 | -0.11 | -0.4 | -2.1 | -2.05 | 2.5 | 0.55 | -1.41 | -1.58 | 1.54 | -0.82 | -0.535 | 0.0525 | -0.24 | 0.745077 | 0.309755 | 0.959857 | 0.989223 | 0.997988 | 0.9971 |
| Zfp219_Q4VA98_S643s_1 | Zfp219_Q4 | 1.18 | 0.78 | -0.44 | -1.64 | 0.44 | -0.43 | -0.44 | -1.33 | -1.16 | -0.06 | 0.8 | -1.66 | 1.25 | 0.7 | 0.43 | 0.2 | -0.41 | -0.49 | 0.675 | 0.28834 | 0.532652 | 0.521885 | 0.989223 | 0.997988 | 0.994666 |
| Zfp362_E9QA68_S145s_1 | Zfp362_E9 | -0.8 | -2.3 | -1.71 | -0.96 | -0.83 | -0.39 | -0.24 | -1.55 | -1.18 | -1.59 | -0.67 | -0.88 | -1.24 | -0.16 | -0.97 | -0.39 | 0.69 | 0.3625 | 0.7525 | 0.10226 | 0.133427 | 0.374801 | 0.989223 | 0.997988 | 0.994666 |
| Zfp37_Q6P3Z4_S586s_1 | Zfp37_Q6P | 3.15 | -0.24 | 2.72 | -1.84 | -0.21 | 3.17 | 1.98 | 1.5 | 0.98 | 2.32 | -0.59 | 2.06 | 0.54 | -1.12 | 3 | 2.86 | 0.6625 | 0.245 | 0.3725 | 0.780753 | 0.581526 | 0.83814 | 0.989223 | 0.997988 | 0.998226 |
| Zfp385b_G5E8Y8_S185s_1 | Zfp385b_G | -0.22 | -1.76 | -1.17 | 0.07 | -1.14 | -0.51 | -0.13 | -0.63 | -0.78 | -0.51 | 0.06 | 0.24 | 0.45 | -0.68 | -0.22 | -2.18 | 0.1675 | 0.5225 | 0.1125 | 0.859152 | 0.705838 | 0.277485 | 0.995437 | 0.997988 | 0.999507 |
| Zfp407_G3UVV3_S221s_1 | Zfp407_G3 | -3.04 | -4.28 | -3.38 | -2.59 | -4.42 | -3.62 | -4.12 | -4.27 | -0.14 | -2.36 | -3.38 | -3 | 0.92 | -2.77 | -2.28 | -2.94 | -0.785 | 1.1025 | 1.555 | 0.094998 | 0.066591 | 0.154093 | 0.989223 | 0.997988 | 0.994666 |
| Zfp532_S4R1H5_S293s_1 | Zfp532_S4 | 3.02 | 1.46 | 2.15 | 2.45 | 0.19 | 3.03 | 1.14 | 2.37 | 3.06 | 2.09 | 1.33 | 1.92 | 2.96 | -1.05 | 2.52 | 1.92 | -0.5875 | -0.17 | -0.6825 | 0.427643 | 0.36451 | 0.71426 | 0.989223 | 0.997988 | 0.994666 |
| Zfp532_S4R1H5_S297s_1 | Zfp532_S4 | 0.22 | 0.28 | 0.51 | -0.04 | 0.1 | -0.28 | -0.01 | 0.17 | -0.01 | 0.06 | 0.52 | 0.29 | 0.93 | 0.75 | 0.59 | 0.26 | -0.2475 | -0.0275 | 0.39 | 0.171614 | 0.292884 | 0.910528 | 0.989223 | 0.997988 | 0.994666 |
| Zfp532_S4R1H5_S306s_1 | Zfp532_S4 | -0.1 | -0.34 | -0.94 | -1.07 | -1.96 | 0.35 | -1.57 | -0.73 | 2 | -0.83 | -3.01 | -0.59 | 2.97 | -1.25 | 1.57 | -0.17 | -0.365 | 0.005 | 1.3925 | 0.124872 | 0.482815 | 0.995711 | 0.989223 | 0.998151 | 0.994666 |
| Zfp532_S4R1H5_S333s_1 | Zfp532_S4 | 0.35 | 0.75 | 0.32 | 0.21 | -0.02 | 0.48 | -0.03 | 0.44 | -0.07 | 0.2 | 2.04 | -0.04 | 1.29 | 0.37 | 1.12 | -0.33 | -0.19 | 0.125 | 0.205 | 0.608855 | 0.444819 | 0.799201 | 0.989223 | 0.997988 | 0.995259 |
| Zfp532_S4R1H5_S349s_1 | Zfp532_S4 | 0.31 | 0.44 | -0.3 | -0.14 | 0.01 | -0.37 | 0.13 | 0.04 | -0.28 | 0.1 | -0.44 | 0.15 | 0.12 | 1.34 | -0.06 | -0.27 | -0.125 | -0.195 | 0.205 | 0.616091 | 0.630897 | 0.491761 | 0.989223 | 0.997988 | 0.995259 |
| Zfp644_Q3UMJ0_S376s_1 | Zfp644_Q3 | -0.19 | -1.35 | -0.48 | -0.4 | -2 | -0.42 | -1.02 | -1.49 | 0.91 | -1.13 | -1.11 | -0.2 | 0.71 | -2.97 | 0.09 | -0.72 | -0.6275 | 0.2225 | -0.1175 | 0.875418 | 0.14591 | 0.665247 | 0.989223 | 0.997988 | 0.999507 |
| Zfp646_Q6NV66_S601s_1 | Zfp646_Q6 | 0.56 | -0.51 | -0.65 | -0.91 | -0.61 | 0.57 | -0.66 | -0.91 | 1.41 | 0.09 | -1.53 | -0.76 | 1.77 | -0.86 | 0.07 | -0.41 | -0.025 | 0.18 | 0.52 | 0.397077 | 0.95416 | 0.779954 | 0.999187 | 0.997988 | 0.994666 |
| Zfp651_E9PZ11_T406t_1 | Zfp651_E9 | -2.06 | -2.17 | -2.78 | -2.51 | -2.94 | -2.27 | -3.82 | -2.48 | -1.49 | -1.68 | -1.33 | -2.28 | -0.6 | -2.51 | -2.81 | -2.52 | -0.4975 | 0.685 | 0.27 | 0.594878 | 0.204713 | 0.045839 | 0.989223 | 0.997988 | 0.995259 |
| Zfp704_Q9ERQ3_S521s_1 | Zfp704_Q9 | 1.14 | 1.03 | 1.25 | -0.77 | 0.92 | 1.27 | 1.31 | 1.3 | -1.77 | 1.67 | -3.51 | 1.22 | 0.26 | -2.12 | 2.33 | 2.27 | 0.5375 | -1.26 | 0.0225 | 0.982132 | 0.255813 | 0.294705 | 0.989223 | 0.997988 | 1 |
| Zfp740_D3Z606_S19s_1 | Zfp740_D3 | -0.32 | -1.57 | -1.1 | -0.42 | -0.31 | -1.15 | -0.47 | -1.01 | -0.32 | -0.5 | 0.14 | -0.57 | -0.32 | -0.58 | -0.04 | -0.23 | 0.1175 | 0.54 | 0.56 | 0.131654 | 0.743211 | 0.148895 | 0.995762 | 0.997988 | 0.994666 |
| Zfp777_G5E8L5_S615s_1 | Zfp777_G5 | 0.23 | -0.05 | 0.18 | 0.15 | -1.26 | 0.89 | -0.5 | -0.28 | 0.68 | 0.03 | -0.86 | 0.13 | 1.66 | -0.91 | -0.13 | 0.17 | -0.415 | -0.1325 | 0.07 | 0.890725 | 0.341689 | 0.70039 | 0.989223 | 0.997988 | 0.999507 |
| Zfpm1_O35615_S681s_1 | Zfpm1_O3 | -1.68 | -1.9 | 0.19 | -0.8 | -1.42 | -2.45 | -0.7 | 0.04 | -0.96 | -2.08 | 1.35 | -1.64 | -0.54 | -2.08 | 0.25 | 1.26 | -0.085 | 0.215 | 0.77 | 0.321491 | 0.893282 | 0.789088 | 0.996493 | 0.997988 | 0.994666 |
| Zfpm2_Q8CCH7_S1057s_1 | Zfpm2_Q8 | -1.58 | -1.87 | -1.47 | -0.96 | -1.26 | -2.2 | -1.92 | -1.18 | -1.22 | -1.9 | -0.77 | -1.34 | -0.98 | -1.59 | -0.77 | -1.19 | -0.17 | 0.1625 | 0.3375 | 0.293624 | 0.604302 | 0.620652 | 0.989223 | 0.997988 | 0.994666 |
| Zfr_O88532_S1054s_1 | Zfr_O8853 | -0.19 | -0.54 | -1.1 | -0.99 | -0.32 | -0.24 | -1.35 | -1.11 | -1.36 | -1.15 | -1.54 | -2.16 | -0.67 | -0.3 | -0.77 | -1.49 | -0.05 | -0.8475 | -0.1025 | 0.772306 | 0.886679 | 0.026846 | 0.996493 | 0.997988 | 0.997453 |
| Zfr2_E9Q5M4_S130s_1 | Zfr2_E9Q5 | 0.96 | 0.39 | 0.34 | 0.62 | 0.43 | 0.84 | 0.2 | 0.81 | -2.39 | -2.46 | -1.48 | -2.74 | -1.49 | -1.74 | -2.5 | -2.77 | -0.0075 | -2.845 | -2.7025 | 3.09E-05 | 0.976896 | 1.31E-05 | 0.999187 | 0.133636 | 0.269064 |
| Zfyve1_Q810J8_S81s_1 | Zfyve1_Q8 | -0.51 | -1.05 | 0.81 | -0.2 | -2.24 | -0.18 | -0.16 | -0.02 | 1.69 | -0.37 | -0.87 | -0.1 | 1.38 | -0.21 | 0.41 | 0.58 | -0.4125 | 0.325 | 0.7775 | 0.134322 | 0.490688 | 0.603479 | 0.989223 | 0.997988 | 0.994666 |
| Zfyve19_Q9DAZ9_S219s_1 | Zfyve19_Q | -1.42 | -1.9 | -3.09 | -1.95 | -1.35 | -1.78 | -2.1 | -2.32 | -2.01 | -1.38 | -1.67 | -2.17 | 0.43 | -1.01 | -2.61 | -2.26 | 0.2025 | 0.2825 | 0.7275 | 0.308821 | 0.612406 | 0.480654 | 0.989223 | 0.997988 | 0.994666 |
| Zfyve19_Q9DAZ9_S94s_1 | Zfyve19_Q | -5.72 | -5.69 | -5.4 | -11.83 | -4.68 | -2.24 | -5.09 | -5.27 | -4.38 | -4.1 | -4.09 | -5.2 | -1.96 | -4.42 | -6.03 | -5.3 | 2.84 | 2.7175 | 2.7325 | 0.101214 | 0.077576 | 0.074993 | 0.989223 | 0.997988 | 0.994666 |
| Zfyve19_Q9DAZ9_T377t_1 | Zfyve19_Q | -1.94 | 0.76 | 1.15 | -0.54 | -0.68 | -1.74 | 1.53 | 1.7 | 2.18 | -1.49 | 0.6 | -0.39 | 1.64 | 0.88 | 2.53 | 1.23 | 0.345 | 0.3675 | 1.7125 | 0.033651 | 0.717289 | 0.69295 | 0.995437 | 0.997988 | 0.994666 |
| Zfyve26_Q5DU37_S605s_2 | Zfyve26_Q | -1.94 | -1.46 | -1.93 | -2.74 | -1.1 | -2.62 | -3.48 | -1.26 | -3.63 | -1.97 | -2.76 | -1.76 | -2.56 | -1.69 | -4.16 | -1.32 | -0.0975 | -0.5125 | -0.415 | 0.509794 | 0.862984 | 0.297793 | 0.996493 | 0.997988 | 0.994666 |
| Zfyve28_Q6ZPK7_S686s_1 | Zfyve28_Q | -0.12 | 0.09 | 0.01 | -0.29 | -0.28 | -0.65 | -0.07 | -0.02 | -0.61 | -0.25 | 0.26 | -0.25 | 0.21 | -0.16 | 0.18 | -0.28 | -0.1775 | -0.135 | 0.065 | 0.798908 | 0.457739 | 0.608747 | 0.989223 | 0.997988 | 0.999507 |
| Zkscan1_Q8BGS3_S13s_1 | Zkscan1_Q | 0.09 | -0.4 | -0.35 | 0.07 | 0.13 | -0.3 | -0.03 | -1.41 | 0.11 | 0.6 | 1.1 | 0.05 | -0.09 | 0.26 | -0.01 | -0.24 | -0.255 | 0.6125 | 0.1275 | 0.630885 | 0.492914 | 0.074761 | 0.989223 | 0.997988 | 0.995259 |
| Zmym2_Q9CU65_S837s_1 | Zmym2_Q9 | 0.68 | -0.76 | -1.21 | -0.27 | -0.45 | 1.16 | -0.69 | -0.25 | 0.01 | -1.09 | -1 | -0.78 | -0.16 | -0.48 | -1.08 | -0.4 | 0.3325 | -0.325 | -0.14 | 0.751153 | 0.532756 | 0.482855 | 0.989223 | 0.997988 | 0.9971 |
| Zmynd8_Q3UH28_S407s_2 | Zmynd8_Q | -1.57 | -1.17 | 0.18 | -0.6 | -0.77 | -1.22 | -0.65 | -0.25 | -0.75 | -0.74 | 1.12 | 0.01 | -1.81 | -0.28 | -0.59 | -0.07 | 0.0675 | 0.7 | 0.1025 | 0.841167 | 0.869134 | 0.216535 | 0.996493 | 0.997988 | 0.999507 |
| Zmynd8_Q3UH28_S419s_2 | Zmynd8_Q | -1.57 | -1.17 | 0.18 | -0.6 | -0.77 | -1.22 | -0.65 | -0.25 | -0.75 | -0.74 | 1.12 | 0.01 | -1.81 | -0.28 | -0.59 | -0.07 | 0.0675 | 0.7 | 0.1025 | 0.841167 | 0.869134 | 0.216535 | 0.996493 | 0.997988 | 0.999507 |
| Zmynd8_Q3UH28_S465s_1 | Zmynd8_Q | -2.43 | -0.56 | -1.73 | -4.47 | -1.35 | -1.66 | -1.95 | -1.69 | -4.18 | -1.37 | -1.24 | -1.9 | -0.06 | -3.98 | -1.24 | -2.26 | 0.635 | 0.125 | 0.4125 | 0.685706 | 0.39366 | 0.894738 | 0.989223 | 0.997988 | 0.996159 |
| Zmynd8_Q3UH28_S524s_1 | Zmynd8_Q | 0.32 | 0.68 | -0.41 | 0.36 | 0.85 | 0.98 | 0.5 | 0.57 | 0.22 | 0.59 | 0.32 | 0.26 | 0.25 | 0.82 | 0.37 | 0.36 | 0.4875 | 0.11 | 0.2125 | 0.503863 | 0.11588 | 0.705587 | 0.989223 | 0.997988 | 0.994666 |
| Zmynd8_Q3UH28_S629s_1 | Zmynd8_Q | -0.01 | 1.04 | 0.34 | 0.38 | 0.94 | -0.38 | 0.56 | 0.31 | -0.18 | -0.12 | 0.59 | 0.65 | 0.39 | 1.61 | 0.24 | 0.45 | -0.08 | -0.2025 | 0.235 | 0.554018 | 0.821432 | 0.549206 | 0.995762 | 0.997988 | 0.994666 |
| Zmynd8_Q3UH28_S734s_1 | Zmynd8_Q | 0.54 | 0.7 | 0.35 | 0.59 | -0.07 | 0.07 | 0.67 | 0.51 | 0.01 | 0.75 | -0.36 | -0.29 | 0.36 | 0.57 | 0.69 | 0.48 | -0.25 | -0.5175 | -0.02 | 0.932982 | 0.327582 | 0.11311 | 0.989223 | 0.997988 | 0.999507 |
| Znf148_Q61624_S665s_1 | Znf148_Q6 | -2.12 | -1.5 | -2.77 | -2.27 | -2.13 | -4.01 | -2.75 | -5.85 | -2.12 | -2.02 | -0.79 | -3.71 | -0.96 | -1.22 | -2.54 | -2.88 | -1.52 | 0.005 | 0.265 | 0.606412 | 0.068757 | 0.993329 | 0.989223 | 0.997988 | 0.995259 |
| Znf148_Q61624_S784s_1 | Znf148_Q6 | 1.63 | 0.25 | 1.49 | 1.07 | 1.07 | 1.56 | -0.31 | 1.45 | 1.07 | 0.98 | 1.27 | 1.5 | 0.02 | 0.31 | 0.16 | 1.09 | -0.1675 | 0.095 | -0.715 | 0.098452 | 0.732169 | 0.784946 | 0.995762 | 0.997988 | 0.994666 |
| Znf318_Q99PP2_S1591s_1 | Znf318_Q9 | -1.59 | -1.69 | -2.47 | -2.44 | -4.81 | -1.94 | -2.96 | -2.06 | -2.02 | -1.68 | -3.44 | -2.19 | -0.43 | -3.31 | -2.05 | -2.76 | -0.895 | -0.285 | -0.09 | 0.882358 | 0.175945 | 0.519278 | 0.989223 | 0.997988 | 0.999507 |
| Znf318_Q99PP2_S2140s_1 | Znf318_Q9 | -1.48 | -0.98 | -0.18 | 0.44 | -0.57 | -1.63 | 0.7 | 0.78 | -1.8 | -1.38 | -0.5 | -0.17 | -1.25 | -1.03 | -0.11 | 0.01 | 0.37 | -0.4125 | -0.045 | 0.928499 | 0.56345 | 0.444402 | 0.989223 | 0.997988 | 0.999507 |
| Znf318_Q99PP2_S2206s_1 | Znf318_Q9 | -2.74 | -2.98 | -1.83 | -0.76 | -2.65 | -3.82 | -0.87 | -1.26 | -4.32 | -3.29 | 0.06 | -2.84 | -1.33 | -1.87 | 0.21 | -0.34 | -0.0725 | -0.52 | 1.245 | 0.07124 | 0.921877 | 0.584287 | 0.997335 | 0.997988 | 0.994666 |
| Znf318_Q99PP2_S41s_1 | Znf318_Q9 | 0.63 | -0.26 | -2.48 | -1.91 | -0.51 | -0.13 | -1.8 | -2.83 | 1.51 | 0.07 | -2.13 | -2.77 | 1.39 | 0.51 | -1.64 | -1.98 | -0.3125 | 0.175 | 0.575 | 0.549792 | 0.706584 | 0.870876 | 0.995437 | 0.997988 | 0.994666 |
| Znf318_Q99PP2_S79s_2 | Znf318_Q9 | -0.61 | -0.94 | -1.73 | -0.82 | -2.59 | 0.02 | -2.59 | -1.26 | 0.29 | -0.36 | -1.44 | -1.63 | -0.17 | -2.02 | -0.79 | -0.78 | -0.58 | 0.24 | 0.085 | 0.849307 | 0.346633 | 0.625165 | 0.989223 | 0.997988 | 0.999507 |
| Znf318_Q99PP2_S81s_2 | Znf318_Q9 | -0.61 | -0.94 | -1.73 | -0.82 | -2.59 | 0.02 | -2.59 | -1.26 | 0.29 | -0.36 | -1.44 | -1.63 | -0.17 | -2.02 | -0.79 | -0.78 | -0.58 | 0.24 | 0.085 | 0.849307 | 0.346633 | 0.625165 | 0.989223 | 0.997988 | 0.999507 |
| Znf365_Q8BG89_S146s_1 | Znf365_Q8 | 0.47 | 0.33 | 1.11 | 0.76 | 0.29 | 1.03 | 1.14 | 0.74 | 0.49 | 0.43 | 1.15 | 0.87 | 1.03 | 1.14 | 0.22 | 0.63 | 0.1325 | 0.0675 | 0.0875 | 0.784243 | 0.646977 | 0.814885 | 0.99033 | 0.997988 | 0.998711 |
| Znf365_Q8BG89_S329s_1 | Znf365_Q8 | 0.42 | -1.52 | -0.17 | -0.75 | 0.07 | 0.62 | -0.56 | -0.35 | -2.67 | -1.72 | -0.65 | -0.91 | -1.39 | -3.04 | -0.82 | -1.07 | 0.45 | -0.9825 | -1.075 | 0.095851 | 0.336443 | 0.110053 | 0.989223 | 0.997988 | 0.994666 |

| | | | | | | | | | | | | | | | | | | | | | | | | | | |
|---|---|---|---|---|---|---|---|---|---|---|---|---|---|---|---|---|---|---|---|---|---|---|---|---|---|---|
| Znf365_Q8BG89_T206t_1 | Znf365_Q8 | 1.08 | 0.55 | 1.54 | 0.62 | 0.88 | 0.66 | 1.12 | 1.12 | 0.53 | 0.64 | 0.82 | 1.02 | 0.87 | 1.06 | 0.73 | 0.89 | -0.0025 | -0.195 | -0.06 | 0.841796 | 0.993041 | 0.514571 | 0.999187 | 0.997988 | 0.999507 |
| Znf385a_Q8VD12_S133s_1 | Znf385a_Q | -0.68 | 0.18 | -1.23 | -1.1 | 0.01 | -0.47 | -1.08 | -2.78 | -0.89 | -0.66 | -0.21 | 0.23 | -0.8 | -0.8 | -1.8 | -1.38 | -0.3725 | 0.325 | -0.4875 | 0.24719 | 0.547329 | 0.426163 | 0.989223 | 0.997988 | 0.994666 |
| Znf385a_Q8VD12_S140s_1 | Znf385a_Q | -0.06 | 0.36 | -0.57 | -0.21 | 0.15 | -0.18 | -0.25 | 0.36 | 0.35 | 0.58 | 0.07 | -0.33 | 0.2 | 0.16 | -0.15 | -0.18 | 0.14 | 0.2875 | 0.1275 | 0.659523 | 0.615983 | 0.364085 | 0.989223 | 0.997988 | 0.995259 |
| Znf451_Q8C0P7_S1032s_1 | Znf451_Q8 | -4.23 | -3.82 | -5.12 | -6.53 | -3.51 | -4.32 | -9.46 | -5.4 | -4.19 | -4.41 | -2.08 | -5.55 | -2 | -3.45 | -4.8 | -3.85 | -0.7475 | 0.8675 | 1.4 | 0.084889 | 0.552356 | 0.314555 | 0.989223 | 0.997988 | 0.994666 |
| Znf592_Q8BHZ4_S332s_1 | Znf592_Q8 | 0.13 | 0.17 | 0.45 | -0.7 | 0.28 | 0.12 | -0.18 | -0.04 | 0.44 | -0.3 | -1.04 | -0.4 | 1.09 | 0.64 | 0.38 | -0.03 | 0.0325 | -0.3375 | 0.5075 | 0.185522 | 0.912112 | 0.398964 | 0.997335 | 0.997988 | 0.994666 |
| Znf592_Q8BHZ4_S346s_1 | Znf592_Q8 | 0.88 | 0.49 | 1.21 | 0.4 | -0.02 | 0.56 | 0.11 | 0.39 | 1.11 | -0.29 | 0.18 | 0.35 | 1.46 | 0.34 | 0.98 | 0.67 | -0.485 | -0.4075 | 0.1175 | 0.729362 | 0.098297 | 0.272357 | 0.989223 | 0.997988 | 0.9971 |
| Znf609_Q8BZ47_S413s_1 | Znf609_Q8 | -4.13 | -5.03 | -2.97 | -3.48 | -2.74 | -6.17 | -4.21 | -2.89 | -3.82 | -4.34 | -2.27 | -0.92 | -0.36 | -1.61 | -3.08 | -4.13 | -0.1 | 1.065 | 1.6075 | 0.076887 | 0.899917 | 0.204201 | 0.996493 | 0.997988 | 0.994666 |
| Znf609_Q8BZ47_S491s_1 | Znf609_Q8 | -0.9 | -0.57 | -0.6 | -0.39 | -0.21 | -0.38 | 0.13 | -0.48 | -0.46 | -0.47 | -0.04 | -0.43 | -0.77 | 0.29 | -0.49 | -0.55 | 0.38 | 0.265 | 0.235 | 0.454108 | 0.13496 | 0.279756 | 0.989223 | 0.997988 | 0.994666 |
| Znf622_Q91VY9_S145s_1 | Znf622_Q9 | -2.05 | -2.11 | -0.39 | -1.14 | -2.45 | -1.06 | -1.17 | -1 | 0.01 | -1.65 | -1.06 | -0.49 | 1.53 | -2.66 | -0.3 | -0.87 | 0.0025 | 0.625 | 0.8475 | 0.32676 | 0.995942 | 0.240379 | 0.999655 | 0.997988 | 0.994666 |
| Znf687_Q9D2D7_S228s_1 | Znf687_Q9 | -0.66 | -0.92 | -0.61 | -0.66 | -2.43 | -1.46 | -1.14 | -1.1 | -0.53 | -0.33 | -1.01 | -1.16 | 1.09 | -1.21 | -0.7 | -0.64 | -0.82 | -0.045 | 0.3475 | 0.478098 | 0.030647 | 0.866644 | 0.989223 | 0.997988 | 0.994666 |
| Znf750_Q8BH05_S373s_1 | Znf750_Q8 | -1.11 | -0.31 | -1.59 | -1.59 | -0.83 | -1.29 | -1.78 | -1.26 | -1.48 | -1.57 | -0.22 | -1.59 | -0.62 | -1.69 | -1.67 | -1.55 | -0.14 | -0.065 | -0.2325 | 0.566122 | 0.696779 | 0.881367 | 0.994893 | 0.997988 | 0.995259 |
| Znf830_Q8R1N0_S216s_1 | Znf830_Q8 | -1.57 | -0.38 | -1.83 | -1.91 | -1.04 | -3.53 | -1.2 | -2.66 | -1.96 | -1.36 | 0.11 | -0.96 | -0.63 | -0.58 | -0.04 | -1.66 | -0.685 | 0.38 | 0.695 | 0.163608 | 0.285022 | 0.475406 | 0.989223 | 0.997988 | 0.994666 |
| Znrf2_Q71FD5_S18s_1 | Znrf2_Q71 | 0.66 | -0.24 | 0.3 | 0.28 | 0.98 | -0.53 | -0.39 | -0.17 | 0.19 | 0.6 | -0.43 | 0.01 | 0.25 | 0.63 | 0.56 | 0.27 | -0.2775 | -0.1575 | 0.1775 | 0.536237 | 0.472905 | 0.618761 | 0.989223 | 0.997988 | 0.994666 |
| Znrf2_Q71FD5_S75s_1 | Znrf2_Q71 | -0.07 | 0.41 | 0.58 | 0.29 | 0.52 | 0.4 | 0.8 | 0.55 | 0.42 | 0.36 | 0.97 | 0.27 | 0.72 | 0.7 | 0.47 | 0.36 | 0.265 | 0.2025 | 0.26 | 0.333249 | 0.272613 | 0.458561 | 0.989223 | 0.997988 | 0.994666 |
| Zranb2_E9PUD0_S106s_1 | Zranb2_E9 | -0.91 | -1.34 | -1.42 | -1.21 | -0.09 | -1.16 | 0.02 | -0.62 | -0.81 | -0.91 | -0.63 | -0.47 | -0.64 | -0.44 | -2.3 | -0.29 | 0.7575 | 0.515 | 0.3025 | 0.518251 | 0.034263 | 0.053134 | 0.989223 | 0.997988 | 0.994666 |
| Zranb2_E9PUD0_S116s_1 | Zranb2_E9 | -1.61 | 0.45 | -1.21 | -0.42 | -1.67 | -1.96 | -1.99 | -2.02 | -0.12 | 0.57 | 1.11 | -1.69 | -2.07 | -0.2 | -3.54 | -3.16 | -1.2125 | 0.665 | -1.545 | 0.071204 | 0.018918 | 0.344983 | 0.989223 | 0.997988 | 0.994666 |
| Zranb2_E9PUD0_S151s_1 | Zranb2_E9 | -0.68 | -0.42 | -0.83 | -1.03 | -0.79 | -0.82 | -2.56 | -2.15 | -0.79 | -0.81 | -0.16 | -1.57 | -0.69 | 0.38 | -3.19 | -3.06 | -0.84 | -0.0925 | -0.9 | 0.271837 | 0.083515 | 0.783953 | 0.989223 | 0.997988 | 0.994666 |
| Zranb2_E9PUD0_S83s_1 | Zranb2_E9 | -0.61 | -0.48 | -0.38 | -0.72 | -0.09 | -0.33 | -0.97 | -0.7 | -0.6 | 0.23 | 0.22 | -0.89 | -0.79 | -0.27 | -1.36 | -0.78 | 0.025 | 0.2875 | -0.2525 | 0.405701 | 0.92296 | 0.384316 | 0.997335 | 0.997988 | 0.994666 |
| Zrsr1_Q64707_S418s_1 | Zrsr1_Q64 | -0.88 | -1.07 | -1.81 | -1.01 | -2.01 | -1.11 | -2.54 | -2.01 | 1.1 | -1.98 | -1.91 | -2.07 | 2.51 | -3.04 | -0.78 | -1.42 | -0.725 | -0.0225 | 0.51 | 0.622443 | 0.069284 | 0.975044 | 0.989223 | 0.997988 | 0.995259 |
| Zswim8_Q3UHH1_S1060s_1 | Zswim8_Q3 | 0.15 | -0.43 | 0.15 | -0.08 | 0.11 | 0.08 | 0.42 | 0.7 | 0.06 | -0.56 | 0.14 | 0.32 | -0.47 | 0.06 | -0.27 | -0.54 | 0.38 | 0.0425 | -0.2525 | 0.369607 | 0.158208 | 0.881071 | 0.989223 | 0.997988 | 0.994666 |
| Zswim8_Q3UHH1_S48s_1 | Zswim8_Q3 | -1.78 | -1.11 | -1.27 | -1.06 | -1.87 | -0.98 | -1.58 | -1.69 | 0.26 | -1.42 | -2.39 | -2.32 | 1.23 | -2.33 | -1.76 | -1.89 | -0.225 | -0.1625 | 0.1175 | 0.873718 | 0.438999 | 0.781966 | 0.989223 | 0.997988 | 0.999507 |
| Zswim8_Q3UHH1_S53s_1 | Zswim8_Q3 | -1.3 | 0.29 | 1.55 | 2.22 | -0.25 | 0.18 | 0.46 | 1.78 | 2.31 | -0.02 | -1.08 | 2.01 | 2.39 | -3.12 | 1.24 | 1.01 | -0.1475 | 0.115 | -0.31 | 0.802146 | 0.849743 | 0.907661 | 0.996349 | 0.997988 | 0.999507 |
| Zswim8_Q3UHH1_S564s_1 | Zswim8_Q3 | 0.15 | -0.54 | 0.33 | -0.09 | -0.72 | -0.22 | -0.59 | -0.36 | 1.19 | -0.15 | -0.6 | 0.2 | 1.15 | -0.55 | -0.15 | -0.59 | -0.435 | 0.1975 | 0.0025 | 0.99546 | 0.125616 | 0.637488 | 0.989223 | 0.997988 | 1 |
| Zwint_Q9CQU5_S216s_1 | Zwint_Q9C | -2.26 | -3.43 | -2.61 | -4.08 | -2.2 | -1.92 | -0.9 | -2.4 | -2.23 | -1.89 | -1.02 | -2.95 | -1.51 | -1.47 | -5.4 | -3.21 | 1.24 | 1.0725 | 0.1975 | 0.824198 | 0.027985 | 0.069895 | 0.989223 | 0.997988 | 0.999507 |
| Zyx_Q7TQE2_S241s_1 | Zyx_Q7TQ | -0.52 | -0.92 | -0.08 | -0.19 | 0.54 | -0.66 | 0.12 | -0.36 | 0.42 | 0.16 | 0.63 | 0.19 | -0.76 | -0.22 | -0.45 | -0.34 | 0.3375 | 0.7775 | -0.015 | 0.958786 | 0.325331 | 0.017752 | 0.989223 | 0.997988 | 0.999507 |
| Zyx_Q7TQE2_S305s_1 | Zyx_Q7TQ | 0.62 | 0.49 | 0.02 | 0.55 | 0.78 | 0.52 | 0.97 | 0.66 | 0.28 | 0.39 | 0.54 | 0.86 | 0 | 0.81 | 0.49 | 0.31 | 0.3125 | 0.0975 | -0.0175 | 0.951531 | 0.205737 | 0.70363 | 0.989223 | 0.997988 | 0.999507 |
| Zzef1_E9Q5W5_S1475s_1 | Zzef1_E9Q | 0.53 | 0.24 | 0.01 | -0.11 | 0.38 | 0.05 | -1.31 | -1.51 | 0.49 | 0.85 | -0.8 | -1.43 | -0.82 | 0.54 | 0.3 | 0.3 | -0.765 | -0.39 | -0.0875 | 0.808447 | 0.122511 | 0.459165 | 0.989223 | 0.997988 | 0.999507 |
| Zzef1_E9Q5W5_S1488s_1 | Zzef1_E9Q | -0.35 | -0.68 | -0.12 | -0.89 | 0.03 | -0.38 | 0.17 | 0.33 | -0.8 | -0.46 | 0.57 | -0.26 | -1.21 | -0.29 | -0.58 | -0.3 | 0.5475 | 0.2725 | -0.085 | 0.791799 | 0.067351 | 0.44701 | 0.989223 | 0.997988 | 0.999507 |
| Zzef1_E9Q5W5_S2407s_1 | Zzef1_E9Q | -0.81 | -0.37 | 0.12 | 0.52 | 0.56 | 0.11 | -0.14 | 0.25 | 0.06 | -0.06 | 0.03 | -0.41 | -0.37 | 0.02 | 0.1 | 0.15 | 0.33 | 0.04 | 0.11 | 0.750395 | 0.334807 | 0.904105 | 0.989223 | 0.997988 | 0.9971 |
| Zzef1_E9Q5W5_T2066t_1 | Zzef1_E9Q | 0.28 | 0.4 | 0.31 | 0.47 | -0.17 | -0.88 | 0.28 | -0.11 | 0.17 | 0.27 | -0.17 | -0.51 | -0.03 | -0.18 | 0.1 | 0.58 | -0.585 | -0.425 | -0.2475 | 0.360968 | 0.06092 | 0.120018 | 0.989223 | 0.997988 | 0.994666 |

**Top altered phosphosites**

| decreased in MNK1KO (logFC>-1.2 | | | |
|---|---|---|---|
| id.query | id.mapped | logFC MNK | logFC MNK2KO |
| Ror1 | Ror1_Q9Z1 | -2.0725 | 0.0625 |
| A0A140LI6 | A0A140LI6 | -2.0025 | 0.3725 |
| Sec62 | Sec62_Q8E | -1.8675 | -0.4975 |
| Kiaa1107 | Kiaa1107_ | -1.82 | -1.1075 |
| Chd9 | Chd9_Q8BY | -1.775 | 0.395 |
| Ubr4 | Ubr4_F6SS | -1.735 | -1.02 |
| Hivep2 | Hivep2_Q3 | -1.575 | 0.7675 |
| Pdap1 | Pdap1_Q3U | -1.56 | 0.43 |
| Poldip3 | Poldip3_F6 | -1.5275 | -0.7075 |
| Znf148 | Znf148_Q6 | -1.52 | 0.005 |
| Hp1bp3 | Hp1bp3_Z4 | -1.5125 | -0.9975 |
| Srrm2 | Srrm2_Q8E | -1.49 | 0.095 |
| Srrm2 | Srrm2_Q8E | -1.49 | 0.095 |
| Prrc2c | Prrc2c_S4R | -1.455 | 0.065 |
| Hdgfrp2 | Hdgfrp2_Q | -1.4525 | 0.4875 |
| Hirip3 | Hirip3_Q8E | -1.4375 | -0.365 |
| Spag9 | Spag9_F8W | -1.4125 | -0.255 |
| Phyhipl | Phyhipl_Q8 | -1.3925 | -0.4325 |
| Usp24 | Usp24_E9P | -1.3625 | 0.615 |
| Dpf2 | Dpf2_F8WI | -1.3425 | 0.975 |
| Bzw1 | Bzw1_Q9C | -1.34 | 0.465 |
| Cdc42bpb | Cdc42bpb_ | -1.3375 | 0.375 |
| Kctd1 | Kctd1_E9Q | -1.3325 | -0.02 |
| Thrap3 | Thrap3_Q5 | -1.3325 | -0.1425 |
| Sec62 | Sec62_Q8E | -1.3275 | 0.1625 |
| Hsp90b1 | Hsp90b1_F | -1.3225 | -0.4125 |
| Macrod2 | Macrod2_ | -1.3225 | -1.8625 |
| Macrod2 | Macrod2_ | -1.3225 | -1.8625 |
| Fnip1 | Fnip1_F6RJ | -1.305 | 0.2325 |
| Eif4b | Eif4b_Q8B | -1.2875 | 0.2675 |
| Arhgap31 | Arhgap31_ | -1.2825 | -0.9475 |
| Pcyt1a | Pcyt1a_P4 | -1.28 | -0.7375 |
| Prrc2c | Prrc2c_S4R | -1.275 | 0.27 |
| Irs2 | Irs2_P8112 | -1.2725 | 0.11 |
| Eif4b | Eif4b_Q8B | -1.2575 | -0.0625 |
| Marchf11 | March11_ | -1.2425 | -0.4425 |
| Srrt | Srrt_Q99M | -1.24 | 0.2675 |
| Kri1 | Kri1_V9GX | -1.24 | -0.3875 |
| U2af1 | U2af1_Q9D | -1.24 | -0.8225 |
| Kiaa1107 | Kiaa1107_ | -1.235 | -0.1575 |
| Fam83h | Fam83h_Q | -1.225 | 0.3875 |
| Gpatch8 | Gpatch8_A | -1.225 | -0.6875 |
| Dnajc8 | Dnajc8_E9I | -1.22 | -0.4875 |
| Ubn1 | Ubn1_Q4G | -1.215 | 0.27 |
| Zranb2 | Zranb2_E9I | -1.2125 | 0.665 |
| Sec62 | Sec62_Q8E | -1.2125 | -0.2225 |
| Tmem169 | Tmem169_ | -1.205 | 0.035 |

| increased in MNK1KO (logFC<1.2 | | | |
|---|---|---|---|
| id.query | id.mapped | logFC MNK | logFC MNK2KO |
| Bcan | Bcan_Q613 | 3.0175 | 2.8425 |
| Gys1 | Gys1_Q9Z1 | 2.9025 | 3.92 |
| Zfyve19 | Zfyve19_Q | 2.84 | 2.7175 |
| Ppp1r10 | Ppp1r10_C | 2.76 | 3.1625 |
| Ccdc85a | Ccdc85a_E | 2.64 | 2.9 |
| Agap2 | Agap2_Q3U | 2.6275 | 2.2225 |
| Epb4 | Epb4.1l1_C | 2.56 | -0.655 |
| Bsn | Bsn_O8873 | 2.5 | 3.5675 |
| Ccar2 | Ccar2_Q8V | 2.485 | 2.8275 |
| Polr3c | Polr3c_F6C | 2.4275 | 2.97 |
| Gabra2 | Gabra2_P2 | 2.415 | 2.5275 |
| Nf1 | Nf1_Q0469 | 2.38 | 0.4175 |
| Q9D9H8 | Q9D9H8_S | 2.375 | 2.52 |
| Atxn2 | Atxn2_F6U | 2.345 | 3.2725 |
| Shroom2 | Shroom2_ | 2.3275 | -0.7625 |
| Cnp | Cnp_P1633 | 2.315 | 3.5275 |
| Ccsap | Ccsap_Q8C | 2.3025 | 1.0825 |
| Asap1 | Asap1_E9C | 2.255 | -0.4325 |
| Ank2 | Ank2_Q8C | 2.245 | -0.63 |
| Rimbp2 | Rimbp2_D3 | 2.1975 | 0.48 |
| Bsn | Bsn_O8873 | 2.1875 | 0.7975 |
| Hcn1 | Hcn1_O887 | 2.18 | -1.565 |
| Cenpv | Cenpv_Q9C | 2.135 | 2.635 |
| Kiaa1671 | Kiaa1671_ | 2.08 | 3.4625 |
| Eml1 | Eml1_D3Z4 | 2.06 | 4.3825 |
| Map1a | Map1a_Q9 | 2.05 | 3.86 |
| Rbm15b | Rbm15b_Q | 2.005 | 0.1275 |
| Map7d2 | Map7d2_A | 1.9975 | 2.7825 |
| Ncbp1 | Ncbp1_Q3U | 1.995 | -0.5475 |
| Dlgap3 | Dlgap3_B1 | 1.9925 | -1.0475 |
| Sptbn1 | Sptbn1_Q6 | 1.96 | 1.095 |
| Rap1gap2 | Rap1gap2_ | 1.9225 | 0.985 |
| Lppr2 | Lppr2_Q8V | 1.9175 | -0.1625 |
| Lppr4 | Lppr4_Q7T | 1.905 | 0.6025 |
| Cobll1 | Cobll1_B1A | 1.905 | -0.715 |
| Dmwd | Dmwd_E9C | 1.9025 | -0.7875 |
| Syn2 | Syn2_Q643 | 1.8775 | 0.9875 |
| Gigyf2 | Gigyf2_G3U | 1.8675 | 2.015 |
| Map2 | Map2_P20 | 1.8425 | 0.3375 |
| Gnaz | Gnaz_O704 | 1.8225 | -0.96 |
| Irs1 | Irs1_Q543V | 1.8175 | -0.83 |
| Ctage5 | Ctage5_H3 | 1.74 | 1.0525 |
| Ank2 | Ank2_Q8C | 1.735 | -0.495 |
| Rasgrf2 | Rasgrf2_P7 | 1.73 | 1.4025 |
| Dlgap2 | Dlgap2_Q0 | 1.73 | -0.3075 |
| Grin2b | Grin2b_G3 | 1.73 | -0.79 |
| Gphn | Gphn_Q8B | 1.7275 | -1.95 |
| Cacna1e | Cacna1e_C | 1.6825 | -1.1175 |
| Mybbp1a | Mybbp1a_ | 1.6675 | 2.4625 |
| Nhsl1 | Nhsl1_Q8C | 1.665 | 3.295 |
| Cit | Cit_E9QL5 | 1.655 | -1.74 |
| Ppp1r3e | Ppp1r3e_H | 1.64 | 1.4025 |
| Clvs1 | Clvs1_Q9D | 1.63 | -2.5075 |
| Cnr1 | Cnr1_P477 | 1.6275 | -1.4225 |
| Ehbp1l1 | Ehbp1l1_G | 1.6225 | 0.015 |
| Ankrd34a | Ankrd34a_ | 1.6125 | -1.2425 |
| Epb4 | Epb4.1l1_C | 1.6075 | -0.2075 |
| Wdr37 | Wdr37_Q8 | 1.605 | 2.8925 |
| Bsn | Bsn_O8873 | 1.5975 | 1.08 |
| Mical3 | Mical3_Q8 | 1.595 | 1.14 |
| Ank2 | Ank2_Q8C | 1.5925 | 2.405 |
| Dlgap2 | Dlgap2_Q0 | 1.57 | 1.3575 |
| Sptbn1 | Sptbn1_Q6 | 1.5675 | 1.6025 |
| Ank3 | Ank3_G5E8 | 1.565 | -0.275 |
| Bsn | Bsn_O8873 | 1.5625 | 2.43 |
| Ank2 | Ank2_Q8C | 1.5525 | 1.15 |
| Ank2 | Ank2_Q8C | 1.5325 | -0.0125 |
| Eif4b | Eif4b_Q8B | 1.5125 | 0.5225 |
| Leo1 | Leo1_Q5XJ | 1.5025 | 2.21 |
| Paip1 | Paip1_G3U | 1.5025 | -0.2 |
| Phactr2 | Phactr2_B1 | 1.4725 | 1.1675 |
| Ifngr1 | Ifngr1_P15 | 1.47 | 0.4075 |
| Ppp1r1a | Ppp1r1a_Q | 1.465 | 1.125 |
| Dlgap2 | Dlgap2_Q0 | 1.46 | -0.1 |
| Lrrc16a | Lrrc16a_D3 | 1.4525 | 2.66 |
| Agap3 | Agap3_Q8 | 1.4475 | 1.4475 |
| Aak1 | Aak1_Q3U | 1.445 | 0.7475 |
| Amer2 | Amer2_Q8 | 1.4425 | -0.3625 |
| Pdha1 | Pdha1_P35 | 1.4425 | -0.5925 |
| Pdha1 | Pdha1_P35 | 1.4425 | -0.5925 |
| Cacna1e | Cacna1e_C | 1.44 | 2.7725 |
| Akap12 | Akap12_Q9 | 1.4375 | 1.2125 |
| Epb4 | Epb4.1_P4 | 1.4375 | 0.1975 |
| Map2 | Map2_P20 | 1.4325 | 2.6475 |
| Scg2 | Scg2_Q4W | 1.4325 | -0.625 |
| Pitpnm2 | Pitpnm2_C | 1.4225 | 2.2525 |
| Lmo7 | Lmo7_E9P | 1.4225 | -0.51 |
| Hcn3 | Hcn3_O887 | 1.41 | 2.805 |
| Gapvd1 | Gapvd1_Q | 1.4075 | 0.765 |
| Inpp4b | Inpp4b_E9 | 1.4 | -0.77 |
| Sgip1 | Sgip1_Q8V | 1.4 | -1.695 |
| Epb4 | Epb4.1l1_C | 1.3925 | 0.4075 |
| Bag2 | Bag2_Q91Y | 1.39 | 0.52 |
| Pex5l | Pex5l_F8SL | 1.385 | -0.775 |
| Fam160a2 | Fam160a2_ | 1.3675 | 1.785 |
| Iqsec1 | Iqsec1_Q8I | 1.3625 | 1.3575 |
| Sgsm1 | Sgsm1_B2F | 1.3625 | -0.8875 |
| Hspa4l | Hspa4l_P48 | 1.35 | -0.985 |
| Sgip1 | Sgip1_Q8V | 1.345 | -0.0975 |
| Tra2a | Tra2a_E9Q | 1.34 | -1.5925 |
| Sptbn1 | Sptbn1_Q6 | 1.3325 | 1.03 |
| Slc30a3 | Slc30a3_P9 | 1.325 | 0.47 |

| decreased in MNK2KO (logFC>-1.2 | | | |
|---|---|---|---|
| id.query | id.concat | logFC MNK | logFC MNK2KO |
| Zfr2 | Zfr2_E9Q5 | -0.0075 | -2.845 |
| Macrod2 | Macrod2_ | -1.3225 | -1.8625 |
| Macrod2 | Macrod2_ | -1.3225 | -1.8625 |
| Kif21b | Kif21b_Q9 | -0.6525 | -1.4775 |
| Atp7a | Atp7a_Q64 | -0.4175 | -1.3875 |
| Plekhf2 | Plekhf2_Q9 | -0.35 | -1.7725 |
| Itsn1 | Itsn1_E9Q | 0.335 | -1.2225 |
| Atp2b2 | Atp2b2_S4 | -0.1175 | -1.2775 |
| Brd8 | Brd8_D3YZ | -0.1825 | -1.6175 |
| Rims1 | Rims1_E9P | -0.0625 | -1.5775 |
| Cacnb1 | Cacnb1_Q8 | -1.0175 | -1.345 |
| Napa | Napa_Q9D | 0.035 | -2.9575 |
| Adgrb1 | Adgrb1_E9 | -0.1025 | -1.295 |
| Arhgef26 | Arhgef26_ | -0.13 | -1.23 |
| Arhgef26 | Arhgef26_ | -0.13 | -1.23 |
| Sf3b1 | Sf3b1_Q99 | -0.2525 | -1.765 |
| Bcas1 | Bcas1_Q80 | -0.2875 | -1.3425 |
| Rbm15 | Rbm15_Q0 | -0.57 | -1.24 |
| Rabgef1 | Rabgef1_Q | -0.5575 | -2.35 |
| Cacng3 | Cacng3_Q9 | -0.055 | -1.3575 |
| Cacng2 | Cacng2_O8 | -0.1725 | -1.2 |
| Atp1a3 | Atp1a3_Q6 | -0.32 | -1.3725 |
| Wdr59 | Wdr59_Q8 | -0.3825 | -1.4925 |
| Rps6ka2 | Rps6ka2_Q | 0.7675 | -1.2025 |
| Gm996 | Gm996_A2 | -0.1325 | -1.9075 |
| Prkag2 | Prkag2_Q9 | 0.4975 | -1.6075 |
| Ank2 | Ank2_Q8C | 0.1525 | -2.21 |
| Atp2a2 | Atp2a2_O5 | 0.39 | -1.715 |
| Ppme1 | Ppme1_Q8 | -0.3 | -1.3375 |
| Cmip | Cmip_Q9D | -0.345 | -1.925 |
| Stmn1 | Stmn1_P54 | 0.0075 | -1.375 |
| Shisa6 | Shisa6_F6V | 0.425 | -1.715 |
| Sptan1 | Sptan1_P1 | 0.0975 | -1.2075 |
| Gphn | Gphn_Q8B | 1.7275 | -1.95 |
| Pcnxl3 | Pcnxl3_Q8 | 0.3525 | -1.2075 |
| Sipa1l1 | Sipa1l1_Q8 | 1.1425 | -1.5225 |
| Ddx3x | Ddx3x_Q62 | 1.08 | -1.5 |
| Psd | Psd_Q5DT | 0.865 | -1.6925 |
| Lysmd2 | Lysmd2_G3 | 0.39 | -1.905 |
| Kcnc1 | Kcnc1_P15 | 0.46 | -1.555 |
| Slc4a10 | Slc4a10_Q | 0.685 | -1.65 |
| Sirt2 | Sirt2_Q8VD | -0.0175 | -1.275 |
| Ncam2 | Ncam2_O3 | 0.23 | -1.305 |
| Shisa7 | Shisa7_F6T | 0.8725 | -2.3025 |
| Pgam1 | Pgam1_Q9 | 0.1325 | -1.52 |
| Efnb3 | Efnb3_O35 | 1.2375 | -1.9325 |
| Sgip1 | Sgip1_Q8V | 1.4 | -1.695 |
| Kcnc2 | Kcnc2_Q14 | 0.7775 | -1.4425 |
| Tbc1d10b | Tbc1d10b_ | 0.4575 | -1.3725 |
| Mta3 | Mta3_H3B | 0.9775 | -1.29 |
| Zfp704 | Zfp704_Q9 | 0.5375 | -1.26 |
| Srsf1 | Srsf1_H7B | 0.95 | -2.2275 |
| Cap2 | Cap2_D3Y | 0.365 | -1.26 |
| Prkcb | Prkcb_P68 | 0.9225 | -1.8875 |
| Mobp | Mobp_Q9D | 0.4175 | -1.61 |
| Ank2 | Ank2_Q8C | 1.07 | -1.8375 |
| Thrap3 | Thrap3_Q5 | 1.0275 | -1.855 |
| Clvs1 | Clvs1_Q9D | 1.63 | -2.5075 |
| Hcn1 | Hcn1_O887 | 2.18 | -1.565 |
| Actr3 | Actr3_Q99 | 1.125 | -1.7175 |
| Bckdha | Bckdha_Q3 | 0.7525 | -1.6875 |
| Ccz1 | Ccz1_Q8C1 | 0.79 | -1.4825 |
| Tra2a | Tra2a_E9Q | 1.34 | -1.5925 |
| Fam129a | Fam129a_ | -0.245 | -1.235 |
| Cap2 | Cap2_D3Y | 0.34 | -1.6 |
| Htt | Htt_G3X9H | 1.0125 | -1.675 |
| Apc | Apc_B2RU | 1.02 | -1.6325 |
| Mbp | Mbp_P043 | 0.6325 | -1.2275 |
| Ppig | Ppig_A2AR | 0.955 | -1.3475 |
| Mapk8ip1 | Mapk8ip1_ | 1.3125 | -1.2125 |
| Grin2b | Grin2b_G3 | 1.155 | -1.25 |
| Fam169a | Fam169a_ | -0.1775 | -1.29 |
| Bnip3l | Bnip3l_Q9 | -0.27 | -1.2175 |
| Ankrd34a | Ankrd34a_ | 1.6125 | -1.2425 |
| Inpp5b | Inpp5b_F2 | 0.585 | -1.23 |
| Kif21b | Kif21b_Q9 | 1.14 | -1.2275 |
| Cnr1 | Cnr1_P477 | 1.6275 | -1.4225 |
| Ranbp2 | Ranbp2_Q | 1.305 | -1.32 |
| Cit | Cit_E9QL5 | 1.655 | -1.74 |

| increased in MNK2KO (logFC<1.2 | | | |
|---|---|---|---|
| id.query | | logFC MNK | logFC MNK2KO |
| Eml1 | Eml1_D3Z4 | 2.06 | 4.3825 |
| Gys1 | Gys1_Q9Z1 | 2.9025 | 3.92 |
| Map1a | Map1a_Q9 | 2.05 | 3.86 |
| Bsn | Bsn_O8873 | 2.5 | 3.5675 |
| Cnp | Cnp_P1633 | 2.315 | 3.5275 |
| Kiaa1671 | Kiaa1671_ | 2.08 | 3.4625 |
| Nhsl1 | Nhsl1_Q8C | 1.665 | 3.295 |
| Dtna | Dtna_Q9D | 0.2825 | 3.2725 |
| Atxn2 | Atxn2_F6U | 2.345 | 3.2725 |
| Ppp1r10 | Ppp1r10_C | 2.76 | 3.1625 |
| Elac2 | Elac2_B1A | 1.2625 | 3.0225 |
| Polr3c | Polr3c_F6C | 2.4275 | 2.97 |
| Ptprd | Ptprd_Q64 | 0.9275 | 2.965 |
| Ccdc85a | Ccdc85a_E | 2.64 | 2.9 |
| Kcnb1 | Kcnb1_Q03 | 0.675 | 2.8925 |
| Wdr37 | Wdr37_Q8 | 1.605 | 2.8925 |
| Bcan | Bcan_Q613 | 3.0175 | 2.8425 |
| Arhgap39 | Arhgap39_ | 1.295 | 2.84 |
| Ccar2 | Ccar2_Q8V | 2.485 | 2.8275 |
| Hcn3 | Hcn3_O887 | 1.41 | 2.805 |
| Map7d2 | Map7d2_A | 1.9975 | 2.7825 |
| Cacna1e | Cacna1e_C | 1.44 | 2.7725 |
| Q3UTZ3 | Q3UTZ3_S | 0.37 | 2.7375 |
| Zfyve19 | Zfyve19_Q | 2.84 | 2.7175 |
| Lrrc16a | Lrrc16a_D3 | 1.4525 | 2.66 |
| Map2 | Map2_P20 | 1.4325 | 2.6475 |
| Cenpv | Cenpv_Q9C | 2.135 | 2.635 |
| Dnm1 | Dnm1_P39 | 0.885 | 2.6075 |
| Mapt | Mapt_P10 | -0.6225 | 2.595 |
| Rptor | Rptor_Q8K | 1.2725 | 2.5775 |
| Gabra2 | Gabra2_P2 | 2.415 | 2.5275 |
| Q9D9H8 | Q9D9H8_S | 2.375 | 2.52 |
| Hivep2 | Hivep2_Q3 | -0.2975 | 2.5 |
| Prpf40a | Prpf40a_Q | 1.245 | 2.475 |
| Mybbp1a | Mybbp1a_ | 1.6675 | 2.4625 |
| Bsn | Bsn_O8873 | 1.5625 | 2.43 |
| Ank2 | Ank2_Q8C | 1.5925 | 2.405 |
| Cdkn2aip | Cdkn2aip_ | 0.3775 | 2.4025 |
| D430042O | D430042O | 1.1575 | 2.395 |
| Alkbh5 | Alkbh5_Q3 | 0.6225 | 2.355 |
| Dtna | Dtna_Q8CF | 0.83 | 2.33 |
| Map2 | Map2_P20 | 1.1975 | 2.295 |
| Pitpnm2 | Pitpnm2_C | 1.4225 | 2.2525 |
| Agap2 | Agap2_Q3U | 2.6275 | 2.2225 |
| Leo1 | Leo1_Q5XJ | 1.5025 | 2.21 |
| Pacs1 | Pacs1_Q8K | 1.01 | 2.1875 |
| Mark4 | Mark4_Q8 | 1.3075 | 2.18 |
| Gas7 | Gas7_B1AT | 1.265 | 2.1175 |
| Zbtb7a | Zbtb7a_D3 | 0.99 | 2.11 |
| Bod1l | Bod1l_E9Q | 0.925 | 2.0925 |
| Cep170 | Cep170_H7 | 0.9 | 2.0625 |
| Dpysl3 | Dpysl3_E9F | 0.55 | 2.0575 |
| Satb2 | Satb2_H3B | 0.8 | 2.055 |
| A0A0U1RP | A0A0U1RP | 0.515 | 2.0425 |
| Dnm1 | Dnm1_P39 | 0.7125 | 2.0375 |
| Gigyf2 | Gigyf2_G3U | 1.8675 | 2.015 |
| Dpysl3 | Dpysl3_E9F | 0.96 | 2.0125 |
| Acin1 | Acin1_F6RJ | 0.65 | 2.0025 |
| Syngap1 | Syngap1_F | 0.7775 | 1.99 |
| Htatsf1 | Htatsf1_Q8 | 0.79 | 1.95 |
| Dnm1 | Dnm1_P39 | 0.7075 | 1.9475 |
| Abl1 | Abl1_P005 | -0.0775 | 1.93 |
| Caskin1 | Caskin1_Q | 0.8375 | 1.9175 |
| Mast1 | Mast1_Q9F | 1.28 | 1.9075 |
| Nasp | Nasp_B1AU | 1.1475 | 1.89 |
| Rbm15 | Rbm15_Q0 | 0.9175 | 1.89 |
| Dmtn | Dmtn_Q9W | 0.3725 | 1.875 |
| Mark3 | Mark3_Q0 | 0.725 | 1.875 |
| Arhgap39 | Arhgap39_ | 0.1275 | 1.87 |
| Prpf4b | Prpf4b_Q6 | 0.3475 | 1.8525 |
| Prrc2c | Prrc2c_S4R | 0.505 | 1.8375 |
| Pja1 | Pja1_O551 | 1.04 | 1.8325 |
| Srrm2 | Srrm2_Q8E | 0.4275 | 1.815 |
| Pak6 | Pak6_Q3UI | 1.175 | 1.795 |
| Pak6 | Pak6_Q3UI | 1.175 | 1.795 |
| Kcnj3 | Kcnj3_P63 | 1.0475 | 1.7875 |
| Fam160a2 | Fam160a2_ | 1.3675 | 1.785 |
| Atxn2l | Atxn2l_Q7 | 0.7175 | 1.7725 |
| Prrc2c | Prrc2c_S4R | 0.6125 | 1.75 |
| Nav1 | Nav1_Q8C | 1.215 | 1.75 |
| Rgs12 | Rgs12_E9Q | 0.8775 | 1.745 |
| Atxn2l | Atxn2l_Q7 | 0.945 | 1.7425 |
| Abl1 | Abl1_P005 | -0.1775 | 1.73 |
| Satb2 | Satb2_H3B | -0.65 | 1.715 |
| Satb2 | Satb2_H3B | -0.65 | 1.715 |
| Wrnip1 | Wrnip1_Q9 | 0.66 | 1.7025 |
| Ppp1r7 | Ppp1r7_Q3 | -0.07 | 1.69 |
| Hecw1 | Hecw1_Q8 | 1.19 | 1.68 |
| Gsk3b | Gsk3b_E9C | 0.2125 | 1.675 |
| Syt17 | Syt17_Q92 | 0.455 | 1.6725 |
| Mdc1 | Mdc1_E9Q | -0.01 | 1.6525 |
| Sorbs2 | Sorbs2_D3 | 0.8475 | 1.6475 |
| Srcin1 | Srcin1_B1A | 0.945 | 1.6475 |
| Vps51 | Vps51_Q3U | 0.73 | 1.6425 |
| Irf2bpl | Irf2bpl_Q8 | 0.615 | 1.6425 |
| Gpr158 | Gpr158_Q8 | 1.2875 | 1.6425 |
| Brf1 | Brf1_G3X8 | 0.425 | 1.6325 |
| Mef2c | Mef2c_Q8 | -0.03 | 1.6275 |
| Fxyd7 | Fxyd7_P59 | 0.5775 | 1.625 |
| Macf1 | Macf1_Q9 | 0.06 | 1.6125 |
| Rnf139 | Rnf139_Q7 | 1.16 | 1.61 |
| Abl2 | Abl2_F8VQ | 0.69 | 1.605 |

| | | | |
|---|---|---|---|
| Srgap3 | Srgap3_E9( | 1.3225 | 0.0325 |
| Map7d1 | Map7d1_F( | 1.32 | 1.37 |
| Aftph | Aftph_H3B | 1.315 | -0.9625 |
| Mapk8ip1 | Mapk8ip1_ | 1.3125 | -1.2125 |
| Mark4 | Mark4_Q8( | 1.3075 | 2.18 |
| Map3k10 | Map3k10_I | 1.3075 | -0.1375 |
| Tnrc6b | Tnrc6b_Q8 | 1.3075 | -0.295 |
| Srrm1 | Srrm1_E9P | 1.305 | -0.0425 |
| Ranbp2 | Ranbp2_Q9 | 1.305 | -1.32 |
| Cbfa2t2 | Cbfa2t2_O | 1.3025 | 0.9425 |
| Ank2 | Ank2_Q8C8 | 1.3 | -0.3175 |
| Kiaa1522 | Kiaa1522_/ | 1.2975 | 0.8025 |
| Arhgap39 | Arhgap39_ | 1.295 | 2.84 |
| Dlgap4 | Dlgap4_B7. | 1.295 | 0.9675 |
| Tex2 | Tex2_B1AT | 1.2925 | 0.4775 |
| Tbc1d10b | Tbc1d10b_ | 1.29 | 0.7975 |
| Shank1 | Shank1_D3 | 1.29 | -0.15 |
| Gpr158 | Gpr158_Q8 | 1.2875 | 1.6425 |
| Fcho2 | Fcho2_Q3U | 1.2875 | -0.805 |
| Gprin1 | Gprin1_Q3 | 1.2825 | 0.6225 |
| Mast1 | Mast1_Q9R | 1.28 | 1.9075 |
| Marcks | Marcks_P2 | 1.28 | 0.7875 |
| Ppp6r2 | Ppp6r2_G3 | 1.275 | 1.55 |
| Rptor | Rptor_Q8K | 1.2725 | 2.5775 |
| Bsn | Bsn_O8873 | 1.2675 | 0.4625 |
| Gas7 | Gas7_B1AT | 1.265 | 2.1175 |
| Slc4a4 | Slc4a4_E9C | 1.265 | 0.945 |
| Elac2 | Elac2_B1A1 | 1.2625 | 3.0225 |
| Shank3 | Shank3_Q4 | 1.255 | 0.7 |
| Cnksr2 | Cnksr2_Q8 | 1.2525 | 0.8275 |
| Prrc2a | Prrc2a_G3U | 1.25 | 1.115 |
| Myh10 | Myh10_Q5 | 1.25 | 0.6425 |
| Tiam2 | Tiam2_Q62 | 1.25 | -1.0025 |
| Prpf40a | Prpf40a_Q9 | 1.245 | 2.475 |
| Zwint | Zwint_Q9C | 1.24 | 1.0725 |
| Efnb3 | Efnb3_O35 | 1.2375 | -1.9325 |
| Mapt | Mapt_P106 | 1.23 | 1.3825 |
| Arhgap32 | Arhgap32_ | 1.23 | 0.2425 |
| Tpd52l2 | Tpd52l2_Q | 1.225 | -0.2125 |
| Chgb | Chgb_P160 | 1.2175 | -0.2325 |
| Mtus2 | Mtus2_Q3U | 1.2175 | -0.2375 |
| Nav1 | Nav1_Q8CI | 1.215 | 1.75 |
| Tmem184b | Tmem184b | 1.2125 | 0.26 |
| Plch2 | Plch2_H3B | 1.21 | 0.6575 |
| Psd3 | Psd3_E9PU | 1.2075 | -0.4775 |
| Scg2 | Scg2_Q4W | 1.2075 | -0.755 |
| Dlg2 | Dlg2_E9Q2 | 1.205 | 0.67 |
| Cacnb1 | Cacnb1_Q8 | 1.205 | 0.1375 |
| Cep170 | Cep170_H7 | 1.2 | 1.0025 |

| | | | |
|---|---|---|---|
| Sptbn1 | Sptbn1_Q6 | 1.5675 | 1.6025 |
| Plcl2 | Plcl2_Q8K3 | 0.875 | 1.5975 |
| Eif4b | Eif4b_Q8B( | 0.915 | 1.565 |
| Set | Set_Q9EQU | 0.3075 | 1.5525 |
| Ppp6r2 | Ppp6r2_G3 | 1.275 | 1.55 |
| Cnksr2 | Cnksr2_Q8 | 0.4675 | 1.545 |
| Rnmt | Rnmt_D3Z4 | 0.7925 | 1.545 |
| Kank3 | Kank3_Q9Z | -0.3975 | 1.5325 |
| Camkk2 | Camkk2_Q. | 0.5375 | 1.5275 |
| Phactr1 | Phactr1_Q2 | 0.8425 | 1.5275 |
| Caskin1 | Caskin1_Q6 | 0.805 | 1.5175 |
| Ccdc85a | Ccdc85a_E( | -0.4175 | 1.5125 |
| Sipa1l1 | Sipa1l1_Q8 | 0.13 | 1.51 |
| Ildr2 | Ildr2_B5TV | 0.94 | 1.51 |
| Cep170 | Cep170_H7 | -0.255 | 1.5025 |
| Tenm4 | Tenm4_Q3 | -0.845 | 1.5025 |
| Tenm4 | Tenm4_Q3 | -0.845 | 1.5025 |
| Tenm4 | Tenm4_Q3 | -0.845 | 1.5025 |
| Eif3a | Eif3a_P231 | 0.7275 | 1.5 |
| Vps51 | Vps51_Q3U | 1.1775 | 1.5 |
| Grb10 | Grb10_B1A | 1.0375 | 1.485 |
| Usp6nl | Usp6nl_Q8 | 0.3925 | 1.475 |
| Tcp11l1 | Tcp11l1_Q8 | 0.4775 | 1.4725 |
| Agfg1 | Agfg1_Q8K | 0.34 | 1.47 |
| Pole4 | Pole4_E0CY | 0.0775 | 1.4675 |
| Nfic | Nfic_P7025 | -0.08 | 1.46 |
| Agap3 | Agap3_Q8V | 1.4475 | 1.4475 |
| Eif4g1 | Eif4g1_E9P | 0.3025 | 1.4375 |
| Mast2 | Mast2_B1A | 0.6975 | 1.435 |
| Psd | Psd_Q5DTT | 0.8225 | 1.42 |
| Ank2 | Ank2_Q8C8 | 0.6525 | 1.415 |
| Tuba4a | Tuba4a_P6 | -0.1125 | 1.4075 |
| Sh3gl3 | Sh3gl3_J3C | 0.525 | 1.405 |
| Tcof1 | Tcof1_H3B | 0.78 | 1.4025 |
| Srrm2 | Srrm2_Q8B | 0.8325 | 1.4025 |
| Ppp1r3e | Ppp1r3e_H | 1.64 | 1.4025 |
| Dnm1 | Dnm1_P39 | 1.1675 | 1.4025 |
| Rasgrf2 | Rasgrf2_P7 | 1.73 | 1.4025 |
| Clasp2 | Clasp2_F7D | 0.445 | 1.3975 |
| Srcin1 | Srcin1_B1A | 0.6125 | 1.3975 |
| Gsk3b | Gsk3b_E9C | -0.2525 | 1.385 |
| Chgb | Chgb_P160 | 0.2775 | 1.385 |
| Rrbp1 | Rrbp1_Q99 | 0.4625 | 1.385 |
| Mapt | Mapt_P106 | 1.23 | 1.3825 |
| Tra2a | Tra2a_E9Q | 0.525 | 1.3725 |
| Eif4b | Eif4b_Q8B( | 0.1475 | 1.37 |
| Map7d1 | Map7d1_F( | 1.32 | 1.37 |
| Nrgn | Nrgn_P607 | 0.3325 | 1.365 |
| Kif1a | Kif1a_Q6TA | 0.98 | 1.3575 |
| Dlgap2 | Dlgap2_Q0 | 1.57 | 1.3575 |
| Iqsec1 | Iqsec1_Q8R | 1.3625 | 1.3575 |
| Aff4 | Aff4_Q9ES( | 0.3575 | 1.3525 |
| Dlg2 | Dlg2_E9Q2 | 0.855 | 1.35 |
| Kiaa1671 | Kiaa1671_( | 0.9375 | 1.3475 |
| Dpp8 | Dpp8_Q80Y | 0.3575 | 1.345 |
| Shank1 | Shank1_D3 | 1.06 | 1.345 |
| Add1 | Add1_F8W | -0.3175 | 1.3425 |
| Syn1 | Syn1_O889 | 0.0975 | 1.3375 |
| Trim2 | Trim2_Q9E | 0.75 | 1.3325 |
| Tom1 | Tom1_Q3U | 1.105 | 1.33 |
| Rbm33 | Rbm33_D3 | -0.7775 | 1.3275 |
| Eif2b4 | Eif2b4_D3Z | 0.4775 | 1.3275 |
| Tmem132c | Tmem132c | -0.12 | 1.325 |
| Srp68 | Srp68_Q8B | 0.275 | 1.3225 |
| Ank2 | Ank2_Q8C8 | -0.095 | 1.3225 |
| Rims1 | Rims1_E9P | 0.99 | 1.3225 |
| Pkp4 | Pkp4_Q68F | 0.64 | 1.3175 |
| Smap | Smap_Q9R | 0.1875 | 1.3175 |
| Prrc2c | Prrc2c_S4R | -0.1875 | 1.3125 |
| Mark2 | Mark2_Q05 | 0.8275 | 1.305 |
| Camkv | Camkv_Q3 | 0.1725 | 1.3 |
| Lmna | Lmna_P48( | -0.185 | 1.3 |
| Uhrf1bp1l | Uhrf1bp1l_ | 0.88 | 1.3 |
| Eef2 | Eef2_P5825 | 0.4825 | 1.2975 |
| Vim | Vim_P2015 | 0.69 | 1.2975 |
| Satb2 | Satb2_H3B | -0.245 | 1.295 |
| Mark4 | Mark4_Q8( | 0.08 | 1.295 |
| Ppp1r16a | Ppp1r16a_ | 0.3425 | 1.2925 |
| Map1a | Map1a_Q9 | 0.5525 | 1.2925 |
| Fsd1 | Fsd1_Q7TP | 0.225 | 1.2875 |
| Eif3f | Eif3f_Q9DC | 1.09 | 1.2875 |
| Sh3gl1 | Sh3gl1_Q6. | 1.1725 | 1.285 |
| Mycbp2 | Mycbp2_F6 | 0.3275 | 1.28 |
| Epb4 | Epb4.1_P48 | 0.925 | 1.28 |
| Phactr2 | Phactr2_B1 | 0.0925 | 1.2775 |
| Pcyt1a | Pcyt1a_P49 | -0.3075 | 1.2775 |
| Sparcl1 | Sparcl1_P7 | 0.4825 | 1.2775 |
| Rgs20 | Rgs20_Q9C | -0.33 | 1.2725 |
| Camkv | Camkv_Q3 | -0.0625 | 1.2675 |
| Bsg | Bsg_K3W4( | -0.0725 | 1.2675 |
| Itpka | Itpka_Q8R( | 1.01 | 1.2675 |
| Csnk1d | Csnk1d_Q9 | 0.2375 | 1.255 |
| Tcf4 | Tcf4_Q607 | 0.31 | 1.2525 |
| Mark3 | Mark3_Q03 | 0.5725 | 1.245 |
| Ank2 | Ank2_Q8C8 | 0.3225 | 1.2425 |
| Golim4 | Golim4_D3 | 0.7725 | 1.2425 |
| Epb4 | Epb4.1l1_C | 0.425 | 1.235 |
| Sh2d5 | Sh2d5_Q8J | 1.1525 | 1.2275 |
| Setd1a | Setd1a_E9I | -0.9425 | 1.225 |
| Ppp2r5d | Ppp2r5d_C | 0.5025 | 1.225 |
| Cttnbp2 | Cttnbp2_B9 | 0.25 | 1.2225 |
| Mtcl1 | Mtcl1_H3B | -0.2375 | 1.215 |
| Mtcl1 | Mtcl1_H3B | -0.2375 | 1.215 |
| Psen2 | Psen2_Q3U | -0.16 | 1.215 |
| Ube2e3 | Ube2e3_B2 | 1.1325 | 1.215 |

| | | | |
|---|---|---|---|
| Camsap1 | Camsap1_/ | 0.66 | 1.2125 |
| Akap12 | Akap12_Q9 | 1.4375 | 1.2125 |
| Ptgdr2 | Ptgdr2_Q9. | 0.255 | 1.21 |
| A0A571BEH | A0A571BEH | 0.3925 | 1.21 |
| Pcyt1a | Pcyt1a_P49 | -0.035 | 1.2075 |
| Plekha1 | Plekha1_Q8 | 0.1 | 1.2075 |
| Dip2c | Dip2c_B7ZI | 0.6125 | 1.2075 |
| Leo1 | Leo1_Q5XJ | 0.2775 | 1.205 |
| Tom1 | Tom1_Q3U | 0.315 | 1.205 |

**Gene Ontology Cellular Component**

**GO performed using Enrichr with total proteome as background list*

**upregulated GO cellular component**

| **SYP MNK1KO / WT** | | | **SYP MNK2KO / WT** | | |
|---|---|---|---|---|---|
| **cellular component** | **NOM p-val** | **Adjusted p** | **cellular component** | **NOM p-val** | **Adjusted p-value** |
| Dendrite (GO:0030425) | 0.000261 | 0.02726 | Microtubule Cytoskeleto | 6.60E-06 | 0.000897 |
| Neuron Projection (GO:004300 | 0.000474 | 0.02726 | Cytoskeleton (GO:00058 | 1.36E-04 | 0.006376 |
| | | | Microtubule (GO:000587 | 0.000141 | 0.006376 |
| | | | Polymeric Cytoskeletal F | 0.000228 | 0.006605 |
| | | | Glutamatergic Synapse ( | 0.000272 | 0.006605 |
| | | | Dendrite (GO:0030425) | 0.000291 | 0.006605 |

downregulated GO cellular component

| **SYP MNK1KO / WT** | | | **SYP MNK2KO / WT** | | |
|---|---|---|---|---|---|
| **cellular component** | **NOM p-val** | **Adjusted p** | **cellular component** | **NOM p-val** | **Adjusted p-value** |
| | | | Dendrite Membrane (GC | 5.5E-06 | 0.000693 |
| | | | Synaptic Membrane (GO | 5.86E-05 | 0.003691 |
| | | | Postsynaptic Density (GC | 0.000169 | 0.006164 |
| | | | Ionotropic Glutamate Re | 0.000196 | 0.006164 |
| | | | AMPA Glutamate Recept | 0.001155 | 0.02911 |
| | | | Asymmetric, Glutamater | 0.001463 | 0.03071 |

**PTM-SEA cortex**

<table>
<tr><th colspan="10">signature</th><th colspan="3">p-value</th><th colspan="3">FDR p.value</th><th colspan="3">NES</th></tr>
<tr><th>id</th><th>Signature.s</th><th>Signature.s</th><th>Signature.s</th><th>Signature.s</th><th>Signature.s</th><th>Signature.s</th><th>Signature.s</th><th>Signature.s</th><th>No.column</th><th>pvalue.MN</th><th>pvalue.MN</th><th>pvalue.MN</th><th>fdr.pvalue.</th><th>fdr.pvalue.</th><th>fdr.pvalue.</th><th>MNK1KO</th><th>MNK2KO</th><th>MNKDKO</th></tr>
<tr><td>PERT-PSP_UV</td><td>n=32|Trp5</td><td>30</td><td>20</td><td>20</td><td>20</td><td>TDDEMTGY</td><td>TDDEMTGY</td><td>TDDEMTGY</td><td>3</td><td>0.141677</td><td>0.220909</td><td>0.181599</td><td>0.220487</td><td>0.358795</td><td>0.399518</td><td>1.838</td><td>1.5195</td><td>1.6896</td></tr>
<tr><td>PERT-PSP_TNF</td><td>n=20|Ikbkb</td><td>18</td><td>27.8</td><td>27.8</td><td>27.8</td><td>TDDEMTGY</td><td>TDDEMTGY</td><td>TDDEMTGY</td><td>3</td><td>0.059984</td><td>0.17374</td><td>0.038932</td><td>0.126613</td><td>0.318523</td><td>0.122359</td><td>2.213</td><td>1.6395</td><td>2.1851</td></tr>
<tr><td>PERT-PSP_PHORBOL_ESTER</td><td>n=41|Mapl</td><td>39</td><td>15.4</td><td>15.4</td><td>15.4</td><td>TDDEMTGY</td><td>TDDEMTGY</td><td>TDDEMTGY</td><td>3</td><td>0.150332</td><td>0.027463</td><td>0.081856</td><td>0.220487</td><td>0.120836</td><td>0.200093</td><td>1.793</td><td>2.1876</td><td>2.1582</td></tr>
<tr><td>PERT-PSP_OKADAIC_ACID</td><td>n=17|Mapl</td><td>15</td><td>40</td><td>40</td><td>40</td><td>TDDEMTGY</td><td>TDDEMTGY</td><td>TDDEMTGY</td><td>3</td><td>0.005528</td><td>0.026825</td><td>0.004605</td><td>0.023975</td><td>0.120836</td><td>0.016884</td><td>3.111</td><td>2.1606</td><td>3.0978</td></tr>
<tr><td>PERT-PSP_LPS</td><td>n=27|Ikbkb</td><td>26</td><td>30.8</td><td>30.8</td><td>30.8</td><td>SGRPRTTSF</td><td>TDDEMTGY</td><td>TDDEMTGY</td><td>3</td><td>0.093556</td><td>0.060529</td><td>0.049338</td><td>0.161642</td><td>0.174409</td><td>0.13568</td><td>2.078</td><td>2.0659</td><td>2.4297</td></tr>
<tr><td>PERT-PSP_INSULIN</td><td>n=25|Eif4e</td><td>25</td><td>32</td><td>32</td><td>32</td><td>IVADQTPTF</td><td>IVADQTPTF</td><td>IVADQTPTF</td><td>3</td><td>0.063306</td><td>0.039656</td><td>0.460168</td><td>0.126613</td><td>0.145404</td><td>0.572332</td><td>-2.283</td><td>-2.5387</td><td>-0.9949</td></tr>
<tr><td>PERT-PSP_EGF</td><td>n=37|Mapl</td><td>35</td><td>22.9</td><td>22.9</td><td>22.9</td><td>TDDEMTGY</td><td>TDDEMTGY</td><td>TDDEMTGY</td><td>3</td><td>0.682486</td><td>0.063422</td><td>0.000798</td><td>0.729044</td><td>0.174409</td><td>0.00585</td><td>0.517</td><td>2.0217</td><td>3.5108</td></tr>
<tr><td>KINASE-PSP_PKCA/Prkca</td><td>n=74|Vim_</td><td>74</td><td>8.1</td><td>8.1</td><td>8.1</td><td>LKSTLSMSP</td><td>TTSTRTYSL</td><td>YSSGPGASF</td><td>3</td><td>0.03864</td><td>0.00326</td><td>0.001582</td><td>0.094452</td><td>0.023907</td><td>0.008699</td><td>2.240</td><td>-3.1413</td><td>-3.2177</td></tr>
<tr><td>KINASE-PSP_PKACA/Prkaca</td><td>n=173|Myl</td><td>172</td><td>17.4</td><td>17.4</td><td>17.4</td><td>MGRVRLRS</td><td>RSAIRRAST</td><td>GGRKKTVSI</td><td>3</td><td>0.038538</td><td>0.715205</td><td>0.271637</td><td>0.094452</td><td>0.749262</td><td>0.459693</td><td>2.520</td><td>0.4814</td><td>-1.3862</td></tr>
<tr><td>KINASE-PSP_P38A/Mapk14</td><td>n=61|Ulk1_</td><td>60</td><td>11.7</td><td>11.7</td><td>11.7</td><td>TDDEMTGY</td><td>TDDEMTGY</td><td>TDDEMTGY</td><td>3</td><td>0.002558</td><td>1.00E-04</td><td>0.000604</td><td>0.021899</td><td>0.0011</td><td>0.00585</td><td>3.439</td><td>3.8524</td><td>3.4295</td></tr>
<tr><td>KINASE-PSP_mTOR/Mtor</td><td>n=88|Dpys</td><td>88</td><td>14.8</td><td>14.8</td><td>14.8</td><td>GPSSSLLSLF</td><td>DVSDERLSF</td><td>DVSDERLSF</td><td>3</td><td>0.91299</td><td>0.432567</td><td>0.832931</td><td>0.91299</td><td>0.559793</td><td>0.832931</td><td>-0.128</td><td>1.0045</td><td>-0.2779</td></tr>
<tr><td>KINASE-PSP_JNK1/Mapk8</td><td>n=37|Jun_</td><td>37</td><td>16.2</td><td>16.2</td><td>16.2</td><td>IVADQTPTF</td><td>GGDSSASSI</td><td>GGDSSASSI</td><td>3</td><td>0.695905</td><td>1.00E-04</td><td>0.000401</td><td>0.729044</td><td>0.0011</td><td>0.00585</td><td>-0.511</td><td>4.1912</td><td>3.895</td></tr>
<tr><td>KINASE-PSP_GSK3B/Gsk3b</td><td>n=70|Cry2_</td><td>70</td><td>12.9</td><td>12.9</td><td>12.9</td><td>SSPGSPGTF</td><td>PVFDLTTTP</td><td>PVFDLTTTP</td><td>3</td><td>0.386516</td><td>0.228324</td><td>0.571252</td><td>0.472409</td><td>0.358795</td><td>0.66145</td><td>-1.100</td><td>-1.5128</td><td>-0.7428</td></tr>
<tr><td>KINASE-PSP_ERK2/Mapk1</td><td>n=187|Irs1</td><td>183</td><td>15.8</td><td>15.8</td><td>15.8</td><td>PTRNEPTTF</td><td>LKHTGPNSI</td><td>KDEILPTTPI</td><td>3</td><td>0.004602</td><td>0.691015</td><td>0.672966</td><td>0.023975</td><td>0.749262</td><td>0.705012</td><td>3.402</td><td>-0.531</td><td>0.5357</td></tr>
<tr><td>KINASE-PSP_ERK1/Mapk3</td><td>n=72|Irf3_</td><td>72</td><td>13.9</td><td>13.9</td><td>13.9</td><td>LKSTLSMSP</td><td>ASLSLPATP</td><td>IVADQTPTF</td><td>3</td><td>0.209539</td><td>0.116944</td><td>0.400199</td><td>0.271168</td><td>0.248514</td><td>0.572332</td><td>1.582</td><td>1.9256</td><td>1.0696</td></tr>
<tr><td>KINASE-PSP_CDK5/Cdk5</td><td>n=71|Pik3c</td><td>70</td><td>21.4</td><td>21.4</td><td>21.4</td><td>VIEPLPVTP</td><td>NVRVSNGS</td><td>STPKSKQSP</td><td>3</td><td>0.031287</td><td>0.124257</td><td>0.30006</td><td>0.094452</td><td>0.248514</td><td>0.471524</td><td>2.571</td><td>1.9261</td><td>-1.3125</td></tr>
<tr><td>KINASE-PSP_CDK1/Cdk1</td><td>n=35|Mcm</td><td>35</td><td>14.3</td><td>14.3</td><td>14.3</td><td>VIEPLPVTP</td><td>NSSDTVTSF</td><td>NPGGRPITF</td><td>3</td><td>0.002986</td><td>0.375372</td><td>0.206999</td><td>0.021899</td><td>0.516137</td><td>0.413999</td><td>3.105</td><td>1.2058</td><td>-1.6123</td></tr>
<tr><td>KINASE-PSP_CAMK2A/Camk2a</td><td>n=48|Gria1</td><td>48</td><td>18.8</td><td>18.8</td><td>18.8</td><td>PIPIMPASP</td><td>RSAIRRAST</td><td>PIPIMPASP</td><td>3</td><td>0.531661</td><td>0.508016</td><td>0.004333</td><td>0.615607</td><td>0.620908</td><td>0.016884</td><td>0.795</td><td>-0.8425</td><td>-2.8207</td></tr>
<tr><td>KINASE-PSP_AMPKA1/Prkaa1</td><td>n=48|Becn</td><td>47</td><td>17</td><td>17</td><td>17</td><td>GRLKRERSN</td><td>KKLMHSSSI</td><td>CGLSRVDST</td><td>3</td><td>1.00E-04</td><td>0.117272</td><td>0.267634</td><td>0.0022</td><td>0.248514</td><td>0.459693</td><td>3.724</td><td>1.95E+00</td><td>-1.416</td></tr>
<tr><td>KINASE-PSP_Akt1</td><td>n=50|Bad_</td><td>50</td><td>16</td><td>16</td><td>16</td><td>SGRPRTTSF</td><td>INRERQKSL</td><td>PFRGRSRSA</td><td>3</td><td>0.006539</td><td>1</td><td>0.468271</td><td>0.023975</td><td>1</td><td>0.572332</td><td>2.826</td><td>3.00E-04</td><td>-0.9336</td></tr>
<tr><td>DISEASE-PSP_Parkinsons_disea</td><td>n=19|Apex</td><td>19</td><td>21.1</td><td>21.1</td><td>21.1</td><td>TPGSRSRTF</td><td>SGYSSPGSF</td><td>SGYSSPGSF</td><td>3</td><td>0.177123</td><td>0.652769</td><td>0.429203</td><td>0.243543</td><td>0.749262</td><td>0.572332</td><td>1.674</td><td>0.5886</td><td>1.0313</td></tr>
<tr><td>DISEASE-PSP_Alzheimers_disea</td><td>n=32|Apex</td><td>31</td><td>19.4</td><td>19.4</td><td>19.4</td><td>TPGSRSRTF</td><td>SGYSSPGSF</td><td>SGYSSPGSF</td><td>3</td><td>0.095516</td><td>0.254261</td><td>0.626068</td><td>0.161642</td><td>0.372916</td><td>0.688675</td><td>2.022</td><td>-1.4422</td><td>0.6263</td></tr>
</table>

**PTM-SEA synaptoneurosomes**

| | signature | | | | | | | | | p-value | | | FDR p.value | | | NES | | |
|---|---|---|---|---|---|---|---|---|---|---|---|---|---|---|---|---|---|---|
| id | Signature.s | Signature.s | Signature.s | Signature.s | Signature.s | Signature.s | Signature.s | Signature.s | No.column | pvalue.MN | pvalue.MN | pvalue.MN | fdr.pvalue. | fdr.pvalue. | fdr.pvalue. | MNK1KO | MNK2KO | MNKDKO |
| PERT-PSP_UV | n=32\|Trp5 | 30 | 20 | 20 | 20 | TDDEMTGY | IVADQTPTF | AGTSFMMT | 3 | 0.094328 | 0.051344 | 0.78322 | 0.173587 | 0.141237 | 0.78322 | 1.9435 | -2.2862 | -0.3665 |
| PERT-PSP_TNF | n=20\|Ikbkb | 18 | 27.8 | 27.8 | 27.8 | TDDEMTGY | AGTSFMMT | AGTSFMMT | 3 | 0.062994 | 0.056909 | 0.270523 | 0.158227 | 0.141237 | 0.59515 | 2.0138 | -2.2254 | -1.4167 |
| PERT-PSP_PHORBOL_ESTER | n=41\|Mapl | 39 | 15.4 | 15.4 | 15.4 | TDDEMTGY | GEKSFRRSV | ______MAS | 3 | 0.016763 | 0.67175 | 0.190081 | 0.061465 | 0.821027 | 0.522723 | 2.5791 | -0.5405 | -1.6517 |
| PERT-PSP_OKADAIC_ACID | n=17\|Mapl | 15 | 40 | 40 | 40 | SFKLSGFSFI | SFKLSGFSFI | SFKLSGFSFI | 3 | 0.02599 | 0.655351 | 0.42728 | 0.081682 | 0.821027 | 0.700516 | 2.5644 | -0.5893 | 1.0901 |
| PERT-PSP_LPS | n=27\|Ikbkb | 26 | 30.8 | 30.8 | 30.8 | TDDEMTGY | IVADQTPTF | AGTSFMMT | 3 | 0.580249 | 0.465296 | 0.219892 | 0.580249 | 0.787423 | 0.537514 | 0.726 | -0.9564 | -1.539 |
| PERT-PSP_INSULIN | n=25\|Eif4e | 25 | 32 | 32 | 32 | TSFMMTPY | IVADQTPTF | AGTSFMMT | 3 | 0.344346 | 0.735453 | 0.036615 | 0.451099 | 0.851577 | 0.225159 | 1.2294 | 0.4354 | 2.3488 |
| PERT-PSP_EGF | n=37\|Mapl | 35 | 22.9 | 22.9 | 22.9 | TDDEMTGY | AGTSFMMT | AGTSFMMT | 3 | 0.148335 | 0.020228 | 0.106742 | 0.233098 | 0.117014 | 0.366176 | 1.7686 | -2.7112 | -1.9869 |
| KINASE-PSP_PKCA/Prkca | n=74\|Vim_ | 74 | 8.1 | 8.1 | 8.1 | LKSTLSMSP | TRSVSSSSY | TTSTRTYSL | 3 | 0.00701 | 0.086782 | 0.669532 | 0.030843 | 0.19092 | 0.701415 | 3.0422 | 2.1386 | 0.5532 |
| KINASE-PSP_PKACA/Prkaca | n=173\|Myl | 172 | 17.4 | 17.4 | 17.4 | SFKLSGFSFI | KVLLRRKSE | SFKLSGFSFI | 3 | 1.00E-04 | 0.031913 | 1.00E-04 | 0.0022 | 0.117014 | 0.0022 | 5.2593 | 2.636 | 4.0322 |
| KINASE-PSP_P38A/Mapk14 | n=61\|Ulk1_ | 60 | 11.7 | 11.7 | 11.7 | TDDEMTGY | IVADQTPTF | LKHTGPNSI | 3 | 0.368295 | 0.187438 | 0.060654 | 0.451099 | 0.374875 | 0.266877 | 1.1817 | -1.6763 | -2.2884 |
| KINASE-PSP_mTOR/Mtor | n=88\|Dpys | 88 | 14.8 | 14.8 | 14.8 | GPSSSLLSLF | EPSTIARSLF | EPSTIARSLF | 3 | 0.50182 | 0.000596 | 0.040938 | 0.525716 | 0.013116 | 0.225159 | 0.8782 | 3.5997 | 2.5447 |
| KINASE-PSP_JNK1/Mapk8 | n=37\|Jun_ | 37 | 16.2 | 16.2 | 16.2 | VGLLKLASP | STPKSKQSP | STPKSKQSP | 3 | 0.480789 | 0.0293 | 0.485312 | 0.525716 | 0.117014 | 0.701415 | -0.9104 | 2.5713 | -0.947 |
| KINASE-PSP_GSK3B/Gsk3b | n=70\|Cry2_ | 70 | 12.9 | 12.9 | 12.9 | SSPGSPGTF | SSPGSPGTF | SVTPKTVTP | 3 | 0.094684 | 0.836824 | 0.539312 | 0.173587 | 0.920507 | 0.701415 | -2.069 | -0.2576 | 0.7777 |
| KINASE-PSP_ERK2/Mapk1 | n=187\|Irs1 | 183 | 15.8 | 15.8 | 15.8 | KEDLPPLTP | KEDLPPLTP | QASSTPLSP | 3 | 0.00339 | 0.438519 | 0.116511 | 0.018644 | 0.787423 | 0.366176 | 3.5489 | 0.9777 | 1.967 |
| KINASE-PSP_ERK1/Mapk3 | n=72\|Irf3_ | 72 | 13.9 | 13.9 | 13.9 | LKSTLSMSP | LKSTLSMSP | LKSTLSMSP | 3 | 0.114229 | 0.963948 | 0.445783 | 0.19331 | 0.971968 | 0.700516 | 1.9293 | 0.065 | -0.9711 |
| KINASE-PSP_CDK5/Cdk5 | n=71\|Pik3c | 70 | 21.4 | 21.4 | 21.4 | LKSTLSMSP | LKSTLSMSP | EQASTPMS | 3 | 0.000609 | 0.001204 | 0.440935 | 0.004465 | 0.013248 | 0.700516 | 3.6099 | 3.9695 | 0.9932 |
| KINASE-PSP_CDK1/Cdk1 | n=35\|Mcm | 35 | 14.3 | 14.3 | 14.3 | LKSTLSMSP | LKSTLSMSP | LKSTLSMSP | 3 | 0.000594 | 0.971968 | 0.665812 | 0.004465 | 0.971968 | 0.701415 | 3.3275 | 0.0455 | -0.5734 |
| KINASE-PSP_CAMK2A/Camk2a | n=48\|Gria1 | 48 | 18.8 | 18.8 | 18.8 | KELEKRASG | KELEKRASG | AGPTRQAS | 3 | 0.369081 | 0.617481 | 0.650547 | 0.451099 | 0.821027 | 0.701415 | -1.1407 | -0.6455 | -0.581 |
| KINASE-PSP_AMPKA1/Prkaa1 | n=48\|Becn | 47 | 17 | 17 | 17 | GRLKRERSN | GHLARECTI | GRLKRERSN | 3 | 0.068952 | 0.057779 | 0.039936 | 0.158227 | 0.141237 | 0.225159 | 2.2267 | 2.3532 | 2.4884 |
| KINASE-PSP_Akt1 | n=50\|Bad_ | 50 | 16 | 16 | 16 | PARNRSASI | PAPSRTASF | PARNRSASI | 3 | 0.426583 | 0.58033 | 0.377995 | 0.493939 | 0.821027 | 0.700516 | -1.0147 | -0.7193 | -1.1327 |
| DISEASE-PSP_Parkinsons_disea | n=19\|Apex | 19 | 21.1 | 21.1 | 21.1 | SSPGSPGTF | TPGSRSRTF | TPGSRSRTF | 3 | 0.193198 | 0.029459 | 0.587106 | 0.283356 | 0.117014 | 0.701415 | -1.6329 | 2.4955 | 0.6909 |
| DISEASE-PSP_Alzheimers_disea | n=32\|Apex | 31 | 19.4 | 19.4 | 19.4 | SSPGSPGTF | TPGSRSRTF | TPGSRSRTF | 3 | 0.071921 | 0.607847 | 0.653434 | 0.158227 | 0.821027 | 0.701415 | -2.262 | 0.6633 | 0.5594 |
