## Supplementary Table 6 for "Distinct roles for MNK1 and MNK2 in social and cognitive behavior through kinase-specific regulation of the synaptic proteome and phosphoproteome"

| Figure | Sample Size | Statistical test | Values |
| --- | --- | --- | --- |
| 1B | wild-type=17<br>MNK1$^{KO}$=14<br>MNK2$^{KO}$=17 | RM two-way ANOVA | trial: $F_{(4, 180}$ = 85.86, P<0.0001<br>genotype: $F_{(2, 45)}$ = 0.7987, P=0.4562<br>trial × genotype: $F_{(8, 180)}$ = 1.933, P=0.0577 |
| 1C | wild-type=17<br>MNK1$^{KO}$=14<br>MNK2$^{KO}$=17 | One-way ANOVA, Tukey's post-hoc test for multiple comparisons | $F_{(2, 45)}$ = 6.565, P=0.0031<br>wild-type vs MNK1$^{KO}$: P=0.0038<br>wild-type vs MNK2$^{KO}$: P= 0.8385<br>MNK1$^{KO}$ vs MNK2$^{KO}$: P= 0.0164 |
| 1D | wild-type=17<br>MNK1$^{KO}$=15<br>MNK2$^{KO}$=17 | RM two-way ANOVA | trials: $F_{(3.882, 178.6)}$ = 41.79, P<0.0001<br>genotype: $F_{(2, 46)}$ = 1.825, P=0.1727<br>trials × genotype: $F_{(22, 506)}$ = 1.265, P=0.1887 |
| 1E | wild-type=17<br>MNK1$^{KO}$=15<br>MNK2$^{KO}$=17 | RM two-way ANOVA, Tukey's post-hoc test for multiple comparisons | trial: $F_{(2.514, 115.6)}$ = 37.21, P<0.0001<br>genotype: $F_{(2, 46)}$ = 10.58, P=0.0002<br>trial × genotype: $F_{(8, 184)}$ = 6.180, P<0.0001<br><br>wild-type vs MNK1$^{KO}$<br>Trial1: P=0.0037<br>Trial2: P=0.0155<br>Trial3: P=0.1663<br>Trial4: P=0.0127<br>Trial5: P=0.0005<br><br>wild-type vs MNK2$^{KO}$<br>Trial1: P=0.8023<br>Trial2: P=0.8813<br>Trial3: P=0.7782<br>Trial4: P=0.9338<br>Trial5: P=0.2891<br><br>MNK1$^{KO}$ vs MNK2$^{KO}$<br>Trial1: P=0.0066<br>Trial2: P=0.0309<br>Trial3: P=0.3216<br>Trial4: P=0.0086<br>Trial5: P<0.0001 |
| 1F | wild-type=17<br>MNK1$^{KO}$=15<br>MNK2$^{KO}$=17 | Kruskal-Wallis test, Dunn's post-hoc test for multiple comparisons | $K_{(3)}$=19.25, P<0.0001<br>wild-type vs MNK1$^{KO}$: P=0.0073<br>wild-type vs MNK2$^{KO}$: P=0.5720<br>MNK1$^{KO}$ vs MNK2$^{KO}$: P<0.0001 |
| 1G | wild-type=17<br>MNK1$^{KO}$=15<br>MNK2$^{KO}$=17 | RM two-way ANOVA, Tukey's post-hoc test for multiple comparisons | familiarity: $F_{(1, 46)}$ = 20.08, P<0.0001<br>genotype: $F_{(2, 46)}$ = 13.48, P=0.0001<br>familiarity × genotype F (2, 46) = 4.183, P=0.0409 |

| | | | |
|---|---|---|---|
| | | | familiar vs novel<br>wild-type: P=0.0002<br>MNK1$^{KO}$: P=0.8610<br>MNK2$^{KO}$: P=0.0006 |
| 1H | wild-type=17<br>MNK1$^{KO}$=15<br>MNK2$^{KO}$=17 | One-way ANOVA, Tukey's post-hoc test for multiple comparisons | $F_{(2, 46)}$ = 4.843, P=0.0123<br>wild-type vs MNK1$^{KO}$: P= 0.0407<br>wild-type vs MNK2$^{KO}$: P= 0.9177<br>MNK1$^{KO}$ vs MNK2$^{KO}$: P= 0.0158 |
| 1I | wild-type=16<br>MNK1$^{KO}$=15<br>MNK2$^{KO}$=17<br><br>*Last three minutes are missing from one MNK1$^{KO}$ mouse* | Mixed-effects model, Tukey's post-hoc test for multiple comparisons | time: $F_{(4.129, 183.0)}$ = 20.74, P<0.0001<br>genotype: $F_{(2, 45)}$ = 3.984, P=0.0255<br>time × genotype: $F_{(18, 399)}$ = 3.815, P<0.0001<br><br>wild-type vs MNK1$^{KO}$<br>Minute 1: P=0.0640<br>Minute 2: P=0.0006<br>Minute 3: P=0.0652<br>Minute 4: P=0.1525<br>Minute 5: P=0.0203<br>Minute 6: P=0.3122<br>Minute 7: P=0.7071<br>Minute 8: P=0.8076<br>Minute 9: P=0.6218<br>Minute 10: P=0.2908<br><br>wild-type vs MNK2$^{KO}$<br>Minute 1: P=0.1390<br>Minute 2: P=0.4599<br>Minute 3: P=0.2032<br>Minute 4: P=0.4710<br>Minute 5: P=0.0012<br>Minute 6: P=0.1773<br>Minute 7: P=0.9988<br>Minute 8: P=0.8445<br>Minute 9: P=0.6583<br>Minute 10: P=0.2778<br><br>MNK1$^{KO}$ vs MNK2$^{KO}$<br>Minute 1: P=0.0003<br>Minute 2: P=0.0266<br>Minute 3: P=0.6507<br>Minute 4: P=0.5080<br>Minute 5: P=0.6591<br>Minute 6: P=0.8818<br>Minute 7: P=0.6743<br>Minute 8: P=0.3860<br>Minute 9: P=0.9824<br>Minute 10: P=0.9967 |
| 1J | wild-type=16<br>MNK1$^{KO}$=15<br>MNK2$^{KO}$=17 | Mixed-effects model | time: $F_{(5.368, 238.0)}$ = 2.066, P=0.0655<br>genotype:<br>$F_{(2, 45)}$ = 0.08049, P=0.9228 |

| | | | |
|---|---|---|---|
| | *Last three minutes are missing from one MNK1$^{KO}$ mouse* | | time × genotype interaction: $F_{(18, 399)} = 1.324$, P=0.1684 |
| 1L | wild-type=17<br>MNK1$^{KO}$=14<br>MNK2$^{KO}$=17<br><br>*Trial 5 is missing from one MNK2$^{KO}$ mouse* | Mixed-effects model, Tukey's post-hoc test for multiple comparisons | trial: $F_{(3.512, 157.2)} = 8.760$, P<0.0001<br>genotype: $F_{(2, 45)} = 4.735$, P=0.0136<br>trial × genotype interaction: $F_{(8, 179)} = 0.8201$, P=0.5858<br><br>wild-type vs MNK1$^{KO}$<br>Trial1: P=0.1996<br>Trial2: P=0.4601<br>Trial3: P=0.8604<br>Trial4: P=0.1793<br>Trial5: P=0.2983<br>wild-type vs MNK2$^{KO}$<br>Trial1: P=0.0500<br>Trial2: P=0.1476<br>Trial3: P=0.2713<br>Trial4: P=0.0609<br>Trial5: P=0.2653<br>MNK1$^{KO}$ vs MNK2$^{KO}$<br>Trial1: P=0.9361<br>Trial2: P=0.7916<br>Trial3: P=0.0967<br>Trial4: P=0.9844<br>Trial5: P=0.9880 |
| 1M | wild-type=17<br>MNK1$^{KO}$=14<br>MNK2$^{KO}$=17<br><br>*Trial 5 is missing from one MNK2$^{KO}$ mouse* | Mixed-effects model, Tukey's post-hoc test for multiple comparisons | trial: $F_{(3.318, 148.5)} = 0.5512$, P=0.6658<br>genotype: $F_{(2, 45)} = 6.569$, P=0.0031<br>trial × genotype interaction: $F_{(8, 179)} = 0.9465$, P=0.4797<br>wild-type vs MNK1$^{KO}$<br>Trial1: P=0.0226<br>Trial2: P=0.0063<br>Trial3: P=0.5086<br>Trial4: P=0.7346<br>Trial5: P=0.066<br>wild-type vs MNK2$^{KO}$<br>Trial1: P=0.2453<br>Trial2: P=0.1319<br>Trial3: P=0.2567<br>Trial4: P=0.1472<br>Trial5: P=0.598<br>MNK1$^{KO}$ vs MNK2$^{KO}$<br>Trial1: P=0.6477<br>Trial2: P=0.3311<br>Trial3: P=0.8572<br>Trial4: P=0.5895 |

| | | | |
|---|---|---|---|
| | | | Trial5: P=0.4593 |
| 1N | wild-type=17<br>MNK1$^{KO}$=14<br>MNK2$^{KO}$=17<br><br>*Trial 5 is missing from one MNK2$^{KO}$ mouse* | Mixed-effects model, Tukey's post-hoc test for multiple comparisons | trial: $F_{(2.703, 121.0)}$ = 3.131, P=0.0328<br>genotype $F_{(2, 45)}$ = 13.24, P<0.0001<br><br>trial $\times$ genotype interaction: $F_{(8, 179)}$ = 1.357, P=0.2183<br><br>wild-type vs MNK1$^{KO}$<br>Trial1: P=0.357<br>Trial2: P=0.0155<br>Trial3: P=0.2736<br>Trial4: P=0.0188<br>Trial5: P=0.0886<br><br>wild-type vs MNK2$^{KO}$<br>Trial1: P=0.0004<br>Trial2: P=0.0001<br>Trial3: P=0.0351<br>Trial4: P=0.3417<br>Trial5: P=0.0667<br><br>MNK1$^{KO}$ vs MNK2$^{KO}$<br>Trial1: P=0.2773<br>Trial2: P=0.6374<br>Trial3: P=0.4472<br>Trial4: P=0.2077<br>Trial5: P=0.9735 |
| 3B | Mean $\log_2$ value relative to wild-type from 4 mice per genotype.<br>Number of pairs: 9114 proteins / genotype | Pearson r | MNK1$^{KO}$ logFC vs MNK2$^{KO}$ logFC: r = 0.5574 (95% confidence interval 0.5430 – 0.5713), P<0.0001<br><br>MNK1$^{KO}$ logFC vs MNK1/2$^{DKO}$ logFC: r = 0.5186 (95% confidence interval 0.5034 – 0.5334), P<0.0001<br><br>MNK2$^{KO}$ logFC vs MNK1/2$^{DKO}$ logFC: r = 0.4770 (95% confidence interval 0.4610 – 0.4927), P<0.0001 |
| 4A | Mean $\log_2$ value relative to wild-type from 4 mice per genotype.<br>Number of pairs: 9114 proteins / genotype | Pearson r | MNK1$^{KO}$ logFC vs MNK2$^{KO}$ logFC: r = 0.05323 (95% confidence interval 0.03273 – 0.07368), P<0.0001<br><br>MNK1$^{KO}$ logFC vs MNK1/2$^{DKO}$ logFC: r = 0.2309 (95% confidence interval 0. 2114 – 0.2502), P<0.0001<br><br>MNK2$^{KO}$ logFC vs MNK1/2$^{DKO}$ logFC: r = 0.6062 (95% confidence interval 0.5930 – 0.6190), P<0.0001 |

| | | | |
|---|---|---|---|
| 4E | Mean $log_2$ value relative to wild-type from 4 mice per genotype.<br>Ribosomes: 73<br>All: 73 randomly selected proteins | Kolmogorov-Smirnov test | $MNK1^{KO}$ CX: D= 0.3699, P<0.0001<br>$MNK2^{KO}$ CX: D= 0.3288, P=0.0007<br>$MNK1/2^{DKO}$ CX: D= 0.4110, P<0.0001<br>$MNK1^{KO}$ SYP: D= 0.6849, P<0.0001<br>$MNK2^{KO}$ SYP: D= 0.3288, P=0.0007<br>$MNK1/2^{DKO}$ SYP: D= 0.6164, P<0.0001 |
| 6B | wild-type=14<br>$MNK1^{KO}$=14<br>$MNK2^{KO}$=16 | One-way ANOVA, Tukey's post-hoc test for multiple comparisons | $F_{(2, 41)}$ = 17.27, P<0.0001<br>wild-type vs $MNK1^{KO}$: P= 0.0026<br>wild-type vs $MNK2^{KO}$: P< 0.0001<br>$MNK1^{KO}$ vs $MNK2^{KO}$: P= 0.0932 |
| 6C | wild-type=14<br>$MNK1^{KO}$=15<br>$MNK2^{KO}$=15 | Kruskal-Wallis test, Dunn's post-hoc test for multiple comparisons | $K_{(3)}$=22.14, P<0.0001<br>wild-type vs $MNK1^{KO}$: P<0.0001<br>wild-type vs $MNK2^{KO}$: P=0.0166<br>$MNK1^{KO}$ vs $MNK2^{KO}$: P= 0.1545 |
| 6D | wild-type=8<br>$MNK1^{KO}$=6<br>$MNK2^{KO}$=9<br>$MNK1/2^{DKO}$=9 | One-way ANOVA, Tukey's post-hoc test for multiple comparisons | F (3, 28) = 4.855, P=0.0076<br>wild-type vs $MNK1^{KO}$: P= 0.0038<br>wild-type vs $MNK2^{KO}$: P= 0.3164<br>wild-type vs $MNK1/2^{DKO}$: P= 0.1924<br>$MNK1^{KO}$ vs $MNK2^{KO}$: P= 0.1274<br>$MNK1^{KO}$ vs $MNK1/2^{DKO}$: P= 0.2121<br>$MNK2^{KO}$ vs $MNK1/2^{DKO}$: P= 0.9893 |
| 6E | wild-type=9<br>$MNK1^{KO}$=9<br>$MNK2^{KO}$=11<br>$MNK1/2^{DKO}$=7 | One-way ANOVA, Tukey's post-hoc test for multiple comparisons | $F_{(3, 32)}$ = 7.421, P=0.0007<br>wild-type vs $MNK1^{KO}$: P= 0.0054<br>wild-type vs $MNK2^{KO}$: P= 0.0022<br>wild-type vs $MNK1/2^{DKO}$: P= 0.8317<br>$MNK1^{KO}$ vs $MNK2^{KO}$: P= 0.9985<br>$MNK1^{KO}$ vs $MNK1/2^{DKO}$: P= 0.0738<br>$MNK2^{KO}$ vs $MNK1/2^{DKO}$: P= 0.0417 |
| 6G | wild-type=18<br>$MNK1^{KO}$=12<br>$MNK2^{KO}$=12 | Kruskal-Wallis test, Dunn's post-hoc test for multiple comparisons | $K_{(3)}$=10.77, P=0.0046<br>wild-type vs $MNK1^{KO}$: P= 0.0142<br>wild-type vs $MNK2^{KO}$: P= 0.0234<br>$MNK1^{KO}$ vs $MNK2^{KO}$: P>0.9999 |
| 6I | Mean $log_2$ value relative to wild-type from 4 mice per genotype.<br>Number of pairs: 10230 phosphosites / genotype | Pearson r | *synaptoneurosomes:*<br>$MNK1^{KO}$ logFC vs $MNK2^{KO}$ logFC: r = 0.2254 (95% confidence interval 0.2069 – 0.2437), P<0.0001<br>*cortex:*<br>$MNK1^{KO}$ logFC vs $MNK2^{KO}$ logFC: r = 0.4523 (95% confidence interval 0.4368 – 0.4676), P<0.0001 |
| **Extended data Figures** | **Sample Size** | **Statistical test** | **Values** |

| | | | |
|---|---|---|---|
| S1A-B | *females:*<br>wild-type=10<br>MNK1$^{KO}$=8<br>MNK2$^{KO}$=12<br><br>*males:*<br>wild-type=7<br>MNK1$^{KO}$=6<br>MNK2$^{KO}$=5 | Three-way Repeated Measures ANOVA | Genotype: $F_{(2, 42)}$=0.28, P=0.760<br>Sex: $F_{(1, 42)}$=1.19, P=0.2820<br>genotype x Sex $F_{(2, 42)}$=1.63, P=0.2080<br>trials: $F_{(3.35, 140.79)}$=81.16, P<0.0001<br>genotype x trials: $F_{(6.7, 140.79)}$=1.887, P=0.0820<br>Sex x trials $_{(3.35, 140.79)}$: F=0.86, P=0.4720<br>Sex x trials x genotype $_{(6.7, 140.79)}$: F=0.67, P=0.6880 |
| S1C | *females:*<br>wild-type=10<br>MNK1$^{KO}$=8<br>MNK2$^{KO}$=12<br><br>*males:*<br>wild-type=7<br>MNK1$^{KO}$=6<br>MNK2$^{KO}$=5 | Two-way ANOVA | Interaction: F $_{(2, 42)}$ = 0.2222, P=0.8017<br>Sex: F $_{(1, 42)}$ = 0.2822, P=0.5980<br>Genotype: F $_{(2, 42)}$ = 6.141, P=0.0046 |
| S1D-E | *females:*<br>wild-type=10<br>MNK1$^{KO}$=8<br>MNK2$^{KO}$=12<br><br>*males:*<br>wild-type=7<br>MNK1$^{KO}$=7<br>MNK2$^{KO}$=5 | Three-way Repeated Measures ANOVA | Genotype: $F_{(2, 43)}$ =2.48, P=0.0950<br>Sex: $F_{(1, 43)}$ =2.23, P=0.1430<br>genotype x sex: $F_{(2, 42)}$ =2.06, P=0.139<br>trials: $F_{(3.87, 166.45)}$ =42.84, P<0.0001<br>genotype x trials: $F_{(7.74, 166.45)}$ =1.47, P=0.175<br>Sex x trials: $F_{(3.87, 166.45)}$ =0.58, P=0.669<br>Sex x trials x genotype: $F_{(7.74, 166.45)}$ =1.62, P=0.125 |
| S1F-G | *females:*<br>wild-type=10<br>MNK1$^{KO}$=8<br>MNK2$^{KO}$=12<br><br>*males:*<br>wild-type=7<br>MNK1$^{KO}$=7<br>MNK2$^{KO}$=5 | Three-way Repeated Measures ANOVA | Genotype: $F_{(2, 43)}$ =9.80, P<0.0001<br>Sex: $F_{(1, 43)}$ =0.54, P=0.465<br>genotype x sex: $F_{(2, 43)}$ =1.74, P=0.188<br>trials: $F_{(2.43, 104.6)}$ =34.96, P<0.0001<br>genotype x trials: $F_{(4.86, 104.6)}$ =5.89, P<0.0001<br>Sex x trials: $F_{(2.43, 104.6)}$ =0.83, P=0.461<br>Sex x trials x genotype: $F_{(4.86, 104.6)}$ =0.61, P=0.689 |
| S1H | *females:*<br>wild-type=10<br>MNK1$^{KO}$=8<br>MNK2$^{KO}$=12<br><br>*males:*<br>wild-type=7<br>MNK1$^{KO}$=7<br>MNK2$^{KO}$=5 | Two-way ANOVA | Interaction: F $_{(2, 43)}$ = 0.9540, P=0.3932<br>Sex: F $_{(1, 43)}$ = 0.6882, P=0.4114<br>Genotype: F $_{(2, 43)}$ = 14.51, P<0.0001 |

| | | | |
|---|---|---|---|
| S1I | *females:*<br>wild-type=10<br>MNK1$^{KO}$=8<br>MNK2$^{KO}$=12<br><br>*males:*<br>wild-type=7<br>MNK1$^{KO}$=7<br>MNK2$^{KO}$=5 | Three-way Repeated Measures ANOVA | Genotype: $F_{(2, 43)}$ =11.66, P<0.0001<br>Sex: $F_{(1, 43)}$ =0.51, P=0.479<br>Novelty: $F_{(1, 43)}$ =21.05, P<0.0001<br>genotype x sex: $F_{(2, 43)}$ =3.23, P=0.049<br>genotype x novelty: $F_{(2, 43)}$ =4.62, P=0.015<br>Sex x novelty: $F_{(1, 43)}$ =0.70, P=0.409<br>Sex x novelty x genotype: $F_{(2, 43)}$ =0.52, P=0.596 |
| S1J | females:<br>wild-type=10<br>MNK1KO=8<br>MNK2KO=12<br><br>males:<br>wild-type=7<br>MNK1KO=7<br>MNK2KO=5 | Two-way ANOVA | Interaction: $F_{(2, 43)}$ = 0.8828, P=0.4210<br>Sex: $F_{(1, 43)}$ = 1.423, P=0.2394<br>Genotype: $F_{(2, 43)}$ = 5.574, P=0.0070 |
| S1K | females:<br>wild-type=10<br>MNK1KO=7<br>MNK2KO=12<br><br>males:<br>wild-type=6<br>MNK1KO=7<br>MNK2KO=5 | Two-way ANOVA | Interaction: $F_{(2, 40)}$ = 2.048, P=0.1424<br>Sex: $F_{(1, 40)}$ = 0.03098, P=0.8612<br>Genotype: $F_{(2, 40)}$ = 2.365, P=0.1069 |
| S1L | females:<br>wild-type=10<br>MNK1KO=7<br>MNK2KO=12<br><br>males:<br>wild-type=6<br>MNK1KO=7<br>MNK2KO=5 | Two-way ANOVA | Interaction: $F_{(2, 40)}$ = 0.1903, P=0.8275<br>Sex: $F_{(1, 40)}$ = 2.104, P=0.1547<br>Genotype: $F_{(2, 40)}$ = 0.06901, P=0.9334 |
| S2A | wild-type=17<br>MNK1$^{KO}$=14<br>MNK2$^{KO}$=17<br><br>*Trial 5 is missing from one MNK2$^{KO}$ mouse* | Mixed-effects model | trial: $F_{(2.235, 100.0)}$ = 5.231, P=0.0052<br>genotype: $F_{(2, 45)}$ = 1.827, P=0.1726<br>trial × genotype: $F_{(8, 179)}$ = 1.186, P=0.3099 |
| S2B | wild-type=17<br>MNK1$^{KO}$=14<br>MNK2$^{KO}$=17<br><br>*Trial 5 is missing from one MNK2$^{KO}$ mouse* | Mixed-effects model | trial: $F_{(2.676, 119.8)}$ = 30.94, P<0.0001<br>genotype: $F_{(2, 45)}$ = 2.411, P=0.1012<br>trial × genotype: $F_{(8, 179)}$ = 0.1849, P=0.9927 |

| | | | |
|---|---|---|---|
| S2C | wild-type=17<br>MNK1$^{KO}$=14<br>MNK2$^{KO}$=17<br><br>*Trial 5 is missing from one MNK2$^{KO}$ mouse* | Mixed-effects model, Tukey's post-hoc test for multiple comparisons | trial: $F_{(3.757, 168.1)}$ = 4.985, P=0.0010<br>genotype: $F_{(2, 45)}$ = 4.599, P=0.0152<br>trial × genotype: $F_{(8, 179)}$ = 0.8569, P=0.5541<br><br>wild-type vs MNK1$^{KO}$<br>Trial1: P= 0.0520<br>Trial2: P= 0.4134<br>Trial3: P= 0.9037<br>Trial4: P= 0.6319<br>Trial5: P= 0.3153<br><br>wild-type vs MNK2$^{KO}$<br>Trial1: P= 0.0155<br>Trial2: P= 0.0752<br>Trial3: P= 0.4734<br>Trial4: P= 0.1237<br>Trial5: P= 0.0995<br><br>MNK1$^{KO}$ vs MNK2$^{KO}$<br>Trial1: P= 0.9999<br>Trial2: P= 0.6942<br>Trial3: P= 0.3563<br>Trial4: P= 0.5535<br>Trial5: P= 0.9831 |
| S2D | wild-type=17<br>MNK1$^{KO}$=14<br>MNK2$^{KO}$=17<br><br>*Trial 5 is missing from one MNK2$^{KO}$ mouse* | Mixed-effects model, Tukey's post-hoc test for multiple comparisons | trial: $F_{(2.492, 111.5)}$ = 3.214, P=0.0336<br>genotype: $F_{(2, 45)}$ = 3.387, P=0.0426<br>trial × genotype: $F_{(8, 179)}$ = 0.7083, P=0.6840<br><br>wild-type vs MNK1$^{KO}$<br>Trial1: P= 0.0714<br>Trial2: P= 0.0610<br>Trial3: P= 0.9046<br>Trial4: P= 0.7391<br>Trial5: P= 0.1718<br><br>wild-type vs MNK2$^{KO}$<br>Trial1: P= 0.2334<br>Trial2: P= 0.4526<br>Trial3: P= 0.5697<br>Trial4: P= 0.3383<br>Trial5: P= 0.9498<br><br>MNK1$^{KO}$ vs MNK2$^{KO}$<br>Trial1: P= 0.7469<br>Trial2: P= 0.6209<br>Trial3: P= 0.7352<br>Trial4: P= 0.7417<br>Trial5: P= 0.2276 |
| S2E | wild-type=17<br>MNK1$^{KO}$=14<br>MNK2$^{KO}$=17 | Mixed-effects model, Tukey's post-hoc test for multiple comparisons | trial: $F_{(3.672, 164.3)}$ = 1.060, P=0.3753<br>genotype: $F_{(2, 45)}$ = 5.983, P=0.0050<br>trial × genotype: $F_{(8, 179)}$ = 0.5915, P=0.7841 |

| | | | |
|---|---|---|---|
| | *Trial 5 is missing from one $MNK2^{KO}$ mouse* | | wild-type vs $MNK1^{KO}$<br>Trial1: P= 0.2486<br>Trial2: P= 0.0274<br>Trial3: P= 0.3846<br>Trial4: P= 0.9537<br>Trial5: P= 0.1072<br><br>wild-type vs $MNK2^{KO}$<br>Trial1: P= 0.3977<br>Trial2: P= 0.1633<br>Trial3: P= 0.2106<br>Trial4: P=0.4885<br>Trial5: P= 0.4039<br><br>$MNK1^{KO}$ vs $MNK2^{KO}$<br>Trial1: P= 0.8071<br>Trial2: P= 0.3965<br>Trial3: P= 0.9997<br>Trial4: P= 0.7163<br>Trial5: P= 0.8183 |
| S2F | wild-type=17<br>$MNK1^{KO}$=14<br>$MNK2^{KO}$=17<br><br>*Trial 5 is missing from one $MNK2^{KO}$ mouse* | Mixed-effects model, Tukey's post-hoc test for multiple comparisons | trial: $F_{(2.962, 132.5)} = 3.233$, P=0.0250<br>genotype: $F_{(2, 45)} = 12.11$, P<0.0001<br>trial $\times$ genotype: $F_{(8, 179)} = 1.633$, P= 0.1182<br><br>wild-type vs $MNK1^{KO}$<br>Trial1: P= 0.5450<br>Trial2: P= 0.0354<br>Trial3: P= 0.3064<br>Trial4: P= 0.0154<br>Trial5: P= 0.2479<br><br>wild-type vs $MNK2^{KO}$<br>Trial1: P= 0.0001<br>Trial2: P= 0.0008<br>Trial3: P= 0.0182<br>Trial4: P= 0.2998<br>Trial5: P= 0.0304<br><br>$MNK1^{KO}$ vs $MNK2^{KO}$<br>Trial1: P= 0.0472<br>Trial2: P= 0.3948<br>Trial3: P= 0.3763<br>Trial4: P= 0.1547<br>Trial5: P= 0.8933 |
| S2G | wild-type=17<br>$MNK1^{KO}$=14<br>$MNK2^{KO}$=17<br><br>*Trial 5 is missing from one $MNK2^{KO}$ mouse* | Mixed-effects model, Tukey's post-hoc test for multiple comparisons | trial: $F_{(3.078, 137.8)} = 1.141$, P=0.3355<br>genotype: $F_{(2, 45)} = 5.391$, P=0.0080<br>trial $\times$ genotype: $F_{(8, 179)} = 0.5523$, P= 0.8157<br><br>wild-type vs $MNK1^{KO}$<br>Trial1: P= 0.3453 |

| | | | |
|---|---|---|---|
| | | | Trial2: P= 0.1734<br>Trial3: P= 0.4833<br>Trial4: P= 0.2026<br>Trial5: P= 0.0955<br><br>wild-type vs MNK2$^{KO}$<br>Trial1: P= 0.0298<br>Trial2: P= 0.0693<br>Trial3: P= 0.2653<br>Trial4: P= 0.6798<br>Trial5: P= 0.7745<br><br>MNK1$^{KO}$ vs MNK2$^{KO}$<br>Trial1: P= 0.9522<br>Trial2: P= 0.9916<br>Trial3: P= 0.8581<br>Trial4: P= 0.5523<br>Trial5: P= 0.3675 |
| S2H | wild-type=17<br>MNK1$^{KO}$=14<br>MNK2$^{KO}$=17<br><br>*Trial 5 is missing from one MNK2$^{KO}$ mouse* | Mixed-effects model, Tukey's post-hoc test for multiple comparisons | trial: $F_{(3.629, 162.4)}$ = 17.83, P<0.0001<br>genotype: $F_{(2, 45)}$ = 1.410, P=0.2548<br>trial × genotype: $F_{(8, 179)}$ = 0.9646, P= 0.4652 |
| S2I | *Trial 1-4*<br>wild-type=17<br>MNK1$^{KO}$=14<br>MNK2$^{KO}$=17<br><br>*Trial 5*<br>wild-type=17<br>MNK1$^{KO}$=14<br>MNK2$^{KO}$=16<br><br>*Trial 5 is missing from one MNK2$^{KO}$ mouse* | *Trial 1-4*<br>RM two-way ANOVA Tukey's post-hoc test for multiple comparisons<br><br>*Trial 5*<br>Kruskal-Wallis test, Dunn's post-hoc test for multiple comparisons | trial: $F_{(2.405, 108.2)}$ = 9.880, P<0.0001<br>genotype: $F_{(2, 45)}$ = 0.1016, P=0.9036<br>trial × genotype: $F_{(6, 135)}$ = 0.2036, P=0.9752<br><br>*Trial 5*<br>$K_{(3)}$= 1.752, P= 0.4164 |
| S2J | *Trial 1-4*<br>wild-type=17<br>MNK1$^{KO}$=14<br>MNK2$^{KO}$=17<br><br>*Trial 5*<br>wild-type=17<br>MNK1$^{KO}$=14<br>MNK2$^{KO}$=16<br><br>*Trial 5 is missing from one MNK2$^{KO}$ mouse* | *Trial 1-4*<br>RM two-way ANOVA Tukey's post-hoc test for multiple comparisons<br><br>*Trial 5*<br>Kruskal-Wallis test, Dunn's post-hoc test for multiple comparisons | trial: $F_{(1.831, 82.39)}$ = 1.237, P=0.2933<br>genotype: $F_{(2, 45)}$ = 1.049, P=0.3587<br>trial × genotype: $F_{(6, 135)}$ = 0.7871, P=0.5815<br><br>*Trial 5*<br>$K_{(3)}$= 1.133, P= 0.5676 |
| S2K | *Trial 1-4*<br>wild-type=17<br>MNK1$^{KO}$=14<br>MNK2$^{KO}$=17<br><br>*Trial 5*<br>wild-type=17<br>MNK1$^{KO}$=14<br>MNK2$^{KO}$=16 | *Trial 1-4*<br>RM two-way ANOVA Tukey's post-hoc test for multiple comparisons<br><br>*Trial 5*<br>Kruskal-Wallis test, Dunn's post-hoc test for multiple comparisons | trial: $F_{(2.726, 122.7)}$ = 0.8485, P=0.4608<br>genotype: $F_{(2, 45)}$ = 4.349, P=0.0188<br>trial × genotype: $F_{(6, 135)}$ = 1.347, P=0.2405<br><br>wild-type vs MNK1$^{KO}$<br>Trial1: P= 0.2406 |

| | | | |
|---|---|---|---|
| | *Trial 5 is missing from one MNK2$^{KO}$ mouse* | | Trial2: P= 0.0848<br>Trial3: P= 0.8704<br>Trial4: P= 0.5397<br><br>wild-type vs MNK2$^{KO}$<br>Trial1: P= 0.0840<br>Trial2: P= 0.0009<br>Trial3: P= 0.1845<br>Trial4: P= 0.7983<br><br>MNK1$^{KO}$ vs MNK2$^{KO}$<br>Trial1: P= 0.8235<br>Trial2: P= 0.5235<br>Trial3: P= 0.4831<br>Trial4: P= 0.8956<br><br>*Trial 5*<br>$K_{(3)}$= 7.106, P= 0.0286<br>wild-type vs MNK1$^{KO}$: P= 0.1021<br>wild-type vs MNK2$^{KO}$: P= 0.0465<br>MNK1$^{KO}$ vs MNK2$^{KO}$: P >0.9999 |
| S2L | *Trial 1-4*<br>wild-type=17<br>MNK1$^{KO}$=14<br>MNK2$^{KO}$=17<br><br>*Trial 5*<br>wild-type=17<br>MNK1$^{KO}$=14<br>MNK2$^{KO}$=16<br><br>*Trial 5 is missing from one MNK2$^{KO}$ mouse* | *Trial 1-4*<br>RM two-way ANOVA<br>Tukey's post-hoc test for multiple comparisons<br><br>*Trial 5*<br>Kruskal-Wallis test,<br>Dunn's post-hoc test for multiple comparisons | trial: $F_{(2.199, 98.94)}$ = 13.31, P<0.0001<br>genotype: $F_{(2, 45)}$ = 9.062, P=0.0005<br>trial × genotype: $F_{(6, 135)}$ = 1.747, P=0.1149<br><br>wild-type vs MNK1$^{KO}$<br>Trial1: P= 0.0098<br>Trial2: P= 0.0122<br>Trial3: P= 0.0029<br>Trial4: P= 0.3759<br><br>wild-type vs MNK2$^{KO}$<br>Trial1: P= 0.1692<br>Trial2: P= 0.1393<br>Trial3: P= 0.0425<br>Trial4: P>0.9999<br><br>MNK1$^{KO}$ vs MNK2$^{KO}$<br>Trial1: P= 0.6840<br>Trial2: P= 0.2173<br>Trial3: P= 0.5598<br>Trial4: P= 0.2081<br><br>*Trial 5*<br>$K_{(3)}$= 8.680, P= 0.0130<br>wild-type vs MNK1$^{KO}$: P= 0.0100<br>wild-type vs MNK2$^{KO}$: P= 0.7436<br>MNK1$^{KO}$ vs MNK2$^{KO}$: P = 0.2177 |
| S2M | *Trial 1-4*<br>wild-type=17<br>MNK1$^{KO}$=14 | *Trial 1-4* | trial: $F_{(2.267, 102.0)}$ = 48.06, P<0.0001<br>genotype: $F_{(2, 45)}$ = 3.162, P=0.0519 |

| | | | |
|---|---|---|---|
| | $MNK2^{KO}$=17<br><br>*Trial 5*<br>wild-type=17<br>$MNK1^{KO}$=14<br>$MNK2^{KO}$=16<br><br>*Trial 5 is missing from one $MNK2^{KO}$ mouse* | RM two-way ANOVA Tukey's post-hoc test for multiple comparisons<br><br>*Trial 5*<br>One-way ANOVA, Tukey's post-hoc test for multiple comparisons | trial × genotype: $F_{(6, 135)}$ = 3.638, P=0.0022<br><br>wild-type vs $MNK1^{KO}$<br>Trial1: P= 0.0014<br>Trial2: P= 0.0247<br>Trial3: P= 0.0725<br>Trial4: P= 0.7854<br><br>wild-type vs $MNK2^{KO}$<br>Trial1: P= 0.9622<br>Trial2: P= 0.9987<br>Trial3: P= 0.6026<br>Trial4: P= 0.8590<br><br>$MNK1^{KO}$ vs $MNK2^{KO}$<br>Trial1: P= 0.0272<br>Trial2: P= 0.0407<br>Trial3: P= 0.4160<br>Trial4: P= 0.9713<br><br>*Trial 5*<br>F (2, 44) = 3.811, P=0.0298<br>wild-type vs $MNK1^{KO}$: P= 0.0577<br>wild-type vs $MNK2^{KO}$: P= 0.9865<br>$MNK1^{KO}$ vs $MNK2^{KO}$: P = 0.0442 |
| S2L | wild-type=17<br>$MNK1^{KO}$=14<br>$MNK2^{KO}$=17 | RM two-way ANOVA, Tukey's post-hoc test for multiple comparisons | PC: $F_{(1, 45)}$ = 0.02775, P=0.8684<br>genotype: $F_{(2, 45)}$ = 16.44, P<0.0001<br>PC × genotype: $F_{(2, 45)}$ = 11.61, P<0.0001<br><br>PC1:<br>wild-type vs $MNK1^{KO}$ : P<0.0001<br>wild-type vs $MNK2^{KO}$ : P=0.0092<br>$MNK1^{KO}$ vs $MNK2^{KO}$ : P=0.0057<br><br>PC1:<br>wild-type vs $MNK1^{KO}$ : P=0.8159<br>wild-type vs $MNK2^{KO}$ : P=0.0021<br>$MNK1^{KO}$ vs $MNK2^{KO}$ : P=0.0005 |
| S4B | Mean $log_2$ value of ETC-168 treated wild-type relative to vehicle-treated wild-type from 4 mice per treatment.<br>Ribosomes: 77<br>All: 77 randomly selected proteins | Kolmogorov-Smirnov test | WT ETC-168: D= 0.3506, P=0.0002 |
| S5C | Mean $log_2$ value relative to wild-type from 3 mice per genotype. | Pearson r | $MNK1^{KO}$ logFC vs $MNK2^{KO}$ logFC: r = 0.1540 (95% confidence interval 0.1398 – 0.1683), P<0.0001 |

| | | | |
|---|---|---|---|
| | Number of pairs: 18019 RNAs / genotype | | |
| S6A | wild-type=6<br>$MNK1^{KO}$=7<br>$MNK2^{KO}$=7 | Kruskal-Wallis test, Dunn's post-hoc test for multiple comparisons | $K_{(3)}$=12.46, P=0.0002<br>wild-type vs $MNK1^{KO}$: P=0.0200<br>wild-type vs $MNK2^{KO}$: P=0.0023<br>$MNK1^{KO}$ vs $MNK2^{KO}$: P>0.9999 |
| S6B | wild-type=6<br>$MNK1^{KO}$=7<br>$MNK2^{KO}$=7 | Kruskal-Wallis test, Dunn's post-hoc test for multiple comparisons | $K_{(3)}$=15.43, P<0.0001<br>wild-type vs $MNK1^{KO}$: P=0.0003<br>wild-type vs $MNK2^{KO}$: P=0.0950<br>$MNK1^{KO}$ vs $MNK2^{KO}$: P=0.1920 |
| S6C | wild-type=14<br>$MNK1^{KO}$=14<br>$MNK2^{KO}$=16<br>$MNK1/2^{DKO}$=8 | One-way ANOVA | F $_{F\ (3,\ 48)}$ = 2.378, P=0.0814 |
| S6D | wild-type=14<br>$MNK1^{KO}$=15<br>$MNK2^{KO}$=15<br>$MNK1/2^{DKO}$=8 | One-way ANOVA | $F_{(3,\ 48)}$ = 1.523, P=0.2204 |
| S6E | wild-type=8<br>$MNK1^{KO}$=8<br>$MNK2^{KO}$=9 | One-way ANOVA | $F_{(2,\ 22)}$ = 0.9563, P=0.3997 |
| S6F | wild-type=9<br>$MNK1^{KO}$=9<br>$MNK2^{KO}$=10<br>$MNK1/2^{DKO}$=7 | One-way ANOVA | F (3, 31) = 0.7150, P=0.5505 |
| S6G | wild-type=6<br>$MNK1^{KO}$=7<br>$MNK2^{KO}$=7<br>$MNK1/2^{DKO}$=4 | One-way ANOVA | F (3, 20) = 1.050, P=0.3922 |
| S6H | wild-type=14<br>$MNK2^{KO}$=16 | Mann-Whitney test | Mann-Whitney U = 52, P=0.0118 |
| S6H | wild-type=8<br>$MNK2^{KO}$=9 | Mann-Whitney test | Mann-Whitney U = 19, P=0.1139 |
